# Aging-dependent dysregulation of EXOSC2 is maintained in cancer as a dependency

**DOI:** 10.1101/2025.04.04.647279

**Authors:** Maria Skamagki, Cheng Zhang, Ezgi Hacisuleyman, Giuseppe Galleti, Chao Wu, Rajasekhar K Vinagolu, HyoKyeong Cha, Deniz Ata, Jongjin Kim, Taylor Weiskittel, Mame Diop, Thiri Aung, Michael Del Latto, Amanda S. Kim, Zhuoning Li, Matthew Miele, Rui Zhao, Laura H. Tang, Ronald C. Hendrickson, Paul B. Romesser, Joshua J. Smith, Paraskevi Giannakakou, Robert B. Darnell, Matthew J. Bott, Hu Li, Kitai Kim

## Abstract

Reprogramming of aged donor tissue cells into induced pluripotent stem cells (A-iPSC) preserved the epigenetic memory of aged-donor tissue, defined as genomic instability and poor tissue differentiation in our previous study. The unbalanced expression of RNA exosome subunits affects the RNA degradation complex function and is associated with geriatric diseases including premature aging and cancer progression. We hypothesized that the age-dependent progressive subtle dysregulation of EXOSC2 (exosome component 2) causes the aging traits (abnormal cell cycle and poor tissue differentiation). We used embryonic stem cells as a tool to study EXOSC2 function as the aging trait epigenetic memory determined in A-iPSC because these aging traits could not be studied in senesced aged cells or immortalized cancer cells. We found that the regulatory subunit of PP2A phosphatase, PPP2R5E, is a key target of EXOSC2 and this regulation is preserved in stem cells and cancer.

## Introduction

Dysregulation of RNA processing and degradation is a hallmark of aging and is often observed in aging-related diseases. However, the underlying mechanism of how dysregulation leads to aging traits remains poorly understood. RNA exosome deficiency is associated with aging-related diseases, including autoimmune dysfunction^1, 2^, neurological disorders^3–5^, premature aging^6^, and cancer therapy resistance^7^. The eukaryotic RNA exosome is an essential and conserved protein complex in all tissue types in the entire animal kingdom that degrade or process RNA substrates to regulate gene expression. In our previous works, we compared the mouse and human induced pluripotent stem cells (iPSC) that originated from young and aged tissue donors (Y-iPSC & A-iPSC), and found that A-iPSC exhibit increased genomic instability and poor tissue differentiation potential as an aging trait^8, 9^. We discovered that EXOSC2, a subunit of RNA exosome, is down-regulated in A-iPSC, which may be preserved as aging trait epigenetic memory of the aged-donor tissue^8–11^and also observed as aging related traits of RNA exosome-deficient diseases^12–14^. Even though there is litterature suggesting that the use of iPSC generated from aged donors is not an ideal model to study aging, because the reprogramming process erases aging signatures^15, 16^, the correlation of EXOSC2 level and specific aging traits became a strong driving force to study the functional significance of subtle dysregulation of EXOSC2 in pluripotency. We hypothesized that the aging-dependent progressive dysregulation of EXOSC2 causes the aging traits (unreliable cell cycle & division and poor tissue differentiation) and to test this hypothesis spherically we combined three approaches.

The first approach is to use the pluripotent stem cells to study the consequences of EXOSC2 donwregulation in a system that is not related to aging like the pluripotent stem cells. The pluripotent stem cells can self-renew as opposed to somatic cells from aged-donors that are readily senescent, rendering them unfit to conduct prolonged tissue culture studies to define the aging traits. The immortalized lines or cancer lines can be used as alternative tools to study the effects of age-dependent dysregulation of EXOSC2, but these cell types are known to have increased genomic instability and poor tissue differentiation potential. In contrast, pluripotent stem cells can become an alternative standard to study the functional impact of deregulation of the aging related genes because genomic stability is well maintained in comparison to immortalized or cancer cell lines, and the tissue differentiation potential is also well preserved. Therefore, we pursued to define the mechanism of the aging traits using pluripotent stem cells. This research tool also allows viewing the impact of EXOSC2 down-regulation to uncouple the concomitant consequences of aging in our functional study. There is a recently published study that reports that the gradual degradation of a reconstituted chimaeric RNA exosome in a mouse ESC Knock Out for EXOSC2 leads to upregulation of senescence signatures^17^.To understand the function of a reconstituted RNA exosme described by the published study^17^, one needs to know the consequences of downregulation of endogenous EXOSC2 in a stem cell context that our paper provides in great detail.

The second approach is to focus on the subtle dysregulation of RNA exosomes that impacts biological function and could be readily ignored in genomic research. We therefore generated embryonic stem cell lines with down-regulated EXOSC2. It is worth mentioning that the homozygous knockout mouse for EXOSC2 is embryonic lethal (International Mouse Phenotyping Consortium) because EXOSC2 is a highly conserved housekeeping gene that is expressed in all tissue types and the entire animal kingdom. Therefore, we decided to down - regulate the levels of EXOSC2 to observe its effects on cell plasticity. We observed that even subtle dysregulation (50%) of the EXOSC2 significantly impacts cellular function and induces aging traits, such as increasing unreliable cell cycle & division and blocking tissue differentiation. Among the top pathways affected by subtle down-regulation of EXOSC2 are microtubule polymerization, differentiation-related proteins, and DNA damage checkpoint.

The third unique research concept is to connect the defined aging traits to cancer because unreliable cell cycle & division and poor tissue differentiation potential are not only aging traits but also cancer-related properties. In addition, the cancer risk by the subtle dysregulation of genes, such as EXOSC2, has been ignored and not documented before. Furthermore, cancer is considered an aging related disease, and it is well-known that cancer incidents increase with aging ^18^. We observed a conserved regulation of PPP2R5E by EXOSC2 in multiple tissue types during embryonic tissue development, including colon, kidney, choroid plexus, dorsal root ganglia, lung, spleen, testis, and liver. Thus, we sought to confirm the aging traits of subtle EXOSC2 modulation in a colon cancer model. Overall, our study indicates that subtle dysregulation of EXOSC2 can be preserved as a dependency in colon cancer. Defining the detailed mechanisms of how the aging related subtle reduction of the EXOSC2 expression may lead to imbalances in tissue homeostasis will contribute to basic cell biology concepts, such as the effect of aging in RNA regulation, but, at the same time, will have a translational impact on unique considerations on the clinical applications of human iPSC and cancer therapies in aged populations.

### The down-regulation of EXOSC2 in ESC leads to a blockade in tissue differentiation, a significant aging trait of stem and progenitor cells

The self-renewal property of pluripotent stem cells and their multipotent differentiation potential involve complex signaling networks of self -renewal, gate-keeping/limiting self-renewal, and house-keeping genes to maintain genomic integrity^19^. However, the mechanisms that ensure faithful cell proliferation and regulated tissue differentiation in normal development remain largely undissected. The eukaryotic RNA exosome is an essential and conserved protein complex that can degrade or process RNA substrates as required during cellular differentiation and development. While the importance of the housekeeping function of the core exosome subunits in RNA processing is extensively studied^20^, the role of the different exosome components in cell fate transitions remains unclear. Recent work has revealed a role for the RNA exosome during erythropoiesis, the process through which hematopoietic stem cells differentiate into erythrocytes^21, 22^. Moreover, RNA exosome deficiency is associated with aging related diseases, including autoimmune dysfunction^1, 2^, neurological disorders^3–5^, premature aging^6^, and cancer therapy resistance^7^.

The RNA exosome complex consists of 10 subunits, and changes in the expression of any of these subunits could alter exosome complex function ^23^ by either missing essential components with the poor expression of few exosome subunits or having dominant-negative effects on the RNA exosome complex formation with excessive expression of few exosome subunits. We also confirmed that the balance of each exosome subunit is important to maintain the exosome complex function. The subtle changes in gene expression analysis might have caused other researchers to overlook or ignore the biological function of the RNA exosome, although increasing or decreasing EXOSC2 expression by 50% can significantly affect downstream RNA targets. Defective functionality or potentially gained function of the RNA exosome complex by the altered exosome subunit expression has not been studied yet. We chose to down-regulate EXOSC2 because our previous work suggested that the core subunit EXOSC2 plays a pivotal role in an age-related differentiation block in the iPSC model and was the most significant down-regulated subunit in iPSC lines generated from aged-donors (A-iPSC)^8, 9^. In other words, we wanted to assess whether the age-dependent down-regulation of EXOSC2 seen in iPSC is also observed in other cell types. To address this question, we looked into RNAseq databases of independently published aging studies such as Tabula Sapiens^24^, Tabula Muris Senis^25^, and human fibroblasts^26–31^. We show that the age-dependent down-regulation of EXOSC2 is observed in scRNA seq data of Tabula Sapiens in the bone marrow **(Figure 1a),** the fat **(Figure 1b),** and in the skin **(Figure 1c)**.

**Figure 1.**
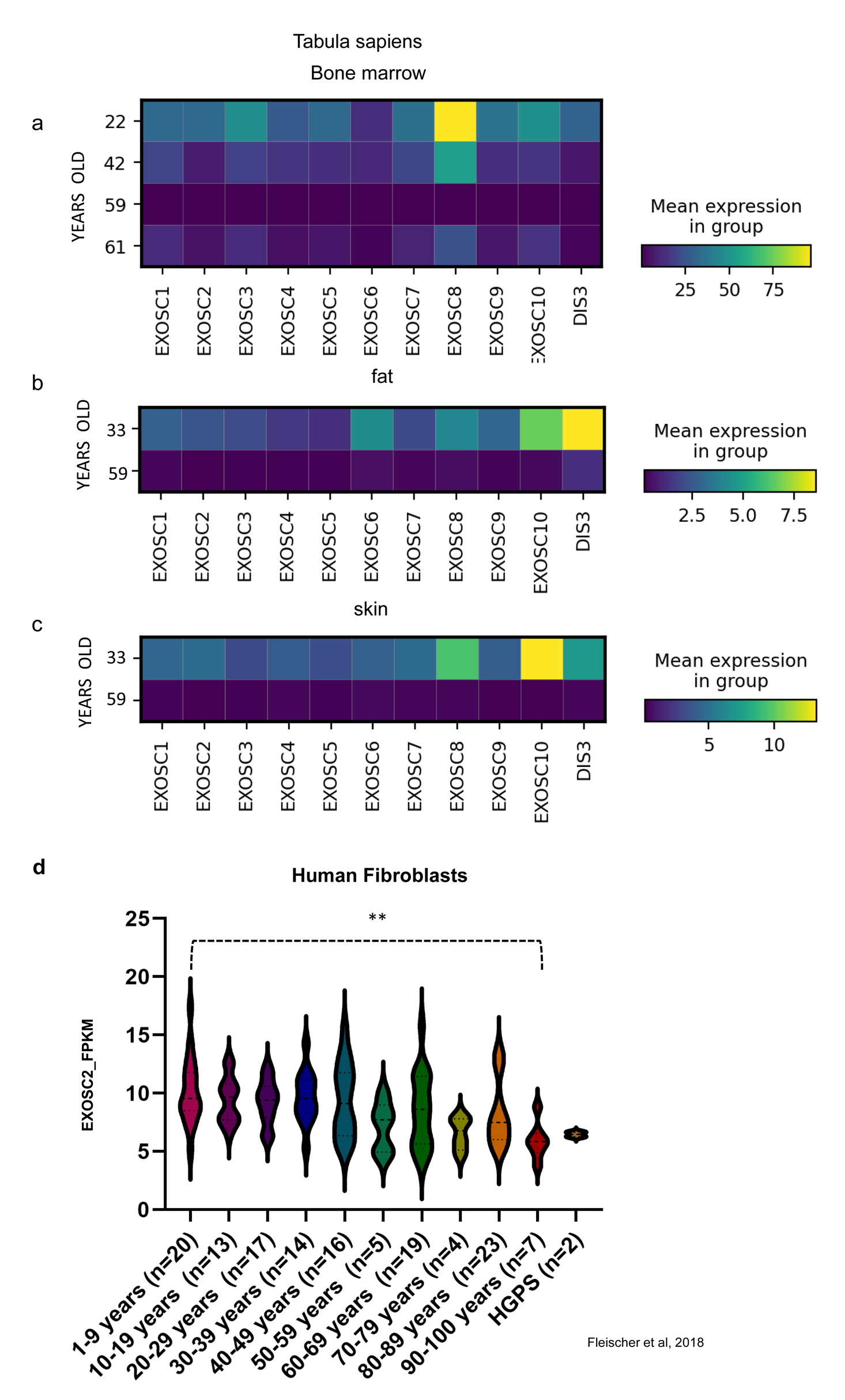
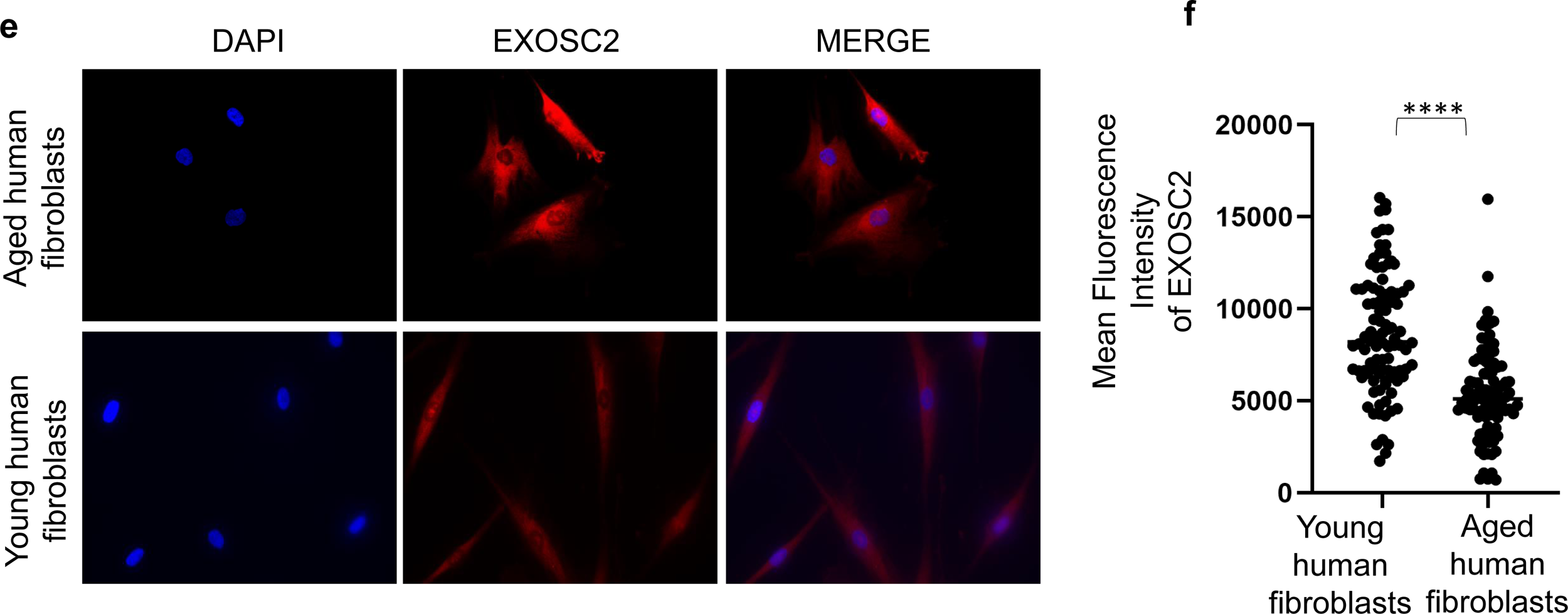
Physiological down-regulation of EXOSC2 in various cell types with aging. (a-c) Heatmaps for the gene expression of RNA exosome subunits from Tabula sapiens ^24^ in aged and young people. Gene expression of RNA exosome subunits from (a) the bone marrow of a 22-year-old, a 42-year-old, a 59-year-old, and a 61-year-old person, (b) from the fat tissue of a 33-year-old person and a 59-year-old person and (c) and from the skin of the 33-year-old person and a 59-year-old. The heatmaps indicate that the RNA exosome subunits drop with age. (d) EXOSC2 RNA levels from an RNAseq dataset of fibroblasts derived from individuals of different ages spanning from 1-year-old to 100 years old and from individuals with Hutchinson-Gilford progeria syndrome (HGPS). (e) IF for EXOSC2 levels in the human aged and young fibroblasts. (f) Quantification of mean fluorescence intensity for EXOSC2 shows a significant drop in EXOS2 levels in aged fibroblasts (n=3) compared to young (n=2) .****Statistical significance by two-sided t-test (p<0.0001) .

Moreover, we checked the EXOSC2 levels in fibroblasts derived from individuals of ages spanning from 1 year old to 100 years old, and from individuals with Hutchinson-Gilford progeria syndrome (HGPS)^31^. Given the importance of the passage number in this type of study, we made sure that cell lines were cultured to a similar mean population doubling^31^ . We interrogated the EXOSC2 RNA levels in an age-dependent manner. We pooled together RNA seq data from individuals whose age range would be no more than ten years apart and ended up with ten age groups, and we included a separate group for people with HPGS. The data analysis confirms a gradual reduction of EXOSC2 with age **(Figure 1 d)**. Moreover, we looked into individuals with Hutchinson-Gilford progeria syndrome (HGPS), a rare genetic disorder characterized by premature aging of various tissues^32^, and we confirmed that the levels of EXOSC2 were low, similar to the oldest group in our analysis **(Figure 1 d)**. In these datasets, we see an overall down-regulation of all the RNA exosome subunits, which is indicative of an RNA exosome-dependent down-regulation of EXOSC2 with aging. In the case of the Tabula Sapiens, we observe a uniform down-regulation of all the RNA exosome components **(Figure 1a, b, and c)** while in the case of the human fibroblasts RNA seq dataset, we see a significant downregulation in all the subunits apart from EXOSC7 and DIS3 (**Figure S1 a-k).** We have also checked the levels of RNA exosome in Tabula Muris Senis, a scRNA seq database from mouse tissue of young and aged mice. In this dataset, we observe that the EXOSC2 down-regulation is observed in different cell types in the bone marrow, the fat, and the skin **(Figure S1 l-n).** Lastly, we performed staining in human fibroblasts from young (n=2) and aged (n=3) donors for EXOSC2 protein and we observed significant decrease in the levels of EXOSC2 protein in the aged donors (**Figure 1e, f**).

EXOSC2 carries an S1 RNA binding domain, and as a peripheral part of the RNA exosome complex, it also stabilizes the hexameric ring of RNase PH-domain subunits through contacts with EXOSC4 and EXOSC7^33–36^. Moreover, it is responsible for tethering the MTR4 helicase to the complex, positioned in a crucial location between the exosome complex and RNA helicase (**Figure S2 a**)^37^. Therefore, we hypothesized that disruption of EXOSC2 has a significant functional impact on the mRNAs bound to the complex and on mRNA degradation (**Figure S2 a**)^9, 23, 38^. To test our hypothesis, we first generated ESC lines that stably express a hairpin for EXOSC2 confirmed by immunofluorescence, qRT-PCR, proteomics, and bulk RNA seq (**Figure 2 a, b, c, and Figure 3 a, b and Figure S3a**). We identified that the perturbation of EXOSC2 by shRNA significantly down-regulated the expression of other exosome subunits (**Figure 2 d and Figure S3 a**) and observed the same auto-regulatory feedback pattern with the perturbation of other exosome subunits, EXOSC3/10 (**Figure S4a, b**) as was suggested earlier^39^. Indeed, we observed more significant and broader down-regulation of the exosome components with EXOSC2 than EXOSC3/10 knockdown as we initially had hypothesized (**Figure 2 d** and **Figure S4a, b**) maybe because of the important position of the protein in the RNA exosome machinery. Thus, these data suggest that there might exist an autoregulatory feedback mechanism in the expression pattern of RNA exosome subunits.

**Figure 2.**
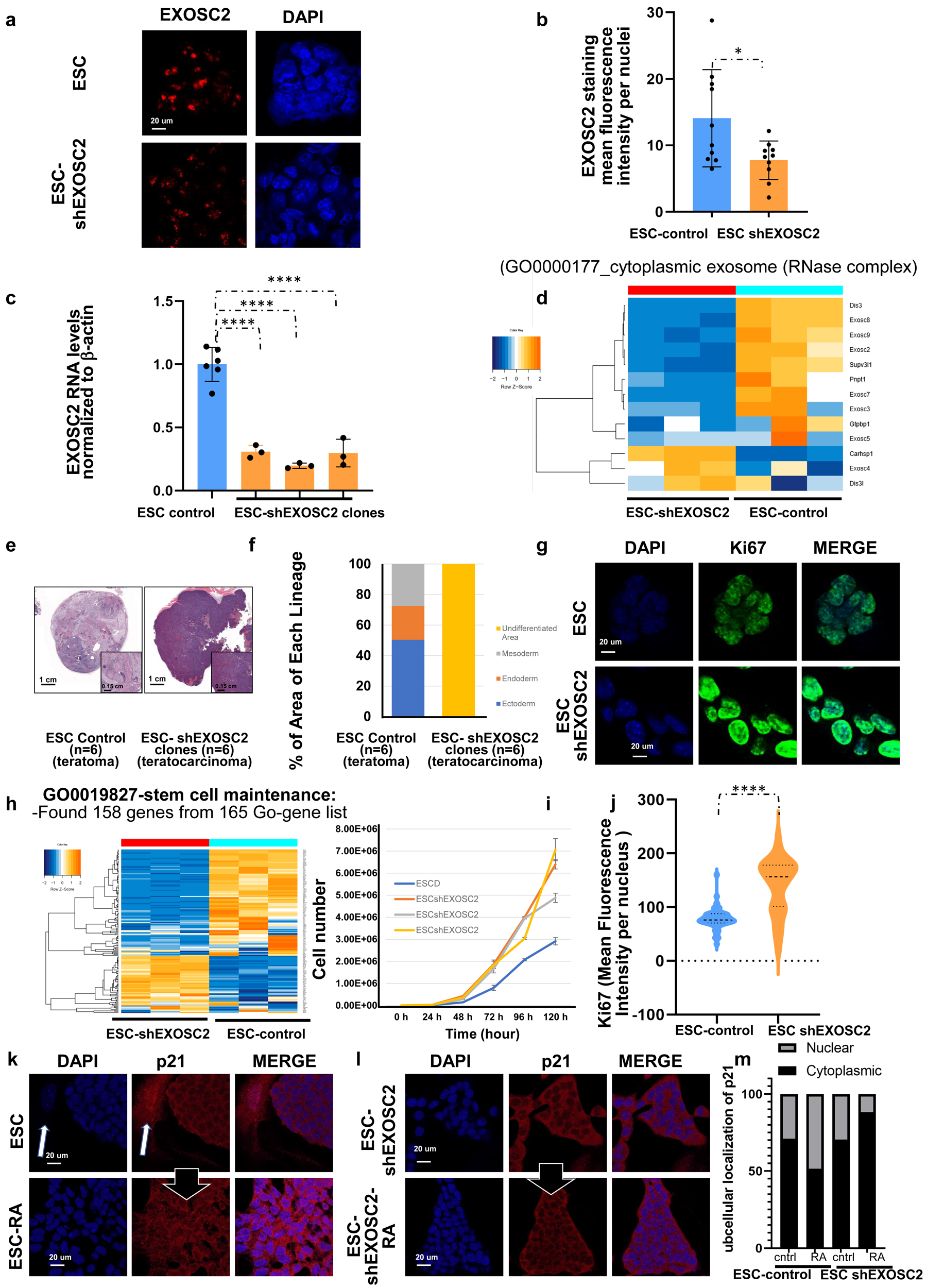
shRNA knockdown of EXOSC2 in ESC. (a) Immunostaining for EXOSC2 in ESC and ESC-shEXOSC2 lines. Nuclei were stained with DAPI. (b) Quantification of EXOSC2 staining in ESC and ESC-shEXOSC2. Mean fluorescence intensity per nucleus (>10) from 10 different colonies. Error bars indicate S.E. *Statistical significance by two -sided t-test (p<0.05). (c) Quantification of EXOSC2 RNA level by qPCR. Error bars indicate SE.****Statistical significance by two-sided t-test (p<0.0001). (d) RNAseq of three independent clones of ESC and ESC-shEXOSC2 showed that down-regulation of EXOSC2 leads to down-regulation of other core components of the RNA degradation complex (GO0000177_cytoplasmic exosome (RNase complex). (e) Teratoma analysis of ESC and ESC-shEXOSC2 by H/E staining. (f) Quantification of the area of undifferentiated teratocarcinoma and benign teratoma with mesoderm endoderm and ectoderm. (g) Ki67 staining in ESC and ESC-shEXOSC2. Nuclei were stained with DAPI. (h) Among the 165 stem cell maintenance genes from GO0019827, 158 genes were differentially regulated by shRNA-EXOSC2 treatment in ESC in RNAseq analysis. (i) Maintenance of self -renewal capacity in ESC-shEXOSC2 with faster proliferation rates than ESC. (j) Quantification of Ki67 staining in ESC and ESC-shEXOSC2. Mean fluorescence intensity per nucleus (>10) from 10 different colonies. Error bars indicate S.E. *Statistical significance by two-sided t-test (p<0.05). (k-l) P21 staining in ESC and ESC-shEXOSC2 before and after R.A. treatment. Nuclei were stained with DAPI. The white arrow shows the irradiated fibroblast that serves as a positive control.(m) Quantification of Immunofluorescence for cytoplasmic and nuclear localization of p21 in embryonic stem cells (ESCs), ESCs treated with retinoic acid, ESCshEXOSC2 and ESCshEXOSC2 treated with RA. Nuclear localization of p21 increases from thirty percent to fifty percent in ESC upon treatment with retinoic acid indicating differentiation of stem cells . Nuclear localization of p21 decreases upon retinoic acid treatment in ESCshEXOSC2 indicating a blockade in differentiation.

**Figure 3.**
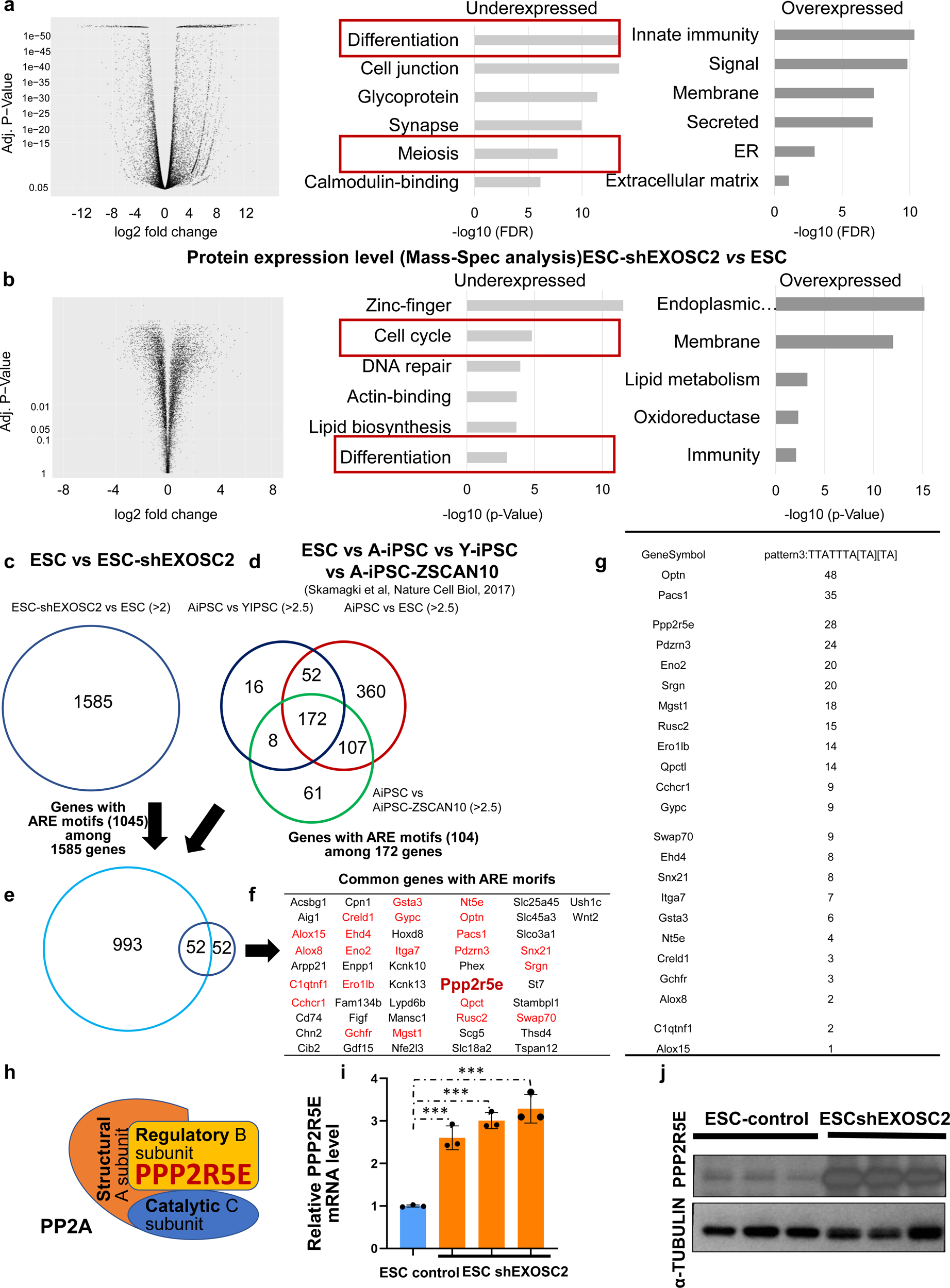
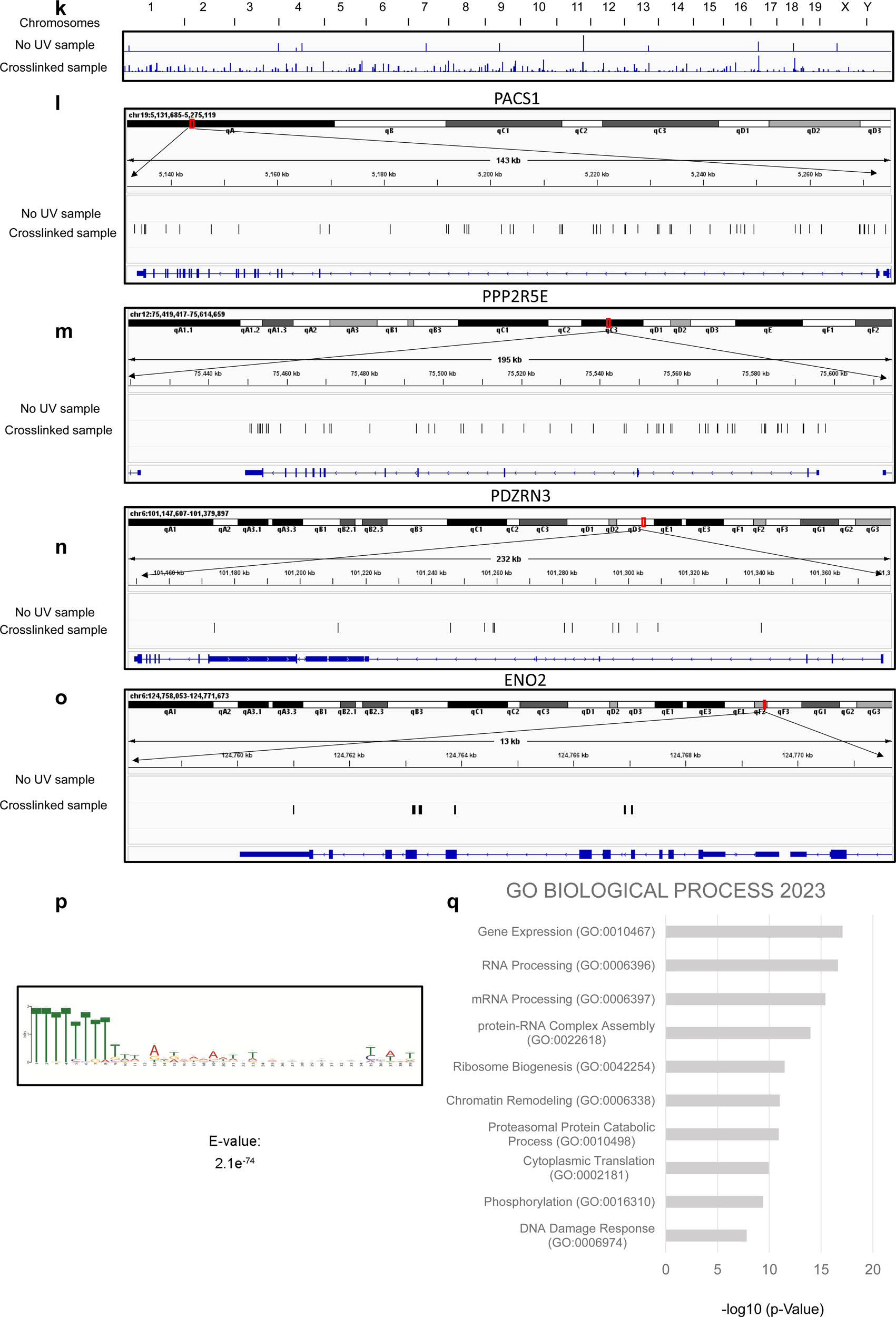
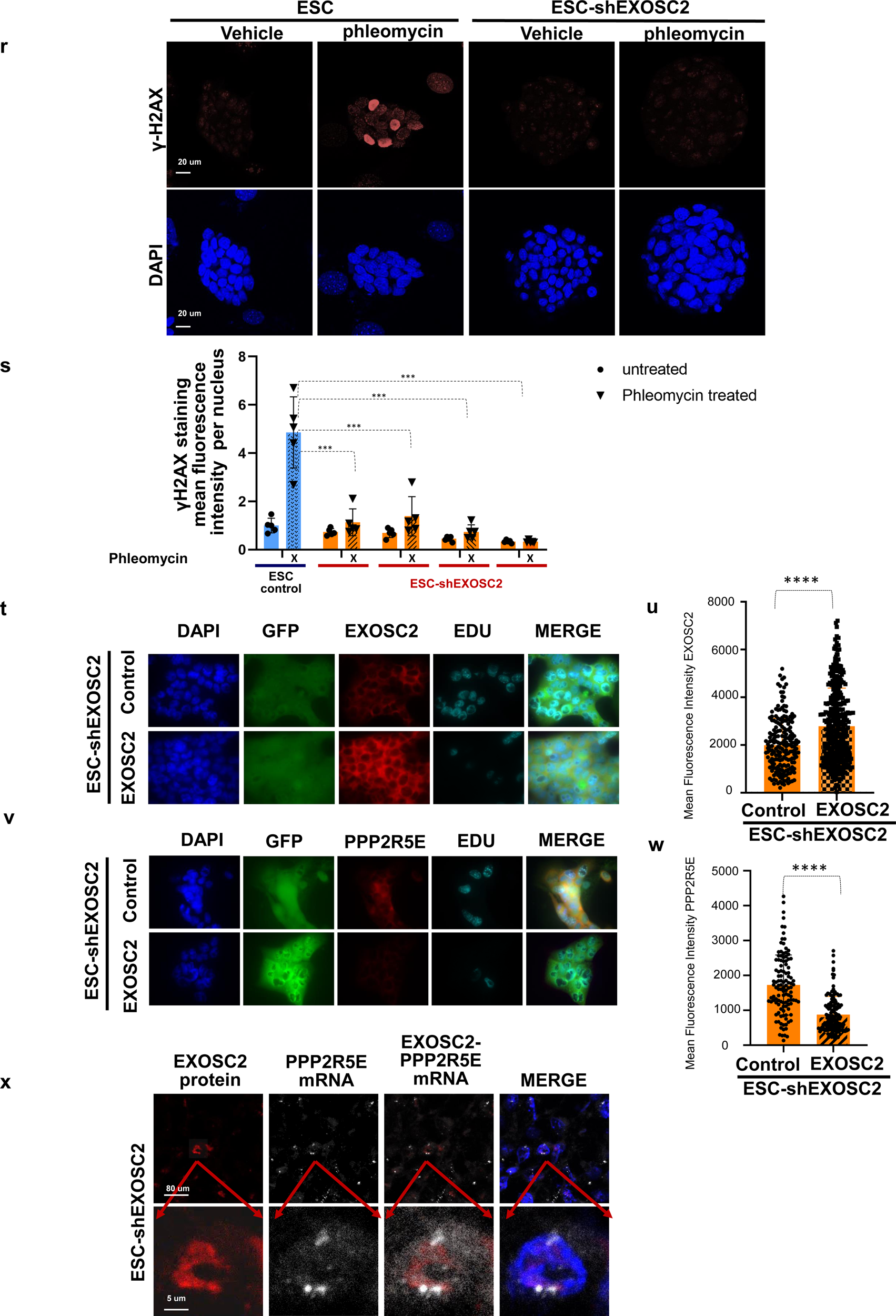
mRNA targets of EXOSC2. (a) Gene expression analysis by RNAseq was performed to look into downstream targets of EXOSC2 and ontology analysis with the significantly down/upregulated group of genes. (b) Protein expression analysis by Mass-spec was performed to look into downstream targets of EXOSC2 with an ontology approach with the significantly down/upregulated group of proteins. (c) We analyzed the genes upregulated more than two folds (log2 scale) in ESC-shEXOSC2. Among 1585 genes, 1046 carry the ARE motif. (d) We cross-referenced the genes that are upregulated more than 2 times in ESC-shEXOSC2 with the genes that are upregulated in A-iPSC over Y-iPSC, ESC, and A-iPSC-ZSCAN10 (172 GENES) and found that 104 gene carries AREs motif. (e) 52 genes carry AREs motif commonly upregulated in two independent pluripotent contexts where EXOSC2 is downregulated. (f) We further confirmed 23 genes (red letters) differentially translated by Mass-spec analysis. (g) The number of AREs consensus sequences in the genes targeted by EXOSC2 in ESC. The AREs consensus sequence is defined as [UUAUUUA[A.U.][A.U.], and PPP2R5E is the third-ranked gene that is commonly identified by the comparative analysis above. (h) PPP2R5E is the structural subunit of the PP2A complex. (i) QPCR analysis also demonstrates the increase in RNA levels. Error bars indicate S.D. Statistical significance by two-sided t-test (p<0.01). (j) immunoblot analysis of PPP2R5E with three independent clones of ESC-shEXOSC2, which shows up-regulation of PPP2R5E expression compared to ESC control. (k) CLIP-sequencing results for EXOSC2 from mouse ESCshEXOSC2 with and without UV crosslinking. (l) The CLIP sequencing results for EXOSC2 demonstrate binding to 15 targets out of the 23 predicted including PACS1 mRNA (m) PPP2R5E mRNA, (n) PDZRN3 mRNA, and (o) ENO2 mRNA. (p) To identify the motifs bound by Exosc2, we generated the unique peaks file using the CTK software. We then grabbed 50 nucleotides around the unique peaks, converted the bed file to fasta (code available upon request), and fed it into MEME (Suite 5.5.5)1 with default settings. (q) Gene ontology analysis of EXOSC2 targets assessed by CLIP-seq. (r) γ-H2AX staining in ESC and ESC-shEXOSC2. (s) Quantification of γ-H2AX staining in undifferentiated ESC. Mean fluorescence intensity per nucleus (>10) from 5 different colonies per clone per condition was measured in ESC control and four independent clones of ESC-shEXOSC2. Nuclei were stained with DAPI. Error bars indicate SE.***Statistical significance by two-sided t-test (p<0.001). (t) Immunostaining for EXOSC2 protein level in ESC-shEXOSC2 line and ESC-shEXOSC2 line after EXOSC2 transient overexpression. Nuclei were stained with DAPI. (u) Quantification of mean fluorescence intensity 18 h after the EXOSC2 overexpression. **Statistical significance by two-sided t-test (p<0.005). n.s (not significant) (v) Immunostaining for PPP2R5E protein level in ESC-shEXOSC2 line and ESC-shEXOSC2 line after EXOSC2 transient overexpression. Nuclei were stained with DAPI. (w) Quantification of mean fluorescence intensity 18 h after the EXOSC2 overexpression. **Statistical significance by two-sided t-test (p<0.005). n.s (not significant) (x) Combined IF for EXOSC2 and FISH for PPP2R5E m RNA shows the interaction of the mRNA with the complex.

To functionally assess the impact of RNA degradation on tissue differentiation potential, we exploited a teratoma formation assay with control ESCs (**Figure 2 e**) and those with down-regulated EXOSC2 (ESC-shEXOSC2; **Figure 2 e** and **Figure S4 d, e**). We quantified the area of undifferentiated teratoma and teratoma with three germ layers (mesoderm, endoderm, and ectoderm structures (**Figure 2 e, f).** The quantification showed that the decrease in EXOSC2 levels impacted pluripotency, as the ESC-shEXOSC2 lost the tissue differentiation capacity to form teratomas, resulting in the formation of exclusively undifferentiated tumors (**Figure 2 e, f)**. These cell lines also had accumulated significantly more structural abnormalities compared to the ESC control **(Figure S3b)**.To further explore these findings, we performed the bulk RNAseq analysis in the embryonic stem cell lines and looked for differential effects in stemness following EXOSC2 down-regulation. We found that 158 related genes involved in stem cell maintenance were differentially regulated by shRNA-EXOSC2 (**Figure 2 h**). These cell lines maintained higher self-renewal capacity in ESC-shEXOSC2 as determined by their faster rate of proliferation (**Figure 2 i**). Moreover, the Ki67 staining in ESC (**Figure 2 g)** and in ESCshEXOSC2 **(Figure 2 g)** confirmed that the down-regulation of EXOSC2 leads to an increase in the levels of Ki67 (**Figure 2 j**) and, therefore to an increase in proliferation. The Edu experiment shows that the ESC shEXOSC2 cells have elongated G1 and G2 phases compared to the ESC control (**Figure S3 c, d**). To confirm in vitro the defect in differentiation potential by EXOSC2 down-regulation, we monitored the nuclear localization of p21 from the cytoplasm, which is a typical characteristic phenomenon of the tissue differentiation of stem cells^40–43^ that is induced by retinoic acid (R.A.) treatment^44^. We observed that the p21 protein in ESC was relocated from the cytoplasm (**Figure 2 k, m**) to the nucleus upon R.A. treatment (**Figure 2 k, m**) to trigger stem cell differentiation as expected^45^. However, in the ESC-shEXOSC2 cells (**Figure 2 l, m**), the p21 protein remained in the cytoplasm upon R.A. treatment (**Figure 2 l, m**), thus preventing the differentiation of stem cells. We also performed a differentiation experiment by leuchemia inhibitory factor (LIF) withdrawal and Embryoid body formation and assessed the defect in differentiation by qPCR for PAX6 (Ectoderm), GATA6 (Endoderm), and FGF8 (Mesoderm) in ESCshEXOSC2 compared to ESC (**Figure S3e**). These findings validate our initial observations that the ESC-shEXOSC2 cell lines have defective differentiation potential as an aging trait.

### Messenger RNA targets of EXOSC2

We further investigated the precise mechanism of how the EXOSC2 down-regulation may affect tissue differentiation as an aging trait^46–49^. We first performed unbiased approaches to investigate the downstream effectors of RNA exosome activity through the shRNA-mediated reduction of EXOSC2 expression. We performed bulk RNAseq to interrogate the genes that will be influenced by RNA exosome complex (**Figure 3 a**) and further confirmed their respective changes in protein levels by mass spectrometry (**Figure 3 b**). The unbiased gene ontology analysis of the gene and protein datasets revealed that EXOSC2 is essential for tissue differentiation and proliferation (**Figure 3 a, b and supplementary table 1-4**). This result corroborates our observed phenotypes with poor tissue differentiation (**Figure 2 e, f**) and increased cell proliferation rates (**Figure 2 i).** To bioinformatically predict the mRNAs targeted to RNA exosome, we employed the in-silico methodology that we have previously developed ^11^. This methodology involves RNA sequencing of different cell lines and mapping specific ARE-motifs ^11, 50^ that mark transcripts for degradation by RNA exosome.

We analyzed the genes that are upregulated more than two-fold (log2) in ESC-shEXOSC2 cells (**Figure 3 c**). Among 1585 genes, 1045 carry the ARE-motifs (**Figure 3 c and supplementary table 5**). We then cross-verified the genes that are upregulated more than two-fold in ESC-shEXOSC2 cells with the genes (172 genes) that are upregulated in our previously published cell lines (A-iPSCs)^8, 9^, which have physiologically low levels of EXOSC2 with shared phenotypes. We identified that 104 genes carried the ARE-motifs (**Figure 3 d**). Fifty-two genes with the ARE-motifs were commonly upregulated in two independent pluripotent cell contexts where EXOSC2 levels are down-regulated (**Figure 3 e**). We further confirmed 23 genes (red letters) that are differentially translated by Mass-spec analysis (**Figure 3 f**). We then counted the number of ARE-consensus sequences in the genes targeted for degradation among the 23 genes (**Figure 3 g**). PPP2R5E, one of the regulatory subunits of the PP2A complex (**Figure 3 h)**, was among the top genes in this 23-gene list. We independently confirmed the high expression levels of this mRNA following EXOSC2 knockdown by qPCR analysis (**Figure 3 i**) and increased protein expression by western blot (**Figure 3 j)**. Therefore, based on the above results, we hypothesized that the candidates identified by our approach might be either degraded by RNA exosome directly or by interacting with the RNA binding protein EXOSC2 might affect the translation of the mRNAs, and thus we see an effect in the protein levels.

### mRNA of the regulatory subunit of PP2A phosphatase, PPP2R5E, is one of the key targets of EXOSC2, a new mechanism for an aging trait

PP2A phosphatase constitutes approximately 1% of the total protein in a cell. The structure of the PP2A complex consists of three parts. The scaffold A, the regulatory subunit B, and the catalytic subunit C. Notably, the scaffold and the catalytic subunits are encoded by two genes, while the regulatory subunits of PP2A comprise four families, each encoded by several genes^51–53^. This trimeric structural arrangement thus generates many distinct combinations of PP2A holoenzymes. Each of these distinct complexes is responsible for the activity and the distinct substrate specificity of protein phosphatase. The fact that there are so many different regulatory subunits implies that their presence or absence controls the specificity. We hypothesized that the mRNA of the regulatory subunit of PP2A phosphatase, PPP2R5E, is a key target of the RNA binding protein (RBP) EXOSC2 and the interaction of EXOSC2 with the PPP2R5E transcript impacts its translation, therefore, leading to dynamic regulation of the subunit’s expression as a key mechanism of the aging trait. To prove that there is direct interaction we performed RIP-qPCR using EXOSC2 antibody and demonstrated the enrichment of PPP2R5E mRNA detected by the pull down of EXOSC2 compared to the IgG control in the (**Figure S3h**). We also performed Cross linking immunoprecipitation (CLIP)^54, 55^ in order to detect all the RNA targets of EXOSC2, and we present the unique positions identified across the whole transcriptome for the non-crosslinked control and the crosslinked sample (**Figure 3k and Figure S3g-o)**. CLIP sequencing confirms EXOSC2 binding to fifteen targets out of the twenty-three predicted, including PACS1 mRNA **(Figure 3l),** PPP2R5E mRNA **(Figure 3m)**, PDZRN3 mRNA **(Figure 3n),** ENO2 mRNA **(Figure 3o),** in the crosslinked sample while no binding is detected in the negative control. We found that fifteen targets that were observed among the 6248 cytoplasmic targets were among the twenty-three that we had predicted, which is statistically significant (**Figure S3o**). The motif analysis of the targets based on the significant peaks shows that the sequence of the transcripts that bind to EXOSC2 is AU rich as well **(Figure 3p)**. We also performed gene ontology analysis on the observed targets of EXOSC2 and found that the targets are involved in a wide range of cellular processes including cytoplasmic translation and phosphorylation (**Figure 3q**).

We interrogated if the high levels of PPP2R5E also led to the over-activation of the PP2A phosphatase. To address this question, we looked into the well-studied downstream target of PP2A phosphatase, such as γ-H2AX^6, 56^. In eukaryotic cells, the induction of DNA double-strand breaks (DSBs) leads to the phosphorylation of H2AX, and this event is critical in maintaining genomic stability^57^. Phosphorylation of histone H2AX (γ-H2AX) in response to DNA double-strand breaks (DSBs) must be eliminated from the sites of DNA damage to facilitate the DNA repair and release cells from their growth arrest phase^58^. Previous studies showed that PP2A interacts with γ-H2AX and leads to its dephosphorylation^6, 56^. Therefore, to investigate whether the PP2A complex was hyperactivated in the present system, we treated ESC and ESC-shEXOSC2 with the radiomimetic drug phleomycin to induce γ-H2AX as we have previously demonstrated^8^. We then quantified the γ-H2AX expression (**Figure 3 r**) and found that ESC-shEXOSC2 lines had significantly lower levels of γ-H2AX expression than ESC after phleomycin treatment (**Figure 3 s**). This result indicates that disruption of EXOSC2 leads to increased PP2A activity, presumably through up-regulation of PPP2R5E activity as a rate-limiting factor that is demonstrated by its ability to remove the phospho-group from H2AX. Thus, upon EXOSC2 downregulation, DNA repair is impaired after phleomycin treatment. It is worth mentioning that the robust DNA double-strand break repair co-evolves with longevity^59^, therefore reinforcing the concept of EXOSC2 down-regulation as an aging trait.

We further investigated how the down-regulation of EXOSC2 protein affects PPP2R5E mRNA by performing a rescue experiment. We explored the two hypotheses mentioned above: that (1) PPP2R5E is a critical transcript targeted by RNA exosome for degradation through its binding to EXOSC2 and alternatively (2) that the EXOSC2 protein directly binds PPP2R5E mRNA affecting its translation. We transiently overexpressed the EXOSC2 component in ESCshEXOSC2 lines and followed the PPP2R5E transcript and protein levels during an initial time course of 24 hours. We observed that at 24 hours post-transfection (**Figure 3 t, u)**, there is no impact on the PPP2R5E transcript levels and the levels of other RNA exosome components (**Figure S4 h, i, j, k, l**). Yet, the levels of PPP2R5E protein significantly dropped (**Figure 3 v, w and Figure S4 f, g**). This intriguing result led us to reason that excess EXOSC2 could bind to PPP2R5E mRNA, but the degradation of the transcript is still rate limited by the availability of the other RNA exosome components (which do not increase following EXOSC2 overexpression) (**Figure S4 f, g, h, I, j, k, l**). However, EXOSC2 binding to PPP2R5E mRNA alone may not suffice to inhibit translation.We performed combined immunofluorescence (IF) imaging for EXOSC2 and fluorescence in situ hybridization (FISH) for PPP2R5E mRNA, which demonstrated the physical proximity of the mRNA with the EXOSC2 subunit (**Figure 3 x**) as we observed spots of PPP2R5E mRNA colocalizing with the EXOSC2 protein and is supplementary to the eCLIP assay **(Figure 3k)**. To assess whether the PPP2R5E transcript specifically depends on EXOSC2 expression, we looked into PPP2R5E expression upon down-regulation of different RNA exosome subunits in sets of stem cell lines. Indeed, we didn’t observe up-regulation of the PPP2R5E transcript upon down-regulation of EXOSC3/10 (**Figure S4c**). These data once again corroborate our original hypothesis that the EXOSC2 directly and specifically regulates the levels of the PPP2R5E transcript by direct binding. The binding of PPP2R5E mRNA by EXOSC2 leads to transcript degradation in Embryonic Stem cell lines that have physiological levels of RNA exosome complex and seems to lead to a decrease in protein levels in ESCshEXOSC2 that lack a functional RNA exosome.

### A conserved functional interaction between EXOSC2 and PPP2R5E prevents defective tissue differentiation potential as an aging trait

We next asked if the above dynamic regulation of PPP2R5E by EXOSC2 is a broadly conserved phenomenon in various tissue types. We hypothesized that the levels of EXOSC2 are essential for cell fate decisions as the down-regulation of EXOSC2 resulted in a blockade of ESC differentiation (**Figure 2 e, f**). Thus, the regulation of PPP2R5E by EXOSC2 is expected to be present in the dynamic surveillance of cell fate regulation as it happens during embryo development. We performed co-staining of mouse embryo tissue with EXOSC2 and PPP2R5E at the E13.5 stage (**Figure 4 a** and **Figure S5 a**), and we observed inverse staining for these two proteins (**Figure 4 b, c, d**) which led us to concur that the regulation of PPP2R5E by EXOSC2 may be crucial for the progression of embryo development. Additionally, we performed co - staining for EXOSC2 and PPP2R5E in adult organs (**Figure 4 e** and **Figure S5 b, c),** and we also observed a similar inverse relationship between EXOSC2 expression and PPP2R5E protein levels in different cell types within the same organ. Interestingly while we observed high levels of EXOSC2 and low levels of PPP2R5E in most tissues, the brain shows the opposite pattern with low levels of EXOSC2 and high levels of PPP2R5E (**Figure 4 e and Figure S5 b, c**). These data suggest that EXOSC2 may regulate PPP2R5E expression dynamically in multiple tissue types. Although the EXOSC2 protein is uniformly expressed in many tissue types, the levels differ in a cell context-dependent manner within each cell type. The pattern of expression of EXOSC2 coincides with the proliferation–differentiation gradient from the crypt compartment to the villus compartment. The crypt invaginations house stem cells^60^, and this is where we observed the higher levels of EXOSC2 (**Figure 4 f**). Among the multiple tissue types, the intestine presents the most significant inverse staining pattern between the expression of EXOSC2 and PPP2R5E as an aging trait.

**Figure 4.**
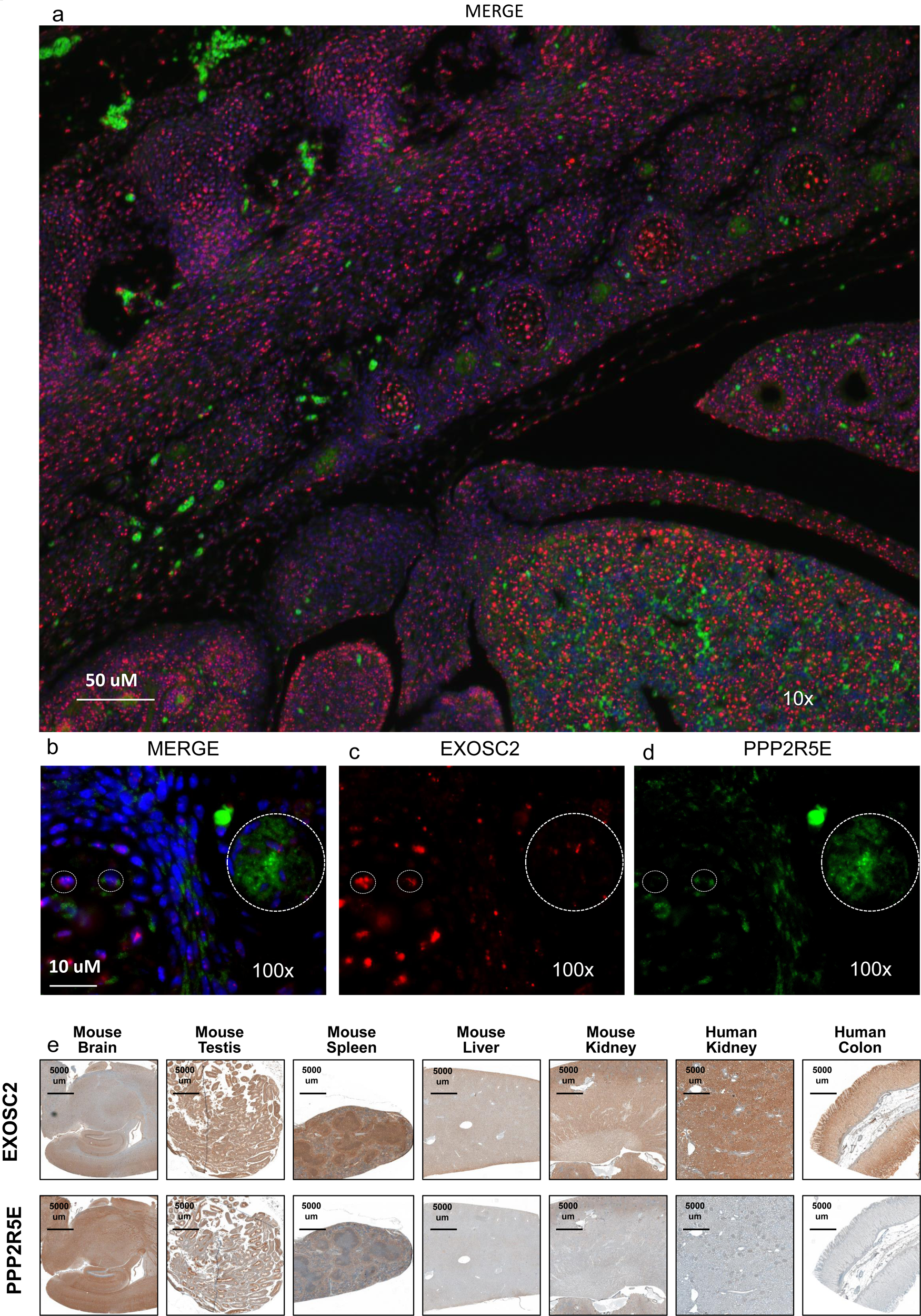
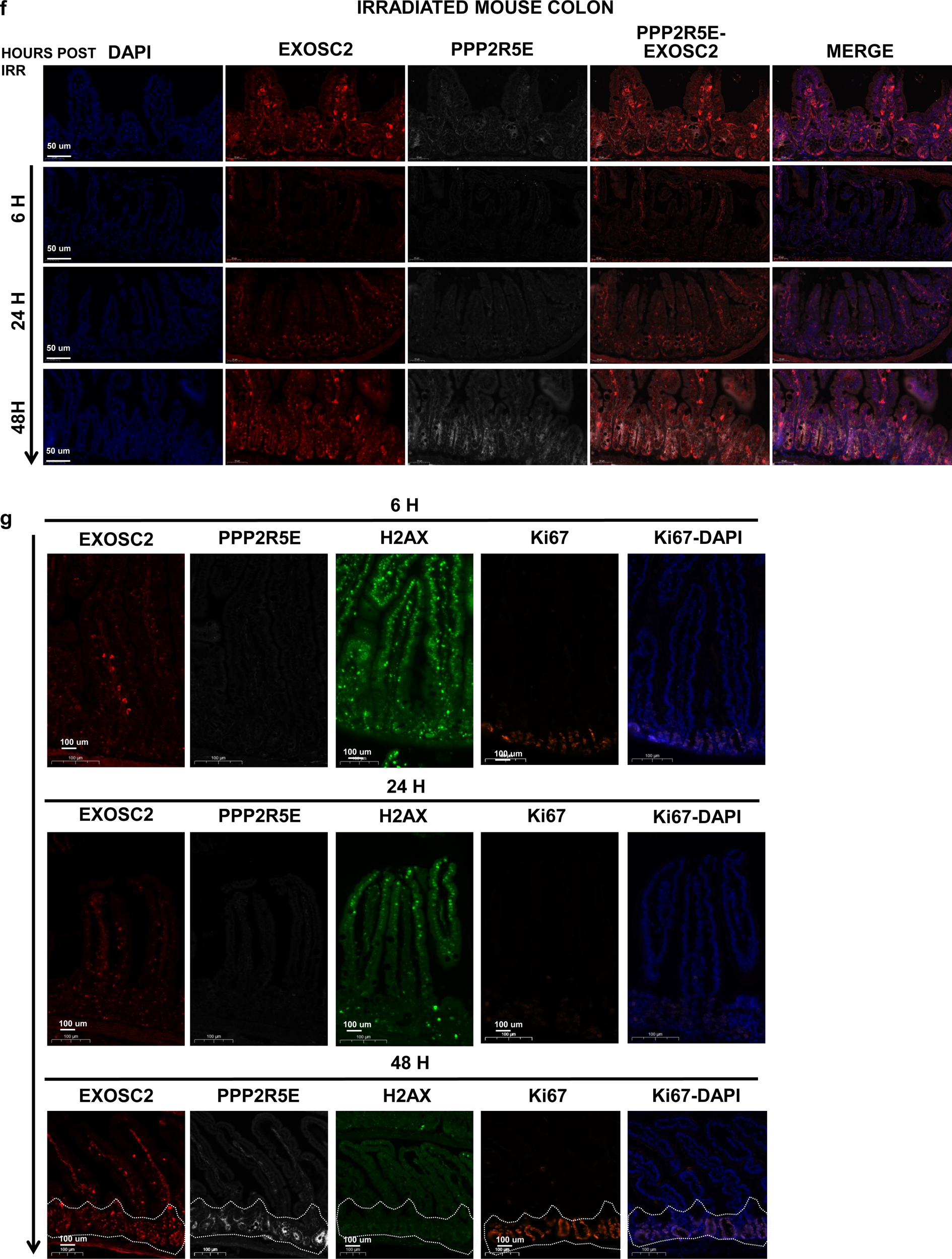
In vivo dynamic regulation of PPP2R5E by EXOSC2. (a) Dynamic regulation of PPP2R5E by EXOSC2 in the developing embryo (E13.5). Co-staining for EXOSC2 and PPP2R5E. Nuclei were stained with DAPI at 10 x magnification and (b, c, d) 100X magnification. The big circle emphasizes areas where levels of PPP2R5E are high and levels of EXOSC2 are low. The two small circles emphasize areas with high EXOSC2 and PPP2R5E and the opposite, an area of high PPP2R5E and low EXOSC2 (e) Dynamic regulation of PPP2R5E by EXOSC2 in adult tissues from mice and humans by immunohistochemistry. Staining for EXOSC2 and PPP2R5E was done independently in consecutive sections of the same tissues. (d) The mouse intestine shows a gradient in EXOSC2 staining as assessed by immunostaining for EXOSC2 and PPP2R5E. Nuclei were stained with DAPI. (f) Co -staining for EXOSC2 and PPP2R5E in mouse intestines of mice subjected to whole abdomen irradiation as previously described 55. Nuclei were stained with DAPI. The staining was performed at 0h, 6h, 24h, and 48h post-irradiation. 6 hours post-irradiation, there was a significant decrease in EXOSC2 and PPP2R5E staining. (g) Co-staining for EXOSC2 and PPP2R5E in mouse colon, and immunostaining for H2AX and Ki67 in consecutive sections of mouse colon 6h post-IRR, 24h post-IRR, and 48h post-IRR. Nuclei were stained with DAPI.

Of its capability for rapid proliferation and differentiation, the intestinal clonogenicity assay is also a routinely employed model to investigate the tissue regenerative capacity following tissue damage induced by radiation treatment^61^. More specifically, to assess the importance of the EXOSC2 in the intestine, we used a preclinical murine platform of abdominopelvic-focused irradiation that enables localized delivery of escalating radiation doses to the intestinal luminal organs while sparing the majority of the mouse bone marrow^62^. Intestinal epithelial regeneration following a radiation injury depends on intestinal stem cell (ISC) repopulation and regeneration of the differentiated cells that populate the functional intestinal villus^63, 64^. Therefore, we reasoned that the intestine is an ideal system to assess the gradient of EXOSC2 levels between the top and basal crypt compartments (**Figure 4 f**). The importance of this gradient in EXOSC2 levels and the regulation of PPP2R5E is manifested when we apply irradiation (IRR) and check the levels of EXOSC2 and PPP2R5E at different times after irradiation as tissue regeneration takes place. We observed that the 6h (**Figure 4 f, g)** and the 24h (**Figure 4 f, g**) post-radiation levels of both the EXOSC2 and the PPP2R5E drop dramatically, and by the 48h timepoint, both proteins increase again (**4 f, g)**. We interrogated why the dynamic regulation of PPP2R5E by EXOSC2 is interrupted in these early time points after irradiation by looking into downstream targets of the phosphatase and, more specifically, H2AX (**Figure 4 g)** at 6h (**Figure 4 g),** at 24h (**Figure 4 g)** and 48h **(Figure 4 g)**. We found that at the 6h post-IRR timepoint, the staining for γ-H2AX is the highest (**Figure 4 g),** and proliferation levels are low as assessed by Ki67 staining (**Figure 4 g**). This increase in γ-H2AX corroborates our contention with the above *in vitro* findings whereby EXOSC2 is a direct regulator of PP2A activity via PPP2R5E. As the γ-H2AX is a target of PPP2R5E, during the early time frame of DNA repair (6h to 24h), the levels of PPP2R5E and its regulator, EXOSC2, decrease dramatically **(Figure 4 f)**. On the contrary, 48h post-IRR (**Figure 4 f, g**), when DNA damage is resolved as assessed by γ-H2AX staining (**Figure 4 g**), both the levels of EXOSC2 and PPP2R5E increase (**Figure 4 f, g**), at the base of the intestinal crypts and demonstrate an inverse pattern of expression. At the same time, the expression of γ-H2AX drops, and proliferation resumes as assessed by Ki67 staining (**Figure 4 g**). Together, all the above data indicates that the dynamic regulation patterns of EXOSC2 and PPP2R5E are conserved not only during normal tissue development but also in the context of tissue regeneration.

### PPP2R5E targets MTCL1 to modulate microtubule dynamics

To find downstream targets of PPP2R5E regulating cell fate decisions, we performed mass spectrometry for the phospho-enriched population of peptides in ESC and ESC-shEXOSC2. We discovered that PP2A governs a series of downstream effectors essential for cytoskeleton assembly and tissue differentiation, such as MTCL1. Phospho-enrichment analysis by mass spectrometry was performed to interrogate downstream targets of PPP2R5E with an unbiased ontology approach with the significantly underphosphorylated/phosphorylated groups of proteins (**Figure 5 a**). The gene ontology analysis shows that the underphosphorylated group of proteins, potential targets of PPP2R5E, are involved in tissue differentiation and cell cycle (**Figure 5 a and supplementary tables 6 -9**). To find the downstream targets of PPP2R5E, we looked into the underphosphorylated proteins that carry the LxxI motif, which is part of a motif that provides binding specificity to the PPP2R5E^65^. One of the downstream targets is Microtubule Crosslinking factor 1 (MTCL1), which has been shown to interact with PPP2R5E (**Figure 5 b**)^66^. Therefore, we measured the protein levels of MTCL1 in ESC (**Figure 5 c**) and ESC treated with shEXOSC2 (**Figure 5 b, c**) and found decreased levels of MTCL1 protein as compared to ESC control. We hypothesized that because the MTCL1 protein decreases by PPP2R5E dephosphorylation, the microtubule reorganization would change in the ESC-shEXOSC2 lines. Indeed, we observed low levels of an α-tubulin network in ESC-shEXOSC2 (**5 b, c**) compared with the ESC (**Figure 5 b, c**) and differentiated cells (i.e., fibroblasts). We also confirmed this result with tyrosinated α-tubulin (**Figure S6a)**, which controls the initiation of processive dynein–dynactin motility^67^. We further performed RNA-seq analysis and confirmed differential regulation of microtubule polymerization genes (G.O. term#: GO031109) following shEXOSC2 treatment in ESC (**Figure 5 d**). To confirm the interaction of MTCL1 with PPP2R5E we performed immunoprecipitation for MTCL1 **(Figure S6b)** and co-immunoprecipitation for PPP2R5E **(Figure S6c)**. Collectively these results are in support of the hypothesis that the defects that we observe upon EXOSC2 dysregulation impact microtubule polymerization and not total levels of alpha-tubulin. Therefore, the alpha tubulin signal by immunofluorescence is dim because there is no proper microtubule network to visualize, but the total levels of alpha-tubulin remain unchanged.

**Figure 5.**
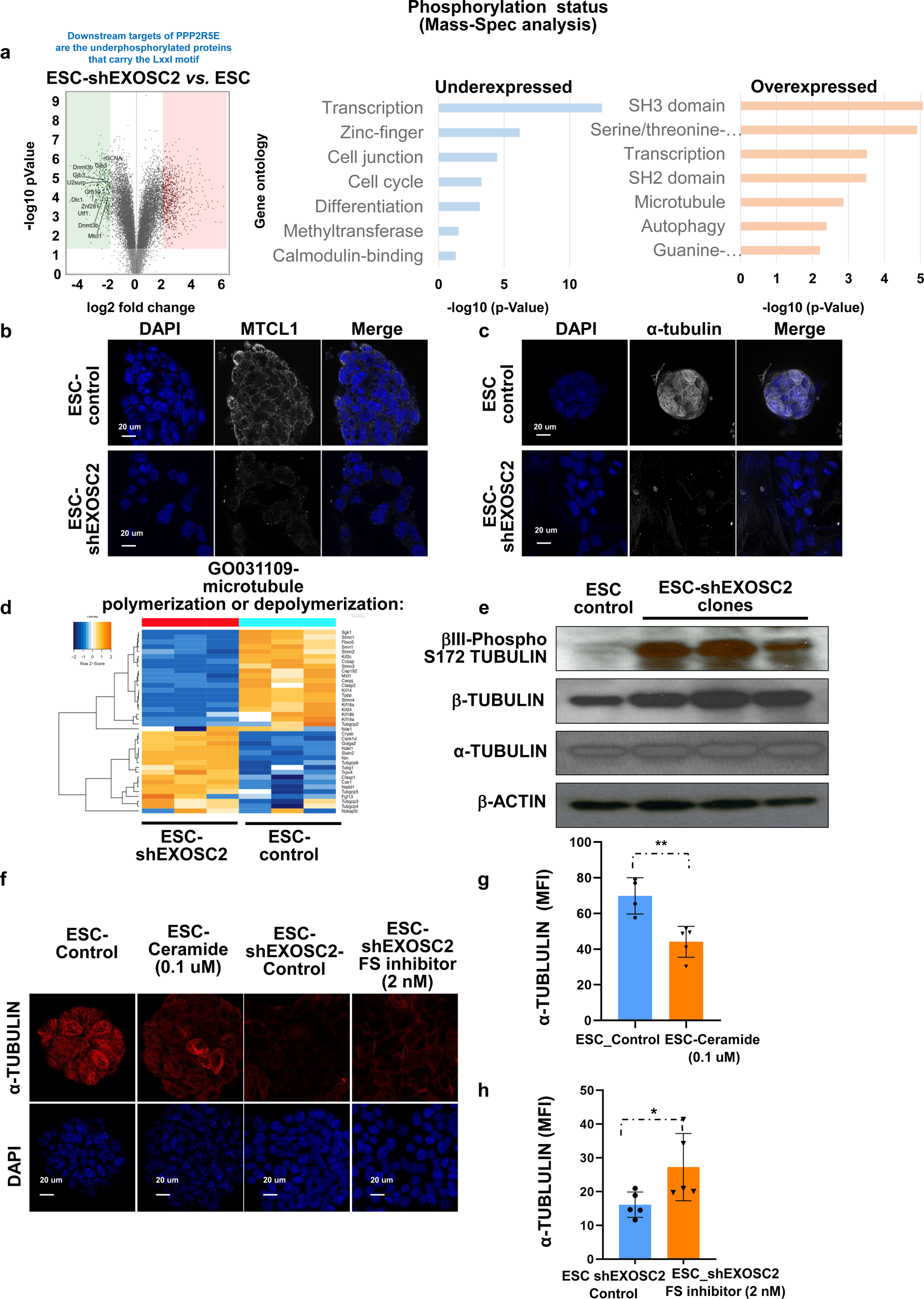
Downstream targets of PPP2R5E. (a) Phosphoenrichment analysis by Mass-spec was performed to look into downstream targets of PPP2R5E with the significantly underphosphorylated group of proteins. Potential downstream targets of PPP2R5E are the underphosphorylated proteins that carry the LxxI motif. One of the downstream targets is MTCL1. Phosphoenrichment analysis by Mass-spec was performed to look into downstream targets of PPP2R5E with an ontology approach with the significantly underphosphorylated/phosphorylated group of proteins. (b) Low levels of MTCL1 in ESC treated with shEXOSC2 in immunostaining. (c) Low levels of α-tubulin in ESC treated with shEXOSC2 in immunostaining compared to fibroblast control and ESC control. Nuclei were stained with DAPI. (d) Among the 42 microtubule polymerization or depolymerization genes from GO031109, 38 genes were differentially regulated by shRNA-EXOSC2 treatment in ESC in RNAseq analysis. (e) Higher phosphorylation of -tubulin on SER172 correlated with increased depolymerization. (f) Activation of PP2A with ceramide (0.1 uM for 2 days) and immunostaining for α-tubulin. (f) Inhibition of PP2A by Fostriecin sodium salt (2nM, 2 days) and immunostaining for α-tubulin. Nuclei were stained with DAPI. (g) Quantification of α-tubulin staining in ESC and ESC treated with ceramide. Mean fluorescence intensity (>10) from 5 different colonies per clone per condition. Error bars indicate SE.**Statistical significance by two-sided t-test (p<0.005). (h) Quantification of α-tubulin staining in ESC-shEXOSC2 and ESC-shEXOSC2 treated with Fostriecin sodium salt. Mean fluorescence intensity (>10) from 5 different colonies per clone per condition. Error bars indicate SE.*Statistical significance by two-sided t-test (p<0.05).

Lastly, we demonstrated that the ESC-shEXOSC2 cells had higher levels of phosphorylated b-tubulin on Ser172, which is known to be present in the soluble tubulin fraction, compared to the controls (**Figure 5 e**)^68^. The data show that ESC-shEXOSC2 cell lines have defects in microtubule polymerization. To confirm that the impact on microtubule polymerization happens through PPP2R5E-mediated dephosphorylation of MTCL1, we used the activator of PP2A, ceramide, in ESC. We observed a decrease in the levels of the α-tubulin network following ceramide treatment (**Figure 5 f, g**). On the contrary, inhibition of PP2A by Fostriecin sodium salt in ESC-shEXOSC2 rescues the staining levels for α-tubulin (**Figure 5 f, h**). Therefore, we suggest that the main target of PPP2R5E, responsible for the observed phenotypes of differentiation blockade, is MTCL1, which impacts tubulin polymerization. In support of this hypothesis, we performed transient overexpression of EXOSC2 in ESCshEXOSC2 and we observed a decrease in the percentage of cells that are in the G1 phase **(Figure S6 d, e)**. Many studies have described how microtubule arrays reorganize as cells undergo differentiation^69–71^; here, we suggest that MTCL1 alters microtubule dynamics to facilitate microtubule reorganization upon embryonic stem cell differentiation.

### Constitutive activation of PP2A by down-regulation of EXOSC2 as a cancer dependency in colorectal cancer

Having previously demonstrated that the RNA binding protein EXOSC2 plays an essential role in stem cell biology, we sought to determine whether the conserved dynamic regulation of PPP2R5E by EXOSC2 is similarly maintained in the context of cancer and more specifically in a model of colorectal cancer. In stem cell models, the subtle down-regulation of EXOSC2 as an aging trait leads to the loss in the dynamic regulation of the PPP2R5E mRNA and consequently to constitutive activation of PPP2R5E which leads to a dependency meaning that the cell lines rely on PPP2R5E for their survival. This type of dependency was also observed in AiPSC that express physiologicaly low levels of EXOSC2. We, therefore, were interested in exploring the role of PPP2R5E/EXOSC2 as aging trait in the context of malignancies. To this end, we chose to study a mouse model of colorectal cancer. We performed orthotopic transplantation of GFP-expressing CT26 colon cancer lines into the colon of syngeneic host mice^72^. Engrafted tumors show low levels of EXOSC2 and high levels of PPP2R5E compared to the surrounding colonic epithelium. This inverse correlation in protein expression suggests that indeed EXOSC2 regulates PPP2R5E also in the context of cancer. We performed an immunostaining experiment for EXOSC2 and PPP2R5E **(Figure 6 a, a’)** and found that the cancer cells had a greater proliferation rate based on higher Ki67 staining of the transplanted region of the CT26 mouse colorectal cancer (circled area) compared to the normal colon region (rectangular area) **(Figure 6 b, b’, c, c’)**. Since we observed that EXOSC2 may regulate PPP2R5E also in the context of colorectal cancer, we wanted to explore whether the down-regulation of EXOSC2 in the colorectal cell line may lead to a PPP2R5E constitutive activation as we have seen in the ESCshEXOSC2 cell lines.

**Figure 6.**
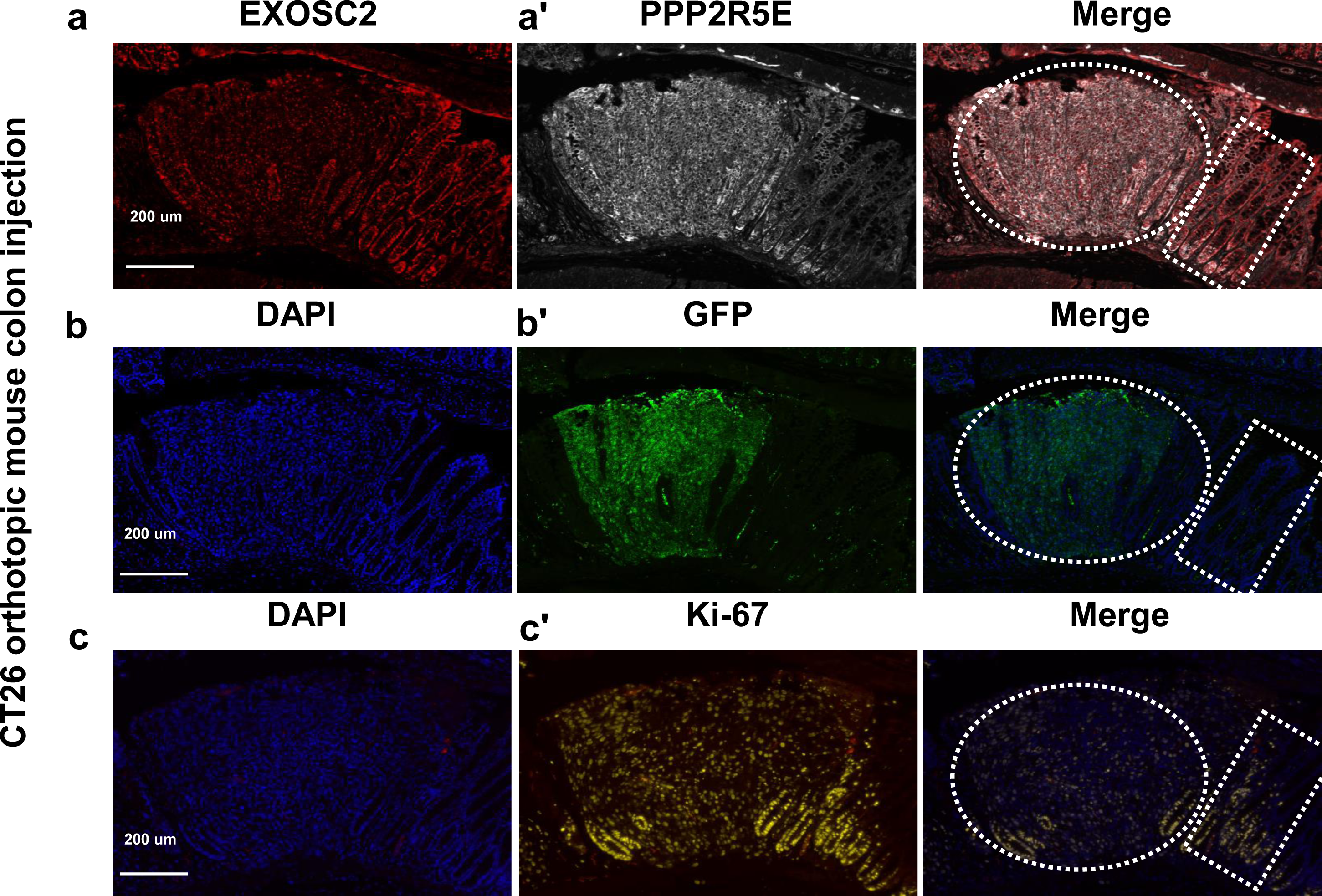
Constitutive activation of PP2A by down-regulation of EXOSC2 in colorectal cancer. (a, a’, b, b’, c, c’) Immunostaining for EXOSC2, PPP2R5E in orthotopically injected CT26 cancer cell line-GFP positive in mouse colon. Up-regulation of PPP2R5E is seen in the transplanted region of the CT26 mouse colorectal cancer (circled area) that expresses low levels of EXOSC2 compared to the normal colon region (rectangular area) as demonstrated by co-immunostaining of PPP2R5E and EXOSC2. Immunostaining for GFP and Ki67 was performed in consecutive sections of the same region. Nuclei were stained with DAPI.

### Cancer cells develop PPP2R5E dependency, thereby blocking microtubule depolymerization

In ESCs, we have observed that the down-regulation of EXOSC2 leads to constitutive up-regulation of PPP2R5E at the transcript and protein levels. Furthermore, we verified that the induction of the PPP2R5E protein correlates with the PP2A phosphatase activity by measuring its downstream effectors, such as H2AX and MTCL1. Here, we hypothesize that CT26 mouse colon cancer cells (marked with GFP+) **(Figure 6)** hijack the EXOSC2-PPP2R5E axis develop PPP2R5E dependency and lead to dysregulation of the microtubule depolymerization process. To test this possibility, we first assessed the levels of alpha-tubulin staining, as a measure of the organized microtubule network, in stem cells and the same cell line containing a constitutively expressed shRNA targeting EXOSC2 (ESC-shEXOSC2) **(Figure 7a, b)**. Similar to what we observed in our parallel study, the levels of alpha-tubulin staining decreased significantly in the ESC-shEXOSC2 cells compared to the parental ESC cells and the stained fibroblasts **(Figure 7 a, b)**. In the same experiment, we also compared alpha-tubulin levels in the CT26 colon cancer cell line **(Figure 7 a, b)**, which were demonstrated to have low levels of EXOSC2 expression in our orthotopic mouse model. We found that levels of alpha-tubulin were relatively low compared to control ESCs and fibroblasts but appeared similar to the ESC-shEXOSC2 cells that express PPP2R5E constitutively.

**Figure 7.**
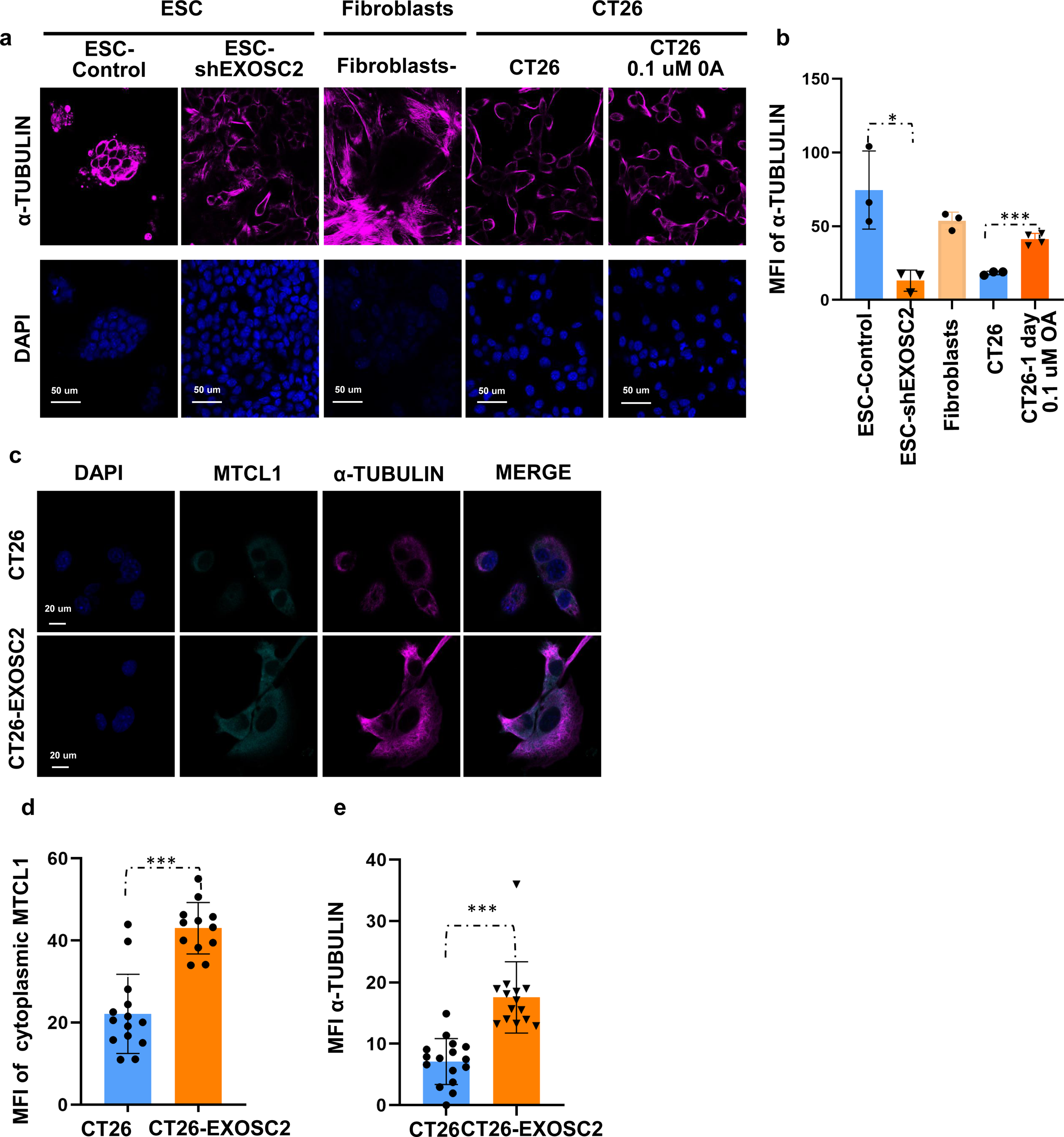
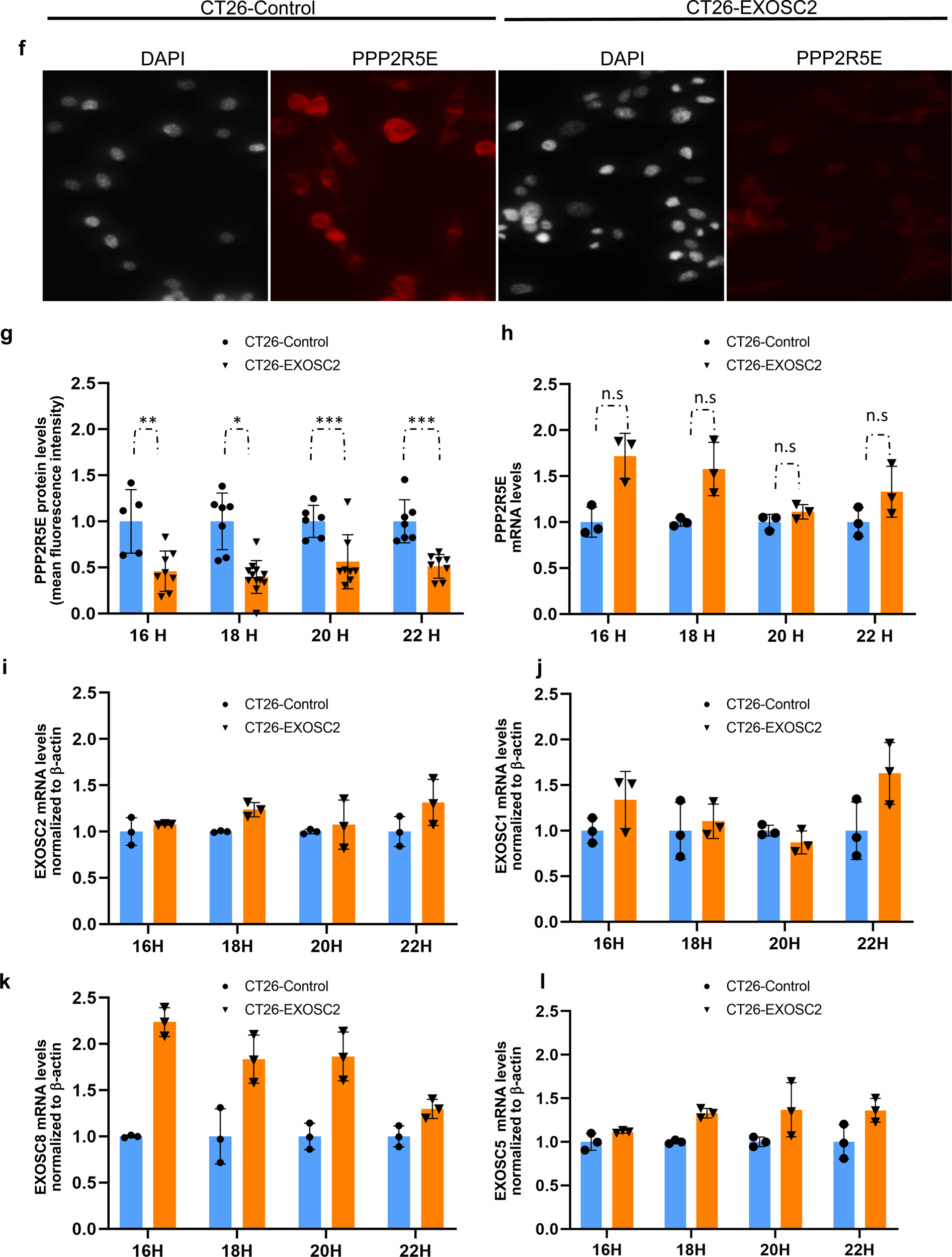
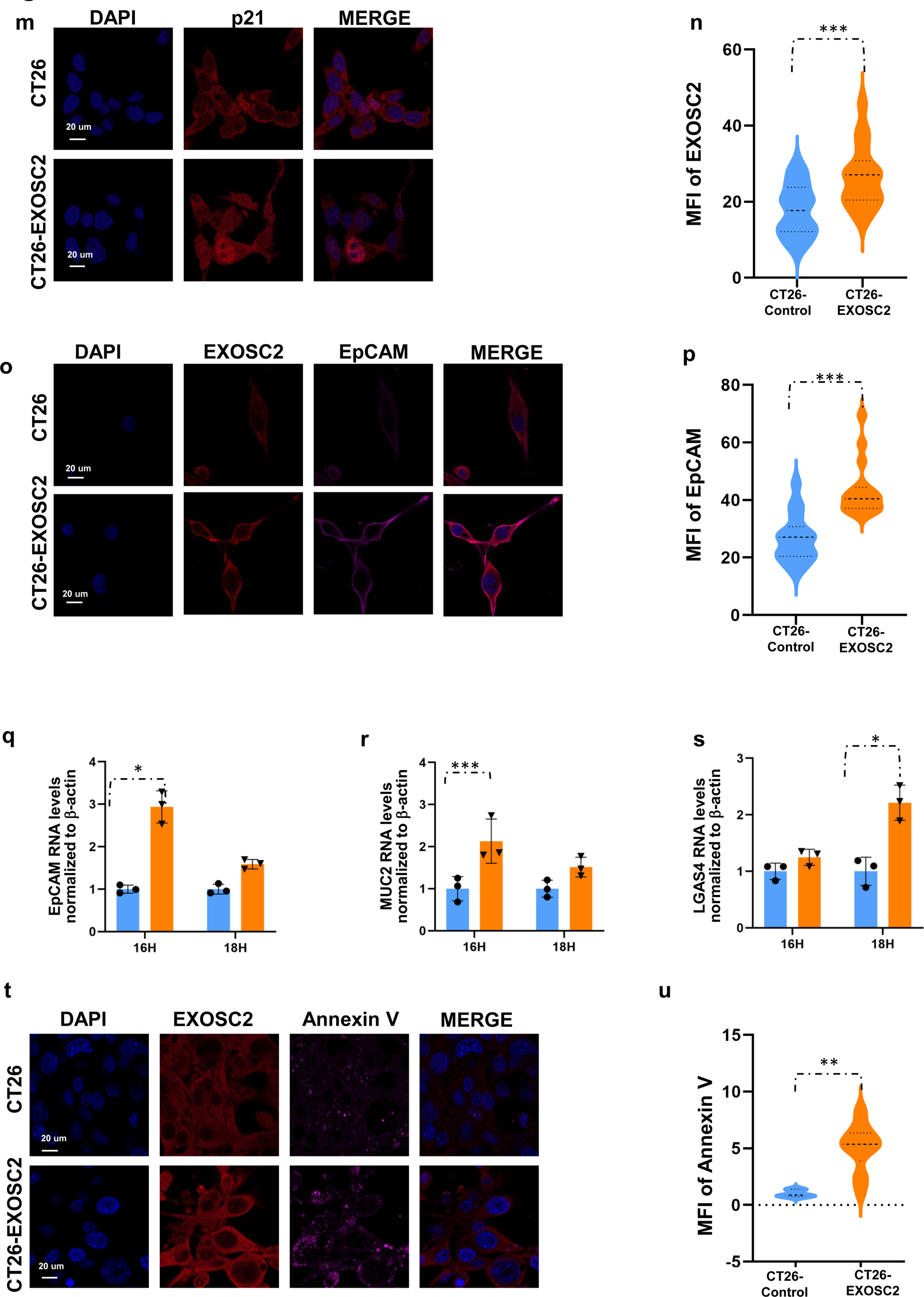
Down-regulation of PPP2R5E through transient EXOSC2 overexpression leads in CT26. (a) Low levels of α-tubulin in CT26 and ESC-sh EXOSC2 lines, compared to CT26 treated with Okadaic acid, PP2A inhibitor, fibroblast control, and ESC control. (b) Quantification of mean fluorescence intensity of α-tubulin in the different cell lines. *Statistical significance by two-sided t-test (p<0.05). ***Statistical significance by two-sided t-test (p<0.0005). (c) MTCL1 and α-tubulin are staining in the CT26 line and CT26 line after EXOSC2 transient overexpression. Nuclei were stained with DAPI. (d, e) Qu antification of mean fluorescence intensity of cytoplasmic MTCL1 and α-tubulin in CT26 and CT26 cell line with transient EXOSC2 overexpression. ***Statistical significance by two -sided t-test (p<0.0005). (f) Immunostaining and (g) quantification for PPP2R5E protein level in CT26 line and CT26 line after EXOSC2 transient overexpression in different time points after the EXOSC2. Nuclei were stained with DAPI. **Statistical significance by two -sided t-test (p<0.005). n.s (not significant). (h) Quantification of PPP2R5E RNA level by qPCR in CT26 line and CT26 line after EXOSC2 transient overexpression in different time points after the EXOSC2 overexpression. Error bars indicate S.E. (i, j, k, l) Quantification of EXOSC2, EXOSC1, EXOSC8, and EXOSC5 RNA level by qPCR in CT26 line and CT26 line after EXOSC2 transient overexpression in different timepoints after the EXOSC2 overexpression. Error bars indicate S.E. (m) p21 staining in the CT26 line and CT26 line after EXOSC2 transient overexpression. Nuclei were stained with DAPI. (n) Quantification of mean fluorescence intensity of EXOSC2 in CT26 and CT26 cell line with transient EXOSC2 overexpression. (o) EXOSC2 and EpCAM staining in CT26 line and CT26 line after EXOSC2 transient overexpression. Nuclei were stained with DAPI. ***Statistical significance by two - sided t-test (p<0.0005). (p) Quantification of mean fluorescence intensity of EpCam in CT26 and CT26 cell line with transient EXOSC2 overexpression. (q) Quantification of EpCAM RNA levels by qPCR in CT26 line and CT26 line after EXOSC2 transient overexpression in different time points after the EXOSC2 overexpression. Error bars indicate SE.*Statistical significance by two-sided t-test (p<0.05). (r) Quantification of MUC2 RNA level by qPCR in CT26 line and CT26 line after EXOSC2 transient overexpression in different time points after the EXOSC2 overexpression. Error bars indicate SE.***Statistical significance by two -sided t-test (p<0.005). (s) Quantification of LGAS4 RNA level by qPCR in CT26 line and CT26 line after EXOSC2 transient overexpression in different time points after the EXOSC2 overexpression. Error bars indicate SE.*Statistical significance by two-sided t-test (p<0.05) (t) Annexin -V staining and (u) quantification of mean fluorescence intensity of Annexin V in CT26 and CT26 cell line with transient EXOSC2 overexpression. **Statistical significance by two-sided t-test (p<0.005)

We previously demonstrated that EXOSC2 contributes to microtubule organization through PPP2R5E-mediated control of PP2A via downstream mediators such as MTCL1. Importantly, we find that levels of alpha-tubulin in the CT26 cell line increase upon treatment with the PP2A inhibitor, okadaic acid, suggesting that PP2A controls microtubule organization in the present model **(Figure 7 a, b)**. To further prove that microtubule polymerization directly results from EXOSC2-mediated control of PPP2R5E, we transiently expressed EXOSC2 in the CT26 cells **(Figure 7 c)**. By immunofluorescence, MTCL1 and alpha-tubulin levels were significantly higher following EXOSC2 expression (**Figure 7 c, d, e**) than parental CT26 cells **(Figure 7 c, d, e)**. This result confirms our hypothesis that the EXOSC2-PPP2R5E, an aging-trait axis, leads to MTCL1 regulation and, thus, the rescue of the tubulin polymerization phenotype.

In our previous work using the embryonic stem cell system, we performed a rescue experiment in the lines that express the hairpins for EXOSC2 by overexpressing EXOSC2 cDNA. Here, we observed that the overexpression of EXOSC2 decreased protein levels of PPP2R5E though the mRNA levels were unchanged. We, therefore, hypothesized that the interaction of EXOSC2 may be directly at the PPP2R5E mRNA to hinder its translation . To determine whether this effect is cell context-dependent and stem cell-specific or if it is also observed in our CT26 colon cancer model, we measured PPP2R5E mRNA levels 24 hours after transient EXOSC2 overexpression **(Figure 7 f)**. Notably, even though the PPP2R5E protein levels were low **(Figure 7 g)**, the corresponding transcript levels were not affected **(Figure 7 h)**, similar to what had been observed with the ESC. Consistent with the hypothesis that the interaction of the RBP EXOSC2 with the PPP2R5E transcript affects its translation is the fact that we did not observe an increase in the levels of RNA exosome components following EXOSC2 transient transfection **(Figure 7 i, j, k, l)** which would mean that the whole complex remains down-regulated after EXOSC2 overexpression. We did observe an increase in the EXOSC2 protein levels as a confirmation that transfection was efficient **(Figure 7 n, t)**. To rationalize this intriguing finding, we reasoned that excess EXOSC2 could bind to PPP2R5E mRNA, but the transcript degradation is still rate limited by the availability of other RNA exosome components (which do not increase following EXOSC2 overexpression). However, EXOSC2 binding to PPP2R5E mRNA alone is enough to inhibit translation. This dataset corroborates our observations in ESCs and shows that in addition to the conserved regulation of PPP2R5E by EXOSC2, the CT26 colon cancer cell line develops a PPP2R5E dependency.

### Down-regulation of PPP2R5E leads to the induction of apoptosis in mouse colon cancer cell lines

As previously described in the embryonic stem cell model that demonstrates the aging traits by EXOSC2, the down-regulation of EXOSC2 led to defects in differentiation which were molecularly manifested by the cytoplasmic localization of p21. Essentially, EXOSC2 knockdown correlated with a significant decrease in nuclear translocation of p21, even after retinoic acid treatment, which typically results in ESC differentiation via p21 translocation under physiologic conditions **(Figure 2 k, l, m)**. Therefore, we examined p21 localization as a differentiation marker in CT26 cancer cell lines after EXOSC2 overexpression. We further reasoned that the restoration of tubulin polymerization that we observed after transient EXOSC2 overexpression in CT26 **(Figure 7 m, n)** might increase p21 nuclear localization as it is the case for p53, which was shown to bind directly to microtubules and be transported to the nucleus by dynein^73^. In CT26 colon cancer cells, p21 is mainly cytoplasmic, reminiscent of the EXOSC2 - downregulated stem cell phenotype **(Figure 2 k, l, m)**. Conversely, CT26 cells transiently expressing EXOSC2 showed an increase in nuclear localization of p21 **(Figure 7 m, n)**. In addition to facilitating p21 translocation, transient expression of EXOSC2 in CT26 cells also led to an increased expression of epithelial differentiation markers such as EpCAM and LGAS4 **(Figure 7 o, p, q, r, s)**^74^. It has been reported that the cytoplasmic accumulation of p21 promotes cell survival by inhibiting cytoplasmically localized apoptosis-related proteins^19^. Indeed, we found that the reduction of cytoplasmic p21 following EXOSC2 expression was associated with cell death in the CT26 cancer cell line, as demonstrated by an increase in Annexin V staining **(Figure 7 t, u)**.

Importantly, we did not observe evidence of cell death after transient EXOSC2 overexpression in control ESC cells **(Figure S7 a, b)**. While transient EXOSC2 expression in the ESC model also leads to a temporal decrease in PPP2R5E protein levels **(Figure S7 c, d)**, there were no significant changes in Annexin V staining, suggesting that cancer cells may have specific vulnerabilities related to PPP2R5E dependency. To further interrogate cellular dependencies on PPP2R5E, we generated cell lines **(Figure 8 a, b, c, d, e, f, g)** that selectively overexpress EXOSC2 upon doxycycline administration (CT26 -EXOSC2). We confirmed that these cell lines significantly down-regulate PPP2R5E one day after doxycycline addition **(Figure 8 a, b)**. Moreover, continuous culture of CT26-EXOSC2 cells in doxycycline media led to a significant decrease in cell proliferation **(Figure 8 c, d, e)**. Interestingly skin fibroblasts containing the same dox-inducible EXOSC2 overexpression vector showed no significant difference in cell growth following doxycycline exposure **(Figure 8 d, g)**. Once again, this data suggests cancer-specific dependencies on PPP2R5E for continued cell proliferation. Moreover, we also generated an additional colon cancer cell line with doxycycline-inducible EXOSC2 overexpression (AKP-EXOSC2) and verified similar defects in cell proliferation in response to the EXOSC2 overexpression **(Figure 8 g)**. Therefore, we conclude that down-regulation of EXOSC2 in cancer cells results in a dependency on PPP2R5E for continued proliferation, suggesting a therapeutic window for targeting PPP2R5E using the concept of conserved aging trait. Indeed the cancer dependency maps show a clear negative gene effect for cancer cells when you knock out PPP2R5E in a set of a thousand cancer cell lines **(Figure S8b)**, further suggesting that cancer cells often rely on this gene for survival. We transiently overexpressed EXOSC2 in the AKP cells and subsequently observed a decrease in PPP2R5E protein **(Figure 8 h, i)** without a change in mRNA transcript levels **(Figure 8 j)**, similar to what we observed in ESC **(Figure 3 t, u, v, w)** and CT26 cells **(Figure 7 f, g, h)**. We also monitored the levels of exosome components after EXOSC2 overexpression. Similarly, we did not see an up-regulation of these components **(Figure 8 k, l, m, n)**, which confirms our above hypothesis that the interaction of PPP2R5E transcript with EXOSC2 is responsible for the change in protein levels.

**Figure 8.**
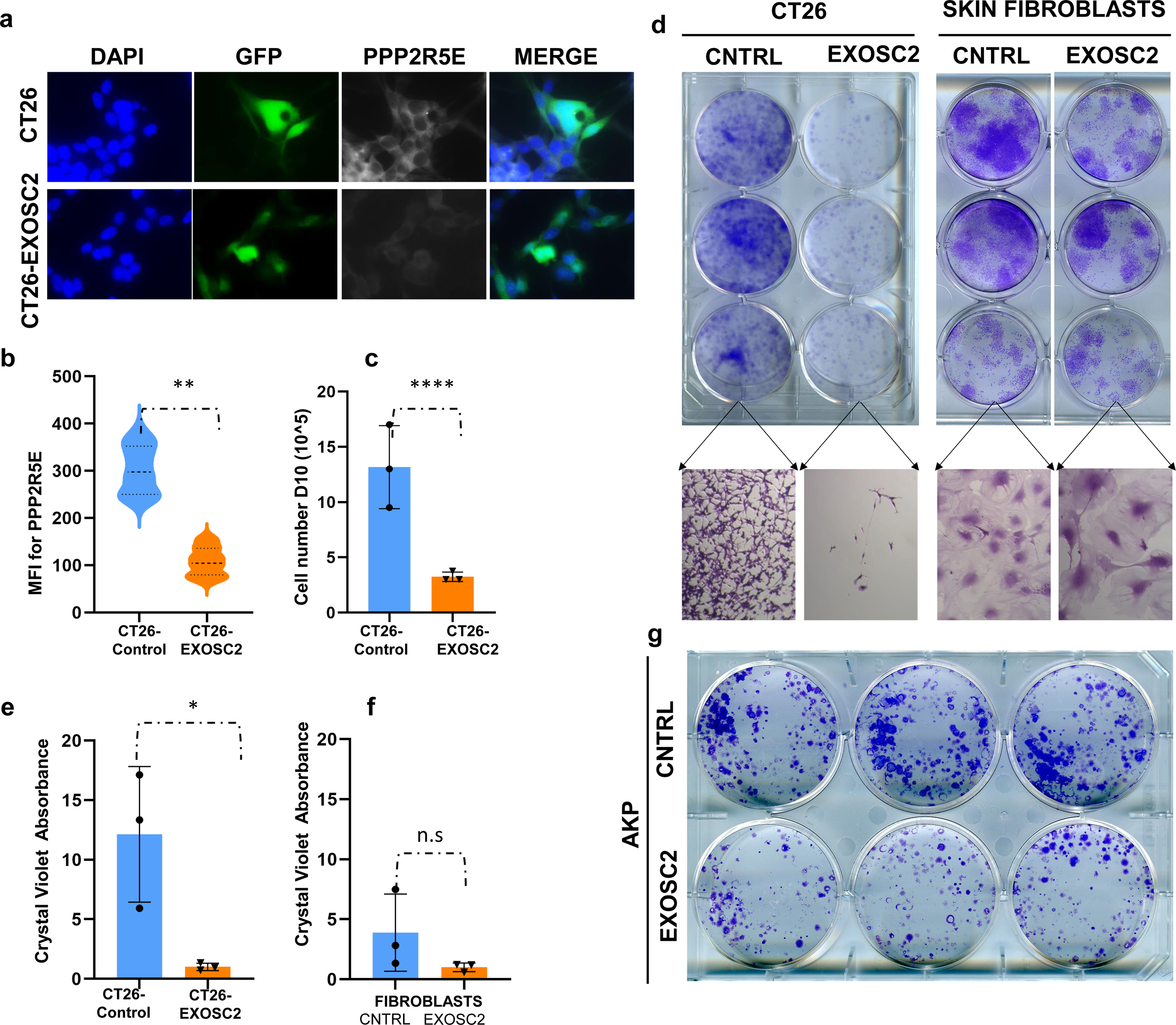
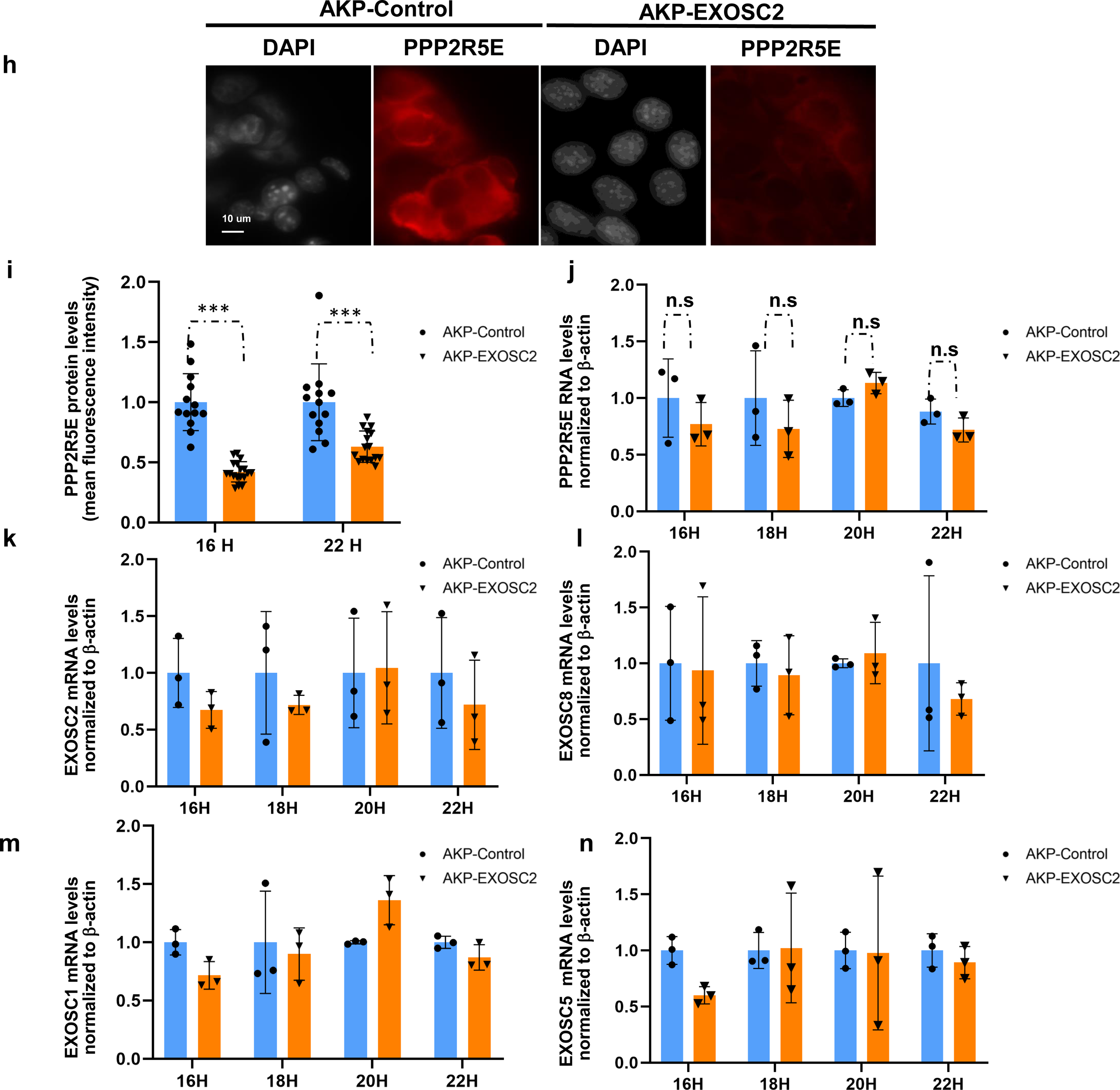
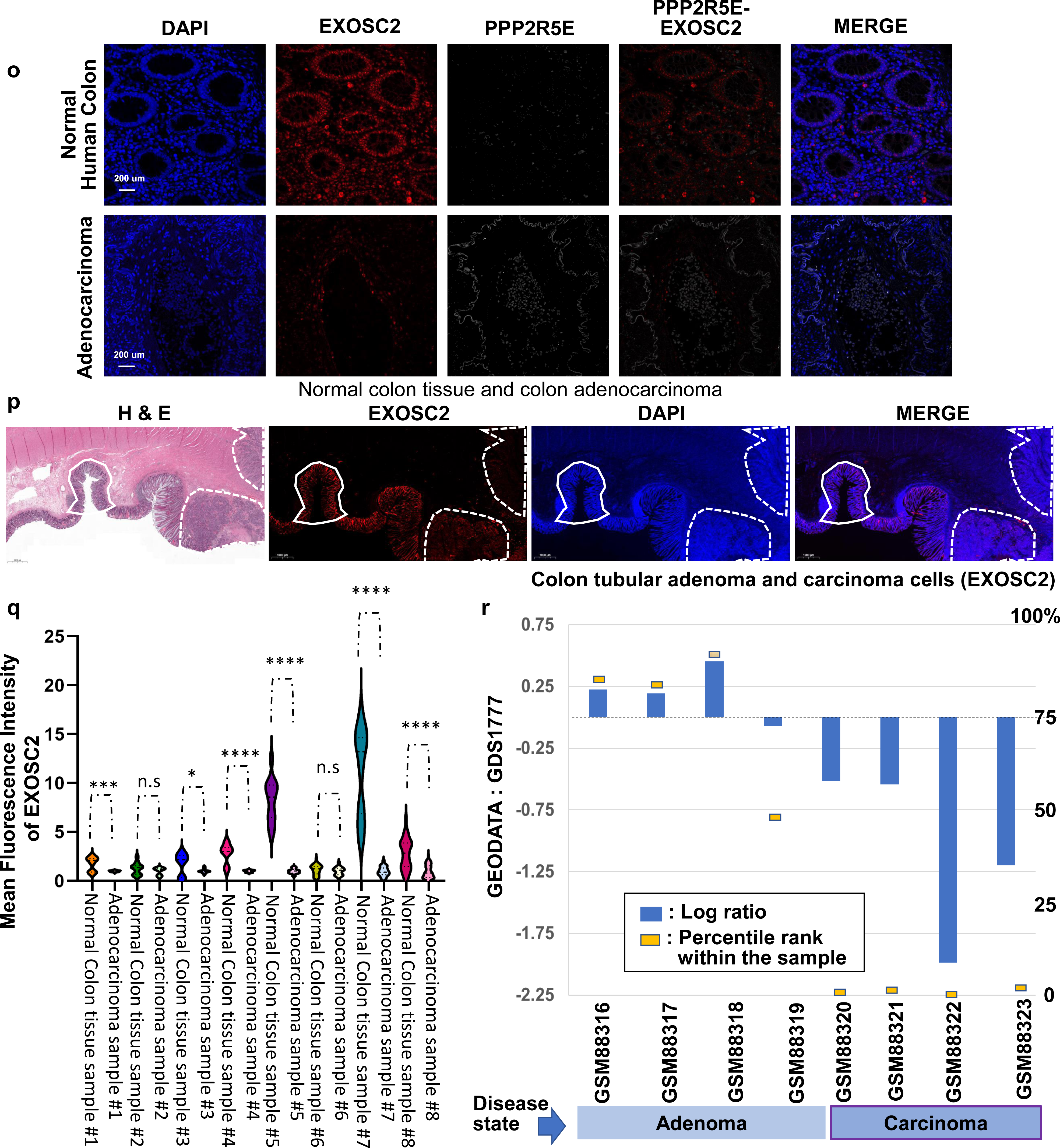
Down-regulation of PPP2R5E through EXOSC2 overexpression leads to a decrease in cell proliferation. (a) PPP2R5E immunostaining in CT26 line and CT26 line after EXOSC2 stable overexpression. (b) Quantification of mean fluorescence intensity of PPP2R5E in the different cell lines. ***Statistical significance by two-sided t-test (p<0.005) (c) Cell number after 12 days of EXOSC2 overexpression in CT26 cell line. (d) Crystal Violet Staining in CT26 and CT26 line after EXOSC2 stable overexpression and skin fibroblasts after EXOSC2 overexpression (e, f) Crystal violet absorbance in CT26 cells and in fibroblasts. (g) Crystal Violet Staining in AKP and AKP cell line after EXOSC2 stable overexpression. (h) Immunostaining and (w) quantification for PPP2R5E protein level in AKP line and AKP line after EXOSC2 transient overexpression in different time points after the EXOSC2 overexpression. Nuclei were stained with DAPI. **Statistical significance by two-sided t-test (p<0.005). n.s (not significant). (j) Quantification of PPP2R5E RNA level by qPCR in AKP line and AKP line after EXOSC2 transient overexpression in different time points after the EXOSC2 overexpression. Error bars indicate S.E. (k, l, m, n) Quantification of EXOSC2, EXOSC1, EXOSC8, and EXOSC5 RNA level by qPCR in AKP line and AKP line after EXOSC2 transient overexpression in different timepoints after the EXOSC2 overexpression. Error bars indicate S.E. (o, p, q) Reduction of EXOSC2 expression during human colorectal cancer progression. (o, p) Immunostaining for EXOSC2 and PPP2R5E in human samples of normal and cancerous isogenic tissue. Straight lines enclose normal colon tissue, and dotted lines colon adenocarcinoma. (q) Quantification of immunostaining for EXOSC2 in normal and cancerous tissue in 8 independent human samples. ***Statistical significance by two-sided t-test (p<0.0005). (r) GEODATA: GDS1777 analysis of colon tubular adenoma and carcinoma cells microdissected from formalin-fixed paraffin-embedded sections of colon tubular adenomas containing focal adenocarcinomas. Results provide insight into the progression of colon cancers from pre-existing adenomas.

### EXOSC2 is down-regulated in human colon adenocarcinoma samples *in vivo*

The hierarchical organization of cells in rapidly renewing tissue, such as the intestinal epithelium in normal physiological conditions, can act as a tumor suppressive mechanism by controlling the number of cell divisions of self -renewing stem cells while reducing proliferation rates on transiently-amplifying cells that are going to terminal differentiation ^75^. We wanted to investigate whether the human colon cancer tissue shows the correlation between the expression levels of EXOSC2 and features of the tumor architecture and progression in the patient population. We sought to understand whether suppression of EXOSC2 and the concomitant increase in PPP2R5E confers an adaptive trait to human colorectal cancer. We wanted to address whether EXOSC2 expression varies during the transition from normal colonic epithelium to pre - invasive adenomas and on to invasive cancer. In this pursuit, we examined the EXOSC2 protein levels in matched human patients’ adjacent healthy colonic epithelium as the negative margin and the adenocarcinoma of the same surgical resection.

We indeed observed that the levels of EXOSC2 are lower in human colon cancer compared with the adjacent normal colonic epithelium, and there is an increase in PPP2R5E levels only in the cancer specimens **(Figure 8 o, p, q)**. Once again, this is consistent with our data from the teratoma formation assay using ESCs. The down-regulation of EXOSC2 in colonic adenocarcinoma was observed in six out of eight independent samples **(Figure 8 o, p, q, r)**. Moreover, we analyzed all available TCGA datasets and noted that PPP2R5E mRNA levels are elevated in numerous cancer types, including colorectal cancer **(Figure S8 a)**. To investigate EXOSC2 expression during the progression from pre-invasive states to invasive cancer, we further mined the publicly available Gene Expression Omnibus (GEO) repository, which contains transcriptional data from colonic tubular adenoma and adenocarcinoma samples which have been micro-dissected from formalin-fixed paraffin-embedded human specimens **(Figure 8 s)**. This result confirms that EXOSC2 levels are lower in invasive cancers than in pre-invasive tubular adenomas. These observations suggest that dysregulation of EXOSC2 as a conserved aging trait is a consistent feature of human colorectal cancer, and levels of EXOSC2 decrease stepwise as normal epithelial cells transition into a pre-malignant state and further progress into invasive cancer. Furthermore, these malignant cells are ultimately dependent upon concomitant increases in PPP2R5E for survival, suggesting potential therapeutic opportunities.

## Discussion

Understanding the mechanisms of cell fate determination is vital for stem cell specification, tissue development, and oncogenesis. Recent work has illuminated a role for the RNA exosome complex during erythropoiesis, a process through which hematopoietic stem cells differentiate into erythrocytes^21, 22^. We showed that the pluripotency factor ZSCAN10 upregulates specific core RNA exosome subunits during reprogramming, and the levels of these core RNA exosome subunits drop upon tissue differentiation ^8, 76^. Recently, mutations in several genes encoding RNA exosome subunits and exosome cofactors have been linked to inherited, tissue-specific diseases^6^. For instance, mutations in the RNA exosome catalytic subunit gene DIS3 have been linked to hematological cancers^77^. However, despite the evident biological and clinical significance, our understanding of how RNA exosome mechanisms contribute to cell fate decisions during tissue development and aging remains limited. In the present study, we report for the first time the aging-dependent subtle down-regulation of EXOSC2 as an aging trait in a series of datasets of aged somatic cells and demonstrate the presence of the significantly conserved aging hallmarks using pluripotent stem cells and cancer cells. We show that the down-regulation of EXOSC2 in stem cell context leads to severe differentiation defects, a known aging hallmark. A recently published study reports that the gradual degradation of a reconstituted chimaeric RNA exosome in a mouse ESC Knock Out for EXOSC2 leads to upregulation of senescence signatures^17^.We compared the signatures that they checked and we also see an upregulation in most of the genes they checked to study senescence in our system (data not shown).

PP2A is a multi-faceted cellular regulator, operating in various signaling mechanisms in any tissue type. Therefore, global PP2A inhibition or over-activation may be detrimental and underscores the importance of PP2A as a significant homeostatic factor. Nevertheless, a substantial part of its physiologic functions remains unclear, presumably due to the highly complex structure and regulation of PP2A. PP2A is comprised of three components: a structural, a regulatory, and a catalytic subunit. Although the prevailing view is that the regulatory subunit provides substrate specificity, how this subunit accomplishes such regulation remains unclear ^65, 78^. Here, we identify that PPP2R5E, the regulatory subunit of Protein Phosphatase 2A (PP2A), is dynamically regulated by EXOSC2 and further show that this interaction controls proliferation in pluripotent stem cells and tissue differentiation. In this study, we also demonstrate that the levels of the RNA binding protein EXOSC2, the regulatory subunit of RNA exosome, are reduced with aging in adult and progenitor cells. Thus, our study shows that the protein levels are crucial for manifesting the functional phenotypes of cell fate decisions by balancing self -renewal, cell proliferation, and tissue differentiation. Therefore, the dysregulation of EXOSC2 may be responsible for functional impairment in adult stem cells as an aging trait. It is known that aging in adult stem cells can impact the cell numbers^79^, and lead to phenotypes related to defective regeneration^26, 80^, self-renewal^81^, or skewed differentiation^82^. We showed that a critical target of the EXOSC2 RNA binding protein is the mRNA of the PPP2R5E subunit of the PP2A. The interaction of EXOSC2 protein with PPP2R5E mRNA impacts its translation. This dynamic regulation is essential in self-renewal and depends, in turn, on the levels of PPP2R5E that impact microtubule polymerization. It has been shown that p53 is associated with cellular microtubules and is transported to the nucleus by dynein^73^. Therefore, one of the possible ways that PPP2R5E overactivation leads to differentiation defects could be via the impairment of microtubule polymerization, leading to defective transport of p21 from the cytoplasm to the nucleus.

We show that the dynamic regulation of PPP2R5E by EXOSC2 is a conserved phenomenon observed during mouse embryonic development where there is a continuum of cell fate decisions. The PPP2R5E regulation by EXOSC2 is also observed in adult tissues, and its importance is further highlighted after focal irradiation in the mouse colon. It is well known that an imbalance towards cell lineage commitment at the expense of self -renewal or vice versa can have detrimental effects on tissue homeostasis. We propose that this misregulation might take place when EXOSC2 levels drop below a threshold during aging.

We also demonstrate key regulators of PPP2R5E mRNA translation and key downstream targets in the context of cell fate decisions. Particularly during self-renewal, it is vital to regulate the levels of the PPP2R5E and the mRNA turnover of this gene to balance tissue differentiation and proliferation, and this balance is often affected by aging. We show that this interaction of EXOSC2 with PPP2R5E mRNA is sufficient to block the PPP2R5E mRNA translation. We then provide evidence that the levels of EXOSC2 are vital for maintaining cell plasticity. Reducing the levels of the exosome complex even by half radically alters the impact on PPP2R5E levels, which again suggests potential therapeutic vulnerability but also demonstrates the fact that the age-related reduction of EXOSC2 in stem cells can potentially lead to imbalances in tissue homeostasis if the levels drop from a developmentally required threshold. We also demonstrate that the high level of the PPP2R5E by reduced EXOSC2 expression leads to overactivation of the PP2A phosphatase and consequently blocks the phosphorylation of γ-H2AX^6, 56^ at Serine 139 ^83^ after DNA double-strand breaks. In eukaryotic cells, the phosphorylation of H2AX is critical for the DNA damage response pathway to maintain the genome by including the DNA repair and release of cells from the growth arrest ^58^. Indeed, we observe severe induction of genomic instability and aneuploidy with knockdown of EXOSC2 (Figure S3b), which has a significant clinical value in maintaining the genomic integrity of stem cells and preventing the induction of additional genomic aberrations during chemotherapy treatment.

In this study, first, we show the dynamic regulation of PPP2R5E by EXOSC2 as a conserved event in different cellular contexts and then present the mechanism of how the constitutively active PP2A can become a dependency and is associated with the RNA exosome deficiency phenotypes, such as aging, poor tissue differentiation, and uncontrolled cellular proliferation. We perform co-staining for EXOSC2 and PPP2R5E in adult organs such as the intestine, liver, and kidney and observed an inverse correlation between EXOSC2 expression and PPP2R5E protein levels, confirming our hypothesis that the dynamic regulation of PPP2R5E by EXOSC2 is highly conserved. Therefore, we hypothesize that it is likely that EXOSC2 regulates PPP2R5E expression in a diverse array of tissue types and biological contexts. Intestinal epithelium represents a suitable model to investigate the developmental and oncologic roles of EXOSC2 for several reasons. First, the intestinal epithelium consists of a proliferation - differentiation gradient, including stem-like progenitor cells in the crypt compartment and more differentiated progeny in the villus. Therefore, the expression and function of EXOSC2 can be explored over a range of cellular phenotypes ^60^. Secondly, emerging data suggests the role of RNA-binding proteins such as LIN28, IGF2BPs/IMPs, Musashi, HuR, Mex3A, CELF1, and RBM3 in intestinal epithelial homeostasis and malignant transformation ^84^. Finally, colon cancer remains a primary cause of cancer-related death worldwide. There is an urgent clinical need to identify new molecular mechanisms underlying the tumorigenic process for early detection and treatment using the concept of the conserved aging traits ^85^.

In cancer, the expression of PP2A is altered through several mechanisms, including somatic mutation, phosphorylation and/or methylation of the C-terminal tail of the catalytic subunit, and the modified expression of cellular endogenous PP2A inhibitors such as CIP2A ^86–88^. Cancer cells generally deregulate PP2A function by overexpressing accessory proteins that mediate posttranslational modifications or block enzymatic activity ^89^. Our study proposes a novel mechanism by which cancer cells deregulate PP2A activity through the down-regulation of EXOSC2 RNA binding protein. EXOSC2 is the non-catalytic component of RNA exosome which has 3’-5’ exoribonuclease activity and participates in many cellular processing and degradation events. EXOSC2 has an S1 RNA binding domain. S1 domains are found in several exoribonucleases and translation initiation factors^90, 91^. S1 domains interact with both ssRNA and dsRNA in the context of the RNA binding channel of exoribonuclease. Therefore, we hypothesize that in colon cancer, this interaction is exploited to lead to hyperactivation of PP2A phosphatase, which gives malignant cells a proliferative advantage compared to normal cells in the tumor microenvironment. This means EXOSC2 is an essential catalyst whose relative increases or decreases in this context may become detrimental to cells. RBPs bind to their target RNA, form ribonucleoprotein complexes, and regulate gene expression post-transcriptionally in many ways. Together, our above studies show that EXOSC2 is a novel addition to previously published RBPs such as LIN28, Musashi, HuR, Mex3A, CELF1, and RBM3 ^92, 93^ that are relevant in the context of intestinal tissue homeostasis and intestinal tumorigenesis, and are derived from the concept of the conserved aging traits. PP2A complex targets pATM and γH2AX and deactivates DNA damage response when the DNA repair is completed ^56^. Consequently, PP2A inhibition has been suggested as a potential cancer treatment, and the knockdown of PP2A in several in vitro cancer models resulted in elevated γH2AX and radio sensitivity ^94–97^. On the other hand, PP2A is a known tumor suppressor; some of its substrates are c-Myc, and ERK ^94, 98, 99^, and in many cases, cancer cells develop a mechanism to evade PP2A-mediated tumor suppression.

The dysregulation of phosphorylation-dependent signaling events is a hallmark of human disease, and targeting these aberrant phosphorylation processes has proven necessary for disease control. Our study shows that increased PP2A activity via up-regulation of PPP2R5E can promote tumor progression and become essential for cancer cell survival. RNAseq data from thirty-two cancer datasets in the TCGA show that PPP2R5E is expressed at very high levels in many cancer types **(Figure S8a)** and cancer cell lines show a negative gene effect when you knock out PPP2R5E **(Figure S8b)** suggesting that the cancer cell lines rely on PPP2R5E for survival. On the contrary, in the absence of tumor suppressor p53, the gene effect is positive because the cells proliferate faster **(Figure S8b)**. Different regulatory subunits confer differential specificity to downstream targets. It has been shown that small molecule inhibitors selectively stabilize the tripartite phosphatase complex that contains PPP2R5A and dephosphorylate of c-Myc, which provides a therapeutic potential in leukemias ^100^. In our system, we observe the opposite situation where PPP2R5E is aberrantly expressed, and the cells cycle faster, as assessed by Ki67 staining and proliferation assays. This disparity can be attributed to the difference in the cancer cell lines and regulatory subunits examined.

Overall, this study shows how a conserved regulatory mechanism for the subunit PPP2R5E is deregulated in colon cancer and opens the road to therapeutic intervention to manage the phosphatase levels. These results will also pave the way for additional avenues toward studying the conserved and significant roles of the aging-traits in a variety of aging-related disease models, including cancer models which show a conserved regulation of the EXOSC2/PPP2R5E axis **(Figure 4e and Figure S5 c).**

## Data availability

The authors declare that all data supporting the findings of this study are available within the paper and its supplementary information files.

## Acknowledgments

The authors wish to acknowledge assistance from the Bioinformatics Core which is funded in part through the NIH/NCI Cancer Center Support Grant P30 CA008748, from the Molecular Cytology Core Facility, which is funded in part by the Core Grant (P30 CA008748), from the Integrated Genomics Operation Core, funded by the NCI Cancer Center Support Grant (CCSG, P30 CA08748), Cycle for Survival, and the Marie-Josée and Henry R. Kravis Center for Molecular Oncology, Molecular Cytogenetics core. Katia Manova for consultation. H.L. is supported by NIH (AG056318, AG61796, and CA208517), the Glenn Foundation for Medical Research, the Mayo Clinic Center for Biomedical Discovery, the Mayo Clinic Cancer Center, and the Margaret T. Grohne Cancer Immunology and Immunotherapy Program. PBR is partly supported by a K12 Paul Calebresi Career Development Award for Clinical Oncology (K12 CA184746).

## Material and methods

### Cell culture

Mouse ESC and iPSC were cultured in ESC media containing 20 % FBS and 1,000 U/ml of LIF (ESGRO^®^ Leukemia Inhibitory Factor [LIF], 1 million units/1 mL). Mouse ESC were generated and their pluripotency was tested as reported in our previous publication ^1^.

### Generation of embryonic stem cells with ESC-shEXOSC2

ESCs were infected with a set of shRNA viruses for EXOSC2 (2 GIPZ Lentiviral shRNA vectors for EXOSC2 from GE DHARMACON: RMM4431-200370629, RMM4431-20033273). Clones were selected with puromycin.

### Lentivirus production

293T cells were seeded overnight at 5×10^6^ cells per 150-mm dish with DMEM supplemented with 10% FBS and penicillin/streptomycin. The cells were transfected with GIPZ Lentiviral shRNA vectors for EXOSC2 with calcium phosphate cell transfection, as previously described ^2^. At 48 hours after transfection, the medium containing the lentivirus was collected and the cellular debris was removed with centrifugation. The supernatant was filtered through a 0.45 -μm filter, and the lentivirus was pelleted with ultracentrifugation at 33,000 rpm in a 45Ti rotor (Beckman) for 90 min at 4°C. The lentivirus particles were resuspended in DMEM medium and stored at -80°C.

### Teratoma analysis of mouse and human iPSC

Mouse teratomas were assessed by injecting 10^6^ undifferentiated cells into the subcutaneous tissue above the rear haunch of Rag2/γC immunodeficient mice (Taconic), and teratoma formation was monitored for 3 months post-injection. Collected tumors were fixed in 10% formalin solution and processed for hematoxylin and eosin (H/E) staining by the Molecular Cytology facility of Memorial Sloan Kettering Cancer Center.

### Cytogenetic analysis and copy number profiling analysis

Cytogenetic analysis was performed by metaphase chromosome preparation, G-band karyotyping, Metaphase chromosome preparation, and the G-band karyotyping were performed by the Molecular Cytogenetics Core Facility of Memorial Sloan Kettering Cancer Center. Copy number profiling analysis was performed according to the published protocol.^3^

### Growth rate

10,000 cells per line were plated in triplicate in 24 -well dishes, and the numbers of cells were counted every 24 hours for 3 days.

### Drug treatments and irradiation

Phleomycin (Sigma) was added at 30 μg/ml for 2 hours. Cells were processed for analysis 30 min after phleomycin treatment unless indicated otherwise. After a 30 -min recovery in ESC media, the cells were collected and processed for following experiments for H2AX immunostaining.

Ceramide activates protein phosphatase PP2A, (Tocris Bioscience, Cat#: 0744/10) and ESC were treated (0.1 uM Ceramide, 2 days) and processed for immunostaining for α-tubulin.

Okadaic Acid is a potent inhibitor of protein phosphatase 2A (IC_50_ = 0.1-1 nM, Okadaic acid, ACROS Organics™, Cat#. AC328370001) treatment and ESCshEXOSC2 and CT26 cell lines were treated with OA (0.5 nM, 2 days and 0.1 nM, 2 days respectively) and were processed for immunostaining for α-tubulin.

Fostriecin sodium salt is a potent inhibitor of protein phosphatase types 2A (IC_50_ values of 1.5 nM, Tocris Bioscience: Cat#: 87860-39-7) and ESCshEXOSC2 were treated with FS (2 nM, 2 days) and were processed for immunostaining for α-tubulin.

### Transfection of EXOSC2-GFP

To introduce DNA (Cat #: MG204028, Origene, pCMV6-AC-GFP, mammalian vector with C-terminal tGFP tag or GFP control only) we used the CalPhos™ Mammalian Transfection Kit (Cat # 631312, TAKARA bio).

### Stable expression of EXOSC2-GFP

To stably express EXOSC2 we introduced EXOSC2 mouse cDNA from (Cat #: MG204028, Origene, pCMV6-AC-GFP) in a *TetON* lentinviral vector ^4^. Briefly on day 1we infect cells, CT26 AKP and skin fibro, with Tet-on lenti EXOSC2 and Tet-on lenti GFP cntr. On day 2 we add doxyxycline (2ug/ml), on day 3 we sort GFP+ cells and plate GFP+ cells in triplicates. On day 7 we replate cells in different concentrations and stain for crystal violet assay at D12 (Crystal violet Assay Kit (Cell viability) (ab232855).

### Immunohistochemistry staining

Immunofluorescence analyses were performed as previously described. ^5^ Briefly, cells were fixed in 3.7% formaldehyde for 20 min at room temperature and washed with PBS. Samples were then permeabilized with 0.1 Triton X-100 in PBS for 20 min and blocked for 1 h with 3% BSA in PBS-T, and primary antibodies were incubated for 2 h at room temperature or overnight at 4°C. Anti-H2AX was purchased from Millipore (05-636),

For the immunostaining of microtubules, the fixation process was performed as previously described^6^.

α-tubulin **(clone DM1A)** from Millipore-Sigma (CP06),

rat anti tyrosinated alpha-tubulin (clone YL 1/2) from Abcam cat #6160

p21 Mouse, Unlabeled, Clone: SXM30, BD, Mouse Monoclonal Antibody, Manufacturer: BD Biosciences 556431

SOGA2 (MTCL1) from mybiosource.com (MBS8530859), Anti-EpCAM antibody (ab71916),

Primary antibodies were used at 1:500 dilution. Alexa 568 -conjugated goat anti-mouse IgM (A-21124) and Alexa 633-conjugated goat anti-rabbit IgG (A-21072) were from Molecular Probes.

Secondary antibodies were used at 1:1000 dilution. The nuclei were stained with 4’,6 -diamidino-2-phenylindole (DAPI, Sigma), Alexa 633-conjugated goat anti-rat IgG (Product # A-21094)

### Protocols for automated IF staining

The immunofluorescent staining was performed at Molecular Cytology Core Facility of Memorial Sloan Kettering Cancer Center using the Discovery XT processor (Ventana Medical Systems).

The tissue sections were deparaffinized with EZPrep buffer (Ventana Medical Systems), antigen retrieval was performed with CC1 buffer (Ventana Medical Systems). Sections were blocked for 30 minutes with Background Buster solution (Innovex), followed by avidin-biotin blocking for 8 minutes (Ventana Medical Systems)

For single staining: First, sections were incubated with Ki67 (AB15580, Abcam, 1 ug/ml), or H2AX (AB11174, Abcam, 0.1 ug/ml) or GFP (AB13970, Abcam,2 ug/ml) for 5 hours, followed by 60 minutes incubation with biotinylated goat anti-rabbit IgG (Vector labs, cat#PK6101) at 1:200 dilution. The detection was performed with Streptavidin-HRP D (part of DABMap kit, Ventana Medical Systems), followed by incubation with Tyramide Alexa 488 (Invitrogen, cat# B40953) prepared according to manufacturer instruction with predetermined dilutions.

Multiplex immunofluorescent stainings were performed as previously described ^7^

### PPPR5E/EXOSC2

#### PPPR5E

First, sections were incubated with anti-PPPR5E (Thermo, cat# PA5-52243, 0.0005ug/ml) for 5 hours, followed by 60 minutes incubation with biotinylated goat anti-rabbit IgG (Vector labs, cat#PK6101) at 1:200 dilution. The detection was performed with Streptavidin -HRP D (part of DABMap kit, Ventana Medical Systems), followed by incubation with Tyramide Alexa 488 (Invitrogen, cat# B40953) prepared according to manufacturer instruction with predetermined dilutions.

#### EXOSC2

Next, sections were incubated with anti-EXOSC2 (Bethyl, cat# A303-886A, 1ug/ml) for 5 hours, followed by 60 minutes incubation with biotinylated goat anti-rabbit IgG (Vector labs, cat#PK6101) at 1:200 dilution. The detection was performed with Streptavidin -HRP D (part of DABMap kit, Ventana Medical Systems), followed by incubation with Tyramide CF594 (Biotum, cat# 92174) prepared according to manufacturer instruction with predetermined dilutions.

After staining, slides were counterstained with DAPI (Sigma Aldrich, cat# D9542, 5 ug/ml) for 10 min and coverslipped with Mowiol.

Imaging + Quantitation Method

Slides were scanned with Pannoramic 250 Flash scanner (3DHistech, Hungary) using a 20x/0.8NA objective lens. Regions of interest around the tissues were then drawn and exported as .tif files from these scans using Caseviewer (3DHistech, Hungary.) These images were then analyzed using ImageJ/FIJI (NIH, USA) where a median filter, thresholding, and water-shedding were used to segment the nuclei in the DAPI channel and get a cell count and area. Next, the TRITC and FITC channels were thresholded, and the total intensity and area of each signal were measured and recorded.

#### Fish-IF

The FISH/ immunofluorescence double detection was performed at the Molecular Cytology Core Facility of Memorial Sloan Kettering Cancer Center.

Cell monolayers were fixed in 4% PFA for 30 min at RT and stained for FISH using BaseScope LS Reagent kit-Red (ACD, cat#323600) on Leica Bond RX instrument following routine manufacturer protocol and IF on Ventana Discovery XT. BaseScope LS-Mm-PPIB-3zz (ACD, cat#701078) and BaseScope LS Mm-Ppp2r5e-2zz-st (ACD, cat# 824928) probes were used with hybridization at 42^0^ C for 2 hours. The sections were pre-treated with Protease III (ACD, cat#322102) for 5 min at RT. After FISH staining, the sections were post-fixed in 4% Paraformaldehyde for 30 min, and slides were processed for immunofluorescence staining with anti-EXOSC2 (Bethyl, cat# A303-886A, 1ug/ml). Tyramide Alexa Fluor 488 (Life Technologies, cat# B40953) was used according to manufacturer instructions for IF. After staining the sections counterstained with 10ug/ml DAPI/PBS for 10 min and mounted with Mowiol mounting media. The slides were digitized using Pannoramic Flash 250 (3DHistech, Budapest, Hungary) with Zeiss 40x/0.95NA air objective. The scans were viewed and exported to tiff images using CaseViewer (3DHistech).

### Probe information

Basescope RNA probe:BA-Mm-Ppp2r5e-2zz-st targeting NM_012024.3 (*Advanced Cell Diagnostics, Bio-Techne Brand), cat number 824928*

RNA scope, Probe NPR-0005240, cat number: 842638, *Advanced Cell Diagnostics, Bio-Techne Brand)*

Species of Target Gene: Mouse(Mus musculus); Species of Sample: Mouse(Mus musculus); Gene Symbol: Ppp2r5e. A bioinformatics service fee for probe design, for a standard custom probe, is typically designed to contain about 40 oligos or 20 signal-generating oligo -pairs. Target sequence spans approximately 1000 bases and the standard design aims to cover as many transcript variants as possible

### Quantitative real time-PCR (Q-PCR) analysis

The expression levels of various genes (ZSCAN10, OCT4, GPX2, EXOSC1/2/5, and β-actin) were quantified by Q-PCR. Total RNA (1 μg) was reverse transcribed in a volume of 20 μl using the M-MuLV Reverse Transcriptase system (New England Biolabs), and the resulting cDNA was diluted into a total volume of 200 μl. 10 μl of this synthesized cDNA solution was used for analysis. For pluripotency genes, each reaction was performed in a 25 μl volume using the Power SYBR Green PCR Mastermix (Applied Biosystems). The conditions were programmed as follows: initial denaturation at 95°C for 10 min followed by 40 cycles of 30 sec at 95°C, 1 min at 55°C, and 1 min at 72°C; then 1 min at 95°C, 30 s at 55°C, and 30 sec at 95°C. All of the samples were duplicated, and the PCR reaction was performed using a Mx3005P reader (Stratagene), which can detect the number of synthesized signals during each PCR cycle. The relative amounts of the mRNAs were determined using the MxPro program (Stratagene). The amount of PCR product was normalized to a percentage of the expression level of β - actin. The PCR products were also evaluated on 1.2% agarose gels after staining with ethidium bromide. The primers used to amplify the cDNA were the following:

β-ACTIN-For: TCGTGGGTGACATCAAAGAGA and

β-ACTIN-Rev: GAACCGCTCGTTGCCAATAGT,

EXOSC2-For: CCCCAAGGAGCATCTGACAA and

EXOSC2-Rev: CCAACCCACCATTACCTCCC

PPP2R5E-For: GCATTGGAATCCGGCCATTG

PPP2R5E-Rev: GTGGCTGTCAGCTCATCAAAC

### Immunoblot analysis

Treated and untreated cells (1 x 10^5^ cells) were collected 30 min after the 2-hour phleomycin treatment (30 μg/ml). To harvest protein, 100-200 ml RIPA buffer (50 mM Tris-HCl [pH 7.4], 150 mM NaCl, 1% NP40, 0.25% Na-deoxycholate, 1 mM PMSF, protease inhibitor cocktail, and phosphatase inhibitor cocktail) was added to floating cell pellets and the remaining adherent cells. The samples were incubated on ice (10 min) and centrifuged (14,000 g, 10 min, 4°C). Protein concentrations were determined using a BCA protein assay kit (Pierce). Samples were adjusted to the same concentration with RIPA buffer (3000 μg/ml) and were combined with Laemmli Sample Buffer (BioRad) and β-Mercaptoethanol (Sigma) then heated at 95°C for 5 min and loaded onto a 4 -15% Mini Protean TGX SDS-PAGE gel (BioRad). Samples on the SDS-PAGE gel were transferred to a 0.2-mm PVDF membrane at 100 V for 1 h, using a wet electro-transfer method (0.2 M glycine, 25 mM Tris, and 20% methanol). The membrane was blocked with 5% BSA in PBS-T (1 h at 4°C), followed by incubation with primary antibodies anti-beta actin (Cell Signaling, #4967) (1:5000) in blocking solution (5% BSA in phosphate-buffered saline containing Tween-20 [1:1000] PBS-T, overnight at 4°C), βIII-Phospho S172 tubulin antibody from Abcam (ab76286), PPP2R5E antibody from NOVUS biological (NB100-845), α-tubulin **(clone DM1A)** from Millipore-Sigma (CP06), anti-beta tubulin (clone TUB 2.1) from sigma cat #T4026

After primary antibody incubation, membranes were washed three times in PBS-T) before the addition of secondary antibody labeled with peroxidase. Secondary antibodies were from Cell Signaling (1:10,000).

### Immunoprecipitation for MTCL1 and Co-Immunoprecipitation for PPP2R5E

For lysate preparation we collected 293T cells by scraping them from one confluent 15 centimeter culture dish (MTCL1 IP and control IgG IP). We lysed them on the dish with 500 ul of 1% TNESV buffer (50 mM Tris-HCl pH 7.5, 2 mM EDTA, 100 mM NaCl, 1% NP-40, H2O) with Protease/Phosphatase Inhibitor Cocktail (Cell signaling, #5872) and RNAse inhibitor (Applied Biosystems, N8080119), rotated in the cold room for 30 min and centrifuged the lysate at 4 degrees, maximum speed for 20 min and collected supernatant. At this step we removed 55 ul to use as input sample during western blot.

For beads preparation, we have added 50 μl of A/G agarose beads (Pierce™ Protein A/G Agarose Catalog number: 20421) per experiment to a fresh Eppendorf and washed beads twice with lysis buffer (without protease inhibitor, TNESV). This and all following washing steps of the beads were performed with 900 μl of the respective buffer. We resuspended the beads in 100 μl lysis buffer with 5 μg antibody per experiment (Rabbit IgG Innovative Research, IRBIGGAP and Rabbit-anti KIAA0802 A302-786A antibodies) and rotated tubes for 30–60 min at room temperature, washed 3 times with 900 μl lysis buffer. Then we added the lysate to the beads and rotated o/n.

Next day we centrifuged and discarded the supernatant and washed twice with high -salt wash (50 mM Tris–HCl, pH 7.4, 1 M NaCl, 1 mM EDTA, 1% IGEPAL CA-630, 0.1% SDS, 0.5% sodium deoxycholate) (rotate the second wash for at least 1 min at 4 °C) and twice with 1% TNESV. To elute protein from the beads, we have added 50 sample buffer (Laemmli buffer). We boiled at 95°C for 5 min, spinned at max speed for 1 minute, moved the supernatant to a clean tube and discarded the beads. We probed for desired proteins MTCL1 (Bethyl Laboratories Rabbit-anti KIAA0802 A302-786A) and PPP2R5E (Invitrogen, PA5_52243) following normal western blot.

### Transcriptome sequencing

After RiboGreen quantification and quality control by Agilent BioAnalyzer, 500ng of total RNA underwent polyA selection and TruSeq library preparation according to instructions provided by Illumina (TruSeq Stranded mRNA LT Kit, catalog # RS-122-2102), with 8 cycles of PCR. Samples were barcoded and run on a HiSeq 4000 in a 50bp/50bp paired**-**end run, using the HiSeq 3000/4000 SBS Kit (Illumina). An average of 31 million paired reads was generated per sample and the percent of mRNA bases averaged 71%.

### Transcriptome sequencing analysis

The analysis was performed by the Bioinformatics core at MSKCC. Briefly, The output data (FASTQ files) are mapped to reference genome mm10 using STAR 2.5.0a^8^ that maps reads genomically and resolves reads across splice junctions. We use the 2 pass mapping method outlined in ^9^ in which the reads are mapped twice. The first mapping pass uses a list of known annotated junctions from Ensemble. Novel junctions found in the first pass are then added to the known junctions and a second mapping pass is done (on the second pass the RemoveNoncanoncial flag is used). After mapping, we post-process the output SAM files using the PICARD tools to add read groups, AddOrReplaceReadGroups which in additional sorts the file and converts it to the compressed BAM format. We then compute the expression count matrix from the mapped reads using HTSeq (www-huber.embl.de/users/anders/HTSeq) against reference genome mm10. The raw count matrix generated by HTSeq is then be processed using the R/Bioconductor package DESeq (www-huber.embl.de/users/anders/DESeq) which is used to both normalize the full dataset and analyze differential expression between sample groups.

### Proteomics

Cell pellets were lysed with 500 μL buffer containing 8M urea and 200mM EPPS (pH at 8.5) with protease inhibitor (Roche) and phosphatase inhibitor cocktails 2 and 3 (Sigma). Samples were then sonicated (Diagenode Bioruptor) for 3 cycles [1 min ON/1 min OFF]. Samples were centrifuged at 4°C, 14,000 g’s for 10 min, and supernatant extracted. The Pierce bicinchoninic acid (BCA) protein concentration assay was used for determining protein concentration. Protein disulfide bonds were reduced with 5 mM tris (2-carboxyethyl) phosphine (room temperature, 15 min), then alkylated with 10 mM iodoacetamide (RT, 30 min, dark). The reaction was quenched with 10 mM dithiothreitol (RT, 15 min). Aliquots of 200ug were taken for each sample and diluted to approximately 100 μL with lysis buffer. Samples were subject to chloroform/methanol precipitation as previously described ^1^. Pellets were reconstituted in 100 μL of 200mM EPPS buffer and digested with Lys-C (1:50 enzyme-to-protein ratio) and incubated at 37°C overnight. Trypsin was then added (1:50 enzyme-to-protein ratio) and digested for 37°C for 7 hours. Anhydrous acetonitrile was added to make a final volume of 30% ACN.

Samples were TMT-labeled as described^10^. Briefly, samples were TMT-tagged by adding 20 μL (20ug/μL) TMT reagents for each respective sample and incubated for 1 hr (RT). A ratio check was performed by taking a 2 μL aliquot from each sample and desalted it by the StageTip method ^11^. TMT-tags were then quenched with hydroxylamine to a final concentration of 0.3% for 15 min (RT). Samples were then pooled 1:1 based on the ratio check and vacuum-centrifuged to dryness. Dried samples were reconstituted in 1mL of 3% ACN/1% TFA, desalted using a 50 mg tC18 SepPak (Waters), and lyophilized overnight.

Phosphopeptides were enriched using the Thermo High-Select Fe-NTA Phosphopeptide Enrichment Kit (Cat. No.: A32992). Unbound peptides and washes were saved for further analysis of the non - phosphorylated (phospho-depleted) peptides. The phosphopeptide elute was vacuum-centrifuged to dryness and reconstituted in 100 μL of 1% ACN/25 mM ammonium bicarbonate (ABC). A StageTip was constructed by placing two plugs with a narrow bore syringe of a C18 disk (3M Empore Solid Phase Extraction Disk, #2315) into a 200 μL tip (VWR, Cat. #89079 -458). StageTips were conditioned with 100 μL of methanol, 70% ACN/25 mM ABC, then 1% ACN/25 mM ABC. Phospho-enriched sample was loaded onto the StageTip and eluted into 7 fractions of 1, 3, 5, 8, 10, 12, and 70% ACN/25 mM ABC with 100 μL each. Fractions were immediately dried down by vacuum-centrifugation and reconstituted in 0.1% formic acid (FA) for LC-MS/MS.

Phospho-depleted peptides were centrifuged to dryness and reconstituted in 1mL of 1% ACN/25 mM ABC. Peptides were fractionated into 48 fractions. Briefly, an Ultimate 3000 HPLC (Dionex) coupled to an Ultimate 3000 Fraction Collector using a Waters XBridge BEH130 C18 column (3.5 um 4.6 x 250 mm) was operated at 1mL/min. Buffer A consisted of 100% water, buffer B consisted of 100% acetonitrile, and buffer C consisted of 25 mM ABC. The fractionation gradient operated as follows: 1% B to 5% B in 1 min, 5% B to 35% B in 61 min, 35% B to 60% B in 5 min, 60% B to 70% B in 3 min, 70% B to 1% B in 10min, with 10% C the entire gradient to maintain pH. The 48 fractions were then concatenated to 12 fractions (i.e. fractions 1, 13, 25, 37 were pooled, followed by fractions 2, 14, 26, 38, etc.) so that every 12^th^ fraction was used to pool. Pooled fractions were vacuum-centrifuged then reconstituted in 1% ACN/0.1% FA for LC-MS/MS.

Phospho-enriched and phospho-depleted fractions were analyzed by LC-MS/MS using a Thermo Easy-nLC 1200 (Thermo Fisher Scientific) with a 50 cm (inner diameter 75µm) EASY-Spray Column (PepMap RSLC, C18, 2µm, 100Å) heated to 60°C coupled to a Orbitrap Fusion Lumos Tribrid Mass Spectrometer (Thermo Fisher Scientific). Peptides were separated at a flow rate of 300nL/min using a linear gradient of 1 to 35% acetonitrile (0.1% FA) in water (0.1% FA) over 4 hours and analyzed by SPS-MS3. MS1 scans were acquired over a range of m/z 375-1500, 120K resolution, AGC target of 4e5, and maximum IT of 50 ms. MS2 scans were acquired on MS1 scans of charge 2 -7 using isolation of 0.7 m/z, collision-induced dissociation with activation of 35%, turbo scan, and max IT of 50 ms. MS3 scans were acquired using specific precursor selection (SPS) of 10 isolation notches, m/z range 100-1000, 50K resolution AGC target of 1e5, HCD activation of 65%, and max IT of 150 ms. The dynamic exclusion was set at 38 s.

### RNA-binding protein immunoprecipitation (RIP)

Briefly the protocol we used is the following: For lysate preparation we collected ESCshEXOSC2 cells by scraping them from one confluent 6 well dish per condition (EXOSC2 IP and control IgG IP). We lysed them on the dish with 500 ul of 1% TNESV buffer (50 mM Tris-HCl pH 7.5, 2 mM EDTA, 100 mM NaCl, 1% NP-40, H2O) with Protease/Phosphatase Inhibitor Cocktail (Cell signaling, #5872) and RNAse inhibitor (Applied Biosystems, N8080119), rotated in the cold room for 30 min and centrifuged the lysate at 4 degrees, maximum speed for 20 min and collected supernatant. At this step we removed 55 ul to use as input sample during RNA isolation and western blot.

For beads preparation, we have added 50 μl of A/G agarose beads (Pierce™ Protein A/G Agarose Catalog number: 20421) per experiment to a fresh Eppendorf and washed beads twice with lysis buffer (without protease inhibitor, TNESV). This and all following washing steps of the beads were performed with 900 μl of the respective buffer. We resuspended the beads in 100 μl lysis buffer with 5 μg antibody per experiment (Rabbit IgG Innovative Research, IRBIGGAP and Rabbit anti-EXOSC2 Bethyl, A303-886A antibodies) and rotated tubes for 30–60 min at room temperature, washed 3 times with 900 μl lysis buffer.

Then we added the lysate to the beads and rotated o/n. Next day we centrifuged and discarded the supernatant and washed twice with high-salt wash (50 mM Tris–HCl, pH 7.4, 1 M NaCl, 1 mM EDTA, 1% IGEPAL CA-630, 0.1% SDS, 0.5% sodium deoxycholate) (rotate the second wash for at least 1 min at 4 °C). We washed twice with PNK buffer (20 mM Tris–HCl, pH 7.4, 10 mM MgCl2, 0.2% Tween-20) and proceeded with IP and RNA isolation.

We have removed 50 μL each out of 500 μL of the bead’s suspension during the last PNK wash to test the efficiency of immunoprecipitation by western blotting. The proteins are eluted off the beads by re - suspending the beads in 1X SDS-PAGE loading buffer followed by heating at 95°C. The beads are centrifuged down and the supernatant directly applied on SDS-PAGE.

For purification of RNA we have prepared the proteinase K buffer. Each immunoprecipitated required 150 μL of proteinase K buffer containing 117 μL of TNESV Buffer, 15 μL of 10% SDS, 18 μL of 10 mg/mL proteinase K (Avantor, IB05406). Order of addition is intended to reduce risk of denaturation of proteinase K by addition of concentrated SDS. We have re-suspended each immunoprecipitate in 150 μL of proteinase K buffer. At this step we made use of the input sample as we isolated RNA from that sample too and added 107 μL of TNESV, 15 μL of 10% SDS, and 18 μL of proteinase K to the tubes and brought up the volume to 150 μL. We incubated all tubes at 55°C for 30 minutes with shaking to digest the protein. After the incubation, we centrifuged the tubes briefly and transfer the supernatant into a new tube. We added 50 μL of TNESV to each tube contacting the supernatant to bring the volume to 200 ul and then added 200 μL of phenol:chloroform:isoamyl alcohol to each tubes. We vortexed for 15 seconds and centrifuged at 14000 rpm for 30 minutes at room temperature to separate the phases. We have removed 200 μL of the aqueous phase carefully and placed it in a new tube. We have added 1 ml of chloroform. Repeated chloroform extraction and removed ∼100 μL of the aqueous phase carefully and place it in a new tube.

To each tube we added 10 ul 3M sodium acetate, 1.5 ul linear acrylamide and 400 ul 100% EtOH, vortex tubes vigorously and put at -20°C overnight to precipitate the RNA, centrifuged and washed the pellet twice with 1 ml cold 80% ethanol. We then removed all EtOH and re-suspended in 7 ul of RNase-free water. We continued with cDNA synthesis. The PPP2R5E primers that I used to amplify the transcript are the following:

M_PPP2R5E F CCAGACAGAAGAGGTCGCAA

M_PPP2R5E R GAACGTCTTTCAGCAGTGGC

### Cross-linking immunoprecipitation (CLIP)

The ESCshEXOSC2 were washed twice with 1x PBS and UV-crosslinked on ice in the same wash buffer with one pulse of 400 mJ/cm2 and one pulse of 200 mJ/cm2. Cells were then immediately scraped in fresh 1x ice-cold PBS with cycloheximide and processed as described previously ^12, 13^ with modifications. One biological replicate was used, prepared from 6 wells of a 6 well culture dish. Pellets were resuspended in 0.5 ml lysis buffer, and immunoprecipitations (IP) were performed overnight at 4°C using the rabbit antibody against EXOSC2.

### Embryonic Stem Cell Differentiation assay

Mouse embryonic stem cells were maintained in an undifferentiated state in EB medium consisting of DMEM F12(Gibco) supplemented with KnockOut™ Serum Replacement (Gibco), MEM Non-Essential Amino Acids Solution(Gibco), GlutaMAX™-I(Gibco), Antibiotic-antimycotic(Gibco) in an incubator under 5% CO2. The cells were cultured on MEF coated with 0.2% gelatin (Sigma) in phosphate buffered saline.

To differentiate EB, differentiation was carried out in three stages.

1. When cells on MEFs showed 70-80% confluency, mESCs were separated from MEF feeder cells using ACCUTASE™ (Stemcelltechnology).
2. We resuspended with EB medium(with 10% FBS) and transfered the mESC suspension from each well into a separate conical tube and centrifuge the tube(s) at 1000rpm for 5 minutes.
3. We plated 10 mL of the cell suspension (at 6×105 cells/mL) in one 100-mm non-adherent dish (i.e., not tissue culture–treated).
4. We incubated the cells for 3 days in a 37°C, 5% CO2 incubator to allow them to form EBs. We replaced spent medium daily. On the 7day after differentiation into EBs, EBs were harvested and frozen. mRNA was isolated and cDNA was synthesized as described above and PAX6, GATA6 and FGF8 were amplified and normalized to GAPDH using the following primers:

msGATA6_F : TTGCTCCGGTAACAGCAGTG

msGATA6_R : GTGGTCGCTTGTGTAGAAGGA

msPAX6_F : TACCAGTGTCTACCAGCCAAT

msPAX6_R : TGCACGAGTATGAGGAGGTCT

FGF8_F CGCTTTAGTTGAGGAACTCGAAGCG

FGF8_R ACATGGCCTTTACCCGCAAG

### Gene Ontology

For gene ontology analysis we have used The **D**atabase for **A**nnotation, **V**isualization, and **I**ntegrated **D**iscovery (**DAVID).** https://david.ncifcrf.gov/content.jsp?file=FAQs.html#1

### AU-rich elements (AREs) mapping

For ARE detection all transcript sequences were first extracted from reference genome GRCm38 and then used as input to the fuzznuc program from the EMBOSS package^14^ to detect occurrence of five AU-rich elements^15^ in all transcripts.

#### AKP cell line generation

(Pancho Barriga generated these; please consider acknowledging him).

Normal colon organoids bearing unrecombined Apc fl/fl, p53 fl/fl, and LSL-KrasG12D were treated with Adeno Cre, then selected for recombination through the removal of Wnt, EGF, and addition of Nutlin-3. These lines were then transduced with retroviral EGFP IRES Luc and experimental liver mets were established by intrasplenic injection. Mice were then sacrificed and macrometastases were dissected and generated 2D lines on collagen-coated plates and using standard DMEM media.

#### Generation of AKP organoids

Colon crypts were isolated from Apcfl/fl; Trp53fl/fl; KrasLSL-G12D/+ C57Bl6 female mice as previously described ^16^. These purified colon crypt fractions were then used to establish organoid cultures by embedding them in Matrigel and culturing in basal organoid media in the presence of EGF (50 ng/mL), Noggin (100 ng/mL), RSPO1 (250 ng/mL), and CHIR99021 (3 M). These established organoids were then treated with an adenoviral Cre to induce the recombination of the aforementioned alleles. Recombined cells were then selected by withdrawing EGF, Rspo1, and CHIR99021 from the media, as well as supplementing with Nutlin-3a (10 M). Organoids growing in this culture conditions were termed AKP and their genotype was confirmed by assessing allele recombination by PCR. AKP organoids were then transduced with a retroviral Luciferase IRES EGFP vector and EGFP+ cells were isolated by FACS.

#### Models of colon cancer metastasis

250.000 AKP LucEGFP cells were injected in the spleens of 6 -8 week old C57Bl6 females. 5 minutes after injection, spleens were removed and mice were then monitored longitudinally for the development of metastasis by in vivo bioluminescence imaging (IVIS). Mice were euthanized when exhibiting overt disease burden and tissues were collected in formalin and embedded in paraffin to assess by immunohistochemistry.

#### Endoluminal Cell Implantation

Colorectal mouse models were performed as previously described ^17^. Briefly, AKP colorectal tumor cells (generated in house from an APC^-/-^, Kras^G12D/+,^ p53^-/-^ mouse) and CT26 cells (obtained from ATCC (CRL2638). Briefly, female C57Bl/6N mice (Jackson Labs) of 6 – 8 weeks of age were pre-treated with 3% DSS drinking water for 5 days then allowed to recover for 2 days. On day 8 mice underwent endorectal cell implantation (1,000,000 cells). Mice there thereafter monitored for tumor development with weekly endoscopy.

#### Mouse Irradiation

Mice were subjected to whole abdomen irradiation as previously described ^18^. Briefly, mice were anesthetized and secured on a X-RAD 225Cx (Precision X-Ray Inc; Northford, CT) orthovoltage small-animal irradiator. Whole abdomen irradiation was performed using a single AP field with a 25mm diameter circular cone. Mouse position was confirmed based on the laser coordinate system and further with live fluoroscopic imaging. A total whole abdomen dose of 14.5Gy was delivered over 245 seconds at a dose rate of 5.9cGy/second.

#### Statistics and Reproducibility

Data were analyzed using GraphPad Prism 8 software (GraphPad Software) and/or Excel software (Microsoft Office). Data are presented as bar graphs that demonstrate individual values with mean and standard deviation to feature information about the distribution of the underlying data (as indicated in figure legends). For samples >10 we represented the data with a violin plot. Statistic tests were performed, and P value thresholds were obtained using GraphPad 8 or Excel. Multiple groups were tested using two-sided t-test followed by post-hoc Holm-Bonferroni correction for a significance level of 0.05 and comparisons between two groups were performed using two-tailed unpaired Student’s t-test. To confirm the reproducibility every experiment was repeated independently. Representative figures are shown main figures and Supplementary Figures.

