## Supplementary Figures for "Aging-dependent dysregulation of EXOSC2 is maintained in cancer as a dependency"

### Supplementary Figure 1

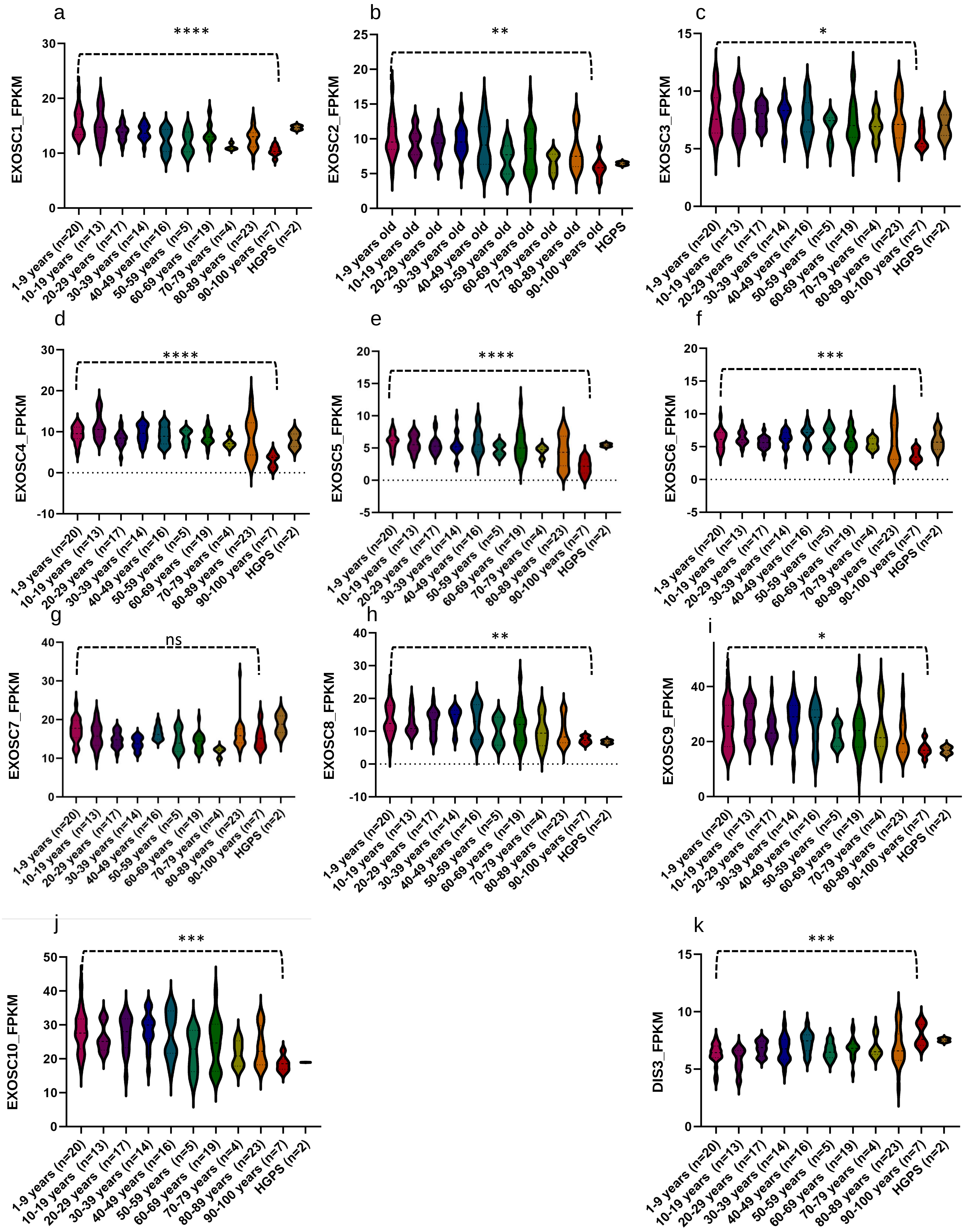

### Supplementary Figure 1

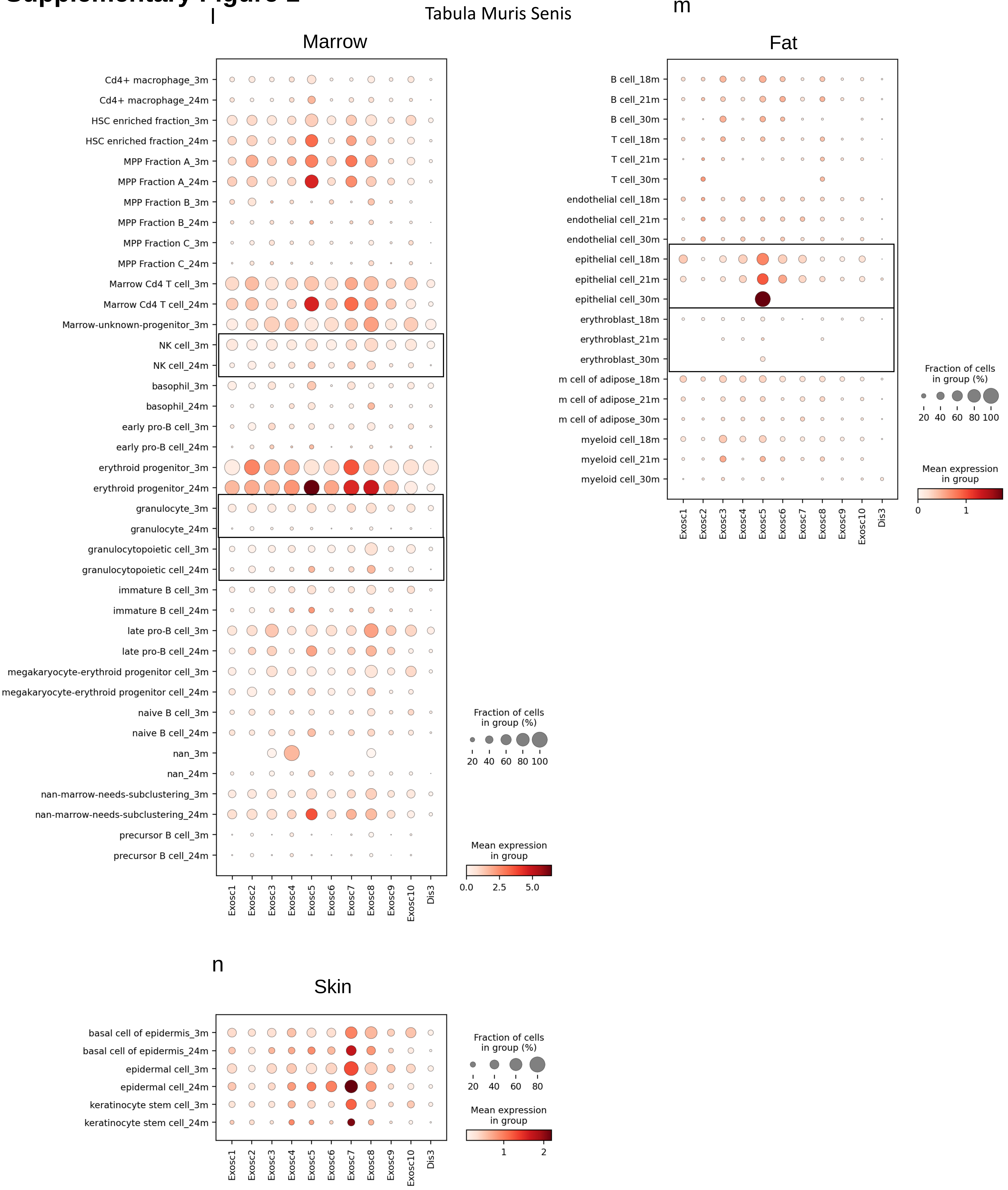

**Figure S1**: Gene expression data for (a-k) RNA exosome subunits from human fibroblasts in young and aged people. (l – n ) Dot plots for gene expression of the RNA exosome subunits from Tabula Muris Senis. The diagrams demonstrate plotted levels of the RNA exosome subunits in the different cell types of young (3 months old) and old (24 months old) mice in (a) the marrow , (b) the fat and (c) the skin.

#### Supplementary Figure 2

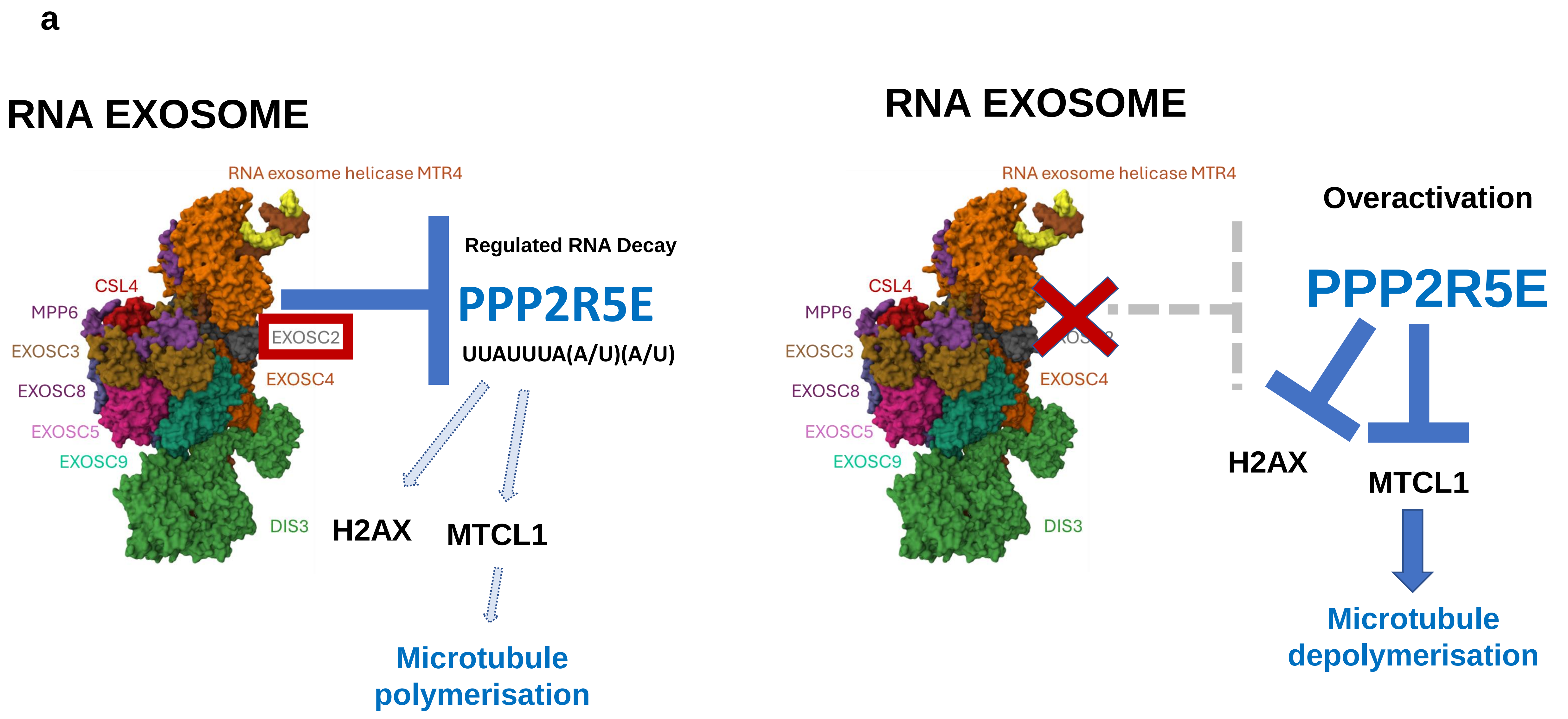

**Figure S2: (a)Schematic summary.** We found that EXOSC2 RNA binding protein regulates the RNA decay of PPP2R5E by directly binding to the PPP2R5E mRNA. In a self renewal context this interaction is crucial to balance proliferation and differentiation as the downstream effector of PPP2R5E is H2AX and the MTCL1 protein that regulates microtubule polymerisation and p21 localisation to the nucleus. We also demonstrate that cancer cells hijack this mechanism , by permanently downregulating the EXOSC2 protein thus they carry an overactivated phosphatase that continuously dephosphorylates H2AX and MTCL1 thus leading to a series of defects in the affected cell, including an increase in cytoplasmic p21. The RNA exosome structure presented here is from https://www.rcsb.org/3d-view/6D6R/1 . Weick, E.M., Puno, M.R., Januszyk, K., Zinder, J.C., DiMattia, M.A., Lima, C.D. (2018) Cell 173: 1663

### Supplementary Figure 3

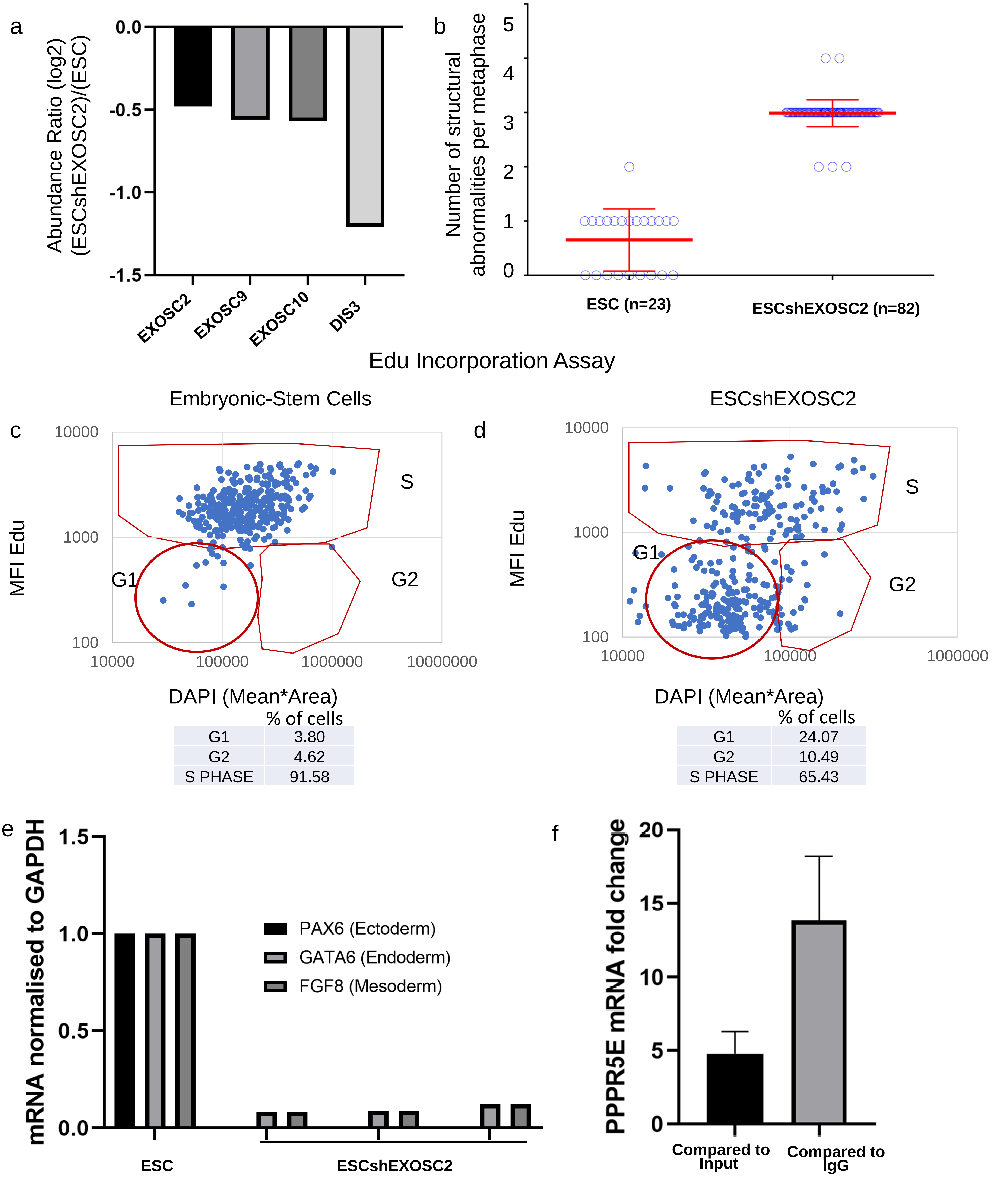

Embryonic-Stem Cells:

| | % of cells |
|---|---|
| G1 | 3.80 |
| G2 | 4.62 |
| S PHASE | 91.58 |

ESCshEXOSC2:

| | % of cells |
|---|---|
| G1 | 24.07 |
| G2 | 10.49 |
| S PHASE | 65.43 |

**Figure S3:** a ) Proteomics data indicating downregulation of EXOSC2 in embryonic stem cell lines with EXOSC2 knock down , accompanied by downregulation in EXOSC9, EXOSC10 and DIS3.
b) Quantification of the number of structural abnormalities per metaphase spread. The number indicates the number of abnormalities counted. c) Edu Incorporation staining with DAPI in ESC and in d)ESCshEXOSC2 to assess cell cycle stage after EXOSC2 downregulation. e)Differentiation assay on ESC cell lines and ESCshEXOSC2 f) RNA immunoprecipitation with EXOSC2 antibody and IgG control and qPCR for PPP2R5E mRNA. The graph shows the fold change of PPP2R5E mRNA compared to the input and the fold change compared to the IgG control confirming that EXOSC2 binds PPP2R5E mRNA directly.

g

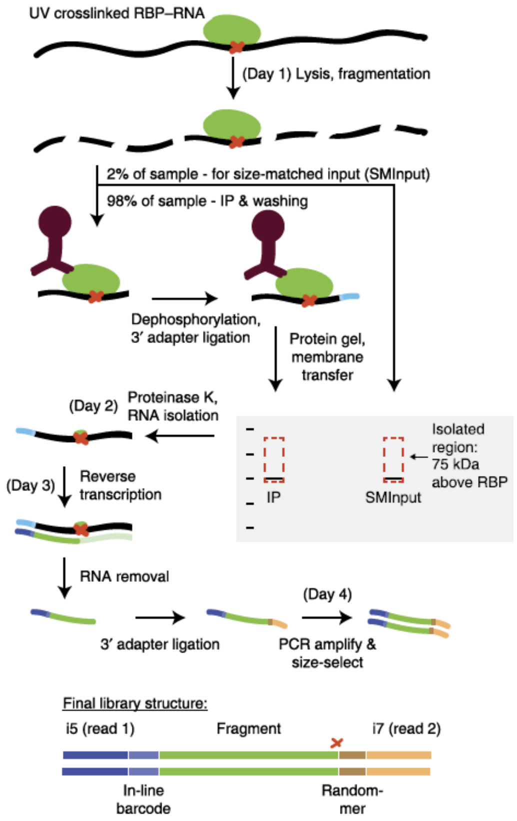

Van Nostrand et al, Nature Methods 2016

h

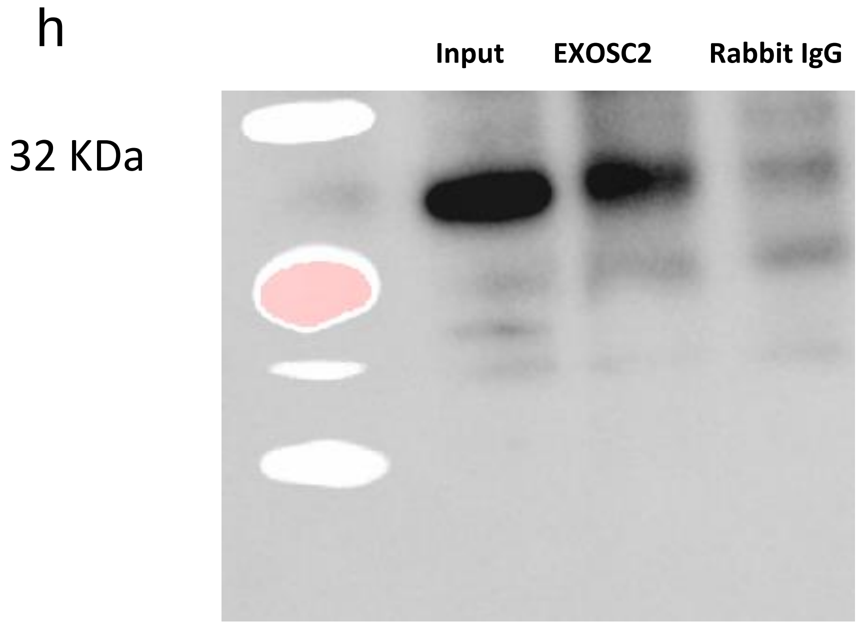

i

Van Nostrand et al, Nature Methods 2016

j

k

Barcodes: 3, 4, 3, 4, 5 (4000-2000), and 6 (4000)

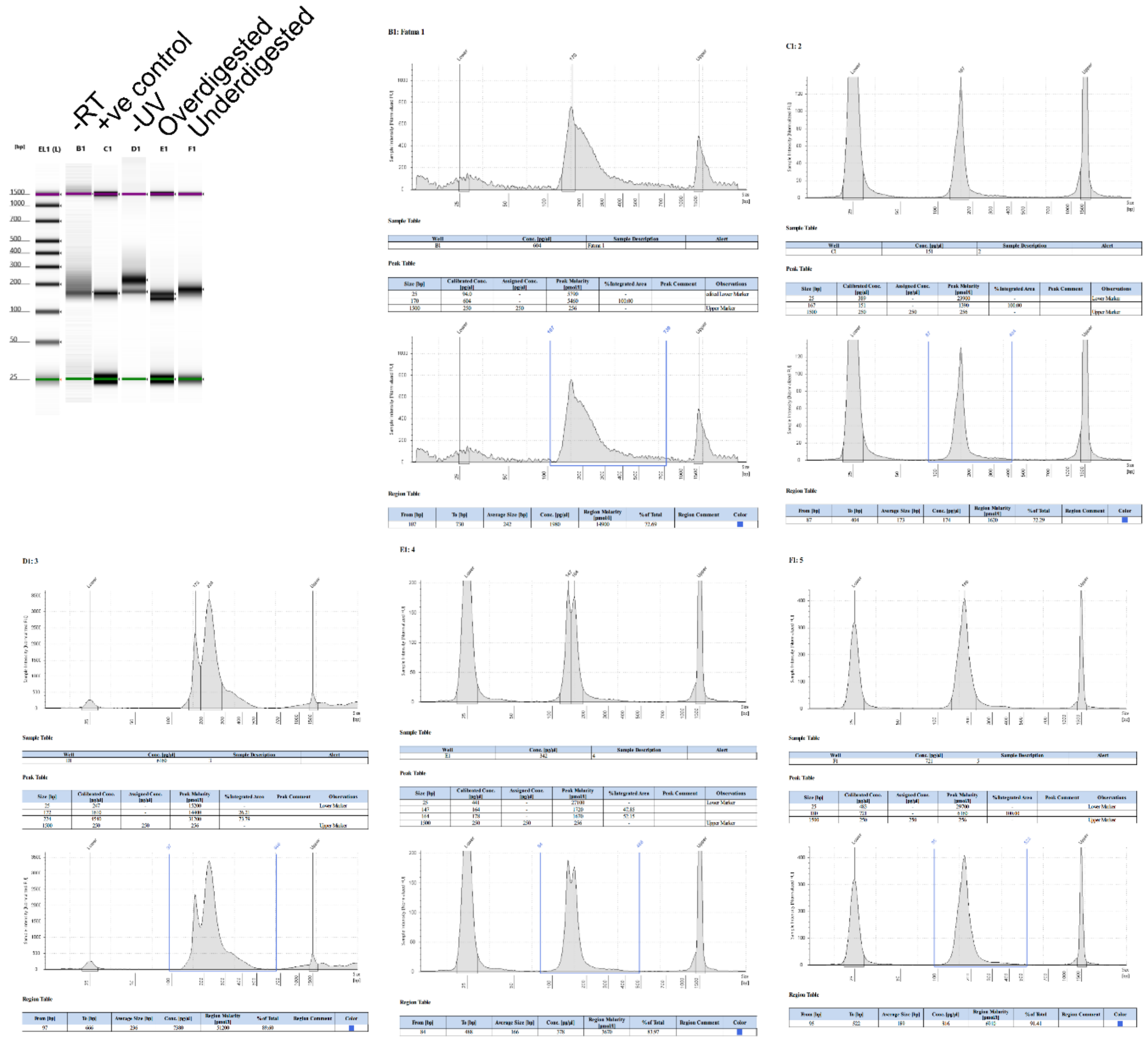

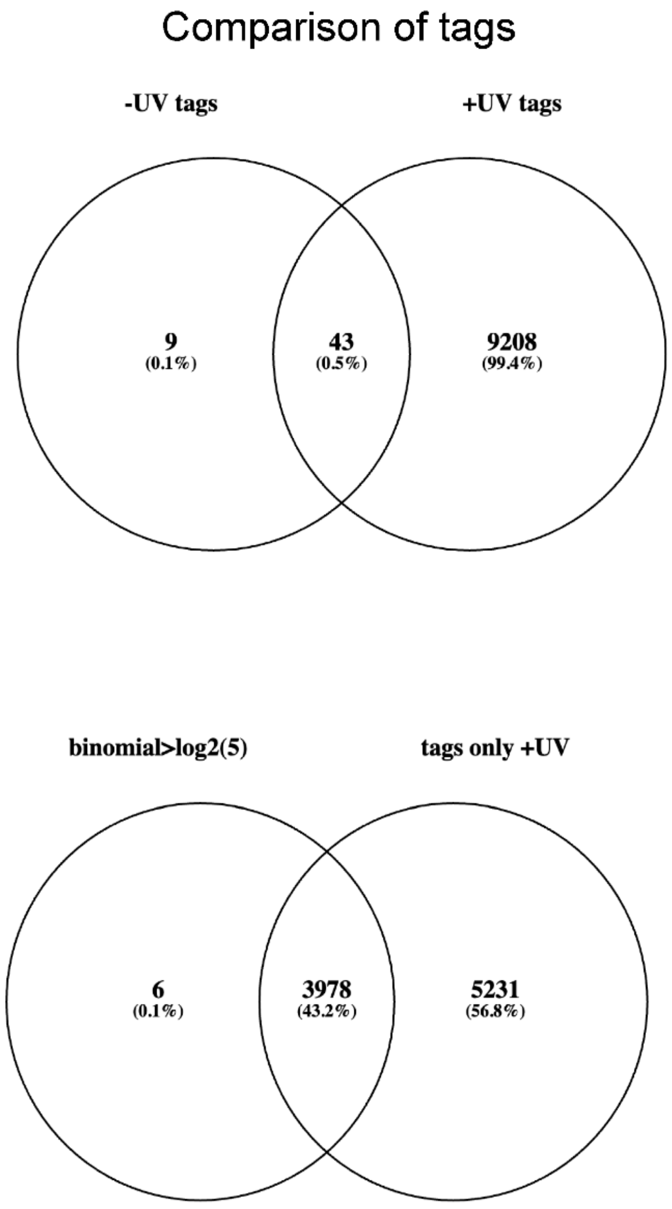

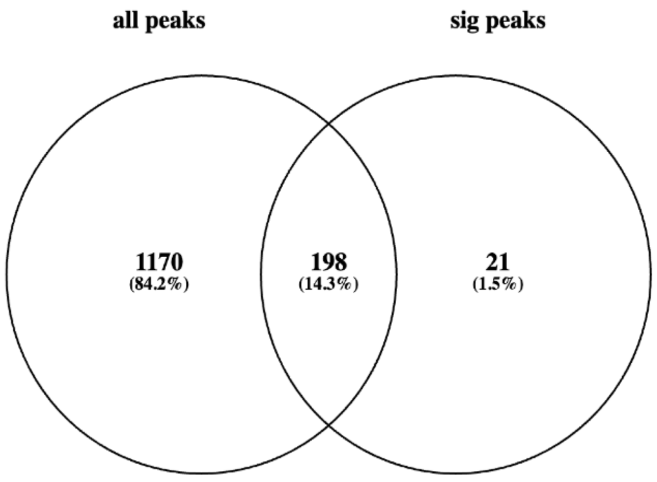

l

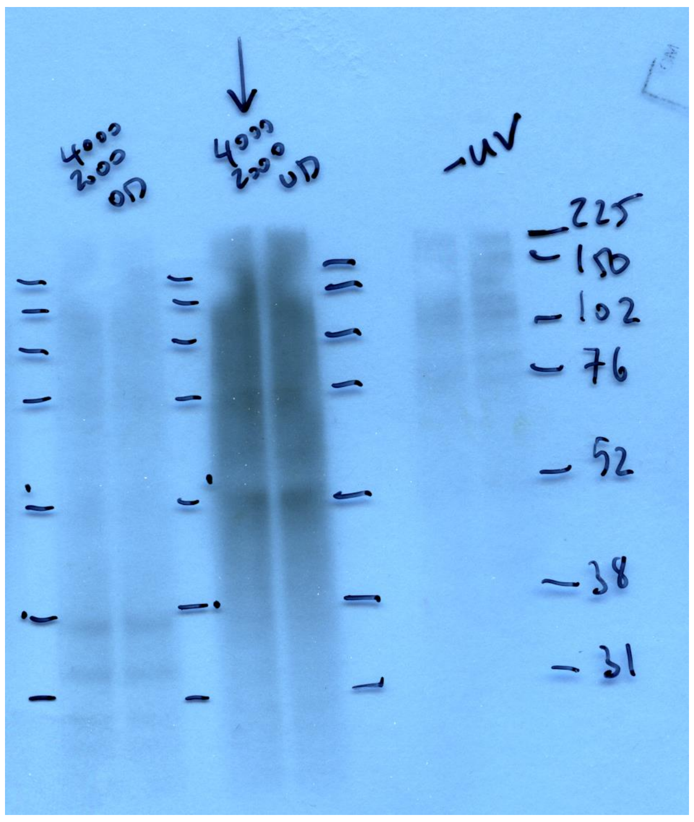

m

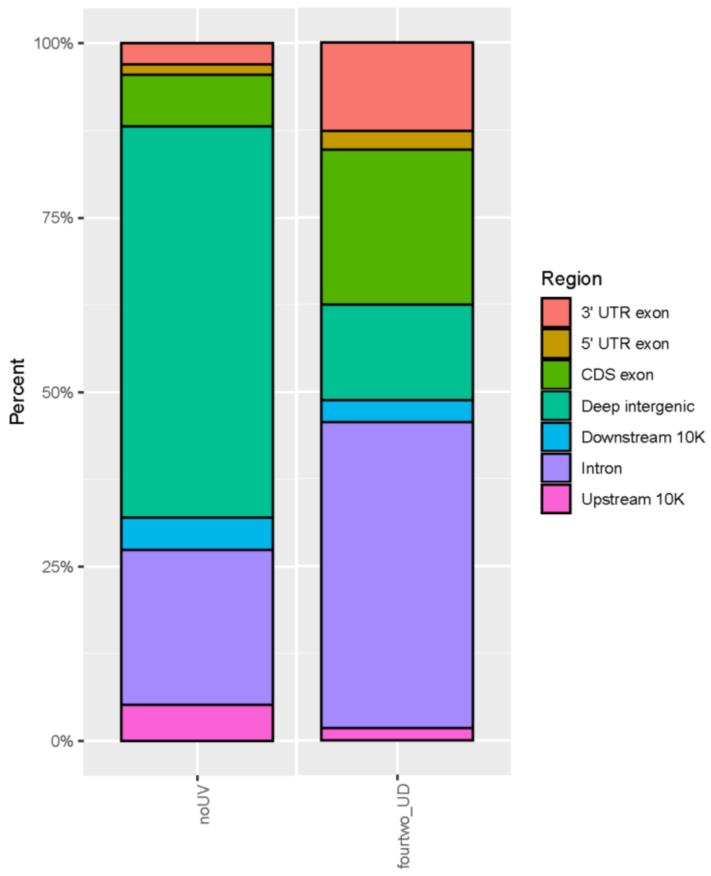

n

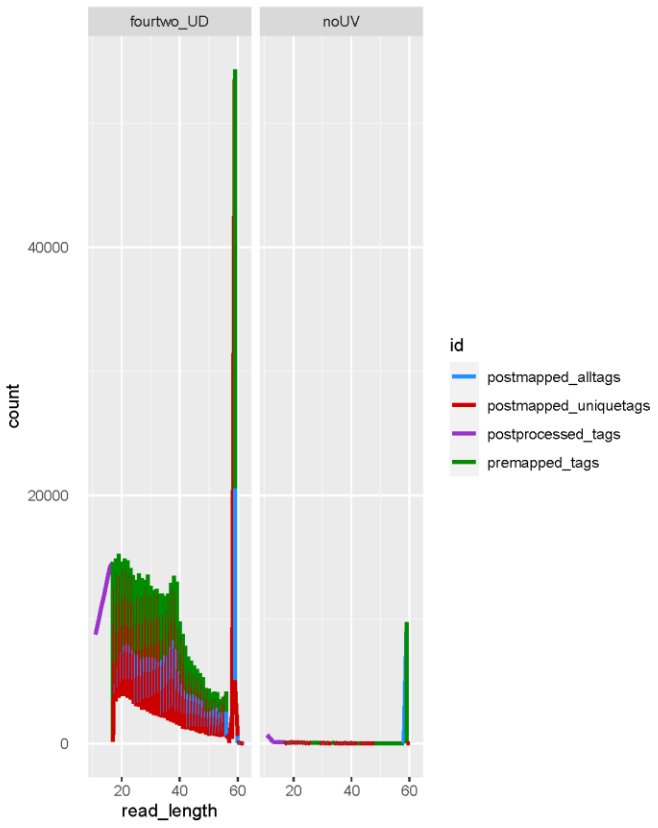

o

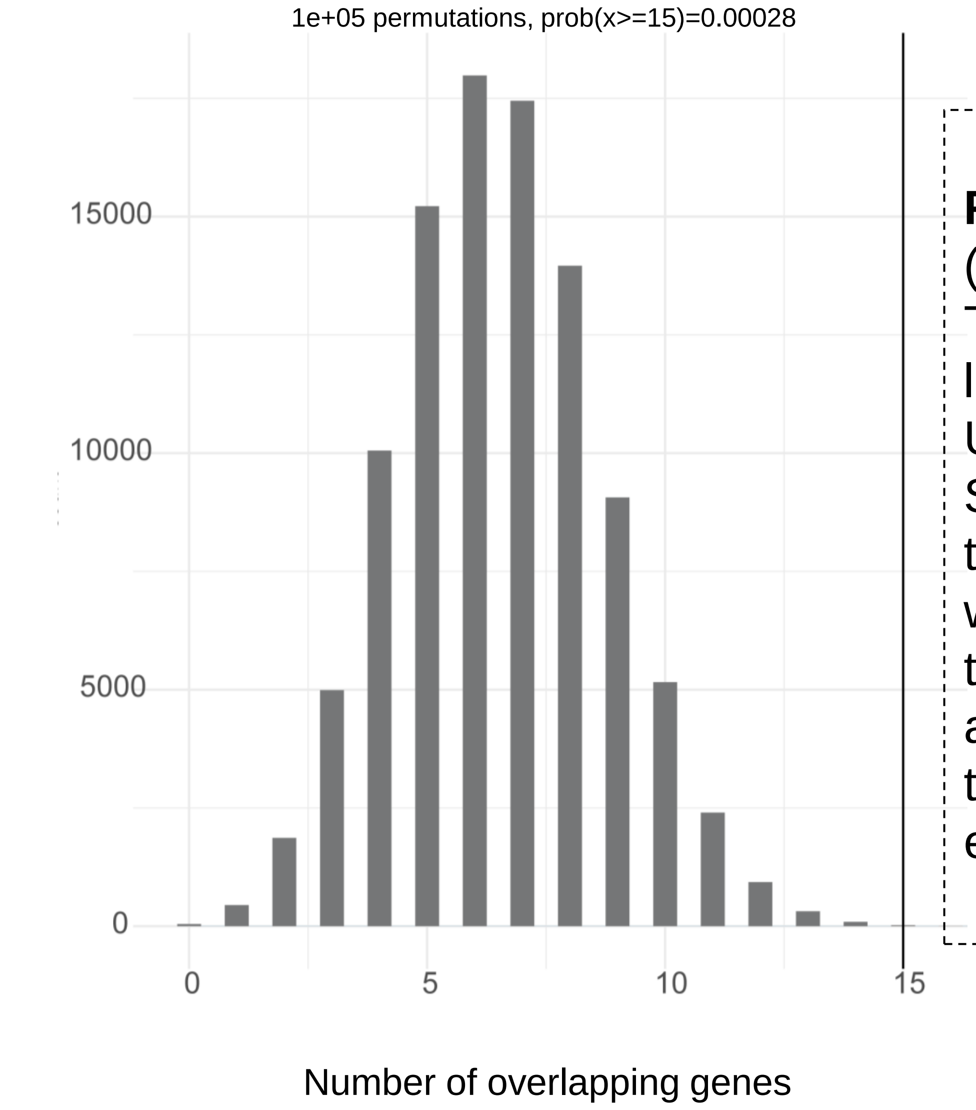

**Figure S3**: (g) Schematic presentation of eCLIP (h) Immunoprecipitation with EXOSC2 and IgG antibodies (i-j) Tape station readouts of the size distribution of the amplified library. (k) comparison of tags in the minus UV condition and the UV treated sample. (l) Protein gel and membrane transfer (m) Summary of mapped tags annotations with and without UV treatment and (n) summary of tag length distribution with and without UV treatment (o) Histogram of permutation show that 15 targets that were observed out of 6248 observed targets were among the 23 mRNAs that we had predicted to be targets out of a total 22018 protein coding genes and this is significant. The empirical p-value from the permutation experiment is 0.00028.

### Supplementary Figure 4

#### GO0000177 cytoplasmic exosome (RNase complex): Data set from Pefanis et al. Cell , 2015

**a**

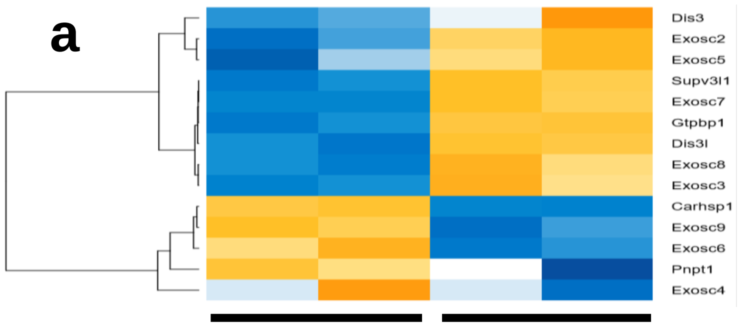

ESC-EXOSC3ko ESC-EXOSC3wt

**b**

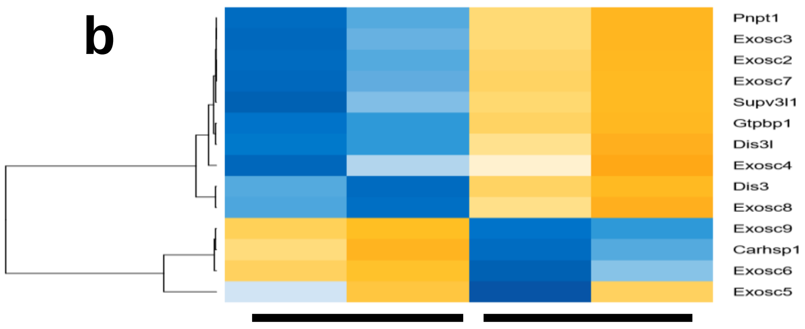

ESC-EXOSC10ko ESC-EXOSC10wt

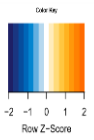

**c**

| Gene | ESC_Exos3ko_vs_ESCExos3wt _log2FoldChange | ESC_Exos3ko_vs_ESCExosc3 _padj |
|---|---|---|
| Ppp2r5e | -0.14 | 0.30115 |
| Exosc2 | -0.49 | 0.00004 |
| Gene | ESC_Exos10ko_vs_ESCExos10wt _log2FoldChange | ESC_Exos10ko_vs_ESCExos10wt _padj |
| Ppp2r5e | -0.14 | 0.30342 |
| Exosc2 | -0.40 | 0.00024 |

**d**

ESC ESC

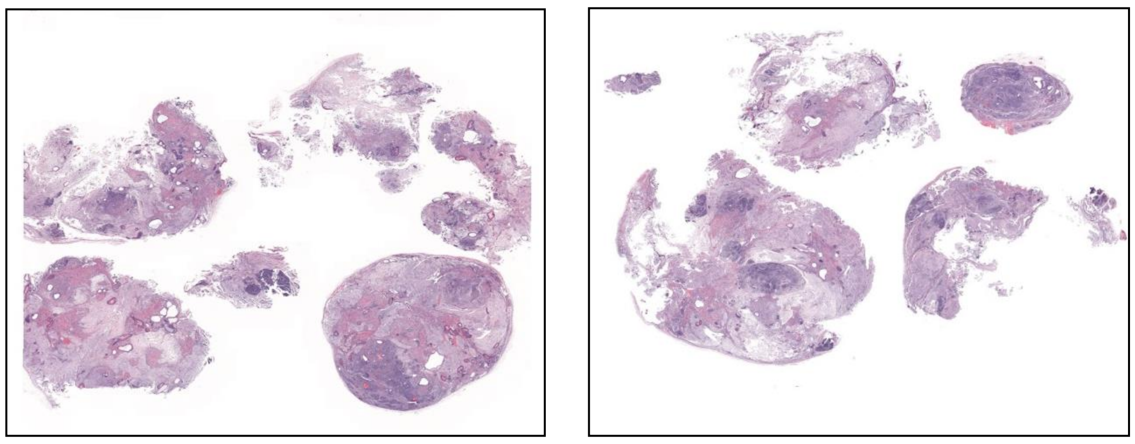

**e**

ESC-shEXOSC2 ESC-shEXOSC2

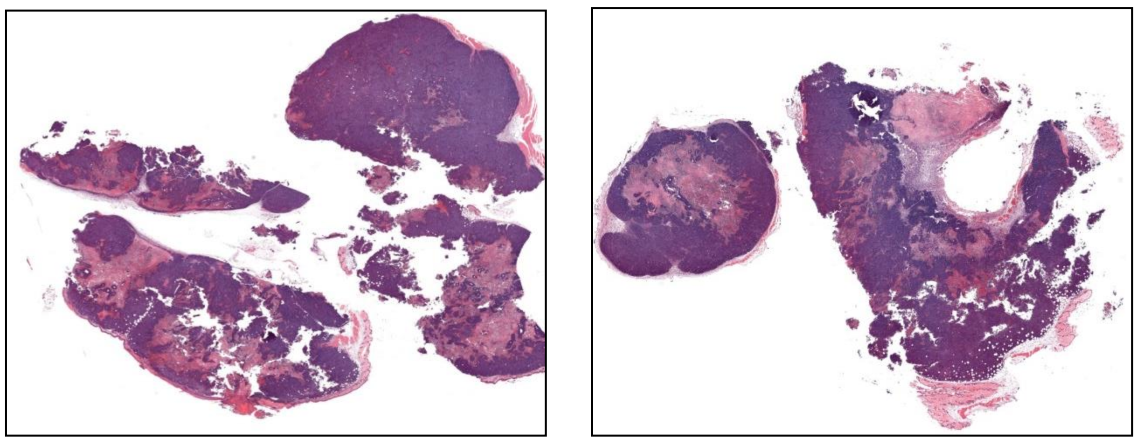

ESC-shEXOSC2 ESC-shEXOSC2 ESCDshEXOSC2 ESC-shEXOSC2

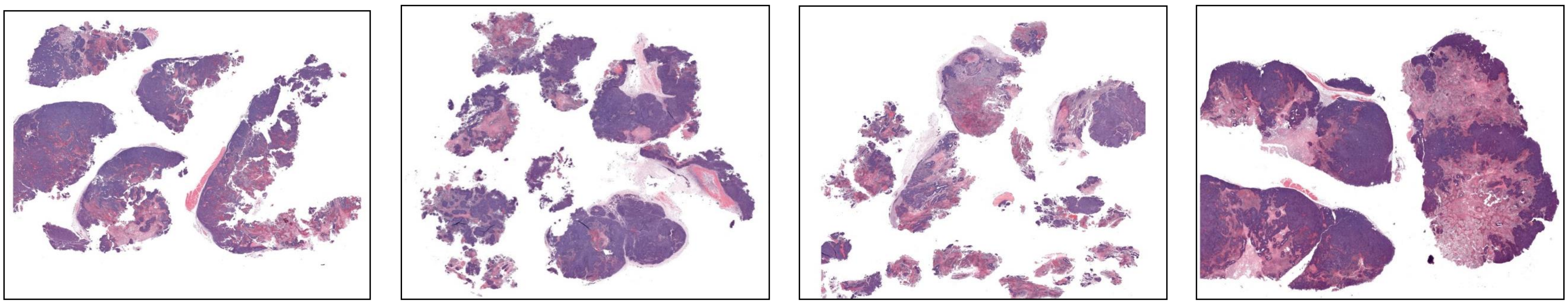

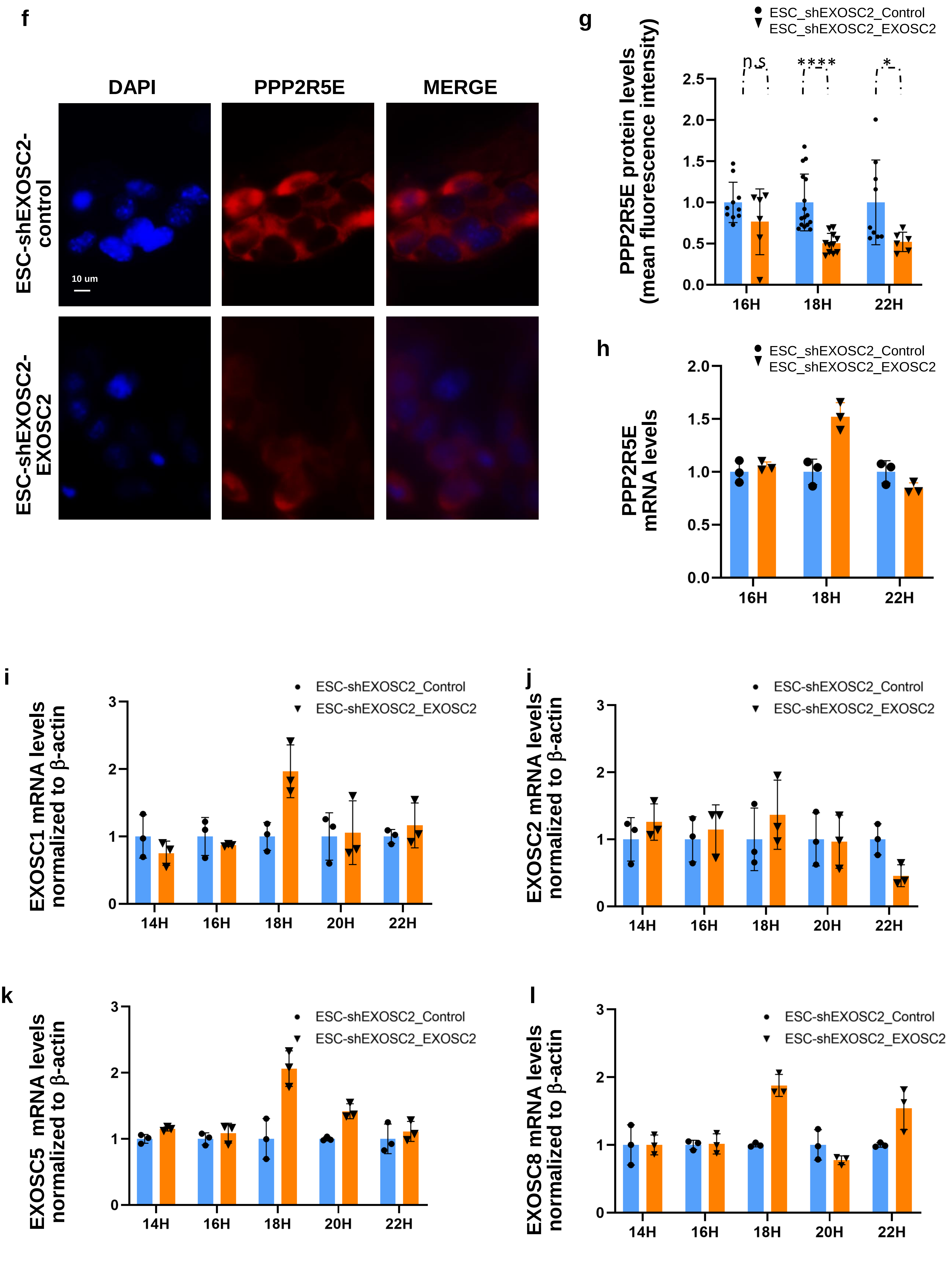

**Figure S4:Downregulation of Exosome subunits in ESC.** RNAseq of two independent clones of ESC and ESC-shEXOSC3 (a) and ESC-shEXOSC10 (b) (Pefanis E. et al, Cell 2015) showed that downregulation of other subunits than EXOSC2 , also lead to downregulation of other components of exosome complex. PPP2R5E levels from RNAseq data in ESC lines with downregulation of EXOSC3 or EXOSC10 do not show upregulation of PPP2R5E transcript (c). Teratoma analysis of ESC (d) and ESC-shEXOSC2 (e) by H/E staining. IF staining for PPP2R5E protein (f) and quantification (g) after EXOSC2 transient overexpression. Transcript levels for PPP2R5E (h), EXOSC1 (i), EXOSC2 (j) , EXOSC5 (k) and EXOSC8 (l) in ESC-shEXOSC2 line and ESC-shEXOSC2 line after EXOSC2 transient overexpression in different timepoints after the EXOSC2 overexpression . Error bars indicate SE.

### Supplementary Figure 5

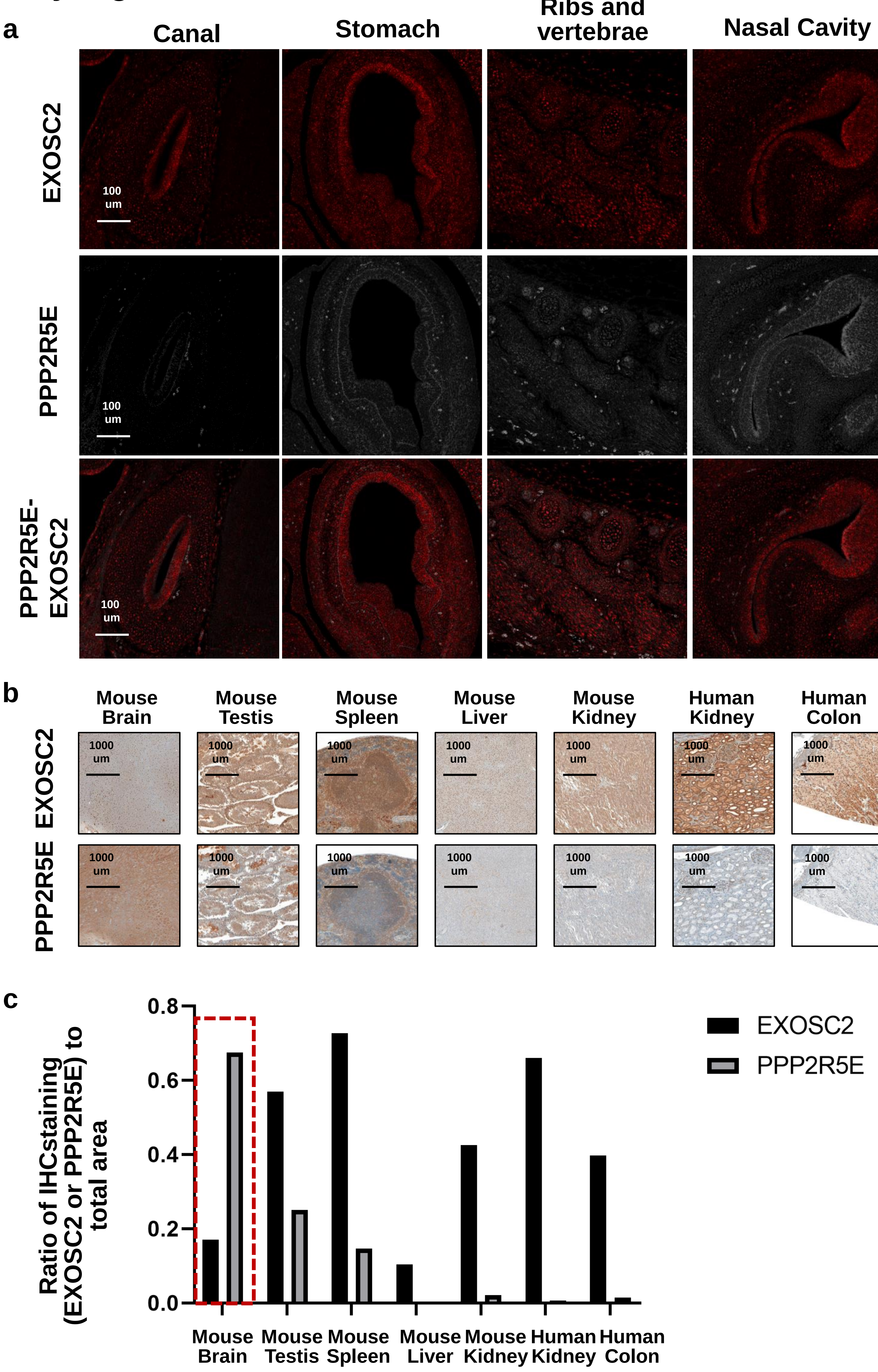

**Figure S5: Broadly conserved reverse-regulatory patterns between EXOSC2 and PPP2R5E in various mouse and human tissues**. Conserved dynamic regulation of PPP2R5E by EXOSC2 in various structures (canal, stomach, ribs and vertebrae and nasal cavity) at the developing embryo (E13.5). Co-staining for EXOSC2 and PPP2R5E and merged images for the two proteins (a). Conserved dynamic regulation of PPP2R5E by EXOSC2 in adult mouse (spleen, testis, kidney) and human (colon, kidney) tissues by immunohistochemistry . Staining for EXOSC2 and PPP2R5E (b) was done independently is consecutive sections of the same tissues.(c) Quantification of the IHC staining for EXOSC2 (c) and for PPP2R5E in adult mouse and human tissues. The mouse brain shows the opposite pattern to all the other organs, low levels of EXOSC2 and high levels of PPP2R5E (red box).

### Supplementary Figure 6

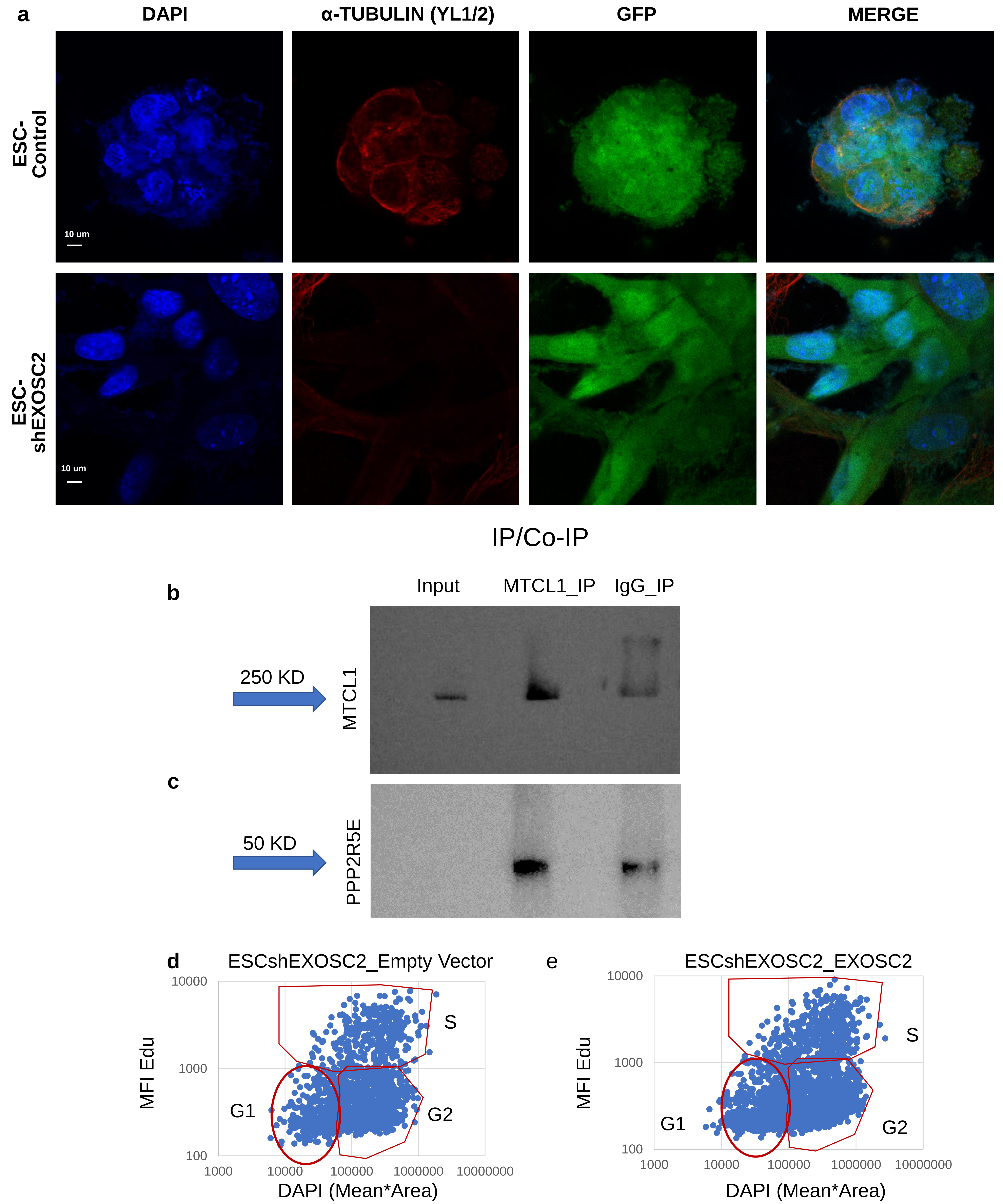

| | |
|---|---|
| G1 | 41.86046512 |
| G2 | 50.46511628 |
| S | 7.674418605 |

| | |
|---|---|
| G1 | 28.4507 |
| G2 | 52.59557 |
| S | 18.95372 |

**Figure S6: Microtubule polymerization defects in ESC-shEXOSC2** Low levels of α-tubulin(YL ½) in ESC treated with shEXOSC2 in immunostaining compared to fibroblast control and ESC control (a). Immunoprecipitation for MTCL1 protein (b) and co-immunoprecipitation for PPP2R5E protein (c) in 293 T cells. Transient overexpression of empty vector (d) and EXOSC2 (e) in ESCshEXOSC2 and effect on cell cycle

#### Supplementary Figure 7

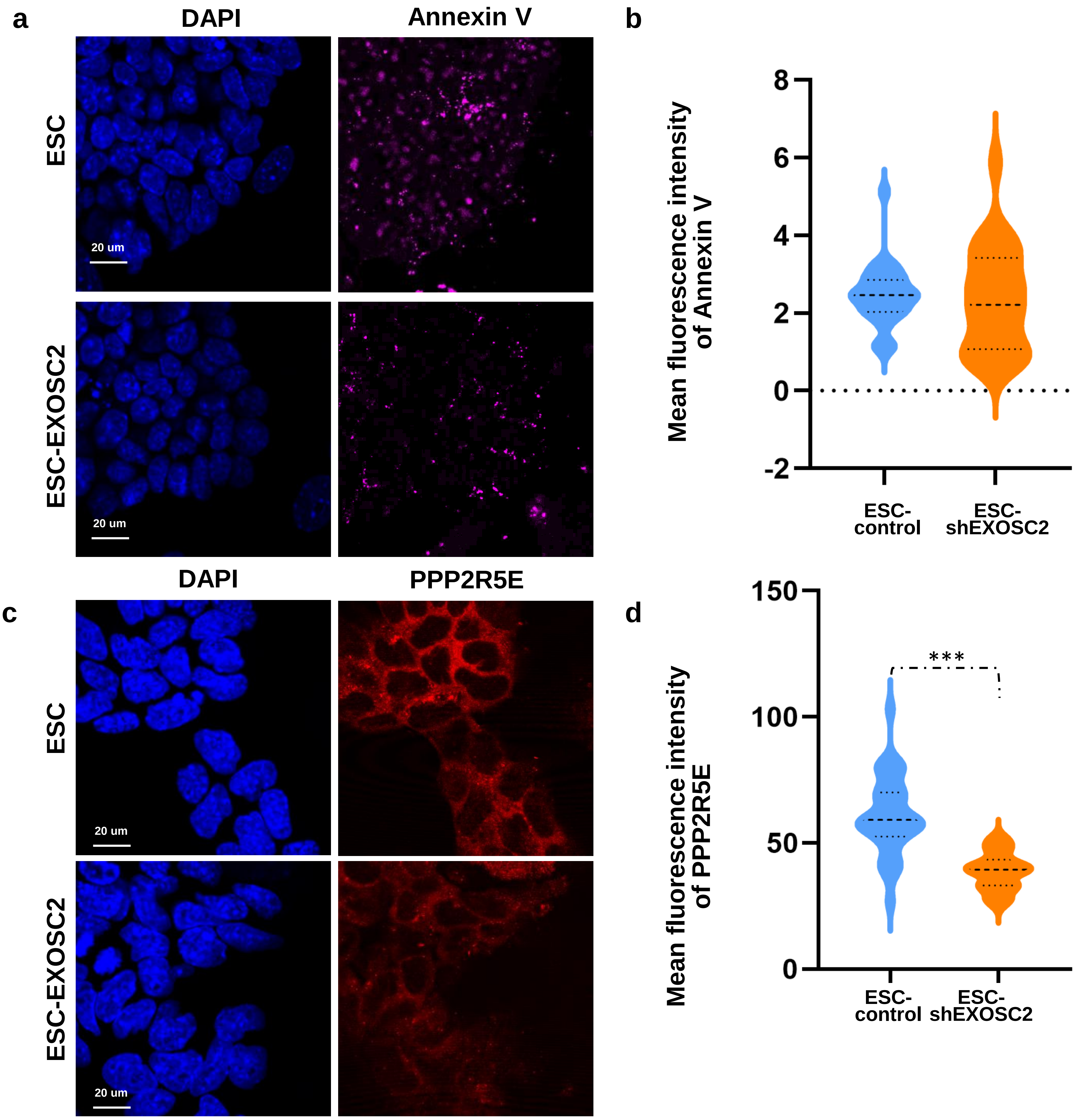

**Figure S7: Conserved inversed regulation of PPP2R5E by EXOSC2.** (a) Annexin V staining and (b) quantification in ESC line and ESC line after EXOSC2 transient overexpression in different timepoints after the EXOSC2 overexpression . Nuclei were stained with DAPI. **Statistical significance by two-sided t-test ($p<0.005$). (c) Immunostaining and (d) quantification for PPP2R5E protein level in ESC line and ESC line after EXOSC2 transient overexpression in different timepoints after the EXOSC2 overexpression . Nuclei were stained with DAPI. **Statistical significance by two-sided t-test ($p<0.005$).

#### Supplementary Figure 8

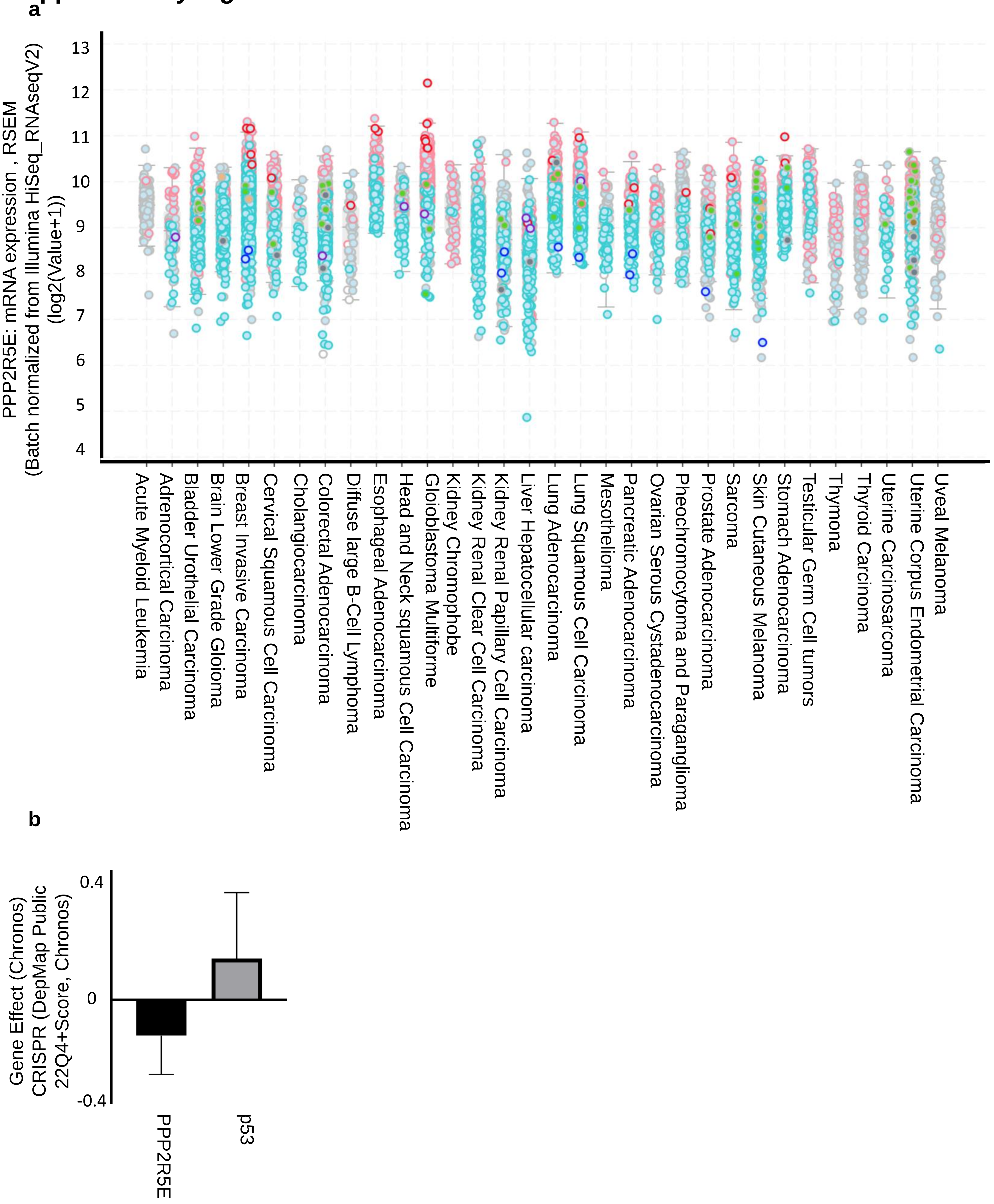

**Figure S8**: (a) RNAseq data for PPP2R5E from TCGA database. (b) Comparative gene effect when you knock out PPP2R5E and the tumor suppressor p53. The graph shows the average gene effect in 1000 cancer cell lines after deleting the different genes. (https://depmap.org/portal/interactive/). The cancer cell lines have on average a positive gene effect when you delete a component of the PP2A complex , which means that they grow faster. On the contrary on average the gene effect when you delete PPP2R5E gene is negative which is indicative of cell death.
