## Supplementary_Table1 for "Aging-dependent dysregulation of EXOSC2 is maintained in cancer as a dependency"

**SUPPLEMENTARY TABLE 1**

Annotation Cluster 1 — Enrichment Score: 19.785838947352495

| Category | Term | Count | % | PValue | | #NAME? | Genes | List Total | Pop Hits | Pop Total | ld Enrichm | Bonferron | Benjamini | FDR | LOG10 | #NAME? |
|---|---|---|---|---|---|---|---|---|---|---|---|---|---|---|---|---|
| UP_KEYWORDS | Differentiation | 137 | 5.53983 | 2.10E-17 | -1.67E+01 | 1.67E+01 | DFI. BMP4. CPNE9. BMP2. SLCO4C1. CPNE5. CSRP2. DDX4. ADGRG1. KDR. NOTCH3. SFRP5. SEMA6A. PLK5. SEMA6C. CLGN. TRIM54. SARM1. DMRTC2. SFRP2. NTRK1. ITGA8. SFRP4. D1PAS1. BMP7. BMP8B. UTP14B. FGF8. ELF3. NR6A1. PAX6. MOV10L1. ANPEP. CBFA2T3. GLI1. ARHGAP22. PRMT7. STRA8. PIWIL1. PIWIL2. PIWI | 2284 | 646 | 22680 | 2.10589 | 8.87E-15 | 4.43E-15 | 2.95E-14 | -13.5297 | 13.5297 |
| UP_KEYWORDS | Cell junction | 139 | 5.6207 | 2.53E-17 | -1.66E+01 | 1.66E+01 | R156. GABRE. PARD6B. GABRA4. KCNB1. SYT11. FSCN1. CRB3. KCNK1. HOMER2. SYNGR3. SHANK3. KDR. TULP1. EPHA4. SARM1. CASS4. CNIH2. RAPSN. CHRNB4. CYFIP2. CHRND. CHRNB2. GRB7. PRX. CLDN7. LZTS1. KCNAB2. CLDN4. GRIP1. TLN2. FERMT3. CLDN5. FERMT1. CPEB1. GJA4. ZFYVE21. EPCAM. SNPH. SNTB1. ESAM. SV | 2284 | 661 | 22680 | 2.08814 | 1.07E-14 | 3.56E-15 | 3.56E-14 | -13.4491 | 13.4491 |
| UP_KEYWORDS | Meiosis | 34 | 1.37485 | 1.33E-11 | -1.09E+01 | 1.09E+01 | TEX19.1, TEX19.2, FKBP6, REC114, MAEL, MOV10L1, SYCP2, TDRD12, SYCP1, MEI4, STRA8, PIWIL1, FBXO43, PIWIL2, STAG3, PIWIL4, CCNA1, TDRD1, HFM1, CCNB1IP1, MSH4, SYCE2, SYCE1, DDX4, REC8, CLGN, TEX15, SYCP3, HORMAD1, HORMAD2, DMC1, SMC1B, UTP14B, TEX11 | 2284 | 90 | 22680 | 3.75131 | 5.61E-09 | 8.01E-10 | 1.87E-08 | -7.72857 | 7.72857 |
| UP_KEYWORDS | Calmodulin-binding | 41 | 1.65791 | 5.23E-10 | -9.28E+00 | 9.28E+00 | KCNH1, SLC8A3, ADCY1, MARCKSL1, MYO7A, CASK, RRAD, TTN, ITPKA, CAMKV, ATP2B2, ENKUR, PDE1C, SNTB1, NOS3, CAMK2B, CNN1, CAMK2A, MYO5B, MYO1A, NOS1, SLC8A2, PHKG1, PNCK, MYH3, MYO1F, MYH6, DAPK2, MYO10, KCNN1, KCNN3, KCNN2, RYR1, AKAP5, IQCG, RYR2, MYH14, NRGN, GAP43, MYH10, SNTA1 | 2284 | 139 | 22680 | 2.92898 | 2.21E-07 | 2.21E-08 | 7.35E-07 | -6.13362 | 6.13362 |
| UP_KEYWORDS | Synapse | 85 | 3.43712 | 7.82E-14 | -1.31E+01 | 1.31E+01 | IRNA4. CHRNA7. SV2A. SV2C. SVOP. MPDZ. SDK2. RIMBP2. CACNG5. PRKCG. PCLO. CBLN4. CBLN1. DOC2A. DACT1. NRGN. ERC2. SEZ6. IGSF9. GAP43. DSCAML1. RIMS1. RIMS4. AMPH. RIMS3. PRR7. ARHGAP44. DMD. RASGRP2. SYN2. SLC30A3. EMB. HTR3A. OLFM2. CAMK2A. GPR156. GABRE. DLGAP1. GABRA4. DLGAP3. COL13 | 2284 | 357 | 22680 | 2.36427 | 3.30E-11 | 6.60E-12 | 1.10E-10 | -9.95913 | 9.95913 |
| UP_KEYWORDS | Glycoprotein | 523 | 21.1484 | 2.70E-15 | -1.46E+01 | 1.46E+01 | PINI1. KCNK1. EMILIN3. TMEM117. CHRND. TREML2. BMP8B. KCNH1. FGF5. FGF8. TRPV2. ST6GALNAC3. SLC23A1. TMEM108. CILP2. XYLT1. NCMAP. PRSS36. DPP4. LAG3. FGF3. HTR5B. MROH7. ZP3. CAR14. CMKLR1. SDK2. PRSS50. MMP17. CACNG4. PKDCC. ST6GALNAC2. C8G. INHBB. SERPINB5. CPXM1. CYBRD1. PRSS44. PRS | 2284 | 3815 | 22680 | 1.3613 | 1.12E-12 | 2.81E-13 | 3.74E-12 | -11.427 | 11.427 |
| UP_KEYWORDS | Cell membrane | 394 | 15.9321 | 0.19684 | -7.06E-01 | 7.06E-01 | CNG5. CACNG4. MMP17. MAST1. CLIC6. RASD1. RASD2. ENOX1. ALPL. NKD1. NKD2. OPRK1. TFR2. GPR68. MREG. HTR3A. TMPRSS2. SLC8A2. GPR75. OPRL1. RTN4R. CDH15. NPHS1. SLC8A3. KCNC3. SRCIN1. GRIK3. GRIK4. SLC7A8. RRAD. SLC7A9. IL17RD. SLC7A7. NRCAM. PACSIN1. SLC2A5. SLC2A4. GPR45. SLC2A3. UNC5A. TDG | 2284 | 3759 | 22680 | 1.04081 | 1 | 0.51731 | 95.4101 | 1.97959 | -1.97959 |
| UP_KEYWORDS | Ion channel | 79 | 3.1945 | 1.26E-12 | -1.19E+01 | 1.19E+01 | 2C. KCNK5. CHRNA5. CHRNA4. CHRNA7. KCNQ2. KCNG1. SCN2B. TRPA1. CACNG6. CACNG5. CACNG4. TRPM1. KCNT1. CLIC5. CLIC6. KCNH6. RYR1. RYR2. KCNH2. KCNH3. KCNH4. CLCN1. KCNMB4. CLCNKA. CACNB2. CLCNKB. KCNA6. FXYD6. KCNJ3. KCNS3. BEST2. KCNS1. BEST1. SCN9A. HTR3A. HCN3. GABRE. GABRA4. KCNB1. TMC | 2284 | 336 | 22680 | 2.33472 | 5.34E-10 | 8.90E-11 | 1.78E-09 | -8.75007 | 8.75007 |
| UP_KEYWORDS | Immunoglobulin domain | 84 | 3.39668 | 7.03E-07 | -6.15E+00 | 6.15E+00 | MO1. KIRREL2. NRG2. LAG3. ISLR2. ICAM1. ICOSL. SCN2B. SDK2. NFAM1. PTPRU. PALLD. TMEM25. VSIG2. SEMA4G. IGSF1. IGSF3. CNTN2. ROR2. GPA33. MADCAM1. NCAN. IGSF9. SEMA4A. KALRN. FGFR2. IL1R2. FGFR4. FGFR3. SIGLECF. IGFBPL1. DSCAML1. CNTFR. KIT. IGSF10. LINGO1. ACAN. OBSL1. CD4. EMB. CSF1R. VSTM2 | 2284 | 481 | 22680 | 1.73413 | 2.97E-04 | 1.85E-05 | 9.89E-04 | -3.00496 | 3.00496 |
| UP_KEYWORDS | Transcription regulation | 222 | 8.97695 | 8.61E-04 | -3.06E+00 | 3.06E+00 | AZF1. RAD54B. KLF2. KAT6B. PARP1. TCF15. HLF. STOX1. HNF1A. ELF3. TBX21. MITF. ZFP42. GLI2. GLI1. KCNIP3. ZFP879. STRA8. HSF5. SPIB. HSF4. BHLHE41. THRSP. SCX. ZFP354B. ZFP239. ZCCHC12. BATF3. TLE3. FOXR1. ZFP37. FOXR2. ZSCAN10. RIPPLY1. RIPPLY3. STAT4. FSBP. TCEA3. BCL11B. BCL11A. GBX2. GBX1. PRKAA2. F | 2284 | 1799 | 22680 | 1.22537 | 0.3049 | 0.00805 | 1.20417 | 0.08069 | -0.08069 |
| UP_KEYWORDS | Lipid metabolism | 70 | 2.83057 | 2.47E-05 | -4.61E+00 | 4.61E+00 | PG. PLA2G1B. THRSP. UGT8A. FA2H. PLD4. CDS1. LPCAT2. LPIN3. LIPO1. PTGDS. DGAT2. SLC27A2. PLA2G5. ALOX12. CHKA. MVD. PTCHD2. HSD17B1. PLBD1. ST8SIA1. CERS6. HMGCS1. CERS4. PLIN5. SULT4A1. ISYNA1. CERS1. DHCR7. PLCH2. PLCD4. PLCD1. ETNK2. PRKAA2. ACAA1B. ACSL3. ACSL6. GAL3ST1. SOAT2. LPL. CPT1B | 2284 | 417 | 22680 | 1.6669 | 0.01037 | 4.74E-04 | 0.03472 | -1.45943 | 1.45943 |
| UP_KEYWORDS | SH3 domain | 41 | 1.65791 | 4.59E-05 | -4.34E+00 | 4.34E+00 | 7A. CASK. CACNB2. STAC3. SKAP1. MCF2L. AMPH. ARHGAP4. DOCK2. PACSIN1. MAP3K9. MYO15B. UBASH3A. ARHGAP9. ARHGEF4. SH3GL3. CASKIN1. HCK. ARHGEF6. ARHGEF37. MPP3. RIMBP2. BZRAP1. ARHGEF16. EFS. MYO1F. VAV1. SHANK3. BC021891. CASS4. LCK. MAPK8IP2. TNK1. MAPK8IP1. TJP3. SRMS. SH3GL2. KALI | 2284 | 208 | 22680 | 1.95735 | 0.01917 | 7.44E-04 | 0.06446 | -1.19073 | 1.19073 |
| UP_KEYWORDS | Wnt signaling pathway | 32 | 1.29397 | 0.00144 | -2.84E+00 | 2.84E+00 | NKD1, NKD2, NPHP3, WNT4, WNT3, RSPO1, FRAT2, PRKAA2, NDRG2, AXIN2, WNT6, WNT8A, FZD9, TCF7, TNIK, DRAXIN, TLE3, LEF1, FZD7, SFRP5, RNF43, WNT7B, FZD10, DACT1, DKK1, KREMEN2, SFRP2, DACT3, SFRP4, PYGO1, ROR2, LRP4 | 2284 | 176 | 22680 | 1.80544 | 0.45517 | 0.01284 | 2.00251 | 0.30157 | -0.30157 |
| UP_KEYWORDS | RNA-mediated gene silencing | 12 | 0.48524 | 0.01131 | -1.95E+00 | 1.95E+00 | FKBP6, MAEL, PIWIL1, PIWIL2, MOV10L1, PIWIL4, TDRD12, LIN28A, DDX4, LIN28B, TRIM71, TDRD1 | 2284 | 51 | 22680 | 2.33646 | 0.99176 | 0.07013 | 14.772 | 1.16944 | -1.16944 |
| UP_KEYWORDS | Neurotransmitter transport | 14 | 0.56611 | 3.67E-05 | -4.44E+00 | 4.44E+00 | STX3, CPLX1, SLC6A1, SLC6A2, SLC6A11, SLC6A15, STX1B, RIMS1, RIMS4, RIMS3, SLC17A7, SLC6A7, SV2A, SV2C | 2284 | 37 | 22680 | 3.75728 | 0.01537 | 6.20E-04 | 0.0516 | -1.28739 | 1.28739 |
| UP_KEYWORDS | Metal-binding | 422 | 17.0643 | 1.01E-06 | -5.99E+00 | 5.99E+00 | ST1. KSR2. ZSCAN10. PPEF2. CPXM1. CYBRD1. GUCY1B3. GM17067. ZSCAN18. ALOX12. ALPL. REPS2. NKD1. NKD2. SORD. ECEL1. FHL1. NUFIP1. TRIM13. NHLRC1. NECAB3. GM13212. NECAB2. TCEA3. PDE1C. GALNT16. C2. GALNT12. CCNB1IP1. SLC8A2. TRIM29. DGKK. DGKH. ZFP2. CELA1. MID1. PHYH. RPS6KA6. RASSF5. PLEK | 2284 | 3395 | 22680 | 1.2343 | 4.28E-04 | 2.52E-05 | 0.00143 | -2.84616 | 2.84616 |
| UP_KEYWORDS | Sarcoplasmic reticulum | 11 | 0.4448 | 0.00353 | -2.45E+00 | 2.45E+00 | JSRP1, SRL, RYR1, RYR2, CAMK2B, FKBP1B, JPH1, CASQ1, ART1, CASQ2, THBS4 | 2284 | 38 | 22680 | 2.87446 | 0.77552 | 0.02832 | 4.85451 | 0.68615 | -0.68615 |
| UP_KEYWORDS | Ligand-gated ion channel | 19 | 0.7683 | 2.57E-04 | -3.59E+00 | 3.59E+00 | GABRB3, GABRA4, GRIK3, GRIK4, GRIA4, TRPM1, CHRNA9, GRIN2C, P2RX3, CHRNB4, CHRNA5, RYR1, CHRNA4, RYR2, CHRNA7, CHRNB2, CHRND, HTR3A, HCN3 | 2284 | 73 | 22680 | 2.58451 | 0.10286 | 0.00328 | 0.36091 | -0.4426 | 0.4426 |
| UP_KEYWORDS | Guanine-nucleotide releasing factor | 27 | 1.09179 | 7.08E-04 | -3.15E+00 | 3.15E+00 | FGD2, FGD1, CYTH4, RASGEF1B, DENND2A, DENND2D, MCF2L, DOCK2, RASGRP4, TIAM1, RASGRP2, FGD5, KNDC1, ARHGEF4, ARHGEF6, ARHGEF37, RAB3IL1, ARHGEF16, PSD4, ARHGEF15, DOCK6, VAV1, ARHGEF10, DENND1C, PSD, DENND4B, KALRN | 2284 | 133 | 22680 | 2.01585 | 0.25821 | 0.00709 | 0.98999 | -0.00437 | 0.00437 |
| UP_KEYWORDS | Chromosome | 35 | 1.41529 | 0.90101 | -4.53E-02 | 4.53E-02 | 2810004N23RIK, HMGB2, FKBP6, SYCP2, SYCP1, MEI4, STAG3, SKA3, SKA1, HIST3H2BA, CCNB1IP1, NUP133, HIST1H1E, HIST1H1D, HIST1H1B, HIST1H1A, SYCE2, SYCE1, CCDC155, ESCO2, FAM208A, REC8, TEX14, SYCP3, RASSF2, H2AFY2, CFDP1, HORMAD1, HORMAD2, ZRANB3, DMC1, KAT6B, MPHOSPH8, SMC1B, TEX11 | 2284 | 409 | 22680 | 0.84975 | 1 | 0.98886 | 100 | 2 | -2 |
| UP_KEYWORDS | Tyrosine-protein kinase | 23 | 0.93004 | 0.00185 | -2.73E+00 | 2.73E+00 | FGFR2, RET, FGFR4, FGFR3, FGR, HCK, FLT4, PKDCC, EPHA10, KIT, EPHA1, EPHB2, KDR, EPHA4, EPHA8, NTRK1, LCK, ZAP70, TNK1, MST1R, SRMS, CSF1R, SYK | 2284 | 113 | 22680 | 2.02114 | 0.54221 | 0.01582 | 2.56908 | 0.40978 | -0.40978 |
| UP_KEYWORDS | SH3-binding | 14 | 0.56611 | 0.00594 | -2.23E+00 | 2.23E+00 | ENAH, KHDRBS3, KHDRBS2, DTX1, EFS, EVL, ELMO3, CBLC, ENKUR, SH3BGR, DNAJC6, INPP5D, ADAM19, LRP2 | 2284 | 60 | 22680 | 2.31699 | 0.91893 | 0.04169 | 8.02828 | 0.90462 | -0.90462 |
| UP_KEYWORDS | Sterol biosynthesis | 9 | 0.36393 | 0.00309 | -2.51E+00 | 2.51E+00 | CYB5R2, MSMO1, MVD, DHCR7, HMGCS1, PRKAA2, IDI1, PMVK, FDFT1 | 2284 | 26 | 22680 | 3.43729 | 0.72908 | 0.02528 | 4.25673 | 0.62908 | -0.62908 |
| UP_KEYWORDS | Sodium | 22 | 0.88961 | 0.00837 | -2.08E+00 | 2.08E+00 | SLC8A3, SLC38A4, SLC38A3, SLC5A5, SLC8A2, SCN2B, SLC6A1, SCN3A, SLC6A2, SCN3B, ATP1B2, ASIC2, ATP1A3, SLC6A15, ASIC1, SLC34A3, SLC17A7, SLC23A1, SCN9A, SLC9B2, HCN3, SLC5A11 | 2284 | 120 | 22680 | 1.82049 | 0.9712 | 0.05561 | 11.1451 | 1.04708 | -1.04708 |
| UP_KEYWORDS | EGF-like domain | 36 | 1.45572 | 0.00581 | -2.24E+00 | 2.24E+00 | MASP2, LTBP4, NELL2, SORL1, BCAN, CSPG5, MEGF11, HMCN2, HMCN1, CRB2, COMP, TDGF1, ACAN, ADAM33, NRG2, THBS4, MATN3, ADAM11, EGFL7, SCUBE2, CRTAC1, SCUBE1, DLL3, TMEM8, CELSR3, NID2, CFC1, NOTCH3, FBLN1, TENM3, ADAM22, ADAM19, NCAN, LRP2, MEGF6, LRP4 | 2284 | 224 | 22680 | 1.59588 | 0.91432 | 0.04148 | 7.85868 | 0.89535 | -0.89535 |
| UP_KEYWORDS | Calcium transport | 17 | 0.68742 | 0.0228 | -1.64E+00 | 1.64E+00 | SLC8A3, SLC8A2, TRPV2, CACNG6, CACNG5, CACNG4, CACNB2, ASIC1, CACNA1S, ATP2B2, BSPRY, ATP2C2, RYR1, RYR2, CACNA1H, CACNA1A, CACNA1B | 2284 | 93 | 22680 | 1.81515 | 0.99994 | 0.12171 | 27.6911 | 1.44234 | -1.44234 |
| UP_KEYWORDS | Vitamin A | 4 | 0.16175 | 0.02613 | -1.58E+00 | 1.58E+00 | RBP4, RBP7, RBP1, CRABP1 | 2284 | 7 | 22680 | 5.67426 | 0.99999 | 0.13506 | 31.0754 | 1.49242 | -1.49242 |
| UP_KEYWORDS | Metal-thiolate cluster | 3 | 0.12131 | 0.05293 | -1.28E+00 | 1.28E+00 | MT2, MT1, MT4 | 2284 | 4 | 22680 | 7.44746 | 1 | 0.23186 | 53.44 | 1.72787 | -1.72787 |
| UP_KEYWORDS | GTPase activation | 23 | 0.93004 | 0.09524 | -1.02E+00 | 1.02E+00 | DLC1, TBC1D10C, TBC1D9, ARHGAP28, ARHGEF15, RGS17, RGS14, ARHGAP30, GARNL3, ARHGAP22, RASAL1, SGSM1, ARHGAP20, RINL, ARHGAP44, STARD8, CHN1, ARHGAP8, ARAP3, TBC1D1, AGAP2, ARHGAP9, ARHGAP10 | 2284 | 163 | 22680 | 1.40116 | 1 | 0.33376 | 75.5085 | 1.878 | -1.878 |
| UP_KEYWORDS | Sexual differentiation | 3 | 0.12131 | 0.15132 | -8.20E-01 | 8.20E-01 | DMRTC2, DMRT3, DMRT1 | 2284 | 7 | 22680 | 4.25569 | 1 | 0.4384 | 90.0366 | 1.95442 | -1.95442 |
| UP_KEYWORDS | Tight junction | 15 | 0.60655 | 0.03692 | -1.43E+00 | 1.43E+00 | CLDN7, PARD6B, CLDN19, CLDN4, MPDZ, CLDN5, CRB3, EPCAM, NPHP4, BVES, CGN, MARVELD3, AMOT, ESAM, TJP3 | 2284 | 83 | 22680 | 1.79457 | 1 | 0.17037 | 41.071 | 1.61354 | -1.61354 |
| UP_KEYWORDS | SH2 domain | 18 | 0.72786 | 0.03307 | -1.48E+00 | 1.48E+00 | PTPN6, SOCS2, FGR, HCK, VAV1, HSH2D, TNS4, SH2D3C, STAT4, LCK, ZAP70, CHN1, INPP5D, SHC2, GRB7, SRMS, SYK, BLNK | 2284 | 105 | 22680 | 1.70228 | 1 | 0.15717 | 37.6709 | 1.57601 | -1.57601 |
| UP_KEYWORDS | Chloride | 12 | 0.48524 | 0.07021 | -1.15E+00 | 1.15E+00 | BEST2, CLCN1, SLC12A8, BEST1, GABRA4, GABRB3, CLIC5, CLIC6, CLCNKA, CLCNKB, NPR1, CTSC | 2284 | 67 | 22680 | 1.7785 | 1 | 0.28133 | 64.0612 | 1.80659 | -1.80659 |
| UP_KEYWORDS | Calcium channel | 11 | 0.4448 | 0.08987 | -1.05E+00 | 1.05E+00 | TRPV2, CACNG6, RYR1, CACNG5, CACNA1H, CACNB2, RYR2, CACNG4, CACNA1S, CACNA1A, CACNA1B | 2284 | 62 | 22680 | 1.76176 | 1 | 0.32794 | 73.386 | 1.86561 | -1.86561 |
| UP_KEYWORDS | Growth factor | 21 | 0.84917 | 0.02901 | -1.54E+00 | 1.54E+00 | MIA, BMP4, GDF3, FGF5, FGF18, FGF8, BMP2, FGF15, NODAL, FGF17, IGF2, MDK, LEFTY1, INHBB, TDGF1, LEFTY2, BMP7, NRG2, BMP8B, FGF3, THBS4 | 2284 | 127 | 22680 | 1.64196 | 1 | 0.14384 | 33.8893 | 1.53006 | -1.53006 |
| UP_KEYWORDS | Sodium channel | 7 | 0.28306 | 0.05638 | -1.25E+00 | 1.25E+00 | SCN2B, SCN3A, SCN3B, SCN9A, ASIC2, ASIC1, HCN3 | 2284 | 28 | 22680 | 2.48249 | 1 | 0.24054 | 55.7671 | 1.74638 | -1.74638 |
| UP_KEYWORDS | Osteogenesis | 8 | 0.32349 | 0.06451 | -1.19E+00 | 1.19E+00 | MSX2, BMP4, BMP2, COL13A1, IGF2, PKDCC, BMP7, BMP8B | 2284 | 36 | 22680 | 2.20665 | 1 | 0.26353 | 60.8355 | 1.78416 | -1.78416 |

| | |
|---|---|
| Calmodulin-binding | 6.13362 |
| Meiosis | 7.72857 |
| Synapse | 9.95913 |
| Glycoprotein | 11.427 |
| Cell junction | 13.4491 |
| Differentiation | 13.5297 |
