## Supplementary_Table2 for "Aging-dependent dysregulation of EXOSC2 is maintained in cancer as a dependency"

**SUPPLEMENTARY TABLE 2**

Annotatior Enrichment Score: 19.068993370061047

| Category | Term | Count | % | PValue | Genes | List Total | Pop Hits | Pop Total | Fold Enrich | Bonferroni | Benjamini | FDR | | | | |
|---|---|---|---|---|---|---|---|---|---|---|---|---|---|---|---|---|
| UP_KEYW( | Signal | 395 | 26.02108 | 1.09E-13 | LTBP1, MA | 1411 | 4543 | 22680 | 1.39756 | 4.27E-11 | 8.54E-12 | 1.51E-10 | -9.82041 | 9.820413 | Extracellul: | 1.066209 |
| UP_KEYW( | Innate immunity | 51 | 3.359684 | 3.29E-14 | MASP1, S1 | 1411 | 241 | 22680 | 3.40149 | 1.29E-11 | 3.23E-12 | 4.57E-11 | -10.3399 | 10.3399 | Endoplasm | 2.965805 |
| UP_KEYW( | Secreted | 173 | 11.39657 | 3.92E-11 | LTBP1, MA | 1411 | 1685 | 22680 | 1.650297 | 1.54E-08 | 1.93E-09 | 5.46E-08 | -7.26305 | 7.263048 | Secreted | 7.263048 |
| UP_KEYW( | Membrane | 658 | 43.34651 | 3.26E-11 | SGMS2, OS | 1411 | 8683 | 22680 | 1.21807 | 1.28E-08 | 1.83E-09 | 4.54E-08 | -7.34276 | 7.342765 | Membrane | 7.342765 |
| UP_KEYW( | Endoplasmic reticulum | 102 | 6.719368 | 7.78E-07 | FAM20A, C | 1411 | 997 | 22680 | 1.644451 | 3.06E-04 | 1.91E-05 | 0.001082 | -2.96581 | 2.965805 | Signal | 9.820413 |
| UP_KEYW( | Extracellular matrix | 32 | 2.108037 | 6.17E-05 | PXDN, COL | 1411 | 235 | 22680 | 2.18876 | 0.023973 | 8.66E-04 | 0.08586 | -1.06621 | 1.066209 | Innate imn | 10.3399 |
| UP_KEYW( | Protease | 64 | 4.216074 | 1.28E-06 | NPEPL1, KI | 1411 | 542 | 22680 | 1.898002 | 5.02E-04 | 2.96E-05 | 0.001778 | -2.74996 | 2.749961 | | |
| UP_KEYW( | Homeobox | 38 | 2.503294 | 1.21E-05 | TSHZ3, HN | 1411 | 280 | 22680 | 2.181432 | 0.004747 | 2.16E-04 | 0.016844 | -1.77355 | 1.773553 | | |
| UP_KEYW( | GPI-anchor | 23 | 1.515152 | 5.30E-05 | TECTA, LY6 | 1411 | 140 | 22680 | 2.64068 | 0.02061 | 7.71E-04 | 0.073695 | -1.13256 | 1.132561 | | |
| UP_KEYW( | Metal-binding | 273 | 17.98419 | 3.24E-06 | S100A4, S1 | 1411 | 3395 | 22680 | 1.292525 | 0.001274 | 6.71E-05 | 0.004512 | -2.34561 | 2.345606 | | |
| UP_KEYW( | Keratin | 12 | 0.790514 | 0.179061 | KRT6A, KR | 1411 | 129 | 22680 | 1.495229 | 1 | 0.502705 | 93.57498 | 1.97116 | -1.97116 | | |
| UP_KEYW( | Oxidoreductase | 61 | 4.018445 | 9.77E-04 | ACOX1, PX | 1411 | 639 | 22680 | 1.534423 | 0.318962 | 0.009801 | 1.350641 | 0.13054 | -0.13054 | | |
| UP_KEYW( | Tyrosine-protein kinase | 15 | 0.988142 | 0.01037 | EGFR, LYN, | 1411 | 113 | 22680 | 2.133678 | 0.98337 | 0.073056 | 13.49929 | 1.130311 | -1.13031 | | |
| UP_KEYW( | Transcription | 139 | 9.156785 | 0.01625 | MEF2A, HN | 1411 | 1859 | 22680 | 1.201853 | 0.998402 | 0.105074 | 20.3824 | 1.309255 | -1.30926 | | |
| UP_KEYW( | Ion channel | 31 | 2.042161 | 0.029953 | ORAI1, CAI | 1411 | 336 | 22680 | 1.482991 | 0.999994 | 0.147304 | 34.4975 | 1.537788 | -1.53779 | | |
| UP_KEYW( | Integrin | 14 | 0.922266 | 1.59E-05 | CD47, ADA | 1411 | 53 | 22680 | 4.245885 | 0.006228 | 2.72E-04 | 0.022114 | -1.65533 | 1.655325 | | |
| UP_KEYW( | DNA replication | 4 | 0.263505 | 0.939854 | NFIX, NFIC, | 1411 | 95 | 22680 | 0.676788 | 1 | 0.994937 | 100 | 2 | -2 | | |
| UP_KEYW( | Laminin EGF-like domain | 8 | 0.527009 | 0.002484 | LAMA4, LA | 1411 | 31 | 22680 | 4.148053 | 0.623659 | 0.021483 | 3.400348 | 0.531523 | -0.53152 | | |
| UP_KEYW( | Cell junction | 63 | 4.150198 | 8.23E-04 | CLDN9, TR | 1411 | 661 | 22680 | 1.531987 | 0.276577 | 0.008712 | 1.139572 | 0.056742 | -0.05674 | | |
| UP_KEYW( | Collagen | 13 | 0.85639 | 0.006193 | FCNA, COL | 1411 | 85 | 22680 | 2.458332 | 0.912948 | 0.048601 | 8.27915 | 0.917986 | -0.91799 | | |
| UP_KEYW( | Mitogen | 5 | 0.329381 | 0.183025 | EREG, BTC, | 1411 | 36 | 22680 | 2.232459 | 1 | 0.508021 | 93.99341 | 1.973097 | -1.9731 | | |
| UP_KEYW( | Sushi | 9 | 0.592885 | 0.007854 | C1RA, C1RI | 1411 | 47 | 22680 | 3.077944 | 0.954894 | 0.060094 | 10.38933 | 1.016588 | -1.01659 | | |
| UP_KEYW( | Molybdenum | 4 | 0.263505 | 0.015171 | XDH, SUO> | 1411 | 9 | 22680 | 7.14387 | 0.997541 | 0.101728 | 19.15831 | 1.282357 | -1.28236 | | |
| UP_KEYW( | Protease inhibitor | 14 | 0.922266 | 0.037862 | SPINK10, F | 1411 | 121 | 22680 | 1.859768 | 1 | 0.170778 | 41.54832 | 1.618554 | -1.61855 | | |
| UP_KEYW( | SH3-binding | 9 | 0.592885 | 0.031556 | SH3BGRL, I | 1411 | 60 | 22680 | 2.411056 | 0.999997 | 0.149179 | 35.98693 | 1.556145 | -1.55614 | | |
| UP_KEYW( | Transferase | 125 | 8.234519 | 0.016163 | SGMS2, RN | 1411 | 1654 | 22680 | 1.21476 | 0.998345 | 0.106267 | 20.28367 | 1.307146 | -1.30715 | | |
| UP_KEYW( | GTPase activation | 18 | 1.185771 | 0.025334 | BCR, ASAP: | 1411 | 163 | 22680 | 1.775011 | 0.999958 | 0.130698 | 30.02249 | 1.477447 | -1.47745 | | |
| UP_KEYW( | Dioxygenase | 7 | 0.461133 | 0.412987 | ALOX15, TI | 1411 | 83 | 22680 | 1.355614 | 1 | 0.756967 | 99.93955 | 1.999737 | -1.99974 | | |
| UP_KEYW( | Keratinization | 6 | 0.395257 | 0.020307 | SPRR1A, SF | 1411 | 26 | 22680 | 3.709317 | 0.999685 | 0.114998 | 24.83077 | 1.39499 | -1.39499 | | |
| UP_KEYW( | Protein phosphatase | 13 | 0.85639 | 0.110301 | PTPRJ, DU! | 1411 | 130 | 22680 | 1.607371 | 1 | 0.374172 | 80.32749 | 1.904864 | -1.90486 | | |
| UP_KEYW( | Metalloprotease | 21 | 1.383399 | 5.55E-04 | STAMBP, C | 1411 | 143 | 22680 | 2.360474 | 0.195861 | 0.006209 | 0.76869 | -0.11425 | 0.114249 | | |
| UP_KEYW( | Molybdenum | 4 | 0.263505 | 0.015171 | XDH, SUO> | 1411 | 9 | 22680 | 7.14387 | 0.997541 | 0.101728 | 19.15831 | 1.282357 | -1.28236 | | |
| UP_KEYW( | Biological rhythms | 13 | 0.85639 | 0.092788 | PROK2, NP | 1411 | 126 | 22680 | 1.658398 | 1 | 0.337348 | 74.19911 | 1.870399 | -1.8704 | | |
| UP_KEYW( | Kringle | 4 | 0.263505 | 0.062258 | KREMEN1, | 1411 | 15 | 22680 | 4.286322 | 1 | 0.252015 | 59.11017 | 1.771662 | -1.77166 | | |
| UP_KEYW( | Anion exchange | 4 | 0.263505 | 0.020684 | SLC4A11, S | 1411 | 10 | 22680 | 6.429483 | 0.999729 | 0.115381 | 25.23197 | 1.401951 | -1.40195 | | |
| UP_KEYW( | Growth factor binding | 6 | 0.395257 | 0.001637 | LTBP1, HTI | 1411 | 15 | 22680 | 6.429483 | 0.474675 | 0.01521 | 2.253066 | 0.352774 | -0.35277 | | |
| UP_KEYW( | Carboxypeptidase | 5 | 0.329381 | 0.158262 | CPA4, ACE, | 1411 | 34 | 22680 | 2.36378 | 1 | 0.468892 | 90.89989 | 1.958563 | -1.95856 | | |
| UP_KEYW( | Lyase | 12 | 0.790514 | 0.299114 | L3HYPDH, | 1411 | 146 | 22680 | 1.321127 | 1 | 0.661341 | 99.28778 | 1.996896 | -1.9969 | | |
| UP_KEYW( | Antiviral defense | 9 | 0.592885 | 0.282434 | IL10RB, TIC | 1411 | 100 | 22680 | 1.446634 | 1 | 0.644838 | 99.01208 | 1.995688 | -1.99569 | | |
| UP_KEYW( | Guanine-nucleotide releasing factor | 12 | 0.790514 | 0.20509 | ARHGEF3, | 1411 | 133 | 22680 | 1.450259 | 1 | 0.537436 | 95.89602 | 1.981801 | -1.9818 | | |
| UP_KEYW( | Protease inhibitor | 14 | 0.922266 | 0.037862 | SPINK10, F | 1411 | 121 | 22680 | 1.859768 | 1 | 0.170778 | 41.54832 | 1.618554 | -1.61855 | | |
| UP_KEYW( | Tight junction | 8 | 0.527009 | 0.257141 | CGNL1, SH | 1411 | 83 | 22680 | 1.549273 | 1 | 0.613162 | 98.40038 | 1.992997 | -1.993 | | |
| UP_KEYW( | Immunoglobulin domain | 36 | 2.371542 | 0.190687 | MPZL3, PS( | 1411 | 481 | 22680 | 1.203022 | 1 | 0.517782 | 94.7314 | 1.976494 | -1.97649 | | |
| UP_KEYW( | Fatty acid metabolism | 13 | 0.85639 | 0.084712 | ACOX1, AL( | 1411 | 124 | 22680 | 1.685147 | 1 | 0.320585 | 70.81355 | 1.850116 | -1.85012 | | |
| UP_KEYW( | SH2 domain | 9 | 0.592885 | 0.328811 | SH2D1B1, I | 1411 | 105 | 22680 | 1.377746 | 1 | 0.684039 | 99.61003 | 1.998303 | -1.9983 | | |
| UP_KEYW( | Prenylation | 14 | 0.922266 | 0.179019 | RHOJ, RAB | 1411 | 157 | 22680 | 1.433324 | 1 | 0.505764 | 93.57045 | 1.971139 | -1.97114 | | |
| UP_KEYW( | Copper | 6 | 0.395257 | 0.292027 | PAM, LOXL | 1411 | 58 | 22680 | 1.662797 | 1 | 0.656538 | 99.18078 | 1.996428 | -1.99643 | | |
| UP_KEYW( | Amino-acid transport | 4 | 0.263505 | 0.406599 | SLC6A9, SL | 1411 | 37 | 22680 | 1.737698 | 1 | 0.752227 | 99.92973 | 1.999695 | -1.99969 | | |
| UP_KEYW( | SH3 domain | 16 | 1.054018 | 0.313748 | SH3PXD2B | 1411 | 208 | 22680 | 1.236439 | 1 | 0.671278 | 99.46896 | 1.997688 | -1.99769 | | |
| UP_KEYW( | Cyclin | 4 | 0.263505 | 0.489395 | CDKN1A, S | 1411 | 42 | 22680 | 1.530829 | 1 | 0.808141 | 99.99131 | 1.999962 | -1.99996 | | |
| UP_KEYW( | Potassium channel | 3 | 0.197628 | 0.926491 | KCNK7, KC | 1411 | 67 | 22680 | 0.719718 | 1 | 0.992788 | 100 | 2 | -2 | | |
| UP_KEYW( | Zinc transport | 3 | 0.197628 | 0.401106 | SLC39A8, S | 1411 | 22 | 22680 | 2.191869 | 1 | 0.748421 | 99.92012 | 1.999653 | -1.99965 | | |
| UP_KEYW( | Mitochondrion inner membrane | 11 | 0.724638 | 0.957788 | MICU1, DU | 1411 | 254 | 22680 | 0.696105 | 1 | 0.996928 | 100 | 2 | -2 | | |
| UP_KEYW( | RNA-binding | 17 | 1.119895 | 0.999985 | RBFOX1, EI | 1411 | 601 | 22680 | 0.454664 | 1 | 1 | 100 | 2 | -2 | | |
| UP_KEYW( | Lectin | 16 | 1.054018 | 0.120801 | GALNT1, E' | 1411 | 173 | 22680 | 1.486586 | 1 | 0.400149 | 83.3225 | 1.920762 | -1.92076 | | |
| UP_KEYW( | Voltage-gated channel | 9 | 0.592885 | 0.615244 | CLIC3, KCN | 1411 | 136 | 22680 | 1.063701 | 1 | 0.884365 | 99.99983 | 1.999999 | -2 | | |
| UP_KEYW( | Synapse | 25 | 1.646904 | 0.378052 | FLRT2, PH/ | 1411 | 357 | 22680 | 1.12561 | 1 | 0.731346 | 99.8649 | 1.999413 | -1.99941 | | |
| UP_KEYW( | Hemostasis | 3 | 0.197628 | 0.767803 | ANXA8, EN | 1411 | 44 | 22680 | 1.095935 | 1 | 0.951211 | 100 | 2 | -2 | | |
| UP_KEYW( | Immunoglobulin domain | 36 | 2.371542 | 0.190687 | MPZL3, PS( | 1411 | 481 | 22680 | 1.203022 | 1 | 0.517782 | 94.7314 | 1.976494 | -1.97649 | | |
| UP_KEYW( | Iron | 26 | 1.71278 | 0.390513 | XDH, PXDN | 1411 | 375 | 22680 | 1.114444 | 1 | 0.741099 | 99.89805 | 1.999557 | -1.99956 | | |
| UP_KEYW( | Hydrogen ion transport | 3 | 0.197628 | 0.778571 | ATP6V1C2, | 1411 | 45 | 22680 | 1.07158 | 1 | 0.954315 | 100 | 2 | -2 | | |
| UP_KEYW( | LIM domain | 5 | 0.329381 | 0.683182 | CRIP1, LIM | 1411 | 74 | 22680 | 1.086061 | 1 | 0.918698 | 99.99999 | 2 | -2 | | |
| UP_KEYW( | Chloride channel | 3 | 0.197628 | 0.7889 | CLIC3, ANC | 1411 | 46 | 22680 | 1.048285 | 1 | 0.95649 | 100 | 2 | -2 | | |
| UP_KEYW( | Peroxisome | 6 | 0.395257 | 0.802412 | HAO1, XDH | 1411 | 107 | 22680 | 0.901329 | 1 | 0.961281 | 100 | 2 | -2 | | |

Annotatior Enrichment Score: 0.17791603524492008

| Category | Term |
|---|---|
| INTERPRO | IPR024156:Small GTPase superfamily, ARF type |
| INTERPRO | IPR006689:Small GTPase superfamily, ARF/SAR type |
| UP_SEQ_F | lipid moiety-binding region:N-myristoyl glycine |

Annotatior Enrichment Score: 0.17363305968109202

| Category | Term |
|---|---|
| UP_SEQ_F | domain:Ig-like C1-type |
| INTERPRO | IPR003597:Immunoglobulin C1-set |
| INTERPRO | IPR003006:Immunoglobulin/major histocompatibility complex, conserved site |
| GOTERM_I | GO:0002474~antigen processing and presentation of peptide antigen via MHC class I |

| | |
|---|---|
| INTERPRO | IPR001039:MHC class I, alpha chain, alpha1/alpha2 |
| SMART | SM00407:IGc1 |
| INTERPRO | IPR011161:MHC class I-like antigen recognition |
| INTERPRO | IPR011162:MHC classes I/II-like antigen recognition protein |
| Annotatior | Enrichment Score: 0.16710023499759513 |
| Category | Term |
| GOTERM_( | GO:0001917~photoreceptor inner segment |
| GOTERM_( | GO:0032420~stereocilium |
| GOTERM_I | GO:0045494~photoreceptor cell maintenance |
| Annotatior | Enrichment Score: 0.15611550761349594 |
| Category | Term |
| INTERPRO | IPR018303:P-type ATPase, phosphorylation site |
| INTERPRO | IPR001757:Cation-transporting P-type ATPase |
| INTERPRO | IPR023299:P-type ATPase, cytoplasmic domain N |
| INTERPRO | IPR008250:P-type ATPase, A domain |
| Annotatior | Enrichment Score: 0.14702969087727702 |
| Category | Term |
| GOTERM_I | GO:0008144~drug binding |
| GOTERM_I | GO:0042220~response to cocaine |
| SMART | SM01381:SM01381 |
| Annotatior | Enrichment Score: 0.1437633581618954 |
| Category | Term |
| UP_SEQ_F | domain:F-box |
| INTERPRO | IPR001810:F-box domain, cyclin-like |
| SMART | SM00256:FBOX |
| Annotatior | Enrichment Score: 0.12466983515768307 |
| Category | Term |
| GOTERM_I | GO:0004867~serine-type endopeptidase inhibitor activity |
| INTERPRO | IPR023796:Serpin domain |
| INTERPRO | IPR000215:Serpin family |
| INTERPRO | IPR023795:Protease inhibitor I4, serpin, conserved site |
| SMART | SM00093:SERPIN |
| Annotatior | Enrichment Score: 0.12169666409389673 |
| Category | Term |
| INTERPRO | IPR011333:BTB/POZ fold |
| UP_SEQ_F | domain:BTB |
| INTERPRO | IPR000210:BTB/POZ-like |
| SMART | SM00225:BTB |
| Annotatior | Enrichment Score: 0.10461835158091745 |
| Category | Term |
| GOTERM_( | GO:0016459~myosin complex |
| INTERPRO | IPR001609:Myosin head, motor domain |
| GOTERM_I | GO:0003774~motor activity |
| UP_KEYW( | Myosin |
| SMART | SM00242:MYSc |
| UP_KEYW( | Motor protein |
| Annotatior | Enrichment Score: 0.10288893333549652 |
| Category | Term |
| GOTERM_I | GO:0008654~phospholipid biosynthetic process |
| UP_KEYW( | Lipid biosynthesis |
| UP_KEYW( | Phospholipid biosynthesis |
| UP_KEYW( | Phospholipid metabolism |
| Annotatior | Enrichment Score: 0.09798767769942301 |
| Category | Term |
| KEGG_PAT | mmu04612:Antigen processing and presentation |
| KEGG_PAT | mmu05332:Graft-versus-host disease |
| KEGG_PAT | mmu05330:Allograft rejection |
| KEGG_PAT | mmu04940:Type I diabetes mellitus |
| Annotatior | Enrichment Score: 0.08771443711055353 |
| Category | Term |
| UP_SEQ_F | zinc finger region:C2H2-type 1 |
| UP_SEQ_F | zinc finger region:C2H2-type 2 |
| UP_SEQ_F | zinc finger region:C2H2-type 3 |
| UP_SEQ_F | zinc finger region:C2H2-type 5 |
| UP_SEQ_F | zinc finger region:C2H2-type 4 |
| UP_SEQ_F | zinc finger region:C2H2-type 6 |
| INTERPRO | IPR015880:Zinc finger, C2H2-like |
| INTERPRO | IPR007087:Zinc finger, C2H2 |
| UP_SEQ_F | zinc finger region:C2H2-type 9 |
| UP_SEQ_F | zinc finger region:C2H2-type 7 |

| | |
|---|---|
| INTERPRO | IPR013087:Zinc finger C2H2-type/integrase DNA-binding domain |
| UP_SEQ_F | zinc finger region:C2H2-type 8 |
| SMART | SM00355:ZnF_C2H2 |
| GOTERM_I | GO:0003676~nucleic acid binding |

| | |
|---|---|
| Annotatior | Enrichment Score: 0.0736668476120031 |
| Category | Term |
| GOTERM_I | GO:0007601~visual perception |
| UP_KEYW( | Vision |
| GOTERM_I | GO:0050896~response to stimulus |
| UP_KEYW( | Sensory transduction |

| | |
|---|---|
| Annotatior | Enrichment Score: 0.07114143744011604 |
| Category | Term |
| KEGG_PAT | mmu04080:Neuroactive ligand-receptor interaction |
| GOTERM_I | GO:0004871~signal transducer activity |
| INTERPRO | IPR000276:G protein-coupled receptor, rhodopsin-like |
| UP_KEYW( | Transducer |
| GOTERM_I | GO:0007186~G-protein coupled receptor signaling pathway |
| INTERPRO | IPR017452:GPCR, rhodopsin-like, 7TM |
| GOTERM_I | GO:0004930~G-protein coupled receptor activity |
| UP_KEYW( | G-protein coupled receptor |

| | |
|---|---|
| Annotatior | Enrichment Score: 0.06318656316440345 |
| Category | Term |
| UP_KEYW( | Lipid biosynthesis |
| UP_KEYW( | Fatty acid biosynthesis |
| GOTERM_I | GO:0006633~fatty acid biosynthetic process |

| | |
|---|---|
| Annotatior | Enrichment Score: 0.06225290592184858 |
| Category | Term |
| INTERPRO | IPR011011:Zinc finger, FYVE/PHD-type |
| INTERPRO | IPR019786:Zinc finger, PHD-type, conserved site |
| UP_SEQ_F | zinc finger region:PHD-type |
| INTERPRO | IPR019787:Zinc finger, PHD-finger |
| INTERPRO | IPR001965:Zinc finger, PHD-type |
| SMART | SM00249:PHD |

| | |
|---|---|
| Annotatior | Enrichment Score: 0.0578828766388336 |
| Category | Term |
| GOTERM_I | GO:0008202~steroid metabolic process |
| GOTERM_I | GO:0008203~cholesterol metabolic process |
| UP_KEYW( | Cholesterol metabolism |
| UP_KEYW( | Sterol metabolism |
| UP_KEYW( | Steroid metabolism |

| | |
|---|---|
| Annotatior | Enrichment Score: 0.05323723411983497 |
| Category | Term |
| INTERPRO | IPR000225:Armadillo |
| SMART | SM00185:ARM |
| INTERPRO | IPR011989:Armadillo-like helical |

| | |
|---|---|
| Annotatior | Enrichment Score: 0.04523589907780113 |
| Category | Term |
| UP_KEYW( | Endonuclease |
| UP_KEYW( | Nuclease |
| GOTERM_I | GO:0004518~nuclease activity |
| GOTERM_I | GO:0004519~endonuclease activity |

| | |
|---|---|
| Annotatior | Enrichment Score: 0.04175260006786172 |
| Category | Term |
| INTERPRO | IPR015915:Kelch-type beta propeller |
| INTERPRO | IPR006652:Kelch repeat type 1 |
| UP_KEYW( | Kelch repeat |

| | |
|---|---|
| Annotatior | Enrichment Score: 0.02142254711233065 |
| Category | Term |
| UP_SEQ_F | repeat:WD 9 |
| UP_SEQ_F | repeat:WD 8 |
| INTERPRO | IPR015943:WD40/YVTN repeat-like-containing domain |
| UP_SEQ_F | repeat:WD 7 |
| UP_SEQ_F | repeat:WD 3 |
| UP_SEQ_F | repeat:WD 1 |
| UP_SEQ_F | repeat:WD 2 |
| UP_KEYW( | WD repeat |
| UP_SEQ_F | repeat:WD 6 |
| UP_SEQ_F | repeat:WD 4 |
| UP_SEQ_F | repeat:WD 5 |
| INTERPRO | IPR019775:WD40 repeat, conserved site |
| INTERPRO | IPR001680:WD40 repeat |

| | |
|---|---|
| INTERPRO | IPR017986:WD40-repeat-containing domain |
| SMART | SM00320:WD40 |
| Annotatior | Enrichment Score: 0.006980563073946442 |
| Category | Term |
| INTERPRO | IPR011990:Tetratricopeptide-like helical |
| INTERPRO | IPR019734:Tetratricopeptide repeat |
| SMART | SM00028:TPR |
| Annotatior | Enrichment Score: 0.006781670027664856 |
| Category | Term |
| INTERPRO | IPR011990:Tetratricopeptide-like helical |
| UP_KEYW( | TPR repeat |
| UP_SEQ_F | repeat:TPR 2 |
| UP_SEQ_F | repeat:TPR 1 |
| INTERPRO | IPR013026:Tetratricopeptide repeat-containing domain |
| UP_SEQ_F | repeat:TPR 3 |
| Annotatior | Enrichment Score: 0.0012152573762811024 |
| Category | Term |
| UP_SEQ_F | domain:RRM |
| INTERPRO | IPR012677:Nucleotide-binding, alpha-beta plait |
| INTERPRO | IPR000504:RNA recognition motif domain |
| SMART | SM00360:RRM |
| Annotatior | Enrichment Score: 2.411303208364301E-4 |
| Category | Term |
| UP_KEYW( | Mitosis |
| UP_KEYW( | Cell cycle |
| GOTERM_I | GO:0007067~mitotic nuclear division |
| UP_KEYW( | Cell division |
| GOTERM_I | GO:0007049~cell cycle |
| GOTERM_I | GO:0051301~cell division |
| Annotatior | Enrichment Score: 1.2340246761732496E-4 |
| Category | Term |
| UP_KEYW( | DNA damage |
| UP_KEYW( | DNA repair |
| GOTERM_I | GO:0006281~DNA repair |
| Annotatior | Enrichment Score: 5.160978785673568E-5 |
| Category | Term |
| UP_KEYW( | Methyltransferase |
| GOTERM_I | GO:0032259~methylation |
| GOTERM_I | GO:0008168~methyltransferase activity |
| Annotatior | Enrichment Score: 8.058640278449858E-6 |
| Category | Term |
| UP_KEYW( | mRNA splicing |
| UP_KEYW( | mRNA processing |
| GOTERM_I | GO:0006397~mRNA processing |
| GOTERM_I | GO:0008380~RNA splicing |
