## Supplementary_Table3 for "Aging-dependent dysregulation of EXOSC2 is maintained in cancer as a dependency"

**SUPPLEMENTARY TABLE 3**

Annotation Cluster 1

Enrichment Score: 12.502183696951953

| Category | Term | Count | % | PValue | Genes | List Total | Pop Hits | Pop Total | ld Enrichm | Bonferroni | Benjamini | FDR |
|---|---|---|---|---|---|---|---|---|---|---|---|---|
| UP_KEYWORDS | Zinc-finger | 104 | 14.11126 | 2.81E-12 | Q8CIP9, E9 | 732 | 1565 | 22680 | 2.058974 | 9.37E-10 | 1.34E-10 | 3.81E-09 |
| UP_KEYWORDS | Lipid biosynthesis | 16 | 2.170963 | 2.16E-04 | 48, Q8K0C4 | 732 | 160 | 22680 | 3.098361 | 0.069752 | 0.001952 | 0.293581 |
| UP_KEYWORDS | DNA repair | 23 | 3.12076 | 1.13E-04 | Z7, B2RSG2 | 732 | 279 | 22680 | 2.554204 | 0.036903 | 0.001105 | 0.152783 |
| UP_KEYWORDS | Differentiation | 37 | 5.020353 | 0.00107 | 5381, Q9Q | 732 | 646 | 22680 | 1.774603 | 0.300586 | 0.007913 | 1.443254 |
| UP_KEYWORDS | Cell cycle | 42 | 5.698779 | 1.57E-05 | 41, Q80YR6 | 732 | 626 | 22680 | 2.078772 | 0.00523 | 2.18E-04 | 0.021319 |
| UP_KEYWORDS | Actin-binding | 21 | 2.849389 | 2.10E-04 | 0A0A6YW | 732 | 252 | 22680 | 2.581967 | 0.067597 | 0.001942 | 0.2842 |

| | | | | | | | | | | | | |
|---|---|---|---|---|---|---|---|---|---|---|---|---|
| GOTERM_BP_DIRECT | GO:0007599~hemostasis | 3 | 0.407056 | 0.504427 | 2, Q3UER8, | 661 | 46 | 18082 | 1.784056 | 1 | 0.999803 | 99.99966 |
| KEGG_PATHWAY | mmu04610:Complement and coagulation cascades | 3 | 0.407056 | 0.763376 | 2, Q3UER8, | 277 | 76 | 7691 | 1.096 | 1 | 0.980542 | 100 |
| Annotation Cluster 74 | Enrichment Score: 0.5197157543925095 | | | | | | | | | | | |
| Category | Term | Count | % | PValue | Genes | List Total | Pop Hits | Pop Total | ld Enrichme | Bonferroni | Benjamini | FDR |
| KEGG_PATHWAY | mmu04520:Adherens junction | 6 | 0.814111 | 0.115677 | R4IZW5, E9 | 277 | 72 | 7691 | 2.313779 | 1 | 0.635071 | 79.66786 |
| KEGG_PATHWAY | mmu05216:Thyroid cancer | 3 | 0.407056 | 0.279448 | A0A0R4IZW | 277 | 29 | 7691 | 2.872277 | 1 | 0.833312 | 98.56904 |
| KEGG_PATHWAY | mmu04390:Hippo signaling pathway | 7 | 0.949796 | 0.459095 | 0A0R4IZW5 | 277 | 151 | 7691 | 1.287135 | 1 | 0.916106 | 99.96518 |
| KEGG_PATHWAY | mmu05213:Endometrial cancer | 3 | 0.407056 | 0.561933 | A0A0R4IZW | 277 | 52 | 7691 | 1.601847 | 1 | 0.951709 | 99.99773 |
| Annotation Cluster 75 | Enrichment Score: 0.5167671061382304 | | | | | | | | | | | |
| Category | Term | Count | % | PValue | Genes | List Total | Pop Hits | Pop Total | ld Enrichme | Bonferroni | Benjamini | FDR |
| INTERPRO | IPR002172:Low-density lipoprotein (LDL) receptor class A repeat | 4 | 0.542741 | 0.232705 | D3K4, O883 | 688 | 50 | 20594 | 2.394651 | 1 | 0.98484 | 98.66615 |
| SMART | SM00192:LDLa | 4 | 0.542741 | 0.286354 | D3K4, O883 | 415 | 47 | 10425 | 2.137913 | 1 | 0.9541 | 98.82733 |
| INTERPRO | IPR023415:Low-density lipoprotein (LDL) receptor class A, conserved site | 3 | 0.407056 | 0.422659 | 4, O88307, | 688 | 43 | 20594 | 2.088359 | 1 | 0.998607 | 99.98706 |
| Annotation Cluster 76 | Enrichment Score: 0.493494868358881 | | | | | | | | | | | |
| Category | Term | Count | % | PValue | Genes | List Total | Pop Hits | Pop Total | ld Enrichme | Bonferroni | Benjamini | FDR |
| BIOCARTA | m_ps1Pathway:Presenilin action in Notch and Wnt signaling | 3 | 0.407056 | 0.146778 | 3, P31266, | 64 | 14 | 1289 | 4.315848 | 1 | 0.981096 | 82.82748 |
| GOTERM_BP_DIRECT | GO:0045596~negative regulation of cell differentiation | 7 | 0.949796 | 0.237492 | A0A0R4J04 | 661 | 114 | 18082 | 1.679725 | 1 | 0.988463 | 99.2251 |
| GOTERM_BP_DIRECT | GO:0030182~neuron differentiation | 3 | 0.407056 | 0.948866 | 3, P31266, | 661 | 127 | 18082 | 0.646193 | 1 | 1 | 100 |
| Annotation Cluster 77 | Enrichment Score: 0.48792090020484385 | | | | | | | | | | | |
| Category | Term | Count | % | PValue | Genes | List Total | Pop Hits | Pop Total | ld Enrichme | Bonferroni | Benjamini | FDR |
| GOTERM_MF_DIRECT | GO:0004843~thiol-dependent ubiquitin-specific protease activity | 6 | 0.814111 | 0.181777 | TCH2, Q6IE | 648 | 81 | 17446 | 1.994284 | 1 | 0.876671 | 95.50262 |
| UP_KEYWORDS | Thiol protease | 7 | 0.949796 | 0.264244 | , Q3TCH2, | 732 | 134 | 22680 | 1.618547 | 1 | 0.602808 | 98.45109 |
| GOTERM_MF_DIRECT | GO:0036459~thiol-dependent ubiquitinyl hydrolase activity | 5 | 0.678426 | 0.319723 | 2, Q6IE21, | 648 | 77 | 17446 | 1.748236 | 1 | 0.952281 | 99.74102 |
| GOTERM_BP_DIRECT | GO:0006511~ubiquitin-dependent protein catabolic process | 8 | 1.085482 | 0.331645 | CH2, Q3US | 661 | 154 | 18082 | 1.421066 | 1 | 0.996615 | 99.92701 |
| GOTERM_MF_DIRECT | GO:0008234~cysteine-type peptidase activity | 7 | 0.949796 | 0.425052 | , Q3TCH2, | 648 | 141 | 17446 | 1.336595 | 1 | 0.976987 | 99.98078 |
| GOTERM_BP_DIRECT | GO:0016579~protein deubiquitination | 4 | 0.542741 | 0.545807 | TCH2, Q99 | 661 | 78 | 18082 | 1.402847 | 1 | 0.999891 | 99.99993 |
| Annotation Cluster 78 | Enrichment Score: 0.48642344720533753 | | | | | | | | | | | |
| Category | Term | Count | % | PValue | Genes | List Total | Pop Hits | Pop Total | ld Enrichme | Bonferroni | Benjamini | FDR |
| GOTERM_BP_DIRECT | GO:0048488~synaptic vesicle endocytosis | 3 | 0.407056 | 0.126196 | A0A0G2JEG | 661 | 17 | 18082 | 4.827445 | 1 | 0.944939 | 91.09045 |
| UP_KEYWORDS | SH3 domain | 9 | 1.221167 | 0.356542 | R4J0P5, Q9 | 732 | 208 | 22680 | 1.340637 | 1 | 0.703891 | 99.74916 |
| UP_SEQ_FEATURE | domain:SH3 | 8 | 1.085482 | 0.376944 | 0A0R4J0P5, | 641 | 166 | 18012 | 1.354209 | 1 | 0.999826 | 99.96868 |
| INTERPRO | IPR001452:Src homology-3 domain | 9 | 1.221167 | 0.414771 | R4J0P5, Q9 | 688 | 212 | 20594 | 1.270746 | 1 | 0.99855 | 99.98386 |
| SMART | SM00326:SH3 | 9 | 1.221167 | 0.525584 | R4J0P5, Q9 | 415 | 197 | 10425 | 1.147636 | 1 | 0.98933 | 99.9946 |
| Annotation Cluster 79 | Enrichment Score: 0.4863318381177524 | | | | | | | | | | | |
| Category | Term | Count | % | PValue | Genes | List Total | Pop Hits | Pop Total | ld Enrichme | Bonferroni | Benjamini | FDR |
| INTERPRO | IPR003347:JmjC domain | 3 | 0.407056 | 0.301903 | 5, Q8BK58, | 688 | 33 | 20594 | 2.721195 | 1 | 0.99329 | 99.71416 |
| UP_SEQ_FEATURE | domain:JmjC | 3 | 0.407056 | 0.328586 | 5, Q8BK58, | 641 | 33 | 18012 | 2.554531 | 1 | 0.999717 | 99.88793 |
| SMART | SM00558:JmjC | 3 | 0.407056 | 0.350339 | 5, Q8BK58, | 415 | 31 | 10425 | 2.431014 | 1 | 0.958992 | 99.65994 |
| Annotation Cluster 80 | Enrichment Score: 0.4849397553954399 | | | | | | | | | | | |
| Category | Term | Count | % | PValue | Genes | List Total | Pop Hits | Pop Total | ld Enrichme | Bonferroni | Benjamini | FDR |
| UP_SEQ_FEATURE | calcium-binding region:5 | 3 | 0.407056 | 0.023508 | 7, Q08331, | 641 | 7 | 18012 | 12.04279 | 1 | 0.903316 | 33.34856 |
| UP_SEQ_FEATURE | domain:EF-hand 6 | 3 | 0.407056 | 0.030615 | 7, Q08331, | 641 | 8 | 18012 | 10.53744 | 1 | 0.945033 | 41.1552 |
| UP_SEQ_FEATURE | domain:EF-hand 5 | 3 | 0.407056 | 0.08653 | 7, Q08331, | 641 | 14 | 18012 | 6.021395 | 1 | 0.988254 | 78.63615 |
| UP_SEQ_FEATURE | calcium-binding region:4 | 3 | 0.407056 | 0.132752 | 7, Q08331, | 641 | 18 | 18012 | 4.683307 | 1 | 0.995085 | 91.18754 |
| INTERPRO | IPR018247:EF-Hand 1, calcium-binding site | 8 | 1.085482 | 0.367076 | 5DDB7, Q9 | 688 | 175 | 20594 | 1.368372 | 1 | 0.997147 | 99.94214 |
| INTERPRO | IPR011992:EF-hand-like domain | 11 | 1.492537 | 0.430147 | 49, G5DDB7 | 688 | 273 | 20594 | 1.206097 | 1 | 0.998732 | 99.98954 |
| UP_SEQ_FEATURE | calcium-binding region:1 | 6 | 0.814111 | 0.465466 | QUG9, E9Q2 | 641 | 126 | 18012 | 1.338088 | 1 | 0.999965 | 99.9977 |
| UP_SEQ_FEATURE | calcium-binding region:2 | 5 | 0.678426 | 0.580663 | G5DDB7, C | 641 | 114 | 18012 | 1.232449 | 1 | 0.999996 | 99.99996 |
| UP_KEYWORDS | Calcium | 27 | 3.663501 | 0.58078 | 8, Q3TJX0, | 732 | 827 | 22680 | 1.011557 | 1 | 0.861277 | 99.99925 |
| UP_SEQ_FEATURE | domain:EF-hand 2 | 7 | 0.949796 | 0.581745 | , Q9QUG9, | 641 | 173 | 18012 | 1.136988 | 1 | 0.999996 | 99.99997 |
| UP_SEQ_FEATURE | domain:EF-hand 1 | 7 | 0.949796 | 0.587392 | , Q9QUG9, | 641 | 174 | 18012 | 1.130453 | 1 | 0.999996 | 99.99997 |
| UP_SEQ_FEATURE | domain:EF-hand 4 | 3 | 0.407056 | 0.625017 | 7, Q08331, | 641 | 59 | 18012 | 1.428806 | 1 | 0.999998 | 99.99999 |
| SMART | SM00054:EFh | 6 | 0.814111 | 0.688211 | QUG9, E9Q2 | 415 | 145 | 10425 | 1.039468 | 1 | 0.997297 | 99.99998 |
| INTERPRO | IPR002048:EF-hand domain | 7 | 0.949796 | 0.757924 | , Q9QUG9, | 688 | 223 | 20594 | 0.939605 | 1 | 0.999999 | 100 |
| UP_SEQ_FEATURE | domain:EF-hand 3 | 3 | 0.407056 | 0.838925 | 7, Q08331, | 641 | 91 | 18012 | 0.926368 | 1 | 1 | 100 |
| GOTERM_MF_DIRECT | GO:0005509~calcium ion binding | 21 | 2.849389 | 0.909457 | 287, O8832 | 648 | 699 | 17446 | 0.808841 | 1 | 0.999992 | 100 |
| Annotation Cluster 81 | Enrichment Score: 0.483815864865665 | | | | | | | | | | | |
| Category | Term | Count | % | PValue | Genes | List Total | Pop Hits | Pop Total | ld Enrichme | Bonferroni | Benjamini | FDR |
| UP_SEQ_FEATURE | domain:EGF-like 5; calcium-binding | 4 | 0.542741 | 0.09639 | 0493, Q088 | 641 | 31 | 18012 | 3.625786 | 1 | 0.988926 | 82.24572 |
| INTERPRO | IPR009030:Insulin-like growth factor binding protein, N-terminal | 8 | 1.085482 | 0.144524 | 8879, Q017 | 688 | 130 | 20594 | 1.842039 | 1 | 0.962298 | 92.14553 |
| UP_SEQ_FEATURE | domain:EGF-like 3; calcium-binding | 4 | 0.542741 | 0.168888 | 0493, Q088 | 641 | 40 | 18012 | 2.809984 | 1 | 0.997656 | 95.73542 |
| INTERPRO | IPR018097:EGF-like calcium-binding, conserved site | 6 | 0.814111 | 0.223089 | 8879, Q3U | 688 | 97 | 20594 | 1.851534 | 1 | 0.984997 | 98.36596 |
| INTERPRO | IPR000742:Epidermal growth factor-like domain | 11 | 1.492537 | 0.270932 | 9137, F8VQ | 688 | 237 | 20594 | 1.389302 | 1 | 0.989946 | 99.42006 |
| SMART | SM00181:EGF | 10 | 1.356852 | 0.292458 | , F8VQJ3, C | 415 | 181 | 10425 | 1.387872 | 1 | 0.95301 | 98.95284 |
| INTERPRO | IPR013032:EGF-like, conserved site | 9 | 1.221167 | 0.33634 | 9137, Q3U | 688 | 197 | 20594 | 1.367504 | 1 | 0.996254 | 99.87467 |
| UP_SEQ_FEATURE | domain:EGF-like 1 | 6 | 0.814111 | 0.359953 | 8879, Q3U | 641 | 111 | 18012 | 1.51891 | 1 | 0.999796 | 99.95044 |
| INTERPRO | IPR001881:EGF-like calcium-binding | 6 | 0.814111 | 0.411568 | 8879, Q3U | 688 | 126 | 20594 | 1.425388 | 1 | 0.998633 | 99.98236 |
| INTERPRO | IPR000152:EGF-type aspartate/asparagine hydroxylation site | 5 | 0.678426 | 0.413976 | 3, Q08879, | 688 | 98 | 20594 | 1.527201 | 1 | 0.998615 | 99.9835 |
| UP_KEYWORDS | EGF-like domain | 9 | 1.221167 | 0.434922 | , O88307, | 732 | 224 | 22680 | 1.244877 | 1 | 0.769268 | 99.95702 |
| UP_SEQ_FEATURE | domain:EGF-like 2; calcium-binding | 3 | 0.407056 | 0.556007 | 9, Q3UTJ7, | 641 | 52 | 18012 | 1.621145 | 1 | 0.999993 | 99.9999 |
| SMART | SM00179:EGF_CA | 6 | 0.814111 | 0.564785 | 8879, Q3U | 415 | 126 | 10425 | 1.196213 | 1 | 0.991338 | 99.99827 |
| UP_SEQ_FEATURE | domain:EGF-like 4 | 3 | 0.407056 | 0.586646 | 2, P10493, | 641 | 55 | 18012 | 1.532719 | 1 | 0.999996 | 99.99997 |
| UP_SEQ_FEATURE | domain:EGF-like 2 | 3 | 0.407056 | 0.804627 | 2, P10493, | 641 | 84 | 18012 | 1.003566 | 1 | 1 | 100 |
| Annotation Cluster 82 | Enrichment Score: 0.4782942542110664 | | | | | | | | | | | |
| Category | Term | Count | % | PValue | Genes | List Total | Pop Hits | Pop Total | ld Enrichme | Bonferroni | Benjamini | FDR |
| INTERPRO | IPR000306:Zinc finger, FYVE-type | 3 | 0.407056 | 0.264488 | 4, Q571N6, | 688 | 30 | 20594 | 2.993314 | 1 | 0.989144 | 99.33062 |
| SMART | SM00064:FYVE | 3 | 0.407056 | 0.335706 | 4, Q571N6, | 415 | 30 | 10425 | 2.512048 | 1 | 0.959324 | 99.54393 |
| INTERPRO | IPR017455:Zinc finger, FYVE-related | 3 | 0.407056 | 0.338976 | 4, Q571N6, | 688 | 36 | 20594 | 2.494428 | 1 | 0.99601 | 99.88254 |
| UP_SEQ_FEATURE | zinc finger region:FYVE-type | 3 | 0.407056 | 0.405779 | 4, Q571N6, | 641 | 39 | 18012 | 2.161526 | 1 | 0.999904 | 99.98604 |
| Annotation Cluster 83 | Enrichment Score: 0.4648824213088968 | | | | | | | | | | | |
| Category | Term | Count | % | PValue | Genes | List Total | Pop Hits | Pop Total | ld Enrichme | Bonferroni | Benjamini | FDR |
| INTERPRO | IPR013026:Tetratricopeptide repeat-containing domain | 7 | 0.949796 | 0.212197 | 5, Q9CX34, | 688 | 120 | 20594 | 1.7461 | 1 | 0.983897 | 97.94974 |
| INTERPRO | IPR011990:Tetratricopeptide-like helical | 10 | 1.356852 | 0.24252 | 16, Q9CX34 | 688 | 204 | 20594 | 1.467311 | 1 | 0.986252 | 98.91861 |
| INTERPRO | IPR019734:Tetratricopeptide repeat | 7 | 0.949796 | 0.244724 | 5, Q9CX34, | 688 | 126 | 20594 | 1.662952 | 1 | 0.986161 | 98.96876 |
| UP_KEYWORDS | TPR repeat | 7 | 0.949796 | 0.320111 | 5, Q9CX34, | 732 | 144 | 22680 | 1.506148 | 1 | 0.6676 | 99.47005 |
| SMART | SM00028:TPR | 7 | 0.949796 | 0.334717 | 5, Q9CX34, | 415 | 119 | 10425 | 1.477675 | 1 | 0.962534 | 99.5349 |
| UP_SEQ_FEATURE | repeat:TPR 1 | 7 | 0.949796 | 0.360311 | 5, Q9CX34, | 641 | 137 | 18012 | 1.435758 | 1 | 0.99978 | 99.95091 |
| UP_SEQ_FEATURE | repeat:TPR 2 | 7 | 0.949796 | 0.360311 | 5, Q9CX34, | 641 | 137 | 18012 | 1.435758 | 1 | 0.99978 | 99.95091 |
| UP_SEQ_FEATURE | repeat:TPR 3 | 6 | 0.814111 | 0.458549 | CX34, Q7TN | 641 | 125 | 18012 | 1.348793 | 1 | 0.99996 | 99.99714 |

| | | | | | | | | | | | | |
|---|---|---|---|---|---|---|---|---|---|---|---|---|
| UP_SEQ_FEATURE | repeat:TPR 4 | 3 | 0.407056 | 0.815038 | ı, Q7TNH6, | 641 | 86 | 18012 | 0.980227 | 1 | 1 | 100 |
| Annotation Cluster 84 | Enrichment Score: 0.44512982680683394 | | | | | | | | | | | |
| Category | Term | Count | % | PValue | Genes | List Total | Pop Hits | Pop Total | ld Enrichme | Bonferroni | Benjamini | FDR |
| GOTERM_BP_DIRECT | GO:0018108~peptidyl-tyrosine phosphorylation | 6 | 0.814111 | 0.08834 | 37, Q3UCJ0 | 661 | 65 | 18082 | 2.525125 | 1 | 0.89737 | 80.94481 |
| INTERPRO | IPR001245:Serine-threonine/tyrosine-protein kinase catalytic domain | 8 | 1.085482 | 0.183119 | 5, Q99JT2, | 688 | 139 | 20594 | 1.722771 | 1 | 0.975311 | 96.29868 |
| GOTERM_MF_DIRECT | GO:0004715~non-membrane spanning protein tyrosine kinase activity | 4 | 0.542741 | 0.232759 | 10, A0A087 | 648 | 45 | 17446 | 2.393141 | 1 | 0.909949 | 98.33658 |
| INTERPRO | IPR008266:Tyrosine-protein kinase, active site | 5 | 0.678426 | 0.421497 | 48025, Q03 | 688 | 99 | 20594 | 1.511775 | 1 | 0.998659 | 99.98663 |
| GOTERM_MF_DIRECT | GO:0004713~protein tyrosine kinase activity | 6 | 0.814111 | 0.46755 | .0, Q3UIF3, | 648 | 121 | 17446 | 1.335017 | 1 | 0.981763 | 99.99414 |
| UP_KEYWORDS | Tyrosine-protein kinase | 5 | 0.678426 | 0.496244 | 47810, Q03 | 732 | 113 | 22680 | 1.370956 | 1 | 0.814353 | 99.99097 |
| GOTERM_BP_DIRECT | GO:0007169~transmembrane receptor protein tyrosine kinase signaling pathway | 5 | 0.678426 | 0.498281 | 48025, Q03 | 661 | 100 | 18082 | 1.367776 | 1 | 0.999794 | 99.99957 |
| INTERPRO | IPR020635:Tyrosine-protein kinase, catalytic domain | 4 | 0.542741 | 0.510135 | 5, Q03137, | 688 | 81 | 20594 | 1.47818 | 1 | 0.999647 | 99.99911 |
| GOTERM_MF_DIRECT | GO:0004714~transmembrane receptor protein tyrosine kinase activity | 3 | 0.407056 | 0.600241 | 9, P97793, ( | 648 | 54 | 17446 | 1.495713 | 1 | 0.994482 | 99.99993 |
| SMART | SM00219:TyrKc | 4 | 0.542741 | 0.629655 | 5, Q03137, | 415 | 81 | 10425 | 1.240518 | 1 | 0.995188 | 99.99979 |
| Annotation Cluster 85 | Enrichment Score: 0.4333607649124296 | | | | | | | | | | | |
| Category | Term | Count | % | PValue | Genes | List Total | Pop Hits | Pop Total | ld Enrichme | Bonferroni | Benjamini | FDR |
| INTERPRO | IPR001723:Steroid hormone receptor | 4 | 0.542741 | 0.215036 | 5E8P8, Q53 | 688 | 48 | 20594 | 2.494428 | 1 | 0.983109 | 98.06692 |
| INTERPRO | IPR000536:Nuclear hormone receptor, ligand-binding, core | 4 | 0.542741 | 0.223836 | 5E8P8, Q53 | 688 | 49 | 20594 | 2.443522 | 1 | 0.984415 | 98.39139 |
| UP_SEQ_FEATURE | region of interest:Ligand-binding | 3 | 0.407056 | 0.275596 | 3, Q53ZY9, | 641 | 29 | 18012 | 2.90688 | 1 | 0.999349 | 99.59062 |
| GOTERM_BP_DIRECT | GO:0043401~steroid hormone mediated signaling pathway | 4 | 0.542741 | 0.305201 | 5E8P8, Q53 | 661 | 53 | 18082 | 2.064568 | 1 | 0.995509 | 99.85366 |
| SMART | SM00430:HOLI | 4 | 0.542741 | 0.308285 | 5E8P8, Q53 | 415 | 49 | 10425 | 2.050652 | 1 | 0.957403 | 99.22279 |
| GOTERM_MF_DIRECT | GO:0003707~steroid hormone receptor activity | 4 | 0.542741 | 0.344486 | 5E8P8, Q53 | 648 | 56 | 17446 | 1.92306 | 1 | 0.957497 | 99.85402 |
| GOTERM_MF_DIRECT | GO:0004879~RNA polymerase II transcription factor activity, ligand-activated sequence-specific DNA binding | 3 | 0.407056 | 0.387879 | 3, Q53ZY9, | 648 | 36 | 17446 | 2.24357 | 1 | 0.968451 | 99.94937 |
| INTERPRO | IPR001628:Zinc finger, nuclear hormone receptor-type | 3 | 0.407056 | 0.467983 | 3, G5E8P8, | 688 | 47 | 20594 | 1.910626 | 1 | 0.999399 | 99.99659 |
| UP_SEQ_FEATURE | DNA-binding region:Nuclear receptor | 3 | 0.407056 | 0.501456 | 3, G5E8P8, | 641 | 47 | 18012 | 1.793607 | 1 | 0.99998 | 99.9993 |
| UP_SEQ_FEATURE | zinc finger region:NR C4-type | 3 | 0.407056 | 0.501456 | 3, G5E8P8, | 641 | 47 | 18012 | 1.793607 | 1 | 0.99998 | 99.9993 |
| SMART | SM00399:ZnF_C4 | 3 | 0.407056 | 0.562269 | 3, G5E8P8, | 415 | 47 | 10425 | 1.603435 | 1 | 0.991903 | 99.99813 |
| INTERPRO | IPR013088:Zinc finger, NHR/GATA-type | 3 | 0.407056 | 0.571362 | 3, G5E8P8, | 688 | 57 | 20594 | 1.575428 | 1 | 0.999853 | 99.9999 |
| Annotation Cluster 86 | Enrichment Score: 0.42545899979955143 | | | | | | | | | | | |
| Category | Term | Count | % | PValue | Genes | List Total | Pop Hits | Pop Total | ld Enrichme | Bonferroni | Benjamini | FDR |
| KEGG_PATHWAY | mmu04142:Lysosome | 9 | 1.221167 | 0.071615 | 29416, Q3 | 277 | 122 | 7691 | 2.048263 | 1 | 0.533112 | 61.8213 |
| GOTERM_CC_DIRECT | GO:0005764~lysosome | 10 | 1.356852 | 0.814961 | 1, Q3TD09, | 683 | 331 | 19662 | 0.869719 | 1 | 0.996292 | 100 |
| UP_KEYWORDS | Lysosome | 6 | 0.814111 | 0.906739 | 9, B7ZWC4 | 732 | 249 | 22680 | 0.746593 | 1 | 0.989473 | 100 |
| Annotation Cluster 87 | Enrichment Score: 0.39634598971641594 | | | | | | | | | | | |
| Category | Term | Count | % | PValue | Genes | List Total | Pop Hits | Pop Total | ld Enrichme | Bonferroni | Benjamini | FDR |
| KEGG_PATHWAY | mmu04915:Estrogen signaling pathway | 7 | 0.949796 | 0.139675 | , B2RSH2, F | 277 | 98 | 7691 | 1.983239 | 1 | 0.67421 | 85.76505 |
| GOTERM_BP_DIRECT | GO:0009408~response to heat | 5 | 0.678426 | 0.15415 | 5, F6W677, | 661 | 57 | 18082 | 2.399607 | 1 | 0.963205 | 95.02559 |
| GOTERM_MF_DIRECT | GO:0051082~unfolded protein binding | 5 | 0.678426 | 0.302856 | , O35685, ( | 648 | 75 | 17446 | 1.794856 | 1 | 0.945157 | 99.62181 |
| UP_KEYWORDS | Chaperone | 8 | 1.085482 | 0.419242 | 0416, Q3TU | 732 | 191 | 22680 | 1.297743 | 1 | 0.755123 | 99.93767 |
| UP_KEYWORDS | Stress response | 4 | 0.542741 | 0.436755 | TU85, P178 | 732 | 75 | 22680 | 1.652459 | 1 | 0.768595 | 99.95888 |
| GOTERM_BP_DIRECT | GO:0006986~response to unfolded protein | 3 | 0.407056 | 0.560196 | , Q3TU85, ( | 661 | 51 | 18082 | 1.609148 | 1 | 0.999919 | 99.99996 |
| GOTERM_MF_DIRECT | GO:0031072~heat shock protein binding | 3 | 0.407056 | 0.59001 | 5, Q3TU85, ( | 648 | 53 | 17446 | 1.523934 | 1 | 0.993846 | 99.9999 |
| KEGG_PATHWAY | mmu04612:Antigen processing and presentation | 3 | 0.407056 | 0.799192 | 3, Q3TU85, | 277 | 82 | 7691 | 1.015805 | 1 | 0.983655 | 100 |
| KEGG_PATHWAY | mmu04141:Protein processing in endoplasmic reticulum | 5 | 0.678426 | 0.859077 | , P17879, C | 277 | 168 | 7691 | 0.826349 | 1 | 0.988948 | 100 |
| Annotation Cluster 88 | Enrichment Score: 0.3836371406168919 | | | | | | | | | | | |
| Category | Term | Count | % | PValue | Genes | List Total | Pop Hits | Pop Total | ld Enrichme | Bonferroni | Benjamini | FDR |
| INTERPRO | IPR000315:Zinc finger, B-box | 6 | 0.814111 | 0.088192 | SW8, B2M | 688 | 71 | 20594 | 2.529561 | 1 | 0.906088 | 77.79253 |
| INTERPRO | IPR006574:SPRY-associated | 3 | 0.407056 | 0.351207 | , B2M0R4, | 688 | 37 | 20594 | 2.427011 | 1 | 0.996501 | 99.91336 |
| SMART | SM00589:PRY | 3 | 0.407056 | 0.393511 | , B2M0R4, | 415 | 34 | 10425 | 2.216513 | 1 | 0.970202 | 99.86259 |
| INTERPRO | IPR003877:SPla/RYanodine receptor SPRY | 4 | 0.542741 | 0.47637 | M0R4, Q7TF | 688 | 77 | 20594 | 1.554968 | 1 | 0.999466 | 99.99737 |
| INTERPRO | IPR001870:B30.2/SPRY domain | 4 | 0.542741 | 0.526582 | M0R4, Q7TF | 688 | 83 | 20594 | 1.442561 | 1 | 0.999741 | 99.99949 |
| INTERPRO | IPR013320:Concanavalin A-like lectin/glucanase, subgroup | 8 | 1.085482 | 0.559451 | 137, B2M0 | 688 | 211 | 20594 | 1.134906 | 1 | 0.99983 | 99.99984 |
| INTERPRO | IPR003879:Butyrophylin-like | 3 | 0.407056 | 0.571362 | , B2M0R4, | 688 | 57 | 20594 | 1.575428 | 1 | 0.999853 | 99.9999 |
| SMART | SM00449:SPRY | 4 | 0.542741 | 0.577216 | M0R4, Q7TF | 415 | 75 | 10425 | 1.339759 | 1 | 0.991884 | 99.99882 |
| UP_SEQ_FEATURE | domain:B30.2/SPRY | 3 | 0.407056 | 0.625017 | , Q7TPM6, | 641 | 59 | 18012 | 1.428806 | 1 | 0.999998 | 99.99999 |
| Annotation Cluster 89 | Enrichment Score: 0.37021841789469456 | | | | | | | | | | | |
| Category | Term | Count | % | PValue | Genes | List Total | Pop Hits | Pop Total | ld Enrichme | Bonferroni | Benjamini | FDR |
| UP_KEYWORDS | Acyltransferase | 8 | 1.085482 | 0.314173 | CAY6, Q8B | 732 | 171 | 22680 | 1.449525 | 1 | 0.662394 | 99.40362 |
| GOTERM_MF_DIRECT | GO:0016746~transferase activity, transferring acyl groups | 8 | 1.085482 | 0.425414 | CAY6, Q8B | 648 | 167 | 17446 | 1.289717 | 1 | 0.97567 | 99.98097 |
| GOTERM_MF_DIRECT | GO:0004402~histone acetyltransferase activity | 3 | 0.407056 | 0.464931 | ), Q8BY71, ( | 648 | 42 | 17446 | 1.92306 | 1 | 0.981744 | 99.99367 |
| INTERPRO | IPR016181:Acyl-CoA N-acyltransferase | 3 | 0.407056 | 0.53181 | ), Q8BY71, ( | 688 | 53 | 20594 | 1.694329 | 1 | 0.999754 | 99.99957 |
| Annotation Cluster 90 | Enrichment Score: 0.3654431582541758 | | | | | | | | | | | |
| Category | Term | Count | % | PValue | Genes | List Total | Pop Hits | Pop Total | ld Enrichme | Bonferroni | Benjamini | FDR |
| GOTERM_BP_DIRECT | GO:0030168~platelet activation | 4 | 0.542741 | 0.169373 | GR2, Q9QU | 661 | 39 | 18082 | 2.805695 | 1 | 0.970286 | 96.40779 |
| UP_KEYWORDS | Adaptive immunity | 5 | 0.678426 | 0.38889 | 7, P48025, ( | 732 | 98 | 22680 | 1.580796 | 1 | 0.734599 | 99.8755 |
| KEGG_PATHWAY | mmu04611:Platelet activation | 5 | 0.678426 | 0.697658 | , Q9QUG9, | 277 | 131 | 7691 | 1.059746 | 1 | 0.975884 | 99.99998 |
| GOTERM_BP_DIRECT | GO:0002250~adaptive immune response | 5 | 0.678426 | 0.751473 | 7, P48025, ( | 661 | 139 | 18082 | 0.984012 | 1 | 0.999999 | 100 |
| Annotation Cluster 91 | Enrichment Score: 0.3525429224986463 | | | | | | | | | | | |
| Category | Term | Count | % | PValue | Genes | List Total | Pop Hits | Pop Total | ld Enrichme | Bonferroni | Benjamini | FDR |
| BIOCARTA | m_caspasePathway:Caspase Cascade in Apoptosis | 3 | 0.407056 | 0.292607 | 3, Q3TX36, | 64 | 22 | 1289 | 2.746449 | 1 | 0.996879 | 97.85546 |
| BIOCARTA | m_fasPathway:FAS signaling pathway ( CD95 ) | 3 | 0.407056 | 0.401695 | 3, Q3TX36, | 64 | 28 | 1289 | 2.157924 | 1 | 0.996678 | 99.66582 |
| BIOCARTA | m_tnfr1Pathway:TNFR1 Signaling Pathway | 3 | 0.407056 | 0.419208 | 3, Q3TX36, | 64 | 29 | 1289 | 2.083513 | 1 | 0.995633 | 99.75969 |
| BIOCARTA | m_hivnefPathway:HIV-I Nef: negative effector of Fas and TNF | 3 | 0.407056 | 0.789255 | 3, Q3TX36, | 64 | 58 | 1289 | 1.041756 | 1 | 0.999998 | 100 |
| Annotation Cluster 92 | Enrichment Score: 0.33951747626602513 | | | | | | | | | | | |
| Category | Term | Count | % | PValue | Genes | List Total | Pop Hits | Pop Total | ld Enrichme | Bonferroni | Benjamini | FDR |
| INTERPRO | IPR013098:Immunoglobulin I-set | 9 | 1.221167 | 0.119168 | 3YUT2, A2A | 688 | 147 | 20594 | 1.832641 | 1 | 0.93639 | 87.35662 |
| UP_SEQ_FEATURE | domain:Ig-like C2-type 3 | 6 | 0.814111 | 0.203277 | 86, A2A8L5 | 641 | 88 | 18012 | 1.915898 | 1 | 0.998216 | 97.92559 |
| INTERPRO | IPR003598:Immunoglobulin subtype 2 | 11 | 1.492537 | 0.292143 | D3YUT2, A | 688 | 242 | 20594 | 1.360597 | 1 | 0.992279 | 99.64157 |
| UP_SEQ_FEATURE | domain:Ig-like C2-type 1 | 7 | 0.949796 | 0.328632 | A2A8L5, A0 | 641 | 132 | 18012 | 1.490143 | 1 | 0.999691 | 99.88806 |
| UP_SEQ_FEATURE | domain:Ig-like C2-type 2 | 7 | 0.949796 | 0.334943 | A2A8L5, A0 | 641 | 133 | 18012 | 1.478939 | 1 | 0.999721 | 99.90471 |
| INTERPRO | IPR013162:CD80-like, immunoglobulin C2-set | 3 | 0.407056 | 0.351207 | 2, Q99K86, | 688 | 37 | 20594 | 2.427011 | 1 | 0.996501 | 99.91336 |
| KEGG_PATHWAY | mmu04514:Cell adhesion molecules (CAMs) | 8 | 1.085482 | 0.362815 | C5, A0A0R4 | 277 | 162 | 7691 | 1.371128 | 1 | 0.891111 | 99.70915 |
| SMART | SM00408:IGc2 | 11 | 1.492537 | 0.491131 | D3YUT2, A | 415 | 242 | 10425 | 1.14184 | 1 | 0.985108 | 99.9864 |
| UP_SEQ_FEATURE | domain:Ig-like C2-type 5 | 3 | 0.407056 | 0.512708 | A0A0A6YX4 | 641 | 48 | 18012 | 1.75624 | 1 | 0.999983 | 99.99953 |
| UP_SEQ_FEATURE | domain:Ig-like C2-type 4 | 3 | 0.407056 | 0.596505 | A0A0A6YX4 | 641 | 56 | 18012 | 1.505349 | 1 | 0.999996 | 99.99998 |
| UP_KEYWORDS | Immunoglobulin domain | 15 | 2.035278 | 0.689231 | X2, F7AGE9 | 732 | 481 | 22680 | 0.966225 | 1 | 0.923317 | 99.99999 |
| INTERPRO | IPR003599:Immunoglobulin subtype | 15 | 2.035278 | 0.824756 | 40, F7AGE9 | 688 | 518 | 20594 | 0.86679 | 1 | 1 | 100 |
| SMART | SM00409:IG | 15 | 2.035278 | 0.955211 | 40, F7AGE9 | 415 | 518 | 10425 | 0.727427 | 1 | 1 | 100 |
| INTERPRO | IPR013783:Immunoglobulin-like fold | 22 | 2.985075 | 0.998639 | AGE9, Q99 | 688 | 1099 | 20594 | 0.599208 | 1 | 1 | 100 |
| INTERPRO | IPR007110:Immunoglobulin-like domain | 17 | 2.306649 | 0.999046 | X2, F7AGE9 | 688 | 920 | 20594 | 0.553112 | 1 | 1 | 100 |
| Annotation Cluster 93 | Enrichment Score: 0.32635195542938006 | | | | | | | | | | | |
| Category | Term | Count | % | PValue | Genes | List Total | Pop Hits | Pop Total | ld Enrichme | Bonferroni | Benjamini | FDR |
| GOTERM_CC_DIRECT | GO:0019005~SCF ubiquitin ligase complex | 4 | 0.542741 | 0.345666 | TJW2, Q56 | 683 | 60 | 19662 | 1.91918 | 1 | 0.857195 | 99.77394 |
| UP_SEQ_FEATURE | domain:F-box | 4 | 0.542741 | 0.427191 | TJW2, Q56 | 641 | 67 | 18012 | 1.677603 | 1 | 0.999928 | 99.99253 |
| INTERPRO | IPR001810:F-box domain, cyclin-like | 4 | 0.542741 | 0.542719 | TJW2, Q56 | 688 | 85 | 20594 | 1.408618 | 1 | 0.999795 | 99.99971 |
| SMART | SM00256:FBOX | 3 | 0.407056 | 0.617645 | 7, Q3TJW2, | 415 | 52 | 10425 | 1.449259 | 1 | 0.994849 | 99.99969 |
| Annotation Cluster 94 | Enrichment Score: 0.32045391326166683 | | | | | | | | | | | |
| Category | Term | Count | % | PValue | Genes | List Total | Pop Hits | Pop Total | ld Enrichme | Bonferroni | Benjamini | FDR |
| INTERPRO | IPR001478:PDZ domain | 7 | 0.949796 | 0.408584 | H3BJD6, E | 688 | 154 | 20594 | 1.360597 | 1 | 0.998635 | 99.98084 |
| UP_SEQ_FEATURE | domain:PDZ | 5 | 0.678426 | 0.484987 | Q61234, E | 641 | 101 | 18012 | 1.391081 | 1 | 0.999975 | 99.99878 |
| SMART | SM00228:PDZ | 7 | 0.949796 | 0.551603 | H3BJD6, E | 415 | 150 | 10425 | 1.172289 | 1 | 0.991584 | 99.99743 |
| Annotation Cluster 95 | Enrichment Score: 0.3150106995730219 | | | | | | | | | | | |
| Category | Term | Count | % | PValue | Genes | List Total | Pop Hits | Pop Total | ld Enrichme | Bonferroni | Benjamini | FDR |
| INTERPRO | IPR003595:Protein-tyrosine phosphatase, catalytic | 4 | 0.542741 | 0.351283 | Q8BWF4, A | 688 | 63 | 20594 | 1.900517 | 1 | 0.996301 | 99.91353 |
| SMART | SM00404:PTPc_motif | 4 | 0.542741 | 0.459635 | Q8BWF4, A | 415 | 63 | 10425 | 1.594951 | 1 | 0.981966 | 99.96999 |
| GOTERM_MF_DIRECT | GO:0004725~protein tyrosine phosphatase activity | 4 | 0.542741 | 0.702907 | A2A8L5, Q3 | 648 | 97 | 17446 | 1.11022 | 1 | 0.998355 | 100 |

| Annotation Cluster 96 | Enrichment Score: 0.29718588677331526 | | | | | | | | | | | |
|---|---|---|---|---|---|---|---|---|---|---|---|---|
| Category | Term | Count | % | PValue | Genes | List Total | Pop Hits | Pop Total | ld Enrichm | Bonferroni | Benjamini | FDR |
| UP_KEYWORDS | Innate immunity | 11 | 1.492537 | 0.250775 | GR2, A0A0R | 732 | 241 | 22680 | 1.41419 | 1 | 0.587158 | 98.01834 |
| UP_KEYWORDS | Immunity | 15 | 2.035278 | 0.41657 | WQ04, O54 | 732 | 401 | 22680 | 1.158988 | 1 | 0.76029 | 99.93366 |
| UP_KEYWORDS | Inflammatory response | 7 | 0.949796 | 0.417829 | Q9EPB4, Q | 732 | 161 | 22680 | 1.347113 | 1 | 0.756258 | 99.93558 |
| GOTERM_BP_DIRECT | GO:0002376~immune system process | 15 | 2.035278 | 0.536426 | WQ04, O54 | 661 | 383 | 18082 | 1.071365 | 1 | 0.999881 | 99.9999 |
| GOTERM_BP_DIRECT | GO:0045087~innate immune response | 13 | 1.763908 | 0.795173 | Q62318, A | 661 | 400 | 18082 | 0.889054 | 1 | 1 | 100 |
| GOTERM_BP_DIRECT | GO:0006954~inflammatory response | 10 | 1.356852 | 0.885 | PB4, Q3TU | 661 | 344 | 18082 | 0.795219 | 1 | 1 | 100 |

| Annotation Cluster 97 | Enrichment Score: 0.2956377490861948 | | | | | | | | | | | |
|---|---|---|---|---|---|---|---|---|---|---|---|---|
| Category | Term | Count | % | PValue | Genes | List Total | Pop Hits | Pop Total | ld Enrichm | Bonferroni | Benjamini | FDR |
| UP_KEYWORDS | Thiol protease | 7 | 0.949796 | 0.264244 | , Q3TCH2, | 732 | 134 | 22680 | 1.618547 | 1 | 0.602808 | 98.45109 |
| GOTERM_MF_DIRECT | GO:0008234~cysteine-type peptidase activity | 7 | 0.949796 | 0.425052 | , Q3TCH2, | 648 | 141 | 17446 | 1.336595 | 1 | 0.976987 | 99.98078 |
| UP_KEYWORDS | Protease | 18 | 2.442334 | 0.580631 | 43E3, Q99K | 732 | 542 | 22680 | 1.028976 | 1 | 0.863031 | 99.99925 |
| GOTERM_MF_DIRECT | GO:0008233~peptidase activity | 19 | 2.578019 | 0.639503 | Q543E3, C | 648 | 516 | 17446 | 0.991345 | 1 | 0.996296 | 99.99999 |
| GOTERM_BP_DIRECT | GO:0006508~proteolysis | 19 | 2.578019 | 0.797297 | 9JKB1, Q80 | 661 | 582 | 18082 | 0.89305 | 1 | 1 | 100 |

| Annotation Cluster 98 | Enrichment Score: 0.27605723494367584 | | | | | | | | | | | |
|---|---|---|---|---|---|---|---|---|---|---|---|---|
| Category | Term | Count | % | PValue | Genes | List Total | Pop Hits | Pop Total | ld Enrichm | Bonferroni | Benjamini | FDR |
| KEGG_PATHWAY | mmu04964:Proximal tubule bicarbonate reclamation | 4 | 0.542741 | 0.042523 | 71F8, P14C | 277 | 22 | 7691 | 5.048244 | 0.999977 | 0.398923 | 43.0545 |
| UP_KEYWORDS | Sodium transport | 5 | 0.678426 | 0.496244 | , Q67BT3, | 732 | 113 | 22680 | 1.370956 | 1 | 0.814353 | 99.99097 |
| UP_KEYWORDS | Sodium | 5 | 0.678426 | 0.543594 | , Q67BT3, | 732 | 120 | 22680 | 1.290984 | 1 | 0.846075 | 99.99764 |
| GOTERM_BP_DIRECT | GO:0006814~sodium ion transport | 5 | 0.678426 | 0.667699 | , Q67BT3, | 661 | 124 | 18082 | 1.103045 | 1 | 0.999989 | 100 |
| UP_KEYWORDS | Symport | 3 | 0.407056 | 0.876896 | , Q8R1S9, | 732 | 111 | 22680 | 0.837395 | 1 | 0.984075 | 100 |
| GOTERM_MF_DIRECT | GO:0015293~symporter activity | 3 | 0.407056 | 0.92352 | , Q8R1S9, | 648 | 112 | 17446 | 0.721147 | 1 | 0.999996 | 100 |
| UP_KEYWORDS | Ion transport | 10 | 1.356852 | 0.998292 | 'BT3, Q3UN | 732 | 619 | 22680 | 0.500543 | 1 | 0.999989 | 100 |
| GOTERM_BP_DIRECT | GO:0006811~ion transport | 10 | 1.356852 | 0.999339 | 'BT3, Q3UN | 661 | 584 | 18082 | 0.468416 | 1 | 1 | 100 |

| Annotation Cluster 99 | Enrichment Score: 0.2677946223824866 | | | | | | | | | | | |
|---|---|---|---|---|---|---|---|---|---|---|---|---|
| Category | Term | Count | % | PValue | Genes | List Total | Pop Hits | Pop Total | ld Enrichm | Bonferroni | Benjamini | FDR |
| UP_KEYWORDS | Translation regulation | 5 | 0.678426 | 0.410804 | , O88477, | 732 | 101 | 22680 | 1.533842 | 1 | 0.756705 | 99.92419 |
| GOTERM_MF_DIRECT | GO:0003730~mRNA 3'-UTR binding | 3 | 0.407056 | 0.59001 | ., O88477, | 648 | 53 | 17446 | 1.523934 | 1 | 0.993846 | 99.9999 |
| GOTERM_BP_DIRECT | GO:0006417~regulation of translation | 5 | 0.678426 | 0.648817 | , O88477, | 661 | 121 | 18082 | 1.130393 | 1 | 0.999984 | 100 |

| Annotation Cluster 100 | Enrichment Score: 0.23640043124325494 | | | | | | | | | | | |
|---|---|---|---|---|---|---|---|---|---|---|---|---|
| Category | Term | Count | % | PValue | Genes | List Total | Pop Hits | Pop Total | ld Enrichm | Bonferroni | Benjamini | FDR |
| UP_KEYWORDS | Prenylation | 9 | 1.221167 | 0.135589 | 39, Q0PD62 | 732 | 157 | 22680 | 1.77613 | 1 | 0.394507 | 86.17841 |
| UP_SEQ_FEATURE | lipid moiety-binding region:S-geranylgeranyl cysteine | 5 | 0.678426 | 0.552285 | 5, Q0PD62, | 641 | 110 | 18012 | 1.277266 | 1 | 0.999993 | 99.99989 |
| GOTERM_BP_DIRECT | GO:0007264~small GTPase mediated signal transduction | 9 | 1.221167 | 0.634169 | Q0PD62, P6 | 661 | 236 | 18082 | 1.043219 | 1 | 0.999979 | 100 |
| UP_SEQ_FEATURE | short sequence motif:Effector region | 4 | 0.542741 | 0.694727 | 'TN89, Q0P | 641 | 100 | 18012 | 1.123994 | 1 | 1 | 100 |
| INTERPRO | IPR001806:Small GTPase superfamily | 4 | 0.542741 | 0.828711 | 'TN89, Q0P | 688 | 134 | 20594 | 0.893527 | 1 | 1 | 100 |
| UP_SEQ_FEATURE | nucleotide phosphate-binding region:GTP | 9 | 1.221167 | 0.883738 | 4, Q7TN89, | 641 | 319 | 18012 | 0.792786 | 1 | 1 | 100 |
| INTERPRO | IPR005225:Small GTP-binding protein domain | 4 | 0.542741 | 0.916354 | 'TN89, Q0P | 688 | 165 | 20594 | 0.725652 | 1 | 1 | 100 |

| Annotation Cluster 101 | Enrichment Score: 0.23505086084190305 | | | | | | | | | | | |
|---|---|---|---|---|---|---|---|---|---|---|---|---|
| Category | Term | Count | % | PValue | Genes | List Total | Pop Hits | Pop Total | ld Enrichm | Bonferroni | Benjamini | FDR |
| UP_KEYWORDS | Mitochondrion | 36 | 4.884668 | 0.452117 | P5, Q60936 | 732 | 1058 | 22680 | 1.054263 | 1 | 0.779315 | 99.97175 |
| UP_KEYWORDS | Transit peptide | 18 | 2.442334 | 0.484036 | 9MR8, Q9D | 732 | 510 | 22680 | 1.093539 | 1 | 0.805464 | 99.9875 |
| UP_SEQ_FEATURE | transit peptide:Mitochondrion | 14 | 1.899593 | 0.900987 | 5N7, Q609 | 641 | 496 | 18012 | 0.793141 | 1 | 1 | 100 |

| Annotation Cluster 102 | Enrichment Score: 0.23416438071688503 | | | | | | | | | | | |
|---|---|---|---|---|---|---|---|---|---|---|---|---|
| Category | Term | Count | % | PValue | Genes | List Total | Pop Hits | Pop Total | ld Enrichm | Bonferroni | Benjamini | FDR |
| INTERPRO | IPR009057:Homeodomain-like | 15 | 2.035278 | 0.263269 | 5, A2RS90, | 688 | 345 | 20594 | 1.301441 | 1 | 0.989443 | 99.31231 |
| UP_KEYWORDS | Homeobox | 10 | 1.356852 | 0.550631 | 238, Q6132 | 732 | 280 | 22680 | 1.106557 | 1 | 0.849655 | 99.99809 |
| INTERPRO | IPR001356:Homeodomain | 10 | 1.356852 | 0.552863 | 238, Q6132 | 688 | 271 | 20594 | 1.104544 | 1 | 0.999814 | 99.9998 |
| SMART | SM00389:HOX | 10 | 1.356852 | 0.735391 | 238, Q6132 | 415 | 266 | 10425 | 0.944379 | 1 | 0.998504 | 100 |
| UP_SEQ_FEATURE | DNA-binding region:Homeobox | 6 | 0.814111 | 0.770857 | 1952, P480 | 641 | 180 | 18012 | 0.936661 | 1 | 1 | 100 |
| INTERPRO | IPR017970:Homeobox, conserved site | 5 | 0.678426 | 0.866252 | 2, P48031, | 688 | 184 | 20594 | 0.813401 | 1 | 1 | 100 |

| Annotation Cluster 103 | Enrichment Score: 0.2250562490284223 | | | | | | | | | | | |
|---|---|---|---|---|---|---|---|---|---|---|---|---|
| Category | Term | Count | % | PValue | Genes | List Total | Pop Hits | Pop Total | ld Enrichm | Bonferroni | Benjamini | FDR |
| UP_KEYWORDS | GTP-binding | 13 | 1.763908 | 0.384169 | 7, Q811U4, | 732 | 332 | 22680 | 1.213214 | 1 | 0.731902 | 99.86179 |
| GOTERM_MF_DIRECT | GO:0005525~GTP binding | 15 | 2.035278 | 0.560756 | 3, Q80YE7, | 648 | 383 | 17446 | 1.054419 | 1 | 0.991995 | 99.9997 |
| GOTERM_MF_DIRECT | GO:0003924~GTPase activity | 8 | 1.085482 | 0.660929 | RSH2, Q8C | 648 | 209 | 17446 | 1.030539 | 1 | 0.99681 | 99.99999 |
| UP_SEQ_FEATURE | nucleotide phosphate-binding region:GTP | 9 | 1.221167 | 0.883738 | 4, Q7TN89, | 641 | 319 | 18012 | 0.792786 | 1 | 1 | 100 |

| Annotation Cluster 104 | Enrichment Score: 0.21657263079990333 | | | | | | | | | | | |
|---|---|---|---|---|---|---|---|---|---|---|---|---|
| Category | Term | Count | % | PValue | Genes | List Total | Pop Hits | Pop Total | ld Enrichm | Bonferroni | Benjamini | FDR |
| INTERPRO | IPR002110:Ankyrin repeat | 9 | 1.221167 | 0.508034 | , Q01705, | 688 | 232 | 20594 | 1.161199 | 1 | 0.999655 | 99.99905 |
| UP_KEYWORDS | ANK repeat | 9 | 1.221167 | 0.512431 | , Q01705, | 732 | 240 | 22680 | 1.161885 | 1 | 0.826441 | 99.99421 |
| UP_SEQ_FEATURE | repeat:ANK 5 | 5 | 0.678426 | 0.537722 | 7, Q9Z1P7, | 641 | 108 | 18012 | 1.300919 | 1 | 0.99999 | 99.99981 |
| INTERPRO | IPR020683:Ankyrin repeat-containing domain | 9 | 1.221167 | 0.56056 | , Q01705, | 688 | 242 | 20594 | 1.113216 | 1 | 0.999822 | 99.99985 |
| UP_SEQ_FEATURE | repeat:ANK 4 | 5 | 0.678426 | 0.665308 | 7, Q9Z1P7, | 641 | 127 | 18012 | 1.106293 | 1 | 0.999999 | 100 |
| SMART | SM00248:ANK | 9 | 1.221167 | 0.675634 | , Q01705, | 415 | 225 | 10425 | 1.004819 | 1 | 0.997034 | 99.99996 |
| UP_SEQ_FEATURE | repeat:ANK 3 | 6 | 0.814111 | 0.676202 | Z1P7, Q017 | 641 | 160 | 18012 | 1.053744 | 1 | 1 | 100 |
| UP_SEQ_FEATURE | repeat:ANK 2 | 7 | 0.949796 | 0.686537 | , Q9Z1P7, | 641 | 193 | 18012 | 1.019165 | 1 | 1 | 100 |
| UP_SEQ_FEATURE | repeat:ANK 1 | 7 | 0.949796 | 0.686537 | , Q9Z1P7, | 641 | 193 | 18012 | 1.019165 | 1 | 1 | 100 |

| Annotation Cluster 105 | Enrichment Score: 0.21090577534507285 | | | | | | | | | | | |
|---|---|---|---|---|---|---|---|---|---|---|---|---|
| Category | Term | Count | % | PValue | Genes | List Total | Pop Hits | Pop Total | ld Enrichm | Bonferroni | Benjamini | FDR |
| KEGG_PATHWAY | mmu04919:Thyroid hormone signaling pathway | 6 | 0.814111 | 0.389324 | 02248, P14 | 277 | 114 | 7691 | 1.461334 | 1 | 0.894257 | 99.8323 |
| KEGG_PATHWAY | mmu04976:Bile secretion | 3 | 0.407056 | 0.729431 | 0, P14094, | 277 | 71 | 7691 | 1.173184 | 1 | 0.977252 | 100 |
| KEGG_PATHWAY | mmu04911:Insulin secretion | 3 | 0.407056 | 0.820329 | 0, P14094, | 277 | 86 | 7691 | 0.968558 | 1 | 0.986554 | 100 |

| Annotation Cluster 106 | Enrichment Score: 0.20621967216538772 | | | | | | | | | | | |
|---|---|---|---|---|---|---|---|---|---|---|---|---|
| Category | Term | Count | % | PValue | Genes | List Total | Pop Hits | Pop Total | ld Enrichm | Bonferroni | Benjamini | FDR |
| GOTERM_BP_DIRECT | GO:0002230~positive regulation of defense response to virus by host | 6 | 0.814111 | 0.460837 | 4094, Q9EP | 661 | 122 | 18082 | 1.345354 | 1 | 0.999632 | 99.99845 |
| GOTERM_BP_DIRECT | GO:0098779~mitophagy in response to mitochondrial depolarization | 5 | 0.678426 | 0.714585 | 4, A2ARP8, | 661 | 132 | 18082 | 1.036194 | 1 | 0.999997 | 100 |
| GOTERM_BP_DIRECT | GO:0098792~xenophagy | 4 | 0.542741 | 0.730701 | ARP8, P14C | 661 | 103 | 18082 | 1.06235 | 1 | 0.999998 | 100 |

| Annotation Cluster 107 | Enrichment Score: 0.20309711782953735 | | | | | | | | | | | |
|---|---|---|---|---|---|---|---|---|---|---|---|---|
| Category | Term | Count | % | PValue | Genes | List Total | Pop Hits | Pop Total | ld Enrichm | Bonferroni | Benjamini | FDR |
| KEGG_PATHWAY | mmu04727:GABAergic synapse | 6 | 0.814111 | 0.201888 | 7445, B2RS | 277 | 87 | 7691 | 1.914851 | 1 | 0.767731 | 94.61802 |
| GOTERM_CC_DIRECT | GO:0005834~heterotrimeric G-protein complex | 3 | 0.407056 | 0.381634 | 5, B2RSH2, | 683 | 38 | 19662 | 2.272713 | 1 | 0.880713 | 99.89965 |
| KEGG_PATHWAY | mmu04724:Glutamatergic synapse | 6 | 0.814111 | 0.396508 | 7445, B2RS | 277 | 115 | 7691 | 1.448627 | 1 | 0.895527 | 99.85614 |
| KEGG_PATHWAY | mmu05032:Morphine addiction | 5 | 0.678426 | 0.429668 | 6, P97445, | 277 | 93 | 7691 | 1.49276 | 1 | 0.911387 | 99.93083 |
| KEGG_PATHWAY | mmu04728:Dopaminergic synapse | 6 | 0.814111 | 0.529279 | ZQK4, Q57 | 277 | 134 | 7691 | 1.243224 | 1 | 0.944769 | 99.99425 |
| KEGG_PATHWAY | mmu04723:Retrograde endocannabinoid signaling | 4 | 0.542741 | 0.721102 | 7445, B2RS | 277 | 103 | 7691 | 1.078266 | 1 | 0.979308 | 99.99999 |
| KEGG_PATHWAY | mmu04725:Cholinergic synapse | 4 | 0.542741 | 0.777342 | 7445, B2RS | 277 | 113 | 7691 | 0.982844 | 1 | 0.981985 | 100 |
| KEGG_PATHWAY | mmu04062:Chemokine signaling pathway | 6 | 0.814111 | 0.838489 | 12, G3UWG | 277 | 196 | 7691 | 0.849959 | 1 | 0.98715 | 100 |
| KEGG_PATHWAY | mmu04726:Serotonergic synapse | 4 | 0.542741 | 0.858542 | 7445, B2RS | 277 | 132 | 7691 | 0.841374 | 1 | 0.989314 | 100 |
| KEGG_PATHWAY | mmu04713:Circadian entrainment | 3 | 0.407056 | 0.872286 | 5, B2RSH2, | 277 | 98 | 7691 | 0.849959 | 1 | 0.989972 | 100 |
| KEGG_PATHWAY | mmu05034:Alcoholism | 5 | 0.678426 | 0.936232 | 5, B2RSH2, | 277 | 202 | 7691 | 0.687261 | 1 | 0.99678 | 100 |
| GOTERM_BP_DIRECT | GO:0007186~G-protein coupled receptor signaling pathway | 8 | 1.085482 | 1 | DDB7, B7ZV | 661 | 1706 | 18082 | 0.128279 | 1 | 1 | 100 |
| UP_KEYWORDS | Transducer | 5 | 0.678426 | 1 | 7, B2RSH2, | 732 | 1801 | 22680 | 0.086018 | 1 | 1 | 100 |

| Annotation Cluster 108 | Enrichment Score: 0.19801480508855065 | | | | | | | | | | | |
|---|---|---|---|---|---|---|---|---|---|---|---|---|
| Category | Term | Count | % | PValue | Genes | List Total | Pop Hits | Pop Total | ld Enrichm | Bonferroni | Benjamini | FDR |
| GOTERM_BP_DIRECT | GO:0016311~dephosphorylation | 7 | 0.949796 | 0.208025 | 5, Q3UCJ0, | 661 | 109 | 18082 | 1.756777 | 1 | 0.982937 | 98.47097 |
| GOTERM_MF_DIRECT | GO:0016791~phosphatase activity | 5 | 0.678426 | 0.64176 | ), Q9D0T2, | 648 | 118 | 17446 | 1.140798 | 1 | 0.996289 | 99.99999 |
| GOTERM_MF_DIRECT | GO:0004725~protein tyrosine phosphatase activity | 4 | 0.542741 | 0.702907 | A2A8L5, Q3 | 648 | 97 | 17446 | 1.11022 | 1 | 0.998355 | 100 |
| GOTERM_BP_DIRECT | GO:0006470~protein dephosphorylation | 5 | 0.678426 | 0.746438 | 5, Q571J7, | 661 | 138 | 18082 | 0.991142 | 1 | 0.999999 | 100 |
| GOTERM_MF_DIRECT | GO:0004721~phosphoprotein phosphatase activity | 5 | 0.678426 | 0.762533 | AK9, A2A8L | 648 | 139 | 17446 | 0.968447 | 1 | 0.99942 | 100 |
| INTERPRO | IPR000387:Protein-tyrosine/Dual specificity phosphatase | 3 | 0.407056 | 0.763513 | 5, Q3UCJ0, | 688 | 82 | 20594 | 1.095115 | 1 | 0.999999 | 100 |
| UP_KEYWORDS | Protein phosphatase | 4 | 0.542741 | 0.794021 | A8L5, Q3U | 732 | 130 | 22680 | 0.953342 | 1 | 0.963051 | 100 |
| UP_SEQ_FEATURE | active site:Phosphocysteine intermediate | 3 | 0.407056 | 0.804627 | 5, Q3UCJ0, | 641 | 84 | 18012 | 1.003566 | 1 | 1 | 100 |

| Annotation Cluster 109 | Enrichment Score: 0.19744899351093445 |
|---|---|

| Category | Term | Count | % | PValue | Genes | List Total | Pop Hits | Pop Total | ld Enrichme | Bonferroni | Benjamini | FDR |
|---|---|---|---|---|---|---|---|---|---|---|---|---|
| UP_SEQ_FEATURE | domain:Fibronectin type-III 6 | 3 | 0.407056 | 0.275596 | , A2A8L5, C | 641 | 29 | 18012 | 2.90688 | 1 | 0.999349 | 99.59062 |
| UP_SEQ_FEATURE | domain:Fibronectin type-III 5 | 3 | 0.407056 | 0.418247 | , A2A8L5, C | 641 | 40 | 18012 | 2.107488 | 1 | 0.999914 | 99.99028 |
| UP_SEQ_FEATURE | domain:Fibronectin type-III 4 | 3 | 0.407056 | 0.61569 | , A2A8L5, C | 641 | 58 | 18012 | 1.45344 | 1 | 0.999997 | 99.99999 |
| UP_SEQ_FEATURE | domain:Fibronectin type-III 3 | 3 | 0.407056 | 0.757619 | , A2A8L5, C | 641 | 76 | 18012 | 1.109204 | 1 | 1 | 100 |
| UP_SEQ_FEATURE | domain:Fibronectin type-III 2 | 4 | 0.542741 | 0.821663 | A8L5, Q031 | 641 | 124 | 18012 | 0.906447 | 1 | 1 | 100 |
| UP_SEQ_FEATURE | domain:Fibronectin type-III 1 | 4 | 0.542741 | 0.825808 | A8L5, Q031 | 641 | 125 | 18012 | 0.899195 | 1 | 1 | 100 |
| INTERPRO | IPR003961:Fibronectin, type III | 6 | 0.814111 | 0.836509 | O88307, F | 688 | 211 | 20594 | 0.851179 | 1 | 1 | 100 |
| SMART | SM00060:FN3 | 5 | 0.678426 | 0.862674 | Q03137, O | 415 | 153 | 10425 | 0.820931 | 1 | 0.999928 | 100 |
| Annotation Cluster 110 | Enrichment Score: 0.18469903440879196 | | | | | | | | | | | |
| Category | Term | Count | % | PValue | Genes | List Total | Pop Hits | Pop Total | ld Enrichme | Bonferroni | Benjamini | FDR |
| GOTERM_MF_DIRECT | GO:0000049~tRNA binding | 4 | 0.542741 | 0.293301 | ARD6, Q8BG | 648 | 51 | 17446 | 2.111595 | 1 | 0.94043 | 99.53322 |
| UP_KEYWORDS | Protein biosynthesis | 3 | 0.407056 | 0.952738 | , Q8BGQ7, | 732 | 147 | 22680 | 0.632319 | 1 | 0.996743 | 100 |
| GOTERM_BP_DIRECT | GO:0006412~translation | 6 | 0.814111 | 0.999117 | 22G0, Q8BG | 661 | 401 | 18082 | 0.40931 | 1 | 1 | 100 |
| Annotation Cluster 111 | Enrichment Score: 0.17277130936276597 | | | | | | | | | | | |
| Category | Term | Count | % | PValue | Genes | List Total | Pop Hits | Pop Total | ld Enrichme | Bonferroni | Benjamini | FDR |
| GOTERM_BP_DIRECT | GO:0010508~positive regulation of autophagy | 3 | 0.407056 | 0.444275 | 7, Q3UFT4, | 661 | 41 | 18082 | 2.001624 | 1 | 0.99951 | 99.99733 |
| UP_KEYWORDS | Autophagy | 4 | 0.542741 | 0.762213 | UKG7, Q3U | 732 | 123 | 22680 | 1.007597 | 1 | 0.954733 | 100 |
| GOTERM_BP_DIRECT | GO:0006914~autophagy | 4 | 0.542741 | 0.895278 | UKG7, Q3U | 661 | 142 | 18082 | 0.770578 | 1 | 1 | 100 |
| Annotation Cluster 112 | Enrichment Score: 0.15991041971100636 | | | | | | | | | | | |
| Category | Term | Count | % | PValue | Genes | List Total | Pop Hits | Pop Total | ld Enrichme | Bonferroni | Benjamini | FDR |
| KEGG_PATHWAY | mmu04728:Dopaminergic synapse | 6 | 0.814111 | 0.529279 | ZQK4, Q57 | 277 | 134 | 7691 | 1.243224 | 1 | 0.944769 | 99.99425 |
| KEGG_PATHWAY | mmu04261:Adrenergic signaling in cardiomyocytes | 5 | 0.678426 | 0.755946 | 4, Q571J7, | 277 | 142 | 7691 | 0.977653 | 1 | 0.980602 | 100 |
| KEGG_PATHWAY | mmu04071:Sphingolipid signaling pathway | 4 | 0.542741 | 0.828121 | 71J7, B2RS | 277 | 124 | 7691 | 0.895656 | 1 | 0.987421 | 100 |
| Annotation Cluster 113 | Enrichment Score: 0.14426527889834181 | | | | | | | | | | | |
| Category | Term | Count | % | PValue | Genes | List Total | Pop Hits | Pop Total | ld Enrichme | Bonferroni | Benjamini | FDR |
| GOTERM_MF_DIRECT | GO:0008237~metallopeptidase activity | 7 | 0.949796 | 0.547173 | C7, Q3UDD | 648 | 160 | 17446 | 1.177874 | 1 | 0.99133 | 99.99952 |
| UP_SEQ_FEATURE | metal ion-binding site:Zinc; catalytic | 4 | 0.542741 | 0.799657 | JDD5, P974 | 641 | 119 | 18012 | 0.944533 | 1 | 1 | 100 |
| UP_KEYWORDS | Metalloprotease | 4 | 0.542741 | 0.843675 | JDD5, P974 | 732 | 143 | 22680 | 0.866674 | 1 | 0.976103 | 100 |
| Annotation Cluster 114 | Enrichment Score: 0.1380664090658601 | | | | | | | | | | | |
| Category | Term | Count | % | PValue | Genes | List Total | Pop Hits | Pop Total | ld Enrichme | Bonferroni | Benjamini | FDR |
| GOTERM_MF_DIRECT | GO:0008237~metallopeptidase activity | 7 | 0.949796 | 0.547173 | C7, Q3UDD | 648 | 160 | 17446 | 1.177874 | 1 | 0.99133 | 99.99952 |
| INTERPRO | IPR024079:Metallopeptidase, catalytic domain | 3 | 0.407056 | 0.838871 | , Q3UG07, | 688 | 97 | 20594 | 0.925767 | 1 | 1 | 100 |
| GOTERM_MF_DIRECT | GO:0004222~metalloendopeptidase activity | 4 | 0.542741 | 0.839423 | 1391, Q3U( | 648 | 123 | 17446 | 0.875539 | 1 | 0.999893 | 100 |
| Annotation Cluster 115 | Enrichment Score: 0.13410544733892663 | | | | | | | | | | | |
| Category | Term | Count | % | PValue | Genes | List Total | Pop Hits | Pop Total | ld Enrichme | Bonferroni | Benjamini | FDR |
| KEGG_PATHWAY | mmu00980:Metabolism of xenobiotics by cytochrome P450 | 3 | 0.407056 | 0.675088 | 2, Q61133, | 277 | 64 | 7691 | 1.3015 | 1 | 0.974963 | 99.99995 |
| KEGG_PATHWAY | mmu00982:Drug metabolism - cytochrome P450 | 3 | 0.407056 | 0.69146 | 2, Q61133, | 277 | 66 | 7691 | 1.262061 | 1 | 0.975489 | 99.99998 |
| KEGG_PATHWAY | mmu05204:Chemical carcinogenesis | 3 | 0.407056 | 0.848313 | 2, Q61133, | 277 | 92 | 7691 | 0.905392 | 1 | 0.988449 | 100 |
| Annotation Cluster 116 | Enrichment Score: 0.1093384678714823 | | | | | | | | | | | |
| Category | Term | Count | % | PValue | Genes | List Total | Pop Hits | Pop Total | ld Enrichme | Bonferroni | Benjamini | FDR |
| INTERPRO | IPR000198:Rho GTPase-activating protein domain | 3 | 0.407056 | 0.651271 | 3, D5M8I5, ( | 688 | 66 | 20594 | 1.360597 | 1 | 0.999976 | 100 |
| UP_SEQ_FEATURE | domain:Rho-GAP | 3 | 0.407056 | 0.660571 | 3, D5M8I5, ( | 641 | 63 | 18012 | 1.338088 | 1 | 0.999999 | 100 |
| SMART | SM00324:RhoGAP | 3 | 0.407056 | 0.728298 | 3, D5M8I5, ( | 415 | 64 | 10425 | 1.177523 | 1 | 0.998484 | 100 |
| INTERPRO | IPR008936:Rho GTPase activation protein | 3 | 0.407056 | 0.801877 | 3, D5M8I5, ( | 688 | 89 | 20594 | 1.008982 | 1 | 1 | 100 |
| UP_KEYWORDS | GTPase activation | 4 | 0.542741 | 0.899785 | M8I5, Q8R3 | 732 | 163 | 22680 | 0.760334 | 1 | 0.98852 | 100 |
| GOTERM_MF_DIRECT | GO:0005096~GTPase activator activity | 5 | 0.678426 | 0.976635 | 5, Q53YX2, ( | 648 | 235 | 17446 | 0.572826 | 1 | 1 | 100 |
| Annotation Cluster 117 | Enrichment Score: 0.09270999426656928 | | | | | | | | | | | |
| Category | Term | Count | % | PValue | Genes | List Total | Pop Hits | Pop Total | ld Enrichme | Bonferroni | Benjamini | FDR |
| UP_SEQ_FEATURE | repeat:LRR 1 | 11 | 1.492537 | 0.512598 | Z0, Q569Z9 | 641 | 274 | 18012 | 1.128096 | 1 | 0.999985 | 99.99952 |
| UP_SEQ_FEATURE | repeat:LRR 2 | 11 | 1.492537 | 0.512598 | Z0, Q569Z9 | 641 | 274 | 18012 | 1.128096 | 1 | 0.999985 | 99.99952 |
| UP_KEYWORDS | Leucine-rich repeat | 8 | 1.085482 | 0.786621 | 9Z9, Q9DAI | 732 | 275 | 22680 | 0.901341 | 1 | 0.961024 | 100 |
| UP_SEQ_FEATURE | repeat:LRR 3 | 8 | 1.085482 | 0.799118 | Q6PB97, Q5 | 641 | 253 | 18012 | 0.888533 | 1 | 1 | 100 |
| UP_SEQ_FEATURE | repeat:LRR 7 | 5 | 0.678426 | 0.817178 | 5K4, Q569Z | 641 | 158 | 18012 | 0.889236 | 1 | 1 | 100 |
| UP_SEQ_FEATURE | repeat:LRR 4 | 7 | 0.949796 | 0.824515 | 5K4, Q6PB9 | 641 | 228 | 18012 | 0.862715 | 1 | 1 | 100 |
| INTERPRO | IPR001611:Leucine-rich repeat | 7 | 0.949796 | 0.866603 | Q9DAM1, ( | 688 | 259 | 20594 | 0.809004 | 1 | 1 | 100 |
| UP_SEQ_FEATURE | repeat:LRR 6 | 5 | 0.678426 | 0.911545 | 5K4, Q569Z | 641 | 191 | 18012 | 0.735598 | 1 | 1 | 100 |
| UP_SEQ_FEATURE | repeat:LRR 5 | 5 | 0.678426 | 0.94351 | 5K4, Q569Z | 641 | 210 | 18012 | 0.669044 | 1 | 1 | 100 |
| UP_SEQ_FEATURE | repeat:LRR 8 | 3 | 0.407056 | 0.953981 | , L7N1W9, | 641 | 134 | 18012 | 0.629101 | 1 | 1 | 100 |
| INTERPRO | IPR003591:Leucine-rich repeat, typical subtype | 3 | 0.407056 | 0.981759 | 4, Q9D3R3, | 688 | 175 | 20594 | 0.51314 | 1 | 1 | 100 |
| SMART | SM00369:LRR_TYP | 3 | 0.407056 | 0.993451 | 4, Q9D3R3, | 415 | 175 | 10425 | 0.430637 | 1 | 1 | 100 |
| Annotation Cluster 118 | Enrichment Score: 0.08982066061601474 | | | | | | | | | | | |
| Category | Term | Count | % | PValue | Genes | List Total | Pop Hits | Pop Total | ld Enrichme | Bonferroni | Benjamini | FDR |
| UP_SEQ_FEATURE | domain:Ig-like V-type | 5 | 0.678426 | 0.537722 | YX2, E9PYN | 641 | 108 | 18012 | 1.300919 | 1 | 0.99999 | 99.99981 |
| INTERPRO | IPR013106:Immunoglobulin V-set | 6 | 0.814111 | 0.99996 | A0A6YX40, | 688 | 554 | 20594 | 0.324186 | 1 | 1 | 100 |
| SMART | SM00406:IGv | 4 | 0.542741 | 0.999995 | A0A6YX40, | 415 | 423 | 10425 | 0.237546 | 1 | 1 | 100 |
| Annotation Cluster 119 | Enrichment Score: 0.07360380756021769 | | | | | | | | | | | |
| Category | Term | Count | % | PValue | Genes | List Total | Pop Hits | Pop Total | ld Enrichme | Bonferroni | Benjamini | FDR |
| KEGG_PATHWAY | mmu04971:Gastric acid secretion | 3 | 0.407056 | 0.736533 | 0, B2RSH2, | 277 | 72 | 7691 | 1.156889 | 1 | 0.97797 | 100 |
| KEGG_PATHWAY | mmu04261:Adrenergic signaling in cardiomyocytes | 5 | 0.678426 | 0.755946 | 4, Q571J7, | 277 | 142 | 7691 | 0.977653 | 1 | 0.980602 | 100 |
| KEGG_PATHWAY | mmu04024:cAMP signaling pathway | 5 | 0.678426 | 0.928016 | 2, B2RSH2, I | 277 | 197 | 7691 | 0.704704 | 1 | 0.996228 | 100 |
| KEGG_PATHWAY | mmu04022:cGMP-PKG signaling pathway | 3 | 0.407056 | 0.982531 | 0, B2RSH2, | 277 | 163 | 7691 | 0.511019 | 1 | 0.999581 | 100 |
| Annotation Cluster 120 | Enrichment Score: 0.04228155067576231 | | | | | | | | | | | |
| Category | Term | Count | % | PValue | Genes | List Total | Pop Hits | Pop Total | ld Enrichme | Bonferroni | Benjamini | FDR |
| GOTERM_CC_DIRECT | GO:0005615~extracellular space | 55 | 7.462687 | 0.416392 | 22W9, Q3U | 683 | 1504 | 19662 | 1.052742 | 1 | 0.89766 | 99.95628 |
| GOTERM_CC_DIRECT | GO:0005576~extracellular region | 36 | 4.884668 | 0.999943 | Q3UER8, Q | 683 | 1753 | 19662 | 0.591191 | 1 | 1 | 100 |
| UP_KEYWORDS | Secreted | 29 | 3.934871 | 0.999986 | H9, Q9D3K4 | 732 | 1685 | 22680 | 0.533249 | 1 | 1 | 100 |
| UP_KEYWORDS | Glycoprotein | 84 | 11.39756 | 0.99999 | 0R4IZW5, Q | 732 | 3815 | 22680 | 0.682208 | 1 | 1 | 100 |
| UP_KEYWORDS | Disulfide bond | 63 | 8.548168 | 0.999997 | O88307, Q | 732 | 3124 | 22680 | 0.624829 | 1 | 1 | 100 |
| UP_SEQ_FEATURE | disulfide bond | 51 | 6.919946 | 1 | 55, O88307, | 641 | 2510 | 18012 | 0.570953 | 1 | 1 | 100 |
| UP_SEQ_FEATURE | signal peptide | 63 | 8.548168 | 1 | 8307, Q3UG | 641 | 3124 | 18012 | 0.566674 | 1 | 1 | 100 |
| UP_SEQ_FEATURE | glycosylation site:N-linked (GlcNAc...) | 74 | 10.04071 | 1 | 0A0R4IZW5, | 641 | 3563 | 18012 | 0.583606 | 1 | 1 | 100 |
| UP_KEYWORDS | Signal | 85 | 11.53324 | 1 | O88307, Q | 732 | 4543 | 22680 | 0.579706 | 1 | 1 | 100 |
| Annotation Cluster 121 | Enrichment Score: 0.021885342473272496 | | | | | | | | | | | |
| Category | Term | Count | % | PValue | Genes | List Total | Pop Hits | Pop Total | ld Enrichme | Bonferroni | Benjamini | FDR |
| UP_KEYWORDS | Endoplasmic reticulum | 27 | 3.663501 | 0.890987 | Q544Z7, D | 732 | 997 | 22680 | 0.839075 | 1 | 0.987149 | 100 |
| GOTERM_CC_DIRECT | GO:0005783~endoplasmic reticulum | 35 | 4.748982 | 0.976893 | 291WB4, Q5 | 683 | 1323 | 19662 | 0.761579 | 1 | 0.999988 | 100 |
| GOTERM_CC_DIRECT | GO:0005789~endoplasmic reticulum membrane | 16 | 2.170963 | 0.987701 | K0C4, Q3TX | 683 | 710 | 19662 | 0.648737 | 1 | 0.999998 | 100 |
| Annotation Cluster 122 | Enrichment Score: 0.006125758599483736 | | | | | | | | | | | |
| Category | Term | Count | % | PValue | Genes | List Total | Pop Hits | Pop Total | ld Enrichme | Bonferroni | Benjamini | FDR |
| INTERPRO | IPR000210:BTB/POZ-like | 4 | 0.542741 | 0.980175 | 9Z45, Q920 | 688 | 222 | 20594 | 0.539336 | 1 | 1 | 100 |
| INTERPRO | IPR011333:BTB/POZ fold | 4 | 0.542741 | 0.984788 | 9Z45, Q920 | 688 | 232 | 20594 | 0.516089 | 1 | 1 | 100 |
| SMART | SM00225:BTB | 4 | 0.542741 | 0.993061 | 9Z45, Q920 | 415 | 218 | 10425 | 0.460926 | 1 | 1 | 100 |
| Annotation Cluster 123 | Enrichment Score: 0.002426528378757891 | | | | | | | | | | | |
| Category | Term | Count | % | PValue | Genes | List Total | Pop Hits | Pop Total | ld Enrichme | Bonferroni | Benjamini | FDR |
| UP_KEYWORDS | mRNA processing | 5 | 0.678426 | 0.990085 | Q8R3W5, ( | 732 | 307 | 22680 | 0.504619 | 1 | 0.999769 | 100 |
| GOTERM_BP_DIRECT | GO:0006397~mRNA processing | 6 | 0.814111 | 0.992239 | 3W5, E9PU | 661 | 322 | 18082 | 0.50973 | 1 | 1 | 100 |
| UP_KEYWORDS | mRNA splicing | 3 | 0.407056 | 0.996651 | 1, A2AFQ9, ( | 732 | 240 | 22680 | 0.387295 | 1 | 0.999966 | 100 |
| GOTERM_BP_DIRECT | GO:0008380~RNA splicing | 3 | 0.407056 | 0.998762 | 1, A2AFQ9, ( | 661 | 241 | 18082 | 0.340525 | 1 | 1 | 100 |
| Annotation Cluster 124 | Enrichment Score: 4.105820882297169E-6 | | | | | | | | | | | |
| Category | Term | Count | % | PValue | Genes | List Total | Pop Hits | Pop Total | ld Enrichme | Bonferroni | Benjamini | FDR |
| GOTERM_CC_DIRECT | GO:0005886~plasma membrane | 126 | 17.09634 | 0.999983 | 36, P63216 | 683 | 4874 | 19662 | 0.744204 | 1 | 1 | 100 |
| GOTERM_CC_DIRECT | GO:0016020~membrane | 194 | 26.32293 | 0.999984 | N7, B7XGA6 | 683 | 6998 | 19662 | 0.798059 | 1 | 1 | 100 |

| | | | | | | | | | | | | |
|---|---|---|---|---|---|---|---|---|---|---|---|---|
| UP_KEYWORDS | Cell membrane | 81 | 10.9905 | 0.999995 | 7, Q61391, | 732 | 3759 | 22680 | 0.667644 | 1 | 1 | 100 |
| UP_KEYWORDS | Membrane | 170 | 23.06649 | 1 | K4, Q8BNE3 | 732 | 8683 | 22680 | 0.606612 | 1 | 1 | 100 |

Annotation Cluster 125 Enrichment Score: 4.960583589669108E-7

| Category | Term | Count | % | PValue | Genes | List Total | Pop Hits | Pop Total | ld Enrichme | Bonferroni | Benjamini | FDR |
|---|---|---|---|---|---|---|---|---|---|---|---|---|
| UP_KEYWORDS | Glycoprotein | 84 | 11.39756 | 0.99999 | )R4IZW5, Q | 732 | 3815 | 22680 | 0.682208 | 1 | 1 | 100 |
| UP_SEQ_FEATURE | topological domain:Extracellular | 41 | 5.563094 | 1 | 3TZP9, Q9C | 641 | 2256 | 18012 | 0.51068 | 1 | 1 | 100 |
| UP_SEQ_FEATURE | glycosylation site:N-linked (GlcNAc...) | 74 | 10.04071 | 1 | )A0R4IZW5 | 641 | 3563 | 18012 | 0.583606 | 1 | 1 | 100 |
| UP_SEQ_FEATURE | topological domain:Cytoplasmic | 51 | 6.919946 | 1 | P13597, O8 | 641 | 2880 | 18012 | 0.497601 | 1 | 1 | 100 |
| UP_SEQ_FEATURE | transmembrane region | 73 | 9.90502 | 1 | 7BT3, O883 | 641 | 4312 | 18012 | 0.475716 | 1 | 1 | 100 |
| UP_KEYWORDS | Membrane | 170 | 23.06649 | 1 | K4, Q8BNE3 | 732 | 8683 | 22680 | 0.606612 | 1 | 1 | 100 |
| UP_KEYWORDS | Transmembrane helix | 94 | 12.75441 | 1 | IK4, Q80X9 | 732 | 6938 | 22680 | 0.419784 | 1 | 1 | 100 |
| UP_KEYWORDS | Transmembrane | 94 | 12.75441 | 1 | IK4, Q80X9 | 732 | 6955 | 22680 | 0.418758 | 1 | 1 | 100 |
| GOTERM_CC_DIRECT | GO:0016021~integral component of membrane | 93 | 12.61872 | 1 | Q8BMK4, Q | 683 | 6878 | 19662 | 0.389249 | 1 | 1 | 100 |
