## Supplementary_Table4 for "Aging-dependent dysregulation of EXOSC2 is maintained in cancer as a dependency"

**SUPPLEMENTARY TABLE 3**

| Annotation Cluster 1 | Enrichment Score: 16.011075451721236 | | | | | | | | | | | |
|---|---|---|---|---|---|---|---|---|---|---|---|---|
| Category | Term | Count | % | PValue | Genes | List Total | Pop Hits | Pop Total | ld Enrichme | Bonferroni | Benjamini | FDR |
| UP_KEYWORDS | Endoplasmic reticulum | 112 | 10.91618 | 4.94E-19 | Q64FW2, Q | 1024 | 997 | 22680 | 2.488089 | 1.77E-16 | 8.87E-17 | 6.78E-16 |
| UP_KEYWORDS | Membrane | 516 | 50.2924 | 7.48E-16 | 64, O55101 | 1024 | 8683 | 22680 | 1.316203 | 2.79E-13 | 6.97E-14 | 1.07E-12 |
| UP_KEYWORDS | Lipid metabolism | 44 | 4.288499 | 4.29E-07 | 8M4, Q9D6 | 1024 | 417 | 22680 | 2.337005 | 1.54E-04 | 1.18E-05 | 5.89E-04 |
| UP_KEYWORDS | Oxidoreductase | | 5.45809 | 3.66E-06 | .7, A3KG36, | 1024 | 639 | 22680 | 1.941021 | 0.001314 | 7.30E-05 | 0.005027 |
| UP_KEYWORDS | Immunity | 40 | 3.898635 | 6.04E-06 | ₹2Q8, Q8VI‹ | 1024 | 401 | 22680 | 2.20932 | 0.002165 | 1.08E-04 | 0.008289 |

| | FDR | LOG10 | |
|---|---|---|---|
| Endoplasmic reticulum | 6.78E-16 | -15.1687 | 15.16871 |
| Membrane | 1.07E-12 | -11.9723 | 11.97232 |
| Lipid metabolism | 5.89E-04 | -3.23003 | 3.23003 |
| Oxidoreductase | 0.005027 | -2.29868 | 2.298683 |
| Immunity | 0.008289 | -2.08151 | 2.081508 |
