## Supplementary_Table5 for "Aging-dependent dysregulation of EXOSC2 is maintained in cancer as a dependency"

**SUPPLEMENTARY TABLE 5**

| GeneID | GeneSymbol | pval | P.adj | log2[ESC/ESCshESXOSC2] | Mean at cond ESCshESXOSC2 | Mean at cond ESC | pattern1:ATTTA | pattern2:TTATTTATT | pattern3:TTATTTA[TA][TA] | pattern4:TATTTAT | pattern5:TAT(2,5)AT | pattern6:TGTANATA | in 3prime utr pattern1:ATTTA | in 3prime utr pattern2:TTATTTATT | in 3prime utr pattern3:TTATTTA[TA][TA] | in 3prime utr pattern4:TATTTAT | in 3prime utr pattern5:TAT(2,5)AT | in 3prime utr pattern6:TGTANATA |
|---|---|---|---|---|---|---|---|---|---|---|---|---|---|---|---|---|---|---|
| SG0000003 | Adgrl3 | 0.001508 | 0.002775 | -1.889294391 | 33.52583919 | 9.049933236 | 29541 | 1049 | 2002 | 4439 | 14737 | 2381 | 19 | 5 | 5 | 5 | 13 | 0 |
| SG0000002 | Nrxn1 | 5.63E-06 | 1.36E-05 | -1.972978036 | 68.08259749 | 17.34245383 | 22638 | 1096 | 1899 | 3306 | 11763 | 1874 | 26 | 0 | 0 | 0 | 11 | 7 |
| SG0000007 | Dtnb | 6.75E-05 | 0.000146 | 0.619716257 | 542.3286729 | 833.3275664 | 6561 | 973 | 1255 | 1645 | 3979 | 425 | 21 | 1 | 1 | 3 | 7 | 1 |
| SG0000003 | 43525 | 7.69E-09 | 2.36E-08 | -4.758676147 | 26.07099758 | 0 | 11913 | 517 | 1052 | 1751 | 6002 | 1175 | 76 | 7 | 7 | 19 | 26 | 4 |
| SG0000005 | Tcf4 | 8.83E-09 | 2.7E-08 | -0.889066797 | 2038.320793 | 1100.618379 | 11879 | 573 | 967 | 1536 | 5057 | 996 | 36 | 0 | 0 | 0 | 26 | 7 |
| SG0000006 | Macrod2 | 0.004597 | 0.007909 | 0.772101792 | 71.39542088 | 121.9259526 | 10750 | 378 | 840 | 1514 | 5337 | 865 | 17 | 2 | 2 | 6 | 10 | 0 |
| SG0000002 | Dach2 | 3.04E-08 | 8.92E-08 | -1.728233241 | 141.3070098 | 42.64947225 | 8939 | 450 | 835 | 1525 | 5024 | 750 | 17 | 0 | 0 | 0 | 11 | 2 |
| SG0000002 | Macf1 | 2.98E-15 | 1.43E-14 | -1.13655437 | 8949.292354 | 4070.5373 | 4059 | 581 | 744 | 977 | 2204 | 392 | 46 | 0 | 11 | 33 | 33 | 11 |
| SG0000000 | Dlgap1 | 1.7E-05 | 3.92E-05 | 2.042538356 | 13.59380322 | 56.00235513 | 7416 | 459 | 739 | 1238 | 3339 | 771 | 34 | 0 | 2 | 6 | 12 | 2 |
| SG0000005 | Lrrc4c | 0.002088 | 0.00377 | -2.489248789 | 24.89667773 | 4.434073069 | 8081 | 401 | 716 | 1348 | 4493 | 687 | 11 | 2 | 2 | 4 | 4 | 0 |
| SG0000005 | Cacna1c | 0.000171 | 0.000353 | -1.327864627 | 98.97353294 | 39.42684178 | 8597 | 452 | 693 | 1069 | 3452 | 888 | 12 | 0 | 1 | 0 | 0 | 1 |
| SG0000002 | Pde4d | 2.06E-66 | 6.75E-65 | -3.541743173 | 2131.053781 | 182.9883655 | 8127 | 371 | 689 | 1252 | 4386 | 769 | 5 | 0 | 0 | 1 | 3 | 0 |
| SG0000000 | Celf2 | 0.005162 | 0.008818 | 1.507902796 | 75.06816485 | 213.4910981 | 10937 | 337 | 675 | 1237 | 5349 | 932 | 65 | 0 | 0 | 3 | 38 | 0 |
| SG0000006 | Fhit | 5.79E-10 | 1.94E-09 | 6.356607729 | 0.34247518 | 28.06458986 | 8378 | 361 | 666 | 1264 | 4277 | 791 | 0 | 0 | 0 | 0 | 3 | 0 |
| SG0000004 | Cacna2d1 | 0.38349 | 0.4523 | 0.333469307 | 143.2125134 | 180.4534671 | 8900 | 381 | 665 | 1386 | 4592 | 692 | 19 | 0 | 0 | 6 | 10 | 2 |
| SG0000003 | Rsrc1 | 0.674603 | 0.731756 | 0.055393891 | 916.3759478 | 952.2454463 | 6471 | 371 | 654 | 1091 | 3413 | 484 | 12 | 0 | 0 | 0 | 10 | 0 |
| SG0000006 | Ank3 | 0.301283 | 0.366093 | -0.169993699 | 1415.407936 | 1258.08048 | 6505 | 544 | 643 | 1244 | 2904 | 679 | 50 | 0 | 0 | 8 | 8 | 14 |
| SG0000003 | Diap2 | 1.51E-11 | 5.7E-11 | -1.198453122 | 1443.88284 | 629.1607434 | 6830 | 354 | 632 | 1098 | 3695 | 614 | 35 | 0 | 2 | 0 | 8 | 8 |
| SG0000002 | Ktn1 | 0.007945 | 0.013198 | -0.378960393 | 3923.259736 | 3016.953496 | 4000 | 398 | 624 | 756 | 2014 | 293 | 26 | 0 | 0 | 20 | 28 | 0 |
| SG0000003 | Etl4 | 0.007224 | 0.012078 | -0.370669411 | 5072.319406 | 3923.051255 | 4883 | 445 | 623 | 923 | 2419 | 495 | 8 | 0 | 0 | 1 | 13 | 12 |
| SG0000003 | Rbms3 | 0.001238 | 0.002299 | -1.328660514 | 115.5099459 | 45.98886713 | 7923 | 372 | 609 | 1174 | 3947 | 1029 | 16 | 0 | 0 | 0 | 37 | 1 |
| SG0000002 | Stau2 | 6.1E-09 | 1.89E-08 | 0.901345075 | 421.7270895 | 787.7046383 | 5813 | 381 | 586 | 1141 | 3418 | 580 | 41 | 0 | 0 | 0 | 10 | 3 |
| SG0000005 | Ptprt | 0.001783 | 0.00325 | -1.84250428 | 63.85458324 | 17.8050427 | 8182 | 362 | 578 | 1234 | 3998 | 740 | 22 | 2 | 2 | 2 | 10 | 0 |
| SG0000004 | Magi2 | 0.041229 | 0.060369 | -0.811487371 | 65.72892081 | 37.45195234 | 6764 | 308 | 572 | 1002 | 3576 | 599 | 8 | 0 | 0 | 1 | 2 | 1 |
| SG0000005 | A830018L16Rik | 6.1E-12 | 2.36E-11 | 5.951400425 | 3.489843332 | 215.9513756 | 6043 | 335 | 567 | 889 | 3259 | 630 | 54 | 0 | 6 | 9 | 34 | 6 |
| SG0000004 | Rims1 | 3.04E-25 | 2.49E-24 | 8.552158217 | 0.34247518 | 128.5538991 | 6311 | 308 | 565 | 938 | 3469 | 535 | 21 | 0 | 5 | 0 | 10 | 0 |
| SG0000001 | Cadps2 | 0.279426 | 0.342673 | 0.569074993 | 18.35952368 | 27.23767616 | 7049 | 250 | 562 | 1074 | 3336 | 718 | 5 | 0 | 0 | 1 | 1 | 0 |
| SG0000004 | Dmd | 1E-131 | 1.2E-129 | 4.086553148 | 72.80632134 | 1236.927271 | 7518 | 285 | 560 | 1095 | 3995 | 640 | 22 | 0 | 0 | 0 | 2 | 1 |
| SG0000003 | Adgrb3 | 2.42E-15 | 1.17E-14 | 7.141587016 | 0.307509278 | 43.42003131 | 8544 | 244 | 557 | 1237 | 4173 | 767 | 11 | 0 | 0 | 6 | 10 | 9 |
| SG0000002 | Lrba | 0.708671 | 0.761471 | 0.049462007 | 1187.316648 | 1228.72899 | 5732 | 274 | 547 | 902 | 2863 | 479 | 5 | 0 | 0 | 0 | 2 | 0 |
| SG0000003 | Stag1 | 5E-11 | 1.81E-10 | 0.836507938 | 1863.562291 | 3327.805192 | 4821 | 345 | 541 | 867 | 2714 | 346 | 27 | 0 | 4 | 4 | 17 | 0 |
| ISG000000 | Eda | 0.649172 | 0.708504 | 0.119202672 | 114.8155341 | 124.7051044 | 7161 | 203 | 511 | 1078 | 3836 | 742 | 28 | 0 | 0 | 7 | 7 | 7 |
| SG0000003 | Bbs9 | 3.72E-16 | 1.89E-15 | 1.507671187 | 155.3249034 | 441.6673983 | 4401 | 304 | 508 | 731 | 2278 | 520 | 6 | 0 | 0 | 0 | 0 | 0 |
| SG0000003 | Fgfr2 | 4.8E-15 | 2.28E-14 | 4.765390992 | 19.58103025 | 532.5509551 | 5494 | 326 | 504 | 865 | 2291 | 447 | 24 | 0 | 0 | 0 | 4 | 8 |
| SG0000003 | Dennd1a | 1.58E-06 | 4.01E-06 | -0.889846596 | 579.7974784 | 312.9001744 | 4153 | 389 | 503 | 801 | 2162 | 250 | 2 | 0 | 0 | 0 | 0 | 0 |
| SG0000002 | Frmd5 | 7.16E-53 | 1.61E-51 | -3.142295524 | 2228.375881 | 252.3848265 | 3222 | 373 | 490 | 654 | 1561 | 254 | 9 | 0 | 0 | 1 | 5 | 0 |
| SG0000005 | Phf21a | 0.631723 | 0.692753 | -0.090932039 | 1295.9811 | 1216.817359 | 3932 | 343 | 483 | 723 | 2130 | 361 | 36 | 5 | 5 | 17 | 26 | 0 |
| SG0000004 | Specc1 | 2.13E-09 | 6.82E-09 | -1.109987415 | 1991.013474 | 922.4327048 | 3159 | 357 | 474 | 653 | 1892 | 252 | 17 | 0 | 0 | 4 | 7 | 2 |
| SG0000003 | Ccser1 | 0.029273 | 0.044023 | -1.768251136 | 31.64831433 | 9.290817974 | 5511 | 252 | 474 | 863 | 3031 | 462 | 14 | 1 | 3 | 3 | 9 | 2 |
| SG0000005 | Prkg1 | 0.196235 | 0.250638 | -0.902180701 | 24.87024237 | 13.30750277 | 5326 | 245 | 461 | 795 | 2597 | 481 | 9 | 1 | 1 | 1 | 3 | 0 |
| SG0000002 | Cux1 | 5.31E-39 | 7.56E-38 | -2.182518718 | 4117.866064 | 907.1280924 | 3485 | 313 | 459 | 671 | 1816 | 288 | 21 | 2 | 4 | 6 | 30 | 0 |
| SG0000003 | Edil3 | 0.030231 | 0.045341 | -0.756190554 | 76.84359499 | 45.49583448 | 4373 | 257 | 451 | 756 | 2512 | 434 | 36 | 0 | 4 | 8 | 27 | 4 |
| SG0000003 | Erc1 | 5.57E-06 | 1.34E-05 | -0.667364165 | 2520.246389 | 1586.888341 | 5807 | 275 | 450 | 857 | 2724 | 562 | 20 | 4 | 5 | 4 | 5 | 2 |
| SG0000002 | Bnc2 | 4.85E-80 | 2.18E-78 | 5.204682158 | 10.3741597 | 382.5765765 | 5256 | 245 | 448 | 752 | 2264 | 501 | 2 | 0 | 0 | 0 | 0 | 7 |
| SG0000002 | Epb41 | 0.038172 | 0.056219 | 0.277994765 | 1610.686881 | 1952.971404 | 2549 | 365 | 447 | 634 | 1451 | 171 | 22 | 0 | 0 | 0 | 11 | 0 |
| SG0000003 | Ank2 | 0.489915 | 0.55798 | -0.13441343 | 355.1118004 | 323.5211143 | 4198 | 311 | 445 | 807 | 2254 | 372 | 13 | 1 | 1 | 4 | 13 | 4 |
| SG0000002 | Cdk14 | 0.539802 | 0.606013 | -0.18332762 | 185.5058343 | 163.3693046 | 4424 | 279 | 435 | 808 | 2213 | 343 | 16 | 0 | 3 | 0 | 5 | 2 |
| SG0000000 | Pde1c | 0.013519 | 0.021563 | 4.591598678 | 0.663202137 | 15.99023503 | 4635 | 241 | 434 | 785 | 2412 | 422 | 25 | 0 | 0 | 0 | 10 | 1 |
| SG0000006 | Clasp1 | 0.008353 | 0.013834 | -0.368534621 | 3163.56169 | 2450.396324 | 4786 | 214 | 432 | 637 | 2415 | 444 | 21 | 0 | 0 | 4 | 11 | 0 |
| SG0000003 | Tcf12 | 0.51522 | 0.5821 | 0.092935469 | 1145.555112 | 1221.778143 | 5822 | 203 | 424 | 801 | 2400 | 489 | 25 | 6 | 6 | 6 | 18 | 4 |
| SG0000002 | Dst | 3.98E-06 | 9.74E-06 | -0.628242564 | 9296.466111 | 6014.479295 | 2543 | 307 | 421 | 569 | 1264 | 170 | 28 | 5 | 5 | 5 | 23 | 0 |
| SG0000006 | Kalrn | 1.41E-09 | 4.6E-09 | 2.107195267 | 82.04013023 | 353.4722489 | 3411 | 337 | 410 | 629 | 1635 | 277 | 15 | 0 | 0 | 4 | 8 | 1 |
| SG0000003 | Grip1 | 3.34E-36 | 4.32E-35 | 6.897274802 | 1.005677317 | 119.8795927 | 5968 | 264 | 408 | 806 | 2541 | 632 | 14 | 0 | 0 | 2 | 2 | 3 |
| SG0000005 | Kcnt2 | 2.98E-39 | 4.29E-38 | 7.555890072 | 0.674076249 | 126.8406855 | 3831 | 150 | 406 | 617 | 1930 | 316 | 19 | 0 | 1 | 3 | 6 | 2 |
| SG0000002 | Pcdh7 | 4.18E-05 | 9.26E-05 | -1.754342249 | 614.6915864 | 182.1996881 | 4817 | 141 | 403 | 718 | 2394 | 397 | 20 | 0 | 0 | 3 | 11 | 2 |
| SG0000002 | St7 | 5.12E-24 | 3.94E-23 | -2.382565119 | 465.1052751 | 89.19236873 | 5796 | 241 | 400 | 842 | 2847 | 381 | 13 | 0 | 0 | 0 | 13 | 0 |
| SG0000005 | Bre | 0.009936 | 0.016258 | -0.37474247 | 1868.590898 | 1441.137785 | 5387 | 150 | 398 | 688 | 2733 | 492 | 0 | 0 | 0 | 0 | 0 | 0 |
| SG0000004 | Ambra1 | 0.407229 | 0.476378 | 0.113146146 | 1455.663113 | 1574.422402 | 2174 | 320 | 397 | 539 | 1244 | 161 | 5 | 0 | 0 | 0 | 0 | 0 |
| SG0000006 | Crem | 0.004985 | 0.008531 | -0.623087347 | 254.3536506 | 165.1467357 | 3991 | 276 | 394 | 670 | 2156 | 319 | 99 | 0 | 7 | 8 | 8 | 0 |
| SG0000003 | Cntnap2 | 0.000224 | 0.000457 | -1.309113013 | 433.3469136 | 174.885347 | 5890 | 171 | 393 | 780 | 3053 | 491 | 23 | 0 | 0 | 3 | 7 | 2 |
| SG0000002 | Pde4b | 1.02E-17 | 5.65E-17 | -3.682952258 | 164.0890355 | 12.77616347 | 4897 | 208 | 390 | 800 | 2377 | 393 | 11 | 0 | 0 | 2 | 7 | 3 |
| SG0000005 | Pard3b | 0.000646 | 0.00124 | -1.465091946 | 60.24328409 | 21.82086772 | 4678 | 206 | 384 | 720 | 2402 | 468 | 10 | 0 | 0 | 0 | 2 | 0 |
| SG0000002 | Arhgef9 | 0.779637 | 0.824129 | 0.103390999 | 52.6659898 | 56.57884045 | 4267 | 175 | 382 | 647 | 2298 | 347 | 64 | 0 | 1 | 19 | 40 | 2 |
| SG0000004 | Adam22 | 3.1E-31 | 3.31E-30 | 4.119539638 | 10.86502903 | 188.8582192 | 3977 | 204 | 371 | 650 | 2519 | 484 | 40 | 0 | 0 | 6 | 39 | 23 |
| SG0000006 | Uty | 0.940948 | 0.957683 | -0.06743342 | 29.87688073 | 28.51253106 | 2659 | 236 | 369 | 485 | 1656 | 205 | 38 | 0 | 0 | 4 | 8 | 21 |
| SG0000000 | Cdkal1 | 3.21E-13 | 1.35E-12 | 1.190455981 | 284.0628421 | 648.302038 | 3608 | 241 | 367 | 646 | 1854 | 300 | 10 | 1 | 2 | 3 | 7 | 0 |
| SG0000001 | Mgat4c | 8.38E-36 | 1.07E-34 | -7.509786999 | 179.565147 | 0.985260105 | 3685 | 197 | 356 | 633 | 2086 | 329 | 35 | 0 | 4 | 20 | 34 | 0 |
| SG0000002 | Cep112 | 4.64E-05 | 0.000102 | 1.532342357 | 41.32280614 | 119.5283209 | 5390 | 148 | 356 | 731 | 2371 | 474 | 0 | 0 | 0 | 0 | 0 | 0 |
| SG0000003 | Ppp1r9a | 3.38E-10 | 1.16E-09 | 2.054426705 | 217.6444278 | 904.0482605 | 3461 | 192 | 353 | 544 | 1618 | 265 | 18 | 1 | 2 | 2 | 4 | 2 |
| SG0000002 | Ralgps2 | 6.46E-15 | 3.04E-14 | 1.023881035 | 1137.528176 | 2313.028939 | 2913 | 223 | 349 | 493 | 1160 | 245 | 24 | 0 | 0 | 0 | 12 | 0 |
| SG0000005 | Robo2 | 0.005506 | 0.009362 | 1.356976129 | 16.22636339 | 41.56346212 | 4459 | 159 | 349 | 677 | 2037 | 385 | 15 | 0 | 0 | 5 | 10 | 2 |
| SG0000005 | Plcb1 | 1.53E-65 | 4.85E-64 | -5.191881138 | 1070.607504 | 29.2899295 | 4511 | 217 | 348 | 638 | 2181 | 452 | 30 | 0 | 2 | 4 | 10 | 4 |
| SG0000002 | Mecom | 5.46E-27 | 4.8E-26 | -3.576641789 | 347.5699132 | 29.13169223 | 3989 | 150 | 347 | 674 | 1933 | 369 | 23 | 0 | 0 | 8 | 16 | 0 |
| SG0000002 | Cnot2 | 0.633507 | 0.694024 | -0.066630808 | 2965.321316 | 2831.482538 | 2470 | 211 | 347 | 444 | 1239 | 112 | 17 | 0 | 1 | 1 | 6 | 4 |
| SG0000006 | Adgrv1 | 0.023854 | 0.036405 | 0.406552139 | 281.1985852 | 372.732721 | 3359 | 210 | 343 | 581 | 1601 | 353 | 17 | 0 | 0 | 2 | 8 | 0 |
| SG0000005 | Camk2d | 5.02E-57 | 1.27E-55 | -2.959540875 | 3336.772507 | 428.9592346 | 6537 | 188 | 342 | 747 | 2645 | 758 | 54 | 1 | 1 | 8 | 14 | 8 |
| SG0000003 | Mast4 | 1.54E-21 | 1.05E-20 | -1.601494425 | 1724.355352 | 568.2362158 | 3811 | 181 | 332 | 525 | 1618 | 391 | 11 | 0 | 2 | 2 | 7 | 4 |
| SG0000001 | Psmd11 | 0.162871 | 0.211955 | 0.179127025 | 4140.663147 | 4688.052954 | 892 | 296 | 328 | 351 | 713 | 59 | 0 | 0 | 0 | 0 | 0 | 0 |
| SG0000005 | Bcas3 | 8.01E-11 | 2.86E-10 | 0.922063199 | 713.7731923 | 1352.473965 | 2757 | 201 | 324 | 453 | 1401 | 225 | 0 | 0 | 0 | 0 | 0 | 0 |
| SG0000003 | Slit2 | 0.968743 | 0.982399 | -0.020918348 | 507.9781414 | 500.6658557 | 5165 | 174 | 324 | 721 | 2692 | 485 | 22 | 0 | 0 | 4 | 19 | 4 |
| SG0000005 | Abi1 | 0.05421 | 0.077731 | -0.270812576 | 2434.349547 | 2017.716893 | 2933 | 190 | 322 | 553 | 1728 | 216 | 51 | 3 | 3 | 3 | 23 | 18 |
| SG0000002 | Tlk2 | 0.038812 | 0.057087 | 0.268672767 | 2263.509613 | 2726.847844 | 1838 | 250 | 320 | 498 | 992 | 130 | 18 | 0 | 0 | 0 | 1 | 0 |
| SG0000003 | Cpeb3 | 0.211024 | 0.267298 | 0.321872573 | 95.84203231 | 119.7979299 | 2654 | 224 | 317 | 446 | 1246 | 274 | 32 | 0 | 0 | 4 | 39 | 4 |
| SG0000003 | Ehmt1 | 0.670973 | 0.728328 | 0.053661932 | 1820.356828 | 1889.341127 | 1948 | 238 | 316 | 455 | 1135 | 166 | 14 | 0 | 0 | 0 | 0 | 0 |
| SG0000002 | Fut8 | 1.77E-09 | 5.72E-09 | -1.001209773 | 1079.968544 | 539.531658 | 2202 | 201 | 315 | 399 | 1123 | 123 | 3 | 0 | 0 | 0 | 4 | 0 |
| SG0000002 | Papola | 0.930438 | 0.949472 | -0.017679959 | 8707.609073 | 8601.550139 | 1314 | 242 | 315 | 351 | 626 | 95 | 9 | 0 | 2 | 3 | 10 | 2 |
| SG0000004 | Sh3kbp1 | 2.32E-62 | 6.84E-61 | -3.606888937 | 3655.199469 | 300.005572 | 3086 | 188 | 313 | 422 | 1358 | 256 | 20 | 2 | 2 | 3 | 8 | 0 |
| SG0000002 | Ptprd | 1.22E-05 | 2.85E-05 | 2.315935389 | 8.055295785 | 40.10952438 | 3629 | 177 | 311 | 542 | 1784 | 291 | 10 | 0 | 0 | 0 | 2 | 6 |
| SG0000003 | Ano4 | 5.2E-122 | 5.4E-120 | -7.428694716 | 1382.871662 | 8.026421061 | 4221 | 139 | 309 | 494 | 1926 | 453 | 11 | 0 | 0 | 0 | 1 | 12 |
| SG0000002 | Exoc4 | 0.001131 | 0.002112 | -0.473501592 | 4817.758463 | 3469.819158 | 3354 | 171 | 307 | 562 | 1571 | 274 | 10 | 0 | 1 | 2 | 4 | 1 |
| SG0000004 | Rfx3 | 0.240709 | 0.300605 | -0.212106866 | 629.9221104 | 543.7963943 | 3479 | 134 | 307 | 455 | 1591 | 345 | 38 | 0 | 2 | 0 | 12 | 6 |
| SG0000002 | Adgrl2 | 3.35E-34 | 4.02E-33 | 1.977139091 | 233.2618592 | 918.3789164 | 3715 | 189 | 303 | 440 | 1247 | 226 | 21 | 0 | 0 | 0 | 12 | 17 |
| SG0000002 | Zbtb20 | 1.7E-105 | 1.4E-103 | -5.380657294 | 4338.416856 | 104.1338768 | 3876 | 178 | 302 | 601 | 1841 | 326 | 46 | 0 | 2 | 4 | 25 | 1 |
| SG0000005 | Gpc6 | 8.64E-06 | 2.05E-05 | -1.317920722 | 283.5641328 | 113.7411535 | 4520 | 154 | 298 | 606 | 2062 | 402 | 2 | 0 | 0 | 0 | 2 | 0 |
| SG0000005 | Dennd1b | 5.93E-08 | 1.7E-07 | -0.892135187 | 1239.626826 | 667.930868 | 2487 | 135 | 294 | 477 | 1291 | 174 | 56 | 0 | 9 | 6 | 23 | 7 |
| SG0000000 | Wwox | 0.000761 | 0.001449 | 0.551857856 | 383.5507987 | 562.2748513 | 2608 | 221 | 293 | 438 | 1131 | 265 | 4 | 0 | 0 | 0 | 1 | 0 |
| SG0000002 | Oxr1 | 1.92E-05 | 4.39E-05 | -0.614333661 | 2925.344095 | 1910.927006 | 3619 | 152 | 291 | 494 | 1576 | 318 | 24 | 0 | 0 | 0 | 6 | 6 |
| SG0000005 | Mta3 | 0.026768 | 0.040548 | -0.376920021 | 2949.26606 | 2271.170407 | 1172 | 229 | 291 | 319 | 808 | 106 | 5 | 1 | 1 | 2 | 4 | 1 |
| SG0000005 | Phactr1 | 1.33E-12 | 5.35E-12 | -2.851319096 | 272.7920961 | 37.80064477 | 3223 | 136 | 289 | 485 | 1341 | 336 | 4 | 0 | 0 | 0 | 0 | 8 |
| SG0000003 | Dysf | 6.39E-08 | 1.82E-07 | 1.357464897 | 140.81949 | 360.8281562 | 1375 | 258 | 289 | 390 | 926 | 171 | 0 | 0 | 0 | 0 | 0 | 0 |
| SG0000002 | Trip12 | 6.54E-20 | 4.08E-19 | 1.104780308 | 10304.52262 | 22161.53756 | 1659 | 230 | 288 | 394 | 748 | 132 | 10 | 0 | 2 | 2 | 2 | 2 |
| SG0000003 | Ctnnd1 | 0.06729 | 0.094861 | -0.271155258 | 1781.463655 | 1476.220053 | 2580 | 240 | 288 | 456 | 1019 | 264 | 62 | 0 | 0 | 0 | 37 | 24 |
| SG0000003 | Zmynd8 | 0.268604 | 0.330964 | 0.14228652 | 3552.911524 | 3921.181122 | 954 | 233 | 287 | 355 | 672 | 18 | 10 | 2 | 2 | 6 | 14 | 0 |
| SG0000001 | Ttll5 | 0.879082 | 0.90693 | -0.034276086 | 507.2097141 | 495.3012597 | 3259 | 140 | 287 | 512 | 1458 | 203 | 6 | 0 | 0 | 0 | 6 | 0 |
| SG0000005 | 9330182L06Rik | 9.27E-06 | 2.19E-05 | 1.867330285 | 36.76688109 | 134.1464605 | 2718 | 125 | 286 | 483 | 1483 | 264 | 39 | 1 | 3 | 6 | 21 | 3 |
| SG0000004 | Pid1 | 1.7E-08 | 5.06E-08 | -1.021815434 | 707.6816711 | 348.5305501 | 2796 | 180 | 285 | 514 | 1488 | 237 | 8 | 1 | 2 | 2 | 2 | 0 |
| SG0000003 | Phc3 | 2.86E-13 | 1.2E-12 | -1.291128674 | 1390.586877 | 568.2372076 | 1258 | 189 | 284 | 294 | 749 | 60 | 20 | 0 | 1 | 3 | 7 | 0 |
| ISG000000 | Gulp1 | 0.054931 | 0.078689 | 0.605501887 | 159.2009495 | 242.2255128 | 2390 | 158 | 284 | 424 | 1227 | 179 | 42 | 0 | 3 | 0 | 15 | 0 |
| SG0000002 | Bbx | 1.02E-12 | 4.14E-12 | -1.180342048 | 4210.476387 | 1857.85953 | 2438 | 154 | 283 | 376 | 1159 | 188 | 20 | 1 | 3 | 4 | 14 | 3 |
| SG0000003 | Sbf2 | 0.000886 | 0.001673 | 0.461254032 | 1487.628642 | 2048.07488 | 2464 | 201 | 282 | 445 | 1265 | 255 | 4 | 0 | 0 | 0 | 0 | 2 |
| SG0000004 | Pbrm1 | 0.028424 | 0.042857 | 0.273095199 | 3776.724125 | 4563.784291 | 2387 | 162 | 282 | 431 | 1276 | 214 | 15 | 2 | 2 | 3 | 8 | 0 |
| SG0000002 | Nfia | 1.35E-19 | 8.26E-19 | -3.832157158 | 295.3831184 | 20.73917224 | 3695 | 195 | 280 | 625 | 1735 | 315 | 35 | 0 | 0 | 0 | 20 | 5 |
| SG0000004 | Fmn1 | 0.588718 | 0.652101 | 0.186101553 | 166.1160476 | 188.9877868 | 2678 | 152 | 277 | 378 | 1111 | 231 | 16 | 2 | 4 | 5 | 8 | 6 |
| SG0000006 | Opcml | 1.92E-18 | 1.1E-17 | -7.697728042 | 69.50667206 | 0.33479538 | 3396 | 152 | 276 | 488 | 1653 | 298 | 10 | 0 | 0 | 1 | 4 | 0 |
| SG0000003 | Cadm1 | 5.4E-258 | 2.7E-255 | 9.182997768 | 2.235714429 | 1299.49349 | 3319 | 195 | 275 | 465 | 1348 | 230 | 18 | 0 | 0 | 0 | 2 | 0 |
| SG0000005 | Dach1 | 0.361679 | 0.430001 | -0.622805268 | 25.37397954 | 16.47803869 | 2718 | 152 | 274 | 382 | 1278 | 212 | 10 | 0 | 0 | 0 | 16 | 4 |
| SG0000004 | Dip2c | 1.38E-22 | 9.93E-22 | 1.808577692 | 136.1231984 | 476.8351693 | 2601 | 168 | 273 | 405 | 1301 | 220 | 9 | 0 | 0 | 0 | 2 | 1 |
| SG0000003 | Zfp280d | 1.38E-10 | 4.86E-10 | 0.917755594 | 652.5192193 | 1232.722351 | 2153 | 178 | 272 | 417 | 1226 | 201 | 38 | 0 | 0 | 0 | 15 | 5 |
| SG0000003 | Sorbs2 | 0.018005 | 0.028082 | -0.767158425 | 95.51980551 | 56.12491963 | 5579 | 151 | 272 | 719 | 2296 | 433 | 35 | 0 | 0 | 0 | 14 | 7 |
| SG0000003 | Arhgap6 | 0.183588 | 0.236086 | 0.758429739 | 13.42147042 | 22.70440889 | 3942 | 126 | 271 | 543 | 1983 | 336 | 15 | 0 | 0 | 4 | 12 | 0 |
| SG0000002 | Kcnq5 | 0.857645 | 0.888858 | 0.076325763 | 23.20470335 | 24.46540248 | 4130 | 112 | 271 | 578 | 2090 | 387 | 38 | 4 | 4 | 8 | 14 | 0 |

| | | | | | | | | | | | | | | | | | | |
|---|---|---|---|---|---|---|---|---|---|---|---|---|---|---|---|---|---|---|
| SG0000005 | Nrf1 | 0.078437 | 0.109131 | 0.24349684 | 1255.453122 | 1486.279042 | 1673 | 185 | 268 | 388 | 1212 | 151 | 8 | 0 | 0 | 0 | 1 | 8 |
| SG0000004 | Madd | 0.001266 | 0.00235 | -0.61934321 | 436.2594757 | 283.9906158 | 1051 | 214 | 267 | 328 | 775 | 148 | 13 | 0 | 13 | 13 | 13 | 13 |
| SG0000004 | Gphn | 2.68E-26 | 2.3E-25 | -1.951234762 | 1238.819236 | 320.3522275 | 2606 | 132 | 266 | 398 | 1374 | 250 | 3 | 0 | 0 | 0 | 0 | 0 |
| SG0000007 | Rbm47 | 8.5E-132 | 1E-129 | 4.031663924 | 78.372261 | 1281.781944 | 1452 | 199 | 265 | 371 | 914 | 113 | 37 | 0 | 0 | 11 | 38 | 4 |
| SG0000000 | Nfib | 2.19E-31 | 2.35E-30 | -2.601836511 | 1295.656345 | 213.4317339 | 3096 | 141 | 264 | 357 | 1221 | 216 | 54 | 3 | 6 | 3 | 15 | 6 |
| SG0000001 | Lmbr1 | 6.26E-05 | 0.000136 | -0.753455631 | 518.3514707 | 307.47626 | 1882 | 166 | 264 | 367 | 880 | 142 | 24 | 0 | 6 | 2 | 5 | 2 |
| SG0000002 | Pdpk1 | 1.27E-23 | 9.59E-23 | -1.581474016 | 6674.034129 | 2230.063898 | 762 | 231 | 262 | 322 | 511 | 37 | 12 | 0 | 4 | 0 | 4 | 0 |
| SG0000001 | Camta1 | 5.1E-101 | 3.7E-99 | 3.498625371 | 98.92331207 | 1118.123638 | 3293 | 171 | 261 | 498 | 1269 | 403 | 53 | 0 | 5 | 5 | 25 | 15 |
| SG0000002 | Gbe1 | 8.04E-12 | 3.09E-11 | -1.566515295 | 673.2198581 | 227.2944469 | 2672 | 174 | 261 | 425 | 1509 | 276 | 4 | 0 | 1 | 2 | 3 | 0 |
| SG0000005 | Unc5c | 0.008042 | 0.013344 | -1.492564474 | 108.0527607 | 38.39982015 | 3577 | 124 | 260 | 527 | 1860 | 308 | 21 | 0 | 0 | 3 | 14 | 0 |
| SG0000002 | Abi2 | 1.64E-22 | 1.18E-21 | 1.57049041 | 971.2407493 | 2884.639969 | 1567 | 190 | 259 | 332 | 832 | 132 | 22 | 3 | 3 | 3 | 11 | 0 |
| SG0000003 | Tenm3 | 1.83E-16 | 9.46E-16 | 4.211501232 | 50.33125837 | 932.4509298 | 2710 | 136 | 259 | 455 | 1274 | 242 | 5 | 0 | 0 | 0 | 0 | 0 |
| SG0000003 | Depdc5 | 1.38E-07 | 3.82E-07 | 0.722317079 | 1002.937378 | 1654.675835 | 1504 | 221 | 259 | 414 | 972 | 151 | 9 | 0 | 0 | 1 | 1 | 10 |
| SG0000002 | Gria4 | 2.64E-09 | 8.4E-09 | 6.406036105 | 0.307509278 | 26.07757748 | 2757 | 101 | 258 | 425 | 1499 | 228 | 10 | 0 | 0 | 2 | 7 | 0 |
| SG0000005 | Whsc1l1 | 0.001662 | 0.003046 | -0.449995945 | 2812.749349 | 2059.058831 | 1483 | 204 | 257 | 367 | 823 | 131 | 34 | 0 | 1 | 7 | 15 | 2 |
| SG0000000 | Uqcc1 | 7.07E-05 | 0.000153 | -0.608994381 | 1386.55339 | 909.0986671 | 1127 | 192 | 256 | 272 | 632 | 119 | 16 | 0 | 0 | 0 | 0 | 0 |
| SG0000001 | Cdh23 | 6.62E-06 | 1.59E-05 | 3.040028808 | 3.559775136 | 29.27941696 | 1591 | 207 | 255 | 365 | 790 | 219 | 0 | 0 | 0 | 0 | 0 | 0 |
| SG0000004 | Sorcs1 | 4.6E-08 | 1.33E-07 | -3.849814597 | 39.15617979 | 2.715755343 | 3603 | 115 | 250 | 461 | 1695 | 340 | 10 | 0 | 0 | 1 | 5 | 2 |
| SG0000004 | Trappc9 | 2.03E-12 | 8.09E-12 | -1.149889956 | 1371.357314 | 618.0153449 | 2055 | 163 | 248 | 360 | 919 | 147 | 2 | 0 | 0 | 2 | 4 | 0 |
| SG0000002 | Fggy | 0.107076 | 0.144882 | -0.374523342 | 223.0886145 | 172.0816836 | 2528 | 156 | 248 | 436 | 1377 | 244 | 2 | 0 | 0 | 1 | 11 | 0 |
| SG0000003 | Psd3 | 5.72E-49 | 1.15E-47 | -3.295346117 | 1064.049304 | 108.383638 | 3249 | 115 | 246 | 479 | 1481 | 280 | 63 | 0 | 3 | 9 | 31 | 9 |
| SG0000003 | Ccdc85a | 0.000351 | 0.000698 | 1.132648611 | 42.73854687 | 93.70892555 | 2882 | 101 | 246 | 418 | 1434 | 229 | 61 | 0 | 4 | 4 | 20 | 0 |
| SG0000007 | Fnbp1 | 0.126025 | 0.167809 | -0.212837769 | 2983.763005 | 2574.505194 | 1124 | 201 | 244 | 305 | 561 | 73 | 8 | 0 | 0 | 0 | 2 | 2 |
| SG0000004 | Klhl32 | 1.65E-15 | 8.09E-15 | -2.746142061 | 153.390205 | 22.86264617 | 2087 | 153 | 243 | 374 | 1226 | 208 | 23 | 0 | 0 | 2 | 10 | 0 |
| SG0000003 | Smarcc1 | 1.56E-10 | 5.48E-10 | 0.774383305 | 5341.364042 | 9136.182819 | 1035 | 224 | 243 | 302 | 639 | 58 | 5 | 0 | 1 | 0 | 2 | 0 |
| SG0000005 | Whsc1 | 9.37E-09 | 2.86E-08 | -0.818248752 | 4820.03088 | 2733.583865 | 1374 | 197 | 243 | 319 | 615 | 101 | 79 | 2 | 3 | 2 | 13 | 4 |
| SG0000002 | Tcf7l2 | 0.237188 | 0.296684 | -0.227667434 | 369.2105343 | 315.3111708 | 3751 | 158 | 242 | 515 | 1650 | 242 | 44 | 16 | 16 | 24 | 37 | 8 |
| SG0000003 | Nr3c2 | 3.79E-06 | 9.31E-06 | 2.15136792 | 11.16400776 | 49.59597193 | 2765 | 92 | 239 | 372 | 1195 | 243 | 27 | 0 | 7 | 0 | 3 | 3 |
| SG0000002 | Plxdc2 | 0.737979 | 0.787645 | 0.134184252 | 243.0555504 | 266.746683 | 3064 | 150 | 238 | 437 | 1603 | 289 | 10 | 0 | 1 | 2 | 7 | 1 |
| SG0000006 | Dpp6 | 3.65E-06 | 8.96E-06 | -1.565095757 | 140.2865714 | 47.41058698 | 4147 | 99 | 237 | 472 | 1977 | 477 | 0 | 0 | 0 | 0 | 0 | 3 |
| SG0000003 | Pphln1 | 4.78E-07 | 1.26E-06 | -0.753205115 | 2132.341367 | 1265.084107 | 954 | 204 | 236 | 284 | 604 | 64 | 6 | 0 | 0 | 0 | 6 | 0 |
| SG0000003 | Trps1 | 0.893127 | 0.918036 | -0.045904258 | 429.289998 | 415.8456965 | 4188 | 122 | 236 | 512 | 2090 | 455 | 24 | 3 | 4 | 7 | 13 | 6 |
| SG0000004 | Sox5 | 0.056111 | 0.080187 | -1.53225636 | 16.47715854 | 5.696750448 | 3272 | 129 | 235 | 430 | 1410 | 377 | 25 | 0 | 2 | 2 | 6 | 4 |
| SG0000002 | Ankhd1 | 4.54E-17 | 2.43E-16 | 1.439845189 | 1294.2167 | 3511.103167 | 1166 | 160 | 234 | 298 | 698 | 73 | 3 | 0 | 0 | 0 | 0 | 0 |
| SG0000005 | Rit2 | 0.000104 | 0.000219 | -5.396338084 | 15.16262524 | 0.360010171 | 2199 | 135 | 234 | 403 | 1214 | 160 | 10 | 0 | 0 | 0 | 3 | 2 |
| SG0000001 | Lama2 | 0.000493 | 0.000963 | -1.973653734 | 40.03271447 | 10.19262541 | 2896 | 102 | 234 | 400 | 1446 | 210 | 0 | 0 | 0 | 0 | 1 | 0 |
| SG0000002 | Helz | 0.342687 | 0.409955 | -0.132768488 | 3220.910539 | 2937.726527 | 1100 | 210 | 234 | 343 | 670 | 84 | 19 | 0 | 1 | 2 | 9 | 2 |
| SG0000003 | Cask | 2.97E-14 | 1.34E-13 | 2.207265089 | 149.6865697 | 691.2511011 | 3085 | 118 | 233 | 470 | 1563 | 221 | 23 | 4 | 4 | 4 | 19 | 0 |
| SG0000001 | Akt3 | 0.001672 | 0.003062 | -0.482109844 | 1633.795121 | 1169.682711 | 2231 | 128 | 233 | 387 | 1082 | 185 | 14 | 0 | 2 | 6 | 8 | 2 |
| SG0000005 | Ccdc171 | 4.69E-06 | 1.14E-05 | 1.256417967 | 56.7594746 | 135.5994293 | 1811 | 161 | 230 | 371 | 1088 | 151 | 3 | 1 | 1 | 1 | 2 | 0 |
| SG0000003 | Pbx3 | 5.28E-08 | 1.52E-07 | 0.93132124 | 266.0985112 | 507.4556281 | 2956 | 70 | 229 | 327 | 1272 | 169 | 9 | 0 | 0 | 2 | 4 | 4 |
| SG0000002 | Sh3gl2 | 2.9E-155 | 4.7E-153 | 9.164557506 | 0 | 572.8609939 | 2158 | 177 | 228 | 385 | 978 | 149 | 13 | 0 | 0 | 2 | 4 | 1 |
| SG0000002 | Mctp1 | 6.63E-10 | 2.22E-09 | -3.660846092 | 54.55403894 | 4.313228102 | 2273 | 116 | 228 | 387 | 1243 | 197 | 5 | 0 | 0 | 1 | 3 | 2 |
| SG0000002 | Eif4g3 | 9.28E-09 | 2.83E-08 | -0.807396508 | 4835.112522 | 2762.8418 | 1431 | 154 | 228 | 289 | 695 | 111 | 30 | 0 | 0 | 5 | 9 | 4 |
| SG0000002 | Dlg1 | 0.025414 | 0.038613 | 0.290088969 | 2857.28976 | 3493.652988 | 2996 | 126 | 227 | 463 | 1587 | 216 | 15 | 0 | 0 | 0 | 0 | 0 |
| SG0000002 | Pdlim5 | 0.040288 | 0.059078 | -0.287755586 | 2314.499671 | 1895.981348 | 1514 | 137 | 226 | 338 | 814 | 102 | 11 | 0 | 0 | 2 | 6 | 1 |
| SG0000001 | Gltp | 1.7E-33 | 1.98E-32 | -1.884743023 | 7581.875446 | 2053.11115 | 285 | 162 | 225 | 180 | 192 | 0 | 0 | 0 | 0 | 0 | 0 | 0 |
| SG0000000 | Homer1 | 0.033453 | 0.049789 | 0.328278643 | 753.855638 | 946.4766534 | 2843 | 82 | 225 | 397 | 1544 | 167 | 60 | 0 | 6 | 13 | 27 | 2 |
| SG0000005 | Ryr3 | 2.13E-23 | 1.59E-22 | -4.798579611 | 121.1025836 | 4.351480172 | 2971 | 134 | 224 | 427 | 1385 | 286 | 0 | 0 | 0 | 0 | 0 | 0 |
| SG0000002 | Kdm2b | 0.010538 | 0.017164 | 0.452879303 | 1274.945369 | 1745.106075 | 949 | 202 | 224 | 297 | 519 | 49 | 16 | 0 | 0 | 4 | 12 | 4 |
| SG0000001 | Esr1 | 4.14E-10 | 1.41E-09 | -4.420235594 | 70.99219146 | 3.315790485 | 2694 | 157 | 223 | 448 | 1686 | 225 | 14 | 0 | 0 | 1 | 3 | 0 |
| SG0000003 | Plch1 | 4.44E-05 | 9.81E-05 | 0.796669826 | 182.258715 | 316.5992037 | 2424 | 124 | 223 | 399 | 1178 | 173 | 16 | 0 | 0 | 0 | 12 | 14 |
| SG0000004 | Tenm4 | 0.000406 | 0.000801 | -0.928644572 | 1201.577611 | 631.250821 | 2792 | 145 | 223 | 379 | 1113 | 209 | 6 | 0 | 0 | 0 | 0 | 3 |
| SG0000003 | Vps8 | 0.002379 | 0.00426 | -0.536792228 | 830.0511509 | 572.1557859 | 2193 | 137 | 221 | 360 | 964 | 207 | 0 | 0 | 0 | 0 | 0 | 0 |
| SG0000005 | Nyap2 | 2.75E-49 | 5.56E-48 | -7.665432101 | 202.0135343 | 0 | 2879 | 122 | 220 | 368 | 1189 | 251 | 63 | 1 | 2 | 3 | 12 | 7 |
| SG0000003 | Smg6 | 0.011119 | 0.018049 | 0.345687319 | 1345.422046 | 1709.703465 | 1016 | 174 | 220 | 276 | 632 | 90 | 2 | 0 | 0 | 0 | 0 | 0 |
| SG0000004 | Atxn1 | 0.318591 | 0.38449 | 0.494658955 | 72.56140689 | 102.2381254 | 1914 | 156 | 219 | 369 | 903 | 216 | 9 | 0 | 0 | 1 | 3 | 1 |
| SG0000003 | Mid1 | 8.3E-14 | 3.63E-13 | 2.368792474 | 24.3803699 | 125.9267163 | 2811 | 146 | 218 | 473 | 1520 | 282 | 0 | 0 | 0 | 0 | 0 | 0 |
| SG0000003 | Mbd5 | 2.27E-05 | 5.17E-05 | -0.85277676 | 508.0397182 | 281.3107198 | 2575 | 92 | 218 | 384 | 1437 | 200 | 34 | 0 | 4 | 7 | 14 | 11 |
| SG0000006 | Fam172a | 0.001405 | 0.002595 | -0.565156866 | 622.4624376 | 420.7111095 | 2198 | 120 | 218 | 334 | 1218 | 208 | 2 | 0 | 0 | 0 | 1 | 0 |
| SG0000002 | Ptpn4 | 0.011586 | 0.018761 | 0.408459431 | 741.8319284 | 984.6095918 | 1853 | 118 | 218 | 317 | 1027 | 122 | 42 | 0 | 0 | 3 | 10 | 4 |
| SG0000002 | Adarb1 | 1.36E-51 | 2.94E-50 | -2.84341267 | 2406.591915 | 335.3126592 | 1152 | 144 | 216 | 258 | 540 | 114 | 27 | 0 | 3 | 0 | 0 | 6 |
| SG0000004 | Poln | 0.368476 | 0.43672 | -0.643548049 | 17.27553715 | 11.05870552 | 1104 | 168 | 216 | 246 | 630 | 81 | 0 | 0 | 0 | 0 | 0 | 0 |
| SG0000007 | Scn9a | 2.25E-06 | 5.63E-06 | 4.027086602 | 0 | 15.30323777 | 2126 | 135 | 215 | 324 | 1143 | 207 | 41 | 0 | 0 | 4 | 8 | 12 |
| SG0000003 | Jarid2 | 1.39E-49 | 2.86E-48 | 1.75870502 | 2628.515301 | 8894.744425 | 2017 | 149 | 214 | 335 | 820 | 133 | 3 | 0 | 0 | 0 | 0 | 1 |
| SG0000002 | Mllt10 | 0.00243 | 0.004348 | 0.415097631 | 1183.864149 | 1578.551325 | 1509 | 140 | 213 | 252 | 854 | 145 | 12 | 0 | 0 | 0 | 9 | 6 |
| SG0000006 | Syne2 | 1.6E-108 | 1.4E-106 | -4.552022617 | 14264.34199 | 608.0739435 | 1323 | 173 | 212 | 297 | 686 | 132 | 20 | 0 | 0 | 0 | 0 | 0 |
| SG0000003 | Cbfa2t2 | 6.92E-22 | 4.83E-21 | 1.203471092 | 1690.728516 | 3893.630857 | 950 | 173 | 212 | 269 | 517 | 71 | 22 | 0 | 3 | 0 | 5 | 3 |
| SG0000000 | Snd1 | 0.007248 | 0.012114 | -0.360082861 | 5901.522083 | 4597.992882 | 2420 | 129 | 212 | 371 | 1011 | 183 | 10 | 0 | 1 | 0 | 1 | 1 |
| SG0000003 | Ralgapa2 | 0.379302 | 0.448098 | -0.198597827 | 953.9898323 | 831.3039478 | 1889 | 124 | 212 | 305 | 949 | 171 | 13 | 1 | 2 | 2 | 3 | 2 |
| SG0000006 | Pclo | 4.94E-26 | 4.18E-25 | 6.407836362 | 1.005677317 | 85.39050993 | 2261 | 97 | 210 | 377 | 1273 | 202 | 22 | 0 | 1 | 1 | 16 | 2 |
| SG0000006 | Slc4a4 | 0.118967 | 0.159191 | -0.559405633 | 134.8495117 | 91.50640694 | 2671 | 130 | 210 | 403 | 1296 | 183 | 6 | 0 | 0 | 0 | 5 | 3 |
| SG0000003 | Map4 | 8.12E-05 | 0.000174 | -0.527230419 | 10093.34673 | 7003.627219 | 1122 | 161 | 209 | 289 | 681 | 41 | 20 | 0 | 1 | 2 | 7 | 0 |
| SG0000005 | Spag16 | 3.98E-31 | 4.22E-30 | 8.200967544 | 0.307509278 | 90.48892895 | 2704 | 93 | 208 | 416 | 1391 | 289 | 4 | 0 | 0 | 0 | 3 | 0 |
| SG0000002 | Atf2 | 0.034762 | 0.05157 | 0.274903236 | 1924.930602 | 2328.998435 | 2011 | 136 | 208 | 354 | 998 | 173 | 40 | 0 | 0 | 0 | 7 | 7 |
| SG0000002 | Park2 | 0.000141 | 0.000294 | 1.664425435 | 15.99966002 | 50.71690421 | 2802 | 86 | 207 | 399 | 1328 | 296 | 1 | 0 | 0 | 0 | 0 | 0 |
| SG0000004 | Wnk1 | 2.31E-09 | 7.39E-09 | -0.821634339 | 15544.57215 | 8795.12855 | 1505 | 133 | 205 | 296 | 788 | 108 | 18 | 0 | 3 | 0 | 8 | 0 |
| SG0000003 | Tmcc1 | 1.44E-08 | 4.34E-08 | -0.973371279 | 940.7657761 | 479.1456593 | 1338 | 137 | 205 | 249 | 759 | 106 | 8 | 0 | 0 | 0 | 3 | 1 |
| SG0000004 | Hecw2 | 4.71E-05 | 0.000104 | -1.353421069 | 176.5542058 | 69.09676725 | 2117 | 130 | 205 | 357 | 1208 | 202 | 42 | 0 | 2 | 8 | 12 | 2 |
| SG0000004 | Magi1 | 0.143376 | 0.188599 | 0.214818788 | 719.8131938 | 835.385055 | 2737 | 122 | 205 | 372 | 1209 | 238 | 11 | 0 | 0 | 2 | 4 | 0 |
| SG0000002 | Unc13b | 5.2E-10 | 1.75E-09 | 0.940794477 | 476.6470829 | 914.9646111 | 1776 | 120 | 204 | 324 | 906 | 124 | 0 | 0 | 0 | 0 | 2 | 0 |
| SG0000003 | Itgbl1 | 0.031051 | 0.046476 | -1.48895111 | 21.5721365 | 7.685536925 | 2249 | 111 | 204 | 423 | 1458 | 168 | 23 | 0 | 0 | 6 | 6 | 3 |
| SG0000002 | Ablim1 | 3.59E-19 | 2.15E-18 | -1.661740877 | 1165.225851 | 368.2784123 | 2843 | 152 | 203 | 492 | 1323 | 342 | 16 | 0 | 0 | 12 | 20 | 0 |
| SG0000005 | Senp7 | 8.56E-10 | 2.84E-09 | -1.46047802 | 535.6778413 | 194.650709 | 1182 | 157 | 203 | 297 | 648 | 99 | 12 | 0 | 0 | 2 | 10 | 1 |
| SG0000006 | Inadl | 0.881277 | 0.908462 | -0.027659657 | 581.3458673 | 570.3063447 | 2669 | 122 | 203 | 411 | 1334 | 249 | 17 | 0 | 2 | 1 | 5 | 0 |
| SG0000003 | Mef2a | 1.27E-27 | 1.15E-26 | -2.070229189 | 1929.061314 | 459.3512978 | 2171 | 109 | 202 | 302 | 982 | 199 | 34 | 0 | 4 | 9 | 23 | 9 |
| SG0000002 | Gtf2ird1 | 0.00123 | 0.002286 | 0.464315927 | 904.5262737 | 1247.941444 | 1533 | 164 | 202 | 339 | 853 | 255 | 9 | 0 | 0 | 0 | 0 | 0 |
| SG0000002 | Shroom3 | 4.94E-06 | 1.2E-05 | -0.959348132 | 1484.83355 | 763.6339144 | 1259 | 156 | 201 | 286 | 740 | 119 | 3 | 0 | 3 | 3 | 3 | 0 |
| SG0000000 | Npepps | 0.000155 | 0.000321 | -0.513852113 | 8266.858352 | 5789.693815 | 1028 | 169 | 201 | 259 | 583 | 107 | 4 | 0 | 0 | 0 | 1 | 1 |
| SG0000003 | Tasp1 | 0.000558 | 0.001082 | 0.636434524 | 235.3452925 | 365.8401762 | 1737 | 117 | 201 | 285 | 978 | 129 | 6 | 0 | 0 | 0 | 6 | 3 |
| SG0000003 | Arid4b | 0.102835 | 0.139554 | -0.243356271 | 1421.162175 | 1200.566173 | 1342 | 129 | 200 | 280 | 717 | 118 | 22 | 0 | 0 | 3 | 19 | 1 |
| SG0000003 | Myo5a | 5.25E-06 | 1.27E-05 | -0.657332954 | 3135.518417 | 1988.073424 | 1406 | 146 | 199 | 298 | 717 | 99 | 19 | 1 | 1 | 3 | 8 | 0 |
| SG0000005 | Arhgap24 | 0.011766 | 0.01902 | -1.030304893 | 77.08017561 | 37.73896783 | 3163 | 90 | 199 | 502 | 1589 | 263 | 12 | 0 | 0 | 0 | 18 | 0 |
| SG0000003 | Phka1 | 3.5E-19 | 2.09E-18 | -1.723385509 | 837.9988465 | 253.777251 | 1728 | 120 | 198 | 378 | 1212 | 192 | 24 | 0 | 0 | 6 | 6 | 6 |
| SG0000004 | Phf8 | 0.018776 | 0.029186 | -0.364176047 | 1477.681394 | 1148.029086 | 1525 | 101 | 198 | 292 | 966 | 196 | 10 | 0 | 0 | 0 | 6 | 0 |
| SG0000002 | Ptprg | 1.21E-46 | 2.27E-45 | 1.95196683 | 500.8288378 | 1937.714968 | 2494 | 127 | 196 | 387 | 1145 | 248 | 12 | 0 | 1 | 4 | 10 | 1 |
| SG0000005 | Ghr | 1.25E-40 | 1.9E-39 | -3.447530328 | 2328.154626 | 213.4036195 | 2132 | 110 | 196 | 323 | 1050 | 224 | 10 | 0 | 0 | 0 | 0 | 0 |
| SG0000002 | Pum2 | 0.000145 | 0.000303 | -0.513421834 | 8901.454117 | 6235.992514 | 2075 | 82 | 196 | 327 | 1013 | 140 | 62 | 0 | 0 | 8 | 31 | 11 |
| SG0000004 | Cdh18 | 0.231087 | 0.289984 | -0.805207278 | 22.51491271 | 12.88483093 | 1936 | 124 | 196 | 318 | 1112 | 170 | 0 | 0 | 0 | 0 | 0 | 0 |
| SG0000003 | Cobll1 | 0.700081 | 0.75405 | -0.227386445 | 183.6544385 | 156.8741147 | 1835 | 140 | 195 | 295 | 805 | 210 | 40 | 0 | 0 | 0 | 15 | 5 |
| SG0000002 | Itsn1 | 0.046778 | 0.067938 | -0.278865856 | 2808.4331 | 2314.819305 | 1996 | 126 | 193 | 326 | 871 | 106 | 20 | 4 | 4 | 5 | 8 | 0 |
| SG0000001 | Kansl1 | 2.74E-06 | 6.81E-06 | -0.661814031 | 3649.919399 | 2307.052083 | 1200 | 142 | 191 | 259 | 651 | 100 | 16 | 0 | 0 | 4 | 4 | 4 |
| SG0000004 | Pcca | 0.058843 | 0.083784 | 0.288955524 | 567.5578524 | 693.4168918 | 1962 | 126 | 191 | 277 | 854 | 163 | 5 | 0 | 0 | 0 | 0 | 5 |
| SG0000003 | Magt1 | 1.36E-12 | 5.47E-12 | -1.170229504 | 2530.093694 | 1124.248772 | 448 | 171 | 190 | 228 | 360 | 33 | 18 | 0 | 3 | 0 | 3 | 0 |
| SG0000002 | Pde10a | 0.036826 | 0.054362 | 1.020871879 | 21.42473921 | 43.47389996 | 2245 | 111 | 189 | 325 | 949 | 351 | 31 | 0 | 3 | 0 | 14 | 2 |
| SG0000002 | Pkp4 | 6.35E-16 | 3.17E-15 | 1.028394451 | 1642.389294 | 3350.068464 | 2301 | 122 | 188 | 326 | 990 | 168 | 3 | 0 | 0 | 0 | 4 | 4 |
| SG0000001 | Map2 | 1.17E-14 | 5.42E-14 | -1.270735191 | 1563.495517 | 647.9883998 | 2968 | 82 | 188 | 424 | 1420 | 261 | 48 | 0 | 0 | 0 | 39 | 0 |
| SG0000002 | Mnat1 | 6.83E-09 | 2.11E-08 | 0.817908209 | 741.9299232 | 1307.909794 | 1193 | 156 | 187 | 303 | 684 | 71 | 12 | 0 | 0 | 1 | 2 | 0 |
| SG0000003 | Rora | 0.000349 | 0.000696 | 0.805904134 | 146.1608732 | 255.5244409 | 2139 | 103 | 187 | 336 | 863 | 203 | 32 | 1 | 2 | 6 | 12 | 4 |
| SG0000002 | Apbb2 | 0.008449 | 0.013984 | -0.416009876 | 984.5929742 | 737.9471862 | 3143 | 130 | 187 | 423 | 1296 | 375 | 35 | 4 | 4 | 4 | 19 | 7 |
| SG0000000 | Tex11 | 7.03E-28 | 6.45E-27 | 8.884194937 | 0.34247518 | 161.8222614 | 1677 | 88 | 186 | 310 | 1005 | 168 | 2 | 0 | 1 | 0 | 1 | 0 |
| SG0000001 | Ncor1 | 0.00022 | 0.000448 | -0.505498682 | 6427.19568 | 4527.424914 | 1380 | 134 | 186 | 313 | 830 | 88 | 25 | 0 | 1 | 4 | 5 | 3 |
| SG0000005 | Glis3 | 0.409324 | 0.478508 | 0.260610174 | 146.1825118 | 175.1244964 | 2419 | 86 | 186 | 277 | 856 | 195 | 10 | 0 | 0 | 0 | 0 | 0 |
| SG0000006 | Nr6a1 | 1.25E-95 | 8.03E-94 | 2.910543147 | 292.4483923 | 2198.923036 | 1179 | 126 | 185 | 222 | 659 | 92 | 15 | 0 | 0 | 3 | 9 | 0 |
| SG0000000 | Nfat5 | 5.38E-11 | 1.95E-10 | -1.22522114 | 4669.255604 | 1997.189776 | 1831 | 112 | 185 | 313 | 937 | 86 | 100 | 0 | 10 | 10 | 37 | 3 |
| SG0000002 | Scn8a | 6.88E-07 | 1.8E-06 | -1.272766901 | 690.7692551 | 285.8854054 | 1233 | 113 | 185 | 195 | 679 | 113 | 22 | 0 | 0 | 6 | 12 | 10 |
| SG0000003 | Runx2 | 1.38E-39 | 2.01E-38 | -5.722909621 | 2366.133839 | 44.79936396 | 3149 | 116 | 184 | 479 | 1239 | 187 | 26 | 0 | 3 | 5 | 11 | 2 |
| SG0000002 | Fbxl17 | 2.18E-26 | 1.88E-25 | -1.916207173 | 1390.028053 | 368.2881354 | 1958 | 102 | 184 | 273 | 929 | 144 | 6 | 0 | 0 | 2 | 5 | 0 |
| SG0000002 | Schip1 | 9.91E-08 | 2.78E-07 | 2.817829242 | 10.20182689 | 71.93314929 | 2612 | 108 | 184 | 414 | 1237 | 211 | 12 | 0 | 0 | 0 | 0 | 0 |
| SG0000004 | Mettl8 | 6.97E-06 | 1.67E-05 | 0.822053736 | 232.4154501 | 410.8921342 | 1078 | 100 | 183 | 298 | 702 | 88 | 1 | 0 | 0 | 1 | 3 | 0 |
| SG0000000 | Tom1l2 | 1.73E-47 | 3.33E-46 | -2.652022182 | 2451.755745 | 390.0667892 | 1137 | 152 | 182 | 314 | 668 | 116 | 13 | 0 | 0 | 8 | 8 | 0 |
| SG0000002 | Cdc42bpa | 6.64E-35 | 8.23E-34 | -2.005649143 | 4700.530219 | 1170.5401 | 1924 | 97 | 182 | 272 | 905 | 134 | 24 | 0 | 0 | 2 | 10 | 6 |
| SG0000003 | Thsd7a | 0.086084 | 0.11883 | -1.171980916 | 103.363662 | 45.87398596 | 2334 | 91 | 182 | 389 | 1267 | 213 | 16 | 1 | 2 | 3 | 11 | 1 |

| | | | | | | | | | | | | | | | | | | |
|---|---|---|---|---|---|---|---|---|---|---|---|---|---|---|---|---|---|---|
| SG0000004 | Wnk3 | 3.86E-38 | 5.36E-37 | 4.166768851 | 33.96583153 | 610.0489166 | 1375 | 112 | 181 | 308 | 973 | 77 | 7 | 0 | 1 | 0 | 1 | 1 |
| SG0000004 | Plekha7 | 5.17E-20 | 3.24E-19 | 1.247357666 | 748.7742458 | 1777.63655 | 845 | 134 | 181 | 203 | 588 | 55 | 9 | 0 | 0 | 0 | 5 | 0 |
| SG0000003 | Lpp | 1.52E-17 | 8.35E-17 | -1.69841209 | 1959.110185 | 603.6509358 | 2805 | 103 | 181 | 384 | 1324 | 262 | 28 | 0 | 0 | 3 | 9 | 0 |
| SG0000004 | Clip1 | 2.45E-13 | 1.04E-12 | -1.14431528 | 2116.812822 | 957.6553379 | 1183 | 160 | 181 | 249 | 607 | 93 | 16 | 0 | 0 | 0 | 4 | 8 |
| SG0000002 | Osbpl9 | 0.000174 | 0.000359 | -0.55579485 | 2238.26779 | 1522.653635 | 1859 | 139 | 181 | 283 | 891 | 143 | 7 | 0 | 0 | 6 | 6 | 6 |
| SG0000005 | Ebf1 | 0.000881 | 0.001664 | -0.917580339 | 145.6404247 | 77.10148298 | 2223 | 120 | 181 | 329 | 958 | 200 | 15 | 0 | 1 | 5 | 7 | 2 |
| SG0000003 | Nox4 | 0.010506 | 0.017121 | 0.884136872 | 45.28177058 | 83.57475629 | 2339 | 85 | 181 | 351 | 1107 | 169 | 36 | 0 | 6 | 6 | 17 | 0 |
| SG0000002 | Asph | 3.86E-17 | 2.08E-16 | 1.148093963 | 971.5718249 | 2153.206255 | 2389 | 93 | 180 | 363 | 1257 | 219 | 39 | 3 | 3 | 8 | 25 | 1 |
| SG0000002 | Kdm4c | 8.5E-16 | 4.22E-15 | 1.00784996 | 1963.961142 | 3949.353058 | 1002 | 134 | 179 | 239 | 605 | 86 | 10 | 0 | 0 | 4 | 6 | 0 |
| SG0000002 | Pou2f1 | 0.000357 | 0.00071 | 0.512190389 | 1532.966375 | 2186.337981 | 2265 | 68 | 179 | 290 | 1002 | 160 | 42 | 2 | 4 | 6 | 28 | 4 |
| SG0000002 | Atp9a | 0.07021 | 0.09862 | -0.487987805 | 1344.537321 | 958.6805026 | 654 | 174 | 179 | 224 | 490 | 35 | 0 | 0 | 0 | 0 | 0 | 0 |
| SG0000005 | Skap1 | 0.000189 | 0.000388 | 3.959919579 | 0.970711415 | 15.10583409 | 2490 | 102 | 178 | 344 | 1131 | 205 | 7 | 0 | 0 | 0 | 1 | 1 |
| SG0000003 | Pds5b | 0.483853 | 0.55225 | -0.104180921 | 2690.961193 | 2503.489951 | 1203 | 144 | 178 | 284 | 721 | 71 | 17 | 0 | 0 | 3 | 16 | 4 |
| SG0000002 | Nf1 | 5.93E-12 | 2.3E-11 | -1.262199586 | 2516.646618 | 1049.209711 | 1391 | 120 | 177 | 267 | 853 | 85 | 13 | 0 | 0 | 2 | 9 | 4 |
| SG0000003 | Rbm6 | 5.58E-05 | 0.000122 | 0.531566221 | 1517.732099 | 2193.878117 | 814 | 128 | 177 | 229 | 493 | 66 | 1 | 0 | 0 | 0 | 2 | 0 |
| SG0000002 | Stk3 | 6.05E-05 | 0.000132 | -0.609045049 | 1784.07929 | 1169.696858 | 1567 | 97 | 177 | 313 | 816 | 246 | 8 | 2 | 2 | 6 | 6 | 0 |
| SG0000002 | Slmap | 4.49E-10 | 1.52E-09 | -0.890890646 | 5084.325228 | 2741.880457 | 1619 | 79 | 176 | 281 | 873 | 132 | 43 | 0 | 5 | 5 | 10 | 3 |
| SG0000007 | Snx29 | 0.013429 | 0.02144 | 1.079464103 | 42.69336298 | 90.22179013 | 1604 | 126 | 176 | 248 | 756 | 104 | 7 | 1 | 1 | 1 | 2 | 1 |
| SG0000004 | Suv420h1 | 0.280532 | 0.343864 | -0.167586881 | 927.0177446 | 825.3523064 | 891 | 158 | 176 | 231 | 558 | 26 | 26 | 5 | 5 | 9 | 16 | 3 |
| SG0000002 | Dync1i2 | 0.981727 | 0.993037 | 0.000803715 | 3473.636227 | 3475.571904 | 1224 | 80 | 176 | 216 | 559 | 104 | 0 | 0 | 0 | 0 | 7 | 0 |
| SG0000004 | Bdnf | 9.49E-21 | 6.23E-20 | -2.22832725 | 684.5731908 | 146.0920371 | 1162 | 138 | 175 | 270 | 443 | 48 | 88 | 0 | 0 | 33 | 44 | 22 |
| JSG000000 | Atf7 | 1.01E-15 | 5E-15 | -1.481612467 | 2047.9942 | 733.3628866 | 1387 | 140 | 174 | 258 | 637 | 103 | 18 | 3 | 4 | 4 | 6 | 0 |
| SG0000002 | Meis2 | 0.947217 | 0.963243 | -0.039801264 | 37.37137538 | 36.35446076 | 2489 | 120 | 174 | 342 | 1008 | 198 | 5 | 0 | 0 | 0 | 6 | 0 |
| SG0000000 | Ssbp2 | 1.27E-95 | 8.13E-94 | -4.589437986 | 3138.219315 | 130.3541295 | 1374 | 105 | 173 | 256 | 782 | 135 | 2 | 0 | 0 | 0 | 1 | 1 |
| SG0000003 | Fgf13 | 0.132678 | 0.175817 | -0.510271636 | 171.6424347 | 120.508477 | 1747 | 102 | 173 | 302 | 979 | 144 | 9 | 0 | 0 | 0 | 3 | 0 |
| SG0000002 | Ptbp3 | 1.46E-09 | 4.73E-09 | -0.928519416 | 6672.964191 | 3505.957108 | 1652 | 69 | 171 | 314 | 1042 | 184 | 32 | 0 | 0 | 0 | 18 | 8 |
| SG0000005 | Ube2e2 | 9.88E-06 | 2.33E-05 | -0.82270907 | 555.9971956 | 314.3492968 | 1715 | 99 | 171 | 264 | 1000 | 171 | 2 | 0 | 0 | 0 | 2 | 0 |
| SG0000006 | Atp11c | 1.41E-05 | 3.29E-05 | -0.81568603 | 801.8417847 | 455.5569974 | 2024 | 113 | 171 | 354 | 1008 | 132 | 39 | 0 | 0 | 3 | 12 | 0 |
| SG0000002 | Mbnl1 | 2.54E-79 | 1.13E-77 | -3.736010462 | 4330.804941 | 325.0254816 | 2652 | 53 | 170 | 368 | 1366 | 169 | 66 | 0 | 0 | 14 | 51 | 15 |
| SG0000005 | Pbx1 | 4.17E-08 | 1.21E-07 | 1.441753257 | 253.495908 | 688.6236438 | 1484 | 133 | 170 | 289 | 839 | 92 | 13 | 1 | 1 | 1 | 16 | 1 |
| SG0000003 | Enox2 | 0.023173 | 0.035428 | -0.424654718 | 464.2851138 | 345.9003102 | 3059 | 70 | 170 | 340 | 1571 | 348 | 4 | 0 | 0 | 0 | 1 | 8 |
| SG0000002 | Cnksr2 | 0.013173 | 0.021071 | 0.806675564 | 44.19159708 | 77.29888666 | 1545 | 96 | 169 | 246 | 824 | 126 | 16 | 1 | 3 | 2 | 7 | 0 |
| SG0000003 | Lace1 | 0.604902 | 0.667468 | 0.096507175 | 283.1130788 | 302.6993492 | 1183 | 134 | 169 | 289 | 703 | 121 | 3 | 0 | 3 | 3 | 3 | 0 |
| SG0000003 | Aebp2 | 7.39E-40 | 1.09E-38 | -2.250417174 | 11102.04077 | 2333.241788 | 534 | 147 | 168 | 185 | 413 | 44 | 23 | 0 | 0 | 2 | 11 | 6 |
| SG0000004 | Pde9a | 5.24E-33 | 5.98E-32 | -2.199731914 | 1528.815828 | 332.7897042 | 796 | 128 | 168 | 208 | 583 | 56 | 0 | 0 | 0 | 0 | 4 | 5 |
| SG0000003 | Dusp16 | 2.72E-08 | 7.99E-08 | -0.852784818 | 1847.822168 | 1023.166619 | 606 | 138 | 168 | 176 | 322 | 34 | 12 | 0 | 1 | 1 | 2 | 1 |
| SG0000005 | Dis3l2 | 1.2E-05 | 2.81E-05 | -0.649712885 | 1875.755344 | 1195.619875 | 1975 | 87 | 168 | 300 | 917 | 167 | 1 | 0 | 0 | 0 | 2 | 0 |
| SG0000000 | Mast2 | 9.5E-05 | 0.000202 | -0.555914142 | 2708.052129 | 1842.087215 | 1336 | 104 | 168 | 260 | 784 | 104 | 0 | 0 | 0 | 0 | 0 | 0 |
| SG0000004 | Ankrd6 | 0.000448 | 0.000879 | 0.551830133 | 481.789204 | 706.2760618 | 907 | 117 | 168 | 207 | 482 | 50 | 4 | 0 | 0 | 0 | 4 | 4 |
| SG0000003 | Kcnk2 | 0.003637 | 0.006342 | -1.47772275 | 41.64972007 | 14.95454534 | 1619 | 107 | 168 | 269 | 810 | 182 | 19 | 0 | 0 | 3 | 3 | 3 |
| SG0000005 | Chn1 | 4.01E-50 | 8.36E-49 | 2.701123115 | 95.86512724 | 623.418567 | 1822 | 72 | 167 | 257 | 940 | 177 | 26 | 0 | 0 | 0 | 3 | 0 |
| SG0000005 | Zgrf1 | 3.56E-11 | 1.31E-10 | 1.150471267 | 363.2831996 | 806.4392825 | 807 | 116 | 167 | 195 | 429 | 63 | 36 | 3 | 6 | 12 | 17 | 0 |
| SG0000002 | Dab2 | 1.08E-25 | 8.96E-25 | -2.750545273 | 1589.301957 | 236.1618741 | 1192 | 127 | 166 | 280 | 546 | 79 | 16 | 0 | 5 | 0 | 6 | 0 |
| SG0000002 | Ryr2 | 1.65E-25 | 1.36E-24 | 6.890597537 | 0.649984458 | 77.12221941 | 2118 | 76 | 166 | 292 | 1090 | 228 | 4 | 0 | 0 | 0 | 2 | 0 |
| SG0000005 | Usp34 | 0.000459 | 0.0009 | -0.483485609 | 5680.801485 | 4063.17888 | 1292 | 123 | 166 | 273 | 709 | 141 | 18 | 3 | 3 | 4 | 10 | 1 |
| SG0000005 | Nebl | 0.057394 | 0.081915 | 1.144891259 | 10.23913636 | 22.64174723 | 1445 | 103 | 166 | 267 | 712 | 104 | 27 | 1 | 1 | 2 | 11 | 0 |
| SG0000005 | Ptprn2 | 0.093871 | 0.128558 | 1.263124787 | 6.526279939 | 15.66405313 | 2074 | 118 | 166 | 304 | 978 | 234 | 0 | 0 | 0 | 0 | 0 | 0 |
| SG0000002 | Tbc1d1 | 2.19E-68 | 7.56E-67 | 3.918569468 | 170.9627271 | 2585.284765 | 1124 | 95 | 164 | 211 | 519 | 67 | 16 | 0 | 4 | 8 | 20 | 0 |
| SG0000002 | Dclk1 | 1.19E-24 | 9.47E-24 | -2.840401675 | 569.9169594 | 79.57295327 | 2412 | 96 | 164 | 264 | 854 | 252 | 46 | 0 | 0 | 0 | 13 | 4 |
| SG0000002 | Brd4 | 2.71E-09 | 8.61E-09 | 0.73720832 | 3666.236376 | 6111.42209 | 593 | 128 | 164 | 185 | 388 | 49 | 1 | 0 | 0 | 0 | 2 | 0 |
| SG0000002 | Tbl1xr1 | 0.33975 | 0.406873 | -0.147890511 | 2065.714108 | 1864.449976 | 1292 | 111 | 164 | 228 | 686 | 121 | 29 | 0 | 0 | 0 | 23 | 7 |
| SG0000003 | 4932438A13Rik | 1.07E-08 | 3.26E-08 | -0.902966206 | 2413.031987 | 1290.455895 | 1943 | 70 | 163 | 286 | 924 | 153 | 20 | 0 | 0 | 3 | 6 | 5 |
| SG0000004 | Tmem164 | 3.24E-08 | 9.47E-08 | -0.918099752 | 1086.367382 | 574.9117082 | 1721 | 125 | 163 | 241 | 894 | 160 | 8 | 0 | 0 | 4 | 4 | 4 |
| SG0000002 | Agtpbp1 | 0.845217 | 0.878928 | 0.026446721 | 453.3533403 | 461.7406142 | 1638 | 93 | 163 | 256 | 849 | 213 | 24 | 0 | 0 | 0 | 3 | 4 |
| SG0000003 | Cnot6l | 2.68E-26 | 2.29E-25 | -1.786400702 | 2826.861172 | 819.4911825 | 1268 | 73 | 162 | 191 | 617 | 53 | 43 | 1 | 2 | 3 | 14 | 1 |
| SG0000002 | Frmd4a | 8.37E-11 | 2.99E-10 | -1.11159269 | 1042.245939 | 482.3335472 | 2248 | 94 | 162 | 315 | 991 | 255 | 8 | 0 | 0 | 0 | 1 | 3 |
| SG0000002 | Dnajc1 | 1.16E-10 | 4.09E-10 | -1.197236005 | 840.7134497 | 366.6435472 | 2407 | 93 | 162 | 296 | 1258 | 205 | 13 | 0 | 1 | 0 | 1 | 0 |
| SG0000002 | Hmbox1 | 1.74E-06 | 4.41E-06 | -0.788830354 | 1198.156723 | 693.5088876 | 1974 | 54 | 162 | 297 | 988 | 241 | 11 | 0 | 2 | 2 | 2 | 0 |
| SG0000002 | B230118H07Rik | 0.001421 | 0.002623 | 0.573529704 | 278.8939358 | 415.0388161 | 995 | 84 | 162 | 160 | 439 | 56 | 1 | 0 | 0 | 1 | 3 | 0 |
| SG0000004 | Gli2 | 4.1E-203 | 1.3E-200 | 6.228086096 | 16.18052337 | 1212.923179 | 1079 | 135 | 161 | 266 | 536 | 136 | 1 | 0 | 0 | 0 | 0 | 1 |
| SG0000004 | Osbpl10 | 8.83E-08 | 2.49E-07 | 1.172801763 | 128.892026 | 290.585915 | 1175 | 121 | 161 | 227 | 621 | 168 | 0 | 0 | 0 | 0 | 1 | 0 |
| SG0000000 | Myo18a | 0.000659 | 0.001264 | -0.501274172 | 2472.572153 | 1746.829075 | 1166 | 133 | 161 | 263 | 534 | 90 | 16 | 0 | 0 | 0 | 8 | 0 |
| SG0000002 | Fam129a | 0.060807 | 0.086292 | -0.264584134 | 2869.148205 | 2388.389874 | 971 | 118 | 161 | 227 | 462 | 66 | 13 | 0 | 0 | 2 | 2 | 0 |
| SG0000002 | Xpr1 | 0.379919 | 0.448678 | 0.117541855 | 1557.045619 | 1689.215124 | 1124 | 120 | 161 | 230 | 563 | 69 | 9 | 0 | 2 | 2 | 5 | 0 |
| SG0000004 | Ubr3 | 0.799632 | 0.841243 | 0.028482396 | 2486.832634 | 2536.416756 | 1031 | 115 | 161 | 231 | 575 | 64 | 16 | 0 | 0 | 2 | 13 | 0 |
| SG0000003 | Mcf2 | 2.2E-154 | 3.6E-152 | 9.194959726 | 0 | 585.0824091 | 787 | 111 | 159 | 178 | 522 | 91 | 15 | 0 | 0 | 0 | 6 | 0 |
| SG0000002 | Odf2 | 9.44E-26 | 7.87E-25 | 1.275863463 | 2622.193672 | 6349.476758 | 684 | 126 | 159 | 186 | 348 | 82 | 6 | 6 | 6 | 6 | 9 | 0 |
| SG0000006 | Msi2 | 1.48E-21 | 1.01E-20 | -1.434042455 | 6674.125091 | 2470.043195 | 1004 | 116 | 158 | 205 | 490 | 69 | 11 | 0 | 1 | 1 | 2 | 4 |
| SG0000005 | Med12l | 1.02E-54 | 2.41E-53 | 6.896432551 | 1.655661775 | 197.2444005 | 2252 | 93 | 157 | 322 | 1059 | 185 | 79 | 0 | 0 | 12 | 33 | 0 |
| SG0000003 | Tfdp2 | 5.96E-30 | 6.03E-29 | 1.507444125 | 763.536404 | 2170.779175 | 1279 | 111 | 157 | 257 | 822 | 164 | 22 | 0 | 0 | 1 | 3 | 0 |
| SG0000002 | Dock7 | 1.44E-09 | 4.66E-09 | -0.991593692 | 2955.510575 | 1486.39102 | 1302 | 83 | 157 | 253 | 694 | 94 | 17 | 0 | 3 | 2 | 2 | 0 |
| SG0000005 | Ankrd17 | 0.000591 | 0.001141 | -0.469939204 | 7708.116195 | 5565.221526 | 1636 | 98 | 157 | 286 | 759 | 175 | 9 | 1 | 2 | 1 | 3 | 0 |
| SG0000003 | Pdss2 | 0.418403 | 0.487538 | -0.1582877 | 362.4674192 | 324.8026994 | 1335 | 105 | 157 | 255 | 647 | 128 | 1 | 0 | 0 | 0 | 0 | 0 |
| SG0000002 | Hira | 0.532686 | 0.599226 | 0.094410306 | 911.2761477 | 972.9047799 | 642 | 128 | 157 | 215 | 381 | 37 | 5 | 0 | 0 | 0 | 0 | 0 |
| SG0000003 | Dhx30 | 0.005409 | 0.009207 | 0.374462618 | 2034.105788 | 2636.926446 | 525 | 115 | 156 | 196 | 339 | 16 | 20 | 0 | 10 | 10 | 10 | 0 |
| SG0000005 | Chd9 | 0.034802 | 0.051617 | -0.295999127 | 3195.324564 | 2602.617599 | 974 | 108 | 156 | 220 | 592 | 88 | 15 | 0 | 0 | 0 | 2 | 1 |
| SG0000006 | Cep128 | 0.049482 | 0.07149 | -0.343972914 | 569.2145981 | 448.4663528 | 1489 | 85 | 156 | 247 | 746 | 113 | 10 | 0 | 0 | 0 | 2 | 0 |
| SG0000003 | Sdk1 | 0.067797 | 0.09547 | 1.2944959 | 7.178607964 | 17.60849879 | 1948 | 85 | 156 | 266 | 875 | 267 | 1 | 0 | 0 | 0 | 0 | 0 |
| SG0000004 | Dync2h1 | 0.243857 | 0.304112 | -0.231676963 | 395.4570084 | 336.7887388 | 2067 | 87 | 156 | 278 | 1129 | 178 | 10 | 0 | 0 | 0 | 2 | 0 |
| SG0000004 | Reps2 | 1.01E-09 | 3.31E-09 | 4.46276139 | 4.460554747 | 98.35895724 | 1447 | 93 | 155 | 257 | 821 | 119 | 54 | 3 | 9 | 9 | 18 | 3 |
| JSG000000 | Cntnap5a | 0.014432 | 0.022936 | -4.303206464 | 18.69596502 | 0.947008034 | 2042 | 80 | 155 | 296 | 1024 | 195 | 7 | 0 | 1 | 1 | 10 | 1 |
| SG0000003 | Ttc28 | 5.99E-27 | 5.26E-26 | -2.598907797 | 984.230186 | 162.4603849 | 1240 | 106 | 154 | 232 | 618 | 114 | 8 | 0 | 2 | 0 | 6 | 0 |
| SG0000000 | Celf1 | 5.87E-11 | 2.12E-10 | 0.792716333 | 5124.797439 | 8877.856047 | 952 | 114 | 154 | 202 | 667 | 72 | 60 | 0 | 4 | 4 | 32 | 0 |
| SG0000000 | Large | 0.003383 | 0.005925 | -0.471679838 | 915.1443103 | 659.9329371 | 1800 | 86 | 154 | 256 | 974 | 198 | 0 | 0 | 0 | 0 | 0 | 0 |
| SG0000006 | Adamtsl1 | 0.572425 | 0.636929 | 0.4396365 | 22.65846659 | 30.73082996 | 2223 | 77 | 154 | 307 | 1043 | 174 | 18 | 6 | 8 | 10 | 10 | 0 |
| SG0000003 | Ephb1 | 5.01E-06 | 1.22E-05 | -3.487954546 | 41.37778925 | 3.687978168 | 1597 | 101 | 153 | 314 | 890 | 152 | 3 | 0 | 0 | 0 | 0 | 0 |
| SG0000003 | Ncam1 | 7.87E-06 | 1.87E-05 | -1.473734412 | 754.1326627 | 271.5247906 | 2022 | 91 | 153 | 262 | 863 | 206 | 5 | 0 | 0 | 0 | 2 | 2 |
| SG0000002 | Rabgap1l | 0.00302 | 0.005325 | 0.462633552 | 524.5359316 | 722.8394047 | 1804 | 84 | 153 | 306 | 933 | 148 | 12 | 0 | 1 | 1 | 4 | 0 |
| SG0000002 | Sgk3 | 4.62E-38 | 6.38E-37 | 3.070758862 | 35.97384574 | 302.257715 | 1166 | 102 | 151 | 173 | 636 | 146 | 27 | 0 | 0 | 12 | 17 | 4 |
| SG0000003 | Myo6 | 2.93E-07 | 7.89E-07 | -0.885070719 | 1672.817821 | 905.7655777 | 1635 | 97 | 151 | 274 | 882 | 180 | 48 | 4 | 4 | 4 | 12 | 0 |
| SG0000005 | Cacnb2 | 0.007214 | 0.012063 | 2.436334802 | 5.723214194 | 30.97774888 | 1406 | 86 | 151 | 240 | 646 | 146 | 1 | 0 | 0 | 0 | 1 | 3 |
| SG0000003 | Slc12a8 | 9.22E-40 | 1.35E-38 | 4.348165898 | 10.06196328 | 204.9326188 | 1126 | 104 | 150 | 236 | 686 | 104 | 6 | 2 | 2 | 2 | 4 | 0 |
| SG0000004 | Zfp407 | 5.04E-14 | 2.24E-13 | -1.387333694 | 1118.0148 | 427.3842337 | 1872 | 65 | 149 | 266 | 886 | 168 | 1 | 0 | 0 | 0 | 0 | 0 |
| SG0000000 | Elavl2 | 1.3E-10 | 4.58E-10 | 6.961363157 | 5.589034583 | 696.4916559 | 1777 | 64 | 149 | 247 | 748 | 145 | 90 | 0 | 6 | 18 | 30 | 18 |
| JSG000000 | Erc2 | 3.53E-10 | 1.21E-09 | 4.022247779 | 15.66556225 | 254.5442073 | 1676 | 82 | 149 | 237 | 810 | 145 | 6 | 0 | 0 | 1 | 3 | 1 |
| SG0000002 | Dock2 | 9.49E-07 | 2.46E-06 | 5.824197801 | 0.331601068 | 18.7877235 | 1801 | 68 | 148 | 269 | 882 | 122 | 6 | 0 | 1 | 0 | 0 | 0 |
| SG0000002 | Ep400 | 0.012289 | 0.01978 | -0.344732039 | 4583.950339 | 3609.651047 | 755 | 116 | 148 | 196 | 466 | 63 | 0 | 0 | 0 | 0 | 2 | 3 |
| SG0000004 | Csrnp3 | 3.89E-97 | 2.6E-95 | 8.303526919 | 0 | 314.9444074 | 1980 | 87 | 147 | 364 | 1041 | 223 | 37 | 1 | 2 | 11 | 18 | 2 |
| SG0000005 | Nav2 | 1.45E-17 | 7.96E-17 | -1.293632841 | 3655.70518 | 1491.244571 | 1037 | 123 | 147 | 218 | 487 | 112 | 9 | 2 | 2 | 2 | 2 | 2 |
| SG0000002 | Zfhx4 | 4.72E-12 | 1.84E-11 | -2.894427121 | 219.5183151 | 29.52306052 | 1769 | 65 | 147 | 252 | 785 | 135 | 12 | 0 | 0 | 0 | 2 | 2 |
| SG0000002 | Ndst3 | 8.71E-05 | 0.000186 | -2.762649853 | 29.87319046 | 4.401909755 | 1545 | 80 | 147 | 216 | 705 | 118 | 21 | 0 | 0 | 0 | 0 | 8 |
| SG0000003 | Trip4 | 0.00085 | 0.001608 | 0.481322218 | 789.0141312 | 1101.481499 | 694 | 104 | 147 | 200 | 431 | 58 | 20 | 0 | 5 | 4 | 9 | 0 |
| SG0000000 | Bicd1 | 0.001604 | 0.002945 | -0.619739657 | 415.7296699 | 270.5520283 | 1311 | 99 | 147 | 289 | 834 | 128 | 43 | 3 | 9 | 7 | 22 | 2 |
| SG0000002 | Rabep1 | 0.006237 | 0.01053 | -0.396541737 | 2235.414306 | 1698.193086 | 1017 | 103 | 147 | 188 | 619 | 72 | 18 | 0 | 1 | 5 | 17 | 0 |
| JSG000000 | Ttbk2 | 0.009678 | 0.015863 | -0.497919391 | 446.3284261 | 316.0573361 | 909 | 93 | 147 | 192 | 474 | 66 | 63 | 3 | 9 | 9 | 39 | 6 |
| SG0000002 | Alcam | 0.05913 | 0.084138 | -0.798984759 | 298.2983831 | 171.4480384 | 2032 | 96 | 147 | 267 | 946 | 165 | 9 | 0 | 0 | 0 | 3 | 1 |
| SG0000005 | Smyd3 | 3.13E-49 | 6.33E-48 | 2.185545786 | 273.6781581 | 1244.958792 | 1337 | 106 | 146 | 232 | 604 | 121 | 9 | 0 | 0 | 0 | 2 | 1 |
| SG0000000 | Pou2f2 | 2.03E-26 | 1.75E-25 | 4.426449989 | 5.215436911 | 112.1462917 | 464 | 138 | 146 | 206 | 310 | 48 | 16 | 2 | 2 | 6 | 6 | 0 |
| SG0000004 | Ino80d | 8.61E-06 | 2.04E-05 | 0.608596193 | 1021.776197 | 1557.978609 | 772 | 105 | 146 | 198 | 492 | 79 | 100 | 0 | 12 | 9 | 45 | 21 |
| SG0000002 | Pfkfb3 | 0.010834 | 0.01762 | 0.752958387 | 652.9222654 | 1100.334015 | 682 | 139 | 146 | 198 | 304 | 46 | 23 | 5 | 10 | 10 | 15 | 0 |
| SG0000004 | Lrrc49 | 0.086873 | 0.119836 | -0.344208929 | 310.7509046 | 244.790855 | 951 | 77 | 146 | 155 | 499 | 100 | 7 | 0 | 2 | 0 | 4 | 0 |
| SG0000003 | Rbpms | 7.31E-60 | 2.02E-58 | 3.411375021 | 609.0821952 | 6480.404457 | 1176 | 90 | 145 | 235 | 494 | 98 | 7 | 0 | 0 | 0 | 10 | 0 |
| SG0000005 | Kdm2a | 2.88E-24 | 2.24E-23 | -1.529200628 | 7358.739682 | 2549.577287 | 692 | 116 | 145 | 164 | 360 | 84 | 9 | 0 | 0 | 0 | 1 | 4 |
| SG0000002 | Pum1 | 3.42E-17 | 1.85E-16 | 1.040211944 | 2549.691214 | 5243.515839 | 1008 | 105 | 145 | 190 | 568 | 85 | 3 | 0 | 0 | 0 | 3 | 5 |
| SG0000002 | Prom1 | 2.5E-14 | 1.13E-13 | -1.646897963 | 1529.346279 | 488.3600665 | 1491 | 115 | 145 | 282 | 686 | 181 | 21 | 0 | 0 | 0 | 1 | 7 |
| SG0000002 | Sgcd | 1.78E-13 | 7.63E-13 | -2.641683871 | 135.2637248 | 21.67480796 | 2086 | 65 | 145 | 261 | 926 | 214 | 2 | 0 | 0 | 0 | 3 | 1 |
| SG0000002 | Katnal2 | 3.54E-22 | 2.5E-21 | 3.3310967 | 16.29010821 | 163.9396465 | 869 | 126 | 144 | 223 | 581 | 58 | 9 | 0 | 0 | 2 | 3 | 0 |
| SG0000002 | Pan3 | 0.000204 | 0.000417 | -0.554069647 | 1872.489203 | 1275.345161 | 1627 | 68 | 144 | 219 | 697 | 125 | 21 | 0 | 0 | 7 | 7 | 3 |
| SG0000002 | Btrc | 0.109206 | 0.147463 | 0.241730852 | 671.7162027 | 794.2442173 | 754 | 114 | 144 | 180 | 436 | 46 | 3 | 0 | 0 | 0 | 0 | 0 |
| SG0000004 | Bptf | 0.294371 | 0.358693 | -0.14716427 | 4130.644954 | 3730.07022 | 637 | 117 | 144 | 156 | 376 | 30 | 18 | 3 | 3 | 6 | 6 | 3 |
| SG0000005 | Cpne8 | 1.9E-34 | 2.31E-33 | -2.439130886 | 1079.448834 | 199.0446467 | 961 | 110 | 143 | 182 | 570 | 87 | 11 | 0 | 0 | 0 | 3 | 2 |
| JSG000000 | Ext1 | 2.01E-14 | 9.16E-14 | -1.178820332 | 2550.420395 | 1126.552649 | 1149 | 122 | 143 | 227 | 478 | 84 | 20 | 0 | 1 | 3 | 5 | 0 |
| SG0000005 | Bmpr1b | 0.001517 | 0.002792 | 1.505174228 | 14.08935969 | 39.99390841 | 2433 | 67 | 143 | 340 | 1104 | 225 | 34 | 2 | 2 | 6 | 14 | 3 |

| | | | | | | | | | | | | | | | | | | |
|---|---|---|---|---|---|---|---|---|---|---|---|---|---|---|---|---|---|---|
| SG0000003 | Tanc1 | 0.062309 | 0.088303 | 0.246736595 | 1810.431082 | 2148.11295 | 1127 | 126 | 143 | 197 | 506 | 131 | 9 | 3 | 3 | 3 | 12 | 3 |
| SG0000000 | Arpp19 | 0.824371 | 0.862127 | -0.034512372 | 3392.549813 | 3312.355748 | 412 | 120 | 143 | 167 | 305 | 32 | 35 | 0 | 0 | 3 | 6 | 11 |
| SG0000001 | Vps53 | 5.44E-15 | 2.57E-14 | -1.235793415 | 2041.033802 | 866.6411584 | 1049 | 125 | 142 | 257 | 664 | 77 | 0 | 0 | 0 | 0 | 0 | 0 |
| SG0000004 | Mkl1 | 0.006569 | 0.011056 | -0.486147789 | 743.3737958 | 530.7160004 | 946 | 117 | 142 | 219 | 518 | 143 | 0 | 0 | 0 | 0 | 8 | 4 |
| SG0000002 | Atxn7l1 | 0.448004 | 0.517118 | -0.147642121 | 372.7130923 | 336.4572892 | 1291 | 106 | 142 | 212 | 617 | 94 | 14 | 0 | 0 | 2 | 4 | 0 |
| SG0000003 | Lin54 | 0.73578 | 0.786026 | 0.042788949 | 1055.198145 | 1086.963037 | 1235 | 109 | 142 | 210 | 676 | 85 | 68 | 0 | 0 | 6 | 26 | 18 |
| SG0000002 | Dab1 | 6.18E-36 | 7.93E-35 | 6.666507089 | 0 | 100.5824306 | 2563 | 62 | 141 | 306 | 1218 | 248 | 2 | 0 | 0 | 0 | 1 | 1 |
| SG0000002 | Kcnj3 | 1.51E-30 | 1.56E-29 | 8.174240843 | 0.307509278 | 88.82800515 | 1570 | 63 | 141 | 275 | 814 | 89 | 34 | 0 | 3 | 2 | 10 | 2 |
| SG0000004 | Mei4 | 4.84E-27 | 4.27E-26 | 7.992904147 | 0.307509278 | 78.33613217 | 1714 | 86 | 141 | 297 | 972 | 181 | 13 | 0 | 0 | 1 | 5 | 1 |
| SG0000006 | Sfmbt2 | 5.63E-21 | 3.73E-20 | 4.769207308 | 45.32023005 | 1235.852274 | 1210 | 104 | 141 | 188 | 558 | 177 | 10 | 0 | 0 | 0 | 14 | 4 |
| SG0000003 | Foxp1 | 5.81E-13 | 2.4E-12 | -1.083036998 | 3042.093518 | 1435.971888 | 1456 | 90 | 141 | 261 | 692 | 129 | 20 | 4 | 4 | 6 | 14 | 2 |
| SG0000002 | Gpd2 | 8.14E-06 | 1.93E-05 | -0.635315723 | 4115.420759 | 2649.507509 | 921 | 102 | 141 | 198 | 489 | 57 | 21 | 3 | 6 | 6 | 12 | 0 |
| SG0000003 | Mphosph9 | 0.00017 | 0.000352 | -0.636781375 | 799.3861876 | 514.1221858 | 561 | 121 | 141 | 161 | 349 | 41 | 18 | 3 | 3 | 3 | 12 | 1 |
| SG0000002 | Dnmt3b | 1E-272 | 6E-270 | 5.519495279 | 65.33441864 | 2996.923595 | 570 | 130 | 140 | 170 | 339 | 20 | 0 | 0 | 0 | 0 | 9 | 0 |
| SG0000002 | Birc6 | 2.7E-33 | 3.12E-32 | -1.853182418 | 13335.29409 | 3690.957993 | 697 | 107 | 140 | 200 | 422 | 61 | 12 | 0 | 0 | 0 | 1 | 0 |
| SG0000003 | Ctps2 | 0.010283 | 0.016781 | 0.385150834 | 641.6772072 | 838.0280727 | 1233 | 89 | 140 | 161 | 673 | 105 | 10 | 8 | 8 | 8 | 12 | 0 |
| SG0000003 | Rere | 0.03474 | 0.051541 | -0.293598665 | 3796.016834 | 3097.035083 | 913 | 100 | 140 | 177 | 421 | 55 | 11 | 3 | 3 | 3 | 3 | 4 |
| SG0000006 | Gtf2i | 0.122717 | 0.163781 | 0.19413374 | 4505.67749 | 5154.661858 | 833 | 118 | 140 | 209 | 488 | 137 | 0 | 0 | 0 | 0 | 0 | 9 |
| SG0000005 | Magi3 | 0.305665 | 0.370764 | -0.172109913 | 570.2778424 | 506.1465155 | 1586 | 78 | 140 | 213 | 795 | 147 | 9 | 0 | 2 | 0 | 3 | 0 |
| SG0000002 | Trim2 | 0.382557 | 0.451437 | 0.175890052 | 858.4540812 | 969.7621984 | 958 | 116 | 140 | 180 | 513 | 81 | 56 | 0 | 0 | 6 | 37 | 0 |
| SG0000001 | Rev3l | 1.37E-07 | 3.81E-07 | 0.747020608 | 766.8947175 | 1287.097233 | 1140 | 98 | 139 | 190 | 629 | 140 | 17 | 0 | 0 | 0 | 3 | 5 |
| SG0000002 | Asb3 | 0.007899 | 0.013129 | -0.499096235 | 445.3198018 | 315.085973 | 954 | 83 | 139 | 186 | 604 | 86 | 10 | 0 | 2 | 0 | 11 | 2 |
| JSG000000 | Adk | 0.017272 | 0.027053 | 0.308932406 | 3490.942187 | 4324.546562 | 1083 | 98 | 139 | 224 | 562 | 78 | 3 | 0 | 0 | 0 | 0 | 0 |
| SG0000002 | Smarca2 | 3.74E-48 | 7.37E-47 | -3.898733807 | 1556.146064 | 104.3312805 | 1347 | 108 | 138 | 237 | 666 | 123 | 4 | 0 | 0 | 0 | 0 | 0 |
| SG0000003 | Cntln | 1.86E-37 | 2.5E-36 | 3.475389304 | 23.42437601 | 260.5340303 | 1629 | 82 | 138 | 258 | 927 | 147 | 15 | 0 | 0 | 0 | 6 | 1 |
| SG0000002 | Epha7 | 4.65E-17 | 2.49E-16 | -2.346816291 | 623.8729882 | 122.6405486 | 1786 | 64 | 138 | 240 | 889 | 213 | 26 | 0 | 2 | 4 | 8 | 5 |
| SG0000003 | Atg7 | 1.82E-09 | 5.85E-09 | -1.041746648 | 885.6991632 | 430.2186621 | 1712 | 96 | 138 | 276 | 686 | 144 | 2 | 0 | 0 | 0 | 2 | 0 |
| SG0000002 | Sgip1 | 0.000158 | 0.000328 | 2.791149926 | 3.276357648 | 22.67833434 | 2141 | 77 | 138 | 260 | 801 | 198 | 33 | 0 | 3 | 0 | 5 | 2 |
| SG0000001 | Apaf1 | 0.000901 | 0.001699 | -0.496759453 | 2301.998992 | 1631.419441 | 608 | 100 | 138 | 177 | 374 | 40 | 10 | 0 | 0 | 1 | 8 | 4 |
| JSG000000 | Vps13b | 0.026734 | 0.040502 | -0.363275383 | 1745.783733 | 1357.16814 | 1359 | 84 | 138 | 228 | 752 | 117 | 2 | 0 | 0 | 0 | 2 | 1 |
| SG0000003 | Senp6 | 0.643525 | 0.703205 | -0.068122763 | 2567.765701 | 2449.336185 | 920 | 105 | 138 | 182 | 445 | 71 | 15 | 0 | 0 | 3 | 4 | 4 |
| SG0000004 | Ehbp1 | 0.59544 | 0.658365 | 0.078352599 | 526.8867272 | 556.2931868 | 2075 | 75 | 137 | 228 | 1005 | 216 | 9 | 6 | 6 | 9 | 9 | 0 |
| SG0000002 | Cfh | 6.68E-84 | 3.33E-82 | -9.931451284 | 11279.29098 | 11.55093297 | 1200 | 80 | 136 | 216 | 680 | 88 | 9 | 0 | 0 | 0 | 6 | 1 |
| SG0000002 | Strbp | 1.08E-50 | 2.28E-49 | 1.882133639 | 999.3435992 | 3683.778564 | 1010 | 69 | 136 | 202 | 548 | 98 | 34 | 0 | 0 | 2 | 11 | 9 |
| SG0000001 | Sulf1 | 2.28E-06 | 5.7E-06 | 1.561883257 | 457.6772054 | 1351.241556 | 1062 | 90 | 135 | 174 | 478 | 101 | 4 | 0 | 0 | 2 | 14 | 0 |
| SG0000004 | Ranbp17 | 0.000261 | 0.000527 | -0.545347226 | 1737.27633 | 1190.427715 | 1316 | 84 | 135 | 225 | 715 | 122 | 17 | 6 | 8 | 9 | 18 | 1 |
| SG0000004 | Reln | 0.367249 | 0.435698 | 0.48817685 | 18.47295193 | 25.91147735 | 2069 | 61 | 135 | 263 | 1001 | 217 | 3 | 0 | 0 | 0 | 1 | 0 |
| SG0000004 | Bach2 | 0.698087 | 0.752173 | -0.128076624 | 215.5397057 | 197.2297299 | 1322 | 87 | 135 | 226 | 649 | 122 | 24 | 0 | 0 | 2 | 2 | 5 |
| SG0000002 | Dnajc6 | 5.1E-130 | 6E-128 | 9.502363332 | 0.649984458 | 471.410623 | 1111 | 89 | 134 | 221 | 623 | 112 | 40 | 10 | 10 | 20 | 35 | 10 |
| SG0000002 | Enah | 1.3E-119 | 1.3E-117 | 2.717490766 | 2503.445463 | 16465.8581 | 1216 | 80 | 134 | 168 | 598 | 148 | 8 | 0 | 2 | 0 | 2 | 0 |
| SG0000002 | Fkbp5 | 5.73E-70 | 2.03E-68 | 2.150471731 | 1314.405571 | 5835.604993 | 609 | 112 | 134 | 176 | 346 | 61 | 6 | 0 | 0 | 0 | 0 | 0 |
| SG0000006 | Armc9 | 2.47E-08 | 7.27E-08 | 1.053424239 | 219.8931063 | 456.3771675 | 626 | 117 | 134 | 182 | 373 | 42 | 3 | 0 | 0 | 2 | 2 | 0 |
| SG0000003 | Fmnl2 | 0.000884 | 0.001669 | -0.724981654 | 2006.308316 | 1213.826037 | 1102 | 108 | 134 | 211 | 542 | 99 | 10 | 0 | 0 | 0 | 0 | 4 |
| SG0000003 | Adgrg2 | 0.008257 | 0.013686 | -1.470541056 | 50.10654873 | 18.08079501 | 1699 | 85 | 134 | 263 | 844 | 210 | 24 | 0 | 6 | 12 | 12 | 12 |
| SG0000002 | Tspan5 | 0.062659 | 0.088765 | 0.274814413 | 750.7925034 | 908.3376359 | 1187 | 85 | 134 | 182 | 520 | 140 | 8 | 0 | 0 | 0 | 5 | 1 |
| SG0000003 | Mtm1 | 0.076571 | 0.106742 | 0.369316262 | 179.9552247 | 232.4554501 | 1207 | 82 | 134 | 176 | 592 | 110 | 42 | 0 | 6 | 6 | 18 | 6 |
| SG0000003 | Mapkap1 | 0.116072 | 0.155678 | -0.2375768 | 1246.82512 | 1057.518078 | 1819 | 80 | 134 | 248 | 896 | 184 | 2 | 0 | 0 | 0 | 4 | 2 |
| SG0000003 | Rapgef6 | 7.85E-19 | 4.61E-18 | -1.419948162 | 2363.176334 | 883.1798252 | 1286 | 96 | 133 | 209 | 684 | 92 | 5 | 0 | 0 | 0 | 12 | 2 |
| SG0000001 | Ppt2 | 2.83E-14 | 1.28E-13 | -1.328869751 | 1011.038882 | 402.4743449 | 255 | 126 | 133 | 164 | 200 | 21 | 3 | 0 | 0 | 3 | 3 | 0 |
| SG0000003 | Foxn3 | 5.77E-08 | 1.65E-07 | 1.395619309 | 140.2186424 | 368.9173093 | 883 | 109 | 133 | 181 | 417 | 76 | 0 | 0 | 0 | 0 | 3 | 0 |
| SG0000003 | Lrig2 | 4.33E-06 | 1.06E-05 | 0.686537911 | 567.2497305 | 912.9411911 | 623 | 101 | 133 | 153 | 279 | 61 | 21 | 0 | 2 | 4 | 9 | 7 |
| SG0000004 | C77370 | 2.8E-16 | 1.43E-15 | -1.732180212 | 531.8870164 | 160.0962603 | 918 | 96 | 132 | 213 | 585 | 76 | 28 | 0 | 0 | 0 | 6 | 4 |
| JSG000000 | Utrn | 2.24E-06 | 5.62E-06 | 0.586230192 | 3148.696111 | 4727.196114 | 1215 | 80 | 132 | 212 | 612 | 96 | 6 | 1 | 2 | 2 | 4 | 0 |
| SG0000000 | Clcn3 | 0.076424 | 0.106568 | -0.242352427 | 5693.472231 | 4813.066439 | 905 | 85 | 132 | 175 | 505 | 62 | 19 | 0 | 7 | 9 | 14 | 0 |
| SG0000002 | Fars2 | 0.68824 | 0.743666 | -0.0749 | 370.6898667 | 351.9358978 | 1512 | 66 | 132 | 222 | 756 | 154 | 0 | 0 | 0 | 0 | 0 | 0 |
| SG0000003 | Thsd4 | 4.24E-76 | 1.77E-74 | -3.939249154 | 2221.489935 | 144.8145484 | 1704 | 81 | 131 | 274 | 831 | 207 | 14 | 0 | 0 | 2 | 6 | 2 |
| SG0000005 | Commd1 | 2.86E-06 | 7.09E-06 | 0.703881421 | 645.9090213 | 1052.109087 | 675 | 103 | 131 | 143 | 355 | 32 | 4 | 0 | 0 | 0 | 4 | 1 |
| SG0000002 | Plce1 | 0.000358 | 0.000712 | -1.019431036 | 1099.283567 | 542.2885262 | 1350 | 81 | 131 | 220 | 698 | 115 | 9 | 1 | 1 | 2 | 3 | 2 |
| SG0000003 | Gas7 | 4.22E-07 | 1.12E-06 | -1.282369886 | 451.035098 | 185.4293762 | 641 | 114 | 130 | 165 | 378 | 71 | 3 | 0 | 0 | 0 | 0 | 0 |
| SG0000004 | Arhgef10l | 0.253913 | 0.314942 | 0.220610442 | 263.2573147 | 306.7544047 | 764 | 126 | 130 | 219 | 362 | 75 | 7 | 0 | 0 | 0 | 7 | 14 |
| SG0000003 | Asxl2 | 0.742979 | 0.791874 | 0.039369695 | 1916.011804 | 1969.017787 | 929 | 103 | 130 | 194 | 474 | 58 | 41 | 0 | 0 | 3 | 12 | 6 |
| SG0000002 | Xrcc4 | 0.800245 | 0.841625 | -0.054237137 | 268.5676625 | 258.6584759 | 1512 | 64 | 130 | 227 | 821 | 155 | 6 | 0 | 0 | 2 | 2 | 0 |
| SG0000003 | Atp2c1 | 4.22E-28 | 3.91E-27 | -1.68904065 | 7712.214028 | 2391.812736 | 982 | 88 | 129 | 207 | 544 | 67 | 17 | 0 | 1 | 4 | 10 | 0 |
| JSG000000 | Ern1 | 1.65E-07 | 4.56E-07 | -1.048016586 | 841.6339656 | 407.0416271 | 587 | 110 | 129 | 154 | 363 | 26 | 6 | 0 | 2 | 4 | 6 | 0 |
| SG0000003 | Stau1 | 4.79E-05 | 0.000105 | -0.590139738 | 2386.743832 | 1585.462762 | 465 | 117 | 129 | 149 | 305 | 30 | 4 | 0 | 0 | 0 | 0 | 0 |
| SG0000002 | Stag2 | 0.000113 | 0.000238 | -0.538691257 | 4708.380906 | 3241.226368 | 1266 | 78 | 129 | 210 | 681 | 102 | 24 | 0 | 0 | 6 | 15 | 3 |
| SG0000000 | Scml2 | 0.214948 | 0.271827 | 0.445941917 | 42.8581621 | 58.38152517 | 1610 | 26 | 129 | 195 | 896 | 241 | 10 | 0 | 0 | 5 | 24 | 0 |
| SG0000000 | Ncoa2 | 0.567713 | 0.632314 | -0.088025829 | 1474.232279 | 1386.971385 | 1220 | 90 | 129 | 163 | 561 | 86 | 26 | 0 | 0 | 4 | 6 | 2 |
| SG0000003 | Rbpj | 2.38E-46 | 4.38E-45 | 1.883801672 | 840.4640582 | 3101.701178 | 918 | 86 | 128 | 185 | 455 | 74 | 29 | 0 | 4 | 5 | 10 | 5 |
| SG0000003 | Csgalnact1 | 5.84E-14 | 2.58E-13 | -2.449027288 | 193.769741 | 35.48585593 | 1513 | 73 | 128 | 226 | 784 | 170 | 7 | 1 | 2 | 2 | 2 | 0 |
| SG0000000 | Etv1 | 8.75E-05 | 0.000187 | -0.926273352 | 323.8567855 | 170.4185783 | 1863 | 42 | 128 | 226 | 790 | 87 | 16 | 0 | 2 | 2 | 4 | 3 |
| SG0000006 | Hnrnpr | 0.004681 | 0.008045 | -0.394235038 | 3278.290777 | 2494.427583 | 476 | 99 | 128 | 133 | 252 | 19 | 35 | 0 | 2 | 3 | 6 | 9 |
| SG0000002 | Rngtt | 0.134866 | 0.178453 | 0.216750851 | 827.131627 | 961.2207445 | 1075 | 90 | 128 | 216 | 631 | 72 | 7 | 0 | 2 | 4 | 9 | 2 |
| SG0000002 | Rsu1 | 0.298449 | 0.363241 | 0.150656165 | 878.8845068 | 975.627051 | 1223 | 94 | 128 | 189 | 655 | 91 | 3 | 0 | 0 | 0 | 4 | 0 |
| SG0000002 | Tpd52 | 4.75E-71 | 1.75E-69 | 2.18123279 | 1206.446318 | 5471.72799 | 723 | 82 | 127 | 165 | 405 | 44 | 27 | 4 | 4 | 6 | 13 | 2 |
| SG0000002 | Slx4ip | 0.043189 | 0.063059 | 0.337697419 | 364.2250976 | 460.2851049 | 1182 | 71 | 127 | 214 | 602 | 97 | 13 | 0 | 0 | 0 | 2 | 1 |
| SG0000002 | Vcan | 0.252744 | 0.313736 | 0.469355025 | 663.3469691 | 918.3974943 | 1052 | 104 | 127 | 176 | 526 | 83 | 15 | 0 | 0 | 1 | 4 | 8 |
| SG0000003 | Ophn1 | 7.2E-69 | 2.49E-67 | -4.046603239 | 1304.788034 | 78.95706745 | 1488 | 90 | 126 | 236 | 847 | 137 | 7 | 0 | 0 | 0 | 0 | 1 |
| SG0000003 | Prdm16 | 4E-52 | 8.82E-51 | 4.160872696 | 15.65952841 | 280.1084231 | 1644 | 72 | 126 | 180 | 480 | 228 | 30 | 0 | 0 | 6 | 18 | 12 |
| SG0000000 | Epn2 | 1.57E-19 | 9.6E-19 | -1.366688173 | 4926.959271 | 1910.578285 | 696 | 108 | 126 | 162 | 390 | 36 | 36 | 0 | 0 | 6 | 18 | 0 |
| SG0000003 | Satb2 | 2.2E-14 | 9.97E-14 | -2.74374633 | 374.7342807 | 55.9465716 | 1132 | 75 | 126 | 180 | 509 | 123 | 27 | 0 | 0 | 9 | 9 | 6 |
| SG0000003 | Zcchc11 | 6.23E-14 | 2.75E-13 | 0.97511855 | 1234.265711 | 2426.322899 | 948 | 82 | 126 | 168 | 523 | 52 | 8 | 0 | 3 | 0 | 6 | 3 |
| SG0000004 | Slc39a11 | 0.000398 | 0.000788 | -0.612023373 | 648.2020399 | 424.1046167 | 1029 | 102 | 126 | 168 | 444 | 102 | 0 | 0 | 0 | 0 | 3 | 0 |
| SG0000003 | Focad | 0.486491 | 0.554751 | 0.094456276 | 1040.401199 | 1110.797814 | 1384 | 71 | 126 | 194 | 687 | 128 | 3 | 0 | 0 | 0 | 0 | 0 |
| SG0000005 | Vwa3b | 7.11E-44 | 1.2E-42 | 6.68679146 | 2.340612135 | 241.1316715 | 538 | 101 | 125 | 135 | 400 | 67 | 0 | 0 | 0 | 0 | 0 | 0 |
| SG0000002 | Parg | 2.29E-19 | 1.38E-18 | 1.167822929 | 1177.956443 | 2646.543052 | 1303 | 65 | 125 | 189 | 653 | 53 | 15 | 0 | 0 | 0 | 3 | 6 |
| SG0000002 | Ppp3cb | 3.9E-09 | 1.22E-08 | 0.740348595 | 2420.847856 | 4044.218675 | 734 | 104 | 125 | 189 | 415 | 40 | 28 | 0 | 1 | 2 | 13 | 1 |
| SG0000003 | Gpatch2 | 0.008827 | 0.014559 | -0.402323347 | 1275.599325 | 965.1679353 | 1499 | 57 | 125 | 168 | 636 | 116 | 27 | 5 | 5 | 6 | 12 | 1 |
| SG0000003 | Rims2 | 7.19E-12 | 2.78E-11 | -1.371416367 | 546.1736535 | 211.1024548 | 1387 | 61 | 124 | 224 | 792 | 109 | 0 | 0 | 0 | 0 | 0 | 0 |
| SG0000002 | Gnb1 | 0.178762 | 0.230474 | -0.174319326 | 14438.40257 | 12795.10342 | 633 | 110 | 124 | 149 | 329 | 36 | 12 | 0 | 0 | 0 | 0 | 0 |
| SG0000003 | Atrx | 0.000947 | 0.001781 | -0.500485934 | 12092.74447 | 8547.981966 | 778 | 88 | 123 | 171 | 451 | 47 | 30 | 1 | 3 | 5 | 12 | 1 |
| SG0000002 | Snx27 | 0.002606 | 0.004645 | 0.382997927 | 2520.939436 | 3287.4282 | 671 | 94 | 123 | 170 | 438 | 55 | 10 | 0 | 0 | 0 | 5 | 0 |
| SG0000003 | Txlng | 0.016196 | 0.025522 | 0.336747496 | 1011.211153 | 1277.06484 | 440 | 101 | 123 | 145 | 274 | 30 | 22 | 2 | 2 | 4 | 6 | 4 |
| SG0000004 | Rnf214 | 0.05912 | 0.084138 | 0.292498501 | 639.8506029 | 783.6631144 | 613 | 87 | 123 | 151 | 343 | 61 | 14 | 0 | 5 | 11 | 17 | 5 |
| SG0000003 | Bckdhb | 6.99E-39 | 9.88E-38 | 2.059001761 | 216.2190157 | 900.980055 | 1263 | 68 | 122 | 175 | 638 | 94 | 0 | 0 | 0 | 0 | 0 | 0 |
| SG0000003 | Gtdc1 | 8.42E-11 | 3.01E-10 | 1.141937226 | 223.3516305 | 492.8860032 | 1517 | 58 | 122 | 205 | 835 | 122 | 6 | 0 | 1 | 1 | 1 | 1 |
| JSG000000 | Ctnnd2 | 0.007285 | 0.012172 | 1.960052631 | 10.84931465 | 42.21209817 | 1747 | 68 | 122 | 241 | 847 | 178 | 3 | 0 | 0 | 0 | 1 | 1 |
| SG0000002 | AU040320 | 3.14E-10 | 1.08E-09 | -1.292113296 | 833.5348073 | 340.3759607 | 598 | 96 | 121 | 166 | 324 | 73 | 1 | 0 | 0 | 0 | 0 | 0 |
| SG0000003 | F8 | 0.000876 | 0.001655 | -1.297038556 | 71.72651897 | 29.18987554 | 905 | 76 | 121 | 184 | 557 | 99 | 2 | 0 | 0 | 0 | 0 | 0 |
| SG0000004 | Lonp2 | 0.10885 | 0.147037 | -0.238121215 | 1576.03874 | 1336.242438 | 753 | 75 | 121 | 131 | 377 | 70 | 10 | 0 | 1 | 1 | 1 | 0 |
| SG0000005 | Chchd3 | 0.140467 | 0.185167 | 0.192411836 | 1857.099762 | 2122.056626 | 1660 | 60 | 121 | 209 | 749 | 107 | 7 | 0 | 0 | 0 | 2 | 0 |
| SG0000005 | Asap2 | 3.02E-72 | 1.15E-70 | -3.330555339 | 7342.914183 | 729.9134955 | 814 | 76 | 120 | 159 | 411 | 66 | 22 | 0 | 0 | 3 | 3 | 6 |
| SG0000002 | Pip5k1b | 8.97E-15 | 4.17E-14 | 2.812781533 | 14.4644572 | 101.6327679 | 928 | 74 | 120 | 180 | 422 | 94 | 8 | 0 | 0 | 2 | 4 | 2 |
| SG0000000 | Sgce | 7.34E-14 | 3.22E-13 | 1.942849235 | 55.60305956 | 213.7738535 | 1351 | 64 | 120 | 184 | 549 | 88 | 9 | 0 | 0 | 0 | 1 | 0 |
| SG0000004 | St7l | 6.52E-09 | 2.01E-08 | -1.155819941 | 492.7788761 | 221.1645983 | 860 | 86 | 120 | 199 | 538 | 36 | 34 | 1 | 7 | 6 | 13 | 0 |
| SG0000002 | Ralgapa1 | 2.65E-08 | 7.8E-08 | -0.83697184 | 2242.901651 | 1255.615307 | 1264 | 60 | 120 | 210 | 678 | 112 | 10 | 2 | 4 | 4 | 6 | 2 |
| SG0000004 | Maob | 0.000393 | 0.000778 | 3.633836946 | 2.580533177 | 32.03342437 | 959 | 85 | 120 | 214 | 663 | 108 | 3 | 0 | 0 | 0 | 0 | 0 |
| SG0000001 | Bicc1 | 0.11771 | 0.157686 | 0.530631242 | 150.4734272 | 217.3680618 | 1148 | 89 | 120 | 212 | 720 | 187 | 9 | 0 | 2 | 1 | 2 | 1 |
| SG0000004 | Ttc3 | 3.7E-99 | 2.57E-97 | 2.770481259 | 448.5218813 | 3060.423981 | 730 | 80 | 119 | 171 | 327 | 57 | 14 | 2 | 2 | 4 | 7 | 0 |
| SG0000004 | Ncmap | 9.84E-20 | 6.08E-19 | 7.412047546 | 0.331601068 | 56.47608222 | 232 | 104 | 119 | 148 | 225 | 12 | 0 | 0 | 0 | 0 | 0 | 0 |
| SG0000000 | Runx1t1 | 4.22E-07 | 1.12E-06 | 1.773831629 | 37.64122535 | 128.7182956 | 1128 | 62 | 119 | 172 | 682 | 117 | 12 | 0 | 0 | 0 | 2 | 2 |
| SG0000002 | Plcd4 | 0.002246 | 0.004038 | 3.773632106 | 1.334934818 | 18.25729855 | 290 | 112 | 119 | 119 | 189 | 21 | 0 | 0 | 0 | 0 | 0 | 0 |
| SG0000003 | Col4a5 | 0.01942 | 0.030105 | -0.936652734 | 240.4269867 | 125.6095457 | 1413 | 57 | 119 | 245 | 831 | 135 | 9 | 0 | 0 | 0 | 9 | 0 |
| SG0000002 | Pard3 | 0.070403 | 0.098845 | 0.246013103 | 1420.825774 | 1684.993191 | 1273 | 62 | 119 | 220 | 639 | 125 | 3 | 0 | 0 | 0 | 1 | 0 |
| SG0000002 | Acvr1 | 1.75E-43 | 2.92E-42 | -2.643110056 | 1680.601932 | 269.0354131 | 948 | 85 | 118 | 187 | 419 | 39 | 18 | 3 | 3 | 3 | 9 | 0 |
| SG0000002 | Micu1 | 7.67E-40 | 1.13E-38 | -2.287924905 | 2824.194838 | 578.3099049 | 988 | 85 | 118 | 182 | 448 | 73 | 4 | 0 | 0 | 0 | 0 | 0 |
| SG0000003 | Limch1 | 1.88E-28 | 1.77E-27 | -3.568155399 | 1057.878342 | 89.18953951 | 1832 | 55 | 118 | 277 | 854 | 196 | 17 | 4 | 4 | 9 | 13 | 0 |
| SG0000002 | Rap1gds1 | 1.76E-25 | 1.45E-24 | -1.595005643 | 5767.555209 | 1909.181505 | 1516 | 70 | 118 | 198 | 721 | 131 | 12 | 0 | 3 | 0 | 2 | 1 |
| SG0000005 | Rpgrip1 | 3.35E-08 | 9.77E-08 | 1.674052453 | 51.06753604 | 162.9614599 | 392 | 90 | 118 | 140 | 283 | 23 | 4 | 0 | 0 | 3 | 3 | 0 |
| SG0000002 | Vps13d | 0.000758 | 0.001443 | -0.513428105 | 1529.178952 | 1071.2752 | 1106 | 89 | 118 | 191 | 536 | 120 | 6 | 0 | 0 | 3 | 3 | 8 |
| JSG000000 | Luzp2 | 2.12E-31 | 2.28E-30 | -10.15012243 | 818.1559187 | 0.720020342 | 1083 | 76 | 117 | 188 | 644 | 82 | 0 | 0 | 0 | 0 | 0 | 0 |
| SG0000002 | Nbea | 1.04E-21 | 7.2E-21 | -1.750117405 | 1084.335748 | 322.3487132 | 1408 | 66 | 117 | 207 | 666 | 139 | 6 | 1 | 2 | 2 | 2 | 1 |
| SG0000006 | Brwd3 | 4.77E-11 | 1.73E-10 | 0.925281312 | 833.3543851 | 1582.585399 | 1000 | 49 | 117 | 188 | 629 | 63 | 35 | 0 | 2 | 5 | 17 | 2 |

| | | | | | | | | | | | | | | | | | | |
|---|---|---|---|---|---|---|---|---|---|---|---|---|---|---|---|---|---|---|
| SG0000002 | Cobl | 1.74E-05 | 4E-05 | -0.600398895 | 4118.946576 | 2716.740033 | 1305 | 81 | 117 | 222 | 615 | 111 | 3 | 0 | 0 | 0 | 3 | 0 |
| SG0000003 | Gab3 | 0.000388 | 0.000769 | 3.354621101 | 1.596604083 | 16.33197893 | 732 | 88 | 117 | 158 | 380 | 50 | 6 | 0 | 0 | 0 | 2 | 0 |
| SG0000002 | Diap3 | 0.311151 | 0.376653 | 0.138072907 | 1835.699851 | 2020.066932 | 1257 | 61 | 117 | 169 | 575 | 95 | 0 | 0 | 0 | 0 | 0 | 0 |
| SG0000002 | Col11a1 | 0.397506 | 0.466404 | 0.357706039 | 89.78542003 | 115.0498525 | 866 | 68 | 117 | 157 | 464 | 81 | 10 | 0 | 1 | 1 | 6 | 1 |
| SG0000002 | Ahcyl2 | 0.706864 | 0.759985 | 0.052889339 | 1001.675265 | 1039.078185 | 998 | 69 | 117 | 134 | 439 | 103 | 16 | 10 | 10 | 10 | 13 | 0 |
| SG0000003 | Itpr2 | 3.44E-26 | 2.93E-25 | -3.248141801 | 1419.724375 | 149.4224753 | 1415 | 60 | 116 | 225 | 720 | 149 | 6 | 0 | 0 | 0 | 0 | 0 |
| SG0000001 | Smurf2 | 2.05E-06 | 5.14E-06 | -0.699635822 | 2470.340527 | 1521.05688 | 1145 | 72 | 116 | 202 | 490 | 68 | 21 | 4 | 4 | 6 | 10 | 0 |
| SG0000003 | Ralgps1 | 0.000149 | 0.000309 | 0.658118226 | 302.1982027 | 476.875712 | 877 | 92 | 116 | 159 | 347 | 88 | 9 | 0 | 0 | 3 | 3 | 0 |
| SG0000003 | Dag1 | 0.173105 | 0.223907 | -0.188495782 | 5105.3455 | 4480.043731 | 513 | 100 | 116 | 132 | 248 | 36 | 5 | 0 | 0 | 0 | 4 | 0 |
| SG0000003 | Pdgfd | 0.961061 | 0.975603 | -0.01879548 | 44.54529621 | 43.96872253 | 966 | 80 | 116 | 173 | 466 | 81 | 8 | 0 | 0 | 3 | 4 | 5 |
| SG0000003 | Lrch2 | 6.34E-13 | 2.61E-12 | 2.735100389 | 80.23172791 | 534.1871094 | 862 | 76 | 115 | 160 | 475 | 92 | 33 | 0 | 3 | 0 | 21 | 0 |
| SG0000002 | 1110008L16Rik | 0.000266 | 0.000537 | 0.533873301 | 686.461887 | 993.8670838 | 695 | 80 | 115 | 137 | 343 | 55 | 0 | 0 | 0 | 0 | 0 | 0 |
| SG000000C | Usp32 | 0.041191 | 0.060319 | -0.296085574 | 2325.26825 | 1893.836239 | 546 | 85 | 115 | 145 | 356 | 44 | 6 | 1 | 1 | 2 | 5 | 3 |
| SG0000002 | Deptor | 2E-48 | 3.98E-47 | -3.068871437 | 13805.5504 | 1645.247995 | 626 | 97 | 114 | 167 | 332 | 37 | 13 | 0 | 0 | 1 | 4 | 0 |
| SG0000002 | Samd4 | 2.98E-32 | 3.31E-31 | -3.241918438 | 1468.140954 | 155.1861862 | 1042 | 96 | 114 | 202 | 479 | 106 | 12 | 0 | 1 | 4 | 10 | 1 |
| SG0000002 | Acsl6 | 2.95E-23 | 2.19E-22 | 5.06502988 | 4.48230297 | 150.0469358 | 501 | 114 | 114 | 134 | 211 | 75 | 10 | 0 | 0 | 0 | 16 | 0 |
| SG000000C | Tek | 1E-19 | 6.18E-19 | -2.458872378 | 390.2456025 | 70.9812555 | 825 | 72 | 114 | 144 | 432 | 84 | 6 | 0 | 0 | 0 | 0 | 6 |
| SG0000003 | Wnk2 | 2.99E-19 | 1.8E-18 | 3.244777761 | 19.23636464 | 182.3469217 | 612 | 84 | 114 | 120 | 327 | 77 | 0 | 0 | 0 | 0 | 0 | 0 |
| SG0000003 | Prickle2 | 3.61E-10 | 1.23E-09 | 1.596058682 | 58.72498922 | 177.5352041 | 1305 | 61 | 114 | 205 | 616 | 114 | 28 | 0 | 3 | 0 | 0 | 0 |
| ISG000000 | Pdgfc | 1.63E-05 | 3.77E-05 | 0.69235325 | 476.8602187 | 770.5666158 | 1313 | 66 | 114 | 207 | 747 | 146 | 9 | 0 | 0 | 2 | 10 | 0 |
| SG0000002 | Rab28 | 0.000838 | 0.001587 | -0.574552894 | 738.3928932 | 495.8266352 | 864 | 57 | 114 | 144 | 492 | 90 | 8 | 0 | 3 | 0 | 4 | 0 |
| SG0000002 | Ppp6r3 | 0.011463 | 0.01858 | 0.31452417 | 4624.932316 | 5751.5724 | 615 | 102 | 114 | 150 | 348 | 72 | 9 | 0 | 0 | 3 | 3 | 3 |
| SG0000001 | Atad5 | 0.012352 | 0.01987 | 0.400821307 | 1414.722936 | 1867.801117 | 328 | 108 | 114 | 142 | 258 | 26 | 2 | 0 | 0 | 0 | 4 | 6 |
| SG0000003 | Fchsd2 | 0.089383 | 0.122967 | 0.245315685 | 727.4666069 | 862.3040822 | 944 | 72 | 114 | 164 | 528 | 94 | 15 | 0 | 0 | 4 | 10 | 0 |
| SG0000002 | Asic2 | 2.46E-11 | 9.16E-11 | 4.901499438 | 0 | 28.88810324 | 1673 | 63 | 113 | 214 | 757 | 189 | 2 | 2 | 2 | 2 | 2 | 0 |
| SG0000005 | Tead1 | 2.47E-05 | 5.6E-05 | 0.603083606 | 3047.27693 | 4628.690891 | 1114 | 78 | 113 | 174 | 472 | 138 | 42 | 0 | 7 | 6 | 6 | 4 |
| SG0000003 | Uggt1 | 0.00664 | 0.011167 | 0.357673973 | 4428.238397 | 5674.16013 | 596 | 81 | 113 | 152 | 358 | 40 | 15 | 0 | 0 | 8 | 8 | 0 |
| SG0000002 | Cdkl3 | 0.080301 | 0.11144 | 0.461358318 | 91.4129499 | 125.8607793 | 967 | 42 | 113 | 146 | 537 | 120 | 14 | 0 | 2 | 2 | 7 | 4 |
| SG0000006 | Casc4 | 0.269834 | 0.332321 | -0.168061158 | 1911.601756 | 1701.398208 | 721 | 72 | 113 | 140 | 377 | 63 | 20 | 0 | 4 | 0 | 20 | 4 |
| SG0000002 | Rps6ka6 | 0 | 0 | 10.63689475 | 2.613155511 | 4160.922603 | 1420 | 52 | 112 | 217 | 712 | 76 | 16 | 0 | 0 | 0 | 20 | 0 |
| SG0000001 | Cbx1 | 1.04E-60 | 2.94E-59 | 1.963123686 | 2293.008113 | 8940.559667 | 236 | 100 | 112 | 123 | 167 | 12 | 16 | 8 | 8 | 8 | 9 | 0 |
| SG0000002 | Fam168a | 4.61E-19 | 2.74E-18 | -1.392735265 | 3090.253945 | 1176.898688 | 605 | 74 | 112 | 144 | 380 | 44 | 11 | 0 | 0 | 0 | 0 | 0 |
| SG0000005 | Samd12 | 1.34E-13 | 5.78E-13 | -6.043774996 | 177.0978018 | 2.684451795 | 1743 | 62 | 112 | 248 | 876 | 188 | 0 | 0 | 0 | 0 | 0 | 0 |
| ISG000000 | Lmbrd1 | 4.29E-11 | 1.57E-10 | -1.143976502 | 1326.98267 | 600.4737266 | 567 | 79 | 112 | 128 | 313 | 54 | 18 | 0 | 0 | 0 | 2 | 4 |
| SG000000C | Luzp1 | 9.46E-09 | 2.89E-08 | 0.731573011 | 2072.211052 | 3440.799814 | 534 | 78 | 112 | 139 | 314 | 20 | 7 | 0 | 0 | 0 | 1 | 0 |
| SG0000004 | Stxbp6 | 1.06E-08 | 3.24E-08 | 4.317161415 | 4.899397089 | 97.66472278 | 1160 | 67 | 112 | 159 | 469 | 81 | 5 | 0 | 0 | 0 | 1 | 1 |
| SG0000002 | Khdrbs2 | 1.88E-06 | 4.75E-06 | 4.056936125 | 0 | 15.64406733 | 1273 | 60 | 112 | 206 | 747 | 121 | 4 | 0 | 0 | 0 | 1 | 1 |
| SG0000003 | Sik2 | 9.23E-05 | 0.000196 | -0.701638605 | 726.86704 | 446.9312372 | 1112 | 84 | 112 | 184 | 472 | 104 | 8 | 0 | 0 | 0 | 2 | 1 |
| SG0000002 | Esco1 | 0.012094 | 0.019497 | -0.393148277 | 1287.590985 | 980.4568907 | 655 | 93 | 112 | 156 | 456 | 43 | 8 | 0 | 0 | 0 | 16 | 4 |
| SG0000006 | Scfd2 | 0.018746 | 0.029143 | -0.552532359 | 306.9686539 | 209.2980664 | 1456 | 62 | 112 | 220 | 710 | 159 | 15 | 0 | 0 | 1 | 3 | 1 |
| SG0000002 | Slc44a1 | 0.040065 | 0.058784 | -0.285299985 | 4988.704717 | 4093.586252 | 1356 | 62 | 112 | 180 | 586 | 95 | 5 | 0 | 0 | 0 | 2 | 4 |
| SG0000002 | Luc7l2 | 0.449114 | 0.5182 | -0.120232583 | 4814.967193 | 4429.959118 | 984 | 75 | 112 | 180 | 607 | 110 | 14 | 0 | 0 | 0 | 6 | 6 |
| SG0000004 | Mdfic | 1.56E-80 | 7.13E-79 | -4.197640368 | 2087.16411 | 113.7472581 | 1120 | 65 | 111 | 180 | 533 | 112 | 34 | 1 | 1 | 4 | 13 | 2 |
| SG0000002 | Spata5 | 9.7E-25 | 7.74E-24 | 1.690259175 | 561.7072701 | 1812.71178 | 1278 | 45 | 111 | 174 | 585 | 123 | 0 | 0 | 0 | 0 | 0 | 0 |
| SG0000006 | Rapgef2 | 7.66E-24 | 5.85E-23 | -1.772213237 | 1453.757184 | 425.6008933 | 888 | 69 | 111 | 155 | 385 | 69 | 12 | 0 | 3 | 0 | 3 | 3 |
| SG0000006 | Kif1b | 1.34E-16 | 6.98E-16 | -1.400937104 | 4556.153786 | 1725.338384 | 742 | 82 | 111 | 153 | 424 | 55 | 9 | 0 | 0 | 1 | 3 | 3 |
| SG0000006 | Arid1b | 3.59E-07 | 9.6E-07 | -0.807386093 | 1316.579903 | 752.3151182 | 1503 | 84 | 111 | 245 | 656 | 140 | 21 | 0 | 0 | 2 | 8 | 2 |
| SG0000005 | Dph6 | 1.01E-06 | 2.61E-06 | 0.681386653 | 887.27171 | 1422.90025 | 1035 | 66 | 111 | 159 | 531 | 90 | 12 | 0 | 0 | 0 | 9 | 3 |
| SG0000001 | Mettl16 | 2.09E-05 | 4.76E-05 | 0.559753729 | 1781.098648 | 2625.370715 | 418 | 89 | 111 | 136 | 249 | 29 | 18 | 2 | 3 | 4 | 9 | 2 |
| SG0000003 | Dzip1l | 0.000266 | 0.000538 | -0.632327289 | 671.6273692 | 433.2902379 | 287 | 99 | 111 | 143 | 220 | 27 | 21 | 7 | 7 | 7 | 22 | 0 |
| SG0000002 | Uimc1 | 0.127675 | 0.169753 | 0.219810576 | 855.6826409 | 996.5114619 | 573 | 90 | 111 | 156 | 408 | 45 | 12 | 0 | 0 | 3 | 9 | 0 |
| SG0000002 | St3gal3 | 0.869566 | 0.899132 | 0.027061455 | 472.5635849 | 481.5113872 | 744 | 99 | 111 | 168 | 441 | 60 | 3 | 0 | 0 | 0 | 3 | 0 |
| SG0000002 | Il1rap | 2.78E-32 | 3.1E-31 | -2.614592483 | 697.8332188 | 113.9412227 | 1401 | 56 | 110 | 244 | 832 | 128 | 24 | 0 | 0 | 6 | 12 | 7 |
| SG0000004 | Mcmdc2 | 1.41E-16 | 7.35E-16 | 7.331743435 | 0.34247518 | 55.17010331 | 500 | 97 | 110 | 135 | 247 | 4 | 12 | 0 | 1 | 1 | 6 | 0 |
| SG0000002 | Gabpb1 | 6.96E-16 | 3.47E-15 | 1.126830297 | 682.2672728 | 1489.92449 | 587 | 95 | 110 | 150 | 334 | 81 | 14 | 0 | 0 | 2 | 20 | 7 |
| SG0000004 | Vprbp | 2.02E-07 | 5.52E-07 | 0.663603379 | 2222.768941 | 3520.941764 | 475 | 76 | 110 | 125 | 252 | 22 | 7 | 0 | 1 | 1 | 3 | 0 |
| SG0000002 | R3hdm2 | 3.37E-05 | 7.55E-05 | -0.585688894 | 3167.80333 | 2110.805833 | 766 | 86 | 110 | 168 | 450 | 66 | 15 | 0 | 5 | 5 | 5 | 0 |
| SG0000004 | Fcho2 | 0.000216 | 0.00044 | -0.53523189 | 2375.945489 | 1639.515847 | 600 | 76 | 110 | 138 | 310 | 56 | 35 | 0 | 0 | 1 | 7 | 2 |
| SG0000003 | Wbscr17 | 0.027426 | 0.041477 | 1.509461326 | 12.13606588 | 34.5518306 | 1174 | 72 | 110 | 184 | 618 | 145 | 4 | 0 | 0 | 0 | 1 | 0 |
| SG0000004 | Sema3d | 0.612827 | 0.675007 | 0.330506886 | 14.45492979 | 18.17642518 | 1195 | 60 | 110 | 204 | 708 | 102 | 20 | 0 | 1 | 4 | 14 | 2 |
| SG0000002 | Tec | 0.917372 | 0.939453 | 0.023424381 | 151.1879566 | 153.6627633 | 1146 | 68 | 110 | 189 | 812 | 107 | 6 | 0 | 0 | 6 | 6 | 12 |
| SG0000002 | Kat6b | 1.4E-201 | 4.4E-199 | 3.848301718 | 439.1829871 | 6325.571332 | 1458 | 70 | 109 | 217 | 686 | 134 | 20 | 0 | 5 | 0 | 4 | 0 |
| SG0000002 | Spag9 | 2.49E-38 | 3.48E-37 | -2.478224104 | 10368.57363 | 1860.797975 | 1166 | 62 | 109 | 150 | 522 | 103 | 48 | 0 | 6 | 12 | 36 | 0 |
| SG0000003 | Sos2 | 2.03E-26 | 1.75E-25 | -1.860791077 | 1631.383064 | 449.1609788 | 809 | 78 | 109 | 178 | 471 | 58 | 5 | 0 | 1 | 1 | 3 | 1 |
| SG0000003 | Csnk1g1 | 4.06E-17 | 2.19E-16 | -1.309996718 | 2740.017777 | 1105.108894 | 831 | 71 | 109 | 161 | 425 | 63 | 11 | 0 | 0 | 1 | 7 | 2 |
| SG0000007 | Txnrd2 | 1.02E-08 | 3.12E-08 | -1.167269659 | 438.5936037 | 195.2895989 | 352 | 93 | 109 | 138 | 238 | 9 | 0 | 0 | 0 | 0 | 0 | 0 |
| SG0000003 | Cdkl5 | 0.013362 | 0.021343 | 0.500659962 | 174.9858867 | 247.5806443 | 1505 | 62 | 109 | 226 | 700 | 157 | 19 | 0 | 2 | 4 | 9 | 3 |
| SG0000004 | Mtus1 | 4.26E-26 | 3.62E-25 | 1.489962157 | 505.2840057 | 1419.249837 | 732 | 89 | 108 | 139 | 343 | 64 | 20 | 0 | 0 | 0 | 4 | 4 |
| SG0000005 | Gpc3 | 3.29E-21 | 2.21E-20 | 6.540085169 | 22.65143589 | 2107.936614 | 1327 | 66 | 108 | 186 | 580 | 86 | 2 | 0 | 0 | 1 | 1 | 2 |
| SG0000001 | Syt14 | 4.8E-17 | 2.57E-16 | 6.576382232 | 4.840339396 | 461.9171953 | 1126 | 69 | 108 | 148 | 528 | 89 | 7 | 0 | 0 | 1 | 6 | 2 |
| SG0000003 | Aff3 | 6.33E-11 | 2.28E-10 | 6.106604383 | 10.48893465 | 722.7739155 | 1441 | 58 | 108 | 216 | 626 | 161 | 8 | 0 | 0 | 2 | 6 | 0 |
| SG0000004 | Alg13 | 9.78E-08 | 2.74E-07 | 0.651693217 | 5258.679532 | 8261.443014 | 710 | 87 | 108 | 138 | 406 | 67 | 31 | 0 | 3 | 3 | 12 | 1 |
| SG0000002 | Tpk1 | 1.15E-06 | 2.96E-06 | 1.024266309 | 130.2003723 | 264.8177606 | 1508 | 70 | 108 | 246 | 907 | 132 | 8 | 0 | 0 | 0 | 3 | 0 |
| SG0000002 | Acaca | 0.000935 | 0.001759 | -0.486506671 | 2145.74164 | 1531.525838 | 985 | 65 | 108 | 177 | 567 | 65 | 4 | 0 | 0 | 2 | 4 | 0 |
| SG0000004 | Setd2 | 0.084062 | 0.116254 | 0.214935996 | 3419.840925 | 3969.246651 | 587 | 88 | 108 | 148 | 352 | 52 | 3 | 0 | 0 | 0 | 4 | 5 |
| SG0000002 | Rprd2 | 0.379093 | 0.44789 | 0.120135388 | 1610.075359 | 1749.889233 | 431 | 91 | 108 | 139 | 255 | 33 | 16 | 0 | 3 | 3 | 7 | 2 |
| SG0000005 | Immt | 0.970469 | 0.983705 | -0.008690452 | 5673.721354 | 5639.646935 | 508 | 84 | 108 | 128 | 292 | 24 | 6 | 0 | 0 | 0 | 0 | 0 |
| SG0000004 | Dmxl2 | 2.84E-13 | 1.19E-12 | -1.190006736 | 2629.453353 | 1152.491546 | 895 | 72 | 107 | 165 | 517 | 76 | 9 | 2 | 2 | 2 | 9 | 0 |
| SG0000004 | Synpo | 3.34E-13 | 1.4E-12 | -1.708881815 | 609.9997507 | 186.5971353 | 206 | 98 | 107 | 111 | 139 | 31 | 6 | 0 | 0 | 0 | 3 | 0 |
| SG0000002 | Sema6d | 3.53E-12 | 1.39E-11 | -3.26618612 | 128.7591931 | 13.38314714 | 1772 | 41 | 107 | 217 | 681 | 167 | 35 | 0 | 0 | 5 | 10 | 10 |
| SG0000005 | Fto | 1.71E-06 | 4.32E-06 | 0.626759177 | 1730.372814 | 2671.855679 | 1458 | 70 | 107 | 205 | 667 | 156 | 18 | 0 | 0 | 1 | 6 | 0 |
| SG0000003 | Sik3 | 0.005129 | 0.008766 | -0.407675196 | 1812.452357 | 1366.293924 | 1029 | 57 | 107 | 153 | 490 | 71 | 0 | 0 | 0 | 0 | 3 | 1 |
| SG0000006 | Phactr4 | 7.72E-14 | 3.38E-13 | -1.149799792 | 2544.880198 | 1146.946382 | 429 | 101 | 106 | 145 | 321 | 22 | 0 | 0 | 0 | 0 | 2 | 2 |
| SG0000002 | Epha4 | 3.44E-11 | 1.27E-10 | 2.738433292 | 69.51793958 | 463.9246554 | 1064 | 72 | 106 | 176 | 507 | 78 | 17 | 0 | 2 | 8 | 11 | 2 |
| SG0000003 | Cnot4 | 0.428809 | 0.498113 | 0.113936839 | 960.649094 | 1039.59261 | 806 | 48 | 106 | 141 | 481 | 64 | 6 | 0 | 0 | 0 | 3 | 0 |
| SG0000002 | Numb | 0.949621 | 0.965413 | -0.011765132 | 953.9066096 | 946.159165 | 783 | 81 | 106 | 158 | 414 | 69 | 9 | 0 | 0 | 0 | 6 | 3 |
| SG0000003 | Dlc1 | 5.14E-45 | 9.03E-44 | 5.440778765 | 188.3286412 | 8180.012757 | 1024 | 65 | 105 | 159 | 593 | 110 | 8 | 0 | 0 | 2 | 6 | 4 |
| SG0000007 | Zfp827 | 4.56E-29 | 4.42E-28 | -2.026498652 | 1459.673823 | 358.2770084 | 853 | 81 | 105 | 161 | 356 | 50 | 14 | 0 | 2 | 2 | 12 | 0 |
| SG0000005 | Gon4l | 1.45E-12 | 5.84E-12 | 0.972769769 | 834.5915056 | 1637.97338 | 474 | 99 | 105 | 162 | 288 | 51 | 0 | 0 | 0 | 0 | 3 | 3 |
| ISG000000 | Tbc1d5 | 2.31E-12 | 9.18E-12 | -1.16803306 | 1272.310287 | 566.2132807 | 1215 | 55 | 105 | 208 | 694 | 108 | 0 | 0 | 0 | 0 | 1 | 0 |
| SG0000002 | Ush2a | 1.46E-06 | 3.71E-06 | -2.944933498 | 60.44708923 | 7.849862952 | 1359 | 57 | 105 | 199 | 663 | 151 | 0 | 0 | 0 | 0 | 0 | 0 |
| SG0000003 | Papd5 | 8.95E-05 | 0.000191 | 0.519790996 | 1424.707946 | 2042.671491 | 402 | 84 | 105 | 118 | 259 | 34 | 25 | 0 | 3 | 6 | 9 | 3 |
| SG0000003 | Camk1d | 0.018409 | 0.028664 | -0.415692139 | 575.9210508 | 431.7448343 | 1196 | 50 | 105 | 193 | 633 | 136 | 6 | 0 | 1 | 2 | 5 | 3 |
| ISG000000 | Zfpm2 | 0.030271 | 0.045397 | -1.025994143 | 68.43345007 | 33.60573691 | 1226 | 41 | 105 | 161 | 603 | 113 | 4 | 0 | 1 | 0 | 2 | 0 |
| SG0000002 | Slc7a11 | 4.3E-153 | 6.8E-151 | -8.895559592 | 18305.11388 | 38.43635268 | 1571 | 50 | 104 | 245 | 776 | 136 | 11 | 0 | 0 | 0 | 4 | 3 |
| SG0000002 | Lrp2 | 2.29E-93 | 1.4E-91 | 7.697970794 | 1.596604083 | 331.5258142 | 579 | 90 | 104 | 143 | 363 | 47 | 5 | 0 | 0 | 1 | 3 | 0 |
| SG0000006 | Tnfaip8 | 9.11E-80 | 4.07E-78 | 3.362816386 | 76.91083339 | 791.2179282 | 651 | 57 | 104 | 127 | 288 | 51 | 9 | 0 | 0 | 0 | 0 | 0 |
| SG0000004 | Shroom2 | 1.9E-46 | 3.51E-45 | -2.42505809 | 5509.061745 | 1025.799336 | 669 | 78 | 104 | 168 | 448 | 62 | 21 | 0 | 2 | 4 | 6 | 4 |
| SG0000002 | Nrcam | 1.03E-26 | 8.95E-26 | 4.857006861 | 3.570649248 | 103.4789337 | 1194 | 56 | 104 | 172 | 550 | 136 | 1 | 0 | 0 | 0 | 2 | 2 |
| SG000000C | Gria3 | 1.84E-15 | 8.95E-15 | -2.243152797 | 570.2117014 | 120.4425242 | 1842 | 36 | 104 | 174 | 681 | 158 | 6 | 0 | 2 | 0 | 0 | 2 |
| SG0000002 | Tmem56 | 9.24E-14 | 4.03E-13 | -3.315191369 | 96.55207133 | 9.700397961 | 543 | 88 | 104 | 104 | 298 | 34 | 49 | 2 | 2 | 4 | 13 | 2 |
| SG0000005 | Pign | 0.000482 | 0.000943 | -0.579992972 | 1869.621672 | 1250.715268 | 1208 | 36 | 104 | 165 | 703 | 104 | 20 | 0 | 0 | 2 | 13 | 0 |
| SG0000005 | Skint4 | 0.001045 | 0.001956 | -3.546392821 | 15.78851791 | 1.351359032 | 929 | 56 | 104 | 150 | 534 | 58 | 0 | 0 | 0 | 0 | 0 | 2 |
| ISG000000 | Atrnl1 | 1.15E-37 | 1.56E-36 | -2.814491988 | 2586.34607 | 367.6547023 | 1176 | 65 | 103 | 196 | 576 | 82 | 1 | 0 | 0 | 1 | 1 | 0 |
| SG0000003 | Dennd4c | 7.36E-24 | 5.63E-23 | -1.948197091 | 2724.545937 | 706.0384825 | 703 | 74 | 103 | 147 | 412 | 65 | 5 | 0 | 0 | 2 | 2 | 1 |
| SG0000002 | Ikzf2 | 0.003163 | 0.00556 | -1.296401084 | 196.6968464 | 80.08326746 | 1111 | 48 | 103 | 144 | 517 | 111 | 27 | 0 | 1 | 3 | 9 | 3 |
| SG0000002 | Dclk2 | 1.3E-13 | 5.62E-13 | 2.093804037 | 249.4276124 | 1064.737008 | 1271 | 54 | 102 | 153 | 486 | 100 | 6 | 0 | 0 | 2 | 2 | 4 |
| SG0000002 | Ppp3ca | 1.56E-11 | 5.87E-11 | -1.142459566 | 1680.747708 | 761.3562717 | 1234 | 58 | 102 | 221 | 622 | 114 | 8 | 0 | 0 | 3 | 4 | 0 |
| SG0000002 | Asap1 | 4.13E-08 | 1.2E-07 | -0.826142213 | 2110.731096 | 1190.527198 | 1230 | 63 | 102 | 159 | 555 | 99 | 3 | 0 | 0 | 3 | 3 | 6 |
| SG0000003 | Wdr44 | 1.09E-07 | 3.04E-07 | -0.983243194 | 636.9334407 | 322.1872551 | 773 | 62 | 102 | 158 | 540 | 89 | 3 | 0 | 0 | 0 | 3 | 4 |
| SG0000004 | Ckap5 | 0.000265 | 0.000536 | 0.462060846 | 4621.079575 | 6365.574738 | 825 | 53 | 102 | 147 | 392 | 81 | 4 | 0 | 0 | 0 | 0 | 0 |
| SG0000002 | Dnm1 | 5.84E-47 | 1.1E-45 | 2.544994297 | 127.7004919 | 745.2675165 | 322 | 101 | 101 | 111 | 254 | 38 | 0 | 0 | 0 | 0 | 0 | 0 |
| SG0000002 | Pikfyve | 2.51E-30 | 2.58E-29 | 1.476893718 | 1221.646584 | 3400.438265 | 730 | 69 | 101 | 153 | 399 | 35 | 15 | 0 | 2 | 1 | 6 | 3 |
| SG0000004 | Dmtf1 | 2.57E-12 | 1.02E-11 | 0.910140236 | 1403.464822 | 2637.430614 | 920 | 48 | 101 | 90 | 475 | 43 | 26 | 7 | 7 | 7 | 21 | 0 |
| SG0000004 | Ipo11 | 2.14E-07 | 5.84E-07 | 0.666838452 | 1875.056484 | 2976.821072 | 967 | 75 | 101 | 172 | 437 | 76 | 15 | 0 | 0 | 3 | 3 | 0 |
| SG0000002 | Rbfox3 | 9.55E-07 | 2.47E-06 | 3.964151808 | 1.631569985 | 25.46445049 | 1611 | 52 | 101 | 215 | 777 | 227 | 0 | 0 | 0 | 0 | 4 | 2 |
| SG0000003 | Rfx7 | 1.16E-06 | 2.99E-06 | -0.706597521 | 3081.063735 | 1887.963673 | 970 | 66 | 101 | 214 | 543 | 110 | 28 | 2 | 3 | 4 | 11 | 4 |
| ISG000000 | Gatad2b | 0.013341 | 0.02132 | 0.33687351 | 1973.060087 | 2492.007486 | 544 | 70 | 101 | 146 | 324 | 58 | 3 | 0 | 0 | 1 | 5 | 1 |
| SG0000003 | Zmat4 | 0.023313 | 0.035624 | 0.546217984 | 316.4056687 | 462.0322101 | 1527 | 54 | 101 | 246 | 818 | 183 | 7 | 1 | 2 | 1 | 3 | 0 |
| SG0000002 | Gapvd1 | 0.033357 | 0.049655 | -0.295998163 | 3146.182124 | 2562.592397 | 590 | 89 | 101 | 134 | 431 | 59 | 19 | 0 | 0 | 10 | 17 | 2 |
| ISG000000 | Scaper | 0.042049 | 0.061514 | -0.411710553 | 660.9193999 | 496.8339605 | 945 | 48 | 101 | 161 | 497 | 79 | 4 | 0 | 0 | 0 | 0 | 1 |
| SG0000003 | Adam32 | 0.077573 | 0.108022 | -1.354331148 | 17.29344196 | 6.763743683 | 883 | 73 | 101 | 170 | 601 | 76 | 0 | 0 | 0 | 0 | 0 | 0 |
| ISG000000 | 2210408I21Rik | 0.536192 | 0.602607 | -0.20836024 | 83.00145828 | 71.83945692 | 1136 | 49 | 101 | 183 | 620 | 97 | 1 | 0 | 0 | 0 | 0 | 0 |

| | | | | | | | | | | | | | | | | | | |
|---|---|---|---|---|---|---|---|---|---|---|---|---|---|---|---|---|---|---|
| SG0000002 | Kif21a | 2.74E-42 | 4.37E-41 | -2.44592825 | 2489.1283 | 456.8246309 | 1345 | 25 | 100 | 185 | 715 | 85 | 15 | 0 | 5 | 0 | 10 | 5 |
| SG0000002 | Invs | 1.29E-19 | 7.95E-19 | -2.171983114 | 393.7442918 | 87.37399703 | 711 | 74 | 100 | 138 | 378 | 38 | 3 | 0 | 0 | 0 | 1 | 0 |
| SG0000004 | Piezo2 | 3.09E-07 | 8.29E-07 | -1.592471782 | 267.4890973 | 88.70014127 | 895 | 61 | 100 | 155 | 484 | 58 | 4 | 0 | 0 | 0 | 2 | 0 |
| SG0000002 | Eya3 | 0.393228 | 0.461929 | 0.112088601 | 2054.760157 | 2220.767877 | 704 | 64 | 100 | 128 | 304 | 72 | 8 | 0 | 0 | 8 | 12 | 0 |
| SG0000002 | Pam | 4.3E-64 | 1.33E-62 | -4.245903071 | 11501.80569 | 606.2082514 | 1797 | 39 | 99 | 272 | 883 | 148 | 7 | 0 | 0 | 0 | 0 | 3 |
| SG0000004 | Prrg1 | 5.08E-18 | 2.86E-17 | -2.188418499 | 412.7630119 | 90.55681961 | 1062 | 57 | 99 | 186 | 660 | 123 | 45 | 0 | 0 | 0 | 15 | 3 |
| SG0000002 | Sfi1 | 7.62E-15 | 3.57E-14 | -1.917682584 | 721.3834061 | 190.9352737 | 593 | 63 | 99 | 149 | 371 | 26 | 2 | 0 | 0 | 0 | 0 | 0 |
| SG0000002 | Nudt3 | 6.4E-14 | 2.82E-13 | -1.149276674 | 2422.769382 | 1092.308528 | 255 | 85 | 99 | 104 | 183 | 12 | 2 | 0 | 1 | 1 | 2 | 0 |
| SG0000003 | Rnf13 | 2.66E-11 | 9.86E-11 | -1.173559449 | 1764.330753 | 782.1740592 | 1321 | 59 | 99 | 163 | 670 | 119 | 10 | 0 | 0 | 0 | 3 | 3 |
| SG0000003 | Gpbp1 | 4.73E-11 | 1.72E-10 | 0.820375684 | 2552.077963 | 4506.627122 | 608 | 57 | 99 | 114 | 321 | 84 | 7 | 0 | 0 | 0 | 0 | 0 |
| SG0000002 | Rufy3 | 1.29E-08 | 3.9E-08 | -0.969452153 | 956.7359169 | 488.604993 | 584 | 49 | 99 | 148 | 392 | 75 | 6 | 0 | 0 | 0 | 0 | 2 |
| SG0000002 | Pcsk5 | 0.002361 | 0.00423 | 2.286695113 | 3.939559785 | 19.22257285 | 1355 | 42 | 99 | 165 | 585 | 122 | 4 | 0 | 1 | 0 | 3 | 0 |
| SG0000002 | Slco3a1 | 2.07E-85 | 1.04E-83 | -4.657028304 | 1636.856061 | 64.87916933 | 1107 | 54 | 98 | 155 | 430 | 117 | 8 | 1 | 2 | 3 | 6 | 0 |
| JSG000000 | Slc7a7 | 4.13E-33 | 4.73E-32 | 3.226257238 | 281.1590508 | 2631.190369 | 254 | 83 | 98 | 107 | 175 | 3 | 3 | 0 | 0 | 0 | 0 | 0 |
| SG0000001 | Myo1b | 1.34E-21 | 9.24E-21 | 1.415176027 | 1075.118643 | 2867.25835 | 1278 | 56 | 98 | 170 | 532 | 80 | 16 | 0 | 4 | 8 | 9 | 4 |
| SG0000006 | Hmcn1 | 1.65E-05 | 3.82E-05 | 2.250677316 | 8.670314341 | 41.2625654 | 1126 | 53 | 98 | 172 | 543 | 112 | 4 | 1 | 1 | 1 | 4 | 1 |
| SG0000002 | Abl1 | 0.000704 | 0.001345 | -0.550970454 | 3227.835435 | 2203.194058 | 572 | 85 | 98 | 127 | 270 | 52 | 0 | 0 | 0 | 0 | 0 | 2 |
| SG0000002 | Mad1l1 | 0.169303 | 0.219639 | -0.204258361 | 964.5004855 | 837.1717349 | 646 | 78 | 98 | 138 | 306 | 50 | 0 | 0 | 0 | 0 | 0 | 0 |
| SG0000002 | Luc7l | 1.67E-11 | 6.3E-11 | 0.848752477 | 2176.604405 | 3919.940349 | 343 | 69 | 97 | 125 | 237 | 44 | 7 | 0 | 0 | 1 | 2 | 3 |
| SG0000002 | Zeb2 | 1.43E-07 | 3.95E-07 | -1.701231489 | 284.1161407 | 87.37238664 | 2032 | 46 | 97 | 200 | 622 | 108 | 51 | 4 | 5 | 9 | 46 | 1 |
| SG0000003 | Frmd4b | 0.124616 | 0.166117 | 0.290792262 | 240.7703717 | 294.5372559 | 784 | 72 | 97 | 162 | 380 | 55 | 2 | 0 | 0 | 0 | 1 | 0 |
| SG0000003 | Mtmr3 | 0.191678 | 0.245449 | 0.169657794 | 2372.720517 | 2668.816234 | 924 | 65 | 97 | 163 | 385 | 23 | 0 | 0 | 0 | 0 | 0 | 3 |
| SG0000002 | Entpd5 | 1.64E-43 | 2.73E-42 | -2.435578816 | 2984.058459 | 551.6010021 | 318 | 84 | 96 | 110 | 262 | 12 | 38 | 0 | 0 | 4 | 16 | 0 |
| SG0000001 | Pax5 | 9.92E-40 | 1.45E-38 | -6.652138371 | 1199.343995 | 11.92478562 | 2418 | 55 | 96 | 422 | 944 | 230 | 6 | 0 | 0 | 0 | 0 | 2 |
| SG0000002 | Ash1l | 2.04E-26 | 1.76E-25 | -1.760801267 | 2514.061538 | 741.859916 | 556 | 87 | 96 | 151 | 361 | 64 | 2 | 0 | 0 | 0 | 4 | 4 |
| SG0000000 | Pou6f1 | 1.3E-22 | 9.38E-22 | -1.960904097 | 740.051948 | 190.0952452 | 224 | 72 | 96 | 127 | 234 | 10 | 5 | 0 | 0 | 0 | 0 | 0 |
| SG0000002 | Rbm26 | 3.13E-17 | 1.69E-16 | 1.044424192 | 2311.553606 | 4767.678694 | 890 | 58 | 96 | 120 | 474 | 66 | 19 | 0 | 2 | 4 | 11 | 1 |
| SG0000002 | Ift80 | 0.063343 | 0.089685 | -0.296685565 | 973.0261874 | 792.1606666 | 859 | 59 | 96 | 139 | 426 | 80 | 3 | 0 | 0 | 0 | 0 | 0 |
| SG0000004 | 1110051M20Rik | 0.232926 | 0.291882 | 0.20727693 | 339.6285922 | 392.1035897 | 946 | 72 | 96 | 163 | 475 | 105 | 2 | 0 | 0 | 0 | 0 | 0 |
| SG0000002 | Rufy2 | 0.898319 | 0.922579 | 0.011454594 | 774.4007424 | 780.5737406 | 304 | 85 | 96 | 100 | 170 | 2 | 22 | 0 | 1 | 2 | 5 | 1 |
| SG0000004 | Osbpl6 | 5.41E-27 | 4.77E-26 | -2.465807706 | 554.5427896 | 100.3813379 | 1209 | 31 | 95 | 146 | 574 | 98 | 11 | 0 | 1 | 0 | 4 | 0 |
| SG0000003 | Pak3 | 3.69E-12 | 1.45E-11 | -1.955474197 | 245.8154695 | 63.38008557 | 2144 | 55 | 95 | 226 | 1107 | 264 | 120 | 6 | 6 | 24 | 48 | 18 |
| SG0000003 | Nptn | 8.91E-12 | 3.42E-11 | -0.980361078 | 5311.005892 | 2691.898572 | 824 | 75 | 95 | 118 | 342 | 62 | 16 | 0 | 0 | 3 | 9 | 1 |
| SG0000002 | Ercc6l2 | 1.84E-05 | 4.22E-05 | 0.57667997 | 1233.88736 | 1840.235839 | 820 | 63 | 95 | 175 | 460 | 43 | 12 | 0 | 2 | 2 | 7 | 0 |
| JSG000000 | Fut9 | 0.015303 | 0.024222 | 0.439511899 | 1186.24228 | 1608.717048 | 1040 | 53 | 95 | 171 | 574 | 91 | 56 | 1 | 1 | 7 | 14 | 5 |
| SG0000000 | Fer | 0.023795 | 0.036332 | -0.358675302 | 1064.809001 | 830.4236116 | 1058 | 41 | 95 | 154 | 516 | 102 | 0 | 0 | 0 | 0 | 0 | 0 |
| JSG000000 | Cog5 | 0.841754 | 0.876291 | 0.029215363 | 477.7730225 | 487.5468157 | 721 | 71 | 95 | 135 | 373 | 64 | 0 | 0 | 0 | 0 | 0 | 0 |
| SG0000000 | Ltbp1 | 2.21E-96 | 1.47E-94 | -5.537175056 | 34526.9078 | 743.5358307 | 1647 | 42 | 94 | 207 | 631 | 142 | 3 | 0 | 0 | 0 | 9 | 0 |
| SG0000003 | Gpr19 | 2.44E-40 | 3.66E-39 | 2.780989428 | 59.17236211 | 406.7055899 | 281 | 90 | 94 | 118 | 170 | 28 | 3 | 0 | 0 | 0 | 0 | 0 |
| SG0000004 | Efna5 | 4.92E-38 | 6.79E-37 | -3.818651311 | 1170.386152 | 82.94698163 | 1152 | 58 | 94 | 159 | 516 | 109 | 4 | 0 | 0 | 1 | 2 | 1 |
| JSG000000 | Faf1 | 1.26E-30 | 1.3E-29 | 1.421017832 | 1760.098258 | 4713.091864 | 835 | 62 | 94 | 140 | 382 | 57 | 5 | 0 | 0 | 1 | 1 | 0 |
| SG0000003 | Ccdc77 | 1.4E-21 | 9.59E-21 | 1.604885796 | 459.3300339 | 1397.151841 | 318 | 60 | 94 | 98 | 212 | 17 | 3 | 0 | 0 | 0 | 2 | 0 |
| SG0000003 | Nubpl | 3.49E-16 | 1.77E-15 | 1.3618875 | 241.3822029 | 620.4034937 | 877 | 66 | 94 | 157 | 553 | 59 | 3 | 0 | 0 | 0 | 1 | 0 |
| SG0000006 | Ica1 | 2.86E-15 | 1.37E-14 | -1.676606258 | 500.9532406 | 156.707007 | 1257 | 39 | 94 | 150 | 571 | 148 | 4 | 0 | 0 | 0 | 0 | 0 |
| SG0000000 | Eif2ak4 | 3.7E-13 | 1.55E-12 | -1.110379127 | 2500.4119 | 1158.121525 | 775 | 45 | 94 | 122 | 297 | 48 | 5 | 0 | 0 | 0 | 1 | 0 |
| SG0000003 | Tesk2 | 6.34E-09 | 1.96E-08 | 0.989575233 | 272.1272409 | 540.335929 | 564 | 81 | 94 | 138 | 291 | 29 | 3 | 0 | 0 | 2 | 2 | 0 |
| SG0000003 | Rapgef1 | 1.02E-08 | 3.1E-08 | -0.805815804 | 4666.333541 | 2669.322519 | 496 | 78 | 94 | 126 | 242 | 33 | 0 | 0 | 0 | 0 | 0 | 4 |
| SG0000006 | Srpk2 | 0.001999 | 0.003619 | 0.401150448 | 1931.149023 | 2550.199207 | 591 | 70 | 94 | 128 | 357 | 43 | 19 | 0 | 0 | 2 | 7 | 0 |
| SG0000003 | Nedd4 | 0.596157 | 0.659077 | -0.074648898 | 54400.69458 | 51657.44089 | 505 | 69 | 94 | 107 | 267 | 24 | 12 | 0 | 1 | 1 | 1 | 0 |
| SG0000002 | 2310035C23Rik | 0.756836 | 0.804107 | 0.037200312 | 1525.669441 | 1565.520855 | 1286 | 46 | 94 | 149 | 543 | 125 | 27 | 2 | 2 | 4 | 20 | 1 |
| SG0000000 | Rad51c | 2.68E-63 | 8.1E-62 | 2.609665362 | 180.7325457 | 1103.122946 | 297 | 72 | 93 | 121 | 232 | 17 | 8 | 0 | 1 | 2 | 6 | 1 |
| SG0000002 | Satb1 | 1.37E-11 | 5.18E-11 | 2.020154761 | 70.55100555 | 286.1741404 | 1392 | 21 | 93 | 170 | 625 | 69 | 23 | 3 | 5 | 6 | 24 | 4 |
| SG0000003 | Pknox2 | 1.38E-08 | 4.15E-08 | 4.003246689 | 4.816247606 | 77.2335753 | 1065 | 61 | 93 | 163 | 556 | 78 | 7 | 0 | 3 | 3 | 3 | 0 |
| SG0000006 | Trmt2b | 2.49E-08 | 7.35E-08 | -1.04297197 | 628.7499465 | 305.1491141 | 437 | 80 | 93 | 143 | 299 | 37 | 7 | 0 | 0 | 0 | 0 | 0 |
| SG0000003 | Kdm5a | 0.000215 | 0.000438 | -0.535265634 | 3277.885156 | 2261.844385 | 633 | 59 | 93 | 134 | 348 | 36 | 16 | 0 | 0 | 1 | 1 | 0 |
| SG0000001 | Fyn | 0.013002 | 0.020818 | -0.400194725 | 960.5522899 | 727.8642605 | 1178 | 49 | 93 | 192 | 522 | 134 | 8 | 0 | 2 | 0 | 4 | 8 |
| SG0000002 | Vps54 | 0.184199 | 0.236785 | -0.195457071 | 2023.963686 | 1767.519749 | 633 | 60 | 93 | 134 | 343 | 79 | 15 | 0 | 3 | 2 | 2 | 2 |
| SG0000004 | Zmym4 | 0.395405 | 0.464152 | 0.102665024 | 2349.278847 | 2522.550214 | 525 | 77 | 93 | 136 | 316 | 52 | 6 | 0 | 0 | 2 | 4 | 0 |
| SG0000002 | Huwe1 | 0.400294 | 0.469246 | 0.101077912 | 13443.17413 | 14418.8063 | 994 | 52 | 93 | 150 | 473 | 70 | 19 | 1 | 3 | 2 | 6 | 1 |
| SG0000004 | Osbpl1a | 0.856626 | 0.888214 | -0.042015989 | 1712.148728 | 1663.004462 | 1733 | 25 | 93 | 297 | 829 | 174 | 21 | 0 | 0 | 14 | 35 | 0 |
| SG0000005 | Phex | 9.2E-163 | 1.7E-160 | -9.464592721 | 2409.639723 | 3.410560894 | 662 | 67 | 92 | 135 | 368 | 43 | 9 | 0 | 0 | 1 | 2 | 0 |
| SG0000002 | Lef1 | 1.61E-98 | 1.1E-96 | 5.834896549 | 8.11200991 | 463.0281077 | 671 | 76 | 92 | 136 | 292 | 92 | 12 | 0 | 0 | 0 | 0 | 0 |
| SG0000000 | Prlr | 5.02E-39 | 7.15E-38 | 5.261110178 | 3.891376204 | 149.2296749 | 1447 | 37 | 92 | 179 | 680 | 142 | 23 | 0 | 1 | 1 | 10 | 4 |
| SG0000002 | Nme7 | 2.14E-22 | 1.53E-21 | 1.366245939 | 567.647276 | 1463.388262 | 872 | 81 | 92 | 165 | 407 | 76 | 1 | 0 | 0 | 0 | 0 | 0 |
| SG0000003 | Dennd5a | 2.74E-18 | 1.56E-17 | -1.324092289 | 3950.694384 | 1577.908948 | 322 | 84 | 92 | 124 | 232 | 18 | 6 | 2 | 2 | 2 | 4 | 0 |
| SG0000003 | Fndc3b | 1.71E-13 | 7.31E-13 | -1.353178282 | 2187.748436 | 856.347688 | 848 | 70 | 92 | 134 | 384 | 114 | 16 | 2 | 4 | 6 | 10 | 2 |
| JSG000000 | Arfip1 | 0.000103 | 0.000217 | -0.782262025 | 397.1914824 | 230.9487376 | 456 | 68 | 92 | 115 | 268 | 45 | 6 | 0 | 0 | 1 | 6 | 1 |
| SG0000003 | Herc2 | 0.000125 | 0.000263 | -0.56248791 | 2223.183714 | 1505.392054 | 842 | 72 | 92 | 164 | 452 | 74 | 3 | 0 | 0 | 0 | 4 | 0 |
| SG0000003 | Gmds | 0.099764 | 0.135743 | -0.317004836 | 361.9502767 | 290.550046 | 1550 | 39 | 92 | 208 | 671 | 132 | 2 | 0 | 0 | 0 | 0 | 0 |
| SG0000001 | Eya4 | 0.200699 | 0.255685 | -0.896625084 | 24.03320759 | 12.90924052 | 1245 | 38 | 92 | 186 | 522 | 102 | 9 | 0 | 0 | 0 | 6 | 0 |
| SG0000003 | Taf1 | 0.261374 | 0.323114 | 0.138964842 | 2919.964103 | 3215.215584 | 492 | 86 | 92 | 131 | 361 | 52 | 10 | 0 | 0 | 1 | 4 | 7 |
| SG0000006 | Hs6st2 | 0.358945 | 0.427062 | 0.376084653 | 114.8311825 | 149.0296216 | 1224 | 56 | 92 | 176 | 588 | 88 | 8 | 0 | 0 | 0 | 2 | 2 |
| SG0000000 | Aire | 1.7E-119 | 1.7E-117 | 7.897126047 | 1.974045165 | 470.5750579 | 273 | 65 | 91 | 78 | 156 | 13 | 4 | 1 | 1 | 1 | 1 | 0 |
| SG0000003 | Smarca1 | 1.53E-61 | 4.45E-60 | 3.84371177 | 27.52170421 | 395.1371476 | 854 | 55 | 91 | 127 | 471 | 69 | 14 | 0 | 1 | 1 | 16 | 1 |
| SG0000000 | Tiam1 | 2.41E-24 | 1.88E-23 | 4.177715841 | 82.49153413 | 1492.887776 | 758 | 66 | 91 | 134 | 361 | 92 | 12 | 3 | 3 | 6 | 10 | 5 |
| SG0000005 | Dcun1d4 | 2.6E-15 | 1.26E-14 | 1.363315876 | 428.5331221 | 1102.512106 | 571 | 59 | 91 | 116 | 283 | 94 | 9 | 3 | 3 | 3 | 6 | 3 |
| SG0000005 | Fry | 1.59E-08 | 4.78E-08 | 1.149605135 | 183.2236988 | 406.4875748 | 576 | 67 | 91 | 127 | 301 | 59 | 12 | 0 | 3 | 3 | 6 | 3 |
| SG0000003 | Cfap61 | 3.59E-05 | 8.01E-05 | 3.478703867 | 1.904113361 | 21.22692134 | 791 | 56 | 91 | 120 | 370 | 94 | 17 | 1 | 3 | 4 | 6 | 1 |
| SG0000001 | Tnpo3 | 0.004087 | 0.007071 | 0.35628669 | 5350.465907 | 6849.274657 | 566 | 69 | 91 | 115 | 274 | 26 | 10 | 0 | 0 | 0 | 0 | 0 |
| SG0000002 | Gpcpd1 | 0.004956 | 0.008483 | -0.488285887 | 729.2450712 | 519.8580868 | 917 | 56 | 91 | 105 | 437 | 96 | 42 | 0 | 0 | 0 | 4 | 12 |
| SG0000006 | Dlgap4 | 0.41901 | 0.48815 | -0.11054077 | 1672.616977 | 1549.246106 | 626 | 80 | 91 | 118 | 212 | 40 | 4 | 0 | 0 | 0 | 6 | 3 |
| SG0000003 | Plcb4 | 0.52497 | 0.591624 | 0.091373661 | 740.6668043 | 789.0946401 | 1444 | 47 | 91 | 199 | 665 | 83 | 9 | 0 | 0 | 4 | 6 | 0 |
| SG0000003 | Gabrb3 | 2.76E-79 | 1.22E-77 | 8.667971803 | 0.639110347 | 259.9533166 | 1755 | 53 | 90 | 201 | 828 | 178 | 23 | 0 | 0 | 3 | 14 | 6 |
| SG0000004 | BC052040 | 3.39E-49 | 6.84E-48 | 2.158996119 | 293.0597641 | 1308.81658 | 902 | 60 | 90 | 134 | 444 | 64 | 4 | 0 | 0 | 0 | 0 | 0 |
| SG0000002 | Sycp1 | 2.64E-41 | 4.1E-40 | 6.886535034 | 0 | 117.3187605 | 759 | 52 | 90 | 166 | 509 | 58 | 1 | 0 | 0 | 0 | 3 | 0 |
| SG0000002 | Rimbp2 | 5.5E-32 | 6.04E-31 | 6.486786283 | 0 | 88.68447012 | 689 | 68 | 90 | 149 | 453 | 95 | 2 | 0 | 0 | 0 | 0 | 0 |
| JSG000000 | 1700101E01Rik | 7.86E-29 | 7.52E-28 | 7.072575092 | 0.649984458 | 87.49036364 | 439 | 75 | 90 | 120 | 219 | 34 | 2 | 0 | 0 | 0 | 0 | 0 |
| SG0000005 | Nrk | 1.07E-26 | 9.34E-26 | 8.992282941 | 1.585729971 | 807.562473 | 927 | 63 | 90 | 156 | 468 | 72 | 24 | 0 | 0 | 0 | 6 | 0 |
| SG0000006 | Fmr1nb | 1.98E-23 | 1.48E-22 | 6.001743681 | 0 | 63.07739894 | 354 | 66 | 90 | 114 | 273 | 12 | 0 | 0 | 0 | 0 | 0 | 0 |
| SG0000004 | Asxl3 | 2.92E-20 | 1.86E-19 | 5.013291051 | 2.25980622 | 72.98307926 | 1000 | 44 | 90 | 158 | 478 | 84 | 26 | 0 | 0 | 4 | 10 | 0 |
| SG0000003 | Slc25a14 | 9.92E-20 | 6.13E-19 | -2.079845813 | 443.8152276 | 104.9799165 | 633 | 55 | 90 | 130 | 375 | 30 | 6 | 0 | 0 | 6 | 6 | 0 |
| SG0000003 | Myo1d | 1.11E-12 | 4.5E-12 | -1.573297293 | 1090.818143 | 366.5579772 | 869 | 49 | 90 | 126 | 453 | 84 | 1 | 0 | 0 | 0 | 0 | 0 |
| SG0000003 | Dmxl1 | 1.39E-09 | 4.52E-09 | -0.935654802 | 2997.000051 | 1566.846997 | 840 | 50 | 90 | 158 | 462 | 66 | 30 | 0 | 0 | 4 | 26 | 6 |
| JSG000000 | Gucy1a2 | 5.57E-09 | 1.73E-08 | 2.960924877 | 6.526279939 | 50.81511368 | 884 | 53 | 90 | 151 | 554 | 61 | 0 | 0 | 0 | 0 | 0 | 0 |
| SG0000000 | Dnah2 | 0.000297 | 0.000596 | -0.847870374 | 227.8423108 | 126.5901638 | 349 | 68 | 90 | 112 | 240 | 28 | 0 | 0 | 0 | 0 | 0 | 0 |
| SG0000005 | Yap1 | 0.011554 | 0.018716 | -0.342396742 | 5791.053749 | 4567.5773 | 660 | 45 | 90 | 115 | 329 | 80 | 10 | 0 | 0 | 0 | 4 | 0 |
| JSG000000 | Ptprm | 0.1072 | 0.145017 | -0.676165347 | 102.8747864 | 64.38176745 | 1215 | 48 | 90 | 158 | 476 | 145 | 0 | 0 | 0 | 0 | 0 | 0 |
| SG0000003 | Zfyve9 | 0.22792 | 0.286351 | 0.169752387 | 815.0171471 | 916.7845932 | 756 | 58 | 90 | 130 | 366 | 52 | 6 | 0 | 0 | 0 | 0 | 0 |
| SG0000003 | Pml | 0.386376 | 0.455134 | 0.106942466 | 4706.073097 | 5068.174489 | 325 | 76 | 90 | 118 | 290 | 26 | 8 | 0 | 0 | 0 | 4 | 0 |
| SG0000003 | Elf2 | 0.890476 | 0.916152 | 0.019308554 | 1279.278121 | 1296.514644 | 998 | 47 | 90 | 138 | 507 | 117 | 15 | 1 | 1 | 3 | 9 | 0 |
| SG0000001 | Abr | 9.14E-36 | 1.16E-34 | -1.973542691 | 8192.522616 | 2086.037451 | 438 | 78 | 89 | 126 | 285 | 24 | 4 | 0 | 4 | 4 | 16 | 0 |
| SG0000002 | Wdr76 | 1.58E-25 | 1.31E-24 | 1.380303359 | 868.1700188 | 2260.047141 | 322 | 74 | 89 | 119 | 199 | 35 | 11 | 0 | 1 | 0 | 0 | 1 |
| SG0000004 | 0610010F05Rik | 4.64E-16 | 2.34E-15 | -1.31786047 | 1810.668882 | 726.3124794 | 736 | 49 | 89 | 132 | 385 | 34 | 19 | 4 | 4 | 8 | 11 | 1 |
| SG0000002 | Zdhhc3 | 2.12E-10 | 7.38E-10 | -0.919221884 | 4057.167014 | 2145.406174 | 482 | 82 | 89 | 116 | 203 | 39 | 7 | 0 | 0 | 1 | 1 | 5 |
| SG0000001 | Pls3 | 1.38E-09 | 4.49E-09 | -0.878355837 | 4351.777433 | 2367.310486 | 661 | 76 | 89 | 122 | 292 | 68 | 12 | 0 | 0 | 0 | 4 | 4 |
| SG0000005 | Mapk14 | 5.55E-09 | 1.72E-08 | -0.821351039 | 5474.548291 | 3098.111125 | 443 | 84 | 89 | 112 | 248 | 20 | 5 | 0 | 0 | 0 | 0 | 0 |
| SG0000003 | Rreb1 | 2.4E-06 | 6.01E-06 | -0.730783378 | 1561.912334 | 941.1718678 | 887 | 68 | 89 | 107 | 408 | 113 | 16 | 0 | 0 | 1 | 31 | 6 |
| SG0000002 | Rprd1b | 0.000314 | 0.000628 | -0.541699234 | 1665.655355 | 1144.240796 | 584 | 71 | 89 | 207 | 369 | 54 | 7 | 0 | 0 | 0 | 0 | 0 |
| SG0000002 | Tnip1 | 0.005686 | 0.009648 | -0.621912059 | 2858.716621 | 1857.620188 | 345 | 83 | 89 | 101 | 191 | 0 | 0 | 0 | 0 | 0 | 0 | 0 |
| JSG000000 | Wdpcp | 0.068451 | 0.096308 | -0.632599254 | 78.17349284 | 50.4229402 | 757 | 55 | 89 | 123 | 393 | 61 | 1 | 0 | 0 | 0 | 0 | 0 |
| SG0000002 | Pld1 | 0.465221 | 0.534683 | -0.215789527 | 129.9259792 | 111.8759866 | 790 | 69 | 89 | 139 | 440 | 66 | 2 | 0 | 0 | 0 | 2 | 0 |
| SG0000001 | Msh3 | 7.17E-36 | 9.16E-35 | 1.804167786 | 376.8376996 | 1316.021441 | 1184 | 46 | 88 | 163 | 532 | 95 | 19 | 0 | 1 | 4 | 6 | 2 |
| SG0000003 | Aamdc | 1E-13 | 4.35E-13 | 1.771425426 | 65.72173697 | 224.3681573 | 655 | 44 | 88 | 110 | 322 | 64 | 7 | 0 | 0 | 0 | 0 | 0 |
| SG0000002 | Creb1 | 7.24E-11 | 2.6E-10 | 0.843891467 | 1458.319628 | 2617.516125 | 1092 | 60 | 88 | 158 | 562 | 92 | 37 | 0 | 1 | 1 | 8 | 5 |
| SG0000002 | Fancc | 5.36E-09 | 1.67E-08 | 0.977593726 | 297.4866997 | 585.8043452 | 787 | 60 | 88 | 136 | 420 | 92 | 0 | 0 | 0 | 0 | 3 | 0 |
| SG0000002 | Raly | 8.3E-07 | 2.16E-06 | 0.612309766 | 4082.400932 | 6240.785572 | 748 | 76 | 88 | 160 | 388 | 44 | 0 | 0 | 0 | 0 | 0 | 0 |
| SG0000003 | Wdr20 | 0.00265 | 0.004714 | -0.503568397 | 743.1921323 | 524.217978 | 387 | 72 | 88 | 103 | 271 | 12 | 6 | 0 | 0 | 0 | 2 | 1 |
| SG0000003 | Polq | 0.011639 | 0.018832 | -0.565960189 | 866.4953185 | 585.3225116 | 791 | 52 | 88 | 136 | 404 | 128 | 19 | 0 | 0 | 0 | 4 | 0 |
| SG0000005 | Sclt1 | 0.025199 | 0.038316 | 0.459881735 | 177.4121886 | 244.0178804 | 922 | 51 | 88 | 138 | 463 | 90 | 6 | 0 | 0 | 0 | 1 | 0 |
| SG0000003 | Dock3 | 0.058094 | 0.082803 | 0.424754518 | 135.4589558 | 181.8325269 | 750 | 69 | 88 | 139 | 422 | 70 | 1 | 0 | 1 | 0 | 2 | 4 |
| SG0000002 | Arhgef2 | 0.067326 | 0.094903 | 0.247104948 | 1835.800418 | 2178.770396 | 305 | 77 | 88 | 89 | 135 | 18 | 19 | 10 | 10 | 10 | 10 | 0 |
| SG0000002 | 9030617O03Rik | 3.9E-103 | 3E-101 | -4.93094048 | 3041.643685 | 99.71198123 | 680 | 68 | 87 | 124 | 422 | 136 | 9 | 0 | 0 | 0 | 0 | 0 |
| SG0000004 | Pitpnc1 | 5.98E-26 | 5.04E-25 | 1.463546482 | 575.8523126 | 1588.117042 | 578 | 70 | 87 | 129 | 315 | 45 | 17 | 2 | 2 | 2 | 4 | 6 |

| | | | | | | | | | | | | | | | | | |
|---|---|---|---|---|---|---|---|---|---|---|---|---|---|---|---|---|---|
| SG0000005 | AI314180 | 6.19E-24 | 4.75E-23 | -1.534360432 | 6057.768535 | 2091.337673 | 818 | 61 | 87 | 124 | 390 | 92 | 10 | 2 | 3 | 2 | 3 | 2 |
| SG0000003 | Rcor1 | 1.67E-14 | 7.66E-14 | -1.185742405 | 2610.003389 | 1147.35296 | 343 | 72 | 87 | 98 | 214 | 44 | 5 | 0 | 1 | 0 | 0 | 0 |
| SG0000003 | Pcsk6 | 3.42E-14 | 1.53E-13 | 1.284079643 | 277.3681163 | 675.4652737 | 888 | 63 | 87 | 180 | 381 | 78 | 0 | 0 | 0 | 0 | 0 | 3 |
| SG0000007 | Ano3 | 1.06E-12 | 4.32E-12 | -2.123940102 | 410.3829435 | 94.14984784 | 982 | 52 | 87 | 157 | 530 | 108 | 11 | 0 | 0 | 1 | 5 | 3 |
| SG0000003 | Zc3hc1 | 4.97E-09 | 1.55E-08 | 0.779054595 | 1193.451544 | 2047.970007 | 298 | 66 | 87 | 101 | 184 | 41 | 0 | 0 | 0 | 0 | 0 | 0 |
| SG0000005 | Ntrk2 | 4.68E-05 | 0.000103 | -1.284624758 | 119.0348647 | 48.86114021 | 709 | 64 | 87 | 148 | 422 | 102 | 30 | 9 | 10 | 15 | 27 | 2 |
| SG0000002 | Itga8 | 8.15E-05 | 0.000175 | 2.038244576 | 9.665117546 | 39.69903006 | 923 | 57 | 87 | 168 | 438 | 89 | 23 | 0 | 1 | 7 | 13 | 2 |
| SG0000002 | 4930402H24Rik | 0.984987 | 0.995772 | -0.007780675 | 604.9109942 | 601.6573981 | 582 | 62 | 87 | 103 | 313 | 37 | 8 | 0 | 0 | 1 | 6 | 0 |
| SG0000007 | Map3k5 | 1.68E-45 | 3E-44 | -2.798388141 | 1498.687446 | 215.4330349 | 729 | 30 | 86 | 114 | 392 | 70 | 1 | 0 | 0 | 0 | 2 | 0 |
| SG0000003 | Gsap | 1.4E-40 | 2.11E-39 | -4.13311911 | 1142.27034 | 65.09926307 | 773 | 33 | 86 | 120 | 392 | 63 | 15 | 0 | 0 | 2 | 2 | 0 |
| SG0000000 | Hmgcll1 | 1.18E-18 | 6.88E-18 | 3.67985386 | 7.200356187 | 92.27846726 | 1290 | 52 | 86 | 226 | 786 | 108 | 14 | 0 | 2 | 2 | 10 | 0 |
| JSG000000 | Cpq | 9.29E-11 | 3.31E-10 | -1.886186433 | 287.0841018 | 77.6623359 | 1051 | 43 | 86 | 169 | 652 | 80 | 2 | 0 | 0 | 0 | 0 | 0 |
| SG0000005 | Sptan1 | 4.5E-08 | 1.3E-07 | -0.747656213 | 16053.82455 | 9561.181575 | 436 | 58 | 86 | 104 | 251 | 39 | 0 | 0 | 0 | 0 | 0 | 0 |
| SG0000003 | Acer3 | 2.17E-06 | 5.44E-06 | -0.846600363 | 710.3975592 | 395.0475357 | 943 | 47 | 86 | 162 | 553 | 69 | 21 | 0 | 0 | 5 | 13 | 0 |
| SG0000005 | Abat | 9.3E-06 | 2.2E-05 | 3.558433677 | 6.992904343 | 82.38589461 | 453 | 62 | 86 | 112 | 245 | 45 | 6 | 0 | 1 | 0 | 2 | 0 |
| SG0000004 | Rnf216 | 6.68E-05 | 0.000145 | -0.581938454 | 2158.698228 | 1442.151894 | 658 | 66 | 86 | 122 | 306 | 67 | 2 | 0 | 0 | 0 | 1 | 1 |
| SG0000005 | Ttn | 9.3E-05 | 0.000198 | 2.713435453 | 4.212103161 | 27.62641058 | 1517 | 41 | 86 | 177 | 647 | 205 | 3 | 0 | 0 | 0 | 3 | 10 |
| SG0000000 | Qrich1 | 0.001141 | 0.002128 | 0.41767923 | 2745.983598 | 3668.021863 | 336 | 66 | 86 | 82 | 196 | 24 | 0 | 0 | 0 | 0 | 0 | 0 |
| SG0000003 | Snx14 | 0.006792 | 0.011402 | -0.438516986 | 934.542081 | 689.5918106 | 756 | 57 | 86 | 136 | 468 | 94 | 4 | 0 | 0 | 0 | 4 | 0 |
| SG0000002 | Phf14 | 0.023437 | 0.035811 | -0.343641091 | 1339.013422 | 1055.209407 | 1012 | 44 | 86 | 187 | 552 | 41 | 7 | 0 | 0 | 0 | 1 | 0 |
| SG0000004 | Mthfd1l | 0.029305 | 0.044061 | -0.321637359 | 3066.236977 | 2453.483965 | 850 | 58 | 86 | 146 | 398 | 107 | 0 | 0 | 0 | 0 | 3 | 0 |
| SG0000002 | Ube3a | 0.035377 | 0.052382 | -0.294044809 | 3858.207594 | 3146.801031 | 888 | 64 | 86 | 150 | 546 | 57 | 30 | 1 | 1 | 5 | 27 | 3 |
| SG0000003 | Ino80 | 0.101395 | 0.137705 | -0.239170172 | 1896.978433 | 1607.18177 | 468 | 70 | 86 | 118 | 242 | 36 | 2 | 0 | 0 | 0 | 1 | 1 |
| SG0000003 | Ccnd3 | 0.170549 | 0.220984 | 0.286042026 | 4360.86471 | 5317.1614 | 438 | 54 | 86 | 114 | 322 | 67 | 5 | 0 | 0 | 0 | 0 | 0 |
| SG0000003 | AW554918 | 0.579921 | 0.643868 | -0.122064627 | 395.5077232 | 363.4208621 | 1320 | 40 | 86 | 198 | 720 | 126 | 16 | 0 | 0 | 4 | 6 | 2 |
| SG0000002 | Mkrn1 | 4.6E-109 | 4E-107 | 2.587883264 | 2924.631942 | 17583.35348 | 222 | 80 | 85 | 85 | 170 | 15 | 12 | 0 | 2 | 0 | 4 | 0 |
| SG0000002 | Mtf2 | 3.32E-92 | 1.96E-90 | 2.446238666 | 1420.319378 | 7740.646375 | 378 | 61 | 85 | 106 | 281 | 24 | 10 | 1 | 1 | 1 | 5 | 6 |
| SG0000002 | Hp1bp3 | 3.74E-41 | 5.75E-40 | -2.231863138 | 4999.563026 | 1064.325082 | 346 | 65 | 85 | 100 | 187 | 25 | 16 | 0 | 0 | 0 | 2 | 5 |
| SG0000006 | Tsga10 | 6.64E-17 | 3.53E-16 | 2.050676618 | 52.34710343 | 216.8741694 | 955 | 49 | 85 | 148 | 459 | 115 | 11 | 0 | 0 | 1 | 7 | 0 |
| SG0000002 | Pspc1 | 3.88E-16 | 1.97E-15 | 1.033491266 | 1719.128001 | 3519.006632 | 400 | 59 | 85 | 100 | 292 | 48 | 12 | 0 | 1 | 1 | 3 | 1 |
| SG0000002 | Txndc11 | 1.36E-12 | 5.47E-12 | -1.190863805 | 1203.137841 | 527.0231111 | 557 | 68 | 85 | 115 | 295 | 65 | 22 | 6 | 9 | 8 | 13 | 1 |
| SG0000002 | Dtna | 0.000105 | 0.000221 | -0.703965343 | 562.4430539 | 345.2739981 | 920 | 55 | 85 | 132 | 427 | 98 | 6 | 0 | 0 | 0 | 1 | 3 |
| SG0000002 | Nedd4l | 0.007469 | 0.012456 | -0.424974692 | 1008.594316 | 751.2532898 | 745 | 49 | 85 | 119 | 347 | 85 | 10 | 0 | 0 | 1 | 3 | 2 |
| SG0000001 | Lcorl | 0.018641 | 0.028999 | -0.423115565 | 886.0810196 | 660.8500954 | 1490 | 49 | 85 | 196 | 675 | 162 | 70 | 7 | 7 | 16 | 25 | 2 |
| SG0000002 | Sdccag8 | 0.066428 | 0.093741 | -0.370681256 | 328.5934208 | 254.1398017 | 946 | 48 | 85 | 141 | 436 | 63 | 7 | 0 | 0 | 0 | 0 | 0 |
| SG0000001 | Rhot1 | 0.132454 | 0.17556 | 0.196364174 | 2053.530417 | 2352.949718 | 561 | 52 | 85 | 125 | 316 | 118 | 14 | 0 | 0 | 10 | 23 | 10 |
| SG0000002 | Myef2 | 0.472839 | 0.541666 | 0.093318947 | 3060.617553 | 3265.132928 | 705 | 56 | 85 | 125 | 311 | 35 | 11 | 3 | 6 | 3 | 10 | 0 |
| SG0000000 | Cd44 | 1.62E-59 | 4.41E-58 | -4.615212333 | 15757.82661 | 642.9526179 | 1126 | 49 | 84 | 157 | 449 | 56 | 20 | 0 | 0 | 10 | 22 | 0 |
| SG0000002 | Itgb5 | 2.82E-32 | 3.13E-31 | -2.291113126 | 6441.64243 | 1316.142277 | 377 | 70 | 84 | 98 | 200 | 26 | 1 | 0 | 0 | 0 | 0 | 0 |
| SG0000002 | Agfg1 | 5.96E-24 | 4.58E-23 | 1.216350253 | 3200.648724 | 7436.968187 | 1106 | 42 | 84 | 169 | 529 | 158 | 28 | 0 | 0 | 8 | 11 | 11 |
| SG0000002 | Kmt2e | 1.23E-18 | 7.17E-18 | -1.348668329 | 3736.456979 | 1467.135971 | 662 | 69 | 84 | 130 | 321 | 59 | 11 | 0 | 0 | 0 | 4 | 4 |
| SG0000004 | Ildr2 | 2.76E-18 | 1.58E-17 | 6.425708726 | 0.639110347 | 54.94231042 | 619 | 63 | 84 | 91 | 260 | 35 | 17 | 0 | 0 | 0 | 1 | 0 |
| SG0000003 | Mga | 1.02E-14 | 4.75E-14 | 1.032996699 | 3078.831156 | 6300.120622 | 814 | 60 | 84 | 132 | 472 | 79 | 18 | 0 | 0 | 4 | 20 | 11 |
| SG0000002 | Banp | 5.14E-13 | 2.13E-12 | 1.072049338 | 565.250759 | 1188.39321 | 379 | 50 | 84 | 69 | 166 | 42 | 5 | 0 | 0 | 2 | 2 | 0 |
| SG0000002 | Adam23 | 2.8E-11 | 1.04E-10 | -1.055246885 | 4724.414555 | 2273.458514 | 1038 | 39 | 84 | 171 | 524 | 114 | 6 | 0 | 0 | 0 | 2 | 6 |
| SG0000002 | Pde8b | 9.25E-09 | 2.83E-08 | -2.010164727 | 125.3933023 | 31.12823243 | 1381 | 60 | 84 | 204 | 733 | 113 | 7 | 0 | 0 | 0 | 11 | 3 |
| SG0000005 | Prkcb | 4.42E-06 | 1.08E-05 | 2.302332069 | 9.866382414 | 48.6663704 | 1313 | 60 | 84 | 171 | 624 | 142 | 0 | 0 | 0 | 0 | 1 | 0 |
| SG0000002 | Tpx2 | 0.000738 | 0.001408 | 0.417702013 | 5188.375732 | 6930.622666 | 420 | 52 | 84 | 116 | 260 | 24 | 12 | 0 | 0 | 0 | 0 | 0 |
| SG0000003 | Smarca4 | 0.019265 | 0.029886 | -0.334959776 | 6655.085283 | 5276.193145 | 380 | 63 | 84 | 111 | 255 | 20 | 2 | 0 | 0 | 0 | 0 | 2 |
| SG0000003 | Foxj3 | 0.098046 | 0.13368 | -0.246790889 | 1526.688652 | 1286.64583 | 748 | 44 | 84 | 124 | 372 | 54 | 26 | 0 | 4 | 6 | 10 | 8 |
| SG0000002 | Adcy3 | 0.155476 | 0.203187 | 0.26058813 | 341.2618587 | 408.8204345 | 363 | 76 | 84 | 96 | 212 | 76 | 4 | 0 | 0 | 0 | 0 | 0 |
| SG0000002 | Slc29a1 | 0.1702 | 0.220605 | 0.250093885 | 3436.2413 | 4086.668539 | 127 | 84 | 84 | 85 | 93 | 1 | 0 | 0 | 0 | 0 | 0 | 0 |
| SG0000004 | Rsbn1 | 0.176927 | 0.228223 | 0.200207499 | 1252.961208 | 1439.4815 | 465 | 69 | 84 | 93 | 220 | 23 | 21 | 0 | 2 | 0 | 2 | 3 |
| SG0000007 | Rasgrp3 | 0.235414 | 0.294609 | -0.736066506 | 22.667994 | 13.60927506 | 385 | 66 | 84 | 97 | 217 | 41 | 16 | 0 | 0 | 4 | 6 | 2 |
| SG0000002 | Dnajb6 | 0.266549 | 0.328726 | 0.148782511 | 1352.45122 | 1499.372733 | 465 | 65 | 84 | 90 | 254 | 39 | 6 | 1 | 2 | 2 | 4 | 0 |
| JSG000000 | Cox16 | 0.306232 | 0.371327 | -0.378394 | 151.6028258 | 116.6270666 | 648 | 64 | 84 | 113 | 435 | 51 | 1 | 0 | 0 | 0 | 1 | 0 |
| SG0000003 | Zfp638 | 0.327222 | 0.39395 | -0.140268726 | 3271.204622 | 2968.127942 | 729 | 54 | 84 | 123 | 339 | 90 | 4 | 0 | 0 | 0 | 0 | 0 |
| SG0000003 | Smg1 | 0.390931 | 0.459562 | 0.10312763 | 3667.187645 | 3938.924376 | 460 | 64 | 84 | 124 | 276 | 46 | 35 | 4 | 6 | 5 | 15 | 4 |
| SG0000003 | Mettl10 | 0.4091 | 0.478379 | -0.158932749 | 401.200471 | 359.3502154 | 294 | 64 | 84 | 98 | 196 | 16 | 4 | 0 | 0 | 0 | 0 | 0 |
| SG0000002 | Axin1 | 1 | 1 | 0.000985762 | 1878.75131 | 1880.035459 | 315 | 63 | 84 | 93 | 180 | 40 | 0 | 0 | 0 | 0 | 0 | 1 |
| SG0000001 | Cyth1 | 9.11E-72 | 3.42E-70 | -3.805966937 | 2042.783672 | 146.0535668 | 447 | 65 | 83 | 124 | 226 | 25 | 7 | 0 | 0 | 3 | 18 | 3 |
| SG0000004 | Klf8 | 1.52E-32 | 1.71E-31 | 3.212634113 | 70.14094214 | 650.2357274 | 527 | 56 | 83 | 121 | 328 | 53 | 10 | 0 | 0 | 0 | 3 | 4 |
| SG0000002 | Mib1 | 2.14E-20 | 1.37E-19 | -1.45414873 | 5780.746289 | 2109.801143 | 729 | 60 | 83 | 120 | 392 | 82 | 19 | 0 | 0 | 0 | 16 | 4 |
| SG0000002 | Lrrc16a | 7.24E-15 | 3.39E-14 | 1.423418256 | 258.3484338 | 692.9428602 | 1016 | 47 | 83 | 155 | 522 | 83 | 12 | 0 | 0 | 2 | 4 | 3 |
| SG0000004 | Arhgap12 | 4.76E-11 | 1.73E-10 | 1.199089358 | 1111.698188 | 2552.400154 | 544 | 62 | 83 | 120 | 321 | 51 | 6 | 0 | 1 | 0 | 1 | 1 |
| SG0000005 | Adam17 | 1.65E-09 | 5.33E-09 | -0.897750415 | 2632.874276 | 1413.124147 | 484 | 74 | 83 | 119 | 252 | 70 | 6 | 0 | 0 | 0 | 1 | 0 |
| SG0000003 | Dnm2 | 7.02E-08 | 2E-07 | -0.924738489 | 4056.223207 | 2136.721019 | 475 | 83 | 83 | 124 | 212 | 31 | 2 | 0 | 0 | 0 | 0 | 4 |
| SG0000002 | Rgl1 | 2.76E-07 | 7.44E-07 | 1.258390761 | 76.36006264 | 182.6752209 | 1117 | 42 | 83 | 171 | 523 | 92 | 9 | 0 | 1 | 3 | 3 | 4 |
| SG0000002 | Fmn2 | 5.94E-07 | 1.56E-06 | -0.923494277 | 593.9477799 | 313.147382 | 755 | 53 | 83 | 131 | 395 | 82 | 9 | 0 | 0 | 1 | 1 | 1 |
| JSG000000 | Tmem117 | 2.3E-05 | 5.22E-05 | 2.501954978 | 6.183804759 | 35.02831652 | 931 | 50 | 83 | 149 | 477 | 98 | 1 | 0 | 0 | 0 | 1 | 1 |
| SG0000004 | Ubap2l | 0.000631 | 0.001214 | 0.422605867 | 6278.232144 | 8415.005661 | 398 | 71 | 83 | 100 | 193 | 29 | 2 | 0 | 0 | 1 | 3 | 1 |
| SG0000003 | Tspan9 | 0.069905 | 0.098207 | 0.641123599 | 815.5307937 | 1271.855577 | 510 | 66 | 83 | 110 | 213 | 39 | 2 | 0 | 0 | 0 | 3 | 0 |
| SG0000002 | Epb41l1 | 0.070808 | 0.099328 | 0.318077175 | 855.4545086 | 1066.46766 | 528 | 48 | 83 | 92 | 187 | 60 | 5 | 0 | 0 | 0 | 0 | 0 |
| SG0000000 | Hltf | 0.56515 | 0.629695 | 0.071512984 | 2259.865836 | 2374.70797 | 550 | 60 | 83 | 106 | 326 | 72 | 11 | 0 | 1 | 3 | 8 | 0 |
| JSG000000 | Pola1 | 0.759387 | 0.806448 | 0.028138518 | 3230.745069 | 3294.376474 | 855 | 52 | 83 | 128 | 431 | 68 | 2 | 0 | 0 | 0 | 0 | 0 |
| SG0000002 | 4930447C04Rik | 7.1E-188 | 1.9E-185 | 9.542952975 | 0 | 744.9592314 | 485 | 57 | 82 | 94 | 244 | 52 | 17 | 0 | 2 | 0 | 7 | 0 |
| SG0000003 | Cdh13 | 3.19E-83 | 1.55E-81 | -4.438948461 | 1833.64112 | 84.53911626 | 1391 | 43 | 82 | 166 | 538 | 172 | 1 | 0 | 0 | 0 | 0 | 0 |
| SG0000004 | Mms22l | 3.95E-34 | 4.73E-33 | 1.580473996 | 881.7233598 | 2636.95323 | 985 | 36 | 82 | 148 | 466 | 104 | 5 | 1 | 3 | 2 | 4 | 0 |
| SG0000003 | Fgfr1 | 4.76E-27 | 4.2E-26 | -1.61182496 | 10476.45674 | 3427.731245 | 516 | 65 | 82 | 107 | 254 | 50 | 15 | 0 | 0 | 0 | 3 | 9 |
| SG0000002 | Rai14 | 1.67E-17 | 9.17E-17 | -1.232156322 | 9652.98299 | 4109.088819 | 518 | 58 | 82 | 108 | 268 | 38 | 7 | 0 | 0 | 2 | 4 | 0 |
| SG0000002 | Inpp5d | 7.03E-16 | 3.5E-15 | 7.613446643 | 10.07887124 | 1973.729838 | 614 | 58 | 82 | 94 | 258 | 80 | 0 | 0 | 0 | 0 | 6 | 0 |
| SG0000006 | Mllt4 | 1.01E-10 | 3.6E-10 | 0.790502586 | 3940.205022 | 6815.281679 | 1277 | 40 | 82 | 214 | 730 | 75 | 13 | 0 | 2 | 0 | 2 | 2 |
| SG0000003 | Ankrd13c | 0.00039 | 0.000773 | -0.514525862 | 2350.255076 | 1645.232505 | 464 | 66 | 82 | 120 | 303 | 51 | 19 | 0 | 4 | 4 | 4 | 0 |
| SG0000003 | Kmt2c | 0.002622 | 0.00467 | -0.513658 | 1643.53334 | 1151.203371 | 810 | 51 | 82 | 132 | 407 | 73 | 8 | 0 | 0 | 2 | 7 | 3 |
| SG0000002 | Atg4c | 0.002917 | 0.005156 | -0.516629445 | 651.6416533 | 455.4994714 | 362 | 66 | 82 | 118 | 268 | 14 | 6 | 0 | 0 | 2 | 2 | 0 |
| SG0000003 | Tmem135 | 0.005962 | 0.010089 | -0.429510557 | 1149.447263 | 853.4802769 | 1067 | 34 | 82 | 135 | 506 | 84 | 6 | 0 | 0 | 0 | 1 | 0 |
| SG0000002 | Ube2g1 | 0.006595 | 0.011096 | 0.352280029 | 1904.910532 | 2431.764078 | 534 | 68 | 82 | 124 | 330 | 40 | 10 | 0 | 0 | 0 | 0 | 4 |
| JSG000000 | Camkmt | 0.046528 | 0.067597 | -0.52507497 | 151.7427239 | 105.4495085 | 731 | 49 | 82 | 114 | 361 | 65 | 0 | 0 | 0 | 0 | 0 | 0 |
| SG0000002 | Anks1 | 0.084941 | 0.117397 | -0.311800083 | 858.219124 | 691.4121551 | 483 | 63 | 82 | 118 | 221 | 47 | 8 | 4 | 8 | 4 | 4 | 4 |
| SG0000004 | Tmtc4 | 0.153524 | 0.200783 | -0.235720384 | 660.6270542 | 561.0446804 | 594 | 47 | 82 | 95 | 244 | 53 | 18 | 2 | 4 | 4 | 10 | 0 |
| SG0000003 | Jmjd1c | 0.685401 | 0.741269 | -0.066368198 | 4150.887896 | 3964.26051 | 863 | 35 | 82 | 135 | 392 | 81 | 21 | 0 | 3 | 3 | 3 | 3 |
| SG0000004 | Fam13c | 3.43E-23 | 2.53E-22 | 4.0637033 | 6.469565814 | 108.186164 | 645 | 69 | 81 | 114 | 312 | 66 | 18 | 0 | 0 | 0 | 6 | 3 |
| SG0000002 | Gsk3b | 1.12E-17 | 6.2E-17 | -1.360140957 | 3967.088953 | 1545.356603 | 792 | 55 | 81 | 112 | 348 | 50 | 12 | 1 | 1 | 2 | 8 | 4 |
| SG0000004 | Prex2 | 1.24E-17 | 6.83E-17 | -1.921884953 | 454.6012386 | 119.9735737 | 1037 | 43 | 81 | 156 | 522 | 92 | 35 | 0 | 1 | 3 | 19 | 5 |
| SG0000004 | Ints8 | 1.18E-15 | 5.81E-15 | 1.04617287 | 1196.086407 | 2469.971655 | 410 | 69 | 81 | 118 | 271 | 43 | 0 | 0 | 0 | 0 | 1 | 2 |
| SG0000002 | Fbxw7 | 1.45E-10 | 5.1E-10 | -1.014400699 | 1649.888936 | 816.7509946 | 911 | 47 | 81 | 126 | 397 | 84 | 9 | 0 | 3 | 6 | 6 | 0 |
| SG0000002 | Hormad2 | 2.14E-10 | 7.44E-10 | 5.156220075 | 0.946619625 | 33.75611132 | 778 | 39 | 81 | 129 | 390 | 39 | 2 | 0 | 0 | 0 | 2 | 1 |
| SG0000002 | Nek7 | 1.05E-08 | 3.18E-08 | -0.849318611 | 2623.870836 | 1456.371175 | 893 | 33 | 81 | 117 | 444 | 72 | 19 | 0 | 6 | 6 | 9 | 0 |
| SG0000002 | Sema3a | 5.73E-08 | 1.64E-07 | -1.897291017 | 169.2836642 | 45.44368536 | 1277 | 20 | 81 | 163 | 619 | 116 | 14 | 0 | 1 | 1 | 5 | 1 |
| SG0000002 | Spopl | 0.000659 | 0.001264 | -0.608740324 | 612.1771114 | 401.4467909 | 618 | 44 | 81 | 115 | 375 | 75 | 28 | 0 | 1 | 1 | 7 | 0 |
| SG0000003 | Tacc2 | 0.002138 | 0.003855 | 0.422572888 | 1433.228524 | 1920.978736 | 593 | 57 | 81 | 140 | 334 | 50 | 0 | 0 | 0 | 0 | 0 | 2 |
| SG0000005 | Kdm5d | 0.002436 | 0.004359 | -1.761105381 | 35.74833594 | 10.54654683 | 517 | 60 | 81 | 85 | 294 | 19 | 8 | 0 | 0 | 0 | 3 | 0 |
| SG0000002 | Wac | 0.003173 | 0.005576 | -0.436482683 | 2453.017225 | 1812.617705 | 765 | 36 | 81 | 142 | 394 | 73 | 19 | 0 | 1 | 2 | 2 | 5 |
| SG0000003 | Ap2b1 | 0.057006 | 0.081394 | -0.260297959 | 7254.795701 | 6057.141143 | 619 | 57 | 81 | 129 | 320 | 42 | 3 | 0 | 0 | 0 | 0 | 0 |
| SG0000002 | Slco5a1 | 0.31996 | 0.386005 | -0.267498865 | 225.2364923 | 187.117141 | 1346 | 48 | 81 | 198 | 783 | 119 | 32 | 0 | 0 | 3 | 19 | 2 |
| SG0000004 | Fam227a | 0.519563 | 0.586638 | -0.301128246 | 36.51040194 | 29.6324786 | 332 | 68 | 81 | 109 | 250 | 32 | 8 | 0 | 1 | 1 | 2 | 0 |
| SG0000002 | Hormad1 | 9.32E-40 | 1.36E-38 | 6.829112058 | 0 | 112.701861 | 409 | 64 | 80 | 96 | 235 | 20 | 4 | 0 | 0 | 0 | 4 | 0 |
| JSG000000 | Xkr6 | 1.9E-31 | 2.04E-30 | 4.12749929 | 8.529104028 | 149.0748925 | 515 | 65 | 80 | 101 | 282 | 43 | 8 | 0 | 0 | 0 | 2 | 0 |
| SG0000004 | Sin3a | 2.36E-28 | 2.22E-27 | 1.335225083 | 2574.472508 | 6495.776287 | 523 | 55 | 80 | 130 | 224 | 79 | 10 | 0 | 0 | 0 | 0 | 15 |
| SG0000007 | Mcc | 2.03E-21 | 1.37E-20 | -2.189962628 | 656.3201315 | 143.8372215 | 681 | 46 | 80 | 115 | 321 | 55 | 8 | 0 | 1 | 1 | 1 | 0 |
| SG0000002 | Prkd3 | 1.56E-17 | 8.57E-17 | -1.297675889 | 3880.306638 | 1578.434917 | 689 | 38 | 80 | 94 | 357 | 72 | 12 | 0 | 0 | 0 | 0 | 2 |
| SG0000006 | Clec16a | 1.96E-15 | 9.51E-15 | -1.381644452 | 1106.866844 | 424.7945681 | 1138 | 46 | 80 | 149 | 455 | 73 | 7 | 0 | 0 | 0 | 1 | 2 |
| SG0000002 | Chm | 2.11E-15 | 1.03E-14 | -1.280582722 | 1810.492058 | 745.2514213 | 790 | 28 | 80 | 128 | 468 | 66 | 18 | 0 | 0 | 2 | 8 | 4 |
| JSG000000 | Cacna2d3 | 5.22E-12 | 2.03E-11 | -3.893044238 | 115.5122023 | 7.775078345 | 1322 | 36 | 80 | 164 | 598 | 148 | 1 | 0 | 0 | 0 | 0 | 0 |
| SG0000004 | Hectd2 | 1.5E-08 | 4.51E-08 | 4.613308175 | 12.39773515 | 303.4495475 | 797 | 36 | 80 | 112 | 389 | 110 | 43 | 2 | 2 | 11 | 16 | 3 |
| SG0000003 | Ccpg1 | 3.55E-08 | 1.03E-07 | 1.055774769 | 171.9536434 | 357.463132 | 318 | 75 | 80 | 110 | 181 | 10 | 5 | 0 | 0 | 0 | 6 | 1 |
| SG0000003 | L3mbtl3 | 5.11E-07 | 1.35E-06 | -0.979616447 | 492.1961392 | 249.59982 | 726 | 48 | 80 | 137 | 359 | 89 | 9 | 0 | 2 | 5 | 11 | 5 |
| SG0000000 | Mid2 | 9E-07 | 2.34E-06 | -2.863431194 | 266.0362226 | 36.55628823 | 736 | 69 | 80 | 137 | 325 | 67 | 26 | 2 | 2 | 4 | 12 | 0 |
| SG0000001 | Reps1 | 0.00172 | 0.003145 | 0.420860762 | 1544.073314 | 2067.091164 | 795 | 50 | 80 | 130 | 400 | 70 | 5 | 0 | 0 | 0 | 0 | 0 |
| SG0000008 | Rps6kc1 | 0.002237 | 0.004024 | 0.47030056 | 592.5867596 | 820.9685475 | 832 | 46 | 80 | 121 | 288 | 70 | 10 | 0 | 0 | 3 | 4 | 0 |
| SG0000003 | Lrrfip2 | 0.068854 | 0.096837 | -0.276928635 | 1146.255411 | 946.0575868 | 1120 | 40 | 80 | 175 | 584 | 75 | 9 | 0 | 0 | 0 | 5 | 0 |
| SG0000002 | Rbm27 | 0.853553 | 0.885694 | 0.015560949 | 2697.377573 | 2726.629032 | 446 | 62 | 80 | 106 | 238 | 30 | 28 | 0 | 4 | 2 | 6 | 6 |
| SG0000002 | Dab2ip | 0.888315 | 0.914244 | 0.02439965 | 1239.723712 | 1260.868905 | 524 | 53 | 80 | 96 | 236 | 45 | 21 | 0 | 0 | 0 | 0 | 7 |

| | | | | | | | | | | | | | | | | | | |
|---|---|---|---|---|---|---|---|---|---|---|---|---|---|---|---|---|---|---|
| SG0000003 | Cdc14a | 4.17E-30 | 4.24E-29 | -2.460140101 | 703.7433829 | 127.8905221 | 790 | 32 | 79 | 119 | 395 | 71 | 16 | 0 | 4 | 6 | 12 | 6 |
| SG0000003 | Ddx26b | 1.05E-19 | 6.46E-19 | 3.852226964 | 40.48342779 | 584.6731416 | 504 | 67 | 79 | 128 | 300 | 51 | 2 | 0 | 0 | 2 | 2 | 4 |
| SG0000002 | Pvrl3 | 7.01E-17 | 3.73E-16 | -1.271985164 | 3142.038926 | 1301.085608 | 808 | 45 | 79 | 149 | 428 | 84 | 13 | 0 | 1 | 0 | 4 | 2 |
| SG0000002 | Ube4b | 3.64E-14 | 1.63E-13 | 0.928511362 | 3117.167176 | 5932.938761 | 440 | 76 | 79 | 115 | 243 | 39 | 6 | 0 | 0 | 2 | 4 | 2 |
| SG0000000 | Fbxo11 | 1.84E-13 | 7.88E-13 | -1.105395905 | 3340.244492 | 1552.461833 | 426 | 61 | 79 | 99 | 241 | 37 | 3 | 0 | 0 | 0 | 2 | 0 |
| JSG000000 | Tmem132d | 4.65E-10 | 1.57E-09 | 6.52563574 | 0.307509278 | 28.33154915 | 941 | 55 | 79 | 165 | 429 | 112 | 1 | 0 | 0 | 1 | 2 | 1 |
| SG0000004 | Dnal1 | 1.09E-07 | 3.04E-07 | -1.049141903 | 497.2098518 | 240.2794022 | 264 | 68 | 79 | 90 | 173 | 13 | 16 | 0 | 3 | 2 | 9 | 1 |
| SG0000003 | Zhx3 | 4.19E-06 | 1.02E-05 | -1.237338685 | 191.2014196 | 81.09891678 | 635 | 53 | 79 | 113 | 307 | 26 | 14 | 0 | 0 | 0 | 4 | 0 |
| SG0000002 | Eif2b3 | 6.03E-06 | 1.45E-05 | 0.630620398 | 968.5932706 | 1499.605771 | 376 | 67 | 79 | 100 | 232 | 34 | 6 | 2 | 2 | 2 | 2 | 0 |
| JSG000000 | Ugt1a7c | 7.38E-05 | 0.000159 | -2.770937814 | 73.16051436 | 10.71868115 | 758 | 55 | 79 | 148 | 351 | 39 | 6 | 0 | 0 | 2 | 2 | 0 |
| SG0000001 | Nbr1 | 0.000131 | 0.000275 | -0.521969734 | 6725.836135 | 4684.009156 | 378 | 79 | 79 | 113 | 240 | 9 | 35 | 0 | 0 | 7 | 14 | 0 |
| SG0000005 | Mdm4 | 0.000154 | 0.00032 | 0.660002672 | 8035.49883 | 12696.77559 | 581 | 60 | 79 | 131 | 405 | 41 | 28 | 0 | 0 | 1 | 7 | 1 |
| SG0000002 | Txndc16 | 0.000606 | 0.001167 | -0.596104718 | 659.5194697 | 436.2969935 | 908 | 51 | 79 | 158 | 441 | 60 | 26 | 0 | 1 | 3 | 9 | 0 |
| SG0000003 | Akap2 | 0.000778 | 0.00148 | -1.657475955 | 47.14035021 | 14.94317303 | 611 | 54 | 79 | 111 | 362 | 43 | 44 | 6 | 6 | 12 | 30 | 0 |
| JSG000000 | Epha3 | 0.002554 | 0.004556 | -3.045147936 | 16.31554671 | 1.976608965 | 1038 | 40 | 79 | 153 | 494 | 93 | 12 | 1 | 1 | 3 | 7 | 1 |
| SG0000003 | Fam193a | 0.283648 | 0.347209 | -0.165675296 | 1216.335777 | 1084.376875 | 314 | 68 | 79 | 95 | 175 | 28 | 6 | 2 | 3 | 3 | 4 | 2 |
| SG0000002 | Gli3 | 0.436895 | 0.506155 | -0.14708382 | 467.5546284 | 422.2364473 | 922 | 43 | 79 | 143 | 389 | 77 | 7 | 0 | 0 | 0 | 3 | 1 |
| SG0000002 | Elavl4 | 0.494941 | 0.562667 | -0.289814216 | 46.12983262 | 37.73454404 | 1248 | 35 | 79 | 195 | 583 | 69 | 63 | 7 | 21 | 14 | 28 | 7 |
| SG0000000 | Arid3b | 1.3E-182 | 3.3E-180 | 4.782643576 | 58.54646944 | 1611.46132 | 409 | 48 | 78 | 96 | 155 | 30 | 10 | 0 | 0 | 0 | 0 | 0 |
| SG0000004 | Gk5 | 1.2E-45 | 2.15E-44 | 3.101873027 | 70.55688625 | 605.7539161 | 375 | 57 | 78 | 123 | 234 | 30 | 6 | 0 | 0 | 0 | 1 | 0 |
| SG0000005 | Agap1 | 4.05E-33 | 4.65E-32 | 1.759388967 | 460.1009985 | 1557.693576 | 1223 | 41 | 78 | 149 | 451 | 124 | 17 | 0 | 0 | 4 | 13 | 2 |
| SG0000003 | Sh3gl3 | 8.55E-30 | 8.58E-29 | 2.51300827 | 56.25319716 | 321.0983541 | 770 | 23 | 78 | 104 | 326 | 72 | 9 | 0 | 0 | 0 | 0 | 0 |
| SG0000000 | Scube2 | 1.16E-29 | 1.15E-28 | 8.022923905 | 0.331601068 | 86.24951655 | 345 | 69 | 78 | 87 | 183 | 18 | 0 | 0 | 0 | 0 | 0 | 0 |
| SG0000003 | Ssh2 | 6.19E-25 | 4.99E-24 | -1.784825976 | 1695.840043 | 492.1514056 | 848 | 50 | 78 | 138 | 385 | 67 | 4 | 0 | 0 | 1 | 5 | 1 |
| SG0000002 | Uso1 | 7.45E-16 | 3.71E-15 | -1.180900322 | 5815.186727 | 2564.940208 | 456 | 63 | 78 | 117 | 273 | 36 | 14 | 0 | 0 | 4 | 7 | 0 |
| SG0000002 | Tmem116 | 1.22E-11 | 4.65E-11 | 4.154951898 | 2.6022814 | 46.35743622 | 492 | 52 | 78 | 103 | 254 | 42 | 0 | 0 | 0 | 0 | 2 | 0 |
| SG0000002 | Cald1 | 3.14E-11 | 1.16E-10 | -1.500781441 | 7759.753427 | 2742.001517 | 1191 | 36 | 78 | 126 | 457 | 105 | 27 | 0 | 0 | 6 | 16 | 0 |
| SG0000003 | Spata6 | 4.28E-11 | 1.56E-10 | -1.217718708 | 985.6768119 | 423.80364 | 867 | 45 | 78 | 153 | 442 | 82 | 11 | 0 | 0 | 3 | 5 | 0 |
| SG0000007 | Ptgr2 | 6.69E-09 | 2.06E-08 | -0.926014185 | 1427.240432 | 751.1714863 | 330 | 59 | 78 | 108 | 193 | 35 | 16 | 0 | 1 | 2 | 4 | 0 |
| SG0000002 | Thrb | 2.11E-08 | 6.24E-08 | -3.693208608 | 107.7585787 | 8.33077266 | 1402 | 31 | 78 | 174 | 611 | 113 | 12 | 0 | 0 | 1 | 2 | 0 |
| SG0000002 | Rbm39 | 1.61E-06 | 4.07E-06 | 0.585396324 | 6051.988932 | 9080.713588 | 752 | 48 | 78 | 108 | 390 | 76 | 35 | 0 | 1 | 3 | 3 | 10 |
| SG0000005 | Frem1 | 1.03E-05 | 2.44E-05 | 2.980120348 | 3.889032637 | 30.68648913 | 660 | 45 | 78 | 102 | 312 | 57 | 12 | 0 | 0 | 0 | 2 | 2 |
| SG0000000 | Abhd3 | 2.01E-05 | 4.59E-05 | -1.336790417 | 122.9587101 | 48.67940768 | 426 | 42 | 78 | 102 | 246 | 54 | 8 | 0 | 0 | 0 | 0 | 0 |
| SG0000003 | Sec24a | 4.06E-05 | 9E-05 | -0.632785555 | 1616.721193 | 1042.67197 | 484 | 58 | 78 | 116 | 265 | 51 | 8 | 0 | 0 | 0 | 7 | 1 |
| SG0000002 | Lrch3 | 0.001901 | 0.003451 | -0.48797812 | 1534.017625 | 1093.790881 | 959 | 45 | 78 | 135 | 475 | 71 | 6 | 0 | 0 | 0 | 6 | 2 |
| SG0000001 | Yes1 | 0.008025 | 0.013317 | -0.385453044 | 2260.818512 | 1730.743751 | 664 | 60 | 78 | 125 | 337 | 57 | 7 | 0 | 0 | 1 | 7 | 0 |
| SG0000002 | Galk2 | 0.099017 | 0.134863 | -0.268940346 | 731.7642994 | 607.3123411 | 529 | 55 | 78 | 95 | 260 | 36 | 5 | 0 | 0 | 0 | 4 | 0 |
| SG0000002 | Robo1 | 0.118728 | 0.158919 | 0.500075289 | 648.5965252 | 917.3018717 | 983 | 43 | 78 | 162 | 465 | 88 | 2 | 1 | 1 | 1 | 2 | 1 |
| SG0000003 | Arhgap26 | 0.838096 | 0.873602 | 0.062737873 | 232.3881093 | 242.716826 | 1088 | 39 | 78 | 153 | 501 | 100 | 6 | 0 | 0 | 0 | 2 | 3 |
| SG0000004 | Gramd1b | 9E-104 | 7E-102 | 5.553055209 | 14.0652679 | 660.365223 | 914 | 65 | 77 | 164 | 435 | 73 | 2 | 0 | 0 | 0 | 2 | 4 |
| JSG000000 | Tmtc2 | 3.77E-45 | 6.65E-44 | -4.78324663 | 897.2060214 | 32.58302993 | 977 | 36 | 77 | 122 | 419 | 83 | 8 | 2 | 3 | 3 | 5 | 1 |
| SG0000005 | Col13a1 | 4.58E-26 | 3.87E-25 | 6.173981392 | 0 | 71.20272553 | 429 | 64 | 77 | 112 | 178 | 43 | 4 | 0 | 0 | 4 | 12 | 4 |
| JSG000000 | Fancl | 6.88E-22 | 4.8E-21 | 1.527828284 | 305.8984885 | 882.0627314 | 462 | 56 | 77 | 102 | 267 | 32 | 2 | 0 | 0 | 0 | 0 | 0 |
| SG0000000 | Mprip | 2.64E-20 | 1.68E-19 | -1.353481515 | 8293.833292 | 3245.762714 | 765 | 69 | 77 | 162 | 410 | 88 | 12 | 0 | 0 | 2 | 4 | 6 |
| SG0000005 | Tanc2 | 2.86E-15 | 1.38E-14 | -1.641765285 | 1026.598837 | 328.9880866 | 858 | 47 | 77 | 143 | 444 | 84 | 9 | 0 | 0 | 1 | 5 | 1 |
| SG0000003 | Farp2 | 9.57E-14 | 4.17E-13 | -2.046925921 | 233.571209 | 56.52404164 | 381 | 54 | 77 | 98 | 207 | 22 | 0 | 0 | 0 | 0 | 0 | 0 |
| SG0000002 | Mtus2 | 1.68E-08 | 5.03E-08 | 1.77290592 | 32.12446614 | 109.7826678 | 763 | 49 | 77 | 131 | 364 | 93 | 2 | 0 | 0 | 1 | 1 | 0 |
| SG0000002 | Baz2b | 0.125187 | 0.166817 | 0.246141176 | 535.981354 | 635.6902622 | 800 | 37 | 77 | 141 | 417 | 69 | 11 | 0 | 1 | 2 | 4 | 6 |
| SG0000000 | Lnp | 0.686101 | 0.741848 | -0.07001499 | 985.2003973 | 938.5295824 | 766 | 45 | 77 | 124 | 427 | 55 | 22 | 0 | 2 | 4 | 16 | 4 |
| SG0000002 | Clcn6 | 9.27E-34 | 1.09E-32 | 1.739034091 | 449.8501261 | 1501.651905 | 196 | 72 | 76 | 80 | 126 | 8 | 1 | 0 | 0 | 0 | 2 | 1 |
| SG0000002 | Manba | 7.83E-30 | 7.88E-29 | 1.724500939 | 1070.61544 | 3538.017792 | 305 | 63 | 76 | 85 | 214 | 37 | 2 | 0 | 0 | 0 | 0 | 1 |
| SG0000003 | Nek1 | 1.22E-22 | 8.85E-22 | 1.727408437 | 716.3367122 | 2372.02337 | 720 | 48 | 76 | 108 | 354 | 66 | 18 | 0 | 0 | 2 | 10 | 0 |
| SG0000004 | Spred2 | 5.32E-20 | 3.33E-19 | -1.383410345 | 4777.117233 | 1831.124566 | 319 | 66 | 76 | 93 | 154 | 33 | 5 | 0 | 0 | 1 | 2 | 3 |
| JSG000000 | Cdc73 | 5.27E-18 | 2.96E-17 | 1.17018227 | 1787.260035 | 4022.052247 | 456 | 59 | 76 | 96 | 243 | 24 | 31 | 3 | 4 | 4 | 14 | 1 |
| SG0000002 | Nav3 | 1.12E-13 | 4.86E-13 | -3.715997226 | 88.13928566 | 6.707225344 | 905 | 37 | 76 | 159 | 499 | 126 | 0 | 0 | 0 | 0 | 0 | 1 |
| SG0000000 | Cul3 | 1.22E-13 | 5.26E-13 | 0.906979586 | 3567.242255 | 6688.991815 | 797 | 45 | 76 | 123 | 367 | 114 | 14 | 0 | 1 | 0 | 1 | 1 |
| SG0000003 | Megf11 | 2.17E-13 | 9.24E-13 | 4.824154891 | 1.666535887 | 47.2095647 | 1188 | 48 | 76 | 144 | 538 | 116 | 3 | 0 | 0 | 0 | 2 | 0 |
| SG0000002 | Mllt3 | 5.6E-12 | 2.17E-11 | -1.225209037 | 901.8822087 | 385.7670666 | 1198 | 34 | 76 | 160 | 550 | 101 | 9 | 0 | 0 | 5 | 7 | 2 |
| SG0000002 | Gabra4 | 9.65E-08 | 2.71E-07 | 4.322048837 | 0 | 19.00167391 | 855 | 56 | 76 | 139 | 575 | 101 | 7 | 0 | 0 | 1 | 2 | 1 |
| SG0000003 | Wrn | 0.000128 | 0.000267 | -0.604325354 | 1305.800842 | 858.9282481 | 770 | 44 | 76 | 124 | 424 | 90 | 7 | 0 | 2 | 0 | 3 | 1 |
| SG0000005 | Arhgap42 | 0.0002 | 0.00041 | -0.636517609 | 767.5088924 | 493.7106809 | 721 | 52 | 76 | 119 | 334 | 80 | 8 | 1 | 2 | 2 | 6 | 0 |
| SG0000002 | Enpp2 | 0.011604 | 0.018782 | 0.73633754 | 63.21482587 | 105.3121872 | 441 | 58 | 76 | 104 | 234 | 36 | 3 | 0 | 0 | 2 | 4 | 0 |
| SG0000003 | Arhgap21 | 0.0323 | 0.048217 | -0.314929352 | 2406.161435 | 1934.289972 | 1065 | 42 | 76 | 141 | 463 | 101 | 6 | 0 | 0 | 0 | 4 | 7 |
| SG0000006 | Cped1 | 0.033263 | 0.049535 | -1.036298147 | 40.66948125 | 19.82950194 | 1126 | 32 | 76 | 140 | 552 | 82 | 13 | 0 | 0 | 0 | 4 | 0 |
| SG0000003 | Snx10 | 0.078659 | 0.109381 | -0.289649844 | 1640.221794 | 1341.866427 | 414 | 71 | 76 | 119 | 290 | 26 | 16 | 0 | 0 | 0 | 0 | 4 |
| SG0000003 | Lrrc28 | 0.079178 | 0.110052 | -0.314833221 | 490.3457403 | 394.2104781 | 806 | 36 | 76 | 131 | 422 | 87 | 6 | 0 | 0 | 0 | 2 | 2 |
| SG0000000 | Xpo6 | 0.862077 | 0.892613 | -0.023750828 | 3772.911046 | 3711.306773 | 562 | 65 | 76 | 111 | 297 | 46 | 2 | 0 | 0 | 0 | 0 | 0 |
| JSG000000 | Arl15 | 2.18E-48 | 4.34E-47 | 2.151153371 | 287.4976679 | 1277.015051 | 693 | 54 | 75 | 121 | 336 | 64 | 9 | 1 | 1 | 3 | 5 | 2 |
| SG0000002 | Mapk4 | 4.59E-46 | 8.34E-45 | 6.122270174 | 2.305646233 | 160.6126087 | 546 | 66 | 75 | 111 | 275 | 41 | 6 | 0 | 0 | 0 | 2 | 2 |
| SG0000003 | Whrn | 2.96E-24 | 2.3E-23 | 3.507859913 | 33.18970413 | 377.5499659 | 784 | 66 | 75 | 179 | 350 | 73 | 36 | 21 | 21 | 35 | 35 | 2 |
| SG0000002 | Gfm2 | 8.28E-21 | 5.45E-20 | 1.24006167 | 899.4974095 | 2124.6905 | 469 | 57 | 75 | 99 | 228 | 21 | 7 | 0 | 2 | 0 | 3 | 0 |
| SG0000002 | Cers6 | 3.08E-20 | 1.96E-19 | 2.752509393 | 122.792258 | 827.482611 | 958 | 37 | 75 | 149 | 470 | 99 | 8 | 1 | 1 | 1 | 4 | 1 |
| JSG000000 | Enox1 | 6.39E-20 | 3.99E-19 | 3.864203097 | 50.3763987 | 733.6149261 | 1011 | 43 | 75 | 144 | 465 | 93 | 0 | 0 | 0 | 0 | 0 | 0 |
| SG0000004 | S100pbp | 3.39E-13 | 1.42E-12 | 0.929323775 | 1616.29653 | 3078.047891 | 436 | 43 | 75 | 92 | 256 | 44 | 13 | 0 | 4 | 0 | 8 | 0 |
| SG0000000 | Ccdc132 | 6.97E-08 | 1.98E-07 | -0.809105689 | 2191.187437 | 1250.588547 | 864 | 45 | 75 | 141 | 384 | 69 | 3 | 0 | 0 | 0 | 3 | 0 |
| SG0000007 | Rasal2 | 2.03E-06 | 5.11E-06 | -0.742337318 | 1422.267363 | 850.1889277 | 768 | 37 | 75 | 119 | 404 | 80 | 9 | 0 | 0 | 2 | 13 | 4 |
| JSG000000 | Adsl | 2.45E-05 | 5.54E-05 | 0.547157399 | 2041.815229 | 2983.508148 | 167 | 70 | 75 | 80 | 124 | 14 | 2 | 0 | 0 | 0 | 0 | 0 |
| SG0000003 | Nup205 | 0.00259 | 0.004617 | 0.372976711 | 5451.976541 | 7060.430072 | 573 | 60 | 75 | 123 | 309 | 66 | 0 | 0 | 0 | 0 | 0 | 0 |
| SG0000003 | Cfap70 | 0.002625 | 0.004673 | 5.341039488 | 0.615018556 | 24.92879655 | 363 | 60 | 75 | 108 | 204 | 24 | 3 | 0 | 0 | 0 | 0 | 0 |
| SG0000005 | Cryzl1 | 0.125353 | 0.167013 | 0.216496155 | 917.089677 | 1065.574065 | 395 | 55 | 75 | 83 | 191 | 34 | 10 | 0 | 2 | 1 | 1 | 0 |
| SG0000005 | Adam10 | 9.94E-25 | 7.91E-24 | -1.640052838 | 3407.595785 | 1093.309127 | 606 | 46 | 74 | 120 | 319 | 38 | 5 | 0 | 0 | 0 | 7 | 2 |
| SG0000003 | Zfp292 | 3.24E-22 | 2.29E-21 | 1.272651832 | 1139.124881 | 2752.18557 | 450 | 52 | 74 | 96 | 228 | 36 | 16 | 2 | 2 | 6 | 12 | 8 |
| SG0000003 | Ibtk | 4.99E-15 | 2.37E-14 | 1.00715413 | 1382.766815 | 2779.281596 | 316 | 62 | 74 | 88 | 210 | 12 | 4 | 0 | 0 | 0 | 0 | 0 |
| SG0000002 | Strn3 | 5.85E-14 | 2.58E-13 | -1.094353507 | 4600.181232 | 2154.476248 | 568 | 36 | 74 | 70 | 256 | 52 | 2 | 0 | 0 | 0 | 0 | 0 |
| SG0000002 | Dapk1 | 5.61E-11 | 2.03E-10 | 1.033071137 | 479.1415674 | 980.503728 | 418 | 58 | 74 | 114 | 198 | 50 | 4 | 0 | 0 | 0 | 0 | 2 |
| SG0000002 | Ptpra | 2.36E-08 | 6.98E-08 | -0.845062036 | 2042.126301 | 1136.8249 | 418 | 64 | 74 | 116 | 248 | 38 | 0 | 0 | 0 | 0 | 0 | 4 |
| SG0000003 | Rbfox2 | 8.96E-08 | 2.52E-07 | -0.751822075 | 5390.194747 | 3200.983683 | 1036 | 42 | 74 | 154 | 412 | 78 | 12 | 0 | 0 | 0 | 5 | 3 |
| SG0000003 | Nrg4 | 1.68E-07 | 4.62E-07 | -1.62782723 | 140.2111523 | 45.36884618 | 697 | 36 | 74 | 113 | 292 | 47 | 2 | 0 | 1 | 1 | 1 | 0 |
| SG0000003 | Atp7a | 5.03E-06 | 1.22E-05 | -0.995575618 | 736.4875771 | 369.3748329 | 474 | 50 | 74 | 124 | 286 | 50 | 0 | 0 | 0 | 0 | 0 | 0 |
| SG0000002 | Nell2 | 9.46E-06 | 2.23E-05 | 3.7607901 | 1.631569985 | 22.1164967 | 1250 | 38 | 74 | 176 | 732 | 98 | 0 | 0 | 0 | 0 | 0 | 0 |
| SG0000002 | Atp8a2 | 1.85E-05 | 4.25E-05 | 4.258320127 | 2.872481209 | 54.97171491 | 1021 | 45 | 74 | 138 | 417 | 101 | 0 | 0 | 0 | 0 | 1 | 0 |
| SG0000006 | Csde1 | 0.000316 | 0.000632 | 0.43919499 | 8677.77801 | 11765.74419 | 773 | 58 | 74 | 104 | 319 | 45 | 58 | 0 | 0 | 0 | 0 | 0 |
| SG0000004 | Prkce | 0.003229 | 0.005669 | -0.660996688 | 316.464845 | 200.1454137 | 1189 | 48 | 74 | 159 | 508 | 138 | 5 | 2 | 2 | 2 | 4 | 1 |
| SG0000004 | Plekha1 | 0.110274 | 0.148748 | -0.223942728 | 2395.678928 | 2051.233659 | 549 | 46 | 74 | 101 | 286 | 68 | 27 | 0 | 0 | 4 | 14 | 5 |
| JSG000000 | Tbc1d32 | 0.240473 | 0.300353 | -0.226856781 | 482.0272318 | 411.8896493 | 550 | 44 | 74 | 90 | 286 | 53 | 12 | 0 | 1 | 0 | 4 | 0 |
| SG0000002 | Gls | 0.476444 | 0.544959 | 0.088368986 | 5533.492897 | 5883.030127 | 803 | 31 | 74 | 102 | 314 | 68 | 47 | 0 | 5 | 3 | 8 | 0 |
| SG0000007 | Nbeal1 | 0.853621 | 0.885712 | -0.038011153 | 1263.163122 | 1230.316761 | 688 | 49 | 74 | 102 | 333 | 58 | 43 | 0 | 1 | 3 | 10 | 4 |
| SG0000002 | Pecam1 | 1.1E-289 | 7.1E-287 | 7.396806481 | 11.12669829 | 1875.110799 | 397 | 56 | 73 | 116 | 260 | 82 | 8 | 4 | 4 | 8 | 12 | 0 |
| SG0000003 | Plcl1 | 8.46E-60 | 2.33E-58 | -5.494668222 | 432.5096941 | 9.592590273 | 783 | 39 | 73 | 115 | 334 | 65 | 17 | 0 | 1 | 3 | 10 | 1 |
| SG0000003 | 2700049A03Rik | 8.76E-15 | 4.08E-14 | 1.070478016 | 784.6524059 | 1647.87135 | 494 | 43 | 73 | 65 | 224 | 55 | 1 | 0 | 0 | 0 | 1 | 0 |
| SG0000004 | Mipol1 | 1.1E-13 | 4.77E-13 | 1.310560169 | 198.930414 | 493.4226807 | 774 | 29 | 73 | 109 | 416 | 73 | 1 | 0 | 0 | 1 | 2 | 0 |
| JSG000000 | Wdr70 | 2.98E-13 | 1.25E-12 | 1.058105488 | 559.2365936 | 1164.439884 | 531 | 54 | 73 | 110 | 294 | 45 | 0 | 0 | 0 | 0 | 0 | 0 |
| SG0000005 | Palld | 8.78E-12 | 3.37E-11 | 2.116970618 | 161.5599694 | 700.818275 | 1215 | 43 | 73 | 149 | 446 | 121 | 0 | 0 | 0 | 0 | 0 | 0 |
| SG0000002 | Tm2d1 | 2.72E-08 | 7.98E-08 | -1.147807923 | 436.5109179 | 197.001937 | 373 | 63 | 73 | 113 | 259 | 34 | 17 | 0 | 0 | 0 | 1 | 0 |
| SG0000003 | Phf20 | 1.82E-07 | 4.99E-07 | 0.663862982 | 2025.921367 | 3209.70601 | 221 | 59 | 73 | 80 | 153 | 19 | 8 | 0 | 1 | 0 | 0 | 1 |
| JSG000000 | Gm7173 | 9.11E-06 | 2.16E-05 | 4.237321212 | 1.016551429 | 19.17300303 | 884 | 38 | 73 | 145 | 537 | 85 | 0 | 0 | 0 | 0 | 0 | 0 |
| SG0000007 | Rc3h2 | 6.41E-05 | 0.000139 | -0.625255814 | 1924.448887 | 1247.629473 | 656 | 39 | 73 | 112 | 306 | 42 | 82 | 4 | 8 | 12 | 27 | 5 |
| SG0000003 | Xrn1 | 0.000471 | 0.000921 | 0.46582575 | 1404.899906 | 1940.317975 | 581 | 56 | 73 | 121 | 348 | 51 | 11 | 1 | 1 | 1 | 5 | 1 |
| SG0000003 | Ubn2 | 0.000978 | 0.001836 | -0.502336806 | 1579.48864 | 1115.059547 | 594 | 40 | 73 | 109 | 301 | 67 | 18 | 0 | 1 | 2 | 5 | 7 |
| SG0000004 | Dnah7b | 0.2625 | 0.324291 | -0.685794362 | 34.74146506 | 21.59749863 | 724 | 46 | 73 | 117 | 400 | 44 | 1 | 0 | 0 | 0 | 0 | 0 |
| SG0000005 | Nudcd3 | 0.561264 | 0.626194 | -0.077268419 | 2931.703184 | 2778.816614 | 308 | 64 | 73 | 98 | 202 | 15 | 4 | 0 | 0 | 0 | 0 | 0 |
| SG0000002 | Pex14 | 0.598451 | 0.661286 | -0.130119058 | 1059.309056 | 967.9500471 | 421 | 51 | 73 | 85 | 241 | 37 | 2 | 0 | 0 | 0 | 0 | 1 |
| SG0000002 | Nfasc | 1.3E-140 | 1.8E-138 | 8.703719049 | 1.289094805 | 537.4837834 | 673 | 45 | 72 | 129 | 352 | 137 | 3 | 0 | 0 | 0 | 0 | 1 |
| SG0000005 | Tln2 | 3.66E-78 | 1.59E-76 | 2.899254612 | 197.5332686 | 1473.678859 | 891 | 30 | 72 | 130 | 397 | 82 | 5 | 0 | 0 | 1 | 2 | 0 |
| SG0000002 | Hdac7 | 7.39E-43 | 1.2E-41 | -2.90376924 | 2429.776046 | 324.6718268 | 317 | 68 | 72 | 88 | 237 | 42 | 0 | 0 | 0 | 0 | 0 | 0 |
| SG0000003 | Mlxip | 3.2E-15 | 1.54E-14 | -1.234517603 | 3165.736488 | 1345.389188 | 223 | 66 | 72 | 85 | 146 | 17 | 3 | 0 | 0 | 0 | 0 | 0 |
| SG0000004 | Zfp423 | 2.49E-12 | 9.85E-12 | 1.31611765 | 179.4748797 | 446.8837485 | 683 | 50 | 72 | 102 | 321 | 56 | 0 | 0 | 0 | 0 | 6 | 0 |
| JSG000000 | Sema5a | 1.67E-11 | 6.3E-11 | 3.016888886 | 19.35432688 | 156.6578349 | 922 | 38 | 72 | 139 | 465 | 95 | 6 | 0 | 0 | 1 | 4 | 1 |
| JSG000000 | Fras1 | 1.27E-10 | 4.47E-10 | 4.814352372 | 1.289094805 | 36.27014833 | 760 | 42 | 72 | 125 | 380 | 53 | 1 | 0 | 0 | 0 | 1 | 0 |
| SG0000003 | Hdx | 1.56E-10 | 5.48E-10 | 3.798894289 | 12.01041682 | 167.1626361 | 826 | 48 | 72 | 144 | 392 | 82 | 36 | 0 | 4 | 2 | 14 | 0 |
| SG0000002 | Gk | 3E-09 | 9.5E-09 | -1.004735432 | 1045.830462 | 521.2016534 | 776 | 60 | 72 | 148 | 500 | 80 | 28 | 0 | 0 | 0 | 16 | 8 |
| SG0000002 | Csnk1a1 | 3.77E-08 | 1.1E-07 | 0.661545682 | 8442.811467 | 13354.64 | 515 | 50 | 72 | 68 | 169 | 49 | 14 | 0 | 1 | 0 | 5 | 2 |
| SG0000003 | Chrdl1 | 1.33E-07 | 3.7E-07 | -3.837504455 | 117.801478 | 8.240371474 | 887 | 32 | 72 | 99 | 399 | 102 | 17 | 0 | 1 | 5 | 11 | 4 |

| | | | | | | | | | | | | | | | | | | |
|---|---|---|---|---|---|---|---|---|---|---|---|---|---|---|---|---|---|---|
| JSG000000 | Slc35f1 | 2.04E-07 | 5.55E-07 | 3.904912731 | 1.939079263 | 29.04634051 | 939 | 43 | 72 | 145 | 516 | 85 | 8 | 0 | 0 | 1 | 2 | 1 |
| SG0000002 | Ncoa3 | 1.05E-06 | 2.72E-06 | 0.61726932 | 2475.900807 | 3797.955066 | 325 | 58 | 72 | 84 | 184 | 21 | 7 | 0 | 0 | 0 | 4 | 5 |
| JSG000000 | Spidr | 5.17E-06 | 1.25E-05 | 0.632479099 | 1189.697451 | 1844.300617 | 523 | 51 | 72 | 90 | 314 | 44 | 2 | 0 | 0 | 0 | 1 | 0 |
| SG0000004 | Golph3l | 0.000475 | 0.000929 | -0.50301884 | 2077.693966 | 1466.080507 | 552 | 40 | 72 | 112 | 268 | 48 | 24 | 0 | 0 | 0 | 12 | 0 |
| SG0000002 | Smoc1 | 0.008629 | 0.014266 | -0.536246031 | 340.7577282 | 234.9738722 | 702 | 51 | 72 | 126 | 408 | 69 | 3 | 0 | 0 | 0 | 0 | 3 |
| SG0000003 | Rnf111 | 0.020288 | 0.03134 | -0.336988263 | 1712.716377 | 1355.944379 | 486 | 58 | 72 | 126 | 284 | 42 | 14 | 0 | 0 | 4 | 6 | 2 |
| SG0000002 | Nup210l | 0.252013 | 0.312932 | 0.483811824 | 26.93312102 | 37.66418322 | 258 | 66 | 72 | 101 | 163 | 22 | 0 | 0 | 0 | 0 | 0 | 0 |
| SG0000003 | D930015E06Rik | 0.642787 | 0.702563 | -0.101954983 | 1191.744756 | 1110.431419 | 684 | 48 | 72 | 105 | 270 | 48 | 0 | 0 | 0 | 0 | 0 | 0 |
| SG0000004 | Zfp532 | 0.747424 | 0.795902 | -0.054675263 | 2048.514042 | 1972.332137 | 701 | 44 | 72 | 84 | 280 | 49 | 9 | 0 | 0 | 4 | 4 | 13 |
| SG0000003 | Fam126b | 0.789551 | 0.832551 | -0.049781268 | 652.8371088 | 630.6946948 | 646 | 43 | 72 | 105 | 309 | 53 | 39 | 0 | 1 | 1 | 10 | 6 |
| JSG000000 | 9130011E15Rik | 0.823016 | 0.861051 | -0.038739623 | 679.7917905 | 661.7807449 | 344 | 56 | 72 | 88 | 195 | 20 | 0 | 0 | 0 | 0 | 0 | 0 |
| SG0000003 | Usp3 | 0.878088 | 0.906218 | 0.016853355 | 861.5458719 | 871.6693416 | 725 | 32 | 72 | 104 | 396 | 68 | 11 | 0 | 0 | 0 | 0 | 0 |
| SG0000003 | Rabgap1 | 0.991771 | 1 | -0.009181054 | 2028.603579 | 2015.734898 | 870 | 46 | 72 | 138 | 394 | 65 | 6 | 0 | 0 | 1 | 2 | 6 |
| SG0000002 | Cpeb1 | 2.81E-63 | 8.5E-62 | 3.665729182 | 35.33089198 | 448.3829002 | 631 | 26 | 71 | 92 | 379 | 60 | 10 | 2 | 4 | 2 | 4 | 2 |
| SG0000002 | Abcc5 | 4.18E-49 | 8.43E-48 | -2.437154878 | 9845.415281 | 1817.930672 | 404 | 65 | 71 | 102 | 300 | 35 | 6 | 0 | 0 | 2 | 2 | 2 |
| SG0000003 | Mecp2 | 7.42E-37 | 9.82E-36 | -2.176355825 | 2935.514472 | 649.4352253 | 536 | 51 | 71 | 124 | 318 | 49 | 22 | 0 | 0 | 2 | 8 | 2 |
| JSG000000 | Nop58 | 3.36E-31 | 3.57E-30 | 1.37814641 | 4374.201509 | 11370.04527 | 351 | 64 | 71 | 103 | 213 | 22 | 18 | 3 | 3 | 3 | 7 | 1 |
| SG0000000 | Kat2b | 1.19E-25 | 9.89E-25 | -1.666338099 | 3491.435825 | 1099.983861 | 322 | 59 | 71 | 87 | 211 | 16 | 10 | 0 | 0 | 0 | 2 | 0 |
| JSG000000 | Dock4 | 2.63E-20 | 1.68E-19 | -1.870035826 | 709.7553893 | 194.1653913 | 814 | 36 | 71 | 113 | 381 | 79 | 2 | 0 | 0 | 1 | 1 | 2 |
| SG0000003 | Pde4dip | 2.59E-09 | 8.24E-09 | -0.941010824 | 1464.379389 | 762.7480802 | 707 | 65 | 71 | 129 | 412 | 58 | 6 | 1 | 1 | 1 | 6 | 0 |
| SG0000003 | Vezt | 5.19E-07 | 1.37E-06 | -0.89901683 | 963.1453937 | 516.4886943 | 863 | 44 | 71 | 125 | 351 | 87 | 35 | 1 | 1 | 2 | 10 | 2 |
| SG0000000 | Fbxo7 | 3.65E-06 | 8.96E-06 | 0.747592717 | 394.3315718 | 662.0783434 | 286 | 58 | 71 | 84 | 194 | 9 | 5 | 1 | 1 | 1 | 1 | 0 |
| SG0000002 | Brwd1 | 6.96E-06 | 1.66E-05 | 0.610844051 | 1708.656953 | 2609.379584 | 440 | 48 | 71 | 92 | 190 | 50 | 12 | 3 | 5 | 5 | 6 | 0 |
| SG0000002 | Spata13 | 1.12E-05 | 2.63E-05 | -0.92868168 | 2920.578976 | 1534.291618 | 412 | 62 | 71 | 97 | 204 | 30 | 18 | 0 | 0 | 0 | 0 | 0 |
| SG0000003 | Col12a1 | 0.00041 | 0.000809 | -1.881131009 | 1123.094359 | 304.8873363 | 627 | 51 | 71 | 125 | 372 | 58 | 7 | 2 | 2 | 3 | 6 | 1 |
| SG0000002 | Antxr2 | 0.012691 | 0.020363 | -1.084884778 | 91.83358115 | 43.29309759 | 710 | 38 | 71 | 115 | 324 | 52 | 11 | 0 | 1 | 2 | 5 | 2 |
| SG0000002 | Kif2a | 0.144986 | 0.190548 | 0.185723107 | 2202.677522 | 2505.296583 | 722 | 43 | 71 | 85 | 310 | 94 | 12 | 0 | 2 | 0 | 1 | 3 |
| JSG000000 | Atg10 | 0.208813 | 0.264872 | -0.323679092 | 156.6148419 | 125.1398992 | 660 | 42 | 71 | 117 | 352 | 81 | 1 | 0 | 0 | 0 | 0 | 0 |
| SG0000001 | Rnf170 | 0.38793 | 0.456628 | 0.147147652 | 357.4646242 | 395.8484295 | 291 | 61 | 71 | 103 | 172 | 31 | 15 | 0 | 0 | 0 | 7 | 0 |
| SG0000003 | Gpm6b | 0.398297 | 0.467179 | -0.328455767 | 60.39152511 | 48.09511408 | 1387 | 39 | 71 | 170 | 621 | 137 | 57 | 0 | 0 | 9 | 9 | 12 |
| SG0000001 | Taok1 | 9.22E-19 | 5.4E-18 | -1.418711203 | 5629.142564 | 2105.559994 | 430 | 41 | 70 | 91 | 231 | 32 | 56 | 0 | 6 | 6 | 20 | 10 |
| SG0000005 | Aak1 | 1.23E-15 | 6.09E-15 | -1.235502164 | 2892.214074 | 1228.307841 | 502 | 54 | 70 | 90 | 268 | 60 | 6 | 0 | 0 | 0 | 4 | 0 |
| SG0000007 | Sms | 6.64E-14 | 2.92E-13 | -1.122594245 | 3783.279937 | 1737.536501 | 268 | 62 | 70 | 81 | 144 | 12 | 8 | 0 | 0 | 1 | 6 | 0 |
| SG0000002 | Mrpl1 | 2.05E-12 | 8.16E-12 | 0.991225691 | 668.4627778 | 1328.8192 | 378 | 60 | 70 | 99 | 198 | 30 | 14 | 2 | 2 | 2 | 8 | 0 |
| SG0000002 | Atp13a3 | 1.51E-11 | 5.71E-11 | -0.950802364 | 9158.094177 | 4737.891095 | 496 | 54 | 70 | 100 | 240 | 30 | 24 | 2 | 2 | 8 | 14 | 4 |
| SG0000002 | Tmem131 | 3.38E-07 | 9.06E-07 | 0.646966549 | 3156.145717 | 4942.121105 | 564 | 54 | 70 | 120 | 261 | 31 | 3 | 0 | 0 | 0 | 3 | 0 |
| JSG000000 | Ptprz1 | 9.97E-07 | 2.58E-06 | 4.317228932 | 2.850732985 | 56.82925301 | 1202 | 30 | 70 | 139 | 549 | 128 | 8 | 0 | 0 | 1 | 6 | 4 |
| SG0000005 | Klhl18 | 1.63E-06 | 4.14E-06 | -0.968554852 | 409.4057043 | 209.2135586 | 317 | 54 | 70 | 84 | 207 | 12 | 9 | 0 | 4 | 4 | 4 | 0 |
| SG0000002 | Pfkfb2 | 1.72E-05 | 3.97E-05 | 1.233382045 | 226.3866108 | 532.2745842 | 471 | 39 | 70 | 125 | 215 | 15 | 13 | 0 | 0 | 3 | 15 | 0 |
| SG0000000 | Cdk12 | 0.0005 | 0.000977 | -0.50065576 | 2554.102835 | 1805.202714 | 481 | 49 | 70 | 98 | 263 | 28 | 7 | 0 | 0 | 0 | 1 | 4 |
| SG0000001 | Rhobtb1 | 0.000513 | 0.001 | 1.555078086 | 19.70298904 | 57.89716058 | 524 | 32 | 70 | 52 | 281 | 73 | 9 | 0 | 0 | 0 | 4 | 0 |
| SG0000005 | Tcf7l1 | 0.000663 | 0.00127 | 0.53146994 | 1104.554898 | 1596.524911 | 418 | 48 | 70 | 84 | 174 | 27 | 2 | 0 | 0 | 0 | 0 | 1 |
| SG0000005 | Ebf4 | 0.001025 | 0.001919 | 1.904751221 | 8.265091197 | 30.94816487 | 615 | 40 | 70 | 120 | 345 | 70 | 0 | 0 | 0 | 0 | 0 | 0 |
| SG0000003 | Pik3c3 | 0.003953 | 0.006856 | 0.400930812 | 1011.684108 | 1335.786742 | 609 | 47 | 70 | 101 | 261 | 55 | 2 | 0 | 0 | 1 | 1 | 3 |
| SG0000007 | Gm11744 | 0.010142 | 0.016571 | 1.735963386 | 6.288702465 | 20.94778453 | 121 | 67 | 70 | 74 | 82 | 4 | 1 | 0 | 0 | 0 | 0 | 0 |
| SG0000003 | Ttll7 | 0.035826 | 0.053012 | -0.637746689 | 105.7890675 | 67.99232715 | 622 | 46 | 70 | 115 | 329 | 62 | 29 | 0 | 0 | 7 | 9 | 4 |
| SG0000004 | Atxn2 | 0.052958 | 0.076082 | 0.250401395 | 2398.966991 | 2853.662467 | 426 | 49 | 70 | 99 | 206 | 34 | 11 | 3 | 6 | 6 | 9 | 3 |
| SG0000003 | Otud5 | 0.265509 | 0.327659 | -0.155779009 | 2383.472127 | 2139.517894 | 291 | 65 | 70 | 85 | 205 | 10 | 6 | 0 | 0 | 0 | 0 | 0 |
| SG0000002 | Nceh1 | 0.33449 | 0.401271 | -0.167268945 | 565.9136457 | 503.9613101 | 346 | 51 | 70 | 80 | 167 | 30 | 9 | 0 | 1 | 1 | 4 | 2 |
| SG0000004 | Uspl1 | 0.706414 | 0.759774 | -0.056695358 | 1866.367158 | 1794.444705 | 525 | 35 | 70 | 84 | 189 | 49 | 1 | 0 | 0 | 0 | 0 | 0 |
| JSG000000 | Ascc3 | 0.990122 | 0.99944 | -0.007625866 | 2267.37283 | 2255.419463 | 796 | 32 | 70 | 109 | 395 | 79 | 3 | 0 | 0 | 0 | 0 | 0 |
| JSG000000 | Gm13225 | 9.6E-276 | 5.9E-273 | 8.103775568 | 5.91360495 | 1626.791268 | 211 | 63 | 69 | 79 | 155 | 18 | 8 | 0 | 0 | 2 | 2 | 2 |
| SG0000002 | Rragd | 3.4E-154 | 5.5E-152 | 4.800441981 | 40.79628032 | 1136.835419 | 267 | 51 | 69 | 87 | 186 | 21 | 9 | 0 | 0 | 3 | 9 | 3 |
| SG0000005 | Prkca | 1.01E-44 | 1.75E-43 | -3.085028582 | 1500.690585 | 176.8499809 | 683 | 44 | 69 | 103 | 267 | 51 | 12 | 0 | 2 | 2 | 7 | 2 |
| SG0000003 | Zwilch | 1.56E-44 | 2.69E-43 | 1.727089567 | 1406.393165 | 4655.99507 | 376 | 52 | 69 | 112 | 216 | 27 | 2 | 0 | 0 | 1 | 4 | 0 |
| SG0000003 | Tmtc1 | 8.86E-37 | 1.17E-35 | 5.192244292 | 5.972662642 | 218.3681333 | 1117 | 18 | 69 | 108 | 413 | 102 | 30 | 2 | 2 | 4 | 8 | 4 |
| SG0000002 | Lats2 | 1.25E-30 | 1.3E-29 | -2.191935494 | 1187.22412 | 259.8330427 | 273 | 58 | 69 | 83 | 149 | 4 | 10 | 2 | 2 | 2 | 7 | 1 |
| SG0000002 | Cdk5rap1 | 8.72E-18 | 4.85E-17 | 1.324127669 | 348.9860473 | 873.7963178 | 239 | 59 | 69 | 69 | 120 | 13 | 0 | 0 | 0 | 0 | 0 | 0 |
| SG0000000 | Mef2d | 4.06E-16 | 2.06E-15 | -1.216637089 | 3869.580039 | 1665.020468 | 134 | 65 | 69 | 78 | 96 | 2 | 2 | 1 | 1 | 2 | 16 | 2 |
| SG0000003 | Gas2 | 3.87E-09 | 1.21E-08 | 2.227797165 | 82.01538232 | 384.1749866 | 776 | 39 | 69 | 123 | 435 | 103 | 10 | 0 | 0 | 3 | 3 | 0 |
| SG0000004 | Elmo1 | 7.17E-09 | 2.2E-08 | -1.507676736 | 2940.365056 | 1034.05903 | 1304 | 30 | 69 | 143 | 515 | 84 | 1 | 0 | 0 | 0 | 1 | 0 |
| SG0000002 | 2010111I01Rik | 1.78E-08 | 5.32E-08 | -0.987024877 | 809.553005 | 408.4333484 | 821 | 45 | 69 | 118 | 344 | 66 | 3 | 0 | 0 | 0 | 0 | 0 |
| SG0000002 | Ola1 | 1.99E-08 | 5.91E-08 | 0.699130719 | 2851.78391 | 4629.946065 | 839 | 31 | 69 | 118 | 469 | 68 | 1 | 0 | 0 | 0 | 2 | 0 |
| SG0000003 | Bbs7 | 0.000424 | 0.000835 | 0.712163579 | 198.1807564 | 324.6714352 | 337 | 60 | 69 | 91 | 223 | 18 | 4 | 0 | 0 | 0 | 4 | 0 |
| JSG000000 | Ctnna1 | 0.002643 | 0.004704 | 0.369067455 | 11294.77697 | 14587.40478 | 372 | 41 | 69 | 85 | 200 | 36 | 1 | 0 | 1 | 0 | 1 | 0 |
| SG0000002 | Tom1l1 | 0.013156 | 0.021049 | 0.36384497 | 776.2506474 | 998.918888 | 330 | 57 | 69 | 93 | 190 | 28 | 14 | 0 | 0 | 0 | 7 | 0 |
| JSG000000 | Wdr7 | 0.01998 | 0.030919 | -0.391006409 | 1027.350173 | 783.4546113 | 675 | 44 | 69 | 131 | 386 | 68 | 4 | 0 | 0 | 0 | 1 | 2 |
| SG0000003 | Lpar1 | 0.099494 | 0.135457 | 0.486621806 | 284.6299609 | 398.812143 | 1095 | 23 | 69 | 150 | 578 | 102 | 40 | 0 | 0 | 0 | 10 | 5 |
| SG0000004 | Kdm5b | 1.3E-141 | 1.8E-139 | 2.970615487 | 2170.136252 | 17011.05971 | 355 | 48 | 68 | 89 | 228 | 14 | 3 | 0 | 0 | 1 | 2 | 0 |
| SG0000002 | Fam3c | 1.62E-36 | 2.12E-35 | 1.679314356 | 813.5074264 | 2605.46663 | 427 | 52 | 68 | 103 | 236 | 34 | 12 | 4 | 4 | 7 | 8 | 6 |
| JSG000000 | Sema3e | 5.04E-29 | 4.87E-28 | -2.293063057 | 2502.596691 | 510.6344414 | 695 | 44 | 68 | 107 | 375 | 56 | 8 | 0 | 0 | 2 | 7 | 0 |
| SG0000004 | Wdfy3 | 5.83E-29 | 5.62E-28 | -1.813839033 | 3466.282782 | 985.9253509 | 640 | 45 | 68 | 113 | 323 | 70 | 11 | 0 | 2 | 0 | 0 | 4 |
| SG0000001 | Sptbn4 | 5.02E-24 | 3.87E-23 | 1.941026649 | 112.5073912 | 432.0045272 | 325 | 61 | 68 | 84 | 272 | 59 | 0 | 0 | 0 | 0 | 0 | 0 |
| SG0000005 | Ankrd44 | 1.87E-14 | 8.53E-14 | -2.770741064 | 1142.327495 | 167.3841994 | 1045 | 36 | 68 | 150 | 496 | 69 | 10 | 0 | 0 | 2 | 10 | 0 |
| SG0000003 | Pccb | 2.26E-14 | 1.03E-13 | 1.259691438 | 271.7511466 | 650.6932377 | 228 | 66 | 68 | 76 | 156 | 18 | 0 | 0 | 0 | 0 | 0 | 0 |
| SG0000002 | Phactr3 | 2.83E-13 | 1.19E-12 | 6.928937541 | 0.307509278 | 37.46935885 | 893 | 43 | 68 | 131 | 355 | 185 | 12 | 1 | 1 | 1 | 6 | 0 |
| SG0000002 | Ogdh | 2.74E-10 | 9.43E-10 | -0.885520499 | 6855.209458 | 3710.671401 | 516 | 64 | 68 | 104 | 256 | 44 | 12 | 4 | 4 | 4 | 4 | 0 |
| SG0000002 | Pigx | 4.96E-08 | 1.43E-07 | 0.939360651 | 272.2293357 | 522.0482642 | 223 | 49 | 68 | 70 | 154 | 28 | 2 | 0 | 0 | 0 | 0 | 0 |
| SG0000005 | Ccser2 | 1.37E-05 | 3.18E-05 | 0.655374023 | 1218.38351 | 1918.98355 | 1171 | 17 | 68 | 128 | 416 | 65 | 24 | 0 | 2 | 8 | 12 | 0 |
| SG0000001 | Pcmt1 | 2.49E-05 | 5.63E-05 | -0.635082888 | 1680.625793 | 1082.161367 | 493 | 53 | 68 | 80 | 175 | 24 | 19 | 2 | 5 | 4 | 4 | 3 |
| SG0000005 | Srcap | 6.84E-05 | 0.000148 | -0.963980769 | 218.6531575 | 112.0904536 | 283 | 53 | 68 | 73 | 132 | 20 | 2 | 0 | 1 | 0 | 5 | 2 |
| SG0000002 | Glrb | 0.000154 | 0.000321 | 1.437442138 | 27.07817474 | 73.33859524 | 786 | 20 | 68 | 100 | 409 | 63 | 10 | 2 | 2 | 3 | 9 | 0 |
| SG0000002 | Ubap2 | 0.000177 | 0.000365 | 0.466897643 | 4421.380784 | 6110.941341 | 356 | 56 | 68 | 110 | 216 | 34 | 4 | 0 | 0 | 2 | 2 | 0 |
| SG0000002 | Papss1 | 0.000718 | 0.00137 | -0.620752519 | 1302.111156 | 846.8040602 | 539 | 37 | 68 | 79 | 240 | 57 | 11 | 0 | 0 | 4 | 4 | 0 |
| SG0000002 | Carf | 0.003481 | 0.006088 | 0.561846904 | 244.5553503 | 361.0022758 | 623 | 48 | 68 | 118 | 440 | 49 | 30 | 0 | 0 | 5 | 19 | 3 |
| SG0000007 | Syndig1 | 0.005816 | 0.009857 | 1.84655276 | 7.833279557 | 28.17159235 | 602 | 44 | 68 | 104 | 304 | 58 | 2 | 0 | 0 | 0 | 0 | 0 |
| SG0000004 | Agbl2 | 0.037307 | 0.055044 | 4.133770585 | 0.922527834 | 16.19453266 | 384 | 67 | 68 | 102 | 225 | 69 | 6 | 0 | 0 | 3 | 6 | 5 |
| SG0000000 | Dlg3 | 0.042848 | 0.062596 | 0.299066052 | 772.6963687 | 950.6851792 | 319 | 64 | 68 | 76 | 148 | 16 | 12 | 0 | 4 | 4 | 12 | 0 |
| SG0000002 | Prdm5 | 0.055695 | 0.079656 | 0.299819664 | 579.8340221 | 713.7701905 | 1408 | 36 | 68 | 212 | 720 | 108 | 0 | 0 | 0 | 0 | 0 | 0 |
| SG0000002 | Slc12a6 | 0.108233 | 0.146337 | 0.246743901 | 728.457807 | 864.3342395 | 797 | 38 | 68 | 113 | 452 | 73 | 6 | 0 | 0 | 0 | 9 | 0 |
| SG0000001 | Suz12 | 0.531303 | 0.597857 | -0.091544679 | 8686.631578 | 8152.55445 | 359 | 52 | 68 | 88 | 197 | 43 | 11 | 0 | 2 | 4 | 6 | 6 |
| SG0000003 | Zfp280c | 0.558272 | 0.623388 | -0.092897067 | 1419.823397 | 1331.28036 | 643 | 26 | 68 | 78 | 293 | 55 | 20 | 0 | 0 | 4 | 16 | 0 |
| SG0000002 | Gm20498 | 0.590545 | 0.653786 | -0.231444777 | 51.28486513 | 43.68349697 | 665 | 48 | 68 | 101 | 380 | 63 | 1 | 0 | 0 | 0 | 1 | 0 |
| SG0000000 | Ikbkg | 1.57E-65 | 4.99E-64 | -3.078348219 | 6672.519251 | 789.9774315 | 483 | 54 | 67 | 111 | 267 | 15 | 82 | 15 | 15 | 15 | 25 | 0 |
| SG0000004 | Hlcs | 1.6E-46 | 2.96E-45 | 2.954336135 | 56.90168177 | 441.0307761 | 501 | 46 | 67 | 93 | 235 | 34 | 6 | 0 | 2 | 0 | 0 | 0 |
| SG0000002 | Hacl1 | 2.74E-24 | 2.14E-23 | 1.831503719 | 145.4650487 | 517.7215148 | 320 | 56 | 67 | 69 | 134 | 19 | 8 | 0 | 0 | 0 | 3 | 0 |
| SG0000004 | Esrp1 | 1.18E-21 | 8.15E-21 | 2.241603497 | 505.1992995 | 2389.200686 | 518 | 49 | 67 | 70 | 239 | 63 | 11 | 1 | 1 | 1 | 8 | 7 |
| SG0000003 | Trim33 | 2.44E-19 | 1.47E-18 | 1.135736046 | 1571.047698 | 3452.073132 | 552 | 44 | 67 | 103 | 293 | 50 | 25 | 0 | 3 | 2 | 9 | 4 |
| SG0000002 | Myh10 | 1.41E-17 | 7.78E-17 | 3.492058416 | 272.2645418 | 3063.412039 | 729 | 37 | 67 | 116 | 340 | 56 | 6 | 0 | 0 | 0 | 3 | 0 |
| SG0000003 | Grik4 | 4.55E-14 | 2.03E-13 | 5.974831704 | 0.674076249 | 42.3947996 | 778 | 46 | 67 | 105 | 298 | 75 | 1 | 0 | 0 | 0 | 0 | 0 |
| SG0000002 | Mpdz | 7.46E-11 | 2.68E-10 | 2.055808411 | 145.6387718 | 605.5318687 | 893 | 35 | 67 | 156 | 492 | 61 | 5 | 0 | 0 | 0 | 7 | 0 |
| JSG000000 | Adamts19 | 2.9E-10 | 9.98E-10 | 4.743050073 | 0 | 25.77936921 | 524 | 35 | 67 | 83 | 294 | 33 | 3 | 0 | 1 | 1 | 4 | 0 |
| SG0000000 | Cttnbp2 | 6.46E-09 | 1.99E-08 | 2.83938443 | 7.453494908 | 53.345752 | 774 | 20 | 67 | 127 | 359 | 59 | 9 | 0 | 0 | 4 | 6 | 9 |
| SG0000007 | Emc1 | 3.41E-05 | 7.63E-05 | -0.614046173 | 1855.099609 | 1212.051064 | 162 | 64 | 67 | 77 | 118 | 0 | 10 | 0 | 2 | 2 | 15 | 0 |
| SG0000002 | Cep97 | 6.53E-05 | 0.000142 | 0.650035438 | 382.9068826 | 600.8600613 | 281 | 46 | 67 | 90 | 167 | 13 | 18 | 1 | 4 | 7 | 18 | 1 |
| SG0000002 | D3Ertd751e | 0.001469 | 0.002706 | 0.759336656 | 97.05750504 | 164.2904176 | 677 | 50 | 67 | 217 | 418 | 60 | 32 | 0 | 11 | 11 | 23 | 8 |
| SG0000002 | Ggnbp2 | 0.183159 | 0.235635 | -0.187920042 | 2686.604098 | 2358.490349 | 410 | 37 | 67 | 88 | 251 | 60 | 18 | 0 | 0 | 3 | 3 | 0 |
| SG0000003 | Shank2 | 0.945112 | 0.961265 | -0.095785917 | 20.18148442 | 18.88507321 | 795 | 54 | 67 | 118 | 283 | 86 | 21 | 3 | 3 | 3 | 3 | 0 |
| SG0000005 | Mtcl1 | 3.5E-110 | 3.1E-108 | 3.743781593 | 122.4196136 | 1639.991364 | 500 | 50 | 66 | 77 | 229 | 59 | 3 | 0 | 0 | 0 | 0 | 3 |
| JSG000000 | Fam169b | 9.37E-18 | 5.2E-17 | 5.88718614 | 0.946619625 | 56.02671016 | 287 | 55 | 66 | 82 | 144 | 27 | 4 | 0 | 0 | 0 | 0 | 2 |
| SG0000001 | Sema6a | 4.54E-17 | 2.44E-16 | 3.134596088 | 14.51398749 | 127.4659512 | 975 | 29 | 66 | 94 | 275 | 68 | 18 | 4 | 10 | 8 | 8 | 0 |
| SG0000005 | 43719 | 3.23E-14 | 1.45E-13 | -1.065717879 | 14991.00659 | 7161.726748 | 559 | 36 | 66 | 101 | 268 | 30 | 8 | 0 | 1 | 1 | 2 | 0 |
| SG0000003 | Dus2 | 3.39E-11 | 1.25E-10 | 1.105887194 | 329.0556988 | 708.230513 | 148 | 56 | 66 | 64 | 114 | 24 | 0 | 0 | 0 | 0 | 0 | 0 |
| SG0000002 | Sufu | 7.46E-11 | 2.68E-10 | -1.092107124 | 1103.262378 | 517.5136461 | 385 | 44 | 66 | 93 | 182 | 23 | 1 | 0 | 0 | 0 | 1 | 1 |
| JSG000000 | Tmem178b | 3.73E-10 | 1.27E-09 | 6.427450007 | 0.331601068 | 28.54113034 | 818 | 31 | 66 | 115 | 426 | 76 | 5 | 1 | 1 | 1 | 3 | 0 |
| SG0000001 | Zc3h18 | 1.1E-08 | 3.35E-08 | 0.738040258 | 1903.2986 | 3174.52816 | 157 | 66 | 66 | 70 | 107 | 0 | 0 | 0 | 0 | 0 | 0 | 0 |
| SG0000007 | Zfx | 1.89E-07 | 5.17E-07 | -0.791655646 | 1892.806563 | 1093.437937 | 504 | 52 | 66 | 110 | 243 | 41 | 39 | 0 | 0 | 3 | 9 | 6 |
| SG0000004 | Tardbp | 2.83E-07 | 7.62E-07 | 0.617989498 | 9129.171382 | 14010.85815 | 381 | 12 | 66 | 73 | 122 | 22 | 66 | 0 | 0 | 0 | 8 | 18 |
| SG0000003 | Shq1 | 1.62E-05 | 3.74E-05 | -0.701996903 | 1045.69296 | 642.8091645 | 543 | 42 | 66 | 72 | 165 | 39 | 3 | 0 | 0 | 0 | 0 | 0 |
| SG0000007 | Ppp1r12b | 0.000103 | 0.000219 | 0.794689988 | 162.9402068 | 282.6531222 | 774 | 36 | 66 | 144 | 399 | 83 | 3 | 0 | 0 | 1 | 2 | 1 |
| SG0000002 | Tmem209 | 0.000104 | 0.00022 | 0.504723855 | 2487.53037 | 3529.436817 | 414 | 42 | 66 | 66 | 182 | 16 | 18 | 0 | 0 | 0 | 2 | 1 |
| SG0000002 | Fbxw11 | 0.000145 | 0.000303 | -0.53327566 | 3756.092388 | 2595.399976 | 742 | 44 | 66 | 118 | 291 | 99 | 6 | 0 | 0 | 0 | 4 | 6 |
| SG0000001 | Med13l | 0.000204 | 0.000417 | 0.501756881 | 1391.007833 | 1969.579191 | 740 | 40 | 66 | 85 | 236 | 66 | 6 | 2 | 2 | 3 | 6 | 2 |

| | | | | | | | | | | | | | | | | | | |
|---|---|---|---|---|---|---|---|---|---|---|---|---|---|---|---|---|---|---|
| SG0000006 | Cnnm2 | 0.004937 | 0.008454 | -0.603277976 | 813.5233963 | 535.5071586 | 408 | 50 | 66 | 80 | 158 | 24 | 4 | 0 | 0 | 0 | 0 | 0 |
| SG0000005 | Rad51b | 0.009845 | 0.01612 | 1.092683959 | 50.83449256 | 108.4149416 | 579 | 47 | 66 | 114 | 301 | 50 | 1 | 0 | 0 | 1 | 2 | 0 |
| SG0000002 | Gfra1 | 0.030081 | 0.04513 | -1.051247583 | 39.51087579 | 19.06600054 | 1478 | 33 | 66 | 203 | 548 | 210 | 10 | 0 | 3 | 0 | 0 | 0 |
| SG0000003 | Pik3cb | 0.043684 | 0.063723 | -0.294659778 | 1449.304456 | 1181.566572 | 432 | 38 | 66 | 86 | 219 | 50 | 12 | 0 | 0 | 8 | 11 | 2 |
| SG0000001 | Frk | 0.05152 | 0.074176 | -0.965374377 | 64.95363713 | 33.26571254 | 704 | 44 | 66 | 128 | 392 | 66 | 4 | 0 | 0 | 2 | 6 | 0 |
| SG0000002 | Col4a3bp | 0.130083 | 0.172737 | -0.224329886 | 1928.775754 | 1651.01764 | 735 | 21 | 66 | 123 | 392 | 73 | 12 | 0 | 0 | 3 | 11 | 2 |
| SG0000003 | Ddx60 | 0.174201 | 0.225064 | 0.573141639 | 182.1738254 | 271.0308973 | 572 | 34 | 66 | 112 | 376 | 44 | 2 | 0 | 0 | 0 | 0 | 2 |
| SG0000006 | Vrk2 | 0.196709 | 0.251137 | -0.239967261 | 416.59084 | 352.7543459 | 602 | 44 | 66 | 100 | 294 | 29 | 0 | 0 | 0 | 0 | 0 | 0 |
| SG0000005 | A430033K04Rik | 0.600207 | 0.6629 | -0.132303934 | 216.573515 | 197.5958834 | 244 | 54 | 66 | 80 | 163 | 16 | 7 | 0 | 0 | 0 | 0 | 0 |
| SG0000002 | Xpnpep3 | 0.665923 | 0.724006 | -0.075652703 | 833.1635785 | 790.5995152 | 348 | 48 | 66 | 66 | 192 | 24 | 8 | 0 | 0 | 0 | 0 | 0 |
| SG0000003 | Ip6k1 | 0.72943 | 0.780732 | -0.043383771 | 1904.681006 | 1848.257318 | 335 | 63 | 66 | 113 | 246 | 26 | 5 | 0 | 0 | 1 | 5 | 0 |
| SG0000003 | Eva1c | 1.6E-90 | 9.16E-89 | -5.224185702 | 1299.111727 | 34.7544087 | 408 | 55 | 65 | 120 | 284 | 60 | 2 | 0 | 0 | 0 | 0 | 0 |
| SG0000003 | Nrxn2 | 2.95E-51 | 6.34E-50 | 6.968896828 | 1.348152497 | 168.8830261 | 353 | 61 | 65 | 93 | 208 | 40 | 5 | 0 | 0 | 0 | 0 | 0 |
| SG0000002 | Rab27b | 5.28E-31 | 5.58E-30 | -5.367103008 | 582.539955 | 14.1145398 | 690 | 38 | 65 | 112 | 282 | 48 | 20 | 0 | 0 | 0 | 9 | 7 |
| SG0000004 | Rbbp8 | 5.27E-28 | 4.87E-27 | 1.644120336 | 335.031208 | 1047.164235 | 588 | 42 | 65 | 109 | 287 | 47 | 8 | 0 | 2 | 2 | 2 | 0 |
| SG0000002 | Dcaf8 | 3.18E-20 | 2.01E-19 | 1.139240063 | 2289.978097 | 5044.019533 | 490 | 38 | 65 | 85 | 217 | 11 | 1 | 0 | 0 | 0 | 0 | 0 |
| SG0000000 | Ulk2 | 3.79E-15 | 1.81E-14 | -1.765847791 | 775.2871543 | 227.9761606 | 249 | 49 | 65 | 76 | 144 | 13 | 4 | 0 | 0 | 1 | 2 | 0 |
| SG0000002 | Slc37a3 | 1.65E-14 | 7.54E-14 | -1.224464295 | 1944.620483 | 832.2127679 | 278 | 50 | 65 | 68 | 167 | 21 | 0 | 0 | 0 | 0 | 3 | 3 |
| SG0000004 | Upf2 | 1.05E-12 | 4.27E-12 | 0.937387442 | 1173.030898 | 2246.421202 | 408 | 57 | 65 | 98 | 236 | 25 | 6 | 0 | 0 | 0 | 0 | 0 |
| SG0000004 | Usp48 | 1E-11 | 3.82E-11 | 0.831260928 | 3884.02439 | 6910.610602 | 256 | 56 | 65 | 85 | 172 | 13 | 10 | 0 | 1 | 2 | 5 | 0 |
| SG0000004 | Dpy19l4 | 2.01E-11 | 7.52E-11 | -1.168376386 | 1716.50321 | 763.7096559 | 366 | 59 | 65 | 96 | 192 | 28 | 30 | 0 | 0 | 2 | 8 | 4 |
| SG0000003 | Sbno1 | 2.23E-05 | 5.08E-05 | 0.520900626 | 4498.484928 | 6454.654236 | 384 | 56 | 65 | 95 | 222 | 28 | 34 | 2 | 2 | 10 | 18 | 4 |
| SG0000003 | Phkb | 0.001889 | 0.003431 | 0.433435985 | 954.5026647 | 1289.004334 | 834 | 23 | 65 | 104 | 427 | 67 | 8 | 0 | 0 | 1 | 3 | 0 |
| SG0000001 | Poc1b | 0.019575 | 0.030331 | 0.349917301 | 596.0040782 | 759.5997882 | 587 | 46 | 65 | 103 | 282 | 64 | 5 | 0 | 0 | 0 | 7 | 0 |
| SG0000002 | Lrp5 | 0.127998 | 0.170157 | -0.280590141 | 2529.724801 | 2082.606548 | 309 | 62 | 65 | 93 | 246 | 34 | 9 | 1 | 1 | 4 | 9 | 4 |
| JSG000000 | Psma3 | 0.131814 | 0.174789 | 0.190610387 | 3447.028568 | 3933.909157 | 324 | 43 | 65 | 73 | 179 | 19 | 9 | 0 | 0 | 0 | 1 | 4 |
| SG0000002 | Eif4enif1 | 0.661995 | 0.72013 | -0.063578349 | 2972.454975 | 2844.305852 | 398 | 45 | 65 | 84 | 184 | 55 | 0 | 0 | 0 | 0 | 0 | 10 |
| SG0000003 | Blm | 1.4E-151 | 2.1E-149 | 3.445504404 | 358.6412876 | 3907.153386 | 364 | 40 | 64 | 68 | 200 | 26 | 2 | 0 | 0 | 0 | 0 | 0 |
| SG0000003 | Ppm1h | 9.32E-43 | 1.51E-41 | 4.492097103 | 8.753463824 | 196.9862501 | 802 | 38 | 64 | 117 | 311 | 83 | 7 | 1 | 2 | 1 | 2 | 5 |
| SG0000005 | Arhgef12 | 3.85E-41 | 5.91E-40 | -2.18278882 | 7245.196466 | 1595.751486 | 612 | 42 | 64 | 84 | 304 | 60 | 14 | 0 | 2 | 4 | 4 | 2 |
| SG0000002 | Zeb1 | 3.78E-33 | 4.35E-32 | -2.37561761 | 1461.635508 | 281.6481649 | 1046 | 19 | 64 | 140 | 562 | 123 | 9 | 1 | 1 | 2 | 8 | 0 |
| SG0000003 | Hectd1 | 3.91E-32 | 4.32E-31 | -1.847096283 | 13689.3828 | 3804.980909 | 448 | 42 | 64 | 88 | 238 | 46 | 4 | 0 | 0 | 0 | 2 | 0 |
| SG0000003 | Cnot1 | 9.89E-32 | 1.07E-30 | 1.419483641 | 6446.120228 | 17242.71126 | 470 | 46 | 64 | 108 | 282 | 42 | 2 | 0 | 0 | 0 | 0 | 2 |
| SG0000002 | Snta1 | 8.98E-25 | 7.17E-24 | 2.810633682 | 73.21773146 | 513.6902394 | 182 | 60 | 64 | 84 | 122 | 10 | 0 | 0 | 0 | 0 | 0 | 0 |
| JSG000000 | Ncoa1 | 3.26E-23 | 2.42E-22 | -1.90885958 | 1616.955957 | 430.6001726 | 530 | 47 | 64 | 91 | 263 | 43 | 4 | 0 | 1 | 0 | 2 | 2 |
| SG0000003 | Carm1 | 2.14E-18 | 1.23E-17 | 1.235293952 | 2392.113257 | 5631.735777 | 240 | 56 | 64 | 84 | 176 | 20 | 1 | 0 | 0 | 0 | 0 | 0 |
| JSG000000 | Ldlrad4 | 1.8E-12 | 7.21E-12 | -1.873053584 | 240.1379513 | 65.55645924 | 500 | 46 | 64 | 102 | 298 | 49 | 0 | 0 | 0 | 0 | 3 | 0 |
| SG0000003 | Kif1bp | 1.87E-11 | 7.02E-11 | -1.043796782 | 1932.504814 | 937.3600579 | 218 | 55 | 64 | 69 | 117 | 14 | 9 | 0 | 0 | 0 | 0 | 0 |
| SG0000000 | Cacna1b | 1.02E-10 | 3.62E-10 | 5.475184568 | 2.556441386 | 113.7183701 | 651 | 48 | 64 | 96 | 315 | 67 | 6 | 0 | 0 | 0 | 2 | 0 |
| SG0000005 | Kcnh1 | 4.2E-10 | 1.43E-09 | 2.750517691 | 11.36661934 | 76.492639 | 878 | 35 | 64 | 129 | 351 | 115 | 9 | 3 | 3 | 3 | 12 | 0 |
| SG0000006 | Tank | 6.13E-08 | 1.75E-07 | -1.004309382 | 721.8298257 | 359.8384556 | 863 | 22 | 64 | 81 | 331 | 77 | 15 | 0 | 0 | 0 | 6 | 0 |
| SG0000002 | Ocln | 1.79E-07 | 4.9E-07 | -0.956185661 | 650.1837193 | 335.116279 | 460 | 52 | 64 | 96 | 227 | 38 | 16 | 0 | 0 | 0 | 3 | 6 |
| SG0000003 | Stox1 | 4.41E-07 | 1.17E-06 | 6.00351796 | 0.307509278 | 19.72864278 | 309 | 61 | 64 | 70 | 202 | 29 | 2 | 0 | 0 | 0 | 2 | 2 |
| SG0000003 | Tmem132c | 1.07E-06 | 2.75E-06 | -1.8676273 | 88.84713425 | 24.34622248 | 416 | 51 | 64 | 91 | 254 | 50 | 3 | 0 | 1 | 2 | 4 | 0 |
| SG0000003 | Abcc4 | 3.01E-06 | 7.43E-06 | -0.685199789 | 5689.496147 | 3538.40971 | 634 | 48 | 64 | 114 | 288 | 58 | 6 | 0 | 0 | 0 | 0 | 4 |
| SG0000004 | Rfwd2 | 0.002264 | 0.004065 | 0.438038336 | 857.9434876 | 1162.308357 | 479 | 36 | 64 | 88 | 255 | 35 | 5 | 0 | 0 | 0 | 1 | 0 |
| SG0000002 | Cubn | 0.165878 | 0.215477 | -0.893011588 | 20.63020401 | 11.1091351 | 524 | 38 | 64 | 89 | 272 | 30 | 1 | 0 | 0 | 0 | 0 | 0 |
| SG0000005 | Pus7 | 0.363963 | 0.432343 | 0.117711237 | 1890.760387 | 2051.498048 | 354 | 48 | 64 | 98 | 172 | 30 | 3 | 0 | 0 | 0 | 0 | 3 |
| SG0000003 | Fam175b | 0.706758 | 0.759916 | -0.063491056 | 983.2589836 | 940.92544 | 255 | 55 | 64 | 74 | 131 | 34 | 16 | 0 | 4 | 0 | 2 | 0 |
| SG0000002 | Grhl2 | 1.98E-47 | 3.8E-46 | 5.409750357 | 13.18858008 | 560.6543127 | 493 | 53 | 63 | 110 | 270 | 43 | 1 | 1 | 1 | 1 | 2 | 1 |
| SG0000000 | Itgb2 | 9.04E-37 | 1.19E-35 | -4.416794118 | 259.1451503 | 12.13264727 | 135 | 54 | 63 | 63 | 95 | 21 | 8 | 0 | 3 | 3 | 11 | 0 |
| SG0000000 | Nfix | 1.56E-33 | 1.82E-32 | -2.815009892 | 586.2134169 | 83.3015991 | 255 | 59 | 63 | 77 | 198 | 8 | 6 | 1 | 1 | 1 | 2 | 0 |
| SG0000003 | Atp8a1 | 9.62E-26 | 8.02E-25 | 4.937379629 | 23.68319872 | 725.6709327 | 1132 | 26 | 63 | 151 | 511 | 136 | 14 | 0 | 1 | 3 | 5 | 2 |
| SG0000003 | Ggta1 | 3.58E-25 | 2.92E-24 | -1.659701536 | 3177.155784 | 1005.584484 | 274 | 46 | 63 | 80 | 164 | 32 | 12 | 0 | 4 | 4 | 8 | 0 |
| SG0000003 | Mcph1 | 3.78E-21 | 2.52E-20 | 1.232234349 | 1037.571755 | 2437.57306 | 596 | 44 | 63 | 112 | 377 | 38 | 3 | 0 | 0 | 0 | 0 | 1 |
| SG0000003 | Arhgap44 | 8.36E-21 | 5.49E-20 | 2.180701749 | 56.84980792 | 257.7422645 | 637 | 36 | 63 | 96 | 292 | 66 | 6 | 0 | 0 | 0 | 4 | 9 |
| SG0000003 | Gtf2e2 | 4.56E-12 | 1.78E-11 | 0.872549713 | 2020.374101 | 3699.094523 | 393 | 45 | 63 | 105 | 273 | 15 | 6 | 0 | 0 | 6 | 9 | 0 |
| SG0000003 | Wwp2 | 7.84E-11 | 2.81E-10 | 0.85756147 | 1527.137364 | 2767.131693 | 191 | 55 | 63 | 69 | 135 | 10 | 0 | 0 | 0 | 0 | 0 | 0 |
| SG0000004 | Tpp2 | 1.63E-06 | 4.13E-06 | 0.595410066 | 3782.701701 | 5715.291479 | 770 | 20 | 63 | 114 | 345 | 56 | 8 | 3 | 3 | 6 | 9 | 0 |
| SG0000002 | Pus10 | 2.02E-06 | 5.07E-06 | -0.708656299 | 2019.955484 | 1235.990111 | 429 | 48 | 63 | 87 | 235 | 33 | 0 | 0 | 0 | 0 | 4 | 0 |
| SG0000003 | Phtf2 | 0.002621 | 0.004669 | -0.449817123 | 1536.625053 | 1125.017979 | 617 | 40 | 63 | 119 | 348 | 60 | 13 | 0 | 2 | 4 | 6 | 1 |
| SG0000003 | Galnt7 | 0.077496 | 0.107932 | -0.27942079 | 1733.543129 | 1428.303866 | 483 | 42 | 63 | 97 | 203 | 20 | 17 | 0 | 1 | 1 | 3 | 0 |
| SG0000003 | Tdg | 0.283822 | 0.347374 | 0.172646203 | 548.6867951 | 618.4380388 | 182 | 56 | 63 | 73 | 99 | 21 | 11 | 0 | 0 | 3 | 3 | 0 |
| SG0000002 | Slc25a12 | 0.338357 | 0.405286 | -0.143816259 | 1415.708042 | 1281.388114 | 374 | 49 | 63 | 80 | 239 | 45 | 8 | 1 | 1 | 2 | 9 | 3 |
| JSG000000 | Pacrg | 0.53885 | 0.605171 | -0.330563106 | 28.66240487 | 22.79309055 | 805 | 21 | 63 | 102 | 376 | 79 | 0 | 0 | 0 | 0 | 0 | 0 |
| SG0000005 | Swt1 | 7.13E-35 | 8.8E-34 | 1.707867727 | 496.3115253 | 1621.338796 | 352 | 44 | 62 | 80 | 207 | 24 | 2 | 0 | 0 | 0 | 1 | 0 |
| JSG000000 | Gab2 | 2.5E-26 | 2.15E-25 | -2.741367109 | 753.0492319 | 112.6132885 | 500 | 36 | 62 | 101 | 279 | 36 | 6 | 0 | 0 | 1 | 3 | 0 |
| SG0000000 | Chn2 | 6.37E-25 | 5.13E-24 | -2.552594011 | 407.5398973 | 69.46447658 | 823 | 25 | 62 | 125 | 390 | 69 | 5 | 0 | 0 | 1 | 5 | 0 |
| SG0000003 | Cdk19 | 3.95E-22 | 2.79E-21 | -1.553323692 | 2769.91555 | 943.7772495 | 622 | 44 | 62 | 116 | 312 | 60 | 12 | 0 | 0 | 3 | 8 | 1 |
| JSG000000 | Kif16b | 7.12E-20 | 4.44E-19 | -1.488415358 | 2069.356022 | 737.5263589 | 581 | 44 | 62 | 98 | 285 | 61 | 11 | 0 | 2 | 0 | 0 | 1 |
| SG0000002 | Trpm7 | 4.75E-18 | 2.68E-17 | -1.277260125 | 11647.86395 | 4805.655996 | 546 | 32 | 62 | 86 | 252 | 42 | 10 | 0 | 0 | 0 | 0 | 0 |
| SG0000001 | Cep57l1 | 4.84E-16 | 2.43E-15 | 1.297604168 | 298.3319151 | 733.3604639 | 826 | 23 | 62 | 145 | 512 | 64 | 10 | 0 | 0 | 0 | 0 | 0 |
| SG0000003 | Atp2b2 | 1.67E-15 | 8.15E-15 | 7.190529895 | 0.307509278 | 44.91830987 | 764 | 29 | 62 | 116 | 306 | 172 | 18 | 0 | 0 | 0 | 3 | 6 |
| SG0000002 | Usp24 | 5.91E-15 | 2.78E-14 | -1.143421374 | 5933.471575 | 2685.992124 | 502 | 50 | 62 | 86 | 232 | 48 | 10 | 0 | 0 | 0 | 0 | 0 |
| SG0000002 | Wwtr1 | 3.54E-11 | 1.3E-10 | -1.077987622 | 2874.987063 | 1361.850033 | 516 | 36 | 62 | 98 | 250 | 31 | 18 | 0 | 2 | 2 | 12 | 1 |
| SG0000007 | Chd2 | 1.91E-10 | 6.66E-10 | 0.817126306 | 1702.684903 | 2999.948598 | 337 | 53 | 62 | 89 | 216 | 34 | 3 | 0 | 0 | 0 | 3 | 2 |
| JSG000000 | Cep192 | 2.12E-10 | 7.36E-10 | 0.953220495 | 2254.580737 | 4365.296563 | 244 | 56 | 62 | 72 | 145 | 12 | 4 | 0 | 0 | 0 | 0 | 1 |
| SG0000006 | Taok3 | 4.67E-10 | 1.58E-09 | 0.952471162 | 445.9415012 | 862.9791069 | 577 | 25 | 62 | 112 | 274 | 69 | 7 | 0 | 0 | 3 | 12 | 5 |
| JSG000000 | Ispd | 6.29E-07 | 1.65E-06 | 2.85310019 | 5.298586394 | 38.28500937 | 734 | 36 | 62 | 113 | 375 | 68 | 2 | 0 | 0 | 0 | 0 | 0 |
| SG0000004 | Ulk4 | 8.22E-07 | 2.14E-06 | 1.196354629 | 94.13349982 | 215.7162364 | 480 | 46 | 62 | 96 | 238 | 54 | 4 | 0 | 0 | 0 | 0 | 2 |
| SG0000007 | Fryl | 1.09E-05 | 2.56E-05 | -0.655847167 | 2094.381203 | 1329.309356 | 714 | 39 | 62 | 109 | 286 | 82 | 0 | 0 | 0 | 0 | 8 | 0 |
| SG0000003 | Gcc2 | 4.01E-05 | 8.9E-05 | -0.633022803 | 1535.361104 | 990.0375859 | 385 | 40 | 62 | 80 | 206 | 21 | 5 | 0 | 0 | 0 | 4 | 3 |
| SG0000002 | Wdr37 | 0.000102 | 0.000217 | 0.557665728 | 770.2669485 | 1133.744974 | 620 | 38 | 62 | 116 | 361 | 51 | 31 | 2 | 5 | 3 | 16 | 2 |
| SG0000001 | Dhdds | 0.004065 | 0.007036 | -0.481464135 | 735.3541566 | 526.6976798 | 357 | 62 | 62 | 134 | 196 | 6 | 2 | 0 | 0 | 0 | 0 | 0 |
| SG0000003 | Fam65b | 0.008345 | 0.013826 | -1.111741231 | 55.69942672 | 25.77408565 | 715 | 42 | 62 | 102 | 389 | 92 | 7 | 0 | 0 | 0 | 2 | 2 |
| JSG000000 | Dock1 | 0.011615 | 0.018796 | 0.334569546 | 1986.367681 | 2504.811851 | 685 | 35 | 62 | 98 | 319 | 82 | 1 | 0 | 0 | 0 | 1 | 1 |
| SG0000004 | Ascc1 | 0.011834 | 0.019123 | 0.360791104 | 848.1134493 | 1089.087804 | 328 | 50 | 62 | 82 | 210 | 34 | 0 | 0 | 0 | 0 | 0 | 0 |
| SG0000004 | 1700019D03Rik | 0.025804 | 0.039169 | 0.316487944 | 909.7480852 | 1132.905055 | 704 | 29 | 62 | 116 | 399 | 41 | 18 | 0 | 0 | 0 | 6 | 0 |
| JSG000000 | 1110059E24Rik | 0.113476 | 0.15255 | 0.278133893 | 848.0138767 | 1028.323128 | 789 | 52 | 62 | 103 | 320 | 40 | 3 | 0 | 2 | 2 | 4 | 4 |
| SG0000004 | Bri3 | 0.220392 | 0.277765 | 0.205799438 | 448.4373506 | 517.194124 | 323 | 49 | 62 | 74 | 171 | 30 | 1 | 0 | 0 | 0 | 1 | 0 |
| SG0000002 | Dip2b | 0.254223 | 0.315306 | 0.16194258 | 1069.173394 | 1196.183251 | 366 | 38 | 62 | 70 | 161 | 36 | 9 | 1 | 1 | 2 | 4 | 2 |
| SG0000001 | Ppp1cb | 0.313297 | 0.378969 | 0.119364994 | 8086.489839 | 8784.003272 | 335 | 41 | 62 | 55 | 135 | 17 | 12 | 0 | 1 | 0 | 1 | 0 |
| SG0000001 | Crlf3 | 0.843494 | 0.877543 | 0.020359632 | 2250.919901 | 2282.91058 | 320 | 48 | 62 | 92 | 175 | 38 | 5 | 0 | 0 | 2 | 2 | 0 |
| SG0000004 | Wbp1l | 4.69E-47 | 8.86E-46 | -2.572839244 | 6613.619417 | 1111.571526 | 220 | 42 | 61 | 67 | 118 | 22 | 8 | 0 | 0 | 0 | 0 | 0 |
| SG0000003 | Snx25 | 4.5E-20 | 2.84E-19 | -1.641353026 | 1200.106302 | 384.7009173 | 695 | 25 | 61 | 74 | 366 | 166 | 4 | 0 | 0 | 4 | 5 | 0 |
| SG0000006 | Chpt1 | 7.38E-18 | 4.12E-17 | 1.387391742 | 288.9655796 | 755.9492569 | 586 | 28 | 61 | 69 | 234 | 32 | 32 | 0 | 1 | 4 | 15 | 2 |
| SG0000002 | Fam135a | 3.26E-16 | 1.66E-15 | -1.335007999 | 3367.472241 | 1334.831885 | 881 | 34 | 61 | 149 | 550 | 94 | 39 | 0 | 0 | 6 | 19 | 0 |
| SG0000002 | Rbms1 | 4.07E-16 | 2.06E-15 | -1.355367491 | 4062.982683 | 1587.956866 | 595 | 35 | 61 | 101 | 248 | 42 | 7 | 0 | 2 | 2 | 3 | 1 |
| SG0000002 | Lnx1 | 1.38E-12 | 5.57E-12 | 5.211791389 | 5.368518198 | 198.9573414 | 1078 | 32 | 61 | 210 | 417 | 128 | 19 | 0 | 0 | 8 | 12 | 0 |
| SG0000004 | Adamts3 | 1.79E-12 | 7.15E-12 | -2.855517422 | 131.1447924 | 18.11985227 | 1056 | 28 | 61 | 134 | 461 | 95 | 1 | 0 | 0 | 0 | 1 | 0 |
| SG0000004 | Klhl15 | 1.24E-11 | 4.69E-11 | -1.275317613 | 689.6894049 | 284.9342781 | 406 | 22 | 61 | 66 | 196 | 21 | 23 | 0 | 1 | 1 | 9 | 0 |
| SG0000002 | Ndufs1 | 2.09E-07 | 5.68E-07 | 0.643690793 | 3491.628916 | 5455.044757 | 211 | 52 | 61 | 82 | 174 | 23 | 2 | 0 | 0 | 0 | 0 | 0 |
| JSG000000 | Epb41l4a | 3.06E-07 | 8.22E-07 | -0.792892269 | 1669.701601 | 963.728067 | 354 | 43 | 61 | 84 | 195 | 36 | 2 | 0 | 0 | 0 | 0 | 0 |
| SG0000001 | Hps5 | 8.66E-07 | 2.25E-06 | -0.778567524 | 1287.891712 | 750.7704494 | 274 | 52 | 61 | 72 | 176 | 13 | 8 | 0 | 0 | 0 | 4 | 0 |
| SG0000003 | Prc1 | 1.02E-06 | 2.64E-06 | 0.608284184 | 3912.373514 | 5964.198586 | 232 | 45 | 61 | 64 | 130 | 20 | 5 | 0 | 0 | 5 | 5 | 0 |
| SG0000002 | Gbf1 | 2.17E-06 | 5.45E-06 | -0.693937631 | 2238.084853 | 1383.504276 | 334 | 46 | 61 | 89 | 217 | 40 | 0 | 0 | 0 | 0 | 0 | 2 |
| SG0000002 | Trim24 | 2.93E-06 | 7.25E-06 | 0.606972032 | 1960.051191 | 2985.274026 | 644 | 40 | 61 | 124 | 332 | 71 | 0 | 0 | 0 | 0 | 0 | 0 |
| SG0000000 | Nf2 | 3.24E-06 | 8E-06 | -0.631133129 | 7605.400968 | 4910.572333 | 566 | 40 | 61 | 101 | 306 | 31 | 7 | 0 | 0 | 0 | 1 | 0 |
| SG0000003 | Spats2l | 3.59E-06 | 8.82E-06 | 1.451662551 | 37.05648556 | 101.3580391 | 1244 | 20 | 61 | 147 | 570 | 162 | 8 | 0 | 0 | 0 | 0 | 0 |
| SG0000002 | Fn1 | 6.77E-06 | 1.62E-05 | 1.082908768 | 22285.07402 | 47206.53267 | 1035 | 31 | 61 | 157 | 415 | 53 | 21 | 0 | 0 | 7 | 21 | 0 |
| JSG000000 | Prkd1 | 0.000171 | 0.000353 | 2.204407998 | 22.35714429 | 103.040809 | 731 | 31 | 61 | 126 | 372 | 76 | 3 | 0 | 0 | 0 | 0 | 0 |
| SG0000002 | Ppm1l | 0.000312 | 0.000624 | -0.692471624 | 473.2184006 | 292.8241514 | 493 | 47 | 61 | 87 | 220 | 41 | 8 | 0 | 0 | 0 | 4 | 0 |
| SG0000002 | Diap1 | 0.031153 | 0.046621 | -0.284122972 | 6887.805936 | 5656.546546 | 764 | 32 | 61 | 103 | 419 | 93 | 9 | 5 | 5 | 5 | 9 | 0 |
| SG0000003 | Rad18 | 0.035286 | 0.052261 | -0.314620364 | 1578.703267 | 1269.376169 | 662 | 38 | 61 | 119 | 316 | 56 | 12 | 0 | 0 | 4 | 4 | 0 |
| JSG000000 | Rad54l2 | 0.073778 | 0.103211 | 0.232268687 | 1859.123337 | 2183.876346 | 286 | 50 | 61 | 88 | 166 | 14 | 7 | 1 | 1 | 3 | 4 | 0 |
| SG0000004 | Ccbe1 | 0.277445 | 0.340523 | -0.347822496 | 96.64145135 | 75.93787484 | 762 | 38 | 61 | 111 | 389 | 100 | 6 | 0 | 1 | 1 | 1 | 0 |
| SG0000002 | Axdnd1 | 0.432774 | 0.501965 | -0.584596634 | 20.57833016 | 13.72236631 | 391 | 43 | 61 | 90 | 239 | 78 | 0 | 0 | 0 | 0 | 0 | 0 |
| SG0000003 | Ttc39b | 0.501721 | 0.569003 | -0.104884862 | 2058.694959 | 1914.33752 | 528 | 46 | 61 | 109 | 263 | 19 | 27 | 2 | 4 | 4 | 9 | 0 |
| SG0000002 | Sybu | 4.1E-75 | 1.68E-73 | -6.232053281 | 559.2091087 | 7.439423198 | 661 | 31 | 60 | 108 | 372 | 110 | 3 | 0 | 0 | 0 | 3 | 0 |
| SG0000003 | Acsl4 | 8.39E-71 | 3.07E-69 | -3.384663922 | 10853.05206 | 1039.121954 | 692 | 36 | 60 | 108 | 392 | 52 | 40 | 0 | 8 | 4 | 8 | 4 |
| SG0000003 | Cd99l2 | 1.36E-56 | 3.42E-55 | -3.128807886 | 2110.758038 | 241.3089486 | 523 | 36 | 60 | 80 | 184 | 52 | 3 | 0 | 0 | 0 | 0 | 0 |
| SG0000002 | Meis1 | 2.26E-53 | 5.17E-52 | -6.216180909 | 2266.08825 | 30.48034701 | 1536 | 12 | 60 | 180 | 639 | 95 | 33 | 0 | 0 | 0 | 16 | 5 |
| SG0000005 | Kcnn2 | 4.8E-49 | 9.66E-48 | 7.16136862 | 0 | 142.1484878 | 940 | 23 | 60 | 114 | 491 | 99 | 0 | 0 | 0 | 0 | 0 | 0 |

| | | | | | | | | | | | | | | | | | | |
|---|---|---|---|---|---|---|---|---|---|---|---|---|---|---|---|---|---|---|
| SG0000006 | D630045J12Rik | 3.82E-47 | 7.24E-46 | -3.033386573 | 4593.136185 | 561.007898 | 404 | 40 | 60 | 72 | 178 | 32 | 20 | 2 | 4 | 4 | 6 | 0 |
| SG0000002 | Ankrd33b | 1.89E-44 | 3.24E-43 | -4.222699987 | 1594.732078 | 85.41386441 | 374 | 43 | 60 | 87 | 212 | 37 | 24 | 0 | 1 | 3 | 6 | 4 |
| SG0000002 | Rffl | 2.26E-20 | 1.45E-19 | 1.392469417 | 375.4414822 | 985.6377264 | 407 | 54 | 60 | 86 | 233 | 27 | 9 | 0 | 0 | 0 | 0 | 0 |
| SG0000002 | Atp2a2 | 2E-16 | 1.03E-15 | -1.159742857 | 22955.60613 | 10274.75269 | 311 | 51 | 60 | 82 | 173 | 31 | 17 | 2 | 9 | 5 | 12 | 3 |
| ISG000000 | Adamts6 | 2.45E-14 | 1.11E-13 | -3.077010489 | 114.9537883 | 13.62231234 | 608 | 29 | 60 | 94 | 316 | 43 | 5 | 0 | 0 | 0 | 4 | 0 |
| SG0000003 | Hk1 | 2.26E-13 | 9.58E-13 | -1.663051921 | 8467.652745 | 2673.834526 | 305 | 55 | 60 | 88 | 157 | 26 | 0 | 0 | 0 | 0 | 0 | 0 |
| SG0000004 | Cux2 | 1.14E-10 | 4.05E-10 | 6.864370809 | 1.359026609 | 158.3468669 | 435 | 44 | 60 | 75 | 148 | 36 | 5 | 0 | 0 | 3 | 6 | 1 |
| SG0000003 | Vav1 | 1.41E-09 | 4.57E-09 | 3.617907473 | 3.524809234 | 43.27472218 | 237 | 54 | 60 | 93 | 174 | 27 | 3 | 0 | 0 | 0 | 0 | 0 |
| SG0000003 | Fntb | 1.67E-09 | 5.41E-09 | 0.87427747 | 641.1371665 | 1175.262006 | 606 | 51 | 60 | 96 | 285 | 55 | 1 | 0 | 0 | 1 | 1 | 0 |
| SG0000002 | Rtn3 | 5.95E-09 | 1.84E-08 | -0.806372814 | 8551.947467 | 4890.154944 | 528 | 28 | 60 | 72 | 239 | 60 | 8 | 0 | 0 | 0 | 3 | 0 |
| SG0000000 | Slc2a9 | 6.05E-09 | 1.87E-08 | -2.422273753 | 86.42770992 | 16.12411727 | 823 | 26 | 60 | 106 | 278 | 80 | 12 | 0 | 0 | 6 | 10 | 0 |
| SG0000002 | Zmynd11 | 6.09E-09 | 1.88E-08 | -0.917327943 | 1454.823353 | 770.312663 | 1098 | 12 | 60 | 142 | 644 | 108 | 40 | 0 | 0 | 12 | 12 | 6 |
| ISG000000 | Bmpr2 | 4.12E-08 | 1.19E-07 | -1.014353611 | 2219.510714 | 1098.768962 | 367 | 43 | 60 | 81 | 233 | 15 | 35 | 1 | 6 | 4 | 14 | 0 |
| SG0000003 | Pigk | 6.16E-08 | 1.76E-07 | -0.895756787 | 1025.117367 | 550.9648964 | 940 | 28 | 60 | 121 | 432 | 102 | 9 | 1 | 1 | 2 | 4 | 4 |
| SG0000003 | Cngb1 | 4.45E-07 | 1.18E-06 | 2.056940413 | 15.70536843 | 65.35049664 | 204 | 52 | 60 | 67 | 92 | 15 | 4 | 0 | 0 | 0 | 1 | 0 |
| SG0000004 | Zfp236 | 4.76E-07 | 1.26E-06 | -0.808219702 | 1202.684119 | 686.8362411 | 509 | 44 | 60 | 94 | 257 | 67 | 14 | 2 | 3 | 4 | 11 | 6 |
| SG0000003 | Slc9a8 | 2.98E-05 | 6.69E-05 | -0.647178468 | 1267.248606 | 809.1735732 | 209 | 60 | 60 | 72 | 117 | 6 | 2 | 0 | 0 | 0 | 0 | 0 |
| SG0000004 | Necab1 | 0.00023 | 0.000468 | -2.522458 | 49.3143571 | 8.582975143 | 850 | 44 | 60 | 118 | 409 | 73 | 7 | 0 | 0 | 0 | 3 | 3 |
| SG0000005 | Synpo2 | 0.000866 | 0.001638 | 1.708789566 | 11.34252754 | 37.07718536 | 867 | 30 | 60 | 117 | 399 | 123 | 35 | 0 | 0 | 2 | 14 | 0 |
| SG0000002 | Scamp1 | 0.0018 | 0.00328 | -0.458464166 | 2027.655416 | 1475.647703 | 398 | 43 | 60 | 85 | 223 | 23 | 8 | 0 | 1 | 1 | 4 | 0 |
| SG0000003 | 2610020H08Rik | 0.006012 | 0.010169 | 0.566706011 | 185.0309856 | 274.0564137 | 544 | 36 | 60 | 92 | 376 | 40 | 2 | 0 | 0 | 0 | 0 | 0 |
| SG0000002 | Rad17 | 0.00799 | 0.013261 | 0.384464915 | 850.3166285 | 1109.982531 | 240 | 48 | 60 | 74 | 166 | 8 | 5 | 0 | 0 | 1 | 1 | 0 |
| SG0000003 | Supt3 | 0.011725 | 0.018957 | -0.461326421 | 485.4516864 | 352.5923809 | 792 | 30 | 60 | 124 | 385 | 63 | 0 | 0 | 0 | 0 | 0 | 0 |
| SG0000002 | Luc7l3 | 0.020328 | 0.031397 | 0.394933954 | 3400.548115 | 4471.321424 | 352 | 55 | 60 | 97 | 244 | 16 | 12 | 0 | 0 | 2 | 20 | 1 |
| SG0000004 | Ythdf2 | 0.035905 | 0.053117 | 0.276708966 | 2181.447634 | 2642.667302 | 201 | 54 | 60 | 67 | 157 | 21 | 4 | 0 | 0 | 1 | 4 | 0 |
| SG0000002 | Pisd | 0.091048 | 0.125065 | 0.25265354 | 880.9012138 | 1049.50256 | 230 | 51 | 60 | 51 | 99 | 28 | 6 | 0 | 0 | 0 | 0 | 0 |
| SG0000002 | Ube2v2 | 0.118681 | 0.158868 | 0.252108904 | 515.5349035 | 613.9746158 | 432 | 52 | 60 | 107 | 226 | 23 | 20 | 0 | 0 | 3 | 10 | 3 |
| SG0000002 | Myom1 | 0.266767 | 0.328928 | 0.687600758 | 13.18758321 | 21.23995861 | 723 | 42 | 60 | 117 | 291 | 96 | 3 | 0 | 0 | 0 | 0 | 0 |
| SG0000003 | Zcchc7 | 0.518159 | 0.585089 | -0.103930426 | 904.1884359 | 841.3423936 | 1054 | 22 | 60 | 156 | 461 | 82 | 6 | 0 | 0 | 0 | 3 | 1 |
| SG0000002 | Lrrig3 | 0.661063 | 0.719378 | -0.292439992 | 23.72200804 | 19.36949237 | 616 | 53 | 60 | 114 | 315 | 40 | 2 | 0 | 1 | 0 | 4 | 1 |
| SG0000002 | Dnmt3a | 1.38E-44 | 2.39E-43 | 2.850859962 | 252.2630995 | 1819.902366 | 308 | 59 | 59 | 76 | 203 | 11 | 8 | 0 | 0 | 3 | 18 | 2 |
| SG0000001 | Aig1 | 2.84E-38 | 3.96E-37 | -2.387499708 | 1766.284516 | 337.5605104 | 1244 | 30 | 59 | 149 | 494 | 118 | 0 | 0 | 0 | 0 | 0 | 1 |
| SG0000004 | Hfm1 | 5.06E-20 | 3.18E-19 | 5.758520941 | 0 | 53.13617081 | 713 | 24 | 59 | 85 | 390 | 79 | 11 | 0 | 0 | 0 | 9 | 1 |
| SG0000000 | Cdh4 | 1.88E-17 | 1.03E-16 | 7.339855914 | 0.307509278 | 49.81670716 | 739 | 40 | 59 | 88 | 292 | 113 | 2 | 1 | 1 | 1 | 1 | 1 |
| SG0000003 | Dock11 | 1.14E-16 | 5.96E-16 | -1.690788585 | 3005.725528 | 931.0462162 | 706 | 33 | 59 | 112 | 416 | 68 | 4 | 0 | 0 | 0 | 0 | 2 |
| SG0000003 | Mitf | 8.52E-16 | 4.23E-15 | 2.197520833 | 37.51607926 | 172.0826683 | 837 | 16 | 59 | 124 | 465 | 121 | 12 | 1 | 1 | 2 | 10 | 0 |
| ISG000000 | Zswim5 | 1.64E-05 | 3.79E-05 | -0.819381306 | 489.8936802 | 277.6153566 | 281 | 43 | 59 | 74 | 149 | 13 | 2 | 0 | 0 | 1 | 1 | 1 |
| SG0000002 | Tnrc6c | 2.21E-05 | 5.04E-05 | 0.599210859 | 910.9027721 | 1379.915415 | 409 | 48 | 59 | 79 | 190 | 30 | 14 | 0 | 2 | 0 | 3 | 0 |
| SG0000006 | Tor3a | 0.000147 | 0.000305 | 0.545892926 | 818.7638377 | 1195.332694 | 187 | 48 | 59 | 59 | 103 | 11 | 3 | 0 | 0 | 0 | 3 | 0 |
| ISG000000 | Syne1 | 0.000204 | 0.000419 | -1.140502219 | 399.6929722 | 181.3013901 | 510 | 37 | 59 | 95 | 311 | 62 | 5 | 0 | 1 | 2 | 5 | 0 |
| ISG000000 | Tjp2 | 0.002797 | 0.004957 | -0.40186425 | 5723.491854 | 4331.99429 | 240 | 44 | 59 | 70 | 160 | 18 | 2 | 0 | 0 | 0 | 1 | 1 |
| ISG000000 | Atad2b | 0.108943 | 0.147131 | 0.292711006 | 408.9673559 | 500.9604454 | 375 | 41 | 59 | 82 | 199 | 23 | 0 | 0 | 0 | 0 | 0 | 0 |
| SG0000003 | Ryk | 0.301494 | 0.366325 | -0.145715287 | 2260.585224 | 2043.413283 | 434 | 38 | 59 | 77 | 218 | 27 | 5 | 2 | 2 | 2 | 2 | 0 |
| SG0000003 | Myo9a | 0.474707 | 0.543493 | -0.132238355 | 956.762112 | 872.9638453 | 521 | 38 | 59 | 91 | 304 | 42 | 10 | 0 | 0 | 2 | 7 | 1 |
| SG0000002 | Tbc1d14 | 0.668032 | 0.725684 | -0.065299053 | 1605.818488 | 1534.756495 | 915 | 37 | 59 | 123 | 328 | 92 | 0 | 0 | 0 | 0 | 9 | 18 |
| SG0000002 | Arih1 | 0.722939 | 0.774849 | -0.054991485 | 5562.537497 | 5354.499288 | 605 | 26 | 59 | 83 | 273 | 52 | 13 | 0 | 0 | 2 | 4 | 1 |
| SG0000002 | Hat1 | 1.7E-135 | 2.2E-133 | 2.921450798 | 1825.513421 | 13830.22836 | 284 | 48 | 58 | 76 | 158 | 14 | 0 | 0 | 0 | 0 | 0 | 0 |
| SG0000003 | Zranb3 | 5.8E-105 | 4.7E-103 | 3.121120301 | 215.0336686 | 1870.928949 | 560 | 22 | 58 | 76 | 342 | 46 | 2 | 0 | 0 | 0 | 0 | 0 |
| SG0000003 | Ryr1 | 6.54E-69 | 2.27E-67 | 4.080169289 | 24.94117104 | 421.8617832 | 270 | 46 | 58 | 68 | 148 | 16 | 0 | 0 | 0 | 0 | 0 | 0 |
| ISG000000 | Lppr1 | 1.34E-46 | 2.5E-45 | 7.081354318 | 0 | 134.425379 | 542 | 40 | 58 | 107 | 288 | 51 | 4 | 0 | 0 | 1 | 2 | 2 |
| SG0000002 | Ntn1 | 1.51E-26 | 1.31E-25 | 1.531832623 | 479.1834635 | 1385.57293 | 401 | 53 | 58 | 82 | 188 | 52 | 2 | 0 | 0 | 0 | 4 | 0 |
| SG0000003 | 1700025G04Rik | 1.75E-25 | 1.45E-24 | 3.009235273 | 64.94226004 | 522.8745206 | 893 | 36 | 58 | 137 | 399 | 94 | 14 | 0 | 0 | 2 | 6 | 7 |
| SG0000002 | Tsc22d1 | 9.75E-24 | 7.43E-23 | 1.210261006 | 3584.87904 | 8294.674955 | 587 | 48 | 58 | 111 | 277 | 42 | 10 | 0 | 0 | 0 | 5 | 0 |
| SG0000002 | Pias2 | 4.64E-21 | 3.08E-20 | 1.545377296 | 1405.391303 | 4102.061645 | 503 | 41 | 58 | 102 | 266 | 45 | 12 | 0 | 1 | 2 | 5 | 1 |
| SG0000002 | Snx13 | 1.73E-19 | 1.05E-18 | -1.64320992 | 2211.495826 | 707.9957471 | 526 | 28 | 58 | 86 | 302 | 41 | 14 | 1 | 2 | 3 | 10 | 2 |
| SG0000001 | Crk | 1.91E-13 | 8.15E-13 | -1.051158338 | 7595.84437 | 3665.606721 | 265 | 48 | 58 | 81 | 145 | 10 | 15 | 0 | 0 | 0 | 3 | 0 |
| SG0000001 | Med15 | 3.39E-13 | 1.42E-12 | -1.175909722 | 3040.904441 | 1345.918226 | 244 | 48 | 58 | 76 | 148 | 38 | 1 | 0 | 0 | 0 | 1 | 0 |
| SG0000002 | Erbb2ip | 1.84E-12 | 7.35E-12 | -1.110550313 | 2040.409197 | 944.9488842 | 832 | 26 | 58 | 126 | 455 | 92 | 12 | 0 | 0 | 0 | 6 | 0 |
| SG0000000 | Krit1 | 2.59E-09 | 8.24E-09 | -0.91330662 | 1898.77547 | 1008.186609 | 570 | 32 | 58 | 124 | 248 | 22 | 9 | 0 | 1 | 0 | 3 | 1 |
| SG0000003 | Ap1g1 | 2.93E-07 | 7.88E-07 | 0.638082264 | 3318.451082 | 5164.369663 | 479 | 46 | 58 | 88 | 251 | 31 | 6 | 0 | 0 | 0 | 3 | 1 |
| SG0000002 | Nt5c2 | 2.74E-06 | 6.79E-06 | 0.657657097 | 863.1423228 | 1361.623145 | 620 | 25 | 58 | 96 | 307 | 59 | 5 | 0 | 0 | 0 | 0 | 0 |
| SG0000004 | Arhgap23 | 5.49E-06 | 1.32E-05 | -0.774307622 | 1604.106071 | 937.8713348 | 221 | 51 | 58 | 81 | 170 | 21 | 8 | 0 | 0 | 0 | 4 | 8 |
| SG0000005 | Ap1s3 | 7.85E-06 | 1.87E-05 | 0.765995602 | 292.4858984 | 497.384887 | 385 | 51 | 58 | 85 | 257 | 25 | 7 | 0 | 1 | 1 | 7 | 1 |
| SG0000003 | Ube2cbp | 1.33E-05 | 3.1E-05 | 1.132589214 | 68.37085525 | 149.9043855 | 656 | 34 | 58 | 106 | 358 | 50 | 0 | 0 | 0 | 0 | 0 | 0 |
| SG0000004 | Filip1l | 1.68E-05 | 3.89E-05 | -1.292564921 | 334.567858 | 136.5788348 | 766 | 34 | 58 | 109 | 340 | 59 | 4 | 0 | 0 | 0 | 0 | 0 |
| SG0000002 | Ptk2 | 4.39E-05 | 9.71E-05 | -0.579422313 | 3221.39973 | 2155.862814 | 448 | 42 | 58 | 84 | 249 | 48 | 2 | 0 | 0 | 0 | 0 | 0 |
| SG0000002 | Acin1 | 5.77E-05 | 0.000126 | 0.505214247 | 4828.470932 | 6853.213534 | 215 | 52 | 58 | 74 | 164 | 18 | 0 | 0 | 0 | 0 | 14 | 0 |
| SG0000006 | Qk | 5.88E-05 | 0.000128 | 0.505349485 | 3734.38846 | 5300.841829 | 666 | 32 | 58 | 98 | 354 | 69 | 34 | 0 | 0 | 2 | 21 | 2 |
| SG0000002 | Cdk15 | 8.49E-05 | 0.000182 | 3.423335939 | 1.91733104 | 20.56950809 | 311 | 50 | 58 | 80 | 199 | 42 | 1 | 0 | 0 | 0 | 1 | 0 |
| SG0000003 | Map2k4 | 9.88E-05 | 0.00021 | -0.558677717 | 2818.245684 | 1913.37502 | 692 | 30 | 58 | 104 | 293 | 82 | 4 | 0 | 0 | 0 | 2 | 2 |
| SG0000003 | Kdm6a | 0.00021 | 0.00043 | -0.551108416 | 2420.772652 | 1652.166607 | 786 | 34 | 58 | 118 | 402 | 72 | 16 | 0 | 0 | 4 | 6 | 6 |
| SG0000002 | Ift74 | 0.00024 | 0.000486 | 0.567473145 | 493.4425722 | 731.2452436 | 396 | 40 | 58 | 93 | 234 | 32 | 4 | 0 | 0 | 0 | 0 | 0 |
| SG0000005 | Zfp426 | 0.005172 | 0.008833 | -0.48520978 | 666.2262196 | 475.9474701 | 291 | 51 | 58 | 70 | 147 | 31 | 8 | 0 | 0 | 0 | 4 | 0 |
| SG0000004 | Cfap69 | 0.007857 | 0.013066 | 0.753886328 | 66.89974709 | 112.8150068 | 838 | 46 | 58 | 123 | 467 | 89 | 20 | 0 | 0 | 4 | 11 | 0 |
| SG0000002 | Lima1 | 0.026308 | 0.039887 | -0.302867071 | 5380.805588 | 4361.89521 | 213 | 49 | 58 | 69 | 114 | 15 | 10 | 0 | 0 | 0 | 2 | 0 |
| SG0000003 | Rbm5 | 0.143633 | 0.188895 | 0.191124569 | 2545.604259 | 2906.197449 | 313 | 52 | 58 | 85 | 192 | 28 | 4 | 0 | 0 | 0 | 7 | 3 |
| SG0000002 | Sema3c | 0.148523 | 0.194812 | -0.745355287 | 425.8068748 | 254.0027216 | 753 | 14 | 58 | 114 | 384 | 60 | 11 | 0 | 3 | 1 | 5 | 2 |
| SG0000002 | Dyrk1a | 0.221878 | 0.279466 | -0.179294854 | 1807.279564 | 1596.071007 | 505 | 42 | 58 | 84 | 242 | 34 | 10 | 0 | 0 | 0 | 2 | 0 |
| SG0000002 | Cfap54 | 0.278553 | 0.34172 | -0.583066481 | 51.44966425 | 34.34488328 | 817 | 22 | 58 | 105 | 385 | 71 | 8 | 2 | 2 | 2 | 4 | 2 |
| SG0000002 | Pter | 0.944747 | 0.961004 | 0.008286475 | 280.9297321 | 282.5479642 | 296 | 44 | 58 | 80 | 222 | 41 | 4 | 0 | 0 | 2 | 4 | 1 |
| SG0000007 | Rad54b | 3.28E-51 | 7.02E-50 | 2.14973416 | 333.7143311 | 1480.844042 | 330 | 36 | 57 | 79 | 177 | 18 | 11 | 1 | 5 | 1 | 2 | 1 |
| SG0000002 | Mgat4a | 1.17E-47 | 2.25E-46 | -2.529142501 | 3928.034932 | 680.4991469 | 543 | 26 | 57 | 97 | 267 | 77 | 7 | 2 | 2 | 2 | 12 | 6 |
| SG0000003 | Usp37 | 4.41E-47 | 8.36E-46 | 1.78994453 | 1187.530584 | 4106.49972 | 412 | 42 | 57 | 80 | 237 | 21 | 4 | 0 | 1 | 0 | 3 | 5 |
| SG0000002 | Traf3 | 3.33E-43 | 5.46E-42 | -2.381258015 | 3345.475343 | 642.1370719 | 438 | 33 | 57 | 114 | 231 | 75 | 9 | 0 | 0 | 6 | 6 | 0 |
| SG0000000 | Pcbp3 | 3.74E-25 | 3.04E-24 | 4.829028829 | 21.22966132 | 603.4281877 | 705 | 37 | 57 | 98 | 384 | 118 | 3 | 0 | 0 | 0 | 0 | 0 |
| SG0000002 | Ptbp2 | 6.26E-21 | 4.14E-20 | 1.345239415 | 501.5305338 | 1274.250444 | 795 | 31 | 57 | 97 | 378 | 53 | 17 | 1 | 1 | 1 | 2 | 0 |
| SG0000003 | Stim2 | 5.78E-18 | 3.24E-17 | -1.492495561 | 1284.917137 | 456.6560219 | 681 | 32 | 57 | 98 | 304 | 37 | 20 | 0 | 0 | 5 | 17 | 0 |
| SG0000006 | Maml3 | 3.27E-16 | 1.67E-15 | -3.04728444 | 477.8184873 | 57.80147583 | 703 | 30 | 57 | 104 | 312 | 74 | 8 | 0 | 0 | 0 | 2 | 2 |
| SG0000000 | Adcy9 | 5.81E-14 | 2.57E-13 | -2.317354905 | 361.5624209 | 72.54207076 | 672 | 30 | 57 | 114 | 318 | 48 | 0 | 0 | 0 | 0 | 0 | 0 |
| SG0000001 | Mapt | 1.88E-13 | 8.05E-13 | -1.254563289 | 1218.831352 | 510.8371057 | 324 | 54 | 57 | 81 | 189 | 30 | 3 | 0 | 0 | 0 | 0 | 0 |
| SG0000003 | Eml4 | 2.62E-13 | 1.11E-12 | 0.913691433 | 2241.057362 | 4221.837382 | 903 | 18 | 57 | 96 | 372 | 60 | 3 | 0 | 0 | 0 | 0 | 0 |
| SG0000002 | Osbpl3 | 5.71E-13 | 2.36E-12 | -1.303335843 | 1932.617467 | 783.0738426 | 605 | 18 | 57 | 87 | 219 | 50 | 9 | 0 | 0 | 0 | 0 | 0 |
| SG0000004 | Trrap | 2.22E-12 | 8.82E-12 | -1.010218858 | 5542.740375 | 2751.809452 | 442 | 45 | 57 | 79 | 190 | 47 | 8 | 0 | 0 | 0 | 3 | 0 |
| SG0000003 | Vav3 | 6.81E-06 | 1.63E-05 | -1.546264039 | 204.0892899 | 69.87926973 | 1041 | 18 | 57 | 127 | 460 | 84 | 8 | 0 | 0 | 0 | 4 | 2 |
| SG0000005 | Pcdh19 | 8.59E-06 | 2.04E-05 | -1.989555261 | 254.4817304 | 64.08269941 | 857 | 35 | 57 | 114 | 437 | 43 | 42 | 6 | 9 | 9 | 35 | 3 |
| ISG000000 | Hydin | 1.48E-05 | 3.43E-05 | 7.073607108 | 1.02742554 | 138.3943227 | 516 | 36 | 57 | 90 | 281 | 41 | 2 | 0 | 0 | 0 | 1 | 0 |
| ISG000000 | Mpp7 | 3.93E-05 | 8.74E-05 | -0.643873517 | 1306.208999 | 835.9637193 | 605 | 37 | 57 | 93 | 320 | 49 | 11 | 0 | 0 | 0 | 4 | 2 |
| SG0000006 | Fkbp15 | 0.000133 | 0.000279 | -0.605654604 | 1107.991821 | 728.1424513 | 431 | 38 | 57 | 73 | 206 | 30 | 3 | 0 | 1 | 1 | 3 | 0 |
| SG0000003 | Fam214a | 0.000226 | 0.000461 | -0.777835569 | 441.0986651 | 257.2668953 | 260 | 44 | 57 | 61 | 130 | 32 | 6 | 0 | 2 | 1 | 1 | 1 |
| SG0000003 | Cyfip1 | 0.000584 | 0.001128 | -0.473695954 | 4476.642572 | 3223.708242 | 481 | 51 | 57 | 99 | 300 | 33 | 4 | 0 | 0 | 0 | 3 | 0 |
| SG0000002 | Mtdh | 0.001 | 0.001875 | -0.45297512 | 4998.436391 | 3651.531668 | 445 | 45 | 57 | 87 | 245 | 19 | 8 | 0 | 0 | 0 | 2 | 1 |
| SG0000002 | Tns3 | 0.002799 | 0.00496 | 0.385928722 | 2392.233644 | 3125.93336 | 330 | 46 | 57 | 84 | 182 | 26 | 2 | 0 | 1 | 1 | 1 | 0 |
| ISG000000 | Rttn | 0.003067 | 0.0054 | -0.468052181 | 1131.692215 | 818.145569 | 506 | 32 | 57 | 95 | 277 | 50 | 1 | 0 | 1 | 0 | 0 | 0 |
| SG0000000 | Prkar2b | 0.005434 | 0.009247 | -0.535140521 | 428.8931613 | 295.9755067 | 621 | 27 | 57 | 105 | 258 | 27 | 15 | 0 | 0 | 0 | 2 | 0 |
| SG0000000 | Arid1a | 0.005746 | 0.009744 | 0.351327173 | 3701.680271 | 4722.358151 | 372 | 39 | 57 | 81 | 187 | 23 | 7 | 0 | 0 | 3 | 3 | 2 |
| ISG000000 | Vkorc1l1 | 0.018706 | 0.029087 | 0.306557533 | 1850.474337 | 2288.5799 | 369 | 44 | 57 | 65 | 194 | 43 | 6 | 0 | 1 | 1 | 5 | 3 |
| SG0000002 | Acly | 0.229023 | 0.287603 | -0.153394667 | 9868.264214 | 8872.875197 | 280 | 36 | 57 | 40 | 99 | 3 | 9 | 0 | 0 | 0 | 3 | 0 |
| SG0000002 | Amph | 1.1E-104 | 8.8E-103 | 9.141484106 | 0.639110347 | 360.9414586 | 898 | 22 | 56 | 118 | 422 | 76 | 4 | 0 | 0 | 4 | 4 | 0 |
| SG0000004 | Rundc3b | 4.49E-74 | 1.82E-72 | 8.480141541 | 0.674076249 | 240.7054902 | 625 | 28 | 56 | 112 | 348 | 55 | 3 | 0 | 0 | 0 | 0 | 3 |
| SG0000004 | Kazn | 3.82E-72 | 1.45E-70 | 4.269964969 | 130.6179694 | 2519.94153 | 329 | 50 | 56 | 78 | 198 | 51 | 8 | 0 | 0 | 4 | 6 | 6 |
| SG0000003 | Me1 | 7.9E-52 | 1.72E-50 | -2.659352814 | 4600.116268 | 728.1550108 | 546 | 37 | 56 | 90 | 256 | 50 | 9 | 0 | 0 | 2 | 2 | 0 |
| SG0000003 | Zfp800 | 3.19E-29 | 3.12E-28 | -2.451873519 | 991.0140411 | 181.1308116 | 516 | 36 | 56 | 81 | 311 | 37 | 18 | 0 | 0 | 2 | 7 | 0 |
| SG0000002 | Pld2 | 9.71E-29 | 9.26E-28 | -2.21359545 | 1166.428374 | 251.4776809 | 123 | 54 | 56 | 67 | 104 | 7 | 14 | 8 | 8 | 9 | 11 | 0 |
| SG0000003 | Sh3glb1 | 1.13E-25 | 9.4E-25 | -1.635561064 | 4984.79555 | 1604.332607 | 372 | 33 | 56 | 74 | 184 | 22 | 24 | 1 | 4 | 7 | 10 | 0 |
| SG0000002 | Pia2 | 1.3E-19 | 7.97E-19 | -1.554444288 | 3578.949323 | 1218.487818 | 446 | 32 | 56 | 86 | 270 | 38 | 28 | 0 | 0 | 0 | 16 | 4 |
| SG0000003 | Kctd1 | 4.25E-18 | 2.4E-17 | -1.865581691 | 523.3617213 | 143.6170186 | 461 | 35 | 56 | 85 | 199 | 44 | 2 | 0 | 0 | 0 | 0 | 0 |
| SG0000004 | Elovl6 | 6.19E-18 | 3.47E-17 | 1.341174683 | 1469.084931 | 3722.037232 | 455 | 41 | 56 | 75 | 178 | 39 | 8 | 2 | 2 | 3 | 7 | 0 |
| SG0000003 | Igf2bp2 | 3.93E-12 | 1.54E-11 | -0.994256093 | 6036.594922 | 3030.338382 | 370 | 52 | 56 | 86 | 172 | 20 | 8 | 0 | 0 | 0 | 0 | 2 |
| SG0000002 | Pitpnm2 | 4.14E-12 | 1.62E-11 | -1.383127696 | 528.7296796 | 202.7079425 | 442 | 44 | 56 | 80 | 202 | 51 | 4 | 0 | 0 | 4 | 7 | 0 |
| SG0000004 | Snx12 | 9.65E-12 | 3.69E-11 | -1.054291621 | 2105.77551 | 1014.001666 | 382 | 41 | 56 | 87 | 228 | 31 | 1 | 0 | 0 | 0 | 0 | 2 |
| SG0000000 | Stat3 | 1.58E-11 | 5.97E-11 | -0.96000145 | 6087.665738 | 3129.403514 | 280 | 44 | 56 | 72 | 130 | 32 | 4 | 0 | 0 | 0 | 2 | 0 |
| SG0000002 | Auts2 | 2.16E-11 | 8.06E-11 | 4.302972814 | 21.64575858 | 427.2643893 | 419 | 41 | 56 | 71 | 207 | 62 | 3 | 0 | 0 | 0 | 0 | 2 |
| SG0000001 | Stx5a | 1.31E-10 | 4.61E-10 | -1.029249673 | 1405.85402 | 688.8191342 | 287 | 56 | 56 | 70 | 112 | 0 | 0 | 0 | 0 | 0 | 0 | 0 |

| | | | | | | | | | | | | | | | | | | |
|---|---|---|---|---|---|---|---|---|---|---|---|---|---|---|---|---|---|---|
| SG0000003 | Alg9 | 2.5E-10 | 8.65E-10 | -1.026940347 | 1378.652204 | 676.5733253 | 382 | 42 | 56 | 80 | 222 | 14 | 5 | 0 | 2 | 0 | 2 | 0 |
| SG0000002 | Eps15 | 1.37E-09 | 4.46E-09 | -0.899634698 | 3004.858969 | 1610.671834 | 717 | 28 | 56 | 89 | 295 | 32 | 11 | 0 | 0 | 0 | 0 | 1 |
| SG0000003 | Ctdspl2 | 5.03E-09 | 1.57E-08 | 0.798976778 | 997.7819864 | 1736.007652 | 639 | 44 | 56 | 112 | 323 | 49 | 19 | 0 | 2 | 6 | 12 | 2 |
| JSG0000004 | Rapgef5 | 3.2E-06 | 7.88E-06 | 1.190403213 | 159.7682003 | 364.617423 | 476 | 40 | 56 | 73 | 208 | 56 | 0 | 0 | 0 | 0 | 0 | 0 |
| SG0000003 | 3110057O12Rik | 0.000214 | 0.000437 | 1.098382429 | 113.0156623 | 241.9829086 | 399 | 38 | 56 | 76 | 200 | 59 | 15 | 0 | 0 | 3 | 3 | 2 |
| SG0000002 | Cit | 0.000238 | 0.000482 | 0.5162346 | 965.0339458 | 1380.208474 | 266 | 40 | 56 | 71 | 132 | 18 | 4 | 0 | 0 | 0 | 0 | 1 |
| SG0000001 | Atp6v0a1 | 0.00031 | 0.00062 | 0.561314504 | 561.5410463 | 828.6173249 | 293 | 40 | 56 | 76 | 192 | 12 | 3 | 0 | 0 | 0 | 0 | 0 |
| JSG000000 | Kctd16 | 0.000334 | 0.000667 | -4.222552917 | 37.97198269 | 2.033987071 | 635 | 34 | 56 | 108 | 377 | 67 | 0 | 0 | 0 | 0 | 0 | 0 |
| SG0000002 | Sass6 | 0.000473 | 0.000927 | 0.612380703 | 284.8555712 | 435.4814679 | 306 | 28 | 56 | 56 | 178 | 48 | 13 | 0 | 0 | 1 | 3 | 1 |
| SG0000003 | Mpp1 | 0.000599 | 0.001156 | -0.508203891 | 1710.972963 | 1202.980346 | 187 | 40 | 56 | 56 | 158 | 6 | 3 | 0 | 0 | 0 | 0 | 0 |
| JSG000000 | Nbas | 0.001327 | 0.002457 | -0.526699743 | 1082.874473 | 751.6673648 | 649 | 38 | 56 | 113 | 351 | 59 | 1 | 0 | 0 | 0 | 0 | 0 |
| SG0000003 | Arhgap39 | 0.020854 | 0.032157 | -0.507062498 | 884.6306599 | 622.4736409 | 336 | 30 | 56 | 66 | 162 | 34 | 4 | 2 | 2 | 2 | 2 | 0 |
| SG0000000 | Dgcr2 | 0.04226 | 0.061803 | -0.294084001 | 1586.284475 | 1293.757762 | 367 | 51 | 56 | 76 | 214 | 36 | 0 | 0 | 0 | 0 | 0 | 4 |
| SG0000002 | Cdh10 | 0.202186 | 0.257359 | -0.731572454 | 28.53206867 | 17.18335679 | 572 | 25 | 56 | 78 | 267 | 45 | 3 | 0 | 1 | 2 | 2 | 0 |
| JSG000000 | Vwa8 | 0.391564 | 0.460189 | -0.129440637 | 1362.006949 | 1245.127454 | 636 | 29 | 56 | 106 | 299 | 81 | 1 | 0 | 0 | 0 | 0 | 0 |
| SG0000003 | Fbxl5 | 0.416977 | 0.486319 | -0.119630388 | 2192.851467 | 2018.351866 | 877 | 29 | 56 | 97 | 272 | 57 | 27 | 0 | 0 | 0 | 1 | 2 |
| JSG000000 | Ubl5 | 0.69537 | 0.749832 | 0.052018202 | 1834.209039 | 1901.550521 | 93 | 49 | 56 | 70 | 77 | 0 | 0 | 0 | 0 | 0 | 0 | 0 |
| SG0000002 | Rab1a | 0.730639 | 0.781606 | 0.03788528 | 8771.445811 | 9004.835643 | 282 | 52 | 56 | 83 | 189 | 12 | 6 | 0 | 0 | 3 | 5 | 0 |
| SG0000002 | Fip1l1 | 0.898665 | 0.922881 | -0.024255738 | 2999.899902 | 2949.88482 | 585 | 28 | 56 | 68 | 286 | 42 | 8 | 0 | 0 | 0 | 4 | 0 |
| SG0000003 | Cul5 | 0.922286 | 0.943622 | 0.007484508 | 2116.004737 | 2127.01081 | 452 | 34 | 56 | 77 | 180 | 30 | 25 | 0 | 0 | 4 | 8 | 2 |
| SG0000002 | Flt1 | 0.991161 | 1 | -0.042738279 | 280.0928069 | 271.9170574 | 374 | 45 | 56 | 81 | 220 | 57 | 13 | 1 | 1 | 2 | 8 | 1 |
| SG0000004 | Tet1 | 2.9E-245 | 1.3E-242 | 4.924893755 | 315.6865617 | 9589.519367 | 325 | 40 | 55 | 77 | 173 | 33 | 20 | 2 | 2 | 6 | 12 | 6 |
| SG0000002 | Myo10 | 3.72E-91 | 2.17E-89 | 2.449587961 | 1253.572529 | 6847.766131 | 338 | 44 | 55 | 70 | 167 | 39 | 2 | 0 | 0 | 2 | 4 | 0 |
| SG0000004 | Ccdc141 | 2.72E-62 | 7.99E-61 | -4.464484604 | 2192.10515 | 99.29276428 | 762 | 29 | 55 | 126 | 376 | 60 | 18 | 1 | 1 | 4 | 7 | 0 |
| SG0000002 | Espn | 8.23E-56 | 2.04E-54 | 4.559161397 | 12.49041204 | 294.4564696 | 98 | 55 | 55 | 55 | 86 | 3 | 0 | 0 | 0 | 0 | 0 | 0 |
| SG0000003 | Klhl13 | 1.08E-47 | 2.1E-46 | 6.871850716 | 26.44978536 | 3097.814038 | 1159 | 14 | 55 | 172 | 628 | 105 | 20 | 0 | 0 | 0 | 0 | 0 |
| SG0000003 | Fam53b | 2.97E-35 | 3.73E-34 | -2.664464935 | 1300.271037 | 205.0926143 | 394 | 31 | 55 | 86 | 196 | 43 | 3 | 0 | 0 | 0 | 6 | 3 |
| SG0000005 | R3hdm1 | 7.76E-28 | 7.12E-27 | 1.392157933 | 1577.425849 | 4140.285119 | 570 | 34 | 55 | 89 | 303 | 74 | 9 | 1 | 1 | 1 | 2 | 0 |
| SG0000003 | Dctd | 7.96E-28 | 7.3E-27 | 1.433801786 | 886.861178 | 2395.923734 | 318 | 45 | 55 | 73 | 162 | 20 | 6 | 0 | 0 | 0 | 0 | 0 |
| JSG0000004 | Tnrc6b | 8.69E-22 | 6.04E-21 | -1.531031354 | 2882.246415 | 997.3434638 | 483 | 33 | 55 | 94 | 228 | 48 | 12 | 0 | 1 | 1 | 11 | 4 |
| SG0000001 | Ndel1 | 7.69E-20 | 4.78E-19 | -1.41180286 | 3206.934884 | 1205.299934 | 242 | 43 | 55 | 59 | 124 | 32 | 18 | 15 | 15 | 15 | 15 | 4 |
| SG0000005 | Ccdc18 | 8.12E-16 | 4.04E-15 | 1.507551697 | 361.9448808 | 1029.107443 | 304 | 39 | 55 | 73 | 221 | 18 | 10 | 0 | 0 | 0 | 4 | 0 |
| SG0000002 | Rptor | 5.91E-12 | 2.29E-11 | -1.034487107 | 2687.129839 | 1311.82832 | 725 | 40 | 55 | 111 | 294 | 55 | 3 | 0 | 0 | 2 | 2 | 2 |
| SG0000003 | Stt3a | 3.69E-10 | 1.26E-09 | -0.924128406 | 10254.04871 | 5403.871414 | 250 | 44 | 55 | 70 | 146 | 12 | 7 | 0 | 0 | 0 | 1 | 2 |
| SG0000003 | Acbd6 | 5.49E-08 | 1.58E-07 | 0.983682738 | 582.5026202 | 1151.902958 | 461 | 30 | 55 | 81 | 270 | 33 | 9 | 0 | 0 | 2 | 4 | 2 |
| SG0000004 | Fut10 | 1.39E-07 | 3.86E-07 | -0.974557043 | 627.8722618 | 319.5217269 | 438 | 36 | 55 | 80 | 239 | 46 | 10 | 0 | 0 | 1 | 7 | 1 |
| SG0000002 | Mkln1 | 2.53E-06 | 6.3E-06 | -0.71583806 | 2549.890935 | 1552.504533 | 342 | 41 | 55 | 65 | 174 | 32 | 22 | 0 | 0 | 1 | 5 | 4 |
| SG0000002 | Snapc3 | 0.00053 | 0.001031 | 0.610022968 | 353.0766856 | 538.8951219 | 479 | 25 | 55 | 69 | 174 | 69 | 23 | 5 | 5 | 5 | 7 | 1 |
| SG0000002 | Trnau1ap | 0.07872 | 0.109458 | -0.300308533 | 636.7855314 | 517.1199713 | 183 | 50 | 55 | 85 | 138 | 5 | 2 | 0 | 0 | 0 | 0 | 0 |
| SG0000003 | Srgap3 | 0.093391 | 0.127959 | 0.478791461 | 222.1587933 | 309.5951148 | 574 | 39 | 55 | 87 | 241 | 68 | 13 | 0 | 0 | 2 | 2 | 6 |
| SG0000000 | Dtx2 | 0.23056 | 0.289383 | -0.330014676 | 644.83643 | 512.9856877 | 215 | 40 | 55 | 95 | 140 | 15 | 0 | 0 | 0 | 0 | 0 | 0 |
| SG0000003 | Pkig | 0.276654 | 0.339692 | -0.348455114 | 388.8512003 | 305.41334 | 378 | 55 | 55 | 94 | 218 | 43 | 4 | 4 | 4 | 4 | 4 | 0 |
| SG0000003 | Slc10a7 | 0.281353 | 0.344706 | -0.170082039 | 1110.635015 | 987.12367 | 417 | 33 | 55 | 75 | 210 | 32 | 5 | 0 | 1 | 1 | 3 | 2 |
| SG0000003 | Senp1 | 0.317545 | 0.383383 | -0.143199959 | 3173.969551 | 2874.056312 | 281 | 36 | 55 | 79 | 160 | 24 | 11 | 0 | 1 | 4 | 4 | 0 |
| SG0000004 | Gng2 | 2.2E-171 | 4.6E-169 | 4.633382988 | 61.63243633 | 1529.66594 | 1006 | 18 | 54 | 98 | 392 | 72 | 10 | 0 | 0 | 2 | 2 | 0 |
| SG0000006 | Shroom4 | 1.57E-51 | 3.39E-50 | -3.015459282 | 2837.917141 | 350.9587031 | 658 | 29 | 54 | 119 | 382 | 73 | 4 | 0 | 0 | 0 | 4 | 2 |
| SG0000001 | Pitpna | 1.8E-43 | 2.99E-42 | -2.259024554 | 7852.804491 | 1640.55439 | 203 | 52 | 54 | 60 | 124 | 9 | 16 | 0 | 2 | 0 | 4 | 1 |
| JSG000000 | St6galnac3 | 3.12E-43 | 5.14E-42 | 4.82011131 | 6.517749395 | 184.1177863 | 845 | 26 | 54 | 107 | 350 | 58 | 9 | 0 | 1 | 2 | 5 | 0 |
| SG0000002 | Ipp | 1.79E-28 | 1.69E-27 | 1.636605778 | 357.3438154 | 1111.101414 | 194 | 50 | 54 | 62 | 156 | 12 | 0 | 0 | 0 | 0 | 0 | 0 |
| SG0000004 | Tmem241 | 1.89E-25 | 1.56E-24 | -2.269957281 | 621.6715491 | 128.8949082 | 434 | 40 | 54 | 74 | 188 | 32 | 11 | 2 | 2 | 2 | 10 | 2 |
| SG0000004 | Kcnk13 | 6.43E-21 | 4.26E-20 | -8.279821801 | 398.3787318 | 1.281803414 | 441 | 42 | 54 | 93 | 207 | 30 | 6 | 0 | 0 | 3 | 3 | 0 |
| SG0000003 | Nploc4 | 1.94E-18 | 1.11E-17 | -1.290813045 | 5823.950453 | 2380.368674 | 216 | 38 | 54 | 70 | 128 | 8 | 4 | 2 | 2 | 2 | 4 | 0 |
| SG0000002 | Ephb2 | 3.42E-16 | 1.74E-15 | 3.226421411 | 107.6022005 | 1007.09578 | 346 | 39 | 54 | 75 | 162 | 37 | 1 | 0 | 0 | 0 | 0 | 1 |
| SG0000002 | Mxi1 | 8.95E-16 | 4.44E-15 | -1.492462665 | 3102.767892 | 1102.740332 | 249 | 42 | 54 | 61 | 112 | 24 | 27 | 0 | 0 | 0 | 3 | 0 |
| SG0000002 | Igsf11 | 2.57E-11 | 9.55E-11 | -2.334733557 | 341.7947773 | 67.75488147 | 465 | 38 | 54 | 92 | 202 | 36 | 7 | 0 | 0 | 0 | 0 | 0 |
| SG0000005 | Ccdc79 | 1.63E-10 | 5.7E-10 | 6.479468354 | 0.331601068 | 29.58899754 | 448 | 36 | 54 | 75 | 234 | 36 | 5 | 0 | 0 | 0 | 0 | 0 |
| SG0000002 | Abcd3 | 3.87E-10 | 1.32E-09 | -0.995531928 | 1635.969316 | 820.5219089 | 434 | 39 | 54 | 87 | 282 | 51 | 6 | 0 | 0 | 0 | 2 | 0 |
| SG0000000 | Bcl11a | 7.06E-09 | 2.17E-08 | 5.085872339 | 1.609821762 | 54.67363159 | 642 | 24 | 54 | 91 | 285 | 54 | 11 | 0 | 3 | 5 | 11 | 1 |
| SG0000001 | Med1 | 1.37E-08 | 4.13E-08 | -0.82150775 | 3526.742864 | 1995.608393 | 302 | 36 | 54 | 75 | 170 | 8 | 8 | 2 | 2 | 2 | 6 | 2 |
| SG0000000 | Col26a1 | 2.3E-07 | 6.23E-07 | 4.245936171 | 0 | 17.97379252 | 294 | 46 | 54 | 74 | 170 | 28 | 4 | 0 | 0 | 0 | 0 | 0 |
| SG0000003 | Pcyt1b | 6.66E-07 | 1.74E-06 | -0.718327913 | 3130.476805 | 1902.708427 | 337 | 37 | 54 | 59 | 202 | 29 | 2 | 0 | 0 | 0 | 0 | 2 |
| SG0000005 | Cntrl | 0.000161 | 0.000333 | 0.668074655 | 914.8370231 | 1453.631222 | 661 | 32 | 54 | 115 | 324 | 49 | 8 | 0 | 1 | 3 | 8 | 0 |
| SG0000004 | Bdp1 | 0.000437 | 0.000859 | -0.583509146 | 2578.535818 | 1720.756476 | 542 | 32 | 54 | 94 | 309 | 42 | 25 | 0 | 2 | 8 | 14 | 4 |
| SG0000002 | Gmeb1 | 0.002928 | 0.005173 | 0.439822483 | 756.2017241 | 1025.740311 | 251 | 50 | 54 | 69 | 147 | 34 | 15 | 2 | 2 | 5 | 17 | 0 |
| SG0000000 | Aven | 0.018637 | 0.028995 | 0.465375798 | 400.378897 | 552.7939143 | 443 | 44 | 54 | 70 | 221 | 44 | 5 | 0 | 0 | 1 | 1 | 0 |
| SG0000001 | Slc25a13 | 0.25169 | 0.312618 | 0.144757897 | 2778.870366 | 3072.166552 | 632 | 32 | 54 | 88 | 288 | 42 | 4 | 0 | 0 | 2 | 6 | 0 |
| SG0000003 | Ate1 | 0.867775 | 0.897591 | 0.017802291 | 1314.956779 | 1331.283354 | 624 | 20 | 54 | 78 | 312 | 54 | 12 | 0 | 2 | 2 | 12 | 4 |
| SG0000002 | Svil | 0.985622 | 0.996301 | -0.004820458 | 952.8377909 | 949.6593994 | 1060 | 35 | 54 | 129 | 407 | 150 | 9 | 0 | 0 | 4 | 8 | 0 |
| JSG000000 | Arhgap10 | 9.86E-49 | 1.97E-47 | 2.753092337 | 87.54666235 | 590.2051136 | 406 | 38 | 53 | 81 | 214 | 35 | 1 | 0 | 0 | 0 | 0 | 0 |
| SG0000002 | Sptbn1 | 3.21E-43 | 5.27E-42 | -2.346392658 | 28286.07948 | 5562.092226 | 702 | 35 | 53 | 98 | 265 | 90 | 12 | 1 | 1 | 1 | 2 | 2 |
| SG0000002 | Tmeff1 | 2.56E-42 | 4.08E-41 | 1.754428159 | 859.2594772 | 2899.077544 | 378 | 36 | 53 | 81 | 215 | 24 | 10 | 0 | 0 | 3 | 5 | 0 |
| JSG000000 | Lemd1 | 4.99E-23 | 3.66E-22 | -1.952918177 | 812.9588207 | 209.9817725 | 340 | 38 | 53 | 88 | 163 | 12 | 4 | 0 | 0 | 0 | 1 | 0 |
| SG0000002 | Itgav | 8.53E-20 | 5.28E-19 | -1.587057108 | 4497.584056 | 1497.019627 | 398 | 34 | 53 | 73 | 187 | 33 | 38 | 0 | 2 | 6 | 12 | 4 |
| SG0000002 | Depdc1a | 8.43E-15 | 3.93E-14 | 1.43804236 | 160.4625685 | 434.7780647 | 386 | 42 | 53 | 68 | 203 | 18 | 5 | 0 | 2 | 2 | 5 | 0 |
| SG0000002 | Nr3c1 | 2.81E-13 | 1.18E-12 | -1.416617855 | 1341.83263 | 502.6363372 | 611 | 21 | 53 | 87 | 297 | 59 | 29 | 0 | 3 | 0 | 8 | 6 |
| SG0000002 | Srrm1 | 7.6E-13 | 3.12E-12 | 0.862829146 | 5670.331385 | 10312.071 | 276 | 48 | 53 | 62 | 172 | 18 | 8 | 0 | 0 | 0 | 7 | 0 |
| SG0000004 | Gigyf2 | 3.27E-09 | 1.03E-08 | 0.761486098 | 2573.117499 | 4362.04132 | 631 | 33 | 53 | 92 | 320 | 74 | 1 | 0 | 0 | 0 | 2 | 1 |
| SG0000003 | Ncoa6 | 4.78E-09 | 1.49E-08 | -0.876187076 | 2498.03811 | 1360.944666 | 333 | 40 | 53 | 77 | 227 | 36 | 5 | 2 | 2 | 2 | 6 | 2 |
| SG0000001 | Arnt | 1.11E-07 | 3.1E-07 | 0.672788213 | 2252.181869 | 3590.317788 | 635 | 53 | 53 | 121 | 385 | 37 | 10 | 0 | 0 | 2 | 5 | 0 |
| SG0000003 | Trpc1 | 2.17E-07 | 5.91E-07 | -1.24859564 | 443.6005392 | 186.6926951 | 513 | 21 | 53 | 101 | 294 | 53 | 9 | 1 | 1 | 5 | 6 | 1 |
| SG0000006 | Phactr2 | 9.75E-06 | 2.3E-05 | 1.86617927 | 42.97981462 | 156.6897096 | 789 | 29 | 53 | 101 | 375 | 64 | 0 | 0 | 0 | 0 | 0 | 0 |
| SG0000003 | Wdr78 | 0.000243 | 0.000492 | 1.668146715 | 14.688817 | 46.68195315 | 655 | 28 | 53 | 111 | 327 | 40 | 13 | 0 | 0 | 0 | 0 | 0 |
| SG0000002 | Kif26b | 0.00267 | 0.004748 | -0.922021037 | 110.853426 | 58.50501983 | 803 | 33 | 53 | 112 | 317 | 106 | 3 | 0 | 0 | 0 | 0 | 3 |
| SG0000004 | Sacs | 0.007287 | 0.012175 | 0.403379754 | 887.8413297 | 1174.261342 | 482 | 28 | 53 | 77 | 220 | 41 | 14 | 0 | 0 | 1 | 7 | 2 |
| SG0000002 | Abcg2 | 0.051213 | 0.07377 | -0.274568291 | 2051.166987 | 1695.694948 | 557 | 30 | 53 | 86 | 298 | 45 | 1 | 0 | 0 | 0 | 6 | 0 |
| JSG000000 | Nalcn | 0.135255 | 0.178942 | 0.653894046 | 29.82016661 | 46.91932843 | 771 | 23 | 53 | 96 | 391 | 76 | 3 | 0 | 0 | 2 | 3 | 0 |
| JSG000000 | Gm28043 | 0.145247 | 0.190863 | 0.570372685 | 40.77303225 | 60.5441655 | 150 | 45 | 53 | 60 | 103 | 6 | 7 | 0 | 1 | 1 | 2 | 0 |
| SG0000002 | Kpna1 | 0.398834 | 0.467717 | -0.118452676 | 4545.027811 | 4186.766461 | 405 | 42 | 53 | 100 | 236 | 47 | 11 | 0 | 1 | 1 | 2 | 0 |
| SG0000003 | Tpm1 | 0.663152 | 0.721344 | 0.107096693 | 4332.884718 | 4666.77058 | 683 | 25 | 53 | 64 | 151 | 18 | 50 | 2 | 12 | 3 | 4 | 0 |
| JSG000000 | Atf6 | 0.89116 | 0.916645 | -0.023129291 | 1865.227839 | 1835.56293 | 355 | 37 | 53 | 76 | 202 | 26 | 7 | 0 | 0 | 0 | 1 | 1 |
| SG0000000 | Mcu | 0.906744 | 0.929909 | 0.011721609 | 850.1812463 | 857.1169366 | 445 | 30 | 53 | 73 | 243 | 24 | 6 | 1 | 2 | 2 | 4 | 1 |
| SG0000004 | Lhfp | 9.38E-59 | 2.5E-57 | -3.309160027 | 1793.798346 | 180.9744188 | 830 | 34 | 52 | 108 | 354 | 70 | 0 | 0 | 0 | 0 | 2 | 0 |
| SG0000003 | Usp10 | 3.49E-47 | 6.65E-46 | 1.741767864 | 2085.45028 | 6974.678807 | 170 | 52 | 52 | 60 | 123 | 4 | 0 | 0 | 0 | 0 | 0 | 0 |
| SG0000006 | Slc4a5 | 9.02E-39 | 1.27E-37 | 5.575724974 | 2.955630691 | 140.9647847 | 262 | 45 | 52 | 61 | 140 | 21 | 4 | 0 | 0 | 0 | 0 | 0 |
| SG0000004 | Urgcp | 5.81E-35 | 7.22E-34 | -2.437354226 | 1907.432522 | 352.1538584 | 327 | 39 | 52 | 67 | 155 | 47 | 3 | 0 | 0 | 0 | 0 | 0 |
| SG0000002 | Prim2 | 6.98E-31 | 7.33E-30 | 1.548605286 | 691.8276022 | 2023.832323 | 456 | 30 | 52 | 82 | 230 | 38 | 1 | 0 | 0 | 0 | 0 | 0 |
| SG0000004 | Lyn | 1.33E-27 | 1.21E-26 | -3.841254569 | 654.1277786 | 45.63833021 | 382 | 42 | 52 | 80 | 230 | 34 | 4 | 0 | 0 | 0 | 0 | 2 |
| SG0000003 | Fam60a | 2.93E-25 | 2.4E-24 | 1.399812318 | 791.6632677 | 2088.940119 | 168 | 52 | 52 | 78 | 123 | 7 | 15 | 4 | 4 | 6 | 6 | 4 |
| SG0000003 | Enpp1 | 7.75E-25 | 6.21E-24 | -3.626814892 | 561.5686225 | 45.45930194 | 346 | 44 | 52 | 78 | 203 | 24 | 2 | 0 | 0 | 0 | 1 | 0 |
| JSG000000 | Ksr2 | 1.15E-23 | 8.75E-23 | 7.67798939 | 0.331601068 | 67.90801472 | 467 | 40 | 52 | 100 | 251 | 65 | 5 | 0 | 1 | 1 | 2 | 2 |
| SG0000002 | Kcnip2 | 1E-22 | 7.27E-22 | 4.260569972 | 8.031203994 | 153.9359363 | 109 | 48 | 52 | 51 | 67 | 12 | 0 | 0 | 0 | 0 | 1 | 0 |
| SG0000003 | Dpp8 | 3.95E-20 | 2.5E-19 | -1.447868788 | 2854.886141 | 1046.494276 | 268 | 40 | 52 | 64 | 156 | 8 | 2 | 0 | 0 | 0 | 0 | 0 |
| SG0000003 | Arhgef6 | 2.46E-19 | 1.48E-18 | 3.171389685 | 19.37373154 | 174.5406948 | 365 | 41 | 52 | 89 | 202 | 21 | 2 | 0 | 0 | 0 | 2 | 1 |
| SG0000003 | Spon1 | 3.08E-19 | 1.85E-18 | 5.629480173 | 1.278220693 | 63.27734317 | 692 | 25 | 52 | 95 | 306 | 61 | 5 | 0 | 0 | 1 | 3 | 1 |
| SG0000002 | Kifap3 | 6.96E-19 | 4.1E-18 | 1.160356437 | 1067.457625 | 2385.902748 | 656 | 18 | 52 | 86 | 354 | 68 | 0 | 0 | 0 | 0 | 0 | 0 |
| SG0000003 | Lrmp | 8.76E-19 | 5.14E-18 | -2.183012676 | 909.0152484 | 200.1791645 | 275 | 48 | 52 | 72 | 229 | 14 | 1 | 0 | 0 | 0 | 0 | 0 |
| SG0000006 | Gak | 1.07E-16 | 5.62E-16 | -1.21827253 | 5452.439941 | 2343.442577 | 483 | 38 | 52 | 72 | 248 | 62 | 11 | 0 | 0 | 0 | 0 | 7 |
| JSG000000 | Grk5 | 4.56E-16 | 2.3E-15 | -2.04638087 | 698.8117611 | 169.1757776 | 360 | 42 | 52 | 78 | 190 | 26 | 1 | 0 | 0 | 0 | 1 | 0 |
| SG0000002 | Ttll11 | 9.04E-13 | 3.69E-12 | -3.339322701 | 84.95191463 | 8.393379755 | 560 | 38 | 52 | 94 | 254 | 44 | 0 | 0 | 0 | 0 | 0 | 0 |
| SG0000007 | Ccdc162 | 1.44E-11 | 5.43E-11 | 3.806133196 | 3.594741038 | 50.28377439 | 373 | 40 | 52 | 78 | 194 | 44 | 0 | 0 | 0 | 0 | 0 | 0 |
| SG0000001 | Plxdc1 | 2.16E-11 | 8.06E-11 | 1.371166015 | 120.7100843 | 312.2523048 | 148 | 46 | 52 | 58 | 112 | 0 | 4 | 0 | 0 | 0 | 0 | 0 |
| SG0000003 | Atp6v1h | 1.02E-09 | 3.35E-09 | 0.784135234 | 1924.910571 | 3314.811095 | 624 | 35 | 52 | 111 | 392 | 52 | 6 | 0 | 0 | 2 | 2 | 0 |
| SG0000003 | Cdk5rap2 | 1.07E-09 | 3.51E-09 | 0.818697254 | 1073.710894 | 1893.82476 | 705 | 34 | 52 | 89 | 295 | 47 | 8 | 0 | 0 | 0 | 5 | 0 |
| SG0000002 | Drosha | 4.58E-08 | 1.32E-07 | 0.672797802 | 4107.022415 | 6547.257059 | 384 | 46 | 52 | 78 | 164 | 24 | 0 | 0 | 0 | 0 | 0 | 0 |
| SG0000004 | Ssh1 | 3.14E-07 | 8.42E-07 | -0.773581596 | 1847.947442 | 1080.981722 | 195 | 46 | 52 | 60 | 118 | 23 | 8 | 0 | 0 | 1 | 4 | 2 |
| SG0000002 | Slk | 3.74E-07 | 9.97E-07 | -0.734128567 | 5850.364188 | 3517.12807 | 356 | 42 | 52 | 80 | 179 | 20 | 4 | 0 | 0 | 0 | 3 | 0 |
| SG0000004 | Sytl3 | 9.92E-05 | 0.00021 | 6.527321442 | 0.663202137 | 61.17380048 | 374 | 33 | 52 | 62 | 130 | 59 | 3 | 0 | 0 | 0 | 0 | 0 |
| SG0000001 | Stard13 | 0.000554 | 0.001075 | -0.884517251 | 204.5773998 | 110.8131831 | 614 | 35 | 52 | 75 | 254 | 46 | 8 | 2 | 2 | 3 | 9 | 5 |
| SG0000004 | Naa25 | 0.002612 | 0.004654 | 0.37983917 | 3468.727982 | 4513.49772 | 197 | 43 | 52 | 67 | 118 | 20 | 3 | 0 | 0 | 0 | 0 | 2 |
| JSG000000 | Uvrag | 0.010212 | 0.016678 | -0.401384742 | 1027.946146 | 778.290117 | 453 | 35 | 52 | 83 | 231 | 26 | 3 | 0 | 0 | 0 | 0 | 0 |
| SG0000007 | Cpne1 | 0.01485 | 0.023547 | -1.380272778 | 31.69884147 | 12.1769881 | 361 | 30 | 52 | 72 | 190 | 52 | 8 | 0 | 0 | 0 | 7 | 0 |
| SG0000004 | 9930021J03Rik | 0.094225 | 0.128995 | -0.29628349 | 1524.700015 | 1241.635688 | 718 | 22 | 52 | 88 | 330 | 79 | 4 | 0 | 0 | 2 | 5 | 2 |
| SG0000001 | Ikzf1 | 0.124415 | 0.165874 | -0.665682058 | 69.09192151 | 43.55489828 | 403 | 36 | 52 | 71 | 204 | 68 | 6 | 0 | 0 | 0 | 2 | 2 |

| | | | | | | | | | | | | | | | | | | |
|---|---|---|---|---|---|---|---|---|---|---|---|---|---|---|---|---|---|---|
| SG0000003 | Larp4b | 0.18731 | 0.240354 | -0.18176433 | 4146.697665 | 3655.828857 | 551 | 27 | 52 | 71 | 199 | 27 | 11 | 0 | 1 | 2 | 4 | 3 |
| SG0000003 | Gramd1c | 0.207434 | 0.263402 | -0.427933525 | 69.89787737 | 51.95689152 | 351 | 39 | 52 | 58 | 157 | 40 | 27 | 0 | 0 | 6 | 15 | 9 |
| SG0000002 | Med27 | 0.396733 | 0.465589 | 0.159351204 | 338.9528721 | 378.5374137 | 659 | 26 | 52 | 68 | 235 | 40 | 1 | 0 | 0 | 0 | 1 | 0 |
| SG0000003 | Eftud1 | 0.507889 | 0.574906 | -0.102432718 | 1086.751941 | 1012.267058 | 538 | 34 | 52 | 76 | 190 | 58 | 0 | 0 | 0 | 0 | 2 | 0 |
| SG0000002 | Map4k3 | 0.529483 | 0.596034 | 0.08881134 | 1385.709649 | 1473.693331 | 552 | 34 | 52 | 96 | 266 | 60 | 16 | 0 | 0 | 0 | 4 | 6 |
| SG0000003 | Dis3l | 0.633216 | 0.693891 | -0.083966192 | 577.2768902 | 544.6378623 | 232 | 48 | 52 | 60 | 100 | 32 | 0 | 0 | 0 | 0 | 0 | 0 |
| SG0000000 | Apc | 0.665549 | 0.723716 | 0.063326083 | 2938.98677 | 3070.864696 | 656 | 32 | 52 | 96 | 321 | 34 | 7 | 0 | 0 | 1 | 3 | 1 |
| SG0000001 | Stk4 | 0.726874 | 0.778616 | 0.042317645 | 2661.884765 | 2741.120521 | 323 | 36 | 52 | 65 | 162 | 17 | 6 | 0 | 0 | 0 | 10 | 2 |
| SG0000005 | Slit3 | 0.811649 | 0.851137 | -0.067765644 | 136.1817622 | 129.9329977 | 756 | 29 | 52 | 92 | 285 | 65 | 0 | 0 | 0 | 0 | 3 | 0 |
| SG0000002 | Chd1 | 0.826712 | 0.863948 | -0.036603905 | 6668.447682 | 6501.384852 | 363 | 34 | 52 | 81 | 178 | 37 | 9 | 1 | 1 | 1 | 6 | 1 |
| SG0000002 | Esrrb | 0 | 0 | 12.83108897 | 2.270680331 | 16546.21468 | 418 | 34 | 51 | 77 | 189 | 40 | 6 | 0 | 1 | 1 | 2 | 2 |
| SG0000003 | Smurf1 | 4.6E-58 | 1.19E-56 | -2.779528984 | 7789.026314 | 1134.387868 | 355 | 48 | 51 | 84 | 237 | 12 | 7 | 0 | 0 | 0 | 3 | 0 |
| SG0000003 | Rps6ka3 | 2.51E-56 | 6.26E-55 | -2.904145023 | 9135.074878 | 1220.330169 | 676 | 28 | 51 | 97 | 344 | 63 | 16 | 0 | 1 | 0 | 5 | 2 |
| SG0000004 | Iqsec2 | 4.26E-30 | 4.32E-29 | -2.558350684 | 713.7734641 | 121.1769739 | 407 | 26 | 51 | 69 | 218 | 44 | 4 | 4 | 4 | 4 | 8 | 4 |
| SG0000004 | Wapl | 1.31E-27 | 1.19E-26 | 1.287754939 | 5039.414931 | 12303.62206 | 661 | 27 | 51 | 99 | 364 | 104 | 8 | 0 | 0 | 1 | 4 | 3 |
| SG0000004 | Plekhg1 | 2.98E-27 | 2.66E-26 | 2.912648632 | 27.90533227 | 210.1269725 | 757 | 28 | 51 | 133 | 314 | 113 | 14 | 0 | 5 | 2 | 2 | 6 |
| SG0000004 | Sipa1l1 | 2.99E-26 | 2.55E-25 | -1.616085072 | 6475.064202 | 2112.292207 | 598 | 18 | 51 | 76 | 313 | 72 | 4 | 0 | 0 | 2 | 2 | 0 |
| SG0000003 | Dennd5b | 2.26E-24 | 1.77E-23 | -1.7526008 | 1717.941748 | 509.827224 | 310 | 42 | 51 | 70 | 164 | 28 | 20 | 2 | 2 | 4 | 6 | 2 |
| SG0000001 | Lrp11 | 3.01E-24 | 2.34E-23 | 1.384201742 | 692.2017464 | 1806.836832 | 238 | 36 | 51 | 54 | 143 | 33 | 7 | 0 | 0 | 0 | 2 | 0 |
| JSG000000 | Vps13a | 9.91E-23 | 7.19E-22 | -1.88212791 | 2807.541453 | 761.6390817 | 558 | 32 | 51 | 89 | 282 | 49 | 0 | 0 | 0 | 0 | 0 | 0 |
| SG0000006 | Ssbp3 | 1.92E-20 | 1.23E-19 | 1.220022686 | 1003.578806 | 2337.840646 | 345 | 39 | 51 | 57 | 139 | 24 | 0 | 0 | 0 | 0 | 1 | 0 |
| SG0000002 | Phf6 | 3.29E-19 | 1.97E-18 | -1.371534094 | 3855.581751 | 1490.105487 | 373 | 16 | 51 | 64 | 170 | 21 | 18 | 2 | 4 | 2 | 6 | 0 |
| SG0000002 | Lrrfip1 | 1.03E-17 | 5.71E-17 | -1.267694345 | 5822.536609 | 2418.233341 | 508 | 36 | 51 | 77 | 236 | 36 | 8 | 0 | 3 | 0 | 5 | 0 |
| SG0000002 | Ttc17 | 6.67E-16 | 3.33E-15 | -1.429932793 | 1402.617569 | 520.5789607 | 645 | 43 | 51 | 101 | 254 | 79 | 7 | 0 | 0 | 3 | 3 | 3 |
| SG0000000 | Nav1 | 1.33E-15 | 6.53E-15 | -1.182473354 | 4626.619502 | 2038.467706 | 496 | 39 | 51 | 87 | 208 | 55 | 25 | 0 | 0 | 5 | 11 | 2 |
| SG0000004 | Tet2 | 1.03E-14 | 4.76E-14 | 1.655848613 | 682.8783426 | 2151.807616 | 347 | 40 | 51 | 83 | 182 | 30 | 11 | 0 | 1 | 3 | 8 | 2 |
| SG0000003 | Lhpp | 3.09E-14 | 1.39E-13 | 1.503822605 | 147.9652789 | 419.6193721 | 285 | 37 | 51 | 72 | 165 | 32 | 0 | 0 | 0 | 0 | 0 | 0 |
| SG0000002 | Cdkl2 | 1.08E-13 | 4.68E-13 | -1.485466502 | 562.4829128 | 200.8812301 | 366 | 37 | 51 | 84 | 196 | 45 | 6 | 0 | 0 | 2 | 4 | 0 |
| SG0000002 | Xiap | 2.28E-13 | 9.67E-13 | -1.113935901 | 3816.294111 | 1763.24929 | 398 | 41 | 51 | 80 | 238 | 25 | 64 | 0 | 0 | 0 | 12 | 4 |
| SG0000002 | Vrk1 | 1.26E-11 | 4.8E-11 | 0.915494642 | 971.6701604 | 1832.779481 | 348 | 39 | 51 | 60 | 165 | 27 | 6 | 0 | 0 | 0 | 2 | 0 |
| JSG000000 | Vti1a | 1.54E-10 | 5.41E-10 | -1.120031001 | 891.5954353 | 410.2087397 | 553 | 28 | 51 | 81 | 208 | 49 | 4 | 0 | 0 | 0 | 2 | 1 |
| SG0000004 | Borcs5 | 2.32E-10 | 8.05E-10 | -1.574566626 | 550.8191432 | 184.9342033 | 313 | 41 | 51 | 90 | 180 | 16 | 1 | 0 | 0 | 0 | 2 | 0 |
| SG0000004 | Gm15800 | 2.59E-10 | 8.95E-10 | -1.018994769 | 1460.651435 | 720.7731477 | 246 | 33 | 51 | 51 | 136 | 22 | 3 | 0 | 0 | 2 | 3 | 0 |
| SG0000004 | Mbtps2 | 2.46E-06 | 6.13E-06 | -0.731384127 | 2463.836608 | 1484.032265 | 293 | 36 | 51 | 71 | 218 | 32 | 11 | 0 | 0 | 2 | 20 | 4 |
| SG0000002 | Srgap2 | 5.18E-06 | 1.25E-05 | 0.595835882 | 1771.551283 | 2677.430472 | 488 | 33 | 51 | 76 | 224 | 41 | 6 | 0 | 0 | 0 | 0 | 1 |
| SG0000002 | Cachd1 | 1.17E-05 | 2.74E-05 | 1.74390054 | 82.62805731 | 276.7540048 | 841 | 32 | 51 | 120 | 349 | 88 | 4 | 0 | 0 | 2 | 3 | 0 |
| SG0000002 | Ociad1 | 1.89E-05 | 4.34E-05 | -0.598181249 | 4405.296347 | 2910.078002 | 523 | 22 | 51 | 72 | 187 | 20 | 11 | 0 | 2 | 0 | 1 | 1 |
| SG0000005 | Adamts17 | 5.59E-05 | 0.000122 | 2.350471507 | 7.10867616 | 36.25360159 | 874 | 25 | 51 | 113 | 407 | 116 | 8 | 0 | 0 | 0 | 0 | 2 |
| SG0000002 | Rtn4 | 7.4E-05 | 0.000159 | -0.526317859 | 12974.3474 | 9008.408251 | 902 | 16 | 51 | 101 | 298 | 44 | 20 | 0 | 0 | 0 | 1 | 0 |
| JSG000000 | Nsmce2 | 0.000109 | 0.000229 | 0.542667725 | 934.5129613 | 1361.270999 | 655 | 34 | 51 | 87 | 292 | 64 | 1 | 0 | 0 | 0 | 0 | 0 |
| SG0000005 | Sv2c | 0.001597 | 0.002933 | 2.493995345 | 3.854066735 | 21.7113405 | 448 | 30 | 51 | 67 | 185 | 26 | 6 | 0 | 0 | 0 | 2 | 1 |
| SG0000002 | Pde6d | 0.002931 | 0.005178 | 0.469470654 | 483.477479 | 669.4235673 | 230 | 48 | 51 | 72 | 145 | 21 | 3 | 0 | 0 | 0 | 4 | 0 |
| SG0000005 | Eml1 | 0.004119 | 0.007123 | -0.945081022 | 429.3485484 | 223.0037998 | 474 | 36 | 51 | 81 | 197 | 28 | 4 | 0 | 0 | 2 | 2 | 2 |
| SG0000005 | Cdyl | 0.005553 | 0.009436 | 0.398750749 | 885.6528638 | 1167.614468 | 536 | 30 | 51 | 77 | 255 | 49 | 8 | 0 | 2 | 2 | 10 | 0 |
| SG0000007 | Scn2a1 | 0.015867 | 0.025042 | 1.633720921 | 6.528623506 | 20.2591223 | 821 | 28 | 51 | 114 | 343 | 73 | 8 | 0 | 0 | 2 | 4 | 10 |
| SG0000000 | Tle1 | 0.022871 | 0.035013 | 0.344633068 | 731.5861294 | 928.9885339 | 677 | 29 | 51 | 101 | 342 | 71 | 11 | 0 | 0 | 0 | 2 | 3 |
| JSG000000 | Tll1 | 0.030306 | 0.04544 | -0.790860623 | 134.4709206 | 77.72408322 | 637 | 24 | 51 | 95 | 335 | 60 | 4 | 0 | 0 | 0 | 4 | 0 |
| SG0000001 | Chfr | 0.031138 | 0.046603 | -0.300288567 | 2844.107789 | 2309.671342 | 408 | 34 | 51 | 73 | 195 | 37 | 22 | 1 | 5 | 3 | 3 | 2 |
| SG0000004 | Zbtb37 | 0.051971 | 0.074742 | 0.312310279 | 490.3673262 | 608.8864182 | 262 | 45 | 51 | 63 | 176 | 9 | 48 | 1 | 1 | 5 | 12 | 1 |
| SG0000002 | Sec11a | 0.074147 | 0.103679 | -0.261695296 | 1454.365071 | 1213.096366 | 251 | 42 | 51 | 50 | 158 | 24 | 0 | 0 | 0 | 0 | 0 | 0 |
| SG0000000 | Ewsr1 | 0.126796 | 0.168694 | 0.18953695 | 9403.23174 | 10723.42322 | 262 | 36 | 51 | 67 | 153 | 9 | 11 | 0 | 0 | 3 | 4 | 0 |
| SG0000002 | Anapc7 | 0.213811 | 0.270503 | -0.177349103 | 2268.864723 | 2006.41702 | 122 | 36 | 51 | 42 | 67 | 9 | 1 | 0 | 0 | 0 | 0 | 0 |
| SG0000003 | Pgm2l1 | 0.305602 | 0.370713 | 0.199452867 | 428.9947095 | 492.5986666 | 431 | 27 | 51 | 61 | 203 | 42 | 15 | 0 | 0 | 2 | 7 | 1 |
| SG0000005 | Pla2g4a | 0.461431 | 0.530838 | 0.294706746 | 340.6404341 | 417.8416885 | 528 | 33 | 51 | 81 | 271 | 62 | 8 | 0 | 2 | 0 | 2 | 0 |
| SG0000000 | Ubp1 | 0.785039 | 0.828763 | 0.034352257 | 2639.71728 | 2703.326334 | 297 | 42 | 51 | 66 | 176 | 21 | 8 | 0 | 0 | 0 | 2 | 0 |
| SG0000002 | Jade1 | 1.3E-130 | 1.6E-128 | 3.184350327 | 742.1943548 | 6746.874192 | 406 | 35 | 50 | 83 | 208 | 14 | 36 | 0 | 0 | 5 | 12 | 0 |
| SG0000003 | Slc22a23 | 4.21E-60 | 1.17E-58 | -5.293270319 | 6100.464894 | 155.5714112 | 633 | 24 | 50 | 100 | 344 | 53 | 4 | 0 | 0 | 2 | 4 | 0 |
| SG0000004 | Acacb | 6.92E-60 | 1.91E-58 | 4.748929093 | 33.4018431 | 898.1327563 | 246 | 47 | 50 | 66 | 144 | 36 | 2 | 0 | 0 | 0 | 0 | 0 |
| SG0000003 | Rap1gap2 | 6.01E-42 | 9.49E-41 | -3.123940321 | 3834.967543 | 439.9080926 | 510 | 40 | 50 | 96 | 252 | 36 | 2 | 0 | 0 | 0 | 0 | 1 |
| SG0000002 | Snx7 | 1.39E-38 | 1.96E-37 | -2.525624157 | 1346.418018 | 233.8251846 | 558 | 29 | 50 | 111 | 322 | 45 | 7 | 0 | 0 | 2 | 8 | 0 |
| SG0000005 | Uxs1 | 3.21E-24 | 2.49E-23 | -1.663925283 | 2473.743249 | 780.6622901 | 418 | 34 | 50 | 87 | 187 | 43 | 9 | 0 | 2 | 4 | 5 | 0 |
| JSG000000 | Chst8 | 1.46E-15 | 7.15E-15 | 7.181764192 | 0.34247518 | 49.72279652 | 292 | 41 | 50 | 71 | 163 | 40 | 0 | 0 | 0 | 0 | 0 | 0 |
| SG0000005 | Tns1 | 3.84E-14 | 1.72E-13 | -1.939969184 | 2049.546895 | 534.1570255 | 616 | 31 | 50 | 107 | 311 | 30 | 12 | 0 | 0 | 6 | 6 | 0 |
| SG0000001 | Eps8 | 1.67E-13 | 7.16E-13 | -1.315330606 | 1927.892579 | 774.6916319 | 826 | 33 | 50 | 113 | 320 | 87 | 28 | 0 | 3 | 0 | 1 | 0 |
| SG0000003 | Ptpn13 | 4.71E-11 | 1.72E-10 | -1.034458163 | 1638.493796 | 799.9113608 | 492 | 26 | 50 | 73 | 254 | 61 | 3 | 0 | 0 | 1 | 2 | 0 |
| SG0000004 | Nmnat2 | 1.71E-10 | 5.98E-10 | 1.035236343 | 327.8668237 | 671.9464519 | 526 | 48 | 50 | 88 | 214 | 34 | 6 | 2 | 2 | 2 | 2 | 0 |
| SG0000005 | Efcab5 | 2.44E-10 | 8.43E-10 | -4.323911255 | 47.30165576 | 2.361833928 | 322 | 41 | 50 | 68 | 160 | 29 | 2 | 0 | 0 | 1 | 1 | 0 |
| SG0000002 | Itch | 4.59E-08 | 1.33E-07 | -0.780510211 | 3796.540515 | 2210.197349 | 454 | 26 | 50 | 108 | 332 | 50 | 14 | 0 | 0 | 0 | 4 | 2 |
| JSG000000 | Angpt1 | 4.45E-07 | 1.18E-06 | -2.888041401 | 92.60163743 | 12.50925875 | 680 | 21 | 50 | 102 | 320 | 44 | 7 | 0 | 1 | 2 | 3 | 1 |
| SG0000004 | Rnf24 | 1.78E-06 | 4.49E-06 | -0.838830722 | 713.018157 | 398.6459725 | 322 | 41 | 50 | 67 | 186 | 30 | 21 | 0 | 0 | 0 | 6 | 9 |
| SG0000002 | Thap4 | 2.27E-06 | 5.69E-06 | 0.662579677 | 966.623144 | 1530.077915 | 308 | 42 | 50 | 73 | 192 | 30 | 2 | 0 | 0 | 1 | 8 | 0 |
| SG0000000 | Lman2l | 8.35E-06 | 1.98E-05 | 0.689608101 | 490.355652 | 790.8678279 | 138 | 32 | 50 | 45 | 60 | 4 | 1 | 0 | 0 | 0 | 1 | 0 |
| JSG000000 | Ube3c | 1.45E-05 | 3.38E-05 | -0.602774609 | 4431.197811 | 2917.883174 | 398 | 39 | 50 | 72 | 194 | 45 | 4 | 1 | 1 | 1 | 3 | 0 |
| SG0000003 | Pigu | 5.51E-05 | 0.00012 | -0.645084351 | 1123.649959 | 718.5240364 | 304 | 44 | 50 | 76 | 192 | 32 | 0 | 0 | 0 | 0 | 0 | 0 |
| SG0000002 | Poc1a | 7.6E-05 | 0.000163 | 0.632040843 | 441.1421271 | 683.6625415 | 144 | 44 | 50 | 50 | 74 | 10 | 0 | 0 | 0 | 0 | 0 | 0 |
| SG0000002 | Klhdc10 | 9.43E-05 | 0.0002 | -0.570924052 | 2455.001648 | 1652.671444 | 248 | 46 | 50 | 62 | 130 | 28 | 6 | 0 | 0 | 2 | 4 | 0 |
| JSG000000 | Rnf150 | 0.000527 | 0.001026 | -1.255343144 | 86.92830336 | 36.41373793 | 420 | 36 | 50 | 63 | 223 | 31 | 11 | 0 | 1 | 1 | 5 | 2 |
| SG0000004 | Pi4ka | 0.001772 | 0.003232 | -0.447664073 | 3759.968149 | 2756.91802 | 239 | 41 | 50 | 64 | 136 | 29 | 0 | 0 | 0 | 0 | 0 | 0 |
| JSG000000 | Wwc2 | 0.006403 | 0.010789 | -0.378415719 | 3252.219471 | 2501.873594 | 293 | 43 | 50 | 69 | 145 | 21 | 13 | 1 | 1 | 3 | 6 | 1 |
| SG0000005 | Usp29 | 0.006511 | 0.01096 | -0.955404779 | 80.12948005 | 41.32252281 | 434 | 42 | 50 | 94 | 228 | 67 | 13 | 0 | 0 | 0 | 2 | 4 |
| JSG000000 | Dnah7a | 0.010294 | 0.016797 | 4.7096289 | 0.68495036 | 17.92250317 | 659 | 26 | 50 | 92 | 332 | 48 | 0 | 0 | 0 | 0 | 0 | 0 |
| SG0000002 | Ttll1 | 0.015561 | 0.024601 | 0.430860659 | 310.7957042 | 418.9642724 | 117 | 44 | 50 | 50 | 91 | 12 | 2 | 0 | 0 | 0 | 2 | 2 |
| SG0000003 | Pnpla8 | 0.064527 | 0.091231 | 0.251567254 | 1212.868304 | 1443.919349 | 453 | 31 | 50 | 82 | 272 | 30 | 7 | 0 | 0 | 0 | 0 | 0 |
| SG0000003 | Hhat | 0.079242 | 0.110123 | 0.798735721 | 34.17113651 | 59.44328942 | 726 | 18 | 50 | 94 | 272 | 82 | 1 | 0 | 0 | 0 | 0 | 0 |
| SG0000002 | Acbd5 | 0.108682 | 0.146855 | -0.247217294 | 1138.453057 | 959.1693809 | 538 | 27 | 50 | 89 | 242 | 18 | 20 | 0 | 0 | 6 | 10 | 3 |
| SG0000000 | Clcn5 | 0.118767 | 0.158959 | 0.229807618 | 1149.001578 | 1347.40952 | 435 | 30 | 50 | 75 | 206 | 44 | 5 | 0 | 0 | 0 | 5 | 0 |
| SG0000000 | BC003331 | 0.232189 | 0.291184 | 0.174488017 | 987.7383305 | 1114.725753 | 508 | 35 | 50 | 100 | 243 | 35 | 8 | 0 | 0 | 0 | 8 | 0 |
| SG0000005 | Setd3 | 0.273035 | 0.33571 | -0.154845252 | 2501.646206 | 2247.050456 | 514 | 16 | 50 | 46 | 178 | 30 | 9 | 0 | 0 | 0 | 4 | 0 |
| SG0000001 | Anapc1 | 0.346072 | 0.413338 | -0.130227191 | 6281.650923 | 5739.466269 | 333 | 33 | 50 | 71 | 179 | 28 | 6 | 0 | 0 | 0 | 1 | 2 |
| SG0000002 | Scaf4 | 0.407213 | 0.476378 | 0.11402057 | 1442.237646 | 1560.847378 | 258 | 42 | 50 | 68 | 136 | 10 | 2 | 0 | 0 | 0 | 0 | 0 |
| SG0000004 | Sgsm3 | 0.41648 | 0.485771 | -0.175326309 | 217.0503664 | 192.212685 | 218 | 43 | 50 | 58 | 144 | 14 | 1 | 0 | 0 | 0 | 0 | 1 |
| SG0000003 | Scarb1 | 0.478426 | 0.547016 | 0.095960038 | 1715.024167 | 1832.977332 | 167 | 42 | 50 | 67 | 181 | 11 | 0 | 0 | 0 | 0 | 0 | 0 |
| SG0000005 | Plekhm3 | 0.487378 | 0.555579 | 0.109262607 | 633.5689311 | 683.416082 | 603 | 32 | 50 | 100 | 258 | 54 | 12 | 2 | 4 | 5 | 11 | 2 |
| SG0000003 | Brip1 | 0.497856 | 0.565227 | 0.109455555 | 677.0994202 | 730.4690965 | 359 | 32 | 50 | 77 | 205 | 29 | 12 | 0 | 0 | 2 | 5 | 1 |
| JSG000000 | Ap3b1 | 0.530319 | 0.596847 | 0.074099807 | 2762.259137 | 2907.841187 | 457 | 24 | 50 | 68 | 214 | 23 | 1 | 0 | 0 | 0 | 0 | 0 |
| SG0000006 | Rp2 | 0.820698 | 0.858816 | -0.043717119 | 1078.586011 | 1046.392503 | 534 | 40 | 50 | 100 | 322 | 59 | 30 | 0 | 0 | 2 | 12 | 4 |
| SG0000005 | Triqk | 0.87858 | 0.906602 | -0.068632474 | 76.37193361 | 72.82380269 | 427 | 22 | 50 | 71 | 298 | 53 | 13 | 0 | 0 | 1 | 6 | 5 |
| SG0000002 | Nr5a2 | 0 | 0 | 10.99505688 | 0.946619625 | 1932.045856 | 619 | 12 | 49 | 75 | 237 | 47 | 5 | 0 | 2 | 1 | 2 | 0 |
| SG0000002 | Naa16 | 1.95E-61 | 5.66E-60 | 2.128054756 | 707.8204627 | 3094.079209 | 362 | 36 | 49 | 72 | 233 | 23 | 5 | 0 | 0 | 0 | 3 | 0 |
| SG0000003 | Caprin2 | 1.17E-57 | 2.99E-56 | 2.341452563 | 270.3466207 | 1370.150899 | 352 | 45 | 49 | 81 | 216 | 36 | 4 | 0 | 0 | 0 | 0 | 0 |
| SG0000001 | Tex14 | 1.25E-56 | 3.15E-55 | 7.690695471 | 5.158722786 | 1065.791408 | 326 | 36 | 49 | 66 | 173 | 22 | 0 | 0 | 0 | 0 | 0 | 0 |
| SG0000002 | C2 | 2.71E-46 | 4.97E-45 | 4.888453116 | 6.6226471 | 196.1564684 | 152 | 38 | 49 | 71 | 102 | 5 | 0 | 0 | 0 | 0 | 0 | 0 |
| SG0000003 | Bcat1 | 1.19E-33 | 1.4E-32 | 2.732224485 | 589.4316386 | 3916.651845 | 356 | 32 | 49 | 64 | 221 | 27 | 8 | 0 | 1 | 0 | 1 | 0 |
| SG0000005 | Usp13 | 2.34E-30 | 2.41E-29 | 3.107372145 | 25.20518387 | 217.2211423 | 571 | 19 | 49 | 81 | 276 | 61 | 13 | 1 | 1 | 3 | 3 | 1 |
| SG0000002 | Stk38 | 7.9E-24 | 6.03E-23 | -1.501754816 | 9961.648153 | 3517.693147 | 211 | 46 | 49 | 68 | 150 | 20 | 5 | 0 | 0 | 0 | 0 | 0 |
| SG0000003 | Gng12 | 2.05E-20 | 1.32E-19 | -1.608130913 | 4390.867608 | 1440.305772 | 1407 | 28 | 49 | 198 | 584 | 217 | 14 | 0 | 0 | 2 | 3 | 0 |
| SG0000003 | Tulp4 | 5.66E-17 | 3.02E-16 | -1.60596119 | 1641.459958 | 539.247081 | 389 | 37 | 49 | 55 | 171 | 28 | 24 | 2 | 2 | 4 | 10 | 2 |
| SG0000003 | Elf1 | 1.5E-16 | 7.79E-16 | -1.315130011 | 2069.168609 | 831.5766755 | 408 | 23 | 49 | 71 | 194 | 28 | 12 | 0 | 0 | 0 | 14 | 2 |
| SG0000003 | Stox2 | 9.33E-16 | 4.63E-15 | 1.644359589 | 149.777406 | 468.2177402 | 493 | 36 | 49 | 97 | 259 | 50 | 8 | 0 | 0 | 1 | 7 | 0 |
| JSG000000 | Hivep1 | 5.41E-15 | 2.56E-14 | -1.688778156 | 1309.842682 | 406.2994711 | 277 | 36 | 49 | 60 | 139 | 19 | 1 | 0 | 0 | 1 | 1 | 0 |
| SG0000000 | Scmh1 | 2.96E-14 | 1.33E-13 | -1.208693185 | 1782.705065 | 771.3057946 | 556 | 40 | 49 | 91 | 322 | 57 | 2 | 0 | 0 | 1 | 1 | 0 |
| SG0000001 | Clvs2 | 4E-14 | 1.79E-13 | 5.234484651 | 0 | 36.647565 | 417 | 29 | 49 | 87 | 321 | 37 | 17 | 1 | 1 | 2 | 8 | 2 |
| SG0000004 | Mbp | 6.49E-14 | 2.85E-13 | -1.42743973 | 1138.359571 | 423.2308216 | 509 | 21 | 49 | 51 | 202 | 95 | 16 | 0 | 1 | 1 | 2 | 2 |
| SG0000004 | Adgrd1 | 6.49E-12 | 2.51E-11 | 5.286930191 | 0.981585527 | 38.32240167 | 285 | 42 | 49 | 67 | 121 | 10 | 3 | 0 | 1 | 1 | 1 | 0 |
| JSG000000 | Plcg2 | 1.91E-10 | 6.65E-10 | 1.35092265 | 675.2941411 | 1722.507896 | 195 | 42 | 49 | 59 | 135 | 33 | 0 | 0 | 0 | 0 | 0 | 0 |
| SG0000002 | Casc5 | 1.16E-09 | 3.81E-09 | 1.010916735 | 1059.774409 | 2135.648076 | 341 | 37 | 49 | 77 | 212 | 39 | 5 | 0 | 0 | 2 | 4 | 1 |
| SG0000007 | Wdfy1 | 1.49E-08 | 4.47E-08 | 0.778937133 | 930.14678 | 1596.007503 | 679 | 35 | 49 | 104 | 276 | 70 | 20 | 0 | 0 | 4 | 12 | 0 |
| SG0000003 | Ppp2r1b | 2.16E-08 | 6.39E-08 | 0.731210674 | 1496.21262 | 2483.760177 | 388 | 17 | 49 | 35 | 162 | 50 | 4 | 0 | 2 | 2 | 2 | 0 |
| JSG000000 | Adamts12 | 2.15E-07 | 5.85E-07 | -2.75056247 | 67.23638514 | 9.990852514 | 568 | 28 | 49 | 78 | 246 | 47 | 4 | 0 | 0 | 0 | 0 | 0 |
| SG0000002 | Ccl25 | 3.32E-05 | 7.45E-05 | 1.321305351 | 38.67149744 | 96.63702091 | 166 | 36 | 49 | 54 | 101 | 16 | 1 | 0 | 0 | 0 | 0 | 0 |
| SG0000004 | Gpr155 | 3.56E-05 | 7.96E-05 | -1.387496879 | 109.1473811 | 41.71912009 | 361 | 28 | 49 | 69 | 222 | 14 | 15 | 0 | 0 | 4 | 8 | 3 |
| SG0000002 | Fbxl20 | 0.000103 | 0.000218 | -0.595206328 | 1932.365741 | 1279.129076 | 175 | 45 | 49 | 60 | 108 | 11 | 8 | 1 | 1 | 1 | 4 | 2 |
| JSG000000 | Ust | 0.009178 | 0.015103 | 1.094561306 | 29.66239819 | 63.34349847 | 547 | 36 | 49 | 90 | 260 | 50 | 3 | 1 | 1 | 1 | 2 | 0 |

| | | | | | | | | | | | | | | | | | |
|---|---|---|---|---|---|---|---|---|---|---|---|---|---|---|---|---|---|
| SG0000005 | Prr14l | 0.024752 | 0.037688 | -0.326506392 | 2273.574505 | 1813.096723 | 187 | 37 | 49 | 52 | 124 | 18 | 5 | 0 | 0 | 0 | 2 | 0 |
| SG0000002 | Map2k6 | 0.035222 | 0.052175 | 0.569590594 | 190.7071227 | 283.0288897 | 417 | 36 | 49 | 86 | 239 | 20 | 6 | 0 | 2 | 2 | 4 | 0 |
| SG0000003 | Bclaf1 | 0.071141 | 0.099755 | 0.231198411 | 6300.172865 | 7395.203383 | 472 | 27 | 49 | 62 | 258 | 23 | 20 | 0 | 4 | 4 | 10 | 0 |
| SG0000003 | Sh3rf3 | 0.137037 | 0.181084 | 0.919391386 | 16.34582548 | 30.91514178 | 800 | 17 | 49 | 105 | 371 | 91 | 1 | 0 | 0 | 0 | 1 | 0 |
| JSG000000 | Hgf | 0.142117 | 0.187136 | -1.534560119 | 17.12579629 | 5.911560627 | 617 | 18 | 49 | 106 | 331 | 73 | 8 | 0 | 1 | 4 | 7 | 1 |
| SG0000003 | Syncrip | 0.148643 | 0.194931 | 0.179420625 | 5043.537952 | 5711.448736 | 608 | 28 | 49 | 74 | 235 | 70 | 58 | 0 | 0 | 5 | 12 | 15 |
| SG0000007 | Tia1 | 0.287203 | 0.351033 | -0.19365383 | 2005.056792 | 1753.198404 | 324 | 33 | 49 | 80 | 159 | 33 | 28 | 0 | 4 | 4 | 11 | 0 |
| SG0000000 | Kpna6 | 0.368232 | 0.436459 | 0.112500486 | 3541.75366 | 3828.991235 | 98 | 47 | 49 | 55 | 79 | 8 | 4 | 0 | 1 | 1 | 1 | 0 |
| SG0000002 | Mapre2 | 0.525743 | 0.592346 | 0.085406872 | 1404.679595 | 1490.346679 | 782 | 26 | 49 | 103 | 309 | 86 | 12 | 0 | 0 | 3 | 8 | 0 |
| SG0000003 | Scai | 0.805204 | 0.845615 | 0.039193456 | 511.0676497 | 525.1420523 | 559 | 31 | 49 | 69 | 274 | 40 | 26 | 1 | 1 | 2 | 8 | 3 |
| SG0000002 | Dmtn | 5.04E-66 | 1.63E-64 | 7.735537611 | 5.983536754 | 1275.224995 | 127 | 45 | 48 | 54 | 86 | 9 | 8 | 0 | 0 | 0 | 0 | 2 |
| SG0000002 | Optn | 2.51E-65 | 7.9E-64 | -3.194226279 | 4343.144523 | 474.5110805 | 188 | 42 | 48 | 62 | 90 | 12 | 0 | 0 | 0 | 0 | 0 | 0 |
| SG0000005 | Spop | 1.9E-42 | 3.04E-41 | -2.249350407 | 6442.720066 | 1355.025033 | 352 | 32 | 48 | 74 | 196 | 22 | 12 | 0 | 0 | 2 | 6 | 0 |
| SG0000000 | Cdh1 | 7.08E-34 | 8.38E-33 | 4.007493041 | 981.9926225 | 15793.69831 | 221 | 44 | 48 | 58 | 100 | 14 | 2 | 0 | 0 | 0 | 0 | 0 |
| SG0000005 | Jam2 | 7.36E-33 | 8.35E-32 | 1.915699473 | 216.199611 | 815.7139928 | 222 | 34 | 48 | 59 | 115 | 11 | 11 | 0 | 1 | 1 | 2 | 0 |
| SG0000002 | Nnt | 2.66E-31 | 2.84E-30 | -2.120533147 | 3785.80593 | 870.5911801 | 479 | 40 | 48 | 63 | 241 | 45 | 3 | 0 | 0 | 0 | 1 | 0 |
| SG0000001 | Bcas1 | 8.54E-30 | 8.58E-29 | 6.37733154 | 0 | 82.1319726 | 419 | 36 | 48 | 71 | 207 | 83 | 10 | 0 | 0 | 0 | 4 | 4 |
| SG0000000 | Pcyt1a | 2.65E-26 | 2.27E-25 | -1.726119179 | 2767.064637 | 836.3838823 | 320 | 36 | 48 | 68 | 204 | 36 | 5 | 0 | 0 | 0 | 0 | 0 |
| SG0000005 | Cspp1 | 1.63E-19 | 9.94E-19 | 1.591528499 | 546.7933937 | 1647.8629 | 670 | 39 | 48 | 76 | 287 | 79 | 27 | 0 | 0 | 1 | 5 | 4 |
| SG0000002 | Rnf6 | 1.88E-18 | 1.08E-17 | -1.378538966 | 2616.252976 | 1006.232074 | 113 | 42 | 48 | 45 | 78 | 9 | 5 | 0 | 0 | 0 | 0 | 0 |
| SG0000002 | Tc2n | 1.01E-14 | 4.71E-14 | 4.747816922 | 1.974045165 | 53.03864157 | 794 | 12 | 48 | 65 | 404 | 85 | 19 | 0 | 2 | 2 | 6 | 3 |
| SG0000000 | Man1a2 | 6.15E-10 | 2.06E-09 | -0.977840061 | 2591.872872 | 1315.995811 | 447 | 24 | 48 | 75 | 259 | 43 | 7 | 0 | 0 | 1 | 4 | 1 |
| SG0000006 | Fnip2 | 9.67E-10 | 3.19E-09 | 0.823788699 | 1018.992362 | 1803.665662 | 412 | 38 | 48 | 84 | 233 | 31 | 4 | 0 | 0 | 0 | 3 | 1 |
| SG0000003 | Mgarp | 4.88E-09 | 1.52E-08 | 2.509596205 | 95.8798012 | 545.9977582 | 102 | 48 | 48 | 54 | 63 | 0 | 9 | 0 | 0 | 0 | 0 | 0 |
| SG0000002 | Setbp1 | 1.38E-07 | 3.83E-07 | 1.940012589 | 37.17963792 | 142.6616154 | 611 | 28 | 48 | 72 | 269 | 52 | 5 | 0 | 0 | 0 | 2 | 2 |
| SG0000002 | Fam188a | 3.41E-05 | 7.64E-05 | -0.627558575 | 1633.42492 | 1057.268327 | 1005 | 15 | 48 | 79 | 408 | 107 | 21 | 5 | 5 | 10 | 27 | 1 |
| SG0000000 | Fmr1 | 7.73E-05 | 0.000166 | 0.501524756 | 2754.238548 | 3899.20032 | 720 | 30 | 48 | 102 | 318 | 96 | 24 | 6 | 6 | 6 | 12 | 12 |
| JSG000000 | Bmper | 8.45E-05 | 0.000181 | -1.776177526 | 94.96185093 | 27.72467463 | 528 | 30 | 48 | 81 | 269 | 39 | 5 | 0 | 0 | 0 | 2 | 1 |
| SG0000002 | Pdxdc1 | 0.000107 | 0.000226 | -0.542650169 | 3332.515374 | 2287.800675 | 365 | 36 | 48 | 69 | 233 | 27 | 3 | 0 | 0 | 0 | 0 | 0 |
| SG0000002 | 2810403A07Rik | 0.000342 | 0.000682 | 0.554036347 | 1858.302006 | 2728.336009 | 254 | 44 | 48 | 63 | 183 | 26 | 11 | 0 | 1 | 3 | 5 | 2 |
| SG0000002 | Actr3 | 0.000426 | 0.000838 | 0.435326112 | 5404.692342 | 7308.313937 | 388 | 39 | 48 | 63 | 161 | 36 | 6 | 0 | 0 | 0 | 3 | 0 |
| SG0000000 | Relb | 0.000585 | 0.001131 | 0.623526818 | 313.1915306 | 482.5144066 | 141 | 42 | 48 | 57 | 78 | 15 | 0 | 0 | 0 | 0 | 0 | 3 |
| JSG000000 | Parp8 | 0.000795 | 0.00151 | -1.218136916 | 118.2032076 | 50.80816516 | 425 | 32 | 48 | 63 | 223 | 30 | 2 | 0 | 0 | 0 | 3 | 0 |
| SG0000003 | Lrrk2 | 0.000841 | 0.001592 | -1.093874349 | 176.9618251 | 82.90688506 | 381 | 37 | 48 | 79 | 173 | 44 | 6 | 0 | 0 | 2 | 2 | 0 |
| SG0000004 | Tex2 | 0.001819 | 0.003312 | 0.44700395 | 898.2507138 | 1224.501067 | 221 | 37 | 48 | 55 | 106 | 10 | 1 | 0 | 0 | 0 | 1 | 0 |
| SG0000004 | 5330417C22Rik | 0.004176 | 0.007218 | -1.448501856 | 52.69541575 | 19.30769048 | 353 | 21 | 48 | 73 | 166 | 30 | 7 | 0 | 0 | 1 | 1 | 0 |
| JSG000000 | Sil1 | 0.01619 | 0.025515 | -0.409181923 | 684.4642439 | 515.4359826 | 410 | 37 | 48 | 82 | 211 | 28 | 0 | 0 | 0 | 0 | 0 | 0 |
| SG0000005 | Tyw1 | 0.0349 | 0.051736 | -0.319958372 | 1222.222887 | 979.1141897 | 214 | 42 | 48 | 71 | 132 | 15 | 6 | 1 | 1 | 1 | 1 | 0 |
| SG0000003 | Rbm41 | 0.085401 | 0.117979 | -0.350261049 | 339.7281467 | 266.4970757 | 637 | 21 | 48 | 100 | 372 | 80 | 17 | 3 | 3 | 11 | 38 | 0 |
| SG0000000 | Adck4 | 0.119646 | 0.16004 | -0.424589884 | 412.8047956 | 307.5604183 | 150 | 48 | 48 | 63 | 150 | 6 | 0 | 0 | 0 | 0 | 0 | 0 |
| SG0000004 | Wdr47 | 0.186321 | 0.239188 | -0.238345128 | 551.1324146 | 467.2043974 | 269 | 36 | 48 | 76 | 165 | 15 | 4 | 1 | 1 | 1 | 2 | 0 |
| SG0000003 | Rab11fip3 | 0.225117 | 0.283241 | -0.200323256 | 938.8404702 | 817.1249914 | 239 | 33 | 48 | 60 | 127 | 18 | 0 | 0 | 0 | 0 | 0 | 0 |
| SG0000003 | Tmem87a | 0.352884 | 0.420746 | 0.130784689 | 1207.7773 | 1322.382146 | 303 | 31 | 48 | 66 | 167 | 23 | 10 | 2 | 2 | 2 | 4 | 0 |
| SG0000000 | Dnaja3 | 0.479598 | 0.548252 | -0.099652436 | 2733.701233 | 2551.247993 | 128 | 44 | 48 | 52 | 90 | 8 | 12 | 0 | 0 | 0 | 0 | 2 |
| SG0000003 | Nhsl1 | 0.777166 | 0.821807 | -0.047872725 | 874.2783954 | 845.7434057 | 345 | 38 | 48 | 63 | 173 | 30 | 8 | 0 | 0 | 4 | 8 | 0 |
| SG0000002 | Cat | 0.846004 | 0.87949 | -0.028180561 | 4002.758028 | 3925.329726 | 155 | 40 | 48 | 58 | 102 | 6 | 3 | 0 | 0 | 0 | 0 | 0 |
| SG0000000 | Rgs20 | 5.18E-33 | 5.91E-32 | -3.976185436 | 416.1720585 | 26.44367641 | 828 | 14 | 47 | 109 | 405 | 57 | 7 | 0 | 0 | 3 | 8 | 2 |
| JSG000000 | Clybl | 8.7E-33 | 9.84E-32 | 1.806311666 | 305.74856 | 1069.346404 | 383 | 30 | 47 | 75 | 186 | 32 | 0 | 0 | 0 | 0 | 0 | 0 |
| SG0000004 | Tmem108 | 8.81E-26 | 7.36E-25 | 5.313597943 | 2.305646233 | 91.69486971 | 1022 | 17 | 47 | 113 | 463 | 106 | 5 | 0 | 0 | 0 | 4 | 1 |
| SG0000003 | Rnf43 | 1.66E-25 | 1.37E-24 | 4.689967639 | 4.247069063 | 109.6251654 | 372 | 44 | 47 | 94 | 233 | 13 | 9 | 0 | 0 | 9 | 9 | 0 |
| SG0000002 | Add1 | 2.57E-22 | 1.83E-21 | -1.447759073 | 7255.921155 | 2659.951068 | 404 | 26 | 47 | 68 | 192 | 35 | 5 | 0 | 0 | 0 | 0 | 0 |
| SG0000002 | Ccdc136 | 8.25E-17 | 4.36E-16 | 1.457131868 | 201.9397682 | 554.4498282 | 190 | 25 | 47 | 67 | 127 | 31 | 0 | 0 | 0 | 0 | 0 | 0 |
| JSG000000 | Mctp2 | 3.71E-14 | 1.66E-13 | 5.301919435 | 3.562118703 | 140.522275 | 556 | 27 | 47 | 81 | 280 | 73 | 14 | 2 | 3 | 3 | 4 | 2 |
| SG0000003 | Snap91 | 5.3E-14 | 2.35E-13 | 4.137763766 | 3.252265858 | 57.25020531 | 912 | 16 | 47 | 114 | 414 | 87 | 15 | 0 | 3 | 6 | 6 | 7 |
| SG0000002 | Zfp346 | 2.26E-12 | 8.97E-12 | 1.030080964 | 526.5650379 | 1075.318922 | 254 | 23 | 47 | 55 | 189 | 32 | 1 | 0 | 0 | 0 | 0 | 0 |
| SG0000002 | Ptpn12 | 9.2E-12 | 3.52E-11 | -1.019419141 | 2860.538079 | 1411.146145 | 499 | 27 | 47 | 81 | 242 | 65 | 5 | 0 | 0 | 0 | 4 | 0 |
| SG0000003 | Ttc7 | 1.12E-10 | 3.97E-10 | -1.548946892 | 1047.858884 | 358.1156691 | 291 | 39 | 47 | 56 | 133 | 17 | 3 | 0 | 0 | 0 | 1 | 1 |
| SG0000003 | Btbd10 | 1.84E-10 | 6.41E-10 | -0.985846196 | 1822.782998 | 920.3768637 | 540 | 8 | 47 | 78 | 276 | 53 | 26 | 0 | 5 | 13 | 13 | 0 |
| SG0000003 | Oxsr1 | 8.66E-10 | 2.87E-09 | 0.763960166 | 2619.763006 | 4448.739034 | 370 | 35 | 47 | 81 | 193 | 38 | 7 | 0 | 0 | 0 | 1 | 0 |
| SG0000003 | Rcan2 | 2.89E-09 | 9.15E-09 | 2.258411627 | 37.41252826 | 179.005634 | 738 | 27 | 47 | 109 | 343 | 67 | 10 | 1 | 1 | 2 | 19 | 1 |
| SG0000003 | Rsf1 | 2.93E-09 | 9.27E-09 | 0.781140695 | 1254.223186 | 2155.368864 | 526 | 23 | 47 | 64 | 203 | 39 | 18 | 0 | 0 | 1 | 11 | 2 |
| JSG000000 | Gtf2ird2 | 1.9E-06 | 4.78E-06 | -1.318409488 | 179.0024617 | 71.77582635 | 127 | 47 | 47 | 59 | 110 | 4 | 2 | 1 | 1 | 1 | 2 | 1 |
| SG0000004 | Arhgap32 | 3.04E-05 | 6.82E-05 | 0.577280612 | 927.0371372 | 1383.17112 | 575 | 25 | 47 | 97 | 324 | 46 | 24 | 0 | 0 | 0 | 4 | 2 |
| JSG000000 | Nmi | 0.000159 | 0.000329 | -0.980995328 | 347.518236 | 176.0631954 | 180 | 40 | 47 | 52 | 103 | 39 | 4 | 0 | 0 | 0 | 0 | 0 |
| SG0000002 | Ncor2 | 0.000181 | 0.000372 | -1.039234193 | 4452.294835 | 2166.422968 | 477 | 47 | 47 | 74 | 184 | 49 | 5 | 0 | 0 | 0 | 0 | 0 |
| SG0000002 | B4galt4 | 0.000402 | 0.000795 | -0.752719373 | 463.2888199 | 274.954423 | 146 | 45 | 47 | 64 | 97 | 13 | 2 | 0 | 0 | 0 | 0 | 0 |
| SG0000004 | Camsap2 | 0.000532 | 0.001034 | -0.497472843 | 2620.894419 | 1856.501388 | 518 | 24 | 47 | 58 | 203 | 41 | 31 | 0 | 3 | 9 | 17 | 3 |
| SG0000002 | Hdac4 | 0.001141 | 0.002128 | -0.478465174 | 1857.765291 | 1333.393935 | 577 | 32 | 47 | 65 | 250 | 50 | 5 | 0 | 0 | 1 | 3 | 1 |
| SG0000003 | Rpap2 | 0.002551 | 0.004552 | 0.436669804 | 832.4649166 | 1126.721674 | 566 | 28 | 47 | 66 | 150 | 91 | 2 | 0 | 0 | 0 | 0 | 0 |
| SG0000002 | Ptn | 0.017703 | 0.027652 | -0.984331086 | 61.74237101 | 31.2083006 | 415 | 31 | 47 | 66 | 189 | 38 | 6 | 0 | 0 | 2 | 3 | 0 |
| SG0000003 | Slc2a13 | 0.02871 | 0.04324 | -1.533750341 | 21.35865082 | 7.376816103 | 579 | 27 | 47 | 99 | 300 | 59 | 14 | 1 | 3 | 3 | 10 | 0 |
| SG0000002 | Sucla2 | 0.082749 | 0.114624 | 0.217373785 | 2769.442917 | 3219.796674 | 210 | 34 | 47 | 56 | 114 | 12 | 2 | 0 | 1 | 2 | 3 | 0 |
| SG0000003 | Zzz3 | 0.099916 | 0.135902 | 0.221644493 | 1773.363069 | 2067.851278 | 720 | 25 | 47 | 75 | 325 | 50 | 30 | 0 | 2 | 5 | 15 | 0 |
| SG0000002 | Eya1 | 0.260504 | 0.322181 | -0.323602915 | 195.4633157 | 156.1892277 | 625 | 28 | 47 | 96 | 286 | 60 | 5 | 0 | 1 | 2 | 12 | 2 |
| SG0000005 | Stam2 | 0.384009 | 0.45278 | -0.129124579 | 1536.449489 | 1404.908098 | 320 | 42 | 47 | 82 | 153 | 181 | 11 | 0 | 0 | 1 | 2 | 2 |
| SG0000002 | Csrp2bp | 0.389695 | 0.45838 | 0.128099072 | 1035.126298 | 1131.240656 | 204 | 42 | 47 | 72 | 136 | 25 | 0 | 0 | 0 | 0 | 0 | 0 |
| SG0000002 | Camkk2 | 0.445129 | 0.51437 | -0.128372923 | 769.1983609 | 703.7108542 | 215 | 43 | 47 | 51 | 95 | 10 | 8 | 0 | 1 | 1 | 1 | 0 |
| SG0000002 | Hdhd2 | 0.706055 | 0.759514 | 0.057294777 | 629.0312516 | 654.5150981 | 486 | 7 | 47 | 48 | 142 | 27 | 39 | 0 | 0 | 0 | 8 | 2 |
| SG0000003 | Cacna1a | 9.94E-59 | 2.65E-57 | 3.931824008 | 30.2014511 | 460.9191732 | 356 | 42 | 46 | 64 | 136 | 58 | 0 | 0 | 0 | 0 | 0 | 0 |
| SG0000002 | Chka | 2.11E-57 | 5.35E-56 | 2.572096535 | 274.0509466 | 1629.706941 | 326 | 38 | 46 | 84 | 175 | 19 | 0 | 0 | 0 | 0 | 3 | 0 |
| SG0000003 | Cul4b | 6.07E-39 | 8.6E-38 | 1.59372406 | 2369.734839 | 7152.510437 | 432 | 34 | 46 | 72 | 216 | 34 | 7 | 0 | 0 | 0 | 0 | 0 |
| JSG000000 | Ugt1a6b | 2.21E-29 | 2.17E-28 | -8.339600427 | 423.400647 | 1.307018205 | 470 | 30 | 46 | 87 | 215 | 22 | 4 | 0 | 0 | 0 | 0 | 0 |
| SG0000006 | Plac1 | 2.01E-28 | 1.9E-27 | 7.186645591 | 0.649984458 | 94.68885464 | 374 | 33 | 46 | 68 | 202 | 47 | 0 | 0 | 0 | 0 | 0 | 0 |
| SG0000003 | Fanci | 7.14E-26 | 6.01E-25 | 1.400226312 | 785.4111659 | 2073.03766 | 274 | 30 | 46 | 64 | 199 | 38 | 2 | 0 | 0 | 0 | 4 | 0 |
| SG0000004 | Tapt1 | 1.59E-25 | 1.32E-24 | -1.823850415 | 1602.366807 | 452.614286 | 292 | 36 | 46 | 59 | 137 | 20 | 9 | 0 | 2 | 4 | 4 | 0 |
| SG0000003 | Wdr59 | 5.44E-25 | 4.4E-24 | 1.602564333 | 605.3835878 | 1838.444726 | 272 | 42 | 46 | 60 | 138 | 20 | 8 | 0 | 0 | 0 | 0 | 0 |
| SG0000002 | Dtwd2 | 2.91E-21 | 1.96E-20 | 1.934556318 | 92.80325214 | 354.7502217 | 344 | 24 | 46 | 69 | 185 | 38 | 16 | 0 | 0 | 5 | 9 | 0 |
| SG0000004 | Dcaf17 | 2.44E-20 | 1.56E-19 | 1.331473487 | 492.3922484 | 1239.152237 | 382 | 21 | 46 | 53 | 143 | 40 | 16 | 0 | 0 | 0 | 1 | 1 |
| SG0000002 | Stk39 | 1E-19 | 6.18E-19 | -2.652633974 | 1050.556962 | 167.0695148 | 449 | 31 | 46 | 81 | 234 | 44 | 8 | 0 | 2 | 2 | 3 | 0 |
| JSG000000 | Klf12 | 8.87E-18 | 4.93E-17 | 3.713287374 | 6.550371729 | 85.91654985 | 582 | 26 | 46 | 84 | 286 | 60 | 0 | 0 | 0 | 0 | 0 | 0 |
| JSG000000 | Parvb | 1.02E-14 | 4.72E-14 | -1.139727813 | 4493.619783 | 2039.407745 | 137 | 43 | 46 | 57 | 95 | 28 | 2 | 0 | 0 | 1 | 1 | 0 |
| SG0000003 | Rbm46 | 2.39E-13 | 1.01E-12 | 5.145486423 | 0 | 34.39531287 | 333 | 32 | 46 | 67 | 186 | 11 | 2 | 0 | 0 | 0 | 0 | 0 |
| JSG000000 | Zmym2 | 6.14E-13 | 2.53E-12 | 0.893066444 | 2560.73091 | 4755.582624 | 249 | 34 | 46 | 66 | 156 | 13 | 10 | 1 | 1 | 1 | 5 | 4 |
| SG0000003 | Pak1 | 1.09E-12 | 4.42E-12 | 0.96893804 | 896.3437539 | 1754.502577 | 372 | 22 | 46 | 64 | 190 | 46 | 2 | 0 | 2 | 2 | 2 | 0 |
| SG0000003 | Dnah10 | 5.59E-12 | 2.17E-11 | 2.658177395 | 13.49608935 | 85.19205114 | 254 | 44 | 46 | 66 | 136 | 24 | 0 | 0 | 0 | 0 | 0 | 0 |
| SG0000005 | Lin9 | 1.8E-10 | 6.3E-10 | 0.986899147 | 407.0602084 | 806.7610019 | 366 | 36 | 46 | 58 | 175 | 41 | 21 | 3 | 3 | 3 | 9 | 0 |
| SG0000004 | E130308A19Rik | 1.46E-09 | 4.74E-09 | 0.978956475 | 329.8851654 | 650.2165952 | 522 | 29 | 46 | 95 | 221 | 42 | 15 | 1 | 1 | 4 | 4 | 2 |
| SG0000002 | Myo5b | 2.29E-08 | 6.78E-08 | 6.397429992 | 1.572512292 | 132.5599623 | 787 | 28 | 46 | 94 | 284 | 96 | 1 | 0 | 0 | 0 | 0 | 2 |
| SG0000001 | Ppfibp1 | 2.45E-08 | 7.22E-08 | -0.809525946 | 3564.958283 | 2034.055789 | 508 | 26 | 46 | 66 | 238 | 42 | 6 | 0 | 0 | 0 | 8 | 2 |
| SG0000002 | Sec23a | 7.12E-08 | 2.02E-07 | -0.784440766 | 3131.304452 | 1817.96323 | 297 | 38 | 46 | 64 | 178 | 14 | 12 | 0 | 2 | 0 | 4 | 0 |
| SG0000003 | Gucd1 | 1.06E-07 | 2.96E-07 | -0.964367111 | 667.7910197 | 342.2450277 | 419 | 35 | 46 | 68 | 190 | 22 | 7 | 0 | 0 | 0 | 1 | 0 |
| SG0000002 | Fgd4 | 3.1E-07 | 8.33E-07 | -1.009584527 | 475.8453548 | 236.3472798 | 485 | 28 | 46 | 76 | 293 | 51 | 23 | 1 | 3 | 2 | 9 | 5 |
| SG0000005 | Mfsd4 | 1.32E-06 | 3.38E-06 | -2.410623083 | 57.0841981 | 10.73608765 | 276 | 33 | 46 | 52 | 242 | 38 | 7 | 0 | 0 | 1 | 7 | 2 |
| SG0000003 | Galm | 1.57E-05 | 3.63E-05 | -0.680406774 | 1328.270418 | 828.8265462 | 153 | 37 | 46 | 55 | 110 | 14 | 1 | 0 | 0 | 0 | 1 | 0 |
| SG0000002 | Pdgfra | 2.95E-05 | 6.63E-05 | -1.938284647 | 561.1845217 | 146.4279105 | 255 | 40 | 46 | 72 | 176 | 14 | 10 | 0 | 0 | 0 | 4 | 4 |
| SG0000003 | Ccdc88a | 4.97E-05 | 0.000109 | -0.680659942 | 1387.949835 | 865.9138984 | 539 | 20 | 46 | 73 | 227 | 54 | 20 | 0 | 0 | 1 | 7 | 2 |
| SG0000003 | Nt5dc1 | 0.000179 | 0.00037 | -0.621698164 | 895.4803933 | 581.9776219 | 570 | 28 | 46 | 91 | 324 | 73 | 4 | 0 | 0 | 0 | 10 | 4 |
| SG0000004 | Ppip5k2 | 0.000268 | 0.000541 | 0.463440394 | 2394.058236 | 3300.98971 | 336 | 20 | 46 | 54 | 188 | 24 | 12 | 0 | 2 | 2 | 6 | 0 |
| SG0000002 | Limk2 | 0.000716 | 0.001367 | 0.440528835 | 2306.436379 | 3130.068721 | 148 | 39 | 46 | 56 | 107 | 10 | 0 | 0 | 0 | 0 | 0 | 4 |
| SG0000003 | C2cd5 | 0.004628 | 0.007959 | 0.370774998 | 1871.865585 | 2420.410652 | 410 | 24 | 46 | 64 | 178 | 26 | 4 | 0 | 0 | 0 | 0 | 0 |
| SG0000004 | Ankrd42 | 0.005456 | 0.009283 | -0.83184348 | 116.8628951 | 65.65484824 | 423 | 24 | 46 | 68 | 199 | 26 | 7 | 0 | 0 | 0 | 0 | 0 |
| SG0000002 | Ica1l | 0.023872 | 0.036428 | 1.454410737 | 7.911741905 | 21.68170192 | 236 | 36 | 46 | 66 | 178 | 30 | 10 | 0 | 0 | 2 | 6 | 0 |
| SG0000002 | Ahrr | 0.025151 | 0.03825 | 0.746968695 | 50.84352608 | 85.32880135 | 332 | 38 | 46 | 74 | 156 | 18 | 2 | 0 | 0 | 0 | 0 | 0 |
| SG0000002 | Copg2 | 0.067404 | 0.094999 | 0.241846108 | 2065.452185 | 2442.407239 | 418 | 32 | 46 | 88 | 213 | 39 | 6 | 0 | 0 | 0 | 0 | 1 |
| SG0000002 | Sepsecs | 0.084981 | 0.117443 | -0.285078472 | 788.7981143 | 647.3642208 | 263 | 23 | 46 | 52 | 123 | 19 | 12 | 0 | 0 | 2 | 4 | 3 |
| SG0000000 | Vav2 | 0.088684 | 0.122109 | 0.291421484 | 643.5800834 | 787.6426119 | 276 | 42 | 46 | 50 | 98 | 24 | 0 | 0 | 0 | 0 | 0 | 2 |
| SG0000003 | Agbl3 | 0.099613 | 0.135552 | -0.872999854 | 50.62051299 | 27.6393229 | 700 | 16 | 46 | 87 | 366 | 109 | 12 | 0 | 0 | 0 | 7 | 0 |
| SG0000003 | Uba6 | 0.272822 | 0.335538 | -0.155537919 | 2881.98153 | 2587.435986 | 444 | 32 | 46 | 67 | 222 | 32 | 14 | 0 | 0 | 1 | 2 | 0 |
| JSG000000 | Abcb7 | 0.410703 | 0.479737 | -0.121817465 | 2454.251181 | 2255.528389 | 354 | 30 | 46 | 70 | 190 | 43 | 13 | 0 | 2 | 1 | 2 | 1 |
| SG0000004 | Aff4 | 0.427136 | 0.496423 | 0.099286349 | 5532.95926 | 5927.146227 | 214 | 37 | 46 | 56 | 121 | 15 | 21 | 2 | 3 | 6 | 11 | 1 |
| SG0000004 | Trp53bp1 | 0.920095 | 0.941871 | -0.023547068 | 2470.412562 | 2430.418786 | 302 | 35 | 46 | 62 | 170 | 32 | 10 | 0 | 0 | 2 | 5 | 2 |
| SG0000002 | Pik3r3 | 0.937484 | 0.9547 | 0.02362492 | 188.8285349 | 191.9461622 | 408 | 32 | 46 | 67 | 195 | 22 | 6 | 0 | 0 | 1 | 1 | 0 |
| SG0000005 | Calcrl | 2.93E-93 | 1.79E-91 | -7.820553462 | 9115.276865 | 40.32261503 | 396 | 19 | 45 | 68 | 239 | 25 | 12 | 0 | 2 | 2 | 14 | 2 |

| | | | | | | | | | | | | | | | | | | |
|---|---|---|---|---|---|---|---|---|---|---|---|---|---|---|---|---|---|---|
| SG0000004 | Fam169a | 7.91E-90 | 4.47E-88 | 2.427376539 | 1293.599287 | 6958.4565 | 306 | 36 | 45 | 68 | 157 | 22 | 20 | 0 | 0 | 0 | 2 | 2 |
| SG0000005 | A630001G21Rik | 5.32E-50 | 1.11E-48 | -5.377715548 | 323.2290868 | 7.774218578 | 189 | 42 | 45 | 62 | 111 | 11 | 0 | 0 | 0 | 0 | 2 | 2 |
| SG0000002 | Lypla1 | 1.22E-46 | 2.27E-45 | 1.776367798 | 1283.593535 | 4397.111836 | 417 | 28 | 45 | 68 | 251 | 37 | 9 | 0 | 1 | 1 | 7 | 0 |
| SG0000002 | Echdc2 | 1.12E-37 | 1.52E-36 | 1.936053174 | 327.9035957 | 1254.747265 | 121 | 45 | 45 | 51 | 75 | 7 | 3 | 0 | 0 | 0 | 0 | 0 |
| SG0000003 | Cgnl1 | 4.47E-32 | 4.93E-31 | -2.032844456 | 4073.268059 | 995.3958687 | 632 | 27 | 45 | 90 | 251 | 45 | 2 | 0 | 0 | 0 | 2 | 0 |
| SG0000001 | Stxbp5 | 3.09E-31 | 3.3E-30 | -2.066816735 | 1767.935209 | 421.9806262 | 649 | 16 | 45 | 107 | 344 | 71 | 11 | 0 | 3 | 1 | 2 | 5 |
| SG0000002 | Col5a2 | 3.42E-30 | 3.48E-29 | -3.051488584 | 4252.879085 | 512.9717399 | 448 | 25 | 45 | 74 | 229 | 41 | 6 | 0 | 0 | 2 | 4 | 1 |
| ISG000000 | Zfp141 | 2.9E-24 | 2.26E-23 | 1.40549116 | 597.6827253 | 1583.309077 | 105 | 42 | 45 | 52 | 85 | 1 | 3 | 0 | 0 | 0 | 1 | 0 |
| ISG000000 | Rb1 | 2.95E-22 | 2.09E-21 | -1.822184818 | 4018.536866 | 1136.411629 | 467 | 23 | 45 | 64 | 223 | 55 | 11 | 0 | 0 | 0 | 1 | 0 |
| SG0000003 | Rnf38 | 6.09E-15 | 2.86E-14 | 1.055907988 | 864.2480231 | 1796.794344 | 443 | 23 | 45 | 47 | 168 | 76 | 20 | 0 | 3 | 0 | 3 | 8 |
| SG0000003 | Slc43a2 | 4.78E-14 | 2.12E-13 | -1.378441109 | 1285.498287 | 494.4465844 | 123 | 36 | 45 | 42 | 69 | 17 | 3 | 0 | 0 | 0 | 3 | 2 |
| SG0000004 | Inpp5f | 1.14E-11 | 4.35E-11 | -1.286790433 | 1002.817763 | 411.0167299 | 265 | 29 | 45 | 57 | 136 | 26 | 14 | 3 | 6 | 3 | 3 | 0 |
| SG0000005 | Klf13 | 1.6E-11 | 6.02E-11 | -1.144560046 | 2907.009887 | 1314.920824 | 183 | 40 | 45 | 53 | 109 | 6 | 3 | 0 | 0 | 0 | 1 | 1 |
| SG0000004 | Ipo9 | 2.44E-11 | 9.1E-11 | 0.810450424 | 5005.039157 | 8777.631962 | 423 | 37 | 45 | 71 | 155 | 22 | 18 | 0 | 0 | 1 | 3 | 0 |
| SG0000003 | Ccdc122 | 9.63E-10 | 3.18E-09 | -1.756465215 | 183.1882391 | 54.21867148 | 474 | 27 | 45 | 81 | 300 | 51 | 2 | 0 | 0 | 0 | 1 | 0 |
| ISG000000 | Pstpip2 | 5.69E-09 | 1.76E-08 | 3.671422205 | 6.068032941 | 77.31364347 | 180 | 33 | 45 | 57 | 129 | 21 | 1 | 0 | 0 | 0 | 0 | 0 |
| SG0000001 | Cmc2 | 3.54E-07 | 9.46E-07 | 0.913796549 | 266.3689077 | 501.8381819 | 290 | 40 | 45 | 75 | 135 | 26 | 0 | 0 | 0 | 0 | 0 | 1 |
| SG0000003 | Cdon | 4.08E-07 | 1.09E-06 | 0.824245961 | 502.8784461 | 890.4012887 | 347 | 26 | 45 | 57 | 170 | 26 | 18 | 2 | 4 | 8 | 14 | 2 |
| SG0000002 | Rgs9 | 1.21E-06 | 3.1E-06 | 2.459661546 | 8.537634573 | 46.96447445 | 223 | 36 | 45 | 44 | 125 | 24 | 5 | 0 | 0 | 0 | 3 | 3 |
| SG0000002 | Zfp142 | 1.34E-06 | 3.42E-06 | 0.739599164 | 761.4123643 | 1271.339258 | 179 | 30 | 45 | 45 | 84 | 23 | 3 | 0 | 0 | 0 | 0 | 0 |
| SG0000003 | Opa1 | 1.48E-06 | 3.76E-06 | -0.688462038 | 3695.022329 | 2292.816734 | 466 | 25 | 45 | 71 | 199 | 35 | 10 | 1 | 1 | 3 | 6 | 0 |
| SG0000002 | Ston2 | 4.17E-06 | 1.02E-05 | 0.755273377 | 367.8624914 | 620.934012 | 344 | 32 | 45 | 61 | 187 | 11 | 1 | 0 | 0 | 0 | 0 | 0 |
| ISG000000 | Frs2 | 4.35E-06 | 1.06E-05 | -0.689296099 | 2426.276184 | 1504.670594 | 201 | 40 | 45 | 57 | 124 | 22 | 7 | 0 | 0 | 2 | 3 | 0 |
| ISG000000 | Rsbn1l | 5.2E-06 | 1.26E-05 | 0.632055982 | 929.3462019 | 1440.27472 | 315 | 30 | 45 | 70 | 182 | 36 | 17 | 0 | 4 | 6 | 10 | 5 |
| SG0000003 | Fam53a | 1.38E-05 | 3.22E-05 | -0.722464936 | 1103.012358 | 668.4928427 | 144 | 45 | 45 | 59 | 113 | 14 | 6 | 0 | 0 | 1 | 2 | 1 |
| ISG000000 | Nsd1 | 1.82E-05 | 4.18E-05 | 0.533115235 | 3723.699139 | 5388.380567 | 276 | 38 | 45 | 67 | 163 | 19 | 6 | 0 | 0 | 1 | 1 | 1 |
| SG0000004 | Pla2g6 | 8.73E-05 | 0.000186 | 0.568098363 | 994.4612814 | 1474.35656 | 185 | 35 | 45 | 55 | 135 | 10 | 0 | 0 | 0 | 0 | 0 | 0 |
| ISG000000 | Dym | 0.000113 | 0.000239 | 0.540902701 | 910.8310566 | 1325.152219 | 393 | 30 | 45 | 67 | 166 | 38 | 0 | 0 | 0 | 0 | 0 | 0 |
| SG0000004 | Mdga1 | 0.000202 | 0.000415 | 1.439538798 | 23.26141747 | 63.09290638 | 249 | 38 | 45 | 59 | 130 | 25 | 2 | 0 | 0 | 0 | 2 | 0 |
| SG0000004 | Ccdc73 | 0.000215 | 0.000439 | 2.383227483 | 6.555058864 | 34.19790919 | 392 | 29 | 45 | 73 | 206 | 26 | 2 | 0 | 0 | 0 | 0 | 0 |
| SG0000003 | Ric1 | 0.000226 | 0.000461 | 0.508419841 | 1111.241087 | 1580.730792 | 536 | 24 | 45 | 78 | 261 | 48 | 21 | 4 | 4 | 4 | 19 | 5 |
| SG0000002 | Grb10 | 0.000668 | 0.001279 | -0.77982016 | 5070.492909 | 2953.254812 | 452 | 28 | 45 | 73 | 179 | 25 | 11 | 0 | 0 | 2 | 6 | 0 |
| SG0000005 | D17Wsu92e | 0.000709 | 0.001355 | 0.426147366 | 3716.467734 | 4993.597185 | 271 | 37 | 45 | 67 | 157 | 18 | 3 | 0 | 0 | 0 | 0 | 0 |
| SG0000002 | Usp7 | 0.00081 | 0.001537 | 0.408401573 | 8604.083053 | 11419.46394 | 333 | 41 | 45 | 73 | 172 | 22 | 5 | 0 | 0 | 0 | 1 | 0 |
| SG0000000 | Rin2 | 0.001183 | 0.002202 | -0.804000892 | 567.4386621 | 325.0053709 | 460 | 35 | 45 | 80 | 256 | 51 | 14 | 0 | 0 | 0 | 0 | 0 |
| SG0000007 | Nhsl2 | 0.002216 | 0.003989 | -1.536709743 | 139.0748985 | 47.93503231 | 336 | 39 | 45 | 78 | 195 | 23 | 11 | 0 | 0 | 2 | 6 | 0 |
| SG0000000 | Rbfox1 | 0.002828 | 0.005007 | -2.326260536 | 21.50454826 | 4.28801331 | 612 | 29 | 45 | 73 | 252 | 64 | 0 | 0 | 0 | 0 | 0 | 0 |
| SG0000002 | Gnl3l | 0.011836 | 0.019125 | 0.316484471 | 3780.72307 | 4708.105501 | 182 | 38 | 45 | 62 | 87 | 7 | 20 | 6 | 8 | 8 | 12 | 0 |
| SG0000003 | Arhgap17 | 0.012131 | 0.019549 | 0.363982415 | 807.5911026 | 1039.348396 | 330 | 39 | 45 | 63 | 144 | 36 | 3 | 0 | 0 | 0 | 0 | 0 |
| SG0000006 | Pla2g16 | 0.027112 | 0.04102 | 0.471164569 | 161.9422072 | 224.4888228 | 168 | 39 | 45 | 66 | 129 | 14 | 23 | 7 | 8 | 9 | 10 | 0 |
| SG0000003 | Phka2 | 0.032843 | 0.048974 | -0.374637658 | 769.8018584 | 593.7474844 | 608 | 10 | 45 | 95 | 344 | 53 | 1 | 0 | 0 | 0 | 0 | 0 |
| ISG000000 | Fam19a4 | 0.081949 | 0.113586 | -0.751364347 | 94.03039914 | 55.85796034 | 413 | 29 | 45 | 64 | 250 | 47 | 2 | 0 | 0 | 0 | 3 | 0 |
| SG0000002 | Prkar1b | 0.089026 | 0.122523 | -0.67112604 | 154.0216376 | 96.72810234 | 318 | 31 | 45 | 94 | 173 | 35 | 3 | 0 | 0 | 3 | 3 | 0 |
| SG0000003 | Rasa2 | 0.184761 | 0.237389 | 0.208309372 | 545.6622295 | 630.4218579 | 427 | 20 | 45 | 62 | 225 | 59 | 7 | 0 | 1 | 0 | 3 | 3 |
| SG0000003 | Zranb1 | 0.46494 | 0.534395 | 0.095311647 | 1203.345581 | 1285.529456 | 261 | 31 | 45 | 67 | 138 | 12 | 10 | 3 | 4 | 5 | 7 | 0 |
| SG0000003 | Brcc3 | 0.469087 | 0.537884 | -0.113011086 | 967.7835494 | 894.867033 | 371 | 30 | 45 | 60 | 241 | 7 | 16 | 0 | 0 | 0 | 8 | 1 |
| SG0000003 | Atm | 0.534165 | 0.600777 | 0.099775221 | 1068.532451 | 1145.046307 | 272 | 30 | 45 | 68 | 152 | 28 | 1 | 0 | 0 | 0 | 0 | 0 |
| SG0000005 | Znrf2 | 0.859968 | 0.890802 | -0.037201962 | 1510.296737 | 1471.849408 | 499 | 11 | 45 | 55 | 191 | 64 | 13 | 0 | 0 | 0 | 0 | 2 |
| SG0000000 | Ap4e1 | 0.925288 | 0.945991 | 0.005791066 | 568.3583027 | 570.6443126 | 294 | 34 | 45 | 77 | 179 | 14 | 9 | 0 | 0 | 3 | 6 | 2 |
| SG0000003 | Lrig1 | 1.66E-61 | 4.83E-60 | -3.434471452 | 4494.108774 | 415.6852873 | 274 | 36 | 44 | 66 | 152 | 24 | 5 | 2 | 4 | 4 | 6 | 2 |
| SG0000003 | Ncoa7 | 9.67E-58 | 2.47E-56 | -3.649767465 | 2980.197315 | 237.4409028 | 425 | 27 | 44 | 69 | 219 | 20 | 20 | 4 | 4 | 6 | 11 | 0 |
| SG0000002 | Dock9 | 5.77E-37 | 7.66E-36 | -2.045556764 | 6307.890888 | 1527.95382 | 561 | 31 | 44 | 90 | 221 | 66 | 5 | 0 | 0 | 0 | 1 | 2 |
| SG0000002 | Eif2a | 3.3E-36 | 4.28E-35 | 1.539593028 | 1753.967774 | 5099.003291 | 250 | 29 | 44 | 50 | 119 | 12 | 6 | 0 | 0 | 0 | 0 | 0 |
| SG0000002 | Ror2 | 7.48E-35 | 9.22E-34 | 3.910240056 | 32.19724449 | 484.0812402 | 422 | 34 | 44 | 66 | 180 | 30 | 4 | 0 | 0 | 0 | 0 | 0 |
| SG0000000 | Nr2c1 | 1.53E-27 | 1.38E-26 | -2.118898487 | 973.4118618 | 224.1014321 | 202 | 38 | 44 | 46 | 107 | 14 | 4 | 0 | 0 | 2 | 5 | 0 |
| SG0000007 | Gm13212 | 4.03E-26 | 3.43E-25 | 3.060330999 | 52.98302649 | 441.9653171 | 234 | 36 | 44 | 66 | 146 | 22 | 4 | 0 | 0 | 2 | 2 | 0 |
| SG0000001 | Arl5b | 2.1E-24 | 1.65E-23 | 1.455213475 | 571.1041471 | 1565.951185 | 325 | 36 | 44 | 64 | 114 | 18 | 52 | 2 | 4 | 10 | 25 | 2 |
| SG0000003 | Tmem63c | 1.24E-23 | 9.38E-23 | -4.764601625 | 1076.962355 | 39.61982166 | 236 | 37 | 44 | 68 | 125 | 20 | 0 | 0 | 0 | 0 | 0 | 0 |
| SG0000006 | Sort1 | 4.7E-22 | 3.3E-21 | 1.380158692 | 511.1241944 | 1330.440952 | 250 | 36 | 44 | 56 | 140 | 14 | 10 | 2 | 2 | 2 | 4 | 4 |
| SG0000002 | Lrp8 | 1.93E-21 | 1.31E-20 | -1.846754632 | 1797.613278 | 499.7671758 | 380 | 27 | 44 | 70 | 127 | 32 | 25 | 2 | 2 | 4 | 4 | 4 |
| ISG000000 | Itga1 | 1.5E-19 | 9.17E-19 | -2.642579705 | 330.9110567 | 52.99263578 | 328 | 28 | 44 | 61 | 192 | 38 | 9 | 0 | 2 | 0 | 1 | 0 |
| SG0000004 | Iffo2 | 3.29E-16 | 1.68E-15 | -1.313632735 | 1883.699231 | 757.8246088 | 190 | 42 | 44 | 65 | 109 | 17 | 2 | 0 | 0 | 1 | 6 | 3 |
| SG0000003 | Slain2 | 1.83E-11 | 6.87E-11 | -1.056246391 | 1806.347203 | 868.6391199 | 314 | 30 | 44 | 58 | 159 | 34 | 8 | 0 | 2 | 0 | 7 | 2 |
| SG0000001 | Lyst | 3.04E-11 | 1.12E-10 | -1.229064051 | 742.0655936 | 316.5608109 | 479 | 26 | 44 | 77 | 267 | 42 | 0 | 0 | 0 | 0 | 0 | 0 |
| SG0000006 | Akap13 | 4.09E-10 | 1.39E-09 | -1.229205039 | 1767.608268 | 753.9775882 | 597 | 28 | 44 | 78 | 293 | 47 | 2 | 1 | 1 | 2 | 4 | 0 |
| SG0000002 | Stxbp1 | 1.1E-09 | 3.59E-09 | -0.960673535 | 1674.494838 | 860.3838762 | 186 | 42 | 44 | 66 | 114 | 12 | 6 | 0 | 2 | 4 | 4 | 2 |
| SG0000001 | Nars2 | 2.39E-08 | 7.06E-08 | 0.944963906 | 285.5774903 | 549.7768557 | 558 | 25 | 44 | 78 | 253 | 46 | 9 | 0 | 0 | 0 | 0 | 1 |
| ISG000000 | Zyg11b | 1.47E-06 | 3.73E-06 | -0.763861265 | 2850.949459 | 1678.975524 | 156 | 38 | 44 | 57 | 98 | 10 | 9 | 0 | 0 | 1 | 1 | 1 |
| SG0000002 | Chid1 | 3.87E-06 | 9.47E-06 | -0.762875236 | 955.3278721 | 562.994458 | 224 | 29 | 44 | 72 | 134 | 30 | 0 | 0 | 0 | 0 | 0 | 0 |
| SG0000006 | Sntb1 | 5.39E-05 | 0.000118 | 2.112297176 | 8.893327432 | 38.45289942 | 739 | 15 | 44 | 100 | 329 | 89 | 0 | 0 | 0 | 0 | 0 | 0 |
| SG0000002 | Ccdc34 | 0.000287 | 0.000578 | 0.565073317 | 506.6584404 | 749.5822608 | 447 | 29 | 44 | 63 | 258 | 31 | 9 | 2 | 2 | 2 | 2 | 1 |
| SG0000002 | Osbpl8 | 0.000515 | 0.001004 | -0.578873936 | 1154.478299 | 772.9072047 | 636 | 26 | 44 | 92 | 304 | 74 | 13 | 0 | 0 | 3 | 10 | 3 |
| SG0000004 | Znrf3 | 0.000621 | 0.001196 | -0.591769846 | 670.658039 | 445.0006657 | 691 | 31 | 44 | 80 | 334 | 74 | 15 | 0 | 3 | 2 | 5 | 1 |
| SG0000006 | Naa15 | 0.002045 | 0.003698 | 0.381133276 | 4743.513774 | 6177.783929 | 210 | 32 | 44 | 57 | 128 | 24 | 7 | 0 | 0 | 2 | 3 | 1 |
| ISG000000 | Suclg2 | 0.0058 | 0.009831 | -0.406191877 | 2431.425909 | 1834.784664 | 463 | 32 | 44 | 73 | 201 | 49 | 1 | 0 | 0 | 0 | 1 | 0 |
| SG0000003 | Etv6 | 0.009229 | 0.015178 | -0.391163429 | 1331.920898 | 1015.608923 | 614 | 34 | 44 | 78 | 308 | 70 | 14 | 6 | 6 | 6 | 12 | 0 |
| SG0000005 | Rnf157 | 0.012142 | 0.019563 | -0.491645721 | 580.470217 | 412.8381586 | 229 | 32 | 44 | 47 | 118 | 20 | 3 | 0 | 0 | 3 | 3 | 0 |
| SG0000005 | Fam73a | 0.015666 | 0.024758 | 0.34677705 | 899.5434553 | 1143.964222 | 252 | 36 | 44 | 56 | 128 | 11 | 8 | 0 | 0 | 2 | 6 | 1 |
| SG0000002 | Tceanc2 | 0.029117 | 0.043808 | -0.394370431 | 476.8755924 | 362.8171151 | 227 | 32 | 44 | 60 | 148 | 12 | 4 | 0 | 0 | 0 | 0 | 0 |
| ISG000000 | Ap4s1 | 0.050315 | 0.072592 | -0.495541375 | 228.2187115 | 161.8744967 | 118 | 36 | 44 | 50 | 94 | 14 | 0 | 0 | 0 | 0 | 0 | 0 |
| SG0000006 | Wdr35 | 0.057538 | 0.082101 | -0.284539589 | 1878.21686 | 1542.022759 | 294 | 32 | 44 | 64 | 172 | 10 | 6 | 0 | 0 | 0 | 2 | 0 |
| SG0000002 | Exoc2 | 0.060046 | 0.08532 | -0.279575111 | 1501.839663 | 1237.26599 | 609 | 10 | 44 | 80 | 297 | 73 | 10 | 0 | 0 | 2 | 4 | 0 |
| SG0000004 | Zfp644 | 0.07078 | 0.099296 | -0.269328556 | 1565.351618 | 1298.781017 | 856 | 20 | 44 | 96 | 352 | 84 | 36 | 0 | 0 | 0 | 4 | 12 |
| SG0000003 | Pbx4 | 0.077987 | 0.108565 | 0.425001091 | 113.5021944 | 152.3850404 | 212 | 40 | 44 | 76 | 188 | 20 | 0 | 0 | 0 | 0 | 0 | 0 |
| SG0000003 | Mpp6 | 0.15825 | 0.206376 | 0.181411959 | 1940.94128 | 2201.014167 | 740 | 12 | 44 | 72 | 245 | 57 | 1 | 0 | 0 | 0 | 0 | 0 |
| SG0000002 | Il6st | 0.226505 | 0.284793 | -0.169139678 | 2585.772226 | 2299.715698 | 431 | 29 | 44 | 83 | 261 | 22 | 17 | 0 | 0 | 0 | 6 | 3 |
| SG0000007 | Rabepk | 0.272264 | 0.334991 | -0.186598169 | 640.1237933 | 562.4608826 | 143 | 31 | 44 | 37 | 110 | 34 | 2 | 0 | 0 | 0 | 0 | 2 |
| SG0000002 | Rnf2 | 0.306114 | 0.371258 | -0.138998776 | 4871.635761 | 4424.172062 | 282 | 41 | 44 | 84 | 173 | 22 | 19 | 0 | 0 | 4 | 10 | 0 |
| SG0000003 | Ythdc1 | 0.317227 | 0.383048 | 0.1298844 | 2597.275096 | 2841.95405 | 452 | 36 | 44 | 80 | 232 | 28 | 11 | 0 | 0 | 0 | 5 | 4 |
| SG0000003 | Kdm4a | 0.708253 | 0.761159 | -0.050764875 | 2095.836686 | 2023.371777 | 167 | 40 | 44 | 55 | 122 | 12 | 6 | 0 | 0 | 3 | 3 | 0 |
| SG0000002 | Foxp4 | 0.731237 | 0.782152 | 0.097131546 | 1410.624431 | 1508.866826 | 217 | 40 | 44 | 44 | 80 | 4 | 1 | 0 | 0 | 0 | 0 | 0 |
| SG0000004 | Wdr72 | 0.799779 | 0.841261 | -0.296619213 | 16.22870695 | 13.21273236 | 398 | 32 | 44 | 79 | 224 | 43 | 3 | 0 | 0 | 0 | 1 | 0 |
| SG0000003 | Pqlc1 | 0.897135 | 0.921574 | -0.027152724 | 406.4927264 | 398.9137395 | 309 | 44 | 44 | 58 | 136 | 14 | 1 | 0 | 0 | 0 | 0 | 0 |
| SG0000002 | Cyfip2 | 1.4E-174 | 3.3E-172 | 5.132383505 | 35.66868003 | 1251.089623 | 475 | 35 | 43 | 75 | 313 | 56 | 15 | 0 | 0 | 3 | 3 | 0 |
| SG0000002 | Clock | 3.71E-46 | 6.76E-45 | -2.624314543 | 3201.551562 | 519.2339791 | 472 | 8 | 43 | 71 | 206 | 74 | 32 | 0 | 2 | 3 | 11 | 2 |
| ISG000000 | Zfhx3 | 9.76E-45 | 1.7E-43 | -2.9239132 | 1460.910188 | 192.5031941 | 301 | 33 | 43 | 60 | 156 | 38 | 11 | 0 | 0 | 1 | 4 | 2 |
| SG0000003 | Pou2f3 | 1.15E-37 | 1.56E-36 | 3.77222749 | 15.43048148 | 210.8303369 | 185 | 38 | 43 | 49 | 113 | 13 | 0 | 0 | 0 | 0 | 2 | 1 |
| SG0000002 | Hipk3 | 4.45E-32 | 4.91E-31 | -2.207494801 | 3164.917406 | 685.2360933 | 366 | 38 | 43 | 72 | 209 | 24 | 16 | 0 | 1 | 0 | 1 | 0 |
| ISG000000 | Adcy2 | 8.18E-26 | 6.86E-25 | -4.32236898 | 554.5744677 | 27.72025084 | 655 | 21 | 43 | 80 | 294 | 63 | 0 | 0 | 0 | 0 | 1 | 0 |
| SG0000005 | Nhs | 8.79E-26 | 7.35E-25 | 4.60099551 | 10.77803614 | 261.5635925 | 629 | 22 | 43 | 84 | 317 | 61 | 12 | 0 | 1 | 3 | 4 | 3 |
| SG0000005 | Aasdh | 1.09E-22 | 7.87E-22 | 1.511575308 | 339.083549 | 966.7990797 | 234 | 34 | 43 | 47 | 103 | 23 | 17 | 0 | 0 | 2 | 2 | 0 |
| SG0000005 | Cep70 | 1.31E-20 | 8.51E-20 | 1.507450972 | 257.5389844 | 732.202046 | 306 | 30 | 43 | 78 | 187 | 38 | 2 | 0 | 1 | 0 | 1 | 1 |
| SG0000003 | Arpp21 | 2.31E-16 | 1.18E-15 | -5.902547493 | 63.09870421 | 1.054815722 | 751 | 9 | 43 | 76 | 295 | 79 | 8 | 0 | 0 | 0 | 0 | 0 |
| ISG000000 | Igf1r | 8.32E-14 | 3.64E-13 | -1.590894449 | 894.9267416 | 297.0848713 | 408 | 36 | 43 | 80 | 180 | 32 | 0 | 0 | 0 | 0 | 0 | 0 |
| SG0000003 | Rpgr | 2.26E-13 | 9.6E-13 | 1.262569257 | 225.1898086 | 540.2813169 | 919 | 17 | 43 | 118 | 599 | 90 | 15 | 0 | 0 | 8 | 12 | 0 |
| SG0000002 | Synj1 | 3.36E-12 | 1.32E-11 | -1.106947879 | 1795.179246 | 833.4572019 | 625 | 24 | 43 | 81 | 259 | 43 | 15 | 0 | 0 | 9 | 12 | 3 |
| ISG000000 | Pwwp2a | 3.48E-10 | 1.19E-09 | 0.883281 | 744.9930263 | 1374.188415 | 226 | 34 | 43 | 52 | 156 | 14 | 4 | 0 | 0 | 0 | 1 | 2 |
| SG0000002 | Slc35a3 | 4.25E-10 | 1.44E-09 | -1.002430569 | 2331.557327 | 1163.816281 | 260 | 30 | 43 | 50 | 140 | 18 | 28 | 0 | 3 | 0 | 4 | 2 |
| SG0000004 | Zfp704 | 8.89E-10 | 2.94E-09 | -1.060824309 | 2491.875939 | 1194.500895 | 469 | 27 | 43 | 76 | 212 | 52 | 8 | 0 | 0 | 0 | 1 | 8 |
| SG0000003 | Glb1l2 | 9.14E-09 | 2.79E-08 | 5.660124033 | 2.947100147 | 149.0259871 | 429 | 33 | 43 | 60 | 169 | 50 | 2 | 0 | 0 | 0 | 2 | 0 |
| ISG000000 | Ltf | 1.03E-08 | 3.15E-08 | -6.798465579 | 258.7382924 | 2.324441624 | 136 | 43 | 43 | 65 | 96 | 20 | 3 | 1 | 1 | 1 | 1 | 0 |
| SG0000002 | Rps6kb1 | 1.95E-06 | 4.92E-06 | -0.691904192 | 2812.724228 | 1741.178268 | 232 | 28 | 43 | 53 | 146 | 15 | 16 | 0 | 0 | 1 | 3 | 3 |
| SG0000002 | Nmd3 | 7.87E-05 | 0.000169 | 0.506861781 | 2322.827873 | 3300.635971 | 193 | 40 | 43 | 57 | 135 | 9 | 16 | 0 | 1 | 1 | 10 | 2 |
| SG0000005 | Lypd6 | 0.000136 | 0.000284 | 1.542774133 | 42.71326152 | 124.4468836 | 551 | 26 | 43 | 86 | 276 | 64 | 21 | 0 | 0 | 6 | 6 | 3 |
| SG0000006 | Col23a1 | 0.000331 | 0.000661 | 1.929356029 | 9.222584933 | 35.12745615 | 327 | 29 | 43 | 58 | 146 | 28 | 1 | 0 | 0 | 0 | 0 | 0 |
| ISG000000 | Peak1 | 0.001149 | 0.002144 | -0.472829035 | 2139.483391 | 1541.605308 | 542 | 14 | 43 | 74 | 253 | 37 | 8 | 0 | 0 | 0 | 4 | 1 |
| SG0000004 | Agps | 0.01107 | 0.017974 | 0.322414435 | 3704.157441 | 4631.757928 | 486 | 26 | 43 | 66 | 229 | 36 | 13 | 0 | 0 | 3 | 7 | 0 |
| SG0000005 | Fam160a1 | 0.01369 | 0.021818 | -0.716590327 | 179.9062319 | 109.479051 | 619 | 23 | 43 | 88 | 275 | 48 | 3 | 0 | 0 | 0 | 3 | 0 |
| SG0000003 | BC026585 | 0.015416 | 0.024386 | 0.63461789 | 138.0351426 | 214.3032163 | 234 | 35 | 43 | 66 | 120 | 23 | 0 | 0 | 0 | 0 | 0 | 0 |
| SG0000003 | Arhgef7 | 0.022385 | 0.034334 | 0.313474646 | 1470.982868 | 1827.986087 | 312 | 27 | 43 | 55 | 139 | 25 | 12 | 3 | 3 | 3 | 5 | 7 |
| SG0000001 | Ints2 | 0.080111 | 0.111202 | -0.309341906 | 2364.772569 | 1908.394632 | 271 | 29 | 43 | 56 | 139 | 50 | 11 | 0 | 0 | 0 | 1 | 2 |
| SG0000002 | Tfpi | 0.272524 | 0.335241 | -0.194918613 | 1288.149364 | 1125.355854 | 804 | 32 | 43 | 102 | 468 | 39 | 46 | 0 | 0 | 6 | 8 | 0 |
| SG0000003 | Nktr | 0.580013 | 0.643868 | 0.076924472 | 2430.759382 | 2563.885016 | 320 | 28 | 43 | 50 | 148 | 40 | 6 | 0 | 0 | 0 | 1 | 7 |

| | | | | | | | | | | | | | | | | | | |
|---|---|---|---|---|---|---|---|---|---|---|---|---|---|---|---|---|---|---|
| SG0000003 | Tbcel | 0.826036 | 0.863393 | 0.025094409 | 1132.321773 | 1152.189805 | 411 | 25 | 43 | 60 | 139 | 33 | 15 | 0 | 0 | 0 | 7 | 3 |
| SG0000002 | Apex2 | 0.99091 | 1 | -0.00447061 | 430.1735815 | 428.8426268 | 375 | 30 | 43 | 73 | 206 | 44 | 13 | 2 | 2 | 3 | 4 | 1 |
| SG0000002 | Dmrt1 | 4.3E-267 | 2.3E-264 | 10.31243914 | 0 | 1270.608758 | 315 | 36 | 42 | 66 | 144 | 34 | 1 | 0 | 0 | 0 | 0 | 0 |
| SG0000004 | Phc1 | 1.06E-85 | 5.36E-84 | 2.32645234 | 2099.787744 | 10531.91473 | 293 | 27 | 42 | 79 | 165 | 71 | 0 | 0 | 0 | 0 | 0 | 0 |
| SG0000003 | Cep85 | 1.02E-83 | 5.02E-82 | 2.517621631 | 543.5148547 | 3112.368755 | 193 | 36 | 42 | 54 | 150 | 22 | 5 | 0 | 2 | 2 | 2 | 0 |
| JSG000000 | Akap12 | 2.43E-72 | 9.34E-71 | 2.082474674 | 4921.151897 | 20842.70647 | 170 | 33 | 42 | 48 | 111 | 14 | 1 | 0 | 0 | 1 | 3 | 0 |
| SG0000002 | Cbs | 4.74E-57 | 1.2E-55 | 3.839549964 | 45.80676209 | 655.7666152 | 105 | 39 | 42 | 45 | 78 | 3 | 0 | 0 | 0 | 0 | 0 | 0 |
| SG0000002 | Ezh2 | 2.09E-54 | 4.91E-53 | 1.86243226 | 1994.458213 | 7252.252507 | 503 | 24 | 42 | 78 | 274 | 52 | 6 | 0 | 0 | 0 | 0 | 0 |
| SG0000004 | Zbtb7c | 1.18E-49 | 2.45E-48 | -3.940626421 | 1176.204161 | 76.60126771 | 415 | 38 | 42 | 80 | 220 | 50 | 2 | 0 | 0 | 2 | 6 | 0 |
| SG0000003 | Porcn | 2.4E-48 | 4.75E-47 | -2.915770251 | 5343.582401 | 708.1058866 | 91 | 42 | 42 | 42 | 78 | 12 | 2 | 0 | 0 | 0 | 0 | 0 |
| SG0000002 | Als2 | 5.23E-41 | 8.01E-40 | 1.704329372 | 1018.142359 | 3317.896005 | 332 | 28 | 42 | 50 | 180 | 18 | 0 | 0 | 0 | 0 | 0 | 0 |
| SG0000004 | Rbm20 | 4.44E-39 | 6.36E-38 | -3.485476799 | 1290.696234 | 115.2367436 | 315 | 35 | 42 | 76 | 174 | 33 | 10 | 0 | 0 | 4 | 8 | 2 |
| JSG000000 | Myo1e | 9.77E-39 | 1.38E-37 | -2.252791008 | 6525.887514 | 1369.247373 | 335 | 35 | 42 | 73 | 173 | 20 | 4 | 1 | 1 | 1 | 1 | 1 |
| SG0000002 | Prkg2 | 9.18E-38 | 1.25E-36 | -3.754261117 | 902.5720367 | 66.88622207 | 467 | 28 | 42 | 72 | 229 | 42 | 1 | 0 | 0 | 0 | 3 | 2 |
| SG0000005 | Ugt1a6a | 5.91E-33 | 6.73E-32 | -6.913665956 | 119.5648841 | 0 | 350 | 26 | 42 | 66 | 164 | 19 | 4 | 0 | 0 | 0 | 0 | 0 |
| SG0000002 | Rcbtb2 | 3.55E-24 | 2.75E-23 | 3.568187855 | 107.8998326 | 1279.830174 | 601 | 20 | 42 | 51 | 234 | 56 | 16 | 0 | 0 | 0 | 1 | 6 |
| SG0000000 | Mlxipl | 1.09E-22 | 7.88E-22 | 3.42295759 | 12.09740971 | 129.7493819 | 115 | 36 | 42 | 39 | 54 | 12 | 0 | 0 | 0 | 0 | 0 | 0 |
| SG0000002 | Sec14l1 | 3.26E-19 | 1.95E-18 | -1.851321428 | 11830.0716 | 3278.567215 | 162 | 42 | 42 | 45 | 72 | 0 | 0 | 0 | 0 | 0 | 0 | 0 |
| SG0000002 | Iqcb1 | 7.58E-19 | 4.46E-18 | 1.305723374 | 438.9089438 | 1085.016474 | 310 | 28 | 42 | 68 | 174 | 14 | 3 | 0 | 0 | 1 | 1 | 0 |
| JSG000000 | Ddx10 | 9.76E-18 | 5.41E-17 | 1.121008369 | 1128.627862 | 2454.753635 | 438 | 24 | 42 | 72 | 251 | 30 | 1 | 0 | 0 | 0 | 0 | 0 |
| JSG000000 | Plag1 | 1.84E-15 | 8.95E-15 | 1.825752131 | 69.51774288 | 246.4346768 | 282 | 27 | 42 | 84 | 165 | 30 | 3 | 0 | 0 | 0 | 3 | 3 |
| SG0000003 | Gripap1 | 4.3E-15 | 2.04E-14 | -1.386566879 | 985.5410937 | 376.9437233 | 133 | 36 | 42 | 51 | 84 | 3 | 3 | 0 | 0 | 0 | 3 | 0 |
| JSG000000 | Stac | 6.34E-15 | 2.98E-14 | -7.253838168 | 54.94619754 | 0.360010171 | 360 | 24 | 42 | 72 | 170 | 48 | 4 | 0 | 0 | 0 | 0 | 0 |
| SG0000002 | Fhl1 | 7.9E-15 | 3.7E-14 | 3.104919486 | 150.8507151 | 1297.840351 | 421 | 27 | 42 | 64 | 178 | 30 | 6 | 0 | 0 | 0 | 0 | 0 |
| SG0000004 | Brca2 | 2.49E-14 | 1.12E-13 | 1.395098831 | 1035.078537 | 2722.324543 | 234 | 26 | 42 | 42 | 98 | 20 | 2 | 0 | 0 | 0 | 0 | 0 |
| SG0000005 | Cd47 | 8.14E-12 | 3.13E-11 | -2.783927931 | 2869.661883 | 416.6629803 | 232 | 35 | 42 | 78 | 176 | 16 | 8 | 1 | 2 | 1 | 4 | 2 |
| SG0000002 | Tor1aip1 | 3.02E-11 | 1.11E-10 | -1.076339324 | 1435.855789 | 680.9267591 | 354 | 34 | 42 | 54 | 168 | 15 | 20 | 0 | 2 | 0 | 12 | 0 |
| SG0000003 | Trpc4ap | 1.06E-10 | 3.75E-10 | -0.932306942 | 3859.845818 | 2022.635392 | 210 | 36 | 42 | 56 | 124 | 32 | 0 | 0 | 0 | 0 | 0 | 0 |
| SG0000002 | Elp4 | 6.01E-10 | 2.02E-09 | 1.103485082 | 210.3074044 | 451.8941019 | 534 | 18 | 42 | 77 | 285 | 34 | 4 | 0 | 0 | 0 | 0 | 0 |
| SG0000003 | Phldb2 | 7.3E-10 | 2.44E-09 | -2.197492975 | 2386.457894 | 520.2864021 | 386 | 22 | 42 | 64 | 244 | 44 | 4 | 0 | 0 | 0 | 4 | 2 |
| SG0000002 | Mbnl2 | 7.83E-10 | 2.61E-09 | -1.012337185 | 4566.111809 | 2263.615626 | 530 | 26 | 42 | 62 | 210 | 32 | 17 | 2 | 4 | 6 | 18 | 2 |
| SG0000000 | Tbc1d8 | 1.02E-09 | 3.36E-09 | 1.035251177 | 329.1334797 | 674.5493351 | 438 | 27 | 42 | 89 | 200 | 57 | 6 | 0 | 0 | 1 | 1 | 0 |
| SG0000002 | Eif4e2 | 6.19E-09 | 1.91E-08 | 0.731332485 | 2619.498451 | 4348.817282 | 145 | 21 | 42 | 35 | 117 | 0 | 8 | 0 | 0 | 0 | 3 | 0 |
| SG0000002 | Cenpl | 1.41E-08 | 4.26E-08 | 0.886726959 | 401.7830204 | 742.8873987 | 247 | 34 | 42 | 49 | 127 | 38 | 20 | 2 | 2 | 9 | 19 | 4 |
| SG0000002 | Scp2 | 1.71E-08 | 5.1E-08 | 0.771978418 | 947.562478 | 1618.067053 | 221 | 32 | 42 | 59 | 116 | 13 | 4 | 0 | 0 | 0 | 0 | 2 |
| SG0000003 | Alg14 | 5.5E-07 | 1.45E-06 | -0.963789593 | 499.1412442 | 255.913903 | 290 | 20 | 42 | 56 | 207 | 42 | 4 | 0 | 0 | 0 | 1 | 0 |
| SG0000002 | Wasf2 | 4.69E-06 | 1.14E-05 | -0.63417601 | 5592.847 | 3603.520004 | 171 | 33 | 42 | 54 | 105 | 8 | 6 | 1 | 1 | 2 | 4 | 0 |
| SG0000007 | Pigb | 4.84E-06 | 1.17E-05 | -0.767850301 | 884.4153619 | 519.4099977 | 169 | 42 | 42 | 54 | 106 | 9 | 12 | 0 | 0 | 1 | 9 | 0 |
| SG0000001 | Gopc | 9.37E-06 | 2.21E-05 | -0.705367913 | 1230.087008 | 754.3952264 | 226 | 26 | 42 | 52 | 126 | 20 | 16 | 0 | 0 | 2 | 5 | 4 |
| SG0000002 | Ascc2 | 3.16E-05 | 7.08E-05 | -0.626159193 | 1694.668307 | 1097.973879 | 123 | 38 | 42 | 66 | 86 | 20 | 1 | 0 | 0 | 0 | 0 | 0 |
| SG0000003 | Ciz1 | 0.0003 | 0.000601 | -0.553892054 | 1241.739944 | 845.8483937 | 120 | 41 | 42 | 43 | 85 | 0 | 0 | 0 | 0 | 0 | 0 | 0 |
| JSG000000 | Nova1 | 0.000443 | 0.00087 | 0.899252825 | 84.30861103 | 157.243973 | 438 | 14 | 42 | 74 | 225 | 39 | 29 | 0 | 2 | 4 | 7 | 1 |
| SG0000004 | Nfrkb | 0.000499 | 0.000974 | -0.532686224 | 1411.532252 | 975.7448642 | 178 | 39 | 42 | 55 | 113 | 28 | 1 | 0 | 0 | 0 | 0 | 0 |
| SG0000002 | Trappc13 | 0.000795 | 0.00151 | -0.468156651 | 3453.135991 | 2496.229363 | 284 | 22 | 42 | 60 | 144 | 55 | 16 | 2 | 6 | 4 | 4 | 3 |
| SG0000002 | Sptb | 0.002224 | 0.004002 | -0.736041275 | 461.474555 | 277.062187 | 269 | 29 | 42 | 55 | 126 | 28 | 1 | 0 | 0 | 0 | 0 | 0 |
| SG0000004 | Rbm33 | 0.004193 | 0.007244 | 0.367671189 | 2395.114331 | 3090.340303 | 591 | 31 | 42 | 63 | 234 | 38 | 24 | 3 | 3 | 4 | 13 | 2 |
| SG0000002 | Tfeb | 0.008216 | 0.013623 | 0.906271061 | 320.972552 | 601.5651502 | 141 | 33 | 42 | 51 | 96 | 12 | 0 | 0 | 0 | 0 | 0 | 0 |
| SG0000003 | Herc1 | 0.008942 | 0.014739 | 0.413994762 | 1857.787994 | 2475.260966 | 463 | 23 | 42 | 71 | 241 | 35 | 2 | 0 | 0 | 1 | 1 | 0 |
| SG0000006 | Dazap1 | 0.014895 | 0.02361 | 0.354889788 | 2702.796386 | 3456.573538 | 100 | 42 | 42 | 42 | 48 | 6 | 4 | 0 | 0 | 0 | 0 | 0 |
| SG0000003 | Nme5 | 0.017703 | 0.027652 | 1.636050546 | 6.471909381 | 20.11558727 | 160 | 39 | 42 | 54 | 105 | 4 | 0 | 0 | 0 | 0 | 0 | 0 |
| SG0000000 | 43715 | 0.018811 | 0.029239 | -0.32245676 | 6488.886894 | 5189.207618 | 382 | 30 | 42 | 72 | 250 | 36 | 12 | 0 | 0 | 0 | 2 | 0 |
| SG0000003 | Galnt18 | 0.019581 | 0.030337 | -1.091701084 | 60.75978862 | 28.50896703 | 726 | 18 | 42 | 84 | 308 | 72 | 2 | 0 | 0 | 2 | 4 | 2 |
| JSG000000 | 2410089E03Rik | 0.02138 | 0.03288 | -0.348931554 | 1186.077708 | 931.2671405 | 245 | 29 | 42 | 55 | 128 | 20 | 2 | 0 | 0 | 0 | 0 | 0 |
| SG0000003 | Cdc14b | 0.027071 | 0.040967 | 0.408714559 | 922.3835581 | 1224.466463 | 402 | 34 | 42 | 77 | 189 | 23 | 16 | 0 | 0 | 4 | 5 | 0 |
| SG0000002 | Kif17 | 0.061163 | 0.086769 | -0.908295977 | 38.51238231 | 20.51993827 | 129 | 39 | 42 | 47 | 72 | 3 | 1 | 0 | 0 | 0 | 0 | 0 |
| JSG000000 | Mei1 | 0.067351 | 0.094932 | 1.802585478 | 6.019849361 | 20.99993364 | 164 | 39 | 42 | 45 | 126 | 12 | 0 | 0 | 0 | 0 | 0 | 0 |
| SG0000002 | Gcnt2 | 0.069053 | 0.097109 | 0.242439958 | 1688.552056 | 1997.543076 | 352 | 35 | 42 | 84 | 197 | 34 | 9 | 3 | 3 | 6 | 9 | 2 |
| SG0000005 | Cds2 | 0.115435 | 0.154928 | 0.2137007 | 2522.486447 | 2925.224232 | 242 | 37 | 42 | 56 | 137 | 20 | 14 | 0 | 2 | 2 | 4 | 4 |
| SG0000003 | Ube2q2 | 0.191725 | 0.245491 | -0.197215577 | 1183.054079 | 1031.898053 | 403 | 33 | 42 | 84 | 201 | 33 | 7 | 0 | 0 | 0 | 0 | 0 |
| SG0000002 | Bmpr1a | 0.423203 | 0.492267 | 0.102457261 | 3657.481139 | 3926.673613 | 343 | 28 | 42 | 59 | 156 | 27 | 10 | 0 | 0 | 0 | 5 | 1 |
| SG0000002 | Ubxn2a | 0.423767 | 0.492795 | -0.12445816 | 954.2031261 | 875.3368174 | 170 | 39 | 42 | 59 | 162 | 6 | 6 | 0 | 0 | 2 | 6 | 0 |
| SG0000003 | Rab6a | 0.49567 | 0.563175 | 0.081590186 | 3883.421489 | 4109.373564 | 315 | 27 | 42 | 66 | 162 | 21 | 21 | 0 | 3 | 6 | 9 | 3 |
| SG0000003 | Cep350 | 0.543414 | 0.60938 | 0.093020713 | 1148.135318 | 1224.602386 | 314 | 25 | 42 | 62 | 177 | 28 | 3 | 0 | 0 | 0 | 1 | 3 |
| SG0000005 | Fam118b | 0.618437 | 0.680432 | 0.078067555 | 494.9833149 | 522.5059444 | 334 | 28 | 42 | 73 | 203 | 34 | 5 | 0 | 0 | 0 | 3 | 0 |
| SG0000000 | Tuft1 | 0.685394 | 0.741269 | 0.069927342 | 396.4021373 | 416.088981 | 306 | 33 | 42 | 94 | 209 | 39 | 9 | 0 | 0 | 3 | 3 | 2 |
| SG0000005 | Zbtb25 | 0.726426 | 0.77826 | 0.055845394 | 459.8100382 | 477.9578369 | 165 | 34 | 42 | 58 | 129 | 16 | 7 | 1 | 2 | 2 | 7 | 1 |
| SG0000003 | Ogt | 0.791467 | 0.833927 | -0.070007448 | 9881.058526 | 9413.023216 | 225 | 38 | 42 | 60 | 134 | 16 | 3 | 0 | 0 | 0 | 2 | 0 |
| SG0000002 | Ube2w | 0.913887 | 0.936365 | 0.013066968 | 786.3924542 | 793.5474256 | 375 | 31 | 42 | 59 | 169 | 26 | 5 | 0 | 1 | 2 | 6 | 1 |
| SG0000002 | Ptprj | 1.08E-97 | 7.39E-96 | -4.50012788 | 4277.332069 | 189.0164019 | 354 | 31 | 41 | 56 | 206 | 11 | 8 | 0 | 2 | 0 | 8 | 2 |
| JSG000000 | Hrh1 | 1.68E-66 | 5.51E-65 | -4.43681766 | 862.2423089 | 39.81201217 | 326 | 31 | 41 | 63 | 141 | 4 | 6 | 0 | 0 | 5 | 5 | 0 |
| SG0000003 | Lmo7 | 2.1E-55 | 5.13E-54 | -3.271333496 | 4065.008961 | 421.0096002 | 448 | 22 | 41 | 73 | 218 | 39 | 2 | 0 | 1 | 1 | 3 | 1 |
| JSG000000 | Rab31 | 5.83E-55 | 1.4E-53 | -2.711630153 | 6488.588423 | 990.5313242 | 196 | 32 | 41 | 57 | 132 | 15 | 2 | 0 | 0 | 0 | 0 | 1 |
| SG0000007 | Zfp568 | 6.52E-51 | 1.39E-49 | 1.882547005 | 1036.023872 | 3820.083704 | 163 | 34 | 41 | 43 | 107 | 8 | 11 | 0 | 0 | 1 | 4 | 0 |
| JSG000000 | Atxn10 | 4.2E-43 | 6.86E-42 | 1.630172818 | 3449.335588 | 10677.42097 | 235 | 32 | 41 | 55 | 132 | 23 | 0 | 0 | 0 | 0 | 0 | 0 |
| SG0000003 | Tead4 | 1.92E-36 | 2.5E-35 | -2.201903452 | 2361.883826 | 513.3570678 | 220 | 38 | 41 | 67 | 119 | 25 | 0 | 0 | 0 | 0 | 2 | 1 |
| SG0000004 | Cdk6 | 8.01E-31 | 8.37E-30 | -2.446198466 | 5452.422568 | 1000.484565 | 599 | 15 | 41 | 80 | 263 | 44 | 17 | 1 | 1 | 2 | 7 | 0 |
| SG0000002 | Trip11 | 1.41E-26 | 1.23E-25 | -1.809470003 | 2004.035021 | 571.7426093 | 253 | 26 | 41 | 49 | 125 | 10 | 12 | 0 | 0 | 1 | 5 | 1 |
| SG0000002 | Atrn | 3.02E-20 | 1.92E-19 | -1.748459882 | 1749.001982 | 520.5367918 | 295 | 30 | 41 | 60 | 176 | 32 | 7 | 0 | 0 | 0 | 1 | 5 |
| SG0000005 | Adap1 | 3.22E-18 | 1.83E-17 | -1.672744341 | 1181.704633 | 370.6489009 | 82 | 39 | 41 | 48 | 62 | 4 | 2 | 0 | 0 | 2 | 2 | 0 |
| SG0000006 | Wtap | 4.28E-17 | 2.3E-16 | 1.078598585 | 1381.532756 | 2917.774428 | 291 | 41 | 41 | 81 | 139 | 6 | 14 | 0 | 0 | 3 | 8 | 2 |
| SG0000005 | Mbtd1 | 8.96E-14 | 3.91E-13 | 0.930719425 | 2095.837831 | 3995.141261 | 603 | 13 | 41 | 66 | 206 | 71 | 29 | 0 | 0 | 2 | 13 | 2 |
| SG0000001 | Rgs17 | 1.8E-12 | 7.2E-12 | 2.647837538 | 26.54615253 | 166.3719504 | 539 | 23 | 41 | 94 | 355 | 71 | 56 | 2 | 2 | 16 | 28 | 2 |
| SG0000002 | Rwdd3 | 2.21E-11 | 8.25E-11 | 1.868426443 | 40.9455182 | 149.5060529 | 143 | 35 | 41 | 45 | 76 | 9 | 2 | 0 | 1 | 1 | 2 | 0 |
| SG0000002 | Map3k3 | 7.15E-11 | 2.57E-10 | -1.01308476 | 1846.233254 | 914.7821217 | 185 | 36 | 41 | 49 | 97 | 8 | 1 | 0 | 0 | 0 | 0 | 0 |
| JSG000000 | Kcnn3 | 8.99E-10 | 2.97E-09 | 3.864236458 | 2.923008357 | 42.56779369 | 170 | 35 | 41 | 49 | 84 | 15 | 0 | 0 | 0 | 0 | 0 | 0 |
| JSG000000 | Efcab11 | 1.62E-08 | 4.84E-08 | 1.324543351 | 83.23669218 | 208.4692924 | 293 | 30 | 41 | 57 | 147 | 28 | 1 | 0 | 0 | 0 | 1 | 0 |
| SG0000003 | Ctnnal1 | 2.72E-07 | 7.34E-07 | -0.75099535 | 2979.536958 | 1770.421397 | 184 | 37 | 41 | 56 | 105 | 10 | 2 | 0 | 0 | 0 | 1 | 0 |
| SG0000003 | Adam5 | 4.66E-07 | 1.23E-06 | -4.153663058 | 40.34975116 | 2.267063519 | 517 | 20 | 41 | 87 | 295 | 54 | 2 | 0 | 0 | 0 | 0 | 0 |
| SG0000004 | Edem3 | 1.57E-06 | 3.98E-06 | -0.701579945 | 2722.721691 | 1674.197324 | 637 | 17 | 41 | 73 | 282 | 63 | 17 | 1 | 1 | 1 | 2 | 3 |
| SG0000002 | Tmem163 | 4.56E-06 | 1.11E-05 | 5.621666973 | 0.34247518 | 16.86240388 | 755 | 20 | 41 | 115 | 420 | 93 | 7 | 2 | 2 | 4 | 4 | 0 |
| SG0000004 | Uggt2 | 0.002476 | 0.004426 | -0.510427868 | 783.1046759 | 549.7506078 | 394 | 24 | 41 | 70 | 159 | 42 | 12 | 0 | 3 | 4 | 4 | 1 |
| SG0000004 | Akap10 | 0.003025 | 0.005332 | -0.463109228 | 1216.53013 | 882.4967218 | 291 | 23 | 41 | 47 | 135 | 44 | 12 | 3 | 5 | 3 | 5 | 4 |
| SG0000002 | Mroh1 | 0.004191 | 0.007241 | -0.658351166 | 2493.842173 | 1580.103431 | 168 | 38 | 41 | 64 | 177 | 6 | 4 | 0 | 0 | 0 | 0 | 0 |
| SG0000002 | Ccdc53 | 0.007976 | 0.01324 | -0.443923029 | 763.4789992 | 561.2585217 | 307 | 20 | 41 | 67 | 164 | 25 | 4 | 0 | 0 | 0 | 0 | 1 |
| SG0000003 | Gatad2a | 0.012387 | 0.019922 | 0.31867824 | 4715.379332 | 5880.961616 | 272 | 35 | 41 | 52 | 159 | 22 | 4 | 0 | 0 | 0 | 3 | 0 |
| SG0000004 | Zfand3 | 0.019311 | 0.029953 | -0.450144484 | 1281.607822 | 938.0978859 | 420 | 30 | 41 | 74 | 189 | 32 | 3 | 0 | 0 | 0 | 0 | 0 |
| SG0000002 | Adap2 | 0.034356 | 0.051006 | 0.98456864 | 18.92032481 | 37.4380553 | 69 | 39 | 41 | 43 | 51 | 3 | 0 | 0 | 0 | 0 | 0 | 0 |
| SG0000004 | Pde3a | 0.039324 | 0.057778 | -1.073601402 | 39.03890813 | 18.54861287 | 585 | 23 | 41 | 91 | 286 | 64 | 12 | 0 | 1 | 1 | 6 | 5 |
| SG0000004 | Rbms2 | 0.043185 | 0.063057 | -0.28464956 | 3548.729795 | 2913.297774 | 153 | 35 | 41 | 52 | 114 | 21 | 5 | 0 | 0 | 0 | 0 | 0 |
| SG0000002 | Pds5a | 0.080019 | 0.1111 | 0.235621844 | 5811.548263 | 6842.599033 | 293 | 21 | 41 | 56 | 166 | 16 | 10 | 0 | 1 | 1 | 10 | 2 |
| JSG000000 | Prkdc | 0.135369 | 0.179079 | 0.202153516 | 1237.853849 | 1424.044771 | 471 | 25 | 41 | 83 | 281 | 53 | 0 | 0 | 0 | 0 | 0 | 0 |
| JSG000000 | Zfp609 | 0.194598 | 0.248833 | -0.180750813 | 3117.472038 | 2750.370178 | 276 | 35 | 41 | 61 | 146 | 19 | 4 | 0 | 0 | 1 | 1 | 2 |
| JSG000000 | Hs2st1 | 0.222411 | 0.280078 | -0.182846098 | 1649.50523 | 1453.153648 | 479 | 25 | 41 | 76 | 198 | 35 | 11 | 0 | 0 | 2 | 7 | 1 |
| SG0000002 | Raph1 | 0.223784 | 0.281727 | 0.16869241 | 2554.885438 | 2871.791504 | 480 | 17 | 41 | 59 | 202 | 31 | 17 | 0 | 0 | 2 | 3 | 3 |
| SG0000002 | Casz1 | 0.429791 | 0.49899 | 0.21289062 | 428.2538816 | 496.3497076 | 281 | 28 | 41 | 65 | 135 | 31 | 6 | 1 | 2 | 2 | 5 | 3 |
| SG0000002 | Mier1 | 0.448479 | 0.517567 | 0.099684457 | 1712.058466 | 1834.53754 | 595 | 24 | 41 | 70 | 211 | 55 | 12 | 0 | 0 | 0 | 8 | 0 |
| JSG000000 | Mettl15 | 0.586653 | 0.650094 | 0.104215802 | 191.3184945 | 205.6501531 | 493 | 17 | 41 | 72 | 298 | 53 | 2 | 0 | 1 | 0 | 1 | 0 |
| SG0000002 | Tmeff2 | 0.79737 | 0.839209 | -0.112788695 | 42.38086029 | 39.19376533 | 786 | 16 | 41 | 96 | 378 | 93 | 7 | 0 | 0 | 1 | 3 | 2 |
| SG0000006 | Cgn | 3.9E-214 | 1.4E-211 | 5.545432326 | 34.47495552 | 1610.071577 | 156 | 40 | 40 | 56 | 82 | 8 | 12 | 2 | 2 | 2 | 6 | 0 |
| SG0000002 | Nin | 6.34E-65 | 1.98E-63 | -3.456915918 | 2218.678369 | 202.0500514 | 488 | 16 | 40 | 61 | 191 | 62 | 14 | 0 | 4 | 4 | 10 | 4 |
| SG0000003 | Igfbp2 | 5.17E-55 | 1.25E-53 | 8.749233204 | 19.62218312 | 8443.625403 | 94 | 36 | 40 | 46 | 64 | 4 | 2 | 0 | 0 | 2 | 2 | 0 |
| SG0000002 | Pdzd2 | 2.72E-53 | 6.17E-52 | -5.109668259 | 981.9378881 | 28.43941141 | 339 | 30 | 40 | 67 | 165 | 35 | 4 | 0 | 0 | 2 | 2 | 0 |
| SG0000006 | Hmgn3 | 1.36E-49 | 2.81E-48 | -3.176872389 | 1142.657652 | 126.3520378 | 361 | 25 | 40 | 45 | 175 | 40 | 0 | 0 | 0 | 0 | 0 | 0 |
| SG0000002 | Itgb4 | 3.69E-46 | 6.73E-45 | -6.237887704 | 12418.79732 | 164.5463415 | 120 | 40 | 40 | 50 | 110 | 5 | 0 | 0 | 0 | 0 | 0 | 0 |
| SG0000006 | Hipk2 | 7.26E-44 | 1.22E-42 | -2.702458639 | 1580.592925 | 242.8280885 | 926 | 10 | 40 | 102 | 415 | 98 | 4 | 0 | 0 | 0 | 4 | 0 |
| JSG000000 | Fhod3 | 1.85E-43 | 3.07E-42 | 3.399205646 | 28.70824488 | 302.8792372 | 637 | 22 | 40 | 89 | 324 | 58 | 0 | 0 | 0 | 0 | 0 | 0 |
| SG0000002 | Synj2 | 6.65E-41 | 1.02E-39 | -2.52346813 | 2191.941091 | 381.231939 | 635 | 19 | 40 | 94 | 185 | 48 | 13 | 0 | 0 | 0 | 2 | 0 |
| SG0000005 | Kctd19 | 1.51E-32 | 1.7E-31 | 7.335202482 | 0.649984458 | 104.958821 | 153 | 32 | 40 | 40 | 101 | 14 | 0 | 0 | 0 | 0 | 2 | 0 |
| SG0000002 | Calu | 2.04E-29 | 2.01E-28 | -1.682200212 | 24232.10309 | 7550.894206 | 217 | 27 | 40 | 51 | 114 | 21 | 12 | 0 | 0 | 0 | 0 | 2 |
| SG0000002 | Senp5 | 1.65E-26 | 1.43E-25 | -2.067307343 | 2228.570814 | 531.7468406 | 179 | 37 | 40 | 49 | 151 | 5 | 5 | 0 | 0 | 1 | 3 | 0 |
| SG0000003 | Garnl3 | 6.91E-26 | 5.82E-25 | 2.424422517 | 52.47609293 | 281.6990565 | 494 | 25 | 40 | 67 | 205 | 31 | 3 | 0 | 0 | 0 | 1 | 0 |
| SG0000002 | Ak2 | 1.55E-24 | 1.22E-23 | 1.278216249 | 1711.012726 | 4149.871714 | 107 | 39 | 40 | 45 | 68 | 5 | 1 | 0 | 0 | 0 | 1 | 0 |
| JSG000000 | Iqgap2 | 1.46E-23 | 1.1E-22 | -2.23122409 | 4694.898697 | 999.9098524 | 421 | 23 | 40 | 57 | 181 | 35 | 4 | 0 | 0 | 0 | 1 | 0 |

| | | | | | | | | | | | | | | | | | | |
|---|---|---|---|---|---|---|---|---|---|---|---|---|---|---|---|---|---|---|
| SG0000003 | Prr5l | 1.46E-23 | 1.1E-22 | 3.773481452 | 8.718497922 | 119.2264941 | 351 | 33 | 40 | 57 | 144 | 23 | 14 | 2 | 2 | 2 | 4 | 0 |
| JSG000000 | Morc1 | 3.87E-23 | 2.85E-22 | 8.706736072 | 4.341092658 | 1813.793235 | 415 | 27 | 40 | 66 | 255 | 36 | 0 | 0 | 0 | 0 | 0 | 0 |
| SG0000002 | Sptlc2 | 1.19E-20 | 7.79E-20 | -1.416948231 | 4626.138851 | 1732.505987 | 170 | 31 | 40 | 43 | 87 | 15 | 7 | 1 | 1 | 1 | 5 | 2 |
| SG0000006 | Hnrnpc | 4.19E-20 | 2.64E-19 | 1.098501632 | 8036.711566 | 17209.1858 | 186 | 30 | 40 | 46 | 110 | 6 | 6 | 0 | 0 | 2 | 2 | 0 |
| SG0000002 | Pip5k1a | 1.99E-19 | 1.21E-18 | -1.409886039 | 2841.550447 | 1069.393112 | 347 | 32 | 40 | 56 | 220 | 28 | 15 | 0 | 0 | 4 | 4 | 0 |
| SG0000005 | Chmp3 | 9.84E-18 | 5.45E-17 | -1.2832155 | 4294.473608 | 1764.507796 | 216 | 24 | 40 | 47 | 121 | 29 | 2 | 0 | 0 | 0 | 1 | 0 |
| SG0000006 | Zfp941 | 3.99E-16 | 2.02E-15 | 1.638662942 | 236.0125256 | 734.8890067 | 95 | 30 | 40 | 45 | 104 | 7 | 14 | 11 | 11 | 11 | 13 | 0 |
| JSG000000 | Pik3c2a | 2.18E-15 | 1.06E-14 | -1.36934818 | 2731.653053 | 1057.330261 | 321 | 29 | 40 | 65 | 175 | 23 | 9 | 0 | 0 | 2 | 5 | 3 |
| SG0000006 | Hvcn1 | 5.22E-15 | 2.47E-14 | 3.141221582 | 38.42319899 | 338.9966755 | 84 | 38 | 40 | 46 | 76 | 4 | 0 | 0 | 0 | 0 | 0 | 0 |
| SG0000003 | Gsr | 9.91E-15 | 4.6E-14 | -1.095397899 | 12794.91106 | 5988.108302 | 104 | 39 | 40 | 48 | 86 | 4 | 3 | 0 | 0 | 1 | 1 | 0 |
| SG0000002 | Lamc1 | 1.92E-14 | 8.74E-14 | 0.917291786 | 5795.411294 | 10945.02115 | 228 | 28 | 40 | 51 | 126 | 18 | 10 | 0 | 2 | 2 | 8 | 2 |
| SG0000002 | Rnpc3 | 3.15E-12 | 1.24E-11 | 1.138886203 | 295.3234137 | 650.3344922 | 353 | 28 | 40 | 64 | 197 | 18 | 23 | 0 | 0 | 0 | 12 | 2 |
| SG0000002 | 3110002H16Rik | 1.3E-11 | 4.94E-11 | -1.012265144 | 2755.240784 | 1365.958127 | 85 | 35 | 40 | 37 | 48 | 5 | 1 | 0 | 0 | 0 | 0 | 0 |
| SG0000002 | Srgap1 | 1.14E-10 | 4.04E-10 | -1.352322275 | 624.6756472 | 244.6611466 | 783 | 14 | 40 | 86 | 335 | 71 | 8 | 0 | 2 | 0 | 0 | 0 |
| SG0000000 | Dnajc5 | 2.13E-10 | 7.4E-10 | -0.916480217 | 5288.869515 | 2802.043135 | 274 | 24 | 40 | 48 | 129 | 19 | 6 | 0 | 0 | 0 | 1 | 3 |
| SG0000003 | Kat7 | 3.57E-10 | 1.22E-09 | 0.769273348 | 4186.965846 | 7136.311046 | 286 | 40 | 40 | 60 | 168 | 40 | 5 | 0 | 0 | 0 | 0 | 0 |
| SG0000003 | Azin1 | 4.44E-10 | 1.51E-09 | -0.872394146 | 8437.632116 | 4608.968926 | 312 | 20 | 40 | 52 | 161 | 36 | 29 | 0 | 3 | 1 | 6 | 0 |
| SG0000004 | Akap9 | 4.49E-10 | 1.52E-09 | -0.969675662 | 1904.713564 | 972.5864488 | 355 | 28 | 40 | 57 | 162 | 22 | 4 | 0 | 0 | 1 | 1 | 1 |
| SG0000002 | Lipo1 | 2.71E-09 | 8.62E-09 | 2.424865042 | 23.87159576 | 128.1854005 | 521 | 26 | 40 | 82 | 373 | 55 | 8 | 0 | 0 | 0 | 4 | 0 |
| SG0000002 | Ext2 | 1.53E-08 | 4.59E-08 | -0.85127281 | 2050.944533 | 1136.829318 | 368 | 32 | 40 | 76 | 162 | 34 | 0 | 0 | 0 | 0 | 0 | 0 |
| SG0000005 | Rnaseh2a | 1.53E-08 | 4.6E-08 | 0.75571652 | 1446.82141 | 2442.914501 | 103 | 40 | 40 | 40 | 62 | 10 | 2 | 0 | 0 | 0 | 0 | 0 |
| SG0000003 | Gpr160 | 2.56E-08 | 7.53E-08 | 5.624499182 | 3.795009043 | 187.2215255 | 300 | 20 | 40 | 55 | 215 | 5 | 0 | 0 | 0 | 0 | 5 | 0 |
| SG0000002 | Tnc | 1.81E-07 | 4.97E-07 | -1.765688283 | 1384.195528 | 407.0730072 | 305 | 30 | 40 | 45 | 159 | 35 | 0 | 0 | 0 | 0 | 3 | 0 |
| SG0000006 | Dgkk | 2.42E-07 | 6.55E-07 | 4.989126441 | 2.011354634 | 63.88006675 | 434 | 18 | 40 | 62 | 198 | 39 | 6 | 0 | 0 | 0 | 4 | 2 |
| SG0000002 | L3mbtl2 | 7.97E-07 | 2.07E-06 | 0.652834893 | 2132.622809 | 3353.026106 | 154 | 40 | 40 | 56 | 100 | 16 | 2 | 0 | 0 | 0 | 0 | 4 |
| SG0000004 | Lix1 | 1.46E-05 | 3.39E-05 | 4.668736522 | 0.674076249 | 17.14505015 | 177 | 31 | 40 | 51 | 94 | 6 | 6 | 0 | 0 | 0 | 0 | 0 |
| SG0000007 | 43526 | 5.05E-05 | 0.000111 | -0.708510864 | 667.333157 | 408.3755021 | 298 | 29 | 40 | 56 | 169 | 53 | 9 | 0 | 0 | 1 | 6 | 2 |
| SG0000000 | Top3a | 0.000941 | 0.00177 | 0.515055246 | 506.0771942 | 723.2090385 | 184 | 36 | 40 | 48 | 108 | 24 | 0 | 0 | 0 | 0 | 0 | 0 |
| SG0000007 | Phyhd1 | 0.001137 | 0.002122 | 1.568881948 | 13.43468809 | 39.8573219 | 96 | 40 | 40 | 52 | 92 | 0 | 0 | 0 | 0 | 0 | 0 | 0 |
| JSG000000 | Stim1 | 0.001591 | 0.002923 | 0.505309283 | 487.6335677 | 692.160554 | 306 | 25 | 40 | 60 | 216 | 40 | 1 | 1 | 1 | 1 | 1 | 1 |
| SG0000002 | Ywhae | 0.001899 | 0.003448 | 0.374152195 | 20716.72146 | 26850.48041 | 160 | 28 | 40 | 43 | 101 | 6 | 4 | 0 | 0 | 1 | 1 | 0 |
| SG0000002 | Anapc5 | 0.003131 | 0.005507 | 0.366376235 | 6606.263192 | 8516.205026 | 366 | 39 | 40 | 54 | 219 | 40 | 9 | 0 | 1 | 0 | 0 | 1 |
| SG0000005 | Supt7l | 0.005073 | 0.008677 | -0.461494828 | 803.1832746 | 583.2985143 | 92 | 36 | 40 | 43 | 57 | 10 | 7 | 2 | 2 | 2 | 3 | 3 |
| JSG000000 | Pappa | 0.006395 | 0.010778 | -1.803324694 | 108.4991276 | 31.0864709 | 450 | 28 | 40 | 74 | 242 | 41 | 14 | 1 | 1 | 2 | 9 | 5 |
| JSG000000 | Dkk2 | 0.00906 | 0.014917 | 0.934573847 | 53.39747076 | 102.0599727 | 268 | 32 | 40 | 58 | 127 | 14 | 6 | 0 | 0 | 0 | 2 | 1 |
| SG0000004 | Rc3h1 | 0.012563 | 0.020176 | -0.362091127 | 2243.96215 | 1745.883404 | 307 | 36 | 40 | 73 | 166 | 24 | 20 | 1 | 3 | 2 | 11 | 2 |
| JSG000000 | Vps13c | 0.017885 | 0.027914 | 0.351006427 | 776.2002552 | 990.0046728 | 360 | 24 | 40 | 59 | 196 | 35 | 0 | 0 | 0 | 0 | 0 | 0 |
| JSG000000 | Ror1 | 0.018612 | 0.02896 | 1.460854639 | 8.456828657 | 23.27922924 | 569 | 21 | 40 | 87 | 259 | 58 | 9 | 0 | 0 | 1 | 3 | 1 |
| SG0000002 | Stx8 | 0.033695 | 0.050107 | -0.354242066 | 706.3825894 | 552.5893351 | 391 | 26 | 40 | 64 | 193 | 53 | 1 | 0 | 0 | 0 | 0 | 0 |
| SG0000002 | Slc4a1ap | 0.03808 | 0.056102 | -0.332793825 | 998.3665041 | 792.7004011 | 246 | 26 | 40 | 52 | 118 | 41 | 6 | 0 | 0 | 0 | 0 | 5 |
| SG0000002 | Bag6 | 0.064201 | 0.090828 | 0.316195013 | 4822.103863 | 6003.724346 | 76 | 40 | 40 | 44 | 61 | 8 | 5 | 0 | 0 | 0 | 0 | 0 |
| SG0000003 | Trim66 | 0.068003 | 0.095731 | 1.263319447 | 10.60086306 | 25.44709856 | 177 | 32 | 40 | 49 | 141 | 15 | 3 | 0 | 0 | 0 | 2 | 0 |
| SG0000002 | Cap2 | 0.087992 | 0.121267 | 0.845104568 | 17.88436873 | 32.12738958 | 408 | 22 | 40 | 74 | 154 | 47 | 2 | 0 | 0 | 0 | 0 | 0 |
| SG0000003 | 9030624J02Rik | 0.136768 | 0.180769 | -0.213370227 | 1917.575307 | 1653.947005 | 402 | 32 | 40 | 53 | 154 | 84 | 7 | 0 | 0 | 1 | 2 | 1 |
| SG0000002 | Khdrbs1 | 0.177638 | 0.229108 | 0.165588384 | 5190.276679 | 5821.535902 | 146 | 34 | 40 | 46 | 107 | 17 | 4 | 0 | 0 | 1 | 1 | 2 |
| SG0000001 | Ebf3 | 0.200625 | 0.255608 | -0.884412206 | 18.16544265 | 9.840368963 | 784 | 16 | 40 | 92 | 239 | 28 | 39 | 0 | 8 | 8 | 18 | 0 |
| SG0000003 | Ddhd1 | 0.279585 | 0.342827 | -0.177369328 | 864.0459399 | 764.087941 | 579 | 22 | 40 | 51 | 187 | 60 | 19 | 0 | 0 | 3 | 9 | 1 |
| SG0000002 | Acad10 | 0.307857 | 0.37312 | 0.256204734 | 115.0712131 | 137.4333085 | 108 | 40 | 40 | 46 | 68 | 14 | 0 | 0 | 0 | 0 | 0 | 0 |
| SG0000006 | Kcnj15 | 0.322242 | 0.388425 | -0.808249872 | 37.77191139 | 21.5705643 | 478 | 16 | 40 | 56 | 205 | 96 | 43 | 0 | 0 | 8 | 19 | 0 |
| SG0000005 | Ep300 | 0.494674 | 0.562399 | -0.096335739 | 3747.234812 | 3505.185132 | 168 | 30 | 40 | 46 | 96 | 17 | 1 | 0 | 0 | 0 | 0 | 0 |
| SG0000003 | Susd1 | 0.552471 | 0.617834 | 0.114143418 | 203.5800128 | 220.3412099 | 382 | 28 | 40 | 64 | 220 | 26 | 0 | 0 | 0 | 0 | 0 | 0 |
| SG0000000 | Rdh13 | 0.656908 | 0.715378 | 0.062453122 | 682.0654178 | 712.2398903 | 98 | 38 | 40 | 56 | 96 | 8 | 5 | 0 | 0 | 0 | 6 | 0 |
| SG0000005 | Klhl26 | 0.691214 | 0.746205 | -0.066959609 | 601.3633274 | 574.0901422 | 76 | 38 | 40 | 42 | 70 | 0 | 0 | 0 | 0 | 0 | 0 | 0 |
| SG0000005 | Pde5a | 0.694099 | 0.748597 | 0.066921516 | 250.3471496 | 262.2334174 | 530 | 22 | 40 | 72 | 242 | 42 | 8 | 0 | 0 | 0 | 14 | 2 |
| SG0000004 | St8sia4 | 0.971127 | 0.984204 | -0.061962483 | 20.13330084 | 19.28689948 | 447 | 22 | 40 | 73 | 218 | 40 | 50 | 1 | 5 | 9 | 25 | 6 |
| JSG000000 | Prtg | 1.9E-167 | 3.9E-165 | 7.611878234 | 3.937216218 | 770.1812422 | 225 | 34 | 39 | 57 | 112 | 21 | 12 | 0 | 0 | 2 | 2 | 0 |
| JSG000000 | Cd109 | 2.14E-63 | 6.5E-62 | -5.795036339 | 15899.10666 | 286.347212 | 277 | 29 | 39 | 64 | 180 | 15 | 3 | 0 | 0 | 0 | 0 | 0 |
| SG0000003 | Adamts9 | 2.4E-61 | 6.93E-60 | 4.558191306 | 13.63964323 | 321.3329934 | 502 | 18 | 39 | 60 | 223 | 50 | 8 | 2 | 4 | 4 | 12 | 2 |
| SG0000000 | Hsf2bp | 5.07E-52 | 1.12E-50 | 8.504910403 | 3.832318512 | 1392.177911 | 292 | 24 | 39 | 56 | 134 | 24 | 2 | 0 | 0 | 0 | 0 | 1 |
| SG0000002 | Cdh2 | 6.63E-38 | 9.09E-37 | -3.21011542 | 2008.09525 | 216.9917778 | 679 | 30 | 39 | 84 | 287 | 84 | 4 | 0 | 0 | 0 | 2 | 2 |
| SG0000002 | Farsb | 1.7E-25 | 1.4E-24 | 1.271488617 | 2478.695204 | 5983.831278 | 428 | 18 | 39 | 72 | 189 | 41 | 4 | 0 | 0 | 1 | 1 | 0 |
| JSG000000 | St14 | 3.16E-25 | 2.58E-24 | 5.589699309 | 51.12813714 | 2462.22155 | 122 | 36 | 39 | 45 | 65 | 8 | 1 | 0 | 0 | 0 | 0 | 0 |
| SG0000005 | Ttc26 | 7.22E-21 | 4.76E-20 | 1.624078759 | 184.4969261 | 568.7030489 | 244 | 27 | 39 | 50 | 124 | 18 | 16 | 0 | 0 | 0 | 1 | 0 |
| SG0000002 | Zbtb46 | 7.73E-20 | 4.81E-19 | 2.531896145 | 30.96336397 | 179.0708204 | 217 | 27 | 39 | 56 | 132 | 11 | 5 | 0 | 0 | 1 | 3 | 0 |
| SG0000003 | Uvssa | 6.12E-18 | 3.43E-17 | -2.354434779 | 468.0780014 | 91.52984763 | 177 | 30 | 39 | 48 | 120 | 12 | 16 | 0 | 0 | 0 | 3 | 3 |
| SG0000004 | Bcl11b | 1.48E-17 | 8.17E-17 | 3.350194264 | 9.521563667 | 97.09955522 | 307 | 24 | 39 | 48 | 155 | 37 | 12 | 0 | 0 | 3 | 11 | 1 |
| SG0000004 | Crebrf | 8.4E-17 | 4.44E-16 | 1.235136136 | 521.7839438 | 1228.297977 | 262 | 32 | 39 | 52 | 121 | 32 | 11 | 0 | 0 | 1 | 4 | 3 |
| SG0000002 | Ankfy1 | 8.99E-17 | 4.74E-16 | -1.230457801 | 5503.118286 | 2345.330985 | 318 | 26 | 39 | 46 | 136 | 14 | 12 | 0 | 1 | 0 | 3 | 0 |
| SG0000003 | Nos1ap | 4.36E-15 | 2.07E-14 | -3.466733976 | 457.8537854 | 41.412924 | 439 | 25 | 39 | 74 | 194 | 38 | 0 | 0 | 0 | 0 | 1 | 0 |
| SG0000003 | Sipa1l3 | 1.39E-14 | 6.4E-14 | -1.483135687 | 2452.586416 | 877.315877 | 379 | 18 | 39 | 67 | 179 | 27 | 9 | 0 | 3 | 3 | 6 | 0 |
| SG0000006 | Med14 | 5.84E-14 | 2.58E-13 | -1.145835041 | 3064.611578 | 1384.983887 | 430 | 22 | 39 | 68 | 201 | 23 | 11 | 0 | 1 | 2 | 6 | 1 |
| SG0000005 | Acbd4 | 4.85E-13 | 2.01E-12 | -1.384887725 | 707.3211035 | 270.8469005 | 69 | 33 | 39 | 39 | 57 | 3 | 0 | 0 | 0 | 0 | 0 | 0 |
| SG0000004 | Catsperg1 | 1.2E-12 | 4.86E-12 | 3.450388814 | 6.903721021 | 75.46642257 | 128 | 29 | 39 | 51 | 87 | 20 | 0 | 0 | 0 | 0 | 0 | 0 |
| SG0000003 | Plekha5 | 2.4E-12 | 9.5E-12 | 0.981772939 | 712.6816286 | 1407.468446 | 327 | 25 | 39 | 48 | 152 | 28 | 0 | 0 | 0 | 0 | 2 | 0 |
| JSG000000 | Alms1 | 6.81E-12 | 2.63E-11 | 0.983177517 | 598.8714128 | 1183.857708 | 290 | 21 | 39 | 55 | 159 | 26 | 2 | 0 | 0 | 0 | 0 | 0 |
| SG0000002 | Cacna1s | 1.24E-11 | 4.72E-11 | 4.944789315 | 0 | 29.79852416 | 220 | 35 | 39 | 46 | 96 | 19 | 0 | 0 | 0 | 0 | 1 | 0 |
| SG0000005 | Lepr | 7.01E-10 | 2.34E-09 | 3.315989299 | 4.932019423 | 49.11758696 | 561 | 22 | 39 | 70 | 298 | 88 | 5 | 0 | 0 | 0 | 0 | 0 |
| SG0000004 | Cyyr1 | 2.48E-09 | 7.89E-09 | 3.406260077 | 4.212103161 | 44.65657956 | 501 | 24 | 39 | 77 | 215 | 67 | 11 | 0 | 1 | 2 | 8 | 0 |
| JSG000000 | Lexm | 3.47E-09 | 1.09E-08 | 6.387004762 | 0.307509278 | 25.73583358 | 191 | 36 | 39 | 59 | 103 | 12 | 0 | 0 | 0 | 0 | 0 | 0 |
| SG0000003 | Mon2 | 1.39E-08 | 4.18E-08 | -0.837872085 | 2736.1309 | 1530.778101 | 483 | 18 | 39 | 69 | 258 | 42 | 7 | 0 | 1 | 2 | 8 | 0 |
| SG0000003 | Zc3h7a | 3.15E-07 | 8.45E-07 | -0.837979498 | 3015.112898 | 1686.734057 | 149 | 30 | 39 | 51 | 107 | 17 | 2 | 0 | 1 | 0 | 1 | 1 |
| SG0000002 | Sec61a2 | 6.49E-07 | 1.7E-06 | 0.674303059 | 1078.733937 | 1721.471309 | 244 | 21 | 39 | 45 | 93 | 15 | 3 | 0 | 0 | 0 | 0 | 0 |
| SG0000003 | Fam222b | 2.94E-05 | 6.62E-05 | -0.701627341 | 845.2466043 | 519.7237451 | 281 | 28 | 39 | 49 | 171 | 15 | 2 | 0 | 0 | 0 | 0 | 0 |
| SG0000002 | Samhd1 | 0.000114 | 0.00024 | -0.566091384 | 3071.351804 | 2074.526935 | 284 | 28 | 39 | 64 | 164 | 35 | 3 | 0 | 1 | 0 | 0 | 0 |
| JSG000000 | Srbd1 | 0.000165 | 0.000342 | 0.509970054 | 1161.404031 | 1653.863308 | 341 | 22 | 39 | 60 | 181 | 30 | 0 | 0 | 0 | 0 | 0 | 0 |
| SG0000005 | Zfp943 | 0.000412 | 0.000813 | -0.79504508 | 266.5934206 | 153.644497 | 368 | 30 | 39 | 81 | 189 | 31 | 5 | 1 | 1 | 1 | 1 | 1 |
| SG0000002 | Cdk8 | 0.000561 | 0.001088 | 0.463771192 | 1511.599027 | 2084.709971 | 467 | 18 | 39 | 60 | 180 | 39 | 5 | 0 | 0 | 0 | 0 | 0 |
| SG0000002 | Mtbp | 0.000832 | 0.001576 | 0.492532758 | 704.9104753 | 991.7474634 | 441 | 21 | 39 | 65 | 187 | 28 | 4 | 0 | 2 | 0 | 0 | 0 |
| SG0000003 | Mboat7 | 0.001282 | 0.002378 | -0.471444772 | 1827.196354 | 1317.850672 | 83 | 39 | 39 | 42 | 50 | 6 | 2 | 0 | 0 | 0 | 2 | 0 |
| SG0000002 | Crcp | 0.002058 | 0.00372 | -0.496805527 | 921.8886009 | 653.3186852 | 144 | 36 | 39 | 51 | 109 | 9 | 0 | 0 | 0 | 0 | 1 | 0 |
| JSG000000 | Efhc2 | 0.003386 | 0.005929 | 3.648649916 | 2.936226035 | 36.82498288 | 483 | 24 | 39 | 80 | 277 | 40 | 4 | 0 | 0 | 0 | 0 | 0 |
| SG0000002 | Gstcd | 0.006161 | 0.010412 | 0.391184407 | 848.9908277 | 1113.425693 | 470 | 12 | 39 | 73 | 304 | 40 | 8 | 0 | 1 | 1 | 4 | 0 |
| SG0000003 | Nufip2 | 0.008955 | 0.014757 | -0.3634005 | 4940.802209 | 3840.634842 | 189 | 28 | 39 | 41 | 112 | 25 | 9 | 0 | 2 | 0 | 2 | 1 |
| SG0000005 | A630033H20Rik | 0.013437 | 0.021451 | -2.201710782 | 21.32602848 | 4.635845969 | 315 | 27 | 39 | 75 | 177 | 27 | 15 | 0 | 0 | 0 | 12 | 0 |
| JSG000000 | Igtp | 0.015656 | 0.024745 | -1.028295732 | 602.065981 | 295.1863214 | 90 | 39 | 39 | 42 | 51 | 3 | 0 | 0 | 0 | 0 | 0 | 0 |
| SG0000002 | Hnrnpll | 0.051446 | 0.074081 | -0.28329806 | 2831.314378 | 2326.520484 | 328 | 24 | 39 | 63 | 123 | 33 | 14 | 0 | 3 | 3 | 9 | 0 |
| SG0000000 | Braf | 0.051717 | 0.07443 | 0.288910265 | 1197.01437 | 1462.412935 | 315 | 22 | 39 | 58 | 173 | 17 | 27 | 1 | 5 | 4 | 8 | 1 |
| SG0000007 | Wipf1 | 0.101845 | 0.138284 | -0.412190214 | 378.9667418 | 284.7865533 | 280 | 27 | 39 | 56 | 128 | 28 | 13 | 0 | 0 | 2 | 6 | 3 |
| SG0000003 | Dner | 0.186676 | 0.239592 | 0.533180228 | 38.70277307 | 56.00738877 | 539 | 23 | 39 | 77 | 262 | 55 | 3 | 0 | 1 | 1 | 2 | 0 |
| SG0000003 | Tspan8 | 0.242001 | 0.30205 | -0.718763126 | 23.13477154 | 14.05710712 | 252 | 33 | 39 | 48 | 111 | 18 | 6 | 0 | 0 | 0 | 1 | 3 |
| SG0000004 | Cept1 | 0.465905 | 0.535195 | 0.10325002 | 947.782544 | 1018.099112 | 413 | 15 | 39 | 69 | 207 | 45 | 8 | 0 | 0 | 0 | 3 | 0 |
| SG0000002 | Cdadc1 | 0.621484 | 0.683155 | -0.10329058 | 304.4054789 | 283.3732833 | 174 | 36 | 39 | 50 | 162 | 7 | 0 | 0 | 0 | 0 | 0 | 0 |
| SG0000002 | Spire1 | 0.73764 | 0.787402 | -0.052772259 | 1463.008779 | 1410.460406 | 302 | 33 | 39 | 66 | 237 | 35 | 4 | 0 | 0 | 0 | 6 | 4 |
| JSG000000 | Chst11 | 0.824826 | 0.86248 | 0.10839343 | 439.6623564 | 473.9679299 | 220 | 28 | 39 | 47 | 95 | 19 | 10 | 0 | 4 | 2 | 3 | 1 |
| JSG000000 | Gldc | 0 | 0 | 11.64000078 | 1.313186595 | 4190.982298 | 164 | 30 | 38 | 45 | 83 | 15 | 1 | 0 | 0 | 0 | 1 | 0 |
| SG0000001 | Cacna2d2 | 1.64E-88 | 8.95E-87 | -4.156730484 | 3843.697733 | 215.5006914 | 212 | 36 | 38 | 48 | 94 | 26 | 2 | 0 | 0 | 0 | 0 | 0 |
| SG0000002 | Rnf144a | 2.09E-68 | 7.21E-67 | 4.983120103 | 9.994375048 | 316.0998324 | 286 | 34 | 38 | 58 | 138 | 32 | 4 | 0 | 0 | 0 | 4 | 0 |
| SG0000000 | Folr1 | 3.67E-43 | 6.01E-42 | 6.769482123 | 6.809697427 | 742.9250285 | 115 | 38 | 38 | 47 | 70 | 5 | 0 | 0 | 0 | 0 | 0 | 0 |
| SG0000003 | Ash2l | 3.43E-38 | 4.77E-37 | 1.733996066 | 3232.896345 | 10754.16285 | 264 | 23 | 38 | 52 | 137 | 7 | 1 | 0 | 0 | 0 | 2 | 0 |
| SG0000005 | Zzef1 | 7.62E-38 | 1.04E-36 | -2.259554657 | 2487.314323 | 519.4418566 | 442 | 24 | 38 | 56 | 250 | 34 | 0 | 0 | 0 | 0 | 0 | 2 |
| SG0000002 | Prkcz | 1.16E-36 | 1.52E-35 | 2.254381649 | 142.8201237 | 681.4373011 | 182 | 34 | 38 | 56 | 118 | 46 | 4 | 1 | 1 | 1 | 2 | 0 |
| SG0000003 | Ctbp2 | 1.8E-29 | 1.77E-28 | 1.3992779 | 1687.11132 | 4450.085544 | 386 | 26 | 38 | 74 | 184 | 24 | 0 | 0 | 0 | 0 | 1 | 0 |
| SG0000004 | Ift88 | 1.37E-27 | 1.24E-26 | 1.67459518 | 291.5944931 | 930.8564183 | 331 | 22 | 38 | 76 | 168 | 32 | 3 | 0 | 0 | 0 | 1 | 0 |
| SG0000003 | Myh14 | 3.18E-22 | 2.25E-21 | 4.515849662 | 3.926342106 | 89.82440347 | 212 | 31 | 38 | 47 | 137 | 15 | 0 | 0 | 0 | 0 | 0 | 0 |
| SG0000005 | Mok | 4.07E-22 | 2.87E-21 | -1.835691213 | 1144.455089 | 320.6274025 | 160 | 35 | 38 | 49 | 84 | 3 | 0 | 0 | 0 | 0 | 0 | 0 |
| JSG000000 | Zfp512 | 7.85E-22 | 5.47E-21 | 1.29840648 | 733.0196787 | 1802.913667 | 128 | 18 | 38 | 36 | 74 | 20 | 4 | 0 | 0 | 0 | 0 | 4 |
| SG0000004 | Spink10 | 6.92E-21 | 4.57E-20 | -7.582868512 | 434.6448754 | 2.267063519 | 447 | 20 | 38 | 78 | 256 | 48 | 5 | 0 | 0 | 2 | 7 | 0 |
| SG0000001 | Serac1 | 3.46E-20 | 2.2E-19 | 1.607621275 | 180.6816687 | 550.6254976 | 133 | 36 | 38 | 42 | 63 | 16 | 3 | 0 | 0 | 0 | 1 | 0 |
| SG0000002 | Gnas | 9.4E-19 | 5.5E-18 | -1.268874425 | 14352.93932 | 5956.231734 | 755 | 24 | 38 | 104 | 256 | 66 | 22 | 1 | 1 | 2 | 2 | 0 |
| SG0000004 | Ptpn11 | 6.86E-18 | 3.83E-17 | -1.265049128 | 6427.683648 | 2674.464024 | 180 | 30 | 38 | 42 | 92 | 2 | 4 | 0 | 0 | 2 | 2 | 0 |
| SG0000002 | Kcnab2 | 1.86E-17 | 1.02E-16 | 3.060398474 | 13.34300807 | 111.3077716 | 237 | 33 | 38 | 49 | 116 | 53 | 6 | 0 | 0 | 3 | 3 | 3 |
| SG0000003 | Rerg | 5.32E-16 | 2.67E-15 | 5.658597703 | 1.005677317 | 50.8003023 | 606 | 13 | 38 | 71 | 275 | 34 | 4 | 0 | 0 | 2 | 10 | 0 |
| SG0000002 | Cdc27 | 3.37E-12 | 1.33E-11 | -1.013862704 | 4870.083287 | 2411.755641 | 328 | 21 | 38 | 57 | 167 | 22 | 12 | 0 | 0 | 0 | 3 | 0 |

| | | | | | | | | | | | | | | | | | | |
|---|---|---|---|---|---|---|---|---|---|---|---|---|---|---|---|---|---|---|
| SG0000004 | Bcor | 2.45E-11 | 9.13E-11 | -1.066988584 | 3002.429068 | 1433.102281 | 297 | 7 | 38 | 46 | 120 | 8 | 15 | 0 | 5 | 5 | 5 | 5 |
| SG0000004 | Zbtb8b | 2.92E-11 | 1.08E-10 | 1.681870684 | 57.57960143 | 184.7405274 | 88 | 30 | 38 | 40 | 44 | 10 | 4 | 0 | 0 | 0 | 0 | 0 |
| SG0000005 | Zfp277 | 3.95E-11 | 1.45E-10 | 0.94221253 | 738.2175262 | 1418.464874 | 530 | 16 | 38 | 68 | 246 | 54 | 2 | 0 | 0 | 0 | 2 | 0 |
| SG0000002 | 43532 | 7.67E-11 | 2.75E-10 | -1.033831106 | 1511.836819 | 738.3983957 | 336 | 28 | 38 | 80 | 174 | 38 | 10 | 0 | 0 | 2 | 2 | 2 |
| SG0000005 | Pcmtd1 | 7.97E-11 | 2.85E-10 | 0.921996611 | 967.3117004 | 1832.800002 | 329 | 26 | 38 | 59 | 139 | 19 | 22 | 0 | 1 | 1 | 5 | 1 |
| SG0000003 | Dcdc2a | 8.65E-11 | 3.09E-10 | 6.630774069 | 0.307509278 | 30.47334391 | 423 | 15 | 38 | 53 | 176 | 37 | 7 | 0 | 0 | 0 | 2 | 2 |
| SG0000004 | Smg7 | 1.01E-09 | 3.31E-09 | 0.75634472 | 3377.331814 | 5705.006964 | 424 | 16 | 38 | 78 | 209 | 30 | 2 | 0 | 0 | 0 | 2 | 0 |
| JSG000000 | Commd10 | 1.28E-09 | 4.17E-09 | 0.844308192 | 814.5746581 | 1462.490272 | 290 | 23 | 38 | 60 | 160 | 26 | 1 | 0 | 0 | 0 | 1 | 0 |
| SG0000005 | Fank1 | 1.71E-09 | 5.53E-09 | 6.442609466 | 0.307509278 | 26.74711367 | 335 | 24 | 38 | 56 | 183 | 30 | 3 | 1 | 1 | 1 | 2 | 0 |
| SG0000002 | Fbn1 | 2.35E-09 | 7.49E-09 | -2.060620208 | 2487.685324 | 596.330403 | 663 | 22 | 38 | 74 | 264 | 62 | 4 | 0 | 0 | 0 | 2 | 0 |
| SG0000002 | Inpp5a | 9.67E-09 | 2.95E-08 | 0.896063179 | 429.2189343 | 798.7682004 | 588 | 23 | 38 | 79 | 230 | 45 | 8 | 0 | 0 | 0 | 0 | 0 |
| SG0000002 | Oxct1 | 1.78E-08 | 5.29E-08 | -0.830150407 | 2876.752276 | 1618.088577 | 534 | 18 | 38 | 70 | 276 | 40 | 6 | 0 | 0 | 0 | 0 | 0 |
| SG0000002 | Fam198b | 2.8E-07 | 7.55E-07 | -1.5454112 | 554.062571 | 189.8207505 | 269 | 32 | 38 | 52 | 116 | 22 | 34 | 0 | 0 | 0 | 8 | 4 |
| SG0000002 | Desi1 | 9.79E-07 | 2.53E-06 | -0.702519746 | 3154.063409 | 1938.165702 | 146 | 25 | 38 | 38 | 80 | 20 | 6 | 0 | 3 | 3 | 10 | 0 |
| SG0000002 | Nckap1 | 1.27E-06 | 3.25E-06 | -0.660633432 | 9065.060769 | 5734.561843 | 412 | 26 | 38 | 66 | 194 | 44 | 4 | 0 | 0 | 0 | 0 | 0 |
| SG0000002 | Arsg | 1.56E-05 | 3.61E-05 | 2.163367243 | 11.29668753 | 50.60454777 | 289 | 25 | 38 | 81 | 162 | 34 | 0 | 0 | 0 | 0 | 3 | 3 |
| SG0000007 | Slc8a3 | 8.26E-05 | 0.000177 | 2.629140366 | 4.306126756 | 26.64016575 | 386 | 16 | 38 | 54 | 166 | 26 | 2 | 0 | 0 | 0 | 0 | 0 |
| SG0000005 | Arhgap35 | 8.64E-05 | 0.000184 | -0.556048689 | 2958.767901 | 2012.442887 | 334 | 24 | 38 | 44 | 169 | 28 | 0 | 0 | 0 | 0 | 1 | 0 |
| SG0000000 | Brd8 | 0.00022 | 0.000449 | 0.618186979 | 2531.383828 | 3885.534996 | 180 | 30 | 38 | 42 | 102 | 30 | 4 | 2 | 2 | 2 | 6 | 2 |
| SG0000003 | Tbcd | 0.000431 | 0.000848 | -0.501268317 | 2470.98137 | 1745.712299 | 324 | 31 | 38 | 57 | 131 | 45 | 0 | 0 | 0 | 0 | 0 | 0 |
| SG0000006 | Dzip3 | 0.00863 | 0.014266 | 0.378764078 | 1139.431547 | 1481.520237 | 376 | 10 | 38 | 50 | 241 | 55 | 7 | 0 | 0 | 0 | 4 | 1 |
| JSG000000 | Tbc1d19 | 0.009545 | 0.015663 | -0.473132121 | 530.6517983 | 382.2809705 | 563 | 16 | 38 | 80 | 286 | 40 | 10 | 0 | 1 | 1 | 4 | 0 |
| JSG000001 | Snrpn | 0.012797 | 0.020517 | -0.437849241 | 3888.583151 | 2870.685627 | 558 | 19 | 38 | 87 | 327 | 66 | 8 | 0 | 0 | 0 | 0 | 0 |
| JSG000000 | Parn | 0.082863 | 0.114764 | 0.229469313 | 1548.98711 | 1816.038077 | 274 | 27 | 38 | 54 | 147 | 11 | 0 | 0 | 0 | 0 | 0 | 0 |
| SG0000003 | Prrc2b | 0.083363 | 0.115394 | 0.213791353 | 8104.048832 | 9398.524098 | 163 | 20 | 38 | 35 | 100 | 23 | 12 | 0 | 4 | 2 | 2 | 0 |
| SG0000007 | Adamtsl3 | 0.096716 | 0.132091 | -0.941498712 | 33.09787096 | 17.23378637 | 467 | 21 | 38 | 75 | 250 | 47 | 5 | 0 | 0 | 2 | 4 | 2 |
| JSG000000 | Rspo2 | 0.103031 | 0.139768 | -1.072263149 | 21.64441187 | 10.29348458 | 401 | 23 | 38 | 66 | 232 | 39 | 6 | 0 | 0 | 3 | 4 | 1 |
| SG0000000 | Raf1 | 0.116421 | 0.1561 | -0.211554742 | 3052.214731 | 2635.911112 | 258 | 30 | 38 | 70 | 128 | 20 | 2 | 0 | 0 | 0 | 0 | 0 |
| SG0000000 | Mark3 | 0.322043 | 0.388238 | 0.134164955 | 2300.16998 | 2524.338593 | 318 | 24 | 38 | 48 | 166 | 32 | 6 | 0 | 0 | 0 | 0 | 0 |
| SG0000001 | Clpx | 0.343201 | 0.41043 | 0.133492986 | 1059.364604 | 1162.066151 | 180 | 30 | 38 | 50 | 100 | 14 | 6 | 0 | 0 | 0 | 0 | 2 |
| JSG000000 | Thada | 0.477929 | 0.546484 | -0.108778269 | 1616.946131 | 1499.51229 | 461 | 16 | 38 | 55 | 191 | 39 | 3 | 0 | 0 | 1 | 2 | 0 |
| JSG000000 | Sash1 | 0.680962 | 0.737493 | -0.064358006 | 894.564677 | 855.5355399 | 271 | 27 | 38 | 57 | 127 | 24 | 6 | 0 | 0 | 1 | 3 | 2 |
| SG0000002 | Aspscr1 | 0.769362 | 0.815239 | 0.059786022 | 1230.601861 | 1282.670049 | 157 | 30 | 38 | 50 | 88 | 5 | 0 | 0 | 0 | 0 | 0 | 0 |
| JSG000000 | Atp10a | 0.833722 | 0.869903 | -0.046247961 | 958.4902114 | 928.2514977 | 348 | 25 | 38 | 69 | 157 | 27 | 2 | 0 | 0 | 1 | 4 | 0 |
| SG0000004 | Tcf20 | 0.84622 | 0.879664 | 0.019064546 | 4013.0335 | 4066.415811 | 406 | 20 | 38 | 60 | 180 | 50 | 6 | 2 | 2 | 4 | 6 | 0 |
| SG0000002 | Plxna4 | 0.857902 | 0.889073 | 0.135118893 | 14.11245461 | 15.49806214 | 563 | 22 | 38 | 83 | 278 | 61 | 8 | 0 | 0 | 1 | 4 | 1 |
| SG0000001 | Fam184a | 6.89E-49 | 1.38E-47 | 6.929362006 | 1.313186595 | 160.0561092 | 505 | 30 | 37 | 92 | 350 | 58 | 6 | 0 | 0 | 3 | 11 | 0 |
| SG0000002 | Snx24 | 1.3E-45 | 2.33E-44 | -2.954888504 | 1179.7121 | 152.1478992 | 255 | 30 | 37 | 56 | 148 | 18 | 2 | 0 | 0 | 0 | 0 | 0 |
| SG0000003 | Nkd1 | 4.39E-39 | 6.29E-38 | 8.556478011 | 0.307509278 | 115.7749927 | 150 | 29 | 37 | 39 | 57 | 12 | 6 | 0 | 2 | 2 | 6 | 0 |
| SG0000003 | Dtl | 2.07E-38 | 2.89E-37 | 1.582222362 | 1772.539922 | 5307.529492 | 244 | 23 | 37 | 44 | 133 | 34 | 9 | 0 | 1 | 0 | 7 | 2 |
| JSG000000 | Rock2 | 5.42E-37 | 7.21E-36 | -2.164715555 | 13352.70759 | 2978.00227 | 301 | 24 | 37 | 50 | 149 | 29 | 8 | 0 | 0 | 1 | 3 | 5 |
| SG0000000 | Rab3b | 1.11E-29 | 1.1E-28 | 3.491057182 | 15.65718484 | 176.0461805 | 265 | 28 | 37 | 59 | 139 | 26 | 10 | 0 | 1 | 0 | 1 | 0 |
| SG0000000 | Agpat3 | 2.73E-29 | 2.67E-28 | -1.857225054 | 2755.284558 | 760.4768682 | 365 | 21 | 37 | 60 | 141 | 20 | 7 | 0 | 0 | 1 | 6 | 0 |
| SG0000002 | Tex10 | 1.74E-26 | 1.51E-25 | 1.326688632 | 1632.53174 | 4094.818971 | 311 | 23 | 37 | 59 | 146 | 38 | 2 | 0 | 0 | 0 | 1 | 0 |
| SG0000000 | Rac1 | 1.38E-23 | 1.04E-22 | -1.473765593 | 13759.72966 | 4954.071284 | 74 | 30 | 37 | 36 | 62 | 6 | 6 | 2 | 3 | 2 | 2 | 0 |
| SG0000002 | Mapk8 | 2.58E-18 | 1.48E-17 | 1.158779452 | 952.5127956 | 2126.660634 | 593 | 17 | 37 | 81 | 306 | 46 | 11 | 0 | 0 | 0 | 7 | 0 |
| SG0000003 | Mtmr14 | 7.98E-18 | 4.45E-17 | 1.172443969 | 771.4492705 | 1738.794171 | 171 | 27 | 37 | 49 | 94 | 8 | 3 | 0 | 0 | 0 | 0 | 0 |
| SG0000002 | Acot7 | 4.27E-17 | 2.29E-16 | -1.653328587 | 4631.675727 | 1472.436751 | 348 | 24 | 37 | 57 | 130 | 19 | 4 | 0 | 0 | 4 | 4 | 0 |
| SG0000004 | Chd7 | 3.52E-16 | 1.79E-15 | 1.148878956 | 618.418474 | 1371.290546 | 692 | 17 | 37 | 86 | 284 | 65 | 6 | 2 | 2 | 2 | 6 | 0 |
| JSG000000 | Megf9 | 2.62E-15 | 1.27E-14 | -1.767510138 | 434.0085408 | 127.4749079 | 498 | 9 | 37 | 50 | 257 | 42 | 18 | 0 | 2 | 3 | 10 | 6 |
| SG0000005 | Msrb3 | 5.05E-14 | 2.24E-13 | -1.971650652 | 606.351133 | 154.5959834 | 412 | 30 | 37 | 70 | 182 | 82 | 5 | 0 | 0 | 0 | 1 | 1 |
| SG0000002 | Nfx1 | 4.23E-13 | 1.77E-12 | 0.893116495 | 2995.781484 | 5563.716429 | 276 | 24 | 37 | 63 | 165 | 33 | 2 | 0 | 0 | 0 | 5 | 1 |
| SG0000002 | 4933427D14Rik | 1.23E-11 | 4.69E-11 | -1.242450572 | 766.9797167 | 324.1671394 | 265 | 28 | 37 | 41 | 128 | 34 | 7 | 0 | 0 | 0 | 1 | 0 |
| SG0000002 | St6gal1 | 4.08E-10 | 1.39E-09 | 2.281594943 | 128.8790395 | 626.6294565 | 457 | 21 | 37 | 59 | 174 | 57 | 2 | 0 | 0 | 0 | 4 | 0 |
| JSG000000 | Neo1 | 5.86E-10 | 1.97E-09 | -0.910881345 | 3256.293108 | 1731.892305 | 434 | 20 | 37 | 58 | 179 | 45 | 4 | 0 | 0 | 0 | 3 | 0 |
| SG0000004 | Eml2 | 1.87E-09 | 6.02E-09 | -1.595885932 | 2379.450361 | 787.1674256 | 125 | 32 | 37 | 48 | 92 | 16 | 3 | 0 | 0 | 3 | 3 | 0 |
| SG0000003 | Otud7a | 2.12E-09 | 6.78E-09 | 6.430976184 | 0.307509278 | 26.5323035 | 489 | 19 | 37 | 71 | 248 | 65 | 0 | 0 | 0 | 0 | 0 | 0 |
| SG0000000 | Akt2 | 8.22E-08 | 2.33E-07 | -0.805727303 | 1954.37517 | 1118.046457 | 181 | 25 | 37 | 50 | 96 | 24 | 2 | 0 | 0 | 0 | 0 | 0 |
| JSG000000 | Kif13a | 2.47E-07 | 6.69E-07 | -0.782656014 | 1756.246237 | 1020.898275 | 270 | 30 | 37 | 60 | 137 | 27 | 2 | 0 | 0 | 0 | 2 | 0 |
| JSG000000 | Gm21985 | 4.3E-07 | 1.14E-06 | 4.202570907 | 0 | 17.41195487 | 332 | 24 | 37 | 60 | 187 | 25 | 1 | 0 | 0 | 0 | 0 | 0 |
| SG0000003 | Acot11 | 6.97E-07 | 1.82E-06 | 1.956030438 | 31.93776565 | 123.9162791 | 187 | 31 | 37 | 43 | 141 | 15 | 5 | 0 | 0 | 0 | 0 | 0 |
| SG0000002 | Neb | 4.28E-05 | 9.48E-05 | 3.111766098 | 2.648121413 | 22.89142498 | 878 | 20 | 37 | 101 | 352 | 69 | 9 | 0 | 0 | 0 | 0 | 0 |
| SG0000002 | Copa | 5.09E-05 | 0.000112 | 0.496962097 | 6541.79274 | 9232.031519 | 256 | 27 | 37 | 50 | 117 | 15 | 3 | 0 | 0 | 0 | 0 | 0 |
| SG0000004 | Usp45 | 0.000149 | 0.00031 | -0.558371036 | 2234.230134 | 1517.195251 | 517 | 18 | 37 | 73 | 266 | 32 | 36 | 0 | 3 | 3 | 13 | 0 |
| SG0000000 | Mov10 | 0.000182 | 0.000375 | 0.769133724 | 2222.2383 | 3787.241067 | 154 | 26 | 37 | 37 | 48 | 5 | 1 | 0 | 0 | 0 | 0 | 0 |
| SG0000004 | 1110037F02Rik | 0.000872 | 0.001648 | -0.478056536 | 2690.077134 | 1931.325061 | 401 | 28 | 37 | 63 | 221 | 29 | 12 | 0 | 2 | 2 | 8 | 2 |
| SG0000003 | Tbc1d24 | 0.002291 | 0.004111 | -0.552323293 | 904.1696093 | 616.5723246 | 348 | 37 | 37 | 100 | 252 | 30 | 71 | 11 | 11 | 22 | 61 | 6 |
| JSG000000 | Dennd4a | 0.002461 | 0.004402 | 0.653797588 | 548.1968264 | 862.4803359 | 276 | 28 | 37 | 50 | 155 | 30 | 2 | 0 | 0 | 0 | 2 | 3 |
| SG0000002 | Zc3h14 | 0.005365 | 0.009142 | 0.350748694 | 3680.852916 | 4693.905496 | 311 | 33 | 37 | 73 | 179 | 16 | 13 | 0 | 0 | 0 | 0 | 0 |
| SG0000002 | Smchd1 | 0.01196 | 0.019307 | -0.369335852 | 4953.178576 | 3834.447677 | 351 | 16 | 37 | 60 | 185 | 39 | 5 | 0 | 0 | 0 | 0 | 2 |
| SG0000003 | Apopt1 | 0.020785 | 0.032067 | 0.391470837 | 393.5848704 | 516.2770978 | 115 | 30 | 37 | 38 | 88 | 10 | 2 | 0 | 0 | 0 | 0 | 0 |
| JSG000000 | Acyp2 | 0.04465 | 0.065023 | -0.921842567 | 50.40333704 | 26.60461794 | 303 | 25 | 37 | 57 | 169 | 33 | 1 | 0 | 0 | 0 | 3 | 0 |
| JSG000000 | D2hgdh | 0.074439 | 0.104014 | 0.335376747 | 256.3303456 | 323.4137528 | 120 | 33 | 37 | 46 | 93 | 6 | 0 | 0 | 0 | 0 | 2 | 0 |
| JSG000000 | Map2k5 | 0.078986 | 0.109802 | -0.314558696 | 481.7759863 | 387.3945853 | 370 | 13 | 37 | 55 | 166 | 20 | 0 | 0 | 0 | 0 | 1 | 0 |
| JSG000000 | Phlpp1 | 0.094546 | 0.129404 | -0.277547884 | 776.9635239 | 640.9887275 | 457 | 19 | 37 | 74 | 210 | 35 | 3 | 0 | 0 | 0 | 0 | 1 |
| SG0000003 | Utp6 | 0.233515 | 0.292519 | 0.15102342 | 2960.584679 | 3287.305503 | 147 | 37 | 37 | 52 | 97 | 6 | 3 | 0 | 0 | 1 | 1 | 0 |
| JSG000000 | Csnk1g3 | 0.285134 | 0.348766 | -0.162913476 | 1335.866863 | 1193.222225 | 269 | 23 | 37 | 43 | 141 | 31 | 8 | 1 | 1 | 1 | 1 | 1 |
| SG0000002 | Tmem180 | 0.401142 | 0.470055 | 0.221699357 | 952.5435835 | 1110.767112 | 106 | 33 | 37 | 41 | 98 | 9 | 0 | 0 | 0 | 0 | 0 | 0 |
| SG0000006 | Mapk1 | 0.597648 | 0.660522 | -0.075238875 | 10127.58766 | 9612.953752 | 268 | 15 | 37 | 34 | 103 | 27 | 11 | 0 | 0 | 0 | 0 | 0 |
| SG0000002 | Pde8a | 0.928166 | 0.947905 | 0.001239908 | 301.4121545 | 301.6713112 | 222 | 28 | 37 | 44 | 110 | 10 | 4 | 1 | 1 | 1 | 2 | 0 |
| SG0000000 | Dnmt3l | 0 | 0 | 9.519147014 | 3.276357648 | 2404.031134 | 97 | 36 | 36 | 45 | 91 | 1 | 0 | 0 | 0 | 0 | 0 | 0 |
| SG0000007 | Cecr2 | 0 | 0 | 10.20827107 | 2.648121413 | 3132.807347 | 448 | 24 | 36 | 67 | 179 | 28 | 8 | 0 | 0 | 1 | 3 | 2 |
| SG0000004 | Sgsm1 | 3.9E-121 | 4E-119 | 6.546636142 | 5.512072078 | 515.2864679 | 153 | 32 | 36 | 47 | 97 | 18 | 7 | 2 | 2 | 4 | 4 | 0 |
| SG0000002 | Irak3 | 4.29E-96 | 2.81E-94 | -4.555560562 | 3529.564843 | 150.0931212 | 341 | 33 | 36 | 45 | 156 | 33 | 2 | 0 | 0 | 0 | 0 | 0 |
| SG0000006 | Syt9 | 3.33E-82 | 1.57E-80 | 3.348319683 | 83.57498321 | 851.1791123 | 570 | 16 | 36 | 80 | 302 | 66 | 5 | 0 | 0 | 0 | 0 | 0 |
| SG0000002 | Cflar | 3.05E-73 | 1.21E-71 | -3.478540253 | 4517.832551 | 405.3079972 | 299 | 24 | 36 | 62 | 146 | 19 | 19 | 2 | 3 | 8 | 10 | 1 |
| SG0000004 | Ldlrad3 | 3.45E-62 | 1.01E-60 | 2.630418304 | 164.2115409 | 1016.806906 | 582 | 16 | 36 | 68 | 244 | 44 | 6 | 2 | 2 | 4 | 4 | 4 |
| SG0000003 | Fez1 | 6.79E-50 | 1.41E-48 | -4.068961163 | 2307.61607 | 137.4941415 | 232 | 26 | 36 | 52 | 149 | 27 | 9 | 1 | 1 | 1 | 1 | 0 |
| SG0000002 | Fubp1 | 6.99E-47 | 1.31E-45 | 2.52162443 | 1743.75626 | 10013.14191 | 506 | 19 | 36 | 86 | 342 | 54 | 39 | 1 | 6 | 8 | 35 | 6 |
| SG0000004 | Eml6 | 3.4E-44 | 5.79E-43 | -4.003350335 | 561.9367329 | 35.0395797 | 535 | 19 | 36 | 80 | 328 | 48 | 6 | 0 | 0 | 0 | 0 | 1 |
| SG0000002 | Ptpn2 | 1.99E-37 | 2.67E-36 | 1.720118179 | 606.1924247 | 1997.182136 | 403 | 24 | 36 | 60 | 193 | 49 | 4 | 0 | 0 | 1 | 6 | 0 |
| SG0000002 | Sirt1 | 5.47E-37 | 7.27E-36 | 1.696356643 | 656.6921724 | 2128.217806 | 251 | 26 | 36 | 58 | 127 | 28 | 21 | 0 | 6 | 9 | 15 | 3 |
| SG0000002 | Dars | 2.81E-34 | 3.4E-33 | 1.4482793 | 3837.37565 | 10471.52241 | 190 | 29 | 36 | 52 | 102 | 20 | 3 | 0 | 0 | 0 | 0 | 0 |
| SG0000002 | Tmcc3 | 1.01E-31 | 1.1E-30 | -3.132578277 | 1612.518194 | 183.8673421 | 192 | 27 | 36 | 45 | 114 | 29 | 18 | 0 | 0 | 0 | 0 | 4 |
| SG0000003 | Bcorl1 | 9.56E-25 | 7.63E-24 | -1.905852723 | 1177.079265 | 314.1136956 | 198 | 33 | 36 | 49 | 117 | 15 | 8 | 0 | 0 | 3 | 6 | 0 |
| SG0000002 | Sf3b1 | 4.03E-24 | 3.11E-23 | 1.196973947 | 9681.935747 | 22196.64102 | 148 | 30 | 36 | 46 | 84 | 11 | 3 | 0 | 2 | 1 | 4 | 0 |
| JSG000000 | Gnaq | 1.04E-23 | 7.94E-23 | -1.640013849 | 2524.729186 | 810.0679289 | 507 | 19 | 36 | 62 | 228 | 43 | 13 | 0 | 1 | 1 | 5 | 1 |
| SG0000002 | Agpat9 | 1.49E-22 | 1.07E-21 | -2.954024739 | 655.7731472 | 84.62595342 | 279 | 18 | 36 | 42 | 177 | 21 | 9 | 0 | 0 | 3 | 15 | 0 |
| SG0000004 | Palb2 | 2.43E-22 | 1.73E-21 | 1.478627496 | 341.1025814 | 950.5965522 | 291 | 36 | 36 | 50 | 131 | 21 | 14 | 0 | 0 | 5 | 5 | 4 |
| SG0000002 | Gbas | 4E-22 | 2.82E-21 | -1.666274937 | 3401.897992 | 1071.821711 | 228 | 24 | 36 | 48 | 144 | 28 | 0 | 0 | 0 | 0 | 0 | 4 |
| SG0000002 | Arhgap19 | 4.44E-22 | 3.12E-21 | 1.216396845 | 1541.237058 | 3581.305583 | 171 | 36 | 36 | 41 | 96 | 23 | 10 | 2 | 2 | 4 | 14 | 0 |
| SG0000003 | Tpst1 | 8.78E-22 | 6.1E-21 | 1.375515455 | 495.4407182 | 1285.473441 | 211 | 32 | 36 | 49 | 108 | 27 | 4 | 0 | 2 | 0 | 2 | 0 |
| SG0000002 | Jak2 | 1.56E-21 | 1.07E-20 | -1.994632088 | 1292.234908 | 324.2629879 | 390 | 10 | 36 | 70 | 198 | 38 | 4 | 0 | 2 | 2 | 2 | 2 |
| SG0000004 | Gm973 | 2.91E-21 | 1.96E-20 | 5.853902655 | 0 | 56.83627192 | 188 | 22 | 36 | 41 | 87 | 14 | 2 | 0 | 0 | 0 | 0 | 0 |
| SG0000001 | Ccdc170 | 3.88E-21 | 2.59E-20 | 3.139403833 | 16.85090935 | 148.4834551 | 181 | 31 | 36 | 48 | 117 | 15 | 3 | 0 | 0 | 0 | 1 | 0 |
| SG0000003 | Iqcg | 7.34E-20 | 4.57E-19 | 3.221271676 | 51.13896768 | 476.9263915 | 111 | 30 | 36 | 39 | 59 | 9 | 5 | 0 | 0 | 1 | 2 | 1 |
| JSG000000 | Hdac9 | 8.24E-20 | 5.11E-19 | -2.889440378 | 205.9564781 | 27.79503544 | 417 | 17 | 36 | 52 | 195 | 35 | 7 | 0 | 0 | 2 | 4 | 3 |
| JSG000000 | Fam35a | 2.56E-18 | 1.47E-17 | 1.366338855 | 316.0633557 | 814.8602381 | 201 | 28 | 36 | 57 | 136 | 19 | 1 | 0 | 0 | 0 | 2 | 0 |
| SG0000002 | Boll | 3.82E-15 | 1.83E-14 | 7.124086201 | 0.307509278 | 42.89650031 | 522 | 16 | 36 | 70 | 268 | 39 | 11 | 1 | 1 | 3 | 7 | 0 |
| SG0000004 | Krba1 | 5.99E-15 | 2.82E-14 | 1.209084966 | 368.5080199 | 851.9572164 | 123 | 24 | 36 | 32 | 40 | 4 | 8 | 0 | 0 | 0 | 0 | 0 |
| SG0000006 | Ppih | 5.01E-13 | 2.08E-12 | 1.170008763 | 306.0430828 | 688.6369168 | 118 | 19 | 36 | 35 | 90 | 12 | 3 | 0 | 0 | 0 | 0 | 3 |
| SG0000004 | Fgfr1op2 | 1.11E-12 | 4.52E-12 | -1.085983774 | 2533.948876 | 1193.669667 | 168 | 24 | 36 | 34 | 112 | 16 | 14 | 0 | 0 | 2 | 6 | 2 |
| SG0000003 | Fam122b | 9.92E-12 | 3.79E-11 | 1.052361083 | 460.6736524 | 955.4008568 | 209 | 27 | 36 | 48 | 109 | 3 | 27 | 0 | 0 | 3 | 16 | 0 |
| JSG000000 | AI481877 | 2E-11 | 7.48E-11 | 4.879335895 | 1.313186595 | 38.6503031 | 261 | 25 | 36 | 55 | 148 | 15 | 1 | 0 | 0 | 0 | 0 | 0 |
| SG0000002 | Adamts20 | 2.53E-11 | 9.4E-11 | 4.899021581 | 0 | 28.83681389 | 438 | 20 | 36 | 80 | 277 | 51 | 6 | 0 | 0 | 3 | 8 | 1 |
| SG0000002 | Tmem68 | 3.1E-11 | 1.14E-10 | -1.19687249 | 801.5149235 | 349.6367643 | 210 | 22 | 36 | 38 | 100 | 14 | 14 | 0 | 0 | 6 | 10 | 4 |
| SG0000002 | Kif24 | 5E-11 | 1.82E-10 | 1.108867964 | 262.3916886 | 565.9168078 | 302 | 28 | 36 | 52 | 158 | 28 | 2 | 0 | 0 | 0 | 0 | 0 |
| SG0000004 | Ppp2r2d | 2.68E-10 | 9.23E-10 | 0.826351132 | 1382.826888 | 2452.021608 | 211 | 32 | 36 | 46 | 94 | 26 | 7 | 0 | 0 | 2 | 2 | 0 |
| SG0000004 | Zmym5 | 3.82E-10 | 1.3E-09 | 0.880082599 | 760.4444559 | 1399.583323 | 158 | 27 | 36 | 36 | 75 | 17 | 2 | 0 | 0 | 0 | 0 | 5 |
| SG0000002 | Pax6 | 5.8E-10 | 1.95E-09 | 4.704015569 | 0 | 25.06452329 | 259 | 27 | 36 | 36 | 92 | 0 | 18 | 0 | 0 | 0 | 9 | 0 |
| SG0000000 | Col18a1 | 1.11E-09 | 3.64E-09 | 1.320879781 | 2037.789234 | 5090.772604 | 172 | 24 | 36 | 46 | 87 | 43 | 2 | 0 | 0 | 0 | 0 | 0 |
| SG0000004 | Ppp2r3a | 3.21E-09 | 1.01E-08 | -1.191901329 | 668.3895055 | 292.5711983 | 482 | 17 | 36 | 63 | 231 | 57 | 10 | 0 | 2 | 0 | 10 | 2 |
| SG0000003 | Ube2h | 4.67E-09 | 1.46E-08 | -0.873148427 | 2505.929681 | 1368.122587 | 317 | 20 | 36 | 52 | 144 | 31 | 5 | 0 | 0 | 0 | 4 | 1 |

| | | | | | | | | | | | | | | | | | | |
|---|---|---|---|---|---|---|---|---|---|---|---|---|---|---|---|---|---|---|
| SG0000002 | Tex26 | 5.98E-09 | 1.85E-08 | 4.550011501 | 1.324060707 | 31.01686072 | 304 | 36 | 36 | 60 | 127 | 30 | 6 | 0 | 0 | 0 | 1 | 0 |
| SG0000002 | Pip4k2a | 1.11E-08 | 3.37E-08 | -1.003444072 | 825.9578102 | 411.9941975 | 268 | 25 | 36 | 46 | 119 | 21 | 3 | 0 | 0 | 1 | 1 | 0 |
| SG0000002 | Ndrg3 | 1.8E-08 | 5.37E-08 | -0.849619024 | 2267.872827 | 1258.513522 | 356 | 27 | 36 | 77 | 228 | 32 | 9 | 1 | 1 | 3 | 6 | 0 |
| SG0000002 | Eaf2 | 2.01E-08 | 5.96E-08 | 2.510511307 | 10.46349616 | 59.62330242 | 414 | 20 | 36 | 82 | 280 | 40 | 3 | 0 | 0 | 0 | 0 | 0 |
| SG0000002 | Mtfr1 | 4.34E-08 | 1.26E-07 | -0.780636524 | 4054.091009 | 2359.926539 | 415 | 20 | 36 | 77 | 171 | 41 | 11 | 0 | 1 | 2 | 6 | 1 |
| SG0000002 | Nsun6 | 1.66E-07 | 4.57E-07 | 1.045766529 | 152.5305782 | 314.8937279 | 586 | 36 | 36 | 135 | 316 | 57 | 19 | 3 | 3 | 5 | 11 | 0 |
| SG0000003 | Phip | 1.93E-07 | 5.27E-07 | -0.753037276 | 3552.450879 | 2107.857627 | 385 | 22 | 36 | 59 | 178 | 22 | 16 | 1 | 1 | 3 | 9 | 1 |
| SG0000003 | Tusc3 | 2.49E-07 | 6.74E-07 | -0.795463341 | 1562.988279 | 900.5283615 | 628 | 16 | 36 | 108 | 314 | 76 | 4 | 0 | 0 | 0 | 2 | 0 |
| SG000000C | Rpa1 | 5.21E-07 | 1.37E-06 | 0.612985438 | 5625.079408 | 8603.113676 | 193 | 28 | 36 | 42 | 95 | 12 | 5 | 0 | 0 | 0 | 1 | 0 |
| SG0000002 | Sgpl1 | 9.91E-07 | 2.56E-06 | -0.693968718 | 7130.651496 | 4407.819592 | 324 | 27 | 36 | 54 | 159 | 36 | 3 | 0 | 0 | 0 | 0 | 0 |
| SG0000003 | Fgf2 | 1.4E-06 | 3.58E-06 | -1.541737643 | 471.6364915 | 161.9936996 | 237 | 28 | 36 | 56 | 93 | 17 | 17 | 0 | 0 | 2 | 3 | 1 |
| SG000000E | Lrrc51 | 2.47E-06 | 6.15E-06 | 1.448816493 | 46.69666759 | 127.4744397 | 181 | 28 | 36 | 40 | 160 | 12 | 0 | 0 | 0 | 0 | 0 | 0 |
| SG0000003 | Wdtc1 | 3.75E-06 | 9.21E-06 | 0.92310392 | 1257.878936 | 2385.178104 | 130 | 24 | 36 | 44 | 92 | 8 | 2 | 0 | 0 | 2 | 2 | 0 |
| SG0000004 | Rhoj | 1.16E-05 | 2.72E-05 | -2.204822504 | 186.6162653 | 40.47918734 | 492 | 28 | 36 | 48 | 160 | 36 | 6 | 0 | 0 | 0 | 1 | 0 |
| SG0000003 | Dzank1 | 1.28E-05 | 2.99E-05 | 1.447324695 | 35.4480105 | 96.66739431 | 242 | 34 | 36 | 54 | 142 | 16 | 10 | 0 | 0 | 0 | 6 | 0 |
| SG0000002 | Pafah1b1 | 2.27E-05 | 5.17E-05 | -0.574317953 | 9116.774691 | 6122.859461 | 310 | 22 | 36 | 50 | 138 | 32 | 14 | 0 | 0 | 2 | 2 | 6 |
| SG0000002 | Il15ra | 3.68E-05 | 8.22E-05 | -1.747319811 | 91.91903064 | 27.37850693 | 445 | 36 | 36 | 68 | 242 | 52 | 26 | 0 | 0 | 6 | 12 | 0 |
| SG0000004 | 9430038I01Rik | 0.000112 | 0.000236 | 0.908179464 | 99.27615838 | 186.3091667 | 191 | 30 | 36 | 39 | 106 | 9 | 9 | 0 | 1 | 1 | 1 | 2 |
| SG000000E | Adgrg5 | 0.000223 | 0.000455 | 2.107797721 | 15.45207657 | 66.60352124 | 81 | 30 | 36 | 36 | 60 | 6 | 2 | 0 | 0 | 0 | 0 | 0 |
| SG0000002 | Clcc1 | 0.00212 | 0.003826 | -0.495865449 | 891.3041869 | 632.0560207 | 165 | 27 | 36 | 39 | 74 | 18 | 5 | 0 | 0 | 0 | 0 | 1 |
| SG000000C | Msh4 | 0.002466 | 0.00441 | 5.289144046 | 0.994803205 | 38.89808179 | 384 | 21 | 36 | 48 | 129 | 12 | 0 | 0 | 0 | 0 | 0 | 0 |
| SG0000003 | Ift122 | 0.00343 | 0.006001 | -0.483882196 | 809.9786388 | 579.1759375 | 255 | 24 | 36 | 39 | 159 | 275 | 5 | 0 | 1 | 4 | 5 | 3 |
| SG0000002 | Atg13 | 0.005382 | 0.009168 | 0.395788837 | 1079.67963 | 1420.493399 | 162 | 24 | 36 | 54 | 88 | 6 | 2 | 0 | 0 | 2 | 4 | 0 |
| SG0000003 | Thoc2 | 0.007075 | 0.011843 | -0.38091143 | 3108.073447 | 2386.852089 | 405 | 18 | 36 | 69 | 224 | 25 | 16 | 0 | 2 | 0 | 2 | 2 |
| SG0000003 | Cacul1 | 0.009263 | 0.015228 | -0.370176168 | 2831.768743 | 2190.90554 | 322 | 20 | 36 | 54 | 140 | 18 | 8 | 0 | 0 | 0 | 0 | 2 |
| JSG000000 | Agpat4 | 0.01481 | 0.023488 | 0.363517486 | 740.0057582 | 952.0609804 | 230 | 24 | 36 | 50 | 114 | 39 | 0 | 0 | 0 | 0 | 0 | 0 |
| SG0000002 | Slit1 | 0.01631 | 0.025681 | 1.480462945 | 8.999571842 | 25.11224861 | 369 | 24 | 36 | 66 | 183 | 27 | 0 | 0 | 0 | 0 | 0 | 0 |
| SG0000004 | D330045A20Rik | 0.017109 | 0.026825 | 3.277042228 | 1.561638181 | 15.1379974 | 384 | 24 | 36 | 50 | 206 | 46 | 6 | 0 | 0 | 0 | 2 | 0 |
| SG0000002 | Inpp4a | 0.038443 | 0.056576 | 0.367200159 | 310.5711913 | 400.5893789 | 675 | 11 | 36 | 81 | 216 | 52 | 18 | 2 | 3 | 5 | 6 | 3 |
| SG0000004 | Slc26a7 | 0.039655 | 0.058249 | -2.822939924 | 16.76157289 | 2.368782451 | 305 | 18 | 36 | 37 | 142 | 24 | 9 | 0 | 2 | 2 | 9 | 1 |
| SG0000003 | 2810474O19Rik | 0.055363 | 0.079244 | 0.256082538 | 1641.904109 | 1960.813621 | 169 | 33 | 36 | 39 | 89 | 17 | 4 | 0 | 0 | 0 | 0 | 2 |
| SG0000004 | Arhgef1 | 0.058277 | 0.083037 | -0.38730165 | 3072.477814 | 2349.088299 | 82 | 36 | 36 | 48 | 84 | 4 | 0 | 0 | 0 | 0 | 4 | 0 |
| SG0000003 | Rpgrip1l | 0.063784 | 0.090267 | 0.320785899 | 344.7900084 | 430.6464284 | 422 | 20 | 36 | 73 | 196 | 36 | 15 | 0 | 0 | 1 | 2 | 0 |
| SG0000002 | Alpk1 | 0.075777 | 0.105727 | 1.100557844 | 14.12567229 | 30.29075162 | 299 | 28 | 36 | 57 | 158 | 23 | 2 | 0 | 0 | 0 | 0 | 0 |
| SG000000C | Smpd4 | 0.083031 | 0.114987 | -0.232515582 | 3294.226526 | 2803.879176 | 140 | 28 | 36 | 53 | 96 | 8 | 16 | 0 | 0 | 9 | 12 | 0 |
| SG0000002 | Pcbd2 | 0.096686 | 0.132061 | 0.419793574 | 102.6818554 | 137.3612281 | 162 | 30 | 36 | 40 | 86 | 22 | 0 | 0 | 0 | 0 | 0 | 0 |
| JSG000000 | Brinp1 | 0.098051 | 0.13368 | -0.737786868 | 66.16930656 | 39.67898969 | 456 | 16 | 36 | 69 | 228 | 44 | 0 | 0 | 0 | 0 | 0 | 0 |
| SG0000002 | Tollip | 0.106184 | 0.143751 | -0.225877623 | 2199.047913 | 1880.35025 | 166 | 35 | 36 | 55 | 89 | 9 | 4 | 0 | 0 | 1 | 1 | 0 |
| SG000000C | Prkacb | 0.125084 | 0.166692 | 0.206841917 | 1378.458975 | 1590.960755 | 462 | 10 | 36 | 33 | 203 | 61 | 24 | 0 | 0 | 0 | 12 | 4 |
| SG0000003 | Atp11b | 0.151988 | 0.198948 | -0.193686134 | 4416.480506 | 3861.632863 | 541 | 21 | 36 | 81 | 295 | 38 | 14 | 0 | 0 | 0 | 0 | 0 |
| SG0000002 | Mme | 0.211322 | 0.267619 | -0.49808334 | 690.6396101 | 489.0051773 | 624 | 18 | 36 | 63 | 247 | 55 | 18 | 0 | 0 | 0 | 7 | 4 |
| SG0000003 | Cenpi | 0.216627 | 0.273737 | 0.189482725 | 664.0936403 | 757.3023512 | 292 | 26 | 36 | 74 | 158 | 22 | 2 | 0 | 0 | 2 | 2 | 2 |
| SG0000002 | Aptx | 0.259927 | 0.321578 | 0.194451959 | 375.8171172 | 430.0435411 | 156 | 36 | 36 | 40 | 96 | 14 | 16 | 4 | 4 | 4 | 6 | 0 |
| SG000000C | Med6 | 0.40619 | 0.475255 | 0.123671678 | 794.4803379 | 865.5896172 | 174 | 30 | 36 | 36 | 87 | 18 | 0 | 0 | 0 | 0 | 0 | 3 |
| SG0000001 | Epb41l2 | 0.545533 | 0.611356 | -0.090484329 | 4109.684934 | 3859.846477 | 455 | 21 | 36 | 51 | 211 | 34 | 8 | 0 | 0 | 0 | 0 | 0 |
| SG0000003 | Susd4 | 0.690095 | 0.745086 | 0.063161624 | 401.7767989 | 419.7574353 | 349 | 24 | 36 | 58 | 166 | 28 | 1 | 0 | 0 | 0 | 0 | 0 |
| SG000000C | Ncaph2 | 0.782097 | 0.826143 | -0.038111166 | 4932.719502 | 4804.119658 | 100 | 24 | 36 | 40 | 60 | 0 | 12 | 0 | 0 | 0 | 0 | 0 |
| JSG000000 | Gm21992 | 0.820452 | 0.858658 | -0.109827383 | 42.1297153 | 39.04156224 | 248 | 25 | 36 | 52 | 124 | 19 | 6 | 0 | 1 | 1 | 3 | 0 |
| SG0000002 | Rnf19b | 0.836149 | 0.871978 | -0.028566444 | 1098.334902 | 1076.800945 | 188 | 33 | 36 | 46 | 80 | 10 | 6 | 0 | 0 | 0 | 0 | 2 |
| JSG000000 | Pacs1 | 4.36E-83 | 2.1E-81 | -3.608934996 | 15084.59675 | 1236.334586 | 272 | 26 | 35 | 56 | 155 | 28 | 0 | 0 | 0 | 0 | 0 | 0 |
| SG0000002 | Igf2bp3 | 1.44E-65 | 4.57E-64 | 3.542232139 | 249.2210569 | 2903.373169 | 253 | 24 | 35 | 46 | 118 | 21 | 7 | 0 | 0 | 0 | 1 | 0 |
| SG0000002 | Ccdc109b | 2.94E-64 | 9.14E-63 | -4.437144019 | 796.2419947 | 36.75628703 | 318 | 21 | 35 | 54 | 158 | 50 | 0 | 0 | 0 | 0 | 0 | 0 |
| SG0000003 | Atf7ip2 | 1.41E-55 | 3.47E-54 | 3.724434142 | 27.24681727 | 360.1489664 | 478 | 15 | 35 | 62 | 224 | 16 | 0 | 0 | 0 | 0 | 1 | 0 |
| SG0000003 | Rasgrp4 | 8.29E-42 | 1.3E-40 | 4.643338033 | 39.93615061 | 998.0452442 | 116 | 25 | 35 | 37 | 114 | 15 | 6 | 0 | 0 | 2 | 4 | 0 |
| SG0000003 | Aktip | 1.07E-38 | 1.5E-37 | 1.791891373 | 502.5656462 | 1740.226792 | 131 | 30 | 35 | 40 | 64 | 12 | 8 | 0 | 0 | 0 | 3 | 2 |
| JSG000000 | Arhgef3 | 4.89E-33 | 5.59E-32 | -2.390971223 | 2597.145117 | 495.1561301 | 256 | 21 | 35 | 54 | 118 | 31 | 3 | 0 | 1 | 1 | 2 | 1 |
| SG0000004 | Camta2 | 2.07E-30 | 2.13E-29 | -2.451687421 | 1321.756955 | 241.6129182 | 91 | 35 | 35 | 35 | 56 | 21 | 14 | 7 | 7 | 7 | 7 | 14 |
| SG0000002 | Rtn1 | 2.43E-30 | 2.49E-29 | 6.405235425 | 0 | 83.75551991 | 411 | 24 | 35 | 66 | 182 | 32 | 2 | 0 | 0 | 0 | 0 | 0 |
| SG0000003 | Poli | 2.42E-24 | 1.89E-23 | 1.500115961 | 395.1966859 | 1117.874875 | 220 | 25 | 35 | 45 | 155 | 25 | 1 | 0 | 0 | 0 | 6 | 0 |
| SG0000007 | Rnf213 | 1.14E-23 | 8.65E-23 | -2.003618732 | 15926.75292 | 3971.713423 | 194 | 27 | 35 | 51 | 101 | 19 | 3 | 0 | 0 | 0 | 0 | 0 |
| JSG000000 | Cenpp | 1.42E-23 | 1.07E-22 | 1.785970405 | 153.6459935 | 529.8487369 | 427 | 16 | 35 | 71 | 217 | 37 | 0 | 0 | 0 | 0 | 0 | 0 |
| SG0000002 | Arhgap28 | 3.03E-21 | 2.03E-20 | 3.307787133 | 42.36010893 | 419.4693322 | 461 | 21 | 35 | 87 | 224 | 77 | 6 | 0 | 0 | 4 | 6 | 0 |
| SG0000002 | Dpysl3 | 3.08E-20 | 1.96E-19 | -1.961485709 | 2881.277509 | 739.8080534 | 418 | 18 | 35 | 59 | 197 | 33 | 4 | 0 | 0 | 0 | 2 | 0 |
| SG0000001 | Efr3a | 1.15E-19 | 7.07E-19 | -1.345826284 | 7369.259605 | 2899.277438 | 537 | 17 | 35 | 82 | 316 | 77 | 8 | 0 | 0 | 0 | 1 | 0 |
| SG0000003 | Itga9 | 3.23E-19 | 1.94E-18 | 3.243351609 | 27.11064393 | 256.7355173 | 580 | 18 | 35 | 73 | 231 | 51 | 8 | 0 | 0 | 2 | 4 | 1 |
| SG0000003 | Cyld | 3.96E-16 | 2.01E-15 | -1.319329885 | 2180.189749 | 873.6478725 | 416 | 16 | 35 | 41 | 202 | 32 | 36 | 0 | 3 | 6 | 27 | 6 |
| SG0000003 | Ubash3b | 1.81E-11 | 6.8E-11 | -1.988341227 | 385.7413202 | 97.21780506 | 251 | 27 | 35 | 46 | 130 | 13 | 8 | 0 | 0 | 1 | 1 | 0 |
| SG0000003 | Rgs22 | 4.2E-11 | 1.53E-10 | 6.550011899 | 0.331601068 | 31.07176866 | 320 | 18 | 35 | 67 | 188 | 48 | 0 | 0 | 0 | 0 | 0 | 0 |
| SG0000002 | Ralgapb | 2.46E-10 | 8.52E-10 | -0.923630342 | 3766.835075 | 1985.803024 | 745 | 18 | 35 | 71 | 315 | 87 | 27 | 0 | 0 | 3 | 6 | 6 |
| SG0000004 | Sgms1 | 9.85E-10 | 3.24E-09 | -0.938016212 | 1986.819767 | 1037.020773 | 446 | 27 | 35 | 71 | 189 | 59 | 5 | 1 | 1 | 1 | 3 | 0 |
| JSG000000 | Ube2d3 | 1.18E-09 | 3.86E-09 | 0.738499478 | 5848.266986 | 9757.480355 | 551 | 14 | 35 | 56 | 242 | 18 | 40 | 0 | 0 | 0 | 11 | 4 |
| SG000000E | Nrg2 | 4.64E-09 | 1.45E-08 | 4.56866627 | 0 | 22.73042888 | 201 | 28 | 35 | 44 | 130 | 23 | 1 | 0 | 0 | 0 | 1 | 0 |
| SG0000003 | St3gal2 | 8.76E-09 | 2.68E-08 | -0.941267855 | 1121.413462 | 584.0041148 | 166 | 22 | 35 | 48 | 96 | 8 | 2 | 0 | 1 | 0 | 2 | 0 |
| SG0000002 | Psd2 | 1.21E-08 | 3.65E-08 | 5.788926048 | 1.265003014 | 69.94101706 | 308 | 30 | 35 | 70 | 129 | 15 | 13 | 0 | 0 | 0 | 4 | 0 |
| SG0000003 | Morf4l2 | 1.53E-06 | 3.88E-06 | 0.582212691 | 7707.074381 | 11538.59776 | 168 | 14 | 35 | 30 | 51 | 21 | 0 | 0 | 0 | 0 | 0 | 7 |
| SG0000002 | 1700016H13Rik | 2.86E-06 | 7.09E-06 | 4.004987108 | 0 | 15.05540451 | 114 | 35 | 35 | 77 | 329 | 21 | 2 | 0 | 0 | 0 | 0 | 0 |
| SG0000002 | Hivep3 | 5.23E-06 | 1.27E-05 | 3.25151927 | 5.206906367 | 49.58891427 | 389 | 24 | 35 | 59 | 166 | 38 | 11 | 0 | 1 | 0 | 2 | 2 |
| SG0000002 | Trappc12 | 1.01E-05 | 2.39E-05 | -0.680965731 | 1390.735085 | 867.4676743 | 180 | 27 | 35 | 34 | 68 | 12 | 1 | 0 | 0 | 0 | 0 | 1 |
| SG0000003 | Far1 | 1.31E-05 | 3.05E-05 | -0.612169238 | 4422.310547 | 2893.1299 | 427 | 24 | 35 | 59 | 204 | 56 | 28 | 0 | 2 | 8 | 24 | 5 |
| JSG000000 | Dapk2 | 1.84E-05 | 4.22E-05 | 2.587727893 | 8.140788835 | 48.93844955 | 168 | 28 | 35 | 42 | 85 | 28 | 1 | 0 | 0 | 0 | 1 | 0 |
| SG0000005 | C77080 | 2.46E-05 | 5.58E-05 | -0.824347534 | 2825.37265 | 1595.594028 | 68 | 34 | 35 | 39 | 51 | 0 | 0 | 0 | 0 | 0 | 0 | 0 |
| JSG000000 | Shprh | 3.65E-05 | 8.15E-05 | -0.673515313 | 1813.504726 | 1137.025959 | 606 | 10 | 35 | 97 | 285 | 69 | 26 | 1 | 1 | 3 | 13 | 3 |
| SG0000003 | Zfp335 | 4.16E-05 | 9.21E-05 | -0.784937991 | 1041.372288 | 604.3884221 | 50 | 33 | 35 | 33 | 35 | 0 | 2 | 0 | 0 | 0 | 0 | 0 |
| SG0000007 | Faap20 | 9.47E-05 | 0.000201 | 0.680574008 | 307.063793 | 492.1548859 | 79 | 35 | 35 | 45 | 50 | 0 | 0 | 0 | 0 | 0 | 0 | 0 |
| SG0000002 | Crebbp | 0.000355 | 0.000707 | -0.548880676 | 1293.888629 | 884.4379079 | 269 | 17 | 35 | 51 | 141 | 19 | 5 | 1 | 2 | 2 | 3 | 1 |
| SG0000005 | Hnrnpm | 0.000469 | 0.000918 | 0.429114742 | 6057.209021 | 8155.469526 | 331 | 13 | 35 | 39 | 168 | 22 | 0 | 0 | 0 | 0 | 0 | 0 |
| SG0000002 | Atat1 | 0.000544 | 0.001057 | 0.711834139 | 164.1715815 | 268.8941846 | 101 | 35 | 35 | 44 | 74 | 12 | 6 | 0 | 0 | 3 | 3 | 2 |
| SG0000007 | Rnf121 | 0.000579 | 0.001119 | -0.560034153 | 884.5619246 | 599.9860638 | 302 | 29 | 35 | 65 | 120 | 26 | 3 | 0 | 0 | 0 | 2 | 0 |
| SG0000002 | Upp2 | 0.003359 | 0.005883 | -1.992089419 | 26.70641765 | 6.7133141 | 478 | 22 | 35 | 66 | 222 | 47 | 7 | 0 | 0 | 0 | 1 | 0 |
| SG000000C | Clpb | 0.01075 | 0.017487 | 0.351915851 | 1357.213771 | 1732.149947 | 341 | 20 | 35 | 59 | 148 | 36 | 1 | 0 | 0 | 0 | 0 | 0 |
| SG0000003 | Nsf | 0.010875 | 0.017679 | -0.359706054 | 2462.323805 | 1918.946434 | 269 | 23 | 35 | 50 | 130 | 15 | 2 | 0 | 0 | 0 | 1 | 0 |
| SG0000004 | Simc1 | 0.012083 | 0.019484 | -0.452960239 | 743.9581016 | 543.4928795 | 264 | 21 | 35 | 46 | 103 | 22 | 4 | 0 | 0 | 4 | 4 | 2 |
| SG0000001 | Aatf | 0.050578 | 0.072932 | 0.283313311 | 1084.207611 | 1319.46616 | 227 | 28 | 35 | 56 | 132 | 19 | 0 | 0 | 0 | 0 | 0 | 0 |
| SG000000E | Exoc5 | 0.061012 | 0.086569 | -0.271025551 | 1938.793542 | 1606.736812 | 404 | 22 | 35 | 61 | 238 | 25 | 9 | 0 | 0 | 2 | 9 | 0 |
| SG0000002 | Zfr | 0.107141 | 0.144953 | -0.219556831 | 3832.326355 | 3291.313825 | 217 | 27 | 35 | 51 | 128 | 14 | 3 | 0 | 0 | 0 | 2 | 0 |
| SG000000C | Prpf6 | 0.113772 | 0.152902 | 0.202858944 | 3228.675379 | 3716.130946 | 125 | 27 | 35 | 40 | 87 | 5 | 0 | 0 | 0 | 0 | 0 | 0 |
| SG0000003 | Col4a6 | 0.125867 | 0.167636 | 1.067986979 | 8.08791812 | 16.9563691 | 647 | 14 | 35 | 72 | 286 | 66 | 3 | 0 | 0 | 0 | 1 | 1 |
| SG0000003 | Prrx2 | 0.195019 | 0.249335 | -0.438886622 | 145.1550954 | 107.081451 | 91 | 30 | 35 | 31 | 57 | 4 | 0 | 0 | 0 | 0 | 0 | 0 |
| SG0000002 | Usp33 | 0.221979 | 0.279573 | -0.180001664 | 1866.818519 | 1647.844399 | 325 | 15 | 35 | 72 | 171 | 27 | 18 | 0 | 0 | 1 | 1 | 3 |
| SG0000002 | Kansl1l | 0.2991 | 0.363885 | 0.249602452 | 128.9158115 | 153.2653608 | 683 | 14 | 35 | 83 | 339 | 81 | 13 | 0 | 0 | 2 | 4 | 0 |
| SG0000004 | Fam117b | 0.334221 | 0.40103 | -0.140225543 | 2266.022983 | 2056.137793 | 219 | 24 | 35 | 33 | 84 | 15 | 8 | 0 | 0 | 0 | 3 | 0 |
| SG0000003 | Lrrc1 | 0.520861 | 0.587917 | 0.10039571 | 462.5325474 | 495.8660998 | 509 | 15 | 35 | 53 | 170 | 40 | 9 | 0 | 0 | 0 | 3 | 2 |
| SG0000005 | Fsd1l | 0.686939 | 0.742574 | 0.062263852 | 349.0074457 | 364.3996736 | 387 | 15 | 35 | 45 | 157 | 28 | 27 | 0 | 0 | 2 | 14 | 0 |
| SG0000004 | Wwp1 | 0.708111 | 0.761051 | -0.062599246 | 2157.232121 | 2065.630452 | 419 | 22 | 35 | 77 | 217 | 35 | 16 | 2 | 2 | 2 | 6 | 1 |
| JSG000000 | Acvr2a | 0.80458 | 0.845059 | -0.062186392 | 403.3111925 | 386.2960836 | 260 | 21 | 35 | 49 | 106 | 17 | 10 | 0 | 0 | 1 | 2 | 1 |
| JSG000000 | Qser1 | 0.825391 | 0.862819 | -0.046902116 | 2237.475002 | 2165.904224 | 180 | 25 | 35 | 42 | 100 | 10 | 12 | 0 | 1 | 2 | 5 | 0 |
| SG000000C | Galnt1 | 1.62E-55 | 3.99E-54 | -2.968372069 | 15106.9942 | 1930.229867 | 194 | 27 | 34 | 45 | 107 | 22 | 4 | 0 | 0 | 0 | 0 | 0 |
| JSG000000 | Trank1 | 5.12E-53 | 1.16E-51 | 9.051270432 | 0.331601068 | 175.9218648 | 169 | 22 | 34 | 42 | 106 | 17 | 1 | 0 | 0 | 0 | 1 | 0 |
| SG0000002 | Ddc | 9.79E-52 | 2.12E-50 | -3.415088771 | 2257.360626 | 211.6200378 | 365 | 23 | 34 | 60 | 191 | 33 | 0 | 0 | 0 | 0 | 0 | 0 |
| SG0000004 | A830080D01Rik | 9.25E-47 | 1.73E-45 | 1.962311648 | 476.4680166 | 1856.728571 | 314 | 22 | 34 | 58 | 139 | 26 | 6 | 0 | 0 | 0 | 1 | 0 |
| SG0000002 | Nos1 | 6.37E-44 | 1.07E-42 | 3.725986227 | 19.76070003 | 261.4784749 | 306 | 23 | 34 | 52 | 167 | 55 | 2 | 0 | 0 | 0 | 0 | 0 |
| JSG000000 | Plcl2 | 7.84E-38 | 1.07E-36 | 5.693739715 | 2.6022814 | 134.6916581 | 388 | 20 | 34 | 56 | 173 | 29 | 2 | 0 | 0 | 1 | 1 | 1 |
| SG0000001 | Psen1 | 8.19E-36 | 1.05E-34 | -2.011241532 | 5126.185684 | 1271.599369 | 188 | 28 | 34 | 48 | 79 | 10 | 2 | 0 | 0 | 0 | 1 | 0 |
| SG000000E | Ppp4r1 | 7.84E-32 | 8.55E-31 | -1.845585479 | 6023.389288 | 1675.96268 | 414 | 21 | 34 | 55 | 153 | 45 | 9 | 0 | 0 | 0 | 3 | 0 |
| SG0000005 | Prx | 4.8E-28 | 4.44E-27 | 2.232551331 | 93.15810127 | 437.8098676 | 76 | 30 | 34 | 34 | 88 | 8 | 1 | 0 | 0 | 0 | 0 | 0 |
| SG000000C | Man1a | 3.88E-24 | 3.01E-23 | -2.32023337 | 1569.796905 | 314.3284038 | 784 | 12 | 34 | 86 | 306 | 90 | 6 | 0 | 0 | 0 | 4 | 0 |
| SG0000002 | Cwc27 | 2.89E-21 | 1.95E-20 | 1.379140182 | 446.468027 | 1161.322729 | 436 | 13 | 34 | 62 | 223 | 39 | 0 | 0 | 0 | 0 | 0 | 0 |
| SG0000003 | Ccdc50 | 4.05E-21 | 2.7E-20 | -1.582332417 | 4749.268996 | 1585.978325 | 368 | 4 | 34 | 58 | 165 | 35 | 18 | 1 | 2 | 5 | 10 | 2 |
| SG0000005 | Adam12 | 7.65E-21 | 5.04E-20 | -3.671273947 | 195.6697047 | 15.35889634 | 567 | 10 | 34 | 68 | 224 | 44 | 9 | 0 | 0 | 0 | 2 | 0 |
| SG0000002 | Ttc39c | 5.38E-20 | 3.37E-19 | -2.490680398 | 439.9068908 | 78.26926499 | 360 | 16 | 34 | 53 | 167 | 36 | 4 | 0 | 0 | 0 | 2 | 0 |
| JSG000000 | Uri1 | 7.74E-18 | 4.32E-17 | 1.082659347 | 1716.961738 | 3636.415964 | 198 | 25 | 34 | 43 | 94 | 14 | 2 | 0 | 1 | 1 | 3 | 0 |
| SG0000002 | Klc1 | 2.75E-17 | 1.49E-16 | -1.315406544 | 2587.566541 | 1039.715936 | 197 | 34 | 34 | 52 | 122 | 12 | 0 | 0 | 0 | 0 | 3 | 0 |
| SG0000003 | Stx11 | 9.14E-17 | 4.82E-16 | -2.171959127 | 292.5954488 | 64.92959891 | 116 | 26 | 34 | 36 | 68 | 8 | 8 | 0 | 2 | 2 | 2 | 2 |

| | | | | | | | | | | | | | | | | | | |
|---|---|---|---|---|---|---|---|---|---|---|---|---|---|---|---|---|---|---|
| SG0000000 | Brinp2 | 1.14E-16 | 5.97E-16 | 5.487776949 | 0 | 43.87303779 | 400 | 16 | 34 | 46 | 176 | 26 | 4 | 0 | 0 | 2 | 2 | 0 |
| SG0000004 | Nrip1 | 1.44E-16 | 7.49E-16 | -1.472339036 | 1438.391309 | 518.3926518 | 340 | 23 | 34 | 48 | 136 | 33 | 7 | 1 | 2 | 1 | 3 | 5 |
| SG0000003 | Tbc1d9 | 2.61E-15 | 1.26E-14 | 4.516263655 | 23.2985738 | 533.1631994 | 357 | 22 | 34 | 68 | 212 | 32 | 7 | 0 | 0 | 4 | 6 | 0 |
| SG0000003 | Me3 | 2.8E-15 | 1.35E-14 | -4.825951902 | 73.96992022 | 2.607947655 | 445 | 12 | 34 | 56 | 221 | 53 | 18 | 1 | 1 | 1 | 2 | 2 |
| SG0000002 | Rnf220 | 2.82E-15 | 1.36E-14 | 1.025631591 | 1352.791063 | 2754.080302 | 370 | 26 | 34 | 58 | 158 | 32 | 0 | 0 | 0 | 0 | 0 | 6 |
| SG0000000 | Ugp2 | 4.31E-14 | 1.92E-13 | -1.223286521 | 1699.27145 | 727.8079991 | 261 | 10 | 34 | 33 | 95 | 34 | 3 | 0 | 0 | 0 | 0 | 0 |
| SG0000002 | Ptpn14 | 9.1E-14 | 3.97E-13 | -1.222307053 | 4179.397668 | 1791.276421 | 253 | 29 | 34 | 65 | 151 | 28 | 8 | 0 | 0 | 0 | 3 | 0 |
| SG0000002 | Zc3hav1 | 8.63E-13 | 3.53E-12 | -1.042197019 | 4891.886619 | 2375.438651 | 337 | 28 | 34 | 64 | 171 | 28 | 14 | 0 | 0 | 0 | 1 | 0 |
| SG0000003 | Chchd6 | 2.76E-12 | 1.09E-11 | 1.518485328 | 133.1594966 | 381.4887861 | 524 | 14 | 34 | 62 | 266 | 44 | 0 | 0 | 0 | 0 | 0 | 0 |
| JSG000000 | Lars | 1.64E-10 | 5.75E-10 | 0.778869482 | 4215.471517 | 7232.846295 | 163 | 30 | 34 | 51 | 109 | 12 | 0 | 0 | 0 | 0 | 0 | 0 |
| SG0000004 | Spen | 4.44E-10 | 1.51E-09 | -0.917949691 | 2971.593907 | 1572.747733 | 188 | 26 | 34 | 40 | 98 | 12 | 2 | 0 | 0 | 0 | 0 | 2 |
| SG0000007 | Fat1 | 2.89E-09 | 9.16E-09 | -0.90779219 | 11070.21713 | 5900.427248 | 453 | 6 | 34 | 50 | 190 | 34 | 4 | 0 | 0 | 2 | 2 | 2 |
| SG0000002 | Mboat2 | 8.28E-08 | 2.34E-07 | -0.868227141 | 1240.605028 | 679.6277833 | 406 | 22 | 34 | 60 | 202 | 32 | 0 | 0 | 0 | 0 | 0 | 0 |
| SG0000000 | C330007P06Rik | 1.59E-07 | 4.4E-07 | -0.866045309 | 1059.87591 | 581.4995802 | 141 | 33 | 34 | 41 | 77 | 22 | 2 | 0 | 0 | 0 | 2 | 5 |
| SG0000003 | Nfyc | 4.65E-07 | 1.23E-06 | 0.807438405 | 405.5059598 | 709.6764946 | 436 | 24 | 34 | 60 | 170 | 42 | 3 | 0 | 0 | 3 | 3 | 0 |
| SG0000002 | Tle4 | 7.03E-07 | 1.84E-06 | -0.708350982 | 4327.860514 | 2648.734097 | 361 | 26 | 34 | 59 | 194 | 37 | 3 | 0 | 0 | 1 | 5 | 2 |
| SG0000005 | Casr | 7.4E-07 | 1.93E-06 | 5.957120068 | 0.307509278 | 19.10425261 | 222 | 28 | 34 | 48 | 100 | 10 | 4 | 0 | 0 | 2 | 2 | 2 |
| SG0000005 | 43717 | 1.22E-06 | 3.13E-06 | -0.744472416 | 9384.295509 | 5601.355632 | 275 | 19 | 34 | 40 | 169 | 57 | 5 | 0 | 0 | 0 | 5 | 5 |
| SG0000003 | Itpr1 | 4.93E-06 | 1.2E-05 | -0.670196836 | 3749.115787 | 2356.022874 | 467 | 23 | 34 | 65 | 191 | 47 | 5 | 0 | 0 | 1 | 2 | 1 |
| SG0000002 | Slc30a6 | 5.47E-06 | 1.32E-05 | -0.794280657 | 743.0134068 | 428.4442784 | 130 | 30 | 34 | 40 | 94 | 12 | 6 | 0 | 0 | 0 | 0 | 0 |
| SG0000004 | Npnt | 7.96E-06 | 1.9E-05 | -0.904698787 | 401.6764788 | 214.5531669 | 506 | 12 | 34 | 36 | 182 | 38 | 14 | 0 | 2 | 0 | 2 | 2 |
| SG0000006 | Rap1a | 8.84E-06 | 2.1E-05 | -0.694569884 | 1482.872056 | 916.2570132 | 437 | 19 | 34 | 55 | 237 | 58 | 5 | 0 | 0 | 0 | 8 | 0 |
| JSG000000 | Rps6ka5 | 2.46E-05 | 5.57E-05 | -0.728339024 | 751.3120076 | 453.4907537 | 405 | 18 | 34 | 64 | 203 | 41 | 5 | 0 | 0 | 2 | 5 | 2 |
| SG0000002 | Rad50 | 4.6E-05 | 0.000101 | 0.521165637 | 2261.442489 | 3245.428436 | 203 | 32 | 34 | 46 | 98 | 11 | 3 | 0 | 0 | 0 | 0 | 0 |
| JSG000000 | Cmss1 | 0.000134 | 0.00028 | 0.667150527 | 289.2105376 | 459.2471112 | 524 | 17 | 34 | 77 | 244 | 31 | 0 | 0 | 0 | 0 | 0 | 0 |
| SG0000003 | Tlr4 | 0.000222 | 0.000452 | -2.136567172 | 54.29356324 | 12.34745745 | 269 | 19 | 34 | 49 | 127 | 20 | 5 | 0 | 0 | 1 | 12 | 1 |
| SG0000003 | Heatr3 | 0.000264 | 0.000533 | 0.483007051 | 1920.681354 | 2684.447598 | 144 | 28 | 34 | 46 | 116 | 16 | 2 | 0 | 0 | 0 | 0 | 2 |
| JSG000000 | Nfatc3 | 0.000455 | 0.000893 | 0.440822147 | 3052.971559 | 4144.035312 | 175 | 21 | 34 | 39 | 96 | 16 | 3 | 0 | 0 | 0 | 0 | 1 |
| SG0000002 | Zcchc8 | 0.00075 | 0.001429 | 0.458646711 | 1343.453047 | 1846.243205 | 199 | 24 | 34 | 53 | 116 | 9 | 12 | 0 | 0 | 3 | 7 | 4 |
| SG0000003 | Bmp2k | 0.003339 | 0.005849 | 0.560285107 | 225.0596691 | 331.864119 | 353 | 15 | 34 | 43 | 175 | 31 | 10 | 0 | 0 | 1 | 2 | 1 |
| JSG000000 | Dok5 | 0.005615 | 0.009534 | 1.748496021 | 8.007112204 | 26.90454575 | 270 | 31 | 34 | 47 | 100 | 30 | 0 | 0 | 0 | 0 | 0 | 0 |
| SG0000003 | Dhx32 | 0.016448 | 0.02588 | -0.341535568 | 2114.671005 | 1668.900174 | 295 | 27 | 34 | 53 | 188 | 15 | 0 | 0 | 0 | 0 | 0 | 0 |
| SG0000002 | Zfat | 0.021216 | 0.032664 | 0.487416938 | 172.4339884 | 241.7408208 | 476 | 14 | 34 | 68 | 198 | 46 | 0 | 0 | 0 | 0 | 0 | 0 |
| JSG000000 | Nipbl | 0.040989 | 0.060038 | -0.301019935 | 3354.071227 | 2722.42705 | 440 | 11 | 34 | 66 | 227 | 49 | 0 | 0 | 0 | 0 | 0 | 0 |
| SG0000003 | Arid2 | 0.089873 | 0.123555 | -0.2400956 | 3376.497509 | 2858.84399 | 413 | 19 | 34 | 57 | 206 | 65 | 8 | 1 | 1 | 1 | 5 | 4 |
| SG0000003 | Zdhhc2 | 0.092729 | 0.127189 | -0.355387274 | 375.4592339 | 293.4813851 | 376 | 26 | 34 | 51 | 178 | 22 | 7 | 0 | 0 | 0 | 1 | 0 |
| SG0000004 | Wdr25 | 0.122521 | 0.163577 | 0.331304817 | 182.5730674 | 229.7044452 | 328 | 18 | 34 | 54 | 122 | 30 | 0 | 0 | 0 | 0 | 0 | 0 |
| SG0000002 | Radil | 0.133598 | 0.17688 | 0.360771949 | 434.2835555 | 557.6690767 | 188 | 22 | 34 | 40 | 99 | 18 | 8 | 0 | 4 | 8 | 8 | 0 |
| SG0000004 | Dpy19l1 | 0.158667 | 0.206842 | -0.198887899 | 2535.160856 | 2208.687621 | 539 | 13 | 34 | 96 | 275 | 44 | 10 | 0 | 0 | 0 | 8 | 2 |
| SG0000003 | Ppip5k1 | 0.183204 | 0.235659 | -0.211622255 | 870.1534818 | 751.4346474 | 315 | 23 | 34 | 65 | 260 | 29 | 3 | 0 | 0 | 0 | 1 | 0 |
| JSG000000 | Pias1 | 0.260651 | 0.322318 | 0.161066248 | 1005.662768 | 1124.444809 | 223 | 22 | 34 | 44 | 122 | 33 | 1 | 0 | 0 | 1 | 3 | 3 |
| SG0000000 | Wars2 | 0.4 | 0.469024 | 0.197629456 | 142.668883 | 163.6144494 | 578 | 16 | 34 | 98 | 250 | 47 | 10 | 2 | 2 | 5 | 7 | 0 |
| SG0000005 | Grb2 | 0.428848 | 0.498121 | -0.114139487 | 2592.688965 | 2395.471509 | 369 | 26 | 34 | 82 | 182 | 31 | 7 | 0 | 4 | 0 | 0 | 7 |
| SG0000003 | Ppfibp2 | 0.491729 | 0.559549 | -0.111714439 | 709.2227981 | 656.3769034 | 428 | 21 | 34 | 69 | 194 | 42 | 0 | 0 | 0 | 0 | 0 | 0 |
| SG0000002 | Matn2 | 0.679914 | 0.73667 | -0.127334384 | 99.70682002 | 91.2837339 | 420 | 24 | 34 | 70 | 200 | 28 | 2 | 0 | 0 | 0 | 0 | 0 |
| SG0000006 | Pym1 | 0.85593 | 0.887645 | 0.029023542 | 838.5902509 | 855.6315024 | 106 | 32 | 34 | 46 | 70 | 0 | 0 | 0 | 0 | 0 | 0 | 0 |
| SG0000002 | Agfg2 | 0.883831 | 0.910518 | 0.040424003 | 538.1033836 | 553.3941456 | 128 | 30 | 34 | 42 | 80 | 6 | 2 | 0 | 0 | 2 | 4 | 0 |
| SG0000003 | Atr | 0.965223 | 0.979329 | -0.009405665 | 2149.441336 | 2135.473612 | 301 | 27 | 34 | 57 | 160 | 28 | 0 | 0 | 0 | 0 | 0 | 0 |
| SG0000005 | Shisa6 | 2.4E-136 | 3.2E-134 | -8.480386856 | 3277.519272 | 9.17686696 | 544 | 21 | 33 | 86 | 273 | 51 | 5 | 0 | 0 | 2 | 6 | 0 |
| JSG000000 | Pced1b | 3.8E-127 | 4.2E-125 | 3.845383643 | 95.23454744 | 1368.895651 | 213 | 26 | 33 | 46 | 90 | 18 | 0 | 0 | 0 | 0 | 0 | 0 |
| SG0000002 | Runx1 | 1.4E-104 | 1.1E-102 | -5.164693511 | 2551.376674 | 71.12903479 | 555 | 18 | 33 | 63 | 210 | 39 | 24 | 0 | 4 | 4 | 14 | 1 |
| SG0000003 | Acsl3 | 3.06E-92 | 1.82E-90 | 2.61071168 | 579.7307683 | 3541.024049 | 243 | 9 | 33 | 30 | 97 | 72 | 9 | 0 | 0 | 0 | 1 | 0 |
| SG0000003 | Ugt8a | 7.39E-67 | 2.45E-65 | 7.667673134 | 0 | 202.3291337 | 348 | 14 | 33 | 34 | 165 | 22 | 15 | 2 | 4 | 2 | 13 | 1 |
| SG0000002 | Zhx1 | 6.8E-63 | 2.05E-61 | -3.88274715 | 1583.07382 | 107.3192945 | 175 | 18 | 33 | 45 | 108 | 30 | 10 | 0 | 0 | 3 | 6 | 3 |
| SG0000003 | Dna2 | 7.64E-51 | 1.62E-49 | 1.993810723 | 574.1628258 | 2286.81959 | 101 | 28 | 33 | 34 | 59 | 8 | 5 | 0 | 1 | 0 | 1 | 0 |
| SG0000002 | Lpin2 | 1.86E-44 | 3.21E-43 | -2.432594188 | 3553.85786 | 658.2884509 | 304 | 12 | 33 | 22 | 113 | 29 | 3 | 0 | 2 | 0 | 0 | 0 |
| SG0000003 | Itgam | 2.24E-37 | 3E-36 | -4.880783038 | 229.2486773 | 7.781167101 | 342 | 20 | 33 | 70 | 214 | 12 | 10 | 0 | 0 | 0 | 6 | 0 |
| SG0000003 | Slc5a11 | 1.49E-33 | 1.73E-32 | 6.562127163 | 0 | 93.49245041 | 111 | 30 | 33 | 35 | 68 | 9 | 0 | 0 | 0 | 0 | 0 | 0 |
| SG0000000 | Homer3 | 6.43E-30 | 6.48E-29 | -2.168177497 | 1696.931542 | 377.552943 | 62 | 27 | 33 | 30 | 42 | 3 | 3 | 0 | 0 | 0 | 0 | 0 |
| SG0000004 | Dok7 | 8.78E-26 | 7.34E-25 | -3.896068327 | 181.367625 | 12.18221709 | 70 | 33 | 33 | 36 | 40 | 6 | 1 | 0 | 0 | 0 | 1 | 0 |
| SG0000002 | Evl | 1.12E-25 | 9.29E-25 | 3.513028113 | 74.71028157 | 852.9178644 | 474 | 25 | 33 | 76 | 194 | 37 | 3 | 0 | 0 | 0 | 3 | 0 |
| SG0000003 | Rpap1 | 1.44E-21 | 9.82E-21 | 1.238332594 | 1177.103821 | 2777.090449 | 87 | 24 | 33 | 30 | 72 | 18 | 0 | 0 | 0 | 0 | 0 | 0 |
| SG0000000 | Tpr | 2.56E-21 | 1.73E-20 | 1.182305835 | 5753.200615 | 13056.26747 | 355 | 15 | 33 | 47 | 133 | 31 | 2 | 0 | 0 | 0 | 0 | 0 |
| SG0000004 | Bend6 | 2.87E-20 | 1.83E-19 | 3.104212115 | 15.04216629 | 129.3514629 | 327 | 27 | 33 | 60 | 192 | 27 | 6 | 0 | 0 | 0 | 0 | 0 |
| SG0000002 | Lypd6b | 5.29E-20 | 3.32E-19 | -2.67466643 | 356.5593542 | 55.84406329 | 290 | 16 | 33 | 40 | 129 | 29 | 1 | 0 | 0 | 0 | 0 | 0 |
| JSG000000 | Espl1 | 1.31E-19 | 8.01E-19 | 1.132180186 | 1906.992493 | 4179.931647 | 64 | 30 | 33 | 35 | 52 | 1 | 1 | 0 | 0 | 0 | 0 | 0 |
| SG0000003 | Mdfi | 8.23E-19 | 4.83E-18 | 3.155099929 | 26.80079109 | 238.7411775 | 72 | 33 | 33 | 42 | 60 | 9 | 0 | 0 | 0 | 0 | 0 | 3 |
| SG0000004 | Glt1d1 | 2.41E-18 | 1.38E-17 | 1.774126107 | 286.770362 | 980.8428688 | 139 | 26 | 33 | 49 | 109 | 13 | 2 | 1 | 2 | 2 | 2 | 0 |
| SG0000002 | Tnks2 | 4.39E-13 | 1.83E-12 | 0.860455322 | 8927.460397 | 16208.79741 | 204 | 28 | 33 | 46 | 103 | 17 | 11 | 0 | 0 | 2 | 4 | 2 |
| SG0000005 | Ptk2b | 1.28E-12 | 5.19E-12 | 1.132488762 | 424.2762375 | 930.1688596 | 519 | 15 | 33 | 63 | 230 | 67 | 12 | 0 | 4 | 0 | 0 | 0 |
| SG0000006 | Armt1 | 2.01E-12 | 7.99E-12 | 0.941055942 | 1013.903418 | 1946.626597 | 153 | 26 | 33 | 36 | 81 | 6 | 9 | 0 | 0 | 0 | 1 | 0 |
| SG0000000 | Tgfbr1 | 6.1E-10 | 2.05E-09 | 0.927929738 | 499.5128046 | 950.3450362 | 435 | 15 | 33 | 46 | 153 | 61 | 7 | 0 | 0 | 0 | 3 | 2 |
| SG0000006 | Arfgef1 | 1.54E-09 | 4.98E-09 | -0.932917719 | 6051.603355 | 3169.817349 | 447 | 23 | 33 | 71 | 198 | 45 | 4 | 0 | 0 | 0 | 0 | 0 |
| SG0000003 | Gnao1 | 2.28E-09 | 7.27E-09 | 3.724032126 | 10.96039933 | 144.8344092 | 486 | 27 | 33 | 78 | 284 | 46 | 8 | 0 | 0 | 0 | 0 | 0 |
| SG0000003 | Cfap53 | 1.22E-08 | 3.69E-08 | 6.143749711 | 0.34247518 | 24.21486496 | 227 | 24 | 33 | 51 | 144 | 27 | 2 | 0 | 0 | 0 | 0 | 0 |
| SG0000002 | Slc38a1 | 3.51E-07 | 9.39E-07 | 0.655309561 | 1984.769628 | 3125.92058 | 297 | 27 | 33 | 60 | 156 | 48 | 7 | 0 | 0 | 5 | 7 | 10 |
| SG0000002 | Marveld2 | 1.24E-05 | 2.89E-05 | 1.615952377 | 65.1467213 | 199.6837391 | 135 | 27 | 33 | 33 | 63 | 11 | 6 | 0 | 0 | 0 | 0 | 2 |
| SG0000003 | Prmt3 | 1.33E-05 | 3.09E-05 | 0.594209983 | 1186.978112 | 1791.91638 | 290 | 15 | 33 | 40 | 95 | 14 | 0 | 0 | 0 | 0 | 1 | 0 |
| SG0000007 | Kifc1 | 3.14E-05 | 7.05E-05 | 0.627882776 | 949.7560699 | 1467.653813 | 75 | 33 | 33 | 41 | 49 | 0 | 3 | 0 | 0 | 3 | 6 | 0 |
| JSG000000 | Epm2a | 3.29E-05 | 7.37E-05 | 1.423307732 | 36.04143397 | 96.66302508 | 237 | 25 | 33 | 44 | 126 | 21 | 3 | 0 | 0 | 0 | 2 | 0 |
| SG0000003 | Sfxn5 | 3.72E-05 | 8.29E-05 | 1.079043579 | 83.57378964 | 176.5609185 | 303 | 19 | 33 | 34 | 121 | 12 | 4 | 0 | 0 | 0 | 0 | 0 |
| JSG000000 | 8030462N17Rik | 4.81E-05 | 0.000106 | 0.576550199 | 837.3830187 | 1248.771716 | 219 | 25 | 33 | 43 | 132 | 11 | 6 | 0 | 0 | 1 | 2 | 0 |
| SG0000001 | Kitl | 7.99E-05 | 0.000172 | 1.818755474 | 33.84422258 | 119.3945478 | 384 | 20 | 33 | 49 | 147 | 18 | 14 | 0 | 1 | 3 | 3 | 4 |
| SG0000007 | Mob3b | 9.4E-05 | 0.0002 | 1.091500919 | 131.4258663 | 280.0627605 | 304 | 26 | 33 | 48 | 153 | 41 | 8 | 1 | 1 | 2 | 7 | 1 |
| JSG000000 | Adamts2 | 0.000304 | 0.00061 | -1.017089348 | 141.7741372 | 70.0523342 | 208 | 27 | 33 | 46 | 106 | 27 | 5 | 0 | 1 | 1 | 1 | 1 |
| SG0000003 | Usp53 | 0.000388 | 0.000769 | 0.716856721 | 438.2330088 | 720.278518 | 293 | 17 | 33 | 66 | 133 | 20 | 18 | 3 | 4 | 4 | 6 | 1 |
| SG0000006 | Fam78b | 0.000397 | 0.000784 | 1.900568056 | 9.462505974 | 35.32922905 | 622 | 21 | 33 | 90 | 238 | 52 | 48 | 15 | 15 | 30 | 35 | 1 |
| SG0000003 | Rnf141 | 0.000442 | 0.000869 | 0.524095491 | 628.4100116 | 903.6735252 | 260 | 13 | 33 | 46 | 108 | 20 | 7 | 0 | 2 | 3 | 4 | 0 |
| SG0000003 | Dctn1 | 0.000554 | 0.001074 | -0.770103326 | 2954.860269 | 1732.657599 | 98 | 33 | 33 | 37 | 63 | 4 | 0 | 0 | 0 | 0 | 0 | 0 |
| SG0000001 | Evi5 | 0.000563 | 0.001091 | -0.493842662 | 2744.569943 | 1949.004529 | 298 | 20 | 33 | 52 | 170 | 29 | 12 | 0 | 1 | 1 | 5 | 2 |
| SG0000002 | Rrm2b | 0.000774 | 0.001473 | -0.64540487 | 639.0990176 | 408.5845351 | 273 | 13 | 33 | 52 | 148 | 35 | 18 | 1 | 4 | 4 | 10 | 0 |
| SG0000002 | Kdm5c | 0.000798 | 0.001514 | -0.453771077 | 5358.306254 | 3912.270066 | 274 | 27 | 33 | 57 | 153 | 21 | 0 | 0 | 0 | 0 | 2 | 0 |
| SG0000002 | Epb41l4b | 0.000813 | 0.001543 | 0.576981508 | 311.6571715 | 464.9067651 | 321 | 26 | 33 | 57 | 171 | 22 | 15 | 1 | 1 | 3 | 9 | 0 |
| SG0000004 | Foxo3 | 0.001731 | 0.003163 | 0.407420225 | 1995.188756 | 2646.242847 | 336 | 10 | 33 | 33 | 104 | 31 | 6 | 0 | 0 | 0 | 1 | 2 |
| SG0000002 | Zdhhc21 | 0.002142 | 0.003861 | -0.47413119 | 1504.036559 | 1082.756209 | 356 | 18 | 33 | 59 | 153 | 33 | 30 | 0 | 1 | 3 | 9 | 3 |
| SG0000002 | Snap23 | 0.006167 | 0.010421 | 0.384994648 | 986.3411071 | 1288.01841 | 219 | 24 | 33 | 39 | 114 | 24 | 15 | 0 | 0 | 0 | 3 | 0 |
| SG0000003 | Fam83b | 0.017402 | 0.027242 | 1.043308392 | 25.51738028 | 52.59000432 | 303 | 20 | 33 | 58 | 185 | 42 | 1 | 0 | 0 | 0 | 1 | 2 |
| SG0000007 | Rnls | 0.032441 | 0.048403 | 1.265291103 | 10.11249043 | 24.30797041 | 590 | 9 | 33 | 74 | 269 | 66 | 1 | 0 | 0 | 0 | 1 | 0 |
| SG0000002 | Nsun4 | 0.128364 | 0.170606 | 0.287386584 | 274.4859546 | 334.990205 | 107 | 25 | 33 | 31 | 56 | 6 | 7 | 0 | 0 | 1 | 3 | 0 |
| SG0000002 | Tiam2 | 0.261941 | 0.323668 | -0.349010352 | 95.72979763 | 75.15961663 | 477 | 17 | 33 | 61 | 180 | 52 | 9 | 0 | 1 | 0 | 0 | 0 |
| SG0000002 | Stil | 0.294022 | 0.358341 | 0.154895817 | 700.2031129 | 779.5649852 | 256 | 19 | 33 | 58 | 121 | 19 | 5 | 0 | 0 | 0 | 0 | 1 |
| SG0000002 | Dpy30 | 0.4622 | 0.531654 | 0.093186767 | 2406.468023 | 2567.036874 | 116 | 25 | 33 | 44 | 104 | 4 | 4 | 0 | 1 | 0 | 0 | 0 |
| SG0000004 | Scml4 | 0.508592 | 0.575519 | -0.39256134 | 27.08535858 | 20.63297495 | 453 | 15 | 33 | 75 | 212 | 43 | 8 | 0 | 0 | 0 | 0 | 0 |
| SG0000003 | Ergic2 | 0.587335 | 0.65077 | -0.078776678 | 2122.985499 | 2010.170451 | 233 | 21 | 33 | 47 | 83 | 31 | 12 | 0 | 0 | 1 | 2 | 0 |
| SG0000003 | Cln3 | 0.670078 | 0.727509 | 0.071998515 | 569.9007076 | 599.0635682 | 99 | 25 | 33 | 44 | 63 | 13 | 6 | 0 | 0 | 3 | 3 | 0 |
| JSG000000 | Mpp5 | 0.829787 | 0.866555 | 0.023355234 | 1298.62798 | 1319.822056 | 278 | 26 | 33 | 51 | 144 | 24 | 7 | 0 | 0 | 0 | 2 | 3 |
| SG0000002 | Ylpm1 | 0.844656 | 0.878447 | -0.039299921 | 2528.476783 | 2460.529154 | 604 | 10 | 33 | 87 | 297 | 78 | 0 | 0 | 0 | 0 | 0 | 0 |
| SG0000006 | Stat4 | 3.2E-151 | 4.9E-149 | 9.113780576 | 0 | 553.0147499 | 411 | 16 | 32 | 45 | 214 | 48 | 2 | 0 | 0 | 0 | 0 | 2 |
| SG0000002 | Dhtkd1 | 5.7E-109 | 5E-107 | 4.314424896 | 39.64554928 | 788.7979023 | 161 | 26 | 32 | 40 | 94 | 16 | 7 | 0 | 0 | 0 | 2 | 0 |
| SG0000001 | Dusp14 | 2.92E-87 | 1.54E-85 | -4.066329782 | 3965.642075 | 236.7152715 | 163 | 32 | 32 | 48 | 100 | 4 | 0 | 0 | 0 | 0 | 0 | 0 |
| SG0000003 | Parva | 1.67E-56 | 4.18E-55 | -3.419265478 | 5933.318045 | 554.6208045 | 646 | 22 | 32 | 75 | 224 | 58 | 11 | 1 | 1 | 1 | 1 | 0 |
| SG0000005 | Slc9b1 | 5.23E-54 | 1.21E-52 | 7.315076787 | 2.316520345 | 368.8872562 | 139 | 25 | 32 | 44 | 94 | 18 | 1 | 0 | 0 | 1 | 1 | 0 |
| SG0000003 | Mobp | 2.08E-43 | 3.45E-42 | 7.063652822 | 0 | 132.7738968 | 206 | 28 | 32 | 33 | 68 | 10 | 5 | 0 | 0 | 0 | 0 | 1 |
| SG0000002 | Klf7 | 1.33E-41 | 2.08E-40 | -2.324079639 | 3602.384248 | 719.4031767 | 346 | 24 | 32 | 65 | 162 | 13 | 19 | 0 | 0 | 1 | 3 | 0 |
| SG0000003 | Map4k5 | 8.08E-41 | 1.23E-39 | -2.214839161 | 5253.96014 | 1131.758462 | 728 | 12 | 32 | 124 | 376 | 88 | 16 | 4 | 4 | 4 | 8 | 0 |
| JSG000000 | Ptpn9 | 1.17E-38 | 1.64E-37 | -2.670822112 | 2061.800314 | 323.7792648 | 138 | 25 | 32 | 38 | 102 | 12 | 5 | 0 | 0 | 0 | 0 | 1 |
| JSG000000 | Dsp | 5.75E-36 | 7.39E-35 | -2.57389557 | 9297.531766 | 1561.521249 | 174 | 22 | 32 | 44 | 74 | 16 | 4 | 0 | 0 | 0 | 0 | 0 |
| SG0000002 | Suv39h2 | 6.37E-33 | 7.24E-32 | 1.706230432 | 415.0470631 | 1354.328085 | 169 | 24 | 32 | 53 | 90 | 12 | 13 | 0 | 0 | 1 | 10 | 0 |
| SG0000001 | Kcnq2 | 4.45E-32 | 4.91E-31 | 6.490795099 | 0 | 88.93402292 | 243 | 24 | 32 | 40 | 141 | 32 | 3 | 0 | 0 | 0 | 0 | 0 |
| SG0000002 | Sumo1 | 1.33E-29 | 1.31E-28 | 1.420958054 | 1361.743592 | 3646.24912 | 168 | 26 | 32 | 36 | 112 | 26 | 0 | 0 | 0 | 0 | 0 | 0 |
| SG0000002 | Wif1 | 2.98E-28 | 2.79E-27 | -5.002692928 | 425.3800354 | 13.26833636 | 252 | 26 | 32 | 60 | 152 | 38 | 2 | 0 | 0 | 0 | 0 | 0 |
| SG0000003 | Greb1 | 5.47E-26 | 4.62E-25 | 3.086504945 | 21.06301251 | 178.9167728 | 349 | 13 | 32 | 40 | 178 | 51 | 14 | 0 | 2 | 0 | 0 | 2 |
| SG0000005 | Aldh7a1 | 9.64E-25 | 7.69E-24 | 1.370610753 | 810.3873119 | 2095.499597 | 235 | 28 | 32 | 48 | 136 | 5 | 9 | 0 | 0 | 0 | 0 | 0 |

| | | | | | | | | | | | | | | | | | | |
|---|---|---|---|---|---|---|---|---|---|---|---|---|---|---|---|---|---|---|
| JSG000000 | Trio | 6.87E-24 | 5.27E-23 | -1.876279882 | 1471.24976 | 400.7466314 | 386 | 21 | 32 | 53 | 163 | 42 | 6 | 1 | 1 | 3 | 8 | 2 |
| SG0000003 | Ets1 | 1.19E-21 | 8.16E-21 | -2.545508024 | 1427.644272 | 244.537652 | 342 | 28 | 32 | 60 | 147 | 25 | 6 | 0 | 0 | 0 | 4 | 1 |
| JSG000000 | Ak7 | 1.91E-20 | 1.23E-19 | 3.546274429 | 9.933970651 | 116.0533947 | 109 | 30 | 32 | 39 | 73 | 1 | 0 | 0 | 0 | 0 | 0 | 0 |
| SG0000005 | Lgr4 | 4.71E-20 | 2.96E-19 | 1.292270197 | 918.1387728 | 2248.642526 | 450 | 26 | 32 | 60 | 168 | 42 | 16 | 0 | 0 | 0 | 0 | 4 |
| SG0000002 | Ubr2 | 3.54E-18 | 2.01E-17 | -1.349265448 | 4512.803797 | 1771.23865 | 240 | 18 | 32 | 40 | 100 | 20 | 3 | 0 | 0 | 0 | 0 | 0 |
| JSG000000 | Pgap1 | 6.9E-17 | 3.67E-16 | -1.726624411 | 4011.175514 | 1212.009228 | 236 | 17 | 32 | 43 | 112 | 14 | 19 | 1 | 3 | 3 | 7 | 3 |
| SG0000002 | Tsc1 | 8.9E-15 | 4.14E-14 | 1.068505234 | 770.3641684 | 1615.653406 | 440 | 21 | 32 | 52 | 196 | 52 | 18 | 5 | 5 | 5 | 9 | 0 |
| SG0000003 | Zyg11a | 2.1E-14 | 9.54E-14 | 6.780221554 | 2.022228746 | 222.2697416 | 138 | 26 | 32 | 40 | 100 | 10 | 1 | 0 | 0 | 0 | 0 | 0 |
| SG0000002 | Anxa3 | 5.85E-14 | 2.58E-13 | -3.048510097 | 4799.676865 | 580.1215764 | 176 | 28 | 32 | 39 | 75 | 8 | 2 | 0 | 0 | 0 | 0 | 0 |
| JSG000000 | Gm5134 | 8.85E-14 | 3.87E-13 | 5.198242456 | 0 | 35.71359425 | 129 | 30 | 32 | 38 | 76 | 7 | 1 | 0 | 0 | 0 | 0 | 0 |
| SG0000005 | Tmem40 | 2.66E-13 | 1.12E-12 | 2.950978581 | 10.9627429 | 84.77197443 | 130 | 32 | 32 | 44 | 76 | 8 | 0 | 0 | 0 | 0 | 0 | 0 |
| SG0000000 | Syn2 | 2.46E-12 | 9.73E-12 | 4.370107046 | 2.270680331 | 46.95580639 | 307 | 16 | 32 | 46 | 132 | 20 | 1 | 0 | 0 | 0 | 0 | 0 |
| SG0000002 | Mark2 | 6.82E-12 | 2.64E-11 | -1.116268723 | 3423.785663 | 1579.342055 | 445 | 26 | 32 | 69 | 316 | 43 | 2 | 0 | 0 | 0 | 3 | 0 |
| SG0000003 | Rnf128 | 1.61E-11 | 6.08E-11 | -1.321695364 | 1490.818887 | 596.4237601 | 383 | 18 | 32 | 58 | 221 | 40 | 8 | 0 | 0 | 0 | 2 | 4 |
| SG0000002 | Tbx15 | 2.11E-11 | 7.87E-11 | -1.867201793 | 207.3872862 | 56.84579974 | 220 | 25 | 32 | 49 | 118 | 14 | 2 | 0 | 0 | 1 | 3 | 1 |
| SG0000004 | Insc | 2.74E-11 | 1.01E-10 | -2.834558689 | 92.12782919 | 12.91527471 | 297 | 19 | 32 | 71 | 142 | 32 | 6 | 0 | 2 | 2 | 2 | 0 |
| SG0000003 | Atcay | 4.1E-11 | 1.5E-10 | 3.838239493 | 4.306126756 | 61.5902586 | 74 | 28 | 32 | 44 | 64 | 6 | 0 | 0 | 0 | 0 | 0 | 0 |
| SG0000002 | Epb41l5 | 7.5E-11 | 2.69E-10 | 1.264388024 | 716.5606968 | 1721.359992 | 639 | 14 | 32 | 55 | 250 | 90 | 13 | 0 | 0 | 1 | 6 | 2 |
| SG0000004 | 1190005I06Rik | 1.55E-10 | 5.44E-10 | 5.264825977 | 0.957493736 | 36.81344685 | 79 | 28 | 32 | 38 | 67 | 3 | 1 | 0 | 0 | 0 | 0 | 0 |
| SG0000002 | Tecpr2 | 2.71E-10 | 9.35E-10 | -1.025920168 | 1299.436323 | 638.1492434 | 276 | 16 | 32 | 28 | 118 | 22 | 6 | 1 | 1 | 1 | 2 | 0 |
| JSG000000 | Vps35 | 1.34E-09 | 4.37E-09 | 0.736825094 | 6229.177308 | 10380.95116 | 123 | 22 | 32 | 32 | 60 | 8 | 2 | 0 | 0 | 0 | 1 | 0 |
| JSG000000 | Slc5a4b | 1.42E-09 | 4.61E-09 | 6.341512093 | 0.331601068 | 26.8906487 | 105 | 28 | 32 | 35 | 72 | 11 | 4 | 0 | 0 | 0 | 0 | 1 |
| SG0000004 | Flt3 | 2.59E-09 | 8.25E-09 | -3.653864985 | 50.79883607 | 4.035810827 | 124 | 29 | 32 | 41 | 81 | 25 | 1 | 0 | 0 | 0 | 1 | 1 |
| SG0000002 | Nusap1 | 4.63E-09 | 1.45E-08 | 0.74246293 | 2223.939902 | 3720.71718 | 126 | 32 | 32 | 46 | 98 | 14 | 1 | 0 | 0 | 0 | 4 | 2 |
| SG0000002 | Spryd7 | 6.08E-09 | 1.88E-08 | 0.906995434 | 398.8223527 | 747.8462542 | 136 | 24 | 32 | 39 | 101 | 12 | 20 | 2 | 6 | 5 | 11 | 4 |
| SG0000002 | Xpo1 | 9.84E-09 | 3E-08 | 0.688785891 | 8143.255352 | 13126.32845 | 425 | 32 | 32 | 87 | 267 | 32 | 9 | 0 | 0 | 3 | 3 | 0 |
| SG0000003 | Iffo1 | 1.1E-08 | 3.33E-08 | -1.228521137 | 367.277642 | 156.7374507 | 84 | 28 | 32 | 32 | 60 | 0 | 0 | 0 | 0 | 0 | 0 | 0 |
| SG0000002 | Fam69a | 1.89E-08 | 5.61E-08 | 1.464370061 | 122.7786469 | 338.7990281 | 179 | 23 | 32 | 36 | 104 | 11 | 7 | 0 | 0 | 0 | 0 | 0 |
| JSG000000 | Gpc4 | 8.91E-07 | 2.31E-06 | 1.745115847 | 205.3681352 | 688.4386992 | 224 | 26 | 32 | 42 | 131 | 18 | 1 | 0 | 0 | 1 | 1 | 0 |
| SG0000002 | Mff | 1.27E-06 | 3.24E-06 | 0.626427745 | 1909.906777 | 2948.395307 | 404 | 7 | 32 | 41 | 164 | 33 | 11 | 0 | 0 | 0 | 0 | 7 |
| SG0000002 | Aff1 | 1.32E-06 | 3.36E-06 | -0.759782263 | 2255.021662 | 1331.782999 | 334 | 24 | 32 | 54 | 162 | 24 | 18 | 0 | 0 | 2 | 10 | 2 |
| SG0000002 | Dlg5 | 5.18E-06 | 1.25E-05 | -0.650333524 | 3620.893707 | 2306.990883 | 227 | 23 | 32 | 41 | 108 | 23 | 4 | 0 | 0 | 0 | 4 | 0 |
| SG0000003 | Plod2 | 6.49E-06 | 1.56E-05 | -0.687465789 | 5007.020118 | 3109.07725 | 337 | 16 | 32 | 56 | 166 | 22 | 11 | 0 | 0 | 4 | 4 | 0 |
| SG0000004 | Fbxl4 | 8.44E-06 | 2E-05 | -0.855666469 | 657.6048981 | 363.3990319 | 699 | 12 | 32 | 79 | 355 | 68 | 17 | 0 | 0 | 3 | 8 | 0 |
| SG0000001 | Calr3 | 1.48E-05 | 3.44E-05 | 1.199900847 | 55.5078424 | 127.5147704 | 73 | 30 | 32 | 31 | 48 | 5 | 0 | 0 | 0 | 0 | 0 | 0 |
| SG0000004 | Prrc2c | 2.36E-05 | 5.36E-05 | 0.701432965 | 8693.61022 | 14136.846 | 594 | 17 | 32 | 76 | 267 | 68 | 13 | 0 | 0 | 0 | 8 | 2 |
| SG0000003 | Smim14 | 5.13E-05 | 0.000113 | -0.644516727 | 1155.378201 | 739.103573 | 186 | 24 | 32 | 38 | 82 | 14 | 6 | 0 | 0 | 2 | 2 | 0 |
| SG0000003 | Tgfb2 | 0.00027 | 0.000544 | 1.04762341 | 73.65507396 | 152.2540103 | 240 | 26 | 32 | 51 | 110 | 20 | 6 | 2 | 2 | 3 | 4 | 0 |
| SG0000000 | Sfmbt1 | 0.00036 | 0.000715 | -0.573987994 | 982.3890293 | 659.927005 | 520 | 16 | 32 | 58 | 237 | 38 | 15 | 4 | 4 | 4 | 7 | 0 |
| JSG000000 | Taf4b | 0.000413 | 0.000814 | 0.511438345 | 734.5514225 | 1047.081501 | 323 | 15 | 32 | 49 | 166 | 29 | 6 | 1 | 1 | 1 | 4 | 1 |
| SG0000003 | Slc18a1 | 0.000651 | 0.00125 | -1.958410782 | 36.2018958 | 9.315172998 | 117 | 30 | 32 | 42 | 71 | 8 | 1 | 0 | 0 | 0 | 1 | 0 |
| SG0000002 | Ccdc150 | 0.000871 | 0.001647 | 1.56987805 | 14.9831086 | 44.48179556 | 470 | 10 | 32 | 58 | 186 | 46 | 2 | 0 | 0 | 0 | 0 | 0 |
| JSG000000 | Col8a1 | 0.002499 | 0.004465 | -1.292466472 | 1001.670432 | 408.9344305 | 282 | 23 | 32 | 58 | 154 | 35 | 11 | 1 | 1 | 2 | 3 | 0 |
| SG0000001 | Wt1 | 0.004083 | 0.007066 | 5.189633415 | 0.615018556 | 22.44520332 | 264 | 24 | 32 | 39 | 163 | 26 | 12 | 0 | 0 | 3 | 9 | 6 |
| SG0000002 | Tnk2 | 0.005474 | 0.009313 | -0.665615997 | 2995.672993 | 1888.530589 | 160 | 28 | 32 | 36 | 87 | 9 | 0 | 0 | 0 | 0 | 5 | 0 |
| JSG000000 | Cep83 | 0.007293 | 0.012184 | 0.430274064 | 989.9119348 | 1333.895969 | 273 | 19 | 32 | 52 | 146 | 30 | 3 | 0 | 0 | 0 | 2 | 0 |
| SG0000006 | Ppm1b | 0.011928 | 0.019265 | 0.322168673 | 2265.024018 | 2831.752196 | 514 | 25 | 32 | 75 | 215 | 64 | 16 | 1 | 1 | 5 | 7 | 0 |
| SG0000003 | Mgat5 | 0.020137 | 0.031137 | -0.355417218 | 1100.069257 | 859.8620552 | 576 | 22 | 32 | 75 | 227 | 75 | 11 | 2 | 3 | 3 | 3 | 1 |
| SG0000003 | Pank1 | 0.028866 | 0.043464 | -0.467186039 | 331.9268424 | 240.1072995 | 258 | 20 | 32 | 56 | 126 | 18 | 8 | 0 | 0 | 0 | 0 | 2 |
| SG0000005 | Uba2 | 0.030438 | 0.045624 | 0.265090613 | 9174.001493 | 11024.50332 | 128 | 29 | 32 | 44 | 73 | 10 | 3 | 0 | 1 | 1 | 2 | 2 |
| SG0000006 | Amn1 | 0.06946 | 0.097628 | -0.395994571 | 335.231476 | 254.7642852 | 162 | 30 | 32 | 47 | 88 | 10 | 12 | 2 | 2 | 5 | 6 | 2 |
| SG0000002 | Cwc22 | 0.076251 | 0.106364 | 0.243571614 | 1204.119793 | 1425.58152 | 567 | 12 | 32 | 83 | 245 | 116 | 5 | 0 | 0 | 0 | 0 | 0 |
| SG0000005 | Pitpnb | 0.080177 | 0.111285 | -0.240272385 | 3916.466396 | 3315.623504 | 188 | 24 | 32 | 42 | 92 | 30 | 8 | 0 | 0 | 4 | 4 | 6 |
| SG0000002 | Soat1 | 0.0803 | 0.11144 | -0.244597913 | 5742.571934 | 4847.023613 | 179 | 28 | 32 | 52 | 140 | 39 | 8 | 0 | 0 | 1 | 7 | 0 |
| SG0000004 | Rhbdl2 | 0.105529 | 0.142972 | 2.017768 | 4.419555006 | 17.89728838 | 117 | 25 | 32 | 32 | 72 | 11 | 0 | 0 | 0 | 0 | 0 | 0 |
| SG0000002 | Kcnab1 | 0.127355 | 0.169353 | -1.140504964 | 24.8399636 | 11.26742694 | 525 | 23 | 32 | 65 | 229 | 58 | 6 | 0 | 0 | 0 | 1 | 2 |
| SG0000003 | Cbfb | 0.13141 | 0.174331 | -0.214461542 | 2552.630618 | 2200.030072 | 448 | 20 | 32 | 44 | 188 | 52 | 16 | 0 | 0 | 0 | 8 | 8 |
| SG0000003 | St5 | 0.138758 | 0.183103 | -0.387197396 | 1006.460894 | 769.5536012 | 452 | 12 | 32 | 56 | 192 | 48 | 0 | 0 | 0 | 0 | 0 | 3 |
| SG0000003 | Slco2a1 | 0.217206 | 0.274314 | -0.81955488 | 308.4119048 | 174.7513469 | 212 | 28 | 32 | 47 | 116 | 9 | 4 | 0 | 0 | 0 | 3 | 1 |
| SG0000002 | Ccdc88c | 0.217775 | 0.274897 | 0.254292811 | 827.6650802 | 987.1982943 | 222 | 24 | 32 | 38 | 104 | 20 | 0 | 0 | 0 | 0 | 0 | 0 |
| SG0000004 | Yeats2 | 0.242888 | 0.30303 | -0.171726205 | 1934.184931 | 1717.130319 | 370 | 14 | 32 | 44 | 174 | 36 | 2 | 0 | 0 | 0 | 2 | 0 |
| SG0000003 | Mier3 | 0.299995 | 0.364752 | 0.155992991 | 683.2145394 | 761.2296038 | 262 | 12 | 32 | 36 | 102 | 22 | 43 | 0 | 4 | 8 | 18 | 6 |
| SG0000003 | Pde4a | 0.360459 | 0.428693 | -0.260824661 | 660.5128615 | 551.2711083 | 118 | 29 | 32 | 39 | 73 | 3 | 1 | 0 | 0 | 0 | 0 | 1 |
| JSG000000 | Vcl | 0.471686 | 0.540518 | 0.093250862 | 4607.194604 | 4914.822789 | 247 | 17 | 32 | 50 | 141 | 24 | 5 | 0 | 0 | 1 | 2 | 0 |
| SG0000007 | Kiz | 0.571395 | 0.636023 | -0.104939014 | 520.154836 | 483.6629826 | 239 | 18 | 32 | 34 | 99 | 16 | 0 | 0 | 0 | 0 | 0 | 0 |
| SG0000002 | Map3k7 | 0.641336 | 0.701233 | 0.059928677 | 2399.286043 | 2501.049902 | 492 | 20 | 32 | 64 | 236 | 36 | 28 | 0 | 0 | 4 | 4 | 0 |
| SG0000004 | Wdr41 | 0.832991 | 0.869394 | -0.036960743 | 614.1482096 | 598.6140387 | 420 | 15 | 32 | 45 | 156 | 55 | 19 | 0 | 3 | 0 | 12 | 6 |
| SG0000000 | Msh5 | 0.925725 | 0.946224 | -0.091150001 | 51.79114257 | 48.62018508 | 103 | 28 | 32 | 37 | 85 | 0 | 6 | 3 | 3 | 3 | 4 | 0 |
| SG0000003 | Sun1 | 1 | 1 | -0.004547039 | 3440.573095 | 3429.746281 | 288 | 27 | 32 | 52 | 139 | 25 | 9 | 0 | 0 | 3 | 3 | 4 |
| SG0000004 | Fam178b | 3.6E-157 | 6.2E-155 | 10.95272585 | 0.307509278 | 609.476883 | 229 | 24 | 31 | 53 | 184 | 50 | 1 | 0 | 0 | 0 | 2 | 0 |
| SG0000004 | Mfsd6 | 1E-72 | 3.9E-71 | -4.60261472 | 3461.500834 | 142.475225 | 371 | 20 | 31 | 64 | 199 | 34 | 9 | 2 | 2 | 4 | 8 | 0 |
| SG0000002 | Lama3 | 1.28E-67 | 4.34E-66 | -4.237796043 | 3163.657527 | 167.6817275 | 496 | 16 | 31 | 66 | 240 | 58 | 0 | 0 | 0 | 0 | 2 | 0 |
| SG0000002 | Nt5c3 | 2.29E-36 | 2.98E-35 | -2.161981967 | 2828.523089 | 632.0308921 | 324 | 3 | 31 | 50 | 121 | 29 | 2 | 0 | 2 | 2 | 6 | 0 |
| JSG000000 | Prune2 | 6.58E-36 | 8.43E-35 | 2.389822386 | 143.7210473 | 753.2317116 | 496 | 17 | 31 | 60 | 226 | 45 | 6 | 0 | 1 | 1 | 4 | 0 |
| JSG000000 | Gna14 | 1.19E-35 | 1.51E-34 | -2.860669715 | 1215.943799 | 167.4041077 | 304 | 21 | 31 | 50 | 106 | 35 | 4 | 0 | 0 | 0 | 1 | 0 |
| SG0000003 | Crtc3 | 3.26E-35 | 4.09E-34 | -3.003530071 | 951.7696192 | 118.6804525 | 234 | 25 | 31 | 52 | 122 | 29 | 1 | 0 | 0 | 0 | 0 | 0 |
| JSG000000 | Tango6 | 2.56E-30 | 2.62E-29 | 1.736236814 | 302.6544124 | 1008.338548 | 266 | 27 | 31 | 53 | 131 | 20 | 2 | 0 | 0 | 0 | 0 | 0 |
| SG0000003 | Hpse | 4.84E-29 | 4.68E-28 | -5.270860248 | 1678.582355 | 43.47660422 | 145 | 22 | 31 | 46 | 112 | 4 | 5 | 0 | 1 | 1 | 3 | 0 |
| SG0000003 | Arap2 | 1.49E-25 | 1.23E-24 | -2.419648738 | 1432.799104 | 267.7925262 | 430 | 10 | 31 | 67 | 222 | 44 | 8 | 0 | 0 | 2 | 3 | 1 |
| SG0000003 | Ppp1r16b | 2.65E-21 | 1.79E-20 | 7.629830159 | 0.307509278 | 60.90682536 | 330 | 31 | 31 | 55 | 208 | 24 | 36 | 12 | 12 | 12 | 24 | 0 |
| SG0000001 | Anp32e | 4.24E-19 | 2.53E-18 | 1.146599617 | 3619.569475 | 8013.417953 | 177 | 14 | 31 | 21 | 65 | 7 | 16 | 0 | 3 | 7 | 9 | 0 |
| SG0000003 | Ints7 | 8.41E-19 | 4.94E-18 | 1.107001084 | 1925.541607 | 4147.567274 | 292 | 24 | 31 | 61 | 145 | 30 | 8 | 0 | 0 | 0 | 2 | 2 |
| SG0000003 | Pgr | 7.49E-17 | 3.97E-16 | 5.495087337 | 0 | 44.10099439 | 451 | 21 | 31 | 57 | 178 | 40 | 6 | 0 | 1 | 0 | 1 | 1 |
| SG0000003 | Armc8 | 3.23E-15 | 1.55E-14 | 1.089810237 | 719.4342965 | 1531.287398 | 222 | 23 | 31 | 51 | 104 | 21 | 5 | 0 | 0 | 1 | 2 | 0 |
| JSG000000 | Atp9b | 6.09E-15 | 2.86E-14 | -1.261563995 | 1707.758448 | 712.2916257 | 418 | 13 | 31 | 57 | 191 | 44 | 0 | 0 | 0 | 0 | 0 | 0 |
| SG0000004 | Abca6 | 8.84E-15 | 4.11E-14 | -5.777298837 | 57.85179497 | 1.054815722 | 224 | 22 | 31 | 42 | 116 | 14 | 0 | 0 | 0 | 0 | 0 | 0 |
| SG0000002 | Ccdc30 | 1.05E-13 | 4.57E-13 | 2.588079958 | 17.43330557 | 104.826103 | 210 | 21 | 31 | 31 | 133 | 15 | 1 | 0 | 0 | 0 | 0 | 0 |
| SG0000002 | Serinc2 | 1.17E-11 | 4.47E-11 | 2.17605956 | 45.99131571 | 207.8428236 | 70 | 28 | 31 | 31 | 59 | 3 | 0 | 0 | 0 | 0 | 0 | 0 |
| SG0000004 | Ermp1 | 1.4E-11 | 5.29E-11 | -0.982686407 | 4428.379074 | 2240.921818 | 266 | 31 | 31 | 55 | 124 | 12 | 11 | 0 | 0 | 2 | 6 | 0 |
| SG0000003 | Itgal | 6.67E-11 | 2.4E-10 | 2.483349203 | 15.69215075 | 87.74958503 | 158 | 24 | 31 | 28 | 131 | 10 | 5 | 0 | 3 | 0 | 0 | 0 |
| SG0000006 | Morf4l1 | 2.19E-10 | 7.6E-10 | 0.787859134 | 3011.642977 | 5199.633316 | 216 | 16 | 31 | 43 | 108 | 25 | 0 | 0 | 0 | 0 | 0 | 2 |
| SG0000002 | Casp8ap2 | 1.89E-08 | 5.62E-08 | 0.732889121 | 1463.172332 | 2431.739723 | 276 | 10 | 31 | 41 | 120 | 12 | 3 | 0 | 0 | 0 | 0 | 3 |
| JSG000001 | Pcdhga1 | 1.94E-07 | 5.31E-07 | 2.3274584 | 11.41245935 | 57.28145429 | 275 | 23 | 31 | 50 | 122 | 23 | 3 | 0 | 0 | 1 | 3 | 1 |
| SG0000001 | Eva2 | 1.97E-06 | 4.95E-06 | -2.819633148 | 73.17304145 | 10.36470516 | 290 | 23 | 31 | 39 | 125 | 29 | 2 | 0 | 0 | 0 | 0 | 0 |
| SG0000002 | Azin2 | 2.13E-06 | 5.35E-06 | 1.390955874 | 119.8598754 | 314.3353777 | 145 | 25 | 31 | 46 | 83 | 18 | 2 | 0 | 0 | 0 | 0 | 4 |
| SG0000004 | Crocc | 2.3E-06 | 5.75E-06 | 1.532749271 | 220.6706556 | 638.4811671 | 97 | 29 | 31 | 39 | 85 | 14 | 0 | 0 | 0 | 0 | 4 | 0 |
| JSG000000 | Ap1ar | 1.48E-05 | 3.45E-05 | 0.602275295 | 1018.480531 | 1546.16437 | 140 | 16 | 31 | 29 | 88 | 10 | 14 | 0 | 2 | 2 | 4 | 2 |
| SG0000006 | Brpf3 | 1.87E-05 | 4.3E-05 | 0.586727641 | 1229.482969 | 1846.482247 | 96 | 25 | 31 | 32 | 60 | 9 | 2 | 0 | 0 | 2 | 2 | 3 |
| SG0000003 | Ppp2r4 | 1.92E-05 | 4.41E-05 | -0.592176082 | 3824.044808 | 2536.648017 | 167 | 27 | 31 | 50 | 81 | 8 | 0 | 0 | 0 | 0 | 0 | 0 |
| SG0000002 | Xrcc6bp1 | 2.13E-05 | 4.86E-05 | 0.962776717 | 106.0992266 | 206.7935026 | 118 | 27 | 31 | 47 | 86 | 5 | 0 | 0 | 0 | 0 | 0 | 1 |
| SG0000000 | Cse1l | 7.47E-05 | 0.000161 | 0.484222345 | 6398.146794 | 8949.930242 | 260 | 27 | 31 | 42 | 138 | 11 | 4 | 0 | 0 | 0 | 1 | 0 |
| SG0000002 | Faf2 | 0.000144 | 0.000299 | -0.543436692 | 2657.323387 | 1823.281245 | 235 | 25 | 31 | 39 | 86 | 9 | 1 | 0 | 0 | 0 | 7 | 0 |
| SG0000002 | Parp9 | 0.000702 | 0.001343 | -0.944470289 | 2569.948782 | 1335.397568 | 158 | 29 | 31 | 49 | 121 | 12 | 0 | 0 | 0 | 0 | 0 | 1 |
| SG0000007 | Tox2 | 0.006 | 0.01015 | 2.875039233 | 2.211622639 | 16.22497644 | 383 | 25 | 31 | 60 | 154 | 56 | 2 | 0 | 0 | 1 | 1 | 0 |
| SG0000002 | Dcun1d1 | 0.006741 | 0.011324 | -0.46828885 | 661.9227863 | 478.4519635 | 379 | 13 | 31 | 44 | 152 | 30 | 45 | 0 | 3 | 2 | 7 | 10 |
| SG0000002 | Ago3 | 0.008747 | 0.014433 | -0.448831682 | 718.5492657 | 526.4349939 | 283 | 17 | 31 | 49 | 154 | 24 | 25 | 1 | 3 | 4 | 11 | 2 |
| SG0000001 | Steap2 | 0.04349 | 0.063457 | -0.360732239 | 851.50236 | 663.123825 | 356 | 6 | 31 | 26 | 152 | 15 | 71 | 0 | 2 | 2 | 24 | 2 |
| SG0000004 | Brox | 0.044487 | 0.064795 | -0.306163494 | 1191.20339 | 963.4330163 | 164 | 21 | 31 | 42 | 86 | 15 | 20 | 3 | 3 | 5 | 5 | 0 |
| SG0000006 | Epc2 | 0.052963 | 0.076083 | -0.304135841 | 950.5662353 | 769.8894579 | 288 | 13 | 31 | 60 | 159 | 21 | 7 | 0 | 0 | 2 | 3 | 1 |
| SG0000003 | Phf12 | 0.056931 | 0.081294 | -0.267002864 | 2214.463249 | 1840.316866 | 161 | 22 | 31 | 38 | 101 | 13 | 4 | 0 | 1 | 0 | 1 | 3 |
| SG0000002 | Kpna4 | 0.060133 | 0.085431 | -0.258540813 | 4141.306786 | 3461.854927 | 302 | 14 | 31 | 38 | 152 | 24 | 16 | 0 | 1 | 0 | 6 | 2 |
| SG0000000 | Trmt1 | 0.081001 | 0.112342 | -0.248381406 | 2699.888571 | 2272.875185 | 62 | 30 | 31 | 36 | 76 | 5 | 0 | 0 | 0 | 0 | 0 | 0 |
| SG0000003 | Lrp6 | 0.31781 | 0.383599 | -0.14883188 | 2198.931974 | 1983.393722 | 306 | 20 | 31 | 50 | 142 | 21 | 9 | 0 | 0 | 2 | 4 | 1 |
| SG0000002 | Nisch | 0.356542 | 0.424636 | 0.118441288 | 7948.21715 | 8628.2775 | 195 | 20 | 31 | 31 | 111 | 14 | 9 | 1 | 3 | 2 | 5 | 1 |
| SG0000003 | Hace1 | 0.504794 | 0.57209 | 0.096835241 | 571.7589628 | 611.4533004 | 597 | 12 | 31 | 76 | 349 | 63 | 14 | 0 | 0 | 1 | 1 | 1 |
| SG0000005 | Kctd18 | 0.643671 | 0.703314 | -0.079910984 | 790.2180392 | 747.6379472 | 230 | 10 | 31 | 48 | 122 | 10 | 9 | 0 | 4 | 0 | 3 | 0 |
| JSG000000 | Arhgef28 | 0.660026 | 0.718381 | -0.104659843 | 154.9026627 | 144.0632245 | 539 | 12 | 31 | 70 | 243 | 42 | 1 | 0 | 0 | 0 | 0 | 0 |
| SG0000002 | App | 0.753117 | 0.801049 | -0.114610928 | 1782.429608 | 1646.307893 | 414 | 13 | 31 | 57 | 193 | 43 | 3 | 0 | 0 | 1 | 2 | 0 |
| SG0000002 | Cblb | 1 | 1 | -0.004497972 | 800.6703641 | 798.1779563 | 401 | 12 | 31 | 45 | 159 | 38 | 4 | 0 | 0 | 0 | 5 | 1 |
| SG0000002 | Dmc1 | 2.1E-102 | 1.5E-100 | 8.397478276 | 0 | 336.2041019 | 117 | 25 | 30 | 40 | 91 | 8 | 4 | 0 | 0 | 1 | 1 | 0 |
| SG0000003 | Tgfbr2 | 4.34E-76 | 1.81E-74 | -5.027781667 | 6505.473224 | 199.4186631 | 262 | 26 | 30 | 46 | 106 | 32 | 1 | 0 | 0 | 0 | 0 | 2 |
| SG0000004 | Myof | 2.28E-53 | 5.2E-52 | -3.833544719 | 13291.97879 | 932.3472398 | 414 | 22 | 30 | 62 | 168 | 60 | 4 | 0 | 0 | 0 | 0 | 2 |
| SG0000002 | Cpeb4 | 8.02E-52 | 1.75E-50 | -3.406886166 | 3900.307433 | 367.7256405 | 577 | 21 | 30 | 99 | 259 | 27 | 34 | 0 | 0 | 6 | 16 | 0 |

| | | | | | | | | | | | | | | | | | |
|---|---|---|---|---|---|---|---|---|---|---|---|---|---|---|---|---|---|
| SG0000001 | Mov10l1 | 8.74E-52 | 1.9E-50 | 9.090593048 | 0.331601068 | 180.7827994 | 217 | 27 | 30 | 38 | 104 | 9 | 0 | 0 | 0 | 0 | 0 | 0 |
| SG0000005 | Syne3 | 2.42E-43 | 4E-42 | -4.715262118 | 1716.176652 | 65.33210542 | 205 | 24 | 30 | 52 | 104 | 31 | 2 | 0 | 0 | 2 | 4 | 2 |
| SG000000C | Itga3 | 4.11E-42 | 6.51E-41 | -4.380597932 | 19655.62539 | 943.6164213 | 141 | 24 | 30 | 30 | 51 | 6 | 3 | 0 | 0 | 3 | 3 | 0 |
| JSG000000 | Cdyl2 | 5.64E-32 | 6.18E-31 | 6.070382522 | 24.84465074 | 1669.552654 | 258 | 23 | 30 | 55 | 149 | 32 | 3 | 0 | 0 | 0 | 1 | 0 |
| SG000000C | Ubl3 | 1.94E-31 | 2.09E-30 | -1.878783408 | 4184.28071 | 1137.759973 | 308 | 21 | 30 | 45 | 120 | 16 | 5 | 0 | 0 | 0 | 3 | 1 |
| SG0000002 | Baiap2 | 5.75E-30 | 5.82E-29 | -2.714022065 | 2009.08352 | 306.1934814 | 175 | 25 | 30 | 35 | 90 | 15 | 6 | 3 | 3 | 3 | 3 | 0 |
| SG0000001 | Sgk1 | 2.09E-29 | 2.05E-28 | 4.086294446 | 461.4894195 | 7838.969013 | 275 | 25 | 30 | 49 | 142 | 12 | 15 | 0 | 0 | 5 | 5 | 0 |
| SG0000001 | Mpped2 | 1.25E-24 | 9.89E-24 | -2.398531265 | 479.9324214 | 91.02280879 | 593 | 12 | 30 | 50 | 239 | 60 | 5 | 0 | 0 | 0 | 1 | 0 |
| SG0000006 | Slc38a10 | 1.26E-24 | 9.99E-24 | -1.620101212 | 7526.895188 | 2448.594151 | 163 | 30 | 30 | 32 | 103 | 5 | 2 | 0 | 0 | 2 | 2 | 0 |
| JSG000000 | Mtr | 4.18E-24 | 3.23E-23 | -1.609706181 | 3367.638032 | 1103.457373 | 140 | 23 | 30 | 40 | 83 | 4 | 0 | 0 | 0 | 0 | 0 | 0 |
| SG0000003 | Pogz | 6.15E-23 | 4.5E-22 | 1.265113445 | 1245.067826 | 2992.472067 | 211 | 15 | 30 | 38 | 118 | 35 | 9 | 0 | 0 | 2 | 3 | 2 |
| SG0000002 | Rab15 | 1.06E-22 | 7.67E-22 | 4.25463153 | 31.66672212 | 604.4700982 | 71 | 27 | 30 | 30 | 79 | 2 | 2 | 0 | 0 | 0 | 0 | 0 |
| SG0000001 | Chp1 | 1.76E-21 | 1.2E-20 | -1.46761847 | 3858.43324 | 1395.127209 | 160 | 22 | 30 | 38 | 106 | 16 | 13 | 0 | 0 | 2 | 2 | 1 |
| JSG000000 | Mlh1 | 6.1E-18 | 3.42E-17 | 1.284663232 | 433.4449458 | 1055.981015 | 220 | 15 | 30 | 34 | 76 | 12 | 5 | 0 | 0 | 0 | 1 | 0 |
| SG0000005 | Tspan7 | 1.4E-17 | 7.73E-17 | -1.908543274 | 2516.101374 | 670.1921885 | 482 | 24 | 30 | 70 | 222 | 32 | 18 | 4 | 4 | 7 | 11 | 1 |
| SG0000002 | Clcn1 | 4.35E-17 | 2.34E-16 | 5.073124641 | 2.992940161 | 100.7536347 | 287 | 24 | 30 | 24 | 104 | 31 | 14 | 0 | 0 | 0 | 0 | 5 |
| JSG000000 | Mfsd7b | 9.7E-17 | 5.1E-16 | -1.611162011 | 1171.433869 | 383.4508311 | 133 | 28 | 30 | 44 | 78 | 2 | 13 | 0 | 0 | 0 | 3 | 0 |
| SG0000004 | Klhdc4 | 1.22E-16 | 6.38E-16 | 1.123001537 | 1144.745262 | 2493.251004 | 79 | 28 | 30 | 32 | 68 | 8 | 0 | 0 | 0 | 0 | 0 | 0 |
| SG0000006 | Zfp462 | 4.83E-16 | 2.43E-15 | 1.04271958 | 1385.156617 | 2853.571453 | 379 | 21 | 30 | 90 | 211 | 28 | 9 | 0 | 3 | 0 | 2 | 1 |
| SG0000001 | Dnase1l1 | 5.96E-15 | 2.81E-14 | -2.495776008 | 383.7239753 | 68.03235332 | 45 | 30 | 30 | 30 | 36 | 3 | 0 | 0 | 0 | 0 | 0 | 0 |
| SG000000C | Cct3 | 1.2E-14 | 5.52E-14 | 0.916064492 | 10915.35642 | 20596.85024 | 141 | 24 | 30 | 36 | 66 | 18 | 3 | 0 | 0 | 0 | 0 | 0 |
| SG0000003 | Gramd4 | 1.88E-14 | 8.6E-14 | -1.248301389 | 2688.479089 | 1131.697876 | 181 | 24 | 30 | 42 | 93 | 12 | 6 | 0 | 3 | 3 | 3 | 3 |
| SG0000004 | Plekha6 | 2.19E-14 | 9.94E-14 | -2.390827805 | 1417.337428 | 270.2479275 | 265 | 24 | 30 | 58 | 121 | 9 | 6 | 0 | 0 | 2 | 2 | 0 |
| SG0000004 | Zmym6 | 9.12E-14 | 3.98E-13 | 1.108075554 | 495.996023 | 1069.158698 | 200 | 20 | 30 | 46 | 114 | 24 | 2 | 0 | 0 | 0 | 0 | 0 |
| SG0000006 | C330021F23Rik | 1.04E-13 | 4.52E-13 | 7.002560403 | 2.839858874 | 364.1476286 | 66 | 28 | 30 | 36 | 51 | 2 | 0 | 0 | 0 | 0 | 0 | 0 |
| SG0000002 | Rassf5 | 1.65E-13 | 7.07E-13 | 3.207445513 | 25.78892679 | 238.2158082 | 172 | 28 | 30 | 35 | 85 | 32 | 12 | 0 | 0 | 0 | 9 | 3 |
| SG0000002 | Rnf130 | 2.37E-13 | 1.01E-12 | 1.520928658 | 106.3957087 | 305.3298461 | 304 | 14 | 30 | 36 | 120 | 26 | 2 | 0 | 0 | 0 | 0 | 0 |
| SG0000005 | Actr3b | 5.29E-13 | 2.19E-12 | 2.596296149 | 69.46770961 | 420.0935357 | 240 | 10 | 30 | 40 | 155 | 22 | 2 | 0 | 0 | 0 | 1 | 4 |
| JSG000000 | Esyt2 | 4.24E-11 | 1.55E-10 | -0.963784008 | 3670.818787 | 1882.066871 | 209 | 21 | 30 | 39 | 111 | 15 | 4 | 0 | 1 | 1 | 7 | 1 |
| JSG000000 | Rock1 | 8.31E-11 | 2.97E-10 | -0.948956 | 4216.948725 | 2184.409716 | 345 | 19 | 30 | 63 | 204 | 30 | 7 | 0 | 0 | 0 | 0 | 0 |
| SG0000003 | Fanca | 2.62E-10 | 9.06E-10 | 0.861171499 | 991.3168197 | 1800.739651 | 137 | 24 | 30 | 48 | 96 | 17 | 2 | 0 | 0 | 0 | 2 | 2 |
| JSG000000 | Baz1b | 3.01E-10 | 1.03E-09 | -0.877300146 | 9947.394144 | 5415.214541 | 164 | 27 | 30 | 41 | 91 | 10 | 4 | 0 | 0 | 1 | 1 | 0 |
| SG0000002 | Itga6 | 3.54E-10 | 1.21E-09 | -0.971097053 | 7098.059284 | 3620.84789 | 244 | 28 | 30 | 46 | 126 | 26 | 8 | 0 | 0 | 2 | 2 | 0 |
| SG0000002 | Cul2 | 5.45E-10 | 1.83E-09 | 0.787128015 | 1904.009744 | 3285.627385 | 332 | 18 | 30 | 47 | 172 | 29 | 10 | 0 | 0 | 2 | 4 | 2 |
| SG0000001 | Prox1 | 5.68E-10 | 1.91E-09 | 4.740973327 | 1.299968916 | 34.76227156 | 244 | 24 | 30 | 42 | 98 | 18 | 17 | 0 | 0 | 0 | 4 | 3 |
| SG0000005 | BC004004 | 1.86E-09 | 5.98E-09 | -0.939425866 | 1605.056969 | 836.9414694 | 138 | 30 | 30 | 39 | 85 | 24 | 0 | 0 | 0 | 0 | 0 | 0 |
| SG0000002 | Fmnl3 | 7.42E-09 | 2.28E-08 | 1.583502079 | 239.3667422 | 717.3736705 | 177 | 21 | 30 | 42 | 72 | 36 | 0 | 0 | 0 | 0 | 0 | 3 |
| JSG000000 | Ccdc61 | 6.06E-08 | 1.73E-07 | 1.11303533 | 266.8430313 | 577.1821428 | 67 | 28 | 30 | 36 | 60 | 7 | 0 | 0 | 0 | 0 | 0 | 3 |
| SG0000002 | Dlg4 | 1.32E-07 | 3.66E-07 | -1.047450862 | 802.0468241 | 388.0481301 | 113 | 21 | 30 | 30 | 68 | 3 | 7 | 4 | 4 | 4 | 4 | 0 |
| SG0000002 | Ppil3 | 4.44E-07 | 1.18E-06 | 0.760668836 | 536.0282345 | 908.179726 | 86 | 19 | 30 | 22 | 46 | 11 | 0 | 0 | 0 | 0 | 0 | 0 |
| SG0000005 | Nrd1 | 9.8E-07 | 2.54E-06 | -0.669616814 | 8366.745396 | 5259.952314 | 421 | 24 | 30 | 63 | 230 | 48 | 4 | 0 | 0 | 0 | 2 | 0 |
| SG0000002 | Dhx35 | 1.11E-06 | 2.86E-06 | 0.694115321 | 764.9353554 | 1237.582804 | 196 | 14 | 30 | 40 | 90 | 18 | 0 | 0 | 0 | 0 | 0 | 2 |
| SG0000003 | Cpeb2 | 1.13E-06 | 2.92E-06 | -1.177809598 | 482.1949737 | 213.140834 | 522 | 9 | 30 | 66 | 240 | 60 | 42 | 0 | 0 | 6 | 39 | 3 |
| JSG000000 | Gemin7 | 3.49E-06 | 8.6E-06 | -0.821887773 | 673.9652971 | 381.2630014 | 79 | 30 | 30 | 30 | 55 | 0 | 0 | 0 | 0 | 0 | 0 | 0 |
| SG0000002 | Mybl1 | 4.17E-06 | 1.02E-05 | -0.802174549 | 1383.281676 | 793.2900787 | 258 | 18 | 30 | 39 | 152 | 19 | 14 | 0 | 4 | 7 | 14 | 1 |
| SG0000002 | Gfpt1 | 7.02E-06 | 1.68E-05 | -0.688368985 | 2390.309911 | 1483.318795 | 243 | 12 | 30 | 37 | 120 | 14 | 21 | 0 | 2 | 1 | 4 | 0 |
| SG0000002 | Zgpat | 2.03E-05 | 4.65E-05 | -0.71409901 | 802.0362774 | 488.909854 | 111 | 21 | 30 | 36 | 69 | 6 | 0 | 0 | 0 | 0 | 0 | 0 |
| SG0000002 | Ndufv3 | 2.06E-05 | 4.7E-05 | 0.656184815 | 1407.985376 | 2218.857726 | 82 | 30 | 30 | 33 | 39 | 0 | 1 | 0 | 0 | 0 | 0 | 0 |
| SG0000006 | Gpatch1 | 8.53E-05 | 0.000182 | 0.638454334 | 396.2482032 | 616.823926 | 140 | 18 | 30 | 37 | 102 | 17 | 4 | 0 | 0 | 2 | 2 | 0 |
| SG0000002 | Dtd1 | 0.000149 | 0.00031 | 0.599409227 | 541.902305 | 821.0340251 | 290 | 19 | 30 | 48 | 155 | 22 | 0 | 0 | 0 | 0 | 1 | 0 |
| JSG000000 | Ubr1 | 0.00016 | 0.000331 | -0.563573383 | 1824.05323 | 1234.198501 | 281 | 18 | 30 | 44 | 146 | 31 | 5 | 0 | 0 | 0 | 0 | 0 |
| SG0000001 | Lims1 | 0.000291 | 0.000584 | -0.525757622 | 3325.928598 | 2310.170888 | 314 | 22 | 30 | 48 | 177 | 39 | 5 | 0 | 0 | 1 | 4 | 0 |
| SG0000004 | Map7d2 | 0.000383 | 0.00076 | 0.528961304 | 605.4343116 | 873.5751048 | 311 | 19 | 30 | 50 | 166 | 19 | 6 | 0 | 0 | 0 | 6 | 0 |
| SG0000006 | Srrm4 | 0.000464 | 0.00091 | 2.253449095 | 5.536163868 | 26.39761605 | 231 | 21 | 30 | 53 | 125 | 16 | 25 | 10 | 10 | 17 | 20 | 3 |
| SG000000C | Spg7 | 0.001032 | 0.001932 | -0.469246951 | 2161.293694 | 1561.192221 | 154 | 18 | 30 | 35 | 122 | 18 | 11 | 0 | 0 | 5 | 5 | 0 |
| SG0000003 | Usp6nl | 0.001119 | 0.00209 | -0.498738274 | 1435.733599 | 1016.105222 | 430 | 17 | 30 | 62 | 177 | 48 | 5 | 0 | 0 | 1 | 3 | 4 |
| SG000000C | Gnb1l | 0.001351 | 0.0025 | -0.644936406 | 347.2233065 | 222.0565893 | 430 | 15 | 30 | 60 | 244 | 30 | 2 | 0 | 0 | 0 | 3 | 0 |
| SG0000004 | Tmem41b | 0.001378 | 0.002548 | 0.513803333 | 443.5823281 | 633.3509863 | 199 | 12 | 30 | 44 | 81 | 7 | 9 | 0 | 0 | 5 | 15 | 1 |
| SG0000002 | Ddx46 | 0.001389 | 0.002566 | 0.400509927 | 4106.346033 | 5420.271554 | 168 | 18 | 30 | 42 | 102 | 20 | 12 | 2 | 2 | 2 | 6 | 4 |
| SG0000002 | Add3 | 0.001552 | 0.002853 | -0.453238984 | 2559.061118 | 1869.141279 | 604 | 6 | 30 | 48 | 289 | 63 | 12 | 0 | 0 | 0 | 3 | 3 |
| JSG000000 | Cdk13 | 0.001764 | 0.003219 | -0.46400045 | 1810.90161 | 1312.855117 | 261 | 19 | 30 | 45 | 152 | 34 | 1 | 0 | 0 | 0 | 1 | 0 |
| SG0000006 | Zfp120 | 0.002011 | 0.003641 | -0.572430596 | 492.9114028 | 331.4744703 | 123 | 24 | 30 | 30 | 69 | 21 | 35 | 3 | 3 | 3 | 6 | 0 |
| SG0000004 | Tlk1 | 0.004377 | 0.007547 | 0.369955196 | 2078.711824 | 2686.345683 | 302 | 22 | 30 | 48 | 131 | 32 | 2 | 0 | 0 | 0 | 0 | 2 |
| SG0000002 | Sfswap | 0.004721 | 0.00811 | -0.411460058 | 1697.869211 | 1276.563472 | 158 | 26 | 30 | 48 | 95 | 15 | 1 | 0 | 0 | 0 | 2 | 2 |
| SG0000002 | Cops2 | 0.007536 | 0.012562 | 0.333703305 | 4075.105597 | 5135.628158 | 240 | 24 | 30 | 49 | 117 | 24 | 15 | 0 | 0 | 1 | 3 | 0 |
| SG000000C | Cbx5 | 0.011168 | 0.018123 | 0.343085857 | 7649.030727 | 9702.542184 | 162 | 26 | 30 | 37 | 100 | 19 | 14 | 0 | 0 | 2 | 9 | 5 |
| SG0000001 | Gfod2 | 0.01226 | 0.01974 | 0.504124578 | 260.3801899 | 369.2874588 | 147 | 22 | 30 | 33 | 84 | 16 | 3 | 0 | 0 | 1 | 4 | 0 |
| SG0000003 | Lmx1b | 0.018098 | 0.028223 | 1.56034015 | 7.405311327 | 21.83999376 | 123 | 25 | 30 | 37 | 71 | 11 | 3 | 1 | 2 | 1 | 1 | 1 |
| SG0000002 | Gbp9 | 0.031894 | 0.047643 | -1.253452637 | 148.0060729 | 62.08014085 | 174 | 23 | 30 | 43 | 97 | 6 | 11 | 1 | 2 | 2 | 8 | 0 |
| JSG000000 | D230025D16Rik | 0.042733 | 0.062444 | -0.280611137 | 2633.852345 | 2168.29843 | 165 | 18 | 30 | 30 | 63 | 18 | 3 | 0 | 3 | 0 | 0 | 3 |
| SG0000003 | Fbxw2 | 0.051535 | 0.07418 | -0.285405192 | 1703.011204 | 1397.339647 | 285 | 24 | 30 | 47 | 95 | 6 | 11 | 0 | 0 | 0 | 1 | 0 |
| SG0000001 | Trim37 | 0.059125 | 0.084138 | 0.237588251 | 2991.074375 | 3526.536657 | 234 | 20 | 30 | 45 | 115 | 16 | 5 | 0 | 0 | 2 | 2 | 0 |
| JSG000000 | Il15 | 0.079735 | 0.110774 | -1.181802757 | 23.02618356 | 10.14994955 | 170 | 26 | 30 | 41 | 120 | 13 | 1 | 0 | 0 | 0 | 1 | 1 |
| SG0000002 | Rftn2 | 0.093011 | 0.127498 | -0.669627244 | 53.65445289 | 33.73089653 | 178 | 21 | 30 | 46 | 104 | 22 | 3 | 0 | 0 | 1 | 2 | 2 |
| SG000000C | Cdc42 | 0.099515 | 0.135459 | -0.217259347 | 11837.25488 | 10182.38277 | 154 | 18 | 30 | 40 | 90 | 18 | 3 | 0 | 0 | 0 | 0 | 0 |
| SG0000002 | Lrrcc1 | 0.100297 | 0.136359 | 0.251448718 | 580.8353095 | 691.427448 | 340 | 12 | 30 | 50 | 187 | 17 | 15 | 0 | 0 | 0 | 4 | 0 |
| SG0000004 | Stk40 | 0.111818 | 0.150605 | -0.226299026 | 1588.185707 | 1357.620816 | 106 | 28 | 30 | 46 | 56 | 4 | 4 | 0 | 0 | 4 | 4 | 0 |
| SG0000003 | Pank2 | 0.149038 | 0.195386 | 0.194380231 | 1403.255433 | 1605.650477 | 374 | 12 | 30 | 62 | 162 | 38 | 12 | 0 | 2 | 2 | 5 | 0 |
| SG0000009 | Rnf8 | 0.248225 | 0.308936 | -0.284712451 | 168.1432698 | 138.0296863 | 72 | 30 | 30 | 34 | 42 | 6 | 0 | 0 | 0 | 0 | 0 | 0 |
| SG0000002 | Wls | 0.332032 | 0.398857 | 0.396011506 | 583.7376312 | 768.1199292 | 268 | 16 | 30 | 40 | 122 | 27 | 3 | 0 | 0 | 1 | 2 | 1 |
| SG0000002 | Usp8 | 0.515767 | 0.582682 | 0.083968763 | 2374.966502 | 2517.297851 | 250 | 20 | 30 | 42 | 118 | 16 | 10 | 0 | 0 | 2 | 4 | 2 |
| JSG000000 | Ttc7b | 0.583188 | 0.646879 | -0.101039487 | 1224.481824 | 1141.659058 | 286 | 19 | 30 | 46 | 133 | 27 | 4 | 0 | 0 | 2 | 3 | 0 |
| SG0000004 | Lemd3 | 0.631962 | 0.69293 | 0.066660211 | 643.310277 | 673.7320575 | 308 | 16 | 30 | 60 | 133 | 28 | 12 | 0 | 0 | 2 | 9 | 8 |
| SG0000003 | Hiatl1 | 0.756277 | 0.803704 | -0.052042741 | 1853.795503 | 1788.114802 | 254 | 16 | 30 | 45 | 163 | 22 | 12 | 0 | 0 | 3 | 5 | 2 |
| SG0000002 | Tbck | 0.782531 | 0.826505 | 0.035629906 | 579.6974554 | 594.1923614 | 359 | 14 | 30 | 53 | 188 | 33 | 5 | 0 | 0 | 1 | 2 | 1 |
| SG0000001 | Jph2 | 0.86866 | 0.898354 | -0.045100635 | 58.42386362 | 56.62570601 | 128 | 28 | 30 | 46 | 88 | 6 | 4 | 0 | 0 | 0 | 0 | 0 |
| SG0000002 | Pnkd | 1.16E-71 | 4.35E-70 | 2.533942001 | 299.3769048 | 1733.847288 | 148 | 22 | 29 | 34 | 68 | 9 | 2 | 0 | 0 | 0 | 0 | 0 |
| SG0000003 | Msn | 3.19E-64 | 9.87E-63 | -2.891507235 | 31684.17621 | 4269.844013 | 176 | 24 | 29 | 49 | 103 | 13 | 6 | 0 | 0 | 0 | 1 | 0 |
| SG0000001 | Slc16a10 | 4.63E-54 | 1.08E-52 | 4.688666297 | 9.724175239 | 250.7736908 | 221 | 21 | 29 | 44 | 96 | 13 | 15 | 4 | 5 | 6 | 9 | 0 |
| SG0000004 | Ankib1 | 2.67E-44 | 4.57E-43 | -2.616255225 | 2389.595622 | 389.7203779 | 313 | 14 | 29 | 43 | 122 | 22 | 7 | 0 | 0 | 0 | 2 | 0 |
| JSG000000 | Map3k15 | 9.18E-33 | 1.04E-31 | 2.531218612 | 63.29044167 | 365.8565663 | 296 | 16 | 29 | 46 | 177 | 27 | 2 | 0 | 0 | 0 | 1 | 0 |
| SG0000002 | Prkch | 1.42E-32 | 1.6E-31 | -3.484904805 | 1275.366042 | 113.9131787 | 265 | 17 | 29 | 42 | 112 | 25 | 2 | 0 | 0 | 1 | 1 | 0 |
| SG0000002 | Rev1 | 8.86E-32 | 9.63E-31 | 1.503635432 | 1185.359283 | 3361.16144 | 260 | 21 | 29 | 34 | 102 | 28 | 1 | 0 | 0 | 0 | 0 | 1 |
| SG0000003 | Rnf144b | 4.7E-22 | 3.29E-21 | 4.349925964 | 4.552234774 | 92.82882345 | 298 | 23 | 29 | 60 | 132 | 30 | 15 | 0 | 0 | 2 | 2 | 4 |
| SG0000004 | Unc5cl | 2.2E-20 | 1.41E-19 | 5.791508861 | 0 | 54.38828107 | 229 | 25 | 29 | 49 | 105 | 14 | 2 | 0 | 0 | 0 | 0 | 0 |
| SG0000004 | Shcbp1l | 8.47E-20 | 5.25E-19 | 7.524196012 | 0.307509278 | 56.60657997 | 94 | 25 | 29 | 38 | 48 | 3 | 1 | 1 | 1 | 1 | 1 | 1 |
| SG0000002 | Mbd2 | 1.16E-18 | 6.74E-18 | -1.454599473 | 1831.424826 | 668.2070048 | 209 | 16 | 29 | 40 | 122 | 46 | 0 | 0 | 0 | 0 | 2 | 2 |
| SG0000002 | Tulp2 | 1.79E-18 | 1.03E-17 | 7.418133224 | 0.307509278 | 52.59431897 | 90 | 25 | 29 | 34 | 69 | 2 | 0 | 0 | 0 | 0 | 0 | 0 |
| SG0000004 | Zfp451 | 3.47E-18 | 1.97E-17 | 1.114987767 | 1379.877828 | 2988.721167 | 294 | 22 | 29 | 38 | 95 | 23 | 36 | 4 | 4 | 5 | 12 | 5 |
| SG0000005 | Ube4a | 4.23E-18 | 2.39E-17 | -1.36890846 | 2475.159608 | 958.342396 | 254 | 24 | 29 | 36 | 106 | 34 | 12 | 0 | 1 | 0 | 3 | 2 |
| SG0000003 | C530008M17Rik | 6.93E-17 | 3.68E-16 | 4.316568473 | 11.23528627 | 223.8724908 | 312 | 27 | 29 | 45 | 141 | 20 | 30 | 6 | 6 | 9 | 16 | 0 |
| JSG000000 | 43530 | 1.76E-16 | 9.09E-16 | -1.213371644 | 5910.224692 | 2548.840922 | 203 | 23 | 29 | 39 | 98 | 16 | 9 | 0 | 0 | 0 | 2 | 2 |
| SG0000002 | Erlec1 | 4.38E-16 | 2.21E-15 | -1.362488772 | 1440.721685 | 560.3122327 | 143 | 21 | 29 | 34 | 77 | 12 | 2 | 0 | 0 | 1 | 2 | 1 |
| JSG000000 | Ttc27 | 1.37E-14 | 6.3E-14 | 1.027588956 | 1006.679363 | 2052.231099 | 300 | 15 | 29 | 44 | 150 | 21 | 0 | 0 | 0 | 0 | 0 | 0 |
| SG0000003 | Ccp110 | 1.55E-13 | 6.68E-13 | 0.995423895 | 1194.829882 | 2382.091977 | 143 | 19 | 29 | 33 | 68 | 6 | 20 | 0 | 4 | 6 | 14 | 0 |
| SG0000003 | Dixdc1 | 2.02E-13 | 8.6E-13 | -1.791390186 | 1696.740508 | 490.1773601 | 311 | 21 | 29 | 46 | 140 | 26 | 18 | 0 | 2 | 0 | 6 | 0 |
| SG000000C | Cp | 5.64E-13 | 2.33E-12 | -4.772696243 | 853.8512503 | 31.23614926 | 439 | 8 | 29 | 54 | 170 | 52 | 9 | 0 | 0 | 1 | 1 | 0 |
| SG0000003 | Trim36 | 6.18E-11 | 2.23E-10 | 3.431444743 | 4.932019423 | 53.20991612 | 120 | 26 | 29 | 36 | 67 | 9 | 8 | 1 | 1 | 3 | 4 | 0 |
| SG0000002 | Pik3ca | 1.85E-10 | 6.46E-10 | -0.982612442 | 2010.186422 | 1017.280023 | 327 | 12 | 29 | 45 | 114 | 38 | 29 | 0 | 3 | 3 | 10 | 4 |
| SG0000005 | Tprkb | 3.19E-09 | 1.01E-08 | -0.919263967 | 1723.470081 | 911.3343093 | 134 | 21 | 29 | 40 | 89 | 8 | 7 | 0 | 0 | 1 | 1 | 0 |
| SG0000003 | Ints10 | 4.24E-08 | 1.23E-07 | 0.719414301 | 1488.79814 | 2451.326169 | 284 | 21 | 29 | 36 | 89 | 31 | 16 | 3 | 3 | 6 | 6 | 3 |
| SG0000006 | Tmem219 | 8.43E-08 | 2.38E-07 | -1.047485611 | 508.7011893 | 246.1150456 | 151 | 20 | 29 | 35 | 115 | 23 | 0 | 0 | 0 | 0 | 0 | 0 |
| SG0000006 | 4931406P16Rik | 6.02E-07 | 1.58E-06 | -0.726861212 | 2780.206281 | 1679.848024 | 210 | 17 | 29 | 52 | 107 | 32 | 9 | 0 | 2 | 0 | 0 | 2 |
| JSG000000 | Tbc1d22a | 3.83E-06 | 9.38E-06 | -0.746223355 | 1098.63595 | 654.9651504 | 341 | 20 | 29 | 54 | 146 | 33 | 0 | 0 | 0 | 0 | 1 | 0 |
| JSG000001 | Gm42715 | 2.47E-05 | 5.6E-05 | 1.5098029 | 28.86132617 | 82.18872503 | 109 | 21 | 29 | 30 | 61 | 8 | 2 | 1 | 1 | 1 | 4 | 0 |
| SG0000003 | Dopey1 | 4.36E-05 | 9.63E-05 | -0.691561189 | 836.9681586 | 518.2368301 | 600 | 26 | 29 | 73 | 343 | 57 | 27 | 0 | 0 | 0 | 5 | 5 |
| SG0000005 | Hadhb | 4.36E-05 | 9.63E-05 | -0.666689721 | 1003.347288 | 632.059084 | 223 | 19 | 29 | 37 | 149 | 21 | 0 | 0 | 0 | 0 | 0 | 0 |
| SG0000005 | Chd8 | 7.14E-05 | 0.000154 | 0.502241805 | 3241.726387 | 4591.622809 | 188 | 25 | 29 | 44 | 143 | 16 | 2 | 0 | 0 | 0 | 0 | 0 |
| JSG000000 | Prcc | 0.000942 | 0.001772 | 0.454108393 | 1248.564246 | 1710.452987 | 61 | 23 | 29 | 27 | 47 | 4 | 0 | 0 | 0 | 0 | 0 | 0 |
| SG0000003 | Ttc23 | 0.009215 | 0.015157 | 0.435570618 | 381.9343306 | 516.5454238 | 351 | 12 | 29 | 48 | 138 | 32 | 0 | 0 | 0 | 0 | 0 | 0 |
| SG000000C | Met | 0.014906 | 0.023626 | -0.571300252 | 320.2035243 | 215.5001591 | 419 | 4 | 29 | 30 | 191 | 39 | 5 | 0 | 0 | 1 | 3 | 3 |
| JSG000000 | Ccdc57 | 0.01705 | 0.026742 | 0.619238224 | 87.84798465 | 134.940117 | 131 | 27 | 29 | 38 | 88 | 12 | 0 | 0 | 0 | 0 | 0 | 0 |
| SG0000002 | Usp16 | 0.029578 | 0.044447 | -0.317658107 | 1511.681779 | 1212.930061 | 234 | 22 | 29 | 34 | 100 | 27 | 4 | 1 | 2 | 1 | 6 | 0 |

| | | | | | | | | | | | | | | | | | | |
|---|---|---|---|---|---|---|---|---|---|---|---|---|---|---|---|---|---|---|
| SG0000002 | Fgf7 | 0.031284 | 0.04679 | -0.941883456 | 375.6047591 | 195.5221272 | 266 | 14 | 29 | 34 | 116 | 24 | 15 | 0 | 3 | 2 | 6 | 2 |
| SG0000003 | Fam175a | 0.033463 | 0.0498 | 0.336897833 | 545.0221223 | 688.38355 | 170 | 23 | 29 | 34 | 99 | 10 | 2 | 0 | 0 | 0 | 2 | 2 |
| USG000000 | Gxylt2 | 0.0362 | 0.053495 | -0.996521505 | 79.35788664 | 39.77472901 | 139 | 25 | 29 | 34 | 90 | 14 | 1 | 0 | 0 | 0 | 1 | 0 |
| SG0000002 | Grb14 | 0.036883 | 0.054441 | 0.753709397 | 76.81231936 | 129.5149838 | 283 | 15 | 29 | 44 | 118 | 19 | 3 | 0 | 0 | 0 | 0 | 0 |
| USG000000 | Acsf2 | 0.055793 | 0.079763 | 0.506938542 | 266.7298911 | 379.031579 | 65 | 26 | 29 | 34 | 60 | 7 | 2 | 0 | 0 | 0 | 0 | 0 |
| USG000000 | Fam120c | 0.070962 | 0.099536 | -0.662746521 | 91.32611015 | 57.68838459 | 288 | 20 | 29 | 59 | 164 | 26 | 6 | 0 | 0 | 2 | 6 | 1 |
| SG0000002 | Dcaf6 | 0.079895 | 0.110954 | 0.263864974 | 660.3653322 | 792.8947415 | 343 | 10 | 29 | 45 | 167 | 33 | 5 | 0 | 1 | 0 | 1 | 0 |
| USG000000 | Pde3b | 0.087914 | 0.121172 | 0.48494559 | 209.8275968 | 293.6606633 | 386 | 14 | 29 | 59 | 197 | 25 | 12 | 1 | 2 | 2 | 5 | 2 |
| USG000000 | Lrch1 | 0.20645 | 0.262284 | -0.239691936 | 829.8231669 | 702.798932 | 328 | 17 | 29 | 62 | 164 | 35 | 6 | 1 | 2 | 3 | 8 | 2 |
| SG0000000 | Pemt | 0.219298 | 0.276488 | -0.369255274 | 96.4871765 | 74.69863815 | 265 | 15 | 29 | 31 | 84 | 11 | 0 | 0 | 0 | 0 | 0 | 0 |
| SG0000002 | Pcgf5 | 0.253786 | 0.314829 | 0.189317603 | 484.559625 | 552.5066401 | 246 | 13 | 29 | 43 | 135 | 21 | 2 | 0 | 0 | 0 | 0 | 0 |
| SG0000002 | Sec16b | 0.329818 | 0.396596 | -0.723332325 | 16.88103498 | 10.22478873 | 311 | 22 | 29 | 52 | 189 | 40 | 2 | 0 | 0 | 0 | 1 | 3 |
| USG000001 | Pcdhga2 | 0.332991 | 0.399794 | 0.598586319 | 16.39635262 | 24.82788282 | 256 | 23 | 29 | 48 | 116 | 20 | 2 | 0 | 0 | 1 | 1 | 0 |
| SG0000002 | Psph | 0.431339 | 0.500625 | -0.13008369 | 802.9986774 | 733.7628796 | 74 | 24 | 29 | 34 | 62 | 4 | 0 | 0 | 0 | 0 | 0 | 0 |
| USG000000 | Tsc22d2 | 0.488921 | 0.557061 | 0.105038712 | 810.2525289 | 871.445433 | 443 | 11 | 29 | 58 | 186 | 51 | 26 | 2 | 2 | 4 | 9 | 0 |
| SG0000002 | Psma6 | 0.495573 | 0.563135 | -0.09783105 | 5059.168325 | 4727.472908 | 147 | 16 | 29 | 29 | 82 | 10 | 0 | 0 | 0 | 0 | 0 | 0 |
| SG0000000 | Trim25 | 0.52265 | 0.589594 | 0.075538308 | 4356.92652 | 4591.129386 | 72 | 27 | 29 | 34 | 56 | 11 | 2 | 0 | 0 | 0 | 4 | 8 |
| USG000000 | Ankrd28 | 0.810016 | 0.849772 | -0.045936936 | 3537.394299 | 3426.534241 | 332 | 9 | 29 | 40 | 165 | 39 | 5 | 0 | 0 | 0 | 0 | 1 |
| SG0000005 | Zfp329 | 0.845626 | 0.879199 | 0.022774525 | 483.9675391 | 491.6681191 | 97 | 23 | 29 | 29 | 67 | 11 | 23 | 2 | 3 | 3 | 9 | 2 |
| SG0000002 | Bfar | 0.891324 | 0.916699 | 0.019350861 | 772.6206687 | 783.0536377 | 172 | 17 | 29 | 31 | 85 | 22 | 4 | 0 | 0 | 0 | 0 | 0 |
| SG0000002 | Stk31 | 1.8E-105 | 1.4E-103 | 10.18080001 | 0.307509278 | 356.9311125 | 218 | 19 | 28 | 34 | 96 | 21 | 0 | 0 | 0 | 0 | 0 | 0 |
| SG0000005 | Slc39a8 | 2.31E-80 | 1.05E-78 | -5.614743179 | 3190.995835 | 65.12085926 | 515 | 12 | 28 | 68 | 290 | 36 | 20 | 0 | 0 | 0 | 14 | 0 |
| USG000000 | Ppp2r5e | 9.17E-48 | 1.78E-46 | -2.418313321 | 7920.623709 | 1481.748625 | 370 | 13 | 28 | 46 | 174 | 37 | 7 | 0 | 0 | 2 | 2 | 2 |
| SG0000003 | Mgll | 9.22E-48 | 1.79E-46 | -4.031539542 | 2113.34156 | 129.227625 | 452 | 0 | 28 | 58 | 196 | 58 | 6 | 0 | 0 | 0 | 0 | 0 |
| SG0000003 | AI661453 | 2.46E-39 | 3.56E-38 | -3.421129932 | 4774.529286 | 445.7258613 | 68 | 28 | 28 | 30 | 52 | 16 | 0 | 0 | 0 | 0 | 1 | 2 |
| SG0000002 | Fam49a | 5.73E-39 | 8.14E-38 | 3.715866908 | 17.09936094 | 224.6814815 | 334 | 12 | 28 | 58 | 146 | 34 | 7 | 0 | 0 | 0 | 3 | 0 |
| SG0000002 | Mfn2 | 6.39E-38 | 8.78E-37 | -2.171325438 | 3285.11755 | 729.3178233 | 120 | 28 | 28 | 36 | 68 | 11 | 7 | 0 | 0 | 0 | 0 | 0 |
| SG0000003 | Tmem33 | 6.74E-31 | 7.09E-30 | -1.83154642 | 5343.691105 | 1501.381066 | 304 | 12 | 28 | 41 | 139 | 29 | 63 | 0 | 0 | 9 | 23 | 9 |
| SG0000000 | Hipk1 | 8.2E-31 | 8.57E-30 | -1.923843443 | 5226.569558 | 1377.470007 | 413 | 28 | 28 | 56 | 232 | 21 | 10 | 0 | 0 | 0 | 1 | 1 |
| SG0000003 | Ankrd35 | 1.13E-30 | 1.18E-29 | 2.460953382 | 64.97304179 | 357.7288654 | 78 | 24 | 28 | 24 | 48 | 2 | 0 | 0 | 0 | 0 | 0 | 0 |
| SG0000003 | Rhpn2 | 7.6E-30 | 7.65E-29 | 2.634529021 | 106.6797823 | 662.4519461 | 158 | 26 | 28 | 46 | 144 | 18 | 6 | 2 | 2 | 2 | 2 | 0 |
| SG0000003 | Ilf3 | 5.41E-28 | 5E-27 | 1.317059403 | 3231.278458 | 8050.980442 | 111 | 22 | 28 | 30 | 68 | 11 | 3 | 0 | 1 | 0 | 4 | 1 |
| SG0000004 | 2410002F23Rik | 7.35E-23 | 5.36E-22 | -1.483558489 | 6526.452338 | 2333.896353 | 75 | 21 | 28 | 28 | 51 | 17 | 8 | 0 | 0 | 0 | 8 | 0 |
| USG000000 | Cast | 2.36E-22 | 1.68E-21 | -1.706518755 | 4073.211657 | 1248.026033 | 257 | 16 | 28 | 39 | 159 | 27 | 0 | 0 | 0 | 0 | 1 | 1 |
| SG0000005 | Tmem62 | 9.78E-22 | 6.77E-21 | -2.198879702 | 465.9961866 | 101.4970957 | 160 | 20 | 28 | 48 | 106 | 22 | 0 | 0 | 0 | 0 | 0 | 0 |
| SG0000003 | Atp6v0a4 | 2.97E-21 | 2E-20 | 2.702474173 | 47.87483085 | 311.6255378 | 166 | 22 | 28 | 46 | 123 | 12 | 2 | 0 | 0 | 0 | 2 | 0 |
| SG0000001 | Kif1a | 9.9E-19 | 5.77E-18 | 1.306772601 | 585.1499759 | 1447.587892 | 319 | 8 | 28 | 39 | 125 | 1 | 3 | 0 | 0 | 3 | 4 | 1 |
| SG0000002 | Supt20 | 3.05E-18 | 1.74E-17 | 1.117090313 | 1380.543374 | 2994.523654 | 373 | 22 | 28 | 53 | 186 | 62 | 13 | 0 | 0 | 4 | 7 | 4 |
| SG0000002 | Ppa2 | 1.74E-17 | 9.55E-17 | 1.296796744 | 447.7940906 | 1100.152972 | 266 | 11 | 28 | 29 | 79 | 39 | 3 | 0 | 0 | 1 | 1 | 3 |
| USG000000 | Tunar | 2.62E-17 | 1.42E-16 | 5.54706978 | 0 | 45.75568154 | 84 | 26 | 28 | 34 | 54 | 14 | 24 | 20 | 20 | 20 | 20 | 0 |
| USG000000 | Eea1 | 4.68E-17 | 2.51E-16 | -1.472247687 | 1734.042138 | 624.9840473 | 275 | 7 | 28 | 44 | 134 | 30 | 12 | 0 | 1 | 2 | 2 | 2 |
| SG0000005 | H2-T3 | 9.51E-17 | 5.01E-16 | 6.306688467 | 0.639110347 | 50.59158088 | 45 | 28 | 28 | 28 | 35 | 0 | 2 | 0 | 0 | 0 | 0 | 0 |
| USG000000 | Mndal | 1.83E-16 | 9.47E-16 | -4.524685806 | 91.13501971 | 3.959306687 | 375 | 19 | 28 | 45 | 153 | 34 | 6 | 0 | 0 | 0 | 0 | 0 |
| USG000000 | Atp6v0d1 | 7.06E-13 | 2.9E-12 | 1.011720164 | 1955.377375 | 3942.654331 | 78 | 23 | 28 | 26 | 52 | 4 | 0 | 0 | 0 | 0 | 0 | 0 |
| SG0000002 | Trim9 | 1.6E-12 | 6.42E-12 | -9.114541178 | 399.1122501 | 0.720020342 | 332 | 18 | 28 | 54 | 158 | 32 | 8 | 0 | 0 | 0 | 0 | 4 |
| SG0000002 | Bad | 1.61E-11 | 6.08E-11 | -1.642520755 | 642.3238258 | 205.7339744 | 65 | 28 | 28 | 28 | 52 | 0 | 0 | 0 | 0 | 0 | 0 | 0 |
| USG000000 | Ubox5 | 1.99E-11 | 7.44E-11 | 1.028816228 | 430.2388262 | 877.837531 | 135 | 21 | 28 | 32 | 73 | 14 | 1 | 0 | 0 | 0 | 0 | 0 |
| SG0000002 | Ctnnbl1 | 3.83E-11 | 1.4E-10 | 0.882364664 | 1162.151673 | 2142.303723 | 194 | 26 | 28 | 46 | 109 | 15 | 0 | 0 | 0 | 0 | 0 | 0 |
| USG000000 | Irs1 | 2.35E-10 | 8.13E-10 | 1.763708688 | 100.8271192 | 342.378573 | 125 | 20 | 28 | 36 | 76 | 13 | 6 | 0 | 0 | 1 | 3 | 4 |
| SG0000002 | Sap130 | 2.87E-10 | 9.86E-10 | 0.82439412 | 1451.998168 | 2571.185599 | 288 | 14 | 28 | 54 | 148 | 24 | 4 | 0 | 0 | 2 | 2 | 0 |
| SG0000007 | Apitd1 | 3.51E-10 | 1.2E-09 | 1.182908338 | 203.49706 | 462.0075056 | 84 | 16 | 28 | 28 | 57 | 7 | 0 | 0 | 0 | 0 | 0 | 0 |
| SG0000005 | Sec31b | 7.15E-10 | 2.39E-09 | 2.592735867 | 12.42048023 | 74.92550089 | 79 | 26 | 28 | 32 | 67 | 2 | 0 | 0 | 0 | 0 | 1 | 0 |
| SG0000000 | Zak | 8.12E-10 | 2.7E-09 | -1.055705418 | 1422.091087 | 684.1139034 | 501 | 15 | 28 | 74 | 226 | 40 | 7 | 0 | 0 | 2 | 2 | 0 |
| SG0000002 | Xrcc6 | 6.73E-09 | 2.08E-08 | 0.752380705 | 1531.883744 | 2580.565982 | 90 | 22 | 28 | 30 | 60 | 10 | 0 | 0 | 0 | 0 | 0 | 0 |
| SG0000002 | Adipor1 | 1.49E-08 | 4.47E-08 | -0.77328957 | 9983.479368 | 5841.152824 | 74 | 26 | 28 | 32 | 38 | 4 | 4 | 0 | 0 | 0 | 0 | 0 |
| SG0000000 | Ddb2 | 2.59E-08 | 7.62E-08 | 1.040573444 | 184.5064535 | 379.5381013 | 114 | 22 | 28 | 34 | 76 | 6 | 0 | 0 | 0 | 0 | 0 | 0 |
| USG000000 | Foxk2 | 2.7E-08 | 7.92E-08 | -0.802811341 | 3369.937817 | 1931.752989 | 106 | 22 | 28 | 35 | 54 | 7 | 9 | 1 | 2 | 1 | 2 | 0 |
| SG0000003 | Ube2f | 2.68E-07 | 7.24E-07 | -0.760470796 | 2254.642246 | 1330.923581 | 286 | 10 | 28 | 40 | 144 | 80 | 6 | 0 | 3 | 0 | 4 | 0 |
| SG0000003 | Thoc5 | 3E-07 | 8.06E-07 | 0.695981585 | 1144.356487 | 1853.841796 | 100 | 24 | 28 | 34 | 78 | 6 | 0 | 0 | 0 | 0 | 0 | 0 |
| USG000000 | As3mt | 3.28E-07 | 8.78E-07 | -0.966446322 | 547.0968374 | 279.9850692 | 76 | 26 | 28 | 29 | 61 | 4 | 1 | 0 | 0 | 0 | 0 | 0 |
| SG0000006 | Sars | 3.3E-07 | 8.83E-07 | 0.634092222 | 4170.336303 | 6472.198932 | 60 | 26 | 28 | 35 | 51 | 4 | 4 | 0 | 0 | 1 | 1 | 0 |
| SG0000003 | Cep250 | 1.24E-06 | 3.18E-06 | -0.719532068 | 1985.416596 | 1205.732348 | 153 | 18 | 28 | 27 | 109 | 17 | 3 | 0 | 0 | 0 | 0 | 0 |
| SG0000002 | Psmd1 | 1.34E-06 | 3.42E-06 | 0.586878677 | 7768.44431 | 11668.15373 | 321 | 18 | 28 | 45 | 133 | 29 | 3 | 0 | 0 | 0 | 0 | 0 |
| SG0000003 | Dcun1d2 | 1.75E-06 | 4.41E-06 | -0.84887356 | 691.0029758 | 383.6573419 | 212 | 24 | 28 | 32 | 64 | 4 | 19 | 0 | 0 | 0 | 4 | 0 |
| SG0000004 | Pxylp1 | 2E-06 | 5.02E-06 | 1.351785096 | 175.5832407 | 448.1371234 | 328 | 12 | 28 | 24 | 124 | 12 | 8 | 0 | 0 | 0 | 4 | 0 |
| USG000000 | Vegfc | 2.47E-06 | 6.16E-06 | 1.883156699 | 29.08937623 | 107.3052725 | 312 | 13 | 28 | 44 | 151 | 42 | 2 | 0 | 0 | 0 | 2 | 0 |
| SG0000003 | Fndc3a | 3.68E-06 | 9.04E-06 | -0.682601399 | 4749.309971 | 2959.013994 | 474 | 7 | 28 | 53 | 225 | 46 | 8 | 0 | 1 | 0 | 1 | 0 |
| SG0000004 | Vstm2a | 4.15E-06 | 1.01E-05 | -2.581951961 | 45.07447187 | 7.528104852 | 394 | 17 | 28 | 41 | 189 | 37 | 4 | 0 | 0 | 1 | 4 | 0 |
| SG0000003 | Cmip | 4.96E-06 | 1.2E-05 | 0.581939557 | 2527.389889 | 3783.14952 | 264 | 24 | 28 | 47 | 128 | 20 | 0 | 0 | 0 | 0 | 0 | 0 |
| SG0000006 | H2-Q7 | 1.69E-05 | 3.89E-05 | 2.380714717 | 15.44988614 | 80.46218539 | 47 | 24 | 28 | 24 | 36 | 4 | 3 | 0 | 0 | 0 | 0 | 0 |
| SG0000003 | Vbp1 | 1.73E-05 | 3.99E-05 | 0.544988578 | 2779.215503 | 4054.899857 | 114 | 26 | 28 | 41 | 103 | 5 | 4 | 1 | 1 | 2 | 5 | 0 |
| SG0000002 | Adrbk1 | 2.69E-05 | 6.07E-05 | -0.648198063 | 2802.141877 | 1787.981663 | 48 | 28 | 28 | 28 | 52 | 4 | 3 | 0 | 0 | 0 | 3 | 0 |
| SG0000000 | Shpk | 8.77E-05 | 0.000187 | -1.06783045 | 165.5176968 | 78.95787265 | 108 | 26 | 28 | 32 | 68 | 0 | 4 | 0 | 0 | 0 | 0 | 0 |
| SG0000002 | Agbl5 | 0.000111 | 0.000233 | 0.657255572 | 362.3334018 | 571.4286533 | 154 | 21 | 28 | 42 | 50 | 0 | 2 | 0 | 0 | 0 | 1 | 0 |
| SG0000001 | Nlk | 0.000144 | 0.0003 | 0.522302225 | 1075.340435 | 1544.452703 | 300 | 14 | 28 | 42 | 147 | 29 | 4 | 0 | 0 | 0 | 4 | 2 |
| SG0000006 | Serpinb5 | 0.000181 | 0.000373 | 2.830275552 | 3.241391746 | 23.05310132 | 153 | 18 | 28 | 35 | 73 | 33 | 9 | 0 | 0 | 0 | 3 | 6 |
| SG0000002 | Pnisr | 0.000427 | 0.00084 | 0.740672579 | 1979.400413 | 3307.488325 | 334 | 26 | 28 | 62 | 146 | 24 | 27 | 0 | 1 | 5 | 10 | 3 |
| USG000001 | Pcdhga3 | 0.001226 | 0.002279 | -1.5306189 | 48.27755729 | 16.71025537 | 249 | 23 | 28 | 46 | 114 | 19 | 2 | 0 | 0 | 1 | 1 | 0 |
| SG0000001 | Zfp207 | 0.001503 | 0.002766 | 0.391884419 | 6016.264475 | 7893.977508 | 348 | 6 | 28 | 62 | 161 | 10 | 27 | 0 | 0 | 2 | 9 | 1 |
| SG0000006 | Zfp27 | 0.001813 | 0.003302 | -0.631501943 | 346.8167711 | 223.871631 | 111 | 24 | 28 | 63 | 106 | 15 | 1 | 0 | 0 | 0 | 4 | 0 |
| USG000000 | Prss32 | 0.002279 | 0.004091 | -2.854174895 | 17.21647946 | 2.380959963 | 40 | 28 | 28 | 28 | 34 | 0 | 1 | 0 | 0 | 0 | 0 | 0 |
| SG0000003 | Mrpl48 | 0.002688 | 0.004778 | 0.507675106 | 368.5552066 | 523.9960169 | 231 | 26 | 28 | 71 | 180 | 22 | 4 | 0 | 0 | 0 | 1 | 0 |
| SG0000004 | 2310022A10Rik | 0.004091 | 0.007076 | 0.402241401 | 1194.043564 | 1577.999638 | 164 | 28 | 28 | 36 | 67 | 10 | 4 | 0 | 0 | 0 | 0 | 0 |
| SG0000006 | Jazf1 | 0.00441 | 0.007598 | 2.291770769 | 3.538026913 | 17.32418756 | 509 | 12 | 28 | 55 | 198 | 41 | 1 | 0 | 0 | 0 | 1 | 0 |
| USG000000 | Cwf19l2 | 0.006213 | 0.010493 | 0.433415196 | 912.4551274 | 1232.203649 | 204 | 22 | 28 | 46 | 133 | 17 | 0 | 0 | 0 | 0 | 2 | 1 |
| SG0000002 | Adck5 | 0.006548 | 0.011021 | -0.51102022 | 486.8345076 | 341.6244452 | 92 | 20 | 28 | 28 | 52 | 15 | 2 | 0 | 0 | 0 | 0 | 3 |
| SG0000006 | Ap2a1 | 0.007358 | 0.012284 | 0.480710635 | 2663.256855 | 3716.390779 | 96 | 24 | 28 | 36 | 76 | 12 | 0 | 0 | 0 | 0 | 8 | 0 |
| USG000000 | Mb21d2 | 0.009829 | 0.016095 | -0.45219964 | 588.8146487 | 430.3808612 | 167 | 23 | 28 | 47 | 78 | 7 | 3 | 0 | 1 | 1 | 3 | 0 |
| SG0000002 | Gtsf1 | 0.013247 | 0.02118 | 1.72978083 | 6.162056536 | 20.43815057 | 387 | 28 | 28 | 54 | 216 | 35 | 0 | 0 | 0 | 0 | 0 | 0 |
| SG0000006 | Ccdc62 | 0.01447 | 0.022986 | 1.083809257 | 20.21645032 | 42.85129972 | 132 | 20 | 28 | 32 | 58 | 8 | 6 | 2 | 4 | 4 | 6 | 0 |
| SG0000002 | Inpp5b | 0.018571 | 0.028897 | -0.373851617 | 1033.612253 | 797.6586854 | 166 | 24 | 28 | 42 | 110 | 10 | 4 | 0 | 2 | 0 | 0 | 2 |
| SG0000002 | Kif6 | 0.027795 | 0.042 | 1.34171106 | 10.95905262 | 27.77590941 | 470 | 14 | 28 | 63 | 223 | 39 | 1 | 1 | 1 | 1 | 1 | 0 |
| USG000000 | Apoo | 0.041407 | 0.060625 | 0.479476279 | 121.6853297 | 169.6582431 | 407 | 20 | 28 | 59 | 211 | 55 | 3 | 0 | 0 | 3 | 3 | 3 |
| SG0000002 | Fam49b | 0.045775 | 0.066568 | 0.255206837 | 2365.72687 | 2823.511205 | 466 | 8 | 28 | 59 | 196 | 46 | 11 | 0 | 0 | 0 | 1 | 3 |
| SG0000002 | Taf15 | 0.097624 | 0.133205 | 0.211936449 | 2878.090885 | 3333.525123 | 175 | 15 | 28 | 32 | 67 | 8 | 0 | 0 | 0 | 0 | 0 | 0 |
| SG0000006 | Prpf40a | 0.100523 | 0.136618 | 0.202901367 | 4971.98623 | 5722.809834 | 382 | 9 | 28 | 50 | 171 | 23 | 18 | 0 | 0 | 3 | 4 | 2 |
| SG0000003 | Uros | 0.114556 | 0.153818 | -0.326392618 | 282.4954104 | 225.2980475 | 147 | 28 | 28 | 31 | 66 | 28 | 4 | 0 | 0 | 0 | 3 | 0 |
| SG0000003 | Ppfia1 | 0.138086 | 0.182323 | -0.208248938 | 2143.975861 | 1855.798052 | 280 | 14 | 28 | 51 | 136 | 36 | 16 | 2 | 2 | 7 | 10 | 0 |
| SG0000002 | Wdr33 | 0.173854 | 0.224712 | 0.181360406 | 1895.560476 | 2149.475835 | 331 | 18 | 28 | 63 | 162 | 29 | 3 | 0 | 0 | 0 | 0 | 0 |
| USG000000 | Cd2ap | 0.24157 | 0.301596 | 0.143546605 | 4049.776656 | 4473.453284 | 249 | 14 | 28 | 42 | 139 | 16 | 7 | 0 | 0 | 0 | 2 | 1 |
| SG0000002 | N4bp2l2 | 0.248757 | 0.309427 | 0.146955148 | 2063.110304 | 2284.337954 | 186 | 22 | 28 | 34 | 94 | 11 | 21 | 0 | 0 | 0 | 9 | 0 |
| SG0000002 | Ndc1 | 0.285503 | 0.349147 | -0.150927789 | 2744.385186 | 2471.788314 | 296 | 12 | 28 | 54 | 153 | 48 | 7 | 0 | 0 | 2 | 5 | 0 |
| USG000000 | Nxn | 0.294089 | 0.358397 | 0.144426312 | 1806.61948 | 1996.840336 | 196 | 23 | 28 | 40 | 83 | 15 | 2 | 0 | 0 | 0 | 0 | 0 |
| SG0000002 | Odf2l | 0.36383 | 0.432242 | -0.177002448 | 607.7291182 | 537.5600668 | 390 | 10 | 28 | 51 | 222 | 39 | 25 | 0 | 0 | 0 | 10 | 0 |
| USG000000 | Ar | 0.418323 | 0.487477 | 0.538452593 | 10.45262205 | 15.18153303 | 436 | 15 | 28 | 57 | 269 | 42 | 10 | 0 | 0 | 0 | 4 | 1 |
| USG000000 | Mamdc2 | 0.437889 | 0.50711 | -0.404308993 | 33.0071878 | 24.94016887 | 272 | 20 | 28 | 45 | 137 | 38 | 2 | 0 | 0 | 0 | 1 | 1 |
| SG0000006 | Clip2 | 0.480965 | 0.549549 | 0.157271391 | 282.3012949 | 314.815623 | 120 | 24 | 28 | 44 | 104 | 10 | 0 | 0 | 0 | 0 | 0 | 0 |
| SG0000005 | Actn2 | 0.483373 | 0.551863 | -0.207014503 | 107.4883789 | 93.12020816 | 232 | 26 | 28 | 48 | 94 | 34 | 0 | 0 | 0 | 0 | 0 | 0 |
| SG0000004 | Rnf10 | 0.508936 | 0.575872 | -0.085199862 | 8515.73528 | 8027.391393 | 152 | 20 | 28 | 52 | 116 | 8 | 0 | 0 | 0 | 0 | 0 | 0 |
| SG0000002 | Gabbr1 | 0.558015 | 0.62314 | -0.158407872 | 1669.703458 | 1496.076584 | 125 | 24 | 28 | 42 | 75 | 11 | 6 | 0 | 2 | 0 | 6 | 4 |
| USG000000 | Fbxo42 | 0.573264 | 0.637704 | 0.074796106 | 1826.165204 | 1923.339429 | 98 | 21 | 28 | 33 | 65 | 3 | 5 | 0 | 0 | 1 | 2 | 0 |
| SG0000007 | Gas2l3 | 0.705925 | 0.759431 | 0.050325815 | 1214.319573 | 1257.426403 | 166 | 18 | 28 | 34 | 88 | 20 | 26 | 4 | 4 | 4 | 11 | 4 |
| SG0000001 | Cnksr3 | 0.746243 | 0.794833 | -0.059655288 | 542.3149087 | 520.3475483 | 187 | 16 | 28 | 38 | 87 | 25 | 6 | 0 | 0 | 2 | 2 | 0 |
| SG0000002 | Nde1 | 0.840294 | 0.875332 | 0.029519084 | 2178.901814 | 2223.943707 | 197 | 24 | 28 | 41 | 105 | 5 | 0 | 0 | 0 | 0 | 0 | 0 |
| SG0000002 | Nasp | 6.3E-192 | 1.7E-189 | 3.699474671 | 1885.119316 | 24490.16368 | 196 | 24 | 27 | 33 | 108 | 18 | 1 | 0 | 0 | 0 | 0 | 3 |
| SG0000000 | Fblim1 | 9.5E-104 | 7.4E-102 | 2.676652633 | 842.101383 | 5384.149508 | 113 | 21 | 27 | 42 | 70 | 9 | 3 | 0 | 0 | 0 | 0 | 0 |
| SG0000002 | Vangl2 | 2.95E-87 | 1.56E-85 | 3.11432718 | 136.9506622 | 1185.95998 | 95 | 21 | 27 | 33 | 90 | 6 | 2 | 0 | 0 | 0 | 2 | 0 |
| USG000000 | Ano6 | 2.46E-60 | 6.92E-59 | -3.494105603 | 10517.7582 | 933.4532832 | 299 | 19 | 27 | 50 | 155 | 51 | 2 | 0 | 0 | 0 | 0 | 0 |
| SG0000002 | Anxa4 | 1.66E-54 | 3.93E-53 | -2.680131659 | 7362.999408 | 1148.827291 | 220 | 22 | 27 | 48 | 120 | 12 | 7 | 0 | 0 | 0 | 3 | 0 |
| USG000000 | Bcl2 | 2.14E-45 | 3.8E-44 | -4.782791449 | 1373.284691 | 49.88809145 | 343 | 16 | 27 | 53 | 133 | 36 | 18 | 2 | 2 | 2 | 9 | 6 |
| SG0000005 | Rgs3 | 8.65E-37 | 1.14E-35 | 3.27356935 | 26.87072289 | 259.8495191 | 280 | 21 | 27 | 39 | 144 | 36 | 9 | 3 | 6 | 3 | 13 | 0 |
| SG0000002 | Egfr | 6.64E-36 | 8.49E-35 | -5.436128152 | 8727.415326 | 201.5802641 | 488 | 10 | 27 | 62 | 197 | 44 | 12 | 0 | 0 | 1 | 5 | 3 |
| SG0000003 | Arhgap18 | 7.93E-34 | 9.37E-33 | -2.680707314 | 1023.203358 | 159.5837426 | 306 | 19 | 27 | 55 | 164 | 25 | 5 | 0 | 0 | 1 | 3 | 2 |

| | | | | | | | | | | | | | | | | | |
|---|---|---|---|---|---|---|---|---|---|---|---|---|---|---|---|---|---|
| SG0000001 | Prpf3 | 2.99E-31 | 3.19E-30 | 1.504646316 | 964.0524383 | 2735.547948 | 161 | 19 | 27 | 40 | 94 | 9 | 1 | 0 | 0 | 0 | 0 |
| SG0000002 | Lmna | 5.81E-30 | 5.88E-29 | -3.129334469 | 7990.070784 | 913.1183807 | 65 | 27 | 27 | 29 | 47 | 0 | 2 | 0 | 0 | 1 | 0 |
| SG0000002 | Casp8 | 4.45E-29 | 4.31E-28 | -2.130788318 | 1141.51742 | 260.6461685 | 192 | 17 | 27 | 36 | 104 | 6 | 6 | 0 | 0 | 0 | 0 |
| JSG000000 | Skap2 | 2.27E-28 | 2.14E-27 | -1.735579829 | 4593.343869 | 1379.327423 | 437 | 16 | 27 | 65 | 203 | 27 | 1 | 0 | 0 | 0 | 0 |
| JSG000000 | Itsn2 | 1.69E-22 | 1.21E-21 | -1.67617259 | 2894.922045 | 905.8549167 | 257 | 14 | 27 | 41 | 125 | 19 | 1 | 0 | 1 | 5 | 0 |
| SG0000006 | Akr1b10 | 2.91E-21 | 1.96E-20 | -1.731638217 | 1089.388139 | 328.0254936 | 101 | 24 | 27 | 33 | 42 | 9 | 2 | 0 | 0 | 0 | 0 |
| JSG000000 | Lnpep | 4.13E-21 | 2.75E-20 | -2.014035235 | 1962.276171 | 485.8196892 | 247 | 17 | 27 | 42 | 133 | 25 | 3 | 0 | 0 | 2 | 1 |
| SG0000002 | Uap1 | 4.62E-20 | 2.91E-19 | 1.190797312 | 1134.360066 | 2589.504314 | 213 | 15 | 27 | 42 | 129 | 27 | 3 | 0 | 3 | 3 | 6 | 3 |
| SG0000003 | Slc16a2 | 1.71E-19 | 1.04E-18 | 2.755746136 | 28.45879643 | 192.2112383 | 218 | 20 | 27 | 49 | 127 | 23 | 4 | 0 | 0 | 0 | 0 |
| SG0000005 | Senp8 | 1.16E-16 | 6.05E-16 | 1.34830302 | 264.6852672 | 673.9216687 | 96 | 21 | 27 | 27 | 63 | 18 | 14 | 0 | 3 | 0 | 8 | 3 |
| SG0000002 | Rnf165 | 3.54E-16 | 1.8E-15 | 8.681085404 | 1.334934818 | 547.9326277 | 253 | 21 | 27 | 43 | 129 | 15 | 9 | 1 | 3 | 3 | 7 | 3 |
| SG0000002 | Fgd2 | 3.57E-16 | 1.82E-15 | 3.399384479 | 9.883443503 | 104.2857525 | 57 | 24 | 27 | 27 | 43 | 0 | 0 | 0 | 0 | 0 | 0 |
| SG0000003 | Wdr17 | 1.27E-13 | 5.48E-13 | 5.133835087 | 1.299968916 | 45.64269943 | 367 | 12 | 27 | 62 | 228 | 36 | 6 | 0 | 1 | 2 | 4 | 1 |
| JSG000000 | B4galnt3 | 4.08E-13 | 1.7E-12 | 4.600547546 | 15.01323423 | 364.2311358 | 105 | 23 | 27 | 37 | 85 | 13 | 0 | 0 | 0 | 0 | 0 |
| SG0000004 | Pacsin1 | 2.93E-12 | 1.16E-11 | 7.459424582 | 2.837515307 | 499.4002583 | 115 | 21 | 27 | 27 | 103 | 15 | 1 | 0 | 0 | 1 | 0 |
| SG0000003 | Pik3cd | 7.85E-10 | 2.61E-09 | 1.785112617 | 275.9890036 | 951.1833113 | 213 | 23 | 27 | 45 | 197 | 22 | 23 | 5 | 5 | 11 | 11 | 0 |
| SG0000004 | Phf3 | 1.66E-09 | 5.37E-09 | 0.744879458 | 3697.057424 | 6195.655425 | 475 | 14 | 27 | 54 | 217 | 59 | 12 | 0 | 0 | 0 | 2 | 4 |
| SG0000005 | Zfp74 | 3.45E-09 | 1.08E-08 | 0.952253113 | 347.2392085 | 671.8704001 | 113 | 15 | 27 | 30 | 64 | 2 | 8 | 1 | 1 | 1 | 4 | 0 |
| SG0000002 | Ifnar1 | 1.81E-08 | 5.39E-08 | -0.880425476 | 1526.91303 | 829.4301612 | 84 | 27 | 27 | 36 | 72 | 12 | 2 | 0 | 0 | 0 | 1 |
| SG0000003 | Tnrc18 | 4.66E-08 | 1.34E-07 | -1.320248465 | 1793.758406 | 718.3391884 | 108 | 26 | 27 | 34 | 75 | 16 | 0 | 0 | 0 | 1 | 0 |
| SG0000005 | Ccdc28a | 2.08E-07 | 5.67E-07 | 0.971977928 | 191.7153403 | 376.0550077 | 154 | 24 | 27 | 40 | 96 | 11 | 1 | 0 | 0 | 3 | 0 |
| SG0000002 | Pqlc2 | 3.06E-07 | 8.21E-07 | -1.193912256 | 393.5594845 | 172.0311061 | 94 | 26 | 27 | 32 | 55 | 1 | 3 | 0 | 0 | 1 | 0 |
| SG0000003 | Ccdc91 | 3.46E-07 | 9.24E-07 | -0.8778097 | 793.6045718 | 431.8740515 | 383 | 8 | 27 | 56 | 171 | 34 | 3 | 0 | 0 | 1 | 0 |
| SG0000002 | Golga3 | 3.51E-07 | 9.38E-07 | -0.753320535 | 2309.858003 | 1370.292262 | 150 | 19 | 27 | 25 | 69 | 17 | 12 | 3 | 5 | 3 | 7 | 1 |
| SG0000002 | Nucks1 | 3.7E-07 | 9.86E-07 | 0.610197727 | 8802.867374 | 13437.2989 | 137 | 20 | 27 | 32 | 66 | 5 | 17 | 0 | 1 | 0 | 6 | 1 |
| JSG000000 | Msra | 8.62E-07 | 2.24E-06 | 1.081578419 | 112.6857578 | 238.4825334 | 457 | 13 | 27 | 60 | 200 | 41 | 0 | 0 | 0 | 0 | 0 |
| SG0000002 | Cfap52 | 1.97E-06 | 4.95E-06 | 5.869185732 | 0.307509278 | 17.97459771 | 138 | 21 | 27 | 30 | 72 | 26 | 0 | 0 | 0 | 0 | 0 |
| SG0000004 | Nup88 | 2.24E-06 | 5.6E-06 | 0.583465673 | 3885.354661 | 5821.988415 | 226 | 19 | 27 | 51 | 110 | 12 | 15 | 3 | 4 | 3 | 9 | 0 |
| SG0000002 | Slc29a3 | 2.27E-06 | 5.68E-06 | -0.914424956 | 672.0268557 | 356.5474117 | 114 | 21 | 27 | 21 | 39 | 15 | 6 | 0 | 0 | 0 | 0 |
| JSG000000 | Nup153 | 2.92E-06 | 7.22E-06 | 0.60168552 | 3779.36314 | 5735.13985 | 166 | 23 | 27 | 43 | 95 | 6 | 2 | 0 | 0 | 0 | 4 | 1 |
| SG0000002 | Snx8 | 3.62E-06 | 8.9E-06 | 0.725118153 | 594.3487753 | 982.4799283 | 142 | 25 | 27 | 49 | 111 | 6 | 0 | 0 | 0 | 0 | 0 |
| JSG000000 | Memo1 | 5.08E-06 | 1.23E-05 | -0.692935983 | 1834.374891 | 1134.732729 | 236 | 14 | 27 | 36 | 145 | 22 | 2 | 0 | 0 | 0 | 0 |
| JSG000000 | Cdh11 | 7.59E-06 | 1.81E-05 | -1.713163939 | 841.784121 | 256.7361334 | 386 | 12 | 27 | 54 | 188 | 21 | 5 | 0 | 1 | 1 | 3 | 0 |
| SG0000003 | Exoc1 | 1.22E-05 | 2.86E-05 | -0.656168781 | 1925.880946 | 1222.089326 | 246 | 9 | 27 | 42 | 117 | 15 | 3 | 0 | 0 | 0 | 0 |
| SG0000003 | Kank1 | 2.14E-05 | 4.88E-05 | -0.692013879 | 1195.98305 | 740.30058 | 299 | 20 | 27 | 49 | 112 | 32 | 1 | 0 | 0 | 1 | 0 |
| SG0000003 | Scn5a | 5.08E-05 | 0.000112 | -1.889813642 | 77.40101214 | 20.8860372 | 198 | 15 | 27 | 31 | 59 | 11 | 2 | 0 | 0 | 5 | 0 |
| SG0000003 | Ube2l3 | 9.73E-05 | 0.000207 | -0.539965466 | 4005.002232 | 2754.589962 | 112 | 27 | 27 | 39 | 75 | 5 | 3 | 0 | 0 | 0 | 0 |
| SG000000C | Drp2 | 0.000127 | 0.000266 | 1.651448583 | 16.7025152 | 52.47076974 | 192 | 24 | 27 | 42 | 120 | 41 | 6 | 0 | 0 | 0 | 2 |
| SG0000005 | Prdm9 | 0.000146 | 0.000304 | -1.236605846 | 111.3759117 | 47.26458179 | 83 | 27 | 27 | 34 | 77 | 14 | 1 | 0 | 0 | 0 | 0 |
| SG0000002 | Parp6 | 0.000167 | 0.000345 | -0.666320072 | 640.1069198 | 403.3389783 | 194 | 21 | 27 | 45 | 126 | 36 | 0 | 0 | 0 | 0 | 3 |
| JSG000000 | Col6a5 | 0.000227 | 0.000462 | -1.656453617 | 56.0603097 | 17.78333736 | 179 | 19 | 27 | 36 | 91 | 19 | 4 | 0 | 0 | 1 | 0 |
| SG0000002 | Lamb3 | 0.000327 | 0.000653 | 2.41147096 | 36.37323175 | 193.5115904 | 235 | 10 | 27 | 25 | 96 | 32 | 0 | 0 | 0 | 0 | 0 |
| SG0000004 | Tox3 | 0.000555 | 0.001077 | 5.53833495 | 1.326404274 | 61.64258724 | 229 | 18 | 27 | 40 | 104 | 31 | 5 | 1 | 1 | 1 | 2 | 1 |
| SG0000002 | Zfp618 | 0.000711 | 0.001359 | 1.284386031 | 60.82921745 | 148.1668168 | 649 | 18 | 27 | 102 | 281 | 45 | 7 | 0 | 0 | 1 | 2 | 0 |
| SG0000002 | Mtl5 | 0.000804 | 0.001525 | 1.568655853 | 14.69734754 | 43.59648028 | 91 | 27 | 27 | 45 | 60 | 7 | 0 | 0 | 0 | 4 | 0 |
| JSG000000 | Pros1 | 0.00145 | 0.002672 | -0.705171868 | 1530.118294 | 938.5277608 | 201 | 21 | 27 | 46 | 109 | 16 | 2 | 0 | 0 | 1 | 1 |
| SG0000006 | Lancl2 | 0.002274 | 0.004082 | 0.462653677 | 616.4449754 | 849.5069799 | 278 | 12 | 27 | 45 | 150 | 21 | 5 | 0 | 0 | 1 | 1 |
| SG0000002 | Lpcat1 | 0.002756 | 0.004889 | -0.423527344 | 2769.7092 | 2065.093592 | 163 | 15 | 27 | 38 | 59 | 32 | 7 | 0 | 2 | 2 | 2 |
| SG0000002 | Spata7 | 0.003665 | 0.006387 | 0.571309421 | 193.8212215 | 287.9934423 | 336 | 6 | 27 | 42 | 150 | 15 | 0 | 0 | 0 | 0 | 0 |
| SG0000002 | Cdk11b | 0.00406 | 0.007029 | 0.373678415 | 2343.68144 | 3036.595838 | 159 | 21 | 27 | 27 | 60 | 3 | 0 | 0 | 0 | 0 | 0 |
| SG0000003 | Triobp | 0.006246 | 0.010541 | -0.692922686 | 2550.79635 | 1577.921065 | 139 | 27 | 27 | 45 | 85 | 15 | 5 | 0 | 1 | 2 | 0 |
| SG0000004 | Dhx9 | 0.011209 | 0.018186 | 0.31642252 | 11404.24364 | 14201.0075 | 382 | 17 | 27 | 42 | 131 | 19 | 10 | 0 | 0 | 2 | 0 |
| SG0000005 | 43714 | 0.015638 | 0.024718 | 0.511621082 | 503.1894145 | 717.3726058 | 375 | 21 | 27 | 74 | 167 | 44 | 4 | 0 | 1 | 1 | 3 |
| SG0000003 | Dph5 | 0.017026 | 0.026709 | 0.40408954 | 388.1501106 | 513.6210174 | 273 | 7 | 27 | 51 | 137 | 31 | 6 | 1 | 1 | 3 | 4 | 1 |
| SG0000003 | Pdpr | 0.025877 | 0.039273 | 0.463653396 | 753.797349 | 1039.508837 | 91 | 19 | 27 | 29 | 58 | 7 | 2 | 0 | 0 | 1 | 0 |
| SG0000005 | Cpsf6 | 0.033108 | 0.049324 | 0.266457594 | 4077.568041 | 4904.705606 | 418 | 21 | 27 | 72 | 201 | 23 | 35 | 1 | 2 | 7 | 16 | 3 |
| JSG000000 | Cdk17 | 0.040786 | 0.059765 | -0.296732771 | 1823.915728 | 1484.838787 | 202 | 21 | 27 | 38 | 90 | 21 | 6 | 2 | 2 | 2 | 5 | 0 |
| JSG000000 | Fancm | 0.048616 | 0.070352 | 0.277426174 | 1026.034499 | 1243.585254 | 139 | 19 | 27 | 37 | 72 | 8 | 5 | 0 | 1 | 1 | 3 | 1 |
| SG0000002 | Hibadh | 0.055619 | 0.079561 | 0.254228034 | 1801.524068 | 2148.67303 | 274 | 19 | 27 | 43 | 110 | 20 | 1 | 0 | 0 | 2 | 0 |
| SG0000005 | Grk4 | 0.067526 | 0.095138 | -1.452242448 | 16.468628 | 6.018508549 | 294 | 18 | 27 | 46 | 166 | 23 | 0 | 0 | 0 | 0 | 0 |
| SG0000003 | Cyb5r4 | 0.07693 | 0.107194 | 0.25098676 | 1031.650975 | 1227.686093 | 160 | 23 | 27 | 40 | 82 | 12 | 2 | 0 | 0 | 2 | 0 |
| SG0000002 | Abca8b | 0.081922 | 0.113558 | 0.752746749 | 22.50284504 | 37.91724547 | 282 | 8 | 27 | 43 | 196 | 28 | 4 | 0 | 1 | 1 | 18 | 0 |
| JSG000000 | Cmpk1 | 0.119575 | 0.159957 | 0.205844959 | 1818.695867 | 2097.614067 | 174 | 21 | 27 | 27 | 60 | 9 | 8 | 1 | 1 | 1 | 5 | 1 |
| SG0000006 | Ipmk | 0.203008 | 0.258332 | -0.179355437 | 2461.872427 | 2174.073277 | 262 | 10 | 27 | 46 | 228 | 29 | 11 | 0 | 0 | 1 | 1 | 0 |
| SG000000C | Mef2c | 0.210489 | 0.266753 | -1.041397415 | 18.96331828 | 9.213454066 | 415 | 12 | 27 | 46 | 174 | 34 | 11 | 0 | 0 | 0 | 1 | 2 |
| JSG000000 | Ccdc15 | 0.218748 | 0.275911 | 0.301648563 | 133.9008548 | 165.039772 | 192 | 24 | 27 | 38 | 110 | 24 | 0 | 0 | 0 | 0 | 0 |
| JSG000001 | Pcdhga4 | 0.245457 | 0.305916 | 0.770103676 | 10.69488665 | 18.23897771 | 231 | 23 | 27 | 43 | 108 | 18 | 2 | 0 | 0 | 1 | 1 | 0 |
| SG0000006 | Nck2 | 0.248586 | 0.309299 | -0.2429093 | 569.5785559 | 481.3163833 | 266 | 11 | 27 | 36 | 116 | 26 | 1 | 0 | 0 | 0 | 4 | 0 |
| JSG000000 | Zfp82 | 0.259217 | 0.320788 | 0.667464891 | 14.23910054 | 22.61567267 | 86 | 21 | 27 | 38 | 64 | 9 | 1 | 0 | 0 | 0 | 0 | 0 |
| JSG000000 | Slc7a14 | 0.260081 | 0.321702 | -0.736080464 | 21.65862641 | 13.00315117 | 213 | 15 | 27 | 49 | 103 | 20 | 9 | 0 | 0 | 3 | 3 | 1 |
| SG0000003 | Ndufab1 | 0.297569 | 0.362342 | 0.157140345 | 689.1244541 | 768.4252355 | 87 | 21 | 27 | 28 | 53 | 6 | 8 | 0 | 0 | 0 | 1 | 0 |
| SG0000006 | Pde7a | 0.333083 | 0.399878 | -0.149819132 | 1469.903379 | 1324.917193 | 540 | 16 | 27 | 47 | 184 | 51 | 29 | 2 | 2 | 2 | 2 | 0 |
| SG0000003 | Rab30 | 0.334404 | 0.401195 | 0.316839407 | 52.41703524 | 65.29059381 | 232 | 17 | 27 | 38 | 107 | 20 | 1 | 0 | 0 | 0 | 0 | 0 |
| JSG000000 | Smad9 | 0.36301 | 0.431383 | -0.488422813 | 28.59247306 | 20.38082704 | 89 | 23 | 27 | 30 | 62 | 10 | 8 | 0 | 0 | 1 | 4 | 3 |
| SG0000004 | Slc38a6 | 0.464815 | 0.534285 | 0.150895191 | 196.3556084 | 218.0054189 | 369 | 22 | 27 | 60 | 179 | 22 | 13 | 0 | 0 | 0 | 5 | 1 |
| JSG000000 | Ube2r2 | 0.492995 | 0.560775 | 0.08695648 | 2237.353701 | 2376.354101 | 139 | 25 | 27 | 41 | 89 | 11 | 4 | 0 | 0 | 1 | 1 | 0 |
| SG0000002 | Psmd14 | 0.526486 | 0.593071 | 0.077036353 | 3538.486308 | 3732.568379 | 272 | 8 | 27 | 39 | 141 | 26 | 2 | 0 | 0 | 0 | 0 | 0 |
| JSG000000 | Colec12 | 0.784849 | 0.828709 | 0.071997573 | 451.815234 | 474.9351349 | 280 | 11 | 27 | 39 | 133 | 36 | 4 | 0 | 0 | 0 | 1 | 0 |
| SG0000005 | Traf7 | 0.787867 | 0.831405 | 0.038065211 | 3048.339691 | 3129.83996 | 137 | 27 | 27 | 36 | 63 | 26 | 4 | 0 | 0 | 0 | 0 | 6 |
| SG0000002 | Mapk8ip3 | 0.905576 | 0.928967 | -0.018740144 | 1951.722565 | 1926.534266 | 217 | 18 | 27 | 18 | 100 | 45 | 1 | 0 | 0 | 0 | 0 | 9 |
| SG0000002 | Gatb | 0.988488 | 0.99841 | 0.004526921 | 376.4609238 | 377.6440465 | 448 | 15 | 27 | 54 | 177 | 28 | 3 | 0 | 0 | 0 | 0 | 0 |
| SG0000004 | Mageb16 | 4.9E-93 | 2.97E-91 | 8.229404442 | 0 | 299.1218284 | 171 | 18 | 26 | 36 | 119 | 14 | 4 | 0 | 0 | 0 | 0 | 0 |
| SG0000005 | Gprc5c | 9.51E-93 | 5.72E-91 | 6.799603773 | 3.252265858 | 362.3020022 | 110 | 22 | 26 | 26 | 79 | 4 | 3 | 0 | 0 | 0 | 0 | 0 |
| JSG000000 | Dennd2a | 3.72E-89 | 2.07E-87 | 3.318509751 | 102.7880998 | 1025.448404 | 145 | 23 | 26 | 40 | 90 | 11 | 1 | 0 | 0 | 0 | 3 | 0 |
| SG0000002 | Trim63 | 7.37E-84 | 3.65E-82 | 8.051231889 | 0 | 264.2542193 | 54 | 24 | 26 | 32 | 40 | 4 | 0 | 0 | 0 | 0 | 0 | 2 |
| SG0000002 | Fam126a | 5.17E-69 | 1.8E-67 | -3.340425751 | 4386.479264 | 433.0596608 | 424 | 15 | 26 | 64 | 218 | 41 | 7 | 0 | 1 | 1 | 2 | 1 |
| SG0000003 | Baz1a | 2.12E-67 | 7.19E-66 | -3.203129545 | 10224.66789 | 1110.225345 | 193 | 17 | 26 | 32 | 94 | 16 | 8 | 0 | 0 | 2 | 2 | 0 |
| SG0000005 | Sema5b | 3.03E-52 | 6.72E-51 | 6.212396845 | 3.99627391 | 296.3282409 | 266 | 22 | 26 | 38 | 104 | 14 | 0 | 0 | 0 | 0 | 0 | 0 |
| JSG000000 | Cep78 | 1.66E-43 | 2.77E-42 | 2.055125267 | 282.7816217 | 1175.183083 | 72 | 24 | 26 | 29 | 52 | 4 | 0 | 0 | 0 | 0 | 0 | 0 |
| SG0000005 | Scn3a | 1.86E-33 | 2.15E-32 | 8.201543372 | 0.331601068 | 97.61723156 | 551 | 6 | 26 | 48 | 238 | 59 | 19 | 0 | 0 | 0 | 4 | 5 |
| SG0000004 | Ccdc60 | 4.43E-33 | 5.07E-32 | 6.541788798 | 0 | 92.16969067 | 284 | 18 | 26 | 49 | 117 | 54 | 3 | 0 | 1 | 1 | 1 | 1 |
| SG0000003 | Kdm1a | 1.78E-29 | 1.75E-28 | 1.347278763 | 3357.723175 | 8543.114134 | 223 | 14 | 26 | 32 | 86 | 18 | 1 | 0 | 0 | 0 | 1 | 0 |
| JSG000000 | Vmp1 | 6.08E-29 | 5.86E-28 | -1.817296727 | 5581.117074 | 1583.654018 | 200 | 18 | 26 | 45 | 138 | 19 | 2 | 0 | 0 | 0 | 0 | 1 |
| SG0000005 | Nup188 | 1.86E-22 | 1.33E-21 | 1.181528841 | 3163.549203 | 7175.467726 | 140 | 24 | 26 | 36 | 82 | 10 | 1 | 0 | 0 | 1 | 2 | 0 |
| JSG000000 | Gm5595 | 3.3E-21 | 2.21E-20 | 1.946433013 | 89.68466291 | 345.6630053 | 78 | 23 | 26 | 30 | 56 | 3 | 3 | 0 | 0 | 0 | 1 | 0 |
| JSG000000 | Lrrc20 | 4.86E-20 | 3.06E-19 | -1.752390212 | 836.4613092 | 248.2697685 | 137 | 24 | 26 | 37 | 106 | 19 | 2 | 0 | 0 | 1 | 1 | 0 |
| SG0000003 | Nmnat3 | 1.78E-19 | 1.08E-18 | 3.446799333 | 9.767671685 | 106.5077087 | 417 | 19 | 26 | 64 | 191 | 74 | 0 | 0 | 0 | 0 | 0 | 0 |
| SG0000003 | Specc1l | 2.75E-19 | 1.66E-18 | -1.366888973 | 4124.207329 | 1599.064228 | 350 | 18 | 26 | 53 | 155 | 44 | 4 | 2 | 2 | 2 | 10 | 0 |
| SG0000002 | Mfsd11 | 5.71E-16 | 2.86E-15 | -1.453750505 | 1066.386253 | 389.306836 | 118 | 23 | 26 | 35 | 63 | 8 | 10 | 2 | 2 | 2 | 6 | 2 |
| SG0000003 | Rbpms2 | 1.37E-15 | 6.72E-15 | 1.023594415 | 1609.922883 | 3272.937377 | 122 | 26 | 26 | 38 | 62 | 2 | 4 | 0 | 2 | 2 | 0 | 0 |
| JSG000000 | Acap2 | 5.47E-15 | 2.58E-14 | -1.308917586 | 4582.028109 | 1849.414652 | 230 | 15 | 26 | 43 | 111 | 19 | 11 | 0 | 0 | 0 | 1 | 1 |
| SG0000005 | Mdn1 | 2.42E-14 | 1.1E-13 | 1.320628015 | 2075.660152 | 5184.4763 | 398 | 18 | 26 | 60 | 168 | 52 | 4 | 0 | 0 | 0 | 2 | 0 |
| JSG000000 | Btbd7 | 1.79E-13 | 7.66E-13 | -1.432431933 | 1494.222663 | 553.6181767 | 220 | 19 | 26 | 48 | 138 | 25 | 1 | 0 | 0 | 0 | 1 | 1 |
| SG0000002 | Slc28a1 | 2.74E-13 | 1.16E-12 | -1.302986564 | 872.611034 | 353.6573272 | 110 | 20 | 26 | 26 | 58 | 22 | 0 | 0 | 0 | 0 | 0 | 0 |
| SG0000004 | Grsf1 | 4.69E-13 | 1.95E-12 | -1.031352092 | 7626.011418 | 3731.036825 | 84 | 24 | 26 | 34 | 54 | 0 | 4 | 0 | 2 | 0 | 0 | 0 |
| SG0000003 | Tcea1 | 1.19E-12 | 4.8E-12 | 0.922940095 | 1378.283687 | 2613.19169 | 218 | 24 | 26 | 52 | 126 | 14 | 0 | 0 | 0 | 0 | 2 | 0 |
| JSG000000 | Eml5 | 1.3E-12 | 5.26E-12 | 1.077778712 | 429.1416565 | 905.825191 | 302 | 10 | 26 | 33 | 141 | 24 | 13 | 0 | 0 | 1 | 2 | 1 |
| SG0000006 | Wdr61 | 1.36E-12 | 5.47E-12 | 0.92607424 | 1338.986843 | 2544.206947 | 137 | 20 | 26 | 40 | 64 | 12 | 5 | 0 | 2 | 0 | 0 | 6 |
| JSG000000 | Slc22a5 | 1.83E-12 | 7.29E-12 | -1.43240685 | 520.5788168 | 192.880829 | 95 | 22 | 26 | 30 | 54 | 4 | 0 | 0 | 0 | 0 | 1 | 0 |
| JSG000000 | Kif11 | 2.68E-12 | 1.06E-11 | 0.854210248 | 3393.481208 | 6134.62974 | 125 | 22 | 26 | 36 | 78 | 18 | 2 | 0 | 0 | 0 | 1 | 3 |
| SG0000007 | Zfp950 | 4.56E-12 | 1.78E-11 | -2.042341393 | 330.1962899 | 80.16156153 | 275 | 14 | 26 | 48 | 143 | 21 | 4 | 0 | 0 | 0 | 2 | 0 |
| JSG000000 | Jakmip2 | 4.81E-12 | 1.88E-11 | 4.991563697 | 0 | 30.81342285 | 339 | 18 | 26 | 51 | 157 | 17 | 2 | 0 | 0 | 0 | 0 | 0 |
| JSG000000 | Lmtk2 | 5.59E-12 | 2.17E-11 | 0.877960995 | 1833.138388 | 3368.897881 | 124 | 21 | 26 | 40 | 72 | 17 | 5 | 0 | 0 | 3 | 3 | 0 |
| SG0000003 | Synrg | 1.01E-11 | 3.87E-11 | -1.080919564 | 1642.72866 | 776.5628717 | 248 | 12 | 26 | 37 | 126 | 26 | 4 | 0 | 0 | 1 | 2 | 2 |
| SG0000003 | 1700021K19Rik | 1.4E-11 | 5.31E-11 | -1.07405016 | 1665.823282 | 791.2388671 | 286 | 20 | 26 | 76 | 162 | 35 | 5 | 0 | 0 | 0 | 0 | 0 |
| SG0000004 | Dcun1d3 | 2.25E-11 | 8.38E-11 | -1.090029176 | 1360.074109 | 638.8973623 | 206 | 14 | 26 | 30 | 131 | 37 | 19 | 2 | 2 | 3 | 5 | 4 |
| SG0000004 | Aftph | 3.03E-11 | 1.12E-10 | -0.991341538 | 3033.986458 | 1526.124973 | 492 | 20 | 26 | 72 | 176 | 55 | 28 | 0 | 0 | 4 | 4 | 6 |
| SG0000002 | Ercc8 | 4.23E-11 | 1.55E-10 | 1.304733165 | 184.0709952 | 454.7253641 | 345 | 6 | 26 | 32 | 131 | 42 | 7 | 0 | 0 | 0 | 4 | 1 |
| JSG000000 | Pten | 4.58E-11 | 1.67E-10 | -0.955194126 | 4561.113039 | 2352.495232 | 204 | 17 | 26 | 39 | 98 | 13 | 20 | 0 | 0 | 1 | 5 | 1 |
| SG0000007 | Cldn34c1 | 1.6E-10 | 5.62E-10 | -5.034897576 | 31.7834908 | 0 | 196 | 10 | 26 | 38 | 102 | 12 | 18 | 0 | 2 | 2 | 4 | 4 |
| SG0000002 | Ddah1 | 1.63E-09 | 5.26E-09 | 1.781415877 | 263.9217884 | 907.2664663 | 396 | 10 | 26 | 48 | 185 | 56 | 7 | 0 | 0 | 0 | 0 | 0 |
| SG0000005 | Fez2 | 2.35E-09 | 7.49E-09 | -1.003002174 | 1054.533723 | 526.1707872 | 156 | 22 | 26 | 46 | 86 | 20 | 2 | 0 | 0 | 0 | 0 | 0 |
| SG0000002 | Caprin1 | 2.57E-09 | 8.19E-09 | 0.713674466 | 12053.33973 | 19767.18465 | 227 | 24 | 26 | 37 | 86 | 10 | 24 | 0 | 0 | 1 | 8 | 0 |

| | | | | | | | | | | | | | | | | | | |
|---|---|---|---|---|---|---|---|---|---|---|---|---|---|---|---|---|---|---|
| SG0000003 | Tmprss5 | 3.51E-09 | 1.1E-08 | -3.145210651 | 92.59076332 | 10.4656189 | 119 | 26 | 26 | 40 | 74 | 11 | 0 | 0 | 0 | 0 | 0 | 0 |
| SG0000003 | 3632451O06Rik | 5.39E-09 | 1.67E-08 | 2.068472952 | 233.0034432 | 977.3154011 | 433 | 10 | 26 | 50 | 174 | 58 | 3 | 0 | 0 | 0 | 6 | 0 |
| SG0000003 | Ttc21b | 1.1E-08 | 3.34E-08 | 0.949289978 | 467.2407354 | 902.2054721 | 302 | 16 | 26 | 54 | 145 | 24 | 2 | 0 | 0 | 0 | 1 | 0 |
| SG0000005 | Lrrc48 | 1.3E-08 | 3.91E-08 | 2.254100694 | 15.56316125 | 74.24201308 | 153 | 20 | 26 | 29 | 58 | 11 | 2 | 0 | 0 | 0 | 1 | 0 |
| SG0000004 | Fkbp6 | 1.64E-08 | 4.92E-08 | 4.468412576 | 0 | 21.13737992 | 212 | 10 | 26 | 28 | 84 | 14 | 0 | 0 | 0 | 0 | 0 | 2 |
| JSG000000 | Rab10 | 1.68E-08 | 5.01E-08 | -0.797208453 | 5045.575074 | 2903.534664 | 152 | 18 | 26 | 32 | 74 | 13 | 6 | 0 | 1 | 0 | 4 | 0 |
| SG0000003 | Syn1 | 1.83E-07 | 5.03E-07 | 1.11935622 | 116.8908303 | 253.9453364 | 216 | 16 | 26 | 34 | 138 | 26 | 2 | 0 | 0 | 0 | 0 | 0 |
| SG0000002 | Mapk9 | 1.91E-07 | 5.23E-07 | -0.754683561 | 3200.014516 | 1896.572981 | 427 | 7 | 26 | 46 | 203 | 60 | 35 | 0 | 5 | 0 | 10 | 0 |
| SG0000003 | Aifm1 | 6.09E-07 | 1.6E-06 | -0.726631787 | 2938.69315 | 1775.890879 | 120 | 22 | 26 | 24 | 74 | 18 | 0 | 0 | 0 | 0 | 0 | 0 |
| SG0000002 | Casp12 | 1.73E-06 | 4.38E-06 | -1.571276716 | 120.6706188 | 40.60692626 | 113 | 22 | 26 | 28 | 82 | 10 | 9 | 0 | 0 | 0 | 4 | 0 |
| SG0000005 | Atp6v1a | 2.87E-06 | 7.11E-06 | -0.635476887 | 8732.28786 | 5621.218142 | 234 | 16 | 26 | 41 | 129 | 18 | 10 | 0 | 0 | 1 | 3 | 0 |
| SG0000004 | Psme4 | 3.74E-06 | 9.18E-06 | 0.568184962 | 5219.329596 | 7738.475993 | 283 | 14 | 26 | 39 | 162 | 21 | 2 | 0 | 0 | 0 | 1 | 0 |
| SG0000002 | Pdcd4 | 4.48E-06 | 1.09E-05 | 0.870693458 | 595.7017245 | 1089.265392 | 149 | 16 | 26 | 24 | 57 | 8 | 0 | 0 | 0 | 0 | 0 | 0 |
| SG0000002 | Soat2 | 7.27E-06 | 1.74E-05 | 2.175910046 | 10.20417046 | 46.10965753 | 48 | 24 | 26 | 30 | 60 | 2 | 0 | 0 | 0 | 0 | 0 | 0 |
| SG0000004 | Lrrc8d | 8.44E-06 | 2E-05 | -0.648583217 | 2460.282681 | 1569.430204 | 324 | 16 | 26 | 54 | 158 | 40 | 4 | 0 | 0 | 0 | 0 | 0 |
| JSG000000 | Bmp6 | 1.06E-05 | 2.49E-05 | -1.468003439 | 110.4506995 | 39.92596318 | 240 | 19 | 26 | 42 | 103 | 27 | 0 | 0 | 0 | 0 | 0 | 0 |
| SG0000004 | Elmsan1 | 1.08E-05 | 2.54E-05 | -0.679606748 | 2460.356881 | 1536.087981 | 126 | 21 | 26 | 32 | 67 | 10 | 19 | 4 | 4 | 6 | 10 | 2 |
| JSG000000 | Pibf1 | 1.6E-05 | 3.71E-05 | 0.751353501 | 277.569722 | 467.2529279 | 328 | 11 | 26 | 42 | 140 | 30 | 0 | 0 | 0 | 0 | 0 | 0 |
| SG0000001 | Ccnt1 | 4.34E-05 | 9.59E-05 | -0.5895484 | 2943.676081 | 1956.222576 | 159 | 26 | 26 | 37 | 72 | 4 | 5 | 0 | 0 | 1 | 4 | 1 |
| SG0000003 | Setd5 | 6.14E-05 | 0.000134 | 0.487586928 | 5902.766585 | 8276.255717 | 455 | 10 | 26 | 53 | 205 | 37 | 3 | 0 | 0 | 3 | 6 | 0 |
| SG0000003 | Bmx | 6.73E-05 | 0.000146 | -5.646731318 | 15.81495327 | 0.315669345 | 245 | 20 | 26 | 41 | 184 | 25 | 4 | 0 | 0 | 0 | 2 | 0 |
| SG0000003 | Man1c1 | 7.43E-05 | 0.00016 | -0.687041297 | 702.898208 | 436.5886061 | 226 | 18 | 26 | 36 | 88 | 26 | 0 | 0 | 0 | 0 | 0 | 0 |
| JSG000000 | Klhl29 | 0.000172 | 0.000355 | 3.430170642 | 4.743972235 | 51.13595745 | 339 | 16 | 26 | 48 | 158 | 54 | 6 | 1 | 1 | 4 | 7 | 1 |
| SG0000004 | Szrd1 | 0.000201 | 0.000412 | -0.517052471 | 3274.080592 | 2287.9212 | 74 | 18 | 26 | 28 | 38 | 2 | 0 | 0 | 0 | 0 | 0 | 0 |
| SG0000002 | Drg1 | 0.000448 | 0.00088 | 0.445109539 | 3072.594613 | 4183.084016 | 181 | 18 | 26 | 36 | 92 | 18 | 0 | 0 | 0 | 0 | 1 | 0 |
| SG0000002 | Slc25a19 | 0.000481 | 0.00094 | 0.535123785 | 662.5714331 | 960.1100552 | 67 | 21 | 26 | 30 | 42 | 0 | 4 | 0 | 2 | 0 | 0 | 0 |
| SG0000002 | Ccnc | 0.000768 | 0.001461 | -0.517302073 | 1342.891556 | 938.2476859 | 257 | 17 | 26 | 26 | 110 | 8 | 19 | 0 | 0 | 0 | 8 | 2 |
| SG0000002 | Trim23 | 0.000838 | 0.001587 | -0.497773304 | 1874.509538 | 1327.525766 | 269 | 15 | 26 | 39 | 118 | 18 | 20 | 0 | 2 | 0 | 10 | 9 |
| SG0000003 | Dhx57 | 0.001219 | 0.002267 | -0.508004394 | 1140.325698 | 801.8707111 | 152 | 20 | 26 | 30 | 80 | 24 | 1 | 0 | 0 | 0 | 1 | 0 |
| SG0000003 | Aplf | 0.001457 | 0.002684 | -0.573908921 | 538.309295 | 361.6330204 | 248 | 12 | 26 | 34 | 116 | 12 | 4 | 0 | 0 | 0 | 0 | 0 |
| JSG000000 | Slc25a16 | 0.002188 | 0.003941 | -0.518374501 | 769.1622106 | 536.9967595 | 84 | 22 | 26 | 37 | 65 | 5 | 4 | 0 | 1 | 1 | 4 | 1 |
| SG0000004 | Arhgef11 | 0.002702 | 0.004802 | -0.455354229 | 1395.391919 | 1017.702688 | 192 | 17 | 26 | 35 | 129 | 22 | 0 | 0 | 0 | 0 | 0 | 2 |
| SG0000002 | Afap1 | 0.002794 | 0.004952 | -0.495147726 | 1048.569932 | 743.9488594 | 127 | 21 | 26 | 36 | 84 | 7 | 10 | 2 | 2 | 3 | 3 | 2 |
| SG0000002 | Avl9 | 0.003091 | 0.005439 | -0.444809631 | 1552.280991 | 1140.431745 | 192 | 19 | 26 | 34 | 102 | 13 | 15 | 0 | 0 | 0 | 4 | 0 |
| SG0000005 | Dtnbp1 | 0.003092 | 0.005441 | -0.509760962 | 630.0827436 | 442.5315564 | 260 | 8 | 26 | 32 | 104 | 24 | 0 | 0 | 0 | 0 | 0 | 0 |
| SG0000002 | Osbp2 | 0.003526 | 0.006162 | 1.363064104 | 16.28307751 | 41.88511108 | 232 | 20 | 26 | 38 | 95 | 17 | 11 | 0 | 2 | 0 | 0 | 0 |
| SG0000003 | Kat6a | 0.004108 | 0.007105 | -0.401609197 | 3452.786271 | 2613.805584 | 364 | 14 | 26 | 50 | 204 | 40 | 4 | 0 | 0 | 0 | 14 | 2 |
| SG0000005 | Wdr91 | 0.005769 | 0.009779 | 0.454487264 | 411.6289668 | 564.0534088 | 82 | 24 | 26 | 36 | 60 | 4 | 2 | 0 | 0 | 0 | 0 | 2 |
| SG0000002 | Rest | 0.006735 | 0.011316 | 0.33375885 | 8248.943449 | 10396.08332 | 76 | 26 | 26 | 34 | 54 | 4 | 10 | 0 | 0 | 2 | 4 | 0 |
| JSG000000 | Zfp608 | 0.00759 | 0.012649 | -0.750059331 | 292.0399727 | 173.6408655 | 202 | 18 | 26 | 35 | 93 | 17 | 1 | 0 | 0 | 0 | 1 | 4 |
| JSG000000 | Gm27029 | 0.019875 | 0.030769 | 1.877829137 | 5.649592118 | 20.7634727 | 69 | 20 | 26 | 30 | 58 | 0 | 1 | 0 | 0 | 0 | 0 | 0 |
| SG0000002 | Ptp4a2 | 0.038797 | 0.057074 | -0.275344817 | 8905.336869 | 7358.059638 | 154 | 22 | 26 | 26 | 68 | 4 | 8 | 0 | 0 | 0 | 0 | 0 |
| SG0000002 | Alkbh8 | 0.045683 | 0.066445 | -0.349989527 | 626.7347485 | 491.7296869 | 159 | 18 | 26 | 43 | 111 | 9 | 3 | 0 | 0 | 0 | 0 | 1 |
| SG0000002 | Gm20517 | 0.052992 | 0.076112 | 1.151599529 | 11.24850395 | 24.98968411 | 129 | 22 | 26 | 31 | 76 | 6 | 2 | 0 | 0 | 0 | 0 | 0 |
| SG0000002 | Depdc1b | 0.06973 | 0.097969 | 0.305582398 | 386.853976 | 478.1195837 | 318 | 20 | 26 | 58 | 177 | 24 | 1 | 0 | 0 | 0 | 0 | 2 |
| JSG000000 | Ppp4r2 | 0.089774 | 0.123438 | -0.238439396 | 2495.806642 | 2115.599842 | 106 | 21 | 26 | 35 | 61 | 5 | 12 | 0 | 0 | 3 | 3 | 0 |
| SG0000003 | Fbxo8 | 0.12274 | 0.1638 | 0.230874563 | 691.5008414 | 811.508142 | 256 | 6 | 26 | 30 | 132 | 24 | 12 | 0 | 0 | 0 | 2 | 0 |
| SG0000002 | R3hcc1l | 0.132354 | 0.175441 | -0.253313238 | 792.163415 | 664.5993289 | 184 | 17 | 26 | 38 | 102 | 18 | 2 | 1 | 1 | 2 | 6 | 0 |
| SG0000006 | Slc9a6 | 0.161278 | 0.210035 | -0.26346744 | 524.945995 | 437.3237106 | 258 | 16 | 26 | 50 | 128 | 22 | 10 | 0 | 0 | 2 | 2 | 2 |
| SG0000006 | Rnasel | 0.17304 | 0.223855 | -0.839590974 | 63.65731492 | 35.57183332 | 50 | 22 | 26 | 29 | 47 | 6 | 6 | 0 | 0 | 0 | 0 | 0 |
| JSG000000 | Ubr5 | 0.182301 | 0.234649 | -0.178838868 | 15615.98089 | 13795.37166 | 304 | 14 | 26 | 51 | 129 | 25 | 2 | 0 | 0 | 0 | 0 | 0 |
| JSG000001 | Pcdhgb4 | 0.222244 | 0.279887 | 0.71841947 | 12.75826828 | 20.99217992 | 186 | 23 | 26 | 40 | 91 | 18 | 2 | 0 | 0 | 1 | 1 | 0 |
| SG0000002 | Atf1 | 0.255761 | 0.316993 | 0.15460891 | 1288.529398 | 1434.287624 | 139 | 15 | 26 | 29 | 63 | 9 | 2 | 0 | 0 | 0 | 2 | 0 |
| JSG000001 | Pcdhgb2 | 0.313569 | 0.379222 | -0.486557669 | 36.02421975 | 25.71142399 | 223 | 23 | 26 | 42 | 106 | 18 | 2 | 0 | 0 | 1 | 1 | 0 |
| JSG000000 | Pcnx | 0.398078 | 0.466953 | -0.145413603 | 785.2753079 | 709.9832214 | 281 | 11 | 26 | 45 | 130 | 34 | 18 | 0 | 2 | 2 | 5 | 1 |
| SG0000003 | Ak5 | 0.492209 | 0.560059 | 0.377139494 | 19.08178351 | 24.78273679 | 346 | 12 | 26 | 38 | 143 | 21 | 2 | 0 | 0 | 0 | 0 | 0 |
| SG0000002 | Abl2 | 0.538023 | 0.604356 | 0.074661383 | 3615.77951 | 3807.827404 | 196 | 18 | 26 | 34 | 106 | 18 | 7 | 1 | 1 | 1 | 1 | 1 |
| JSG000000 | Wdr43 | 0.558895 | 0.623967 | -0.082333752 | 6510.086188 | 6148.962004 | 95 | 21 | 26 | 29 | 51 | 5 | 5 | 0 | 1 | 0 | 1 | 1 |
| SG0000004 | Tcaim | 0.722617 | 0.774572 | 0.065496325 | 295.6423001 | 309.3733719 | 134 | 6 | 26 | 20 | 86 | 12 | 10 | 0 | 2 | 0 | 3 | 2 |
| SG0000002 | Mfn1 | 0.73189 | 0.782654 | -0.05816195 | 1701.169367 | 1633.951127 | 201 | 24 | 26 | 46 | 122 | 28 | 4 | 0 | 0 | 1 | 2 | 1 |
| JSG000001 | Pcdhga7 | 0.798583 | 0.840288 | -0.114368019 | 44.41730358 | 41.03212282 | 192 | 23 | 26 | 41 | 94 | 18 | 3 | 0 | 0 | 1 | 1 | 0 |
| JSG000000 | Mcm3ap | 0.878673 | 0.906612 | -0.020388831 | 2336.635871 | 2303.845705 | 96 | 23 | 26 | 35 | 68 | 10 | 0 | 0 | 0 | 0 | 0 | 0 |
| JSG000001 | Pcdhga5 | 0.881195 | 0.90843 | -0.120261855 | 17.36706404 | 15.97805752 | 214 | 23 | 26 | 42 | 105 | 18 | 2 | 0 | 0 | 1 | 1 | 0 |
| SG0000002 | Rasa1 | 0.8889 | 0.914793 | 0.016132631 | 1450.684019 | 1466.997023 | 284 | 15 | 26 | 44 | 158 | 30 | 8 | 0 | 0 | 0 | 3 | 0 |
| JSG000001 | Pcdhgb5 | 0.953922 | 0.96918 | -0.070718105 | 15.54141303 | 14.79797303 | 167 | 23 | 26 | 40 | 87 | 16 | 2 | 0 | 0 | 1 | 1 | 0 |
| JSG000000 | Sorl1 | 2E-216 | 7.1E-214 | 5.298203033 | 48.55962807 | 1910.703889 | 248 | 16 | 25 | 35 | 124 | 20 | 6 | 0 | 1 | 1 | 6 | 2 |
| JSG000000 | Lin28b | 2.35E-86 | 1.22E-84 | 5.467969394 | 25.05928642 | 1109.153115 | 264 | 10 | 25 | 45 | 120 | 25 | 10 | 0 | 1 | 1 | 2 | 2 |
| SG0000003 | Rcbtb1 | 1.29E-82 | 6.2E-81 | 2.411670777 | 795.1134053 | 4230.720345 | 288 | 4 | 25 | 42 | 111 | 21 | 9 | 0 | 0 | 0 | 3 | 0 |
| JSG000000 | Tipin | 1.51E-55 | 3.71E-54 | 1.938159466 | 1220.663355 | 4677.78269 | 79 | 23 | 25 | 30 | 54 | 9 | 2 | 0 | 0 | 0 | 1 | 1 |
| SG0000004 | Lrp1 | 1.92E-41 | 2.99E-40 | -3.018269472 | 19299.45712 | 2382.075085 | 102 | 22 | 25 | 31 | 53 | 9 | 1 | 0 | 0 | 0 | 1 | 0 |
| SG0000003 | Wdr13 | 2.53E-35 | 3.19E-34 | -2.289599666 | 1608.643945 | 329.0195855 | 61 | 20 | 25 | 33 | 48 | 18 | 6 | 0 | 0 | 3 | 3 | 3 |
| SG0000002 | 4930452B06Rik | 3.71E-32 | 4.11E-31 | 4.309299597 | 37.94035722 | 752.1939821 | 461 | 15 | 25 | 70 | 227 | 44 | 1 | 0 | 0 | 0 | 0 | 1 |
| SG0000003 | Jade3 | 7.93E-29 | 7.57E-28 | 1.480841965 | 758.4887659 | 2117.029954 | 257 | 16 | 25 | 48 | 167 | 20 | 2 | 0 | 0 | 0 | 0 | 0 |
| SG0000003 | Ddx19b | 1.35E-28 | 1.28E-27 | 1.465614684 | 809.6041792 | 2235.973866 | 102 | 20 | 25 | 32 | 74 | 17 | 5 | 0 | 0 | 1 | 3 | 0 |
| JSG000000 | Txnrd1 | 7.12E-25 | 5.72E-24 | -1.496503624 | 34275.18956 | 12147.51338 | 69 | 23 | 25 | 25 | 46 | 3 | 4 | 2 | 2 | 2 | 6 | 0 |
| SG0000003 | Frrs1 | 1.37E-23 | 1.04E-22 | -1.51268579 | 6421.087915 | 2250.322795 | 218 | 18 | 25 | 25 | 107 | 18 | 10 | 0 | 2 | 1 | 6 | 1 |
| JSG000000 | Ppm1e | 4.24E-23 | 3.12E-22 | 7.698223086 | 0.331601068 | 68.86712987 | 250 | 14 | 25 | 39 | 121 | 25 | 7 | 1 | 1 | 3 | 6 | 2 |
| SG0000002 | 3110009E18Rik | 5.22E-22 | 3.66E-21 | 3.254675883 | 14.03883254 | 133.9942187 | 370 | 11 | 25 | 46 | 153 | 42 | 1 | 0 | 0 | 0 | 0 | 0 |
| SG0000003 | Idh2 | 1.1E-20 | 7.18E-20 | 1.495697584 | 1263.911473 | 3564.236337 | 89 | 21 | 25 | 35 | 68 | 4 | 4 | 0 | 0 | 0 | 0 | 0 |
| SG0000006 | Serpinb6a | 1.74E-20 | 1.12E-19 | -1.385004607 | 6098.713927 | 2335.126066 | 243 | 15 | 25 | 43 | 101 | 32 | 0 | 0 | 0 | 0 | 0 | 0 |
| SG0000002 | Msh2 | 5.32E-20 | 3.33E-19 | 1.241192905 | 4159.311262 | 9832.358769 | 153 | 22 | 25 | 41 | 84 | 5 | 1 | 0 | 0 | 0 | 0 | 0 |
| SG0000002 | Lcor | 1.26E-19 | 7.74E-19 | -2.657178132 | 570.8954581 | 90.50368577 | 237 | 13 | 25 | 42 | 114 | 23 | 4 | 0 | 0 | 0 | 1 | 1 |
| SG0000004 | Mob1a | 3.55E-18 | 2.01E-17 | -1.284508498 | 6408.774424 | 2630.870056 | 181 | 21 | 25 | 42 | 82 | 11 | 11 | 3 | 3 | 3 | 3 | 0 |
| SG0000005 | Lclat1 | 1.02E-16 | 5.37E-16 | -1.473046418 | 1079.790313 | 388.962981 | 309 | 14 | 25 | 48 | 132 | 20 | 5 | 0 | 2 | 1 | 1 | 0 |
| SG0000006 | Bbs5 | 5.13E-16 | 2.58E-15 | 1.847956772 | 70.08791828 | 252.3094866 | 126 | 11 | 25 | 25 | 56 | 4 | 11 | 0 | 0 | 0 | 6 | 0 |
| SG0000005 | Mmel1 | 6.8E-15 | 3.19E-14 | 3.590826955 | 6.127090634 | 73.82457023 | 118 | 25 | 25 | 40 | 65 | 10 | 0 | 0 | 0 | 0 | 0 | 0 |
| SG0000002 | Acss2 | 1.07E-12 | 4.37E-12 | -1.232024136 | 1065.144599 | 453.4530727 | 332 | 15 | 25 | 73 | 179 | 36 | 0 | 0 | 0 | 0 | 0 | 0 |
| JSG000000 | Anapc10 | 5.56E-11 | 2.01E-10 | 1.317275091 | 345.3323926 | 860.5510377 | 152 | 14 | 25 | 33 | 100 | 10 | 7 | 0 | 0 | 1 | 2 | 0 |
| JSG000000 | Zfp521 | 1.09E-10 | 3.87E-10 | 3.296363381 | 27.5725812 | 270.8829552 | 529 | 10 | 25 | 60 | 235 | 50 | 1 | 0 | 0 | 0 | 2 | 1 |
| JSG000000 | Rab7 | 3.98E-10 | 1.35E-09 | -0.868199834 | 10181.65415 | 5577.815549 | 274 | 14 | 25 | 59 | 166 | 16 | 6 | 0 | 0 | 0 | 4 | 0 |
| SG0000002 | Tti1 | 5.62E-10 | 1.89E-09 | 0.813821451 | 1381.950209 | 2429.277142 | 112 | 16 | 25 | 29 | 62 | 9 | 2 | 0 | 2 | 2 | 2 | 0 |
| JSG000000 | Ints6 | 6.25E-10 | 2.09E-09 | 0.834146381 | 1027.993477 | 1832.708696 | 229 | 11 | 25 | 38 | 109 | 26 | 4 | 0 | 0 | 0 | 2 | 2 |
| SG0000003 | Med13 | 8.9E-10 | 2.94E-09 | -0.925552362 | 3475.013476 | 1829.521298 | 262 | 18 | 25 | 48 | 141 | 18 | 10 | 2 | 2 | 2 | 5 | 2 |
| JSG000000 | Dscaml1 | 2.33E-08 | 6.87E-08 | 4.032788991 | 1.974045165 | 32.31078707 | 272 | 13 | 25 | 34 | 109 | 23 | 0 | 0 | 0 | 0 | 0 | 0 |
| SG0000006 | Gm608 | 3.43E-08 | 1E-07 | -1.172987576 | 1459.239672 | 647.1756718 | 228 | 15 | 25 | 56 | 127 | 28 | 18 | 0 | 4 | 8 | 10 | 4 |
| SG0000003 | Tbc1d2b | 8.32E-08 | 2.35E-07 | 1.156760666 | 236.9026369 | 528.1892439 | 160 | 18 | 25 | 30 | 86 | 18 | 8 | 0 | 0 | 1 | 4 | 1 |
| SG0000003 | 6430573F11Rik | 9.4E-08 | 2.64E-07 | 5.926583648 | 1.302312483 | 79.21267025 | 293 | 7 | 25 | 47 | 144 | 41 | 5 | 0 | 0 | 1 | 7 | 0 |
| JSG000000 | 43534 | 1.19E-06 | 3.05E-06 | -5.099403479 | 23.8197219 | 0.694805551 | 343 | 11 | 25 | 42 | 116 | 21 | 0 | 0 | 0 | 0 | 0 | 0 |
| SG0000000 | Srpk1 | 1.33E-06 | 3.4E-06 | 0.609228511 | 3066.467994 | 4677.722908 | 91 | 19 | 25 | 29 | 50 | 4 | 3 | 0 | 0 | 0 | 0 | 0 |
| SG0000001 | Top2b | 2.14E-06 | 5.36E-06 | -0.671045157 | 4714.263565 | 2960.800454 | 221 | 17 | 25 | 45 | 130 | 24 | 1 | 0 | 0 | 0 | 1 | 0 |
| SG0000002 | Ankra2 | 2.48E-06 | 6.18E-06 | 0.772945185 | 365.5659567 | 624.6624941 | 101 | 15 | 25 | 30 | 45 | 25 | 1 | 0 | 0 | 0 | 0 | 0 |
| SG0000001 | Mpp2 | 2.58E-06 | 6.41E-06 | 1.310621483 | 166.9299004 | 414.0668986 | 82 | 23 | 25 | 32 | 52 | 3 | 6 | 0 | 0 | 5 | 7 | 0 |
| SG0000003 | Foxn2 | 1.22E-05 | 2.86E-05 | -0.649799587 | 2260.946811 | 1441.057065 | 214 | 9 | 25 | 18 | 96 | 16 | 10 | 1 | 1 | 2 | 4 | 0 |
| SG0000002 | Dnah12 | 1.71E-05 | 3.93E-05 | 5.475692803 | 0.34247518 | 15.23971633 | 271 | 14 | 25 | 46 | 158 | 22 | 0 | 0 | 0 | 0 | 0 | 0 |
| SG0000005 | Ide | 2.09E-05 | 4.78E-05 | 0.51968555 | 5853.73542 | 8392.165268 | 136 | 18 | 25 | 34 | 71 | 8 | 3 | 0 | 0 | 0 | 1 | 0 |
| SG0000007 | Armc2 | 4.05E-05 | 8.99E-05 | 2.577349954 | 5.16725333 | 30.84035718 | 507 | 10 | 25 | 67 | 199 | 125 | 8 | 0 | 2 | 3 | 6 | 2 |
| SG0000003 | Clasp2 | 6.65E-05 | 0.000144 | 0.537591039 | 1300.510389 | 1887.75163 | 228 | 15 | 25 | 45 | 119 | 20 | 3 | 0 | 1 | 2 | 3 | 1 |
| SG0000000 | Txn2 | 7.29E-05 | 0.000157 | -0.583721976 | 3643.450578 | 2431.056538 | 96 | 22 | 25 | 42 | 61 | 0 | 1 | 0 | 0 | 0 | 0 | 0 |
| SG0000004 | Dpy19l3 | 8.56E-05 | 0.000183 | 0.717250774 | 235.1992419 | 386.6784019 | 164 | 18 | 25 | 33 | 70 | 19 | 13 | 0 | 1 | 0 | 1 | 0 |
| SG0000005 | Inca1 | 0.000126 | 0.000264 | -1.660979827 | 74.65821101 | 23.60874107 | 90 | 20 | 25 | 30 | 70 | 15 | 0 | 0 | 0 | 0 | 0 | 0 |
| SG0000002 | Tgfbr3 | 0.000168 | 0.000348 | -0.893400145 | 402.1031003 | 216.4697322 | 304 | 17 | 25 | 37 | 138 | 31 | 6 | 0 | 0 | 1 | 3 | 0 |
| JSG000000 | Klhl2 | 0.000216 | 0.000441 | 0.528795279 | 873.7063885 | 1260.51713 | 166 | 17 | 25 | 37 | 97 | 10 | 4 | 1 | 1 | 2 | 3 | 0 |
| SG0000002 | Stxbp4 | 0.000328 | 0.000654 | 0.698898437 | 314.66172 | 510.7793202 | 273 | 11 | 25 | 29 | 135 | 37 | 13 | 0 | 2 | 1 | 5 | 2 |
| SG0000006 | Hdac8 | 0.000617 | 0.001188 | -0.696641868 | 368.5983532 | 227.4276648 | 576 | 9 | 25 | 76 | 282 | 72 | 1 | 0 | 0 | 0 | 0 | 0 |
| SG0000003 | Hook3 | 0.001862 | 0.003384 | -0.476565239 | 1706.990858 | 1226.791514 | 264 | 13 | 25 | 41 | 116 | 18 | 37 | 0 | 3 | 3 | 13 | 2 |
| SG0000002 | Rhbdd1 | 0.002043 | 0.003696 | 0.468182687 | 850.7605423 | 1176.913032 | 425 | 13 | 25 | 65 | 217 | 29 | 8 | 0 | 0 | 2 | 6 | 0 |
| SG0000003 | Trim14 | 0.003727 | 0.006486 | -0.959236402 | 100.3037371 | 51.58912763 | 117 | 20 | 25 | 28 | 57 | 4 | 4 | 2 | 2 | 2 | 6 | 0 |
| SG0000003 | Tmem260 | 0.005852 | 0.009914 | -0.55512118 | 345.3372856 | 235.036518 | 343 | 9 | 25 | 29 | 137 | 34 | 8 | 0 | 1 | 2 | 6 | 1 |
| SG0000002 | Fubp3 | 0.006877 | 0.011534 | 0.349448864 | 2313.418466 | 2947.465889 | 182 | 12 | 25 | 33 | 69 | 12 | 9 | 0 | 0 | 0 | 2 | 0 |
| JSG000001 | K230010J24Rik | 0.007803 | 0.01298 | 1.139820741 | 21.30058999 | 46.93662579 | 54 | 21 | 25 | 25 | 33 | 3 | 0 | 0 | 0 | 0 | 0 | 0 |
| SG0000002 | Saxo1 | 0.013936 | 0.022189 | -1.736346258 | 21.67553436 | 6.505452443 | 205 | 17 | 25 | 38 | 98 | 19 | 0 | 0 | 0 | 0 | 0 | 0 |
| JSG000000 | Kcnk10 | 0.015492 | 0.024501 | -4.233336603 | 17.81209336 | 0.947008034 | 264 | 16 | 25 | 46 | 152 | 26 | 2 | 0 | 1 | 0 | 0 | 1 |
| SG0000002 | Pcx | 0.017366 | 0.027195 | 0.665293306 | 310.0000281 | 491.6261488 | 163 | 24 | 25 | 36 | 105 | 12 | 0 | 0 | 0 | 0 | 0 | 0 |

| | | | | | | | | | | | | | | | | | | |
|---|---|---|---|---|---|---|---|---|---|---|---|---|---|---|---|---|---|---|
| SG0000000 | Def8 | 0.022618 | 0.034661 | -0.31094521 | 2971.507448 | 2395.372106 | 94 | 20 | 25 | 35 | 135 | 5 | 7 | 2 | 2 | 3 | 12 | 1 |
| JSG000000 | Snx4 | 0.023241 | 0.03552 | -0.325807692 | 2194.816956 | 1751.138187 | 96 | 17 | 25 | 27 | 60 | 15 | 3 | 0 | 0 | 0 | 4 | 0 |
| SG0000003 | Rtel1 | 0.027552 | 0.041654 | 0.294860337 | 1958.814992 | 2403.008048 | 205 | 25 | 25 | 45 | 90 | 20 | 0 | 0 | 0 | 0 | 0 | 0 |
| SG0000003 | Plekha2 | 0.032935 | 0.049095 | 0.506969139 | 2299.176996 | 3267.272244 | 189 | 17 | 25 | 33 | 94 | 26 | 11 | 1 | 1 | 1 | 3 | 0 |
| SG0000005 | Tdrd9 | 0.07677 | 0.107003 | 1.387031858 | 5.771397775 | 15.09451634 | 321 | 18 | 25 | 43 | 198 | 40 | 3 | 0 | 0 | 0 | 0 | 0 |
| SG0000002 | Uhrf2 | 0.087113 | 0.12013 | 0.228023259 | 1883.164891 | 2205.617452 | 229 | 10 | 25 | 33 | 109 | 20 | 6 | 0 | 0 | 0 | 2 | 0 |
| SG0000004 | Gpr1 | 0.117221 | 0.157067 | 0.75143994 | 19.82579156 | 33.37616991 | 85 | 22 | 25 | 30 | 55 | 6 | 1 | 0 | 0 | 0 | 0 | 0 |
| JSG000000 | Glis1 | 0.137413 | 0.181542 | 0.96458143 | 10.09074221 | 19.69205567 | 340 | 12 | 25 | 52 | 154 | 30 | 0 | 0 | 0 | 0 | 0 | 2 |
| SG0000003 | Dnajc12 | 0.176819 | 0.2281 | 0.307763746 | 142.0066777 | 175.7741014 | 62 | 21 | 25 | 26 | 48 | 6 | 3 | 0 | 0 | 0 | 1 | 1 |
| SG0000003 | Atf7ip | 0.200444 | 0.255395 | -0.186137776 | 2096.346854 | 1842.595844 | 259 | 14 | 25 | 43 | 139 | 21 | 9 | 0 | 0 | 1 | 6 | 2 |
| SG0000007 | Safb | 0.214144 | 0.270867 | 0.167961374 | 2613.257644 | 2935.916105 | 39 | 23 | 25 | 23 | 35 | 6 | 0 | 0 | 0 | 0 | 0 | 0 |
| JSG000000 | Nxpe3 | 0.237864 | 0.297466 | -0.256712979 | 239.9639655 | 200.8480979 | 129 | 19 | 25 | 30 | 80 | 12 | 8 | 2 | 2 | 2 | 8 | 1 |
| JSG000000 | Scaf8 | 0.28294 | 0.346508 | -0.157304758 | 2134.082632 | 1913.629163 | 237 | 12 | 25 | 26 | 107 | 16 | 0 | 0 | 0 | 0 | 0 | 0 |
| SG0000004 | C2cd3 | 0.484776 | 0.553041 | -0.109074823 | 1213.517641 | 1125.15227 | 333 | 13 | 25 | 37 | 156 | 41 | 5 | 0 | 0 | 0 | 4 | 2 |
| SG0000001 | Nxf1 | 0.618991 | 0.680959 | -0.064661229 | 4871.6467 | 4658.121535 | 86 | 21 | 25 | 21 | 21 | 11 | 3 | 0 | 0 | 0 | 0 | 3 |
| SG0000004 | Ttc41 | 0.661465 | 0.719684 | 0.173708451 | 35.70364593 | 40.27207631 | 145 | 17 | 25 | 45 | 107 | 14 | 0 | 0 | 0 | 0 | 0 | 0 |
| SG0000002 | G3bp2 | 0.729636 | 0.780889 | -0.054817882 | 13701.98928 | 13191.12381 | 364 | 20 | 25 | 52 | 120 | 27 | 14 | 0 | 0 | 2 | 10 | 2 |
| SG0000000 | Tnpo1 | 0.825085 | 0.86265 | -0.034321514 | 8765.897848 | 8559.819367 | 292 | 15 | 25 | 47 | 173 | 21 | 22 | 0 | 0 | 0 | 16 | 2 |
| SG0000001 | Cuedc1 | 0.862115 | 0.892613 | -0.041718365 | 171.2554753 | 166.3742023 | 154 | 23 | 25 | 32 | 77 | 3 | 7 | 0 | 0 | 0 | 0 | 0 |
| JSG000000 | Ubxn7 | 0.903941 | 0.927501 | 0.005478024 | 3100.717747 | 3112.513792 | 144 | 15 | 25 | 29 | 91 | 10 | 27 | 0 | 3 | 3 | 10 | 1 |
| SG0000002 | Sppl3 | 0.953139 | 0.968649 | 0.002241777 | 1767.915129 | 1770.664395 | 288 | 16 | 25 | 39 | 140 | 21 | 3 | 0 | 1 | 0 | 2 | 1 |
| JSG000000 | Trim71 | 1.2E-133 | 1.5E-131 | 4.171471424 | 348.0828806 | 6272.211424 | 94 | 19 | 24 | 33 | 64 | 4 | 8 | 0 | 1 | 1 | 7 | 0 |
| SG0000002 | Acss1 | 4.3E-130 | 5E-128 | 9.504521838 | 0.649984458 | 472.1164576 | 88 | 20 | 24 | 29 | 47 | 4 | 2 | 0 | 0 | 0 | 1 | 0 |
| JSG000000 | Slco4c1 | 2.2E-111 | 2E-109 | 8.272424794 | 1.299968916 | 401.9582693 | 173 | 12 | 24 | 34 | 96 | 16 | 11 | 1 | 1 | 2 | 3 | 2 |
| SG0000003 | Nphp3 | 4.44E-87 | 2.32E-85 | 3.696731638 | 56.2998809 | 730.0197086 | 240 | 12 | 24 | 46 | 111 | 19 | 10 | 0 | 0 | 0 | 2 | 1 |
| SG0000004 | Gprasp1 | 1.69E-83 | 8.3E-82 | 2.586851982 | 419.4765389 | 2520.157695 | 408 | 15 | 24 | 39 | 139 | 32 | 0 | 0 | 0 | 0 | 4 | 4 |
| SG0000003 | Rasgrp2 | 1.08E-75 | 4.47E-74 | 5.988413339 | 9.591495471 | 608.9454188 | 96 | 24 | 24 | 36 | 60 | 0 | 0 | 0 | 0 | 0 | 0 | 0 |
| SG0000002 | Hook1 | 2.87E-71 | 1.06E-69 | 6.946179198 | 31.30199574 | 3859.937936 | 265 | 8 | 24 | 39 | 123 | 23 | 16 | 0 | 0 | 1 | 3 | 0 |
| SG0000003 | Wdhd1 | 3.09E-65 | 9.72E-64 | 2.110478192 | 1058.244602 | 4569.864289 | 254 | 13 | 24 | 32 | 133 | 21 | 6 | 0 | 0 | 0 | 7 | 2 |
| SG0000003 | Pdzrn3 | 3.97E-46 | 7.22E-45 | -3.389563889 | 745.6312149 | 71.14816083 | 347 | 9 | 24 | 40 | 151 | 30 | 0 | 0 | 0 | 0 | 1 | 2 |
| SG0000002 | Terf1 | 1.9E-41 | 2.96E-40 | 1.696043244 | 1114.886583 | 3612.356298 | 174 | 20 | 24 | 35 | 97 | 16 | 6 | 0 | 0 | 1 | 1 | 0 |
| JSG000000 | Gpr39 | 1.71E-40 | 2.57E-39 | -5.323787449 | 844.8785284 | 21.09475862 | 321 | 15 | 24 | 44 | 136 | 31 | 0 | 0 | 0 | 0 | 0 | 0 |
| SG0000002 | Lrriq4 | 6.84E-40 | 1.01E-38 | 6.888716135 | 0 | 117.4977729 | 183 | 12 | 24 | 38 | 98 | 21 | 0 | 0 | 0 | 0 | 0 | 0 |
| JSG000000 | Aoah | 1.86E-37 | 2.5E-36 | -3.458609741 | 2005.590885 | 182.4303418 | 352 | 7 | 24 | 52 | 182 | 41 | 2 | 0 | 0 | 0 | 0 | 0 |
| SG0000001 | Lpl | 7.84E-37 | 1.04E-35 | 2.987092272 | 64.31886408 | 509.9677819 | 138 | 24 | 24 | 30 | 72 | 10 | 5 | 1 | 1 | 2 | 4 | 1 |
| SG0000002 | Rnf138 | 1.72E-33 | 2E-32 | 1.591242182 | 742.5830859 | 2237.467213 | 192 | 14 | 24 | 36 | 80 | 8 | 11 | 0 | 0 | 5 | 6 | 4 |
| SG0000004 | 2900011O08Rik | 5.81E-31 | 6.13E-30 | 3.338476643 | 33.28706815 | 336.7108838 | 384 | 12 | 24 | 55 | 165 | 32 | 4 | 0 | 0 | 1 | 3 | 0 |
| SG0000003 | Mtmr7 | 2.99E-30 | 3.06E-29 | 1.615818087 | 471.816927 | 1446.049732 | 314 | 12 | 24 | 40 | 118 | 30 | 3 | 0 | 0 | 0 | 0 | 0 |
| JSG000000 | Zdhhc15 | 4.46E-29 | 4.33E-28 | -2.576350516 | 2074.983271 | 347.9011105 | 285 | 16 | 24 | 41 | 183 | 35 | 12 | 0 | 1 | 1 | 5 | 1 |
| SG0000002 | Psip1 | 1.34E-28 | 1.27E-27 | 1.323208327 | 3642.647562 | 9114.703607 | 171 | 9 | 24 | 22 | 80 | 13 | 2 | 0 | 0 | 0 | 0 | 0 |
| SG0000003 | Dcps | 1.86E-28 | 1.76E-27 | 1.615818785 | 413.3719622 | 1266.925093 | 199 | 15 | 24 | 36 | 108 | 24 | 0 | 0 | 0 | 0 | 0 | 0 |
| SG0000002 | Rad51 | 3.18E-27 | 2.83E-26 | 1.37445119 | 1240.221103 | 3215.512132 | 137 | 18 | 24 | 30 | 83 | 17 | 4 | 0 | 0 | 2 | 3 | 0 |
| SG0000003 | Kars | 3.14E-26 | 2.68E-25 | 1.251377542 | 5851.061644 | 13929.5424 | 93 | 21 | 24 | 30 | 51 | 6 | 0 | 0 | 0 | 0 | 0 | 0 |
| SG0000002 | Afap1l2 | 4.67E-24 | 3.6E-23 | -2.629358624 | 801.2282498 | 129.4912544 | 267 | 18 | 24 | 36 | 105 | 63 | 3 | 0 | 0 | 0 | 6 | 0 |
| JSG000000 | Mipep | 2.04E-21 | 1.38E-20 | 1.31150382 | 630.5690607 | 1565.073173 | 200 | 16 | 24 | 38 | 98 | 9 | 0 | 0 | 0 | 0 | 0 | 0 |
| SG0000003 | Pdcd6ip | 3.81E-19 | 2.27E-18 | -1.306892507 | 8268.409073 | 3342.012802 | 217 | 20 | 24 | 36 | 136 | 12 | 13 | 0 | 0 | 0 | 4 | 0 |
| SG0000006 | Zbtb34 | 5.11E-19 | 3.03E-18 | 1.328181873 | 432.0939032 | 1084.927511 | 104 | 24 | 24 | 36 | 57 | 8 | 14 | 2 | 2 | 4 | 10 | 6 |
| SG0000001 | Piezo1 | 7.68E-19 | 4.52E-18 | -1.363492317 | 7778.776006 | 3023.146405 | 103 | 21 | 24 | 34 | 78 | 1 | 2 | 0 | 1 | 1 | 1 | 0 |
| SG0000003 | Caps2 | 4.17E-18 | 2.36E-17 | 5.049537141 | 1.939079263 | 64.21813749 | 262 | 12 | 24 | 47 | 192 | 52 | 5 | 2 | 2 | 2 | 6 | 2 |
| SG0000003 | Usp9x | 1.13E-17 | 6.22E-17 | -1.379466613 | 19967.50013 | 7674.725787 | 292 | 13 | 24 | 41 | 153 | 34 | 10 | 0 | 0 | 0 | 1 | 2 |
| SG0000001 | Slc6a15 | 3.61E-17 | 1.95E-16 | 5.894250814 | 3.854066735 | 229.2268717 | 298 | 10 | 24 | 56 | 166 | 16 | 6 | 0 | 0 | 4 | 4 | 2 |
| SG0000002 | Zfp326 | 4.89E-16 | 2.46E-15 | -1.261598885 | 2587.217408 | 1079.080434 | 220 | 10 | 24 | 48 | 88 | 18 | 6 | 0 | 0 | 1 | 2 | 0 |
| SG0000002 | Prmt2 | 1.24E-15 | 6.1E-15 | -1.413524596 | 1024.308027 | 384.51853 | 188 | 20 | 24 | 36 | 116 | 20 | 0 | 0 | 0 | 0 | 0 | 4 |
| SG0000002 | Mtch2 | 1.25E-15 | 6.15E-15 | 0.987301236 | 2591.179269 | 5136.943006 | 179 | 16 | 24 | 24 | 91 | 20 | 7 | 0 | 0 | 0 | 3 | 0 |
| JSG000000 | Chek2 | 2.27E-13 | 9.65E-13 | 0.986290682 | 937.3148111 | 1856.900184 | 126 | 18 | 24 | 30 | 74 | 12 | 0 | 0 | 0 | 0 | 0 | 2 |
| SG0000005 | Pcbp2 | 1.5E-12 | 6.03E-12 | 0.840154631 | 11317.6967 | 20261.41612 | 180 | 15 | 24 | 39 | 93 | 9 | 17 | 1 | 2 | 2 | 9 | 0 |
| SG0000002 | Brap | 1.83E-12 | 7.29E-12 | -1.09748634 | 1945.195581 | 909.0483151 | 152 | 24 | 24 | 34 | 82 | 17 | 9 | 0 | 0 | 3 | 6 | 3 |
| SG0000003 | Ric8b | 2.42E-12 | 9.58E-12 | -1.119887532 | 1749.47635 | 804.9860179 | 364 | 12 | 24 | 40 | 140 | 38 | 10 | 2 | 4 | 4 | 4 | 2 |
| SG0000004 | Ccbl2 | 2.94E-11 | 1.09E-10 | 1.285606985 | 149.5751007 | 364.641059 | 263 | 15 | 24 | 36 | 117 | 27 | 7 | 0 | 0 | 0 | 0 | 0 |
| SG0000002 | Mgrn1 | 7.79E-11 | 2.79E-10 | -0.975803108 | 2762.614741 | 1404.67004 | 120 | 22 | 24 | 44 | 64 | 6 | 0 | 0 | 0 | 0 | 0 | 0 |
| SG0000001 | Rad51d | 4.26E-10 | 1.45E-09 | -1.049815415 | 1055.453942 | 509.815877 | 66 | 12 | 24 | 24 | 36 | 24 | 6 | 0 | 0 | 0 | 0 | 0 |
| JSG000000 | Rab8b | 7.54E-10 | 2.51E-09 | -1.337482674 | 1582.600672 | 626.2517354 | 144 | 17 | 24 | 31 | 91 | 15 | 11 | 0 | 0 | 0 | 3 | 2 |
| SG0000003 | Ldah | 1.06E-09 | 3.49E-09 | -0.943543875 | 1743.632563 | 906.6088673 | 280 | 16 | 24 | 54 | 154 | 24 | 3 | 0 | 0 | 0 | 2 | 0 |
| JSG000000 | Ell2 | 1.31E-09 | 4.28E-09 | 0.841213666 | 882.635919 | 1581.292254 | 185 | 19 | 24 | 35 | 90 | 12 | 4 | 0 | 0 | 1 | 1 | 0 |
| SG0000002 | Eif2d | 1.38E-09 | 4.48E-09 | 0.765808763 | 2332.115071 | 3965.348704 | 161 | 24 | 24 | 46 | 97 | 18 | 3 | 0 | 0 | 1 | 1 | 0 |
| JSG000000 | Epha10 | 1.61E-09 | 5.2E-09 | 4.395304738 | 1.904113361 | 40.06926411 | 49 | 23 | 24 | 26 | 42 | 4 | 1 | 0 | 0 | 0 | 1 | 0 |
| SG0000004 | Setd8 | 4.71E-09 | 1.47E-08 | -0.819962824 | 5592.443082 | 3167.875953 | 80 | 18 | 24 | 27 | 51 | 5 | 7 | 0 | 0 | 1 | 1 | 1 |
| SG0000003 | Pnpla7 | 8.05E-09 | 2.47E-08 | 1.002488236 | 249.0322756 | 498.924311 | 110 | 16 | 24 | 27 | 68 | 3 | 0 | 0 | 0 | 0 | 0 | 0 |
| SG0000001 | Wasf1 | 8.94E-09 | 2.74E-08 | 0.78124317 | 1029.415567 | 1769.165083 | 268 | 16 | 24 | 36 | 116 | 30 | 0 | 0 | 0 | 0 | 0 | 0 |
| SG0000003 | Hmgxb4 | 1.25E-08 | 3.79E-08 | -0.956072243 | 6118.158485 | 3153.655861 | 189 | 16 | 24 | 26 | 69 | 8 | 13 | 0 | 0 | 1 | 1 | 3 |
| SG0000002 | Ifih1 | 6.11E-08 | 1.75E-07 | -1.272248483 | 285.6710109 | 118.2718027 | 278 | 14 | 24 | 43 | 136 | 18 | 2 | 0 | 0 | 1 | 4 | 0 |
| JSG000000 | 1700019A02Rik | 8.06E-08 | 2.28E-07 | 6.154616211 | 0.307509278 | 21.90697007 | 168 | 12 | 24 | 36 | 130 | 24 | 2 | 0 | 0 | 0 | 0 | 0 |
| JSG000000 | Ttc37 | 8.09E-08 | 2.29E-07 | -0.88571095 | 2240.335061 | 1212.515816 | 233 | 14 | 24 | 39 | 116 | 26 | 1 | 0 | 0 | 0 | 0 | 0 |
| JSG000000 | Tmem8b | 2.53E-07 | 6.85E-07 | 1.234535992 | 88.65639365 | 208.6135076 | 96 | 16 | 24 | 32 | 52 | 16 | 0 | 0 | 0 | 0 | 0 | 8 |
| SG0000004 | Greb1l | 5.99E-07 | 1.57E-06 | -1.434837219 | 840.1601114 | 310.7656556 | 539 | 10 | 24 | 54 | 238 | 62 | 6 | 0 | 0 | 1 | 1 | 1 |
| JSG000000 | Pcdh17 | 7.67E-07 | 2E-06 | -3.661733657 | 33.56699207 | 2.652288481 | 327 | 9 | 24 | 46 | 171 | 25 | 14 | 1 | 1 | 1 | 5 | 1 |
| SG0000004 | Rasef | 1.01E-06 | 2.61E-06 | 4.104249127 | 0 | 16.19895645 | 310 | 18 | 24 | 42 | 186 | 40 | 14 | 2 | 2 | 4 | 14 | 2 |
| JSG000000 | Pon3 | 1.24E-06 | 3.18E-06 | -1.471927062 | 141.0900306 | 50.86301854 | 156 | 18 | 24 | 34 | 68 | 12 | 4 | 0 | 0 | 0 | 0 | 0 |
| SG0000003 | Larp1 | 2.5E-06 | 6.24E-06 | -0.634583584 | 15332.27747 | 9875.929395 | 154 | 22 | 24 | 30 | 74 | 18 | 4 | 0 | 0 | 0 | 8 | 0 |
| JSG000000 | Gtf2f2 | 2.71E-06 | 6.72E-06 | 0.639078289 | 1038.030111 | 1616.559471 | 249 | 12 | 24 | 32 | 117 | 17 | 1 | 0 | 1 | 1 | 1 | 0 |
| SG0000003 | Celf6 | 3.21E-06 | 7.93E-06 | 4.004512264 | 0 | 15.05012095 | 170 | 18 | 24 | 36 | 74 | 11 | 0 | 0 | 0 | 0 | 0 | 5 |
| SG0000004 | C87436 | 5.27E-06 | 1.28E-05 | -0.928504369 | 424.2932895 | 222.9248571 | 132 | 18 | 24 | 24 | 66 | 21 | 0 | 0 | 0 | 0 | 0 | 3 |
| JSG000000 | Mrps28 | 6.57E-06 | 1.58E-05 | 0.796000827 | 282.6576347 | 490.7732081 | 218 | 11 | 24 | 39 | 148 | 26 | 1 | 0 | 0 | 0 | 0 | 0 |
| SG0000004 | Fbxo30 | 1.06E-05 | 2.48E-05 | 0.600886819 | 1096.180819 | 1662.521058 | 138 | 14 | 24 | 26 | 47 | 3 | 8 | 0 | 0 | 0 | 3 | 1 |
| SG0000006 | Itgb1bp1 | 1.51E-05 | 3.5E-05 | -0.777831163 | 625.4130749 | 364.7677586 | 103 | 15 | 24 | 21 | 54 | 15 | 4 | 0 | 0 | 0 | 0 | 0 |
| SG0000003 | Zfp384 | 2.07E-05 | 4.73E-05 | -0.63434264 | 1737.752271 | 1119.519604 | 298 | 0 | 24 | 0 | 96 | 8 | 10 | 0 | 0 | 0 | 0 | 0 |
| SG0000003 | Ap4b1 | 3.8E-05 | 8.46E-05 | -0.979177931 | 237.0111813 | 120.2283555 | 139 | 24 | 24 | 59 | 109 | 30 | 3 | 0 | 0 | 3 | 3 | 0 |
| SG0000003 | Wdr26 | 3.82E-05 | 8.51E-05 | -0.558353707 | 10108.15509 | 6864.211918 | 314 | 24 | 24 | 46 | 175 | 37 | 37 | 0 | 0 | 5 | 14 | 4 |
| SG0000006 | Pepd | 5.55E-05 | 0.000121 | 0.619883872 | 536.7054635 | 824.7829137 | 220 | 9 | 24 | 34 | 86 | 21 | 0 | 0 | 0 | 0 | 0 | 0 |
| SG0000002 | Cmtr1 | 6.52E-05 | 0.000141 | -0.780251609 | 434.1196346 | 252.7727715 | 76 | 17 | 24 | 26 | 38 | 3 | 1 | 0 | 0 | 0 | 0 | 2 |
| JSG000001 | Pcdhga11 | 6.8E-05 | 0.000147 | 2.115105067 | 15.77045996 | 68.32103377 | 138 | 21 | 24 | 37 | 77 | 12 | 2 | 0 | 0 | 1 | 1 | 0 |
| JSG000000 | Pcm1 | 7.19E-05 | 0.000155 | 0.493389994 | 4600.549854 | 6476.418864 | 302 | 15 | 24 | 42 | 163 | 23 | 1 | 0 | 0 | 0 | 0 | 0 |
| JSG000000 | Uhrf1bp1l | 9.55E-05 | 0.000203 | -0.553052794 | 2900.510295 | 1976.919254 | 176 | 12 | 24 | 27 | 90 | 9 | 6 | 0 | 1 | 0 | 1 | 1 |
| JSG000000 | Plekhm1 | 0.000105 | 0.000223 | -0.57507753 | 1736.679703 | 1165.746559 | 101 | 17 | 24 | 28 | 57 | 5 | 0 | 0 | 0 | 0 | 1 | 0 |
| SG0000005 | B3glct | 0.000107 | 0.000226 | 0.584631836 | 588.6103842 | 882.7132357 | 180 | 19 | 24 | 38 | 84 | 21 | 5 | 0 | 0 | 0 | 2 | 0 |
| SG0000004 | Lgr6 | 0.000118 | 0.000247 | -1.921768538 | 542.2425013 | 143.1144811 | 133 | 19 | 24 | 31 | 85 | 19 | 6 | 0 | 1 | 1 | 4 | 3 |
| SG0000002 | Hmgcr | 0.00018 | 0.000372 | 0.466166976 | 4149.112146 | 5731.725936 | 108 | 23 | 24 | 25 | 46 | 4 | 6 | 1 | 1 | 1 | 2 | 0 |
| SG0000005 | Pgbd5 | 0.00026 | 0.000526 | 1.031324758 | 59.89128151 | 122.4118026 | 237 | 21 | 24 | 45 | 145 | 79 | 3 | 0 | 0 | 0 | 0 | 0 |
| SG0000004 | Entpd1 | 0.000294 | 0.00059 | 3.189177397 | 1.963171053 | 17.90590186 | 498 | 18 | 24 | 61 | 209 | 36 | 3 | 0 | 0 | 1 | 5 | 0 |
| SG0000001 | Qdpr | 0.000356 | 0.000708 | 0.464178945 | 2077.711072 | 2866.268955 | 112 | 6 | 24 | 12 | 142 | 0 | 0 | 0 | 0 | 0 | 0 | 0 |
| SG0000002 | Efhd1 | 0.000394 | 0.00078 | 6.275035505 | 0.615018556 | 47.62797645 | 166 | 24 | 24 | 38 | 78 | 4 | 4 | 0 | 0 | 4 | 4 | 0 |
| SG0000003 | Yif1b | 0.000503 | 0.000983 | 0.966347241 | 526.3584704 | 1028.445121 | 51 | 24 | 24 | 28 | 28 | 0 | 0 | 0 | 0 | 0 | 0 | 0 |
| SG0000003 | Sspn | 0.000626 | 0.001204 | -1.467834693 | 94.63443401 | 34.212666 | 134 | 19 | 24 | 30 | 57 | 4 | 6 | 0 | 0 | 3 | 9 | 0 |
| SG0000002 | Svep1 | 0.000736 | 0.001403 | -1.821983083 | 132.9593157 | 37.60514015 | 303 | 10 | 24 | 46 | 158 | 36 | 2 | 0 | 0 | 0 | 0 | 1 |
| SG0000002 | Gnal | 0.000881 | 0.001663 | 1.127394125 | 38.79898709 | 84.76173472 | 339 | 2 | 24 | 51 | 173 | 46 | 10 | 0 | 0 | 2 | 4 | 2 |
| SG0000003 | Ythdc2 | 0.001184 | 0.002206 | 0.600078446 | 535.2024145 | 811.2592766 | 217 | 10 | 24 | 32 | 98 | 20 | 9 | 0 | 0 | 0 | 3 | 4 |
| JSG000000 | 2210404O09Rik | 0.002281 | 0.004094 | -0.7558762 | 214.4932254 | 127.0200182 | 113 | 14 | 24 | 32 | 85 | 18 | 0 | 0 | 0 | 0 | 4 | 0 |
| SG0000002 | Ccdc163 | 0.002946 | 0.005204 | 0.620444922 | 202.2038155 | 310.8578784 | 104 | 6 | 24 | 24 | 118 | 21 | 0 | 0 | 0 | 0 | 0 | 0 |
| SG0000003 | Nipa2 | 0.004651 | 0.007997 | -0.425730535 | 1346.751428 | 1002.60481 | 252 | 16 | 24 | 32 | 120 | 36 | 20 | 0 | 0 | 4 | 12 | 4 |
| JSG000001 | Pcdhga9 | 0.005405 | 0.009202 | 2.122851312 | 4.212103161 | 18.3459802 | 159 | 21 | 24 | 38 | 84 | 14 | 2 | 0 | 0 | 1 | 1 | 0 |
| SG0000004 | Oacyl | 0.00557 | 0.009464 | 1.966657408 | 7.383563104 | 28.85950395 | 158 | 16 | 24 | 32 | 88 | 38 | 10 | 4 | 4 | 4 | 6 | 0 |
| SG0000004 | Cage1 | 0.006096 | 0.010305 | -1.194073425 | 52.55590198 | 22.97045385 | 204 | 16 | 24 | 35 | 89 | 12 | 5 | 1 | 1 | 1 | 2 | 0 |
| SG0000000 | Kmt2a | 0.007995 | 0.013268 | -0.459797218 | 2644.248815 | 1922.602771 | 210 | 14 | 24 | 28 | 110 | 34 | 10 | 0 | 2 | 0 | 2 | 4 |
| SG0000007 | Ube2v1 | 0.008537 | 0.014124 | -0.871594146 | 92.59211002 | 50.60553249 | 66 | 21 | 24 | 24 | 48 | 15 | 0 | 0 | 0 | 0 | 0 | 0 |
| SG0000002 | Taf3 | 0.008627 | 0.014263 | 0.4100339 | 543.8621701 | 722.6389306 | 335 | 6 | 24 | 45 | 145 | 30 | 10 | 0 | 2 | 6 | 12 | 2 |
| SG0000004 | Zfp444 | 0.008661 | 0.014312 | 0.622345604 | 332.568415 | 511.9478166 | 112 | 8 | 24 | 24 | 51 | 5 | 0 | 0 | 0 | 0 | 0 | 1 |
| SG0000002 | Vps26a | 0.009778 | 0.016019 | -0.360616967 | 3593.511769 | 2798.739952 | 166 | 14 | 24 | 32 | 74 | 10 | 6 | 0 | 0 | 0 | 0 | 4 |
| SG0000003 | Kdm3b | 0.010021 | 0.016391 | -0.352978475 | 5093.537616 | 3988.066662 | 132 | 17 | 24 | 34 | 67 | 7 | 2 | 0 | 0 | 0 | 0 | 0 |
| SG0000002 | Fam131b | 0.010905 | 0.017723 | 1.097301334 | 23.326509 | 49.90802268 | 48 | 24 | 24 | 24 | 42 | 3 | 0 | 0 | 0 | 0 | 0 | 0 |
| SG0000005 | Spink2 | 0.011202 | 0.018176 | 1.390583184 | 11.39071113 | 29.86469529 | 57 | 21 | 24 | 21 | 30 | 9 | 0 | 0 | 0 | 0 | 0 | 0 |
| SG0000002 | Lias | 0.01353 | 0.021577 | 0.356292786 | 903.1755403 | 1156.184075 | 88 | 22 | 24 | 24 | 39 | 2 | 8 | 0 | 0 | 0 | 3 | 0 |
| SG0000002 | Ift81 | 0.016115 | 0.025408 | 0.399296274 | 415.5805851 | 548.0944519 | 193 | 16 | 24 | 33 | 90 | 17 | 4 | 0 | 0 | 0 | 2 | 0 |
| SG0000005 | Atl2 | 0.020371 | 0.031461 | -0.32802435 | 2886.512023 | 2299.472404 | 258 | 14 | 24 | 32 | 112 | 22 | 10 | 0 | 0 | 2 | 2 | 4 |

| | | | | | | | | | | | | | | | | | | |
|---|---|---|---|---|---|---|---|---|---|---|---|---|---|---|---|---|---|---|
| ISG000000 | Lin52 | 0.021007 | 0.032357 | -0.408148534 | 543.4961061 | 409.5731568 | 175 | 17 | 24 | 28 | 82 | 15 | 2 | 0 | 0 | 1 | 3 | 0 |
| SG0000003 | Agl | 0.023184 | 0.035442 | -0.389028352 | 1060.239364 | 809.6451551 | 163 | 19 | 24 | 34 | 87 | 13 | 7 | 0 | 0 | 1 | 1 | 0 |
| SG0000001 | P4ha1 | 0.024001 | 0.036618 | -0.332969093 | 4631.209138 | 3676.721281 | 304 | 9 | 24 | 36 | 151 | 24 | 13 | 0 | 3 | 3 | 7 | 0 |
| SG0000003 | Alg12 | 0.026223 | 0.039772 | -0.434786979 | 361.363381 | 267.3377069 | 62 | 22 | 24 | 28 | 38 | 2 | 2 | 0 | 0 | 0 | 0 | 0 |
| SG0000004 | Pkd1l1 | 0.036392 | 0.053765 | -1.568311462 | 20.48914683 | 6.908998244 | 268 | 17 | 24 | 37 | 143 | 27 | 0 | 0 | 0 | 0 | 0 | 0 |
| ISG000001 | Pcdhga12 | 0.039564 | 0.058121 | 0.627985073 | 58.95637972 | 91.11149045 | 125 | 21 | 24 | 37 | 70 | 12 | 2 | 0 | 0 | 1 | 1 | 0 |
| SG0000002 | Tbl1x | 0.040971 | 0.060025 | -0.281767394 | 4635.085166 | 3812.740793 | 294 | 17 | 24 | 39 | 132 | 40 | 2 | 0 | 0 | 0 | 3 | 0 |
| ISG000001 | Pcdhga10 | 0.047036 | 0.068285 | 1.103712779 | 17.66135564 | 37.95549754 | 153 | 21 | 24 | 38 | 81 | 14 | 2 | 0 | 0 | 1 | 1 | 0 |
| SG0000003 | Wdr36 | 0.047962 | 0.069501 | 0.247276615 | 3946.373384 | 4684.204551 | 198 | 8 | 24 | 30 | 110 | 8 | 0 | 0 | 0 | 0 | 2 | 0 |
| SG0000003 | Thsd1 | 0.048098 | 0.069664 | 0.578437474 | 78.88607212 | 117.7951373 | 174 | 21 | 24 | 31 | 88 | 18 | 9 | 0 | 0 | 1 | 1 | 0 |
| ISG000000 | Ubac2 | 0.054245 | 0.077775 | 0.274662185 | 1108.474287 | 1340.933312 | 199 | 16 | 24 | 30 | 89 | 23 | 0 | 0 | 0 | 0 | 0 | 0 |
| SG0000002 | Ttc39a | 0.055126 | 0.078936 | -0.726495744 | 60.76562576 | 36.72492891 | 93 | 20 | 24 | 29 | 48 | 9 | 2 | 0 | 0 | 1 | 1 | 1 |
| SG0000002 | Slf1 | 0.059333 | 0.084401 | -0.322870532 | 662.8906856 | 529.9662361 | 429 | 8 | 24 | 49 | 224 | 46 | 10 | 0 | 1 | 3 | 4 | 0 |
| SG0000005 | Ndfip2 | 0.072095 | 0.101006 | -0.268345173 | 1377.82429 | 1143.968037 | 255 | 10 | 24 | 32 | 130 | 18 | 16 | 0 | 0 | 2 | 4 | 2 |
| SG0000002 | Epb41l3 | 0.092213 | 0.12654 | 0.522198972 | 517.2163311 | 742.7963886 | 414 | 10 | 24 | 32 | 176 | 52 | 10 | 0 | 0 | 0 | 4 | 0 |
| SG0000002 | Klhl20 | 0.104034 | 0.141085 | 0.229690211 | 1055.992033 | 1238.238447 | 229 | 18 | 24 | 46 | 122 | 10 | 15 | 0 | 0 | 3 | 3 | 0 |
| SG0000002 | Rab40c | 0.108535 | 0.146701 | -0.254550292 | 841.8838572 | 705.7077859 | 208 | 16 | 24 | 44 | 68 | 16 | 9 | 0 | 4 | 4 | 4 | 0 |
| ISG000000 | Acbd3 | 0.125592 | 0.167318 | 0.208596025 | 1546.822158 | 1787.450588 | 80 | 18 | 24 | 27 | 49 | 9 | 3 | 0 | 0 | 0 | 1 | 2 |
| ISG000000 | Mocos | 0.199786 | 0.254682 | -0.321138625 | 149.3382138 | 119.5359655 | 88 | 18 | 24 | 29 | 55 | 11 | 0 | 0 | 0 | 0 | 0 | 0 |
| SG0000003 | Cab39 | 0.218615 | 0.275782 | 0.155508372 | 3687.90089 | 4107.635756 | 204 | 18 | 24 | 33 | 107 | 26 | 8 | 0 | 0 | 1 | 1 | 4 |
| SG0000002 | Apool | 0.261533 | 0.323253 | -0.215053663 | 465.6089778 | 401.128706 | 331 | 10 | 24 | 36 | 140 | 34 | 1 | 0 | 0 | 0 | 2 | 0 |
| SG0000003 | Ss18 | 0.290177 | 0.35428 | -0.141226489 | 3923.942063 | 3558.026488 | 315 | 15 | 24 | 39 | 124 | 21 | 6 | 0 | 0 | 0 | 4 | 3 |
| ISG000001 | Pcdhgb6 | 0.334746 | 0.401498 | -0.724302925 | 15.75589557 | 9.536877131 | 157 | 21 | 24 | 38 | 84 | 14 | 2 | 0 | 0 | 1 | 1 | 0 |
| SG0000003 | Vwa9 | 0.359463 | 0.427621 | -0.14154828 | 982.1482645 | 890.3623263 | 98 | 14 | 24 | 24 | 50 | 8 | 2 | 0 | 0 | 0 | 0 | 0 |
| SG0000002 | Snx16 | 0.545337 | 0.61123 | -0.119302661 | 355.2401337 | 327.0456262 | 187 | 24 | 24 | 45 | 73 | 6 | 7 | 0 | 0 | 0 | 4 | 0 |
| ISG000000 | Nol10 | 0.553016 | 0.618328 | -0.080601604 | 2176.179576 | 2057.932992 | 157 | 17 | 24 | 30 | 65 | 8 | 0 | 0 | 0 | 0 | 1 | 0 |
| SG0000002 | Dnm1l | 0.561056 | 0.626068 | 0.066377074 | 4936.947727 | 5169.398579 | 411 | 12 | 24 | 54 | 186 | 36 | 9 | 0 | 0 | 3 | 6 | 0 |
| SG0000003 | Ttll3 | 0.564406 | 0.628983 | -0.212529184 | 59.92624741 | 51.71772632 | 48 | 20 | 24 | 26 | 72 | 10 | 0 | 0 | 0 | 0 | 0 | 0 |
| ISG000000 | Nemf | 0.571536 | 0.636058 | -0.085808037 | 2007.787903 | 1891.851485 | 130 | 14 | 24 | 30 | 76 | 9 | 1 | 0 | 0 | 0 | 0 | 1 |
| SG0000000 | Arvcf | 0.612669 | 0.674881 | 0.105066178 | 1075.518502 | 1156.767176 | 200 | 18 | 24 | 50 | 120 | 9 | 7 | 0 | 0 | 0 | 1 | 0 |
| ISG000001 | Pcdhgb7 | 0.612668 | 0.674881 | 0.336898076 | 14.38130772 | 18.16413853 | 146 | 21 | 24 | 37 | 79 | 12 | 2 | 0 | 0 | 1 | 1 | 0 |
| SG0000003 | Rdx | 0.63217 | 0.693116 | 0.056323399 | 4997.772048 | 5196.745832 | 300 | 9 | 24 | 30 | 101 | 26 | 7 | 0 | 0 | 0 | 4 | 0 |
| SG0000006 | Chd4 | 0.633478 | 0.694024 | -0.067046214 | 13896.62514 | 13265.58545 | 129 | 21 | 24 | 33 | 83 | 6 | 0 | 0 | 0 | 0 | 0 | 0 |
| SG0000002 | Csnk2b | 0.661829 | 0.719993 | 0.057412347 | 2623.353984 | 2729.855976 | 54 | 24 | 24 | 24 | 47 | 4 | 0 | 0 | 0 | 0 | 0 | 0 |
| SG0000006 | Btbd9 | 0.668079 | 0.725691 | 0.066740989 | 669.4494094 | 701.1465513 | 539 | 14 | 24 | 56 | 211 | 40 | 4 | 0 | 0 | 0 | 0 | 0 |
| SG0000004 | Ikzf5 | 0.706087 | 0.759514 | 0.060731942 | 466.0381832 | 486.075394 | 133 | 15 | 24 | 32 | 83 | 12 | 26 | 0 | 0 | 5 | 12 | 0 |
| SG0000002 | Snx30 | 0.747039 | 0.795586 | 0.045436739 | 1114.410553 | 1150.066723 | 159 | 18 | 24 | 35 | 87 | 11 | 6 | 0 | 0 | 2 | 2 | 1 |
| SG0000000 | Scin | 0.751204 | 0.799261 | -0.199664006 | 22.7238644 | 19.78688065 | 336 | 16 | 24 | 52 | 190 | 42 | 0 | 0 | 0 | 0 | 0 | 0 |
| SG0000003 | Gpatch8 | 0.850004 | 0.882727 | 0.017006869 | 1918.221293 | 1940.967597 | 182 | 18 | 24 | 34 | 102 | 15 | 6 | 0 | 0 | 0 | 1 | 2 |
| ISG000000 | Dram1 | 0.860535 | 0.891286 | -0.044874063 | 37.31282067 | 36.17009436 | 136 | 20 | 24 | 32 | 68 | 8 | 12 | 0 | 0 | 0 | 2 | 0 |
| ISG000000 | Cep290 | 0.933385 | 0.951445 | -0.02492012 | 593.7458498 | 583.5779627 | 276 | 13 | 24 | 44 | 137 | 15 | 6 | 0 | 0 | 1 | 4 | 0 |
| SG0000002 | Larp1b | 0.990487 | 0.999753 | -0.005870057 | 544.6566709 | 542.4450669 | 363 | 14 | 24 | 39 | 177 | 47 | 6 | 2 | 2 | 2 | 4 | 6 |
| ISG000000 | Gm2381 | 6.5E-113 | 6E-111 | 8.570363932 | 0 | 379.133914 | 102 | 20 | 23 | 30 | 67 | 7 | 8 | 0 | 0 | 1 | 2 | 1 |
| ISG000000 | Gnai1 | 5.4E-105 | 4.3E-103 | 5.416355203 | 11.9575461 | 510.6548337 | 210 | 15 | 23 | 36 | 120 | 12 | 13 | 0 | 1 | 2 | 5 | 1 |
| SG0000002 | Pecr | 2.26E-81 | 1.05E-79 | 2.974115947 | 151.5161737 | 1190.575949 | 153 | 20 | 23 | 39 | 90 | 30 | 1 | 0 | 0 | 0 | 0 | 0 |
| ISG000000 | Cfdp1 | 3.3E-63 | 9.96E-62 | 2.001973929 | 2076.899426 | 8318.97213 | 140 | 10 | 23 | 28 | 72 | 9 | 0 | 0 | 0 | 0 | 0 | 0 |
| SG0000004 | Slc25a24 | 2.36E-53 | 5.38E-52 | -4.685207878 | 7254.524682 | 281.9816553 | 175 | 18 | 23 | 36 | 117 | 4 | 7 | 0 | 0 | 1 | 2 | 0 |
| SG0000004 | BC055324 | 4.73E-50 | 9.85E-49 | 1.883038705 | 925.5866273 | 3414.03685 | 304 | 8 | 23 | 44 | 123 | 17 | 5 | 0 | 0 | 0 | 0 | 0 |
| SG0000004 | Fam110b | 3.82E-41 | 5.87E-40 | 2.97475435 | 46.09486671 | 362.3621708 | 466 | 8 | 23 | 79 | 262 | 55 | 2 | 0 | 0 | 2 | 2 | 2 |
| ISG000000 | Arid5b | 1.39E-40 | 2.1E-39 | -2.216567133 | 4969.551569 | 1069.21245 | 342 | 15 | 23 | 54 | 189 | 42 | 4 | 0 | 0 | 1 | 1 | 1 |
| ISG000000 | Aldh1a2 | 9.21E-37 | 1.21E-35 | 7.438504401 | 0.649984458 | 112.7498363 | 184 | 19 | 23 | 45 | 101 | 8 | 6 | 0 | 1 | 1 | 1 | 0 |
| SG0000002 | Ing5 | 2.39E-36 | 3.11E-35 | 1.732421351 | 658.4935785 | 2188.075664 | 60 | 20 | 23 | 26 | 51 | 15 | 12 | 8 | 8 | 8 | 12 | 0 |
| SG0000003 | Vopp1 | 6.82E-35 | 8.43E-34 | -2.931295287 | 2185.648107 | 286.5315397 | 492 | 7 | 23 | 44 | 238 | 29 | 20 | 1 | 2 | 3 | 8 | 1 |
| SG0000003 | Chek1 | 2.16E-34 | 2.63E-33 | 1.622269092 | 874.1579273 | 2691.172636 | 129 | 12 | 23 | 15 | 58 | 15 | 12 | 0 | 2 | 0 | 2 | 0 |
| ISG000000 | Fam120a | 2.78E-28 | 2.61E-27 | -1.659064059 | 12750.2165 | 4037.285973 | 148 | 16 | 23 | 25 | 78 | 14 | 2 | 0 | 0 | 0 | 1 | 0 |
| SG0000007 | Arhgef10 | 4.55E-22 | 3.2E-21 | 2.955460062 | 101.2338421 | 785.2498272 | 185 | 15 | 23 | 32 | 85 | 10 | 9 | 0 | 3 | 3 | 6 | 0 |
| SG0000002 | Abcc1 | 1.32E-21 | 9.05E-21 | -1.602250043 | 7855.203318 | 2587.212536 | 135 | 19 | 23 | 37 | 81 | 20 | 2 | 0 | 0 | 1 | 2 | 0 |
| SG0000002 | Mdm2 | 2.93E-21 | 1.97E-20 | -1.438368356 | 10265.64941 | 3787.864266 | 149 | 18 | 23 | 24 | 78 | 9 | 24 | 3 | 3 | 3 | 12 | 0 |
| SG0000009 | Synj2bp | 3.81E-21 | 2.54E-20 | -1.533863165 | 2257.660222 | 779.6860413 | 253 | 12 | 23 | 37 | 121 | 28 | 7 | 0 | 0 | 0 | 2 | 0 |
| SG0000003 | Fancd2 | 3.47E-20 | 2.2E-19 | 1.190085448 | 1185.759247 | 2705.502485 | 136 | 13 | 23 | 24 | 46 | 9 | 4 | 0 | 2 | 0 | 0 | 3 |
| SG0000003 | Ston1 | 2.71E-19 | 1.63E-18 | -1.690829837 | 959.2906709 | 297.1390444 | 106 | 19 | 23 | 34 | 86 | 2 | 2 | 0 | 0 | 0 | 1 | 0 |
| SG0000002 | Entpd7 | 3.95E-18 | 2.23E-17 | -1.881272716 | 505.423563 | 137.1942682 | 97 | 20 | 23 | 25 | 51 | 3 | 6 | 0 | 0 | 0 | 2 | 0 |
| ISG000000 | Exoc6 | 1.27E-17 | 6.99E-17 | -1.423354223 | 1605.632869 | 598.651306 | 300 | 11 | 23 | 46 | 131 | 17 | 0 | 0 | 0 | 0 | 1 | 0 |
| ISG000000 | Slc4a7 | 2.33E-17 | 1.27E-16 | -1.379786835 | 7790.421626 | 2993.668708 | 294 | 10 | 23 | 39 | 124 | 22 | 23 | 3 | 5 | 8 | 11 | 3 |
| SG0000002 | Smap1 | 5.7E-16 | 2.86E-15 | 1.043318666 | 1390.271674 | 2865.298601 | 313 | 16 | 23 | 44 | 155 | 25 | 2 | 0 | 0 | 0 | 0 | 3 |
| SG0000005 | Lrguk | 8.4E-15 | 3.92E-14 | 4.214116536 | 3.239048179 | 60.11638964 | 256 | 16 | 23 | 39 | 107 | 13 | 2 | 0 | 0 | 0 | 0 | 0 |
| ISG000000 | Cpne3 | 2.79E-14 | 1.26E-13 | -1.240064585 | 4506.362136 | 1907.7851 | 137 | 20 | 23 | 34 | 78 | 10 | 19 | 1 | 1 | 2 | 4 | 2 |
| SG0000000 | Dennd2c | 6.68E-13 | 2.75E-12 | 0.984125138 | 830.0502981 | 1641.933622 | 233 | 17 | 23 | 40 | 96 | 11 | 3 | 0 | 0 | 0 | 0 | 0 |
| SG0000003 | Rftn1 | 1.79E-11 | 6.72E-11 | 1.593836412 | 71.41566926 | 215.5688866 | 369 | 13 | 23 | 38 | 142 | 25 | 4 | 0 | 2 | 0 | 1 | 0 |
| SG0000002 | Rnf115 | 3.87E-11 | 1.42E-10 | 0.882897126 | 1082.608389 | 1996.410779 | 148 | 20 | 23 | 41 | 95 | 9 | 3 | 0 | 0 | 0 | 1 | 0 |
| ISG000000 | Cstf3 | 9.01E-11 | 3.21E-10 | 0.873808913 | 1017.008023 | 1863.661667 | 226 | 15 | 23 | 38 | 113 | 16 | 1 | 0 | 0 | 1 | 1 | 0 |
| SG0000000 | Gpr107 | 1.47E-09 | 4.75E-09 | -0.920010052 | 2025.209858 | 1070.33422 | 88 | 23 | 23 | 28 | 62 | 11 | 6 | 0 | 0 | 0 | 1 | 1 |
| SG0000003 | Hip1 | 1.68E-09 | 5.42E-09 | -1.399492402 | 4113.399949 | 1559.235617 | 195 | 18 | 23 | 40 | 115 | 13 | 12 | 0 | 0 | 2 | 4 | 0 |
| ISG000000 | Creb3l2 | 3.92E-09 | 1.23E-08 | -1.200057464 | 697.9975676 | 303.8089866 | 170 | 19 | 23 | 36 | 100 | 23 | 3 | 0 | 0 | 2 | 3 | 1 |
| SG0000003 | Ttll4 | 7.45E-08 | 2.11E-07 | 0.675746011 | 2829.005775 | 4519.117699 | 117 | 17 | 23 | 26 | 53 | 12 | 12 | 0 | 1 | 3 | 4 | 3 |
| ISG000000 | Slc25a17 | 4.94E-07 | 1.31E-06 | -0.768653944 | 1799.116293 | 1056.018055 | 89 | 16 | 23 | 26 | 50 | 8 | 2 | 0 | 0 | 2 | 2 | 0 |
| SG0000002 | Smn1 | 1.27E-06 | 3.25E-06 | 0.667190853 | 976.4754904 | 1550.621518 | 52 | 23 | 23 | 25 | 40 | 2 | 0 | 0 | 0 | 0 | 0 | 0 |
| ISG000001 | Mrpl33 | 4.95E-06 | 1.2E-05 | -0.71132013 | 1454.372194 | 888.2738291 | 102 | 15 | 23 | 33 | 70 | 4 | 5 | 2 | 3 | 5 | 12 | 0 |
| SG0000003 | Fnbp1l | 2.06E-05 | 4.7E-05 | 0.544619459 | 2341.373944 | 3415.211266 | 308 | 9 | 23 | 42 | 154 | 25 | 10 | 0 | 2 | 4 | 8 | 2 |
| ISG000000 | Mtss1 | 2.97E-05 | 6.68E-05 | 0.719869574 | 1098.629522 | 1809.479219 | 177 | 19 | 23 | 35 | 79 | 17 | 8 | 0 | 0 | 0 | 0 | 1 |
| ISG000000 | Dock5 | 3.93E-05 | 8.74E-05 | -0.586949122 | 2997.930581 | 1995.87014 | 261 | 15 | 23 | 38 | 102 | 34 | 3 | 1 | 1 | 1 | 2 | 0 |
| SG0000002 | Fn3k | 5.07E-05 | 0.000111 | 6.53649094 | 0.615018556 | 57.0910537 | 56 | 21 | 23 | 28 | 57 | 7 | 2 | 0 | 0 | 0 | 3 | 0 |
| ISG000000 | Anln | 6.55E-05 | 0.000142 | -0.553714799 | 9511.970517 | 6480.160044 | 143 | 16 | 23 | 36 | 88 | 10 | 4 | 0 | 0 | 1 | 4 | 0 |
| SG0000002 | Postn | 8.92E-05 | 0.00019 | -1.325555576 | 196.7981845 | 78.52159243 | 307 | 9 | 23 | 44 | 153 | 26 | 3 | 0 | 0 | 0 | 0 | 0 |
| ISG000000 | Slc15a1 | 0.000111 | 0.000234 | 7.804039749 | 0.663202137 | 148.2163004 | 76 | 19 | 23 | 24 | 44 | 7 | 4 | 0 | 2 | 1 | 2 | 0 |
| ISG000000 | Chmp4c | 0.000124 | 0.00026 | 0.600246195 | 487.16944 | 738.5368109 | 206 | 17 | 23 | 34 | 104 | 14 | 16 | 0 | 1 | 2 | 6 | 5 |
| SG0000004 | Tmem194 | 0.000137 | 0.000287 | 0.604022131 | 1194.399177 | 1815.424845 | 162 | 10 | 23 | 33 | 81 | 17 | 12 | 1 | 1 | 2 | 3 | 2 |
| ISG000000 | Ddr2 | 0.000274 | 0.000552 | -1.392137424 | 411.456646 | 156.7649466 | 213 | 17 | 23 | 38 | 148 | 18 | 0 | 0 | 0 | 0 | 0 | 0 |
| SG0000001 | Tmem167 | 0.000336 | 0.00067 | -0.519221923 | 2142.67332 | 1495.046115 | 265 | 4 | 23 | 32 | 93 | 24 | 20 | 2 | 3 | 2 | 8 | 4 |
| SG0000003 | Disp1 | 0.000566 | 0.001096 | -0.608831105 | 575.8367169 | 377.5921252 | 548 | 9 | 23 | 72 | 225 | 52 | 0 | 0 | 0 | 0 | 0 | 0 |
| ISG000000 | Cep85l | 0.000569 | 0.001102 | -1.449905336 | 125.335285 | 45.87840975 | 297 | 13 | 23 | 49 | 142 | 29 | 0 | 0 | 0 | 0 | 0 | 0 |
| SG0000002 | Lrsam1 | 0.000572 | 0.001108 | -0.600573369 | 654.3153663 | 431.5156201 | 101 | 18 | 23 | 25 | 58 | 4 | 4 | 2 | 4 | 4 | 4 | 0 |
| ISG000000 | Noc3l | 0.000686 | 0.001312 | 0.461304961 | 1287.215258 | 1772.220719 | 80 | 21 | 23 | 27 | 48 | 11 | 3 | 0 | 0 | 0 | 2 | 1 |
| SG0000002 | Tmem50b | 0.000933 | 0.001757 | 0.511754514 | 560.7604284 | 799.5227355 | 116 | 17 | 23 | 41 | 68 | 0 | 2 | 0 | 0 | 0 | 0 | 0 |
| SG0000003 | Mical2 | 0.001391 | 0.002571 | -1.424825401 | 750.3261839 | 279.4698218 | 366 | 12 | 23 | 53 | 190 | 17 | 11 | 0 | 0 | 2 | 2 | 0 |
| ISG000000 | Pak2 | 0.001824 | 0.003319 | -0.422319798 | 5865.433066 | 4376.925518 | 131 | 15 | 23 | 29 | 77 | 8 | 9 | 1 | 1 | 3 | 4 | 0 |
| SG0000003 | Nck1 | 0.004659 | 0.008009 | -0.465735928 | 883.5494702 | 639.7795976 | 182 | 18 | 23 | 35 | 103 | 18 | 8 | 0 | 0 | 1 | 2 | 2 |
| SG0000002 | Fktn | 0.00686 | 0.011509 | -0.446952518 | 793.5802833 | 582.1632072 | 310 | 21 | 23 | 47 | 141 | 45 | 25 | 3 | 3 | 6 | 18 | 0 |
| ISG000000 | Senp2 | 0.017842 | 0.027856 | -0.329600585 | 3422.570461 | 2723.533586 | 94 | 18 | 23 | 32 | 54 | 7 | 4 | 0 | 0 | 1 | 1 | 0 |
| SG0000000 | Fnbp4 | 0.022259 | 0.034156 | 0.294498198 | 2762.877468 | 3388.554039 | 91 | 21 | 23 | 28 | 54 | 7 | 5 | 0 | 0 | 1 | 1 | 2 |
| ISG000000 | Mms19 | 0.02531 | 0.038469 | 0.289434874 | 2390.135874 | 2921.131913 | 300 | 8 | 23 | 29 | 134 | 87 | 8 | 0 | 0 | 0 | 0 | 0 |
| ISG000000 | Trpm6 | 0.078518 | 0.109226 | 1.02268333 | 12.85463544 | 26.11668932 | 299 | 16 | 23 | 43 | 163 | 31 | 1 | 0 | 0 | 0 | 0 | 0 |
| ISG000000 | Glrx3 | 0.078654 | 0.109381 | 0.223308455 | 3237.686616 | 3779.700195 | 76 | 20 | 23 | 24 | 44 | 3 | 1 | 0 | 0 | 0 | 0 | 0 |
| ISG000000 | Atad1 | 0.082158 | 0.113858 | -0.242632601 | 3201.226061 | 2705.68138 | 126 | 19 | 23 | 34 | 87 | 13 | 7 | 0 | 0 | 0 | 1 | 3 |
| SG0000000 | Uba1 | 0.090201 | 0.12395 | 0.207166882 | 14647.6345 | 16909.50677 | 91 | 21 | 23 | 27 | 48 | 2 | 2 | 0 | 0 | 0 | 0 | 0 |
| ISG000000 | Fbn2 | 0.092194 | 0.126523 | -0.722920684 | 59.45697316 | 36.02317484 | 389 | 6 | 23 | 39 | 142 | 33 | 5 | 1 | 1 | 1 | 4 | 0 |
| SG0000002 | Naa50 | 0.118463 | 0.1586 | 0.189342585 | 7828.278719 | 8926.14788 | 230 | 16 | 23 | 28 | 99 | 12 | 26 | 0 | 3 | 0 | 3 | 0 |
| SG0000002 | Zfp64 | 0.130222 | 0.172896 | 0.424708928 | 767.8853863 | 1030.733828 | 133 | 21 | 23 | 36 | 94 | 14 | 1 | 0 | 0 | 0 | 1 | 0 |
| SG0000000 | Naa20 | 0.130545 | 0.173311 | -0.22740715 | 1534.015304 | 1310.307777 | 128 | 15 | 23 | 33 | 62 | 22 | 6 | 0 | 0 | 0 | 3 | 0 |
| SG0000003 | Parp16 | 0.131908 | 0.174901 | -0.294254729 | 417.4928792 | 340.4627266 | 52 | 20 | 23 | 25 | 39 | 1 | 4 | 0 | 0 | 0 | 0 | 0 |
| SG0000005 | Zbtb8os | 0.184038 | 0.236613 | -0.224334505 | 833.9323345 | 713.8375646 | 134 | 18 | 23 | 32 | 67 | 12 | 9 | 1 | 1 | 2 | 5 | 0 |
| SG0000003 | Mbnl3 | 0.21592 | 0.272979 | 0.308268535 | 151.8093153 | 187.9734434 | 660 | 6 | 23 | 35 | 251 | 45 | 7 | 0 | 0 | 0 | 2 | 2 |
| SG0000002 | Tubd1 | 0.302788 | 0.367726 | -0.243322734 | 206.5527481 | 174.4952205 | 148 | 19 | 23 | 46 | 130 | 13 | 0 | 0 | 0 | 0 | 0 | 0 |
| SG0000002 | Mtor | 0.508548 | 0.575506 | 0.080681313 | 2637.600307 | 2789.308048 | 156 | 20 | 23 | 38 | 85 | 12 | 0 | 0 | 0 | 0 | 2 | 0 |
| ISG000000 | Ube2n | 0.523624 | 0.590367 | 0.080569586 | 1865.026137 | 1972.144735 | 91 | 20 | 23 | 27 | 58 | 4 | 5 | 0 | 0 | 1 | 3 | 1 |
| SG0000004 | Ankrd16 | 0.616957 | 0.679013 | -0.11471356 | 602.8085539 | 556.733281 | 64 | 19 | 23 | 22 | 33 | 4 | 0 | 0 | 0 | 0 | 0 | 0 |
| SG0000003 | Sec24d | 0.638364 | 0.69841 | -0.073528199 | 680.8177597 | 646.9886811 | 150 | 18 | 23 | 37 | 91 | 20 | 0 | 0 | 0 | 0 | 0 | 1 |
| SG0000000 | Pxmp4 | 0.669779 | 0.72736 | -0.093187382 | 251.0536606 | 235.3500858 | 55 | 16 | 23 | 19 | 31 | 3 | 1 | 0 | 0 | 0 | 0 | 0 |
| ISG000000 | Lmln | 0.925849 | 0.946294 | 0.015277163 | 315.1107458 | 318.4652849 | 132 | 19 | 23 | 30 | 71 | 14 | 10 | 1 | 1 | 3 | 3 | 2 |
| SG0000003 | Acer2 | 2.7E-161 | 5.1E-159 | 5.717428614 | 17.87115105 | 940.307795 | 150 | 12 | 22 | 28 | 106 | 6 | 12 | 0 | 0 | 2 | 6 | 0 |
| SG0000004 | Dlgap3 | 2.1E-106 | 1.8E-104 | 4.819131919 | 23.54717853 | 664.7251904 | 188 | 14 | 22 | 42 | 81 | 10 | 0 | 0 | 0 | 0 | 0 | 0 |
| ISG000000 | Lmnb1 | 1.14E-95 | 7.38E-94 | 2.48218993 | 1500.905365 | 8386.233184 | 93 | 19 | 22 | 23 | 51 | 4 | 0 | 0 | 0 | 0 | 0 | 0 |
| SG0000002 | Trim44 | 2.8E-92 | 1.67E-90 | 2.870999294 | 260.7328395 | 1907.447702 | 217 | 18 | 22 | 33 | 117 | 22 | 8 | 0 | 0 | 1 | 1 | 0 |

| | | | | | | | | | | | | | | | | | | |
|---|---|---|---|---|---|---|---|---|---|---|---|---|---|---|---|---|---|---|
| SG0000002 | Homer2 | 1.12E-87 | 5.99E-86 | 6.16965564 | 4.883835843 | 351.5705337 | 252 | 12 | 22 | 42 | 142 | 20 | 11 | 1 | 1 | 2 | 13 | 0 |
| SG0000006 | Prmt7 | 9.69E-63 | 2.9E-61 | 2.067728702 | 1123.528128 | 4710.123687 | 111 | 18 | 22 | 25 | 65 | 13 | 0 | 0 | 0 | 0 | 0 | 0 |
| SG0000002 | Dpysl5 | 3.95E-61 | 1.13E-59 | 9.814154803 | 0.992459638 | 893.4450183 | 260 | 16 | 22 | 29 | 136 | 65 | 0 | 0 | 0 | 0 | 0 | 0 |
| SG0000006 | Ahnak | 7.83E-59 | 2.09E-57 | -3.720177863 | 26274.42773 | 1993.646431 | 160 | 19 | 22 | 24 | 82 | 12 | 0 | 0 | 0 | 0 | 0 | 2 |
| JSG000000 | Bard1 | 5.8E-58 | 1.5E-56 | 2.0617089 | 730.0068745 | 3047.636534 | 180 | 10 | 22 | 28 | 74 | 10 | 10 | 1 | 2 | 1 | 3 | 0 |
| SG0000002 | Ptch1 | 7.37E-54 | 1.7E-52 | 1.9187635 | 1240.18235 | 4689.11533 | 327 | 8 | 22 | 37 | 72 | 28 | 31 | 0 | 3 | 6 | 9 | 5 |
| SG0000002 | Ppap2a | 1.34E-46 | 2.5E-45 | -2.641559766 | 2321.568844 | 372.0427467 | 260 | 16 | 22 | 34 | 134 | 18 | 0 | 0 | 0 | 0 | 0 | 0 |
| SG0000003 | Ctsc | 3.18E-40 | 4.74E-39 | 2.427102069 | 105.9675873 | 569.9064492 | 129 | 20 | 22 | 28 | 62 | 7 | 6 | 0 | 0 | 0 | 1 | 0 |
| SG000000C | Arhgef18 | 5.57E-37 | 7.4E-36 | -2.029904203 | 7326.241179 | 1793.986449 | 96 | 18 | 22 | 26 | 45 | 8 | 3 | 0 | 0 | 0 | 1 | 2 |
| SG0000002 | Fam134b | 3.22E-34 | 3.87E-33 | -2.596950284 | 2424.015685 | 400.6595443 | 253 | 12 | 22 | 46 | 125 | 26 | 4 | 1 | 1 | 3 | 3 | 0 |
| SG0000005 | Capg | 6.53E-34 | 7.75E-33 | -2.356608418 | 6162.877487 | 1203.299434 | 91 | 20 | 22 | 32 | 93 | 10 | 0 | 0 | 0 | 0 | 0 | 0 |
| SG000000C | Sqrdl | 4.95E-32 | 5.44E-31 | -2.92409078 | 802.3240757 | 105.7087141 | 158 | 18 | 22 | 38 | 150 | 29 | 0 | 0 | 0 | 0 | 0 | 0 |
| JSG000000 | 43718 | 9.28E-30 | 9.28E-29 | -2.057417805 | 1789.570663 | 429.9365615 | 148 | 16 | 22 | 32 | 79 | 13 | 0 | 0 | 0 | 0 | 3 | 0 |
| JSG000000 | Inpp5k | 1.07E-29 | 1.07E-28 | -2.131944906 | 1623.682999 | 370.4434136 | 49 | 20 | 22 | 20 | 30 | 2 | 0 | 0 | 0 | 0 | 0 | 0 |
| JSG000000 | Flrt2 | 9.04E-27 | 7.89E-26 | -2.615363649 | 1136.44203 | 185.4574906 | 383 | 12 | 22 | 56 | 181 | 33 | 14 | 0 | 0 | 0 | 0 | 0 |
| JSG000000 | Cacng7 | 1.26E-26 | 1.09E-25 | 1.79227182 | 197.3484178 | 683.5357398 | 68 | 21 | 22 | 31 | 57 | 10 | 2 | 0 | 0 | 0 | 0 | 1 |
| SG0000002 | Zfp148 | 1.38E-24 | 1.09E-23 | -1.639717434 | 3106.746642 | 997.0150301 | 230 | 11 | 22 | 38 | 136 | 15 | 2 | 0 | 0 | 0 | 1 | 1 |
| SG0000002 | Lrp12 | 1.95E-23 | 1.46E-22 | -1.938088423 | 881.576934 | 230.0580684 | 342 | 8 | 22 | 36 | 140 | 20 | 14 | 2 | 2 | 6 | 12 | 0 |
| SG0000003 | G2e3 | 3.82E-23 | 2.82E-22 | 1.221699907 | 2105.248003 | 4909.88796 | 205 | 12 | 22 | 36 | 118 | 21 | 16 | 0 | 4 | 3 | 10 | 3 |
| SG0000002 | Cdk18 | 1.12E-21 | 7.72E-21 | 1.450351523 | 356.9797543 | 975.5350973 | 47 | 22 | 22 | 24 | 36 | 0 | 1 | 0 | 0 | 0 | 0 | 0 |
| SG0000002 | Gata4 | 1.39E-21 | 9.53E-21 | 8.069371147 | 0.34247518 | 91.99238193 | 160 | 16 | 22 | 24 | 72 | 13 | 6 | 0 | 0 | 0 | 0 | 0 |
| SG0000005 | Tmem245 | 1.5E-20 | 9.72E-20 | -1.552641925 | 4732.764377 | 1613.329561 | 210 | 9 | 22 | 29 | 91 | 27 | 22 | 4 | 6 | 9 | 15 | 0 |
| SG0000003 | U2surp | 3.53E-20 | 2.24E-19 | 1.191761881 | 2745.834178 | 6272.351388 | 393 | 9 | 22 | 47 | 204 | 39 | 12 | 0 | 1 | 2 | 3 | 4 |
| SG0000004 | Mmgt2 | 5.73E-20 | 3.58E-19 | -1.827381836 | 698.6231286 | 196.8548925 | 77 | 20 | 22 | 34 | 56 | 0 | 1 | 0 | 0 | 1 | 1 | 0 |
| SG0000002 | Cenpk | 1.59E-19 | 9.71E-19 | 1.58612547 | 181.0556162 | 543.6048769 | 150 | 20 | 22 | 36 | 100 | 11 | 4 | 0 | 0 | 0 | 1 | 1 |
| SG0000005 | Sh3rf2 | 1.28E-17 | 7.07E-17 | -3.828921903 | 250.0588483 | 17.59632127 | 266 | 12 | 22 | 50 | 176 | 34 | 0 | 0 | 0 | 0 | 2 | 0 |
| SG0000003 | Srrm3 | 2E-17 | 1.09E-16 | -3.282574065 | 187.8645795 | 19.30597094 | 114 | 16 | 22 | 24 | 60 | 18 | 0 | 0 | 0 | 0 | 0 | 0 |
| SG0000003 | Prmt10 | 4.13E-17 | 2.22E-16 | 1.204575552 | 531.0178623 | 1223.833969 | 126 | 22 | 22 | 28 | 74 | 20 | 0 | 0 | 0 | 0 | 0 | 0 |
| SG0000003 | Spag1 | 8.47E-17 | 4.47E-16 | 1.769319395 | 88.52605745 | 301.7792209 | 163 | 12 | 22 | 32 | 82 | 18 | 7 | 2 | 2 | 2 | 4 | 2 |
| SG0000002 | Fam107b | 2.94E-15 | 1.41E-14 | -1.380528977 | 1776.513359 | 682.3195981 | 511 | 14 | 22 | 72 | 222 | 47 | 6 | 0 | 0 | 0 | 0 | 2 |
| SG0000001 | Tnfaip1 | 1.07E-14 | 4.97E-14 | -1.145297845 | 4006.01915 | 1811.106611 | 46 | 18 | 22 | 22 | 30 | 6 | 2 | 0 | 0 | 0 | 0 | 0 |
| JSG000000 | Spats2 | 9.92E-14 | 4.31E-13 | 0.946250459 | 1669.056047 | 3216.034448 | 133 | 16 | 22 | 31 | 74 | 13 | 2 | 0 | 0 | 0 | 0 | 0 |
| SG0000003 | Mvp | 2.11E-13 | 8.99E-13 | -1.794510353 | 3419.855533 | 985.8397297 | 49 | 20 | 22 | 26 | 43 | 10 | 1 | 0 | 0 | 0 | 0 | 0 |
| SG0000002 | D5Ertd579e | 1.69E-12 | 6.79E-12 | -1.082023411 | 2379.983153 | 1124.222777 | 198 | 15 | 22 | 31 | 81 | 11 | 3 | 2 | 2 | 2 | 2 | 0 |
| JSG000000 | Morc3 | 9.06E-12 | 3.47E-11 | 0.865633238 | 1855.968351 | 3381.832832 | 183 | 12 | 22 | 30 | 112 | 5 | 6 | 0 | 0 | 2 | 6 | 2 |
| JSG000000 | Stk10 | 2.49E-11 | 9.26E-11 | 1.143925827 | 250.8967795 | 554.4355554 | 94 | 19 | 22 | 28 | 46 | 8 | 1 | 0 | 0 | 0 | 0 | 1 |
| SG0000002 | Rpn2 | 4.72E-11 | 1.72E-10 | -0.910603734 | 12712.46249 | 6762.552449 | 146 | 16 | 22 | 28 | 64 | 12 | 2 | 0 | 0 | 0 | 0 | 0 |
| SG0000002 | Hsd17b12 | 9.13E-11 | 3.26E-10 | -0.934133801 | 4310.355737 | 2255.853126 | 269 | 12 | 22 | 39 | 119 | 18 | 2 | 0 | 0 | 1 | 1 | 1 |
| JSG000000 | Sgpp2 | 1.16E-10 | 4.1E-10 | 6.679072985 | 0.307509278 | 31.5108077 | 168 | 13 | 22 | 28 | 75 | 15 | 3 | 0 | 0 | 0 | 0 | 0 |
| SG0000002 | Nmt2 | 1.66E-10 | 5.81E-10 | 0.828791389 | 2414.593092 | 4288.792375 | 268 | 10 | 22 | 28 | 128 | 26 | 10 | 1 | 1 | 1 | 5 | 2 |
| SG0000002 | Pot1a | 9.07E-10 | 3E-09 | 0.888404929 | 598.7450823 | 1108.353996 | 295 | 15 | 22 | 37 | 159 | 23 | 6 | 0 | 0 | 1 | 2 | 1 |
| SG0000007 | Heg1 | 1.25E-09 | 4.09E-09 | -3.047952127 | 1503.627963 | 181.809 | 131 | 12 | 22 | 31 | 61 | 10 | 4 | 0 | 0 | 0 | 0 | 0 |
| SG0000003 | Iars | 1.37E-09 | 4.47E-09 | 0.738688484 | 4930.939838 | 8228.053274 | 147 | 9 | 22 | 22 | 70 | 11 | 2 | 0 | 0 | 0 | 0 | 0 |
| SG0000002 | Xpo7 | 2.8E-09 | 8.89E-09 | 0.730354801 | 3757.926655 | 6234.577129 | 222 | 14 | 22 | 31 | 113 | 22 | 4 | 0 | 0 | 1 | 1 | 0 |
| SG0000007 | Gm13157 | 6.65E-09 | 2.05E-08 | 0.858181796 | 541.476365 | 981.5625205 | 203 | 14 | 22 | 42 | 114 | 22 | 2 | 0 | 0 | 2 | 4 | 0 |
| SG0000004 | Sdk2 | 1.02E-08 | 3.1E-08 | 5.320715576 | 2.696304994 | 107.7615931 | 243 | 15 | 22 | 31 | 108 | 31 | 3 | 0 | 0 | 0 | 1 | 1 |
| SG0000002 | Snx15 | 1.57E-08 | 4.69E-08 | -1.140536362 | 572.9937143 | 259.9047411 | 65 | 18 | 22 | 22 | 30 | 2 | 1 | 0 | 0 | 0 | 0 | 0 |
| SG0000005 | Nup107 | 2.15E-08 | 6.37E-08 | 0.71349983 | 2113.092441 | 3465.000863 | 112 | 18 | 22 | 28 | 92 | 14 | 0 | 0 | 0 | 0 | 2 | 0 |
| SG0000002 | Son | 1.13E-07 | 3.14E-07 | -0.729034101 | 10603.46919 | 6397.154601 | 321 | 5 | 22 | 39 | 100 | 23 | 24 | 0 | 0 | 1 | 3 | 7 |
| SG0000002 | Zc3h7b | 1.88E-07 | 5.14E-07 | -0.737021329 | 4242.643972 | 2545.488042 | 73 | 19 | 22 | 22 | 39 | 8 | 1 | 0 | 0 | 0 | 1 | 1 |
| SG0000004 | Bend7 | 3.18E-07 | 8.52E-07 | 2.076797322 | 15.50510042 | 65.41131382 | 333 | 14 | 22 | 50 | 140 | 21 | 5 | 0 | 1 | 1 | 2 | 0 |
| SG0000006 | Phf11d | 4.69E-07 | 1.24E-06 | 0.69011027 | 947.8312966 | 1529.237489 | 91 | 18 | 22 | 18 | 46 | 3 | 4 | 0 | 1 | 0 | 1 | 0 |
| SG0000003 | Zfp949 | 7.14E-07 | 1.86E-06 | 0.834464363 | 303.6905561 | 541.5394384 | 100 | 20 | 22 | 30 | 66 | 4 | 0 | 0 | 0 | 0 | 2 | 0 |
| JSG000000 | 2700097O09Rik | 7.69E-07 | 2E-06 | 0.714069504 | 674.8523814 | 1107.044617 | 84 | 17 | 22 | 27 | 60 | 10 | 0 | 0 | 0 | 0 | 0 | 0 |
| SG0000002 | Htt | 1.49E-06 | 3.78E-06 | -0.689019742 | 3634.74153 | 2254.539887 | 275 | 11 | 22 | 38 | 137 | 21 | 5 | 0 | 1 | 0 | 0 | 0 |
| JSG000000 | Plekhh2 | 2.76E-06 | 6.86E-06 | 0.95432674 | 152.7042578 | 295.8912488 | 226 | 13 | 22 | 45 | 128 | 24 | 7 | 0 | 0 | 3 | 6 | 0 |
| SG0000002 | Snx5 | 2.83E-06 | 7.01E-06 | 0.56646473 | 7693.601674 | 11393.37974 | 92 | 12 | 22 | 18 | 32 | 6 | 8 | 0 | 0 | 0 | 0 | 0 |
| SG0000002 | Ncf2 | 3.06E-06 | 7.55E-06 | -1.412118529 | 152.6429661 | 57.35704409 | 83 | 20 | 22 | 31 | 70 | 4 | 2 | 0 | 0 | 0 | 0 | 0 |
| SG0000002 | Rfc1 | 8E-06 | 1.9E-05 | 0.548631515 | 4715.815135 | 6897.811383 | 172 | 17 | 22 | 50 | 136 | 11 | 0 | 0 | 0 | 0 | 3 | 1 |
| JSG000000 | Sos1 | 8.04E-06 | 1.91E-05 | -0.647315996 | 3086.554791 | 1970.663438 | 213 | 17 | 22 | 35 | 102 | 13 | 10 | 0 | 1 | 2 | 7 | 2 |
| JSG000000 | Insr | 9E-06 | 2.13E-05 | -0.713839072 | 2014.254985 | 1228.082307 | 202 | 13 | 22 | 39 | 116 | 26 | 7 | 0 | 0 | 2 | 9 | 1 |
| JSG000000 | Caap1 | 2.12E-05 | 4.85E-05 | 0.704018683 | 346.4617253 | 564.3987001 | 157 | 11 | 22 | 27 | 104 | 20 | 2 | 0 | 0 | 0 | 2 | 1 |
| SG0000003 | Sergef | 2.15E-05 | 4.91E-05 | -0.957848102 | 278.5138104 | 143.3856617 | 239 | 15 | 22 | 42 | 125 | 20 | 0 | 0 | 0 | 0 | 0 | 0 |
| JSG000000 | Coq2 | 3.3E-05 | 7.39E-05 | -0.649728184 | 1240.518709 | 790.7071142 | 50 | 15 | 22 | 28 | 44 | 20 | 2 | 0 | 1 | 1 | 2 | 0 |
| JSG000000 | Sstr2 | 4.77E-05 | 0.000105 | 2.95228147 | 3.217299956 | 24.90100246 | 32 | 22 | 22 | 22 | 24 | 0 | 2 | 0 | 0 | 0 | 0 | 0 |
| SG0000002 | Hacd4 | 0.000179 | 0.000369 | -1.42911274 | 292.6548037 | 108.6800564 | 174 | 2 | 22 | 23 | 89 | 13 | 11 | 0 | 0 | 0 | 3 | 0 |
| SG0000003 | Apbb1 | 0.000216 | 0.000441 | 1.179595775 | 150.8106342 | 341.6061473 | 149 | 22 | 22 | 26 | 95 | 32 | 0 | 0 | 0 | 0 | 0 | 0 |
| SG0000005 | Eapp | 0.000276 | 0.000556 | -0.564631721 | 1330.294846 | 899.4495815 | 114 | 16 | 22 | 34 | 168 | 8 | 0 | 0 | 0 | 0 | 0 | 0 |
| SG000000C | Ppard | 0.000305 | 0.000611 | -0.682400387 | 1403.528088 | 874.5771429 | 125 | 20 | 22 | 28 | 52 | 8 | 1 | 0 | 0 | 0 | 0 | 0 |
| SG0000003 | Usp4 | 0.000376 | 0.000746 | -0.49835499 | 3193.647447 | 2260.826168 | 189 | 18 | 22 | 33 | 134 | 34 | 0 | 0 | 0 | 0 | 0 | 0 |
| JSG000000 | Ccar1 | 0.000404 | 0.000798 | -0.490861181 | 4507.440656 | 3207.495666 | 139 | 18 | 22 | 36 | 76 | 24 | 2 | 0 | 1 | 1 | 2 | 0 |
| JSG000000 | Taf2 | 0.000414 | 0.000815 | -0.503417495 | 2876.501312 | 2029.181107 | 152 | 13 | 22 | 29 | 68 | 3 | 4 | 0 | 0 | 0 | 1 | 1 |
| SG0000004 | Zfp410 | 0.000418 | 0.000823 | -0.562395356 | 1007.955249 | 682.564001 | 83 | 17 | 22 | 29 | 48 | 5 | 0 | 0 | 0 | 0 | 0 | 0 |
| SG0000003 | Tjp1 | 0.000503 | 0.000983 | 0.428531415 | 4331.012045 | 5828.948276 | 347 | 14 | 22 | 42 | 151 | 31 | 7 | 0 | 0 | 0 | 9 | 1 |
| SG0000001 | Kif3a | 0.000511 | 0.000997 | 0.539736365 | 483.6159959 | 703.035884 | 179 | 19 | 22 | 28 | 132 | 15 | 1 | 0 | 0 | 0 | 0 | 0 |
| SG0000005 | Trmt1l | 0.000878 | 0.001659 | 0.462582736 | 1043.089826 | 1437.384603 | 150 | 20 | 22 | 34 | 96 | 8 | 9 | 0 | 0 | 0 | 2 | 0 |
| SG0000001 | Baiap2l2 | 0.001343 | 0.002486 | 1.94983861 | 7.202699755 | 27.82628442 | 55 | 12 | 22 | 15 | 36 | 2 | 1 | 0 | 1 | 0 | 0 | 0 |
| SG0000003 | Gemin5 | 0.001449 | 0.00267 | 0.405349487 | 2822.668867 | 3738.370041 | 184 | 19 | 22 | 34 | 62 | 13 | 2 | 0 | 0 | 0 | 1 | 1 |
| SG0000003 | Helq | 0.00188 | 0.003416 | 0.472808846 | 610.9178451 | 847.8371768 | 201 | 7 | 22 | 32 | 78 | 17 | 1 | 1 | 1 | 1 | 4 | 0 |
| JSG000000 | Usp49 | 0.002088 | 0.00377 | 0.525410964 | 324.3194134 | 466.8070495 | 99 | 19 | 22 | 26 | 64 | 4 | 4 | 0 | 0 | 0 | 0 | 0 |
| SG0000003 | N4bp2 | 0.002385 | 0.004271 | 0.436500151 | 1705.286553 | 2307.79382 | 165 | 17 | 22 | 32 | 90 | 19 | 11 | 0 | 0 | 2 | 5 | 0 |
| SG0000002 | Traf2 | 0.002453 | 0.004387 | -0.526030239 | 626.1538957 | 434.8407201 | 69 | 18 | 22 | 18 | 40 | 6 | 1 | 0 | 0 | 0 | 0 | 0 |
| SG0000006 | Vsig10 | 0.003046 | 0.005366 | -0.510847139 | 616.9646328 | 432.9920842 | 81 | 14 | 22 | 26 | 53 | 12 | 1 | 0 | 0 | 0 | 3 | 0 |
| JSG000000 | P3h2 | 0.003246 | 0.005697 | -1.424002854 | 45.8557894 | 17.08939158 | 299 | 11 | 22 | 38 | 163 | 25 | 3 | 0 | 1 | 1 | 1 | 0 |
| SG0000003 | Dennd3 | 0.006659 | 0.011194 | -0.616938113 | 374.0533359 | 243.9027263 | 67 | 18 | 22 | 24 | 40 | 7 | 1 | 0 | 1 | 0 | 0 | 1 |
| SG0000007 | Zfp946 | 0.008851 | 0.014595 | 0.566499012 | 145.8942105 | 216.0584485 | 164 | 18 | 22 | 30 | 114 | 16 | 2 | 0 | 0 | 0 | 2 | 0 |
| SG0000007 | Setdb2 | 0.010293 | 0.016795 | 0.417983374 | 415.4382248 | 555.0498467 | 236 | 12 | 22 | 43 | 142 | 28 | 2 | 0 | 0 | 0 | 4 | 3 |
| JSG000000 | Cab39l | 0.013362 | 0.021343 | 0.327803669 | 1503.308179 | 1886.803865 | 250 | 5 | 22 | 41 | 120 | 22 | 1 | 0 | 0 | 0 | 0 | 0 |
| JSG000000 | Slc5a1 | 0.015731 | 0.024853 | -1.309612251 | 37.5836675 | 15.16235243 | 106 | 15 | 22 | 28 | 84 | 8 | 3 | 0 | 0 | 0 | 6 | 1 |
| SG0000004 | Ago1 | 0.017922 | 0.027961 | -0.34141117 | 2520.012433 | 1988.96747 | 91 | 14 | 22 | 25 | 63 | 4 | 3 | 0 | 0 | 1 | 3 | 0 |
| SG0000007 | Fign | 0.029652 | 0.044553 | -0.698462153 | 142.5055746 | 87.816029 | 305 | 14 | 22 | 29 | 127 | 21 | 4 | 0 | 0 | 0 | 0 | 0 |
| JSG000000 | Usp15 | 0.041472 | 0.060715 | 0.261293134 | 2938.326789 | 3521.738991 | 233 | 7 | 22 | 34 | 96 | 15 | 2 | 0 | 0 | 0 | 0 | 0 |
| SG000000C | Ankrd13d | 0.049288 | 0.071255 | -0.44690352 | 222.5905269 | 163.2959121 | 69 | 7 | 22 | 34 | 62 | 4 | 6 | 2 | 4 | 4 | 4 | 0 |
| JSG000000 | Appbp2 | 0.054846 | 0.07858 | 0.24337167 | 3054.093145 | 3615.300913 | 128 | 16 | 22 | 22 | 57 | 9 | 13 | 0 | 1 | 1 | 4 | 0 |
| JSG000000 | Actr2 | 0.078087 | 0.108687 | -0.23302306 | 14059.60001 | 11962.61249 | 119 | 16 | 22 | 30 | 64 | 9 | 8 | 0 | 0 | 1 | 1 | 0 |
| SG0000002 | Zcchc4 | 0.083342 | 0.115374 | 0.27773074 | 500.1032284 | 606.2684128 | 155 | 8 | 22 | 29 | 70 | 2 | 16 | 0 | 0 | 3 | 7 | 0 |
| SG0000001 | Anxa6 | 0.093214 | 0.127737 | 0.221633027 | 2248.011262 | 2621.299664 | 128 | 18 | 22 | 34 | 66 | 16 | 0 | 0 | 0 | 0 | 0 | 0 |
| JSG000000 | Sdhaf3 | 0.093829 | 0.128511 | 0.332980122 | 209.6733311 | 264.1071748 | 179 | 18 | 22 | 35 | 106 | 16 | 9 | 0 | 0 | 2 | 4 | 0 |
| SG0000003 | Ankrd12 | 0.113462 | 0.152543 | -0.280107629 | 605.3779818 | 498.5466735 | 232 | 16 | 22 | 37 | 124 | 15 | 7 | 0 | 2 | 2 | 7 | 1 |
| SG0000005 | Tor1aip2 | 0.113896 | 0.153057 | 0.190080376 | 6650.851412 | 7587.472677 | 221 | 10 | 22 | 20 | 92 | 11 | 18 | 0 | 0 | 0 | 4 | 0 |
| SG0000003 | Kras | 0.173905 | 0.224746 | -0.191305614 | 3846.907251 | 3369.170485 | 327 | 12 | 22 | 54 | 144 | 13 | 12 | 0 | 1 | 0 | 0 | 1 |
| SG000000C | Pex1 | 0.185856 | 0.238693 | -0.215977533 | 849.7304985 | 731.586176 | 228 | 19 | 22 | 40 | 120 | 14 | 2 | 0 | 0 | 0 | 4 | 0 |
| SG0000002 | Mapkapk5 | 0.261083 | 0.322796 | 0.316517057 | 111.2718577 | 138.5691225 | 79 | 22 | 22 | 29 | 49 | 5 | 0 | 0 | 0 | 0 | 0 | 0 |
| JSG000000 | Btaf1 | 0.3418 | 0.409137 | -0.136564673 | 2662.238505 | 2421.792319 | 253 | 16 | 22 | 40 | 164 | 16 | 3 | 0 | 0 | 0 | 0 | 3 |
| SG0000002 | Il1r1 | 0.362595 | 0.430946 | -0.366070095 | 96.61970313 | 74.96656635 | 220 | 11 | 22 | 26 | 98 | 25 | 16 | 0 | 2 | 4 | 8 | 0 |
| SG0000002 | Golga5 | 0.409048 | 0.478348 | 0.118169027 | 1063.338295 | 1154.101157 | 158 | 16 | 22 | 20 | 72 | 12 | 1 | 0 | 0 | 0 | 0 | 0 |
| SG0000003 | Atp13a2 | 0.480081 | 0.548734 | -0.303098872 | 773.0683722 | 626.5793152 | 46 | 18 | 22 | 22 | 38 | 2 | 0 | 0 | 0 | 0 | 0 | 0 |
| SG0000002 | Unk | 0.622135 | 0.683745 | -0.081098676 | 841.1705873 | 795.1900635 | 102 | 18 | 22 | 32 | 78 | 12 | 2 | 0 | 0 | 0 | 0 | 0 |
| SG0000002 | Stam | 0.632998 | 0.693769 | -0.078076608 | 1651.03393 | 1564.057076 | 311 | 13 | 22 | 36 | 153 | 22 | 18 | 1 | 2 | 4 | 10 | 0 |
| SG0000003 | Prkar2a | 0.682778 | 0.739058 | -0.055673706 | 2805.503986 | 2699.301739 | 136 | 17 | 22 | 31 | 90 | 7 | 6 | 0 | 0 | 0 | 0 | 0 |
| SG0000002 | Usp43 | 0.709393 | 0.762018 | 0.070282297 | 371.8484377 | 390.4118947 | 198 | 12 | 22 | 52 | 118 | 16 | 0 | 0 | 0 | 0 | 0 | 0 |
| SG0000004 | Zfand4 | 0.775508 | 0.820393 | -0.159807879 | 29.10409376 | 26.05236269 | 284 | 6 | 22 | 26 | 172 | 14 | 4 | 0 | 0 | 0 | 0 | 0 |
| SG0000003 | Znrf1 | 0.813634 | 0.852968 | 0.037860039 | 1138.968574 | 1169.253689 | 151 | 17 | 22 | 29 | 65 | 10 | 4 | 1 | 1 | 2 | 3 | 0 |
| SG0000003 | Cog7 | 0.819811 | 0.858238 | 0.028586621 | 943.8181238 | 962.7061432 | 95 | 17 | 22 | 22 | 63 | 7 | 1 | 0 | 0 | 1 | 2 | 1 |
| SG0000002 | Lamc2 | 0.906143 | 0.929442 | -0.013343786 | 1159.346373 | 1148.672775 | 179 | 16 | 22 | 38 | 102 | 24 | 7 | 0 | 0 | 4 | 4 | 0 |
| JSG000000 | St3gal4 | 0.962395 | 0.976847 | -0.03735949 | 172.4635675 | 168.0548453 | 109 | 16 | 22 | 34 | 69 | 3 | 3 | 0 | 0 | 0 | 0 | 0 |
| SG0000002 | Celf4 | 0.969076 | 0.98262 | -0.004370495 | 349.970086 | 348.9114921 | 494 | 8 | 22 | 54 | 172 | 42 | 10 | 0 | 0 | 4 | 7 | 1 |
| JSG000000 | Cd38 | 0.98559 | 0.996301 | -0.007908068 | 218.7988583 | 217.6028032 | 104 | 17 | 22 | 27 | 54 | 10 | 6 | 1 | 2 | 1 | 1 | 0 |
| SG0000003 | Morc4 | 1 | 1 | -0.013539227 | 431.1392124 | 427.1120367 | 232 | 8 | 22 | 40 | 104 | 32 | 6 | 0 | 0 | 0 | 0 | 0 |
| JSG000000 | Bin2 | 1.4E-170 | 3E-168 | 10.04317777 | 0.639110347 | 674.3317519 | 93 | 18 | 21 | 30 | 54 | 3 | 0 | 0 | 0 | 0 | 0 | 0 |
| JSG000000 | Gab1 | 1E-161 | 1.9E-159 | 3.292115761 | 921.4172639 | 9025.721703 | 186 | 12 | 21 | 27 | 85 | 11 | 3 | 0 | 0 | 0 | 2 | 2 |
| SG0000003 | Sult6b1 | 1.87E-81 | 8.7E-80 | 5.719586253 | 6.515405827 | 343.3274255 | 55 | 17 | 21 | 21 | 37 | 5 | 0 | 0 | 0 | 0 | 0 | 0 |
| SG0000004 | Sestd1 | 1.81E-66 | 5.93E-65 | -3.129114316 | 7226.22899 | 825.9513181 | 425 | 11 | 21 | 37 | 163 | 29 | 13 | 0 | 1 | 0 | 2 | 4 |

| | | | | | | | | | | | | | | | | | | |
|---|---|---|---|---|---|---|---|---|---|---|---|---|---|---|---|---|---|---|
| SG0000001 | Hivep2 | 2.4E-59 | 6.5E-58 | -4.447422147 | 2429.048765 | 111.3342439 | 380 | 13 | 21 | 53 | 182 | 40 | 3 | 0 | 0 | 1 | 4 | 3 |
| SG000000 | Reep3 | 1.28E-51 | 2.77E-50 | -3.326455481 | 5205.634683 | 518.9324251 | 123 | 20 | 21 | 29 | 74 | 7 | 6 | 0 | 1 | 0 | 9 | 1 |
| SG0000003 | Cerk | 1.2E-45 | 2.15E-44 | -2.751639813 | 2216.244347 | 329.0724465 | 63 | 19 | 21 | 25 | 37 | 3 | 7 | 0 | 0 | 0 | 0 | 0 |
| SG0000003 | Dlgap5 | 2.21E-43 | 3.66E-42 | 1.799317957 | 1550.858495 | 5397.850474 | 189 | 18 | 21 | 47 | 97 | 10 | 8 | 0 | 0 | 1 | 1 | 1 |
| SG0000003 | Slc19a3 | 4.42E-42 | 6.98E-41 | 6.944670999 | 0 | 122.1839939 | 105 | 14 | 21 | 25 | 61 | 20 | 3 | 0 | 1 | 1 | 1 | 0 |
| SG0000005 | Gda | 1.08E-41 | 1.69E-40 | -5.376222777 | 3339.911854 | 80.41381858 | 321 | 14 | 21 | 57 | 139 | 16 | 15 | 0 | 1 | 1 | 1 | 0 |
| SG0000003 | Clmp | 2.58E-39 | 3.72E-38 | -3.097525432 | 1515.287688 | 177.0300643 | 134 | 19 | 21 | 28 | 63 | 13 | 1 | 0 | 0 | 0 | 0 | 0 |
| SG0000003 | Frem2 | 7.92E-38 | 1.08E-36 | 7.304735579 | 16.89056239 | 2670.475696 | 253 | 11 | 21 | 39 | 132 | 15 | 4 | 0 | 0 | 1 | 1 | 0 |
| SG0000005 | Fam171a1 | 5.91E-35 | 7.34E-34 | -2.232996235 | 1833.561726 | 390.02881 | 540 | 12 | 21 | 53 | 307 | 46 | 2 | 0 | 0 | 0 | 0 | 2 |
| SG000000 | Susd6 | 5.53E-33 | 6.3E-32 | -1.971457197 | 7043.659471 | 1796.100361 | 164 | 15 | 21 | 34 | 83 | 24 | 3 | 0 | 0 | 1 | 4 | 1 |
| SG0000002 | Btbd11 | 4.72E-32 | 5.19E-31 | 3.073750363 | 43.40928269 | 365.4884592 | 398 | 15 | 21 | 45 | 148 | 45 | 0 | 0 | 0 | 0 | 0 | 2 |
| SG0000002 | Lyrm7 | 2.26E-29 | 2.21E-28 | -2.643063414 | 604.7305376 | 96.81006957 | 104 | 21 | 21 | 25 | 79 | 3 | 7 | 0 | 0 | 0 | 0 | 0 |
| SG0000002 | Tex30 | 3.98E-28 | 3.7E-27 | 1.562158611 | 474.5140666 | 1401.217947 | 212 | 7 | 21 | 28 | 133 | 28 | 2 | 0 | 0 | 0 | 0 | 0 |
| SG000000 | Gm13051 | 1.35E-27 | 1.22E-26 | 6.256626027 | 0 | 75.45961483 | 102 | 19 | 21 | 31 | 57 | 9 | 2 | 0 | 0 | 1 | 2 | 0 |
| SG0000002 | Ppp4r4 | 5.21E-25 | 4.21E-24 | 5.460760904 | 1.963171053 | 86.45915231 | 238 | 9 | 21 | 30 | 101 | 22 | 3 | 0 | 0 | 0 | 0 | 0 |
| SG0000002 | Herc3 | 6.69E-24 | 5.13E-23 | -1.772400072 | 2680.152611 | 784.5379172 | 362 | 15 | 21 | 69 | 172 | 38 | 6 | 0 | 0 | 2 | 4 | 1 |
| SG0000004 | Zc3h6 | 3.73E-19 | 2.23E-18 | 1.612980619 | 201.2189991 | 615.4950055 | 205 | 10 | 21 | 39 | 96 | 31 | 2 | 1 | 1 | 1 | 6 | 0 |
| SG000000 | Sntb2 | 9.71E-19 | 5.67E-18 | 1.27780083 | 504.0516686 | 1222.169482 | 157 | 15 | 21 | 29 | 104 | 14 | 1 | 0 | 0 | 0 | 0 | 0 |
| SG0000002 | Rab22a | 1.26E-17 | 6.97E-17 | -1.366474201 | 2172.591307 | 842.6133095 | 183 | 18 | 21 | 29 | 78 | 16 | 18 | 0 | 0 | 2 | 6 | 4 |
| SG0000002 | Cap1 | 6.46E-17 | 3.44E-16 | -1.18735478 | 14310.24794 | 6283.733448 | 153 | 18 | 21 | 27 | 123 | 3 | 0 | 0 | 0 | 0 | 0 | 0 |
| SG0000002 | Cryl1 | 2.77E-16 | 1.42E-15 | 1.4956615 | 164.8750928 | 464.9369131 | 168 | 19 | 21 | 36 | 88 | 19 | 0 | 0 | 0 | 0 | 0 | 0 |
| SG0000004 | Cltc | 3.3E-16 | 1.68E-15 | -1.383646692 | 40263.93708 | 15431.10695 | 371 | 19 | 21 | 53 | 185 | 12 | 8 | 0 | 0 | 0 | 0 | 0 |
| SG0000003 | Slc18b1 | 3.56E-16 | 1.81E-15 | -1.664239462 | 584.294227 | 184.3510423 | 138 | 15 | 21 | 27 | 96 | 16 | 6 | 0 | 0 | 0 | 0 | 0 |
| SG000000 | Dennd6a | 1.23E-15 | 6.07E-15 | 1.04603439 | 1169.725924 | 2415.304236 | 130 | 14 | 21 | 23 | 70 | 10 | 6 | 1 | 1 | 1 | 7 | 1 |
| SG000000 | Arsb | 2.07E-15 | 1.01E-14 | -2.189682758 | 452.3796292 | 99.16144552 | 260 | 14 | 21 | 38 | 157 | 29 | 3 | 0 | 0 | 0 | 3 | 0 |
| SG0000002 | Adgra3 | 4.76E-15 | 2.26E-14 | -1.173386673 | 3702.171954 | 1641.466416 | 204 | 13 | 21 | 40 | 95 | 17 | 2 | 0 | 0 | 0 | 1 | 0 |
| SG000000 | Picalm | 8.43E-15 | 3.93E-14 | -1.107289868 | 12448.8271 | 5778.311777 | 261 | 8 | 21 | 34 | 123 | 15 | 6 | 0 | 0 | 2 | 3 | 1 |
| SG0000002 | Tmem57 | 3.84E-14 | 1.72E-13 | 0.979259892 | 1360.537297 | 2682.236406 | 109 | 12 | 21 | 27 | 65 | 9 | 3 | 0 | 1 | 1 | 1 | 0 |
| SG0000003 | Fkbp14 | 5E-14 | 2.22E-13 | -1.610503131 | 464.8964857 | 152.2462178 | 130 | 18 | 21 | 30 | 71 | 8 | 14 | 0 | 3 | 3 | 6 | 5 |
| SG0000002 | Il10rb | 5.66E-14 | 2.5E-13 | -2.359129157 | 299.9875769 | 58.47020683 | 105 | 21 | 21 | 33 | 54 | 17 | 11 | 0 | 0 | 0 | 0 | 2 |
| SG0000005 | Cep170 | 7.53E-14 | 3.3E-13 | -1.144495615 | 2860.298909 | 1293.849941 | 235 | 13 | 21 | 30 | 114 | 18 | 9 | 3 | 3 | 3 | 7 | 1 |
| SG000000 | Vat1l | 1.04E-13 | 4.51E-13 | 5.141129449 | 1.299968916 | 45.87405635 | 259 | 12 | 21 | 32 | 122 | 43 | 2 | 0 | 1 | 1 | 1 | 0 |
| SG000000 | Rictor | 1.39E-13 | 5.96E-13 | -1.114885985 | 4003.520552 | 1848.536112 | 230 | 11 | 21 | 42 | 116 | 21 | 11 | 0 | 1 | 2 | 3 | 1 |
| SG0000004 | Fndc7 | 2.16E-13 | 9.21E-13 | -7.174782117 | 45.60968138 | 0.315669345 | 178 | 6 | 21 | 12 | 84 | 10 | 1 | 0 | 0 | 0 | 0 | 0 |
| SG0000002 | Kif20b | 5.77E-13 | 2.38E-12 | 0.985410824 | 2203.073821 | 4361.815243 | 150 | 13 | 21 | 24 | 77 | 16 | 1 | 0 | 0 | 0 | 1 | 2 |
| SG0000004 | Zfp710 | 6.52E-13 | 2.69E-12 | 1.563678674 | 1303.072516 | 3851.968877 | 201 | 18 | 21 | 33 | 105 | 6 | 5 | 0 | 0 | 0 | 0 | 0 |
| SG000000 | Znhit6 | 3.15E-12 | 1.24E-11 | 1.025944234 | 516.625221 | 1051.999643 | 110 | 17 | 21 | 32 | 80 | 10 | 4 | 0 | 0 | 1 | 1 | 0 |
| SG0000003 | Parl | 1.09E-11 | 4.14E-11 | -1.018475736 | 2954.561792 | 1458.482864 | 119 | 17 | 21 | 35 | 97 | 13 | 2 | 0 | 0 | 0 | 0 | 0 |
| SG000000 | Rilpl2 | 1.16E-11 | 4.4E-11 | -1.967097232 | 876.5512288 | 224.1929913 | 45 | 20 | 21 | 25 | 36 | 1 | 0 | 0 | 0 | 0 | 0 | 0 |
| SG0000004 | Arid4a | 2.77E-11 | 1.03E-10 | -1.085458246 | 1425.856495 | 671.9242687 | 155 | 12 | 21 | 27 | 85 | 15 | 2 | 0 | 0 | 1 | 7 | 0 |
| SG000000 | Stard4 | 3.14E-11 | 1.16E-10 | 0.908467795 | 874.3183547 | 1641.140081 | 101 | 18 | 21 | 27 | 56 | 5 | 8 | 0 | 0 | 0 | 2 | 2 |
| SG0000002 | Pfkfb1 | 1.44E-10 | 5.07E-10 | 1.203879733 | 171.6466371 | 395.4023169 | 186 | 19 | 21 | 39 | 126 | 26 | 2 | 0 | 0 | 0 | 0 | 0 |
| SG0000004 | Arel1 | 1.85E-09 | 5.94E-09 | -0.923042189 | 1887.953459 | 995.6986028 | 146 | 15 | 21 | 28 | 88 | 23 | 4 | 0 | 0 | 0 | 0 | 2 |
| SG0000002 | Ercc5 | 6E-09 | 1.86E-08 | 0.999370416 | 986.0810593 | 1971.301666 | 90 | 15 | 21 | 25 | 52 | 6 | 2 | 0 | 0 | 0 | 0 | 0 |
| SG000000 | Atxn7 | 6.79E-09 | 2.09E-08 | -1.205677953 | 964.037952 | 417.9736488 | 195 | 13 | 21 | 34 | 100 | 13 | 15 | 1 | 1 | 3 | 9 | 2 |
| SG0000004 | Mpped1 | 8.13E-09 | 2.49E-08 | 4.528080911 | 0 | 22.07215587 | 465 | 7 | 21 | 48 | 194 | 76 | 3 | 0 | 0 | 0 | 6 | 0 |
| SG0000002 | Plekhb2 | 9.03E-09 | 2.76E-08 | -0.891364603 | 2841.000689 | 1531.594705 | 86 | 11 | 21 | 17 | 47 | 26 | 18 | 1 | 5 | 2 | 2 | 0 |
| SG0000002 | Ect2 | 1.23E-08 | 3.72E-08 | 0.705746271 | 3566.513996 | 5816.941948 | 507 | 0 | 21 | 68 | 255 | 52 | 20 | 0 | 0 | 0 | 12 | 4 |
| SG0000002 | Lgals12 | 2.22E-08 | 6.56E-08 | 4.444468333 | 0 | 20.77300052 | 51 | 21 | 21 | 24 | 29 | 0 | 9 | 0 | 0 | 0 | 2 | 0 |
| SG000000 | Abcb1b | 3.45E-08 | 1.01E-07 | -1.187360361 | 508.4016852 | 223.2419961 | 315 | 10 | 21 | 28 | 123 | 23 | 9 | 0 | 0 | 0 | 0 | 0 |
| SG0000002 | Slc11a2 | 8.55E-08 | 2.41E-07 | -0.816900438 | 1750.509993 | 993.6948958 | 112 | 21 | 21 | 27 | 65 | 18 | 6 | 0 | 0 | 1 | 2 | 0 |
| SG0000000 | Usp40 | 8.57E-08 | 2.42E-07 | -0.843865926 | 1466.874026 | 817.2668106 | 407 | 15 | 21 | 68 | 145 | 42 | 5 | 0 | 0 | 3 | 6 | 2 |
| SG0000002 | Man2a1 | 9.17E-08 | 2.58E-07 | -0.884793894 | 2041.32602 | 1105.510493 | 396 | 6 | 21 | 46 | 151 | 38 | 5 | 0 | 0 | 0 | 0 | 0 |
| SG0000006 | l7Rn6 | 1.98E-07 | 5.4E-07 | 0.777705749 | 902.2575689 | 1546.832514 | 142 | 17 | 21 | 23 | 117 | 6 | 1 | 0 | 1 | 0 | 0 | 0 |
| SG0000002 | Fbxo3 | 2.44E-07 | 6.62E-07 | -0.762582412 | 2496.644118 | 1471.622639 | 313 | 13 | 21 | 85 | 163 | 26 | 31 | 1 | 3 | 7 | 11 | 4 |
| SG000000 | Map3k2 | 8.57E-07 | 2.23E-06 | -1.149366959 | 1272.51115 | 573.6773002 | 190 | 14 | 21 | 40 | 97 | 25 | 29 | 0 | 0 | 2 | 10 | 3 |
| SG0000002 | Stx18 | 1.4E-06 | 3.58E-06 | -0.932080886 | 489.5960482 | 256.598196 | 384 | 18 | 21 | 42 | 153 | 36 | 3 | 0 | 0 | 0 | 0 | 0 |
| SG0000006 | Nmral1 | 1.93E-06 | 4.85E-06 | 0.814781624 | 345.5617549 | 607.8541409 | 51 | 13 | 21 | 17 | 49 | 0 | 3 | 1 | 1 | 1 | 1 | 0 |
| SG0000001 | Pacsin2 | 1.94E-06 | 4.88E-06 | -0.682742157 | 3081.237111 | 1919.549215 | 282 | 11 | 21 | 38 | 95 | 23 | 3 | 0 | 0 | 0 | 3 | 3 |
| SG0000001 | Anxa9 | 9.93E-06 | 2.34E-05 | -1.829621459 | 73.56135665 | 20.69563661 | 66 | 18 | 21 | 24 | 51 | 3 | 0 | 0 | 0 | 0 | 0 | 0 |
| SG0000004 | Ccdc63 | 1.86E-05 | 4.28E-05 | 3.498002758 | 1.963171053 | 22.18001813 | 60 | 21 | 21 | 26 | 58 | 8 | 1 | 0 | 0 | 0 | 0 | 0 |
| SG0000000 | Ezh1 | 2.59E-05 | 5.87E-05 | -0.626437595 | 2412.325604 | 1562.641737 | 154 | 18 | 21 | 36 | 93 | 14 | 4 | 0 | 0 | 0 | 1 | 0 |
| SG0000000 | Cd244 | 3.49E-05 | 7.79E-05 | -6.081680201 | 42.75929823 | 0.631338689 | 110 | 17 | 21 | 30 | 85 | 5 | 3 | 0 | 0 | 0 | 6 | 0 |
| SG000000 | Cnst | 5.05E-05 | 0.000111 | 0.707539775 | 540.0523746 | 881.9146872 | 140 | 14 | 21 | 25 | 58 | 12 | 9 | 2 | 2 | 3 | 3 | 0 |
| SG000000 | Zdhhc14 | 6.49E-05 | 0.000141 | 1.622751031 | 19.35916716 | 59.61882405 | 334 | 6 | 21 | 43 | 138 | 40 | 0 | 0 | 0 | 0 | 0 | 0 |
| SG0000000 | Ltbp2 | 7.39E-05 | 0.000159 | -1.734644657 | 3188.371224 | 958.0513941 | 243 | 18 | 21 | 36 | 108 | 21 | 0 | 0 | 0 | 0 | 0 | 0 |
| SG0000003 | Akap7 | 9.1E-05 | 0.000194 | -0.665878401 | 829.9018259 | 523.0911207 | 331 | 11 | 21 | 50 | 150 | 36 | 7 | 0 | 0 | 0 | 4 | 1 |
| SG000000 | Jmy | 9.92E-05 | 0.00021 | -0.600521506 | 1599.99742 | 1055.223115 | 179 | 9 | 21 | 30 | 87 | 15 | 15 | 0 | 1 | 3 | 4 | 0 |
| SG000000 | Slc20a2 | 0.000205 | 0.00042 | -0.559637242 | 3254.856522 | 2208.331424 | 157 | 16 | 21 | 34 | 76 | 14 | 1 | 0 | 0 | 0 | 1 | 3 |
| SG0000007 | Ccnyl1 | 0.000629 | 0.001211 | -0.546809968 | 1054.354617 | 721.739545 | 260 | 15 | 21 | 42 | 84 | 3 | 2 | 0 | 0 | 0 | 0 | 0 |
| SG0000005 | Tmem150c | 0.000706 | 0.00135 | 4.107816645 | 1.302312483 | 22.45387137 | 129 | 17 | 21 | 29 | 70 | 20 | 1 | 0 | 0 | 0 | 0 | 0 |
| SG000000 | Tssc1 | 0.000779 | 0.001481 | -0.727925751 | 868.8469689 | 524.5849234 | 143 | 18 | 21 | 37 | 80 | 14 | 0 | 0 | 0 | 0 | 0 | 0 |
| SG0000001 | Acot8 | 0.001587 | 0.002915 | -0.815828112 | 409.0683412 | 232.384489 | 52 | 12 | 21 | 21 | 39 | 0 | 0 | 0 | 0 | 0 | 0 | 0 |
| SG0000000 | Arpc2 | 0.001626 | 0.002983 | -0.422031309 | 8147.193937 | 6080.845532 | 153 | 15 | 21 | 18 | 45 | 18 | 0 | 0 | 0 | 0 | 0 | 0 |
| SG000000 | Tnfrsf21 | 0.002521 | 0.004501 | 0.903284177 | 316.0890436 | 591.1872708 | 119 | 18 | 21 | 25 | 82 | 11 | 2 | 0 | 0 | 0 | 1 | 0 |
| SG000000 | Msl2 | 0.002769 | 0.00491 | 0.402879183 | 1352.187591 | 1787.786547 | 109 | 11 | 21 | 22 | 71 | 20 | 11 | 1 | 3 | 2 | 3 | 0 |
| SG0000002 | Cep89 | 0.005597 | 0.009505 | 0.468485197 | 519.113274 | 718.2741403 | 72 | 15 | 21 | 27 | 60 | 5 | 0 | 0 | 0 | 0 | 0 | 0 |
| SG0000002 | Stard7 | 0.00566 | 0.009608 | 0.367219774 | 1686.262158 | 2175.05008 | 161 | 18 | 21 | 33 | 83 | 5 | 2 | 0 | 0 | 0 | 0 | 0 |
| SG0000004 | Ptger3 | 0.007778 | 0.012945 | 4.889287562 | 0.615018556 | 18.2268002 | 313 | 14 | 21 | 46 | 122 | 22 | 5 | 0 | 1 | 0 | 0 | 0 |
| SG0000003 | Ift57 | 0.026673 | 0.040417 | -0.470923041 | 681.5774955 | 491.7600919 | 409 | 5 | 21 | 63 | 245 | 69 | 4 | 0 | 0 | 0 | 1 | 0 |
| SG000000 | Glb1 | 0.031319 | 0.046834 | 0.296027262 | 1289.241746 | 1582.878007 | 87 | 17 | 21 | 21 | 65 | 12 | 0 | 0 | 0 | 0 | 0 | 0 |
| SG0000005 | Usp46 | 0.033581 | 0.049955 | -0.350676185 | 837.0871928 | 656.4575489 | 219 | 10 | 21 | 35 | 136 | 32 | 26 | 0 | 1 | 8 | 12 | 0 |
| SG000000 | Vps41 | 0.03798 | 0.055969 | -0.289289532 | 3151.508083 | 2578.894634 | 327 | 11 | 21 | 56 | 158 | 36 | 2 | 0 | 0 | 0 | 0 | 0 |
| SG000000 | Cdh6 | 0.038969 | 0.057298 | -1.250685471 | 27.10929723 | 11.39264113 | 261 | 11 | 21 | 37 | 111 | 16 | 0 | 0 | 0 | 0 | 0 | 0 |
| SG000000 | Creb5 | 0.051933 | 0.074704 | 1.470096049 | 13.09924362 | 36.2901887 | 197 | 10 | 21 | 32 | 111 | 9 | 0 | 0 | 0 | 0 | 0 | 0 |
| SG0000003 | Gabpb2 | 0.119425 | 0.159769 | 0.206853497 | 2091.046259 | 2413.419113 | 179 | 18 | 21 | 36 | 94 | 19 | 35 | 0 | 0 | 3 | 9 | 6 |
| SG000000 | Fbxo28 | 0.147399 | 0.193521 | -0.204422683 | 1993.692146 | 1730.297334 | 156 | 4 | 21 | 41 | 67 | 20 | 5 | 0 | 1 | 1 | 3 | 0 |
| SG0000004 | Blvrb | 0.166123 | 0.215764 | -0.508709861 | 1008.591091 | 708.8889398 | 114 | 15 | 21 | 18 | 79 | 2 | 0 | 0 | 0 | 0 | 0 | 0 |
| SG0000002 | Prkcd | 0.25101 | 0.311903 | -0.167285761 | 1778.111547 | 1583.437662 | 118 | 21 | 21 | 28 | 51 | 32 | 0 | 0 | 0 | 0 | 5 | 9 |
| SG000000 | Ssu72 | 0.255628 | 0.316873 | -0.160014792 | 2191.191946 | 1961.151618 | 118 | 15 | 21 | 29 | 56 | 15 | 0 | 0 | 0 | 0 | 0 | 1 |
| SG000000 | Rab18 | 0.25679 | 0.318071 | -0.155665357 | 4352.249279 | 3907.093642 | 87 | 13 | 21 | 28 | 58 | 2 | 1 | 0 | 0 | 0 | 0 | 1 |
| SG0000003 | Dhx8 | 0.264715 | 0.326757 | -0.15599565 | 2532.311316 | 2272.781711 | 84 | 19 | 21 | 37 | 66 | 5 | 2 | 0 | 0 | 1 | 1 | 0 |
| SG000000 | Cpe | 0.282987 | 0.346542 | -0.30338144 | 1312.234431 | 1063.370273 | 198 | 15 | 21 | 39 | 109 | 14 | 3 | 0 | 0 | 0 | 0 | 0 |
| SG0000001 | Tmpo | 0.29889 | 0.363687 | -0.143370604 | 7940.236144 | 7189.10069 | 276 | 13 | 21 | 31 | 94 | 13 | 15 | 0 | 0 | 0 | 1 | 0 |
| SG0000001 | Mpv17 | 0.313348 | 0.37898 | 0.15164284 | 790.5741255 | 878.1963744 | 69 | 21 | 21 | 21 | 56 | 12 | 6 | 0 | 0 | 0 | 0 | 5 |
| SG000000 | Zfp438 | 0.411124 | 0.480118 | 0.223068796 | 106.6582216 | 124.4929439 | 275 | 9 | 21 | 34 | 133 | 26 | 4 | 0 | 1 | 2 | 2 | 0 |
| SG0000005 | Tmem120b | 0.454501 | 0.523964 | 0.180309424 | 162.9689858 | 184.6645398 | 143 | 12 | 21 | 35 | 89 | 18 | 6 | 0 | 0 | 2 | 2 | 0 |
| SG000000 | Hs3st1 | 0.526102 | 0.592712 | -0.392510753 | 25.44410805 | 19.38338942 | 319 | 5 | 21 | 42 | 157 | 32 | 6 | 0 | 0 | 0 | 1 | 0 |
| SG0000005 | Tmco1 | 0.560163 | 0.625188 | 0.074214993 | 1874.997929 | 1973.975379 | 176 | 21 | 21 | 34 | 111 | 42 | 5 | 0 | 0 | 1 | 3 | 0 |
| SG0000003 | Birc3 | 0.591016 | 0.654201 | 0.106399048 | 201.9978199 | 217.4582993 | 77 | 12 | 21 | 16 | 51 | 11 | 11 | 4 | 5 | 4 | 6 | 1 |
| SG0000002 | Racgap1 | 0.737539 | 0.787388 | -0.046372586 | 4319.880589 | 4183.234426 | 183 | 19 | 21 | 45 | 115 | 4 | 2 | 0 | 0 | 0 | 2 | 0 |
| SG0000004 | Tmem29 | 0.744799 | 0.793578 | -0.07574531 | 262.2811069 | 248.8659026 | 266 | 11 | 21 | 43 | 151 | 12 | 4 | 0 | 0 | 0 | 1 | 2 |
| SG0000000 | Rtca | 0.795311 | 0.837386 | 0.034946719 | 1057.616725 | 1083.548414 | 115 | 19 | 21 | 30 | 66 | 0 | 5 | 0 | 0 | 0 | 0 | 0 |
| SG0000000 | Ap1m2 | 4.9E-278 | 3.1E-275 | 10.40793029 | 0 | 1357.623778 | 58 | 20 | 20 | 28 | 62 | 4 | 0 | 0 | 0 | 0 | 0 | 0 |
| SG0000002 | Srgn | 3.1E-269 | 1.7E-266 | -12.33725418 | 27579.97155 | 5.329791754 | 100 | 17 | 20 | 28 | 84 | 9 | 8 | 0 | 0 | 0 | 3 | 0 |
| SG0000002 | Il6ra | 2.4E-104 | 1.9E-102 | -5.955279081 | 3977.724841 | 64.1087194 | 105 | 18 | 20 | 28 | 93 | 6 | 5 | 0 | 0 | 0 | 7 | 0 |
| SG000000 | Cry1 | 6.19E-96 | 4.03E-94 | 3.00555005 | 219.9735624 | 1766.571447 | 134 | 14 | 20 | 27 | 80 | 6 | 3 | 0 | 1 | 0 | 0 | 2 |
| SG000000 | Mgat3 | 6.8E-84 | 3.38E-82 | -3.791390606 | 6857.463104 | 495.2690418 | 62 | 19 | 20 | 29 | 50 | 3 | 2 | 0 | 0 | 1 | 2 | 1 |
| SG0000006 | Zfp936 | 1.69E-80 | 7.72E-79 | 8.069876593 | 0 | 267.7044852 | 67 | 20 | 20 | 26 | 44 | 4 | 5 | 0 | 0 | 0 | 0 | 0 |
| SG000000 | Abca1 | 2.5E-71 | 9.26E-70 | -4.203231559 | 10837.53021 | 588.3442906 | 200 | 11 | 20 | 28 | 98 | 13 | 6 | 0 | 0 | 0 | 2 | 1 |
| SG0000001 | Lrrk1 | 7.5E-68 | 2.57E-66 | -3.721045591 | 1771.316842 | 134.3228549 | 169 | 12 | 20 | 24 | 62 | 15 | 1 | 0 | 0 | 0 | 0 | 0 |
| SG0000002 | Kremen1 | 2.5E-62 | 7.37E-61 | -3.318951996 | 8430.71662 | 844.8126082 | 118 | 16 | 20 | 28 | 75 | 7 | 5 | 0 | 0 | 0 | 1 | 1 |
| SG0000002 | Tspan12 | 4.87E-58 | 1.26E-56 | -4.049721014 | 775.9664643 | 46.85494723 | 336 | 6 | 20 | 46 | 154 | 32 | 8 | 0 | 0 | 2 | 4 | 2 |
| SG0000002 | Cpne5 | 2.76E-52 | 6.15E-51 | 3.580259188 | 59.84659149 | 715.8216482 | 81 | 17 | 20 | 25 | 42 | 6 | 0 | 0 | 0 | 0 | 0 | 0 |
| SG0000001 | Ptgis | 8.86E-51 | 1.88E-49 | 3.314973433 | 39.86990908 | 396.7817454 | 108 | 17 | 20 | 31 | 45 | 0 | 0 | 0 | 0 | 0 | 0 | 0 |
| SG0000005 | Rab7b | 3.36E-47 | 6.42E-46 | -5.555553032 | 1746.612615 | 37.13714277 | 118 | 12 | 20 | 24 | 60 | 12 | 6 | 0 | 0 | 0 | 0 | 0 |
| SG0000003 | Adam9 | 3.48E-46 | 6.36E-45 | -2.76227193 | 9449.525768 | 1392.782517 | 299 | 12 | 20 | 54 | 154 | 32 | 11 | 0 | 0 | 0 | 0 | 2 |
| SG000000 | Il17rd | 2.39E-44 | 4.11E-43 | 4.232236872 | 143.6519248 | 2699.862809 | 121 | 16 | 20 | 26 | 58 | 12 | 5 | 0 | 1 | 2 | 5 | 3 |
| SG0000002 | Pxdn | 1.34E-42 | 2.16E-41 | -2.481824028 | 5149.44567 | 921.8430889 | 169 | 16 | 20 | 30 | 84 | 14 | 9 | 0 | 0 | 1 | 1 | 2 |
| SG0000002 | Mink1 | 1.42E-42 | 2.29E-41 | -2.223788144 | 7434.709819 | 1591.611569 | 216 | 16 | 20 | 40 | 112 | 23 | 0 | 0 | 0 | 0 | 0 | 3 |
| SG0000000 | Eno2 | 1.61E-39 | 2.34E-38 | -2.787940087 | 9930.964689 | 1437.930334 | 33 | 20 | 20 | 23 | 41 | 0 | 20 | 20 | 20 | 20 | 20 | 0 |
| SG0000003 | Agpat5 | 3.99E-39 | 5.73E-38 | 1.77187131 | 569.6504408 | 1945.337015 | 200 | 8 | 20 | 27 | 79 | 12 | 14 | 0 | 0 | 3 | 7 | 0 |
| SG0000006 | Tacc1 | 1.74E-36 | 2.26E-35 | 1.724493424 | 545.7405042 | 1803.476285 | 124 | 16 | 20 | 26 | 64 | 12 | 22 | 4 | 4 | 6 | 12 | 0 |

| | | | | | | | | | | | | | | | | | | |
|---|---|---|---|---|---|---|---|---|---|---|---|---|---|---|---|---|---|---|
| SG0000003 | Arhgef26 | 2.15E-35 | 2.72E-34 | -2.907765523 | 621.160397 | 82.77117414 | 256 | 16 | 20 | 38 | 120 | 30 | 4 | 0 | 0 | 0 | 2 | 1 |
| SG0000003 | Rtkn2 | 3.59E-35 | 4.5E-34 | 2.208191473 | 142.1545689 | 656.8900626 | 559 | 5 | 20 | 72 | 243 | 59 | 19 | 0 | 0 | 0 | 0 | 0 |
| JSG000000 | Napsa | 3.74E-35 | 4.68E-34 | 8.28648216 | 0.331601068 | 103.5370079 | 38 | 18 | 20 | 23 | 32 | 3 | 0 | 0 | 0 | 0 | 0 | 0 |
| SG0000001 | Snx31 | 4.93E-35 | 6.14E-34 | 8.385260405 | 0.307509278 | 102.8189412 | 162 | 12 | 20 | 30 | 78 | 10 | 2 | 0 | 0 | 0 | 0 | 0 |
| SG0000002 | Tmem2 | 1.9E-34 | 2.31E-33 | 1.837452552 | 571.0944139 | 2040.968237 | 340 | 12 | 20 | 46 | 164 | 44 | 8 | 0 | 2 | 0 | 0 | 0 |
| SG0000003 | Coil | 5.51E-27 | 4.84E-26 | 1.641678109 | 311.6894876 | 972.5602359 | 81 | 20 | 20 | 30 | 58 | 8 | 1 | 0 | 0 | 0 | 0 | 0 |
| SG0000002 | Golga1 | 1.27E-26 | 1.1E-25 | -2.095824512 | 1379.060932 | 322.6096928 | 265 | 16 | 20 | 28 | 191 | 26 | 11 | 0 | 0 | 0 | 11 | 3 |
| SG0000001 | Map7 | 7.5E-26 | 6.3E-25 | 1.404532844 | 791.4783733 | 2095.29689 | 425 | 10 | 20 | 43 | 125 | 28 | 4 | 0 | 0 | 1 | 2 | 0 |
| SG0000002 | Eprs | 7.31E-25 | 5.87E-24 | 1.211113258 | 7294.931858 | 16888.95032 | 158 | 14 | 20 | 31 | 81 | 6 | 0 | 0 | 0 | 0 | 0 | 0 |
| SG0000005 | Sh3pxd2a | 4.03E-24 | 3.11E-23 | -1.5532609 | 4930.978787 | 1680.176911 | 366 | 14 | 20 | 52 | 156 | 48 | 20 | 0 | 0 | 2 | 8 | 0 |
| SG0000001 | Myo1c | 4.2E-23 | 3.1E-22 | -1.626110881 | 6806.689142 | 2205.097341 | 82 | 20 | 20 | 24 | 24 | 8 | 4 | 0 | 0 | 0 | 0 | 0 |
| SG0000002 | Xpo4 | 4.48E-21 | 2.98E-20 | 1.289593159 | 1361.224804 | 3327.638017 | 419 | 14 | 20 | 54 | 216 | 60 | 4 | 0 | 0 | 1 | 1 | 0 |
| SG0000002 | Yars | 2.79E-19 | 1.68E-18 | 1.139302648 | 1600.844337 | 3526.252242 | 76 | 13 | 20 | 18 | 34 | 6 | 1 | 0 | 0 | 0 | 0 | 0 |
| SG0000003 | Apba2 | 3.58E-19 | 2.14E-18 | 7.741622717 | 0.34247518 | 73.29760017 | 322 | 13 | 20 | 45 | 131 | 37 | 1 | 0 | 0 | 0 | 0 | 0 |
| SG0000003 | Trpm1 | 3.95E-19 | 2.36E-18 | 6.004121998 | 0.992459638 | 63.69915524 | 199 | 13 | 20 | 34 | 104 | 26 | 4 | 0 | 0 | 0 | 0 | 0 |
| SG0000003 | Sgsm2 | 4.77E-19 | 2.84E-18 | -1.656961975 | 1745.129366 | 553.3913552 | 104 | 18 | 20 | 22 | 50 | 4 | 0 | 0 | 0 | 0 | 0 | 0 |
| SG0000002 | Cd74 | 9.76E-19 | 5.7E-18 | -2.26375296 | 316.2102591 | 65.84433449 | 32 | 20 | 20 | 22 | 24 | 2 | 0 | 0 | 0 | 0 | 0 | 0 |
| SG0000002 | Paip1 | 4.3E-18 | 2.43E-17 | -1.313653517 | 4141.632572 | 1666.182063 | 295 | 8 | 20 | 40 | 115 | 18 | 8 | 0 | 0 | 0 | 3 | 6 |
| SG0000002 | Cers5 | 4.65E-18 | 2.62E-17 | -1.346778803 | 2919.25542 | 1147.760509 | 186 | 13 | 20 | 29 | 87 | 14 | 6 | 0 | 3 | 0 | 2 | 2 |
| JSG000000 | Asah2 | 2.66E-17 | 1.45E-16 | -2.570737791 | 467.7416442 | 78.7293837 | 173 | 12 | 20 | 32 | 101 | 20 | 12 | 0 | 2 | 2 | 2 | 0 |
| SG0000005 | Camk2b | 3.86E-17 | 2.08E-16 | 8.463801341 | 2.22249675 | 784.691983 | 600 | 10 | 20 | 30 | 160 | 50 | 0 | 0 | 0 | 0 | 0 | 0 |
| SG0000002 | Uhmk1 | 1.15E-16 | 6.03E-16 | -1.32191106 | 4327.531462 | 1731.033024 | 104 | 16 | 20 | 22 | 46 | 12 | 26 | 4 | 6 | 4 | 8 | 2 |
| SG0000001 | Cyth3 | 1.43E-16 | 7.43E-16 | -1.377685309 | 1571.492333 | 604.7664137 | 181 | 4 | 20 | 21 | 73 | 19 | 3 | 0 | 0 | 1 | 2 | 3 |
| SG0000003 | Ccsap | 4.65E-16 | 2.34E-15 | 1.66670785 | 106.0186174 | 336.5977379 | 66 | 20 | 20 | 22 | 68 | 10 | 6 | 0 | 0 | 0 | 0 | 0 |
| SG0000002 | Phc2 | 5.62E-16 | 2.82E-15 | -1.331930991 | 2240.792363 | 890.1238447 | 172 | 16 | 20 | 22 | 68 | 32 | 0 | 0 | 0 | 0 | 0 | 3 |
| SG0000004 | Napepld | 7.41E-16 | 3.69E-15 | 1.86466786 | 168.3633928 | 613.1526741 | 158 | 12 | 20 | 34 | 96 | 10 | 8 | 0 | 0 | 2 | 14 | 0 |
| JSG000000 | Lrrc3b | 2.48E-15 | 1.2E-14 | 7.139824642 | 0.307509278 | 43.36702243 | 256 | 14 | 20 | 32 | 108 | 40 | 8 | 0 | 0 | 0 | 0 | 0 |
| JSG000000 | M1ap | 3.92E-15 | 1.87E-14 | -1.895260941 | 344.5731387 | 92.62986395 | 115 | 12 | 20 | 22 | 61 | 12 | 0 | 0 | 0 | 0 | 0 | 0 |
| SG0000001 | 43716 | 1.25E-14 | 5.78E-14 | -1.163781269 | 3091.950645 | 1380.064812 | 104 | 20 | 20 | 21 | 66 | 8 | 12 | 0 | 0 | 1 | 3 | 2 |
| JSG000000 | Npl | 2.52E-13 | 1.07E-12 | 7.690661069 | 2.674556771 | 552.54988 | 86 | 18 | 20 | 27 | 61 | 5 | 0 | 0 | 0 | 0 | 2 | 0 |
| SG0000005 | Sdhc | 2.9E-13 | 1.22E-12 | 0.91992965 | 2010.516871 | 3803.944223 | 106 | 18 | 20 | 32 | 61 | 14 | 8 | 2 | 2 | 2 | 3 | 0 |
| SG0000003 | Gpt2 | 4.41E-13 | 1.83E-12 | 1.379853251 | 760.569087 | 1979.319429 | 67 | 20 | 20 | 27 | 47 | 4 | 2 | 0 | 0 | 1 | 2 | 0 |
| SG0000002 | Strn | 7.56E-13 | 3.11E-12 | -1.131925927 | 2420.180151 | 1104.34314 | 297 | 10 | 20 | 25 | 117 | 25 | 10 | 0 | 0 | 2 | 13 | 1 |
| SG0000003 | Ptpn3 | 1.02E-12 | 4.17E-12 | -1.066523042 | 3094.32722 | 1477.443259 | 127 | 15 | 20 | 29 | 74 | 12 | 5 | 0 | 0 | 1 | 3 | 2 |
| SG0000002 | Cacnb1 | 1.25E-12 | 5.03E-12 | -1.608617711 | 584.6354215 | 191.7091943 | 51 | 20 | 20 | 20 | 39 | 4 | 0 | 0 | 0 | 0 | 0 | 0 |
| JSG000000 | Col22a1 | 3.31E-12 | 1.3E-11 | 6.70273067 | 0.331601068 | 34.54131805 | 392 | 9 | 20 | 48 | 169 | 28 | 9 | 0 | 0 | 0 | 6 | 0 |
| SG0000002 | Ahctf1 | 7.86E-12 | 3.03E-11 | 0.827560266 | 4971.02514 | 8821.987191 | 184 | 10 | 20 | 28 | 78 | 21 | 6 | 0 | 1 | 0 | 0 | 2 |
| SG0000002 | Tra2b | 1.02E-11 | 3.88E-11 | 0.836331666 | 3295.291707 | 5883.757748 | 167 | 12 | 20 | 32 | 51 | 12 | 8 | 0 | 0 | 0 | 3 | 1 |
| SG0000004 | Agrn | 1.69E-11 | 6.35E-11 | 0.828828081 | 3411.368356 | 6059.415455 | 71 | 20 | 20 | 25 | 35 | 4 | 0 | 0 | 0 | 0 | 0 | 0 |
| SG0000006 | Mocs1 | 2.19E-11 | 8.17E-11 | -1.300735365 | 719.263788 | 291.9630116 | 191 | 8 | 20 | 20 | 55 | 27 | 2 | 0 | 0 | 0 | 0 | 0 |
| SG0000002 | Aass | 2.68E-11 | 9.92E-11 | 2.089401125 | 30.24963468 | 128.7337872 | 161 | 12 | 20 | 26 | 95 | 6 | 4 | 0 | 0 | 0 | 1 | 0 |
| SG0000002 | Tcerg1 | 4.11E-11 | 1.5E-10 | 0.814350243 | 3062.353762 | 5385.16741 | 173 | 11 | 20 | 32 | 96 | 17 | 3 | 0 | 0 | 0 | 1 | 0 |
| SG0000002 | Shank3 | 5E-11 | 1.81E-10 | 6.561031852 | 2.22249675 | 209.849783 | 118 | 14 | 20 | 30 | 80 | 10 | 4 | 0 | 0 | 0 | 10 | 0 |
| SG0000003 | AI118078 | 5.65E-11 | 2.04E-10 | 3.931875448 | 2.990596593 | 45.64259029 | 286 | 14 | 20 | 30 | 84 | 32 | 1 | 0 | 0 | 0 | 0 | 1 |
| SG0000004 | Tmem229b | 2.97E-10 | 1.02E-09 | -1.447022176 | 329.9870726 | 121.0318989 | 206 | 18 | 20 | 30 | 81 | 14 | 8 | 0 | 0 | 4 | 12 | 0 |
| JSG000000 | Wdr93 | 3.12E-10 | 1.07E-09 | 2.751993618 | 9.826729378 | 66.19750528 | 87 | 11 | 20 | 25 | 43 | 5 | 1 | 0 | 0 | 0 | 0 | 0 |
| JSG000000 | Herc4 | 3.45E-10 | 1.18E-09 | 0.79047067 | 2172.471393 | 3757.590495 | 204 | 16 | 20 | 36 | 106 | 21 | 0 | 0 | 0 | 0 | 0 | 0 |
| SG0000003 | Pir | 1.91E-09 | 6.15E-09 | -1.016708074 | 1512.442971 | 747.5140855 | 200 | 13 | 20 | 34 | 96 | 20 | 1 | 0 | 0 | 0 | 0 | 0 |
| SG0000003 | Eda2r | 2.64E-09 | 8.41E-09 | -0.966699088 | 2782.970324 | 1423.977585 | 183 | 10 | 20 | 31 | 125 | 19 | 5 | 0 | 0 | 1 | 5 | 1 |
| SG0000005 | Cdc42se2 | 3.56E-09 | 1.12E-08 | 0.774862306 | 1406.3421 | 2406.288938 | 167 | 17 | 20 | 37 | 76 | 16 | 6 | 0 | 0 | 1 | 3 | 1 |
| SG0000001 | Scube1 | 5.35E-09 | 1.66E-08 | 3.680210266 | 3.003814272 | 38.50585373 | 286 | 12 | 20 | 28 | 124 | 24 | 0 | 0 | 0 | 0 | 0 | 0 |
| SG0000003 | Anxa10 | 5.45E-09 | 1.69E-08 | -5.332099977 | 38.91879902 | 0.966134069 | 349 | 2 | 20 | 42 | 153 | 14 | 1 | 0 | 0 | 0 | 1 | 0 |
| SG0000002 | Kctd3 | 9.15E-09 | 2.8E-08 | 0.74143783 | 1786.025283 | 2985.950673 | 128 | 10 | 20 | 32 | 66 | 20 | 6 | 0 | 0 | 2 | 2 | 2 |
| SG0000003 | Fam149b | 1.11E-08 | 3.38E-08 | 0.863610685 | 486.5371637 | 885.2964851 | 220 | 12 | 20 | 38 | 122 | 4 | 0 | 0 | 0 | 0 | 0 | 0 |
| SG0000002 | Maats1 | 1.64E-08 | 4.91E-08 | 2.502840262 | 10.74925721 | 60.92681116 | 129 | 12 | 20 | 26 | 68 | 6 | 4 | 0 | 0 | 0 | 1 | 0 |
| JSG000000 | Spast | 2.06E-08 | 6.11E-08 | -0.843046015 | 2322.963959 | 1294.971867 | 119 | 17 | 20 | 32 | 75 | 12 | 4 | 0 | 0 | 0 | 1 | 3 |
| SG0000003 | Slc35a2 | 2.29E-08 | 6.75E-08 | -0.977993471 | 809.3452875 | 410.8927441 | 55 | 15 | 20 | 26 | 31 | 6 | 3 | 0 | 0 | 2 | 2 | 2 |
| SG0000005 | Arhgap20 | 2.85E-08 | 8.36E-08 | 3.825470583 | 4.020365701 | 56.99633786 | 293 | 4 | 20 | 39 | 104 | 20 | 7 | 0 | 0 | 1 | 2 | 0 |
| SG0000000 | Fbxo9 | 1.05E-07 | 2.93E-07 | -1.017776127 | 521.4694565 | 257.5418037 | 96 | 14 | 20 | 24 | 56 | 12 | 2 | 0 | 2 | 0 | 0 | 4 |
| JSG000000 | Gm1305 | 1.65E-07 | 4.56E-07 | 5.989504005 | 0.331601068 | 21.0686295 | 83 | 18 | 20 | 24 | 53 | 7 | 0 | 0 | 0 | 0 | 0 | 0 |
| SG0000003 | Mto1 | 2.53E-07 | 6.85E-07 | 0.799810949 | 461.4578601 | 803.3395235 | 130 | 20 | 20 | 29 | 68 | 12 | 2 | 0 | 0 | 0 | 0 | 2 |
| SG0000002 | Nfs1 | 2.64E-07 | 7.14E-07 | -0.802918667 | 1488.733192 | 853.3246056 | 62 | 16 | 20 | 22 | 41 | 10 | 2 | 0 | 0 | 0 | 0 | 0 |
| SG0000002 | Eif4e | 2.78E-07 | 7.5E-07 | 0.646657913 | 2557.237214 | 4003.45033 | 308 | 16 | 20 | 32 | 96 | 21 | 12 | 0 | 0 | 0 | 3 | 1 |
| SG0000002 | Ralbp1 | 2.94E-07 | 7.9E-07 | -0.799217949 | 4274.970916 | 2456.657365 | 118 | 14 | 20 | 24 | 50 | 12 | 2 | 0 | 0 | 2 | 3 | 0 |
| SG0000002 | Crispld1 | 3.61E-07 | 9.65E-07 | 5.657130254 | 1.302312483 | 65.71750991 | 230 | 8 | 20 | 32 | 99 | 32 | 8 | 0 | 0 | 2 | 11 | 0 |
| SG0000005 | Kcnt1 | 5.56E-07 | 1.46E-06 | 5.826878127 | 0.34247518 | 19.43990776 | 77 | 15 | 20 | 20 | 33 | 4 | 0 | 0 | 0 | 0 | 0 | 0 |
| JSG000000 | Yaf2 | 1.13E-06 | 2.9E-06 | -0.894511777 | 629.193979 | 338.4618172 | 226 | 14 | 20 | 34 | 98 | 16 | 8 | 0 | 0 | 0 | 2 | 0 |
| SG0000003 | Tcte2 | 1.16E-06 | 2.97E-06 | 1.853917124 | 20.09599137 | 72.64305489 | 375 | 20 | 20 | 44 | 110 | 30 | 14 | 2 | 2 | 2 | 5 | 0 |
| SG0000002 | Ptdss2 | 1.38E-06 | 3.52E-06 | 0.65367413 | 1230.710575 | 1936.116352 | 100 | 14 | 20 | 23 | 45 | 5 | 0 | 0 | 0 | 0 | 0 | 0 |
| JSG000000 | St8sia1 | 1.61E-06 | 4.08E-06 | 2.668413027 | 6.633521212 | 42.17125098 | 240 | 15 | 20 | 39 | 98 | 18 | 11 | 0 | 0 | 0 | 2 | 1 |
| JSG000000 | Zfp652 | 1.65E-06 | 4.18E-06 | 0.643860767 | 1482.90704 | 2317.0496 | 163 | 9 | 20 | 33 | 95 | 13 | 11 | 1 | 1 | 2 | 8 | 0 |
| SG0000002 | Rasgrp1 | 1.81E-06 | 4.57E-06 | 1.092424084 | 99.92264927 | 213.0670816 | 424 | 0 | 20 | 37 | 230 | 54 | 14 | 0 | 0 | 2 | 10 | 4 |
| SG0000003 | Pdzk1 | 5.41E-06 | 1.31E-05 | 8.368561636 | 0.663202137 | 219.196689 | 139 | 16 | 20 | 20 | 48 | 16 | 4 | 0 | 0 | 0 | 0 | 0 |
| SG0000002 | Stradb | 5.44E-06 | 1.31E-05 | 0.847480898 | 231.4410484 | 416.4449106 | 193 | 12 | 20 | 28 | 80 | 20 | 8 | 0 | 0 | 0 | 0 | 0 |
| SG0000002 | Shoc2 | 5.57E-06 | 1.34E-05 | -0.659852383 | 2704.918582 | 1712.059435 | 256 | 9 | 20 | 45 | 153 | 16 | 8 | 0 | 0 | 1 | 2 | 1 |
| SG0000005 | Pdlim1 | 6.28E-06 | 1.51E-05 | 0.940287773 | 700.6975557 | 1344.576222 | 103 | 16 | 20 | 22 | 60 | 8 | 2 | 0 | 0 | 0 | 2 | 0 |
| SG0000002 | B4galt1 | 1.44E-05 | 3.36E-05 | 0.568188456 | 1713.158242 | 2540.032352 | 103 | 19 | 20 | 26 | 66 | 8 | 1 | 0 | 0 | 0 | 1 | 0 |
| SG0000002 | Cd4 | 1.53E-05 | 3.55E-05 | 4.738689273 | 0.639110347 | 17.06331702 | 49 | 19 | 20 | 26 | 39 | 5 | 1 | 1 | 1 | 1 | 1 | 0 |
| SG0000002 | Myocd | 1.71E-05 | 3.94E-05 | -3.813896039 | 24.33702659 | 1.730495238 | 327 | 2 | 20 | 22 | 123 | 22 | 12 | 0 | 0 | 0 | 0 | 0 |
| SG0000004 | Ccdc6 | 1.88E-05 | 4.32E-05 | -0.603302411 | 4351.278834 | 2864.209573 | 165 | 11 | 20 | 25 | 79 | 11 | 5 | 0 | 1 | 1 | 2 | 1 |
| SG0000005 | Pgbd1 | 2.15E-05 | 4.9E-05 | 0.964413484 | 107.349862 | 209.4685745 | 164 | 12 | 20 | 30 | 136 | 22 | 5 | 0 | 4 | 2 | 12 | 2 |
| SG0000002 | Zmat3 | 2.18E-05 | 4.96E-05 | -0.590540804 | 5254.803267 | 3489.682901 | 128 | 18 | 20 | 24 | 66 | 10 | 16 | 0 | 2 | 0 | 2 | 8 |
| SG0000000 | Ubqln1 | 2.41E-05 | 5.46E-05 | 0.52465051 | 4630.927644 | 6661.983245 | 172 | 16 | 20 | 38 | 90 | 12 | 14 | 0 | 0 | 0 | 0 | 0 |
| SG0000000 | Por | 2.41E-05 | 5.46E-05 | -0.748940462 | 2078.038275 | 1236.516735 | 83 | 18 | 20 | 32 | 60 | 14 | 0 | 0 | 0 | 0 | 0 | 2 |
| SG0000001 | Mrpl45 | 3.32E-05 | 7.45E-05 | 0.527922649 | 2495.297506 | 3597.84861 | 58 | 19 | 20 | 26 | 44 | 2 | 3 | 0 | 0 | 1 | 2 | 0 |
| SG0000002 | Sumo2 | 3.67E-05 | 8.19E-05 | 0.700410815 | 332.936857 | 541.011554 | 130 | 20 | 20 | 35 | 85 | 20 | 5 | 0 | 0 | 0 | 0 | 5 |
| SG0000005 | Zfp217 | 5.31E-05 | 0.000116 | -0.59582263 | 4050.222292 | 2679.898687 | 100 | 17 | 20 | 33 | 77 | 12 | 10 | 0 | 0 | 4 | 8 | 8 |
| SG0000001 | Pmp22 | 6.42E-05 | 0.000139 | 0.857799263 | 171.9538401 | 311.6270778 | 141 | 20 | 20 | 42 | 74 | 6 | 6 | 0 | 0 | 2 | 2 | 6 |
| SG0000004 | Cyp20a1 | 6.45E-05 | 0.00014 | -0.645842965 | 1025.114162 | 655.1701891 | 210 | 12 | 20 | 36 | 90 | 18 | 0 | 0 | 0 | 0 | 2 | 0 |
| SG0000002 | Smad4 | 8.64E-05 | 0.000185 | -0.565874591 | 2525.107108 | 1705.825407 | 215 | 9 | 20 | 35 | 80 | 23 | 2 | 0 | 1 | 0 | 0 | 0 |
| SG0000003 | Wbp11 | 0.000253 | 0.000511 | 0.462488652 | 3649.700585 | 5028.983308 | 44 | 18 | 20 | 20 | 28 | 2 | 0 | 0 | 0 | 0 | 0 | 1 |
| SG0000002 | Chuk | 0.000262 | 0.000529 | -0.522853941 | 2638.573579 | 1836.430469 | 192 | 2 | 20 | 36 | 112 | 12 | 14 | 0 | 0 | 0 | 0 | 0 |
| SG0000000 | Rasa4 | 0.000282 | 0.000568 | 1.174682749 | 119.9778376 | 270.841851 | 84 | 12 | 20 | 26 | 39 | 11 | 2 | 0 | 0 | 0 | 1 | 0 |
| SG0000003 | Pxk | 0.00039 | 0.000771 | -0.542341834 | 1548.480483 | 1063.272487 | 192 | 18 | 20 | 30 | 118 | 68 | 2 | 0 | 0 | 0 | 4 | 0 |
| JSG000000 | Fig4 | 0.000537 | 0.001043 | 0.47166087 | 1178.884908 | 1634.765552 | 212 | 9 | 20 | 30 | 93 | 14 | 0 | 0 | 0 | 0 | 0 | 0 |
| SG0000002 | Ptpn21 | 0.000558 | 0.001082 | -0.530329219 | 1209.990619 | 837.7935275 | 220 | 10 | 20 | 28 | 93 | 18 | 8 | 0 | 2 | 0 | 0 | 0 |
| SG0000003 | Xkr5 | 0.00064 | 0.001229 | 0.672734478 | 229.0589336 | 365.1409081 | 99 | 8 | 20 | 27 | 99 | 12 | 5 | 2 | 2 | 5 | 5 | 0 |
| SG0000004 | Rcsd1 | 0.000825 | 0.001563 | 2.17678715 | 5.592877993 | 25.28794695 | 152 | 14 | 20 | 26 | 61 | 26 | 8 | 0 | 0 | 2 | 3 | 0 |
| SG0000000 | Tmem161a | 0.000837 | 0.001585 | -0.504597932 | 1345.636513 | 948.4810307 | 78 | 14 | 20 | 19 | 34 | 4 | 7 | 0 | 0 | 5 | 5 | 0 |
| SG0000003 | Rabep2 | 0.000902 | 0.0017 | -0.852194368 | 417.4920537 | 231.2661906 | 70 | 20 | 20 | 28 | 40 | 4 | 2 | 0 | 0 | 0 | 0 | 0 |
| SG0000004 | Zfp367 | 0.000969 | 0.001819 | 0.492927006 | 646.5565526 | 909.8972069 | 100 | 14 | 20 | 28 | 46 | 16 | 12 | 1 | 2 | 4 | 7 | 1 |
| SG0000006 | Bod1l | 0.001202 | 0.002237 | -0.648725427 | 2691.483917 | 1716.745732 | 252 | 14 | 20 | 26 | 88 | 22 | 6 | 2 | 2 | 2 | 8 | 0 |
| JSG000000 | Haus3 | 0.001531 | 0.002815 | 0.44998692 | 910.9316515 | 1244.358025 | 68 | 18 | 20 | 19 | 38 | 3 | 1 | 0 | 0 | 0 | 0 | 0 |
| JSG000000 | Ubtd2 | 0.001571 | 0.002887 | 0.726835537 | 110.9480966 | 183.6196497 | 149 | 15 | 20 | 29 | 75 | 10 | 2 | 0 | 0 | 0 | 0 | 0 |
| SG0000003 | Fam20b | 0.001751 | 0.003197 | -0.450208937 | 2168.506697 | 1587.209936 | 148 | 16 | 20 | 26 | 61 | 16 | 8 | 0 | 0 | 0 | 1 | 0 |
| SG0000003 | Mapkbp1 | 0.002628 | 0.004679 | -0.519781513 | 633.2216157 | 441.6578001 | 79 | 15 | 20 | 24 | 50 | 5 | 4 | 0 | 1 | 2 | 2 | 0 |
| JSG000000 | Cyb5b | 0.002958 | 0.005224 | -0.399665027 | 7350.01751 | 5571.56514 | 83 | 15 | 20 | 24 | 57 | 8 | 7 | 0 | 0 | 1 | 2 | 0 |
| SG0000003 | Sh3bgrl2 | 0.003748 | 0.00652 | 0.396676471 | 1133.178357 | 1491.797198 | 339 | 7 | 20 | 32 | 112 | 50 | 6 | 0 | 0 | 0 | 4 | 2 |
| SG0000003 | Eef2k | 0.004436 | 0.007642 | -0.456677498 | 930.7296357 | 678.1877134 | 315 | 16 | 20 | 79 | 208 | 55 | 4 | 0 | 0 | 0 | 3 | 0 |
| JSG000000 | Nvl | 0.006777 | 0.011378 | 0.375833095 | 4680.33212 | 6073.145796 | 135 | 15 | 20 | 23 | 85 | 11 | 10 | 0 | 1 | 1 | 5 | 0 |
| JSG000000 | Tex21 | 0.007386 | 0.01233 | 0.990281183 | 45.7222659 | 90.83057954 | 101 | 13 | 20 | 22 | 57 | 8 | 0 | 0 | 0 | 0 | 0 | 0 |
| JSG000000 | Leo1 | 0.009539 | 0.015653 | 0.363426785 | 1121.219832 | 1442.424646 | 57 | 19 | 20 | 25 | 43 | 4 | 1 | 0 | 0 | 1 | 1 | 0 |
| SG0000000 | Nup214 | 0.010308 | 0.016817 | 0.340095218 | 2082.178013 | 2635.704501 | 177 | 17 | 20 | 39 | 76 | 11 | 1 | 0 | 0 | 1 | 1 | 0 |
| SG0000002 | Podn | 0.012866 | 0.020617 | -1.593172565 | 25.78289296 | 8.54552827 | 141 | 15 | 20 | 32 | 64 | 13 | 0 | 0 | 0 | 0 | 3 | 0 |
| SG0000002 | Galnt10 | 0.014563 | 0.023121 | 0.651010246 | 313.1897772 | 491.7916937 | 191 | 14 | 20 | 24 | 64 | 21 | 0 | 0 | 0 | 0 | 1 | 0 |
| SG0000002 | Sp3 | 0.014692 | 0.023309 | 0.30587081 | 3445.554901 | 4259.27286 | 230 | 16 | 20 | 46 | 132 | 15 | 12 | 0 | 0 | 2 | 10 | 1 |
| JSG000000 | Myh11 | 0.016492 | 0.025942 | 1.586502808 | 7.145985629 | 21.46085755 | 140 | 15 | 20 | 26 | 57 | 11 | 0 | 0 | 0 | 0 | 0 | 0 |
| JSG000000 | Parm1 | 0.017772 | 0.027751 | -0.92126147 | 470.1546715 | 248.2638118 | 198 | 12 | 20 | 33 | 85 | 25 | 6 | 0 | 1 | 3 | 4 | 0 |
| SG0000003 | Zdhhc17 | 0.019077 | 0.029624 | -0.391879601 | 737.0777909 | 561.7535309 | 154 | 13 | 20 | 30 | 78 | 19 | 7 | 0 | 1 | 1 | 4 | 4 |
| JSG000000 | Npr3 | 0.019668 | 0.03047 | 0.767941054 | 47.44286607 | 80.78750752 | 139 | 14 | 20 | 30 | 100 | 12 | 14 | 0 | 0 | 0 | 9 | 1 |
| SG0000002 | Sec23b | 0.021265 | 0.032728 | 0.303712954 | 2116.1204 | 2611.963376 | 91 | 17 | 20 | 30 | 50 | 5 | 1 | 0 | 0 | 0 | 0 | 0 |
| SG0000002 | Serf1 | 0.022692 | 0.034761 | 0.653863837 | 64.52297549 | 101.518926 | 44 | 20 | 20 | 20 | 22 | 0 | 1 | 0 | 0 | 0 | 0 | 0 |
| SG0000002 | Ppp2r5a | 0.027285 | 0.041275 | -0.308102202 | 2887.080419 | 2331.905125 | 119 | 17 | 20 | 27 | 76 | 7 | 3 | 0 | 1 | 0 | 0 | 0 |

| | | | | | | | | | | | | | | | | | | |
|---|---|---|---|---|---|---|---|---|---|---|---|---|---|---|---|---|---|---|
| JSG000000 | Tnks | 0.047713 | 0.069178 | -0.309670882 | 2630.626732 | 2122.45751 | 296 | 14 | 20 | 41 | 117 | 25 | 3 | 1 | 1 | 1 | 2 | 0 |
| JSG000000 | Nim1k | 0.0488 | 0.070601 | -0.932558613 | 65.57095569 | 34.3544815 | 95 | 17 | 20 | 25 | 53 | 4 | 3 | 0 | 1 | 0 | 0 | 0 |
| SG0000002 | Camsap1 | 0.053435 | 0.076705 | -0.276921175 | 1746.751493 | 1441.682222 | 224 | 20 | 20 | 43 | 130 | 22 | 13 | 0 | 0 | 8 | 12 | 0 |
| SG0000005 | Klhdc1 | 0.055809 | 0.07978 | -0.64727989 | 130.9500643 | 83.60938976 | 192 | 14 | 20 | 28 | 84 | 32 | 7 | 0 | 0 | 2 | 4 | 2 |
| JSG000000 | Smap2 | 0.064704 | 0.091475 | 0.271556294 | 797.4028433 | 962.5523747 | 98 | 16 | 20 | 31 | 54 | 6 | 1 | 0 | 0 | 0 | 1 | 0 |
| JSG000000 | Mrc1 | 0.071058 | 0.099662 | 2.16553369 | 5.324024889 | 23.88529857 | 236 | 10 | 20 | 43 | 148 | 20 | 5 | 0 | 0 | 2 | 3 | 1 |
| SG0000005 | Ccdc157 | 0.071797 | 0.100604 | -0.392443812 | 235.9969735 | 179.7914664 | 48 | 16 | 20 | 20 | 30 | 6 | 10 | 2 | 2 | 2 | 2 | 0 |
| JSG000000 | Nup98 | 0.074682 | 0.104312 | 0.240067592 | 2212.648514 | 2613.244088 | 212 | 13 | 20 | 30 | 98 | 26 | 2 | 0 | 0 | 0 | 0 | 1 |
| JSG000000 | Hspbp1 | 0.084912 | 0.117365 | 0.469742831 | 592.1416613 | 820.0348325 | 46 | 16 | 20 | 23 | 36 | 4 | 0 | 0 | 0 | 0 | 0 | 0 |
| SG0000003 | Phrf1 | 0.086817 | 0.119768 | 0.227696231 | 2653.200954 | 3106.801742 | 104 | 14 | 20 | 14 | 35 | 9 | 3 | 0 | 0 | 0 | 3 | 0 |
| SG0000001 | Ehmt2 | 0.096405 | 0.131717 | -0.364935315 | 5248.765628 | 4075.685876 | 68 | 16 | 20 | 20 | 36 | 4 | 0 | 0 | 0 | 0 | 0 | 0 |
| JSG000000 | Nup155 | 0.097332 | 0.132862 | 0.207148541 | 3596.708149 | 4152.055245 | 109 | 12 | 20 | 23 | 60 | 13 | 6 | 0 | 1 | 2 | 6 | 1 |
| SG0000005 | Lins1 | 0.14609 | 0.191859 | -0.277094367 | 407.7152207 | 336.4675676 | 203 | 20 | 20 | 34 | 73 | 12 | 13 | 0 | 0 | 0 | 1 | 0 |
| SG0000002 | Cutc | 0.155228 | 0.202907 | 0.320566448 | 146.0874478 | 182.4370404 | 96 | 18 | 20 | 32 | 80 | 14 | 4 | 0 | 0 | 0 | 0 | 0 |
| SG0000005 | Rlim | 0.180842 | 0.232938 | 0.162435928 | 7088.669215 | 7933.462945 | 112 | 18 | 20 | 24 | 54 | 12 | 22 | 0 | 0 | 2 | 4 | 4 |
| SG0000005 | Raet1e | 0.191659 | 0.245449 | 0.283244857 | 316.6226225 | 385.3072136 | 327 | 2 | 20 | 33 | 159 | 37 | 2 | 0 | 0 | 0 | 6 | 0 |
| SG0000000 | Gm38393 | 0.195529 | 0.249826 | -0.354596105 | 984.6173837 | 770.0579966 | 270 | 8 | 20 | 44 | 138 | 30 | 3 | 0 | 0 | 0 | 1 | 0 |
| SG0000002 | Sumf2 | 0.20385 | 0.259219 | -0.211673905 | 1169.617669 | 1010.00547 | 55 | 18 | 20 | 20 | 39 | 0 | 7 | 2 | 2 | 2 | 3 | 0 |
| SG0000004 | Il7 | 0.249427 | 0.310174 | -0.629286264 | 30.38850143 | 19.64604988 | 392 | 4 | 20 | 43 | 251 | 39 | 8 | 0 | 0 | 0 | 2 | 0 |
| JSG000000 | Atl1 | 0.255165 | 0.316408 | 0.3214773 | 81.8599139 | 102.2929242 | 150 | 10 | 20 | 21 | 79 | 12 | 1 | 0 | 0 | 0 | 2 | 0 |
| JSG000000 | Eif3a | 0.298309 | 0.363096 | 0.123658836 | 17801.15258 | 19394.25679 | 95 | 14 | 20 | 22 | 46 | 2 | 2 | 0 | 0 | 1 | 2 | 0 |
| SG0000003 | Zfand6 | 0.311044 | 0.37655 | 0.134487628 | 1995.057435 | 2189.980325 | 326 | 12 | 20 | 40 | 138 | 28 | 6 | 0 | 0 | 4 | 6 | 2 |
| SG0000003 | Fam81a | 0.342211 | 0.409519 | 0.164540438 | 388.3843386 | 435.3046599 | 83 | 15 | 20 | 21 | 50 | 5 | 7 | 0 | 0 | 1 | 1 | 0 |
| SG0000002 | Hsdl2 | 0.343599 | 0.410769 | -0.182570772 | 505.9146951 | 445.7773681 | 182 | 16 | 20 | 32 | 96 | 22 | 4 | 0 | 0 | 0 | 6 | 0 |
| JSG000000 | Ubxn4 | 0.402835 | 0.471785 | -0.115947974 | 2775.254707 | 2560.937797 | 87 | 18 | 20 | 26 | 63 | 5 | 3 | 0 | 0 | 0 | 0 | 0 |
| SG0000001 | Lamp2 | 0.409344 | 0.478508 | -0.115808993 | 4373.693809 | 4036.327487 | 276 | 9 | 20 | 56 | 149 | 21 | 6 | 0 | 1 | 0 | 1 | 2 |
| JSG000000 | Ipo7 | 0.427104 | 0.496417 | 0.093275375 | 7080.316495 | 7553.20648 | 115 | 11 | 20 | 23 | 64 | 6 | 0 | 0 | 0 | 0 | 0 | 0 |
| SG0000004 | Usp14 | 0.473713 | 0.542529 | 0.082043426 | 4901.463836 | 5188.279152 | 193 | 4 | 20 | 28 | 95 | 22 | 5 | 0 | 0 | 2 | 3 | 0 |
| SG0000004 | Gng7 | 0.474025 | 0.542816 | 0.174072877 | 111.1781404 | 125.4355282 | 234 | 20 | 20 | 42 | 121 | 60 | 0 | 0 | 0 | 0 | 0 | 0 |
| JSG000000 | Dock8 | 0.505004 | 0.572255 | 0.288707133 | 31.48670249 | 38.46242724 | 286 | 12 | 20 | 43 | 131 | 22 | 2 | 0 | 1 | 1 | 1 | 0 |
| SG0000005 | Cacng8 | 0.528645 | 0.595241 | -0.355319741 | 26.71879161 | 20.88598263 | 68 | 19 | 20 | 24 | 42 | 2 | 0 | 0 | 0 | 0 | 0 | 0 |
| JSG000000 | Zfp131 | 0.706728 | 0.759916 | -0.063385641 | 1069.194359 | 1023.235689 | 124 | 12 | 20 | 20 | 60 | 10 | 8 | 0 | 0 | 0 | 4 | 4 |
| SG0000002 | Nphp1 | 0.767489 | 0.813517 | 0.036384179 | 556.4165897 | 570.6276337 | 160 | 14 | 20 | 24 | 74 | 10 | 0 | 0 | 0 | 0 | 0 | 0 |
| SG0000007 | Arfgef2 | 0.797863 | 0.839629 | -0.041161302 | 1921.652491 | 1867.600865 | 124 | 15 | 20 | 25 | 64 | 9 | 4 | 0 | 0 | 1 | 4 | 0 |
| SG0000002 | Atad3a | 0.847349 | 0.880685 | 0.028128015 | 2430.261262 | 2478.108633 | 77 | 20 | 20 | 20 | 38 | 3 | 1 | 0 | 0 | 0 | 0 | 0 |
| SG0000004 | Slc17a5 | 0.84776 | 0.880958 | 0.031858899 | 298.5079038 | 305.1731488 | 154 | 16 | 20 | 26 | 84 | 16 | 6 | 0 | 0 | 2 | 2 | 4 |
| SG0000002 | Fam76a | 0.93202 | 0.950396 | 0.003655419 | 1213.966623 | 1217.046403 | 61 | 12 | 20 | 14 | 46 | 2 | 3 | 2 | 2 | 2 | 4 | 0 |
| SG0000002 | Rnf185 | 0.947655 | 0.963579 | -0.013793665 | 1214.453933 | 1202.897823 | 160 | 18 | 20 | 34 | 84 | 6 | 4 | 0 | 0 | 0 | 0 | 0 |
| SG0000002 | St3gal6 | 5.1E-120 | 5.1E-118 | -5.904977297 | 2634.840556 | 43.97228656 | 306 | 9 | 19 | 18 | 134 | 16 | 12 | 0 | 1 | 3 | 5 | 1 |
| SG0000002 | Plac8 | 3.38E-91 | 1.97E-89 | 3.596722146 | 75.98730869 | 919.3107002 | 82 | 13 | 19 | 24 | 47 | 1 | 6 | 0 | 0 | 0 | 0 | 0 |
| JSG000000 | Plxnc1 | 1.64E-70 | 5.92E-69 | -5.187833032 | 1375.190956 | 37.72850985 | 279 | 13 | 19 | 40 | 103 | 36 | 6 | 0 | 0 | 0 | 1 | 0 |
| SG0000003 | Barx2 | 4.19E-67 | 1.4E-65 | -5.933023903 | 912.5548815 | 14.9362245 | 127 | 15 | 19 | 30 | 81 | 10 | 0 | 0 | 0 | 0 | 0 | 0 |
| JSG000000 | Gm13152 | 7.8E-58 | 2E-56 | 6.912532038 | 1.620695873 | 195.2454922 | 87 | 19 | 19 | 24 | 69 | 6 | 0 | 0 | 0 | 0 | 0 | 0 |
| SG0000003 | Mboat1 | 1.2E-54 | 2.84E-53 | -4.005140157 | 2191.495041 | 136.4813055 | 195 | 7 | 19 | 27 | 85 | 24 | 1 | 0 | 0 | 0 | 1 | 0 |
| SG0000003 | Usp54 | 1.43E-43 | 2.39E-42 | 1.978366766 | 408.6743674 | 1610.368023 | 264 | 14 | 19 | 47 | 156 | 21 | 2 | 0 | 0 | 1 | 2 | 0 |
| SG0000003 | Sh3bgrl | 5.99E-40 | 8.86E-39 | -2.199851258 | 4920.711852 | 1071.042537 | 198 | 12 | 19 | 32 | 121 | 23 | 4 | 0 | 0 | 0 | 2 | 0 |
| JSG000000 | Ccdc138 | 1.47E-38 | 2.07E-37 | 2.566127102 | 76.63744629 | 453.8603621 | 196 | 10 | 19 | 34 | 105 | 12 | 1 | 0 | 0 | 0 | 0 | 0 |
| SG0000003 | Maml2 | 1.53E-30 | 1.58E-29 | -3.248718325 | 354.1706589 | 37.26069199 | 363 | 11 | 19 | 44 | 144 | 26 | 0 | 0 | 0 | 0 | 0 | 0 |
| SG0000002 | Capn1 | 2.88E-28 | 2.7E-27 | -1.882259701 | 4432.379407 | 1202.320629 | 49 | 16 | 19 | 20 | 38 | 4 | 0 | 0 | 0 | 0 | 0 | 1 |
| SG0000002 | Cxadr | 6.02E-28 | 5.53E-27 | -1.791175755 | 3055.303112 | 882.7885114 | 166 | 14 | 19 | 42 | 92 | 20 | 16 | 0 | 1 | 2 | 5 | 3 |
| SG0000002 | Igf2r | 2.81E-27 | 2.51E-26 | 1.385155126 | 1924.317232 | 5026.317409 | 105 | 15 | 19 | 24 | 56 | 12 | 3 | 0 | 0 | 1 | 1 | 1 |
| SG0000002 | Rgs12 | 3.96E-27 | 3.51E-26 | -1.795529692 | 2379.178262 | 685.3598958 | 214 | 14 | 19 | 15 | 142 | 25 | 1 | 0 | 1 | 0 | 0 | 0 |
| JSG000000 | Arfgef3 | 8.13E-25 | 6.51E-24 | -3.120803768 | 1081.659072 | 124.3468683 | 284 | 7 | 19 | 40 | 125 | 25 | 0 | 0 | 0 | 0 | 0 | 0 |
| JSG000000 | Prkaa2 | 2.39E-22 | 1.71E-21 | 5.985389943 | 8.667970774 | 549.1605696 | 185 | 10 | 19 | 30 | 100 | 18 | 23 | 1 | 3 | 7 | 11 | 2 |
| JSG000000 | Uchl3 | 2.88E-22 | 2.05E-21 | 1.243467467 | 1231.257075 | 2915.209329 | 120 | 15 | 19 | 21 | 62 | 15 | 1 | 0 | 0 | 0 | 1 | 0 |
| SG0000002 | Mettl3 | 1.88E-21 | 1.27E-20 | 1.308055804 | 672.5148125 | 1665.198026 | 134 | 15 | 19 | 24 | 58 | 10 | 8 | 0 | 1 | 2 | 6 | 4 |
| JSG000000 | Tbc1d12 | 3.67E-21 | 2.45E-20 | -1.992158044 | 610.3884291 | 153.428827 | 209 | 13 | 19 | 32 | 97 | 17 | 6 | 0 | 0 | 0 | 2 | 3 |
| JSG000000 | Clip4 | 4.48E-21 | 2.98E-20 | 4.237094374 | 12.21921537 | 230.4283024 | 151 | 13 | 19 | 32 | 77 | 11 | 0 | 0 | 0 | 0 | 1 | 0 |
| JSG000000 | Plekho2 | 1.66E-18 | 9.57E-18 | -1.510000997 | 1332.194632 | 467.7481579 | 46 | 18 | 19 | 24 | 31 | 3 | 1 | 0 | 0 | 0 | 0 | 0 |
| SG0000004 | Zc2hc1a | 7.96E-18 | 4.44E-17 | 1.387719454 | 297.1029185 | 777.4135377 | 250 | 9 | 19 | 31 | 125 | 50 | 12 | 3 | 3 | 3 | 3 | 4 |
| JSG000000 | Snupn | 1.62E-17 | 8.9E-17 | 1.376828994 | 268.9385232 | 698.425105 | 66 | 14 | 19 | 24 | 57 | 7 | 0 | 0 | 0 | 0 | 0 | 1 |
| SG0000000 | Araf | 1.87E-17 | 1.02E-16 | -1.353579445 | 3318.074215 | 1298.428624 | 137 | 18 | 19 | 34 | 101 | 13 | 4 | 0 | 0 | 0 | 6 | 2 |
| SG0000003 | Sumf1 | 7.52E-17 | 3.99E-16 | -1.426553545 | 1288.41771 | 479.3152847 | 176 | 9 | 19 | 26 | 97 | 8 | 5 | 0 | 0 | 0 | 1 | 0 |
| SG0000005 | Commd7 | 1.41E-16 | 7.35E-16 | -1.536717404 | 831.4403467 | 286.5715429 | 63 | 17 | 19 | 22 | 44 | 2 | 5 | 1 | 1 | 2 | 4 | 0 |
| JSG000000 | Cradd | 2.4E-16 | 1.23E-15 | 1.771904907 | 93.35686944 | 318.8179804 | 231 | 16 | 19 | 44 | 113 | 28 | 1 | 0 | 0 | 0 | 1 | 0 |
| SG0000000 | Nr2c2 | 1.46E-15 | 7.15E-15 | -1.264590905 | 2253.475411 | 937.9356193 | 309 | 8 | 19 | 40 | 186 | 46 | 8 | 0 | 0 | 1 | 2 | 4 |
| SG0000002 | Aifm2 | 1.42E-14 | 6.52E-14 | -1.388818079 | 899.4977684 | 343.4979819 | 95 | 11 | 19 | 19 | 74 | 8 | 0 | 0 | 0 | 0 | 0 | 0 |
| JSG000000 | Cenpj | 9.86E-14 | 4.29E-13 | 1.163603075 | 344.1058582 | 770.8528719 | 126 | 17 | 19 | 20 | 43 | 13 | 1 | 0 | 0 | 0 | 0 | 0 |
| JSG000000 | Cep126 | 2.12E-13 | 9.02E-13 | 1.621264994 | 85.15233578 | 261.9666218 | 132 | 11 | 19 | 29 | 78 | 19 | 4 | 0 | 0 | 1 | 2 | 1 |
| JSG000000 | Ndufs4 | 3.75E-13 | 1.57E-12 | 0.986897472 | 886.8666744 | 1757.697267 | 219 | 12 | 19 | 35 | 108 | 24 | 3 | 0 | 0 | 0 | 0 | 0 |
| JSG000000 | Wwc1 | 1.13E-12 | 4.57E-12 | 1.200531925 | 469.3602023 | 1078.704232 | 169 | 13 | 19 | 33 | 94 | 15 | 4 | 0 | 1 | 1 | 2 | 0 |
| SG0000002 | Tmem59 | 9.52E-12 | 3.65E-11 | -0.976605979 | 5536.64972 | 2813.580562 | 174 | 15 | 19 | 36 | 79 | 9 | 4 | 0 | 1 | 0 | 1 | 0 |
| SG0000002 | Nif3l1 | 8.31E-11 | 2.97E-10 | 0.909434015 | 874.062269 | 1641.758566 | 123 | 17 | 19 | 34 | 81 | 9 | 8 | 0 | 0 | 2 | 2 | 0 |
| JSG000000 | Cyp39a1 | 2.04E-10 | 7.09E-10 | 2.744130899 | 33.7438153 | 226.0788305 | 164 | 6 | 19 | 23 | 83 | 10 | 0 | 0 | 0 | 0 | 1 | 1 |
| SG0000004 | Git2 | 3.36E-10 | 1.15E-09 | -0.949993448 | 2175.28459 | 1126.003031 | 237 | 13 | 19 | 26 | 107 | 7 | 26 | 0 | 6 | 0 | 13 | 0 |
| SG0000002 | Gtf2h2 | 2.63E-09 | 8.36E-09 | 0.872247567 | 588.6458875 | 1077.523612 | 235 | 16 | 19 | 43 | 129 | 31 | 5 | 0 | 0 | 0 | 1 | 2 |
| SG0000004 | Ccdc64 | 4.74E-09 | 1.48E-08 | -1.612108072 | 1187.252507 | 388.3740549 | 150 | 15 | 19 | 26 | 65 | 21 | 1 | 0 | 0 | 0 | 0 | 6 |
| SG0000002 | Recql5 | 6.62E-09 | 2.04E-08 | -0.937022924 | 1215.116147 | 634.6668071 | 104 | 12 | 19 | 15 | 44 | 16 | 2 | 0 | 0 | 0 | 0 | 0 |
| JSG000000 | Clic4 | 1.58E-08 | 4.73E-08 | 0.699592037 | 4079.248135 | 6624.884505 | 103 | 15 | 19 | 20 | 58 | 8 | 8 | 0 | 0 | 0 | 1 | 1 |
| JSG000000 | Polr3b | 3.86E-08 | 1.12E-07 | 0.721999439 | 1425.961777 | 2352.076119 | 177 | 10 | 19 | 27 | 78 | 14 | 0 | 0 | 0 | 0 | 1 | 0 |
| SG0000004 | Ctdspl | 3.87E-08 | 1.12E-07 | -1.091067561 | 496.2966111 | 232.9685231 | 129 | 13 | 19 | 22 | 55 | 9 | 1 | 0 | 0 | 0 | 1 | 1 |
| SG0000005 | Map6 | 7.27E-08 | 2.06E-07 | -0.987561802 | 860.6677024 | 434.0600147 | 327 | 12 | 19 | 24 | 118 | 10 | 4 | 0 | 0 | 0 | 0 | 0 |
| SG0000002 | Rrbp1 | 8.13E-08 | 2.3E-07 | 0.657377358 | 5257.011699 | 8291.426338 | 134 | 10 | 19 | 20 | 54 | 11 | 1 | 0 | 1 | 0 | 2 | 0 |
| SG0000002 | Itgb1 | 8.17E-08 | 2.31E-07 | -0.756507464 | 15961.33662 | 9447.955042 | 250 | 15 | 19 | 35 | 117 | 24 | 5 | 1 | 1 | 1 | 3 | 1 |
| SG0000000 | Fap | 1.6E-07 | 4.43E-07 | -4.103404254 | 33.97570878 | 1.976608965 | 227 | 8 | 19 | 34 | 109 | 9 | 2 | 0 | 0 | 0 | 0 | 0 |
| JSG000000 | Rybp | 1.7E-07 | 4.67E-07 | 0.708811881 | 1054.778851 | 1723.991224 | 155 | 15 | 19 | 32 | 82 | 7 | 14 | 1 | 1 | 4 | 4 | 0 |
| JSG000000 | Katnbl1 | 5.92E-07 | 1.56E-06 | 0.734988939 | 625.0605784 | 1040.341269 | 114 | 15 | 19 | 30 | 61 | 3 | 3 | 0 | 0 | 0 | 2 | 1 |
| JSG000000 | Heatr5a | 1.32E-06 | 3.36E-06 | -0.690553993 | 3290.554317 | 2038.879684 | 182 | 16 | 19 | 29 | 81 | 18 | 5 | 0 | 0 | 0 | 2 | 1 |
| SG0000002 | Cops7b | 7.16E-06 | 1.71E-05 | 0.648445725 | 760.8978626 | 1192.691098 | 117 | 14 | 19 | 35 | 55 | 8 | 8 | 0 | 0 | 0 | 0 | 0 |
| SG0000003 | Fgd3 | 1.09E-05 | 2.56E-05 | -2.015796829 | 112.4604012 | 27.80893249 | 105 | 16 | 19 | 28 | 52 | 16 | 1 | 0 | 0 | 0 | 0 | 0 |
| JSG000000 | Fgd6 | 1.1E-05 | 2.59E-05 | -0.758005055 | 803.4604869 | 475.0969749 | 182 | 12 | 19 | 33 | 94 | 16 | 8 | 1 | 2 | 2 | 4 | 1 |
| SG0000000 | Mkl2 | 1.28E-05 | 2.99E-05 | -0.696997773 | 1174.962138 | 724.7807254 | 622 | 10 | 19 | 71 | 267 | 38 | 6 | 0 | 0 | 0 | 1 | 5 |
| SG0000002 | Rint1 | 2.26E-05 | 5.14E-05 | -0.732419576 | 719.8748789 | 433.2880951 | 126 | 9 | 19 | 23 | 109 | 11 | 4 | 1 | 1 | 2 | 4 | 0 |
| SG0000004 | Myrip | 4.21E-05 | 9.32E-05 | 3.987635068 | 2.211622639 | 35.08397509 | 181 | 12 | 19 | 32 | 92 | 26 | 6 | 0 | 0 | 1 | 1 | 0 |
| SG0000002 | Kntc1 | 0.00011 | 0.000233 | 0.49288327 | 2338.919448 | 3291.454937 | 140 | 16 | 19 | 23 | 77 | 8 | 1 | 1 | 1 | 1 | 1 | 0 |
| SG0000001 | Ahi1 | 0.000161 | 0.000333 | 0.599000142 | 445.9659338 | 675.4896445 | 368 | 8 | 19 | 39 | 172 | 43 | 14 | 0 | 0 | 4 | 6 | 0 |
| SG0000002 | Oxnad1 | 0.0004 | 0.00079 | 0.503503011 | 898.9291273 | 1274.368306 | 99 | 9 | 19 | 24 | 59 | 9 | 5 | 0 | 0 | 0 | 0 | 0 |
| JSG000000 | Itga2 | 0.000411 | 0.00081 | -2.046931526 | 42.43472787 | 10.26912956 | 170 | 9 | 19 | 30 | 94 | 25 | 2 | 0 | 0 | 1 | 3 | 1 |
| SG0000003 | Spg20 | 0.000773 | 0.001472 | -0.522415846 | 1238.322835 | 862.126594 | 199 | 12 | 19 | 23 | 83 | 4 | 8 | 0 | 0 | 0 | 3 | 0 |
| SG0000006 | Myo1h | 0.001458 | 0.002686 | 1.086547081 | 35.54068739 | 75.4760366 | 133 | 12 | 19 | 25 | 69 | 41 | 6 | 0 | 0 | 1 | 1 | 0 |
| SG0000003 | Fnip1 | 0.001499 | 0.002761 | 0.455667775 | 1312.200529 | 1799.574691 | 233 | 9 | 19 | 24 | 112 | 27 | 13 | 0 | 0 | 0 | 3 | 5 |
| SG0000007 | Rec114 | 0.002092 | 0.003776 | 4.968873266 | 1.7123759 | 53.62644463 | 171 | 11 | 19 | 22 | 78 | 15 | 0 | 0 | 0 | 0 | 2 | 0 |
| SG0000004 | Exog | 0.004702 | 0.00808 | 0.532467155 | 237.4569012 | 343.4577983 | 69 | 15 | 19 | 26 | 54 | 0 | 5 | 3 | 3 | 4 | 4 | 0 |
| SG0000003 | Nexn | 0.007923 | 0.013165 | 0.719272037 | 75.90265937 | 124.9624267 | 274 | 15 | 19 | 43 | 122 | 32 | 8 | 0 | 0 | 2 | 4 | 2 |
| SG0000002 | Slc28a3 | 0.009943 | 0.016269 | -1.75831191 | 54.37198202 | 16.07202273 | 150 | 11 | 19 | 23 | 53 | 19 | 9 | 0 | 0 | 1 | 2 | 3 |
| SG0000002 | Ece2 | 0.012762 | 0.020465 | 0.493977649 | 278.8250444 | 392.6755579 | 72 | 15 | 19 | 19 | 37 | 4 | 2 | 0 | 0 | 0 | 0 | 0 |
| JSG000000 | Usp47 | 0.015455 | 0.024445 | -0.345879445 | 3170.412294 | 2494.569772 | 202 | 7 | 19 | 33 | 103 | 17 | 2 | 0 | 0 | 0 | 1 | 1 |
| SG0000000 | Ick | 0.04605 | 0.066951 | 0.342691972 | 372.3784916 | 472.220842 | 229 | 7 | 19 | 58 | 178 | 23 | 7 | 0 | 0 | 1 | 10 | 4 |
| JSG000000 | Sh3tc2 | 0.04907 | 0.070969 | -0.669582568 | 79.54339356 | 50.00802208 | 102 | 14 | 19 | 22 | 54 | 6 | 1 | 0 | 0 | 0 | 1 | 1 |
| SG0000002 | Zfp523 | 0.049599 | 0.071641 | -0.510417443 | 513.872645 | 360.7485041 | 57 | 17 | 19 | 24 | 38 | 12 | 3 | 0 | 0 | 0 | 3 | 0 |
| SG0000002 | Kif5c | 0.050327 | 0.072605 | -0.320125788 | 897.9752367 | 719.2781966 | 217 | 12 | 19 | 37 | 120 | 23 | 7 | 0 | 0 | 1 | 1 | 0 |
| SG0000007 | Csnk2a1 | 0.059659 | 0.08483 | 0.233040612 | 5015.097974 | 5894.291857 | 137 | 12 | 19 | 24 | 72 | 6 | 9 | 0 | 0 | 2 | 2 | 2 |
| JSG000000 | Fkbp9 | 0.097908 | 0.133526 | -0.22195556 | 4941.749093 | 4237.067767 | 102 | 18 | 19 | 32 | 57 | 7 | 0 | 0 | 0 | 0 | 0 | 1 |
| JSG000000 | Lars2 | 0.11035 | 0.148817 | 0.20185725 | 17662.10469 | 20314.56569 | 162 | 9 | 19 | 31 | 68 | 10 | 0 | 0 | 0 | 0 | 0 | 0 |
| SG0000002 | Dhx15 | 0.188954 | 0.242272 | 0.159789027 | 8559.000622 | 9561.462986 | 371 | 2 | 19 | 38 | 159 | 22 | 6 | 0 | 0 | 2 | 5 | 0 |
| JSG000000 | Grem2 | 0.208162 | 0.264196 | -0.732500477 | 554.1763309 | 333.5364927 | 181 | 14 | 19 | 35 | 87 | 17 | 5 | 0 | 0 | 0 | 3 | 0 |
| SG0000003 | Eri3 | 0.210409 | 0.266669 | 0.18645635 | 847.0942571 | 963.9637261 | 168 | 8 | 19 | 22 | 63 | 13 | 0 | 0 | 0 | 0 | 0 | 0 |
| SG0000003 | Polg | 0.215708 | 0.272749 | -0.172939071 | 1767.328945 | 1567.680451 | 74 | 9 | 19 | 27 | 41 | 0 | 3 | 0 | 0 | 0 | 4 | 0 |
| JSG000000 | Eps15l1 | 0.298144 | 0.362944 | -0.151349909 | 2146.346765 | 1932.586875 | 91 | 17 | 19 | 32 | 69 | 12 | 0 | 0 | 0 | 0 | 0 | 0 |
| SG0000003 | Trub2 | 0.397726 | 0.466601 | -0.133444515 | 754.8922369 | 688.1991907 | 78 | 18 | 19 | 25 | 68 | 5 | 10 | 0 | 0 | 0 | 0 | 1 |
| SG0000005 | Prdm2 | 0.400867 | 0.469824 | -0.122971806 | 1550.552766 | 1423.863469 | 181 | 7 | 19 | 27 | 70 | 11 | 4 | 0 | 1 | 1 | 1 | 1 |
| JSG000000 | Rapgefl1 | 0.562534 | 0.627326 | -0.160739331 | 110.5927536 | 98.93257458 | 38 | 16 | 19 | 20 | 27 | 1 | 0 | 0 | 0 | 0 | 1 | 1 |
| JSG000000 | Phldb3 | 0.742123 | 0.791197 | 0.105426702 | 91.26620873 | 98.18533125 | 36 | 19 | 19 | 23 | 34 | 1 | 1 | 0 | 0 | 1 | 1 | 0 |

| | | | | | | | | | | | | | | | | | | |
|---|---|---|---|---|---|---|---|---|---|---|---|---|---|---|---|---|---|---|
| SG0000003 | Rab3gap1 | 0.807381 | 0.847454 | -0.039367927 | 1607.618666 | 1564.343468 | 179 | 11 | 19 | 24 | 97 | 18 | 2 | 0 | 0 | 1 | 2 | 0 |
| JSG000000 | Mvb12b | 0.860249 | 0.891041 | -0.044454437 | 272.9876769 | 264.7042558 | 208 | 15 | 19 | 32 | 78 | 9 | 2 | 0 | 0 | 1 | 1 | 0 |
| SG0000001 | Zfp318 | 0.898861 | 0.923029 | -0.030701391 | 1462.184831 | 1431.397435 | 146 | 10 | 19 | 22 | 65 | 27 | 1 | 0 | 0 | 0 | 1 | 0 |
| JSG000000 | Rnf17 | 0 | 0 | 11.57201835 | 0.639110347 | 1945.810225 | 286 | 7 | 18 | 32 | 132 | 21 | 1 | 0 | 0 | 0 | 0 | 0 |
| SG0000000 | Pdzd4 | 0 | 0 | 7.131018976 | 18.14135086 | 2542.847025 | 94 | 12 | 18 | 22 | 86 | 2 | 0 | 0 | 0 | 0 | 0 | 0 |
| SG0000001 | Nanog | 0 | 0 | 13.81251681 | 0.674076249 | 9698.194726 | 63 | 18 | 18 | 21 | 39 | 3 | 9 | 3 | 3 | 3 | 9 | 0 |
| SG0000001 | Ehf | 6.3E-260 | 3.2E-257 | -13.2992414 | 16996.82978 | 1.686154412 | 149 | 6 | 18 | 19 | 63 | 12 | 14 | 0 | 0 | 3 | 6 | 0 |
| SG0000003 | Gga2 | 2.8E-189 | 7.7E-187 | 3.658769201 | 586.8652419 | 7412.038479 | 90 | 12 | 18 | 26 | 60 | 12 | 6 | 0 | 0 | 0 | 0 | 0 |
| SG0000002 | Tnfrsf8 | 9.6E-124 | 1E-121 | -6.795344735 | 2681.568989 | 24.14273005 | 108 | 16 | 18 | 32 | 56 | 8 | 0 | 0 | 0 | 0 | 0 | 0 |
| SG0000003 | Il34 | 9.5E-123 | 1E-120 | -7.379381887 | 3009.436847 | 18.07465168 | 132 | 14 | 18 | 34 | 110 | 10 | 2 | 0 | 0 | 0 | 0 | 0 |
| SG0000006 | Aox3 | 7.9E-119 | 7.7E-117 | -7.29466374 | 1323.805645 | 8.431631826 | 165 | 13 | 18 | 31 | 103 | 22 | 2 | 0 | 0 | 0 | 1 | 0 |
| JSG000000 | Crtac1 | 5.24E-73 | 2.06E-71 | 9.652675071 | 0.307509278 | 247.5158179 | 179 | 15 | 18 | 35 | 76 | 13 | 0 | 0 | 0 | 0 | 0 | 0 |
| JSG000000 | Smc1b | 3.84E-70 | 1.36E-68 | 9.59469164 | 0.307509278 | 237.765154 | 155 | 7 | 18 | 25 | 110 | 19 | 0 | 0 | 0 | 0 | 0 | 0 |
| SG0000000 | Mgst1 | 3.21E-67 | 1.08E-65 | -5.479626778 | 3451.732806 | 77.3579843 | 166 | 0 | 18 | 24 | 66 | 18 | 4 | 0 | 0 | 0 | 0 | 0 |
| SG0000000 | Syce2 | 5.29E-64 | 1.63E-62 | 4.984649056 | 94.4071932 | 2989.055436 | 121 | 18 | 18 | 18 | 45 | 9 | 2 | 0 | 0 | 0 | 0 | 0 |
| SG0000002 | Lcp1 | 1E-62 | 2.99E-61 | 4.732273163 | 12.41813667 | 330.0749794 | 398 | 2 | 18 | 47 | 168 | 46 | 0 | 0 | 0 | 0 | 4 | 0 |
| JSG000000 | Golph3 | 1.55E-62 | 4.59E-61 | -2.854984528 | 21615.18942 | 2987.604229 | 84 | 15 | 18 | 22 | 50 | 5 | 1 | 0 | 0 | 0 | 2 | 2 |
| SG0000005 | Hmga2 | 1.08E-57 | 2.76E-56 | -3.018136752 | 6854.638214 | 846.1256185 | 420 | 6 | 18 | 56 | 204 | 48 | 6 | 0 | 0 | 0 | 0 | 0 |
| SG0000003 | Syde2 | 5.99E-55 | 1.44E-53 | 3.234830657 | 70.86505165 | 667.1341411 | 128 | 12 | 18 | 32 | 85 | 2 | 8 | 0 | 2 | 6 | 18 | 0 |
| SG0000002 | Gclm | 7.39E-53 | 1.67E-51 | -2.565094996 | 11507.67087 | 1944.540069 | 140 | 18 | 18 | 31 | 89 | 27 | 7 | 0 | 0 | 1 | 3 | 6 |
| SG0000004 | Kirrel | 2.77E-52 | 6.17E-51 | -2.613644492 | 6056.473158 | 989.542398 | 309 | 15 | 18 | 36 | 122 | 26 | 9 | 0 | 0 | 0 | 2 | 0 |
| SG0000000 | Elk3 | 8.75E-51 | 1.86E-49 | -2.923441006 | 3780.326627 | 498.2942686 | 275 | 15 | 18 | 54 | 117 | 25 | 11 | 0 | 0 | 0 | 0 | 1 |
| SG0000002 | Tes | 1.25E-47 | 2.42E-46 | -2.519923176 | 6446.897263 | 1124.030994 | 172 | 14 | 18 | 32 | 88 | 16 | 10 | 0 | 0 | 0 | 2 | 0 |
| SG0000002 | Klhl12 | 4.75E-44 | 8.05E-43 | 1.79148039 | 850.0065039 | 2942.466878 | 198 | 12 | 18 | 39 | 96 | 9 | 3 | 0 | 0 | 3 | 3 | 0 |
| SG0000003 | Zdhhc9 | 4.82E-44 | 8.14E-43 | -2.482652832 | 2727.204825 | 487.9381611 | 166 | 10 | 18 | 24 | 74 | 6 | 2 | 0 | 0 | 2 | 4 | 0 |
| SG0000007 | H2-Bl | 1.49E-42 | 2.39E-41 | 4.993479228 | 5.487980287 | 174.8234043 | 36 | 18 | 18 | 18 | 36 | 0 | 0 | 0 | 0 | 0 | 0 | 0 |
| SG0000003 | Ttll6 | 2.44E-42 | 3.89E-41 | 3.048221906 | 43.26438211 | 357.8794264 | 44 | 16 | 18 | 24 | 36 | 10 | 0 | 0 | 0 | 0 | 0 | 0 |
| SG0000000 | Trpv1 | 9.33E-41 | 1.42E-39 | -9.01422746 | 163.224468 | 0.315669345 | 89 | 16 | 18 | 28 | 48 | 4 | 0 | 0 | 0 | 0 | 0 | 0 |
| JSG000000 | Pcolce2 | 1.28E-40 | 1.94E-39 | -2.969233142 | 2124.601801 | 271.2996949 | 105 | 14 | 18 | 17 | 46 | 7 | 5 | 0 | 0 | 1 | 2 | 1 |
| JSG000000 | Plekha3 | 1.23E-39 | 1.79E-38 | -2.288066664 | 2968.193169 | 607.7367 | 55 | 14 | 18 | 21 | 32 | 6 | 8 | 2 | 3 | 3 | 4 | 1 |
| SG0000002 | Cnot6 | 3.97E-37 | 5.29E-36 | -2.033787471 | 8009.497433 | 1956.024242 | 174 | 14 | 18 | 24 | 87 | 12 | 16 | 0 | 0 | 2 | 3 | 0 |
| SG0000002 | Prepl | 1.86E-34 | 2.27E-33 | -2.581432668 | 6280.272595 | 1049.276219 | 102 | 12 | 18 | 22 | 68 | 8 | 2 | 0 | 0 | 0 | 1 | 0 |
| SG0000003 | Igfbp7 | 2.82E-34 | 3.41E-33 | -2.117005758 | 2352.959214 | 542.415592 | 176 | 6 | 18 | 22 | 86 | 12 | 4 | 0 | 0 | 0 | 2 | 0 |
| SG0000003 | Stag3 | 3.39E-34 | 4.08E-33 | 7.656524507 | 4.306126756 | 868.8210787 | 158 | 14 | 18 | 28 | 70 | 9 | 0 | 0 | 0 | 0 | 0 | 0 |
| SG0000005 | Nup160 | 5.67E-34 | 6.75E-33 | 1.485436205 | 1855.03794 | 5194.139993 | 141 | 10 | 18 | 20 | 50 | 13 | 4 | 1 | 1 | 1 | 1 | 0 |
| SG0000002 | Fastkd1 | 7.83E-34 | 9.25E-33 | 1.823688888 | 299.9271725 | 1061.697687 | 98 | 14 | 18 | 28 | 100 | 10 | 2 | 0 | 0 | 0 | 0 | 0 |
| SG0000003 | Ramp1 | 2.46E-31 | 2.63E-30 | -5.292115725 | 160.3452624 | 4.092329166 | 94 | 18 | 18 | 25 | 49 | 6 | 1 | 0 | 0 | 0 | 0 | 0 |
| SG0000002 | Dip2a | 1.9E-30 | 1.96E-29 | -3.597852393 | 5893.54196 | 486.7600082 | 372 | 16 | 18 | 49 | 150 | 46 | 10 | 3 | 3 | 3 | 10 | 0 |
| JSG000000 | Eed | 6.31E-29 | 6.07E-28 | 1.372079503 | 1944.609705 | 5033.493772 | 84 | 12 | 18 | 16 | 34 | 5 | 2 | 0 | 0 | 0 | 1 | 0 |
| SG0000002 | Sall4 | 3.56E-28 | 3.31E-27 | 3.493104404 | 477.9960671 | 5382.121863 | 102 | 18 | 18 | 21 | 45 | 12 | 3 | 0 | 0 | 0 | 0 | 0 |
| SG0000004 | Atp10d | 1.47E-27 | 1.33E-26 | -1.981382311 | 1570.071853 | 397.6161533 | 223 | 15 | 18 | 24 | 95 | 21 | 9 | 0 | 2 | 2 | 2 | 0 |
| SG0000004 | Rnpep | 3.56E-25 | 2.9E-24 | 1.419048813 | 643.4839565 | 1720.734956 | 52 | 12 | 18 | 18 | 30 | 4 | 0 | 0 | 0 | 0 | 0 | 0 |
| SG0000004 | Stx3 | 7.58E-25 | 6.08E-24 | 3.308031617 | 237.6647465 | 2353.864775 | 188 | 12 | 18 | 27 | 113 | 9 | 7 | 0 | 0 | 0 | 1 | 1 |
| SG0000004 | Liph | 8.58E-25 | 6.86E-24 | -1.675090089 | 2621.672149 | 820.9676007 | 164 | 16 | 18 | 28 | 92 | 8 | 4 | 0 | 0 | 0 | 0 | 0 |
| SG0000002 | 43531 | 1.93E-24 | 1.51E-23 | 1.261107071 | 1943.736767 | 4658.740347 | 295 | 7 | 18 | 38 | 94 | 29 | 8 | 0 | 0 | 1 | 1 | 1 |
| SG0000003 | Snrpa1 | 1.32E-23 | 9.98E-23 | 1.275181039 | 1285.788853 | 3111.984314 | 72 | 18 | 18 | 24 | 38 | 6 | 0 | 0 | 0 | 0 | 0 | 0 |
| SG0000000 | Bcl2l1 | 6.02E-23 | 4.41E-22 | -1.817363811 | 3077.817535 | 873.2967608 | 121 | 14 | 18 | 28 | 78 | 10 | 3 | 0 | 0 | 2 | 2 | 2 |
| SG0000001 | Rnf145 | 4.14E-22 | 2.91E-21 | -1.438110221 | 9012.914705 | 3326.219823 | 168 | 13 | 18 | 31 | 70 | 12 | 9 | 1 | 2 | 3 | 4 | 0 |
| SG0000001 | Dcbld1 | 1.1E-21 | 7.57E-21 | 3.677859591 | 29.66474175 | 379.6528188 | 170 | 12 | 18 | 29 | 96 | 14 | 3 | 0 | 0 | 0 | 1 | 0 |
| JSG000000 | Slc35f2 | 1.37E-21 | 9.39E-21 | 1.292715359 | 738.3618051 | 1808.903178 | 91 | 15 | 18 | 21 | 57 | 7 | 1 | 0 | 0 | 0 | 1 | 0 |
| SG0000003 | Myo7a | 3.51E-21 | 2.35E-20 | 3.799142242 | 16.8607866 | 234.7110868 | 188 | 16 | 18 | 36 | 84 | 25 | 0 | 0 | 0 | 0 | 0 | 1 |
| SG0000000 | Dbt | 6.62E-21 | 4.38E-20 | 1.356103618 | 473.8691192 | 1213.071149 | 135 | 14 | 18 | 23 | 65 | 8 | 4 | 0 | 0 | 0 | 3 | 0 |
| JSG000000 | Hacd2 | 1.15E-20 | 7.51E-20 | -1.529504736 | 2087.478034 | 723.0945868 | 146 | 9 | 18 | 24 | 67 | 14 | 4 | 0 | 0 | 0 | 0 | 0 |
| SG0000002 | Ly75 | 3.07E-20 | 1.95E-19 | 2.08876465 | 211.9751775 | 901.7077887 | 186 | 9 | 18 | 29 | 93 | 23 | 5 | 0 | 0 | 1 | 2 | 1 |
| SG0000003 | Ccdc68 | 1E-19 | 6.18E-19 | -3.103685333 | 600.5339177 | 69.86105803 | 146 | 6 | 18 | 22 | 138 | 16 | 6 | 0 | 0 | 2 | 2 | 2 |
| JSG000000 | Chrna7 | 3.72E-19 | 2.22E-18 | 6.496811671 | 0.639110347 | 57.7179686 | 260 | 10 | 18 | 32 | 100 | 12 | 2 | 0 | 0 | 0 | 0 | 0 |
| SG0000003 | Kidins220 | 5.57E-19 | 3.3E-18 | -1.506955421 | 3940.148984 | 1386.353106 | 205 | 11 | 18 | 25 | 95 | 13 | 6 | 0 | 0 | 0 | 1 | 0 |
| JSG000000 | Ammecr1 | 2.11E-18 | 1.21E-17 | -1.377907401 | 2798.341015 | 1076.735865 | 304 | 8 | 18 | 42 | 166 | 25 | 8 | 0 | 0 | 0 | 2 | 0 |
| SG0000002 | Lpgat1 | 4.41E-17 | 2.37E-16 | -1.451146939 | 4088.985722 | 1495.467012 | 285 | 4 | 18 | 35 | 112 | 45 | 17 | 0 | 0 | 1 | 2 | 1 |
| SG0000004 | Appl1 | 2.05E-15 | 9.95E-15 | 1.054258328 | 2027.543001 | 4210.497884 | 139 | 12 | 18 | 25 | 64 | 12 | 14 | 0 | 1 | 0 | 1 | 2 |
| SG0000004 | Fhdc1 | 3.23E-15 | 1.55E-14 | -1.953924371 | 466.3017457 | 120.3586191 | 141 | 12 | 18 | 18 | 56 | 12 | 0 | 0 | 0 | 0 | 2 | 0 |
| SG0000006 | Tmed5 | 6.89E-15 | 3.23E-14 | -1.350270052 | 2411.609315 | 945.8780879 | 161 | 12 | 18 | 22 | 86 | 25 | 30 | 2 | 2 | 4 | 17 | 2 |
| SG0000002 | Canx | 7.22E-15 | 3.39E-14 | -1.087990963 | 32053.18396 | 15078.3306 | 143 | 16 | 18 | 29 | 67 | 22 | 13 | 0 | 0 | 3 | 5 | 0 |
| JSG000000 | Dcaf5 | 1.1E-14 | 5.07E-14 | -1.269965068 | 1571.658104 | 651.7191726 | 200 | 14 | 18 | 31 | 85 | 22 | 4 | 0 | 0 | 0 | 0 | 1 |
| SG0000004 | Agr | 1.2E-14 | 5.53E-14 | 0.950595307 | 2776.778777 | 5366.596986 | 289 | 4 | 18 | 36 | 101 | 24 | 1 | 0 | 0 | 0 | 0 | 1 |
| JSG000000 | Skiv2l2 | 2.08E-14 | 9.48E-14 | 0.951856645 | 2331.012337 | 4509.018048 | 169 | 9 | 18 | 35 | 94 | 9 | 1 | 0 | 0 | 0 | 1 | 1 |
| SG0000003 | Tmem30a | 5.79E-14 | 2.56E-13 | -1.15253248 | 4454.264854 | 2003.683812 | 150 | 6 | 18 | 30 | 84 | 8 | 10 | 2 | 2 | 4 | 8 | 2 |
| SG0000006 | Grm4 | 1.24E-13 | 5.36E-13 | 2.739538177 | 15.45457327 | 103.2143742 | 111 | 12 | 18 | 20 | 45 | 6 | 0 | 0 | 0 | 0 | 0 | 0 |
| SG0000004 | Sertad2 | 1.36E-13 | 5.86E-13 | -1.161638833 | 2049.914943 | 916.3211445 | 245 | 7 | 18 | 25 | 90 | 22 | 26 | 2 | 4 | 10 | 20 | 2 |
| SG0000002 | Tpm3 | 2.22E-13 | 9.43E-13 | 0.896163143 | 3462.045446 | 6443.246577 | 136 | 18 | 18 | 36 | 84 | 12 | 8 | 0 | 0 | 0 | 6 | 0 |
| JSG000000 | Adora1 | 2.81E-13 | 1.18E-12 | -2.492766763 | 163.2004858 | 28.99505116 | 117 | 18 | 18 | 36 | 69 | 18 | 5 | 0 | 0 | 0 | 0 | 0 |
| JSG000000 | Parp4 | 4.39E-13 | 1.83E-12 | -1.473323801 | 1481.870606 | 533.6980624 | 135 | 13 | 18 | 30 | 77 | 10 | 1 | 0 | 0 | 0 | 0 | 0 |
| SG0000002 | Stat5b | 7.89E-13 | 3.24E-12 | -1.228587075 | 1108.573908 | 473.0674758 | 170 | 13 | 18 | 31 | 91 | 29 | 6 | 2 | 2 | 2 | 6 | 0 |
| SG0000001 | Timm23 | 7.9E-13 | 3.24E-12 | 1.070138877 | 464.0699407 | 974.3777091 | 160 | 12 | 18 | 18 | 63 | 9 | 1 | 0 | 0 | 0 | 0 | 0 |
| SG0000006 | Tdrd5 | 8.91E-13 | 3.64E-12 | 7.433376622 | 3.392129466 | 586.3295086 | 200 | 14 | 18 | 38 | 126 | 26 | 2 | 0 | 0 | 0 | 0 | 2 |
| SG0000002 | Pxn | 5.26E-12 | 2.05E-11 | -1.326594051 | 3631.310116 | 1447.83389 | 90 | 18 | 18 | 24 | 32 | 2 | 2 | 0 | 0 | 0 | 0 | 0 |
| SG0000004 | Slc7a1 | 5.97E-12 | 2.31E-11 | -1.020481756 | 3394.296062 | 1673.224063 | 122 | 15 | 18 | 23 | 64 | 23 | 6 | 0 | 0 | 0 | 6 | 4 |
| SG0000000 | Jup | 8.89E-12 | 3.41E-11 | -1.508348695 | 6584.875537 | 2314.671528 | 67 | 18 | 18 | 28 | 62 | 2 | 0 | 0 | 0 | 0 | 1 | 0 |
| SG0000001 | Zwint | 3.88E-11 | 1.42E-10 | -0.93323219 | 6408.36563 | 3355.957099 | 104 | 9 | 18 | 12 | 57 | 25 | 14 | 0 | 0 | 0 | 1 | 1 |
| SG0000003 | Aspm | 4.1E-11 | 1.5E-10 | 0.842289853 | 1801.090288 | 3229.163206 | 198 | 12 | 18 | 32 | 110 | 12 | 4 | 0 | 0 | 0 | 0 | 4 |
| SG0000000 | Lpcat3 | 7.24E-11 | 2.6E-10 | -0.984420234 | 2600.303372 | 1314.268202 | 93 | 12 | 18 | 18 | 46 | 4 | 0 | 0 | 0 | 0 | 0 | 0 |
| SG0000002 | Dapp1 | 1.11E-10 | 3.93E-10 | -1.259909863 | 870.4485391 | 363.4732383 | 160 | 8 | 18 | 26 | 60 | 10 | 2 | 0 | 0 | 0 | 2 | 0 |
| SG0000002 | Trit1 | 1.74E-10 | 6.09E-10 | 0.993089912 | 400.6440163 | 797.4592735 | 75 | 14 | 18 | 20 | 45 | 6 | 0 | 0 | 0 | 0 | 0 | 1 |
| JSG000000 | Fam63b | 2.46E-10 | 8.51E-10 | -1.352497539 | 835.2318755 | 327.0880909 | 137 | 11 | 18 | 25 | 76 | 14 | 18 | 0 | 1 | 3 | 9 | 3 |
| JSG000000 | Map3k4 | 3.31E-10 | 1.13E-09 | 1.097897568 | 953.6297423 | 2041.173342 | 117 | 13 | 18 | 29 | 47 | 18 | 0 | 0 | 0 | 0 | 0 | 0 |
| JSG000000 | Gosr1 | 5.53E-10 | 1.86E-09 | -0.975141897 | 1632.931135 | 830.6554305 | 88 | 15 | 18 | 19 | 38 | 5 | 6 | 1 | 2 | 1 | 4 | 0 |
| JSG000000 | Fam13b | 6.75E-10 | 2.26E-09 | -1.26464786 | 1553.67848 | 646.6424493 | 158 | 14 | 18 | 31 | 91 | 10 | 0 | 0 | 0 | 0 | 0 | 0 |
| SG0000003 | Ppargc1b | 6.8E-10 | 2.27E-09 | 1.973666395 | 30.63660317 | 120.3298561 | 216 | 6 | 18 | 18 | 74 | 12 | 0 | 0 | 0 | 0 | 0 | 0 |
| JSG000000 | Zdhhc20 | 9.28E-10 | 3.07E-09 | -0.880919857 | 4604.131576 | 2500.14056 | 123 | 9 | 18 | 23 | 55 | 10 | 12 | 0 | 0 | 2 | 6 | 0 |
| JSG000000 | Rnf152 | 1.14E-09 | 3.73E-09 | 2.937859448 | 7.261757447 | 55.64492426 | 230 | 10 | 18 | 28 | 102 | 32 | 48 | 0 | 4 | 4 | 12 | 4 |
| JSG000000 | Casc3 | 1.24E-09 | 4.05E-09 | 0.786779424 | 1582.826068 | 2730.721442 | 99 | 16 | 18 | 19 | 54 | 11 | 4 | 0 | 0 | 0 | 0 | 0 |
| SG0000002 | Slc33a1 | 1.33E-09 | 4.34E-09 | -0.997742848 | 1222.417496 | 612.1657561 | 218 | 11 | 18 | 32 | 91 | 14 | 8 | 0 | 0 | 0 | 0 | 0 |
| SG0000002 | Eln | 1.34E-09 | 4.37E-09 | 4.293433295 | 1.952296942 | 38.28230511 | 85 | 16 | 18 | 22 | 52 | 34 | 2 | 0 | 0 | 0 | 2 | 0 |
| SG0000002 | Notch2 | 2.03E-09 | 6.53E-09 | -0.928890177 | 1773.577405 | 931.5932679 | 179 | 17 | 18 | 27 | 59 | 24 | 4 | 0 | 0 | 0 | 1 | 0 |
| SG0000003 | Reep6 | 2.49E-09 | 7.93E-09 | 1.014600905 | 306.0540755 | 618.3344921 | 30 | 18 | 18 | 24 | 36 | 0 | 0 | 0 | 0 | 0 | 0 | 0 |
| SG0000002 | Fam219a | 7.03E-09 | 2.16E-08 | -1.4470948 | 442.3715172 | 162.2438393 | 159 | 9 | 18 | 21 | 57 | 27 | 6 | 0 | 0 | 0 | 0 | 3 |
| SG0000002 | Fyco1 | 9.89E-09 | 3.01E-08 | -0.935438231 | 1250.610736 | 653.9238631 | 180 | 16 | 18 | 28 | 86 | 24 | 6 | 0 | 0 | 0 | 0 | 2 |
| SG0000002 | E2f7 | 1.19E-08 | 3.6E-08 | 0.870164762 | 468.3834824 | 856.1448837 | 150 | 9 | 18 | 21 | 55 | 22 | 16 | 3 | 5 | 3 | 8 | 2 |
| SG0000006 | Stx17 | 1.5E-08 | 4.5E-08 | -1.196253148 | 699.4606047 | 305.249653 | 355 | 9 | 18 | 40 | 240 | 18 | 8 | 0 | 0 | 2 | 3 | 0 |
| JSG000000 | Rrn3 | 1.78E-08 | 5.3E-08 | 0.697539297 | 4137.957554 | 6710.676187 | 67 | 11 | 18 | 20 | 42 | 10 | 4 | 0 | 1 | 1 | 1 | 0 |
| SG0000003 | Abcd1 | 2.21E-08 | 6.53E-08 | -1.156423581 | 545.0432895 | 244.5191129 | 59 | 14 | 18 | 18 | 39 | 9 | 1 | 0 | 0 | 0 | 0 | 0 |
| JSG000000 | Megf8 | 1.65E-07 | 4.55E-07 | -1.308128507 | 2732.726052 | 1103.596154 | 51 | 15 | 18 | 20 | 29 | 4 | 3 | 0 | 1 | 1 | 1 | 0 |
| SG0000001 | Setdb1 | 2.27E-07 | 6.16E-07 | 0.6508964 | 2668.399028 | 4189.769337 | 132 | 12 | 18 | 27 | 87 | 18 | 3 | 0 | 0 | 0 | 0 | 0 |
| SG0000005 | Slc25a40 | 4.68E-07 | 1.24E-06 | -0.766715576 | 1745.926483 | 1026.175322 | 252 | 9 | 18 | 39 | 126 | 18 | 31 | 1 | 4 | 7 | 16 | 0 |
| SG0000001 | Rbm25 | 7.66E-07 | 2E-06 | 0.605804405 | 5316.45498 | 8090.724964 | 156 | 13 | 18 | 28 | 78 | 23 | 2 | 0 | 0 | 0 | 0 | 4 |
| SG0000003 | Abhd12 | 1.1E-06 | 2.83E-06 | -0.769058783 | 1247.848601 | 732.237783 | 330 | 9 | 18 | 52 | 126 | 17 | 8 | 0 | 0 | 1 | 1 | 0 |
| SG0000000 | Myo9b | 1.23E-06 | 3.16E-06 | -0.691238767 | 3290.537671 | 2037.901853 | 231 | 12 | 18 | 42 | 111 | 30 | 0 | 0 | 0 | 0 | 0 | 0 |
| JSG000000 | Dstn | 1.27E-06 | 3.26E-06 | 0.597927118 | 6009.784456 | 9096.051151 | 71 | 17 | 18 | 21 | 44 | 2 | 0 | 0 | 0 | 0 | 0 | 0 |
| SG0000003 | Ovol2 | 1.58E-06 | 4.01E-06 | 3.390000151 | 13.09590319 | 137.2862956 | 89 | 14 | 18 | 30 | 44 | 16 | 0 | 0 | 0 | 0 | 0 | 0 |
| JSG000000 | Serbp1 | 1.89E-06 | 4.75E-06 | 0.569431272 | 16517.87177 | 24511.50533 | 92 | 11 | 18 | 16 | 30 | 6 | 19 | 0 | 0 | 0 | 3 | 1 |
| JSG000000 | G3bp1 | 2.29E-06 | 5.72E-06 | 0.573112056 | 8466.313982 | 12595.58348 | 77 | 15 | 18 | 26 | 53 | 4 | 3 | 0 | 0 | 1 | 2 | 0 |
| SG0000002 | Glipr2 | 4.29E-06 | 1.05E-05 | 1.8406232 | 299.6822012 | 1073.35589 | 62 | 16 | 18 | 22 | 30 | 4 | 4 | 0 | 0 | 2 | 2 | 0 |
| SG0000003 | Loxl2 | 4.42E-06 | 1.08E-05 | -0.860101475 | 2125.613289 | 1171.029709 | 147 | 8 | 18 | 13 | 66 | 12 | 3 | 0 | 0 | 1 | 2 | 0 |
| SG0000005 | 5730455P16Rik | 5.59E-06 | 1.35E-05 | -0.755528813 | 975.7462343 | 577.9630157 | 66 | 12 | 18 | 18 | 38 | 6 | 12 | 0 | 2 | 4 | 4 | 4 |
| JSG000000 | Wdr89 | 5.94E-06 | 1.43E-05 | 0.914957197 | 648.3522057 | 1222.476597 | 153 | 14 | 18 | 40 | 85 | 10 | 5 | 0 | 0 | 2 | 3 | 0 |
| SG0000007 | Rab11b | 6.07E-06 | 1.46E-05 | -0.705830039 | 1272.351092 | 780.0652892 | 44 | 15 | 18 | 18 | 37 | 2 | 6 | 0 | 0 | 0 | 1 | 2 |
| SG0000002 | Grhl1 | 1.19E-05 | 2.79E-05 | -1.198677773 | 728.0444424 | 317.1903209 | 107 | 15 | 18 | 27 | 62 | 9 | 4 | 0 | 0 | 0 | 2 | 4 |
| SG0000003 | Nup210 | 1.49E-05 | 3.47E-05 | 0.558559194 | 1956.64623 | 2881.743898 | 269 | 16 | 18 | 39 | 124 | 26 | 0 | 0 | 0 | 0 | 0 | 0 |
| SG0000002 | Gga3 | 3.2E-05 | 7.17E-05 | 0.564053913 | 1271.247905 | 1879.435376 | 69 | 12 | 18 | 24 | 48 | 0 | 4 | 0 | 0 | 0 | 0 | 0 |
| SG0000003 | Metap2 | 0.00018 | 0.00037 | -0.51066549 | 8614.904268 | 6046.789154 | 268 | 18 | 18 | 52 | 144 | 21 | 11 | 0 | 0 | 4 | 4 | 1 |
| SG0000002 | Hps3 | 0.000183 | 0.000377 | -0.616036886 | 901.0696591 | 587.9126233 | 188 | 10 | 18 | 24 | 88 | 6 | 6 | 2 | 2 | 2 | 4 | 2 |
| SG0000001 | Tmem87b | 0.000189 | 0.000388 | -0.574206947 | 1590.43086 | 1068.221413 | 152 | 10 | 18 | 29 | 71 | 13 | 8 | 0 | 0 | 1 | 1 | 1 |
| SG0000002 | Pmf1 | 0.000189 | 0.000389 | 0.730175886 | 605.7223909 | 1004.797328 | 60 | 12 | 18 | 12 | 30 | 3 | 0 | 0 | 0 | 0 | 0 | 0 |

| | | | | | | | | | | | | | | | | | | |
|---|---|---|---|---|---|---|---|---|---|---|---|---|---|---|---|---|---|---|
| SG0000000 | Il11 | 0.000243 | 0.000492 | -1.757766744 | 62.00019687 | 18.33380269 | 25 | 14 | 18 | 16 | 22 | 0 | 14 | 10 | 14 | 12 | 12 | 0 |
| SG0000005 | Kcnmb4 | 0.000306 | 0.000613 | 3.665158299 | 1.893239249 | 24.01751586 | 95 | 12 | 18 | 17 | 48 | 4 | 1 | 0 | 0 | 0 | 0 | 0 |
| SG0000002 | Vamp4 | 0.000625 | 0.001202 | -0.607365631 | 1030.932607 | 676.6981708 | 386 | 0 | 18 | 30 | 154 | 46 | 20 | 0 | 0 | 0 | 4 | 2 |
| SG0000003 | Tmem184a | 0.000631 | 0.001214 | 1.055154855 | 54.15385277 | 112.5285237 | 24 | 14 | 18 | 16 | 20 | 4 | 6 | 0 | 2 | 2 | 2 | 0 |
| JSG0000000 | Gramd3 | 0.00066 | 0.001266 | 0.604562847 | 264.4647508 | 402.1234006 | 117 | 14 | 18 | 26 | 60 | 10 | 2 | 0 | 0 | 0 | 0 | 0 |
| SG0000001 | Mlkl | 0.000851 | 0.00161 | -1.08098223 | 181.3092052 | 85.70610889 | 94 | 14 | 18 | 22 | 46 | 6 | 0 | 0 | 0 | 0 | 0 | 0 |
| SG0000002 | Pigg | 0.000861 | 0.001629 | -0.580795402 | 917.0465213 | 613.1327729 | 206 | 12 | 18 | 21 | 69 | 10 | 8 | 0 | 0 | 0 | 3 | 1 |
| SG0000007 | H60b | 0.000919 | 0.001731 | 1.744978865 | 10.63348539 | 35.64237367 | 209 | 2 | 18 | 26 | 100 | 28 | 0 | 0 | 0 | 0 | 0 | 0 |
| SG0000003 | Coro7 | 0.000948 | 0.001783 | -0.549424365 | 805.3874731 | 550.3153872 | 125 | 18 | 18 | 30 | 60 | 19 | 1 | 0 | 0 | 0 | 0 | 1 |
| SG0000000 | Ccm2 | 0.001263 | 0.002345 | -0.507995296 | 1007.808133 | 708.6895633 | 198 | 8 | 18 | 33 | 134 | 20 | 1 | 0 | 0 | 0 | 0 | 0 |
| JSG0000000 | Smc5 | 0.001272 | 0.00236 | 0.41240308 | 2425.966798 | 3228.721235 | 170 | 11 | 18 | 26 | 81 | 17 | 7 | 1 | 1 | 1 | 4 | 0 |
| SG0000000 | Snap47 | 0.001287 | 0.002386 | -0.454607517 | 2406.839476 | 1756.291612 | 98 | 12 | 18 | 28 | 58 | 10 | 0 | 0 | 0 | 0 | 0 | 0 |
| SG0000002 | Ankle2 | 0.001776 | 0.003239 | 0.412775598 | 1790.058432 | 2383.005496 | 126 | 9 | 18 | 24 | 90 | 15 | 12 | 0 | 0 | 0 | 0 | 0 |
| SG0000002 | Abcb9 | 0.00182 | 0.003313 | -1.393071986 | 51.53601012 | 19.62250005 | 49 | 16 | 18 | 19 | 37 | 5 | 1 | 0 | 0 | 0 | 0 | 0 |
| SG0000002 | Erp44 | 0.002666 | 0.004742 | -0.441585111 | 1815.072041 | 1336.483197 | 211 | 10 | 18 | 36 | 108 | 11 | 5 | 0 | 0 | 1 | 4 | 0 |
| SG0000001 | Glod4 | 0.002852 | 0.005049 | -0.431595951 | 2433.988634 | 1804.659277 | 193 | 14 | 18 | 27 | 65 | 21 | 3 | 0 | 0 | 0 | 3 | 2 |
| SG0000002 | Gne | 0.003409 | 0.005968 | -0.520280686 | 523.1557033 | 364.7630778 | 231 | 17 | 18 | 61 | 145 | 24 | 4 | 0 | 0 | 1 | 4 | 5 |
| SG0000002 | Sorcs2 | 0.003727 | 0.006487 | -0.780238831 | 145.2018887 | 84.54676084 | 451 | 9 | 18 | 42 | 146 | 46 | 2 | 0 | 0 | 0 | 0 | 1 |
| SG0000002 | Hoga1 | 0.003873 | 0.006724 | -1.364388332 | 47.72508999 | 18.53638078 | 71 | 16 | 18 | 18 | 47 | 10 | 1 | 0 | 0 | 0 | 1 | 0 |
| SG0000003 | 6430548M08Rik | 0.003991 | 0.006916 | 1.535687149 | 11.37514988 | 32.97957264 | 116 | 12 | 18 | 19 | 78 | 9 | 3 | 0 | 0 | 0 | 0 | 2 |
| SG0000003 | Ccbl1 | 0.004727 | 0.008119 | 0.628167474 | 155.2213524 | 239.9102012 | 79 | 18 | 18 | 24 | 93 | 0 | 0 | 0 | 0 | 0 | 0 | 0 |
| SG0000005 | Zfp182 | 0.00486 | 0.008332 | -0.686409266 | 342.1779938 | 212.6289024 | 274 | 6 | 18 | 36 | 123 | 20 | 16 | 0 | 0 | 0 | 0 | 2 |
| SG0000002 | Ubap1 | 0.006083 | 0.010285 | -0.389244857 | 2321.541949 | 1772.564952 | 163 | 14 | 18 | 24 | 112 | 20 | 1 | 0 | 0 | 0 | 2 | 0 |
| SG0000002 | Tnnt2 | 0.007394 | 0.012339 | -1.617222013 | 52.43274962 | 17.09111111 | 88 | 8 | 18 | 22 | 40 | 10 | 0 | 0 | 0 | 0 | 0 | 0 |
| SG0000005 | Zmat1 | 0.007622 | 0.012694 | 2.335010671 | 5.230998157 | 26.39324683 | 228 | 12 | 18 | 34 | 112 | 30 | 2 | 0 | 0 | 0 | 0 | 4 |
| JSG0000004 | Cox10 | 0.008878 | 0.014638 | 0.450866592 | 440.8592471 | 602.5933336 | 219 | 12 | 18 | 30 | 82 | 16 | 0 | 0 | 0 | 0 | 0 | 0 |
| SG0000002 | Sema4d | 0.009628 | 0.015786 | 0.354025602 | 1280.991023 | 1637.262865 | 199 | 15 | 18 | 24 | 60 | 27 | 6 | 0 | 0 | 0 | 3 | 0 |
| SG0000003 | Smim19 | 0.01166 | 0.018859 | -0.485732987 | 428.1768918 | 305.7757403 | 58 | 16 | 18 | 22 | 48 | 0 | 2 | 0 | 0 | 0 | 0 | 0 |
| SG0000003 | Golim4 | 0.013849 | 0.022055 | -0.489842061 | 600.7068751 | 427.7651986 | 414 | 9 | 18 | 49 | 174 | 44 | 18 | 0 | 0 | 4 | 9 | 2 |
| SG0000003 | Cep135 | 0.0154 | 0.024368 | -0.410718306 | 746.4316562 | 561.5022744 | 206 | 4 | 18 | 28 | 113 | 26 | 10 | 0 | 2 | 2 | 4 | 0 |
| SG0000002 | Larp4 | 0.016293 | 0.025658 | 0.299109619 | 3738.97703 | 4600.380612 | 217 | 10 | 18 | 24 | 91 | 20 | 17 | 0 | 0 | 1 | 6 | 0 |
| SG0000006 | Fbxl18 | 0.01973 | 0.030558 | -0.356025281 | 1208.158813 | 943.9516223 | 52 | 18 | 18 | 24 | 28 | 4 | 4 | 0 | 0 | 0 | 0 | 0 |
| JSG0000002 | Kif4 | 0.020017 | 0.030973 | 0.315556062 | 1554.196507 | 1934.183921 | 190 | 7 | 18 | 36 | 116 | 26 | 0 | 0 | 0 | 0 | 0 | 0 |
| SG0000003 | Cntrob | 0.021797 | 0.033495 | 0.343086509 | 745.5388879 | 945.6918187 | 112 | 18 | 18 | 30 | 82 | 16 | 3 | 0 | 0 | 0 | 4 | 1 |
| JSG0000002 | Dirc2 | 0.02928 | 0.044029 | -0.382982367 | 604.6078102 | 463.643924 | 290 | 10 | 18 | 32 | 103 | 17 | 8 | 0 | 0 | 0 | 1 | 1 |
| SG0000002 | Nedd9 | 0.03097 | 0.046371 | -0.656555152 | 106.1532909 | 67.34272219 | 337 | 8 | 18 | 42 | 146 | 26 | 3 | 0 | 0 | 0 | 1 | 1 |
| JSG000000 | Arnt2 | 0.032162 | 0.048027 | -0.45191155 | 297.7767635 | 217.6967139 | 204 | 8 | 18 | 33 | 93 | 24 | 3 | 0 | 0 | 1 | 1 | 0 |
| SG0000002 | Crls1 | 0.033283 | 0.049556 | 0.344740719 | 461.6604372 | 586.2730836 | 141 | 6 | 18 | 24 | 59 | 14 | 1 | 0 | 0 | 0 | 1 | 2 |
| JSG0000002 | Sh3rf1 | 0.035201 | 0.052148 | -0.36390534 | 574.6050268 | 446.5015781 | 272 | 15 | 18 | 40 | 139 | 23 | 2 | 0 | 0 | 0 | 2 | 0 |
| SG0000002 | Rad23b | 0.035717 | 0.05286 | 0.25870143 | 8569.760158 | 10252.87253 | 103 | 14 | 18 | 31 | 50 | 7 | 5 | 0 | 0 | 0 | 2 | 0 |
| SG0000003 | Otud4 | 0.036221 | 0.053517 | 0.27930351 | 5509.588775 | 6686.48569 | 190 | 10 | 18 | 30 | 80 | 22 | 16 | 0 | 0 | 2 | 10 | 6 |
| JSG0000002 | Cog3 | 0.039702 | 0.058315 | 0.282782979 | 1303.462867 | 1585.713861 | 138 | 15 | 18 | 24 | 75 | 4 | 2 | 0 | 0 | 0 | 2 | 1 |
| SG0000002 | Tmem47 | 0.050693 | 0.07308 | 0.278167469 | 979.0666005 | 1187.268615 | 186 | 16 | 18 | 31 | 99 | 6 | 12 | 0 | 0 | 1 | 7 | 0 |
| JSG0000000 | Crkl | 0.057105 | 0.08153 | -0.271850015 | 2166.703219 | 1794.586597 | 77 | 15 | 18 | 24 | 39 | 7 | 5 | 0 | 0 | 1 | 1 | 0 |
| JSG0000007 | Top1 | 0.059073 | 0.084077 | 0.2382593 | 4725.63485 | 5574.210395 | 234 | 9 | 18 | 30 | 87 | 23 | 5 | 0 | 2 | 0 | 1 | 0 |
| SG0000003 | Sema3f | 0.077476 | 0.107912 | 0.54294334 | 167.5284479 | 244.0792141 | 71 | 18 | 18 | 23 | 36 | 13 | 3 | 0 | 0 | 2 | 2 | 1 |
| JSG0000000 | Pcnt | 0.093712 | 0.12837 | 0.222959045 | 1694.004139 | 1977.114528 | 101 | 14 | 18 | 18 | 69 | 13 | 1 | 0 | 0 | 0 | 0 | 0 |
| JSG0000002 | B4galt6 | 0.126587 | 0.168444 | 0.201991408 | 1444.114103 | 1661.142852 | 121 | 15 | 18 | 29 | 88 | 8 | 10 | 1 | 1 | 2 | 4 | 1 |
| SG0000002 | Eif3h | 0.148076 | 0.194239 | 0.177984257 | 6047.702816 | 6841.779004 | 203 | 10 | 18 | 28 | 76 | 10 | 0 | 0 | 0 | 0 | 0 | 0 |
| JSG0000004 | Pskh1 | 0.172955 | 0.223777 | 0.198546032 | 1021.236481 | 1171.911002 | 57 | 15 | 18 | 20 | 29 | 1 | 2 | 1 | 1 | 1 | 1 | 0 |
| SG0000003 | Zfc3h1 | 0.173319 | 0.224134 | -0.240025585 | 2211.390654 | 1872.451463 | 153 | 13 | 18 | 25 | 92 | 11 | 2 | 0 | 0 | 0 | 1 | 0 |
| JSG0000007 | Nfxl1 | 0.181694 | 0.233884 | 0.188741077 | 941.5220739 | 1073.117388 | 260 | 10 | 18 | 32 | 138 | 22 | 6 | 0 | 0 | 2 | 2 | 0 |
| JSG000000 | Ppp1r12a | 0.18821 | 0.241352 | 0.190026888 | 1201.859655 | 1371.063438 | 256 | 6 | 18 | 39 | 131 | 35 | 0 | 0 | 0 | 0 | 0 | 0 |
| SG0000003 | Rcor3 | 0.227551 | 0.285948 | -0.234365982 | 363.1170629 | 308.6707422 | 553 | 12 | 18 | 42 | 170 | 44 | 29 | 0 | 0 | 3 | 10 | 6 |
| JSG0000002 | Neto2 | 0.252747 | 0.313736 | -0.28617219 | 178.8001473 | 146.6295356 | 157 | 12 | 18 | 23 | 77 | 15 | 14 | 0 | 1 | 2 | 10 | 1 |
| SG0000004 | Rusc1 | 0.255725 | 0.316971 | -0.254469954 | 229.0238145 | 191.9895342 | 56 | 18 | 18 | 27 | 36 | 0 | 3 | 0 | 0 | 0 | 0 | 0 |
| SG0000002 | Pdcd10 | 0.30079 | 0.365568 | 0.158434775 | 638.8056133 | 712.9553681 | 209 | 8 | 18 | 30 | 105 | 22 | 5 | 0 | 0 | 0 | 1 | 2 |
| SG0000003 | Zdhhc18 | 0.358079 | 0.426174 | -0.128491967 | 2055.184109 | 1880.056051 | 41 | 17 | 18 | 22 | 40 | 0 | 2 | 0 | 0 | 1 | 1 | 0 |
| JSG0000002 | Sfxn2 | 0.361859 | 0.430158 | 0.188837975 | 187.8731881 | 214.1463615 | 44 | 16 | 18 | 20 | 29 | 2 | 3 | 0 | 0 | 0 | 0 | 0 |
| SG0000006 | Serpinb2 | 0.37103 | 0.439485 | -0.481644849 | 74.9742944 | 53.69363923 | 144 | 8 | 18 | 27 | 84 | 22 | 12 | 2 | 2 | 2 | 2 | 0 |
| SG0000002 | Itgb8 | 0.391178 | 0.459823 | 0.645429087 | 9.750610596 | 15.25194842 | 230 | 8 | 18 | 27 | 83 | 17 | 20 | 0 | 0 | 3 | 5 | 2 |
| SG0000004 | Hnrnpf | 0.417955 | 0.487143 | 0.099973938 | 4096.683399 | 4390.637233 | 160 | 18 | 18 | 26 | 73 | 7 | 5 | 0 | 0 | 0 | 0 | 0 |
| SG0000003 | Afg3l1 | 0.44196 | 0.51123 | 0.101628427 | 2433.170956 | 2610.753259 | 72 | 12 | 18 | 26 | 44 | 4 | 2 | 0 | 0 | 0 | 0 | 0 |
| SG0000006 | Hspb11 | 0.445221 | 0.51437 | 0.150832555 | 316.9876806 | 351.9228835 | 136 | 10 | 18 | 30 | 76 | 22 | 2 | 0 | 0 | 2 | 2 | 0 |
| SG0000004 | Slc25a53 | 0.459325 | 0.528721 | 0.168050847 | 125.170136 | 140.6335921 | 264 | 14 | 18 | 24 | 114 | 18 | 18 | 0 | 0 | 0 | 6 | 0 |
| SG0000004 | Slc19a2 | 0.481193 | 0.54969 | 0.096441735 | 2308.654975 | 2468.259903 | 84 | 18 | 18 | 20 | 40 | 14 | 4 | 0 | 0 | 0 | 4 | 0 |
| SG0000005 | Cklf | 0.496276 | 0.563756 | -0.185425633 | 140.2076152 | 123.2971179 | 65 | 12 | 18 | 16 | 26 | 2 | 1 | 0 | 1 | 1 | 1 | 0 |
| JSG0000004 | Garem | 0.499094 | 0.566453 | -0.158545072 | 347.0895772 | 310.9673194 | 312 | 10 | 18 | 49 | 127 | 39 | 2 | 0 | 0 | 1 | 1 | 2 |
| JSG0000002 | Nmt1 | 0.554176 | 0.619432 | 0.072780554 | 4168.584085 | 4384.27408 | 59 | 14 | 18 | 21 | 40 | 6 | 1 | 0 | 0 | 0 | 3 | 0 |
| SG0000004 | Tifa | 0.562839 | 0.627627 | -0.437642641 | 16.62789626 | 12.27704207 | 98 | 16 | 18 | 22 | 34 | 4 | 18 | 0 | 2 | 6 | 6 | 0 |
| JSG0000004 | Oscp1 | 0.598387 | 0.661257 | 0.161463503 | 72.02600068 | 80.55539997 | 92 | 16 | 18 | 19 | 62 | 7 | 4 | 0 | 0 | 0 | 0 | 1 |
| SG0000002 | Tsr2 | 0.600973 | 0.663705 | 0.07737261 | 812.5352242 | 857.3016779 | 57 | 15 | 18 | 21 | 41 | 0 | 9 | 0 | 0 | 0 | 2 | 0 |
| SG0000002 | Vdac1 | 0.690854 | 0.745861 | 0.044538543 | 15665.01846 | 16156.16755 | 79 | 14 | 18 | 24 | 74 | 8 | 1 | 0 | 0 | 0 | 0 | 0 |
| SG0000003 | Eif2ak3 | 0.692256 | 0.747059 | 0.057167498 | 823.2641986 | 856.5414043 | 135 | 16 | 18 | 25 | 58 | 14 | 3 | 0 | 1 | 1 | 2 | 1 |
| SG0000005 | Rcan3 | 0.71824 | 0.770643 | 0.04432759 | 1058.043609 | 1091.057157 | 53 | 16 | 18 | 24 | 39 | 0 | 3 | 0 | 0 | 0 | 1 | 0 |
| JSG0000002 | Rras2 | 0.7205 | 0.772697 | -0.051860355 | 2206.213285 | 2128.31533 | 129 | 13 | 18 | 25 | 69 | 9 | 2 | 0 | 0 | 1 | 2 | 0 |
| SG0000002 | Brd1 | 0.740547 | 0.789959 | 0.04558504 | 2190.32984 | 2260.643001 | 111 | 15 | 18 | 19 | 58 | 15 | 5 | 1 | 2 | 1 | 2 | 1 |
| SG0000002 | Rb1cc1 | 0.758265 | 0.805578 | -0.052562696 | 1569.008408 | 1512.872479 | 369 | 8 | 18 | 37 | 172 | 31 | 15 | 0 | 0 | 0 | 2 | 0 |
| SG0000005 | Fam133b | 0.780989 | 0.825071 | -0.049774422 | 846.8082152 | 818.0907243 | 182 | 12 | 18 | 42 | 96 | 18 | 5 | 0 | 0 | 0 | 0 | 3 |
| JSG0000002 | Exph5 | 0.841287 | 0.876182 | 0.112624125 | 18.76824039 | 20.29209081 | 134 | 14 | 18 | 26 | 66 | 16 | 0 | 0 | 0 | 0 | 0 | 0 |
| SG0000002 | Kdm4b | 0.849438 | 0.882303 | -0.035106438 | 1320.624794 | 1288.87665 | 96 | 16 | 18 | 25 | 51 | 9 | 3 | 0 | 0 | 3 | 3 | 0 |
| JSG0000002 | Dopey2 | 0.86776 | 0.897591 | -0.034456305 | 721.5578858 | 704.5288728 | 91 | 10 | 18 | 18 | 53 | 7 | 0 | 0 | 0 | 0 | 0 | 0 |
| SG0000002 | Samd8 | 0.869287 | 0.898895 | 0.013160406 | 1651.068852 | 1666.198969 | 242 | 15 | 18 | 36 | 134 | 14 | 6 | 0 | 0 | 0 | 2 | 3 |
| JSG000000 | Trmt11 | 0.927811 | 0.947759 | 0.006538301 | 598.7017734 | 601.4212503 | 167 | 13 | 18 | 32 | 87 | 9 | 3 | 0 | 0 | 0 | 0 | 0 |
| SG0000004 | Cnpy1 | 3.6E-175 | 8.2E-173 | 9.11824515 | 1.289094805 | 716.3910576 | 136 | 6 | 17 | 20 | 46 | 6 | 14 | 0 | 3 | 4 | 5 | 0 |
| JSG0000002 | Gca | 4.8E-173 | 1E-170 | 6.614613228 | 8.753463824 | 857.7820077 | 80 | 15 | 17 | 27 | 56 | 8 | 8 | 0 | 1 | 2 | 5 | 0 |
| SG0000004 | Trim16 | 2.08E-95 | 1.33E-93 | -5.093769025 | 1745.379709 | 51.11079723 | 120 | 13 | 17 | 33 | 76 | 3 | 7 | 2 | 2 | 4 | 4 | 0 |
| SG0000000 | Vwf | 1.7E-90 | 9.67E-89 | 7.021959034 | 2.578189609 | 335.0696958 | 262 | 16 | 17 | 34 | 146 | 27 | 2 | 0 | 0 | 0 | 0 | 0 |
| JSG0000000 | Zfp534 | 6.06E-77 | 2.58E-75 | 10.41251119 | 0.331601068 | 451.9538789 | 96 | 16 | 17 | 25 | 49 | 9 | 2 | 0 | 0 | 1 | 2 | 0 |
| SG0000002 | Wdr31 | 3.25E-73 | 1.29E-71 | 3.531661546 | 52.78729248 | 610.4715751 | 105 | 17 | 17 | 24 | 53 | 7 | 1 | 1 | 1 | 1 | 1 | 1 |
| SG0000007 | Naip2 | 4.02E-60 | 1.12E-58 | -8.155613438 | 370.9693446 | 1.300929449 | 300 | 8 | 17 | 40 | 163 | 26 | 0 | 0 | 0 | 0 | 0 | 0 |
| SG0000004 | Pls1 | 9.51E-51 | 2.01E-49 | -2.646164696 | 4067.219219 | 649.714612 | 181 | 10 | 17 | 29 | 84 | 15 | 6 | 0 | 1 | 1 | 2 | 1 |
| SG0000003 | Sesn1 | 2.65E-41 | 4.11E-40 | 2.424710679 | 111.2723607 | 597.445022 | 224 | 10 | 17 | 30 | 89 | 18 | 3 | 0 | 0 | 0 | 0 | 0 |
| JSG0000002 | Slc41a2 | 6.95E-34 | 8.23E-33 | -5.344886031 | 1388.126257 | 34.1552879 | 210 | 11 | 17 | 35 | 108 | 15 | 10 | 0 | 1 | 1 | 6 | 1 |
| SG0000006 | Cpped1 | 1.41E-31 | 1.52E-30 | -2.069726686 | 1802.958082 | 429.4729492 | 301 | 8 | 17 | 37 | 160 | 33 | 2 | 0 | 0 | 0 | 2 | 0 |
| SG0000001 | Uchl5 | 1.64E-31 | 1.77E-30 | 1.498937654 | 1021.351391 | 2886.691544 | 233 | 10 | 17 | 27 | 111 | 11 | 5 | 2 | 2 | 2 | 4 | 0 |
| SG0000003 | Eogt | 5.65E-28 | 5.21E-27 | -2.165938901 | 2107.727526 | 469.6798074 | 144 | 11 | 17 | 25 | 79 | 19 | 7 | 1 | 1 | 3 | 3 | 1 |
| SG0000002 | Mgme1 | 8.23E-26 | 6.9E-25 | 1.491546712 | 491.508036 | 1382.07281 | 63 | 13 | 17 | 19 | 28 | 6 | 8 | 1 | 2 | 1 | 1 | 0 |
| SG0000004 | Rlf | 2.12E-25 | 1.74E-24 | 1.301594725 | 1535.963858 | 3786.169487 | 274 | 13 | 17 | 33 | 107 | 13 | 0 | 0 | 0 | 0 | 0 | 0 |
| JSG0000007 | Hsf5 | 4.69E-23 | 3.45E-22 | 5.879365285 | 1.278220693 | 75.24387449 | 95 | 9 | 17 | 22 | 56 | 8 | 5 | 0 | 0 | 1 | 1 | 0 |
| SG0000002 | Atl3 | 4.93E-23 | 3.63E-22 | -1.696634638 | 5334.29136 | 1645.655083 | 99 | 13 | 17 | 24 | 68 | 8 | 14 | 1 | 1 | 3 | 6 | 1 |
| SG0000002 | Efemp1 | 1.12E-21 | 7.71E-21 | -2.896778437 | 1939.885616 | 260.4707272 | 195 | 5 | 17 | 24 | 105 | 18 | 6 | 0 | 2 | 2 | 6 | 0 |
| SG0000002 | Sel1l3 | 1.26E-18 | 7.3E-18 | 5.653914247 | 0 | 49.34980364 | 167 | 10 | 17 | 20 | 83 | 20 | 1 | 0 | 0 | 0 | 0 | 0 |
| SG0000003 | Iqgap1 | 3.36E-18 | 1.91E-17 | -1.437291787 | 15684.58069 | 5791.685523 | 149 | 15 | 17 | 34 | 72 | 9 | 6 | 1 | 1 | 1 | 1 | 0 |
| SG0000003 | Rif1 | 1.36E-16 | 7.1E-16 | 0.999129333 | 3810.837269 | 7617.076244 | 246 | 6 | 17 | 29 | 78 | 25 | 26 | 0 | 0 | 4 | 9 | 4 |
| JSG0000002 | Wee1 | 3.43E-16 | 1.74E-15 | 1.238690054 | 385.3453617 | 909.3540334 | 49 | 15 | 17 | 20 | 39 | 0 | 0 | 0 | 0 | 0 | 0 | 0 |
| SG0000002 | St13 | 5.29E-16 | 2.66E-15 | 0.963984722 | 7270.003493 | 14181.52504 | 95 | 14 | 17 | 21 | 51 | 8 | 2 | 0 | 0 | 0 | 0 | 1 |
| SG0000002 | Hdac1 | 2.27E-15 | 1.1E-14 | 0.97346333 | 2829.052155 | 5554.981489 | 45 | 17 | 17 | 20 | 32 | 3 | 1 | 0 | 0 | 1 | 1 | 0 |
| JSG0000002 | 2610002M06Rik | 1.49E-14 | 6.81E-14 | -1.334604375 | 1122.9936 | 445.2678433 | 107 | 16 | 17 | 25 | 64 | 9 | 17 | 0 | 0 | 2 | 5 | 1 |
| SG0000006 | Sycp2 | 8.98E-13 | 3.67E-12 | 5.080713987 | 0 | 32.84132117 | 191 | 5 | 17 | 26 | 81 | 16 | 3 | 0 | 0 | 1 | 4 | 1 |
| SG0000007 | Ywhaq | 2.79E-12 | 1.1E-11 | 0.883324383 | 2072.631939 | 3823.220282 | 87 | 14 | 17 | 20 | 43 | 7 | 2 | 0 | 0 | 0 | 0 | 0 |
| JSG0000002 | Zc3h12c | 7.22E-12 | 2.78E-11 | -1.555725604 | 361.5584589 | 122.9867867 | 118 | 11 | 17 | 20 | 64 | 16 | 12 | 0 | 2 | 2 | 5 | 1 |
| SG0000002 | Sdcbp | 1.84E-11 | 6.89E-11 | -0.967345759 | 4610.03752 | 2357.785923 | 224 | 3 | 17 | 36 | 118 | 29 | 9 | 0 | 1 | 0 | 8 | 0 |
| JSG0000007 | Myo18b | 5.21E-11 | 1.89E-10 | 6.65918805 | 0.307509278 | 31.07946781 | 209 | 9 | 17 | 33 | 80 | 54 | 1 | 0 | 0 | 0 | 0 | 0 |
| SG0000000 | Pex19 | 5.82E-11 | 2.1E-10 | 0.848556502 | 1556.189097 | 2802.227098 | 35 | 15 | 17 | 19 | 28 | 0 | 7 | 3 | 3 | 3 | 6 | 0 |
| SG0000002 | Faxc | 2.45E-10 | 8.49E-10 | -1.447840052 | 337.7228259 | 123.7989981 | 126 | 14 | 17 | 24 | 64 | 7 | 0 | 0 | 0 | 0 | 0 | 0 |
| SG0000002 | Cd55 | 3.3E-10 | 1.13E-09 | 2.276208621 | 82.3508268 | 398.9101026 | 81 | 13 | 17 | 24 | 40 | 12 | 10 | 0 | 1 | 3 | 3 | 1 |
| SG0000003 | Tyk2 | 2.06E-09 | 6.61E-09 | -1.228227781 | 1390.086173 | 593.3464112 | 50 | 15 | 17 | 20 | 28 | 2 | 1 | 0 | 0 | 0 | 0 | 0 |
| SG0000006 | Zfp13 | 2.91E-09 | 9.21E-09 | 1.215714447 | 162.2754613 | 376.8941243 | 58 | 13 | 17 | 19 | 32 | 5 | 0 | 0 | 0 | 0 | 0 | 0 |
| SG0000004 | Wipi1 | 4.54E-09 | 1.42E-08 | -1.02208849 | 1398.144247 | 688.4504472 | 113 | 12 | 17 | 23 | 55 | 8 | 6 | 0 | 0 | 0 | 0 | 0 |
| JSG0000009 | Tcerg1l | 7.4E-09 | 2.27E-08 | 3.403365416 | 4.244725496 | 44.91223693 | 250 | 14 | 17 | 35 | 120 | 23 | 0 | 0 | 0 | 0 | 1 | 0 |
| SG0000003 | Cog4 | 2.97E-08 | 8.69E-08 | 0.716031367 | 1893.991247 | 3111.178199 | 164 | 14 | 17 | 22 | 74 | 7 | 0 | 0 | 0 | 0 | 0 | 0 |
| JSG0000002 | Efcab2 | 7.07E-08 | 2.01E-07 | 1.216524441 | 95.0702422 | 220.9301297 | 180 | 8 | 17 | 28 | 62 | 19 | 7 | 0 | 0 | 0 | 1 | 3 |

| | | | | | | | | | | | | | | | | | | |
|---|---|---|---|---|---|---|---|---|---|---|---|---|---|---|---|---|---|---|
| SG0000002 | Nadk | 7.33E-08 | 2.08E-07 | -0.75212218 | 5235.219014 | 3108.304223 | 82 | 10 | 17 | 12 | 36 | 14 | 3 | 0 | 0 | 0 | 0 | 3 |
| SG0000003 | Ap5z1 | 8.03E-08 | 2.27E-07 | -1.330063877 | 767.2718016 | 305.1828428 | 44 | 17 | 17 | 21 | 30 | 3 | 2 | 0 | 0 | 2 | 2 | 0 |
| SG0000003 | Lca5 | 8.25E-08 | 2.33E-07 | 1.232859014 | 179.4342207 | 421.7283692 | 264 | 8 | 17 | 24 | 120 | 36 | 8 | 0 | 0 | 2 | 3 | 1 |
| SG0000002 | 5830415F09Rik | 2.62E-07 | 7.07E-07 | 1.111267054 | 117.2512103 | 253.3040017 | 82 | 11 | 17 | 14 | 34 | 9 | 5 | 3 | 3 | 3 | 5 | 0 |
| SG0000000 | Sez6 | 1.69E-06 | 4.27E-06 | 4.056338438 | 0 | 15.63717338 | 88 | 15 | 17 | 19 | 36 | 30 | 3 | 0 | 0 | 0 | 3 | 6 |
| SG0000004 | Suco | 9.37E-06 | 2.22E-05 | 0.589071307 | 1647.338169 | 2478.054732 | 217 | 11 | 17 | 38 | 106 | 15 | 12 | 0 | 0 | 2 | 5 | 1 |
| JSG000000 | Etfb | 1.63E-05 | 3.76E-05 | 0.726888064 | 361.2254766 | 597.8517207 | 31 | 17 | 17 | 18 | 31 | 0 | 0 | 0 | 0 | 0 | 0 | 0 |
| JSG000000 | Irgm2 | 4.89E-05 | 0.000108 | -1.586629685 | 740.0837218 | 246.4096574 | 64 | 17 | 17 | 21 | 34 | 7 | 4 | 0 | 0 | 0 | 7 | 4 |
| SG0000000 | Blvra | 0.000111 | 0.000234 | -0.709759062 | 561.9740424 | 343.6034286 | 75 | 14 | 17 | 20 | 47 | 9 | 0 | 0 | 0 | 0 | 0 | 0 |
| SG0000004 | Pik3r1 | 0.000112 | 0.000236 | -0.767414645 | 419.8773467 | 246.665049 | 180 | 11 | 17 | 20 | 54 | 18 | 9 | 0 | 0 | 1 | 3 | 3 |
| SG0000002 | Bche | 0.000138 | 0.000288 | 1.461850469 | 30.00721693 | 82.65831699 | 242 | 7 | 17 | 42 | 125 | 15 | 11 | 0 | 0 | 2 | 5 | 2 |
| SG0000005 | Txndc9 | 0.000184 | 0.000378 | 0.499330278 | 1489.244235 | 2105.131932 | 121 | 10 | 17 | 24 | 57 | 5 | 19 | 2 | 5 | 3 | 9 | 0 |
| SG0000002 | Mterf3 | 0.000254 | 0.000515 | 0.593160116 | 406.5678392 | 613.3268941 | 191 | 9 | 17 | 24 | 63 | 25 | 1 | 0 | 0 | 0 | 0 | 2 |
| SG0000003 | Pcid2 | 0.000349 | 0.000694 | 0.484359613 | 1399.635163 | 1958.040257 | 148 | 9 | 17 | 25 | 49 | 6 | 7 | 2 | 2 | 2 | 2 | 0 |
| SG0000003 | Cnbd2 | 0.000429 | 0.000843 | 1.40757881 | 22.57881068 | 59.89968039 | 182 | 14 | 17 | 35 | 100 | 17 | 0 | 0 | 0 | 0 | 1 | 0 |
| SG0000007 | Capza1 | 0.000618 | 0.00119 | -0.502264252 | 2124.209065 | 1499.687087 | 102 | 12 | 17 | 22 | 53 | 7 | 4 | 0 | 0 | 0 | 1 | 0 |
| SG0000003 | Cnnm4 | 0.000802 | 0.001522 | -0.568703292 | 780.9420026 | 526.5286801 | 59 | 15 | 17 | 19 | 31 | 4 | 4 | 0 | 0 | 1 | 1 | 1 |
| JSG000000 | Tgm3 | 0.001788 | 0.003259 | -1.142411729 | 105.1944505 | 47.65326164 | 53 | 13 | 17 | 19 | 29 | 4 | 1 | 0 | 0 | 0 | 0 | 0 |
| SG0000006 | Scyl2 | 0.001981 | 0.003589 | -0.445908785 | 2401.278203 | 1762.830512 | 197 | 6 | 17 | 25 | 97 | 23 | 3 | 0 | 0 | 0 | 1 | 0 |
| SG0000000 | Pknox1 | 0.003074 | 0.005412 | -0.46085086 | 1083.068529 | 786.9117032 | 130 | 11 | 17 | 27 | 73 | 18 | 4 | 0 | 0 | 0 | 5 | 3 |
| SG0000005 | Kxd1 | 0.005267 | 0.008984 | 0.394020275 | 1053.385533 | 1384.201358 | 87 | 12 | 17 | 18 | 39 | 1 | 0 | 0 | 0 | 0 | 0 | 0 |
| SG0000004 | Sft2d2 | 0.007152 | 0.011965 | -0.408795586 | 2530.828863 | 1906.351784 | 54 | 9 | 17 | 14 | 26 | 3 | 6 | 0 | 2 | 2 | 2 | 0 |
| JSG000000 | Tfcp2 | 0.008848 | 0.014591 | 0.461071027 | 311.4845325 | 428.7781982 | 83 | 14 | 17 | 21 | 45 | 9 | 2 | 0 | 0 | 0 | 0 | 0 |
| SG0000003 | Gmeb2 | 0.014687 | 0.023303 | 0.355452091 | 835.3694827 | 1068.760383 | 72 | 14 | 17 | 23 | 39 | 6 | 4 | 0 | 0 | 0 | 0 | 1 |
| JSG000000 | Zfp953 | 0.015184 | 0.024054 | 0.505289533 | 159.8742916 | 226.9268784 | 66 | 14 | 17 | 19 | 42 | 2 | 11 | 0 | 0 | 2 | 5 | 0 |
| SG0000000 | Zfp592 | 0.020956 | 0.032284 | -0.353826084 | 1124.536074 | 879.9563445 | 81 | 15 | 17 | 20 | 50 | 14 | 6 | 0 | 0 | 2 | 2 | 0 |
| JSG000000 | Ranbp9 | 0.022236 | 0.034126 | -0.336765572 | 1997.586107 | 1581.717684 | 209 | 10 | 17 | 28 | 80 | 13 | 5 | 0 | 1 | 1 | 2 | 0 |
| SG0000003 | Prkci | 0.024172 | 0.036864 | -0.30871836 | 7345.811377 | 5930.704503 | 158 | 12 | 17 | 24 | 80 | 10 | 9 | 1 | 2 | 1 | 6 | 1 |
| SG0000003 | Rnf146 | 0.028376 | 0.042794 | -0.341261678 | 1097.662495 | 866.4406589 | 208 | 3 | 17 | 31 | 72 | 7 | 18 | 0 | 3 | 0 | 0 | 0 |
| SG0000003 | Fbxo34 | 0.036496 | 0.05391 | 0.295633166 | 1093.120759 | 1341.72216 | 343 | 13 | 17 | 34 | 194 | 14 | 5 | 0 | 0 | 0 | 5 | 5 |
| SG0000003 | Ccdc55 | 0.037774 | 0.055687 | -0.319121271 | 1207.96965 | 968.2576745 | 94 | 12 | 17 | 25 | 45 | 11 | 5 | 1 | 1 | 1 | 3 | 1 |
| SG0000002 | Brdt | 0.046337 | 0.067348 | -0.746245936 | 66.47527243 | 39.62941987 | 173 | 10 | 17 | 35 | 75 | 17 | 1 | 0 | 0 | 0 | 1 | 0 |
| JSG000000 | Baiap2l1 | 0.055091 | 0.078892 | -0.312997124 | 819.9394606 | 660.0248166 | 155 | 13 | 17 | 24 | 65 | 18 | 1 | 0 | 0 | 0 | 2 | 1 |
| SG0000002 | Fam45a | 0.059445 | 0.08454 | -0.287701188 | 1235.109907 | 1011.809885 | 142 | 14 | 17 | 35 | 86 | 7 | 6 | 0 | 0 | 0 | 2 | 0 |
| JSG000000 | Ranbp10 | 0.065767 | 0.092882 | -0.266408497 | 1583.295097 | 1316.330422 | 65 | 15 | 17 | 24 | 63 | 9 | 2 | 0 | 0 | 1 | 3 | 0 |
| SG0000003 | Trappc8 | 0.066656 | 0.094048 | 0.236183143 | 2196.134003 | 2586.76542 | 145 | 9 | 17 | 30 | 76 | 9 | 3 | 0 | 0 | 0 | 0 | 0 |
| SG0000003 | Kdm1b | 0.066878 | 0.094331 | 0.3087348 | 547.4894464 | 678.1319203 | 103 | 14 | 17 | 18 | 46 | 9 | 12 | 0 | 0 | 0 | 2 | 0 |
| SG0000002 | Uck2 | 0.081625 | 0.113172 | 0.220657182 | 6336.045344 | 7383.16726 | 234 | 9 | 17 | 31 | 118 | 20 | 6 | 0 | 0 | 1 | 1 | 4 |
| JSG000000 | Mrpl32 | 0.093726 | 0.128379 | -0.295941747 | 569.9515319 | 464.2485783 | 26 | 16 | 17 | 19 | 25 | 0 | 1 | 0 | 1 | 0 | 0 | 0 |
| JSG000000 | Ptprb | 0.133396 | 0.176664 | -1.036353776 | 19.85841389 | 9.682131692 | 157 | 10 | 17 | 28 | 85 | 18 | 11 | 0 | 0 | 1 | 2 | 8 |
| JSG000000 | Snx2 | 0.220239 | 0.277596 | 0.151501635 | 3196.058564 | 3549.942063 | 121 | 13 | 17 | 23 | 78 | 6 | 0 | 0 | 0 | 0 | 1 | 0 |
| JSG000000 | Ddx50 | 0.263055 | 0.32491 | 0.145437729 | 1738.430161 | 1922.818833 | 104 | 13 | 17 | 22 | 48 | 3 | 0 | 0 | 0 | 0 | 1 | 0 |
| JSG000000 | Efhb | 0.282073 | 0.345517 | -0.407585693 | 75.48337482 | 56.90570258 | 133 | 12 | 17 | 26 | 85 | 8 | 1 | 0 | 0 | 0 | 0 | 0 |
| JSG000000 | Kif3c | 0.366995 | 0.435514 | 0.152306967 | 402.9739074 | 447.8431049 | 71 | 13 | 17 | 18 | 52 | 6 | 6 | 2 | 3 | 2 | 6 | 1 |
| JSG000000 | Mgmt | 0.376027 | 0.444844 | -0.175252328 | 318.6303305 | 282.1830391 | 315 | 9 | 17 | 34 | 123 | 29 | 0 | 0 | 0 | 0 | 0 | 0 |
| SG0000005 | Pla2r1 | 0.398697 | 0.467618 | -0.464022388 | 46.7749768 | 33.9100885 | 219 | 10 | 17 | 34 | 100 | 33 | 6 | 0 | 0 | 1 | 1 | 0 |
| SG0000002 | Wasl | 0.439447 | 0.50875 | -0.113671204 | 2667.164847 | 2465.08226 | 239 | 3 | 17 | 23 | 79 | 23 | 9 | 0 | 0 | 2 | 7 | 0 |
| SG0000000 | Tubgcp3 | 0.490322 | 0.558254 | -0.100179228 | 1754.165742 | 1636.491194 | 183 | 12 | 17 | 29 | 80 | 19 | 5 | 0 | 0 | 2 | 2 | 2 |
| JSG000000 | Immp1l | 0.495289 | 0.562991 | 0.148218969 | 164.1381155 | 181.8979474 | 166 | 12 | 17 | 28 | 97 | 15 | 3 | 0 | 0 | 1 | 1 | 0 |
| JSG000000 | Exd2 | 0.575817 | 0.64008 | 0.088266743 | 632.6648111 | 672.5810343 | 94 | 14 | 17 | 26 | 54 | 8 | 2 | 0 | 0 | 0 | 1 | 0 |
| JSG000000 | Abcb1a | 0.590568 | 0.653786 | -0.176458884 | 99.26782454 | 87.83934473 | 199 | 11 | 17 | 35 | 117 | 13 | 3 | 0 | 1 | 1 | 2 | 2 |
| SG0000002 | Gna13 | 0.604501 | 0.667127 | 0.058819062 | 3373.05958 | 3513.421983 | 117 | 9 | 17 | 17 | 49 | 9 | 9 | 0 | 1 | 0 | 0 | 1 |
| SG0000003 | Pole4 | 0.798137 | 0.839867 | -0.038701256 | 898.036428 | 874.266253 | 84 | 14 | 17 | 22 | 38 | 5 | 6 | 1 | 2 | 2 | 3 | 0 |
| SG0000000 | Pkn2 | 0.857308 | 0.888609 | 0.015253851 | 3279.830894 | 3314.693061 | 276 | 7 | 17 | 36 | 150 | 27 | 5 | 0 | 1 | 1 | 5 | 1 |
| SG0000002 | Ccnl1 | 0.901456 | 0.92527 | -0.024244797 | 1658.967917 | 1631.32156 | 191 | 6 | 17 | 28 | 42 | 22 | 17 | 0 | 0 | 0 | 9 | 0 |
| SG0000003 | Tmod2 | 0.96806 | 0.981818 | -0.004117919 | 250.404673 | 249.690956 | 151 | 7 | 17 | 22 | 56 | 16 | 14 | 0 | 1 | 1 | 3 | 3 |
| JSG000000 | Egln1 | 1 | 1 | -0.005511136 | 2029.423147 | 2021.685482 | 62 | 17 | 17 | 21 | 32 | 9 | 3 | 0 | 0 | 0 | 1 | 0 |
| SG0000003 | Mcf2l | 4E-125 | 4.3E-123 | 4.410832436 | 43.95920972 | 935.0670323 | 371 | 6 | 16 | 25 | 99 | 50 | 8 | 0 | 0 | 4 | 16 | 8 |
| SG0000004 | Foxn4 | 2.6E-124 | 2.8E-122 | 8.830918734 | 0 | 454.3773528 | 39 | 14 | 16 | 18 | 24 | 0 | 3 | 0 | 1 | 2 | 2 | 0 |
| SG0000003 | Fgd5 | 2.1E-122 | 2.2E-120 | 7.677927247 | 2.294772122 | 469.9222092 | 154 | 8 | 16 | 20 | 64 | 8 | 6 | 0 | 0 | 2 | 2 | 0 |
| SG0000002 | Osmr | 2.6E-105 | 2.1E-103 | -4.980311003 | 3277.291194 | 103.8226312 | 193 | 4 | 16 | 26 | 82 | 14 | 5 | 0 | 0 | 0 | 2 | 0 |
| JSG000000 | Uprt | 2.14E-97 | 1.45E-95 | -5.367486919 | 1569.80025 | 38.02505316 | 58 | 13 | 16 | 15 | 44 | 3 | 3 | 1 | 1 | 1 | 2 | 1 |
| SG0000002 | Clmn | 1.95E-89 | 1.09E-87 | 4.915189428 | 15.27121322 | 460.7793343 | 152 | 6 | 16 | 28 | 74 | 38 | 6 | 1 | 2 | 1 | 2 | 0 |
| SG0000006 | Tnfrsf19 | 5.12E-86 | 2.62E-84 | 8.214969197 | 0.981585527 | 291.6622944 | 242 | 4 | 16 | 30 | 127 | 28 | 12 | 0 | 0 | 1 | 4 | 0 |
| SG0000004 | Tmem194b | 6.5E-73 | 2.54E-71 | 2.59525651 | 265.1000613 | 1601.990307 | 201 | 12 | 16 | 20 | 63 | 16 | 9 | 0 | 0 | 0 | 3 | 1 |
| SG0000002 | Fgr | 1.56E-64 | 4.85E-63 | 8.374812961 | 0.649984458 | 215.7609679 | 54 | 14 | 16 | 20 | 38 | 2 | 3 | 0 | 2 | 2 | 4 | 0 |
| SG0000002 | Nabp1 | 1.39E-62 | 4.12E-61 | 2.494628786 | 475.5444828 | 2680.089146 | 155 | 8 | 16 | 8 | 44 | 16 | 32 | 8 | 8 | 8 | 12 | 0 |
| JSG000000 | Ntn4 | 3.95E-61 | 1.13E-59 | -6.530072583 | 2397.695054 | 25.944555 | 176 | 13 | 16 | 33 | 87 | 14 | 3 | 0 | 0 | 0 | 0 | 0 |
| SG0000000 | Atp7b | 1.64E-59 | 4.46E-58 | 3.98059107 | 27.99316889 | 441.9054697 | 105 | 13 | 16 | 25 | 59 | 12 | 2 | 0 | 0 | 1 | 1 | 1 |
| SG0000001 | Cpsf4l | 7.82E-54 | 1.8E-52 | 4.98858836 | 17.77478389 | 564.3117088 | 55 | 4 | 16 | 16 | 60 | 16 | 8 | 0 | 4 | 4 | 4 | 0 |
| SG0000000 | Nab1 | 3.54E-52 | 7.84E-51 | -2.708067986 | 3964.602421 | 606.7222862 | 176 | 4 | 16 | 24 | 92 | 8 | 12 | 0 | 0 | 2 | 6 | 0 |
| SG0000005 | Cbx7 | 9.28E-52 | 2.02E-50 | 1.888031662 | 1139.266043 | 4216.764202 | 68 | 16 | 16 | 27 | 59 | 4 | 6 | 0 | 0 | 2 | 5 | 0 |
| SG0000003 | Atp6v0a2 | 1.16E-50 | 2.43E-49 | -2.609752701 | 4346.493171 | 712.0740472 | 100 | 16 | 16 | 25 | 69 | 11 | 6 | 0 | 0 | 0 | 3 | 0 |
| SG0000002 | Mpzl1 | 2.74E-50 | 5.75E-49 | 1.967228823 | 621.4266438 | 2429.879594 | 148 | 12 | 16 | 22 | 54 | 22 | 2 | 0 | 0 | 0 | 0 | 0 |
| SG0000000 | Tmbim1 | 1.66E-49 | 3.41E-48 | -2.596579138 | 8538.231733 | 1411.626267 | 31 | 16 | 16 | 22 | 28 | 2 | 2 | 0 | 0 | 2 | 2 | 0 |
| SG0000002 | Syk | 8.57E-49 | 1.72E-47 | 3.195627008 | 111.5492504 | 1021.989046 | 168 | 10 | 16 | 40 | 117 | 61 | 6 | 0 | 0 | 0 | 0 | 4 |
| SG0000002 | Serpini1 | 4.21E-44 | 7.15E-43 | 4.06211475 | 13.84709508 | 231.30076 | 257 | 9 | 16 | 31 | 113 | 26 | 18 | 1 | 2 | 2 | 5 | 1 |
| JSG000000 | Grik3 | 2.53E-43 | 4.18E-42 | 10.7580632 | 0.34247518 | 593.1012252 | 207 | 10 | 16 | 32 | 84 | 17 | 1 | 0 | 0 | 1 | 1 | 2 |
| JSG000000 | Nt5dc3 | 3.86E-43 | 6.32E-42 | -2.780431681 | 2719.934681 | 395.8814367 | 76 | 14 | 16 | 22 | 41 | 2 | 9 | 2 | 2 | 4 | 4 | 0 |
| SG0000005 | Skp2 | 1.11E-41 | 1.74E-40 | 1.798362183 | 638.1465953 | 2219.634245 | 188 | 13 | 16 | 29 | 76 | 11 | 11 | 0 | 0 | 1 | 6 | 0 |
| JSG000000 | Wipf3 | 8.74E-40 | 1.28E-38 | 6.117050719 | 1.963171053 | 136.2617283 | 260 | 8 | 16 | 30 | 168 | 40 | 20 | 0 | 0 | 0 | 4 | 12 |
| SG0000002 | Herc6 | 6.41E-39 | 9.07E-38 | -3.297394051 | 541.5179773 | 55.08056189 | 205 | 6 | 16 | 30 | 116 | 23 | 8 | 0 | 1 | 0 | 3 | 1 |
| SG0000000 | Tex15 | 1.2E-33 | 1.4E-32 | 5.295882927 | 19.1154027 | 750.9362441 | 179 | 9 | 16 | 26 | 90 | 19 | 0 | 0 | 0 | 0 | 0 | 0 |
| SG0000004 | Tmem104 | 4.6E-33 | 5.26E-32 | -2.497534549 | 831.996623 | 147.3291723 | 98 | 14 | 16 | 22 | 40 | 18 | 10 | 0 | 0 | 2 | 2 | 0 |
| JSG000000 | Atp2b1 | 7.6E-32 | 8.29E-31 | -2.124785846 | 7461.980165 | 1710.920446 | 225 | 9 | 16 | 28 | 89 | 29 | 7 | 0 | 0 | 1 | 2 | 3 |
| SG0000002 | Slc39a14 | 9.14E-32 | 9.93E-31 | -1.799120144 | 10765.0227 | 3093.326919 | 106 | 14 | 16 | 20 | 44 | 9 | 4 | 0 | 0 | 0 | 2 | 1 |
| SG0000004 | Rell1 | 3.88E-31 | 4.12E-30 | 1.65992232 | 423.2072566 | 1337.332856 | 198 | 10 | 16 | 36 | 82 | 24 | 4 | 0 | 0 | 2 | 4 | 0 |
| SG0000002 | Steap3 | 1.47E-28 | 1.39E-27 | 2.374111697 | 75.25552146 | 390.1370567 | 147 | 8 | 16 | 24 | 78 | 11 | 2 | 0 | 0 | 0 | 5 | 0 |
| SG0000002 | Tcp11l1 | 4.01E-28 | 3.72E-27 | -2.524032151 | 809.5470281 | 140.7449093 | 96 | 14 | 16 | 26 | 76 | 10 | 8 | 0 | 0 | 0 | 4 | 0 |
| SG0000002 | Kif18a | 2.04E-27 | 1.83E-26 | 1.481125246 | 639.6217526 | 1785.608939 | 183 | 3 | 16 | 23 | 90 | 14 | 3 | 0 | 0 | 1 | 3 | 2 |
| JSG000000 | Pald1 | 2.88E-26 | 2.46E-25 | 1.86959386 | 1029.057409 | 3760.481318 | 69 | 14 | 16 | 24 | 39 | 4 | 0 | 0 | 0 | 0 | 0 | 0 |
| SG0000003 | Tbx20 | 2.54E-25 | 2.09E-24 | -3.204948832 | 592.7924937 | 64.28608271 | 213 | 3 | 16 | 30 | 111 | 21 | 17 | 0 | 1 | 3 | 7 | 0 |
| SG0000004 | Kif14 | 2.04E-24 | 1.6E-23 | 1.334621023 | 952.6546529 | 2402.68288 | 285 | 14 | 16 | 60 | 138 | 39 | 21 | 2 | 4 | 6 | 12 | 6 |
| SG0000002 | Blmh | 2.11E-24 | 1.65E-23 | -1.523051482 | 9955.685807 | 3464.072727 | 164 | 12 | 16 | 24 | 56 | 12 | 0 | 0 | 0 | 0 | 0 | 0 |
| JSG000000 | Tpd52l1 | 3.79E-24 | 2.94E-23 | 2.258686474 | 62.04115304 | 296.9013963 | 222 | 9 | 16 | 41 | 163 | 22 | 5 | 0 | 0 | 1 | 1 | 0 |
| SG0000001 | Sgk2 | 5.26E-24 | 4.04E-23 | 6.043386053 | 0 | 64.95389937 | 76 | 14 | 16 | 16 | 42 | 10 | 6 | 0 | 0 | 0 | 0 | 0 |
| JSG000000 | Kif15 | 1.06E-23 | 8.08E-23 | 1.299190718 | 1059.118444 | 2606.39304 | 144 | 10 | 16 | 30 | 86 | 12 | 1 | 0 | 0 | 0 | 1 | 0 |
| SG0000003 | Cc2d2a | 2.93E-23 | 2.18E-22 | -1.85498144 | 1662.950186 | 459.6996628 | 132 | 12 | 16 | 22 | 50 | 13 | 0 | 0 | 0 | 0 | 0 | 0 |
| SG0000002 | Hells | 9.14E-23 | 6.65E-22 | 1.170864645 | 4607.036724 | 10372.58621 | 122 | 10 | 16 | 23 | 70 | 14 | 17 | 2 | 4 | 3 | 7 | 1 |
| JSG000000 | Fbxo36 | 1.82E-22 | 1.31E-21 | 2.978153125 | 22.50653531 | 177.3462731 | 101 | 14 | 16 | 22 | 57 | 8 | 0 | 0 | 0 | 0 | 0 | 0 |
| SG0000002 | Eif5b | 3.35E-22 | 2.37E-21 | 1.155682862 | 4819.327509 | 10736.96773 | 134 | 8 | 16 | 22 | 53 | 15 | 4 | 0 | 0 | 0 | 5 | 1 |
| JSG000000 | 4930539E08Rik | 9.12E-22 | 6.34E-21 | 4.445125243 | 6.4237258 | 139.9274846 | 39 | 15 | 16 | 23 | 41 | 5 | 5 | 2 | 3 | 3 | 6 | 3 |
| SG0000003 | Wdr62 | 1.21E-21 | 8.32E-21 | 1.395356519 | 439.7456288 | 1156.766487 | 119 | 13 | 16 | 22 | 56 | 5 | 0 | 0 | 0 | 0 | 0 | 0 |
| JSG000000 | Dnajb14 | 1.49E-21 | 1.02E-20 | -1.828885307 | 1231.603725 | 346.6743083 | 117 | 10 | 16 | 20 | 50 | 11 | 17 | 1 | 3 | 3 | 7 | 3 |
| SG0000003 | Naa30 | 5.33E-21 | 3.54E-20 | 1.202049809 | 1307.093186 | 3007.181213 | 202 | 9 | 16 | 38 | 90 | 8 | 26 | 0 | 4 | 2 | 6 | 2 |
| JSG000000 | P3h3 | 2.34E-20 | 1.49E-19 | -1.947402161 | 591.8435517 | 153.4547925 | 56 | 14 | 16 | 18 | 48 | 0 | 1 | 0 | 0 | 0 | 0 | 0 |
| SG0000003 | Ankrd27 | 8.57E-20 | 5.3E-19 | 1.205835061 | 940.2374787 | 2168.852822 | 154 | 16 | 16 | 24 | 71 | 25 | 3 | 0 | 0 | 0 | 3 | 0 |
| SG0000003 | Aplp2 | 1.16E-19 | 7.16E-19 | 1.084219229 | 4648.843696 | 9856.607309 | 236 | 12 | 16 | 26 | 122 | 22 | 2 | 0 | 0 | 0 | 2 | 0 |
| SG0000004 | Hibch | 1.19E-18 | 6.89E-18 | 1.314994248 | 404.8738275 | 1007.331503 | 254 | 4 | 16 | 32 | 110 | 26 | 0 | 0 | 0 | 0 | 0 | 0 |
| SG0000003 | Slc9b2 | 1.43E-18 | 8.26E-18 | 5.658114748 | 0 | 49.49661402 | 137 | 4 | 16 | 22 | 58 | 25 | 6 | 0 | 0 | 1 | 3 | 1 |
| SG0000005 | Dnajc8 | 1.8E-18 | 1.04E-17 | 1.071926265 | 2811.588758 | 5910.629775 | 189 | 8 | 16 | 31 | 111 | 21 | 3 | 0 | 0 | 0 | 0 | 0 |
| SG0000005 | 1600014C10Rik | 2.38E-18 | 1.37E-17 | -1.73350481 | 691.5470313 | 207.9623785 | 83 | 12 | 16 | 24 | 73 | 7 | 25 | 12 | 16 | 16 | 21 | 0 |
| SG0000002 | Gtf2a1 | 2.86E-18 | 1.63E-17 | 1.173518523 | 833.928269 | 1881.018089 | 139 | 8 | 16 | 14 | 68 | 6 | 17 | 0 | 0 | 2 | 4 | 0 |
| SG0000002 | Ldb3 | 6.32E-18 | 3.54E-17 | 3.120250212 | 20.66516991 | 179.6916465 | 229 | 12 | 16 | 42 | 134 | 21 | 12 | 0 | 0 | 2 | 5 | 1 |
| JSG000000 | Kdm7a | 8.46E-18 | 4.71E-17 | -1.553824464 | 2732.506562 | 930.7079889 | 192 | 11 | 16 | 28 | 88 | 20 | 19 | 0 | 0 | 2 | 13 | 1 |
| SG0000004 | Mum1l1 | 2.42E-17 | 1.32E-16 | 4.293662806 | 17.66135564 | 346.3740373 | 232 | 4 | 16 | 26 | 94 | 16 | 24 | 0 | 0 | 4 | 4 | 0 |
| JSG000000 | Khdrbs3 | 4.04E-17 | 2.18E-16 | 5.523943247 | 15.07463549 | 693.6165244 | 299 | 10 | 16 | 31 | 139 | 31 | 1 | 0 | 0 | 0 | 1 | 0 |
| SG0000000 | Rps6ka1 | 5.15E-17 | 2.75E-16 | 1.057105242 | 1875.486754 | 3902.423405 | 130 | 13 | 16 | 26 | 47 | 29 | 0 | 0 | 0 | 0 | 0 | 0 |
| JSG000000 | Prkaa1 | 7.88E-17 | 4.18E-16 | -1.32128174 | 4121.055621 | 1649.160938 | 90 | 14 | 16 | 22 | 51 | 8 | 16 | 0 | 0 | 1 | 3 | 2 |
| JSG000000 | Rassf3 | 1.42E-16 | 7.36E-16 | -1.29109385 | 2627.290947 | 1073.619012 | 72 | 12 | 16 | 21 | 61 | 14 | 3 | 0 | 0 | 1 | 3 | 0 |
| SG0000002 | Rfx2 | 1.51E-16 | 7.82E-16 | 1.839788048 | 254.4293851 | 910.7489022 | 114 | 12 | 16 | 34 | 62 | 6 | 0 | 0 | 0 | 0 | 0 | 0 |
| SG0000004 | Proca1 | 1.04E-15 | 5.15E-15 | 4.538143063 | 2.578189609 | 59.90071969 | 59 | 13 | 16 | 20 | 29 | 0 | 2 | 0 | 0 | 0 | 0 | 0 |
| SG0000001 | Tada2a | 1.07E-15 | 5.27E-15 | 1.058347296 | 1062.075115 | 2211.818646 | 108 | 10 | 16 | 25 | 68 | 3 | 1 | 0 | 0 | 0 | 1 | 0 |

| | | | | | | | | | | | | | | | | | | |
|---|---|---|---|---|---|---|---|---|---|---|---|---|---|---|---|---|---|---|
| SG0000002 | Fcgr2b | 1.1E-15 | 5.42E-15 | -4.444220361 | 192.8841728 | 8.860392418 | 102 | 8 | 16 | 16 | 25 | 8 | 2 | 0 | 0 | 0 | 1 | 4 |
| SG0000005 | Nrros | 2.8E-15 | 1.35E-14 | 4.230693418 | 3.298105871 | 61.91989538 | 153 | 12 | 16 | 30 | 103 | 16 | 10 | 0 | 0 | 5 | 5 | 0 |
| SG0000002 | Rab12 | 2.28E-14 | 1.03E-13 | -1.201697486 | 2186.742006 | 950.7154656 | 72 | 12 | 16 | 22 | 38 | 14 | 4 | 0 | 0 | 4 | 4 | 2 |
| SG0000005 | 3110082I17Rik | 5.29E-14 | 2.34E-13 | 1.19027479 | 358.7386172 | 818.6279049 | 142 | 10 | 16 | 20 | 52 | 20 | 1 | 0 | 0 | 0 | 0 | 0 |
| JSG000000 | Babam1 | 1.78E-13 | 7.61E-13 | -1.132448715 | 2499.342315 | 1140.05213 | 22 | 16 | 16 | 17 | 20 | 0 | 0 | 0 | 0 | 0 | 1 | 0 |
| SG0000002 | Dfna5 | 2.57E-13 | 1.09E-12 | -2.319126223 | 172.0464734 | 34.47618623 | 289 | 5 | 16 | 46 | 119 | 41 | 13 | 0 | 1 | 2 | 9 | 0 |
| SG0000002 | Pkia | 1.7E-12 | 6.8E-12 | 3.514958129 | 12.53994232 | 143.3518722 | 245 | 4 | 16 | 29 | 95 | 20 | 11 | 0 | 1 | 0 | 2 | 0 |
| SG0000005 | Usp26 | 1.88E-12 | 7.52E-12 | 5.036296025 | 0 | 31.81528426 | 258 | 12 | 16 | 38 | 147 | 14 | 6 | 0 | 0 | 0 | 2 | 0 |
| SG0000004 | Cul9 | 2.55E-12 | 1.01E-11 | -2.199701151 | 827.6773536 | 180.1710646 | 114 | 10 | 16 | 22 | 64 | 16 | 0 | 0 | 0 | 0 | 0 | 0 |
| SG0000002 | Hcfc2 | 5.09E-12 | 1.98E-11 | -1.298829686 | 712.5683879 | 289.6275409 | 151 | 6 | 16 | 20 | 92 | 17 | 7 | 1 | 2 | 1 | 3 | 0 |
| JSG000000 | Nek9 | 6.18E-12 | 2.39E-11 | -1.087226865 | 4911.631827 | 2311.734245 | 73 | 12 | 16 | 18 | 53 | 5 | 2 | 0 | 0 | 0 | 0 | 0 |
| SG0000004 | Eif4g1 | 2.02E-11 | 7.56E-11 | -0.91607466 | 30901.05209 | 16375.9806 | 136 | 8 | 16 | 16 | 32 | 8 | 0 | 0 | 0 | 0 | 0 | 0 |
| SG0000003 | Rnft2 | 2.75E-11 | 1.02E-10 | 2.249192115 | 53.02856023 | 252.1065495 | 111 | 9 | 16 | 24 | 47 | 18 | 6 | 0 | 0 | 0 | 0 | 2 |
| SG0000003 | Hykk | 5.6E-11 | 2.03E-10 | 1.681116775 | 54.83610972 | 175.8462908 | 84 | 9 | 16 | 18 | 35 | 3 | 10 | 0 | 1 | 0 | 1 | 0 |
| SG0000002 | Ctc1 | 7.24E-11 | 2.6E-10 | 0.81340584 | 2731.466559 | 4800.157023 | 134 | 12 | 16 | 32 | 64 | 16 | 0 | 0 | 0 | 0 | 0 | 0 |
| SG0000000 | Ift46 | 1.56E-10 | 5.49E-10 | 0.844105819 | 1312.17199 | 2355.547868 | 74 | 12 | 16 | 15 | 31 | 4 | 2 | 0 | 0 | 0 | 0 | 0 |
| SG0000003 | Mtmr12 | 2.33E-10 | 8.07E-10 | -0.96675777 | 2256.173644 | 1154.381702 | 178 | 10 | 16 | 26 | 94 | 16 | 4 | 0 | 0 | 2 | 8 | 0 |
| SG0000003 | Mre11a | 6.96E-10 | 2.33E-09 | 0.824199034 | 1299.945102 | 2301.620111 | 188 | 10 | 16 | 18 | 74 | 12 | 4 | 0 | 0 | 0 | 2 | 0 |
| SG0000004 | Orc3 | 1.74E-09 | 5.61E-09 | 0.778842898 | 1485.640977 | 2548.994766 | 218 | 14 | 16 | 34 | 114 | 24 | 0 | 0 | 0 | 0 | 0 | 0 |
| SG0000003 | Chic1 | 2.41E-09 | 7.68E-09 | -1.250176978 | 908.0784534 | 381.7531248 | 107 | 12 | 16 | 20 | 65 | 7 | 19 | 0 | 1 | 2 | 12 | 3 |
| JSG000000 | Kcnq3 | 2.49E-09 | 7.94E-09 | 6.412611498 | 0.307509278 | 26.19670292 | 465 | 11 | 16 | 46 | 218 | 56 | 17 | 1 | 2 | 1 | 5 | 0 |
| SG0000004 | Elmod1 | 2.98E-09 | 9.43E-09 | -4.191205209 | 197.883824 | 10.83257759 | 180 | 11 | 16 | 29 | 87 | 9 | 8 | 0 | 2 | 2 | 6 | 0 |
| SG0000006 | Lsm14a | 3.14E-09 | 9.92E-09 | 0.737432941 | 2886.767811 | 4812.838505 | 101 | 11 | 16 | 20 | 49 | 14 | 1 | 0 | 0 | 0 | 0 | 1 |
| SG0000003 | Edc3 | 4.18E-09 | 1.31E-08 | 0.793724861 | 1041.061564 | 1804.726535 | 132 | 10 | 16 | 34 | 70 | 12 | 2 | 2 | 2 | 2 | 4 | 2 |
| JSG000000 | Kif3b | 6.03E-09 | 1.87E-08 | -0.946345104 | 1665.183763 | 864.1395947 | 72 | 11 | 16 | 20 | 44 | 3 | 3 | 0 | 0 | 1 | 2 | 0 |
| SG0000003 | Tirap | 7.42E-09 | 2.28E-08 | -1.11240928 | 559.9243905 | 258.9767791 | 79 | 12 | 16 | 12 | 53 | 0 | 3 | 0 | 0 | 0 | 1 | 0 |
| JSG000000 | Ywhag | 1.8E-08 | 5.37E-08 | -0.850837939 | 16678.42222 | 9247.561386 | 82 | 14 | 16 | 20 | 36 | 2 | 6 | 0 | 0 | 1 | 1 | 0 |
| JSG000000 | Wdr60 | 2.02E-08 | 6E-08 | -1.104670757 | 515.8077095 | 239.8550109 | 119 | 8 | 16 | 23 | 68 | 11 | 2 | 0 | 0 | 0 | 0 | 3 |
| SG0000002 | Spred1 | 3.69E-08 | 1.07E-07 | -0.973171172 | 4499.519337 | 2291.988366 | 349 | 6 | 16 | 44 | 176 | 30 | 13 | 0 | 0 | 3 | 10 | 4 |
| SG0000002 | Jak1 | 5.13E-08 | 1.48E-07 | -0.834923519 | 4198.347282 | 2353.647047 | 197 | 7 | 16 | 28 | 93 | 14 | 4 | 0 | 0 | 0 | 0 | 0 |
| SG0000003 | Magohb | 5.38E-08 | 1.54E-07 | -1.365302589 | 389.3270712 | 151.1184778 | 106 | 14 | 16 | 21 | 57 | 1 | 5 | 0 | 2 | 2 | 3 | 1 |
| SG0000002 | Myo1f | 6.07E-08 | 1.73E-07 | 4.41750114 | 1.91733104 | 40.97297062 | 122 | 14 | 16 | 30 | 60 | 16 | 0 | 0 | 0 | 0 | 0 | 0 |
| SG0000003 | Tial1 | 6.71E-08 | 1.91E-07 | 0.691419903 | 1982.604152 | 3201.652124 | 360 | 0 | 16 | 45 | 168 | 31 | 16 | 0 | 1 | 7 | 10 | 2 |
| SG0000004 | Adck2 | 9.7E-08 | 2.72E-07 | -1.193475938 | 326.8199499 | 142.9014063 | 55 | 16 | 16 | 18 | 31 | 7 | 5 | 0 | 0 | 0 | 0 | 1 |
| SG0000004 | Prkd2 | 1.43E-07 | 3.97E-07 | -1.260507791 | 671.3753552 | 280.2300732 | 74 | 14 | 16 | 16 | 42 | 6 | 0 | 0 | 0 | 0 | 0 | 0 |
| SG0000004 | Kif21b | 1.55E-07 | 4.29E-07 | 1.223052965 | 700.8082857 | 1635.968196 | 77 | 12 | 16 | 14 | 27 | 18 | 10 | 0 | 0 | 0 | 0 | 2 |
| JSG000000 | Zfp677 | 1.69E-07 | 4.66E-07 | 2.131613559 | 15.29296144 | 67.01487533 | 96 | 8 | 16 | 20 | 40 | 0 | 10 | 0 | 0 | 0 | 0 | 0 |
| JSG000000 | Fbxo16 | 2.61E-07 | 7.06E-07 | 3.950908205 | 7.388403376 | 114.2595417 | 80 | 14 | 16 | 22 | 46 | 8 | 0 | 0 | 0 | 0 | 0 | 0 |
| SG0000002 | 1200014J11Rik | 2.72E-07 | 7.34E-07 | 0.688870048 | 1222.36835 | 1970.48278 | 71 | 13 | 16 | 20 | 38 | 6 | 6 | 0 | 0 | 2 | 8 | 1 |
| SG0000002 | Zfp871 | 4.05E-07 | 1.08E-06 | -1.355786518 | 989.5629228 | 386.6437597 | 75 | 10 | 16 | 17 | 39 | 6 | 6 | 0 | 0 | 3 | 11 | 0 |
| SG0000004 | Cblc | 5.65E-07 | 1.49E-06 | 3.356416253 | 15.01089066 | 153.7405504 | 44 | 8 | 16 | 16 | 34 | 8 | 0 | 0 | 0 | 0 | 0 | 0 |
| SG0000000 | Ap2a2 | 7.52E-07 | 1.96E-06 | 0.622809399 | 3076.124365 | 4736.834381 | 105 | 11 | 16 | 26 | 56 | 9 | 1 | 0 | 0 | 0 | 0 | 0 |
| JSG000000 | Tfap2e | 8.39E-07 | 2.18E-06 | 4.133631195 | 0 | 16.5528233 | 38 | 14 | 16 | 20 | 39 | 3 | 2 | 1 | 2 | 2 | 2 | 0 |
| JSG000000 | Nipa1 | 1.19E-06 | 3.07E-06 | -1.170321833 | 251.3652009 | 111.6871419 | 74 | 13 | 16 | 17 | 41 | 13 | 3 | 0 | 0 | 0 | 1 | 0 |
| SG0000003 | Mb21d1 | 1.31E-06 | 3.36E-06 | 2.001568866 | 15.63778019 | 62.61917929 | 70 | 9 | 16 | 15 | 46 | 0 | 9 | 0 | 1 | 0 | 3 | 0 |
| SG0000003 | Cdc123 | 1.44E-06 | 3.66E-06 | 0.633346251 | 1572.946324 | 2439.889298 | 122 | 9 | 16 | 20 | 71 | 22 | 2 | 0 | 0 | 0 | 0 | 0 |
| SG0000003 | Prex1 | 2.09E-06 | 5.24E-06 | -1.080144368 | 1939.584212 | 917.3874317 | 156 | 14 | 16 | 26 | 62 | 7 | 3 | 0 | 0 | 0 | 2 | 0 |
| SG0000002 | Lmo4 | 2.11E-06 | 5.29E-06 | -1.022935857 | 323.9537428 | 159.4221366 | 120 | 8 | 16 | 12 | 32 | 8 | 10 | 0 | 0 | 0 | 0 | 0 |
| SG0000001 | Capza2 | 3.51E-06 | 8.63E-06 | -0.64227308 | 6210.905454 | 3979.343742 | 213 | 14 | 16 | 51 | 99 | 26 | 4 | 0 | 0 | 1 | 4 | 0 |
| SG0000001 | Tsacc | 3.68E-06 | 9.04E-06 | 1.961756992 | 17.7926887 | 69.30894356 | 46 | 14 | 16 | 16 | 34 | 2 | 0 | 0 | 0 | 0 | 0 | 0 |
| SG0000002 | Ifi44 | 3.82E-06 | 9.37E-06 | -1.783639622 | 409.0157271 | 118.7985062 | 67 | 12 | 16 | 15 | 39 | 4 | 7 | 0 | 0 | 1 | 5 | 1 |
| SG0000005 | Cnn3 | 4.76E-06 | 1.16E-05 | 0.558002351 | 5670.238065 | 8347.890398 | 101 | 14 | 16 | 23 | 43 | 6 | 3 | 0 | 0 | 1 | 3 | 0 |
| SG0000000 | Rhoa | 7.09E-06 | 1.69E-05 | 0.551518514 | 5826.747813 | 8539.842043 | 72 | 16 | 16 | 21 | 50 | 6 | 3 | 0 | 0 | 0 | 0 | 0 |
| SG0000003 | Zer1 | 7.1E-06 | 1.7E-05 | 0.660643312 | 707.0858035 | 1117.752298 | 54 | 16 | 16 | 16 | 28 | 4 | 0 | 0 | 0 | 0 | 2 | 0 |
| SG0000000 | Sp1 | 9.38E-06 | 2.22E-05 | 0.541022515 | 4797.25698 | 6980.025172 | 112 | 14 | 16 | 24 | 77 | 9 | 10 | 0 | 0 | 0 | 5 | 1 |
| SG0000003 | Dera | 1.29E-05 | 3.01E-05 | -0.663778205 | 1675.212582 | 1057.432797 | 133 | 12 | 16 | 25 | 72 | 18 | 2 | 0 | 0 | 0 | 0 | 0 |
| JSG000000 | Ptk7 | 1.31E-05 | 3.04E-05 | -1.08781107 | 7264.813857 | 3417.91084 | 81 | 13 | 16 | 17 | 50 | 5 | 0 | 0 | 0 | 0 | 1 | 0 |
| JSG000000 | Tceb1 | 1.64E-05 | 3.8E-05 | 0.557445877 | 2265.756149 | 3334.426105 | 186 | 15 | 16 | 25 | 75 | 30 | 6 | 0 | 1 | 0 | 0 | 0 |
| SG0000000 | Srr | 2.06E-05 | 4.72E-05 | -1.211993042 | 263.2077844 | 113.6193942 | 137 | 12 | 16 | 20 | 96 | 20 | 17 | 0 | 0 | 0 | 4 | 0 |
| SG0000002 | Fam234a | 3.36E-05 | 7.53E-05 | -0.71957748 | 1866.07809 | 1133.223073 | 84 | 16 | 16 | 22 | 53 | 4 | 2 | 0 | 0 | 0 | 0 | 0 |
| JSG000000 | Crim1 | 4.42E-05 | 9.75E-05 | -1.571123987 | 811.8554506 | 273.2267799 | 262 | 7 | 16 | 29 | 116 | 19 | 9 | 0 | 0 | 2 | 10 | 2 |
| JSG000000 | Bcar3 | 5.62E-05 | 0.000123 | 0.658168678 | 407.9342554 | 643.7521465 | 128 | 8 | 16 | 26 | 73 | 14 | 3 | 0 | 0 | 1 | 2 | 2 |
| SG0000002 | Fopnl | 8.23E-05 | 0.000176 | -0.644646605 | 927.0551727 | 592.9903375 | 92 | 16 | 16 | 20 | 50 | 6 | 4 | 0 | 0 | 0 | 4 | 0 |
| SG0000001 | Fam134c | 8.57E-05 | 0.000183 | -0.574027008 | 2008.717676 | 1349.334279 | 78 | 12 | 16 | 18 | 74 | 6 | 0 | 0 | 0 | 0 | 0 | 0 |
| SG0000002 | Srek1ip1 | 9.22E-05 | 0.000196 | -0.658989461 | 777.7828941 | 492.5868261 | 97 | 10 | 16 | 20 | 42 | 10 | 9 | 0 | 0 | 2 | 2 | 2 |
| SG0000002 | Ube2e3 | 0.000137 | 0.000287 | 0.562117406 | 655.5005603 | 967.8036762 | 274 | 8 | 16 | 46 | 119 | 22 | 2 | 0 | 0 | 0 | 1 | 0 |
| SG0000003 | Sec31a | 0.000138 | 0.000287 | -0.517384939 | 6174.889001 | 4314.006243 | 99 | 11 | 16 | 19 | 76 | 13 | 0 | 0 | 0 | 0 | 0 | 0 |
| JSG000000 | Fuk | 0.00015 | 0.000312 | 0.649602951 | 488.4215407 | 766.2046493 | 39 | 16 | 16 | 20 | 35 | 2 | 0 | 0 | 0 | 0 | 0 | 0 |
| SG0000003 | Mau2 | 0.000169 | 0.000349 | -0.536560649 | 2488.677914 | 1715.725649 | 60 | 16 | 16 | 20 | 46 | 14 | 8 | 2 | 2 | 2 | 4 | 4 |
| SG0000003 | Aim2 | 0.000171 | 0.000354 | -0.778649365 | 968.2382654 | 564.3979494 | 234 | 7 | 16 | 29 | 121 | 17 | 2 | 0 | 0 | 0 | 2 | 0 |
| SG0000005 | Ece1 | 0.000235 | 0.000476 | -0.526848003 | 2619.401731 | 1818.046798 | 103 | 16 | 16 | 27 | 84 | 17 | 3 | 0 | 0 | 1 | 4 | 1 |
| SG0000002 | Ccdc39 | 0.000286 | 0.000576 | 1.314198001 | 28.6250954 | 71.18032414 | 92 | 10 | 16 | 20 | 47 | 8 | 6 | 0 | 1 | 1 | 1 | 1 |
| SG0000001 | Cwc25 | 0.000512 | 0.000997 | -0.558865219 | 981.8347214 | 666.5046627 | 76 | 9 | 16 | 26 | 47 | 3 | 7 | 1 | 1 | 2 | 3 | 1 |
| SG0000003 | Srprb | 0.000537 | 0.001043 | -0.558321751 | 886.8515157 | 602.2534801 | 41 | 15 | 16 | 19 | 34 | 2 | 4 | 2 | 3 | 4 | 4 | 0 |
| SG0000008 | Rbm12 | 0.001013 | 0.0019 | 0.474160181 | 902.8955819 | 1254.220702 | 90 | 12 | 16 | 22 | 54 | 10 | 16 | 0 | 0 | 0 | 2 | 0 |
| SG0000002 | Morn1 | 0.001124 | 0.0021 | 1.279863872 | 24.47823691 | 59.43709152 | 78 | 11 | 16 | 15 | 29 | 5 | 0 | 0 | 0 | 0 | 0 | 0 |
| JSG000000 | Mastl | 0.001813 | 0.003301 | 0.508455178 | 984.1997043 | 1400.04982 | 102 | 8 | 16 | 20 | 50 | 12 | 11 | 1 | 1 | 3 | 3 | 1 |
| SG0000000 | Snf8 | 0.001881 | 0.003418 | -0.46428365 | 1401.204753 | 1015.636492 | 62 | 14 | 16 | 24 | 44 | 2 | 0 | 0 | 0 | 0 | 0 | 0 |
| SG0000002 | Wipi2 | 0.001955 | 0.003543 | -0.428316871 | 3536.556862 | 2628.115329 | 88 | 14 | 16 | 22 | 36 | 8 | 4 | 0 | 0 | 0 | 0 | 0 |
| SG0000002 | Tmem234 | 0.002258 | 0.004055 | -0.439253643 | 2153.189028 | 1588.011473 | 152 | 0 | 16 | 43 | 129 | 6 | 18 | 0 | 0 | 0 | 0 | 0 |
| SG0000001 | Pex3 | 0.002696 | 0.004793 | -0.524631504 | 646.2244485 | 449.2142922 | 334 | 4 | 16 | 60 | 168 | 52 | 6 | 0 | 0 | 2 | 2 | 2 |
| SG0000003 | Arhgap29 | 0.004876 | 0.008356 | 0.397089541 | 1232.293738 | 1622.744345 | 207 | 7 | 16 | 27 | 87 | 13 | 19 | 0 | 0 | 3 | 3 | 2 |
| SG0000002 | Sec16a | 0.006496 | 0.010936 | 0.350103965 | 3085.909807 | 3933.462587 | 70 | 10 | 16 | 18 | 38 | 28 | 4 | 0 | 0 | 0 | 0 | 0 |
| SG0000002 | Pbld2 | 0.007128 | 0.011925 | 1.676811059 | 7.785095976 | 24.89054448 | 66 | 13 | 16 | 19 | 36 | 4 | 5 | 0 | 0 | 0 | 0 | 0 |
| SG0000000 | Phax | 0.01044 | 0.017024 | 0.36096089 | 1043.082743 | 1339.611341 | 88 | 14 | 16 | 22 | 41 | 2 | 1 | 0 | 0 | 0 | 0 | 0 |
| SG0000003 | Iqsec1 | 0.012226 | 0.019689 | -0.422347524 | 1058.935024 | 790.1872889 | 167 | 14 | 16 | 24 | 63 | 17 | 4 | 0 | 0 | 0 | 2 | 0 |
| SG0000006 | Myh13 | 0.012427 | 0.019982 | 1.849020709 | 10.54799234 | 37.99972923 | 135 | 6 | 16 | 14 | 56 | 12 | 0 | 0 | 0 | 0 | 0 | 0 |
| SG0000002 | Rnf20 | 0.012453 | 0.020016 | -0.357780173 | 2067.612527 | 1613.491162 | 112 | 12 | 16 | 20 | 56 | 4 | 2 | 0 | 0 | 2 | 4 | 0 |
| SG0000003 | 5031439G07Rik | 0.014444 | 0.022952 | -0.364696331 | 1538.699756 | 1195.003975 | 67 | 16 | 16 | 16 | 36 | 6 | 5 | 0 | 0 | 0 | 2 | 2 |
| SG0000002 | Hspa4 | 0.016198 | 0.025523 | 0.291767366 | 18592.77002 | 22760.13462 | 79 | 13 | 16 | 24 | 55 | 14 | 0 | 0 | 0 | 0 | 1 | 1 |
| JSG000000 | Apba1 | 0.02166 | 0.033297 | 2.371222846 | 3.171459942 | 16.40848307 | 281 | 6 | 16 | 27 | 116 | 38 | 4 | 0 | 0 | 1 | 2 | 2 |
| SG0000002 | Btf3l4 | 0.026861 | 0.040674 | 0.343517931 | 581.0685497 | 737.2869912 | 190 | 12 | 16 | 28 | 72 | 16 | 8 | 0 | 0 | 0 | 5 | 0 |
| SG0000002 | Meaf6 | 0.028668 | 0.043187 | 0.346047093 | 547.2322106 | 695.5721858 | 86 | 12 | 16 | 16 | 52 | 13 | 0 | 0 | 0 | 0 | 0 | 1 |
| SG0000002 | Papln | 0.033118 | 0.049336 | 2.05396668 | 4.301439621 | 17.86156104 | 52 | 16 | 16 | 20 | 28 | 0 | 0 | 0 | 0 | 0 | 0 | 0 |
| JSG000000 | 2210018M11Rik | 0.033393 | 0.049703 | 0.358342632 | 1272.988735 | 1631.910894 | 183 | 7 | 16 | 21 | 81 | 16 | 0 | 0 | 0 | 0 | 0 | 0 |
| JSG000000 | Sobp | 0.034128 | 0.050694 | 1.553706662 | 6.113872955 | 17.94852315 | 272 | 9 | 16 | 35 | 122 | 28 | 3 | 1 | 1 | 1 | 2 | 0 |
| SG0000002 | Gpr143 | 0.037743 | 0.055651 | -1.772543019 | 16.05253074 | 4.698453064 | 66 | 15 | 16 | 23 | 45 | 6 | 0 | 0 | 0 | 0 | 0 | 0 |
| SG0000003 | Ankrd11 | 0.038128 | 0.056159 | 0.258357058 | 5245.575561 | 6274.315763 | 257 | 6 | 16 | 26 | 78 | 25 | 0 | 0 | 0 | 0 | 0 | 0 |
| SG0000002 | Tmem50a | 0.043577 | 0.063573 | -0.29706731 | 1441.12408 | 1172.938399 | 55 | 14 | 16 | 22 | 54 | 2 | 1 | 0 | 0 | 0 | 0 | 0 |
| JSG000000 | Gbp4 | 0.052335 | 0.075229 | -0.769212969 | 89.93510822 | 52.76829778 | 108 | 8 | 16 | 22 | 70 | 6 | 25 | 2 | 2 | 4 | 14 | 0 |
| JSG000000 | Zswim3 | 0.063175 | 0.089468 | 0.317255453 | 400.5679066 | 499.0907931 | 43 | 15 | 16 | 17 | 27 | 0 | 0 | 0 | 0 | 0 | 2 | 0 |
| JSG000000 | Txnl1 | 0.067715 | 0.09537 | 0.228348017 | 3170.804225 | 3714.574139 | 100 | 8 | 16 | 20 | 69 | 4 | 8 | 0 | 1 | 0 | 3 | 1 |
| SG0000004 | Ccdc125 | 0.107213 | 0.145024 | -0.418696121 | 161.5393277 | 120.8476416 | 144 | 12 | 16 | 30 | 68 | 6 | 1 | 0 | 0 | 0 | 0 | 0 |
| SG0000002 | Grin1 | 0.115848 | 0.155436 | 1.033561775 | 81.4841167 | 166.8038456 | 96 | 8 | 16 | 16 | 40 | 24 | 0 | 0 | 0 | 0 | 0 | 0 |
| JSG000000 | Snx32 | 0.133675 | 0.176968 | 0.448111939 | 61.76007912 | 84.25641542 | 36 | 14 | 16 | 18 | 44 | 3 | 0 | 0 | 0 | 0 | 0 | 0 |
| SG0000002 | Gpr137b | 0.147823 | 0.193964 | -0.329617026 | 312.0801464 | 248.3370564 | 50 | 15 | 16 | 18 | 31 | 6 | 4 | 0 | 0 | 0 | 1 | 0 |
| SG0000004 | Stat2 | 0.164914 | 0.214395 | 0.304454557 | 1318.482502 | 1628.262139 | 66 | 16 | 16 | 24 | 66 | 16 | 0 | 0 | 0 | 0 | 6 | 0 |
| JSG000001 | Gtf3c2 | 0.173074 | 0.223882 | 0.187185749 | 1494.313818 | 1701.336963 | 108 | 16 | 16 | 24 | 44 | 10 | 3 | 0 | 0 | 0 | 0 | 0 |
| SG0000003 | Gxylt1 | 0.201704 | 0.256818 | 0.176486673 | 1138.474677 | 1286.622546 | 128 | 12 | 16 | 28 | 71 | 14 | 9 | 0 | 0 | 1 | 2 | 0 |
| SG0000002 | Mttp | 0.203794 | 0.259166 | -0.356745138 | 169.9737175 | 132.7366295 | 83 | 13 | 16 | 20 | 42 | 3 | 8 | 0 | 0 | 2 | 3 | 0 |
| JSG000000 | Zufsp | 0.206759 | 0.262601 | -0.250533228 | 336.9301403 | 283.2186488 | 91 | 10 | 16 | 19 | 55 | 7 | 2 | 0 | 0 | 1 | 1 | 0 |
| SG0000004 | Smek1 | 0.233223 | 0.292213 | 0.15638889 | 2831.050544 | 3155.188935 | 210 | 6 | 16 | 30 | 106 | 12 | 2 | 0 | 0 | 0 | 1 | 0 |
| SG0000004 | 1110034G24Rik | 0.250379 | 0.311249 | 0.282610617 | 124.1129347 | 150.9702303 | 232 | 11 | 16 | 42 | 128 | 27 | 0 | 0 | 0 | 0 | 0 | 0 |
| SG0000000 | Wbscr22 | 0.283356 | 0.346923 | 0.170550997 | 1028.723254 | 1157.815937 | 52 | 4 | 16 | 12 | 24 | 0 | 0 | 0 | 0 | 0 | 0 | 0 |
| SG0000000 | Tfe3 | 0.300529 | 0.365326 | -0.141348002 | 3668.078079 | 3325.742216 | 56 | 16 | 16 | 16 | 40 | 8 | 0 | 0 | 0 | 0 | 0 | 0 |
| JSG000000 | Isy1 | 0.323822 | 0.390198 | -0.157810658 | 981.4269489 | 879.735743 | 59 | 11 | 16 | 20 | 40 | 4 | 2 | 0 | 0 | 0 | 1 | 0 |
| JSG000000 | Gkap1 | 0.361109 | 0.429381 | 0.158602163 | 368.2252585 | 411.014917 | 108 | 14 | 16 | 20 | 43 | 6 | 2 | 0 | 0 | 0 | 0 | 0 |
| SG0000002 | Mob4 | 0.375717 | 0.444507 | 0.120111804 | 1781.043045 | 1935.671571 | 215 | 4 | 16 | 22 | 112 | 20 | 20 | 0 | 0 | 2 | 12 | 4 |
| JSG000000 | Slc16a14 | 0.379906 | 0.448678 | 0.508226642 | 36.54486487 | 51.97779166 | 56 | 14 | 16 | 21 | 37 | 3 | 0 | 0 | 0 | 0 | 1 | 0 |
| JSG000000 | Vcpip1 | 0.444028 | 0.513323 | 0.094867901 | 2101.410854 | 2244.23874 | 87 | 11 | 16 | 20 | 64 | 9 | 19 | 1 | 2 | 1 | 12 | 1 |
| SG0000003 | Rab27a | 0.49844 | 0.565819 | 0.130449033 | 850.6592702 | 931.1608562 | 70 | 11 | 16 | 21 | 37 | 9 | 5 | 0 | 0 | 0 | 0 | 1 |
| SG0000002 | Smad2 | 0.506566 | 0.573698 | 0.093710675 | 958.5077439 | 1022.834457 | 222 | 7 | 16 | 30 | 119 | 24 | 2 | 0 | 0 | 0 | 0 | 0 |
| JSG000000 | Adgrg6 | 0.513118 | 0.580011 | 0.240159239 | 117.1659139 | 138.3873583 | 326 | 6 | 16 | 42 | 165 | 38 | 6 | 1 | 1 | 2 | 3 | 1 |

| | | | | | | | | | | | | | | | | | |
|---|---|---|---|---|---|---|---|---|---|---|---|---|---|---|---|---|---|
| SG0000002 | Megf10 | 0.55539 | 0.62045 | -0.382045669 | 20.51074192 | 15.73889231 | 258 | 13 | 16 | 35 | 131 | 41 | 8 | 0 | 0 | 0 | 2 | 0 |
| SG0000002 | Nmnat1 | 0.609115 | 0.671663 | -0.114318062 | 215.2635592 | 198.8645404 | 77 | 13 | 16 | 16 | 37 | 6 | 7 | 2 | 2 | 2 | 3 | 0 |
| SG0000002 | Ndufv2 | 0.62926 | 0.69039 | -0.073036779 | 3109.996902 | 2956.471488 | 138 | 9 | 16 | 15 | 48 | 6 | 6 | 1 | 2 | 3 | 4 | 0 |
| SG0000001 | Brca1 | 0.649568 | 0.708764 | 0.04857187 | 4128.293669 | 4269.648993 | 128 | 15 | 16 | 23 | 62 | 7 | 1 | 0 | 0 | 0 | 0 | 0 |
| SG0000003 | Tsr1 | 0.687447 | 0.742933 | -0.053643595 | 4050.590075 | 3902.743086 | 45 | 9 | 16 | 18 | 33 | 6 | 4 | 0 | 0 | 0 | 0 | 2 |
| SG0000003 | Clstn1 | 0.701936 | 0.755775 | -0.058386045 | 3634.708396 | 3490.548072 | 172 | 12 | 16 | 40 | 118 | 19 | 0 | 0 | 0 | 0 | 0 | 1 |
| SG0000004 | Setx | 0.711753 | 0.764461 | 0.038552567 | 5519.084986 | 5668.557557 | 127 | 8 | 16 | 21 | 57 | 15 | 7 | 1 | 1 | 1 | 3 | 0 |
| JSG000000 | Lmod1 | 0.736326 | 0.786455 | 0.260689189 | 29.14125009 | 34.91270054 | 61 | 14 | 16 | 19 | 40 | 4 | 1 | 0 | 0 | 0 | 0 | 0 |
| JSG000000 | Scfd1 | 0.775155 | 0.820213 | 0.037370848 | 1213.834227 | 1245.687555 | 184 | 7 | 16 | 32 | 80 | 15 | 0 | 0 | 0 | 0 | 0 | 0 |
| JSG000000 | Mkx | 0.829579 | 0.866489 | -0.094124535 | 154.8933889 | 145.1104114 | 139 | 13 | 16 | 25 | 58 | 13 | 1 | 0 | 0 | 0 | 3 | 1 |
| SG0000002 | Iars2 | 0.853804 | 0.885851 | -0.031515141 | 4352.159947 | 4258.119498 | 162 | 8 | 16 | 32 | 61 | 18 | 7 | 0 | 1 | 0 | 1 | 1 |
| SG0000002 | Slirp | 0.864012 | 0.894371 | 0.017080633 | 1188.073149 | 1202.22281 | 64 | 16 | 16 | 20 | 28 | 0 | 4 | 0 | 0 | 0 | 1 | 0 |
| SG0000002 | Taf12 | 0.939424 | 0.956295 | -0.013254942 | 726.7012876 | 720.0552059 | 59 | 12 | 16 | 20 | 35 | 2 | 1 | 0 | 0 | 0 | 1 | 0 |
| SG0000002 | Coq10b | 0.958156 | 0.972985 | 0.011971838 | 710.5107865 | 716.4313107 | 83 | 16 | 16 | 25 | 41 | 5 | 3 | 0 | 0 | 1 | 1 | 1 |
| JSG000000 | Gm13151 | 1.9E-137 | 2.5E-135 | 9.591232281 | 0.674076249 | 519.9452759 | 83 | 14 | 15 | 23 | 51 | 8 | 2 | 0 | 0 | 1 | 4 | 0 |
| JSG000000 | Pawr | 7.2E-104 | 5.6E-102 | 3.776155925 | 69.92530959 | 958.0112288 | 157 | 10 | 15 | 22 | 55 | 11 | 1 | 0 | 0 | 1 | 2 | 0 |
| JSG000000 | Hunk | 8.85E-96 | 5.74E-94 | 6.023838116 | 6.183804759 | 402.3571573 | 150 | 11 | 15 | 27 | 67 | 12 | 3 | 0 | 0 | 2 | 3 | 0 |
| SG0000001 | Lama4 | 3.71E-88 | 2E-86 | -6.423232777 | 9638.675704 | 112.3134153 | 294 | 7 | 15 | 40 | 150 | 20 | 0 | 0 | 0 | 0 | 0 | 0 |
| SG0000002 | Tspan13 | 8.6E-71 | 3.14E-69 | 3.056443886 | 99.0149921 | 823.7249997 | 89 | 11 | 15 | 20 | 39 | 7 | 2 | 0 | 0 | 0 | 0 | 0 |
| SG0000004 | Pla2g5 | 2.72E-70 | 9.76E-69 | 8.512111417 | 0.970711415 | 354.3977857 | 185 | 15 | 15 | 33 | 102 | 17 | 3 | 0 | 0 | 0 | 0 | 3 |
| SG0000002 | Pms1 | 1.48E-60 | 4.19E-59 | 2.367888929 | 286.5070488 | 1478.907138 | 223 | 9 | 15 | 26 | 122 | 20 | 0 | 0 | 0 | 0 | 0 | 0 |
| JSG000000 | Fam208a | 8.69E-59 | 2.32E-57 | 2.009525892 | 1051.936308 | 4235.620246 | 173 | 8 | 15 | 25 | 90 | 14 | 9 | 0 | 2 | 2 | 3 | 2 |
| SG0000002 | Rnaseh2b | 3.9E-56 | 9.74E-55 | 2.133206827 | 472.9028982 | 2074.584923 | 147 | 7 | 15 | 13 | 74 | 8 | 1 | 0 | 0 | 0 | 0 | 0 |
| SG0000002 | Nfatc2 | 1.48E-55 | 3.65E-54 | -3.316331193 | 1397.30285 | 140.2734182 | 375 | 15 | 15 | 62 | 137 | 77 | 2 | 0 | 0 | 2 | 6 | 2 |
| SG0000002 | Eng | 2.41E-53 | 5.48E-52 | -3.195040393 | 3969.14083 | 433.4045963 | 118 | 12 | 15 | 18 | 108 | 12 | 4 | 0 | 0 | 0 | 0 | 0 |
| SG0000001 | Adgrl1 | 1.5E-52 | 3.36E-51 | 1.910733517 | 1232.669125 | 4634.838659 | 121 | 10 | 15 | 15 | 30 | 11 | 1 | 0 | 0 | 0 | 0 | 6 |
| SG0000002 | Slc38a4 | 1.47E-37 | 1.99E-36 | 6.794729136 | 78.26905984 | 8689.75235 | 145 | 14 | 15 | 32 | 74 | 16 | 4 | 0 | 0 | 0 | 2 | 2 |
| JSG000000 | Hs3st2 | 6.39E-37 | 8.47E-36 | -7.376243422 | 164.7013038 | 0.991348861 | 159 | 8 | 15 | 18 | 61 | 14 | 0 | 0 | 0 | 0 | 0 | 0 |
| SG0000003 | Antxr1 | 1.9E-34 | 2.31E-33 | -3.376522412 | 2981.7879 | 287.1059684 | 217 | 9 | 15 | 25 | 94 | 18 | 5 | 0 | 0 | 0 | 1 | 1 |
| JSG000000 | Cdc5l | 8.66E-33 | 9.8E-32 | 1.48857981 | 1432.775021 | 4020.547295 | 97 | 12 | 15 | 17 | 41 | 8 | 2 | 0 | 0 | 0 | 0 | 0 |
| JSG000000 | Cfap74 | 3.16E-32 | 3.51E-31 | 4.834080342 | 4.541360663 | 129.535611 | 74 | 14 | 15 | 17 | 42 | 2 | 0 | 0 | 0 | 0 | 1 | 0 |
| SG0000004 | D630023F18Rik | 4.18E-31 | 4.43E-30 | 9.286791483 | 0.34247518 | 213.9104013 | 98 | 12 | 15 | 24 | 50 | 11 | 5 | 0 | 0 | 0 | 1 | 2 |
| JSG000000 | Mertk | 8.04E-31 | 8.4E-30 | -3.023452628 | 1668.147157 | 205.156097 | 160 | 11 | 15 | 21 | 59 | 19 | 2 | 0 | 1 | 1 | 1 | 0 |
| SG0000002 | Fxr1 | 1.75E-29 | 1.72E-28 | 1.338368415 | 4261.53741 | 10775.94443 | 149 | 9 | 15 | 23 | 64 | 12 | 0 | 0 | 0 | 0 | 0 | 0 |
| SG0000006 | Dmkn | 9.1E-28 | 8.32E-27 | -6.718929501 | 3743.8006 | 35.53967001 | 45 | 15 | 15 | 20 | 67 | 10 | 0 | 0 | 0 | 0 | 0 | 0 |
| SG0000002 | Hif1a | 1.03E-27 | 9.39E-27 | -1.664416513 | 8711.132258 | 2748.11759 | 343 | 10 | 15 | 44 | 155 | 23 | 18 | 4 | 4 | 6 | 8 | 0 |
| SG0000002 | Orc2 | 8.59E-27 | 7.51E-26 | 1.325461985 | 1836.491156 | 4602.487916 | 117 | 10 | 15 | 26 | 69 | 8 | 4 | 0 | 0 | 0 | 1 | 0 |
| SG0000002 | Plxna2 | 2.28E-26 | 1.96E-25 | -3.17542571 | 2319.384965 | 256.7287167 | 277 | 7 | 15 | 34 | 129 | 30 | 7 | 0 | 1 | 1 | 5 | 2 |
| SG0000002 | Ppt1 | 8.49E-25 | 6.79E-24 | -1.76421941 | 2391.562073 | 704.042208 | 126 | 6 | 15 | 24 | 82 | 18 | 5 | 0 | 0 | 0 | 2 | 2 |
| SG0000005 | Rnf103 | 1.96E-24 | 1.54E-23 | -2.206245384 | 978.6826661 | 212.0781167 | 156 | 6 | 15 | 15 | 36 | 9 | 16 | 1 | 1 | 2 | 2 | 0 |
| SG0000003 | Dap | 5.49E-24 | 4.22E-23 | -1.724877539 | 3244.242052 | 981.4617334 | 140 | 15 | 15 | 21 | 36 | 18 | 5 | 0 | 0 | 0 | 0 | 0 |
| SG0000002 | Acox3 | 2.19E-23 | 1.63E-22 | -1.846172365 | 1070.412237 | 297.713005 | 118 | 10 | 15 | 26 | 56 | 10 | 5 | 0 | 0 | 1 | 1 | 0 |
| SG0000004 | Zbtb38 | 2.87E-22 | 2.04E-21 | -1.595064782 | 2140.785865 | 708.6158859 | 176 | 11 | 15 | 23 | 74 | 17 | 9 | 0 | 0 | 1 | 3 | 1 |
| SG0000000 | St8sia6 | 1.04E-21 | 7.22E-21 | 7.551238206 | 0.331601068 | 62.1963825 | 298 | 8 | 15 | 43 | 176 | 30 | 14 | 1 | 2 | 2 | 8 | 0 |
| SG0000004 | Pofut1 | 1.38E-21 | 9.45E-21 | -1.601934496 | 1781.842146 | 587.0010697 | 181 | 10 | 15 | 32 | 85 | 4 | 7 | 0 | 0 | 0 | 0 | 4 |
| JSG000000 | Golga4 | 3.26E-21 | 2.19E-20 | -1.480439668 | 4531.484934 | 1623.991656 | 156 | 11 | 15 | 22 | 48 | 18 | 1 | 1 | 1 | 1 | 2 | 1 |
| SG0000002 | Cdc7 | 3.55E-21 | 2.37E-20 | 1.284582762 | 734.5131597 | 1789.359174 | 202 | 10 | 15 | 59 | 93 | 4 | 8 | 0 | 0 | 4 | 8 | 4 |
| JSG000000 | Rcn2 | 1.28E-20 | 8.33E-20 | -1.426503023 | 4284.790463 | 1594.077346 | 95 | 8 | 15 | 19 | 51 | 10 | 13 | 0 | 0 | 0 | 0 | 0 |
| JSG000000 | 2610021A01Rik | 8.69E-20 | 5.38E-19 | 1.463677202 | 628.3275436 | 1732.993125 | 76 | 11 | 15 | 20 | 53 | 5 | 5 | 0 | 0 | 1 | 3 | 0 |
| SG0000002 | Klhl8 | 3.58E-19 | 2.14E-18 | 1.394837592 | 315.742935 | 830.2743019 | 205 | 6 | 15 | 24 | 102 | 26 | 14 | 4 | 4 | 6 | 10 | 0 |
| SG0000003 | Timeless | 3.92E-19 | 2.34E-18 | 1.470176809 | 796.0669685 | 2205.550075 | 86 | 15 | 15 | 21 | 59 | 0 | 3 | 0 | 0 | 0 | 0 | 0 |
| SG0000002 | Mark1 | 5.69E-19 | 3.37E-18 | -1.51283801 | 1496.696227 | 524.4740974 | 163 | 10 | 15 | 24 | 76 | 17 | 4 | 0 | 1 | 0 | 0 | 0 |
| JSG000000 | Uaca | 1.94E-18 | 1.11E-17 | 1.188293399 | 812.6148457 | 1851.811211 | 170 | 8 | 15 | 26 | 67 | 21 | 0 | 0 | 0 | 0 | 0 | 0 |
| SG0000003 | Edrf1 | 1.08E-17 | 5.98E-17 | 1.197847433 | 638.1763802 | 1463.958393 | 130 | 11 | 15 | 25 | 83 | 19 | 4 | 0 | 2 | 2 | 2 | 0 |
| SG0000002 | Myh9 | 3.02E-17 | 1.64E-16 | -1.404910454 | 26817.99739 | 10127.59105 | 104 | 15 | 15 | 22 | 48 | 6 | 1 | 0 | 0 | 0 | 0 | 0 |
| SG0000002 | P2rx4 | 4.28E-17 | 2.3E-16 | -1.927283983 | 889.4647808 | 233.861358 | 135 | 15 | 15 | 28 | 83 | 5 | 1 | 0 | 0 | 0 | 0 | 0 |
| JSG000000 | Mthfd1 | 4.42E-17 | 2.37E-16 | 1.014454358 | 4701.882441 | 9498.454734 | 105 | 9 | 15 | 28 | 57 | 16 | 0 | 0 | 0 | 0 | 0 | 0 |
| SG0000003 | Adamts7 | 1.44E-16 | 7.47E-16 | -2.160348191 | 529.3421143 | 118.4150332 | 116 | 10 | 15 | 30 | 64 | 21 | 1 | 0 | 0 | 0 | 4 | 1 |
| SG0000002 | Rab2b | 3.82E-16 | 1.94E-15 | 1.249172292 | 372.8442723 | 886.2695035 | 150 | 15 | 15 | 43 | 92 | 10 | 10 | 2 | 2 | 8 | 10 | 0 |
| JSG000000 | Cnot10 | 1.46E-15 | 7.14E-15 | 1.032352967 | 1344.751976 | 2750.498422 | 96 | 9 | 15 | 21 | 39 | 13 | 1 | 0 | 0 | 0 | 0 | 0 |
| SG0000006 | Ecsit | 7.32E-15 | 3.43E-14 | 1.162196342 | 567.8263992 | 1270.783418 | 57 | 12 | 15 | 15 | 21 | 6 | 0 | 0 | 0 | 0 | 0 | 0 |
| JSG000000 | Chst7 | 1.46E-14 | 6.72E-14 | -2.508209554 | 188.0602039 | 33.05602221 | 79 | 10 | 15 | 23 | 60 | 4 | 0 | 0 | 0 | 0 | 0 | 0 |
| SG0000002 | Zkscan1 | 1.74E-14 | 7.95E-14 | -1.281429864 | 1385.827549 | 570.1122385 | 111 | 9 | 15 | 9 | 35 | 13 | 24 | 0 | 0 | 0 | 8 | 6 |
| JSG000000 | Smarcad1 | 6.46E-14 | 2.85E-13 | 0.947037384 | 1874.044007 | 3612.987381 | 197 | 10 | 15 | 32 | 86 | 19 | 7 | 0 | 1 | 1 | 5 | 1 |
| SG0000003 | Rusc2 | 1.24E-13 | 5.35E-13 | -3.046322509 | 2233.676737 | 270.3870388 | 117 | 15 | 15 | 28 | 51 | 7 | 15 | 12 | 12 | 15 | 18 | 3 |
| SG0000003 | Tmtc3 | 1.39E-13 | 5.97E-13 | -1.181053297 | 3341.606836 | 1473.746823 | 289 | 2 | 15 | 32 | 105 | 19 | 19 | 0 | 2 | 3 | 6 | 1 |
| JSG000000 | Ndfip1 | 1.62E-13 | 6.95E-13 | -1.130597647 | 2652.849278 | 1211.626522 | 109 | 12 | 15 | 25 | 61 | 9 | 0 | 0 | 0 | 0 | 0 | 0 |
| JSG000000 | Mmrn2 | 2.16E-13 | 9.18E-13 | 4.652928636 | 1.939079263 | 48.78279158 | 46 | 12 | 15 | 20 | 33 | 3 | 1 | 0 | 1 | 0 | 0 | 0 |
| JSG000000 | Frrs1l | 1.26E-12 | 5.1E-12 | 3.655073344 | 4.611292466 | 58.09107062 | 104 | 15 | 15 | 33 | 57 | 22 | 23 | 1 | 1 | 7 | 11 | 6 |
| SG0000004 | Sh3bgr | 2.34E-12 | 9.28E-12 | 6.71467124 | 0.331601068 | 34.82838811 | 125 | 9 | 15 | 21 | 56 | 13 | 0 | 0 | 0 | 0 | 0 | 0 |
| SG0000004 | Hjurp | 6.83E-12 | 2.64E-11 | 0.843949029 | 5582.036479 | 10019.51369 | 54 | 8 | 15 | 8 | 14 | 0 | 4 | 0 | 0 | 0 | 0 | 0 |
| SG0000003 | Skp1a | 7.86E-12 | 3.03E-11 | 0.81991076 | 6371.232492 | 11247.11629 | 120 | 15 | 15 | 19 | 60 | 2 | 5 | 0 | 0 | 0 | 1 | 2 |
| SG0000006 | Zfp229 | 9E-12 | 3.45E-11 | 1.186849086 | 222.1707738 | 505.7828542 | 153 | 10 | 15 | 30 | 98 | 17 | 18 | 0 | 0 | 2 | 6 | 0 |
| SG0000001 | Slc38a3 | 9.1E-12 | 3.49E-11 | 9.556480822 | 0.68495036 | 515.7586004 | 26 | 15 | 15 | 17 | 20 | 2 | 0 | 0 | 0 | 0 | 1 | 1 |
| JSG000000 | Atg4a | 1.02E-11 | 3.88E-11 | -1.613542844 | 315.303974 | 103.0397151 | 112 | 11 | 15 | 16 | 54 | 13 | 0 | 0 | 0 | 0 | 0 | 0 |
| SG0000002 | Atp5c1 | 4.73E-11 | 1.72E-10 | 0.802359715 | 4028.968288 | 7026.324299 | 121 | 12 | 15 | 21 | 63 | 15 | 0 | 0 | 0 | 0 | 0 | 0 |
| SG0000002 | Polr1d | 5.97E-11 | 2.15E-10 | -0.957090783 | 3478.124008 | 1791.562766 | 103 | 3 | 15 | 18 | 51 | 15 | 0 | 0 | 0 | 0 | 0 | 0 |
| SG0000002 | Wsb2 | 6.45E-11 | 2.32E-10 | -0.939031564 | 4185.11924 | 2182.886568 | 47 | 11 | 15 | 23 | 36 | 0 | 2 | 0 | 0 | 1 | 1 | 0 |
| SG0000000 | Cav1 | 1.09E-10 | 3.87E-10 | -1.15110008 | 1309.577465 | 589.6788363 | 379 | 10 | 15 | 35 | 160 | 20 | 12 | 0 | 0 | 0 | 0 | 0 |
| SG0000003 | Igsf8 | 2.14E-10 | 7.45E-10 | 1.99444098 | 469.6023889 | 1871.185559 | 70 | 15 | 15 | 20 | 36 | 2 | 5 | 0 | 0 | 0 | 0 | 0 |
| SG0000002 | Pgm1 | 3.3E-10 | 1.13E-09 | 0.910524334 | 584.4507583 | 1098.608773 | 103 | 10 | 15 | 25 | 56 | 13 | 7 | 0 | 0 | 1 | 5 | 0 |
| SG0000002 | Fyb | 4.41E-10 | 1.5E-09 | -4.691564449 | 42.4253536 | 1.641813585 | 203 | 10 | 15 | 34 | 112 | 16 | 0 | 0 | 0 | 0 | 1 | 0 |
| SG0000002 | Auh | 5.71E-10 | 1.92E-09 | 0.994319635 | 349.0031084 | 695.26334 | 317 | 4 | 15 | 32 | 114 | 28 | 15 | 0 | 0 | 3 | 3 | 1 |
| SG0000002 | Anapc16 | 9.05E-10 | 2.99E-09 | -1.016436488 | 1082.369867 | 535.0542542 | 130 | 15 | 15 | 20 | 50 | 0 | 5 | 0 | 0 | 0 | 0 | 0 |
| JSG000000 | Ints9 | 1.21E-09 | 3.94E-09 | 0.807536787 | 1231.289588 | 2155.028422 | 130 | 9 | 15 | 27 | 79 | 20 | 0 | 0 | 0 | 0 | 1 | 0 |
| SG0000002 | Rbl1 | 1.64E-09 | 5.3E-09 | -0.950286953 | 1626.111201 | 841.5605408 | 114 | 10 | 15 | 18 | 52 | 12 | 4 | 0 | 1 | 0 | 1 | 3 |
| JSG000000 | Cpt1a | 2.49E-09 | 7.94E-09 | -0.922868467 | 5567.656066 | 2936.711668 | 81 | 13 | 15 | 20 | 42 | 10 | 3 | 0 | 0 | 1 | 1 | 1 |
| SG0000003 | Xylt1 | 2.85E-09 | 9.02E-09 | 3.160327147 | 15.86682712 | 141.8545784 | 324 | 6 | 15 | 33 | 127 | 35 | 0 | 0 | 0 | 0 | 0 | 0 |
| JSG000000 | B4galnt2 | 6.16E-09 | 1.9E-08 | 4.548220406 | 0 | 22.39649327 | 67 | 11 | 15 | 23 | 45 | 4 | 0 | 0 | 0 | 0 | 0 | 0 |
| JSG000000 | Ppp1r14c | 1.2E-08 | 3.63E-08 | -1.691502346 | 166.145933 | 51.43950385 | 177 | 10 | 15 | 21 | 76 | 21 | 7 | 0 | 0 | 0 | 2 | 1 |
| JSG000000 | Celsr1 | 1.33E-08 | 4E-08 | 1.275610038 | 237.4637008 | 574.9023924 | 122 | 11 | 15 | 25 | 66 | 19 | 1 | 0 | 0 | 1 | 1 | 1 |
| JSG000000 | Cenph | 1.69E-08 | 5.04E-08 | 0.883858347 | 396.0848948 | 730.8969585 | 40 | 13 | 15 | 16 | 20 | 3 | 1 | 0 | 0 | 0 | 0 | 0 |
| JSG000000 | Npat | 2.24E-08 | 6.62E-08 | 0.744852163 | 1278.188542 | 2141.991602 | 104 | 10 | 15 | 19 | 43 | 7 | 8 | 0 | 0 | 0 | 2 | 2 |
| JSG000000 | Rab2a | 3.23E-08 | 9.44E-08 | -0.793420035 | 4036.11851 | 2328.738269 | 158 | 8 | 15 | 19 | 65 | 8 | 1 | 0 | 0 | 1 | 2 | 1 |
| JSG000000 | Gpr156 | 6.04E-08 | 1.73E-07 | 6.179087958 | 0.307509278 | 22.28173706 | 129 | 11 | 15 | 27 | 60 | 11 | 1 | 0 | 0 | 0 | 0 | 0 |
| SG0000002 | Fmo5 | 1.02E-07 | 2.86E-07 | 2.196489589 | 15.40254628 | 70.59952417 | 131 | 9 | 15 | 21 | 49 | 9 | 2 | 0 | 0 | 0 | 1 | 0 |
| JSG000000 | Acer1 | 1.22E-07 | 3.4E-07 | 4.304694193 | 0 | 18.76250871 | 69 | 12 | 15 | 16 | 37 | 6 | 1 | 0 | 0 | 0 | 0 | 1 |
| SG0000002 | Nfkb1 | 1.36E-07 | 3.76E-07 | -0.804778952 | 1716.872038 | 982.8230268 | 192 | 8 | 15 | 23 | 60 | 12 | 4 | 0 | 0 | 0 | 0 | 1 |
| SG0000005 | Tnip2 | 1.36E-07 | 3.78E-07 | -1.051856517 | 455.6002786 | 219.7574557 | 80 | 12 | 15 | 15 | 56 | 3 | 2 | 0 | 0 | 0 | 2 | 0 |
| JSG000000 | Cars2 | 1.39E-07 | 3.84E-07 | 0.906031862 | 320.8968611 | 601.3235828 | 41 | 14 | 15 | 16 | 24 | 2 | 0 | 0 | 0 | 0 | 0 | 0 |
| SG0000006 | Ninl | 1.43E-07 | 3.97E-07 | 0.970821053 | 381.4621317 | 747.648883 | 113 | 11 | 15 | 22 | 59 | 11 | 1 | 0 | 0 | 0 | 0 | 0 |
| SG0000002 | Vps33a | 2.87E-07 | 7.73E-07 | -0.78051258 | 1684.707656 | 980.7692722 | 87 | 10 | 15 | 14 | 40 | 10 | 2 | 0 | 0 | 0 | 0 | 1 |
| SG0000002 | Klhl7 | 2.95E-07 | 7.92E-07 | 0.739058432 | 667.3141682 | 1113.804844 | 155 | 8 | 15 | 22 | 58 | 8 | 1 | 0 | 0 | 0 | 3 | 0 |
| SG0000002 | Slc15a4 | 4.27E-07 | 1.13E-06 | -0.852260263 | 910.341893 | 504.2531567 | 112 | 12 | 15 | 18 | 41 | 9 | 0 | 0 | 0 | 0 | 0 | 0 |
| JSG000000 | Ncs1 | 5.04E-07 | 1.33E-06 | -1.196747041 | 543.3104896 | 237.02346 | 68 | 11 | 15 | 17 | 37 | 2 | 3 | 1 | 1 | 1 | 1 | 0 |
| SG0000002 | Slc35f5 | 5.15E-07 | 1.36E-06 | -0.853767067 | 929.0193975 | 514.0617444 | 85 | 12 | 15 | 17 | 43 | 4 | 2 | 0 | 0 | 0 | 1 | 0 |
| SG0000003 | Tmem237 | 8.52E-07 | 2.21E-06 | 0.734568613 | 578.6049819 | 962.7407513 | 112 | 6 | 15 | 12 | 53 | 9 | 7 | 0 | 0 | 0 | 11 | 0 |
| JSG000000 | Snx9 | 1.15E-06 | 2.96E-06 | -0.768804493 | 1257.787984 | 738.2003189 | 112 | 9 | 15 | 23 | 48 | 9 | 3 | 0 | 0 | 0 | 0 | 0 |
| JSG000000 | Bud13 | 1.49E-06 | 3.8E-06 | 0.741952711 | 482.017748 | 806.1447886 | 54 | 13 | 15 | 23 | 40 | 4 | 2 | 0 | 0 | 2 | 2 | 0 |
| SG0000004 | Esf1 | 1.96E-06 | 4.94E-06 | 0.631753657 | 1369.263713 | 2121.601919 | 137 | 10 | 15 | 26 | 72 | 17 | 4 | 0 | 0 | 0 | 1 | 1 |
| JSG000000 | Pigl | 2.58E-06 | 6.42E-06 | 0.802005418 | 301.7197073 | 526.0552575 | 89 | 10 | 15 | 17 | 37 | 10 | 3 | 0 | 0 | 0 | 0 | 0 |
| SG0000004 | Dgkz | 3.23E-06 | 7.97E-06 | -0.694956453 | 1843.644665 | 1138.870866 | 104 | 14 | 15 | 18 | 36 | 1 | 2 | 0 | 0 | 0 | 0 | 0 |
| SG0000002 | Smad5 | 3.24E-06 | 7.98E-06 | -0.697917125 | 1982.979905 | 1222.430914 | 262 | 9 | 15 | 30 | 69 | 27 | 16 | 3 | 3 | 3 | 9 | 0 |
| SG0000002 | Vipas39 | 1.06E-05 | 2.5E-05 | -0.680985648 | 1359.101907 | 847.724852 | 132 | 15 | 15 | 37 | 120 | 9 | 0 | 0 | 0 | 0 | 0 | 0 |
| JSG000000 | Exd1 | 1.18E-05 | 2.76E-05 | -1.474222235 | 110.0846264 | 39.62247135 | 98 | 13 | 15 | 21 | 48 | 11 | 12 | 4 | 4 | 4 | 10 | 0 |
| SG0000004 | Svop | 1.96E-05 | 4.49E-05 | 2.676201549 | 10.5744277 | 67.58865639 | 69 | 7 | 15 | 16 | 27 | 8 | 1 | 0 | 0 | 0 | 0 | 0 |
| SG0000001 | Atraid | 1.97E-05 | 4.51E-05 | -0.709023263 | 863.3260418 | 528.1260279 | 38 | 15 | 15 | 15 | 41 | 0 | 2 | 0 | 0 | 0 | 3 | 0 |
| SG0000000 | Fkbp7 | 2.54E-05 | 5.75E-05 | -1.138445169 | 171.0758055 | 77.71099138 | 49 | 7 | 15 | 14 | 29 | 10 | 1 | 0 | 0 | 0 | 0 | 0 |
| SG0000000 | Fance | 2.59E-05 | 5.86E-05 | 0.696047714 | 463.8594547 | 751.4803924 | 57 | 15 | 15 | 23 | 90 | 27 | 0 | 0 | 0 | 0 | 0 | 0 |
| SG0000003 | Slc7a2 | 2.95E-05 | 6.64E-05 | 1.033568587 | 96.32556467 | 197.1862876 | 308 | 0 | 15 | 40 | 127 | 21 | 31 | 0 | 0 | 2 | 4 | 2 |
| SG0000003 | Ncoa5 | 3.34E-05 | 7.49E-05 | -0.659606741 | 1213.787622 | 768.3892669 | 189 | 9 | 15 | 27 | 51 | 9 | 0 | 0 | 0 | 0 | 0 | 0 |
| SG0000002 | Arpc3 | 3.42E-05 | 7.65E-05 | -0.577757175 | 4023.914943 | 2696.041371 | 63 | 13 | 15 | 17 | 23 | 7 | 0 | 0 | 0 | 0 | 0 | 0 |
| SG0000002 | Ctage5 | 4.78E-05 | 0.000105 | -0.573486594 | 3077.54818 | 2068.084091 | 135 | 9 | 15 | 27 | 66 | 16 | 2 | 0 | 0 | 0 | 1 | 1 |

| | | | | | | | | | | | | | | | | | |
|---|---|---|---|---|---|---|---|---|---|---|---|---|---|---|---|---|---|
| ISG000000 | Itfg1 | 6.25E-05 | 0.000136 | -0.581985679 | 2461.406687 | 1644.327157 | 294 | 3 | 15 | 39 | 141 | 26 | 6 | 0 | 1 | 1 | 2 | 2 |
| SG0000002 | Tmem38b | 0.000169 | 0.000349 | 0.646621554 | 315.9877399 | 494.6781367 | 91 | 12 | 15 | 21 | 40 | 8 | 5 | 1 | 1 | 1 | 4 | 1 |
| SG0000005 | Elmod3 | 0.000211 | 0.000431 | -0.734960073 | 406.2207956 | 244.0715307 | 104 | 9 | 15 | 15 | 36 | 3 | 1 | 0 | 0 | 0 | 0 | 0 |
| ISG000000 | Pgpep1 | 0.000226 | 0.00046 | -0.563961216 | 1793.556471 | 1213.237465 | 28 | 15 | 15 | 16 | 20 | 0 | 3 | 1 | 1 | 1 | 1 | 0 |
| SG0000002 | P2rx3 | 0.000287 | 0.000577 | 2.576843048 | 3.915467995 | 23.36096237 | 107 | 6 | 15 | 18 | 45 | 15 | 0 | 0 | 0 | 0 | 0 | 0 |
| ISG000000 | Car8 | 0.00029 | 0.000583 | 2.72945625 | 7.651419345 | 50.74464373 | 197 | 12 | 15 | 26 | 106 | 15 | 8 | 0 | 0 | 0 | 3 | 0 |
| SG0000004 | Mat2b | 0.000292 | 0.000586 | 0.474366264 | 1825.739136 | 2536.513625 | 73 | 6 | 15 | 18 | 45 | 7 | 1 | 0 | 0 | 0 | 0 | 0 |
| SG0000001 | Arrb1 | 0.000305 | 0.000611 | -0.695399757 | 2511.976537 | 1551.241421 | 177 | 15 | 15 | 28 | 140 | 9 | 2 | 0 | 0 | 0 | 0 | 0 |
| ISG000000 | Cenpa | 0.000336 | 0.00067 | 0.454857776 | 3228.842838 | 4425.605935 | 76 | 6 | 15 | 16 | 37 | 10 | 6 | 0 | 4 | 0 | 0 | 0 |
| SG0000000 | Dnmt1 | 0.000379 | 0.000753 | 0.434368633 | 6903.027333 | 9328.194239 | 116 | 12 | 15 | 21 | 45 | 9 | 0 | 0 | 0 | 0 | 0 | 0 |
| SG0000003 | Psmf1 | 0.000434 | 0.000854 | -0.531103539 | 1397.88079 | 967.368721 | 143 | 6 | 15 | 24 | 54 | 8 | 6 | 0 | 0 | 0 | 3 | 2 |
| SG0000001 | Snx3 | 0.000449 | 0.000881 | 0.447655633 | 2559.618569 | 3490.864772 | 174 | 9 | 15 | 36 | 96 | 27 | 6 | 0 | 0 | 0 | 0 | 0 |
| SG0000007 | Trappc2 | 0.000488 | 0.000954 | -0.781051001 | 276.5245447 | 160.9214387 | 89 | 9 | 15 | 18 | 45 | 9 | 21 | 0 | 6 | 6 | 6 | 0 |
| SG0000002 | Polr3gl | 0.001158 | 0.002159 | 0.780947669 | 254.9093712 | 438.0003678 | 40 | 14 | 15 | 17 | 31 | 1 | 1 | 0 | 0 | 0 | 0 | 0 |
| ISG000000 | Ap3d1 | 0.001446 | 0.002666 | -0.442139916 | 3203.4764 | 2357.89332 | 34 | 14 | 15 | 15 | 23 | 3 | 1 | 0 | 0 | 0 | 1 | 0 |
| ISG000000 | Fat4 | 0.001506 | 0.002772 | -1.412405233 | 81.94779408 | 30.7865431 | 268 | 8 | 15 | 34 | 116 | 31 | 9 | 1 | 2 | 2 | 2 | 0 |
| ISG000000 | Bcl2l13 | 0.003429 | 0.006001 | -0.414822482 | 2703.798376 | 2028.151032 | 145 | 12 | 15 | 23 | 71 | 10 | 12 | 0 | 0 | 2 | 3 | 0 |
| SG0000003 | Ice2 | 0.004129 | 0.007138 | 0.425726389 | 687.2061262 | 923.0887022 | 135 | 11 | 15 | 23 | 80 | 14 | 8 | 2 | 2 | 2 | 7 | 0 |
| SG0000004 | Zfp758 | 0.008676 | 0.014333 | -0.731510398 | 138.1114581 | 83.1808087 | 123 | 9 | 15 | 21 | 63 | 6 | 6 | 0 | 3 | 3 | 3 | 0 |
| SG0000003 | Ttc29 | 0.010574 | 0.017215 | 3.434764983 | 1.561638181 | 16.88681348 | 342 | 2 | 15 | 48 | 152 | 35 | 1 | 0 | 0 | 0 | 0 | 0 |
| ISG000000 | Elavl1 | 0.011158 | 0.018109 | 0.317278208 | 3848.387821 | 4795.005273 | 70 | 13 | 15 | 19 | 36 | 4 | 17 | 8 | 8 | 8 | 9 | 0 |
| ISG000000 | Gm364 | 0.013093 | 0.020952 | 2.011150187 | 3.937216218 | 15.87105502 | 183 | 8 | 15 | 28 | 128 | 10 | 0 | 0 | 0 | 0 | 0 | 0 |
| SG0000004 | Tsfm | 0.013605 | 0.021691 | 0.382253986 | 760.1565795 | 990.7702894 | 61 | 13 | 15 | 24 | 44 | 1 | 3 | 0 | 0 | 0 | 0 | 0 |
| SG0000000 | Ap1b1 | 0.014052 | 0.02236 | -0.34224158 | 2516.90163 | 1985.369111 | 161 | 15 | 15 | 25 | 47 | 19 | 3 | 0 | 0 | 0 | 0 | 0 |
| SG0000002 | Aco2 | 0.018687 | 0.029066 | -0.310371432 | 8252.131785 | 6654.800556 | 119 | 12 | 15 | 21 | 52 | 7 | 1 | 0 | 0 | 1 | 1 | 0 |
| ISG000000 | Supt6 | 0.020841 | 0.032142 | -0.307600166 | 7256.712033 | 5863.31211 | 57 | 14 | 15 | 18 | 36 | 9 | 1 | 0 | 0 | 1 | 1 | 0 |
| ISG000000 | Rrp12 | 0.023723 | 0.036225 | 0.325529657 | 1180.062411 | 1478.765017 | 49 | 13 | 15 | 17 | 40 | 2 | 0 | 0 | 0 | 0 | 0 | 0 |
| ISG000000 | Fra10ac1 | 0.044513 | 0.064828 | 0.330956196 | 410.0820806 | 515.8204244 | 113 | 8 | 15 | 20 | 49 | 13 | 1 | 0 | 0 | 0 | 0 | 1 |
| SG0000002 | Rhof | 0.050597 | 0.072954 | 0.851706542 | 52.79232946 | 95.2708505 | 69 | 13 | 15 | 19 | 37 | 3 | 9 | 2 | 2 | 2 | 2 | 3 |
| ISG000000 | Pdgfrl | 0.053233 | 0.07644 | 0.438738103 | 147.7156773 | 200.2165251 | 102 | 7 | 15 | 17 | 57 | 11 | 0 | 0 | 0 | 0 | 0 | 0 |
| SG0000007 | Dgkd | 0.055465 | 0.079377 | 0.247220729 | 2568.5474 | 3048.656118 | 143 | 9 | 15 | 26 | 62 | 16 | 7 | 0 | 0 | 3 | 3 | 4 |
| SG0000005 | Chd6 | 0.05679 | 0.081105 | 0.259614101 | 1605.799822 | 1922.396809 | 374 | 8 | 15 | 34 | 164 | 37 | 8 | 1 | 1 | 3 | 3 | 0 |
| SG0000005 | Elp6 | 0.057919 | 0.08258 | 0.362867627 | 294.468177 | 378.6800344 | 50 | 15 | 15 | 19 | 40 | 2 | 1 | 0 | 0 | 1 | 1 | 2 |
| ISG000000 | Sec23ip | 0.066193 | 0.093432 | -0.257580295 | 2432.821034 | 2035.029771 | 103 | 12 | 15 | 21 | 54 | 7 | 7 | 2 | 3 | 2 | 4 | 0 |
| SG0000002 | Mta1 | 0.068791 | 0.09677 | -0.321185842 | 3721.377808 | 2978.63433 | 135 | 0 | 15 | 20 | 50 | 5 | 0 | 0 | 0 | 0 | 5 | 0 |
| SG0000002 | Wiz | 0.07461 | 0.10422 | 0.244069715 | 1606.404284 | 1902.510922 | 176 | 10 | 15 | 29 | 68 | 5 | 2 | 0 | 0 | 0 | 0 | 0 |
| SG0000002 | Wdsub1 | 0.084732 | 0.117126 | 0.337115416 | 237.4660879 | 299.9738865 | 128 | 7 | 15 | 25 | 83 | 8 | 1 | 0 | 0 | 0 | 0 | 0 |
| SG0000003 | Rad52 | 0.085336 | 0.117906 | -0.319671355 | 442.2835183 | 354.3807225 | 84 | 9 | 15 | 18 | 39 | 12 | 3 | 0 | 0 | 0 | 0 | 1 |
| SG0000003 | BC027231 | 0.086626 | 0.119533 | 0.249420371 | 818.6353765 | 973.135961 | 81 | 15 | 15 | 25 | 37 | 6 | 1 | 0 | 0 | 0 | 0 | 0 |
| ISG000000 | Ccny | 0.090039 | 0.123755 | -0.249476479 | 2021.568157 | 1700.546394 | 252 | 3 | 15 | 34 | 132 | 21 | 3 | 0 | 0 | 0 | 1 | 1 |
| SG0000003 | Dnajc13 | 0.097472 | 0.133039 | 0.206041206 | 3306.154261 | 3813.710273 | 292 | 10 | 15 | 49 | 107 | 17 | 4 | 0 | 0 | 1 | 1 | 1 |
| SG0000003 | Tbkbp1 | 0.112022 | 0.150856 | -0.465677308 | 362.3014925 | 262.3537554 | 50 | 10 | 15 | 10 | 20 | 5 | 0 | 0 | 0 | 0 | 0 | 0 |
| SG0000005 | Nod2 | 0.123493 | 0.164681 | 0.408086743 | 85.36781518 | 113.2766814 | 135 | 3 | 15 | 36 | 69 | 9 | 12 | 0 | 0 | 6 | 6 | 0 |
| SG0000000 | Dgke | 0.155661 | 0.20337 | -0.253337989 | 576.7239544 | 483.8443307 | 151 | 14 | 15 | 41 | 104 | 15 | 32 | 4 | 5 | 14 | 24 | 3 |
| ISG000000 | Mrps21 | 0.157099 | 0.205024 | 0.235646319 | 582.4153646 | 685.7557124 | 47 | 12 | 15 | 17 | 24 | 2 | 0 | 0 | 0 | 0 | 0 | 0 |
| ISG000000 | Atrip | 0.168125 | 0.218158 | 0.958100022 | 8.181941714 | 15.89546462 | 78 | 12 | 15 | 15 | 27 | 6 | 0 | 0 | 0 | 0 | 0 | 0 |
| SG0000004 | Slc35e2 | 0.177765 | 0.229239 | 0.204246738 | 668.4099506 | 770.0648518 | 77 | 15 | 15 | 18 | 53 | 0 | 5 | 0 | 0 | 0 | 2 | 0 |
| SG0000002 | Cyp4f13 | 0.182906 | 0.235343 | -0.291482448 | 227.6156075 | 185.9760751 | 165 | 10 | 15 | 20 | 58 | 0 | 0 | 0 | 0 | 0 | 0 | 0 |
| SG0000001 | Ppp1r12c | 0.192754 | 0.246706 | 0.195212041 | 1007.634147 | 1153.632697 | 57 | 12 | 15 | 17 | 23 | 7 | 2 | 0 | 0 | 2 | 2 | 2 |
| SG0000003 | Cep19 | 0.246294 | 0.306788 | 0.241466316 | 197.4809009 | 233.4606573 | 52 | 12 | 15 | 25 | 52 | 0 | 0 | 0 | 0 | 0 | 6 | 0 |
| ISG000000 | Churc1 | 0.362771 | 0.431128 | 0.162240054 | 363.1569126 | 406.3810405 | 120 | 15 | 15 | 27 | 51 | 3 | 0 | 0 | 0 | 0 | 0 | 0 |
| SG0000002 | Cryz | 0.373742 | 0.442434 | 0.162605191 | 1106.629892 | 1238.658072 | 163 | 0 | 15 | 11 | 46 | 4 | 3 | 0 | 0 | 0 | 0 | 0 |
| SG0000002 | Mtf1 | 0.37452 | 0.443267 | -0.138613565 | 744.4854549 | 676.2844107 | 112 | 12 | 15 | 26 | 62 | 12 | 10 | 0 | 1 | 0 | 4 | 2 |
| SG0000002 | Skil | 0.394801 | 0.463504 | -0.122356999 | 4849.241478 | 4454.927949 | 244 | 4 | 15 | 30 | 92 | 50 | 28 | 0 | 0 | 2 | 4 | 6 |
| SG0000006 | Cnih4 | 0.400472 | 0.469423 | -0.120488648 | 2109.888871 | 1940.8362 | 94 | 12 | 15 | 22 | 54 | 18 | 6 | 0 | 0 | 2 | 4 | 0 |
| SG0000000 | Dgcr6 | 0.42916 | 0.498419 | 0.147963995 | 459.9138611 | 509.5867174 | 57 | 13 | 15 | 16 | 26 | 2 | 0 | 0 | 0 | 0 | 0 | 0 |
| ISG000000 | Atg2b | 0.496032 | 0.563547 | -0.098626672 | 2381.989164 | 2224.591097 | 125 | 10 | 15 | 29 | 69 | 13 | 2 | 0 | 0 | 0 | 0 | 0 |
| SG0000002 | Mrps18b | 0.612676 | 0.674881 | 0.072946845 | 1011.303686 | 1063.752953 | 33 | 15 | 15 | 20 | 35 | 0 | 3 | 0 | 0 | 0 | 0 | 0 |
| SG0000003 | Tbce | 0.708616 | 0.761457 | 0.074860253 | 751.7730169 | 791.8117013 | 180 | 12 | 15 | 33 | 114 | 30 | 2 | 0 | 0 | 0 | 0 | 0 |
| SG0000004 | Ipo8 | 0.722637 | 0.774572 | 0.037288628 | 2855.307679 | 2930.069414 | 151 | 5 | 15 | 20 | 59 | 15 | 7 | 0 | 1 | 0 | 0 | 0 |
| ISG000000 | Smim13 | 0.739266 | 0.788762 | -0.070648396 | 497.2328596 | 473.4700748 | 62 | 13 | 15 | 15 | 27 | 2 | 13 | 2 | 2 | 3 | 3 | 0 |
| SG0000003 | Adipor2 | 0.810582 | 0.850316 | 0.024065143 | 5047.214486 | 5132.111588 | 245 | 6 | 15 | 40 | 125 | 20 | 9 | 0 | 0 | 3 | 4 | 2 |
| SG0000000 | Slc22a18 | 0.862971 | 0.893448 | 0.04292884 | 124.7846674 | 128.5535559 | 55 | 10 | 15 | 15 | 39 | 0 | 0 | 0 | 0 | 0 | 0 | 0 |
| SG0000003 | Rbbp6 | 0.887552 | 0.913563 | 0.018248527 | 2708.849666 | 2743.331294 | 186 | 8 | 15 | 26 | 66 | 15 | 3 | 0 | 0 | 0 | 2 | 3 |
| ISG000000 | Tmem45a | 1.4E-116 | 1.3E-114 | -5.30532071 | 13092.30762 | 331.0972173 | 155 | 10 | 14 | 28 | 81 | 8 | 2 | 0 | 0 | 0 | 0 | 0 |
| SG0000000 | Slc2a3 | 5.7E-108 | 4.9E-106 | 10.67733096 | 30.71356567 | 50295.26835 | 101 | 11 | 14 | 21 | 73 | 9 | 4 | 1 | 2 | 2 | 3 | 0 |
| SG0000004 | Tph1 | 3.6E-100 | 2.55E-98 | -11.7065211 | 42780.07658 | 12.8005185 | 135 | 9 | 14 | 18 | 110 | 11 | 2 | 0 | 0 | 0 | 1 | 0 |
| SG0000002 | Shmt1 | 4.7E-99 | 3.26E-97 | 2.631894972 | 844.322746 | 5233.446891 | 57 | 12 | 14 | 18 | 41 | 7 | 0 | 0 | 0 | 0 | 0 | 0 |
| ISG000000 | Jph1 | 3.58E-78 | 1.56E-76 | 4.824937632 | 13.76863273 | 390.2489741 | 207 | 3 | 14 | 25 | 98 | 21 | 13 | 0 | 3 | 1 | 3 | 1 |
| ISG000000 | Trim6 | 1.41E-77 | 6.08E-76 | 4.515093663 | 249.1686801 | 5697.33921 | 36 | 12 | 14 | 14 | 34 | 0 | 4 | 0 | 0 | 0 | 3 | 0 |
| SG0000005 | Ankrd29 | 3.74E-76 | 1.57E-74 | -5.617979813 | 832.7474286 | 16.9563691 | 172 | 10 | 14 | 30 | 79 | 4 | 7 | 0 | 0 | 0 | 1 | 0 |
| SG0000004 | Fam221a | 9.99E-74 | 3.99E-72 | 6.150301891 | 3.974525687 | 282.299858 | 111 | 10 | 14 | 19 | 58 | 8 | 7 | 2 | 2 | 5 | 10 | 2 |
| SG0000000 | Clgn | 9.95E-66 | 3.18E-64 | 6.697487765 | 2.235714429 | 232.0390307 | 114 | 14 | 14 | 25 | 66 | 18 | 6 | 0 | 0 | 1 | 2 | 0 |
| SG0000003 | Anpep | 2.32E-61 | 6.71E-60 | 2.972360384 | 87.68168569 | 688.142727 | 51 | 10 | 14 | 12 | 18 | 5 | 2 | 0 | 0 | 0 | 0 | 2 |
| SG0000004 | Mlf1 | 5.71E-58 | 1.47E-56 | 4.848966381 | 9.128561339 | 263.0792934 | 194 | 0 | 14 | 27 | 100 | 7 | 0 | 0 | 0 | 0 | 0 | 0 |
| ISG000000 | Gap43 | 7.51E-57 | 1.89E-55 | 5.079274935 | 7.747786507 | 261.9339286 | 218 | 6 | 14 | 26 | 98 | 17 | 1 | 0 | 0 | 0 | 0 | 0 |
| SG0000007 | Gad1 | 1.82E-55 | 4.44E-54 | 7.361049329 | 0 | 163.3980446 | 148 | 4 | 14 | 20 | 60 | 7 | 4 | 0 | 2 | 2 | 2 | 1 |
| SG0000004 | Os9 | 2.02E-55 | 4.93E-54 | -2.835487017 | 3864.684645 | 541.4363599 | 84 | 12 | 14 | 26 | 38 | 2 | 0 | 0 | 0 | 0 | 0 | 0 |
| SG0000000 | Clstn3 | 5.3E-55 | 1.28E-53 | 5.962827726 | 12.56887439 | 783.9464363 | 72 | 12 | 14 | 18 | 58 | 2 | 0 | 0 | 0 | 0 | 2 | 0 |
| SG0000002 | Ivns1abp | 8.18E-55 | 1.95E-53 | 1.829251429 | 3612.836117 | 12838.30862 | 165 | 8 | 14 | 18 | 36 | 18 | 21 | 0 | 2 | 2 | 4 | 5 |
| SG0000002 | Lrrc9 | 1.75E-53 | 4E-52 | 7.300874765 | 0 | 156.682065 | 280 | 8 | 14 | 43 | 154 | 21 | 1 | 0 | 0 | 0 | 0 | 0 |
| ISG000000 | Fem1b | 3.61E-52 | 7.99E-51 | 1.803792561 | 2550.502355 | 8904.743122 | 48 | 12 | 14 | 14 | 20 | 6 | 4 | 0 | 1 | 1 | 2 | 1 |
| SG0000003 | Rab11fip1 | 5.56E-52 | 1.22E-50 | 1.988224473 | 646.5132093 | 2565.030934 | 62 | 8 | 14 | 20 | 52 | 6 | 5 | 0 | 2 | 1 | 2 | 3 |
| SG0000000 | Kit | 6.67E-50 | 1.38E-48 | 4.142192323 | 102.1193668 | 1803.151636 | 121 | 7 | 14 | 21 | 74 | 27 | 3 | 0 | 0 | 0 | 2 | 5 |
| SG0000002 | Kif1c | 6.49E-48 | 1.26E-46 | -2.408156975 | 8094.148616 | 1524.908167 | 93 | 14 | 14 | 31 | 61 | 6 | 9 | 0 | 0 | 0 | 0 | 2 |
| ISG000000 | Ero1lb | 1.81E-45 | 3.22E-44 | -2.798660166 | 1552.282838 | 223.0951857 | 97 | 9 | 14 | 19 | 37 | 7 | 8 | 0 | 3 | 3 | 5 | 0 |
| SG0000003 | 9530068E07Rik | 3.65E-41 | 5.63E-40 | -2.182942299 | 7324.000863 | 1612.936523 | 50 | 12 | 14 | 14 | 42 | 2 | 6 | 0 | 2 | 0 | 4 | 0 |
| SG0000000 | Tmem39a | 2.64E-39 | 3.8E-38 | 1.686264527 | 914.7533017 | 2943.880582 | 79 | 12 | 14 | 19 | 44 | 6 | 1 | 0 | 0 | 0 | 3 | 0 |
| SG0000003 | Stk26 | 3.22E-39 | 4.63E-38 | 2.399891186 | 106.3433409 | 561.2411856 | 280 | 8 | 14 | 54 | 140 | 16 | 22 | 0 | 0 | 6 | 6 | 0 |
| SG0000002 | Acox1 | 4.83E-39 | 6.9E-38 | -2.179001671 | 4248.087222 | 938.098759 | 112 | 14 | 14 | 18 | 59 | 3 | 6 | 0 | 0 | 0 | 0 | 0 |
| SG0000006 | Syt11 | 2.22E-38 | 3.1E-37 | 2.576142861 | 74.57271796 | 444.7093356 | 90 | 10 | 14 | 24 | 60 | 10 | 0 | 0 | 0 | 0 | 0 | 3 |
| SG0000003 | Celf5 | 4.44E-38 | 6.15E-37 | 2.801894551 | 75.89697537 | 529.2718221 | 46 | 12 | 14 | 16 | 16 | 2 | 2 | 0 | 2 | 0 | 0 | 2 |
| ISG000000 | Dgkh | 6.54E-38 | 8.99E-37 | 4.162915495 | 11.32077932 | 202.7863686 | 298 | 5 | 14 | 32 | 116 | 40 | 1 | 0 | 0 | 0 | 0 | 0 |
| ISG000000 | Gprin3 | 1.73E-37 | 2.34E-36 | -6.457263546 | 173.6803869 | 1.976608965 | 144 | 7 | 14 | 27 | 103 | 19 | 2 | 0 | 1 | 1 | 1 | 0 |
| SG0000003 | Mfap3l | 7.06E-37 | 9.36E-36 | -2.564062424 | 1073.37638 | 181.5065633 | 238 | 10 | 14 | 36 | 77 | 27 | 18 | 0 | 0 | 2 | 7 | 1 |
| SG0000003 | Ttk | 5.5E-34 | 6.56E-33 | 1.611308182 | 726.8903317 | 2220.859004 | 164 | 8 | 14 | 26 | 78 | 18 | 0 | 0 | 0 | 0 | 0 | 2 |
| ISG000000 | Elovl7 | 6.89E-34 | 8.16E-33 | -2.569430245 | 2745.296765 | 462.5021145 | 129 | 8 | 14 | 19 | 71 | 14 | 6 | 0 | 0 | 0 | 6 | 0 |
| ISG000000 | Nkain1 | 6.77E-30 | 6.82E-29 | 3.649337035 | 37.21610367 | 466.971972 | 76 | 14 | 14 | 20 | 48 | 8 | 1 | 0 | 0 | 0 | 0 | 0 |
| SG0000005 | St3gal5 | 8.77E-30 | 8.79E-29 | 2.353767526 | 73.18944642 | 374.1132505 | 152 | 8 | 14 | 16 | 44 | 12 | 6 | 0 | 0 | 2 | 2 | 0 |
| ISG000000 | Tbc1d8b | 3.26E-29 | 3.18E-28 | -2.834736173 | 825.3399886 | 115.689054 | 211 | 7 | 14 | 35 | 111 | 20 | 6 | 0 | 0 | 1 | 2 | 0 |
| SG0000003 | Prickle1 | 1.24E-28 | 1.17E-27 | 2.945280681 | 28.76880241 | 221.5846205 | 186 | 7 | 14 | 25 | 83 | 18 | 12 | 0 | 0 | 4 | 6 | 4 |
| ISG000000 | Kpna3 | 2.3E-27 | 2.06E-26 | 1.303188494 | 2967.133282 | 7322.103807 | 174 | 8 | 14 | 17 | 83 | 13 | 6 | 0 | 1 | 1 | 2 | 0 |
| SG0000003 | Slc25a36 | 2.76E-27 | 2.46E-26 | 1.925906026 | 1773.328876 | 6738.213519 | 148 | 13 | 14 | 25 | 78 | 27 | 18 | 0 | 0 | 0 | 12 | 0 |
| ISG000000 | Zbtb10 | 8.57E-27 | 7.5E-26 | 1.798037753 | 237.5412007 | 826.0422184 | 85 | 8 | 14 | 15 | 57 | 11 | 13 | 1 | 2 | 2 | 7 | 0 |
| SG0000002 | Rad54l | 8.94E-27 | 7.81E-26 | 1.427532172 | 760.7152501 | 2046.219062 | 92 | 10 | 14 | 16 | 48 | 6 | 2 | 0 | 0 | 0 | 2 | 0 |
| SG0000005 | Mettl5 | 1.29E-25 | 1.07E-24 | 1.672791085 | 244.3025179 | 778.9117158 | 80 | 12 | 14 | 18 | 66 | 8 | 0 | 0 | 0 | 0 | 0 | 0 |
| SG0000002 | Tfrc | 9.73E-24 | 7.41E-23 | 1.427524024 | 9901.822885 | 26634.38693 | 103 | 7 | 14 | 16 | 54 | 7 | 5 | 0 | 0 | 2 | 2 | 0 |
| ISG000000 | Kcng1 | 2.11E-23 | 1.58E-22 | 6.425951644 | 0.992459638 | 85.33300687 | 34 | 12 | 14 | 14 | 22 | 4 | 1 | 0 | 0 | 0 | 1 | 0 |
| SG0000004 | Ccdc69 | 2.45E-23 | 1.82E-22 | -3.507052953 | 187.4798014 | 16.4902162 | 62 | 14 | 14 | 19 | 49 | 6 | 2 | 0 | 0 | 1 | 1 | 0 |
| SG0000004 | Mis18bp1 | 2.46E-23 | 1.83E-22 | 1.277118453 | 1199.275925 | 2906.498331 | 161 | 7 | 14 | 28 | 78 | 7 | 1 | 0 | 0 | 0 | 2 | 0 |
| SG0000006 | Irak2 | 9.04E-23 | 6.58E-22 | -1.912174615 | 850.3684244 | 225.9358824 | 212 | 12 | 14 | 40 | 154 | 30 | 8 | 0 | 2 | 4 | 4 | 0 |
| ISG000000 | Snapc1 | 1.03E-22 | 7.48E-22 | -1.673254728 | 1696.480815 | 531.9233916 | 50 | 11 | 14 | 17 | 34 | 2 | 1 | 0 | 0 | 0 | 0 | 0 |
| SG0000000 | Pdcd2l | 2.4E-22 | 1.71E-21 | 1.539531824 | 298.6966415 | 868.3115896 | 34 | 12 | 14 | 16 | 30 | 4 | 3 | 0 | 2 | 2 | 2 | 0 |
| SG0000002 | Pomgnt1 | 1.04E-21 | 7.22E-21 | -1.582887791 | 1930.353067 | 644.3770216 | 114 | 14 | 14 | 22 | 44 | 4 | 18 | 0 | 0 | 7 | 7 | 0 |
| SG0000002 | Ppp2r2c | 3.66E-21 | 2.44E-20 | 8.21514891 | 2.248932108 | 668.3171189 | 125 | 10 | 14 | 17 | 48 | 17 | 2 | 0 | 0 | 0 | 2 | 1 |
| ISG000000 | Nrg1 | 5.26E-21 | 3.49E-20 | -3.790733582 | 384.8423242 | 27.80726753 | 247 | 4 | 14 | 25 | 96 | 10 | 0 | 0 | 0 | 0 | 0 | 0 |
| SG0000000 | Ttc25 | 7.95E-20 | 4.94E-19 | 3.849965511 | 6.517749395 | 93.98374771 | 66 | 12 | 14 | 20 | 39 | 2 | 0 | 0 | 0 | 0 | 1 | 0 |
| SG0000005 | Zfp667 | 3.2E-19 | 1.92E-18 | 1.671177729 | 138.9713911 | 442.5884022 | 137 | 7 | 14 | 30 | 96 | 10 | 12 | 0 | 0 | 3 | 7 | 0 |
| SG0000002 | Sp100 | 2.74E-17 | 1.49E-16 | -3.628803033 | 174.1265572 | 14.07623316 | 429 | 9 | 14 | 63 | 234 | 53 | 4 | 0 | 0 | 1 | 3 | 2 |
| SG0000003 | Zdhhc5 | 4.56E-17 | 2.45E-16 | -1.248959241 | 4346.480461 | 1828.78873 | 66 | 9 | 14 | 15 | 38 | 8 | 1 | 0 | 0 | 0 | 2 | 1 |
| ISG000000 | Nek5 | 7.13E-17 | 3.78E-16 | 5.504372493 | 0 | 44.39219957 | 94 | 10 | 14 | 21 | 67 | 14 | 0 | 0 | 0 | 0 | 0 | 0 |
| ISG000000 | Galnt16 | 2.63E-16 | 1.35E-15 | 4.118344157 | 3.961308008 | 68.79925504 | 98 | 10 | 14 | 19 | 50 | 9 | 2 | 0 | 0 | 0 | 0 | 0 |
| ISG000000 | Tmco3 | 4.58E-16 | 2.31E-15 | -1.495731975 | 864.9309797 | 306.7052873 | 59 | 14 | 14 | 19 | 33 | 4 | 10 | 1 | 1 | 2 | 2 | 0 |

| | | | | | | | | | | | | | | | | | |
|---|---|---|---|---|---|---|---|---|---|---|---|---|---|---|---|---|---|
| SG0000003 | Gnptab | 7.37E-16 | 3.67E-15 | -1.212153144 | 3776.051126 | 1629.834138 | 218 | 8 | 14 | 44 | 152 | 20 | 6 | 0 | 2 | 1 | 7 | 4 |
| JSG000000 | Sord | 1.31E-15 | 6.46E-15 | 2.391892019 | 137.4162806 | 721.2227768 | 46 | 13 | 14 | 17 | 39 | 9 | 1 | 0 | 0 | 0 | 0 | 0 |
| SG0000005 | Pkn1 | 1.8E-15 | 8.74E-15 | -1.586001072 | 2737.078031 | 911.7027884 | 73 | 11 | 14 | 15 | 42 | 9 | 1 | 0 | 0 | 1 | 1 | 0 |
| SG0000005 | Adamts14 | 2.49E-15 | 1.2E-14 | 3.054385083 | 46.15746154 | 383.445323 | 143 | 12 | 14 | 23 | 75 | 18 | 1 | 0 | 0 | 1 | 1 | 0 |
| SG0000002 | B4galt7 | 2.56E-15 | 1.24E-14 | -1.532474638 | 674.5584143 | 233.1839906 | 34 | 14 | 14 | 16 | 26 | 2 | 2 | 0 | 0 | 0 | 4 | 0 |
| SG0000000 | Slc1a3 | 7.59E-15 | 3.55E-14 | 6.791659446 | 3.810570289 | 422.1659386 | 156 | 10 | 14 | 25 | 83 | 11 | 2 | 0 | 0 | 0 | 1 | 0 |
| SG0000001 | Ksr1 | 1.26E-14 | 5.79E-14 | 1.472606862 | 144.6098028 | 401.3253298 | 198 | 8 | 14 | 30 | 87 | 14 | 2 | 0 | 0 | 0 | 0 | 0 |
| SG0000002 | Ppat | 1.76E-14 | 8.05E-14 | 0.949292176 | 2463.035929 | 4755.937807 | 114 | 10 | 14 | 20 | 48 | 8 | 10 | 0 | 0 | 2 | 6 | 2 |
| SG0000002 | Enoph1 | 4.83E-14 | 2.15E-13 | 1.095464379 | 690.5835128 | 1475.651634 | 94 | 8 | 14 | 16 | 50 | 4 | 0 | 0 | 0 | 0 | 1 | 0 |
| SG0000003 | Pold3 | 1.04E-13 | 4.49E-13 | 0.942834308 | 1841.971765 | 3540.824293 | 181 | 10 | 14 | 24 | 77 | 23 | 1 | 0 | 0 | 0 | 0 | 1 |
| SG0000004 | Ppp1r2 | 1.65E-13 | 7.07E-13 | -1.114312606 | 3006.646973 | 1388.803873 | 94 | 10 | 14 | 18 | 57 | 11 | 9 | 0 | 1 | 1 | 4 | 2 |
| SG0000003 | Afap1l1 | 2.12E-13 | 9.02E-13 | 3.990516754 | 10.5744277 | 168.0823507 | 100 | 12 | 14 | 18 | 42 | 16 | 0 | 0 | 0 | 0 | 0 | 0 |
| SG0000004 | Maff | 6.77E-13 | 2.78E-12 | -1.200278408 | 1265.121777 | 550.5699799 | 34 | 12 | 14 | 22 | 30 | 0 | 8 | 4 | 4 | 4 | 6 | 0 |
| SG0000002 | Rabgef1 | 7.74E-13 | 3.18E-12 | -1.173825008 | 1418.899113 | 628.9193834 | 105 | 12 | 14 | 28 | 61 | 7 | 7 | 0 | 0 | 2 | 2 | 2 |
| SG0000001 | Slc13a3 | 1.17E-12 | 4.73E-12 | -4.120074847 | 64.46157423 | 3.707104204 | 104 | 10 | 14 | 22 | 48 | 16 | 0 | 0 | 0 | 0 | 0 | 0 |
| SG0000004 | Amot | 1.18E-12 | 4.78E-12 | 3.714086806 | 67.30247353 | 883.2477933 | 331 | 9 | 14 | 26 | 114 | 27 | 15 | 0 | 0 | 0 | 6 | 3 |
| JSG000000 | Foxo1 | 1.53E-12 | 6.14E-12 | 0.953969412 | 949.6791237 | 1839.71394 | 133 | 7 | 14 | 14 | 58 | 12 | 13 | 0 | 0 | 0 | 2 | 4 |
| SG0000002 | Knstrn | 3.62E-12 | 1.42E-11 | -1.066669063 | 2305.748717 | 1100.810573 | 180 | 2 | 14 | 24 | 70 | 16 | 2 | 0 | 0 | 0 | 0 | 0 |
| JSG000000 | Ptdss1 | 7.85E-12 | 3.02E-11 | -1.059034019 | 2029.396089 | 974.0153767 | 111 | 10 | 14 | 19 | 62 | 11 | 0 | 0 | 0 | 0 | 1 | 0 |
| SG0000001 | Mrpl24 | 9.9E-12 | 3.78E-11 | 0.864839998 | 1877.319818 | 3418.857864 | 32 | 14 | 14 | 17 | 24 | 0 | 8 | 0 | 0 | 3 | 3 | 0 |
| JSG000000 | Abtb2 | 1.3E-11 | 4.94E-11 | 1.915500436 | 39.67248762 | 149.6623365 | 195 | 8 | 14 | 20 | 56 | 16 | 1 | 0 | 0 | 1 | 2 | 0 |
| SG0000001 | Dync1h1 | 2.04E-11 | 7.64E-11 | -0.926013717 | 18865.05541 | 9928.878128 | 88 | 12 | 14 | 16 | 28 | 3 | 1 | 0 | 0 | 0 | 0 | 0 |
| SG0000003 | Cul4a | 2.25E-11 | 8.41E-11 | -0.967749374 | 4400.075773 | 2249.772301 | 95 | 12 | 14 | 14 | 48 | 10 | 4 | 0 | 0 | 0 | 0 | 0 |
| SG0000004 | Dsg2 | 2.37E-11 | 8.84E-11 | 1.456667684 | 678.5507879 | 1862.443152 | 178 | 11 | 14 | 21 | 68 | 25 | 3 | 0 | 0 | 0 | 2 | 3 |
| JSG000000 | Sgpp1 | 3.78E-11 | 1.39E-10 | -1.112433615 | 1138.689603 | 526.65895 | 58 | 14 | 14 | 15 | 32 | 3 | 5 | 0 | 0 | 0 | 2 | 0 |
| SG0000005 | Colq | 1.49E-10 | 5.25E-10 | -4.586996232 | 47.96520774 | 1.995735 | 101 | 14 | 14 | 27 | 57 | 11 | 0 | 0 | 0 | 0 | 0 | 0 |
| SG0000003 | Zbtb41 | 1.56E-10 | 5.48E-10 | -1.114424 | 1557.791139 | 719.5055979 | 193 | 8 | 14 | 21 | 65 | 12 | 23 | 0 | 2 | 1 | 9 | 0 |
| SG0000003 | Zc3h12b | 2.08E-10 | 7.23E-10 | -2.183085729 | 137.0880291 | 30.18736772 | 114 | 10 | 14 | 18 | 61 | 9 | 8 | 0 | 0 | 0 | 2 | 0 |
| SG0000005 | Zkscan5 | 2.61E-10 | 9.01E-10 | 0.948819051 | 497.8104591 | 960.9195116 | 44 | 12 | 14 | 16 | 40 | 4 | 0 | 0 | 0 | 0 | 0 | 0 |
| SG0000002 | Itgb3bp | 3.7E-10 | 1.26E-09 | 1.76235223 | 43.14257645 | 146.361537 | 174 | 8 | 14 | 24 | 82 | 13 | 9 | 0 | 0 | 0 | 1 | 1 |
| SG0000002 | Dnajc27 | 4.01E-10 | 1.37E-09 | 1.001781705 | 355.4052827 | 711.6889472 | 64 | 10 | 14 | 18 | 33 | 6 | 14 | 6 | 6 | 8 | 9 | 0 |
| JSG000000 | Fzd3 | 5.17E-10 | 1.74E-09 | 1.403448733 | 88.48240786 | 234.0653345 | 154 | 4 | 14 | 23 | 72 | 10 | 1 | 0 | 0 | 0 | 1 | 1 |
| SG0000001 | Nol11 | 5.71E-10 | 1.92E-09 | 0.79779071 | 1658.644922 | 2883.449551 | 78 | 10 | 14 | 26 | 58 | 8 | 0 | 0 | 0 | 0 | 0 | 0 |
| JSG000000 | Qpctl | 5.75E-10 | 1.93E-09 | -1.316066863 | 401.7912537 | 161.370772 | 22 | 13 | 14 | 14 | 19 | 1 | 1 | 0 | 0 | 0 | 0 | 0 |
| JSG000000 | Chrna5 | 1.87E-09 | 6E-09 | 4.6201944 | 1.324060707 | 32.56304413 | 71 | 10 | 14 | 17 | 33 | 1 | 2 | 0 | 0 | 0 | 0 | 0 |
| SG0000002 | Plekhm2 | 2.31E-09 | 7.39E-09 | -1.33077025 | 822.6315997 | 327.0420718 | 52 | 8 | 14 | 14 | 40 | 2 | 2 | 0 | 0 | 0 | 0 | 0 |
| SG0000001 | Rxrg | 2.64E-09 | 8.39E-09 | 4.608759568 | 0 | 23.39915988 | 196 | 7 | 14 | 36 | 93 | 20 | 3 | 0 | 0 | 0 | 0 | 3 |
| JSG000000 | A2m | 1.16E-08 | 3.51E-08 | 4.412233187 | 19.32519812 | 411.4702987 | 104 | 9 | 14 | 25 | 76 | 12 | 0 | 0 | 0 | 0 | 0 | 0 |
| SG0000003 | Synm | 1.47E-08 | 4.41E-08 | 2.762793667 | 35.92451216 | 243.8229856 | 86 | 12 | 14 | 22 | 44 | 8 | 16 | 0 | 0 | 2 | 4 | 0 |
| SG0000004 | Myzap | 1.81E-08 | 5.38E-08 | 3.885252848 | 2.294772122 | 33.90917417 | 259 | 8 | 14 | 33 | 117 | 30 | 3 | 0 | 0 | 1 | 1 | 0 |
| JSG000000 | Dmgdh | 2.04E-08 | 6.07E-08 | 4.454745651 | 0 | 20.92865849 | 152 | 8 | 14 | 20 | 70 | 19 | 0 | 0 | 0 | 0 | 0 | 0 |
| JSG000000 | Rbp7 | 5.18E-08 | 1.49E-07 | 6.18504834 | 0.307509278 | 22.37398274 | 28 | 12 | 14 | 14 | 16 | 0 | 0 | 0 | 0 | 0 | 0 | 0 |
| SG0000003 | Mtmr2 | 6.35E-08 | 1.81E-07 | -0.809293304 | 2367.310172 | 1350.932358 | 494 | 11 | 14 | 69 | 260 | 37 | 12 | 0 | 0 | 4 | 10 | 0 |
| SG0000003 | Cttn | 6.37E-08 | 1.82E-07 | -0.749619614 | 6245.679062 | 3714.682287 | 50 | 10 | 14 | 14 | 26 | 10 | 2 | 0 | 0 | 0 | 0 | 0 |
| SG0000003 | Nlrp4c | 8.88E-08 | 2.5E-07 | 4.337837208 | 0 | 19.22176765 | 169 | 14 | 14 | 32 | 120 | 30 | 1 | 0 | 0 | 0 | 0 | 0 |
| SG0000003 | Ifi203 | 9.87E-08 | 2.77E-07 | -2.012772368 | 206.0829709 | 51.06663593 | 251 | 5 | 14 | 29 | 83 | 34 | 16 | 0 | 4 | 4 | 13 | 4 |
| SG0000003 | Trpm8 | 1.16E-07 | 3.24E-07 | -6.234517022 | 25.20902728 | 0.33479538 | 317 | 5 | 14 | 43 | 159 | 27 | 0 | 0 | 0 | 0 | 0 | 0 |
| JSG000000 | Chic2 | 1.56E-07 | 4.32E-07 | -0.877927089 | 979.9037042 | 533.2133466 | 115 | 8 | 14 | 20 | 61 | 7 | 16 | 0 | 3 | 2 | 4 | 3 |
| SG0000003 | Rab43 | 1.69E-07 | 4.64E-07 | -3.939418 | 60.85042822 | 3.966255209 | 57 | 11 | 14 | 24 | 63 | 3 | 5 | 1 | 1 | 3 | 3 | 0 |
| JSG000000 | Sfxn4 | 1.76E-07 | 4.83E-07 | 2.356040889 | 11.0326747 | 56.48326484 | 109 | 14 | 14 | 26 | 58 | 21 | 4 | 2 | 2 | 2 | 4 | 0 |
| JSG000000 | Cidea | 1.87E-07 | 5.11E-07 | 4.275965045 | 0 | 18.37285995 | 51 | 12 | 14 | 17 | 28 | 3 | 0 | 0 | 0 | 0 | 0 | 0 |
| SG0000002 | Smek2 | 1.87E-07 | 5.12E-07 | -0.758051136 | 3003.348185 | 1775.863388 | 279 | 8 | 14 | 38 | 110 | 28 | 2 | 0 | 0 | 0 | 1 | 2 |
| JSG000000 | Tm6sf1 | 2.37E-07 | 6.43E-07 | 1.675263277 | 31.64212735 | 101.0578772 | 123 | 11 | 14 | 23 | 40 | 15 | 5 | 2 | 2 | 2 | 4 | 0 |
| JSG000000 | Kcna6 | 2.44E-07 | 6.61E-07 | 4.663084105 | 1.005677317 | 25.47926188 | 64 | 10 | 14 | 12 | 26 | 14 | 0 | 0 | 0 | 0 | 0 | 0 |
| SG0000002 | Glt8d2 | 2.6E-07 | 7.02E-07 | -2.315820377 | 69.64657924 | 13.98841127 | 94 | 4 | 14 | 21 | 67 | 19 | 0 | 0 | 0 | 0 | 0 | 0 |
| SG0000004 | Pitpnm3 | 5.55E-07 | 1.46E-06 | -2.339368204 | 64.40251654 | 12.72573389 | 146 | 8 | 14 | 32 | 80 | 12 | 0 | 0 | 0 | 0 | 0 | 0 |
| SG0000003 | Uba3 | 7.75E-07 | 2.02E-06 | -0.693306011 | 5838.841873 | 3610.94446 | 150 | 6 | 14 | 28 | 78 | 12 | 8 | 0 | 0 | 2 | 6 | 0 |
| JSG000000 | Ppp2r5d | 1.23E-06 | 3.15E-06 | -0.687428623 | 3611.694718 | 2242.716606 | 37 | 13 | 14 | 15 | 23 | 3 | 0 | 0 | 0 | 0 | 0 | 0 |
| SG0000002 | Rnf41 | 1.37E-06 | 3.49E-06 | -0.781786585 | 1062.187683 | 617.8172768 | 96 | 4 | 14 | 20 | 54 | 6 | 4 | 0 | 0 | 1 | 1 | 1 |
| JSG000000 | Prdx3 | 1.38E-06 | 3.53E-06 | -0.696944767 | 3067.201141 | 1892.086456 | 31 | 12 | 14 | 16 | 37 | 2 | 1 | 0 | 1 | 1 | 1 | 0 |
| SG0000005 | Kcmf1 | 1.58E-06 | 4E-06 | -0.691788072 | 3056.399728 | 1892.174529 | 124 | 8 | 14 | 16 | 44 | 20 | 6 | 0 | 1 | 2 | 2 | 0 |
| JSG000000 | Mblac2 | 1.82E-06 | 4.59E-06 | -1.446466016 | 144.0277129 | 52.84652145 | 89 | 11 | 14 | 16 | 49 | 6 | 13 | 3 | 5 | 3 | 8 | 0 |
| SG0000003 | Rab3gap2 | 2.31E-06 | 5.78E-06 | 0.612041573 | 1935.54327 | 2958.324136 | 272 | 4 | 14 | 29 | 135 | 19 | 4 | 0 | 0 | 1 | 3 | 1 |
| SG0000002 | Cul1 | 3.26E-06 | 8.04E-06 | -0.650402444 | 4564.178132 | 2907.849605 | 228 | 4 | 14 | 26 | 112 | 17 | 2 | 0 | 0 | 0 | 0 | 1 |
| SG0000006 | Cd302 | 3.63E-06 | 8.91E-06 | 1.399538057 | 42.75929823 | 112.8063388 | 134 | 12 | 14 | 24 | 56 | 12 | 0 | 0 | 0 | 0 | 0 | 0 |
| JSG000000 | Rarb | 4.68E-06 | 1.14E-05 | -1.91796865 | 122.0045568 | 32.28568142 | 208 | 5 | 14 | 25 | 68 | 20 | 2 | 0 | 0 | 1 | 1 | 1 |
| SG0000004 | Vwa5b2 | 6.56E-06 | 1.57E-05 | 4.104721222 | 1.642444096 | 28.25756974 | 62 | 10 | 14 | 20 | 32 | 8 | 0 | 0 | 0 | 0 | 0 | 0 |
| SG0000004 | Ddx58 | 7.03E-06 | 1.68E-05 | -1.230495719 | 4680.78913 | 1994.816517 | 75 | 12 | 14 | 17 | 34 | 3 | 2 | 0 | 0 | 0 | 0 | 0 |
| SG0000002 | Top2a | 7.52E-06 | 1.79E-05 | 0.620506385 | 15586.37757 | 23962.72628 | 74 | 11 | 14 | 23 | 42 | 7 | 2 | 0 | 0 | 0 | 0 | 1 |
| SG0000002 | Dnajc2 | 9.3E-06 | 2.2E-05 | 0.560256421 | 2471.518528 | 3644.331362 | 155 | 10 | 14 | 24 | 61 | 16 | 0 | 0 | 0 | 0 | 1 | 0 |
| SG0000003 | Ncapd3 | 1.15E-05 | 2.69E-05 | 0.56274213 | 1945.922095 | 2874.270982 | 208 | 8 | 14 | 20 | 88 | 20 | 6 | 0 | 2 | 4 | 8 | 2 |
| SG0000006 | Tnnt1 | 1.18E-05 | 2.75E-05 | 1.43553972 | 58.2320826 | 157.5080802 | 28 | 14 | 14 | 14 | 22 | 2 | 0 | 0 | 0 | 0 | 2 | 0 |
| SG0000002 | Xpnpep1 | 1.25E-05 | 2.92E-05 | -0.767120907 | 6145.435631 | 3610.989877 | 293 | 9 | 14 | 42 | 126 | 17 | 1 | 0 | 0 | 0 | 0 | 0 |
| SG0000004 | Ripk2 | 1.34E-05 | 3.12E-05 | 0.794854423 | 237.2921021 | 411.6786144 | 164 | 10 | 14 | 32 | 84 | 26 | 3 | 0 | 2 | 2 | 2 | 0 |
| SG0000002 | Fas | 1.34E-05 | 3.12E-05 | -1.372133701 | 126.1229426 | 48.72374851 | 143 | 2 | 14 | 27 | 72 | 8 | 0 | 0 | 0 | 0 | 0 | 0 |
| SG0000002 | Desi2 | 1.44E-05 | 3.35E-05 | 0.545703623 | 2510.285393 | 3664.344175 | 252 | 10 | 14 | 27 | 101 | 14 | 13 | 0 | 0 | 1 | 3 | 0 |
| SG0000007 | Zhx2 | 1.49E-05 | 3.46E-05 | -1.275137544 | 215.9700266 | 89.23574065 | 160 | 9 | 14 | 23 | 57 | 11 | 1 | 0 | 0 | 0 | 0 | 0 |
| SG0000002 | Pik3cg | 3.44E-05 | 7.69E-05 | -4.612171845 | 31.34898575 | 1.281803414 | 196 | 9 | 14 | 18 | 71 | 12 | 7 | 0 | 1 | 0 | 3 | 0 |
| SG0000003 | Tmed7 | 3.66E-05 | 8.17E-05 | -0.577085622 | 4662.786234 | 3125.542605 | 72 | 11 | 14 | 14 | 34 | 6 | 0 | 0 | 0 | 0 | 0 | 0 |
| JSG000000 | Tmem243 | 4.28E-05 | 9.47E-05 | 1.004375625 | 83.53498033 | 167.577445 | 165 | 9 | 14 | 19 | 60 | 16 | 8 | 0 | 0 | 0 | 0 | 1 |
| JSG000000 | Mrpl39 | 4.64E-05 | 0.000102 | 0.573869531 | 908.942907 | 1352.971185 | 45 | 13 | 14 | 14 | 24 | 6 | 2 | 0 | 1 | 0 | 1 | 5 |
| SG0000004 | Obfc1 | 4.73E-05 | 0.000104 | -0.652521786 | 1035.951768 | 659.0386809 | 89 | 7 | 14 | 14 | 35 | 9 | 0 | 0 | 0 | 0 | 0 | 0 |
| JSG000000 | Gltscr1l | 5.05E-05 | 0.000111 | 0.551402582 | 1163.724635 | 1705.449818 | 38 | 13 | 14 | 17 | 42 | 4 | 2 | 0 | 0 | 1 | 4 | 0 |
| JSG000000 | Tmem55a | 5.48E-05 | 0.00012 | -0.60059429 | 1795.944461 | 1184.393474 | 124 | 8 | 14 | 26 | 57 | 6 | 7 | 0 | 0 | 1 | 1 | 0 |
| SG0000002 | Stat1 | 6.56E-05 | 0.000142 | 0.613456317 | 772.361487 | 1181.651369 | 334 | 0 | 14 | 43 | 109 | 42 | 13 | 0 | 3 | 2 | 14 | 0 |
| SG0000004 | Fbxw4 | 8.3E-05 | 0.000178 | -0.813247476 | 343.1793429 | 195.3031527 | 128 | 13 | 14 | 17 | 61 | 8 | 1 | 0 | 0 | 0 | 0 | 0 |
| SG0000002 | Paqr8 | 9.2E-05 | 0.000196 | 0.703186281 | 254.9657882 | 415.108927 | 165 | 5 | 14 | 27 | 63 | 7 | 9 | 0 | 0 | 2 | 4 | 0 |
| JSG000000 | Ero1l | 9.87E-05 | 0.000209 | -1.162670292 | 17669.99495 | 7892.921663 | 90 | 8 | 14 | 21 | 69 | 6 | 12 | 0 | 0 | 3 | 7 | 0 |
| JSG000000 | Rala | 0.000111 | 0.000233 | -0.571101185 | 2059.265459 | 1386.097396 | 137 | 12 | 14 | 28 | 84 | 15 | 8 | 0 | 0 | 1 | 4 | 0 |
| JSG000000 | Ccdc126 | 0.000112 | 0.000236 | -0.827430112 | 314.223337 | 177.0749604 | 94 | 10 | 14 | 14 | 40 | 12 | 10 | 0 | 2 | 2 | 6 | 4 |
| SG0000001 | Cd40 | 0.00023 | 0.000468 | 3.14973723 | 6.076563486 | 53.92913127 | 76 | 5 | 14 | 17 | 32 | 0 | 11 | 0 | 4 | 2 | 3 | 0 |
| SG0000005 | Smagp | 0.000244 | 0.000494 | -0.601544603 | 857.6371029 | 565.2239971 | 46 | 8 | 14 | 14 | 30 | 6 | 0 | 0 | 0 | 0 | 0 | 0 |
| SG0000005 | Gan | 0.000248 | 0.000502 | -0.605273279 | 878.3719237 | 577.395019 | 123 | 12 | 14 | 34 | 80 | 16 | 11 | 0 | 0 | 0 | 2 | 0 |
| SG0000002 | Pag1 | 0.000259 | 0.000523 | 1.137304171 | 54.57094689 | 120.0394949 | 393 | 6 | 14 | 35 | 177 | 49 | 13 | 0 | 0 | 1 | 7 | 1 |
| JSG000000 | Stx12 | 0.000312 | 0.000625 | -0.519756638 | 2499.080381 | 1743.082275 | 75 | 14 | 14 | 21 | 40 | 6 | 7 | 0 | 0 | 2 | 4 | 3 |
| JSG000000 | Olfml2b | 0.000579 | 0.00112 | -1.043971789 | 119.2667582 | 57.84323736 | 48 | 13 | 14 | 20 | 42 | 4 | 1 | 0 | 0 | 0 | 1 | 0 |
| SG0000002 | Stx16 | 0.000639 | 0.001228 | -0.48759304 | 2593.256834 | 1849.54712 | 106 | 4 | 14 | 14 | 28 | 4 | 30 | 2 | 6 | 6 | 10 | 2 |
| SG0000002 | Sh3yl1 | 0.000666 | 0.001276 | 0.925390795 | 92.07546145 | 174.8695897 | 224 | 8 | 14 | 38 | 126 | 26 | 0 | 0 | 0 | 0 | 0 | 0 |
| JSG000000 | Cbl | 0.000713 | 0.001363 | -0.499660586 | 3213.303926 | 2272.683613 | 159 | 9 | 14 | 30 | 90 | 12 | 3 | 0 | 0 | 0 | 0 | 0 |
| JSG000000 | Cep152 | 0.001154 | 0.002151 | 0.540149865 | 378.1197637 | 549.8328575 | 128 | 11 | 14 | 26 | 80 | 13 | 3 | 0 | 0 | 1 | 1 | 0 |
| SG0000000 | Stx1a | 0.001829 | 0.003328 | 0.744104662 | 109.3525329 | 183.1583279 | 48 | 8 | 14 | 14 | 32 | 6 | 0 | 0 | 0 | 0 | 0 | 0 |
| SG0000002 | Hadha | 0.001969 | 0.003567 | -0.431808948 | 3173.127262 | 2352.339735 | 64 | 13 | 14 | 19 | 37 | 14 | 0 | 0 | 0 | 0 | 0 | 0 |
| SG0000003 | Ybx3 | 0.002146 | 0.003869 | -0.432003122 | 9953.329078 | 7377.725031 | 78 | 12 | 14 | 22 | 36 | 6 | 4 | 2 | 2 | 2 | 2 | 0 |
| SG0000002 | Sympk | 0.002258 | 0.004056 | 0.423474214 | 3450.974323 | 4628.285795 | 72 | 12 | 14 | 16 | 34 | 14 | 1 | 0 | 0 | 0 | 0 | 0 |
| SG0000004 | Eif4h | 0.002291 | 0.004111 | -0.404316392 | 10368.47955 | 7834.363416 | 71 | 14 | 14 | 18 | 32 | 0 | 3 | 0 | 0 | 0 | 0 | 0 |
| SG0000003 | Brsk1 | 0.002386 | 0.004273 | 0.633714935 | 173.0286055 | 268.4634158 | 64 | 10 | 14 | 16 | 37 | 6 | 8 | 6 | 6 | 6 | 10 | 2 |
| SG0000002 | Sephs1 | 0.003611 | 0.006299 | 0.385809541 | 1690.67187 | 2209.019611 | 95 | 14 | 14 | 22 | 47 | 8 | 9 | 0 | 0 | 0 | 3 | 0 |
| SG0000006 | Nuggc | 0.003829 | 0.006654 | 1.680252101 | 8.644875846 | 27.70543945 | 107 | 8 | 14 | 16 | 58 | 6 | 0 | 0 | 0 | 0 | 0 | 0 |
| SG0000000 | Slc25a42 | 0.004042 | 0.006999 | -0.527021013 | 501.4405068 | 347.9928176 | 42 | 12 | 14 | 20 | 38 | 8 | 4 | 0 | 0 | 0 | 0 | 0 |
| SG0000005 | Nsun3 | 0.004757 | 0.008168 | -0.624920905 | 284.9916005 | 184.8043216 | 240 | 6 | 14 | 24 | 94 | 18 | 14 | 2 | 2 | 4 | 10 | 1 |
| SG0000000 | Hddc2 | 0.005293 | 0.009026 | 0.445266767 | 518.666964 | 706.1992302 | 54 | 8 | 14 | 14 | 30 | 2 | 0 | 0 | 0 | 0 | 0 | 0 |
| SG0000003 | Tcaf1 | 0.006363 | 0.010727 | 0.958203611 | 282.0333635 | 547.9595692 | 150 | 8 | 14 | 26 | 77 | 21 | 15 | 0 | 2 | 3 | 6 | 2 |
| JSG000000 | Chmp5 | 0.009246 | 0.015204 | -0.372859386 | 2237.410655 | 1727.841269 | 43 | 12 | 14 | 16 | 27 | 1 | 4 | 0 | 1 | 1 | 2 | 0 |
| SG0000006 | Cdc26 | 0.011503 | 0.018639 | 0.411307528 | 456.8784993 | 607.5984035 | 94 | 14 | 14 | 23 | 45 | 5 | 8 | 1 | 1 | 1 | 4 | 1 |
| SG0000006 | Fbxl12 | 0.01155 | 0.018711 | 0.462194859 | 301.5654325 | 415.447428 | 110 | 13 | 14 | 18 | 41 | 7 | 13 | 0 | 0 | 0 | 0 | 0 |
| JSG000000 | Afg3l2 | 0.016247 | 0.025595 | 0.31495275 | 2332.808724 | 2901.946194 | 89 | 10 | 14 | 16 | 39 | 4 | 1 | 0 | 0 | 0 | 1 | 0 |
| SG0000002 | Xpa | 0.016948 | 0.026592 | -0.570194264 | 206.6044253 | 139.1534707 | 102 | 13 | 14 | 24 | 54 | 36 | 4 | 0 | 1 | 0 | 0 | 0 |
| SG0000002 | Ngef | 0.018166 | 0.028315 | -0.690519161 | 118.2826324 | 73.29156598 | 134 | 11 | 14 | 23 | 70 | 10 | 0 | 0 | 0 | 0 | 0 | 0 |
| SG0000003 | Nlrp3 | 0.01847 | 0.028752 | 4.39931868 | 1.02742554 | 21.68089672 | 67 | 12 | 14 | 26 | 75 | 4 | 10 | 0 | 0 | 4 | 4 | 0 |
| SG0000004 | Zfp866 | 0.019154 | 0.029731 | 0.352808242 | 685.1422507 | 874.9568056 | 71 | 4 | 14 | 14 | 42 | 5 | 3 | 0 | 0 | 0 | 0 | 0 |
| SG0000000 | Ankrd26 | 0.02084 | 0.032142 | -0.384473847 | 778.1009186 | 596.0706905 | 134 | 8 | 14 | 28 | 78 | 12 | 8 | 0 | 1 | 2 | 9 | 1 |
| SG0000000 | Cdc45 | 0.020893 | 0.032209 | -0.340058346 | 1769.552478 | 1397.963023 | 93 | 10 | 14 | 16 | 68 | 4 | 2 | 0 | 2 | 2 | 2 | 0 |
| JSG000000 | Ptges3l | 0.026194 | 0.039732 | -0.884804143 | 59.29801117 | 32.11349254 | 42 | 10 | 14 | 18 | 41 | 0 | 0 | 0 | 0 | 0 | 1 | 0 |

| | | | | | | | | | | | | | | | | | | |
|---|---|---|---|---|---|---|---|---|---|---|---|---|---|---|---|---|---|---|
| SG0000002 | 2010107G12Rik | 0.035074 | 0.051968 | -0.688249076 | 110.1239823 | 68.34366926 | 57 | 11 | 14 | 17 | 38 | 3 | 1 | 0 | 0 | 0 | 0 | 0 |
| SG0000005 | Tonsl | 0.037538 | 0.055359 | -0.292728884 | 2223.915953 | 1815.50812 | 24 | 12 | 14 | 14 | 14 | 0 | 0 | 0 | 0 | 0 | 0 | 0 |
| SG0000003 | Parpbp | 0.04083 | 0.059824 | -0.330299689 | 797.3309958 | 634.1741469 | 137 | 8 | 14 | 22 | 80 | 9 | 6 | 0 | 0 | 1 | 2 | 0 |
| SG000000C | Stk38l | 0.042093 | 0.061568 | -0.317051174 | 1225.481168 | 983.7046593 | 182 | 8 | 14 | 24 | 73 | 8 | 2 | 0 | 0 | 0 | 3 | 0 |
| SG0000004 | Tmco4 | 0.046923 | 0.068133 | -0.404364117 | 287.4546744 | 217.1919111 | 138 | 2 | 14 | 20 | 68 | 38 | 2 | 0 | 0 | 0 | 0 | 0 |
| SG0000004 | Xxylt1 | 0.047289 | 0.068625 | -0.313051128 | 962.5071362 | 774.7581946 | 113 | 12 | 14 | 22 | 54 | 16 | 2 | 0 | 0 | 0 | 0 | 1 |
| SG0000005 | Kdm3a | 0.047572 | 0.068996 | 0.24890975 | 3866.74896 | 4594.891686 | 180 | 10 | 14 | 22 | 58 | 22 | 4 | 0 | 0 | 0 | 2 | 0 |
| JSG000000 | Rnf169 | 0.053822 | 0.077217 | -0.287618468 | 1421.376623 | 1164.467528 | 115 | 7 | 14 | 18 | 53 | 13 | 6 | 0 | 1 | 0 | 1 | 0 |
| SG0000005 | Max | 0.055402 | 0.079294 | -0.264246745 | 3032.720622 | 2525.14434 | 76 | 10 | 14 | 16 | 28 | 12 | 2 | 0 | 0 | 2 | 2 | 4 |
| SG0000004 | Gemin8 | 0.065172 | 0.092099 | 0.38407815 | 177.7329156 | 231.9460115 | 144 | 12 | 14 | 30 | 78 | 4 | 8 | 0 | 2 | 2 | 2 | 0 |
| SG0000002 | Pdss1 | 0.067529 | 0.095138 | -0.328863603 | 552.311959 | 439.7305489 | 146 | 10 | 14 | 23 | 81 | 15 | 3 | 0 | 0 | 0 | 1 | 1 |
| SG0000002 | Pfkp | 0.07857 | 0.109274 | 0.222415893 | 6186.731266 | 7217.970576 | 226 | 0 | 14 | 33 | 108 | 6 | 10 | 0 | 0 | 3 | 3 | 1 |
| JSG000000 | Hprt | 0.080672 | 0.111929 | 0.219153386 | 4109.935599 | 4784.17172 | 94 | 11 | 14 | 24 | 61 | 1 | 2 | 0 | 0 | 0 | 0 | 0 |
| SG0000004 | Rnf149 | 0.089432 | 0.123025 | -0.264900722 | 988.7710648 | 822.9106551 | 134 | 4 | 14 | 15 | 45 | 5 | 3 | 0 | 0 | 1 | 3 | 0 |
| SG0000002 | Galnt14 | 0.115598 | 0.155124 | 1.073348961 | 8.478576881 | 17.84157523 | 509 | 2 | 14 | 59 | 187 | 41 | 0 | 0 | 0 | 0 | 1 | 1 |
| SG0000003 | Car12 | 0.133168 | 0.176428 | 0.382064854 | 313.9129721 | 409.0931843 | 124 | 12 | 14 | 38 | 58 | 4 | 8 | 0 | 0 | 2 | 2 | 0 |
| SG0000003 | Morc2a | 0.144571 | 0.190045 | 0.190899084 | 2411.385131 | 2752.535502 | 171 | 10 | 14 | 34 | 100 | 14 | 2 | 0 | 0 | 2 | 2 | 0 |
| SG0000006 | Cttnbp2nl | 0.211563 | 0.267886 | -0.190977744 | 1201.09233 | 1052.171088 | 152 | 8 | 14 | 26 | 48 | 16 | 8 | 2 | 2 | 2 | 6 | 0 |
| JSG000000 | Sltm | 0.220236 | 0.277596 | 0.16334486 | 2023.945597 | 2266.577311 | 134 | 10 | 14 | 19 | 54 | 6 | 1 | 0 | 0 | 0 | 1 | 0 |
| SG0000002 | Gdap2 | 0.221397 | 0.278918 | 0.196065483 | 456.6778815 | 523.1564321 | 211 | 9 | 14 | 20 | 62 | 11 | 9 | 0 | 0 | 0 | 0 | 0 |
| SG0000005 | Trim32 | 0.238784 | 0.298514 | 0.173577765 | 855.6710853 | 965.0703444 | 40 | 14 | 14 | 14 | 24 | 0 | 2 | 0 | 0 | 0 | 2 | 0 |
| SG0000004 | Lhfpl2 | 0.239496 | 0.299236 | 0.305250497 | 246.8238929 | 304.9837875 | 217 | 6 | 14 | 25 | 78 | 16 | 11 | 0 | 0 | 0 | 0 | 0 |
| SG0000002 | Atg4b | 0.250228 | 0.311104 | 0.158658362 | 1551.721521 | 1732.10697 | 112 | 10 | 14 | 25 | 45 | 2 | 4 | 0 | 0 | 1 | 1 | 0 |
| SG0000002 | Gin1 | 0.254711 | 0.315867 | 0.229975057 | 285.452182 | 334.7825071 | 85 | 12 | 14 | 21 | 65 | 13 | 3 | 0 | 0 | 0 | 3 | 2 |
| SG0000007 | Nrbf2 | 0.256197 | 0.317433 | -0.225163292 | 316.1978852 | 270.5067186 | 78 | 10 | 14 | 16 | 44 | 4 | 2 | 0 | 0 | 0 | 0 | 0 |
| SG0000003 | Dchs1 | 0.300016 | 0.364752 | 0.547377182 | 19.37373154 | 28.31328289 | 38 | 13 | 14 | 14 | 23 | 0 | 1 | 1 | 1 | 1 | 1 | 0 |
| JSG000000 | Cyp7b1 | 0.316443 | 0.382259 | 0.43827053 | 30.97558478 | 41.971268 | 417 | 5 | 14 | 47 | 204 | 45 | 2 | 0 | 0 | 0 | 0 | 0 |
| SG0000004 | Ago4 | 0.389194 | 0.457882 | -0.324666942 | 76.84574186 | 61.35999555 | 91 | 12 | 14 | 18 | 43 | 11 | 3 | 0 | 0 | 0 | 2 | 2 |
| JSG000000 | Herpud2 | 0.435513 | 0.504782 | -0.130172034 | 817.5491651 | 747.0130564 | 128 | 8 | 14 | 19 | 57 | 8 | 2 | 0 | 0 | 0 | 1 | 0 |
| SG0000003 | Tbc1d10a | 0.463834 | 0.533328 | -0.138070507 | 607.5709837 | 552.1202305 | 68 | 12 | 14 | 22 | 36 | 0 | 2 | 2 | 2 | 2 | 2 | 0 |
| SG0000003 | Frmpd1 | 0.467565 | 0.536516 | -0.265514769 | 89.92004996 | 74.80465592 | 233 | 9 | 14 | 32 | 96 | 29 | 0 | 0 | 0 | 0 | 0 | 0 |
| SG0000002 | Cenpc1 | 0.469334 | 0.538098 | 0.100140333 | 1126.666378 | 1207.648589 | 142 | 6 | 14 | 22 | 79 | 16 | 0 | 0 | 0 | 0 | 0 | 2 |
| JSG000000 | Mcm9 | 0.48961 | 0.557704 | 0.121213913 | 292.6229155 | 318.2712112 | 178 | 6 | 14 | 29 | 93 | 23 | 0 | 0 | 0 | 0 | 0 | 0 |
| JSG000000 | S100a13 | 0.494481 | 0.562251 | -0.137625239 | 322.9754542 | 293.589279 | 29 | 14 | 14 | 16 | 23 | 0 | 0 | 0 | 0 | 0 | 0 | 0 |
| SG0000002 | Cyb5rl | 0.495529 | 0.563135 | 0.19523948 | 87.31511872 | 99.96831884 | 96 | 7 | 14 | 14 | 54 | 21 | 6 | 0 | 0 | 0 | 0 | 0 |
| JSG000000 | Eefsec | 0.736678 | 0.786704 | 0.059514785 | 942.4523159 | 982.1438949 | 190 | 11 | 14 | 28 | 119 | 14 | 0 | 0 | 0 | 0 | 0 | 0 |
| JSG000000 | Arhgap5 | 0.75961 | 0.806543 | 0.031532802 | 2236.764374 | 2286.191239 | 160 | 8 | 14 | 23 | 83 | 11 | 1 | 0 | 0 | 0 | 0 | 0 |
| SG0000004 | Tmem67 | 0.772861 | 0.818319 | 0.051058946 | 193.4611822 | 200.4306392 | 388 | 6 | 14 | 35 | 201 | 18 | 13 | 0 | 2 | 2 | 10 | 0 |
| JSG000000 | Slc7a6 | 0.801482 | 0.842596 | -0.034858392 | 2322.629764 | 2267.182942 | 69 | 13 | 14 | 22 | 33 | 3 | 5 | 1 | 1 | 1 | 2 | 0 |
| SG0000002 | Arl13b | 0.836787 | 0.872491 | -0.041515203 | 792.8723476 | 770.3816862 | 116 | 9 | 14 | 22 | 75 | 6 | 5 | 0 | 1 | 0 | 1 | 1 |
| SG0000003 | Cdr2 | 0.911888 | 0.934478 | -0.016041526 | 731.0173352 | 722.9340747 | 102 | 12 | 14 | 14 | 36 | 8 | 2 | 0 | 0 | 0 | 0 | 0 |
| SG0000003 | Trim3 | 0.921722 | 0.943099 | -0.017089851 | 516.4269649 | 510.3455743 | 56 | 14 | 14 | 19 | 45 | 3 | 3 | 2 | 2 | 3 | 3 | 0 |
| SG000000C | Nosip | 0.957336 | 0.972208 | -0.01869347 | 943.5321843 | 931.3853953 | 40 | 10 | 14 | 13 | 29 | 0 | 1 | 0 | 0 | 0 | 0 | 0 |
| SG0000003 | C330027C09Rik | 0.971786 | 0.984761 | -0.010702845 | 3014.050547 | 2991.773107 | 178 | 2 | 14 | 30 | 86 | 14 | 12 | 2 | 2 | 4 | 4 | 0 |
| SG0000003 | Strip2 | 1 | 1 | -0.023436387 | 38.63368499 | 38.01115612 | 189 | 9 | 14 | 28 | 105 | 18 | 16 | 0 | 2 | 3 | 10 | 0 |
| SG0000001 | Rab11fip4 | 9.9E-227 | 3.7E-224 | 5.242437437 | 54.97482333 | 2081.108684 | 96 | 10 | 13 | 18 | 45 | 10 | 3 | 0 | 0 | 1 | 1 | 1 |
| SG0000004 | Ankrd45 | 3.6E-139 | 4.7E-137 | 7.73415657 | 2.613155511 | 556.3887941 | 62 | 10 | 13 | 16 | 32 | 4 | 5 | 0 | 0 | 0 | 0 | 0 |
| SG0000005 | Cemip | 1.3E-121 | 1.4E-119 | -6.859365575 | 8171.145865 | 70.37334168 | 183 | 5 | 13 | 34 | 98 | 16 | 2 | 1 | 1 | 2 | 4 | 1 |
| SG0000002 | BC067074 | 1.5E-105 | 1.2E-103 | 5.30284526 | 13.3406645 | 526.6146637 | 197 | 4 | 13 | 27 | 78 | 21 | 2 | 0 | 0 | 1 | 2 | 0 |
| JSG000000 | Aspa | 2.53E-92 | 1.51E-90 | -12.70242456 | 13006.36131 | 1.951394174 | 67 | 12 | 13 | 17 | 38 | 9 | 3 | 0 | 0 | 0 | 2 | 0 |
| JSG000000 | Hspbap1 | 5.88E-88 | 3.17E-86 | 2.771294796 | 322.1830658 | 2199.609416 | 110 | 9 | 13 | 21 | 48 | 11 | 5 | 1 | 1 | 2 | 3 | 1 |
| JSG000000 | Nrp1 | 1.36E-87 | 7.27E-86 | -4.478617986 | 2228.538077 | 99.95895473 | 247 | 10 | 13 | 34 | 85 | 28 | 0 | 0 | 0 | 0 | 0 | 0 |
| SG0000002 | Cth | 1.41E-70 | 5.13E-69 | -3.241487837 | 7748.875327 | 819.3200899 | 58 | 11 | 13 | 17 | 41 | 5 | 1 | 0 | 0 | 0 | 0 | 0 |
| SG0000002 | Wasf3 | 7.81E-68 | 2.67E-66 | 6.40374982 | 28.12953893 | 2381.679915 | 132 | 6 | 13 | 18 | 67 | 15 | 9 | 0 | 1 | 0 | 3 | 0 |
| SG0000007 | 2410141K09Rik | 1.35E-62 | 4.02E-61 | 9.511812303 | 0.331601068 | 242.0787887 | 73 | 11 | 13 | 16 | 40 | 8 | 1 | 0 | 0 | 0 | 0 | 0 |
| SG0000003 | Nxf7 | 3.97E-48 | 7.81E-47 | 7.261847765 | 0 | 152.4737133 | 41 | 11 | 13 | 16 | 33 | 0 | 0 | 0 | 0 | 0 | 0 | 0 |
| JSG000000 | Gprc5a | 3.14E-43 | 5.16E-42 | -2.713763439 | 2300.601229 | 350.6849685 | 28 | 11 | 13 | 11 | 18 | 2 | 0 | 0 | 0 | 0 | 0 | 0 |
| JSG000000 | Eif4e3 | 1.06E-42 | 1.72E-41 | -3.380026788 | 1341.524471 | 128.8573364 | 57 | 9 | 13 | 18 | 30 | 9 | 3 | 0 | 1 | 0 | 1 | 1 |
| SG0000002 | Bub1 | 1.14E-40 | 1.73E-39 | 1.602984911 | 2383.213465 | 7239.515208 | 94 | 9 | 13 | 15 | 45 | 5 | 2 | 0 | 0 | 1 | 1 | 0 |
| SG0000002 | Pmvk | 2.64E-39 | 3.8E-38 | 2.008024497 | 258.3878901 | 1039.316354 | 62 | 9 | 13 | 14 | 54 | 0 | 0 | 0 | 0 | 0 | 0 | 0 |
| JSG000000 | Oip5 | 7.87E-35 | 9.7E-34 | 1.993451431 | 199.0711213 | 792.6782434 | 37 | 12 | 13 | 14 | 25 | 1 | 2 | 0 | 0 | 0 | 0 | 0 |
| SG0000004 | Dzip1 | 1.65E-33 | 1.92E-32 | 1.90177269 | 269.3708378 | 1006.563555 | 144 | 10 | 13 | 25 | 57 | 22 | 3 | 0 | 0 | 0 | 0 | 0 |
| SG0000002 | Rph3al | 2.52E-32 | 2.81E-31 | -2.855660416 | 522.7358813 | 72.21756965 | 196 | 9 | 13 | 35 | 105 | 15 | 3 | 1 | 1 | 2 | 2 | 0 |
| SG0000003 | Nup93 | 1.04E-30 | 1.08E-29 | 1.383632259 | 2977.457411 | 7768.915023 | 356 | 6 | 13 | 36 | 149 | 28 | 1 | 0 | 0 | 0 | 2 | 0 |
| JSG000000 | Fbxo4 | 3.15E-30 | 3.22E-29 | -3.116097678 | 365.7774051 | 42.18693795 | 41 | 13 | 13 | 15 | 35 | 2 | 0 | 0 | 0 | 0 | 0 | 0 |
| JSG000000 | Phf20l1 | 1.96E-29 | 1.93E-28 | -2.18197468 | 1955.284031 | 430.8935814 | 193 | 2 | 13 | 23 | 75 | 18 | 13 | 0 | 2 | 2 | 5 | 2 |
| SG0000001 | Cr1l | 2.38E-29 | 2.33E-28 | -1.939275316 | 1999.482249 | 521.3597657 | 331 | 7 | 13 | 49 | 130 | 25 | 2 | 0 | 0 | 0 | 0 | 0 |
| SG0000002 | Slc30a9 | 3.35E-29 | 3.26E-28 | -1.76796039 | 5321.629653 | 1562.555813 | 142 | 11 | 13 | 32 | 74 | 6 | 4 | 0 | 0 | 1 | 3 | 0 |
| SG0000002 | Amhr2 | 1.15E-23 | 8.76E-23 | 5.083904954 | 2.567315498 | 87.07372599 | 21 | 12 | 13 | 16 | 18 | 0 | 0 | 0 | 0 | 0 | 0 | 0 |
| SG0000003 | Emid1 | 2.64E-21 | 1.78E-20 | 5.452036704 | 1.655661775 | 72.47666609 | 166 | 13 | 13 | 30 | 102 | 17 | 4 | 0 | 0 | 4 | 4 | 0 |
| SG0000007 | Ept1 | 2.7E-20 | 1.72E-19 | -1.495683515 | 3281.133202 | 1163.531806 | 193 | 7 | 13 | 27 | 92 | 20 | 20 | 0 | 0 | 4 | 15 | 6 |
| SG0000002 | Hsd17b7 | 3.74E-20 | 2.36E-19 | 1.617947457 | 176.8938871 | 542.9546305 | 68 | 9 | 13 | 19 | 29 | 6 | 14 | 1 | 2 | 5 | 8 | 0 |
| SG0000004 | Pigs | 1.49E-19 | 9.1E-19 | -1.440330738 | 3153.267535 | 1161.924913 | 35 | 12 | 13 | 16 | 20 | 1 | 1 | 0 | 0 | 0 | 0 | 0 |
| SG0000004 | Rgs8 | 1.6E-19 | 9.75E-19 | -4.052937096 | 116.4506412 | 7.015946166 | 237 | 12 | 13 | 30 | 156 | 13 | 8 | 0 | 0 | 2 | 6 | 3 |
| SG0000002 | Smc3 | 1.88E-19 | 1.14E-18 | 1.08230334 | 3922.402094 | 8305.348907 | 112 | 10 | 13 | 20 | 51 | 14 | 4 | 0 | 0 | 0 | 3 | 0 |
| JSG000000 | Myom2 | 2.22E-19 | 1.34E-18 | 7.494232672 | 0.307509278 | 55.44304222 | 103 | 8 | 13 | 15 | 57 | 17 | 0 | 0 | 0 | 0 | 0 | 0 |
| SG0000003 | Rims3 | 2.29E-19 | 1.39E-18 | 7.534789752 | 3.675546953 | 681.5844935 | 78 | 8 | 13 | 14 | 42 | 7 | 0 | 0 | 0 | 0 | 0 | 2 |
| SG0000002 | Atf3 | 2.59E-18 | 1.48E-17 | 3.360110036 | 13.58677252 | 139.511566 | 85 | 8 | 13 | 26 | 46 | 3 | 6 | 0 | 0 | 2 | 2 | 0 |
| JSG000000 | Arhgap31 | 3.23E-18 | 1.84E-17 | -2.242995404 | 348.2934537 | 73.57604092 | 169 | 9 | 13 | 25 | 67 | 11 | 4 | 0 | 0 | 1 | 3 | 0 |
| SG0000001 | Sec63 | 2.04E-17 | 1.11E-16 | -1.271472577 | 4723.715875 | 1956.736666 | 187 | 7 | 13 | 32 | 91 | 20 | 11 | 0 | 1 | 0 | 3 | 1 |
| SG0000002 | Ccnh | 5.14E-17 | 2.75E-16 | 1.187309322 | 584.7900898 | 1331.728593 | 276 | 12 | 13 | 31 | 116 | 39 | 3 | 0 | 0 | 0 | 0 | 0 |
| JSG000000 | Tmem130 | 6.17E-16 | 3.09E-15 | 5.425236684 | 0 | 41.96936894 | 52 | 11 | 13 | 15 | 30 | 1 | 0 | 0 | 0 | 0 | 0 | 0 |
| JSG000000 | Pfdn1 | 8.4E-16 | 4.17E-15 | 1.063955707 | 1070.890414 | 2238.863483 | 106 | 8 | 13 | 23 | 64 | 10 | 1 | 0 | 0 | 0 | 0 | 0 |
| JSG000000 | Gtf3c3 | 1.27E-15 | 6.23E-15 | 1.035005181 | 1271.429022 | 2605.311915 | 134 | 9 | 13 | 17 | 50 | 12 | 6 | 0 | 0 | 2 | 3 | 2 |
| SG0000003 | Nfatc1 | 1.34E-15 | 6.56E-15 | -1.99380409 | 413.6558167 | 103.8590388 | 119 | 10 | 13 | 20 | 61 | 5 | 5 | 0 | 0 | 1 | 6 | 2 |
| JSG000000 | Clptm1 | 5.73E-15 | 2.7E-14 | -1.14527093 | 4534.67531 | 2050.148374 | 48 | 9 | 13 | 16 | 26 | 6 | 5 | 0 | 0 | 0 | 0 | 0 |
| JSG000000 | Extl3 | 3.94E-14 | 1.76E-13 | -1.101564292 | 4856.928732 | 2263.381892 | 66 | 8 | 13 | 14 | 31 | 2 | 3 | 0 | 2 | 1 | 4 | 1 |
| SG0000002 | Stx6 | 6.73E-14 | 2.96E-13 | 1.11630437 | 466.0672684 | 1010.391545 | 218 | 6 | 13 | 16 | 57 | 30 | 1 | 0 | 0 | 0 | 2 | 0 |
| JSG000000 | Tcaf2 | 1.34E-13 | 5.78E-13 | 2.867943578 | 39.0355677 | 284.9689503 | 42 | 8 | 13 | 12 | 22 | 2 | 4 | 0 | 0 | 0 | 0 | 0 |
| SG0000003 | Nucb2 | 8.96E-13 | 3.66E-12 | -1.427354424 | 568.3768637 | 211.3293879 | 215 | 11 | 13 | 31 | 75 | 19 | 4 | 0 | 0 | 0 | 0 | 0 |
| SG0000002 | Trub1 | 2.76E-11 | 1.02E-10 | 1.044103426 | 596.0041562 | 1229.01091 | 100 | 13 | 13 | 23 | 47 | 10 | 19 | 1 | 1 | 2 | 7 | 2 |
| SG0000002 | Trdmt1 | 4.18E-11 | 1.53E-10 | 1.381393301 | 114.337926 | 297.8729935 | 99 | 4 | 13 | 16 | 53 | 14 | 11 | 0 | 2 | 0 | 5 | 0 |
| SG0000005 | Phtf1 | 8.69E-11 | 3.1E-10 | -1.075846752 | 1223.10249 | 580.2307049 | 450 | 0 | 13 | 43 | 147 | 86 | 12 | 0 | 0 | 0 | 3 | 6 |
| SG0000002 | Speg | 1.41E-10 | 4.95E-10 | 1.334677523 | 120.4557955 | 303.8125215 | 83 | 10 | 13 | 18 | 34 | 3 | 3 | 2 | 2 | 3 | 3 | 0 |
| JSG000000 | Syap1 | 1.98E-10 | 6.9E-10 | -1.043515963 | 1312.681556 | 636.839184 | 58 | 10 | 13 | 16 | 34 | 1 | 1 | 0 | 0 | 0 | 0 | 0 |
| SG0000006 | Nol4l | 3.1E-10 | 1.06E-09 | 1.720680017 | 152.0103834 | 501.0136434 | 133 | 11 | 13 | 29 | 66 | 5 | 12 | 0 | 0 | 4 | 4 | 2 |
| JSG000000 | Ippk | 7.14E-10 | 2.38E-09 | -1.0025746 | 1233.89534 | 615.8476613 | 67 | 11 | 13 | 18 | 37 | 9 | 0 | 0 | 0 | 0 | 1 | 0 |
| JSG000000 | Prmt8 | 1.1E-09 | 3.62E-09 | 4.657848265 | 0 | 24.24364378 | 91 | 11 | 13 | 19 | 41 | 7 | 1 | 0 | 0 | 0 | 0 | 0 |
| SG0000004 | Pdp1 | 2.79E-09 | 8.85E-09 | 1.099349943 | 187.6660425 | 402.0897518 | 108 | 0 | 13 | 20 | 40 | 0 | 21 | 0 | 1 | 2 | 4 | 0 |
| SG0000002 | Tbc1d31 | 3.77E-09 | 1.18E-08 | 0.847503286 | 688.3190012 | 1238.550357 | 140 | 8 | 13 | 24 | 55 | 14 | 2 | 0 | 0 | 2 | 2 | 0 |
| JSG000000 | Adss | 5.66E-09 | 1.76E-08 | 0.71402739 | 4211.867966 | 6909.051194 | 87 | 6 | 13 | 16 | 43 | 2 | 2 | 0 | 1 | 0 | 1 | 0 |
| JSG000000 | Sdhaf2 | 6.45E-09 | 1.99E-08 | -0.917339994 | 1453.702866 | 769.7129481 | 47 | 10 | 13 | 17 | 36 | 3 | 1 | 0 | 0 | 0 | 1 | 0 |
| SG0000006 | Strada | 8.7E-09 | 2.66E-08 | -0.91824032 | 1356.348298 | 717.7171885 | 85 | 10 | 13 | 16 | 67 | 15 | 8 | 0 | 1 | 2 | 12 | 1 |
| JSG000000 | Dpp4 | 2.59E-08 | 7.63E-08 | 8.09408859 | 0.663202137 | 181.2213536 | 171 | 9 | 13 | 26 | 76 | 14 | 10 | 0 | 1 | 2 | 3 | 0 |
| SG0000007 | Kcnip3 | 2.64E-08 | 7.77E-08 | 3.624068864 | 2.920664789 | 36.01099732 | 115 | 11 | 13 | 18 | 44 | 7 | 0 | 0 | 0 | 0 | 0 | 0 |
| SG000000C | Tfam | 9.04E-08 | 2.55E-07 | 0.690693829 | 1693.956484 | 2734.146682 | 72 | 12 | 13 | 18 | 31 | 10 | 6 | 0 | 1 | 0 | 1 | 1 |
| SG0000007 | Spef2 | 1.24E-07 | 3.47E-07 | 4.302024791 | 0 | 18.72597617 | 216 | 5 | 13 | 20 | 106 | 19 | 7 | 0 | 0 | 1 | 1 | 0 |
| SG0000001 | Klhl3 | 1.59E-07 | 4.38E-07 | 4.087313795 | 1.644787664 | 27.9585017 | 164 | 7 | 13 | 21 | 87 | 14 | 0 | 0 | 0 | 0 | 0 | 0 |
| SG0000002 | Trp53inp1 | 1.8E-07 | 4.95E-07 | 0.677537794 | 3690.203361 | 5902.139896 | 41 | 11 | 13 | 12 | 30 | 10 | 8 | 1 | 2 | 2 | 9 | 4 |
| SG0000003 | Bcl7c | 2.91E-07 | 7.82E-07 | 1.162407135 | 385.5382238 | 862.9524321 | 91 | 10 | 13 | 18 | 40 | 8 | 5 | 0 | 1 | 0 | 0 | 0 |
| JSG000000 | Slc5a8 | 3.33E-07 | 8.93E-07 | -5.297401365 | 66.82194086 | 1.699191691 | 99 | 10 | 13 | 24 | 65 | 8 | 11 | 1 | 1 | 3 | 6 | 1 |
| SG0000002 | Elk4 | 6.3E-07 | 1.65E-06 | -0.886482976 | 710.3874283 | 384.2707259 | 91 | 10 | 13 | 12 | 33 | 6 | 7 | 0 | 0 | 0 | 2 | 0 |
| SG0000002 | Rcc1 | 8.32E-07 | 2.16E-06 | 0.653221653 | 2597.362423 | 4084.810037 | 47 | 13 | 13 | 13 | 39 | 0 | 0 | 0 | 0 | 0 | 0 | 0 |
| SG0000001 | Mocs2 | 1.08E-06 | 2.77E-06 | -0.850881651 | 765.8519274 | 424.6233869 | 237 | 0 | 13 | 14 | 50 | 0 | 25 | 0 | 1 | 3 | 4 | 0 |
| SG0000002 | Papolg | 1.45E-06 | 3.7E-06 | 0.709675207 | 609.8793487 | 997.4185759 | 194 | 4 | 13 | 40 | 98 | 3 | 13 | 0 | 3 | 6 | 11 | 1 |
| JSG000000 | Gpr176 | 2.33E-06 | 5.82E-06 | -1.667599615 | 115.2003993 | 36.26239461 | 169 | 7 | 13 | 30 | 85 | 15 | 2 | 1 | 1 | 1 | 4 | 0 |
| SG0000001 | Prep | 3.67E-06 | 9.01E-06 | -0.651138968 | 4000.311306 | 2547.307821 | 147 | 10 | 13 | 21 | 44 | 15 | 1 | 0 | 0 | 0 | 0 | 0 |
| JSG000000 | Ist1 | 4E-06 | 9.79E-06 | 0.597000729 | 2136.653327 | 3231.835076 | 43 | 10 | 13 | 18 | 35 | 2 | 1 | 0 | 0 | 0 | 1 | 1 |
| JSG000000 | Hadh | 5.13E-06 | 1.24E-05 | 0.899538796 | 1341.351107 | 2502.24962 | 55 | 10 | 13 | 17 | 31 | 3 | 0 | 0 | 0 | 0 | 0 | 0 |
| SG0000002 | Zfp54 | 9.89E-06 | 2.33E-05 | -1.165733978 | 187.5757092 | 83.60951472 | 61 | 10 | 13 | 13 | 36 | 8 | 3 | 0 | 1 | 1 | 2 | 0 |
| JSG000000 | Tep1 | 1.59E-05 | 3.67E-05 | -0.707965511 | 1415.065679 | 866.2789131 | 61 | 12 | 13 | 21 | 51 | 6 | 0 | 0 | 0 | 0 | 0 | 0 |
| SG0000002 | Spata1 | 1.79E-05 | 4.13E-05 | 1.734828369 | 18.60662856 | 61.93028297 | 275 | 8 | 13 | 33 | 110 | 26 | 7 | 0 | 0 | 1 | 4 | 0 |
| SG0000002 | Clta | 3.57E-05 | 7.98E-05 | 0.52862202 | 2673.734541 | 3856.997622 | 155 | 7 | 13 | 31 | 52 | 1 | 0 | 0 | 0 | 0 | 0 | 0 |

| | | | | | | | | | | | | | | | | | | |
|---|---|---|---|---|---|---|---|---|---|---|---|---|---|---|---|---|---|---|
| ISG000000 | Gpr45 | 3.75E-05 | 8.36E-05 | 3.144197422 | 2.637247302 | 23.31570721 | 123 | 8 | 13 | 20 | 65 | 14 | 3 | 0 | 0 | 0 | 0 | 1 |
| ISG000000 | 6720489N17Rik | 6.55E-05 | 0.000142 | -0.871378043 | 304.1115372 | 166.2348086 | 38 | 11 | 13 | 13 | 27 | 1 | 4 | 1 | 1 | 1 | 1 | 0 |
| SG0000003 | Kifc3 | 6.72E-05 | 0.000145 | -1.403592809 | 1689.26759 | 638.5205992 | 82 | 11 | 13 | 18 | 42 | 17 | 4 | 0 | 0 | 2 | 2 | 4 |
| ISG000000 | Prelid2 | 7.46E-05 | 0.00016 | 0.740198817 | 281.0948375 | 469.542509 | 141 | 9 | 13 | 21 | 82 | 11 | 0 | 0 | 0 | 0 | 0 | 0 |
| SG0000001 | Lhx9 | 0.000192 | 0.000394 | -1.402901929 | 92.16668206 | 34.85446267 | 197 | 2 | 13 | 15 | 117 | 9 | 36 | 0 | 0 | 4 | 25 | 0 |
| SG0000002 | Paxbp1 | 0.000205 | 0.00042 | 0.574009395 | 1212.672682 | 1805.25119 | 148 | 9 | 13 | 26 | 47 | 14 | 5 | 0 | 0 | 2 | 3 | 3 |
| SG0000002 | Fgf10 | 0.000228 | 0.000463 | 1.460285959 | 53.37806611 | 146.876634 | 210 | 7 | 13 | 25 | 107 | 24 | 10 | 1 | 2 | 2 | 5 | 0 |
| ISG000000 | Letm1 | 0.000242 | 0.000491 | -0.534382892 | 1880.187155 | 1298.182653 | 89 | 13 | 13 | 26 | 62 | 10 | 6 | 2 | 2 | 3 | 7 | 1 |
| ISG000000 | Fbln5 | 0.000289 | 0.000581 | -1.18766809 | 266.2635597 | 116.8928679 | 117 | 10 | 13 | 21 | 47 | 8 | 7 | 0 | 0 | 0 | 0 | 0 |
| ISG000000 | Hgsnat | 0.000338 | 0.000673 | -0.616469084 | 1090.350789 | 711.1980485 | 52 | 13 | 13 | 19 | 30 | 7 | 3 | 1 | 1 | 1 | 1 | 0 |
| SG0000003 | Col4a2 | 0.000543 | 0.001054 | -1.104450602 | 1627.437066 | 756.8876919 | 122 | 7 | 13 | 16 | 46 | 16 | 0 | 0 | 0 | 0 | 0 | 0 |
| SG0000002 | Trim26 | 0.000546 | 0.001059 | -0.489802499 | 3070.563757 | 2186.617781 | 77 | 8 | 13 | 20 | 62 | 4 | 0 | 0 | 0 | 0 | 0 | 0 |
| SG0000002 | Polr1b | 0.0006 | 0.001158 | 0.456339151 | 1774.923864 | 2435.294574 | 93 | 7 | 13 | 18 | 34 | 4 | 1 | 0 | 0 | 0 | 1 | 0 |
| ISG000000 | Lyrm4 | 0.000658 | 0.001263 | 0.656052599 | 204.3193773 | 321.9593689 | 245 | 11 | 13 | 37 | 118 | 24 | 2 | 0 | 0 | 0 | 0 | 0 |
| SG0000002 | Gnpda2 | 0.000896 | 0.00169 | 0.541753095 | 398.0554464 | 579.4653907 | 245 | 0 | 13 | 28 | 99 | 28 | 11 | 0 | 2 | 0 | 4 | 0 |
| SG0000002 | Trak2 | 0.001275 | 0.002365 | -0.475839393 | 1707.040997 | 1227.444943 | 170 | 2 | 13 | 20 | 80 | 29 | 7 | 0 | 1 | 2 | 2 | 1 |
| SG0000005 | 1700030K09Rik | 0.001686 | 0.003086 | 0.583949281 | 253.3083203 | 379.6957225 | 51 | 12 | 13 | 13 | 23 | 0 | 1 | 0 | 1 | 1 | 1 | 0 |
| ISG000000 | Slc30a7 | 0.002299 | 0.004123 | -0.489159037 | 1092.276335 | 778.1816424 | 127 | 4 | 13 | 16 | 58 | 10 | 11 | 0 | 0 | 0 | 4 | 2 |
| SG0000003 | Plekhg5 | 0.003462 | 0.006055 | 0.619621754 | 576.5572457 | 885.8642353 | 77 | 5 | 13 | 13 | 33 | 28 | 4 | 0 | 0 | 4 | 4 | 4 |
| SG0000003 | Fam13a | 0.004393 | 0.007571 | 1.558001379 | 10.68166897 | 31.45171006 | 208 | 5 | 13 | 22 | 84 | 15 | 4 | 0 | 0 | 0 | 2 | 0 |
| ISG000000 | Ranbp3 | 0.005309 | 0.00905 | -0.418501899 | 2705.21724 | 2024.046667 | 48 | 13 | 13 | 15 | 21 | 1 | 0 | 0 | 0 | 0 | 0 | 0 |
| ISG000000 | Lactb | 0.007834 | 0.01303 | 0.493618485 | 249.5569608 | 351.3691744 | 47 | 9 | 13 | 14 | 27 | 2 | 0 | 0 | 0 | 0 | 0 | 0 |
| SG0000002 | Arf4 | 0.009947 | 0.016274 | 0.323691203 | 5255.986775 | 6578.017509 | 214 | 12 | 13 | 41 | 90 | 8 | 11 | 0 | 0 | 3 | 7 | 0 |
| SG0000001 | Tbp | 0.011988 | 0.019344 | 0.356559715 | 935.3578705 | 1197.603274 | 203 | 9 | 13 | 25 | 95 | 22 | 35 | 1 | 5 | 5 | 15 | 6 |
| ISG000000 | Fxr2 | 0.013361 | 0.021343 | 0.315026138 | 4534.601372 | 5641.199153 | 47 | 12 | 13 | 17 | 31 | 6 | 2 | 0 | 0 | 1 | 1 | 0 |
| ISG000000 | Hhip | 0.014468 | 0.022984 | -1.199831153 | 41.43389082 | 18.03725938 | 174 | 6 | 13 | 28 | 82 | 14 | 5 | 0 | 0 | 1 | 5 | 1 |
| SG0000002 | Hdgfrp3 | 0.01489 | 0.023605 | -0.452830643 | 648.1349202 | 473.5325253 | 267 | 5 | 13 | 22 | 110 | 10 | 16 | 1 | 1 | 1 | 7 | 2 |
| ISG000000 | Sacm1l | 0.015778 | 0.024923 | -0.351412819 | 2098.448917 | 1644.798125 | 137 | 10 | 13 | 23 | 75 | 15 | 6 | 0 | 1 | 1 | 2 | 1 |
| SG0000002 | Farp1 | 0.0166 | 0.026098 | -0.376859695 | 1327.639637 | 1022.431304 | 258 | 6 | 13 | 30 | 107 | 36 | 1 | 0 | 0 | 0 | 0 | 0 |
| SG0000002 | Adck1 | 0.020258 | 0.031305 | 0.433514252 | 262.8566256 | 354.9929447 | 124 | 11 | 13 | 25 | 55 | 12 | 0 | 0 | 0 | 0 | 0 | 1 |
| SG0000002 | Ttl | 0.028883 | 0.043479 | -0.400951895 | 479.2975043 | 362.9999961 | 85 | 10 | 13 | 19 | 42 | 5 | 3 | 0 | 0 | 0 | 3 | 0 |
| ISG000000 | Stk32a | 0.073846 | 0.10329 | -0.788444346 | 43.96908697 | 25.45669677 | 237 | 6 | 13 | 28 | 107 | 21 | 5 | 0 | 0 | 0 | 0 | 1 |
| ISG000000 | Rab5a | 0.088808 | 0.122257 | -0.244145986 | 2270.550155 | 1917.060569 | 83 | 11 | 13 | 21 | 46 | 8 | 4 | 0 | 0 | 0 | 1 | 1 |
| ISG000000 | Arsj | 0.097628 | 0.133205 | -1.03754522 | 28.53241851 | 13.89972962 | 178 | 6 | 13 | 23 | 95 | 13 | 6 | 1 | 1 | 3 | 5 | 0 |
| ISG000000 | Peli2 | 0.114729 | 0.154039 | -0.277428042 | 629.7902399 | 519.6151006 | 189 | 7 | 13 | 22 | 76 | 16 | 7 | 0 | 1 | 3 | 4 | 2 |
| ISG000000 | Pcsk1 | 0.122821 | 0.163895 | 1.125442851 | 13.40341246 | 29.24202465 | 111 | 9 | 13 | 23 | 57 | 8 | 0 | 0 | 0 | 0 | 0 | 0 |
| SG0000002 | Rbm18 | 0.126451 | 0.168276 | 0.225327212 | 830.9379381 | 971.4016518 | 141 | 4 | 13 | 29 | 54 | 12 | 5 | 0 | 1 | 1 | 2 | 0 |
| ISG000000 | Arl8b | 0.156853 | 0.204733 | -0.19923051 | 3237.28388 | 2819.72286 | 85 | 10 | 13 | 23 | 53 | 7 | 5 | 0 | 0 | 0 | 0 | 1 |
| SG0000004 | Zbtb44 | 0.172942 | 0.223776 | 0.177462864 | 1697.122098 | 1919.264072 | 113 | 7 | 13 | 16 | 50 | 12 | 21 | 0 | 1 | 3 | 9 | 3 |
| SG0000006 | Gpn1 | 0.18314 | 0.235627 | 0.190128549 | 1314.56313 | 1499.739546 | 58 | 6 | 13 | 12 | 24 | 3 | 4 | 0 | 1 | 2 | 2 | 0 |
| ISG000000 | Spag5 | 0.185696 | 0.238504 | 0.174758425 | 1962.119198 | 2214.791841 | 43 | 11 | 13 | 19 | 32 | 1 | 2 | 0 | 1 | 0 | 0 | 0 |
| SG0000003 | Acpp | 0.195694 | 0.250019 | 0.422031935 | 98.54386817 | 132.0303812 | 114 | 7 | 13 | 16 | 56 | 22 | 6 | 1 | 1 | 2 | 3 | 0 |
| SG0000003 | Ufsp2 | 0.217624 | 0.274725 | 0.169010563 | 1325.429156 | 1490.162975 | 137 | 5 | 13 | 25 | 46 | 6 | 1 | 0 | 0 | 0 | 0 | 0 |
| ISG000001 | Zc3h11a | 0.258259 | 0.319736 | 0.196517747 | 528.2289148 | 605.3128688 | 102 | 8 | 13 | 20 | 49 | 7 | 6 | 0 | 0 | 3 | 5 | 1 |
| ISG000000 | Wbscr16 | 0.282275 | 0.34574 | 0.174769768 | 613.2890422 | 692.2710138 | 39 | 12 | 13 | 15 | 27 | 3 | 0 | 0 | 0 | 0 | 0 | 0 |
| SG0000002 | Nln | 0.289208 | 0.35317 | 0.148631197 | 1074.449074 | 1191.045313 | 129 | 9 | 13 | 17 | 54 | 12 | 2 | 0 | 0 | 0 | 3 | 0 |
| SG0000003 | Wtip | 0.308026 | 0.373274 | -0.183607691 | 815.2536679 | 717.8295607 | 71 | 13 | 13 | 20 | 44 | 5 | 2 | 0 | 0 | 1 | 1 | 0 |
| SG0000002 | Rhobtb3 | 0.31693 | 0.382795 | 0.147740483 | 634.1454375 | 702.5272736 | 183 | 4 | 13 | 27 | 113 | 8 | 10 | 0 | 1 | 1 | 3 | 2 |
| ISG000000 | Dync1li1 | 0.357247 | 0.425377 | -0.127595558 | 2472.58608 | 2263.295864 | 69 | 8 | 13 | 14 | 38 | 7 | 2 | 0 | 0 | 1 | 1 | 1 |
| SG0000001 | Actn1 | 0.358307 | 0.42636 | -0.146683614 | 8063.523511 | 7283.979066 | 87 | 8 | 13 | 17 | 60 | 6 | 0 | 0 | 0 | 0 | 3 | 0 |
| ISG000000 | Mall | 0.360736 | 0.428993 | -0.604633867 | 62.60982861 | 41.17461856 | 33 | 12 | 13 | 13 | 21 | 3 | 2 | 0 | 0 | 0 | 0 | 1 |
| SG0000002 | Zdbf2 | 0.385666 | 0.454388 | 0.180602906 | 1272.580723 | 1442.288804 | 168 | 12 | 13 | 23 | 87 | 6 | 48 | 2 | 2 | 6 | 18 | 2 |
| SG0000002 | Ift140 | 0.42002 | 0.489072 | 0.113913278 | 987.7027491 | 1068.852004 | 163 | 11 | 13 | 22 | 65 | 15 | 0 | 0 | 0 | 0 | 0 | 0 |
| SG0000004 | Proser2 | 0.427518 | 0.496834 | 0.248606842 | 150.5589862 | 178.8730031 | 82 | 7 | 13 | 21 | 51 | 4 | 8 | 1 | 1 | 1 | 5 | 0 |
| SG0000002 | Pi4k2b | 0.451234 | 0.520411 | -0.120107396 | 1047.799523 | 964.1004097 | 135 | 5 | 13 | 16 | 71 | 8 | 9 | 0 | 2 | 2 | 8 | 0 |
| SG0000002 | Lrpprc | 0.502689 | 0.569956 | 0.080184255 | 3899.775862 | 4122.659935 | 226 | 10 | 13 | 28 | 106 | 20 | 0 | 0 | 0 | 0 | 0 | 0 |
| SG0000000 | Trap1 | 0.509133 | 0.575986 | 0.081281799 | 4735.97856 | 5010.464474 | 61 | 12 | 13 | 13 | 31 | 6 | 2 | 0 | 0 | 0 | 0 | 0 |
| SG0000002 | Polr1e | 0.51359 | 0.580442 | 0.09310741 | 810.1468879 | 864.1554857 | 76 | 12 | 13 | 17 | 61 | 0 | 1 | 0 | 0 | 0 | 0 | 0 |
| SG0000002 | Fyttd1 | 0.554846 | 0.620064 | -0.08746629 | 3684.091483 | 3467.371931 | 92 | 9 | 13 | 19 | 44 | 6 | 13 | 0 | 1 | 1 | 2 | 1 |
| SG0000002 | Rmdn1 | 0.590094 | 0.653463 | -0.218691227 | 70.17256761 | 60.302421 | 149 | 6 | 13 | 24 | 67 | 7 | 3 | 0 | 0 | 0 | 0 | 0 |
| SG0000002 | Tm7sf2 | 0.590539 | 0.653786 | -0.097385278 | 347.4329488 | 324.7544513 | 32 | 8 | 13 | 17 | 18 | 5 | 0 | 0 | 0 | 0 | 0 | 0 |
| SG0000003 | Pcdh18 | 0.636292 | 0.696468 | 0.278461843 | 44.08939278 | 53.47607022 | 60 | 10 | 13 | 16 | 46 | 4 | 16 | 2 | 2 | 4 | 6 | 0 |
| SG0000002 | Nr1d2 | 0.682457 | 0.738812 | -0.070367425 | 1175.100117 | 1119.15996 | 68 | 11 | 13 | 18 | 41 | 7 | 6 | 0 | 1 | 1 | 2 | 2 |
| SG0000002 | Dgcr8 | 0.717796 | 0.770213 | 0.052262372 | 1634.317347 | 1694.606751 | 66 | 10 | 13 | 17 | 46 | 1 | 0 | 0 | 0 | 0 | 0 | 0 |
| SG0000004 | Smc1a | 0.788613 | 0.831947 | 0.03036682 | 5477.72288 | 5594.243695 | 109 | 10 | 13 | 19 | 62 | 5 | 4 | 0 | 0 | 0 | 1 | 0 |
| ISG000000 | Maoa | 0.802075 | 0.842972 | 0.027889768 | 1570.870222 | 1601.533266 | 180 | 6 | 13 | 23 | 101 | 13 | 7 | 0 | 0 | 0 | 1 | 0 |
| SG0000004 | Slc25a26 | 0.810819 | 0.850465 | -0.057776199 | 279.2251961 | 268.263901 | 187 | 4 | 13 | 23 | 64 | 19 | 1 | 0 | 0 | 1 | 1 | 0 |
| ISG000000 | Atad2 | 0.853938 | 0.885928 | -0.040850455 | 5349.478679 | 5200.130572 | 100 | 7 | 13 | 19 | 53 | 7 | 4 | 0 | 0 | 1 | 2 | 1 |
| ISG000000 | Map2k1 | 0.906754 | 0.929909 | -0.019412052 | 4593.841805 | 4532.443779 | 104 | 11 | 13 | 17 | 47 | 8 | 1 | 0 | 0 | 1 | 2 | 0 |
| ISG000000 | Ell | 0.962798 | 0.977034 | -0.010004847 | 987.6267925 | 980.8014609 | 44 | 10 | 13 | 11 | 21 | 2 | 0 | 0 | 0 | 0 | 0 | 0 |
| ISG000000 | Fam83c | 1 | 1 | -0.039106317 | 19.44600691 | 18.92597497 | 18 | 13 | 13 | 13 | 15 | 1 | 1 | 0 | 0 | 0 | 1 | 1 |
| SG0000002 | Tfcp2l1 | 0 | 0 | 5.985813272 | 191.6310408 | 12144.37591 | 87 | 10 | 12 | 19 | 45 | 10 | 7 | 0 | 0 | 2 | 8 | 1 |
| SG0000002 | Chst10 | 7.2E-197 | 2E-194 | 8.803969521 | 1.928205151 | 861.8112229 | 172 | 8 | 12 | 20 | 64 | 12 | 3 | 0 | 0 | 0 | 0 | 0 |
| SG0000004 | Rimklb | 3.7E-178 | 8.6E-176 | 10.04889611 | 0.674076249 | 714.049264 | 114 | 7 | 12 | 15 | 46 | 7 | 13 | 0 | 2 | 0 | 0 | 1 |
| SG0000004 | Ltbp4 | 4.1E-114 | 3.8E-112 | 4.087508085 | 60.833214 | 1034.19707 | 122 | 12 | 12 | 28 | 48 | 16 | 0 | 0 | 0 | 0 | 0 | 0 |
| SG0000000 | Lck | 2.8E-106 | 2.3E-104 | 3.833725335 | 67.46673521 | 961.9577893 | 37 | 11 | 12 | 12 | 23 | 0 | 0 | 0 | 0 | 0 | 0 | 0 |
| ISG000000 | Slc22a4 | 5.39E-92 | 3.16E-90 | -6.754114083 | 766.2313369 | 7.098539062 | 72 | 10 | 12 | 18 | 35 | 7 | 0 | 0 | 0 | 0 | 0 | 0 |
| ISG000000 | Sycp3 | 6.91E-84 | 3.43E-82 | 8.73919478 | 0.649984458 | 277.7555459 | 151 | 3 | 12 | 24 | 75 | 9 | 8 | 0 | 0 | 0 | 0 | 0 |
| SG0000003 | Acsbg1 | 4.48E-82 | 2.11E-80 | -5.673769946 | 796.3278903 | 15.59972651 | 81 | 11 | 12 | 19 | 48 | 2 | 1 | 0 | 0 | 0 | 0 | 0 |
| SG0000003 | Zfp185 | 4.26E-74 | 1.73E-72 | 4.639406231 | 15.36289325 | 382.8894942 | 116 | 10 | 12 | 20 | 60 | 4 | 10 | 0 | 0 | 4 | 6 | 0 |
| SG0000002 | Acat2 | 4.03E-73 | 1.58E-71 | 2.535050262 | 315.0603718 | 1826.080634 | 48 | 8 | 12 | 14 | 24 | 22 | 0 | 0 | 0 | 4 | 0 | 0 |
| ISG000000 | Sugt1 | 1.23E-71 | 4.59E-70 | 2.143047454 | 1738.904975 | 7680.640107 | 111 | 7 | 12 | 17 | 50 | 8 | 2 | 0 | 0 | 0 | 0 | 0 |
| SG0000000 | Hck | 5.51E-71 | 2.02E-69 | 4.58508463 | 16.64345751 | 399.4767959 | 114 | 12 | 12 | 18 | 57 | 9 | 6 | 0 | 0 | 0 | 0 | 0 |
| SG0000002 | Rab3il1 | 3.32E-67 | 1.11E-65 | 6.075496216 | 3.867284414 | 260.8030778 | 62 | 12 | 12 | 13 | 35 | 3 | 3 | 0 | 0 | 0 | 0 | 0 |
| ISG000000 | Mael | 2.05E-62 | 6.07E-61 | 7.555920167 | 0 | 187.1735661 | 160 | 4 | 12 | 18 | 81 | 12 | 0 | 0 | 0 | 0 | 1 | 0 |
| SG0000002 | P2rx7 | 4.14E-61 | 1.19E-59 | -3.650916376 | 1327.502156 | 105.6817252 | 172 | 9 | 12 | 19 | 60 | 4 | 5 | 0 | 0 | 0 | 4 | 1 |
| SG0000007 | Vrtn | 1.41E-55 | 3.47E-54 | 7.454319435 | 0 | 174.3774484 | 43 | 9 | 12 | 18 | 45 | 0 | 0 | 0 | 0 | 0 | 0 | 0 |
| ISG000000 | Rrp15 | 5.54E-52 | 1.22E-50 | 1.988096381 | 645.8416642 | 2562.139096 | 48 | 8 | 12 | 14 | 29 | 1 | 2 | 0 | 0 | 0 | 0 | 0 |
| SG0000000 | Tspan32 | 2.65E-48 | 5.25E-47 | -6.951358163 | 921.4290133 | 7.445511955 | 84 | 6 | 12 | 12 | 18 | 0 | 6 | 0 | 0 | 0 | 0 | 0 |
| ISG000000 | Zbtb8a | 4.97E-47 | 9.39E-46 | 2.294856727 | 189.1376536 | 928.1084088 | 48 | 9 | 12 | 12 | 37 | 1 | 2 | 0 | 0 | 0 | 0 | 0 |
| ISG000000 | Gm17067 | 2.09E-44 | 3.6E-43 | 10.86961169 | 0.34247518 | 640.7789889 | 44 | 12 | 12 | 18 | 35 | 3 | 3 | 0 | 0 | 1 | 2 | 1 |
| ISG000000 | Scarb2 | 1.7E-43 | 2.83E-42 | -2.629608105 | 19606.93823 | 3168.245774 | 104 | 8 | 12 | 17 | 58 | 14 | 6 | 0 | 0 | 0 | 0 | 2 |
| SG0000002 | Tmbim4 | 7.13E-42 | 1.12E-40 | -2.430299324 | 2422.790447 | 449.4929221 | 102 | 3 | 12 | 10 | 24 | 6 | 0 | 0 | 0 | 0 | 1 | 0 |
| SG0000003 | Melk | 2.88E-40 | 4.31E-39 | 1.71533585 | 848.0559695 | 2784.787488 | 132 | 9 | 12 | 23 | 57 | 5 | 4 | 0 | 0 | 0 | 1 | 0 |
| ISG000000 | Oas3 | 1.41E-37 | 1.91E-36 | -3.594918774 | 2189.997339 | 181.2446535 | 29 | 11 | 12 | 14 | 21 | 3 | 2 | 0 | 0 | 2 | 3 | 2 |
| SG0000000 | Spc25 | 1.51E-37 | 2.05E-36 | 1.720761446 | 610.8836793 | 2013.535695 | 93 | 12 | 12 | 21 | 48 | 18 | 3 | 0 | 0 | 0 | 0 | 6 |
| SG0000002 | Prpsap2 | 5.88E-36 | 7.55E-35 | 1.786411756 | 398.6475232 | 1375.15798 | 96 | 8 | 12 | 30 | 60 | 16 | 2 | 0 | 0 | 2 | 2 | 0 |
| ISG000000 | Cspg5 | 2.89E-35 | 3.63E-34 | 5.108005335 | 3.950433897 | 136.2409373 | 64 | 8 | 12 | 12 | 34 | 4 | 3 | 0 | 0 | 0 | 0 | 0 |
| SG0000000 | Dusp3 | 6.76E-35 | 8.35E-34 | -2.121626144 | 2468.109454 | 567.1413423 | 91 | 12 | 12 | 20 | 47 | 0 | 16 | 0 | 0 | 0 | 0 | 0 |
| SG0000003 | Capn5 | 8.7E-34 | 1.03E-32 | -2.104866454 | 5294.226781 | 1230.76348 | 73 | 12 | 12 | 17 | 33 | 18 | 4 | 0 | 0 | 0 | 0 | 0 |
| SG0000002 | Tnn | 3.73E-33 | 4.3E-32 | -5.811854667 | 389.1386397 | 6.927264512 | 189 | 10 | 12 | 32 | 107 | 19 | 1 | 0 | 0 | 0 | 0 | 1 |
| ISG000000 | Mllt11 | 2.42E-31 | 2.59E-30 | -2.232057044 | 1157.765124 | 246.4361077 | 53 | 12 | 12 | 15 | 28 | 0 | 8 | 3 | 3 | 6 | 7 | 0 |
| SG0000006 | Gm13154 | 3.58E-31 | 3.81E-30 | 7.540945301 | 4.29056551 | 799.0340166 | 94 | 10 | 12 | 25 | 58 | 9 | 5 | 0 | 0 | 2 | 2 | 0 |
| SG0000003 | Chst15 | 1.16E-30 | 1.2E-29 | 4.152523719 | 8.127571156 | 144.5423055 | 164 | 2 | 12 | 20 | 60 | 18 | 6 | 0 | 0 | 2 | 6 | 0 |
| SG0000000 | Sult5a1 | 1.32E-30 | 1.37E-29 | 6.422785008 | 0 | 84.79281999 | 51 | 12 | 12 | 16 | 23 | 0 | 0 | 0 | 0 | 0 | 0 | 0 |
| SG0000003 | Mapkapk3 | 4.34E-30 | 4.41E-29 | -1.775551972 | 6033.292458 | 1762.21939 | 56 | 8 | 12 | 20 | 44 | 2 | 0 | 0 | 0 | 0 | 0 | 0 |
| ISG000000 | Zfp345 | 6.94E-30 | 6.99E-29 | 3.514819314 | 15.30383555 | 174.9308205 | 42 | 11 | 12 | 12 | 20 | 2 | 2 | 0 | 0 | 0 | 0 | 0 |
| SG0000002 | Smarcc2 | 8.49E-30 | 8.53E-29 | -1.897866686 | 2509.885281 | 673.5022818 | 96 | 6 | 12 | 24 | 81 | 12 | 0 | 0 | 0 | 0 | 11 | 0 |
| ISG000000 | Irf2 | 1.19E-29 | 1.18E-28 | -3.742179044 | 859.2838843 | 64.21382283 | 171 | 4 | 12 | 23 | 70 | 26 | 1 | 0 | 0 | 0 | 4 | 1 |
| SG0000003 | Abhd5 | 1.77E-29 | 1.75E-28 | -1.981787709 | 1766.480678 | 447.2304302 | 213 | 8 | 12 | 28 | 80 | 16 | 16 | 0 | 0 | 0 | 8 | 0 |
| SG0000003 | Ccdc40 | 1.75E-28 | 1.65E-27 | 5.435971168 | 6.124747066 | 265.1420047 | 62 | 10 | 12 | 12 | 40 | 8 | 0 | 0 | 0 | 0 | 0 | 0 |
| SG0000002 | Rbbp5 | 3.07E-28 | 2.87E-27 | 1.39634677 | 1267.490043 | 3336.466884 | 205 | 12 | 12 | 21 | 129 | 37 | 1 | 0 | 0 | 1 | 1 | 1 |
| SG0000001 | Pcgf2 | 3.95E-28 | 3.67E-27 | 1.526377033 | 551.9963877 | 1590.089271 | 40 | 12 | 12 | 12 | 16 | 7 | 4 | 0 | 0 | 0 | 0 | 7 |
| SG0000000 | Slc30a4 | 1.95E-27 | 1.76E-26 | -2.20904543 | 2314.283926 | 500.5272223 | 116 | 2 | 12 | 12 | 49 | 13 | 10 | 0 | 0 | 0 | 1 | 1 |
| ISG000000 | Msantd2 | 2.7E-27 | 2.42E-26 | 1.652947787 | 360.7622271 | 1134.508945 | 75 | 10 | 12 | 15 | 36 | 5 | 3 | 0 | 0 | 0 | 1 | 1 |
| SG0000005 | Rbbp4 | 4.27E-27 | 3.78E-26 | 1.268433748 | 6731.943378 | 16217.24435 | 69 | 9 | 12 | 16 | 35 | 3 | 6 | 0 | 0 | 0 | 1 | 0 |
| ISG000000 | Sae1 | 5.05E-27 | 4.46E-26 | 1.286167787 | 3845.839203 | 9379.208827 | 86 | 8 | 12 | 15 | 41 | 4 | 2 | 0 | 0 | 0 | 0 | 0 |
| SG0000001 | Tnfrsf22 | 6.16E-27 | 5.41E-26 | -2.035806282 | 1099.840114 | 268.2197784 | 32 | 10 | 12 | 14 | 24 | 2 | 8 | 2 | 2 | 4 | 12 | 0 |
| SG0000003 | Polr3g | 2.09E-26 | 1.8E-25 | 1.8789106 | 469.8096379 | 1727.946817 | 103 | 9 | 12 | 20 | 51 | 22 | 3 | 0 | 0 | 0 | 0 | 0 |
| SG0000003 | Nrip3 | 4.67E-26 | 3.96E-25 | 5.32811769 | 2.294772122 | 92.18554134 | 84 | 6 | 12 | 18 | 58 | 14 | 8 | 0 | 0 | 0 | 4 | 0 |
| ISG000000 | Sdc2 | 2.12E-25 | 1.74E-24 | -1.718395604 | 2443.758268 | 742.6250982 | 222 | 4 | 12 | 24 | 107 | 16 | 11 | 0 | 0 | 1 | 3 | 0 |
| SG0000004 | Phldb1 | 2.94E-25 | 2.4E-24 | -1.768649268 | 2285.856114 | 670.8607883 | 170 | 12 | 12 | 55 | 90 | 18 | 0 | 0 | 0 | 0 | 0 | 0 |
| SG0000000 | Ube2g2 | 3.1E-25 | 2.54E-24 | -1.681540749 | 2595.648215 | 809.1920894 | 66 | 10 | 12 | 13 | 32 | 6 | 2 | 0 | 0 | 1 | 2 | 0 |

| | | | | | | | | | | | | | | | | | | |
|---|---|---|---|---|---|---|---|---|---|---|---|---|---|---|---|---|---|---|
| SG0000002 | Dusp12 | 8.35E-25 | 6.68E-24 | 1.65345678 | 246.5841159 | 775.7203501 | 125 | 12 | 12 | 36 | 95 | 6 | 12 | 0 | 0 | 0 | 5 | 0 |
| SG0000002 | Rfc4 | 9.87E-25 | 7.86E-24 | 1.82100396 | 576.198032 | 2035.863165 | 98 | 12 | 12 | 17 | 50 | 13 | 0 | 0 | 0 | 0 | 0 | 0 |
| SG0000003 | Cep57 | 1.16E-24 | 9.2E-24 | 1.341741478 | 921.5709396 | 2335.786815 | 253 | 12 | 12 | 31 | 70 | 25 | 4 | 0 | 0 | 0 | 0 | 0 |
| SG0000002 | Tmem44 | 1.32E-24 | 1.05E-23 | 3.233101234 | 16.61936572 | 156.269742 | 180 | 8 | 12 | 23 | 116 | 7 | 3 | 0 | 0 | 0 | 2 | 0 |
| SG0000001 | Ncapg | 1.59E-24 | 1.26E-23 | 1.257295892 | 2024.184968 | 4838.758779 | 94 | 2 | 12 | 17 | 44 | 5 | 5 | 0 | 0 | 2 | 3 | 0 |
| JSG000000 | Rnf217 | 1.27E-23 | 9.65E-23 | -2.860448761 | 977.0995587 | 134.5420185 | 282 | 4 | 12 | 30 | 101 | 36 | 32 | 1 | 3 | 2 | 10 | 5 |
| SG0000003 | Rbl2 | 1.33E-23 | 1E-22 | -2.065156056 | 1372.144485 | 327.8883044 | 101 | 7 | 12 | 19 | 47 | 9 | 5 | 0 | 0 | 1 | 1 | 0 |
| SG0000001 | Hint3 | 1.97E-23 | 1.47E-22 | -2.162143015 | 588.2516825 | 131.4296271 | 101 | 9 | 12 | 13 | 43 | 3 | 2 | 0 | 0 | 1 | 1 | 0 |
| SG0000002 | Got1 | 2.7E-23 | 2.01E-22 | 1.223390275 | 2196.13114 | 5127.851494 | 68 | 6 | 12 | 13 | 21 | 1 | 5 | 0 | 1 | 1 | 1 | 0 |
| SG0000003 | Kpna7 | 6.72E-23 | 4.91E-22 | -8.871013108 | 470.2640161 | 1.00438614 | 55 | 10 | 12 | 14 | 28 | 14 | 0 | 0 | 0 | 0 | 2 | 0 |
| SG0000003 | Dmpk | 2.15E-22 | 1.54E-21 | -2.898965962 | 617.3369981 | 82.76497625 | 48 | 8 | 12 | 24 | 36 | 0 | 12 | 0 | 0 | 8 | 8 | 0 |
| SG0000005 | Idi1 | 4.83E-22 | 3.39E-21 | 2.375790292 | 83.5430079 | 433.6049675 | 68 | 8 | 12 | 16 | 51 | 4 | 8 | 0 | 0 | 0 | 3 | 0 |
| JSG000000 | Tex9 | 6.44E-22 | 4.5E-21 | 1.474025473 | 331.7438231 | 921.5708452 | 67 | 7 | 12 | 12 | 38 | 10 | 2 | 0 | 0 | 0 | 0 | 0 |
| SG0000002 | Gins1 | 1.36E-21 | 9.37E-21 | 1.453069766 | 345.4378024 | 945.774167 | 50 | 10 | 12 | 14 | 29 | 5 | 0 | 0 | 0 | 0 | 0 | 0 |
| SG0000002 | Oprk1 | 4.89E-21 | 3.25E-20 | 5.837847264 | 0 | 56.19619479 | 113 | 3 | 12 | 13 | 56 | 20 | 11 | 0 | 0 | 1 | 5 | 5 |
| SG0000005 | Prokr2 | 1.16E-20 | 7.58E-20 | -3.886101262 | 179.1932683 | 12.11960999 | 145 | 11 | 12 | 18 | 58 | 3 | 18 | 0 | 0 | 0 | 6 | 0 |
| SG0000003 | Smc4 | 1.8E-20 | 1.16E-19 | 1.112411668 | 3986.773738 | 8619.674545 | 323 | 4 | 12 | 4 | 101 | 12 | 5 | 0 | 0 | 0 | 1 | 0 |
| SG000000C | Psap | 1.44E-19 | 8.8E-19 | -1.403497337 | 25390.12878 | 9597.764871 | 34 | 12 | 12 | 14 | 28 | 6 | 0 | 0 | 0 | 0 | 0 | 0 |
| JSG000000 | Ap3s1 | 2.23E-19 | 1.35E-18 | -1.514089711 | 1672.134549 | 585.4432474 | 142 | 5 | 12 | 25 | 78 | 5 | 1 | 0 | 1 | 0 | 0 | 0 |
| SG0000002 | Wdr5 | 5.21E-19 | 3.09E-18 | 1.097079985 | 2530.091394 | 5412.403828 | 41 | 12 | 12 | 13 | 30 | 0 | 1 | 0 | 0 | 0 | 0 | 0 |
| SG000000C | Zfp40 | 1.04E-18 | 6.08E-18 | 3.262614447 | 20.03927724 | 192.32107 | 102 | 10 | 12 | 28 | 84 | 17 | 6 | 0 | 0 | 1 | 1 | 0 |
| SG0000004 | Arl4c | 1.16E-18 | 6.75E-18 | -1.390512607 | 3741.601211 | 1427.15615 | 93 | 10 | 12 | 17 | 27 | 8 | 11 | 0 | 0 | 1 | 3 | 0 |
| JSG000000 | Eif4ebp1 | 2E-18 | 1.15E-17 | 1.093337188 | 2269.634358 | 4842.651162 | 36 | 10 | 12 | 14 | 20 | 1 | 3 | 0 | 0 | 0 | 1 | 0 |
| SG0000006 | Flot2 | 7.64E-18 | 4.27E-17 | -1.30071917 | 3655.645689 | 1483.913584 | 88 | 12 | 12 | 32 | 64 | 4 | 1 | 0 | 0 | 0 | 0 | 0 |
| SG0000003 | Tbc1d9b | 1.87E-17 | 1.02E-16 | -1.454035242 | 2316.921188 | 845.6741167 | 75 | 8 | 12 | 10 | 23 | 2 | 1 | 0 | 0 | 0 | 1 | 0 |
| SG0000002 | Mxra7 | 1.17E-15 | 5.8E-15 | 2.812809094 | 49.39182258 | 347.0523278 | 34 | 12 | 12 | 19 | 43 | 2 | 3 | 0 | 0 | 0 | 0 | 0 |
| SG0000003 | Mrps10 | 1.35E-15 | 6.64E-15 | -1.243561917 | 2425.075683 | 1024.17898 | 72 | 8 | 12 | 20 | 20 | 4 | 12 | 4 | 4 | 8 | 8 | 0 |
| SG0000004 | Fbxo41 | 1.46E-15 | 7.15E-15 | 4.414746137 | 2.874824776 | 61.31721055 | 117 | 12 | 12 | 15 | 27 | 12 | 3 | 0 | 0 | 0 | 0 | 3 |
| SG0000003 | Sccpdh | 1.71E-15 | 8.33E-15 | 1.164987994 | 494.1737124 | 1108.092384 | 86 | 10 | 12 | 16 | 40 | 6 | 6 | 2 | 2 | 2 | 3 | 0 |
| SG0000002 | Pex7 | 2.31E-15 | 1.12E-14 | -1.34336589 | 1322.549788 | 521.2170359 | 123 | 5 | 12 | 17 | 42 | 10 | 5 | 0 | 0 | 0 | 1 | 0 |
| SG0000001 | Cep95 | 3E-15 | 1.44E-14 | 1.399421275 | 191.5912346 | 505.4095181 | 109 | 6 | 12 | 23 | 49 | 2 | 4 | 0 | 0 | 2 | 2 | 0 |
| SG0000003 | Nipsnap1 | 4.4E-15 | 2.09E-14 | 1.037685753 | 1038.78484 | 2132.55451 | 42 | 12 | 12 | 18 | 24 | 10 | 2 | 0 | 0 | 0 | 0 | 0 |
| SG0000001 | Gid4 | 5.62E-15 | 2.65E-14 | 1.019277167 | 1198.397156 | 2429.034991 | 76 | 8 | 12 | 14 | 40 | 9 | 4 | 0 | 0 | 0 | 2 | 0 |
| SG0000002 | Mdm1 | 1.29E-14 | 5.96E-14 | 1.204948874 | 350.9320042 | 809.00039 | 159 | 6 | 12 | 21 | 69 | 12 | 4 | 0 | 0 | 2 | 4 | 2 |
| JSG000000 | Rab21 | 3E-14 | 1.35E-13 | -1.18950238 | 2128.168219 | 933.1040126 | 60 | 10 | 12 | 20 | 35 | 3 | 2 | 0 | 1 | 0 | 1 | 0 |
| JSG000000 | Esyt1 | 3.07E-14 | 1.38E-13 | -1.143460857 | 3112.081737 | 1408.753403 | 24 | 11 | 12 | 11 | 17 | 2 | 0 | 0 | 0 | 0 | 0 | 0 |
| SG0000005 | Zscan22 | 3.2E-14 | 1.44E-13 | -1.398718639 | 803.1338223 | 304.6012281 | 63 | 12 | 12 | 24 | 72 | 8 | 3 | 0 | 0 | 0 | 0 | 0 |
| SG0000002 | P4hb | 4.46E-14 | 1.98E-13 | -1.047012109 | 25536.94561 | 12359.10155 | 28 | 12 | 12 | 14 | 24 | 2 | 1 | 0 | 0 | 0 | 0 | 0 |
| SG000000C | Pdk1 | 4.89E-14 | 2.17E-13 | 1.347724187 | 3826.92744 | 9739.925479 | 49 | 10 | 12 | 18 | 32 | 3 | 5 | 0 | 0 | 1 | 1 | 1 |
| SG0000002 | Spag6 | 8.82E-14 | 3.86E-13 | 5.523179891 | 0.957493736 | 44.03304916 | 145 | 5 | 12 | 22 | 110 | 11 | 0 | 0 | 0 | 0 | 1 | 0 |
| JSG000000 | Bcap31 | 9.09E-14 | 3.96E-13 | -1.087699817 | 4683.295183 | 2203.541634 | 56 | 11 | 12 | 13 | 29 | 4 | 0 | 0 | 0 | 0 | 0 | 0 |
| SG000000C | Prg4 | 9.93E-14 | 4.32E-13 | 1.996920486 | 83.79499662 | 334.465289 | 266 | 0 | 12 | 40 | 104 | 18 | 14 | 0 | 0 | 4 | 8 | 0 |
| SG0000003 | Ogfod1 | 2.53E-13 | 1.07E-12 | 1.021034969 | 727.4190479 | 1476.205508 | 129 | 12 | 12 | 24 | 54 | 21 | 9 | 0 | 0 | 0 | 0 | 3 |
| SG0000002 | Pik3c2b | 3.34E-13 | 1.4E-12 | 1.20326356 | 273.9339467 | 630.7601952 | 56 | 10 | 12 | 15 | 32 | 12 | 1 | 0 | 0 | 0 | 2 | 1 |
| SG0000002 | Cnot8 | 4.14E-13 | 1.73E-12 | 0.924249921 | 1710.32791 | 3245.684731 | 56 | 10 | 12 | 18 | 34 | 8 | 6 | 0 | 0 | 2 | 4 | 2 |
| JSG000000 | Chrnb2 | 7.87E-13 | 3.23E-12 | 3.090859205 | 8.788429726 | 74.87772099 | 29 | 11 | 12 | 14 | 18 | 0 | 6 | 0 | 0 | 1 | 1 | 0 |
| SG0000006 | Pvrl2 | 9.94E-13 | 4.05E-12 | 1.572560319 | 916.5732749 | 2726.183068 | 54 | 12 | 12 | 19 | 34 | 3 | 0 | 0 | 0 | 0 | 0 | 0 |
| SG0000003 | Scap | 1.59E-12 | 6.39E-12 | -1.112445675 | 1739.774362 | 804.6618476 | 78 | 7 | 12 | 17 | 45 | 2 | 0 | 0 | 0 | 0 | 0 | 0 |
| SG0000006 | Pcnxl2 | 2.07E-12 | 8.25E-12 | 5.037017814 | 0 | 31.83170604 | 211 | 9 | 12 | 30 | 84 | 31 | 0 | 0 | 0 | 0 | 0 | 0 |
| SG0000002 | Ttc8 | 2.16E-12 | 8.58E-12 | -1.378963724 | 574.796961 | 221.006486 | 206 | 6 | 12 | 24 | 118 | 20 | 2 | 0 | 0 | 0 | 6 | 2 |
| SG0000004 | Cystm1 | 2.25E-12 | 8.95E-12 | 1.662215581 | 115.1307738 | 364.3914454 | 163 | 4 | 12 | 26 | 84 | 16 | 3 | 0 | 0 | 1 | 1 | 0 |
| SG0000005 | Tmem123 | 4.24E-12 | 1.66E-11 | -1.027406225 | 3392.361076 | 1664.263083 | 113 | 6 | 12 | 15 | 67 | 11 | 7 | 0 | 0 | 0 | 9 | 2 |
| JSG000000 | Cyp27a1 | 5.96E-12 | 2.31E-11 | 4.026643122 | 6.293389599 | 102.5710921 | 47 | 10 | 12 | 13 | 44 | 2 | 2 | 0 | 0 | 0 | 0 | 0 |
| SG0000006 | Dap3 | 6.38E-12 | 2.47E-11 | 0.884582776 | 1623.693226 | 2997.71215 | 120 | 2 | 12 | 15 | 33 | 6 | 6 | 0 | 2 | 3 | 3 | 0 |
| JSG000000 | Cmtm4 | 8.33E-12 | 3.2E-11 | 0.860479299 | 2118.004178 | 3845.536094 | 74 | 6 | 12 | 20 | 48 | 5 | 10 | 0 | 1 | 2 | 3 | 0 |
| SG0000002 | Ptpn22 | 2.04E-11 | 7.62E-11 | -2.429835766 | 121.9592197 | 22.63399351 | 230 | 8 | 12 | 34 | 112 | 26 | 2 | 0 | 0 | 0 | 0 | 0 |
| SG0000003 | Otud7b | 3.14E-11 | 1.16E-10 | -0.993243388 | 2928.374188 | 1471.060443 | 305 | 3 | 12 | 33 | 120 | 36 | 21 | 0 | 3 | 3 | 6 | 0 |
| JSG000000 | Pinx1 | 4.51E-11 | 1.64E-10 | 1.064610424 | 361.9203387 | 756.9943898 | 93 | 8 | 12 | 16 | 37 | 9 | 0 | 0 | 0 | 0 | 0 | 0 |
| JSG000000 | Rab3ip | 1.29E-10 | 4.55E-10 | 0.921470853 | 657.7978727 | 1245.898983 | 78 | 11 | 12 | 18 | 37 | 9 | 3 | 1 | 1 | 2 | 4 | 0 |
| JSG000000 | Dvx1c1 | 2.17E-10 | 7.54E-10 | 3.886349722 | 2.933882468 | 43.38609389 | 44 | 10 | 12 | 16 | 34 | 6 | 0 | 0 | 0 | 0 | 2 | 0 |
| SG0000002 | Gtf2a1l | 2.6E-10 | 8.99E-10 | 4.75424809 | 0 | 25.98803606 | 105 | 8 | 12 | 21 | 64 | 11 | 0 | 0 | 0 | 0 | 0 | 0 |
| SG0000002 | Cdh17 | 3.69E-10 | 1.26E-09 | -4.107993916 | 86.1468327 | 4.99585614 | 188 | 6 | 12 | 26 | 83 | 12 | 2 | 0 | 0 | 1 | 2 | 0 |
| SG0000001 | Echdc1 | 4.04E-10 | 1.37E-09 | -1.050763255 | 1088.66612 | 525.5129824 | 188 | 4 | 12 | 25 | 91 | 36 | 16 | 0 | 0 | 1 | 1 | 2 |
| SG0000002 | Tmem144 | 4.26E-10 | 1.45E-09 | 3.933174255 | 11.83090017 | 180.7262423 | 120 | 8 | 12 | 23 | 44 | 10 | 6 | 0 | 0 | 0 | 0 | 0 |
| JSG000000 | Dnajc18 | 5.3E-10 | 1.79E-09 | -1.081324841 | 905.327943 | 427.8531296 | 53 | 7 | 12 | 15 | 33 | 7 | 7 | 1 | 1 | 2 | 2 | 0 |
| SG0000002 | Il6 | 9.85E-10 | 3.24E-09 | -2.954551114 | 70.64622271 | 9.113400098 | 72 | 4 | 12 | 7 | 21 | 4 | 20 | 0 | 0 | 3 | 13 | 0 |
| JSG000000 | Stk24 | 1.02E-09 | 3.36E-09 | -0.873957096 | 5165.089508 | 2818.321613 | 112 | 11 | 12 | 19 | 46 | 11 | 1 | 0 | 0 | 0 | 1 | 2 |
| SG0000003 | Stk36 | 1.35E-09 | 4.41E-09 | 1.326420626 | 103.4598325 | 259.4562675 | 85 | 2 | 12 | 8 | 59 | 17 | 3 | 0 | 0 | 0 | 0 | 0 |
| SG0000001 | Dguok | 1.44E-09 | 4.67E-09 | 0.962151313 | 362.2796783 | 705.7979276 | 136 | 8 | 12 | 20 | 68 | 8 | 0 | 0 | 0 | 0 | 0 | 0 |
| SG000000C | Pole | 1.57E-09 | 5.1E-09 | 0.757724638 | 2484.95799 | 4201.621201 | 99 | 9 | 12 | 16 | 42 | 10 | 0 | 0 | 0 | 0 | 0 | 1 |
| JSG000000 | Gm14139 | 1.92E-09 | 6.17E-09 | 5.802516261 | 2.033102857 | 113.472545 | 40 | 10 | 12 | 11 | 16 | 3 | 0 | 0 | 0 | 0 | 0 | 0 |
| SG0000004 | Tyw5 | 3.8E-09 | 1.19E-08 | 1.124180488 | 272.0238775 | 592.9513664 | 133 | 6 | 12 | 18 | 55 | 5 | 9 | 0 | 0 | 4 | 4 | 2 |
| SG0000004 | Ssr2 | 9.43E-09 | 2.88E-08 | 0.711415486 | 3647.210752 | 5971.978838 | 64 | 12 | 12 | 12 | 74 | 4 | 0 | 0 | 0 | 0 | 2 | 0 |
| SG000000C | Mettl17 | 1.04E-08 | 3.16E-08 | 0.923782603 | 402.4211774 | 763.4262702 | 57 | 12 | 12 | 16 | 31 | 0 | 1 | 0 | 0 | 0 | 0 | 0 |
| SG0000001 | Cdc6 | 1.6E-08 | 4.81E-08 | 0.745808582 | 1307.029458 | 2191.775813 | 76 | 3 | 12 | 24 | 42 | 0 | 10 | 0 | 0 | 3 | 9 | 0 |
| SG0000007 | Med12 | 1.65E-08 | 4.94E-08 | -0.977536103 | 2290.349047 | 1163.145339 | 72 | 4 | 12 | 8 | 28 | 8 | 4 | 0 | 4 | 4 | 8 | 0 |
| JSG000000 | Hao1 | 2.15E-08 | 6.37E-08 | -3.017908699 | 55.45481855 | 6.846336579 | 131 | 7 | 12 | 22 | 80 | 10 | 1 | 0 | 0 | 0 | 1 | 0 |
| SG0000003 | Mbtps1 | 2.44E-08 | 7.21E-08 | 0.694199752 | 3594.733714 | 5816.231353 | 120 | 10 | 12 | 22 | 48 | 24 | 2 | 0 | 0 | 2 | 2 | 0 |
| SG0000001 | Rbsn | 3.32E-08 | 9.7E-08 | -0.844901149 | 1841.985604 | 1025.523561 | 70 | 7 | 12 | 13 | 43 | 8 | 1 | 0 | 0 | 0 | 1 | 0 |
| SG0000004 | Tiprl | 4.04E-08 | 1.17E-07 | 0.73781794 | 1139.310987 | 1899.973821 | 112 | 2 | 12 | 16 | 78 | 10 | 12 | 0 | 0 | 2 | 6 | 0 |
| SG0000002 | Denr | 4.47E-08 | 1.29E-07 | 0.697123889 | 2000.373471 | 3243.144413 | 40 | 11 | 12 | 12 | 20 | 8 | 3 | 0 | 0 | 0 | 0 | 2 |
| SG0000006 | Zfp398 | 6.16E-08 | 1.76E-07 | -1.166742013 | 372.1085981 | 165.74688 | 136 | 8 | 12 | 20 | 74 | 14 | 4 | 0 | 0 | 2 | 2 | 0 |
| JSG000000 | Yme1l1 | 8.42E-08 | 2.38E-07 | 0.661137655 | 3855.486144 | 6096.792469 | 132 | 11 | 12 | 19 | 59 | 12 | 5 | 0 | 0 | 0 | 2 | 3 |
| SG0000002 | Brd3 | 8.69E-08 | 2.45E-07 | 0.698336351 | 1661.696715 | 2696.32321 | 52 | 9 | 12 | 20 | 42 | 7 | 16 | 3 | 3 | 3 | 3 | 3 |
| SG0000002 | Rit1 | 8.7E-08 | 2.45E-07 | -1.03537826 | 512.1196791 | 249.8570174 | 100 | 8 | 12 | 20 | 32 | 4 | 2 | 0 | 0 | 0 | 0 | 0 |
| SG0000001 | H2afy | 9.09E-08 | 2.56E-07 | 0.659255534 | 3924.993487 | 6198.614544 | 122 | 4 | 12 | 12 | 60 | 6 | 1 | 0 | 0 | 0 | 1 | 0 |
| SG0000004 | Kcns3 | 1.54E-07 | 4.27E-07 | 3.066200252 | 12.00203941 | 100.5248184 | 198 | 12 | 12 | 36 | 122 | 22 | 6 | 0 | 0 | 0 | 1 | 2 |
| SG0000002 | Rhoq | 1.63E-07 | 4.49E-07 | -0.799061586 | 1895.890543 | 1089.611691 | 55 | 10 | 12 | 12 | 27 | 3 | 6 | 0 | 0 | 0 | 3 | 1 |
| JSG000000 | 4933430I17Rik | 1.83E-07 | 5.03E-07 | 2.796518181 | 6.23198834 | 43.29746681 | 48 | 11 | 12 | 15 | 30 | 0 | 1 | 0 | 0 | 0 | 0 | 0 |
| SG0000007 | Dnajc25 | 2.03E-07 | 5.55E-07 | -1.163386062 | 314.4881936 | 140.40748 | 98 | 9 | 12 | 9 | 58 | 12 | 14 | 0 | 0 | 0 | 1 | 0 |
| SG0000005 | Pdik1l | 2.68E-07 | 7.23E-07 | -0.929839052 | 667.8066499 | 350.5429956 | 87 | 9 | 12 | 24 | 51 | 6 | 27 | 0 | 3 | 6 | 12 | 3 |
| SG0000007 | Cass4 | 3.27E-07 | 8.77E-07 | 3.849469125 | 1.963171053 | 28.29852607 | 128 | 9 | 12 | 25 | 89 | 15 | 2 | 0 | 0 | 1 | 2 | 0 |
| JSG000000 | Ints4 | 3.32E-07 | 8.9E-07 | 0.657878194 | 1750.7442 | 2762.254985 | 114 | 10 | 12 | 21 | 61 | 13 | 0 | 0 | 0 | 0 | 0 | 0 |
| JSG000000 | Best1 | 3.63E-07 | 9.69E-07 | 3.679354145 | 2.305646233 | 29.53851339 | 28 | 12 | 12 | 15 | 20 | 0 | 0 | 0 | 0 | 0 | 0 | 0 |
| JSG000000 | Fam19a5 | 3.67E-07 | 9.79E-07 | 1.902774418 | 20.70632279 | 77.42748534 | 126 | 11 | 12 | 19 | 45 | 25 | 5 | 0 | 0 | 0 | 0 | 0 |
| SG0000004 | Ppig | 3.72E-07 | 9.92E-07 | 0.650865247 | 2134.358152 | 3351.176174 | 156 | 4 | 12 | 20 | 60 | 6 | 18 | 0 | 0 | 0 | 0 | 0 |
| SG000000C | Gabpa | 3.94E-07 | 1.05E-06 | 0.642345305 | 2305.39563 | 3598.410206 | 156 | 8 | 12 | 30 | 83 | 7 | 14 | 0 | 0 | 0 | 3 | 1 |
| SG0000004 | Tdrkh | 6.68E-07 | 1.75E-06 | 1.797007999 | 61.47681477 | 213.6311941 | 198 | 6 | 12 | 26 | 100 | 6 | 14 | 0 | 0 | 0 | 3 | 0 |
| JSG000000 | Clic5 | 1.07E-06 | 2.76E-06 | 3.772227764 | 1.939079263 | 26.49410599 | 113 | 6 | 12 | 20 | 67 | 15 | 7 | 0 | 1 | 1 | 5 | 1 |
| SG0000004 | D1Ertd622e | 1.12E-06 | 2.9E-06 | -0.8590595 | 797.1815176 | 439.4956122 | 194 | 9 | 12 | 26 | 90 | 33 | 92 | 6 | 9 | 17 | 48 | 24 |
| SG0000003 | Zc4h2 | 1.31E-06 | 3.36E-06 | 0.959487288 | 157.3660338 | 306.0169296 | 126 | 3 | 12 | 18 | 87 | 9 | 6 | 0 | 0 | 3 | 9 | 0 |
| SG0000002 | Agk | 1.53E-06 | 3.88E-06 | 0.783241475 | 358.3779309 | 616.7660155 | 217 | 3 | 12 | 26 | 74 | 16 | 7 | 0 | 0 | 0 | 3 | 1 |
| SG0000002 | Tmed2 | 1.88E-06 | 4.75E-06 | -0.735993148 | 1578.370013 | 947.6605614 | 42 | 10 | 12 | 18 | 28 | 0 | 6 | 0 | 0 | 0 | 2 | 0 |
| SG0000003 | Ccdc174 | 2.74E-06 | 6.79E-06 | -0.828921365 | 732.7759234 | 412.5161885 | 72 | 4 | 12 | 12 | 38 | 8 | 6 | 0 | 0 | 0 | 2 | 0 |
| SG000000C | Ncstn | 3.27E-06 | 8.06E-06 | -0.654002847 | 3378.595875 | 2147.146974 | 48 | 9 | 12 | 12 | 42 | 6 | 4 | 0 | 0 | 0 | 3 | 0 |
| JSG000000 | Gm20390 | 4.09E-06 | 1E-05 | 3.721809426 | 5.908917816 | 77.96222495 | 24 | 12 | 12 | 14 | 23 | 1 | 0 | 0 | 0 | 0 | 0 | 0 |
| SG0000005 | Maml1 | 4.09E-06 | 1E-05 | -0.735125353 | 1211.944932 | 728.0950946 | 118 | 10 | 12 | 16 | 42 | 6 | 4 | 0 | 0 | 2 | 2 | 2 |
| SG0000002 | Ywhaz | 4.3E-06 | 1.05E-05 | -0.620638093 | 12193.08692 | 7930.178949 | 151 | 9 | 12 | 24 | 39 | 3 | 16 | 0 | 0 | 0 | 0 | 0 |
| SG0000002 | Sypl | 5.52E-06 | 1.33E-05 | -0.658022844 | 2642.67591 | 1674.785885 | 204 | 3 | 12 | 26 | 87 | 12 | 15 | 0 | 0 | 2 | 4 | 1 |
| SG0000002 | Tm9sf2 | 5.67E-06 | 1.37E-05 | -0.620746694 | 7517.686347 | 4889.008895 | 198 | 2 | 12 | 27 | 83 | 17 | 4 | 0 | 0 | 0 | 3 | 2 |
| SG0000003 | Cnep1r1 | 8.42E-06 | 2E-05 | 0.623094665 | 896.6746553 | 1381.036251 | 93 | 6 | 12 | 6 | 58 | 6 | 10 | 0 | 0 | 0 | 6 | 2 |
| SG0000002 | Arpc1a | 1.26E-05 | 2.95E-05 | -0.617649098 | 3395.223611 | 2212.776125 | 74 | 10 | 12 | 22 | 46 | 4 | 3 | 0 | 0 | 1 | 3 | 0 |
| SG0000002 | Npr2 | 1.64E-05 | 3.79E-05 | -1.131292849 | 334.2897058 | 152.605407 | 49 | 9 | 12 | 21 | 33 | 6 | 0 | 0 | 0 | 0 | 0 | 6 |
| JSG000000 | Mrrf | 1.73E-05 | 3.99E-05 | 0.703863925 | 374.7095419 | 610.3499408 | 123 | 6 | 12 | 15 | 43 | 14 | 3 | 0 | 0 | 1 | 2 | 0 |
| SG0000004 | Cox19 | 3.14E-05 | 7.05E-05 | -0.776693116 | 512.311885 | 299.038087 | 31 | 12 | 12 | 13 | 18 | 7 | 9 | 8 | 8 | 9 | 9 | 0 |
| SG0000003 | Rpf2 | 3.5E-05 | 7.82E-05 | 0.581604792 | 1134.828863 | 1698.286123 | 70 | 10 | 12 | 22 | 67 | 8 | 2 | 2 | 2 | 2 | 3 | 0 |
| SG0000007 | Prdm11 | 3.55E-05 | 7.93E-05 | 2.793820003 | 11.36043236 | 78.78045477 | 112 | 6 | 12 | 11 | 34 | 7 | 5 | 0 | 0 | 1 | 8 | 1 |
| JSG000000 | Smoc2 | 3.69E-05 | 8.23E-05 | -1.806605303 | 870.0591253 | 248.7175206 | 164 | 8 | 12 | 24 | 61 | 29 | 2 | 0 | 0 | 0 | 0 | 0 |
| SG0000003 | Fbxo22 | 3.85E-05 | 8.57E-05 | 0.527319938 | 2285.972002 | 3294.655617 | 96 | 0 | 12 | 12 | 32 | 4 | 5 | 0 | 0 | 0 | 1 | 0 |
| SG0000004 | Neurl4 | 3.99E-05 | 8.85E-05 | -0.600022956 | 2060.852375 | 1359.633871 | 47 | 12 | 12 | 12 | 19 | 0 | 5 | 0 | 0 | 0 | 1 | 0 |
| JSG000000 | Smyd4 | 5.83E-05 | 0.000127 | 0.681714014 | 328.0419939 | 526.1939005 | 109 | 8 | 12 | 20 | 56 | 7 | 0 | 0 | 0 | 0 | 1 | 0 |
| SG0000001 | Pltp | 7.46E-05 | 0.000161 | 1.02849085 | 714.7141159 | 1457.937665 | 81 | 6 | 12 | 24 | 75 | 9 | 3 | 0 | 0 | 0 | 0 | 0 |
| SG0000002 | Col5a1 | 8.81E-05 | 0.000188 | -1.314726432 | 2915.032977 | 1171.848261 | 100 | 10 | 12 | 22 | 62 | 27 | 0 | 0 | 0 | 0 | 0 | 1 |

| | | | | | | | | | | | | | | | | | | |
|---|---|---|---|---|---|---|---|---|---|---|---|---|---|---|---|---|---|---|
| ISG000000 | Tarsl2 | 9.09E-05 | 0.000193 | 0.692924084 | 278.1686854 | 449.675444 | 99 | 6 | 12 | 21 | 63 | 5 | 2 | 0 | 0 | 1 | 2 | 0 |
| SG0000002 | Per3 | 9.61E-05 | 0.000204 | 0.752699585 | 188.7262435 | 317.9929183 | 132 | 6 | 12 | 24 | 48 | 15 | 5 | 0 | 0 | 1 | 3 | 2 |
| SG0000002 | Polk | 0.000124 | 0.000261 | -0.666642524 | 1244.709474 | 784.1309552 | 153 | 3 | 12 | 25 | 72 | 9 | 10 | 0 | 0 | 2 | 8 | 0 |
| ISG000000 | Fbxo24 | 0.000148 | 0.000307 | -1.491641396 | 74.02194721 | 26.32277688 | 44 | 9 | 12 | 9 | 37 | 6 | 0 | 0 | 0 | 0 | 0 | 0 |
| SG0000002 | Tmem80 | 0.000182 | 0.000375 | -0.995500259 | 166.3340237 | 83.42681329 | 34 | 8 | 12 | 12 | 22 | 5 | 2 | 0 | 0 | 0 | 2 | 5 |
| SG0000002 | Hnrnpk | 0.000202 | 0.000414 | 0.447013378 | 17117.6209 | 23335.00182 | 163 | 0 | 12 | 12 | 55 | 6 | 5 | 0 | 0 | 0 | 1 | 0 |
| SG0000002 | Glud1 | 0.000203 | 0.000415 | 0.452565098 | 9278.741855 | 12697.69122 | 75 | 8 | 12 | 17 | 39 | 7 | 2 | 0 | 1 | 1 | 2 | 0 |
| SG000000C | Psmd2 | 0.000204 | 0.000417 | -0.490399209 | 16648.65212 | 11850.97841 | 50 | 10 | 12 | 12 | 34 | 0 | 1 | 0 | 0 | 0 | 0 | 0 |
| SG0000003 | Pygm | 0.000224 | 0.000457 | 2.887064065 | 7.68019827 | 56.81530139 | 38 | 8 | 12 | 12 | 22 | 0 | 0 | 0 | 0 | 0 | 0 | 0 |
| ISG000000 | B630005N14Rik | 0.00025 | 0.000507 | -0.551922356 | 1683.053123 | 1148.028421 | 149 | 6 | 12 | 21 | 70 | 11 | 11 | 0 | 0 | 0 | 0 | 0 |
| SG0000001 | Cad | 0.000259 | 0.000524 | 0.455651187 | 5847.97115 | 8019.917419 | 50 | 8 | 12 | 8 | 16 | 12 | 0 | 0 | 0 | 0 | 0 | 0 |
| ISG000000 | Psat1 | 0.000278 | 0.000559 | 0.443986067 | 6549.472547 | 8909.625506 | 83 | 8 | 12 | 20 | 54 | 6 | 1 | 0 | 0 | 0 | 2 | 0 |
| SG0000001 | Tmem175 | 0.000325 | 0.000649 | -0.693139493 | 512.9782745 | 317.280363 | 134 | 12 | 12 | 12 | 47 | 9 | 9 | 0 | 0 | 0 | 0 | 2 |
| SG0000002 | Stra6l | 0.000332 | 0.000663 | 6.051514024 | 0.615018556 | 40.79204328 | 87 | 9 | 12 | 18 | 45 | 3 | 3 | 0 | 0 | 3 | 3 | 0 |
| ISG000000 | A230046K03Rik | 0.000334 | 0.000667 | 0.457238851 | 2517.661068 | 3456.526364 | 135 | 6 | 12 | 18 | 57 | 8 | 9 | 0 | 0 | 0 | 2 | 0 |
| SG000000C | Gna12 | 0.0004 | 0.000791 | -0.534336769 | 1686.905564 | 1164.767852 | 163 | 6 | 12 | 20 | 42 | 10 | 1 | 0 | 0 | 0 | 0 | 0 |
| SG0000002 | Arrdc1 | 0.000443 | 0.000871 | -0.684998842 | 573.9967418 | 357.0296044 | 56 | 8 | 12 | 12 | 36 | 4 | 0 | 0 | 0 | 0 | 0 | 0 |
| ISG000000 | Ak8 | 0.000495 | 0.000966 | 5.931952198 | 0.663202137 | 40.48941122 | 125 | 11 | 12 | 16 | 51 | 12 | 0 | 0 | 0 | 0 | 0 | 0 |
| SG0000003 | Acad8 | 0.000517 | 0.001006 | -0.564835551 | 913.0402352 | 617.2448466 | 136 | 6 | 12 | 14 | 76 | 8 | 10 | 0 | 0 | 0 | 2 | 0 |
| SG0000002 | Mknk1 | 0.000577 | 0.001116 | 0.5903815 | 336.2768155 | 506.3134525 | 98 | 6 | 12 | 20 | 50 | 6 | 6 | 0 | 2 | 0 | 2 | 0 |
| SG0000001 | Exosc10 | 0.000605 | 0.001167 | 0.46270441 | 1502.665374 | 2070.857359 | 82 | 9 | 12 | 16 | 38 | 3 | 0 | 0 | 0 | 0 | 0 | 0 |
| ISG000000 | Micu2 | 0.000654 | 0.001255 | -0.519663444 | 1404.991263 | 980.0299305 | 158 | 6 | 12 | 23 | 64 | 20 | 1 | 0 | 0 | 0 | 0 | 1 |
| SG0000003 | Ikbkb | 0.000746 | 0.001422 | -0.493625064 | 1895.933845 | 1346.56471 | 146 | 3 | 12 | 15 | 44 | 27 | 5 | 0 | 0 | 1 | 3 | 0 |
| SG0000003 | Nme1 | 0.001277 | 0.002369 | -0.489493754 | 2893.719438 | 2061.123993 | 26 | 12 | 12 | 12 | 24 | 2 | 2 | 0 | 0 | 0 | 0 | 0 |
| SG0000005 | Zfp341 | 0.001443 | 0.002662 | 0.703354674 | 136.7251524 | 222.6277364 | 81 | 9 | 12 | 21 | 45 | 9 | 0 | 0 | 0 | 0 | 0 | 0 |
| ISG000000 | Cmc1 | 0.001473 | 0.002714 | -0.584487882 | 529.251278 | 352.95028 | 162 | 9 | 12 | 17 | 65 | 55 | 2 | 0 | 0 | 1 | 1 | 0 |
| SG0000003 | Phlpp2 | 0.001717 | 0.003139 | 0.620135735 | 347.5953517 | 534.2609322 | 264 | 8 | 12 | 40 | 152 | 32 | 6 | 0 | 0 | 2 | 8 | 0 |
| SG000000C | Cdk4 | 0.00175 | 0.003195 | 0.387628797 | 6829.335298 | 8934.419309 | 22 | 12 | 12 | 15 | 18 | 0 | 1 | 0 | 0 | 0 | 0 | 0 |
| SG0000002 | Ing3 | 0.002026 | 0.003666 | 0.478414147 | 535.7857822 | 746.4628409 | 248 | 8 | 12 | 37 | 107 | 32 | 19 | 0 | 0 | 5 | 11 | 4 |
| SG0000006 | Trim12a | 0.002175 | 0.003919 | 6.352275658 | 1.326404274 | 108.3680918 | 91 | 9 | 12 | 18 | 50 | 3 | 1 | 0 | 0 | 0 | 1 | 0 |
| ISG000000 | Fermt3 | 0.002282 | 0.004095 | 2.08719161 | 5.298586394 | 22.51475894 | 34 | 11 | 12 | 14 | 34 | 0 | 0 | 0 | 0 | 0 | 1 | 0 |
| SG0000004 | Zfp560 | 0.002389 | 0.004276 | -0.849003951 | 144.9586181 | 80.4763711 | 159 | 6 | 12 | 28 | 84 | 8 | 5 | 0 | 1 | 0 | 3 | 0 |
| SG0000002 | Aatk | 0.0025 | 0.004466 | 2.359858739 | 10.28263281 | 52.78282049 | 60 | 12 | 12 | 15 | 27 | 3 | 0 | 0 | 0 | 0 | 0 | 0 |
| SG000000C | Hmox2 | 0.003487 | 0.006096 | -0.450939677 | 1304.81861 | 954.5611899 | 128 | 10 | 12 | 23 | 83 | 9 | 1 | 0 | 0 | 0 | 0 | 0 |
| SG0000002 | Cox7a2l | 0.003776 | 0.006566 | -0.40841947 | 2803.737505 | 2112.47141 | 40 | 10 | 12 | 12 | 30 | 0 | 0 | 0 | 0 | 0 | 0 | 0 |
| ISG000000 | F11r | 0.003824 | 0.006645 | -0.398111347 | 4436.561565 | 3366.68943 | 43 | 8 | 12 | 13 | 32 | 1 | 2 | 0 | 0 | 0 | 0 | 0 |
| SG0000002 | Casq2 | 0.004346 | 0.007495 | 5.164739387 | 0.663202137 | 23.78961383 | 166 | 6 | 12 | 24 | 70 | 14 | 0 | 0 | 0 | 0 | 0 | 0 |
| SG0000004 | Zfp62 | 0.005107 | 0.008733 | -0.461857786 | 1509.690292 | 1096.111704 | 66 | 7 | 12 | 11 | 28 | 5 | 8 | 0 | 0 | 0 | 0 | 3 |
| ISG000000 | Itga11 | 0.005468 | 0.009303 | -1.010140481 | 78.07363211 | 38.76339434 | 81 | 11 | 12 | 19 | 50 | 8 | 3 | 0 | 0 | 0 | 0 | 0 |
| ISG000000 | Siah1a | 0.00676 | 0.011353 | 0.409244559 | 611.4369123 | 811.9822607 | 62 | 7 | 12 | 11 | 21 | 2 | 7 | 0 | 0 | 1 | 1 | 0 |
| SG0000002 | Snph | 0.006957 | 0.011658 | 4.980287251 | 0.922527834 | 29.12026535 | 131 | 0 | 12 | 8 | 52 | 16 | 3 | 0 | 0 | 0 | 0 | 0 |
| SG0000007 | Pcnp | 0.007878 | 0.013096 | 0.353918759 | 1518.892322 | 1941.185995 | 134 | 9 | 12 | 27 | 71 | 15 | 11 | 0 | 0 | 0 | 2 | 4 |
| ISG000000 | Col4a1 | 0.007902 | 0.013133 | -0.682438298 | 2380.665154 | 1483.419273 | 133 | 9 | 12 | 15 | 55 | 16 | 1 | 0 | 0 | 0 | 0 | 0 |
| SG0000003 | Rbbp7 | 0.008436 | 0.013963 | 0.318564815 | 8283.723393 | 10330.54309 | 140 | 6 | 12 | 24 | 65 | 9 | 6 | 0 | 0 | 0 | 0 | 0 |
| SG0000004 | Asun | 0.008815 | 0.014541 | 0.347099333 | 1719.421303 | 2187.104901 | 97 | 9 | 12 | 17 | 52 | 5 | 2 | 0 | 0 | 0 | 2 | 0 |
| SG0000003 | Azi2 | 0.00943 | 0.015486 | -0.387015448 | 1486.086155 | 1136.42487 | 292 | 0 | 12 | 23 | 50 | 9 | 10 | 0 | 0 | 0 | 2 | 3 |
| SG0000003 | Prkrir | 0.010353 | 0.01689 | -0.367041231 | 1875.302972 | 1454.055628 | 95 | 6 | 12 | 24 | 43 | 0 | 0 | 0 | 0 | 0 | 1 | 0 |
| SG0000003 | Ppp1r16a | 0.012951 | 0.020751 | -0.81652779 | 875.168019 | 496.9264136 | 51 | 12 | 12 | 19 | 45 | 2 | 0 | 0 | 0 | 0 | 0 | 0 |
| SG0000004 | Ttc19 | 0.013649 | 0.021757 | 0.428030669 | 324.8921291 | 437.1085089 | 119 | 10 | 12 | 16 | 61 | 24 | 11 | 0 | 0 | 0 | 3 | 4 |
| SG0000002 | Anapc4 | 0.014349 | 0.022812 | -0.351308953 | 2190.655108 | 1717.19445 | 144 | 10 | 12 | 22 | 46 | 10 | 2 | 0 | 0 | 0 | 0 | 0 |
| SG0000002 | Erlin1 | 0.014877 | 0.023586 | -0.363067367 | 1514.438195 | 1177.49043 | 208 | 3 | 12 | 32 | 124 | 11 | 15 | 0 | 0 | 0 | 3 | 0 |
| SG0000002 | Cfap36 | 0.016167 | 0.025482 | -0.397388325 | 779.680777 | 591.9581731 | 63 | 9 | 12 | 11 | 35 | 0 | 3 | 0 | 1 | 0 | 1 | 0 |
| SG0000002 | Dnajc24 | 0.01692 | 0.02656 | 0.499577781 | 165.1231946 | 233.4511295 | 147 | 4 | 12 | 24 | 62 | 16 | 0 | 0 | 0 | 0 | 0 | 0 |
| ISG000000 | Nkap | 0.017099 | 0.026813 | -0.369808403 | 1221.931314 | 945.6346397 | 71 | 10 | 12 | 17 | 39 | 1 | 8 | 0 | 1 | 1 | 3 | 0 |
| SG0000006 | Rdh11 | 0.017184 | 0.026929 | 0.342212499 | 959.9467763 | 1216.92379 | 77 | 12 | 12 | 27 | 69 | 3 | 2 | 0 | 0 | 0 | 1 | 0 |
| ISG000000 | Dctn5 | 0.017637 | 0.027567 | -0.34695938 | 1584.737658 | 1245.983231 | 50 | 8 | 12 | 13 | 37 | 10 | 5 | 0 | 0 | 0 | 0 | 0 |
| ISG000000 | Fastkd5 | 0.019152 | 0.029731 | 0.46693429 | 287.6456026 | 397.5749074 | 45 | 9 | 12 | 14 | 25 | 4 | 1 | 0 | 0 | 0 | 0 | 1 |
| SG0000003 | Tacc3 | 0.019367 | 0.030027 | 0.305366908 | 2451.871065 | 3029.857669 | 68 | 10 | 12 | 10 | 26 | 3 | 9 | 0 | 0 | 0 | 0 | 0 |
| SG0000005 | Srsf11 | 0.022062 | 0.033883 | 0.366250167 | 5328.055012 | 6867.85293 | 190 | 0 | 12 | 3 | 68 | 30 | 5 | 0 | 0 | 0 | 0 | 2 |
| SG0000002 | Tubgcp4 | 0.023444 | 0.035818 | -0.333951871 | 1664.779086 | 1320.769651 | 146 | 6 | 12 | 14 | 65 | 13 | 16 | 0 | 0 | 0 | 6 | 2 |
| SG0000006 | Pick1 | 0.024047 | 0.03668 | 0.337634535 | 840.2845282 | 1061.853223 | 50 | 12 | 12 | 17 | 38 | 9 | 2 | 0 | 0 | 2 | 2 | 0 |
| SG0000002 | Pde6a | 0.024542 | 0.037385 | 0.967838576 | 23.16505031 | 45.30870925 | 94 | 12 | 12 | 27 | 68 | 8 | 2 | 0 | 0 | 2 | 4 | 0 |
| SG0000005 | Smim4 | 0.031179 | 0.046657 | 0.632279788 | 72.5929888 | 112.5200352 | 96 | 7 | 12 | 17 | 42 | 4 | 2 | 0 | 1 | 1 | 1 | 0 |
| SG0000002 | Nsun2 | 0.035293 | 0.052268 | 0.257726725 | 8348.454715 | 9981.356574 | 96 | 9 | 12 | 21 | 38 | 3 | 0 | 0 | 0 | 0 | 2 | 0 |
| SG0000005 | Tmem248 | 0.036817 | 0.054353 | -0.287427169 | 3081.506144 | 2524.868932 | 77 | 6 | 12 | 21 | 45 | 6 | 8 | 0 | 0 | 2 | 2 | 0 |
| SG0000004 | Dtx3 | 0.037501 | 0.055312 | -0.294267151 | 1872.125484 | 1526.693006 | 30 | 12 | 12 | 20 | 24 | 3 | 2 | 0 | 0 | 0 | 0 | 3 |
| SG0000005 | Rab11fip5 | 0.037516 | 0.055331 | -0.364904026 | 670.7384697 | 520.8421923 | 46 | 11 | 12 | 18 | 29 | 3 | 0 | 0 | 0 | 0 | 0 | 0 |
| ISG000000 | Taf1b | 0.038783 | 0.057057 | -0.326273939 | 983.5786792 | 784.4962347 | 111 | 7 | 12 | 13 | 40 | 9 | 0 | 0 | 0 | 0 | 0 | 0 |
| SG0000002 | 3110043O21Rik | 0.039116 | 0.057496 | -0.352414041 | 826.375167 | 647.2768292 | 216 | 9 | 12 | 36 | 90 | 18 | 13 | 0 | 0 | 3 | 4 | 0 |
| SG0000002 | Impa1 | 0.041083 | 0.060171 | 0.286187037 | 1212.042623 | 1477.980742 | 75 | 9 | 12 | 12 | 43 | 6 | 12 | 0 | 0 | 0 | 1 | 0 |
| ISG000000 | Rtn4rl1 | 0.047639 | 0.069088 | 0.930116645 | 19.76673387 | 37.66412865 | 67 | 6 | 12 | 15 | 33 | 2 | 1 | 0 | 0 | 0 | 0 | 0 |
| SG0000001 | Zfp790 | 0.047729 | 0.069197 | -0.41116869 | 298.1739936 | 224.2309997 | 60 | 12 | 12 | 24 | 28 | 12 | 6 | 0 | 0 | 0 | 0 | 0 |
| ISG000000 | Snx6 | 0.052811 | 0.075882 | 0.254107797 | 2230.134802 | 2659.654257 | 88 | 11 | 12 | 18 | 55 | 5 | 1 | 0 | 0 | 0 | 0 | 0 |
| SG0000003 | Papd7 | 0.061464 | 0.087169 | -0.292078764 | 1226.682854 | 1001.861819 | 86 | 6 | 12 | 14 | 30 | 4 | 16 | 0 | 4 | 2 | 4 | 2 |
| SG0000002 | Patz1 | 0.066081 | 0.093289 | 0.258098352 | 1332.311453 | 1593.313027 | 56 | 4 | 12 | 8 | 21 | 0 | 0 | 0 | 0 | 0 | 4 | 0 |
| SG0000006 | Gspt1 | 0.073211 | 0.10249 | 0.219538693 | 6776.451093 | 7890.236851 | 172 | 9 | 12 | 24 | 70 | 14 | 3 | 0 | 0 | 0 | 0 | 0 |
| SG0000002 | Golm1 | 0.090728 | 0.124654 | 0.216730318 | 2115.979687 | 2458.973345 | 92 | 6 | 12 | 12 | 38 | 10 | 2 | 0 | 0 | 0 | 0 | 2 |
| SG0000004 | Acp1 | 0.09485 | 0.12977 | -0.246194558 | 1559.949447 | 1315.220525 | 68 | 10 | 12 | 16 | 42 | 10 | 14 | 0 | 0 | 2 | 2 | 2 |
| ISG000000 | Atg5 | 0.096744 | 0.13212 | -0.246985067 | 1346.277192 | 1134.447949 | 192 | 5 | 12 | 22 | 77 | 14 | 2 | 0 | 0 | 0 | 0 | 0 |
| ISG000000 | Cbwd1 | 0.097764 | 0.13336 | 0.246232424 | 753.0909749 | 893.2454006 | 137 | 6 | 12 | 16 | 77 | 6 | 3 | 0 | 0 | 1 | 4 | 0 |
| ISG000000 | Timm10b | 0.102384 | 0.138963 | -0.340870508 | 254.8217749 | 201.1982916 | 24 | 12 | 12 | 14 | 16 | 0 | 13 | 6 | 6 | 7 | 8 | 0 |
| ISG000000 | Polr2d | 0.119872 | 0.160319 | 0.231460113 | 774.3087561 | 909.0559367 | 20 | 12 | 12 | 13 | 29 | 1 | 0 | 0 | 0 | 0 | 0 | 0 |
| ISG000000 | Nid1 | 0.124682 | 0.166193 | 0.62811887 | 2625.402691 | 4057.68703 | 124 | 8 | 12 | 21 | 65 | 12 | 1 | 0 | 0 | 0 | 2 | 0 |
| ISG000000 | Hsd17b4 | 0.126282 | 0.168089 | 0.204266999 | 1722.507201 | 1984.501995 | 162 | 9 | 12 | 30 | 77 | 2 | 0 | 0 | 0 | 0 | 0 | 0 |
| SG0000003 | Znrd1 | 0.126803 | 0.168694 | 0.224585001 | 793.0861832 | 926.6744861 | 26 | 12 | 12 | 12 | 30 | 0 | 2 | 0 | 0 | 0 | 0 | 0 |
| ISG000000 | Fbxo47 | 0.138007 | 0.182242 | -0.756074839 | 56.52424069 | 33.46829063 | 88 | 6 | 12 | 13 | 53 | 6 | 1 | 0 | 0 | 0 | 0 | 0 |
| SG0000002 | Angel2 | 0.141641 | 0.186604 | 0.205621574 | 1151.444403 | 1327.826203 | 168 | 4 | 12 | 14 | 27 | 15 | 13 | 0 | 0 | 1 | 1 | 0 |
| SG0000001 | Mtmr4 | 0.15811 | 0.206208 | -0.200041129 | 2143.873307 | 1866.296909 | 63 | 9 | 12 | 18 | 51 | 3 | 6 | 0 | 0 | 0 | 0 | 0 |
| ISG000000 | Papd4 | 0.172462 | 0.223285 | 0.197937959 | 778.0993843 | 892.5248867 | 160 | 9 | 12 | 27 | 89 | 10 | 1 | 0 | 0 | 0 | 2 | 0 |
| SG0000004 | Sall2 | 0.186629 | 0.239549 | -0.241570237 | 470.0939742 | 397.6168652 | 34 | 11 | 12 | 13 | 28 | 0 | 6 | 0 | 0 | 0 | 2 | 0 |
| SG0000001 | Med23 | 0.196061 | 0.250452 | -0.190614571 | 1608.585722 | 1409.494891 | 170 | 12 | 12 | 32 | 90 | 19 | 0 | 0 | 0 | 0 | 0 | 3 |
| SG0000002 | Cnga3 | 0.201539 | 0.256663 | 0.402343858 | 72.7024205 | 96.08739954 | 176 | 3 | 12 | 22 | 64 | 9 | 1 | 0 | 0 | 1 | 1 | 0 |
| SG0000003 | Prpf39 | 0.205839 | 0.261581 | -0.233588298 | 1748.764749 | 1487.353863 | 178 | 4 | 12 | 30 | 84 | 12 | 11 | 0 | 0 | 0 | 7 | 1 |
| SG0000003 | Ing4 | 0.239145 | 0.298881 | -0.394958958 | 94.55128453 | 71.90729301 | 36 | 12 | 12 | 12 | 30 | 3 | 3 | 0 | 0 | 0 | 0 | 0 |
| SG0000003 | Secisbp2 | 0.240673 | 0.300582 | 0.174256441 | 799.401824 | 902.0311817 | 75 | 10 | 12 | 16 | 31 | 3 | 2 | 0 | 1 | 1 | 1 | 1 |
| ISG000000 | Gnpnat1 | 0.24855 | 0.309275 | -0.178747384 | 1385.480803 | 1224.030356 | 41 | 10 | 12 | 15 | 27 | 4 | 5 | 0 | 1 | 0 | 2 | 2 |
| SG0000005 | Fat2 | 0.261385 | 0.323114 | -0.536442801 | 36.47558918 | 25.14878115 | 160 | 6 | 12 | 22 | 66 | 12 | 0 | 0 | 0 | 0 | 0 | 0 |
| SG0000002 | Nudt5 | 0.275312 | 0.338206 | 0.147086384 | 1916.746456 | 2122.472551 | 81 | 6 | 12 | 16 | 45 | 6 | 11 | 0 | 0 | 2 | 3 | 0 |
| SG0000002 | Camk2g | 0.288778 | 0.352693 | 0.149989266 | 1101.148998 | 1221.792222 | 162 | 9 | 12 | 21 | 84 | 21 | 13 | 0 | 0 | 0 | 0 | 0 |
| SG0000001 | Hbs1l | 0.290311 | 0.35442 | 0.138557697 | 2538.490171 | 2794.380289 | 175 | 6 | 12 | 15 | 70 | 25 | 2 | 0 | 0 | 0 | 4 | 1 |
| SG0000002 | Ccdc43 | 0.297401 | 0.362163 | 0.133992344 | 1849.09 | 2029.054744 | 44 | 12 | 12 | 14 | 26 | 4 | 10 | 0 | 0 | 0 | 2 | 0 |
| SG0000003 | Micu3 | 0.299191 | 0.363971 | -0.192876921 | 451.8574497 | 395.3117362 | 215 | 7 | 12 | 29 | 105 | 13 | 0 | 0 | 0 | 0 | 0 | 0 |
| ISG000000 | Dse | 0.301054 | 0.365865 | 0.206397325 | 1508.270997 | 1740.24809 | 120 | 5 | 12 | 17 | 59 | 8 | 1 | 0 | 0 | 0 | 3 | 0 |
| ISG000000 | Topbp1 | 0.309806 | 0.375178 | 0.124301952 | 6266.96978 | 6830.873089 | 101 | 10 | 12 | 13 | 39 | 9 | 2 | 0 | 0 | 0 | 0 | 1 |
| SG0000004 | Uevld | 0.321744 | 0.387903 | 0.145569462 | 734.5768265 | 812.5647873 | 71 | 10 | 12 | 19 | 51 | 4 | 10 | 0 | 1 | 2 | 2 | 1 |
| SG0000005 | Zfp595 | 0.326366 | 0.393078 | 0.21601 | 153.6638984 | 178.4831968 | 154 | 0 | 12 | 12 | 83 | 1 | 9 | 0 | 0 | 0 | 3 | 1 |
| ISG000000 | Dcun1d5 | 0.329668 | 0.396469 | -0.145847529 | 1717.701018 | 1552.54105 | 74 | 4 | 12 | 9 | 27 | 8 | 1 | 0 | 0 | 0 | 0 | 0 |
| SG0000003 | Sec22a | 0.355753 | 0.423771 | 0.137184394 | 635.9834625 | 699.4270755 | 189 | 6 | 12 | 29 | 81 | 20 | 4 | 0 | 1 | 0 | 1 | 0 |
| SG0000002 | Nme2 | 0.366131 | 0.434717 | -0.550115333 | 1937.016511 | 1322.915504 | 20 | 12 | 12 | 16 | 20 | 0 | 0 | 0 | 0 | 0 | 0 | 0 |
| ISG000000 | Bbs1 | 0.366229 | 0.434804 | 0.20441357 | 152.7806828 | 176.0366369 | 32 | 10 | 12 | 13 | 22 | 3 | 1 | 0 | 0 | 0 | 2 | 0 |
| SG0000002 | Nfu1 | 0.377872 | 0.446674 | -0.135523368 | 1618.061936 | 1472.985741 | 97 | 3 | 12 | 15 | 64 | 3 | 0 | 0 | 0 | 0 | 0 | 0 |
| SG0000003 | Efcab12 | 0.402765 | 0.47176 | -0.308623563 | 62.93474881 | 50.81425392 | 126 | 9 | 12 | 12 | 51 | 0 | 1 | 0 | 0 | 0 | 2 | 0 |
| SG0000003 | Rgs10 | 0.404067 | 0.473111 | 0.226348996 | 85.4964984 | 100.0198423 | 62 | 11 | 12 | 18 | 38 | 3 | 0 | 0 | 0 | 0 | 1 | 0 |
| SG0000002 | Ssfa2 | 0.40929 | 0.478508 | 0.124645811 | 780.9016365 | 851.3702318 | 237 | 3 | 12 | 24 | 99 | 18 | 15 | 0 | 0 | 3 | 6 | 0 |
| SG0000002 | Cnpy3 | 0.409548 | 0.478683 | -0.122633876 | 1183.339644 | 1086.908385 | 52 | 9 | 12 | 20 | 42 | 10 | 0 | 0 | 0 | 0 | 0 | 1 |
| ISG000000 | Traf6 | 0.417009 | 0.486325 | -0.125665363 | 1054.916117 | 966.9163052 | 57 | 9 | 12 | 18 | 35 | 5 | 11 | 0 | 0 | 1 | 3 | 1 |
| SG0000002 | Rangap1 | 0.445063 | 0.514354 | -0.09662628 | 7455.554958 | 6972.564933 | 93 | 9 | 12 | 21 | 57 | 9 | 2 | 0 | 0 | 0 | 0 | 0 |
| SG000000C | Uhrf1 | 0.452457 | 0.521788 | -0.098112282 | 9297.853669 | 8686.562864 | 64 | 8 | 12 | 12 | 24 | 0 | 8 | 0 | 0 | 0 | 8 | 0 |
| SG0000003 | Sugp2 | 0.465678 | 0.535036 | -0.101780323 | 1878.858423 | 1750.874937 | 56 | 12 | 12 | 12 | 37 | 0 | 1 | 0 | 0 | 0 | 0 | 0 |
| SG0000003 | Med21 | 0.480702 | 0.549374 | 0.097178403 | 1039.693385 | 1112.13855 | 44 | 12 | 12 | 18 | 30 | 4 | 1 | 0 | 0 | 0 | 2 | 0 |
| SG0000002 | Bag1 | 0.490639 | 0.558485 | 0.089065621 | 2464.337963 | 2621.2696 | 78 | 9 | 12 | 18 | 30 | 3 | 0 | 0 | 0 | 0 | 0 | 0 |
| SG0000002 | Ube2b | 0.496883 | 0.564307 | -0.102642751 | 2619.282208 | 2439.40408 | 77 | 10 | 12 | 20 | 42 | 6 | 3 | 0 | 0 | 0 | 0 | 0 |
| SG0000005 | Rexo4 | 0.559735 | 0.624788 | -0.084542159 | 1367.169756 | 1289.355617 | 26 | 12 | 12 | 12 | 18 | 0 | 0 | 0 | 0 | 0 | 0 | 0 |

| | | | | | | | | | | | | | | | | | | |
|---|---|---|---|---|---|---|---|---|---|---|---|---|---|---|---|---|---|---|
| SG0000002 | Bnip3l | 0.578823 | 0.642792 | 0.061598325 | 4735.399849 | 4941.964251 | 78 | 6 | 12 | 10 | 24 | 2 | 8 | 0 | 0 | 0 | 0 | 0 |
| SG000000C | Kctd20 | 0.58321 | 0.646879 | 0.065448934 | 2835.375227 | 2966.966432 | 86 | 6 | 12 | 12 | 30 | 6 | 0 | 0 | 0 | 0 | 0 | 0 |
| JSG000000 | Zfp787 | 0.593818 | 0.656775 | 0.177964385 | 177.2833178 | 200.5582303 | 38 | 10 | 12 | 11 | 15 | 1 | 0 | 0 | 0 | 0 | 0 | 0 |
| SG0000004 | Zfp260 | 0.626841 | 0.688031 | 0.061688087 | 1435.125151 | 1497.820396 | 58 | 10 | 12 | 12 | 56 | 4 | 6 | 0 | 0 | 0 | 4 | 4 |
| SG000000C | Rbm14 | 0.681429 | 0.737909 | -0.053997764 | 2375.787125 | 2288.508752 | 66 | 6 | 12 | 24 | 39 | 6 | 1 | 0 | 0 | 0 | 0 | 0 |
| SG0000002 | Tmem216 | 0.6918 | 0.746657 | -0.093035244 | 207.5523916 | 194.5903696 | 69 | 10 | 12 | 11 | 48 | 19 | 2 | 0 | 0 | 0 | 0 | 0 |
| SG0000007 | Pak6 | 0.720355 | 0.772588 | 0.120490097 | 66.81909432 | 72.63931133 | 43 | 12 | 12 | 15 | 30 | 3 | 0 | 0 | 0 | 0 | 0 | 0 |
| JSG000000 | Snrnp70 | 0.755164 | 0.802617 | 0.042827634 | 7806.333172 | 8041.544796 | 40 | 12 | 12 | 14 | 21 | 1 | 1 | 0 | 0 | 0 | 0 | 0 |
| SG0000003 | Vps33b | 0.804043 | 0.84459 | 0.040191637 | 523.3753669 | 538.1609317 | 84 | 12 | 12 | 15 | 33 | 24 | 2 | 0 | 0 | 0 | 3 | 6 |
| SG0000003 | 43527 | 0.823027 | 0.861051 | -0.105392738 | 47.72294313 | 44.36095059 | 278 | 6 | 12 | 32 | 132 | 25 | 1 | 0 | 0 | 0 | 0 | 0 |
| JSG000000 | Mfhas1 | 0.88332 | 0.910061 | 0.04094488 | 188.1537681 | 193.5702261 | 138 | 8 | 12 | 23 | 81 | 10 | 2 | 0 | 0 | 0 | 1 | 0 |
| SG0000002 | Fam21 | 0.893094 | 0.918036 | -0.023147749 | 2975.063128 | 2927.709747 | 97 | 10 | 12 | 16 | 46 | 7 | 0 | 0 | 0 | 0 | 0 | 0 |
| SG0000002 | Epc1 | 0.909517 | 0.932315 | 0.025285936 | 460.6847668 | 468.8303057 | 226 | 6 | 12 | 28 | 79 | 18 | 2 | 0 | 0 | 2 | 2 | 2 |
| SG0000003 | Gpbp1l1 | 0.932038 | 0.950396 | 0.006281977 | 2770.807411 | 2782.898738 | 186 | 6 | 12 | 36 | 82 | 16 | 2 | 0 | 0 | 0 | 2 | 0 |
| SG0000002 | Sirt4 | 0.943624 | 0.960025 | -0.019279856 | 197.0830583 | 194.4668045 | 26 | 10 | 12 | 14 | 32 | 0 | 2 | 0 | 0 | 0 | 0 | 0 |
| JSG000000 | Ckap2 | 0.950933 | 0.966582 | 0.010515388 | 2976.148169 | 2997.919701 | 61 | 11 | 12 | 15 | 31 | 2 | 4 | 0 | 0 | 0 | 0 | 0 |
| SG0000002 | Zbtb11 | 0.986388 | 0.99685 | -0.010038193 | 1084.754639 | 1077.233173 | 82 | 7 | 12 | 18 | 45 | 5 | 8 | 0 | 0 | 2 | 4 | 1 |
| SG0000007 | Ndufb6 | 1 | 1 | 0.003383169 | 1281.112213 | 1284.11999 | 52 | 10 | 12 | 14 | 30 | 2 | 2 | 0 | 0 | 0 | 0 | 0 |
| JSG000000 | Mreg | 0 | 0 | 6.248372808 | 59.0953996 | 4492.63683 | 88 | 7 | 11 | 18 | 56 | 5 | 6 | 0 | 0 | 0 | 2 | 0 |
| SG0000003 | Raver2 | 7.3E-148 | 1.1E-145 | 6.693078091 | 6.480439925 | 670.5354724 | 235 | 5 | 11 | 32 | 90 | 30 | 10 | 0 | 0 | 0 | 3 | 1 |
| SG000000C | Scpep1 | 1.08E-94 | 6.78E-93 | 2.510621942 | 1139.686128 | 6494.680251 | 61 | 9 | 11 | 13 | 36 | 3 | 3 | 0 | 0 | 0 | 3 | 0 |
| JSG000000 | Adcy1 | 1.48E-94 | 9.23E-93 | 6.003717524 | 6.186148326 | 396.9349949 | 120 | 5 | 11 | 15 | 59 | 17 | 10 | 2 | 3 | 2 | 6 | 2 |
| JSG000000 | Nufip1 | 2.03E-65 | 6.42E-64 | 2.202883414 | 657.7212074 | 3028.139163 | 53 | 10 | 11 | 16 | 29 | 2 | 12 | 8 | 8 | 8 | 8 | 0 |
| JSG000000 | Stx1b | 7.72E-61 | 2.2E-59 | 5.882915151 | 3.915467995 | 231.0561862 | 32 | 10 | 11 | 12 | 17 | 4 | 0 | 0 | 0 | 0 | 0 | 1 |
| JSG000000 | Igf2bp1 | 7.26E-59 | 1.94E-57 | 2.397422522 | 1871.346796 | 9859.397336 | 70 | 9 | 11 | 16 | 45 | 0 | 10 | 0 | 0 | 0 | 4 | 0 |
| JSG000000 | Impa2 | 1.06E-58 | 2.81E-57 | 2.578356545 | 233.2566255 | 1393.14605 | 45 | 10 | 11 | 11 | 29 | 5 | 1 | 0 | 0 | 0 | 0 | 0 |
| SG0000002 | Fstl1 | 1.39E-58 | 3.67E-57 | -2.712437419 | 27202.78693 | 4150.38491 | 80 | 10 | 11 | 14 | 38 | 6 | 3 | 0 | 0 | 0 | 1 | 0 |
| JSG000000 | Shcbp1 | 5.12E-54 | 1.19E-52 | 2.047357419 | 580.1753038 | 2398.143866 | 93 | 2 | 11 | 9 | 51 | 8 | 0 | 0 | 0 | 0 | 1 | 0 |
| SG0000002 | Apbb1ip | 5.39E-52 | 1.18E-50 | -3.538180173 | 896.8185154 | 77.19802749 | 193 | 5 | 11 | 19 | 103 | 19 | 0 | 0 | 0 | 0 | 0 | 0 |
| SG0000005 | Sh2d3c | 1.3E-50 | 2.73E-49 | 4.304264247 | 13.35153861 | 263.7812753 | 57 | 7 | 11 | 9 | 29 | 6 | 1 | 0 | 0 | 0 | 0 | 2 |
| SG0000002 | Serpinb8 | 4.83E-50 | 1.01E-48 | -5.500416885 | 2014.188215 | 44.49483283 | 69 | 8 | 11 | 14 | 35 | 13 | 8 | 0 | 1 | 2 | 9 | 5 |
| SG0000003 | Gcnt1 | 2.79E-46 | 5.12E-45 | 3.878383714 | 18.55844498 | 272.9300887 | 92 | 9 | 11 | 26 | 63 | 5 | 12 | 0 | 2 | 2 | 8 | 2 |
| JSG000000 | Usp28 | 3.48E-45 | 6.14E-44 | 2.961933937 | 590.3079463 | 4599.489055 | 92 | 10 | 11 | 17 | 42 | 12 | 1 | 0 | 0 | 0 | 0 | 0 |
| JSG000000 | Erbb3 | 3.19E-43 | 5.24E-42 | -2.307806122 | 5563.122622 | 1123.569046 | 25 | 9 | 11 | 10 | 22 | 1 | 4 | 1 | 1 | 1 | 2 | 0 |
| SG0000005 | Gm13247 | 2.43E-42 | 3.88E-41 | 6.150126715 | 4.870618164 | 345.9048906 | 74 | 11 | 11 | 20 | 57 | 19 | 5 | 0 | 0 | 0 | 3 | 1 |
| JSG000000 | Ppp1r21 | 4.2E-42 | 6.65E-41 | -2.376324014 | 3082.984406 | 593.7813056 | 96 | 3 | 11 | 14 | 43 | 2 | 1 | 0 | 0 | 0 | 0 | 0 |
| SG0000006 | Arhgap22 | 2.17E-40 | 3.25E-39 | 3.371092843 | 27.21554164 | 281.5898821 | 175 | 5 | 11 | 29 | 74 | 17 | 0 | 0 | 0 | 0 | 0 | 0 |
| SG0000002 | Tdrd1 | 4.62E-38 | 6.38E-37 | 6.554166052 | 1.313186595 | 123.4033698 | 318 | 7 | 11 | 27 | 65 | 12 | 4 | 0 | 0 | 0 | 0 | 0 |
| SG0000004 | Arl4a | 2.38E-36 | 3.09E-35 | -2.381383957 | 3765.884063 | 722.768045 | 139 | 4 | 11 | 26 | 60 | 7 | 11 | 0 | 1 | 2 | 3 | 1 |
| JSG000000 | Nme4 | 7.5E-34 | 8.88E-33 | 1.92085612 | 230.6210747 | 873.2413679 | 11 | 10 | 11 | 11 | 13 | 0 | 0 | 0 | 0 | 0 | 0 | 0 |
| SG0000002 | Prrg4 | 1.46E-33 | 1.71E-32 | -2.849491803 | 2523.836695 | 350.1698461 | 63 | 10 | 11 | 15 | 34 | 4 | 4 | 0 | 0 | 0 | 0 | 0 |
| SG0000006 | Nlrp9b | 4.11E-33 | 4.72E-32 | 6.538664671 | 0 | 91.96815186 | 108 | 6 | 11 | 20 | 77 | 13 | 1 | 0 | 0 | 0 | 0 | 0 |
| JSG000000 | Lurap1l | 1.9E-32 | 2.12E-31 | -2.767059758 | 572.2941905 | 84.07199445 | 103 | 11 | 11 | 19 | 55 | 6 | 9 | 1 | 1 | 3 | 6 | 0 |
| SG0000005 | Nfic | 5.94E-31 | 6.25E-30 | -2.546390512 | 4370.646171 | 748.1793625 | 61 | 10 | 11 | 19 | 33 | 1 | 4 | 0 | 0 | 0 | 0 | 1 |
| SG0000002 | Sppl2a | 1.72E-29 | 1.7E-28 | -1.885897728 | 2615.802979 | 707.771784 | 85 | 8 | 11 | 18 | 48 | 6 | 9 | 0 | 0 | 1 | 3 | 1 |
| SG0000005 | Ptp4a3 | 2.5E-29 | 2.45E-28 | -2.554544284 | 4899.066698 | 833.909454 | 70 | 11 | 11 | 14 | 44 | 6 | 20 | 5 | 5 | 5 | 14 | 0 |
| JSG000000 | AU018091 | 9.01E-28 | 8.23E-27 | 4.590659469 | 106.681273 | 2570.480949 | 30 | 11 | 11 | 14 | 27 | 1 | 6 | 0 | 0 | 1 | 5 | 0 |
| JSG000000 | Parp12 | 1.66E-27 | 1.49E-26 | -2.105325498 | 7154.12371 | 1662.60974 | 60 | 7 | 11 | 10 | 36 | 8 | 1 | 0 | 0 | 0 | 1 | 0 |
| SG0000002 | Hspa4l | 2.85E-27 | 2.55E-26 | -2.397683302 | 1680.092808 | 318.8296327 | 120 | 5 | 11 | 13 | 57 | 9 | 0 | 0 | 0 | 0 | 0 | 0 |
| SG0000003 | Ciapin1 | 1.76E-24 | 1.39E-23 | 1.300444077 | 1484.048077 | 3655.279959 | 56 | 11 | 11 | 26 | 49 | 5 | 2 | 0 | 0 | 1 | 1 | 0 |
| SG0000002 | Tm9sf3 | 3.83E-24 | 2.96E-23 | -1.512564277 | 9439.203533 | 3308.324741 | 135 | 6 | 11 | 22 | 61 | 13 | 13 | 0 | 1 | 1 | 3 | 2 |
| SG0000002 | Stambpl1 | 9.97E-23 | 7.24E-22 | -2.214805232 | 592.4814593 | 127.6297607 | 156 | 9 | 11 | 32 | 64 | 18 | 4 | 0 | 0 | 4 | 6 | 0 |
| JSG000000 | Mthfd2l | 8.28E-22 | 5.76E-21 | -2.525084529 | 328.8403725 | 57.12930577 | 237 | 9 | 11 | 39 | 118 | 16 | 10 | 1 | 1 | 5 | 6 | 0 |
| JSG000000 | Daam1 | 2.29E-21 | 1.55E-20 | -1.642894215 | 2682.934463 | 859.1116844 | 214 | 6 | 11 | 18 | 89 | 24 | 6 | 1 | 1 | 3 | 5 | 2 |
| JSG000000 | Zfp1 | 4.63E-21 | 3.08E-20 | 1.550783977 | 255.2023597 | 747.6824375 | 44 | 10 | 11 | 14 | 23 | 3 | 0 | 0 | 0 | 0 | 1 | 0 |
| JSG000000 | Myo5c | 8.14E-21 | 5.36E-20 | 7.229304524 | 4.260286742 | 639.2582481 | 84 | 6 | 11 | 18 | 53 | 5 | 5 | 0 | 0 | 1 | 3 | 0 |
| SG0000001 | Pgs1 | 1.17E-20 | 7.65E-20 | -1.678585767 | 4179.882871 | 1305.74823 | 87 | 11 | 11 | 15 | 36 | 9 | 0 | 0 | 0 | 0 | 0 | 3 |
| JSG000000 | Akap8l | 4.9E-20 | 3.08E-19 | 1.325716767 | 487.0290735 | 1220.774363 | 50 | 6 | 11 | 10 | 25 | 3 | 0 | 0 | 0 | 0 | 0 | 0 |
| JSG000000 | Catsperd | 1.86E-19 | 1.13E-18 | 3.775995948 | 6.849350463 | 93.82907446 | 33 | 10 | 11 | 14 | 22 | 4 | 0 | 0 | 0 | 0 | 0 | 0 |
| SG0000001 | Acsl1 | 2.56E-19 | 1.54E-18 | 1.239328446 | 698.9149235 | 1650.058534 | 143 | 4 | 11 | 23 | 51 | 23 | 3 | 0 | 0 | 0 | 1 | 0 |
| JSG000000 | Seh1l | 3.99E-19 | 2.38E-18 | 1.328045876 | 2373.924995 | 5960.032632 | 53 | 10 | 11 | 17 | 39 | 5 | 3 | 0 | 0 | 0 | 5 | 1 |
| JSG000000 | Sarnp | 8.12E-19 | 4.77E-18 | 1.14928486 | 1253.378517 | 2780.042677 | 141 | 7 | 11 | 24 | 56 | 13 | 0 | 0 | 0 | 0 | 1 | 0 |
| JSG000000 | Clic6 | 1.43E-18 | 8.27E-18 | 6.45096687 | 0.639110347 | 55.91268876 | 61 | 10 | 11 | 14 | 39 | 3 | 2 | 0 | 0 | 0 | 0 | 0 |
| JSG000000 | Mrps27 | 1.52E-18 | 8.81E-18 | 1.214287636 | 770.731929 | 1788.299901 | 104 | 5 | 11 | 18 | 40 | 11 | 0 | 0 | 0 | 0 | 0 | 0 |
| SG0000005 | Aph1c | 1.53E-18 | 8.85E-18 | 4.470168899 | 3.298105871 | 73.1003602 | 83 | 11 | 11 | 19 | 36 | 8 | 7 | 1 | 1 | 1 | 2 | 2 |
| JSG000000 | Kif23 | 1.03E-17 | 5.68E-17 | 1.081672047 | 1797.21861 | 3803.790931 | 67 | 9 | 11 | 15 | 24 | 3 | 3 | 0 | 0 | 0 | 0 | 1 |
| SG0000001 | Rab4a | 1.89E-17 | 1.03E-16 | 1.429621316 | 249.7694496 | 672.8189506 | 65 | 10 | 11 | 16 | 42 | 4 | 2 | 0 | 1 | 0 | 0 | 0 |
| JSG000000 | Atp6v1c1 | 2.03E-17 | 1.11E-16 | -1.266441654 | 4785.327382 | 1989.182955 | 66 | 4 | 11 | 18 | 34 | 3 | 4 | 0 | 2 | 2 | 2 | 0 |
| JSG000000 | Tshz3 | 2.13E-17 | 1.16E-16 | -2.352578901 | 253.952424 | 49.72279652 | 124 | 6 | 11 | 16 | 57 | 6 | 9 | 0 | 1 | 3 | 5 | 1 |
| JSG000000 | B3galnt2 | 2.68E-17 | 1.45E-16 | -1.343399897 | 2302.933981 | 907.5651753 | 108 | 6 | 11 | 17 | 43 | 10 | 3 | 0 | 0 | 0 | 1 | 0 |
| SG0000001 | Etv5 | 5.02E-17 | 2.68E-16 | -1.331815003 | 8321.570421 | 3305.89479 | 97 | 8 | 11 | 12 | 41 | 10 | 5 | 0 | 0 | 1 | 3 | 1 |
| SG0000003 | Tdrd12 | 1.12E-16 | 5.85E-16 | 9.958885012 | 1.265003014 | 1258.967945 | 78 | 8 | 11 | 15 | 46 | 7 | 0 | 0 | 0 | 0 | 0 | 0 |
| SG0000006 | Fgfr1op | 2.02E-16 | 1.04E-15 | 1.100752096 | 936.1002692 | 2007.62118 | 124 | 2 | 11 | 13 | 66 | 6 | 4 | 0 | 0 | 0 | 4 | 2 |
| SG0000003 | Rasa3 | 1.06E-15 | 5.25E-15 | -1.335686804 | 1480.746482 | 586.6768027 | 182 | 6 | 11 | 20 | 77 | 30 | 0 | 0 | 0 | 0 | 0 | 0 |
| SG0000002 | Zfp106 | 3.5E-15 | 1.67E-14 | 0.935190937 | 8694.560613 | 16625.24918 | 114 | 7 | 11 | 21 | 58 | 14 | 4 | 0 | 0 | 0 | 0 | 0 |
| SG0000005 | Zfp420 | 6.27E-15 | 2.95E-14 | 1.693744924 | 87.67699855 | 283.6310034 | 70 | 11 | 11 | 19 | 39 | 2 | 6 | 1 | 1 | 1 | 3 | 0 |
| JSG000000 | Kcnb1 | 8.78E-15 | 4.09E-14 | 3.840122719 | 4.624510145 | 66.23047379 | 81 | 9 | 11 | 14 | 34 | 4 | 0 | 0 | 0 | 0 | 0 | 0 |
| JSG000000 | Prune | 5.32E-14 | 2.36E-13 | 0.997872955 | 1037.905621 | 2072.753014 | 53 | 7 | 11 | 11 | 28 | 5 | 3 | 0 | 2 | 0 | 1 | 0 |
| SG0000002 | Prkag2 | 1.84E-13 | 7.88E-13 | -1.275022907 | 1080.525287 | 446.4931442 | 336 | 9 | 11 | 38 | 128 | 55 | 3 | 3 | 3 | 3 | 3 | 3 |
| JSG000000 | Heatr1 | 2.08E-13 | 8.88E-13 | 1.098212424 | 2404.561686 | 5147.908321 | 87 | 10 | 11 | 19 | 39 | 6 | 0 | 0 | 0 | 0 | 0 | 0 |
| SG0000002 | Zc3h13 | 4.38E-13 | 1.82E-12 | 1.092455582 | 1323.901153 | 2823.042779 | 164 | 5 | 11 | 20 | 65 | 14 | 2 | 0 | 0 | 0 | 1 | 0 |
| SG0000002 | Sh3bp5 | 5.26E-13 | 2.18E-12 | -1.336279789 | 764.0936426 | 302.6120984 | 122 | 8 | 11 | 24 | 64 | 6 | 12 | 0 | 0 | 2 | 4 | 0 |
| SG0000004 | Samd14 | 1.92E-12 | 7.65E-12 | -2.299007244 | 196.5898018 | 39.94761395 | 20 | 11 | 11 | 11 | 14 | 1 | 0 | 0 | 0 | 0 | 0 | 0 |
| SG0000003 | Pcgf3 | 6.93E-12 | 2.68E-11 | -1.013774139 | 4186.922126 | 2073.568843 | 136 | 4 | 11 | 18 | 54 | 19 | 17 | 1 | 1 | 3 | 7 | 0 |
| JSG000000 | Cnnm1 | 1.34E-11 | 5.08E-11 | 5.251267787 | 0.981585527 | 37.38671138 | 65 | 9 | 11 | 15 | 35 | 5 | 1 | 0 | 0 | 1 | 1 | 1 |
| SG0000004 | Chchd7 | 2.89E-11 | 1.07E-10 | 1.455997188 | 95.31854065 | 261.5027208 | 99 | 1 | 11 | 3 | 51 | 30 | 2 | 0 | 0 | 0 | 1 | 1 |
| SG0000004 | Tm7sf3 | 2.97E-11 | 1.1E-10 | 1.230851083 | 1446.16043 | 3394.216733 | 89 | 6 | 11 | 15 | 71 | 2 | 6 | 0 | 1 | 2 | 2 | 0 |
| SG000000C | Ufd1l | 4.32E-11 | 1.58E-10 | -0.995446388 | 2400.876044 | 1204.232979 | 189 | 2 | 11 | 16 | 70 | 15 | 15 | 2 | 2 | 5 | 11 | 0 |
| SG0000002 | Cenpf | 1.07E-10 | 3.81E-10 | 0.866836751 | 4942.93134 | 9014.225981 | 95 | 7 | 11 | 19 | 48 | 16 | 1 | 0 | 0 | 1 | 1 | 1 |
| SG0000003 | Tdrd7 | 1.93E-10 | 6.71E-10 | -1.148169894 | 802.4126598 | 362.0463669 | 188 | 4 | 11 | 25 | 92 | 24 | 2 | 0 | 0 | 0 | 0 | 0 |
| JSG000000 | Ltn1 | 5.48E-10 | 1.84E-09 | -0.906870694 | 3723.669198 | 1985.984302 | 111 | 7 | 11 | 16 | 56 | 11 | 8 | 1 | 2 | 2 | 5 | 3 |
| JSG000000 | Nfe2l2 | 8.26E-10 | 2.74E-09 | -0.881370052 | 4725.742671 | 2565.377293 | 57 | 10 | 11 | 13 | 26 | 2 | 1 | 0 | 1 | 1 | 1 | 1 |
| SG0000002 | Csnk1d | 9.09E-10 | 3E-09 | -0.859877173 | 5918.28937 | 3260.974285 | 99 | 8 | 11 | 20 | 42 | 16 | 0 | 0 | 0 | 0 | 0 | 3 |
| SG0000003 | Arhgap27 | 1.18E-09 | 3.86E-09 | -1.596954649 | 619.9310686 | 204.9331195 | 42 | 11 | 11 | 19 | 25 | 2 | 6 | 0 | 0 | 4 | 4 | 0 |
| SG0000002 | Col15a1 | 1.67E-09 | 5.39E-09 | -3.099673704 | 86.01680278 | 10.03433357 | 213 | 0 | 11 | 19 | 70 | 29 | 0 | 0 | 0 | 0 | 0 | 4 |
| SG0000003 | Cops4 | 2.19E-09 | 7.02E-09 | -0.851442135 | 5344.546032 | 2962.110119 | 95 | 3 | 11 | 21 | 61 | 16 | 1 | 0 | 0 | 1 | 1 | 0 |
| JSG000000 | Zfp799 | 5.4E-09 | 1.68E-08 | 0.913597921 | 386.6456369 | 728.3388765 | 92 | 2 | 11 | 12 | 39 | 17 | 16 | 2 | 5 | 4 | 11 | 2 |
| JSG000000 | Polr3h | 8.73E-09 | 2.67E-08 | -0.973997002 | 939.8097507 | 478.4511837 | 22 | 11 | 11 | 13 | 16 | 2 | 3 | 0 | 0 | 0 | 0 | 0 |
| JSG000000 | Ugcg | 9.72E-09 | 2.96E-08 | -0.838205866 | 3291.393606 | 1841.003919 | 71 | 6 | 11 | 19 | 45 | 5 | 6 | 0 | 2 | 1 | 4 | 2 |
| JSG000000 | Lmbrd2 | 1.39E-08 | 4.19E-08 | -1.172116174 | 981.6847312 | 435.6421666 | 177 | 2 | 11 | 25 | 87 | 12 | 6 | 0 | 0 | 0 | 2 | 0 |
| JSG000000 | Tbc1d15 | 1.62E-08 | 4.85E-08 | 0.715050195 | 2279.213601 | 3741.421046 | 110 | 6 | 11 | 17 | 51 | 10 | 9 | 0 | 1 | 2 | 4 | 2 |
| SG0000002 | Nprl3 | 1.77E-08 | 5.28E-08 | -0.939991825 | 990.8866701 | 516.4856397 | 255 | 5 | 11 | 38 | 77 | 49 | 14 | 0 | 0 | 11 | 14 | 0 |
| JSG000000 | Pi4k2a | 1.87E-08 | 5.58E-08 | -0.831690857 | 2438.746935 | 1370.259563 | 59 | 9 | 11 | 14 | 36 | 3 | 3 | 0 | 0 | 0 | 0 | 0 |
| SG0000006 | Rai1 | 2.94E-08 | 8.63E-08 | -1.22713426 | 1570.197666 | 670.7336679 | 162 | 11 | 11 | 26 | 65 | 16 | 0 | 0 | 0 | 0 | 3 | 9 |
| JSG000000 | Mtmr6 | 4.18E-08 | 1.21E-07 | -0.83177505 | 2012.575721 | 1130.740609 | 94 | 6 | 11 | 15 | 33 | 11 | 5 | 0 | 1 | 0 | 1 | 1 |
| SG000000C | Rab11a | 5.97E-08 | 1.71E-07 | -0.788943381 | 3292.810197 | 1905.772586 | 176 | 11 | 11 | 33 | 58 | 13 | 6 | 0 | 0 | 1 | 3 | 0 |
| JSG000000 | Dnajc17 | 7.89E-08 | 2.23E-07 | 1.177325825 | 109.9512996 | 248.662763 | 51 | 8 | 11 | 16 | 35 | 5 | 0 | 0 | 0 | 0 | 0 | 0 |
| JSG000000 | Fam163b | 2.67E-07 | 7.2E-07 | 4.240234432 | 0 | 17.89895334 | 28 | 11 | 11 | 13 | 16 | 1 | 0 | 0 | 0 | 0 | 0 | 0 |
| JSG000000 | Taf8 | 3.35E-07 | 8.96E-07 | 0.72189652 | 816.7550142 | 1347.113895 | 30 | 11 | 11 | 13 | 20 | 4 | 2 | 0 | 0 | 1 | 1 | 0 |
| SG0000004 | B3gntl1 | 3.7E-07 | 9.88E-07 | 0.981464005 | 175.761717 | 347.0358902 | 149 | 8 | 11 | 20 | 50 | 17 | 2 | 0 | 0 | 0 | 0 | 0 |
| SG0000002 | Dennd2d | 4.05E-07 | 1.08E-06 | 2.554747485 | 8.042078106 | 47.25240427 | 31 | 9 | 11 | 16 | 23 | 2 | 4 | 0 | 1 | 2 | 2 | 0 |
| JSG000000 | Slc39a10 | 1.41E-06 | 3.6E-06 | 0.788694246 | 2505.118853 | 4327.618422 | 105 | 10 | 11 | 16 | 48 | 5 | 6 | 0 | 0 | 0 | 2 | 2 |
| SG0000002 | Hspg2 | 2.11E-06 | 5.29E-06 | -0.951961077 | 7979.312555 | 4124.740568 | 56 | 11 | 11 | 17 | 41 | 7 | 0 | 0 | 0 | 0 | 4 | 0 |
| SG0000002 | Nus1 | 2.62E-06 | 6.52E-06 | -0.663119014 | 4053.723002 | 2559.972826 | 68 | 11 | 11 | 14 | 31 | 7 | 13 | 0 | 0 | 0 | 1 | 2 |
| JSG000000 | Hpcal1 | 3.22E-06 | 7.95E-06 | -0.884452725 | 1514.432842 | 820.357625 | 88 | 10 | 11 | 19 | 40 | 11 | 3 | 0 | 0 | 1 | 1 | 1 |
| SG0000002 | Peli1 | 5.51E-06 | 1.33E-05 | -0.717208035 | 1361.814818 | 828.355812 | 203 | 8 | 11 | 26 | 87 | 14 | 10 | 0 | 0 | 4 | 10 | 0 |
| SG0000002 | Arhgap1 | 6.09E-06 | 1.47E-05 | -0.651258003 | 2578.976967 | 1642.098746 | 89 | 11 | 11 | 19 | 35 | 0 | 12 | 0 | 0 | 0 | 0 | 0 |
| JSG000000 | Ikzf3 | 6.38E-06 | 1.53E-05 | 4.390048151 | 0.946619625 | 19.84776821 | 132 | 7 | 11 | 23 | 74 | 10 | 2 | 0 | 0 | 0 | 1 | 1 |
| SG0000002 | Gpr89 | 7.47E-06 | 1.78E-05 | -0.747476301 | 898.549871 | 535.2163829 | 112 | 6 | 11 | 15 | 51 | 10 | 1 | 0 | 0 | 0 | 0 | 1 |
| SG0000003 | Slc22a15 | 9.42E-06 | 2.23E-05 | -1.800431603 | 76.46496035 | 21.95222524 | 266 | 3 | 11 | 22 | 104 | 11 | 11 | 0 | 2 | 1 | 3 | 2 |
| JSG000000 | Eif3k | 1.44E-05 | 3.35E-05 | 0.541860868 | 4130.986178 | 6014.092681 | 21 | 10 | 11 | 11 | 18 | 2 | 0 | 0 | 0 | 0 | 0 | 0 |
| JSG000000 | Fam204a | 1.56E-05 | 3.62E-05 | -0.777079376 | 641.4155379 | 374.2960769 | 88 | 7 | 11 | 15 | 38 | 6 | 6 | 1 | 2 | 3 | 3 | 0 |
| JSG000000 | Eif3j1 | 1.89E-05 | 4.33E-05 | 0.657599591 | 521.8312927 | 823.1658047 | 61 | 10 | 11 | 14 | 49 | 9 | 6 | 0 | 0 | 2 | 4 | 1 |

| | | | | | | | | | | | | | | | | | | |
|---|---|---|---|---|---|---|---|---|---|---|---|---|---|---|---|---|---|---|
| SG0000002 | Lhx6 | 2.19E-05 | 4.98E-05 | 6.73851823 | 0.663202137 | 70.81773467 | 109 | 0 | 11 | 12 | 38 | 6 | 20 | 0 | 5 | 5 | 5 | 0 |
| JSG000000 | Heca | 2.66E-05 | 6.02E-05 | -0.667861778 | 1152.003373 | 725.1157232 | 110 | 4 | 11 | 15 | 71 | 11 | 5 | 0 | 0 | 1 | 3 | 1 |
| SG0000002 | Spin1 | 2.88E-05 | 6.49E-05 | 0.550159475 | 1704.625603 | 2495.993854 | 101 | 6 | 11 | 16 | 42 | 6 | 9 | 0 | 0 | 1 | 4 | 1 |
| SG0000004 | Gcn1l1 | 4.51E-05 | 9.94E-05 | -0.741063557 | 4248.219994 | 2541.702051 | 56 | 8 | 11 | 11 | 17 | 6 | 0 | 0 | 0 | 0 | 0 | 0 |
| SG0000003 | Zcwpw1 | 4.52E-05 | 9.98E-05 | 0.857643195 | 237.8665713 | 431.0322244 | 68 | 9 | 11 | 16 | 36 | 5 | 0 | 0 | 0 | 0 | 0 | 0 |
| SG0000000 | Slc1a5 | 6.33E-05 | 0.000137 | -0.552592162 | 3937.015177 | 2684.233381 | 24 | 11 | 11 | 12 | 15 | 0 | 0 | 0 | 0 | 0 | 0 | 0 |
| SG0000003 | Chmp4b | 8.49E-05 | 0.000182 | -0.555163803 | 3799.101994 | 2585.592003 | 57 | 9 | 11 | 13 | 40 | 6 | 2 | 0 | 0 | 0 | 0 | 0 |
| JSG000000 | Clcn7 | 9.87E-05 | 0.000209 | -0.58811838 | 1397.461405 | 929.6052099 | 28 | 11 | 11 | 14 | 18 | 4 | 0 | 0 | 0 | 0 | 0 | 0 |
| SG0000001 | Gle1 | 0.000122 | 0.000256 | 0.530168951 | 1173.292629 | 1694.349893 | 45 | 9 | 11 | 14 | 24 | 0 | 2 | 0 | 0 | 0 | 0 | 0 |
| JSG000000 | Psma5 | 0.000123 | 0.000258 | 0.497842149 | 1984.859883 | 2802.820427 | 67 | 8 | 11 | 12 | 25 | 3 | 3 | 0 | 0 | 0 | 0 | 1 |
| SG0000002 | Zbtb43 | 0.000182 | 0.000374 | 0.52454742 | 972.9900715 | 1399.628948 | 70 | 9 | 11 | 14 | 39 | 0 | 15 | 0 | 0 | 2 | 11 | 0 |
| JSG000000 | Dnajc3 | 0.000184 | 0.000378 | -0.528612314 | 2859.194188 | 1982.054034 | 82 | 6 | 11 | 13 | 34 | 9 | 5 | 0 | 0 | 0 | 3 | 0 |
| JSG000000 | Papl | 0.000206 | 0.000422 | 2.161749141 | 7.143642062 | 31.96472852 | 36 | 9 | 11 | 11 | 38 | 2 | 0 | 0 | 0 | 0 | 0 | 0 |
| SG0000002 | Zfp711 | 0.000227 | 0.000461 | 1.711001915 | 13.51783758 | 44.25579259 | 226 | 8 | 11 | 32 | 123 | 10 | 20 | 0 | 0 | 2 | 8 | 0 |
| JSG000000 | Epg5 | 0.000473 | 0.000926 | -0.61492412 | 782.0506619 | 510.6511131 | 177 | 9 | 11 | 22 | 78 | 14 | 1 | 0 | 0 | 0 | 0 | 0 |
| SG0000002 | Tfg | 0.000499 | 0.000974 | -0.496308536 | 2821.720663 | 2000.369674 | 156 | 7 | 11 | 25 | 63 | 7 | 3 | 0 | 0 | 0 | 0 | 0 |
| SG0000001 | Casd1 | 0.001148 | 0.002141 | -0.648061961 | 487.6815546 | 311.207633 | 211 | 4 | 11 | 29 | 113 | 21 | 6 | 0 | 0 | 3 | 6 | 3 |
| SG0000002 | Vangl1 | 0.001426 | 0.002633 | -0.482856549 | 1277.151192 | 913.8775483 | 176 | 4 | 11 | 25 | 89 | 40 | 4 | 0 | 0 | 0 | 0 | 0 |
| SG0000002 | Usp20 | 0.001663 | 0.003047 | -0.604574848 | 427.6783447 | 281.2691474 | 62 | 8 | 11 | 13 | 65 | 0 | 0 | 0 | 0 | 0 | 2 | 0 |
| SG0000003 | Ntng2 | 0.002401 | 0.004298 | -2.617294763 | 19.02756609 | 3.100980305 | 116 | 11 | 11 | 16 | 27 | 9 | 0 | 0 | 0 | 0 | 0 | 0 |
| SG0000001 | Sp2 | 0.002659 | 0.004731 | 0.483561851 | 746.7011927 | 1044.031175 | 70 | 9 | 11 | 9 | 32 | 9 | 3 | 0 | 2 | 0 | 0 | 0 |
| SG0000003 | Rassf8 | 0.003016 | 0.00532 | -0.569859915 | 659.2573635 | 444.1299506 | 107 | 10 | 11 | 22 | 56 | 8 | 11 | 0 | 0 | 2 | 5 | 0 |
| SG0000006 | Nr2f1 | 0.00304 | 0.005356 | -2.222195887 | 23.04309152 | 4.938478035 | 67 | 3 | 11 | 5 | 14 | 6 | 10 | 0 | 0 | 0 | 0 | 0 |
| SG0000003 | Spg11 | 0.004341 | 0.007489 | 0.415343405 | 1121.788148 | 1496.034713 | 112 | 8 | 11 | 24 | 59 | 8 | 2 | 0 | 0 | 0 | 0 | 0 |
| SG0000003 | Ndrg4 | 0.005492 | 0.009341 | 0.520592332 | 686.9585021 | 985.4722749 | 48 | 6 | 11 | 6 | 22 | 6 | 9 | 0 | 5 | 0 | 0 | 4 |
| SG0000000 | Slc19a1 | 0.006641 | 0.011168 | -0.485819341 | 501.3205508 | 357.9886966 | 30 | 7 | 11 | 10 | 23 | 22 | 0 | 0 | 0 | 0 | 0 | 0 |
| SG0000003 | Cstf2 | 0.011162 | 0.018116 | 0.331073418 | 2368.4426 | 2979.379968 | 143 | 5 | 11 | 19 | 57 | 15 | 10 | 0 | 0 | 1 | 6 | 1 |
| SG0000004 | Asxl1 | 0.011376 | 0.018442 | 0.322829438 | 3056.138359 | 3822.560398 | 115 | 9 | 11 | 22 | 63 | 8 | 4 | 0 | 0 | 1 | 2 | 0 |
| SG0000002 | Ppm1d | 0.01468 | 0.023295 | 0.385970791 | 534.7384926 | 698.7635348 | 79 | 9 | 11 | 17 | 43 | 10 | 2 | 0 | 0 | 0 | 1 | 0 |
| SG0000003 | Wipf2 | 0.016931 | 0.026575 | -0.356841019 | 1494.422328 | 1166.953356 | 61 | 8 | 11 | 17 | 39 | 9 | 9 | 1 | 1 | 4 | 12 | 4 |
| JSG000000 | Dpysl2 | 0.017972 | 0.028035 | 0.528305286 | 618.6852118 | 892.2888647 | 88 | 7 | 11 | 19 | 35 | 10 | 3 | 0 | 0 | 0 | 1 | 0 |
| JSG000000 | Plekhg6 | 0.024295 | 0.037036 | 1.087777141 | 43.08302488 | 91.57137506 | 18 | 10 | 11 | 10 | 18 | 1 | 0 | 0 | 0 | 0 | 0 | 0 |
| JSG000000 | Ofd1 | 0.029739 | 0.044658 | 0.323855886 | 702.7282877 | 879.5849137 | 124 | 7 | 11 | 21 | 65 | 9 | 2 | 0 | 1 | 1 | 1 | 0 |
| SG0000003 | Cpxm2 | 0.039846 | 0.058497 | 1.45436384 | 8.472389903 | 23.21737278 | 150 | 5 | 11 | 22 | 66 | 21 | 2 | 0 | 0 | 0 | 0 | 0 |
| JSG000000 | Arl3 | 0.052159 | 0.074988 | 0.300738994 | 583.1697347 | 718.3340203 | 64 | 5 | 11 | 11 | 30 | 5 | 0 | 0 | 0 | 0 | 1 | 0 |
| SG0000005 | Fbxo31 | 0.052694 | 0.075733 | 0.446826494 | 553.2113321 | 754.0484389 | 29 | 9 | 11 | 10 | 22 | 4 | 0 | 0 | 0 | 0 | 0 | 0 |
| JSG000001 | Gm42878 | 0.052916 | 0.076027 | -1.476215812 | 17.9566441 | 6.454163094 | 44 | 11 | 11 | 14 | 24 | 6 | 2 | 0 | 0 | 0 | 0 | 0 |
| JSG000000 | Marf1 | 0.054691 | 0.078363 | -0.273405874 | 2980.437024 | 2465.906359 | 96 | 6 | 11 | 19 | 35 | 5 | 6 | 0 | 0 | 0 | 0 | 0 |
| JSG000000 | Copb1 | 0.058948 | 0.083913 | -0.260013921 | 4769.591862 | 3982.990112 | 92 | 5 | 11 | 16 | 41 | 4 | 2 | 0 | 0 | 0 | 1 | 0 |
| JSG000000 | Scaf11 | 0.079475 | 0.11043 | -0.236728863 | 6470.072303 | 5490.939333 | 113 | 3 | 11 | 14 | 50 | 10 | 3 | 0 | 1 | 2 | 2 | 0 |
| JSG000000 | Itpkc | 0.111011 | 0.149619 | 0.275477219 | 379.9873769 | 459.9345933 | 42 | 9 | 11 | 17 | 26 | 2 | 1 | 0 | 0 | 1 | 1 | 0 |
| SG0000002 | Gaa | 0.11168 | 0.150464 | -0.41909614 | 2599.626475 | 1944.242543 | 20 | 11 | 11 | 11 | 16 | 0 | 0 | 0 | 0 | 0 | 0 | 0 |
| SG0000002 | Noct | 0.130675 | 0.173472 | -0.288142447 | 1766.093714 | 1446.352748 | 70 | 7 | 11 | 18 | 39 | 3 | 12 | 3 | 3 | 6 | 9 | 0 |
| JSG000000 | Sync | 0.133585 | 0.17688 | -0.886437802 | 36.73241817 | 19.87033331 | 35 | 9 | 11 | 13 | 43 | 2 | 1 | 0 | 0 | 0 | 0 | 0 |
| JSG000000 | Atp6v1d | 0.149741 | 0.196236 | -0.201421625 | 2102.214077 | 1828.281182 | 49 | 7 | 11 | 14 | 30 | 4 | 4 | 0 | 0 | 1 | 2 | 0 |
| SG0000002 | Zcchc17 | 0.174199 | 0.225064 | -0.228228877 | 618.0228309 | 527.5951339 | 61 | 8 | 11 | 13 | 38 | 2 | 0 | 0 | 0 | 0 | 0 | 0 |
| SG0000003 | Acadsb | 0.17671 | 0.227976 | -0.217095544 | 984.8363271 | 847.2504131 | 169 | 6 | 11 | 17 | 79 | 10 | 11 | 0 | 0 | 0 | 1 | 0 |
| SG0000000 | Plekha8 | 0.182539 | 0.234905 | 0.185374098 | 875.4882501 | 995.5280147 | 154 | 9 | 11 | 26 | 83 | 8 | 12 | 2 | 2 | 2 | 6 | 0 |
| SG0000002 | Dnaaf5 | 0.203145 | 0.258451 | -0.183115948 | 1383.694041 | 1218.75575 | 80 | 7 | 11 | 17 | 34 | 6 | 2 | 1 | 1 | 1 | 2 | 0 |
| JSG000000 | Ubl7 | 0.256533 | 0.317774 | -0.169856625 | 1009.552356 | 897.4224046 | 39 | 11 | 11 | 12 | 23 | 7 | 0 | 0 | 0 | 0 | 0 | 0 |
| SG0000003 | Mettl25 | 0.278198 | 0.341354 | -0.294508402 | 153.7276432 | 125.3418516 | 188 | 8 | 11 | 24 | 93 | 21 | 2 | 0 | 0 | 0 | 0 | 1 |
| JSG000000 | Adam1a | 0.294899 | 0.359311 | -0.825067416 | 18.6151591 | 10.50743499 | 47 | 11 | 11 | 14 | 24 | 7 | 3 | 0 | 0 | 0 | 0 | 0 |
| JSG000000 | Naa40 | 0.301221 | 0.366043 | -0.143969343 | 2353.051235 | 2129.571828 | 20 | 10 | 11 | 11 | 14 | 1 | 0 | 0 | 0 | 0 | 0 | 0 |
| JSG000000 | Ank | 0.333439 | 0.400267 | 0.299449552 | 359.0667022 | 441.8943315 | 160 | 8 | 11 | 20 | 70 | 14 | 4 | 0 | 0 | 0 | 1 | 1 |
| SG0000003 | Mrpl3 | 0.401972 | 0.470935 | -0.132389869 | 1279.381073 | 1167.20355 | 163 | 7 | 11 | 14 | 61 | 20 | 15 | 2 | 3 | 3 | 7 | 0 |
| JSG000000 | Dus3l | 0.448822 | 0.51793 | -0.246868137 | 932.8921712 | 786.1704832 | 16 | 11 | 11 | 11 | 14 | 0 | 0 | 0 | 0 | 0 | 0 | 0 |
| SG0000002 | Rilpl1 | 0.45017 | 0.519351 | -0.159670387 | 402.5893921 | 360.4099327 | 46 | 8 | 11 | 12 | 29 | 1 | 2 | 0 | 0 | 1 | 1 | 0 |
| JSG000000 | Nckap1l | 0.456016 | 0.525554 | -0.373031919 | 44.97456758 | 34.72747438 | 103 | 9 | 11 | 22 | 43 | 5 | 1 | 0 | 0 | 0 | 0 | 0 |
| JSG000000 | Rad21 | 0.480398 | 0.549061 | 0.082510352 | 10761.37234 | 11394.77487 | 74 | 9 | 11 | 18 | 33 | 7 | 3 | 0 | 0 | 1 | 5 | 1 |
| SG0000004 | Wdr11 | 0.483128 | 0.551618 | -0.101259775 | 1621.749572 | 1511.825138 | 175 | 2 | 11 | 17 | 50 | 17 | 3 | 0 | 0 | 1 | 2 | 0 |
| JSG000000 | Hmg20a | 0.493754 | 0.561567 | 0.091543614 | 1114.66841 | 1187.689911 | 143 | 7 | 11 | 27 | 77 | 10 | 0 | 0 | 0 | 0 | 0 | 0 |
| JSG000000 | Pdk3 | 0.503746 | 0.570938 | 0.083383465 | 1390.915484 | 1473.67483 | 139 | 7 | 11 | 23 | 76 | 22 | 0 | 0 | 0 | 0 | 0 | 0 |
| SG0000002 | Mrpl13 | 0.520558 | 0.587613 | 0.081276405 | 1444.42849 | 1528.138374 | 135 | 5 | 11 | 13 | 56 | 12 | 2 | 0 | 0 | 1 | 1 | 0 |
| SG0000002 | Cand1 | 0.633791 | 0.694171 | -0.067518658 | 6747.119215 | 6438.626643 | 97 | 6 | 11 | 15 | 41 | 8 | 12 | 0 | 2 | 1 | 4 | 0 |
| JSG000000 | Aox2 | 0.641095 | 0.701086 | -0.336801537 | 19.12627682 | 15.14408616 | 128 | 8 | 11 | 18 | 77 | 12 | 1 | 0 | 0 | 0 | 0 | 0 |
| SG0000007 | Smarcd2 | 0.672495 | 0.729823 | 0.053544678 | 2203.60922 | 2286.931387 | 16 | 10 | 11 | 10 | 13 | 0 | 0 | 0 | 0 | 0 | 0 | 0 |
| SG0000002 | Tcf3 | 0.70493 | 0.758407 | 0.121474831 | 2575.693916 | 2801.959283 | 130 | 0 | 11 | 22 | 33 | 22 | 4 | 0 | 0 | 0 | 0 | 0 |
| JSG000000 | Copb2 | 0.766196 | 0.812365 | 0.034724699 | 4993.406278 | 5115.052303 | 70 | 10 | 11 | 13 | 32 | 5 | 1 | 0 | 0 | 0 | 1 | 0 |
| SG0000004 | Ccdc9 | 0.773534 | 0.818885 | -0.04535546 | 805.8651683 | 780.9244691 | 27 | 11 | 11 | 11 | 15 | 2 | 0 | 0 | 0 | 0 | 0 | 0 |
| SG0000003 | Uba5 | 0.793353 | 0.835619 | 0.038504537 | 785.1497369 | 806.3869855 | 77 | 6 | 11 | 13 | 54 | 12 | 2 | 0 | 0 | 1 | 1 | 0 |
| SG0000002 | Toe1 | 0.79652 | 0.838461 | -0.048568386 | 558.2367534 | 539.7565054 | 24 | 11 | 11 | 15 | 27 | 2 | 1 | 0 | 0 | 0 | 1 | 0 |
| JSG000000 | Ppp6c | 0.798836 | 0.84046 | 0.028493262 | 1833.506012 | 1870.077758 | 88 | 8 | 11 | 20 | 56 | 8 | 8 | 0 | 1 | 0 | 4 | 0 |
| JSG000000 | Xrn2 | 0.938262 | 0.955437 | -0.016555264 | 6263.024704 | 6191.565815 | 157 | 7 | 11 | 21 | 74 | 10 | 1 | 0 | 0 | 0 | 1 | 0 |
| SG0000006 | 2610305D13Rik | 0 | 0 | 6.684115939 | 41.28699651 | 4245.537514 | 128 | 8 | 10 | 22 | 82 | 22 | 12 | 0 | 0 | 4 | 6 | 4 |
| SG0000007 | Nup62cl | 2.1E-147 | 3E-145 | 8.339397327 | 4.622166578 | 1497.112147 | 154 | 2 | 10 | 21 | 98 | 5 | 6 | 0 | 0 | 0 | 0 | 0 |
| JSG000000 | Lama1 | 2.1E-132 | 2.6E-130 | 8.050386441 | 1.952296942 | 517.5516166 | 170 | 6 | 10 | 25 | 78 | 18 | 1 | 0 | 0 | 1 | 1 | 0 |
| SG0000004 | Sptssb | 9.6E-113 | 8.7E-111 | -10.35604127 | 1309.62684 | 0 | 124 | 8 | 10 | 24 | 92 | 6 | 4 | 0 | 0 | 0 | 2 | 0 |
| SG0000002 | Crmp1 | 3.2E-92 | 1.89E-90 | 6.422097615 | 4.257943175 | 365.1269407 | 102 | 6 | 10 | 12 | 61 | 16 | 0 | 0 | 0 | 0 | 0 | 0 |
| SG0000002 | Fgd1 | 2.95E-81 | 1.37E-79 | 2.447958793 | 622.332067 | 3395.714776 | 90 | 4 | 10 | 8 | 38 | 14 | 0 | 0 | 0 | 0 | 2 | 0 |
| JSG000000 | Plekhd1 | 4.74E-78 | 2.06E-76 | 8.620634405 | 0.649984458 | 255.8423937 | 43 | 9 | 10 | 14 | 24 | 4 | 0 | 0 | 0 | 0 | 0 | 2 |
| JSG000000 | Tbc1d16 | 2.02E-77 | 8.66E-76 | 2.936165975 | 147.4985014 | 1128.916027 | 81 | 5 | 10 | 12 | 41 | 7 | 3 | 0 | 0 | 0 | 0 | 0 |
| JSG000000 | Slc18a2 | 1.71E-75 | 7.06E-74 | -6.683037324 | 800.164244 | 7.787255857 | 64 | 8 | 10 | 12 | 29 | 1 | 4 | 0 | 0 | 1 | 4 | 0 |
| SG0000002 | Fetub | 3.1E-74 | 1.26E-72 | -3.799745438 | 2451.016891 | 175.9984939 | 72 | 4 | 10 | 14 | 44 | 2 | 0 | 0 | 0 | 0 | 0 | 0 |
| SG0000004 | Tshz1 | 1.27E-72 | 4.94E-71 | -4.8461232 | 2798.389801 | 97.29258967 | 118 | 8 | 10 | 21 | 57 | 11 | 7 | 0 | 2 | 1 | 3 | 1 |
| SG0000003 | Cpne9 | 7.81E-60 | 2.15E-58 | 9.38661876 | 0.331601068 | 221.9574496 | 60 | 8 | 10 | 12 | 16 | 4 | 0 | 0 | 0 | 0 | 0 | 0 |
| SG0000004 | Igsf3 | 1.18E-55 | 2.92E-54 | 2.479607951 | 286.9883472 | 1600.665682 | 180 | 2 | 10 | 22 | 68 | 10 | 2 | 0 | 0 | 0 | 0 | 0 |
| SG0000002 | Ptpru | 7.31E-55 | 1.75E-53 | 3.725304519 | 28.97475442 | 383.2200003 | 104 | 2 | 10 | 12 | 54 | 11 | 1 | 0 | 0 | 0 | 0 | 2 |
| SG0000005 | Slain1 | 3.44E-54 | 8.04E-53 | 3.335318061 | 56.75613417 | 572.8535227 | 109 | 7 | 10 | 16 | 52 | 10 | 4 | 0 | 0 | 1 | 1 | 0 |
| SG0000002 | Slc23a2 | 1.05E-53 | 2.41E-52 | -2.609812489 | 9132.876986 | 1496.152166 | 159 | 6 | 10 | 26 | 80 | 21 | 8 | 0 | 0 | 1 | 3 | 1 |
| JSG000000 | Asic1 | 1.28E-52 | 2.86E-51 | 5.332598903 | 5.239528701 | 211.1370496 | 34 | 8 | 10 | 10 | 20 | 1 | 0 | 0 | 0 | 0 | 0 | 1 |
| SG0000002 | Orc1 | 3.91E-51 | 8.37E-50 | 1.991695707 | 601.3762206 | 2391.698336 | 65 | 7 | 10 | 15 | 36 | 11 | 1 | 1 | 1 | 1 | 2 | 1 |
| JSG000000 | Xylb | 1.18E-48 | 2.36E-47 | 2.812113001 | 75.4082529 | 529.6015451 | 40 | 8 | 10 | 13 | 47 | 2 | 1 | 0 | 0 | 0 | 0 | 0 |
| SG0000002 | Asb1 | 1.21E-47 | 2.33E-46 | -2.613253713 | 2851.34296 | 465.9954795 | 54 | 4 | 10 | 15 | 46 | 4 | 16 | 2 | 2 | 7 | 8 | 0 |
| JSG000000 | Lix1l | 3.68E-42 | 5.84E-41 | -2.502269365 | 2125.081907 | 375.0744988 | 49 | 8 | 10 | 11 | 26 | 5 | 2 | 0 | 0 | 0 | 2 | 1 |
| SG0000003 | Galnt12 | 2.47E-40 | 3.69E-39 | 4.895644814 | 8.459172224 | 251.8046838 | 62 | 6 | 10 | 18 | 36 | 6 | 0 | 0 | 0 | 0 | 0 | 0 |
| SG0000003 | Emc8 | 6.01E-39 | 8.53E-38 | 1.746206952 | 771.527383 | 2588.284524 | 63 | 5 | 10 | 10 | 30 | 0 | 8 | 0 | 0 | 0 | 0 | 0 |
| SG0000003 | Fgf1 | 3.22E-38 | 4.48E-37 | -3.48224037 | 437.719303 | 39.16849597 | 217 | 3 | 10 | 32 | 104 | 37 | 5 | 0 | 0 | 0 | 0 | 1 |
| SG0000003 | Bcat2 | 1.26E-37 | 1.71E-36 | 1.771697254 | 582.7450815 | 1989.814692 | 50 | 8 | 10 | 12 | 14 | 6 | 0 | 0 | 0 | 0 | 0 | 0 |
| JSG000000 | Bcr | 1.48E-36 | 1.94E-35 | -2.103519324 | 3704.684617 | 862.0428005 | 111 | 6 | 10 | 20 | 60 | 15 | 3 | 1 | 1 | 1 | 1 | 0 |
| SG0000002 | Lrp4 | 6.65E-35 | 8.23E-34 | 2.76035066 | 48.84404242 | 330.9481417 | 66 | 9 | 10 | 13 | 40 | 8 | 2 | 0 | 0 | 0 | 0 | 0 |
| SG0000002 | Bin1 | 1.11E-34 | 1.36E-33 | -2.308952488 | 1448.894319 | 292.396892 | 78 | 0 | 10 | 8 | 44 | 8 | 0 | 0 | 0 | 0 | 0 | 0 |
| JSG000000 | Wscd2 | 4.09E-33 | 4.7E-32 | 8.190067869 | 0.331601068 | 96.84384328 | 83 | 9 | 10 | 11 | 37 | 12 | 0 | 0 | 0 | 0 | 0 | 0 |
| SG0000002 | Fzd6 | 4.74E-32 | 5.21E-31 | -2.213344983 | 1337.54862 | 288.420647 | 96 | 4 | 10 | 12 | 50 | 14 | 6 | 0 | 0 | 0 | 4 | 4 |
| SG0000003 | Nup37 | 7.89E-31 | 8.26E-30 | 1.595498596 | 540.69336 | 1633.969598 | 160 | 2 | 10 | 14 | 92 | 24 | 0 | 0 | 0 | 0 | 2 | 0 |
| SG0000006 | Blnk | 9.26E-31 | 9.64E-30 | 4.944827478 | 3.891376204 | 119.8518144 | 170 | 6 | 10 | 26 | 76 | 14 | 2 | 0 | 0 | 0 | 0 | 0 |
| SG0000006 | Mrps9 | 2.57E-30 | 2.64E-29 | 1.59182549 | 523.3401698 | 1577.506986 | 144 | 1 | 10 | 24 | 56 | 15 | 1 | 0 | 0 | 0 | 0 | 0 |
| SG0000003 | Abhd2 | 2.58E-30 | 2.64E-29 | -2.233529695 | 5461.589383 | 1161.340606 | 115 | 9 | 10 | 15 | 47 | 9 | 4 | 0 | 0 | 0 | 1 | 0 |
| SG0000004 | Lat2 | 2.91E-30 | 2.98E-29 | -6.909735723 | 585.4240627 | 4.868922417 | 58 | 10 | 10 | 13 | 21 | 7 | 4 | 0 | 0 | 0 | 0 | 0 |
| SG0000004 | Slc25a51 | 1.25E-29 | 1.24E-28 | -1.888379569 | 5769.711501 | 1558.458365 | 44 | 8 | 10 | 8 | 14 | 4 | 8 | 0 | 0 | 0 | 4 | 2 |
| SG0000003 | Ppp1r26 | 4.57E-29 | 4.42E-28 | 2.89980649 | 31.14906759 | 232.4736776 | 22 | 8 | 10 | 12 | 14 | 2 | 4 | 0 | 2 | 0 | 0 | 2 |
| SG0000004 | Arhgap30 | 6.72E-29 | 6.45E-28 | 3.369567365 | 19.14568147 | 197.8844547 | 35 | 10 | 10 | 13 | 26 | 7 | 4 | 0 | 0 | 0 | 0 | 0 |
| SG0000003 | Bcl2l14 | 8.78E-29 | 8.39E-28 | 8.086500864 | 0.307509278 | 83.58677009 | 148 | 8 | 10 | 27 | 38 | 18 | 0 | 0 | 0 | 0 | 0 | 0 |
| SG0000002 | Nat10 | 1.55E-28 | 1.47E-27 | 1.430073392 | 1260.295774 | 3395.998322 | 49 | 8 | 10 | 14 | 34 | 3 | 0 | 0 | 0 | 0 | 0 | 0 |
| SG0000002 | Slc43a3 | 1.68E-28 | 1.58E-27 | 3.28737249 | 18.81777068 | 183.7240025 | 41 | 7 | 10 | 12 | 24 | 4 | 1 | 0 | 0 | 0 | 0 | 0 |
| JSG000000 | Drc1 | 4.97E-28 | 4.6E-27 | 4.55730928 | 4.932019423 | 116.1212149 | 44 | 10 | 10 | 11 | 30 | 1 | 0 | 0 | 0 | 0 | 0 | 0 |
| JSG000000 | Emb | 5.53E-28 | 5.11E-27 | 5.828105205 | 64.19575528 | 3647.042275 | 113 | 8 | 10 | 22 | 54 | 15 | 1 | 0 | 0 | 1 | 1 | 0 |
| JSG000000 | Mnd1 | 1.26E-27 | 1.14E-26 | 1.90195204 | 156.5964341 | 585.2298681 | 133 | 5 | 10 | 24 | 78 | 11 | 0 | 0 | 0 | 0 | 0 | 0 |
| SG0000002 | Tspan2 | 2.45E-27 | 2.2E-26 | -2.575425234 | 1623.157911 | 272.3206191 | 184 | 1 | 10 | 14 | 42 | 16 | 11 | 0 | 1 | 1 | 2 | 2 |
| JSG000000 | Diexf | 4.1E-27 | 3.63E-26 | 1.821376112 | 487.5471123 | 1723.07996 | 150 | 6 | 10 | 19 | 57 | 7 | 8 | 0 | 0 | 0 | 4 | 0 |
| JSG000000 | Wdfy2 | 4.73E-26 | 4E-25 | -2.341911732 | 861.0129188 | 169.8337461 | 199 | 2 | 10 | 25 | 81 | 22 | 0 | 0 | 0 | 0 | 0 | 0 |
| SG0000003 | Dcbld2 | 6.53E-26 | 5.5E-25 | -1.775570871 | 5982.594215 | 1747.388427 | 114 | 2 | 10 | 18 | 43 | 15 | 11 | 0 | 1 | 1 | 4 | 0 |
| SG0000003 | Pptc7 | 7.67E-26 | 6.44E-25 | -1.734442977 | 2426.300098 | 729.1638911 | 82 | 4 | 10 | 18 | 57 | 9 | 14 | 0 | 2 | 4 | 6 | 1 |
| SG0000005 | Lifr | 3.59E-25 | 2.93E-24 | -1.626460416 | 4332.036989 | 1403.068208 | 206 | 5 | 10 | 12 | 81 | 19 | 14 | 0 | 0 | 0 | 6 | 5 |

| | | | | | | | | | | | | | | | | | | |
|---|---|---|---|---|---|---|---|---|---|---|---|---|---|---|---|---|---|---|
| SG0000002 | Pop1 | 4.59E-25 | 3.73E-24 | 1.397703754 | 855.0756499 | 2252.969405 | 136 | 4 | 10 | 16 | 72 | 14 | 4 | 0 | 0 | 0 | 0 | 0 |
| JSG000000 | Tmem231 | 7.55E-25 | 6.05E-24 | 1.538668193 | 360.833318 | 1048.315269 | 40 | 8 | 10 | 15 | 30 | 3 | 0 | 0 | 0 | 0 | 0 | 0 |
| JSG000000 | Mccc2 | 1.41E-24 | 1.12E-23 | 1.805536647 | 158.7178766 | 554.8128788 | 111 | 6 | 10 | 16 | 69 | 17 | 1 | 0 | 0 | 0 | 0 | 0 |
| SG0000002 | Orc5 | 2.8E-24 | 2.18E-23 | 1.342741273 | 865.5906701 | 2195.42177 | 169 | 2 | 10 | 27 | 64 | 11 | 1 | 0 | 0 | 0 | 0 | 0 |
| SG0000002 | Mak | 1.43E-23 | 1.08E-22 | 8.333197251 | 0.34247518 | 110.4514534 | 175 | 6 | 10 | 23 | 101 | 12 | 2 | 0 | 0 | 1 | 2 | 0 |
| SG0000006 | Zfp600 | 2.65E-23 | 1.97E-22 | 8.30032932 | 0.34247518 | 107.9635613 | 162 | 8 | 10 | 20 | 103 | 30 | 10 | 0 | 0 | 0 | 8 | 2 |
| SG0000001 | Cacfd1 | 6.13E-23 | 4.49E-22 | -1.71959248 | 1421.898174 | 431.7373084 | 32 | 10 | 10 | 10 | 34 | 4 | 1 | 0 | 0 | 0 | 2 | 1 |
| JSG000000 | Spred3 | 2.17E-22 | 1.55E-21 | -1.849301959 | 961.466781 | 266.8326217 | 18 | 10 | 10 | 11 | 16 | 1 | 3 | 0 | 0 | 1 | 2 | 0 |
| SG0000002 | Mcm10 | 3.73E-22 | 2.63E-21 | 1.211033507 | 1663.621578 | 3851.34085 | 55 | 8 | 10 | 12 | 34 | 2 | 2 | 0 | 2 | 0 | 2 | 0 |
| SG0000000 | Adgre5 | 9.11E-22 | 6.33E-21 | -1.636224808 | 3772.890813 | 1213.728333 | 74 | 5 | 10 | 43 | 53 | 0 | 0 | 0 | 0 | 0 | 0 | 0 |
| JSG000000 | Irg1 | 1.8E-21 | 1.22E-20 | -6.453614438 | 84.67767471 | 0.966134069 | 20 | 8 | 10 | 11 | 16 | 1 | 1 | 1 | 1 | 1 | 1 | 0 |
| JSG000000 | Smim3 | 3.46E-21 | 2.32E-20 | -1.92644282 | 671.1842241 | 176.5731506 | 51 | 7 | 10 | 14 | 20 | 5 | 4 | 0 | 0 | 1 | 3 | 0 |
| SG0000002 | Stx2 | 9.61E-21 | 6.31E-20 | -1.7020697 | 1057.642207 | 325.0609045 | 49 | 10 | 10 | 10 | 32 | 2 | 6 | 0 | 0 | 0 | 0 | 0 |
| SG0000003 | Iqce | 1.02E-20 | 6.66E-20 | -1.787075531 | 856.0294625 | 248.0420848 | 52 | 8 | 10 | 16 | 30 | 10 | 2 | 0 | 0 | 0 | 0 | 0 |
| JSG000000 | Frmd6 | 1.37E-20 | 8.93E-20 | -1.663480399 | 2503.675593 | 790.3520031 | 121 | 6 | 10 | 19 | 50 | 11 | 3 | 0 | 0 | 0 | 2 | 0 |
| SG0000002 | Gsn | 1.78E-20 | 1.15E-19 | -2.658495129 | 7580.038194 | 1200.562065 | 72 | 4 | 10 | 12 | 31 | 18 | 0 | 0 | 0 | 0 | 0 | 0 |
| JSG000000 | Tnfrsf11a | 2E-20 | 1.29E-19 | -2.615385992 | 335.533061 | 54.75523976 | 98 | 7 | 10 | 14 | 50 | 11 | 5 | 0 | 0 | 1 | 3 | 0 |
| SG0000002 | Acot9 | 2.15E-20 | 1.38E-19 | -1.406255708 | 4290.881039 | 1618.904852 | 101 | 8 | 10 | 22 | 51 | 13 | 0 | 0 | 0 | 0 | 0 | 0 |
| SG0000005 | Mast1 | 2.53E-20 | 1.62E-19 | 5.204502155 | 1.939079263 | 71.50012861 | 32 | 8 | 10 | 12 | 36 | 2 | 0 | 0 | 0 | 0 | 0 | 0 |
| SG0000006 | Zfp143 | 3.06E-20 | 1.94E-19 | 1.28675372 | 610.4209074 | 1489.295465 | 128 | 8 | 10 | 10 | 48 | 10 | 8 | 2 | 2 | 2 | 4 | 2 |
| SG0000000 | Ocrl | 4.09E-20 | 2.58E-19 | -1.433477812 | 3226.170835 | 1194.448808 | 184 | 8 | 10 | 17 | 82 | 15 | 4 | 0 | 0 | 0 | 1 | 0 |
| JSG000000 | Ccdc66 | 7.02E-20 | 4.37E-19 | 1.344803062 | 425.1461827 | 1079.852266 | 53 | 7 | 10 | 18 | 36 | 7 | 0 | 0 | 0 | 0 | 0 | 0 |
| SG0000003 | Myo15b | 8.2E-20 | 5.1E-19 | 3.798350439 | 6.860224575 | 95.44556412 | 23 | 10 | 10 | 10 | 14 | 2 | 0 | 0 | 0 | 0 | 0 | 0 |
| SG0000003 | Zbed4 | 1.9E-19 | 1.15E-18 | 1.525581136 | 225.4004913 | 648.9340182 | 59 | 7 | 10 | 14 | 29 | 4 | 2 | 0 | 0 | 1 | 2 | 1 |
| SG0000004 | Selplg | 2.42E-19 | 1.46E-18 | -2.627436092 | 292.2909301 | 47.30186495 | 19 | 10 | 10 | 11 | 27 | 0 | 0 | 0 | 0 | 0 | 4 | 0 |
| SG0000003 | Rhbdd2 | 3.17E-19 | 1.9E-18 | -1.551305984 | 1307.337191 | 446.0648297 | 39 | 6 | 10 | 17 | 24 | 5 | 6 | 2 | 2 | 2 | 2 | 2 |
| SG0000003 | Unc93b1 | 6.32E-19 | 3.73E-18 | -2.358436939 | 1160.637385 | 226.326962 | 46 | 6 | 10 | 8 | 24 | 2 | 0 | 0 | 0 | 0 | 2 | 0 |
| JSG000000 | Gpr37 | 3.42E-18 | 1.94E-17 | -6.154235058 | 70.17141761 | 0.985260105 | 140 | 4 | 10 | 14 | 44 | 10 | 7 | 0 | 1 | 0 | 2 | 1 |
| SG0000007 | Spint2 | 5.31E-18 | 2.99E-17 | 1.556041379 | 1204.719414 | 3542.428053 | 60 | 4 | 10 | 14 | 26 | 6 | 2 | 0 | 0 | 0 | 0 | 0 |
| SG0000004 | Rhod | 4.46E-17 | 2.39E-16 | -1.680562516 | 664.5238832 | 207.3055205 | 32 | 8 | 10 | 12 | 22 | 2 | 2 | 0 | 0 | 2 | 2 | 0 |
| SG0000000 | Ctcf | 8.52E-17 | 4.5E-16 | 1.034069027 | 2316.023912 | 4742.734722 | 104 | 6 | 10 | 16 | 50 | 13 | 3 | 0 | 0 | 0 | 1 | 1 |
| SG0000003 | Trf | 1.02E-16 | 5.35E-16 | 5.494887319 | 0 | 44.09474192 | 66 | 8 | 10 | 9 | 32 | 2 | 0 | 0 | 0 | 0 | 1 | 0 |
| SG0000003 | Primpol | 1.45E-16 | 7.53E-16 | 1.237216075 | 477.7171521 | 1126.185637 | 121 | 3 | 10 | 14 | 53 | 11 | 5 | 0 | 0 | 0 | 0 | 0 |
| SG0000002 | Myl12a | 1.94E-16 | 1E-15 | -1.176772752 | 9457.881653 | 4183.59831 | 50 | 10 | 10 | 12 | 35 | 0 | 6 | 0 | 0 | 2 | 3 | 0 |
| JSG000000 | Heatr5b | 1.95E-16 | 1.01E-15 | -1.49622397 | 2068.325956 | 733.1801306 | 202 | 6 | 10 | 28 | 94 | 19 | 3 | 1 | 1 | 2 | 2 | 0 |
| JSG000000 | Igdcc3 | 4.4E-16 | 2.22E-15 | 7.110631082 | 0.331601068 | 45.82781646 | 54 | 7 | 10 | 12 | 27 | 1 | 1 | 0 | 0 | 0 | 0 | 0 |
| JSG000000 | Psmb7 | 1.82E-15 | 8.87E-15 | 0.972504226 | 3276.472992 | 6429.23848 | 109 | 7 | 10 | 17 | 47 | 5 | 0 | 0 | 0 | 0 | 0 | 0 |
| SG0000007 | Naip6 | 2.11E-15 | 1.03E-14 | -4.893298919 | 69.44127425 | 2.336619136 | 122 | 6 | 10 | 22 | 81 | 10 | 12 | 0 | 0 | 0 | 2 | 0 |
| JSG000000 | Ebag9 | 3.77E-15 | 1.8E-14 | -1.414987987 | 954.190665 | 357.8337797 | 68 | 6 | 10 | 10 | 35 | 7 | 1 | 0 | 0 | 1 | 2 | 1 |
| JSG000000 | Pole2 | 4.32E-15 | 2.05E-14 | 1.177369647 | 435.7890515 | 985.5981843 | 59 | 9 | 10 | 13 | 33 | 3 | 0 | 0 | 0 | 0 | 1 | 0 |
| SG0000002 | Pcmtd2 | 5.2E-15 | 2.46E-14 | -1.246612366 | 2034.806583 | 857.5420373 | 115 | 2 | 10 | 10 | 62 | 11 | 12 | 0 | 2 | 2 | 6 | 2 |
| SG0000000 | Stat5a | 5.76E-15 | 2.72E-14 | -1.685131572 | 585.7609199 | 182.1567394 | 58 | 10 | 10 | 19 | 48 | 21 | 3 | 0 | 0 | 0 | 3 | 3 |
| JSG000000 | Ccnf | 1.05E-14 | 4.88E-14 | 0.968045919 | 2369.162382 | 4634.529883 | 36 | 8 | 10 | 15 | 27 | 1 | 0 | 0 | 0 | 0 | 0 | 0 |
| SG0000001 | Atp6v1e1 | 1.37E-14 | 6.29E-14 | -1.129634799 | 4541.734097 | 2075.715104 | 48 | 6 | 10 | 9 | 23 | 3 | 2 | 1 | 1 | 1 | 1 | 0 |
| SG0000007 | St6galnac4 | 3.22E-14 | 1.45E-13 | -1.402691211 | 786.1852742 | 297.3533089 | 34 | 10 | 10 | 10 | 29 | 0 | 6 | 0 | 0 | 0 | 0 | 0 |
| JSG000000 | Mypn | 4.39E-14 | 1.96E-13 | 6.929465004 | 0.34247518 | 41.74514008 | 137 | 7 | 10 | 18 | 78 | 18 | 2 | 0 | 0 | 0 | 0 | 0 |
| JSG000000 | Notch3 | 4.63E-14 | 2.06E-13 | 2.318696132 | 303.6983391 | 1515.093739 | 58 | 6 | 10 | 16 | 32 | 4 | 2 | 0 | 1 | 1 | 6 | 1 |
| SG0000002 | Ddx4 | 8.98E-14 | 3.92E-13 | 8.085513911 | 2.912134245 | 791.0311921 | 278 | 0 | 10 | 26 | 114 | 26 | 0 | 0 | 0 | 0 | 0 | 0 |
| JSG000000 | Ccdc47 | 1.62E-13 | 6.97E-13 | -1.166891443 | 4902.775137 | 2183.597805 | 50 | 7 | 10 | 10 | 23 | 3 | 5 | 0 | 0 | 0 | 3 | 0 |
| JSG000000 | Rnd3 | 1.63E-13 | 7.01E-13 | -1.197009619 | 1588.399287 | 692.8255309 | 56 | 4 | 10 | 9 | 31 | 8 | 6 | 0 | 1 | 1 | 2 | 1 |
| SG0000002 | Fuca1 | 8.11E-13 | 3.33E-12 | -1.189241833 | 1266.505254 | 555.4046762 | 26 | 9 | 10 | 14 | 28 | 1 | 2 | 0 | 0 | 0 | 1 | 0 |
| SG0000001 | Slc35b4 | 1.2E-12 | 4.86E-12 | -1.136816105 | 1684.615718 | 766.099363 | 69 | 8 | 10 | 14 | 34 | 12 | 6 | 1 | 1 | 3 | 4 | 1 |
| JSG000000 | Flnb | 1.45E-12 | 5.84E-12 | -1.101142802 | 25165.67545 | 11730.90664 | 156 | 9 | 10 | 28 | 78 | 18 | 0 | 0 | 0 | 0 | 0 | 0 |
| JSG000000 | Bcl7a | 1.75E-12 | 7.02E-12 | 1.149386207 | 307.3318368 | 681.7219485 | 64 | 9 | 10 | 12 | 45 | 6 | 16 | 3 | 4 | 4 | 7 | 0 |
| JSG000000 | Spats1 | 2.06E-12 | 8.19E-12 | 3.378355976 | 6.116216522 | 63.60178971 | 38 | 8 | 10 | 13 | 26 | 7 | 0 | 0 | 0 | 0 | 0 | 0 |
| SG0000002 | Sh3glb2 | 4.05E-12 | 1.58E-11 | -1.450635907 | 1822.066369 | 666.6214305 | 30 | 10 | 10 | 10 | 25 | 5 | 0 | 0 | 0 | 0 | 0 | 0 |
| SG0000002 | Ndufa10 | 4.96E-12 | 1.93E-11 | 0.87145192 | 2196.057145 | 4017.693522 | 86 | 10 | 10 | 14 | 48 | 8 | 2 | 0 | 0 | 0 | 0 | 0 |
| SG0000007 | Ptges3 | 9.2E-12 | 3.52E-11 | 0.83213123 | 5543.72948 | 9869.575879 | 75 | 6 | 10 | 8 | 50 | 4 | 7 | 0 | 0 | 0 | 0 | 0 |
| SG0000003 | Klra2 | 1.02E-11 | 3.89E-11 | -4.937418274 | 162.5669063 | 5.305436729 | 72 | 4 | 10 | 12 | 43 | 3 | 2 | 0 | 0 | 1 | 2 | 0 |
| SG0000001 | Evi5l | 1.45E-11 | 5.46E-11 | -1.313767445 | 598.3582669 | 240.700996 | 207 | 10 | 10 | 23 | 57 | 10 | 8 | 0 | 0 | 0 | 0 | 0 |
| SG0000002 | Rpe | 1.55E-11 | 5.86E-11 | 0.871470636 | 1568.016504 | 2868.728389 | 197 | 0 | 10 | 29 | 104 | 20 | 15 | 0 | 0 | 0 | 0 | 0 |
| JSG000000 | Txndc12 | 1.99E-11 | 7.45E-11 | 0.904986622 | 996.6416102 | 1866.238456 | 67 | 8 | 10 | 13 | 31 | 5 | 0 | 0 | 0 | 0 | 0 | 0 |
| SG0000002 | Eif3m | 2.33E-11 | 8.69E-11 | 0.830926793 | 2637.170414 | 4691.071836 | 79 | 0 | 10 | 2 | 21 | 2 | 0 | 0 | 0 | 0 | 0 | 0 |
| JSG000000 | Cdc42bpb | 2.39E-11 | 8.89E-11 | -1.333292735 | 5027.703198 | 1995.301485 | 74 | 7 | 10 | 9 | 29 | 9 | 1 | 0 | 0 | 0 | 1 | 1 |
| SG0000005 | Zfp169 | 3.85E-11 | 1.41E-10 | -1.785295019 | 225.3097645 | 65.36611322 | 64 | 8 | 10 | 20 | 46 | 8 | 4 | 0 | 0 | 4 | 4 | 4 |
| SG0000003 | Tbc1d2 | 3.9E-11 | 1.43E-10 | -1.735208659 | 997.6373371 | 299.6559044 | 55 | 8 | 10 | 14 | 23 | 3 | 0 | 0 | 0 | 0 | 0 | 0 |
| JSG000000 | Cep120 | 4.08E-11 | 1.49E-10 | 0.897291301 | 911.306749 | 1697.368675 | 126 | 5 | 10 | 16 | 64 | 6 | 4 | 0 | 0 | 1 | 1 | 2 |
| SG0000003 | Srp72 | 8.31E-10 | 2.76E-09 | -0.854740278 | 8712.0967 | 4817.48338 | 130 | 8 | 10 | 19 | 57 | 2 | 4 | 0 | 0 | 1 | 1 | 0 |
| JSG000000 | Sp4 | 1.03E-09 | 3.39E-09 | 1.202980635 | 144.335069 | 332.2807021 | 153 | 3 | 10 | 12 | 63 | 27 | 6 | 0 | 2 | 2 | 7 | 1 |
| JSG000000 | Cmtm8 | 1.2E-09 | 3.91E-09 | 5.139395442 | 4.301439621 | 151.6093615 | 58 | 9 | 10 | 15 | 39 | 9 | 2 | 0 | 0 | 0 | 0 | 0 |
| SG0000006 | Ei24 | 1.28E-09 | 4.18E-09 | -0.851884307 | 7118.489907 | 3944.074815 | 62 | 0 | 10 | 6 | 26 | 6 | 6 | 0 | 0 | 0 | 0 | 0 |
| SG0000007 | Eif2s2 | 1.79E-09 | 5.77E-09 | 0.730579694 | 5124.909111 | 8503.790843 | 96 | 4 | 10 | 16 | 40 | 4 | 0 | 0 | 0 | 0 | 2 | 0 |
| JSG000000 | Scnm1 | 2.8E-09 | 8.89E-09 | 1.005929866 | 276.4629903 | 555.2033327 | 24 | 10 | 10 | 12 | 16 | 0 | 0 | 0 | 0 | 0 | 0 | 0 |
| SG0000004 | Lonrf2 | 3.41E-09 | 1.07E-08 | 6.27886853 | 0.331601068 | 25.74801109 | 113 | 8 | 10 | 23 | 62 | 14 | 15 | 0 | 0 | 0 | 2 | 0 |
| SG0000002 | Slc35b3 | 3.71E-09 | 1.17E-08 | -1.275007957 | 383.5650568 | 158.4978574 | 82 | 2 | 10 | 10 | 36 | 10 | 6 | 0 | 2 | 0 | 0 | 0 |
| SG0000003 | Map9 | 4.92E-09 | 1.53E-08 | -1.141252788 | 936.5492768 | 424.5993363 | 130 | 10 | 10 | 24 | 44 | 8 | 38 | 10 | 10 | 12 | 18 | 2 |
| SG0000002 | Snrnp48 | 5.2E-09 | 1.62E-08 | 0.866148439 | 551.2435428 | 1004.801272 | 50 | 6 | 10 | 14 | 29 | 2 | 0 | 0 | 0 | 0 | 1 | 1 |
| SG0000003 | Pik3r4 | 9.19E-09 | 2.81E-08 | 0.779806849 | 1021.957529 | 1754.599899 | 156 | 6 | 10 | 14 | 66 | 12 | 0 | 0 | 0 | 0 | 1 | 0 |
| SG0000002 | Pik3r5 | 9.24E-09 | 2.82E-08 | 2.018235048 | 22.53297067 | 91.2783412 | 63 | 4 | 10 | 9 | 40 | 4 | 1 | 0 | 0 | 0 | 2 | 0 |
| SG0000002 | Zfp91 | 1.07E-08 | 3.25E-08 | -0.817912408 | 4225.344388 | 2396.878078 | 200 | 6 | 10 | 21 | 51 | 13 | 7 | 1 | 1 | 1 | 2 | 0 |
| SG0000002 | P3h1 | 1.62E-08 | 4.84E-08 | -0.897560449 | 1302.822275 | 699.3467441 | 68 | 10 | 10 | 19 | 33 | 6 | 13 | 0 | 0 | 4 | 8 | 1 |
| JSG000000 | Gbgt1 | 2.23E-08 | 6.61E-08 | 3.912545755 | 2.235714429 | 33.66742966 | 22 | 10 | 10 | 12 | 46 | 2 | 0 | 0 | 0 | 0 | 0 | 0 |
| SG0000002 | Atp2b4 | 2.37E-08 | 6.98E-08 | -0.968927648 | 2074.449763 | 1059.80664 | 96 | 10 | 10 | 19 | 51 | 11 | 10 | 0 | 0 | 1 | 3 | 3 |
| SG0000003 | Nod1 | 3.45E-08 | 1E-07 | -1.601581211 | 797.6953167 | 262.8530722 | 98 | 6 | 10 | 12 | 46 | 6 | 2 | 0 | 0 | 0 | 0 | 0 |
| SG0000002 | 3010026O09Rik | 4.29E-08 | 1.24E-07 | 1.075447392 | 160.4086574 | 338.0412723 | 72 | 8 | 10 | 14 | 24 | 2 | 0 | 0 | 0 | 0 | 0 | 0 |
| SG0000003 | Fam179b | 5E-08 | 1.44E-07 | -0.825558844 | 1979.128703 | 1116.75026 | 167 | 4 | 10 | 17 | 64 | 18 | 4 | 0 | 0 | 0 | 1 | 0 |
| JSG000000 | Prl7d1 | 5.03E-08 | 1.45E-07 | 6.312474238 | 0.307509278 | 24.44006273 | 33 | 9 | 10 | 15 | 24 | 4 | 1 | 0 | 0 | 0 | 0 | 0 |
| JSG000000 | Eepd1 | 5.11E-08 | 1.47E-07 | 2.01713528 | 19.95862447 | 80.78836728 | 180 | 2 | 10 | 16 | 69 | 18 | 0 | 0 | 0 | 0 | 0 | 0 |
| JSG000000 | Fech | 5.77E-08 | 1.65E-07 | -0.853784021 | 1489.778395 | 824.3413246 | 59 | 7 | 10 | 16 | 40 | 5 | 2 | 0 | 0 | 0 | 3 | 1 |
| JSG000000 | Car5b | 9.88E-08 | 2.77E-07 | -1.419532787 | 311.0114899 | 116.2664695 | 115 | 8 | 10 | 23 | 74 | 13 | 4 | 0 | 0 | 0 | 3 | 0 |
| SG0000002 | Tnf | 1.11E-07 | 3.1E-07 | -2.823881811 | 54.33237255 | 7.673359413 | 18 | 10 | 10 | 16 | 22 | 0 | 16 | 10 | 10 | 16 | 22 | 0 |
| SG0000004 | Ginm1 | 1.16E-07 | 3.23E-07 | -0.942445064 | 737.9075638 | 383.9703844 | 56 | 2 | 10 | 4 | 10 | 4 | 8 | 0 | 2 | 0 | 0 | 0 |
| SG0000000 | Zfp287 | 2.47E-07 | 6.68E-07 | 1.683762074 | 30.442369 | 97.80055866 | 126 | 6 | 10 | 20 | 123 | 25 | 8 | 0 | 0 | 0 | 2 | 2 |
| SG0000000 | Paxip1 | 3.63E-07 | 9.69E-07 | 0.678201307 | 1338.454939 | 2141.719996 | 82 | 5 | 10 | 17 | 49 | 7 | 2 | 0 | 1 | 1 | 3 | 1 |
| SG0000003 | Hmces | 4.51E-07 | 1.19E-06 | 0.645540863 | 2406.344551 | 3764.306475 | 108 | 4 | 10 | 18 | 36 | 16 | 0 | 0 | 0 | 0 | 0 | 0 |
| SG0000002 | Sdf4 | 5.29E-07 | 1.39E-06 | -0.714638511 | 3734.526864 | 2275.66306 | 49 | 3 | 10 | 6 | 26 | 3 | 1 | 0 | 1 | 0 | 1 | 0 |
| SG0000002 | Ssrp1 | 7.26E-07 | 1.89E-06 | 0.600935197 | 7915.641461 | 12005.64878 | 30 | 10 | 10 | 12 | 20 | 0 | 2 | 0 | 0 | 0 | 0 | 0 |
| SG0000004 | Rap1gap | 7.7E-07 | 2.01E-06 | -0.957137052 | 812.787 | 418.6487376 | 71 | 10 | 10 | 19 | 45 | 3 | 3 | 0 | 0 | 3 | 3 | 0 |
| SG0000000 | Coro1c | 1.02E-06 | 2.63E-06 | -0.661804184 | 9235.592117 | 5837.700819 | 142 | 6 | 10 | 20 | 72 | 4 | 2 | 0 | 0 | 0 | 0 | 0 |
| SG0000000 | Kif5b | 1.39E-06 | 3.54E-06 | -0.710961264 | 6571.867872 | 4014.839211 | 103 | 8 | 10 | 17 | 45 | 10 | 12 | 1 | 1 | 3 | 5 | 0 |
| SG0000002 | Gars | 1.55E-06 | 3.93E-06 | 0.583304661 | 7419.545446 | 11116.53651 | 91 | 2 | 10 | 13 | 31 | 3 | 1 | 0 | 0 | 0 | 0 | 0 |
| SG0000002 | Lipa | 1.66E-06 | 4.19E-06 | -0.874751122 | 874.4001666 | 476.8523259 | 68 | 9 | 10 | 19 | 55 | 3 | 2 | 0 | 0 | 0 | 0 | 0 |
| SG0000002 | Fam213a | 2.51E-06 | 6.25E-06 | -0.722866116 | 1593.263878 | 965.3467224 | 62 | 10 | 10 | 12 | 20 | 0 | 6 | 0 | 0 | 0 | 0 | 0 |
| SG0000000 | Gtf2h1 | 3.33E-06 | 8.2E-06 | 0.59712832 | 2102.479895 | 3180.426672 | 173 | 6 | 10 | 52 | 102 | 10 | 8 | 0 | 0 | 0 | 0 | 0 |
| SG0000003 | Proser3 | 4.74E-06 | 1.15E-05 | 0.964691517 | 154.0361584 | 300.6240783 | 25 | 9 | 10 | 10 | 14 | 2 | 0 | 0 | 0 | 0 | 0 | 0 |
| SG0000006 | Naca | 6.76E-06 | 1.62E-05 | 0.544493215 | 9444.23727 | 13774.4948 | 42 | 2 | 10 | 14 | 20 | 0 | 0 | 0 | 0 | 0 | 1 | 0 |
| JSG000000 | Tgfbrap1 | 7.36E-06 | 1.75E-05 | -0.740397074 | 925.3950114 | 553.9179337 | 140 | 6 | 10 | 17 | 64 | 8 | 0 | 0 | 0 | 0 | 0 | 0 |
| JSG000000 | AU041133 | 7.36E-06 | 1.75E-05 | -2.273662491 | 103.0422354 | 21.3095688 | 54 | 5 | 10 | 15 | 39 | 4 | 1 | 0 | 0 | 0 | 2 | 0 |
| JSG000000 | Tab3 | 9.41E-06 | 2.22E-05 | 0.618190755 | 857.0854612 | 1315.582502 | 156 | 6 | 10 | 16 | 69 | 19 | 5 | 0 | 0 | 1 | 3 | 1 |
| SG0000002 | Gucy1b3 | 1.25E-05 | 2.91E-05 | 5.284281142 | 1.668879454 | 65.03574164 | 187 | 6 | 10 | 28 | 82 | 6 | 7 | 0 | 0 | 0 | 2 | 2 |
| SG0000003 | Nudt1 | 1.28E-05 | 2.99E-05 | -0.725326021 | 905.7734408 | 547.8664337 | 58 | 10 | 10 | 20 | 25 | 10 | 4 | 2 | 2 | 2 | 3 | 0 |
| SG0000003 | Trmt13 | 1.32E-05 | 3.08E-05 | 0.870299822 | 251.1960989 | 459.1972115 | 158 | 2 | 10 | 18 | 79 | 19 | 8 | 0 | 0 | 0 | 2 | 0 |
| SG0000004 | Katnal1 | 1.6E-05 | 3.69E-05 | -0.984315327 | 428.6344917 | 216.6599691 | 86 | 7 | 10 | 12 | 30 | 8 | 11 | 2 | 3 | 3 | 4 | 0 |
| SG0000001 | Mdh2 | 1.67E-05 | 3.86E-05 | -0.60969845 | 10235.76583 | 6707.841927 | 26 | 10 | 10 | 16 | 18 | 2 | 0 | 0 | 0 | 0 | 0 | 0 |
| JSG000000 | Ncapg2 | 1.72E-05 | 3.98E-05 | -0.618588825 | 3129.635056 | 2038.355393 | 137 | 7 | 10 | 21 | 84 | 13 | 1 | 1 | 1 | 1 | 3 | 0 |
| SG0000000 | Cdip1 | 2.04E-05 | 4.66E-05 | 0.617621514 | 710.823696 | 1090.647727 | 137 | 10 | 10 | 26 | 47 | 9 | 13 | 0 | 0 | 0 | 0 | 0 |
| SG0000003 | 1700088E04Rik | 2.05E-05 | 4.69E-05 | -1.775601802 | 69.32735212 | 20.24860975 | 24 | 10 | 10 | 17 | 27 | 10 | 0 | 0 | 0 | 0 | 0 | 0 |
| SG0000002 | Eif4ebp2 | 2.44E-05 | 5.54E-05 | 0.543006887 | 2181.904763 | 3179.048482 | 78 | 4 | 10 | 4 | 18 | 4 | 0 | 0 | 0 | 0 | 0 | 0 |
| SG0000001 | Slc47a1 | 2.45E-05 | 5.55E-05 | 2.419852887 | 9.318952095 | 49.86724588 | 108 | 10 | 10 | 18 | 50 | 4 | 2 | 0 | 0 | 0 | 1 | 0 |
| JSG000000 | Tbc1d23 | 4.01E-05 | 8.91E-05 | -0.636786699 | 1424.229425 | 915.9843568 | 126 | 7 | 10 | 20 | 68 | 14 | 4 | 0 | 0 | 1 | 2 | 1 |
| SG0000002 | Mina | 4.81E-05 | 0.000106 | -0.604006388 | 2017.672112 | 1327.475611 | 78 | 4 | 10 | 10 | 30 | 6 | 2 | 0 | 0 | 0 | 0 | 0 |
| SG0000002 | Pdia3 | 5.22E-05 | 0.000114 | -0.537477865 | 18857.41568 | 12992.27531 | 97 | 4 | 10 | 16 | 39 | 3 | 3 | 0 | 0 | 0 | 0 | 0 |
| SG0000003 | Mtg2 | 5.38E-05 | 0.000118 | -0.791834595 | 473.8035337 | 273.6732381 | 38 | 10 | 10 | 12 | 17 | 2 | 0 | 0 | 0 | 0 | 0 | 0 |

| | | | | | | | | | | | | | | | | | | |
|---|---|---|---|---|---|---|---|---|---|---|---|---|---|---|---|---|---|---|
| ISG000000 | Lrrc40 | 6.81E-05 | 0.000147 | 0.520938866 | 1705.398422 | 2447.056976 | 358 | 0 | 10 | 19 | 127 | 33 | 21 | 0 | 1 | 1 | 10 | 0 |
| SG0000001 | Abca5 | 7.38E-05 | 0.000159 | -1.022395863 | 334.8423171 | 164.8422434 | 199 | 3 | 10 | 26 | 102 | 15 | 14 | 0 | 1 | 0 | 1 | 0 |
| ISG000000 | Snrpd3 | 7.51E-05 | 0.000162 | 0.498262035 | 3020.92079 | 4267.083645 | 41 | 9 | 10 | 17 | 26 | 4 | 3 | 0 | 0 | 1 | 4 | 0 |
| ISG000000 | Hiat1 | 8.41E-05 | 0.00018 | 0.528859916 | 1414.041628 | 2040.163122 | 92 | 3 | 10 | 19 | 39 | 6 | 4 | 0 | 2 | 0 | 1 | 3 |
| SG0000005 | Gpd1l | 9.3E-05 | 0.000198 | 0.558212869 | 819.061776 | 1206.022683 | 105 | 4 | 10 | 27 | 42 | 15 | 8 | 0 | 2 | 2 | 3 | 3 |
| SG0000002 | Btd | 9.83E-05 | 0.000209 | -0.732657135 | 482.3172297 | 290.2558637 | 51 | 7 | 10 | 12 | 30 | 1 | 0 | 0 | 0 | 0 | 0 | 0 |
| SG0000000 | Snrnp200 | 0.000145 | 0.000302 | 0.466959143 | 5899.249042 | 8153.901099 | 51 | 9 | 10 | 13 | 35 | 3 | 0 | 0 | 0 | 0 | 0 | 0 |
| ISG000000 | Stt3b | 0.000169 | 0.00035 | -0.513370966 | 6829.459139 | 4784.606779 | 142 | 6 | 10 | 19 | 67 | 5 | 4 | 0 | 0 | 1 | 1 | 1 |
| SG0000002 | Dpf2 | 0.000203 | 0.000416 | 0.491016915 | 1787.63381 | 2512.403444 | 49 | 10 | 10 | 10 | 20 | 0 | 7 | 1 | 1 | 1 | 1 | 0 |
| ISG000000 | Cd80 | 0.000234 | 0.000476 | -1.231819801 | 97.48232955 | 41.50602944 | 49 | 7 | 10 | 11 | 24 | 0 | 0 | 0 | 0 | 0 | 1 | 0 |
| ISG000000 | Nek4 | 0.000294 | 0.00059 | 0.565978942 | 488.2832024 | 722.8503951 | 78 | 7 | 10 | 14 | 36 | 6 | 0 | 0 | 0 | 0 | 0 | 0 |
| ISG000000 | Adamts5 | 0.000295 | 0.000591 | -1.550223684 | 91.17840658 | 31.13351599 | 90 | 2 | 10 | 8 | 40 | 5 | 15 | 1 | 1 | 1 | 7 | 1 |
| SG0000003 | Ahdc1 | 0.000304 | 0.000609 | 1.625033835 | 468.0520407 | 1443.703706 | 112 | 10 | 10 | 31 | 68 | 13 | 8 | 0 | 0 | 4 | 8 | 7 |
| SG0000002 | Limk1 | 0.000318 | 0.000636 | -0.627357011 | 1179.309629 | 763.4393945 | 50 | 10 | 10 | 16 | 39 | 1 | 0 | 0 | 0 | 0 | 3 | 0 |
| ISG000000 | Gfod1 | 0.000328 | 0.000655 | -0.743632644 | 335.0188431 | 200.0845261 | 115 | 8 | 10 | 17 | 64 | 30 | 7 | 0 | 0 | 2 | 4 | 3 |
| SG0000002 | Pla2g12a | 0.000338 | 0.000673 | 0.579650548 | 411.7807358 | 615.4010419 | 103 | 7 | 10 | 34 | 46 | 3 | 4 | 1 | 1 | 1 | 4 | 0 |
| SG0000005 | Nfam1 | 0.000394 | 0.00078 | 6.359414946 | 0.615018556 | 50.49668551 | 63 | 10 | 10 | 13 | 39 | 8 | 5 | 0 | 0 | 0 | 0 | 0 |
| SG0000003 | Suv39h1 | 0.000471 | 0.000922 | -0.510999214 | 2008.768876 | 1409.625928 | 33 | 5 | 10 | 8 | 31 | 5 | 0 | 0 | 0 | 0 | 3 | 3 |
| SG0000003 | Tecta | 0.000575 | 0.001113 | -3.417803743 | 17.7506921 | 1.66093962 | 203 | 4 | 10 | 12 | 54 | 28 | 0 | 0 | 0 | 0 | 0 | 0 |
| ISG000000 | Dnajc15 | 0.000589 | 0.001138 | 0.65556632 | 456.182247 | 718.5938673 | 112 | 3 | 10 | 13 | 56 | 7 | 0 | 0 | 0 | 0 | 0 | 0 |
| SG0000000 | Gcdh | 0.000612 | 0.001178 | 0.50493336 | 729.14511 | 1034.699061 | 26 | 10 | 10 | 12 | 20 | 2 | 0 | 0 | 0 | 0 | 0 | 0 |
| ISG000000 | Gm4070 | 0.000699 | 0.001336 | -3.637450246 | 40.07367065 | 3.220160309 | 232 | 6 | 10 | 30 | 90 | 20 | 4 | 0 | 0 | 0 | 2 | 0 |
| SG0000002 | Plaa | 0.000965 | 0.001813 | 0.40993638 | 3966.516953 | 5270.022406 | 98 | 6 | 10 | 23 | 57 | 3 | 0 | 0 | 0 | 0 | 0 | 0 |
| SG0000002 | Sdsl | 0.001174 | 0.002188 | -3.463583081 | 32.04146979 | 2.904490964 | 28 | 10 | 10 | 12 | 30 | 14 | 0 | 0 | 0 | 0 | 0 | 0 |
| ISG000000 | Sesn2 | 0.002225 | 0.004003 | 0.452540966 | 1541.884155 | 2109.988802 | 29 | 9 | 10 | 10 | 18 | 0 | 1 | 0 | 0 | 0 | 0 | 0 |
| ISG000000 | Ubn1 | 0.002354 | 0.004218 | -0.4428053 | 1990.725195 | 1464.581808 | 72 | 6 | 10 | 12 | 38 | 6 | 2 | 0 | 1 | 0 | 0 | 0 |
| SG0000003 | Epn1 | 0.002545 | 0.004542 | -0.724349797 | 3414.65176 | 2066.785486 | 44 | 6 | 10 | 8 | 14 | 4 | 0 | 0 | 0 | 0 | 0 | 0 |
| SG0000004 | Ggact | 0.002718 | 0.004827 | -0.618286251 | 351.982337 | 229.2968892 | 184 | 6 | 10 | 28 | 86 | 8 | 2 | 0 | 0 | 0 | 0 | 0 |
| SG0000004 | Mcmbp | 0.002874 | 0.005083 | 0.383419966 | 2372.484771 | 3094.741254 | 149 | 4 | 10 | 24 | 90 | 19 | 3 | 0 | 0 | 0 | 2 | 1 |
| SG0000002 | Npc1 | 0.003194 | 0.005612 | -0.446402308 | 3134.650269 | 2300.427809 | 56 | 7 | 10 | 9 | 24 | 7 | 0 | 0 | 0 | 0 | 0 | 0 |
| ISG000000 | Atg3 | 0.003996 | 0.006924 | -0.458614067 | 984.1522386 | 716.1527996 | 97 | 3 | 10 | 19 | 46 | 9 | 2 | 0 | 0 | 0 | 0 | 0 |
| ISG000000 | Fam185a | 0.004339 | 0.007485 | -0.493937403 | 906.7086398 | 643.8397405 | 122 | 4 | 10 | 11 | 52 | 13 | 5 | 0 | 0 | 1 | 1 | 1 |
| ISG000000 | Slc1a6 | 0.00466 | 0.00801 | 2.156464736 | 6.696269176 | 29.85337754 | 81 | 5 | 10 | 14 | 43 | 6 | 0 | 0 | 0 | 0 | 0 | 0 |
| SG0000002 | Nrp2 | 0.004728 | 0.008119 | -0.428704627 | 3899.783886 | 2897.260786 | 802 | 4 | 10 | 102 | 393 | 68 | 22 | 0 | 0 | 4 | 15 | 4 |
| SG0000001 | Ube2a | 0.004874 | 0.008353 | 0.397527961 | 1039.477036 | 1369.24997 | 139 | 0 | 10 | 60 | 84 | 6 | 14 | 0 | 0 | 0 | 4 | 1 |
| ISG000000 | Bnip3 | 0.005864 | 0.009933 | 0.752641842 | 2255.640792 | 3800.473516 | 71 | 8 | 10 | 22 | 46 | 5 | 5 | 0 | 0 | 0 | 0 | 1 |
| ISG000000 | Zcchc14 | 0.00596 | 0.010086 | 0.380558332 | 1250.344748 | 1627.755784 | 49 | 7 | 10 | 10 | 24 | 4 | 5 | 0 | 0 | 0 | 1 | 1 |
| ISG000000 | Enpp3 | 0.006183 | 0.010445 | -0.684563878 | 1394.625533 | 867.7274793 | 130 | 6 | 10 | 15 | 54 | 15 | 0 | 0 | 0 | 0 | 0 | 0 |
| SG0000005 | Spock2 | 0.006647 | 0.011177 | 0.669509416 | 404.8448174 | 643.9186453 | 39 | 10 | 10 | 14 | 18 | 6 | 3 | 0 | 0 | 0 | 0 | 0 |
| SG0000003 | Neil3 | 0.006909 | 0.011583 | 0.526049264 | 259.195787 | 373.2368761 | 121 | 7 | 10 | 19 | 58 | 11 | 0 | 0 | 0 | 0 | 0 | 1 |
| SG0000003 | Casp4 | 0.007303 | 0.012198 | -1.232472265 | 54.62346776 | 23.24706593 | 155 | 8 | 10 | 22 | 97 | 13 | 3 | 1 | 1 | 1 | 3 | 2 |
| SG0000002 | Ppp3cc | 0.007485 | 0.012481 | 0.641950027 | 112.5242991 | 175.5870695 | 130 | 6 | 10 | 17 | 74 | 11 | 4 | 0 | 1 | 1 | 1 | 0 |
| ISG000000 | Hcn2 | 0.008081 | 0.013405 | 0.643565177 | 167.400718 | 261.5108661 | 32 | 10 | 10 | 14 | 20 | 2 | 0 | 0 | 0 | 0 | 0 | 0 |
| ISG000000 | Tmod3 | 0.009297 | 0.01528 | -0.370013468 | 3046.398404 | 2357.227758 | 106 | 6 | 10 | 17 | 56 | 10 | 2 | 0 | 0 | 0 | 2 | 1 |
| SG0000002 | Tnfrsf1b | 0.009576 | 0.015708 | -0.442989267 | 693.2081803 | 509.9300693 | 61 | 4 | 10 | 12 | 22 | 3 | 2 | 0 | 0 | 0 | 0 | 0 |
| ISG000000 | Ift43 | 0.009804 | 0.016056 | -0.478326175 | 440.5530845 | 316.2333872 | 70 | 8 | 10 | 17 | 36 | 8 | 1 | 0 | 0 | 1 | 1 | 0 |
| ISG000000 | Map3k14 | 0.009846 | 0.01612 | -0.743546049 | 125.388462 | 74.8907037 | 57 | 5 | 10 | 13 | 29 | 7 | 0 | 0 | 0 | 0 | 0 | 0 |
| SG0000002 | Eif2b4 | 0.012277 | 0.019763 | 0.376325252 | 799.915994 | 1038.316055 | 45 | 10 | 10 | 15 | 25 | 6 | 0 | 0 | 0 | 0 | 0 | 1 |
| SG0000002 | Zfyve16 | 0.012445 | 0.020007 | -0.436843525 | 944.0472488 | 697.4141048 | 137 | 7 | 10 | 22 | 68 | 9 | 10 | 0 | 1 | 1 | 3 | 0 |
| ISG000000 | Slc38a9 | 0.012662 | 0.02032 | 0.487819868 | 260.9187308 | 365.8926369 | 127 | 2 | 10 | 18 | 79 | 10 | 9 | 0 | 0 | 2 | 4 | 0 |
| ISG000000 | Zfp160 | 0.012806 | 0.02053 | -0.556327537 | 516.6234149 | 351.3199479 | 54 | 9 | 10 | 15 | 51 | 6 | 6 | 0 | 0 | 1 | 1 | 0 |
| SG0000003 | Commd2 | 0.013372 | 0.021355 | 0.410307614 | 435.5566641 | 578.8413691 | 46 | 6 | 10 | 13 | 15 | 2 | 6 | 0 | 0 | 1 | 1 | 0 |
| SG0000002 | Itga4 | 0.014498 | 0.023021 | -1.282668492 | 37.32199823 | 15.34063007 | 173 | 4 | 10 | 22 | 90 | 13 | 16 | 0 | 1 | 2 | 8 | 3 |
| SG0000003 | Ip6k2 | 0.014653 | 0.023256 | -0.375519463 | 1118.037157 | 861.8141392 | 102 | 6 | 10 | 14 | 30 | 16 | 2 | 2 | 2 | 2 | 2 | 0 |
| SG0000003 | Zfp653 | 0.016517 | 0.025977 | 0.630377328 | 93.82749039 | 145.2421218 | 42 | 8 | 10 | 12 | 20 | 8 | 1 | 0 | 0 | 1 | 1 | 0 |
| ISG000000 | Abcd2 | 0.017395 | 0.027236 | -1.347954281 | 31.87985796 | 12.52396099 | 120 | 5 | 10 | 17 | 71 | 6 | 11 | 0 | 1 | 0 | 1 | 0 |
| ISG000000 | Pdia5 | 0.018831 | 0.029264 | 0.379030235 | 502.4574081 | 653.4294795 | 98 | 6 | 10 | 12 | 45 | 12 | 0 | 0 | 0 | 0 | 0 | 0 |
| SG0000003 | Jam3 | 0.019177 | 0.029758 | 0.430021923 | 257.0847248 | 346.3584365 | 101 | 5 | 10 | 15 | 46 | 14 | 1 | 0 | 0 | 0 | 1 | 1 |
| SG0000006 | Zbtb18 | 0.019881 | 0.030773 | -0.37466471 | 935.9771919 | 721.9048337 | 40 | 4 | 10 | 4 | 28 | 0 | 16 | 0 | 4 | 0 | 12 | 0 |
| SG0000006 | Ifi27 | 0.022447 | 0.034426 | -1.124428145 | 2040.695171 | 936.0338124 | 40 | 5 | 10 | 5 | 10 | 0 | 0 | 0 | 0 | 0 | 0 | 0 |
| ISG000000 | Ttc5 | 0.022524 | 0.034533 | 0.340244768 | 1729.059879 | 2188.940287 | 37 | 7 | 10 | 12 | 27 | 2 | 2 | 0 | 0 | 1 | 1 | 1 |
| SG0000000 | Zfp276 | 0.02531 | 0.038469 | -0.432874511 | 331.7023386 | 245.719895 | 20 | 10 | 10 | 10 | 17 | 0 | 1 | 0 | 0 | 0 | 0 | 0 |
| SG0000005 | Atxn7l3 | 0.02567 | 0.038973 | -0.309334566 | 2553.063463 | 2060.357712 | 24 | 10 | 10 | 15 | 20 | 5 | 10 | 5 | 5 | 10 | 15 | 5 |
| SG0000002 | Eftud2 | 0.02691 | 0.040742 | 0.277938849 | 6532.572492 | 7920.492155 | 109 | 10 | 10 | 12 | 34 | 9 | 0 | 0 | 0 | 0 | 0 | 0 |
| SG0000002 | Smim8 | 0.027562 | 0.041665 | 0.525186153 | 115.324505 | 165.9657162 | 95 | 5 | 10 | 25 | 95 | 15 | 0 | 0 | 0 | 0 | 5 | 0 |
| ISG000000 | Dcaf13 | 0.028295 | 0.042676 | -0.316022305 | 2379.26799 | 1911.222144 | 75 | 7 | 10 | 13 | 46 | 3 | 0 | 0 | 0 | 0 | 0 | 1 |
| SG0000002 | Atp2a3 | 0.029424 | 0.044233 | -0.960380053 | 615.8989893 | 316.5237398 | 130 | 5 | 10 | 5 | 55 | 15 | 0 | 0 | 0 | 0 | 5 | 0 |
| ISG000000 | Ranbp2 | 0.031217 | 0.046706 | -0.299678233 | 7236.081761 | 5878.835771 | 136 | 5 | 10 | 19 | 61 | 14 | 0 | 0 | 0 | 0 | 0 | 0 |
| SG0000007 | Tbpl1 | 0.032317 | 0.048238 | -0.326699194 | 1329.729358 | 1060.271286 | 102 | 6 | 10 | 18 | 44 | 17 | 4 | 0 | 0 | 2 | 2 | 3 |
| SG0000003 | Fbxl2 | 0.032321 | 0.04824 | -0.522846123 | 257.873576 | 179.4793453 | 123 | 5 | 10 | 22 | 54 | 11 | 4 | 0 | 0 | 0 | 0 | 0 |
| SG0000001 | Trnt1 | 0.034383 | 0.051042 | -0.303779934 | 2085.617992 | 1689.615529 | 110 | 3 | 10 | 20 | 54 | 6 | 2 | 0 | 0 | 0 | 6 | 3 |
| SG0000000 | Arhgef40 | 0.035775 | 0.052942 | -0.425938179 | 389.3167437 | 289.7896792 | 37 | 8 | 10 | 13 | 29 | 6 | 3 | 0 | 0 | 1 | 3 | 0 |
| ISG000000 | Ppp2ca | 0.041693 | 0.061018 | -0.27079814 | 9305.947726 | 7713.335926 | 62 | 7 | 10 | 12 | 26 | 8 | 6 | 0 | 0 | 0 | 1 | 2 |
| SG0000001 | Dhx40 | 0.042023 | 0.061481 | -0.280928114 | 3126.464419 | 2573.272043 | 83 | 8 | 10 | 14 | 33 | 3 | 4 | 0 | 0 | 0 | 1 | 0 |
| SG0000003 | Plscr1 | 0.04935 | 0.071333 | -0.317928344 | 821.4380882 | 658.9748929 | 104 | 10 | 10 | 21 | 49 | 14 | 10 | 0 | 0 | 1 | 3 | 0 |
| ISG000000 | Fam228b | 0.049883 | 0.072027 | 1.100343076 | 11.87070635 | 25.45146778 | 67 | 7 | 10 | 16 | 42 | 6 | 7 | 1 | 2 | 2 | 3 | 0 |
| SG0000002 | Prdx1 | 0.05034 | 0.072617 | 0.241861328 | 26327.89501 | 31133.19214 | 65 | 8 | 10 | 12 | 30 | 4 | 3 | 0 | 0 | 0 | 0 | 0 |
| SG0000005 | Tmem39b | 0.064339 | 0.091008 | -0.381152679 | 353.2910332 | 271.2652893 | 37 | 8 | 10 | 10 | 50 | 8 | 0 | 0 | 0 | 0 | 2 | 0 |
| SG0000007 | Ttc9c | 0.067246 | 0.094806 | -0.256538946 | 2540.399584 | 2126.552531 | 84 | 9 | 10 | 15 | 31 | 3 | 9 | 0 | 1 | 0 | 1 | 0 |
| SG0000004 | Mars | 0.068059 | 0.095801 | 0.235213611 | 4147.631165 | 4882.097871 | 36 | 4 | 10 | 10 | 30 | 9 | 0 | 0 | 0 | 0 | 0 | 0 |
| ISG000000 | Rspo3 | 0.070399 | 0.098845 | 1.045926779 | 31.97407826 | 66.01663252 | 211 | 5 | 10 | 25 | 78 | 18 | 0 | 0 | 0 | 0 | 1 | 1 |
| ISG000000 | Tbca | 0.073873 | 0.103319 | 0.246577253 | 1056.927988 | 1253.927849 | 146 | 3 | 10 | 15 | 54 | 13 | 1 | 0 | 0 | 0 | 0 | 0 |
| SG0000002 | Slc1a4 | 0.083381 | 0.115409 | 0.265874133 | 662.234786 | 796.2474948 | 54 | 4 | 10 | 6 | 22 | 6 | 2 | 0 | 0 | 0 | 2 | 0 |
| ISG000000 | Gpatch2l | 0.09434 | 0.129141 | -0.273722237 | 715.7520778 | 592.0576726 | 78 | 5 | 10 | 18 | 55 | 11 | 7 | 0 | 1 | 2 | 2 | 1 |
| SG0000002 | Akr1a1 | 0.096885 | 0.132301 | -0.217334134 | 8259.900514 | 7104.78145 | 36 | 8 | 10 | 12 | 20 | 3 | 0 | 0 | 0 | 0 | 0 | 0 |
| SG0000003 | Trappc11 | 0.099229 | 0.135142 | -0.229767753 | 2738.122681 | 2334.994795 | 93 | 6 | 10 | 19 | 56 | 12 | 3 | 0 | 0 | 1 | 3 | 1 |
| SG0000002 | Psmd9 | 0.099818 | 0.135789 | -0.288608996 | 665.8670514 | 545.139386 | 42 | 8 | 10 | 11 | 38 | 7 | 2 | 0 | 0 | 0 | 2 | 0 |
| ISG000000 | Aggf1 | 0.100416 | 0.1365 | 0.2223096 | 1789.23493 | 2087.320921 | 63 | 6 | 10 | 11 | 26 | 1 | 2 | 0 | 0 | 0 | 0 | 0 |
| SG0000003 | Plscr2 | 0.100416 | 0.1365 | -1.265872857 | 16.9027832 | 7.028983445 | 157 | 4 | 10 | 20 | 74 | 18 | 6 | 0 | 0 | 0 | 6 | 0 |
| SG0000004 | Ppp1r15a | 0.108774 | 0.146947 | -0.256119169 | 868.1286439 | 726.9165114 | 16 | 10 | 10 | 10 | 12 | 0 | 8 | 8 | 8 | 8 | 10 | 0 |
| ISG000000 | Pabpc1l | 0.115829 | 0.155422 | 0.88396957 | 13.67226557 | 25.23142862 | 39 | 8 | 10 | 12 | 26 | 2 | 0 | 0 | 0 | 0 | 0 | 0 |
| SG0000002 | Slc12a2 | 0.119021 | 0.159252 | 0.263239737 | 924.08889 | 1109.064504 | 198 | 4 | 10 | 27 | 83 | 17 | 9 | 0 | 1 | 1 | 5 | 0 |
| SG0000002 | Tnfrsf9 | 0.121475 | 0.162305 | 0.701493459 | 30.75706212 | 50.01674471 | 98 | 0 | 10 | 17 | 52 | 55 | 8 | 0 | 0 | 0 | 0 | 0 |
| SG0000003 | Pgap3 | 0.126723 | 0.168612 | -0.394704575 | 140.712052 | 107.0319358 | 24 | 8 | 10 | 10 | 18 | 0 | 0 | 0 | 0 | 0 | 0 | 0 |
| ISG000000 | Zw10 | 0.138097 | 0.182324 | 0.206908526 | 1148.688725 | 1325.830577 | 51 | 8 | 10 | 11 | 23 | 2 | 1 | 0 | 0 | 0 | 0 | 0 |
| SG0000002 | Ap3m1 | 0.138385 | 0.182651 | 0.193619875 | 2444.124743 | 2795.17342 | 79 | 9 | 10 | 20 | 56 | 3 | 14 | 0 | 1 | 2 | 5 | 0 |
| ISG000000 | Wdr46 | 0.147503 | 0.193606 | -0.219244916 | 1514.572708 | 1301.040544 | 13 | 10 | 10 | 11 | 15 | 1 | 0 | 0 | 0 | 0 | 0 | 0 |
| SG0000006 | Zfp467 | 0.152218 | 0.199221 | -0.782144094 | 46.88486792 | 27.26364158 | 88 | 10 | 10 | 34 | 59 | 9 | 12 | 0 | 0 | 4 | 4 | 0 |
| ISG000000 | Vps36 | 0.160905 | 0.20961 | 0.182917414 | 2198.735612 | 2495.954354 | 58 | 7 | 10 | 10 | 36 | 7 | 3 | 0 | 0 | 0 | 0 | 2 |
| ISG000000 | Grem1 | 0.166213 | 0.215865 | 0.311180507 | 830.8200012 | 1030.817089 | 28 | 6 | 10 | 11 | 15 | 3 | 7 | 0 | 1 | 0 | 0 | 1 |
| SG0000000 | Pex5 | 0.17266 | 0.223492 | 0.185252314 | 1722.132665 | 1958.092066 | 120 | 10 | 10 | 25 | 80 | 30 | 10 | 0 | 0 | 0 | 0 | 0 |
| SG0000003 | Gtf2a2 | 0.197242 | 0.251709 | 0.183035996 | 985.9786898 | 1119.352448 | 115 | 10 | 10 | 15 | 55 | 5 | 5 | 5 | 5 | 5 | 5 | 0 |
| SG0000006 | Arf2 | 0.208735 | 0.264827 | 0.208695037 | 462.3469218 | 534.3077044 | 70 | 6 | 10 | 10 | 28 | 6 | 2 | 0 | 0 | 0 | 0 | 0 |
| SG0000002 | Hdac3 | 0.213832 | 0.27051 | 0.161984824 | 1938.982149 | 2169.382346 | 46 | 7 | 10 | 14 | 30 | 2 | 0 | 0 | 0 | 0 | 0 | 0 |
| ISG000000 | Emp2 | 0.253909 | 0.314942 | -0.293461858 | 278.758812 | 227.4509647 | 44 | 8 | 10 | 12 | 40 | 4 | 1 | 0 | 0 | 0 | 0 | 1 |
| SG0000003 | Arcn1 | 0.257233 | 0.318576 | 0.140280657 | 7286.009889 | 8030.053556 | 66 | 10 | 10 | 20 | 35 | 5 | 3 | 0 | 0 | 0 | 1 | 0 |
| ISG000000 | Cdr2l | 0.283502 | 0.347077 | 0.166614278 | 595.6017451 | 668.5160784 | 19 | 10 | 10 | 11 | 19 | 1 | 1 | 0 | 0 | 0 | 0 | 0 |
| ISG000000 | Cep76 | 0.304068 | 0.369127 | 0.151857695 | 706.0836016 | 784.4582713 | 64 | 6 | 10 | 14 | 36 | 8 | 10 | 4 | 4 | 5 | 7 | 1 |
| SG0000002 | Arl6 | 0.315407 | 0.381135 | 0.159122023 | 556.3457046 | 621.2197312 | 171 | 7 | 10 | 22 | 80 | 16 | 8 | 0 | 0 | 0 | 3 | 0 |
| SG0000002 | Fzr1 | 0.34037 | 0.407507 | -0.223060915 | 2392.501867 | 2049.765877 | 24 | 10 | 10 | 10 | 24 | 1 | 0 | 0 | 0 | 0 | 0 | 1 |
| SG0000002 | Slc25a25 | 0.371986 | 0.44053 | 0.150226165 | 469.7377874 | 521.2884226 | 144 | 10 | 10 | 34 | 74 | 8 | 14 | 0 | 0 | 14 | 19 | 0 |
| SG0000002 | Dclre1c | 0.377154 | 0.445913 | 0.170107335 | 276.5879833 | 311.2008095 | 278 | 5 | 10 | 41 | 80 | 21 | 24 | 1 | 3 | 6 | 17 | 7 |
| SG0000002 | Tyro3 | 0.379685 | 0.448491 | -0.18109141 | 1151.477412 | 1015.643928 | 37 | 6 | 10 | 10 | 16 | 2 | 0 | 0 | 0 | 0 | 0 | 0 |
| ISG000000 | Etnk1 | 0.43825 | 0.507463 | -0.136531771 | 8510.346132 | 7741.891232 | 88 | 7 | 10 | 12 | 24 | 9 | 21 | 2 | 2 | 3 | 4 | 3 |
| ISG000000 | Ppme1 | 0.462786 | 0.53226 | -0.108073923 | 1903.12924 | 1765.772665 | 111 | 6 | 10 | 18 | 47 | 7 | 2 | 0 | 0 | 1 | 1 | 0 |
| SG0000002 | Amz2 | 0.467388 | 0.536383 | 0.099829618 | 999.3056336 | 1070.902778 | 73 | 10 | 10 | 11 | 32 | 8 | 3 | 1 | 1 | 1 | 2 | 0 |
| ISG000000 | Ddx21 | 0.473135 | 0.541971 | 0.085818587 | 10842.35167 | 11506.87667 | 43 | 10 | 10 | 13 | 25 | 2 | 5 | 0 | 0 | 1 | 2 | 1 |
| ISG000000 | Bloc1s5 | 0.495521 | 0.563135 | -0.153496675 | 247.764929 | 222.7577018 | 46 | 6 | 10 | 7 | 20 | 7 | 0 | 0 | 0 | 0 | 0 | 0 |
| SG0000002 | Cdk2ap1 | 0.529844 | 0.596366 | 0.114598181 | 372.3645833 | 403.1492508 | 38 | 10 | 10 | 13 | 28 | 7 | 1 | 0 | 0 | 1 | 1 | 0 |
| ISG000000 | D6Wsu163e | 0.532268 | 0.598834 | -0.11391597 | 444.6565997 | 410.8966672 | 69 | 5 | 10 | 16 | 40 | 10 | 2 | 0 | 0 | 0 | 0 | 0 |
| ISG000000 | Arntl | 0.547493 | 0.613302 | 0.098026503 | 387.3525322 | 414.5866535 | 135 | 6 | 10 | 21 | 55 | 13 | 0 | 0 | 0 | 0 | 0 | 0 |
| SG0000004 | Ythdf3 | 0.550515 | 0.61607 | 0.073272974 | 3155.522952 | 3319.928316 | 204 | 6 | 10 | 20 | 92 | 10 | 18 | 0 | 0 | 2 | 10 | 0 |
| SG0000005 | Zfp516 | 0.567538 | 0.632159 | -0.134536694 | 1996.469496 | 1818.708508 | 114 | 5 | 10 | 17 | 45 | 16 | 9 | 2 | 2 | 2 | 11 | 1 |

| | | | | | | | | | | | | | | | | | | |
|---|---|---|---|---|---|---|---|---|---|---|---|---|---|---|---|---|---|---|
| SG0000000 | Smarcb1 | 0.647936 | 0.707457 | -0.05980959 | 2690.97878 | 2581.700105 | 40 | 10 | 10 | 10 | 28 | 8 | 0 | 0 | 0 | 0 | 0 | 0 |
| SG0000002 | Aldh1l2 | 0.660414 | 0.718738 | 0.160627343 | 99.24941675 | 110.9383269 | 136 | 4 | 10 | 32 | 80 | 10 | 8 | 0 | 0 | 0 | 2 | 0 |
| SG0000004 | E130311K13Rik | 0.697328 | 0.751582 | -0.124359329 | 90.97894783 | 83.46511993 | 42 | 8 | 10 | 11 | 25 | 3 | 1 | 0 | 0 | 0 | 0 | 1 |
| SG0000003 | Smad1 | 0.717533 | 0.770022 | -0.066639058 | 632.0918987 | 603.5591924 | 187 | 6 | 10 | 18 | 78 | 9 | 5 | 0 | 0 | 0 | 0 | 2 |
| SG0000002 | Xrcc2 | 0.74657 | 0.795134 | -0.061783316 | 577.8328419 | 553.6095719 | 60 | 6 | 10 | 12 | 18 | 6 | 5 | 0 | 0 | 0 | 1 | 0 |
| JSG000000 | Ecscr | 0.75071 | 0.798878 | -0.124820056 | 84.25120632 | 77.26833374 | 38 | 10 | 10 | 14 | 24 | 7 | 0 | 0 | 0 | 0 | 0 | 0 |
| SG0000002 | Mospd1 | 0.789967 | 0.832739 | 0.048851931 | 272.7675448 | 282.1620431 | 82 | 10 | 10 | 16 | 56 | 6 | 8 | 2 | 2 | 2 | 2 | 0 |
| JSG000000 | 2510009E07Rik | 0.827643 | 0.864668 | 0.082541868 | 42.61105722 | 45.12008277 | 88 | 7 | 10 | 12 | 34 | 8 | 5 | 0 | 0 | 0 | 2 | 0 |
| SG0000001 | Kpna2 | 0.832861 | 0.869359 | 0.024624181 | 4896.584954 | 4980.878074 | 49 | 7 | 10 | 10 | 29 | 14 | 2 | 0 | 0 | 0 | 2 | 4 |
| SG0000001 | Kansl3 | 0.941218 | 0.957849 | 0.009632865 | 2464.391227 | 2480.901007 | 229 | 10 | 10 | 25 | 112 | 22 | 5 | 0 | 0 | 0 | 0 | 0 |
| SG0000004 | Meis3 | 0.957107 | 0.972031 | -0.005268388 | 116.8609014 | 116.4349306 | 17 | 10 | 10 | 12 | 13 | 0 | 0 | 0 | 0 | 0 | 0 | 0 |
| SG0000002 | Ppp1r13b | 0.973491 | 0.986209 | -0.012837085 | 388.8038949 | 385.3596681 | 108 | 5 | 10 | 15 | 50 | 9 | 0 | 0 | 0 | 0 | 0 | 0 |
| JSG000000 | Smc6 | 0.982881 | 0.99398 | -0.013243703 | 2971.942908 | 2944.785801 | 139 | 5 | 10 | 23 | 58 | 14 | 3 | 0 | 0 | 1 | 2 | 3 |
| SG0000002 | Il33 | 0.98857 | 0.998436 | -0.019281542 | 48.91666764 | 48.26724839 | 99 | 6 | 10 | 13 | 54 | 8 | 6 | 0 | 0 | 0 | 2 | 0 |
| SG0000002 | Zscan10 | 4.2E-271 | 2.4E-268 | 7.664074904 | 8.090261687 | 1640.888248 | 60 | 6 | 9 | 12 | 24 | 6 | 0 | 0 | 0 | 0 | 0 | 2 |
| SG0000005 | Phgdh | 3.3E-199 | 1E-196 | 4.852698401 | 95.1923977 | 2750.489836 | 30 | 8 | 9 | 9 | 15 | 1 | 0 | 0 | 0 | 0 | 0 | 0 |
| SG0000003 | Ano1 | 7.6E-181 | 1.8E-178 | -7.577295692 | 12617.31602 | 66.06534257 | 372 | 6 | 9 | 36 | 129 | 24 | 0 | 0 | 0 | 0 | 0 | 3 |
| SG0000002 | Ildr1 | 8.4E-166 | 1.7E-163 | 9.439821051 | 0.946619625 | 657.4228486 | 120 | 3 | 9 | 6 | 27 | 12 | 16 | 0 | 0 | 0 | 3 | 0 |
| SG0000003 | Usp11 | 3.2E-157 | 5.5E-155 | 3.689239211 | 218.8657904 | 2823.251376 | 33 | 7 | 9 | 13 | 21 | 5 | 1 | 0 | 0 | 1 | 1 | 0 |
| SG0000002 | Pkp1 | 1E-103 | 7.7E-102 | -8.317890705 | 2742.762463 | 8.595152656 | 109 | 6 | 9 | 9 | 24 | 18 | 1 | 0 | 0 | 0 | 0 | 0 |
| SG0000003 | Tead2 | 2.79E-90 | 1.58E-88 | 2.8998884 | 470.2930577 | 3510.119971 | 36 | 9 | 9 | 15 | 21 | 0 | 6 | 0 | 0 | 3 | 3 | 0 |
| JSG000000 | Klf9 | 2.07E-85 | 1.04E-83 | 2.571733282 | 478.0716513 | 2842.24722 | 48 | 6 | 9 | 12 | 28 | 4 | 5 | 0 | 0 | 0 | 0 | 1 |
| SG0000006 | Jakmip1 | 1.2E-74 | 4.89E-73 | 7.858003724 | 0 | 231.0036551 | 76 | 4 | 9 | 14 | 34 | 6 | 1 | 0 | 0 | 1 | 1 | 0 |
| SG0000000 | Lbr | 1.45E-72 | 5.6E-71 | 2.148641141 | 1928.650082 | 8551.826344 | 51 | 9 | 9 | 13 | 24 | 1 | 3 | 0 | 0 | 0 | 1 | 0 |
| SG0000003 | Slc7a3 | 9.63E-72 | 3.61E-70 | -3.196007579 | 17109.82111 | 1867.030107 | 15 | 9 | 9 | 9 | 15 | 0 | 0 | 0 | 0 | 0 | 0 | 0 |
| JSG000000 | Cpn1 | 3.39E-68 | 1.16E-66 | -4.505406233 | 3433.890298 | 151.1903241 | 33 | 5 | 9 | 10 | 16 | 5 | 1 | 0 | 0 | 0 | 1 | 0 |
| SG0000002 | Wscd1 | 5.28E-66 | 1.7E-64 | 7.649485088 | 0 | 199.7818553 | 126 | 6 | 9 | 24 | 45 | 24 | 0 | 0 | 0 | 0 | 0 | 0 |
| JSG000000 | Il13ra1 | 8.13E-63 | 2.44E-61 | -4.377672712 | 4625.894188 | 222.528119 | 135 | 3 | 9 | 16 | 61 | 10 | 5 | 0 | 0 | 0 | 1 | 0 |
| SG0000003 | Urb2 | 2.87E-61 | 8.28E-60 | 2.035104382 | 1202.528738 | 4928.592674 | 78 | 6 | 9 | 18 | 43 | 12 | 3 | 0 | 0 | 0 | 4 | 2 |
| SG0000002 | Prr13 | 2.68E-58 | 7.04E-57 | -3.588439722 | 11196.52575 | 930.7976744 | 23 | 9 | 9 | 9 | 16 | 0 | 4 | 0 | 0 | 0 | 0 | 0 |
| JSG000000 | Gypc | 4.99E-54 | 1.16E-52 | -3.725324967 | 835.3770418 | 63.1608516 | 63 | 0 | 9 | 9 | 67 | 4 | 8 | 0 | 0 | 0 | 0 | 0 |
| JSG000000 | Patl2 | 1.85E-49 | 3.78E-48 | 7.479774637 | 0.970711415 | 173.2713117 | 124 | 6 | 9 | 19 | 62 | 18 | 1 | 0 | 0 | 0 | 0 | 0 |
| SG0000000 | Fbln1 | 2.27E-48 | 4.5E-47 | 4.751633622 | 82.89456685 | 2233.111023 | 96 | 5 | 9 | 19 | 45 | 15 | 1 | 0 | 0 | 1 | 1 | 0 |
| SG0000004 | Cchcr1 | 4.45E-48 | 8.73E-47 | -2.615574635 | 2995.018343 | 488.689565 | 36 | 6 | 9 | 15 | 18 | 0 | 0 | 0 | 0 | 0 | 0 | 0 |
| SG0000000 | Zfp184 | 1.26E-46 | 2.35E-45 | 2.86105108 | 65.77695126 | 477.8987939 | 83 | 9 | 9 | 26 | 62 | 0 | 2 | 0 | 0 | 2 | 2 | 0 |
| JSG000000 | Efr3b | 2.78E-45 | 4.92E-44 | 3.385515137 | 39.39859754 | 411.7393453 | 104 | 8 | 9 | 13 | 36 | 13 | 6 | 0 | 0 | 1 | 1 | 1 |
| JSG000000 | Swap70 | 6.51E-44 | 1.1E-42 | -2.278621514 | 8660.091428 | 1784.797926 | 119 | 5 | 9 | 22 | 61 | 8 | 2 | 0 | 1 | 1 | 1 | 0 |
| JSG000000 | Xrcc5 | 3.67E-42 | 5.83E-41 | 1.643177388 | 2027.460141 | 6332.832585 | 134 | 6 | 9 | 19 | 67 | 15 | 1 | 0 | 0 | 0 | 0 | 0 |
| SG0000002 | Pask | 1.31E-40 | 1.98E-39 | 1.948197085 | 624.2049301 | 2408.756805 | 61 | 7 | 9 | 15 | 30 | 7 | 3 | 1 | 1 | 2 | 2 | 0 |
| JSG000000 | Fkbp3 | 4.88E-39 | 6.97E-38 | 1.729845545 | 1878.529569 | 6230.938035 | 45 | 7 | 9 | 10 | 20 | 1 | 1 | 0 | 0 | 0 | 0 | 0 |
| SG0000004 | Kank3 | 6.78E-39 | 9.59E-38 | 2.281250335 | 227.9266194 | 1107.949148 | 40 | 6 | 9 | 12 | 42 | 3 | 1 | 0 | 0 | 0 | 0 | 0 |
| SG0000002 | Atp6v1c2 | 1.03E-38 | 1.45E-37 | -6.272207879 | 438.3523886 | 5.671535656 | 121 | 9 | 9 | 17 | 53 | 14 | 4 | 0 | 0 | 4 | 4 | 0 |
| SG0000002 | Tpmt | 1.08E-36 | 1.42E-35 | 2.689823962 | 58.87941721 | 379.9104139 | 90 | 9 | 9 | 9 | 60 | 6 | 0 | 0 | 0 | 0 | 0 | 0 |
| JSG000000 | Nid2 | 8.95E-35 | 1.1E-33 | 6.350806085 | 138.9830084 | 11343.44152 | 82 | 3 | 9 | 11 | 25 | 6 | 2 | 0 | 2 | 1 | 1 | 0 |
| SG0000004 | Spice1 | 2.79E-34 | 3.38E-33 | 1.738489268 | 419.3456653 | 1399.295943 | 97 | 4 | 9 | 18 | 44 | 6 | 2 | 0 | 0 | 1 | 3 | 0 |
| SG0000004 | Nsmce4a | 9.95E-34 | 1.17E-32 | 1.551672518 | 1014.414579 | 2973.82531 | 30 | 7 | 9 | 8 | 25 | 5 | 0 | 0 | 0 | 0 | 0 | 1 |
| JSG000000 | Hs3st3a1 | 4.57E-33 | 5.23E-32 | -7.735580809 | 1764.778437 | 8.280343078 | 144 | 6 | 9 | 21 | 74 | 9 | 3 | 0 | 0 | 0 | 1 | 0 |
| SG0000000 | Fkbp10 | 7.13E-33 | 8.09E-32 | -2.063316454 | 2359.025307 | 564.4330774 | 24 | 9 | 9 | 12 | 24 | 0 | 0 | 0 | 0 | 0 | 0 | 0 |
| SG0000004 | Zmpste24 | 7.5E-33 | 8.49E-32 | -1.961460363 | 3640.638445 | 934.8009524 | 72 | 6 | 9 | 13 | 36 | 5 | 7 | 0 | 0 | 1 | 2 | 0 |
| JSG000000 | Trib2 | 1.27E-32 | 1.43E-31 | -2.419754819 | 1138.232419 | 212.7218828 | 43 | 9 | 9 | 12 | 21 | 2 | 6 | 0 | 0 | 1 | 2 | 0 |
| SG0000003 | Elf4 | 3.74E-32 | 4.14E-31 | -2.843811638 | 3065.411532 | 426.9885033 | 98 | 9 | 9 | 18 | 39 | 12 | 2 | 0 | 0 | 0 | 0 | 2 |
| JSG000000 | Rufy1 | 2.58E-31 | 2.76E-30 | -2.122327667 | 1476.339756 | 339.0798845 | 46 | 6 | 9 | 9 | 19 | 5 | 0 | 0 | 0 | 0 | 0 | 0 |
| JSG000000 | Phf2 | 5.01E-31 | 5.29E-30 | -2.1813176 | 1271.045837 | 280.2329412 | 74 | 7 | 9 | 15 | 31 | 14 | 1 | 0 | 0 | 0 | 2 | 1 |
| JSG000000 | Plbd1 | 6.76E-31 | 7.1E-30 | 6.50099616 | 0 | 89.57218517 | 67 | 5 | 9 | 12 | 44 | 8 | 0 | 0 | 0 | 0 | 0 | 0 |
| SG0000003 | Nek3 | 8.28E-31 | 8.64E-30 | 2.243741862 | 93.81830373 | 444.3459021 | 221 | 6 | 9 | 24 | 78 | 30 | 0 | 0 | 0 | 0 | 0 | 0 |
| SG0000002 | Tapbp | 1.41E-30 | 1.46E-29 | -2.483232923 | 7133.870541 | 1275.844154 | 15 | 8 | 9 | 9 | 11 | 1 | 0 | 0 | 0 | 0 | 0 | 0 |
| SG0000007 | Ttpa | 1.56E-30 | 1.61E-29 | 2.645568321 | 53.2927262 | 333.4750585 | 168 | 3 | 9 | 15 | 71 | 3 | 21 | 0 | 3 | 3 | 8 | 0 |
| SG0000004 | Zfp454 | 9.3E-30 | 9.29E-29 | 4.746539375 | 4.495520649 | 120.6787281 | 81 | 3 | 9 | 3 | 18 | 0 | 3 | 0 | 0 | 0 | 0 | 0 |
| SG0000004 | Ggnbp1 | 2.68E-29 | 2.62E-28 | 2.583478409 | 49.90074987 | 299.0966619 | 79 | 7 | 9 | 13 | 48 | 3 | 0 | 0 | 0 | 0 | 0 | 0 |
| SG0000002 | Fh1 | 3.09E-28 | 2.88E-27 | 1.35979575 | 1854.112746 | 4758.558887 | 82 | 3 | 9 | 12 | 40 | 12 | 6 | 0 | 0 | 0 | 2 | 0 |
| SG0000003 | Arhgef4 | 1.12E-27 | 1.02E-26 | 2.623645711 | 42.72313877 | 263.3050623 | 162 | 6 | 9 | 24 | 88 | 26 | 4 | 0 | 0 | 0 | 2 | 0 |
| SG0000004 | Atp1a3 | 1.74E-27 | 1.57E-26 | 6.908178172 | 11.47770402 | 1378.554266 | 30 | 6 | 9 | 9 | 45 | 3 | 0 | 0 | 0 | 0 | 0 | 0 |
| JSG000000 | Mfsd7c | 2.7E-27 | 2.42E-26 | 4.442309063 | 5.50972851 | 119.7839079 | 96 | 6 | 9 | 14 | 48 | 8 | 2 | 0 | 1 | 0 | 0 | 0 |
| SG0000003 | Amfr | 1.77E-26 | 1.53E-25 | -1.876469232 | 4528.328243 | 1233.287629 | 88 | 6 | 9 | 15 | 38 | 7 | 2 | 0 | 0 | 0 | 1 | 0 |
| SG0000003 | Dusp11 | 3.71E-25 | 3.02E-24 | -1.582511349 | 6741.675073 | 2251.046325 | 72 | 5 | 9 | 12 | 34 | 10 | 18 | 3 | 5 | 4 | 6 | 0 |
| JSG000000 | Slc44a3 | 1.74E-24 | 1.37E-23 | 5.883865908 | 1.348152497 | 79.60844651 | 111 | 4 | 9 | 12 | 43 | 8 | 1 | 0 | 0 | 0 | 0 | 0 |
| SG0000003 | Igsf10 | 2.04E-24 | 1.6E-23 | 3.495110072 | 17.91080409 | 201.9519511 | 168 | 3 | 9 | 34 | 82 | 18 | 12 | 0 | 0 | 1 | 1 | 3 |
| SG0000003 | Gmip | 5.12E-24 | 3.94E-23 | -2.089256553 | 849.547685 | 199.6451984 | 28 | 7 | 9 | 9 | 11 | 0 | 6 | 2 | 2 | 2 | 2 | 0 |
| SG0000006 | Tmprss11d | 1.57E-22 | 1.13E-21 | -6.529141854 | 180.2239682 | 1.951394174 | 226 | 4 | 9 | 25 | 113 | 17 | 8 | 0 | 0 | 0 | 2 | 0 |
| JSG000000 | Neurod4 | 1.59E-21 | 1.09E-20 | -8.211106127 | 106.68516 | 0.360010171 | 38 | 4 | 9 | 11 | 25 | 1 | 15 | 2 | 5 | 6 | 8 | 0 |
| JSG000000 | Bzw2 | 1.71E-21 | 1.17E-20 | 1.175770523 | 2154.902392 | 4868.215542 | 120 | 7 | 9 | 10 | 38 | 11 | 0 | 0 | 0 | 0 | 0 | 0 |
| SG0000003 | Smarca5 | 1.9E-21 | 1.29E-20 | 1.136881272 | 4465.58865 | 9820.058606 | 112 | 2 | 9 | 17 | 43 | 15 | 12 | 0 | 0 | 0 | 4 | 1 |
| SG0000002 | Nadk2 | 1.03E-20 | 6.73E-20 | 1.597357946 | 734.3674469 | 2222.112656 | 207 | 6 | 9 | 36 | 120 | 15 | 33 | 0 | 0 | 3 | 15 | 0 |
| SG0000006 | Mdc1 | 1.39E-20 | 9.03E-20 | 1.121940677 | 3356.464094 | 7304.992979 | 31 | 9 | 9 | 15 | 25 | 6 | 2 | 0 | 0 | 0 | 3 | 1 |
| SG0000003 | Rad9b | 1.75E-20 | 1.13E-19 | 2.469396184 | 79.04733411 | 437.7735825 | 102 | 6 | 9 | 13 | 40 | 6 | 2 | 0 | 0 | 0 | 1 | 0 |
| JSG000000 | Themis2 | 1.87E-20 | 1.2E-19 | 5.799864012 | 0 | 54.7099846 | 25 | 9 | 9 | 9 | 23 | 2 | 4 | 1 | 1 | 1 | 1 | 0 |
| SG0000002 | Ostf1 | 2.02E-20 | 1.3E-19 | -1.434345558 | 3345.737733 | 1237.9721 | 84 | 5 | 9 | 13 | 32 | 5 | 3 | 0 | 0 | 1 | 3 | 0 |
| JSG000000 | Adgrf5 | 2.11E-20 | 1.35E-19 | 5.522061423 | 1.655661775 | 76.08126909 | 105 | 6 | 9 | 15 | 44 | 16 | 7 | 0 | 1 | 1 | 2 | 3 |
| SG0000000 | Dbf4 | 4.14E-20 | 2.61E-19 | 1.120144956 | 2991.470333 | 6502.523516 | 206 | 6 | 9 | 30 | 78 | 36 | 0 | 0 | 0 | 0 | 2 | 2 |
| SG0000003 | Kdf1 | 1.12E-19 | 6.91E-19 | 3.945649927 | 12.8824175 | 198.4980982 | 57 | 9 | 9 | 12 | 21 | 3 | 0 | 0 | 0 | 0 | 0 | 0 |
| SG0000003 | Sidt2 | 1.31E-19 | 8.05E-19 | -1.362539036 | 4365.011567 | 1697.541025 | 22 | 6 | 9 | 6 | 12 | 4 | 2 | 0 | 0 | 0 | 0 | 0 |
| SG0000004 | Rin3 | 1.8E-19 | 1.09E-18 | -4.120422273 | 1919.820354 | 110.3798737 | 261 | 2 | 9 | 6 | 68 | 23 | 0 | 0 | 0 | 0 | 0 | 0 |
| SG0000002 | Abca3 | 1.61E-18 | 9.31E-18 | -1.7416023 | 1918.690172 | 573.7600664 | 139 | 9 | 9 | 18 | 61 | 22 | 0 | 0 | 0 | 0 | 0 | 0 |
| JSG000000 | Phyhipl | 4.49E-18 | 2.53E-17 | 6.940524357 | 1.609821762 | 197.7351196 | 187 | 4 | 9 | 22 | 60 | 21 | 5 | 0 | 1 | 2 | 3 | 0 |
| SG0000003 | Lasp1 | 5.52E-18 | 3.1E-17 | -1.228268356 | 16988.43058 | 7251.162571 | 39 | 9 | 9 | 14 | 20 | 1 | 1 | 0 | 0 | 0 | 0 | 0 |
| SG0000003 | Arhgdib | 6.59E-18 | 3.68E-17 | -1.623827401 | 866.9814409 | 281.3125098 | 69 | 3 | 9 | 12 | 18 | 3 | 0 | 0 | 0 | 0 | 0 | 0 |
| SG0000002 | Trp53bp2 | 3.09E-17 | 1.67E-16 | 1.172171678 | 689.3095107 | 1553.363535 | 99 | 6 | 9 | 16 | 52 | 8 | 4 | 0 | 0 | 1 | 1 | 0 |
| JSG000000 | Fam184b | 4.74E-17 | 2.54E-16 | 7.193174107 | 0.331601068 | 48.52629026 | 207 | 6 | 9 | 24 | 84 | 17 | 0 | 0 | 0 | 0 | 0 | 2 |
| JSG000000 | Ube2ql1 | 1.59E-16 | 8.23E-16 | -2.991457145 | 200.7364727 | 25.2410814 | 45 | 6 | 9 | 13 | 23 | 5 | 3 | 0 | 1 | 1 | 1 | 0 |
| SG0000001 | Vac14 | 1.91E-16 | 9.83E-16 | 1.083932319 | 1154.578266 | 2447.482072 | 78 | 9 | 9 | 14 | 35 | 5 | 0 | 0 | 0 | 0 | 0 | 0 |
| SG0000006 | Dusp22 | 1.98E-16 | 1.02E-15 | 2.895936902 | 14.83955472 | 110.4548379 | 169 | 0 | 9 | 18 | 96 | 11 | 0 | 0 | 0 | 0 | 0 | 0 |
| JSG000000 | 0610040J01Rik | 3.25E-16 | 1.66E-15 | 5.394872642 | 1.278220693 | 53.78043764 | 115 | 3 | 9 | 11 | 63 | 28 | 1 | 0 | 0 | 0 | 0 | 0 |
| JSG000000 | Zfp7 | 3.94E-16 | 2E-15 | 1.077951735 | 1126.601877 | 2378.298031 | 30 | 7 | 9 | 9 | 15 | 3 | 3 | 0 | 1 | 0 | 0 | 0 |
| JSG000000 | Grik5 | 4.9E-16 | 2.46E-15 | -2.240589009 | 1353.742664 | 286.4518401 | 62 | 7 | 9 | 11 | 38 | 6 | 0 | 0 | 0 | 0 | 0 | 0 |
| SG0000002 | Prim1 | 5.87E-16 | 2.94E-15 | 1.057019792 | 1177.880809 | 2450.733058 | 36 | 8 | 9 | 10 | 19 | 1 | 2 | 0 | 0 | 0 | 1 | 0 |
| SG0000002 | Donson | 7.78E-16 | 3.87E-15 | 1.33935753 | 249.4592469 | 631.2282084 | 145 | 9 | 9 | 18 | 45 | 27 | 1 | 0 | 0 | 0 | 8 | 0 |
| SG0000000 | Fhl2 | 9.59E-16 | 4.75E-15 | -1.97467362 | 603.4152793 | 153.5254261 | 171 | 5 | 9 | 15 | 48 | 18 | 1 | 0 | 0 | 0 | 0 | 0 |
| SG0000004 | Coro2b | 1.24E-15 | 6.1E-15 | 1.591705077 | 117.9829975 | 355.6070794 | 93 | 9 | 9 | 17 | 45 | 18 | 2 | 0 | 0 | 1 | 1 | 0 |
| SG0000002 | Nuf2 | 1.67E-15 | 8.17E-15 | 1.064143988 | 948.9647292 | 1984.217753 | 118 | 4 | 9 | 17 | 41 | 14 | 2 | 0 | 1 | 1 | 1 | 0 |
| JSG000000 | Nos2 | 1.7E-15 | 8.28E-15 | -5.807623442 | 3566.885956 | 63.68253811 | 49 | 8 | 9 | 15 | 26 | 4 | 3 | 0 | 0 | 2 | 2 | 0 |
| SG0000004 | Gtpbp10 | 3.79E-15 | 1.81E-14 | 1.121131805 | 587.7221661 | 1278.398855 | 233 | 0 | 9 | 42 | 117 | 39 | 7 | 0 | 1 | 1 | 2 | 1 |
| JSG000000 | Naf1 | 4.08E-15 | 1.94E-14 | 1.040875122 | 1209.970231 | 2489.483883 | 94 | 7 | 9 | 12 | 37 | 3 | 4 | 0 | 1 | 0 | 0 | 0 |
| SG0000000 | Farsa | 5.16E-15 | 2.45E-14 | 1.011178558 | 1487.883011 | 2998.912966 | 39 | 3 | 9 | 9 | 24 | 6 | 3 | 0 | 0 | 3 | 3 | 0 |
| SG0000003 | Glce | 8.45E-15 | 3.94E-14 | 1.110805119 | 580.2608313 | 1253.166902 | 154 | 2 | 9 | 20 | 84 | 30 | 4 | 0 | 2 | 4 | 4 | 2 |
| SG0000005 | Trim34a | 2.08E-14 | 9.44E-14 | 1.755735842 | 86.31527853 | 291.4854151 | 74 | 9 | 9 | 18 | 50 | 13 | 2 | 0 | 0 | 0 | 2 | 4 |
| JSG000000 | Zcrb1 | 2.39E-14 | 1.08E-13 | -1.286351993 | 1283.628732 | 526.2703405 | 79 | 6 | 9 | 11 | 23 | 8 | 3 | 0 | 0 | 2 | 2 | 2 |
| SG0000006 | Dnaic1 | 2.44E-14 | 1.1E-13 | 7.043959819 | 0.307509278 | 40.57900721 | 87 | 7 | 9 | 19 | 46 | 7 | 0 | 0 | 0 | 0 | 0 | 0 |
| SG0000003 | Ostm1 | 2.59E-14 | 1.17E-13 | -1.190656625 | 2219.525928 | 972.381895 | 47 | 7 | 9 | 10 | 30 | 9 | 8 | 0 | 0 | 0 | 0 | 3 |
| SG0000003 | Sema4b | 3.44E-14 | 1.54E-13 | -1.104325385 | 4395.519555 | 2044.443677 | 46 | 9 | 9 | 12 | 23 | 1 | 1 | 0 | 0 | 0 | 0 | 0 |
| JSG000000 | Asap3 | 4.86E-14 | 2.16E-13 | 2.521169598 | 48.85292281 | 280.4388809 | 49 | 7 | 9 | 15 | 31 | 4 | 0 | 0 | 0 | 0 | 0 | 0 |
| JSG000000 | Mtpn | 8.6E-14 | 3.76E-13 | -1.076495145 | 10880.63982 | 5159.375125 | 91 | 3 | 9 | 13 | 44 | 5 | 9 | 0 | 2 | 3 | 3 | 0 |
| JSG000000 | Gpat4 | 1.9E-13 | 8.12E-13 | -1.079089802 | 3939.771699 | 1864.801706 | 62 | 4 | 9 | 11 | 27 | 7 | 1 | 0 | 0 | 0 | 0 | 0 |
| JSG000000 | Zglp1 | 3.54E-13 | 1.48E-12 | 4.486032101 | 2.235714429 | 50.10101839 | 17 | 8 | 9 | 10 | 16 | 0 | 0 | 0 | 0 | 0 | 0 | 0 |
| SG0000002 | Mad2l2 | 4.28E-13 | 1.78E-12 | 1.227054302 | 252.5162818 | 591.1116747 | 44 | 6 | 9 | 13 | 68 | 4 | 0 | 0 | 0 | 0 | 0 | 0 |
| SG0000003 | Ftsj1 | 4.34E-13 | 1.81E-12 | -1.159254604 | 1664.491425 | 745.2657344 | 42 | 9 | 9 | 9 | 18 | 0 | 0 | 0 | 0 | 0 | 0 | 0 |
| SG0000003 | Sema6c | 4.65E-13 | 1.93E-12 | 2.531783411 | 55.75026015 | 322.3960019 | 42 | 5 | 9 | 20 | 20 | 0 | 5 | 5 | 5 | 5 | 5 | 0 |
| SG0000005 | Gm6792 | 1.84E-12 | 7.35E-12 | 1.613298223 | 77.14003346 | 236.0103125 | 74 | 6 | 9 | 12 | 51 | 12 | 2 | 0 | 0 | 0 | 0 | 0 |
| SG0000000 | Sec24b | 2.13E-12 | 8.46E-12 | -1.063034586 | 2676.295542 | 1280.93996 | 145 | 7 | 9 | 23 | 68 | 16 | 4 | 0 | 0 | 1 | 2 | 1 |
| JSG000000 | Carnmt1 | 2.5E-12 | 9.89E-12 | 1.383646466 | 131.369502 | 342.7785547 | 71 | 8 | 9 | 13 | 44 | 12 | 4 | 0 | 0 | 0 | 1 | 1 |
| SG0000002 | Clk1 | 7.1E-12 | 2.74E-11 | 0.971991316 | 2355.788524 | 4620.988031 | 84 | 9 | 9 | 12 | 27 | 3 | 10 | 0 | 0 | 0 | 2 | 0 |
| SG0000002 | Ttc33 | 9.01E-12 | 3.45E-11 | -1.129370676 | 1259.778016 | 575.8635307 | 180 | 9 | 9 | 30 | 87 | 42 | 3 | 0 | 0 | 0 | 0 | 0 |
| SG0000004 | Gbp7 | 1.29E-11 | 4.89E-11 | -1.912116369 | 320.5255544 | 85.16443658 | 87 | 8 | 9 | 22 | 57 | 1 | 9 | 0 | 1 | 0 | 3 | 1 |
| JSG000000 | Ccdc113 | 1.37E-11 | 5.18E-11 | 4.214607721 | 3.677890521 | 68.28450123 | 43 | 9 | 9 | 11 | 28 | 3 | 0 | 0 | 0 | 0 | 1 | 0 |

| | | | | | | | | | | | | | | | | | | |
|---|---|---|---|---|---|---|---|---|---|---|---|---|---|---|---|---|---|---|
| JSG000000 | Gmps | 1.41E-11 | 5.33E-11 | 0.835535245 | 2884.617974 | 5147.655523 | 121 | 4 | 9 | 18 | 67 | 10 | 22 | 0 | 0 | 1 | 8 | 1 |
| JSG000000 | 1190002N15Rik | 3.02E-11 | 1.11E-10 | 1.009163168 | 433.5021102 | 872.5284565 | 60 | 7 | 9 | 12 | 27 | 5 | 13 | 0 | 0 | 2 | 5 | 1 |
| SG0000003 | Dido1 | 5.46E-11 | 1.98E-10 | -0.956735264 | 3613.236767 | 1861.617309 | 221 | 7 | 9 | 30 | 78 | 9 | 6 | 0 | 0 | 0 | 2 | 1 |
| JSG000000 | Fbxo25 | 1.56E-10 | 5.47E-10 | -1.052395086 | 1222.818238 | 589.6026399 | 51 | 8 | 9 | 14 | 27 | 8 | 1 | 0 | 0 | 1 | 1 | 1 |
| JSG000000 | Cyp3a13 | 2.63E-10 | 9.08E-10 | -5.283181888 | 39.34791725 | 1.010474896 | 70 | 8 | 9 | 12 | 47 | 6 | 3 | 0 | 0 | 0 | 1 | 0 |
| SG0000002 | Reck | 3E-10 | 1.03E-09 | 1.234659341 | 146.7375854 | 345.3113587 | 130 | 8 | 9 | 19 | 66 | 14 | 2 | 0 | 0 | 0 | 4 | 1 |
| JSG000000 | Ccdc65 | 4.59E-10 | 1.55E-09 | 4.245836006 | 1.987262844 | 37.70329506 | 19 | 8 | 9 | 9 | 31 | 0 | 0 | 0 | 0 | 0 | 0 | 0 |
| SG0000002 | Smu1 | 4.88E-10 | 1.65E-09 | 0.773297508 | 2876.915022 | 4917.142329 | 40 | 8 | 9 | 13 | 39 | 7 | 1 | 0 | 0 | 0 | 0 | 2 |
| SG0000007 | Rab9 | 5.37E-10 | 1.81E-09 | 0.864766743 | 786.6674508 | 1432.556996 | 48 | 9 | 9 | 14 | 30 | 4 | 2 | 0 | 0 | 0 | 1 | 0 |
| SG0000002 | Npas2 | 7.83E-10 | 2.6E-09 | -2.619472041 | 86.15467266 | 14.01971482 | 190 | 4 | 9 | 17 | 59 | 24 | 0 | 0 | 0 | 0 | 0 | 0 |
| JSG000000 | Slc25a30 | 2.05E-09 | 6.57E-09 | 0.963022413 | 369.5047823 | 720.3087602 | 45 | 9 | 9 | 16 | 24 | 4 | 0 | 0 | 0 | 0 | 1 | 1 |
| JSG000000 | Olfm2 | 2.67E-09 | 8.49E-09 | 3.156956762 | 11.58244859 | 103.309215 | 71 | 8 | 9 | 20 | 43 | 2 | 0 | 0 | 0 | 0 | 0 | 0 |
| JSG000000 | Fem1c | 2.99E-09 | 9.47E-09 | 0.789050294 | 1154.645936 | 1995.154986 | 59 | 4 | 9 | 8 | 28 | 10 | 5 | 0 | 0 | 0 | 3 | 0 |
| SG0000002 | Rsrc2 | 5.52E-09 | 1.71E-08 | 0.734056902 | 2334.236274 | 3882.558066 | 174 | 0 | 9 | 19 | 69 | 10 | 16 | 0 | 1 | 1 | 4 | 3 |
| JSG000000 | Zbtb1 | 1.03E-08 | 3.12E-08 | -0.97027862 | 1342.682913 | 685.3154037 | 56 | 5 | 9 | 6 | 18 | 4 | 19 | 2 | 4 | 2 | 6 | 2 |
| SG000000C | Gstt3 | 1.08E-08 | 3.27E-08 | -1.353257287 | 735.8425607 | 288.0141787 | 28 | 6 | 9 | 9 | 15 | 5 | 0 | 0 | 0 | 0 | 0 | 3 |
| JSG000000 | Fam189a2 | 1.8E-08 | 5.36E-08 | 3.426218154 | 3.632050507 | 39.04328178 | 80 | 7 | 9 | 16 | 41 | 8 | 1 | 0 | 0 | 1 | 1 | 0 |
| SG0000003 | Txndc5 | 2.64E-08 | 7.76E-08 | -0.776097658 | 5950.745248 | 3474.902983 | 72 | 6 | 9 | 12 | 33 | 15 | 0 | 0 | 0 | 0 | 3 | 0 |
| JSG000000 | Sfrp1 | 2.68E-08 | 7.89E-08 | 1.142658814 | 1668.954665 | 3684.844094 | 46 | 9 | 9 | 10 | 28 | 6 | 2 | 0 | 0 | 0 | 2 | 1 |
| JSG000000 | Slc2a12 | 3.42E-08 | 9.97E-08 | 4.245911023 | 1.644787664 | 31.20731587 | 99 | 4 | 9 | 22 | 53 | 7 | 9 | 0 | 1 | 3 | 3 | 1 |
| JSG000000 | Egfl6 | 3.71E-08 | 1.08E-07 | -1.853265502 | 123.1042577 | 34.07097547 | 113 | 4 | 9 | 12 | 33 | 7 | 1 | 0 | 0 | 0 | 1 | 0 |
| JSG000000 | Vit | 4.42E-08 | 1.28E-07 | -4.365141962 | 34.74849576 | 1.686154412 | 139 | 7 | 9 | 18 | 47 | 17 | 0 | 0 | 0 | 0 | 0 | 0 |
| SG0000003 | Zfp157 | 4.81E-08 | 1.38E-07 | 0.845008007 | 437.4258025 | 785.7369085 | 132 | 9 | 9 | 26 | 72 | 11 | 12 | 0 | 0 | 2 | 4 | 1 |
| SG0000002 | Ikbkap | 6.34E-08 | 1.81E-07 | 0.696858089 | 1898.329366 | 3077.136442 | 90 | 5 | 9 | 15 | 50 | 15 | 0 | 0 | 0 | 0 | 1 | 0 |
| SG0000002 | Pigg | 8.55E-08 | 2.41E-07 | -0.780447825 | 2708.072361 | 1576.601949 | 46 | 9 | 9 | 19 | 25 | 3 | 1 | 0 | 0 | 0 | 0 | 0 |
| SG0000003 | Golgb1 | 9.16E-08 | 2.58E-07 | -0.767317839 | 3960.59214 | 2326.882404 | 124 | 6 | 9 | 16 | 80 | 25 | 0 | 0 | 0 | 0 | 0 | 0 |
| JSG000000 | Klhl36 | 1.28E-07 | 3.56E-07 | 2.433583253 | 29.54308924 | 159.6016814 | 23 | 8 | 9 | 10 | 11 | 1 | 0 | 0 | 0 | 0 | 0 | 0 |
| JSG000000 | Crtap | 1.44E-07 | 3.98E-07 | -0.79798932 | 1766.099411 | 1015.772437 | 33 | 7 | 9 | 11 | 27 | 5 | 0 | 0 | 0 | 0 | 1 | 0 |
| JSG000000 | Ptpdc1 | 1.56E-07 | 4.31E-07 | -1.095130963 | 548.0040007 | 256.5171818 | 82 | 7 | 9 | 15 | 46 | 9 | 5 | 2 | 3 | 2 | 3 | 0 |
| SG0000005 | 9430015G10Rik | 1.69E-07 | 4.64E-07 | 0.907482729 | 714.9652892 | 1341.110243 | 54 | 9 | 9 | 12 | 60 | 6 | 3 | 0 | 0 | 0 | 0 | 0 |
| SG0000003 | Nlrx1 | 1.98E-07 | 5.4E-07 | 1.772506664 | 55.26592764 | 188.8144107 | 51 | 3 | 9 | 12 | 15 | 0 | 0 | 0 | 0 | 0 | 0 | 0 |
| SG000000C | Golga2 | 2.02E-07 | 5.52E-07 | -0.781773368 | 1816.883692 | 1056.792916 | 33 | 6 | 9 | 12 | 27 | 12 | 9 | 3 | 3 | 3 | 3 | 6 |
| SG0000004 | Fam161a | 2.09E-07 | 5.68E-07 | 2.15305834 | 23.55705578 | 104.7746342 | 114 | 1 | 9 | 20 | 38 | 13 | 7 | 1 | 1 | 2 | 4 | 1 |
| JSG000000 | Patl1 | 3E-07 | 8.06E-07 | 0.661861518 | 1916.81101 | 3032.630267 | 77 | 4 | 9 | 13 | 36 | 6 | 5 | 0 | 0 | 0 | 1 | 1 |
| JSG000000 | Slc9a1 | 3.75E-07 | 9.99E-07 | -0.78941086 | 1439.332176 | 832.7692612 | 47 | 7 | 9 | 12 | 27 | 8 | 0 | 0 | 0 | 0 | 0 | 0 |
| SG0000002 | Zmynd19 | 4.76E-07 | 1.26E-06 | 0.677619153 | 1414.350747 | 2262.251135 | 39 | 6 | 9 | 19 | 27 | 3 | 3 | 0 | 1 | 1 | 1 | 1 |
| JSG000000 | Nomo1 | 4.88E-07 | 1.29E-06 | -0.71428043 | 6752.226787 | 4115.5434 | 53 | 4 | 9 | 8 | 27 | 4 | 3 | 0 | 0 | 0 | 3 | 0 |
| SG0000005 | Zscan5b | 5.33E-07 | 1.41E-06 | 4.169424893 | 0 | 16.9937614 | 38 | 8 | 9 | 9 | 23 | 3 | 1 | 0 | 0 | 0 | 0 | 0 |
| SG0000002 | Tsc22d4 | 5.47E-07 | 1.44E-06 | -1.294450817 | 1926.085022 | 785.2481647 | 97 | 9 | 9 | 30 | 55 | 8 | 3 | 0 | 0 | 2 | 2 | 0 |
| SG0000002 | Adcyap1r1 | 5.52E-07 | 1.45E-06 | 3.277695804 | 3.677890521 | 35.66839365 | 95 | 9 | 9 | 20 | 47 | 11 | 3 | 0 | 0 | 0 | 0 | 3 |
| SG0000002 | Kif2c | 6.22E-07 | 1.63E-06 | 0.63927503 | 2211.686736 | 3444.804444 | 54 | 7 | 9 | 15 | 30 | 4 | 1 | 0 | 0 | 1 | 1 | 0 |
| SG0000003 | Pgap2 | 6.42E-07 | 1.68E-06 | 0.726496213 | 647.8154873 | 1071.885703 | 122 | 8 | 9 | 23 | 53 | 7 | 0 | 0 | 0 | 0 | 2 | 0 |
| SG0000002 | Ssr3 | 7.05E-07 | 1.84E-06 | -0.67473535 | 11539.5512 | 7228.918674 | 67 | 4 | 9 | 7 | 38 | 4 | 9 | 0 | 1 | 1 | 4 | 0 |
| JSG000000 | Nudt13 | 7.79E-07 | 2.03E-06 | 1.126973087 | 99.00496171 | 216.2268709 | 51 | 8 | 9 | 12 | 34 | 4 | 4 | 0 | 0 | 0 | 2 | 0 |
| SG0000002 | Stxbp3 | 9.89E-07 | 2.56E-06 | -0.797393488 | 1123.489659 | 646.4422322 | 130 | 2 | 9 | 21 | 90 | 12 | 7 | 0 | 1 | 2 | 6 | 3 |
| JSG000000 | Tmed10 | 1.03E-06 | 2.66E-06 | -0.67947822 | 4819.501524 | 3009.253653 | 69 | 6 | 9 | 12 | 33 | 2 | 0 | 0 | 0 | 0 | 0 | 1 |
| SG0000006 | Fiz1 | 1.04E-06 | 2.67E-06 | 0.96531278 | 693.6778662 | 1354.396863 | 32 | 9 | 9 | 12 | 18 | 0 | 0 | 0 | 0 | 0 | 0 | 0 |
| SG0000002 | Pear1 | 2.06E-06 | 5.17E-06 | -1.591930755 | 177.4132381 | 58.85279792 | 77 | 9 | 9 | 17 | 63 | 11 | 0 | 0 | 0 | 0 | 0 | 0 |
| JSG000000 | Spta1 | 2.48E-06 | 6.19E-06 | 1.168742466 | 99.50570829 | 223.7044045 | 185 | 4 | 9 | 22 | 88 | 11 | 5 | 0 | 0 | 0 | 1 | 0 |
| SG0000003 | Rbm43 | 2.57E-06 | 6.38E-06 | -1.027477226 | 510.7968779 | 250.5802102 | 42 | 6 | 9 | 9 | 24 | 0 | 3 | 0 | 0 | 0 | 0 | 0 |
| JSG000000 | Col6a4 | 3.79E-06 | 9.29E-06 | 5.7936822 | 0.307509278 | 17.05808803 | 159 | 5 | 9 | 23 | 96 | 45 | 0 | 0 | 0 | 0 | 0 | 0 |
| JSG000000 | Macrod1 | 3.86E-06 | 9.46E-06 | -1.862431914 | 195.6222432 | 53.79852438 | 95 | 7 | 9 | 13 | 26 | 4 | 0 | 0 | 0 | 0 | 0 | 0 |
| SG0000001 | Ptprs | 3.87E-06 | 9.47E-06 | -1.105225083 | 6052.446274 | 2813.358268 | 53 | 8 | 9 | 11 | 15 | 6 | 0 | 0 | 0 | 0 | 0 | 2 |
| JSG000000 | 2410131K14Rik | 4.71E-06 | 1.14E-05 | -0.772050866 | 827.4184376 | 484.5233655 | 26 | 9 | 9 | 11 | 14 | 4 | 5 | 1 | 1 | 1 | 2 | 2 |
| SG0000001 | Plgrkt | 5.44E-06 | 1.31E-05 | -0.866636876 | 541.9042897 | 297.1932176 | 60 | 6 | 9 | 15 | 30 | 0 | 0 | 0 | 0 | 0 | 0 | 0 |
| SG0000002 | Rnf25 | 8.04E-06 | 1.91E-05 | 0.671143201 | 591.8034392 | 942.3488818 | 42 | 9 | 9 | 9 | 24 | 0 | 0 | 0 | 0 | 0 | 0 | 0 |
| JSG000000 | B3gat3 | 8.06E-06 | 1.92E-05 | -1.141313094 | 979.5139135 | 444.0594673 | 18 | 9 | 9 | 13 | 16 | 1 | 0 | 0 | 0 | 0 | 0 | 0 |
| JSG000000 | Supt5 | 8.56E-06 | 2.03E-05 | 0.561066627 | 4155.758451 | 6131.238091 | 33 | 8 | 9 | 8 | 15 | 3 | 0 | 0 | 0 | 0 | 0 | 0 |
| SG0000005 | Kcp | 9.34E-06 | 2.21E-05 | 2.081806465 | 11.48004759 | 48.59928494 | 123 | 4 | 9 | 13 | 42 | 17 | 0 | 0 | 0 | 0 | 0 | 1 |
| SG0000002 | Aldh18a1 | 9.91E-06 | 2.34E-05 | 0.623209721 | 1718.14231 | 2646.45158 | 54 | 8 | 9 | 12 | 26 | 8 | 0 | 0 | 0 | 0 | 0 | 0 |
| JSG000000 | Trabd2b | 1.09E-05 | 2.56E-05 | -2.145626529 | 53.87746598 | 12.17612833 | 238 | 6 | 9 | 31 | 97 | 19 | 5 | 0 | 0 | 2 | 3 | 0 |
| SG0000004 | Cep164 | 1.77E-05 | 4.07E-05 | 0.594199534 | 1029.458373 | 1554.10613 | 60 | 6 | 9 | 16 | 40 | 5 | 1 | 0 | 0 | 0 | 0 | 0 |
| JSG000000 | Exoc3l4 | 2.57E-05 | 5.81E-05 | -1.830144742 | 156.7888277 | 44.0947261 | 18 | 8 | 9 | 9 | 11 | 0 | 0 | 0 | 0 | 0 | 0 | 0 |
| SG0000003 | Bri3bp | 2.7E-05 | 6.11E-05 | -0.598165555 | 3279.532705 | 2166.437636 | 31 | 9 | 9 | 11 | 27 | 4 | 5 | 1 | 1 | 1 | 5 | 2 |
| JSG000000 | Drd1 | 3.5E-05 | 7.83E-05 | -2.08481089 | 49.29979271 | 11.62129378 | 15 | 8 | 9 | 10 | 11 | 1 | 14 | 8 | 9 | 9 | 10 | 1 |
| SG0000004 | Apobec1 | 4.12E-05 | 9.14E-05 | -0.768161397 | 560.9853826 | 329.3911475 | 144 | 9 | 9 | 30 | 109 | 24 | 3 | 0 | 0 | 0 | 3 | 0 |
| JSG000000 | Ankmy2 | 4.2E-05 | 9.3E-05 | 0.62612567 | 559.6170779 | 863.7211758 | 84 | 8 | 9 | 13 | 35 | 5 | 1 | 0 | 0 | 0 | 0 | 0 |
| JSG000000 | Cdkn2a | 4.27E-05 | 9.45E-05 | -1.303131704 | 610.9893173 | 247.6007101 | 46 | 7 | 9 | 10 | 30 | 4 | 0 | 0 | 0 | 0 | 0 | 0 |
| JSG000000 | Ndufaf1 | 4.84E-05 | 0.000106 | 0.642958826 | 432.64533 | 675.5879781 | 54 | 3 | 9 | 12 | 30 | 3 | 0 | 0 | 0 | 0 | 0 | 0 |
| SG0000005 | Btn2a2 | 6.73E-05 | 0.000146 | 1.980279232 | 10.53342796 | 41.56168801 | 44 | 9 | 9 | 15 | 18 | 0 | 2 | 0 | 0 | 0 | 0 | 0 |
| SG0000005 | Utp18 | 6.79E-05 | 0.000147 | 0.518283056 | 1865.434872 | 2671.768573 | 74 | 4 | 9 | 10 | 20 | 2 | 4 | 0 | 1 | 0 | 0 | 1 |
| JSG000001 | Gbp5 | 8.93E-05 | 0.00019 | -2.158671419 | 42.32629303 | 9.479499025 | 90 | 6 | 9 | 18 | 66 | 21 | 6 | 0 | 0 | 0 | 3 | 0 |
| SG0000003 | Stvk1 | 0.000103 | 0.000217 | -2.295365671 | 37.88030266 | 7.716840473 | 71 | 4 | 9 | 19 | 60 | 6 | 1 | 0 | 0 | 1 | 2 | 0 |
| JSG000000 | Aaed1 | 0.000104 | 0.000221 | -0.769095249 | 418.5613136 | 245.6056458 | 37 | 5 | 9 | 9 | 20 | 2 | 0 | 0 | 0 | 0 | 0 | 0 |
| SG0000002 | Orc4 | 0.000155 | 0.000321 | 0.506508972 | 1401.074015 | 1990.377584 | 186 | 2 | 9 | 27 | 105 | 8 | 11 | 0 | 1 | 0 | 0 | 1 |
| JSG000000 | Fam208b | 0.000182 | 0.000375 | 0.488681969 | 1813.471312 | 2544.594714 | 112 | 7 | 9 | 17 | 55 | 10 | 1 | 0 | 0 | 1 | 1 | 0 |
| SG0000002 | Ptcd1 | 0.000183 | 0.000376 | -0.613515941 | 876.6009242 | 572.9480202 | 23 | 5 | 9 | 9 | 22 | 1 | 0 | 0 | 0 | 0 | 0 | 0 |
| SG0000006 | Mthfs | 0.000198 | 0.000406 | 1.229496934 | 37.09664158 | 86.9861382 | 128 | 3 | 9 | 20 | 62 | 2 | 2 | 0 | 0 | 0 | 0 | 0 |
| SG0000001 | Unkl | 0.000203 | 0.000415 | -0.898036104 | 212.7843957 | 114.1836632 | 128 | 4 | 9 | 19 | 65 | 13 | 33 | 0 | 0 | 9 | 21 | 0 |
| JSG000000 | Polr2b | 0.000217 | 0.000442 | 0.459944738 | 3901.79169 | 5366.872057 | 75 | 6 | 9 | 11 | 49 | 9 | 1 | 0 | 0 | 0 | 0 | 2 |
| SG0000002 | Klhdc2 | 0.000228 | 0.000464 | -0.549136193 | 1902.485561 | 1300.214197 | 39 | 8 | 9 | 13 | 21 | 1 | 3 | 0 | 0 | 1 | 1 | 1 |
| SG0000002 | Akap11 | 0.000266 | 0.000537 | 0.499042771 | 2593.131267 | 3664.808997 | 130 | 5 | 9 | 24 | 53 | 9 | 19 | 3 | 3 | 5 | 7 | 1 |
| SG0000005 | Paqr3 | 0.000289 | 0.00058 | 0.923231131 | 80.15222514 | 151.9972907 | 84 | 0 | 9 | 12 | 38 | 7 | 1 | 0 | 0 | 0 | 0 | 0 |
| JSG000000 | Tbc1d25 | 0.000339 | 0.000676 | -0.649346752 | 685.1925902 | 436.8575119 | 90 | 9 | 9 | 24 | 117 | 12 | 0 | 0 | 0 | 0 | 0 | 0 |
| JSG000000 | Cisd2 | 0.000353 | 0.000702 | -0.559937327 | 1195.178444 | 810.7273433 | 74 | 6 | 9 | 10 | 12 | 8 | 6 | 0 | 0 | 0 | 0 | 2 |
| SG0000005 | Bzw1 | 0.000377 | 0.000748 | 0.427858026 | 13898.39471 | 18696.60449 | 71 | 6 | 9 | 15 | 37 | 6 | 5 | 0 | 0 | 0 | 1 | 3 |
| SG0000002 | Mapre1 | 0.000422 | 0.000832 | -0.478728174 | 6584.703449 | 4725.248802 | 59 | 6 | 9 | 18 | 39 | 7 | 10 | 1 | 1 | 1 | 5 | 1 |
| SG0000001 | Plagl1 | 0.000558 | 0.001082 | 1.652098582 | 74.5173941 | 234.201077 | 218 | 3 | 9 | 21 | 81 | 21 | 20 | 0 | 0 | 0 | 6 | 1 |
| SG0000002 | Nuak1 | 0.000565 | 0.001095 | 1.672570918 | 121.4452991 | 387.145954 | 93 | 5 | 9 | 11 | 36 | 6 | 4 | 1 | 2 | 1 | 3 | 0 |
| JSG000000 | Qrsl1 | 0.000624 | 0.001201 | 0.620291025 | 336.706318 | 517.5799809 | 58 | 6 | 9 | 14 | 30 | 8 | 0 | 0 | 0 | 0 | 0 | 0 |
| SG0000003 | Qsox1 | 0.000738 | 0.001407 | -0.91285069 | 1454.923944 | 772.7603881 | 117 | 9 | 9 | 9 | 36 | 6 | 0 | 0 | 0 | 0 | 0 | 0 |
| JSG000000 | Rspry1 | 0.000745 | 0.00142 | 0.49047492 | 710.1348586 | 997.673633 | 180 | 2 | 9 | 29 | 88 | 16 | 6 | 0 | 1 | 0 | 5 | 0 |
| SG0000004 | Prdm10 | 0.000926 | 0.001743 | 0.619487408 | 231.8278638 | 356.1639221 | 85 | 6 | 9 | 23 | 42 | 6 | 4 | 0 | 0 | 2 | 2 | 1 |
| SG0000002 | Prpf40b | 0.00136 | 0.002515 | 0.643168682 | 776.0241632 | 1211.960065 | 80 | 9 | 9 | 22 | 49 | 6 | 4 | 0 | 0 | 4 | 4 | 0 |
| SG0000002 | Atg16l1 | 0.001418 | 0.002618 | 0.453097357 | 1158.534955 | 1586.006772 | 141 | 6 | 9 | 15 | 69 | 18 | 6 | 3 | 6 | 3 | 3 | 3 |
| JSG000000 | Ap3s2 | 0.001637 | 0.003002 | 0.418689281 | 1677.533792 | 2242.380509 | 67 | 8 | 9 | 15 | 41 | 4 | 2 | 0 | 0 | 0 | 3 | 0 |
| JSG000000 | Mtrr | 0.002054 | 0.003713 | 0.460898135 | 704.4668018 | 969.6269891 | 33 | 6 | 9 | 10 | 21 | 1 | 3 | 0 | 1 | 1 | 2 | 0 |
| SG0000005 | Lrif1 | 0.002211 | 0.00398 | -0.539725863 | 595.3739353 | 409.5586886 | 237 | 1 | 9 | 19 | 116 | 21 | 4 | 0 | 0 | 0 | 10 | 2 |
| JSG000000 | Emc2 | 0.002301 | 0.004125 | 0.456138294 | 969.268863 | 1329.705822 | 145 | 7 | 9 | 26 | 88 | 11 | 2 | 0 | 0 | 0 | 0 | 0 |
| SG0000002 | Trmu | 0.004624 | 0.007953 | -0.629381451 | 244.6177575 | 158.1340104 | 45 | 6 | 9 | 17 | 37 | 0 | 0 | 0 | 0 | 0 | 0 | 0 |
| SG0000003 | Cnot7 | 0.00504 | 0.008622 | 0.35941255 | 2458.743908 | 3154.330935 | 53 | 8 | 9 | 13 | 36 | 5 | 5 | 0 | 0 | 1 | 7 | 2 |
| JSG000000 | Galnt2 | 0.005173 | 0.008833 | -1.327452067 | 44.96488703 | 17.91721961 | 104 | 6 | 9 | 13 | 25 | 11 | 0 | 0 | 0 | 0 | 0 | 0 |
| SG0000003 | Ttc13 | 0.005543 | 0.009421 | 0.385103988 | 1289.82043 | 1684.446041 | 174 | 3 | 9 | 27 | 66 | 27 | 3 | 0 | 0 | 0 | 3 | 0 |
| SG0000002 | Rab14 | 0.007215 | 0.012063 | -0.37253533 | 3826.731224 | 2955.858575 | 69 | 2 | 9 | 11 | 33 | 7 | 5 | 0 | 0 | 0 | 4 | 2 |
| SG0000006 | Hacd1 | 0.007454 | 0.012432 | 0.92759691 | 36.94455715 | 70.27242794 | 138 | 3 | 9 | 30 | 66 | 12 | 0 | 0 | 0 | 0 | 0 | 0 |
| SG0000007 | Alg6 | 0.007737 | 0.012877 | 0.512608792 | 618.1133428 | 881.8175558 | 112 | 2 | 9 | 18 | 45 | 13 | 5 | 0 | 0 | 0 | 1 | 1 |
| SG0000003 | Lrrc56 | 0.008606 | 0.014229 | 0.694202164 | 93.04617287 | 150.5477222 | 33 | 3 | 9 | 9 | 12 | 0 | 0 | 0 | 0 | 0 | 0 | 0 |
| SG0000004 | Nupl2 | 0.009265 | 0.01523 | -0.470745614 | 545.8763277 | 393.8997554 | 79 | 6 | 9 | 12 | 38 | 2 | 2 | 0 | 1 | 0 | 0 | 0 |
| SG0000006 | Mospd2 | 0.012163 | 0.019592 | 0.382747573 | 589.6101808 | 768.7470711 | 249 | 3 | 9 | 30 | 131 | 36 | 6 | 0 | 0 | 0 | 1 | 1 |
| SG0000001 | Gorasp2 | 0.013379 | 0.021363 | -0.341589754 | 2857.292384 | 2254.892911 | 228 | 0 | 9 | 24 | 94 | 12 | 7 | 0 | 0 | 0 | 2 | 2 |
| SG0000002 | Ak4 | 0.014379 | 0.022857 | -0.383968438 | 2270.306491 | 1739.796584 | 189 | 6 | 9 | 32 | 79 | 14 | 6 | 0 | 0 | 3 | 3 | 0 |
| SG0000004 | Flywch1 | 0.01603 | 0.025285 | -0.408069298 | 890.2682715 | 670.9339942 | 31 | 9 | 9 | 9 | 14 | 2 | 1 | 0 | 0 | 0 | 0 | 0 |
| SG0000002 | Fsip1 | 0.018203 | 0.028369 | 1.623796107 | 6.469565814 | 19.93822397 | 131 | 3 | 9 | 11 | 51 | 8 | 0 | 0 | 0 | 0 | 1 | 0 |
| SG0000005 | Mtfmt | 0.018385 | 0.028632 | -0.446162392 | 436.788192 | 320.5993418 | 43 | 8 | 9 | 14 | 23 | 2 | 3 | 0 | 0 | 1 | 2 | 0 |
| SG0000001 | Shroom1 | 0.020982 | 0.032321 | 1.348939692 | 12.92357038 | 32.91954484 | 45 | 9 | 9 | 21 | 39 | 0 | 0 | 0 | 0 | 0 | 0 | 0 |
| SG0000002 | Fam92a | 0.021264 | 0.032728 | -0.377217603 | 828.6710576 | 638.0112806 | 153 | 9 | 9 | 12 | 70 | 24 | 5 | 0 | 0 | 0 | 4 | 0 |
| SG0000002 | Tada1 | 0.022913 | 0.035067 | -0.414854188 | 488.8609114 | 366.6922731 | 37 | 9 | 9 | 10 | 20 | 1 | 2 | 0 | 0 | 0 | 0 | 0 |
| SG0000004 | Tnip3 | 0.026055 | 0.039534 | -1.833518902 | 17.62269947 | 4.944566791 | 100 | 2 | 9 | 19 | 59 | 10 | 3 | 0 | 0 | 1 | 2 | 0 |
| SG0000003 | Lap3 | 0.027525 | 0.041616 | -0.293569845 | 8934.435766 | 7289.433891 | 36 | 8 | 9 | 12 | 22 | 2 | 2 | 0 | 0 | 0 | 0 | 0 |
| SG000000C | Sulf2 | 0.027693 | 0.041853 | -0.300767105 | 4393.024539 | 3566.347919 | 132 | 0 | 9 | 12 | 38 | 18 | 4 | 0 | 0 | 0 | 2 | 0 |
| SG0000004 | Tmcc2 | 0.027988 | 0.042259 | -0.425239861 | 359.7218993 | 267.8902604 | 82 | 9 | 9 | 19 | 30 | 4 | 18 | 9 | 9 | 15 | 18 | 0 |
| SG0000003 | Tspan3 | 0.030315 | 0.045451 | -0.319922264 | 3001.621478 | 2404.638113 | 65 | 6 | 9 | 10 | 21 | 7 | 2 | 0 | 0 | 0 | 0 | 1 |
| SG000000C | Lmf1 | 0.032118 | 0.047966 | -0.408226417 | 376.9372287 | 284.0407409 | 144 | 3 | 9 | 24 | 51 | 9 | 0 | 0 | 0 | 0 | 0 | 0 |

| | | | | | | | | | | | | | | | | | | |
|---|---|---|---|---|---|---|---|---|---|---|---|---|---|---|---|---|---|---|
| SG0000002 | Scyl3 | 0.034843 | 0.051665 | -0.336534973 | 910.5023893 | 721.0642571 | 191 | 5 | 9 | 27 | 119 | 17 | 16 | 0 | 0 | 0 | 8 | 0 |
| ISG000000 | Erich1 | 0.03978 | 0.058414 | -0.355053823 | 587.6883593 | 459.4785456 | 114 | 5 | 9 | 17 | 66 | 12 | 3 | 1 | 1 | 2 | 6 | 2 |
| ISG000000 | Frzb | 0.04257 | 0.062221 | -1.750759694 | 15.72711665 | 4.673238273 | 77 | 6 | 9 | 11 | 41 | 10 | 4 | 0 | 0 | 0 | 0 | 0 |
| SG0000000 | Stxbp2 | 0.046474 | 0.067531 | -0.315700733 | 2662.361947 | 2139.103051 | 33 | 9 | 9 | 13 | 18 | 4 | 2 | 0 | 0 | 0 | 0 | 0 |
| SG0000005 | Actn4 | 0.04726 | 0.068589 | -0.273558913 | 15094.15902 | 12487.03942 | 74 | 8 | 9 | 14 | 27 | 9 | 3 | 0 | 0 | 0 | 1 | 0 |
| SG0000000 | Rnf114 | 0.048031 | 0.069579 | -0.277167187 | 3267.82257 | 2696.639123 | 46 | 6 | 9 | 6 | 28 | 0 | 3 | 0 | 0 | 0 | 0 | 0 |
| SG0000003 | Idua | 0.04945 | 0.07146 | 0.359465637 | 354.6165411 | 454.9555093 | 68 | 9 | 9 | 14 | 21 | 5 | 11 | 0 | 0 | 0 | 0 | 0 |
| ISG000000 | Kdsr | 0.051951 | 0.074719 | -0.289665017 | 1650.237511 | 1350.046091 | 52 | 5 | 9 | 10 | 32 | 4 | 9 | 1 | 3 | 2 | 6 | 0 |
| SG0000000 | Psmd4 | 0.052105 | 0.074922 | 0.250548181 | 3606.51244 | 4290.520214 | 43 | 9 | 9 | 16 | 31 | 6 | 0 | 0 | 0 | 0 | 0 | 0 |
| SG0000002 | Pmm1 | 0.066814 | 0.094248 | -0.301962628 | 8852.264991 | 7180.498527 | 30 | 6 | 9 | 12 | 21 | 0 | 0 | 0 | 0 | 0 | 0 | 0 |
| ISG000000 | Usp38 | 0.067516 | 0.095135 | -0.268008806 | 1751.858399 | 1454.857007 | 75 | 7 | 9 | 11 | 35 | 6 | 3 | 0 | 0 | 0 | 0 | 0 |
| SG0000000 | Tmem143 | 0.069372 | 0.097527 | 0.557101751 | 355.2419054 | 522.6711494 | 36 | 9 | 9 | 11 | 21 | 2 | 1 | 0 | 0 | 0 | 2 | 0 |
| ISG000000 | Zfyve1 | 0.073233 | 0.102504 | 0.283890411 | 930.023152 | 1132.278573 | 74 | 9 | 9 | 19 | 43 | 7 | 2 | 0 | 0 | 0 | 0 | 0 |
| ISG000000 | Vps26b | 0.073844 | 0.10329 | -0.261750207 | 2027.630156 | 1691.196527 | 34 | 9 | 9 | 12 | 20 | 5 | 2 | 0 | 0 | 0 | 0 | 0 |
| SG0000002 | Rars2 | 0.075337 | 0.105153 | 0.257755415 | 867.8036486 | 1037.560866 | 111 | 5 | 9 | 14 | 49 | 6 | 1 | 0 | 0 | 0 | 1 | 0 |
| SG0000002 | Yars2 | 0.078929 | 0.109739 | 0.27455751 | 742.2256831 | 897.8132784 | 51 | 9 | 9 | 18 | 36 | 17 | 8 | 3 | 3 | 4 | 5 | 5 |
| SG0000000 | Sh2b2 | 0.080769 | 0.112045 | -0.49338011 | 122.1291654 | 86.75553191 | 79 | 9 | 9 | 21 | 35 | 6 | 0 | 0 | 0 | 0 | 2 | 0 |
| SG0000002 | Adamts10 | 0.091807 | 0.12604 | 0.396252913 | 168.7969445 | 222.1511643 | 41 | 8 | 9 | 10 | 19 | 4 | 3 | 1 | 1 | 1 | 1 | 0 |
| SG0000003 | Psmd8 | 0.097045 | 0.132491 | 0.211908423 | 3976.737192 | 4605.933705 | 18 | 9 | 9 | 9 | 15 | 0 | 0 | 0 | 0 | 0 | 0 | 0 |
| ISG000000 | Fbxo45 | 0.097963 | 0.133581 | -0.258711158 | 1206.122351 | 1008.118413 | 31 | 5 | 9 | 6 | 16 | 5 | 6 | 0 | 1 | 1 | 1 | 1 |
| SG0000002 | Vcam1 | 0.101012 | 0.137205 | -0.786946204 | 303.7328365 | 176.0343146 | 79 | 4 | 9 | 17 | 28 | 7 | 7 | 0 | 1 | 2 | 3 | 3 |
| SG0000001 | Wsb1 | 0.104724 | 0.141967 | -0.232353271 | 4627.439749 | 3939.086055 | 93 | 0 | 9 | 7 | 26 | 8 | 7 | 0 | 2 | 2 | 5 | 3 |
| ISG000000 | Rer1 | 0.112068 | 0.150879 | -0.225539012 | 2520.275024 | 2155.529286 | 30 | 8 | 9 | 11 | 20 | 2 | 2 | 0 | 0 | 0 | 1 | 0 |
| SG0000006 | Slc31a1 | 0.11235 | 0.151172 | 0.219664505 | 1764.27738 | 2054.435311 | 76 | 9 | 9 | 15 | 38 | 8 | 12 | 0 | 0 | 0 | 2 | 0 |
| SG0000003 | Rara | 0.116517 | 0.156182 | 0.409630851 | 778.5133844 | 1034.135047 | 128 | 9 | 9 | 24 | 56 | 10 | 6 | 0 | 0 | 3 | 3 | 5 |
| SG0000005 | Ccdc158 | 0.124155 | 0.16554 | -0.44221031 | 112.3819388 | 82.71379604 | 117 | 7 | 9 | 14 | 69 | 15 | 0 | 0 | 0 | 0 | 0 | 0 |
| SG0000007 | Dpm1 | 0.12645 | 0.168276 | 0.39946293 | 207.3374496 | 273.4815775 | 104 | 6 | 9 | 16 | 47 | 17 | 8 | 0 | 0 | 1 | 2 | 2 |
| SG0000006 | Eif2s3y | 0.128252 | 0.170483 | 0.668993975 | 54.1210773 | 86.05055705 | 66 | 4 | 9 | 12 | 30 | 7 | 1 | 0 | 0 | 0 | 3 | 0 |
| SG0000004 | Myo15 | 0.150744 | 0.197464 | 0.939056413 | 9.259894402 | 17.75375335 | 99 | 6 | 9 | 24 | 63 | 0 | 0 | 0 | 0 | 0 | 0 | 0 |
| ISG000000 | Fam160b1 | 0.173563 | 0.224401 | 0.198957197 | 974.038948 | 1118.068487 | 45 | 7 | 9 | 11 | 23 | 3 | 7 | 0 | 0 | 1 | 3 | 2 |
| SG0000006 | 3300002I08Rik | 0.173817 | 0.224682 | -0.781564519 | 27.37949704 | 15.92762793 | 94 | 4 | 9 | 14 | 51 | 10 | 0 | 0 | 0 | 0 | 0 | 0 |
| ISG000000 | Med19 | 0.173978 | 0.224809 | -0.203389403 | 1607.550191 | 1396.16977 | 30 | 8 | 9 | 12 | 20 | 0 | 2 | 0 | 0 | 0 | 1 | 0 |
| SG0000003 | Slc35d2 | 0.208482 | 0.264546 | 0.291200732 | 133.5777842 | 163.4535624 | 60 | 6 | 9 | 14 | 30 | 4 | 2 | 2 | 2 | 2 | 3 | 0 |
| SG0000001 | Fhod1 | 0.213219 | 0.269773 | 0.239917182 | 737.056783 | 870.4086845 | 22 | 9 | 9 | 9 | 11 | 1 | 0 | 0 | 0 | 0 | 0 | 0 |
| SG0000000 | Tmem242 | 0.239109 | 0.298857 | 0.181955155 | 698.1051243 | 791.9445697 | 75 | 6 | 9 | 11 | 21 | 2 | 2 | 0 | 0 | 0 | 0 | 0 |
| ISG000000 | Oma1 | 0.241754 | 0.301783 | -0.211851899 | 486.0862972 | 419.7006116 | 110 | 3 | 9 | 13 | 48 | 9 | 0 | 0 | 0 | 0 | 3 | 0 |
| ISG000000 | Fam104a | 0.24642 | 0.306924 | -0.183202064 | 891.3291631 | 785.0346025 | 47 | 8 | 9 | 14 | 22 | 3 | 2 | 0 | 0 | 1 | 1 | 0 |
| SG0000002 | Capzb | 0.255561 | 0.316811 | -0.147665756 | 5629.837977 | 5082.11016 | 273 | 9 | 9 | 26 | 81 | 25 | 4 | 0 | 0 | 0 | 0 | 0 |
| SG0000004 | Bdh1 | 0.271432 | 0.334082 | 0.174646316 | 528.605137 | 596.6300992 | 66 | 6 | 9 | 12 | 39 | 2 | 2 | 0 | 0 | 0 | 0 | 0 |
| SG0000002 | Atxn3 | 0.279198 | 0.34244 | 0.14267092 | 1690.018852 | 1865.691254 | 80 | 6 | 9 | 16 | 32 | 9 | 11 | 0 | 0 | 2 | 3 | 2 |
| ISG000000 | Pot1b | 0.290965 | 0.355121 | -0.190634735 | 520.9872182 | 456.4994883 | 120 | 5 | 9 | 19 | 74 | 13 | 1 | 0 | 1 | 1 | 1 | 1 |
| ISG000000 | Tnfrsf11b | 0.311902 | 0.377384 | -0.585555031 | 24.82908949 | 16.54592934 | 65 | 7 | 9 | 16 | 54 | 6 | 3 | 0 | 1 | 1 | 4 | 1 |
| SG0000003 | Ago2 | 0.318736 | 0.384639 | -0.145306599 | 4864.176132 | 4398.125802 | 93 | 3 | 9 | 16 | 41 | 8 | 20 | 2 | 4 | 5 | 12 | 0 |
| ISG000000 | Bub3 | 0.329908 | 0.396678 | 0.132468192 | 2697.524618 | 2956.938664 | 42 | 7 | 9 | 9 | 13 | 3 | 2 | 1 | 1 | 1 | 2 | 0 |
| ISG000000 | Tomm70a | 0.343293 | 0.410458 | 0.1131671 | 7202.054355 | 7789.742225 | 76 | 9 | 9 | 15 | 35 | 7 | 5 | 0 | 0 | 0 | 1 | 1 |
| SG0000002 | 2010300C02Rik | 0.374458 | 0.443223 | 0.232401986 | 109.9303951 | 129.145048 | 154 | 2 | 9 | 25 | 94 | 17 | 0 | 0 | 0 | 0 | 0 | 0 |
| SG0000000 | Scrn3 | 0.434468 | 0.503701 | 0.113969882 | 749.5619724 | 811.1775047 | 85 | 5 | 9 | 10 | 62 | 5 | 11 | 0 | 0 | 0 | 5 | 1 |
| SG0000002 | Dram2 | 0.456342 | 0.525896 | -0.126233521 | 645.0372609 | 590.9962516 | 167 | 3 | 9 | 18 | 106 | 19 | 17 | 0 | 0 | 3 | 13 | 4 |
| SG0000002 | Ttc14 | 0.52421 | 0.590916 | 0.130498353 | 2007.551503 | 2197.609959 | 219 | 7 | 9 | 15 | 68 | 14 | 37 | 4 | 6 | 5 | 21 | 1 |
| ISG000000 | Fam178a | 0.524551 | 0.591213 | -0.095307238 | 2853.175628 | 2670.780326 | 134 | 7 | 9 | 14 | 39 | 11 | 10 | 0 | 0 | 0 | 0 | 2 |
| ISG000000 | Vgll3 | 0.526633 | 0.593199 | -0.352231608 | 213.3950019 | 167.1675447 | 140 | 2 | 9 | 18 | 43 | 9 | 6 | 0 | 1 | 1 | 2 | 0 |
| SG0000002 | Ube2i2 | 0.535416 | 0.601919 | 0.085028575 | 1146.028433 | 1215.602386 | 127 | 9 | 9 | 30 | 48 | 9 | 8 | 2 | 2 | 2 | 6 | 2 |
| SG0000005 | Anks1b | 0.553645 | 0.618916 | -0.407754278 | 21.67553436 | 16.33892745 | 119 | 6 | 9 | 18 | 58 | 6 | 5 | 0 | 0 | 0 | 3 | 0 |
| SG0000002 | Adhfe1 | 0.62667 | 0.687922 | 0.198179653 | 69.8560339 | 80.14232635 | 143 | 9 | 9 | 33 | 102 | 20 | 7 | 0 | 0 | 0 | 6 | 0 |
| SG0000000 | 1700109H08Rik | 0.658305 | 0.716769 | -0.265207873 | 44.2387838 | 36.81015568 | 58 | 6 | 9 | 9 | 22 | 0 | 0 | 0 | 0 | 0 | 0 | 0 |
| ISG000000 | Socs7 | 0.66752 | 0.725421 | -0.071293502 | 1188.751137 | 1131.434619 | 74 | 8 | 9 | 13 | 36 | 6 | 5 | 0 | 0 | 0 | 0 | 1 |
| SG0000003 | Myrf | 0.684846 | 0.740804 | 0.097327643 | 1082.642446 | 1158.200079 | 48 | 9 | 9 | 10 | 53 | 3 | 2 | 0 | 0 | 1 | 1 | 1 |
| SG0000000 | Tead3 | 0.727273 | 0.778935 | 0.076860001 | 239.4367926 | 252.5387649 | 42 | 6 | 9 | 12 | 21 | 0 | 3 | 0 | 0 | 3 | 6 | 0 |
| SG0000003 | Msantd3 | 0.759959 | 0.806851 | 0.041525715 | 744.0307674 | 765.7577149 | 99 | 6 | 9 | 15 | 63 | 15 | 3 | 0 | 0 | 0 | 0 | 0 |
| SG0000004 | Thrap3 | 0.779817 | 0.824221 | -0.040040258 | 4699.625845 | 4570.986776 | 117 | 9 | 9 | 22 | 74 | 11 | 0 | 0 | 0 | 0 | 2 | 0 |
| ISG000000 | Brms1l | 0.811317 | 0.850887 | 0.031551371 | 721.4953345 | 737.4480577 | 71 | 9 | 9 | 20 | 44 | 6 | 4 | 0 | 0 | 0 | 0 | 0 |
| SG0000003 | 1110004F10Rik | 0.8523 | 0.884496 | 0.023044683 | 2116.646004 | 2150.727419 | 58 | 6 | 9 | 9 | 39 | 9 | 4 | 0 | 0 | 0 | 3 | 0 |
| SG0000002 | Bcl2l11 | 0.87791 | 0.906098 | 0.013156772 | 1060.601166 | 1070.317641 | 173 | 4 | 9 | 31 | 79 | 18 | 15 | 0 | 1 | 3 | 6 | 10 |
| SG0000002 | Gid8 | 0.886757 | 0.91306 | -0.021960453 | 1691.941741 | 1666.382322 | 23 | 7 | 9 | 13 | 16 | 4 | 0 | 0 | 0 | 0 | 0 | 0 |
| ISG000000 | Tbc1d22b | 0.905022 | 0.928451 | -0.023278593 | 968.6002142 | 953.0967879 | 78 | 8 | 9 | 15 | 40 | 5 | 2 | 0 | 0 | 1 | 3 | 1 |
| SG0000003 | Podxl2 | 0.934957 | 0.95256 | -0.037314103 | 46.92955793 | 45.73132652 | 77 | 9 | 9 | 18 | 51 | 9 | 0 | 0 | 0 | 0 | 0 | 0 |
| SG0000004 | Metap1d | 0.938481 | 0.955606 | 0.010402835 | 413.7712295 | 416.7655907 | 120 | 7 | 9 | 15 | 80 | 8 | 0 | 0 | 0 | 0 | 0 | 0 |
| ISG000000 | Mtx2 | 1 | 1 | -0.007501821 | 1233.680905 | 1227.282579 | 157 | 3 | 9 | 22 | 81 | 19 | 5 | 0 | 0 | 0 | 0 | 1 |
| SG0000002 | Tdh | 0 | 0 | 9.992352191 | 11.53057473 | 11744.88291 | 34 | 8 | 8 | 12 | 27 | 13 | 2 | 0 | 0 | 0 | 0 | 0 |
| ISG000000 | Zfp42 | 0 | 0 | 12.6169139 | 0 | 6280.589294 | 28 | 6 | 8 | 9 | 17 | 1 | 0 | 0 | 0 | 0 | 0 | 0 |
| ISG000000 | Kcnk1 | 8.8E-224 | 3.2E-221 | 9.913168709 | 0 | 963.186693 | 42 | 7 | 8 | 12 | 18 | 6 | 3 | 0 | 0 | 2 | 3 | 2 |
| SG0000002 | Srl | 2.3E-128 | 2.6E-126 | 4.604072369 | 36.73710531 | 893.4470485 | 108 | 6 | 8 | 14 | 50 | 4 | 2 | 0 | 0 | 0 | 0 | 0 |
| SG0000001 | Aldh1a3 | 2.4E-128 | 2.6E-126 | -7.804380614 | 5833.238011 | 26.09498399 | 91 | 7 | 8 | 15 | 29 | 2 | 4 | 0 | 1 | 1 | 1 | 0 |
| SG0000005 | Hic2 | 4.5E-103 | 3.4E-101 | 2.839606024 | 421.1129147 | 3014.429627 | 38 | 6 | 8 | 12 | 28 | 6 | 8 | 0 | 0 | 3 | 5 | 0 |
| ISG000000 | Akr1b8 | 3.6E-101 | 2.6E-99 | -4.40067864 | 7312.591476 | 346.2063584 | 29 | 7 | 8 | 13 | 16 | 3 | 0 | 0 | 0 | 0 | 0 | 0 |
| SG0000003 | Ush1c | 3.83E-89 | 2.12E-87 | -5.213541204 | 1760.136719 | 47.43666153 | 56 | 4 | 8 | 12 | 26 | 10 | 0 | 0 | 0 | 0 | 0 | 0 |
| ISG000000 | H2afy2 | 3.15E-86 | 1.63E-84 | 3.888626421 | 43.70457115 | 647.3215387 | 55 | 7 | 8 | 11 | 30 | 11 | 1 | 0 | 0 | 0 | 0 | 0 |
| SG0000002 | Asns | 3.62E-83 | 1.75E-81 | 2.372877473 | 1189.470756 | 6161.139819 | 66 | 6 | 8 | 12 | 38 | 14 | 0 | 0 | 0 | 0 | 0 | 0 |
| ISG000000 | Tbc1d30 | 1.58E-72 | 6.11E-71 | -4.979270831 | 789.6624809 | 25.03407951 | 87 | 8 | 8 | 15 | 40 | 6 | 4 | 0 | 0 | 0 | 0 | 1 |
| SG0000002 | Elovl2 | 2.3E-67 | 7.79E-66 | 9.74125585 | 0.331601068 | 283.8089766 | 92 | 0 | 8 | 6 | 30 | 8 | 8 | 0 | 0 | 2 | 4 | 0 |
| SG0000003 | Gjc1 | 3.65E-67 | 1.22E-65 | 4.626611756 | 67.09113471 | 1657.349319 | 73 | 6 | 8 | 10 | 33 | 14 | 3 | 0 | 0 | 0 | 3 | 3 |
| SG0000002 | Srd5a1 | 1.08E-66 | 3.58E-65 | -3.5755102 | 2089.034414 | 175.23053 | 49 | 3 | 8 | 11 | 20 | 10 | 1 | 0 | 0 | 0 | 1 | 0 |
| SG0000003 | Fndc3c1 | 2.32E-60 | 6.53E-59 | 7.007118413 | 1.596604083 | 205.3761749 | 149 | 1 | 8 | 9 | 61 | 9 | 1 | 0 | 0 | 0 | 0 | 0 |
| SG0000000 | Ggt1 | 1.49E-59 | 4.07E-58 | 7.224095281 | 1.324060707 | 197.9599196 | 18 | 6 | 8 | 8 | 11 | 0 | 0 | 0 | 0 | 0 | 0 | 0 |
| SG0000002 | Pla2g10 | 1.25E-58 | 3.32E-57 | 7.455012263 | 0 | 174.4616905 | 54 | 4 | 8 | 6 | 26 | 0 | 0 | 0 | 0 | 0 | 0 | 0 |
| ISG000000 | Snrpe | 6.65E-58 | 1.71E-56 | 1.957041621 | 1405.052577 | 5455.327234 | 77 | 8 | 8 | 16 | 40 | 8 | 4 | 0 | 0 | 0 | 5 | 0 |
| SG0000005 | Serpinb6c | 1.6E-56 | 4.01E-55 | 7.393837499 | 0 | 167.1771042 | 80 | 6 | 8 | 18 | 58 | 0 | 0 | 0 | 0 | 0 | 0 | 0 |
| ISG000000 | Fam78a | 5.97E-55 | 1.43E-53 | 7.765320216 | 0.970711415 | 211.1953966 | 47 | 8 | 8 | 15 | 31 | 4 | 1 | 0 | 0 | 0 | 0 | 0 |
| ISG000000 | Itm2b | 1.5E-54 | 3.56E-53 | -2.656603541 | 19840.84687 | 3146.609524 | 51 | 7 | 8 | 10 | 22 | 3 | 1 | 0 | 0 | 0 | 1 | 0 |
| SG0000002 | Mcm6 | 1.08E-52 | 2.42E-51 | 1.772251707 | 6014.972928 | 20546.34616 | 100 | 6 | 8 | 22 | 46 | 10 | 4 | 0 | 0 | 0 | 0 | 0 |
| SG0000002 | Usp12 | 1.58E-52 | 3.53E-51 | -2.643221571 | 5721.638383 | 915.8649268 | 88 | 6 | 8 | 12 | 32 | 6 | 5 | 1 | 2 | 2 | 4 | 0 |
| SG0000002 | Ccnt2 | 1.38E-51 | 2.97E-50 | 1.855893515 | 1534.98155 | 5556.26277 | 165 | 4 | 8 | 22 | 52 | 16 | 25 | 0 | 0 | 8 | 8 | 4 |
| ISG000000 | Bub1b | 4.71E-48 | 9.21E-47 | 1.749906481 | 2114.490574 | 7111.809157 | 71 | 5 | 8 | 17 | 39 | 5 | 0 | 0 | 0 | 0 | 1 | 0 |
| ISG000000 | Lrrc8b | 1E-47 | 1.94E-46 | -2.579044499 | 3321.771687 | 555.9042887 | 143 | 3 | 8 | 15 | 48 | 10 | 7 | 0 | 0 | 0 | 0 | 0 |
| ISG000000 | Il17ra | 1.16E-46 | 2.17E-45 | -2.822549589 | 1570.688708 | 222.0332577 | 29 | 7 | 8 | 9 | 16 | 1 | 1 | 0 | 0 | 1 | 1 | 0 |
| SG0000002 | Loxl4 | 2.45E-46 | 4.5E-45 | -4.600782962 | 2710.169323 | 111.6921755 | 20 | 7 | 8 | 9 | 10 | 0 | 1 | 0 | 0 | 0 | 0 | 0 |
| SG0000002 | Slc30a2 | 1.1E-45 | 1.99E-44 | 8.484877301 | 0.992459638 | 355.5620741 | 32 | 4 | 8 | 4 | 11 | 4 | 4 | 0 | 0 | 0 | 3 | 0 |
| ISG000000 | 2700060E02Rik | 1.74E-45 | 3.1E-44 | 1.722061145 | 1714.528636 | 5656.356871 | 32 | 7 | 8 | 9 | 18 | 1 | 4 | 0 | 0 | 0 | 1 | 0 |
| ISG000000 | Supt16 | 1.36E-43 | 2.27E-42 | 1.614387458 | 5783.456011 | 17707.87657 | 70 | 2 | 8 | 13 | 37 | 4 | 0 | 0 | 0 | 0 | 0 | 0 |
| SG0000000 | Kcnn1 | 1.59E-43 | 2.66E-42 | 3.518781781 | 24.92560979 | 285.6963265 | 24 | 6 | 8 | 12 | 18 | 0 | 2 | 0 | 0 | 0 | 0 | 0 |
| SG0000000 | Hnrnpd | 5.94E-43 | 9.65E-42 | 1.611969834 | 4234.952052 | 12944.9319 | 315 | 0 | 8 | 33 | 127 | 45 | 11 | 0 | 0 | 0 | 3 | 0 |
| SG0000002 | Adgrb2 | 1.23E-42 | 1.99E-41 | 4.84394587 | 6.927812811 | 198.9615469 | 91 | 8 | 8 | 8 | 20 | 11 | 0 | 0 | 0 | 0 | 0 | 0 |
| SG0000002 | Slc25a45 | 7.9E-42 | 1.24E-40 | -4.136805142 | 365.3976205 | 20.77128099 | 31 | 8 | 8 | 8 | 20 | 0 | 3 | 0 | 0 | 0 | 0 | 0 |
| SG0000002 | Ehd4 | 2.34E-41 | 3.64E-40 | -2.215577103 | 5785.591511 | 1245.64016 | 117 | 7 | 8 | 19 | 64 | 10 | 3 | 0 | 0 | 0 | 0 | 1 |
| SG0000002 | Shisa5 | 4.37E-40 | 6.5E-39 | -2.192402394 | 4906.113222 | 1073.392813 | 42 | 8 | 8 | 8 | 30 | 2 | 2 | 0 | 0 | 0 | 0 | 0 |
| SG0000000 | Poldip2 | 5.19E-40 | 7.71E-39 | -2.103053423 | 10495.30713 | 2442.941062 | 36 | 8 | 8 | 8 | 16 | 0 | 0 | 0 | 0 | 0 | 0 | 0 |
| ISG000000 | Lyar | 9.89E-40 | 1.44E-38 | 1.652154069 | 1352.714782 | 4251.617325 | 90 | 8 | 8 | 14 | 30 | 4 | 0 | 0 | 0 | 0 | 0 | 0 |
| SG0000002 | Plec | 2.09E-37 | 2.8E-36 | -2.740416124 | 13105.49318 | 1961.127884 | 132 | 6 | 8 | 24 | 51 | 0 | 13 | 0 | 0 | 0 | 0 | 0 |
| SG0000000 | Tmem184b | 2.45E-36 | 3.18E-35 | -2.346710263 | 5922.44375 | 1164.315728 | 64 | 6 | 8 | 12 | 24 | 12 | 3 | 1 | 1 | 1 | 1 | 2 |
| SG0000002 | Wdr75 | 2.75E-35 | 3.46E-34 | 1.512484493 | 2007.320565 | 5726.904413 | 77 | 6 | 8 | 23 | 42 | 9 | 0 | 0 | 0 | 0 | 2 | 0 |
| SG0000003 | Trim67 | 1.01E-33 | 1.19E-32 | 6.568605261 | 0 | 93.91770155 | 66 | 6 | 8 | 10 | 26 | 10 | 3 | 0 | 0 | 0 | 3 | 0 |
| ISG000000 | Tceb3 | 1.21E-33 | 1.41E-32 | -2.006449752 | 3420.994871 | 851.4337533 | 35 | 7 | 8 | 9 | 19 | 2 | 6 | 0 | 0 | 1 | 1 | 1 |
| ISG000000 | Prps1 | 5.51E-33 | 6.28E-32 | 1.505085872 | 1225.706175 | 3479.063596 | 49 | 7 | 8 | 9 | 28 | 4 | 3 | 0 | 0 | 0 | 0 | 0 |
| SG0000003 | Fam234b | 3.43E-32 | 3.8E-31 | 2.114872747 | 130.9584417 | 567.2488613 | 125 | 2 | 8 | 13 | 33 | 12 | 1 | 0 | 0 | 0 | 0 | 1 |
| ISG000000 | Acot13 | 4.85E-31 | 5.13E-30 | 1.802043254 | 262.1623009 | 914.1959923 | 29 | 5 | 8 | 10 | 17 | 1 | 0 | 0 | 0 | 0 | 0 | 0 |
| SG0000003 | Mras | 7.71E-31 | 8.08E-30 | -3.12618839 | 5213.267301 | 597.0813632 | 166 | 6 | 8 | 30 | 78 | 0 | 1 | 0 | 0 | 0 | 4 | 0 |
| ISG000000 | Inpp1 | 1.15E-29 | 1.15E-28 | -2.217280214 | 1036.474192 | 222.8900283 | 69 | 4 | 8 | 13 | 43 | 1 | 15 | 1 | 2 | 3 | 12 | 0 |
| SG0000005 | Pla2g4e | 4.08E-29 | 3.97E-28 | -3.964592913 | 214.1390759 | 13.71622299 | 90 | 5 | 8 | 11 | 58 | 7 | 6 | 1 | 2 | 3 | 3 | 0 |
| SG0000004 | Zfp473 | 1.39E-27 | 1.26E-26 | 3.59568104 | 109.471492 | 1323.454105 | 96 | 8 | 8 | 8 | 32 | 20 | 0 | 0 | 0 | 0 | 0 | 0 |
| ISG000000 | Lrrc2 | 1.87E-26 | 1.62E-25 | 3.201833655 | 68.88497265 | 633.8295192 | 73 | 8 | 8 | 10 | 59 | 6 | 7 | 0 | 0 | 0 | 2 | 0 |
| SG0000005 | Plagl2 | 1.36E-25 | 1.13E-24 | -1.753224275 | 2166.310829 | 642.6102887 | 31 | 6 | 8 | 9 | 23 | 2 | 10 | 0 | 2 | 2 | 6 | 0 |
| SG0000003 | Nae1 | 2.13E-25 | 1.75E-24 | 1.344598928 | 1091.925263 | 2773.048872 | 94 | 4 | 8 | 26 | 70 | 16 | 0 | 0 | 0 | 0 | 0 | 0 |
| ISG000000 | Taf5 | 3.4E-25 | 2.78E-24 | 1.564447985 | 337.0318417 | 996.8199099 | 33 | 6 | 8 | 7 | 20 | 2 | 6 | 0 | 0 | 0 | 1 | 1 |

| | | | | | | | | | | | | | | | | | | |
|---|---|---|---|---|---|---|---|---|---|---|---|---|---|---|---|---|---|---|
| SG0000002 | Stk17b | 4.17E-25 | 3.38E-24 | 1.439558698 | 558.9558169 | 1516.099934 | 93 | 3 | 8 | 10 | 39 | 7 | 6 | 0 | 0 | 0 | 6 | 1 |
| SG0000004 | Cmklr1 | 8.25E-25 | 6.6E-24 | 6.079192273 | 2.992940161 | 202.3565458 | 26 | 7 | 8 | 8 | 28 | 1 | 1 | 0 | 0 | 0 | 0 | 0 |
| SG0000002 | Gusb | 1.15E-24 | 9.12E-24 | -1.663158028 | 2695.321871 | 851.0404084 | 51 | 4 | 8 | 10 | 20 | 0 | 5 | 0 | 0 | 0 | 0 | 0 |
| SG0000004 | Rragb | 1.39E-24 | 1.1E-23 | 1.728574783 | 214.6885 | 711.4782712 | 85 | 6 | 8 | 10 | 39 | 7 | 1 | 0 | 0 | 1 | 1 | 1 |
| SG0000003 | Rasgef1a | 7.16E-24 | 5.48E-23 | -2.576604022 | 526.5840582 | 88.27395998 | 33 | 8 | 8 | 15 | 21 | 15 | 0 | 0 | 0 | 0 | 0 | 0 |
| JSG000000 | Tmco6 | 8.44E-24 | 6.44E-23 | 1.856594721 | 132.7261851 | 480.6703265 | 13 | 8 | 8 | 10 | 13 | 1 | 1 | 0 | 0 | 0 | 0 | 0 |
| SG0000007 | Prss36 | 1.16E-23 | 8.84E-23 | 2.15814343 | 73.28012959 | 327.0786043 | 32 | 6 | 8 | 12 | 26 | 4 | 0 | 0 | 0 | 0 | 0 | 2 |
| JSG000000 | Zfp850 | 1.19E-23 | 9.03E-23 | 1.572598069 | 385.9491563 | 1147.966688 | 49 | 5 | 8 | 9 | 34 | 7 | 8 | 0 | 1 | 0 | 1 | 2 |
| SG0000002 | Vdac2 | 1.42E-23 | 1.07E-22 | 1.188625648 | 5180.959415 | 11809.24585 | 31 | 5 | 8 | 6 | 13 | 1 | 3 | 0 | 1 | 0 | 0 | 1 |
| SG0000003 | Spryd3 | 1.87E-23 | 1.4E-22 | -1.702405952 | 1693.769461 | 520.4500346 | 25 | 7 | 8 | 12 | 21 | 3 | 1 | 0 | 0 | 1 | 1 | 0 |
| JSG000000 | Nutf2 | 2.51E-23 | 1.86E-22 | 1.333227844 | 875.5888541 | 2206.184335 | 29 | 6 | 8 | 10 | 20 | 2 | 0 | 0 | 0 | 0 | 0 | 0 |
| SG0000002 | Mettl14 | 1.43E-22 | 1.03E-21 | 1.366744143 | 569.0112869 | 1467.411318 | 130 | 0 | 8 | 12 | 66 | 14 | 35 | 0 | 0 | 0 | 18 | 1 |
| SG0000004 | Cyp4f16 | 4.17E-22 | 2.94E-21 | -1.880310628 | 818.1547434 | 222.2314666 | 66 | 4 | 8 | 12 | 30 | 8 | 0 | 0 | 0 | 0 | 0 | 0 |
| JSG000000 | Tpm4 | 9.18E-22 | 6.37E-21 | -1.549018078 | 13550.31677 | 4630.720223 | 31 | 7 | 8 | 10 | 23 | 2 | 2 | 0 | 0 | 1 | 1 | 0 |
| SG0000002 | Mlh3 | 1.22E-21 | 8.4E-21 | 1.38666242 | 460.1670778 | 1203.21303 | 128 | 0 | 8 | 16 | 50 | 20 | 4 | 0 | 2 | 2 | 2 | 0 |
| SG0000002 | Rnf32 | 1.44E-21 | 9.85E-21 | 3.26864253 | 13.42381398 | 129.3705344 | 184 | 0 | 8 | 0 | 55 | 8 | 5 | 0 | 0 | 0 | 0 | 0 |
| JSG000000 | 2610318N02Rik | 1.9E-21 | 1.29E-20 | 1.566757636 | 253.1612292 | 749.9602415 | 26 | 7 | 8 | 10 | 17 | 1 | 0 | 0 | 0 | 0 | 0 | 0 |
| SG0000002 | Ppil1 | 2.92E-21 | 1.97E-20 | 1.266488953 | 1124.357707 | 2704.927724 | 36 | 2 | 8 | 16 | 26 | 4 | 0 | 0 | 0 | 0 | 0 | 0 |
| SG0000003 | Paqr5 | 3.42E-21 | 2.29E-20 | 5.850622285 | 0 | 56.70491441 | 89 | 5 | 8 | 13 | 22 | 7 | 6 | 0 | 1 | 1 | 1 | 0 |
| JSG000000 | Poc5 | 3.59E-21 | 2.4E-20 | -1.570791267 | 1890.895963 | 636.5204284 | 51 | 2 | 8 | 5 | 23 | 7 | 2 | 0 | 1 | 0 | 1 | 1 |
| JSG000000 | Hspa12a | 8.27E-21 | 5.44E-20 | 4.386068998 | 12.85832571 | 268.8578703 | 63 | 3 | 8 | 9 | 20 | 7 | 4 | 0 | 0 | 0 | 0 | 1 |
| SG0000005 | Dppa4 | 1.36E-20 | 8.86E-20 | 7.815022829 | 12.15262399 | 2736.694687 | 38 | 6 | 8 | 8 | 20 | 4 | 0 | 0 | 0 | 0 | 0 | 0 |
| SG0000002 | Smad7 | 5.98E-20 | 3.74E-19 | 1.277303785 | 632.1913527 | 1532.340584 | 64 | 6 | 8 | 11 | 43 | 5 | 4 | 0 | 0 | 1 | 3 | 1 |
| SG0000002 | Slc43a1 | 9.23E-20 | 5.71E-19 | 7.162451469 | 3.489843332 | 499.9408975 | 87 | 4 | 8 | 12 | 41 | 5 | 4 | 0 | 0 | 0 | 0 | 0 |
| SG0000000 | Gpa33 | 1.48E-19 | 9.05E-19 | 3.965179142 | 155.7163054 | 2432.046928 | 84 | 4 | 8 | 12 | 48 | 12 | 4 | 0 | 0 | 0 | 2 | 0 |
| JSG000000 | Cdh3 | 1.67E-19 | 1.02E-18 | 6.218626009 | 24.37552963 | 1815.294206 | 63 | 8 | 8 | 14 | 28 | 6 | 1 | 0 | 0 | 0 | 0 | 0 |
| SG0000001 | Ikbip | 4.08E-19 | 2.43E-18 | -1.656456617 | 1343.05129 | 426.0391113 | 59 | 1 | 8 | 6 | 28 | 0 | 5 | 0 | 0 | 0 | 2 | 0 |
| SG0000002 | Ppef2 | 6.05E-19 | 3.58E-18 | 3.346782036 | 10.51167974 | 106.9433791 | 43 | 5 | 8 | 12 | 30 | 4 | 1 | 0 | 0 | 0 | 0 | 0 |
| SG0000000 | Keap1 | 6.1E-19 | 3.6E-18 | -1.965374262 | 1889.144247 | 483.7583945 | 36 | 8 | 8 | 20 | 54 | 0 | 0 | 0 | 0 | 0 | 2 | 0 |
| SG0000004 | E2f8 | 7.59E-19 | 4.47E-18 | 1.179249448 | 930.5787793 | 2107.378771 | 52 | 6 | 8 | 8 | 20 | 6 | 6 | 0 | 2 | 2 | 6 | 2 |
| SG0000005 | Hopx | 8.86E-19 | 5.19E-18 | -2.62885749 | 525.1747103 | 84.90609077 | 87 | 5 | 8 | 13 | 35 | 11 | 12 | 0 | 3 | 0 | 0 | 3 |
| SG0000002 | Slc11a1 | 9.45E-19 | 5.52E-18 | 3.966132165 | 8.400114532 | 131.2834424 | 26 | 6 | 8 | 8 | 12 | 2 | 0 | 0 | 0 | 0 | 0 | 0 |
| SG0000000 | Pttg1ip | 1.62E-18 | 9.35E-18 | -1.314300529 | 5342.207214 | 2148.210559 | 52 | 8 | 8 | 13 | 32 | 10 | 2 | 0 | 0 | 0 | 1 | 0 |
| JSG000000 | Serinc5 | 2.52E-18 | 1.44E-17 | 1.457126324 | 281.9334157 | 774.0789969 | 132 | 6 | 8 | 18 | 60 | 25 | 4 | 0 | 0 | 2 | 2 | 0 |
| JSG000000 | Me2 | 2.54E-18 | 1.45E-17 | 1.195500071 | 1423.417735 | 3259.971129 | 74 | 6 | 8 | 15 | 48 | 6 | 1 | 0 | 0 | 0 | 0 | 0 |
| SG0000004 | Mab21l3 | 1.72E-17 | 9.45E-17 | 2.841650884 | 29.54912307 | 211.8198959 | 46 | 3 | 8 | 8 | 29 | 6 | 4 | 0 | 0 | 0 | 2 | 0 |
| JSG000000 | Moxd1 | 1.8E-17 | 9.86E-17 | 2.815167474 | 18.10872852 | 127.4494045 | 180 | 2 | 8 | 19 | 70 | 19 | 1 | 0 | 0 | 0 | 1 | 0 |
| SG0000002 | Unc5a | 2.16E-17 | 1.18E-16 | 7.245343879 | 0.331601068 | 50.31317888 | 80 | 6 | 8 | 10 | 24 | 6 | 0 | 0 | 0 | 0 | 0 | 2 |
| JSG000000 | Usp25 | 2.53E-17 | 1.38E-16 | -1.367945482 | 2055.856871 | 796.5265445 | 251 | 1 | 8 | 25 | 122 | 21 | 4 | 0 | 0 | 1 | 2 | 0 |
| JSG000000 | Soga1 | 3.74E-17 | 2.02E-16 | -1.940907486 | 1158.219941 | 301.6613468 | 56 | 7 | 8 | 9 | 24 | 4 | 5 | 0 | 0 | 0 | 0 | 1 |
| JSG000000 | Slc39a9 | 4.4E-17 | 2.36E-16 | -1.347332706 | 2000.651353 | 786.2919939 | 85 | 7 | 8 | 19 | 41 | 5 | 4 | 0 | 0 | 2 | 3 | 0 |
| SG0000001 | Tmc4 | 6.8E-17 | 3.62E-16 | 2.056797988 | 54.7457764 | 227.7762868 | 32 | 4 | 8 | 10 | 14 | 4 | 1 | 0 | 0 | 0 | 0 | 0 |
| SG0000003 | Ankrd10 | 8.53E-17 | 4.5E-16 | 1.115606819 | 1147.223013 | 2485.873165 | 96 | 4 | 8 | 16 | 36 | 2 | 3 | 0 | 0 | 1 | 2 | 0 |
| JSG000000 | Slc7a8 | 2.38E-16 | 1.22E-15 | 2.526542288 | 24.39124401 | 140.5396815 | 65 | 7 | 8 | 16 | 34 | 10 | 2 | 0 | 0 | 0 | 0 | 0 |
| SG0000002 | Slc45a3 | 2.67E-16 | 1.37E-15 | -2.074943185 | 513.637258 | 121.9091718 | 27 | 8 | 8 | 12 | 23 | 2 | 12 | 4 | 4 | 4 | 8 | 0 |
| SG0000002 | Nacc2 | 2.68E-16 | 1.37E-15 | -1.614377488 | 867.4064712 | 283.3000229 | 119 | 7 | 8 | 17 | 59 | 8 | 8 | 0 | 0 | 0 | 0 | 0 |
| SG0000002 | Helb | 1.46E-15 | 7.14E-15 | 1.247415356 | 971.9202032 | 2307.491176 | 111 | 4 | 8 | 9 | 43 | 6 | 6 | 0 | 0 | 1 | 3 | 0 |
| SG0000004 | Nxt2 | 2.17E-15 | 1.05E-14 | -1.507304153 | 762.72058 | 268.300636 | 121 | 4 | 8 | 12 | 56 | 8 | 28 | 0 | 4 | 4 | 20 | 0 |
| JSG000000 | Itpr3 | 3.01E-15 | 1.45E-14 | -1.403263575 | 8127.120448 | 3072.644164 | 47 | 5 | 8 | 9 | 25 | 6 | 0 | 0 | 0 | 0 | 0 | 0 |
| SG0000001 | Katna1 | 4.11E-15 | 1.96E-14 | 1.05758963 | 900.7458635 | 1874.858422 | 67 | 3 | 8 | 12 | 34 | 6 | 0 | 0 | 0 | 0 | 0 | 0 |
| JSG000000 | Xndc1 | 5.36E-15 | 2.54E-14 | 1.45945727 | 155.5476102 | 427.7635336 | 55 | 8 | 8 | 18 | 28 | 6 | 0 | 0 | 0 | 0 | 0 | 0 |
| SG0000004 | Dynlrb1 | 6.55E-15 | 3.08E-14 | -1.182990713 | 3039.819853 | 1338.850724 | 58 | 6 | 8 | 8 | 16 | 6 | 2 | 0 | 0 | 0 | 0 | 0 |
| SG0000006 | Dhrs3 | 1.3E-14 | 5.99E-14 | -2.406875809 | 347.6711824 | 65.55817877 | 140 | 0 | 8 | 8 | 48 | 16 | 0 | 0 | 0 | 0 | 0 | 0 |
| JSG000000 | Lonrf1 | 2.59E-14 | 1.17E-13 | 1.151066103 | 418.7383335 | 929.925473 | 58 | 3 | 8 | 10 | 28 | 5 | 5 | 0 | 1 | 1 | 1 | 2 |
| JSG000000 | Ddx43 | 3.8E-14 | 1.7E-13 | 7.403689 | 1.254128903 | 212.3609425 | 61 | 4 | 8 | 9 | 35 | 4 | 0 | 0 | 0 | 0 | 0 | 0 |
| JSG000000 | Abcb10 | 4.2E-14 | 1.87E-13 | 1.108581005 | 514.472512 | 1109.374836 | 41 | 8 | 8 | 10 | 19 | 1 | 2 | 0 | 0 | 0 | 0 | 0 |
| JSG000000 | Edem1 | 1.89E-13 | 8.09E-13 | -1.124986939 | 2663.888883 | 1221.409496 | 51 | 6 | 8 | 11 | 23 | 5 | 5 | 2 | 2 | 2 | 3 | 1 |
| SG0000003 | Fam71f1 | 2.27E-13 | 9.64E-13 | -5.699275226 | 51.19218824 | 0.985260105 | 82 | 8 | 8 | 21 | 25 | 4 | 2 | 0 | 0 | 1 | 1 | 0 |
| JSG000000 | Btc | 2.48E-13 | 1.05E-12 | -5.054652487 | 56.67582214 | 1.705280447 | 161 | 4 | 8 | 20 | 89 | 15 | 5 | 0 | 0 | 0 | 5 | 1 |
| SG0000002 | Pef1 | 2.55E-13 | 1.08E-12 | -1.155937429 | 1762.310061 | 790.8797967 | 51 | 2 | 8 | 4 | 18 | 5 | 0 | 0 | 0 | 0 | 0 | 0 |
| SG0000004 | Mios | 4.72E-13 | 1.96E-12 | 1.009560778 | 727.0601333 | 1463.788761 | 60 | 4 | 8 | 14 | 35 | 4 | 3 | 0 | 0 | 0 | 0 | 1 |
| SG0000004 | Rcc2 | 5.07E-13 | 2.1E-12 | 0.895858654 | 7683.615373 | 14297.03379 | 34 | 8 | 8 | 14 | 26 | 2 | 2 | 0 | 0 | 0 | 0 | 0 |
| SG0000002 | Hsd17b11 | 7.59E-13 | 3.12E-12 | -1.118626613 | 2132.457289 | 982.0648722 | 93 | 6 | 8 | 13 | 40 | 7 | 4 | 0 | 0 | 0 | 1 | 0 |
| SG0000004 | Shb | 9.18E-13 | 3.75E-12 | -1.395485764 | 1608.454063 | 611.4002173 | 65 | 4 | 8 | 8 | 24 | 5 | 2 | 0 | 0 | 1 | 1 | 1 |
| JSG000000 | Prcp | 1.03E-12 | 4.21E-12 | 1.046132804 | 505.6753115 | 1044.21302 | 113 | 7 | 8 | 17 | 71 | 10 | 2 | 0 | 0 | 0 | 2 | 0 |
| JSG000000 | Tbx21 | 1.06E-12 | 4.33E-12 | 5.075290145 | 0 | 32.71433287 | 30 | 6 | 8 | 10 | 23 | 0 | 1 | 0 | 0 | 1 | 1 | 0 |
| SG0000000 | Capns1 | 1.55E-12 | 6.21E-12 | -0.978784094 | 14175.81454 | 7192.910411 | 30 | 8 | 8 | 11 | 29 | 0 | 0 | 0 | 0 | 0 | 0 | 0 |
| SG0000005 | Mpp3 | 1.66E-12 | 6.65E-12 | 2.090031896 | 32.57568243 | 138.6934065 | 83 | 0 | 8 | 7 | 39 | 26 | 5 | 0 | 0 | 0 | 0 | 0 |
| SG0000002 | Gtf3c5 | 1.91E-12 | 7.62E-12 | 1.029574391 | 545.7901877 | 1114.188054 | 33 | 8 | 8 | 8 | 15 | 0 | 0 | 0 | 0 | 0 | 0 | 0 |
| JSG000000 | Atp2c2 | 2.29E-12 | 9.1E-12 | 3.601912942 | 8.300057099 | 100.7778102 | 50 | 8 | 8 | 13 | 23 | 14 | 2 | 0 | 0 | 1 | 2 | 0 |
| SG0000000 | Sri | 2.58E-12 | 1.02E-11 | -1.119741276 | 1664.567338 | 765.9945104 | 55 | 6 | 8 | 8 | 23 | 2 | 1 | 0 | 0 | 1 | 2 | 0 |
| JSG000000 | Map1b | 2.87E-12 | 1.13E-11 | 0.95669639 | 1028.202923 | 1995.598386 | 122 | 6 | 8 | 16 | 53 | 8 | 13 | 2 | 2 | 3 | 8 | 1 |
| SG0000002 | Coro2a | 7.78E-12 | 3E-11 | 1.367045234 | 153.4013854 | 395.6861987 | 78 | 4 | 8 | 5 | 35 | 14 | 3 | 0 | 3 | 0 | 3 | 0 |
| SG0000004 | Sys1 | 1.07E-11 | 4.1E-11 | -1.268881222 | 690.6144386 | 286.592186 | 30 | 3 | 8 | 6 | 25 | 0 | 0 | 0 | 0 | 0 | 6 | 0 |
| SG0000002 | Spp1 | 1.09E-11 | 4.16E-11 | 0.811596321 | 10731.23777 | 18834.95794 | 68 | 0 | 8 | 0 | 35 | 0 | 8 | 0 | 0 | 0 | 0 | 0 |
| JSG000000 | Adcy5 | 1.1E-11 | 4.2E-11 | 4.679086822 | 1.620695873 | 41.51901215 | 107 | 5 | 8 | 18 | 55 | 12 | 3 | 0 | 0 | 1 | 2 | 0 |
| JSG000000 | Ube2d1 | 1.24E-11 | 4.71E-11 | -1.103241884 | 1431.980595 | 666.5430943 | 55 | 7 | 8 | 11 | 18 | 4 | 1 | 0 | 0 | 0 | 1 | 0 |
| SG0000001 | Cars | 2.07E-11 | 7.75E-11 | 0.856809935 | 1880.141306 | 3404.991106 | 48 | 6 | 8 | 16 | 38 | 10 | 0 | 0 | 0 | 0 | 0 | 0 |
| SG0000001 | Slc3a2 | 2.53E-11 | 9.39E-11 | -0.924313151 | 13726.49259 | 7232.918907 | 36 | 8 | 8 | 13 | 22 | 1 | 1 | 0 | 0 | 0 | 0 | 0 |
| SG0000003 | Fam3a | 3.38E-11 | 1.24E-10 | -1.284735773 | 577.3754478 | 236.9814256 | 48 | 8 | 8 | 16 | 16 | 13 | 2 | 0 | 0 | 0 | 0 | 0 |
| SG0000005 | B3gnt2 | 4.49E-11 | 1.64E-10 | -2.079626205 | 157.4962169 | 37.25977766 | 100 | 6 | 8 | 8 | 24 | 12 | 4 | 2 | 2 | 2 | 8 | 4 |
| SG0000003 | Rab3a | 5.87E-11 | 2.12E-10 | -1.347258273 | 475.0054045 | 186.695306 | 25 | 8 | 8 | 16 | 20 | 4 | 17 | 8 | 8 | 16 | 20 | 4 |
| SG0000002 | Fam73b | 7.27E-11 | 2.61E-10 | -1.189014364 | 715.3417172 | 313.7505992 | 52 | 8 | 8 | 16 | 25 | 0 | 3 | 2 | 2 | 2 | 2 | 0 |
| SG0000006 | Tm9sf4 | 8.6E-11 | 3.07E-10 | -0.948496713 | 3572.098546 | 1850.961861 | 74 | 5 | 8 | 15 | 35 | 8 | 1 | 0 | 0 | 0 | 0 | 2 |
| SG0000003 | Tmf1 | 1.13E-10 | 4.01E-10 | -0.996370181 | 2013.660858 | 1009.366802 | 108 | 8 | 8 | 16 | 54 | 34 | 1 | 0 | 0 | 0 | 0 | 0 |
| SG0000004 | Agtr1a | 1.41E-10 | 4.98E-10 | 6.446378183 | 0.34247518 | 29.86636025 | 89 | 5 | 8 | 12 | 28 | 6 | 2 | 0 | 0 | 1 | 1 | 2 |
| SG0000006 | Tcp1 | 1.67E-10 | 5.83E-10 | 0.762234165 | 11960.37321 | 20286.17086 | 37 | 2 | 8 | 8 | 10 | 0 | 5 | 0 | 2 | 2 | 2 | 0 |
| JSG000000 | Kif27 | 2.16E-10 | 7.49E-10 | 3.018259612 | 7.250883335 | 58.74590457 | 98 | 3 | 8 | 15 | 43 | 9 | 2 | 0 | 0 | 0 | 1 | 0 |
| JSG000000 | Mettl7a1 | 2.34E-10 | 8.1E-10 | -1.036052595 | 1251.931528 | 610.516869 | 22 | 7 | 8 | 8 | 11 | 1 | 1 | 0 | 0 | 0 | 0 | 0 |
| SG0000002 | Map3k8 | 2.48E-10 | 8.56E-10 | -1.895846563 | 252.4866936 | 67.84718172 | 100 | 0 | 8 | 12 | 26 | 10 | 8 | 0 | 2 | 0 | 0 | 2 |
| SG0000003 | Crebl2 | 3.19E-10 | 1.1E-09 | -1.177138391 | 666.4613004 | 294.7277111 | 79 | 4 | 8 | 11 | 39 | 4 | 3 | 0 | 0 | 1 | 3 | 0 |
| SG0000002 | Eif6 | 3.34E-10 | 1.15E-09 | -1.233753538 | 5713.285213 | 2429.344373 | 41 | 8 | 8 | 11 | 23 | 3 | 3 | 0 | 0 | 0 | 0 | 0 |
| JSG000000 | Rprd1a | 3.56E-10 | 1.22E-09 | 0.997867449 | 442.2053532 | 883.1043612 | 97 | 4 | 8 | 12 | 34 | 9 | 5 | 0 | 0 | 2 | 4 | 1 |
| SG0000001 | Ubxn11 | 4.79E-10 | 1.62E-09 | 2.386320397 | 15.70068129 | 82.08664704 | 45 | 8 | 8 | 10 | 30 | 2 | 1 | 0 | 0 | 0 | 0 | 0 |
| JSG000000 | Lpcat2 | 6.35E-10 | 2.13E-09 | 3.884192628 | 5.14316154 | 75.9431584 | 144 | 3 | 8 | 14 | 65 | 13 | 3 | 0 | 0 | 0 | 0 | 0 |
| JSG000000 | Rmi1 | 7.03E-10 | 2.35E-09 | 1.07554426 | 234.6409375 | 494.5097617 | 23 | 5 | 8 | 5 | 10 | 2 | 4 | 0 | 0 | 0 | 0 | 0 |
| SG0000002 | Nos3 | 7.75E-10 | 2.58E-09 | 2.68875568 | 10.19095278 | 65.70688822 | 26 | 6 | 8 | 10 | 12 | 0 | 4 | 0 | 0 | 2 | 2 | 0 |
| JSG000000 | BC021891 | 7.96E-10 | 2.65E-09 | 6.491219228 | 0.307509278 | 27.66367793 | 44 | 6 | 8 | 12 | 20 | 7 | 7 | 0 | 0 | 2 | 3 | 0 |
| SG0000003 | Fam76b | 9.3E-10 | 3.07E-09 | 0.823132792 | 1041.332948 | 1842.37177 | 118 | 2 | 8 | 20 | 46 | 14 | 20 | 0 | 2 | 6 | 12 | 8 |
| SG0000003 | Gcfc2 | 9.63E-10 | 3.18E-09 | 0.963095857 | 380.0417819 | 740.8871925 | 151 | 4 | 8 | 26 | 70 | 10 | 6 | 0 | 0 | 2 | 5 | 2 |
| SG0000000 | Rnmt | 1.23E-09 | 4.02E-09 | 0.853146678 | 1814.618444 | 3277.993302 | 119 | 6 | 8 | 14 | 36 | 4 | 5 | 0 | 0 | 0 | 0 | 2 |
| SG0000001 | Mpdu1 | 1.37E-09 | 4.46E-09 | 0.830231532 | 1021.709217 | 1816.56903 | 32 | 8 | 8 | 16 | 32 | 4 | 0 | 0 | 0 | 0 | 0 | 0 |
| SG0000005 | Snx21 | 1.61E-09 | 5.22E-09 | -2.447108323 | 673.9964947 | 123.596061 | 51 | 8 | 8 | 12 | 20 | 20 | 3 | 0 | 0 | 0 | 4 | 0 |
| JSG000000 | Fam207a | 1.69E-09 | 5.45E-09 | -1.030834778 | 1496.090807 | 732.227045 | 30 | 7 | 8 | 11 | 19 | 3 | 1 | 0 | 0 | 0 | 0 | 0 |
| SG0000002 | Gtf2e1 | 1.88E-09 | 6.04E-09 | 0.81049771 | 1076.982429 | 1888.829427 | 71 | 6 | 8 | 13 | 30 | 7 | 1 | 0 | 0 | 1 | 1 | 0 |
| SG0000002 | Il1rl1 | 1.98E-09 | 6.36E-09 | -3.29546614 | 984.3866529 | 100.2608907 | 123 | 7 | 8 | 23 | 86 | 6 | 7 | 0 | 0 | 4 | 8 | 2 |
| SG0000003 | Igdcc4 | 2.02E-09 | 6.48E-09 | 1.788360831 | 70.87711933 | 244.825309 | 99 | 6 | 8 | 21 | 99 | 6 | 6 | 0 | 2 | 0 | 12 | 0 |
| SG0000004 | Cfap43 | 2.35E-09 | 7.51E-09 | -1.210485247 | 699.8745641 | 302.4320696 | 157 | 5 | 8 | 22 | 86 | 15 | 2 | 0 | 0 | 0 | 0 | 0 |
| SG0000003 | Ubac1 | 2.83E-09 | 8.97E-09 | 0.816212945 | 963.3661504 | 1696.27383 | 31 | 4 | 8 | 9 | 17 | 0 | 2 | 0 | 0 | 0 | 0 | 0 |
| JSG000000 | Cpsf3l | 3.31E-09 | 1.04E-08 | 0.781762543 | 1373.315901 | 2361.045864 | 60 | 8 | 8 | 16 | 45 | 6 | 0 | 0 | 0 | 0 | 0 | 0 |
| JSG000000 | Zbtb2 | 4.72E-09 | 1.47E-08 | -1.165666574 | 490.1174366 | 218.473894 | 47 | 6 | 8 | 6 | 16 | 8 | 3 | 0 | 0 | 0 | 0 | 0 |
| SG0000004 | Cacng5 | 9.39E-09 | 2.87E-08 | 4.510577997 | 0 | 21.7939334 | 86 | 8 | 8 | 12 | 36 | 12 | 2 | 0 | 0 | 0 | 0 | 2 |
| SG0000002 | Prrx1 | 1.22E-08 | 3.68E-08 | -1.843041111 | 1573.943928 | 438.7110761 | 298 | 0 | 8 | 22 | 114 | 20 | 8 | 0 | 0 | 2 | 6 | 4 |
| SG0000003 | Secisbp2l | 1.22E-08 | 3.69E-08 | -0.888679041 | 1653.714752 | 893.1852708 | 85 | 3 | 8 | 7 | 29 | 5 | 12 | 0 | 1 | 1 | 3 | 1 |
| JSG000000 | Fgl1 | 1.36E-08 | 4.09E-08 | 4.492237993 | 0 | 21.50600357 | 59 | 4 | 8 | 11 | 35 | 5 | 0 | 0 | 0 | 0 | 0 | 0 |
| SG0000004 | Kcnj12 | 1.77E-08 | 5.29E-08 | 2.773420759 | 11.90083198 | 81.36922182 | 55 | 6 | 8 | 9 | 23 | 8 | 0 | 0 | 0 | 0 | 0 | 0 |
| JSG000000 | Gbx1 | 1.79E-08 | 5.33E-08 | 4.471859492 | 0 | 21.19033423 | 34 | 7 | 8 | 8 | 34 | 1 | 1 | 1 | 1 | 1 | 1 | 1 |
| SG0000001 | Tsen15 | 1.81E-08 | 5.39E-08 | 0.945032818 | 292.2218856 | 562.5951239 | 78 | 6 | 8 | 6 | 32 | 6 | 0 | 0 | 0 | 0 | 0 | 0 |
| JSG000000 | Tbx18 | 2.12E-08 | 6.29E-08 | -2.046321985 | 111.4100774 | 26.9724364 | 73 | 6 | 8 | 10 | 36 | 3 | 6 | 1 | 1 | 1 | 3 | 2 |
| JSG000000 | BC030336 | 2.51E-08 | 7.41E-08 | -0.920883371 | 1069.749988 | 565.0264431 | 113 | 3 | 8 | 14 | 30 | 15 | 10 | 0 | 0 | 2 | 4 | 2 |
| SG0000003 | Pbdc1 | 2.7E-08 | 7.93E-08 | 0.743732675 | 1185.635898 | 1985.350474 | 104 | 2 | 8 | 8 | 67 | 9 | 9 | 0 | 1 | 1 | 8 | 1 |
| SG0000003 | Chsy1 | 3.17E-08 | 9.28E-08 | 0.782988938 | 749.5845643 | 1289.804213 | 111 | 5 | 8 | 16 | 51 | 22 | 5 | 0 | 1 | 1 | 3 | 1 |
| JSG000000 | Abce1 | 3.43E-08 | 1E-07 | 0.665061263 | 9619.243701 | 15252.6153 | 71 | 4 | 8 | 8 | 33 | 3 | 5 | 0 | 0 | 0 | 1 | 0 |

| | | | | | | | | | | | | | | | | | | |
|---|---|---|---|---|---|---|---|---|---|---|---|---|---|---|---|---|---|---|
| SG0000007 | Arpc4 | 3.79E-08 | 1.1E-07 | -0.779221012 | 4151.032172 | 2418.728942 | 28 | 4 | 8 | 8 | 14 | 2 | 0 | 0 | 0 | 0 | 0 | 0 |
| SG0000006 | Myl4 | 3.98E-08 | 1.16E-07 | 4.804732036 | 2.863950664 | 80.04496082 | 97 | 6 | 8 | 18 | 47 | 7 | 0 | 0 | 0 | 0 | 0 | 0 |
| SG0000002 | Dmbx1 | 4.87E-08 | 1.4E-07 | 4.385064335 | 0 | 19.89468834 | 69 | 8 | 8 | 16 | 33 | 0 | 9 | 3 | 3 | 6 | 9 | 0 |
| SG0000003 | Fermt2 | 5.15E-08 | 1.48E-07 | 0.683832393 | 2587.990901 | 4157.352095 | 123 | 2 | 8 | 9 | 46 | 8 | 3 | 0 | 1 | 1 | 3 | 0 |
| SG0000000 | Pparg | 5.36E-08 | 1.54E-07 | -2.869531204 | 166.088725 | 22.72611423 | 143 | 5 | 8 | 23 | 69 | 12 | 4 | 0 | 0 | 0 | 2 | 0 |
| SG0000002 | Nbn | 7.37E-08 | 2.09E-07 | 1.044475254 | 494.5649612 | 1020.097599 | 84 | 4 | 8 | 15 | 41 | 17 | 0 | 0 | 0 | 0 | 0 | 0 |
| SG0000003 | Rad51ap1 | 8.62E-08 | 2.43E-07 | 0.766896362 | 690.8827796 | 1175.609743 | 58 | 2 | 8 | 8 | 32 | 5 | 1 | 0 | 0 | 0 | 2 | 2 |
| SG0000003 | Mettl20 | 9.25E-08 | 2.6E-07 | 1.100686818 | 131.644389 | 282.3202962 | 54 | 6 | 8 | 10 | 44 | 2 | 6 | 0 | 0 | 2 | 2 | 0 |
| SG0000002 | Cct8 | 1.05E-07 | 2.94E-07 | 0.634831876 | 14595.78217 | 22663.69987 | 52 | 8 | 8 | 12 | 34 | 4 | 3 | 0 | 0 | 1 | 1 | 0 |
| SG0000001 | Rnf168 | 2.06E-07 | 5.61E-07 | -0.777521032 | 2267.529892 | 1322.805123 | 94 | 6 | 8 | 12 | 43 | 6 | 10 | 0 | 0 | 0 | 5 | 0 |
| SG0000002 | Mpc1 | 2.22E-07 | 6.03E-07 | -1.376217887 | 197.5670501 | 76.10823506 | 116 | 8 | 8 | 12 | 24 | 12 | 4 | 0 | 0 | 0 | 4 | 0 |
| JSG000000 | 43526 | 2.45E-07 | 6.65E-07 | -0.765741368 | 2299.060534 | 1352.194875 | 48 | 5 | 8 | 12 | 24 | 7 | 0 | 0 | 0 | 0 | 0 | 0 |
| SG0000006 | Clk2 | 2.84E-07 | 7.65E-07 | 0.697403715 | 1141.227059 | 1850.595482 | 60 | 8 | 8 | 16 | 28 | 8 | 4 | 0 | 0 | 0 | 1 | 8 |
| SG0000002 | Tmem230 | 6.32E-07 | 1.66E-06 | -0.827422248 | 1008.468667 | 568.3076388 | 60 | 4 | 8 | 16 | 68 | 16 | 4 | 0 | 0 | 0 | 4 | 4 |
| JSG000000 | Slc25a33 | 7.06E-07 | 1.84E-06 | 0.910559782 | 428.8337284 | 806.1108062 | 46 | 7 | 8 | 13 | 25 | 9 | 1 | 0 | 0 | 0 | 0 | 0 |
| SG0000002 | Rragc | 7.88E-07 | 2.05E-06 | -0.720114866 | 2597.816858 | 1577.002406 | 92 | 6 | 8 | 16 | 44 | 4 | 12 | 0 | 1 | 2 | 2 | 1 |
| JSG000000 | Slc45a4 | 9.82E-07 | 2.54E-06 | 0.813632029 | 948.4408461 | 1667.008811 | 58 | 5 | 8 | 14 | 29 | 2 | 4 | 0 | 0 | 0 | 0 | 0 |
| SG0000003 | Rbm11 | 1.14E-06 | 2.92E-06 | 5.004817538 | 0.639110347 | 20.51993827 | 61 | 3 | 8 | 3 | 13 | 3 | 7 | 0 | 2 | 0 | 4 | 0 |
| JSG000000 | Syt17 | 1.16E-06 | 2.97E-06 | -1.296257456 | 368.6680974 | 150.1146857 | 56 | 3 | 8 | 13 | 31 | 17 | 0 | 0 | 0 | 0 | 0 | 0 |
| SG0000003 | Ssx2ip | 1.2E-06 | 3.08E-06 | 0.720875483 | 574.7595897 | 947.3083595 | 250 | 0 | 8 | 11 | 72 | 35 | 6 | 0 | 0 | 3 | 12 | 3 |
| SG0000002 | Fundc1 | 1.24E-06 | 3.17E-06 | -0.782826627 | 1131.237992 | 657.5059484 | 154 | 0 | 8 | 28 | 80 | 8 | 3 | 0 | 0 | 0 | 0 | 0 |
| JSG000000 | Zdhhc6 | 1.32E-06 | 3.38E-06 | -0.720683106 | 2081.917135 | 1263.328249 | 45 | 7 | 8 | 10 | 19 | 1 | 0 | 0 | 0 | 0 | 0 | 0 |
| SG0000002 | Gosr2 | 1.75E-06 | 4.42E-06 | -0.692086125 | 2524.474018 | 1562.54388 | 59 | 2 | 8 | 6 | 42 | 4 | 5 | 0 | 0 | 0 | 0 | 0 |
| SG0000001 | Mb | 2.35E-06 | 5.87E-06 | 3.912553349 | 1.607478194 | 24.2070021 | 40 | 3 | 8 | 10 | 16 | 3 | 0 | 0 | 0 | 0 | 0 | 0 |
| JSG000000 | BC023829 | 2.55E-06 | 6.34E-06 | -0.753406408 | 1216.94558 | 721.8936639 | 102 | 4 | 8 | 19 | 49 | 6 | 2 | 0 | 0 | 1 | 1 | 0 |
| JSG000000 | Actr10 | 2.93E-06 | 7.25E-06 | -0.68203932 | 2573.432132 | 1603.978069 | 55 | 4 | 8 | 9 | 42 | 7 | 9 | 2 | 2 | 3 | 6 | 1 |
| SG0000002 | Dscc1 | 2.95E-06 | 7.3E-06 | 0.887423431 | 188.4125473 | 348.5386083 | 58 | 6 | 8 | 14 | 30 | 2 | 0 | 0 | 0 | 0 | 0 | 0 |
| SG0000002 | Vti1b | 2.96E-06 | 7.33E-06 | -0.735741786 | 1324.36523 | 795.2934828 | 123 | 8 | 8 | 20 | 56 | 10 | 2 | 0 | 0 | 0 | 0 | 0 |
| SG0000003 | Galnt11 | 3.83E-06 | 9.4E-06 | -0.709003393 | 1557.171142 | 952.587895 | 134 | 6 | 8 | 12 | 39 | 15 | 12 | 0 | 0 | 3 | 6 | 0 |
| SG0000004 | Dhx33 | 4.2E-06 | 1.03E-05 | -0.765852207 | 2040.140129 | 1199.818384 | 81 | 3 | 8 | 9 | 28 | 7 | 7 | 0 | 1 | 2 | 2 | 0 |
| SG0000002 | Slc46a3 | 4.66E-06 | 1.13E-05 | -1.541061233 | 205.6172653 | 70.65679314 | 42 | 8 | 8 | 10 | 28 | 4 | 0 | 0 | 0 | 0 | 0 | 0 |
| SG0000004 | Arglu1 | 4.97E-06 | 1.21E-05 | 0.627230965 | 1743.974005 | 2693.737932 | 57 | 5 | 8 | 8 | 13 | 7 | 1 | 0 | 0 | 0 | 0 | 1 |
| JSG000000 | Mtrf1l | 5.73E-06 | 1.38E-05 | -1.072805666 | 247.9802988 | 117.8882427 | 35 | 8 | 8 | 10 | 18 | 0 | 4 | 0 | 0 | 0 | 0 | 0 |
| SG0000004 | Dhrs7b | 6.1E-06 | 1.47E-05 | -0.737522503 | 959.1254305 | 575.2531563 | 88 | 4 | 8 | 11 | 35 | 4 | 2 | 0 | 0 | 1 | 3 | 0 |
| SG0000004 | Ddx24 | 6.31E-06 | 1.51E-05 | -0.627294453 | 5360.81621 | 3470.535345 | 56 | 8 | 8 | 20 | 30 | 4 | 0 | 0 | 0 | 0 | 0 | 0 |
| JSG000000 | Enkur | 1.07E-05 | 2.52E-05 | 2.427037006 | 7.097802049 | 38.17111292 | 60 | 8 | 8 | 13 | 29 | 4 | 0 | 0 | 0 | 0 | 0 | 0 |
| SG0000002 | Dnajc10 | 1.35E-05 | 3.15E-05 | -0.608997341 | 3953.973072 | 2592.431204 | 104 | 4 | 8 | 13 | 48 | 6 | 4 | 0 | 0 | 0 | 1 | 0 |
| JSG000000 | Ecd | 1.57E-05 | 3.65E-05 | 0.569596032 | 1693.450006 | 2513.262613 | 60 | 5 | 8 | 15 | 30 | 4 | 2 | 0 | 0 | 0 | 1 | 1 |
| JSG000000 | Rmdn2 | 1.72E-05 | 3.97E-05 | -1.25073591 | 145.4821533 | 61.13651731 | 120 | 7 | 8 | 18 | 70 | 12 | 1 | 0 | 0 | 0 | 0 | 0 |
| JSG000000 | Wbp4 | 2.86E-05 | 6.46E-05 | 0.58315545 | 907.7827491 | 1359.969623 | 52 | 4 | 8 | 11 | 29 | 1 | 6 | 0 | 1 | 1 | 2 | 0 |
| SG0000003 | Oas2 | 3.46E-05 | 7.75E-05 | -2.786965726 | 3108.854479 | 450.4432812 | 44 | 5 | 8 | 9 | 23 | 0 | 3 | 0 | 0 | 0 | 0 | 0 |
| SG0000000 | Ifrd1 | 4.16E-05 | 9.23E-05 | -0.595375895 | 2281.804789 | 1510.262657 | 118 | 0 | 8 | 14 | 46 | 10 | 5 | 0 | 0 | 0 | 4 | 2 |
| SG0000003 | Poglut1 | 4.37E-05 | 9.65E-05 | 0.525762001 | 2245.544592 | 3232.896615 | 51 | 6 | 8 | 11 | 26 | 3 | 1 | 0 | 0 | 0 | 1 | 0 |
| SG0000006 | Pex26 | 4.41E-05 | 9.73E-05 | -0.709922981 | 680.8718991 | 416.252925 | 53 | 4 | 8 | 4 | 31 | 1 | 10 | 0 | 0 | 0 | 1 | 1 |
| JSG000000 | Galc | 4.59E-05 | 0.000101 | -0.818486313 | 638.5752287 | 362.0955073 | 107 | 2 | 8 | 17 | 73 | 15 | 5 | 0 | 0 | 0 | 1 | 0 |
| JSG000000 | Pacs2 | 5.37E-05 | 0.000118 | -0.600300983 | 1734.323529 | 1143.988119 | 50 | 4 | 8 | 6 | 17 | 5 | 1 | 0 | 0 | 0 | 0 | 2 |
| SG0000003 | Gnai2 | 7.26E-05 | 0.000156 | -0.621246397 | 9685.890753 | 6296.885063 | 30 | 5 | 8 | 5 | 22 | 12 | 2 | 0 | 0 | 0 | 0 | 0 |
| JSG000000 | Dcp2 | 7.37E-05 | 0.000159 | 0.7477054 | 1968.316207 | 3305.039251 | 90 | 6 | 8 | 16 | 39 | 8 | 18 | 0 | 0 | 2 | 15 | 1 |
| SG0000004 | Cenpe | 7.42E-05 | 0.00016 | 0.562084324 | 2546.444564 | 3759.57253 | 96 | 3 | 8 | 16 | 50 | 17 | 1 | 1 | 1 | 1 | 2 | 0 |
| JSG000000 | Rtf1 | 7.7E-05 | 0.000165 | 0.520806614 | 1608.975843 | 2308.489675 | 110 | 5 | 8 | 13 | 43 | 12 | 4 | 0 | 0 | 0 | 1 | 3 |
| SG0000003 | Wnk4 | 7.76E-05 | 0.000167 | -1.228701425 | 120.3637657 | 51.35938111 | 49 | 8 | 8 | 8 | 43 | 16 | 0 | 0 | 0 | 0 | 0 | 0 |
| JSG000000 | Itih5 | 8.14E-05 | 0.000174 | 1.475607392 | 25.88310353 | 71.98109289 | 164 | 4 | 8 | 20 | 64 | 16 | 10 | 0 | 0 | 2 | 3 | 1 |
| JSG000001 | Gm28230 | 8.4E-05 | 0.00018 | -5.529843742 | 15.46779095 | 0.33479538 | 39 | 7 | 8 | 15 | 22 | 1 | 5 | 0 | 0 | 3 | 4 | 0 |
| SG0000004 | Tmem183a | 8.82E-05 | 0.000188 | 0.49764113 | 2750.31713 | 3883.18142 | 68 | 4 | 8 | 11 | 23 | 6 | 9 | 0 | 0 | 1 | 1 | 0 |
| SG0000003 | 3830406C13Rik | 0.000102 | 0.000217 | -0.82386232 | 338.4768643 | 191.2148945 | 80 | 6 | 8 | 14 | 42 | 8 | 6 | 0 | 0 | 0 | 4 | 0 |
| SG0000002 | Cutal | 0.000103 | 0.000218 | 2.868391521 | 3.562118703 | 26.01239109 | 69 | 0 | 8 | 12 | 36 | 8 | 7 | 0 | 0 | 0 | 0 | 0 |
| SG0000002 | Slc36a1 | 0.000112 | 0.000237 | -0.675965944 | 644.3479732 | 403.3057757 | 108 | 3 | 8 | 3 | 15 | 3 | 5 | 0 | 1 | 0 | 1 | 0 |
| SG0000002 | 1700037H04Rik | 0.000128 | 0.000268 | 1.022440161 | 803.5110729 | 1632.213717 | 32 | 4 | 8 | 4 | 8 | 4 | 0 | 0 | 0 | 0 | 0 | 0 |
| SG0000002 | Eci2 | 0.000149 | 0.000309 | -0.594494744 | 1259.316366 | 834.0154526 | 92 | 0 | 8 | 12 | 68 | 20 | 0 | 0 | 0 | 0 | 0 | 0 |
| JSG000000 | Rrp1 | 0.000176 | 0.000364 | -0.509125266 | 6945.809516 | 4880.461514 | 17 | 5 | 8 | 9 | 15 | 2 | 0 | 0 | 0 | 0 | 0 | 1 |
| JSG000000 | Txlnb | 0.000243 | 0.000493 | 2.678230771 | 7.218261001 | 46.20195777 | 98 | 2 | 8 | 11 | 41 | 1 | 6 | 0 | 0 | 0 | 3 | 0 |
| JSG000000 | Zfp960 | 0.000245 | 0.000495 | 0.85635367 | 106.1211716 | 192.1277856 | 59 | 8 | 8 | 15 | 44 | 5 | 2 | 0 | 0 | 0 | 0 | 0 |
| JSG000000 | Pomt2 | 0.000258 | 0.000522 | -0.628414283 | 857.3982568 | 554.6398214 | 58 | 8 | 8 | 12 | 34 | 4 | 3 | 0 | 0 | 0 | 0 | 0 |
| SG0000000 | Brat1 | 0.000262 | 0.000529 | -0.689173622 | 596.9524034 | 370.2352562 | 48 | 4 | 8 | 12 | 28 | 4 | 0 | 0 | 0 | 0 | 0 | 0 |
| SG0000000 | Polr2a | 0.000325 | 0.000649 | 0.489544051 | 7941.626062 | 11150.05161 | 36 | 6 | 8 | 9 | 21 | 2 | 1 | 0 | 0 | 0 | 0 | 1 |
| SG0000004 | Rab11fip2 | 0.000341 | 0.00068 | -0.767975318 | 450.2832845 | 264.4248207 | 197 | 6 | 8 | 18 | 72 | 8 | 22 | 0 | 0 | 0 | 7 | 3 |
| SG0000001 | Birc5 | 0.000382 | 0.000758 | 0.457590671 | 2401.992666 | 3298.52815 | 26 | 8 | 8 | 8 | 12 | 2 | 6 | 3 | 3 | 3 | 3 | 1 |
| SG0000002 | Fam114a1 | 0.000448 | 0.000879 | -0.706251197 | 669.0565855 | 410.0719421 | 124 | 4 | 8 | 14 | 61 | 9 | 3 | 0 | 0 | 0 | 0 | 0 |
| JSG000000 | Usp39 | 0.000506 | 0.000987 | 0.458150727 | 1935.226439 | 2658.5749 | 50 | 8 | 8 | 15 | 29 | 1 | 0 | 0 | 0 | 0 | 1 | 0 |
| JSG000000 | Gltscr1 | 0.000513 | 0.001001 | -0.779026327 | 377.7596991 | 220.1432289 | 27 | 7 | 8 | 11 | 22 | 3 | 0 | 0 | 0 | 0 | 0 | 0 |
| SG0000002 | Napb | 0.000626 | 0.001204 | -0.714475115 | 334.5716143 | 203.8969133 | 176 | 8 | 8 | 20 | 56 | 10 | 9 | 0 | 0 | 0 | 2 | 2 |
| SG0000006 | Mrps36 | 0.000661 | 0.001267 | 0.719117632 | 151.8325633 | 249.942956 | 40 | 6 | 8 | 6 | 18 | 10 | 0 | 0 | 0 | 0 | 0 | 0 |
| SG0000003 | Wfs1 | 0.000873 | 0.00165 | -0.829001276 | 675.2479828 | 380.1097808 | 40 | 8 | 8 | 14 | 22 | 2 | 2 | 2 | 2 | 2 | 2 | 0 |
| SG0000000 | Metap1 | 0.00093 | 0.001751 | 0.437053151 | 1880.592591 | 2546.014243 | 111 | 8 | 8 | 21 | 63 | 8 | 12 | 0 | 0 | 1 | 11 | 1 |
| SG0000001 | Sirt2 | 0.001031 | 0.00193 | -0.515414623 | 1034.696957 | 723.8655533 | 72 | 8 | 8 | 12 | 24 | 4 | 0 | 0 | 0 | 0 | 0 | 0 |
| SG0000003 | Flcn | 0.00117 | 0.002181 | -0.581011525 | 1819.015555 | 1216.002635 | 28 | 6 | 8 | 8 | 14 | 0 | 2 | 0 | 0 | 0 | 0 | 0 |
| SG0000002 | Col17a1 | 0.001181 | 0.0022 | 1.40850052 | 18.60428499 | 49.38714137 | 92 | 6 | 8 | 12 | 22 | 4 | 4 | 0 | 0 | 0 | 0 | 0 |
| JSG000000 | Fam162a | 0.001259 | 0.002338 | -0.65109783 | 3747.360073 | 2386.302237 | 46 | 6 | 8 | 11 | 25 | 2 | 0 | 0 | 0 | 0 | 0 | 0 |
| SG0000003 | Ruvbl1 | 0.001332 | 0.002466 | 0.420585241 | 1993.406723 | 2668.115905 | 44 | 7 | 8 | 9 | 15 | 4 | 0 | 0 | 0 | 0 | 0 | 0 |
| SG0000001 | Trpv4 | 0.001591 | 0.002923 | -0.811164192 | 174.4441495 | 99.41951848 | 64 | 8 | 8 | 20 | 32 | 2 | 4 | 2 | 2 | 2 | 4 | 0 |
| JSG000000 | Gm4944 | 0.001648 | 0.00302 | -0.707240499 | 262.2120188 | 160.6023848 | 66 | 4 | 8 | 14 | 44 | 9 | 4 | 0 | 1 | 1 | 2 | 0 |
| SG0000002 | Slc16a7 | 0.001711 | 0.003131 | 3.363461454 | 2.3079898 | 23.75399563 | 415 | 0 | 8 | 28 | 175 | 37 | 3 | 0 | 0 | 0 | 0 | 1 |
| SG0000005 | Tatdn2 | 0.001762 | 0.003216 | -0.473791175 | 1696.284258 | 1221.443201 | 65 | 8 | 8 | 8 | 26 | 0 | 2 | 0 | 0 | 0 | 0 | 0 |
| SG0000005 | Zfp961 | 0.001905 | 0.003457 | -0.459674764 | 2090.215404 | 1519.900478 | 148 | 8 | 8 | 20 | 87 | 12 | 7 | 1 | 1 | 1 | 1 | 0 |
| JSG000000 | Tmem106c | 0.002402 | 0.004299 | -0.486696816 | 961.7263125 | 686.3429198 | 15 | 8 | 8 | 8 | 8 | 0 | 1 | 0 | 0 | 0 | 0 | 0 |
| SG0000003 | Zfp273 | 0.002625 | 0.004673 | 0.919944874 | 58.9986826 | 111.6280442 | 68 | 2 | 8 | 8 | 26 | 6 | 2 | 0 | 0 | 0 | 0 | 2 |
| JSG000000 | Eif3l | 0.003 | 0.005294 | 0.365102968 | 8569.60363 | 11037.42283 | 33 | 8 | 8 | 11 | 28 | 3 | 0 | 0 | 0 | 0 | 0 | 0 |
| JSG000000 | Ppp1r37 | 0.003682 | 0.006414 | -0.980816886 | 902.4214257 | 457.250401 | 23 | 8 | 8 | 10 | 16 | 2 | 0 | 0 | 0 | 0 | 0 | 0 |
| SG0000007 | Stard6 | 0.00387 | 0.006719 | -1.869239267 | 28.26571227 | 7.736826275 | 200 | 4 | 8 | 28 | 120 | 20 | 0 | 0 | 0 | 0 | 0 | 0 |
| SG0000003 | Asb7 | 0.003883 | 0.00674 | -0.485102468 | 803.9461316 | 574.3763587 | 156 | 2 | 8 | 18 | 54 | 20 | 23 | 2 | 4 | 4 | 6 | 2 |
| SG0000002 | Prrc2a | 0.004222 | 0.007289 | -0.428215874 | 11954.48994 | 8884.341181 | 21 | 8 | 8 | 10 | 16 | 6 | 0 | 0 | 0 | 0 | 0 | 2 |
| SG0000000 | Gstt1 | 0.004343 | 0.00749 | 1.081370198 | 98.94771014 | 209.3777913 | 47 | 2 | 8 | 12 | 50 | 12 | 0 | 0 | 0 | 0 | 0 | 0 |
| JSG000000 | Pggt1b | 0.004393 | 0.007571 | -0.531537978 | 464.4820758 | 321.3365187 | 89 | 6 | 8 | 11 | 41 | 16 | 4 | 0 | 0 | 0 | 2 | 2 |
| JSG000000 | Chst5 | 0.004666 | 0.00802 | 5.064031322 | 0.68495036 | 22.91313032 | 30 | 6 | 8 | 11 | 38 | 1 | 1 | 0 | 0 | 0 | 0 | 0 |
| JSG000000 | Ttyh2 | 0.005838 | 0.009893 | -0.478758151 | 1072.734923 | 769.7892736 | 39 | 7 | 8 | 12 | 27 | 6 | 0 | 0 | 0 | 0 | 0 | 0 |
| SG0000002 | Mfsd8 | 0.006459 | 0.010876 | -0.723618573 | 176.9166321 | 107.1365772 | 71 | 5 | 8 | 11 | 35 | 7 | 5 | 0 | 0 | 0 | 1 | 0 |
| SG0000004 | Nlrp4a | 0.006648 | 0.011177 | 4.953636937 | 0.663202137 | 20.55129639 | 178 | 5 | 8 | 30 | 124 | 17 | 5 | 0 | 0 | 0 | 3 | 0 |
| SG0000003 | Abcc10 | 0.007956 | 0.013212 | -0.567945759 | 280.0873196 | 188.9403597 | 59 | 8 | 8 | 8 | 32 | 0 | 0 | 0 | 0 | 0 | 0 | 0 |
| SG0000003 | Ano7 | 0.009388 | 0.015425 | 3.48092036 | 1.572512292 | 17.55720944 | 61 | 8 | 8 | 10 | 27 | 6 | 0 | 0 | 0 | 0 | 1 | 0 |
| JSG000000 | Zfp97 | 0.009562 | 0.015688 | 0.705549802 | 72.20701717 | 117.7527501 | 59 | 8 | 8 | 15 | 44 | 6 | 2 | 0 | 0 | 0 | 0 | 1 |
| JSG000000 | Acad11 | 0.011474 | 0.018597 | 0.373903476 | 834.3386072 | 1081.181241 | 187 | 6 | 8 | 25 | 102 | 24 | 2 | 0 | 0 | 0 | 0 | 0 |
| SG0000002 | Syt7 | 0.011968 | 0.019318 | -0.475749204 | 430.51586 | 309.581054 | 74 | 6 | 8 | 14 | 28 | 6 | 0 | 0 | 0 | 0 | 0 | 0 |
| SG0000001 | Dcn | 0.012021 | 0.01939 | -1.665387795 | 28.00304614 | 8.828229104 | 169 | 2 | 8 | 27 | 116 | 18 | 0 | 0 | 0 | 0 | 3 | 0 |
| SG0000004 | Zfp30 | 0.012298 | 0.019791 | 0.665341551 | 80.04882727 | 126.9529328 | 38 | 6 | 8 | 10 | 22 | 2 | 2 | 0 | 0 | 0 | 0 | 0 |
| SG0000002 | Commd8 | 0.012596 | 0.020228 | -0.369585041 | 1720.943363 | 1332.018923 | 63 | 4 | 8 | 14 | 22 | 8 | 15 | 4 | 4 | 4 | 6 | 2 |
| SG0000004 | Trafd1 | 0.013136 | 0.02102 | -0.400059665 | 837.8166546 | 634.9200331 | 46 | 8 | 8 | 16 | 24 | 0 | 5 | 0 | 0 | 1 | 1 | 0 |
| JSG000000 | Zfyve26 | 0.01353 | 0.021577 | -0.374549156 | 1218.40153 | 939.8096601 | 65 | 7 | 8 | 10 | 31 | 8 | 1 | 0 | 0 | 0 | 0 | 0 |
| SG0000002 | Tmem192 | 0.016787 | 0.026366 | 0.382715471 | 508.8342534 | 663.4148302 | 56 | 6 | 8 | 8 | 26 | 4 | 0 | 0 | 0 | 0 | 0 | 0 |
| SG0000005 | Zfp455 | 0.017174 | 0.026916 | 0.806538791 | 42.790727 | 74.84142255 | 70 | 4 | 8 | 10 | 28 | 6 | 16 | 0 | 0 | 0 | 4 | 0 |
| JSG000000 | Tas1r1 | 0.017918 | 0.027957 | 1.8599879 | 4.530486551 | 16.44587537 | 17 | 7 | 8 | 10 | 16 | 0 | 0 | 0 | 0 | 0 | 0 | 0 |
| JSG000000 | BC068281 | 0.019077 | 0.029624 | 0.447411001 | 233.5929573 | 318.5252582 | 26 | 8 | 8 | 10 | 17 | 0 | 3 | 0 | 0 | 0 | 0 | 0 |
| SG0000003 | Adgra2 | 0.020269 | 0.031316 | -1.114521506 | 492.9635696 | 227.6724189 | 53 | 2 | 8 | 16 | 26 | 8 | 4 | 0 | 0 | 2 | 2 | 0 |
| SG0000002 | Rdh12 | 0.02383 | 0.036377 | 0.742026939 | 44.06276072 | 73.69602612 | 46 | 4 | 8 | 8 | 28 | 4 | 2 | 0 | 0 | 0 | 0 | 0 |
| SG0000004 | Zfp668 | 0.024779 | 0.037723 | -0.650835822 | 330.593373 | 210.5586264 | 23 | 8 | 8 | 8 | 20 | 0 | 0 | 0 | 0 | 0 | 0 | 0 |
| SG0000002 | Sh3d19 | 0.024873 | 0.037856 | -0.357923078 | 1336.187634 | 1042.60995 | 89 | 4 | 8 | 11 | 37 | 5 | 9 | 0 | 0 | 0 | 3 | 1 |
| JSG000000 | Tatdn1 | 0.02678 | 0.040561 | -0.468001249 | 304.9811507 | 220.4908566 | 70 | 4 | 8 | 6 | 36 | 59 | 0 | 0 | 0 | 0 | 0 | 0 |
| JSG000000 | 1810009A15Rik | 0.033099 | 0.049317 | -0.510571299 | 325.5688207 | 228.5312159 | 14 | 8 | 8 | 8 | 10 | 0 | 0 | 0 | 0 | 0 | 0 | 0 |
| SG0000002 | Glmn | 0.033179 | 0.049419 | 0.337220965 | 450.6646785 | 569.3340348 | 386 | 4 | 8 | 24 | 120 | 32 | 7 | 0 | 0 | 0 | 1 | 0 |
| SG0000002 | Zkscan3 | 0.036905 | 0.05447 | -0.377743097 | 1306.241034 | 1005.336197 | 99 | 0 | 8 | 4 | 21 | 0 | 8 | 0 | 0 | 0 | 1 | 0 |
| SG0000005 | Eif3b | 0.044858 | 0.065304 | -0.257915966 | 12176.62129 | 10183.24885 | 25 | 8 | 8 | 10 | 23 | 1 | 0 | 0 | 0 | 0 | 0 | 0 |
| SG0000000 | Itfg2 | 0.045186 | 0.065765 | -0.32547245 | 728.7695186 | 581.5850435 | 62 | 8 | 8 | 12 | 20 | 2 | 5 | 1 | 1 | 3 | 3 | 0 |
| SG0000003 | Tubgcp5 | 0.048412 | 0.070078 | -0.315138506 | 991.7790196 | 797.1660118 | 102 | 5 | 8 | 16 | 40 | 10 | 3 | 1 | 1 | 1 | 5 | 5 |
| JSG000000 | Cdk7 | 0.049285 | 0.071255 | 0.269196323 | 1419.740869 | 1710.981258 | 63 | 4 | 8 | 10 | 38 | 5 | 3 | 0 | 0 | 0 | 1 | 1 |

| | | | | | | | | | | | | | | | | | | |
|---|---|---|---|---|---|---|---|---|---|---|---|---|---|---|---|---|---|---|
| SG0000004 | Shtn1 | 0.049922 | 0.072078 | 0.794848302 | 110.3466891 | 191.4399128 | 165 | 3 | 8 | 25 | 71 | 12 | 3 | 0 | 1 | 0 | 0 | 0 |
| JSG000000 | Tspan15 | 0.050172 | 0.072404 | 0.419778052 | 167.2846399 | 223.7802997 | 37 | 8 | 8 | 13 | 27 | 10 | 1 | 0 | 0 | 0 | 0 | 0 |
| SG0000002 | Rbm19 | 0.053506 | 0.076789 | -0.309053118 | 1155.597611 | 932.7652895 | 94 | 6 | 8 | 12 | 44 | 4 | 0 | 0 | 0 | 0 | 0 | 0 |
| JSG000000 | Fbxw10 | 0.056569 | 0.080802 | 2.538840664 | 3.145024585 | 18.27642458 | 184 | 0 | 8 | 40 | 92 | 20 | 0 | 0 | 0 | 0 | 0 | 0 |
| JSG000000 | 1810024B03Rik | 0.058267 | 0.083031 | -0.89201526 | 38.74511951 | 20.87822891 | 42 | 6 | 8 | 9 | 19 | 6 | 1 | 0 | 0 | 0 | 0 | 0 |
| SG0000003 | Uhrf1bp1 | 0.058931 | 0.083896 | 0.260891807 | 1325.60027 | 1588.35964 | 77 | 7 | 8 | 13 | 64 | 3 | 2 | 0 | 0 | 0 | 0 | 0 |
| JSG000000 | Malt1 | 0.071345 | 0.100018 | 0.281322527 | 570.7992877 | 693.6972 | 99 | 7 | 8 | 21 | 61 | 19 | 4 | 0 | 0 | 2 | 5 | 4 |
| JSG000000 | Erp29 | 0.073364 | 0.10268 | 0.246224213 | 1531.406622 | 1816.399587 | 58 | 8 | 8 | 10 | 17 | 5 | 12 | 0 | 0 | 0 | 0 | 0 |
| JSG000000 | Cs | 0.074309 | 0.103856 | -0.234114332 | 11363.10072 | 9660.985263 | 46 | 7 | 8 | 11 | 31 | 3 | 3 | 0 | 0 | 2 | 2 | 0 |
| SG0000004 | A4galt | 0.078983 | 0.109802 | 0.500558447 | 558.9118399 | 790.7267243 | 52 | 8 | 8 | 16 | 40 | 0 | 0 | 0 | 0 | 0 | 2 | 0 |
| JSG000000 | Sparcl1 | 0.080356 | 0.111508 | -1.387846154 | 16.50978087 | 6.308963102 | 63 | 5 | 8 | 13 | 26 | 10 | 0 | 0 | 0 | 0 | 0 | 0 |
| JSG000000 | Itgb3 | 0.089061 | 0.122552 | -0.418267201 | 311.5181517 | 233.1161617 | 79 | 5 | 8 | 9 | 46 | 6 | 2 | 0 | 0 | 1 | 3 | 0 |
| SG0000003 | D430042O09Rik | 0.097661 | 0.13324 | -0.305379724 | 599.6007124 | 485.2144021 | 212 | 6 | 8 | 19 | 96 | 24 | 0 | 0 | 0 | 0 | 0 | 0 |
| SG0000002 | Col2a1 | 0.108124 | 0.146201 | -0.591058005 | 141.2081986 | 93.74189406 | 44 | 6 | 8 | 10 | 18 | 0 | 4 | 2 | 2 | 4 | 4 | 0 |
| SG0000002 | Tmem222 | 0.111012 | 0.149619 | -0.242404829 | 990.4432938 | 837.2564293 | 52 | 8 | 8 | 16 | 40 | 4 | 3 | 0 | 0 | 0 | 0 | 0 |
| SG0000002 | Iqgap3 | 0.114118 | 0.153332 | 0.222417554 | 1321.711381 | 1542.023474 | 62 | 8 | 8 | 16 | 44 | 2 | 2 | 0 | 0 | 2 | 2 | 0 |
| SG0000004 | Atpaf2 | 0.116451 | 0.156116 | -0.249538802 | 851.7640292 | 716.4743236 | 68 | 8 | 8 | 12 | 32 | 6 | 0 | 0 | 0 | 0 | 0 | 0 |
| SG0000007 | Zfp58 | 0.12461 | 0.166117 | 0.384973319 | 104.6356522 | 136.6369635 | 40 | 4 | 8 | 6 | 16 | 0 | 2 | 0 | 0 | 0 | 0 | 0 |
| SG0000003 | Pkm | 0.126858 | 0.168754 | 0.190136169 | 72070.77784 | 82223.48868 | 48 | 2 | 8 | 14 | 38 | 2 | 0 | 0 | 0 | 0 | 1 | 0 |
| SG0000000 | Zmat5 | 0.127119 | 0.169064 | -0.34445558 | 421.1741284 | 331.7189057 | 49 | 6 | 8 | 7 | 28 | 3 | 0 | 0 | 0 | 0 | 0 | 0 |
| SG0000002 | Cdan1 | 0.137611 | 0.181777 | 0.239176136 | 531.8211338 | 627.7182896 | 52 | 6 | 8 | 10 | 18 | 6 | 8 | 0 | 0 | 0 | 2 | 2 |
| SG0000003 | Exosc1 | 0.141722 | 0.186697 | -0.239892877 | 953.1597459 | 807.1434767 | 34 | 4 | 8 | 8 | 18 | 4 | 4 | 0 | 0 | 0 | 0 | 2 |
| SG0000002 | Dusp19 | 0.160978 | 0.209674 | -0.309153392 | 255.3081011 | 206.0630631 | 43 | 2 | 8 | 10 | 21 | 4 | 2 | 0 | 0 | 0 | 1 | 0 |
| JSG000000 | Prdm4 | 0.165153 | 0.214659 | -0.202247573 | 1871.650643 | 1626.830109 | 48 | 7 | 8 | 11 | 23 | 8 | 3 | 0 | 0 | 2 | 2 | 0 |
| SG0000002 | Serpine2 | 0.170382 | 0.220802 | 0.508545933 | 1241.371185 | 1765.994067 | 104 | 3 | 8 | 15 | 46 | 9 | 0 | 0 | 0 | 0 | 1 | 0 |
| SG0000001 | Ppp2r5c | 0.174464 | 0.225371 | -0.185902624 | 4009.958647 | 3525.150213 | 190 | 4 | 8 | 12 | 52 | 34 | 2 | 0 | 0 | 1 | 2 | 1 |
| JSG000000 | Mob2 | 0.183179 | 0.235644 | -0.232270199 | 468.2295665 | 398.6011405 | 67 | 6 | 8 | 12 | 36 | 3 | 1 | 0 | 0 | 0 | 0 | 0 |
| SG0000002 | Dffa | 0.187887 | 0.24099 | -0.210614158 | 733.5252087 | 633.8899473 | 40 | 6 | 8 | 12 | 22 | 2 | 0 | 0 | 0 | 0 | 0 | 0 |
| SG0000004 | Fam160a2 | 0.193992 | 0.248146 | 0.185430413 | 1104.722962 | 1256.242545 | 210 | 2 | 8 | 32 | 126 | 20 | 18 | 1 | 1 | 1 | 6 | 1 |
| JSG000000 | Rnf7 | 0.208926 | 0.264996 | 0.172739305 | 1300.540812 | 1465.965323 | 42 | 8 | 8 | 10 | 22 | 4 | 0 | 0 | 0 | 0 | 0 | 0 |
| SG0000002 | Vldlr | 0.211774 | 0.268116 | -0.475597162 | 563.1693236 | 405.013947 | 209 | 0 | 8 | 25 | 52 | 25 | 13 | 0 | 0 | 1 | 4 | 13 |
| SG0000003 | Thumpd3 | 0.213058 | 0.269588 | 0.161019171 | 1845.355548 | 2063.249057 | 52 | 7 | 8 | 10 | 27 | 2 | 0 | 0 | 0 | 0 | 1 | 0 |
| SG0000003 | Crispld2 | 0.239319 | 0.299057 | -0.374680275 | 592.348313 | 456.8642276 | 119 | 5 | 8 | 32 | 57 | 59 | 6 | 0 | 0 | 0 | 0 | 0 |
| SG0000000 | Rrnad1 | 0.270244 | 0.33271 | 0.2068407 | 449.819397 | 519.1626423 | 25 | 5 | 8 | 8 | 13 | 20 | 3 | 0 | 3 | 3 | 3 | 2 |
| SG0000002 | Tatdn3 | 0.278888 | 0.342083 | 0.220166827 | 183.6931698 | 213.9783465 | 92 | 8 | 8 | 16 | 52 | 8 | 0 | 0 | 0 | 0 | 0 | 0 |
| SG0000003 | Lamtor1 | 0.331989 | 0.398832 | -0.133568409 | 1395.452897 | 1272.0585 | 35 | 8 | 8 | 11 | 17 | 4 | 1 | 0 | 0 | 0 | 0 | 0 |
| SG0000002 | Msrb2 | 0.355306 | 0.423295 | -0.278439572 | 128.6107989 | 106.0373274 | 40 | 8 | 8 | 10 | 24 | 6 | 0 | 0 | 0 | 0 | 0 | 0 |
| SG0000000 | Dhx34 | 0.359807 | 0.427975 | -0.231790567 | 481.511292 | 410.0440959 | 52 | 8 | 8 | 8 | 8 | 8 | 0 | 0 | 0 | 0 | 0 | 0 |
| JSG000000 | Tubgcp2 | 0.363034 | 0.431383 | 0.12810827 | 1254.171758 | 1370.633836 | 48 | 7 | 8 | 9 | 32 | 4 | 0 | 0 | 0 | 0 | 0 | 0 |
| SG0000004 | Atg16l2 | 0.368751 | 0.436992 | -0.184642099 | 268.2405519 | 236.0160968 | 26 | 8 | 8 | 12 | 21 | 0 | 0 | 0 | 0 | 0 | 0 | 0 |
| SG0000003 | Nadsyn1 | 0.382923 | 0.451787 | -0.218653148 | 196.0054028 | 168.4406414 | 72 | 8 | 8 | 12 | 61 | 10 | 3 | 0 | 0 | 0 | 3 | 0 |
| SG0000005 | Zfp846 | 0.395484 | 0.464215 | 0.168154116 | 195.1196905 | 219.2403726 | 38 | 8 | 8 | 14 | 24 | 4 | 2 | 0 | 0 | 2 | 2 | 0 |
| SG0000007 | Apcdd1 | 0.404481 | 0.473502 | -0.202716041 | 168.5180175 | 146.4275286 | 43 | 8 | 8 | 12 | 18 | 6 | 0 | 0 | 0 | 0 | 0 | 0 |
| SG0000000 | Tor2a | 0.40976 | 0.4789 | 0.190301113 | 716.962838 | 818.0559146 | 20 | 4 | 8 | 12 | 12 | 0 | 0 | 0 | 0 | 0 | 0 | 0 |
| JSG000000 | Kank2 | 0.422547 | 0.4916 | -0.144286315 | 3407.893415 | 3083.553547 | 39 | 8 | 8 | 12 | 25 | 5 | 1 | 0 | 0 | 0 | 0 | 0 |
| JSG000000 | Cdkn2aipnl | 0.445333 | 0.514466 | 0.092886048 | 2137.813415 | 2279.981298 | 32 | 8 | 8 | 12 | 18 | 3 | 6 | 0 | 0 | 4 | 4 | 1 |
| JSG000000 | Aoc2 | 0.457256 | 0.526746 | 0.206706338 | 94.00151064 | 108.482489 | 26 | 8 | 8 | 10 | 14 | 0 | 0 | 0 | 0 | 0 | 0 | 0 |
| SG0000000 | Hdgfrp2 | 0.483992 | 0.552324 | -0.112406996 | 2346.237622 | 2170.371695 | 23 | 8 | 8 | 10 | 14 | 3 | 0 | 0 | 0 | 0 | 0 | 0 |
| SG0000003 | Mfap5 | 0.493986 | 0.561724 | 0.265399744 | 99.6362412 | 119.7596779 | 88 | 8 | 8 | 20 | 48 | 12 | 0 | 0 | 0 | 0 | 0 | 0 |
| SG0000003 | Stk35 | 0.496061 | 0.563547 | -0.100787796 | 1524.947659 | 1422.049742 | 88 | 6 | 8 | 14 | 26 | 7 | 2 | 0 | 0 | 0 | 0 | 1 |
| SG0000001 | Slc12a4 | 0.499466 | 0.566769 | 0.096399294 | 1688.595033 | 1805.280065 | 52 | 8 | 8 | 22 | 22 | 8 | 6 | 0 | 0 | 4 | 4 | 0 |
| SG0000002 | Pex11b | 0.510902 | 0.577768 | -0.123705498 | 443.8403071 | 407.3687162 | 61 | 8 | 8 | 12 | 40 | 6 | 3 | 0 | 0 | 0 | 0 | 6 |
| SG0000000 | Apobec3 | 0.513144 | 0.580011 | 0.078048253 | 2960.518825 | 3125.091172 | 116 | 8 | 8 | 9 | 44 | 17 | 11 | 0 | 0 | 0 | 9 | 6 |
| SG0000007 | Aarsd1 | 0.517579 | 0.584471 | 0.182980705 | 102.7099437 | 116.599093 | 21 | 7 | 8 | 9 | 15 | 0 | 0 | 0 | 0 | 0 | 0 | 0 |
| SG0000002 | Nckap5l | 0.525727 | 0.592346 | -0.285216362 | 298.0981121 | 244.6248349 | 33 | 7 | 8 | 9 | 15 | 1 | 1 | 0 | 0 | 0 | 0 | 0 |
| JSG000000 | Rnaset2a | 0.534629 | 0.601222 | 0.252545919 | 34.2567827 | 40.81036412 | 37 | 7 | 8 | 13 | 21 | 2 | 0 | 0 | 0 | 0 | 0 | 0 |
| JSG000000 | Ppp2r3c | 0.535912 | 0.602363 | 0.094829399 | 482.4818321 | 515.2612206 | 55 | 3 | 8 | 13 | 30 | 3 | 0 | 0 | 0 | 0 | 0 | 0 |
| SG0000003 | Eif2s3x | 0.545551 | 0.611356 | 0.076223574 | 2014.648433 | 2123.952757 | 74 | 8 | 8 | 14 | 47 | 9 | 4 | 0 | 0 | 1 | 2 | 0 |
| JSG000000 | Mtap | 0.561371 | 0.626224 | 0.07978392 | 2318.603058 | 2450.438219 | 65 | 6 | 8 | 10 | 37 | 5 | 0 | 0 | 0 | 0 | 0 | 0 |
| SG0000003 | Gtf3c4 | 0.646271 | 0.705941 | 0.056613501 | 2105.775119 | 2190.051624 | 50 | 2 | 8 | 10 | 24 | 2 | 14 | 0 | 2 | 2 | 8 | 0 |
| JSG000000 | Egf | 0.651967 | 0.710777 | 0.153093081 | 37.2639374 | 41.43566863 | 127 | 5 | 8 | 14 | 66 | 11 | 5 | 0 | 1 | 0 | 1 | 0 |
| SG0000003 | Palm | 0.654313 | 0.712899 | 0.143435181 | 1230.407988 | 1359.024962 | 48 | 8 | 8 | 12 | 46 | 10 | 0 | 0 | 0 | 0 | 0 | 0 |
| SG0000003 | Sytl2 | 0.678737 | 0.735572 | 0.253888788 | 24.10917323 | 28.7481868 | 424 | 0 | 8 | 40 | 184 | 40 | 4 | 0 | 0 | 0 | 4 | 0 |
| SG0000002 | Taf11 | 0.728245 | 0.779743 | 0.052576875 | 649.1635211 | 673.2576914 | 12 | 4 | 8 | 4 | 6 | 2 | 0 | 0 | 0 | 0 | 2 | 0 |
| SG0000000 | Crbn | 0.746176 | 0.794809 | -0.05584607 | 1020.846001 | 982.084574 | 104 | 0 | 8 | 10 | 30 | 16 | 4 | 0 | 0 | 0 | 0 | 0 |
| SG0000000 | Ndufb2 | 0.759444 | 0.806448 | 0.043126342 | 490.1625242 | 505.0361045 | 28 | 6 | 8 | 6 | 17 | 4 | 4 | 0 | 0 | 0 | 3 | 2 |
| SG0000005 | Txlna | 0.776748 | 0.821463 | -0.038124183 | 2338.850646 | 2277.854434 | 40 | 8 | 8 | 20 | 28 | 2 | 6 | 0 | 0 | 2 | 2 | 0 |
| SG0000002 | Bcap29 | 0.79884 | 0.84046 | 0.026152278 | 888.9141014 | 905.1747192 | 188 | 2 | 8 | 14 | 94 | 8 | 8 | 0 | 0 | 0 | 0 | 0 |
| SG0000006 | Pex11g | 0.832616 | 0.869154 | 0.052928369 | 143.1923746 | 148.5432488 | 30 | 8 | 8 | 12 | 18 | 0 | 2 | 0 | 0 | 0 | 0 | 0 |
| JSG000000 | Siae | 0.837979 | 0.873531 | -0.075566082 | 718.3413424 | 681.684163 | 60 | 4 | 8 | 15 | 39 | 4 | 5 | 1 | 1 | 2 | 4 | 0 |
| SG0000003 | Pla2g4c | 0.840369 | 0.87536 | -0.184346659 | 17.24291482 | 15.17458451 | 128 | 4 | 8 | 20 | 63 | 8 | 2 | 0 | 0 | 0 | 2 | 0 |
| SG0000000 | Zbtb17 | 0.841766 | 0.876291 | 0.048879808 | 470.6748888 | 486.8950026 | 29 | 6 | 8 | 10 | 13 | 2 | 1 | 0 | 0 | 0 | 0 | 0 |
| JSG000000 | Ddhd2 | 0.843484 | 0.877543 | -0.031981265 | 1129.393968 | 1104.633332 | 71 | 5 | 8 | 14 | 42 | 5 | 2 | 0 | 0 | 0 | 0 | 0 |
| SG0000002 | Smc2 | 0.866821 | 0.896915 | 0.014741079 | 5675.175015 | 5733.459724 | 254 | 2 | 8 | 28 | 98 | 16 | 2 | 0 | 0 | 0 | 4 | 0 |
| JSG000000 | Rab38 | 0.926414 | 0.94671 | -0.076864413 | 23.23966925 | 22.0339038 | 149 | 4 | 8 | 20 | 83 | 12 | 0 | 0 | 0 | 0 | 1 | 0 |
| JSG000000 | Ctsb | 0.96706 | 0.980915 | 0.004121872 | 8102.796921 | 8125.980234 | 32 | 7 | 8 | 8 | 12 | 2 | 2 | 0 | 0 | 1 | 1 | 0 |
| JSG000000 | Cds1 | 5.3E-209 | 1.7E-206 | 7.780312163 | 4.587200676 | 1008.45164 | 101 | 6 | 7 | 12 | 31 | 6 | 3 | 0 | 0 | 0 | 1 | 0 |
| SG0000002 | Alpl | 2.4E-207 | 7.8E-205 | 7.03614536 | 40.66444428 | 5337.103793 | 48 | 5 | 7 | 13 | 30 | 5 | 1 | 0 | 0 | 0 | 0 | 0 |
| SG0000000 | Nphs1 | 8.4E-199 | 2.6E-196 | 9.663196765 | 0 | 809.7969011 | 43 | 7 | 7 | 12 | 18 | 6 | 1 | 0 | 0 | 0 | 0 | 0 |
| SG0000002 | Dtx1 | 3.1E-159 | 5.7E-157 | 6.805105079 | 6.871098686 | 768.363876 | 26 | 7 | 7 | 9 | 9 | 0 | 0 | 0 | 0 | 0 | 0 | 0 |
| JSG000000 | Elavl3 | 2.7E-148 | 4E-146 | 9.145875988 | 0 | 565.4779492 | 42 | 6 | 7 | 15 | 27 | 2 | 8 | 0 | 1 | 3 | 3 | 0 |
| JSG000000 | Gcnt3 | 1.1E-115 | 1.1E-113 | -8.040651375 | 4300.683378 | 16.33278413 | 14 | 6 | 7 | 8 | 12 | 2 | 3 | 0 | 0 | 0 | 1 | 1 |
| JSG000000 | Rab6b | 3.02E-95 | 1.92E-93 | 7.100429654 | 2.6022814 | 357.1055216 | 56 | 6 | 7 | 13 | 25 | 7 | 2 | 0 | 0 | 0 | 0 | 1 |
| SG0000006 | Ap3b2 | 1.62E-89 | 9.07E-88 | 9.611808499 | 0.970711415 | 759.5087954 | 45 | 4 | 7 | 5 | 12 | 8 | 1 | 0 | 0 | 0 | 1 | 0 |
| JSG000000 | Qpct | 4.05E-87 | 2.13E-85 | -4.93547546 | 5085.845349 | 166.2022863 | 90 | 4 | 7 | 12 | 48 | 6 | 5 | 0 | 0 | 0 | 1 | 0 |
| SG0000004 | Cables1 | 2.2E-72 | 8.47E-71 | 6.159388325 | 3.913124428 | 279.6947396 | 113 | 3 | 7 | 15 | 42 | 17 | 0 | 0 | 0 | 0 | 1 | 0 |
| JSG000000 | Vax2 | 4.51E-67 | 1.5E-65 | -6.270942136 | 1347.546076 | 17.45026151 | 48 | 7 | 7 | 12 | 41 | 2 | 0 | 0 | 0 | 0 | 0 | 0 |
| JSG000000 | Itga7 | 4.58E-64 | 1.41E-62 | -7.13765413 | 12750.71221 | 90.54958242 | 38 | 5 | 7 | 11 | 24 | 1 | 1 | 0 | 0 | 1 | 1 | 0 |
| SG0000002 | Chrna9 | 1.1E-63 | 3.35E-62 | 7.638207927 | 0 | 198.2185153 | 30 | 7 | 7 | 8 | 21 | 2 | 0 | 0 | 0 | 0 | 0 | 0 |
| SG0000005 | Megf6 | 1.39E-62 | 4.12E-61 | 9.30898537 | 0.307509278 | 195.048246 | 64 | 5 | 7 | 13 | 31 | 17 | 2 | 0 | 0 | 2 | 2 | 0 |
| JSG000000 | Ccnjl | 2.46E-55 | 5.99E-54 | 5.411698836 | 5.204562799 | 221.5480492 | 66 | 4 | 7 | 10 | 17 | 3 | 1 | 0 | 0 | 0 | 0 | 0 |
| SG0000001 | Aim1l | 3.65E-50 | 7.62E-49 | 5.168328751 | 14.37662058 | 516.9860124 | 33 | 6 | 7 | 10 | 16 | 1 | 0 | 0 | 0 | 0 | 0 | 0 |
| SG0000003 | Plekhb1 | 1.15E-48 | 2.3E-47 | 4.115383903 | 19.42675539 | 336.7086477 | 97 | 7 | 7 | 14 | 26 | 0 | 7 | 0 | 0 | 0 | 0 | 0 |
| JSG000000 | Pmel | 4.74E-48 | 9.27E-47 | 4.490571989 | 10.7383831 | 241.3991631 | 22 | 6 | 7 | 6 | 10 | 1 | 0 | 0 | 0 | 0 | 0 | 0 |
| SG0000003 | Slc26a2 | 5.93E-48 | 1.16E-46 | -3.059677557 | 8142.878835 | 976.6146863 | 40 | 3 | 7 | 8 | 19 | 2 | 15 | 0 | 1 | 2 | 4 | 2 |
| SG0000001 | H13 | 3.57E-47 | 6.79E-46 | -2.434583537 | 5486.01056 | 1014.784833 | 63 | 5 | 7 | 15 | 40 | 5 | 0 | 0 | 0 | 0 | 0 | 0 |
| JSG000000 | Laptm4b | 1.06E-44 | 1.84E-43 | 1.872609881 | 596.5748092 | 2184.623773 | 69 | 5 | 7 | 11 | 39 | 4 | 1 | 0 | 0 | 0 | 0 | 0 |
| SG0000004 | Ino80c | 6.62E-44 | 1.12E-42 | -2.506347594 | 2513.740207 | 442.4198319 | 87 | 3 | 7 | 13 | 28 | 13 | 14 | 0 | 0 | 4 | 4 | 0 |
| SG0000002 | Npr1 | 1.71E-42 | 2.75E-41 | 3.370816485 | 40.0499287 | 414.3035161 | 22 | 5 | 7 | 8 | 16 | 0 | 0 | 0 | 0 | 0 | 0 | 0 |
| SG0000003 | Ap1s2 | 1.18E-41 | 1.85E-40 | -2.410082141 | 2586.006545 | 486.5445398 | 286 | 7 | 7 | 31 | 142 | 4 | 20 | 1 | 1 | 5 | 11 | 0 |
| JSG000000 | Acadl | 2.33E-41 | 3.63E-40 | -2.202468428 | 6254.666528 | 1358.923781 | 57 | 4 | 7 | 11 | 53 | 5 | 1 | 0 | 0 | 0 | 0 | 1 |
| SG0000002 | Klf3 | 1.37E-40 | 2.07E-39 | -2.153731884 | 8117.335765 | 1824.213271 | 74 | 3 | 7 | 4 | 32 | 7 | 11 | 0 | 0 | 0 | 3 | 1 |
| SG0000002 | Ube2j1 | 1.96E-39 | 2.84E-38 | -2.292084376 | 2667.797 | 544.7116076 | 80 | 5 | 7 | 15 | 38 | 6 | 12 | 0 | 0 | 0 | 0 | 0 |
| SG0000002 | Pla2g7 | 9.47E-39 | 1.34E-37 | -2.703055394 | 1501.602269 | 230.5972705 | 82 | 0 | 7 | 9 | 38 | 12 | 1 | 0 | 0 | 0 | 0 | 0 |
| JSG000000 | Ptpre | 3.62E-37 | 4.84E-36 | -5.280219491 | 1019.561056 | 26.23667452 | 161 | 5 | 7 | 18 | 60 | 16 | 7 | 0 | 0 | 0 | 1 | 1 |
| JSG000000 | Cpd | 4.61E-37 | 6.14E-36 | -2.695945224 | 5560.844959 | 858.183989 | 155 | 3 | 7 | 25 | 75 | 13 | 6 | 0 | 0 | 1 | 4 | 1 |
| SG0000002 | Crybg3 | 8.28E-37 | 1.09E-35 | -2.461225848 | 1895.597502 | 344.2257235 | 188 | 4 | 7 | 22 | 99 | 19 | 2 | 0 | 1 | 1 | 1 | 1 |
| JSG000000 | Psma8 | 1.23E-36 | 1.61E-35 | 10.56076721 | 0.34247518 | 517.293247 | 104 | 2 | 7 | 13 | 55 | 6 | 1 | 0 | 0 | 0 | 0 | 0 |
| SG0000002 | Gart | 5.52E-36 | 7.1E-35 | 1.479229441 | 4438.893584 | 12375.62571 | 76 | 7 | 7 | 21 | 38 | 3 | 3 | 0 | 0 | 0 | 0 | 0 |
| SG0000006 | Atp8b2 | 1.13E-35 | 1.44E-34 | -2.0261351 | 4960.831621 | 1217.943232 | 69 | 7 | 7 | 7 | 37 | 0 | 2 | 0 | 0 | 0 | 6 | 0 |
| JSG000000 | Prnp | 1.65E-35 | 2.1E-34 | -2.423438234 | 4953.084696 | 923.3113057 | 51 | 4 | 7 | 11 | 26 | 7 | 0 | 0 | 0 | 0 | 0 | 0 |
| SG0000002 | Slc27a2 | 3.97E-35 | 4.96E-34 | 4.434318663 | 7.734568828 | 167.2241651 | 49 | 5 | 7 | 11 | 20 | 10 | 1 | 0 | 0 | 0 | 0 | 0 |
| SG0000000 | Zmiz1 | 9.99E-35 | 1.23E-33 | -1.989607406 | 4923.805789 | 1239.850741 | 141 | 6 | 7 | 23 | 47 | 15 | 1 | 0 | 0 | 0 | 0 | 0 |
| JSG000000 | Igsf21 | 2.23E-34 | 2.71E-33 | 6.606559813 | 0 | 96.44794206 | 151 | 4 | 7 | 19 | 54 | 17 | 0 | 0 | 0 | 0 | 0 | 0 |
| SG0000002 | Cabp1 | 1.33E-33 | 1.56E-32 | 2.753356168 | 47.08817008 | 317.5079535 | 77 | 6 | 7 | 11 | 35 | 4 | 0 | 0 | 0 | 0 | 0 | 0 |
| SG0000002 | Zfp37 | 1.25E-31 | 1.35E-30 | 2.413329586 | 72.86303547 | 388.1430863 | 35 | 4 | 7 | 6 | 27 | 4 | 9 | 0 | 1 | 1 | 5 | 1 |
| SG0000002 | Tmem8 | 8.38E-31 | 8.73E-30 | 2.190469092 | 104.6294652 | 477.5852515 | 13 | 7 | 7 | 7 | 12 | 0 | 2 | 0 | 0 | 0 | 2 | 0 |
| JSG000000 | Fam171b | 1.13E-29 | 1.13E-28 | 3.574549883 | 38.46404559 | 458.24918 | 147 | 5 | 7 | 19 | 67 | 12 | 6 | 0 | 0 | 1 | 1 | 0 |
| SG0000002 | Col3a1 | 1.02E-28 | 9.74E-28 | -3.549972613 | 4121.292795 | 351.8724214 | 108 | 3 | 7 | 14 | 48 | 8 | 2 | 1 | 1 | 1 | 1 | 0 |
| JSG000000 | Nudt21 | 1.18E-28 | 1.12E-27 | 1.561641454 | 492.7263462 | 1454.476487 | 34 | 4 | 7 | 8 | 16 | 4 | 1 | 0 | 0 | 0 | 1 | 0 |
| JSG000000 | D15Ertd621e | 2.38E-28 | 2.24E-27 | -1.669317327 | 12703.14655 | 3993.895715 | 92 | 3 | 7 | 19 | 55 | 4 | 8 | 0 | 1 | 3 | 7 | 0 |
| SG0000002 | Ly96 | 5.29E-28 | 4.89E-27 | -4.074261925 | 194.7824055 | 11.56311048 | 47 | 4 | 7 | 6 | 29 | 6 | 0 | 0 | 0 | 0 | 0 | 1 |
| SG0000000 | Lamb1 | 3.59E-27 | 3.19E-26 | -1.648139336 | 9175.89645 | 2927.582394 | 88 | 4 | 7 | 12 | 37 | 4 | 1 | 0 | 0 | 0 | 0 | 0 |

| | | | | | | | | | | | | | | | | | | |
|---|---|---|---|---|---|---|---|---|---|---|---|---|---|---|---|---|---|---|
| SG0000003 | Dcaf10 | 4.06E-27 | 3.6E-26 | -1.879007188 | 1749.718846 | 475.6973629 | 90 | 6 | 7 | 15 | 46 | 10 | 21 | 1 | 1 | 1 | 9 | 2 |
| SG0000007 | Foxr2 | 7.01E-27 | 6.14E-26 | 8.009689539 | 0.307509278 | 79.25287595 | 33 | 6 | 7 | 10 | 16 | 2 | 3 | 0 | 0 | 1 | 1 | 0 |
| SG0000002 | Cpm | 2.28E-26 | 1.96E-25 | -2.368895126 | 781.6999534 | 151.3322875 | 81 | 5 | 7 | 13 | 33 | 9 | 3 | 0 | 1 | 0 | 2 | 0 |
| SG0000007 | Fam149a | 3.31E-26 | 2.82E-25 | -2.6710819 | 617.1209238 | 96.89346766 | 88 | 2 | 7 | 10 | 22 | 14 | 9 | 0 | 0 | 1 | 1 | 1 |
| SG0000003 | Kat8 | 4.28E-26 | 3.63E-25 | 1.636950627 | 289.5831385 | 900.6260632 | 20 | 7 | 7 | 9 | 17 | 3 | 1 | 0 | 0 | 0 | 0 | 1 |
| SG0000004 | Gpr161 | 4.56E-26 | 3.87E-25 | -1.965959436 | 1117.952314 | 286.1610486 | 49 | 4 | 7 | 13 | 40 | 7 | 5 | 0 | 1 | 2 | 6 | 1 |
| SG000000 | Ccdc14 | 1.99E-25 | 1.64E-24 | 1.522282541 | 409.4978783 | 1176.2628 | 75 | 3 | 7 | 10 | 39 | 9 | 4 | 0 | 1 | 1 | 2 | 0 |
| SG000000 | Gfpt2 | 2.56E-25 | 2.1E-24 | 1.312420382 | 1386.956607 | 3444.615691 | 55 | 4 | 7 | 7 | 46 | 6 | 2 | 0 | 0 | 1 | 1 | 0 |
| SG0000004 | Creg1 | 3.51E-24 | 2.73E-23 | -1.702712791 | 1941.213001 | 596.3559644 | 21 | 4 | 7 | 9 | 16 | 6 | 2 | 0 | 0 | 2 | 2 | 0 |
| SG000000 | Foxk1 | 3.89E-24 | 3.01E-23 | -1.65441079 | 2516.226432 | 799.3232532 | 47 | 5 | 7 | 10 | 29 | 3 | 2 | 0 | 0 | 0 | 1 | 0 |
| SG000000 | Lactb2 | 7.29E-24 | 5.58E-23 | 1.281909663 | 1327.644119 | 3228.307806 | 65 | 1 | 7 | 6 | 22 | 4 | 6 | 0 | 0 | 1 | 3 | 1 |
| SG000000 | Lrig3 | 3.75E-23 | 2.77E-22 | 1.740187016 | 169.4551097 | 566.1130401 | 98 | 4 | 7 | 18 | 48 | 20 | 2 | 0 | 1 | 1 | 4 | 0 |
| SG000000 | 4921524J17Rik | 3.79E-23 | 2.8E-22 | 1.439352952 | 445.1280643 | 1207.183774 | 74 | 5 | 7 | 19 | 38 | 7 | 6 | 1 | 2 | 2 | 4 | 0 |
| SG0000002 | Ercc4 | 1.71E-22 | 1.23E-21 | -1.665999646 | 1616.103601 | 509.2761425 | 126 | 4 | 7 | 11 | 44 | 6 | 2 | 0 | 0 | 2 | 2 | 0 |
| SG0000004 | Gyltl1b | 8.26E-22 | 5.75E-21 | 4.9962305 | 21.24756613 | 678.1479222 | 42 | 0 | 7 | 0 | 0 | 0 | 16 | 0 | 7 | 0 | 0 | 0 |
| SG000000 | Gdpd5 | 1.68E-21 | 1.14E-20 | -2.679799733 | 510.3990535 | 79.65439773 | 57 | 7 | 7 | 13 | 28 | 3 | 1 | 0 | 0 | 1 | 1 | 0 |
| SG0000002 | Phykpl | 3.82E-21 | 2.55E-20 | -2.261985087 | 494.5649177 | 103.1093253 | 52 | 6 | 7 | 9 | 15 | 0 | 3 | 0 | 1 | 0 | 0 | 0 |
| SG000000 | Smo | 2.33E-20 | 1.49E-19 | -1.520182599 | 1999.023365 | 696.9430783 | 43 | 7 | 7 | 9 | 21 | 5 | 0 | 0 | 0 | 0 | 0 | 0 |
| SG0000003 | Adat1 | 1.93E-19 | 1.17E-18 | 1.629297944 | 227.216265 | 702.9217306 | 89 | 6 | 7 | 15 | 37 | 12 | 4 | 0 | 0 | 0 | 1 | 0 |
| SG0000003 | Sirpa | 4.87E-19 | 2.89E-18 | -2.184525965 | 1117.24577 | 245.7767954 | 146 | 7 | 7 | 7 | 63 | 41 | 6 | 0 | 0 | 0 | 0 | 6 |
| SG000000 | Plekhg3 | 1.78E-18 | 1.03E-17 | -1.395838953 | 2225.215175 | 845.6343562 | 44 | 6 | 7 | 11 | 31 | 4 | 1 | 0 | 0 | 0 | 0 | 1 |
| SG0000002 | Dek | 3.97E-18 | 2.24E-17 | 1.096080353 | 4035.585284 | 8626.995937 | 55 | 1 | 7 | 6 | 22 | 5 | 5 | 0 | 0 | 1 | 2 | 2 |
| SG000000 | Rsph1 | 8.54E-18 | 4.75E-17 | 1.764207638 | 98.494994 | 334.5750686 | 24 | 6 | 7 | 8 | 14 | 0 | 0 | 0 | 0 | 0 | 0 | 0 |
| SG000000 | Ap5m1 | 1.08E-17 | 5.98E-17 | 1.223486672 | 538.1707254 | 1256.684497 | 51 | 4 | 7 | 11 | 24 | 9 | 8 | 0 | 1 | 1 | 2 | 0 |
| SG000000 | Eaf1 | 2.42E-17 | 1.32E-16 | -1.239531918 | 6094.408599 | 2581.043239 | 40 | 6 | 7 | 13 | 16 | 2 | 8 | 0 | 0 | 1 | 2 | 1 |
| SG000000 | Def6 | 1.06E-16 | 5.55E-16 | -2.080610966 | 1480.38052 | 349.9830824 | 27 | 7 | 7 | 8 | 16 | 3 | 0 | 0 | 0 | 0 | 0 | 0 |
| SG000000C | Neurl1a | 1.31E-16 | 6.83E-16 | 1.700051629 | 141.7095486 | 460.4321589 | 96 | 7 | 7 | 10 | 43 | 10 | 4 | 2 | 2 | 2 | 2 | 0 |
| SG000000 | Stx7 | 1.8E-16 | 9.31E-16 | -1.256175129 | 3311.244895 | 1386.26069 | 82 | 5 | 7 | 16 | 49 | 9 | 5 | 1 | 2 | 1 | 2 | 1 |
| SG000000 | Dhfr | 3.05E-16 | 1.56E-15 | 1.249242508 | 815.9573171 | 1939.665799 | 65 | 3 | 7 | 10 | 34 | 0 | 9 | 0 | 1 | 0 | 0 | 0 |
| SG000000C | Ralb | 5.59E-16 | 2.8E-15 | 1.074553409 | 1011.437787 | 2130.158811 | 49 | 6 | 7 | 9 | 25 | 3 | 4 | 0 | 0 | 0 | 1 | 0 |
| SG000000 | Snrnp40 | 5.66E-16 | 2.84E-15 | 1.055237661 | 1258.602068 | 2615.45125 | 49 | 2 | 7 | 12 | 23 | 6 | 0 | 0 | 0 | 0 | 0 | 0 |
| SG000000C | Rarg | 1.35E-15 | 6.65E-15 | -1.513705098 | 4192.026808 | 1468.092474 | 27 | 7 | 7 | 9 | 14 | 2 | 6 | 2 | 2 | 4 | 4 | 0 |
| SG0000005 | Birc2 | 1.38E-14 | 6.35E-14 | 1.026242108 | 978.4559446 | 1992.833124 | 68 | 3 | 7 | 7 | 30 | 10 | 2 | 0 | 0 | 0 | 0 | 2 |
| SG0000003 | Fbxl21 | 1.75E-14 | 8.02E-14 | -2.36929576 | 189.5473672 | 36.68501188 | 148 | 3 | 7 | 22 | 46 | 15 | 17 | 0 | 1 | 2 | 5 | 2 |
| SG0000003 | Pon2 | 1.97E-14 | 8.98E-14 | -1.780830572 | 439.2172968 | 127.8191766 | 77 | 4 | 7 | 11 | 37 | 4 | 2 | 0 | 0 | 0 | 1 | 0 |
| SG000000 | Exosc7 | 4.39E-14 | 1.96E-13 | 1.05958358 | 814.5788604 | 1697.851004 | 35 | 6 | 7 | 8 | 24 | 4 | 1 | 0 | 0 | 0 | 0 | 1 |
| SG000000 | Gtf3c1 | 6.29E-14 | 2.77E-13 | -1.084497523 | 5355.083311 | 2525.224225 | 61 | 5 | 7 | 12 | 27 | 7 | 2 | 1 | 1 | 2 | 2 | 0 |
| SG000000 | Maea | 9.65E-14 | 4.2E-13 | -1.109808572 | 5634.457078 | 2610.756725 | 68 | 6 | 7 | 12 | 45 | 8 | 1 | 0 | 0 | 0 | 0 | 0 |
| SG000000 | Zfp251 | 1.21E-13 | 5.24E-13 | -1.519513456 | 529.0148941 | 184.5222623 | 36 | 6 | 7 | 9 | 17 | 2 | 1 | 0 | 0 | 0 | 0 | 0 |
| SG000000 | Apmap | 1.3E-13 | 5.59E-13 | -1.305385168 | 991.4784224 | 401.1651294 | 41 | 4 | 7 | 8 | 23 | 3 | 1 | 0 | 0 | 0 | 0 | 0 |
| SG0000001 | Kif19a | 1.54E-13 | 6.64E-13 | 6.860068261 | 0.331601068 | 38.52141574 | 18 | 7 | 7 | 8 | 12 | 2 | 0 | 0 | 0 | 0 | 2 | 0 |
| SG000000 | Ammecr1l | 2.6E-13 | 1.1E-12 | -1.128204963 | 2383.653732 | 1090.484838 | 42 | 6 | 7 | 11 | 17 | 1 | 1 | 0 | 0 | 0 | 0 | 0 |
| SG000000 | Polh | 2.9E-13 | 1.22E-12 | -1.200850217 | 1400.493852 | 609.241209 | 59 | 6 | 7 | 12 | 33 | 6 | 0 | 0 | 0 | 0 | 0 | 0 |
| SG000000 | Gpi1 | 3.39E-13 | 1.42E-12 | 0.894595514 | 15947.10782 | 29647.08608 | 20 | 6 | 7 | 7 | 13 | 4 | 0 | 0 | 0 | 0 | 0 | 0 |
| SG000000 | Plvap | 9.08E-13 | 3.71E-12 | 5.22290831 | 9.241989589 | 345.1576798 | 19 | 7 | 7 | 9 | 16 | 0 | 1 | 0 | 0 | 0 | 0 | 0 |
| SG0000003 | Lzts1 | 9.56E-13 | 3.9E-12 | 6.707773864 | 0.34247518 | 35.79894597 | 58 | 6 | 7 | 10 | 27 | 13 | 3 | 0 | 0 | 1 | 1 | 0 |
| SG000000 | Mup6 | 1.36E-12 | 5.47E-12 | 5.819601182 | 1.609821762 | 90.91851056 | 81 | 3 | 7 | 15 | 58 | 7 | 1 | 0 | 0 | 0 | 0 | 0 |
| SG000000 | Ccdc82 | 1.38E-12 | 5.57E-12 | -1.987973611 | 309.4468865 | 78.00930879 | 104 | 7 | 7 | 12 | 41 | 4 | 0 | 0 | 0 | 0 | 0 | 0 |
| SG000000 | Bbs2 | 3.53E-12 | 1.39E-11 | 1.132532093 | 293.5169707 | 643.5159972 | 59 | 5 | 7 | 12 | 28 | 5 | 3 | 0 | 0 | 0 | 0 | 1 |
| SG000000 | Art1 | 4.23E-12 | 1.65E-11 | 5.002953182 | 0 | 31.06557077 | 15 | 6 | 7 | 7 | 7 | 1 | 0 | 0 | 0 | 0 | 0 | 0 |
| SG0000003 | Gse1 | 5.21E-12 | 2.03E-11 | 1.325663147 | 1060.00247 | 2656.875774 | 88 | 7 | 7 | 14 | 38 | 6 | 6 | 0 | 0 | 1 | 8 | 3 |
| SG0000003 | Eva1a | 1.39E-11 | 5.26E-11 | 3.351201867 | 5.830455467 | 59.49969862 | 59 | 6 | 7 | 13 | 33 | 4 | 2 | 0 | 1 | 0 | 2 | 0 |
| SG000000 | Zdhhc16 | 1.4E-11 | 5.29E-11 | -1.115388571 | 1289.86083 | 595.3569639 | 25 | 7 | 7 | 9 | 14 | 2 | 0 | 0 | 0 | 0 | 0 | 0 |
| SG0000001 | Adam19 | 2.87E-11 | 1.06E-10 | 2.412038269 | 666.1848778 | 3545.607356 | 128 | 5 | 7 | 13 | 52 | 18 | 3 | 0 | 0 | 0 | 0 | 1 |
| SG000000 | Sod2 | 3.51E-11 | 1.29E-10 | 0.849956959 | 1991.083434 | 3588.822664 | 28 | 7 | 7 | 7 | 25 | 1 | 8 | 0 | 0 | 0 | 3 | 0 |
| SG0000002 | Map3k1 | 3.98E-11 | 1.46E-10 | 1.392582555 | 628.3645784 | 1649.760176 | 75 | 3 | 7 | 8 | 28 | 6 | 4 | 0 | 0 | 0 | 1 | 1 |
| SG000000 | Nhlrc2 | 9.65E-11 | 3.44E-10 | -1.084311706 | 1144.079895 | 539.5677769 | 104 | 5 | 7 | 17 | 45 | 14 | 9 | 0 | 0 | 1 | 5 | 0 |
| SG000000 | Ppp6r2 | 1.33E-10 | 4.7E-10 | -1.493884535 | 520.8826014 | 184.9421049 | 34 | 5 | 7 | 8 | 19 | 3 | 1 | 0 | 0 | 0 | 0 | 0 |
| SG0000002 | 43710 | 1.58E-10 | 5.54E-10 | 0.768315825 | 7536.756051 | 12837.2077 | 72 | 5 | 7 | 15 | 44 | 8 | 7 | 0 | 0 | 0 | 2 | 0 |
| SG0000003 | Mtmr10 | 2.71E-10 | 9.36E-10 | -1.091978904 | 901.629516 | 422.9701464 | 85 | 6 | 7 | 16 | 44 | 9 | 8 | 0 | 0 | 0 | 0 | 0 |
| SG000000 | Tmem65 | 7.98E-10 | 2.65E-09 | -0.981678463 | 1478.937126 | 748.9193433 | 101 | 3 | 7 | 14 | 54 | 5 | 10 | 0 | 0 | 1 | 4 | 0 |
| SG000000 | Fsbp | 1.44E-09 | 4.68E-09 | 2.130248751 | 21.16691336 | 92.66725625 | 37 | 2 | 7 | 11 | 18 | 3 | 16 | 1 | 4 | 1 | 3 | 2 |
| SG0000002 | Arsk | 1.5E-09 | 4.87E-09 | -1.351711374 | 355.0491964 | 139.1178525 | 69 | 5 | 7 | 10 | 57 | 8 | 4 | 0 | 0 | 1 | 2 | 0 |
| SG000000 | Dnajc22 | 2.12E-09 | 6.79E-09 | 2.684971889 | 23.16136004 | 148.9433555 | 16 | 7 | 7 | 8 | 11 | 1 | 1 | 0 | 0 | 1 | 2 | 0 |
| SG0000002 | Col1a2 | 3.25E-09 | 1.02E-08 | -2.217024877 | 8690.332695 | 1869.155417 | 79 | 2 | 7 | 11 | 44 | 10 | 0 | 0 | 0 | 0 | 0 | 0 |
| SG0000003 | Prkx | 3.82E-09 | 1.2E-08 | -0.949186955 | 1350.590878 | 699.5037821 | 128 | 1 | 7 | 21 | 61 | 11 | 15 | 0 | 2 | 2 | 4 | 0 |
| SG000000 | Tnrc6a | 4.07E-09 | 1.27E-08 | -0.843603657 | 4062.674443 | 2263.924903 | 132 | 2 | 7 | 19 | 57 | 6 | 3 | 0 | 0 | 0 | 3 | 0 |
| SG0000004 | Ccdc74a | 4.21E-09 | 1.32E-08 | -2.241305159 | 99.53129993 | 21.05036323 | 18 | 0 | 7 | 0 | 4 | 0 | 4 | 0 | 0 | 0 | 4 | 0 |
| SG000000 | Fbxw8 | 4.57E-09 | 1.43E-08 | -0.841362846 | 3712.71359 | 2072.125208 | 78 | 5 | 7 | 8 | 25 | 4 | 0 | 0 | 0 | 0 | 0 | 0 |
| SG0000005 | Fhad1 | 8.93E-09 | 2.73E-08 | 3.368674564 | 8.291526554 | 85.64590156 | 109 | 3 | 7 | 9 | 48 | 11 | 2 | 0 | 0 | 0 | 0 | 1 |
| SG000000 | Zfp947 | 1.07E-08 | 3.26E-08 | 2.494211298 | 11.13138543 | 62.71647443 | 59 | 5 | 7 | 11 | 28 | 5 | 3 | 0 | 1 | 1 | 3 | 0 |
| SG000000 | Recql4 | 1.24E-08 | 3.75E-08 | 0.93657844 | 342.7045723 | 655.9309183 | 13 | 7 | 7 | 7 | 8 | 0 | 0 | 0 | 0 | 0 | 0 | 0 |
| SG0000004 | Pogk | 1.27E-08 | 3.83E-08 | 2.205129245 | 49.05637811 | 226.2067331 | 115 | 3 | 7 | 12 | 29 | 1 | 16 | 2 | 5 | 4 | 10 | 0 |
| SG0000006 | Slc22a21 | 1.9E-08 | 5.65E-08 | -1.84941074 | 129.9264822 | 36.05533815 | 89 | 4 | 7 | 11 | 28 | 6 | 0 | 0 | 0 | 0 | 0 | 0 |
| SG0000002 | Naa35 | 2.78E-08 | 8.14E-08 | 0.737046759 | 1283.809364 | 2139.802654 | 117 | 6 | 7 | 13 | 59 | 13 | 1 | 0 | 0 | 0 | 0 | 0 |
| SG0000003 | Cnot3 | 8.27E-08 | 2.34E-07 | 0.702148933 | 1817.750665 | 2957.346434 | 22 | 7 | 7 | 9 | 9 | 0 | 2 | 0 | 0 | 2 | 2 | 0 |
| SG000000 | Pygb | 8.99E-08 | 2.53E-07 | -0.776867594 | 3292.017109 | 1921.328545 | 45 | 7 | 7 | 11 | 21 | 2 | 0 | 0 | 0 | 0 | 0 | 0 |
| SG000000 | Ddx6 | 9.51E-08 | 2.67E-07 | -0.753901306 | 7137.609326 | 4232.586734 | 79 | 6 | 7 | 10 | 37 | 4 | 7 | 0 | 0 | 0 | 1 | 2 |
| SG0000001 | Tab2 | 1.7E-07 | 4.68E-07 | -0.750012621 | 3707.390188 | 2204.40811 | 115 | 3 | 7 | 16 | 52 | 11 | 5 | 0 | 1 | 2 | 3 | 1 |
| SG000000 | Zc3h3 | 4.24E-07 | 1.13E-06 | -1.067132686 | 366.4016103 | 174.8712546 | 76 | 6 | 7 | 13 | 43 | 8 | 0 | 0 | 0 | 0 | 1 | 0 |
| SG000000C | Chtop | 5.02E-07 | 1.33E-06 | 0.635947405 | 2516.909654 | 3911.171603 | 171 | 0 | 7 | 7 | 42 | 0 | 10 | 0 | 0 | 0 | 0 | 0 |
| SG000000C | Tax1bp1 | 5.19E-07 | 1.37E-06 | -0.695754662 | 6619.326736 | 4086.681538 | 143 | 5 | 7 | 23 | 83 | 15 | 4 | 0 | 0 | 1 | 1 | 0 |
| SG000000 | L2hgdh | 6.49E-07 | 1.7E-06 | 0.736403794 | 592.275057 | 986.7407736 | 87 | 6 | 7 | 17 | 34 | 4 | 8 | 0 | 0 | 1 | 2 | 0 |
| SG000000 | Dlst | 6.58E-07 | 1.72E-06 | -0.688056748 | 5544.003988 | 3441.104125 | 42 | 5 | 7 | 10 | 14 | 3 | 1 | 0 | 0 | 0 | 0 | 0 |
| SG0000001 | Stk32c | 8.72E-07 | 2.26E-06 | 4.127400426 | 0 | 16.47717892 | 167 | 6 | 7 | 28 | 76 | 12 | 0 | 0 | 0 | 0 | 0 | 2 |
| SG000000 | Dcakd | 9.76E-07 | 2.53E-06 | 0.904008584 | 1312.970572 | 2456.906857 | 33 | 5 | 7 | 7 | 14 | 1 | 0 | 0 | 0 | 0 | 0 | 0 |
| SG000000 | Tmem136 | 1.02E-06 | 2.64E-06 | -0.904321242 | 609.4582145 | 325.6237814 | 19 | 6 | 7 | 8 | 13 | 0 | 8 | 3 | 3 | 3 | 5 | 0 |
| SG0000003 | Plp1 | 1.31E-06 | 3.35E-06 | 1.460335715 | 41.01060973 | 112.8498744 | 49 | 5 | 7 | 5 | 21 | 2 | 9 | 1 | 1 | 1 | 7 | 0 |
| SG000000 | Slc6a11 | 1.37E-06 | 3.49E-06 | 4.091123431 | 0 | 16.04318934 | 137 | 5 | 7 | 13 | 46 | 21 | 3 | 0 | 0 | 0 | 3 | 0 |
| SG000000 | Bace2 | 1.49E-06 | 3.78E-06 | -2.680883883 | 51.55291808 | 8.03945834 | 105 | 2 | 7 | 8 | 44 | 8 | 1 | 0 | 0 | 0 | 0 | 0 |
| SG000000 | Actr6 | 1.5E-06 | 3.82E-06 | 0.776332973 | 377.8627816 | 647.192728 | 51 | 3 | 7 | 11 | 24 | 6 | 1 | 0 | 1 | 0 | 1 | 2 |
| SG0000001 | E2f3 | 2.32E-06 | 5.8E-06 | 0.668857757 | 869.7365844 | 1382.719177 | 109 | 5 | 7 | 26 | 56 | 11 | 6 | 0 | 0 | 2 | 2 | 3 |
| SG000000 | Trim40 | 2.43E-06 | 6.06E-06 | -2.743225229 | 67.06988947 | 10.01692707 | 23 | 4 | 7 | 6 | 13 | 2 | 1 | 0 | 1 | 0 | 1 | 0 |
| SG0000002 | Tmod1 | 4.92E-06 | 1.19E-05 | 4.78913262 | 0.674076249 | 18.63723995 | 85 | 4 | 7 | 14 | 34 | 4 | 3 | 0 | 1 | 0 | 1 | 0 |
| SG000000 | Crocc2 | 5.7E-06 | 1.37E-05 | -2.433551112 | 49.02590263 | 9.075148028 | 44 | 4 | 7 | 7 | 17 | 3 | 2 | 0 | 0 | 0 | 0 | 1 |
| SG000000 | Cpsf3 | 6.42E-06 | 1.54E-05 | 0.572021149 | 2562.448288 | 3809.347867 | 45 | 2 | 7 | 6 | 13 | 3 | 0 | 0 | 0 | 0 | 0 | 0 |
| SG000000 | Smpd3 | 6.51E-06 | 1.56E-05 | 3.441047573 | 2.305646233 | 25.04097346 | 96 | 5 | 7 | 11 | 34 | 6 | 2 | 0 | 0 | 0 | 0 | 0 |
| SG000000 | Cgref1 | 6.85E-06 | 1.64E-05 | -0.957272163 | 430.3477985 | 221.641944 | 17 | 6 | 7 | 7 | 14 | 2 | 0 | 0 | 0 | 0 | 0 | 0 |
| SG000000 | Dthd1 | 7.48E-06 | 1.78E-05 | -5.828020659 | 19.01903554 | 0.33479538 | 195 | 1 | 7 | 25 | 86 | 18 | 0 | 0 | 0 | 0 | 0 | 0 |
| SG000000 | Trmt10b | 7.81E-06 | 1.86E-05 | 0.861012799 | 211.7782063 | 384.6554985 | 48 | 7 | 7 | 12 | 34 | 7 | 1 | 0 | 0 | 0 | 1 | 1 |
| SG000000 | Nedd1 | 8.92E-06 | 2.11E-05 | -0.674670619 | 1763.121918 | 1104.552215 | 93 | 4 | 7 | 13 | 41 | 9 | 7 | 1 | 1 | 2 | 4 | 2 |
| SG0000005 | Cml1 | 9.22E-06 | 2.18E-05 | -1.991764222 | 63.19891478 | 15.89018106 | 13 | 4 | 7 | 6 | 7 | 0 | 0 | 0 | 0 | 0 | 0 | 0 |
| SG000000 | Kctd5 | 2.03E-05 | 4.64E-05 | 0.576666121 | 1304.002773 | 1944.788217 | 31 | 7 | 7 | 10 | 20 | 2 | 5 | 0 | 0 | 2 | 3 | 0 |
| SG0000003 | Traf3ip1 | 2.17E-05 | 4.96E-05 | 0.65174259 | 510.7482349 | 802.4185196 | 43 | 4 | 7 | 6 | 12 | 5 | 2 | 0 | 0 | 0 | 1 | 3 |
| SG000000 | Lrr1 | 2.46E-05 | 5.57E-05 | 0.858572866 | 154.7783168 | 280.6508135 | 24 | 5 | 7 | 5 | 13 | 0 | 3 | 0 | 0 | 0 | 1 | 0 |
| SG000000 | Morn4 | 2.61E-05 | 5.9E-05 | -0.71293103 | 799.6089695 | 487.8249765 | 28 | 7 | 7 | 9 | 16 | 0 | 1 | 0 | 0 | 1 | 2 | 0 |
| SG000000 | Cpsf2 | 5.57E-05 | 0.000122 | -0.563871305 | 3907.068515 | 2643.071128 | 56 | 4 | 7 | 11 | 32 | 5 | 4 | 0 | 0 | 0 | 1 | 2 |
| SG000000 | Copz1 | 6.08E-05 | 0.000132 | 0.508528941 | 2884.622526 | 4103.660797 | 47 | 3 | 7 | 6 | 23 | 4 | 1 | 0 | 0 | 0 | 0 | 0 |
| SG0000003 | Tgfb1i1 | 7.8E-05 | 0.000168 | -1.240441422 | 114.4356399 | 48.43409915 | 28 | 0 | 7 | 14 | 35 | 0 | 0 | 0 | 0 | 0 | 0 | 0 |
| SG0000003 | Parp11 | 8.24E-05 | 0.000177 | -0.701565656 | 640.61635 | 393.9178967 | 177 | 6 | 7 | 25 | 96 | 20 | 6 | 2 | 2 | 3 | 6 | 1 |
| SG000000 | Ubxn2b | 8.28E-05 | 0.000177 | 0.632433049 | 424.0128717 | 657.2950455 | 55 | 4 | 7 | 9 | 24 | 6 | 6 | 0 | 0 | 2 | 6 | 0 |
| SG000000 | Srebf2 | 8.3E-05 | 0.000178 | 0.759829016 | 3472.272825 | 5879.564604 | 47 | 2 | 7 | 8 | 29 | 5 | 2 | 0 | 1 | 0 | 0 | 1 |
| SG0000001 | Tango2 | 8.37E-05 | 0.000179 | -0.767906807 | 428.4411104 | 251.6101481 | 70 | 4 | 7 | 8 | 35 | 4 | 0 | 0 | 0 | 0 | 0 | 0 |
| SG0000002 | Tyms | 9.29E-05 | 0.000198 | 0.52923858 | 1257.226934 | 1814.388831 | 96 | 4 | 7 | 18 | 47 | 3 | 9 | 0 | 0 | 0 | 1 | 0 |
| SG0000004 | Eno4 | 0.000115 | 0.000243 | 1.783595983 | 13.87487714 | 47.76887761 | 43 | 6 | 7 | 14 | 20 | 5 | 0 | 0 | 0 | 0 | 1 | 0 |
| SG000000 | Abcd4 | 0.000155 | 0.000322 | 0.596502045 | 460.284962 | 695.9720426 | 26 | 6 | 7 | 10 | 15 | 0 | 0 | 0 | 0 | 0 | 0 | 0 |
| SG0000005 | Amz1 | 0.000234 | 0.000475 | 1.61558736 | 16.66286216 | 51.0610637 | 48 | 4 | 7 | 8 | 32 | 3 | 1 | 0 | 0 | 0 | 1 | 0 |
| SG000000 | F5 | 0.000278 | 0.000561 | 1.468851385 | 33.72575735 | 93.35337792 | 135 | 4 | 7 | 19 | 61 | 13 | 1 | 0 | 0 | 0 | 1 | 0 |
| SG0000003 | Slc12a9 | 0.000282 | 0.000568 | -0.731228496 | 444.3132372 | 267.6501808 | 23 | 6 | 7 | 7 | 13 | 3 | 0 | 0 | 0 | 0 | 0 | 0 |
| SG000000 | Lrrc17 | 0.00029 | 0.000583 | -2.629262485 | 37.74278262 | 6.100241679 | 63 | 6 | 7 | 10 | 33 | 4 | 1 | 0 | 0 | 0 | 1 | 0 |
| SG0000005 | Tubgcp6 | 0.000327 | 0.000653 | -0.548849355 | 1355.30538 | 926.4394569 | 60 | 7 | 7 | 11 | 34 | 2 | 2 | 0 | 0 | 0 | 0 | 0 |
| SG000000C | Tulp3 | 0.000351 | 0.000699 | -0.505794344 | 2807.079767 | 1976.949085 | 87 | 5 | 7 | 10 | 32 | 9 | 7 | 0 | 0 | 2 | 3 | 0 |
| SG0000002 | Mrc2 | 0.000377 | 0.000749 | -1.101474597 | 732.1181383 | 341.1962691 | 103 | 4 | 7 | 10 | 35 | 12 | 0 | 0 | 0 | 0 | 1 | 0 |

| | | | | | | | | | | | | | | | | | | |
|---|---|---|---|---|---|---|---|---|---|---|---|---|---|---|---|---|---|---|
| ISG000000 | Rnf34 | 0.000387 | 0.000767 | 0.492237273 | 1107.030952 | 1557.176884 | 39 | 5 | 7 | 12 | 18 | 3 | 8 | 0 | 0 | 1 | 2 | 0 |
| ISG000000 | C2cd2 | 0.000412 | 0.000811 | -0.815967579 | 353.1643346 | 200.6070178 | 96 | 6 | 7 | 13 | 36 | 17 | 7 | 0 | 0 | 0 | 0 | 0 |
| SG0000005 | Ncoa4 | 0.00047 | 0.00092 | 0.505934252 | 785.5040486 | 1115.449244 | 111 | 4 | 7 | 5 | 14 | 5 | 14 | 0 | 0 | 0 | 6 | 0 |
| SG0000004 | Pcf11 | 0.000474 | 0.000927 | 0.463729629 | 1644.570332 | 2268.030965 | 110 | 5 | 7 | 19 | 49 | 26 | 4 | 0 | 1 | 1 | 1 | 2 |
| SG0000002 | Bend5 | 0.000477 | 0.000933 | 3.133680491 | 5.390266421 | 47.30881347 | 116 | 5 | 7 | 19 | 50 | 12 | 0 | 0 | 0 | 0 | 0 | 0 |
| ISG000000 | Cpox | 0.000499 | 0.000975 | -0.506955163 | 1993.699405 | 1402.978349 | 28 | 5 | 7 | 7 | 14 | 1 | 6 | 0 | 2 | 0 | 5 | 1 |
| ISG000000 | Zbtb6 | 0.000561 | 0.001087 | -0.578218031 | 845.0777378 | 566.0250995 | 68 | 4 | 7 | 6 | 21 | 1 | 26 | 2 | 4 | 4 | 10 | 0 |
| SG0000004 | Coq5 | 0.000612 | 0.00118 | 0.525142042 | 579.9295456 | 834.5622606 | 28 | 4 | 7 | 10 | 16 | 4 | 0 | 0 | 0 | 0 | 0 | 0 |
| ISG000000 | Psmb2 | 0.00063 | 0.001213 | 0.449253278 | 2624.659458 | 3583.535205 | 74 | 2 | 7 | 8 | 27 | 6 | 0 | 0 | 0 | 0 | 0 | 0 |
| SG0000007 | H2-Q6 | 0.000647 | 0.001243 | 1.534799948 | 16.06106128 | 46.53664402 | 11 | 6 | 7 | 6 | 9 | 1 | 0 | 0 | 0 | 0 | 0 | 0 |
| ISG000000 | Gmpr2 | 0.00073 | 0.001393 | 0.525144504 | 538.5719611 | 775.0469067 | 17 | 6 | 7 | 7 | 12 | 1 | 0 | 0 | 0 | 0 | 0 | 0 |
| ISG000000 | 1700027J19Rik | 0.000731 | 0.001395 | -1.21396389 | 365.292448 | 157.4712044 | 26 | 6 | 7 | 9 | 25 | 1 | 0 | 0 | 0 | 0 | 0 | 0 |
| ISG000000 | Ccdc3 | 0.00075 | 0.001429 | 1.969063269 | 16.21933269 | 63.50093054 | 139 | 2 | 7 | 15 | 40 | 12 | 0 | 0 | 0 | 0 | 0 | 0 |
| ISG000000 | Rbks | 0.000882 | 0.001664 | -1.006900205 | 119.3376868 | 59.38413721 | 111 | 5 | 7 | 14 | 52 | 12 | 0 | 0 | 0 | 0 | 0 | 0 |
| ISG000000 | Pvrl1 | 0.001426 | 0.002633 | 0.951157438 | 289.9702032 | 560.63511 | 45 | 6 | 7 | 9 | 21 | 2 | 1 | 0 | 0 | 0 | 0 | 0 |
| SG0000003 | Zcchc2 | 0.001456 | 0.002684 | -0.478000148 | 1703.492306 | 1223.060087 | 148 | 2 | 7 | 7 | 57 | 9 | 4 | 0 | 0 | 2 | 6 | 4 |
| ISG000000 | Zc3h4 | 0.001874 | 0.003405 | -0.449298854 | 2208.025519 | 1617.155031 | 29 | 5 | 7 | 5 | 14 | 7 | 1 | 0 | 0 | 0 | 0 | 2 |
| ISG000000 | Ddx42 | 0.001935 | 0.003509 | -0.426994668 | 4438.879848 | 3301.681579 | 64 | 5 | 7 | 13 | 35 | 7 | 2 | 0 | 0 | 0 | 0 | 0 |
| SG0000000 | Stard3nl | 0.002402 | 0.004299 | -0.487964887 | 949.8887315 | 677.299364 | 69 | 5 | 7 | 10 | 37 | 7 | 1 | 0 | 0 | 1 | 1 | 0 |
| SG0000002 | Srd5a3 | 0.002485 | 0.00444 | -0.534118807 | 553.4321232 | 382.1893505 | 39 | 4 | 7 | 7 | 32 | 3 | 0 | 0 | 0 | 0 | 1 | 0 |
| SG0000001 | Ift52 | 0.002979 | 0.005259 | -0.442703896 | 1650.590597 | 1214.429239 | 53 | 5 | 7 | 9 | 24 | 2 | 2 | 0 | 0 | 0 | 0 | 0 |
| ISG000000 | Pigf | 0.003065 | 0.005397 | -0.462506564 | 1033.02461 | 749.6910374 | 73 | 4 | 7 | 9 | 24 | 5 | 2 | 0 | 0 | 1 | 1 | 0 |
| ISG000000 | Kifc5b | 0.003136 | 0.005515 | 0.499285448 | 454.8217732 | 642.8966211 | 20 | 7 | 7 | 10 | 14 | 2 | 2 | 0 | 0 | 2 | 3 | 1 |
| SG0000001 | Plod1 | 0.003605 | 0.006289 | -0.626525162 | 1949.773082 | 1262.935595 | 23 | 6 | 7 | 7 | 17 | 3 | 0 | 0 | 0 | 0 | 0 | 0 |
| SG0000003 | Lif | 0.003981 | 0.006902 | -0.670180831 | 294.8707067 | 185.3049831 | 20 | 4 | 7 | 10 | 29 | 0 | 14 | 4 | 6 | 10 | 28 | 0 |
| ISG000000 | Zfp395 | 0.003989 | 0.006914 | 0.765190812 | 914.2132877 | 1553.792125 | 64 | 5 | 7 | 9 | 25 | 5 | 0 | 0 | 0 | 0 | 3 | 0 |
| ISG000000 | Ptgfr | 0.005843 | 0.009899 | -2.370029805 | 18.60662856 | 3.599296515 | 174 | 2 | 7 | 25 | 91 | 20 | 14 | 0 | 1 | 1 | 5 | 2 |
| SG0000002 | Dck | 0.006189 | 0.010454 | -0.405831338 | 1844.618562 | 1392.320343 | 51 | 5 | 7 | 8 | 16 | 4 | 6 | 0 | 1 | 1 | 1 | 0 |
| ISG000000 | Dhx36 | 0.0071 | 0.011882 | 0.340471036 | 2985.543948 | 3780.205966 | 96 | 3 | 7 | 11 | 46 | 22 | 5 | 0 | 0 | 1 | 4 | 2 |
| ISG000000 | Ctps | 0.00712 | 0.011913 | 0.339961728 | 3870.640582 | 4899.158873 | 53 | 6 | 7 | 12 | 31 | 2 | 1 | 0 | 0 | 0 | 2 | 0 |
| SG0000007 | Nlrc5 | 0.007609 | 0.012675 | 1.309152879 | 14.66941234 | 36.3502165 | 80 | 6 | 7 | 12 | 39 | 13 | 6 | 0 | 0 | 2 | 8 | 0 |
| SG0000004 | Fbxo40 | 0.008288 | 0.013735 | -4.579040399 | 16.00434716 | 0.66959076 | 43 | 4 | 7 | 12 | 44 | 4 | 1 | 1 | 1 | 1 | 2 | 0 |
| ISG000000 | Gm13139 | 0.009396 | 0.015436 | 1.101243429 | 21.62131694 | 46.3862696 | 91 | 6 | 7 | 19 | 70 | 14 | 2 | 0 | 0 | 1 | 2 | 0 |
| ISG000000 | Zfp11 | 0.010168 | 0.016611 | -0.599767528 | 223.1948589 | 147.277421 | 27 | 4 | 7 | 8 | 15 | 2 | 9 | 4 | 4 | 5 | 8 | 1 |
| SG0000007 | Il11ra1 | 0.011738 | 0.018975 | -0.638948326 | 164.5705832 | 105.6840862 | 79 | 7 | 7 | 20 | 42 | 4 | 0 | 0 | 0 | 0 | 0 | 0 |
| SG0000001 | Rtn4ip1 | 0.018684 | 0.029063 | 0.362712986 | 589.0614473 | 757.4397429 | 116 | 3 | 7 | 11 | 45 | 17 | 4 | 0 | 2 | 0 | 0 | 0 |
| ISG000000 | Zfp825 | 0.019154 | 0.029731 | 0.483915705 | 176.505725 | 246.8493221 | 30 | 7 | 7 | 13 | 18 | 4 | 0 | 0 | 0 | 0 | 0 | 0 |
| SG0000002 | Ppil2 | 0.022251 | 0.034146 | -0.324552983 | 2104.493423 | 1680.534335 | 29 | 6 | 7 | 7 | 19 | 3 | 0 | 0 | 0 | 0 | 1 | 0 |
| ISG000000 | Sh3gl1 | 0.022317 | 0.034239 | -0.335142293 | 1698.444967 | 1346.367427 | 14 | 7 | 7 | 7 | 9 | 0 | 0 | 0 | 0 | 0 | 0 | 0 |
| ISG000000 | Asf1b | 0.024583 | 0.037443 | 0.313751705 | 1289.837141 | 1603.184674 | 25 | 5 | 7 | 11 | 21 | 3 | 0 | 0 | 0 | 0 | 1 | 0 |
| ISG000000 | Osbpl11 | 0.027516 | 0.041606 | 0.32955255 | 794.5361332 | 998.4328355 | 95 | 1 | 7 | 15 | 47 | 6 | 3 | 0 | 0 | 1 | 5 | 0 |
| SG0000003 | Snx1 | 0.027959 | 0.042223 | -0.323539462 | 1436.155289 | 1147.641697 | 56 | 4 | 7 | 11 | 45 | 5 | 1 | 0 | 0 | 0 | 0 | 0 |
| SG0000004 | Zfp354c | 0.030553 | 0.045789 | 0.517638169 | 388.5394887 | 556.2368797 | 40 | 7 | 7 | 10 | 27 | 4 | 8 | 0 | 0 | 1 | 3 | 3 |
| SG0000002 | Sike1 | 0.033947 | 0.050453 | 0.302832924 | 917.1636834 | 1131.380381 | 31 | 7 | 7 | 7 | 18 | 3 | 11 | 1 | 1 | 1 | 6 | 0 |
| SG0000005 | C030039L03Rik | 0.036146 | 0.053433 | 0.685878081 | 46.52986565 | 74.851701 | 29 | 7 | 7 | 8 | 15 | 2 | 5 | 0 | 0 | 1 | 1 | 1 |
| SG0000003 | Mthfsd | 0.037432 | 0.05522 | -0.366868701 | 489.8286323 | 379.8443615 | 56 | 7 | 7 | 7 | 14 | 0 | 2 | 0 | 0 | 0 | 0 | 0 |
| ISG000000 | Tldc1 | 0.040312 | 0.059103 | 0.377823442 | 385.535574 | 500.9572271 | 24 | 7 | 7 | 11 | 18 | 2 | 1 | 0 | 0 | 0 | 0 | 0 |
| SG0000003 | Pomt1 | 0.043224 | 0.063099 | -0.319908639 | 919.9155181 | 736.9632795 | 27 | 7 | 7 | 8 | 13 | 3 | 1 | 0 | 0 | 0 | 0 | 1 |
| SG0000002 | Ppm1a | 0.044749 | 0.065161 | 0.264486389 | 2444.475955 | 2936.324888 | 84 | 5 | 7 | 15 | 43 | 13 | 2 | 0 | 0 | 0 | 0 | 0 |
| ISG000000 | Trak1 | 0.048255 | 0.069868 | 0.279412324 | 1289.801843 | 1565.432998 | 105 | 6 | 7 | 14 | 36 | 9 | 5 | 2 | 2 | 3 | 5 | 0 |
| ISG000000 | Csad | 0.051137 | 0.073672 | 0.354152078 | 311.9570031 | 398.7540784 | 17 | 7 | 7 | 9 | 14 | 0 | 0 | 0 | 0 | 0 | 0 | 0 |
| SG0000002 | Tmem63a | 0.052765 | 0.075822 | 0.274330399 | 994.1940195 | 1202.41068 | 64 | 5 | 7 | 10 | 50 | 0 | 2 | 0 | 0 | 0 | 0 | 0 |
| ISG000000 | Zfp809 | 0.053747 | 0.077123 | -0.421620013 | 249.7197662 | 186.4372331 | 34 | 4 | 7 | 9 | 22 | 3 | 0 | 0 | 0 | 0 | 0 | 0 |
| ISG000000 | Gnai3 | 0.055624 | 0.079561 | 0.23691944 | 4172.183477 | 4916.808081 | 88 | 3 | 7 | 13 | 41 | 5 | 4 | 0 | 0 | 0 | 1 | 0 |
| SG0000004 | Pkp2 | 0.057776 | 0.082383 | 0.249757384 | 1655.203302 | 1968.048553 | 108 | 5 | 7 | 12 | 52 | 17 | 0 | 0 | 0 | 0 | 0 | 0 |
| ISG000000 | Tmem200a | 0.065392 | 0.092381 | -0.622224249 | 78.25314875 | 50.83860894 | 158 | 3 | 7 | 15 | 86 | 26 | 3 | 0 | 0 | 0 | 5 | 1 |
| ISG000000 | Xpot | 0.066027 | 0.093219 | -0.246320289 | 5442.15464 | 4587.975422 | 59 | 5 | 7 | 11 | 24 | 11 | 5 | 0 | 0 | 0 | 2 | 0 |
| SG0000003 | Man2c1 | 0.069401 | 0.097559 | -0.282373132 | 1460.940822 | 1201.240156 | 53 | 7 | 7 | 13 | 20 | 0 | 0 | 0 | 0 | 0 | 0 | 0 |
| SG0000002 | Gfm1 | 0.070632 | 0.099112 | 0.235952427 | 1903.026558 | 2241.163843 | 78 | 2 | 7 | 10 | 50 | 13 | 7 | 0 | 0 | 0 | 3 | 2 |
| ISG000000 | Cdc40 | 0.079212 | 0.110091 | -0.258873536 | 1637.703186 | 1368.694414 | 86 | 0 | 7 | 11 | 43 | 9 | 1 | 0 | 1 | 1 | 2 | 1 |
| SG0000003 | Vps18 | 0.081387 | 0.112868 | -0.383521105 | 867.8937253 | 665.2964284 | 13 | 7 | 7 | 8 | 12 | 2 | 0 | 0 | 0 | 0 | 0 | 0 |
| ISG000000 | Sugp1 | 0.083621 | 0.115698 | 0.247377962 | 1001.604752 | 1188.952661 | 37 | 7 | 7 | 8 | 17 | 3 | 0 | 0 | 0 | 0 | 0 | 0 |
| ISG000000 | Hmox1 | 0.092811 | 0.127273 | -0.226967088 | 5710.330783 | 4879.07355 | 17 | 4 | 7 | 6 | 9 | 0 | 2 | 0 | 1 | 0 | 1 | 0 |
| SG0000003 | Slc26a11 | 0.094894 | 0.129821 | 0.360404713 | 181.7603211 | 233.3413523 | 42 | 7 | 7 | 12 | 19 | 0 | 6 | 2 | 2 | 2 | 2 | 0 |
| ISG000000 | 6030458C11Rik | 0.100986 | 0.13718 | 0.224865483 | 1587.078611 | 1854.768425 | 30 | 4 | 7 | 6 | 16 | 3 | 8 | 2 | 3 | 2 | 3 | 1 |
| SG0000003 | Cfap97 | 0.132498 | 0.175592 | -0.243906699 | 998.7600439 | 843.4084103 | 181 | 2 | 7 | 17 | 101 | 17 | 15 | 0 | 0 | 4 | 4 | 2 |
| ISG000000 | Pi15 | 0.139393 | 0.183895 | -1.036150859 | 19.16727656 | 9.346476546 | 79 | 3 | 7 | 12 | 45 | 8 | 27 | 0 | 2 | 5 | 15 | 1 |
| ISG000000 | Bckdha | 0.147794 | 0.19394 | 0.430976846 | 792.174282 | 1067.966665 | 21 | 6 | 7 | 8 | 19 | 6 | 0 | 0 | 0 | 0 | 0 | 0 |
| SG0000002 | Boc | 0.169997 | 0.220411 | 0.430242014 | 75.10478374 | 101.2006616 | 84 | 2 | 7 | 16 | 41 | 8 | 3 | 0 | 0 | 3 | 4 | 1 |
| SG0000002 | Zbtb49 | 0.175562 | 0.226627 | -0.341885754 | 159.0523242 | 125.493766 | 129 | 0 | 7 | 31 | 96 | 7 | 10 | 0 | 0 | 3 | 5 | 0 |
| SG0000004 | Zfp507 | 0.203865 | 0.25922 | 0.174110656 | 1267.44779 | 1430.021637 | 61 | 3 | 7 | 8 | 27 | 2 | 12 | 0 | 1 | 1 | 1 | 0 |
| SG0000001 | Cd164 | 0.209375 | 0.26551 | -0.168020552 | 9519.375087 | 8472.844061 | 43 | 5 | 7 | 8 | 14 | 8 | 9 | 0 | 0 | 0 | 2 | 2 |
| ISG000000 | Daam2 | 0.238037 | 0.297642 | -0.656487454 | 28.50094617 | 18.0816002 | 100 | 4 | 7 | 13 | 53 | 20 | 0 | 0 | 0 | 0 | 0 | 0 |
| SG0000002 | Zfp605 | 0.248636 | 0.30934 | 0.214630036 | 269.9040881 | 313.1984048 | 48 | 5 | 7 | 8 | 32 | 6 | 0 | 0 | 0 | 0 | 0 | 0 |
| ISG000000 | Vps45 | 0.251215 | 0.312136 | -0.221317387 | 419.0776123 | 359.4771491 | 126 | 3 | 7 | 13 | 46 | 9 | 0 | 0 | 0 | 0 | 0 | 0 |
| ISG000000 | Zc3hav1l | 0.270651 | 0.333166 | 0.183281385 | 486.2520932 | 552.1215243 | 31 | 4 | 7 | 5 | 14 | 1 | 15 | 0 | 2 | 0 | 3 | 0 |
| SG0000002 | Mylk | 0.272925 | 0.335619 | 0.619517304 | 13.79038095 | 21.1870043 | 252 | 5 | 7 | 23 | 107 | 27 | 2 | 0 | 0 | 0 | 2 | 0 |
| ISG000000 | Sesn3 | 0.280705 | 0.34403 | -0.198572189 | 403.5636842 | 351.6704627 | 102 | 3 | 7 | 10 | 46 | 16 | 0 | 0 | 0 | 0 | 0 | 0 |
| SG0000006 | Tmem151a | 0.284309 | 0.347923 | 0.301622901 | 98.45803437 | 121.3524927 | 21 | 7 | 7 | 8 | 19 | 1 | 1 | 1 | 1 | 1 | 3 | 0 |
| SG0000002 | Vps29 | 0.314292 | 0.379916 | -0.14323152 | 2558.068037 | 2316.3016 | 53 | 7 | 7 | 7 | 26 | 15 | 4 | 0 | 0 | 0 | 0 | 3 |
| ISG000000 | Tmem206 | 0.32073 | 0.386785 | 0.18637342 | 288.8638783 | 328.6981256 | 33 | 7 | 7 | 9 | 18 | 2 | 1 | 0 | 0 | 0 | 0 | 0 |
| SG0000002 | Rab37 | 0.342378 | 0.409681 | -0.510269806 | 28.68565293 | 20.1399423 | 63 | 4 | 7 | 8 | 48 | 4 | 2 | 0 | 0 | 0 | 0 | 0 |
| SG0000001 | Pdhx | 0.344672 | 0.411914 | 0.143133369 | 641.4259181 | 708.3272371 | 177 | 3 | 7 | 27 | 86 | 19 | 9 | 0 | 0 | 1 | 1 | 2 |
| SG0000002 | Tcn2 | 0.375143 | 0.443917 | 0.139970797 | 759.5538253 | 836.9393046 | 105 | 7 | 7 | 14 | 63 | 14 | 7 | 0 | 0 | 0 | 0 | 0 |
| ISG000000 | Gm10320 | 0.377845 | 0.446672 | -0.591955771 | 19.32420125 | 12.8205043 | 21 | 7 | 7 | 8 | 12 | 1 | 0 | 0 | 0 | 0 | 1 | 0 |
| SG0000001 | Zbtb24 | 0.424637 | 0.493774 | -0.13833032 | 480.2443206 | 436.3356142 | 25 | 5 | 7 | 7 | 11 | 2 | 0 | 0 | 0 | 0 | 0 | 0 |
| ISG000000 | Zfp213 | 0.439588 | 0.508815 | 0.308261184 | 215.511526 | 266.8494825 | 16 | 7 | 7 | 8 | 11 | 1 | 1 | 0 | 0 | 0 | 0 | 0 |
| ISG000000 | Cpsf7 | 0.450889 | 0.520047 | 0.100918454 | 2086.486015 | 2237.664438 | 38 | 5 | 7 | 13 | 29 | 5 | 2 | 0 | 0 | 0 | 0 | 0 |
| SG0000006 | Zfp26 | 0.489096 | 0.557189 | 0.092001198 | 1035.502134 | 1103.687486 | 57 | 5 | 7 | 12 | 35 | 3 | 8 | 0 | 0 | 1 | 4 | 0 |
| SG0000000 | Smg5 | 0.49228 | 0.560104 | -0.086235915 | 3419.786692 | 3221.36118 | 26 | 1 | 7 | 7 | 18 | 1 | 0 | 0 | 0 | 0 | 0 | 0 |
| ISG000000 | Nudcd1 | 0.524573 | 0.591213 | 0.078079215 | 2147.811074 | 2267.254451 | 145 | 2 | 7 | 23 | 75 | 11 | 1 | 0 | 0 | 1 | 1 | 0 |
| SG0000003 | Catsper2 | 0.537929 | 0.604289 | -0.316527261 | 33.45087041 | 26.86106469 | 86 | 6 | 7 | 12 | 54 | 4 | 3 | 0 | 1 | 0 | 2 | 0 |
| SG0000002 | Lrrc42 | 0.575861 | 0.64008 | 0.069400592 | 1590.671293 | 1669.060592 | 54 | 5 | 7 | 10 | 27 | 1 | 1 | 0 | 0 | 0 | 0 | 0 |
| SG0000001 | Ddx27 | 0.626703 | 0.687922 | 0.06348121 | 3042.615153 | 3179.48494 | 41 | 7 | 7 | 8 | 28 | 4 | 1 | 0 | 0 | 0 | 2 | 0 |
| SG0000003 | Cep63 | 0.760456 | 0.807235 | 0.047961526 | 465.8012963 | 481.546855 | 94 | 6 | 7 | 11 | 60 | 6 | 0 | 0 | 0 | 0 | 0 | 0 |
| ISG000000 | Cramp1l | 0.771971 | 0.817666 | 0.034333193 | 1049.052296 | 1074.317028 | 79 | 4 | 7 | 10 | 36 | 7 | 6 | 1 | 2 | 1 | 2 | 1 |
| ISG000000 | Rnf182 | 0.784957 | 0.828726 | 0.143629938 | 44.27190911 | 48.9063408 | 94 | 6 | 7 | 19 | 52 | 11 | 7 | 0 | 0 | 3 | 4 | 1 |
| ISG000000 | Fam174a | 0.809965 | 0.849768 | -0.058865148 | 292.1853854 | 280.5035191 | 109 | 3 | 7 | 22 | 51 | 9 | 17 | 1 | 1 | 4 | 5 | 1 |
| ISG000000 | Fam98b | 0.814492 | 0.853767 | -0.037113439 | 1391.151771 | 1355.820682 | 76 | 4 | 7 | 12 | 22 | 5 | 2 | 0 | 0 | 1 | 2 | 1 |
| ISG000000 | Nhp2l1 | 0.848812 | 0.881846 | -0.032639831 | 2023.233186 | 1977.973058 | 15 | 7 | 7 | 7 | 11 | 0 | 2 | 0 | 0 | 0 | 2 | 0 |
| ISG000000 | Rrm1 | 0.874636 | 0.903592 | 0.012815836 | 7985.857862 | 8057.114343 | 74 | 7 | 7 | 15 | 37 | 8 | 4 | 0 | 0 | 0 | 1 | 1 |
| SG0000002 | Errfi1 | 0.892972 | 0.917981 | -0.033837638 | 1033.84944 | 1009.883212 | 39 | 3 | 7 | 11 | 16 | 4 | 8 | 0 | 0 | 2 | 2 | 0 |
| ISG000000 | Rbm4 | 0.902778 | 0.926361 | -0.04526233 | 113.8855162 | 110.3680007 | 73 | 7 | 7 | 11 | 23 | 3 | 7 | 1 | 1 | 2 | 2 | 0 |
| SG0000005 | Rft1 | 0.910902 | 0.933521 | 0.013361292 | 582.5532254 | 587.9735108 | 52 | 4 | 7 | 11 | 27 | 4 | 1 | 0 | 0 | 0 | 0 | 0 |
| SG0000004 | Mvk | 0.913035 | 0.93557 | -0.023864928 | 319.6179589 | 314.3743575 | 40 | 0 | 7 | 7 | 14 | 8 | 0 | 0 | 0 | 0 | 0 | 0 |
| ISG000000 | Trim34b | 0.925727 | 0.946224 | -0.11105199 | 16.19239435 | 14.99274284 | 66 | 4 | 7 | 12 | 37 | 4 | 0 | 0 | 0 | 0 | 0 | 0 |
| SG0000003 | Ldhb | 1.6E-303 | 1.1E-300 | 5.33510179 | 153.1422564 | 6181.881992 | 50 | 2 | 6 | 10 | 20 | 8 | 0 | 0 | 0 | 0 | 0 | 0 |
| SG0000003 | Prok2 | 6E-194 | 1.7E-191 | -8.746037883 | 7651.845286 | 17.82158943 | 60 | 0 | 6 | 6 | 21 | 6 | 6 | 0 | 0 | 0 | 1 | 0 |
| SG0000002 | Tmigd1 | 6.1E-186 | 1.6E-183 | -11.51393794 | 2923.42634 | 0 | 39 | 4 | 6 | 6 | 27 | 4 | 6 | 0 | 0 | 2 | 2 | 0 |
| ISG000000 | Padi2 | 1E-165 | 2.1E-163 | 10.01530771 | 0.649984458 | 672.6838637 | 36 | 4 | 6 | 5 | 15 | 3 | 6 | 2 | 2 | 2 | 4 | 0 |
| SG0000002 | 43711 | 1.8E-144 | 2.6E-142 | 6.534522486 | 7.132767951 | 661.219071 | 59 | 6 | 6 | 11 | 22 | 8 | 9 | 0 | 0 | 1 | 2 | 2 |
| ISG000000 | Peg10 | 1.8E-138 | 2.3E-136 | 7.530035999 | 54.62616117 | 10096.41947 | 65 | 4 | 6 | 6 | 21 | 0 | 13 | 0 | 0 | 0 | 2 | 0 |
| ISG000000 | Slc15a2 | 3.8E-129 | 4.3E-127 | 8.838183837 | 0 | 456.6763193 | 202 | 4 | 6 | 14 | 80 | 16 | 3 | 0 | 0 | 0 | 5 | 0 |
| SG0000002 | Gsta3 | 3.8E-114 | 3.6E-112 | -5.892871175 | 3201.208 | 53.87445742 | 79 | 2 | 6 | 6 | 34 | 2 | 1 | 0 | 0 | 0 | 0 | 0 |
| SG0000004 | Bmf | 5E-99 | 3.46E-97 | 4.176442262 | 38.63518483 | 698.5819985 | 33 | 4 | 6 | 8 | 14 | 8 | 1 | 0 | 0 | 0 | 0 | 4 |
| SG0000002 | Slc30a3 | 6.1E-99 | 4.23E-97 | 8.455422646 | 0.981585527 | 344.5593096 | 29 | 5 | 6 | 8 | 18 | 1 | 0 | 0 | 0 | 0 | 0 | 0 |
| ISG000000 | Col9a2 | 2.56E-96 | 1.69E-94 | 8.322837148 | 0 | 319.2016979 | 15 | 6 | 6 | 6 | 11 | 3 | 1 | 0 | 0 | 0 | 1 | 1 |
| SG0000005 | Cdca7 | 2.01E-92 | 1.2E-90 | 2.514888674 | 918.7692775 | 5251.258421 | 23 | 4 | 6 | 4 | 9 | 3 | 1 | 0 | 0 | 0 | 0 | 1 |
| ISG000000 | Ccdc112 | 2.41E-84 | 1.21E-82 | 3.795677139 | 47.1028876 | 654.1241822 | 59 | 3 | 6 | 11 | 38 | 6 | 6 | 0 | 1 | 0 | 1 | 0 |
| SG0000002 | Adck3 | 3.4E-83 | 1.65E-81 | 3.966222523 | 37.72253424 | 589.5936674 | 129 | 0 | 6 | 12 | 66 | 33 | 4 | 0 | 0 | 0 | 2 | 0 |
| SG0000003 | Pde4c | 2.47E-82 | 1.17E-80 | 8.768830674 | 0.639110347 | 278.7769795 | 52 | 6 | 6 | 6 | 24 | 2 | 0 | 0 | 0 | 0 | 0 | 0 |
| SG0000007 | Mphosph8 | 2.43E-78 | 1.07E-76 | 2.28139838 | 1239.634719 | 6026.471064 | 70 | 3 | 6 | 7 | 29 | 10 | 2 | 0 | 0 | 0 | 0 | 0 |
| SG0000002 | Gss | 2.65E-77 | 1.13E-75 | -3.525879491 | 6604.105432 | 573.3482748 | 93 | 0 | 6 | 5 | 26 | 16 | 0 | 0 | 0 | 0 | 0 | 0 |

| | | | | | | | | | | | | | | | | | |
|---|---|---|---|---|---|---|---|---|---|---|---|---|---|---|---|---|---|
| ISG000000 | Esco2 | 4.91E-74 | 1.98E-72 | 3.146862136 | 770.2420383 | 6822.241878 | 48 | 4 | 6 | 9 | 24 | 2 | 2 | 0 | 0 | 0 | 1 | 1 |
| ISG000000 | St6galnac2 | 2.78E-73 | 1.11E-71 | 5.899340538 | 4.872961731 | 290.8516 | 23 | 6 | 6 | 8 | 17 | 7 | 1 | 0 | 0 | 0 | 0 | 0 |
| SG0000002 | Scara5 | 4.16E-72 | 1.58E-70 | -5.537590487 | 1667.761763 | 35.9048546 | 166 | 6 | 6 | 16 | 90 | 26 | 3 | 0 | 0 | 0 | 0 | 0 |
| ISG000000 | Nlrp4f | 4.55E-72 | 1.73E-70 | 7.276611925 | 1.596604083 | 247.557274 | 69 | 5 | 6 | 18 | 43 | 8 | 1 | 1 | 1 | 1 | 1 | 0 |
| SG0000006 | Mtap7d3 | 3.22E-70 | 1.15E-68 | 6.767107781 | 2.28389801 | 248.7591888 | 53 | 5 | 6 | 9 | 26 | 6 | 0 | 0 | 0 | 0 | 0 | 0 |
| SG0000002 | Tm4sf1 | 6.65E-70 | 2.35E-68 | -4.807377195 | 5857.596885 | 209.1966591 | 74 | 6 | 6 | 11 | 40 | 9 | 10 | 4 | 4 | 4 | 6 | 0 |
| ISG000000 | Cnr1 | 3.76E-67 | 1.26E-65 | -4.55221634 | 824.8341706 | 35.15709473 | 123 | 2 | 6 | 8 | 28 | 8 | 20 | 2 | 2 | 2 | 4 | 2 |
| SG0000002 | Stra8 | 6.52E-67 | 2.17E-65 | 7.671621234 | 0 | 202.8863292 | 60 | 3 | 6 | 9 | 81 | 9 | 0 | 0 | 0 | 0 | 0 | 0 |
| SG0000002 | Ehbp1l1 | 1.44E-66 | 4.74E-65 | -3.488418569 | 3346.72821 | 298.1960029 | 26 | 4 | 6 | 6 | 8 | 0 | 4 | 0 | 0 | 2 | 2 | 0 |
| SG0000002 | Cmbl | 2.04E-63 | 6.19E-62 | -5.991411687 | 512.3009147 | 8.052495619 | 31 | 4 | 6 | 7 | 13 | 0 | 0 | 0 | 0 | 0 | 0 | 0 |
| SG0000000 | Zdhhc4 | 4.35E-54 | 1.01E-52 | -2.936449873 | 2494.340284 | 325.8338562 | 37 | 0 | 6 | 3 | 9 | 3 | 1 | 0 | 0 | 0 | 0 | 0 |
| SG0000005 | Socs6 | 1.16E-49 | 2.4E-48 | -2.547157947 | 5199.997488 | 889.6766412 | 178 | 0 | 6 | 15 | 75 | 13 | 13 | 0 | 0 | 0 | 3 | 1 |
| SG0000002 | Jade2 | 8.4E-49 | 1.68E-47 | -2.680573624 | 2688.385708 | 419.3324887 | 99 | 3 | 6 | 9 | 27 | 3 | 3 | 0 | 0 | 3 | 3 | 0 |
| SG0000004 | Mgat5b | 2.63E-48 | 5.21E-47 | 5.484889122 | 4.174793692 | 186.9610623 | 44 | 4 | 6 | 8 | 25 | 4 | 1 | 0 | 0 | 0 | 1 | 0 |
| SG0000002 | Slc4a8 | 2.64E-45 | 4.68E-44 | -2.496966044 | 3169.791432 | 561.5248879 | 55 | 4 | 6 | 8 | 19 | 2 | 5 | 0 | 0 | 0 | 2 | 1 |
| SG0000000 | Mmd | 2.87E-45 | 5.08E-44 | -2.767754268 | 4526.133254 | 664.5846108 | 66 | 6 | 6 | 14 | 38 | 5 | 3 | 0 | 0 | 1 | 6 | 1 |
| ISG000000 | Rims4 | 3.17E-45 | 5.59E-44 | 7.197153935 | 0.981585527 | 144.04141 | 62 | 4 | 6 | 7 | 24 | 5 | 3 | 1 | 1 | 1 | 1 | 0 |
| ISG000000 | Pard6b | 5.63E-45 | 9.86E-44 | 2.143508747 | 246.7997575 | 1090.448495 | 26 | 4 | 6 | 7 | 14 | 2 | 2 | 1 | 1 | 2 | 4 | 0 |
| SG0000003 | Aars | 1.23E-44 | 2.13E-43 | 1.653804536 | 4110.41058 | 12933.91424 | 31 | 5 | 6 | 9 | 17 | 0 | 2 | 0 | 0 | 0 | 1 | 0 |
| SG0000002 | Pxdc1 | 4.36E-43 | 7.1E-42 | -5.067780534 | 3057.328952 | 91.15662065 | 90 | 6 | 6 | 21 | 48 | 3 | 2 | 0 | 0 | 0 | 0 | 3 |
| SG0000004 | Pyhin1 | 2.4E-42 | 3.83E-41 | -9.632178254 | 1287.684542 | 1.62268755 | 68 | 2 | 6 | 10 | 56 | 4 | 5 | 0 | 0 | 0 | 2 | 0 |
| ISG000000 | Parp2 | 1.51E-41 | 2.35E-40 | 1.74382357 | 836.7720058 | 2802.530478 | 27 | 4 | 6 | 7 | 20 | 1 | 0 | 0 | 0 | 0 | 0 | 0 |
| SG0000002 | Dut | 6.88E-41 | 1.05E-39 | 1.682515095 | 1416.555549 | 4546.959869 | 26 | 4 | 6 | 6 | 18 | 2 | 4 | 0 | 2 | 0 | 2 | 0 |
| ISG000000 | Casp1 | 3.15E-40 | 4.7E-39 | -7.530872247 | 183.334818 | 0.991348861 | 35 | 5 | 6 | 11 | 24 | 2 | 0 | 0 | 0 | 0 | 0 | 0 |
| SG0000004 | Als2cl | 4.76E-40 | 7.07E-39 | -2.246713853 | 3484.725993 | 734.2439443 | 21 | 6 | 6 | 9 | 12 | 0 | 6 | 3 | 3 | 3 | 3 | 0 |
| SG0000005 | Sema3b | 2.05E-39 | 2.97E-38 | -3.530259619 | 937.0662996 | 81.10661593 | 20 | 6 | 6 | 6 | 6 | 0 | 6 | 6 | 6 | 6 | 6 | 0 |
| SG0000006 | Zic3 | 6.75E-39 | 9.55E-38 | 4.25092377 | 205.4300618 | 3911.285888 | 42 | 6 | 6 | 17 | 31 | 6 | 18 | 0 | 0 | 6 | 8 | 4 |
| SG0000000 | 4930432K21Rik | 1.51E-38 | 2.12E-37 | 5.224971746 | 3.950433897 | 147.7467788 | 42 | 6 | 6 | 20 | 36 | 4 | 0 | 0 | 0 | 0 | 0 | 0 |
| SG0000002 | Capn2 | 9.41E-37 | 1.24E-35 | -2.484225734 | 8680.093765 | 1551.307723 | 51 | 4 | 6 | 5 | 17 | 5 | 1 | 0 | 0 | 0 | 1 | 1 |
| ISG000000 | Sh3pxd2b | 1.36E-36 | 1.78E-35 | -2.240905618 | 2232.447093 | 472.2819708 | 83 | 3 | 6 | 13 | 41 | 10 | 4 | 0 | 0 | 0 | 0 | 0 |
| SG0000003 | Ppp1r11 | 2.08E-36 | 2.7E-35 | -2.148830135 | 2934.043919 | 661.6133872 | 15 | 6 | 6 | 10 | 14 | 0 | 1 | 0 | 0 | 0 | 0 | 0 |
| SG0000002 | E2f5 | 6.29E-36 | 8.06E-35 | 1.849545606 | 326.5024538 | 1176.671285 | 112 | 6 | 6 | 16 | 66 | 10 | 4 | 0 | 0 | 0 | 2 | 0 |
| ISG000000 | Bora | 7.87E-36 | 1.01E-34 | 1.701905181 | 572.5220783 | 1862.587776 | 52 | 3 | 6 | 10 | 30 | 10 | 3 | 0 | 0 | 0 | 0 | 0 |
| SG0000003 | Dagla | 1.9E-35 | 2.41E-34 | -3.253976339 | 630.2889674 | 66.06861792 | 60 | 2 | 6 | 8 | 24 | 6 | 2 | 0 | 0 | 0 | 0 | 3 |
| ISG000000 | Dgka | 2.8E-34 | 3.39E-33 | -1.969454126 | 4694.417659 | 1198.717813 | 37 | 4 | 6 | 8 | 29 | 6 | 2 | 0 | 0 | 1 | 2 | 0 |
| SG0000002 | Chaf1b | 4.13E-34 | 4.94E-33 | 1.563496719 | 984.4556987 | 2909.749404 | 53 | 6 | 6 | 13 | 29 | 3 | 0 | 0 | 0 | 0 | 0 | 0 |
| ISG000000 | Adad2 | 4.2E-34 | 5.02E-33 | 6.670410895 | 0 | 100.8576759 | 7 | 6 | 6 | 6 | 6 | 0 | 0 | 0 | 0 | 0 | 0 | 0 |
| SG0000002 | Cacna1h | 1.67E-33 | 1.94E-32 | 3.101975396 | 30.82350036 | 264.6485972 | 59 | 0 | 6 | 3 | 30 | 15 | 2 | 0 | 0 | 0 | 0 | 0 |
| SG0000000 | Mob1b | 1.82E-33 | 2.11E-32 | -2.145042101 | 2831.324416 | 640.1291727 | 163 | 2 | 6 | 18 | 104 | 10 | 17 | 0 | 0 | 4 | 14 | 2 |
| SG0000003 | Mtss1l | 6.21E-33 | 7.06E-32 | 2.913943795 | 64.19022442 | 483.7860953 | 28 | 6 | 6 | 8 | 33 | 7 | 2 | 0 | 0 | 0 | 4 | 1 |
| ISG000000 | Scg5 | 7.04E-33 | 7.99E-32 | -3.715476169 | 294.8454122 | 22.44525789 | 73 | 2 | 6 | 11 | 32 | 5 | 1 | 0 | 0 | 1 | 1 | 0 |
| SG0000002 | Naaa | 5.55E-32 | 6.09E-31 | -2.164229787 | 1502.397506 | 335.1866943 | 56 | 2 | 6 | 4 | 18 | 10 | 4 | 0 | 0 | 0 | 0 | 0 |
| SG0000002 | Rasal1 | 8.14E-32 | 8.87E-31 | 3.54546673 | 16.09368362 | 187.9088756 | 26 | 4 | 6 | 8 | 34 | 8 | 0 | 0 | 0 | 0 | 0 | 0 |
| SG0000002 | Dars2 | 9.65E-32 | 1.05E-30 | 1.525099317 | 900.0422335 | 2590.380305 | 58 | 1 | 6 | 6 | 25 | 7 | 3 | 0 | 0 | 0 | 0 | 0 |
| SG0000005 | Lrrc8c | 3.62E-31 | 3.84E-30 | -2.068569563 | 1792.93097 | 427.4271278 | 147 | 5 | 6 | 22 | 60 | 13 | 6 | 0 | 0 | 0 | 2 | 2 |
| ISG000000 | Capsl | 4.11E-31 | 4.36E-30 | 9.405893478 | 0.34247518 | 232.3192385 | 58 | 3 | 6 | 8 | 28 | 6 | 0 | 0 | 0 | 0 | 0 | 0 |
| SG0000003 | Adgrg1 | 5.55E-31 | 5.85E-30 | 2.80839195 | 65.79750591 | 460.913733 | 31 | 6 | 6 | 16 | 25 | 12 | 0 | 0 | 0 | 0 | 0 | 0 |
| SG0000003 | Cox4i1 | 1.23E-30 | 1.28E-29 | 1.469349214 | 4522.615291 | 12522.98425 | 30 | 6 | 6 | 6 | 24 | 0 | 0 | 0 | 0 | 0 | 0 | 0 |
| SG0000004 | Ticrr | 3.16E-30 | 3.22E-29 | 1.470021951 | 1030.272885 | 2854.12488 | 72 | 3 | 6 | 12 | 33 | 11 | 1 | 0 | 0 | 0 | 1 | 0 |
| SG0000001 | Tnfaip3 | 1.02E-29 | 1.01E-28 | -2.545478099 | 1247.142344 | 213.6243547 | 40 | 2 | 6 | 8 | 29 | 4 | 10 | 2 | 6 | 4 | 12 | 2 |
| SG0000003 | Vgll4 | 4.37E-29 | 4.24E-28 | -2.090283308 | 1243.744112 | 292.0741105 | 69 | 5 | 6 | 11 | 36 | 6 | 0 | 0 | 0 | 0 | 0 | 0 |
| ISG000000 | Pkd1 | 4.44E-29 | 4.31E-28 | -1.993670311 | 2598.838862 | 652.5665182 | 43 | 5 | 6 | 6 | 17 | 3 | 2 | 0 | 0 | 0 | 0 | 1 |
| ISG000000 | Mgst2 | 2.57E-28 | 2.41E-27 | -3.967771432 | 205.7396084 | 13.14921093 | 63 | 3 | 6 | 12 | 18 | 6 | 0 | 0 | 0 | 0 | 0 | 0 |
| SG0000000 | Peg3 | 3.5E-28 | 3.26E-27 | -1.790900667 | 12931.41405 | 3737.069697 | 101 | 3 | 6 | 9 | 42 | 8 | 12 | 0 | 0 | 0 | 1 | 2 |
| SG0000002 | Pdlim2 | 4.49E-28 | 4.17E-27 | 2.610344211 | 49.05284097 | 299.5408596 | 14 | 6 | 6 | 6 | 14 | 4 | 0 | 0 | 0 | 0 | 0 | 0 |
| SG0000002 | Mcm8 | 5.86E-28 | 5.4E-27 | 1.627943558 | 353.5740917 | 1092.798969 | 103 | 6 | 6 | 24 | 56 | 18 | 3 | 0 | 0 | 0 | 2 | 0 |
| SG0000003 | Calhm2 | 6.39E-28 | 5.88E-27 | -3.005652443 | 349.0168382 | 43.45650928 | 48 | 1 | 6 | 6 | 8 | 2 | 5 | 0 | 0 | 1 | 1 | 0 |
| SG0000001 | Sqstm1 | 1.38E-27 | 1.25E-26 | -1.757903385 | 24133.92086 | 7135.858603 | 33 | 3 | 6 | 9 | 21 | 6 | 1 | 0 | 0 | 0 | 0 | 0 |
| ISG000000 | Epcam | 1.69E-27 | 1.53E-26 | 2.557567061 | 1005.898204 | 5921.864062 | 27 | 6 | 6 | 8 | 15 | 4 | 0 | 0 | 0 | 0 | 1 | 1 |
| SG0000004 | Kctd12b | 3.22E-27 | 2.86E-26 | -4.340948168 | 1156.781094 | 57.08152588 | 52 | 0 | 6 | 4 | 26 | 6 | 20 | 0 | 4 | 2 | 6 | 4 |
| ISG000000 | Lag3 | 5.36E-27 | 4.73E-26 | 5.421483492 | 2.235714429 | 95.81764266 | 11 | 5 | 6 | 5 | 8 | 0 | 0 | 0 | 0 | 0 | 0 | 0 |
| SG0000003 | Tnfsf10 | 6.74E-27 | 5.91E-26 | -5.857481867 | 135.1240052 | 2.33053038 | 62 | 4 | 6 | 12 | 39 | 2 | 8 | 0 | 0 | 3 | 5 | 0 |
| SG0000003 | Zfp365 | 7.48E-27 | 6.55E-26 | -1.79265601 | 2357.362677 | 680.4295597 | 58 | 4 | 6 | 15 | 49 | 7 | 8 | 2 | 3 | 4 | 6 | 0 |
| SG0000005 | Lig1 | 9.05E-27 | 7.9E-26 | 1.292917176 | 2910.561395 | 7131.544828 | 115 | 6 | 6 | 21 | 78 | 6 | 0 | 0 | 0 | 0 | 0 | 0 |
| SG0000004 | Fam102b | 1.32E-26 | 1.15E-25 | -2.361858853 | 1819.632147 | 353.9918315 | 142 | 2 | 6 | 14 | 78 | 12 | 12 | 0 | 0 | 2 | 10 | 0 |
| ISG000000 | Gipr | 2.43E-26 | 2.08E-25 | -2.832458821 | 494.5136382 | 69.42616994 | 17 | 5 | 6 | 6 | 8 | 1 | 0 | 0 | 0 | 0 | 0 | 1 |
| SG0000002 | Helz2 | 3.79E-26 | 3.22E-25 | -2.347190796 | 2654.062971 | 521.5985647 | 12 | 3 | 6 | 6 | 12 | 3 | 0 | 0 | 0 | 0 | 3 | 0 |
| SG0000000 | Tcf7 | 4.38E-26 | 3.71E-25 | 4.182318547 | 110.9049064 | 2013.511404 | 30 | 6 | 6 | 6 | 14 | 7 | 0 | 0 | 0 | 0 | 0 | 1 |
| ISG000000 | Gpr4 | 8.47E-26 | 7.09E-25 | 4.699188389 | 3.939559785 | 102.339751 | 14 | 5 | 6 | 6 | 10 | 2 | 1 | 0 | 0 | 0 | 0 | 1 |
| SG0000002 | Rnf11 | 9.05E-26 | 7.56E-25 | -1.68227282 | 3205.148177 | 998.6965108 | 116 | 2 | 6 | 16 | 53 | 16 | 8 | 0 | 2 | 2 | 9 | 0 |
| ISG000000 | Mvd | 9.25E-26 | 7.72E-25 | 2.893520048 | 179.8640477 | 1336.536089 | 18 | 4 | 6 | 4 | 8 | 0 | 0 | 0 | 0 | 0 | 0 | 0 |
| SG0000003 | Eomes | 1.61E-25 | 1.33E-24 | 2.072535203 | 95.85190956 | 403.1772632 | 44 | 6 | 6 | 10 | 14 | 0 | 4 | 0 | 0 | 0 | 0 | 0 |
| ISG000000 | Smarcd1 | 1.79E-25 | 1.48E-24 | 1.296495791 | 1839.615531 | 4518.675869 | 18 | 6 | 6 | 7 | 9 | 1 | 0 | 0 | 0 | 0 | 0 | 0 |
| ISG000000 | Card6 | 2.91E-25 | 2.38E-24 | -2.634368495 | 577.9660247 | 93.08471492 | 29 | 4 | 6 | 8 | 13 | 3 | 9 | 0 | 0 | 0 | 2 | 1 |
| SG0000002 | Dvl2 | 4.83E-25 | 3.91E-24 | 1.772397193 | 702.4395107 | 2399.681763 | 28 | 6 | 6 | 9 | 18 | 3 | 7 | 0 | 0 | 3 | 3 | 0 |
| SG0000003 | Calml4 | 4.91E-25 | 3.98E-24 | 6.000592166 | 2.923008357 | 187.149336 | 28 | 4 | 6 | 10 | 12 | 4 | 0 | 0 | 0 | 0 | 0 | 0 |
| SG0000003 | C1s1 | 6.86E-25 | 5.52E-24 | -3.395522876 | 3545.599823 | 336.9267403 | 32 | 6 | 6 | 10 | 16 | 6 | 4 | 0 | 0 | 0 | 0 | 0 |
| SG0000005 | Lgals8 | 8.72E-25 | 6.97E-24 | -2.065724709 | 1870.43643 | 446.7842473 | 89 | 0 | 6 | 3 | 19 | 9 | 12 | 0 | 0 | 0 | 3 | 0 |
| SG0000003 | Armcx1 | 9.68E-25 | 7.72E-24 | 3.443399154 | 70.53982517 | 767.3629231 | 66 | 0 | 6 | 6 | 24 | 0 | 18 | 0 | 0 | 0 | 6 | 0 |
| ISG000000 | Ubxn8 | 9.92E-25 | 7.89E-24 | -1.826260755 | 1392.387654 | 392.645746 | 43 | 6 | 6 | 10 | 31 | 2 | 4 | 0 | 0 | 0 | 0 | 0 |
| SG0000003 | Ppp1r18 | 1.03E-24 | 8.2E-24 | -1.623919099 | 3331.562864 | 1080.935184 | 36 | 3 | 6 | 12 | 15 | 0 | 7 | 0 | 3 | 0 | 0 | 0 |
| ISG000000 | Fermt1 | 1.48E-24 | 1.17E-23 | 8.464601693 | 0.34247518 | 120.9840486 | 69 | 3 | 6 | 6 | 21 | 3 | 0 | 0 | 0 | 0 | 0 | 0 |
| SG0000004 | Tsku | 1.94E-24 | 1.53E-23 | -1.77485504 | 1528.095198 | 446.5455915 | 18 | 0 | 6 | 0 | 0 | 0 | 0 | 0 | 0 | 0 | 0 | 0 |
| ISG000000 | Ireb2 | 2.12E-24 | 1.66E-23 | 1.251216419 | 2596.654023 | 6181.128346 | 134 | 4 | 6 | 27 | 70 | 17 | 4 | 0 | 1 | 0 | 3 | 2 |
| SG0000003 | Fam63a | 4.16E-24 | 3.21E-23 | -1.572261149 | 4175.770995 | 1404.232133 | 27 | 6 | 6 | 11 | 23 | 3 | 0 | 0 | 0 | 0 | 3 | 0 |
| ISG000000 | Rqcd1 | 4.21E-24 | 3.25E-23 | 1.240569629 | 2325.469117 | 5494.893078 | 66 | 6 | 6 | 10 | 25 | 4 | 4 | 0 | 0 | 0 | 0 | 0 |
| SG0000001 | Tox4 | 4.23E-24 | 3.26E-23 | 1.249178041 | 2016.818553 | 4794.097785 | 27 | 3 | 6 | 8 | 16 | 1 | 6 | 1 | 2 | 2 | 3 | 0 |
| SG0000003 | BC017158 | 1.26E-23 | 9.52E-23 | -2.040793694 | 722.1399848 | 175.501679 | 71 | 6 | 6 | 8 | 32 | 10 | 0 | 0 | 0 | 0 | 0 | 0 |
| SG0000002 | Pdpn | 1.32E-23 | 9.98E-23 | 4.020063325 | 113.8412196 | 1846.967209 | 52 | 2 | 6 | 10 | 52 | 4 | 0 | 0 | 0 | 0 | 0 | 0 |
| SG0000003 | Arhgap4 | 1.46E-23 | 1.1E-22 | 3.2986286 | 14.59713697 | 143.6328693 | 64 | 6 | 6 | 12 | 51 | 16 | 3 | 1 | 1 | 2 | 4 | 1 |
| ISG000000 | Cda | 1.73E-23 | 1.29E-22 | -4.152492528 | 711.6730267 | 40.01802933 | 26 | 5 | 6 | 6 | 21 | 7 | 0 | 0 | 0 | 0 | 0 | 0 |
| SG0000002 | Gba | 9.24E-23 | 6.72E-22 | -1.604423077 | 2286.169017 | 751.8459469 | 9 | 6 | 6 | 9 | 27 | 0 | 0 | 0 | 0 | 0 | 0 | 0 |
| ISG000000 | 2900026A02Rik | 1.03E-22 | 7.48E-22 | -1.641161162 | 1985.392125 | 636.5134095 | 51 | 3 | 6 | 8 | 35 | 16 | 1 | 0 | 0 | 0 | 0 | 1 |
| SG0000004 | Mtmr11 | 2.21E-22 | 1.58E-21 | -2.049452994 | 629.5013351 | 152.0722003 | 26 | 4 | 6 | 6 | 18 | 0 | 3 | 0 | 0 | 0 | 0 | 0 |
| SG0000002 | Cldnd1 | 5.61E-22 | 3.93E-21 | -1.533284051 | 2711.410692 | 936.7653389 | 31 | 1 | 6 | 6 | 12 | 3 | 7 | 1 | 3 | 3 | 3 | 0 |
| SG0000003 | Vash2 | 6.33E-22 | 4.42E-21 | 3.784286088 | 38.08426094 | 524.7218594 | 52 | 6 | 6 | 8 | 22 | 6 | 8 | 0 | 0 | 0 | 2 | 0 |
| SG0000002 | Rbck1 | 1.05E-21 | 7.27E-21 | -1.856187968 | 4427.927952 | 1223.016426 | 36 | 6 | 6 | 8 | 16 | 2 | 0 | 0 | 0 | 0 | 0 | 0 |
| ISG000000 | Asb16 | 3E-21 | 2.01E-20 | 3.525599204 | 9.88578707 | 113.8472029 | 16 | 3 | 6 | 4 | 7 | 1 | 1 | 0 | 0 | 0 | 0 | 0 |
| SG0000004 | Fbrsl1 | 1.23E-20 | 8.03E-20 | 1.406299298 | 763.9992783 | 2025.028874 | 72 | 5 | 6 | 10 | 45 | 14 | 4 | 0 | 0 | 1 | 2 | 1 |
| ISG000000 | Nampt | 1.26E-20 | 8.2E-20 | 1.520659114 | 3405.947113 | 9772.415107 | 66 | 0 | 6 | 5 | 18 | 6 | 12 | 0 | 0 | 0 | 0 | 0 |
| ISG000000 | Tmem64 | 1.31E-20 | 8.5E-20 | -1.534733411 | 2074.649969 | 716.0511423 | 60 | 0 | 6 | 8 | 15 | 2 | 14 | 0 | 1 | 3 | 6 | 0 |
| ISG000000 | Epha2 | 1.54E-20 | 9.96E-20 | -2.123093334 | 12751.59104 | 2927.181143 | 31 | 3 | 6 | 13 | 19 | 3 | 2 | 1 | 1 | 2 | 4 | 0 |
| ISG000000 | Srfbp1 | 1.6E-20 | 1.04E-19 | 1.318398101 | 547.2757415 | 1364.845754 | 66 | 5 | 6 | 16 | 38 | 3 | 1 | 0 | 1 | 0 | 0 | 0 |
| SG0000003 | Galnt9 | 6.26E-20 | 3.91E-19 | 7.856196529 | 0.34247518 | 79.35602575 | 68 | 5 | 6 | 14 | 33 | 22 | 0 | 0 | 0 | 0 | 0 | 1 |
| ISG000000 | Sbspon | 1.13E-19 | 6.97E-19 | -7.512761041 | 324.1340596 | 1.774836065 | 85 | 4 | 6 | 10 | 32 | 8 | 4 | 0 | 0 | 1 | 2 | 0 |
| ISG000000 | Gtpbp4 | 2.28E-19 | 1.38E-18 | 1.10344228 | 2618.876205 | 5627.094316 | 57 | 1 | 6 | 8 | 27 | 8 | 4 | 0 | 0 | 0 | 0 | 0 |
| ISG000000 | Serpinb9e | 2.97E-19 | 1.79E-18 | 4.937285232 | 2.589063721 | 79.32565947 | 31 | 4 | 6 | 7 | 8 | 1 | 1 | 0 | 0 | 0 | 0 | 0 |
| SG0000003 | Fam214b | 3.67E-19 | 2.19E-18 | -1.641200855 | 954.2799101 | 305.9321331 | 56 | 6 | 6 | 6 | 30 | 0 | 19 | 6 | 6 | 6 | 18 | 0 |
| ISG000000 | Clns1a | 4.71E-19 | 2.8E-18 | 1.125123152 | 1629.58579 | 3554.455213 | 47 | 5 | 6 | 10 | 19 | 5 | 0 | 0 | 0 | 0 | 0 | 0 |
| ISG000000 | Msh6 | 4.78E-19 | 2.84E-18 | 1.053021294 | 8329.16679 | 17281.94244 | 27 | 3 | 6 | 7 | 14 | 2 | 0 | 0 | 0 | 0 | 0 | 0 |
| ISG000000 | 9530053A07Rik | 4.8E-19 | 2.85E-18 | -3.184871765 | 172.3604099 | 18.95376906 | 48 | 4 | 6 | 8 | 24 | 6 | 0 | 0 | 0 | 0 | 0 | 0 |
| ISG000000 | Reep1 | 5.06E-19 | 3.01E-18 | -1.978860224 | 823.6596449 | 208.9543918 | 117 | 4 | 6 | 20 | 63 | 13 | 3 | 0 | 0 | 0 | 0 | 1 |
| SG0000002 | Mapk1ip1l | 5.94E-19 | 3.51E-18 | 1.088336719 | 2811.342278 | 5977.722106 | 68 | 4 | 6 | 9 | 23 | 8 | 8 | 0 | 2 | 2 | 3 | 2 |
| ISG000000 | Itgb7 | 1.31E-18 | 7.61E-18 | -1.928978026 | 2041.142544 | 536.0349268 | 25 | 5 | 6 | 8 | 17 | 3 | 0 | 0 | 0 | 0 | 0 | 0 |
| ISG000000 | Lancl3 | 2.77E-18 | 1.58E-17 | -4.245589233 | 221.5201423 | 11.67786669 | 155 | 3 | 6 | 20 | 65 | 11 | 6 | 0 | 0 | 1 | 5 | 1 |
| ISG000000 | Etaa1 | 4.05E-18 | 2.29E-17 | 1.21929831 | 710.012484 | 1653.14653 | 35 | 3 | 6 | 6 | 31 | 2 | 3 | 0 | 1 | 0 | 6 | 2 |
| SG0000004 | Dennd4b | 8.47E-18 | 4.72E-17 | 2.407799757 | 103.0222933 | 546.7026302 | 27 | 6 | 6 | 6 | 8 | 6 | 4 | 2 | 2 | 2 | 2 | 2 |
| ISG000000 | Ercc1 | 2.43E-17 | 1.32E-16 | -1.41053636 | 1442.387526 | 542.5855253 | 31 | 6 | 6 | 6 | 12 | 6 | 0 | 0 | 0 | 0 | 1 | 0 |
| SG0000007 | Mpzl3 | 3.94E-17 | 2.13E-16 | -2.333332695 | 729.3940857 | 144.7301814 | 83 | 5 | 6 | 12 | 26 | 3 | 10 | 1 | 2 | 2 | 2 | 0 |
| ISG000000 | Ercc6 | 5.51E-17 | 2.94E-16 | 1.337756363 | 517.3863337 | 1307.734994 | 127 | 3 | 6 | 12 | 66 | 8 | 11 | 1 | 2 | 2 | 4 | 0 |
| SG0000007 | 2010315B03Rik | 6.46E-17 | 3.44E-16 | -2.332201571 | 274.4545918 | 54.50142689 | 53 | 3 | 6 | 11 | 28 | 3 | 4 | 0 | 0 | 0 | 1 | 0 |
| SG0000004 | Tgif1 | 7.28E-17 | 3.86E-16 | 1.046730265 | 1974.345428 | 4078.686689 | 67 | 0 | 6 | 6 | 11 | 1 | 0 | 0 | 0 | 0 | 5 | 0 |
| ISG000000 | Amica1 | 9.63E-17 | 5.07E-16 | -5.36440248 | 69.4613786 | 1.686154412 | 29 | 4 | 6 | 9 | 19 | 5 | 1 | 0 | 1 | 0 | 0 | 0 |

| | | | | | | | | | | | | | | | | | | |
|---|---|---|---|---|---|---|---|---|---|---|---|---|---|---|---|---|---|---|
| ISG000000 | Polb | 1.21E-16 | 6.32E-16 | 1.280145136 | 360.8218313 | 876.3041957 | 45 | 4 | 6 | 8 | 25 | 7 | 1 | 0 | 0 | 0 | 0 | 0 |
| SG0000002 | Arpc1b | 1.56E-16 | 8.09E-16 | -1.792030555 | 7918.37922 | 2286.553391 | 33 | 4 | 6 | 10 | 18 | 2 | 2 | 0 | 0 | 0 | 0 | 0 |
| ISG000000 | Chd1l | 1.64E-16 | 8.52E-16 | 1.141066876 | 716.5467661 | 1580.301299 | 83 | 4 | 6 | 17 | 37 | 7 | 0 | 0 | 0 | 0 | 0 | 0 |
| SG0000002 | Slc35d1 | 1.64E-16 | 8.52E-16 | -1.611850603 | 1075.492182 | 351.8778079 | 70 | 2 | 6 | 11 | 40 | 9 | 5 | 0 | 0 | 0 | 1 | 0 |
| ISG000000 | Gdpd1 | 1.97E-16 | 1.01E-15 | -1.739240923 | 757.0900517 | 226.7690873 | 67 | 4 | 6 | 9 | 28 | 5 | 2 | 0 | 0 | 1 | 1 | 0 |
| ISG000000 | Mrpl42 | 2.62E-16 | 1.34E-15 | 1.062998155 | 1362.668336 | 2846.980512 | 57 | 3 | 6 | 18 | 33 | 4 | 0 | 0 | 0 | 0 | 0 | 0 |
| SG0000002 | Slc1a1 | 3.06E-16 | 1.56E-15 | 7.279978624 | 0.34247518 | 53.22565766 | 136 | 2 | 6 | 17 | 57 | 10 | 5 | 0 | 0 | 1 | 6 | 1 |
| SG0000002 | Rpl22 | 4.28E-16 | 2.16E-15 | 1.139781138 | 7033.629724 | 15498.43381 | 24 | 6 | 6 | 6 | 9 | 5 | 3 | 0 | 0 | 0 | 0 | 2 |
| SG000000C | Ptpn6 | 1.03E-15 | 5.11E-15 | 2.506861301 | 194.43854 | 1105.153995 | 51 | 6 | 6 | 12 | 19 | 16 | 4 | 0 | 0 | 0 | 0 | 8 |
| SG0000003 | Setd1b | 1.19E-15 | 5.87E-15 | -1.496235409 | 824.4139763 | 292.2359276 | 65 | 6 | 6 | 15 | 22 | 11 | 5 | 0 | 0 | 3 | 7 | 5 |
| SG0000003 | Znfx1 | 1.72E-15 | 8.38E-15 | -1.24262898 | 2414.819373 | 1020.507157 | 38 | 5 | 6 | 8 | 21 | 2 | 2 | 0 | 0 | 0 | 0 | 0 |
| SG0000003 | Ccdc151 | 2.13E-15 | 1.04E-14 | 3.329568001 | 8.408645077 | 84.53291836 | 26 | 6 | 6 | 6 | 16 | 2 | 0 | 0 | 0 | 0 | 0 | 0 |
| SG0000004 | Ghitm | 2.6E-15 | 1.26E-14 | 0.945254513 | 6807.785594 | 13108.58551 | 92 | 6 | 6 | 16 | 42 | 8 | 9 | 0 | 0 | 3 | 3 | 0 |
| SG0000005 | Nbeal2 | 3.54E-15 | 1.69E-14 | -1.446586919 | 977.8024158 | 358.7443352 | 55 | 0 | 6 | 3 | 6 | 6 | 0 | 0 | 0 | 0 | 0 | 0 |
| ISG000000 | Rnf19a | 6.69E-15 | 3.14E-14 | -1.326518915 | 3618.782883 | 1442.914322 | 105 | 1 | 6 | 11 | 43 | 6 | 3 | 0 | 0 | 0 | 0 | 1 |
| SG000000C | Lsr | 9.7E-15 | 4.51E-14 | 2.596278086 | 199.5910189 | 1206.975838 | 33 | 0 | 6 | 0 | 18 | 9 | 0 | 0 | 0 | 0 | 0 | 0 |
| SG0000006 | Myadm | 1.16E-14 | 5.38E-14 | -1.200339909 | 7568.278746 | 3293.508597 | 25 | 6 | 6 | 6 | 18 | 4 | 7 | 3 | 3 | 3 | 8 | 1 |
| SG0000006 | Cd59b | 1.27E-14 | 5.84E-14 | 6.959303083 | 0.331601068 | 41.26433951 | 76 | 2 | 6 | 6 | 28 | 4 | 2 | 0 | 0 | 0 | 2 | 0 |
| SG0000003 | Unc13a | 2.36E-14 | 1.07E-13 | 1.64634296 | 105.152607 | 329.1687956 | 61 | 6 | 6 | 10 | 20 | 6 | 5 | 0 | 0 | 0 | 1 | 0 |
| SG0000003 | Cep44 | 3.04E-14 | 1.37E-13 | 1.130712611 | 475.3527108 | 1040.864957 | 92 | 2 | 6 | 20 | 64 | 12 | 2 | 0 | 0 | 0 | 0 | 2 |
| ISG000000 | Tbk1 | 3.61E-14 | 1.62E-13 | -1.185866714 | 2216.00699 | 974.0688925 | 56 | 4 | 6 | 12 | 25 | 5 | 0 | 0 | 0 | 0 | 0 | 2 |
| ISG000000 | Man2b2 | 4.17E-14 | 1.86E-13 | -1.891676798 | 1273.37705 | 343.167033 | 21 | 6 | 6 | 6 | 18 | 1 | 0 | 0 | 0 | 0 | 1 | 0 |
| SG0000002 | Tcof1 | 5.42E-14 | 2.4E-13 | 0.95968543 | 2921.588036 | 5682.155025 | 77 | 6 | 6 | 9 | 30 | 3 | 0 | 0 | 0 | 0 | 0 | 0 |
| ISG000000 | Zfp180 | 1.08E-13 | 4.68E-13 | -1.194233225 | 1629.91534 | 712.3034117 | 37 | 3 | 6 | 9 | 25 | 2 | 3 | 0 | 0 | 1 | 1 | 0 |
| ISG000000 | Dpp9 | 1.96E-13 | 8.36E-13 | -1.146735856 | 1927.311809 | 870.4625531 | 25 | 5 | 6 | 6 | 12 | 1 | 1 | 0 | 1 | 0 | 0 | 0 |
| ISG000000 | Tiparp | 2E-13 | 8.53E-13 | -1.062402368 | 6292.213027 | 3012.925806 | 87 | 2 | 6 | 16 | 37 | 7 | 9 | 1 | 3 | 2 | 3 | 1 |
| SG0000003 | Cdv3 | 3.6E-13 | 1.51E-12 | 0.88644442 | 4033.006464 | 7455.474309 | 150 | 6 | 6 | 6 | 46 | 0 | 32 | 0 | 0 | 0 | 2 | 0 |
| SG0000002 | Lbh | 3.6E-13 | 1.51E-12 | -1.795466682 | 609.3393649 | 175.5374996 | 55 | 0 | 6 | 6 | 17 | 7 | 3 | 0 | 0 | 0 | 0 | 1 |
| SG0000002 | Abcb8 | 3.6E-13 | 1.51E-12 | -1.205053306 | 1366.589949 | 592.7629228 | 36 | 4 | 6 | 4 | 16 | 6 | 0 | 0 | 0 | 0 | 0 | 0 |
| SG0000002 | Nefm | 4.03E-13 | 1.68E-12 | 4.925663304 | 3.182334054 | 96.72041901 | 91 | 2 | 6 | 18 | 58 | 8 | 2 | 0 | 0 | 1 | 3 | 0 |
| SG0000001 | Sec14l4 | 5.68E-13 | 2.35E-12 | 5.072698336 | 5.355300519 | 180.2263157 | 18 | 4 | 6 | 7 | 13 | 5 | 0 | 0 | 0 | 0 | 0 | 0 |
| ISG000000 | Smndc1 | 8.11E-13 | 3.33E-12 | 0.930370939 | 1336.217332 | 2546.517473 | 33 | 5 | 6 | 7 | 9 | 5 | 3 | 0 | 0 | 0 | 1 | 1 |
| SG0000002 | Hmmr | 9.43E-13 | 3.85E-12 | 0.880055179 | 2984.685618 | 5493.151788 | 61 | 3 | 6 | 8 | 31 | 5 | 4 | 0 | 0 | 0 | 2 | 0 |
| SG0000003 | Man2a2 | 1.01E-12 | 4.11E-12 | 1.127188765 | 338.8149586 | 740.0826194 | 21 | 5 | 6 | 8 | 20 | 0 | 1 | 0 | 0 | 0 | 7 | 0 |
| SG000000C | Chd5 | 1.3E-12 | 5.26E-12 | 1.562114907 | 92.79991171 | 274.0255175 | 84 | 6 | 6 | 9 | 21 | 9 | 9 | 0 | 0 | 0 | 0 | 0 |
| SG0000002 | Cdc23 | 1.36E-12 | 5.46E-12 | 0.909453611 | 1553.913702 | 2918.768919 | 73 | 4 | 6 | 12 | 42 | 6 | 5 | 0 | 0 | 1 | 1 | 0 |
| SG0000002 | Sub1 | 1.39E-12 | 5.6E-12 | 0.85887504 | 4231.071884 | 7673.571663 | 60 | 4 | 6 | 4 | 14 | 4 | 5 | 0 | 0 | 0 | 1 | 0 |
| SG0000003 | Strap | 1.46E-12 | 5.88E-12 | -0.994945531 | 12088.13793 | 6065.281445 | 51 | 2 | 6 | 8 | 20 | 1 | 7 | 0 | 1 | 1 | 1 | 0 |
| SG0000003 | Copg1 | 1.81E-12 | 7.22E-12 | -0.995970795 | 7651.895156 | 3836.647742 | 35 | 5 | 6 | 7 | 16 | 1 | 2 | 0 | 0 | 1 | 1 | 0 |
| SG0000004 | Rrp9 | 1.99E-12 | 7.95E-12 | 1.059075996 | 752.1756491 | 1567.230634 | 11 | 6 | 6 | 8 | 21 | 1 | 2 | 0 | 0 | 0 | 0 | 0 |
| ISG000000 | Scamp5 | 2.14E-12 | 8.53E-12 | 0.955837626 | 904.6755805 | 1754.804196 | 26 | 4 | 6 | 9 | 17 | 3 | 0 | 0 | 0 | 0 | 0 | 0 |
| ISG000000 | Arl6ip6 | 2.18E-12 | 8.66E-12 | 0.974349107 | 753.4655258 | 1480.374835 | 86 | 2 | 6 | 10 | 32 | 7 | 9 | 0 | 0 | 2 | 4 | 2 |
| SG0000003 | Ccdc90b | 2.38E-12 | 9.45E-12 | 1.169401551 | 271.9236325 | 611.6062249 | 92 | 0 | 6 | 10 | 52 | 4 | 0 | 0 | 0 | 0 | 4 | 0 |
| SG000000C | Clint1 | 2.52E-12 | 9.98E-12 | -1.009438357 | 5392.238239 | 2678.538221 | 214 | 0 | 6 | 26 | 74 | 22 | 6 | 0 | 0 | 2 | 6 | 0 |
| SG0000003 | Mcee | 2.79E-12 | 1.1E-11 | 1.41029802 | 125.4080633 | 333.3246779 | 39 | 5 | 6 | 10 | 26 | 4 | 1 | 0 | 0 | 0 | 1 | 1 |
| SG0000003 | Ears2 | 3.34E-12 | 1.31E-11 | 1.18885012 | 273.2637574 | 622.9620377 | 33 | 5 | 6 | 7 | 22 | 6 | 0 | 0 | 0 | 0 | 0 | 0 |
| SG0000003 | Tmem5 | 4.59E-12 | 1.79E-11 | -1.204790825 | 992.7644831 | 430.6932393 | 77 | 2 | 6 | 16 | 24 | 8 | 8 | 0 | 0 | 0 | 2 | 2 |
| SG0000002 | Lrrc41 | 5.1E-12 | 1.99E-11 | -1.218276668 | 1734.845539 | 745.6293482 | 40 | 6 | 6 | 12 | 21 | 2 | 0 | 0 | 0 | 0 | 0 | 0 |
| SG0000005 | Akap17b | 6.15E-12 | 2.38E-11 | 1.487518805 | 160.3320356 | 449.5811418 | 80 | 4 | 6 | 8 | 35 | 6 | 7 | 0 | 1 | 0 | 0 | 0 |
| SG0000003 | Nphp4 | 6.91E-12 | 2.67E-11 | 5.262686866 | 6.696269176 | 257.0748006 | 92 | 0 | 6 | 8 | 34 | 10 | 0 | 0 | 0 | 0 | 2 | 0 |
| SG0000003 | Prdm1 | 7.74E-12 | 2.98E-11 | -1.212050033 | 1851.085828 | 799.0300998 | 60 | 2 | 6 | 12 | 40 | 8 | 8 | 0 | 0 | 4 | 10 | 4 |
| SG0000002 | Capn10 | 8.77E-12 | 3.37E-11 | 1.063218683 | 396.3439233 | 828.1956016 | 26 | 0 | 6 | 5 | 14 | 3 | 2 | 0 | 0 | 2 | 2 | 0 |
| SG0000004 | Serpinb9 | 1.01E-11 | 3.86E-11 | -1.990424092 | 240.6644771 | 60.56680099 | 59 | 2 | 6 | 8 | 16 | 4 | 10 | 0 | 2 | 2 | 2 | 0 |
| SG0000006 | Acvr2b | 1.08E-11 | 4.13E-11 | 2.018776523 | 36.90724768 | 149.5629241 | 62 | 4 | 6 | 12 | 38 | 12 | 0 | 0 | 0 | 0 | 0 | 0 |
| ISG000000 | Bmp7 | 1.21E-11 | 4.6E-11 | 9.503845056 | 0.615018556 | 446.5094506 | 60 | 5 | 6 | 8 | 20 | 6 | 4 | 1 | 1 | 2 | 2 | 1 |
| SG0000003 | AW551984 | 1.45E-11 | 5.49E-11 | 4.590862112 | 7.666980591 | 184.7615366 | 58 | 0 | 6 | 6 | 18 | 6 | 6 | 0 | 2 | 0 | 2 | 2 |
| SG0000002 | Src | 1.56E-11 | 5.9E-11 | -1.353593861 | 2969.972793 | 1162.197981 | 120 | 3 | 6 | 12 | 38 | 29 | 4 | 0 | 0 | 0 | 0 | 4 |
| ISG000000 | Setd7 | 2.18E-11 | 8.15E-11 | -1.120448973 | 2276.389084 | 1047.026672 | 59 | 2 | 6 | 13 | 25 | 5 | 10 | 0 | 1 | 1 | 2 | 2 |
| SG0000001 | Pea15a | 3.27E-11 | 1.2E-10 | 1.727674437 | 437.202698 | 1447.986996 | 22 | 6 | 6 | 8 | 10 | 8 | 8 | 0 | 0 | 0 | 0 | 2 |
| ISG000000 | Nol8 | 3.55E-11 | 1.3E-10 | 0.876634799 | 1243.266507 | 2282.74607 | 66 | 4 | 6 | 9 | 37 | 3 | 1 | 0 | 0 | 0 | 0 | 1 |
| ISG000000 | Impad1 | 3.7E-11 | 1.36E-10 | -0.959031321 | 4629.296904 | 2381.320448 | 74 | 4 | 6 | 11 | 32 | 7 | 14 | 0 | 1 | 3 | 6 | 2 |
| ISG000000 | Hrk | 5.07E-11 | 1.84E-10 | 6.794093564 | 1.265003014 | 140.3839689 | 31 | 4 | 6 | 7 | 18 | 2 | 7 | 0 | 1 | 2 | 8 | 2 |
| SG0000002 | Rassf2 | 5.29E-11 | 1.92E-10 | 2.304821565 | 46.20929183 | 228.3230402 | 86 | 4 | 6 | 6 | 28 | 9 | 10 | 0 | 2 | 0 | 4 | 1 |
| SG0000004 | Plekha4 | 6.31E-11 | 2.27E-10 | 2.277319848 | 36.86624794 | 178.7189167 | 38 | 4 | 6 | 7 | 14 | 1 | 0 | 0 | 0 | 0 | 0 | 0 |
| ISG000000 | Fgf5 | 6.46E-11 | 2.33E-10 | 5.694190126 | 0.639110347 | 33.09008458 | 92 | 2 | 6 | 12 | 42 | 6 | 4 | 0 | 0 | 0 | 2 | 0 |
| SG000000C | Tm9sf1 | 1.1E-10 | 3.9E-10 | -0.99514101 | 1891.342623 | 948.8616973 | 33 | 6 | 6 | 6 | 30 | 0 | 3 | 0 | 0 | 0 | 6 | 0 |
| ISG000000 | Ulbp1 | 1.97E-10 | 6.86E-10 | 3.314960409 | 5.819581356 | 57.91542685 | 74 | 0 | 6 | 6 | 26 | 4 | 2 | 0 | 0 | 0 | 0 | 0 |
| SG0000003 | Awat2 | 2.05E-10 | 7.15E-10 | -2.735769978 | 160.0541581 | 24.0280284 | 84 | 4 | 6 | 11 | 60 | 7 | 0 | 0 | 0 | 0 | 0 | 0 |
| SG0000003 | E330009J07Rik | 2.18E-10 | 7.57E-10 | 1.159342218 | 190.5214626 | 425.5402944 | 68 | 2 | 6 | 4 | 30 | 18 | 3 | 0 | 0 | 0 | 5 | 2 |
| SG0000003 | Smarcal1 | 2.52E-10 | 8.71E-10 | 0.910579409 | 645.3962941 | 1213.21615 | 124 | 6 | 6 | 10 | 58 | 16 | 0 | 0 | 0 | 0 | 0 | 0 |
| SG0000004 | Spata2 | 2.64E-10 | 9.12E-10 | -1.00437449 | 1497.182812 | 746.3249906 | 20 | 6 | 6 | 8 | 18 | 4 | 5 | 0 | 0 | 1 | 2 | 2 |
| SG0000004 | Csgalnact2 | 4.46E-10 | 1.51E-09 | -1.098458623 | 863.7780927 | 403.397489 | 81 | 4 | 6 | 16 | 42 | 4 | 6 | 1 | 1 | 3 | 6 | 0 |
| ISG000000 | Ubap1l | 7.33E-10 | 2.44E-09 | 6.500034743 | 0.307509278 | 27.83323294 | 36 | 6 | 6 | 9 | 18 | 0 | 0 | 0 | 0 | 0 | 0 | 0 |
| SG0000005 | Chst3 | 7.42E-10 | 2.48E-09 | -2.161850927 | 194.891103 | 43.55219402 | 37 | 3 | 6 | 6 | 14 | 4 | 6 | 2 | 4 | 2 | 6 | 0 |
| ISG000000 | Rgcc | 7.58E-10 | 2.52E-09 | 4.770219784 | 8.269778331 | 225.6696262 | 23 | 6 | 6 | 7 | 13 | 3 | 1 | 0 | 0 | 0 | 0 | 0 |
| SG0000004 | Fcgbp | 8.4E-10 | 2.79E-09 | 4.687648887 | 0 | 24.77050471 | 76 | 4 | 6 | 16 | 58 | 6 | 0 | 0 | 0 | 0 | 0 | 0 |
| ISG000000 | Osbpl2 | 8.83E-10 | 2.92E-09 | -0.962434205 | 1548.451403 | 794.6502332 | 49 | 4 | 6 | 9 | 20 | 5 | 1 | 0 | 0 | 0 | 0 | 0 |
| ISG000000 | Phf7 | 8.95E-10 | 2.96E-09 | 1.022104542 | 280.9589704 | 570.5937738 | 30 | 4 | 6 | 9 | 15 | 1 | 0 | 0 | 0 | 0 | 0 | 0 |
| SG0000003 | Rbm17 | 9.21E-10 | 3.04E-09 | 0.777173802 | 2159.342095 | 3700.61647 | 33 | 4 | 6 | 11 | 19 | 2 | 0 | 0 | 0 | 0 | 0 | 0 |
| ISG000000 | Zfp449 | 9.44E-10 | 3.11E-09 | 2.113915265 | 60.37546088 | 261.3438966 | 36 | 4 | 6 | 8 | 21 | 4 | 4 | 0 | 0 | 1 | 2 | 0 |
| ISG000000 | Zfp821 | 1.07E-09 | 3.51E-09 | 1.180719227 | 159.9034204 | 362.4846807 | 31 | 3 | 6 | 8 | 19 | 2 | 0 | 0 | 0 | 0 | 0 | 0 |
| SG0000002 | Hgs | 1.37E-09 | 4.47E-09 | -1.190599729 | 2416.072262 | 1058.531261 | 22 | 6 | 6 | 6 | 10 | 0 | 0 | 0 | 0 | 0 | 0 | 0 |
| SG0000004 | Mapk6 | 1.81E-09 | 5.84E-09 | -0.835601009 | 8254.432564 | 4625.367618 | 78 | 6 | 6 | 22 | 30 | 8 | 8 | 2 | 2 | 2 | 4 | 2 |
| ISG000000 | Il12rb1 | 2.32E-09 | 7.42E-09 | 2.806903335 | 8.232468862 | 57.60924658 | 15 | 5 | 6 | 7 | 13 | 0 | 0 | 0 | 0 | 0 | 1 | 0 |
| ISG000000 | A130010J15Rik | 3.49E-09 | 1.1E-08 | 0.986403543 | 291.9704343 | 578.4634589 | 29 | 4 | 6 | 4 | 8 | 4 | 16 | 4 | 6 | 4 | 8 | 2 |
| SG0000002 | Mitd1 | 3.67E-09 | 1.15E-08 | 1.045518742 | 548.2392164 | 1131.625175 | 64 | 4 | 6 | 10 | 20 | 1 | 2 | 0 | 0 | 0 | 0 | 0 |
| SG000000C | Ppp1cc | 5E-09 | 1.56E-08 | 0.759316476 | 1599.974243 | 2708.257951 | 87 | 0 | 6 | 8 | 23 | 6 | 5 | 0 | 0 | 0 | 0 | 0 |
| SG0000007 | Htr1d | 6.05E-09 | 1.87E-08 | 6.467809847 | 0.307509278 | 27.21842517 | 60 | 3 | 6 | 9 | 66 | 6 | 0 | 0 | 0 | 0 | 0 | 0 |
| ISG000000 | Cdc42ep4 | 6.85E-09 | 2.11E-08 | 0.832895061 | 991.7612426 | 1766.580894 | 42 | 6 | 6 | 8 | 28 | 0 | 0 | 0 | 0 | 0 | 0 | 0 |
| SG0000004 | Snhg11 | 6.88E-09 | 2.12E-08 | 3.040902208 | 181.5729463 | 1494.355475 | 29 | 3 | 6 | 6 | 11 | 3 | 11 | 3 | 3 | 3 | 8 | 0 |
| SG000000C | Trmt10a | 7.08E-09 | 2.18E-08 | 0.881804034 | 474.5276342 | 874.4016595 | 64 | 0 | 6 | 2 | 26 | 6 | 2 | 0 | 0 | 0 | 4 | 2 |
| ISG000000 | Il17f | 7.82E-09 | 2.4E-08 | 4.302863839 | 1.631569985 | 32.20303395 | 19 | 4 | 6 | 8 | 13 | 0 | 3 | 0 | 0 | 2 | 3 | 0 |
| ISG000000 | Med26 | 7.87E-09 | 2.42E-08 | 0.958841739 | 426.4824389 | 828.9747029 | 64 | 5 | 6 | 11 | 27 | 10 | 1 | 0 | 0 | 0 | 0 | 0 |
| SG0000002 | Mr1 | 8.02E-09 | 2.46E-08 | -2.917063129 | 216.9926028 | 28.72906077 | 67 | 6 | 6 | 9 | 43 | 3 | 10 | 0 | 0 | 0 | 1 | 0 |
| ISG000000 | Qtrt1 | 1.07E-08 | 3.24E-08 | 0.960545027 | 349.5295998 | 680.2003092 | 14 | 6 | 6 | 7 | 15 | 0 | 0 | 0 | 0 | 0 | 0 | 0 |
| SG0000005 | Evc2 | 1.95E-08 | 5.78E-08 | -0.936861717 | 1018.521068 | 532.0427607 | 106 | 5 | 6 | 13 | 43 | 5 | 0 | 0 | 0 | 0 | 0 | 0 |
| SG0000005 | Smco4 | 1.95E-08 | 5.79E-08 | -1.471860847 | 542.669977 | 195.6417462 | 57 | 6 | 6 | 12 | 27 | 6 | 4 | 0 | 0 | 2 | 2 | 0 |
| SG0000002 | Dclre1b | 1.98E-08 | 5.88E-08 | -1.04320584 | 628.1418146 | 304.8045568 | 55 | 0 | 6 | 6 | 32 | 0 | 15 | 0 | 1 | 1 | 6 | 0 |
| SG0000004 | Ufl1 | 2.14E-08 | 6.35E-08 | 0.81531804 | 637.0706454 | 1121.044314 | 177 | 2 | 6 | 20 | 85 | 30 | 9 | 0 | 2 | 0 | 3 | 0 |
| SG0000005 | Thnsl2 | 2.25E-08 | 6.67E-08 | 2.158709911 | 17.90846052 | 79.96403289 | 40 | 4 | 6 | 8 | 18 | 4 | 2 | 0 | 0 | 0 | 0 | 0 |
| ISG000000 | Tma16 | 2.28E-08 | 6.73E-08 | 0.807533547 | 666.0229955 | 1165.684561 | 46 | 4 | 6 | 6 | 16 | 0 | 7 | 0 | 0 | 0 | 0 | 0 |
| SG0000002 | 2810021J22Rik | 3.21E-08 | 9.39E-08 | 1.18993458 | 112.9653319 | 257.7218167 | 28 | 5 | 6 | 9 | 12 | 3 | 3 | 0 | 0 | 0 | 0 | 1 |
| ISG000000 | Nudt4 | 3.44E-08 | 1E-07 | 0.690060977 | 3143.009891 | 5070.780373 | 24 | 5 | 6 | 6 | 11 | 3 | 5 | 0 | 0 | 0 | 1 | 1 |
| SG0000002 | Ccne2 | 3.56E-08 | 1.04E-07 | -0.898681162 | 1176.779805 | 631.1974277 | 100 | 3 | 6 | 6 | 30 | 18 | 19 | 3 | 6 | 3 | 9 | 6 |
| SG0000002 | Tsn | 4.78E-08 | 1.38E-07 | 0.680983703 | 2902.12324 | 4652.778437 | 41 | 6 | 6 | 8 | 18 | 2 | 5 | 1 | 1 | 1 | 1 | 0 |
| SG0000002 | Etfdh | 5.92E-08 | 1.7E-07 | -0.898134027 | 1343.114047 | 720.6885283 | 123 | 4 | 6 | 10 | 60 | 14 | 1 | 0 | 0 | 0 | 0 | 0 |
| SG0000003 | Tbc1d17 | 6.42E-08 | 1.83E-07 | 0.844373814 | 458.6825644 | 823.5577861 | 18 | 6 | 6 | 8 | 10 | 0 | 0 | 0 | 0 | 0 | 0 | 0 |
| SG0000002 | Dctn3 | 7.31E-08 | 2.08E-07 | -0.875138829 | 1104.962355 | 602.4270223 | 24 | 6 | 6 | 10 | 20 | 0 | 2 | 0 | 0 | 2 | 2 | 0 |
| SG0000002 | Ccdc28b | 7.66E-08 | 2.17E-07 | -1.442377726 | 203.7732501 | 74.98047922 | 13 | 6 | 6 | 6 | 8 | 0 | 0 | 0 | 0 | 0 | 0 | 0 |
| ISG000000 | Manbal | 8.55E-08 | 2.41E-07 | -0.898406024 | 963.2700114 | 516.7742656 | 29 | 6 | 6 | 9 | 14 | 4 | 0 | 0 | 0 | 0 | 0 | 0 |
| SG0000002 | Il1r2 | 8.68E-08 | 2.45E-07 | 4.336709816 | 0 | 19.20597154 | 110 | 4 | 6 | 20 | 50 | 18 | 1 | 0 | 0 | 0 | 0 | 0 |
| SG0000004 | Bod1 | 9.76E-08 | 2.74E-07 | -0.810844819 | 1879.869016 | 1071.615477 | 13 | 4 | 6 | 6 | 6 | 2 | 0 | 0 | 0 | 0 | 0 | 0 |
| SG0000002 | Flywch2 | 1.06E-07 | 2.97E-07 | 1.788080999 | 27.50848654 | 95.00199214 | 20 | 4 | 6 | 8 | 14 | 2 | 0 | 0 | 0 | 0 | 0 | 0 |
| ISG000000 | Smpdl3b | 1.06E-07 | 2.98E-07 | 1.069814911 | 261.2770735 | 548.4634371 | 31 | 3 | 6 | 8 | 18 | 0 | 0 | 0 | 0 | 0 | 0 | 0 |
| ISG000000 | Ptgfrn | 1.3E-07 | 3.62E-07 | 0.672514112 | 2225.255294 | 3546.71885 | 101 | 3 | 6 | 10 | 29 | 9 | 5 | 0 | 0 | 0 | 2 | 1 |
| SG0000002 | Pdcl2 | 1.37E-07 | 3.8E-07 | 4.304371213 | 0 | 18.75808492 | 78 | 4 | 6 | 14 | 46 | 0 | 4 | 0 | 0 | 0 | 0 | 0 |
| ISG000000 | Spire2 | 1.51E-07 | 4.18E-07 | 1.639409569 | 141.9919693 | 442.3593771 | 49 | 2 | 6 | 9 | 20 | 7 | 1 | 0 | 0 | 0 | 0 | 1 |
| SG0000001 | Notch4 | 1.91E-07 | 5.22E-07 | -1.280094854 | 478.5992491 | 197.0720637 | 27 | 4 | 6 | 7 | 19 | 6 | 1 | 0 | 0 | 1 | 1 | 0 |
| SG0000003 | Jmjd4 | 2.03E-07 | 5.54E-07 | -0.913194822 | 734.7067559 | 390.1350777 | 45 | 3 | 6 | 9 | 15 | 3 | 18 | 3 | 3 | 6 | 9 | 3 |
| SG0000004 | Tmem201 | 2.16E-07 | 5.88E-07 | 0.720536512 | 928.3008026 | 1529.649148 | 29 | 6 | 6 | 9 | 22 | 0 | 3 | 0 | 0 | 1 | 3 | 0 |
| SG0000003 | Pold1 | 2.17E-07 | 5.9E-07 | 1.428553863 | 1036.538363 | 2790.120395 | 20 | 4 | 6 | 6 | 16 | 0 | 0 | 0 | 0 | 0 | 0 | 0 |
| SG000000C | Vdac3 | 2.22E-07 | 6.04E-07 | 0.67181334 | 1817.917977 | 2896.077977 | 72 | 2 | 6 | 10 | 40 | 14 | 2 | 0 | 0 | 0 | 0 | 0 |
| SG000000C | Ctr9 | 2.47E-07 | 6.7E-07 | 0.690214262 | 1264.822142 | 2040.819786 | 44 | 4 | 6 | 8 | 23 | 2 | 1 | 0 | 0 | 0 | 0 | 0 |

| | | | | | | | | | | | | | | | | | | |
|---|---|---|---|---|---|---|---|---|---|---|---|---|---|---|---|---|---|---|
| ISG000000 | Tspan14 | 2.47E-07 | 6.7E-07 | 0.688142301 | 1360.886327 | 2192.670253 | 57 | 4 | 6 | 12 | 29 | 6 | 1 | 0 | 1 | 1 | 1 | 1 |
| SG0000003 | Pth1r | 2.54E-07 | 6.87E-07 | 3.184148075 | 6.82760224 | 62.05721669 | 41 | 3 | 6 | 9 | 15 | 0 | 0 | 0 | 0 | 0 | 0 | 0 |
| ISG000000 | Btbd1 | 2.98E-07 | 8E-07 | -0.795463206 | 1560.847226 | 899.2948608 | 83 | 3 | 6 | 11 | 43 | 6 | 4 | 0 | 0 | 1 | 1 | 0 |
| ISG000000 | Cand2 | 3.58E-07 | 9.56E-07 | 1.755420364 | 166.4950546 | 562.1283937 | 34 | 6 | 6 | 8 | 22 | 5 | 4 | 0 | 0 | 0 | 0 | 0 |
| SG0000003 | Tmx4 | 3.85E-07 | 1.02E-06 | -0.983793431 | 1159.159965 | 586.1274225 | 222 | 0 | 6 | 19 | 104 | 17 | 16 | 0 | 1 | 5 | 15 | 4 |
| SG0000003 | Inip | 4E-07 | 1.06E-06 | 0.71671029 | 801.5646069 | 1317.315579 | 48 | 4 | 6 | 8 | 26 | 4 | 4 | 0 | 2 | 2 | 4 | 0 |
| SG0000000 | Jkamp | 4.05E-07 | 1.08E-06 | -0.805146001 | 1255.448488 | 718.4983921 | 38 | 4 | 6 | 7 | 18 | 4 | 4 | 0 | 0 | 1 | 2 | 0 |
| ISG000000 | Ppa1 | 5.32E-07 | 1.4E-06 | 0.649873146 | 3447.583116 | 5409.36212 | 45 | 2 | 6 | 6 | 18 | 3 | 0 | 0 | 0 | 0 | 0 | 0 |
| ISG000000 | Pkd2 | 5.52E-07 | 1.45E-06 | -0.753264303 | 1954.465172 | 1159.505428 | 86 | 2 | 6 | 13 | 44 | 8 | 10 | 0 | 0 | 2 | 5 | 1 |
| SG0000002 | Ccndbp1 | 6.04E-07 | 1.59E-06 | -0.817005486 | 1045.989324 | 593.7233018 | 58 | 6 | 6 | 20 | 45 | 3 | 2 | 0 | 0 | 2 | 2 | 0 |
| SG0000002 | Aldh4a1 | 6.3E-07 | 1.65E-06 | 0.829591909 | 340.2501911 | 604.6867141 | 23 | 6 | 6 | 7 | 12 | 2 | 0 | 0 | 0 | 0 | 0 | 0 |
| ISG000000 | Limd1 | 6.6E-07 | 1.73E-06 | -0.760990956 | 1586.625519 | 936.2532348 | 72 | 5 | 6 | 12 | 32 | 3 | 1 | 0 | 0 | 0 | 1 | 0 |
| ISG000000 | Mmachc | 8.03E-07 | 2.09E-06 | 0.771241649 | 437.6695669 | 746.9873021 | 19 | 6 | 6 | 6 | 10 | 0 | 6 | 0 | 0 | 0 | 0 | 0 |
| SG0000001 | Med28 | 9.61E-07 | 2.49E-06 | -0.771225213 | 1331.480552 | 780.1406449 | 20 | 2 | 6 | 8 | 20 | 4 | 2 | 0 | 0 | 2 | 2 | 0 |
| ISG000000 | Zfp820 | 1.27E-06 | 3.26E-06 | 1.111425752 | 95.63239004 | 206.6224464 | 69 | 4 | 6 | 9 | 38 | 9 | 5 | 1 | 1 | 1 | 4 | 1 |
| ISG000000 | Gm8206 | 1.28E-06 | 3.27E-06 | 4.084046507 | 0 | 15.95979125 | 35 | 4 | 6 | 7 | 21 | 3 | 6 | 2 | 2 | 2 | 2 | 0 |
| SG0000002 | Abhd6 | 1.62E-06 | 4.1E-06 | -0.862286032 | 666.331314 | 366.5356849 | 132 | 2 | 6 | 24 | 68 | 6 | 2 | 0 | 0 | 0 | 0 | 0 |
| ISG000000 | Ndc80 | 1.99E-06 | 5.01E-06 | 0.664543148 | 906.5614828 | 1436.960017 | 61 | 4 | 6 | 6 | 19 | 6 | 0 | 0 | 0 | 0 | 1 | 0 |
| SG0000004 | Fbxo10 | 2.03E-06 | 5.1E-06 | -1.061064122 | 743.4660358 | 356.3272247 | 95 | 4 | 6 | 14 | 45 | 6 | 2 | 0 | 0 | 0 | 0 | 0 |
| SG0000002 | Lig3 | 2.1E-06 | 5.26E-06 | -0.683087122 | 2991.244079 | 1863.039825 | 157 | 6 | 6 | 13 | 138 | 14 | 23 | 0 | 0 | 0 | 5 | 0 |
| ISG000000 | Deaf1 | 2.41E-06 | 6.01E-06 | 0.768656564 | 381.7923201 | 650.452924 | 34 | 4 | 6 | 7 | 19 | 9 | 0 | 0 | 0 | 0 | 0 | 1 |
| SG0000001 | Cyp2j9 | 3.11E-06 | 7.68E-06 | 3.647751738 | 1.987262844 | 24.90795098 | 61 | 3 | 6 | 7 | 24 | 4 | 1 | 0 | 1 | 0 | 0 | 0 |
| ISG000000 | Glod5 | 4.42E-06 | 1.08E-05 | 3.512463187 | 5.154035651 | 58.81717972 | 21 | 3 | 6 | 4 | 16 | 5 | 0 | 0 | 0 | 0 | 0 | 0 |
| SG0000002 | Vwa5a | 5.01E-06 | 1.21E-05 | -0.864969818 | 1516.825751 | 832.8251514 | 131 | 4 | 6 | 18 | 69 | 6 | 9 | 0 | 0 | 1 | 5 | 2 |
| SG0000002 | Cldn10 | 5.39E-06 | 1.3E-05 | -3.140213224 | 38.47237943 | 4.363657684 | 269 | 2 | 6 | 32 | 122 | 17 | 3 | 0 | 0 | 0 | 1 | 0 |
| ISG000000 | Zfp874b | 7.13E-06 | 1.7E-05 | 0.85210858 | 192.1654066 | 346.8849446 | 35 | 3 | 6 | 12 | 20 | 0 | 3 | 0 | 0 | 1 | 2 | 0 |
| SG0000003 | Haus6 | 7.75E-06 | 1.85E-05 | 0.578714576 | 1910.718024 | 2853.691663 | 90 | 4 | 6 | 14 | 57 | 6 | 16 | 1 | 1 | 2 | 4 | 0 |
| ISG000000 | Cript | 8.92E-06 | 2.11E-05 | -0.678793432 | 1643.104327 | 1026.426735 | 26 | 4 | 6 | 8 | 15 | 2 | 3 | 0 | 1 | 1 | 1 | 1 |
| SG0000005 | Hmcn2 | 9.54E-06 | 2.25E-05 | 2.144929027 | 11.53910528 | 51.03405899 | 96 | 4 | 6 | 15 | 38 | 12 | 0 | 0 | 0 | 0 | 0 | 0 |
| SG0000001 | Ddx39b | 9.77E-06 | 2.3E-05 | 0.549881826 | 4795.737279 | 7020.795239 | 36 | 6 | 6 | 9 | 15 | 3 | 3 | 0 | 0 | 0 | 0 | 0 |
| SG0000002 | Rcan1 | 9.82E-06 | 2.31E-05 | -0.679435909 | 1521.189728 | 949.8451191 | 72 | 3 | 6 | 12 | 28 | 13 | 0 | 0 | 0 | 0 | 0 | 2 |
| SG0000002 | Nabp2 | 1.05E-05 | 2.46E-05 | -0.686442661 | 1374.249181 | 853.9366546 | 33 | 6 | 6 | 9 | 21 | 9 | 0 | 0 | 0 | 0 | 0 | 0 |
| SG0000000 | Prkcsh | 1.12E-05 | 2.63E-05 | -0.73872771 | 5386.293591 | 3227.831254 | 20 | 6 | 6 | 8 | 14 | 2 | 0 | 0 | 0 | 0 | 0 | 0 |
| SG0000000 | Ccdc130 | 1.23E-05 | 2.87E-05 | 0.815732766 | 218.8347114 | 385.1910978 | 26 | 4 | 6 | 4 | 8 | 6 | 2 | 2 | 2 | 2 | 4 | 0 |
| ISG000000 | Sar1a | 1.36E-05 | 3.16E-05 | -0.608089465 | 4372.702607 | 2868.776882 | 25 | 5 | 6 | 8 | 25 | 2 | 3 | 0 | 0 | 0 | 0 | 1 |
| SG0000004 | Inhba | 1.69E-05 | 3.89E-05 | -2.40858324 | 497.2367495 | 93.64992123 | 50 | 2 | 6 | 11 | 26 | 0 | 1 | 0 | 0 | 0 | 0 | 0 |
| SG0000006 | Pgk1 | 1.75E-05 | 4.03E-05 | -0.578152544 | 18672.9646 | 12507.54333 | 56 | 0 | 6 | 6 | 28 | 4 | 4 | 0 | 0 | 2 | 2 | 0 |
| SG0000002 | Ifitm1 | 2.03E-05 | 4.64E-05 | -0.831538902 | 6847.120556 | 3847.59917 | 9 | 6 | 6 | 6 | 9 | 0 | 0 | 0 | 0 | 0 | 0 | 0 |
| ISG000000 | Sf3b3 | 2.15E-05 | 4.9E-05 | 0.518203749 | 8434.389818 | 12079.48842 | 76 | 4 | 6 | 10 | 31 | 4 | 1 | 0 | 0 | 0 | 0 | 1 |
| SG0000004 | Hmga1 | 2.35E-05 | 5.33E-05 | 1.071690515 | 2229.950842 | 4687.122245 | 29 | 6 | 6 | 6 | 18 | 0 | 5 | 0 | 0 | 0 | 0 | 0 |
| ISG000000 | Mkrn2 | 2.51E-05 | 5.69E-05 | 0.600594656 | 820.5483267 | 1244.231439 | 30 | 6 | 6 | 8 | 18 | 1 | 4 | 0 | 0 | 0 | 2 | 0 |
| SG0000002 | Ablim2 | 2.77E-05 | 6.26E-05 | -1.236322902 | 223.2544287 | 94.76103702 | 373 | 0 | 6 | 46 | 145 | 81 | 0 | 0 | 0 | 0 | 6 | 5 |
| SG0000002 | Arhgef19 | 2.96E-05 | 6.67E-05 | -1.398512392 | 398.485239 | 151.1534483 | 14 | 4 | 6 | 7 | 11 | 1 | 0 | 0 | 0 | 0 | 0 | 1 |
| ISG000000 | E030030I06Rik | 3.02E-05 | 6.79E-05 | 2.112396454 | 12.26136511 | 53.01924269 | 83 | 3 | 6 | 12 | 52 | 10 | 3 | 0 | 1 | 0 | 0 | 0 |
| SG0000003 | Tbc1d13 | 3.95E-05 | 8.78E-05 | 0.527803522 | 2709.198234 | 3905.939174 | 27 | 5 | 6 | 7 | 16 | 1 | 2 | 0 | 0 | 2 | 2 | 0 |
| SG0000002 | Cyb5a | 5.26E-05 | 0.000115 | 0.554122888 | 1133.551504 | 1664.366126 | 163 | 0 | 6 | 24 | 40 | 9 | 5 | 0 | 0 | 0 | 1 | 0 |
| SG0000002 | Ccz1 | 5.29E-05 | 0.000116 | -0.593004083 | 2413.082205 | 1599.779401 | 43 | 3 | 6 | 14 | 25 | 7 | 1 | 0 | 0 | 1 | 1 | 0 |
| SG0000002 | Dclre1a | 5.35E-05 | 0.000117 | -0.705664265 | 700.0326297 | 429.2320802 | 48 | 6 | 6 | 8 | 38 | 2 | 3 | 0 | 0 | 0 | 1 | 0 |
| ISG000000 | Masp1 | 5.41E-05 | 0.000118 | -2.138001817 | 320.7922382 | 72.8821497 | 62 | 4 | 6 | 12 | 33 | 12 | 0 | 0 | 0 | 0 | 0 | 0 |
| SG0000003 | Card10 | 5.5E-05 | 0.00012 | -1.270654351 | 1542.404736 | 639.2831776 | 30 | 4 | 6 | 6 | 16 | 3 | 0 | 0 | 0 | 0 | 0 | 1 |
| SG0000005 | Pkp3 | 5.69E-05 | 0.000124 | -1.176710674 | 867.8724759 | 383.9111327 | 22 | 4 | 6 | 8 | 25 | 4 | 0 | 0 | 0 | 0 | 0 | 0 |
| ISG000000 | Fundc2 | 5.78E-05 | 0.000126 | -0.604986942 | 1852.426565 | 1217.928474 | 45 | 4 | 6 | 12 | 28 | 2 | 7 | 0 | 0 | 2 | 5 | 0 |
| SG0000002 | Supv3l1 | 5.79E-05 | 0.000126 | 0.554485154 | 1075.914685 | 1580.136113 | 62 | 4 | 6 | 10 | 33 | 0 | 3 | 0 | 0 | 0 | 1 | 0 |
| ISG000000 | Iws1 | 6.74E-05 | 0.000146 | 0.525874144 | 1558.760119 | 2244.311719 | 109 | 2 | 6 | 16 | 42 | 4 | 24 | 1 | 1 | 4 | 6 | 1 |
| SG0000004 | 1520401A03Rik | 8.3E-05 | 0.000178 | 2.299689096 | 6.585337631 | 32.42301855 | 28 | 2 | 6 | 6 | 12 | 0 | 2 | 0 | 0 | 0 | 0 | 0 |
| ISG000000 | Ptcd3 | 0.000103 | 0.000218 | 0.492519681 | 2623.873136 | 3691.526841 | 56 | 2 | 6 | 5 | 30 | 10 | 0 | 0 | 0 | 0 | 0 | 0 |
| ISG000000 | Mgea5 | 0.000106 | 0.000223 | 0.479236237 | 5181.907037 | 7223.606826 | 61 | 6 | 6 | 10 | 35 | 3 | 6 | 0 | 0 | 1 | 3 | 1 |
| SG0000001 | Cnih1 | 0.000121 | 0.000254 | 0.50202492 | 1877.359302 | 2658.716056 | 48 | 4 | 6 | 4 | 14 | 10 | 3 | 0 | 0 | 0 | 1 | 1 |
| SG0000001 | Myb | 0.000139 | 0.00029 | 1.000152842 | 108.1246427 | 216.2721965 | 101 | 2 | 6 | 12 | 54 | 16 | 9 | 0 | 0 | 0 | 6 | 0 |
| SG0000002 | Ahcyl1 | 0.00015 | 0.000312 | -0.518552 | 6944.767024 | 4847.948245 | 102 | 3 | 6 | 17 | 45 | 5 | 4 | 0 | 0 | 0 | 0 | 0 |
| SG0000006 | Ubr4 | 0.000191 | 0.000393 | -0.50539206 | 7758.625781 | 5465.710635 | 212 | 2 | 6 | 24 | 86 | 26 | 0 | 0 | 0 | 0 | 0 | 0 |
| ISG000000 | Dicer1 | 0.000201 | 0.000412 | 0.472218769 | 2533.916849 | 3515.154138 | 114 | 4 | 6 | 13 | 41 | 8 | 11 | 0 | 0 | 0 | 5 | 1 |
| ISG000000 | Eif3e | 0.000202 | 0.000413 | 0.453021567 | 5607.407249 | 7676.003763 | 96 | 4 | 6 | 13 | 37 | 7 | 0 | 0 | 0 | 0 | 0 | 0 |
| SG0000003 | Zfp60 | 0.000207 | 0.000424 | 0.549241949 | 640.5350411 | 937.3055636 | 87 | 3 | 6 | 13 | 62 | 2 | 10 | 0 | 0 | 1 | 8 | 2 |
| SG0000004 | Accs | 0.000209 | 0.000427 | -0.992886742 | 170.1328326 | 85.48687492 | 51 | 3 | 6 | 18 | 33 | 6 | 0 | 0 | 0 | 0 | 0 | 0 |
| ISG000000 | Plscr4 | 0.000224 | 0.000457 | -2.741055398 | 26.59817952 | 3.978432722 | 73 | 0 | 6 | 3 | 30 | 8 | 5 | 0 | 2 | 0 | 1 | 0 |
| SG0000004 | Poldip3 | 0.000242 | 0.000491 | -0.502578285 | 4971.884846 | 3509.376167 | 52 | 2 | 6 | 8 | 26 | 2 | 0 | 0 | 0 | 0 | 0 | 0 |
| SG0000002 | Rapgef3 | 0.000336 | 0.00067 | -2.072278226 | 279.8814427 | 66.55124717 | 83 | 0 | 6 | 6 | 18 | 7 | 1 | 0 | 0 | 0 | 0 | 0 |
| ISG000000 | 2810417H13Rik | 0.000343 | 0.000682 | 0.616264134 | 2095.937161 | 3212.863297 | 43 | 6 | 6 | 15 | 25 | 3 | 4 | 0 | 0 | 1 | 1 | 0 |
| SG0000005 | Trim30d | 0.000365 | 0.000725 | -2.056555565 | 50.51791529 | 12.14396502 | 149 | 2 | 6 | 23 | 69 | 3 | 3 | 0 | 0 | 1 | 3 | 1 |
| ISG000000 | Cdkl4 | 0.000401 | 0.000792 | 2.310326815 | 10.31994228 | 51.1864416 | 74 | 2 | 6 | 7 | 23 | 5 | 1 | 0 | 0 | 0 | 0 | 2 |
| ISG000000 | Wdyhv1 | 0.000439 | 0.000863 | -0.633771754 | 777.2608152 | 500.9362557 | 41 | 5 | 6 | 10 | 26 | 2 | 0 | 0 | 0 | 0 | 0 | 0 |
| ISG000000 | Dimt1 | 0.000559 | 0.001083 | 0.551110734 | 429.1464532 | 628.791105 | 36 | 3 | 6 | 7 | 17 | 4 | 9 | 0 | 2 | 2 | 6 | 0 |
| ISG000000 | Tcp11l2 | 0.000566 | 0.001096 | -0.793877878 | 238.4848298 | 137.5560525 | 52 | 5 | 6 | 8 | 21 | 5 | 5 | 0 | 0 | 0 | 0 | 0 |
| SG0000003 | Bgn | 0.00058 | 0.001122 | -1.094652763 | 8749.373728 | 4096.883846 | 24 | 6 | 6 | 10 | 10 | 0 | 2 | 0 | 0 | 0 | 0 | 0 |
| ISG000000 | Zfp456 | 0.000618 | 0.00119 | 1.172031878 | 33.98423933 | 76.57628702 | 57 | 2 | 6 | 10 | 28 | 2 | 9 | 0 | 0 | 1 | 5 | 0 |
| ISG000000 | Elp2 | 0.000637 | 0.001225 | 0.433837754 | 2741.192522 | 3702.863608 | 74 | 2 | 6 | 9 | 20 | 6 | 0 | 0 | 0 | 0 | 0 | 1 |
| SG0000006 | Arih2 | 0.000742 | 0.001416 | 0.451076398 | 1608.509306 | 2198.928479 | 72 | 2 | 6 | 7 | 30 | 6 | 1 | 0 | 0 | 0 | 1 | 0 |
| ISG000000 | Degs1 | 0.000791 | 0.001503 | -0.459235695 | 4488.849347 | 3265.061085 | 14 | 6 | 6 | 6 | 12 | 1 | 1 | 0 | 0 | 0 | 2 | 0 |
| SG0000002 | Phospho2 | 0.000807 | 0.001531 | -0.610291516 | 525.1963307 | 344.0373794 | 47 | 6 | 6 | 9 | 15 | 0 | 0 | 0 | 0 | 0 | 0 | 0 |
| SG0000003 | N4bp1 | 0.000841 | 0.001593 | 0.414948501 | 3902.355395 | 5202.819559 | 65 | 5 | 6 | 10 | 27 | 6 | 3 | 0 | 0 | 0 | 0 | 0 |
| SG0000006 | Gmfb | 0.000862 | 0.00163 | 0.417019805 | 3859.705006 | 5153.349333 | 56 | 4 | 6 | 10 | 26 | 0 | 18 | 0 | 2 | 6 | 8 | 0 |
| SG0000000 | Lrrc8a | 0.000869 | 0.001644 | -0.619946602 | 482.2101415 | 313.7717334 | 34 | 6 | 6 | 8 | 16 | 2 | 4 | 0 | 0 | 2 | 2 | 0 |
| ISG000000 | Rsph10b | 0.001094 | 0.002045 | 2.976686786 | 1.987262844 | 15.64326214 | 70 | 5 | 6 | 12 | 36 | 5 | 0 | 0 | 0 | 0 | 0 | 0 |
| ISG000000 | Slc38a7 | 0.001129 | 0.002109 | 0.454333454 | 1074.788437 | 1472.620979 | 20 | 6 | 6 | 8 | 13 | 1 | 0 | 0 | 0 | 0 | 0 | 0 |
| SG0000003 | Ccdc101 | 0.001258 | 0.002335 | 0.565165177 | 384.0012584 | 568.1517105 | 43 | 4 | 6 | 7 | 23 | 0 | 1 | 0 | 0 | 0 | 0 | 0 |
| SG0000000 | Tmem79 | 0.0013 | 0.002408 | -0.821636188 | 342.3313335 | 193.6910323 | 18 | 6 | 6 | 6 | 12 | 3 | 0 | 0 | 0 | 0 | 0 | 0 |
| SG0000003 | Slx4 | 0.001349 | 0.002497 | -0.495034331 | 1149.598744 | 815.6918034 | 28 | 3 | 6 | 7 | 13 | 2 | 1 | 0 | 1 | 0 | 0 | 0 |
| SG0000000 | Hnrnpa2b1 | 0.001358 | 0.002513 | 0.384524533 | 29355.05291 | 38320.95167 | 116 | 3 | 6 | 6 | 21 | 3 | 9 | 0 | 1 | 0 | 0 | 0 |
| ISG000000 | Sfxn1 | 0.001634 | 0.002997 | -0.438368825 | 3600.11561 | 2656.771978 | 58 | 2 | 6 | 3 | 13 | 5 | 3 | 0 | 0 | 0 | 0 | 0 |
| SG0000003 | Prkrip1 | 0.001764 | 0.003219 | -0.544049413 | 617.6637632 | 423.6204632 | 41 | 6 | 6 | 11 | 27 | 8 | 5 | 0 | 0 | 1 | 1 | 0 |
| ISG000000 | Ddx17 | 0.001859 | 0.00338 | 0.378537161 | 10110.60945 | 13144.02495 | 25 | 3 | 6 | 4 | 12 | 2 | 2 | 0 | 0 | 0 | 0 | 0 |
| ISG000000 | Draxin | 0.001874 | 0.003405 | 4.050375711 | 1.561638181 | 25.87408505 | 25 | 4 | 6 | 9 | 17 | 3 | 1 | 0 | 0 | 0 | 0 | 1 |
| SG0000004 | Fancb | 0.002165 | 0.003902 | 0.556806755 | 255.5197462 | 375.8720238 | 99 | 3 | 6 | 12 | 69 | 3 | 3 | 0 | 0 | 0 | 3 | 0 |
| SG0000005 | Clca3a1 | 0.002446 | 0.004374 | -1.888984261 | 29.00622675 | 7.831596684 | 110 | 2 | 6 | 14 | 70 | 10 | 5 | 0 | 0 | 0 | 0 | 2 |
| SG0000004 | Ankrd50 | 0.003062 | 0.005394 | 0.443851409 | 711.009146 | 967.1365019 | 122 | 2 | 6 | 13 | 54 | 31 | 19 | 0 | 2 | 5 | 11 | 2 |
| SG0000003 | Ogg1 | 0.003294 | 0.005775 | -0.558402033 | 427.6452285 | 290.3941539 | 24 | 4 | 6 | 8 | 16 | 2 | 0 | 0 | 0 | 0 | 0 | 0 |
| ISG000000 | Zfp429 | 0.003427 | 0.005999 | 0.635380997 | 140.8107628 | 218.7281047 | 46 | 2 | 6 | 8 | 18 | 0 | 2 | 0 | 0 | 0 | 0 | 0 |
| SG0000005 | Zscan29 | 0.00343 | 0.006001 | -0.625265135 | 289.0081228 | 187.3641462 | 37 | 2 | 6 | 4 | 46 | 5 | 7 | 0 | 0 | 0 | 2 | 1 |
| ISG000000 | Gtf3c6 | 0.003452 | 0.006039 | -0.497060178 | 674.1536099 | 477.6709622 | 16 | 6 | 6 | 6 | 13 | 3 | 0 | 0 | 0 | 0 | 0 | 0 |
| SG0000003 | Fbxo17 | 0.003719 | 0.006474 | -0.655970371 | 240.6852285 | 152.7505137 | 66 | 3 | 6 | 12 | 30 | 3 | 0 | 0 | 0 | 0 | 0 | 0 |
| ISG000000 | Psmb1 | 0.004081 | 0.007061 | 0.361069402 | 3781.229694 | 4856.526695 | 35 | 5 | 6 | 7 | 17 | 2 | 0 | 0 | 0 | 0 | 0 | 0 |
| SG0000002 | Mx2 | 0.004271 | 0.00737 | -1.054611297 | 129.1757842 | 62.18869917 | 64 | 4 | 6 | 12 | 30 | 6 | 2 | 0 | 0 | 0 | 0 | 0 |
| SG0000006 | Slc31a2 | 0.004486 | 0.007725 | -0.540001604 | 447.9508186 | 308.0871993 | 68 | 3 | 6 | 20 | 67 | 7 | 0 | 0 | 0 | 0 | 0 | 0 |
| SG0000003 | Pfkm | 0.004748 | 0.008152 | -0.405455832 | 2443.825819 | 1845.082889 | 64 | 4 | 6 | 12 | 30 | 19 | 2 | 2 | 2 | 2 | 2 | 0 |
| SG0000002 | Ntan1 | 0.00481 | 0.008253 | -0.439639276 | 1248.012792 | 920.1834701 | 40 | 4 | 6 | 8 | 18 | 2 | 0 | 0 | 0 | 0 | 0 | 0 |
| ISG000000 | Thoc1 | 0.005015 | 0.008579 | 0.528706638 | 980.6424035 | 1414.709355 | 88 | 3 | 6 | 11 | 41 | 12 | 3 | 0 | 1 | 0 | 3 | 0 |
| ISG000000 | AW209491 | 0.005384 | 0.009169 | -0.47291396 | 685.8485994 | 494.1592952 | 32 | 2 | 6 | 4 | 6 | 0 | 5 | 0 | 1 | 1 | 1 | 0 |
| SG0000003 | Aox4 | 0.005424 | 0.009233 | 2.883446353 | 2.211622639 | 16.31980142 | 78 | 6 | 6 | 9 | 41 | 4 | 2 | 0 | 0 | 0 | 0 | 0 |
| SG0000002 | 3110040N11Rik | 0.005572 | 0.009467 | -0.613342402 | 250.8823682 | 163.9969155 | 19 | 5 | 6 | 6 | 7 | 4 | 0 | 0 | 0 | 0 | 0 | 0 |
| ISG000000 | Tefm | 0.00575 | 0.009749 | 0.439420141 | 491.9846908 | 667.1603564 | 9 | 4 | 6 | 4 | 14 | 3 | 2 | 2 | 2 | 2 | 2 | 0 |
| SG0000001 | Srp19 | 0.005853 | 0.009914 | 0.387803863 | 1117.457955 | 1462.08219 | 30 | 3 | 6 | 6 | 9 | 6 | 3 | 0 | 0 | 0 | 0 | 0 |
| ISG000000 | Sdcbp2 | 0.005918 | 0.01002 | -0.954934741 | 142.3652607 | 73.44124433 | 25 | 5 | 6 | 5 | 7 | 0 | 1 | 0 | 0 | 0 | 0 | 0 |
| ISG000000 | Cabin1 | 0.006075 | 0.010272 | -0.385030843 | 2363.228638 | 1809.672171 | 114 | 2 | 6 | 18 | 48 | 5 | 0 | 0 | 0 | 0 | 0 | 0 |
| ISG000000 | Ifi205 | 0.006119 | 0.010344 | -2.471792727 | 23.46871618 | 4.230635205 | 51 | 1 | 6 | 6 | 30 | 4 | 0 | 0 | 0 | 0 | 0 | 0 |
| ISG000000 | Crot | 0.006408 | 0.010797 | -0.419146416 | 1352.548865 | 1011.526629 | 61 | 3 | 6 | 12 | 32 | 7 | 0 | 0 | 0 | 0 | 0 | 0 |
| ISG000000 | Zfp839 | 0.006475 | 0.010903 | -0.472284555 | 626.0856513 | 451.2964666 | 15 | 4 | 6 | 6 | 23 | 1 | 2 | 0 | 1 | 1 | 1 | 0 |
| ISG000000 | Ostc | 0.008357 | 0.01384 | 0.349152887 | 1603.057533 | 2041.994656 | 39 | 6 | 6 | 12 | 22 | 3 | 0 | 0 | 0 | 0 | 0 | 0 |
| SG0000000 | Cbfa2t3 | 0.00841 | 0.013922 | 2.148013977 | 3.607958717 | 15.99109479 | 85 | 2 | 6 | 20 | 40 | 15 | 2 | 0 | 0 | 0 | 2 | 2 |
| SG0000002 | Ndufa12 | 0.008491 | 0.01405 | 0.411547988 | 529.6109674 | 704.4420285 | 108 | 0 | 6 | 18 | 57 | 9 | 0 | 0 | 0 | 0 | 0 | 0 |
| SG0000002 | Vps16 | 0.008685 | 0.014342 | 0.359670285 | 1391.440156 | 1785.402247 | 29 | 5 | 6 | 9 | 18 | 1 | 0 | 0 | 0 | 0 | 1 | 1 |
| ISG000000 | Cep162 | 0.009099 | 0.014979 | 0.450587625 | 486.785453 | 665.239429 | 131 | 1 | 6 | 19 | 53 | 15 | 5 | 0 | 0 | 2 | 4 | 0 |
| SG0000002 | Smg8 | 0.009754 | 0.01598 | -0.391818333 | 1230.051107 | 937.5060631 | 44 | 6 | 6 | 12 | 28 | 3 | 2 | 0 | 0 | 2 | 2 | 0 |

| | | | | | | | | | | | | | | | | | | |
|---|---|---|---|---|---|---|---|---|---|---|---|---|---|---|---|---|---|---|
| SG0000004 | Nqo2 | 0.00995 | 0.016278 | -0.399135835 | 1154.621241 | 875.563571 | 88 | 4 | 6 | 16 | 56 | 22 | 3 | 0 | 0 | 0 | 4 | 0 |
| SG0000003 | Cnppd1 | 0.010873 | 0.017679 | 0.373972619 | 932.3740166 | 1208.278751 | 27 | 6 | 6 | 6 | 12 | 0 | 4 | 0 | 0 | 0 | 0 | 0 |
| JSG000000 | Gclc | 0.011586 | 0.018761 | 0.335364508 | 1823.11919 | 2300.222456 | 83 | 4 | 6 | 13 | 31 | 5 | 4 | 0 | 0 | 0 | 1 | 2 |
| SG0000007 | Sft2d1 | 0.011602 | 0.018781 | -0.518362493 | 323.2573718 | 225.6866024 | 65 | 6 | 6 | 9 | 27 | 8 | 4 | 0 | 0 | 0 | 2 | 2 |
| SG0000007 | Pms2 | 0.013453 | 0.021473 | 0.364129954 | 757.5386527 | 975.0319245 | 56 | 5 | 6 | 14 | 35 | 4 | 5 | 0 | 0 | 0 | 1 | 0 |
| SG0000003 | Panx1 | 0.013837 | 0.022039 | -0.499292009 | 319.2902267 | 225.8831076 | 101 | 0 | 6 | 18 | 34 | 6 | 2 | 0 | 0 | 0 | 0 | 0 |
| SG0000003 | Zrsr2 | 0.014618 | 0.023203 | 0.341398617 | 1109.544578 | 1405.775532 | 59 | 4 | 6 | 15 | 28 | 11 | 7 | 0 | 0 | 2 | 4 | 0 |
| JSG000000 | Exoc3 | 0.015233 | 0.024124 | 0.337531228 | 1099.356429 | 1389.138489 | 65 | 4 | 6 | 9 | 37 | 6 | 2 | 0 | 0 | 0 | 0 | 0 |
| JSG000000 | Gpr3 | 0.0165 | 0.025952 | 1.216664286 | 14.60566751 | 33.94484694 | 7 | 5 | 6 | 6 | 7 | 0 | 7 | 5 | 6 | 6 | 7 | 0 |
| SG0000002 | Extl2 | 0.016547 | 0.026022 | -0.44113805 | 483.4334887 | 356.0745602 | 87 | 0 | 6 | 12 | 45 | 9 | 3 | 0 | 0 | 0 | 3 | 0 |
| JSG000000 | Rel | 0.016626 | 0.026137 | -0.5718205 | 209.8597253 | 141.1867617 | 104 | 3 | 6 | 15 | 45 | 9 | 12 | 1 | 1 | 2 | 6 | 0 |
| SG0000004 | Oaz2 | 0.017208 | 0.02696 | 0.338385348 | 942.348047 | 1191.449048 | 38 | 6 | 6 | 9 | 45 | 6 | 2 | 0 | 0 | 0 | 3 | 0 |
| JSG000000 | Ak6 | 0.017397 | 0.027236 | 0.334183484 | 1105.881252 | 1394.144367 | 53 | 6 | 6 | 7 | 23 | 0 | 3 | 0 | 0 | 0 | 0 | 0 |
| SG0000003 | Mbd3 | 0.017597 | 0.027519 | -0.594222567 | 2871.128785 | 1901.83947 | 21 | 3 | 6 | 6 | 6 | 0 | 0 | 0 | 0 | 0 | 0 | 0 |
| SG0000002 | Ints3 | 0.018521 | 0.028825 | 0.306734696 | 3265.589441 | 4039.223898 | 120 | 6 | 6 | 18 | 117 | 24 | 0 | 0 | 0 | 0 | 0 | 0 |
| SG000000C | Lgals9 | 0.018886 | 0.029345 | -0.637704826 | 1505.525114 | 967.6531697 | 54 | 4 | 6 | 12 | 24 | 2 | 0 | 0 | 0 | 0 | 0 | 0 |
| SG0000002 | Ssc4d | 0.019219 | 0.029821 | 0.651285681 | 76.49039884 | 120.1333122 | 27 | 6 | 6 | 8 | 8 | 0 | 0 | 0 | 0 | 0 | 0 | 0 |
| SG0000002 | Kansl2 | 0.019408 | 0.030088 | -0.35934316 | 1130.043424 | 880.8907764 | 56 | 6 | 6 | 8 | 20 | 7 | 2 | 0 | 0 | 0 | 0 | 1 |
| JSG000000 | Rsph9 | 0.019758 | 0.030598 | -0.976703964 | 53.04811802 | 26.9558351 | 30 | 4 | 6 | 8 | 40 | 11 | 1 | 0 | 0 | 1 | 1 | 1 |
| JSG000000 | Lamtor5 | 0.020023 | 0.03098 | 0.352669922 | 595.9540885 | 760.9866333 | 24 | 2 | 6 | 6 | 8 | 2 | 2 | 0 | 2 | 2 | 2 | 2 |
| SG0000002 | Malsu1 | 0.020122 | 0.031117 | 0.390867333 | 368.5528195 | 483.2396147 | 44 | 0 | 6 | 10 | 24 | 6 | 6 | 0 | 0 | 2 | 2 | 1 |
| SG0000003 | Asah1 | 0.02108 | 0.032464 | 0.29106726 | 3855.592267 | 4717.49096 | 118 | 2 | 6 | 8 | 33 | 9 | 4 | 0 | 0 | 1 | 3 | 0 |
| SG0000002 | Nsmaf | 0.021286 | 0.032755 | -0.348268336 | 1177.133206 | 924.6692074 | 107 | 2 | 6 | 16 | 60 | 9 | 0 | 0 | 0 | 0 | 0 | 0 |
| JSG000000 | Lims2 | 0.021412 | 0.032927 | 1.736584081 | 5.771397775 | 19.23290587 | 21 | 6 | 6 | 7 | 13 | 0 | 0 | 0 | 0 | 0 | 0 | 0 |
| SG0000002 | Gnb4 | 0.02178 | 0.033472 | -0.666224844 | 480.6284559 | 302.8696936 | 160 | 6 | 6 | 10 | 54 | 31 | 5 | 0 | 0 | 0 | 0 | 6 |
| SG0000003 | Prpf18 | 0.023095 | 0.035311 | -0.335344375 | 1715.450739 | 1359.657541 | 38 | 3 | 6 | 14 | 35 | 4 | 2 | 0 | 0 | 1 | 3 | 0 |
| JSG000000 | Polr3a | 0.02463 | 0.037508 | 0.308423644 | 1420.515506 | 1759.100884 | 35 | 4 | 6 | 9 | 17 | 6 | 2 | 1 | 1 | 1 | 1 | 1 |
| JSG000000 | Zfp599 | 0.026794 | 0.04058 | 0.417462688 | 241.0698534 | 321.9672476 | 20 | 4 | 6 | 4 | 10 | 2 | 3 | 2 | 3 | 2 | 5 | 1 |
| SG0000002 | Ccdc93 | 0.030023 | 0.045062 | -0.324474884 | 1562.08362 | 1247.462806 | 155 | 4 | 6 | 32 | 72 | 20 | 3 | 0 | 0 | 0 | 0 | 0 |
| SG0000002 | Psmd13 | 0.031212 | 0.046701 | 0.280032222 | 3205.521487 | 3892.21472 | 70 | 6 | 6 | 6 | 25 | 0 | 1 | 0 | 0 | 0 | 0 | 0 |
| SG0000002 | D16Ertd472e | 0.03133 | 0.046847 | 0.468449966 | 143.4863164 | 198.5308389 | 146 | 3 | 6 | 23 | 45 | 12 | 2 | 0 | 0 | 2 | 3 | 0 |
| JSG000000 | Pip4k2b | 0.0366 | 0.054055 | -0.28915509 | 2667.445798 | 2182.987494 | 45 | 5 | 6 | 12 | 15 | 2 | 7 | 0 | 0 | 1 | 1 | 1 |
| JSG000000 | 4833420G17Rik | 0.038808 | 0.057085 | 0.301401965 | 756.1156845 | 931.7926472 | 56 | 5 | 6 | 12 | 35 | 10 | 0 | 0 | 0 | 0 | 1 | 0 |
| SG0000002 | Cep55 | 0.039241 | 0.057661 | -0.28883338 | 2956.139616 | 2419.788622 | 56 | 3 | 6 | 8 | 20 | 1 | 2 | 0 | 1 | 0 | 2 | 0 |
| SG000000C | Rapsn | 0.040626 | 0.059539 | 4.522900143 | 1.02742554 | 23.61994967 | 45 | 3 | 6 | 15 | 33 | 6 | 0 | 0 | 0 | 0 | 0 | 0 |
| JSG000000 | Pla1a | 0.042535 | 0.062183 | -0.858846136 | 118.4934246 | 65.33649046 | 54 | 6 | 6 | 10 | 22 | 2 | 1 | 0 | 0 | 0 | 0 | 0 |
| JSG000000 | Plekhh1 | 0.045234 | 0.065824 | 0.297011139 | 1074.393203 | 1319.995693 | 49 | 5 | 6 | 7 | 25 | 11 | 2 | 0 | 0 | 0 | 3 | 1 |
| SG0000007 | Vps25 | 0.047681 | 0.069144 | -0.678898249 | 75.02044069 | 46.86092685 | 60 | 6 | 6 | 17 | 23 | 0 | 3 | 0 | 0 | 0 | 0 | 0 |
| SG0000003 | Pafah2 | 0.048305 | 0.069929 | 0.308152995 | 617.6013995 | 764.6656231 | 51 | 6 | 6 | 10 | 42 | 8 | 1 | 0 | 0 | 0 | 0 | 0 |
| SG0000002 | Gpld1 | 0.050004 | 0.07218 | 1.344540054 | 7.701946493 | 19.55903319 | 70 | 3 | 6 | 11 | 38 | 3 | 0 | 0 | 0 | 0 | 0 | 0 |
| JSG000000 | Siah2 | 0.052752 | 0.075809 | 0.417164604 | 151.8203425 | 202.7258013 | 29 | 5 | 6 | 6 | 12 | 4 | 2 | 0 | 1 | 0 | 0 | 0 |
| SG0000002 | Nans | 0.055347 | 0.079228 | 0.266146779 | 1678.740972 | 2018.83978 | 18 | 4 | 6 | 7 | 12 | 1 | 1 | 0 | 1 | 0 | 0 | 0 |
| SG0000003 | Faim | 0.056377 | 0.080541 | 0.299961051 | 494.4241232 | 608.6910639 | 63 | 6 | 6 | 10 | 41 | 5 | 1 | 0 | 0 | 0 | 1 | 1 |
| JSG000000 | Gata6 | 0.057752 | 0.082357 | 0.874819885 | 20.08995753 | 36.84054489 | 59 | 2 | 6 | 7 | 29 | 9 | 4 | 0 | 0 | 0 | 1 | 1 |
| JSG000000 | Gucy2g | 0.058044 | 0.082744 | -1.118102526 | 25.99169151 | 11.97435543 | 63 | 5 | 6 | 12 | 25 | 11 | 0 | 0 | 0 | 0 | 0 | 0 |
| JSG000000 | Trappc6b | 0.060585 | 0.086018 | -0.297019065 | 910.5242997 | 741.1052551 | 45 | 4 | 6 | 11 | 24 | 4 | 4 | 0 | 1 | 0 | 0 | 2 |
| SG000000C | Ndrg1 | 0.060776 | 0.086255 | -1.244916205 | 5523.703642 | 2330.629558 | 116 | 6 | 6 | 13 | 36 | 15 | 2 | 0 | 0 | 1 | 2 | 0 |
| SG0000003 | Oser1 | 0.061891 | 0.087739 | 0.364042598 | 215.6530244 | 277.5513186 | 55 | 4 | 6 | 8 | 23 | 4 | 2 | 0 | 0 | 0 | 0 | 0 |
| SG0000005 | P4ha3 | 0.063913 | 0.090435 | -0.668627446 | 104.3291833 | 65.63400267 | 115 | 3 | 6 | 16 | 63 | 30 | 0 | 0 | 0 | 0 | 0 | 0 |
| JSG000000 | Tomm22 | 0.066772 | 0.094197 | -0.262394258 | 3286.915528 | 2740.311926 | 9 | 6 | 6 | 6 | 8 | 0 | 1 | 0 | 0 | 0 | 0 | 0 |
| JSG000000 | Ppapdc3 | 0.068416 | 0.096273 | 1.135829406 | 9.80029402 | 21.53564216 | 25 | 5 | 6 | 11 | 19 | 0 | 1 | 0 | 0 | 0 | 0 | 0 |
| JSG000000 | Il1rl2 | 0.07025 | 0.098664 | 0.808315135 | 21.91326497 | 38.37374559 | 134 | 2 | 6 | 14 | 91 | 24 | 2 | 0 | 0 | 0 | 3 | 0 |
| SG0000002 | Rita1 | 0.070262 | 0.098669 | 0.35280575 | 259.5635131 | 331.4734564 | 45 | 6 | 6 | 13 | 27 | 6 | 3 | 0 | 0 | 2 | 3 | 0 |
| SG0000001 | Ube2i | 0.071409 | 0.100092 | 0.238082868 | 2001.122637 | 2360.172738 | 36 | 0 | 6 | 0 | 15 | 0 | 0 | 0 | 0 | 0 | 0 | 0 |
| JSG000000 | Tyw3 | 0.071666 | 0.100437 | -0.418820061 | 209.6577698 | 156.8315955 | 53 | 2 | 6 | 14 | 22 | 8 | 7 | 0 | 0 | 2 | 4 | 0 |
| SG0000006 | Lmnb2 | 0.073231 | 0.102504 | 0.257131344 | 1646.855395 | 1968.156887 | 20 | 6 | 6 | 10 | 14 | 2 | 2 | 0 | 0 | 2 | 2 | 0 |
| SG0000002 | Ctbs | 0.080061 | 0.111149 | -0.365724229 | 461.5640264 | 358.2102346 | 106 | 0 | 6 | 20 | 51 | 15 | 20 | 0 | 0 | 4 | 12 | 2 |
| JSG000000 | Whamm | 0.083237 | 0.115246 | 0.287564133 | 437.8621136 | 534.44473 | 49 | 5 | 6 | 8 | 30 | 3 | 1 | 0 | 0 | 1 | 2 | 1 |
| JSG000000 | Usp31 | 0.085924 | 0.118618 | -0.286856773 | 1406.127743 | 1152.583198 | 127 | 3 | 6 | 12 | 57 | 12 | 15 | 3 | 4 | 4 | 8 | 1 |
| SG0000002 | Rab36 | 0.087704 | 0.120902 | -0.863888756 | 41.08153839 | 22.57305138 | 35 | 6 | 6 | 8 | 30 | 2 | 2 | 0 | 0 | 0 | 0 | 0 |
| JSG000000 | Tsc2 | 0.091954 | 0.126232 | 0.233458397 | 1788.633772 | 2102.806965 | 30 | 5 | 6 | 8 | 17 | 1 | 0 | 0 | 0 | 0 | 0 | 0 |
| SG000000C | Bax | 0.092476 | 0.12688 | -0.235544437 | 3938.178956 | 3344.949045 | 16 | 5 | 6 | 7 | 13 | 2 | 0 | 0 | 0 | 0 | 0 | 0 |
| SG0000002 | Col11a2 | 0.097712 | 0.1333 | -0.972697221 | 27.68950302 | 14.10925624 | 36 | 0 | 6 | 6 | 12 | 3 | 9 | 0 | 6 | 0 | 0 | 0 |
| SG0000003 | Atp11a | 0.09853 | 0.134242 | -0.251308186 | 3434.658947 | 2885.57467 | 184 | 2 | 6 | 18 | 56 | 26 | 6 | 0 | 0 | 0 | 2 | 0 |
| SG0000002 | Krt7 | 0.099383 | 0.135321 | 1.102242309 | 252.4514832 | 541.9833237 | 31 | 5 | 6 | 9 | 18 | 3 | 13 | 4 | 4 | 4 | 6 | 0 |
| SG0000003 | Strn4 | 0.100053 | 0.136068 | -0.23250267 | 2620.072591 | 2230.093368 | 49 | 6 | 6 | 15 | 29 | 0 | 2 | 0 | 0 | 2 | 2 | 0 |
| SG0000002 | Fam114a2 | 0.100561 | 0.136655 | 0.217862494 | 1729.523812 | 2011.452085 | 106 | 6 | 6 | 10 | 54 | 8 | 0 | 0 | 0 | 0 | 0 | 0 |
| SG0000005 | Mia3 | 0.111375 | 0.150075 | 0.255835133 | 1282.678203 | 1531.55206 | 58 | 4 | 6 | 10 | 23 | 4 | 2 | 0 | 0 | 0 | 2 | 0 |
| SG0000003 | Klhdc8b | 0.113189 | 0.152199 | 0.497814668 | 66.33540882 | 93.67043938 | 16 | 6 | 6 | 7 | 15 | 0 | 0 | 0 | 0 | 0 | 4 | 0 |
| SG0000007 | Lyrm9 | 0.114173 | 0.153384 | -0.286043533 | 489.1911658 | 401.2092361 | 37 | 6 | 6 | 7 | 16 | 2 | 16 | 4 | 4 | 5 | 8 | 1 |
| SG0000004 | Tmem53 | 0.123316 | 0.164471 | -0.636991208 | 48.77326689 | 31.36377904 | 87 | 6 | 6 | 15 | 36 | 0 | 0 | 0 | 0 | 0 | 0 | 0 |
| JSG000000 | Dnmbp | 0.124818 | 0.166362 | -0.259525305 | 680.3822893 | 568.3660113 | 84 | 5 | 6 | 14 | 44 | 7 | 0 | 0 | 0 | 0 | 0 | 0 |
| SG0000001 | Zfp687 | 0.136662 | 0.180643 | 0.201932734 | 1353.546999 | 1556.901548 | 14 | 4 | 6 | 4 | 4 | 0 | 4 | 0 | 0 | 0 | 0 | 0 |
| SG0000003 | Nono | 0.137571 | 0.181736 | 0.181195432 | 9637.531083 | 10927.25435 | 43 | 6 | 6 | 8 | 26 | 3 | 0 | 0 | 0 | 0 | 0 | 0 |
| SG000000C | Wdr83 | 0.139784 | 0.184361 | -0.27253829 | 458.2870218 | 379.3982807 | 16 | 6 | 6 | 6 | 8 | 0 | 2 | 0 | 0 | 0 | 0 | 0 |
| SG0000003 | Hexdc | 0.139972 | 0.184569 | 0.327977374 | 152.6737823 | 191.6441171 | 65 | 6 | 6 | 9 | 18 | 9 | 0 | 0 | 0 | 0 | 0 | 0 |
| SG0000002 | Wdr53 | 0.141525 | 0.186479 | 0.264821611 | 324.7264862 | 390.1548065 | 34 | 4 | 6 | 4 | 26 | 2 | 0 | 0 | 0 | 0 | 0 | 0 |
| JSG000000 | 5730508B09Rik | 0.14574 | 0.191427 | -0.434273545 | 101.8821301 | 75.39953246 | 87 | 2 | 6 | 10 | 24 | 15 | 8 | 0 | 0 | 3 | 7 | 1 |
| JSG000000 | Fbxo46 | 0.148811 | 0.195118 | -0.323011055 | 289.7751913 | 231.6462024 | 28 | 5 | 6 | 9 | 20 | 3 | 1 | 0 | 0 | 1 | 1 | 0 |
| JSG000000 | Smad6 | 0.158603 | 0.206777 | 0.365833182 | 113.2843714 | 145.9811432 | 55 | 4 | 6 | 9 | 41 | 7 | 2 | 0 | 1 | 1 | 3 | 0 |
| SG0000002 | Pgam5 | 0.159637 | 0.208004 | -0.196619932 | 2649.643648 | 2312.059318 | 48 | 6 | 6 | 12 | 22 | 2 | 6 | 2 | 2 | 2 | 2 | 2 |
| JSG000000 | Pum3 | 0.164267 | 0.213569 | 0.180317319 | 3311.568934 | 3752.448336 | 72 | 5 | 6 | 11 | 34 | 8 | 3 | 0 | 0 | 1 | 1 | 0 |
| SG0000001 | Etv4 | 0.170758 | 0.221207 | 0.178943486 | 1953.692798 | 2211.686833 | 45 | 6 | 6 | 12 | 45 | 3 | 0 | 0 | 0 | 0 | 3 | 3 |
| JSG000000 | Sart3 | 0.172806 | 0.223648 | 0.182962183 | 2205.314174 | 2503.499874 | 50 | 4 | 6 | 8 | 23 | 0 | 0 | 0 | 0 | 0 | 2 | 0 |
| SG0000004 | D11Wsu47e | 0.179933 | 0.231868 | 0.2127529 | 595.8329916 | 690.5093304 | 32 | 6 | 6 | 6 | 12 | 2 | 0 | 0 | 0 | 0 | 0 | 0 |
| JSG000000 | Txnl4b | 0.184319 | 0.236906 | 0.289794437 | 171.8291879 | 210.0553541 | 13 | 5 | 6 | 6 | 7 | 1 | 0 | 0 | 0 | 0 | 0 | 0 |
| SG0000005 | Ech1 | 0.185071 | 0.237754 | 0.18807963 | 1124.643202 | 1281.245558 | 17 | 6 | 6 | 6 | 32 | 2 | 0 | 0 | 0 | 0 | 0 | 0 |
| SG000000C | Vars | 0.192463 | 0.246401 | 0.226998083 | 6185.74903 | 7239.782641 | 20 | 6 | 6 | 8 | 17 | 1 | 0 | 0 | 0 | 0 | 0 | 0 |
| JSG000000 | Tram2 | 0.196474 | 0.250873 | 0.202702846 | 488.4284001 | 562.1090082 | 99 | 2 | 6 | 12 | 43 | 16 | 3 | 0 | 0 | 1 | 1 | 0 |
| SG0000004 | Hps4 | 0.196495 | 0.250882 | 0.212637591 | 539.2634138 | 624.9010432 | 38 | 6 | 6 | 14 | 44 | 4 | 0 | 0 | 0 | 0 | 0 | 0 |
| JSG000000 | BC048403 | 0.211251 | 0.267547 | -0.208084674 | 630.3014194 | 545.6429603 | 30 | 2 | 6 | 10 | 18 | 9 | 8 | 0 | 0 | 2 | 4 | 5 |
| SG0000003 | Nova2 | 0.214723 | 0.271561 | -0.27775471 | 212.2759623 | 175.1008762 | 43 | 5 | 6 | 8 | 16 | 5 | 3 | 0 | 0 | 0 | 0 | 3 |
| SG000000C | Map2k7 | 0.219495 | 0.276717 | -0.168924463 | 2058.49398 | 1831.041851 | 31 | 0 | 6 | 7 | 19 | 0 | 7 | 0 | 0 | 3 | 7 | 0 |
| SG0000003 | Bsn | 0.226298 | 0.284592 | 0.389316346 | 53.64103851 | 70.25756199 | 95 | 6 | 6 | 15 | 57 | 11 | 8 | 0 | 0 | 0 | 0 | 2 |
| SG0000003 | Tstd2 | 0.243 | 0.30315 | 0.167569447 | 1074.184423 | 1206.485981 | 88 | 4 | 6 | 8 | 24 | 2 | 2 | 0 | 0 | 0 | 2 | 0 |
| JSG000000 | Apol8 | 0.243063 | 0.303207 | 0.631502665 | 17.1621089 | 26.58715687 | 18 | 6 | 6 | 8 | 14 | 2 | 0 | 0 | 0 | 0 | 0 | 0 |
| SG0000003 | Osbpl7 | 0.24487 | 0.305227 | -0.214473086 | 423.2279735 | 364.7636331 | 38 | 4 | 6 | 12 | 20 | 2 | 0 | 0 | 0 | 0 | 0 | 0 |
| SG000000C | Tspan33 | 0.26825 | 0.330573 | -0.282225587 | 169.5512801 | 139.425659 | 54 | 2 | 6 | 6 | 18 | 4 | 0 | 0 | 0 | 0 | 0 | 0 |
| JSG000000 | 9530077C05Rik | 0.273217 | 0.335904 | 0.3349392 | 59.72228913 | 75.32917165 | 47 | 5 | 6 | 13 | 22 | 7 | 0 | 0 | 0 | 0 | 0 | 0 |
| SG000000C | Vps4b | 0.276176 | 0.339174 | 0.134805022 | 3081.358164 | 3383.159998 | 125 | 0 | 6 | 11 | 48 | 28 | 15 | 0 | 1 | 4 | 5 | 3 |
| SG0000002 | Guf1 | 0.276392 | 0.339417 | 0.147004511 | 1179.428529 | 1305.943524 | 211 | 0 | 6 | 10 | 82 | 24 | 7 | 0 | 0 | 0 | 1 | 0 |
| SG0000002 | Csnk1e | 0.279624 | 0.342846 | 0.141170894 | 2496.591118 | 2753.240154 | 62 | 6 | 6 | 24 | 33 | 1 | 5 | 0 | 0 | 0 | 6 | 1 |
| JSG000000 | Dlat | 0.28864 | 0.352596 | 0.135831968 | 2388.912937 | 2624.76133 | 48 | 4 | 6 | 11 | 27 | 5 | 8 | 0 | 1 | 3 | 6 | 1 |
| SG0000002 | Mtif2 | 0.290509 | 0.354638 | 0.154810352 | 971.7821578 | 1081.861038 | 88 | 4 | 6 | 12 | 42 | 12 | 0 | 0 | 0 | 0 | 0 | 0 |
| SG0000003 | Rbm10 | 0.298304 | 0.363096 | 0.144473663 | 1599.832079 | 1768.338111 | 132 | 3 | 6 | 18 | 66 | 12 | 0 | 0 | 0 | 0 | 0 | 0 |
| SG0000004 | Zfp738 | 0.304422 | 0.369456 | 0.189490859 | 271.1403122 | 309.1978918 | 72 | 3 | 6 | 19 | 41 | 7 | 3 | 0 | 0 | 0 | 0 | 2 |
| JSG000000 | Akirin2 | 0.306146 | 0.371272 | 0.135035074 | 1827.594257 | 2006.91691 | 45 | 5 | 6 | 8 | 18 | 8 | 3 | 1 | 1 | 1 | 3 | 1 |
| SG0000002 | Fam111a | 0.308801 | 0.374112 | -0.266035368 | 2058.845742 | 1712.139643 | 112 | 3 | 6 | 12 | 44 | 12 | 3 | 0 | 0 | 3 | 6 | 0 |
| JSG000000 | Ckap2l | 0.31397 | 0.379579 | 0.13741058 | 1543.728692 | 1697.992167 | 60 | 6 | 6 | 17 | 49 | 4 | 1 | 0 | 0 | 0 | 0 | 0 |
| SG0000006 | Sco1 | 0.33135 | 0.398154 | -0.165548977 | 685.8286573 | 611.4774142 | 38 | 2 | 6 | 6 | 20 | 4 | 2 | 0 | 0 | 2 | 2 | 0 |
| JSG000000 | Fam151b | 0.341607 | 0.408933 | -0.541914504 | 25.24768345 | 17.34159406 | 70 | 2 | 6 | 8 | 35 | 1 | 0 | 0 | 0 | 0 | 0 | 0 |
| SG0000002 | Qtrtd1 | 0.349291 | 0.416934 | 0.137102021 | 955.5758987 | 1050.840998 | 56 | 3 | 6 | 15 | 35 | 6 | 6 | 0 | 0 | 2 | 2 | 1 |
| SG0000001 | Igfbp4 | 0.352849 | 0.420731 | -0.218580337 | 1322.012002 | 1136.151275 | 22 | 6 | 6 | 7 | 14 | 0 | 4 | 0 | 0 | 1 | 2 | 0 |
| JSG000000 | Mtfr2 | 0.378454 | 0.447302 | 0.136892801 | 618.4914491 | 680.0526524 | 26 | 3 | 6 | 6 | 15 | 6 | 2 | 0 | 0 | 0 | 1 | 1 |
| JSG000000 | Vwa7 | 0.383182 | 0.452025 | -0.589492107 | 17.9146475 | 11.90565958 | 12 | 4 | 6 | 6 | 10 | 0 | 6 | 4 | 4 | 4 | 6 | 0 |
| JSG000000 | Mob3a | 0.383433 | 0.452263 | -0.146756968 | 639.0300917 | 577.2222456 | 18 | 5 | 6 | 5 | 16 | 1 | 0 | 0 | 0 | 0 | 0 | 0 |
| SG0000002 | Wnt10b | 0.387365 | 0.45618 | -0.678005443 | 23.8801263 | 14.92576652 | 21 | 6 | 6 | 10 | 14 | 0 | 1 | 0 | 0 | 0 | 0 | 0 |
| JSG000000 | Mpc2 | 0.394532 | 0.463249 | -0.163226781 | 395.9676323 | 353.6091787 | 90 | 2 | 6 | 12 | 48 | 6 | 0 | 0 | 0 | 0 | 0 | 0 |
| SG0000002 | Psmd12 | 0.41105 | 0.480063 | 0.10131712 | 4296.012946 | 4608.558169 | 60 | 2 | 6 | 5 | 23 | 9 | 0 | 0 | 0 | 0 | 0 | 0 |
| SG0000006 | Ddx3y | 0.436679 | 0.505938 | -0.315303998 | 74.96576385 | 60.24860692 | 126 | 4 | 6 | 17 | 62 | 13 | 20 | 0 | 2 | 3 | 6 | 1 |
| SG0000002 | Mrpl9 | 0.450391 | 0.519541 | 0.109146572 | 1161.148842 | 1252.403537 | 18 | 6 | 6 | 7 | 13 | 5 | 1 | 0 | 0 | 0 | 0 | 0 |
| JSG000000 | Ddi2 | 0.468645 | 0.537411 | -0.109065483 | 1646.529607 | 1526.643252 | 107 | 4 | 6 | 24 | 63 | 10 | 9 | 0 | 0 | 0 | 1 | 2 |
| SG0000001 | Galns | 0.476784 | 0.545278 | -0.138889784 | 391.8319541 | 355.8687551 | 22 | 4 | 6 | 6 | 18 | 1 | 0 | 0 | 0 | 0 | 0 | 0 |

| | | | | | | | | | | | | | | | | | |
|---|---|---|---|---|---|---|---|---|---|---|---|---|---|---|---|---|---|
| ISG000000 | Ndufaf7 | 0.522679 | 0.589594 | -0.110125801 | 928.3263851 | 860.1011429 | 27 | 4 | 6 | 5 | 9 | 1 | 1 | 0 | 0 | 0 | 0 | 0 |
| ISG000000 | Rgl3 | 0.524099 | 0.590865 | 0.220106315 | 90.7344928 | 105.6892994 | 29 | 5 | 6 | 9 | 17 | 1 | 0 | 0 | 0 | 0 | 0 | 0 |
| SG0000002 | Jtb | 0.52617 | 0.592752 | -0.111808798 | 532.1584189 | 492.4738052 | 12 | 6 | 6 | 6 | 13 | 2 | 0 | 0 | 0 | 0 | 1 | 0 |
| SG0000005 | Flot1 | 0.532271 | 0.598834 | -0.122675736 | 896.7364997 | 823.6368538 | 19 | 6 | 6 | 6 | 9 | 3 | 1 | 0 | 0 | 0 | 0 | 0 |
| SG0000002 | Rabl3 | 0.537692 | 0.604098 | 0.092515249 | 536.9711293 | 572.5333654 | 106 | 2 | 6 | 18 | 64 | 12 | 2 | 0 | 0 | 2 | 4 | 0 |
| SG0000002 | Adprm | 0.541539 | 0.60762 | -0.113806379 | 489.0576423 | 451.9609533 | 52 | 4 | 6 | 4 | 18 | 2 | 1 | 0 | 0 | 0 | 0 | 0 |
| SG0000002 | Srp68 | 0.545523 | 0.611356 | 0.077717042 | 2962.971624 | 3126.962357 | 66 | 6 | 6 | 14 | 42 | 4 | 0 | 0 | 0 | 0 | 0 | 0 |
| ISG000000 | Dkk3 | 0.556989 | 0.62211 | 0.131978607 | 182.6676284 | 200.1663684 | 39 | 6 | 6 | 8 | 14 | 7 | 4 | 1 | 1 | 1 | 2 | 0 |
| SG0000000 | Ubqln4 | 0.579173 | 0.643078 | -0.071279139 | 5178.415014 | 4928.782959 | 29 | 6 | 6 | 7 | 13 | 2 | 0 | 0 | 0 | 0 | 0 | 1 |
| SG0000003 | Igsf9b | 0.580028 | 0.643868 | -0.201745383 | 114.9433736 | 99.94303366 | 58 | 4 | 6 | 13 | 33 | 6 | 0 | 0 | 0 | 0 | 0 | 0 |
| SG0000002 | Cyp2u1 | 0.599008 | 0.661779 | 0.211934576 | 35.6049352 | 41.23907015 | 25 | 2 | 6 | 5 | 12 | 1 | 1 | 0 | 0 | 0 | 1 | 0 |
| SG0000000 | Nsun5 | 0.626193 | 0.687488 | -0.087930313 | 541.5428123 | 509.5222271 | 20 | 6 | 6 | 9 | 12 | 0 | 2 | 0 | 0 | 0 | 0 | 0 |
| SG0000003 | Renbp | 0.63047 | 0.691548 | 0.071055612 | 1412.198362 | 1483.493248 | 18 | 6 | 6 | 8 | 20 | 0 | 0 | 0 | 0 | 0 | 0 | 0 |
| SG0000005 | B4galt3 | 0.641123 | 0.701086 | -0.073593319 | 951.7030054 | 904.3731169 | 21 | 6 | 6 | 15 | 15 | 0 | 6 | 0 | 0 | 6 | 6 | 0 |
| SG0000003 | Hdlbp | 0.641464 | 0.701288 | 0.058051159 | 13183.24911 | 13724.53373 | 47 | 3 | 6 | 5 | 23 | 8 | 0 | 0 | 0 | 0 | 0 | 0 |
| SG0000002 | Lsm12 | 0.652811 | 0.711565 | 0.053940473 | 2016.694454 | 2093.523326 | 66 | 6 | 6 | 10 | 22 | 4 | 2 | 0 | 0 | 0 | 0 | 0 |
| SG0000002 | Arpc5l | 0.660435 | 0.718738 | -0.06758359 | 1030.186745 | 983.0401813 | 20 | 4 | 6 | 4 | 8 | 2 | 2 | 0 | 1 | 0 | 0 | 0 |
| SG0000002 | Usp1 | 0.667241 | 0.725176 | -0.063452747 | 3501.213169 | 3350.559796 | 60 | 2 | 6 | 6 | 24 | 12 | 8 | 0 | 0 | 2 | 4 | 4 |
| SG0000002 | Pcsk4 | 0.684164 | 0.740226 | -0.145169892 | 88.93762071 | 80.42387875 | 12 | 6 | 6 | 6 | 9 | 0 | 0 | 0 | 0 | 0 | 0 | 0 |
| ISG000000 | Them4 | 0.688834 | 0.744039 | -0.094239804 | 171.2919846 | 160.4604602 | 43 | 2 | 6 | 7 | 24 | 3 | 1 | 1 | 1 | 1 | 1 | 0 |
| SG0000002 | Hnrnpab | 0.703987 | 0.757755 | 0.045804384 | 17926.10009 | 18504.3703 | 18 | 0 | 6 | 0 | 3 | 0 | 7 | 0 | 3 | 0 | 3 | 0 |
| SG0000004 | Dcp1b | 0.704927 | 0.758407 | 0.080146429 | 253.4785722 | 267.9586192 | 127 | 4 | 6 | 13 | 57 | 14 | 3 | 0 | 0 | 1 | 1 | 0 |
| SG0000002 | Echs1 | 0.709583 | 0.762176 | 0.045310994 | 1901.873261 | 1962.553664 | 15 | 6 | 6 | 6 | 13 | 0 | 0 | 0 | 0 | 0 | 1 | 0 |
| ISG000000 | Klhl28 | 0.715085 | 0.767534 | -0.100981828 | 218.1522578 | 203.4047791 | 63 | 1 | 6 | 12 | 41 | 10 | 0 | 0 | 0 | 0 | 0 | 0 |
| ISG000000 | Zfp963 | 0.728644 | 0.78003 | 0.150584021 | 34.23972161 | 38.00673232 | 43 | 4 | 6 | 13 | 27 | 5 | 3 | 0 | 0 | 1 | 1 | 1 |
| SG0000002 | Tmem14a | 0.749027 | 0.797371 | 0.075002224 | 116.3939271 | 122.6050166 | 58 | 4 | 6 | 12 | 26 | 0 | 10 | 0 | 2 | 0 | 2 | 0 |
| ISG000000 | Pfkl | 0.754523 | 0.802031 | -0.058785628 | 9204.188149 | 8836.682485 | 17 | 6 | 6 | 6 | 10 | 3 | 3 | 0 | 0 | 0 | 0 | 1 |
| SG0000002 | Ndufa5 | 0.762958 | 0.80946 | 0.042162538 | 661.5753086 | 681.1950392 | 34 | 6 | 6 | 10 | 16 | 4 | 0 | 0 | 0 | 0 | 0 | 0 |
| SG0000003 | Gnptg | 0.764463 | 0.810865 | -0.056860912 | 348.4545281 | 334.9880173 | 20 | 0 | 6 | 10 | 15 | 2 | 0 | 0 | 0 | 0 | 1 | 0 |
| SG0000004 | Sgtb | 0.769641 | 0.815393 | -0.081871053 | 248.3966588 | 234.6929842 | 63 | 4 | 6 | 10 | 30 | 1 | 8 | 0 | 0 | 2 | 2 | 1 |
| SG0000002 | Ccl20 | 0.789213 | 0.832433 | -0.163256913 | 22.95609862 | 20.49995247 | 30 | 0 | 6 | 3 | 18 | 0 | 19 | 0 | 6 | 3 | 11 | 0 |
| SG0000006 | Suds3 | 0.809811 | 0.849656 | -0.035688088 | 2503.752734 | 2442.576935 | 39 | 6 | 6 | 6 | 16 | 2 | 1 | 0 | 0 | 0 | 0 | 0 |
| SG0000002 | Dnajb2 | 0.83214 | 0.868708 | -0.040124564 | 413.2580927 | 401.9228244 | 67 | 0 | 6 | 6 | 6 | 12 | 7 | 0 | 0 | 0 | 0 | 0 |
| SG0000001 | Mrpl52 | 0.863035 | 0.893462 | 0.02152033 | 759.1394815 | 770.5482595 | 18 | 3 | 6 | 6 | 9 | 0 | 0 | 0 | 0 | 0 | 0 | 0 |
| ISG000000 | Zfp433 | 0.871822 | 0.9011 | 0.085919641 | 19.85706719 | 21.07557802 | 38 | 6 | 6 | 10 | 26 | 4 | 1 | 0 | 0 | 0 | 1 | 0 |
| SG0000002 | Syngr1 | 0.934232 | 0.951984 | 0.024839934 | 1031.342628 | 1049.25376 | 47 | 6 | 6 | 8 | 18 | 0 | 2 | 0 | 0 | 0 | 0 | 0 |
| SG0000004 | 6330416G13Rik | 0.944025 | 0.960378 | -0.011034124 | 261.8355837 | 259.8406327 | 55 | 4 | 6 | 14 | 24 | 6 | 2 | 0 | 0 | 0 | 4 | 2 |
| ISG000000 | Champ1 | 0.978471 | 0.990358 | -0.000315079 | 2822.846033 | 2822.229602 | 31 | 6 | 6 | 6 | 17 | 3 | 1 | 0 | 0 | 0 | 1 | 1 |
| ISG000000 | Nsg1 | 0.985914 | 0.996533 | -0.003462637 | 60.26522902 | 60.12075886 | 47 | 3 | 6 | 9 | 15 | 0 | 9 | 0 | 0 | 0 | 0 | 0 |
| ISG000000 | Riok1 | 0.989693 | 0.999176 | -0.004504602 | 892.6277446 | 889.8449932 | 48 | 4 | 6 | 9 | 31 | 4 | 0 | 0 | 0 | 0 | 4 | 0 |
| SG0000002 | Dsc2 | 1 | 1 | -0.010058144 | 309.0420223 | 306.8949444 | 257 | 6 | 6 | 26 | 84 | 9 | 27 | 0 | 0 | 1 | 9 | 0 |
| SG0000002 | Eif3i | 1 | 1 | 0.002226053 | 4766.057839 | 4773.41746 | 10 | 6 | 6 | 6 | 11 | 1 | 0 | 0 | 0 | 0 | 0 | 0 |
| ISG000000 | Vasp | 1 | 1 | 0.003701568 | 3252.597469 | 3260.953475 | 18 | 5 | 6 | 6 | 9 | 0 | 0 | 0 | 0 | 0 | 0 | 0 |
| ISG000000 | Icam1 | 8.2E-252 | 4E-249 | 5.285213178 | 67.71384009 | 2640.493748 | 25 | 4 | 5 | 8 | 13 | 1 | 4 | 0 | 0 | 2 | 2 | 1 |
| ISG000000 | Lrrc34 | 1E-248 | 4.9E-246 | 11.86761651 | 0.307509278 | 1149.12367 | 58 | 4 | 5 | 11 | 30 | 1 | 0 | 0 | 0 | 0 | 1 | 0 |
| SG0000002 | Otx2 | 2.7E-198 | 7.9E-196 | 9.65383946 | 0 | 804.5550988 | 29 | 2 | 5 | 2 | 9 | 0 | 4 | 0 | 3 | 0 | 0 | 0 |
| SG0000005 | Gm1564 | 1.9E-183 | 4.9E-181 | 9.520706349 | 0 | 733.544635 | 69 | 3 | 5 | 12 | 33 | 6 | 10 | 0 | 0 | 1 | 3 | 0 |
| ISG000000 | Ogdhl | 7.4E-140 | 9.9E-138 | 3.311560121 | 520.2008383 | 5164.757994 | 19 | 4 | 5 | 4 | 11 | 1 | 0 | 0 | 0 | 0 | 0 | 0 |
| SG0000002 | Ppfia4 | 1E-120 | 1E-118 | 5.82027347 | 10.41765614 | 588.636131 | 75 | 5 | 5 | 14 | 59 | 6 | 3 | 0 | 0 | 0 | 0 | 0 |
| SG0000003 | Tcfl5 | 1.5E-106 | 1.3E-104 | 5.911159423 | 7.852684213 | 472.5572478 | 44 | 3 | 5 | 8 | 25 | 3 | 1 | 0 | 0 | 1 | 1 | 0 |
| SG0000000 | Tcea3 | 6.6E-103 | 4.9E-101 | 5.741926372 | 9.071847213 | 485.4983946 | 27 | 4 | 5 | 7 | 18 | 4 | 0 | 0 | 0 | 0 | 0 | 0 |
| SG0000002 | Tgfa | 1.55E-95 | 9.94E-94 | -4.524245538 | 3377.080524 | 146.7600333 | 88 | 2 | 5 | 12 | 36 | 9 | 4 | 1 | 1 | 2 | 5 | 0 |
| SG0000000 | Osgep | 4.32E-95 | 2.72E-93 | 2.8517957 | 302.8843031 | 2186.516777 | 82 | 5 | 5 | 5 | 28 | 14 | 9 | 0 | 0 | 0 | 1 | 1 |
| ISG000000 | Dock6 | 2.07E-94 | 1.28E-92 | 3.065198979 | 627.9948859 | 5256.212833 | 41 | 5 | 5 | 7 | 24 | 0 | 1 | 0 | 0 | 0 | 0 | 0 |
| ISG000000 | Acsl5 | 9.67E-91 | 5.57E-89 | -3.935354083 | 10006.27932 | 654.0530866 | 73 | 4 | 5 | 13 | 35 | 6 | 0 | 0 | 0 | 0 | 0 | 0 |
| SG0000003 | Zbed5 | 4.64E-89 | 2.56E-87 | 4.033928262 | 38.39142038 | 628.8797163 | 17 | 4 | 5 | 4 | 8 | 1 | 0 | 0 | 0 | 0 | 0 | 0 |
| SG0000002 | Parp1 | 1.81E-87 | 9.6E-86 | 2.283879373 | 5886.300729 | 28665.44209 | 54 | 2 | 5 | 13 | 37 | 4 | 1 | 0 | 0 | 0 | 0 | 0 |
| SG0000002 | Obsl1 | 1.08E-81 | 5.04E-80 | 3.735834336 | 199.2120344 | 2654.078504 | 42 | 5 | 5 | 6 | 20 | 0 | 0 | 0 | 0 | 0 | 0 | 0 |
| ISG000000 | Fdft1 | 1.02E-79 | 4.53E-78 | 2.412721033 | 667.4089351 | 3553.803566 | 43 | 1 | 5 | 3 | 16 | 5 | 3 | 0 | 1 | 0 | 0 | 0 |
| ISG000000 | Piwil2 | 6.25E-79 | 2.75E-77 | 6.695164825 | 6.951904602 | 720.3594239 | 89 | 2 | 5 | 16 | 61 | 14 | 1 | 0 | 0 | 1 | 1 | 0 |
| ISG000000 | Ncan | 1.38E-78 | 6.05E-77 | 7.941964196 | 0 | 244.9061823 | 21 | 5 | 5 | 6 | 13 | 0 | 0 | 0 | 0 | 0 | 0 | 0 |
| ISG000000 | Clec4a1 | 1.21E-77 | 5.23E-76 | -9.654579116 | 559.9911828 | 0.694805551 | 29 | 1 | 5 | 3 | 17 | 1 | 1 | 0 | 0 | 0 | 0 | 0 |
| SG0000003 | Dis3 | 3.74E-74 | 1.52E-72 | 2.214783035 | 1317.156106 | 6114.393388 | 60 | 4 | 5 | 11 | 25 | 3 | 2 | 0 | 0 | 0 | 1 | 0 |
| SG0000002 | Anxa1 | 3.77E-73 | 1.49E-71 | -4.611751294 | 29384.94229 | 1201.847516 | 40 | 3 | 5 | 6 | 22 | 2 | 0 | 0 | 0 | 0 | 0 | 0 |
| ISG000000 | Ypel2 | 4.23E-73 | 1.66E-71 | -3.36833345 | 6914.994985 | 669.6108246 | 63 | 4 | 5 | 5 | 34 | 8 | 4 | 0 | 0 | 0 | 1 | 3 |
| SG0000004 | Adrbk2 | 2.29E-70 | 8.24E-69 | 8.468891519 | 0.649984458 | 230.2996879 | 114 | 3 | 5 | 8 | 44 | 22 | 8 | 3 | 4 | 3 | 7 | 0 |
| SG0000002 | Ociad2 | 6.54E-70 | 2.32E-68 | -3.957173963 | 1499.940784 | 96.57084979 | 148 | 5 | 5 | 15 | 40 | 0 | 18 | 0 | 0 | 0 | 0 | 0 |
| SG0000000 | Htra1 | 2.74E-67 | 9.23E-66 | -4.03693956 | 6953.000449 | 423.5769893 | 58 | 4 | 5 | 9 | 25 | 7 | 0 | 0 | 0 | 0 | 0 | 0 |
| ISG000000 | C1rl | 4.72E-66 | 1.53E-64 | -5.005848224 | 1813.346413 | 56.43783015 | 26 | 1 | 5 | 4 | 13 | 2 | 4 | 0 | 0 | 0 | 0 | 0 |
| SG0000002 | Gucy2e | 2.91E-60 | 8.14E-59 | -7.356995602 | 2060.274709 | 12.56749662 | 52 | 0 | 5 | 0 | 2 | 5 | 7 | 0 | 1 | 0 | 1 | 2 |
| ISG000000 | Abcg1 | 3.08E-60 | 8.61E-59 | 3.157382526 | 64.54138328 | 575.8443597 | 47 | 3 | 5 | 9 | 19 | 5 | 0 | 0 | 0 | 0 | 1 | 0 |
| ISG000000 | Atp12a | 4.27E-60 | 1.18E-58 | -4.460154699 | 662.0997971 | 30.08036523 | 33 | 4 | 5 | 7 | 17 | 3 | 3 | 0 | 0 | 2 | 2 | 0 |
| ISG000000 | Erap1 | 2.64E-53 | 6.01E-52 | -3.042191792 | 1881.016038 | 228.3502632 | 104 | 3 | 5 | 13 | 65 | 11 | 0 | 0 | 0 | 0 | 0 | 0 |
| ISG000000 | Ipo5 | 5.39E-51 | 1.15E-49 | 1.72915265 | 11397.3842 | 37786.09424 | 64 | 3 | 5 | 8 | 26 | 7 | 2 | 0 | 0 | 0 | 1 | 0 |
| ISG000000 | Gnpat | 3.61E-47 | 6.86E-46 | 1.766454829 | 1522.167189 | 5178.6701 | 30 | 3 | 5 | 4 | 9 | 5 | 1 | 0 | 0 | 0 | 0 | 0 |
| ISG000000 | Dcaf4 | 6.65E-45 | 1.16E-43 | 1.930878227 | 482.0669375 | 1838.05969 | 21 | 5 | 5 | 6 | 15 | 1 | 0 | 0 | 0 | 0 | 0 | 0 |
| SG0000002 | Plin2 | 6.87E-45 | 1.2E-43 | -3.491691251 | 16578.35254 | 1473.796662 | 23 | 4 | 5 | 6 | 13 | 2 | 0 | 0 | 0 | 0 | 0 | 0 |
| ISG000000 | Kank4 | 2.54E-44 | 4.34E-43 | 4.152522739 | 12.76061185 | 226.9370477 | 69 | 4 | 5 | 8 | 28 | 4 | 1 | 0 | 0 | 0 | 0 | 0 |
| SG0000002 | Hnf1a | 1.19E-40 | 1.8E-39 | 6.867168204 | 0 | 115.7410552 | 29 | 5 | 5 | 7 | 18 | 4 | 2 | 0 | 0 | 0 | 0 | 0 |
| SG0000003 | Fam102a | 4.57E-39 | 6.54E-38 | -2.19999482 | 3664.551535 | 797.5472141 | 26 | 5 | 5 | 8 | 14 | 4 | 0 | 0 | 0 | 0 | 0 | 0 |
| SG0000000 | Comt | 5.51E-39 | 7.84E-38 | -2.352543424 | 2082.125455 | 407.6812676 | 33 | 5 | 5 | 7 | 17 | 8 | 4 | 0 | 0 | 0 | 1 | 0 |
| ISG000000 | Rnf219 | 5.35E-37 | 7.13E-36 | 2.204351996 | 139.1895639 | 641.4794466 | 86 | 2 | 5 | 11 | 39 | 6 | 3 | 0 | 0 | 0 | 1 | 0 |
| SG0000000 | Ampd3 | 2.25E-35 | 2.84E-34 | -2.560805434 | 2188.076308 | 370.8371333 | 80 | 0 | 5 | 6 | 50 | 11 | 1 | 0 | 0 | 0 | 2 | 0 |
| ISG000000 | Slc24a1 | 2.67E-35 | 3.37E-34 | 6.63637016 | 0 | 98.48245189 | 42 | 5 | 5 | 6 | 15 | 0 | 2 | 0 | 0 | 1 | 1 | 0 |
| ISG000000 | Rhd | 1.11E-34 | 1.37E-33 | 6.815598203 | 0.970711415 | 109.3426599 | 28 | 4 | 5 | 8 | 23 | 2 | 0 | 0 | 0 | 0 | 0 | 0 |
| ISG000000 | Tdp2 | 2.11E-34 | 2.57E-33 | 1.596972164 | 818.7633256 | 2476.822618 | 21 | 5 | 5 | 5 | 11 | 2 | 0 | 0 | 0 | 0 | 0 | 0 |
| SG0000002 | Dtymk | 3.57E-34 | 4.29E-33 | 1.53340682 | 1260.938947 | 3650.022212 | 29 | 5 | 5 | 7 | 19 | 4 | 3 | 0 | 0 | 0 | 0 | 0 |
| ISG000000 | Slc25a27 | 6.42E-34 | 7.62E-33 | 4.684850859 | 5.512072078 | 141.7736504 | 36 | 2 | 5 | 6 | 19 | 4 | 2 | 1 | 1 | 1 | 1 | 0 |
| SG0000002 | Rpa2 | 1.14E-33 | 1.33E-32 | 1.575649855 | 930.1387434 | 2772.462009 | 15 | 5 | 5 | 5 | 13 | 1 | 0 | 0 | 0 | 0 | 0 | 0 |
| SG0000003 | Flna | 9.28E-33 | 1.05E-31 | -1.788886909 | 36224.69296 | 10483.25367 | 63 | 0 | 5 | 6 | 32 | 5 | 0 | 0 | 0 | 0 | 2 | 1 |
| ISG000000 | Ano9 | 1.31E-31 | 1.42E-30 | 4.594320133 | 67.11457948 | 1621.231537 | 18 | 4 | 5 | 8 | 8 | 2 | 2 | 0 | 0 | 1 | 1 | 0 |
| ISG000000 | Ephx2 | 2.58E-31 | 2.76E-30 | 6.475292652 | 25.72518197 | 2288.841791 | 46 | 2 | 5 | 10 | 25 | 5 | 0 | 0 | 0 | 0 | 0 | 0 |
| SG0000003 | Esyt3 | 6.27E-31 | 6.6E-30 | -2.148946155 | 1373.435504 | 309.6784741 | 49 | 4 | 5 | 9 | 25 | 8 | 4 | 2 | 2 | 2 | 5 | 1 |
| SG0000003 | Brd7 | 1.21E-29 | 1.2E-28 | 1.437420934 | 1184.580043 | 3208.272355 | 77 | 2 | 5 | 6 | 32 | 10 | 2 | 0 | 0 | 0 | 7 | 2 |
| ISG000000 | Zfp280b | 1.35E-29 | 1.34E-28 | 1.48531931 | 821.0695935 | 2298.823535 | 20 | 3 | 5 | 10 | 14 | 0 | 8 | 1 | 2 | 3 | 4 | 0 |
| SG0000003 | Ncapd2 | 6.4E-29 | 6.15E-28 | 1.333024463 | 3878.045603 | 9769.970672 | 30 | 1 | 5 | 3 | 22 | 6 | 0 | 0 | 0 | 0 | 0 | 0 |
| ISG000000 | Traf3ip2 | 7.01E-29 | 6.72E-28 | -2.081185727 | 1262.696193 | 298.4004959 | 56 | 1 | 5 | 6 | 19 | 4 | 2 | 0 | 0 | 0 | 0 | 0 |
| SG0000002 | Tdp1 | 5.66E-28 | 5.22E-27 | 1.527486555 | 541.8624989 | 1562.098324 | 178 | 1 | 5 | 23 | 64 | 11 | 1 | 0 | 0 | 0 | 0 | 0 |
| ISG000000 | Tcf24 | 7.8E-28 | 7.15E-27 | 3.480443545 | 21.90857784 | 244.5299687 | 35 | 2 | 5 | 5 | 10 | 1 | 19 | 1 | 3 | 3 | 3 | 0 |
| SG0000002 | Arfgap1 | 9.6E-28 | 8.76E-27 | -1.809373563 | 2498.571651 | 712.8794446 | 50 | 0 | 5 | 0 | 15 | 0 | 0 | 0 | 0 | 0 | 0 | 0 |
| ISG000000 | Micall1 | 1.15E-27 | 1.05E-26 | -2.321445086 | 1734.16583 | 346.9493038 | 24 | 5 | 5 | 8 | 12 | 3 | 3 | 1 | 1 | 3 | 3 | 0 |
| SG0000002 | Nsg2 | 3.18E-27 | 2.83E-26 | 7.009899946 | 0.639110347 | 82.36941827 | 147 | 2 | 5 | 21 | 46 | 14 | 6 | 0 | 0 | 2 | 4 | 2 |
| SG0000002 | Ifnar2 | 6.9E-27 | 6.05E-26 | -2.128300899 | 882.5829611 | 201.8706254 | 114 | 0 | 5 | 16 | 78 | 12 | 0 | 0 | 0 | 0 | 0 | 0 |
| ISG000000 | Slc12a7 | 1.27E-26 | 1.11E-25 | 1.428754658 | 1096.082337 | 2950.809599 | 41 | 5 | 5 | 9 | 29 | 6 | 3 | 1 | 1 | 2 | 2 | 0 |
| ISG000000 | Galnt3 | 1.86E-25 | 1.54E-24 | 6.223999822 | 3.513935122 | 262.6663131 | 91 | 0 | 5 | 14 | 43 | 8 | 4 | 0 | 1 | 2 | 4 | 0 |
| ISG000000 | E2f2 | 1.3E-24 | 1.03E-23 | -2.114828675 | 1388.249939 | 320.5094184 | 25 | 4 | 5 | 8 | 23 | 2 | 3 | 0 | 0 | 1 | 1 | 1 |
| SG0000002 | Plcd3 | 7.38E-24 | 5.64E-23 | -2.446095837 | 2385.54705 | 437.763721 | 34 | 4 | 5 | 6 | 12 | 1 | 2 | 0 | 0 | 0 | 0 | 0 |
| ISG000000 | Ppwd1 | 1.8E-23 | 1.35E-22 | 1.381050611 | 614.503365 | 1600.523019 | 70 | 4 | 5 | 13 | 48 | 2 | 0 | 0 | 0 | 0 | 0 | 0 |
| ISG000000 | Ngdn | 4.86E-23 | 3.58E-22 | 1.358413349 | 663.4379374 | 1701.075018 | 22 | 4 | 5 | 8 | 12 | 4 | 0 | 0 | 0 | 0 | 0 | 0 |
| SG0000002 | Hcn3 | 4.88E-23 | 3.59E-22 | 5.578473297 | 1.631569985 | 77.96388992 | 12 | 5 | 5 | 5 | 11 | 3 | 2 | 2 | 2 | 2 | 2 | 0 |
| SG0000002 | Actl6a | 5.12E-23 | 3.76E-22 | 1.232638732 | 1746.085117 | 4103.237385 | 89 | 5 | 5 | 9 | 20 | 12 | 0 | 0 | 0 | 0 | 0 | 0 |
| ISG000000 | Mtrf1 | 1.17E-22 | 8.49E-22 | 1.729196477 | 161.93233 | 536.8754843 | 43 | 4 | 5 | 8 | 23 | 7 | 0 | 0 | 0 | 0 | 0 | 0 |
| SG0000000 | Stambp | 1.65E-22 | 1.18E-21 | -2.02243894 | 898.6213118 | 221.1881869 | 36 | 3 | 5 | 7 | 18 | 1 | 2 | 0 | 1 | 0 | 1 | 0 |
| ISG000000 | Cmtm6 | 3.05E-22 | 2.16E-21 | 1.240108902 | 1296.332399 | 3062.149237 | 29 | 5 | 5 | 5 | 15 | 2 | 4 | 0 | 0 | 0 | 0 | 1 |
| SG0000004 | Sned1 | 4.4E-22 | 3.09E-21 | -3.245130515 | 1226.491942 | 129.3549566 | 77 | 3 | 5 | 11 | 23 | 8 | 7 | 0 | 0 | 1 | 1 | 1 |
| SG0000002 | Fastkd2 | 1.71E-21 | 1.17E-20 | 1.260459315 | 914.3537626 | 2190.535614 | 57 | 3 | 5 | 11 | 27 | 7 | 4 | 0 | 0 | 1 | 1 | 0 |
| ISG000000 | Lcmt1 | 2.9E-21 | 1.96E-20 | 1.401369491 | 404.2750263 | 1067.901424 | 70 | 4 | 5 | 10 | 32 | 11 | 0 | 0 | 0 | 0 | 0 | 0 |
| ISG000000 | Mrps6 | 6.42E-21 | 4.25E-20 | -1.733350722 | 1332.467878 | 400.743224 | 80 | 5 | 5 | 7 | 38 | 7 | 1 | 0 | 0 | 0 | 0 | 0 |
| ISG000000 | Ngly1 | 7.44E-21 | 4.91E-20 | -1.672747645 | 1197.937778 | 375.7396657 | 149 | 2 | 5 | 15 | 73 | 12 | 3 | 0 | 1 | 0 | 0 | 1 |
| ISG000000 | Abracl | 1.07E-20 | 6.99E-20 | -1.608156117 | 1543.11328 | 506.1678767 | 26 | 3 | 5 | 3 | 8 | 0 | 0 | 0 | 0 | 0 | 2 | 0 |
| SG0000002 | Ephb4 | 1.38E-20 | 8.96E-20 | -1.870171522 | 4337.162853 | 1186.391439 | 105 | 5 | 5 | 15 | 70 | 30 | 3 | 0 | 0 | 0 | 0 | 0 |
| SG0000000 | Atp6v1b2 | 1.85E-20 | 1.19E-19 | 1.129242458 | 3137.130383 | 6862.279791 | 48 | 4 | 5 | 14 | 33 | 6 | 1 | 0 | 0 | 0 | 0 | 0 |
| ISG000000 | Has2 | 8.52E-20 | 5.28E-19 | -3.001481215 | 1443.147061 | 180.2082677 | 64 | 1 | 5 | 14 | 35 | 3 | 4 | 0 | 0 | 2 | 2 | 1 |

| | | | | | | | | | | | | | | | | | | |
|---|---|---|---|---|---|---|---|---|---|---|---|---|---|---|---|---|---|---|
| JSG000000 | Mllt1 | 1.61E-19 | 9.84E-19 | -1.945089077 | 3393.196527 | 881.2088646 | 42 | 5 | 5 | 11 | 24 | 3 | 0 | 0 | 0 | 0 | 0 | 0 |
| SG0000005 | Yipf1 | 1.13E-18 | 6.58E-18 | -1.575645461 | 1084.926181 | 363.9851312 | 46 | 5 | 5 | 8 | 24 | 5 | 0 | 0 | 0 | 0 | 0 | 0 |
| SG0000003 | Paip2 | 1.54E-18 | 8.92E-18 | 1.06950684 | 2950.136347 | 6191.497867 | 41 | 4 | 5 | 8 | 21 | 0 | 2 | 0 | 0 | 0 | 1 | 0 |
| SG0000002 | Cdca7l | 2.63E-18 | 1.5E-17 | 1.289445405 | 434.0309796 | 1060.919602 | 47 | 0 | 5 | 4 | 14 | 11 | 2 | 0 | 0 | 0 | 0 | 0 |
| SG0000004 | Pex2 | 3.28E-18 | 1.87E-17 | -1.614255594 | 908.4080383 | 296.716443 | 205 | 0 | 5 | 31 | 148 | 0 | 2 | 0 | 0 | 0 | 0 | 0 |
| SG0000007 | Hoxa3 | 2.76E-17 | 1.5E-16 | -5.044588482 | 231.3848373 | 7.010717176 | 55 | 1 | 5 | 6 | 16 | 4 | 6 | 0 | 1 | 3 | 4 | 0 |
| SG0000003 | Hspa13 | 2.08E-16 | 1.07E-15 | -1.337410606 | 1857.454636 | 735.051058 | 51 | 2 | 5 | 6 | 23 | 7 | 15 | 0 | 1 | 0 | 3 | 1 |
| SG0000006 | Tgif2 | 2.1E-16 | 1.08E-15 | 1.127573538 | 746.506556 | 1631.049282 | 36 | 0 | 5 | 2 | 12 | 5 | 4 | 0 | 0 | 0 | 2 | 2 |
| JSG000000 | Aldh3b1 | 2.53E-16 | 1.3E-15 | -2.849628661 | 737.5534368 | 102.3221808 | 13 | 4 | 5 | 4 | 7 | 0 | 0 | 0 | 0 | 0 | 0 | 0 |
| SG0000000 | Atp5f1 | 2.8E-16 | 1.43E-15 | 1.008865659 | 2609.36271 | 5250.894307 | 40 | 2 | 5 | 6 | 19 | 3 | 1 | 0 | 0 | 0 | 0 | 0 |
| JSG000000 | Arfgap3 | 3.28E-16 | 1.67E-15 | -1.423765729 | 1183.975155 | 441.3126717 | 49 | 5 | 5 | 8 | 31 | 3 | 1 | 0 | 0 | 0 | 0 | 0 |
| JSG000000 | Zfp712 | 4.13E-16 | 2.09E-15 | 2.2517244 | 34.84017579 | 165.9269476 | 56 | 5 | 5 | 15 | 42 | 7 | 8 | 1 | 1 | 4 | 11 | 2 |
| JSG000000 | Cops5 | 4.55E-16 | 2.3E-15 | 1.008666775 | 2294.206109 | 4616.059379 | 83 | 0 | 5 | 10 | 42 | 2 | 1 | 0 | 0 | 0 | 1 | 0 |
| JSG000000 | Ticam2 | 5.28E-16 | 2.65E-15 | 6.231959305 | 0.639110347 | 48.03773597 | 29 | 4 | 5 | 5 | 18 | 2 | 0 | 0 | 0 | 0 | 0 | 0 |
| JSG000000 | Ggcx | 6.87E-16 | 3.43E-15 | -1.539903117 | 743.9362935 | 255.8460511 | 32 | 4 | 5 | 5 | 33 | 1 | 0 | 0 | 0 | 0 | 0 | 0 |
| JSG000000 | Slc6a14 | 8.87E-16 | 4.4E-15 | 6.139801823 | 0.674076249 | 47.53062675 | 80 | 1 | 5 | 8 | 48 | 14 | 2 | 0 | 0 | 0 | 1 | 1 |
| SG0000003 | Tamm41 | 2.56E-15 | 1.24E-14 | 1.14707145 | 547.0510319 | 1211.520467 | 37 | 3 | 5 | 9 | 24 | 3 | 0 | 0 | 0 | 0 | 0 | 0 |
| SG0000003 | Piwil4 | 2.62E-15 | 1.26E-14 | 3.880156199 | 4.624510145 | 68.09404608 | 52 | 3 | 5 | 9 | 22 | 6 | 0 | 0 | 0 | 0 | 0 | 0 |
| SG0000004 | Arntl2 | 2.81E-15 | 1.35E-14 | -2.263990835 | 574.4129304 | 119.5900524 | 92 | 4 | 5 | 11 | 20 | 6 | 4 | 0 | 1 | 0 | 0 | 0 |
| JSG000000 | Aacs | 3.89E-15 | 1.85E-14 | -1.811708138 | 1777.373575 | 506.2910219 | 26 | 5 | 5 | 6 | 9 | 3 | 0 | 0 | 0 | 0 | 0 | 0 |
| JSG000000 | Lingo1 | 4.92E-15 | 2.33E-14 | 3.269368632 | 9.613243694 | 92.69322167 | 57 | 1 | 5 | 9 | 16 | 7 | 1 | 0 | 0 | 0 | 0 | 0 |
| JSG000000 | Ppp2r2a | 5.03E-15 | 2.38E-14 | 0.956434642 | 3249.94586 | 6306.547081 | 118 | 3 | 5 | 10 | 55 | 14 | 7 | 0 | 0 | 0 | 1 | 1 |
| JSG000000 | Gm8281 | 5.32E-15 | 2.52E-14 | 5.320700288 | 0 | 38.96597244 | 36 | 2 | 5 | 5 | 14 | 2 | 5 | 2 | 2 | 2 | 2 | 0 |
| SG0000002 | Hn1 | 9.53E-15 | 4.43E-14 | -1.22594336 | 4421.715414 | 1890.362369 | 27 | 3 | 5 | 4 | 7 | 1 | 0 | 0 | 0 | 0 | 0 | 0 |
| SG0000002 | Dnajb12 | 2.32E-14 | 1.05E-13 | -1.31945575 | 1669.686597 | 669.020156 | 100 | 5 | 5 | 14 | 26 | 13 | 15 | 0 | 0 | 4 | 6 | 8 |
| JSG000000 | Fam89a | 3.67E-14 | 1.64E-13 | 3.518240995 | 6.159712968 | 70.57591978 | 15 | 5 | 5 | 5 | 6 | 0 | 2 | 1 | 1 | 1 | 2 | 0 |
| SG0000002 | Hspd1 | 4.24E-14 | 1.89E-13 | 0.915064712 | 14254.08338 | 26878.26437 | 19 | 5 | 5 | 5 | 9 | 3 | 1 | 0 | 0 | 0 | 0 | 0 |
| SG0000002 | Bspry | 4.45E-14 | 1.98E-13 | 2.678349881 | 122.3142656 | 782.9621451 | 32 | 5 | 5 | 9 | 22 | 6 | 0 | 0 | 0 | 0 | 0 | 0 |
| JSG000000 | Kcnd1 | 5.19E-14 | 2.3E-13 | 1.650626221 | 87.04173161 | 273.2848311 | 23 | 5 | 5 | 6 | 13 | 2 | 2 | 0 | 0 | 0 | 0 | 0 |
| SG0000002 | Emilin2 | 5.66E-14 | 2.5E-13 | -2.283196018 | 370.5920657 | 76.13522395 | 57 | 4 | 5 | 11 | 37 | 10 | 2 | 0 | 0 | 1 | 1 | 0 |
| JSG000000 | Hspa9 | 6.3E-14 | 2.77E-13 | 0.884599722 | 14603.168 | 26961.13258 | 51 | 4 | 5 | 11 | 21 | 4 | 1 | 0 | 0 | 0 | 0 | 0 |
| JSG000000 | Fbxo33 | 1.12E-13 | 4.85E-13 | -1.205779982 | 1527.917822 | 662.4056895 | 47 | 1 | 5 | 9 | 23 | 3 | 7 | 0 | 0 | 1 | 2 | 0 |
| JSG000000 | Zfp282 | 2.19E-13 | 9.33E-13 | -1.542189418 | 504.8177402 | 173.3362253 | 43 | 4 | 5 | 7 | 23 | 3 | 2 | 0 | 0 | 0 | 2 | 0 |
| SG0000005 | Mical3 | 2.28E-13 | 9.66E-13 | 1.044190304 | 624.8058963 | 1288.480119 | 52 | 5 | 5 | 7 | 21 | 7 | 0 | 0 | 0 | 0 | 0 | 0 |
| SG0000007 | Sox13 | 2.36E-13 | 1E-12 | -1.26724894 | 2698.854444 | 1121.242419 | 38 | 5 | 5 | 6 | 17 | 5 | 0 | 0 | 0 | 0 | 0 | 2 |
| SG0000007 | Zfp951 | 5.74E-13 | 2.37E-12 | 4.769525061 | 4.009491589 | 109.3602301 | 72 | 4 | 5 | 11 | 42 | 15 | 0 | 0 | 0 | 0 | 0 | 0 |
| JSG000000 | Mccc1 | 6.66E-13 | 2.74E-12 | 1.112350881 | 375.2407112 | 811.2616218 | 161 | 2 | 5 | 22 | 95 | 18 | 2 | 0 | 0 | 0 | 0 | 0 |
| SG0000003 | Cep104 | 7.21E-13 | 2.96E-12 | -1.179351989 | 1317.502946 | 581.74314 | 28 | 3 | 5 | 9 | 13 | 4 | 5 | 0 | 1 | 2 | 5 | 1 |
| JSG000000 | Fads2 | 7.56E-13 | 3.1E-12 | 1.135454068 | 343.344142 | 754.2848332 | 43 | 4 | 5 | 9 | 17 | 9 | 0 | 0 | 0 | 0 | 0 | 0 |
| JSG000000 | Dcaf7 | 8.55E-13 | 3.5E-12 | -1.031865506 | 5389.150741 | 2635.711448 | 36 | 5 | 5 | 10 | 22 | 3 | 5 | 0 | 0 | 0 | 1 | 0 |
| SG0000005 | Zfp637 | 9.89E-13 | 4.03E-12 | -1.431629566 | 539.5354542 | 200.0122275 | 15 | 5 | 5 | 10 | 10 | 0 | 0 | 0 | 0 | 0 | 0 | 0 |
| JSG000000 | Zfp870 | 2.01E-12 | 8E-12 | 1.236253603 | 209.0857447 | 492.5766726 | 16 | 5 | 5 | 6 | 10 | 0 | 7 | 1 | 1 | 2 | 5 | 0 |
| SG0000003 | Dlec1 | 2.46E-12 | 9.73E-12 | 3.992111788 | 3.241391746 | 51.57947485 | 28 | 4 | 5 | 6 | 23 | 5 | 1 | 0 | 0 | 0 | 0 | 0 |
| SG0000002 | Mfsd2a | 2.82E-12 | 1.11E-11 | -2.40700667 | 203.04385 | 38.28323526 | 13 | 5 | 5 | 6 | 7 | 3 | 0 | 0 | 0 | 0 | 0 | 1 |
| SG0000000 | Tbl2 | 4.54E-12 | 1.78E-11 | 1.047865112 | 1333.383939 | 2756.72884 | 18 | 4 | 5 | 5 | 12 | 0 | 0 | 0 | 0 | 0 | 0 | 0 |
| JSG000000 | Cetn3 | 5.25E-12 | 2.04E-11 | 0.939866013 | 883.6424969 | 1695.135644 | 41 | 3 | 5 | 9 | 22 | 2 | 1 | 0 | 0 | 0 | 1 | 0 |
| SG0000002 | Lancl1 | 7.07E-12 | 2.73E-11 | 0.941467536 | 876.2584835 | 1682.837588 | 139 | 0 | 5 | 21 | 67 | 17 | 10 | 0 | 2 | 6 | 10 | 2 |
| JSG000000 | Sar1b | 7.1E-12 | 2.74E-11 | -1.11293809 | 1538.40918 | 711.2857604 | 58 | 5 | 5 | 11 | 28 | 1 | 1 | 0 | 0 | 0 | 0 | 0 |
| JSG000000 | Mansc1 | 1.07E-11 | 4.08E-11 | -2.049116751 | 284.7200449 | 68.79746512 | 38 | 5 | 5 | 8 | 26 | 5 | 0 | 0 | 0 | 0 | 2 | 0 |
| JSG000000 | Ggh | 1.5E-11 | 5.66E-11 | -1.23903314 | 750.9076282 | 318.1268868 | 54 | 3 | 5 | 10 | 24 | 12 | 2 | 0 | 0 | 0 | 1 | 0 |
| SG0000003 | Mbd4 | 2.36E-11 | 8.78E-11 | 1.016821667 | 453.1455386 | 916.9201791 | 45 | 5 | 5 | 10 | 28 | 7 | 3 | 0 | 0 | 0 | 2 | 1 |
| SG0000005 | Zfp93 | 3.2E-11 | 1.18E-10 | -1.452937666 | 388.09055 | 141.760504 | 13 | 4 | 5 | 5 | 8 | 6 | 0 | 0 | 0 | 0 | 0 | 0 |
| SG0000006 | Pstk | 3.41E-11 | 1.26E-10 | 1.177135227 | 209.1538359 | 472.9539384 | 86 | 2 | 5 | 9 | 43 | 11 | 0 | 0 | 0 | 0 | 1 | 0 |
| SG0000004 | Ccdc114 | 4.94E-11 | 1.8E-10 | 6.050549809 | 5.290055849 | 350.6365946 | 18 | 5 | 5 | 7 | 15 | 0 | 0 | 0 | 0 | 0 | 0 | 0 |
| JSG000000 | Emc7 | 1.24E-10 | 4.37E-10 | -1.072814495 | 1170.510122 | 556.4495954 | 36 | 4 | 5 | 7 | 13 | 1 | 8 | 0 | 0 | 0 | 0 | 0 |
| JSG000000 | Zfp472 | 1.38E-10 | 4.86E-10 | 1.201132584 | 173.4065404 | 398.6964883 | 36 | 2 | 5 | 4 | 14 | 5 | 2 | 0 | 1 | 0 | 0 | 1 |
| JSG000000 | Rbm34 | 1.38E-10 | 4.87E-10 | 0.940900531 | 566.3590249 | 1087.254235 | 37 | 3 | 5 | 9 | 20 | 5 | 4 | 0 | 1 | 0 | 0 | 0 |
| JSG000000 | Mef2b | 1.48E-10 | 5.2E-10 | 4.789923036 | 0 | 26.66371558 | 46 | 5 | 5 | 11 | 32 | 2 | 0 | 0 | 0 | 0 | 0 | 0 |
| JSG000000 | Car13 | 1.52E-10 | 5.36E-10 | -1.296005381 | 1595.965062 | 649.9603825 | 55 | 1 | 5 | 11 | 29 | 5 | 2 | 0 | 1 | 0 | 2 | 0 |
| SG0000004 | Baz2a | 1.7E-10 | 5.97E-10 | 0.770104571 | 6328.410757 | 10792.42988 | 51 | 3 | 5 | 8 | 25 | 3 | 4 | 0 | 0 | 1 | 3 | 0 |
| JSG000000 | Tubb6 | 2.1E-10 | 7.3E-10 | -1.033014501 | 4055.036482 | 1981.647551 | 25 | 4 | 5 | 8 | 13 | 3 | 0 | 0 | 0 | 0 | 0 | 0 |
| JSG000000 | Fam131c | 2.69E-10 | 9.27E-10 | 3.036857636 | 28.01876052 | 229.9503952 | 13 | 4 | 5 | 6 | 10 | 0 | 0 | 0 | 0 | 0 | 0 | 0 |
| JSG000000 | Colgalt1 | 3.04E-10 | 1.04E-09 | -0.877230408 | 7751.837978 | 4220.190164 | 17 | 5 | 5 | 6 | 20 | 0 | 3 | 0 | 0 | 1 | 1 | 0 |
| JSG000000 | Gm17296 | 4.28E-10 | 1.45E-09 | 1.035316652 | 301.89354 | 618.7499646 | 65 | 3 | 5 | 11 | 48 | 0 | 2 | 0 | 0 | 1 | 1 | 0 |
| JSG000000 | Osgin2 | 4.84E-10 | 1.63E-09 | 1.098076411 | 229.5545336 | 491.405272 | 57 | 3 | 5 | 5 | 20 | 4 | 9 | 1 | 1 | 1 | 2 | 0 |
| JSG000000 | Sgol2a | 5.47E-10 | 1.84E-09 | 0.75184289 | 4553.16013 | 7667.259923 | 73 | 4 | 5 | 9 | 38 | 6 | 7 | 1 | 1 | 1 | 2 | 0 |
| JSG000000 | Manea | 6.24E-10 | 2.09E-09 | -1.206555964 | 986.0803476 | 427.2703602 | 55 | 2 | 5 | 11 | 34 | 9 | 6 | 0 | 0 | 3 | 9 | 1 |
| JSG000000 | Ccdc173 | 7.83E-10 | 2.61E-09 | 2.049000276 | 26.67145175 | 110.3715647 | 61 | 2 | 5 | 7 | 29 | 5 | 1 | 0 | 0 | 0 | 0 | 0 |
| JSG000000 | Ubtd1 | 8.38E-10 | 2.78E-09 | -1.93825745 | 1207.143173 | 314.9815752 | 59 | 4 | 5 | 15 | 33 | 2 | 0 | 0 | 0 | 0 | 0 | 0 |
| JSG000000 | Srsf12 | 9.51E-10 | 3.14E-09 | 2.870933527 | 13.46496686 | 98.5013984 | 59 | 3 | 5 | 13 | 24 | 7 | 8 | 1 | 1 | 3 | 4 | 0 |
| JSG000000 | Abhd17b | 1.42E-09 | 4.6E-09 | -0.971920338 | 1400.520525 | 714.0231665 | 74 | 2 | 5 | 9 | 30 | 6 | 4 | 0 | 1 | 0 | 1 | 0 |
| SG0000001 | Atp6v0e | 1.91E-09 | 6.15E-09 | -0.914607165 | 1918.517371 | 1017.751017 | 31 | 4 | 5 | 10 | 26 | 4 | 0 | 0 | 0 | 0 | 0 | 0 |
| SG0000002 | Ilkap | 1.96E-09 | 6.3E-09 | 0.806036887 | 1127.040785 | 1970.5203 | 157 | 5 | 5 | 21 | 51 | 8 | 4 | 0 | 0 | 0 | 0 | 1 |
| JSG000000 | Cpt1c | 2.44E-09 | 7.8E-09 | -0.982797946 | 1524.636478 | 771.4621883 | 15 | 5 | 5 | 6 | 21 | 0 | 0 | 0 | 0 | 0 | 0 | 0 |
| SG0000006 | Zfp69 | 3.13E-09 | 9.88E-09 | 1.839332101 | 32.13768381 | 115.0028778 | 26 | 2 | 5 | 5 | 18 | 3 | 0 | 0 | 0 | 0 | 0 | 1 |
| JSG000000 | Vps37b | 3.14E-09 | 9.92E-09 | -1.137577425 | 1023.566381 | 465.2336139 | 19 | 4 | 5 | 4 | 4 | 5 | 2 | 0 | 1 | 0 | 0 | 0 |
| JSG000000 | Slc16a1 | 5.15E-09 | 1.6E-08 | -0.813732278 | 9463.65923 | 5383.953118 | 41 | 3 | 5 | 4 | 15 | 4 | 11 | 0 | 0 | 0 | 2 | 0 |
| SG0000001 | Phf21b | 7.11E-09 | 2.19E-08 | 5.309605334 | 3.158242264 | 125.2552423 | 49 | 3 | 5 | 8 | 22 | 6 | 1 | 0 | 0 | 0 | 0 | 0 |
| JSG000000 | Xrcc1 | 7.27E-09 | 2.23E-08 | 0.788348271 | 1254.434705 | 2166.529152 | 24 | 1 | 5 | 3 | 7 | 1 | 0 | 0 | 0 | 0 | 0 | 0 |
| SG0000002 | Pnpt1 | 8.06E-09 | 2.47E-08 | 0.7522188 | 1641.985741 | 2765.730146 | 96 | 4 | 5 | 12 | 47 | 6 | 2 | 0 | 0 | 1 | 3 | 0 |
| SG0000002 | Tmbim6 | 9.34E-09 | 2.85E-08 | -0.85069801 | 9495.831581 | 5265.59418 | 120 | 5 | 5 | 5 | 45 | 5 | 0 | 0 | 0 | 0 | 0 | 0 |
| JSG000000 | Ganab | 9.62E-09 | 2.93E-08 | -0.782315764 | 13781.97673 | 8013.292608 | 30 | 5 | 5 | 10 | 15 | 1 | 1 | 0 | 0 | 1 | 1 | 0 |
| SG0000003 | Coq9 | 1.02E-08 | 3.1E-08 | 0.804869011 | 788.9436275 | 1378.274393 | 18 | 5 | 5 | 5 | 11 | 1 | 1 | 0 | 0 | 0 | 0 | 0 |
| SG0000002 | Oprl1 | 1.16E-08 | 3.52E-08 | 4.860971262 | 0.970711415 | 28.20903922 | 36 | 5 | 5 | 10 | 10 | 5 | 6 | 0 | 0 | 0 | 0 | 0 |
| SG0000000 | Ramp2 | 1.25E-08 | 3.78E-08 | 3.121685468 | 5.22865459 | 45.51042758 | 20 | 4 | 5 | 6 | 7 | 1 | 2 | 0 | 0 | 0 | 0 | 0 |
| JSG000000 | Hacd3 | 1.26E-08 | 3.81E-08 | -0.866622338 | 1970.355673 | 1080.600863 | 52 | 5 | 5 | 10 | 37 | 3 | 3 | 0 | 0 | 0 | 0 | 0 |
| JSG000000 | Sptlc1 | 1.66E-08 | 4.96E-08 | -0.908518965 | 1270.733106 | 676.9598078 | 54 | 1 | 5 | 11 | 21 | 3 | 1 | 0 | 0 | 0 | 2 | 0 |
| SG0000003 | Ankrd13b | 2.7E-08 | 7.92E-08 | -1.380429122 | 371.6003269 | 142.7333683 | 32 | 5 | 5 | 14 | 29 | 0 | 0 | 0 | 0 | 0 | 0 | 0 |
| JSG000000 | Slc13a4 | 3.42E-08 | 9.97E-08 | 6.225733926 | 0.307509278 | 23.01393491 | 46 | 2 | 5 | 5 | 27 | 3 | 1 | 0 | 0 | 0 | 0 | 0 |
| JSG000000 | Tsnax | 3.56E-08 | 1.04E-07 | 0.727549257 | 1410.264105 | 2335.149025 | 34 | 2 | 5 | 9 | 14 | 3 | 2 | 0 | 0 | 0 | 0 | 1 |
| SG0000002 | Tbc1d20 | 4.29E-08 | 1.24E-07 | -0.809036425 | 2412.865878 | 1377.174429 | 23 | 4 | 5 | 5 | 9 | 1 | 2 | 0 | 0 | 0 | 0 | 0 |
| JSG000000 | Rps6ka2 | 4.82E-08 | 1.39E-07 | 1.128618767 | 321.3822432 | 702.7000431 | 136 | 2 | 5 | 24 | 64 | 6 | 1 | 0 | 0 | 1 | 2 | 0 |
| JSG000000 | Pcp4l1 | 5.12E-08 | 1.47E-07 | 2.43034902 | 10.98683469 | 59.22165568 | 31 | 4 | 5 | 5 | 20 | 1 | 1 | 0 | 0 | 0 | 0 | 0 |
| SG0000002 | Osbp | 8.97E-08 | 2.53E-07 | -0.804084758 | 2109.848212 | 1208.363451 | 43 | 3 | 5 | 6 | 20 | 2 | 1 | 0 | 0 | 0 | 1 | 1 |
| SG0000002 | Rpf1 | 9.77E-08 | 2.74E-07 | 0.753031655 | 788.6502576 | 1329.136441 | 30 | 3 | 5 | 4 | 14 | 1 | 1 | 0 | 0 | 0 | 0 | 0 |
| SG0000002 | Psd4 | 9.88E-08 | 2.77E-07 | 5.232649558 | 5.575816904 | 209.6490882 | 81 | 1 | 5 | 18 | 47 | 15 | 8 | 1 | 1 | 1 | 4 | 0 |
| JSG000000 | Cybrd1 | 1.29E-07 | 3.59E-07 | 4.328728754 | 1.324060707 | 26.60633748 | 32 | 4 | 5 | 7 | 17 | 0 | 1 | 0 | 0 | 1 | 4 | 0 |
| SG0000003 | Edem2 | 1.61E-07 | 4.45E-07 | -0.822959988 | 1450.980114 | 820.2114332 | 42 | 1 | 5 | 10 | 19 | 3 | 0 | 0 | 0 | 0 | 0 | 0 |
| JSG000000 | B3galt1 | 1.68E-07 | 4.62E-07 | 3.799825878 | 4.792155815 | 66.74096751 | 50 | 3 | 5 | 10 | 29 | 8 | 18 | 1 | 1 | 4 | 7 | 7 |
| JSG000000 | Rhbdl3 | 1.68E-07 | 4.64E-07 | 5.175467129 | 0.639110347 | 23.09652781 | 52 | 4 | 5 | 8 | 29 | 6 | 1 | 0 | 0 | 0 | 1 | 0 |
| JSG000000 | Arl6ip1 | 1.83E-07 | 5.02E-07 | 0.640323416 | 5553.438893 | 8656.026006 | 22 | 4 | 5 | 5 | 13 | 2 | 1 | 0 | 0 | 0 | 0 | 1 |
| SG0000002 | Setd4 | 1.98E-07 | 5.4E-07 | 1.091043001 | 127.556435 | 271.7309098 | 30 | 4 | 5 | 6 | 17 | 2 | 0 | 0 | 0 | 0 | 0 | 0 |
| SG0000000 | Fpgs | 2.39E-07 | 6.48E-07 | 0.782325282 | 907.0327345 | 1560.006246 | 24 | 4 | 5 | 6 | 11 | 2 | 0 | 0 | 0 | 0 | 0 | 1 |
| JSG000000 | Fam192a | 2.82E-07 | 7.6E-07 | 0.704160884 | 1021.716348 | 1664.577003 | 45 | 4 | 5 | 9 | 20 | 4 | 1 | 0 | 0 | 0 | 0 | 0 |
| JSG000000 | Spsb4 | 3.03E-07 | 8.16E-07 | 3.737823066 | 2.235714429 | 29.82724841 | 86 | 4 | 5 | 13 | 22 | 3 | 1 | 0 | 0 | 1 | 1 | 0 |
| JSG000000 | Ms4a4d | 3.29E-07 | 8.81E-07 | 2.978589319 | 4.872961731 | 38.40941836 | 52 | 3 | 5 | 9 | 30 | 4 | 2 | 0 | 0 | 0 | 0 | 0 |
| SG0000002 | Fbxl3 | 7.04E-07 | 1.84E-06 | 0.654213039 | 1497.922389 | 2357.366245 | 37 | 0 | 5 | 3 | 16 | 2 | 4 | 0 | 2 | 0 | 4 | 0 |
| SG0000002 | Casp9 | 8.3E-07 | 2.16E-06 | 0.743861747 | 534.7384146 | 895.5010152 | 56 | 5 | 5 | 11 | 41 | 6 | 2 | 1 | 1 | 1 | 1 | 0 |
| SG0000003 | Sh3bp4 | 1.09E-06 | 2.82E-06 | 0.890052838 | 1828.361891 | 3388.400699 | 89 | 2 | 5 | 12 | 30 | 8 | 2 | 0 | 1 | 0 | 0 | 2 |
| JSG000000 | Celf3 | 1.45E-06 | 3.69E-06 | 5.901825832 | 0.307509278 | 18.38589723 | 35 | 5 | 5 | 15 | 20 | 5 | 5 | 5 | 5 | 5 | 5 | 5 |
| SG0000003 | Gde1 | 1.69E-06 | 4.28E-06 | -0.744266283 | 1377.79956 | 822.5069298 | 19 | 4 | 5 | 9 | 17 | 1 | 0 | 0 | 0 | 0 | 0 | 0 |
| JSG000000 | Scamp2 | 1.88E-06 | 4.75E-06 | -0.743415951 | 1252.38555 | 748.0791448 | 33 | 4 | 5 | 9 | 18 | 4 | 0 | 0 | 0 | 0 | 1 | 0 |
| SG0000001 | Slc12a5 | 2.57E-06 | 6.39E-06 | 2.339271405 | 9.11534366 | 46.12792379 | 82 | 5 | 5 | 18 | 51 | 10 | 4 | 0 | 0 | 3 | 9 | 3 |
| JSG000000 | Syt13 | 2.93E-06 | 7.24E-06 | 4.886325264 | 3.762386708 | 111.2740524 | 42 | 4 | 5 | 5 | 10 | 10 | 3 | 0 | 0 | 0 | 0 | 0 |
| SG0000004 | Crb3 | 3.35E-06 | 8.25E-06 | 2.055498458 | 118.356801 | 491.9941177 | 22 | 5 | 5 | 7 | 7 | 1 | 4 | 0 | 0 | 2 | 2 | 0 |
| JSG000000 | Tmem19 | 4.8E-06 | 1.17E-05 | -0.731490567 | 1155.25148 | 695.7863932 | 40 | 1 | 5 | 1 | 16 | 2 | 4 | 0 | 0 | 0 | 1 | 1 |
| SG0000002 | Ncdn | 4.88E-06 | 1.19E-05 | -1.152875837 | 1180.282296 | 530.8058305 | 21 | 1 | 5 | 3 | 3 | 0 | 1 | 1 | 1 | 1 | 1 | 0 |
| JSG000000 | Casp7 | 5.58E-06 | 1.35E-05 | -0.857900706 | 522.4588384 | 288.2691946 | 62 | 3 | 5 | 10 | 20 | 2 | 3 | 0 | 0 | 1 | 2 | 0 |
| SG0000002 | Nkiras1 | 6.93E-06 | 1.66E-05 | 0.899169863 | 162.9912279 | 303.9774248 | 44 | 4 | 5 | 7 | 15 | 2 | 14 | 2 | 3 | 3 | 5 | 2 |
| SG0000003 | Abcg4 | 7.22E-06 | 1.72E-05 | 3.594636296 | 1.963171053 | 23.71654875 | 17 | 4 | 5 | 7 | 10 | 0 | 1 | 0 | 0 | 1 | 1 | 0 |
| JSG000000 | Mnda | 8.22E-06 | 1.95E-05 | -3.490050053 | 102.3789897 | 9.111735134 | 59 | 2 | 5 | 6 | 25 | 3 | 0 | 0 | 0 | 0 | 0 | 0 |
| JSG000000 | Atp6ap2 | 1.03E-05 | 2.42E-05 | -0.62184682 | 3927.721565 | 2552.384788 | 60 | 2 | 5 | 9 | 27 | 5 | 5 | 0 | 0 | 0 | 4 | 1 |
| JSG000000 | Gm10644 | 1.12E-05 | 2.63E-05 | 2.842355057 | 4.578670132 | 32.83775714 | 27 | 4 | 5 | 6 | 19 | 1 | 3 | 1 | 1 | 1 | 1 | 0 |

| | | | | | | | | | | | | | | | | | | |
|---|---|---|---|---|---|---|---|---|---|---|---|---|---|---|---|---|---|---|
| ISG000000 | Gm9918 | 1.17E-05 | 2.75E-05 | 3.129921103 | 2.990596593 | 26.17929642 | 10 | 4 | 5 | 5 | 7 | 1 | 7 | 4 | 5 | 5 | 6 | 0 |
| ISG000000 | Cd274 | 1.22E-05 | 2.85E-05 | -1.283294998 | 148.1061394 | 60.85030702 | 35 | 2 | 5 | 9 | 19 | 4 | 11 | 1 | 2 | 4 | 6 | 1 |
| SG0000001 | Slc35c2 | 1.82E-05 | 4.19E-05 | -0.767247567 | 1025.638044 | 602.6006421 | 16 | 0 | 5 | 5 | 5 | 0 | 10 | 0 | 5 | 5 | 5 | 0 |
| ISG000000 | Spdl1 | 2.31E-05 | 5.25E-05 | 0.623192981 | 655.1587757 | 1009.128201 | 45 | 1 | 5 | 5 | 16 | 0 | 4 | 0 | 1 | 1 | 1 | 0 |
| SG0000003 | Vps37a | 2.34E-05 | 5.32E-05 | -0.668907815 | 1943.826554 | 1222.633226 | 100 | 1 | 5 | 9 | 37 | 5 | 16 | 0 | 0 | 2 | 5 | 0 |
| SG0000002 | Hps1 | 2.61E-05 | 5.9E-05 | -0.654534966 | 1260.213286 | 800.5885899 | 98 | 0 | 5 | 15 | 33 | 5 | 2 | 0 | 0 | 0 | 2 | 0 |
| SG0000006 | Utp14a | 2.75E-05 | 6.21E-05 | 0.5478537 | 1690.751602 | 2471.725309 | 36 | 3 | 5 | 8 | 16 | 3 | 0 | 0 | 0 | 0 | 0 | 0 |
| ISG000000 | Slc35g1 | 3.03E-05 | 6.82E-05 | -0.676369234 | 1091.090485 | 682.7368447 | 16 | 5 | 5 | 8 | 19 | 1 | 3 | 0 | 0 | 1 | 1 | 0 |
| SG0000004 | Paip2b | 3.48E-05 | 7.79E-05 | 0.532368631 | 2003.725641 | 2897.992 | 56 | 5 | 5 | 10 | 17 | 2 | 4 | 0 | 0 | 0 | 1 | 0 |
| SG0000004 | Swi5 | 3.75E-05 | 8.36E-05 | 0.519806313 | 3147.861424 | 4513.286444 | 47 | 5 | 5 | 5 | 20 | 5 | 1 | 0 | 0 | 0 | 0 | 0 |
| ISG000000 | Gdi2 | 4.03E-05 | 8.95E-05 | 0.495435705 | 10962.52103 | 15454.37505 | 71 | 2 | 5 | 6 | 28 | 4 | 2 | 0 | 0 | 0 | 1 | 0 |
| SG0000003 | Wnt5b | 4.27E-05 | 9.45E-05 | 1.561472851 | 23.12774084 | 68.26268676 | 300 | 0 | 5 | 46 | 147 | 19 | 0 | 0 | 0 | 0 | 8 | 0 |
| ISG000000 | Tgs1 | 4.82E-05 | 0.000106 | 0.555617485 | 1059.608504 | 1557.410018 | 97 | 5 | 5 | 12 | 28 | 6 | 1 | 0 | 0 | 0 | 0 | 0 |
| SG0000006 | Btbd3 | 6.52E-05 | 0.000141 | 0.635204306 | 450.0412166 | 698.9849732 | 81 | 4 | 5 | 9 | 26 | 6 | 4 | 0 | 0 | 2 | 4 | 0 |
| SG0000003 | Itpkb | 6.56E-05 | 0.000142 | -1.011802122 | 620.0660302 | 307.5071025 | 86 | 1 | 5 | 7 | 26 | 13 | 2 | 0 | 0 | 1 | 1 | 0 |
| SG0000000 | Itgae | 0.000102 | 0.000216 | 2.821662702 | 6.750639734 | 47.72545112 | 48 | 3 | 5 | 10 | 21 | 2 | 1 | 0 | 0 | 0 | 0 | 0 |
| SG0000003 | Fam168b | 0.000106 | 0.000223 | 0.48927407 | 3286.079214 | 4612.795481 | 164 | 0 | 5 | 5 | 67 | 7 | 4 | 0 | 0 | 0 | 2 | 2 |
| ISG000000 | Trappc3 | 0.000126 | 0.000264 | -0.575832413 | 1544.281781 | 1036.057166 | 19 | 4 | 5 | 5 | 13 | 1 | 0 | 0 | 0 | 0 | 0 | 0 |
| SG0000002 | Lgmn | 0.000132 | 0.000276 | 0.517336543 | 1290.578334 | 1847.21815 | 80 | 1 | 5 | 18 | 46 | 1 | 0 | 0 | 0 | 0 | 2 | 0 |
| ISG000000 | Nenf | 0.000144 | 0.000301 | -0.737009836 | 483.5058827 | 290.0946335 | 17 | 4 | 5 | 5 | 11 | 3 | 0 | 0 | 0 | 0 | 0 | 0 |
| ISG000000 | Cyp46a1 | 0.000153 | 0.000318 | 2.154769882 | 7.496991354 | 33.38392363 | 23 | 5 | 5 | 6 | 11 | 6 | 0 | 0 | 0 | 0 | 0 | 0 |
| SG0000003 | Gipc2 | 0.000154 | 0.00032 | 1.082359806 | 55.70092657 | 117.9465351 | 104 | 4 | 5 | 14 | 28 | 10 | 1 | 0 | 0 | 0 | 0 | 0 |
| ISG000000 | Cdc25c | 0.000193 | 0.000397 | 0.58342958 | 486.9164889 | 729.5990933 | 30 | 3 | 5 | 10 | 26 | 3 | 0 | 0 | 0 | 0 | 0 | 1 |
| SG0000004 | Rbm48 | 0.000222 | 0.000452 | 0.653754733 | 276.2755993 | 434.6526322 | 55 | 5 | 5 | 10 | 22 | 5 | 3 | 0 | 0 | 1 | 1 | 0 |
| ISG000000 | Tkt | 0.000239 | 0.000485 | 0.66361073 | 9559.020159 | 15141.8909 | 40 | 3 | 5 | 7 | 20 | 6 | 2 | 0 | 0 | 0 | 0 | 0 |
| SG0000002 | Stmn2 | 0.000279 | 0.000562 | 0.508212858 | 1867.610264 | 2656.278252 | 108 | 2 | 5 | 11 | 42 | 10 | 2 | 0 | 0 | 0 | 2 | 1 |
| SG0000001 | Glrx2 | 0.000304 | 0.000609 | 0.53959996 | 629.2557645 | 914.666928 | 77 | 0 | 5 | 5 | 12 | 5 | 17 | 0 | 1 | 0 | 2 | 0 |
| SG0000002 | Rassf6 | 0.000327 | 0.000653 | 2.008063128 | 15.07328878 | 60.63107305 | 73 | 0 | 5 | 6 | 24 | 3 | 1 | 0 | 0 | 0 | 1 | 0 |
| SG0000002 | Gpsm2 | 0.00034 | 0.000677 | -0.528441606 | 1946.059602 | 1349.209414 | 67 | 1 | 5 | 4 | 27 | 5 | 1 | 0 | 0 | 0 | 0 | 0 |
| ISG000000 | Sh2d4a | 0.000396 | 0.000782 | 1.454241264 | 20.20557621 | 55.36578746 | 119 | 1 | 5 | 11 | 65 | 10 | 0 | 0 | 0 | 0 | 0 | 1 |
| ISG000000 | Nfatc2ip | 0.000405 | 0.0008 | 0.641208105 | 1446.918772 | 2256.664642 | 31 | 5 | 5 | 5 | 22 | 4 | 1 | 1 | 1 | 1 | 4 | 0 |
| ISG000000 | Wnt7a | 0.000508 | 0.00099 | -1.680279879 | 99.85456716 | 31.15683172 | 29 | 2 | 5 | 7 | 18 | 4 | 4 | 0 | 2 | 1 | 2 | 1 |
| ISG000000 | Vwa2 | 0.000561 | 0.001088 | 1.151059986 | 38.28318224 | 85.01814272 | 42 | 0 | 5 | 2 | 9 | 4 | 3 | 0 | 0 | 0 | 0 | 0 |
| SG0000007 | Rgma | 0.00074 | 0.001411 | 1.273145692 | 26.75744778 | 64.66953357 | 93 | 5 | 5 | 12 | 40 | 21 | 3 | 3 | 3 | 3 | 3 | 3 |
| SG0000001 | Nt5c3b | 0.000991 | 0.00186 | -0.551702212 | 798.7551019 | 544.9224876 | 45 | 0 | 5 | 0 | 20 | 30 | 0 | 0 | 0 | 0 | 0 | 0 |
| SG0000003 | Zscan21 | 0.00102 | 0.001911 | 0.486000253 | 694.8180457 | 973.1319763 | 105 | 0 | 5 | 10 | 25 | 25 | 0 | 0 | 0 | 0 | 0 | 0 |
| SG0000000 | Usp19 | 0.001024 | 0.001918 | -0.445001887 | 4342.685586 | 3190.06455 | 22 | 5 | 5 | 5 | 16 | 11 | 2 | 0 | 0 | 0 | 0 | 0 |
| ISG000000 | Bms1 | 0.001289 | 0.002389 | 0.410004378 | 3205.865385 | 4259.600783 | 65 | 3 | 5 | 8 | 36 | 5 | 0 | 0 | 0 | 0 | 0 | 0 |
| SG0000006 | Dot1l | 0.001338 | 0.002477 | 0.436678439 | 1646.095225 | 2227.964485 | 51 | 5 | 5 | 12 | 28 | 2 | 17 | 1 | 1 | 6 | 10 | 0 |
| ISG000000 | Rab44 | 0.001434 | 0.002647 | -1.736269669 | 39.03203057 | 11.715259 | 12 | 3 | 5 | 5 | 13 | 0 | 1 | 0 | 0 | 0 | 0 | 0 |
| ISG000000 | Zfp619 | 0.00149 | 0.002743 | 1.201897374 | 42.60152981 | 98.00141723 | 48 | 5 | 5 | 7 | 30 | 4 | 1 | 0 | 0 | 0 | 3 | 0 |
| ISG000000 | Zkscan16 | 0.001727 | 0.003156 | 2.572986593 | 4.295252644 | 25.55847028 | 39 | 2 | 5 | 5 | 17 | 2 | 1 | 0 | 1 | 0 | 1 | 0 |
| SG0000003 | Zxdc | 0.001825 | 0.003321 | -0.5235005 | 762.3981746 | 530.3865407 | 100 | 5 | 5 | 15 | 44 | 6 | 9 | 0 | 0 | 1 | 3 | 0 |
| SG0000002 | Brix1 | 0.001871 | 0.0034 | 0.392460501 | 3013.685477 | 3955.854487 | 37 | 4 | 5 | 8 | 16 | 4 | 3 | 0 | 0 | 1 | 2 | 2 |
| ISG000000 | Zbtb40 | 0.002035 | 0.003681 | -0.543613751 | 550.3667454 | 377.5792675 | 85 | 2 | 5 | 9 | 30 | 12 | 1 | 0 | 0 | 0 | 0 | 0 |
| SG0000004 | Clspn | 0.002035 | 0.003681 | 0.422446872 | 1167.905641 | 1565.225579 | 54 | 3 | 5 | 7 | 26 | 5 | 2 | 0 | 0 | 1 | 1 | 0 |
| SG0000002 | Smtn | 0.00208 | 0.003758 | 0.552450846 | 501.9875092 | 736.2023282 | 47 | 5 | 5 | 15 | 24 | 0 | 0 | 0 | 0 | 0 | 0 | 0 |
| ISG000000 | Lcat | 0.002671 | 0.00475 | 1.484191083 | 14.06292433 | 39.34247478 | 7 | 4 | 5 | 5 | 8 | 0 | 0 | 0 | 0 | 0 | 0 | 0 |
| ISG000000 | Slc25a46 | 0.002727 | 0.004843 | -0.435337435 | 2113.366016 | 1562.878055 | 88 | 3 | 5 | 8 | 39 | 13 | 13 | 2 | 3 | 3 | 7 | 1 |
| ISG000000 | Katnb1 | 0.002909 | 0.005143 | 0.45512282 | 766.9216741 | 1051.372546 | 14 | 4 | 5 | 5 | 9 | 0 | 2 | 0 | 0 | 0 | 0 | 0 |
| ISG000000 | Aaas | 0.003117 | 0.005482 | 0.38365037 | 2496.556956 | 3257.104912 | 24 | 4 | 5 | 7 | 15 | 3 | 1 | 0 | 0 | 0 | 0 | 1 |
| SG0000006 | Mtx1 | 0.003234 | 0.005677 | -0.439407622 | 1359.301672 | 1002.399812 | 45 | 5 | 5 | 5 | 10 | 5 | 0 | 0 | 0 | 0 | 0 | 0 |
| ISG000000 | Clca3b | 0.004044 | 0.007002 | 5.137857294 | 0.615018556 | 21.65396239 | 70 | 2 | 5 | 6 | 32 | 3 | 1 | 0 | 0 | 0 | 0 | 0 |
| ISG000000 | Ppm1k | 0.004496 | 0.007741 | -0.544462917 | 446.4126161 | 306.0812949 | 83 | 3 | 5 | 14 | 31 | 5 | 14 | 0 | 1 | 2 | 4 | 1 |
| SG0000003 | Itih2 | 0.004805 | 0.008244 | -2.04235568 | 44.77199957 | 10.8691647 | 56 | 4 | 5 | 11 | 41 | 5 | 0 | 0 | 0 | 0 | 1 | 0 |
| ISG000000 | Tmem170b | 0.005659 | 0.009607 | -0.507516621 | 518.3554673 | 364.6279513 | 65 | 2 | 5 | 6 | 21 | 11 | 16 | 0 | 1 | 2 | 3 | 2 |
| SG0000003 | Fam188b | 0.00638 | 0.010754 | -0.742361764 | 149.4348872 | 89.3261964 | 106 | 2 | 5 | 10 | 40 | 7 | 0 | 0 | 0 | 0 | 0 | 0 |
| SG0000000 | Sult2b1 | 0.006651 | 0.011183 | -0.824864533 | 115.6408946 | 65.28335662 | 23 | 3 | 5 | 5 | 15 | 3 | 0 | 0 | 0 | 0 | 0 | 0 |
| SG0000004 | Tmem205 | 0.008101 | 0.013437 | -0.572201018 | 272.0000576 | 182.9444867 | 13 | 5 | 5 | 5 | 10 | 1 | 0 | 0 | 0 | 0 | 0 | 0 |
| ISG000000 | Tpcn1 | 0.008114 | 0.013456 | -0.465428373 | 783.4744048 | 567.4360348 | 45 | 4 | 5 | 10 | 21 | 1 | 1 | 0 | 0 | 0 | 1 | 0 |
| ISG000000 | Fam199x | 0.008224 | 0.013635 | -0.466377845 | 872.2676878 | 631.3294725 | 88 | 2 | 5 | 8 | 31 | 14 | 22 | 0 | 0 | 1 | 3 | 3 |
| ISG000000 | Chordc1 | 0.008584 | 0.014195 | -0.373312743 | 2898.139484 | 2237.386175 | 48 | 2 | 5 | 9 | 30 | 4 | 3 | 0 | 1 | 0 | 2 | 0 |
| ISG000000 | Lbp | 0.008777 | 0.014479 | 1.712594733 | 6.860224575 | 22.48436973 | 33 | 3 | 5 | 7 | 22 | 8 | 1 | 0 | 0 | 1 | 1 | 1 |
| ISG000000 | Intu | 0.013157 | 0.021049 | 0.469667639 | 483.9177025 | 670.1245936 | 144 | 1 | 5 | 10 | 59 | 17 | 11 | 0 | 0 | 1 | 4 | 0 |
| SG0000003 | Rbmx | 0.013252 | 0.021186 | -0.448674871 | 604.0212988 | 442.5757952 | 63 | 5 | 5 | 5 | 25 | 6 | 9 | 0 | 0 | 0 | 0 | 1 |
| ISG000000 | Zswim4 | 0.014068 | 0.022383 | 0.674001085 | 538.2743697 | 858.812278 | 17 | 5 | 5 | 6 | 13 | 3 | 1 | 1 | 1 | 1 | 1 | 0 |
| ISG000000 | Rsl1 | 0.014268 | 0.022689 | 0.62922955 | 87.03888507 | 134.6264171 | 22 | 3 | 5 | 9 | 16 | 0 | 0 | 0 | 0 | 0 | 0 | 0 |
| SG0000002 | H6pd | 0.015535 | 0.024561 | -0.379420659 | 1058.526613 | 813.7383465 | 55 | 5 | 5 | 12 | 30 | 5 | 2 | 0 | 0 | 0 | 0 | 0 |
| SG0000003 | Qsox2 | 0.015796 | 0.024944 | 0.362585829 | 640.3325482 | 823.2936923 | 47 | 4 | 5 | 9 | 21 | 4 | 10 | 2 | 2 | 2 | 3 | 0 |
| SG0000000 | Gmcl1 | 0.016034 | 0.025288 | -0.373852424 | 1131.385193 | 873.1114767 | 91 | 2 | 5 | 11 | 41 | 12 | 5 | 0 | 0 | 1 | 1 | 0 |
| SG0000005 | Suclg1 | 0.01645 | 0.025881 | 0.313457698 | 1935.144043 | 2404.769566 | 75 | 0 | 5 | 6 | 17 | 5 | 0 | 0 | 0 | 0 | 0 | 0 |
| ISG000000 | Ppap2b | 0.017683 | 0.027629 | -0.539055017 | 767.1757781 | 527.9869071 | 88 | 2 | 5 | 9 | 40 | 6 | 2 | 0 | 0 | 0 | 2 | 0 |
| SG0000002 | Wnt5a | 0.019239 | 0.02985 | -0.715342402 | 440.9610791 | 268.5720032 | 33 | 1 | 5 | 6 | 11 | 6 | 16 | 1 | 3 | 6 | 10 | 4 |
| ISG000000 | Zfp944 | 0.019932 | 0.030848 | 0.433874818 | 251.3561765 | 339.5462169 | 57 | 2 | 5 | 9 | 29 | 8 | 1 | 0 | 0 | 0 | 0 | 1 |
| SG0000005 | Isoc2b | 0.020906 | 0.032221 | -1.343627154 | 45.20261765 | 17.81113145 | 27 | 5 | 5 | 6 | 18 | 2 | 1 | 0 | 0 | 0 | 0 | 0 |
| ISG000000 | Timm17a | 0.021682 | 0.033327 | 0.304058139 | 2280.058019 | 2814.987811 | 29 | 2 | 5 | 4 | 21 | 5 | 11 | 1 | 2 | 2 | 8 | 5 |
| SG0000004 | Syt12 | 0.023301 | 0.035609 | -1.393024514 | 38.29505322 | 14.58144332 | 33 | 5 | 5 | 10 | 19 | 2 | 1 | 0 | 0 | 0 | 0 | 0 |
| SG0000004 | Chtf8 | 0.02426 | 0.036992 | 0.290286294 | 4178.839343 | 5110.231287 | 66 | 0 | 5 | 5 | 25 | 5 | 6 | 0 | 0 | 0 | 0 | 0 |
| SG0000003 | Dnajc11 | 0.02645 | 0.040092 | 0.294837733 | 2323.196002 | 2849.973704 | 36 | 4 | 5 | 9 | 13 | 6 | 1 | 0 | 0 | 1 | 1 | 0 |
| SG0000002 | Ddx5 | 0.028073 | 0.042373 | -0.28390257 | 25988.46251 | 21346.04389 | 21 | 2 | 5 | 5 | 13 | 1 | 10 | 0 | 1 | 1 | 1 | 1 |
| SG0000001 | B4galt5 | 0.029687 | 0.044597 | 0.299655126 | 1314.974777 | 1618.536896 | 54 | 4 | 5 | 11 | 23 | 4 | 1 | 0 | 0 | 0 | 0 | 2 |
| SG0000006 | Ccni | 0.029858 | 0.04483 | 0.373413194 | 5005.152817 | 6483.744819 | 78 | 3 | 5 | 8 | 15 | 4 | 5 | 0 | 0 | 0 | 2 | 0 |
| SG0000000 | Vps72 | 0.030601 | 0.045854 | 0.346140083 | 884.0246089 | 1123.732399 | 26 | 5 | 5 | 6 | 20 | 3 | 1 | 0 | 0 | 0 | 0 | 0 |
| SG0000002 | Csrp1 | 0.033001 | 0.049185 | 0.291411154 | 2738.859046 | 3351.916416 | 62 | 4 | 5 | 5 | 12 | 7 | 6 | 0 | 0 | 0 | 0 | 0 |
| ISG000000 | Coq3 | 0.036704 | 0.0542 | 0.337451697 | 466.3926825 | 589.2977876 | 72 | 3 | 5 | 13 | 41 | 8 | 3 | 0 | 0 | 2 | 2 | 0 |
| ISG000000 | Zfp623 | 0.036747 | 0.054259 | -0.38085644 | 557.6562414 | 428.2697055 | 16 | 5 | 5 | 5 | 8 | 0 | 0 | 0 | 0 | 0 | 0 | 0 |
| SG0000000 | Insig2 | 0.044106 | 0.064295 | -0.316189843 | 1185.251032 | 951.9797604 | 193 | 0 | 5 | 27 | 103 | 15 | 7 | 0 | 0 | 2 | 3 | 0 |
| SG0000002 | Tarbp2 | 0.045091 | 0.065633 | 0.300794334 | 769.3322089 | 947.6806914 | 15 | 0 | 5 | 5 | 5 | 0 | 0 | 0 | 0 | 0 | 0 | 0 |
| ISG000000 | Cyp2j6 | 0.045304 | 0.065915 | -0.780631663 | 60.19260381 | 35.03882907 | 91 | 3 | 5 | 11 | 45 | 2 | 3 | 0 | 1 | 0 | 1 | 0 |
| SG0000005 | Zfp697 | 0.051653 | 0.074343 | 0.784247149 | 31.79889891 | 54.76385325 | 138 | 4 | 5 | 22 | 67 | 14 | 14 | 0 | 1 | 2 | 5 | 2 |
| SG0000004 | Polr1a | 0.054475 | 0.078073 | -0.259303068 | 3610.236678 | 3016.321798 | 93 | 2 | 5 | 10 | 47 | 4 | 0 | 0 | 0 | 0 | 0 | 0 |
| SG0000006 | Cdpf1 | 0.054959 | 0.078722 | 0.377877756 | 236.0272522 | 306.7006452 | 21 | 0 | 5 | 15 | 25 | 5 | 1 | 0 | 0 | 0 | 0 | 0 |
| ISG000000 | Arl5a | 0.055591 | 0.079538 | -0.280696475 | 1866.287118 | 1536.315456 | 65 | 3 | 5 | 6 | 21 | 7 | 13 | 0 | 0 | 1 | 3 | 2 |
| ISG000000 | Prrc1 | 0.060459 | 0.085859 | -0.300348332 | 891.3505959 | 723.8268717 | 70 | 1 | 5 | 5 | 31 | 8 | 8 | 0 | 0 | 0 | 2 | 0 |
| SG0000003 | Aph1b | 0.063805 | 0.090289 | 0.508221668 | 93.49807975 | 132.981938 | 149 | 5 | 5 | 22 | 55 | 15 | 15 | 2 | 2 | 4 | 11 | 0 |
| SG0000003 | Tmem161b | 0.065025 | 0.091907 | 0.305093878 | 394.8428336 | 487.8279378 | 191 | 1 | 5 | 23 | 108 | 19 | 2 | 0 | 0 | 1 | 3 | 1 |
| ISG000000 | Fdx1 | 0.065282 | 0.092241 | 0.390051429 | 164.0311805 | 214.9530395 | 53 | 3 | 5 | 9 | 20 | 5 | 0 | 0 | 0 | 0 | 1 | 0 |
| SG0000002 | Adal | 0.065337 | 0.092311 | 0.308667966 | 427.0983356 | 528.9884569 | 170 | 0 | 5 | 10 | 39 | 5 | 4 | 0 | 0 | 0 | 0 | 0 |
| SG0000005 | Fbxo18 | 0.070545 | 0.09902 | -0.267368736 | 1782.57828 | 1481.025721 | 50 | 3 | 5 | 10 | 18 | 2 | 1 | 0 | 0 | 0 | 0 | 0 |
| SG0000002 | Capn7 | 0.071135 | 0.099755 | -0.263181658 | 1897.95119 | 1581.464568 | 96 | 2 | 5 | 11 | 34 | 10 | 4 | 0 | 0 | 1 | 1 | 0 |
| SG0000001 | Vta1 | 0.073529 | 0.10287 | -0.266471807 | 1259.317622 | 1046.933927 | 97 | 2 | 5 | 19 | 70 | 15 | 2 | 0 | 0 | 0 | 1 | 0 |
| ISG000000 | Icam5 | 0.089163 | 0.122674 | 1.020059764 | 11.46682991 | 23.25476508 | 12 | 5 | 5 | 7 | 8 | 0 | 1 | 0 | 0 | 1 | 1 | 0 |
| ISG000000 | Itpk1 | 0.089738 | 0.123398 | -0.299578961 | 4824.038413 | 3919.480464 | 100 | 5 | 5 | 13 | 40 | 8 | 1 | 0 | 0 | 0 | 0 | 0 |
| SG0000003 | Etfa | 0.095181 | 0.130184 | 0.214257434 | 2304.40439 | 2673.354845 | 115 | 3 | 5 | 13 | 67 | 14 | 0 | 0 | 0 | 0 | 1 | 0 |
| ISG000000 | Rpap3 | 0.098259 | 0.133922 | 0.231006396 | 1038.980081 | 1219.402485 | 61 | 4 | 5 | 7 | 25 | 4 | 0 | 0 | 0 | 0 | 0 | 0 |
| SG0000003 | Zfp113 | 0.108281 | 0.146391 | -0.348923208 | 259.7915976 | 203.9805455 | 26 | 5 | 5 | 9 | 17 | 2 | 11 | 2 | 2 | 4 | 8 | 0 |
| ISG000000 | Nipal1 | 0.114226 | 0.153432 | 0.887556788 | 25.52107055 | 47.21501197 | 37 | 4 | 5 | 9 | 17 | 4 | 4 | 0 | 0 | 1 | 1 | 0 |
| SG0000002 | Dnajc4 | 0.132483 | 0.175586 | -0.340275639 | 213.4039047 | 168.5656918 | 6 | 5 | 5 | 5 | 6 | 0 | 0 | 0 | 0 | 0 | 0 | 0 |
| ISG000000 | 2610008E11Rik | 0.14244 | 0.187478 | -0.400366281 | 188.1991051 | 142.5920439 | 58 | 2 | 5 | 5 | 42 | 7 | 2 | 0 | 0 | 0 | 0 | 0 |
| ISG000000 | Acat1 | 0.145464 | 0.191121 | 0.267610695 | 770.7046977 | 927.783944 | 42 | 4 | 5 | 9 | 32 | 5 | 1 | 0 | 0 | 0 | 4 | 1 |
| ISG000000 | Utp20 | 0.152472 | 0.199509 | -0.231335436 | 4184.455013 | 3564.511317 | 119 | 3 | 5 | 12 | 45 | 5 | 0 | 0 | 0 | 0 | 1 | 0 |
| SG0000003 | Pde2a | 0.170208 | 0.220605 | 0.322072099 | 127.6410844 | 159.567282 | 55 | 4 | 5 | 9 | 20 | 5 | 0 | 0 | 0 | 0 | 0 | 0 |
| ISG000000 | Nap1l4 | 0.180761 | 0.232851 | 0.169797164 | 3782.457321 | 4254.887443 | 43 | 4 | 5 | 6 | 21 | 1 | 2 | 1 | 1 | 1 | 1 | 0 |
| SG0000003 | Idh3a | 0.194018 | 0.248161 | -0.178783319 | 4072.41547 | 3597.766194 | 29 | 5 | 5 | 9 | 14 | 3 | 4 | 0 | 0 | 1 | 1 | 1 |
| ISG000000 | Atp5l | 0.195498 | 0.249805 | -0.47333872 | 129.44814 | 93.24094399 | 18 | 5 | 5 | 6 | 10 | 1 | 0 | 0 | 0 | 0 | 0 | 0 |
| ISG000000 | BC024978 | 0.210158 | 0.266427 | 0.24863186 | 252.4452 | 299.9250673 | 20 | 4 | 5 | 9 | 9 | 0 | 8 | 2 | 3 | 3 | 3 | 0 |
| ISG000000 | Tmed8 | 0.22633 | 0.284612 | 0.17186706 | 924.413723 | 1041.366312 | 54 | 3 | 5 | 8 | 44 | 4 | 11 | 1 | 1 | 3 | 10 | 0 |
| ISG000000 | Ube2z | 0.234921 | 0.294074 | -0.157955429 | 5058.754213 | 4534.13302 | 25 | 5 | 5 | 5 | 23 | 3 | 2 | 0 | 0 | 0 | 0 | 2 |
| SG0000002 | Stk25 | 0.248412 | 0.309126 | 0.153335526 | 2519.11788 | 2801.606157 | 14 | 4 | 5 | 7 | 10 | 2 | 1 | 0 | 0 | 0 | 0 | 0 |
| SG0000002 | Ttf1 | 0.270429 | 0.332916 | 0.161684062 | 832.5160999 | 931.2459421 | 35 | 4 | 5 | 9 | 19 | 4 | 0 | 0 | 0 | 0 | 0 | 0 |
| ISG000000 | Ttc9 | 0.283315 | 0.346895 | 0.405795696 | 42.3903877 | 56.15960768 | 44 | 5 | 5 | 5 | 16 | 3 | 0 | 0 | 0 | 0 | 1 | 1 |
| ISG000000 | Slc25a37 | 0.284349 | 0.347947 | 0.150282636 | 1946.823171 | 2160.558788 | 44 | 2 | 5 | 8 | 16 | 3 | 5 | 0 | 0 | 1 | 3 | 1 |
| ISG000000 | Rab33b | 0.285611 | 0.349253 | -0.218911581 | 267.7451921 | 230.0502601 | 31 | 4 | 5 | 7 | 14 | 4 | 6 | 0 | 0 | 1 | 1 | 0 |

| | | | | | | | | | | | | | | | | | | |
|---|---|---|---|---|---|---|---|---|---|---|---|---|---|---|---|---|---|---|
| ISG000000 | Ddx1 | 0.291675 | 0.355818 | 0.131591237 | 6059.571273 | 6638.268397 | 69 | 2 | 5 | 7 | 26 | 4 | 1 | 0 | 0 | 0 | 0 | 0 |
| ISG000000 | Gtpbp1 | 0.331358 | 0.398154 | 0.129716483 | 1901.705284 | 2080.615193 | 30 | 5 | 5 | 7 | 16 | 1 | 1 | 0 | 0 | 1 | 2 | 0 |
| ISG000000 | Pdcd11 | 0.347023 | 0.414419 | -0.131078738 | 3353.820637 | 3062.535988 | 47 | 3 | 5 | 8 | 20 | 4 | 0 | 0 | 0 | 0 | 1 | 0 |
| SG0000003 | Mtg1 | 0.360066 | 0.428254 | 0.154066241 | 473.3553171 | 526.7030338 | 26 | 3 | 5 | 3 | 9 | 3 | 2 | 1 | 1 | 1 | 1 | 0 |
| ISG000000 | Unc5b | 0.37676 | 0.445653 | -0.198646 | 1638.061573 | 1427.3544 | 51 | 3 | 5 | 5 | 15 | 4 | 1 | 0 | 0 | 0 | 1 | 2 |
| ISG000000 | Timmdc1 | 0.390864 | 0.459514 | -0.141616987 | 679.1063026 | 615.6116078 | 56 | 5 | 5 | 13 | 33 | 3 | 2 | 1 | 1 | 1 | 1 | 0 |
| ISG000000 | Tmem17 | 0.394524 | 0.463249 | -0.286698841 | 72.76297803 | 59.6493224 | 19 | 4 | 5 | 7 | 16 | 1 | 0 | 0 | 0 | 0 | 1 | 0 |
| ISG000000 | Pus7l | 0.400221 | 0.469199 | 0.160054017 | 257.4441079 | 287.6497606 | 50 | 3 | 5 | 12 | 30 | 4 | 2 | 0 | 1 | 1 | 1 | 0 |
| SG0000003 | Nudt7 | 0.40561 | 0.474669 | -0.219427796 | 146.625154 | 125.9372289 | 151 | 0 | 5 | 26 | 80 | 5 | 7 | 0 | 0 | 2 | 6 | 1 |
| ISG000000 | Sirt5 | 0.413348 | 0.482526 | 0.196616494 | 134.2316121 | 153.830435 | 27 | 3 | 5 | 6 | 17 | 9 | 0 | 0 | 0 | 0 | 1 | 0 |
| ISG000000 | Arhgef17 | 0.445713 | 0.514772 | 0.181334472 | 572.7786192 | 649.4921606 | 40 | 3 | 5 | 9 | 19 | 5 | 4 | 0 | 0 | 2 | 2 | 0 |
| ISG000000 | Snw1 | 0.465372 | 0.534822 | -0.100756166 | 3599.792567 | 3356.965266 | 39 | 5 | 5 | 6 | 16 | 4 | 0 | 0 | 0 | 0 | 0 | 0 |
| SG0000002 | Lztfl1 | 0.465417 | 0.53484 | 0.113299962 | 449.4822124 | 486.2047496 | 176 | 0 | 5 | 13 | 100 | 35 | 60 | 0 | 0 | 0 | 14 | 0 |
| ISG000000 | Gabarapl2 | 0.468487 | 0.537326 | 0.117096117 | 671.9151767 | 728.7253782 | 24 | 5 | 5 | 7 | 11 | 4 | 3 | 1 | 1 | 1 | 1 | 1 |
| ISG000001 | Gm28557 | 0.480793 | 0.549442 | 0.381228631 | 18.61047197 | 24.23921999 | 90 | 2 | 5 | 12 | 36 | 5 | 6 | 0 | 0 | 0 | 0 | 1 |
| SG0000002 | Mib2 | 0.481355 | 0.54984 | 0.191591474 | 471.3962613 | 538.3452856 | 17 | 5 | 5 | 5 | 10 | 2 | 0 | 0 | 0 | 0 | 1 | 0 |
| ISG000000 | Polr3c | 0.48346 | 0.551927 | 0.098384506 | 1402.794752 | 1501.795573 | 53 | 3 | 5 | 9 | 24 | 2 | 0 | 0 | 0 | 0 | 0 | 0 |
| SG0000003 | Wbp2 | 0.486544 | 0.554776 | -0.087604484 | 3808.01678 | 3583.663934 | 14 | 4 | 5 | 9 | 9 | 0 | 2 | 0 | 1 | 1 | 1 | 0 |
| SG0000002 | Ska2 | 0.512293 | 0.579195 | 0.090821888 | 1200.119003 | 1278.098794 | 36 | 4 | 5 | 9 | 15 | 5 | 8 | 4 | 4 | 6 | 8 | 2 |
| ISG000000 | Papss2 | 0.581524 | 0.64521 | -0.238432416 | 41.81267861 | 35.44318007 | 65 | 2 | 5 | 7 | 45 | 8 | 4 | 0 | 0 | 1 | 3 | 0 |
| SG0000003 | Bend3 | 0.598757 | 0.661543 | 0.081794719 | 958.3143626 | 1014.216475 | 52 | 3 | 5 | 6 | 21 | 2 | 7 | 0 | 0 | 2 | 4 | 0 |
| ISG000000 | Mterf2 | 0.603893 | 0.666559 | 0.109417371 | 147.5849913 | 159.2135789 | 17 | 5 | 5 | 6 | 6 | 1 | 0 | 0 | 0 | 0 | 0 | 0 |
| ISG000000 | Bcar1 | 0.637729 | 0.697843 | -0.10977738 | 1567.447564 | 1452.602333 | 21 | 1 | 5 | 3 | 11 | 5 | 1 | 0 | 0 | 1 | 1 | 1 |
| SG0000007 | Zfp65 | 0.642328 | 0.702104 | -0.094181642 | 482.1831941 | 451.7108998 | 51 | 1 | 5 | 8 | 29 | 4 | 7 | 0 | 1 | 1 | 2 | 1 |
| SG0000001 | Nipsnap3b | 0.646779 | 0.706399 | -0.074900757 | 731.30624 | 694.3075768 | 43 | 3 | 5 | 7 | 22 | 10 | 5 | 1 | 1 | 1 | 2 | 0 |
| ISG000000 | Habp4 | 0.678399 | 0.735295 | -0.096916755 | 331.2015393 | 309.6830933 | 45 | 3 | 5 | 7 | 19 | 6 | 1 | 0 | 0 | 0 | 0 | 1 |
| SG0000002 | Slc35a1 | 0.755349 | 0.802765 | 0.044876064 | 676.563948 | 697.9396853 | 63 | 4 | 5 | 11 | 30 | 5 | 2 | 0 | 0 | 0 | 1 | 0 |
| SG0000002 | Tfb2m | 0.792877 | 0.835216 | -0.064288059 | 250.672223 | 239.747219 | 39 | 5 | 5 | 7 | 15 | 6 | 4 | 0 | 0 | 0 | 2 | 1 |
| SG0000004 | Lsm11 | 0.828616 | 0.865585 | 0.033528474 | 421.5975971 | 431.5103365 | 40 | 5 | 5 | 7 | 19 | 0 | 12 | 1 | 1 | 1 | 1 | 0 |
| ISG000000 | Vapa | 0.849537 | 0.882344 | -0.033720788 | 5457.08913 | 5331.017134 | 82 | 3 | 5 | 14 | 51 | 6 | 3 | 0 | 0 | 0 | 1 | 0 |
| SG0000003 | Micall2 | 0.864589 | 0.894916 | -0.039153802 | 134.4304238 | 130.831132 | 36 | 4 | 5 | 5 | 15 | 3 | 0 | 0 | 0 | 0 | 0 | 0 |
| SG0000001 | Psen2 | 0.87718 | 0.905594 | -0.032195197 | 386.537058 | 378.0066297 | 62 | 0 | 5 | 0 | 10 | 13 | 0 | 0 | 0 | 0 | 0 | 0 |
| SG0000005 | Ccdc176 | 0.880221 | 0.907739 | 0.063961109 | 23.35778463 | 24.41663786 | 60 | 2 | 5 | 6 | 30 | 2 | 1 | 0 | 0 | 1 | 2 | 0 |
| SG0000003 | Mrpl55 | 0.890655 | 0.916284 | 0.024573965 | 390.9290277 | 397.6449029 | 22 | 5 | 5 | 5 | 10 | 5 | 7 | 0 | 0 | 0 | 0 | 0 |
| SG0000003 | Noa1 | 0.893335 | 0.918196 | 0.01927037 | 537.4928149 | 544.7203777 | 24 | 2 | 5 | 4 | 10 | 1 | 1 | 0 | 1 | 1 | 1 | 0 |
| SG0000001 | Fuca2 | 0.91891 | 0.940867 | 0.010929673 | 637.6485076 | 642.4975961 | 66 | 3 | 5 | 10 | 37 | 6 | 4 | 1 | 1 | 2 | 5 | 2 |
| SG0000004 | Arhgap11a | 0.941475 | 0.958055 | 0.003467942 | 2546.849096 | 2552.978561 | 137 | 2 | 5 | 17 | 57 | 49 | 14 | 2 | 5 | 5 | 5 | 32 |
| ISG000000 | Chaf1a | 0.96274 | 0.977031 | -0.007292653 | 3220.942907 | 3204.702502 | 36 | 5 | 5 | 10 | 20 | 4 | 0 | 0 | 0 | 0 | 0 | 0 |
| SG0000005 | Rnd1 | 0.969533 | 0.982978 | 0.024524388 | 131.3772324 | 133.6296052 | 21 | 5 | 5 | 7 | 15 | 1 | 2 | 1 | 1 | 2 | 4 | 0 |
| SG0000003 | Tsen34 | 0.969784 | 0.983121 | -0.011025309 | 1867.347199 | 1853.131021 | 31 | 5 | 5 | 5 | 30 | 0 | 0 | 0 | 0 | 0 | 0 | 0 |
| SG0000005 | Tmem154 | 0.989431 | 0.999097 | 0.026857115 | 23.89718739 | 24.34622248 | 68 | 3 | 5 | 12 | 26 | 3 | 4 | 0 | 0 | 1 | 2 | 0 |
| ISG000000 | Akr7a5 | 0.991312 | 1 | -0.006569913 | 533.2733058 | 530.8503446 | 11 | 4 | 5 | 5 | 7 | 0 | 0 | 0 | 0 | 0 | 0 | 0 |
| ISG000000 | Prpf4 | 0.993412 | 1 | -0.006601541 | 2014.227343 | 2005.031617 | 23 | 4 | 5 | 4 | 12 | 3 | 4 | 0 | 1 | 0 | 1 | 0 |
| ISG000000 | Pim1 | 1.8E-157 | 3.2E-155 | 3.430156552 | 434.3436445 | 4681.806745 | 8 | 4 | 4 | 5 | 7 | 1 | 5 | 3 | 3 | 4 | 5 | 1 |
| SG0000002 | Ptch2 | 7.5E-143 | 1.1E-140 | 9.77046022 | 0.639110347 | 558.1840804 | 16 | 3 | 4 | 6 | 15 | 1 | 1 | 0 | 1 | 1 | 1 | 1 |
| SG0000004 | Slc16a6 | 2.5E-141 | 3.5E-139 | 5.600156244 | 16.6364268 | 807.0028406 | 49 | 2 | 4 | 7 | 28 | 12 | 7 | 0 | 0 | 1 | 4 | 0 |
| SG0000003 | Camkv | 4.1E-141 | 5.6E-139 | 6.735649671 | 5.854547258 | 623.9156898 | 34 | 4 | 4 | 8 | 10 | 6 | 0 | 0 | 0 | 0 | 0 | 2 |
| ISG000000 | Cdcp1 | 1.3E-129 | 1.5E-127 | 8.656904083 | 1.278220693 | 515.9333987 | 38 | 3 | 4 | 8 | 27 | 4 | 5 | 1 | 2 | 2 | 2 | 0 |
| SG0000003 | Galnt6 | 7.7E-115 | 7.3E-113 | 10.54041561 | 0.970711415 | 1445.677439 | 25 | 4 | 4 | 4 | 7 | 11 | 0 | 0 | 0 | 0 | 0 | 0 |
| ISG000000 | Hmgcs1 | 7.9E-107 | 6.7E-105 | 2.66522329 | 1075.765216 | 6823.85285 | 49 | 3 | 4 | 7 | 14 | 3 | 4 | 0 | 0 | 0 | 0 | 0 |
| ISG000000 | Nt5e | 1.8E-105 | 1.5E-103 | -5.424242253 | 8348.727952 | 194.4288411 | 60 | 3 | 4 | 8 | 19 | 6 | 2 | 0 | 0 | 0 | 2 | 0 |
| SG0000002 | Mal | 5.5E-102 | 4E-100 | -8.142760218 | 18939.025 | 67.01045154 | 22 | 2 | 4 | 2 | 17 | 4 | 2 | 0 | 0 | 0 | 1 | 0 |
| SG0000004 | Stab1 | 3.6E-100 | 2.54E-98 | -6.763938748 | 1404.383412 | 12.92222323 | 22 | 3 | 4 | 6 | 8 | 0 | 0 | 0 | 0 | 0 | 0 | 0 |
| SG0000002 | Gem | 3.99E-95 | 2.52E-93 | -5.24298763 | 1555.535488 | 41.07554932 | 31 | 2 | 4 | 12 | 14 | 1 | 8 | 0 | 0 | 6 | 8 | 0 |
| ISG000000 | Slc25a48 | 7.91E-93 | 4.79E-91 | -6.050505314 | 1321.101617 | 19.93208064 | 46 | 3 | 4 | 10 | 28 | 8 | 1 | 0 | 0 | 0 | 0 | 0 |
| SG0000002 | Stom | 3.73E-89 | 2.07E-87 | -5.636939609 | 40599.61184 | 815.8944432 | 25 | 2 | 4 | 4 | 21 | 5 | 1 | 0 | 0 | 0 | 0 | 2 |
| ISG000000 | Kif5a | 4.3E-86 | 2.21E-84 | 5.015957458 | 14.03648897 | 454.1634086 | 38 | 2 | 4 | 8 | 14 | 8 | 2 | 0 | 0 | 0 | 0 | 1 |
| SG0000003 | Msmo1 | 5.57E-86 | 2.84E-84 | 2.468466153 | 768.3170477 | 4252.289509 | 22 | 2 | 4 | 4 | 15 | 4 | 1 | 0 | 0 | 0 | 1 | 1 |
| SG0000004 | Ric3 | 1.83E-85 | 9.25E-84 | 6.783464917 | 2.885698887 | 317.89041 | 273 | 2 | 4 | 22 | 77 | 27 | 14 | 0 | 0 | 0 | 0 | 2 |
| ISG000000 | Dusp27 | 1.84E-83 | 9E-82 | 5.037993342 | 13.06577756 | 429.26193 | 66 | 2 | 4 | 12 | 28 | 8 | 2 | 0 | 0 | 0 | 0 | 0 |
| ISG000000 | Tmem170 | 1.33E-72 | 5.14E-71 | 3.120933065 | 93.55194733 | 813.8554929 | 19 | 1 | 4 | 5 | 16 | 0 | 4 | 0 | 1 | 0 | 0 | 0 |
| ISG000000 | Ska3 | 8E-71 | 2.93E-69 | 2.367848602 | 468.8293119 | 2419.960131 | 23 | 2 | 4 | 6 | 17 | 2 | 0 | 0 | 0 | 0 | 0 | 1 |
| SG0000004 | Dact1 | 2.03E-69 | 7.1E-68 | 4.900277053 | 11.01327004 | 328.8869704 | 20 | 1 | 4 | 3 | 10 | 1 | 2 | 0 | 0 | 0 | 0 | 0 |
| SG000000C | Timp1 | 5.89E-66 | 1.9E-64 | -3.017652039 | 10886.39738 | 1344.251108 | 16 | 2 | 4 | 4 | 8 | 2 | 0 | 0 | 0 | 0 | 0 | 0 |
| ISG000000 | Socs5 | 1.69E-65 | 5.37E-64 | -3.27681061 | 3394.948548 | 350.2796549 | 47 | 4 | 4 | 6 | 21 | 7 | 3 | 0 | 0 | 1 | 2 | 1 |
| SG0000003 | Tnfrsf1a | 1.25E-61 | 3.64E-60 | -3.43281626 | 3303.948115 | 305.9514774 | 9 | 4 | 4 | 5 | 5 | 3 | 0 | 0 | 0 | 0 | 0 | 2 |
| ISG000000 | Gm3604 | 9.23E-60 | 2.53E-58 | 5.509561401 | 5.239528701 | 238.6906837 | 35 | 4 | 4 | 4 | 18 | 4 | 0 | 0 | 0 | 0 | 0 | 0 |
| ISG000000 | Cav2 | 1.05E-59 | 2.86E-58 | -4.501063571 | 2519.960003 | 111.2854793 | 20 | 2 | 4 | 4 | 16 | 0 | 4 | 0 | 0 | 0 | 6 | 0 |
| ISG000000 | St8sia2 | 1.08E-59 | 2.95E-58 | -6.821129926 | 2095.109244 | 18.5285725 | 76 | 1 | 4 | 15 | 31 | 10 | 10 | 0 | 1 | 6 | 8 | 0 |
| SG0000002 | Cdkn1a | 4.15E-59 | 1.12E-57 | -2.745226083 | 15730.7719 | 2346.143857 | 17 | 0 | 4 | 4 | 14 | 3 | 4 | 0 | 1 | 0 | 5 | 2 |
| ISG000000 | Calb2 | 5.43E-57 | 1.37E-55 | 7.488036105 | 0 | 178.5243969 | 29 | 3 | 4 | 5 | 9 | 1 | 1 | 0 | 0 | 0 | 0 | 0 |
| SG0000002 | Socs2 | 5.52E-56 | 1.37E-54 | 3.612285182 | 133.9662961 | 1638.331355 | 74 | 2 | 4 | 15 | 44 | 2 | 11 | 0 | 0 | 5 | 12 | 0 |
| SG0000001 | Mybl2 | 2.33E-55 | 5.68E-54 | 1.848961238 | 3253.034775 | 11718.75567 | 24 | 3 | 4 | 3 | 10 | 2 | 0 | 0 | 0 | 0 | 0 | 0 |
| SG0000002 | Idh1 | 1.02E-54 | 2.43E-53 | 1.936794133 | 1117.61806 | 4278.844863 | 78 | 2 | 4 | 10 | 37 | 2 | 0 | 0 | 0 | 0 | 4 | 0 |
| ISG000000 | Tnfsf11 | 2.72E-54 | 6.36E-53 | -4.850295296 | 2293.79631 | 79.51892091 | 69 | 1 | 4 | 13 | 37 | 3 | 6 | 0 | 0 | 1 | 4 | 0 |
| SG0000003 | Mrpl15 | 4.32E-52 | 9.51E-51 | 1.847628082 | 1616.440256 | 5817.697438 | 86 | 4 | 4 | 8 | 46 | 14 | 6 | 0 | 0 | 0 | 4 | 1 |
| ISG000000 | Mrps31 | 5.63E-52 | 1.23E-50 | 2.130637038 | 375.0046772 | 1642.18599 | 23 | 0 | 4 | 4 | 17 | 2 | 1 | 0 | 1 | 0 | 1 | 0 |
| SG0000002 | Klhl22 | 2.26E-46 | 4.18E-45 | 3.291209297 | 139.4542764 | 1365.16315 | 72 | 2 | 4 | 10 | 44 | 10 | 0 | 0 | 0 | 0 | 0 | 0 |
| SG0000004 | Zmym1 | 3.61E-46 | 6.6E-45 | 1.910498482 | 588.9223839 | 2213.988757 | 53 | 2 | 4 | 6 | 36 | 0 | 2 | 0 | 0 | 0 | 0 | 0 |
| ISG000000 | Hebp1 | 9.99E-45 | 1.74E-43 | 2.669284864 | 83.86308783 | 533.4646672 | 30 | 1 | 4 | 5 | 18 | 3 | 0 | 0 | 0 | 0 | 0 | 0 |
| SG0000002 | Abcc3 | 2.3E-44 | 3.94E-43 | -6.053270638 | 1162.231303 | 17.50155086 | 84 | 4 | 4 | 14 | 46 | 12 | 2 | 0 | 0 | 0 | 0 | 0 |
| ISG000000 | Rela | 3.4E-43 | 5.57E-42 | -2.455670103 | 2543.785529 | 463.7139187 | 9 | 3 | 4 | 3 | 4 | 2 | 0 | 0 | 0 | 0 | 0 | 0 |
| ISG000000 | Gcsh | 2.68E-42 | 4.27E-41 | 1.782223226 | 988.9024323 | 3401.387373 | 18 | 3 | 4 | 4 | 6 | 0 | 1 | 0 | 0 | 0 | 0 | 0 |
| SG0000001 | Slc2a4 | 2.83E-42 | 4.49E-41 | 7.737398474 | 0.639110347 | 136.3844565 | 17 | 0 | 4 | 9 | 9 | 1 | 3 | 0 | 0 | 1 | 1 | 1 |
| SG0000002 | Mpv17l | 3.58E-42 | 5.68E-41 | 2.708708681 | 70.74159301 | 462.4634321 | 112 | 3 | 4 | 12 | 58 | 8 | 37 | 0 | 0 | 0 | 6 | 0 |
| ISG000000 | Six1 | 6.11E-42 | 9.64E-41 | 2.693433888 | 74.49809902 | 481.8918513 | 14 | 4 | 4 | 6 | 6 | 1 | 13 | 3 | 3 | 5 | 5 | 1 |
| SG0000002 | Jph3 | 1.13E-41 | 1.77E-40 | 8.659338072 | 0.307509278 | 124.3307994 | 79 | 0 | 4 | 14 | 46 | 24 | 1 | 0 | 0 | 0 | 0 | 0 |
| SG0000002 | Ripk1 | 1.74E-41 | 2.71E-40 | -2.946985796 | 897.3249805 | 116.3639987 | 99 | 2 | 4 | 6 | 46 | 16 | 1 | 0 | 0 | 0 | 0 | 0 |
| SG0000004 | Ccnb1 | 1.91E-40 | 2.86E-39 | 1.593500382 | 2817.888325 | 8503.842161 | 26 | 2 | 4 | 2 | 14 | 4 | 8 | 2 | 2 | 2 | 4 | 4 |
| SG0000002 | Nkd2 | 5.79E-39 | 8.23E-38 | 4.488686504 | 7.841810101 | 176.0538797 | 45 | 0 | 4 | 10 | 20 | 1 | 1 | 0 | 0 | 0 | 0 | 1 |
| SG0000002 | Tnfaip2 | 2.45E-38 | 3.43E-37 | -3.862084331 | 3873.123063 | 266.3532907 | 18 | 4 | 4 | 4 | 10 | 0 | 4 | 0 | 0 | 0 | 4 | 0 |
| SG0000006 | Tcp11 | 3.88E-38 | 5.38E-37 | 3.787317411 | 15.46779095 | 213.5622254 | 47 | 4 | 4 | 4 | 28 | 4 | 0 | 0 | 0 | 0 | 0 | 0 |
| ISG000000 | Ccnb1ip1 | 4.46E-38 | 6.17E-37 | 10.12385662 | 0.34247518 | 382.1321835 | 15 | 4 | 4 | 5 | 17 | 3 | 1 | 0 | 0 | 0 | 0 | 0 |
| ISG000000 | Atg14 | 5.3E-38 | 7.3E-37 | 1.849872923 | 376.5458703 | 1357.328996 | 49 | 2 | 4 | 8 | 24 | 7 | 2 | 0 | 0 | 0 | 0 | 1 |
| SG0000005 | Zfp819 | 5.74E-38 | 7.9E-37 | 3.765947593 | 28.38901777 | 386.2011424 | 26 | 4 | 4 | 6 | 11 | 2 | 0 | 0 | 0 | 0 | 1 | 0 |
| SG0000003 | Ephx1 | 1.03E-37 | 1.4E-36 | -2.511212639 | 5511.243187 | 966.7167535 | 56 | 4 | 4 | 10 | 26 | 6 | 0 | 0 | 0 | 0 | 0 | 0 |
| SG0000003 | Npepl1 | 1.77E-37 | 2.39E-36 | -2.242598392 | 2779.615622 | 587.3478181 | 17 | 0 | 4 | 2 | 8 | 0 | 0 | 0 | 0 | 0 | 0 | 0 |
| SG0000003 | Lgi2 | 3.33E-36 | 4.3E-35 | 6.860192192 | 0.981585527 | 114.0385724 | 66 | 2 | 4 | 6 | 14 | 6 | 0 | 0 | 0 | 0 | 0 | 0 |
| SG0000004 | Gjb1 | 5.03E-36 | 6.48E-35 | 6.67484173 | 0 | 101.1709842 | 10 | 4 | 4 | 6 | 22 | 0 | 0 | 0 | 0 | 0 | 0 | 0 |
| SG000000C | Nes | 6.66E-35 | 8.23E-34 | -2.401626336 | 5084.244924 | 962.1989936 | 22 | 4 | 4 | 6 | 12 | 2 | 0 | 0 | 0 | 0 | 0 | 0 |
| SG0000004 | Rnf123 | 1.29E-34 | 1.58E-33 | -2.181493566 | 1910.304766 | 421.1217383 | 58 | 4 | 4 | 8 | 33 | 0 | 0 | 0 | 0 | 0 | 4 | 0 |
| SG0000003 | Orai2 | 3.75E-34 | 4.5E-33 | -2.897314378 | 570.3269793 | 76.55003293 | 36 | 4 | 4 | 4 | 16 | 6 | 6 | 0 | 0 | 0 | 0 | 4 |
| ISG000000 | Aldh5a1 | 3.92E-34 | 4.7E-33 | 2.010033306 | 183.9004685 | 740.7354901 | 40 | 4 | 4 | 7 | 17 | 4 | 3 | 0 | 0 | 0 | 3 | 0 |
| SG0000001 | Zfp385c | 1.31E-33 | 1.53E-32 | 6.563308691 | 0 | 93.56986889 | 18 | 4 | 4 | 4 | 13 | 2 | 2 | 0 | 0 | 0 | 0 | 0 |
| SG0000003 | Cotl1 | 1.69E-33 | 1.97E-32 | 5.465730813 | 80.30147211 | 3548.725685 | 50 | 2 | 4 | 6 | 16 | 4 | 2 | 0 | 0 | 0 | 0 | 0 |
| ISG000000 | Trpa1 | 4.92E-33 | 5.62E-32 | 6.535746233 | 0 | 91.78027601 | 102 | 2 | 4 | 11 | 56 | 10 | 2 | 0 | 0 | 0 | 3 | 0 |
| ISG000000 | Chrnb1 | 1.29E-32 | 1.45E-31 | -3.031609782 | 895.1921313 | 109.473947 | 14 | 4 | 4 | 4 | 7 | 2 | 1 | 0 | 0 | 0 | 1 | 0 |
| ISG000000 | Mcm5 | 1.94E-32 | 2.17E-31 | 1.40034744 | 5347.809323 | 14116.3526 | 28 | 3 | 4 | 6 | 15 | 3 | 4 | 0 | 1 | 1 | 1 | 0 |
| SG0000002 | Ppm1f | 5.54E-32 | 6.07E-31 | -1.853514241 | 6153.929852 | 1702.900047 | 34 | 4 | 4 | 6 | 13 | 3 | 4 | 0 | 0 | 0 | 2 | 0 |
| ISG000000 | Ass1 | 5.6E-32 | 6.13E-31 | 2.690419096 | 532.0919237 | 3434.65813 | 41 | 4 | 4 | 6 | 19 | 2 | 0 | 0 | 0 | 0 | 0 | 0 |
| SG0000002 | Slc52a3 | 4.88E-31 | 5.16E-30 | -3.123130809 | 1202.727281 | 138.0419571 | 26 | 4 | 4 | 8 | 16 | 6 | 0 | 0 | 0 | 0 | 4 | 0 |
| SG0000003 | Figf | 2.55E-30 | 2.62E-29 | -2.440030422 | 757.6269051 | 139.6153089 | 60 | 1 | 4 | 5 | 15 | 1 | 1 | 0 | 0 | 0 | 0 | 0 |
| ISG000000 | Mgst3 | 6.07E-30 | 6.14E-29 | -2.056693659 | 2545.018851 | 611.7366777 | 26 | 2 | 4 | 5 | 10 | 1 | 0 | 0 | 0 | 0 | 0 | 0 |
| ISG000000 | Elp3 | 6.43E-30 | 6.49E-29 | 1.392970679 | 1984.350922 | 5211.280315 | 94 | 1 | 4 | 8 | 29 | 4 | 2 | 0 | 1 | 1 | 1 | 0 |
| SG0000003 | Rnh1 | 9.24E-30 | 9.25E-29 | -2.174428206 | 6118.118124 | 1355.344646 | 15 | 4 | 4 | 4 | 9 | 2 | 0 | 0 | 0 | 0 | 0 | 0 |
| SG000000C | Axin2 | 1.18E-29 | 1.17E-28 | 5.800427704 | 5.121413317 | 285.4253571 | 37 | 2 | 4 | 6 | 25 | 2 | 3 | 0 | 0 | 2 | 3 | 0 |
| SG0000005 | Irak4 | 4.47E-29 | 4.33E-28 | -2.585817193 | 550.7303878 | 91.73403612 | 79 | 2 | 4 | 8 | 26 | 14 | 1 | 0 | 0 | 0 | 1 | 1 |
| ISG000000 | Gm2237 | 6.14E-29 | 5.91E-28 | 6.174954804 | 1.299968916 | 93.92465007 | 32 | 2 | 4 | 4 | 14 | 1 | 6 | 2 | 2 | 2 | 2 | 0 |
| SG0000002 | Ddx20 | 1.39E-28 | 1.32E-27 | 1.469265651 | 799.0598862 | 2212.44438 | 44 | 4 | 4 | 10 | 23 | 4 | 2 | 0 | 0 | 0 | 0 | 0 |
| SG0000001 | Rassf1 | 3.69E-28 | 3.43E-27 | -1.917871405 | 1739.52413 | 460.3558185 | 23 | 4 | 4 | 4 | 8 | 4 | 0 | 0 | 0 | 0 | 0 | 0 |
| SG0000002 | Prkar1a | 1.02E-27 | 9.3E-27 | -1.640692945 | 11053.7105 | 3544.951484 | 61 | 1 | 4 | 7 | 21 | 13 | 16 | 0 | 0 | 2 | 4 | 2 |
| SG0000002 | Ssbp1 | 2.31E-27 | 2.07E-26 | 1.423159618 | 898.393209 | 2409.240716 | 76 | 4 | 4 | 16 | 61 | 10 | 9 | 1 | 1 | 1 | 2 | 2 |
| ISG000000 | Map2k3 | 3.12E-27 | 2.78E-26 | -1.722138061 | 3496.423233 | 1059.763077 | 24 | 2 | 4 | 6 | 20 | 4 | 0 | 0 | 0 | 0 | 2 | 2 |

| | | | | | | | | | | | | | | | | | | |
|---|---|---|---|---|---|---|---|---|---|---|---|---|---|---|---|---|---|---|
| SG0000001 | Mtmr1 | 4.26E-27 | 3.77E-26 | -2.061045142 | 1064.488121 | 255.0964467 | 186 | 4 | 4 | 16 | 112 | 32 | 18 | 0 | 0 | 0 | 4 | 2 |
| JSG000000 | Kcnk5 | 1.13E-26 | 9.84E-26 | 6.555970178 | 18.31483367 | 1723.243516 | 35 | 4 | 4 | 6 | 13 | 0 | 1 | 0 | 0 | 0 | 1 | 0 |
| SG0000004 | Klhl42 | 1.69E-26 | 1.46E-25 | -1.95562673 | 1230.719149 | 317.2901953 | 31 | 2 | 4 | 2 | 22 | 3 | 8 | 0 | 0 | 0 | 8 | 1 |
| JSG000000 | Cplx1 | 1.8E-26 | 1.56E-25 | 5.542405969 | 1.963171053 | 91.49315138 | 45 | 3 | 4 | 7 | 16 | 5 | 1 | 0 | 0 | 0 | 0 | 0 |
| JSG000000 | Taf7l | 7.95E-26 | 6.67E-25 | 6.251068313 | 0 | 75.165635 | 109 | 0 | 4 | 6 | 42 | 5 | 3 | 0 | 0 | 0 | 0 | 1 |
| SG0000002 | Pitpnm1 | 1.06E-25 | 8.86E-25 | -2.694704133 | 2407.43339 | 371.8497605 | 12 | 2 | 4 | 2 | 6 | 0 | 2 | 2 | 2 | 2 | 2 | 0 |
| JSG000000 | Actn3 | 1.25E-25 | 1.04E-24 | 5.579422388 | 14.71759592 | 703.7369786 | 14 | 3 | 4 | 4 | 8 | 2 | 0 | 0 | 0 | 0 | 0 | 0 |
| SG0000002 | Cfap45 | 5.88E-25 | 4.75E-24 | 6.334653268 | 0.970711415 | 78.34490936 | 29 | 2 | 4 | 5 | 11 | 3 | 0 | 0 | 0 | 0 | 0 | 0 |
| SG0000002 | Sema4a | 7E-25 | 5.62E-24 | 3.469612455 | 18.81642397 | 208.4464544 | 73 | 4 | 4 | 10 | 35 | 51 | 5 | 0 | 0 | 0 | 5 | 1 |
| SG0000002 | Zbtb7b | 2.23E-24 | 1.74E-23 | -2.319110296 | 2311.704541 | 463.2448116 | 32 | 0 | 4 | 0 | 0 | 8 | 12 | 0 | 0 | 0 | 0 | 8 |
| SG0000000 | Rmnd5a | 3.18E-24 | 2.47E-23 | 1.261675404 | 1714.388379 | 4110.658095 | 110 | 3 | 4 | 8 | 37 | 12 | 11 | 0 | 0 | 0 | 1 | 0 |
| SG0000003 | Dnaic2 | 8.23E-24 | 6.28E-23 | 4.713767887 | 3.642924619 | 95.59511751 | 49 | 2 | 4 | 6 | 24 | 16 | 0 | 0 | 0 | 0 | 2 | 0 |
| SG0000002 | Xbp1 | 1.49E-23 | 1.12E-22 | -1.596804158 | 3027.666114 | 1000.972235 | 18 | 0 | 4 | 0 | 0 | 0 | 3 | 0 | 1 | 0 | 0 | 0 |
| SG0000002 | Slc6a9 | 1.62E-23 | 1.21E-22 | -2.204082189 | 547.1214758 | 118.7377761 | 52 | 4 | 4 | 9 | 22 | 5 | 0 | 0 | 0 | 0 | 0 | 0 |
| SG0000001 | Xab2 | 1.67E-23 | 1.25E-22 | 1.265606322 | 1468.453322 | 3530.576381 | 19 | 4 | 4 | 6 | 12 | 5 | 0 | 0 | 0 | 0 | 0 | 0 |
| SG0000003 | Aldh1l1 | 2.25E-23 | 1.67E-22 | 4.208123094 | 5.57112977 | 102.9709101 | 71 | 2 | 4 | 9 | 36 | 14 | 1 | 0 | 0 | 1 | 2 | 0 |
| SG0000000 | Rab23 | 3.05E-23 | 2.26E-22 | 1.392937019 | 567.5174427 | 1490.373223 | 74 | 4 | 4 | 8 | 30 | 6 | 6 | 0 | 0 | 2 | 2 | 0 |
| JSG000000 | Mthfd2 | 3.26E-23 | 2.41E-22 | 1.277564015 | 1158.489972 | 2808.519016 | 18 | 3 | 4 | 8 | 13 | 1 | 3 | 0 | 0 | 1 | 2 | 0 |
| SG0000003 | Anp32a | 6.74E-23 | 4.93E-22 | 1.181089102 | 3988.651162 | 9044.182198 | 28 | 2 | 4 | 8 | 17 | 3 | 0 | 0 | 0 | 0 | 0 | 1 |
| JSG000000 | 5730507C01Rik | 7.34E-23 | 5.36E-22 | 8.233049809 | 0.34247518 | 103.0443185 | 19 | 3 | 4 | 4 | 12 | 1 | 0 | 0 | 0 | 0 | 0 | 0 |
| JSG000000 | Zfp808 | 9.58E-23 | 6.96E-22 | 2.332164663 | 82.09868494 | 413.4164172 | 47 | 2 | 4 | 4 | 38 | 4 | 1 | 0 | 0 | 0 | 0 | 0 |
| JSG000000 | Crisp1 | 1.26E-22 | 9.09E-22 | 6.710520843 | 0.649984458 | 68.07239531 | 83 | 1 | 4 | 15 | 43 | 7 | 4 | 0 | 0 | 0 | 0 | 0 |
| SG0000003 | Nnmt | 1.61E-22 | 1.16E-21 | -5.777867467 | 178.7793959 | 3.258412379 | 56 | 2 | 4 | 8 | 14 | 6 | 0 | 0 | 0 | 0 | 0 | 0 |
| SG0000003 | Snrk | 2.39E-22 | 1.71E-21 | -1.881015108 | 864.8286974 | 234.7946098 | 116 | 0 | 4 | 8 | 46 | 17 | 2 | 0 | 0 | 0 | 0 | 3 |
| SG0000003 | Ablim3 | 2.97E-22 | 2.11E-21 | 8.167336149 | 0.34247518 | 98.45601828 | 116 | 1 | 4 | 15 | 57 | 9 | 2 | 0 | 0 | 0 | 2 | 0 |
| JSG000000 | Dnajc14 | 1.18E-21 | 8.1E-21 | -1.517512794 | 2811.88399 | 982.1561245 | 22 | 4 | 4 | 6 | 10 | 1 | 4 | 0 | 0 | 0 | 1 | 1 |
| SG0000000 | Pfdn2 | 1.43E-21 | 9.77E-21 | 1.218036742 | 1373.424105 | 3194.99557 | 48 | 0 | 4 | 5 | 18 | 8 | 4 | 0 | 0 | 1 | 6 | 0 |
| JSG000000 | Yipf4 | 1.65E-21 | 1.13E-20 | -1.508168563 | 2955.129677 | 1038.89718 | 24 | 2 | 4 | 7 | 12 | 4 | 7 | 0 | 0 | 4 | 6 | 2 |
| JSG000000 | Sipa1l2 | 2.36E-21 | 1.6E-20 | 1.33304438 | 717.3282368 | 1807.191897 | 59 | 1 | 4 | 13 | 20 | 7 | 4 | 0 | 0 | 1 | 1 | 0 |
| SG0000007 | Slc4a11 | 2.72E-21 | 1.83E-20 | -2.554200919 | 2788.38646 | 474.7466113 | 15 | 4 | 4 | 4 | 8 | 0 | 0 | 0 | 0 | 0 | 0 | 0 |
| SG0000002 | Syvn1 | 3.12E-21 | 2.09E-20 | -1.609700356 | 2069.732887 | 678.1819229 | 20 | 0 | 4 | 0 | 0 | 8 | 4 | 0 | 4 | 0 | 0 | 8 |
| JSG000000 | Kcnq4 | 5.7E-21 | 3.78E-20 | 5.736831144 | 1.289094805 | 68.745316 | 27 | 2 | 4 | 5 | 13 | 2 | 0 | 0 | 0 | 0 | 0 | 0 |
| SG0000001 | Dnttip1 | 9.75E-21 | 6.39E-20 | -1.678141571 | 1158.574784 | 362.0371373 | 23 | 2 | 4 | 5 | 16 | 1 | 1 | 0 | 0 | 0 | 0 | 0 |
| JSG000000 | Gna15 | 1.01E-20 | 6.6E-20 | 3.099317116 | 15.68127664 | 134.3905819 | 28 | 3 | 4 | 5 | 17 | 0 | 2 | 0 | 0 | 0 | 0 | 0 |
| JSG000000 | L3hypdh | 1.71E-20 | 1.11E-19 | -2.477886573 | 393.2852011 | 70.59751597 | 25 | 4 | 4 | 6 | 16 | 0 | 0 | 0 | 0 | 0 | 3 | 0 |
| JSG000000 | Prpf38a | 1.91E-20 | 1.23E-19 | 1.21936151 | 992.5722862 | 2311.141495 | 26 | 3 | 4 | 5 | 13 | 6 | 0 | 0 | 0 | 0 | 0 | 0 |
| SG0000002 | Nfe2l3 | 2.94E-20 | 1.87E-19 | -2.193888307 | 551.5931238 | 120.5570621 | 60 | 4 | 4 | 9 | 17 | 10 | 2 | 0 | 0 | 0 | 0 | 0 |
| SG0000003 | Sh3tc1 | 3.1E-20 | 1.97E-19 | -3.267857834 | 1859.419311 | 193.0430219 | 76 | 4 | 4 | 10 | 53 | 7 | 3 | 0 | 0 | 0 | 20 | 0 |
| JSG000000 | Pan2 | 4E-20 | 2.53E-19 | 1.247634786 | 758.2135726 | 1800.391892 | 18 | 4 | 4 | 4 | 9 | 1 | 0 | 0 | 0 | 0 | 1 | 0 |
| SG0000006 | Scoc | 4.14E-20 | 2.61E-19 | -1.603177006 | 1323.653376 | 435.6822877 | 45 | 2 | 4 | 7 | 20 | 3 | 5 | 0 | 0 | 2 | 3 | 0 |
| SG0000000 | Slbp | 8.93E-20 | 5.53E-19 | 1.150553138 | 1511.827099 | 3356.240949 | 82 | 4 | 4 | 19 | 49 | 0 | 5 | 0 | 0 | 0 | 6 | 0 |
| JSG000000 | Ehhadh | 1.37E-19 | 8.4E-19 | 2.325445099 | 41.05275947 | 205.7648002 | 34 | 3 | 4 | 5 | 38 | 3 | 1 | 0 | 0 | 0 | 0 | 1 |
| SG0000002 | Cntfr | 1.53E-19 | 9.38E-19 | 5.27555898 | 4.493177082 | 174.0423873 | 45 | 0 | 4 | 6 | 14 | 4 | 2 | 0 | 0 | 0 | 0 | 2 |
| JSG000000 | Nudc | 2.49E-19 | 1.5E-18 | 1.139923796 | 2517.025309 | 5546.753139 | 11 | 2 | 4 | 2 | 11 | 1 | 1 | 0 | 0 | 0 | 0 | 0 |
| SG0000003 | Daglb | 2.71E-19 | 1.64E-18 | -1.760775089 | 731.9834346 | 216.0006884 | 36 | 3 | 4 | 6 | 13 | 9 | 6 | 0 | 1 | 0 | 0 | 0 |
| SG0000003 | Furin | 4.1E-19 | 2.44E-18 | -2.058999474 | 3707.97555 | 889.8488818 | 21 | 3 | 4 | 6 | 19 | 10 | 6 | 0 | 0 | 0 | 0 | 0 |
| JSG000000 | Dhrs7 | 5.45E-19 | 3.23E-18 | 1.726130145 | 291.5373947 | 964.5199574 | 32 | 3 | 4 | 4 | 13 | 1 | 1 | 0 | 0 | 0 | 0 | 0 |
| SG0000003 | Ttc21a | 7.19E-19 | 4.23E-18 | 6.477312053 | 0.639110347 | 56.94309488 | 24 | 4 | 4 | 7 | 14 | 3 | 0 | 0 | 0 | 0 | 0 | 0 |
| JSG000000 | Pdcd7 | 7.94E-19 | 4.67E-18 | 1.316049405 | 421.0834797 | 1048.427852 | 21 | 2 | 4 | 3 | 7 | 0 | 1 | 0 | 1 | 0 | 1 | 0 |
| SG0000001 | Scrn1 | 2.52E-18 | 1.44E-17 | -1.715700278 | 729.519158 | 222.1055722 | 98 | 1 | 4 | 13 | 37 | 10 | 1 | 0 | 0 | 0 | 2 | 0 |
| SG0000003 | Cbarp | 3.54E-18 | 2.01E-17 | -2.971308477 | 1566.34399 | 199.7257989 | 28 | 4 | 4 | 10 | 10 | 0 | 8 | 4 | 4 | 4 | 4 | 0 |
| SG0000003 | Exosc2 | 3.56E-18 | 2.02E-17 | 1.143933905 | 1038.43491 | 2294.762256 | 18 | 4 | 4 | 6 | 12 | 0 | 1 | 0 | 0 | 0 | 0 | 0 |
| JSG000000 | Hpgd | 4.27E-18 | 2.41E-17 | 4.935107444 | 2.246588541 | 68.72882383 | 62 | 0 | 4 | 8 | 32 | 7 | 4 | 0 | 0 | 1 | 2 | 0 |
| SG0000001 | Ctdnep1 | 4.76E-18 | 2.68E-17 | -1.356382879 | 2536.556222 | 990.6780958 | 11 | 4 | 4 | 6 | 10 | 1 | 0 | 0 | 0 | 0 | 0 | 0 |
| SG0000004 | Lrrc75a | 6.08E-18 | 3.41E-17 | -2.652264697 | 739.4180938 | 117.6193685 | 36 | 3 | 4 | 6 | 19 | 3 | 1 | 0 | 0 | 0 | 0 | 0 |
| JSG000000 | Sppl2b | 6.46E-18 | 3.61E-17 | -2.106812021 | 686.1473997 | 159.2956103 | 9 | 4 | 4 | 4 | 6 | 1 | 0 | 0 | 0 | 0 | 0 | 0 |
| SG0000004 | Adamtsl5 | 7.13E-18 | 3.98E-17 | -2.19215645 | 357.3347565 | 78.19345691 | 10 | 4 | 4 | 4 | 6 | 0 | 1 | 0 | 0 | 0 | 0 | 0 |
| SG0000002 | Dnajb4 | 9.44E-18 | 5.24E-17 | -1.547199575 | 1048.981989 | 358.9339597 | 149 | 2 | 4 | 16 | 66 | 10 | 6 | 0 | 0 | 0 | 0 | 2 |
| SG0000001 | Gata2 | 9.45E-18 | 5.24E-17 | -2.915043647 | 1179.194863 | 156.339962 | 25 | 2 | 4 | 2 | 4 | 6 | 5 | 0 | 0 | 0 | 0 | 4 |
| JSG000000 | Tmem38a | 1.34E-17 | 7.4E-17 | 1.512099596 | 182.1072867 | 519.4152047 | 24 | 3 | 4 | 5 | 10 | 1 | 2 | 0 | 0 | 0 | 0 | 0 |
| SG0000003 | Nme6 | 2.96E-17 | 1.61E-16 | 1.437992326 | 216.9397756 | 587.7843386 | 32 | 0 | 4 | 8 | 28 | 4 | 0 | 0 | 0 | 0 | 0 | 0 |
| JSG000000 | Ubr7 | 4.99E-17 | 2.67E-16 | 1.02086227 | 3157.531148 | 6407.0455 | 24 | 2 | 4 | 5 | 11 | 3 | 1 | 0 | 0 | 0 | 0 | 0 |
| SG0000000 | Nuak2 | 6.87E-17 | 3.65E-16 | 2.866980175 | 70.08108429 | 511.2670947 | 24 | 4 | 4 | 6 | 12 | 2 | 4 | 2 | 2 | 2 | 2 | 0 |
| SG0000002 | Bdh2 | 7.08E-17 | 3.76E-16 | 4.377856309 | 12.57356152 | 261.4113119 | 64 | 2 | 4 | 10 | 44 | 2 | 0 | 0 | 0 | 0 | 0 | 0 |
| SG0000003 | Fignl1 | 8.68E-17 | 4.58E-16 | 1.043011846 | 1886.800169 | 3887.79859 | 68 | 4 | 4 | 20 | 59 | 8 | 4 | 0 | 0 | 0 | 11 | 0 |
| SG0000006 | Acadm | 8.86E-17 | 4.67E-16 | 1.097902965 | 1020.029511 | 2183.305263 | 68 | 3 | 4 | 8 | 24 | 7 | 0 | 0 | 0 | 0 | 0 | 0 |
| JSG000000 | She | 9.24E-17 | 4.87E-16 | -2.4786899 | 757.9035455 | 135.9733911 | 44 | 3 | 4 | 5 | 17 | 2 | 7 | 0 | 0 | 0 | 2 | 0 |
| JSG000000 | Pard6g | 1.66E-16 | 8.6E-16 | -1.513189715 | 998.5078422 | 349.8130338 | 40 | 2 | 4 | 6 | 23 | 13 | 3 | 0 | 1 | 0 | 2 | 1 |
| SG0000003 | Zfp275 | 1.88E-16 | 9.71E-16 | -1.352525972 | 1710.654474 | 669.9022093 | 70 | 2 | 4 | 6 | 32 | 2 | 24 | 0 | 0 | 0 | 8 | 2 |
| JSG000000 | Arv1 | 1.97E-16 | 1.02E-15 | 1.887252391 | 68.34910703 | 252.8438672 | 19 | 4 | 4 | 4 | 11 | 0 | 0 | 0 | 0 | 0 | 1 | 0 |
| JSG000000 | Podxl | 2.9E-16 | 1.48E-15 | -1.952235639 | 5856.485104 | 1513.406368 | 43 | 4 | 4 | 9 | 24 | 7 | 5 | 0 | 0 | 0 | 2 | 2 |
| SG0000002 | Rpp40 | 3.1E-16 | 1.58E-15 | 2.371480063 | 172.2479259 | 891.3343309 | 43 | 2 | 4 | 5 | 12 | 7 | 3 | 0 | 0 | 0 | 1 | 0 |
| JSG000000 | Wdr34 | 3.39E-16 | 1.73E-15 | 1.463593432 | 249.310512 | 687.5845587 | 22 | 4 | 4 | 4 | 10 | 1 | 0 | 0 | 0 | 0 | 0 | 0 |
| JSG000000 | Ccnb2 | 3.65E-16 | 1.85E-15 | 1.020946654 | 1963.029699 | 3983.478128 | 20 | 2 | 4 | 5 | 13 | 0 | 0 | 0 | 0 | 0 | 1 | 0 |
| SG0000002 | Polm | 3.98E-16 | 2.02E-15 | -1.446271147 | 1040.952473 | 381.9969575 | 10 | 4 | 4 | 4 | 6 | 0 | 0 | 0 | 0 | 0 | 0 | 0 |
| JSG000000 | Dusp4 | 5.26E-16 | 2.64E-15 | -1.426638296 | 3961.485668 | 1473.659571 | 12 | 4 | 4 | 6 | 12 | 2 | 2 | 0 | 0 | 0 | 4 | 0 |
| SG0000003 | Fa2h | 6.36E-16 | 3.18E-15 | 5.416454956 | 0 | 41.70860754 | 57 | 3 | 4 | 10 | 19 | 5 | 1 | 0 | 1 | 1 | 1 | 1 |
| SG0000000 | Ergic3 | 7.08E-16 | 3.53E-15 | -1.234874097 | 2860.421648 | 1215.334739 | 6 | 4 | 4 | 4 | 6 | 0 | 0 | 0 | 0 | 0 | 0 | 0 |
| SG0000005 | Npm1 | 1.41E-15 | 6.9E-15 | 0.932482811 | 42495.06772 | 81104.28916 | 120 | 0 | 4 | 12 | 38 | 4 | 3 | 0 | 0 | 0 | 0 | 0 |
| SG0000004 | Slc39a3 | 1.77E-15 | 8.61E-15 | -2.176059237 | 334.4476364 | 74.00635733 | 24 | 2 | 4 | 4 | 15 | 0 | 3 | 0 | 0 | 0 | 0 | 0 |
| JSG000000 | Ap3m2 | 2.51E-15 | 1.21E-14 | 1.338323474 | 240.5698065 | 608.298336 | 29 | 3 | 4 | 6 | 13 | 2 | 2 | 0 | 0 | 0 | 0 | 0 |
| SG0000003 | Nt5m | 2.71E-15 | 1.31E-14 | -2.095007717 | 264.3386514 | 61.87290486 | 34 | 4 | 4 | 6 | 18 | 1 | 0 | 0 | 0 | 0 | 0 | 0 |
| JSG000000 | 4931414P19Rik | 2.82E-15 | 1.36E-14 | 1.67463982 | 94.85107252 | 302.8022261 | 35 | 2 | 4 | 11 | 22 | 2 | 0 | 0 | 0 | 0 | 1 | 1 |
| SG0000003 | Cmtm7 | 3.41E-15 | 1.63E-14 | -1.344966547 | 1182.792726 | 465.6218705 | 40 | 4 | 4 | 6 | 16 | 2 | 1 | 0 | 0 | 0 | 0 | 0 |
| JSG000000 | Gm14124 | 3.52E-15 | 1.68E-14 | 6.699506046 | 2.353829814 | 244.6399253 | 30 | 3 | 4 | 5 | 11 | 2 | 1 | 0 | 0 | 0 | 0 | 0 |
| SG0000002 | Krt19 | 5.67E-15 | 2.67E-14 | -3.344745217 | 11093.45086 | 1091.938115 | 9 | 3 | 4 | 4 | 6 | 0 | 0 | 0 | 0 | 0 | 0 | 0 |
| JSG000000 | Txndc15 | 8.27E-15 | 3.86E-14 | -1.399116103 | 928.7393731 | 352.1420945 | 12 | 3 | 4 | 3 | 9 | 0 | 0 | 0 | 0 | 0 | 0 | 0 |
| SG0000000 | Cct4 | 8.66E-15 | 4.03E-14 | 0.919022397 | 9840.045438 | 18605.88841 | 44 | 4 | 4 | 5 | 17 | 0 | 2 | 0 | 0 | 0 | 0 | 0 |
| SG0000002 | Vegfa | 1.05E-14 | 4.87E-14 | -1.347151875 | 5510.733582 | 2166.088978 | 32 | 2 | 4 | 6 | 8 | 8 | 18 | 2 | 4 | 6 | 8 | 4 |
| SG0000001 | Tctex1d2 | 1.32E-14 | 6.07E-14 | 1.238774895 | 302.5143957 | 713.9280529 | 26 | 4 | 4 | 8 | 14 | 2 | 2 | 0 | 0 | 2 | 4 | 0 |
| JSG000000 | Zfp383 | 1.32E-14 | 6.09E-14 | 1.342560682 | 207.0066487 | 524.9710214 | 26 | 4 | 4 | 4 | 32 | 2 | 10 | 0 | 0 | 0 | 7 | 0 |
| SG0000005 | Ndufaf6 | 2.04E-14 | 9.27E-14 | 1.583583813 | 105.8760604 | 317.3247901 | 35 | 3 | 4 | 4 | 18 | 3 | 1 | 0 | 0 | 0 | 0 | 0 |
| SG0000003 | Arl5c | 2.6E-14 | 1.18E-13 | 1.906609343 | 52.88849991 | 198.2932137 | 13 | 2 | 4 | 4 | 4 | 0 | 4 | 1 | 2 | 2 | 2 | 0 |
| JSG000000 | Impact | 5.83E-14 | 2.58E-13 | -1.097159834 | 4761.57983 | 2225.732922 | 49 | 1 | 4 | 7 | 19 | 5 | 2 | 0 | 1 | 0 | 2 | 1 |
| JSG000000 | Scg3 | 6.58E-14 | 2.9E-13 | 5.212827728 | 0 | 36.0866417 | 75 | 1 | 4 | 10 | 25 | 5 | 2 | 0 | 0 | 0 | 0 | 0 |
| SG0000005 | Zfp933 | 7.44E-14 | 3.26E-13 | -1.710480048 | 383.6530811 | 117.2284139 | 44 | 3 | 4 | 6 | 23 | 3 | 6 | 2 | 3 | 3 | 4 | 0 |
| SG0000001 | Med22 | 9.08E-14 | 3.96E-13 | -1.149333728 | 2200.302623 | 991.969889 | 14 | 2 | 4 | 2 | 2 | 0 | 2 | 0 | 0 | 0 | 0 | 0 |
| JSG000000 | Dhrs11 | 1.03E-13 | 4.45E-13 | -1.799011521 | 319.211883 | 91.73237116 | 20 | 4 | 4 | 4 | 10 | 0 | 0 | 0 | 0 | 0 | 0 | 0 |
| SG0000003 | Zfand1 | 1.12E-13 | 4.86E-13 | 1.152694212 | 363.3561311 | 807.8449722 | 42 | 0 | 4 | 3 | 27 | 9 | 3 | 0 | 1 | 0 | 3 | 0 |
| SG0000007 | Fam217b | 1.24E-13 | 5.34E-13 | 3.532375543 | 19.84135281 | 229.5737742 | 14 | 4 | 4 | 7 | 10 | 1 | 4 | 1 | 1 | 2 | 2 | 0 |
| JSG000000 | Apela | 1.47E-13 | 6.33E-13 | 5.205407892 | 20.3261883 | 749.9630382 | 26 | 0 | 4 | 4 | 30 | 4 | 2 | 0 | 0 | 0 | 2 | 0 |
| SG0000002 | Zc3h8 | 2.1E-13 | 8.93E-13 | 1.626786733 | 130.8656117 | 404.1449373 | 40 | 3 | 4 | 8 | 25 | 1 | 2 | 1 | 1 | 2 | 2 | 0 |
| JSG000000 | Ankrd1 | 2.61E-13 | 1.1E-12 | -2.693830533 | 412.838724 | 63.80528214 | 13 | 3 | 4 | 6 | 10 | 0 | 4 | 2 | 2 | 4 | 5 | 0 |
| JSG000000 | Cish | 2.76E-13 | 1.16E-12 | 1.570069466 | 100.3723657 | 298.024635 | 16 | 4 | 4 | 8 | 10 | 2 | 6 | 4 | 4 | 6 | 6 | 0 |
| SG0000002 | Map7d1 | 3.3E-13 | 1.39E-12 | -1.609350737 | 3836.142108 | 1257.279521 | 38 | 0 | 4 | 4 | 20 | 10 | 4 | 0 | 0 | 0 | 0 | 4 |
| SG0000003 | Sec61a1 | 3.72E-13 | 1.56E-12 | -1.019392179 | 9776.089998 | 4822.781293 | 17 | 3 | 4 | 3 | 6 | 1 | 1 | 0 | 1 | 0 | 0 | 0 |
| JSG000000 | Lnx2 | 4.24E-13 | 1.77E-12 | -1.423325872 | 1039.444839 | 387.5588568 | 71 | 2 | 4 | 12 | 35 | 7 | 3 | 0 | 0 | 2 | 3 | 0 |
| SG0000004 | Zfp518b | 4.65E-13 | 1.93E-12 | 1.002853155 | 772.6942908 | 1548.447853 | 42 | 0 | 4 | 4 | 10 | 10 | 10 | 0 | 0 | 2 | 8 | 6 |
| JSG000000 | Chmp2b | 4.96E-13 | 2.06E-12 | -1.12752863 | 2094.707028 | 958.7455281 | 53 | 2 | 4 | 11 | 33 | 7 | 4 | 0 | 0 | 0 | 0 | 0 |
| JSG000000 | Lman2 | 6.08E-13 | 2.51E-12 | -1.051164849 | 3892.639026 | 1878.503387 | 17 | 4 | 4 | 6 | 11 | 1 | 2 | 0 | 0 | 0 | 0 | 0 |
| JSG000000 | Pdzd8 | 6.66E-13 | 2.74E-12 | -1.09143043 | 2800.497904 | 1314.26201 | 138 | 3 | 4 | 14 | 58 | 10 | 11 | 0 | 0 | 2 | 3 | 1 |
| SG0000002 | Mmp28 | 1.13E-12 | 4.59E-12 | -2.792816849 | 234.8633035 | 33.8917131 | 81 | 4 | 4 | 8 | 24 | 4 | 1 | 0 | 0 | 0 | 0 | 0 |
| SG0000004 | Pklr | 1.22E-12 | 4.92E-12 | 5.064160135 | 0 | 32.45523644 | 18 | 4 | 4 | 4 | 16 | 0 | 0 | 0 | 0 | 0 | 0 | 0 |
| SG0000001 | Thg1l | 1.3E-12 | 5.23E-12 | 1.264555334 | 408.3137247 | 980.9865771 | 20 | 0 | 4 | 4 | 10 | 4 | 2 | 0 | 0 | 0 | 2 | 0 |
| SG0000005 | Arhgef15 | 1.34E-12 | 5.39E-12 | 6.384515591 | 3.001470705 | 250.763781 | 37 | 4 | 4 | 12 | 26 | 8 | 2 | 0 | 0 | 0 | 2 | 2 |
| JSG000000 | Irs2 | 1.74E-12 | 6.97E-12 | 1.830557524 | 56.54848562 | 201.1285406 | 27 | 3 | 4 | 7 | 10 | 3 | 0 | 0 | 0 | 0 | 0 | 0 |
| JSG000000 | Nudt19 | 1.75E-12 | 7.02E-12 | 1.079089026 | 406.4806496 | 858.772326 | 16 | 3 | 4 | 7 | 16 | 1 | 0 | 0 | 0 | 0 | 0 | 0 |
| SG0000002 | Chrnd | 1.78E-12 | 7.12E-12 | 5.041065601 | 0 | 31.92395172 | 26 | 2 | 4 | 16 | 36 | 0 | 4 | 0 | 0 | 0 | 0 | 0 |
| SG0000000 | Hpn | 1.91E-12 | 7.64E-12 | 6.277756031 | 2.556441386 | 198.3484041 | 59 | 0 | 4 | 4 | 26 | 26 | 1 | 0 | 0 | 0 | 0 | 6 |
| SG0000002 | Mapre3 | 1.96E-12 | 7.8E-12 | -1.552990403 | 664.9826332 | 226.6280224 | 52 | 2 | 4 | 12 | 21 | 3 | 3 | 1 | 2 | 2 | 2 | 0 |
| JSG000000 | Prkcg | 2E-12 | 7.97E-12 | 2.115872456 | 30.76075239 | 133.3331164 | 54 | 2 | 4 | 6 | 20 | 2 | 0 | 0 | 0 | 0 | 0 | 0 |
| SG0000001 | Trpv2 | 2.7E-12 | 1.07E-11 | 2.479307788 | 19.76204673 | 110.1990714 | 45 | 2 | 4 | 10 | 37 | 2 | 0 | 0 | 0 | 0 | 1 | 0 |
| SG0000001 | Strip1 | 4.32E-12 | 1.69E-11 | -1.046915621 | 2671.50392 | 1293.012761 | 12 | 3 | 4 | 3 | 13 | 4 | 0 | 0 | 0 | 0 | 0 | 1 |

| | | | | | | | | | | | | | | | | | | |
|---|---|---|---|---|---|---|---|---|---|---|---|---|---|---|---|---|---|---|
| SG0000003 | Prkab2 | 4.97E-12 | 1.94E-11 | 1.092078997 | 408.3742823 | 870.5764604 | 57 | 4 | 4 | 4 | 10 | 9 | 9 | 1 | 1 | 1 | 1 | 3 |
| SG0000002 | Car6 | 5.32E-12 | 2.07E-11 | 6.624048329 | 0.34247518 | 33.78052091 | 25 | 2 | 4 | 8 | 24 | 8 | 2 | 0 | 0 | 0 | 0 | 0 |
| JSG000000 | Taldo1 | 1.03E-11 | 3.94E-11 | 0.847332646 | 3009.015753 | 5413.735132 | 12 | 3 | 4 | 3 | 6 | 0 | 0 | 0 | 0 | 0 | 0 | 0 |
| JSG000000 | Tmem168 | 1.08E-11 | 4.13E-11 | -1.096494008 | 1503.798838 | 703.253915 | 124 | 0 | 4 | 8 | 68 | 2 | 12 | 0 | 0 | 2 | 10 | 0 |
| JSG000000 | Sf3a3 | 1.25E-11 | 4.74E-11 | 0.867661315 | 1707.852353 | 3116.322565 | 20 | 4 | 4 | 6 | 13 | 1 | 0 | 0 | 0 | 0 | 0 | 0 |
| SG0000003 | Fam124a | 1.28E-11 | 4.86E-11 | -2.094806686 | 165.7517807 | 38.80239704 | 64 | 0 | 4 | 8 | 29 | 6 | 0 | 0 | 0 | 0 | 0 | 1 |
| JSG000000 | Oprd1 | 1.52E-11 | 5.72E-11 | 6.724998006 | 0.307509278 | 32.53002105 | 43 | 3 | 4 | 8 | 19 | 4 | 3 | 1 | 1 | 1 | 1 | 0 |
| SG0000002 | Atic | 1.77E-11 | 6.66E-11 | 0.825060243 | 5206.769349 | 9224.359515 | 43 | 4 | 4 | 10 | 26 | 3 | 0 | 0 | 0 | 0 | 0 | 0 |
| SG0000000 | Acp5 | 1.87E-11 | 7E-11 | -1.512482039 | 801.6874936 | 280.9976236 | 18 | 4 | 4 | 7 | 8 | 0 | 0 | 0 | 0 | 0 | 0 | 0 |
| JSG000000 | Nsdhl | 2.37E-11 | 8.84E-11 | 1.008612294 | 471.4579282 | 948.5615091 | 51 | 1 | 4 | 9 | 33 | 4 | 1 | 0 | 0 | 0 | 0 | 0 |
| JSG000000 | C1galt1 | 2.45E-11 | 9.12E-11 | -1.065165899 | 2039.986048 | 974.9454377 | 71 | 0 | 4 | 8 | 23 | 4 | 4 | 0 | 1 | 1 | 2 | 0 |
| JSG000000 | Clcf1 | 2.67E-11 | 9.91E-11 | -1.958602558 | 235.4343662 | 60.57197541 | 16 | 4 | 4 | 8 | 8 | 0 | 8 | 4 | 4 | 8 | 8 | 0 |
| JSG000000 | Areg | 2.94E-11 | 1.09E-10 | -5.435885074 | 255.0699111 | 5.892434592 | 20 | 3 | 4 | 5 | 18 | 4 | 4 | 1 | 2 | 1 | 2 | 2 |
| JSG000000 | Rcl1 | 4.5E-11 | 1.64E-10 | 0.898303826 | 942.9189566 | 1757.481506 | 58 | 2 | 4 | 8 | 20 | 8 | 0 | 0 | 0 | 0 | 0 | 0 |
| SG0000003 | 2700029M09Rik | 4.98E-11 | 1.81E-10 | 0.877586268 | 1093.676252 | 2009.410061 | 122 | 4 | 4 | 8 | 55 | 16 | 0 | 0 | 0 | 0 | 0 | 0 |
| SG0000003 | Tpbg | 5.41E-11 | 1.96E-10 | -1.293926402 | 547.9892487 | 223.4916896 | 9 | 0 | 4 | 2 | 6 | 2 | 8 | 0 | 4 | 2 | 6 | 2 |
| JSG000000 | Eif2ak2 | 5.55E-11 | 2.01E-10 | -1.355112852 | 3349.492405 | 1309.330791 | 42 | 2 | 4 | 7 | 22 | 8 | 3 | 0 | 1 | 2 | 3 | 1 |
| SG0000002 | Paics | 5.77E-11 | 2.09E-10 | 0.776576377 | 13910.15978 | 23828.94979 | 140 | 0 | 4 | 19 | 37 | 8 | 12 | 0 | 0 | 0 | 0 | 0 |
| JSG000000 | Pla2g15 | 5.86E-11 | 2.12E-10 | 0.944962216 | 616.3745061 | 1186.606271 | 10 | 4 | 4 | 4 | 11 | 2 | 1 | 0 | 0 | 0 | 0 | 0 |
| SG0000002 | Thbs3 | 6.18E-11 | 2.23E-10 | -1.248300495 | 668.6812133 | 281.4772044 | 16 | 2 | 4 | 6 | 10 | 0 | 0 | 0 | 0 | 0 | 2 | 0 |
| SG0000002 | Anxa7 | 6.18E-11 | 2.23E-10 | -0.91334241 | 8181.003026 | 4343.732693 | 88 | 4 | 4 | 8 | 34 | 8 | 2 | 0 | 0 | 0 | 0 | 0 |
| SG0000004 | Oasl1 | 8.09E-11 | 2.89E-10 | -3.872667842 | 1128.174107 | 77.01706141 | 26 | 4 | 4 | 10 | 20 | 2 | 0 | 0 | 0 | 0 | 0 | 0 |
| SG0000000 | Marveld3 | 8.5E-11 | 3.04E-10 | 7.27714623 | 1.561638181 | 242.2254195 | 38 | 1 | 4 | 2 | 5 | 0 | 0 | 0 | 0 | 0 | 0 | 0 |
| JSG000000 | Ubd | 8.88E-11 | 3.17E-10 | 5.31466752 | 1.949953374 | 77.606584 | 7 | 2 | 4 | 2 | 5 | 0 | 1 | 0 | 1 | 0 | 0 | 0 |
| SG0000003 | Parp14 | 1.05E-10 | 3.72E-10 | -2.268805236 | 2480.923731 | 514.7957858 | 60 | 4 | 4 | 10 | 36 | 5 | 3 | 0 | 0 | 0 | 0 | 0 |
| JSG000000 | Rpn1 | 1.16E-10 | 4.12E-10 | -0.894220326 | 9743.791895 | 5242.52921 | 26 | 3 | 4 | 4 | 14 | 1 | 1 | 0 | 0 | 0 | 0 | 0 |
| JSG000000 | Oxa1l | 1.23E-10 | 4.34E-10 | 0.833602192 | 1566.788851 | 2792.220663 | 13 | 4 | 4 | 5 | 9 | 1 | 1 | 0 | 0 | 0 | 0 | 0 |
| SG0000001 | Trabd | 1.44E-10 | 5.07E-10 | -1.444684412 | 1640.859009 | 602.8064956 | 36 | 4 | 4 | 4 | 12 | 4 | 4 | 0 | 0 | 0 | 0 | 0 |
| SG0000002 | Timm22 | 1.72E-10 | 6E-10 | -1.08821489 | 1022.527057 | 480.9384342 | 25 | 4 | 4 | 9 | 18 | 2 | 2 | 0 | 0 | 1 | 2 | 0 |
| JSG000000 | Mppe1 | 1.89E-10 | 6.61E-10 | -1.663487703 | 234.350686 | 73.97867238 | 30 | 3 | 4 | 4 | 15 | 4 | 0 | 0 | 0 | 0 | 0 | 0 |
| SG0000001 | Nlrp14 | 1.92E-10 | 6.69E-10 | 6.5729463 | 0.307509278 | 29.27603246 | 99 | 1 | 4 | 9 | 47 | 5 | 2 | 0 | 0 | 0 | 0 | 0 |
| JSG000000 | Ebpl | 2.08E-10 | 7.23E-10 | 1.249417282 | 215.0864957 | 511.3581982 | 33 | 2 | 4 | 6 | 16 | 1 | 1 | 0 | 0 | 0 | 0 | 0 |
| SG0000005 | Pcdh1 | 2.21E-10 | 7.66E-10 | 2.7704657 | 46.31573294 | 316.0252757 | 61 | 3 | 4 | 13 | 29 | 3 | 1 | 0 | 1 | 0 | 0 | 3 |
| SG0000002 | Otub1 | 2.86E-10 | 9.85E-10 | -0.958095704 | 2988.815854 | 1538.450802 | 14 | 2 | 4 | 6 | 6 | 0 | 0 | 0 | 0 | 0 | 0 | 0 |
| JSG000000 | Tmem238 | 4.13E-10 | 1.4E-09 | -2.835409431 | 138.6518577 | 19.42595614 | 13 | 4 | 4 | 5 | 6 | 0 | 1 | 0 | 0 | 0 | 0 | 0 |
| SG0000002 | Tspan4 | 4.45E-10 | 1.51E-09 | -1.037041437 | 1257.500764 | 612.8125889 | 16 | 4 | 4 | 4 | 10 | 0 | 0 | 0 | 0 | 0 | 0 | 0 |
| SG0000002 | Zfp263 | 4.58E-10 | 1.55E-09 | -0.963508789 | 1876.431278 | 962.249329 | 27 | 4 | 4 | 6 | 14 | 0 | 10 | 0 | 0 | 0 | 0 | 0 |
| SG0000003 | Osbpl5 | 5.2E-10 | 1.75E-09 | -1.79453837 | 495.0251766 | 142.6978434 | 75 | 0 | 4 | 8 | 22 | 6 | 0 | 0 | 0 | 0 | 0 | 0 |
| SG0000001 | Rpain | 6.03E-10 | 2.02E-09 | 1.159414244 | 176.6224937 | 394.515869 | 33 | 4 | 4 | 4 | 9 | 5 | 0 | 0 | 0 | 0 | 0 | 1 |
| SG0000004 | Cyb5d1 | 7.08E-10 | 2.37E-09 | -1.498949385 | 275.5257541 | 97.48402955 | 9 | 4 | 4 | 4 | 14 | 0 | 1 | 0 | 0 | 0 | 0 | 0 |
| SG0000002 | Tpst2 | 7.31E-10 | 2.44E-09 | -1.783942042 | 1914.438641 | 555.9316329 | 44 | 4 | 4 | 7 | 16 | 15 | 2 | 0 | 0 | 0 | 0 | 0 |
| SG0000003 | Fam173b | 7.32E-10 | 2.44E-09 | -1.224521302 | 502.5576186 | 215.0642475 | 24 | 2 | 4 | 6 | 14 | 0 | 2 | 0 | 0 | 0 | 0 | 0 |
| SG0000003 | Mns1 | 9.08E-10 | 3E-09 | 1.05561728 | 285.4654087 | 593.3705059 | 47 | 3 | 4 | 9 | 32 | 0 | 7 | 0 | 0 | 1 | 4 | 0 |
| SG0000003 | Hmbs | 9.4E-10 | 3.1E-09 | 0.851461648 | 833.3315051 | 1503.603394 | 13 | 4 | 4 | 4 | 8 | 0 | 0 | 0 | 0 | 0 | 0 | 0 |
| SG0000002 | Polg2 | 9.87E-10 | 3.25E-09 | 1.291475953 | 112.7976862 | 276.1043136 | 39 | 4 | 4 | 4 | 13 | 5 | 0 | 0 | 0 | 0 | 0 | 0 |
| SG0000004 | Ptchd2 | 9.96E-10 | 3.28E-09 | 4.959179203 | 1.016551429 | 31.62212485 | 38 | 2 | 4 | 8 | 18 | 4 | 0 | 0 | 0 | 0 | 0 | 0 |
| SG0000004 | Skiv2l | 1.05E-09 | 3.45E-09 | -0.95361272 | 1652.541794 | 853.2698868 | 15 | 4 | 4 | 4 | 6 | 3 | 0 | 0 | 0 | 0 | 0 | 0 |
| SG0000002 | Cdca3 | 1.32E-09 | 4.3E-09 | 0.781888811 | 1743.866276 | 2998.369589 | 5 | 4 | 4 | 4 | 6 | 0 | 0 | 0 | 0 | 0 | 0 | 0 |
| SG0000003 | Haus7 | 1.75E-09 | 5.65E-09 | 1.14366608 | 297.9126651 | 658.2134639 | 66 | 4 | 4 | 8 | 30 | 6 | 0 | 0 | 0 | 0 | 0 | 0 |
| SG0000000 | Tpd52l2 | 2.35E-09 | 7.49E-09 | -0.853272296 | 4230.058849 | 2341.455206 | 55 | 0 | 4 | 12 | 27 | 4 | 3 | 0 | 0 | 0 | 3 | 4 |
| SG0000005 | Adnp2 | 4.09E-09 | 1.28E-08 | 0.77917489 | 1516.749666 | 2602.968556 | 45 | 2 | 4 | 5 | 14 | 3 | 2 | 0 | 0 | 0 | 0 | 0 |
| SG0000001 | Aldh3a2 | 4.11E-09 | 1.29E-08 | -0.935052797 | 2319.606354 | 1213.208237 | 98 | 0 | 4 | 12 | 32 | 8 | 4 | 0 | 0 | 0 | 1 | 0 |
| SG0000000 | Etv3 | 5.94E-09 | 1.84E-08 | -0.964832728 | 1127.145111 | 577.4790585 | 26 | 2 | 4 | 4 | 12 | 7 | 15 | 2 | 4 | 4 | 6 | 2 |
| SG0000004 | Pvr | 9.56E-09 | 2.92E-08 | -0.959469218 | 2157.275425 | 1109.37042 | 30 | 4 | 4 | 6 | 11 | 0 | 5 | 0 | 0 | 1 | 1 | 0 |
| SG0000000 | P2rx5 | 1.23E-08 | 3.73E-08 | 3.363865411 | 3.950433897 | 40.66953335 | 13 | 2 | 4 | 5 | 13 | 0 | 2 | 0 | 1 | 1 | 1 | 0 |
| SG0000003 | Elovl5 | 1.24E-08 | 3.74E-08 | 0.727760614 | 1942.244735 | 3216.486459 | 91 | 2 | 4 | 10 | 42 | 17 | 6 | 0 | 0 | 1 | 3 | 1 |
| SG0000002 | Uqcrc1 | 1.41E-08 | 4.23E-08 | 0.722441721 | 3826.754218 | 6314.038091 | 26 | 2 | 4 | 4 | 4 | 4 | 1 | 0 | 0 | 0 | 0 | 0 |
| SG0000003 | Entpd6 | 1.42E-08 | 4.26E-08 | 0.834521545 | 601.3661558 | 1072.395542 | 20 | 4 | 4 | 6 | 8 | 2 | 0 | 0 | 0 | 0 | 0 | 0 |
| SG0000003 | Oraov1 | 2.22E-08 | 6.58E-08 | -1.044350717 | 594.9315649 | 288.4603458 | 34 | 4 | 4 | 6 | 15 | 7 | 1 | 0 | 0 | 0 | 1 | 2 |
| SG0000002 | Nol9 | 2.23E-08 | 6.59E-08 | 0.718245815 | 2190.526924 | 3603.812024 | 43 | 4 | 4 | 6 | 36 | 8 | 0 | 0 | 0 | 0 | 0 | 1 |
| JSG000000 | Atp1b3 | 2.66E-08 | 7.83E-08 | -0.784344113 | 5032.568951 | 2921.989352 | 62 | 2 | 4 | 8 | 35 | 8 | 5 | 0 | 0 | 0 | 1 | 1 |
| SG0000003 | Piga | 2.77E-08 | 8.13E-08 | 0.768782434 | 865.9627679 | 1475.454402 | 145 | 0 | 4 | 14 | 37 | 5 | 24 | 0 | 4 | 4 | 14 | 0 |
| JSG000000 | 1810013L24Rik | 2.77E-08 | 8.14E-08 | -0.81317884 | 2842.678261 | 1617.843415 | 61 | 3 | 4 | 8 | 23 | 5 | 5 | 0 | 1 | 2 | 7 | 0 |
| JSG000000 | Gm3696 | 4.11E-08 | 1.19E-07 | 6.201717404 | 0.307509278 | 22.63399351 | 24 | 2 | 4 | 5 | 14 | 2 | 5 | 2 | 2 | 3 | 3 | 0 |
| JSG000000 | 1700030J22Rik | 5.39E-08 | 1.55E-07 | 1.587255168 | 41.69186982 | 125.2745321 | 9 | 4 | 4 | 4 | 7 | 2 | 1 | 0 | 0 | 0 | 0 | 0 |
| SG0000002 | Nab2 | 7.4E-08 | 2.1E-07 | -1.29361449 | 656.4479547 | 267.7833283 | 10 | 2 | 4 | 2 | 6 | 0 | 6 | 2 | 2 | 2 | 4 | 0 |
| SG0000006 | Slc25a3 | 9.95E-08 | 2.79E-07 | -0.719124583 | 19474.79875 | 11830.27686 | 61 | 0 | 4 | 4 | 40 | 4 | 3 | 0 | 0 | 0 | 0 | 0 |
| SG0000002 | Cpt2 | 1E-07 | 2.8E-07 | -0.843716795 | 1410.942747 | 786.1860403 | 46 | 2 | 4 | 6 | 28 | 8 | 2 | 0 | 0 | 2 | 2 | 0 |
| SG0000002 | Fam20c | 1.1E-07 | 3.07E-07 | -1.628359481 | 186.681576 | 60.38323979 | 78 | 2 | 4 | 10 | 48 | 18 | 0 | 0 | 0 | 0 | 0 | 0 |
| SG0000002 | Tra2a | 1.15E-07 | 3.2E-07 | 0.658131352 | 3598.317504 | 5678.279418 | 56 | 2 | 4 | 7 | 16 | 5 | 4 | 1 | 2 | 1 | 1 | 0 |
| SG0000007 | Srsf3 | 1.25E-07 | 3.47E-07 | 0.660273444 | 6069.674683 | 9592.405443 | 43 | 4 | 4 | 8 | 16 | 4 | 6 | 0 | 0 | 0 | 3 | 0 |
| SG0000002 | Wars | 1.28E-07 | 3.56E-07 | -0.74520514 | 4033.083647 | 2406.069287 | 123 | 0 | 4 | 10 | 55 | 10 | 0 | 0 | 0 | 0 | 0 | 0 |
| JSG000000 | U90926 | 1.52E-07 | 4.2E-07 | -2.946432116 | 134.9559528 | 17.50763962 | 18 | 4 | 4 | 8 | 10 | 2 | 0 | 0 | 0 | 0 | 0 | 0 |
| SG0000002 | Plbd2 | 1.62E-07 | 4.48E-07 | -0.768783278 | 2353.471771 | 1381.281409 | 21 | 2 | 4 | 5 | 10 | 0 | 3 | 0 | 0 | 1 | 1 | 0 |
| SG0000004 | N4bp2l1 | 1.66E-07 | 4.58E-07 | 4.160204329 | 8.343553545 | 149.1754472 | 59 | 2 | 4 | 13 | 36 | 6 | 9 | 0 | 0 | 7 | 14 | 4 |
| SG0000002 | Ten1 | 1.92E-07 | 5.24E-07 | -1.086985398 | 372.0452122 | 175.1380502 | 25 | 2 | 4 | 2 | 4 | 0 | 0 | 0 | 0 | 0 | 0 | 0 |
| SG0000003 | Prickle3 | 2.51E-07 | 6.8E-07 | -1.200232456 | 283.1659059 | 123.2352614 | 18 | 4 | 4 | 4 | 11 | 5 | 1 | 0 | 0 | 0 | 1 | 0 |
| JSG000000 | Lbhd1 | 2.76E-07 | 7.44E-07 | -1.109149566 | 487.9302029 | 226.1884439 | 22 | 4 | 4 | 6 | 11 | 1 | 3 | 0 | 0 | 0 | 1 | 0 |
| SG0000003 | Zfp39 | 2.82E-07 | 7.61E-07 | 1.025731055 | 149.3867037 | 304.149957 | 32 | 1 | 4 | 6 | 12 | 1 | 1 | 0 | 0 | 0 | 1 | 0 |
| SG0000002 | Rasl10b | 3.15E-07 | 8.45E-07 | 2.324559115 | 11.29668753 | 56.5865396 | 54 | 0 | 4 | 4 | 16 | 8 | 6 | 0 | 0 | 0 | 0 | 0 |
| SG0000000 | Hk2 | 3.79E-07 | 1.01E-06 | -0.861697155 | 15840.29277 | 8716.989313 | 60 | 3 | 4 | 9 | 31 | 5 | 7 | 1 | 1 | 2 | 3 | 1 |
| SG0000001 | Mtif3 | 4.36E-07 | 1.16E-06 | -0.966992161 | 512.6145449 | 262.2389992 | 47 | 4 | 4 | 12 | 49 | 0 | 3 | 0 | 0 | 0 | 1 | 0 |
| SG0000002 | Zfp202 | 4.66E-07 | 1.23E-06 | 1.498384632 | 41.78958368 | 118.0665203 | 118 | 0 | 4 | 12 | 30 | 6 | 1 | 0 | 1 | 0 | 0 | 0 |
| SG0000001 | Psmc3ip | 4.91E-07 | 1.3E-06 | 0.885700285 | 244.0176005 | 450.8618671 | 8 | 3 | 4 | 5 | 7 | 0 | 2 | 0 | 0 | 1 | 1 | 0 |
| JSG000000 | Gprin1 | 5.15E-07 | 1.36E-06 | 4.568483355 | 0.981585527 | 23.29049242 | 10 | 0 | 4 | 0 | 2 | 4 | 0 | 0 | 0 | 0 | 0 | 0 |
| SG0000002 | Gpam | 5.4E-07 | 1.42E-06 | -0.908544432 | 651.855786 | 347.2581108 | 53 | 1 | 4 | 4 | 13 | 2 | 1 | 0 | 0 | 0 | 0 | 0 |
| SG0000002 | Acp6 | 7.86E-07 | 2.05E-06 | 0.691945651 | 1229.291288 | 1985.871761 | 26 | 3 | 4 | 7 | 13 | 16 | 0 | 0 | 0 | 0 | 1 | 0 |
| SG0000004 | Ndufa7 | 8.2E-07 | 2.13E-06 | 1.092701604 | 1289.332872 | 2749.79941 | 60 | 0 | 4 | 8 | 20 | 0 | 0 | 0 | 0 | 0 | 0 | 0 |
| SG0000003 | Sin3b | 9.63E-07 | 2.49E-06 | 0.632232571 | 2212.720954 | 3429.632742 | 48 | 2 | 4 | 5 | 9 | 6 | 2 | 0 | 0 | 2 | 2 | 0 |
| JSG000000 | Gimap9 | 1.18E-06 | 3.03E-06 | -1.30644479 | 215.9185806 | 87.29933738 | 8 | 4 | 4 | 6 | 11 | 0 | 3 | 2 | 2 | 2 | 3 | 0 |
| SG0000005 | Zfp729b | 1.4E-06 | 3.57E-06 | 0.757234058 | 420.8105864 | 711.273817 | 34 | 1 | 4 | 7 | 24 | 2 | 0 | 0 | 0 | 0 | 0 | 0 |
| SG0000002 | Mzt2 | 1.75E-06 | 4.42E-06 | 1.203434949 | 170.6728824 | 393.0379995 | 37 | 0 | 4 | 0 | 14 | 6 | 3 | 0 | 0 | 0 | 0 | 0 |
| JSG000000 | Agmat | 1.85E-06 | 4.66E-06 | 4.060173626 | 0 | 15.68145964 | 16 | 4 | 4 | 4 | 9 | 2 | 0 | 0 | 0 | 0 | 0 | 0 |
| SG0000002 | Frmd8 | 1.96E-06 | 4.94E-06 | -1.213999046 | 2397.821008 | 1033.633635 | 19 | 3 | 4 | 5 | 11 | 2 | 4 | 1 | 1 | 1 | 1 | 0 |
| SG0000002 | Vps39 | 2.48E-06 | 6.18E-06 | -0.710128957 | 1875.435399 | 1146.388953 | 66 | 2 | 4 | 12 | 48 | 4 | 2 | 0 | 0 | 2 | 2 | 0 |
| JSG000000 | Slc47a2 | 2.53E-06 | 6.29E-06 | 4.026466756 | 0 | 15.29623467 | 49 | 3 | 4 | 6 | 19 | 12 | 0 | 0 | 0 | 0 | 0 | 0 |
| JSG000000 | Etf1 | 2.57E-06 | 6.39E-06 | 0.569260968 | 7649.083439 | 11349.42934 | 51 | 4 | 4 | 6 | 23 | 8 | 7 | 0 | 0 | 0 | 1 | 0 |
| JSG000000 | Dnaja2 | 2.63E-06 | 6.52E-06 | 0.573380367 | 5678.852299 | 8450.166433 | 42 | 2 | 4 | 6 | 17 | 2 | 2 | 0 | 0 | 0 | 0 | 0 |
| SG0000002 | 4833439L19Rik | 3E-06 | 7.42E-06 | 0.598117298 | 2416.898227 | 3658.55519 | 30 | 0 | 4 | 4 | 12 | 8 | 8 | 0 | 0 | 0 | 0 | 0 |
| SG0000002 | Zc3h15 | 3.32E-06 | 8.19E-06 | 0.592744918 | 2275.267316 | 3431.361139 | 44 | 1 | 4 | 5 | 17 | 5 | 1 | 0 | 0 | 1 | 1 | 1 |
| JSG000000 | Snrpc | 3.62E-06 | 8.91E-06 | 0.91844767 | 780.4373872 | 1475.089476 | 23 | 3 | 4 | 9 | 12 | 3 | 0 | 0 | 0 | 0 | 0 | 0 |
| SG0000000 | Abcf3 | 3.91E-06 | 9.58E-06 | -0.711845561 | 1374.434556 | 839.1453643 | 13 | 3 | 4 | 3 | 7 | 1 | 1 | 0 | 0 | 0 | 0 | 0 |
| SG0000003 | Psma1 | 4.49E-06 | 1.09E-05 | 0.570452499 | 3626.029486 | 5384.614851 | 58 | 0 | 4 | 6 | 26 | 2 | 0 | 0 | 0 | 0 | 0 | 0 |
| JSG000000 | AI846148 | 4.95E-06 | 1.2E-05 | -0.757976279 | 1231.694173 | 728.3318309 | 42 | 1 | 4 | 5 | 17 | 3 | 2 | 0 | 0 | 0 | 1 | 1 |
| JSG000000 | Otulin | 5.18E-06 | 1.25E-05 | -0.835247417 | 776.2851539 | 435.0976667 | 45 | 2 | 4 | 8 | 29 | 2 | 0 | 0 | 0 | 0 | 0 | 0 |
| JSG000000 | Sall3 | 5.72E-06 | 1.38E-05 | 4.73118386 | 1.984919276 | 52.71947859 | 40 | 3 | 4 | 7 | 13 | 5 | 9 | 0 | 0 | 2 | 6 | 1 |
| SG0000002 | 2810459M11Rik | 5.98E-06 | 1.44E-05 | 5.750909044 | 0.307509278 | 16.55977182 | 25 | 2 | 4 | 4 | 11 | 2 | 3 | 0 | 0 | 0 | 1 | 0 |
| JSG000000 | Anks3 | 7.17E-06 | 1.71E-05 | -0.820259311 | 560.1616617 | 317.2420335 | 22 | 4 | 4 | 4 | 20 | 2 | 1 | 0 | 0 | 0 | 0 | 0 |
| SG0000003 | Carkd | 8.39E-06 | 1.99E-05 | -0.742009806 | 884.3647314 | 528.7668327 | 52 | 0 | 4 | 0 | 8 | 16 | 4 | 0 | 0 | 0 | 0 | 0 |
| SG0000003 | Tctn1 | 9.07E-06 | 2.15E-05 | -0.76518893 | 783.7337741 | 461.1303877 | 51 | 4 | 4 | 11 | 15 | 6 | 1 | 0 | 0 | 1 | 1 | 0 |
| SG0000006 | Trim54 | 9.54E-06 | 2.25E-05 | 4.906129158 | 1.337278385 | 40.09718316 | 58 | 2 | 4 | 6 | 20 | 4 | 0 | 0 | 0 | 0 | 0 | 0 |
| SG0000005 | Mpz | 9.58E-06 | 2.26E-05 | 3.544300049 | 1.987262844 | 23.18440427 | 26 | 1 | 4 | 1 | 2 | 2 | 1 | 0 | 0 | 0 | 0 | 0 |
| SG0000002 | Aip | 1.04E-05 | 2.45E-05 | -0.893417239 | 1299.580574 | 699.6129236 | 16 | 4 | 4 | 7 | 17 | 2 | 2 | 0 | 0 | 2 | 2 | 2 |
| SG0000003 | Uqcrc2 | 1.11E-05 | 2.61E-05 | 0.54738337 | 3688.140068 | 5389.968409 | 51 | 1 | 4 | 8 | 20 | 5 | 2 | 0 | 0 | 1 | 1 | 0 |
| SG0000000 | Saal1 | 1.25E-05 | 2.92E-05 | 0.629973644 | 754.9145886 | 1168.258048 | 45 | 2 | 4 | 6 | 17 | 4 | 5 | 0 | 0 | 0 | 1 | 1 |
| JSG000000 | Mut | 1.26E-05 | 2.94E-05 | -0.65236791 | 1943.398604 | 1236.458594 | 62 | 1 | 4 | 9 | 32 | 4 | 9 | 0 | 0 | 2 | 3 | 0 |
| SG0000002 | Erh | 1.47E-05 | 3.42E-05 | 0.583119498 | 1380.409388 | 2067.970619 | 35 | 0 | 4 | 0 | 16 | 2 | 1 | 0 | 0 | 0 | 0 | 0 |
| SG0000003 | Pi4kb | 1.53E-05 | 3.56E-05 | -0.671995507 | 1294.53996 | 812.5024072 | 123 | 0 | 4 | 16 | 51 | 13 | 0 | 0 | 0 | 0 | 0 | 0 |
| SG0000003 | Eri2 | 2.23E-05 | 5.08E-05 | 0.658520843 | 543.6165996 | 858.0789259 | 146 | 4 | 4 | 17 | 63 | 6 | 12 | 0 | 0 | 0 | 0 | 0 |
| SG0000001 | Slc25a22 | 2.25E-05 | 5.11E-05 | -0.680352547 | 1002.007531 | 625.265493 | 18 | 4 | 4 | 8 | 8 | 0 | 12 | 4 | 4 | 8 | 8 | 0 |
| SG0000002 | Tor1b | 2.29E-05 | 5.22E-05 | -0.653832753 | 1345.184787 | 854.9853561 | 23 | 0 | 4 | 8 | 17 | 0 | 4 | 0 | 0 | 0 | 4 | 0 |
| JSG000000 | Dhx29 | 2.38E-05 | 5.39E-05 | 0.585271103 | 1045.081514 | 1567.957631 | 69 | 1 | 4 | 7 | 26 | 4 | 0 | 0 | 0 | 0 | 0 | 0 |
| JSG000000 | Nkrf | 2.38E-05 | 5.41E-05 | 0.689686054 | 483.2889724 | 779.5124849 | 30 | 4 | 4 | 4 | 20 | 6 | 3 | 0 | 0 | 0 | 0 | 0 |

| | | | | | | | | | | | | | | | | | | |
|---|---|---|---|---|---|---|---|---|---|---|---|---|---|---|---|---|---|---|
| SG0000002 | Dtwd1 | 2.44E-05 | 5.54E-05 | 0.779259133 | 223.4153753 | 383.4364817 | 28 | 1 | 4 | 3 | 10 | 1 | 1 | 0 | 0 | 0 | 0 | 0 |
| JSG000000 | Cela1 | 2.86E-05 | 6.46E-05 | 2.043586856 | 10.78422311 | 44.46003565 | 15 | 4 | 4 | 6 | 9 | 1 | 0 | 0 | 0 | 0 | 0 | 0 |
| JSG000000 | Ptar1 | 2.94E-05 | 6.62E-05 | -1.128376641 | 172.5817265 | 78.94408474 | 68 | 1 | 4 | 6 | 31 | 3 | 5 | 1 | 2 | 1 | 2 | 0 |
| SG0000003 | Tle2 | 3.29E-05 | 7.37E-05 | -1.105340249 | 177.5296134 | 82.5144933 | 33 | 0 | 4 | 0 | 7 | 0 | 3 | 0 | 0 | 0 | 0 | 0 |
| SG0000001 | Ap4m1 | 3.86E-05 | 8.58E-05 | -0.694500826 | 799.8557337 | 494.2493215 | 14 | 2 | 4 | 4 | 4 | 2 | 0 | 0 | 0 | 0 | 0 | 0 |
| JSG000000 | Casp3 | 4.09E-05 | 9.08E-05 | -0.661987204 | 1990.479534 | 1257.997307 | 37 | 2 | 4 | 8 | 22 | 5 | 4 | 0 | 0 | 0 | 0 | 0 |
| JSG000000 | Ypel5 | 4.11E-05 | 9.12E-05 | -0.634845471 | 1419.440934 | 914.1338605 | 39 | 2 | 4 | 6 | 13 | 1 | 8 | 0 | 1 | 2 | 2 | 0 |
| SG0000003 | Mst1r | 4.16E-05 | 9.22E-05 | 2.661228645 | 4.835652262 | 30.58895989 | 20 | 4 | 4 | 6 | 10 | 0 | 2 | 0 | 0 | 2 | 2 | 0 |
| JSG000000 | Thap2 | 4.36E-05 | 9.63E-05 | -0.918430212 | 296.5198317 | 156.8840174 | 40 | 2 | 4 | 4 | 7 | 3 | 5 | 0 | 0 | 1 | 2 | 0 |
| SG0000002 | Ahcy | 4.49E-05 | 9.9E-05 | 0.535467713 | 1729.12152 | 2506.209428 | 16 | 2 | 4 | 3 | 14 | 2 | 2 | 0 | 0 | 0 | 0 | 0 |
| SG0000002 | Pkn3 | 4.91E-05 | 0.000108 | 0.695996867 | 328.9551566 | 532.9084818 | 14 | 4 | 4 | 5 | 8 | 4 | 1 | 0 | 0 | 0 | 0 | 0 |
| JSG000000 | Ndufaf5 | 4.93E-05 | 0.000108 | 0.782472371 | 196.5776246 | 338.1284685 | 79 | 0 | 4 | 7 | 36 | 5 | 1 | 0 | 0 | 0 | 0 | 0 |
| SG0000002 | Pes1 | 4.96E-05 | 0.000109 | 0.5080886 | 4148.874802 | 5900.383483 | 24 | 4 | 4 | 6 | 16 | 4 | 0 | 0 | 0 | 0 | 2 | 0 |
| SG0000003 | Plekhh3 | 5.59E-05 | 0.000122 | -1.506060071 | 379.1292155 | 133.4805525 | 25 | 0 | 4 | 0 | 0 | 0 | 8 | 0 | 0 | 0 | 0 | 0 |
| SG0000002 | Gtf2h3 | 6.44E-05 | 0.00014 | 0.534940392 | 1400.597103 | 2029.300337 | 27 | 1 | 4 | 3 | 7 | 0 | 2 | 0 | 1 | 0 | 1 | 0 |
| SG0000007 | Ccdc13 | 6.62E-05 | 0.000143 | 4.032502851 | 1.016551429 | 16.63541619 | 44 | 3 | 4 | 9 | 18 | 1 | 0 | 0 | 0 | 0 | 0 | 0 |
| SG0000001 | Fis1 | 6.97E-05 | 0.00015 | -0.593386654 | 1561.296349 | 1034.804151 | 24 | 4 | 4 | 4 | 22 | 4 | 0 | 0 | 0 | 0 | 0 | 2 |
| SG0000002 | Aar2 | 7.42E-05 | 0.00016 | -0.620373794 | 1178.542495 | 766.6446793 | 45 | 2 | 4 | 2 | 22 | 8 | 3 | 0 | 0 | 0 | 0 | 0 |
| JSG000000 | Fbxo43 | 7.86E-05 | 0.000169 | 2.578976011 | 4.613636034 | 27.56718799 | 30 | 1 | 4 | 1 | 10 | 2 | 9 | 1 | 3 | 1 | 2 | 1 |
| JSG000000 | Vapb | 7.9E-05 | 0.00017 | -0.562191971 | 2712.484816 | 1837.090999 | 48 | 2 | 4 | 8 | 28 | 3 | 7 | 0 | 1 | 0 | 6 | 0 |
| SG0000002 | Ldb1 | 8.2E-05 | 0.000176 | 0.513563943 | 2979.56775 | 4253.54877 | 32 | 4 | 4 | 12 | 15 | 6 | 1 | 0 | 0 | 0 | 3 | 0 |
| SG0000001 | Afmid | 8.33E-05 | 0.000178 | -1.382671662 | 119.8730495 | 45.97226582 | 23 | 4 | 4 | 7 | 12 | 2 | 1 | 0 | 0 | 0 | 0 | 0 |
| SG0000002 | Llgl2 | 9.27E-05 | 0.000197 | 1.101712623 | 823.3938279 | 1767.079763 | 52 | 0 | 4 | 8 | 20 | 2 | 0 | 0 | 0 | 0 | 0 | 0 |
| SG0000002 | Polr3f | 0.000109 | 0.00023 | 0.553950246 | 783.1649272 | 1149.764433 | 47 | 4 | 4 | 6 | 22 | 5 | 12 | 0 | 0 | 2 | 4 | 1 |
| JSG000000 | Tnfrsf26 | 0.00011 | 0.000231 | -3.899874994 | 19.79566593 | 1.326144241 | 32 | 2 | 4 | 6 | 19 | 6 | 2 | 0 | 0 | 0 | 5 | 1 |
| JSG000000 | Mrps35 | 0.000116 | 0.000244 | 0.561108366 | 746.4580916 | 1101.325967 | 48 | 4 | 4 | 7 | 21 | 1 | 1 | 0 | 0 | 0 | 0 | 0 |
| SG0000004 | Coa7 | 0.000122 | 0.000256 | 0.663189543 | 347.2966132 | 549.9718912 | 19 | 4 | 4 | 5 | 11 | 3 | 3 | 0 | 0 | 0 | 0 | 0 |
| SG0000003 | Trim30a | 0.000123 | 0.000258 | -2.692816867 | 466.5153016 | 72.15181216 | 91 | 3 | 4 | 10 | 50 | 7 | 1 | 0 | 0 | 0 | 0 | 0 |
| SG0000005 | Kbtbd2 | 0.00014 | 0.000292 | -0.55767457 | 2132.503294 | 1448.815012 | 106 | 2 | 4 | 8 | 32 | 8 | 6 | 0 | 0 | 0 | 2 | 2 |
| SG0000002 | Ttc4 | 0.000178 | 0.000367 | 0.525814199 | 957.4320912 | 1378.458914 | 30 | 2 | 4 | 4 | 10 | 4 | 0 | 0 | 0 | 0 | 0 | 0 |
| JSG000000 | Dcp1a | 0.000187 | 0.000385 | -0.534860345 | 3161.440352 | 2182.10671 | 85 | 0 | 4 | 8 | 30 | 10 | 6 | 0 | 0 | 1 | 1 | 2 |
| SG0000003 | Efcab14 | 0.00021 | 0.000429 | -0.56102367 | 1583.529426 | 1073.349566 | 261 | 0 | 4 | 27 | 125 | 20 | 6 | 0 | 0 | 0 | 0 | 2 |
| JSG000000 | Minpp1 | 0.000223 | 0.000455 | -0.550668256 | 1646.882821 | 1124.333204 | 47 | 2 | 4 | 7 | 20 | 1 | 2 | 0 | 1 | 1 | 1 | 0 |
| SG0000002 | Znhit3 | 0.000238 | 0.000482 | 0.625192163 | 345.4047207 | 532.7578566 | 12 | 4 | 4 | 6 | 7 | 0 | 3 | 0 | 0 | 1 | 1 | 0 |
| SG0000004 | Taf13 | 0.000263 | 0.000532 | 0.583205868 | 436.2401055 | 653.5638908 | 17 | 2 | 4 | 5 | 22 | 3 | 5 | 0 | 1 | 1 | 4 | 1 |
| SG0000002 | Gnl2 | 0.00027 | 0.000545 | 0.472925587 | 2147.186999 | 2980.126204 | 41 | 4 | 4 | 5 | 16 | 1 | 0 | 0 | 0 | 0 | 0 | 0 |
| SG0000003 | Urb1 | 0.000271 | 0.000546 | 0.498695458 | 1259.879445 | 1780.128208 | 66 | 2 | 4 | 5 | 26 | 6 | 1 | 0 | 0 | 0 | 0 | 0 |
| JSG000000 | Urm1 | 0.000316 | 0.000632 | -0.606641166 | 1055.372021 | 693.0880144 | 14 | 3 | 4 | 4 | 17 | 1 | 1 | 0 | 0 | 0 | 0 | 0 |
| SG0000002 | Psmc5 | 0.000353 | 0.000702 | 0.451735392 | 3725.836282 | 5095.768275 | 14 | 0 | 4 | 2 | 12 | 4 | 0 | 0 | 0 | 0 | 0 | 0 |
| SG0000003 | Znrd1as | 0.000406 | 0.000802 | 0.614207059 | 298.9430903 | 457.5971049 | 26 | 4 | 4 | 6 | 20 | 2 | 0 | 0 | 0 | 0 | 0 | 0 |
| JSG000000 | Tmem28 | 0.00046 | 0.000901 | -1.076338802 | 301.1057516 | 142.7936144 | 38 | 3 | 4 | 7 | 19 | 3 | 1 | 0 | 0 | 0 | 0 | 0 |
| JSG000000 | Cpa4 | 0.000551 | 0.00107 | -3.991429229 | 15.36658352 | 0.966134069 | 37 | 3 | 4 | 10 | 22 | 2 | 2 | 2 | 2 | 2 | 3 | 1 |
| SG0000005 | Iigp1 | 0.000563 | 0.001091 | -1.747934919 | 45.68714686 | 13.60232654 | 74 | 2 | 4 | 6 | 30 | 2 | 10 | 0 | 0 | 0 | 0 | 0 |
| JSG000000 | Csdc2 | 0.000572 | 0.001108 | 1.319147207 | 41.65087008 | 103.9266566 | 13 | 4 | 4 | 4 | 8 | 1 | 0 | 0 | 0 | 0 | 0 | 0 |
| SG0000002 | Dhx37 | 0.000605 | 0.001167 | -0.502628619 | 1691.186143 | 1193.672313 | 13 | 2 | 4 | 5 | 8 | 4 | 1 | 0 | 0 | 0 | 0 | 0 |
| SG0000002 | Tagln3 | 0.000635 | 0.001221 | 4.07724542 | 2.604624967 | 43.96614323 | 20 | 3 | 4 | 4 | 7 | 0 | 3 | 3 | 3 | 3 | 3 | 0 |
| SG0000003 | Tnpo2 | 0.000649 | 0.001247 | 0.423360441 | 6644.983015 | 8911.238493 | 34 | 2 | 4 | 2 | 18 | 2 | 4 | 0 | 2 | 0 | 8 | 0 |
| SG0000002 | Anapc11 | 0.000769 | 0.001464 | 0.606072394 | 611.9077391 | 931.3907406 | 17 | 4 | 4 | 6 | 14 | 0 | 1 | 0 | 0 | 0 | 2 | 0 |
| JSG000000 | Slc35f3 | 0.00078 | 0.001482 | 3.808881245 | 1.254128903 | 17.57639004 | 83 | 2 | 4 | 8 | 30 | 10 | 2 | 0 | 0 | 1 | 1 | 1 |
| JSG000000 | S100a1 | 0.000873 | 0.00165 | -0.946720397 | 219.7102057 | 113.9879633 | 8 | 4 | 4 | 4 | 4 | 0 | 0 | 0 | 0 | 0 | 0 | 0 |
| JSG000000 | Ddx52 | 0.00091 | 0.001715 | -0.48271694 | 2140.757052 | 1531.987088 | 47 | 0 | 4 | 1 | 14 | 4 | 4 | 0 | 0 | 0 | 0 | 0 |
| SG0000005 | Tvp23a | 0.000953 | 0.001791 | 1.419520536 | 30.07096176 | 80.43879927 | 46 | 4 | 4 | 8 | 18 | 4 | 0 | 0 | 0 | 0 | 0 | 0 |
| JSG000000 | Tk2 | 0.000968 | 0.001818 | 0.551488744 | 381.1267744 | 558.5783687 | 27 | 2 | 4 | 3 | 6 | 2 | 1 | 0 | 0 | 0 | 0 | 0 |
| SG0000004 | Cirbp | 0.00097 | 0.001822 | -0.665355024 | 1237.640576 | 780.3738701 | 8 | 0 | 4 | 4 | 4 | 0 | 1 | 0 | 1 | 1 | 1 | 0 |
| JSG000000 | Ptrh2 | 0.001002 | 0.001879 | -0.533844624 | 972.3514459 | 671.6144311 | 42 | 4 | 4 | 12 | 30 | 0 | 8 | 0 | 0 | 0 | 4 | 0 |
| JSG000000 | Ssc5d | 0.001018 | 0.001908 | -0.973920234 | 126.4908563 | 64.39911939 | 22 | 3 | 4 | 7 | 20 | 2 | 0 | 0 | 0 | 0 | 0 | 0 |
| SG0000003 | Recql | 0.001068 | 0.001999 | -0.521413121 | 990.1259762 | 689.8099665 | 192 | 3 | 4 | 16 | 36 | 27 | 8 | 0 | 0 | 0 | 1 | 1 |
| JSG000000 | 4930579G24Rik | 0.001076 | 0.002012 | 0.558373367 | 328.1297961 | 483.2065371 | 14 | 3 | 4 | 4 | 9 | 1 | 5 | 0 | 0 | 0 | 1 | 1 |
| JSG000000 | Gm3468 | 0.001144 | 0.002134 | 5.628773809 | 0.663202137 | 32.81524661 | 32 | 2 | 4 | 4 | 14 | 2 | 6 | 2 | 2 | 2 | 2 | 0 |
| SG0000001 | Psma2 | 0.001215 | 0.00226 | 0.401634794 | 4429.340914 | 5851.176899 | 31 | 0 | 4 | 8 | 18 | 6 | 0 | 0 | 0 | 0 | 0 | 0 |
| SG0000002 | Ppp1r8 | 0.001298 | 0.002406 | 0.419048879 | 2133.126503 | 2852.087818 | 68 | 2 | 4 | 12 | 26 | 2 | 2 | 0 | 0 | 0 | 0 | 0 |
| JSG000000 | Rdm1 | 0.001393 | 0.002574 | -0.531741514 | 752.9487677 | 520.8291479 | 30 | 2 | 4 | 4 | 12 | 0 | 1 | 0 | 0 | 0 | 0 | 0 |
| JSG000000 | Zfp53 | 0.001404 | 0.002593 | -0.667670629 | 322.3694691 | 202.938767 | 35 | 4 | 4 | 6 | 21 | 5 | 1 | 0 | 0 | 0 | 0 | 0 |
| SG0000000 | Lhx2 | 0.001599 | 0.002937 | 2.707814438 | 2.6022814 | 17.00151512 | 24 | 4 | 4 | 11 | 15 | 1 | 2 | 0 | 0 | 1 | 1 | 0 |
| SG0000002 | Agap3 | 0.001701 | 0.003114 | -0.790561731 | 1613.156048 | 932.5960811 | 34 | 4 | 4 | 8 | 22 | 3 | 2 | 0 | 0 | 2 | 2 | 2 |
| JSG000000 | Ifit3b | 0.001718 | 0.003142 | -1.359430017 | 152.0517766 | 59.26008728 | 10 | 3 | 4 | 3 | 4 | 1 | 0 | 0 | 0 | 0 | 0 | 0 |
| SG0000003 | Rps27l | 0.001756 | 0.003206 | 0.395558569 | 2785.196878 | 3663.792707 | 49 | 0 | 4 | 4 | 12 | 0 | 2 | 0 | 0 | 0 | 0 | 0 |
| JSG000000 | Fam210a | 0.001772 | 0.003233 | -0.46509783 | 1884.157595 | 1364.925123 | 73 | 1 | 4 | 10 | 27 | 5 | 19 | 1 | 2 | 4 | 6 | 0 |
| SG0000001 | Zkscan6 | 0.001815 | 0.003305 | -0.590827173 | 432.1466298 | 286.9289896 | 96 | 2 | 4 | 20 | 62 | 10 | 0 | 0 | 0 | 0 | 0 | 2 |
| JSG000000 | Exosc5 | 0.00193 | 0.003501 | 0.469396375 | 1006.944007 | 1394.144244 | 12 | 3 | 4 | 4 | 6 | 1 | 0 | 0 | 0 | 0 | 0 | 0 |
| JSG000000 | Ints12 | 0.001935 | 0.003509 | 0.473073593 | 571.1846032 | 792.8405129 | 51 | 1 | 4 | 7 | 26 | 6 | 7 | 1 | 1 | 3 | 5 | 2 |
| SG0000004 | Bivm | 0.002016 | 0.003647 | 0.431272607 | 1010.548454 | 1362.646274 | 81 | 0 | 4 | 2 | 31 | 2 | 7 | 0 | 0 | 0 | 0 | 0 |
| SG0000000 | Armc6 | 0.002033 | 0.003678 | 0.602231966 | 275.389887 | 418.0592854 | 15 | 3 | 4 | 3 | 11 | 0 | 0 | 0 | 0 | 0 | 0 | 0 |
| SG0000002 | Ngf | 0.002082 | 0.003761 | -1.215340589 | 117.7684054 | 50.71948351 | 129 | 2 | 4 | 11 | 35 | 7 | 4 | 2 | 2 | 2 | 2 | 0 |
| JSG000000 | Gm5141 | 0.002116 | 0.003818 | 0.764747414 | 89.27694306 | 151.6879988 | 16 | 3 | 4 | 3 | 12 | 1 | 1 | 0 | 0 | 0 | 1 | 1 |
| SG0000003 | Shank1 | 0.002166 | 0.003903 | 1.521636254 | 13.60233376 | 39.05454495 | 71 | 4 | 4 | 12 | 32 | 12 | 0 | 0 | 0 | 0 | 0 | 0 |
| JSG000000 | Ice1 | 0.002235 | 0.00402 | 0.403244201 | 1870.891655 | 2474.213873 | 65 | 3 | 4 | 10 | 31 | 5 | 0 | 0 | 0 | 0 | 1 | 1 |
| SG0000005 | Tdrp | 0.002249 | 0.004041 | -0.470320746 | 1136.601058 | 820.4033096 | 25 | 4 | 4 | 9 | 15 | 2 | 0 | 0 | 0 | 0 | 0 | 0 |
| JSG000000 | Chchd2 | 0.002263 | 0.004065 | 0.432838786 | 2295.796616 | 3099.066314 | 13 | 4 | 4 | 5 | 7 | 0 | 0 | 0 | 0 | 0 | 0 | 0 |
| SG0000002 | Pnpla2 | 0.002301 | 0.004126 | -0.465735676 | 1280.944741 | 927.5343171 | 12 | 4 | 4 | 4 | 4 | 0 | 6 | 4 | 4 | 4 | 4 | 0 |
| JSG000000 | Rpl7l1 | 0.002353 | 0.004217 | -0.407486567 | 6659.599973 | 5020.91031 | 14 | 3 | 4 | 4 | 8 | 2 | 3 | 0 | 0 | 0 | 0 | 0 |
| SG0000003 | Surf6 | 0.002366 | 0.004239 | 0.42786889 | 1074.111491 | 1444.943059 | 47 | 4 | 4 | 10 | 20 | 4 | 2 | 0 | 0 | 1 | 1 | 0 |
| JSG000000 | Cct7 | 0.002524 | 0.004506 | 0.369915329 | 15194.23765 | 19635.16363 | 24 | 1 | 4 | 1 | 6 | 2 | 1 | 0 | 1 | 0 | 0 | 0 |
| JSG000000 | Ptgs2 | 0.002578 | 0.004597 | -1.200148796 | 2129.957146 | 927.0220806 | 26 | 1 | 4 | 6 | 12 | 2 | 14 | 1 | 4 | 6 | 9 | 0 |
| SG0000004 | Mis12 | 0.002751 | 0.00488 | -0.449317602 | 1553.674184 | 1137.894174 | 27 | 4 | 4 | 8 | 12 | 2 | 5 | 0 | 0 | 0 | 0 | 0 |
| JSG000000 | Utp11l | 0.002845 | 0.005037 | -0.440466597 | 1905.23045 | 1403.95716 | 41 | 4 | 4 | 5 | 18 | 1 | 1 | 0 | 0 | 1 | 3 | 0 |
| SG0000001 | Cluap1 | 0.002912 | 0.005148 | -0.576468234 | 400.879097 | 268.8308972 | 86 | 0 | 4 | 4 | 22 | 9 | 0 | 0 | 0 | 0 | 0 | 0 |
| SG0000005 | Taok2 | 0.002926 | 0.005171 | -0.424604959 | 2067.160329 | 1540.122739 | 29 | 4 | 4 | 4 | 14 | 6 | 1 | 0 | 0 | 0 | 1 | 0 |
| SG0000004 | Col27a1 | 0.00328 | 0.005756 | 1.091131293 | 33.59108386 | 71.56279027 | 140 | 2 | 4 | 8 | 63 | 4 | 4 | 0 | 2 | 0 | 1 | 0 |
| JSG000000 | Zfpl1 | 0.003291 | 0.00577 | -0.590521301 | 631.9002799 | 419.6468071 | 8 | 4 | 4 | 6 | 6 | 0 | 0 | 0 | 0 | 0 | 0 | 0 |
| SG0000002 | Eps8l2 | 0.003296 | 0.005777 | 1.101572708 | 375.3784897 | 805.5190397 | 28 | 2 | 4 | 6 | 16 | 4 | 0 | 0 | 0 | 0 | 4 | 0 |
| SG0000000 | Ndufa9 | 0.003302 | 0.005787 | -0.412016029 | 3070.442788 | 2307.660021 | 34 | 2 | 4 | 3 | 13 | 4 | 0 | 0 | 0 | 0 | 0 | 0 |
| JSG000000 | Ik | 0.003366 | 0.005895 | 0.375952864 | 2626.169111 | 3407.970117 | 26 | 3 | 4 | 8 | 17 | 4 | 0 | 0 | 0 | 0 | 0 | 0 |
| SG0000004 | Ttc1 | 0.003574 | 0.006241 | 0.407575342 | 1079.359038 | 1431.720806 | 44 | 4 | 4 | 6 | 20 | 2 | 2 | 0 | 0 | 0 | 0 | 0 |
| SG0000002 | Il1rn | 0.003969 | 0.006881 | -1.2439532 | 54.40080451 | 22.96878889 | 50 | 0 | 4 | 12 | 24 | 19 | 0 | 0 | 0 | 0 | 2 | 1 |
| SG0000003 | Zscan2 | 0.00402 | 0.006964 | 0.990816921 | 43.01493366 | 85.48400695 | 17 | 4 | 4 | 8 | 22 | 2 | 0 | 0 | 0 | 0 | 0 | 0 |
| JSG000000 | Spcs2 | 0.004229 | 0.007302 | -0.410673681 | 2392.098927 | 1799.509069 | 31 | 3 | 4 | 6 | 19 | 3 | 1 | 0 | 0 | 0 | 0 | 1 |
| JSG000000 | Rps19bp1 | 0.004615 | 0.007938 | -0.486339643 | 674.4532103 | 481.4475771 | 6 | 4 | 4 | 4 | 4 | 0 | 0 | 0 | 0 | 0 | 0 | 0 |
| SG0000003 | R3hcc1 | 0.004708 | 0.008088 | 0.434415599 | 605.9722236 | 818.8886347 | 22 | 4 | 4 | 6 | 18 | 0 | 2 | 0 | 0 | 0 | 0 | 0 |
| SG0000003 | Jak3 | 0.004889 | 0.008377 | 0.566852732 | 781.0155902 | 1156.909477 | 26 | 4 | 4 | 4 | 8 | 0 | 0 | 0 | 0 | 0 | 0 | 0 |
| JSG000000 | Gch1 | 0.004937 | 0.008454 | -0.553612249 | 400.7095581 | 273.0082736 | 69 | 1 | 4 | 12 | 23 | 4 | 7 | 0 | 1 | 1 | 1 | 0 |
| JSG000000 | Wdr92 | 0.005116 | 0.008746 | 0.418279975 | 699.777694 | 935.136296 | 39 | 3 | 4 | 9 | 19 | 4 | 2 | 0 | 0 | 0 | 0 | 0 |
| JSG000000 | Fbln7 | 0.005252 | 0.00896 | 1.699866719 | 7.914085472 | 25.71056422 | 56 | 4 | 4 | 12 | 44 | 10 | 0 | 0 | 0 | 0 | 0 | 0 |
| JSG000000 | Dcaf12 | 0.005536 | 0.009411 | 0.365279582 | 2034.050034 | 2620.123381 | 32 | 2 | 4 | 4 | 7 | 2 | 3 | 0 | 0 | 1 | 1 | 0 |
| SG0000002 | Snrpb | 0.006721 | 0.011295 | 0.567635325 | 3295.337863 | 4883.994967 | 34 | 2 | 4 | 4 | 14 | 1 | 0 | 0 | 0 | 0 | 0 | 1 |
| JSG000000 | Pacrgl | 0.006996 | 0.011717 | -0.500259812 | 463.5699282 | 327.7344134 | 152 | 4 | 4 | 13 | 57 | 11 | 3 | 0 | 0 | 3 | 6 | 0 |
| SG0000002 | Mkks | 0.008405 | 0.013917 | 0.438516816 | 409.9571312 | 555.5781543 | 84 | 0 | 4 | 6 | 22 | 0 | 4 | 0 | 0 | 0 | 0 | 0 |
| SG0000003 | 2410004B18Rik | 0.008712 | 0.01438 | 0.475328769 | 282.3903159 | 392.5889003 | 31 | 0 | 4 | 8 | 23 | 7 | 12 | 0 | 4 | 4 | 4 | 2 |
| SG0000002 | Csf1r | 0.009254 | 0.015216 | 2.134811594 | 16.92587813 | 74.33503232 | 43 | 2 | 4 | 10 | 16 | 5 | 5 | 0 | 0 | 0 | 0 | 1 |
| JSG000000 | Kpnb1 | 0.009567 | 0.015696 | 0.312340816 | 22072.91577 | 27408.39854 | 41 | 4 | 4 | 7 | 41 | 2 | 5 | 0 | 0 | 0 | 1 | 0 |
| SG0000000 | Taf6l | 0.010408 | 0.016973 | -0.409597441 | 883.6257603 | 665.2229938 | 54 | 4 | 4 | 9 | 29 | 0 | 0 | 0 | 0 | 0 | 0 | 0 |
| SG0000004 | Krtcap2 | 0.01098 | 0.017834 | -0.474565219 | 1309.998968 | 942.7848346 | 8 | 4 | 4 | 4 | 6 | 0 | 0 | 0 | 0 | 0 | 0 | 0 |
| SG0000000 | Mcts1 | 0.011276 | 0.018289 | -0.403738702 | 1001.044948 | 756.6867312 | 39 | 3 | 4 | 11 | 26 | 1 | 0 | 0 | 0 | 0 | 0 | 0 |
| JSG000000 | Kin | 0.011642 | 0.018834 | 0.45635096 | 281.3567034 | 386.0402554 | 23 | 3 | 4 | 4 | 14 | 0 | 2 | 0 | 0 | 0 | 1 | 0 |
| SG0000002 | Aloxe3 | 0.011705 | 0.018929 | 0.74698616 | 62.67307955 | 105.1831749 | 41 | 3 | 4 | 8 | 22 | 2 | 0 | 0 | 0 | 0 | 0 | 0 |
| JSG000000 | Gen1 | 0.011951 | 0.019294 | -0.443501092 | 608.0997253 | 447.1648681 | 68 | 3 | 4 | 10 | 26 | 2 | 0 | 0 | 0 | 0 | 1 | 0 |
| JSG000000 | Taco1 | 0.012758 | 0.02046 | -0.523552053 | 285.5957449 | 198.6766646 | 14 | 4 | 4 | 6 | 6 | 2 | 0 | 0 | 0 | 0 | 0 | 0 |
| SG0000002 | Ctnnbip1 | 0.012956 | 0.020758 | 0.408930762 | 431.4765066 | 572.871978 | 50 | 4 | 4 | 13 | 41 | 12 | 3 | 1 | 1 | 2 | 3 | 0 |
| JSG000000 | Mysm1 | 0.0133 | 0.021259 | 0.365233301 | 994.0263332 | 1280.395292 | 100 | 2 | 4 | 10 | 44 | 17 | 15 | 0 | 0 | 0 | 4 | 4 |
| JSG000000 | Tmem263 | 0.013498 | 0.021538 | -0.385576625 | 1210.098717 | 926.2978817 | 39 | 1 | 4 | 5 | 21 | 4 | 14 | 0 | 0 | 1 | 1 | 0 |
| JSG000000 | Rnmtl1 | 0.014604 | 0.023183 | -0.53660963 | 242.4257272 | 167.1256512 | 9 | 2 | 4 | 3 | 6 | 0 | 0 | 0 | 0 | 0 | 0 | 0 |
| SG0000004 | N6amt1 | 0.014831 | 0.02352 | -0.465984173 | 423.7589328 | 306.7917329 | 74 | 0 | 4 | 6 | 14 | 17 | 12 | 0 | 0 | 2 | 6 | 3 |
| SG0000005 | Trp53 | 0.014967 | 0.023719 | 0.30365876 | 7558.636547 | 9329.403102 | 44 | 0 | 4 | 0 | 0 | 4 | 4 | 0 | 0 | 0 | 0 | 0 |

| | | | | | | | | | | | | | | | | | | |
|---|---|---|---|---|---|---|---|---|---|---|---|---|---|---|---|---|---|---|
| SG0000001 | Adssl1 | 0.015116 | 0.023948 | 0.921273336 | 121.4159391 | 229.9358092 | 28 | 4 | 4 | 6 | 8 | 2 | 2 | 0 | 0 | 0 | 0 | 0 |
| SG0000002 | Aurkb | 0.01634 | 0.025721 | 0.315681416 | 2067.865816 | 2573.664482 | 14 | 4 | 4 | 6 | 8 | 6 | 2 | 0 | 0 | 0 | 2 | 0 |
| SG0000002 | Pygl | 0.016374 | 0.025772 | -1.090046268 | 712.2735813 | 334.587886 | 92 | 2 | 4 | 18 | 74 | 6 | 0 | 0 | 0 | 0 | 0 | 0 |
| SG0000007 | Mthfsl | 0.017155 | 0.026894 | -0.754033957 | 104.0674705 | 61.70610867 | 108 | 0 | 4 | 12 | 46 | 4 | 0 | 0 | 0 | 0 | 0 | 0 |
| SG0000003 | Grip2 | 0.017485 | 0.027366 | -0.898016942 | 100.0254001 | 53.67601445 | 73 | 3 | 4 | 6 | 64 | 9 | 0 | 0 | 0 | 0 | 0 | 0 |
| SG0000002 | Srrd | 0.017615 | 0.027542 | -0.4728315 | 341.0396894 | 245.7358161 | 8 | 2 | 4 | 4 | 8 | 0 | 0 | 0 | 0 | 0 | 0 | 0 |
| SG0000003 | Cipc | 0.018502 | 0.028798 | -0.348628236 | 1526.874714 | 1199.101225 | 37 | 4 | 4 | 4 | 15 | 3 | 11 | 0 | 0 | 0 | 0 | 0 |
| SG0000002 | Tesc | 0.018977 | 0.029478 | 0.961648137 | 69.20819631 | 134.7852801 | 50 | 2 | 4 | 8 | 30 | 0 | 0 | 0 | 0 | 0 | 2 | 0 |
| SG0000007 | Lekr1 | 0.019292 | 0.029926 | -0.674559059 | 126.4629211 | 79.23201456 | 64 | 3 | 4 | 11 | 31 | 3 | 11 | 0 | 0 | 3 | 9 | 0 |
| SG0000003 | Abcf1 | 0.020055 | 0.031015 | 0.299039711 | 4469.670517 | 5499.148315 | 15 | 2 | 4 | 2 | 8 | 5 | 0 | 0 | 0 | 0 | 0 | 0 |
| SG0000002 | Klc2 | 0.02023 | 0.031265 | -0.354234725 | 1076.718885 | 842.3004886 | 12 | 4 | 4 | 4 | 8 | 4 | 4 | 4 | 4 | 4 | 4 | 4 |
| SG000000C | Ppp5c | 0.020875 | 0.032186 | 0.305140607 | 2406.971241 | 2973.906957 | 30 | 2 | 4 | 4 | 9 | 2 | 2 | 0 | 0 | 2 | 3 | 0 |
| SG0000001 | Hars2 | 0.021225 | 0.032673 | -0.423147566 | 814.1097647 | 607.1595828 | 31 | 4 | 4 | 6 | 18 | 8 | 3 | 0 | 0 | 0 | 0 | 0 |
| JSG000000 | Sp6 | 0.021273 | 0.032739 | 0.477999865 | 186.3980489 | 259.6173182 | 14 | 2 | 4 | 4 | 6 | 0 | 6 | 2 | 4 | 2 | 4 | 0 |
| JSG000000 | Fn3krp | 0.022305 | 0.034224 | 0.398088052 | 324.5046231 | 427.6193341 | 19 | 4 | 4 | 7 | 11 | 0 | 2 | 0 | 0 | 0 | 0 | 0 |
| JSG000000 | Gpalpp1 | 0.022877 | 0.03502 | -0.337423269 | 1892.304996 | 1497.671656 | 38 | 1 | 4 | 6 | 19 | 2 | 11 | 0 | 0 | 2 | 4 | 1 |
| SG000000C | Tcf25 | 0.024276 | 0.037013 | 0.285947238 | 5359.932927 | 6534.886528 | 47 | 0 | 4 | 14 | 28 | 8 | 0 | 0 | 0 | 0 | 0 | 0 |
| JSG000000 | Lpar4 | 0.025816 | 0.039184 | 0.562713244 | 93.50610731 | 138.112677 | 46 | 2 | 4 | 4 | 20 | 6 | 17 | 0 | 1 | 1 | 4 | 2 |
| SG0000004 | Tom1 | 0.027816 | 0.042024 | 0.418915261 | 243.5059889 | 325.5485057 | 40 | 4 | 4 | 6 | 16 | 6 | 0 | 0 | 0 | 0 | 0 | 0 |
| SG000000C | Cnn2 | 0.028448 | 0.042889 | -0.322968143 | 2232.128247 | 1784.415738 | 11 | 2 | 4 | 4 | 8 | 0 | 1 | 0 | 0 | 0 | 0 | 0 |
| SG0000003 | Nup54 | 0.028985 | 0.043625 | 0.287446425 | 1897.199018 | 2315.489881 | 60 | 4 | 4 | 9 | 24 | 2 | 3 | 0 | 0 | 0 | 2 | 0 |
| SG0000002 | C330018D20Rik | 0.029521 | 0.044368 | -0.45686785 | 306.1002563 | 223.0143281 | 83 | 0 | 4 | 8 | 24 | 4 | 5 | 0 | 0 | 0 | 2 | 0 |
| SG0000003 | Wdr3 | 0.03077 | 0.046083 | 0.281941771 | 2150.799448 | 2615.006945 | 62 | 3 | 4 | 11 | 34 | 7 | 2 | 0 | 0 | 1 | 1 | 0 |
| JSG000000 | Kbtbd3 | 0.031749 | 0.047442 | -0.814204613 | 73.17403831 | 41.61568162 | 47 | 2 | 4 | 5 | 26 | 3 | 0 | 0 | 0 | 0 | 0 | 0 |
| SG0000007 | AI607873 | 0.032216 | 0.0481 | -0.976038355 | 67.81005411 | 34.4728563 | 98 | 2 | 4 | 10 | 41 | 8 | 0 | 0 | 0 | 0 | 1 | 0 |
| JSG000000 | Fst | 0.034791 | 0.051605 | -0.64024873 | 245.4257507 | 157.4657317 | 12 | 3 | 4 | 4 | 6 | 4 | 7 | 2 | 3 | 3 | 3 | 1 |
| SG0000006 | Zfp266 | 0.03548 | 0.05253 | 0.291590387 | 1702.988241 | 2084.438577 | 49 | 2 | 4 | 5 | 29 | 5 | 13 | 0 | 0 | 0 | 2 | 1 |
| SG0000003 | Dctn6 | 0.037822 | 0.05575 | -0.327442501 | 989.364829 | 788.4723223 | 101 | 4 | 4 | 16 | 33 | 16 | 4 | 0 | 0 | 0 | 2 | 0 |
| SG0000001 | Chdh | 0.038086 | 0.056102 | 1.530888881 | 24.26125765 | 70.10625742 | 55 | 4 | 4 | 5 | 25 | 3 | 9 | 0 | 0 | 0 | 0 | 0 |
| SG0000001 | Zfp715 | 0.039817 | 0.058465 | 0.320835707 | 536.7700521 | 670.4547144 | 151 | 4 | 4 | 28 | 60 | 8 | 25 | 0 | 0 | 0 | 12 | 3 |
| JSG000000 | Lysmd2 | 0.040199 | 0.058968 | -0.558896805 | 139.1778461 | 94.47685075 | 30 | 4 | 4 | 6 | 14 | 2 | 2 | 0 | 0 | 1 | 1 | 1 |
| SG0000002 | Mum1 | 0.041188 | 0.060319 | 0.282062212 | 1389.461603 | 1689.49043 | 18 | 4 | 4 | 5 | 7 | 1 | 0 | 0 | 0 | 0 | 0 | 0 |
| SG000000C | Rab8a | 0.041518 | 0.060777 | -0.288032876 | 2307.016157 | 1889.487837 | 23 | 4 | 4 | 5 | 13 | 1 | 3 | 0 | 0 | 0 | 1 | 0 |
| SG000000C | Tmem110 | 0.041815 | 0.061192 | 0.324157703 | 538.3912165 | 674.0298934 | 53 | 0 | 4 | 5 | 19 | 3 | 1 | 0 | 0 | 0 | 0 | 0 |
| SG0000003 | Atxn2l | 0.043694 | 0.063733 | 0.255562058 | 5800.28131 | 6924.380141 | 28 | 2 | 4 | 4 | 12 | 6 | 2 | 0 | 2 | 2 | 4 | 0 |
| SG0000002 | E2f1 | 0.044111 | 0.064295 | 0.280483346 | 1570.548565 | 1907.591027 | 14 | 4 | 4 | 6 | 6 | 2 | 6 | 4 | 4 | 6 | 6 | 0 |
| SG0000004 | Dstyk | 0.04432 | 0.064573 | -0.354767476 | 913.1725216 | 714.0969577 | 167 | 4 | 4 | 12 | 50 | 30 | 3 | 0 | 0 | 0 | 2 | 0 |
| SG0000003 | Arfrp1 | 0.048021 | 0.06957 | -0.303876013 | 1044.912871 | 846.4557972 | 10 | 4 | 4 | 4 | 10 | 2 | 2 | 0 | 0 | 0 | 0 | 0 |
| JSG000000 | Ccdc12 | 0.049287 | 0.071255 | 0.34762894 | 437.4339578 | 556.6205451 | 57 | 1 | 4 | 5 | 27 | 6 | 1 | 0 | 0 | 0 | 0 | 2 |
| SG0000003 | Tfdp1 | 0.050933 | 0.073392 | 0.248051499 | 4142.36324 | 4919.479111 | 41 | 3 | 4 | 8 | 15 | 3 | 0 | 0 | 0 | 0 | 0 | 0 |
| JSG000000 | 4930533O14Rik | 0.051825 | 0.074567 | 0.262887411 | 2318.582384 | 2782.015388 | 23 | 4 | 4 | 4 | 9 | 0 | 0 | 0 | 0 | 0 | 0 | 0 |
| SG0000007 | Psme2 | 0.054062 | 0.077543 | -0.28933591 | 2854.275422 | 2335.592598 | 6 | 4 | 4 | 4 | 6 | 2 | 2 | 0 | 0 | 0 | 0 | 0 |
| JSG000000 | Clec2l | 0.055163 | 0.078977 | -1.554599104 | 23.35793777 | 7.951581884 | 22 | 3 | 4 | 6 | 11 | 1 | 0 | 0 | 0 | 0 | 0 | 0 |
| SG0000001 | Pnpo | 0.059826 | 0.085035 | -0.290614242 | 1092.029742 | 892.7931773 | 24 | 4 | 4 | 4 | 8 | 2 | 0 | 0 | 0 | 0 | 0 | 0 |
| SG0000003 | Polr3e | 0.06338 | 0.08973 | 0.268658568 | 1084.996743 | 1307.081664 | 40 | 0 | 4 | 6 | 24 | 2 | 2 | 0 | 0 | 0 | 0 | 0 |
| JSG000000 | 1700029J07Rik | 0.067907 | 0.095609 | 0.726853649 | 30.70403827 | 50.81597345 | 90 | 0 | 4 | 4 | 36 | 6 | 2 | 0 | 0 | 0 | 0 | 0 |
| SG0000002 | Prpf19 | 0.071227 | 0.099861 | 0.227266553 | 5582.029998 | 6534.407517 | 22 | 2 | 4 | 4 | 9 | 0 | 4 | 0 | 0 | 0 | 1 | 0 |
| SG0000003 | Lsm6 | 0.072563 | 0.101606 | 0.24699214 | 1427.800502 | 1694.414164 | 70 | 2 | 4 | 3 | 14 | 11 | 10 | 0 | 0 | 1 | 3 | 2 |
| SG0000002 | Snx11 | 0.075297 | 0.105105 | -0.323635051 | 773.8171393 | 618.3216976 | 31 | 4 | 4 | 8 | 14 | 0 | 7 | 0 | 0 | 0 | 0 | 0 |
| JSG000000 | Sprtn | 0.076299 | 0.106414 | 0.339514742 | 256.5232766 | 324.5868341 | 5 | 4 | 4 | 4 | 5 | 0 | 0 | 0 | 0 | 0 | 0 | 0 |
| JSG000000 | Hist1h2be | 0.077505 | 0.107935 | 1.034316506 | 11.93579788 | 24.44622188 | 132 | 2 | 4 | 12 | 44 | 8 | 12 | 0 | 0 | 0 | 0 | 0 |
| SG0000003 | Elmod2 | 0.07839 | 0.109084 | 0.291108701 | 608.0507416 | 743.9988657 | 36 | 2 | 4 | 5 | 17 | 5 | 10 | 0 | 0 | 1 | 6 | 1 |
| JSG000000 | Necap1 | 0.081701 | 0.113269 | -0.245627616 | 2561.732906 | 2160.690521 | 30 | 4 | 4 | 6 | 19 | 4 | 3 | 0 | 0 | 0 | 1 | 1 |
| SG0000007 | Ppia | 0.082779 | 0.114656 | 0.21418882 | 14637.94934 | 16980.77468 | 10 | 4 | 4 | 8 | 12 | 6 | 0 | 0 | 0 | 0 | 0 | 0 |
| SG000000C | Ctnnb1 | 0.083056 | 0.115014 | 0.211633908 | 20848.15285 | 24142.13849 | 100 | 0 | 4 | 4 | 29 | 9 | 0 | 0 | 0 | 0 | 0 | 0 |
| SG0000001 | Tsg101 | 0.084079 | 0.116268 | -0.252739294 | 1594.739292 | 1338.466747 | 57 | 4 | 4 | 7 | 23 | 3 | 1 | 0 | 0 | 0 | 0 | 2 |
| SG0000002 | Pfas | 0.085584 | 0.118194 | 0.216463064 | 3777.902544 | 4389.475463 | 35 | 3 | 4 | 8 | 19 | 3 | 5 | 0 | 0 | 1 | 2 | 0 |
| JSG000000 | Dpep1 | 0.086957 | 0.119933 | -1.149685251 | 21.46589209 | 9.67518317 | 13 | 2 | 4 | 5 | 12 | 0 | 1 | 0 | 0 | 0 | 0 | 0 |
| SG000000C | Cyth2 | 0.08883 | 0.122272 | -0.250606143 | 1398.895995 | 1175.832502 | 14 | 2 | 4 | 2 | 2 | 2 | 10 | 2 | 4 | 2 | 2 | 0 |
| SG0000001 | Copz2 | 0.089042 | 0.122536 | -0.392441205 | 192.8955499 | 146.9554288 | 20 | 2 | 4 | 4 | 8 | 0 | 0 | 0 | 0 | 0 | 1 | 0 |
| SG0000002 | Map1a | 0.090123 | 0.123862 | 0.470265025 | 458.0793508 | 634.6066131 | 28 | 4 | 4 | 4 | 5 | 1 | 13 | 4 | 4 | 4 | 4 | 1 |
| JSG000000 | Crnkl1 | 0.100292 | 0.136359 | 0.229394844 | 1236.293235 | 1449.359835 | 31 | 2 | 4 | 4 | 11 | 2 | 3 | 0 | 0 | 0 | 1 | 0 |
| JSG000000 | Pcnxl4 | 0.100582 | 0.136674 | 0.255207766 | 554.6281041 | 661.9528634 | 78 | 1 | 4 | 11 | 41 | 11 | 2 | 0 | 0 | 1 | 1 | 1 |
| SG0000001 | Ppil4 | 0.105966 | 0.14351 | 0.219132889 | 1383.852464 | 1610.850974 | 67 | 2 | 4 | 10 | 38 | 12 | 4 | 0 | 0 | 0 | 3 | 1 |
| JSG000000 | Slc25a20 | 0.106259 | 0.143841 | -0.28227598 | 647.0884217 | 532.09612 | 38 | 3 | 4 | 13 | 21 | 2 | 1 | 0 | 0 | 0 | 0 | 0 |
| SG0000007 | Itpa | 0.108495 | 0.146657 | 0.232930876 | 1043.010751 | 1225.767115 | 12 | 2 | 4 | 4 | 11 | 3 | 0 | 0 | 0 | 0 | 0 | 0 |
| SG0000003 | Ddx47 | 0.109274 | 0.147511 | 0.21604554 | 2048.23093 | 2379.113207 | 13 | 2 | 4 | 3 | 11 | 4 | 1 | 0 | 1 | 0 | 2 | 0 |
| JSG000000 | Mon1b | 0.109935 | 0.148324 | 0.258268789 | 500.485169 | 598.6015883 | 24 | 2 | 4 | 4 | 14 | 4 | 4 | 0 | 0 | 1 | 4 | 0 |
| JSG000000 | Eci1 | 0.112658 | 0.151535 | 0.264895497 | 489.0983449 | 587.6756324 | 18 | 4 | 4 | 6 | 10 | 2 | 0 | 0 | 0 | 0 | 0 | 0 |
| SG0000006 | Zfp442 | 0.116033 | 0.155638 | -0.535243795 | 98.05215511 | 67.66011107 | 71 | 3 | 4 | 7 | 57 | 7 | 4 | 0 | 0 | 0 | 0 | 0 |
| SG0000005 | Zfp945 | 0.117041 | 0.156836 | -0.303185393 | 628.8073857 | 509.6238407 | 96 | 2 | 4 | 17 | 49 | 6 | 8 | 0 | 0 | 1 | 3 | 4 |
| SG0000001 | Rmnd1 | 0.120948 | 0.161625 | 0.249752998 | 499.5373903 | 593.9517203 | 45 | 3 | 4 | 6 | 28 | 4 | 1 | 0 | 0 | 0 | 2 | 0 |
| JSG000000 | Pgrmc2 | 0.123121 | 0.164243 | -0.226626227 | 1750.050116 | 1495.647308 | 36 | 3 | 4 | 6 | 16 | 2 | 2 | 0 | 0 | 0 | 1 | 1 |
| SG0000002 | Glyr1 | 0.123477 | 0.164673 | 0.196879353 | 4009.019361 | 4595.20345 | 68 | 4 | 4 | 4 | 36 | 8 | 2 | 0 | 0 | 0 | 0 | 0 |
| SG000000C | Ergic1 | 0.126073 | 0.167828 | -0.2022274 | 5912.860276 | 5139.502755 | 91 | 3 | 4 | 10 | 50 | 19 | 1 | 0 | 0 | 0 | 0 | 0 |
| SG000000C | Atp5d | 0.130181 | 0.172855 | -0.493203075 | 4113.021714 | 2922.079872 | 12 | 4 | 4 | 4 | 4 | 0 | 0 | 0 | 0 | 0 | 0 | 0 |
| SG0000002 | Cog6 | 0.131627 | 0.174567 | -0.231411642 | 1157.10337 | 985.6218274 | 100 | 4 | 4 | 13 | 40 | 11 | 4 | 0 | 0 | 1 | 2 | 1 |
| SG0000007 | Zfp942 | 0.135753 | 0.179547 | 0.234816322 | 487.245209 | 573.3691654 | 69 | 2 | 4 | 7 | 39 | 14 | 1 | 0 | 0 | 0 | 0 | 1 |
| SG0000006 | Anks6 | 0.13666 | 0.180643 | 0.853968339 | 20.86309435 | 37.70932925 | 70 | 0 | 4 | 4 | 28 | 4 | 0 | 0 | 0 | 0 | 0 | 0 |
| JSG000000 | Bach1 | 0.13801 | 0.182242 | 0.203203234 | 1438.060337 | 1655.569347 | 56 | 3 | 4 | 7 | 18 | 10 | 5 | 0 | 0 | 0 | 1 | 5 |
| SG0000002 | Timm9 | 0.139874 | 0.184467 | 0.248360431 | 453.5121582 | 538.7073164 | 70 | 4 | 4 | 10 | 20 | 4 | 0 | 0 | 0 | 0 | 0 | 0 |
| JSG000000 | Mmp2 | 0.1418 | 0.186786 | 0.443288787 | 574.4104639 | 781.0261309 | 27 | 3 | 4 | 8 | 15 | 4 | 1 | 0 | 0 | 0 | 0 | 0 |
| JSG000000 | Vcp | 0.143247 | 0.188457 | -0.191697153 | 10788.80306 | 9446.407417 | 38 | 2 | 4 | 5 | 12 | 4 | 0 | 0 | 0 | 0 | 0 | 0 |
| SG0000002 | Btg3 | 0.144395 | 0.189828 | 0.628735869 | 27.81499895 | 43.00781745 | 39 | 3 | 4 | 8 | 26 | 2 | 4 | 0 | 1 | 2 | 4 | 0 |
| JSG000000 | Twistnb | 0.14795 | 0.194088 | 0.202563575 | 1078.933873 | 1241.573802 | 25 | 1 | 4 | 3 | 13 | 1 | 5 | 0 | 0 | 0 | 3 | 0 |
| SG0000001 | Matn4 | 0.151311 | 0.198149 | 1.094880236 | 7.357127746 | 15.71448272 | 52 | 0 | 4 | 0 | 16 | 12 | 4 | 0 | 0 | 0 | 0 | 0 |
| JSG000000 | Xkr8 | 0.161481 | 0.210268 | 0.269629526 | 239.125584 | 288.2654669 | 11 | 4 | 4 | 4 | 6 | 1 | 6 | 3 | 3 | 3 | 4 | 0 |
| JSG000000 | Ndnf | 0.163138 | 0.212287 | -0.817676803 | 28.6205614 | 16.23801372 | 73 | 1 | 4 | 5 | 25 | 13 | 9 | 0 | 0 | 0 | 3 | 2 |
| SG0000005 | Pfdn4 | 0.171604 | 0.222206 | 0.193802006 | 935.5850678 | 1070.097868 | 66 | 4 | 4 | 14 | 45 | 0 | 6 | 3 | 3 | 3 | 10 | 0 |
| SG0000003 | Manf | 0.173663 | 0.224514 | 0.180663234 | 1765.692745 | 2001.244656 | 128 | 1 | 4 | 24 | 51 | 9 | 6 | 0 | 0 | 0 | 0 | 0 |
| SG0000002 | Ddx55 | 0.175329 | 0.226341 | -0.23556584 | 565.1199198 | 479.9856261 | 44 | 2 | 4 | 4 | 6 | 172 | 4 | 0 | 0 | 2 | 2 | 2 |
| SG0000003 | Kif9 | 0.179284 | 0.231098 | 0.543441124 | 33.91181082 | 49.42458824 | 210 | 4 | 4 | 24 | 79 | 12 | 3 | 0 | 0 | 0 | 0 | 0 |
| SG0000002 | Brd2 | 0.181002 | 0.233094 | 0.171765197 | 5737.743531 | 6463.200198 | 42 | 4 | 4 | 8 | 15 | 3 | 7 | 0 | 0 | 0 | 3 | 3 |
| SG000000C | Acvp1 | 0.183689 | 0.236198 | 0.281612587 | 183.2961709 | 222.8061778 | 59 | 4 | 4 | 14 | 21 | 5 | 0 | 0 | 0 | 0 | 0 | 0 |
| JSG000000 | Cnrip1 | 0.184733 | 0.23737 | -0.787095418 | 24.345404 | 14.10839647 | 59 | 3 | 4 | 8 | 31 | 6 | 1 | 0 | 0 | 1 | 1 | 0 |
| SG000000C | Sec14l2 | 0.185503 | 0.238291 | -0.31079647 | 195.7072769 | 157.7785331 | 17 | 4 | 4 | 4 | 7 | 1 | 0 | 0 | 0 | 0 | 0 | 0 |
| JSG000000 | Glt28d2 | 0.192336 | 0.246257 | -1.071395828 | 20.56042534 | 9.783850624 | 39 | 4 | 4 | 8 | 15 | 4 | 3 | 0 | 0 | 0 | 1 | 0 |
| JSG000000 | Decr1 | 0.197116 | 0.251584 | 0.199670325 | 595.6069697 | 684.0164216 | 56 | 2 | 4 | 6 | 33 | 10 | 3 | 0 | 0 | 0 | 0 | 2 |
| SG0000002 | Glt8d1 | 0.198521 | 0.253215 | 0.218931953 | 396.3912977 | 461.3485886 | 34 | 4 | 4 | 4 | 8 | 8 | 4 | 0 | 0 | 0 | 0 | 2 |
| SG0000004 | Safb2 | 0.199265 | 0.25411 | -0.178300932 | 1908.791593 | 1686.881537 | 56 | 0 | 4 | 0 | 7 | 0 | 1 | 0 | 0 | 0 | 0 | 0 |
| SG0000003 | Det1 | 0.200048 | 0.254931 | 0.240402558 | 336.6542565 | 397.6971612 | 47 | 4 | 4 | 6 | 21 | 0 | 2 | 0 | 0 | 0 | 0 | 0 |
| SG0000005 | Rbm12b2 | 0.206742 | 0.262599 | 0.228164947 | 317.6921227 | 372.1267925 | 48 | 0 | 4 | 2 | 8 | 0 | 10 | 0 | 2 | 0 | 0 | 0 |
| SG0000005 | Usf2 | 0.216792 | 0.273868 | 0.199084552 | 564.3628127 | 647.8714038 | 21 | 0 | 4 | 4 | 16 | 0 | 1 | 0 | 0 | 0 | 0 | 0 |
| JSG000000 | Ccar2 | 0.220404 | 0.277765 | 0.166624085 | 1563.946145 | 1755.418381 | 20 | 4 | 4 | 6 | 11 | 3 | 0 | 0 | 0 | 0 | 0 | 1 |
| JSG000000 | Xpc | 0.223273 | 0.281104 | -0.188645928 | 1024.05926 | 898.5391543 | 56 | 2 | 4 | 10 | 28 | 8 | 3 | 0 | 0 | 0 | 0 | 2 |
| JSG000000 | Akirin1 | 0.22452 | 0.282575 | -0.180579386 | 1387.593964 | 1224.341556 | 31 | 3 | 4 | 8 | 22 | 4 | 3 | 0 | 0 | 0 | 3 | 3 |
| SG0000006 | Nr1h2 | 0.227243 | 0.285621 | -0.36802834 | 1410.782243 | 1093.131515 | 4 | 0 | 4 | 0 | 5 | 5 | 4 | 0 | 4 | 0 | 0 | 0 |
| SG0000001 | Cops3 | 0.231804 | 0.290782 | 0.15250729 | 3301.318111 | 3669.413405 | 34 | 4 | 4 | 8 | 25 | 7 | 2 | 0 | 0 | 1 | 1 | 0 |
| JSG000000 | Ddx11 | 0.233551 | 0.292543 | 0.173178457 | 838.4674737 | 945.4055117 | 23 | 4 | 4 | 5 | 14 | 5 | 1 | 0 | 0 | 0 | 0 | 0 |
| JSG000000 | Mettl9 | 0.248755 | 0.309427 | -0.170328032 | 1905.196993 | 1693.035407 | 99 | 1 | 4 | 11 | 37 | 6 | 1 | 0 | 0 | 0 | 0 | 0 |
| JSG000000 | Gldn | 0.248908 | 0.309592 | 0.514134265 | 29.41629017 | 42.01048898 | 72 | 2 | 4 | 8 | 37 | 3 | 7 | 0 | 0 | 1 | 2 | 1 |
| SG0000002 | Pmpcb | 0.253567 | 0.314623 | -0.159665302 | 2176.632921 | 1948.593046 | 44 | 4 | 4 | 11 | 30 | 0 | 2 | 0 | 0 | 2 | 2 | 0 |
| JSG000000 | Tmem45b | 0.25594 | 0.317172 | 0.635255374 | 15.43901203 | 23.98006898 | 49 | 2 | 4 | 7 | 17 | 5 | 1 | 0 | 0 | 0 | 1 | 0 |
| SG0000006 | Zfp938 | 0.280943 | 0.344274 | 0.211442771 | 233.1504246 | 269.9521727 | 54 | 0 | 4 | 8 | 28 | 10 | 3 | 0 | 0 | 1 | 1 | 0 |
| JSG000000 | Brd9 | 0.29144 | 0.355604 | -0.179725727 | 512.537123 | 452.5040723 | 26 | 3 | 4 | 5 | 18 | 3 | 1 | 0 | 0 | 1 | 2 | 0 |
| SG0000006 | Cenpm | 0.292756 | 0.357016 | 0.394606421 | 267.2444201 | 351.3152608 | 20 | 4 | 4 | 6 | 14 | 2 | 0 | 0 | 0 | 0 | 0 | 0 |
| SG0000002 | Rsad2 | 0.299528 | 0.364344 | 0.32218846 | 62.19308433 | 77.75538677 | 44 | 0 | 4 | 18 | 36 | 4 | 5 | 0 | 0 | 0 | 6 | 0 |
| SG0000004 | Mfsd9 | 0.30283 | 0.367748 | -0.310470468 | 196.8601548 | 158.7438462 | 58 | 0 | 4 | 8 | 26 | 2 | 8 | 0 | 2 | 4 | 4 | 0 |
| SG000000C | Pnpla6 | 0.312985 | 0.378617 | -0.136770748 | 1885.762946 | 1715.200918 | 66 | 2 | 4 | 10 | 28 | 8 | 0 | 0 | 0 | 0 | 2 | 0 |
| SG0000002 | Rbm28 | 0.31763 | 0.383459 | -0.135639331 | 3266.495271 | 2973.380758 | 69 | 4 | 4 | 11 | 37 | 5 | 3 | 0 | 0 | 0 | 0 | 0 |
| JSG000000 | Coa4 | 0.319701 | 0.385726 | -0.228797989 | 192.4563577 | 164.2317819 | 19 | 4 | 4 | 4 | 6 | 2 | 1 | 0 | 0 | 0 | 0 | 0 |
| SG0000002 | Mmadhc | 0.321089 | 0.387192 | -0.143895171 | 1704.671374 | 1542.850713 | 57 | 2 | 4 | 9 | 32 | 4 | 2 | 0 | 0 | 1 | 1 | 0 |
| JSG000000 | Tmem70 | 0.329008 | 0.395782 | -0.156165959 | 1010.551947 | 906.8762015 | 54 | 2 | 4 | 6 | 14 | 6 | 4 | 0 | 0 | 0 | 0 | 0 |

| | | | | | | | | | | | | | | | | | | |
|---|---|---|---|---|---|---|---|---|---|---|---|---|---|---|---|---|---|---|
| JSG000000 | Lamtor3 | 0.333589 | 0.400405 | 0.143389372 | 800.844 | 884.5297171 | 44 | 2 | 4 | 6 | 13 | 1 | 4 | 0 | 0 | 2 | 3 | 1 |
| JSG000000 | Cdh5 | 0.338417 | 0.405331 | 0.473914372 | 35.16593972 | 48.84101363 | 55 | 3 | 4 | 6 | 27 | 4 | 1 | 0 | 0 | 0 | 1 | 0 |
| SG0000001 | Cdk1 | 0.367172 | 0.435635 | 0.116491704 | 4905.076657 | 5317.570987 | 49 | 2 | 4 | 6 | 20 | 4 | 5 | 0 | 0 | 0 | 2 | 2 |
| SG0000001 | Dnajc7 | 0.368072 | 0.436327 | 0.112602677 | 3758.157502 | 4063.233312 | 46 | 3 | 4 | 6 | 22 | 7 | 0 | 0 | 0 | 0 | 0 | 0 |
| SG0000001 | Ager | 0.375416 | 0.44421 | -0.562632095 | 22.59202836 | 15.29628924 | 12 | 4 | 4 | 4 | 8 | 0 | 0 | 0 | 0 | 0 | 0 | 0 |
| SG0000003 | Myom3 | 0.378647 | 0.447471 | 0.572670428 | 12.07331792 | 17.95633144 | 39 | 3 | 4 | 5 | 9 | 9 | 2 | 0 | 0 | 0 | 0 | 0 |
| SG0000007 | Fndc1 | 0.405379 | 0.47443 | -0.482641079 | 48.90010043 | 34.9961532 | 101 | 1 | 4 | 11 | 40 | 9 | 2 | 0 | 0 | 0 | 0 | 1 |
| SG0000002 | Wdr4 | 0.414127 | 0.483342 | -0.13514249 | 970.5801492 | 883.7907694 | 61 | 4 | 4 | 6 | 14 | 3 | 6 | 0 | 0 | 0 | 0 | 0 |
| SG0000002 | Wash1 | 0.424809 | 0.493911 | -0.14943075 | 751.7011067 | 677.7383351 | 10 | 0 | 4 | 4 | 10 | 2 | 0 | 0 | 0 | 0 | 0 | 0 |
| SG0000004 | Zfpm1 | 0.429531 | 0.498753 | 0.275686173 | 307.8556003 | 372.6806686 | 21 | 4 | 4 | 4 | 7 | 2 | 0 | 0 | 0 | 0 | 0 | 0 |
| SG0000003 | Apol6 | 0.448189 | 0.517299 | -0.585593103 | 16.40353646 | 10.93091203 | 18 | 4 | 4 | 4 | 10 | 3 | 0 | 0 | 0 | 0 | 0 | 0 |
| SG0000000 | Elk1 | 0.467855 | 0.536746 | 0.109407555 | 828.8985793 | 894.2034702 | 22 | 4 | 4 | 6 | 11 | 4 | 5 | 4 | 4 | 4 | 5 | 1 |
| SG0000005 | Rab4b | 0.47072 | 0.539515 | -0.274043378 | 85.43221613 | 70.65231478 | 24 | 2 | 4 | 10 | 16 | 2 | 2 | 2 | 2 | 2 | 2 | 0 |
| SG0000002 | Sf3a2 | 0.480978 | 0.549549 | -0.140363767 | 2165.669683 | 1964.891224 | 13 | 2 | 4 | 4 | 6 | 0 | 0 | 0 | 0 | 0 | 0 | 0 |
| SG0000003 | Hccs | 0.485381 | 0.553697 | 0.100964964 | 962.8914984 | 1032.692053 | 187 | 2 | 4 | 11 | 72 | 11 | 21 | 0 | 1 | 1 | 5 | 0 |
| SG0000003 | Plet1 | 0.494672 | 0.562399 | 0.109534107 | 538.9459838 | 581.4579216 | 20 | 3 | 4 | 5 | 12 | 4 | 3 | 0 | 0 | 1 | 1 | 1 |
| SG0000003 | H2afz | 0.499373 | 0.566699 | 0.157306234 | 5242.671758 | 5846.642812 | 10 | 4 | 4 | 4 | 7 | 0 | 2 | 0 | 0 | 0 | 3 | 0 |
| JSG000000 | Aagab | 0.522709 | 0.589594 | 0.095477782 | 852.7594885 | 911.1045944 | 57 | 4 | 4 | 8 | 30 | 5 | 0 | 0 | 0 | 0 | 0 | 0 |
| SG0000002 | Rhbdf2 | 0.527403 | 0.594029 | -0.136368902 | 962.2008702 | 875.4162728 | 17 | 3 | 4 | 4 | 8 | 0 | 2 | 0 | 0 | 0 | 2 | 0 |
| SG0000002 | Idh3b | 0.536196 | 0.602607 | -0.084364324 | 2689.40001 | 2536.642181 | 13 | 4 | 4 | 4 | 4 | 2 | 0 | 0 | 0 | 0 | 0 | 0 |
| JSG000000 | Itga10 | 0.540921 | 0.606983 | 0.553848277 | 16.18421365 | 23.75836485 | 30 | 4 | 4 | 6 | 16 | 4 | 2 | 0 | 0 | 0 | 0 | 0 |
| JSG000000 | Cenpv | 0.543542 | 0.609485 | 0.096893038 | 684.9474791 | 732.5292697 | 23 | 4 | 4 | 4 | 16 | 5 | 0 | 0 | 0 | 0 | 0 | 0 |
| SG0000003 | Serpine1 | 0.545752 | 0.611542 | -0.326368376 | 3516.840236 | 2804.826286 | 16 | 0 | 4 | 2 | 4 | 0 | 8 | 0 | 4 | 2 | 2 | 0 |
| JSG000000 | Zfp664 | 0.546592 | 0.612369 | 0.077880705 | 3248.418552 | 3428.59676 | 40 | 3 | 4 | 6 | 9 | 1 | 3 | 0 | 0 | 1 | 3 | 1 |
| SG0000002 | Fen1 | 0.551121 | 0.616631 | 0.078535171 | 1743.910957 | 1841.474667 | 18 | 4 | 4 | 4 | 6 | 0 | 2 | 0 | 0 | 0 | 0 | 0 |
| JSG000000 | Styx | 0.551542 | 0.616988 | 0.193437312 | 58.21871178 | 66.57221769 | 62 | 1 | 4 | 4 | 28 | 3 | 0 | 0 | 0 | 0 | 2 | 0 |
| JSG000000 | Socs4 | 0.561275 | 0.626194 | 0.079685814 | 1434.714681 | 1516.189074 | 46 | 3 | 4 | 7 | 21 | 1 | 17 | 0 | 1 | 1 | 4 | 0 |
| JSG000000 | Ccdc84 | 0.567383 | 0.632025 | -0.165342353 | 105.4636534 | 94.04372093 | 15 | 4 | 4 | 4 | 7 | 1 | 0 | 0 | 0 | 0 | 0 | 0 |
| SG0000005 | Slc25a44 | 0.577619 | 0.641791 | -0.086150282 | 792.2053547 | 746.2836973 | 44 | 4 | 4 | 12 | 20 | 4 | 4 | 0 | 0 | 4 | 4 | 0 |
| JSG000000 | Polr2f | 0.589124 | 0.652469 | -0.165528681 | 1378.489475 | 1229.063599 | 13 | 4 | 4 | 4 | 10 | 1 | 0 | 0 | 0 | 0 | 0 | 0 |
| JSG000000 | 1810030O07Rik | 0.591582 | 0.654732 | -0.087000855 | 973.4011317 | 916.4356595 | 31 | 2 | 4 | 4 | 14 | 5 | 3 | 0 | 1 | 0 | 1 | 0 |
| SG0000003 | Nfkbib | 0.599913 | 0.662738 | 0.160855416 | 425.5428155 | 475.7355083 | 17 | 4 | 4 | 8 | 20 | 2 | 0 | 0 | 0 | 0 | 0 | 0 |
| SG0000002 | Wrb | 0.619945 | 0.681799 | -0.077583173 | 1108.649645 | 1050.604992 | 15 | 3 | 4 | 6 | 7 | 0 | 4 | 0 | 1 | 1 | 2 | 0 |
| SG0000000 | Slc44a4 | 0.651417 | 0.710435 | 0.167970407 | 174.2114994 | 195.722588 | 12 | 2 | 4 | 4 | 12 | 2 | 0 | 0 | 0 | 0 | 1 | 0 |
| SG0000007 | Lrrc61 | 0.652879 | 0.711596 | -0.089221119 | 334.8714114 | 314.7892114 | 44 | 4 | 4 | 12 | 32 | 2 | 2 | 0 | 0 | 2 | 6 | 0 |
| JSG000000 | AA415398 | 0.655911 | 0.714466 | 0.249139371 | 24.14179556 | 28.69247366 | 14 | 2 | 4 | 2 | 3 | 0 | 1 | 0 | 0 | 0 | 0 | 0 |
| JSG000000 | Angpt2 | 0.65928 | 0.717699 | -0.297164143 | 17.80121925 | 14.48753268 | 73 | 3 | 4 | 9 | 28 | 7 | 6 | 0 | 0 | 0 | 1 | 0 |
| JSG000000 | Rmi2 | 0.681031 | 0.737523 | 0.078209954 | 260.3924016 | 274.8981704 | 39 | 3 | 4 | 7 | 20 | 0 | 3 | 1 | 1 | 2 | 4 | 0 |
| JSG000000 | Gpr63 | 0.692752 | 0.747415 | -0.196589769 | 34.37005781 | 29.99168358 | 73 | 2 | 4 | 6 | 30 | 4 | 1 | 0 | 1 | 0 | 0 | 0 |
| SG0000003 | Cc2d1a | 0.699432 | 0.753441 | -0.096332211 | 754.0895616 | 705.3814657 | 16 | 4 | 4 | 8 | 8 | 0 | 0 | 0 | 0 | 0 | 0 | 0 |
| SG0000004 | Ube2q1 | 0.703586 | 0.757415 | -0.053000756 | 2727.957585 | 2629.558252 | 18 | 2 | 4 | 2 | 9 | 0 | 4 | 0 | 2 | 0 | 1 | 0 |
| SG0000002 | Spata24 | 0.721347 | 0.773467 | 0.109379826 | 84.48978066 | 91.1445681 | 12 | 4 | 4 | 4 | 5 | 0 | 0 | 0 | 0 | 0 | 0 | 0 |
| SG0000002 | Chit1 | 0.724091 | 0.775944 | 0.20891111 | 18.10269469 | 20.92337493 | 104 | 2 | 4 | 11 | 74 | 10 | 0 | 0 | 0 | 0 | 0 | 0 |
| JSG000000 | Zfp28 | 0.725882 | 0.777725 | 0.082855868 | 119.3130921 | 126.3659895 | 23 | 4 | 4 | 4 | 5 | 1 | 6 | 0 | 0 | 0 | 0 | 1 |
| SG0000003 | Paqr7 | 0.729938 | 0.781136 | 0.113895193 | 74.60818684 | 80.73695297 | 16 | 4 | 4 | 6 | 15 | 2 | 6 | 0 | 0 | 0 | 1 | 0 |
| JSG000000 | Tbccd1 | 0.73549 | 0.785837 | 0.052488915 | 513.7303101 | 532.7652962 | 59 | 0 | 4 | 7 | 29 | 7 | 0 | 0 | 0 | 0 | 2 | 0 |
| SG0000003 | Taf4a | 0.754028 | 0.801552 | -0.054967224 | 1186.779554 | 1142.413358 | 59 | 3 | 4 | 8 | 22 | 16 | 3 | 0 | 0 | 1 | 1 | 3 |
| SG0000002 | Plk3 | 0.773782 | 0.8191 | 0.090183939 | 526.598225 | 560.5669149 | 10 | 4 | 4 | 6 | 6 | 0 | 8 | 4 | 4 | 6 | 6 | 0 |
| JSG000000 | Nudt15 | 0.776633 | 0.821389 | -0.142348164 | 34.18919447 | 30.97688911 | 42 | 2 | 4 | 5 | 15 | 1 | 2 | 0 | 0 | 0 | 0 | 0 |
| SG0000003 | Robo3 | 0.793012 | 0.835309 | -0.12594909 | 27.19529326 | 24.92179346 | 35 | 4 | 4 | 6 | 11 | 0 | 0 | 0 | 0 | 0 | 0 | 0 |
| JSG000000 | Lcmt2 | 0.802895 | 0.843536 | 0.059503429 | 182.8625187 | 190.5623125 | 23 | 4 | 4 | 7 | 19 | 3 | 4 | 1 | 1 | 2 | 4 | 0 |
| SG0000005 | Ptges | 0.806634 | 0.846819 | -0.086101872 | 425.1191966 | 400.4898098 | 10 | 1 | 4 | 5 | 13 | 2 | 0 | 0 | 0 | 0 | 0 | 0 |
| SG0000001 | Med24 | 0.815051 | 0.854103 | 0.035477072 | 1873.285555 | 1919.922275 | 50 | 2 | 4 | 4 | 19 | 2 | 0 | 0 | 0 | 0 | 1 | 2 |
| SG0000002 | Tsen54 | 0.817091 | 0.855891 | 0.045675693 | 453.4455324 | 468.0312819 | 17 | 4 | 4 | 8 | 15 | 0 | 0 | 0 | 0 | 0 | 0 | 0 |
| JSG000000 | Nrde2 | 0.818281 | 0.856886 | 0.0416198 | 595.9790513 | 613.422643 | 43 | 3 | 4 | 9 | 19 | 2 | 0 | 0 | 0 | 0 | 1 | 0 |
| JSG000000 | Timp2 | 0.828118 | 0.865114 | -0.082640003 | 2275.628984 | 2148.940233 | 46 | 3 | 4 | 4 | 29 | 10 | 0 | 0 | 0 | 0 | 4 | 0 |
| SG0000003 | Mesdc2 | 0.841801 | 0.876291 | 0.025601028 | 2180.909885 | 2219.956161 | 27 | 2 | 4 | 4 | 10 | 0 | 3 | 0 | 0 | 0 | 0 | 0 |
| JSG000000 | Zfp759 | 0.842663 | 0.877075 | 0.038714809 | 179.7755549 | 184.6651558 | 30 | 3 | 4 | 4 | 19 | 2 | 2 | 0 | 0 | 0 | 0 | 0 |
| JSG000000 | Ssr1 | 0.849115 | 0.88206 | 0.017489434 | 5974.824989 | 6047.697124 | 36 | 2 | 4 | 6 | 13 | 6 | 9 | 0 | 0 | 1 | 4 | 1 |
| SG0000002 | Nfya | 0.859713 | 0.890692 | -0.029096663 | 1891.945249 | 1854.170196 | 185 | 0 | 4 | 19 | 48 | 11 | 12 | 0 | 0 | 3 | 3 | 6 |
| SG0000002 | Ccnl2 | 0.867623 | 0.897537 | -0.026717495 | 3308.325389 | 3247.621819 | 46 | 4 | 4 | 8 | 24 | 4 | 2 | 0 | 0 | 0 | 2 | 4 |
| SG0000003 | Wdr19 | 0.880589 | 0.907909 | -0.035263724 | 722.765331 | 705.3129761 | 90 | 3 | 4 | 15 | 37 | 6 | 1 | 0 | 0 | 0 | 0 | 1 |
| SG0000002 | Mrpl20 | 0.881568 | 0.908553 | -0.025314578 | 1073.597799 | 1054.923982 | 20 | 4 | 4 | 7 | 16 | 0 | 5 | 0 | 0 | 2 | 2 | 0 |
| SG0000001 | H3f3b | 0.895849 | 0.920411 | 0.011307332 | 10288.91713 | 10369.87485 | 15 | 2 | 4 | 5 | 12 | 0 | 8 | 0 | 2 | 2 | 4 | 0 |
| SG0000003 | Fkbp1a | 0.899231 | 0.923356 | 0.011660623 | 7877.761414 | 7941.691649 | 48 | 0 | 4 | 4 | 22 | 8 | 0 | 0 | 0 | 0 | 8 | 0 |
| SG0000001 | Srsf6 | 0.907922 | 0.930893 | -0.015041522 | 8097.863356 | 8013.873726 | 28 | 2 | 4 | 6 | 9 | 0 | 11 | 0 | 0 | 0 | 1 | 0 |
| JSG000000 | Dok4 | 0.921344 | 0.942862 | -0.0192524 | 256.3260083 | 252.928125 | 14 | 4 | 4 | 5 | 7 | 2 | 1 | 0 | 0 | 1 | 1 | 1 |
| SG0000002 | Nr4a2 | 0.924515 | 0.945417 | -0.093108143 | 24.80983797 | 23.25924344 | 85 | 0 | 4 | 10 | 39 | 4 | 12 | 0 | 0 | 4 | 17 | 0 |
| JSG000000 | Eif1a | 0.931972 | 0.950396 | -0.017555946 | 5999.033378 | 5926.47439 | 22 | 3 | 4 | 4 | 13 | 1 | 3 | 0 | 0 | 0 | 0 | 0 |
| SG0000004 | Lacc1 | 0.935235 | 0.952789 | -0.023481336 | 422.3548573 | 415.5362409 | 30 | 3 | 4 | 8 | 15 | 1 | 4 | 0 | 0 | 1 | 1 | 0 |
| JSG000000 | Neil2 | 1E-210 | 3.5E-208 | 9.922072107 | 0.946619625 | 918.3681785 | 16 | 1 | 3 | 6 | 8 | 0 | 0 | 0 | 0 | 0 | 1 | 0 |
| SG0000005 | Mest | 1.9E-185 | 4.9E-183 | 4.317703284 | 122.2088213 | 2437.029332 | 78 | 0 | 3 | 3 | 13 | 3 | 15 | 0 | 0 | 0 | 0 | 3 |
| SG0000001 | Aim1 | 3.9E-181 | 9.6E-179 | -7.363045077 | 20893.99492 | 126.9182447 | 90 | 2 | 3 | 14 | 34 | 7 | 2 | 0 | 0 | 0 | 1 | 0 |
| SG0000003 | Hsd17b14 | 1.1E-142 | 1.5E-140 | 7.443905781 | 3.60561515 | 627.7956393 | 11 | 1 | 3 | 4 | 10 | 0 | 0 | 0 | 0 | 0 | 0 | 0 |
| JSG000000 | Qprt | 5.3E-140 | 7.1E-138 | 8.970030071 | 0 | 500.4736448 | 11 | 2 | 3 | 3 | 7 | 1 | 0 | 0 | 0 | 0 | 0 | 0 |
| JSG000000 | Lyplal1 | 5.2E-134 | 6.6E-132 | 5.952939482 | 10.4790574 | 649.1358069 | 60 | 0 | 3 | 4 | 26 | 2 | 5 | 0 | 0 | 0 | 1 | 0 |
| SG0000007 | Ceacam1 | 1.2E-130 | 1.5E-128 | -6.751900233 | 2606.329477 | 24.18270165 | 60 | 0 | 3 | 6 | 30 | 0 | 1 | 0 | 1 | 0 | 0 | 0 |
| SG0000000 | Calcoco2 | 2.8E-130 | 3.4E-128 | 5.041466684 | 23.56273977 | 775.9942105 | 23 | 2 | 3 | 5 | 17 | 2 | 0 | 0 | 0 | 0 | 0 | 0 |
| JSG000000 | Pnldc1 | 6.3E-129 | 7.1E-127 | 8.917477315 | 0.981585527 | 474.6312223 | 22 | 3 | 3 | 4 | 21 | 1 | 0 | 0 | 0 | 0 | 0 | 0 |
| JSG000000 | Gsdmd | 1.1E-127 | 1.2E-125 | -5.563435749 | 6139.936112 | 129.8382977 | 7 | 3 | 3 | 3 | 4 | 1 | 0 | 0 | 0 | 0 | 0 | 0 |
| JSG000000 | Padi3 | 4.9E-106 | 4.1E-104 | 8.461628708 | 0 | 351.5364713 | 30 | 1 | 3 | 6 | 17 | 2 | 0 | 0 | 0 | 0 | 0 | 0 |
| JSG000000 | Ube2t | 3.3E-104 | 2.6E-102 | 3.054274343 | 292.1020737 | 2426.402431 | 25 | 3 | 3 | 8 | 26 | 8 | 4 | 0 | 0 | 2 | 2 | 2 |
| SG0000002 | Wnt7b | 7.6E-101 | 5.5E-99 | 6.07862398 | 6.836132784 | 462.0177279 | 74 | 3 | 3 | 9 | 78 | 9 | 6 | 0 | 0 | 2 | 4 | 0 |
| JSG000000 | E130309D14Rik | 2.93E-97 | 1.97E-95 | 6.491577531 | 4.282034965 | 385.309543 | 13 | 2 | 3 | 4 | 7 | 0 | 0 | 0 | 0 | 0 | 0 | 0 |
| JSG000000 | Utp14b | 3.66E-97 | 2.45E-95 | 3.213554285 | 146.734245 | 1361.155412 | 33 | 0 | 3 | 2 | 9 | 10 | 13 | 0 | 1 | 0 | 1 | 0 |
| JSG000000 | Six4 | 2.23E-88 | 1.21E-86 | 2.953292369 | 197.6130777 | 1530.542069 | 32 | 0 | 3 | 7 | 9 | 4 | 22 | 0 | 2 | 5 | 5 | 3 |
| SG0000002 | Slc5a9 | 1.2E-86 | 6.26E-85 | -10.60195406 | 490.612481 | 0.315669345 | 57 | 0 | 3 | 6 | 17 | 7 | 2 | 0 | 0 | 0 | 0 | 0 |
| SG0000001 | Apol7a | 2.34E-83 | 1.14E-81 | -7.489470501 | 549.2872148 | 3.056639479 | 19 | 3 | 3 | 8 | 17 | 3 | 0 | 0 | 0 | 0 | 0 | 0 |
| JSG000000 | Igfbpl1 | 1.15E-82 | 5.54E-81 | 8.132904576 | 1.005677317 | 282.2974037 | 17 | 3 | 3 | 4 | 10 | 2 | 0 | 0 | 0 | 0 | 0 | 0 |
| SG0000005 | Sh3bp2 | 1.47E-82 | 7.03E-81 | -4.173510549 | 2367.367828 | 131.1939714 | 40 | 3 | 3 | 3 | 26 | 9 | 0 | 0 | 0 | 0 | 4 | 0 |
| SG0000002 | Myc | 1.33E-81 | 6.2E-80 | -5.486339892 | 11858.8757 | 264.5395698 | 20 | 0 | 3 | 0 | 4 | 8 | 7 | 0 | 3 | 0 | 0 | 8 |
| JSG000000 | Pnp2 | 3.14E-76 | 1.32E-74 | 6.73978365 | 2.648121413 | 283.0185725 | 14 | 3 | 3 | 4 | 12 | 1 | 0 | 0 | 0 | 0 | 0 | 0 |
| JSG000000 | Ehd2 | 2.8E-75 | 1.15E-73 | -3.685592279 | 3251.529104 | 252.70499 | 21 | 3 | 3 | 6 | 11 | 4 | 2 | 0 | 0 | 1 | 1 | 0 |
| SG0000003 | Necab2 | 1.81E-73 | 7.23E-72 | 7.959970625 | 0.981585527 | 244.4094923 | 21 | 2 | 3 | 3 | 14 | 0 | 0 | 0 | 0 | 0 | 0 | 0 |
| JSG000000 | Hmgb2 | 3.58E-70 | 1.27E-68 | 2.232055034 | 1901.829481 | 8934.846432 | 11 | 2 | 3 | 4 | 6 | 0 | 11 | 2 | 3 | 4 | 6 | 0 |
| SG0000005 | Cntn2 | 1.42E-67 | 4.82E-66 | 7.754857215 | 0 | 214.9954654 | 22 | 2 | 3 | 3 | 5 | 5 | 1 | 0 | 0 | 0 | 0 | 1 |
| SG0000003 | Zfyve28 | 2.53E-67 | 8.54E-66 | 7.150558458 | 1.620695873 | 230.2682752 | 93 | 0 | 3 | 13 | 29 | 7 | 1 | 0 | 0 | 0 | 1 | 0 |
| JSG000000 | Endod1 | 1.13E-66 | 3.73E-65 | -3.620244166 | 2532.70028 | 205.9594451 | 38 | 1 | 3 | 5 | 13 | 7 | 3 | 1 | 1 | 1 | 3 | 1 |
| JSG000000 | Entpd3 | 2.7E-66 | 8.79E-65 | -5.651092395 | 2305.14094 | 45.87215729 | 31 | 2 | 3 | 4 | 13 | 4 | 4 | 2 | 2 | 2 | 3 | 0 |
| JSG000000 | Soga3 | 6.03E-66 | 1.93E-64 | -4.911186322 | 646.2952423 | 21.47912382 | 108 | 0 | 3 | 9 | 68 | 7 | 1 | 0 | 0 | 0 | 1 | 0 |
| JSG000000 | Map3k9 | 1.22E-65 | 3.9E-64 | 8.119898863 | 2.340612135 | 651.1223036 | 43 | 1 | 3 | 2 | 22 | 9 | 0 | 0 | 0 | 0 | 0 | 0 |
| SG0000007 | Vma21 | 5.31E-64 | 1.63E-62 | -3.941389461 | 9664.987566 | 629.1074775 | 61 | 3 | 3 | 9 | 39 | 3 | 45 | 3 | 3 | 6 | 21 | 0 |
| SG0000001 | Aoc3 | 2.6E-61 | 7.52E-60 | 5.723960033 | 4.519612439 | 238.882788 | 19 | 3 | 3 | 3 | 8 | 2 | 1 | 0 | 0 | 0 | 1 | 0 |
| JSG000000 | Nqo1 | 1E-59 | 2.74E-58 | -2.870019799 | 6844.517199 | 936.2261588 | 29 | 3 | 3 | 7 | 11 | 0 | 1 | 0 | 0 | 0 | 0 | 0 |
| JSG000000 | Haus4 | 7.37E-56 | 1.83E-54 | 2.10015818 | 555.3221316 | 2380.979137 | 14 | 3 | 3 | 4 | 6 | 0 | 1 | 0 | 0 | 0 | 0 | 0 |
| SG0000002 | Stmn1 | 3.87E-55 | 9.37E-54 | 2.042325585 | 745.4380212 | 3070.526163 | 27 | 0 | 3 | 0 | 3 | 0 | 0 | 0 | 0 | 0 | 0 | 0 |
| SG0000004 | Xaf1 | 5.78E-55 | 1.39E-53 | -4.412588421 | 4064.370479 | 190.8410577 | 20 | 3 | 3 | 3 | 17 | 0 | 1 | 0 | 0 | 0 | 0 | 0 |
| SG0000004 | Pm20d1 | 8.84E-55 | 2.11E-53 | 5.024845782 | 9.719488104 | 316.4263901 | 53 | 3 | 3 | 8 | 23 | 6 | 3 | 0 | 0 | 0 | 2 | 0 |
| SG0000002 | Urah | 7.37E-54 | 1.7E-52 | -3.386682602 | 3419.797199 | 326.9695453 | 6 | 3 | 3 | 3 | 3 | 0 | 0 | 0 | 0 | 0 | 0 | 0 |
| JSG000000 | Rfwd3 | 5.84E-53 | 1.32E-51 | 1.878247559 | 1401.495652 | 5152.293674 | 50 | 2 | 3 | 3 | 22 | 2 | 4 | 0 | 0 | 0 | 0 | 0 |
| SG0000001 | Tbx3 | 8.6E-52 | 1.87E-50 | 3.489529437 | 106.3780005 | 1194.826523 | 51 | 3 | 3 | 9 | 27 | 18 | 6 | 0 | 0 | 3 | 3 | 9 |
| JSG000000 | Tmprss11b | 3.64E-50 | 7.6E-49 | -8.127992885 | 278.7497216 | 0 | 40 | 0 | 3 | 3 | 31 | 4 | 11 | 0 | 1 | 1 | 1 | 2 |
| SG0000003 | Mks1 | 1.77E-49 | 3.62E-48 | 3.253636735 | 51.23568469 | 488.6690081 | 13 | 2 | 3 | 3 | 4 | 0 | 1 | 0 | 0 | 1 | 1 | 0 |
| SG0000002 | Extl1 | 1.85E-49 | 3.79E-48 | 7.88483155 | 0.674076249 | 159.3234977 | 9 | 1 | 3 | 3 | 9 | 3 | 0 | 0 | 0 | 0 | 0 | 1 |
| JSG000000 | Zfp518a | 2.64E-49 | 5.36E-48 | 2.011043237 | 475.691233 | 1917.385702 | 58 | 2 | 3 | 8 | 29 | 5 | 5 | 0 | 0 | 0 | 2 | 2 |
| SG0000003 | Arhgef5 | 3.68E-49 | 7.42E-48 | -3.775881707 | 6429.558839 | 469.3831779 | 45 | 1 | 3 | 3 | 9 | 2 | 0 | 0 | 0 | 0 | 0 | 0 |
| JSG000000 | Gcnt4 | 1.67E-47 | 3.22E-46 | -3.03259198 | 1209.302113 | 147.7861248 | 31 | 2 | 3 | 5 | 13 | 3 | 5 | 0 | 1 | 1 | 2 | 1 |
| SG0000001 | Cers2 | 2.23E-46 | 4.12E-45 | -2.417909718 | 6198.890537 | 1159.980305 | 5 | 3 | 3 | 3 | 6 | 2 | 1 | 1 | 1 | 1 | 1 | 0 |
| SG0000000 | Gmpr | 3.39E-45 | 5.98E-44 | 4.122224548 | 14.3005018 | 249.0373663 | 39 | 2 | 3 | 8 | 18 | 4 | 0 | 0 | 0 | 0 | 0 | 0 |
| SG0000002 | Ptma | 1.76E-44 | 3.02E-43 | 1.614387669 | 14440.23432 | 44213.34303 | 11 | 3 | 3 | 3 | 3 | 2 | 5 | 0 | 0 | 0 | 0 | 2 |
| SG0000004 | Lats1 | 2.77E-43 | 4.56E-42 | -2.437679433 | 2897.748531 | 534.8673218 | 45 | 3 | 3 | 9 | 19 | 5 | 2 | 0 | 0 | 0 | 2 | 1 |
| JSG000000 | Rimkla | 1.44E-42 | 2.33E-41 | 3.491819519 | 25.0388849 | 281.6808993 | 26 | 1 | 3 | 3 | 15 | 7 | 4 | 0 | 0 | 0 | 1 | 1 |
| JSG000000 | Nppb | 1.29E-41 | 2.02E-40 | -5.758392326 | 1269.827027 | 23.45825751 | 6 | 2 | 3 | 5 | 7 | 0 | 6 | 2 | 3 | 5 | 7 | 0 |

| | | | | | | | | | | | | | | | | | | |
|---|---|---|---|---|---|---|---|---|---|---|---|---|---|---|---|---|---|---|
| JSG000000 | Adh5 | 6.41E-41 | 9.82E-40 | 1.592667771 | 3185.222053 | 9606.838421 | 52 | 3 | 3 | 11 | 16 | 10 | 2 | 0 | 0 | 0 | 0 | 0 |
| JSG000000 | Cyp51 | 1.73E-40 | 2.6E-39 | 1.74969864 | 719.5169521 | 2419.651417 | 40 | 1 | 3 | 7 | 26 | 2 | 3 | 0 | 0 | 0 | 2 | 0 |
| JSG000000 | Gal3st3 | 5.88E-40 | 8.7E-39 | 6.836794115 | 0 | 113.3089151 | 13 | 2 | 3 | 4 | 8 | 1 | 0 | 0 | 0 | 0 | 0 | 0 |
| JSG000000 | Acrbp | 1.77E-39 | 2.57E-38 | 3.148797843 | 34.37724165 | 304.8973348 | 15 | 3 | 3 | 3 | 5 | 1 | 0 | 0 | 0 | 0 | 0 | 0 |
| SG0000001 | Bnip2 | 6.2E-39 | 8.79E-38 | -2.19324468 | 3849.334214 | 841.6919687 | 127 | 0 | 3 | 9 | 21 | 22 | 4 | 0 | 0 | 2 | 4 | 3 |
| JSG000000 | Cenpn | 2.22E-38 | 3.11E-37 | 1.88389983 | 347.0478118 | 1280.854116 | 19 | 2 | 3 | 3 | 7 | 6 | 0 | 0 | 0 | 0 | 1 | 0 |
| JSG000000 | D430019H16Rik | 1.06E-36 | 1.4E-35 | 6.239385381 | 1.620695873 | 122.44574 | 31 | 0 | 3 | 4 | 13 | 3 | 3 | 0 | 0 | 0 | 0 | 0 |
| SG0000002 | Pycr1 | 2.91E-36 | 3.77E-35 | 3.029067218 | 103.5228867 | 845.0384404 | 7 | 2 | 3 | 4 | 5 | 0 | 1 | 0 | 0 | 0 | 0 | 0 |
| SG0000003 | Tet3 | 3.44E-36 | 4.45E-35 | -3.488778297 | 1543.818095 | 137.52106 | 101 | 1 | 3 | 14 | 38 | 11 | 17 | 0 | 0 | 4 | 8 | 3 |
| JSG000000 | Ezr | 4.88E-36 | 6.28E-35 | 1.510615951 | 2424.309293 | 6907.624789 | 56 | 1 | 3 | 8 | 26 | 4 | 3 | 0 | 0 | 1 | 1 | 0 |
| SG0000001 | Amotl1 | 5.98E-36 | 7.68E-35 | -2.554475124 | 5388.950006 | 917.3403975 | 87 | 0 | 3 | 9 | 31 | 9 | 5 | 0 | 0 | 1 | 2 | 1 |
| JSG000000 | Mcoln3 | 2.92E-35 | 3.67E-34 | 6.638773547 | 0 | 98.64831792 | 41 | 3 | 3 | 8 | 15 | 1 | 5 | 0 | 0 | 1 | 1 | 0 |
| SG0000004 | Ikbke | 5.56E-35 | 6.91E-34 | -3.653811877 | 1119.898652 | 88.97576862 | 35 | 3 | 3 | 5 | 27 | 4 | 0 | 0 | 0 | 0 | 0 | 0 |
| SG0000003 | Cdc25a | 1.88E-34 | 2.3E-33 | 1.542215732 | 1251.544925 | 3645.017676 | 73 | 3 | 3 | 6 | 22 | 0 | 3 | 0 | 0 | 0 | 2 | 0 |
| SG0000002 | Wdr12 | 3.87E-34 | 4.64E-33 | 1.70635628 | 1221.669613 | 3986.742586 | 132 | 0 | 3 | 11 | 68 | 29 | 30 | 0 | 0 | 2 | 8 | 5 |
| SG0000003 | Il17rc | 8.08E-34 | 9.54E-33 | -3.026071608 | 980.0870795 | 120.3168189 | 14 | 3 | 3 | 4 | 4 | 2 | 0 | 0 | 0 | 0 | 0 | 0 |
| SG000000C | Bcas2 | 1.78E-32 | 2E-31 | 1.468096855 | 1660.47474 | 4593.814594 | 33 | 3 | 3 | 6 | 6 | 3 | 0 | 0 | 0 | 0 | 0 | 0 |
| JSG000000 | Rex2 | 4.71E-32 | 5.18E-31 | 5.668254789 | 2.305646233 | 117.2485246 | 72 | 3 | 3 | 6 | 43 | 12 | 5 | 0 | 0 | 0 | 3 | 1 |
| SG0000006 | H3f3a | 2.23E-31 | 2.39E-30 | 1.468307083 | 1495.205417 | 4137.188764 | 50 | 0 | 3 | 0 | 7 | 1 | 2 | 0 | 0 | 0 | 1 | 1 |
| JSG000000 | Wnt3 | 2.42E-31 | 2.59E-30 | 6.458673929 | 0 | 86.95379536 | 27 | 1 | 3 | 1 | 11 | 4 | 4 | 1 | 2 | 1 | 4 | 2 |
| SG000000C | Sgsh | 9.36E-31 | 9.74E-30 | -2.475796264 | 725.6229161 | 130.4433823 | 25 | 3 | 3 | 6 | 27 | 0 | 4 | 0 | 0 | 0 | 0 | 0 |
| JSG000000 | Epas1 | 1.78E-30 | 1.83E-29 | -3.890639825 | 23991.80414 | 1617.571989 | 67 | 1 | 3 | 7 | 30 | 8 | 2 | 0 | 0 | 0 | 2 | 0 |
| JSG000000 | Gm21411 | 3.02E-30 | 3.08E-29 | 6.397742749 | 0 | 83.31648087 | 48 | 3 | 3 | 11 | 29 | 6 | 0 | 0 | 0 | 0 | 0 | 0 |
| SG0000002 | Ak1 | 4.72E-30 | 4.78E-29 | -1.913151353 | 3625.508235 | 962.6157262 | 19 | 3 | 3 | 5 | 6 | 1 | 3 | 0 | 0 | 0 | 0 | 0 |
| SG000000C | Pax9 | 5.1E-30 | 5.16E-29 | -6.572648317 | 479.7464866 | 5.040196967 | 46 | 1 | 3 | 7 | 26 | 6 | 6 | 0 | 1 | 2 | 4 | 2 |
| SG0000003 | Creld1 | 1.78E-29 | 1.76E-28 | -2.025491885 | 3011.446268 | 739.6756212 | 17 | 3 | 3 | 3 | 4 | 1 | 1 | 1 | 1 | 1 | 1 | 0 |
| SG0000002 | Slc13a5 | 3.14E-29 | 3.07E-28 | -2.93796529 | 392.4567404 | 51.21251616 | 28 | 3 | 3 | 5 | 15 | 1 | 0 | 0 | 0 | 0 | 0 | 0 |
| SG0000003 | Bicd2 | 3.36E-29 | 3.27E-28 | -1.86765371 | 2542.067297 | 696.5739926 | 132 | 0 | 3 | 12 | 78 | 12 | 7 | 0 | 0 | 3 | 19 | 0 |
| JSG000000 | Bag4 | 4.8E-29 | 4.65E-28 | 1.436948566 | 1059.155575 | 2867.638331 | 46 | 0 | 3 | 4 | 13 | 5 | 13 | 0 | 1 | 1 | 2 | 1 |
| SG0000002 | Notch1 | 5.05E-29 | 4.88E-28 | 1.805308572 | 254.1757869 | 888.3542993 | 21 | 1 | 3 | 5 | 11 | 5 | 4 | 1 | 2 | 4 | 4 | 1 |
| SG0000003 | Scube3 | 5.89E-29 | 5.68E-28 | -2.485416132 | 2858.818316 | 510.5070553 | 19 | 3 | 3 | 7 | 10 | 0 | 3 | 0 | 0 | 2 | 4 | 0 |
| SG0000003 | Srcin1 | 6.18E-29 | 5.95E-28 | 3.344232862 | 20.00062107 | 203.1224311 | 88 | 0 | 3 | 12 | 38 | 6 | 4 | 0 | 0 | 0 | 3 | 0 |
| JSG000000 | Cdipt | 1.04E-28 | 9.92E-28 | -1.885378361 | 3495.33733 | 946.0926215 | 7 | 1 | 3 | 2 | 2 | 1 | 0 | 0 | 0 | 0 | 0 | 0 |
| JSG000000 | Gchfr | 3.3E-28 | 3.08E-27 | -3.236730565 | 537.3506392 | 57.00392788 | 4 | 3 | 3 | 3 | 6 | 0 | 0 | 0 | 0 | 0 | 0 | 0 |
| JSG000000 | Gdf11 | 2.27E-27 | 2.04E-26 | -2.408054733 | 600.9807967 | 113.230621 | 9 | 2 | 3 | 2 | 5 | 0 | 4 | 2 | 2 | 2 | 3 | 0 |
| SG0000002 | Fam129b | 2.01E-26 | 1.73E-25 | -2.228987392 | 10091.45497 | 2152.591999 | 36 | 2 | 3 | 6 | 17 | 4 | 1 | 0 | 1 | 0 | 0 | 0 |
| SG0000001 | Rab3d | 2.03E-26 | 1.75E-25 | -1.946914742 | 1275.836109 | 330.9140089 | 11 | 3 | 3 | 5 | 5 | 0 | 2 | 0 | 0 | 2 | 2 | 0 |
| JSG000000 | Dynlt3 | 2.26E-26 | 1.94E-25 | -2.063501766 | 2365.468969 | 565.9021285 | 22 | 0 | 3 | 0 | 3 | 1 | 8 | 0 | 1 | 0 | 0 | 0 |
| SG0000005 | Gnpda1 | 4.29E-26 | 3.64E-25 | 1.719607381 | 1019.475279 | 3357.608855 | 11 | 1 | 3 | 4 | 9 | 2 | 1 | 0 | 0 | 0 | 0 | 0 |
| SG000000C | Upk3bl | 5.18E-26 | 4.38E-25 | -3.854096259 | 299.3670366 | 20.7016708 | 19 | 0 | 3 | 3 | 10 | 7 | 0 | 0 | 0 | 0 | 0 | 0 |
| JSG000000 | Psmc6 | 3.95E-25 | 3.21E-24 | 1.252410277 | 2913.546932 | 6941.208316 | 42 | 2 | 3 | 4 | 23 | 5 | 2 | 0 | 0 | 0 | 1 | 0 |
| JSG000000 | Slc16a9 | 4.3E-25 | 3.49E-24 | 5.333896397 | 2.305646233 | 92.99411838 | 56 | 2 | 3 | 11 | 35 | 7 | 3 | 0 | 0 | 0 | 1 | 0 |
| SG0000005 | Lgals3 | 1.47E-24 | 1.16E-23 | -4.379656686 | 14911.70082 | 716.3399107 | 66 | 3 | 3 | 12 | 27 | 3 | 3 | 0 | 0 | 0 | 0 | 0 |
| SG0000002 | Ltbp3 | 1.53E-24 | 1.21E-23 | -3.961370882 | 5900.033826 | 378.7590704 | 9 | 2 | 3 | 2 | 3 | 0 | 2 | 0 | 0 | 0 | 0 | 0 |
| SG0000003 | AF529169 | 1.54E-23 | 1.16E-22 | 6.008735596 | 0 | 63.38869906 | 96 | 1 | 3 | 10 | 29 | 7 | 12 | 0 | 0 | 3 | 5 | 0 |
| SG0000002 | Nek6 | 1.74E-23 | 1.3E-22 | -1.61406502 | 2570.570297 | 839.7451399 | 196 | 0 | 3 | 14 | 58 | 15 | 10 | 0 | 0 | 2 | 4 | 0 |
| JSG000000 | Fbp2 | 2.32E-23 | 1.73E-22 | -3.966418097 | 398.0711699 | 25.46536483 | 27 | 3 | 3 | 4 | 17 | 1 | 0 | 0 | 0 | 0 | 0 | 0 |
| JSG000000 | P2ry2 | 3.32E-23 | 2.45E-22 | -2.302127345 | 466.6223946 | 94.61417206 | 18 | 2 | 3 | 5 | 11 | 4 | 2 | 0 | 1 | 1 | 1 | 0 |
| SG0000001 | Gdap1l1 | 7.29E-23 | 5.32E-22 | 7.634074509 | 0.331601068 | 65.87207401 | 36 | 0 | 3 | 3 | 9 | 0 | 0 | 0 | 0 | 0 | 0 | 0 |
| JSG000000 | Cacng4 | 8.11E-23 | 5.91E-22 | 5.964053694 | 0 | 61.4250728 | 43 | 1 | 3 | 4 | 15 | 5 | 1 | 0 | 0 | 0 | 0 | 1 |
| JSG000000 | Samd5 | 9.54E-23 | 6.94E-22 | -4.068461476 | 182.6645852 | 10.88743097 | 116 | 1 | 3 | 12 | 32 | 8 | 5 | 0 | 0 | 0 | 1 | 0 |
| JSG000000 | Aard | 1.73E-22 | 1.24E-21 | 5.938079066 | 0 | 60.31121401 | 9 | 3 | 3 | 4 | 12 | 0 | 2 | 1 | 1 | 1 | 1 | 0 |
| JSG000000 | Bnc1 | 3.38E-22 | 2.39E-21 | -5.142675374 | 250.120643 | 7.080272794 | 51 | 1 | 3 | 6 | 22 | 5 | 10 | 0 | 0 | 1 | 4 | 1 |
| JSG000000 | Fgfr4 | 3.46E-22 | 2.45E-21 | 6.714887849 | 0.639110347 | 67.13647093 | 15 | 3 | 3 | 4 | 12 | 3 | 1 | 0 | 0 | 0 | 0 | 0 |
| SG0000002 | Aldh9a1 | 1.27E-21 | 8.75E-21 | 1.191328199 | 1718.602279 | 3924.649762 | 19 | 2 | 3 | 7 | 18 | 2 | 1 | 0 | 0 | 0 | 0 | 0 |
| SG0000002 | Pmm2 | 2.99E-21 | 2.01E-20 | -1.892159292 | 1667.629312 | 449.2652383 | 25 | 3 | 3 | 5 | 22 | 0 | 0 | 0 | 0 | 0 | 0 | 0 |
| JSG000000 | Rassf4 | 5.37E-21 | 3.56E-20 | -2.651321148 | 2199.269623 | 350.0670879 | 37 | 3 | 3 | 7 | 14 | 3 | 5 | 0 | 0 | 0 | 1 | 1 |
| SG0000003 | Nfkbiz | 6.92E-21 | 4.57E-20 | -1.822987396 | 1554.963057 | 439.4871562 | 167 | 3 | 3 | 7 | 76 | 18 | 16 | 0 | 0 | 0 | 8 | 12 |
| JSG000000 | Tgfb3 | 1.15E-20 | 7.53E-20 | -2.267193916 | 544.2478881 | 113.0585413 | 15 | 3 | 3 | 5 | 7 | 3 | 0 | 0 | 0 | 0 | 0 | 0 |
| SG0000007 | Cpa2 | 1.27E-20 | 8.27E-20 | 5.80947173 | 0 | 55.08222686 | 51 | 2 | 3 | 8 | 39 | 2 | 0 | 0 | 0 | 0 | 0 | 0 |
| JSG000000 | Lrp10 | 1.38E-20 | 8.97E-20 | -1.760325962 | 4717.472503 | 1392.510442 | 8 | 2 | 3 | 2 | 2 | 0 | 0 | 0 | 0 | 0 | 0 | 0 |
| JSG000000 | Tnfrsf10b | 1.64E-20 | 1.06E-19 | -1.406866832 | 4347.783251 | 1639.678752 | 23 | 2 | 3 | 7 | 17 | 3 | 4 | 1 | 2 | 3 | 8 | 0 |
| SG0000001 | Smpdl3a | 5.47E-20 | 3.42E-19 | -2.908525643 | 482.5159452 | 64.26258745 | 38 | 1 | 3 | 5 | 21 | 7 | 1 | 0 | 0 | 0 | 0 | 0 |
| SG0000001 | Ctsz | 8.65E-20 | 5.35E-19 | -1.358964534 | 7239.606816 | 2822.44763 | 10 | 3 | 3 | 3 | 12 | 1 | 0 | 0 | 0 | 0 | 0 | 0 |
| JSG000000 | Zfp459 | 1.26E-19 | 7.74E-19 | 8.175621023 | 2.532349596 | 732.2018498 | 35 | 2 | 3 | 6 | 10 | 4 | 5 | 0 | 0 | 0 | 0 | 0 |
| SG0000003 | Tmc3 | 1.81E-19 | 1.1E-18 | -3.108385651 | 238.7268541 | 27.681139 | 61 | 1 | 3 | 6 | 16 | 8 | 2 | 0 | 0 | 0 | 0 | 0 |
| SG0000005 | Zfp811 | 1.86E-19 | 1.13E-18 | 3.275913895 | 24.69740659 | 239.2212178 | 24 | 2 | 3 | 7 | 16 | 3 | 2 | 0 | 0 | 1 | 1 | 0 |
| JSG000000 | Tgoln1 | 2.12E-19 | 1.29E-18 | -1.493798659 | 5477.440125 | 1944.909668 | 15 | 3 | 3 | 3 | 5 | 0 | 5 | 0 | 0 | 0 | 0 | 0 |
| SG000000C | Stat6 | 3.12E-19 | 1.87E-18 | -1.389868412 | 3127.973825 | 1193.63358 | 17 | 3 | 3 | 3 | 8 | 0 | 0 | 0 | 0 | 0 | 0 | 0 |
| JSG000000 | Dgat2 | 3.19E-19 | 1.92E-18 | 2.318243191 | 40.65975714 | 202.7801865 | 16 | 0 | 3 | 2 | 3 | 2 | 2 | 0 | 1 | 1 | 1 | 0 |
| SG0000007 | Il4i1 | 3.83E-19 | 2.29E-18 | 1.221054713 | 2017.972942 | 4704.239601 | 21 | 3 | 3 | 6 | 9 | 2 | 0 | 0 | 0 | 0 | 0 | 0 |
| JSG000000 | Ehd1 | 5.54E-19 | 3.28E-18 | -1.643601183 | 7180.365946 | 2298.123276 | 13 | 3 | 3 | 3 | 12 | 1 | 0 | 0 | 0 | 0 | 0 | 0 |
| JSG000000 | Gm3739 | 9.41E-19 | 5.5E-18 | 5.664563935 | 0 | 49.72285109 | 35 | 2 | 3 | 3 | 12 | 4 | 6 | 2 | 2 | 2 | 2 | 0 |
| JSG000000 | Dppa2 | 1.02E-18 | 5.93E-18 | 5.599800027 | 17.15457522 | 831.9318403 | 15 | 3 | 3 | 3 | 5 | 2 | 1 | 0 | 0 | 0 | 0 | 0 |
| SG0000003 | L1cam | 3.28E-18 | 1.87E-17 | 3.38320594 | 9.580621359 | 99.96319899 | 34 | 1 | 3 | 7 | 10 | 7 | 3 | 0 | 0 | 0 | 0 | 3 |
| JSG000000 | Mmp13 | 3.55E-18 | 2.01E-17 | -2.632346604 | 1528.28493 | 246.4841762 | 49 | 2 | 3 | 12 | 17 | 0 | 6 | 0 | 0 | 4 | 5 | 0 |
| SG0000001 | Lonrf3 | 3.76E-18 | 2.13E-17 | 5.253692696 | 1.620695873 | 61.8330424 | 49 | 2 | 3 | 5 | 18 | 8 | 0 | 0 | 0 | 0 | 0 | 0 |
| JSG000000 | Nelfa | 2.46E-17 | 1.34E-16 | 1.120575182 | 1005.492744 | 2186.279503 | 67 | 2 | 3 | 9 | 36 | 3 | 1 | 0 | 0 | 0 | 1 | 0 |
| SG0000001 | Dhx58 | 2.57E-17 | 1.4E-16 | -2.584829583 | 521.2271181 | 86.87919028 | 10 | 3 | 3 | 4 | 9 | 0 | 0 | 0 | 0 | 0 | 1 | 0 |
| SG0000002 | Pcyox1 | 6.04E-17 | 3.23E-16 | -1.268952448 | 3643.545395 | 1511.929131 | 19 | 2 | 3 | 3 | 5 | 0 | 6 | 0 | 0 | 0 | 0 | 0 |
| JSG000000 | Pdgfb | 6.96E-17 | 3.7E-16 | 1.829273096 | 91.22739942 | 324.1838436 | 15 | 1 | 3 | 5 | 8 | 0 | 3 | 0 | 2 | 1 | 2 | 0 |
| JSG000000 | Slc17a7 | 1.06E-16 | 5.57E-16 | 3.633434043 | 6.609429422 | 82.02318018 | 10 | 3 | 3 | 3 | 6 | 1 | 0 | 0 | 0 | 0 | 0 | 0 |
| JSG000000 | Cox5a | 1.18E-16 | 6.17E-16 | 1.024390921 | 2327.591579 | 4734.555061 | 20 | 2 | 3 | 2 | 7 | 4 | 0 | 0 | 0 | 0 | 0 | 0 |
| JSG000000 | Trappc10 | 1.23E-16 | 6.41E-16 | -1.303603799 | 2385.9507 | 966.5795887 | 63 | 2 | 3 | 4 | 19 | 6 | 3 | 0 | 0 | 1 | 3 | 2 |
| JSG000000 | Atp6v0e2 | 1.4E-16 | 7.31E-16 | -1.596505453 | 745.1284564 | 246.3968321 | 9 | 3 | 3 | 4 | 5 | 1 | 2 | 0 | 0 | 1 | 1 | 1 |
| SG0000003 | Ilk | 2.08E-16 | 1.07E-15 | -1.258412388 | 2912.087228 | 1217.263231 | 23 | 0 | 3 | 0 | 3 | 0 | 0 | 0 | 0 | 0 | 3 | 0 |
| SG0000002 | Pdrg1 | 2.4E-16 | 1.23E-15 | -1.342197952 | 1594.994249 | 629.0965126 | 3 | 0 | 3 | 3 | 6 | 0 | 0 | 0 | 0 | 0 | 0 | 0 |
| SG0000006 | Gemin2 | 5.2E-16 | 2.61E-15 | 1.232440491 | 396.0387576 | 930.5489963 | 34 | 3 | 3 | 8 | 13 | 1 | 1 | 0 | 0 | 1 | 1 | 0 |
| JSG000000 | Haus1 | 6.54E-16 | 3.27E-15 | 1.153327811 | 570.3566019 | 1268.623462 | 24 | 1 | 3 | 4 | 10 | 1 | 0 | 0 | 0 | 0 | 0 | 0 |
| JSG000000 | Acsf3 | 1.52E-15 | 7.45E-15 | 1.419698151 | 189.8952292 | 508.025816 | 26 | 3 | 3 | 6 | 7 | 2 | 0 | 0 | 0 | 0 | 0 | 0 |
| SG0000003 | Cul7 | 3.34E-15 | 1.6E-14 | -1.750770627 | 6479.943134 | 1925.46984 | 19 | 0 | 3 | 0 | 9 | 0 | 0 | 0 | 0 | 0 | 2 | 0 |
| SG0000001 | Irf1 | 3.99E-15 | 1.9E-14 | -1.284628774 | 3869.93593 | 1588.517417 | 20 | 0 | 3 | 3 | 3 | 7 | 8 | 0 | 2 | 2 | 2 | 2 |
| SG0000003 | Ctdp1 | 4.46E-15 | 2.12E-14 | 1.099543088 | 697.4573393 | 1494.559122 | 102 | 2 | 3 | 9 | 24 | 5 | 6 | 0 | 0 | 0 | 0 | 2 |
| JSG000000 | Rinl | 7.96E-15 | 3.72E-14 | 4.268390142 | 39.83893972 | 767.7524968 | 9 | 3 | 3 | 4 | 11 | 4 | 1 | 0 | 0 | 0 | 3 | 0 |
| JSG000000 | Ddx19a | 9.19E-15 | 4.27E-14 | 1.015832299 | 1142.905182 | 2311.033251 | 31 | 3 | 3 | 4 | 9 | 1 | 2 | 0 | 0 | 0 | 0 | 0 |
| JSG000000 | Srp54b | 1.11E-14 | 5.15E-14 | -1.586067009 | 541.6445227 | 180.4100019 | 60 | 3 | 3 | 6 | 35 | 9 | 5 | 0 | 0 | 0 | 0 | 0 |
| SG0000002 | Rtfdc1 | 1.55E-14 | 7.08E-14 | -1.159431275 | 2913.966612 | 1304.550753 | 31 | 2 | 3 | 7 | 12 | 1 | 1 | 0 | 0 | 0 | 0 | 0 |
| SG0000003 | Slc8b1 | 1.62E-14 | 7.41E-14 | -1.248657817 | 1804.092943 | 759.2336538 | 39 | 3 | 3 | 6 | 30 | 3 | 0 | 0 | 0 | 0 | 0 | 0 |
| JSG000000 | Dact3 | 1.94E-14 | 8.87E-14 | 2.959946502 | 73.49396511 | 571.8529442 | 13 | 3 | 3 | 5 | 10 | 1 | 1 | 0 | 0 | 1 | 2 | 1 |
| SG0000004 | Zfp386 | 2.53E-14 | 1.15E-13 | -1.396639406 | 846.7519787 | 321.6072761 | 52 | 2 | 3 | 9 | 37 | 6 | 6 | 0 | 1 | 1 | 1 | 0 |
| SG000000C | Dvl3 | 3.93E-14 | 1.76E-13 | -1.725829504 | 872.3006092 | 263.7179722 | 34 | 0 | 3 | 0 | 12 | 3 | 0 | 0 | 0 | 0 | 0 | 0 |
| SG0000004 | Zfp322a | 4.8E-14 | 2.13E-13 | 1.04779325 | 724.226649 | 1497.240957 | 45 | 1 | 3 | 4 | 17 | 4 | 12 | 1 | 1 | 2 | 8 | 1 |
| JSG000000 | 2610507B11Rik | 7.27E-14 | 3.19E-13 | -1.049681499 | 16495.63128 | 7968.623785 | 45 | 2 | 3 | 9 | 27 | 1 | 1 | 0 | 0 | 0 | 0 | 0 |
| JSG000000 | AI597479 | 8.07E-14 | 3.53E-13 | 1.002805985 | 977.5821215 | 1958.970661 | 27 | 1 | 3 | 1 | 15 | 4 | 3 | 0 | 1 | 0 | 1 | 0 |
| JSG000000 | Tjp3 | 8.3E-14 | 3.63E-13 | 3.552531481 | 79.26681916 | 930.0578598 | 13 | 3 | 3 | 4 | 12 | 0 | 0 | 0 | 0 | 0 | 0 | 0 |
| SG0000003 | Rtkn | 8.64E-14 | 3.78E-13 | -2.044091332 | 786.3515478 | 190.6706913 | 27 | 0 | 3 | 0 | 6 | 6 | 3 | 0 | 0 | 0 | 0 | 0 |
| SG0000002 | Tekt1 | 9.4E-14 | 4.09E-13 | 5.518673677 | 0.957493736 | 43.89572785 | 60 | 3 | 3 | 6 | 33 | 6 | 0 | 0 | 0 | 0 | 0 | 0 |
| JSG000000 | Ak3 | 1.16E-13 | 5.02E-13 | -1.339361071 | 1235.493133 | 488.2614971 | 41 | 1 | 3 | 2 | 28 | 1 | 3 | 0 | 0 | 0 | 2 | 0 |
| JSG000000 | Lama5 | 1.19E-13 | 5.16E-13 | -1.565206321 | 12455.08022 | 4208.937502 | 29 | 1 | 3 | 7 | 14 | 1 | 1 | 0 | 0 | 1 | 2 | 0 |
| JSG000000 | Ifi202b | 1.27E-13 | 5.49E-13 | -2.746598979 | 991.5868615 | 147.7481614 | 51 | 0 | 3 | 6 | 29 | 3 | 0 | 0 | 0 | 0 | 0 | 0 |
| JSG000000 | Sall1 | 1.49E-13 | 6.39E-13 | 0.925332023 | 2724.109099 | 5173.413496 | 27 | 2 | 3 | 2 | 10 | 2 | 4 | 1 | 1 | 1 | 1 | 2 |
| JSG000000 | Foxr1 | 1.75E-13 | 7.49E-13 | 5.169943751 | 0 | 35.00046786 | 11 | 2 | 3 | 3 | 6 | 0 | 0 | 0 | 0 | 0 | 1 | 0 |
| JSG000000 | Ano8 | 1.97E-13 | 8.4E-13 | -1.471750378 | 555.8109636 | 200.394639 | 9 | 3 | 3 | 3 | 4 | 0 | 0 | 0 | 0 | 0 | 0 | 0 |
| SG0000004 | Wdr81 | 2.17E-13 | 9.24E-13 | -1.333713462 | 1339.573482 | 531.4700357 | 12 | 3 | 3 | 3 | 5 | 2 | 0 | 0 | 0 | 0 | 0 | 1 |
| JSG000000 | Fsd1 | 3.7E-13 | 1.55E-12 | 2.618646642 | 92.49036514 | 568.0514612 | 6 | 2 | 3 | 2 | 2 | 1 | 0 | 0 | 0 | 0 | 0 | 0 |
| SG0000002 | Hmgcl | 4.94E-13 | 2.05E-12 | 1.144509314 | 347.8354785 | 768.9631193 | 17 | 2 | 3 | 4 | 22 | 2 | 0 | 0 | 0 | 0 | 0 | 0 |
| SG0000004 | Aida | 5.07E-13 | 2.1E-12 | -1.112117836 | 2445.835327 | 1131.478709 | 151 | 0 | 3 | 12 | 53 | 12 | 12 | 0 | 0 | 0 | 5 | 0 |
| SG0000002 | Nfyb | 1.04E-12 | 4.23E-12 | 0.916515121 | 1503.755442 | 2838.414096 | 32 | 1 | 3 | 4 | 15 | 0 | 4 | 1 | 2 | 1 | 4 | 0 |
| SG0000003 | Crb2 | 1.31E-12 | 5.27E-12 | 6.748946368 | 0.34247518 | 36.8353159 | 16 | 3 | 3 | 4 | 28 | 1 | 1 | 0 | 0 | 0 | 1 | 0 |
| JSG000000 | Angptl2 | 1.65E-12 | 6.6E-12 | -1.701349193 | 1463.828001 | 450.1247677 | 24 | 1 | 3 | 8 | 15 | 4 | 0 | 0 | 0 | 0 | 0 | 1 |
| SG000000C | Hip1r | 1.67E-12 | 6.71E-12 | -1.090037564 | 2078.866508 | 976.5457613 | 21 | 3 | 3 | 5 | 7 | 1 | 3 | 0 | 0 | 1 | 1 | 0 |
| JSG000000 | Lrrc58 | 2.24E-12 | 8.91E-12 | -1.062440906 | 2909.599717 | 1393.178182 | 50 | 1 | 3 | 5 | 22 | 8 | 22 | 0 | 2 | 1 | 9 | 6 |
| SG0000002 | Nudt12 | 2.93E-12 | 1.15E-11 | -1.59797952 | 543.4079106 | 179.5089838 | 41 | 1 | 3 | 6 | 27 | 8 | 8 | 0 | 0 | 1 | 2 | 1 |

| | | | | | | | | | | | | | | | | | | |
|---|---|---|---|---|---|---|---|---|---|---|---|---|---|---|---|---|---|---|
| ISG000000 | Sdhaf4 | 3.1E-12 | 1.22E-11 | 1.466375067 | 189.8627509 | 524.6415096 | 16 | 1 | 3 | 1 | 5 | 3 | 2 | 0 | 1 | 0 | 0 | 1 |
| ISG000000 | Rps25 | 3.46E-12 | 1.36E-11 | 0.832407894 | 6587.073476 | 11729.30428 | 4 | 3 | 3 | 3 | 3 | 0 | 0 | 0 | 0 | 0 | 0 | 0 |
| SG0000001 | Plcg1 | 4.59E-12 | 1.79E-11 | 0.888865275 | 1563.284835 | 2894.764412 | 97 | 0 | 3 | 1 | 9 | 8 | 1 | 0 | 0 | 0 | 0 | 0 |
| ISG000000 | Cyp4v3 | 5.06E-12 | 1.97E-11 | -2.549271152 | 226.1688538 | 38.63898535 | 62 | 1 | 3 | 8 | 30 | 13 | 8 | 0 | 0 | 0 | 0 | 0 |
| ISG000000 | Ccdc36 | 9.17E-12 | 3.51E-11 | 6.797384324 | 1.254128903 | 139.4950351 | 48 | 3 | 3 | 7 | 28 | 2 | 4 | 0 | 0 | 0 | 2 | 0 |
| SG0000001 | Ttpal | 9.83E-12 | 3.76E-11 | -1.201162564 | 905.0032721 | 393.6082457 | 74 | 0 | 3 | 6 | 34 | 12 | 11 | 0 | 0 | 0 | 4 | 0 |
| ISG000000 | Tab1 | 1.67E-11 | 6.3E-11 | -1.253383723 | 1169.765967 | 490.6738185 | 31 | 3 | 3 | 5 | 10 | 2 | 0 | 0 | 0 | 0 | 0 | 0 |
| ISG000000 | Mapk1ip1 | 1.84E-11 | 6.91E-11 | 1.229423334 | 185.6461573 | 435.2905762 | 41 | 3 | 3 | 9 | 27 | 15 | 2 | 0 | 0 | 0 | 0 | 0 |
| ISG000000 | Wnt2 | 2.13E-11 | 7.96E-11 | -3.242101758 | 75.88610126 | 8.020332305 | 76 | 3 | 3 | 8 | 33 | 6 | 3 | 0 | 0 | 1 | 2 | 0 |
| SG0000003 | Ttf2 | 2.55E-11 | 9.48E-11 | 0.874818358 | 1356.570733 | 2487.648469 | 39 | 0 | 3 | 4 | 17 | 3 | 4 | 0 | 1 | 0 | 0 | 0 |
| ISG000000 | Zfp119a | 2.74E-11 | 1.01E-10 | -1.915857312 | 192.614476 | 51.04561083 | 35 | 3 | 3 | 6 | 19 | 2 | 2 | 0 | 0 | 0 | 2 | 0 |
| ISG000000 | Slc39a1 | 3.04E-11 | 1.12E-10 | -0.988084318 | 3116.743673 | 1571.29623 | 6 | 3 | 3 | 3 | 7 | 0 | 2 | 0 | 0 | 0 | 0 | 0 |
| SG0000003 | Fhl3 | 3.75E-11 | 1.38E-10 | -1.761633784 | 394.2352707 | 116.2655006 | 18 | 3 | 3 | 6 | 9 | 0 | 0 | 0 | 0 | 0 | 0 | 0 |
| SG0000002 | Slc40a1 | 3.99E-11 | 1.46E-10 | 1.311942084 | 135.9060224 | 337.4213992 | 50 | 0 | 3 | 7 | 20 | 5 | 2 | 0 | 0 | 1 | 2 | 1 |
| ISG000000 | Pdhb | 4.19E-11 | 1.53E-10 | 0.8179819 | 2802.634874 | 4940.872075 | 15 | 1 | 3 | 3 | 6 | 0 | 1 | 0 | 0 | 0 | 1 | 0 |
| SG0000002 | Prdx4 | 4.24E-11 | 1.55E-10 | 0.923484264 | 720.2803121 | 1366.14881 | 35 | 0 | 3 | 4 | 31 | 7 | 0 | 0 | 0 | 0 | 0 | 0 |
| ISG000000 | Fut2 | 5.7E-11 | 2.06E-10 | 3.282789413 | 5.830455467 | 56.74408081 | 29 | 3 | 3 | 9 | 13 | 5 | 0 | 0 | 0 | 0 | 0 | 0 |
| ISG000000 | C1s2 | 5.82E-11 | 2.1E-10 | -3.128211561 | 81.00632101 | 9.264743416 | 16 | 3 | 3 | 5 | 8 | 3 | 2 | 0 | 0 | 0 | 0 | 0 |
| SG0000002 | Rab29 | 5.9E-11 | 2.13E-10 | -1.352869957 | 470.661806 | 184.2699507 | 36 | 3 | 3 | 6 | 24 | 0 | 0 | 0 | 0 | 0 | 1 | 0 |
| SG0000002 | Stpg1 | 7.23E-11 | 2.6E-10 | 5.165512907 | 0.981585527 | 35.22917508 | 42 | 3 | 3 | 4 | 9 | 7 | 0 | 0 | 0 | 0 | 0 | 0 |
| ISG000000 | Rnf125 | 7.84E-11 | 2.81E-10 | -1.458249264 | 409.4292932 | 149.0054461 | 70 | 3 | 3 | 9 | 37 | 3 | 3 | 0 | 0 | 0 | 0 | 0 |
| ISG000000 | Eif5a2 | 8.12E-11 | 2.9E-10 | 0.998949837 | 430.1532805 | 859.6805567 | 39 | 3 | 3 | 6 | 17 | 8 | 10 | 0 | 0 | 1 | 6 | 4 |
| ISG000000 | Tmx3 | 9.93E-11 | 3.53E-10 | -0.955767097 | 3616.196478 | 1864.392958 | 114 | 1 | 3 | 8 | 36 | 9 | 14 | 0 | 1 | 1 | 5 | 1 |
| SG000000C | Hlf | 1.05E-10 | 3.72E-10 | 3.888910179 | 3.003814272 | 44.49914749 | 74 | 0 | 3 | 5 | 27 | 7 | 8 | 0 | 0 | 2 | 4 | 0 |
| SG0000002 | Lztr1 | 1.18E-10 | 4.19E-10 | -0.984845994 | 2072.010817 | 1046.944918 | 14 | 3 | 3 | 4 | 10 | 1 | 0 | 0 | 0 | 0 | 1 | 0 |
| ISG000000 | Dus4l | 1.25E-10 | 4.42E-10 | 1.192887874 | 178.2711248 | 407.545438 | 38 | 2 | 3 | 4 | 11 | 2 | 0 | 0 | 0 | 0 | 0 | 0 |
| ISG000000 | A730046J19Rik | 1.37E-10 | 4.82E-10 | -5.036737797 | 31.82533426 | 0 | 42 | 3 | 3 | 8 | 19 | 7 | 15 | 0 | 0 | 1 | 6 | 0 |
| SG0000002 | Myo19 | 1.38E-10 | 4.86E-10 | -1.03506069 | 1351.817311 | 659.6805638 | 50 | 0 | 3 | 4 | 19 | 1 | 6 | 0 | 1 | 1 | 2 | 1 |
| SG0000003 | Slc7a9 | 1.46E-10 | 5.12E-10 | 4.791509842 | 0 | 26.69415936 | 38 | 3 | 3 | 6 | 15 | 13 | 2 | 0 | 0 | 0 | 2 | 0 |
| SG0000002 | Rnase4 | 1.8E-10 | 6.3E-10 | -1.167001948 | 704.6808249 | 313.8266801 | 37 | 2 | 3 | 6 | 18 | 2 | 3 | 0 | 0 | 0 | 1 | 1 |
| SG0000007 | Sp110 | 1.95E-10 | 6.79E-10 | -3.589798456 | 60.2248763 | 5.001944897 | 21 | 1 | 3 | 4 | 7 | 3 | 0 | 0 | 0 | 0 | 0 | 0 |
| ISG000000 | Lsm5 | 3.32E-10 | 1.14E-09 | 1.235821398 | 413.0659437 | 972.8339088 | 23 | 3 | 3 | 3 | 12 | 3 | 2 | 0 | 0 | 0 | 0 | 0 |
| ISG000000 | 9430020K01Rik | 3.93E-10 | 1.34E-09 | -1.111802782 | 1331.149274 | 615.9433532 | 67 | 2 | 3 | 7 | 25 | 6 | 5 | 0 | 0 | 0 | 2 | 1 |
| SG0000002 | Kdelc1 | 5.71E-10 | 1.92E-09 | 0.873150614 | 750.0600691 | 1373.853964 | 58 | 3 | 3 | 6 | 23 | 9 | 2 | 0 | 0 | 0 | 0 | 0 |
| SG0000002 | Zfp354a | 5.96E-10 | 2E-09 | 1.890288325 | 45.17333574 | 167.4617587 | 36 | 0 | 3 | 6 | 16 | 3 | 6 | 0 | 0 | 0 | 7 | 0 |
| SG000000C | Gys1 | 7.5E-10 | 2.5E-09 | -1.01668753 | 3111.284359 | 1537.751867 | 10 | 3 | 3 | 5 | 13 | 1 | 1 | 0 | 0 | 0 | 0 | 0 |
| ISG000000 | Mrpl44 | 9.23E-10 | 3.05E-09 | 0.909305238 | 615.0584821 | 1155.166442 | 18 | 3 | 3 | 3 | 7 | 0 | 0 | 0 | 0 | 0 | 1 | 0 |
| SG0000003 | Tnks1bp1 | 9.62E-10 | 3.17E-09 | -0.903911273 | 2555.345342 | 1365.668192 | 32 | 2 | 3 | 5 | 22 | 4 | 4 | 0 | 0 | 0 | 0 | 0 |
| ISG000000 | Harbi1 | 1.05E-09 | 3.44E-09 | 1.013898727 | 291.881251 | 589.4135673 | 46 | 3 | 3 | 3 | 39 | 0 | 13 | 1 | 1 | 1 | 11 | 0 |
| SG0000004 | Spata21 | 1.06E-09 | 3.48E-09 | 6.502197769 | 0.307509278 | 27.87499447 | 12 | 2 | 3 | 2 | 7 | 1 | 0 | 0 | 0 | 0 | 0 | 0 |
| SG0000005 | AI987944 | 1.21E-09 | 3.95E-09 | -1.181089248 | 573.6919737 | 253.008716 | 40 | 2 | 3 | 8 | 23 | 3 | 3 | 1 | 1 | 1 | 1 | 0 |
| SG0000004 | Tmprss12 | 1.24E-09 | 4.06E-09 | 4.656452562 | 0 | 24.21923419 | 18 | 2 | 3 | 3 | 7 | 0 | 1 | 0 | 0 | 0 | 0 | 0 |
| SG0000002 | Rgp1 | 1.55E-09 | 5.04E-09 | -0.923775568 | 1852.777882 | 976.6505578 | 10 | 0 | 3 | 0 | 2 | 0 | 2 | 0 | 0 | 0 | 1 | 0 |
| SG0000003 | Bhlhe40 | 1.66E-09 | 5.35E-09 | -0.906432103 | 15838.54178 | 8449.90726 | 8 | 0 | 3 | 0 | 3 | 1 | 0 | 0 | 0 | 0 | 2 | 1 |
| SG000000C | Ddx23 | 1.69E-09 | 5.44E-09 | -0.849443572 | 5341.155519 | 2964.334639 | 13 | 2 | 3 | 2 | 6 | 0 | 1 | 0 | 0 | 0 | 0 | 0 |
| SG0000002 | Atp5a1 | 1.91E-09 | 6.13E-09 | 0.714590197 | 17419.31663 | 28585.39495 | 51 | 0 | 3 | 0 | 9 | 3 | 9 | 0 | 3 | 0 | 0 | 0 |
| SG0000005 | Zfp105 | 2.09E-09 | 6.69E-09 | 1.000873946 | 288.7281989 | 577.8063114 | 15 | 0 | 3 | 2 | 3 | 0 | 1 | 0 | 0 | 0 | 0 | 0 |
| ISG000000 | Sox30 | 2.09E-09 | 6.7E-09 | 4.622882622 | 0 | 23.63918485 | 68 | 2 | 3 | 11 | 50 | 6 | 3 | 0 | 0 | 1 | 2 | 0 |
| SG0000004 | Zfp672 | 3.83E-09 | 1.2E-08 | -1.971056462 | 197.5456426 | 50.38721286 | 21 | 3 | 3 | 6 | 6 | 7 | 2 | 0 | 0 | 0 | 0 | 1 |
| ISG000000 | 1700102P08Rik | 3.9E-09 | 1.22E-08 | 3.749536583 | 2.896572999 | 38.95896935 | 37 | 2 | 3 | 6 | 11 | 1 | 0 | 0 | 0 | 0 | 0 | 0 |
| SG0000003 | Ccdc186 | 4.39E-09 | 1.37E-08 | -0.940540746 | 2065.248348 | 1076.07194 | 85 | 2 | 3 | 8 | 24 | 5 | 9 | 0 | 0 | 1 | 3 | 1 |
| ISG000000 | Thumpd1 | 4.85E-09 | 1.51E-08 | 0.766052935 | 1443.089577 | 2454.1336 | 10 | 3 | 3 | 4 | 9 | 0 | 0 | 0 | 0 | 0 | 0 | 0 |
| SG0000002 | Tmem169 | 5.29E-09 | 1.64E-08 | 3.678473476 | 2.990596593 | 38.29029292 | 34 | 1 | 3 | 6 | 21 | 4 | 9 | 0 | 2 | 2 | 12 | 0 |
| ISG000000 | L1td1 | 5.3E-09 | 1.65E-08 | 0.741574403 | 5023.446664 | 8399.200236 | 24 | 2 | 3 | 5 | 12 | 1 | 1 | 0 | 0 | 0 | 1 | 0 |
| SG0000004 | Slitrk4 | 5.9E-09 | 1.83E-08 | 4.540268768 | 0 | 22.26789458 | 54 | 3 | 3 | 6 | 15 | 1 | 28 | 1 | 1 | 4 | 7 | 1 |
| ISG000000 | Alg10b | 6.76E-09 | 2.09E-08 | -0.964542321 | 2222.379896 | 1138.838469 | 24 | 2 | 3 | 6 | 10 | 1 | 7 | 0 | 0 | 0 | 2 | 1 |
| ISG000000 | Ptchd3 | 7E-09 | 2.16E-08 | 4.532806595 | 0 | 22.14785481 | 20 | 2 | 3 | 2 | 9 | 4 | 0 | 0 | 0 | 0 | 0 | 0 |
| ISG000000 | 1110008F13Rik | 8.9E-09 | 2.72E-08 | -0.840926948 | 3023.855009 | 1688.172136 | 16 | 1 | 3 | 2 | 6 | 1 | 0 | 0 | 0 | 0 | 0 | 0 |
| ISG000000 | Map3k13 | 9.19E-09 | 2.81E-08 | 1.643799862 | 52.24320259 | 163.253634 | 59 | 2 | 3 | 8 | 22 | 4 | 2 | 1 | 1 | 1 | 4 | 0 |
| ISG000000 | Ccdc181 | 1.2E-08 | 3.62E-08 | 0.94434889 | 307.6952074 | 592.1041079 | 32 | 2 | 3 | 9 | 18 | 0 | 1 | 0 | 0 | 0 | 0 | 0 |
| ISG000000 | Rap1b | 1.22E-08 | 3.7E-08 | -0.803806686 | 6061.338753 | 3472.151234 | 56 | 0 | 3 | 3 | 16 | 5 | 4 | 0 | 0 | 0 | 2 | 0 |
| SG0000002 | Ell3 | 1.56E-08 | 4.68E-08 | 4.848231893 | 1.609821762 | 46.37041893 | 5 | 3 | 3 | 4 | 6 | 0 | 0 | 0 | 0 | 0 | 0 | 0 |
| ISG000000 | Naglu | 2.05E-08 | 6.07E-08 | -2.234649518 | 1043.807099 | 221.7806803 | 11 | 2 | 3 | 2 | 7 | 1 | 0 | 0 | 0 | 0 | 0 | 0 |
| SG0000005 | Nrgn | 2.42E-08 | 7.15E-08 | 5.915487102 | 1.337278385 | 80.71623237 | 6 | 3 | 3 | 3 | 8 | 0 | 0 | 0 | 0 | 0 | 0 | 0 |
| ISG000000 | Sap30bp | 2.67E-08 | 7.85E-08 | -0.846125001 | 1842.045174 | 1024.687106 | 30 | 2 | 3 | 5 | 11 | 5 | 0 | 0 | 0 | 0 | 0 | 0 |
| SG000000C | Acvr1b | 3.12E-08 | 9.14E-08 | 0.755856016 | 969.8387383 | 1637.701928 | 32 | 1 | 3 | 3 | 12 | 1 | 5 | 0 | 0 | 1 | 3 | 1 |
| SG0000003 | Aen | 3.42E-08 | 9.97E-08 | -0.812716365 | 2197.993611 | 1251.337391 | 18 | 0 | 3 | 3 | 9 | 0 | 0 | 0 | 0 | 0 | 0 | 0 |
| ISG000000 | Opa3 | 3.62E-08 | 1.05E-07 | -0.858778199 | 1445.961857 | 797.331348 | 48 | 2 | 3 | 10 | 39 | 0 | 4 | 0 | 0 | 0 | 0 | 0 |
| SG0000002 | Col9a3 | 4.14E-08 | 1.2E-07 | 2.564156174 | 9.473380086 | 56.02642149 | 19 | 2 | 3 | 2 | 8 | 1 | 2 | 0 | 0 | 0 | 0 | 0 |
| ISG000000 | Exoc3l | 4.3E-08 | 1.25E-07 | 2.736691161 | 18.11122523 | 120.718536 | 9 | 1 | 3 | 4 | 6 | 0 | 0 | 0 | 0 | 0 | 0 | 0 |
| ISG000000 | Myl12b | 4.64E-08 | 1.34E-07 | -0.782535046 | 6329.402181 | 3679.562388 | 18 | 2 | 3 | 3 | 7 | 1 | 1 | 0 | 0 | 1 | 1 | 0 |
| SG0000006 | Fam196b | 4.73E-08 | 1.36E-07 | 4.388843021 | 0 | 19.94948714 | 140 | 1 | 3 | 10 | 43 | 11 | 6 | 0 | 0 | 0 | 2 | 1 |
| SG0000004 | Mroh7 | 5.42E-08 | 1.55E-07 | 4.783046877 | 0.957493736 | 26.36188871 | 38 | 2 | 3 | 7 | 23 | 6 | 1 | 0 | 0 | 0 | 0 | 0 |
| ISG000000 | Opn3 | 6.68E-08 | 1.9E-07 | 2.240844838 | 16.12630595 | 76.22481994 | 66 | 1 | 3 | 4 | 22 | 8 | 2 | 0 | 0 | 0 | 0 | 0 |
| SG0000001 | Slmo2 | 6.71E-08 | 1.91E-07 | -0.781212268 | 3074.429647 | 1788.94189 | 32 | 0 | 3 | 3 | 7 | 0 | 5 | 0 | 3 | 0 | 4 | 0 |
| SG000000C | Kdelr1 | 6.93E-08 | 1.97E-07 | -0.775281158 | 3570.288685 | 2086.029519 | 10 | 3 | 3 | 5 | 8 | 1 | 1 | 0 | 0 | 0 | 0 | 0 |
| ISG000000 | Hexb | 7.14E-08 | 2.03E-07 | 0.735044132 | 1392.071733 | 2317.031546 | 38 | 2 | 3 | 8 | 16 | 4 | 0 | 0 | 0 | 0 | 0 | 0 |
| SG0000005 | St8sia3 | 7.36E-08 | 2.09E-07 | 4.344207444 | 0 | 19.3112545 | 52 | 2 | 3 | 9 | 17 | 8 | 7 | 0 | 0 | 0 | 2 | 2 |
| ISG000000 | Narf | 8.91E-08 | 2.51E-07 | 0.807623234 | 622.8270109 | 1090.149996 | 23 | 3 | 3 | 4 | 21 | 3 | 5 | 0 | 0 | 0 | 1 | 0 |
| ISG000000 | Bysl | 9.82E-08 | 2.76E-07 | -0.82849838 | 2105.117834 | 1185.423539 | 17 | 3 | 3 | 6 | 23 | 0 | 2 | 0 | 0 | 1 | 2 | 0 |
| SG0000005 | Ism2 | 1.15E-07 | 3.22E-07 | 3.105725747 | 4.565452453 | 39.30071325 | 16 | 1 | 3 | 2 | 9 | 4 | 1 | 0 | 0 | 0 | 0 | 0 |
| ISG000000 | Rpia | 1.26E-07 | 3.51E-07 | -1.086321161 | 423.0342423 | 199.2325384 | 38 | 2 | 3 | 3 | 19 | 7 | 1 | 0 | 0 | 0 | 0 | 0 |
| SG0000002 | Ptpn1 | 1.26E-07 | 3.52E-07 | -0.822892602 | 1944.580372 | 1099.285606 | 31 | 2 | 3 | 6 | 16 | 2 | 3 | 1 | 1 | 1 | 1 | 0 |
| SG0000001 | Cox6c | 1.33E-07 | 3.69E-07 | 0.679571024 | 2507.042625 | 4015.438516 | 41 | 3 | 3 | 6 | 20 | 11 | 0 | 0 | 0 | 0 | 0 | 0 |
| SG0000003 | Dnajb13 | 1.39E-07 | 3.86E-07 | 4.286168552 | 0 | 18.5103608 | 24 | 2 | 3 | 5 | 13 | 0 | 0 | 0 | 0 | 0 | 0 | 0 |
| SG0000004 | Mgat4e | 1.55E-07 | 4.28E-07 | 4.286678236 | 0 | 18.51725475 | 25 | 3 | 3 | 9 | 20 | 3 | 0 | 0 | 0 | 0 | 0 | 0 |
| SG0000002 | Peli3 | 1.82E-07 | 4.99E-07 | 1.84044755 | 24.3383733 | 87.16085182 | 20 | 0 | 3 | 0 | 2 | 0 | 2 | 0 | 1 | 0 | 0 | 0 |
| ISG000000 | H2-DMb1 | 2.07E-07 | 5.65E-07 | -1.770172904 | 116.7956131 | 34.24139025 | 13 | 2 | 3 | 4 | 4 | 1 | 1 | 0 | 0 | 1 | 1 | 0 |
| SG0000001 | Tm4sf5 | 2.15E-07 | 5.85E-07 | 4.169249413 | 4.325531412 | 77.82311372 | 8 | 3 | 3 | 3 | 3 | 1 | 0 | 0 | 0 | 0 | 0 | 0 |
| SG0000000 | Stoml1 | 2.19E-07 | 5.94E-07 | 1.220913695 | 400.9419201 | 934.5727421 | 10 | 3 | 3 | 3 | 7 | 0 | 1 | 0 | 0 | 0 | 0 | 0 |
| ISG000000 | Xdh | 2.42E-07 | 6.56E-07 | -2.756034801 | 154.3820085 | 22.85322748 | 73 | 3 | 3 | 7 | 44 | 8 | 0 | 0 | 0 | 0 | 0 | 0 |
| SG0000007 | Zfp788 | 2.71E-07 | 7.3E-07 | 0.956044026 | 195.9202596 | 380.0819772 | 114 | 3 | 3 | 24 | 58 | 12 | 3 | 0 | 0 | 0 | 1 | 0 |
| SG000000C | Pvrl4 | 2.92E-07 | 7.86E-07 | 1.298317848 | 70.47892689 | 173.3372487 | 33 | 0 | 3 | 0 | 9 | 12 | 3 | 0 | 0 | 0 | 0 | 6 |
| ISG000000 | Mrpl18 | 3.1E-07 | 8.31E-07 | 0.702312112 | 1184.740658 | 1927.703806 | 10 | 1 | 3 | 2 | 6 | 0 | 2 | 0 | 1 | 0 | 0 | 0 |
| SG0000005 | Capn11 | 4.41E-07 | 1.17E-06 | 4.198614003 | 0 | 17.36152529 | 24 | 2 | 3 | 4 | 9 | 2 | 1 | 0 | 0 | 0 | 0 | 0 |
| SG0000001 | Tnfrsf13b | 4.56E-07 | 1.21E-06 | 5.840690828 | 0.34247518 | 19.62692385 | 30 | 3 | 3 | 8 | 15 | 3 | 2 | 0 | 0 | 0 | 0 | 1 |
| ISG000000 | Padi1 | 5.74E-07 | 1.51E-06 | 5.979858709 | 0.307509278 | 19.40774444 | 22 | 2 | 3 | 3 | 4 | 1 | 0 | 0 | 0 | 0 | 0 | 0 |
| SG0000003 | Vamp1 | 7.25E-07 | 1.89E-06 | -1.138097148 | 388.7165867 | 176.6166704 | 41 | 2 | 3 | 3 | 16 | 9 | 6 | 0 | 0 | 0 | 0 | 2 |
| SG0000003 | Cdc16 | 7.41E-07 | 1.93E-06 | 0.640905334 | 1888.61234 | 2944.927129 | 55 | 1 | 3 | 10 | 25 | 3 | 2 | 0 | 0 | 0 | 0 | 0 |
| SG0000003 | Abhd10 | 1.04E-06 | 2.67E-06 | 0.768894272 | 446.5493795 | 760.9037589 | 45 | 2 | 3 | 7 | 19 | 10 | 5 | 0 | 1 | 3 | 3 | 2 |
| ISG000000 | Efhc1 | 1.04E-06 | 2.67E-06 | 1.331184904 | 65.73745136 | 165.4015017 | 58 | 3 | 3 | 9 | 31 | 7 | 0 | 0 | 0 | 0 | 0 | 0 |
| SG0000002 | Sbds | 1.24E-06 | 3.18E-06 | -0.715414047 | 2281.706722 | 1389.62857 | 10 | 0 | 3 | 0 | 10 | 0 | 3 | 0 | 0 | 0 | 1 | 0 |
| ISG000000 | Btbd18 | 1.41E-06 | 3.6E-06 | 4.517355245 | 0.957493736 | 21.92781564 | 6 | 2 | 3 | 2 | 9 | 0 | 0 | 0 | 0 | 0 | 0 | 0 |
| ISG000000 | Armc1 | 1.71E-06 | 4.33E-06 | 0.650201813 | 1127.615341 | 1769.665665 | 66 | 0 | 3 | 8 | 27 | 5 | 6 | 0 | 1 | 2 | 4 | 0 |
| ISG000000 | Gm648 | 1.72E-06 | 4.36E-06 | 4.14990532 | 0 | 16.75194651 | 5 | 3 | 3 | 3 | 4 | 0 | 0 | 0 | 0 | 0 | 0 | 0 |
| SG0000003 | Muc20 | 2.29E-06 | 5.72E-06 | -3.920448534 | 35.0906211 | 2.317493101 | 34 | 2 | 3 | 8 | 18 | 4 | 2 | 0 | 0 | 0 | 2 | 0 |
| ISG000000 | Trim56 | 2.95E-06 | 7.29E-06 | -0.733079572 | 1409.632016 | 848.0604589 | 13 | 2 | 3 | 4 | 7 | 2 | 13 | 2 | 3 | 4 | 7 | 2 |
| ISG000000 | Eny2 | 3.03E-06 | 7.49E-06 | -0.714814353 | 1657.247204 | 1009.733426 | 27 | 0 | 3 | 1 | 6 | 1 | 4 | 0 | 0 | 1 | 2 | 0 |
| SG0000002 | Smarcd3 | 3.18E-06 | 7.84E-06 | -1.051242079 | 355.4805051 | 171.5380189 | 28 | 2 | 3 | 5 | 9 | 1 | 0 | 0 | 0 | 0 | 0 | 0 |
| SG0000002 | Mief1 | 3.93E-06 | 9.62E-06 | -0.67309362 | 2528.88284 | 1586.015181 | 23 | 3 | 3 | 3 | 8 | 2 | 11 | 3 | 3 | 3 | 4 | 0 |
| ISG000000 | Plek2 | 4.01E-06 | 9.8E-06 | -1.142378221 | 225.2129035 | 102.0241908 | 29 | 1 | 3 | 3 | 10 | 3 | 0 | 0 | 0 | 0 | 0 | 0 |
| SG0000001 | Cyb5r3 | 4.36E-06 | 1.06E-05 | -0.621163811 | 9051.097725 | 5884.537226 | 20 | 2 | 3 | 6 | 11 | 3 | 2 | 0 | 0 | 1 | 1 | 0 |
| SG0000003 | Dexi | 4.59E-06 | 1.12E-05 | -1.206358804 | 192.35261 | 83.35811744 | 28 | 3 | 3 | 5 | 34 | 5 | 1 | 0 | 0 | 0 | 0 | 0 |
| SG0000002 | Srsf4 | 5.29E-06 | 1.28E-05 | 0.700725517 | 1178.586766 | 1915.582936 | 52 | 2 | 3 | 4 | 20 | 3 | 0 | 0 | 0 | 0 | 0 | 0 |
| ISG000000 | Caml | 5.52E-06 | 1.33E-05 | -0.8205356 | 625.5442204 | 354.2029614 | 18 | 3 | 3 | 4 | 9 | 1 | 1 | 0 | 0 | 0 | 0 | 0 |
| ISG000000 | Brf1 | 7.37E-06 | 1.76E-05 | -0.795116506 | 650.8651851 | 375.0914142 | 41 | 3 | 3 | 7 | 21 | 3 | 0 | 0 | 0 | 0 | 0 | 0 |
| SG000000C | Kcnh6 | 7.5E-06 | 1.79E-05 | 2.139341584 | 12.60618385 | 55.53792177 | 68 | 0 | 3 | 3 | 24 | 6 | 2 | 0 | 0 | 0 | 0 | 0 |
| ISG000000 | Ttc12 | 9.45E-06 | 2.23E-05 | 0.801596447 | 237.2965926 | 413.6148065 | 74 | 1 | 3 | 11 | 38 | 1 | 1 | 0 | 0 | 0 | 0 | 1 |
| ISG000000 | Med20 | 1.04E-05 | 2.44E-05 | -0.674876443 | 1601.537244 | 1003.180288 | 20 | 2 | 3 | 4 | 9 | 1 | 3 | 0 | 0 | 0 | 0 | 0 |
| SG0000005 | Prl2c5 | 1.04E-05 | 2.45E-05 | 4.209257654 | 1.609821762 | 29.77766278 | 83 | 0 | 3 | 8 | 31 | 12 | 2 | 0 | 0 | 0 | 0 | 0 |
| SG0000003 | Nfe2l1 | 1.06E-05 | 2.49E-05 | -0.593412215 | 9071.042947 | 6012.046718 | 44 | 0 | 3 | 0 | 5 | 0 | 5 | 0 | 0 | 0 | 0 | 0 |
| SG0000002 | Gm16286 | 1.06E-05 | 2.5E-05 | -0.613943037 | 3849.812693 | 2515.50005 | 45 | 3 | 3 | 12 | 21 | 3 | 3 | 0 | 0 | 3 | 3 | 0 |
| SG0000002 | Pabpc1 | 1.16E-05 | 2.73E-05 | 0.522180224 | 47660.14674 | 68445.87591 | 37 | 0 | 3 | 1 | 10 | 4 | 4 | 0 | 0 | 0 | 2 | 0 |

| | | | | | | | | | | | | | | | | | | |
|---|---|---|---|---|---|---|---|---|---|---|---|---|---|---|---|---|---|---|
| ISG000000 | Aplp1 | 1.23E-05 | 2.87E-05 | 1.297239186 | 869.7161939 | 2137.398317 | 10 | 3 | 3 | 3 | 4 | 0 | 2 | 0 | 0 | 0 | 0 | 0 |
| SG0000002 | Wdr82 | 1.53E-05 | 3.55E-05 | 0.542012697 | 3032.732823 | 4415.666155 | 50 | 3 | 3 | 10 | 22 | 10 | 7 | 0 | 0 | 1 | 2 | 2 |
| SG0000002 | Eif4a3 | 1.65E-05 | 3.82E-05 | 0.543635284 | 3011.663572 | 4389.923814 | 13 | 2 | 3 | 4 | 10 | 1 | 1 | 0 | 0 | 0 | 0 | 1 |
| SG0000003 | Vstm2b | 1.69E-05 | 3.9E-05 | 5.636402812 | 0.307509278 | 15.29623467 | 36 | 1 | 3 | 5 | 11 | 3 | 0 | 0 | 0 | 0 | 0 | 0 |
| SG000000 | Bbc3 | 1.79E-05 | 4.12E-05 | 1.712219105 | 147.4887864 | 483.2683036 | 7 | 2 | 3 | 4 | 6 | 2 | 0 | 0 | 0 | 0 | 0 | 1 |
| SG0000004 | Serpinb6b | 1.95E-05 | 4.46E-05 | 2.2606766 | 9.794107043 | 46.93494501 | 73 | 0 | 3 | 12 | 36 | 9 | 12 | 0 | 0 | 0 | 0 | 0 |
| SG0000002 | Mif4gd | 2.16E-05 | 4.92E-05 | 0.637277818 | 602.2064965 | 936.666904 | 21 | 3 | 3 | 6 | 11 | 0 | 0 | 0 | 0 | 0 | 2 | 0 |
| SG0000003 | Prr3 | 2.53E-05 | 5.72E-05 | 0.698468023 | 367.850664 | 596.940947 | 32 | 3 | 3 | 6 | 12 | 4 | 0 | 0 | 0 | 0 | 0 | 1 |
| ISG000000 | Zfp867 | 2.9E-05 | 6.54E-05 | -0.832700088 | 436.1296334 | 244.8769415 | 32 | 3 | 3 | 4 | 15 | 3 | 3 | 0 | 0 | 0 | 4 | 2 |
| SG0000002 | Pgm2 | 2.92E-05 | 6.58E-05 | -0.780672925 | 5754.320584 | 3349.562504 | 112 | 2 | 3 | 15 | 67 | 16 | 0 | 0 | 0 | 0 | 0 | 0 |
| ISG000000 | Ywhab | 3.54E-05 | 7.9E-05 | -0.555559114 | 9133.673262 | 6214.490292 | 29 | 2 | 3 | 2 | 6 | 1 | 3 | 0 | 0 | 0 | 0 | 0 |
| SG0000004 | Flad1 | 3.66E-05 | 8.17E-05 | -0.687737877 | 807.3870439 | 501.2472997 | 48 | 3 | 3 | 6 | 18 | 0 | 0 | 0 | 0 | 0 | 0 | 0 |
| SG0000004 | Ankrd13a | 3.77E-05 | 8.4E-05 | -0.570944107 | 4249.637305 | 2860.754328 | 31 | 3 | 3 | 4 | 13 | 3 | 0 | 0 | 0 | 0 | 1 | 1 |
| ISG000000 | Depdc7 | 3.78E-05 | 8.43E-05 | -0.73717911 | 662.5150071 | 397.4501781 | 40 | 2 | 3 | 5 | 14 | 3 | 1 | 0 | 0 | 0 | 0 | 0 |
| SG0000004 | Tmie | 3.93E-05 | 8.74E-05 | 2.673953376 | 4.530486551 | 28.91245826 | 12 | 1 | 3 | 4 | 5 | 3 | 0 | 0 | 0 | 0 | 0 | 1 |
| ISG000000 | Dusp1 | 4.23E-05 | 9.36E-05 | 0.645951186 | 1047.801001 | 1639.568219 | 5 | 2 | 3 | 4 | 4 | 2 | 4 | 2 | 2 | 3 | 3 | 1 |
| SG0000001 | Card14 | 4.51E-05 | 9.94E-05 | 2.394750023 | 6.421382233 | 33.7691486 | 43 | 2 | 3 | 5 | 23 | 4 | 0 | 0 | 0 | 0 | 0 | 0 |
| SG0000002 | Rhbdf1 | 4.96E-05 | 0.000109 | -0.749667734 | 603.1168471 | 358.6980249 | 14 | 0 | 3 | 3 | 3 | 0 | 2 | 0 | 0 | 0 | 0 | 0 |
| ISG000000 | Rbm24 | 5.45E-05 | 0.000119 | 4.292318902 | 1.984919276 | 38.89193846 | 25 | 2 | 3 | 6 | 22 | 3 | 9 | 0 | 1 | 3 | 3 | 0 |
| ISG000000 | Lypd1 | 5.83E-05 | 0.000127 | -2.018375472 | 49.22667362 | 12.15091354 | 58 | 1 | 3 | 7 | 21 | 6 | 0 | 0 | 0 | 0 | 0 | 0 |
| SG0000003 | Tsc22d3 | 5.86E-05 | 0.000128 | 1.161964932 | 76.57469832 | 171.3450619 | 88 | 0 | 3 | 8 | 38 | 5 | 1 | 0 | 0 | 0 | 0 | 0 |
| SG0000002 | Phkg1 | 6.04E-05 | 0.000131 | 6.169780911 | 1.326404274 | 95.49156991 | 19 | 2 | 3 | 6 | 12 | 1 | 2 | 0 | 0 | 0 | 0 | 0 |
| ISG000000 | Gm1141 | 6.51E-05 | 0.000141 | -2.017062346 | 49.05518454 | 12.11960999 | 23 | 3 | 3 | 5 | 14 | 4 | 0 | 0 | 0 | 0 | 0 | 0 |
| ISG000000 | Hapln4 | 6.55E-05 | 0.000142 | 1.265643824 | 74.8812241 | 180.0402844 | 7 | 2 | 3 | 3 | 4 | 0 | 2 | 2 | 2 | 2 | 3 | 0 |
| ISG000000 | Thumpd2 | 6.57E-05 | 0.000142 | 0.768796537 | 193.279825 | 329.31942 | 72 | 0 | 3 | 5 | 15 | 2 | 0 | 0 | 0 | 0 | 0 | 0 |
| ISG000000 | Vmn2r1 | 6.7E-05 | 0.000145 | 4.567289332 | 0.639110347 | 15.15189445 | 61 | 1 | 3 | 10 | 34 | 4 | 0 | 0 | 0 | 0 | 0 | 0 |
| SG0000002 | Haus2 | 6.89E-05 | 0.000149 | 0.559847831 | 890.6817442 | 1312.966185 | 81 | 3 | 3 | 24 | 51 | 5 | 1 | 0 | 0 | 0 | 9 | 2 |
| SG0000002 | Sdha | 6.96E-05 | 0.00015 | 0.490627871 | 5916.672594 | 8313.257707 | 57 | 2 | 3 | 6 | 15 | 3 | 1 | 0 | 0 | 0 | 0 | 0 |
| SG0000002 | Blzf1 | 7.03E-05 | 0.000152 | 0.597051467 | 604.9204345 | 915.0159321 | 82 | 0 | 3 | 10 | 29 | 5 | 10 | 0 | 0 | 4 | 8 | 2 |
| SG0000003 | Ephx4 | 7.48E-05 | 0.000161 | -1.704513495 | 65.1015283 | 19.97475651 | 47 | 1 | 3 | 8 | 24 | 4 | 0 | 0 | 0 | 0 | 0 | 0 |
| ISG000000 | Ifit3 | 8.01E-05 | 0.000172 | -1.163780665 | 279.0074076 | 124.5325398 | 10 | 2 | 3 | 3 | 5 | 1 | 0 | 0 | 0 | 0 | 0 | 0 |
| SG000000 | Nob1 | 9.34E-05 | 0.000198 | 0.524583974 | 1721.655813 | 2476.63408 | 39 | 3 | 3 | 3 | 9 | 0 | 0 | 0 | 0 | 0 | 0 | 0 |
| SG0000002 | Hr | 0.000105 | 0.000222 | 0.848476491 | 131.6981125 | 237.1354186 | 10 | 2 | 3 | 4 | 5 | 2 | 0 | 0 | 0 | 0 | 1 | 0 |
| ISG000000 | N6amt2 | 0.000133 | 0.000277 | 0.628236704 | 509.8974886 | 788.1356365 | 22 | 1 | 3 | 5 | 10 | 4 | 0 | 0 | 0 | 0 | 0 | 0 |
| SG0000002 | Per1 | 0.000145 | 0.000302 | 0.558033301 | 761.3366269 | 1120.886106 | 19 | 3 | 3 | 3 | 12 | 0 | 3 | 3 | 3 | 3 | 3 | 0 |
| ISG000000 | Tmprss2 | 0.000198 | 0.000407 | 7.551515621 | 0.615018556 | 115.3774494 | 56 | 2 | 3 | 5 | 20 | 4 | 0 | 0 | 0 | 0 | 0 | 0 |
| SG0000002 | Mis18a | 0.000205 | 0.000419 | 0.581675618 | 473.5436615 | 708.699027 | 14 | 2 | 3 | 4 | 9 | 1 | 4 | 0 | 0 | 0 | 1 | 1 |
| SG000000 | Ssb | 0.000215 | 0.000439 | 0.456005794 | 5430.387483 | 7449.073195 | 90 | 3 | 3 | 12 | 40 | 6 | 12 | 0 | 0 | 0 | 0 | 0 |
| SG0000003 | Bahcc1 | 0.000217 | 0.000443 | -0.630779412 | 1341.166396 | 866.1620262 | 83 | 0 | 3 | 12 | 61 | 6 | 3 | 0 | 0 | 3 | 3 | 3 |
| SG0000003 | Phb | 0.000226 | 0.000461 | 0.494318252 | 1746.745329 | 2460.561428 | 13 | 2 | 3 | 3 | 5 | 0 | 1 | 0 | 0 | 0 | 0 | 0 |
| ISG000000 | Ethe1 | 0.000233 | 0.000474 | 0.687446911 | 237.178827 | 381.9601521 | 23 | 2 | 3 | 6 | 15 | 3 | 0 | 0 | 0 | 0 | 0 | 0 |
| ISG000000 | Serinc3 | 0.000234 | 0.000476 | -0.500078597 | 5548.260818 | 3922.999121 | 36 | 3 | 3 | 4 | 14 | 4 | 4 | 0 | 0 | 0 | 1 | 1 |
| SG0000002 | Atp1b1 | 0.000236 | 0.00048 | 0.742653224 | 2061.336972 | 3449.132589 | 31 | 2 | 3 | 7 | 26 | 1 | 6 | 0 | 0 | 2 | 5 | 1 |
| SG0000002 | Ripk3 | 0.00028 | 0.000564 | 0.587343955 | 425.3738576 | 639.1149022 | 3 | 0 | 3 | 0 | 0 | 0 | 0 | 0 | 0 | 0 | 0 | 0 |
| ISG000000 | Zbtb26 | 0.000327 | 0.000653 | -0.598748251 | 824.3557533 | 544.344062 | 43 | 2 | 3 | 2 | 14 | 2 | 15 | 2 | 3 | 2 | 8 | 0 |
| ISG000000 | Tmppe | 0.000328 | 0.000655 | -0.580120714 | 958.5373412 | 641.1731097 | 11 | 3 | 3 | 3 | 19 | 0 | 1 | 0 | 0 | 0 | 0 | 0 |
| ISG000000 | Tarm1 | 0.000373 | 0.000741 | -1.652448188 | 52.13496445 | 16.58412684 | 24 | 3 | 3 | 8 | 32 | 1 | 0 | 0 | 0 | 0 | 0 | 0 |
| SG000000 | H2-Eb1 | 0.000391 | 0.000774 | 3.335354986 | 2.982066049 | 30.09949127 | 17 | 2 | 3 | 2 | 5 | 2 | 0 | 0 | 0 | 0 | 0 | 0 |
| SG0000003 | Ankrd54 | 0.000428 | 0.000842 | -0.640715857 | 649.0457211 | 416.2944295 | 6 | 3 | 3 | 3 | 6 | 0 | 0 | 0 | 0 | 0 | 0 | 0 |
| ISG000000 | Ly6g6d | 0.000491 | 0.00096 | 2.17830362 | 6.456348135 | 29.22278948 | 9 | 2 | 3 | 3 | 7 | 0 | 0 | 0 | 0 | 0 | 0 | 0 |
| ISG000000 | Gatsl2 | 0.000529 | 0.001028 | -0.620094598 | 609.9582705 | 396.856091 | 37 | 2 | 3 | 5 | 18 | 6 | 3 | 0 | 0 | 0 | 5 | 1 |
| ISG000000 | Ngrn | 0.000528 | 0.001028 | 0.508020145 | 710.0688138 | 1009.78692 | 10 | 3 | 3 | 4 | 7 | 1 | 0 | 0 | 0 | 0 | 0 | 0 |
| ISG000000 | Rab5b | 0.00053 | 0.001031 | -0.548235546 | 1937.285492 | 1324.824291 | 35 | 3 | 3 | 7 | 16 | 3 | 5 | 0 | 0 | 1 | 1 | 1 |
| ISG000000 | Fam8a1 | 0.00053 | 0.001032 | -0.516278281 | 1773.470036 | 1239.962658 | 20 | 1 | 3 | 4 | 9 | 0 | 12 | 0 | 2 | 3 | 5 | 0 |
| ISG000000 | Cox15 | 0.00058 | 0.001122 | -0.509684603 | 1858.545603 | 1305.397775 | 44 | 3 | 3 | 6 | 20 | 3 | 7 | 0 | 0 | 0 | 2 | 0 |
| SG0000002 | Fam107a | 0.000582 | 0.001125 | 3.277335562 | 1.620695873 | 15.71367752 | 33 | 3 | 3 | 7 | 29 | 9 | 0 | 0 | 0 | 0 | 0 | 4 |
| SG0000002 | Cyb5r1 | 0.000601 | 0.00116 | 0.439014466 | 2997.067745 | 4063.058568 | 9 | 0 | 3 | 0 | 4 | 0 | 0 | 0 | 0 | 0 | 0 | 0 |
| ISG000000 | Yipf5 | 0.000613 | 0.001181 | -0.545743803 | 1095.029397 | 750.1368904 | 37 | 1 | 3 | 4 | 11 | 3 | 1 | 0 | 0 | 0 | 0 | 0 |
| SG000000 | Rnd2 | 0.000661 | 0.001268 | 0.884061716 | 240.8474528 | 444.4993724 | 6 | 3 | 3 | 3 | 3 | 0 | 0 | 0 | 0 | 0 | 0 | 0 |
| SG0000003 | Ccdc32 | 0.000749 | 0.001427 | -0.6464121 | 449.074952 | 286.8992419 | 36 | 3 | 3 | 10 | 23 | 3 | 1 | 0 | 0 | 0 | 0 | 2 |
| ISG000000 | Lsm1 | 0.000778 | 0.001479 | 0.501200259 | 708.1319685 | 1002.283343 | 57 | 1 | 3 | 9 | 14 | 4 | 3 | 0 | 0 | 0 | 0 | 1 |
| SG000000 | Sardh | 0.000823 | 0.001561 | -2.024683583 | 40.06199638 | 9.845597953 | 34 | 3 | 3 | 6 | 29 | 2 | 0 | 0 | 0 | 0 | 0 | 0 |
| ISG000000 | Pdcd6 | 0.00088 | 0.001661 | -0.484716309 | 1918.799117 | 1371.245962 | 30 | 1 | 3 | 3 | 13 | 2 | 0 | 0 | 0 | 0 | 0 | 1 |
| ISG000000 | Ogfod3 | 0.001075 | 0.002011 | -0.816089358 | 185.6433199 | 105.4415911 | 26 | 3 | 3 | 5 | 20 | 3 | 0 | 0 | 0 | 0 | 0 | 0 |
| SG0000003 | Mettl22 | 0.001079 | 0.002018 | 0.733542183 | 129.7444689 | 215.728266 | 16 | 2 | 3 | 3 | 13 | 2 | 0 | 0 | 0 | 0 | 0 | 0 |
| ISG000000 | Prl2c3 | 0.001233 | 0.002292 | 1.387066917 | 45.60150068 | 119.2690837 | 32 | 2 | 3 | 5 | 17 | 3 | 1 | 0 | 0 | 0 | 0 | 0 |
| ISG000000 | Purg | 0.001309 | 0.002425 | 0.685175441 | 142.1498817 | 228.5624395 | 78 | 2 | 3 | 12 | 32 | 10 | 0 | 0 | 0 | 0 | 0 | 0 |
| ISG000000 | Slc9a3r1 | 0.001413 | 0.00261 | -0.623702603 | 2409.652249 | 1563.871912 | 7 | 2 | 3 | 2 | 2 | 1 | 0 | 0 | 0 | 0 | 0 | 0 |
| ISG000000 | Stc1 | 0.00142 | 0.002622 | 0.927585866 | 101.9661233 | 193.9487912 | 11 | 1 | 3 | 3 | 6 | 1 | 7 | 1 | 2 | 2 | 5 | 1 |
| ISG000000 | Bmp1 | 0.001708 | 0.003126 | -0.824997466 | 1117.498628 | 630.8091948 | 38 | 2 | 3 | 5 | 24 | 3 | 0 | 0 | 0 | 0 | 0 | 0 |
| SG000000 | Gng5 | 0.001913 | 0.003471 | 0.477733123 | 595.7838111 | 829.6610428 | 27 | 0 | 3 | 6 | 27 | 3 | 5 | 0 | 1 | 2 | 9 | 0 |
| ISG000000 | Glrx | 0.001969 | 0.003567 | -0.870703621 | 172.9789129 | 94.59871919 | 13 | 1 | 3 | 2 | 3 | 2 | 0 | 0 | 0 | 0 | 0 | 0 |
| ISG000000 | Rars | 0.002591 | 0.00462 | -0.404563199 | 7559.811069 | 5711.17258 | 50 | 0 | 3 | 7 | 18 | 3 | 0 | 0 | 0 | 0 | 0 | 0 |
| ISG000000 | Ndufa8 | 0.002646 | 0.004708 | -0.455807496 | 1263.136142 | 920.9550489 | 21 | 3 | 3 | 6 | 9 | 1 | 1 | 0 | 0 | 0 | 0 | 0 |
| SG0000002 | Il18rap | 0.002707 | 0.00481 | -1.995650281 | 27.4002484 | 6.870746174 | 59 | 0 | 3 | 7 | 20 | 5 | 4 | 0 | 0 | 1 | 1 | 1 |
| ISG000000 | Rheb | 0.002792 | 0.004949 | -0.42777051 | 2512.399791 | 1867.742469 | 65 | 2 | 3 | 7 | 25 | 8 | 1 | 0 | 0 | 0 | 1 | 1 |
| ISG000000 | Atp5s | 0.0028 | 0.00496 | 0.636224595 | 152.8621793 | 237.587033 | 36 | 2 | 3 | 4 | 12 | 5 | 1 | 0 | 0 | 0 | 0 | 0 |
| SG0000004 | Shfm1 | 0.002977 | 0.005255 | -0.423751244 | 2356.537581 | 1756.760408 | 69 | 2 | 3 | 5 | 27 | 7 | 6 | 0 | 0 | 0 | 1 | 0 |
| SG0000003 | Ttc16 | 0.002981 | 0.005261 | 1.403879635 | 15.52819535 | 41.08950093 | 45 | 0 | 3 | 12 | 12 | 0 | 3 | 0 | 2 | 0 | 0 | 0 |
| ISG000000 | 1700001C19Rik | 0.003001 | 0.005294 | -2.233850318 | 36.54437098 | 7.768989589 | 35 | 2 | 3 | 10 | 29 | 4 | 0 | 0 | 0 | 0 | 0 | 0 |
| SG0000002 | 2310067B10Rik | 0.003077 | 0.005416 | -0.697623327 | 2019.061766 | 1244.927492 | 28 | 3 | 3 | 3 | 15 | 4 | 1 | 0 | 0 | 0 | 0 | 0 |
| ISG000000 | Rpp14 | 0.003079 | 0.005419 | 0.400987261 | 1395.549543 | 1842.699221 | 23 | 2 | 3 | 2 | 7 | 5 | 6 | 0 | 0 | 0 | 2 | 0 |
| ISG000000 | Ppp1r36 | 0.003286 | 0.005764 | 2.446724238 | 3.182334054 | 17.34934778 | 22 | 2 | 3 | 5 | 23 | 1 | 1 | 0 | 0 | 0 | 0 | 0 |
| ISG000000 | Nxpe4 | 0.003523 | 0.006157 | 1.24833872 | 20.1876714 | 47.95938734 | 52 | 2 | 3 | 4 | 33 | 3 | 3 | 0 | 0 | 0 | 2 | 1 |
| SG0000002 | Sfpq | 0.003594 | 0.006273 | 0.360212389 | 7883.143842 | 10118.92053 | 17 | 2 | 3 | 3 | 10 | 3 | 3 | 0 | 0 | 0 | 0 | 0 |
| ISG000000 | Syt4 | 0.003979 | 0.006898 | -1.100933194 | 68.13696805 | 31.76646507 | 31 | 2 | 3 | 4 | 15 | 1 | 7 | 1 | 2 | 1 | 5 | 1 |
| ISG000000 | Ttbk1 | 0.004503 | 0.007751 | 2.872391148 | 3.001470705 | 21.97910499 | 20 | 2 | 3 | 2 | 10 | 6 | 1 | 0 | 0 | 0 | 0 | 0 |
| ISG000000 | Dhdh | 0.004863 | 0.008337 | 0.606034113 | 317.5867971 | 483.3891365 | 26 | 2 | 3 | 3 | 16 | 3 | 5 | 1 | 1 | 1 | 5 | 0 |
| SG0000002 | Zfp729a | 0.004899 | 0.008393 | 0.447676149 | 469.0871813 | 639.7606353 | 49 | 3 | 3 | 10 | 31 | 5 | 3 | 0 | 0 | 1 | 3 | 2 |
| ISG000000 | BC003965 | 0.005086 | 0.008698 | -0.463895186 | 727.2880302 | 527.3027961 | 6 | 3 | 3 | 3 | 4 | 0 | 6 | 3 | 3 | 3 | 4 | 0 |
| SG0000001 | Col20a1 | 0.005157 | 0.008812 | -1.580821281 | 36.54068071 | 12.21524017 | 19 | 1 | 3 | 3 | 8 | 5 | 0 | 0 | 0 | 0 | 0 | 0 |
| ISG000000 | Was | 0.005369 | 0.009148 | 5.039066303 | 0.68495036 | 22.5200425 | 12 | 3 | 3 | 4 | 10 | 0 | 1 | 0 | 0 | 0 | 0 | 0 |
| SG000000 | Hnrnph1 | 0.00625 | 0.010547 | 0.542042849 | 9512.378479 | 13850.33501 | 74 | 0 | 3 | 0 | 15 | 15 | 9 | 0 | 0 | 0 | 0 | 0 |
| SG0000003 | Eif3c | 0.006825 | 0.011454 | 0.330538063 | 11251.3092 | 14148.32187 | 34 | 2 | 3 | 5 | 16 | 3 | 2 | 0 | 0 | 0 | 0 | 0 |
| ISG000000 | Itga5 | 0.007167 | 0.011988 | -0.370762502 | 2665.181623 | 2061.181215 | 20 | 1 | 3 | 6 | 12 | 0 | 2 | 0 | 1 | 1 | 1 | 0 |
| ISG000000 | Dctn4 | 0.007686 | 0.012796 | -0.38786168 | 2009.930087 | 1536.112246 | 55 | 2 | 3 | 6 | 18 | 0 | 1 | 0 | 0 | 0 | 0 | 0 |
| SG0000003 | Zfp661 | 0.007794 | 0.012967 | 0.556556292 | 164.2460129 | 241.5655361 | 23 | 0 | 3 | 3 | 11 | 1 | 4 | 0 | 0 | 0 | 2 | 1 |
| ISG000000 | Zfp746 | 0.007793 | 0.012967 | -0.43077326 | 876.0979872 | 649.9456028 | 30 | 1 | 3 | 7 | 19 | 3 | 1 | 0 | 0 | 0 | 0 | 0 |
| SG000000 | Ddx3x | 0.008963 | 0.014769 | 0.319109877 | 15221.9579 | 18990.31469 | 41 | 0 | 3 | 8 | 14 | 3 | 7 | 0 | 0 | 3 | 3 | 1 |
| ISG000000 | Tmed9 | 0.00899 | 0.014811 | -0.358237796 | 3366.532352 | 2626.288731 | 6 | 3 | 3 | 4 | 6 | 0 | 0 | 0 | 0 | 0 | 0 | 0 |
| ISG000000 | Rfc2 | 0.00997 | 0.016309 | 0.344035985 | 1814.294379 | 2302.888474 | 21 | 2 | 3 | 6 | 11 | 1 | 1 | 0 | 0 | 0 | 0 | 0 |
| SG0000003 | Aste1 | 0.010057 | 0.01644 | 0.514507307 | 200.5496536 | 286.4864253 | 109 | 0 | 3 | 8 | 27 | 13 | 13 | 0 | 2 | 4 | 6 | 0 |
| SG0000004 | Rhno1 | 0.010556 | 0.01719 | -0.371811089 | 1787.057686 | 1381.059155 | 21 | 0 | 3 | 0 | 0 | 4 | 3 | 0 | 0 | 0 | 0 | 4 |
| ISG000000 | Mpzl2 | 0.011699 | 0.018922 | 0.352366155 | 1658.045954 | 2116.748912 | 21 | 2 | 3 | 4 | 8 | 2 | 6 | 2 | 3 | 3 | 4 | 1 |
| ISG000000 | Bves | 0.01187 | 0.019177 | 2.453121613 | 4.923488879 | 26.96100952 | 53 | 1 | 3 | 6 | 26 | 9 | 1 | 0 | 0 | 0 | 1 | 0 |
| SG0000003 | Acaa1a | 0.012038 | 0.019414 | -0.40841308 | 773.6557121 | 582.912205 | 16 | 3 | 3 | 6 | 9 | 6 | 0 | 0 | 0 | 0 | 0 | 0 |
| ISG000000 | Prr12 | 0.012408 | 0.019954 | -0.615417552 | 1511.051049 | 986.3248924 | 15 | 1 | 3 | 7 | 13 | 4 | 3 | 1 | 2 | 2 | 2 | 2 |
| SG0000003 | Map1lc3b | 0.012661 | 0.02032 | 0.326069632 | 2269.033956 | 2844.446273 | 29 | 3 | 3 | 3 | 9 | 3 | 5 | 0 | 0 | 0 | 0 | 0 |
| SG0000005 | D8Ertd82e | 0.012969 | 0.020776 | -0.567698967 | 1214.199799 | 819.2109322 | 72 | 1 | 3 | 9 | 30 | 8 | 1 | 0 | 1 | 0 | 1 | 0 |
| SG000000 | Csn3 | 0.013971 | 0.022237 | -2.066313649 | 17.85324624 | 4.262798519 | 59 | 0 | 3 | 6 | 32 | 3 | 4 | 0 | 0 | 0 | 1 | 0 |
| ISG000000 | Insm1 | 0.014491 | 0.023013 | -1.585459727 | 37.31611753 | 12.43441957 | 5 | 2 | 3 | 4 | 6 | 0 | 5 | 2 | 3 | 4 | 6 | 0 |
| SG0000002 | Coq4 | 0.0148 | 0.023474 | -0.462494606 | 375.645987 | 272.6176718 | 21 | 3 | 3 | 3 | 3 | 0 | 0 | 0 | 0 | 0 | 0 | 0 |
| ISG000000 | Fcgrt | 0.014865 | 0.02357 | 0.668797755 | 91.16480459 | 144.9290317 | 16 | 3 | 3 | 7 | 11 | 3 | 0 | 0 | 0 | 0 | 0 | 0 |
| ISG000000 | Dld | 0.015018 | 0.023794 | -0.336275638 | 3648.910131 | 2890.241372 | 57 | 2 | 3 | 9 | 26 | 2 | 3 | 0 | 0 | 0 | 0 | 1 |
| SG000000 | Lipe | 0.015249 | 0.024146 | 0.707269462 | 104.5359446 | 170.6769882 | 38 | 0 | 3 | 5 | 8 | 3 | 2 | 0 | 0 | 0 | 0 | 2 |
| ISG000000 | Zfp955b | 0.015669 | 0.024761 | 0.554855416 | 124.1944312 | 182.4442776 | 23 | 2 | 3 | 6 | 26 | 5 | 0 | 0 | 0 | 0 | 0 | 0 |
| ISG000000 | Saraf | 0.01626 | 0.025612 | -0.337160808 | 2423.600596 | 1918.51648 | 27 | 3 | 3 | 7 | 17 | 4 | 1 | 0 | 0 | 0 | 0 | 0 |
| ISG000000 | Rsph3a | 0.016376 | 0.025774 | -0.869700882 | 87.14109846 | 47.68886402 | 46 | 1 | 3 | 3 | 14 | 3 | 2 | 0 | 0 | 0 | 0 | 0 |
| SG0000001 | Rabl6 | 0.016661 | 0.026188 | -0.33630107 | 2524.301407 | 1999.422154 | 22 | 3 | 3 | 3 | 14 | 1 | 0 | 0 | 0 | 0 | 0 | 0 |
| ISG000000 | Trim62 | 0.017537 | 0.027437 | 0.766127985 | 46.87618424 | 79.72228838 | 21 | 1 | 3 | 2 | 12 | 2 | 1 | 0 | 1 | 0 | 1 | 0 |
| SG0000002 | Zcchc9 | 0.017885 | 0.027914 | 0.411051993 | 651.0456145 | 865.6660739 | 17 | 0 | 3 | 2 | 7 | 1 | 4 | 0 | 2 | 2 | 2 | 0 |
| ISG000000 | Fbxo38 | 0.018127 | 0.028259 | 0.309310106 | 1997.037264 | 2474.558861 | 88 | 1 | 3 | 6 | 33 | 6 | 4 | 1 | 1 | 1 | 1 | 0 |
| SG0000003 | Zfp707 | 0.018252 | 0.028438 | -0.564696307 | 286.3935861 | 193.6300514 | 33 | 3 | 3 | 9 | 24 | 3 | 0 | 0 | 0 | 0 | 0 | 0 |

| | | | | | | | | | | | | | | | | | | |
|---|---|---|---|---|---|---|---|---|---|---|---|---|---|---|---|---|---|---|
| SG0000002 | Mycbp | 0.018833 | 0.029265 | 0.383531745 | 425.0267206 | 554.4607315 | 22 | 3 | 3 | 3 | 9 | 4 | 6 | 1 | 1 | 1 | 4 | 0 |
| SG0000002 | Sdc3 | 0.019326 | 0.029974 | -0.649031764 | 616.8111974 | 393.3455307 | 21 | 3 | 3 | 4 | 8 | 3 | 3 | 0 | 0 | 0 | 0 | 2 |
| JSG000000 | Jag1 | 0.020303 | 0.031361 | -0.429363266 | 442.1605971 | 328.3437968 | 53 | 1 | 3 | 6 | 12 | 4 | 10 | 1 | 3 | 5 | 6 | 1 |
| JSG000000 | Grasp | 0.021076 | 0.032461 | 0.394328887 | 593.4235632 | 779.9551142 | 6 | 3 | 3 | 4 | 4 | 0 | 3 | 2 | 2 | 3 | 3 | 0 |
| SG0000005 | Klhl5 | 0.021577 | 0.033176 | -0.563812364 | 640.0105751 | 432.974896 | 72 | 1 | 3 | 3 | 33 | 4 | 3 | 0 | 0 | 0 | 2 | 0 |
| SG0000003 | Cep131 | 0.022158 | 0.034021 | 0.696929164 | 302.6066882 | 490.5407669 | 27 | 0 | 3 | 0 | 9 | 3 | 0 | 0 | 0 | 0 | 0 | 0 |
| JSG000000 | Bcdin3d | 0.023643 | 0.036109 | -0.533972777 | 246.5534964 | 170.2822355 | 12 | 2 | 3 | 2 | 5 | 1 | 1 | 1 | 1 | 1 | 1 | 0 |
| SG0000003 | Has3 | 0.024471 | 0.037289 | 4.342233617 | 0.922527834 | 18.71207913 | 23 | 3 | 3 | 3 | 12 | 9 | 4 | 0 | 0 | 0 | 4 | 4 |
| JSG000000 | Coq10a | 0.024481 | 0.0373 | -0.436902019 | 367.0681874 | 271.1603179 | 11 | 3 | 3 | 3 | 4 | 0 | 5 | 2 | 2 | 2 | 3 | 0 |
| JSG000000 | Ntmt1 | 0.024624 | 0.037502 | 0.6208653 | 444.315743 | 683.2675254 | 37 | 3 | 3 | 6 | 9 | 0 | 0 | 0 | 0 | 0 | 0 | 0 |
| JSG000000 | Serpinb9b | 0.025406 | 0.038604 | -0.559660666 | 302.0608794 | 204.9367452 | 34 | 1 | 3 | 5 | 17 | 1 | 5 | 0 | 1 | 0 | 2 | 0 |
| SG0000006 | Zfp563 | 0.025582 | 0.038849 | -0.653003223 | 120.4974423 | 76.63096086 | 41 | 3 | 3 | 8 | 16 | 5 | 1 | 0 | 0 | 0 | 0 | 1 |
| SG0000003 | Zfp57 | 0.026029 | 0.039497 | 0.329457757 | 3214.381541 | 4039.002226 | 53 | 2 | 3 | 2 | 8 | 1 | 0 | 0 | 0 | 0 | 0 | 0 |
| JSG000000 | Mcat | 0.027158 | 0.041086 | -0.361626145 | 757.4365579 | 589.5029001 | 8 | 2 | 3 | 3 | 6 | 0 | 0 | 0 | 0 | 0 | 0 | 0 |
| JSG000000 | Neu3 | 0.027672 | 0.041825 | 0.510186283 | 122.874367 | 175.0018615 | 26 | 2 | 3 | 4 | 8 | 3 | 6 | 1 | 1 | 1 | 1 | 0 |
| SG0000002 | Nup35 | 0.029271 | 0.044023 | 0.398352049 | 333.1746312 | 439.1246749 | 53 | 1 | 3 | 5 | 17 | 4 | 2 | 0 | 0 | 0 | 0 | 0 |
| SG0000004 | Mtfr1l | 0.029665 | 0.04457 | -0.320495832 | 1629.136891 | 1304.604041 | 48 | 3 | 3 | 6 | 15 | 0 | 6 | 0 | 0 | 0 | 3 | 0 |
| SG0000004 | Ublcp1 | 0.032064 | 0.047889 | -0.333694924 | 1121.839222 | 890.1812415 | 33 | 2 | 3 | 3 | 13 | 4 | 1 | 0 | 0 | 0 | 0 | 1 |
| SG0000004 | Ppp1r13l | 0.032331 | 0.048247 | -0.613479644 | 794.1644349 | 519.0804347 | 16 | 2 | 3 | 3 | 7 | 2 | 0 | 0 | 0 | 0 | 0 | 0 |
| JSG000000 | Ppp1r3b | 0.032658 | 0.04871 | -0.671637463 | 758.215239 | 476.0027767 | 25 | 3 | 3 | 9 | 12 | 2 | 4 | 0 | 0 | 2 | 4 | 0 |
| SG0000004 | Zfp740 | 0.032697 | 0.048766 | 0.273010914 | 2695.742602 | 3257.33853 | 38 | 3 | 3 | 3 | 21 | 0 | 15 | 0 | 0 | 0 | 0 | 0 |
| SG0000002 | Nudt9 | 0.032953 | 0.049118 | 0.308437779 | 965.3542225 | 1195.461857 | 31 | 2 | 3 | 8 | 15 | 4 | 2 | 0 | 0 | 0 | 0 | 0 |
| JSG000000 | Zbtb5 | 0.033057 | 0.049261 | 0.371959071 | 322.2832764 | 417.0696694 | 66 | 3 | 3 | 15 | 69 | 6 | 6 | 0 | 0 | 0 | 0 | 0 |
| JSG000000 | Zfp955a | 0.034352 | 0.051003 | 0.470566551 | 148.7175113 | 206.0709417 | 23 | 2 | 3 | 6 | 30 | 5 | 0 | 0 | 0 | 0 | 0 | 0 |
| JSG000000 | Tstd3 | 0.034818 | 0.051632 | -0.372647901 | 605.9638371 | 468.0244171 | 18 | 1 | 3 | 3 | 8 | 0 | 0 | 0 | 0 | 0 | 0 | 0 |
| SG0000004 | Dnajc21 | 0.035987 | 0.053224 | -0.290043054 | 4125.884295 | 3374.468552 | 34 | 1 | 3 | 3 | 16 | 3 | 0 | 0 | 0 | 0 | 2 | 0 |
| SG0000002 | Gpatch4 | 0.040262 | 0.05905 | -0.343636286 | 981.1750473 | 773.217483 | 54 | 0 | 3 | 3 | 24 | 0 | 0 | 0 | 0 | 0 | 0 | 0 |
| SG0000006 | Zfp868 | 0.043176 | 0.06305 | 0.339577718 | 403.7035478 | 510.8408809 | 38 | 3 | 3 | 4 | 11 | 4 | 2 | 0 | 0 | 0 | 0 | 0 |
| JSG000000 | Tmc7 | 0.043208 | 0.063081 | 0.451693252 | 184.5539465 | 252.4041866 | 67 | 2 | 3 | 13 | 24 | 3 | 3 | 1 | 1 | 2 | 3 | 0 |
| SG0000001 | Tvp23b | 0.04326 | 0.063142 | -0.319565194 | 972.9846937 | 779.6636724 | 30 | 2 | 3 | 6 | 10 | 1 | 3 | 0 | 1 | 1 | 2 | 0 |
| SG0000001 | Fkbp8 | 0.045269 | 0.065869 | -0.5430725 | 5190.045598 | 3561.968391 | 12 | 0 | 3 | 0 | 0 | 0 | 0 | 0 | 0 | 0 | 0 | 0 |
| SG0000002 | Psmd5 | 0.04752 | 0.068932 | 0.256082272 | 2409.153676 | 2877.086698 | 44 | 2 | 3 | 7 | 18 | 5 | 4 | 0 | 0 | 0 | 4 | 3 |
| SG0000002 | Net1 | 0.048564 | 0.070293 | 0.265298246 | 1348.622548 | 1620.888519 | 80 | 0 | 3 | 14 | 30 | 11 | 7 | 0 | 2 | 2 | 4 | 1 |
| SG0000001 | Csf1 | 0.048588 | 0.070322 | -0.82568845 | 2402.286983 | 1355.401298 | 52 | 0 | 3 | 5 | 18 | 3 | 8 | 0 | 0 | 2 | 6 | 0 |
| JSG000000 | Zc2hc1c | 0.04917 | 0.071102 | -0.625142191 | 89.79994085 | 58.222319 | 22 | 1 | 3 | 4 | 9 | 0 | 4 | 0 | 0 | 1 | 1 | 0 |
| SG0000002 | Ube2e1 | 0.049483 | 0.07149 | 0.260839708 | 1678.855116 | 2011.563722 | 66 | 2 | 3 | 8 | 29 | 9 | 2 | 0 | 0 | 0 | 2 | 1 |
| SG0000002 | Eral1 | 0.051269 | 0.073845 | 0.302522079 | 585.2424652 | 721.7786823 | 12 | 3 | 3 | 3 | 11 | 0 | 0 | 0 | 0 | 0 | 4 | 0 |
| JSG000000 | Psmb3 | 0.054071 | 0.07755 | 0.258807933 | 2441.833649 | 2921.62872 | 14 | 3 | 3 | 5 | 17 | 1 | 0 | 0 | 0 | 0 | 0 | 0 |
| JSG000000 | Chmp7 | 0.055144 | 0.078956 | 0.262310579 | 1400.623198 | 1679.904786 | 20 | 2 | 3 | 3 | 8 | 3 | 2 | 0 | 1 | 0 | 0 | 1 |
| SG0000002 | Appl2 | 0.057372 | 0.08189 | -0.409050925 | 971.2461179 | 731.4635641 | 54 | 0 | 3 | 3 | 9 | 8 | 5 | 0 | 0 | 1 | 1 | 0 |
| JSG000000 | Spg21 | 0.06333 | 0.089673 | 0.25371444 | 1485.383759 | 1770.982737 | 41 | 0 | 3 | 4 | 14 | 2 | 1 | 0 | 0 | 0 | 0 | 0 |
| JSG000000 | B230219D22Rik | 0.064964 | 0.091828 | -0.252344483 | 3819.119673 | 3206.269389 | 27 | 3 | 3 | 5 | 10 | 2 | 16 | 3 | 3 | 4 | 8 | 0 |
| JSG000000 | Csnk2a2 | 0.070175 | 0.098578 | 0.235293879 | 2100.45186 | 2472.539555 | 79 | 2 | 3 | 9 | 34 | 8 | 2 | 0 | 0 | 0 | 3 | 1 |
| SG0000005 | Klhl25 | 0.074164 | 0.103694 | 0.323664991 | 361.7493181 | 452.7313741 | 15 | 3 | 3 | 3 | 13 | 0 | 0 | 0 | 0 | 0 | 1 | 0 |
| JSG000000 | Rsph3b | 0.076895 | 0.107153 | 1.716162044 | 4.951424079 | 16.2684575 | 55 | 1 | 3 | 3 | 14 | 4 | 1 | 0 | 0 | 0 | 0 | 0 |
| SG0000002 | Slc6a17 | 0.078523 | 0.109226 | 1.176944487 | 18.92870222 | 42.79730611 | 167 | 0 | 3 | 18 | 69 | 14 | 7 | 0 | 0 | 0 | 2 | 0 |
| JSG000000 | Cyp2c44 | 0.08007 | 0.111153 | 1.157373143 | 8.912732088 | 19.87993153 | 42 | 3 | 3 | 7 | 19 | 8 | 1 | 0 | 0 | 0 | 0 | 0 |
| SG0000004 | Cir1 | 0.083498 | 0.115564 | 0.34873918 | 194.9774834 | 248.2935366 | 66 | 2 | 3 | 9 | 29 | 1 | 5 | 0 | 0 | 1 | 1 | 0 |
| SG0000002 | Commd9 | 0.083817 | 0.11596 | -0.318566927 | 379.7682508 | 304.5232481 | 20 | 3 | 3 | 4 | 13 | 3 | 0 | 0 | 0 | 0 | 0 | 0 |
| JSG000000 | Otog | 0.0865 | 0.119376 | -1.183277032 | 20.56745604 | 9.056881759 | 54 | 1 | 3 | 5 | 23 | 8 | 1 | 0 | 0 | 0 | 0 | 0 |
| JSG000000 | 1700066M21Rik | 0.088454 | 0.121829 | 0.281732181 | 429.0075955 | 521.5246241 | 22 | 1 | 3 | 7 | 12 | 0 | 6 | 0 | 1 | 3 | 3 | 0 |
| SG0000003 | Ino80e | 0.093028 | 0.127512 | 0.250054399 | 1409.333261 | 1676.052339 | 22 | 0 | 3 | 6 | 12 | 3 | 9 | 0 | 3 | 0 | 0 | 0 |
| SG0000005 | Zfp68 | 0.094443 | 0.129272 | -0.273330106 | 890.9923972 | 737.2137687 | 108 | 3 | 3 | 13 | 52 | 13 | 9 | 0 | 0 | 1 | 13 | 1 |
| SG0000002 | Dtd2 | 0.095145 | 0.130144 | 0.268390501 | 532.7851644 | 641.7202064 | 57 | 1 | 3 | 11 | 18 | 2 | 3 | 0 | 0 | 1 | 1 | 0 |
| SG0000003 | Egr2 | 0.098771 | 0.13456 | 0.745111305 | 32.3175503 | 54.16756166 | 9 | 0 | 3 | 0 | 0 | 3 | 9 | 0 | 3 | 0 | 0 | 3 |
| SG0000004 | Trp53i13 | 0.100936 | 0.137123 | 0.50297058 | 98.57280933 | 139.6903372 | 8 | 3 | 3 | 5 | 9 | 0 | 0 | 0 | 0 | 0 | 0 | 0 |
| SG0000003 | Tmem184c | 0.109771 | 0.148139 | -0.238389877 | 1448.531948 | 1227.907281 | 36 | 1 | 3 | 3 | 13 | 0 | 4 | 0 | 0 | 0 | 0 | 0 |
| JSG000000 | Timm8a1 | 0.109888 | 0.148272 | 0.334255054 | 183.1738623 | 230.9320975 | 10 | 3 | 3 | 4 | 4 | 0 | 0 | 0 | 0 | 0 | 0 | 0 |
| SG0000003 | Chst12 | 0.115277 | 0.15474 | -0.397906912 | 570.3145364 | 432.8451197 | 24 | 2 | 3 | 7 | 14 | 1 | 1 | 0 | 0 | 0 | 0 | 0 |
| SG0000004 | Prox2 | 0.115567 | 0.155094 | -0.978670133 | 23.78241243 | 12.06832064 | 30 | 3 | 3 | 5 | 18 | 0 | 0 | 0 | 0 | 0 | 0 | 0 |
| JSG000000 | Prl8a9 | 0.116464 | 0.156122 | -2.862360772 | 26.64202581 | 3.663623144 | 16 | 2 | 3 | 5 | 17 | 0 | 0 | 0 | 0 | 0 | 0 | 0 |
| JSG000000 | Lin7c | 0.117424 | 0.157326 | -0.214039733 | 4330.069717 | 3733.038843 | 26 | 0 | 3 | 3 | 10 | 2 | 14 | 0 | 3 | 3 | 4 | 0 |
| SG0000003 | Dbndd1 | 0.125295 | 0.166948 | 0.944105457 | 11.13757241 | 21.42863967 | 15 | 3 | 3 | 6 | 9 | 0 | 2 | 0 | 0 | 2 | 2 | 0 |
| JSG000000 | Murc | 0.130778 | 0.173596 | 0.60489282 | 30.99448646 | 47.13845326 | 23 | 2 | 3 | 5 | 13 | 1 | 2 | 0 | 0 | 0 | 1 | 0 |
| JSG000000 | Efna1 | 0.142049 | 0.187074 | -0.290678933 | 335.4273632 | 274.2176535 | 31 | 0 | 3 | 3 | 9 | 6 | 0 | 0 | 0 | 0 | 0 | 3 |
| JSG000000 | Mrm1 | 0.152951 | 0.200107 | 0.249684731 | 368.2895063 | 437.8768027 | 13 | 2 | 3 | 5 | 6 | 0 | 2 | 0 | 0 | 0 | 0 | 0 |
| JSG000000 | Slc30a5 | 0.163481 | 0.212718 | -0.204311775 | 1657.305471 | 1438.462655 | 77 | 2 | 3 | 10 | 36 | 8 | 4 | 0 | 0 | 1 | 1 | 2 |
| JSG000000 | Dpm3 | 0.164147 | 0.213461 | -1.043247314 | 220.29203 | 106.8931932 | 22 | 2 | 3 | 4 | 9 | 0 | 0 | 0 | 0 | 0 | 0 | 0 |
| JSG000000 | Hoxc4 | 0.171858 | 0.222518 | 0.831522065 | 11.4629865 | 20.39909331 | 41 | 0 | 3 | 7 | 26 | 2 | 8 | 0 | 2 | 4 | 4 | 0 |
| SG0000003 | Slc41a3 | 0.18128 | 0.233385 | -0.373610989 | 133.7541136 | 103.2378694 | 59 | 1 | 3 | 9 | 21 | 7 | 2 | 0 | 0 | 0 | 0 | 0 |
| SG0000002 | Tspan18 | 0.187492 | 0.240536 | 0.583514408 | 33.75119585 | 50.57600305 | 110 | 2 | 3 | 12 | 30 | 7 | 2 | 0 | 0 | 0 | 0 | 0 |
| JSG000000 | Ptgr1 | 0.187977 | 0.241088 | -0.198518029 | 1685.384615 | 1468.720456 | 35 | 2 | 3 | 4 | 11 | 2 | 1 | 0 | 0 | 0 | 0 | 0 |
| JSG000000 | Dr1 | 0.188662 | 0.241915 | 0.174728528 | 2016.255846 | 2275.852792 | 37 | 1 | 3 | 3 | 10 | 2 | 11 | 0 | 0 | 0 | 2 | 0 |
| JSG000000 | Fam174b | 0.189843 | 0.243307 | 0.807803018 | 22.99106452 | 40.24686152 | 47 | 3 | 3 | 8 | 31 | 5 | 3 | 0 | 0 | 1 | 1 | 1 |
| SG0000002 | Irf5 | 0.194418 | 0.248655 | 0.464849236 | 45.08300242 | 62.22234792 | 44 | 3 | 3 | 10 | 16 | 1 | 4 | 0 | 0 | 0 | 0 | 0 |
| SG0000002 | Oasl2 | 0.195444 | 0.249767 | 0.543441834 | 962.0738768 | 1402.170143 | 26 | 1 | 3 | 2 | 6 | 2 | 1 | 0 | 0 | 0 | 0 | 0 |
| SG0000004 | Smim7 | 0.195454 | 0.249767 | -0.185199311 | 1983.918003 | 1744.910559 | 9 | 3 | 3 | 3 | 10 | 0 | 3 | 3 | 3 | 3 | 6 | 0 |
| JSG000000 | Trnp1 | 0.196022 | 0.25042 | 0.446350444 | 46.08867974 | 62.79992709 | 10 | 2 | 3 | 3 | 5 | 0 | 1 | 0 | 0 | 0 | 0 | 0 |
| SG000000C | Kmt2b | 0.197091 | 0.251571 | -0.174064691 | 2903.054003 | 2573.098558 | 35 | 3 | 3 | 3 | 14 | 6 | 3 | 0 | 0 | 0 | 0 | 3 |
| SG0000003 | Jdp2 | 0.21787 | 0.27497 | -0.37017911 | 144.5321497 | 111.8225641 | 99 | 0 | 3 | 18 | 30 | 3 | 0 | 0 | 0 | 0 | 0 | 0 |
| JSG000000 | Nfkbil1 | 0.21815 | 0.275254 | -0.45178099 | 221.2959012 | 161.7982204 | 24 | 3 | 3 | 7 | 16 | 2 | 0 | 0 | 0 | 0 | 0 | 0 |
| JSG000000 | Rbx1 | 0.22256 | 0.280245 | -0.169078558 | 2018.473052 | 1795.251251 | 21 | 3 | 3 | 4 | 6 | 2 | 1 | 0 | 0 | 1 | 1 | 0 |
| SG0000006 | Lmtk3 | 0.230477 | 0.289319 | -0.302723148 | 195.5895113 | 158.5684619 | 24 | 0 | 3 | 0 | 2 | 0 | 1 | 0 | 1 | 0 | 0 | 0 |
| SG0000002 | Nub1 | 0.236321 | 0.295662 | -0.164046602 | 4610.193471 | 4114.681306 | 24 | 2 | 3 | 4 | 11 | 1 | 1 | 0 | 0 | 0 | 0 | 0 |
| JSG000000 | Tchh | 0.23948 | 0.299236 | 0.525551196 | 60.55248992 | 87.1643067 | 8 | 2 | 3 | 2 | 5 | 2 | 3 | 0 | 0 | 0 | 1 | 0 |
| SG0000006 | Rplp0 | 0.242216 | 0.302277 | -0.167364487 | 51131.1393 | 45530.63832 | 10 | 0 | 3 | 2 | 3 | 0 | 2 | 0 | 0 | 0 | 0 | 0 |
| SG0000002 | Lrwd1 | 0.24587 | 0.306346 | 0.259659725 | 1107.563911 | 1325.971416 | 13 | 2 | 3 | 4 | 4 | 0 | 0 | 0 | 0 | 0 | 0 | 0 |
| SG0000002 | Zfp12 | 0.248037 | 0.308723 | -0.212313516 | 418.8993328 | 361.573782 | 45 | 3 | 3 | 6 | 9 | 12 | 10 | 0 | 0 | 0 | 0 | 3 |
| JSG000000 | Dnase1l3 | 0.251979 | 0.312913 | 0.6320548 | 22.53546737 | 34.92480766 | 33 | 3 | 3 | 4 | 11 | 3 | 2 | 0 | 0 | 0 | 0 | 0 |
| SG0000002 | Srsf9 | 0.258503 | 0.319971 | 0.204535133 | 1959.88134 | 2258.40064 | 7 | 1 | 3 | 3 | 12 | 0 | 0 | 0 | 0 | 0 | 6 | 0 |
| SG0000002 | Prpf38b | 0.260479 | 0.322172 | 0.188476787 | 3168.727094 | 3610.954397 | 58 | 3 | 3 | 6 | 18 | 15 | 9 | 0 | 0 | 0 | 0 | 4 |
| JSG000000 | Drg2 | 0.263081 | 0.324919 | -0.151372706 | 2093.330474 | 1884.820822 | 9 | 2 | 3 | 3 | 5 | 0 | 0 | 0 | 0 | 0 | 0 | 0 |
| SG0000002 | Tmem165 | 0.269985 | 0.332461 | -0.155406645 | 2505.758951 | 2249.868985 | 48 | 1 | 3 | 5 | 17 | 7 | 0 | 0 | 0 | 0 | 1 | 2 |
| SG0000002 | Tpm2 | 0.272633 | 0.33535 | -0.196321577 | 1052.021303 | 918.175818 | 21 | 0 | 3 | 3 | 6 | 0 | 0 | 0 | 0 | 0 | 0 | 0 |
| SG0000002 | Rplp2 | 0.273231 | 0.335904 | 0.440974657 | 7958.578148 | 10803.93803 | 9 | 3 | 3 | 3 | 6 | 6 | 0 | 0 | 0 | 0 | 0 | 0 |
| SG0000005 | Nrm | 0.274771 | 0.337657 | -0.175934145 | 799.797323 | 707.9759275 | 8 | 3 | 3 | 3 | 6 | 1 | 1 | 0 | 0 | 0 | 0 | 0 |
| SG0000002 | Elac2 | 0.280757 | 0.34407 | -0.154191916 | 1571.562217 | 1412.26172 | 78 | 3 | 3 | 6 | 21 | 6 | 0 | 0 | 0 | 0 | 0 | 0 |
| SG0000001 | Nek8 | 0.287003 | 0.350836 | -0.203592459 | 378.9601045 | 329.0834597 | 12 | 3 | 3 | 5 | 8 | 0 | 0 | 0 | 0 | 0 | 0 | 0 |
| JSG000000 | Phf11c | 0.302664 | 0.367647 | -0.397693131 | 60.4505828 | 45.88628843 | 21 | 0 | 3 | 2 | 9 | 3 | 1 | 0 | 0 | 0 | 0 | 0 |
| SG0000002 | Ggt7 | 0.304776 | 0.369836 | -0.377506785 | 62.93958909 | 48.44880139 | 31 | 3 | 3 | 5 | 19 | 3 | 0 | 0 | 0 | 0 | 0 | 0 |
| SG000000C | Grin2d | 0.323276 | 0.389593 | 0.435001879 | 28.58544236 | 38.64501954 | 32 | 3 | 3 | 10 | 17 | 6 | 0 | 0 | 0 | 0 | 0 | 0 |
| SG0000002 | Nop16 | 0.323773 | 0.390165 | -0.146086367 | 1623.733223 | 1467.365481 | 27 | 3 | 3 | 6 | 13 | 0 | 0 | 0 | 0 | 0 | 1 | 0 |
| SG0000001 | Tnfsfm13 | 0.326819 | 0.393502 | 0.647351013 | 10.54430207 | 16.51543099 | 27 | 0 | 3 | 0 | 24 | 6 | 0 | 0 | 0 | 0 | 0 | 0 |
| JSG000000 | Zfp382 | 0.328222 | 0.395022 | 0.261932889 | 96.18166095 | 115.3299036 | 24 | 3 | 3 | 5 | 8 | 2 | 0 | 0 | 0 | 0 | 0 | 0 |
| JSG000000 | Alkbh5 | 0.334184 | 0.401012 | 0.114321848 | 6905.672624 | 7475.156532 | 33 | 2 | 3 | 6 | 22 | 4 | 5 | 0 | 1 | 0 | 0 | 1 |
| SG0000002 | Mrps24 | 0.335112 | 0.401857 | 0.143702766 | 663.381486 | 732.8619769 | 6 | 1 | 3 | 2 | 3 | 0 | 0 | 0 | 0 | 0 | 0 | 0 |
| JSG000000 | Hs6st1 | 0.342593 | 0.409894 | -0.1226551 | 4370.044112 | 4013.866803 | 33 | 2 | 3 | 4 | 10 | 5 | 5 | 1 | 2 | 2 | 2 | 0 |
| SG000000C | Tbx4 | 0.342972 | 0.410211 | -0.401272107 | 45.58224916 | 34.5144383 | 60 | 0 | 3 | 9 | 26 | 10 | 6 | 0 | 0 | 3 | 9 | 0 |
| JSG000000 | Fam96a | 0.357029 | 0.425169 | -0.150663478 | 1016.885943 | 916.0475508 | 18 | 3 | 3 | 4 | 13 | 2 | 1 | 0 | 0 | 0 | 1 | 0 |
| SG0000002 | Ftsj3 | 0.357986 | 0.426091 | 0.117494247 | 4642.687941 | 5036.615366 | 7 | 2 | 3 | 2 | 3 | 0 | 0 | 0 | 0 | 0 | 0 | 0 |
| JSG000000 | Aco1 | 0.367982 | 0.436249 | 0.120206673 | 2054.077865 | 2232.557829 | 89 | 2 | 3 | 11 | 30 | 8 | 0 | 0 | 0 | 0 | 0 | 1 |
| SG0000002 | Supt4a | 0.380359 | 0.449124 | 0.13076606 | 787.7431034 | 862.480165 | 15 | 3 | 3 | 9 | 27 | 0 | 0 | 0 | 0 | 0 | 0 | 0 |
| JSG000000 | Rtcb | 0.383312 | 0.452149 | -0.117683653 | 4781.263258 | 4406.729037 | 42 | 1 | 3 | 5 | 14 | 1 | 0 | 0 | 0 | 0 | 0 | 0 |
| JSG000000 | Oat | 0.385564 | 0.454297 | 0.11580268 | 1768.102677 | 1915.876727 | 37 | 2 | 3 | 6 | 17 | 3 | 3 | 0 | 0 | 0 | 2 | 0 |
| JSG000000 | Lysmd3 | 0.385798 | 0.454513 | -0.174478959 | 427.6574402 | 378.941903 | 27 | 2 | 3 | 2 | 8 | 2 | 2 | 0 | 0 | 0 | 1 | 0 |
| SG0000005 | Zfp191 | 0.396378 | 0.465233 | 0.117901735 | 1133.527796 | 1230.053874 | 41 | 3 | 3 | 5 | 17 | 2 | 17 | 1 | 1 | 1 | 9 | 0 |
| JSG000000 | Zfp35 | 0.415481 | 0.48489 | 0.129657051 | 822.3203413 | 899.6460857 | 26 | 3 | 3 | 6 | 16 | 6 | 1 | 0 | 0 | 0 | 1 | 0 |
| SG0000003 | Bphl | 0.416295 | 0.485619 | -0.172685732 | 279.2927844 | 247.785638 | 49 | 2 | 3 | 4 | 21 | 2 | 1 | 0 | 0 | 0 | 0 | 0 |
| JSG000000 | Rnaseh1 | 0.4195 | 0.488594 | -0.143468849 | 503.4582106 | 455.8007138 | 17 | 3 | 3 | 4 | 8 | 1 | 2 | 1 | 1 | 1 | 1 | 0 |
| JSG000000 | Fam222a | 0.4288 | 0.498113 | 0.30276265 | 77.64597015 | 95.77663176 | 31 | 3 | 3 | 5 | 9 | 2 | 2 | 0 | 0 | 1 | 1 | 0 |
| JSG000000 | Uxt | 0.43754 | 0.506771 | -0.162606321 | 293.1243182 | 261.8800829 | 28 | 1 | 3 | 4 | 10 | 5 | 1 | 0 | 0 | 0 | 0 | 0 |
| SG0000002 | Wrap73 | 0.446924 | 0.515905 | 0.133657343 | 352.7308355 | 386.97089 | 12 | 2 | 3 | 3 | 5 | 1 | 0 | 0 | 0 | 0 | 0 | 1 |
| JSG000000 | Prl2c2 | 0.456486 | 0.526028 | 0.379272712 | 29.35957604 | 38.18758927 | 31 | 2 | 3 | 5 | 17 | 3 | 1 | 0 | 0 | 0 | 0 | 0 |

| | | | | | | | | | | | | | | | | | | |
|---|---|---|---|---|---|---|---|---|---|---|---|---|---|---|---|---|---|---|
| JSG000000 | Zfp51 | 0.467646 | 0.536576 | 0.119920104 | 386.1271822 | 419.5946326 | 38 | 3 | 3 | 7 | 24 | 1 | 1 | 0 | 0 | 1 | 1 | 0 |
| SG0000005 | Rap2a | 0.470554 | 0.539359 | 0.113133288 | 731.2405014 | 790.891214 | 38 | 2 | 3 | 4 | 15 | 3 | 4 | 0 | 1 | 0 | 2 | 0 |
| JSG000000 | Abhd17c | 0.486417 | 0.554702 | 0.096621774 | 1340.626568 | 1433.48738 | 66 | 2 | 3 | 8 | 24 | 6 | 6 | 0 | 0 | 0 | 0 | 1 |
| SG0000002 | Uqcc2 | 0.497047 | 0.564452 | 0.1157404 | 966.5142588 | 1047.248139 | 21 | 0 | 3 | 3 | 9 | 3 | 0 | 0 | 0 | 0 | 0 | 0 |
| SG0000002 | Scamp3 | 0.497427 | 0.564777 | -0.113394963 | 1537.027004 | 1420.843393 | 6 | 0 | 3 | 0 | 9 | 0 | 3 | 0 | 3 | 0 | 0 | 0 |
| SG0000002 | Vps51 | 0.515154 | 0.582063 | -0.142104714 | 617.6471706 | 559.7094957 | 14 | 0 | 3 | 3 | 5 | 2 | 2 | 0 | 1 | 1 | 1 | 0 |
| SG0000004 | Rfesd | 0.520975 | 0.587972 | -0.149802805 | 264.3031389 | 238.2358873 | 34 | 0 | 3 | 6 | 19 | 3 | 5 | 0 | 0 | 0 | 1 | 0 |
| SG0000007 | Ang | 0.549351 | 0.614964 | -0.332118953 | 29.78151043 | 23.65750568 | 21 | 2 | 3 | 4 | 10 | 0 | 0 | 0 | 0 | 0 | 0 | 0 |
| SG0000002 | Pithd1 | 0.566049 | 0.630579 | 0.090711743 | 787.028574 | 838.1031168 | 16 | 3 | 3 | 3 | 13 | 1 | 0 | 0 | 0 | 0 | 0 | 0 |
| SG0000002 | Ugdh | 0.574531 | 0.638949 | -0.083891146 | 2060.654536 | 1944.247002 | 32 | 3 | 3 | 5 | 17 | 8 | 1 | 0 | 0 | 0 | 2 | 0 |
| JSG000000 | Zcchc24 | 0.578744 | 0.642762 | -0.088050534 | 709.6944595 | 667.6756884 | 36 | 3 | 3 | 11 | 28 | 4 | 4 | 0 | 0 | 3 | 7 | 1 |
| SG0000004 | Ttll13 | 0.592414 | 0.655424 | -0.321978156 | 28.68046282 | 22.9435741 | 13 | 3 | 3 | 5 | 11 | 2 | 0 | 0 | 0 | 0 | 0 | 0 |
| JSG000000 | Tfap4 | 0.602556 | 0.665371 | 0.105936792 | 556.0158752 | 598.3803778 | 18 | 2 | 3 | 3 | 9 | 1 | 1 | 0 | 0 | 0 | 1 | 0 |
| SG0000002 | Clk4 | 0.612323 | 0.6747 | -0.103085704 | 1427.18389 | 1328.764589 | 147 | 3 | 3 | 4 | 76 | 13 | 0 | 0 | 0 | 0 | 0 | 3 |
| SG0000003 | Zfp551 | 0.61884 | 0.680834 | -0.185385422 | 63.92955202 | 56.22054981 | 15 | 0 | 3 | 0 | 11 | 3 | 2 | 0 | 0 | 0 | 0 | 0 |
| JSG000000 | Samm50 | 0.629398 | 0.690499 | 0.061748289 | 3527.642593 | 3681.905729 | 19 | 2 | 3 | 4 | 8 | 0 | 0 | 0 | 0 | 0 | 0 | 0 |
| SG0000000 | Metrn | 0.647287 | 0.706792 | -0.215403355 | 253.3863547 | 218.2430149 | 4 | 3 | 3 | 3 | 5 | 0 | 0 | 0 | 0 | 0 | 0 | 0 |
| SG0000004 | Tsen2 | 0.668829 | 0.726461 | -0.074805034 | 534.7295778 | 507.7099188 | 47 | 0 | 3 | 10 | 12 | 0 | 0 | 0 | 0 | 0 | 0 | 0 |
| JSG000000 | Dync2li1 | 0.67573 | 0.732846 | 0.067687359 | 542.282037 | 568.3307363 | 48 | 0 | 3 | 3 | 18 | 10 | 0 | 0 | 0 | 0 | 0 | 0 |
| SG0000002 | Scrib | 0.682743 | 0.739058 | -0.098859659 | 2502.664855 | 2336.915301 | 30 | 3 | 3 | 9 | 9 | 0 | 0 | 0 | 0 | 0 | 0 | 0 |
| JSG000000 | Zfp385a | 0.694299 | 0.748728 | -0.105729216 | 959.2613473 | 891.4752283 | 24 | 3 | 3 | 5 | 12 | 3 | 0 | 0 | 0 | 0 | 0 | 0 |
| JSG000000 | Selk | 0.698769 | 0.752863 | -0.065209474 | 880.0770033 | 841.1833448 | 16 | 3 | 3 | 3 | 7 | 1 | 2 | 0 | 0 | 0 | 0 | 1 |
| JSG000000 | Med18 | 0.698893 | 0.752951 | -0.085504629 | 353.0369981 | 332.721422 | 11 | 1 | 3 | 4 | 7 | 0 | 1 | 1 | 1 | 1 | 1 | 0 |
| JSG000000 | Prkra | 0.718918 | 0.771232 | -0.086785557 | 514.2706226 | 484.2466893 | 38 | 1 | 3 | 3 | 9 | 5 | 1 | 0 | 0 | 0 | 0 | 0 |
| JSG000000 | Spef1 | 0.735968 | 0.78618 | -0.190902266 | 32.69130111 | 28.63946478 | 29 | 0 | 3 | 3 | 9 | 5 | 2 | 0 | 0 | 0 | 2 | 0 |
| SG0000001 | Prpsap1 | 0.737999 | 0.787645 | 0.043309091 | 2081.570406 | 2145.005649 | 24 | 1 | 3 | 5 | 10 | 0 | 0 | 0 | 0 | 0 | 0 | 0 |
| JSG000000 | Kdm8 | 0.743499 | 0.792287 | 0.091225362 | 125.3550395 | 133.537539 | 14 | 0 | 3 | 3 | 10 | 0 | 0 | 0 | 0 | 1 | 0 | 0 |
| JSG000000 | Tagap | 0.759236 | 0.806448 | -0.184019737 | 28.03182506 | 24.6749291 | 12 | 1 | 3 | 2 | 6 | 1 | 1 | 1 | 1 | 1 | 3 | 0 |
| SG0000002 | Asl | 0.762521 | 0.809092 | -0.05117464 | 897.1328333 | 865.8679205 | 46 | 3 | 3 | 6 | 12 | 0 | 0 | 0 | 0 | 0 | 0 | 0 |
| SG0000002 | Lsg1 | 0.784945 | 0.828726 | -0.040888752 | 1940.19603 | 1885.97908 | 43 | 3 | 3 | 6 | 19 | 3 | 2 | 0 | 0 | 0 | 0 | 0 |
| SG0000004 | Rexo1 | 0.785274 | 0.828913 | 0.057212633 | 1188.26741 | 1236.337109 | 15 | 2 | 3 | 2 | 7 | 5 | 2 | 0 | 0 | 0 | 2 | 2 |
| JSG000000 | Calcoco1 | 0.789943 | 0.832739 | 0.051026946 | 966.4376462 | 1001.231444 | 17 | 2 | 3 | 5 | 17 | 2 | 3 | 0 | 0 | 1 | 1 | 0 |
| JSG000000 | Tmem35 | 0.791787 | 0.834215 | -0.161591304 | 25.23196907 | 22.55834914 | 19 | 1 | 3 | 4 | 13 | 0 | 2 | 0 | 0 | 1 | 1 | 0 |
| JSG000000 | Troap | 0.806991 | 0.847095 | -0.033650936 | 564.4898865 | 551.4755017 | 4 | 3 | 3 | 3 | 6 | 3 | 1 | 1 | 1 | 1 | 1 | 0 |
| JSG000000 | Map1s | 0.818638 | 0.85721 | -0.126988809 | 922.9420764 | 845.1757058 | 13 | 2 | 3 | 3 | 8 | 0 | 0 | 0 | 0 | 0 | 0 | 0 |
| SG0000005 | Txnl4a | 0.877118 | 0.905581 | -0.029852577 | 1132.726331 | 1109.528533 | 27 | 3 | 3 | 8 | 15 | 3 | 0 | 0 | 0 | 0 | 0 | 0 |
| JSG000000 | AU022252 | 0.895553 | 0.920248 | -0.028435531 | 746.6374297 | 732.06528 | 26 | 3 | 3 | 5 | 11 | 0 | 10 | 1 | 1 | 3 | 3 | 0 |
| JSG000000 | Faim2 | 0.896661 | 0.921192 | 0.145380759 | 43.75110175 | 48.38970375 | 15 | 2 | 3 | 5 | 17 | 2 | 1 | 0 | 0 | 0 | 0 | 1 |
| JSG000000 | Lta4h | 0.906202 | 0.929449 | -0.025787586 | 2201.511901 | 2162.51038 | 48 | 2 | 3 | 4 | 17 | 4 | 0 | 0 | 0 | 0 | 0 | 0 |
| SG0000000 | Stk11 | 0.914306 | 0.936741 | -0.018095712 | 1926.663159 | 1902.647965 | 25 | 0 | 3 | 6 | 12 | 3 | 2 | 0 | 0 | 0 | 0 | 0 |
| SG0000002 | Ercc3 | 0.922691 | 0.943898 | -0.017482555 | 1515.235542 | 1496.984748 | 31 | 0 | 3 | 3 | 13 | 7 | 0 | 0 | 0 | 0 | 0 | 1 |
| JSG000000 | Cecr5 | 0.964469 | 0.978619 | -0.015717594 | 571.3059901 | 565.1156183 | 18 | 2 | 3 | 2 | 6 | 2 | 0 | 0 | 0 | 0 | 0 | 0 |
| JSG000000 | Zfp174 | 0.975101 | 0.98745 | -0.02577559 | 149.6453732 | 146.9955095 | 21 | 3 | 3 | 6 | 14 | 0 | 10 | 1 | 1 | 2 | 2 | 0 |
| JSG000000 | Cmtr2 | 0.985963 | 0.996533 | 0.005578183 | 484.0117443 | 485.8867992 | 16 | 2 | 3 | 3 | 5 | 0 | 6 | 1 | 2 | 2 | 2 | 0 |
| SG0000007 | Zfp558 | 1 | 1 | -0.036553687 | 37.13494791 | 36.20587627 | 90 | 3 | 3 | 8 | 39 | 6 | 1 | 0 | 0 | 0 | 0 | 0 |
| SG0000003 | Tdgf1 | 0 | 0 | 13.65189891 | 1.939079263 | 24958.96846 | 17 | 2 | 2 | 2 | 16 | 2 | 1 | 0 | 0 | 0 | 4 | 0 |
| JSG000000 | Triml1 | 6.7E-302 | 4.6E-299 | 10.6371873 | 0 | 1591.621031 | 20 | 1 | 2 | 2 | 11 | 1 | 0 | 0 | 0 | 0 | 0 | 0 |
| JSG000000 | Dppa3 | 1.2E-248 | 5.4E-246 | 10.14664377 | 0 | 1132.558999 | 10 | 2 | 2 | 4 | 8 | 4 | 2 | 0 | 0 | 0 | 0 | 2 |
| SG0000002 | Alox8 | 3.7E-244 | 1.6E-241 | -11.4312746 | 14737.7984 | 5.336740277 | 17 | 2 | 2 | 6 | 15 | 0 | 0 | 0 | 0 | 0 | 1 | 0 |
| SG0000004 | Camsap3 | 4E-236 | 1.6E-233 | 5.428187914 | 49.39297259 | 2126.730934 | 22 | 2 | 2 | 4 | 34 | 4 | 0 | 0 | 0 | 0 | 0 | 0 |
| JSG000000 | Gli1 | 3.6E-234 | 1.4E-231 | 7.108269185 | 9.853164735 | 1359.496161 | 13 | 2 | 2 | 2 | 5 | 1 | 0 | 0 | 0 | 0 | 0 | 0 |
| SG0000004 | Pnma2 | 5.1E-187 | 1.3E-184 | 7.462050729 | 4.942893535 | 871.5300279 | 15 | 0 | 2 | 0 | 0 | 2 | 5 | 0 | 2 | 0 | 0 | 0 |
| JSG000000 | Prdm14 | 2.2E-178 | 5.1E-176 | 10.10455764 | 0.649984458 | 715.6124949 | 21 | 2 | 2 | 2 | 6 | 1 | 1 | 0 | 0 | 0 | 0 | 0 |
| SG0000002 | Arhgef16 | 5.9E-163 | 1.1E-160 | 9.90393902 | 0.674076249 | 645.7906382 | 23 | 2 | 2 | 4 | 7 | 2 | 0 | 0 | 0 | 0 | 0 | 0 |
| SG0000003 | Alpk3 | 3.3E-121 | 3.3E-119 | 9.447247103 | 0.639110347 | 446.1496974 | 42 | 2 | 2 | 6 | 21 | 3 | 1 | 0 | 0 | 0 | 0 | 0 |
| SG0000000 | Acap1 | 5.7E-119 | 5.6E-117 | -5.13590962 | 6769.912855 | 192.5396721 | 13 | 1 | 2 | 1 | 5 | 2 | 0 | 0 | 0 | 0 | 0 | 0 |
| JSG000000 | Trpv3 | 1.3E-118 | 1.2E-116 | -7.875913347 | 1160.056888 | 4.938478035 | 43 | 2 | 2 | 7 | 23 | 7 | 6 | 0 | 0 | 1 | 3 | 0 |
| SG0000001 | C1qtnf1 | 5.9E-103 | 4.4E-101 | -5.08540296 | 2530.99169 | 74.54729484 | 24 | 2 | 2 | 4 | 10 | 5 | 0 | 0 | 0 | 0 | 0 | 0 |
| SG0000006 | Rnf183 | 6.63E-95 | 4.17E-93 | -8.646529872 | 918.61217 | 2.29227831 | 28 | 2 | 2 | 4 | 16 | 2 | 0 | 0 | 0 | 0 | 0 | 0 |
| SG0000000 | Axl | 7.2E-91 | 4.16E-89 | -3.959311561 | 10644.69311 | 684.3237345 | 40 | 2 | 2 | 6 | 16 | 4 | 0 | 0 | 0 | 0 | 0 | 0 |
| JSG000000 | Nup133 | 6.77E-89 | 3.72E-87 | 2.439283052 | 1104.623927 | 5991.172342 | 60 | 2 | 2 | 8 | 28 | 14 | 4 | 0 | 0 | 1 | 1 | 3 |
| SG0000000 | Cers4 | 2.72E-86 | 1.41E-84 | 4.260083661 | 48.48585286 | 929.0262924 | 95 | 0 | 2 | 3 | 19 | 5 | 14 | 0 | 0 | 1 | 3 | 1 |
| SG0000004 | Mcm3 | 7.64E-86 | 3.89E-84 | 2.277504351 | 3941.077804 | 19107.86364 | 32 | 1 | 2 | 4 | 8 | 0 | 2 | 0 | 0 | 0 | 0 | 0 |
| SG0000004 | Dusp18 | 1.13E-83 | 5.57E-82 | -3.950304223 | 3931.593274 | 254.3364002 | 8 | 1 | 2 | 3 | 3 | 1 | 6 | 1 | 2 | 3 | 3 | 1 |
| SG0000002 | Tfap2a | 7.57E-80 | 3.4E-78 | -5.421750349 | 1096.695983 | 25.58449026 | 83 | 0 | 2 | 8 | 18 | 6 | 16 | 0 | 0 | 6 | 6 | 4 |
| JSG000000 | Dnah1 | 1.32E-79 | 5.89E-78 | 6.184787094 | 4.255599608 | 309.5759342 | 25 | 0 | 2 | 4 | 11 | 4 | 0 | 0 | 0 | 0 | 0 | 0 |
| SG0000004 | Emp3 | 3.5E-79 | 1.54E-77 | -5.321619815 | 3883.94507 | 97.11937731 | 7 | 2 | 2 | 2 | 4 | 0 | 1 | 0 | 0 | 0 | 0 | 0 |
| SG0000003 | Ptger4 | 1.37E-76 | 5.8E-75 | -4.548572181 | 1189.591465 | 50.83246562 | 42 | 2 | 2 | 8 | 12 | 6 | 10 | 0 | 0 | 4 | 6 | 0 |
| SG0000005 | Gm13242 | 2.18E-75 | 9.01E-74 | 9.686884416 | 0.307509278 | 253.4550784 | 67 | 2 | 2 | 8 | 45 | 14 | 5 | 0 | 0 | 0 | 3 | 1 |
| JSG000000 | Csrp2 | 1.3E-74 | 5.31E-73 | 2.332258602 | 703.894436 | 3544.764023 | 28 | 2 | 2 | 6 | 16 | 2 | 0 | 0 | 0 | 0 | 0 | 0 |
| SG0000003 | Amt | 2.09E-70 | 7.56E-69 | 7.726338191 | 11.08554541 | 2347.559549 | 5 | 2 | 2 | 2 | 6 | 0 | 0 | 0 | 0 | 0 | 0 | 0 |
| JSG000000 | Sptbn2 | 7.52E-70 | 2.65E-68 | 7.222861699 | 8.042078106 | 1201.341037 | 41 | 2 | 2 | 2 | 9 | 3 | 1 | 0 | 0 | 0 | 0 | 0 |
| JSG000000 | Plekhf2 | 1.22E-66 | 4.04E-65 | 2.154164089 | 927.8185327 | 4129.8193 | 31 | 1 | 2 | 4 | 15 | 4 | 7 | 0 | 0 | 1 | 1 | 0 |
| JSG000000 | Gnl3 | 1.25E-64 | 3.91E-63 | 2.000095024 | 2840.764134 | 11363.805 | 8 | 2 | 2 | 2 | 4 | 1 | 0 | 0 | 0 | 0 | 0 | 1 |
| JSG000000 | Tigit | 9.07E-63 | 2.72E-61 | -8.147940055 | 282.6444951 | 0 | 26 | 2 | 2 | 6 | 20 | 6 | 0 | 0 | 0 | 0 | 0 | 0 |
| SG0000003 | Plk5 | 7.98E-60 | 2.2E-58 | 7.363719908 | 1.289094805 | 212.3173231 | 6 | 0 | 2 | 2 | 2 | 0 | 2 | 0 | 2 | 2 | 2 | 0 |
| JSG000000 | Nudt11 | 1.08E-58 | 2.86E-57 | 6.958299896 | 1.596604083 | 198.5428369 | 20 | 1 | 2 | 1 | 3 | 3 | 6 | 0 | 0 | 0 | 2 | 1 |
| JSG000000 | Zscan18 | 3.06E-58 | 7.99E-57 | 6.898347012 | 3.206425844 | 382.4992356 | 10 | 1 | 2 | 2 | 3 | 1 | 1 | 0 | 0 | 1 | 1 | 0 |
| JSG000000 | Ppp1r1a | 3.97E-58 | 1.03E-56 | 5.077980763 | 7.45115134 | 251.67954 | 8 | 2 | 2 | 2 | 2 | 0 | 0 | 0 | 0 | 0 | 0 | 0 |
| JSG000000 | 2810004N23Rik | 5.62E-58 | 1.45E-56 | 2.117955215 | 572.6599046 | 2485.792228 | 23 | 2 | 2 | 3 | 16 | 2 | 0 | 0 | 0 | 0 | 0 | 0 |
| JSG000000 | Tmem171 | 9.92E-58 | 2.53E-56 | -7.723632556 | 285.6378947 | 1.351359032 | 10 | 1 | 2 | 1 | 6 | 0 | 1 | 0 | 1 | 0 | 1 | 0 |
| JSG000000 | Tor4a | 2.08E-56 | 5.21E-55 | -3.533797583 | 2597.214665 | 224.2481017 | 2 | 1 | 2 | 2 | 2 | 0 | 2 | 1 | 2 | 2 | 2 | 0 |
| SG0000002 | Ogfrl1 | 1.64E-55 | 4.02E-54 | 4.317269545 | 17.127143 | 341.4385522 | 38 | 0 | 2 | 3 | 7 | 5 | 11 | 0 | 1 | 2 | 5 | 2 |
| SG0000003 | Anxa2 | 2.89E-55 | 7.01E-54 | -4.100887785 | 52669.2043 | 3069.492136 | 50 | 2 | 2 | 8 | 21 | 7 | 0 | 0 | 0 | 0 | 0 | 0 |
| SG0000003 | Cd276 | 5.82E-55 | 1.4E-53 | 2.277797109 | 284.4102011 | 1379.210083 | 35 | 0 | 2 | 2 | 10 | 9 | 4 | 0 | 0 | 0 | 2 | 0 |
| JSG000000 | Sohlh2 | 2.08E-54 | 4.89E-53 | 7.33032014 | 0 | 159.9334197 | 40 | 1 | 2 | 4 | 22 | 4 | 0 | 0 | 0 | 0 | 0 | 0 |
| SG0000003 | Igsf1 | 3.6E-54 | 8.4E-53 | 9.067389666 | 0.307509278 | 164.9736167 | 37 | 0 | 2 | 6 | 38 | 2 | 8 | 0 | 0 | 1 | 1 | 0 |
| JSG000000 | Cdc42ep3 | 1.22E-53 | 2.8E-52 | -3.281743616 | 1309.460037 | 134.644613 | 30 | 2 | 2 | 2 | 18 | 3 | 0 | 0 | 0 | 0 | 1 | 0 |
| JSG000000 | 2310030G06Rik | 1.09E-52 | 2.45E-51 | -2.993593199 | 1901.050087 | 238.6888938 | 14 | 1 | 2 | 4 | 7 | 1 | 2 | 0 | 1 | 1 | 1 | 0 |
| JSG000000 | Htr1b | 3.15E-51 | 6.74E-50 | -6.169143789 | 613.0686039 | 8.519453712 | 13 | 1 | 2 | 1 | 3 | 0 | 7 | 1 | 2 | 1 | 3 | 0 |
| JSG000000 | Gnaz | 4.83E-51 | 1.03E-49 | 5.515821619 | 4.303783188 | 196.9147021 | 33 | 0 | 2 | 6 | 30 | 6 | 6 | 0 | 0 | 0 | 4 | 2 |
| JSG000000 | Srxn1 | 9.76E-51 | 2.06E-49 | -2.716930496 | 5812.715235 | 884.1001655 | 7 | 2 | 2 | 2 | 4 | 0 | 2 | 0 | 0 | 0 | 0 | 0 |
| SG0000000 | Trim28 | 7.21E-50 | 1.49E-48 | 1.734529497 | 8497.24216 | 28276.35348 | 5 | 2 | 2 | 3 | 5 | 1 | 1 | 0 | 0 | 0 | 0 | 0 |
| JSG000000 | Acan | 1.42E-49 | 2.91E-48 | 6.091620509 | 2.591407288 | 176.7242598 | 52 | 1 | 2 | 8 | 20 | 4 | 2 | 0 | 0 | 1 | 1 | 0 |
| SG0000005 | Zfp940 | 2.32E-49 | 4.72E-48 | 4.288239215 | 28.17637581 | 550.5208395 | 58 | 2 | 2 | 8 | 22 | 5 | 7 | 1 | 1 | 1 | 2 | 0 |
| JSG000000 | Kif22 | 2.6E-49 | 5.27E-48 | 1.843301545 | 1146.843303 | 4115.220773 | 18 | 1 | 2 | 4 | 8 | 4 | 0 | 0 | 0 | 0 | 0 | 0 |
| SG0000000 | P3h4 | 1.79E-47 | 3.44E-46 | -3.306516567 | 2521.421787 | 254.8501147 | 3 | 1 | 2 | 2 | 14 | 0 | 1 | 0 | 0 | 1 | 1 | 0 |
| SG0000005 | Zfand2a | 3.34E-47 | 6.39E-46 | -2.681821101 | 2184.043834 | 340.3711611 | 22 | 2 | 2 | 4 | 6 | 0 | 2 | 2 | 2 | 2 | 2 | 0 |
| JSG000000 | Tmem71 | 4.69E-46 | 8.51E-45 | -7.529368425 | 183.742044 | 0 | 88 | 0 | 2 | 4 | 22 | 11 | 2 | 0 | 0 | 0 | 2 | 1 |
| SG0000002 | Slc2a5 | 4.73E-46 | 8.57E-45 | 7.781809394 | 0.649984458 | 143.041159 | 21 | 2 | 2 | 4 | 17 | 2 | 0 | 0 | 0 | 0 | 0 | 0 |
| SG0000003 | Lss | 4.34E-45 | 7.63E-44 | 1.875386657 | 623.3942393 | 2287.233071 | 21 | 0 | 2 | 2 | 11 | 3 | 2 | 0 | 1 | 0 | 1 | 0 |
| SG0000002 | Pla2g1b | 1.41E-44 | 2.44E-43 | 7.85128346 | 0.639110347 | 147.5868537 | 28 | 2 | 2 | 2 | 21 | 7 | 0 | 0 | 0 | 0 | 0 | 0 |
| SG0000004 | Trim7 | 2.78E-44 | 4.75E-43 | -3.571099295 | 1063.802064 | 89.50613901 | 66 | 0 | 2 | 10 | 29 | 3 | 7 | 0 | 0 | 0 | 3 | 1 |
| JSG000000 | Arhgap8 | 3.46E-44 | 5.89E-43 | 6.528855377 | 4.54370423 | 419.5573749 | 43 | 2 | 2 | 9 | 27 | 1 | 0 | 0 | 0 | 0 | 0 | 0 |
| SG0000001 | Cxcl16 | 3.52E-44 | 5.99E-43 | -2.86169183 | 2322.117683 | 319.469312 | 15 | 2 | 2 | 4 | 8 | 0 | 2 | 0 | 0 | 0 | 0 | 0 |
| JSG000000 | Svip | 1.92E-42 | 3.07E-41 | 3.650013438 | 20.63973142 | 259.1000942 | 24 | 1 | 2 | 3 | 12 | 0 | 9 | 0 | 0 | 1 | 5 | 0 |
| SG0000002 | Prmt5 | 1.94E-41 | 3.02E-40 | 1.617113401 | 2627.097078 | 8058.899901 | 18 | 2 | 2 | 5 | 8 | 3 | 1 | 0 | 0 | 1 | 1 | 0 |
| JSG000000 | Kcnn4 | 2.42E-41 | 3.76E-40 | -4.435496101 | 1097.107249 | 50.70277306 | 9 | 2 | 2 | 2 | 3 | 0 | 1 | 0 | 0 | 0 | 0 | 0 |
| JSG000000 | Cfap20 | 2.78E-41 | 4.3E-40 | 1.819392672 | 574.2249928 | 2026.627163 | 21 | 1 | 2 | 1 | 6 | 1 | 1 | 0 | 0 | 0 | 1 | 0 |
| SG0000004 | Nup43 | 5.05E-41 | 7.75E-40 | 1.759165564 | 711.0192677 | 2406.816453 | 28 | 1 | 2 | 2 | 9 | 2 | 0 | 0 | 0 | 0 | 0 | 0 |
| SG0000007 | Etnk2 | 9.52E-41 | 1.45E-39 | 3.653601947 | 19.60527517 | 246.7270242 | 16 | 2 | 2 | 2 | 8 | 2 | 0 | 0 | 0 | 0 | 0 | 0 |
| JSG000000 | Fam212b | 3.22E-40 | 4.79E-39 | -3.382778881 | 534.1705076 | 51.21085119 | 31 | 2 | 2 | 6 | 8 | 3 | 5 | 2 | 2 | 2 | 2 | 1 |
| JSG000000 | Pdlim3 | 2.87E-39 | 4.13E-38 | 6.810578298 | 0 | 111.2505184 | 69 | 1 | 2 | 5 | 37 | 2 | 2 | 0 | 0 | 0 | 1 | 1 |
| SG0000003 | Mturn | 5.24E-38 | 7.23E-37 | 2.739421268 | 57.30705806 | 382.6979064 | 56 | 0 | 2 | 4 | 13 | 6 | 7 | 0 | 1 | 0 | 1 | 0 |
| JSG000001 | Pcdhgc3 | 5.5E-38 | 7.57E-37 | -3.048785172 | 1564.194325 | 189.0231163 | 58 | 2 | 2 | 14 | 26 | 8 | 4 | 0 | 0 | 2 | 2 | 0 |
| SG0000000 | Cacnb3 | 6.91E-38 | 9.47E-37 | -2.734307304 | 4479.605111 | 673.1800168 | 16 | 2 | 2 | 2 | 4 | 0 | 0 | 0 | 0 | 0 | 0 | 0 |
| SG0000005 | Sarm1 | 8.26E-38 | 1.13E-36 | 5.701765275 | 2.589063721 | 134.7550704 | 34 | 2 | 2 | 2 | 16 | 2 | 2 | 0 | 0 | 0 | 0 | 0 |
| JSG000000 | Ccdc160 | 1.33E-37 | 1.81E-36 | 10.07013534 | 0.34247518 | 368.1644929 | 13 | 1 | 2 | 2 | 4 | 1 | 1 | 0 | 0 | 1 | 1 | 0 |
| SG0000002 | Aifm3 | 1.68E-37 | 2.27E-36 | 4.620616113 | 6.587681198 | 162.060294 | 20 | 2 | 2 | 4 | 22 | 4 | 0 | 0 | 0 | 0 | 0 | 0 |
| SG0000002 | Efemp2 | 1.79E-37 | 2.41E-36 | -2.281822662 | 2185.587958 | 449.4394837 | 7 | 2 | 2 | 3 | 3 | 0 | 0 | 0 | 0 | 0 | 0 | 0 |
| SG0000004 | Cltb | 3.86E-37 | 5.16E-36 | -2.317262263 | 3643.289316 | 731.0182363 | 37 | 2 | 2 | 4 | 16 | 0 | 2 | 2 | 2 | 2 | 2 | 0 |
| JSG000000 | Efcab10 | 1.55E-36 | 2.02E-35 | 6.69159376 | 0 | 102.3642697 | 15 | 2 | 2 | 4 | 10 | 4 | 0 | 0 | 0 | 0 | 0 | 0 |
| JSG000000 | Sgms2 | 2.75E-36 | 3.57E-35 | -2.496539454 | 1145.440115 | 202.9734005 | 41 | 0 | 2 | 4 | 13 | 7 | 9 | 0 | 1 | 2 | 5 | 1 |

| | | | | | | | | | | | | | | | | | | |
|---|---|---|---|---|---|---|---|---|---|---|---|---|---|---|---|---|---|---|
| ISG000000 | Pnpla3 | 1.38E-35 | 1.76E-34 | -3.40012794 | 633.1418937 | 59.9736598 | 15 | 2 | 2 | 4 | 7 | 3 | 0 | 0 | 0 | 0 | 0 | 0 |
| SG0000006 | Sepp1 | 2.07E-35 | 2.62E-34 | 2.372087552 | 101.7532937 | 526.7662982 | 64 | 0 | 2 | 4 | 26 | 2 | 8 | 0 | 2 | 0 | 0 | 0 |
| ISG000000 | Naa11 | 2.3E-35 | 2.9E-34 | 6.643248666 | 0 | 98.95789851 | 12 | 0 | 2 | 2 | 4 | 2 | 5 | 0 | 1 | 2 | 3 | 2 |
| SG0000002 | Esd | 5.41E-35 | 6.73E-34 | -1.890610472 | 18358.77935 | 4951.232723 | 213 | 0 | 2 | 37 | 92 | 13 | 1 | 0 | 0 | 0 | 0 | 1 |
| ISG000000 | Abcc8 | 9.5E-35 | 1.17E-33 | 4.393488968 | 7.806844199 | 164.0769133 | 69 | 1 | 2 | 8 | 32 | 7 | 0 | 0 | 0 | 0 | 0 | 0 |
| SG0000002 | Dhx16 | 1.22E-34 | 1.5E-33 | 1.534702205 | 1378.837585 | 3994.88711 | 15 | 2 | 2 | 4 | 12 | 17 | 0 | 0 | 0 | 0 | 0 | 0 |
| SG0000002 | Dmrtb1 | 1.56E-34 | 1.91E-33 | 6.609430175 | 0 | 96.64201581 | 18 | 2 | 2 | 6 | 12 | 0 | 4 | 0 | 0 | 0 | 0 | 0 |
| ISG000000 | Slc25a15 | 5.61E-34 | 6.68E-33 | 2.127437999 | 403.3272131 | 1762.301363 | 34 | 0 | 2 | 4 | 8 | 2 | 2 | 0 | 1 | 0 | 1 | 0 |
| SG0000005 | Fam57b | 9.32E-34 | 1.1E-32 | 6.815239918 | 0.946619625 | 106.6024404 | 17 | 2 | 2 | 2 | 7 | 0 | 4 | 2 | 2 | 2 | 2 | 0 |
| SG0000002 | Hal | 9.79E-34 | 1.15E-32 | 4.275313724 | 8.103479366 | 156.9167131 | 74 | 2 | 2 | 4 | 16 | 12 | 5 | 0 | 0 | 2 | 2 | 0 |
| SG0000004 | Atmin | 1.13E-33 | 1.32E-32 | 1.530494127 | 1181.864649 | 3414.226925 | 20 | 1 | 2 | 2 | 18 | 3 | 2 | 0 | 0 | 0 | 1 | 1 |
| SG0000001 | Chd3 | 1.75E-33 | 2.04E-32 | -1.949870924 | 4661.677711 | 1206.625805 | 19 | 1 | 2 | 2 | 8 | 4 | 0 | 0 | 0 | 0 | 0 | 1 |
| SG0000004 | Mgat2 | 4.45E-33 | 5.1E-32 | -2.004932805 | 2919.6295 | 727.4159664 | 3 | 0 | 2 | 0 | 0 | 0 | 3 | 0 | 2 | 0 | 0 | 0 |
| ISG000000 | Gm13251 | 6.44E-33 | 7.32E-32 | 2.76234374 | 44.97191774 | 305.1333884 | 46 | 2 | 2 | 7 | 19 | 2 | 4 | 0 | 0 | 1 | 4 | 0 |
| SG0000000 | Ilf2 | 1.83E-32 | 2.05E-31 | 1.43558733 | 2490.597111 | 6736.872494 | 22 | 2 | 2 | 5 | 19 | 3 | 1 | 0 | 0 | 0 | 1 | 0 |
| SG0000006 | Kcnc3 | 1.89E-32 | 2.11E-31 | 3.936776559 | 10.81918902 | 165.6847507 | 12 | 2 | 2 | 2 | 6 | 2 | 6 | 0 | 0 | 0 | 2 | 2 |
| SG0000003 | Tmem63b | 1.96E-32 | 2.19E-31 | -2.223278007 | 7321.844085 | 1568.003789 | 28 | 1 | 2 | 4 | 10 | 0 | 3 | 0 | 0 | 0 | 0 | 0 |
| SG0000003 | Ppp1r9b | 8.37E-32 | 9.11E-31 | -2.08266164 | 3517.443811 | 830.3927408 | 2 | 2 | 2 | 2 | 2 | 6 | 0 | 0 | 0 | 0 | 0 | 4 |
| SG0000004 | Zfp2 | 8.84E-32 | 9.61E-31 | 2.282876182 | 91.08718597 | 443.273218 | 81 | 2 | 2 | 5 | 22 | 12 | 9 | 0 | 0 | 0 | 5 | 3 |
| SG0000002 | Aldh2 | 1.24E-31 | 1.34E-30 | -2.863682514 | 27317.64669 | 3753.08974 | 20 | 2 | 2 | 4 | 10 | 1 | 0 | 0 | 0 | 0 | 0 | 0 |
| ISG000000 | Omd | 6.63E-31 | 6.97E-30 | -6.802979448 | 110.6608357 | 0 | 27 | 2 | 2 | 4 | 14 | 0 | 2 | 0 | 0 | 0 | 0 | 0 |
| ISG000000 | Slc6a6 | 7.24E-31 | 7.6E-30 | -1.935286459 | 21670.26618 | 5666.109608 | 67 | 0 | 2 | 9 | 39 | 8 | 1 | 0 | 0 | 0 | 1 | 1 |
| ISG000000 | Tnfaip8l1 | 8.25E-31 | 8.62E-30 | -3.27241509 | 466.6724188 | 48.29665288 | 14 | 2 | 2 | 5 | 6 | 0 | 1 | 0 | 0 | 0 | 0 | 0 |
| SG0000003 | Siglecf | 8.49E-31 | 8.85E-30 | 6.428697351 | 0 | 85.14513101 | 20 | 0 | 2 | 4 | 6 | 2 | 4 | 0 | 0 | 0 | 0 | 2 |
| SG0000004 | Slc26a9 | 1.32E-30 | 1.37E-29 | 6.340477887 | 1.324060707 | 107.2956039 | 35 | 0 | 2 | 2 | 8 | 1 | 2 | 0 | 1 | 0 | 0 | 0 |
| SG0000002 | Necab3 | 1.99E-30 | 2.05E-29 | 3.690585659 | 12.87638366 | 166.2535369 | 16 | 2 | 2 | 4 | 6 | 0 | 2 | 2 | 2 | 2 | 2 | 0 |
| SG0000005 | Set | 2.57E-30 | 2.63E-29 | 1.334382104 | 10937.70488 | 27581.33225 | 26 | 2 | 2 | 4 | 8 | 2 | 4 | 0 | 0 | 0 | 0 | 0 |
| SG0000002 | Sqle | 2.67E-30 | 2.73E-29 | 1.508347966 | 801.1990023 | 2279.283683 | 66 | 2 | 2 | 8 | 28 | 8 | 0 | 0 | 0 | 0 | 0 | 4 |
| ISG000000 | Clic1 | 4.06E-30 | 4.13E-29 | -1.759506305 | 7269.812521 | 2147.13344 | 7 | 2 | 2 | 2 | 2 | 1 | 0 | 0 | 0 | 0 | 0 | 0 |
| SG0000001 | Atp6ap1 | 1.31E-29 | 1.3E-28 | -1.779873265 | 6821.468805 | 1986.472953 | 17 | 2 | 2 | 2 | 7 | 3 | 2 | 2 | 2 | 2 | 4 | 0 |
| ISG000000 | Sde2 | 7.42E-29 | 7.1E-28 | 1.400498227 | 1408.241373 | 3717.654916 | 27 | 1 | 2 | 4 | 13 | 3 | 2 | 0 | 0 | 1 | 5 | 0 |
| ISG000000 | F2rl1 | 2.42E-28 | 2.27E-27 | 3.050479991 | 174.092291 | 1442.332929 | 20 | 0 | 2 | 1 | 7 | 1 | 1 | 0 | 0 | 1 | 3 | 1 |
| ISG000000 | Pou4f1 | 3.46E-28 | 3.23E-27 | 7.941052079 | 0.331601068 | 81.49121522 | 12 | 1 | 2 | 1 | 5 | 1 | 10 | 1 | 2 | 1 | 5 | 1 |
| SG0000003 | Cpne7 | 4.6E-28 | 4.26E-27 | 5.859179202 | 1.620695873 | 94.07846355 | 20 | 2 | 2 | 4 | 7 | 1 | 0 | 0 | 0 | 0 | 0 | 0 |
| SG0000003 | Msl3 | 6.49E-28 | 5.96E-27 | -2.083462179 | 1099.36485 | 259.3923933 | 64 | 0 | 2 | 8 | 38 | 4 | 2 | 0 | 0 | 0 | 2 | 0 |
| SG0000003 | Tlr3 | 6.54E-28 | 6E-27 | -3.584316921 | 351.7910408 | 29.32904134 | 37 | 1 | 2 | 4 | 16 | 1 | 4 | 0 | 1 | 3 | 6 | 0 |
| ISG000000 | Dusp10 | 1.51E-27 | 1.36E-26 | -2.309051163 | 774.8240853 | 156.3541636 | 47 | 0 | 2 | 1 | 16 | 6 | 3 | 0 | 1 | 0 | 1 | 0 |
| ISG000000 | Lect2 | 2.1E-27 | 1.89E-26 | -6.615388751 | 97.04612795 | 0 | 8 | 0 | 2 | 1 | 7 | 0 | 0 | 0 | 0 | 0 | 0 | 0 |
| SG0000003 | Camk1 | 2.78E-27 | 2.48E-26 | -2.094378572 | 1000.707816 | 234.3346548 | 11 | 2 | 2 | 4 | 8 | 0 | 0 | 0 | 0 | 0 | 1 | 0 |
| ISG000000 | Sftpd | 3.17E-27 | 2.82E-26 | -6.610582011 | 133.9238056 | 1.370485067 | 21 | 0 | 2 | 1 | 8 | 1 | 0 | 0 | 0 | 0 | 0 | 0 |
| ISG000000 | 9330159F19Rik | 3.17E-27 | 2.82E-26 | -7.102382288 | 444.2857208 | 3.233197588 | 49 | 0 | 2 | 10 | 25 | 2 | 8 | 0 | 0 | 1 | 5 | 0 |
| ISG000000 | Slc5a3 | 3.88E-27 | 3.44E-26 | -2.104010408 | 1957.895608 | 455.4275389 | 49 | 2 | 2 | 3 | 23 | 6 | 24 | 1 | 1 | 1 | 8 | 4 |
| SG0000003 | Bhlhe41 | 4.3E-27 | 3.8E-26 | 3.636167309 | 26.40743892 | 328.3384271 | 18 | 0 | 2 | 0 | 5 | 3 | 10 | 0 | 0 | 0 | 5 | 1 |
| SG0000004 | Pqlc3 | 5.48E-27 | 4.82E-26 | -2.3705432 | 1844.711362 | 356.7170055 | 28 | 2 | 2 | 4 | 7 | 2 | 5 | 0 | 0 | 0 | 1 | 0 |
| ISG000000 | Snx19 | 8.56E-27 | 7.49E-26 | -1.913943022 | 1474.603064 | 391.3099332 | 44 | 1 | 2 | 5 | 24 | 5 | 3 | 0 | 0 | 1 | 1 | 1 |
| SG0000003 | Ptprf | 1.71E-26 | 1.48E-25 | 1.293067487 | 2850.314331 | 6984.653344 | 60 | 1 | 2 | 2 | 17 | 2 | 3 | 0 | 0 | 1 | 0 | 0 |
| SG0000002 | Tmx1 | 2.1E-26 | 1.81E-25 | -1.650171487 | 5053.088282 | 1609.925467 | 57 | 1 | 2 | 10 | 34 | 3 | 5 | 0 | 1 | 0 | 3 | 0 |
| ISG000000 | Sult4a1 | 2.17E-26 | 1.87E-25 | 3.63132723 | 94.08082581 | 1165.841582 | 17 | 2 | 2 | 4 | 9 | 2 | 0 | 0 | 0 | 0 | 1 | 0 |
| SG0000002 | Slc39a6 | 3.58E-26 | 3.04E-25 | -1.738254929 | 2563.088408 | 768.2396407 | 29 | 2 | 2 | 4 | 11 | 6 | 3 | 0 | 0 | 0 | 0 | 2 |
| SG0000003 | Rbm3 | 5.44E-26 | 4.59E-25 | -1.590574636 | 8745.403049 | 2903.8164 | 33 | 2 | 2 | 8 | 37 | 0 | 15 | 0 | 0 | 4 | 5 | 0 |
| SG0000001 | Ifi35 | 7.66E-26 | 6.43E-25 | -2.506849485 | 1144.793402 | 201.414266 | 26 | 2 | 2 | 8 | 8 | 0 | 0 | 0 | 0 | 0 | 0 | 0 |
| SG0000002 | Cdca8 | 2.24E-25 | 1.84E-24 | 1.329499286 | 1298.535246 | 3263.419517 | 40 | 0 | 2 | 6 | 14 | 8 | 0 | 0 | 0 | 0 | 0 | 0 |
| SG0000004 | Zfp239 | 3.39E-25 | 2.77E-24 | 2.1545344 | 79.5239889 | 354.0606293 | 12 | 2 | 2 | 2 | 4 | 3 | 2 | 0 | 0 | 0 | 0 | 1 |
| SG0000000 | Calm1 | 4.01E-25 | 3.26E-24 | -1.522194798 | 19398.34334 | 6753.646864 | 32 | 0 | 2 | 6 | 18 | 6 | 20 | 0 | 2 | 4 | 12 | 4 |
| ISG000000 | Cpne2 | 1.85E-24 | 1.46E-23 | -2.338956495 | 501.7112096 | 99.16484583 | 42 | 2 | 2 | 6 | 30 | 8 | 0 | 0 | 0 | 0 | 0 | 0 |
| ISG000000 | Chchd10 | 5.37E-24 | 4.13E-23 | 3.940136326 | 193.0878999 | 2963.83676 | 4 | 2 | 2 | 3 | 4 | 0 | 0 | 0 | 0 | 0 | 0 | 0 |
| ISG000000 | Klk13 | 5.64E-24 | 4.33E-23 | 7.700588005 | 0.331601068 | 68.98011198 | 17 | 1 | 2 | 3 | 12 | 5 | 0 | 0 | 0 | 0 | 0 | 0 |
| ISG000000 | Slc7a6os | 6.63E-24 | 5.09E-23 | 1.467378567 | 434.8820849 | 1202.531553 | 24 | 2 | 2 | 4 | 18 | 1 | 0 | 0 | 0 | 0 | 0 | 0 |
| SG0000005 | Msi1 | 1.24E-23 | 9.38E-23 | 5.209201628 | 31.81311345 | 1176.879824 | 41 | 2 | 2 | 18 | 26 | 7 | 3 | 2 | 2 | 2 | 2 | 1 |
| ISG000000 | Rmnd5b | 1.43E-23 | 1.07E-22 | 1.384183472 | 668.1234645 | 1743.963812 | 10 | 2 | 2 | 2 | 8 | 4 | 0 | 0 | 0 | 0 | 0 | 2 |
| SG0000000 | Ndrg2 | 1.9E-23 | 1.42E-22 | 4.364422059 | 90.73214013 | 1868.887281 | 8 | 2 | 2 | 2 | 4 | 0 | 2 | 0 | 0 | 0 | 0 | 0 |
| SG0000004 | H2afv | 2.94E-23 | 2.18E-22 | 1.68882485 | 638.573948 | 2058.72403 | 67 | 0 | 2 | 3 | 19 | 10 | 1 | 0 | 0 | 1 | 3 | 0 |
| ISG000000 | Ifngr2 | 3.78E-23 | 2.79E-22 | -1.93812522 | 847.8123673 | 221.241157 | 27 | 0 | 2 | 2 | 5 | 3 | 5 | 0 | 0 | 0 | 0 | 1 |
| ISG000000 | Pabpc4l | 5.14E-23 | 3.78E-22 | 4.177562637 | 5.582003881 | 101.0094012 | 36 | 2 | 2 | 6 | 14 | 2 | 36 | 2 | 2 | 6 | 14 | 2 |
| SG0000007 | Accsl | 5.5E-23 | 4.03E-22 | 6.060856786 | 0 | 65.75744276 | 15 | 0 | 2 | 1 | 3 | 3 | 1 | 0 | 0 | 0 | 0 | 0 |
| SG0000002 | Bmp4 | 1.02E-22 | 7.38E-22 | 3.363270166 | 52.97334594 | 545.1331698 | 14 | 2 | 2 | 4 | 8 | 0 | 2 | 0 | 0 | 2 | 2 | 0 |
| ISG000000 | Chac2 | 2.12E-22 | 1.52E-21 | -2.022901065 | 643.7955495 | 158.4141865 | 48 | 0 | 2 | 5 | 14 | 12 | 8 | 0 | 0 | 1 | 2 | 0 |
| SG0000003 | Fam167a | 2.82E-22 | 2E-21 | 2.645635799 | 32.35970004 | 202.4977585 | 15 | 0 | 2 | 3 | 8 | 2 | 1 | 0 | 0 | 0 | 1 | 1 |
| SG0000003 | Spsb1 | 3.49E-22 | 2.47E-21 | 1.555131266 | 266.2059583 | 782.2740611 | 68 | 2 | 2 | 9 | 18 | 6 | 4 | 1 | 1 | 2 | 2 | 0 |
| SG0000004 | Oard1 | 4.32E-22 | 3.03E-21 | 1.307328076 | 745.8706766 | 1845.901397 | 8 | 0 | 2 | 0 | 4 | 0 | 0 | 0 | 0 | 0 | 0 | 0 |
| ISG000000 | Tomm20 | 6.77E-22 | 4.73E-21 | 1.179466147 | 2393.696178 | 5421.553099 | 36 | 2 | 2 | 6 | 15 | 2 | 10 | 1 | 1 | 4 | 4 | 1 |
| ISG000000 | Lrrn1 | 1.07E-21 | 7.35E-21 | 5.888534927 | 0 | 58.24144503 | 82 | 1 | 2 | 8 | 32 | 9 | 0 | 0 | 0 | 0 | 7 | 0 |
| SG0000001 | Nedd8 | 1.08E-21 | 7.47E-21 | 1.213402393 | 1487.856276 | 3450.098532 | 30 | 0 | 2 | 6 | 16 | 4 | 0 | 0 | 0 | 0 | 0 | 0 |
| ISG000000 | Dnajc9 | 1.65E-21 | 1.12E-20 | 1.289921907 | 751.1080838 | 1836.570576 | 8 | 0 | 2 | 0 | 1 | 1 | 4 | 0 | 2 | 0 | 1 | 0 |
| SG0000004 | Zfp286 | 1.76E-21 | 1.2E-20 | 3.941322733 | 6.517749395 | 100.12765 | 28 | 0 | 2 | 4 | 13 | 6 | 0 | 0 | 0 | 0 | 5 | 0 |
| SG0000003 | Nucb1 | 2.06E-21 | 1.4E-20 | -1.73527018 | 6069.237245 | 1822.91215 | 10 | 1 | 2 | 2 | 6 | 0 | 1 | 0 | 0 | 0 | 0 | 0 |
| ISG000000 | Ficd | 2.17E-21 | 1.46E-20 | -2.259254698 | 544.1494057 | 113.6618518 | 5 | 1 | 2 | 2 | 4 | 0 | 5 | 1 | 2 | 2 | 4 | 0 |
| ISG000000 | Erbb2 | 3.31E-21 | 2.22E-20 | -1.790331911 | 897.2098394 | 259.3883223 | 11 | 1 | 2 | 4 | 6 | 3 | 0 | 0 | 0 | 0 | 0 | 0 |
| ISG000000 | Irs4 | 4.46E-21 | 2.97E-20 | 5.842359094 | 0 | 56.37534801 | 37 | 0 | 2 | 8 | 21 | 4 | 4 | 0 | 0 | 0 | 1 | 2 |
| SG0000006 | Cry2 | 5.28E-21 | 3.51E-20 | 1.475781814 | 301.8361697 | 839.5100007 | 45 | 0 | 2 | 2 | 8 | 2 | 2 | 0 | 0 | 0 | 0 | 1 |
| ISG000000 | Rspo1 | 8.19E-21 | 5.39E-20 | 6.062515094 | 0.981585527 | 65.60350432 | 22 | 1 | 2 | 3 | 7 | 0 | 0 | 0 | 0 | 0 | 0 | 0 |
| SG0000003 | Cd59a | 1.06E-20 | 6.93E-20 | -2.438353051 | 327.5644518 | 60.43372394 | 44 | 0 | 2 | 4 | 46 | 2 | 2 | 0 | 0 | 0 | 2 | 0 |
| ISG000000 | Klhl23 | 1.09E-20 | 7.16E-20 | 2.568286363 | 30.76543953 | 182.471212 | 26 | 2 | 2 | 3 | 7 | 1 | 7 | 0 | 0 | 0 | 0 | 0 |
| SG0000004 | Rnf31 | 1.59E-20 | 1.03E-19 | 1.261332872 | 763.896571 | 1831.190766 | 16 | 2 | 2 | 4 | 14 | 2 | 0 | 0 | 0 | 0 | 0 | 0 |
| ISG000000 | Plcd1 | 3.98E-20 | 2.51E-19 | 2.836562579 | 101.222968 | 723.0519276 | 16 | 1 | 2 | 3 | 12 | 0 | 0 | 0 | 0 | 0 | 0 | 0 |
| SG0000002 | Sirt3 | 4.51E-20 | 2.84E-19 | 1.556620972 | 210.8021607 | 620.1041853 | 46 | 2 | 2 | 4 | 26 | 4 | 0 | 0 | 0 | 0 | 0 | 0 |
| SG0000002 | Chrna4 | 5.24E-20 | 3.29E-19 | 7.544991331 | 0.307509278 | 57.42842838 | 20 | 0 | 2 | 0 | 1 | 2 | 2 | 0 | 0 | 0 | 0 | 0 |
| SG0000002 | Ppie | 6.02E-20 | 3.76E-19 | 1.315216954 | 506.5364816 | 1260.464108 | 26 | 0 | 2 | 1 | 8 | 2 | 0 | 0 | 0 | 0 | 0 | 0 |
| SG0000000 | Lrp3 | 1.05E-19 | 6.46E-19 | 3.249939237 | 20.33122528 | 193.4161556 | 31 | 2 | 2 | 6 | 8 | 0 | 1 | 0 | 0 | 0 | 0 | 0 |
| SG0000000 | Ap1s1 | 1.84E-19 | 1.12E-18 | -1.391037053 | 3269.748855 | 1246.72457 | 11 | 1 | 2 | 3 | 4 | 2 | 0 | 0 | 0 | 0 | 0 | 1 |
| ISG000000 | Pbxip1 | 5.39E-19 | 3.19E-18 | -1.603969126 | 1629.58228 | 536.0847766 | 9 | 1 | 2 | 1 | 2 | 2 | 1 | 0 | 0 | 0 | 0 | 0 |
| SG0000002 | Artn | 5.95E-19 | 3.52E-18 | -2.600085242 | 242.5540352 | 40.00413229 | 6 | 2 | 2 | 4 | 6 | 0 | 6 | 2 | 2 | 4 | 6 | 0 |
| ISG000000 | Gdf15 | 6.1E-19 | 3.6E-18 | -4.134204126 | 804.4053683 | 45.80949562 | 10 | 0 | 2 | 6 | 8 | 0 | 10 | 0 | 2 | 6 | 8 | 0 |
| SG0000000 | Tnk1 | 8.78E-19 | 5.15E-18 | 7.980083983 | 1.914987472 | 483.515704 | 11 | 0 | 2 | 0 | 2 | 0 | 2 | 0 | 0 | 0 | 0 | 0 |
| ISG000000 | Cks2 | 1.5E-18 | 8.65E-18 | 1.223518385 | 690.1650629 | 1611.642477 | 14 | 1 | 2 | 3 | 4 | 2 | 2 | 0 | 0 | 0 | 0 | 1 |
| ISG000000 | Fam193b | 1.51E-18 | 8.74E-18 | 1.284465269 | 540.9582257 | 1317.729605 | 32 | 2 | 2 | 3 | 9 | 4 | 0 | 0 | 0 | 0 | 0 | 0 |
| SG0000002 | Aldh6a1 | 1.66E-18 | 9.56E-18 | -1.43498666 | 2419.50699 | 894.8553792 | 43 | 1 | 2 | 10 | 20 | 7 | 5 | 0 | 1 | 1 | 3 | 0 |
| ISG000000 | Basp1 | 2.58E-18 | 1.47E-17 | 3.189796659 | 186.1821101 | 1698.8789 | 61 | 0 | 2 | 8 | 38 | 13 | 1 | 0 | 0 | 0 | 0 | 0 |
| ISG000000 | Epha1 | 2.8E-18 | 1.6E-17 | 5.165387289 | 12.47854106 | 447.8167416 | 23 | 1 | 2 | 5 | 11 | 1 | 0 | 0 | 0 | 0 | 0 | 1 |
| ISG000000 | Alg5 | 2.8E-18 | 1.6E-17 | -1.544007335 | 1128.160437 | 386.8818397 | 63 | 2 | 2 | 6 | 22 | 2 | 10 | 2 | 2 | 3 | 4 | 1 |
| SG0000004 | Smtnl2 | 3.66E-18 | 2.07E-17 | 2.768356878 | 296.0218354 | 2016.889969 | 31 | 0 | 2 | 2 | 10 | 4 | 9 | 0 | 0 | 2 | 4 | 0 |
| SG0000003 | Ido1 | 3.71E-18 | 2.1E-17 | -3.301031957 | 141.9179974 | 14.39885103 | 32 | 0 | 2 | 7 | 15 | 6 | 4 | 0 | 0 | 2 | 4 | 0 |
| SG0000003 | C2cd2l | 4.08E-18 | 2.31E-17 | -1.994774355 | 740.7456347 | 185.8583963 | 13 | 1 | 2 | 3 | 4 | 0 | 6 | 1 | 2 | 2 | 2 | 0 |
| SG0000003 | Plekhs1 | 4.52E-18 | 2.55E-17 | -4.413665931 | 94.09603722 | 4.414947034 | 56 | 2 | 2 | 4 | 26 | 14 | 2 | 0 | 0 | 0 | 0 | 2 |
| ISG000000 | Hyls1 | 5.85E-18 | 3.28E-17 | 1.430166658 | 240.7770616 | 648.8408328 | 22 | 2 | 2 | 4 | 9 | 3 | 0 | 0 | 0 | 0 | 0 | 0 |
| SG0000003 | Becn1 | 6.86E-18 | 3.83E-17 | -1.290059055 | 4483.973914 | 1833.650687 | 121 | 2 | 2 | 20 | 65 | 6 | 13 | 2 | 2 | 2 | 5 | 0 |
| ISG000000 | Egln3 | 9.26E-18 | 5.14E-17 | -1.947028898 | 13706.82791 | 3554.862929 | 37 | 1 | 2 | 4 | 11 | 6 | 4 | 1 | 1 | 1 | 2 | 0 |
| SG0000002 | Tmc6 | 9.87E-18 | 5.47E-17 | -1.60876897 | 1448.518908 | 474.9374897 | 34 | 2 | 2 | 2 | 4 | 4 | 2 | 0 | 0 | 0 | 0 | 0 |
| SG0000002 | Sel1l | 1.13E-17 | 6.27E-17 | -1.299687337 | 3455.282715 | 1403.584987 | 69 | 0 | 2 | 8 | 27 | 5 | 7 | 0 | 0 | 1 | 4 | 1 |
| SG0000001 | Tbrg1 | 1.6E-17 | 8.81E-17 | 1.036475696 | 3212.705661 | 6589.936241 | 20 | 2 | 2 | 4 | 12 | 2 | 0 | 0 | 0 | 0 | 0 | 0 |
| SG0000005 | Slc29a4 | 2.09E-17 | 1.14E-16 | 5.881303295 | 6.100655276 | 359.6045606 | 16 | 2 | 2 | 2 | 5 | 2 | 1 | 0 | 0 | 0 | 0 | 0 |
| ISG000000 | Arf3 | 2.56E-17 | 1.39E-16 | -1.225893498 | 8120.58072 | 3471.813386 | 21 | 0 | 2 | 0 | 5 | 4 | 2 | 0 | 0 | 0 | 0 | 1 |
| ISG000000 | 1700028J19Rik | 2.94E-17 | 1.59E-16 | -3.878491077 | 104.5943026 | 7.111576341 | 6 | 1 | 2 | 2 | 3 | 0 | 0 | 0 | 0 | 0 | 0 | 0 |
| ISG000000 | Tspo | 5.28E-17 | 2.82E-16 | -2.80566804 | 2562.088814 | 366.4409012 | 5 | 2 | 2 | 2 | 3 | 0 | 0 | 0 | 0 | 0 | 0 | 0 |
| SG0000002 | Ecm1 | 7.9E-17 | 4.18E-16 | -1.784116375 | 2304.727138 | 669.186192 | 2 | 2 | 2 | 2 | 2 | 2 | 0 | 0 | 0 | 0 | 0 | 0 |
| SG0000002 | Ppid | 1.07E-16 | 5.63E-16 | 1.059495807 | 2981.058692 | 6213.131491 | 48 | 1 | 2 | 11 | 13 | 2 | 1 | 0 | 0 | 0 | 1 | 0 |
| SG0000002 | Anp32b | 1.1E-16 | 5.78E-16 | 0.98969452 | 6266.756054 | 12444.30155 | 28 | 0 | 2 | 2 | 10 | 5 | 0 | 0 | 0 | 0 | 0 | 1 |
| SG0000006 | Dhps | 1.16E-16 | 6.08E-16 | 1.142639729 | 747.7157758 | 1650.841494 | 4 | 2 | 2 | 2 | 4 | 0 | 2 | 2 | 2 | 2 | 3 | 0 |
| SG0000003 | 1110007C09Rik | 1.17E-16 | 6.1E-16 | -2.478681134 | 1209.610456 | 217.0142046 | 21 | 1 | 2 | 6 | 14 | 1 | 1 | 0 | 1 | 1 | 1 | 0 |
| SG0000002 | Rcor2 | 1.37E-16 | 7.16E-16 | 2.181387185 | 550.775944 | 2498.261748 | 6 | 0 | 2 | 2 | 2 | 2 | 0 | 0 | 0 | 0 | 0 | 0 |
| ISG000000 | Scd3 | 1.39E-16 | 7.23E-16 | 4.925583971 | 1.952296942 | 59.33273872 | 59 | 2 | 2 | 6 | 29 | 2 | 1 | 0 | 0 | 0 | 0 | 0 |
| ISG000000 | Psmg2 | 1.39E-16 | 7.25E-16 | 1.189458073 | 570.1052602 | 1300.222289 | 20 | 1 | 2 | 3 | 12 | 2 | 0 | 0 | 0 | 0 | 0 | 0 |
| ISG000000 | Fgf4 | 2.08E-16 | 1.07E-15 | 1.047552192 | 1662.594408 | 3436.615522 | 6 | 0 | 2 | 1 | 1 | 1 | 6 | 0 | 2 | 1 | 1 | 1 |

| | | | | | | | | | | | | | | | | | | |
|---|---|---|---|---|---|---|---|---|---|---|---|---|---|---|---|---|---|---|
| JSG000000 | Flii | 2.27E-16 | 1.16E-15 | -1.217879344 | 4381.719335 | 1883.763346 | 5 | 1 | 2 | 2 | 3 | 0 | 0 | 0 | 0 | 0 | 0 | 0 |
| SG0000004 | Ankrd34b | 2.69E-16 | 1.38E-15 | 7.232399317 | 0.307509278 | 46.24101504 | 23 | 0 | 2 | 3 | 8 | 3 | 6 | 0 | 2 | 1 | 2 | 2 |
| JSG000000 | Ifi204 | 3.4E-16 | 1.73E-15 | -2.816634868 | 343.469926 | 48.75252732 | 61 | 0 | 2 | 6 | 25 | 3 | 0 | 0 | 0 | 0 | 0 | 0 |
| SG0000006 | Fam57a | 4.07E-16 | 2.06E-15 | -1.60745248 | 667.1423982 | 218.9410159 | 31 | 2 | 2 | 6 | 6 | 4 | 1 | 0 | 0 | 0 | 0 | 0 |
| JSG000000 | Lin28a | 4.29E-16 | 2.17E-15 | 12.88929044 | 0.922527834 | 6999.105246 | 23 | 2 | 2 | 2 | 9 | 5 | 5 | 0 | 0 | 0 | 0 | 1 |
| JSG000000 | Cdh15 | 4.56E-16 | 2.3E-15 | 6.228637457 | 0.649984458 | 48.74271083 | 12 | 2 | 2 | 3 | 6 | 0 | 0 | 0 | 0 | 0 | 0 | 0 |
| SG0000003 | BC017643 | 4.94E-16 | 2.48E-15 | -1.374481183 | 1307.382716 | 504.246247 | 4 | 0 | 2 | 0 | 2 | 0 | 0 | 0 | 0 | 0 | 0 | 0 |
| SG0000005 | Cyb5d2 | 5.63E-16 | 2.82E-15 | -2.200593355 | 358.0286217 | 77.88846382 | 46 | 2 | 2 | 8 | 28 | 6 | 0 | 0 | 0 | 0 | 0 | 0 |
| SG0000006 | Ddrgk1 | 6.25E-16 | 3.12E-15 | -1.528476087 | 1036.742784 | 359.3799727 | 10 | 0 | 2 | 0 | 8 | 0 | 0 | 0 | 0 | 0 | 0 | 0 |
| JSG000000 | Gpr179 | 7.72E-16 | 3.84E-15 | 4.019472409 | 6.304263711 | 102.2388918 | 7 | 1 | 2 | 1 | 1 | 0 | 2 | 0 | 0 | 0 | 0 | 0 |
| JSG000000 | Htr5b | 1E-15 | 4.96E-15 | 5.399939034 | 0 | 41.22246884 | 29 | 1 | 2 | 4 | 11 | 1 | 1 | 0 | 0 | 0 | 0 | 0 |
| JSG000000 | Fam118a | 1.09E-15 | 5.4E-15 | 1.248880115 | 345.8353388 | 821.9014465 | 27 | 2 | 2 | 6 | 18 | 1 | 2 | 0 | 0 | 0 | 1 | 0 |
| SG0000002 | Adam11 | 1.15E-15 | 5.68E-15 | 3.339969857 | 9.036881311 | 91.50602495 | 20 | 2 | 2 | 4 | 8 | 6 | 2 | 0 | 0 | 0 | 0 | 2 |
| JSG000000 | Tmem189 | 1.42E-15 | 6.95E-15 | -1.273578997 | 1882.793892 | 778.784503 | 12 | 1 | 2 | 2 | 4 | 2 | 3 | 1 | 1 | 2 | 3 | 2 |
| SG0000000 | Clcnkb | 1.95E-15 | 9.49E-15 | 7.159158755 | 0.34247518 | 48.94976729 | 12 | 0 | 2 | 4 | 14 | 2 | 0 | 0 | 0 | 0 | 0 | 0 |
| SG0000006 | Pwwp2b | 2.09E-15 | 1.02E-14 | -1.657779067 | 888.8038533 | 281.6856372 | 12 | 2 | 2 | 2 | 6 | 2 | 0 | 0 | 0 | 0 | 0 | 0 |
| JSG000000 | Dmrt3 | 2.47E-15 | 1.2E-14 | 5.359626082 | 0 | 40.05898566 | 21 | 0 | 2 | 1 | 6 | 1 | 2 | 0 | 1 | 0 | 0 | 0 |
| JSG000000 | Actr1a | 3.12E-15 | 1.5E-14 | -1.1502033 | 5007.149714 | 2256.030064 | 22 | 2 | 2 | 2 | 7 | 1 | 1 | 1 | 1 | 1 | 1 | 0 |
| JSG000000 | Matn3 | 3.72E-15 | 1.78E-14 | 5.360393924 | 6.240518884 | 256.365784 | 45 | 1 | 2 | 8 | 19 | 4 | 7 | 0 | 0 | 2 | 6 | 0 |
| JSG000000 | Cyct | 3.93E-15 | 1.87E-14 | 7.110522658 | 0.34247518 | 47.32707974 | 17 | 0 | 2 | 1 | 7 | 1 | 1 | 0 | 0 | 0 | 0 | 0 |
| SG0000002 | Cxcl1 | 4.11E-15 | 1.96E-14 | -1.973006057 | 715.3502386 | 182.2152501 | 9 | 2 | 2 | 6 | 8 | 0 | 7 | 2 | 2 | 6 | 7 | 0 |
| SG0000003 | Fbxl8 | 4.28E-15 | 2.04E-14 | -2.372418041 | 218.7172086 | 42.23903249 | 18 | 0 | 2 | 0 | 2 | 0 | 0 | 0 | 0 | 0 | 0 | 0 |
| SG0000001 | Derl2 | 4.89E-15 | 2.32E-14 | -1.259171236 | 1823.24863 | 761.7239036 | 83 | 2 | 2 | 14 | 48 | 6 | 8 | 1 | 1 | 3 | 6 | 0 |
| SG0000004 | Rbp1 | 5.87E-15 | 2.77E-14 | 2.597269264 | 43.10646055 | 260.8544921 | 19 | 0 | 2 | 2 | 7 | 1 | 0 | 0 | 0 | 0 | 0 | 0 |
| SG0000003 | Usb1 | 6.61E-15 | 3.11E-14 | 1.117280988 | 571.7359956 | 1240.311089 | 19 | 2 | 2 | 4 | 9 | 0 | 2 | 0 | 0 | 1 | 1 | 0 |
| SG0000000 | Kifc2 | 7.32E-15 | 3.43E-14 | 2.010745022 | 46.70853857 | 188.2308678 | 6 | 2 | 2 | 3 | 3 | 1 | 0 | 0 | 0 | 0 | 0 | 0 |
| SG0000003 | Icmt | 7.79E-15 | 3.64E-14 | -1.200556325 | 2373.036203 | 1032.525767 | 20 | 0 | 2 | 2 | 8 | 2 | 12 | 0 | 2 | 2 | 4 | 2 |
| JSG000000 | Tigar | 8.28E-15 | 3.87E-14 | -1.252600875 | 3741.416859 | 1570.23865 | 76 | 0 | 2 | 11 | 42 | 2 | 4 | 0 | 0 | 1 | 14 | 0 |
| SG0000002 | Ybx1 | 8.35E-15 | 3.89E-14 | 0.92028602 | 14846.26083 | 28096.40684 | 32 | 2 | 2 | 4 | 21 | 1 | 0 | 0 | 0 | 0 | 2 | 0 |
| JSG000000 | Ctso | 1.14E-14 | 5.26E-14 | -3.057741968 | 327.6357395 | 39.34768795 | 43 | 0 | 2 | 5 | 23 | 6 | 7 | 0 | 0 | 2 | 5 | 1 |
| SG0000002 | Camk2a | 2.92E-14 | 1.31E-13 | 2.960158872 | 11.6632545 | 90.76457212 | 130 | 0 | 2 | 13 | 43 | 2 | 5 | 0 | 0 | 0 | 1 | 0 |
| JSG000000 | Mcm4 | 3.12E-14 | 1.4E-13 | 0.912726513 | 5396.45428 | 10159.36492 | 34 | 2 | 2 | 7 | 13 | 2 | 0 | 0 | 0 | 0 | 1 | 1 |
| JSG000000 | Gns | 3.19E-14 | 1.43E-13 | -1.125847933 | 3887.425931 | 1781.345368 | 30 | 1 | 2 | 2 | 12 | 1 | 0 | 0 | 0 | 0 | 0 | 0 |
| SG0000006 | Kndc1 | 4.15E-14 | 1.86E-13 | 7.339674111 | 1.265003014 | 204.9055049 | 44 | 2 | 2 | 7 | 16 | 5 | 4 | 0 | 0 | 1 | 1 | 0 |
| SG0000003 | Gins2 | 4.16E-14 | 1.86E-13 | 1.15119803 | 443.3107472 | 984.585415 | 11 | 2 | 2 | 2 | 4 | 0 | 4 | 0 | 0 | 0 | 0 | 0 |
| JSG000000 | Cyp1b1 | 4.93E-14 | 2.19E-13 | -1.534932866 | 1192.055535 | 411.3728778 | 16 | 0 | 2 | 1 | 6 | 1 | 11 | 0 | 2 | 1 | 2 | 1 |
| SG0000003 | Tmem9b | 5.89E-14 | 2.6E-13 | -1.390888753 | 800.2139141 | 305.1454251 | 62 | 0 | 2 | 2 | 16 | 0 | 4 | 0 | 0 | 0 | 0 | 0 |
| SG0000004 | Myh6 | 7.16E-14 | 3.14E-13 | 5.206454727 | 0 | 35.92317544 | 20 | 2 | 2 | 3 | 4 | 3 | 0 | 0 | 0 | 0 | 0 | 0 |
| SG0000003 | Hid1 | 8.15E-14 | 3.57E-13 | 2.122952256 | 40.84546076 | 177.9164514 | 36 | 2 | 2 | 6 | 20 | 4 | 2 | 0 | 0 | 0 | 0 | 0 |
| JSG000000 | Got2 | 9.3E-14 | 4.05E-13 | 0.911591914 | 3529.866011 | 6640.101688 | 28 | 2 | 2 | 4 | 10 | 2 | 0 | 0 | 0 | 0 | 0 | 0 |
| SG0000002 | Dnajc19 | 1E-13 | 4.36E-13 | 1.275985112 | 226.8596849 | 549.3727546 | 92 | 0 | 2 | 5 | 19 | 2 | 7 | 0 | 0 | 1 | 2 | 0 |
| JSG000000 | Klf17 | 1.39E-13 | 5.96E-13 | 5.171758367 | 0 | 35.04577759 | 15 | 2 | 2 | 2 | 11 | 1 | 1 | 0 | 0 | 0 | 2 | 0 |
| SG0000003 | BC030867 | 1.7E-13 | 7.3E-13 | 1.025879415 | 762.0503998 | 1551.687177 | 12 | 2 | 2 | 4 | 8 | 10 | 0 | 0 | 0 | 0 | 0 | 0 |
| JSG000000 | Mrpl14 | 1.9E-13 | 8.13E-13 | -2.280727359 | 1878.80543 | 386.6468351 | 12 | 2 | 2 | 2 | 15 | 1 | 0 | 0 | 0 | 0 | 0 | 0 |
| SG0000007 | Ccdc58 | 2.45E-13 | 1.04E-12 | 1.294872205 | 879.6767209 | 2158.333113 | 81 | 2 | 2 | 4 | 14 | 2 | 1 | 0 | 0 | 0 | 2 | 0 |
| SG0000000 | Cnot11 | 2.7E-13 | 1.14E-12 | 0.964890726 | 1159.78696 | 2263.806343 | 38 | 0 | 2 | 0 | 10 | 6 | 12 | 0 | 2 | 0 | 0 | 2 |
| SG0000003 | Kctd9 | 2.78E-13 | 1.17E-12 | -1.130286258 | 2304.289361 | 1052.657157 | 72 | 1 | 2 | 6 | 31 | 5 | 6 | 0 | 0 | 1 | 4 | 0 |
| JSG000000 | Snap29 | 3.09E-13 | 1.3E-12 | -1.194059199 | 1442.764565 | 630.5910926 | 38 | 2 | 2 | 2 | 14 | 2 | 3 | 0 | 0 | 0 | 0 | 0 |
| JSG000000 | Pura | 4.02E-13 | 1.68E-12 | 0.981991728 | 888.8373701 | 1755.623039 | 57 | 0 | 2 | 4 | 29 | 3 | 2 | 0 | 0 | 0 | 6 | 1 |
| SG0000002 | Mrps5 | 4.9E-13 | 2.03E-12 | 0.975599115 | 958.5736102 | 1884.994407 | 27 | 0 | 2 | 1 | 9 | 3 | 5 | 0 | 1 | 0 | 2 | 0 |
| SG0000005 | Fmnl1 | 5.11E-13 | 2.12E-12 | -1.771366037 | 561.5515179 | 164.4959823 | 30 | 2 | 2 | 4 | 14 | 8 | 0 | 0 | 0 | 0 | 2 | 0 |
| JSG000000 | Spa17 | 5.24E-13 | 2.17E-12 | 2.773568558 | 12.87872723 | 88.06437879 | 31 | 0 | 2 | 0 | 5 | 3 | 0 | 0 | 0 | 0 | 0 | 0 |
| SG0000002 | Ptpn18 | 5.37E-13 | 2.22E-12 | 2.998189504 | 9.883443503 | 78.96838519 | 16 | 1 | 2 | 2 | 10 | 2 | 0 | 0 | 0 | 0 | 0 | 0 |
| JSG000000 | Aadac | 6.17E-13 | 2.54E-12 | -5.577538745 | 113.9902607 | 2.387048719 | 23 | 1 | 2 | 4 | 16 | 2 | 1 | 0 | 0 | 0 | 0 | 0 |
| SG0000001 | Exoc3l2 | 6.6E-13 | 2.72E-12 | 4.176562473 | 2.885698887 | 52.18210511 | 20 | 0 | 2 | 4 | 20 | 0 | 2 | 0 | 0 | 0 | 1 | 0 |
| JSG000000 | Gm1045 | 7.37E-13 | 3.03E-12 | -3.046918946 | 99.68772164 | 12.06223189 | 11 | 0 | 2 | 2 | 6 | 0 | 0 | 0 | 0 | 0 | 0 | 0 |
| SG0000003 | R3hdm4 | 7.58E-13 | 3.11E-12 | -1.090796442 | 2443.910329 | 1147.421065 | 6 | 2 | 2 | 2 | 8 | 2 | 0 | 0 | 0 | 0 | 0 | 0 |
| JSG000000 | Aqp11 | 8.48E-13 | 3.47E-12 | 2.508980531 | 17.96283108 | 102.2476144 | 20 | 1 | 2 | 2 | 5 | 0 | 0 | 0 | 0 | 0 | 0 | 0 |
| SG0000004 | Lppr4 | 9.14E-13 | 3.73E-12 | 6.868143325 | 0.307509278 | 35.92323001 | 92 | 0 | 2 | 11 | 47 | 10 | 6 | 0 | 0 | 1 | 4 | 0 |
| SG0000004 | Bcl2l15 | 1.07E-12 | 4.34E-12 | -5.025788658 | 53.48561366 | 1.641813585 | 44 | 2 | 2 | 7 | 27 | 2 | 6 | 0 | 0 | 0 | 0 | 1 |
| SG0000004 | Col16a1 | 1.18E-12 | 4.78E-12 | -1.762991652 | 301.7398117 | 88.90357914 | 82 | 2 | 2 | 2 | 58 | 6 | 0 | 0 | 0 | 0 | 0 | 0 |
| JSG000000 | Pigw | 1.24E-12 | 5.03E-12 | -1.443477409 | 523.465547 | 192.46781 | 14 | 1 | 2 | 3 | 5 | 0 | 2 | 1 | 2 | 1 | 3 | 0 |
| SG0000002 | Sec11c | 2.1E-12 | 8.37E-12 | -1.326110706 | 698.6265696 | 278.6416276 | 42 | 0 | 2 | 2 | 2 | 2 | 0 | 0 | 0 | 0 | 0 | 0 |
| JSG000000 | Ccdc117 | 2.15E-12 | 8.56E-12 | 0.947908492 | 963.004227 | 1857.706423 | 23 | 1 | 2 | 3 | 11 | 0 | 5 | 0 | 0 | 1 | 2 | 0 |
| JSG000000 | Trim60 | 2.36E-12 | 9.35E-12 | 6.667004327 | 0.34247518 | 34.80145379 | 33 | 2 | 2 | 6 | 13 | 2 | 6 | 0 | 0 | 0 | 0 | 0 |
| SG0000002 | 1810043G02Rik | 2.48E-12 | 9.82E-12 | -1.516580901 | 405.2847873 | 141.6524076 | 9 | 2 | 2 | 4 | 8 | 0 | 0 | 0 | 0 | 0 | 0 | 0 |
| JSG000000 | Hsf2 | 2.73E-12 | 1.08E-11 | 0.951969285 | 914.7139985 | 1769.524725 | 71 | 0 | 2 | 7 | 21 | 8 | 4 | 0 | 0 | 1 | 1 | 1 |
| SG0000002 | Psmg1 | 2.99E-12 | 1.18E-11 | 1.133917543 | 629.8350486 | 1382.197227 | 19 | 1 | 2 | 3 | 14 | 2 | 0 | 0 | 0 | 0 | 1 | 0 |
| SG0000002 | Slc35a5 | 3.23E-12 | 1.27E-11 | -1.291303393 | 742.618292 | 303.420268 | 108 | 0 | 2 | 17 | 39 | 7 | 5 | 0 | 0 | 1 | 5 | 1 |
| SG0000000 | Hap1 | 3.36E-12 | 1.32E-11 | 0.955003615 | 833.8698492 | 1616.527157 | 19 | 2 | 2 | 4 | 10 | 0 | 4 | 0 | 0 | 1 | 1 | 0 |
| SG0000002 | Trp73 | 3.97E-12 | 1.55E-11 | 3.695517144 | 4.268817286 | 55.30559595 | 141 | 1 | 2 | 10 | 33 | 55 | 0 | 0 | 0 | 0 | 0 | 0 |
| JSG000000 | Hook2 | 4.99E-12 | 1.95E-11 | 0.959221028 | 815.3703706 | 1585.291853 | 9 | 2 | 2 | 3 | 10 | 0 | 1 | 0 | 0 | 0 | 0 | 0 |
| SG0000005 | Las1l | 5.27E-12 | 2.05E-11 | 0.867999216 | 2220.735982 | 4053.131829 | 58 | 2 | 2 | 10 | 20 | 10 | 0 | 0 | 0 | 0 | 0 | 0 |
| SG0000004 | Gjb3 | 5.36E-12 | 2.08E-11 | 0.895590807 | 1558.369275 | 2899.145912 | 11 | 2 | 2 | 2 | 16 | 0 | 4 | 0 | 0 | 0 | 0 | 0 |
| SG0000003 | Cercam | 5.8E-12 | 2.25E-11 | 1.164203018 | 269.6316978 | 604.2699199 | 20 | 2 | 2 | 3 | 7 | 2 | 1 | 0 | 0 | 0 | 0 | 0 |
| JSG000000 | Trip13 | 8.22E-12 | 3.16E-11 | 0.948202969 | 788.6383867 | 1521.652375 | 42 | 1 | 2 | 6 | 19 | 5 | 2 | 0 | 0 | 0 | 1 | 1 |
| SG0000003 | AA467197 | 9.2E-12 | 3.52E-11 | -1.574954182 | 368.9659697 | 123.8448244 | 20 | 2 | 2 | 2 | 8 | 0 | 0 | 0 | 0 | 0 | 0 | 0 |
| JSG000000 | Tmprss13 | 1.18E-11 | 4.5E-11 | 4.949425854 | 0 | 29.8976638 | 19 | 1 | 2 | 3 | 15 | 1 | 1 | 0 | 0 | 0 | 3 | 0 |
| JSG000000 | Atp8b3 | 1.85E-11 | 6.95E-11 | 3.343676433 | 7.270287991 | 73.80716373 | 7 | 2 | 2 | 2 | 5 | 0 | 0 | 0 | 0 | 0 | 0 | 0 |
| SG0000000 | Txnrd3 | 2.63E-11 | 9.77E-11 | 1.841208096 | 68.83628609 | 246.6472605 | 70 | 0 | 2 | 0 | 18 | 2 | 0 | 0 | 0 | 0 | 0 | 0 |
| JSG000000 | Bbs4 | 2.8E-11 | 1.04E-10 | 1.093826917 | 292.2740657 | 623.8281858 | 54 | 0 | 2 | 5 | 29 | 5 | 0 | 0 | 0 | 0 | 0 | 0 |
| SG0000002 | Cdc25b | 3.29E-11 | 1.21E-10 | -1.043334263 | 1749.489818 | 848.8608623 | 14 | 0 | 2 | 4 | 8 | 2 | 0 | 0 | 0 | 0 | 0 | 0 |
| SG0000001 | Zfyve27 | 3.45E-11 | 1.27E-10 | -1.054432227 | 1543.151264 | 743.0067687 | 25 | 2 | 2 | 6 | 12 | 3 | 4 | 2 | 2 | 4 | 4 | 0 |
| SG0000002 | Exosc9 | 3.53E-11 | 1.3E-10 | 0.928690406 | 749.1059249 | 1425.958548 | 20 | 2 | 2 | 2 | 10 | 3 | 0 | 0 | 0 | 0 | 0 | 0 |
| SG0000002 | Mcm7 | 3.59E-11 | 1.32E-10 | 0.798626902 | 6038.887979 | 10504.31233 | 8 | 2 | 2 | 2 | 6 | 2 | 0 | 0 | 0 | 0 | 0 | 0 |
| SG0000002 | Zfp362 | 3.63E-11 | 1.33E-10 | -1.403907359 | 421.5059261 | 159.28888 | 56 | 0 | 2 | 6 | 30 | 10 | 6 | 0 | 0 | 2 | 4 | 4 |
| JSG000000 | Gm4924 | 4.15E-11 | 1.52E-10 | 2.469538786 | 31.35651943 | 173.6733175 | 6 | 0 | 2 | 1 | 2 | 0 | 6 | 0 | 2 | 1 | 2 | 0 |
| SG0000004 | Samd9l | 4.41E-11 | 1.61E-10 | -4.679833539 | 2162.99915 | 84.38904633 | 46 | 0 | 2 | 3 | 21 | 6 | 0 | 0 | 0 | 0 | 0 | 0 |
| JSG000000 | Mbd1 | 4.44E-11 | 1.62E-10 | -1.189566886 | 770.3168377 | 337.7334499 | 12 | 1 | 2 | 2 | 7 | 3 | 1 | 0 | 0 | 0 | 0 | 0 |
| JSG000000 | Tob2 | 4.7E-11 | 1.71E-10 | -1.112974369 | 1105.762334 | 511.2379947 | 11 | 1 | 2 | 2 | 6 | 1 | 4 | 0 | 1 | 1 | 2 | 0 |
| SG0000002 | Phf10 | 4.94E-11 | 1.8E-10 | 0.896842075 | 1423.080494 | 2649.75568 | 93 | 0 | 2 | 6 | 49 | 2 | 7 | 0 | 0 | 2 | 5 | 1 |
| JSG000000 | Bbs12 | 5.15E-11 | 1.87E-10 | -1.42040188 | 410.5960793 | 153.4020723 | 38 | 0 | 2 | 0 | 6 | 2 | 0 | 0 | 0 | 0 | 0 | 0 |
| JSG000000 | Trib3 | 6.25E-11 | 2.25E-10 | 1.376653702 | 182.8708525 | 474.8523017 | 5 | 2 | 2 | 3 | 5 | 0 | 0 | 0 | 0 | 0 | 0 | 0 |
| JSG000000 | Hs3st3b1 | 7.99E-11 | 2.86E-10 | 5.843746245 | 5.358990791 | 307.76974 | 57 | 1 | 2 | 3 | 17 | 2 | 5 | 0 | 0 | 0 | 5 | 0 |
| JSG000000 | Gm3636 | 9.44E-11 | 3.37E-10 | 4.365173934 | 1.939079263 | 39.96169052 | 10 | 2 | 2 | 2 | 6 | 1 | 6 | 2 | 2 | 2 | 2 | 0 |
| SG0000003 | Zdhhc13 | 9.65E-11 | 3.44E-10 | -0.966376039 | 2571.106678 | 1315.866775 | 83 | 2 | 2 | 6 | 32 | 7 | 1 | 0 | 0 | 0 | 0 | 0 |
| SG0000000 | Mmp11 | 1.31E-10 | 4.63E-10 | 0.991624456 | 420.7712832 | 836.6711581 | 10 | 2 | 2 | 4 | 4 | 0 | 0 | 0 | 0 | 0 | 0 | 0 |
| SG0000003 | Dnaja4 | 1.33E-10 | 4.71E-10 | -1.644947616 | 399.4296935 | 127.7208421 | 34 | 2 | 2 | 6 | 11 | 2 | 2 | 0 | 0 | 0 | 1 | 0 |
| SG0000002 | Arl14ep | 1.59E-10 | 5.59E-10 | 0.887038434 | 825.3413353 | 1526.366311 | 45 | 0 | 2 | 2 | 6 | 8 | 5 | 0 | 0 | 0 | 0 | 2 |
| JSG000000 | Tgfbi | 1.73E-10 | 6.04E-10 | -1.32508317 | 445.9681333 | 177.9975589 | 25 | 1 | 2 | 2 | 6 | 2 | 0 | 0 | 0 | 0 | 0 | 0 |
| SG0000000 | Irf3 | 1.75E-10 | 6.13E-10 | -1.133869071 | 1059.701139 | 482.8973251 | 8 | 2 | 2 | 2 | 6 | 0 | 0 | 0 | 0 | 0 | 0 | 0 |
| SG0000002 | Wdr73 | 1.94E-10 | 6.75E-10 | 0.918009871 | 644.4948675 | 1217.77757 | 10 | 2 | 2 | 2 | 7 | 1 | 1 | 0 | 0 | 0 | 0 | 0 |
| SG0000002 | Irf6 | 1.95E-10 | 6.81E-10 | 1.736045519 | 322.0358834 | 1072.768571 | 28 | 2 | 2 | 3 | 5 | 4 | 4 | 0 | 0 | 0 | 0 | 2 |
| JSG000000 | Pank3 | 2.15E-10 | 7.48E-10 | -1.040553991 | 2528.756245 | 1229.331485 | 50 | 1 | 2 | 6 | 18 | 3 | 16 | 0 | 0 | 1 | 2 | 0 |
| JSG000000 | Ntrk1 | 2.42E-10 | 8.39E-10 | 6.413750467 | 0.34247518 | 29.19848902 | 14 | 2 | 2 | 4 | 6 | 1 | 1 | 0 | 0 | 1 | 2 | 0 |
| JSG000000 | Col5a3 | 2.89E-10 | 9.93E-10 | -1.658058006 | 892.5544997 | 282.8196297 | 40 | 1 | 2 | 3 | 13 | 2 | 1 | 0 | 0 | 0 | 0 | 0 |
| JSG000000 | Serpine3 | 2.9E-10 | 9.97E-10 | -6.565845989 | 34.10604498 | 0.360010171 | 50 | 0 | 2 | 6 | 25 | 2 | 2 | 0 | 0 | 0 | 1 | 0 |
| JSG000000 | Cisd1 | 3.36E-10 | 1.15E-09 | -0.928363871 | 2722.199014 | 1430.390115 | 21 | 2 | 2 | 6 | 20 | 0 | 1 | 0 | 0 | 0 | 0 | 0 |
| JSG000000 | Zfhx2 | 4.15E-10 | 1.41E-09 | -1.139208304 | 732.1006536 | 332.3800309 | 21 | 2 | 2 | 4 | 15 | 4 | 0 | 0 | 0 | 0 | 0 | 0 |
| JSG000000 | Fbxo32 | 5.71E-10 | 1.92E-09 | -1.754297835 | 312.1662955 | 92.53152951 | 43 | 1 | 2 | 4 | 20 | 8 | 6 | 0 | 0 | 0 | 2 | 4 |
| JSG000000 | Tmem213 | 6.51E-10 | 2.18E-09 | 4.703385963 | 0 | 25.05315097 | 9 | 2 | 2 | 2 | 6 | 2 | 0 | 0 | 0 | 0 | 0 | 0 |
| JSG000000 | Pcyox1l | 7.64E-10 | 2.54E-09 | 1.089966227 | 262.3890043 | 558.5449891 | 15 | 1 | 2 | 1 | 4 | 2 | 0 | 0 | 0 | 0 | 1 | 0 |
| JSG000000 | Psmd6 | 8.37E-10 | 2.78E-09 | 0.768739209 | 2391.701338 | 4074.933327 | 18 | 1 | 2 | 3 | 9 | 2 | 0 | 0 | 0 | 0 | 0 | 0 |
| JSG000000 | Ednra | 8.47E-10 | 2.81E-09 | -2.770837409 | 79.65447824 | 11.67091817 | 87 | 0 | 2 | 7 | 26 | 11 | 0 | 0 | 0 | 0 | 4 | 1 |
| JSG000000 | Zcchc6 | 9.16E-10 | 3.03E-09 | -0.937422888 | 2255.101856 | 1177.53481 | 78 | 0 | 2 | 11 | 32 | 7 | 1 | 0 | 0 | 1 | 1 | 1 |
| JSG000000 | Cfl2 | 9.57E-10 | 3.16E-09 | -0.912212798 | 2631.794739 | 1398.455334 | 16 | 1 | 2 | 5 | 10 | 0 | 12 | 1 | 2 | 4 | 7 | 0 |
| JSG000000 | Utp15 | 1.01E-09 | 3.32E-09 | 0.798186988 | 1413.865211 | 2458.590698 | 28 | 0 | 2 | 0 | 17 | 7 | 6 | 0 | 1 | 0 | 3 | 0 |
| JSG000000 | Chst14 | 1.02E-09 | 3.34E-09 | -1.752796896 | 373.4234812 | 110.8044446 | 6 | 0 | 2 | 0 | 0 | 0 | 6 | 0 | 2 | 0 | 0 | 0 |
| JSG000000 | Pyurf | 1.14E-09 | 3.74E-09 | -1.314156413 | 383.5486954 | 154.2481666 | 11 | 0 | 2 | 3 | 7 | 0 | 10 | 0 | 2 | 3 | 6 | 0 |
| JSG000000 | Cxcl11 | 1.17E-09 | 3.82E-09 | 6.351323863 | 0.331601068 | 27.07415533 | 10 | 0 | 2 | 0 | 4 | 2 | 2 | 0 | 0 | 0 | 2 | 0 |
| JSG000000 | Trim68 | 1.38E-09 | 4.5E-09 | 3.01508587 | 15.17953319 | 122.7127539 | 16 | 2 | 2 | 5 | 12 | 0 | 0 | 0 | 0 | 0 | 0 | 0 |
| SG0000005 | Pgm3 | 1.63E-09 | 5.27E-09 | 0.90965418 | 502.2171808 | 943.462754 | 57 | 0 | 2 | 6 | 30 | 0 | 3 | 0 | 0 | 0 | 0 | 0 |
| SG0000000 | M6pr | 1.71E-09 | 5.52E-09 | -0.838451572 | 7938.081076 | 4439.320059 | 18 | 0 | 2 | 4 | 6 | 2 | 0 | 0 | 0 | 0 | 0 | 0 |

| | | | | | | | | | | | | | | | | | | |
|---|---|---|---|---|---|---|---|---|---|---|---|---|---|---|---|---|---|---|
| JSG000000 | Gcc1 | 2.1E-09 | 6.73E-09 | -1.079261776 | 718.2459679 | 339.9249395 | 11 | 2 | 2 | 2 | 8 | 0 | 3 | 2 | 2 | 2 | 4 | 0 |
| SG0000006 | Htatsf1 | 2.65E-09 | 8.43E-09 | 0.781364627 | 1352.47654 | 2324.576969 | 43 | 2 | 2 | 9 | 14 | 4 | 1 | 0 | 0 | 0 | 0 | 0 |
| JSG000000 | Zfp36l1 | 2.67E-09 | 8.49E-09 | 0.813614108 | 957.1591817 | 1682.311528 | 13 | 1 | 2 | 3 | 6 | 0 | 10 | 1 | 2 | 3 | 6 | 0 |
| SG0000001 | Rcn3 | 2.76E-09 | 8.77E-09 | 2.118238023 | 350.6772742 | 1522.512283 | 7 | 1 | 2 | 2 | 23 | 0 | 0 | 0 | 0 | 0 | 0 | 0 |
| JSG000000 | Enc1 | 2.77E-09 | 8.78E-09 | 1.514897005 | 781.7040328 | 2233.941475 | 19 | 1 | 2 | 6 | 8 | 2 | 10 | 1 | 1 | 5 | 5 | 1 |
| SG0000003 | Zkscan2 | 3.02E-09 | 9.54E-09 | 5.871785995 | 2.248932108 | 131.6922014 | 34 | 0 | 2 | 4 | 20 | 4 | 3 | 0 | 0 | 1 | 1 | 0 |
| JSG000000 | Tgds | 3.08E-09 | 9.73E-09 | 0.871762143 | 572.5346054 | 1047.679137 | 31 | 2 | 2 | 6 | 13 | 4 | 2 | 0 | 0 | 1 | 1 | 0 |
| SG0000003 | Nlgn3 | 3.24E-09 | 1.02E-08 | 4.957839127 | 0.970711415 | 30.16813255 | 40 | 2 | 2 | 2 | 12 | 8 | 8 | 2 | 2 | 2 | 8 | 2 |
| SG0000002 | Alox5 | 3.37E-09 | 1.06E-08 | 3.630133864 | 3.276357648 | 40.56677513 | 65 | 1 | 2 | 11 | 34 | 5 | 2 | 0 | 0 | 1 | 1 | 0 |
| SG0000005 | Armc7 | 3.48E-09 | 1.1E-08 | -1.506307522 | 231.6963776 | 81.55967698 | 14 | 0 | 2 | 2 | 4 | 0 | 0 | 0 | 0 | 0 | 1 | 0 |
| SG0000002 | Ulk1 | 3.82E-09 | 1.2E-08 | 0.882007416 | 2280.977975 | 4203.700635 | 36 | 0 | 2 | 5 | 7 | 4 | 1 | 0 | 0 | 1 | 1 | 0 |
| JSG000000 | Ceacam9 | 3.97E-09 | 1.24E-08 | 4.574050941 | 0 | 22.81916511 | 8 | 0 | 2 | 2 | 7 | 0 | 0 | 0 | 0 | 0 | 0 | 0 |
| SG0000002 | Rgs11 | 4.29E-09 | 1.34E-08 | -1.196373002 | 460.123188 | 200.7843969 | 2 | 2 | 2 | 2 | 2 | 2 | 0 | 0 | 0 | 0 | 0 | 0 |
| JSG000000 | Riok3 | 4.68E-09 | 1.46E-08 | -0.837442209 | 4638.501618 | 2595.867301 | 52 | 1 | 2 | 5 | 19 | 10 | 7 | 0 | 0 | 1 | 3 | 1 |
| SG0000001 | Slc35e1 | 4.77E-09 | 1.49E-08 | -0.856561759 | 2889.719519 | 1595.897186 | 13 | 1 | 2 | 5 | 7 | 0 | 7 | 1 | 1 | 1 | 1 | 0 |
| SG0000002 | Nup85 | 5.11E-09 | 1.59E-08 | 0.733442353 | 2738.673511 | 4553.322481 | 18 | 1 | 2 | 5 | 9 | 3 | 0 | 0 | 0 | 0 | 0 | 1 |
| SG0000001 | Rab5c | 5.16E-09 | 1.61E-08 | -0.899890943 | 5700.848218 | 3055.239862 | 60 | 2 | 2 | 4 | 22 | 12 | 2 | 0 | 0 | 0 | 0 | 0 |
| JSG000000 | Paqr9 | 5.22E-09 | 1.62E-08 | -1.923998569 | 132.9828609 | 35.04405806 | 14 | 1 | 2 | 1 | 5 | 2 | 14 | 1 | 2 | 1 | 5 | 2 |
| SG000000C | Rpl3l | 5.35E-09 | 1.66E-08 | 6.251369109 | 0.331601068 | 25.2618724 | 14 | 2 | 2 | 2 | 4 | 0 | 0 | 0 | 0 | 0 | 0 | 0 |
| SG0000002 | Grin2c | 7.06E-09 | 2.17E-08 | 4.576786086 | 0 | 22.8643657 | 10 | 2 | 2 | 6 | 12 | 4 | 0 | 0 | 0 | 0 | 0 | 0 |
| SG0000003 | Armc10 | 7.11E-09 | 2.19E-08 | 0.774324744 | 1096.512639 | 1875.463499 | 38 | 2 | 2 | 4 | 20 | 7 | 6 | 0 | 0 | 0 | 2 | 1 |
| SG0000002 | Mrpl37 | 8.31E-09 | 2.55E-08 | 0.805379927 | 859.8552969 | 1502.688237 | 12 | 1 | 2 | 5 | 6 | 0 | 0 | 0 | 0 | 0 | 0 | 0 |
| JSG000000 | Selt | 9.11E-09 | 2.79E-08 | -0.808013331 | 5418.050132 | 3094.616021 | 45 | 1 | 2 | 7 | 13 | 6 | 8 | 0 | 0 | 1 | 4 | 1 |
| SG0000002 | Lmbr1l | 1.12E-08 | 3.39E-08 | 0.975975438 | 277.5257316 | 545.8849715 | 8 | 1 | 2 | 1 | 4 | 1 | 0 | 0 | 0 | 0 | 0 | 0 |
| SG0000002 | Arfgap2 | 1.42E-08 | 4.28E-08 | -0.879114958 | 1596.25913 | 867.8874782 | 22 | 0 | 2 | 0 | 2 | 6 | 0 | 0 | 0 | 0 | 0 | 0 |
| JSG000000 | Rundc1 | 1.5E-08 | 4.51E-08 | -1.033191797 | 642.63795 | 314.0108441 | 10 | 2 | 2 | 3 | 11 | 1 | 0 | 0 | 0 | 0 | 0 | 0 |
| SG0000003 | Fah | 1.66E-08 | 4.95E-08 | -1.04140633 | 634.5767644 | 308.3114281 | 19 | 1 | 2 | 3 | 7 | 1 | 0 | 0 | 0 | 0 | 0 | 0 |
| SG0000002 | F3 | 1.72E-08 | 5.13E-08 | 2.187032087 | 64.88589575 | 295.4694368 | 35 | 0 | 2 | 4 | 9 | 3 | 14 | 0 | 2 | 2 | 4 | 2 |
| JSG000000 | Dennd6b | 1.85E-08 | 5.5E-08 | 1.086404516 | 167.3796604 | 355.4209547 | 9 | 2 | 2 | 3 | 8 | 0 | 2 | 0 | 0 | 0 | 0 | 0 |
| SG0000004 | Usp22 | 2.3E-08 | 6.78E-08 | 0.691089505 | 3250.310215 | 5247.633351 | 26 | 0 | 2 | 2 | 6 | 1 | 4 | 0 | 0 | 0 | 1 | 0 |
| JSG000000 | AW549877 | 2.33E-08 | 6.88E-08 | 0.746128869 | 2024.640687 | 3395.901855 | 45 | 1 | 2 | 6 | 20 | 3 | 13 | 1 | 2 | 2 | 8 | 0 |
| JSG000000 | Gsx1 | 2.46E-08 | 7.24E-08 | -4.664623069 | 24.36246509 | 0 | 4 | 2 | 2 | 4 | 4 | 0 | 4 | 2 | 2 | 4 | 4 | 0 |
| JSG000000 | Fam159a | 2.49E-08 | 7.34E-08 | 5.248709006 | 0.674076249 | 25.62877652 | 13 | 2 | 2 | 3 | 16 | 1 | 0 | 0 | 0 | 0 | 0 | 0 |
| SG0000001 | Srsf1 | 2.58E-08 | 7.6E-08 | 0.666913809 | 10817.86851 | 17175.23783 | 38 | 0 | 2 | 0 | 8 | 8 | 19 | 0 | 2 | 0 | 2 | 4 |
| SG0000002 | Thpo | 2.62E-08 | 7.71E-08 | 1.880277905 | 26.41965974 | 97.26291234 | 12 | 2 | 2 | 6 | 8 | 6 | 0 | 0 | 0 | 0 | 0 | 0 |
| JSG000000 | Laptm4a | 2.68E-08 | 7.87E-08 | -0.766720876 | 8233.962658 | 4839.526934 | 30 | 1 | 2 | 2 | 8 | 0 | 2 | 0 | 0 | 0 | 0 | 0 |
| SG000000C | Nom1 | 2.79E-08 | 8.19E-08 | -0.844768967 | 1995.326576 | 1110.997773 | 24 | 0 | 2 | 3 | 7 | 5 | 4 | 0 | 0 | 2 | 3 | 4 |
| SG0000002 | Actb | 3.07E-08 | 8.99E-08 | -0.893424527 | 91307.00301 | 49153.73305 | 4 | 2 | 2 | 2 | 2 | 0 | 2 | 0 | 0 | 0 | 0 | 0 |
| SG0000002 | Pdgfrb | 3.3E-08 | 9.64E-08 | -1.67022117 | 776.6405272 | 244.0244698 | 38 | 2 | 2 | 2 | 12 | 4 | 2 | 0 | 0 | 0 | 2 | 0 |
| SG0000005 | Tmem199 | 3.48E-08 | 1.01E-07 | -0.855625645 | 1560.208356 | 862.2111113 | 10 | 2 | 2 | 4 | 8 | 2 | 2 | 1 | 1 | 2 | 2 | 0 |
| JSG000000 | Vps37c | 3.53E-08 | 1.03E-07 | -1.045689419 | 562.9695108 | 272.7099562 | 22 | 1 | 2 | 4 | 12 | 3 | 1 | 0 | 0 | 0 | 0 | 0 |
| JSG000000 | Zfp882 | 3.84E-08 | 1.12E-07 | 1.296970394 | 94.41207704 | 231.9821367 | 13 | 0 | 2 | 2 | 10 | 1 | 1 | 0 | 0 | 0 | 0 | 0 |
| SG0000002 | Tcam1 | 4.09E-08 | 1.19E-07 | 3.007355608 | 5.498854399 | 44.21569603 | 22 | 0 | 2 | 4 | 8 | 0 | 4 | 0 | 0 | 0 | 0 | 0 |
| JSG000000 | Ocel1 | 4.11E-08 | 1.19E-07 | -1.48578396 | 200.4157893 | 71.55935121 | 2 | 0 | 2 | 0 | 2 | 0 | 1 | 0 | 1 | 0 | 1 | 0 |
| SG0000005 | Atpif1 | 4.14E-08 | 1.2E-07 | 0.815743258 | 1907.094782 | 3356.877301 | 8 | 0 | 2 | 0 | 2 | 0 | 0 | 0 | 0 | 0 | 0 | 0 |
| SG0000002 | Mrpl30 | 4.29E-08 | 1.24E-07 | 0.727658647 | 1303.455758 | 2158.456867 | 27 | 0 | 2 | 2 | 22 | 6 | 0 | 0 | 0 | 0 | 0 | 0 |
| JSG000000 | Adprh | 4.41E-08 | 1.28E-07 | 0.868799196 | 1045.911558 | 1909.983346 | 10 | 0 | 2 | 1 | 2 | 2 | 1 | 0 | 0 | 0 | 0 | 0 |
| JSG000000 | Amacr | 4.68E-08 | 1.35E-07 | 0.937163225 | 277.1056034 | 530.5905971 | 24 | 1 | 2 | 4 | 11 | 3 | 3 | 0 | 0 | 1 | 3 | 0 |
| SG0000003 | Ppp3r1 | 5.36E-08 | 1.54E-07 | -0.773711999 | 4593.769093 | 2686.944165 | 90 | 1 | 2 | 8 | 48 | 8 | 11 | 0 | 1 | 2 | 5 | 3 |
| JSG000000 | Tti2 | 6.6E-08 | 1.88E-07 | 0.862568854 | 388.7010345 | 706.7646098 | 20 | 1 | 2 | 6 | 19 | 2 | 4 | 0 | 0 | 2 | 5 | 0 |
| JSG000000 | Pcsk7 | 6.79E-08 | 1.93E-07 | -1.014735127 | 717.8197307 | 355.2627562 | 18 | 1 | 2 | 3 | 7 | 3 | 0 | 0 | 0 | 0 | 0 | 0 |
| JSG000000 | Brk1 | 6.89E-08 | 1.96E-07 | -0.786672966 | 2857.158227 | 1656.235664 | 11 | 2 | 2 | 2 | 7 | 1 | 0 | 0 | 0 | 0 | 0 | 0 |
| JSG000000 | Slc44a2 | 7.04E-08 | 2E-07 | -0.78882711 | 3162.795619 | 1830.671866 | 24 | 1 | 2 | 7 | 16 | 3 | 0 | 0 | 0 | 0 | 0 | 0 |
| JSG000000 | Bpgm | 7.41E-08 | 2.1E-07 | 0.910072114 | 303.3926269 | 570.1170467 | 42 | 1 | 2 | 3 | 16 | 11 | 2 | 0 | 0 | 0 | 0 | 0 |
| SG0000006 | Zfp874a | 7.73E-08 | 2.19E-07 | 0.979786121 | 212.4237003 | 418.9362892 | 88 | 0 | 2 | 8 | 31 | 10 | 5 | 0 | 0 | 2 | 4 | 0 |
| JSG000000 | Wbp2nl | 7.91E-08 | 2.24E-07 | 6.046768986 | 0.331601068 | 21.92172689 | 19 | 2 | 2 | 3 | 9 | 2 | 0 | 0 | 0 | 0 | 0 | 0 |
| JSG000000 | Anks4b | 9.48E-08 | 2.66E-07 | -4.962256446 | 30.11795177 | 0.966134069 | 21 | 2 | 2 | 2 | 7 | 0 | 1 | 0 | 0 | 0 | 0 | 0 |
| JSG000000 | Isg20l2 | 1.1E-07 | 3.07E-07 | 0.6861757 | 1786.955089 | 2875.233405 | 15 | 1 | 2 | 3 | 8 | 0 | 1 | 0 | 0 | 0 | 1 | 0 |
| JSG000000 | Hint1 | 1.17E-07 | 3.26E-07 | -0.790241536 | 6070.821835 | 3510.436176 | 13 | 0 | 2 | 0 | 2 | 0 | 3 | 0 | 0 | 0 | 0 | 0 |
| JSG000000 | Nop9 | 1.27E-07 | 3.54E-07 | 0.680791478 | 1980.319078 | 3174.489004 | 7 | 2 | 2 | 4 | 10 | 0 | 0 | 0 | 0 | 0 | 2 | 0 |
| JSG000000 | BC031181 | 1.33E-07 | 3.7E-07 | -0.85995552 | 1130.657768 | 622.9579959 | 13 | 2 | 2 | 4 | 8 | 1 | 5 | 0 | 0 | 1 | 2 | 0 |
| SG0000005 | Slc6a2 | 1.5E-07 | 4.14E-07 | 6.209231591 | 0.307509278 | 22.75218879 | 124 | 2 | 2 | 12 | 46 | 4 | 9 | 0 | 0 | 0 | 0 | 2 |
| SG0000002 | Pdyn | 1.5E-07 | 4.15E-07 | 3.706245762 | 8.070857031 | 105.3442413 | 12 | 0 | 2 | 0 | 5 | 1 | 2 | 0 | 0 | 0 | 0 | 0 |
| SG0000004 | Yipf6 | 1.54E-07 | 4.24E-07 | -0.851777688 | 1218.283824 | 675.0529569 | 48 | 2 | 2 | 7 | 21 | 5 | 16 | 0 | 0 | 1 | 1 | 1 |
| JSG000000 | Tut1 | 1.61E-07 | 4.43E-07 | -0.802456943 | 1595.096034 | 914.5832121 | 11 | 2 | 2 | 2 | 11 | 4 | 0 | 0 | 0 | 0 | 0 | 0 |
| SG000000C | Timm44 | 1.65E-07 | 4.56E-07 | 0.680589351 | 1816.972063 | 2912.232602 | 16 | 2 | 2 | 4 | 8 | 4 | 0 | 0 | 0 | 0 | 0 | 1 |
| SG000000C | Ap1m1 | 2.04E-07 | 5.58E-07 | -0.750674958 | 2543.005448 | 1511.372832 | 15 | 2 | 2 | 4 | 12 | 0 | 0 | 0 | 0 | 0 | 0 | 0 |
| SG0000002 | Larp7 | 2.06E-07 | 5.6E-07 | 0.702326824 | 1199.864508 | 1952.331889 | 40 | 1 | 2 | 7 | 18 | 4 | 1 | 0 | 0 | 0 | 0 | 0 |
| SG0000001 | 4931428F04Rik | 2.22E-07 | 6.04E-07 | 0.833812332 | 379.7136835 | 676.7975383 | 10 | 1 | 2 | 2 | 6 | 2 | 2 | 1 | 1 | 1 | 2 | 0 |
| SG0000002 | Rbp4 | 2.23E-07 | 6.07E-07 | 2.727907645 | 6.798823315 | 45.04180452 | 22 | 0 | 2 | 2 | 4 | 0 | 2 | 0 | 0 | 2 | 2 | 0 |
| SG0000003 | Fancd2os | 2.5E-07 | 6.78E-07 | 4.233255129 | 0 | 17.80774696 | 10 | 0 | 2 | 0 | 6 | 2 | 1 | 0 | 0 | 0 | 2 | 0 |
| JSG000000 | 1810055G02Rik | 2.86E-07 | 7.71E-07 | -0.847601632 | 1323.28583 | 735.3602432 | 11 | 2 | 2 | 3 | 5 | 4 | 2 | 0 | 0 | 1 | 1 | 0 |
| JSG000000 | Atp5g3 | 3.45E-07 | 9.24E-07 | 0.636852871 | 4131.171951 | 6423.697613 | 14 | 2 | 2 | 4 | 4 | 0 | 0 | 0 | 0 | 0 | 0 | 0 |
| SG0000002 | Cnksr1 | 3.82E-07 | 1.02E-06 | 2.437592475 | 31.48336206 | 170.557003 | 9 | 1 | 2 | 3 | 5 | 1 | 0 | 0 | 0 | 0 | 0 | 0 |
| SG0000003 | Ppp1r10 | 4.15E-07 | 1.1E-06 | -0.97679928 | 2033.550648 | 1033.258795 | 36 | 0 | 2 | 2 | 2 | 6 | 1 | 0 | 0 | 0 | 0 | 2 |
| SG0000003 | Ubfd1 | 4.3E-07 | 1.14E-06 | 0.658262794 | 1744.615758 | 2753.319667 | 23 | 2 | 2 | 5 | 10 | 2 | 2 | 0 | 0 | 1 | 2 | 0 |
| JSG000000 | Gm527 | 4.3E-07 | 1.14E-06 | 1.565718025 | 36.31397735 | 107.4983773 | 23 | 0 | 2 | 2 | 5 | 2 | 0 | 0 | 0 | 0 | 1 | 1 |
| SG0000004 | Vps37d | 4.41E-07 | 1.17E-06 | 1.948280241 | 20.5169289 | 79.17774811 | 4 | 0 | 2 | 0 | 4 | 2 | 0 | 0 | 0 | 0 | 2 | 0 |
| SG0000003 | Smyd5 | 4.7E-07 | 1.24E-06 | -0.709030599 | 4288.881499 | 2623.641966 | 14 | 1 | 2 | 4 | 10 | 0 | 1 | 0 | 0 | 0 | 0 | 0 |
| JSG000000 | Prr11 | 5.79E-07 | 1.52E-06 | -0.74986344 | 2050.700504 | 1219.46924 | 35 | 2 | 2 | 3 | 17 | 1 | 7 | 0 | 0 | 0 | 1 | 1 |
| JSG000000 | Epdr1 | 5.83E-07 | 1.53E-06 | 1.194608401 | 85.78041887 | 196.336536 | 54 | 0 | 2 | 5 | 23 | 2 | 3 | 0 | 0 | 0 | 1 | 1 |
| JSG000001 | 9130401M01Rik | 6.29E-07 | 1.65E-06 | 0.866777403 | 318.9500515 | 581.6325118 | 24 | 2 | 2 | 2 | 6 | 2 | 0 | 0 | 0 | 0 | 0 | 0 |
| JSG000000 | Rbm44 | 6.38E-07 | 1.67E-06 | 9.097081886 | 0.68495036 | 375.1056615 | 48 | 2 | 2 | 5 | 10 | 6 | 2 | 0 | 0 | 0 | 0 | 0 |
| SG0000002 | Fstl3 | 9.89E-07 | 2.56E-06 | 1.249920904 | 107.6321294 | 255.9797538 | 6 | 2 | 2 | 2 | 2 | 0 | 3 | 1 | 1 | 1 | 1 | 0 |
| JSG000000 | Cct2 | 9.89E-07 | 2.56E-06 | 0.584501228 | 15998.74268 | 23990.44232 | 27 | 2 | 2 | 3 | 11 | 3 | 3 | 0 | 0 | 0 | 0 | 0 |
| SG0000003 | Fam103a1 | 1.01E-06 | 2.6E-06 | 0.775018816 | 406.7799782 | 696.08693 | 24 | 2 | 2 | 4 | 24 | 4 | 6 | 0 | 0 | 0 | 4 | 2 |
| JSG000000 | Ky | 1.21E-06 | 3.1E-06 | 5.75530314 | 0.34247518 | 18.49898848 | 36 | 1 | 2 | 4 | 18 | 3 | 1 | 0 | 0 | 0 | 1 | 0 |
| SG0000006 | Uqcrh | 1.24E-06 | 3.18E-06 | 0.633449265 | 2220.564284 | 3444.693454 | 32 | 2 | 2 | 8 | 18 | 6 | 0 | 0 | 0 | 0 | 0 | 0 |
| SG0000003 | Zfp606 | 1.26E-06 | 3.22E-06 | 0.869159783 | 240.0034654 | 438.3900394 | 61 | 0 | 2 | 3 | 22 | 4 | 17 | 0 | 0 | 1 | 3 | 2 |
| JSG000000 | Frg1 | 1.44E-06 | 3.66E-06 | 0.659521894 | 1088.715965 | 1719.691182 | 39 | 0 | 2 | 3 | 14 | 8 | 0 | 0 | 0 | 0 | 0 | 0 |
| SG0000002 | Api5 | 1.47E-06 | 3.74E-06 | 0.593840342 | 4014.003137 | 6058.170101 | 43 | 0 | 2 | 3 | 15 | 11 | 2 | 0 | 0 | 1 | 3 | 2 |
| SG0000004 | Arf1 | 1.51E-06 | 3.83E-06 | -0.648770469 | 10599.5371 | 6760.635598 | 28 | 0 | 2 | 2 | 12 | 6 | 4 | 0 | 2 | 0 | 0 | 2 |
| JSG000000 | Krt8 | 1.66E-06 | 4.2E-06 | -1.708820246 | 7525.590145 | 2302.154208 | 6 | 1 | 2 | 1 | 10 | 0 | 0 | 0 | 0 | 0 | 0 | 0 |
| JSG000000 | 2310022B05Rik | 1.66E-06 | 4.21E-06 | -0.702027827 | 2464.540196 | 1514.971532 | 31 | 2 | 2 | 6 | 20 | 6 | 0 | 0 | 0 | 0 | 1 | 2 |
| SG000000C | Cadm3 | 1.71E-06 | 4.32E-06 | 3.433355564 | 2.613155511 | 28.22983022 | 79 | 2 | 2 | 14 | 32 | 5 | 1 | 0 | 0 | 0 | 4 | 1 |
| JSG000000 | Slc9a5 | 1.82E-06 | 4.59E-06 | 0.780489506 | 503.961173 | 865.6609574 | 25 | 0 | 2 | 3 | 8 | 2 | 1 | 0 | 0 | 0 | 0 | 0 |
| SG0000003 | Smad3 | 1.83E-06 | 4.61E-06 | -0.747116244 | 1385.892922 | 825.7056892 | 99 | 1 | 2 | 13 | 31 | 10 | 4 | 0 | 0 | 2 | 2 | 0 |
| JSG000000 | Acta2 | 1.86E-06 | 4.7E-06 | -1.649579503 | 1304.904599 | 415.9162139 | 21 | 1 | 2 | 3 | 12 | 1 | 2 | 0 | 0 | 0 | 1 | 0 |
| SG0000004 | Dtx3l | 2.13E-06 | 5.34E-06 | -1.101809554 | 2452.781098 | 1142.828508 | 26 | 1 | 2 | 1 | 3 | 2 | 2 | 0 | 0 | 0 | 1 | 0 |
| SG0000002 | Ski | 2.15E-06 | 5.38E-06 | 0.616233538 | 1993.936769 | 3056.442018 | 116 | 2 | 2 | 4 | 20 | 10 | 20 | 0 | 0 | 2 | 6 | 2 |
| JSG000000 | Bmp2 | 2.16E-06 | 5.41E-06 | 2.840886347 | 4.88617941 | 35.00752552 | 22 | 1 | 2 | 7 | 11 | 3 | 8 | 0 | 0 | 4 | 5 | 1 |
| JSG000000 | Scd1 | 2.41E-06 | 6.01E-06 | -0.652463521 | 6039.441966 | 3842.250904 | 16 | 2 | 2 | 3 | 8 | 1 | 5 | 0 | 0 | 0 | 1 | 0 |
| JSG000000 | Srp9 | 2.41E-06 | 6.02E-06 | 0.66616073 | 789.4933923 | 1252.803221 | 16 | 2 | 2 | 2 | 6 | 2 | 0 | 0 | 0 | 0 | 0 | 0 |
| JSG000000 | C1d | 2.51E-06 | 6.24E-06 | -0.752530882 | 1263.984679 | 750.2524835 | 43 | 1 | 2 | 4 | 19 | 2 | 7 | 0 | 0 | 0 | 3 | 0 |
| JSG000000 | Nipal4 | 2.53E-06 | 6.29E-06 | 2.245416511 | 10.3741597 | 49.19156637 | 32 | 1 | 2 | 4 | 9 | 4 | 3 | 0 | 0 | 0 | 0 | 0 |
| JSG000000 | Vnn1 | 2.59E-06 | 6.45E-06 | 2.717099955 | 6.57446352 | 43.23036553 | 20 | 0 | 2 | 1 | 11 | 1 | 4 | 0 | 0 | 1 | 3 | 0 |
| SG0000002 | Hsph1 | 2.6E-06 | 6.47E-06 | -0.697731693 | 7489.677887 | 4617.6921 | 34 | 0 | 2 | 4 | 9 | 0 | 1 | 0 | 0 | 1 | 2 | 0 |
| SG0000003 | Lppr3 | 3.01E-06 | 7.43E-06 | 2.022391605 | 14.10642078 | 57.30827947 | 27 | 0 | 2 | 0 | 3 | 2 | 0 | 0 | 0 | 0 | 0 | 0 |
| JSG000000 | Thoc3 | 3.6E-06 | 8.86E-06 | 0.615832762 | 1402.072055 | 2148.594535 | 11 | 1 | 2 | 2 | 3 | 3 | 1 | 0 | 0 | 0 | 0 | 1 |
| JSG000000 | Batf3 | 4.01E-06 | 9.81E-06 | 3.631133185 | 1.998136955 | 24.75741286 | 8 | 2 | 2 | 4 | 9 | 1 | 0 | 0 | 0 | 0 | 0 | 0 |
| SG000000C | Abhd16a | 4.06E-06 | 9.92E-06 | -0.92487508 | 648.9306054 | 341.8087025 | 18 | 2 | 2 | 4 | 10 | 0 | 3 | 0 | 0 | 0 | 0 | 0 |
| JSG000000 | Pdk4 | 4.09E-06 | 9.99E-06 | 1.852718139 | 140.3184371 | 506.8021776 | 29 | 2 | 2 | 3 | 15 | 3 | 4 | 0 | 0 | 1 | 5 | 1 |
| JSG000000 | Nicn1 | 4.22E-06 | 1.03E-05 | 0.832563233 | 239.7290814 | 426.9207146 | 8 | 1 | 2 | 1 | 5 | 0 | 0 | 0 | 0 | 0 | 1 | 0 |
| SG0000002 | Paox | 4.37E-06 | 1.07E-05 | 1.031814148 | 133.2185542 | 272.3778105 | 12 | 2 | 2 | 6 | 14 | 4 | 0 | 0 | 0 | 0 | 0 | 0 |
| JSG000000 | Ndufs6 | 4.58E-06 | 1.12E-05 | 0.64818145 | 940.3442352 | 1473.699464 | 16 | 2 | 2 | 4 | 10 | 4 | 0 | 0 | 0 | 0 | 0 | 1 |
| JSG000000 | Snrpg | 4.71E-06 | 1.14E-05 | 0.754877169 | 1072.118855 | 1809.187609 | 19 | 1 | 2 | 2 | 7 | 3 | 0 | 0 | 0 | 0 | 0 | 0 |
| JSG000000 | Plin3 | 5.14E-06 | 1.25E-05 | -1.053128116 | 3430.468839 | 1653.218767 | 7 | 2 | 2 | 2 | 4 | 1 | 0 | 0 | 0 | 0 | 0 | 0 |
| JSG000000 | Tubb3 | 5.16E-06 | 1.25E-05 | -1.116410117 | 1610.955026 | 743.0370742 | 19 | 1 | 2 | 4 | 7 | 1 | 2 | 0 | 0 | 2 | 2 | 0 |
| SG0000006 | Tmem128 | 5.77E-06 | 1.39E-05 | -0.77797885 | 837.5708347 | 488.4572683 | 16 | 0 | 2 | 4 | 13 | 4 | 1 | 0 | 0 | 0 | 1 | 0 |
| SG0000004 | Mypop | 5.78E-06 | 1.39E-05 | -1.874136234 | 246.0081784 | 67.10862226 | 10 | 0 | 2 | 4 | 6 | 4 | 0 | 0 | 0 | 0 | 0 | 0 |
| JSG000000 | Zfp958 | 6.34E-06 | 1.52E-05 | 0.683154158 | 574.3398113 | 922.1866148 | 28 | 1 | 2 | 5 | 16 | 0 | 1 | 0 | 0 | 0 | 0 | 0 |
| SG0000003 | Pdcd5 | 6.88E-06 | 1.65E-05 | 0.663857691 | 622.5578079 | 986.3266377 | 10 | 0 | 2 | 0 | 2 | 4 | 0 | 0 | 0 | 0 | 0 | 0 |
| JSG000000 | Lipg | 6.9E-06 | 1.65E-05 | 2.280579094 | 41.80165136 | 203.1029135 | 24 | 1 | 2 | 4 | 11 | 10 | 5 | 1 | 1 | 2 | 2 | 1 |

| | | | | | | | | | | | | | | | | | |
|---|---|---|---|---|---|---|---|---|---|---|---|---|---|---|---|---|---|
| SG0000002 | Scly | 7.13E-06 | 1.7E-05 | 0.711717069 | 438.6283729 | 718.364574 | 20 | 0 | 2 | 3 | 3 | 1 | 1 | 0 | 0 | 0 | 0 | 0 |
| ISG000000 | Zic5 | 7.34E-06 | 1.75E-05 | 0.693959088 | 499.6137149 | 808.2335103 | 20 | 1 | 2 | 4 | 13 | 1 | 11 | 1 | 2 | 3 | 9 | 0 |
| SG0000002 | Tpgs2 | 7.6E-06 | 1.81E-05 | 0.709324332 | 433.765979 | 709.2239405 | 63 | 1 | 2 | 7 | 25 | 4 | 3 | 0 | 0 | 0 | 0 | 0 |
| ISG000000 | Fam161b | 8E-06 | 1.9E-05 | 1.411593667 | 38.37101886 | 102.0787555 | 34 | 2 | 2 | 4 | 10 | 2 | 11 | 2 | 2 | 2 | 2 | 0 |
| SG0000002 | Apod | 8.01E-06 | 1.91E-05 | 3.040239906 | 8.234812429 | 67.74186002 | 21 | 2 | 2 | 4 | 13 | 3 | 0 | 0 | 0 | 0 | 0 | 0 |
| ISG000000 | Ogn | 8.2E-06 | 1.95E-05 | -2.742715048 | 159.2607069 | 23.79409219 | 42 | 1 | 2 | 5 | 20 | 1 | 3 | 0 | 0 | 0 | 3 | 0 |
| SG0000005 | Slx1b | 9.47E-06 | 2.24E-05 | 0.73107036 | 360.3027956 | 598.055845 | 10 | 0 | 2 | 2 | 6 | 1 | 0 | 0 | 0 | 0 | 0 | 1 |
| ISG000000 | Snrpf | 9.53E-06 | 2.25E-05 | 0.651454233 | 839.8817055 | 1319.244786 | 21 | 2 | 2 | 4 | 11 | 0 | 2 | 0 | 0 | 0 | 1 | 0 |
| SG0000004 | Gal3st1 | 1.04E-05 | 2.46E-05 | 4.28453892 | 0.981585527 | 19.1294674 | 24 | 0 | 2 | 7 | 15 | 0 | 3 | 0 | 0 | 0 | 0 | 0 |
| ISG000000 | Rpusd2 | 1.05E-05 | 2.47E-05 | 0.779482997 | 268.6439344 | 461.1315132 | 9 | 2 | 2 | 2 | 5 | 1 | 2 | 1 | 1 | 1 | 2 | 1 |
| SG0000003 | Lrfn1 | 1.15E-05 | 2.7E-05 | 1.483593388 | 94.23101699 | 263.5117526 | 11 | 2 | 2 | 3 | 16 | 2 | 0 | 0 | 0 | 0 | 0 | 0 |
| ISG000000 | Grtp1 | 1.15E-05 | 2.7E-05 | 1.045275593 | 405.4539897 | 836.7599505 | 20 | 1 | 2 | 3 | 7 | 2 | 0 | 0 | 0 | 0 | 0 | 1 |
| ISG000000 | Slc25a31 | 1.24E-05 | 2.9E-05 | 4.823161807 | 1.337278385 | 37.85630334 | 44 | 0 | 2 | 2 | 11 | 4 | 1 | 0 | 1 | 0 | 2 | 1 |
| SG0000000 | Zp3 | 1.33E-05 | 3.1E-05 | 2.992371287 | 3.546557457 | 28.22282713 | 4 | 2 | 2 | 2 | 5 | 0 | 0 | 0 | 0 | 0 | 0 | 0 |
| SG0000001 | Akap1 | 1.57E-05 | 3.64E-05 | 0.561896542 | 2219.530457 | 3276.489817 | 54 | 2 | 2 | 5 | 24 | 14 | 6 | 0 | 0 | 0 | 0 | 6 |
| SG0000002 | Lipn | 1.67E-05 | 3.85E-05 | -4.025421397 | 15.2844309 | 0 | 86 | 0 | 2 | 6 | 48 | 21 | 2 | 0 | 0 | 0 | 0 | 0 |
| SG0000000 | Aif1l | 1.95E-05 | 4.48E-05 | 0.728125728 | 318.1230906 | 526.9663428 | 29 | 2 | 2 | 6 | 14 | 0 | 5 | 0 | 0 | 2 | 2 | 0 |
| SG0000003 | Mllt6 | 2.08E-05 | 4.76E-05 | -0.608693818 | 2823.228136 | 1851.445259 | 18 | 0 | 2 | 2 | 4 | 2 | 0 | 0 | 0 | 0 | 0 | 0 |
| ISG000000 | Stip1 | 2.65E-05 | 6E-05 | 0.51032715 | 10898.80981 | 15523.97211 | 29 | 2 | 2 | 6 | 10 | 2 | 1 | 0 | 0 | 1 | 1 | 0 |
| SG0000002 | Atg2a | 2.7E-05 | 6.11E-05 | -0.895264017 | 2382.064784 | 1280.714261 | 12 | 2 | 2 | 2 | 4 | 2 | 0 | 0 | 0 | 0 | 0 | 0 |
| ISG000000 | Pex11a | 2.95E-05 | 6.64E-05 | 0.86412964 | 146.8269218 | 267.2604134 | 12 | 1 | 2 | 1 | 3 | 1 | 2 | 0 | 0 | 0 | 1 | 1 |
| SG0000003 | Ido2 | 3.25E-05 | 7.29E-05 | 1.587841588 | 22.76586101 | 68.43401588 | 45 | 1 | 2 | 7 | 32 | 5 | 0 | 0 | 0 | 0 | 0 | 0 |
| SG0000002 | Col7a1 | 3.49E-05 | 7.8E-05 | -1.832089123 | 1020.204136 | 286.5321108 | 16 | 0 | 2 | 0 | 0 | 6 | 0 | 0 | 0 | 0 | 0 | 0 |
| ISG000000 | Tmem51 | 3.66E-05 | 8.16E-05 | 1.171657904 | 159.255242 | 358.7549569 | 34 | 1 | 2 | 5 | 22 | 6 | 0 | 0 | 0 | 0 | 0 | 0 |
| SG0000000 | Cox4i2 | 3.82E-05 | 8.5E-05 | 4.155107398 | 2.011354634 | 35.83443921 | 15 | 2 | 2 | 2 | 12 | 2 | 0 | 0 | 0 | 0 | 0 | 0 |
| SG0000003 | Mospd3 | 4.14E-05 | 9.17E-05 | -1.07520276 | 600.9154171 | 285.1970731 | 6 | 2 | 2 | 2 | 4 | 2 | 0 | 0 | 0 | 0 | 0 | 0 |
| SG0000004 | Zfp219 | 4.24E-05 | 9.4E-05 | 1.265046314 | 763.2923049 | 1834.457989 | 10 | 0 | 2 | 3 | 6 | 1 | 0 | 0 | 0 | 0 | 0 | 0 |
| SG0000002 | Pmpca | 4.31E-05 | 9.53E-05 | -0.560360406 | 5854.937739 | 3970.424943 | 8 | 1 | 2 | 1 | 4 | 0 | 4 | 1 | 2 | 1 | 2 | 0 |
| SG0000002 | Gbp2 | 4.47E-05 | 9.86E-05 | -1.134930603 | 711.2574233 | 323.8759184 | 69 | 2 | 2 | 11 | 48 | 4 | 0 | 0 | 0 | 0 | 0 | 0 |
| ISG000000 | Gas8 | 4.48E-05 | 9.88E-05 | -0.839760828 | 360.7849813 | 201.5836257 | 19 | 2 | 2 | 3 | 9 | 1 | 1 | 0 | 0 | 0 | 0 | 0 |
| SG0000002 | Nkain4 | 4.84E-05 | 0.000106 | 4.654284815 | 0.639110347 | 16.09367349 | 14 | 2 | 2 | 2 | 8 | 6 | 0 | 0 | 0 | 0 | 0 | 0 |
| SG0000003 | Fbxl19 | 5.45E-05 | 0.000119 | 0.624416538 | 1470.891067 | 2267.506904 | 40 | 0 | 2 | 0 | 12 | 16 | 0 | 0 | 0 | 0 | 0 | 0 |
| ISG000000 | Igbp1 | 5.54E-05 | 0.000121 | -0.586437934 | 2427.805163 | 1616.882356 | 36 | 0 | 2 | 3 | 15 | 6 | 0 | 0 | 0 | 0 | 1 | 0 |
| SG0000003 | Gng10 | 5.69E-05 | 0.000124 | -2.13537801 | 70.81467764 | 16.11797395 | 10 | 2 | 2 | 2 | 6 | 0 | 3 | 0 | 0 | 0 | 2 | 0 |
| SG0000007 | 1700049G17Rik | 5.79E-05 | 0.000126 | -0.922780132 | 265.1559317 | 139.8675114 | 39 | 0 | 2 | 3 | 31 | 5 | 14 | 0 | 0 | 1 | 3 | 2 |
| SG0000003 | Ptpn23 | 5.82E-05 | 0.000127 | -0.560988585 | 3302.816313 | 2238.77284 | 18 | 1 | 2 | 3 | 5 | 1 | 0 | 0 | 0 | 0 | 0 | 0 |
| SG0000002 | Dvl1 | 6.18E-05 | 0.000134 | -0.767526892 | 1435.587013 | 843.2976763 | 8 | 2 | 2 | 4 | 6 | 4 | 4 | 2 | 2 | 2 | 2 | 0 |
| SG0000002 | Zbtb48 | 6.5E-05 | 0.000141 | 0.816384208 | 181.772192 | 320.0984559 | 5 | 2 | 2 | 3 | 3 | 0 | 0 | 0 | 0 | 0 | 0 | 0 |
| ISG000000 | Yeats4 | 6.62E-05 | 0.000143 | 0.550937683 | 1031.376726 | 1511.005674 | 14 | 2 | 2 | 3 | 8 | 0 | 0 | 0 | 0 | 0 | 0 | 0 |
| SG0000002 | Timm10 | 7.01E-05 | 0.000151 | 0.721238296 | 256.8823099 | 423.4952649 | 12 | 2 | 2 | 4 | 6 | 3 | 0 | 0 | 0 | 0 | 0 | 1 |
| ISG000000 | Cxcr2 | 7.49E-05 | 0.000161 | 4.627836898 | 0.649984458 | 16.07017823 | 20 | 2 | 2 | 6 | 20 | 17 | 4 | 0 | 0 | 0 | 0 | 1 |
| ISG000000 | Ppil6 | 7.55E-05 | 0.000162 | 3.730097755 | 1.324060707 | 17.57030128 | 32 | 2 | 2 | 4 | 20 | 3 | 1 | 0 | 0 | 0 | 0 | 0 |
| SG0000004 | Pramef8 | 7.99E-05 | 0.000171 | -0.637931385 | 962.8730997 | 618.7747332 | 16 | 2 | 2 | 2 | 12 | 0 | 4 | 2 | 2 | 2 | 6 | 0 |
| SG0000005 | Pm20d2 | 8.79E-05 | 0.000187 | 2.104091351 | 8.491794559 | 36.50850834 | 38 | 2 | 2 | 5 | 11 | 0 | 0 | 0 | 0 | 0 | 0 | 0 |
| SG0000002 | Angel1 | 9.81E-05 | 0.000208 | 0.630789175 | 405.9007592 | 628.5015111 | 38 | 1 | 2 | 7 | 38 | 3 | 0 | 0 | 0 | 0 | 0 | 0 |
| SG0000002 | Oxsm | 9.89E-05 | 0.00021 | -0.690329353 | 663.6744654 | 411.287269 | 29 | 0 | 2 | 4 | 21 | 2 | 10 | 0 | 0 | 2 | 6 | 2 |
| SG0000002 | 2610015P09Rik | 0.000103 | 0.000218 | 0.632893144 | 385.5983128 | 597.9363351 | 38 | 0 | 2 | 4 | 17 | 3 | 1 | 0 | 0 | 0 | 1 | 0 |
| SG0000000 | Pwp1 | 0.000109 | 0.000229 | 0.50208158 | 1966.642763 | 2785.268672 | 25 | 2 | 2 | 3 | 10 | 1 | 0 | 0 | 0 | 0 | 0 | 0 |
| ISG000000 | Traip | 0.000114 | 0.00024 | 0.612204515 | 670.9411411 | 1025.596064 | 10 | 2 | 2 | 2 | 18 | 2 | 0 | 0 | 0 | 0 | 0 | 0 |
| ISG000000 | Mrs2 | 0.000115 | 0.000241 | -0.827907497 | 425.6760581 | 239.8027913 | 35 | 1 | 2 | 3 | 19 | 3 | 2 | 0 | 0 | 0 | 1 | 0 |
| SG0000002 | Prdx6 | 0.000115 | 0.000241 | -0.513815877 | 13305.70254 | 9318.882134 | 26 | 2 | 2 | 4 | 6 | 4 | 2 | 0 | 0 | 0 | 0 | 0 |
| ISG000000 | Gm20604 | 0.000116 | 0.000244 | 0.76431782 | 180.9185992 | 307.302297 | 36 | 0 | 2 | 2 | 6 | 4 | 4 | 0 | 1 | 0 | 0 | 0 |
| SG0000003 | Cmas | 0.000117 | 0.000246 | -0.554071029 | 2561.465544 | 1744.602613 | 25 | 2 | 2 | 3 | 6 | 1 | 1 | 1 | 1 | 1 | 1 | 0 |
| SG0000003 | Mphosph10 | 0.000119 | 0.00025 | 0.505895297 | 1705.479118 | 2421.787679 | 30 | 0 | 2 | 2 | 14 | 1 | 1 | 0 | 0 | 0 | 0 | 0 |
| ISG000000 | Cinp | 0.00012 | 0.000252 | 0.544416696 | 876.1221044 | 1277.76325 | 27 | 0 | 2 | 8 | 12 | 3 | 1 | 0 | 0 | 0 | 0 | 0 |
| SG0000001 | Pelp1 | 0.000124 | 0.00026 | 0.49244873 | 2927.92088 | 4119.089002 | 30 | 2 | 2 | 3 | 9 | 2 | 0 | 0 | 0 | 0 | 0 | 0 |
| ISG000000 | Tysnd1 | 0.000139 | 0.00029 | -0.761356462 | 513.6774049 | 303.0395626 | 10 | 2 | 2 | 2 | 5 | 2 | 0 | 0 | 0 | 0 | 0 | 0 |
| ISG000000 | Ss18l1 | 0.000149 | 0.000309 | 0.740999044 | 184.8961154 | 309.0229441 | 22 | 0 | 2 | 3 | 11 | 6 | 8 | 0 | 1 | 0 | 1 | 2 |
| SG0000003 | Alg1 | 0.000149 | 0.000311 | -0.779409727 | 625.1649914 | 364.2243218 | 20 | 2 | 2 | 6 | 20 | 4 | 2 | 0 | 0 | 2 | 2 | 2 |
| ISG000000 | Fam109b | 0.000154 | 0.000319 | 1.788901256 | 13.211675 | 45.65315741 | 7 | 1 | 2 | 2 | 2 | 0 | 3 | 0 | 0 | 1 | 1 | 0 |
| SG0000002 | Vil1 | 0.000154 | 0.00032 | 6.287686152 | 0.68495036 | 53.51077408 | 25 | 1 | 2 | 2 | 21 | 3 | 0 | 0 | 0 | 0 | 0 | 0 |
| ISG000000 | Zfp764 | 0.000158 | 0.000327 | -0.9916901 | 231.6846597 | 116.5115052 | 3 | 2 | 2 | 2 | 2 | 0 | 3 | 2 | 2 | 2 | 2 | 0 |
| ISG000000 | Cox14 | 0.000158 | 0.000328 | -0.952472468 | 782.1151126 | 404.1549429 | 4 | 2 | 2 | 2 | 3 | 0 | 0 | 0 | 0 | 0 | 0 | 0 |
| ISG000000 | Snn | 0.000161 | 0.000333 | 0.580560319 | 514.0658986 | 768.7495341 | 9 | 2 | 2 | 2 | 6 | 1 | 3 | 0 | 0 | 0 | 0 | 1 |
| ISG000000 | Ndufa1 | 0.000167 | 0.000346 | -0.66258712 | 959.3080473 | 606.0374887 | 16 | 1 | 2 | 2 | 3 | 0 | 0 | 0 | 0 | 0 | 1 | 0 |
| SG0000003 | Nsmce1 | 0.000176 | 0.000364 | -0.590656918 | 1200.314147 | 797.0571744 | 31 | 1 | 2 | 2 | 7 | 4 | 1 | 0 | 0 | 0 | 0 | 0 |
| SG0000006 | H2-Q10 | 0.000188 | 0.000387 | 1.679851366 | 38.36179773 | 122.9092591 | 8 | 2 | 2 | 2 | 6 | 2 | 0 | 0 | 0 | 0 | 0 | 0 |
| SG0000004 | Scn3b | 0.00021 | 0.000429 | 5.128602737 | 1.668879454 | 58.38324471 | 76 | 2 | 2 | 10 | 26 | 6 | 2 | 0 | 0 | 0 | 2 | 0 |
| SG0000005 | Thra | 0.000214 | 0.000438 | -0.950166125 | 703.0833124 | 363.8968411 | 42 | 0 | 2 | 3 | 15 | 0 | 0 | 0 | 0 | 0 | 0 | 0 |
| SG0000002 | Srebf1 | 0.000217 | 0.000442 | -0.940241723 | 2593.082718 | 1351.373534 | 13 | 0 | 2 | 4 | 5 | 5 | 2 | 0 | 2 | 2 | 2 | 2 |
| SG0000004 | Wrap53 | 0.000234 | 0.000476 | 0.632843179 | 423.1930947 | 656.210767 | 19 | 2 | 2 | 3 | 7 | 0 | 0 | 0 | 0 | 0 | 0 | 0 |
| SG0000002 | Zfand5 | 0.000238 | 0.000484 | -0.525658462 | 2695.998803 | 1872.753988 | 50 | 0 | 2 | 3 | 14 | 2 | 4 | 0 | 0 | 1 | 2 | 2 |
| ISG000000 | Csrnp1 | 0.000241 | 0.000489 | 0.613742999 | 379.0258521 | 579.9945131 | 9 | 2 | 2 | 2 | 4 | 0 | 0 | 0 | 0 | 0 | 0 | 0 |
| SG0000002 | Pla2g2e | 0.00027 | 0.000544 | -2.08491218 | 44.74706406 | 10.54735203 | 9 | 0 | 2 | 0 | 2 | 0 | 0 | 0 | 0 | 0 | 0 | 0 |
| SG0000006 | Oas1g | 0.000284 | 0.000571 | -1.32575992 | 368.1766724 | 146.8800343 | 30 | 0 | 2 | 0 | 14 | 2 | 2 | 0 | 0 | 0 | 1 | 0 |
| SG0000003 | Ccdc92 | 0.000289 | 0.000581 | -0.772101563 | 572.5718895 | 335.2774096 | 42 | 2 | 2 | 8 | 24 | 8 | 0 | 0 | 0 | 0 | 0 | 0 |
| ISG000000 | Ndufs3 | 0.000294 | 0.00059 | 0.53021639 | 731.2523288 | 1056.03497 | 17 | 1 | 2 | 3 | 6 | 5 | 0 | 0 | 0 | 0 | 0 | 1 |
| SG0000005 | Cyhr1 | 0.000298 | 0.000598 | -0.537444188 | 1557.89976 | 1073.377998 | 13 | 2 | 2 | 2 | 3 | 2 | 3 | 0 | 0 | 0 | 0 | 0 |
| SG0000003 | Rsl24d1 | 0.00031 | 0.00062 | 0.484849395 | 1362.840481 | 1907.213167 | 25 | 2 | 2 | 4 | 14 | 3 | 2 | 0 | 0 | 0 | 0 | 1 |
| ISG000000 | Rac2 | 0.000316 | 0.000633 | -2.150151916 | 34.67756709 | 7.812470649 | 7 | 1 | 2 | 1 | 2 | 1 | 0 | 0 | 0 | 0 | 1 | 0 |
| SG0000001 | Ndufs2 | 0.000335 | 0.000668 | 0.447188009 | 4879.398971 | 6652.476281 | 30 | 0 | 2 | 10 | 14 | 8 | 0 | 0 | 0 | 0 | 0 | 0 |
| ISG000000 | Agap2 | 0.000337 | 0.000671 | 3.836463673 | 2.351486247 | 33.59178529 | 8 | 1 | 2 | 2 | 5 | 0 | 1 | 0 | 0 | 0 | 0 | 0 |
| SG0000002 | Tmem9 | 0.000393 | 0.000778 | 0.626866223 | 811.7798888 | 1253.556243 | 21 | 2 | 2 | 2 | 4 | 0 | 1 | 0 | 0 | 0 | 0 | 0 |
| ISG000000 | Rpp30 | 0.000395 | 0.00078 | 0.559114046 | 462.1899536 | 680.9741087 | 38 | 0 | 2 | 2 | 19 | 5 | 1 | 0 | 0 | 0 | 1 | 0 |
| SG0000002 | Nifk | 0.000471 | 0.000922 | 0.494778095 | 2028.715904 | 2858.671695 | 11 | 2 | 2 | 4 | 8 | 0 | 0 | 0 | 0 | 0 | 0 | 0 |
| SG0000004 | Arrdc4 | 0.000477 | 0.000933 | 0.525607866 | 620.8683738 | 893.7648519 | 34 | 0 | 2 | 4 | 12 | 6 | 4 | 0 | 0 | 0 | 0 | 1 |
| ISG000000 | Eif2s1 | 0.000482 | 0.000942 | 0.433008127 | 5048.022484 | 6815.060492 | 64 | 1 | 2 | 7 | 28 | 8 | 13 | 0 | 0 | 1 | 5 | 1 |
| SG0000003 | Smarce1 | 0.000487 | 0.000952 | -0.505398907 | 2186.103027 | 1540.034304 | 37 | 1 | 2 | 6 | 19 | 3 | 1 | 0 | 0 | 0 | 1 | 0 |
| ISG000000 | BC034090 | 0.000507 | 0.00099 | 0.93153624 | 76.31641306 | 145.5587442 | 34 | 0 | 2 | 6 | 18 | 5 | 2 | 0 | 0 | 0 | 0 | 0 |
| SG0000002 | Slc20a1 | 0.000508 | 0.00099 | -0.538294052 | 6660.49077 | 4586.311776 | 40 | 0 | 2 | 4 | 14 | 2 | 4 | 0 | 0 | 0 | 0 | 2 |
| SG0000002 | Frs3 | 0.000514 | 0.001001 | 0.89860603 | 90.82701655 | 169.3255201 | 6 | 2 | 2 | 5 | 7 | 0 | 3 | 0 | 0 | 3 | 4 | 0 |
| SG0000000 | Heatr6 | 0.000538 | 0.001046 | -0.538208308 | 1172.926522 | 807.7072143 | 44 | 0 | 2 | 4 | 12 | 3 | 8 | 0 | 0 | 0 | 0 | 0 |
| ISG000000 | Ggps1 | 0.000542 | 0.001053 | -0.565827249 | 950.7036682 | 642.2649104 | 29 | 1 | 2 | 3 | 8 | 3 | 4 | 1 | 1 | 1 | 2 | 0 |
| SG0000003 | Tgm2 | 0.000582 | 0.001125 | -2.401268736 | 3889.958089 | 736.3613833 | 19 | 0 | 2 | 4 | 10 | 5 | 1 | 0 | 0 | 0 | 0 | 0 |
| ISG000000 | Sertm1 | 0.000638 | 0.001227 | 2.825479643 | 2.613155511 | 18.52334351 | 66 | 0 | 2 | 2 | 22 | 8 | 8 | 0 | 2 | 0 | 0 | 2 |
| SG0000006 | Numa1 | 0.000661 | 0.001268 | -0.458345993 | 5269.446567 | 3835.20968 | 60 | 1 | 2 | 9 | 25 | 9 | 0 | 0 | 0 | 0 | 0 | 2 |
| SG0000003 | Cops7a | 0.000702 | 0.001343 | -0.460149945 | 5181.441049 | 3766.444961 | 10 | 0 | 2 | 0 | 0 | 0 | 1 | 0 | 0 | 0 | 0 | 0 |
| ISG000000 | Pom121 | 0.000709 | 0.001355 | -0.466062691 | 3705.056643 | 2682.229916 | 19 | 1 | 2 | 3 | 10 | 2 | 3 | 0 | 0 | 0 | 0 | 1 |
| SG0000003 | Dpf1 | 0.00075 | 0.001429 | 0.644979363 | 216.433848 | 338.4412699 | 50 | 2 | 2 | 8 | 12 | 4 | 0 | 0 | 0 | 0 | 0 | 0 |
| SG0000002 | Masp2 | 0.000759 | 0.001445 | 2.169296898 | 5.819581356 | 26.17671712 | 15 | 0 | 2 | 3 | 6 | 1 | 2 | 0 | 0 | 0 | 1 | 0 |
| ISG000000 | Eif4b | 0.000776 | 0.001476 | 0.403374786 | 22680.15329 | 29996.72874 | 31 | 1 | 2 | 4 | 10 | 3 | 1 | 0 | 0 | 0 | 2 | 0 |
| SG0000006 | Kdr | 0.000778 | 0.001479 | 3.818129749 | 1.254128903 | 17.68942672 | 58 | 2 | 2 | 4 | 24 | 3 | 2 | 0 | 0 | 1 | 3 | 0 |
| ISG000000 | Ifi47 | 0.000874 | 0.001652 | -2.283840149 | 43.63463935 | 8.960391817 | 42 | 0 | 2 | 3 | 22 | 4 | 2 | 0 | 0 | 0 | 2 | 0 |
| ISG000000 | Thoc7 | 0.000879 | 0.00166 | 0.456141959 | 1101.795708 | 1511.518574 | 33 | 1 | 2 | 3 | 12 | 1 | 1 | 0 | 0 | 0 | 0 | 0 |
| SG0000004 | Cyp2s1 | 0.000886 | 0.001672 | 1.098477785 | 197.3173958 | 422.5130602 | 14 | 2 | 2 | 4 | 12 | 2 | 0 | 0 | 0 | 0 | 0 | 0 |
| ISG000000 | Pctp | 0.000888 | 0.001675 | -0.585827526 | 641.1150593 | 427.1538457 | 24 | 2 | 2 | 3 | 10 | 3 | 3 | 0 | 0 | 0 | 0 | 0 |
| SG0000000 | Cyba | 0.000899 | 0.001696 | 0.961809583 | 482.8533671 | 940.4783423 | 5 | 2 | 2 | 2 | 5 | 0 | 0 | 0 | 0 | 0 | 0 | 0 |
| SG0000002 | Traf5 | 0.000919 | 0.001731 | 1.166823692 | 33.53686644 | 75.29591447 | 5 | 1 | 2 | 3 | 3 | 0 | 1 | 0 | 1 | 1 | 1 | 0 |
| SG0000004 | Alkbh3 | 0.000941 | 0.001771 | -0.639122084 | 425.4339466 | 273.1726542 | 94 | 0 | 2 | 20 | 38 | 2 | 0 | 0 | 0 | 0 | 0 | 0 |
| SG0000001 | Gata3 | 0.000965 | 0.001813 | -1.422445504 | 125.3330946 | 46.75919209 | 34 | 1 | 2 | 3 | 11 | 1 | 6 | 0 | 0 | 1 | 4 | 0 |
| SG0000002 | Aimp1 | 0.000991 | 0.001859 | 0.426004277 | 2088.52548 | 2805.949456 | 64 | 1 | 2 | 12 | 37 | 4 | 0 | 0 | 0 | 0 | 0 | 0 |
| SG0000003 | 43723 | 0.001208 | 0.002247 | -0.445626334 | 4986.033831 | 3661.072486 | 132 | 0 | 2 | 5 | 44 | 14 | 5 | 0 | 0 | 0 | 4 | 0 |
| SG0000000 | Tbcb | 0.001246 | 0.002314 | 0.48468352 | 856.4577552 | 1198.423245 | 14 | 2 | 2 | 3 | 5 | 0 | 0 | 0 | 0 | 0 | 0 | 0 |
| ISG000000 | AA986860 | 0.001259 | 0.002338 | 1.934868761 | 7.45115134 | 28.48898123 | 16 | 0 | 2 | 2 | 5 | 3 | 2 | 0 | 0 | 1 | 1 | 0 |
| SG0000003 | Tmem126a | 0.001587 | 0.002916 | 0.501394794 | 471.9258122 | 668.0494407 | 32 | 0 | 2 | 0 | 4 | 4 | 0 | 0 | 0 | 0 | 0 | 0 |
| ISG000000 | Polr2g | 0.001612 | 0.002958 | -0.456750871 | 1925.347529 | 1402.85705 | 15 | 2 | 2 | 3 | 8 | 1 | 2 | 1 | 1 | 1 | 1 | 0 |
| SG0000003 | Nr2f2 | 0.001644 | 0.003014 | -1.324259513 | 84.43111637 | 33.71791382 | 26 | 0 | 2 | 0 | 9 | 4 | 8 | 0 | 0 | 0 | 3 | 4 |
| ISG000000 | Mrpl50 | 0.001665 | 0.003049 | 0.425969015 | 1399.542323 | 1880.249524 | 26 | 0 | 2 | 3 | 10 | 1 | 1 | 0 | 0 | 0 | 2 | 0 |
| ISG000000 | Elof1 | 0.001685 | 0.003085 | -0.495959525 | 1148.236793 | 814.2031256 | 6 | 2 | 2 | 2 | 6 | 0 | 0 | 0 | 0 | 0 | 0 | 0 |
| SG0000002 | Cdk2 | 0.001739 | 0.003178 | 0.413877218 | 1660.773351 | 2212.584291 | 22 | 0 | 2 | 2 | 6 | 0 | 3 | 0 | 0 | 2 | 4 | 0 |
| ISG000000 | Cd34 | 0.001795 | 0.003272 | 1.200228897 | 25.49094492 | 58.57210529 | 90 | 2 | 2 | 8 | 34 | 10 | 2 | 0 | 0 | 0 | 4 | 0 |
| SG0000004 | Zfp956 | 0.001816 | 0.003306 | 0.630054996 | 178.1881284 | 275.7682219 | 26 | 0 | 2 | 0 | 4 | 2 | 0 | 0 | 0 | 0 | 0 | 0 |
| SG0000003 | Sdr42e1 | 0.00189 | 0.003433 | -1.617808364 | 40.70559716 | 13.26310737 | 32 | 2 | 2 | 8 | 10 | 0 | 2 | 0 | 0 | 0 | 0 | 0 |

| | | | | | | | | | | | | | | | | | |
|---|---|---|---|---|---|---|---|---|---|---|---|---|---|---|---|---|---|
| ISG000000 | Peo1 | 0.002042 | 0.003694 | -0.474554927 | 1199.795242 | 863.4791743 | 7 | 2 | 2 | 3 | 6 | 0 | 0 | 0 | 0 | 0 | 0 |
| SG0000005 | Zfp85 | 0.002078 | 0.003754 | 0.800098094 | 79.70250868 | 138.7795635 | 50 | 2 | 2 | 4 | 12 | 4 | 4 | 0 | 0 | 0 | 1 |
| ISG000000 | Il20ra | 0.002124 | 0.003832 | -5.127330121 | 23.40397449 | 0.66959076 | 66 | 1 | 2 | 7 | 28 | 11 | 1 | 0 | 0 | 2 | 1 |
| SG000000C | Crtc1 | 0.002136 | 0.003852 | -0.611255281 | 340.6400062 | 222.9921 | 29 | 1 | 2 | 4 | 10 | 7 | 2 | 0 | 0 | 0 | 1 |
| SG0000002 | Ivd | 0.002194 | 0.003952 | -0.475432122 | 1165.052889 | 837.9658342 | 16 | 2 | 2 | 4 | 9 | 3 | 0 | 0 | 0 | 0 | 0 |
| SG0000002 | Dnaja1 | 0.002223 | 0.004 | 0.415204626 | 2257.808646 | 3010.760284 | 51 | 0 | 2 | 0 | 2 | 4 | 15 | 0 | 0 | 0 | 2 |
| SG0000002 | Eif4a2 | 0.002247 | 0.004039 | 0.370342658 | 14301.63723 | 18487.15176 | 125 | 0 | 2 | 22 | 29 | 0 | 37 | 0 | 2 | 9 | 0 |
| ISG000000 | Zfp9 | 0.002588 | 0.004615 | -0.970163112 | 138.1297127 | 70.50808369 | 25 | 0 | 2 | 6 | 13 | 4 | 3 | 0 | 0 | 2 | 0 |
| ISG000000 | Terf2ip | 0.002618 | 0.004665 | 0.472776079 | 494.2290798 | 685.8799006 | 10 | 2 | 2 | 2 | 6 | 2 | 1 | 0 | 0 | 0 | 1 |
| SG0000004 | Gpr153 | 0.00286 | 0.005061 | -0.816097149 | 213.5339782 | 121.2822569 | 14 | 2 | 2 | 2 | 8 | 0 | 3 | 0 | 0 | 0 | 0 |
| SG0000005 | 2810428I15Rik | 0.002869 | 0.005075 | 0.816167293 | 290.5570118 | 511.590177 | 2 | 0 | 2 | 2 | 2 | 0 | 0 | 0 | 0 | 0 | 0 |
| SG000000C | Atp5g1 | 0.00302 | 0.005325 | 0.417014977 | 1670.399245 | 2230.253871 | 6 | 0 | 2 | 0 | 0 | 2 | 6 | 0 | 2 | 0 | 2 |
| ISG000000 | Zscan25 | 0.003048 | 0.005369 | -0.605101414 | 309.6083634 | 203.5443294 | 8 | 2 | 2 | 3 | 4 | 0 | 0 | 0 | 0 | 0 | 0 |
| ISG000000 | Lfng | 0.003175 | 0.005579 | -0.701318685 | 203.4999065 | 125.1544377 | 7 | 2 | 2 | 4 | 6 | 1 | 1 | 0 | 1 | 2 | 1 |
| SG0000003 | U2af2 | 0.003252 | 0.005707 | 0.373559554 | 5032.637775 | 6520.010632 | 16 | 0 | 2 | 0 | 12 | 10 | 0 | 0 | 0 | 2 | 0 |
| ISG000000 | Ttc30a1 | 0.003328 | 0.005831 | 0.865817328 | 59.40993958 | 108.2669986 | 6 | 0 | 2 | 0 | 2 | 2 | 5 | 0 | 2 | 0 | 1 | 1 |
| ISG000000 | Snrnp35 | 0.003346 | 0.005861 | -0.573382589 | 372.8427815 | 250.5649852 | 8 | 2 | 2 | 2 | 8 | 0 | 0 | 0 | 0 | 0 | 0 |
| SG0000001 | Sh3bp5l | 0.003593 | 0.006271 | -0.470687339 | 836.2819548 | 603.4783832 | 36 | 0 | 2 | 0 | 8 | 10 | 2 | 0 | 2 | 0 | 0 | 0 |
| ISG000000 | Pcif1 | 0.003753 | 0.006528 | -0.504075822 | 1360.068916 | 959.0008069 | 7 | 2 | 2 | 2 | 4 | 1 | 0 | 0 | 0 | 0 | 1 |
| SG0000002 | Nfkb2 | 0.00385 | 0.006689 | -0.430998191 | 1336.600331 | 991.4211528 | 12 | 0 | 2 | 0 | 0 | 0 | 2 | 0 | 2 | 0 | 0 | 0 |
| SG0000004 | Zfp84 | 0.00399 | 0.006916 | -0.569514863 | 366.6879088 | 247.0902068 | 19 | 1 | 2 | 3 | 5 | 1 | 6 | 1 | 1 | 1 | 2 | 0 |
| SG0000002 | Selp | 0.004197 | 0.007248 | -1.310654668 | 76.79426141 | 30.95867741 | 59 | 0 | 2 | 4 | 16 | 4 | 3 | 0 | 0 | 0 | 0 |
| ISG000000 | 9230110C19Rik | 0.004337 | 0.007483 | 0.865861847 | 51.09447437 | 93.11600264 | 35 | 1 | 2 | 2 | 17 | 3 | 1 | 0 | 0 | 1 | 0 |
| SG0000003 | Zpr1 | 0.004343 | 0.00749 | 0.39072257 | 1359.333442 | 1782.15394 | 17 | 2 | 2 | 3 | 11 | 0 | 2 | 0 | 1 | 2 | 0 |
| SG0000003 | Ttc38 | 0.004486 | 0.007724 | -0.632988836 | 251.0306092 | 161.8743559 | 13 | 2 | 2 | 3 | 9 | 0 | 0 | 0 | 0 | 0 | 0 |
| ISG000000 | Ctdsp2 | 0.004528 | 0.007793 | -0.399417296 | 3637.913107 | 2758.136367 | 12 | 1 | 2 | 3 | 5 | 0 | 0 | 0 | 0 | 0 | 0 |
| SG0000003 | Gtf2h5 | 0.004906 | 0.008405 | 0.403881971 | 894.071377 | 1182.912931 | 20 | 2 | 2 | 5 | 8 | 0 | 2 | 0 | 1 | 2 | 0 |
| SG0000006 | Zscan20 | 0.004919 | 0.008424 | 0.599345608 | 179.9383603 | 272.6118717 | 67 | 2 | 2 | 8 | 28 | 0 | 1 | 0 | 0 | 0 | 0 |
| SG0000003 | Rwdd4a | 0.005268 | 0.008986 | -0.421916328 | 1364.079936 | 1018.193571 | 48 | 2 | 2 | 8 | 20 | 6 | 2 | 0 | 0 | 0 | 0 |
| ISG000000 | Mrpl12 | 0.005299 | 0.009035 | -0.437712694 | 2138.718486 | 1579.02488 | 6 | 2 | 2 | 2 | 5 | 0 | 0 | 0 | 0 | 0 | 0 |
| ISG000000 | Pcsk9 | 0.005405 | 0.009202 | 1.588421765 | 34.88517208 | 104.9067581 | 22 | 1 | 2 | 3 | 7 | 2 | 1 | 0 | 1 | 2 | 0 |
| SG0000005 | Trim12c | 0.005431 | 0.009244 | -0.514231047 | 808.1194691 | 565.8177923 | 40 | 2 | 2 | 6 | 18 | 6 | 3 | 0 | 0 | 0 | 2 |
| SG0000006 | Hmgb1 | 0.005477 | 0.009317 | 0.34937414 | 3857.250254 | 4914.167012 | 18 | 0 | 2 | 0 | 9 | 4 | 8 | 0 | 0 | 2 | 0 |
| ISG000000 | Adat2 | 0.006219 | 0.010502 | -0.555138932 | 330.4647587 | 224.9115104 | 13 | 2 | 2 | 3 | 8 | 2 | 0 | 0 | 0 | 0 | 0 |
| SG0000003 | Lrrc23 | 0.006263 | 0.010568 | -1.583172138 | 31.5845695 | 10.54126327 | 12 | 2 | 2 | 2 | 8 | 2 | 0 | 0 | 0 | 0 | 0 |
| SG0000002 | Tef | 0.006426 | 0.010826 | -0.43130478 | 1416.538387 | 1050.491813 | 29 | 0 | 2 | 4 | 13 | 0 | 2 | 0 | 0 | 0 | 0 |
| ISG000000 | Cdca4 | 0.006435 | 0.010839 | 0.372756673 | 1389.101853 | 1798.643242 | 20 | 1 | 2 | 4 | 5 | 1 | 4 | 0 | 2 | 3 | 0 |
| SG0000002 | Acads | 0.006941 | 0.011633 | -0.698898321 | 1691.755922 | 1042.193468 | 9 | 2 | 2 | 2 | 5 | 1 | 0 | 0 | 0 | 0 | 0 |
| ISG000000 | Srf | 0.007163 | 0.011982 | -0.549047112 | 1386.594773 | 947.6978796 | 7 | 2 | 2 | 3 | 6 | 0 | 2 | 1 | 1 | 1 | 1 | 0 |
| SG0000005 | Cope | 0.007639 | 0.012721 | -0.512076008 | 2551.914649 | 1789.434914 | 15 | 0 | 2 | 4 | 7 | 6 | 0 | 0 | 0 | 0 | 0 |
| SG0000006 | Zfp617 | 0.008111 | 0.013452 | -0.509316866 | 394.9683386 | 277.486992 | 25 | 0 | 2 | 3 | 7 | 1 | 1 | 0 | 0 | 1 | 0 |
| ISG000000 | Cebpz | 0.008312 | 0.013771 | 0.334383901 | 3416.425662 | 4307.562224 | 32 | 2 | 2 | 2 | 14 | 4 | 3 | 0 | 0 | 0 | 0 |
| SG0000002 | Gpn3 | 0.008545 | 0.014135 | 0.383144011 | 839.2449225 | 1094.527209 | 34 | 0 | 2 | 7 | 9 | 14 | 6 | 0 | 2 | 2 | 4 | 1 |
| ISG000000 | Slc41a1 | 0.008653 | 0.014301 | -0.383399651 | 1764.315635 | 1352.57541 | 11 | 1 | 2 | 4 | 6 | 1 | 1 | 0 | 1 | 1 | 1 | 0 |
| ISG000000 | Slmo1 | 0.00868 | 0.014337 | 0.569902304 | 147.9211264 | 219.577529 | 22 | 2 | 2 | 5 | 17 | 3 | 0 | 0 | 0 | 0 | 0 |
| ISG000000 | Kctd7 | 0.008752 | 0.014441 | -0.76629141 | 121.3227592 | 71.32890863 | 36 | 0 | 2 | 3 | 4 | 0 | 1 | 0 | 1 | 2 | 0 |
| ISG000000 | Trove2 | 0.009082 | 0.014951 | 0.387520482 | 857.6471151 | 1121.925244 | 56 | 0 | 2 | 3 | 17 | 10 | 20 | 0 | 0 | 7 | 6 |
| SG0000003 | Mrpl23 | 0.009232 | 0.015183 | 0.554556211 | 156.5061008 | 229.8631349 | 5 | 2 | 2 | 2 | 2 | 1 | 0 | 0 | 0 | 0 | 0 |
| ISG000000 | Pllp | 0.009625 | 0.015784 | 1.871288815 | 14.41023978 | 52.72119812 | 20 | 2 | 2 | 6 | 13 | 3 | 1 | 0 | 0 | 1 | 0 |
| SG000000C | Zfp81 | 0.009686 | 0.015874 | 0.410073534 | 477.1278783 | 633.9853567 | 57 | 1 | 2 | 11 | 41 | 7 | 14 | 1 | 1 | 2 | 6 | 3 |
| ISG000000 | Rbm7 | 0.010358 | 0.016896 | -0.374603201 | 2043.42591 | 1576.13022 | 14 | 1 | 2 | 2 | 2 | 0 | 3 | 0 | 1 | 0 | 0 | 0 |
| ISG000000 | Padi4 | 0.010471 | 0.017068 | -0.716884247 | 408.4220126 | 248.4880336 | 13 | 0 | 2 | 2 | 8 | 3 | 0 | 0 | 0 | 0 | 0 |
| SG0000001 | Hdac2 | 0.010941 | 0.01778 | -0.347066458 | 5567.87084 | 4377.354685 | 110 | 0 | 2 | 10 | 43 | 9 | 7 | 0 | 0 | 3 | 5 | 0 |
| ISG000000 | Myh7b | 0.010941 | 0.01778 | -0.744622859 | 158.8001823 | 94.77573926 | 9 | 2 | 2 | 3 | 6 | 1 | 0 | 0 | 0 | 0 | 0 |
| SG0000004 | Rwdd2b | 0.011949 | 0.019292 | -0.553099665 | 253.4769628 | 172.7582933 | 6 | 1 | 2 | 3 | 4 | 1 | 0 | 0 | 0 | 0 | 1 |
| ISG000000 | Mettl4 | 0.012151 | 0.019576 | 0.413947959 | 409.6708672 | 545.8154608 | 58 | 2 | 2 | 7 | 16 | 4 | 0 | 0 | 0 | 0 | 0 |
| ISG000000 | Sgta | 0.012327 | 0.019835 | -0.455718462 | 3500.109794 | 2552.09444 | 15 | 2 | 2 | 5 | 7 | 0 | 0 | 0 | 0 | 0 | 0 |
| SG0000005 | Kif18b | 0.012504 | 0.020089 | 0.346011342 | 1470.269567 | 1868.773903 | 22 | 0 | 2 | 3 | 9 | 1 | 1 | 0 | 0 | 1 | 1 | 0 |
| SG0000003 | Srpx2 | 0.012849 | 0.020591 | -0.837365498 | 93.55229717 | 52.3579126 | 62 | 0 | 2 | 0 | 18 | 6 | 0 | 0 | 0 | 0 | 0 |
| ISG000000 | Gpatch3 | 0.012984 | 0.020796 | -0.624243495 | 182.7601177 | 118.5674317 | 16 | 2 | 2 | 2 | 2 | 0 | 0 | 0 | 0 | 0 | 0 |
| SG000000C | Rsl1d1 | 0.013272 | 0.021217 | 0.306165635 | 6407.689887 | 7922.578021 | 17 | 1 | 2 | 5 | 10 | 0 | 3 | 0 | 0 | 0 | 0 |
| SG000000C | Ppp2r1a | 0.013835 | 0.022038 | 0.306566409 | 7450.794544 | 9214.850056 | 50 | 0 | 2 | 8 | 22 | 4 | 0 | 0 | 0 | 1 | 0 |
| SG0000002 | Hspa5 | 0.013894 | 0.022124 | -0.323098942 | 17551.73296 | 14029.99544 | 25 | 2 | 2 | 2 | 13 | 2 | 13 | 0 | 0 | 1 | 2 |
| ISG000000 | Col8a2 | 0.0142 | 0.022587 | 4.275753105 | 1.845055668 | 35.73875447 | 26 | 2 | 2 | 3 | 7 | 1 | 1 | 0 | 0 | 0 | 0 |
| ISG000000 | Dph1 | 0.014267 | 0.022689 | -0.456008176 | 425.226657 | 309.9904703 | 12 | 2 | 2 | 3 | 6 | 1 | 1 | 0 | 0 | 1 | 0 |
| ISG000000 | Ada | 0.014678 | 0.023293 | 0.708436288 | 169.7251219 | 277.3362902 | 17 | 0 | 2 | 1 | 8 | 2 | 2 | 0 | 0 | 2 | 0 |
| SG0000002 | Aimp2 | 0.014998 | 0.023765 | 0.353773132 | 1139.212242 | 1455.797495 | 12 | 2 | 2 | 4 | 6 | 0 | 0 | 0 | 0 | 0 | 0 |
| ISG000000 | Pts | 0.015263 | 0.024164 | 0.450954886 | 263.0411701 | 359.5627349 | 18 | 1 | 2 | 2 | 6 | 2 | 4 | 0 | 0 | 0 | 0 |
| SG0000003 | Sc5d | 0.015347 | 0.024289 | -0.377851527 | 1053.626835 | 810.8530969 | 38 | 0 | 2 | 4 | 12 | 0 | 2 | 0 | 0 | 0 | 0 |
| SG0000002 | Ralgds | 0.016292 | 0.025658 | 0.524104446 | 887.8001788 | 1276.692736 | 31 | 2 | 2 | 4 | 17 | 3 | 2 | 0 | 0 | 2 | 4 | 2 |
| ISG000000 | 4930503L19Rik | 0.016589 | 0.026083 | -0.458006446 | 417.2411308 | 303.7480057 | 48 | 0 | 2 | 0 | 12 | 2 | 1 | 0 | 0 | 0 | 0 |
| SG0000003 | Itga2b | 0.016939 | 0.026581 | 0.882662508 | 37.48595363 | 69.11565919 | 13 | 1 | 2 | 1 | 2 | 1 | 0 | 0 | 0 | 0 | 0 |
| ISG000000 | Gm11992 | 0.017158 | 0.026896 | 1.380937774 | 12.30954869 | 32.05869373 | 30 | 1 | 2 | 4 | 16 | 2 | 4 | 0 | 0 | 5 | 0 |
| SG0000004 | Ap1g2 | 0.01747 | 0.027344 | 0.38593018 | 507.634351 | 663.3268288 | 4 | 0 | 2 | 0 | 2 | 0 | 2 | 0 | 2 | 0 | 0 | 0 |
| SG000000C | Myg1 | 0.017576 | 0.027488 | -0.379298774 | 888.253261 | 682.89904 | 10 | 2 | 2 | 2 | 4 | 1 | 0 | 0 | 0 | 0 | 0 |
| SG0000002 | Cc2d1b | 0.017626 | 0.027554 | 0.329707661 | 1507.796333 | 1894.936139 | 22 | 0 | 2 | 2 | 9 | 5 | 0 | 0 | 0 | 0 | 0 |
| SG0000002 | Vegfb | 0.017653 | 0.027589 | 0.381929805 | 475.6262602 | 619.7808485 | 10 | 2 | 2 | 2 | 6 | 2 | 0 | 0 | 0 | 0 | 0 |
| ISG000000 | Trpm4 | 0.017795 | 0.027785 | 0.459843952 | 248.4102263 | 341.6616968 | 29 | 1 | 2 | 3 | 14 | 1 | 0 | 0 | 0 | 0 | 0 |
| SG0000005 | H2-T24 | 0.01791 | 0.027948 | -2.056305447 | 16.54709034 | 3.978432722 | 39 | 2 | 2 | 6 | 20 | 6 | 7 | 2 | 2 | 4 | 4 | 2 |
| SG0000002 | Nubp1 | 0.01839 | 0.028636 | -0.346989767 | 1342.762116 | 1055.710283 | 15 | 0 | 2 | 0 | 6 | 0 | 0 | 0 | 0 | 0 | 0 |
| ISG000000 | Ccdc51 | 0.018702 | 0.029084 | 0.526172967 | 157.4120705 | 226.6897468 | 7 | 0 | 2 | 0 | 4 | 1 | 2 | 0 | 1 | 0 | 2 | 0 |
| SG000000C | Ranbp1 | 0.020199 | 0.031222 | 0.290899409 | 5192.308008 | 6352.283828 | 18 | 0 | 2 | 4 | 8 | 2 | 2 | 0 | 2 | 2 | 2 | 0 |
| SG000000C | Timm50 | 0.020413 | 0.031517 | 0.331949483 | 1266.294222 | 1593.901122 | 5 | 1 | 2 | 1 | 1 | 0 | 0 | 0 | 0 | 0 | 0 |
| SG0000003 | Pced1a | 0.020761 | 0.032036 | -0.405977101 | 504.4394899 | 380.7130755 | 7 | 2 | 2 | 2 | 2 | 2 | 1 | 0 | 0 | 0 | 0 |
| ISG000000 | Klhdc7a | 0.02093 | 0.03225 | 3.3627875 | 2.495040127 | 25.66713773 | 5 | 0 | 2 | 0 | 1 | 1 | 5 | 0 | 2 | 0 | 0 | 1 |
| SG0000002 | Cfap126 | 0.021319 | 0.032798 | 0.846483747 | 55.61658352 | 100.0049059 | 47 | 1 | 2 | 9 | 18 | 0 | 3 | 0 | 0 | 1 | 1 | 0 |
| SG0000002 | Get4 | 0.021872 | 0.033602 | 0.332623598 | 944.9147246 | 1189.93242 | 35 | 0 | 2 | 5 | 9 | 0 | 5 | 0 | 0 | 3 | 3 | 0 |
| ISG000000 | Pip4k2c | 0.023816 | 0.036358 | -0.313504157 | 2539.881113 | 2043.803948 | 14 | 1 | 2 | 1 | 8 | 1 | 0 | 0 | 0 | 0 | 0 |
| SG0000003 | Usp5 | 0.024291 | 0.037033 | -0.299522835 | 4655.190488 | 3782.440448 | 14 | 2 | 2 | 2 | 4 | 4 | 0 | 0 | 0 | 0 | 0 |
| SG0000005 | Ddx49 | 0.026166 | 0.039693 | 0.363388496 | 1243.366527 | 1599.52123 | 7 | 1 | 2 | 3 | 4 | 0 | 0 | 0 | 0 | 0 | 0 |
| ISG000000 | Isoc2a | 0.026758 | 0.040536 | 0.544210881 | 399.4558977 | 582.4955832 | 22 | 2 | 2 | 3 | 14 | 1 | 0 | 0 | 0 | 0 | 0 |
| ISG000000 | Hars | 0.026981 | 0.040846 | -0.307956218 | 3038.397032 | 2454.372432 | 25 | 1 | 2 | 2 | 18 | 4 | 0 | 0 | 0 | 2 | 0 |
| ISG000000 | Ctns | 0.028483 | 0.042938 | -0.374031238 | 681.2690765 | 525.6831473 | 14 | 0 | 2 | 2 | 10 | 2 | 0 | 0 | 0 | 2 | 0 |
| SG0000003 | Fam53c | 0.028492 | 0.042948 | -0.324567357 | 1505.34488 | 1202.074819 | 26 | 2 | 2 | 6 | 10 | 2 | 2 | 0 | 0 | 0 | 0 |
| SG0000003 | Tns2 | 0.028498 | 0.042953 | -0.561014605 | 444.8159339 | 301.5074946 | 14 | 0 | 2 | 4 | 8 | 0 | 0 | 0 | 0 | 0 | 0 |
| ISG000000 | Tmem97 | 0.028883 | 0.043479 | 0.315861862 | 947.1534985 | 1178.974095 | 13 | 2 | 2 | 5 | 6 | 1 | 0 | 0 | 0 | 1 | 0 |
| ISG000000 | Pin4 | 0.031561 | 0.047173 | -0.474330893 | 265.8199775 | 191.3373657 | 16 | 1 | 2 | 4 | 14 | 0 | 0 | 0 | 0 | 0 | 0 |
| ISG000000 | Hey2 | 0.031804 | 0.047516 | 1.250773034 | 10.58161154 | 25.18094446 | 23 | 2 | 2 | 3 | 10 | 0 | 4 | 0 | 0 | 2 | 0 |
| ISG000000 | Trappc2l | 0.034289 | 0.050919 | 0.353763049 | 441.6603535 | 564.393112 | 5 | 2 | 2 | 2 | 2 | 0 | 1 | 1 | 1 | 1 | 1 | 0 |
| ISG000000 | Mrpl54 | 0.03537 | 0.052376 | -0.384856503 | 721.7805672 | 552.7793791 | 4 | 0 | 2 | 0 | 0 | 0 | 0 | 0 | 0 | 0 | 0 |
| ISG000000 | Wdr74 | 0.035504 | 0.052562 | 0.307181254 | 1293.189608 | 1600.047876 | 4 | 2 | 2 | 2 | 2 | 0 | 0 | 0 | 0 | 0 | 0 |
| SG0000003 | Sec13 | 0.03774 | 0.055651 | -0.279948604 | 3292.084732 | 2711.428007 | 22 | 1 | 2 | 1 | 6 | 1 | 1 | 0 | 0 | 1 | 0 |
| ISG000000 | Nup50 | 0.038354 | 0.056478 | 0.268228978 | 2576.419602 | 3102.855584 | 22 | 1 | 2 | 2 | 15 | 1 | 2 | 0 | 0 | 2 | 0 |
| SG0000002 | Ran | 0.038579 | 0.056772 | 0.251243008 | 11117.17988 | 13232.02505 | 20 | 2 | 2 | 2 | 4 | 2 | 18 | 2 | 2 | 2 | 4 | 0 |
| SG0000005 | Fgf18 | 0.039739 | 0.058358 | 2.295756386 | 3.786478498 | 18.59203936 | 56 | 2 | 2 | 6 | 14 | 2 | 0 | 0 | 0 | 0 | 0 |
| SG0000003 | Dapk3 | 0.040239 | 0.05902 | -0.615337346 | 652.2079255 | 425.7464873 | 22 | 2 | 2 | 2 | 14 | 4 | 0 | 0 | 0 | 0 | 0 |
| SG0000005 | Taf9 | 0.041691 | 0.061018 | 0.759610201 | 36.20658294 | 61.29894428 | 16 | 2 | 2 | 2 | 10 | 0 | 2 | 0 | 0 | 0 | 0 |
| ISG000000 | Npc2 | 0.041907 | 0.061322 | 0.262941684 | 4036.860866 | 4843.921712 | 22 | 0 | 2 | 2 | 6 | 5 | 2 | 0 | 1 | 1 | 1 | 0 |
| ISG000000 | Mterf1a | 0.043422 | 0.063363 | 0.502069005 | 98.58267748 | 139.6170443 | 18 | 0 | 2 | 0 | 4 | 6 | 4 | 0 | 2 | 0 | 0 | 0 |
| ISG000000 | Ddb1 | 0.04383 | 0.063921 | -0.260844234 | 32926.42384 | 27480.37314 | 34 | 2 | 2 | 2 | 12 | 1 | 0 | 0 | 0 | 0 | 0 |
| SG0000002 | Rabl2 | 0.043904 | 0.064019 | 0.410384208 | 249.8101431 | 332.0075995 | 16 | 0 | 2 | 0 | 2 | 4 | 2 | 0 | 0 | 0 | 0 |
| SG0000005 | Tmem220 | 0.044246 | 0.064475 | -0.671837295 | 83.8711154 | 52.64646809 | 24 | 0 | 2 | 2 | 19 | 0 | 2 | 0 | 0 | 1 | 0 |
| ISG000000 | Mex3c | 0.046538 | 0.067606 | 0.264179142 | 1765.268808 | 2120.004063 | 38 | 0 | 2 | 4 | 17 | 2 | 1 | 0 | 0 | 4 | 1 |
| SG0000002 | Cbll1 | 0.046614 | 0.067712 | 0.279800554 | 941.8583621 | 1143.441518 | 126 | 0 | 2 | 16 | 48 | 16 | 14 | 0 | 2 | 4 | 12 | 0 |
| SG0000003 | Cdk16 | 0.047212 | 0.068531 | -0.269972836 | 3576.834022 | 2966.39422 | 18 | 0 | 2 | 10 | 18 | 2 | 0 | 0 | 0 | 2 | 0 |
| SG0000002 | Eif2ak1 | 0.048011 | 0.069566 | -0.287769151 | 1711.08657 | 1401.66696 | 41 | 2 | 2 | 4 | 20 | 7 | 1 | 0 | 0 | 0 | 0 |
| ISG000000 | Mon1a | 0.051535 | 0.07418 | -0.551696644 | 163.1901583 | 111.3311577 | 26 | 2 | 2 | 2 | 34 | 4 | 1 | 0 | 0 | 0 | 0 |
| ISG000000 | Alyref | 0.053352 | 0.076598 | 0.255615202 | 1996.335721 | 2383.314861 | 8 | 2 | 2 | 2 | 5 | 0 | 1 | 0 | 0 | 0 | 0 |
| ISG000000 | Gm2115 | 0.053454 | 0.076725 | -0.822485824 | 222.4683489 | 125.7984609 | 59 | 1 | 2 | 7 | 27 | 7 | 0 | 0 | 0 | 1 | 0 |
| ISG000000 | Zfp786 | 0.054576 | 0.078206 | 1.070666076 | 12.95000574 | 27.20022929 | 25 | 1 | 2 | 3 | 10 | 1 | 1 | 0 | 0 | 0 | 0 |
| SG0000002 | Tbc1d7 | 0.055728 | 0.079696 | 0.330107212 | 385.6287883 | 484.7765685 | 70 | 0 | 2 | 12 | 34 | 4 | 1 | 0 | 0 | 0 | 0 |
| SG0000002 | Trim35 | 0.058055 | 0.082754 | -0.262076175 | 2619.418192 | 2184.298811 | 29 | 2 | 2 | 4 | 7 | 4 | 1 | 0 | 0 | 1 | 0 |
| SG0000004 | Polr2k | 0.059365 | 0.084439 | 0.357264099 | 224.0675066 | 287.0292073 | 24 | 2 | 2 | 6 | 14 | 0 | 4 | 0 | 0 | 0 | 0 |

| | | | | | | | | | | | | | | | | | | |
|---|---|---|---|---|---|---|---|---|---|---|---|---|---|---|---|---|---|---|
| SG0000001 | Git1 | 0.060535 | 0.08596 | -0.437514801 | 3808.643433 | 2812.323248 | 12 | 2 | 2 | 2 | 4 | 2 | 0 | 0 | 0 | 0 | 0 | 2 |
| JSG000000 | Pde12 | 0.061703 | 0.087495 | 0.261001355 | 1127.90134 | 1351.57562 | 24 | 0 | 2 | 1 | 9 | 0 | 20 | 0 | 2 | 1 | 7 | 0 |
| SG0000002 | Coch | 0.070574 | 0.099054 | -0.692335792 | 487.4579156 | 301.6638619 | 42 | 0 | 2 | 10 | 26 | 2 | 5 | 0 | 0 | 3 | 5 | 0 |
| SG0000002 | Bcl10 | 0.070693 | 0.099181 | 0.262568866 | 928.7972899 | 1114.197152 | 17 | 0 | 2 | 0 | 6 | 3 | 2 | 0 | 1 | 0 | 2 | 1 |
| SG0000003 | Zfp189 | 0.071364 | 0.100038 | 0.372032098 | 181.5909789 | 235.0104047 | 32 | 0 | 2 | 0 | 14 | 0 | 4 | 0 | 0 | 0 | 2 | 0 |
| SG0000002 | Dnal4 | 0.073508 | 0.102849 | -0.354060326 | 465.5454388 | 364.2330057 | 16 | 0 | 2 | 2 | 5 | 0 | 2 | 0 | 2 | 2 | 2 | 0 |
| JSG000000 | Ccl7 | 0.075677 | 0.105603 | -0.602183061 | 590.4244916 | 388.945902 | 2 | 0 | 2 | 0 | 2 | 0 | 2 | 0 | 2 | 0 | 2 | 0 |
| JSG000000 | Clk3 | 0.075707 | 0.105637 | 0.256080952 | 1002.04875 | 1196.676874 | 10 | 2 | 2 | 3 | 4 | 3 | 1 | 0 | 0 | 1 | 1 | 0 |
| SG0000003 | Upf3b | 0.07672 | 0.106942 | 0.271179519 | 1261.166069 | 1521.967739 | 35 | 1 | 2 | 5 | 18 | 3 | 1 | 0 | 0 | 0 | 1 | 0 |
| JSG000000 | Gpr180 | 0.077163 | 0.107501 | 0.284846163 | 534.9417603 | 651.7090349 | 38 | 0 | 2 | 4 | 13 | 5 | 2 | 0 | 1 | 1 | 1 | 0 |
| SG0000003 | Atp1a1 | 0.078081 | 0.108687 | -0.234006562 | 13481.68489 | 11463.07616 | 26 | 0 | 2 | 5 | 12 | 0 | 0 | 0 | 0 | 0 | 1 | 0 |
| SG0000002 | Ampd2 | 0.078397 | 0.109084 | -0.243946897 | 1807.109033 | 1525.980639 | 14 | 0 | 2 | 0 | 5 | 15 | 5 | 0 | 2 | 0 | 0 | 0 |
| SG0000007 | Eef1g | 0.07968 | 0.110706 | 0.213859201 | 19444.1584 | 22551.07165 | 22 | 0 | 2 | 3 | 8 | 1 | 0 | 0 | 0 | 0 | 0 | 0 |
| SG0000002 | Wdr45b | 0.080563 | 0.111786 | 0.236491099 | 1893.635061 | 2230.936456 | 35 | 2 | 2 | 4 | 16 | 3 | 2 | 0 | 0 | 0 | 0 | 0 |
| SG0000002 | Bst1 | 0.087371 | 0.120476 | 1.379358148 | 9.030847472 | 23.49398485 | 33 | 0 | 2 | 3 | 17 | 8 | 5 | 0 | 0 | 1 | 2 | 0 |
| SG0000007 | 6430550D23Rik | 0.087667 | 0.120866 | 1.189094468 | 10.25704117 | 23.38703693 | 64 | 0 | 2 | 7 | 29 | 3 | 9 | 0 | 0 | 0 | 2 | 0 |
| SG0000006 | Sec22c | 0.088511 | 0.121889 | -0.31789749 | 512.9029994 | 411.470362 | 57 | 0 | 2 | 8 | 35 | 12 | 6 | 0 | 2 | 2 | 2 | 0 |
| SG0000002 | Ypel1 | 0.092355 | 0.126724 | 0.404189564 | 121.8568187 | 161.2586496 | 21 | 2 | 2 | 5 | 14 | 1 | 2 | 0 | 0 | 0 | 3 | 1 |
| SG0000002 | Pdlim7 | 0.092895 | 0.127358 | -0.389751676 | 1414.910518 | 1079.945974 | 21 | 2 | 2 | 2 | 5 | 2 | 1 | 0 | 0 | 0 | 0 | 0 |
| SG0000003 | Rxrb | 0.095193 | 0.13019 | 0.340390438 | 899.1085841 | 1138.360653 | 4 | 2 | 2 | 2 | 6 | 0 | 0 | 0 | 0 | 0 | 0 | 0 |
| SG0000003 | Zfp248 | 0.095269 | 0.130274 | 0.329723285 | 217.4962395 | 273.3432485 | 82 | 2 | 2 | 4 | 43 | 10 | 6 | 0 | 0 | 0 | 3 | 2 |
| SG0000003 | Lmo2 | 0.098508 | 0.134221 | 0.734574911 | 22.43056966 | 37.32238475 | 69 | 1 | 2 | 9 | 39 | 13 | 8 | 0 | 0 | 4 | 12 | 0 |
| SG0000002 | Glo1 | 0.100526 | 0.136618 | -0.217611584 | 3572.87076 | 3072.62594 | 32 | 0 | 2 | 1 | 12 | 2 | 3 | 0 | 0 | 0 | 0 | 1 |
| SG0000002 | Casp2 | 0.100617 | 0.13671 | 0.228629364 | 1126.24925 | 1319.650149 | 72 | 0 | 2 | 12 | 27 | 0 | 2 | 0 | 0 | 0 | 3 | 0 |
| JSG000000 | Cox6b1 | 0.102973 | 0.139721 | 0.213802081 | 3191.77398 | 3701.629686 | 13 | 1 | 2 | 2 | 3 | 1 | 0 | 0 | 0 | 0 | 0 | 0 |
| JSG000000 | Prorsd1 | 0.104126 | 0.141189 | -0.365216762 | 221.4816393 | 171.9477626 | 17 | 0 | 2 | 1 | 5 | 2 | 6 | 0 | 0 | 1 | 3 | 0 |
| SG0000006 | Zfp932 | 0.104949 | 0.142231 | 0.308892149 | 370.8332584 | 459.3719567 | 49 | 0 | 2 | 2 | 24 | 4 | 0 | 0 | 0 | 0 | 0 | 0 |
| JSG000000 | Zfp937 | 0.105672 | 0.143155 | 0.442823203 | 79.11023521 | 107.5315095 | 53 | 0 | 2 | 6 | 25 | 1 | 3 | 0 | 0 | 0 | 0 | 0 |
| SG0000006 | Krr1 | 0.105889 | 0.143428 | 0.211126569 | 2238.445018 | 2591.205528 | 25 | 1 | 2 | 3 | 14 | 4 | 6 | 0 | 0 | 0 | 5 | 2 |
| JSG000000 | Lrat | 0.106064 | 0.143621 | -0.908271963 | 29.55011994 | 15.74498107 | 29 | 1 | 2 | 5 | 18 | 2 | 13 | 0 | 1 | 2 | 7 | 1 |
| JSG000000 | St3gal1 | 0.10617 | 0.143743 | 0.643741397 | 105.9232907 | 165.4919733 | 12 | 2 | 2 | 2 | 5 | 0 | 6 | 2 | 2 | 2 | 3 | 0 |
| SG0000003 | Rrp8 | 0.10773 | 0.145678 | -0.244431707 | 1095.970736 | 925.1617781 | 8 | 0 | 2 | 0 | 0 | 0 | 4 | 0 | 2 | 0 | 0 | 0 |
| JSG000000 | Amer1 | 0.108732 | 0.1469 | 0.284129708 | 495.0858164 | 602.8538647 | 36 | 1 | 2 | 2 | 12 | 2 | 7 | 0 | 0 | 1 | 3 | 0 |
| SG0000004 | Cbln3 | 0.109774 | 0.148139 | -0.52986452 | 122.9118296 | 85.13116358 | 5 | 2 | 2 | 2 | 4 | 0 | 5 | 2 | 2 | 2 | 4 | 0 |
| SG0000002 | Gsto2 | 0.112894 | 0.151824 | -0.748599338 | 36.95912154 | 21.99737126 | 42 | 0 | 2 | 4 | 10 | 6 | 0 | 0 | 0 | 0 | 0 | 0 |
| JSG000000 | Pcbd1 | 0.116643 | 0.156339 | 0.418599225 | 87.26574157 | 116.6419325 | 4 | 1 | 2 | 1 | 3 | 1 | 0 | 0 | 0 | 0 | 1 | 0 |
| SG0000001 | Golga7 | 0.117479 | 0.157388 | 0.204029139 | 2051.597753 | 2363.257831 | 52 | 2 | 2 | 8 | 20 | 2 | 10 | 0 | 0 | 2 | 4 | 0 |
| SG0000005 | Slfn5 | 0.118501 | 0.158639 | 0.93952632 | 11.13991597 | 21.36522737 | 18 | 1 | 2 | 1 | 3 | 0 | 5 | 1 | 1 | 1 | 1 | 0 |
| SG0000003 | Dbr1 | 0.121763 | 0.162678 | 0.240661722 | 717.449583 | 847.6914145 | 29 | 2 | 2 | 6 | 17 | 5 | 2 | 0 | 0 | 0 | 1 | 1 |
| JSG000000 | Isca1 | 0.122068 | 0.163048 | 0.26287602 | 416.1486228 | 499.3235263 | 15 | 2 | 2 | 4 | 8 | 0 | 1 | 0 | 0 | 0 | 1 | 0 |
| JSG000000 | Rgs16 | 0.122629 | 0.163688 | -0.335369396 | 366.6827623 | 290.6258541 | 10 | 2 | 2 | 3 | 5 | 1 | 4 | 2 | 2 | 3 | 3 | 0 |
| JSG000000 | Tmem101 | 0.125768 | 0.167529 | 0.260411101 | 411.484416 | 492.8842545 | 6 | 2 | 2 | 3 | 3 | 0 | 1 | 0 | 0 | 1 | 1 | 0 |
| SG0000002 | Ccng2 | 0.129912 | 0.172535 | 0.390152963 | 1207.10186 | 1581.946033 | 28 | 0 | 2 | 2 | 14 | 6 | 12 | 0 | 2 | 2 | 8 | 4 |
| SG0000002 | Ccdc127 | 0.129996 | 0.172635 | -0.22183873 | 1699.046215 | 1456.884357 | 26 | 2 | 2 | 4 | 13 | 4 | 14 | 1 | 1 | 2 | 5 | 1 |
| SG0000003 | Pik3ip1 | 0.130913 | 0.173749 | 0.473808634 | 180.1214801 | 250.147472 | 18 | 2 | 2 | 2 | 4 | 0 | 1 | 0 | 0 | 0 | 0 | 0 |
| SG0000003 | Atp6v0b | 0.13153 | 0.174464 | -0.200941249 | 2513.995072 | 2187.132423 | 4 | 2 | 2 | 2 | 4 | 0 | 2 | 2 | 2 | 2 | 2 | 0 |
| JSG000000 | Dlx2 | 0.14514 | 0.190737 | -0.861281572 | 25.70109018 | 14.14750831 | 3 | 1 | 2 | 1 | 3 | 1 | 3 | 1 | 2 | 1 | 3 | 1 |
| SG0000004 | Bsdc1 | 0.147507 | 0.193606 | 0.21313715 | 997.9545432 | 1156.835041 | 26 | 2 | 2 | 6 | 11 | 3 | 1 | 0 | 0 | 0 | 1 | 0 |
| JSG000000 | Gpc1 | 0.148888 | 0.195205 | 0.343470884 | 1330.543934 | 1688.201407 | 16 | 1 | 2 | 4 | 10 | 1 | 2 | 0 | 0 | 0 | 0 | 0 |
| SG0000004 | Slc25a23 | 0.149585 | 0.196075 | -0.222571461 | 1297.531064 | 1112.031471 | 10 | 2 | 2 | 2 | 7 | 2 | 0 | 0 | 0 | 0 | 0 | 0 |
| JSG000000 | B3galnt1 | 0.151152 | 0.19797 | 0.285334107 | 203.7106988 | 248.2606931 | 36 | 1 | 2 | 4 | 21 | 1 | 3 | 0 | 1 | 1 | 4 | 0 |
| SG0000003 | Tnxb | 0.152972 | 0.20012 | -0.868598084 | 31.99078951 | 17.5206769 | 34 | 1 | 2 | 5 | 13 | 3 | 0 | 0 | 0 | 0 | 0 | 0 |
| SG0000003 | 4931406C07Rik | 0.15828 | 0.2064 | 0.203866423 | 721.1722204 | 830.632458 | 30 | 1 | 2 | 4 | 15 | 6 | 2 | 0 | 0 | 0 | 4 | 0 |
| SG0000007 | 2510002D24Rik | 0.16112 | 0.209843 | -0.285848976 | 299.4911767 | 245.660265 | 4 | 2 | 2 | 4 | 10 | 2 | 0 | 0 | 0 | 0 | 0 | 0 |
| JSG000000 | Dand5 | 0.161706 | 0.21053 | 0.515770356 | 41.93698097 | 59.95970819 | 11 | 1 | 2 | 5 | 12 | 1 | 1 | 0 | 0 | 0 | 0 | 0 |
| JSG000000 | Sec22b | 0.16194 | 0.210789 | 0.197997886 | 1117.663441 | 1282.077704 | 153 | 2 | 2 | 14 | 57 | 6 | 7 | 0 | 0 | 0 | 4 | 0 |
| SG0000002 | Mettl2 | 0.16556 | 0.215112 | 0.212563365 | 753.1942287 | 872.7601987 | 16 | 1 | 2 | 1 | 6 | 1 | 0 | 0 | 0 | 0 | 0 | 0 |
| JSG000000 | Mrps33 | 0.170577 | 0.221003 | 0.301806149 | 151.9090229 | 187.2562293 | 41 | 0 | 2 | 2 | 11 | 3 | 1 | 0 | 0 | 0 | 0 | 0 |
| JSG000000 | Sec62 | 0.171239 | 0.221782 | -0.189121521 | 2642.80744 | 2318.111226 | 91 | 1 | 2 | 13 | 47 | 4 | 8 | 0 | 0 | 1 | 2 | 1 |
| SG0000003 | Pus3 | 0.173468 | 0.224311 | 0.226428463 | 474.0799114 | 554.6427139 | 44 | 2 | 2 | 2 | 6 | 4 | 4 | 0 | 0 | 0 | 0 | 0 |
| SG0000006 | Rbak | 0.174808 | 0.225734 | 0.353483846 | 225.612971 | 288.2526479 | 22 | 0 | 2 | 4 | 8 | 2 | 8 | 0 | 2 | 2 | 4 | 0 |
| JSG000000 | Mri1 | 0.175933 | 0.227072 | -0.209022439 | 883.0378332 | 763.9365465 | 8 | 2 | 2 | 3 | 4 | 6 | 2 | 0 | 0 | 1 | 1 | 0 |
| SG0000001 | Sparc | 0.184274 | 0.236865 | 0.362566982 | 4786.827647 | 6154.479475 | 28 | 2 | 2 | 4 | 4 | 14 | 2 | 0 | 0 | 0 | 0 | 0 |
| SG0000006 | Flnc | 0.195133 | 0.249444 | 0.571115927 | 720.0176193 | 1069.71023 | 16 | 0 | 2 | 0 | 4 | 0 | 0 | 0 | 0 | 0 | 0 | 0 |
| JSG000000 | Syf2 | 0.195321 | 0.249631 | -0.200200377 | 1182.378407 | 1029.177235 | 12 | 2 | 2 | 2 | 4 | 1 | 0 | 0 | 0 | 0 | 1 | 0 |
| JSG000000 | Rbmx2 | 0.199689 | 0.254579 | 0.207883802 | 453.9977939 | 524.3641627 | 21 | 2 | 2 | 2 | 18 | 7 | 5 | 1 | 1 | 1 | 1 | 1 |
| JSG000000 | Mettl21b | 0.20351 | 0.258861 | 1.212209812 | 6.781762226 | 15.71281775 | 12 | 2 | 2 | 4 | 11 | 1 | 3 | 1 | 1 | 1 | 2 | 0 |
| SG0000004 | Zfp574 | 0.210521 | 0.266774 | -0.329484763 | 623.7909725 | 496.4257364 | 6 | 2 | 2 | 2 | 2 | 2 | 2 | 0 | 0 | 0 | 0 | 2 |
| SG0000007 | Rmdn3 | 0.216852 | 0.273924 | 0.18385413 | 787.0739861 | 894.0486807 | 33 | 1 | 2 | 5 | 13 | 4 | 4 | 0 | 0 | 1 | 1 | 1 |
| JSG000000 | Pomk | 0.217292 | 0.274383 | 0.181494136 | 752.4735692 | 853.3484987 | 15 | 2 | 2 | 3 | 7 | 1 | 0 | 0 | 0 | 0 | 0 | 0 |
| SG0000002 | Vdr | 0.223992 | 0.28197 | 0.631768253 | 21.35865082 | 33.09443799 | 47 | 1 | 2 | 6 | 19 | 9 | 2 | 0 | 0 | 0 | 2 | 1 |
| JSG000000 | Tmem218 | 0.224651 | 0.282699 | 0.297975812 | 154.9022693 | 190.4396776 | 28 | 2 | 2 | 4 | 14 | 3 | 0 | 0 | 0 | 0 | 0 | 0 |
| JSG000000 | Zfp935 | 0.227298 | 0.28567 | -0.368600042 | 105.678136 | 81.85127996 | 20 | 2 | 2 | 3 | 13 | 2 | 0 | 0 | 0 | 0 | 0 | 0 |
| JSG000000 | Mbip | 0.23097 | 0.289858 | 0.189007629 | 568.186985 | 647.7214018 | 47 | 0 | 2 | 6 | 20 | 1 | 3 | 0 | 0 | 0 | 1 | 1 |
| JSG000000 | Ovgp1 | 0.23501 | 0.294165 | 0.481821143 | 36.82259836 | 51.42295711 | 10 | 1 | 2 | 4 | 10 | 2 | 0 | 0 | 0 | 0 | 0 | 0 |
| JSG000000 | Ndufc2 | 0.238924 | 0.298667 | 0.297092743 | 1058.304878 | 1300.303182 | 14 | 0 | 2 | 1 | 2 | 1 | 0 | 0 | 0 | 0 | 0 | 0 |
| SG0000005 | Zfp101 | 0.238992 | 0.298732 | -0.262181069 | 514.7818528 | 429.2386918 | 34 | 2 | 2 | 8 | 18 | 0 | 4 | 0 | 0 | 1 | 1 | 0 |
| SG0000002 | Cep41 | 0.23964 | 0.299375 | 0.224295817 | 243.2180719 | 284.129031 | 276 | 0 | 2 | 9 | 68 | 38 | 11 | 0 | 2 | 0 | 3 | 5 |
| SG0000002 | Zkscan17 | 0.247048 | 0.307599 | 0.151568545 | 1970.204701 | 2188.457058 | 40 | 2 | 2 | 6 | 8 | 4 | 6 | 0 | 0 | 0 | 0 | 0 |
| JSG000000 | Tuba8 | 0.25163 | 0.312566 | 0.755122799 | 12.83189035 | 21.65734689 | 11 | 2 | 2 | 3 | 10 | 2 | 1 | 0 | 0 | 0 | 0 | 0 |
| SG0000003 | Mfge8 | 0.261988 | 0.323703 | 0.420687404 | 1872.341792 | 2506.25155 | 22 | 2 | 2 | 4 | 10 | 2 | 0 | 0 | 0 | 0 | 0 | 0 |
| SG0000003 | Adcy7 | 0.262227 | 0.323977 | 0.294591949 | 133.7986505 | 164.1091312 | 23 | 2 | 2 | 5 | 9 | 14 | 2 | 1 | 1 | 2 | 2 | 0 |
| JSG000000 | Map6d1 | 0.26545 | 0.327619 | 0.379353389 | 45.75104483 | 59.51107093 | 9 | 2 | 2 | 3 | 4 | 1 | 5 | 2 | 2 | 2 | 3 | 1 |
| JSG000000 | 2310009B15Rik | 0.265571 | 0.327701 | -0.346394248 | 96.20072486 | 75.66643719 | 14 | 0 | 2 | 2 | 6 | 0 | 0 | 0 | 0 | 0 | 1 | 0 |
| JSG000000 | Mfap1a | 0.267883 | 0.330144 | -0.208246367 | 467.3253938 | 404.5115977 | 20 | 0 | 2 | 2 | 8 | 1 | 4 | 0 | 0 | 0 | 0 | 1 |
| SG0000000 | Mid1ip1 | 0.268668 | 0.331021 | 0.253600193 | 612.8688795 | 730.6490669 | 4 | 0 | 2 | 0 | 2 | 0 | 4 | 0 | 2 | 0 | 2 | 0 |
| JSG000000 | Tmem41a | 0.275117 | 0.33799 | -0.20460213 | 334.6069391 | 290.3645312 | 13 | 2 | 2 | 3 | 13 | 1 | 1 | 0 | 0 | 0 | 0 | 0 |
| SG0000003 | Bcl9 | 0.280329 | 0.343639 | 0.176113902 | 793.8270699 | 896.8947289 | 36 | 0 | 2 | 7 | 9 | 5 | 6 | 0 | 0 | 0 | 0 | 0 |
| SG0000000 | Ppan | 0.288667 | 0.352605 | 0.268139043 | 1513.048045 | 1822.093311 | 5 | 2 | 2 | 2 | 3 | 0 | 0 | 0 | 0 | 0 | 0 | 0 |
| SG0000002 | Pdap1 | 0.298131 | 0.362944 | 0.129882934 | 3897.815153 | 4265.008478 | 5 | 2 | 2 | 2 | 4 | 0 | 0 | 0 | 0 | 0 | 0 | 0 |
| JSG000000 | Gdnf | 0.301828 | 0.366706 | -0.668551222 | 20.88099916 | 13.13703341 | 27 | 2 | 2 | 5 | 26 | 2 | 3 | 2 | 2 | 3 | 5 | 0 |
| JSG000000 | Gm166 | 0.311324 | 0.376812 | 0.358929029 | 50.02958623 | 64.16161915 | 4 | 2 | 2 | 2 | 3 | 2 | 0 | 0 | 0 | 0 | 0 | 0 |
| SG0000004 | Lgalsl | 0.311539 | 0.377021 | -0.164227513 | 832.4565047 | 742.8893691 | 18 | 0 | 2 | 3 | 5 | 0 | 12 | 0 | 1 | 2 | 2 | 0 |
| JSG000000 | Mfap1b | 0.313896 | 0.379514 | 0.197160666 | 227.5364546 | 260.8568594 | 22 | 0 | 2 | 3 | 9 | 0 | 6 | 0 | 0 | 1 | 1 | 0 |
| JSG000000 | Rwdd1 | 0.314143 | 0.379761 | -0.157961978 | 1079.143166 | 967.2255742 | 42 | 2 | 2 | 9 | 20 | 4 | 0 | 0 | 0 | 0 | 0 | 0 |
| JSG000001 | Pcdhgc5 | 0.322111 | 0.388293 | -0.451706194 | 58.00776637 | 42.41398021 | 32 | 2 | 2 | 9 | 11 | 6 | 2 | 0 | 0 | 1 | 1 | 0 |
| JSG000000 | Lurap1 | 0.328908 | 0.395688 | 0.286641823 | 70.77152194 | 86.32693503 | 15 | 2 | 2 | 3 | 6 | 0 | 5 | 0 | 0 | 0 | 0 | 0 |
| SG0000004 | Setd1a | 0.32974 | 0.396529 | 0.145048316 | 1005.113244 | 1111.421745 | 34 | 2 | 2 | 12 | 20 | 10 | 2 | 0 | 0 | 2 | 2 | 2 |
| JSG000000 | Sf3b2 | 0.335047 | 0.401832 | 0.11855084 | 8910.233568 | 9673.339884 | 29 | 1 | 2 | 2 | 12 | 4 | 0 | 0 | 0 | 0 | 0 | 1 |
| JSG000000 | Ndufb4 | 0.342745 | 0.409993 | 0.217745254 | 170.8490586 | 198.6828783 | 17 | 1 | 2 | 2 | 8 | 0 | 1 | 0 | 0 | 0 | 0 | 0 |
| JSG000000 | Zfp865 | 0.344357 | 0.411592 | -0.279699146 | 97.38726553 | 80.22400491 | 24 | 2 | 2 | 4 | 4 | 0 | 0 | 0 | 0 | 0 | 0 | 0 |
| SG0000004 | Acad12 | 0.35249 | 0.420359 | -0.225594763 | 168.1949469 | 143.8474454 | 35 | 2 | 2 | 2 | 11 | 5 | 1 | 0 | 0 | 0 | 0 | 0 |
| SG0000004 | Nyap1 | 0.353301 | 0.421159 | 0.296532412 | 232.2786958 | 285.2821045 | 18 | 0 | 2 | 8 | 8 | 1 | 13 | 0 | 2 | 8 | 8 | 0 |
| SG0000006 | Numbl | 0.354942 | 0.422917 | 0.14947167 | 574.8562287 | 637.6093802 | 12 | 1 | 2 | 2 | 11 | 3 | 0 | 0 | 0 | 0 | 1 | 0 |
| JSG000000 | Igfbp6 | 0.355173 | 0.423165 | -0.514623561 | 39.58549473 | 27.70887852 | 4 | 2 | 2 | 2 | 4 | 0 | 0 | 0 | 0 | 0 | 0 | 0 |
| JSG000000 | Fam65a | 0.357611 | 0.42573 | 0.138224845 | 827.5653319 | 910.7771233 | 12 | 1 | 2 | 3 | 9 | 2 | 1 | 0 | 0 | 0 | 1 | 0 |
| SG0000002 | 2210016L21Rik | 0.364763 | 0.433207 | 0.141426646 | 746.3751134 | 823.2482434 | 8 | 2 | 2 | 2 | 5 | 0 | 0 | 0 | 0 | 0 | 0 | 0 |
| SG0000002 | Ppm1g | 0.367083 | 0.435588 | -0.11813951 | 7056.919436 | 6502.069737 | 59 | 0 | 2 | 8 | 26 | 9 | 2 | 0 | 0 | 0 | 0 | 0 |
| SG0000004 | Hscb | 0.372709 | 0.441269 | 0.25036439 | 252.9704542 | 300.910257 | 47 | 2 | 2 | 5 | 8 | 2 | 2 | 0 | 0 | 0 | 0 | 0 |
| JSG000000 | Med30 | 0.380638 | 0.449379 | 0.130172834 | 628.0052824 | 687.3045588 | 31 | 1 | 2 | 5 | 17 | 3 | 4 | 0 | 0 | 1 | 1 | 0 |
| SG0000004 | Nrep | 0.383863 | 0.452679 | 0.167151391 | 220.9614881 | 248.1042299 | 46 | 1 | 2 | 3 | 20 | 4 | 2 | 0 | 0 | 0 | 0 | 1 |
| SG0000003 | Hemk1 | 0.38422 | 0.452952 | 0.187860235 | 167.0380289 | 190.2685122 | 11 | 2 | 2 | 2 | 4 | 0 | 1 | 0 | 0 | 0 | 0 | 0 |
| SG0000002 | Top3b | 0.398808 | 0.467717 | -0.112017973 | 3263.019048 | 3019.248359 | 22 | 2 | 2 | 4 | 16 | 1 | 0 | 0 | 0 | 0 | 0 | 0 |
| SG0000003 | Fam46a | 0.417889 | 0.487098 | -0.270861047 | 83.64206847 | 69.32461471 | 24 | 2 | 2 | 2 | 22 | 6 | 18 | 0 | 0 | 0 | 16 | 4 |
| SG0000002 | Prpf8 | 0.430721 | 0.500037 | -0.1062993 | 17016.06718 | 15807.38073 | 56 | 0 | 2 | 4 | 14 | 8 | 0 | 0 | 0 | 0 | 0 | 0 |
| SG0000002 | Lox | 0.431482 | 0.500694 | -0.378334185 | 2813.595962 | 2164.570792 | 31 | 1 | 2 | 2 | 9 | 3 | 4 | 0 | 0 | 0 | 0 | 0 |
| JSG000000 | Rnft1 | 0.448989 | 0.518089 | 0.102587285 | 1018.015848 | 1093.040839 | 36 | 0 | 2 | 3 | 14 | 1 | 8 | 0 | 0 | 0 | 5 | 0 |
| SG0000002 | Dsn1 | 0.457641 | 0.527155 | 0.109708062 | 803.4454722 | 866.9255946 | 34 | 0 | 2 | 0 | 8 | 6 | 6 | 0 | 2 | 0 | 0 | 0 |
| SG0000003 | Mrps11 | 0.45803 | 0.527535 | 0.147402863 | 334.6243065 | 370.6211226 | 27 | 0 | 2 | 0 | 10 | 2 | 0 | 0 | 0 | 0 | 0 | 0 |
| JSG000000 | Kantr | 0.46604 | 0.535315 | -0.410170193 | 29.24729779 | 22.00960334 | 2 | 2 | 2 | 2 | 4 | 1 | 0 | 0 | 0 | 0 | 0 | 0 |
| JSG000000 | Spry2 | 0.466503 | 0.535675 | 0.102807067 | 2555.4548 | 2744.202874 | 6 | 1 | 2 | 4 | 7 | 1 | 2 | 0 | 0 | 2 | 5 | 1 |
| JSG000000 | Lage3 | 0.471376 | 0.540232 | 0.109066941 | 673.5885375 | 726.4857646 | 6 | 2 | 2 | 2 | 4 | 0 | 6 | 2 | 2 | 2 | 4 | 0 |
| SG0000003 | Srrm2 | 0.497414 | 0.564777 | 0.126984967 | 17008.45793 | 18573.39198 | 54 | 2 | 2 | 9 | 29 | 5 | 0 | 0 | 0 | 0 | 3 | 0 |

| | | | | | | | | | | | | | | | | | | |
|---|---|---|---|---|---|---|---|---|---|---|---|---|---|---|---|---|---|---|
| JSG000000 | Cln8 | 0.501839 | 0.569101 | 0.096807489 | 1055.786353 | 1129.062548 | 17 | 1 | 2 | 3 | 11 | 1 | 5 | 0 | 0 | 2 | 5 | 1 |
| SG0000004 | Rgl2 | 0.539939 | 0.60609 | 0.124172857 | 1038.573888 | 1131.923701 | 8 | 2 | 2 | 4 | 5 | 0 | 0 | 0 | 0 | 0 | 0 | 0 |
| SG0000003 | Slc37a2 | 0.546567 | 0.612369 | 0.213251732 | 51.60678565 | 59.82765462 | 45 | 2 | 2 | 4 | 12 | 0 | 4 | 0 | 0 | 0 | 0 | 0 |
| SG0000002 | Hagh | 0.548957 | 0.614635 | 0.100086054 | 580.3874518 | 622.0809738 | 16 | 0 | 2 | 4 | 8 | 0 | 4 | 0 | 0 | 4 | 4 | 0 |
| JSG000000 | Med4 | 0.549067 | 0.614719 | -0.091714475 | 2043.098612 | 1917.257834 | 21 | 0 | 2 | 3 | 8 | 2 | 0 | 0 | 0 | 0 | 0 | 2 |
| JSG000000 | Acot6 | 0.564019 | 0.62859 | 0.252040745 | 27.8991453 | 33.2248266 | 19 | 1 | 2 | 3 | 11 | 1 | 2 | 0 | 0 | 1 | 5 | 0 |
| JSG000000 | Creb3l1 | 0.571483 | 0.636039 | -0.151944935 | 312.3495987 | 281.1259715 | 28 | 1 | 2 | 4 | 26 | 3 | 0 | 0 | 0 | 0 | 2 | 1 |
| JSG000000 | Klf10 | 0.575865 | 0.64008 | 0.065769012 | 3247.504172 | 3398.97651 | 9 | 1 | 2 | 4 | 6 | 0 | 4 | 0 | 1 | 1 | 2 | 0 |
| JSG000000 | Pgf | 0.576024 | 0.640177 | 0.273829933 | 116.0517491 | 140.3080905 | 8 | 2 | 2 | 5 | 6 | 0 | 2 | 0 | 0 | 2 | 2 | 0 |
| SG0000007 | Efcab7 | 0.591768 | 0.654872 | 0.158745561 | 120.3837078 | 134.3862672 | 10 | 1 | 2 | 3 | 11 | 0 | 2 | 1 | 1 | 1 | 5 | 0 |
| SG0000002 | Lpin1 | 0.604518 | 0.667127 | 0.072986866 | 845.5892895 | 889.4687804 | 130 | 2 | 2 | 28 | 66 | 16 | 4 | 1 | 1 | 2 | 5 | 0 |
| SG0000005 | Homez | 0.62313 | 0.684713 | -0.102219864 | 460.0510657 | 428.5828573 | 29 | 2 | 2 | 8 | 13 | 0 | 13 | 0 | 0 | 5 | 7 | 0 |
| JSG000000 | Rarres1 | 0.625787 | 0.687085 | -0.399058252 | 22.50169504 | 17.06423135 | 56 | 0 | 2 | 6 | 26 | 4 | 0 | 0 | 0 | 0 | 2 | 0 |
| SG0000003 | Phkg2 | 0.635906 | 0.696146 | 0.071873146 | 662.7747262 | 696.6295808 | 36 | 2 | 2 | 8 | 14 | 2 | 0 | 0 | 0 | 0 | 0 | 0 |
| SG0000000 | Tchp | 0.639952 | 0.700019 | 0.077592553 | 446.8446244 | 471.5353144 | 7 | 0 | 2 | 3 | 5 | 0 | 0 | 0 | 0 | 0 | 0 | 0 |
| SG0000003 | Ift172 | 0.649528 | 0.708763 | -0.084958792 | 755.8772633 | 712.6497502 | 52 | 1 | 2 | 11 | 28 | 9 | 0 | 0 | 0 | 0 | 0 | 0 |
| SG0000002 | Hus1 | 0.655226 | 0.713806 | -0.076300577 | 1850.435441 | 1755.113373 | 54 | 0 | 2 | 4 | 22 | 6 | 18 | 0 | 0 | 2 | 4 | 2 |
| JSG000000 | 0610009B22Rik | 0.661118 | 0.719387 | -0.08149725 | 568.8735792 | 537.6289509 | 18 | 0 | 2 | 6 | 18 | 0 | 4 | 0 | 0 | 0 | 2 | 0 |
| JSG000000 | Lysmd1 | 0.661576 | 0.719762 | 0.261903072 | 15.74267789 | 18.87640515 | 7 | 1 | 2 | 2 | 6 | 0 | 2 | 0 | 0 | 1 | 1 | 0 |
| SG0000002 | Ddx54 | 0.666848 | 0.724793 | -0.126323156 | 3410.314258 | 3124.405127 | 12 | 2 | 2 | 3 | 4 | 2 | 0 | 0 | 0 | 0 | 0 | 1 |
| SG0000002 | 4933434E20Rik | 0.670336 | 0.727745 | 0.055460908 | 961.7201165 | 999.410936 | 18 | 2 | 2 | 3 | 16 | 0 | 3 | 0 | 0 | 0 | 0 | 0 |
| JSG000000 | Ccdc134 | 0.676106 | 0.733164 | -0.080730311 | 313.2035983 | 296.1587017 | 16 | 2 | 2 | 2 | 5 | 2 | 0 | 0 | 0 | 0 | 0 | 0 |
| JSG000000 | Letmd1 | 0.681501 | 0.737943 | -0.103260067 | 189.0775991 | 176.0174404 | 11 | 1 | 2 | 3 | 10 | 3 | 1 | 0 | 0 | 0 | 0 | 0 |
| JSG000000 | Eri1 | 0.688502 | 0.743859 | -0.066299584 | 2530.781299 | 2417.110211 | 52 | 0 | 2 | 6 | 29 | 7 | 6 | 0 | 0 | 0 | 3 | 3 |
| JSG000000 | Hmgxb3 | 0.694089 | 0.748597 | 0.056764436 | 1651.336857 | 1717.605698 | 52 | 2 | 2 | 8 | 21 | 2 | 3 | 0 | 0 | 0 | 0 | 0 |
| JSG000000 | Bdkrb2 | 0.704726 | 0.758407 | -0.25399994 | 17.45171337 | 14.6344522 | 20 | 2 | 2 | 3 | 8 | 2 | 1 | 0 | 0 | 0 | 0 | 0 |
| JSG000000 | Twsg1 | 0.708953 | 0.761718 | -0.056637856 | 2399.413845 | 2307.041818 | 51 | 0 | 2 | 5 | 18 | 3 | 6 | 0 | 0 | 0 | 2 | 0 |
| JSG000000 | Arl2 | 0.720027 | 0.772282 | 0.053894768 | 844.1782795 | 876.3107014 | 12 | 2 | 2 | 4 | 6 | 0 | 0 | 0 | 0 | 0 | 0 | 0 |
| SG0000005 | Otop1 | 0.729664 | 0.780889 | -0.29294266 | 24.36730536 | 19.88945935 | 51 | 0 | 2 | 7 | 13 | 0 | 4 | 0 | 2 | 0 | 0 | 0 |
| SG0000003 | Dmwd | 0.742041 | 0.791171 | 0.073855303 | 1105.138803 | 1163.186865 | 2 | 0 | 2 | 2 | 2 | 2 | 2 | 0 | 2 | 2 | 2 | 0 |
| JSG000000 | Prkaca | 0.745492 | 0.794174 | -0.04514684 | 2369.232883 | 2296.23958 | 18 | 2 | 2 | 4 | 12 | 2 | 3 | 0 | 0 | 0 | 0 | 0 |
| SG0000003 | Ythdf1 | 0.750079 | 0.798349 | -0.040904308 | 3968.959848 | 3858.009402 | 67 | 0 | 2 | 12 | 26 | 4 | 5 | 0 | 0 | 0 | 4 | 0 |
| JSG000000 | Gtf2b | 0.753279 | 0.801136 | 0.039737789 | 1084.142586 | 1114.419418 | 49 | 2 | 2 | 4 | 16 | 3 | 2 | 0 | 0 | 0 | 1 | 2 |
| SG0000002 | Dscr3 | 0.753416 | 0.801234 | -0.050560938 | 1563.923426 | 1510.063207 | 32 | 1 | 2 | 6 | 15 | 3 | 1 | 0 | 0 | 0 | 0 | 0 |
| SG0000003 | Srpr | 0.761371 | 0.808111 | 0.038416813 | 3050.028919 | 3132.337648 | 53 | 1 | 2 | 4 | 23 | 9 | 1 | 0 | 0 | 0 | 0 | 0 |
| SG0000003 | Mpi | 0.765653 | 0.811934 | -0.043230162 | 853.2734964 | 828.0845477 | 7 | 0 | 2 | 0 | 10 | 2 | 0 | 0 | 0 | 0 | 0 | 0 |
| SG0000002 | Oas1b | 0.779952 | 0.824315 | 0.058326983 | 114.7922861 | 119.5283368 | 16 | 0 | 2 | 0 | 6 | 4 | 0 | 0 | 0 | 0 | 3 | 0 |
| JSG000000 | Col10a1 | 0.791233 | 0.833779 | -0.14901884 | 206.8508171 | 186.5512228 | 21 | 1 | 2 | 2 | 2 | 0 | 4 | 1 | 1 | 2 | 2 | 0 |
| SG0000005 | Sipa1 | 0.799732 | 0.841261 | -0.067904524 | 573.4765694 | 547.1096563 | 16 | 0 | 2 | 2 | 4 | 4 | 0 | 0 | 0 | 0 | 0 | 2 |
| JSG000000 | Dpcd | 0.813315 | 0.852733 | 0.039616221 | 476.4339035 | 489.6979991 | 17 | 1 | 2 | 1 | 7 | 0 | 0 | 0 | 0 | 0 | 0 | 0 |
| SG0000002 | Gpn2 | 0.817987 | 0.856728 | -0.044571467 | 410.9010665 | 398.4005415 | 26 | 2 | 2 | 4 | 4 | 0 | 2 | 0 | 0 | 0 | 0 | 0 |
| JSG000000 | Ppp6r1 | 0.826502 | 0.863778 | -0.028601898 | 2177.13422 | 2134.396867 | 19 | 2 | 2 | 5 | 13 | 6 | 0 | 0 | 0 | 0 | 2 | 0 |
| SG0000000 | Eif4g2 | 0.829775 | 0.866555 | -0.029092591 | 30835.12446 | 30219.54792 | 37 | 0 | 2 | 2 | 6 | 5 | 7 | 0 | 0 | 0 | 2 | 1 |
| SG0000000 | Mcoln1 | 0.839985 | 0.875062 | 0.030447168 | 1330.398 | 1358.773581 | 10 | 2 | 2 | 2 | 5 | 0 | 2 | 2 | 2 | 2 | 2 | 0 |
| SG0000000 | Pih1d2 | 0.84104 | 0.876008 | -0.110389492 | 58.93109436 | 54.59010854 | 24 | 2 | 2 | 4 | 18 | 0 | 1 | 0 | 0 | 0 | 0 | 0 |
| SG0000001 | Rpl19 | 0.844535 | 0.878371 | -0.02956657 | 17376.00961 | 17023.53012 | 12 | 0 | 2 | 0 | 2 | 4 | 0 | 0 | 0 | 0 | 0 | 2 |
| SG0000002 | Rab35 | 0.876365 | 0.905103 | -0.016129636 | 2270.500954 | 2245.257648 | 14 | 0 | 2 | 0 | 4 | 4 | 2 | 0 | 2 | 0 | 1 | 2 |
| JSG000000 | Eif1ax | 0.876401 | 0.905103 | -0.030798678 | 4385.188437 | 4292.565535 | 33 | 1 | 2 | 7 | 15 | 6 | 7 | 0 | 1 | 0 | 3 | 1 |
| JSG000000 | Usp42 | 0.880912 | 0.90819 | -0.030292374 | 834.5446252 | 817.2043152 | 29 | 1 | 2 | 4 | 11 | 7 | 5 | 0 | 1 | 0 | 1 | 2 |
| JSG000000 | Ppib | 0.882062 | 0.908905 | -0.021894509 | 4379.14588 | 4313.189198 | 8 | 2 | 2 | 2 | 4 | 0 | 0 | 0 | 0 | 0 | 0 | 0 |
| SG0000002 | Pex16 | 0.884145 | 0.91079 | 0.037372055 | 178.2304749 | 182.9077359 | 4 | 2 | 2 | 2 | 2 | 0 | 0 | 0 | 0 | 0 | 0 | 0 |
| JSG000000 | Mfsd1 | 0.890335 | 0.91606 | 0.014260942 | 1626.828783 | 1642.989618 | 37 | 0 | 2 | 5 | 10 | 2 | 2 | 0 | 0 | 1 | 2 | 0 |
| SG0000002 | Mdh1 | 0.930089 | 0.949327 | 0.005730547 | 4196.977431 | 4213.68145 | 41 | 0 | 2 | 3 | 9 | 6 | 1 | 0 | 0 | 0 | 1 | 0 |
| SG0000007 | Zfp94 | 0.931252 | 0.949851 | 0.022392335 | 138.4460588 | 140.6116685 | 18 | 2 | 2 | 4 | 12 | 8 | 2 | 0 | 0 | 0 | 0 | 0 |
| SG0000002 | Telo2 | 0.936548 | 0.953881 | 0.015349558 | 570.5018179 | 576.6040788 | 14 | 2 | 2 | 2 | 8 | 2 | 0 | 0 | 0 | 0 | 0 | 0 |
| SG0000002 | Mrpl40 | 0.939062 | 0.956035 | -0.017169774 | 1404.552062 | 1387.93531 | 17 | 2 | 2 | 2 | 2 | 0 | 1 | 0 | 0 | 0 | 0 | 0 |
| SG0000001 | Apip | 0.95351 | 0.968817 | -0.01555476 | 344.878348 | 341.1799335 | 41 | 2 | 2 | 7 | 17 | 4 | 0 | 0 | 0 | 0 | 0 | 0 |
| JSG000000 | Cwc15 | 0.959375 | 0.974113 | -0.0139288 | 2052.690807 | 2032.968039 | 15 | 1 | 2 | 4 | 10 | 2 | 0 | 0 | 0 | 0 | 0 | 0 |
| JSG000000 | Mark4 | 0.960132 | 0.974781 | -0.014049055 | 542.4649123 | 537.2079924 | 28 | 1 | 2 | 4 | 11 | 4 | 0 | 0 | 0 | 0 | 0 | 0 |
| SG0000004 | Abhd13 | 0.961787 | 0.976285 | 0.001570142 | 1321.35476 | 1322.793625 | 42 | 0 | 2 | 2 | 24 | 4 | 20 | 0 | 0 | 2 | 6 | 2 |
| JSG000000 | Zfp810 | 0.974962 | 0.987365 | -0.019709737 | 174.2725073 | 171.9078297 | 49 | 2 | 2 | 6 | 21 | 7 | 4 | 0 | 0 | 0 | 0 | 0 |
| SG0000006 | Gmfg | 0.9822 | 0.993403 | -0.008066662 | 225.7852693 | 224.52634 | 53 | 0 | 2 | 7 | 24 | 2 | 0 | 0 | 0 | 0 | 0 | 0 |
| JSG000000 | Serp2 | 0.989607 | 0.999176 | -0.033313914 | 41.37029913 | 40.42594436 | 28 | 1 | 2 | 7 | 21 | 3 | 0 | 0 | 0 | 0 | 0 | 0 |
| SG0000000 | Snrpb2 | 0.994107 | 1 | -0.00870742 | 2132.769633 | 2119.936019 | 25 | 1 | 2 | 5 | 11 | 1 | 3 | 0 | 1 | 1 | 1 | 0 |
| JSG000000 | Slc5a7 | 1 | 1 | -0.008613583 | 25.71764829 | 25.56455903 | 54 | 0 | 2 | 7 | 33 | 5 | 12 | 0 | 0 | 3 | 7 | 1 |
| JSG000000 | Adat3 | 0.999581 | 1 | -0.001045698 | 180.5270967 | 180.396294 | 7 | 2 | 2 | 2 | 6 | 1 | 0 | 0 | 0 | 0 | 0 | 0 |
| JSG000000 | Ndufa13 | 1 | 1 | 0.016099177 | 1794.653966 | 1814.792845 | 10 | 1 | 2 | 4 | 8 | 0 | 0 | 0 | 0 | 0 | 0 | 0 |
| JSG000000 | Trappc5 | 1 | 1 | -0.002058043 | 822.7684545 | 821.5955899 | 10 | 1 | 2 | 2 | 4 | 0 | 4 | 0 | 0 | 0 | 2 | 0 |
| JSG000000 | Fxyd6 | 5.6E-159 | 1E-156 | 8.680112084 | 4.622166578 | 1895.918637 | 24 | 1 | 1 | 1 | 5 | 2 | 1 | 0 | 0 | 0 | 0 | 0 |
| SG0000005 | C1ra | 9.5E-158 | 1.7E-155 | -7.531646905 | 3845.806412 | 20.78431827 | 10 | 0 | 1 | 2 | 3 | 0 | 1 | 0 | 0 | 0 | 0 | 0 |
| JSG000000 | Taf7 | 1.4E-151 | 2.2E-149 | 4.743331433 | 153.3993825 | 4108.739925 | 6 | 0 | 1 | 0 | 2 | 0 | 5 | 0 | 1 | 0 | 1 | 0 |
| JSG000000 | Des | 2.5E-151 | 3.8E-149 | 6.95575383 | 5.592877993 | 694.2659895 | 6 | 0 | 1 | 0 | 0 | 0 | 1 | 0 | 0 | 0 | 0 | 0 |
| SG0000002 | Igfbp3 | 1.6E-129 | 1.8E-127 | -7.113402532 | 119052.5512 | 859.7875918 | 12 | 0 | 1 | 1 | 4 | 1 | 4 | 0 | 0 | 1 | 2 | 1 |
| JSG000000 | Efs | 6.5E-121 | 6.6E-119 | 5.209173387 | 18.07007235 | 668.4629384 | 10 | 0 | 1 | 2 | 3 | 0 | 1 | 0 | 0 | 0 | 0 | 0 |
| SG0000005 | Dhcr7 | 2E-119 | 2E-117 | 3.313211755 | 205.7740804 | 2045.34602 | 74 | 0 | 1 | 11 | 30 | 6 | 3 | 0 | 0 | 0 | 0 | 0 |
| JSG000000 | Krt20 | 1.5E-117 | 1.4E-115 | -8.662184234 | 2742.902395 | 6.770692205 | 10 | 0 | 1 | 1 | 2 | 0 | 2 | 0 | 0 | 1 | 1 | 0 |
| SG0000002 | Sema4g | 1.4E-113 | 1.3E-111 | 4.114017369 | 57.18509926 | 990.2059009 | 11 | 1 | 1 | 5 | 7 | 1 | 2 | 0 | 0 | 1 | 1 | 0 |
| SG0000005 | Wdr86 | 9.6E-112 | 8.7E-110 | 8.55680215 | 0 | 375.57728 | 12 | 1 | 1 | 3 | 6 | 2 | 1 | 0 | 0 | 0 | 0 | 0 |
| JSG000000 | Mmp17 | 9.7E-112 | 8.8E-110 | 4.381822641 | 37.30025001 | 777.6278275 | 18 | 1 | 1 | 3 | 8 | 11 | 1 | 0 | 0 | 0 | 0 | 1 |
| SG0000003 | Tub | 2.8E-99 | 1.95E-97 | 8.342195535 | 0 | 323.5271872 | 17 | 1 | 1 | 2 | 3 | 4 | 4 | 1 | 1 | 2 | 2 | 0 |
| JSG000000 | Chga | 1.68E-97 | 1.14E-95 | 8.34758158 | 0 | 324.7410163 | 5 | 1 | 1 | 1 | 1 | 0 | 0 | 0 | 0 | 0 | 0 | 0 |
| JSG000000 | Pou3f1 | 2.84E-96 | 1.87E-94 | 8.344426157 | 0 | 324.0293428 | 6 | 1 | 1 | 4 | 6 | 1 | 6 | 1 | 1 | 4 | 6 | 1 |
| SG0000000 | Zbtb32 | 5.17E-91 | 2.99E-89 | -4.22639598 | 3926.538825 | 209.7673697 | 16 | 1 | 1 | 1 | 2 | 2 | 2 | 0 | 0 | 0 | 0 | 0 |
| JSG000000 | Cldn1 | 2.19E-86 | 1.14E-84 | -7.758887246 | 1433.57439 | 6.618543691 | 27 | 1 | 1 | 3 | 17 | 3 | 3 | 0 | 0 | 0 | 3 | 0 |
| JSG000000 | Tmem30b | 8.11E-82 | 3.8E-80 | 10.41034642 | 0.331601068 | 451.2762308 | 4 | 1 | 1 | 1 | 6 | 0 | 4 | 1 | 1 | 1 | 6 | 0 |
| JSG000000 | Arpin | 3.78E-81 | 1.74E-79 | -4.63182887 | 3316.843928 | 133.7844421 | 11 | 1 | 1 | 1 | 2 | 2 | 0 | 0 | 0 | 0 | 0 | 1 |
| JSG000000 | Alox15 | 3.02E-76 | 1.27E-74 | -10.44532982 | 58753.51544 | 42.1382279 | 5 | 0 | 1 | 0 | 1 | 0 | 1 | 0 | 1 | 0 | 0 | 0 |
| JSG000000 | Gstt2 | 4.81E-74 | 1.94E-72 | 4.406024162 | 20.3865927 | 432.2052925 | 3 | 0 | 1 | 1 | 1 | 1 | 0 | 0 | 0 | 0 | 0 | 0 |
| JSG000000 | Cd24a | 2.5E-72 | 9.57E-71 | -4.209510111 | 33488.83009 | 1810.135833 | 12 | 1 | 1 | 3 | 7 | 2 | 3 | 0 | 0 | 1 | 1 | 2 |
| JSG000000 | Nefh | 3.38E-71 | 1.25E-69 | 9.921595812 | 0.331601068 | 321.5983898 | 6 | 0 | 1 | 0 | 1 | 0 | 1 | 0 | 0 | 0 | 0 | 0 |
| JSG000000 | Fosl1 | 9.36E-71 | 3.41E-69 | -4.139352427 | 2009.881508 | 114.0515551 | 7 | 1 | 1 | 3 | 6 | 0 | 1 | 0 | 0 | 1 | 2 | 0 |
| JSG000000 | Kcnj10 | 1.11E-70 | 4.04E-69 | 7.76160964 | 0 | 216.0087854 | 26 | 0 | 1 | 3 | 15 | 5 | 4 | 0 | 0 | 0 | 1 | 2 |
| SG0000002 | Avil | 2.97E-69 | 1.04E-67 | -11.42275215 | 17785.67553 | 6.478572688 | 18 | 0 | 1 | 0 | 4 | 0 | 1 | 0 | 1 | 0 | 0 | 0 |
| JSG000000 | Cdsn | 1.48E-67 | 5.01E-66 | -4.776465014 | 728.8608579 | 26.59410539 | 4 | 1 | 1 | 2 | 2 | 0 | 0 | 0 | 0 | 0 | 0 | 0 |
| JSG000000 | Tubb2b | 1.39E-66 | 4.58E-65 | 3.075737555 | 352.389563 | 2971.066003 | 3 | 1 | 1 | 1 | 2 | 0 | 0 | 0 | 0 | 0 | 0 | 0 |
| JSG000000 | Ltbr | 2.18E-66 | 7.1E-65 | -3.826872179 | 2156.834648 | 151.9894824 | 4 | 1 | 1 | 2 | 5 | 0 | 1 | 0 | 0 | 1 | 1 | 0 |
| JSG000000 | Phlda1 | 2.31E-65 | 7.3E-64 | -4.215385089 | 3385.769411 | 182.2636327 | 3 | 1 | 1 | 2 | 2 | 0 | 3 | 1 | 1 | 2 | 2 | 0 |
| JSG000000 | Plk2 | 2.28E-64 | 7.09E-63 | -3.688645503 | 10656.88024 | 826.489129 | 9 | 1 | 1 | 2 | 10 | 0 | 2 | 1 | 1 | 2 | 8 | 0 |
| SG0000003 | Cx3cl1 | 8.05E-64 | 2.46E-62 | -5.4078641 | 5412.682691 | 127.4920962 | 19 | 1 | 1 | 1 | 19 | 0 | 1 | 1 | 1 | 1 | 1 | 0 |
| JSG000000 | Il4ra | 1.63E-63 | 4.97E-62 | -4.494823766 | 5684.910352 | 252.142957 | 14 | 1 | 1 | 1 | 4 | 0 | 0 | 0 | 0 | 0 | 0 | 0 |
| SG0000003 | Dhodh | 6.31E-62 | 1.85E-60 | 2.47427757 | 385.7445166 | 2143.539681 | 17 | 1 | 1 | 3 | 9 | 3 | 1 | 0 | 0 | 0 | 0 | 0 |
| SG0000000 | S100a3 | 4.01E-61 | 1.15E-59 | -5.221509918 | 505.2037899 | 13.54052464 | 32 | 1 | 1 | 1 | 8 | 4 | 0 | 0 | 0 | 0 | 0 | 0 |
| JSG000000 | Vim | 1.75E-60 | 4.93E-59 | -2.770064516 | 27646.71518 | 4052.947976 | 11 | 0 | 1 | 1 | 3 | 1 | 0 | 0 | 0 | 0 | 0 | 0 |
| SG0000004 | Nrsn1 | 3.43E-59 | 9.26E-58 | 7.46903033 | 0 | 176.1748883 | 42 | 0 | 1 | 1 | 6 | 3 | 12 | 0 | 0 | 0 | 0 | 0 |
| SG0000002 | Celsr3 | 5.65E-59 | 1.52E-57 | 4.474706687 | 13.38884808 | 297.689933 | 8 | 1 | 1 | 1 | 2 | 1 | 1 | 1 | 1 | 1 | 1 | 0 |
| SG0000007 | Ly6c1 | 1.39E-58 | 3.67E-57 | -8.440270533 | 8337.218783 | 24.00195385 | 10 | 1 | 1 | 2 | 5 | 0 | 2 | 0 | 0 | 0 | 2 | 0 |
| JSG000000 | Gja1 | 3.2E-58 | 8.37E-57 | -3.275930807 | 29938.06631 | 3090.795771 | 22 | 0 | 1 | 1 | 5 | 1 | 4 | 0 | 0 | 0 | 2 | 0 |
| JSG000000 | Phyh | 6.95E-58 | 1.78E-56 | 2.468850877 | 200.4209703 | 1109.535825 | 24 | 1 | 1 | 2 | 14 | 2 | 1 | 0 | 0 | 0 | 0 | 0 |
| JSG000000 | Shisa2 | 9.28E-56 | 2.3E-54 | 6.861032251 | 1.620695873 | 188.3987578 | 8 | 1 | 1 | 1 | 7 | 0 | 1 | 0 | 0 | 0 | 3 | 0 |
| JSG000000 | Sox11 | 2.69E-55 | 6.52E-54 | 5.522946171 | 25.26793183 | 1161.828855 | 10 | 1 | 1 | 1 | 4 | 1 | 10 | 1 | 1 | 1 | 4 | 1 |
| SG0000001 | Phox2b | 4.03E-55 | 9.73E-54 | -12.56147819 | 4352.376848 | 0.720020342 | 15 | 0 | 1 | 1 | 2 | 0 | 4 | 0 | 1 | 1 | 1 | 0 |
| SG0000000 | Ccne1 | 1.02E-54 | 2.41E-53 | 1.936495224 | 1364.809345 | 5224.143963 | 9 | 1 | 1 | 3 | 4 | 0 | 0 | 0 | 0 | 0 | 0 | 0 |
| JSG000000 | Rgs9bp | 3.68E-52 | 8.13E-51 | 6.334412319 | 2.329738024 | 187.9988535 | 10 | 1 | 1 | 1 | 3 | 2 | 10 | 1 | 1 | 1 | 3 | 2 |
| JSG000000 | Hexa | 1.91E-50 | 4E-49 | 1.945664247 | 701.1555878 | 2700.957181 | 35 | 0 | 1 | 2 | 7 | 2 | 0 | 0 | 0 | 0 | 0 | 0 |
| SG0000003 | Elovl4 | 2.02E-50 | 4.23E-49 | 3.213773276 | 44.9951658 | 417.453409 | 47 | 0 | 1 | 5 | 23 | 7 | 9 | 0 | 0 | 2 | 4 | 0 |
| SG0000003 | Ebp | 2.21E-49 | 4.5E-48 | -2.635444728 | 3428.830221 | 551.8208 | 12 | 0 | 1 | 1 | 12 | 1 | 0 | 0 | 0 | 0 | 0 | 0 |
| SG0000002 | Rps6ka4 | 8E-49 | 1.6E-47 | -3.598071111 | 3335.092734 | 275.4105569 | 3 | 1 | 1 | 1 | 3 | 1 | 0 | 0 | 0 | 0 | 1 | 0 |
| SG0000004 | Palm3 | 2.21E-48 | 4.39E-47 | 8.302217796 | 3.274014081 | 1033.468229 | 3 | 1 | 1 | 1 | 2 | 2 | 0 | 0 | 0 | 0 | 0 | 0 |
| JSG000000 | Adamts15 | 4.18E-48 | 8.2E-47 | 7.133312773 | 0 | 139.3915966 | 20 | 1 | 1 | 1 | 3 | 3 | 2 | 0 | 0 | 0 | 0 | 1 |
| JSG000000 | Cxxc5 | 3.55E-47 | 6.75E-46 | 2.083060352 | 330.3487993 | 1399.704572 | 18 | 1 | 1 | 3 | 5 | 2 | 1 | 0 | 0 | 0 | 0 | 0 |
| JSG000000 | Rdh10 | 1.24E-46 | 2.32E-45 | -2.887556815 | 1423.469575 | 192.3565474 | 45 | 1 | 1 | 2 | 13 | 1 | 4 | 0 | 0 | 0 | 1 | 0 |
| JSG000000 | Mal2 | 1.55E-46 | 2.87E-45 | 7.817256822 | 0.639110347 | 144.1466771 | 57 | 1 | 1 | 7 | 22 | 9 | 5 | 0 | 0 | 2 | 4 | 1 |
| SG0000004 | Dnd1 | 2.44E-46 | 4.48E-45 | 4.204093036 | 14.04736308 | 258.9124391 | 2 | 0 | 1 | 1 | 1 | 0 | 1 | 0 | 1 | 1 | 1 | 0 |

| | | | | | | | | | | | | | | | | | | |
|---|---|---|---|---|---|---|---|---|---|---|---|---|---|---|---|---|---|---|
| SG0000000 | Arrdc2 | 3.76E-46 | 6.85E-45 | 2.575023362 | 103.5672269 | 617.1370485 | 3 | 1 | 1 | 1 | 2 | 1 | 0 | 0 | 0 | 0 | 0 | 0 |
| JSG000000 | Hoxa7 | 1.23E-45 | 2.21E-44 | -4.17132498 | 449.8276964 | 24.96618885 | 15 | 1 | 1 | 2 | 6 | 0 | 4 | 0 | 0 | 1 | 1 | 0 |
| SG0000002 | Zap70 | 1.49E-45 | 2.67E-44 | 7.048605794 | 0 | 131.385912 | 24 | 1 | 1 | 5 | 10 | 3 | 0 | 0 | 0 | 0 | 0 | 0 |
| SG0000001 | Serinc1 | 1.5E-45 | 2.69E-44 | -2.485187223 | 11614.4516 | 2074.353697 | 37 | 0 | 1 | 6 | 24 | 6 | 6 | 0 | 0 | 2 | 2 | 2 |
| JSG000000 | Mcur1 | 1.62E-45 | 2.9E-44 | -3.005536472 | 1077.708567 | 134.1975861 | 22 | 1 | 1 | 1 | 11 | 3 | 6 | 1 | 1 | 1 | 6 | 0 |
| JSG000000 | Prr15l | 2.39E-45 | 4.25E-44 | 7.045078719 | 0 | 131.0626526 | 16 | 0 | 1 | 5 | 8 | 0 | 9 | 0 | 1 | 5 | 5 | 0 |
| JSG000000 | 1810011O10Rik | 5.06E-45 | 8.9E-44 | -4.724641178 | 969.3243897 | 36.66151662 | 3 | 1 | 1 | 1 | 4 | 0 | 3 | 1 | 1 | 1 | 4 | 0 |
| JSG000000 | Reep5 | 6.33E-45 | 1.11E-43 | -2.768027499 | 5654.756712 | 830.1462656 | 43 | 1 | 1 | 8 | 22 | 4 | 4 | 0 | 0 | 0 | 0 | 1 |
| SG0000002 | Mutyh | 8.05E-45 | 1.4E-43 | 2.708063651 | 80.37808477 | 525.2257899 | 6 | 0 | 1 | 0 | 2 | 2 | 0 | 0 | 0 | 0 | 0 | 0 |
| JSG000000 | Twist1 | 4.24E-44 | 7.19E-43 | -3.913862138 | 1028.638728 | 68.24533483 | 2 | 1 | 1 | 2 | 5 | 1 | 2 | 1 | 1 | 2 | 5 | 1 |
| JSG000000 | Sv2a | 5.92E-44 | 9.99E-43 | 4.369388954 | 10.82387615 | 223.7174972 | 8 | 0 | 1 | 2 | 2 | 2 | 3 | 0 | 1 | 1 | 1 | 0 |
| JSG000000 | Hoxa5 | 1.94E-43 | 3.21E-42 | -5.519403128 | 336.3229526 | 7.332475277 | 4 | 0 | 1 | 1 | 2 | 0 | 2 | 0 | 1 | 1 | 2 | 0 |
| SG0000002 | Slc17a9 | 1.29E-41 | 2.02E-40 | -2.489269604 | 1979.353598 | 352.5157835 | 6 | 1 | 1 | 1 | 7 | 0 | 2 | 0 | 0 | 0 | 0 | 0 |
| SG0000002 | Shc2 | 2.84E-41 | 4.39E-40 | 3.279345451 | 32.84104197 | 318.8587027 | 12 | 1 | 1 | 1 | 9 | 2 | 0 | 0 | 0 | 0 | 0 | 0 |
| JSG000000 | Cst13 | 1.8E-40 | 2.71E-39 | -7.367777379 | 164.1665101 | 0 | 20 | 1 | 1 | 3 | 7 | 4 | 0 | 0 | 0 | 0 | 0 | 0 |
| JSG000000 | Arhgap9 | 4.69E-40 | 6.98E-39 | 3.91615474 | 14.38983826 | 217.2380806 | 11 | 1 | 1 | 3 | 5 | 0 | 0 | 0 | 0 | 0 | 0 | 0 |
| JSG000000 | Fem1a | 7.12E-40 | 1.05E-38 | -2.375170745 | 2146.562105 | 413.757443 | 5 | 0 | 1 | 1 | 1 | 1 | 5 | 0 | 1 | 1 | 1 | 1 |
| SG0000002 | Pgc | 7.59E-40 | 1.12E-38 | 6.905242651 | 0 | 118.8630061 | 4 | 1 | 1 | 1 | 2 | 0 | 0 | 0 | 0 | 0 | 0 | 0 |
| JSG000000 | C1rb | 7.84E-40 | 1.15E-38 | -5.934922545 | 203.223323 | 3.321879241 | 7 | 0 | 1 | 2 | 4 | 0 | 0 | 0 | 0 | 0 | 0 | 0 |
| SG0000003 | Samd10 | 8.79E-40 | 1.28E-38 | -2.534863004 | 1438.365805 | 248.1987275 | 3 | 1 | 1 | 1 | 2 | 0 | 1 | 1 | 1 | 1 | 1 | 0 |
| JSG000000 | Rab39 | 1.39E-39 | 2.01E-38 | 4.365583153 | 8.928293334 | 184.0517402 | 41 | 1 | 1 | 6 | 25 | 2 | 7 | 0 | 0 | 1 | 4 | 1 |
| JSG000000 | Gulo | 5E-39 | 7.13E-38 | 7.550707532 | 0.639110347 | 119.8299295 | 12 | 1 | 1 | 5 | 12 | 5 | 0 | 0 | 0 | 0 | 0 | 0 |
| JSG000000 | Adcy6 | 5.27E-39 | 7.51E-38 | -2.363707777 | 2009.819677 | 390.4902117 | 16 | 1 | 1 | 1 | 4 | 0 | 3 | 0 | 0 | 0 | 0 | 0 |
| SG0000000 | Ccnd2 | 1.99E-38 | 2.79E-37 | -2.734525237 | 8155.003955 | 1225.321505 | 31 | 0 | 1 | 2 | 14 | 1 | 2 | 0 | 0 | 0 | 5 | 0 |
| SG0000003 | Kcnh2 | 1.14E-37 | 1.55E-36 | 3.83506324 | 14.49707954 | 206.8948079 | 15 | 0 | 1 | 0 | 7 | 1 | 2 | 0 | 0 | 0 | 0 | 0 |
| SG0000000 | Hoxa4 | 1.56E-36 | 2.04E-35 | -5.099148582 | 208.4689292 | 6.08197541 | 6 | 0 | 1 | 0 | 1 | 0 | 4 | 0 | 1 | 0 | 0 | 0 |
| JSG000000 | Stfa3 | 1.65E-36 | 2.15E-35 | -7.171524191 | 143.1597087 | 0 | 5 | 0 | 1 | 0 | 0 | 2 | 0 | 0 | 0 | 0 | 0 | 0 |
| SG0000002 | Fosl2 | 2.04E-35 | 2.58E-34 | -2.643682853 | 2999.089284 | 479.9119425 | 17 | 1 | 1 | 3 | 16 | 1 | 8 | 1 | 1 | 3 | 6 | 1 |
| JSG000000 | Slc35d3 | 5.09E-35 | 6.34E-34 | -4.396565064 | 240.4819699 | 11.41785592 | 5 | 1 | 1 | 3 | 5 | 1 | 4 | 1 | 1 | 3 | 5 | 1 |
| SG0000002 | Aoc1 | 6.26E-35 | 7.77E-34 | -5.802510546 | 170.2581754 | 3.050550723 | 21 | 0 | 1 | 2 | 9 | 1 | 0 | 0 | 0 | 0 | 0 | 0 |
| JSG000000 | Ticam1 | 6.58E-35 | 8.16E-34 | -2.572556217 | 901.4026475 | 151.5312628 | 3 | 1 | 1 | 1 | 3 | 2 | 0 | 0 | 0 | 0 | 0 | 1 |
| JSG000000 | Inafm2 | 1.64E-34 | 2E-33 | -2.534352617 | 900.7374201 | 155.4826908 | 6 | 0 | 1 | 2 | 2 | 1 | 6 | 0 | 1 | 2 | 2 | 1 |
| JSG000000 | Tmem173 | 1.89E-34 | 2.3E-33 | -3.85677464 | 1784.109822 | 123.1449852 | 6 | 0 | 1 | 2 | 2 | 0 | 1 | 0 | 0 | 0 | 0 | 0 |
| SG0000003 | Gngt2 | 3.11E-34 | 3.75E-33 | -2.883616497 | 967.4579266 | 131.0922524 | 5 | 1 | 1 | 1 | 1 | 0 | 0 | 0 | 0 | 0 | 0 | 0 |
| JSG000000 | Aunip | 4.13E-34 | 4.94E-33 | 1.920652785 | 234.2062884 | 886.6917159 | 21 | 1 | 1 | 2 | 8 | 2 | 0 | 0 | 0 | 0 | 0 | 1 |
| JSG000000 | Aqp2 | 4.32E-34 | 5.16E-33 | -6.969868315 | 124.3543556 | 0 | 2 | 1 | 1 | 2 | 4 | 0 | 0 | 0 | 0 | 0 | 0 | 0 |
| JSG000000 | Emc3 | 5.93E-34 | 7.05E-33 | -2.026159821 | 3201.949123 | 786.1031649 | 21 | 0 | 1 | 1 | 6 | 3 | 1 | 0 | 0 | 0 | 0 | 0 |
| SG0000002 | Pgd | 9.89E-34 | 1.16E-32 | -2.082661624 | 16624.9175 | 3924.785075 | 20 | 0 | 1 | 2 | 12 | 0 | 0 | 0 | 0 | 0 | 0 | 0 |
| SG0000000 | Ddx18 | 1.9E-33 | 2.2E-32 | 1.49804629 | 1515.883721 | 4281.764303 | 19 | 0 | 1 | 1 | 4 | 2 | 0 | 0 | 0 | 0 | 1 | 0 |
| JSG000000 | Mogat2 | 2.8E-33 | 3.23E-32 | 4.264415597 | 8.077044008 | 155.227784 | 13 | 0 | 1 | 1 | 6 | 2 | 0 | 0 | 0 | 0 | 0 | 0 |
| JSG000000 | Camk2n1 | 3.76E-33 | 4.33E-32 | -2.996022995 | 1970.720693 | 247.0200968 | 2 | 0 | 1 | 1 | 1 | 1 | 2 | 0 | 1 | 1 | 1 | 1 |
| JSG000000 | Sepn1 | 1.63E-32 | 1.83E-31 | -2.561136354 | 2755.172376 | 466.8419883 | 7 | 0 | 1 | 0 | 1 | 1 | 0 | 0 | 0 | 0 | 0 | 0 |
| JSG000000 | Psmb5 | 1.78E-32 | 1.99E-31 | 1.582878057 | 956.4203638 | 2865.118501 | 6 | 1 | 1 | 2 | 2 | 1 | 0 | 0 | 0 | 0 | 0 | 0 |
| JSG000000 | Itpripl2 | 2.26E-32 | 2.52E-31 | -2.988167007 | 4216.268893 | 531.3741206 | 7 | 0 | 1 | 1 | 4 | 0 | 7 | 0 | 1 | 1 | 4 | 0 |
| JSG000000 | Bambi | 3.47E-32 | 3.84E-31 | 3.212742523 | 24.08373473 | 223.2830228 | 9 | 1 | 1 | 3 | 4 | 0 | 4 | 0 | 0 | 0 | 1 | 0 |
| JSG000000 | Cep72 | 6.14E-32 | 6.71E-31 | 2.008008648 | 168.1879162 | 676.49661 | 33 | 1 | 1 | 3 | 16 | 2 | 2 | 0 | 0 | 0 | 0 | 0 |
| JSG000000 | Aox1 | 2.13E-31 | 2.28E-30 | -3.594341825 | 849.2222527 | 70.30992939 | 122 | 0 | 1 | 13 | 54 | 7 | 2 | 0 | 0 | 0 | 0 | 0 |
| JSG000000 | Dennd1c | 2.36E-31 | 2.53E-30 | 7.214033799 | 0.992459638 | 147.3511117 | 6 | 0 | 1 | 0 | 2 | 0 | 0 | 0 | 0 | 0 | 0 | 0 |
| JSG000000 | Fads3 | 3.04E-31 | 3.25E-30 | -3.171743444 | 885.0292922 | 98.21271795 | 13 | 1 | 1 | 1 | 7 | 2 | 1 | 0 | 0 | 0 | 1 | 0 |
| JSG000000 | Ppm1n | 3.48E-31 | 3.7E-30 | 6.664050078 | 0.970711415 | 98.43943279 | 5 | 0 | 1 | 0 | 1 | 1 | 1 | 0 | 0 | 0 | 0 | 0 |
| JSG000000 | Adh7 | 7.45E-31 | 7.82E-30 | -5.019593864 | 434.1167789 | 13.38314714 | 32 | 0 | 1 | 5 | 11 | 3 | 7 | 0 | 0 | 1 | 2 | 1 |
| SG0000001 | Mafk | 2.26E-30 | 2.32E-29 | -2.047737537 | 1572.337332 | 380.2903491 | 13 | 0 | 1 | 1 | 3 | 0 | 8 | 0 | 1 | 1 | 3 | 0 |
| JSG000000 | Fads1 | 2.61E-30 | 2.67E-29 | 1.514246219 | 768.9594114 | 2196.529041 | 15 | 1 | 1 | 2 | 5 | 0 | 3 | 0 | 0 | 0 | 0 | 0 |
| SG0000005 | 2700081O15Rik | 1.39E-29 | 1.37E-28 | -2.201629104 | 1242.576997 | 270.1263248 | 5 | 1 | 1 | 2 | 5 | 0 | 3 | 1 | 1 | 2 | 5 | 0 |
| JSG000000 | S100a7a | 3.36E-29 | 3.27E-28 | -4.616789981 | 223.2805922 | 9.100362819 | 2 | 0 | 1 | 1 | 2 | 0 | 0 | 0 | 0 | 0 | 1 | 0 |
| SG0000000 | Tmed4 | 3.73E-29 | 3.63E-28 | -1.831541156 | 3019.054459 | 848.2465869 | 7 | 1 | 1 | 5 | 10 | 2 | 2 | 0 | 0 | 2 | 4 | 1 |
| SG0000000 | Ripk4 | 4.1E-29 | 3.99E-28 | 4.618241449 | 29.34167123 | 720.6328406 | 12 | 0 | 1 | 1 | 2 | 2 | 0 | 0 | 0 | 0 | 0 | 0 |
| JSG000000 | Gareml | 6.78E-29 | 6.5E-28 | 3.594731005 | 13.59145965 | 164.2055982 | 10 | 1 | 1 | 2 | 8 | 0 | 1 | 0 | 0 | 1 | 1 | 0 |
| SG0000002 | Nek2 | 6.88E-29 | 6.6E-28 | 1.379099804 | 1838.987561 | 4783.317398 | 25 | 0 | 1 | 4 | 10 | 2 | 5 | 0 | 1 | 1 | 1 | 1 |
| SG0000003 | G6pdx | 7.57E-29 | 7.25E-28 | -1.716425453 | 7684.375356 | 2338.368886 | 23 | 0 | 1 | 3 | 8 | 2 | 1 | 0 | 0 | 0 | 0 | 0 |
| JSG000000 | Zfyve21 | 1.02E-28 | 9.68E-28 | 2.017359751 | 131.1083357 | 530.7818962 | 11 | 1 | 1 | 4 | 6 | 2 | 0 | 0 | 0 | 0 | 0 | 1 |
| SG0000004 | Slc7a5 | 1.17E-28 | 1.11E-27 | 1.329800773 | 3549.93802 | 8923.40679 | 19 | 1 | 1 | 1 | 1 | 0 | 0 | 0 | 0 | 0 | 0 | 0 |
| JSG000000 | Hn1l | 1.4E-28 | 1.32E-27 | 1.34633256 | 2658.213107 | 6758.904027 | 27 | 1 | 1 | 3 | 6 | 0 | 3 | 0 | 0 | 0 | 0 | 0 |
| SG0000002 | Fbf1 | 1.9E-28 | 1.79E-27 | -1.836277336 | 2525.70487 | 707.3071419 | 25 | 1 | 1 | 7 | 11 | 0 | 0 | 0 | 0 | 0 | 0 | 0 |
| JSG000000 | Polr2h | 3.52E-28 | 3.28E-27 | 1.544097448 | 557.6449829 | 1626.21307 | 7 | 0 | 1 | 5 | 12 | 1 | 0 | 0 | 0 | 0 | 0 | 0 |
| JSG000000 | Ccng1 | 4.15E-28 | 3.85E-27 | -1.723708278 | 17758.35111 | 5376.686849 | 20 | 0 | 1 | 3 | 8 | 0 | 8 | 0 | 1 | 1 | 2 | 0 |
| JSG000000 | Magoh | 5.11E-28 | 4.73E-27 | 1.504214801 | 703.7217351 | 1996.24913 | 13 | 0 | 1 | 0 | 4 | 2 | 0 | 0 | 0 | 0 | 0 | 0 |
| JSG000000 | Rasd1 | 8.56E-28 | 7.83E-27 | 4.937526622 | 3.583866926 | 109.8235538 | 2 | 0 | 1 | 0 | 2 | 0 | 2 | 0 | 1 | 0 | 2 | 0 |
| JSG000000 | B3gnt3 | 1.12E-27 | 1.02E-26 | -5.437678368 | 2729.001638 | 62.9650425 | 9 | 0 | 1 | 1 | 2 | 1 | 1 | 0 | 0 | 0 | 0 | 0 |
| SG0000006 | Cyp26b1 | 1.87E-27 | 1.68E-26 | -3.998878622 | 723.0444983 | 45.2254203 | 18 | 1 | 1 | 3 | 8 | 1 | 1 | 0 | 0 | 1 | 1 | 0 |
| SG0000000 | Alox12 | 2.56E-27 | 2.29E-26 | 4.859830072 | 3.62970694 | 105.3964996 | 25 | 0 | 1 | 2 | 3 | 0 | 6 | 0 | 0 | 0 | 0 | 0 |
| JSG000000 | Rnf122 | 2.86E-27 | 2.55E-26 | 3.724931284 | 22.30995757 | 294.9950936 | 26 | 1 | 1 | 3 | 13 | 4 | 3 | 0 | 0 | 0 | 1 | 1 |
| SG0000002 | Anxa5 | 3.28E-27 | 2.91E-26 | -2.072673489 | 4836.260073 | 1149.668828 | 46 | 0 | 1 | 6 | 23 | 7 | 1 | 0 | 0 | 0 | 1 | 0 |
| JSG000000 | Plau | 2.65E-26 | 2.27E-25 | -2.435357548 | 707.6081585 | 130.8209081 | 4 | 0 | 1 | 1 | 4 | 0 | 2 | 0 | 1 | 1 | 3 | 0 |
| SG0000007 | Art5 | 2.69E-26 | 2.3E-25 | 7.846078987 | 0.331601068 | 76.29938627 | 2 | 0 | 1 | 1 | 2 | 0 | 0 | 0 | 0 | 0 | 0 | 0 |
| JSG000000 | Gins3 | 2.73E-26 | 2.33E-25 | 1.645938563 | 290.5259238 | 909.2049032 | 13 | 1 | 1 | 2 | 7 | 3 | 0 | 0 | 0 | 0 | 1 | 0 |
| SG0000000 | Ephb3 | 3.05E-26 | 2.6E-25 | 1.974474893 | 337.5824775 | 1326.649093 | 8 | 0 | 1 | 0 | 2 | 1 | 0 | 0 | 0 | 0 | 0 | 0 |
| JSG000000 | Klf5 | 4.21E-26 | 3.58E-25 | 2.276274746 | 510.7867997 | 2474.381399 | 26 | 1 | 1 | 3 | 12 | 3 | 0 | 0 | 0 | 0 | 0 | 0 |
| JSG000000 | Rab20 | 8.26E-26 | 6.92E-25 | 3.629983359 | 11.35340166 | 140.5593944 | 24 | 1 | 1 | 2 | 7 | 5 | 1 | 0 | 0 | 0 | 0 | 0 |
| SG0000007 | Lrrc31 | 1.9E-25 | 1.56E-24 | 6.131668472 | 0 | 69.11583872 | 32 | 0 | 1 | 4 | 10 | 4 | 7 | 0 | 1 | 0 | 3 | 0 |
| JSG000000 | Hspb8 | 2.4E-25 | 1.97E-24 | -1.598315441 | 5330.779579 | 1760.555965 | 16 | 0 | 1 | 4 | 10 | 2 | 2 | 0 | 0 | 0 | 0 | 0 |
| JSG000000 | Klhl24 | 2.61E-25 | 2.14E-24 | 1.483714753 | 853.1896696 | 2386.097766 | 67 | 0 | 1 | 5 | 32 | 4 | 5 | 0 | 0 | 0 | 2 | 2 |
| JSG000000 | Ano10 | 3.38E-25 | 2.76E-24 | -2.526802617 | 1321.813322 | 229.364787 | 130 | 0 | 1 | 13 | 65 | 13 | 0 | 0 | 0 | 0 | 1 | 0 |
| JSG000000 | C130074G19Rik | 4.75E-25 | 3.85E-24 | 3.458150728 | 36.88300275 | 405.3529796 | 7 | 0 | 1 | 0 | 1 | 1 | 2 | 0 | 0 | 0 | 0 | 1 |
| JSG000000 | Tuba3a | 5.72E-25 | 4.62E-24 | 6.863119836 | 0.649984458 | 75.66724238 | 7 | 1 | 1 | 2 | 6 | 0 | 1 | 0 | 0 | 0 | 1 | 0 |
| JSG000000 | Gsta4 | 5.83E-25 | 4.7E-24 | 1.674209812 | 2336.638279 | 7457.252692 | 30 | 0 | 1 | 2 | 12 | 5 | 0 | 0 | 0 | 0 | 0 | 0 |
| JSG000000 | Clec4d | 5.95E-25 | 4.8E-24 | -7.462701238 | 294.062682 | 1.667028377 | 41 | 0 | 1 | 3 | 12 | 3 | 3 | 0 | 0 | 0 | 0 | 0 |
| JSG000000 | Naip5 | 5.96E-25 | 4.81E-24 | -6.625191496 | 99.74861992 | 1.010474896 | 53 | 1 | 1 | 10 | 47 | 4 | 0 | 0 | 0 | 0 | 0 | 0 |
| SG0000000 | Dll3 | 9.84E-25 | 7.84E-24 | 5.111793716 | 2.613155511 | 90.35839245 | 11 | 0 | 1 | 1 | 2 | 1 | 2 | 0 | 1 | 1 | 1 | 1 |
| SG0000000 | Foxm1 | 3.9E-24 | 3.02E-23 | -1.535866832 | 5638.702854 | 1944.631083 | 9 | 1 | 1 | 1 | 5 | 1 | 1 | 0 | 0 | 0 | 0 | 0 |
| JSG000000 | Carhsp1 | 1.41E-23 | 1.07E-22 | -1.537527412 | 4688.018232 | 1614.906646 | 5 | 0 | 1 | 0 | 1 | 0 | 1 | 0 | 1 | 0 | 0 | 0 |
| SG0000001 | Alox12e | 1.48E-23 | 1.12E-22 | -4.399999504 | 200.2790125 | 9.486447548 | 7 | 1 | 1 | 3 | 4 | 0 | 0 | 0 | 0 | 0 | 0 | 0 |
| SG0000003 | 2310007B03Rik | 2.85E-23 | 2.12E-22 | -4.064865238 | 168.2670347 | 10.05431938 | 11 | 0 | 1 | 0 | 0 | 2 | 1 | 0 | 1 | 0 | 0 | 0 |
| JSG000000 | Zswim7 | 5.56E-23 | 4.07E-22 | 1.984431732 | 96.74565847 | 382.829116 | 17 | 1 | 1 | 4 | 14 | 1 | 0 | 0 | 0 | 0 | 0 | 0 |
| JSG000000 | Hoxd8 | 5.62E-23 | 4.12E-22 | -4.801129268 | 456.5864642 | 16.37717952 | 4 | 0 | 1 | 1 | 7 | 0 | 4 | 0 | 1 | 1 | 7 | 0 |
| JSG000000 | Rasl11b | 1.21E-22 | 8.74E-22 | 3.243908759 | 14.75640523 | 139.795893 | 5 | 1 | 1 | 2 | 3 | 0 | 1 | 0 | 0 | 0 | 1 | 0 |
| JSG000000 | Nfkbia | 1.69E-22 | 1.21E-21 | -2.092965574 | 4324.196167 | 1013.584521 | 3 | 1 | 1 | 2 | 2 | 0 | 2 | 1 | 1 | 2 | 2 | 0 |
| JSG000000 | Slc27a4 | 1.7E-22 | 1.22E-21 | -1.740663041 | 1219.888558 | 365.0298418 | 4 | 0 | 1 | 1 | 3 | 0 | 0 | 0 | 0 | 0 | 0 | 0 |
| JSG000000 | 4930523C07Rik | 2.31E-22 | 1.65E-21 | -3.570066437 | 333.1921517 | 28.05418645 | 43 | 0 | 1 | 6 | 21 | 5 | 8 | 0 | 1 | 3 | 4 | 1 |
| SG0000002 | Ddx59 | 2.46E-22 | 1.75E-21 | 1.548554815 | 271.8162381 | 795.1276335 | 41 | 1 | 1 | 4 | 27 | 6 | 2 | 0 | 0 | 0 | 0 | 0 |
| SG0000006 | Nsl1 | 3.6E-22 | 2.54E-21 | 1.354900448 | 570.5607043 | 1459.377122 | 33 | 1 | 1 | 4 | 12 | 3 | 4 | 0 | 0 | 0 | 3 | 0 |
| SG0000002 | 2010107G23Rik | 4.1E-22 | 2.89E-21 | 6.01873975 | 5.428922595 | 351.9936675 | 45 | 1 | 1 | 14 | 21 | 4 | 5 | 0 | 0 | 5 | 5 | 0 |
| SG0000003 | Gpkow | 6.95E-22 | 4.84E-21 | -1.597091411 | 1924.584354 | 636.1573224 | 17 | 1 | 1 | 2 | 6 | 5 | 4 | 0 | 0 | 0 | 1 | 1 |
| JSG000000 | Ppp2r5b | 9.75E-22 | 6.76E-21 | -1.841433204 | 1604.915677 | 447.8427933 | 3 | 1 | 1 | 2 | 2 | 0 | 3 | 1 | 1 | 2 | 2 | 0 |
| JSG000000 | H2-Ab1 | 1.04E-21 | 7.19E-21 | 5.494815455 | 1.631569985 | 73.57156255 | 7 | 1 | 1 | 1 | 6 | 0 | 0 | 0 | 0 | 0 | 0 | 0 |
| JSG000000 | Slc25a10 | 1.52E-21 | 1.04E-20 | -2.78917248 | 966.8488596 | 139.8730386 | 3 | 1 | 1 | 1 | 2 | 1 | 0 | 0 | 0 | 0 | 0 | 0 |
| JSG000000 | Ccdc25 | 1.66E-21 | 1.13E-20 | 1.252497663 | 961.9997341 | 2291.998442 | 53 | 0 | 1 | 3 | 16 | 1 | 2 | 0 | 0 | 1 | 1 | 0 |
| JSG000000 | Fabp3 | 1.71E-21 | 1.17E-20 | 9.279154826 | 2.224840318 | 1382.301512 | 8 | 1 | 1 | 2 | 2 | 0 | 0 | 0 | 0 | 0 | 0 | 0 |
| JSG000000 | BC049352 | 1.96E-21 | 1.33E-20 | -6.171638547 | 71.08556804 | 0 | 63 | 1 | 1 | 7 | 23 | 8 | 3 | 0 | 0 | 0 | 0 | 0 |
| JSG000000 | Isoc1 | 3.88E-21 | 2.59E-20 | 1.406846774 | 904.2122439 | 2397.581958 | 30 | 1 | 1 | 4 | 10 | 2 | 0 | 0 | 0 | 0 | 1 | 0 |
| JSG000000 | Stap2 | 4.8E-21 | 3.19E-20 | -1.934755331 | 653.4717947 | 170.9257358 | 8 | 1 | 1 | 2 | 6 | 0 | 0 | 0 | 0 | 0 | 0 | 0 |
| JSG000000 | Bccip | 5.5E-21 | 3.65E-20 | 1.257876357 | 1280.180177 | 3061.467209 | 13 | 0 | 1 | 0 | 3 | 0 | 1 | 0 | 0 | 0 | 1 | 0 |
| SG0000000 | Bcam | 6.61E-21 | 4.37E-20 | 3.635114401 | 117.902004 | 1464.871937 | 7 | 1 | 1 | 2 | 4 | 0 | 0 | 0 | 0 | 0 | 0 | 0 |
| SG0000004 | Atp1b2 | 7.13E-21 | 4.71E-20 | 2.753628219 | 87.28330564 | 588.648276 | 2 | 0 | 1 | 0 | 1 | 0 | 1 | 0 | 1 | 0 | 0 | 0 |
| JSG000000 | Lipt1 | 7.78E-21 | 5.13E-20 | 1.888490807 | 98.22848446 | 363.6887129 | 8 | 0 | 1 | 1 | 1 | 0 | 0 | 0 | 0 | 0 | 0 | 0 |
| JSG000000 | E2f6 | 8.33E-21 | 5.48E-20 | -1.524865413 | 2133.333845 | 741.3590522 | 18 | 0 | 1 | 2 | 4 | 3 | 2 | 0 | 0 | 0 | 0 | 1 |
| SG0000002 | Litaf | 9.63E-21 | 6.32E-20 | -1.394911121 | 6082.266269 | 2312.891925 | 58 | 1 | 1 | 9 | 20 | 2 | 0 | 0 | 0 | 0 | 0 | 0 |
| JSG000000 | Ggct | 1.53E-20 | 9.91E-20 | 1.65607036 | 182.260368 | 574.4061697 | 15 | 1 | 1 | 2 | 7 | 0 | 3 | 0 | 0 | 1 | 2 | 0 |
| SG0000002 | Oplah | 2.1E-20 | 1.35E-19 | 2.238104041 | 50.25509602 | 237.0918126 | 3 | 1 | 1 | 1 | 1 | 0 | 0 | 0 | 0 | 0 | 0 | 0 |
| JSG000000 | Tmem181a | 3.63E-20 | 2.3E-19 | 1.610673031 | 181.3366901 | 553.7918755 | 36 | 0 | 1 | 6 | 20 | 4 | 3 | 0 | 0 | 1 | 3 | 1 |
| SG0000002 | Slc23a1 | 4.08E-20 | 2.57E-19 | 5.769760984 | 0 | 53.55959327 | 19 | 0 | 1 | 1 | 2 | 0 | 2 | 0 | 0 | 0 | 0 | 0 |
| SG0000002 | Adamts1 | 4.16E-20 | 2.62E-19 | -2.117364775 | 554.4422727 | 127.7809245 | 14 | 0 | 1 | 3 | 6 | 2 | 6 | 0 | 0 | 3 | 4 | 1 |
| SG0000000 | Acp2 | 4.23E-20 | 2.67E-19 | -1.691737161 | 1058.481092 | 327.6570178 | 12 | 1 | 1 | 2 | 4 | 3 | 2 | 0 | 0 | 0 | 0 | 3 |

| | | | | | | | | | | | | | | | | | | |
|---|---|---|---|---|---|---|---|---|---|---|---|---|---|---|---|---|---|---|
| JSG00000 | Cenpb | 5.34E-20 | 3.35E-19 | -1.942024466 | 3306.785174 | 860.5941014 | 2 | 0 | 1 | 1 | 1 | 0 | 2 | 0 | 1 | 1 | 1 | 0 |
| SG0000002 | Fos | 7.98E-20 | 4.96E-19 | 1.593369118 | 181.6406623 | 548.106529 | 5 | 1 | 1 | 3 | 5 | 0 | 4 | 1 | 1 | 2 | 4 | 0 |
| SG0000003 | Exo1 | 9.01E-20 | 5.57E-19 | 1.173555802 | 1236.206574 | 2788.473662 | 38 | 1 | 1 | 5 | 19 | 6 | 5 | 0 | 0 | 1 | 3 | 0 |
| SG0000002 | Cdk5 | 1.3E-19 | 7.99E-19 | -1.563900285 | 1331.519755 | 450.3671633 | 5 | 0 | 1 | 3 | 3 | 0 | 1 | 0 | 1 | 0 | 0 | 0 |
| JSG000000 | Map3k6 | 1.79E-19 | 1.09E-18 | -1.948700569 | 515.0299818 | 133.4182341 | 8 | 1 | 1 | 1 | 6 | 1 | 0 | 0 | 0 | 0 | 0 | 0 |
| JSG000000 | Cib2 | 2.09E-19 | 1.26E-18 | -2.191003172 | 371.6978532 | 81.40149427 | 13 | 1 | 1 | 4 | 6 | 2 | 0 | 0 | 0 | 0 | 0 | 0 |
| SG0000002 | Ifngr1 | 2.43E-19 | 1.47E-18 | -1.746269796 | 837.0374749 | 249.4969914 | 31 | 0 | 1 | 2 | 11 | 3 | 0 | 0 | 0 | 0 | 0 | 2 |
| JSG00000 | Clcn4 | 2.61E-19 | 1.57E-18 | -1.9106799 | 1224.035069 | 325.5531477 | 24 | 0 | 1 | 1 | 3 | 3 | 3 | 0 | 1 | 1 | 1 | 1 |
| JSG000000 | AI464131 | 4.52E-19 | 2.69E-18 | 1.790295258 | 104.2399564 | 360.551075 | 6 | 1 | 1 | 1 | 3 | 1 | 2 | 0 | 0 | 0 | 0 | 1 |
| JSG000000 | Osgin1 | 6.22E-19 | 3.67E-18 | -3.93381103 | 142.5586075 | 9.328210277 | 4 | 1 | 1 | 1 | 6 | 0 | 0 | 0 | 0 | 0 | 0 | 0 |
| JSG00000 | Gnmt | 1.17E-18 | 6.83E-18 | 5.550355167 | 1.313186595 | 61.53891468 | 6 | 1 | 1 | 2 | 4 | 0 | 0 | 0 | 0 | 0 | 0 | 0 |
| JSG000000 | Chrnb4 | 1.22E-18 | 7.1E-18 | 5.655508175 | 0 | 49.40546221 | 19 | 1 | 1 | 4 | 10 | 3 | 3 | 0 | 0 | 1 | 4 | 0 |
| SG0000003 | Stard9 | 1.31E-18 | 7.6E-18 | 2.578010242 | 51.66918377 | 308.5247529 | 11 | 1 | 1 | 1 | 5 | 1 | 3 | 0 | 0 | 0 | 0 | 0 |
| SG0000003 | E130309D02Rik | 1.42E-18 | 8.2E-18 | -1.889706061 | 2395.362336 | 646.4173569 | 4 | 1 | 1 | 1 | 3 | 0 | 0 | 0 | 0 | 0 | 0 | 0 |
| JSG000000 | Cbr3 | 1.47E-18 | 8.53E-18 | -2.00883875 | 900.4073849 | 223.7269634 | 10 | 1 | 1 | 2 | 7 | 0 | 0 | 0 | 0 | 0 | 1 | 0 |
| JSG000000 | Fam120b | 1.8E-18 | 1.03E-17 | -1.583501361 | 1016.581914 | 339.2040048 | 62 | 1 | 1 | 9 | 24 | 3 | 1 | 0 | 0 | 1 | 1 | 0 |
| JSG00000 | Slc13a2 | 2.33E-18 | 1.34E-17 | -5.506074199 | 76.35188194 | 1.680065656 | 12 | 1 | 1 | 2 | 5 | 0 | 2 | 0 | 0 | 1 | 1 | 0 |
| JSG00000 | Gprc5b | 2.34E-18 | 1.34E-17 | 3.401541008 | 113.1293309 | 1195.476637 | 23 | 0 | 1 | 2 | 5 | 2 | 3 | 0 | 0 | 1 | 3 | 1 |
| JSG000000 | Neurod1 | 2.76E-18 | 1.58E-17 | 5.620646873 | 0 | 48.202062 | 11 | 0 | 1 | 1 | 3 | 1 | 6 | 0 | 0 | 1 | 3 | 1 |
| SG0000002 | Klf11 | 4.91E-18 | 2.76E-17 | 1.572769865 | 381.093562 | 1133.659202 | 9 | 0 | 1 | 1 | 5 | 2 | 4 | 0 | 1 | 1 | 1 | 1 |
| JSG00000 | Mmgt1 | 5.54E-18 | 3.11E-17 | -1.426028752 | 1715.019474 | 638.2511806 | 28 | 1 | 1 | 6 | 14 | 1 | 11 | 1 | 1 | 2 | 4 | 0 |
| SG0000003 | Slc6a1 | 7.1E-18 | 3.97E-17 | 5.588961162 | 0 | 47.13322427 | 24 | 1 | 1 | 2 | 5 | 1 | 6 | 0 | 0 | 1 | 3 | 0 |
| SG0000006 | Prr36 | 8.01E-18 | 4.46E-17 | 2.772448636 | 37.19784901 | 254.1604836 | 4 | 1 | 1 | 1 | 2 | 0 | 0 | 0 | 0 | 0 | 0 | 0 |
| JSG000000 | Snx33 | 8.49E-18 | 4.72E-17 | -1.993236762 | 622.1485973 | 156.2680067 | 11 | 1 | 1 | 5 | 7 | 0 | 0 | 0 | 0 | 0 | 0 | 0 |
| SG000000C | Sdf2 | 1.33E-17 | 7.36E-17 | -1.353311265 | 2211.848672 | 865.701413 | 16 | 1 | 1 | 4 | 10 | 0 | 0 | 0 | 0 | 0 | 0 | 0 |
| SG0000003 | Anapc15 | 1.58E-17 | 8.68E-17 | 1.213523416 | 572.2689396 | 1327.110616 | 48 | 0 | 1 | 9 | 14 | 4 | 6 | 0 | 0 | 2 | 2 | 0 |
| JSG000000 | Wnt4 | 1.72E-17 | 9.44E-17 | 3.301598702 | 11.76346507 | 115.9886544 | 4 | 1 | 1 | 1 | 1 | 1 | 2 | 1 | 1 | 1 | 1 | 0 |
| JSG000000 | Zfp612 | 1.72E-17 | 9.45E-17 | 5.007623505 | 1.928205151 | 62.02947717 | 9 | 0 | 1 | 2 | 5 | 0 | 0 | 0 | 0 | 0 | 0 | 0 |
| JSG000000 | Hoxb4 | 2E-17 | 1.09E-16 | -3.432932472 | 124.3433283 | 11.5134861 | 3 | 0 | 1 | 1 | 18 | 0 | 1 | 0 | 1 | 1 | 18 | 0 |
| SG0000003 | Ipo13 | 4.17E-17 | 2.25E-16 | -1.514958311 | 1748.783352 | 611.9107973 | 12 | 1 | 1 | 2 | 5 | 0 | 0 | 0 | 0 | 0 | 0 | 0 |
| JSG000000 | E2f4 | 4.19E-17 | 2.26E-16 | 1.047777669 | 2307.777169 | 4770.966638 | 13 | 1 | 1 | 2 | 8 | 1 | 3 | 0 | 0 | 0 | 1 | 0 |
| SG0000003 | Tmem127 | 4.7E-17 | 2.52E-16 | -1.3733674 | 1931.459802 | 745.522734 | 10 | 0 | 1 | 0 | 1 | 0 | 1 | 0 | 0 | 0 | 0 | 0 |
| JSG000000 | Ribc1 | 6.16E-17 | 3.29E-16 | 2.343104275 | 48.06756517 | 243.8915723 | 17 | 0 | 1 | 0 | 10 | 0 | 0 | 0 | 0 | 0 | 0 | 0 |
| SG000000C | Spib | 6.23E-17 | 3.32E-16 | 7.1826649 | 0.331601068 | 48.17408838 | 4 | 0 | 1 | 1 | 2 | 0 | 0 | 0 | 0 | 0 | 0 | 0 |
| SG0000004 | Asb8 | 6.6E-17 | 3.51E-16 | -1.373961048 | 1668.840195 | 643.8893816 | 57 | 1 | 1 | 7 | 27 | 2 | 0 | 0 | 0 | 0 | 0 | 0 |
| SG000000C | Esam | 1.03E-16 | 5.44E-16 | 3.421984654 | 8.831926172 | 94.66195195 | 6 | 0 | 1 | 0 | 4 | 2 | 0 | 0 | 0 | 0 | 0 | 0 |
| JSG000000 | Rbbp9 | 1.28E-16 | 6.68E-16 | 1.30028516 | 324.5332489 | 799.2525554 | 9 | 0 | 1 | 0 | 3 | 2 | 0 | 0 | 0 | 0 | 0 | 2 |
| SG0000004 | Col6a3 | 1.35E-16 | 7.04E-16 | -2.304683013 | 1047.72471 | 212.0647424 | 72 | 0 | 1 | 7 | 31 | 5 | 1 | 0 | 0 | 0 | 0 | 0 |
| JSG000000 | Galnt4 | 1.71E-16 | 8.83E-16 | -1.953783362 | 408.1801577 | 105.3669702 | 8 | 0 | 1 | 0 | 4 | 0 | 6 | 0 | 1 | 0 | 4 | 0 |
| JSG000000 | Pycr2 | 1.86E-16 | 9.59E-16 | 1.168743445 | 1581.378966 | 3555.189809 | 1 | 1 | 1 | 1 | 1 | 0 | 1 | 1 | 1 | 1 | 1 | 0 |
| SG0000003 | Cyp11a1 | 1.86E-16 | 9.62E-16 | -5.359377196 | 2593.473819 | 63.17548345 | 12 | 1 | 1 | 2 | 3 | 2 | 0 | 0 | 0 | 0 | 0 | 0 |
| JSG000000 | Sav1 | 1.88E-16 | 9.7E-16 | -1.294522818 | 2338.575741 | 953.3694736 | 45 | 0 | 1 | 5 | 12 | 3 | 1 | 0 | 0 | 0 | 0 | 1 |
| JSG000000 | C1galt1c1 | 1.97E-16 | 1.02E-15 | -1.639886147 | 623.830738 | 200.1759278 | 9 | 1 | 1 | 3 | 3 | 0 | 4 | 0 | 0 | 2 | 2 | 0 |
| JSG000000 | Pcgf6 | 2.02E-16 | 1.04E-15 | 1.111770179 | 861.6277618 | 1862.069368 | 38 | 0 | 1 | 4 | 12 | 3 | 4 | 0 | 0 | 0 | 1 | 1 |
| JSG00000 | Prkag1 | 2.42E-16 | 1.24E-15 | -1.325014019 | 1832.828698 | 731.5649872 | 25 | 1 | 1 | 1 | 9 | 0 | 0 | 0 | 0 | 0 | 1 | 0 |
| JSG000000 | Tfb1m | 2.88E-16 | 1.47E-15 | 1.322107219 | 278.922374 | 697.3925174 | 64 | 0 | 1 | 4 | 22 | 6 | 1 | 0 | 0 | 0 | 0 | 1 |
| JSG000000 | Itm2a | 4.3E-16 | 2.17E-15 | 2.776600702 | 59.75042104 | 409.4313255 | 23 | 0 | 1 | 0 | 8 | 0 | 6 | 0 | 1 | 0 | 1 | 0 |
| SG0000004 | Mlec | 8.25E-16 | 4.1E-15 | -1.139666328 | 15590.55763 | 7076.001221 | 17 | 1 | 1 | 2 | 4 | 0 | 2 | 0 | 0 | 0 | 1 | 0 |
| JSG000000 | Phf19 | 9.64E-16 | 4.78E-15 | 1.41841321 | 192.9776934 | 515.8127032 | 12 | 0 | 1 | 1 | 5 | 2 | 0 | 0 | 0 | 0 | 0 | 0 |
| SG0000003 | Faah | 1.3E-15 | 6.38E-15 | 3.451715694 | 7.453494908 | 81.55111806 | 18 | 1 | 1 | 1 | 9 | 4 | 3 | 0 | 0 | 0 | 0 | 0 |
| JSG000000 | Oaf | 1.45E-15 | 7.11E-15 | -1.669308751 | 1016.31269 | 319.5327243 | 14 | 1 | 1 | 2 | 5 | 1 | 3 | 0 | 0 | 1 | 1 | 0 |
| SG0000002 | Creb3 | 1.47E-15 | 7.18E-15 | -1.594583117 | 982.2402944 | 325.2373376 | 5 | 0 | 1 | 1 | 2 | 1 | 2 | 0 | 1 | 1 | 2 | 0 |
| SG0000005 | Krcc1 | 1.72E-15 | 8.4E-15 | -1.295001781 | 1724.349538 | 702.7339322 | 14 | 0 | 1 | 0 | 3 | 0 | 5 | 0 | 1 | 0 | 1 | 0 |
| SG000000C | Ppfia3 | 3.29E-15 | 1.58E-14 | 2.184391463 | 40.14629586 | 182.4789753 | 19 | 1 | 1 | 4 | 13 | 0 | 1 | 0 | 0 | 0 | 0 | 0 |
| SG0000002 | Gar1 | 3.44E-15 | 1.65E-14 | 1.107297388 | 862.1945187 | 1857.52636 | 9 | 0 | 1 | 1 | 5 | 3 | 0 | 0 | 0 | 0 | 0 | 0 |
| JSG00000 | Mtfp1 | 4.81E-15 | 2.28E-14 | 1.873245639 | 68.43510306 | 250.7160011 | 3 | 1 | 1 | 2 | 5 | 0 | 1 | 0 | 0 | 1 | 2 | 0 |
| JSG000000 | S1pr2 | 5.59E-15 | 2.64E-14 | 1.126408408 | 544.5368589 | 1188.803295 | 11 | 1 | 1 | 5 | 10 | 2 | 1 | 0 | 0 | 1 | 1 | 0 |
| SG0000002 | Pola2 | 6.09E-15 | 2.87E-14 | 1.002468818 | 1420.619223 | 2846.10469 | 19 | 0 | 1 | 2 | 8 | 2 | 0 | 0 | 0 | 0 | 0 | 0 |
| JSG00000 | Crabp2 | 6.94E-15 | 3.25E-14 | -2.800520873 | 2024.559431 | 290.5960747 | 7 | 0 | 1 | 1 | 2 | 1 | 2 | 0 | 0 | 1 | 1 | 0 |
| SG0000004 | Gls2 | 7.32E-15 | 3.43E-14 | 2.490190268 | 136.3013247 | 765.8117798 | 20 | 0 | 1 | 2 | 6 | 3 | 1 | 0 | 0 | 0 | 0 | 0 |
| JSG000000 | Mphosph6 | 8.11E-15 | 3.79E-14 | 1.545426542 | 514.8276492 | 1502.732254 | 14 | 0 | 1 | 3 | 5 | 4 | 2 | 0 | 0 | 2 | 2 | 1 |
| SG0000003 | Inppl1 | 8.28E-15 | 3.87E-14 | -1.121649435 | 6067.145011 | 2788.266306 | 15 | 1 | 1 | 2 | 4 | 0 | 1 | 0 | 0 | 0 | 0 | 0 |
| JSG000000 | Sp5 | 1.06E-14 | 4.92E-14 | 5.301067596 | 0 | 38.42578556 | 2 | 0 | 1 | 0 | 0 | 0 | 2 | 0 | 1 | 0 | 0 | 0 |
| SG0000004 | Alkbh2 | 1.09E-14 | 5.03E-14 | 1.581692336 | 115.5513615 | 345.8692116 | 7 | 1 | 1 | 1 | 4 | 0 | 1 | 1 | 1 | 1 | 2 | 0 |
| JSG000000 | Plk1 | 1.12E-14 | 5.18E-14 | 1.004186454 | 1772.419871 | 3555.141199 | 9 | 0 | 1 | 0 | 1 | 0 | 0 | 0 | 0 | 0 | 0 | 0 |
| SG0000003 | Rfc3 | 1.26E-14 | 5.8E-14 | 1.082148025 | 689.2562902 | 1459.283774 | 16 | 0 | 1 | 1 | 4 | 1 | 1 | 0 | 0 | 0 | 1 | 0 |
| JSG000000 | Trim8 | 1.46E-14 | 6.69E-14 | -2.054184518 | 2461.879847 | 592.7829403 | 10 | 1 | 1 | 3 | 6 | 0 | 1 | 0 | 0 | 0 | 0 | 0 |
| SG0000003 | Lgals3bp | 1.48E-14 | 6.79E-14 | -2.011701368 | 3163.136962 | 784.396312 | 12 | 0 | 1 | 0 | 4 | 2 | 2 | 0 | 0 | 0 | 0 | 0 |
| JSG00000 | Il27ra | 1.56E-14 | 7.15E-14 | 3.410636219 | 7.167733853 | 76.2229367 | 8 | 0 | 1 | 0 | 3 | 0 | 0 | 0 | 0 | 0 | 0 | 0 |
| JSG000000 | Rpl4 | 1.83E-14 | 8.35E-14 | 0.898779032 | 47473.45453 | 88513.65685 | 8 | 1 | 1 | 1 | 4 | 2 | 0 | 0 | 0 | 0 | 0 | 0 |
| JSG000000 | Cptp | 1.87E-14 | 8.54E-14 | -1.645774499 | 454.768937 | 145.3326867 | 4 | 0 | 1 | 0 | 1 | 1 | 0 | 0 | 0 | 0 | 0 | 0 |
| JSG000000 | Igfbp5 | 1.9E-14 | 8.67E-14 | 3.141808707 | 17.08983353 | 150.8399896 | 25 | 0 | 1 | 4 | 28 | 1 | 7 | 0 | 1 | 1 | 1 | 0 |
| JSG000000 | Mmp9 | 2.48E-14 | 1.12E-13 | 2.032791454 | 48.28239756 | 197.5695589 | 7 | 1 | 1 | 1 | 5 | 0 | 4 | 1 | 1 | 1 | 3 | 0 |
| SG0000003 | Akna | 2.81E-14 | 1.27E-13 | 4.987851994 | 12.7775198 | 405.4521675 | 25 | 1 | 1 | 2 | 22 | 1 | 1 | 0 | 0 | 0 | 0 | 0 |
| JSG000000 | Myd88 | 2.94E-14 | 1.33E-13 | -1.388681439 | 828.1991081 | 316.3005818 | 3 | 1 | 1 | 1 | 3 | 0 | 1 | 0 | 0 | 0 | 0 | 0 |
| JSG000000 | Ubiad1 | 2.97E-14 | 1.34E-13 | -1.351632159 | 975.2079906 | 382.1337781 | 7 | 0 | 1 | 1 | 3 | 1 | 3 | 0 | 1 | 1 | 3 | 0 |
| SG0000001 | Ahr | 4.76E-14 | 2.11E-13 | -1.326428278 | 1033.805993 | 412.2345111 | 121 | 0 | 1 | 16 | 62 | 6 | 9 | 0 | 1 | 1 | 3 | 0 |
| JSG000000 | Saysd1 | 4.86E-14 | 2.16E-13 | 1.815013166 | 61.59143659 | 216.7173788 | 9 | 1 | 1 | 2 | 6 | 2 | 5 | 1 | 1 | 2 | 3 | 2 |
| JSG000000 | Vstm5 | 5.96E-14 | 2.63E-13 | -2.271901541 | 316.3718619 | 65.50694399 | 14 | 1 | 1 | 2 | 5 | 1 | 0 | 0 | 0 | 0 | 0 | 0 |
| SG0000005 | Shisa7 | 6.95E-14 | 3.05E-13 | 3.909882425 | 6.326011934 | 95.08716434 | 23 | 0 | 1 | 0 | 0 | 2 | 4 | 0 | 0 | 0 | 0 | 2 |
| JSG00000 | Izumo1 | 9.58E-14 | 4.17E-13 | 5.566269766 | 1.607478194 | 76.16566773 | 7 | 0 | 1 | 1 | 2 | 0 | 0 | 0 | 0 | 0 | 0 | 0 |
| JSG000000 | Prps2 | 9.63E-14 | 4.19E-13 | -1.152357716 | 2418.144861 | 1087.897743 | 56 | 0 | 1 | 2 | 17 | 8 | 5 | 0 | 0 | 0 | 3 | 2 |
| JSG000000 | Kdelr3 | 1.15E-13 | 4.98E-13 | 1.199149351 | 399.7128047 | 917.7575904 | 10 | 1 | 1 | 4 | 9 | 0 | 0 | 0 | 0 | 0 | 0 | 0 |
| SG0000003 | Cenpu | 1.28E-13 | 5.51E-13 | 1.161176477 | 358.4346886 | 801.6023526 | 45 | 0 | 1 | 6 | 16 | 4 | 2 | 0 | 0 | 0 | 0 | 0 |
| JSG000000 | Tagap1 | 1.7E-13 | 7.3E-13 | 1.260205474 | 240.5348406 | 576.1527506 | 6 | 1 | 1 | 1 | 5 | 0 | 1 | 1 | 1 | 1 | 3 | 0 |
| SG0000005 | Mafg | 1.92E-13 | 8.2E-13 | -1.078740549 | 3892.335267 | 1842.794801 | 7 | 0 | 1 | 4 | 4 | 1 | 4 | 0 | 1 | 3 | 3 | 1 |
| SG0000006 | Hsh2d | 2.13E-13 | 9.05E-13 | 5.23739677 | 3.333071773 | 125.7355809 | 10 | 1 | 1 | 2 | 9 | 0 | 0 | 0 | 0 | 0 | 0 | 0 |
| SG0000003 | Taf1d | 2.24E-13 | 9.51E-13 | 1.021119956 | 725.7678766 | 1472.941429 | 29 | 0 | 1 | 3 | 5 | 0 | 0 | 0 | 0 | 0 | 0 | 0 |
| SG0000003 | Dusp8 | 2.83E-13 | 1.19E-12 | -2.617291779 | 320.8066809 | 52.28294846 | 10 | 1 | 1 | 2 | 10 | 0 | 5 | 1 | 1 | 2 | 9 | 0 |
| JSG000000 | Tert | 2.93E-13 | 1.23E-12 | 1.44902935 | 153.7878944 | 419.878319 | 17 | 1 | 1 | 2 | 5 | 2 | 0 | 0 | 0 | 0 | 0 | 0 |
| JSG000000 | Rgmb | 3.98E-13 | 1.66E-12 | -1.513023823 | 479.275612 | 167.9267086 | 26 | 0 | 1 | 5 | 14 | 3 | 2 | 0 | 0 | 1 | 1 | 1 |
| JSG000000 | Inhbb | 4.05E-13 | 1.69E-12 | 3.001119827 | 81.0980446 | 649.2881421 | 5 | 0 | 1 | 0 | 3 | 0 | 2 | 0 | 1 | 0 | 3 | 0 |
| JSG000000 | H2-DMa | 6E-13 | 2.48E-12 | 2.304679479 | 24.92076952 | 123.1229754 | 3 | 0 | 1 | 0 | 0 | 0 | 0 | 0 | 0 | 0 | 0 | 0 |
| SG000000C | Gatad1 | 6.22E-13 | 2.56E-12 | -1.087442688 | 2924.478006 | 1376.244169 | 66 | 0 | 1 | 7 | 28 | 16 | 7 | 0 | 0 | 1 | 3 | 2 |
| SG0000003 | Plxna1 | 6.63E-13 | 2.73E-12 | -1.151824947 | 5417.089911 | 2437.991488 | 38 | 1 | 1 | 3 | 5 | 10 | 3 | 1 | 1 | 1 | 1 | 4 |
| SG0000005 | Zfp709 | 8.66E-13 | 3.54E-12 | 1.965018127 | 40.79393675 | 159.2667128 | 19 | 1 | 1 | 4 | 13 | 0 | 4 | 0 | 0 | 1 | 1 | 0 |
| JSG000000 | Fam72a | 8.88E-13 | 3.63E-12 | 1.508421303 | 102.8784331 | 292.6876511 | 27 | 0 | 1 | 1 | 7 | 6 | 2 | 0 | 0 | 0 | 1 | 0 |
| SG0000003 | Syp | 8.96E-13 | 3.66E-12 | 2.115380026 | 31.86279687 | 138.0628185 | 19 | 0 | 1 | 3 | 12 | 2 | 1 | 0 | 0 | 1 | 2 | 0 |
| JSG00000 | Cbln4 | 1.08E-12 | 4.38E-12 | 5.076704902 | 0 | 32.74741052 | 15 | 0 | 1 | 3 | 6 | 1 | 4 | 0 | 0 | 2 | 3 | 1 |
| JSG000000 | Ppp1r15b | 1.1E-12 | 4.45E-12 | 0.870461339 | 3709.81753 | 6782.46439 | 7 | 1 | 1 | 2 | 7 | 1 | 3 | 0 | 0 | 0 | 1 | 1 |
| JSG000000 | Slc29a2 | 1.14E-12 | 4.62E-12 | 1.390098529 | 141.024939 | 369.6215423 | 4 | 1 | 1 | 1 | 5 | 0 | 0 | 0 | 0 | 0 | 0 | 0 |
| JSG000000 | Zfp872 | 1.19E-12 | 4.8E-12 | 5.06633318 | 0 | 32.50566602 | 26 | 1 | 1 | 3 | 8 | 0 | 8 | 1 | 1 | 1 | 3 | 0 |
| SG0000002 | Tmem106b | 1.76E-12 | 7.02E-12 | -1.198527098 | 3007.220064 | 1310.305618 | 44 | 0 | 1 | 3 | 17 | 5 | 11 | 0 | 1 | 1 | 5 | 1 |
| SG0000005 | Fes | 2.89E-12 | 1.14E-11 | -2.218956771 | 269.0454237 | 57.79004895 | 9 | 1 | 1 | 2 | 5 | 2 | 0 | 0 | 0 | 0 | 0 | 0 |
| JSG000000 | Gpr157 | 3E-12 | 1.18E-11 | 3.382367987 | 6.550371729 | 68.30620657 | 17 | 1 | 1 | 2 | 8 | 1 | 1 | 0 | 0 | 0 | 2 | 0 |
| SG0000006 | Tspan6 | 3.91E-12 | 1.53E-11 | 2.928085667 | 61.85798969 | 470.8010442 | 13 | 0 | 1 | 2 | 3 | 1 | 1 | 0 | 0 | 0 | 1 | 0 |
| SG0000004 | Rbm15 | 4.76E-12 | 1.86E-11 | 0.950296904 | 834.3698392 | 1612.228039 | 22 | 0 | 1 | 2 | 6 | 1 | 2 | 0 | 0 | 0 | 0 | 0 |
| JSG000000 | Slc51a | 5.73E-12 | 2.23E-11 | 5.816347456 | 0.649984458 | 36.62671943 | 21 | 1 | 1 | 2 | 12 | 0 | 0 | 0 | 0 | 0 | 0 | 0 |
| SG0000004 | Proser1 | 8.57E-12 | 3.29E-11 | 0.98929007 | 574.5081131 | 1140.51802 | 24 | 0 | 1 | 2 | 6 | 2 | 4 | 0 | 0 | 0 | 0 | 1 |
| SG0000003 | Lpar2 | 9.38E-12 | 3.59E-11 | 1.047816976 | 414.4777405 | 856.8909407 | 10 | 1 | 1 | 1 | 1 | 0 | 3 | 0 | 0 | 0 | 0 | 0 |
| JSG000000 | Insl6 | 9.58E-12 | 3.67E-11 | 2.481847145 | 16.87400428 | 94.26025065 | 4 | 0 | 1 | 0 | 0 | 0 | 0 | 0 | 0 | 0 | 0 | 0 |
| JSG000000 | Ereg | 1.16E-11 | 4.41E-11 | -3.204095295 | 1270.715868 | 137.8858372 | 40 | 1 | 1 | 8 | 19 | 1 | 9 | 1 | 1 | 3 | 7 | 0 |
| SG0000002 | Sgol1 | 1.29E-11 | 4.91E-11 | 0.887903713 | 1250.50088 | 2314.033556 | 25 | 0 | 1 | 4 | 12 | 2 | 6 | 0 | 0 | 2 | 6 | 1 |
| SG0000002 | Smyd2 | 1.3E-11 | 4.92E-11 | -1.031061096 | 2180.650621 | 1067.101602 | 35 | 1 | 1 | 5 | 12 | 1 | 0 | 0 | 0 | 0 | 0 | 0 |
| JSG000000 | Mylip | 1.5E-11 | 5.66E-11 | 2.247430306 | 33.6657028 | 159.856986 | 28 | 1 | 1 | 4 | 8 | 4 | 2 | 0 | 0 | 1 | 1 | 0 |
| JSG000000 | Bin3 | 1.76E-11 | 6.61E-11 | -1.169883343 | 973.39237 | 432.6313234 | 30 | 1 | 1 | 5 | 13 | 6 | 0 | 0 | 0 | 0 | 0 | 0 |
| JSG000000 | Kdelc2 | 1.78E-11 | 6.7E-11 | -1.208327438 | 840.1992452 | 363.6130685 | 31 | 1 | 1 | 7 | 20 | 2 | 2 | 0 | 0 | 2 | 4 | 0 |
| JSG000000 | Loxl1 | 2.39E-11 | 8.89E-11 | -1.941614034 | 547.2037337 | 142.4508312 | 16 | 1 | 1 | 5 | 8 | 1 | 0 | 0 | 0 | 0 | 0 | 0 |
| JSG000000 | Nlrc4 | 2.57E-11 | 9.53E-11 | -2.550287682 | 302.1993872 | 51.59177732 | 46 | 1 | 1 | 3 | 12 | 3 | 1 | 0 | 0 | 0 | 0 | 0 |
| SG0000002 | Inpp5e | 2.69E-11 | 9.96E-11 | -1.281798882 | 607.99349 | 250.0570004 | 8 | 1 | 1 | 2 | 7 | 0 | 2 | 1 | 1 | 1 | 1 | 0 |
| JSG000000 | Prss22 | 2.8E-11 | 1.04E-10 | -3.296429061 | 99.27720791 | 10.10474896 | 3 | 1 | 1 | 2 | 2 | 1 | 2 | 1 | 1 | 2 | 2 | 1 |
| JSG00000 | Hoxc13 | 2.8E-11 | 1.04E-10 | 5.214012345 | 0.981585527 | 36.43361459 | 8 | 0 | 1 | 1 | 5 | 1 | 2 | 0 | 0 | 1 | 1 | 1 |
| SG000000C | Gcat | 3.83E-11 | 1.4E-10 | 2.278926121 | 280.7171652 | 1362.36692 | 4 | 1 | 1 | 1 | 3 | 1 | 0 | 0 | 0 | 0 | 0 | 0 |

| | | | | | | | | | | | | | | | | | | |
|---|---|---|---|---|---|---|---|---|---|---|---|---|---|---|---|---|---|---|
| SG0000007 | Etohi1 | 4.72E-11 | 1.72E-10 | -1.934010642 | 183.3599157 | 47.98540733 | 19 | 0 | 1 | 1 | 11 | 1 | 1 | 0 | 0 | 0 | 1 | 0 |
| ISG000000 | Rap2c | 5.22E-11 | 1.89E-10 | -1.039711439 | 2463.113089 | 1198.119184 | 31 | 0 | 1 | 8 | 23 | 2 | 7 | 0 | 0 | 1 | 6 | 0 |
| ISG000000 | Msl3l2 | 5.79E-11 | 2.09E-10 | 2.928842458 | 8.609909944 | 65.56439249 | 23 | 0 | 1 | 0 | 3 | 4 | 0 | 0 | 0 | 0 | 0 | 0 |
| ISG000000 | Hoxd13 | 6.14E-11 | 2.21E-10 | -2.277296325 | 395.1493278 | 81.51297513 | 6 | 1 | 1 | 2 | 4 | 3 | 6 | 1 | 1 | 2 | 4 | 1 |
| SG0000002 | Eef2kmt | 8.45E-11 | 3.02E-10 | -1.098352775 | 1080.667613 | 504.7252189 | 8 | 0 | 1 | 1 | 3 | 1 | 1 | 0 | 0 | 0 | 0 | 1 |
| ISG000000 | Cbx8 | 9.54E-11 | 3.4E-10 | -2.172893028 | 137.1850524 | 30.42291433 | 3 | 0 | 1 | 0 | 2 | 2 | 3 | 0 | 1 | 0 | 1 | 2 |
| ISG000000 | Sprr2a3 | 1.15E-10 | 4.08E-10 | 4.851314823 | 0 | 27.86631059 | 10 | 0 | 1 | 2 | 8 | 0 | 8 | 0 | 1 | 2 | 4 | 0 |
| ISG000000 | Rexo2 | 1.16E-10 | 4.12E-10 | -1.043217004 | 1890.849733 | 917.5239436 | 11 | 1 | 1 | 2 | 7 | 3 | 0 | 0 | 0 | 0 | 0 | 0 |
| ISG000000 | Rcn1 | 1.19E-10 | 4.2E-10 | 2.284459587 | 488.0128705 | 2377.508813 | 16 | 1 | 1 | 2 | 6 | 1 | 4 | 1 | 1 | 1 | 2 | 0 |
| ISG000000 | Tmem151b | 2.12E-10 | 7.38E-10 | 5.32459288 | 2.257462652 | 90.46544952 | 2 | 0 | 1 | 0 | 1 | 0 | 1 | 0 | 0 | 0 | 1 | 0 |
| ISG000000 | Xk | 2.27E-10 | 7.85E-10 | -1.251715378 | 554.4654863 | 232.8469979 | 81 | 1 | 1 | 10 | 42 | 13 | 4 | 0 | 0 | 1 | 4 | 1 |
| SG0000002 | Fam20a | 2.47E-10 | 8.54E-10 | -2.417100542 | 103.3539379 | 19.35117154 | 60 | 1 | 1 | 4 | 15 | 14 | 0 | 0 | 0 | 0 | 0 | 0 |
| ISG000000 | Acbd7 | 2.81E-10 | 9.67E-10 | 5.126044249 | 1.976388732 | 69.01848901 | 6 | 0 | 1 | 1 | 3 | 1 | 0 | 0 | 0 | 0 | 0 | 0 |
| ISG000000 | Pygo1 | 3E-10 | 1.03E-09 | 2.118593717 | 23.70629365 | 102.9494231 | 44 | 0 | 1 | 2 | 13 | 8 | 11 | 0 | 1 | 1 | 1 | 2 |
| ISG000000 | Arap3 | 3.4E-10 | 1.16E-09 | 6.189933888 | 3.710512855 | 270.8879659 | 20 | 0 | 1 | 1 | 9 | 4 | 0 | 0 | 0 | 0 | 0 | 0 |
| SG0000000 | Sema4f | 3.51E-10 | 1.2E-09 | 1.81269583 | 39.23932927 | 137.8470236 | 32 | 0 | 1 | 8 | 20 | 1 | 1 | 0 | 0 | 0 | 0 | 0 |
| SG0000003 | Pcsk1n | 3.6E-10 | 1.23E-09 | 4.828970433 | 1.289094805 | 36.63952261 | 8 | 0 | 1 | 0 | 0 | 0 | 4 | 0 | 1 | 0 | 0 | 0 |
| SG0000002 | Psmb8 | 5.17E-10 | 1.74E-09 | -1.678652819 | 435.0139778 | 135.8871409 | 5 | 1 | 1 | 1 | 3 | 0 | 0 | 0 | 0 | 0 | 0 | 0 |
| ISG000000 | Tars | 5.82E-10 | 1.96E-09 | 0.772922863 | 3019.758039 | 5159.945471 | 30 | 1 | 1 | 5 | 10 | 1 | 2 | 0 | 0 | 0 | 0 | 0 |
| SG0000000 | Ruvbl2 | 6.56E-10 | 2.2E-09 | 1.645879268 | 2130.816344 | 6668.145202 | 12 | 0 | 1 | 3 | 4 | 2 | 0 | 0 | 0 | 0 | 0 | 0 |
| ISG000000 | Cby1 | 6.64E-10 | 2.22E-09 | -1.069673558 | 889.5263696 | 423.7941921 | 6 | 0 | 1 | 1 | 4 | 3 | 1 | 0 | 0 | 0 | 0 | 0 |
| SG0000002 | Usp30 | 6.65E-10 | 2.22E-09 | -1.034856786 | 1036.470419 | 505.8642379 | 12 | 0 | 1 | 3 | 6 | 1 | 1 | 0 | 0 | 1 | 2 | 0 |
| ISG000000 | Dynlt1f | 7.27E-10 | 2.42E-09 | 1.170651933 | 164.0982657 | 369.4071845 | 7 | 0 | 1 | 0 | 1 | 0 | 0 | 0 | 0 | 0 | 0 | 0 |
| ISG000000 | C1qbp | 7.48E-10 | 2.49E-09 | 0.749932899 | 4629.560107 | 7785.598874 | 5 | 1 | 1 | 1 | 3 | 0 | 2 | 0 | 0 | 0 | 1 | 0 |
| ISG000000 | Alox5ap | 7.61E-10 | 2.54E-09 | -2.760406085 | 172.1491807 | 25.40621262 | 45 | 0 | 1 | 5 | 16 | 2 | 2 | 0 | 0 | 0 | 0 | 0 |
| ISG000000 | Dynlt1a | 8.4E-10 | 2.79E-09 | 1.546438664 | 62.45241003 | 182.4204937 | 7 | 0 | 1 | 0 | 1 | 0 | 0 | 0 | 0 | 0 | 0 | 0 |
| SG0000005 | Nap1l1 | 9.89E-10 | 3.26E-09 | 0.729524697 | 12478.73761 | 20690.90417 | 57 | 0 | 1 | 6 | 20 | 8 | 0 | 0 | 0 | 0 | 0 | 0 |
| SG0000002 | Glul | 1.4E-09 | 4.55E-09 | -1.179199723 | 13572.38688 | 5993.517024 | 8 | 0 | 1 | 0 | 3 | 3 | 2 | 0 | 0 | 0 | 1 | 0 |
| SG0000002 | Mterf4 | 1.46E-09 | 4.74E-09 | 0.967292003 | 361.7438217 | 707.2696625 | 14 | 0 | 1 | 0 | 3 | 3 | 5 | 0 | 1 | 0 | 1 | 0 |
| SG0000003 | Tecr | 1.57E-09 | 5.07E-09 | 0.749836502 | 3441.155153 | 5786.654237 | 11 | 0 | 1 | 4 | 8 | 2 | 0 | 0 | 0 | 0 | 0 | 0 |
| SG0000005 | Mat2a | 1.59E-09 | 5.13E-09 | -0.99954586 | 9728.565166 | 4865.814032 | 12 | 1 | 1 | 2 | 6 | 0 | 3 | 1 | 1 | 2 | 2 | 0 |
| ISG000000 | Cd9 | 1.71E-09 | 5.51E-09 | -0.826536238 | 11274.47707 | 6357.468362 | 35 | 1 | 1 | 2 | 13 | 3 | 1 | 0 | 0 | 0 | 0 | 1 |
| ISG000000 | Fam50a | 2.63E-09 | 8.36E-09 | 0.820490426 | 891.6422721 | 1574.645801 | 9 | 1 | 1 | 2 | 6 | 0 | 0 | 0 | 0 | 0 | 0 | 0 |
| ISG000000 | Ccdc8 | 2.96E-09 | 9.36E-09 | 2.694034411 | 42.43138744 | 274.5822124 | 4 | 0 | 1 | 1 | 2 | 2 | 1 | 0 | 0 | 0 | 0 | 0 |
| ISG000000 | Vimp | 2.96E-09 | 9.37E-09 | -1.053309908 | 754.3874938 | 363.5102011 | 14 | 0 | 1 | 1 | 10 | 0 | 1 | 0 | 0 | 1 | 2 | 0 |
| SG0000002 | Car2 | 3.09E-09 | 9.75E-09 | 0.92441261 | 437.6911711 | 830.6990436 | 41 | 1 | 1 | 7 | 15 | 2 | 0 | 0 | 0 | 0 | 0 | 0 |
| ISG000000 | Pitrm1 | 3.31E-09 | 1.04E-08 | 0.741889003 | 2659.726891 | 4448.031843 | 45 | 0 | 1 | 3 | 12 | 10 | 1 | 0 | 0 | 0 | 0 | 0 |
| SG0000002 | Plk4 | 3.63E-09 | 1.14E-08 | 0.799824316 | 993.1151389 | 1728.903337 | 52 | 0 | 1 | 7 | 21 | 8 | 4 | 0 | 0 | 1 | 4 | 1 |
| ISG000000 | Dleu7 | 3.93E-09 | 1.23E-08 | 4.575473316 | 0 | 22.84266036 | 25 | 0 | 1 | 1 | 6 | 3 | 4 | 0 | 0 | 1 | 2 | 2 |
| ISG000000 | Lrrc59 | 5.14E-09 | 1.6E-08 | -0.803118202 | 10361.9494 | 5938.527858 | 15 | 1 | 1 | 3 | 8 | 1 | 1 | 0 | 0 | 0 | 0 | 0 |
| SG0000004 | Dnajc16 | 5.54E-09 | 1.72E-08 | -0.982365317 | 978.1953339 | 495.1128065 | 29 | 1 | 1 | 4 | 9 | 1 | 4 | 0 | 0 | 1 | 2 | 0 |
| ISG000000 | Ccdc97 | 6.64E-09 | 2.05E-08 | -0.927547106 | 1318.747928 | 693.333667 | 8 | 1 | 1 | 3 | 5 | 0 | 1 | 0 | 0 | 0 | 1 | 0 |
| SG0000003 | Rsph4a | 6.92E-09 | 2.13E-08 | 5.325820955 | 0.674076249 | 27.03590326 | 20 | 1 | 1 | 2 | 8 | 2 | 2 | 0 | 0 | 0 | 0 | 1 |
| ISG000000 | Lat | 7.28E-09 | 2.24E-08 | 2.888722217 | 7.097802049 | 52.56733008 | 12 | 1 | 1 | 2 | 3 | 0 | 0 | 0 | 0 | 0 | 0 | 0 |
| SG0000003 | Neurl1b | 8.17E-09 | 2.51E-08 | -1.75585851 | 350.2048605 | 103.6945878 | 21 | 1 | 1 | 2 | 10 | 0 | 4 | 0 | 0 | 0 | 1 | 0 |
| ISG000000 | Lemd2 | 1.01E-08 | 3.09E-08 | -0.936131473 | 1110.170608 | 580.2111519 | 13 | 1 | 1 | 2 | 5 | 0 | 1 | 0 | 0 | 1 | 1 | 0 |
| ISG000000 | Nrsn2 | 1.15E-08 | 3.49E-08 | 4.499026031 | 0 | 21.6121463 | 5 | 0 | 1 | 0 | 1 | 1 | 0 | 0 | 0 | 0 | 0 | 0 |
| ISG000000 | Mmp15 | 1.22E-08 | 3.7E-08 | 1.653023354 | 117.7790483 | 370.4056939 | 14 | 0 | 1 | 3 | 10 | 2 | 1 | 0 | 0 | 1 | 1 | 1 |
| ISG000000 | Cyth4 | 1.26E-08 | 3.8E-08 | 3.376691328 | 3.913124428 | 40.64517833 | 11 | 1 | 1 | 2 | 4 | 0 | 0 | 0 | 0 | 0 | 0 | 0 |
| ISG000000 | Sorbs3 | 1.29E-08 | 3.9E-08 | 1.680725893 | 130.8933938 | 419.6301038 | 16 | 1 | 1 | 1 | 9 | 1 | 1 | 0 | 0 | 0 | 0 | 0 |
| ISG000000 | Dynlt1b | 1.63E-08 | 4.87E-08 | 2.021405673 | 21.71434367 | 88.15571013 | 6 | 0 | 1 | 0 | 1 | 0 | 0 | 0 | 0 | 0 | 0 | 0 |
| ISG000000 | 1700016K19Rik | 1.81E-08 | 5.38E-08 | 4.47050359 | 0 | 21.16948866 | 4 | 1 | 1 | 1 | 1 | 0 | 1 | 0 | 0 | 0 | 0 | 0 |
| ISG000000 | Vash1 | 2.07E-08 | 6.13E-08 | 2.761149557 | 12.14090616 | 82.30760055 | 16 | 0 | 1 | 1 | 4 | 0 | 0 | 0 | 0 | 0 | 0 | 0 |
| ISG000000 | Fam43a | 2.15E-08 | 6.36E-08 | -1.454674845 | 734.1909046 | 267.8602302 | 2 | 1 | 1 | 1 | 1 | 2 | 0 | 0 | 0 | 0 | 0 | 2 |
| ISG000000 | Fam105a | 2.88E-08 | 8.44E-08 | 2.980979387 | 5.808707244 | 45.86101907 | 29 | 0 | 1 | 3 | 12 | 5 | 5 | 0 | 1 | 0 | 3 | 1 |
| SG0000002 | Evc | 3.25E-08 | 9.5E-08 | -0.911113149 | 1096.024354 | 582.8379432 | 95 | 0 | 1 | 9 | 21 | 30 | 0 | 0 | 0 | 0 | 0 | 0 |
| SG0000005 | Myl3 | 3.35E-08 | 9.79E-08 | 4.419532049 | 0 | 20.3998985 | 8 | 1 | 1 | 1 | 3 | 0 | 0 | 0 | 0 | 0 | 0 | 0 |
| SG0000007 | Trpc5os | 3.57E-08 | 1.04E-07 | 3.710115775 | 2.613155511 | 34.19962872 | 70 | 0 | 1 | 6 | 34 | 4 | 10 | 0 | 0 | 0 | 6 | 0 |
| ISG000000 | Mrps22 | 3.92E-08 | 1.14E-07 | 0.843755179 | 450.4211108 | 808.3777573 | 15 | 1 | 1 | 1 | 4 | 5 | 1 | 0 | 0 | 0 | 0 | 0 |
| ISG000000 | Atg12 | 4.85E-08 | 1.4E-07 | 0.731677272 | 1138.608779 | 1890.737918 | 23 | 0 | 1 | 4 | 20 | 1 | 7 | 0 | 0 | 1 | 3 | 0 |
| ISG000000 | Sdpr | 5.74E-08 | 1.65E-07 | -1.717659451 | 467.4411208 | 142.1215376 | 29 | 0 | 1 | 4 | 13 | 1 | 3 | 0 | 1 | 1 | 4 | 0 |
| SG0000003 | Pdxk | 5.75E-08 | 1.65E-07 | 0.786463595 | 660.0986351 | 1138.565362 | 19 | 0 | 1 | 1 | 7 | 6 | 0 | 0 | 0 | 0 | 0 | 0 |
| ISG000000 | Csrnp2 | 5.79E-08 | 1.66E-07 | 1.034496794 | 183.4362312 | 375.7506015 | 16 | 1 | 1 | 3 | 6 | 0 | 6 | 1 | 1 | 2 | 2 | 0 |
| ISG000000 | Fzd9 | 5.79E-08 | 1.66E-07 | 2.396548886 | 20.46505504 | 107.7571305 | 2 | 0 | 1 | 0 | 1 | 0 | 2 | 0 | 1 | 0 | 0 | 0 |
| SG0000002 | Esrra | 5.99E-08 | 1.71E-07 | -2.29867476 | 1644.59142 | 334.2627388 | 6 | 1 | 1 | 2 | 3 | 2 | 1 | 0 | 0 | 1 | 1 | 0 |
| ISG000000 | Zfp770 | 7.22E-08 | 2.05E-07 | 0.818044589 | 597.0477991 | 1052.604108 | 22 | 0 | 1 | 1 | 5 | 0 | 8 | 0 | 0 | 0 | 0 | 0 |
| SG0000003 | Cog2 | 7.85E-08 | 2.22E-07 | 0.793812049 | 598.1175802 | 1036.926098 | 66 | 1 | 1 | 3 | 17 | 2 | 0 | 0 | 0 | 0 | 1 | 0 |
| SG0000000 | N4bp3 | 8.83E-08 | 2.49E-07 | 1.700144779 | 219.5363386 | 713.3472516 | 2 | 1 | 1 | 1 | 1 | 1 | 0 | 0 | 0 | 0 | 0 | 0 |
| ISG000000 | Leprotl1 | 9.68E-08 | 2.72E-07 | -0.813954275 | 1732.681273 | 985.5848742 | 14 | 0 | 1 | 2 | 5 | 5 | 4 | 0 | 0 | 1 | 2 | 2 |
| SG0000002 | Plch2 | 9.86E-08 | 2.76E-07 | 3.017432011 | 5.178127442 | 41.92859214 | 15 | 0 | 1 | 0 | 4 | 2 | 2 | 0 | 0 | 0 | 0 | 0 |
| ISG000000 | Eid2b | 1.14E-07 | 3.18E-07 | 1.204274204 | 94.71689291 | 218.247914 | 4 | 0 | 1 | 0 | 1 | 1 | 2 | 0 | 1 | 0 | 0 | 1 |
| SG0000006 | Rnf14 | 1.17E-07 | 3.26E-07 | -0.778189717 | 2592.81281 | 1511.863967 | 43 | 1 | 1 | 8 | 27 | 3 | 5 | 0 | 0 | 2 | 3 | 0 |
| ISG000000 | Ncbp1 | 1.33E-07 | 3.69E-07 | 0.668419644 | 2551.908804 | 4055.827848 | 69 | 0 | 1 | 4 | 26 | 8 | 0 | 0 | 0 | 0 | 1 | 1 |
| SG0000005 | Zfp61 | 1.42E-07 | 3.94E-07 | 1.030497678 | 184.4824053 | 376.847522 | 8 | 1 | 1 | 4 | 8 | 0 | 0 | 0 | 0 | 0 | 0 | 0 |
| ISG000000 | Them6 | 1.5E-07 | 4.14E-07 | -1.478163248 | 789.7974922 | 283.4942628 | 2 | 0 | 1 | 0 | 0 | 0 | 1 | 0 | 1 | 0 | 0 | 0 |
| SG0000002 | Adpgk | 1.55E-07 | 4.28E-07 | 0.713646625 | 1137.655447 | 1865.69132 | 39 | 1 | 1 | 6 | 12 | 0 | 2 | 0 | 0 | 1 | 1 | 0 |
| SG0000004 | Kmt2d | 2.06E-07 | 5.62E-07 | -0.815051813 | 2584.252412 | 1468.858248 | 20 | 1 | 1 | 4 | 8 | 3 | 5 | 0 | 0 | 2 | 2 | 2 |
| SG0000002 | Piwil1 | 2.07E-07 | 5.64E-07 | 4.26112842 | 0 | 18.17465108 | 10 | 0 | 1 | 0 | 2 | 2 | 2 | 0 | 1 | 0 | 0 | 0 |
| ISG000000 | Cyp26a1 | 2.4E-07 | 6.5E-07 | -1.485501166 | 162.8314657 | 58.15109842 | 8 | 0 | 1 | 1 | 1 | 1 | 2 | 0 | 1 | 1 | 1 | 0 |
| ISG000000 | Kcnj4 | 2.45E-07 | 6.63E-07 | 1.924588192 | 22.24017891 | 84.43005099 | 4 | 0 | 1 | 1 | 4 | 2 | 0 | 0 | 0 | 0 | 0 | 1 |
| ISG000000 | 2610001J05Rik | 2.64E-07 | 7.13E-07 | -0.833057981 | 1191.010646 | 668.5597136 | 11 | 0 | 1 | 1 | 12 | 0 | 7 | 0 | 1 | 1 | 8 | 0 |
| SG0000002 | Nol6 | 2.7E-07 | 7.3E-07 | 0.66405365 | 2116.886105 | 3354.26646 | 4 | 1 | 1 | 2 | 3 | 0 | 1 | 0 | 0 | 0 | 0 | 0 |
| ISG000000 | Lsm3 | 2.71E-07 | 7.33E-07 | 0.698372027 | 1074.517624 | 1743.590409 | 10 | 0 | 1 | 2 | 2 | 1 | 0 | 0 | 0 | 0 | 0 | 0 |
| SG0000007 | Haus5 | 3.06E-07 | 8.21E-07 | 1.277559987 | 73.58295173 | 178.3861296 | 10 | 1 | 1 | 1 | 6 | 0 | 2 | 0 | 0 | 0 | 0 | 0 |
| ISG000000 | Gcm1 | 3.61E-07 | 9.64E-07 | 4.211879332 | 0 | 17.53113488 | 24 | 1 | 1 | 1 | 6 | 1 | 0 | 0 | 0 | 0 | 0 | 0 |
| SG0000002 | Tfip11 | 3.66E-07 | 9.78E-07 | -1.066079877 | 1892.32282 | 903.8018085 | 8 | 0 | 1 | 0 | 2 | 0 | 0 | 0 | 0 | 0 | 0 | 0 |
| SG0000003 | Zfp300 | 4.21E-07 | 1.12E-06 | -2.340786736 | 65.83600895 | 12.99620264 | 23 | 1 | 1 | 4 | 15 | 1 | 4 | 0 | 0 | 0 | 1 | 0 |
| ISG000000 | B430305J03Rik | 4.59E-07 | 1.22E-06 | -4.679908636 | 50.50958145 | 1.970520209 | 3 | 0 | 1 | 1 | 1 | 0 | 2 | 0 | 1 | 1 | 1 | 0 |
| ISG000000 | Fitm2 | 4.73E-07 | 1.25E-06 | -0.928325327 | 588.6433908 | 309.313235 | 6 | 0 | 1 | 0 | 2 | 0 | 4 | 0 | 0 | 0 | 1 | 0 |
| ISG000000 | Fthl17a | 4.81E-07 | 1.27E-06 | 4.279911273 | 0 | 18.42592341 | 32 | 1 | 1 | 4 | 16 | 1 | 0 | 0 | 0 | 0 | 0 | 0 |
| SG0000005 | Rpl5 | 6.09E-07 | 1.6E-06 | 0.596157945 | 10580.56953 | 15994.49266 | 20 | 0 | 1 | 0 | 2 | 2 | 2 | 0 | 1 | 0 | 0 | 1 |
| ISG000000 | Gabarapl1 | 7.78E-07 | 2.03E-06 | -0.712113135 | 2853.525001 | 1741.864099 | 16 | 0 | 1 | 1 | 3 | 1 | 2 | 0 | 0 | 0 | 0 | 0 |
| ISG000000 | Tmem211 | 7.92E-07 | 2.06E-06 | 5.835838273 | 0.331601068 | 18.93992659 | 20 | 0 | 1 | 3 | 3 | 1 | 1 | 0 | 0 | 0 | 0 | 0 |
| SG0000005 | Rrp1b | 7.98E-07 | 2.08E-06 | 0.676780158 | 2142.580616 | 3425.060911 | 44 | 0 | 1 | 1 | 4 | 5 | 3 | 0 | 1 | 0 | 1 | 0 |
| SG0000000 | Hif3a | 8.03E-07 | 2.09E-06 | 3.180901129 | 64.76838383 | 587.3672303 | 44 | 1 | 1 | 5 | 14 | 1 | 2 | 0 | 0 | 0 | 0 | 1 |
| SG0000005 | Cgrrf1 | 8.29E-07 | 2.16E-06 | -0.998889499 | 415.9100394 | 208.1151527 | 76 | 0 | 1 | 3 | 18 | 1 | 1 | 0 | 0 | 0 | 0 | 0 |
| ISG000000 | Asf1a | 9.27E-07 | 2.4E-06 | 0.715200655 | 666.2536518 | 1093.796307 | 31 | 0 | 1 | 2 | 10 | 4 | 5 | 0 | 0 | 0 | 2 | 1 |
| ISG000000 | E430025E21Rik | 9.41E-07 | 2.44E-06 | -0.693996362 | 4269.428834 | 2639.101278 | 57 | 0 | 1 | 6 | 17 | 3 | 0 | 0 | 0 | 0 | 0 | 0 |
| SG0000007 | Runx3 | 1.06E-06 | 2.73E-06 | 2.088016336 | 14.20647821 | 60.40070086 | 47 | 0 | 1 | 6 | 15 | 2 | 1 | 0 | 0 | 0 | 0 | 0 |
| ISG000000 | Zfp524 | 1.07E-06 | 2.77E-06 | -1.433071143 | 204.2139421 | 75.62893574 | 3 | 1 | 1 | 1 | 1 | 0 | 0 | 0 | 0 | 0 | 0 | 0 |
| SG0000003 | Zfr2 | 1.11E-06 | 2.86E-06 | 2.116582386 | 15.63778019 | 67.81564408 | 10 | 1 | 1 | 2 | 3 | 0 | 2 | 0 | 0 | 0 | 0 | 0 |
| ISG000000 | Alox12b | 1.44E-06 | 3.66E-06 | -2.661156725 | 186.6067379 | 29.50117566 | 10 | 0 | 1 | 1 | 4 | 4 | 0 | 0 | 0 | 0 | 0 | 0 |
| ISG000000 | Bhmt2 | 1.6E-06 | 4.05E-06 | 4.065508683 | 0 | 15.74326153 | 17 | 1 | 1 | 2 | 6 | 3 | 0 | 0 | 0 | 0 | 0 | 0 |
| ISG000000 | Zfp948 | 1.73E-06 | 4.36E-06 | 0.877353582 | 372.1149726 | 683.5761313 | 58 | 0 | 1 | 7 | 31 | 5 | 0 | 0 | 0 | 0 | 0 | 0 |
| ISG000000 | Dhrs4 | 2.05E-06 | 5.14E-06 | 0.773988354 | 377.717737 | 645.8937617 | 19 | 1 | 1 | 2 | 6 | 0 | 1 | 0 | 0 | 0 | 0 | 0 |
| SG0000006 | Zkscan8 | 2.13E-06 | 5.34E-06 | -0.853019306 | 1268.412493 | 702.224719 | 29 | 0 | 1 | 3 | 14 | 10 | 11 | 0 | 0 | 1 | 5 | 3 |
| ISG000000 | Rnf135 | 2.16E-06 | 5.41E-06 | -1.139284518 | 243.9291169 | 110.7400793 | 13 | 0 | 1 | 6 | 14 | 2 | 0 | 0 | 0 | 0 | 0 | 0 |
| ISG000000 | Best2 | 2.61E-06 | 6.5E-06 | 5.839357276 | 0.307509278 | 17.60677925 | 4 | 1 | 1 | 1 | 2 | 0 | 0 | 0 | 0 | 0 | 0 | 0 |
| ISG000000 | Fzd4 | 2.66E-06 | 6.62E-06 | -1.453860212 | 175.9649755 | 64.23484793 | 7 | 1 | 1 | 3 | 4 | 0 | 3 | 1 | 1 | 2 | 3 | 0 |
| ISG000000 | Mplkip | 2.68E-06 | 6.65E-06 | 0.806664966 | 286.5317967 | 501.1908967 | 9 | 0 | 1 | 2 | 4 | 1 | 6 | 0 | 1 | 1 | 2 | 1 |
| ISG000000 | Thbs4 | 2.77E-06 | 6.86E-06 | 4.439910203 | 0.970711415 | 21.0686295 | 49 | 0 | 1 | 3 | 17 | 4 | 0 | 0 | 0 | 0 | 1 | 0 |
| ISG000000 | Fam83f | 2.78E-06 | 6.89E-06 | 7.217273869 | 0.663202137 | 98.68742976 | 24 | 1 | 1 | 1 | 9 | 3 | 0 | 0 | 0 | 0 | 1 | 0 |
| SG0000005 | Rps10 | 2.78E-06 | 6.89E-06 | 0.600042784 | 6573.061836 | 9963.194178 | 13 | 1 | 1 | 1 | 12 | 0 | 0 | 0 | 0 | 0 | 0 | 0 |
| ISG000000 | Npy4r | 2.8E-06 | 6.95E-06 | 4.008420247 | 0 | 15.09365658 | 6 | 0 | 1 | 0 | 3 | 0 | 1 | 0 | 0 | 0 | 1 | 0 |
| ISG000000 | Topors | 3.06E-06 | 7.56E-06 | 0.609299735 | 1786.916307 | 2725.973996 | 17 | 0 | 1 | 0 | 10 | 1 | 2 | 0 | 1 | 0 | 1 | 0 |
| ISG000000 | Comp | 3.15E-06 | 7.77E-06 | 4.086011333 | 1.313186595 | 22.30172286 | 5 | 1 | 1 | 3 | 3 | 0 | 0 | 0 | 0 | 0 | 0 | 0 |
| ISG000000 | Tkfc | 3.41E-06 | 8.41E-06 | -0.994471653 | 874.6050619 | 438.9814707 | 5 | 0 | 1 | 0 | 2 | 0 | 0 | 0 | 0 | 0 | 0 | 0 |
| ISG000000 | Aga | 3.46E-06 | 8.52E-06 | -0.81900106 | 842.4892388 | 477.5517893 | 21 | 1 | 1 | 4 | 8 | 2 | 0 | 0 | 0 | 0 | 0 | 0 |
| SG0000002 | Pigh | 3.72E-06 | 9.14E-06 | -0.860722383 | 568.2806588 | 312.9389493 | 17 | 0 | 1 | 0 | 7 | 1 | 1 | 0 | 0 | 0 | 0 | 0 |
| SG0000002 | Gpnmb | 3.8E-06 | 9.31E-06 | -0.945093171 | 390.5094713 | 202.8290603 | 20 | 0 | 1 | 0 | 15 | 4 | 2 | 0 | 0 | 0 | 2 | 1 |
| SG0000000 | Efnb2 | 4.29E-06 | 1.05E-05 | -0.967157252 | 1003.062101 | 513.0792818 | 71 | 0 | 1 | 9 | 26 | 9 | 6 | 0 | 1 | 2 | 5 | 3 |
| ISG000000 | Mn1 | 4.42E-06 | 1.08E-05 | 2.179341249 | 20.59523811 | 93.28544852 | 34 | 1 | 1 | 2 | 13 | 9 | 1 | 0 | 0 | 0 | 0 | 2 |
| ISG000000 | Aars2 | 4.42E-06 | 1.08E-05 | 1.127364364 | 296.3016244 | 647.2983829 | 16 | 1 | 1 | 1 | 3 | 1 | 0 | 0 | 0 | 0 | 0 | 0 |

| | | | | | | | | | | | | | | | | | | |
|---|---|---|---|---|---|---|---|---|---|---|---|---|---|---|---|---|---|---|
| JSG000000 | Spry4 | 4.76E-06 | 1.16E-05 | -0.752528495 | 2791.286847 | 1656.802799 | 22 | 0 | 1 | 5 | 7 | 2 | 7 | 0 | 0 | 4 | 4 | 2 |
| JSG000000 | Zfp763 | 4.97E-06 | 1.21E-05 | 1.030203201 | 256.1384459 | 523.114593 | 20 | 0 | 1 | 0 | 4 | 6 | 4 | 0 | 1 | 0 | 0 | 2 |
| JSG000000 | Sox7 | 6.17E-06 | 1.48E-05 | 3.302307413 | 3.182334054 | 31.39347219 | 8 | 1 | 1 | 3 | 6 | 4 | 2 | 0 | 0 | 0 | 3 | 1 |
| JSG000000 | Cnn1 | 6.25E-06 | 1.5E-05 | 2.126251694 | 25.39592447 | 110.8739615 | 4 | 1 | 1 | 1 | 4 | 0 | 1 | 0 | 0 | 0 | 0 | 0 |
| JSG000000 | Mad2l1bp | 6.95E-06 | 1.66E-05 | -0.847612807 | 503.0157125 | 279.5275844 | 10 | 1 | 1 | 1 | 8 | 0 | 0 | 0 | 0 | 0 | 0 | 0 |
| JSG000000 | Dsg3 | 8.07E-06 | 1.92E-05 | -4.237975215 | 25.8602053 | 1.370485067 | 51 | 1 | 1 | 7 | 23 | 3 | 1 | 0 | 0 | 0 | 1 | 1 |
| JSG000000 | Psmd7 | 8.15E-06 | 1.94E-05 | 0.550730155 | 4941.81212 | 7238.89915 | 8 | 1 | 1 | 2 | 2 | 1 | 0 | 0 | 0 | 0 | 0 | 0 |
| JSG000000 | Borcs7 | 9.79E-06 | 2.31E-05 | -1.102418162 | 212.0325477 | 98.75102158 | 13 | 0 | 1 | 1 | 5 | 3 | 3 | 0 | 1 | 0 | 1 | 0 |
| JSG000000 | Map3k11 | 1.11E-05 | 2.6E-05 | -1.568320063 | 1089.361824 | 367.3336973 | 5 | 1 | 1 | 1 | 3 | 1 | 0 | 0 | 0 | 0 | 0 | 0 |
| JSG000000 | Slc30a10 | 1.52E-05 | 3.53E-05 | 3.537332001 | 4.705316061 | 54.63008014 | 25 | 0 | 1 | 4 | 16 | 2 | 8 | 0 | 1 | 2 | 5 | 2 |
| SG0000005 | Sphk2 | 1.53E-05 | 3.54E-05 | -1.267217351 | 787.4434179 | 327.1515057 | 15 | 0 | 1 | 4 | 8 | 1 | 3 | 0 | 0 | 0 | 0 | 0 |
| JSG000000 | Zfp3 | 1.77E-05 | 4.08E-05 | -1.17595707 | 170.5270285 | 75.47356644 | 10 | 1 | 1 | 3 | 5 | 0 | 1 | 0 | 0 | 1 | 1 | 0 |
| JSG000000 | Rln1 | 1.86E-05 | 4.27E-05 | 5.505937771 | 0.331601068 | 15.06844178 | 2 | 1 | 1 | 1 | 5 | 1 | 1 | 1 | 1 | 1 | 1 | 0 |
| SG0000003 | Atp5j2 | 1.89E-05 | 4.33E-05 | 0.556655212 | 2270.294254 | 3339.274076 | 11 | 1 | 1 | 4 | 11 | 1 | 0 | 0 | 0 | 0 | 0 | 0 |
| SG0000002 | Pdcl3 | 2.05E-05 | 4.69E-05 | 0.57281043 | 1365.893529 | 2031.655034 | 29 | 0 | 1 | 2 | 4 | 0 | 4 | 0 | 1 | 0 | 0 | 0 |
| JSG000000 | Tspan11 | 2.11E-05 | 4.82E-05 | -2.633581308 | 37.50520515 | 6.04372334 | 41 | 1 | 1 | 3 | 18 | 7 | 0 | 0 | 0 | 0 | 2 | 1 |
| SG0000002 | Cpsf4 | 2.27E-05 | 5.16E-05 | 0.598299116 | 933.3524057 | 1413.03081 | 10 | 1 | 1 | 2 | 5 | 0 | 1 | 0 | 0 | 0 | 0 | 0 |
| JSG000000 | Hsf1 | 2.67E-05 | 6.04E-05 | -0.687491924 | 1184.74891 | 735.6489624 | 16 | 0 | 1 | 2 | 13 | 4 | 0 | 0 | 0 | 0 | 0 | 0 |
| JSG000000 | Gpr75 | 2.87E-05 | 6.48E-05 | 4.028195368 | 2.532349596 | 41.31723925 | 17 | 1 | 1 | 1 | 6 | 0 | 4 | 1 | 1 | 1 | 3 | 0 |
| JSG000000 | Tlx3 | 2.89E-05 | 6.51E-05 | -2.937256933 | 37.1533557 | 4.850601579 | 3 | 0 | 1 | 0 | 1 | 1 | 2 | 0 | 1 | 0 | 1 | 1 |
| JSG000000 | 2510039O18Rik | 3.02E-05 | 6.79E-05 | -0.959912209 | 1666.993214 | 856.9815335 | 6 | 1 | 1 | 1 | 1 | 0 | 2 | 0 | 0 | 0 | 0 | 0 |
| SG0000004 | Zfp646 | 3.08E-05 | 6.92E-05 | 0.560804483 | 1377.961316 | 2032.619074 | 5 | 1 | 1 | 1 | 3 | 0 | 1 | 0 | 0 | 0 | 0 | 0 |
| SG0000001 | Gdi1 | 3.1E-05 | 6.96E-05 | -0.603074278 | 5545.750178 | 3651.042215 | 9 | 0 | 1 | 1 | 3 | 2 | 3 | 0 | 0 | 0 | 0 | 0 |
| JSG000000 | Mmp24 | 3.13E-05 | 7.02E-05 | 2.884129891 | 3.597084605 | 26.55585332 | 42 | 0 | 1 | 4 | 14 | 1 | 2 | 0 | 0 | 0 | 1 | 0 |
| JSG000000 | Gm4353 | 3.22E-05 | 7.23E-05 | 1.580122479 | 22.77054814 | 68.08285329 | 4 | 1 | 1 | 1 | 2 | 0 | 2 | 0 | 0 | 0 | 0 | 0 |
| JSG000000 | Gpx2 | 3.26E-05 | 7.31E-05 | 3.456077773 | 15.92020081 | 174.7155737 | 2 | 0 | 1 | 0 | 1 | 0 | 0 | 0 | 0 | 0 | 0 | 0 |
| SG0000002 | Slu7 | 3.28E-05 | 7.36E-05 | -0.617914166 | 1900.367462 | 1238.302892 | 37 | 1 | 1 | 1 | 13 | 0 | 6 | 0 | 0 | 0 | 0 | 0 |
| SG0000003 | Rtn2 | 3.72E-05 | 8.3E-05 | 1.284853892 | 98.67366692 | 240.4256669 | 19 | 1 | 1 | 3 | 11 | 3 | 2 | 0 | 0 | 0 | 0 | 0 |
| JSG000000 | Pde1b | 4.1E-05 | 9.1E-05 | 1.276559122 | 164.6583763 | 398.9021851 | 26 | 0 | 1 | 3 | 8 | 3 | 1 | 0 | 0 | 0 | 0 | 1 |
| JSG000000 | Esm1 | 4.25E-05 | 9.4E-05 | -1.705196801 | 98.49254086 | 30.20563399 | 15 | 0 | 1 | 2 | 8 | 0 | 5 | 0 | 0 | 1 | 6 | 0 |
| JSG000000 | Klhl9 | 4.29E-05 | 9.48E-05 | -0.59313514 | 2549.675574 | 1690.181933 | 7 | 0 | 1 | 1 | 2 | 0 | 4 | 0 | 0 | 0 | 1 | 0 |
| JSG000000 | Gorab | 4.29E-05 | 9.49E-05 | -0.883505752 | 940.6807243 | 509.8947168 | 40 | 1 | 1 | 5 | 13 | 3 | 4 | 0 | 0 | 2 | 2 | 0 |
| SG0000004 | Fam46c | 4.62E-05 | 0.000102 | 1.487536874 | 25.85048119 | 72.48728778 | 26 | 1 | 1 | 2 | 4 | 3 | 2 | 0 | 0 | 0 | 0 | 0 |
| JSG000000 | Tmem120a | 4.66E-05 | 0.000103 | -0.685655235 | 833.7503963 | 518.3621243 | 5 | 1 | 1 | 1 | 3 | 0 | 0 | 0 | 0 | 0 | 0 | 0 |
| JSG000000 | Dctn2 | 4.67E-05 | 0.000103 | -0.556843722 | 4834.306741 | 3286.302522 | 11 | 1 | 1 | 2 | 11 | 1 | 0 | 0 | 0 | 0 | 0 | 0 |
| JSG000000 | Ccdc94 | 5.36E-05 | 0.000117 | -1.035694521 | 240.1205494 | 117.1262426 | 6 | 1 | 1 | 1 | 1 | 0 | 0 | 0 | 0 | 0 | 0 | 0 |
| JSG000000 | Rhobtb2 | 5.4E-05 | 0.000118 | 0.605709617 | 593.1990411 | 902.6870329 | 26 | 1 | 1 | 2 | 11 | 2 | 2 | 0 | 0 | 0 | 2 | 0 |
| JSG000000 | Hapln3 | 5.41E-05 | 0.000118 | 3.573331206 | 1.644787664 | 19.57896442 | 10 | 0 | 1 | 3 | 11 | 0 | 2 | 0 | 1 | 0 | 8 | 0 |
| SG0000003 | Ndufc1 | 5.61E-05 | 0.000122 | 0.696833961 | 506.5014377 | 821.010302 | 11 | 0 | 1 | 0 | 2 | 0 | 2 | 0 | 0 | 0 | 1 | 0 |
| JSG000000 | Mrpl51 | 5.62E-05 | 0.000123 | -0.633758905 | 1762.059479 | 1135.638543 | 8 | 1 | 1 | 1 | 4 | 0 | 8 | 1 | 1 | 1 | 4 | 0 |
| JSG000000 | Cmtm3 | 5.94E-05 | 0.000129 | -0.854956788 | 372.428678 | 205.90907 | 3 | 1 | 1 | 1 | 2 | 0 | 0 | 0 | 0 | 0 | 0 | 0 |
| JSG000000 | Atox1 | 6.22E-05 | 0.000135 | -0.722315182 | 1577.014328 | 955.8661961 | 19 | 1 | 1 | 2 | 7 | 2 | 0 | 0 | 0 | 0 | 0 | 0 |
| SG0000005 | Minos1 | 6.86E-05 | 0.000148 | 0.538940821 | 1334.746464 | 1939.260412 | 39 | 1 | 1 | 4 | 16 | 6 | 3 | 0 | 0 | 0 | 1 | 0 |
| JSG000000 | Sptssa | 7.02E-05 | 0.000152 | -0.61242154 | 1461.225165 | 955.7842935 | 24 | 1 | 1 | 4 | 8 | 0 | 1 | 0 | 0 | 0 | 1 | 0 |
| JSG000000 | Steap4 | 7.16E-05 | 0.000154 | -1.844359413 | 57.55416294 | 16.02762733 | 44 | 0 | 1 | 4 | 14 | 4 | 2 | 0 | 0 | 0 | 1 | 0 |
| JSG000000 | Rfk | 7.3E-05 | 0.000157 | -0.589986691 | 1856.177365 | 1233.149642 | 18 | 1 | 1 | 1 | 5 | 1 | 6 | 0 | 0 | 0 | 1 | 0 |
| SG0000002 | Mthfr | 7.46E-05 | 0.000161 | -0.65518157 | 870.10327 | 552.5117212 | 34 | 0 | 1 | 0 | 3 | 0 | 3 | 0 | 0 | 0 | 0 | 0 |
| JSG000000 | Eif3g | 7.53E-05 | 0.000162 | 0.513566608 | 2728.509684 | 3895.152389 | 2 | 1 | 1 | 1 | 1 | 2 | 0 | 0 | 0 | 0 | 0 | 0 |
| JSG000000 | Cox7b | 7.85E-05 | 0.000169 | 0.497562528 | 3093.796642 | 4367.903242 | 17 | 1 | 1 | 1 | 3 | 1 | 3 | 0 | 0 | 0 | 0 | 0 |
| JSG000000 | Zfp397 | 8.11E-05 | 0.000174 | -0.627148988 | 1111.718279 | 719.7871503 | 22 | 0 | 1 | 0 | 7 | 1 | 8 | 0 | 0 | 0 | 2 | 1 |
| JSG000000 | Acot2 | 8.13E-05 | 0.000174 | 1.4230946 | 27.76546867 | 74.45590892 | 2 | 0 | 1 | 1 | 3 | 1 | 0 | 0 | 0 | 0 | 1 | 1 |
| JSG000000 | Tubg1 | 8.81E-05 | 0.000188 | -0.590234324 | 2091.978322 | 1389.565248 | 6 | 1 | 1 | 3 | 4 | 0 | 1 | 0 | 0 | 1 | 1 | 0 |
| SG0000002 | Rab13 | 9.66E-05 | 0.000205 | -0.842056226 | 318.3479008 | 177.5897142 | 11 | 1 | 1 | 1 | 4 | 1 | 0 | 0 | 0 | 0 | 0 | 0 |
| JSG000000 | Grpel1 | 0.000113 | 0.000239 | -0.545464309 | 2999.636038 | 2055.26319 | 13 | 0 | 1 | 1 | 1 | 1 | 4 | 0 | 0 | 0 | 0 | 0 |
| JSG000000 | Dtx4 | 0.000127 | 0.000265 | 0.959165674 | 80.63925106 | 156.7776406 | 47 | 1 | 1 | 5 | 10 | 6 | 2 | 0 | 0 | 0 | 1 | 1 |
| SG0000002 | Vamp2 | 0.000129 | 0.000269 | -0.918939856 | 1496.106863 | 791.2872252 | 6 | 0 | 1 | 1 | 3 | 1 | 2 | 0 | 1 | 1 | 3 | 1 |
| JSG000000 | Gsta1 | 0.000131 | 0.000273 | -4.195216432 | 18.04832413 | 0.985260105 | 12 | 0 | 1 | 2 | 7 | 2 | 0 | 0 | 0 | 0 | 0 | 0 |
| JSG000000 | Echdc3 | 0.000131 | 0.000275 | -1.579306567 | 84.64405551 | 28.32551497 | 24 | 0 | 1 | 1 | 10 | 2 | 2 | 0 | 0 | 1 | 1 | 0 |
| JSG000000 | Grpel2 | 0.000133 | 0.000277 | -0.564533884 | 2021.404595 | 1366.820916 | 19 | 1 | 1 | 4 | 11 | 2 | 6 | 1 | 1 | 3 | 6 | 2 |
| JSG000000 | Ormdl1 | 0.000136 | 0.000284 | 0.616391125 | 452.6884507 | 693.987475 | 23 | 0 | 1 | 1 | 15 | 2 | 4 | 0 | 0 | 0 | 0 | 0 |
| JSG000000 | Emc4 | 0.000155 | 0.000321 | -0.615888514 | 1091.385647 | 712.1595809 | 12 | 0 | 1 | 1 | 5 | 1 | 1 | 0 | 0 | 1 | 1 | 0 |
| JSG000000 | Clasrp | 0.000159 | 0.00033 | 0.589943478 | 519.0668965 | 781.2931473 | 11 | 0 | 1 | 1 | 4 | 2 | 1 | 0 | 1 | 0 | 0 | 0 |
| JSG000000 | Zfp771 | 0.000163 | 0.000337 | -1.556973974 | 124.4100728 | 42.28240441 | 12 | 1 | 1 | 2 | 7 | 0 | 0 | 0 | 0 | 0 | 0 | 0 |
| JSG000000 | Ccnk | 0.000168 | 0.000347 | 0.504100941 | 1502.191453 | 2130.466899 | 27 | 0 | 1 | 1 | 3 | 6 | 0 | 0 | 0 | 0 | 0 | 1 |
| JSG000000 | Cdc37l1 | 0.000177 | 0.000365 | 0.541296965 | 776.8829492 | 1130.58238 | 68 | 0 | 1 | 8 | 25 | 3 | 3 | 0 | 0 | 0 | 1 | 0 |
| JSG000000 | Lamp1 | 0.00018 | 0.00037 | -0.494668081 | 18460.44348 | 13101.83724 | 17 | 0 | 1 | 1 | 8 | 2 | 3 | 0 | 0 | 0 | 1 | 0 |
| JSG000000 | Mtpap | 0.000181 | 0.000372 | 0.52933582 | 860.2840562 | 1241.617524 | 36 | 0 | 1 | 4 | 11 | 2 | 0 | 0 | 0 | 0 | 1 | 0 |
| JSG000000 | Fam32a | 0.000187 | 0.000386 | -0.529861118 | 2859.791137 | 1980.75256 | 5 | 1 | 1 | 2 | 3 | 0 | 1 | 0 | 0 | 0 | 0 | 0 |
| JSG000000 | Guca1a | 0.000193 | 0.000396 | -1.747791587 | 113.5224083 | 33.80211711 | 7 | 0 | 1 | 0 | 3 | 0 | 0 | 0 | 0 | 0 | 0 | 0 |
| SG0000007 | Tmsb15l | 0.000201 | 0.000413 | 1.978885099 | 9.176744919 | 36.17365839 | 34 | 1 | 1 | 2 | 15 | 6 | 1 | 0 | 0 | 0 | 0 | 0 |
| JSG000000 | Zfp36 | 0.000246 | 0.000498 | 0.75335516 | 422.4967764 | 712.2064464 | 3 | 1 | 1 | 3 | 4 | 0 | 0 | 0 | 0 | 0 | 0 | 0 |
| JSG000000 | Spcs3 | 0.000246 | 0.000498 | 0.459731067 | 3043.962806 | 4186.317687 | 14 | 0 | 1 | 5 | 11 | 2 | 5 | 0 | 0 | 1 | 3 | 0 |
| JSG000000 | Fkbp11 | 0.000262 | 0.000529 | 0.697310754 | 204.9186379 | 332.2713634 | 2 | 1 | 1 | 1 | 4 | 0 | 0 | 0 | 0 | 0 | 0 | 0 |
| JSG000000 | Ppp4c | 0.000272 | 0.000549 | 0.479029739 | 2667.211642 | 3717.57551 | 5 | 1 | 1 | 1 | 3 | 0 | 0 | 0 | 0 | 0 | 0 | 0 |
| SG0000001 | Pdcd2 | 0.000284 | 0.000571 | 0.56197741 | 527.5625 | 778.8359235 | 27 | 1 | 1 | 4 | 14 | 2 | 3 | 0 | 0 | 0 | 1 | 0 |
| JSG000000 | Ppp1r3d | 0.000301 | 0.000604 | 2.729260665 | 3.276357648 | 21.72604274 | 1 | 1 | 1 | 1 | 2 | 0 | 1 | 1 | 1 | 1 | 2 | 0 |
| JSG000000 | Dhcr24 | 0.000314 | 0.000629 | 0.68860076 | 2220.100753 | 3578.17987 | 29 | 0 | 1 | 2 | 8 | 3 | 1 | 0 | 0 | 0 | 0 | 0 |
| JSG000000 | Alg2 | 0.000322 | 0.000643 | -0.620796345 | 716.2969587 | 465.8164125 | 5 | 1 | 1 | 1 | 2 | 1 | 1 | 1 | 1 | 1 | 2 | 0 |
| JSG000000 | Rgs2 | 0.000373 | 0.00074 | -1.526266163 | 94.02007157 | 32.64139276 | 14 | 0 | 1 | 0 | 3 | 0 | 6 | 0 | 1 | 0 | 1 | 0 |
| JSG000000 | A930004D18Rik | 0.000403 | 0.000795 | -1.528611256 | 61.21883578 | 21.21911305 | 9 | 0 | 1 | 0 | 4 | 2 | 2 | 0 | 1 | 0 | 0 | 0 |
| JSG000000 | Mtmr9 | 0.000403 | 0.000795 | 0.49930631 | 1241.21588 | 1754.500514 | 23 | 0 | 1 | 2 | 6 | 1 | 0 | 0 | 0 | 0 | 0 | 0 |
| JSG000000 | Fam25c | 0.000407 | 0.000804 | -1.881512752 | 258.9288018 | 70.27301486 | 3 | 1 | 1 | 1 | 3 | 0 | 0 | 0 | 0 | 0 | 0 | 0 |
| SG0000002 | Guk1 | 0.000434 | 0.000854 | 0.51439831 | 833.9205731 | 1191.170719 | 6 | 1 | 1 | 1 | 3 | 0 | 0 | 0 | 0 | 0 | 0 | 0 |
| JSG000000 | Fbxo48 | 0.000443 | 0.000871 | -1.090141712 | 159.2186576 | 74.78742895 | 21 | 0 | 1 | 3 | 6 | 2 | 11 | 0 | 1 | 1 | 2 | 0 |
| SG0000003 | Lzts2 | 0.000446 | 0.000876 | 0.842977852 | 265.1428762 | 475.5997112 | 7 | 1 | 1 | 1 | 4 | 0 | 1 | 0 | 0 | 0 | 0 | 0 |
| JSG000000 | Foxq1 | 0.00046 | 0.000901 | 4.151821509 | 1.359026609 | 24.15743229 | 2 | 0 | 1 | 1 | 2 | 2 | 0 | 0 | 0 | 0 | 0 | 0 |
| SG0000000 | Psmc3 | 0.000492 | 0.000962 | 0.444849121 | 3603.825437 | 4905.4254 | 18 | 0 | 1 | 1 | 4 | 4 | 0 | 0 | 0 | 0 | 0 | 0 |
| JSG000000 | Hsdl1 | 0.000566 | 0.001095 | 0.524479789 | 606.3973544 | 872.2509405 | 13 | 0 | 1 | 2 | 3 | 0 | 0 | 0 | 0 | 0 | 0 | 0 |
| SG0000002 | Twf1 | 0.000576 | 0.001115 | -0.472620305 | 4255.922176 | 3067.04953 | 23 | 1 | 1 | 3 | 11 | 2 | 1 | 0 | 0 | 0 | 1 | 0 |
| SG0000002 | Necap2 | 0.000581 | 0.001122 | -0.607383908 | 1265.698736 | 830.7867684 | 10 | 1 | 1 | 1 | 1 | 0 | 1 | 0 | 0 | 0 | 0 | 0 |
| SG0000002 | Khk | 0.000582 | 0.001125 | -1.033930667 | 388.8870009 | 189.9237524 | 10 | 1 | 1 | 1 | 6 | 0 | 0 | 0 | 0 | 0 | 0 | 0 |
| SG0000005 | Il17d | 0.000584 | 0.001128 | 2.634154616 | 3.276357648 | 20.33999567 | 22 | 0 | 1 | 2 | 3 | 1 | 1 | 0 | 0 | 0 | 0 | 0 |
| JSG000000 | Fibin | 0.0006 | 0.001157 | -2.510194421 | 25.07754108 | 4.401909755 | 5 | 1 | 1 | 2 | 3 | 0 | 5 | 1 | 1 | 2 | 3 | 0 |
| SG0000003 | Zc3h10 | 0.000611 | 0.001177 | 0.528927613 | 696.4819063 | 1004.9233 | 7 | 1 | 1 | 1 | 1 | 0 | 3 | 1 | 1 | 1 | 1 | 0 |
| JSG000000 | Dusp5 | 0.000628 | 0.001208 | -0.615095246 | 632.3381029 | 412.8451934 | 11 | 1 | 1 | 2 | 10 | 1 | 1 | 0 | 0 | 0 | 0 | 0 |
| JSG000000 | Gsdma | 0.000631 | 0.001214 | -1.380864251 | 67.67902733 | 25.9879815 | 10 | 1 | 1 | 3 | 6 | 14 | 3 | 0 | 0 | 0 | 2 | 0 |
| JSG000000 | D10Jhu81e | 0.000646 | 0.00124 | -0.653776212 | 725.6784367 | 461.2516875 | 4 | 1 | 1 | 1 | 1 | 1 | 0 | 0 | 0 | 0 | 0 | 0 |
| SG0000002 | Anapc2 | 0.00065 | 0.001248 | -0.476767985 | 3061.313214 | 2199.815721 | 6 | 0 | 1 | 0 | 3 | 1 | 0 | 0 | 0 | 0 | 0 | 0 |
| JSG000000 | Fpgt | 0.000662 | 0.001268 | -0.594532317 | 703.4108165 | 465.8402193 | 19 | 0 | 1 | 3 | 9 | 2 | 3 | 0 | 1 | 1 | 4 | 0 |
| JSG000000 | Zfp747 | 0.000671 | 0.001286 | -1.004170372 | 241.5641158 | 120.4334196 | 1 | 1 | 1 | 1 | 1 | 0 | 1 | 1 | 1 | 1 | 1 | 0 |
| JSG000000 | Il23a | 0.000674 | 0.00129 | -1.070603717 | 108.6379599 | 51.72467484 | 7 | 1 | 1 | 5 | 6 | 0 | 6 | 1 | 1 | 5 | 6 | 0 |
| SG0000003 | Szt2 | 0.000719 | 0.001373 | -0.533300523 | 1211.638926 | 837.2087586 | 27 | 1 | 1 | 5 | 10 | 4 | 1 | 0 | 0 | 0 | 0 | 0 |
| JSG000000 | Kctd13 | 0.000724 | 0.001382 | 0.745123432 | 282.3974653 | 473.3313805 | 22 | 1 | 1 | 3 | 15 | 4 | 1 | 1 | 1 | 1 | 3 | 0 |
| SG0000006 | Selenbp1 | 0.000749 | 0.001427 | -1.187495233 | 163.949828 | 71.98454777 | 10 | 1 | 1 | 1 | 2 | 1 | 0 | 0 | 0 | 0 | 0 | 0 |
| JSG000000 | Pomgnt2 | 0.000789 | 0.001499 | -0.942717438 | 270.061678 | 140.5001331 | 10 | 1 | 1 | 3 | 5 | 1 | 1 | 0 | 0 | 1 | 1 | 0 |
| JSG000000 | Ndufb3 | 0.000816 | 0.001548 | 0.481946562 | 858.9131241 | 1199.581088 | 33 | 1 | 1 | 5 | 16 | 3 | 0 | 0 | 0 | 0 | 1 | 0 |
| JSG000000 | Steap1 | 0.000872 | 0.001648 | 0.800977333 | 101.8716058 | 177.4889642 | 42 | 0 | 1 | 0 | 13 | 7 | 0 | 0 | 0 | 0 | 0 | 0 |
| JSG000000 | Ptcd2 | 0.000911 | 0.001716 | 0.492483753 | 667.5055994 | 939.0901712 | 32 | 0 | 1 | 4 | 15 | 3 | 0 | 0 | 0 | 0 | 0 | 0 |
| JSG000000 | Chac1 | 0.000936 | 0.001761 | 1.070269043 | 69.04074734 | 144.9734817 | 1 | 1 | 1 | 1 | 1 | 0 | 1 | 1 | 1 | 1 | 1 | 0 |
| JSG000000 | Foxa3 | 0.000944 | 0.001775 | 1.756834904 | 21.87226523 | 73.91860584 | 13 | 1 | 1 | 3 | 5 | 0 | 0 | 0 | 0 | 0 | 0 | 0 |
| SG0000003 | Arap1 | 0.001023 | 0.001918 | -0.600176362 | 1437.40673 | 948.2188542 | 82 | 1 | 1 | 16 | 53 | 6 | 1 | 0 | 0 | 0 | 0 | 0 |
| JSG000000 | Cherp | 0.001114 | 0.002081 | 0.452211807 | 3259.637945 | 4459.628508 | 9 | 1 | 1 | 2 | 5 | 1 | 1 | 0 | 0 | 0 | 0 | 0 |
| SG0000002 | Atp5b | 0.001186 | 0.002208 | 0.390768613 | 36960.03488 | 48457.99561 | 12 | 1 | 1 | 1 | 7 | 0 | 4 | 0 | 0 | 0 | 1 | 0 |
| JSG000000 | Ceacam10 | 0.001314 | 0.002434 | 1.80090115 | 8.917419222 | 31.07165952 | 15 | 1 | 1 | 1 | 11 | 0 | 1 | 0 | 0 | 0 | 0 | 0 |
| JSG000000 | Ercc6l | 0.001398 | 0.002582 | 0.43653393 | 1344.703612 | 1819.853291 | 27 | 1 | 1 | 3 | 12 | 1 | 2 | 0 | 0 | 0 | 0 | 0 |
| JSG000000 | Ccdc64b | 0.001583 | 0.002909 | 3.35710011 | 5.334899 | 54.66558921 | 6 | 1 | 1 | 2 | 2 | 0 | 1 | 0 | 0 | 1 | 1 | 0 |
| JSG000000 | Ing2 | 0.00176 | 0.003212 | 0.465355347 | 626.9960679 | 865.666747 | 12 | 1 | 1 | 2 | 3 | 0 | 2 | 0 | 0 | 0 | 0 | 0 |
| SG0000003 | Pak1ip1 | 0.001762 | 0.003216 | 0.40413622 | 2152.745402 | 2848.720199 | 18 | 1 | 1 | 3 | 10 | 0 | 0 | 0 | 0 | 0 | 1 | 0 |
| SG0000002 | Sapcd2 | 0.001765 | 0.00322 | 0.570578363 | 269.8316251 | 400.7320249 | 9 | 1 | 1 | 1 | 3 | 3 | 3 | 1 | 1 | 1 | 3 | 0 |
| JSG000000 | Shkbp1 | 0.001893 | 0.003437 | 0.550835251 | 616.6568487 | 903.3613228 | 10 | 1 | 1 | 2 | 9 | 0 | 0 | 0 | 0 | 0 | 0 | 0 |
| SG0000004 | Mier2 | 0.002176 | 0.00392 | 0.50096707 | 619.3495942 | 876.4799228 | 5 | 1 | 1 | 3 | 6 | 0 | 2 | 1 | 1 | 2 | 3 | 0 |
| JSG000000 | Caskin1 | 0.00232 | 0.004158 | 2.059792375 | 12.31189226 | 51.33151663 | 14 | 1 | 1 | 2 | 5 | 3 | 3 | 0 | 0 | 0 | 1 | 0 |
| SG0000004 | Msantd4 | 0.002398 | 0.004292 | -0.44913557 | 1700.993952 | 1245.946776 | 3 | 0 | 1 | 1 | 1 | 0 | 2 | 0 | 0 | 1 | 1 | 0 |

| | | | | | | | | | | | | | | | | | | |
|---|---|---|---|---|---|---|---|---|---|---|---|---|---|---|---|---|---|---|
| SG0000002 | Abcb6 | 0.002445 | 0.004374 | 0.729852287 | 101.2602339 | 167.936987 | 13 | 1 | 1 | 2 | 4 | 0 | 0 | 0 | 0 | 0 | 0 | 0 |
| SG0000003 | Sh2d4b | 0.002561 | 0.004568 | 1.465012727 | 14.00386664 | 38.65990131 | 52 | 1 | 1 | 5 | 26 | 4 | 3 | 0 | 0 | 0 | 1 | 1 |
| JSG000000 | Hsd17b1 | 0.002623 | 0.004672 | 2.267398621 | 3.97218212 | 19.12423841 | 2 | 1 | 1 | 1 | 4 | 0 | 2 | 1 | 1 | 1 | 4 | 0 |
| JSG000000 | Thbd | 0.002679 | 0.004763 | -1.405603808 | 438.3982976 | 165.4778809 | 9 | 1 | 1 | 3 | 5 | 1 | 7 | 1 | 1 | 3 | 5 | 1 |
| JSG000000 | Nolc1 | 0.002788 | 0.004942 | 0.371813117 | 6496.262206 | 8406.020539 | 7 | 0 | 1 | 1 | 3 | 0 | 0 | 0 | 0 | 0 | 1 | 0 |
| JSG000000 | Cmpk2 | 0.003013 | 0.005314 | 1.224758404 | 32.23774125 | 75.34485861 | 14 | 1 | 1 | 2 | 5 | 3 | 7 | 0 | 0 | 0 | 1 | 1 |
| JSG000000 | Mterf1b | 0.003034 | 0.005346 | 1.198457743 | 23.47490316 | 53.87354308 | 14 | 0 | 1 | 2 | 6 | 3 | 1 | 0 | 0 | 0 | 0 | 0 |
| JSG000000 | Cdk10 | 0.003063 | 0.005395 | 0.479381108 | 460.4374308 | 641.9167599 | 3 | 1 | 1 | 1 | 1 | 0 | 0 | 0 | 0 | 0 | 0 | 0 |
| JSG000000 | Amd1 | 0.003116 | 0.005481 | -0.423665555 | 2410.039816 | 1796.752167 | 28 | 1 | 1 | 5 | 10 | 0 | 7 | 0 | 0 | 2 | 4 | 0 |
| JSG000000 | Cnpy2 | 0.003144 | 0.005526 | -0.440251146 | 1603.960937 | 1182.129313 | 6 | 0 | 1 | 0 | 1 | 1 | 0 | 0 | 0 | 0 | 0 | 0 |
| SG0000001 | Surf2 | 0.003221 | 0.005657 | 0.443750054 | 708.1204474 | 963.1395388 | 1 | 1 | 1 | 1 | 1 | 0 | 1 | 1 | 1 | 1 | 1 | 0 |
| JSG000000 | Vipr1 | 0.003282 | 0.005758 | 3.489829795 | 2.826641195 | 31.75514733 | 11 | 0 | 1 | 0 | 2 | 6 | 1 | 0 | 0 | 0 | 1 | 0 |
| JSG000000 | Glp1r | 0.003427 | 0.005998 | 2.071896046 | 4.921145312 | 20.69040763 | 27 | 1 | 1 | 4 | 10 | 14 | 0 | 0 | 0 | 0 | 0 | 0 |
| JSG000000 | Thbs2 | 0.003547 | 0.006195 | -1.470492823 | 767.5512375 | 276.9777788 | 35 | 1 | 1 | 10 | 30 | 8 | 10 | 1 | 1 | 5 | 8 | 2 |
| SG0000004 | Gltscr2 | 0.003734 | 0.006498 | 0.700599735 | 2326.979065 | 3781.760412 | 5 | 0 | 1 | 3 | 7 | 0 | 3 | 0 | 1 | 3 | 6 | 0 |
| SG0000002 | Utp23 | 0.003779 | 0.00657 | -0.509807773 | 592.9315347 | 416.4253093 | 47 | 1 | 1 | 14 | 28 | 0 | 3 | 1 | 1 | 2 | 4 | 0 |
| JSG000000 | Mettl6 | 0.00419 | 0.007239 | 0.415721938 | 809.1236901 | 1079.343524 | 23 | 1 | 1 | 2 | 15 | 1 | 1 | 0 | 0 | 0 | 0 | 0 |
| JSG000000 | Pnrc1 | 0.004401 | 0.007582 | -0.432339696 | 1537.302576 | 1139.231902 | 6 | 0 | 1 | 1 | 4 | 2 | 1 | 0 | 0 | 0 | 3 | 2 |
| SG0000007 | Alkbh1 | 0.004481 | 0.007718 | 0.398646909 | 1009.852817 | 1331.259618 | 166 | 0 | 1 | 13 | 42 | 22 | 9 | 0 | 1 | 1 | 2 | 0 |
| SG000000C | Sra1 | 0.004515 | 0.007772 | -0.417625869 | 1743.076838 | 1304.964263 | 31 | 0 | 1 | 3 | 6 | 1 | 5 | 0 | 0 | 2 | 2 | 0 |
| JSG000000 | Nefl | 0.004707 | 0.008088 | 0.854741034 | 288.5016668 | 521.7362547 | 7 | 0 | 1 | 2 | 3 | 1 | 1 | 0 | 0 | 0 | 1 | 1 |
| JSG000000 | Yod1 | 0.004794 | 0.008226 | 0.479658474 | 342.237598 | 477.2207375 | 23 | 0 | 1 | 6 | 11 | 0 | 22 | 0 | 1 | 6 | 11 | 0 |
| JSG000000 | 2210016F16Rik | 0.005118 | 0.008748 | 0.516727606 | 262.5643712 | 375.652517 | 8 | 1 | 1 | 1 | 2 | 0 | 2 | 0 | 0 | 0 | 0 | 0 |
| JSG000000 | Zfp458 | 0.00527 | 0.008988 | -1.014809119 | 79.26850661 | 39.22949267 | 24 | 0 | 1 | 5 | 16 | 1 | 2 | 0 | 0 | 0 | 0 | 0 |
| JSG000000 | Twist2 | 0.005497 | 0.009349 | -1.037206074 | 70.91073853 | 34.55269037 | 47 | 1 | 1 | 5 | 20 | 6 | 1 | 1 | 1 | 1 | 1 | 0 |
| SG000000C | Bloc1s6 | 0.005741 | 0.009737 | -0.423031322 | 1296.409208 | 966.9343532 | 15 | 1 | 1 | 2 | 7 | 3 | 1 | 0 | 0 | 0 | 1 | 1 |
| SG0000003 | Vgf | 0.006606 | 0.011114 | 2.826161553 | 3.917811562 | 27.7845229 | 8 | 1 | 1 | 1 | 3 | 2 | 2 | 0 | 0 | 0 | 2 | 2 |
| JSG000000 | Atoh8 | 0.006973 | 0.011682 | -1.18260666 | 67.96248838 | 29.94119943 | 21 | 1 | 1 | 2 | 9 | 3 | 0 | 0 | 0 | 0 | 0 | 0 |
| JSG000000 | Gas6 | 0.0071 | 0.011882 | -0.564003714 | 955.0002991 | 645.9835698 | 21 | 1 | 1 | 3 | 11 | 4 | 0 | 0 | 0 | 0 | 2 | 1 |
| JSG000000 | Nars | 0.007271 | 0.012151 | 0.329892549 | 7519.238599 | 9451.079551 | 21 | 0 | 1 | 3 | 10 | 0 | 0 | 0 | 0 | 0 | 1 | 0 |
| SG0000003 | Cuedc2 | 0.00742 | 0.012379 | -0.407885036 | 1966.415079 | 1482.14115 | 6 | 0 | 1 | 0 | 1 | 0 | 0 | 0 | 0 | 0 | 0 | 0 |
| JSG000000 | Zfp748 | 0.007605 | 0.012671 | 0.452916755 | 353.8293864 | 484.323373 | 31 | 1 | 1 | 5 | 24 | 2 | 2 | 0 | 0 | 0 | 1 | 0 |
| SG0000001 | Trip10 | 0.0077 | 0.012819 | -0.391146594 | 2058.666698 | 1569.78141 | 8 | 1 | 1 | 2 | 18 | 1 | 1 | 0 | 0 | 0 | 0 | 1 |
| SG0000003 | Usp18 | 0.007782 | 0.012949 | 0.532595441 | 263.8469292 | 381.6622862 | 46 | 1 | 1 | 3 | 15 | 3 | 2 | 0 | 0 | 1 | 2 | 0 |
| SG0000002 | Tor1a | 0.007912 | 0.013148 | -0.44673541 | 697.4708542 | 511.735217 | 3 | 0 | 1 | 0 | 2 | 0 | 1 | 0 | 1 | 0 | 0 | 0 |
| SG0000002 | Sypl2 | 0.007976 | 0.01324 | 3.066432492 | 2.342955702 | 19.62692385 | 20 | 0 | 1 | 4 | 17 | 0 | 9 | 0 | 0 | 1 | 2 | 0 |
| JSG000000 | Ehd3 | 0.008084 | 0.01341 | -0.588199456 | 804.8346215 | 535.3538996 | 20 | 1 | 1 | 2 | 9 | 2 | 1 | 0 | 0 | 0 | 0 | 0 |
| JSG000000 | Ccno | 0.00847 | 0.014016 | 2.57916267 | 2.556441386 | 15.27710864 | 3 | 1 | 1 | 1 | 3 | 2 | 1 | 1 | 1 | 1 | 2 | 1 |
| JSG000000 | Mmp25 | 0.008539 | 0.014126 | 1.778042856 | 13.60483047 | 46.65920852 | 10 | 0 | 1 | 1 | 2 | 0 | 0 | 0 | 0 | 0 | 0 | 0 |
| JSG000000 | Mrpl46 | 0.008719 | 0.01439 | 0.431579898 | 421.1634419 | 568.0272374 | 9 | 1 | 1 | 1 | 2 | 0 | 1 | 0 | 0 | 0 | 0 | 0 |
| JSG000000 | Rdh14 | 0.008932 | 0.014724 | -0.589999482 | 240.6122625 | 159.8491073 | 12 | 1 | 1 | 2 | 5 | 2 | 2 | 1 | 1 | 1 | 2 | 1 |
| JSG000000 | Spin4 | 0.009213 | 0.015156 | 0.675911209 | 81.44147308 | 130.1113457 | 15 | 0 | 1 | 0 | 10 | 1 | 14 | 0 | 1 | 0 | 10 | 1 |
| JSG000000 | Ccdc59 | 0.009497 | 0.015587 | -0.402254117 | 1177.974798 | 891.3442019 | 19 | 1 | 1 | 2 | 4 | 3 | 4 | 0 | 0 | 1 | 1 | 1 |
| JSG000000 | Zfp114 | 0.009572 | 0.015703 | 2.160875634 | 3.548901024 | 15.87019526 | 6 | 0 | 1 | 0 | 6 | 0 | 0 | 0 | 0 | 0 | 0 | 0 |
| SG0000001 | Stard3 | 0.009636 | 0.015798 | -0.465882946 | 476.4560016 | 344.9674042 | 17 | 1 | 1 | 4 | 8 | 4 | 0 | 0 | 0 | 0 | 0 | 0 |
| JSG000000 | Mmp10 | 0.009687 | 0.015874 | -1.651160992 | 28.79892804 | 9.16911324 | 17 | 0 | 1 | 1 | 14 | 0 | 1 | 0 | 1 | 0 | 3 | 0 |
| JSG000000 | Zfp784 | 0.00989 | 0.016188 | -1.359591203 | 35.75083265 | 13.93183836 | 6 | 0 | 1 | 0 | 2 | 0 | 0 | 0 | 0 | 0 | 0 | 0 |
| SG0000007 | Gm14326 | 0.010533 | 0.017159 | -0.720959176 | 134.0562796 | 81.33113346 | 51 | 1 | 1 | 2 | 22 | 4 | 2 | 0 | 0 | 0 | 0 | 0 |
| JSG000000 | 9430016H08Rik | 0.010728 | 0.017453 | 0.475881209 | 422.7920466 | 588.0054171 | 13 | 0 | 1 | 0 | 10 | 2 | 2 | 0 | 0 | 0 | 1 | 1 |
| JSG000000 | Zfp503 | 0.010887 | 0.017698 | -0.917509707 | 80.10354767 | 42.40858751 | 7 | 0 | 1 | 2 | 2 | 0 | 5 | 0 | 1 | 2 | 2 | 0 |
| JSG000000 | Tmem138 | 0.011124 | 0.018056 | -0.442624653 | 595.39449 | 438.088158 | 11 | 1 | 1 | 3 | 11 | 1 | 1 | 0 | 0 | 0 | 0 | 0 |
| JSG000000 | Ccl2 | 0.011214 | 0.018192 | -0.771152409 | 896.8946329 | 525.5347265 | 3 | 0 | 1 | 0 | 4 | 0 | 1 | 0 | 0 | 0 | 4 | 0 |
| JSG000000 | Ndufa4 | 0.011652 | 0.018848 | 0.317736583 | 2926.28511 | 3647.244512 | 17 | 0 | 1 | 3 | 7 | 3 | 0 | 0 | 0 | 0 | 0 | 0 |
| SG000000C | Prrg2 | 0.012158 | 0.019586 | -0.904542086 | 185.2713752 | 98.97238248 | 5 | 0 | 1 | 0 | 1 | 1 | 0 | 0 | 0 | 0 | 0 | 0 |
| SG0000002 | Lzic | 0.012329 | 0.019836 | -0.400804634 | 902.0214688 | 683.2232817 | 11 | 1 | 1 | 1 | 2 | 1 | 0 | 0 | 0 | 0 | 0 | 0 |
| JSG000000 | Gzmm | 0.012439 | 0.019999 | 1.266244815 | 15.14222372 | 36.42229684 | 3 | 0 | 1 | 1 | 1 | 1 | 0 | 0 | 0 | 0 | 0 | 0 |
| SG000000C | Emg1 | 0.012468 | 0.020039 | 0.327041797 | 2510.615687 | 3149.413094 | 14 | 0 | 1 | 4 | 7 | 1 | 0 | 0 | 0 | 0 | 0 | 0 |
| SG0000003 | Pdha1 | 0.012634 | 0.020283 | -0.332468288 | 9755.438372 | 7747.540631 | 30 | 0 | 1 | 1 | 10 | 1 | 5 | 0 | 0 | 0 | 0 | 0 |
| JSG000000 | Stac3 | 0.012725 | 0.02041 | 2.58391602 | 6.155869558 | 36.90843554 | 10 | 1 | 1 | 3 | 4 | 0 | 1 | 0 | 0 | 0 | 0 | 0 |
| JSG000000 | Gm561 | 0.013719 | 0.02186 | -0.70136222 | 187.9171784 | 115.5674197 | 5 | 1 | 1 | 1 | 3 | 0 | 3 | 0 | 0 | 0 | 0 | 0 |
| JSG000000 | Per2 | 0.013959 | 0.022221 | -0.730209776 | 113.3191842 | 68.31064618 | 27 | 0 | 1 | 5 | 14 | 6 | 1 | 0 | 1 | 0 | 2 | 2 |
| SG0000003 | Trmt5 | 0.014426 | 0.022929 | 0.388427581 | 514.1072918 | 672.9489452 | 12 | 0 | 1 | 0 | 4 | 0 | 0 | 0 | 0 | 0 | 0 | 0 |
| JSG000000 | Rbm22 | 0.014605 | 0.023183 | 0.330680724 | 1445.597366 | 1817.992825 | 18 | 0 | 1 | 4 | 7 | 2 | 3 | 0 | 0 | 0 | 1 | 0 |
| JSG000000 | Fam26e | 0.015208 | 0.024089 | 1.35997631 | 10.9977088 | 28.22902503 | 8 | 1 | 1 | 1 | 2 | 1 | 2 | 0 | 0 | 0 | 0 | 0 |
| JSG000000 | Rbm45 | 0.015267 | 0.024169 | -0.376559322 | 1157.220104 | 891.3747319 | 30 | 0 | 1 | 4 | 14 | 1 | 0 | 0 | 0 | 0 | 0 | 0 |
| SG0000002 | Cct5 | 0.015358 | 0.024304 | 0.296164521 | 16065.60605 | 19726.56726 | 14 | 1 | 1 | 2 | 4 | 2 | 1 | 0 | 0 | 0 | 0 | 0 |
| JSG000000 | Kctd21 | 0.015813 | 0.024969 | 0.601621239 | 100.9855437 | 153.2375667 | 11 | 1 | 1 | 1 | 6 | 2 | 0 | 0 | 0 | 0 | 0 | 0 |
| SG0000005 | Tcea2 | 0.016184 | 0.025507 | -0.538743596 | 230.8377477 | 158.9018106 | 11 | 1 | 1 | 1 | 1 | 0 | 0 | 0 | 0 | 0 | 0 | 0 |
| JSG000000 | Lrrc75b | 0.016933 | 0.026575 | 1.717876991 | 5.771397775 | 18.98512718 | 11 | 0 | 1 | 1 | 4 | 0 | 4 | 0 | 0 | 0 | 0 | 0 |
| JSG000000 | Slc38a2 | 0.018493 | 0.028787 | -0.317838441 | 7628.185632 | 6119.872139 | 11 | 0 | 1 | 3 | 8 | 5 | 5 | 0 | 1 | 2 | 4 | 5 |
| JSG000000 | Mmaa | 0.019036 | 0.029565 | -0.560617276 | 193.8751326 | 131.4496675 | 52 | 0 | 1 | 8 | 23 | 4 | 4 | 0 | 1 | 1 | 1 | 0 |
| JSG000000 | Prrt3 | 0.021351 | 0.032839 | -1.15403159 | 40.04040129 | 17.99286398 | 6 | 0 | 1 | 0 | 4 | 0 | 1 | 0 | 0 | 0 | 0 | 0 |
| SG0000005 | Gemin6 | 0.021828 | 0.033538 | 0.340723995 | 783.652447 | 992.4111537 | 11 | 1 | 1 | 4 | 4 | 0 | 2 | 0 | 0 | 0 | 0 | 0 |
| JSG000000 | Mmp3 | 0.022115 | 0.033958 | -0.814251807 | 394.69053 | 224.4618243 | 19 | 0 | 1 | 4 | 7 | 0 | 1 | 0 | 1 | 0 | 0 | 0 |
| JSG000000 | Zfp760 | 0.022207 | 0.034094 | 0.373651963 | 398.2647388 | 516.0030942 | 31 | 0 | 1 | 3 | 18 | 9 | 4 | 0 | 0 | 0 | 0 | 1 |
| JSG000000 | 4930550C14Rik | 0.022683 | 0.034751 | 1.219344445 | 13.02228111 | 30.3211954 | 64 | 1 | 1 | 6 | 23 | 3 | 0 | 0 | 0 | 0 | 0 | 0 |
| SG0000001 | Psmd3 | 0.022708 | 0.034782 | -0.397154663 | 6890.289166 | 5232.171633 | 9 | 1 | 1 | 2 | 3 | 1 | 0 | 0 | 0 | 0 | 0 | 0 |
| JSG000000 | Klhl10 | 0.023 | 0.035187 | 1.439332774 | 8.232468862 | 22.32607788 | 28 | 1 | 1 | 7 | 15 | 1 | 1 | 0 | 0 | 0 | 0 | 0 |
| JSG000000 | Capn6 | 0.02407 | 0.036712 | 1.255529149 | 27.85365512 | 66.50191145 | 60 | 1 | 1 | 6 | 26 | 3 | 4 | 0 | 0 | 0 | 0 | 0 |
| JSG000000 | Akap8 | 0.024408 | 0.037196 | 0.341369588 | 2674.016906 | 3387.869198 | 20 | 1 | 1 | 4 | 11 | 2 | 2 | 0 | 0 | 0 | 2 | 0 |
| SG0000004 | Gck | 0.024882 | 0.037866 | 1.39469698 | 8.594348698 | 22.5974064 | 55 | 0 | 1 | 2 | 28 | 9 | 3 | 0 | 0 | 0 | 0 | 0 |
| SG000000C | Dynlt1c | 0.025014 | 0.038054 | 1.323401891 | 10.30188433 | 25.78103417 | 8 | 0 | 1 | 0 | 1 | 0 | 2 | 0 | 0 | 0 | 0 | 0 |
| JSG000000 | Zfp36l2 | 0.02538 | 0.038568 | -0.336018752 | 1540.588889 | 1220.492283 | 5 | 1 | 1 | 3 | 14 | 0 | 5 | 1 | 1 | 3 | 14 | 0 |
| JSG000000 | Rfx1 | 0.025533 | 0.038781 | 0.508430507 | 483.3608817 | 687.5817382 | 11 | 1 | 1 | 3 | 6 | 2 | 0 | 0 | 0 | 0 | 0 | 0 |
| JSG000000 | Hes1 | 0.026133 | 0.039646 | -0.344766563 | 987.9363482 | 777.9360318 | 1 | 0 | 1 | 1 | 3 | 1 | 0 | 0 | 0 | 0 | 2 | 1 |
| JSG000000 | Slc30a1 | 0.0269 | 0.04073 | 0.319242218 | 870.5551678 | 1086.169945 | 6 | 1 | 1 | 1 | 2 | 0 | 6 | 1 | 1 | 1 | 1 | 0 |
| SG0000003 | Pwp2 | 0.028207 | 0.042557 | -0.310508958 | 1943.484508 | 1567.142804 | 6 | 0 | 1 | 1 | 2 | 0 | 1 | 0 | 1 | 1 | 1 | 0 |
| SG0000005 | Sec61b | 0.028254 | 0.042618 | 0.526552122 | 631.09418 | 909.0801147 | 13 | 0 | 1 | 4 | 7 | 0 | 0 | 0 | 0 | 0 | 1 | 0 |
| JSG000000 | Gm9958 | 0.028402 | 0.04283 | -1.019997398 | 42.93178418 | 21.170403 | 2 | 1 | 1 | 1 | 3 | 0 | 0 | 0 | 0 | 0 | 0 | 0 |
| JSG000000 | Erlin2 | 0.029429 | 0.044237 | -0.304962255 | 2763.795861 | 2237.191578 | 13 | 1 | 1 | 1 | 7 | 1 | 2 | 0 | 0 | 0 | 0 | 0 |
| JSG000000 | Gm21972 | 0.030005 | 0.045042 | 1.612982275 | 5.514415645 | 16.86768744 | 40 | 0 | 1 | 10 | 22 | 3 | 1 | 0 | 0 | 0 | 0 | 0 |
| JSG000000 | Ptx3 | 0.031248 | 0.046747 | 1.043514116 | 54.94255084 | 113.2499108 | 10 | 0 | 1 | 1 | 2 | 2 | 2 | 0 | 0 | 1 | 1 | 1 |
| SG0000003 | Rsad1 | 0.031532 | 0.047134 | -0.489803492 | 230.1715459 | 163.9102421 | 10 | 1 | 1 | 1 | 19 | 2 | 6 | 1 | 1 | 1 | 7 | 1 |
| SG0000002 | Aasdhppt | 0.032234 | 0.048123 | 0.30169988 | 1132.517407 | 1395.936295 | 47 | 0 | 1 | 4 | 22 | 6 | 10 | 0 | 0 | 1 | 2 | 0 |
| JSG000000 | Trpt1 | 0.03264 | 0.048688 | -0.709558883 | 84.99790778 | 51.97682276 | 2 | 1 | 1 | 2 | 2 | 0 | 2 | 1 | 1 | 2 | 2 | 0 |
| SG0000004 | Map3k10 | 0.033042 | 0.049242 | -0.674419678 | 307.8169877 | 192.8732644 | 18 | 1 | 1 | 3 | 7 | 0 | 3 | 0 | 0 | 1 | 1 | 0 |
| JSG000000 | Thbs1 | 0.035536 | 0.052604 | -0.926224594 | 9729.555639 | 5120.019564 | 13 | 1 | 1 | 3 | 7 | 3 | 6 | 1 | 1 | 3 | 4 | 3 |
| JSG000000 | Rhob | 0.035641 | 0.052752 | 0.36446978 | 497.3957086 | 640.3514101 | 3 | 0 | 1 | 0 | 0 | 0 | 3 | 0 | 1 | 0 | 0 | 0 |
| JSG000000 | Maf | 0.036604 | 0.054057 | 1.30924962 | 13.29951162 | 32.9578673 | 43 | 0 | 1 | 2 | 29 | 5 | 10 | 0 | 0 | 0 | 9 | 1 |
| SG0000002 | Nr1d1 | 0.037168 | 0.054845 | -0.351650756 | 717.9541203 | 562.6512224 | 6 | 0 | 1 | 1 | 1 | 1 | 2 | 0 | 1 | 1 | 1 | 1 |
| JSG000000 | Rps6kl1 | 0.03838 | 0.056502 | 0.466705804 | 152.5225507 | 210.7785952 | 7 | 0 | 1 | 1 | 3 | 0 | 0 | 0 | 0 | 0 | 0 | 0 |
| SG0000001 | Epn3 | 0.038652 | 0.056874 | 1.410692176 | 9.985844503 | 26.54883441 | 2 | 0 | 1 | 0 | 12 | 0 | 1 | 0 | 1 | 0 | 12 | 0 |
| JSG000000 | Rbm8a2 | 0.040521 | 0.059395 | 1.200097604 | 15.89810275 | 36.52672004 | 4 | 0 | 1 | 0 | 2 | 0 | 4 | 0 | 1 | 0 | 2 | 0 |
| SG0000004 | Hnrnpul1 | 0.044205 | 0.064427 | 0.252520283 | 4818.230482 | 5739.89239 | 58 | 1 | 1 | 9 | 31 | 6 | 1 | 0 | 0 | 0 | 0 | 0 |
| JSG000000 | Slitrk6 | 0.044229 | 0.064456 | 1.362170863 | 21.46439225 | 55.17888051 | 24 | 0 | 1 | 2 | 9 | 2 | 4 | 0 | 0 | 1 | 3 | 2 |
| SG000000C | P4htm | 0.046461 | 0.067517 | -0.577377276 | 110.3131795 | 73.92979862 | 7 | 1 | 1 | 2 | 5 | 3 | 1 | 1 | 1 | 1 | 1 | 0 |
| SG0000003 | Timm17b | 0.047843 | 0.06935 | 0.31452832 | 592.4781371 | 736.8087724 | 13 | 1 | 1 | 3 | 5 | 1 | 0 | 0 | 0 | 0 | 1 | 0 |
| SG0000002 | Pcna | 0.047863 | 0.069368 | 0.248225191 | 5388.62528 | 6400.312986 | 6 | 1 | 1 | 2 | 3 | 2 | 1 | 0 | 0 | 0 | 0 | 2 |
| JSG000000 | B230307C23Rik | 0.048067 | 0.069624 | -0.700335415 | 73.07601817 | 44.97310867 | 26 | 0 | 1 | 1 | 12 | 1 | 3 | 0 | 0 | 0 | 0 | 0 |
| SG0000003 | Golt1b | 0.049316 | 0.071289 | -0.282049573 | 2132.234629 | 1753.59626 | 24 | 1 | 1 | 7 | 9 | 3 | 5 | 0 | 0 | 4 | 4 | 1 |
| JSG000000 | Ube2o | 0.049478 | 0.07149 | 0.261163582 | 1979.241635 | 2372.012046 | 33 | 1 | 1 | 4 | 12 | 2 | 1 | 0 | 0 | 0 | 0 | 0 |
| JSG000000 | Eif1b | 0.050983 | 0.073456 | 0.327665369 | 421.4505496 | 528.9123759 | 3 | 0 | 1 | 0 | 1 | 0 | 0 | 0 | 0 | 0 | 0 | 0 |
| JSG000000 | Lpar3 | 0.051816 | 0.074561 | 1.294073103 | 8.151662946 | 19.98945875 | 93 | 0 | 1 | 10 | 25 | 7 | 3 | 0 | 0 | 1 | 2 | 0 |
| JSG000000 | Tube1 | 0.052974 | 0.076092 | 0.326805889 | 387.6739497 | 486.2336313 | 30 | 0 | 1 | 4 | 10 | 2 | 4 | 0 | 0 | 0 | 2 | 1 |
| JSG000000 | Slfn2 | 0.053458 | 0.076725 | -1.255711828 | 23.64489239 | 9.902170861 | 6 | 1 | 1 | 1 | 2 | 0 | 0 | 0 | 0 | 0 | 0 | 0 |
| JSG000000 | Sap30l | 0.05503 | 0.078811 | -0.358153704 | 432.1211478 | 337.1246534 | 8 | 1 | 1 | 1 | 4 | 2 | 0 | 0 | 0 | 0 | 0 | 1 |
| JSG000000 | Cyr61 | 0.057407 | 0.081927 | 0.901307896 | 970.0709741 | 1811.858268 | 6 | 0 | 1 | 2 | 2 | 0 | 5 | 0 | 1 | 2 | 2 | 0 |
| SG000000C | Irx2 | 0.0583 | 0.083063 | 1.029602846 | 21.21025666 | 43.29993697 | 15 | 0 | 1 | 2 | 5 | 3 | 7 | 0 | 0 | 2 | 4 | 3 |
| JSG000000 | Klc4 | 0.059009 | 0.083993 | -0.417941827 | 557.8832256 | 417.5709494 | 15 | 1 | 1 | 1 | 6 | 2 | 0 | 0 | 0 | 0 | 0 | 0 |
| JSG000000 | Tram1l1 | 0.059139 | 0.084144 | 0.907470191 | 18.85508014 | 35.36748112 | 10 | 0 | 1 | 0 | 2 | 2 | 10 | 0 | 1 | 0 | 2 | 1 |
| SG0000001 | Traf4 | 0.060413 | 0.085801 | -0.2861868 | 2257.15479 | 1851.01753 | 3 | 1 | 1 | 2 | 2 | 0 | 2 | 1 | 1 | 2 | 2 | 0 |
| JSG000000 | Ccs | 0.060726 | 0.086205 | -0.296657029 | 823.9627165 | 670.818292 | 7 | 0 | 1 | 0 | 4 | 1 | 0 | 0 | 0 | 0 | 0 | 0 |
| JSG000000 | Zdhhc1 | 0.061823 | 0.087658 | -0.460334074 | 161.1193961 | 117.1044668 | 5 | 0 | 1 | 1 | 2 | 1 | 0 | 0 | 0 | 0 | 0 | 0 |

| | | | | | | | | | | | | | | | | | | |
|---|---|---|---|---|---|---|---|---|---|---|---|---|---|---|---|---|---|---|
| SG0000005 | Zfp658 | 0.062457 | 0.088486 | 0.473401687 | 99.41367842 | 138.0238862 | 15 | 0 | 1 | 5 | 10 | 2 | 1 | 0 | 0 | 0 | 0 | 0 |
| JSG000000 | Cdkn3 | 0.065189 | 0.092116 | -0.418048727 | 221.882516 | 166.0649104 | 25 | 1 | 1 | 3 | 9 | 2 | 0 | 0 | 0 | 0 | 0 | 0 |
| SG0000005 | Gdpgp1 | 0.068506 | 0.096378 | 0.358823603 | 207.2006953 | 265.7099857 | 13 | 0 | 1 | 2 | 3 | 0 | 0 | 0 | 0 | 0 | 0 | 0 |
| SG0000004 | Cdk5r1 | 0.069605 | 0.097808 | 1.202087269 | 10.60086306 | 24.38964897 | 4 | 1 | 1 | 1 | 2 | 0 | 4 | 1 | 1 | 1 | 2 | 0 |
| SG0000002 | Atp6v0c | 0.070453 | 0.098907 | -0.871430648 | 583.6243502 | 319.0117205 | 5 | 0 | 1 | 1 | 1 | 0 | 1 | 0 | 1 | 1 | 1 | 0 |
| JSG000000 | 2410004P03Rik | 0.074477 | 0.104051 | 1.190835015 | 11.87923689 | 27.11849616 | 15 | 0 | 1 | 3 | 3 | 2 | 3 | 0 | 1 | 1 | 1 | 0 |
| JSG000000 | Map3k12 | 0.075611 | 0.10552 | 0.269129606 | 637.733804 | 768.5206158 | 23 | 0 | 1 | 4 | 11 | 2 | 4 | 0 | 0 | 1 | 2 | 1 |
| JSG000000 | Plekhf1 | 0.080012 | 0.111099 | 0.385476545 | 168.7293127 | 220.4096171 | 4 | 0 | 1 | 1 | 4 | 0 | 1 | 0 | 0 | 0 | 0 | 0 |
| JSG000000 | Rce1 | 0.08185 | 0.113466 | -0.298932114 | 527.131882 | 428.4811796 | 2 | 0 | 1 | 1 | 1 | 0 | 2 | 0 | 1 | 1 | 1 | 0 |
| JSG000000 | Thyn1 | 0.083546 | 0.115621 | 0.229891335 | 1468.367758 | 1722.023316 | 12 | 0 | 1 | 1 | 3 | 0 | 0 | 0 | 0 | 0 | 0 | 0 |
| SG0000006 | Chadl | 0.085538 | 0.118141 | -0.904731698 | 31.44101562 | 16.79365346 | 37 | 0 | 1 | 0 | 3 | 1 | 2 | 0 | 0 | 0 | 0 | 0 |
| JSG000000 | Klf16 | 0.088182 | 0.121491 | -0.249166897 | 1428.070882 | 1201.553338 | 11 | 1 | 1 | 1 | 1 | 1 | 4 | 1 | 1 | 1 | 1 | 1 |
| SG0000004 | Tbc1d10b | 0.089986 | 0.123691 | -0.270200433 | 2871.914115 | 2381.403638 | 10 | 1 | 1 | 2 | 5 | 0 | 1 | 0 | 0 | 0 | 0 | 0 |
| JSG000000 | Hmgn5 | 0.091167 | 0.125209 | 0.222287859 | 1606.694311 | 1874.340845 | 25 | 1 | 1 | 1 | 9 | 0 | 1 | 0 | 0 | 0 | 2 | 0 |
| SG0000003 | Vhl | 0.096064 | 0.131261 | -0.264697207 | 989.6470438 | 823.7558896 | 8 | 1 | 1 | 3 | 4 | 1 | 1 | 1 | 1 | 1 | 2 | 1 |
| JSG000000 | Rhcg | 0.096507 | 0.131836 | -1.231915787 | 17.53486285 | 7.465497756 | 16 | 1 | 1 | 3 | 9 | 0 | 0 | 0 | 0 | 0 | 0 | 0 |
| JSG000000 | Mkrn3 | 0.100665 | 0.136765 | 0.385208814 | 124.0152208 | 161.9699315 | 2 | 0 | 1 | 1 | 2 | 0 | 2 | 0 | 1 | 1 | 1 | 0 |
| JSG000000 | Gm2a | 0.102159 | 0.138668 | 0.377423342 | 744.6084148 | 967.2610209 | 9 | 1 | 1 | 1 | 8 | 4 | 0 | 0 | 0 | 0 | 1 | 3 |
| SG0000003 | Eif1 | 0.103024 | 0.139768 | -0.210581797 | 8063.920627 | 6968.748752 | 6 | 0 | 1 | 0 | 0 | 0 | 5 | 0 | 1 | 0 | 0 | 0 |
| JSG000000 | Ier3ip1 | 0.103519 | 0.140409 | -0.263138948 | 897.5039597 | 747.8657718 | 23 | 1 | 1 | 4 | 11 | 3 | 4 | 1 | 1 | 1 | 2 | 0 |
| JSG000000 | Kptn | 0.107421 | 0.145282 | 0.350367096 | 173.0946939 | 220.6758257 | 13 | 1 | 1 | 2 | 4 | 1 | 1 | 0 | 0 | 0 | 0 | 0 |
| SG0000003 | Plxnd1 | 0.108908 | 0.147094 | -0.453755302 | 242.3655946 | 176.9607753 | 14 | 0 | 1 | 1 | 3 | 4 | 0 | 0 | 0 | 0 | 0 | 0 |
| JSG000000 | Cycs | 0.112045 | 0.150865 | -0.251007143 | 1686.232636 | 1416.957458 | 8 | 1 | 1 | 2 | 2 | 0 | 8 | 1 | 1 | 2 | 2 | 0 |
| JSG000000 | Gm15446 | 0.112661 | 0.151535 | 0.312076703 | 232.2742398 | 288.3669579 | 31 | 0 | 1 | 1 | 13 | 3 | 0 | 0 | 0 | 0 | 0 | 0 |
| SG0000002 | Napg | 0.1135 | 0.152571 | -0.248540779 | 1156.373301 | 973.3741898 | 42 | 0 | 1 | 6 | 14 | 6 | 9 | 0 | 1 | 2 | 3 | 1 |
| JSG000000 | Pkmyt1 | 0.1142 | 0.153409 | 0.375639503 | 458.5424696 | 594.9196635 | 9 | 0 | 1 | 0 | 4 | 0 | 0 | 0 | 0 | 0 | 0 | 0 |
| SG0000003 | Slco4a1 | 0.117893 | 0.15786 | 0.61420485 | 120.7462873 | 184.8280448 | 13 | 0 | 1 | 3 | 3 | 2 | 2 | 0 | 0 | 0 | 0 | 2 |
| SG0000001 | Mapkapk2 | 0.119908 | 0.160355 | -0.219912226 | 1873.473628 | 1608.597568 | 51 | 1 | 1 | 5 | 20 | 0 | 1 | 0 | 0 | 0 | 0 | 0 |
| JSG000000 | Chmp1a | 0.120786 | 0.161421 | -0.206675833 | 2686.041742 | 2327.539887 | 5 | 1 | 1 | 1 | 3 | 0 | 0 | 0 | 0 | 0 | 0 | 0 |
| JSG000000 | Vpreb3 | 0.122488 | 0.16356 | -1.171281659 | 17.23822768 | 7.654233378 | 5 | 1 | 1 | 1 | 2 | 0 | 0 | 0 | 0 | 0 | 0 | 0 |
| JSG000000 | Cdnf | 0.131541 | 0.174466 | -0.766347841 | 45.02394473 | 26.46975097 | 24 | 0 | 1 | 5 | 16 | 2 | 2 | 0 | 0 | 1 | 1 | 0 |
| JSG000000 | Ndufb9 | 0.137673 | 0.181845 | -0.214006281 | 2159.015322 | 1861.37302 | 4 | 1 | 1 | 1 | 2 | 1 | 0 | 0 | 0 | 0 | 0 | 0 |
| JSG000000 | 4930453N24Rik | 0.138286 | 0.182534 | -0.232901963 | 1254.022458 | 1067.074735 | 22 | 1 | 1 | 3 | 15 | 1 | 12 | 1 | 1 | 3 | 11 | 1 |
| SG0000003 | Fchsd1 | 0.142357 | 0.187395 | 0.307487989 | 201.8559626 | 249.8070498 | 14 | 1 | 1 | 3 | 12 | 0 | 1 | 0 | 0 | 0 | 0 | 0 |
| SG0000006 | Celsr2 | 0.142394 | 0.187431 | -0.276610321 | 416.3801229 | 343.7335989 | 6 | 1 | 1 | 1 | 3 | 2 | 3 | 1 | 1 | 1 | 1 | 1 |
| JSG000000 | Edn1 | 0.142656 | 0.187735 | -0.811291918 | 30.78469105 | 17.54331239 | 17 | 1 | 1 | 3 | 5 | 0 | 1 | 0 | 0 | 0 | 0 | 0 |
| JSG000000 | Ndufs8 | 0.150362 | 0.196978 | -0.236389915 | 851.3439761 | 722.6776313 | 1 | 0 | 1 | 0 | 0 | 0 | 0 | 0 | 0 | 0 | 0 | 0 |
| SG0000000 | Btn1a1 | 0.153115 | 0.200292 | 0.901938825 | 11.00858291 | 20.57036786 | 12 | 0 | 1 | 0 | 1 | 3 | 1 | 0 | 0 | 0 | 0 | 1 |
| SG0000004 | Lman1 | 0.15558 | 0.203293 | 0.179332096 | 3446.4128 | 3902.578384 | 72 | 0 | 1 | 3 | 21 | 2 | 10 | 0 | 1 | 2 | 5 | 1 |
| JSG000000 | Fam83d | 0.156339 | 0.204226 | 0.222853762 | 691.7394684 | 807.2874904 | 18 | 1 | 1 | 4 | 6 | 3 | 1 | 0 | 0 | 0 | 0 | 0 |
| JSG000000 | Gm9833 | 0.156832 | 0.204733 | -0.305391911 | 241.0625164 | 195.0731783 | 16 | 0 | 1 | 1 | 3 | 1 | 1 | 0 | 0 | 1 | 1 | 0 |
| JSG000000 | Rbfa | 0.158846 | 0.207048 | 0.236991098 | 530.6014243 | 625.3308298 | 12 | 1 | 1 | 1 | 2 | 1 | 1 | 0 | 0 | 0 | 0 | 0 |
| SG0000000 | Sf3a1 | 0.160068 | 0.20855 | 0.180624873 | 4712.201065 | 5340.689357 | 19 | 0 | 1 | 2 | 5 | 1 | 1 | 0 | 0 | 0 | 2 | 0 |
| JSG000000 | Rpl23a | 0.162749 | 0.211812 | 0.227516537 | 12087.88137 | 14152.70621 | 5 | 1 | 1 | 1 | 1 | 0 | 0 | 0 | 0 | 0 | 0 | 0 |
| SG0000005 | Oas1a | 0.164956 | 0.214435 | -0.506950034 | 767.4060108 | 540.0301782 | 13 | 0 | 1 | 0 | 8 | 1 | 1 | 0 | 0 | 0 | 0 | 0 |
| JSG000000 | Dnaaf2 | 0.166375 | 0.21606 | 0.21494094 | 644.9393938 | 748.5530348 | 22 | 0 | 1 | 3 | 8 | 2 | 4 | 0 | 0 | 1 | 1 | 0 |
| SG0000000 | Ddx56 | 0.167118 | 0.216961 | -0.196116638 | 2066.35746 | 1803.717263 | 6 | 1 | 1 | 2 | 3 | 0 | 0 | 0 | 0 | 0 | 0 | 0 |
| SG0000000 | Paf1 | 0.172524 | 0.223333 | 0.185705748 | 1652.234772 | 1879.207592 | 6 | 1 | 1 | 1 | 5 | 1 | 0 | 0 | 0 | 0 | 0 | 0 |
| JSG000000 | Rnf225 | 0.172676 | 0.223497 | 0.642690957 | 22.38223295 | 34.94400409 | 4 | 0 | 1 | 1 | 1 | 0 | 2 | 0 | 1 | 1 | 1 | 0 |
| JSG000000 | Mars2 | 0.174759 | 0.22572 | -0.219851217 | 845.3856314 | 725.8937402 | 31 | 0 | 1 | 4 | 13 | 2 | 30 | 0 | 1 | 3 | 11 | 2 |
| JSG000000 | Hspa1b | 0.183304 | 0.235771 | -0.386624313 | 102.6341748 | 78.50665609 | 2 | 0 | 1 | 0 | 0 | 0 | 2 | 0 | 1 | 0 | 0 | 0 |
| JSG000000 | Dlx1 | 0.188012 | 0.241117 | -1.529583871 | 18.32705448 | 6.348074939 | 9 | 1 | 1 | 2 | 8 | 2 | 2 | 0 | 0 | 0 | 6 | 2 |
| SG0000000 | Pex6 | 0.190634 | 0.244251 | -0.421359368 | 832.8597839 | 621.9136415 | 9 | 0 | 1 | 1 | 5 | 5 | 0 | 0 | 0 | 0 | 0 | 0 |
| JSG000000 | Isca2 | 0.193035 | 0.246992 | -0.237073845 | 475.2920093 | 403.2683834 | 3 | 0 | 1 | 0 | 0 | 0 | 2 | 0 | 1 | 0 | 0 | 0 |
| SG0000003 | Mrpl19 | 0.193296 | 0.247308 | -0.196176464 | 1205.341121 | 1052.095039 | 14 | 1 | 1 | 3 | 4 | 1 | 6 | 0 | 0 | 0 | 0 | 0 |
| JSG000000 | Xrra1 | 0.196293 | 0.250678 | 0.854821156 | 8.939167445 | 16.16679314 | 59 | 1 | 1 | 5 | 26 | 10 | 0 | 0 | 0 | 0 | 0 | 0 |
| JSG000000 | Scara3 | 0.19911 | 0.25393 | 0.4459751 | 59.58277537 | 81.16571357 | 22 | 0 | 1 | 1 | 7 | 8 | 0 | 0 | 0 | 0 | 0 | 0 |
| JSG000000 | Ptrhd1 | 0.200341 | 0.255282 | -0.27272465 | 242.096785 | 200.3967722 | 9 | 0 | 1 | 0 | 2 | 0 | 4 | 0 | 0 | 0 | 2 | 0 |
| JSG000000 | Gla | 0.201634 | 0.256756 | -0.211349995 | 740.6962394 | 639.7605428 | 32 | 0 | 1 | 3 | 20 | 0 | 4 | 0 | 0 | 0 | 0 | 0 |
| SG0000003 | Alg8 | 0.202157 | 0.25734 | 0.193173636 | 684.888609 | 783.016659 | 25 | 1 | 1 | 5 | 18 | 1 | 3 | 0 | 0 | 2 | 2 | 0 |
| JSG000000 | Ftsj2 | 0.206401 | 0.26224 | -0.300981379 | 210.549525 | 170.9030616 | 3 | 1 | 1 | 1 | 2 | 1 | 0 | 0 | 0 | 0 | 0 | 0 |
| JSG000000 | Nfil3 | 0.206496 | 0.262304 | 0.278042828 | 201.6185818 | 244.4723723 | 15 | 0 | 1 | 1 | 8 | 3 | 0 | 0 | 0 | 0 | 1 | 1 |
| SG0000002 | Gnb2l1 | 0.207546 | 0.263489 | 0.152421208 | 42018.0744 | 46700.28182 | 9 | 0 | 1 | 2 | 4 | 1 | 1 | 0 | 0 | 0 | 0 | 0 |
| JSG000000 | Mak16 | 0.210672 | 0.266946 | 0.158540836 | 3810.391017 | 4252.99715 | 15 | 1 | 1 | 3 | 8 | 2 | 2 | 0 | 0 | 2 | 3 | 1 |
| JSG000000 | Ssbp4 | 0.21633 | 0.273439 | -0.604605359 | 1771.430586 | 1164.983543 | 5 | 1 | 1 | 2 | 2 | 1 | 0 | 0 | 0 | 0 | 0 | 0 |
| SG0000005 | Bcl3 | 0.220591 | 0.277982 | -0.296951145 | 1031.911894 | 839.9460968 | 8 | 0 | 1 | 1 | 2 | 0 | 0 | 0 | 0 | 0 | 0 | 0 |
| SG0000001 | Vezf1 | 0.222977 | 0.280751 | 0.168846006 | 1282.422931 | 1441.647182 | 46 | 0 | 1 | 1 | 23 | 8 | 8 | 0 | 0 | 0 | 8 | 8 |
| SG0000002 | Ccdc167 | 0.225032 | 0.283159 | -0.328928272 | 121.5812757 | 96.79421889 | 75 | 0 | 1 | 1 | 12 | 0 | 0 | 0 | 0 | 0 | 0 | 0 |
| SG0000002 | Yrdc | 0.22813 | 0.286554 | -0.201714756 | 1248.96701 | 1085.997374 | 13 | 1 | 1 | 2 | 6 | 0 | 7 | 1 | 1 | 2 | 3 | 0 |
| JSG000000 | Trp53rkb | 0.22852 | 0.287007 | -0.380487486 | 86.6341649 | 66.55045779 | 7 | 1 | 1 | 3 | 5 | 0 | 6 | 1 | 1 | 3 | 5 | 0 |
| SG0000002 | Psma7 | 0.23052 | 0.289353 | -0.163878827 | 5546.698099 | 4951.104419 | 6 | 0 | 1 | 0 | 3 | 1 | 0 | 0 | 0 | 0 | 0 | 0 |
| SG0000000 | Ap2s1 | 0.237144 | 0.29665 | -0.325860809 | 1352.44695 | 1079.012214 | 5 | 1 | 1 | 2 | 7 | 1 | 0 | 0 | 0 | 0 | 0 | 0 |
| JSG000000 | 1500011B03Rik | 0.241245 | 0.301211 | 0.239438319 | 212.6472164 | 251.0370473 | 10 | 1 | 1 | 3 | 4 | 0 | 1 | 0 | 0 | 1 | 1 | 0 |
| JSG000000 | Zbtb45 | 0.241933 | 0.301987 | -0.279402699 | 418.2707214 | 344.6266608 | 5 | 1 | 1 | 1 | 3 | 1 | 1 | 0 | 0 | 0 | 0 | 1 |
| JSG000000 | Ankrd49 | 0.248744 | 0.309427 | -0.245050549 | 723.6191024 | 610.5798352 | 4 | 0 | 1 | 0 | 1 | 0 | 1 | 0 | 0 | 0 | 1 | 0 |
| JSG000000 | Ppcdc | 0.253222 | 0.314238 | -0.200596837 | 503.7638702 | 438.3705314 | 30 | 1 | 1 | 3 | 16 | 2 | 2 | 0 | 0 | 0 | 0 | 0 |
| JSG000000 | Tmem18 | 0.256204 | 0.317433 | -0.184055553 | 737.0262669 | 648.7490332 | 7 | 0 | 1 | 1 | 5 | 1 | 4 | 0 | 1 | 0 | 1 | 1 |
| SG0000002 | Gba2 | 0.25995 | 0.321584 | 0.214927729 | 292.8387538 | 339.8820612 | 9 | 1 | 1 | 2 | 4 | 0 | 1 | 1 | 1 | 1 | 1 | 0 |
| JSG000000 | Dpcr1 | 0.261091 | 0.322796 | 0.751396134 | 10.39590792 | 17.5006911 | 8 | 1 | 1 | 1 | 1 | 1 | 2 | 0 | 0 | 0 | 0 | 0 |
| JSG000000 | Mrpl17 | 0.273255 | 0.33591 | -0.153240837 | 1766.746821 | 1588.708539 | 4 | 0 | 1 | 1 | 1 | 1 | 4 | 0 | 1 | 1 | 1 | 1 |
| JSG000000 | Hist3h2a | 0.275024 | 0.337899 | -0.497890227 | 36.02337602 | 25.50965109 | 8 | 0 | 1 | 0 | 1 | 0 | 8 | 0 | 1 | 0 | 1 | 0 |
| JSG000000 | Morn2 | 0.282595 | 0.346109 | -0.281522615 | 147.4645324 | 121.3223534 | 17 | 0 | 1 | 2 | 12 | 5 | 0 | 0 | 0 | 0 | 0 | 0 |
| JSG000000 | Cdc37 | 0.289584 | 0.353581 | -0.209026316 | 5847.903852 | 5059.143901 | 11 | 1 | 1 | 2 | 5 | 1 | 2 | 0 | 0 | 0 | 0 | 0 |
| JSG000000 | 3110001I22Rik | 0.295973 | 0.360497 | -0.284432082 | 187.6536685 | 154.075814 | 15 | 0 | 1 | 1 | 4 | 0 | 1 | 0 | 0 | 0 | 0 | 0 |
| JSG000000 | Tpgs1 | 0.298896 | 0.363687 | -0.620117043 | 163.1764031 | 106.1655282 | 4 | 0 | 1 | 1 | 1 | 0 | 0 | 0 | 0 | 0 | 0 | 0 |
| JSG000000 | Bcl2l2 | 0.300686 | 0.365491 | -0.156906701 | 1012.184295 | 907.87482 | 11 | 0 | 1 | 3 | 4 | 0 | 6 | 0 | 1 | 0 | 1 | 0 |
| JSG000000 | Hectd3 | 0.304967 | 0.369993 | -0.16259695 | 830.4816159 | 741.9650765 | 10 | 0 | 1 | 1 | 6 | 0 | 3 | 0 | 0 | 0 | 1 | 0 |
| JSG000000 | Ccdc124 | 0.315243 | 0.380963 | 0.29191683 | 942.7655407 | 1154.195621 | 3 | 1 | 1 | 1 | 2 | 0 | 0 | 0 | 0 | 0 | 0 | 0 |
| SG0000002 | Mrpl38 | 0.315629 | 0.381378 | 0.144697193 | 1418.343039 | 1567.976239 | 10 | 0 | 1 | 2 | 4 | 0 | 0 | 0 | 0 | 0 | 0 | 0 |
| JSG000000 | BC005561 | 0.31611 | 0.381907 | 0.20541442 | 202.1771834 | 233.1138323 | 40 | 0 | 1 | 0 | 5 | 4 | 0 | 0 | 0 | 0 | 0 | 0 |
| SG0000003 | Olr1 | 0.330045 | 0.39679 | -1.125103393 | 31.39871274 | 14.39534157 | 50 | 1 | 1 | 3 | 17 | 3 | 5 | 0 | 0 | 0 | 2 | 0 |
| JSG000000 | Psmc2 | 0.33097 | 0.397768 | -0.133231618 | 4527.279199 | 4127.913281 | 37 | 1 | 1 | 6 | 22 | 3 | 1 | 0 | 0 | 0 | 0 | 0 |
| JSG000000 | Casp6 | 0.332147 | 0.398941 | -0.19139614 | 376.656955 | 329.8602713 | 19 | 1 | 1 | 4 | 10 | 1 | 3 | 0 | 0 | 0 | 0 | 0 |
| JSG000000 | Mlycd | 0.336546 | 0.403305 | 0.215381115 | 182.0089258 | 211.3142879 | 7 | 0 | 1 | 1 | 3 | 1 | 0 | 0 | 0 | 0 | 0 | 0 |
| JSG000000 | Neurl3 | 0.336861 | 0.40363 | -0.659842153 | 20.36250091 | 12.88839496 | 8 | 1 | 1 | 2 | 3 | 5 | 1 | 0 | 0 | 0 | 0 | 1 |
| JSG000000 | Rpl23 | 0.337735 | 0.404595 | 0.111701626 | 16979.68251 | 18346.57943 | 8 | 1 | 1 | 1 | 3 | 0 | 0 | 0 | 0 | 0 | 0 | 0 |
| SG0000002 | Ptprn | 0.342995 | 0.410211 | 0.228942575 | 395.8137944 | 463.8841223 | 13 | 1 | 1 | 1 | 5 | 1 | 0 | 0 | 0 | 0 | 0 | 0 |
| JSG000000 | Med9 | 0.34344 | 0.410606 | -0.158430472 | 636.1738279 | 570.0113114 | 13 | 1 | 1 | 2 | 7 | 5 | 1 | 0 | 0 | 0 | 1 | 0 |
| JSG000000 | Zfp334 | 0.348986 | 0.416625 | 0.14514359 | 475.8807365 | 526.2482919 | 13 | 0 | 1 | 2 | 5 | 2 | 7 | 0 | 0 | 1 | 4 | 2 |
| JSG000000 | Top1mt | 0.357045 | 0.425169 | 0.185455811 | 302.1449386 | 343.5920333 | 14 | 1 | 1 | 3 | 7 | 0 | 0 | 0 | 0 | 0 | 0 | 0 |
| JSG000000 | Dhrs9 | 0.374978 | 0.44375 | -0.685662468 | 22.54284791 | 14.0153456 | 45 | 0 | 1 | 3 | 17 | 5 | 4 | 0 | 0 | 0 | 2 | 3 |
| JSG000000 | Sart1 | 0.377095 | 0.445873 | 0.12108757 | 2105.403968 | 2289.741354 | 11 | 1 | 1 | 2 | 3 | 0 | 4 | 0 | 0 | 0 | 0 | 0 |
| JSG000000 | Unc119b | 0.378838 | 0.447667 | 0.121404123 | 1814.731577 | 1974.052411 | 7 | 1 | 1 | 1 | 6 | 2 | 1 | 0 | 0 | 0 | 1 | 1 |
| SG0000003 | Calm2 | 0.380399 | 0.449128 | -0.120018755 | 14319.05815 | 13176.04919 | 30 | 1 | 1 | 2 | 10 | 4 | 1 | 0 | 0 | 0 | 0 | 0 |
| JSG000000 | Trmt10c | 0.388727 | 0.457392 | -0.138457353 | 1196.236948 | 1086.769386 | 12 | 1 | 1 | 2 | 8 | 1 | 2 | 1 | 1 | 2 | 3 | 1 |
| JSG000000 | Fosb | 0.393465 | 0.462178 | -0.303024328 | 118.6306384 | 96.15623617 | 4 | 1 | 1 | 3 | 8 | 0 | 1 | 0 | 0 | 1 | 4 | 0 |
| JSG000000 | Gzf1 | 0.412008 | 0.481088 | 0.104604129 | 1287.885504 | 1384.733617 | 20 | 0 | 1 | 1 | 3 | 2 | 6 | 0 | 0 | 0 | 0 | 0 |
| JSG000000 | Atg4d | 0.416194 | 0.485532 | -0.205585923 | 557.8992898 | 483.8026903 | 6 | 0 | 1 | 3 | 4 | 1 | 2 | 0 | 0 | 2 | 2 | 0 |
| JSG000000 | Relt | 0.423448 | 0.492487 | 0.174157932 | 201.9551763 | 227.8671658 | 5 | 1 | 1 | 2 | 3 | 0 | 2 | 1 | 1 | 2 | 3 | 0 |
| JSG000000 | Rpl24 | 0.436522 | 0.505788 | 0.105819832 | 1565.530257 | 1684.676079 | 12 | 1 | 1 | 3 | 6 | 2 | 0 | 0 | 0 | 0 | 0 | 0 |
| JSG000000 | Fdx1l | 0.44644 | 0.515546 | 0.14181564 | 406.6431233 | 448.646407 | 14 | 1 | 1 | 3 | 5 | 0 | 0 | 0 | 0 | 0 | 0 | 0 |
| JSG000000 | Apln | 0.449178 | 0.51824 | 0.332231672 | 64.40785069 | 81.08686422 | 5 | 0 | 1 | 0 | 0 | 1 | 4 | 0 | 1 | 0 | 0 | 1 |
| JSG000000 | Layn | 0.45697 | 0.526497 | -0.228365509 | 111.8753551 | 95.49696261 | 29 | 0 | 1 | 3 | 9 | 1 | 0 | 0 | 0 | 0 | 0 | 0 |
| JSG000000 | Ppp1r3e | 0.458439 | 0.527905 | -0.441010172 | 29.21217875 | 21.51823566 | 6 | 1 | 1 | 2 | 4 | 0 | 0 | 0 | 0 | 0 | 0 | 0 |
| SG0000007 | Ubxn1 | 0.458832 | 0.52829 | -0.099124022 | 3702.891991 | 3457.018792 | 2 | 0 | 1 | 0 | 1 | 0 | 0 | 0 | 0 | 0 | 0 | 0 |
| JSG000000 | P2ry6 | 0.462374 | 0.531821 | 0.467054137 | 13.57355484 | 18.76250871 | 16 | 1 | 1 | 1 | 5 | 6 | 0 | 0 | 0 | 0 | 0 | 0 |
| JSG000000 | Wrnip1 | 0.466897 | 0.535957 | 0.105016914 | 887.8050789 | 954.8405766 | 29 | 0 | 1 | 3 | 19 | 0 | 2 | 0 | 0 | 0 | 0 | 0 |
| JSG000000 | Sdr39u1 | 0.473664 | 0.542508 | -0.134024344 | 508.8311188 | 463.6906259 | 1 | 0 | 1 | 0 | 2 | 0 | 0 | 0 | 0 | 0 | 0 | 0 |
| JSG000000 | Tnfsf12 | 0.477131 | 0.54564 | 0.242013351 | 54.95791538 | 64.99553593 | 6 | 0 | 1 | 0 | 7 | 2 | 1 | 0 | 0 | 0 | 3 | 0 |
| JSG000000 | Ctu1 | 0.478969 | 0.547567 | -0.149707663 | 327.2931111 | 295.032845 | 6 | 0 | 1 | 0 | 0 | 1 | 1 | 0 | 1 | 0 | 0 | 0 |
| JSG000000 | Akip1 | 0.502252 | 0.569496 | -0.128468424 | 446.1072627 | 408.0998589 | 10 | 1 | 1 | 2 | 4 | 0 | 0 | 0 | 0 | 0 | 0 | 0 |
| SG0000005 | Pddc1 | 0.503561 | 0.570764 | 0.102704021 | 736.2010341 | 790.5210053 | 3 | 0 | 1 | 1 | 2 | 0 | 0 | 0 | 0 | 0 | 1 | 0 |
| JSG000000 | Lig4 | 0.505892 | 0.573062 | 0.105303058 | 560.3185773 | 602.7460887 | 22 | 1 | 1 | 1 | 16 | 4 | 3 | 0 | 0 | 0 | 1 | 0 |

| | | | | | | | | | | | | | | | | | | |
|---|---|---|---|---|---|---|---|---|---|---|---|---|---|---|---|---|---|---|
| JSG000000 | Otud3 | 0.50755 | 0.57474 | -0.145893119 | 212.2689316 | 191.8528385 | 24 | 1 | 1 | 4 | 9 | 1 | 0 | 0 | 0 | 0 | 0 | 0 |
| JSG000000 | Sec24c | 0.508322 | 0.575323 | 0.084658052 | 4597.029138 | 4874.856875 | 13 | 1 | 1 | 1 | 8 | 1 | 0 | 0 | 0 | 0 | 0 | 0 |
| JSG000000 | Cadm4 | 0.510292 | 0.577261 | -0.104985019 | 719.721532 | 669.2075866 | 24 | 1 | 1 | 2 | 12 | 2 | 0 | 0 | 0 | 0 | 0 | 0 |
| JSG000000 | Slitrk5 | 0.513649 | 0.580472 | -0.290903958 | 43.54545603 | 35.59360905 | 12 | 0 | 1 | 0 | 4 | 0 | 4 | 0 | 0 | 0 | 3 | 0 |
| SG0000001 | Scn1b | 0.528025 | 0.594666 | 0.214148812 | 172.5733582 | 200.1884265 | 14 | 0 | 1 | 2 | 3 | 1 | 0 | 0 | 0 | 0 | 0 | 0 |
| JSG000000 | Dact2 | 0.529422 | 0.596003 | 0.241209538 | 463.1426568 | 547.4268427 | 6 | 0 | 1 | 0 | 1 | 2 | 1 | 0 | 0 | 0 | 0 | 0 |
| JSG000000 | Zfp597 | 0.533351 | 0.599936 | -0.122079454 | 329.8940894 | 303.1272342 | 25 | 1 | 1 | 3 | 14 | 7 | 14 | 0 | 0 | 1 | 3 | 0 |
| JSG000000 | 2310057M21Rik | 0.535037 | 0.601568 | 0.11536164 | 290.9160795 | 315.1338391 | 41 | 0 | 1 | 6 | 26 | 6 | 1 | 0 | 1 | 1 | 1 | 0 |
| JSG000000 | Kcnk6 | 0.544027 | 0.609991 | 0.175834814 | 212.3435941 | 239.867118 | 6 | 0 | 1 | 1 | 2 | 1 | 2 | 0 | 0 | 1 | 1 | 0 |
| SG0000003 | Pim2 | 0.544498 | 0.610467 | -0.100691144 | 842.943896 | 786.1177739 | 10 | 0 | 1 | 1 | 2 | 1 | 2 | 0 | 1 | 0 | 0 | 1 |
| JSG000000 | Tmem126b | 0.563652 | 0.628266 | 0.095210624 | 348.3201518 | 372.0830228 | 7 | 0 | 1 | 1 | 7 | 0 | 0 | 0 | 0 | 0 | 0 | 0 |
| SG0000003 | Pak4 | 0.573757 | 0.638212 | 0.08808792 | 5203.654666 | 5531.279108 | 23 | 1 | 1 | 5 | 6 | 0 | 2 | 0 | 0 | 2 | 2 | 0 |
| JSG000000 | Plekhn1 | 0.584237 | 0.647898 | -0.163458478 | 144.5606224 | 129.0755857 | 9 | 0 | 1 | 3 | 3 | 0 | 4 | 0 | 0 | 1 | 1 | 0 |
| SG0000003 | Pstpip1 | 0.60459 | 0.667164 | -0.153805596 | 124.757729 | 112.1417587 | 31 | 1 | 1 | 4 | 8 | 1 | 0 | 0 | 0 | 0 | 0 | 0 |
| JSG000000 | F2r | 0.607969 | 0.670646 | 0.128445373 | 1042.185609 | 1139.228869 | 24 | 1 | 1 | 2 | 6 | 2 | 7 | 0 | 0 | 1 | 1 | 0 |
| SG0000002 | Pdia6 | 0.608868 | 0.671513 | -0.066923669 | 9181.383091 | 8765.204978 | 25 | 0 | 1 | 2 | 10 | 2 | 0 | 0 | 0 | 0 | 0 | 0 |
| JSG000000 | Asb13 | 0.615649 | 0.67774 | -0.106617994 | 306.055763 | 284.25319 | 33 | 1 | 1 | 7 | 13 | 2 | 0 | 0 | 0 | 0 | 0 | 1 |
| JSG000000 | Vps4a | 0.616922 | 0.679013 | 0.06992447 | 1188.413402 | 1247.432066 | 20 | 0 | 1 | 2 | 4 | 1 | 0 | 0 | 0 | 0 | 0 | 0 |
| SG0000002 | Hrsp12 | 0.624189 | 0.685751 | -0.103124488 | 467.657148 | 435.3955767 | 20 | 0 | 1 | 2 | 10 | 4 | 0 | 0 | 0 | 0 | 0 | 0 |
| SG0000002 | Eif5 | 0.624427 | 0.685928 | -0.066681757 | 9053.716707 | 8644.774961 | 12 | 0 | 1 | 2 | 7 | 1 | 2 | 0 | 1 | 1 | 1 | 0 |
| JSG000000 | Rsg1 | 0.625499 | 0.686853 | -0.262830294 | 25.73470938 | 21.44862547 | 7 | 1 | 1 | 3 | 6 | 2 | 0 | 0 | 0 | 0 | 0 | 0 |
| JSG000000 | 0610030E20Rik | 0.62703 | 0.688154 | 0.075429175 | 662.9082497 | 698.4894775 | 4 | 0 | 1 | 1 | 2 | 0 | 3 | 0 | 1 | 1 | 2 | 0 |
| JSG000000 | Gm12258 | 0.638179 | 0.698286 | 0.196130372 | 91.53409944 | 104.863425 | 19 | 0 | 1 | 1 | 5 | 5 | 0 | 0 | 0 | 0 | 0 | 4 |
| JSG000000 | 4930430F08Rik | 0.64075 | 0.700764 | 0.065914095 | 558.4522237 | 584.5587215 | 34 | 0 | 1 | 4 | 13 | 4 | 7 | 0 | 0 | 2 | 3 | 1 |
| SG0000004 | Tspyl2 | 0.645913 | 0.705592 | -0.080351292 | 786.8354716 | 744.2104797 | 5 | 1 | 1 | 1 | 2 | 0 | 1 | 1 | 1 | 1 | 2 | 0 |
| SG0000004 | Ifit2 | 0.65146 | 0.710439 | 0.113923913 | 103.8018047 | 112.3309467 | 42 | 1 | 1 | 5 | 20 | 5 | 3 | 0 | 0 | 0 | 0 | 2 |
| JSG000000 | Ppic | 0.652411 | 0.711216 | 0.059327995 | 783.1535065 | 816.0305219 | 13 | 0 | 1 | 1 | 7 | 5 | 1 | 0 | 0 | 0 | 0 | 0 |
| SG0000004 | Mybbp1a | 0.655322 | 0.713867 | -0.057793466 | 14715.73932 | 14137.88742 | 9 | 1 | 1 | 1 | 1 | 0 | 3 | 1 | 1 | 1 | 1 | 0 |
| JSG000000 | Tlr5 | 0.669835 | 0.727377 | -0.586779285 | 16.66770243 | 11.09781735 | 29 | 1 | 1 | 1 | 7 | 6 | 0 | 0 | 0 | 0 | 0 | 0 |
| JSG000000 | Angptl7 | 0.673184 | 0.730482 | -0.237938979 | 26.5498428 | 22.51309397 | 6 | 1 | 1 | 1 | 5 | 1 | 1 | 0 | 0 | 0 | 0 | 1 |
| JSG000000 | Pck2 | 0.680575 | 0.737207 | -0.084209397 | 918.4328768 | 866.3589521 | 7 | 0 | 1 | 0 | 2 | 0 | 1 | 0 | 1 | 0 | 0 | 0 |
| SG0000001 | Raver1 | 0.689166 | 0.744263 | -0.075292928 | 3376.15886 | 3204.479101 | 18 | 0 | 1 | 2 | 8 | 0 | 0 | 0 | 0 | 0 | 3 | 0 |
| JSG000000 | Umps | 0.701316 | 0.755199 | -0.056538156 | 3632.18181 | 3492.592349 | 18 | 1 | 1 | 3 | 8 | 5 | 2 | 0 | 0 | 1 | 1 | 1 |
| JSG000000 | Gm10053 | 0.703909 | 0.757717 | -0.25633687 | 27.9761078 | 23.42190451 | 6 | 1 | 1 | 1 | 1 | 0 | 6 | 1 | 1 | 1 | 1 | 0 |
| JSG000000 | Mrpl49 | 0.707308 | 0.76029 | -0.057877051 | 1787.365118 | 1717.080078 | 7 | 1 | 1 | 3 | 4 | 1 | 3 | 0 | 0 | 0 | 0 | 1 |
| JSG000000 | Kcnj2 | 0.714158 | 0.766676 | 0.211893446 | 35.39498665 | 40.99473052 | 10 | 1 | 1 | 3 | 9 | 2 | 4 | 1 | 1 | 1 | 6 | 2 |
| JSG000000 | Zfp622 | 0.717503 | 0.770022 | 0.049552907 | 1325.99565 | 1372.331432 | 37 | 0 | 1 | 5 | 13 | 2 | 3 | 0 | 0 | 0 | 1 | 0 |
| SG0000002 | Ndufb8 | 0.726602 | 0.778402 | 0.043563258 | 2324.78182 | 2396.050947 | 22 | 0 | 1 | 4 | 4 | 0 | 5 | 0 | 1 | 1 | 1 | 0 |
| JSG000000 | Zkscan7 | 0.730132 | 0.78125 | 0.086167421 | 129.444834 | 137.4116964 | 12 | 0 | 1 | 0 | 5 | 1 | 0 | 0 | 0 | 0 | 0 | 0 |
| JSG000000 | Mki67 | 0.749841 | 0.798143 | -0.077968525 | 9066.051956 | 8589.093667 | 42 | 0 | 1 | 8 | 28 | 3 | 1 | 0 | 0 | 0 | 0 | 1 |
| JSG000000 | Lrrc29 | 0.750898 | 0.799022 | 0.147120337 | 22.58734122 | 25.01224921 | 9 | 0 | 1 | 0 | 6 | 1 | 0 | 0 | 0 | 0 | 0 | 0 |
| SG0000003 | Pdk2 | 0.756543 | 0.803891 | 0.060874645 | 234.4838252 | 244.5895986 | 8 | 1 | 1 | 3 | 3 | 3 | 0 | 0 | 0 | 0 | 0 | 1 |
| JSG000000 | 43533 | 0.775146 | 0.820213 | -0.062054127 | 303.2859891 | 290.5174303 | 4 | 1 | 1 | 1 | 3 | 0 | 3 | 1 | 1 | 1 | 2 | 0 |
| SG0000003 | Man1b1 | 0.788443 | 0.831856 | -0.03929757 | 2375.332439 | 2311.504016 | 8 | 1 | 1 | 1 | 6 | 0 | 0 | 0 | 0 | 0 | 0 | 0 |
| JSG000000 | Uqcr10 | 0.79076 | 0.833427 | 0.04599234 | 1479.099553 | 1527.012105 | 3 | 1 | 1 | 1 | 1 | 0 | 0 | 0 | 0 | 0 | 0 | 0 |
| SG0000001 | Nelfb | 0.791334 | 0.833836 | -0.034900508 | 1796.032281 | 1753.105444 | 11 | 1 | 1 | 2 | 3 | 2 | 0 | 0 | 0 | 0 | 0 | 1 |
| JSG000000 | Plat | 0.79415 | 0.836311 | -0.077234606 | 160.2333685 | 151.8808537 | 30 | 0 | 1 | 1 | 9 | 4 | 2 | 0 | 0 | 1 | 2 | 0 |
| JSG000000 | Ldlr | 0.801271 | 0.842424 | 0.038783722 | 1696.380225 | 1742.602228 | 21 | 1 | 1 | 4 | 6 | 0 | 3 | 0 | 0 | 3 | 3 | 0 |
| JSG000000 | Zfp772 | 0.801535 | 0.842602 | 0.051880582 | 216.2829572 | 224.2022209 | 11 | 1 | 1 | 1 | 6 | 1 | 4 | 0 | 0 | 0 | 0 | 0 |
| JSG000000 | Nacc1 | 0.807872 | 0.84792 | 0.036209475 | 6061.138253 | 6215.188837 | 12 | 1 | 1 | 1 | 5 | 2 | 0 | 0 | 0 | 0 | 0 | 1 |
| JSG000000 | Ddias | 0.812013 | 0.851468 | -0.035986761 | 1322.477293 | 1289.897287 | 36 | 1 | 1 | 2 | 10 | 6 | 1 | 0 | 0 | 0 | 0 | 0 |
| SG0000000 | Vrk3 | 0.816902 | 0.855743 | 0.036163381 | 905.5511274 | 928.5370516 | 22 | 0 | 1 | 5 | 15 | 2 | 0 | 0 | 0 | 0 | 0 | 0 |
| SG0000000 | Gtpbp3 | 0.819883 | 0.858263 | 0.042968335 | 672.8479976 | 693.1890951 | 22 | 0 | 1 | 8 | 11 | 0 | 4 | 0 | 0 | 4 | 4 | 0 |
| SG0000005 | Gpr173 | 0.82331 | 0.861297 | 0.084218566 | 40.65272644 | 43.09649911 | 51 | 1 | 1 | 6 | 21 | 14 | 5 | 0 | 0 | 1 | 2 | 0 |
| SG0000000 | Smg9 | 0.839429 | 0.874635 | -0.040753912 | 607.941897 | 591.0087686 | 22 | 1 | 1 | 3 | 11 | 0 | 0 | 0 | 0 | 0 | 0 | 0 |
| SG0000003 | Amotl2 | 0.850419 | 0.883004 | -0.039413058 | 1300.015903 | 1264.981436 | 39 | 1 | 1 | 4 | 7 | 10 | 4 | 0 | 0 | 0 | 0 | 4 |
| JSG000000 | Upf3a | 0.852216 | 0.884496 | 0.032912016 | 643.793359 | 658.6489389 | 11 | 1 | 1 | 1 | 9 | 2 | 0 | 0 | 0 | 0 | 0 | 0 |
| JSG000000 | Rhou | 0.879177 | 0.906955 | 0.01897431 | 1266.758349 | 1283.528782 | 13 | 0 | 1 | 2 | 2 | 0 | 5 | 0 | 1 | 1 | 1 | 0 |
| SG0000002 | Car9 | 0.879327 | 0.907026 | 0.069314047 | 24.93882747 | 26.16625914 | 6 | 0 | 1 | 1 | 1 | 1 | 1 | 0 | 1 | 0 | 0 | 0 |
| JSG000000 | Nov | 0.886312 | 0.912812 | 0.070134164 | 29.11112446 | 30.56127494 | 21 | 0 | 1 | 1 | 2 | 5 | 6 | 0 | 0 | 1 | 1 | 3 |
| SG0000002 | Yy1 | 0.886406 | 0.912853 | 0.014056335 | 2244.186375 | 2266.158592 | 29 | 0 | 1 | 3 | 10 | 3 | 3 | 0 | 0 | 0 | 1 | 0 |
| SG0000000 | Eif2b2 | 0.889901 | 0.915665 | 0.028076862 | 1218.583133 | 1242.530735 | 11 | 0 | 1 | 1 | 6 | 1 | 0 | 0 | 0 | 0 | 0 | 0 |
| JSG000000 | Dnajb9 | 0.900955 | 0.924967 | -0.024260611 | 478.193313 | 470.2191662 | 15 | 0 | 1 | 1 | 4 | 0 | 4 | 0 | 0 | 0 | 1 | 0 |
| SG0000003 | Add2 | 0.901373 | 0.92527 | -0.073722182 | 25.60337631 | 24.32790164 | 38 | 1 | 1 | 8 | 18 | 3 | 0 | 0 | 0 | 0 | 0 | 0 |
| JSG000000 | Flrt1 | 0.915216 | 0.937566 | -0.062644327 | 38.51457274 | 36.87799176 | 12 | 1 | 1 | 1 | 3 | 0 | 4 | 0 | 0 | 0 | 0 | 0 |
| JSG000000 | Mtx3 | 0.921553 | 0.942979 | 0.010746616 | 379.9228137 | 382.7634202 | 30 | 0 | 1 | 3 | 21 | 5 | 13 | 0 | 0 | 0 | 3 | 2 |
| SG0000004 | Alkbh6 | 0.944805 | 0.961008 | 0.017244946 | 260.054426 | 263.1815834 | 9 | 1 | 1 | 1 | 3 | 0 | 1 | 1 | 1 | 1 | 1 | 0 |
| SG0000000 | Ube2c | 0.95051 | 0.966207 | -0.008029166 | 3893.592592 | 3871.983403 | 11 | 0 | 1 | 0 | 2 | 0 | 1 | 0 | 0 | 0 | 0 | 0 |
| JSG000000 | Higd1a | 0.958364 | 0.973141 | -0.01867378 | 1271.23655 | 1254.888106 | 13 | 0 | 1 | 1 | 5 | 1 | 0 | 0 | 0 | 0 | 0 | 0 |
| JSG000000 | Neurl2 | 0.986908 | 0.997208 | -0.030587493 | 47.07276197 | 46.08524793 | 4 | 1 | 1 | 3 | 4 | 0 | 0 | 0 | 0 | 0 | 0 | 0 |
| JSG000000 | Dad1 | 1 | 1 | 0.005073467 | 1672.481298 | 1678.373198 | 62 | 0 | 1 | 1 | 6 | 3 | 6 | 0 | 0 | 0 | 1 | 0 |
| SG0000003 | Dnhd1 | 1 | 1 | -0.017648462 | 46.21063853 | 45.64878819 | 11 | 1 | 1 | 5 | 11 | 0 | 0 | 0 | 0 | 0 | 0 | 0 |
