## Supplementary_Table6 for "Aging-dependent dysregulation of EXOSC2 is maintained in cancer as a dependency"

| Annotation Cluster 1 | Enrichment Score: 9.214187076220206 | | | | | | | | | | | | | |
|---|---|---|---|---|---|---|---|---|---|---|---|---|---|---|
| Category | Term | Count | % | PValue | | | Genes | List Total | Pop Hits | Pop Total | ld Enrichme | Bonferroni | Benjamini | FDR |
| UP_KEYWORDS | Transcription | 55 | 24.44444 | 3.34E-13 | -1.25E+01 | 12 | 3, G3UZP1, I | 225 | 1859 | 22680 | 2.982249 | 6.07E-11 | 2.02E-11 | 4.12E-10 |
| UP_KEYWORDS | Differentiation | 17 | 7.555556 | 7.29E-04 | -3.14E+00 | 3 | JEJ9, Q99M | 225 | 646 | 22680 | 2.652632 | 0.124315 | 0.006301 | 0.896046 |
| UP_KEYWORDS | Cell junction | 20 | 8.888889 | 3.36E-05 | -4.47E+00 | 4 | A6YXC8, S4I | 225 | 661 | 22680 | 3.049924 | 0.00609 | 4.07E-04 | 0.041408 |
| UP_KEYWORDS | Zinc-finger | 38 | 16.88889 | 6.43E-07 | -6.19E+00 | 6 | , A6H5X4, E | 225 | 1565 | 22680 | 2.44754 | 1.17E-04 | 9.76E-06 | 7.94E-04 |
| UP_KEYWORDS | Cell cycle | 17 | 7.555556 | 5.24E-04 | -3.28E+00 | 3 | 'M6, Q8CIB! | 225 | 626 | 22680 | 2.73738 | 0.09091 | 0.004754 | 0.644161 |
| UP_KEYWORDS | Calmodulin-binding | 5 | 2.222222 | 0.049282 | -1.31E+00 | 1 | ), Q3UVD6, | 225 | 139 | 22680 | 3.625899 | 0.999899 | 0.264054 | 46.40168 |
| UP_KEYWORDS | Methyltransferase | 6 | 2.666667 | 0.030616 | -1.51E+00 | 2 | CWR8, B9EI | 225 | 176 | 22680 | 3.436364 | 0.996515 | 0.189094 | 31.86766 |

| | |
|---|---|
| Calmodulin-binding | 1.307308 |
| Methyltransferase | 1.514045 |
| Differentiation | 3.137198 |
| Cell cycle | 3.281039 |
| Cell junction | 4.474159 |
| Zinc-finger | 6.191602 |
| Transcription | 12.47661 |
