## Supplementary_Table7 for "Aging-dependent dysregulation of EXOSC2 is maintained in cancer as a dependency"

| Annotation Cluster 1 | Enrichment Score: 9.262024237091895 | | | | | | | | | | | | | |
|---|---|---|---|---|---|---|---|---|---|---|---|---|---|---|
| Category | Term | Count | % | PValue | | | Genes | List Total | Pop Hits | Pop Total | ld Enrichme | Bonferroni | Benjamini | FDR |
| UP_KEYWORDS | Keratin | 7 | 1.891892 | 0.019085 | -1.71931 | 1.719308 | , Q0VDR7, ( | 370 | 129 | 22680 | 3.32621 | 0.991094 | 0.11979 | 22.08354 |
| UP_KEYWORDS | SH3 domain | 15 | 4.054054 | 8.55E-06 | -5.06819 | 5.068185 | (C4, D3Z3U! | 370 | 208 | 22680 | 4.420478 | 0.002092 | 2.09E-04 | 0.011068 |
| UP_KEYWORDS | Serine/threonine-protein kinase | 21 | 5.675676 | 1.23E-05 | -4.91148 | 4.911484 | '8, A0A0G2J | 370 | 405 | 22680 | 3.178378 | 0.002999 | 2.50E-04 | 0.015876 |
| UP_KEYWORDS | SH2 domain | 9 | 2.432432 | 3.22E-04 | -3.49226 | 3.492259 | Q3U4K7, Q | 370 | 105 | 22680 | 5.254054 | 0.075851 | 0.004373 | 0.416068 |
| UP_KEYWORDS | Transcription | 51 | 13.78378 | 3.07E-04 | -3.51329 | 3.513292 | i564, Q505( | 370 | 1859 | 22680 | 1.681636 | 0.072398 | 0.004411 | 0.396434 |
| UP_KEYWORDS | Microtubule | 13 | 3.513514 | 0.001362 | -2.86588 | 2.865885 | Q3UU92, ( | 370 | 264 | 22680 | 3.018428 | 0.283853 | 0.014411 | 1.749214 |
| UP_KEYWORDS | Calmodulin-binding | 8 | 2.162162 | 0.0078 | -2.10792 | 2.107922 | Q3TSK3, Q | 370 | 139 | 22680 | 3.527902 | 0.853161 | 0.058189 | 9.642802 |
| UP_KEYWORDS | Autophagy | 8 | 2.162162 | 0.004041 | -2.39351 | 2.393508 | 325, Q6EB\ | 370 | 123 | 22680 | 3.986816 | 0.629187 | 0.037437 | 5.108495 |
| UP_KEYWORDS | Guanine-nucleotide releasing factor | 8 | 2.162162 | 0.006172 | -2.20954 | 2.209543 | 5YWG7, F8\ | 370 | 133 | 22680 | 3.687055 | 0.780617 | 0.054634 | 7.704869 |

| | |
|---|---|
| Guanine-nucleotide releasing factor | 2.209543 |
| Autophagy | 2.393508 |
| Microtubule | 2.865885 |
| SH2 domain | 3.492259 |
| Transcription | 3.513292 |
| Serine/threonine-protein kinase | 4.911484 |
| SH3 domain | 5.068185 |
