## Supplementary_Table8 for "Aging-dependent dysregulation of EXOSC2 is maintained in cancer as a dependency"

| Sequence | Annotated Sequence | Modifications | Modifications in Master Proteins | Qvality PEP | Qvality q-value | # Protein Groups | # Proteins | # PSMs | Master Protein Accessions | Master Protein Descriptions | Positions in Master Proteins | # Missed Cleavages | Theo. MH+ [Da] | # Razor Quan Results | Abundance Ratio (log2): (ESCDshEXOSC2) / (ESCD) | Abundance Ratio Adj. P-Value: (ESCDshEXOSC2) / (ESCD) | Abundance Ratio Variability [%]: (ESCDshEXOSC2) / (ESCD) | Abundances (Grouped): ESCD | Abundances (Grouped): ESCDshEXOSC2 | Abundances (Grouped) CV [%]: ESCD | Abundances (Grouped) CV [%]: ESCDshEXOSC2 | Quan Info | Found in Sample: [S11] F2: 127N, Sample, 1. ESCD | Found in Sample: [S12] F2: 127C, Sample, 2. ESCD | Found in Sample: [S13] F2: 128N, Sample, 3. ESCD | Found in Sample: [S14] F2: 128C, Sample, 4. ESCD | Found in Sample: [S15] F2: 129N, Sample, 5. ESCD | Found in Sample: [S16] F2: 129C, Sample, 1, ESCDshEXOSC2 | Found in Sample: [S17] F2: 130N, Sample, 2, ESCDshEXOSC2 | Found in Sample: [S18] F2: 130C, Sample, 3, ESCDshEXOSC2 | Found in Sample: [S19] F2: 131N, Sample, 4, ESCDshEXOSC2 | Found in Sample: [S20] F2: 131C, Sample, 5, ESCDshEXOSC2 | Confidence (by Search Engine): Sequest HT | Percolator q-Value (by Search Engine): Sequest HT | Percolator PEP (by Search Engine): Sequest HT | XCorr (by Search Engine): Sequest HT |
|---|---|---|---|---|---|---|---|---|---|---|---|---|---|---|---|---|---|---|---|---|---|---|---|---|---|---|---|---|---|---|---|---|---|---|---|---|
| DAPSPPVLEAICTEPVCTPETR | [K].DAPSPPVLEAICTEPVCTPETR.[G] | 2xCarbami | Q3KR45 2x | 0.00185 | 5.7E-05 | 1 | 5 | 2 | Q3KR45 | DNA (cytosine-5)-methyltransferase 3B OS=Mus musculus GN=Dnmt3b PE=1 SV=1 | Q3KR45 [3 | 0 | 2828.25 | 0 | -4.22 | 0.00019 | 0.05 | 190.2 | 9.8 | 4.28 | 50.08 | | High | High | High | High | High | High | High | High | High | High | High | 2.6E-05 | 0.00047 | 4.53 |
| LSIYDNVPGSILYSSSGELADLENI | [R].LSIYDNVPGSILYSSSGELADLENEDIFPELDDI | 1xPhospho | E9PXD2 1x | 0.00219 | 0.00011 | 1 | 7 | 3 | E9PXD2 | Rho GTPase-activating protein 7 OS=Mus musculus GN=Dlc1 PE=1 SV=1 | E9PXD2 [8 | 0 | 4921.44 | 0 | -4.08 | 0.00083 | 0.14 | 188.8 | 11.2 | 4.7 | 95.33 | | High | High | High | High | High | High | High | High | High | High | High | 4.9E-05 | 0.00062 | 3.01 |
| ASSPQVAPQSLNTALLQTLTHLG | [R].ASSPQVAPQSLNTALLQTLTHLGDISTVLGPL | 1xPhospho | Q6J1H4 1x | 1.1E-06 | 0 | 1 | 2 | 5 | Q6J1H4 | Undifferentiated embryonic cell transcription factor 1 OS=Mus musculus GN=Utf1 PE=1 SV=2 | Q6J1H4 [2 | 0 | 3707.99 | 0 | -3.64 | 0.0021 | 0.36 | 183.3 | 16.7 | 17.69 | 53.22 | | High | High | High | High | High | High | High | High | High | High | High | 0 | 1.2E-07 | 4.37 |
| TASLPAIPNPFPELTGAAPGSPPS | [R].TASLPAIPNPFPELTGAAPGSPPSVAPSSLPPF | 1xPhospho | Q60760 1x | 0.00869 | 0.00074 | 1 | 6 | 1 | Q60760 | Growth factor receptor-bound protein 10 OS=Mus musculus GN=Grb10 PE=1 SV=2 | Q60760 [7 | 0 | 4378.31 | 0 | -3.34 | 0.00078 | 0.24 | 181.8 | 18.2 | 12.77 | 33.1 | | High | High | High | High | High | High | High | High | High | High | High | 0.00033 | 0.00269 | 4.82 |
| TSWGFLQSLVSIK | [R].TSWGFLQSLVSIK.[Q] | 1xPhospho | Q99LI5 1x | 0.00809 | 0.00074 | 1 | 1 | 1 | Q99LI5 | Zinc finger protein 281 OS=Mus musculus GN=Znf281 PE=1 SV=1 | Q99LI5 [11 | 0 | 2004.1 | 0 | -3.06 | 0.00075 | 0.65 | 179 | 21 | 6.35 | 50.8 | | High | High | High | High | High | High | High | High | High | High | High | 0.00033 | 0.00247 | 2.06 |
| LQASAPSLTPI | [K].LQASAPSLTPI.[-] | 1xPhospho | P28231 1x | 7.7E-05 | 0 | 1 | 1 | 4 | P28231 | Gap junction beta-3 protein OS=Mus musculus GN=Gjb3 PE=1 SV=1 | P28231 [2 | 0 | 1406.75 | 0 | -2.71 | 0.00018 | 0.23 | 173.5 | 26.5 | 4.92 | 9.32 | | High | High | High | High | High | High | High | High | High | High | High | 0.00018 | 0.00126 | 3.07 |
| LQASAPSLTPI | [K].LQASAPSLTPI.[-] | 2xPhospho | P28231 2x | 0.00217 | 5.7E-05 | 1 | 1 | 3 | P28231 | Gap junction beta-3 protein OS=Mus musculus GN=Gjb3 PE=1 SV=1 | P28231 [2 | 0 | 1486.72 | 0 | -2.48 | 0.00025 | 0.2 | 170.6 | 29.4 | 7.16 | 17.71 | | High | High | High | High | High | High | High | High | High | High | High | 4.9E-05 | 0.00057 | 3.01 |
| LPLIVIDDDDDGSPDLK | [K].LPLIVIDDDDDGSPDLK.[N] | 1xPhospho | A0A1D9BZ | 2.9E-09 | 0 | 1 | 1 | 12 | A0A1D9BZ | GCNA OS=Mus musculus PE=2 SV=1 | A0A1D9BZ | 0 | 2378.21 | 0 | -2.46 | 0.00015 | 0.51 | 169.3 | 30.7 | 6.25 | 16.55 | | High | High | High | High | High | High | High | High | High | High | High | 0 | 1.4E-10 | 4.46 |
| DAPSPPVLEAICTEPVCTPETR | [K].DAPSPPVLEAICTEPVCTPETR.[G] | 2xCarbami | Q3KR45 1x | 1.9E-07 | 0 | 1 | 5 | 6 | Q3KR45 | DNA (cytosine-5)-methyltransferase 3B OS=Mus musculus GN=Dnmt3b PE=1 SV=1 | Q3KR45 [3 | 0 | 2748.28 | 0 | -2.42 | 0.00046 | 0.58 | 168.5 | 31.5 | 5.91 | 23.03 | | High | High | High | High | High | High | High | High | High | High | High | 0 | 2.6E-08 | 6.02 |
| LYSILQGDSPTK | [K].LYSILQGDSPTK.[W] | 1xPhospho | Q6NV83 1 | 0.00081 | 0 | 1 | 3 | 1 | Q6NV83 | U2 snRNP-associated SURP motif-containing protein OS=Mus musculus GN=U2surp PE=1 SV=3 | Q6NV83 [4 | 0 | 1859.99 | 0 | -2.19 | 0.00027 | 0.52 | 163.4 | 36.6 | 8.05 | 10.83 | | High | High | High | High | High | High | High | High | High | High | High | 0 | 0.00019 | 3.15 |
| GNGLASPGVQGGGGEGNSPSEI | [R].GNGLASPGVQGGGGEGNSPSEPHLLETINVF | 1xPhospho | Q3UHU5 1 | 0.00017 | 0 | 1 | 5 | 1 | Q3UHU5 | Microtubule cross-linking factor 1 OS=Mus musculus GN=Mtcl1 PE=1 SV=1 | Q3UHU5 [ | 0 | 3309.61 | 0 | -2.15 | 0.00067 | 1.85 | 164.3 | 35.7 | 6.13 | 28.49 | | High | High | High | High | High | High | High | High | High | High | High | 0 | 3.2E-05 | 3.54 |
| ELGNIHETSQDLSPEK | [R].ELGNIHETSQDLSPEK.[A] | 1xPhospho | A0A0R4J0 | 6.1E-05 | 0 | 1 | 6 | 2 | A0A0R4J0 | Disks large-associated protein 5 OS=Mus musculus GN=Dlgap5 PE=1 SV=1 | A0A0R4J0 | 0 | 2335.16 | 0 | -1.94 | 0.00245 | 0.96 | 157.3 | 42.7 | 13.7 | 24.96 | | High | High | High | High | High | High | High | High | High | High | High | 0 | 1E-05 | 3.24 |
| QLSNANIGQSPHTDDHSDTDHE | [K].QLSNANIGQSPHTDDHSDTDHEEDRDNQQ | 1xPhospho | O88622 1x | 0.07286 | 0.0056 | 1 | 5 | 1 | O88622 | Poly(ADP-ribose) glycohydrolase OS=Mus musculus GN=Parg PE=1 SV=2 | O88622 [1 | 1 | 4541.1 | 0 | -1.87 | 0.00308 | 0.49 | 150.9 | 49.1 | 17.14 | 24.94 | | High | High | High | High | High | High | High | High | High | High | High | 0.00287 | 0.02974 | 3.36 |
| YHHCTDEPLHAIYDSVEK | [K].YHHCTDEPLHAIYDSVEK.[L] | 1xCarbami | Q3TIA9 1x | 0.00152 | 5.7E-05 | 1 | 4 | 1 | Q3TIA9 | Putative uncharacterized protein OS=Mus musculus GN=Asns PE=2 SV=1 | Q3TIA9 [2 | 0 | 2752.28 | 0 | -1.64 | 0.00204 | 0.22 | 148 | 52 | 12.83 | 15.16 | | High | High | High | High | High | High | High | High | High | High | High | 2.6E-05 | 0.00038 | 2.46 |
| SPLPPILGSPTK | [R].SPLPPILGSPTK.[A] | 1xPhospho | Q3UFT4 1x | 0.00057 | 0 | 1 | 7 | 1 | Q3UFT4 | Putative uncharacterized protein OS=Mus musculus GN=Ulk1 PE=2 SV=1 | Q3UFT4 [6 | 0 | 1745 | 0 | -1.6 | 0.0005 | 1.63 | 150.4 | 49.6 | 1.38 | 18.5 | | High | High | High | High | High | High | High | High | High | High | High | 0 | 0.00013 | 2.52 |
| LTYIPENTPIDTVVFK | [K].LTYIPENTPIDTVVFK.[A] | 1xDeamida | Q2PZL6 2x | 0.02464 | 0.00161 | 1 | 1 | 1 | Q2PZL6 | Protocadherin Fat 4 OS=Mus musculus GN=Fat4 PE=1 SV=2 | Q2PZL6 [2 | 0 | 2240.07 | 0 | -1.59 | 0.00062 | 0.53 | 149.1 | 50.9 | 8.46 | 4.02 | | High | High | High | High | High | High | High | High | High | High | High | 0.00087 | 0.00865 | 2.86 |
| LVSFHDDSDEDLLHI | [K].LVSFHDDSDEDLLHI.[-] | 1xPhospho | B7ZWC4 1 | 3E-05 | 0 | 1 | 8 | 6 | B7ZWC4 | Insulin-like growth factor 2 receptor OS=Mus musculus GN=Igf2r PE=2 SV=1 | B7ZWC4 [2 | 0 | 2063.95 | 0 | -1.54 | 0.00025 | 1.24 | 149 | 51 | 3.5 | 10.68 | | High | High | High | High | High | High | High | High | High | High | High | 0 | 0.0002 | 3.06 |
| TESTYLAEIHRQSPK | [K].TESTYLAEIHRQSPK.[E] | 1xPhospho | Q80TI0-4 1 | 6.1E-05 | 0 | 1 | 8 | 2 | Q80TI0-4 | Isoform 4 of GRAM domain-containing protein 1B OS=Mus musculus GN=Gramd1b | Q80TI0-4 [ | 1 | 2298.19 | 0 | -1.49 | 0.00049 | 0.71 | 147.7 | 52.3 | 6.47 | 10.22 | | High | High | High | High | High | High | High | High | High | High | High | 0.00196 | 0.01996 | 3.19 |
| VLTPIK | [R].VLTPIK.[E] | 1xPhospho | Q7TSG1 1x | 0.05769 | 0.00423 | 1 | 2 | 1 | Q7TSG1 | Centrosomal protein of 120 kDa OS=Mus musculus GN=Cep120 PE=1 SV=1 | Q7TSG1 [3 | 0 | 1208.74 | 0 | -1.45 | 0.00041 | 0.22 | 144.4 | 55.6 | 5.91 | 5.28 | | High | High | High | High | High | High | High | High | High | High | High | 0.00216 | 0.02268 | 1.98 |
| TLQAISPK | [R].TLQAISPK.[N] | 1xDeamida | B0V2N1-3 | 0.03518 | 0.00227 | 1 | 2 | 1 | B0V2N1-3 | Isoform 3 of Receptor-type tyrosine-protein phosphatase S OS=Mus musculus GN=Ptprs | B0V2N1-3 | 0 | 1167.62 | 0 | -1.39 | 0.00171 | 1.11 | 145.4 | 54.6 | 9.94 | 8.98 | | High | High | High | High | High | High | High | High | High | High | High | 0.00121 | 0.01294 | 1.87 |
| SPGLYAIGYEECIERPLSPDVER | [K].SPGLYAIGYEECIERPLSPDVER.[H] | 1xCarbami | Q6NZR2 1 | 0.00359 | 0.00021 | 1 | 3 | 4 | Q6NZR2 | Myb/SANT-like DNA-binding domain-containing protein 2 OS=Mus musculus GN=Msantd2 PE=1 SV=1 | Q6NZR2 [4 | 0 | 2959.41 | 0 | -1.38 | 0.00014 | 1.05 | 145 | 55 | 2.39 | 8.62 | | High | High | High | High | High | High | High | High | High | High | High | 9.4E-05 | 0.001 | 2.55 |
| LSALSQVTPLLTNIEK | [K].LSALSQVTPLLTNIEK.[G] | 1xPhospho | G3UYN0 1 | 3.6E-06 | 0 | 1 | 2 | 1 | G3UYN0 | L1TD1 OS=Mus musculus GN=L1td1 PE=1 SV=1 | G3UYN0 [5 | 0 | 2265.29 | 0 | -1.35 | 0.00037 | 0.77 | 145.3 | 54.7 | 5.85 | 5.6 | | High | High | High | High | High | High | High | High | High | High | High | 0 | 4.3E-07 | 2.21 |
| NQLVLIVNESPASPAK | [K].NQLVLIVNESPASPAK.[TA] | 1xPhospho | Q6P5E3 1x | 9.7E-09 | 0 | 1 | 3 | 1 | Q6P5E3 | Sal-like 1 (Drosophila) OS=Mus musculus GN=Sall1 PE=1 SV=1 | Q6P5E3 [6 | 0 | 1989.06 | 0 | -1.35 | 0.1098 | 15.9 | 143.8 | 56.2 | 35.88 | 41.77 | | High | High | High | High | High | High | High | High | High | High | High | 0 | 5.6E-10 | 4.82 |
| GNSPSCQSPDLPGINR | [K].GNSPSCQSPDLPGINR.[G] | 1xCarbami | O88466 2x | 6E-06 | 0 | 2 | 4 | 3 | O88466; R | Zinc finger protein 106 OS=Mus musculus GN=Znf106 PE=1 SV=3Zinc finger protein 106 OS=Mus musculus GN=Zfp106 PE=1 SV=1 | O88466 [1 | 0 | 2087.88 | 0 | -1.31 | 0.00024 | 0.93 | 143.9 | 56.1 | 4.28 | 8.7 | Shared | High | High | High | High | High | High | High | High | High | High | High | 0 | 7.6E-07 | 4.25 |
| TPSPPPPILEDIILGK | [R].TPSPPPPILEDIILGK.[K] | 1xPhospho | E9Q784 1x | 0.06432 | 0.00466 | 1 | 3 | 1 | E9Q784 | Zinc finger CCCH domain-containing protein 13 OS=Mus musculus GN=Zc3h13 PE=1 SV=1 | E9Q784 [2 | 0 | 2225.26 | 0 | -1.3 | 0.006 | 0.9 | 140.4 | 59.6 | 11.1 | 16.38 | | High | High | High | High | High | High | High | High | High | High | High | 0.0024 | 0.02575 | 1.72 |
| YPNQDHLTNQNLPPIHTLHQQT | [R].YPNQDHLTNQNLPPIHTLHQQTFGDSPSK.[Y | 1xDeamida | D0VLQ5 1 | 0.00984 | 0.00074 | 1 | 6 | 4 | D0VLQ5 | Tet oncogene 2 OS=Mus musculus GN=Tet2 PE=2 SV=1 | D0VLQ5 [1 | 0 | 3866.89 | 0 | -1.3 | 0.01875 | 10.27 | 146.1 | 53.9 | 24.16 | 43.86 | | High | High | High | High | High | High | High | High | High | High | High | 0.00045 | 0.00407 | 3.77 |
| RLSTIFEECDEELER | [K].RLSTIFEECDEELER.[M] | 1xCarbami | E9QAU4 1 | 1.1E-06 | 0 | 1 | 3 | 3 | E9QAU4 | Enhancer trap locus 4 OS=Mus musculus GN=Etl4 PE=1 SV=1 | E9QAU4 [1 | 1 | 2235.02 | 0 | -1.28 | 0.00076 | 1.44 | 141.7 | 58.3 | 6.18 | 5.65 | | High | High | High | High | High | High | High | High | High | High | High | 0 | 6.4E-07 | 4.66 |
| NQLVLIVNESPASPAK | [K].NQLVLIVNESPASPAK.[TA] | 1xPhospho | Q6P5E3 1x | 1.5E-07 | 0 | 1 | 3 | 1 | Q6P5E3 | Sal-like 1 (Drosophila) OS=Mus musculus GN=Sall1 PE=1 SV=1 | Q6P5E3 [6 | 0 | 2218.22 | 0 | -1.27 | 0.00074 | 2.43 | 141.2 | 58.8 | 5.2 | 13.81 | | High | High | High | High | High | High | High | High | High | High | High | 0 | 1.2E-08 | 4.19 |
| APPGAPGPGPGSGAPGSQEEEE | [R].APPGAPGPGPGSGAPGSQEEEEEPGLVEADF | 1xDeamida | Q8CCS6 1x | 1.1E-06 | 0 | 1 | 3 | 2 | Q8CCS6 | Polyadenylate-binding protein 2 OS=Mus musculus GN=Pabpn1 PE=1 SV=3 | Q8CCS6 [7 | 0 | 4859.27 | 0 | -1.26 | 0.00207 | 5 | 141.1 | 58.9 | 16.87 | 29.99 | | High | High | High | High | High | High | High | High | High | High | High | 0 | 1.2E-07 | 5.4 |
| YMDDDGSLSPIEEPLTEDEATNP | [K].YMDDDGSLSPIEEPLTEDEATNPQAECDIK.[L | 1xCarbami | Q9D4V4 1 | 0.0003 | 0 | 1 | 5 | 3 | Q9D4V4 | TATA box-binding protein-associated factor RNA polymerase I subunit D OS=Mus musculus GN=Taf1d PE=1 SV=2 | Q9D4V4 [1 | 0 | 3936.73 | 0 | -1.25 | 0.00236 | 3.42 | 138.6 | 61.4 | 7.65 | 13.46 | | High | High | High | High | High | High | High | High | High | High | High | 0 | 6.1E-05 | 4.22 |
| SSTGPELPAPTPLLVEQHVDSPR | [R].SSTGPELPAPTPLLVEQHVDSPRPLAAIPSSQE | 1xPhospho | Q8BTI8 1x | 0.01307 | 0.00089 | 1 | 3 | 3 | Q8BTI8 | Serine/arginine repetitive matrix protein 2 OS=Mus musculus GN=Srrm2 PE=1 SV=3 | Q8BTI8 [35 | 0 | 4952.49 | 0 | -1.23 | 0.00037 | 2.51 | 140.8 | 59.2 | 10.02 | 17.81 | | High | High | High | High | High | High | High | High | High | High | High | 0.00047 | 0.00424 | 5.14 |
| VVSISSEHLEPITPTK | [K].VVSISSEHLEPITPTK.[N] | 1xPhospho | O55201 1x | 0.00457 | 0.00046 | 1 | 2 | 1 | O55201 | Transcription elongation factor SPT5 OS=Mus musculus GN=Supt5h PE=1 SV=1 | O55201 [1 | 0 | 2275.24 | 0 | -1.23 | 0.01511 | 4.06 | 143.5 | 56.5 | 18.71 | 20.1 | | High | High | High | High | High | High | High | High | High | High | High | 0.0002 | 0.00132 | 2.64 |
| EELADISTDK | [K].EELADISTDK.[D] | 1xPhospho | E9Q784 1x | 0.00022 | 0 | 1 | 1 | 2 | E9Q784 | Zinc finger CCCH domain-containing protein 13 OS=Mus musculus GN=Zc3h13 PE=1 SV=1 | E9Q784 [1 | 0 | 1658.83 | 0 | -1.21 | 0.00044 | 1.94 | 141.8 | 58.2 | 5.08 | 6.3 | | High | High | High | High | High | High | High | High | High | High | High | 0 | 4.3E-05 | 2.82 |
| GFGSLPCILNACSGDIK | [R].GFGSLPCILNACSGDIK.[S] | 2xCarbami | Q8K1S6 1x | 3.6E-06 | 0 | 1 | 2 | 1 | Q8K1S6 | Protein spire homolog 2 OS=Mus musculus GN=Spire2 PE=1 SV=1 | Q8K1S6 [3 | 0 | 2347.16 | 0 | -1.17 | 0.00165 | 5.03 | 141.3 | 58.7 | 7.45 | 16.22 | | High | High | High | High | High | High | High | High | High | High | High | 0 | 4.3E-07 | 3.58 |
| APPGAPGPGPGSGAPGSQEEEE | [R].APPGAPGPGPGSGAPGSQEEEEEPGLVEADF | 1xPhospho | Q8CCS6 1x | 4.2E-06 | 0 | 1 | 3 | 4 | Q8CCS6 | Polyadenylate-binding protein 2 OS=Mus musculus GN=Pabpn1 PE=1 SV=3 | Q8CCS6 [7 | 0 | 4858.29 | 0 | -1.15 | 0.00041 | 2.6 | 138.1 | 61.9 | 4.11 | 9.39 | | High | High | High | High | High | High | High | High | High | High | High | 0 | 1.4E-06 | 4.3 |
| TGSYGALAEISASK | [K].TGSYGALAEISASK.[E] | 1xPhospho | Q9DBR7 1 | 5E-05 | 0 | 1 | 2 | 7 | Q9DBR7 | Protein phosphatase 1 regulatory subunit 12A OS=Mus musculus GN=Ppp1r12a PE=1 SV=2 | Q9DBR7 [4 | 0 | 1892.98 | 0 | -1.14 | 0.00262 | 5.84 | 137.7 | 62.3 | 8.14 | 23.07 | | High | High | High | High | High | High | High | High | High | High | High | 0 | 7.2E-05 | 3.24 |
| AHLDHIPDYTPPLLTTISPEQESD | [R].AHLDHIPDYTPPLLTTISPEQESDER.[K] | 1xPhospho | Q8CFC7 1x | 2E-06 | 0 | 1 | 5 | 2 | Q8CFC7 | CLK4-associating serine/arginine rich protein OS=Mus musculus GN=Clasrp PE=1 SV=3 | Q8CFC7 [8 | 0 | 3283.57 | 0 | -1.13 | 0.00015 | 0.49 | 137.4 | 62.6 | 5.94 | 12.43 | | High | High | High | High | High | High | High | High | High | High | High | 0 | 2.2E-07 | 3.72 |
| GNSPSCQSPDLPGINR | [K].GNSPSCQSPDLPGINR.[G] | 1xCarbami | O88466 1x | 2.9E-05 | 0 | 2 | 4 | 4 | O88466; R | Zinc finger protein 106 OS=Mus musculus GN=Znf106 PE=1 SV=3Zinc finger protein 106 OS=Mus musculus GN=Zfp106 PE=1 SV=1 | O88466 [1 | 0 | 2007.92 | 0 | -1.07 | 0.00086 | 2.07 | 132.5 | 67.5 | 6.05 | 6.48 | Shared | High | High | High | High | High | High | High | High | High | High | High | 0 | 4.5E-06 | 3.7 |
| RNSPMAQTPPCHLPNIPGVTSP | [R].RNSPMAQTPPCHLPNIPGVTSPGALIEDSK.[N | 1xCarbami | P70445 1x | 0.09696 | 0.00778 | 1 | 2 | 1 | P70445 | Eukaryotic translation initiation factor 4E-binding protein 2 OS=Mus musculus GN=Eif4ebp2 PE=1 SV=1 | P70445 [6 | 1 | 3723.86 | 0 | -1.07 | 0.00572 | 6.37 | 135 | 65 | 10.97 | 15.49 | | High | High | High | High | High | High | High | High | High | High | High | 0.00392 | 0.0418 | 2.71 |
| SPLSAEELTAIEDEGVLDK | [R].SPLSAEELTAIEDEGVLDK.[M] | 1xPhospho | D3Z3Q3 1 | 2E-06 | 0 | 1 | 7 | 1 | D3Z3Q3 | Smoothelin OS=Mus musculus GN=Smtn PE=1 SV=1 | D3Z3Q3 [5 | 0 | 2554.29 | 0 | -1.07 | 0.00256 | 4.33 | 138.5 | 61.5 | 11.83 | 8.08 | | High | High | High | High | High | High | High | High | High | High | High | 0 | 2.2E-07 | 3.37 |
| GELIPISPSTEFGGSGIGTPPSVLK | [R].GELIPISPSTEFGGSGIGTPPSVLK.[R] | 1xPhospho | P48972 1x | 0.00016 | 0 | 1 | 1 | 4 | P48972 | Myb-related protein B OS=Mus musculus GN=Mybl2 PE=1 SV=1 | P48972 [3 | 0 | 2978.59 | 0 | -1.04 | 0.00047 | 1.35 | 134.8 | 65.2 | 6.22 | 3.68 | | High | High | High | High | High | High | High | High | High | High | High | 0.00304 | 0.03181 | 2.61 |
| TGSYGALAEISASK | [K].TGSYGALAEISASK.[E] | 1xPhospho | Q9DBR7 1 | 1.1E-05 | 0 | 1 | 2 | 6 | Q9DBR7 | Protein phosphatase 1 regulatory subunit 12A OS=Mus musculus GN=Ppp1r12a PE=1 SV=2 | Q9DBR7 [4 | 0 | 1663.81 | 0 | -1.04 | 0.18389 | 4.43 | 135 | 65 | 36.59 | 49.3 | | High | High | High | High | High | High | High | High | High | High | High | 0 | 1.6E-06 | 3.7 |
| VGSPERPLSDLGLISYR | [K].VGSPERPLSDLGLISYR.[S] | 1xPhospho | Q5SVQ0 1 | 0.00244 | 0.00011 | 1 | 8 | 1 | Q5SVQ0 | Histone acetyltransferase KAT7 OS=Mus musculus GN=Kat7 PE=1 SV=1 | Q5SVQ0 [5 | 0 | 2168.13 | 0 | -1.03 | 0.00476 | 2.72 | 131 | 69 | 8.32 | 14.46 | | High | High | High | High | High | High | High | High | High | High | High | 4.9E-05 | 0.00065 | 1.49 |
| ATENDIANFFSPLNPIR | [R].ATENDIANFFSPLNPIR.[V] | 1xPhospho | D3Z3N4 1 | 0.00041 | 0 | 1 | 2 | 1 | D3Z3N4 | Heterogeneous nuclear ribonucleoprotein H3 OS=Mus musculus GN=Hnrnph3 PE=1 SV=1 | D3Z3N4 [2 | 0 | 2228.1 | 0 | -1.01 | 0.00061 | 2.12 | 133.7 | 66.3 | 6.62 | 5.92 | | High | High | High | High | High | High | High | High | High | High | High | 0 | 8.9E-05 | 3.64 |
| IGFPETAEEELEEIASENSDSIYSS | [K].IGFPETAEEELEEIASENSDSIYSSTPEVK.[A] | 2xPhospho | Q9Z1Z2 2x | 0.02442 | 0.00161 | 1 | 1 | 2 | Q9Z1Z2 | Serine-threonine kinase receptor-associated protein OS=Mus musculus GN=Strap PE=1 SV=2 | Q9Z1Z2 [3 | 0 | 3918.78 | 0 | -1.01 | 0.08733 | 12.9 | 123.6 | 76.4 | 15.17 | 39.66 | | High | High | High | High | High | High | High | High | High | High | High | 0.0024 | 0.02576 | 2.86 |
| NQLVLIVNESPASPAK | [K].NQLVLIVNESPASPAK.[TA] | 1xDeamida | Q6P5E3 1x | 1.5E-06 | 0 | 1 | 3 | 1 | Q6P5E3 | Sal-like 1 (Drosophila) OS=Mus musculus GN=Sall1 PE=1 SV=1 | Q6P5E3 [6 | 0 | 2219.21 | 0 | -1 | 0.00052 | 2 | 134.9 | 65.1 | 12.9 | 15.1 | | High | High | High | High | High | High | High | High | High | High | High | 0 | 1.6E-07 | 3.66 |
| EGSQGADADAALELEYRPVSPLF | [R].EGSQGADADAALELEYRPVSPLPEIGHK.[Q] | 1xPhospho | I3PQW4 1 | 0.00279 | 0.00011 | 1 | 1 | 1 | I3PQW4 | Uncharacterized protein OS=Mus musculus PE=4 SV=1 | I3PQW4 [4 | 0 | 3487.75 | 0 | -0.99 | 0.00093 | 2.24 | 133.1 | 66.9 | 10.77 | 5.48 | | High | High | High | High | High | High | High | High | High | High | High | 4.9E-05 | 0.00076 | 2.93 |
| IILDLISESPIK | [K].IILDLISESPIK.[G] | 1xPhospho | Q5FWJ5 1 | 7E-07 | 0 | 2 | 14 | 2 | Q5FWJ5; C | Hnrpk protein OS=Mus musculus GN=Hnrnpk PE=2 SV=1Putative uncharacterized protein OS=Mus musculus GN=Hnrnpk PE=2 SV=1 | Q5FWJ5 [2 | 0 | 1879.1 | 0 | -0.99 | 0.00121 | 2.58 | 132.9 | 67.1 | 3.83 | 10.2 | Shared | High | High | High | High | High | High | High | High | High | High | High | 0 | 6.8E-08 | 3.93 |
| SRLSAIEIDIPVVSHST | [R].SRLSAIEIDIPVVSHST.[-] | 1xPhospho | Q9ET43 1x | 0.00051 | 0 | 1 | 1 | 1 | Q9ET43 | Claudin-12 OS=Mus musculus GN=Cldn12 PE=1 SV=2 | Q9ET43 [2 | 1 | 2133.12 | 0 | -0.99 | 0.00151 | 1.37 | 132.8 | 67.2 | 7.55 | 6.01 | | High | High | High | High | High | High | High | High | High | High | High | 0 | 0.00011 | 4.57 |
| NEPTTPSWLAEIPPWVPK | [R].NEPTTPSWLAEIPPWVPK.[D] | 1xPhospho | A2ARP8 1x | 0.00084 | 0 | 1 | 3 | 2 | A2ARP8 | Microtubule-associated protein 1A OS=Mus musculus GN=Map1a PE=1 SV=1 | A2ARP8 [2 | 0 | 2600.36 | 0 | -0.98 | 0.0113 | 3.15 | 147.3 | 52.7 | 18.42 | 38.42 | | High | High | High | High | High | High | High | High | High | High | High | 0 | 0.00033 | 2.26 |
| EASAEDLEDAACDSALFVIDK | [K].EASAEDLEDAACDSALFVIDK.[T] | 1xCarbami | Q8R2M2 1 | 2.3E-07 | 0 | 1 | 1 | 4 | Q8R2M2 | Deoxynucleotidyltransferase terminal-interacting protein 2 OS=Mus musculus GN=Dnttip2 PE=1 SV=1 | Q8R2M2 [ | 0 | 2807.31 | 0 | -0.96 | 0.00094 | 2.14 | 132.9 | 67.1 | 5.16 | 10.88 | | High | High | High | High | High | High | High | High | High | High | High | 0 | 2E-08 | 3.9 |
| TLLFGALSEIPSSSK | [K].TLLFGALSEIPSSSK.[K] | 1xPhospho | Q8BQ33 1 | 0.001 | 0 | 1 | 1 | 2 | Q8BQ33 | Treslin OS=Mus musculus GN=Ticrr PE=1 SV=2 | Q8BQ33 [1 | 0 | 2088.14 | 0 | -0.96 | 0.00851 | 9.06 | 134.6 | 65.4 | 14.21 | 13.07 | | High | High | High | High | High | High | High | High | High | High | High | 0 | 0.00024 | 3.01 |
| ILPQINPELLYQTSVSSQSFSPVK | [R].ILPQINPELLYQTSVSSQSFSPVK.[E] | 1xPhospho | F7BJB9 1x | 1.6E-06 | 0 | 1 | 1 | 3 | F7BJB9 | MORC family CW-type zinc finger protein 3 OS=Mus musculus GN=Morc3 PE=1 SV=1 | F7BJB9 [48 | 0 | 3213.72 | 0 | -0.95 | 0.00024 | 0.28 | 133 | 67 | 5.56 | 5.22 | | High | High | High | High | High | High | High | High | High | High | High | 0 | 1.7E-07 | 4.91 |
| LIPITGGNAGSPEDQHGK | [K].LIPITGGNAGSPEDQHGK.[H] | 1xPhospho | Q69ZR9-2 | 1.2E-07 | 0 | 1 | 2 | 2 | Q69ZR9-2 | Isoform 2 of Protein TASOR OS=Mus musculus GN=Fam208a | Q69ZR9-2 | 0 | 2329.2 | 0 | -0.94 | 0.00093 | 4.22 | 131.6 | 68.4 | 6.54 | 6.31 | | High | High | High | High | High | High | High | High | High | High | High | 0 | 8.3E-07 | 3.62 |
| LVSFHDDSDEDLLHI | [K].LVSFHDDSDEDLLHI.[-] | 2xPhospho | B7ZWC4 2 | 0.02068 | 0.00147 | 1 | 8 | 1 | B7ZWC4 | Insulin-like growth factor 2 receptor OS=Mus musculus GN=Igf2r PE=2 SV=1 | B7ZWC4 [2 | 0 | 2143.92 | 0 | -0.93 | 0.00541 | 3.76 | 131.3 | 68.7 | 12.34 | 13.8 | | High | High | High | High | High | High | High | High | High | High | High | 0.00079 | 0.0071 | 3.05 |
| LVATSPGSSLPGTASASSLLQGLS | [R].LVATSPGSSLPGTASASSLLQGLSFSLQDISSK | 1xPhospho | A2RSY1 1x | 7.1E-05 | 0 | 1 | 3 | 2 | A2RSY1 | KAT8 regulatory NSL complex subunit 3 OS=Mus musculus GN=Kansl3 PE=1 SV=1 | A2RSY1 [7 | 0 | 3744.97 | 0 | -0.9 | 0.01463 | 3.17 | 129.7 | 70.3 | 17.77 | 13.36 | | High | High | High | High | High | High | High | High | High | High | High | 0 | 1.9E-05 | 3.88 |
| CCSSDTNGSPLANISK | [K].CCSSDTNGSPLANISK.[S] | 2xCarbami | E9QAE4 1x | 0.00308 | 0.00016 | 1 | 2 | 1 | E9QAE4 | Histone-lysine N-methyltransferase OS=Mus musculus GN=Nsd1 PE=1 SV=1 | E9QAE4 [8 | 0 | 2249.03 | 0 | -0.89 | 0.00101 | 1.65 | 129.9 | 70.1 | 7.82 | 9.82 | | High | High | High | High | High | High | High | High | High | High | High | 7.2E-05 | 0.00084 | 2.21 |
| SPIIADLSLSLPSIPEVASDDERVD | [K].SPIIADLSLSLPSIPEVASDDERVDEVGDGGK.[ | 1xPhospho | E9Q8L9 1x | 9.2E-05 | 0 | 1 | 2 | 1 | E9Q8L9 | Rab11 family-interacting protein 1 OS=Mus musculus GN=Rab11fip1 PE=1 SV=1 | E9Q8L9 [8 | 1 | 3818.94 | 0 | -0.89 | 0.00298 | 5.37 | 130.1 | 69.9 | 10 | 13.22 | | High | High | High | High | High | High | High | High | High | High | High | 0 | 1.6E-05 | 4.19 |
| LYSLLGIDLTAPSNNSSPRDSPCK | [K].LYSLLGIDLTAPSNNSSPRDSPCK.[E] | 1xCarbami | B2RRI2 1x | 5.5E-05 | 0 | 1 | 7 | 1 | B2RRI2 | Protein strawberry notch homolog 1 OS=Mus musculus GN=Sbno1 PE=1 SV=1 | B2RRI2 [67 | 1 | 3143.58 | 0 | -0.87 | 0.003 | 4.4 | 131.2 | 68.8 | 7.93 | 10.34 | | High | High | High | High | High | High | High | High | High | High | High | 0 | 9.2E-06 | 3.2 |
| DLQYITISK | [K].DLQYITISK.[E] | 1xPhospho | E9Q9C3 1x | 0.00403 | 0.00031 | 1 | 6 | 1 | E9Q9C3 | Afadin OS=Mus musculus GN=Afdn PE=1 SV=1 | E9Q9C3 [1 | 0 | 1618.89 | 0 | -0.86 | 0.07378 | 11.95 | 129.2 | 70.8 | 4.84 | 47.12 | | High | High | High | High | High | High | High | High | High | High | High | 0.00016 | 0.00114 | 2.43 |
| LSTIDESGSILSDISFDK | [K].LSTIDESGSILSDISFDK.[T] | 2xPhospho | Q9WVM1 | 0.00011 | 0 | 1 | 3 | 2 | Q9WVM1 | Rac GTPase-activating protein 1 OS=Mus musculus GN=Racgap1 PE=1 SV=1 | Q9WVM1 | 0 | 2545.21 | 0 | -0.86 | 0.02443 | 8.97 | 137.4 | 62.6 | 19.95 | 21.27 | | High | High | High | High | High | High | High | High | High | High | High | 9.4E-05 | 0.00094 | 3.2 |
| QLSFISPPAPQPK | [K].QLSFISPPAPQPK.[T] | 1xPhospho | Q3UT19 1 | 0.00238 | 0.00011 | 1 | 2 | 1 | Q3UT19 | Putative uncharacterized protein OS=Mus musculus GN=Prpf3 PE=2 SV=1 | Q3UT19 [1 | 0 | 1948.07 | 0 | -0.86 | 0.00104 | 2.62 | 129.1 | 70.9 | 4.49 | 6.42 | | High | High | High | High | High | High | High | High | High | High | High | 4.9E-05 | 0.00063 | 1.43 |
| TFSLDTILSSYLLGQWPR | [R].TFSLDTILSSYLLGQWPR.[D] | 1xPhospho | Q7TNF9 1 | 8.6E-05 | 0 | 1 | 2 | 2 | Q7TNF9 | Protein FAM117A OS=Mus musculus GN=Fam117a PE=1 SV=1 | Q7TNF9 [8 | 0 | 2406.23 | 0 | -0.86 | 0.00149 | 5.89 | 128.4 | 71.6 | 8.25 | 9.69 | | High | High | High | High | High | High | High | High | High | High | High | 0 | 1.5E-05 | 3.23 |
| VFTPPPNSASQAGDAETLTFI | [K].VFTPPPNSASQAGDAETLTFI.[-] | 1xPhospho | Q9DB07 1 | 0.00082 | 0 | 1 | 1 | 3 | Q9DB07 | Intraflagellar transport protein 46 homolog OS=Mus musculus GN=Ift46 PE=1 SV=1 | Q9DB07 [2 | 0 | 2472.19 | 0 | -0.86 | 0.03291 | 14.89 | 128.8 | 71.2 | 14.21 | 22.3 | | High | High | High | High | High | High | High | High | High | High | High | 0 | 0.00019 | 3.25 |
| LGVDNTLSPIR | [KR].LGVDNTLSPIR.[N] | 1xPhospho | Q8C9G3 1 | 0.00042 | 0 | 1 | 6 | 2 | Q8C9G3 | Putative uncharacterized protein OS=Mus musculus GN=Orc1 PE=2 SV=1 | Q8C9G3 [3 | 0 | 1493.79 | 0 | -0.85 | 0.00049 | 3.49 | 127.5 | 72.5 | 3.92 | 5.3 | | High | High | High | High | High | High | High | High | High | High | High | 0.00033 | 0.00279 | 2.01 |
| DLLSDLQDISDSER | [R].DLLSDLQDISDSER.[K] | 2xPhospho | P24788 2x | 6E-05 | 0 | 1 | 4 | 3 | P24788 | Cyclin-dependent kinase 11B OS=Mus musculus GN=Cdk11b PE=1 SV=2 | P24788 [2 | 0 | 1994.86 | 0 | -0.84 | 0.00569 | 8.64 | 128.6 | 71.4 | 10.48 | 9.2 | | High | High | High | High | High | High | High | High | High | High | High | 0 | 3.3E-05 | 3.97 |
| CSSLEGIVDRPVNSSEK | [R].CSSLEGIVDRPVNSSEK.[K] | 1xCarbami | D3Z6J1 1x | 0.02358 | 0.00158 | 1 | 2 | 2 | D3Z6J1 | DEP domain containing 1a, isoform CRA_c OS=Mus musculus GN=Depdc1a PE=4 SV=1 | D3Z6J1 [4 | 0 | 2415.2 | 0 | -0.8 | 0.05931 | 13.16 | 121.6 | 78.4 | 13.3 | 35.18 | | High | High | High | High | High | High | High | High | High | High | High | 0.00086 | 0.0082 | 1.75 |
| LEDIFQQEFSTPSR | [K].LEDIFQQEFSTPSR.[K] | 1xPhospho | Q9EP97 1x | 7.4E-06 | 0 | 1 | 2 | 3 | Q9EP97 | Sentrin-specific protease 3 OS=Mus musculus GN=Senp3 PE=1 SV=1 | Q9EP97 [3 | 0 | 2005.95 | 0 | -0.8 | 0.00564 | 4.61 | 129.5 | 70.5 | 9.67 | 11.89 | | High | High | High | High | High | High | High | High | High | High | High | 0 | 9.6E-07 | 3.7 |
| EADLVFISVNTPTK | [R].EADLVFISVNTPTK.[T] | 1xPhospho | Q3TJE8 1x | 5.8E-06 | 0 | 1 | 6 | 2 | Q3TJE8 | UDP-glucose 6-dehydrogenase OS=Mus musculus GN=Ugdh PE=2 SV=1 | Q3TJE8 [8 | 0 | 2072.11 | 0 | -0.79 | 0.00114 | 3.64 | 126.7 | 73.3 | 4.79 | 8.78 | | High | High | High | High | High | High | High | High | High | High | High | 0 | 1.2E-05 | 3.87 |
| LSPLPSLMVVTPLAQIK | [R].LSPLPSLMVVTPLAQIK.[Q] | 1xPhospho | Q8CCN5 1 | 3.9E-05 | 0 | 1 | 7 | 2 | Q8CCN5 | Breast carcinoma-amplified sequence 3 homolog OS=Mus musculus GN=Bcas3 PE=1 SV=2 | Q8CCN5 [5 | 0 | 2345.37 | 0 | -0.79 | 0.00494 | 6.67 | 130.4 | 69.6 | 10.96 | 11.64 | | High | High | High | High | High | High | High | High | High | High | High | 0 | 6.2E-06 | 4.25 |
| ILPQINPELLYQTSVSSQSFSPVK | [R].ILPQINPELLYQTSVSSQSFSPVK.[E] | 1xDeamida | F7BJB9 1x | 6.8E-06 | 0 | 1 | 1 | 8 | F7BJB9 | MORC family CW-type zinc finger protein 3 OS=Mus musculus GN=Morc3 PE=1 SV=1 | F7BJB9 [48 | 0 | 3214.71 | 0 | -0.77 | 0.06045 | 6.86 | 115.7 | 84.3 | 26.04 | 26.21 | | High | High | High | High | High | High | High | High | High | High | High | 0 | 1.7E-06 | 3.21 |
| LQDISELLATGVGLSDSEVEPDGI | [R].LQDISELLATGVGLSDSEVEPDGEHNTTELPQ | 1xPhospho | Q91YU8 1 | 0.00609 | 0.00062 | 1 | 1 | 1 | Q91YU8 | Suppressor of SWI4 1 homolog OS=Mus musculus GN=Ppan PE=1 SV=2 | Q91YU8 [2 | 0 | 4157.01 | 0 | -0.72 | 0.01539 | 9.18 | 124.6 | 75.4 | 14.21 | 19.83 | | High | High | High | High | High | High | High | High | High | High | High | 0.00027 | 0.00179 | 3.32 |
| DPLEEIQPLNDLDESQPPAPGSP | [K].DPLEEIQPLNDLDESQPPAPGSPESR.[G] | 1xPhospho | E9Q8S5 1x | 4.6E-06 | 0 | 1 | 2 | 3 | E9Q8S5 | Microtubule-associated serine/threonine-protein kinase 3 OS=Mus musculus GN=Mast3 PE=1 SV=2 | E9Q8S5 [3 | 0 | 3139.47 | 0 | -0.71 | 0.00346 | 2.39 | 121.2 | 78.8 | 5.87 | 7.12 | | High | High | High | High | High | High | High | High | High | High | High | 0 | 5.7E-07 | 4.81 |
| LGVDNTLSPIRNGLR | [KR].LGVDNTLSPIRNGLR.[S] | 1xDeamida | Q8C9G3 1 | 0.0095 | 0.00074 | 1 | 6 | 2 | Q8C9G3 | Putative uncharacterized protein OS=Mus musculus GN=Orc1 PE=2 SV=1 | Q8C9G3 [3 | 1 | 1935.03 | 0 | -0.7 | 0.00242 | 4.85 | 123.8 | 76.2 | 6.99 | 4.66 | | High | High | High | High | High | High | High | High | High | High | High | 0.00033 | 0.00295 | 2.5 |
| ALETIPITPIER | [K].ALETIPITPIER.[N] | 1xPhospho | A2AAE1-3 | 0.00014 | 0 | 1 | 8 | 1 | A2AAE1-3 | Isoform 3 of Uncharacterized protein KIAA1109 OS=Mus musculus GN=Kiaa1109 | A2AAE1-3 | 0 | 1661.91 | 0 | -0.69 | 0.00166 | 4.87 | 124.6 | 75.4 | 5.95 | 6.04 | | High | High | High | High | High | High | High | High | High | High | High | 0 | 2.7E-05 | 2.31 |
| ANSPLESIVEETCIDLTSESPGSCE | [R].ANSPLESIVEETCIDLTSESPGSCEIK.[R] | 2xCarbami | Q9WUF3 1 | 7.2E-09 | 0 | 1 | 1 | 5 | Q9WUF3 | CASP8-associated protein 2 OS=Mus musculus GN=Casp8ap2 PE=1 SV=2 | Q9WUF3 [ | 0 | 3503.66 | 0 | -0.69 | 0.01155 | 2.9 | 123 | 77 | 10.15 | 13.98 | | High | High | High | High | High | High | High | High | High | High | High | 0 | 4E-10 | 5.39 |
| ASPHDILETIFVR | [R].ASPHDILETIFVR.[K] | 1xPhospho | Q91YI1-2 | 0.00186 | 5.7E-05 | 1 | 2 | 1 | Q91YI1-2 | Isoform 2 of Autophagy-related protein 13 OS=Mus musculus GN=Atg13 | Q91YI1-2 [ | 0 | 1806.94 | 0 | -0.69 | 0.00125 | 1.54 | 123.5 | 76.5 | 4.56 | 7.31 | | High | High | High | High | High | High | High | High | High | High | High | 2.6E-05 | 0.00048 | 2.8 |
| DNPSPEPQLDDIKR | [K].DNPSPEPQLDDIKR.[E] | 1xPhospho | Q9D4C5 1 | 0.00715 | 0.00062 | 1 | 1 | 1 | Q9D4C5 | ELL-associated factor 1 OS=Mus musculus GN=Eaf1 PE=1 SV=2 | Q9D4C5 [1 | 1 | 2162.09 | 0 | -0.69 | 0.01055 | 8.3 | 129 | 71 | 11.37 | 16.21 | | High | High | High | High | High | High | High | High | High | High | High | 0.00027 | 0.00216 | 2.14 |
| SLSPIIGK | [R].SLSPIIGK.[D] | 1xPhospho | A2AUK5 1 | 0.02442 | 0.00161 | 2 | 8 | 1 | A2AUK5; C | Band 4.1-like protein 1 OS=Mus musculus GN=Epb41l1 PE=1 SV=1Band 4.1-like protein 1 OS=Mus musculus GN=Epb41l1 PE=1 SV=2 | A2AUK5 [7 | 0 | 1352.8 | 0 | -0.69 | 0.00099 | 5.91 | 123.2 | 76.8 | 3.4 | 8.87 | Shared | High | High | High | High | High | High | High | High | High | High | High | 0.00086 | 0.00855 | 1.32 |
| LSPLPSLMVVTPLAQIK | [R].LSPLPSLMVVTPLAQIK.[Q] | 1xPhospho | Q8CCN5 1 | 0.0013 | 0 | 1 | 7 | 1 | Q8CCN5 | Breast carcinoma-amplified sequence 3 homolog OS=Mus musculus GN=Bcas3 PE=1 SV=2 | Q8CCN5 [5 | 0 | 2361.36 | 0 | -0.67 | 0.05712 | 6.86 | 124.8 | 75.2 | 16.88 | 15.16 | | High | High | High | High | High | High | High | High | High | High | High | 0 | 0.00032 | 2.86 |
| DLPAIPGVTSPTSDEPPMQASQ | [K].DLPAIPGVTSPTSDEPPMQASQSQLPSSPEDI | 1xPhospho | Q3TII9 1xP | 9.7E-06 | 0 | 1 | 2 | 7 | Q3TII9 | Putative uncharacterized protein OS=Mus musculus GN=Eif4ebp1 PE=2 SV=1 | Q3TII9 [73 | 0 | 3860.86 | 0 | -0.66 | 0.32905 | 6.77 | 117.3 | 82.7 | 12.09 | 38.69 | | High | High | High | High | High | High | High | High | High | High | High | 0 | 1.8E-05 | 3.67 |
| ELGNIHETSQDLSPEK | [R].ELGNIHETSQDLSPEK.[A] | 2xPhospho | A0A0R4J0 | 0.00177 | 5.7E-05 | 1 | 6 | 1 | A0A0R4J0 | Disks large-associated protein 5 OS=Mus musculus GN=Dlgap5 PE=1 SV=1 | A0A0R4J0 | 0 | 2415.12 | 0 | -0.66 | 0.00386 | 5.32 | 122.6 | 77.4 | 3.36 | 11.55 | | High | High | High | High | High | High | High | High | High | High | High | 2.6E-05 | 0.00045 | 1.6 |
| LELGLHEGSLPTSPLGSIATR | [K].LELGLHEGSLPTSPLGSIATR.[F] | 2xPhospho | U5KVR9 2x | 0.00389 | 0.00026 | 1 | 2 | 1 | U5KVR9 | Centrosomal protein 63 OS=Mus musculus GN=Cep63 PE=1 SV=1 | U5KVR9 [5 | 0 | 2537.26 | 0 | -0.66 | 0.0031 | 2.29 | 118.4 | 81.6 | 7.91 | 7.33 | | High | High | High | High | High | High | High | High | High | High | High | 0.00014 | 0.00109 | 2.86 |
| RVSLSEIGFGK | [R].RVSLSEIGFGK.[L] | 1xPhospho | Q3UPI3 1x | 0.0076 | 0.00066 | 1 | 4 | 1 | Q3UPI3 | Putative uncharacterized protein OS=Mus musculus GN=Cdk16 PE=2 SV=1 | Q3UPI3 [1 | 1 | 1730.96 | 0 | -0.66 | 0.00189 | 3.26 | 124.9 | 75.1 | 6.86 | 7.92 | | High | High | High | High | High | High | High | High | High | High | High | 0.00029 | 0.00231 | 2.96 |
| DNPSPEPQLDDIK | [K].DNPSPEPQLDDIK.[R] | 1xPhospho | Q9D4C5 1 | 2.9E-06 | 0 | 1 | 1 | 4 | Q9D4C5 | ELL-associated factor 1 OS=Mus musculus GN=Eaf1 PE=1 SV=2 | Q9D4C5 [1 | 0 | 2005.99 | 0 | -0.64 | 0.00024 | 1.21 | 122.2 | 77.8 | 1.98 | 2.15 | | High | High | High | High | High | High | High | High | High | High | High | 0 | 1.7E-06 | 2.16 |
| VASSVEQLNIIEDEVSQPLAAPPS | [R].VASSVEQLNIIEDEVSQPLAAPPSK.[I] | 1xPhospho | Q6ZPQ6 1 | 0.0276 | 0.00172 | 1 | 3 | 2 | Q6ZPQ6 | Membrane-associated phosphatidylinositol transfer protein 2 OS=Mus musculus GN=Pitpnm2 PE=1 SV=2 | Q6ZPQ6 [3 | 0 | 3159.66 | 0 | -0.6 | 0.04369 | 13.02 | 123.2 | 76.8 | 19.87 | 16.33 | | High | High | High | High | High | High | High | High | High | High | High | 0.001 | 0.01042 | 2.72 |
| APPGAPGPGPGSGAPGSQEEEE | [R].APPGAPGPGPGSGAPGSQEEEEEPGLVEADF | 2xPhospho | Q8CCS6 2x | 0.00038 | 0 | 1 | 3 | 1 | Q8CCS6 | Polyadenylate-binding protein 2 OS=Mus musculus GN=Pabpn1 PE=1 SV=3 | Q8CCS6 [7 | 0 | 4938.25 | 0 | -0.59 | 0.1363 | 2.79 | 117.7 | 82.3 | 17.09 | 14.99 | | High | High | High | High | High | High | High | High | High | High | High | 0 | 8E-05 | 4.27 |
| LLDLVVHPVPQPSQCLEFIQQSP | [R].LLDLVVHPVPQPSQCLEFIQQSPTR.[S] | 1xCarbami | Q8CJF7 1x | 5.1E-06 | 0 | 1 | 1 | 1 | Q8CJF7 | Protein ELYS OS=Mus musculus GN=Ahctf1 PE=1 SV=1 | Q8CJF7 [1 | 0 | 3210.66 | 0 | -0.59 | 0.001 | 2.8 | 120.1 | 79.9 | 3.44 | 6.05 | | High | High | High | High | High | High | High | High | High | High | High | 0 | 6.4E-07 | 3.57 |
| ADQSISEEPGWEEEEEELEGIVPS | [R].ADQSISEEPGWEEEEEELEGIVPSPK.[E] | 1xPhospho | Q80Y55 1 | 0.002 | 5.7E-05 | 1 | 1 | 3 | Q80Y55 | BSD domain-containing protein 1 OS=Mus musculus GN=Bsdc1 PE=1 SV=1 | Q80Y55 [2 | 0 | 3451.61 | 0 | -0.58 | 0.03559 | 7.47 | 118.8 | 81.2 | 11.41 | 21.01 | | High | High | High | High | High | High | High | High | High | High | High | 2.6E-05 | 0.00052 | 3.21 |
| AEDPEGTTEQETGGSSAPEPLEV | [K].AEDPEGTTEQETGGSSAPEPLEVIK.[E] | 1xPhospho | G3X961 1x | 4.7E-05 | 0 | 1 | 5 | 2 | G3X961 | Kelch domain containing 4, isoform CRA_a OS=Mus musculus GN=Klhdc4 PE=1 SV=1 | G3X961 [3 | 0 | 3109.49 | 0 | -0.58 | 0.01576 | 9.04 | 120 | 80 | 14.13 | 10.63 | | High | High | High | High | High | High | High | High | High | High | High | 0 | 7.7E-06 | 3.61 |
| HLPSPPTLDSIITEYLR | [K].HLPSPPTLDSIITEYLR.[E] | 1xPhospho | Q80TR8-2 | 1.2E-05 | 0 | 1 | 3 | 2 | Q80TR8-2 | Isoform 2 of DDB1- and CUL4-associated factor 1 OS=Mus musculus GN=Dcaf1 | Q80TR8-2 | 0 | 2261.18 | 0 | -0.57 | 0.0107 | 2.35 | 121.2 | 78.8 | 6.94 | 10.9 | | High | High | High | High | High | High | High | High | High | High | High | 0 | 1.6E-06 | 4.27 |
| RASLSEIGFGK | [R].RASLSEIGFGK.[M] | 1xPhospho | Q8K0D0 1 | 4.1E-06 | 0 | 1 | 4 | 1 | Q8K0D0 | Cyclin-dependent kinase 17 OS=Mus musculus GN=Cdk17 PE=1 SV=2 | Q8K0D0 [1 | 1 | 1702.93 | 0 | -0.57 | 0.01928 | 9.05 | 119.4 | 80.6 | 4.97 | 17.48 | | High | High | High | High | High | High | High | High | High | High | High | 0 | 5E-07 | 3.34 |
| GTGLQPGEEELPDIAPPLVTPDEI | [R].GTGLQPGEEELPDIAPPLVTPDEPK.[GA] | 1xPhospho | P56399 1x | 6.3E-07 | 0 | 1 | 2 | 2 | P56399 | Ubiquitin carboxyl-terminal hydrolase 5 OS=Mus musculus GN=Usp5 PE=1 SV=1 | P56399 [6 | 0 | 3137.61 | 0 | -0.55 | 0.02453 | 8.08 | 124.2 | 75.8 | 25.27 | 12.07 | | High | High | High | High | High | High | High | High | High | High | High | 0 | 6.1E-08 | 3.8 |
| RVSLSSQLNDIK | [K].RVSLSSQLNDIK.[I] | 1xPhospho | Z4YKQ9 1x | 0.0006 | 0 | 1 | 4 | 1 | Z4YKQ9 | ATPase family AAA domain-containing protein 5 OS=Mus musculus GN=Atad5 PE=1 SV=1 | Z4YKQ9 [1 | 1 | 1898.05 | 0 | -0.54 | 0.00143 | 1.04 | 120 | 80 | 2.68 | 8.06 | | High | High | High | High | High | High | High | High | High | High | High | 0 | 0.00014 | 1.77 |
| STSSAMGGSHQDLSVIQPIVK | [R].STSSAMGGSHQDLSVIQPIVK.[D] | 1xPhospho | A0A1W2P | 0.00022 | 0 | 1 | 2 | 2 | A0A1W2P | RAB3A interacting protein, isoform CRA_b OS=Mus musculus GN=Rab3ip PE=1 SV=1 | A0A1W2P | 0 | 2696.37 | 0 | -0.54 | 0.1852 | 3.17 | 118.4 | 81.6 | 9.21 | 27.75 | | High | High | High | High | High | High | High | High | High | High | High | 0 | 4.4E-05 | 2.99 |
| ASFETLPNISDLCLK | [R].ASFETLPNISDLCLK.[D] | 1xCarbami | Q9CWE0 1 | 0.00058 | 0 | 1 | 5 | 2 | Q9CWE0 | Mitochondrial fission regulator 1-like OS=Mus musculus GN=Mtfr1l PE=1 SV=1 | Q9CWE0 [ | 0 | 2246.15 | 0 | -0.53 | 0.00841 | 8.44 | 117.8 | 82.2 | 8.77 | 5.26 | | High | High | High | High | High | High | High | High | High | High | High | 0 | 0.00013 | 3.51 |
| DLLSDLQDISDSER | [R].DLLSDLQDISDSER.[K] | 1xPhospho | P24788 1x | 2.4E-07 | 0 | 1 | 4 | 6 | P24788 | Cyclin-dependent kinase 11B OS=Mus musculus GN=Cdk11b PE=1 SV=2 | P24788 [2 | 0 | 1914.89 | 0 | -0.51 | 0.01125 | 7.15 | 118.1 | 81.9 | 9.15 | 4.98 | | High | High | High | High | High | High | High | High | High | High | High | 0 | 4.2E-07 | 3.78 |
| GPRTPSPPPPILEDIILGK | [K].GPRTPSPPPPILEDIILGK.[K] | 2xPhospho | E9Q784 2x | 0.00019 | 0 | 1 | 3 | 5 | E9Q784 | Zinc finger CCCH domain-containing protein 13 OS=Mus musculus GN=Zc3h13 PE=1 SV=1 | E9Q784 [2 | 1 | 2615.4 | 0 | -0.5 | 0.00162 | 4.65 | 119.9 | 80.1 | 7.27 | 5.68 | | High | High | High | High | High | High | High | High | High | High | High | 0 | 3.6E-05 | 3.64 |
| RASSCSLAVISPFLVEK | [R].RASSCSLAVISPFLVEK.[G] | 1xCarbami | P58801 1x | 2.2E-05 | 0 | 1 | 1 | 2 | P58801 | Receptor-interacting serine/threonine-protein kinase 2 OS=Mus musculus GN=Ripk2 PE=1 SV=1 | P58801 [4 | 1 | 2402.29 | 0 | -0.49 | 0.00923 | 8.3 | 118.2 | 81.8 | 14.4 | 13.04 | | High | High | High | High | High | High | High | High | High | High | High | 0 | 5.9E-06 | 4.78 |
| RNSLPQIPTLNLEAR | [K].RNSLPQIPTLNLEAR.[R] | 1xPhospho | Q9QVP9 1 | 6.5E-06 | 0 | 1 | 6 | 1 | Q9QVP9 | Protein-tyrosine kinase 2-beta OS=Mus musculus GN=Ptk2b PE=1 SV=2 | Q9QVP9 [3 | 1 | 2031.09 | 0 | -0.48 | 0.00695 | 3.57 | 117.4 | 82.6 | 10.63 | 4.4 | | High | High | High | High | High | High | High | High | High | High | High | 0 | 8.4E-07 | 3.76 |
| RPSLPALHFIK | [R].RPSLPALHFIK.[G] | 1xPhospho | Q3SYK5 1x | 0.06595 | 0.00477 | 1 | 4 | 1 | Q3SYK5 | Tyrosine-protein kinase OS=Mus musculus GN=Abl1 PE=2 SV=1 | Q3SYK5 [1 | 0 | 1817.06 | 0 | -0.48 | 0.16364 | 15.78 | 115 | 85 | 12.64 | 25.1 | | High | High | High | High | High | High | High | High | High | High | High | 0.00246 | 0.02653 | 2.03 |
| AASTAPSSTSTPAASSAGLIYIDPS | [R].AASTAPSSTSTPAASSAGLIYIDPSNLR.[R] | 1xPhospho | E9Q2H1 1 | 0.00131 | 0 | 1 | 2 | 2 | E9Q2H1 | E3 ubiquitin-protein ligase UBR5 OS=Mus musculus GN=Ubr5 PE=1 SV=1 | E9Q2H1 [1 | 0 | 3015.49 | 0 | -0.47 | 0.00832 | 3.98 | 116.3 | 83.7 | 8.85 | 4.15 | | High | High | High | High | High | High | High | High | High | High | High | 0 | 0.00032 | 3.15 |
| GALSPITDTTEVDTGIHVPSEDIPI | [K].GALSPITDTTEVDTGIHVPSEDIPETMDEDSSL | 1xPhospho | H7BX08 1x | 0.04003 | 0.00287 | 1 | 5 | 3 | H7BX08 | Calmodulin-regulated spectrin-associated protein 2 OS=Mus musculus GN=Camsap2 PE=1 SV=1 | H7BX08 [5 | 0 | 3936.81 | 0 | -0.45 | 0.06325 | 9.88 | 116.1 | 83.9 | 16.41 | 13.91 | | High | High | High | High | High | High | High | High | High | High | High | 0.00444 | 0.04793 | 3.85 |
| VSTSPVGLPSIDPSGNSPSAAAPI | [R].VSTSPVGLPSIDPSGNSPSAAAPLTSFSGIPGT | 1xPhospho | E9PUR0 1x | 6.2E-06 | 0 | 1 | 7 | 5 | E9PUR0 | Ankyrin repeat and KH domain-containing 1 OS=Mus musculus GN=Ankhd1 PE=1 SV=1 | E9PUR0 [2 | 0 | 3533.77 | 0 | -0.45 | 0.02566 | 4.6 | 110 | 90 | 10.44 | 5.84 | | High | High | High | High | High | High | High | High | High | High | High | 0 | 1.5E-06 | 5.4 |
| ADLSEPSVEFGTNDNLGEIPLYPA | [K].ADLSEPSVEFGTNDNLGEIPLYPAHEAYWGT | 1xDeamida | F8VQG7 1 | 0.00025 | 0 | 1 | 4 | 1 | F8VQG7 | NEDD4-binding protein 2 OS=Mus musculus GN=N4bp2 PE=1 SV=1 | F8VQG7 [8 | 0 | 4615.18 | 0 | -0.44 | 0.04625 | 3.49 | 116.1 | 83.9 | 16.51 | 14.81 | | High | High | High | High | High | High | High | High | High | High | High | 0 | 5.1E-05 | 4.4 |
| FDGSETRPFSPSPLNNISSTVTVR | [K].FDGSETRPFSPSPLNNISSTVTVR.[N] | 1xPhospho | Q6PR54-3 | 0.00014 | 0 | 1 | 3 | 2 | Q6PR54-3 | Isoform 3 of Telomere-associated protein RIF1 OS=Mus musculus GN=Rif1 | Q6PR54-3 | 0 | 2917.43 | 0 | -0.44 | 0.00293 | 3.48 | 114.6 | 85.4 | 8.03 | 3.05 | | High | High | High | High | High | High | High | High | High | High | High | 0 | 2.6E-05 | 2.32 |
| HFGSGDLMDPELLGLTYIK | [K].HFGSGDLMDPELLGLTYIK.[G] | 1xPhospho | B9EK76 1x | 0.00033 | 0 | 1 | 5 | 1 | B9EK76 | Signal-induced proliferation-associated 1 like 2 OS=Mus musculus GN=Sipa1l2 PE=2 SV=1 | B9EK76 [1 | 0 | 2644.35 | 0 | -0.44 | 0.04581 | 4.69 | 116.3 | 83.7 | 10.02 | 14.9 | | High | High | High | High | High | High | High | High | High | High | High | 0 | 6.9E-05 | 2.65 |
| HFGSGDLMDPELLGLTYIK | [K].HFGSGDLMDPELLGLTYIK.[G] | 1xPhospho | B9EK76 1x | 0.00014 | 0 | 1 | 5 | 1 | B9EK76 | Signal-induced proliferation-associated 1 like 2 OS=Mus musculus GN=Sipa1l2 PE=2 SV=1 | B9EK76 [1 | 0 | 2660.34 | 0 | -0.44 | 0.02578 | 5.2 | 115.9 | 84.1 | 7.17 | 30.79 | | High | High | High | High | High | High | High | High | High | High | High | 0 | 2.7E-05 | 2.97 |
| EEPPEPVPSPIIPILPSISGK | [K].EEPPEPVPSPIIPILPSISGK.[N] | 1xPhospho | F8WI37 1x | 0.00231 | 0.00011 | 1 | 6 | 1 | F8WI37 | Histone-lysine N-methyltransferase OS=Mus musculus GN=Kmt2c PE=1 SV=1 | F8WI37 [4 | 0 | 2734.51 | 0 | -0.41 | 0.00915 | 7.84 | 114.4 | 85.6 | 3.02 | 9.1 | | High | High | High | High | High | High | High | High | High | High | High | 4.9E-05 | 0.00061 | 2.95 |
| LELGLHEGSLPTSPLGSIATR | [K].LELGLHEGSLPTSPLGSIATR.[F] | 1xPhospho | U5KVR9 1x | 0.00034 | 0 | 1 | 2 | 3 | U5KVR9 | Centrosomal protein 63 OS=Mus musculus GN=Cep63 PE=1 SV=1 | U5KVR9 [5 | 0 | 2457.3 | 0 | -0.39 | 0.03345 | 4.68 | 116.8 | 83.2 | 12.08 | 12.09 | | High | High | High | High | High | High | High | High | High | High | High | 0 | 7.2E-05 | 2.63 |
| LSGIMVPAPIQDLEALR | [K].LSGIMVPAPIQDLEALR.[A] | 1xPhospho | P28658 1x | 0.08378 | 0.00634 | 1 | 4 | 1 | P28658 | Ataxin-10 OS=Mus musculus GN=Atxn10 PE=1 SV=2 | P28658 [1 | 0 | 2148.13 | 0 | -0.39 | 0.16925 | 3.75 | 114.1 | 85.9 | 17.97 | 6.84 | | High | High | High | High | High | High | High | High | High | High | High | 0.00327 | 0.03507 | 2.63 |
| SLEEIYLFSLPIK | [K].SLEEIYLFSLPIK.[E] | 1xPhospho | P25444 1x | 0.01307 | 0.00089 | 1 | 14 | 1 | P25444 | 40S ribosomal protein S2 OS=Mus musculus GN=Rps2 PE=1 SV=3 | P25444 [7 | 0 | 2090.16 | 0 | -0.37 | 0.09007 | 11.61 | 112.7 | 87.3 | 13.09 | 12.04 | | High | High | High | High | High | High | High | High | High | High | High | 0.00047 | 0.00422 | 2.28 |
| SQSSHSYDDSTLPLIDR | [R].SQSSHSYDDSTLPLIDR.[N] | 1xPhospho | E9Q8Z8 1x | 1E-05 | 0 | 1 | 19 | 3 | E9Q8Z8 | Catenin delta-1 OS=Mus musculus GN=Ctnnd1 PE=1 SV=1 | E9Q8Z8 [8 | 0 | 2230.02 | 0 | -0.37 | 0.17897 | 7.77 | 111.6 | 88.4 | 12.67 | 19.81 | | High | High | High | High | High | High | High | High | High | High | High | 0 | 5.1E-05 | 3.66 |
| DSSAGSQSLTLGCDNLPGPGSPI | [K].DSSAGSQSLTLGCDNLPGPGSPLDGILTSETG | 1xCarbami | Q8VCI7 1x | 0.00267 | 0.00011 | 1 | 1 | 2 | Q8VCI7 | ATPase SWSAP1 OS=Mus musculus GN=Swsap1 PE=2 SV=2 | Q8VCI7 [2 | 0 | 4042.92 | 0 | -0.35 | 0.30575 | 11.44 | 109.5 | 90.5 | 8.92 | 14.84 | | High | High | High | High | High | High | High | High | High | High | High | 0.00058 | 0.0055 | 3.99 |
| HLPSPPLTQEMDLHSIAGPPVPF | [R].HLPSPPLTQEMDLHSIAGPPVPPR.[Q] | 1xPhospho | Q62245 1x | 0.00028 | 0 | 1 | 2 | 1 | Q62245 | Son of sevenless homolog 1 OS=Mus musculus GN=Sos1 PE=1 SV=2 | Q62245 [1 | 0 | 2911.47 | 0 | -0.35 | 0.39588 | 1.24 | 107 | 93 | 10.07 | 22.2 | | High | High | High | High | High | High | High | High | High | High | High | 0 | 5.7E-05 | 2.77 |
| IGFPETAEEELEEIASENSDSIYSS | [K].IGFPETAEEELEEIASENSDSIYSSTPEVK.[A] | 1xPhospho | Q9Z1Z2 1x | 8.1E-09 | 0 | 1 | 1 | 16 | Q9Z1Z2 | Serine-threonine kinase receptor-associated protein OS=Mus musculus GN=Strap PE=1 SV=2 | Q9Z1Z2 [3 | 0 | 3838.81 | 0 | -0.35 | 0.01958 | 6.61 | 108.7 | 91.3 | 6.43 | 7.66 | | High | High | High | High | High | High | High | High | High | High | High | 0 | 4.5E-10 | 4.89 |
| EISPLAAITPFDFK | [K].EISPLAAITPFDFK.[S] | 1xPhospho | L7N209 1x | 1.4E-05 | 0 | 1 | 2 | 1 | L7N209 | HBS1-like protein OS=Mus musculus GN=Hbs1l PE=1 SV=1 | L7N209 [5 | 0 | 2087.12 | 0 | -0.34 | 0.06916 | 13.2 | 111.8 | 88.2 | 6.88 | 12.22 | | High | High | High | High | High | High | High | High | High | High | High | 0 | 2E-06 | 2.93 |
| GSDSPEDLGAITMLNQQETVAM | [K].GSDSPEDLGAITMLNQQETVAMDMDR.[N] | 1xPhospho | D0VLQ4 1 | 0.11833 | 0.00962 | 1 | 3 | 1 | D0VLQ4 | Tet oncogene 1 OS=Mus musculus GN=Tet1 PE=2 SV=1 | D0VLQ4 [3 | 0 | 3149.37 | 0 | -0.34 | 0.15554 | 6.11 | 110.2 | 89.8 | 4.57 | 19.39 | | High | High | High | High | High | High | High | High | High | High | High | 0.00486 | 0.05261 | 2.34 |
| NKPLSPIK | [K].NKPLSPIK.[L] | 1xPhospho | G3UWX1 1 | 0.005 | 0.0005 | 2 | 9 | 2 | G3UWX1; | Replication factor C subunit 1 OS=Mus musculus GN=Rfc1 PE=1 SV=2Replication factor C subunit 1 OS=Mus musculus GN=Rfc1 PE=1 SV=2 | G3UWX1 [ | 0 | 1664.01 | 0 | -0.34 | 0.00837 | 7.11 | 111.9 | 88.1 | 3.74 | 7.66 | Shared | High | High | High | High | High | High | High | High | High | High | High | 0.00269 | 0.02878 | 2.11 |
| TPLATIADQQQLQLSPLK | [R].TPLATIADQQQLQLSPLK.[R] | 1xPhospho | P11157 1x | 6.9E-07 | 0 | 1 | 2 | 4 | P11157 | Ribonucleoside-diphosphate reductase subunit M2 OS=Mus musculus GN=Rrm2 PE=1 SV=1 | P11157 [6 | 0 | 2503.39 | 0 | -0.34 | 0.02089 | 4.87 | 111.4 | 88.6 | 3.6 | 7.69 | | High | High | High | High | High | High | High | High | High | High | High | 0 | 6.7E-08 | 3.51 |
| EGGIDVSDSDDEDDNLPAIFDTC | [K].EGGIDVSDSDDEDDNLPAIFDTCHR.[A] | 1xCarbami | B2RS91 2x | 8.5E-06 | 0 | 1 | 1 | 4 | B2RS91 | RNA polymerase I-specific transcription initiation factor RRN3 OS=Mus musculus GN=Rrn3 PE=1 SV=1 | B2RS91 [1 | 0 | 3181.25 | 0 | -0.33 | 0.02077 | 10.83 | 116 | 84 | 6.52 | 10.38 | | High | High | High | High | High | High | High | High | High | High | High | 0 | 1.1E-06 | 3.49 |
| GKLEAIITPPPAK | [K].GKLEAIITPPPAK.[K] | 1xPhospho | Q9QZQ8 1 | 0.0166 | 0.00112 | 1 | 3 | 1 | Q9QZQ8 | Core histone macro-H2A.1 OS=Mus musculus GN=H2afy PE=1 SV=3 | Q9QZQ8 [ | 1 | 2102.26 | 0 | -0.33 | 0.19716 | 33.85 | 114.4 | 85.6 | 27.92 | 25.08 | | High | High | High | High | High | High | High | High | High | High | High | 0.00058 | 0.00552 | 2.52 |
| RLSPIMLGK | [R].RLSPIMLGK.[Q] | 1xPhospho | Q9CQT7 1 | 0.05769 | 0.00423 | 1 | 3 | 1 | Q9CQT7 | Desumoylating isopeptidase 1 OS=Mus musculus GN=Desi1 PE=1 SV=1 | Q9CQT7 [2 | 1 | 1568.9 | 0 | -0.33 | 0.05879 | 12.56 | 111.3 | 88.7 | 9.1 | 6.32 | | High | High | High | High | High | High | High | High | High | High | High | 0.00216 | 0.02265 | 1.24 |
| AGSPQLDDIR | [K].AGSPQLDDIR.[V] | 1xPhospho | Q8CHW4 1 | 0.00437 | 0.00041 | 1 | 1 | 7 | Q8CHW4 | Translation initiation factor eIF-2B subunit epsilon OS=Mus musculus GN=Eif2b5 PE=1 SV=1 | Q8CHW4 [ | 0 | 1380.67 | 0 | -0.32 | 0.01133 | 5.63 | 110.2 | 89.8 | 1.38 | 8.82 | | High | High | High | High | High | High | High | High | High | High | High | 0.00168 | 0.01684 | 2.22 |
| DEPQTVPDVPSFGDSPPLSPIDM | [K].DEPQTVPDVPSFGDSPPLSPIDMDTQER.[I] | 1xPhospho | P15066 1x | 2.5E-05 | 0 | 1 | 2 | 2 | P15066 | Transcription factor jun-D OS=Mus musculus GN=Jund PE=1 SV=1 | P15066 [2 | 0 | 3378.53 | 0 | -0.32 | 0.12841 | 15.21 | 111.2 | 88.8 | 6.82 | 21.13 | | High | High | High | High | High | High | High | High | High | High | High | 0 | 3.8E-06 | 3.72 |
| DEPQTVPDVPSFGDSPPLSPIDM | [K].DEPQTVPDVPSFGDSPPLSPIDMDTQER.[I] | 1xPhospho | P15066 1x | 0.01115 | 0.00077 | 1 | 2 | 3 | P15066 | Transcription factor jun-D OS=Mus musculus GN=Jund PE=1 SV=1 | P15066 [2 | 0 | 3394.52 | 0 | -0.32 | 0.43656 | 5.86 | 111.3 | 88.7 | 9.21 | 22.25 | | High | High | High | High | High | High | High | High | High | High | High | 0.00057 | 0.00507 | 3.15 |
| GSAEGTSDLEQEGLGEIEPTEIR | [K].GSAEGTSDLEQEGLGEIEPTEIR.[G] | 1xPhospho | A6H619 1x | 2.2E-06 | 0 | 1 | 2 | 8 | A6H619 | PHD and RING finger domain-containing protein 1 OS=Mus musculus GN=Phrf1 PE=1 SV=2 | A6H619 [9 | 0 | 2726.26 | 0 | -0.32 | 0.04652 | 1.04 | 109.4 | 90.6 | 9.28 | 5.1 | | High | High | High | High | High | High | High | High | High | High | High | 0 | 3E-07 | 4.06 |
| EDLSPICDHSQNR | [K].EDLSPICDHSQNR.[V] | 1xCarbami | Q06180 1x | 0.01719 | 0.0012 | 1 | 2 | 3 | Q06180 | Tyrosine-protein phosphatase non-receptor type 2 OS=Mus musculus GN=Ptpn2 PE=1 SV=2 | Q06180 [2 | 0 | 1879.82 | 0 | -0.31 | 0.00416 | 3.35 | 111.5 | 88.5 | 4.25 | 3.92 | | High | High | High | High | High | High | High | High | High | High | High | 0.00085 | 0.00798 | 2.52 |
| SQELEPIPDSLDDSETISER | [K].SQELEPIPDSLDDSETISER.[L] | 1xPhospho | Q62187 1x | 0.0016 | 5.7E-05 | 1 | 2 | 3 | Q62187 | Transcription termination factor 1 OS=Mus musculus GN=Ttf1 PE=1 SV=2 | Q62187 [3 | 0 | 2569.18 | 0 | -0.31 | 0.00636 | 0.75 | 110.8 | 89.2 | 7.22 | 7.07 | | High | High | High | High | High | High | High | High | High | High | High | 0.0002 | 0.00134 | 3.82 |
| AVEYLLTGIPGSPEPEHGSVQES | [R].AVEYLLTGIPGSPEPEHGSVQESQAPEQPATE | 1xDeamida | P54726 1x | 2.7E-05 | 0 | 1 | 3 | 1 | P54726 | UV excision repair protein RAD23 homolog A OS=Mus musculus GN=Rad23a PE=1 SV=2 | P54726 [1 | 0 | 4855.32 | 0 | -0.3 | 0.187 | 6.52 | 109.6 | 90.4 | 10.97 | 14.07 | | High | High | High | High | High | High | High | High | High | High | High | 0 | 4.1E-06 | 5.25 |
| DLPAIPGVTSPTSDEPPMQASQ | [K].DLPAIPGVTSPTSDEPPMQASQSQLPSSPEDI | 2xPhospho | Q3TII9 2xP | 0.01296 | 0.00089 | 1 | 2 | 2 | Q3TII9 | Putative uncharacterized protein OS=Mus musculus GN=Eif4ebp1 PE=2 SV=1 | Q3TII9 [73 | 0 | 3924.83 | 0 | -0.3 | 0.18329 | 18.91 | 119.8 | 80.2 | 20.55 | 32.95 | | High | High | High | High | High | High | High | High | High | High | High | 0.00045 | 0.00419 | 3.03 |
| TLSDISNASSSFGWLSLDGDPTN | [R].TLSDISNASSSFGWLSLDGDPTNFNEGSQVP | 1xDeamida | P22682 1x | 0.0002 | 0 | 1 | 5 | 2 | P22682 | E3 ubiquitin-protein ligase CBL OS=Mus musculus GN=Cbl PE=1 SV=3 | P22682 [7 | 0 | 4886.37 | 0 | -0.3 | 0.96183 | 15.43 | 100.5 | 99.5 | 15.65 | 25.05 | | High | High | High | High | High | High | High | High | High | High | High | 0 | 4E-05 | 4.99 |
| TPSSCSSSLDSIK | [R].TPSSCSSSLDSIK.[S] | 1xCarbami | A3KGI3 1x | 0.00659 | 0.00062 | 1 | 1 | 4 | A3KGI3 | Kinetochore scaffold 1 OS=Mus musculus GN=Knl1 PE=1 SV=1 | A3KGI3 [1 | 0 | 1906.92 | 0 | -0.3 | 0.01015 | 6.62 | 108.7 | 91.3 | 5.62 | 3.65 | | High | High | High | High | High | High | High | High | High | High | High | 0.00027 | 0.00196 | 3.21 |
| AGFDSDYDQALADIR | [K].AGFDSDYDQALADIR.[E] | 1xPhospho | P54276 1x | 9.7E-05 | 0 | 1 | 3 | 3 | P54276 | DNA mismatch repair protein Msh6 OS=Mus musculus GN=Msh6 PE=1 SV=3 | P54276 [9 | 0 | 1965.88 | 0 | -0.29 | 0.02168 | 7.23 | 109.9 | 90.1 | 4.98 | 5.63 | | High | High | High | High | High | High | High | High | High | High | High | 0 | 1.7E-05 | 3.49 |
| ELQPIVYTPEDCK | [K].ELQPIVYTPEDCK.[K] | 1xCarbami | Q91WD7 1 | 0.00043 | 0 | 1 | 1 | 1 | Q91WD7 | Kinesin-like protein KIF18A OS=Mus musculus GN=Kif18a PE=2 SV=1 | Q91WD7 [ | 0 | 2130.06 | 0 | -0.27 | 0.1771 | 11.23 | 105.1 | 94.9 | 12.65 | 11.81 | | High | High | High | High | High | High | High | High | High | High | High | 0 | 9.3E-05 | 3.48 |
| SSPAEPELNTIVIPR | [R].SSPAEPELNTIVIPR.[K] | 1xPhospho | B7ZWN9 1 | 0.00022 | 0 | 2 | 6 | 2 | B7ZWN9; | SUMO/sentrin specific peptidase 6, isoform CRA_a OS=Mus musculus GN=Senp6 PE=2 SV=1Sentrin-specific protease 6 OS=Mus musculus GN=Senp6 PE=1 SV=2 | B7ZWN9 [ | 0 | 1932 | 0 | -0.26 | 0.02711 | 6.98 | 107.7 | 92.3 | 5.47 | 5.97 | Shared | High | High | High | High | High | High | High | High | High | High | High | 0 | 4.3E-05 | 2.99 |
| RCSSSLPPIQAPSHPPPQPPTQPI | [R].RCSSSLPPIQAPSHPPPQPPTQPR.[L] | 1xCarbami | Q3UIA2 1x | 0.00314 | 0.00016 | 1 | 5 | 1 | Q3UIA2 | Rho GTPase-activating protein 17 OS=Mus musculus GN=Arhgap17 PE=1 SV=1 | Q3UIA2 [6 | 1 | 2941.47 | 0 | -0.25 | 0.30177 | 12.38 | 108.8 | 91.2 | 6.88 | 19.29 | | High | High | High | High | High | High | High | High | High | High | High | 7.2E-05 | 0.00086 | 2.39 |
| TASPLEHILQTLFGK | [K].TASPLEHILQTLFGK.[K] | 1xPhospho | Q8C9B9 1x | 0.01574 | 0.00112 | 1 | 1 | 1 | Q8C9B9 | Death-inducer obliterator 1 OS=Mus musculus GN=Dido1 PE=1 SV=4 | Q8C9B9 [1 | 0 | 2193.21 | 0 | -0.25 | 0.2088 | 10.19 | 108.3 | 91.7 | 4.58 | 12.08 | | High | High | High | High | High | High | High | High | High | High | High | 0.00058 | 0.00522 | 2.3 |
| ASSCSLAVISPFLVEK | [R].ASSCSLAVISPFLVEK.[G] | 1xCarbami | P58801 1x | 7.3E-05 | 0 | 1 | 1 | 5 | P58801 | Receptor-interacting serine/threonine-protein kinase 2 OS=Mus musculus GN=Ripk2 PE=1 SV=1 | P58801 [4 | 0 | 2246.19 | 0 | -0.24 | 0.00967 | 7.34 | 107.5 | 92.5 | 8.41 | 6.31 | | High | High | High | High | High | High | High | High | High | High | High | 0 | 1.5E-05 | 3.22 |
| DLPAIPGVTSPTSDEPPMQASQ | [K].DLPAIPGVTSPTSDEPPMQASQSQLPSSPEDI | 1xPhospho | Q3TII9 1xP | 3.6E-05 | 0 | 1 | 2 | 7 | Q3TII9 | Putative uncharacterized protein OS=Mus musculus GN=Eif4ebp1 PE=2 SV=1 | Q3TII9 [73 | 0 | 3844.86 | 0 | -0.24 | 0.24654 | 19.06 | 117.9 | 82.1 | 24.76 | 30.31 | | High | High | High | High | High | High | High | High | High | High | High | 0.0015 | 0.01497 | 3.78 |
| RASLSDIGFGK | [R].RASLSDIGFGK.[L] | 1xPhospho | Q0VDL6 1 | 0.00067 | 0 | 1 | 1 | 1 | Q0VDL6 | PCTAIRE-motif protein kinase 3 OS=Mus musculus GN=Cdk18 PE=1 SV=1 | Q0VDL6 [1 | 1 | 1688.91 | 0 | -0.24 | 0.18338 | 16.39 | 106.2 | 93.8 | 6.25 | 14.83 | | High | High | High | High | High | High | High | High | High | High | High | 0 | 0.00015 | 2.03 |

| | | | | | | | | | | | | | | | | | | | | | | | | | | | | | | | | | | | | |
|---|---|---|---|---|---|---|---|---|---|---|---|---|---|---|---|---|---|---|---|---|---|---|---|---|---|---|---|---|---|---|---|---|---|---|---|---|
| EEPEPLSPELEYIPR | [K].EEPEPLSPELEYIPR.[K] | 1xPhosph | Q69ZP3-3 | 0.0205 | 0.00147 | 1 | 4 | 1 | Q69ZP3-3 | Isoform 3 of Probable hydrolase PNKD OS=Mus musculus GN=Pnkd | Q69ZP3-3 | 0 | 2107.02 | 0 | -0.23 | 0.3003 | 16.2 | 106.6 | 93.4 | 14.77 | 21.66 | | High | High | High | High | High | High | High | High | High | High | High | 0.00079 | 0.00698 | 2.13 |
| STTPQCQHFSPSMLAAIER | [R].STTPQCQHFSPSMLAAIER.[N] | 1xCarbami | Q80WQ8 1 | 0.00015 | 0 | 1 | 2 | 2 | Q80WQ8 | Mis18-binding protein 1 OS=Mus musculus GN=Mis18bp1 PE=1 SV=1 | Q80WQ8 [ | 0 | 2486.14 | 0 | -0.23 | 0.01207 | 5.3 | 108.5 | 91.5 | 3.54 | 6.45 | | High | High | High | High | High | High | High | High | High | High | High | 0.00335 | 0.03657 | 2.77 |
| TPGRPLSSYGMDSRPPMAIFELL | [R].TPGRPLSSYGMDSRPPMAIFELLDYIVNEPPP | 1xPhosph | P31938 1x | 0.00089 | 0 | 1 | 4 | 4 | P31938 | Dual specificity mitogen-activated protein kinase kinase 1 OS=Mus musculus GN=Map2k1 PE=1 SV=2 | P31938 [2 | 0 | 4226.15 | 0 | -0.23 | 0.03646 | 5.17 | 103.3 | 96.7 | 9.21 | 16.03 | | High | High | High | High | High | High | High | High | High | High | High | 4.9E-05 | 0.00074 | 3.39 |
| ASAVLCQIVASIQPPQTPPEAPQ | [K].ASAVLCQIVASIQPPQTPPEAPQSSPK.[A] | 1xCarbami | Q69Z38 1x | 0.06595 | 0.00477 | 1 | 1 | 2 | Q69Z38 | Pseudopodium-enriched atypical kinase 1 OS=Mus musculus GN=Peak1 PE=1 SV=4 | Q69Z38 [7 | 0 | 3339.74 | 0 | -0.22 | 0.1388 | 6.95 | 105.4 | 94.6 | 16.58 | 11.26 | | High | High | High | High | High | High | High | High | High | High | High | 0.00246 | 0.02657 | 2.66 |
| DSSGDGELDLSGIDDLEIDR | [K].DSSGDGELDLSGIDDLEIDR.[Y] | 1xPhosph | G3X8S2 1x | 0.00019 | 0 | 1 | 2 | 2 | G3X8S2 | BRF1 homolog, subunit of RNA polymerase III transcription initiation factor IIIB (S. cerevisiae) OS=Mus musculus GN=Brf1 PE=1 SV=1 | G3X8S2 [4 | 0 | 2430.08 | 0 | -0.22 | 0.73437 | 18.95 | 99.5 | 100.5 | 23.21 | 15.57 | | High | High | High | High | High | High | High | High | High | High | High | 0 | 3.7E-05 | 2.88 |
| RASLSDLTHLEDIEGLTVR | [R].RASLSDLTHLEDIEGLTVR.[Q] | 1xPhosph | Q3UCG9 1 | 0.00659 | 0.00062 | 1 | 6 | 4 | Q3UCG9 | E3 ubiquitin-protein ligase rififylin OS=Mus musculus GN=Rffl PE=1 SV=1 | Q3UCG9 [2 | 1 | 2434.25 | 0 | -0.22 | 0.0057 | 2.44 | 107.9 | 92.1 | 3.28 | 2.87 | | High | High | High | High | High | High | High | High | High | High | High | 0.00058 | 0.00549 | 2.64 |
| TLPGISPPVVK | [R].TLPGISPPVVK.[E] | 1xPhosph | Q9CPY3 1x | 0.03835 | 0.00256 | 1 | 2 | 1 | Q9CPY3 | Sororin OS=Mus musculus GN=Cdca5 PE=1 SV=1 | Q9CPY3 [2 | 0 | 1645.97 | 0 | -0.21 | 0.01727 | 4.23 | 108.6 | 91.4 | 5.43 | 6.15 | | High | High | High | High | High | High | High | High | High | High | High | 0.00136 | 0.01428 | 1.49 |
| TSFSVGSDDELGPIR | [R].TSFSVGSDDELGPIR.[K-] | 1xPhosph | B2RTN5 1x | 8.8E-06 | 0 | 1 | 4 | 2 | B2RTN5 | Glucocorticoid receptor DNA binding factor 1 OS=Mus musculus GN=Arhgap35 PE=1 SV=1 | B2RTN5 [1 | 0 | 1888.89 | 0 | -0.21 | 0.55725 | 0.48 | 103.7 | 96.3 | 6.09 | 11.17 | | High | High | High | High | High | High | High | High | High | High | High | 0 | 1.2E-06 | 3.13 |
| VLSPILLNNILPTTAK | [R].VLSPILLNNILPTTAK.[S] | 1xPhosph | Q9WU62 1 | 0.00081 | 0 | 1 | 2 | 1 | Q9WU62 | Inner centromere protein OS=Mus musculus GN=Incenp PE=1 SV=2 | Q9WU62 [ | 0 | 2245.33 | 0 | -0.21 | 0.02263 | 9.13 | 107.2 | 92.8 | 6.03 | 7.65 | | High | High | High | High | High | High | High | High | High | High | High | 0 | 0.00019 | 2.71 |
| VTIAQGGVLPNIQAVLLPK | [KR].VTIAQGGVLPNIQAVLLPK.[K] | 1xPhosph | C0HKE6 1x | 0.09015 | 0.00698 | 2 | 13 | 1 | C0HKE6; B | Histone H2A type 1-I OS=Mus musculus GN=Hist1h2ai PE=1 SV=1Histone H2A OS=Mus musculus GN=Hist2h2aa1 PE=2 SV=1 | C0HKE6 [1 | 0 | 2469.46 | 0 | -0.21 | 0.47361 | 10.47 | 98.3 | 101.7 | 18.44 | 9.5 | Shared | High | High | High | High | High | High | High | High | High | High | High | 0.00361 | 0.03831 | 2.08 |
| HSPTLVPTATKPDAVQPDSLPYP | [K].HSPTLVPTATKPDAVQPDSLPYPQYLAVIK.[A | 1xPhosph | Q9DB91 1 | 0.00349 | 0.00021 | 1 | 1 | 3 | Q9DB91 | Mediator of RNA polymerase II transcription subunit 29 OS=Mus musculus GN=Med29 PE=2 SV=1 | Q9DB91 [1 | 0 | 4014.2 | 0 | -0.2 | 0.35183 | 17.89 | 106.7 | 93.3 | 16.45 | 38.87 | | High | High | High | High | High | High | High | High | High | High | High | 9.4E-05 | 0.00097 | 3.59 |
| ACSDSLESIPAGQVASDDLR | [R].ACSDSLESIPAGQVASDDLR.[D] | 1xCarbami | Q80TM9 1 | 4E-08 | 0 | 1 | 5 | 7 | Q80TM9 | Nischarin OS=Mus musculus GN=Nisch PE=1 SV=2 | Q80TM9 [ | 0 | 2400.1 | 0 | -0.18 | 0.15278 | 14.13 | 107.3 | 92.7 | 6.85 | 9.07 | | High | High | High | High | High | High | High | High | High | High | High | 0 | 2.7E-09 | 4.02 |
| SHSFADPASNLGLEDIIR | [R].SHSFADPASNLGLEDIIR.[K] | 1xPhosph | Q5RIM6 1 | 0.00238 | 0.00011 | 1 | 11 | 1 | Q5RIM6 | Nuclear receptor co-repressor 1, isoform CRA_a OS=Mus musculus GN=Ncor1 PE=1 SV=1 | Q5RIM6 [2 | 0 | 2251.1 | 0 | -0.17 | 0.20069 | 6.5 | 108.8 | 91.2 | 15.62 | 14.01 | | High | High | High | High | High | High | High | High | High | High | High | 4.9E-05 | 0.00063 | 2.45 |
| LQDISELLATGVGLSDSEVEPDGI | [R].LQDISELLATGVGLSDSEVEPDGEHNTTELPQ | 1xDeamid | Q91YU8 1 | 2.3E-06 | 0 | 1 | 1 | 4 | Q91YU8 | Suppressor of SWI4 1 homolog OS=Mus musculus GN=Ppan PE=1 SV=2 | Q91YU8 [2 | 0 | 4158 | 0 | -0.16 | 0.21082 | 2.79 | 115.5 | 84.5 | 33.9 | 23.09 | | High | High | High | High | High | High | High | High | High | High | High | 0 | 7E-06 | 4.71 |
| SQSSHSYDDSTLPLIDR | [R].SQSSHSYDDSTLPLIDR.[N] | 2xPhosph | E9Q8Z8 2x | 0.00015 | 0 | 1 | 19 | 3 | E9Q8Z8 | Catenin delta-1 OS=Mus musculus GN=Ctnnd1 PE=1 SV=1 | E9Q8Z8 [8 | 0 | 2309.99 | 0 | -0.16 | 0.01023 | 4.57 | 107.1 | 92.9 | 5.71 | 4.77 | | High | High | High | High | High | High | High | High | High | High | High | 0 | 2.8E-05 | 3.72 |
| EMTHSWPPPLTAIHTPCK | [K].EMTHSWPPPLTAIHTPCK.[T] | 1xCarbami | Q9ESC8 1x | 0.00065 | 0 | 1 | 1 | 1 | Q9ESC8 | AF4/FMR2 family member 4 OS=Mus musculus GN=Aff4 PE=1 SV=1 | Q9ESC8 [3 | 0 | 2657.3 | 0 | -0.15 | 0.09674 | 10.57 | 107.7 | 92.3 | 5.16 | 9.69 | | High | High | High | High | High | High | High | High | High | High | High | 0 | 0.00015 | 3.18 |
| SCSITGVDDLEQILADAADLTQLE | [R].SCSITGVDDLEQILADAADLTQLETQPVCSK.[I | 2xCarbami | A3KGI3 1x | 0.00487 | 0.0005 | 1 | 2 | 2 | A3KGI3 | Kinetochore scaffold 1 OS=Mus musculus GN=Knl1 PE=1 SV=1 | A3KGI3 [14 | 0 | 3915.9 | 0 | -0.15 | 0.43253 | 14.09 | 105.5 | 94.5 | 6.07 | 9.3 | | High | High | High | High | High | High | High | High | High | High | High | 0.00022 | 0.0014 | 4.31 |
| DLPAIPGVTSPTSDEPPMQASQ | [K].DLPAIPGVTSPTSDEPPMQASQSQLPSSPEDI | 1xDeamid | Q3TII9 1xP | 4.8E-07 | 0 | 1 | 2 | 4 | Q3TII9 | Putative uncharacterized protein OS=Mus musculus GN=Eif4ebp1 PE=2 SV=1 | Q3TII9 [73 | 0 | 3845.84 | 0 | -0.14 | 0.2587 | 24.29 | 119.8 | 80.2 | 25.51 | 28.71 | | High | High | High | High | High | High | High | High | High | High | High | 0 | 4.5E-08 | 4.19 |
| LDFIGNSTGHK | [K].LDFIGNSTGHK.[R] | 1xPhosph | E9PVX6 1x | 0.00567 | 0.00062 | 1 | 2 | 1 | E9PVX6 | Proliferation marker protein Ki-67 OS=Mus musculus GN=Mki67 PE=1 SV=1 | E9PVX6 [1 | 0 | 1726.89 | 0 | -0.14 | 0.13089 | 9.88 | 105.5 | 94.5 | 11.44 | 7.7 | | High | High | High | High | High | High | High | High | High | High | High | 0.00027 | 0.00166 | 2.89 |
| NKPLSPIK | [K].NKPLSPIK.[L] | 1xPhosph | G3UWX1 1 | 0.09237 | 0.00721 | 2 | 9 | 3 | G3UWX1: | Replication factor C subunit 1 OS=Mus musculus GN=Rfc1 PE=1 SV=2Replication factor C subunit 1 OS=Mus musculus GN=Rfc1 PE=1 SV=2 | G3UWX1 [ | 0 | 1434.85 | 0 | -0.14 | 0.73957 | 32.37 | 104.7 | 95.3 | 48.17 | 49.51 | Shared | High | High | High | High | High | High | High | High | High | High | High | 0.00553 | 0.05995 | 2.48 |
| TPLATIADQQQLQLSPLK | [R].TPLATIADQQQLQLSPLK.[R] | 1xDeamid | P11157 1x | 0.00609 | 0.00062 | 1 | 2 | 2 | P11157 | Ribonucleoside-diphosphate reductase subunit M2 OS=Mus musculus GN=Rrm2 PE=1 SV=1 | P11157 [6 | 0 | 2504.38 | 0 | -0.14 | 0.57537 | 18.11 | 106 | 94 | 12.93 | 11.87 | | High | High | High | High | High | High | High | High | High | High | High | 0.00027 | 0.00179 | 2.42 |
| VHGEPASDLSDID | [R].VHGEPASDLSDID.[-] | 1xPhosph | E9PZ26 1x | 0.00045 | 0 | 1 | 5 | 3 | E9PZ26 | Bromodomain-containing 1 OS=Mus musculus GN=Brd1 PE=1 SV=1 | E9PZ26 [1 | 0 | 1663.74 | 0 | -0.14 | 0.14495 | 9.12 | 106.1 | 93.9 | 7.21 | 8.44 | | High | High | High | High | High | High | High | High | High | High | High | 0 | 9.6E-05 | 3.06 |
| LGLAVIHGEAQCTELDMDDGRH | [R].LGLAVIHGEAQCTELDMDDGRHSPPMVK.[N | 1xCarbami | B1AT82 1x | 0.00067 | 0 | 1 | 3 | 2 | B1AT82 | MCG6846, isoform CRA_c OS=Mus musculus GN=Prpsap1 PE=1 SV=1 | B1AT82 [2 | 1 | 3630.75 | 0 | -0.13 | 0.42678 | 1.94 | 104.8 | 95.2 | 2.19 | 12.63 | | High | High | High | High | High | High | High | High | High | High | High | 0 | 0.00015 | 4.22 |
| QLTEMLPSILNQLGADSLTSLR | [K].QLTEMLPSILNQLGADSLTSLR.[R] | 1xPhosph | Q64152 1 | 6.8E-09 | 0 | 1 | 2 | 1 | Q64152 | Transcription factor BTF3 OS=Mus musculus GN=Btf3 PE=1 SV=3 | Q64152 [1 | 0 | 2709.41 | 0 | -0.12 | 0.93842 | 17.51 | 98.7 | 101.3 | 7.37 | 19.71 | | High | High | High | High | High | High | High | High | High | High | High | 0 | 3.7E-10 | 3.81 |
| LSSTSSLAGIPSPSLVSLPSFFER | [R].LSSTSSLAGIPSPSLVSLPSFFER.[R] | 1xPhosph | Q07139 1x | 0.00025 | 0 | 1 | 2 | 5 | Q07139 | Protein ECT2 OS=Mus musculus GN=Ect2 PE=1 SV=2 | Q07139 [8 | 0 | 2788.44 | 0 | -0.11 | 0.85173 | 6.73 | 100.6 | 99.4 | 11.21 | 9.72 | | High | High | High | High | High | High | High | High | High | High | High | 0 | 6.9E-05 | 3.88 |
| VLSPILLNNILPTTAK | [R].VLSPILLNNILPTTAK.[S] | 1xDeamid | Q9WU62 1 | 0.00066 | 0 | 1 | 2 | 3 | Q9WU62 | Inner centromere protein OS=Mus musculus GN=Incenp PE=1 SV=2 | Q9WU62 [ | 0 | 2246.32 | 0 | -0.1 | 0.30804 | 13.62 | 102.2 | 97.8 | 10.25 | 14.8 | | High | High | High | High | High | High | High | High | High | High | High | 0.00027 | 0.0016 | 3.74 |
| ANFDEENAYFEDEEEDSSNVDLI | [R].ANFDEENAYFEDEEEDSSNVDLPYIPAENSPT | 1xPhosph | Q810A7 1 | 2.8E-06 | 0 | 1 | 2 | 57 | Q810A7 | ATP-dependent RNA helicase DDX42 OS=Mus musculus GN=Ddx42 PE=1 SV=3 | Q810A7 [6 | 0 | 4115.74 | 0 | -0.09 | 0.44896 | 10.63 | 103.5 | 96.5 | 7.25 | 4.72 | | High | High | High | High | High | High | High | High | High | High | High | 0 | 1.5E-06 | 4.83 |
| TPAPATTPSHNPLANILSK | [K].TPAPATTPSHNPLANILSK.[V] | 1xPhosph | Q6NXI6 1x | 0.00036 | 0 | 1 | 4 | 2 | Q6NXI6 | Regulation of nuclear pre-mRNA domain-containing protein 2 OS=Mus musculus GN=Rprd2 PE=1 SV=1 | Q6NXI6 [5 | 0 | 2468.33 | 0 | -0.09 | 0.24933 | 17.43 | 113.8 | 86.2 | 14.01 | 16.09 | | High | High | High | High | High | High | High | High | High | High | High | 0 | 7.6E-05 | 2.65 |
| LNLSCIHSPVVNELMR | [K].LNLSCIHSPVVNELMR.[G] | 1xCarbami | Q6DFW4 1 | 0.00023 | 0 | 1 | 3 | 1 | Q6DFW4 | Nucleolar protein 58 OS=Mus musculus GN=Nop58 PE=1 SV=1 | Q6DFW4 [ | 0 | 2191.1 | 0 | -0.08 | 0.78337 | 6.19 | 102.1 | 97.9 | 9.81 | 8.6 | | High | High | High | High | High | High | High | High | High | High | High | 0 | 4.6E-05 | 2.69 |
| SAVLDGILEQLEDEDNSDGEQAL | [R].SAVLDGILEQLEDEDNSDGEQALQL.[-] | 1xPhosph | Q8R3C6 1 | 1.4E-05 | 0 | 1 | 1 | 5 | Q8R3C6 | Probable RNA-binding protein 19 OS=Mus musculus GN=Rbm19 PE=1 SV=1 | Q8R3C6 [9 | 0 | 3010.4 | 0 | -0.08 | 0.0737 | 2.2 | 103.4 | 96.6 | 8.7 | 9.54 | | High | High | High | High | High | High | High | High | High | High | High | 0 | 2E-06 | 4.17 |
| SSPPPGYIPDELHQIAR | [R].SSPPPGYIPDELHQIAR.[N] | 1xPhosph | G5E8L8 1x | 0.0001 | 0 | 1 | 1 | 1 | G5E8L8 | Mitogen activated protein kinase kinase kinase 2 OS=Mus musculus GN=Map3k2 PE=1 SV=1 | G5E8L8 [1 | 0 | 2186.08 | 0 | -0.07 | 0.82624 | 4.53 | 102.5 | 97.5 | 15.44 | 8.4 | | High | High | High | High | High | High | High | High | High | High | High | 0 | 1.8E-05 | 3.57 |
| ELAQIAGRPTEDEDDK | [K].ELAQIAGRPTEDEDDK.[D] | 1xPhosph | Q61074 1 | 0.00909 | 0.00074 | 1 | 3 | 1 | Q61074 | Protein phosphatase 1G OS=Mus musculus GN=Ppm1g PE=1 SV=3 | Q61074 [1 | 0 | 2325.14 | 0 | -0.06 | 0.51261 | 13.56 | 102.4 | 97.6 | 6.87 | 9.91 | | High | High | High | High | High | High | High | High | High | High | High | 0.00033 | 0.00282 | 1.86 |
| SAVDLTPIVVEDK | [K].SAVDLTPIVVEDK.[E] | 1xPhosph | Q6PDQ2 1 | 3.2E-05 | 0 | 1 | 5 | 3 | Q6PDQ2 | Chromodomain-helicase-DNA-binding protein 4 OS=Mus musculus GN=Chd4 PE=1 SV=1 | Q6PDQ2 [ | 0 | 1924.04 | 0 | -0.06 | 0.21452 | 5.51 | 102.2 | 97.8 | 2.26 | 8.46 | | High | High | High | High | High | High | High | High | High | High | High | 0 | 5E-06 | 3.06 |
| LEAIITPPPAK | [K].LEAIITPPPAK.[K] | 1xPhosph | Q9QZQ8 1 | 0.0014 | 5.7E-05 | 1 | 3 | 3 | Q9QZQ8 | Core histone macro-H2A.1 OS=Mus musculus GN=H2afy PE=1 SV=3 | Q9QZQ8 [ | 0 | 1687.98 | 0 | -0.05 | 0.85825 | 7.71 | 102.6 | 97.4 | 4.89 | 10.31 | | High | High | High | High | High | High | High | High | High | High | High | 2.6E-05 | 0.00035 | 2.06 |
| LGEELVAIVESPPGPVGLLAAGD | [R].LGEELVAIVESPPGPVGLLAAGDGR.[G] | 1xPhosph | A0A140LI3 | 4E-05 | 0 | 1 | 1 | 1 | A0A140LI3 | Zinc finger protein 654 OS=Mus musculus GN=Zfp654 PE=1 SV=1 | A0A140LI3 | 0 | 2725.44 | 0 | -0.04 | 0.57644 | 9.55 | 104 | 96 | 9.63 | 12.48 | | High | High | High | High | High | High | High | High | High | High | High | 0 | 6.4E-06 | 3.68 |
| SPPSGSQSLLEGIMTASSSK | [R].SPPSGSQSLLEGIMTASSSK.[G] | 1xPhosph | B9EKJ7 1x | 0.00337 | 0.00016 | 1 | 3 | 2 | B9EKJ7 | Tankyrase 1 binding protein 1 OS=Mus musculus GN=Tnks1bp1 PE=2 SV=1 | B9EKJ7 [14 | 0 | 2502.26 | 0 | -0.04 | 0.95911 | 29.33 | 101.5 | 98.5 | 9.11 | 18.53 | | High | High | High | High | High | High | High | High | High | High | High | 9.4E-05 | 0.00093 | 2.43 |
| TSVLDGIVPIESR | [K].TSVLDGIVPIESR.[T] | 1xPhosph | Q8CJF7 1x | 0.04072 | 0.00306 | 1 | 1 | 1 | Q8CJF7 | Protein ELYS OS=Mus musculus GN=Ahctf1 PE=1 SV=1 | Q8CJF7 [14 | 0 | 1694.89 | 0 | -0.04 | 0.95242 | 26.25 | 101.5 | 98.5 | 12.48 | 29.73 | | High | High | High | High | High | High | High | High | High | High | High | 0.00158 | 0.01524 | 1.73 |
| GPLVGIGPTSTPR | [R].GPLVGIGPTSTPR.[A] | 1xPhosph | Q8CDZ5 1 | 0.01076 | 0.00077 | 1 | 2 | 2 | Q8CDZ5 | Putative uncharacterized protein OS=Mus musculus GN=Nup133 PE=2 SV=1 | Q8CDZ5 [1 | 0 | 1560.83 | 0 | -0.03 | 0.97743 | 22.44 | 101 | 99 | 9.13 | 5.68 | | High | High | High | High | High | High | High | High | High | High | High | 0.0004 | 0.00341 | 2.9 |
| HLDVDLDRQSLSSIDR | [R].HLDVDLDRQSLSSIDR.[N] | 1xPhosph | A2A5R2 1x | 0.00223 | 0.00011 | 1 | 2 | 1 | A2A5R2 | Brefeldin A-inhibited guanine nucleotide-exchange protein 2 OS=Mus musculus GN=Arfgef2 PE=1 SV=1 | A2A5R2 [1 | 1 | 2178.08 | 0 | -0.03 | 0.25831 | 5.4 | 105.2 | 94.8 | 9.25 | 10.51 | | High | High | High | High | High | High | High | High | High | High | High | 4.9E-05 | 0.00059 | 2.46 |
| SLSELCLISVAPGVR | [R].SLSELCLISVAPGVR.[T] | 1xCarbami | Q66T02 1x | 0.00223 | 0.00011 | 1 | 4 | 1 | Q66T02 | Pleckstrin homology domain-containing family G member 5 OS=Mus musculus GN=Plekhg5 PE=1 SV=1 | Q66T02 [9 | 0 | 1910 | 0 | -0.03 | 0.79425 | 5.33 | 100.6 | 99.4 | 10.65 | 10.93 | | High | High | High | High | High | High | High | High | High | High | High | 4.9E-05 | 0.00059 | 2.65 |
| VFGALGPIGPSSPGLTLGGLAVN | [R].VFGALGPIGPSSPGLTLGGLAVNEHR.[L] | 1xPhosph | Q91VU8 1 | 0.00076 | 0 | 1 | 2 | 2 | Q91VU8 | Prostate tumor-overexpressed gene 1 protein homolog OS=Mus musculus GN=Ptov1 PE=1 SV=1 | Q91VU8 [6 | 0 | 2825.49 | 0 | -0.03 | 0.91885 | 8.78 | 101.7 | 98.3 | 19.93 | 6.12 | | High | High | High | High | High | High | High | High | High | High | High | 0 | 0.00018 | 4.13 |
| DRCPLPCIPPLVGTPVK | [R].DRCPLPCIPPLVGTPVK.[I] | 2xCarbami | Q9WVM1 | 0.0887 | 0.00667 | 1 | 1 | 1 | Q9WVM1 | Rac GTPase-activating protein 1 OS=Mus musculus GN=Racgap1 PE=1 SV=1 | Q9WVM1 | 1 | 2457.32 | 0 | -0.01 | 0.47733 | 17.62 | 99.5 | 100.5 | 12.94 | 9.57 | | High | High | High | High | High | High | High | High | High | High | High | 0.00346 | 0.03759 | 2.32 |
| GGRSPDEVTLTSIVPTR | [K].GGRSPDEVTLTSIVPTR.[D] | 1xPhosph | Q543N7 1 | 0.00149 | 5.7E-05 | 1 | 2 | 2 | Q543N7 | Protein kinase C and casein kinase substrate in neurons 3, isoform CRA_a OS=Mus musculus GN=Pacsin3 PE=1 SV=1 | Q543N7 [3 | 1 | 2094.08 | 0 | -0.01 | 0.4019 | 8.22 | 99.7 | 100.3 | 10.95 | 3.08 | | High | High | High | High | High | High | High | High | High | High | High | 2.6E-05 | 0.00037 | 3.12 |
| SSAELRKILANGQMNEQDIR | [K].SSAELRKILANGQMNEQDIR.[Y] | 1xDeamid | Q9WVS7 1 | 0.05169 | 0.00374 | 1 | 4 | 2 | Q9WVS7 | Dual specificity mitogen-activated protein kinase kinase 5 OS=Mus musculus GN=Map2k5 PE=1 SV=1 | Q9WVS7 [ | 2 | 2599.27 | 0 | -0.01 | 0.6853 | 6.74 | 97.1 | 102.9 | 7.41 | 6.27 | | High | High | High | High | High | High | High | High | High | High | High | 0.00772 | 0.0836 | 2.94 |
| AALEDSNGSSELQEIMR | [K].AALEDSNGSSELQEIMR.[R] | 1xDeamid | Q3U2Z5 1 | 3.1E-05 | 0 | 1 | 3 | 1 | Q3U2Z5 | Putative uncharacterized protein OS=Mus musculus GN=Eps8 PE=2 SV=1 | Q3U2Z5 [8 | 0 | 2175.97 | 0 | 0 | 0.84853 | 24.87 | 99.9 | 100.1 | 6.64 | 12.96 | | High | High | High | High | High | High | High | High | High | High | High | 0 | 4.8E-06 | 3.42 |
| AVEYLLTGIPGSPEPEHGSVQESQ | [R].AVEYLLTGIPGSPEPEHGSVQESQAPEQPATE | 1xPhosph | Q8CAP3 1 | 0.00051 | 0 | 1 | 2 | 1 | Q8CAP3 | Putative uncharacterized protein OS=Mus musculus GN=Rad23a PE=1 SV=1 | Q8CAP3 [1 | 0 | 4783.3 | 0 | 0 | 0.80033 | 56.07 | 102.6 | 97.4 | 19.15 | 28.79 | | High | High | High | High | High | High | High | High | High | High | High | 0 | 0.00011 | 4.52 |
| QLTEMLPSILNQLGADSLTSLR | [K].QLTEMLPSILNQLGADSLTSLR.[R] | 1xPhosph | Q64152 1 | 2.6E-07 | 0 | 1 | 2 | 1 | Q64152 | Transcription factor BTF3 OS=Mus musculus GN=Btf3 PE=1 SV=3 | Q64152 [1 | 0 | 2725.4 | 0 | 0 | 0.62858 | 8.92 | 95.1 | 104.9 | 18.27 | 19.51 | | High | High | High | High | High | High | High | High | High | High | High | 0 | 2.2E-08 | 3.92 |
| RESLSYIPK | [R].RESLSYIPK.[G] | 1xPhosph | Q4VAC9 1 | 0.0047 | 0.00046 | 1 | 3 | 1 | Q4VAC9 | Pleckstrin homology domain-containing family G member 3 OS=Mus musculus GN=Plekhg3 PE=1 SV=2 | Q4VAC9 [8 | 1 | 1630.9 | 0 | 0 | 0.86871 | 23.99 | 99.9 | 100.1 | 5.91 | 13.21 | | High | High | High | High | High | High | High | High | High | High | High | 0.0002 | 0.00135 | 1.74 |
| YTPQTLPVGELDAISPIVSETLQL | [R].YTPQTLPVGELDAISPIVSETLQLSDALK.[R] | 1xPhosph | Q8CFT2 1x | 0.00021 | 0 | 1 | 3 | 2 | Q8CFT2 | Histone-lysine N-methyltransferase SETD1B OS=Mus musculus GN=Setd1b PE=1 SV=2 | Q8CFT2 [1 | 0 | 3636.94 | 0 | 0 | 0.6781 | 4.67 | 100.1 | 99.9 | 10.29 | 7.01 | | High | High | High | High | High | High | High | High | High | High | High | 0 | 4.1E-05 | 3.73 |
| ILLSQTTGMAIPLQASSLDDISLL | [R].ILLSQTTGMAIPLQASSLDDISLLSTPK.[R] | 1xPhosph | F6ZDS4 1x | 0.00033 | 0 | 1 | 7 | 3 | F6ZDS4 | Nucleoprotein TPR OS=Mus musculus GN=Tpr PE=1 SV=1 | F6ZDS4 [6 | 0 | 3467.87 | 0 | 0.01 | 0.67537 | 26 | 94.9 | 105.1 | 7.31 | 16.82 | | High | High | High | High | High | High | High | High | High | High | High | 0 | 6.8E-05 | 4.29 |
| SPQPPAEEDEDDFDDTLVAIDTY | [R].SPQPPAEEDEDDFDDTLVAIDTYNCDLHFK.[ | 1xCarbami | Q8VDM6 1 | 1.3E-05 | 0 | 1 | 3 | 1 | Q8VDM6 | Heterogeneous nuclear ribonucleoprotein U-like protein 1 OS=Mus musculus GN=Hnrnpul1 PE=1 SV=1 | Q8VDM6 [ | 0 | 4034.79 | 0 | 0.02 | 0.35183 | 19.37 | 85.4 | 114.6 | 31.14 | 12.7 | | High | High | High | High | High | High | High | High | High | High | High | 0 | 1.9E-06 | 3.58 |
| APVSVACGGESPLDGICLSEADK | [K].APVSVACGGESPLDGICLSEADK.[T] | 2xCarbami | F8VQ95 1 | 1.4E-07 | 0 | 1 | 5 | 1 | F8VQ95 | Transforming acidic coiled-coil-containing protein 1 OS=Mus musculus GN=Tacc1 PE=1 SV=1 | F8VQ95 [5 | 0 | 2870.37 | 0 | 0.04 | 0.74057 | 28.88 | 99.9 | 100.1 | 18.85 | 19.82 | | High | High | High | High | High | High | High | High | High | High | High | 0 | 1.1E-08 | 3.52 |
| CPLPCIPPLVGTPVK | [R].CPLPCIPPLVGTPVK.[I] | 2xCarbami | Q9WVM1 | 1.3E-05 | 0 | 1 | 1 | 1 | Q9WVM1 | Rac GTPase-activating protein 1 OS=Mus musculus GN=Racgap1 PE=1 SV=1 | Q9WVM1 | 0 | 2186.19 | 0 | 0.04 | 0.56052 | 22.19 | 99 | 101 | 14.91 | 0.59 | | High | High | High | High | High | High | High | High | High | High | High | 0 | 1.8E-06 | 3.21 |
| LSNIFVIGK | [R].LSNIFVIGK.[G] | 1xPhosph | Q3UXQ6 1 | 0.00182 | 5.7E-05 | 1 | 5 | 1 | Q3UXQ6 | 40S ribosomal protein S4 OS=Mus musculus GN=Rps4x PE=2 SV=1 | Q3UXQ6 [ | 0 | 1528.89 | 0 | 0.04 | 0.95065 | 14.59 | 98.3 | 101.7 | 14.85 | 12.39 | | High | High | High | High | High | High | High | High | High | High | High | 2.6E-05 | 0.00047 | 2.49 |
| LSPIMTEQDDLGDGDVHSLVYPI | [R].LSPIMTEQDDLGDGDVHSLVYPPSAAK.[M] | 1xPhosph | Q9R1E0 1x | 0.00044 | 0 | 1 | 3 | 2 | Q9R1E0 | Forkhead box protein O1 OS=Mus musculus GN=Foxo1 PE=1 SV=2 | Q9R1E0 [3 | 0 | 3409.66 | 0 | 0.04 | 0.83198 | 3.83 | 99.8 | 100.2 | 13.42 | 23.88 | | High | High | High | High | High | High | High | High | High | High | High | 0 | 9.4E-05 | 3.68 |
| LLSPIASNR | [R].LLSPIASNR.[L] | 1xPhosph | Q9CW46 1 | 0.01845 | 0.00132 | 1 | 2 | 2 | Q9CW46 | Ribonucleoprotein PTB-binding 1 OS=Mus musculus GN=Raver1 PE=1 SV=2 | Q9CW46 [ | 0 | 1279.7 | 0 | 0.05 | 0.56402 | 6.53 | 98.7 | 101.3 | 5.85 | 3.87 | | High | High | High | High | High | High | High | High | High | High | High | 0.00381 | 0.04107 | 2.06 |
| DDLLGDILQDLNTETAQITPPPVI | [K].DDLLGDILQDLNTETAQITPPPVLIPK.[K] | 1xPhosph | P33609 1x | 8E-05 | 0 | 1 | 3 | 1 | P33609 | DNA polymerase alpha catalytic subunit OS=Mus musculus GN=Pola1 PE=1 SV=2 | P33609 [1 | 0 | 3467.87 | 0 | 0.06 | 0.5672 | 17.84 | 97.9 | 102.1 | 17.21 | 15.68 | | High | High | High | High | High | High | High | High | High | High | High | 0 | 1.4E-05 | 4.05 |
| RDSICSSVSMESSLAEPQDELLQI | [R].RDSICSSVSMESSLAEPQDELLQILK.[D] | 1xCarbami | E9QP99 1x | 0.09541 | 0.00744 | 1 | 5 | 1 | E9QP99 | Golgin subfamily A member 3 OS=Mus musculus GN=Golga3 PE=1 SV=1 | E9QP99 [3 | 1 | 3489.73 | 0 | 0.08 | 0.19048 | 5.94 | 94.4 | 105.6 | 6.58 | 17.81 | | High | High | High | High | High | High | High | High | High | High | High | 0.00377 | 0.04071 | 4.65 |
| SAFSSLQNIPESLR | [R].SAFSSLQNIPESLR.[R] | 1xPhosph | B7ZNA0 1 | 0.00102 | 0 | 2 | 8 | 4 | B7ZNA0; C | Shroom3 protein OS=Mus musculus GN=Shroom3 PE=2 SV=1Protein Shroom3 OS=Mus musculus GN=Shroom3 PE=1 SV=2 | B7ZNA0 [6 | 0 | 1857.93 | 0 | 0.08 | 0.02585 | 3.84 | 98.2 | 101.8 | 3.2 | 1.48 | Shared | High | High | High | High | High | High | High | High | High | High | High | 0.00391 | 0.04148 | 2.52 |
| TPLATIADQQQLQLSPLK | [R].TPLATIADQQQLQLSPLK.[R] | 1xPhosph | P11157 1x | 1.4E-06 | 0 | 1 | 2 | 2 | P11157 | Ribonucleoside-diphosphate reductase subunit M2 OS=Mus musculus GN=Rrm2 PE=1 SV=1 | P11157 [6 | 0 | 2274.23 | 0 | 0.08 | 0.84878 | 31.09 | 106 | 94 | 71.78 | 82.52 | | High | High | High | High | High | High | High | High | High | High | High | 0 | 1.5E-07 | 4.26 |
| FSLCSDNLEGISEGPSNR | [R].FSLCSDNLEGISEGPSNR.[S] | 1xCarbami | Q6PAR5 1 | 3.2E-05 | 0 | 1 | 4 | 2 | Q6PAR5 | GTPase-activating protein and VPS9 domain-containing protein 1 OS=Mus musculus GN=Gapvd1 PE=1 SV=2 | Q6PAR5 [5 | 0 | 2291.02 | 0 | 0.11 | 0.98683 | 22.6 | 101.8 | 98.2 | 17.85 | 10.42 | | High | High | High | High | High | High | High | High | High | High | High | 0 | 5E-06 | 3.34 |
| YLPAITSTPTVNENDSSTSK | [K].YLPAITSTPTVNENDSSTSK.[C] | 1xPhosph | Q9R0L6-2 | 9.8E-08 | 0 | 1 | 3 | 12 | Q9R0L6-2 | Isoform 2 of Pericentriolar material 1 protein OS=Mus musculus GN=Pcm1 | Q9R0L6-2 | 0 | 2663.32 | 0 | 0.11 | 0.25049 | 2.24 | 96.6 | 103.4 | 4.62 | 10.97 | | High | High | High | High | High | High | High | High | High | High | High | 0 | 7.3E-08 | 3.99 |
| NLGSINTELQDVQR | [R].NLGSINTELQDVQR.[I] | 1xPhosph | O08547 1 | 3.2E-06 | 0 | 1 | 3 | 3 | O08547 | Vesicle-trafficking protein SEC22b OS=Mus musculus GN=Sec22b PE=1 SV=3 | O08547 [1 | 0 | 1895.94 | 0 | 0.12 | 0.96371 | 14.8 | 98.1 | 101.9 | 12.4 | 12.11 | | High | High | High | High | High | High | High | High | High | High | High | 0 | 3.7E-07 | 4.11 |
| QLSSSVTGLTNIEEENCQR | [K].QLSSSVTGLTNIEEENCQR.[Y] | 1xCarbami | Q6ZQK2 1 | 9E-07 | 0 | 1 | 7 | 4 | Q6ZQK2 | MKIAA0051 protein (Fragment) OS=Mus musculus GN=Iqgap1 PE=2 SV=1 | Q6ZQK2 [5 | 0 | 2474.14 | 0 | 0.12 | 0.38009 | 16.08 | 94.5 | 105.5 | 3.59 | 14.47 | | High | High | High | High | High | High | High | High | High | High | High | 0 | 9E-08 | 3.62 |
| EEALQNISADDLPDSASQAAHP | [R].EEALQNISADDLPDSASQAAHPQDSAFSYR.[ | 1xPhosph | Q6PAR5 1 | 0.00017 | 0 | 1 | 7 | 1 | Q6PAR5 | GTPase-activating protein and VPS9 domain-containing protein 1 OS=Mus musculus GN=Gapvd1 PE=1 SV=2 | Q6PAR5 [1 | 0 | 3542.59 | 0 | 0.13 | 0.20967 | 19.18 | 92.5 | 107.5 | 11.92 | 18.99 | | High | High | High | High | High | High | High | High | High | High | High | 0 | 3.3E-05 | 3.66 |
| RSDSMTFLGCIPPNPFEVPLAEA | [R].RSDSMTFLGCIPPNPFEVPLAEAIPLADQPHL | 1xCarbami | E9Q2H1 1 | 0.11466 | 0.00941 | 1 | 2 | 2 | E9Q2H1 | E3 ubiquitin-protein ligase UBR5 OS=Mus musculus GN=Ubr5 PE=1 SV=1 | E9Q2H1 [2 | 1 | 4537.26 | 0 | 0.14 | 0.53769 | 26.22 | 99 | 101 | 7.72 | 16.82 | | High | High | High | High | High | High | High | High | High | High | High | 0.00473 | 0.05097 | 3.22 |
| VTEECEEAIPQLVPIGETPDK | [K].VTEECEEAIPQLVPIGETPDK.[E] | 1xCarbami | Q3UL54 1 | 1.2E-06 | 0 | 1 | 8 | 3 | Q3UL54 | Putative uncharacterized protein OS=Mus musculus GN=Rsl1d1 PE=2 SV=1 | Q3UL54 [3 | 0 | 2892.44 | 0 | 0.14 | 0.12142 | 11.32 | 95.6 | 104.4 | 3.84 | 7.17 | | High | High | High | High | High | High | High | High | High | High | High | 0 | 1.3E-07 | 3.46 |
| IMETSPTLLQISPGPPK | [R].IMETSPTLLQISPGPPK.[-] | 1xPhosph | Q6ZQ33 1 | 0.07719 | 0.00589 | 1 | 4 | 1 | Q6ZQ33 | MKIAA0857 protein (Fragment) OS=Mus musculus GN=Rab11fip5 PE=2 SV=1 | Q6ZQ33 [1 | 0 | 2363.27 | 0 | 0.15 | 0.39611 | 18.25 | 93.6 | 106.4 | 10.75 | 6.85 | | High | High | High | High | High | High | High | High | High | High | High | 0.00306 | 0.0319 | 2.85 |
| NSSPSISPNTSFASDGSPSPLGGI | [R].NSSPSISPNTSFASDGSPSPLGGIK.[R] | 1xPhosph | Q9DBR1 1 | 1.9E-05 | 0 | 1 | 2 | 8 | Q9DBR1 | 5'-3' exoribonuclease 2 OS=Mus musculus GN=Xrn2 PE=1 SV=1 | Q9DBR1 [4 | 0 | 2713.29 | 0 | 0.15 | 0.25096 | 24.39 | 94.7 | 105.3 | 16.39 | 25.53 | | High | High | High | High | High | High | High | High | High | High | High | 0 | 2.9E-05 | 4.1 |
| HHGLNVLAALCSIGK | [R].HHGLNVLAALCSIGK.[Q] | 1xCarbami | Q9D8T2 1 | 0.01877 | 0.00136 | 1 | 1 | 1 | Q9D8T2 | Gasdermin-D OS=Mus musculus GN=Gsdmdc1 PE=1 SV=1 | Q9D8T2 [2 | 0 | 2128.15 | 0 | 0.17 | 0.18795 | 27.41 | 92.2 | 107.8 | 14.63 | 23.97 | | High | High | High | High | High | High | High | High | High | High | High | 0.00072 | 0.00635 | 1.58 |
| LLPIDGANDLFFQPPPLTPTSK | [R].LLPIDGANDLFFQPPPLTPTSK.[V] | 1xPhosph | Q9EQQ9-3 | 0.0011 | 0 | 1 | 2 | 1 | Q9EQQ9-3 | Isoform 3 of Protein O-GlcNAcase OS=Mus musculus GN=Mgea5 | Q9EQQ9-3 | 0 | 2919.57 | 0 | 0.17 | 0.16887 | 16.22 | 89.2 | 110.8 | 11.34 | 13.23 | | High | High | High | High | High | High | High | High | High | High | High | 0 | 0.00026 | 3.66 |
| SLQQIAVEETDCPYQTTEPSPAV | [K].SLQQIAVEETDCPYQTTEPSPAVTPETQGCDF | 2xCarbami | Q9CXH7 2 | 8.4E-06 | 0 | 1 | 1 | 1 | Q9CXH7 | Shugoshin 1 OS=Mus musculus GN=Sgo1 PE=2 SV=1 | Q9CXH7 [1 | 0 | 4604.02 | 0 | 0.17 | 0.11115 | 12.21 | 92.7 | 107.3 | 10.14 | 4.93 | | High | High | High | High | High | High | High | High | High | High | High | 0 | 1.1E-06 | 5.52 |
| SPQPPAEEDEDDFDDTLVAIDTY | [R].SPQPPAEEDEDDFDDTLVAIDTYNCDLHFK.[ | 1xCarbami | Q8VDM6 1 | 0.00137 | 5.7E-05 | 1 | 3 | 2 | Q8VDM6 | Heterogeneous nuclear ribonucleoprotein U-like protein 1 OS=Mus musculus GN=Hnrnpul1 PE=1 SV=1 | Q8VDM6 [ | 0 | 3805.63 | 0 | 0.17 | 0.57959 | 9.15 | 94.4 | 105.6 | 21.45 | 12.46 | | High | High | High | High | High | High | High | High | High | High | High | 2.6E-05 | 0.00034 | 3.16 |
| LLCELQSIMSEDEDEK | [R].LLCELQSIMSEDEDEK.[L] | 1xCarbami | Q8BP27 1 | 2.2E-07 | 0 | 1 | 3 | 2 | Q8BP27 | Swi5-dependent recombination DNA repair protein 1 homolog OS=Mus musculus GN=Sfr1 PE=1 SV=2 | Q8BP27 [2 | 0 | 2493.15 | 0 | 0.18 | 0.75606 | 17.05 | 91.6 | 108.4 | 4.83 | 21.56 | | High | High | High | High | High | High | High | High | High | High | High | 0.00033 | 0.0029 | 2.93 |
| SLQQIAVEETDCPYQTTEPSPAV | [K].SLQQIAVEETDCPYQTTEPSPAVTPETQGCDF | 2xCarbami | Q9CXH7 1 | 0.00013 | 0 | 1 | 1 | 1 | Q9CXH7 | Shugoshin 1 OS=Mus musculus GN=Sgo1 PE=2 SV=1 | Q9CXH7 [1 | 0 | 4524.05 | 0 | 0.18 | 0.19301 | 12.45 | 94.7 | 105.3 | 7.72 | 15.34 | | High | High | High | High | High | High | High | High | High | High | High | 0 | 2.3E-05 | 4.64 |
| NSLESISSIDR | [R].NSLESISSIDR.[E] | 2xPhosph | Q80U72-4 | 0.00491 | 0.0005 | 2 | 5 | 2 | Q80U72-4 | Isoform 4 of Protein scribble homolog OS=Mus musculus GN=ScribProtein scribble homolog OS=Mus musculus GN=Scrib PE=1 SV=2 | Q80U72-4 | 0 | 1609.71 | 0 | 0.19 | 0.04329 | 13.04 | 94.3 | 105.7 | 2.58 | 8.31 | Shared | High | High | High | High | High | High | High | High | High | High | High | 0.00027 | 0.00196 | 3.21 |
| QFSPTMSPTLSSITQGVPLDTSK | [K].QFSPTMSPTLSSITQGVPLDTSK.[L] | 1xPhosph | Q3U182 1 | 1.6E-07 | 0 | 2 | 4 | 2 | Q3U182; C | CREB-regulated transcription coactivator 2 OS=Mus musculus GN=Crtc2 PE=1 SV=2Putative uncharacterized protein OS=Mus musculus GN=Crtc2 PE=2 SV=1 | Q3U182 [4 | 0 | 2960.51 | 0 | 0.19 | 0.89335 | 9.24 | 95.8 | 104.2 | 8.13 | 21.33 | Shared | High | High | High | High | High | High | High | High | High | High | High | 0 | 1.3E-08 | 5.65 |
| LSPIMTEQDDLGDGDVHSLVYPI | [R].LSPIMTEQDDLGDGDVHSLVYPPSAAK.[M] | 1xPhosph | Q9R1E0 1x | 0.01296 | 0.00089 | 1 | 3 | 2 | Q9R1E0 | Forkhead box protein O1 OS=Mus musculus GN=Foxo1 PE=1 SV=2 | Q9R1E0 [3 | 0 | 3393.67 | 0 | 0.2 | 0.18073 | 19.84 | 95.9 | 104.1 | 11.1 | 11.02 | | High | High | High | High | High | High | High | High | High | High | High | 0.00045 | 0.00421 | 2.91 |
| EGSQGADADAALEPEYRPVSPLF | [R].EGSQGADADAALEPEYRPVSPLPEIGHK.[Q] | 1xPhosph | Q0VGT4 1 | 3E-07 | 0 | 1 | 3 | 1 | Q0VGT4 | Protein ZGRF1 OS=Mus musculus GN=Zgrf1 PE=1 SV=2 | Q0VGT4 [4 | 0 | 3471.72 | 0 | 0.21 | 0.45237 | 29.7 | 97.2 | 102.8 | 15.52 | 14.82 | | High | High | High | High | High | High | High | High | High | High | High | 0 | 2.6E-08 | 3.97 |
| NLEQILNGGESPK | [K].NLEQILNGGESPK.[Q-] | 1xDeamid | Q9ERG2 1 | 7E-09 | 0 | 2 | 5 | 9 | Q9ERG2: B | Striatin-3 OS=Mus musculus GN=Strn3 PE=1 SV=1Striatin-3 OS=Mus musculus GN=Strn3 PE=1 SV=1 | Q9ERG2 [2 | 0 | 1938 | 0 | 0.22 | 0.10124 | 2.89 | 92.2 | 107.8 | 8.36 | 9.54 | Shared | High | High | High | High | High | High | High | High | High | High | High | 0 | 4.2E-06 | 4.17 |
| NSLESISSIDR | [R].NSLESISSIDR.[E] | 1xPhosph | Q80U72-4 | 0.00246 | 0.00011 | 2 | 5 | 1 | Q80U72-4 | Isoform 4 of Protein scribble homolog OS=Mus musculus GN=ScribProtein scribble homolog OS=Mus musculus GN=Scrib PE=1 SV=2 | Q80U72-4 | 0 | 1529.74 | 0 | 0.22 | 0.05606 | 12.63 | 88.2 | 111.8 | 6.85 | 12.46 | Shared | High | High | High | High | High | High | High | High | High | High | High | 4.9E-05 | 0.00065 | 2.85 |
| NLEQILNGGESPK | [K].NLEQILNGGESPK.[Q-] | 1xPhosph | Q9ERG2 1 | 1.4E-08 | 0 | 2 | 5 | 2 | Q9ERG2: B | Striatin-3 OS=Mus musculus GN=Strn3 PE=1 SV=1Striatin-3 OS=Mus musculus GN=Strn3 PE=1 SV=1 | Q9ERG2 [2 | 0 | 1937.01 | 0 | 0.23 | 0.07272 | 10.77 | 91.6 | 108.4 | 7.17 | 7.27 | Shared | High | High | High | High | High | High | High | High | High | High | High | 0 | 8.4E-10 | 3.69 |
| SQSWLSNIGPTHR | [R].SQSWLSNIGPTHR.[D] | 1xPhosph | E9QPA1 1 | 0.00074 | 0 | 1 | 4 | 1 | E9QPA1 | RAB6A-GEF complex partner protein 1 OS=Mus musculus GN=Ric1 PE=1 SV=1 | E9QPA1 [1 | 0 | 1791.87 | 0 | 0.23 | 0.23676 | 18.28 | 94.8 | 105.2 | 6.14 | 11.34 | | High | High | High | High | High | High | High | High | High | High | High | 0 | 0.00017 | 2.42 |
| RSSYLLAITTER | [R].RSSYLLAITTER.[S] | 1xPhosph | B1AQY2 1 | 0.03769 | 0.00249 | 1 | 4 | 2 | B1AQY2 | Rho GTPase-activating protein 23 OS=Mus musculus GN=Arhgap23 PE=1 SV=1 | B1AQY2 [6 | 1 | 1718.9 | 0 | 0.24 | 0.09288 | 10.31 | 92.8 | 107.2 | 8.64 | 4.87 | | High | High | High | High | High | High | High | High | High | High | High | 0.00246 | 0.02655 | 1.99 |
| STEDGSHDDLFLIK | [K].STEDGSHDDLFLIK.[Q] | 1xPhosph | B1AX39 1x | 0.04003 | 0.00287 | 1 | 2 | 2 | B1AX39 | Zinc finger CCHC domain-containing protein 7 OS=Mus musculus GN=Zcchc7 PE=2 SV=1 | B1AX39 [5 | 0 | 2115.04 | 0 | 0.24 | 0.03999 | 9.12 | 88.6 | 111.4 | 8.81 | 9.5 | | High | High | High | High | High | High | High | High | High | High | High | 0.0015 | 0.01494 | 2.38 |
| CPLPCIPPLVGTPVK | [R].CPLPCIPPLVGTPVK.[I] | 2xCarbami | Q9WVM1 | 0.00072 | 0 | 1 | 1 | 1 | Q9WVM1 | Rac GTPase-activating protein 1 OS=Mus musculus GN=Racgap1 PE=1 SV=1 | Q9WVM1 | 0 | 1957.03 | 0 | 0.25 | 0.22416 | 14.9 | 90.6 | 109.4 | 11.45 | 13.38 | | High | High | High | High | High | High | High | High | High | High | High | 0 | 0.00016 | 3.06 |
| LDHINFPVFEPSTPDPAPAK | [K].LDHINFPVFEPSTPDPAPAK.[N] | 1xPhosph | Q8VE18 1 | 3.5E-05 | 0 | 1 | 2 | 1 | Q8VE18 | Protein SMG8 OS=Mus musculus GN=Smg8 PE=1 SV=1 | Q8VE18 [6 | 0 | 2730.39 | 0 | 0.26 | 0.09022 | 12.11 | 95.2 | 104.8 | 7.23 | 8.15 | | High | High | High | High | High | High | High | High | High | High | High | 0 | 5.5E-06 | 2.84 |
| TSSTDLSDIPALPANPIPVIK | [K].TSSTDLSDIPALPANPIPVIK.[N] | 1xPhosph | Q3V141 1 | 9.8E-07 | 0 | 1 | 7 | 7 | Q3V141 | Putative uncharacterized protein (Fragment) OS=Mus musculus GN=Papola PE=2 SV=1 | Q3V141 [7 | 0 | 2687.47 | 0 | 0.26 | 0.08305 | 8.46 | 92.5 | 107.5 | 4.88 | 8.44 | | High | High | High | High | High | High | High | High | High | High | High | 0 | 2.1E-07 | 4.13 |
| QEPVISESYNGSTPSILSLIK | [K].QEPVISESYNGSTPSILSLIK.[Q] | 1xDeamid | F8WGX5 1 | 0.00422 | 0.00036 | 1 | 2 | 4 | F8WGX5 | DmX-like protein 1 OS=Mus musculus GN=Dmxl1 PE=1 SV=1 | F8WGX5 [ | 0 | 2801.46 | 0 | 0.27 | 0.03849 | 4.66 | 86 | 114 | 4.89 | 14.55 | | High | High | High | High | High | High | High | High | High | High | High | 0.0008 | 0.00717 | 3.09 |
| QNSLGCIGECGVDSAFEAPR | [R].QNSLGCIGECGVDSAFEAPR.[L] | 2xCarbami | G3UYU0 1 | 1.5E-08 | 0 | 1 | 1 | 2 | G3UYU0 | Mex3 RNA-binding family member A OS=Mus musculus GN=Mex3a PE=1 SV=1 | G3UYU0 [3 | 0 | 2476.08 | 0 | 0.27 | 0.00143 | 1.57 | 88.8 | 111.2 | 2.86 | 4.94 | | High | High | High | High | High | High | High | High | High | High | High | 0 | 8.8E-10 | 5.79 |
| VHGEPASDLSDID | [R].VHGEPASDLSDID.[-] | 2xPhosph | E9PZ26 2x | 0.01029 | 0.00077 | 1 | 5 | 3 | E9PZ26 | Bromodomain-containing 1 OS=Mus musculus GN=Brd1 PE=1 SV=1 | E9PZ26 [1 | 0 | 1743.71 | 0 | 0.27 | 0.02101 | 6.91 | 87.7 | 112.3 | 5.49 | 7.72 | | High | High | High | High | High | High | High | High | High | High | High | 0.00038 | 0.00323 | 2.3 |
| ACSDSLESIPAGQVASDDLR | [R].ACSDSLESIPAGQVASDDLR.[D] | 1xCarbami | Q80TM9 1 | 1.1E-05 | 0 | 1 | 5 | 3 | Q80TM9 | Nischarin OS=Mus musculus GN=Nisch PE=1 SV=2 | Q80TM9 [ | 0 | 2480.06 | 0 | 0.28 | 0.24045 | 37.97 | 88.3 | 111.7 | 15.61 | 13.77 | | High | High | High | High | High | High | High | High | High | High | High | 0 | 1.6E-06 | 4.1 |
| GRSPQPPAEEDEDDFDDTLVAII | [R].GRSPQPPAEEDEDDFDDTLVAIDTYNCDLHF | 1xCarbami | Q8VDM6 1 | 1.2E-06 | 0 | 1 | 3 | 6 | Q8VDM6 | Heterogeneous nuclear ribonucleoprotein U-like protein 1 OS=Mus musculus GN=Hnrnpul1 PE=1 SV=1 | Q8VDM6 [ | 1 | 4247.92 | 0 | 0.28 | 0.0876 | 7.92 | 92.1 | 107.9 | 12.74 | 8.3 | | High | High | High | High | High | High | High | High | High | High | High | 0 | 2.2E-07 | 5.74 |
| SLLECIPAQSDEEK | [K].SLLECIPAQSDEEK.[E] | 1xCarbami | A2CG63 1 | 2.4E-06 | 0 | 1 | 4 | 3 | A2CG63 | AT-rich interactive domain-containing protein 4B OS=Mus musculus GN=Arid4b PE=1 SV=1 | A2CG63 [4 | 0 | 2157.06 | 0 | 0.28 | 0.01757 | 7.51 | 92.9 | 107.1 | 4.31 | 6.13 | | High | High | High | High | High | High | High | High | High | High | High | 0 | 2.8E-07 | 3.09 |
| VASGSDLHLTDIDSDSNR | [K].VASGSDLHLTDIDSDSNR.[G] | 1xPhosph | Q9JMH9-6 | 2.2E-05 | 0 | 1 | 8 | 1 | Q9JMH9-6 | Isoform 6 of Unconventional myosin-XVIIIa OS=Mus musculus GN=Myo18a | Q9JMH9-6 | 0 | 2211.01 | 0 | 0.28 | 0.49094 | 25.74 | 90.3 | 109.7 | 18.11 | 18.4 | | High | High | High | High | High | High | High | High | High | High | High | 0 | 3.3E-06 | 2.47 |
| LLLDIPLQTPHK | [K].LLLDIPLQTPHK.[L] | 1xPhosph | Q9ERU9 1 | 9E-05 | 0 | 1 | 1 | 1 | Q9ERU9 | E3 SUMO-protein ligase RanBP2 OS=Mus musculus GN=Ranbp2 PE=1 SV=2 | Q9ERU9 [1 | 0 | 1926.12 | 0 | 0.29 | 0.32534 | 32.85 | 86.4 | 113.6 | 10.22 | 26.37 | | High | High | High | High | High | High | High | High | High | High | High | 0 | 1.6E-05 | 2.88 |
| QTLINIPALNDSDSEVEDISEIESN | [K].QTLINIPALNDSDSEVEDISEIESNR.[S] | 1xPhosph | Q9Z0U1 1 | 1.5E-06 | 0 | 1 | 1 | 7 | Q9Z0U1 | Tight junction protein ZO-2 OS=Mus musculus GN=Tjp2 PE=1 SV=2 | Q9Z0U1 [3 | 0 | 3210.53 | 0 | 0.29 | 0.17263 | 16.21 | 97.2 | 102.8 | 8.71 | 14.4 | | High | High | High | High | High | High | High | High | High | High | High | 0 | 1.6E-07 | 4.16 |
| VTEETEEPIVECQECETEVSPSQT | [K].VTEETEEPIVECQECETEVSPSQTGGSSGDLGI | 2xCarbami | P70399 1x | 2E-06 | 0 | 1 | 4 | 5 | P70399 | TP53-binding protein 1 OS=Mus musculus GN=Tp53bp1 PE=1 SV=3 | P70399 [1 | 0 | 4831.13 | 0 | 0.3 | 0.08216 | 9.95 | 93.8 | 106.2 | 9.42 | 12.3 | | High | High | High | High | High | High | High | High | High | High | High | 0 | 2.2E-07 | 5.15 |
| NEPLLTPIPLK | [R].NEPLLTPIPLK.[R] | 1xPhosph | P33609 1x | 0.00213 | 5.7E-05 | 1 | 3 | 2 | P33609 | DNA polymerase alpha catalytic subunit OS=Mus musculus GN=Pola1 PE=1 SV=2 | P33609 [2 | 0 | 1773.03 | 0 | 0.33 | 0.01272 | 9.38 | 86.6 | 113.4 | 10.37 | 9.07 | | High | High | High | High | High | High | High | High | High | High | High | 2.6E-05 | 0.00056 | 2.27 |
| QGSIEDLLHSQVTLHSIRPR | [R].QGSIEDLLHSQVTLHSIRPR.[D] | 1xPhosph | Q9D6X5 1 | 0.03428 | 0.0022 | 1 | 1 | 1 | Q9D6X5 | Solute carrier family 52, riboflavin transporter, member 3 OS=Mus musculus GN=Slc52a3 PE=1 SV=3 | Q9D6X5 [2 | 0 | 2595.36 | 0 | 0.33 | 0.70393 | 14.32 | 90.7 | 109.3 | 18.06 | 12.32 | | High | High | High | High | High | High | High | High | High | High | High | 0.00115 | 0.01247 | 2.06 |
| SPPSGSQSLLEGIMTASSSK | [R].SPPSGSQSLLEGIMTASSSK.[G] | 1xPhosph | B9EKJ7 1x | 2.8E-05 | 0 | 1 | 3 | 5 | B9EKJ7 | Tankyrase 1 binding protein 1 OS=Mus musculus GN=Tnks1bp1 PE=2 SV=1 | B9EKJ7 [14 | 0 | 2518.25 | 0 | 0.33 | 0.11868 | 25.82 | 90.5 | 109.5 | 8.87 | 20.28 | | High | High | High | High | High | High | High | High | High | High | High | 0 | 7.9E-06 | 3.07 |
| NEPLLTPIPLK | [R].NEPLLTPIPLK.[R] | 1xPhosph | P33609 1x | 0.00727 | 0.00062 | 1 | 3 | 1 | P33609 | DNA polymerase alpha catalytic subunit OS=Mus musculus GN=Pola1 PE=1 SV=2 | P33609 [2 | 0 | 1543.87 | 0 | 0.34 | 0.73928 | 18.71 | 88.4 | 111.6 | 25.97 | 38.18 | | High | High | High | High | High | High | High | High | High | High | High | 0.00027 | 0.0022 | 2.41 |
| VTEETEEPIVECQECETEVSPSQT | [K].VTEETEEPIVECQECETEVSPSQTGGSSGDLGI | 2xCarbami | P70399 2x | 6.2E-05 | 0 | 1 | 4 | 2 | P70399 | TP53-binding protein 1 OS=Mus musculus GN=Tp53bp1 PE=1 SV=3 | P70399 [1 | 0 | 4910.11 | 0 | 0.34 | 0.7088 | 19.6 | 96.3 | 103.7 | 12.62 | 29.99 | | High | High | High | High | High | High | High | High | High | High | High | 0 | 1.1E-05 | 4.76 |
| RLSSLGEVDATALTPELLLYIR | [R].RLSSLGEVDATALTPELLLYIR.[C] | 1xPhosph | Q3UPD0 1 | 0.00665 | 0.00062 | 1 | 5 | 1 | Q3UPD0 | Putative uncharacterized protein OS=Mus musculus GN=Scfd2 PE=2 SV=1 | Q3UPD0 [1 | 1 | 2739.49 | 0 | 0.35 | 0.03022 | 8.61 | 90.4 | 109.6 | 15.72 | 8.09 | | High | High | High | High | High | High | High | High | High | High | High | 0.00027 | 0.00198 | 2.93 |
| EPIYTLPAIISK | [K].EPIYTLPAIISK.[K] | 1xPhosph | B2RQE8 1x | 0.01441 | 0.00101 | 1 | 1 | 2 | B2RQE8 | Rho GTPase-activating protein 42 OS=Mus musculus GN=Arhgap42 PE=1 SV=1 | B2RQE8 [3 | 0 | 1883.07 | 0 | 0.36 | 0.02583 | 20.54 | 88.9 | 111.1 | 15.64 | 4.31 | | High | High | High | High | High | High | High | High | High | High | High | 0.001 | 0.01032 | 2.34 |
| SLTLGPSLSLGNISGVSMK | [R].SLTLGPSLSLGNISGVSMK.[S] | 1xPhosph | B2RWU4 1 | 0.00015 | 0 | 2 | 11 | 2 | B2RWU4: ( | Cordon-bleu OS=Mus musculus GN=Cobl PE=2 SV=1Protein cordon-bleu OS=Mus musculus GN=Cobl PE=1 SV=1 | B2RWU4 [ | 0 | 2399.3 | 0 | 0.36 | 0.02368 | 9.46 | 86 | 114 | 4.57 | 12.29 | Shared | High | High | High | High | High | High | High | High | High | High | High | 0 | 2.8E-05 | 2.29 |
| VTEETEEPIVECQECETEVSPSQT | [K].VTEETEEPIVECQECETEVSPSQTGGSSGDLGI | 2xCarbami | P70399 1x | 1.4E-06 | 0 | 1 | 4 | 6 | P70399 | TP53-binding protein 1 OS=Mus musculus GN=Tp53bp1 PE=1 SV=3 | P70399 [1 | 0 | 4830.14 | 0 | 0.36 | 0.06305 | 7.24 | 85.8 | 114.2 | 4.92 | 16.55 | | High | High | High | High | High | High | High | High | High | High | High | 0 | 1.4E-07 | 6.63 |
| YHSLGNISR | [R].YHSLGNISR.[G] | 1xPhosph | Q08274 1 | 0.00478 | 0.00046 | 1 | 4 | 2 | Q08274 | Dystrophia myotonica WD repeat-containing protein OS=Mus musculus GN=Dmwd PE=1 SV=3 | Q08274 [5 | 0 | 1355.67 | 0 | 0.36 | 0.00303 | 7.21 | 87.8 | 112.2 | 3.22 | 5.22 | | High | High | High | High | High | High | High | High | High | High | High | 0.00027 | 0.00214 | 1.8 |
| HSETTLVNIYDLSDEDTGR | [R].HSETTLVNIYDLSDEDTGR.[R] | 1xPhosph | Q3UQI9 1x | 5.8E-08 | 0 | 1 | 4 | 1 | Q3UQI9 | Probable ubiquitin carboxyl-terminal hydrolase MINDY-4 OS=Mus musculus GN=Mindy4 PE=1 SV=1 | Q3UQI9 [1 | 0 | 2474.13 | 0 | 0.37 | 0.01081 | 8.68 | 91.4 | 108.6 | 9.45 | 8.72 | | High | High | High | High | High | High | High | High | High | High | High | 0 | 4.1E-09 | 3.7 |
| ILLSQTTGMAIPLQASSLDDISLL | [R].ILLSQTTGMAIPLQASSLDDISLLSTPK.[R] | 1xPhosph | F6ZDS4 1x | 6.8E-05 | 0 | 1 | 7 | 1 | F6ZDS4 | Nucleoprotein TPR OS=Mus musculus GN=Tpr PE=1 SV=1 | F6ZDS4 [6 | 0 | 3451.88 | 0 | 0.38 | 0.02885 | 22.32 | 86.2 | 113.8 | 9.98 | 8.55 | | High | High | High | High | High | High | High | High | High | High | High | 0 | 1.2E-05 | 3.71 |
| RQNSSDSISSLNSITSHSSIGSSK | [K].RQNSSDSISSLNSITSHSSIGSSK.[D] | 1xPhosph | Q8CH77 1 | 6E-06 | 0 | 1 | 5 | 1 | Q8CH77 | Neuron navigator 1 OS=Mus musculus GN=Nav1 PE=1 SV=2 | Q8CH77 [1 | 1 | 3017.49 | 0 | 0.38 | 0.26965 | 30.73 | 86.5 | 113.5 | 21.72 | 21.16 | | High | High | High | High | High | High | High | High | High | High | High | 0 | 7.6E-07 | 3.81 |
| GVGDVLLQLGCISLGSGVSPK | [R].GVGDVLLQLGCISLGSGVSPK.[R] | 1xCarbami | Q7TSV9 1 | 3.9E-08 | 0 | 1 | 2 | 1 | Q7TSV9 | Enkurin domain-containing protein 1 OS=Mus musculus GN=Enkd1 PE=1 SV=1 | Q7TSV9 [7 | 0 | 2594.4 | 0 | 0.39 | 0.02074 | 0.91 | 86.6 | 113.4 | 9.98 | 6.73 | | High | High | High | High | High | High | High | High | High | High | High | 0 | 2.7E-09 | 3.77 |
| NLEQILNGGESPK | [K].NLEQILNGGESPK.[Q-] | 1xPhosph | Q9ERG2 1 | 1.3E-05 | 0 | 2 | 5 | 2 | Q9ERG2; B | Striatin-3 OS=Mus musculus GN=Strn3 PE=1 SV=1Striatin-3 OS=Mus musculus GN=Strn3 PE=1 SV=1 | Q9ERG2 [2 | 0 | 1707.85 | 0 | 0.4 | 0.69909 | 10.74 | 89.9 | 110.1 | 36.59 | 42.07 | Shared | High | High | High | High | High | High | High | High | High | High | High | 0 | 1.8E-06 | 3.76 |
| GYALVPIIVSPSTPK | [K].GYALVPIIVSPSTPK.[T] | 1xPhosph | B9EKI3 1xP | 0.00019 | 0 | 1 | 2 | 2 | B9EKI3 | TATA element modulatory factor OS=Mus musculus GN=Tmf1 PE=1 SV=2 | B9EKI3 [34 | 0 | 2080.19 | 0 | 0.41 | 0.07466 | 4.23 | 86.1 | 113.9 | 10.22 | 13.15 | | High | High | High | High | High | High | High | High | High | High | High | 0 | 3.7E-05 | 3.22 |
| QNSSDSISSLNSITSHSSIGSSK | [R].QNSSDSISSLNSITSHSSIGSSK.[D] | 1xPhosph | Q8CH77 1 | 5.7E-07 | 0 | 1 | 5 | 3 | Q8CH77 | Neuron navigator 1 OS=Mus musculus GN=Nav1 PE=1 SV=2 | Q8CH77 [1 | 0 | 2861.39 | 0 | 0.41 | 0.30332 | 17.77 | 84.9 | 115.1 | 23.12 | 24.98 | | High | High | High | High | High | High | High | High | High | High | High | 0 | 1.2E-07 | 4.49 |
| HLLPISTSDLSSR | [R].HLLPISTSDLSSR.[E] | 1xPhosph | Q8CD26 1 | 0.00839 | 0.00074 | 1 | 2 | 1 | Q8CD26 | Solute carrier family 35 member E1 OS=Mus musculus GN=Slc35e1 PE=1 SV=2 | Q8CD26 [3 | 0 | 1734.9 | 0 | 0.42 | 0.01874 | 14.02 | 82.6 | 117.4 | 11.2 | 19.51 | | High | High | High | High | High | High | High | High | High | High | High | 0.00033 | 0.00258 | 1.94 |
| LSSIGIQVDCIQPVPK | [K].LSSIGIQVDCIQPVPK.[E] | 1xCarbami | B1AZP2 1x | 0.00013 | 0 | 1 | 4 | 2 | B1AZP2 | Disks large-associated protein 4 OS=Mus musculus GN=Dlgap4 PE=1 SV=1 | B1AZP2 [6 | 0 | 2292.24 | 0 | 0.42 | 0.02038 | 13.62 | 81.8 | 118.2 | 11.29 | 11.72 | | High | High | High | High | High | High | High | High | High | High | High | 0 | 2.4E-05 | 3.52 |
| EALPSIWDSPTK | [K].EALPSIWDSPTK.[Q] | 1xPhosph | Q9CTE8 1x | 0.00648 | 0.00062 | 1 | 4 | 2 | Q9CTE8 | Putative uncharacterized protein (Fragment) OS=Mus musculus GN=Alg5 PE=2 SV=3 | Q9CTE8 [5 | 0 | 1881.98 | 0 | 0.43 | 0.03301 | 24.52 | 81.8 | 118.2 | 12.91 | 12.35 | | High | High | High | High | High | High | High | High | High | High | High | 0.00027 | 0.00193 | 3.31 |
| ELYEITPPEVSEEMIGSPIPEPR | [K].ELYEITPPEVSEEMIGSPIPEPR.[Q] | 1xPhosph | Q7TPS5 1x | 1.4E-06 | 0 | 2 | 6 | 1 | Q7TPS5; A | C2 domain-containing protein 5 OS=Mus musculus GN=C2cd5 PE=1 SV=2C2 domain-containing protein 5 OS=Mus musculus GN=C2cd5 PE=1 SV=1 | Q7TPS5 [6 | 0 | 2921.41 | 0 | 0.43 | 0.00067 | 4.38 | 86.6 | 113.4 | 11.18 | 4.59 | Shared | High | High | High | High | High | High | High | High | High | High | High | 0 | 1.5E-07 | 4.53 |
| NSLGEGHLELWGTPVPVFPLTV | [R].NSLGEGHLELWGTPVPVFPLTVSPGPLIPIPVI | 1xPhosph | Q5SUQ9 1 | 0.00893 | 0.00074 | 1 | 2 | 2 | Q5SUQ9 | CST complex subunit CTC1 OS=Mus musculus GN=Ctc1 PE=1 SV=2 | Q5SUQ9 [1 | 0 | 4501.46 | 0 | 0.43 | 0.05635 | 22.07 | 92.5 | 107.5 | 7.66 | 15.15 | | High | High | High | High | High | High | High | High | High | High | High | 0.00033 | 0.00278 | 3.31 |
| GPSPSLLSTISLDVMPK | [R].GPSPSLLSTISLDVMPK.[F] | 1xPhosph | E9Q8S5 1x | 0.00024 | 0 | 1 | 3 | 1 | E9Q8S5 | Microtubule-associated serine/threonine-protein kinase 3 OS=Mus musculus GN=Mast3 PE=1 SV=2 | E9Q8S5 [7 | 0 | 2296.23 | 0 | 0.44 | 0.0093 | 12.21 | 88.3 | 111.7 | 3.61 | 10.37 | | High | High | High | High | High | High | High | High | High | High | High | 0 | 4.8E-05 | 2.27 |
| LSGQNTMGSQDSLLEITFR | [K].LSGQNTMGSQDSLLEITFR.[S] | 1xPhosph | Q9D3A8 1 | 0.00043 | 0 | 1 | 2 | 1 | Q9D3A8 | Carboxyl-terminal PDZ ligand of neuronal nitric oxide synthase protein OS=Mus musculus GN=Nos1ap PE=1 SV=3 | Q9D3A8 [3 | 0 | 2422.15 | 0 | 0.44 | 0.10194 | 14.88 | 89.9 | 110.1 | 20.47 | 12.74 | | High | High | High | High | High | High | High | High | High | High | High | 0 | 9.1E-05 | 3.29 |
| LWLISEGYAAGPGDISPDWSPEF | [R].LWLISEGYAAGPGDISPDWSPER.[C] | 1xPhosph | Q5EBP3 1x | 0.00011 | 0 | 1 | 2 | 1 | Q5EBP3 | Armadillo repeat-containing protein 5 OS=Mus musculus GN=Armc5 PE=2 SV=1 | Q5EBP3 [4 | 0 | 2825.34 | 0 | 0.45 | 0.04282 | 26.67 | 89.8 | 110.2 | 11.62 | 13.66 | | High | High | High | High | High | High | High | High | High | High | High | 0 | 1.9E-05 | 2.89 |
| SLSEIAIDLTETGTLK | [R].SLSEIAIDLTETGTLK.[T] | 1xPhosph | Q8BIE6-2 | 0.09237 | 0.00721 | 1 | 7 | 1 | Q8BIE6-2 | Isoform 2 of FERM domain-containing protein 4A OS=Mus musculus GN=Frmd4a | Q8BIE6-2 [ | 0 | 2229.2 | 0 | 0.48 | 0.0221 | 10.15 | 83.5 | 116.5 | 7.86 | 16.87 | | High | High | High | High | High | High | High | High | High | High | High | 0.00367 | 0.03948 | 2.45 |
| SRPALSPLGDIDFCPPNPGPDGP | [R].SRPALSPLGDIDFCPPNPGPDGPR.[R] | 1xCarbami | E9PYL2 1x | 1.4E-07 | 0 | 1 | 2 | 1 | E9PYL2 | Proline-rich protein 12 OS=Mus musculus GN=Prr12 PE=1 SV=1 | E9PYL2 [11 | 0 | 2841.36 | 0 | 0.49 | 0.02413 | 9.57 | 83.3 | 116.7 | 8.03 | 9.97 | | High | High | High | High | High | High | High | High | High | High | High | 0 | 1.1E-08 | 3.42 |
| VDLSDGNTASCPLSPIK | [K].VDLSDGNTASCPLSPIK.[M] | 1xCarbami | Q3U285-3 | 0.00031 | 0 | 1 | 3 | 1 | Q3U285-3 | Isoform 3 of Ligand-dependent nuclear receptor corepressor-like protein OS=Mus musculus GN=Lcorl | Q3U285-3 | 0 | 2312.16 | 0 | 0.49 | 0.01263 | 9.7 | 83 | 117 | 2.75 | 10.58 | | High | High | High | High | High | High | High | High | High | High | High | 0 | 6.4E-05 | 2.94 |
| LGSIAEIDLGVPPPIMK | [R].LGSIAEIDLGVPPPIMK.[S] | 1xPhosph | Q99MR6 1 | 0.01588 | 0.00112 | 1 | 5 | 1 | Q99MR6 | Serrate RNA effector molecule homolog OS=Mus musculus GN=Srrt PE=1 SV=1 | Q99MR6 [ | 0 | 2288.27 | 0 | 0.51 | 0.00811 | 14.85 | 79.2 | 120.8 | 10.85 | 14.04 | | High | High | High | High | High | High | High | High | High | High | High | 0.00058 | 0.00527 | 2.84 |
| TSSIADEGTYTLDSILR | [R].TSSIADEGTYTLDSILR.[Q] | 1xPhosph | D3Z4J3 1x | 1E-07 | 0 | 1 | 4 | 8 | D3Z4J3 | Unconventional myosin-Va OS=Mus musculus GN=Myo5a PE=1 SV=1 | D3Z4J3 [16 | 0 | 2151.04 | 0 | 0.51 | 0.00878 | 19.88 | 82.4 | 117.6 | 9.78 | 4.43 | | High | High | High | High | High | High | High | High | High | High | High | 0 | 1.8E-08 | 4.27 |
| AGTQVPVLGPEDDLAGIFLQIFPI | [R].AGTQVPVLGPEDDLAGIFLQIFPLSPDPR.[W] | 1xPhosph | K3W4P2 1 | 0.00115 | 0 | 1 | 4 | 1 | K3W4P2 | Integrator complex subunit 1 OS=Mus musculus GN=Ints1 PE=1 SV=2 | K3W4P2 [ | 0 | 3371.75 | 0 | 0.52 | 0.00751 | 8.29 | 83.4 | 116.6 | 8.52 | 6.67 | | High | High | High | High | High | High | High | High | High | High | High | 0 | 0.00028 | 3.69 |
| QSTSFLVLQEILESDGK | [K].QSTSFLVLQEILESDGK.[G] | 1xPhosph | O70400 1 | 0.00033 | 0 | 1 | 2 | 3 | O70400 | PDZ and LIM domain protein 1 OS=Mus musculus GN=Pdlim1 PE=1 SV=4 | O70400 [2 | 0 | 2432.27 | 0 | 0.52 | 0.03297 | 30.19 | 78.6 | 121.4 | 4.33 | 17.73 | | High | High | High | High | High | High | High | High | High | High | High | 0 | 0.00011 | 2.81 |
| TLSMIEEEIR | [R].TLSMIEEEIR.[A] | 1xPhosph | F7AA26 1x | 0.00325 | 0.00016 | 1 | 10 | 5 | F7AA26 | Paralemmin A kinase anchor protein (Fragment) OS=Mus musculus GN=Pakap PE=1 SV=1 | F7AA26 [9 | 0 | 1545.74 | 0 | 0.52 | 0.01123 | 7.54 | 81.5 | 118.5 | 4.34 | 13.97 | | High | High | High | High | High | High | High | High | High | High | High | 7.2E-05 | 0.00089 | 3.02 |
| TLSMIEEEIR | [R].TLSMIEEEIR.[A] | 1xPhosph | F7AA26 1x | 0.00282 | 0.00011 | 1 | 10 | 2 | F7AA26 | Paralemmin A kinase anchor protein (Fragment) OS=Mus musculus GN=Pakap PE=1 SV=1 | F7AA26 [9 | 0 | 1529.75 | 0 | 0.52 | 0.00453 | 15.34 | 82.9 | 117.1 | 7.36 | 8.79 | | High | High | High | High | High | High | High | High | High | High | High | 7.2E-05 | 0.00076 | 3.62 |
| LSPPPPVTTLLPSIK | [R].LSPPPPVTTLLPSIK.[C] | 1xPhosph | Q3TTA7 1 | 1.7E-05 | 0 | 1 | 3 | 3 | Q3TTA7 | E3 ubiquitin-protein ligase CBL-B OS=Mus musculus GN=Cblb PE=1 SV=3 | Q3TTA7 [6 | 0 | 2098.23 | 0 | 0.53 | 0.03866 | 19.83 | 81.8 | 118.2 | 10.82 | 14.92 | | High | High | High | High | High | High | High | High | High | High | High | 0 | 2.5E-06 | 3.07 |
| RLSQISAGETEYNTQDCK | [R].RLSQISAGETEYNTQDCK.[-] | 1xCarbami | E9PYP2 1x | 0.00064 | 0 | 1 | 3 | 1 | E9PYP2 | Neurobeachin-like 1 OS=Mus musculus GN=Nbeal1 PE=1 SV=1 | E9PYP2 [2 | 1 | 2638.26 | 0 | 0.54 | 0.05315 | 21.61 | 82.9 | 117.1 | 14.39 | 19.34 | | High | High | High | High | High | High | High | High | High | High | High | 0 | 0.00015 | 4.58 |
| RPSLPCISR | [R].RPSLPCISR.[E] | 1xCarbami | Q61409 1 | 0.01962 | 0.00144 | 1 | 2 | 1 | Q61409 | cGMP-inhibited 3',5'-cyclic phosphodiesterase B OS=Mus musculus GN=Pde3b PE=1 SV=2 | Q61409 [2 | 0 | 1394.72 | 0 | 0.54 | 0.00519 | 7.84 | 76.9 | 123.1 | 10.13 | 12.45 | | High | High | High | High | High | High | High | High | High | High | High | 0.00076 | 0.00669 | 1.67 |
| LANISVPASESPR | [R].LANISVPASESPR.[L] | 1xPhosph | B9EKJ7 1x | 0.00206 | 5.7E-05 | 1 | 2 | 3 | B9EKJ7 | Tankyrase 1 binding protein 1 OS=Mus musculus GN=Tnks1bp1 PE=2 SV=1 | B9EKJ7 [29 | 0 | 1649.85 | 0 | 0.55 | 0.00312 | 10.14 | 80 | 120 | 4.38 | 6.88 | | High | High | High | High | High | High | High | High | High | High | High | 2.6E-05 | 0.00054 | 2.27 |
| LQSIGTENTEENR | [R].LQSIGTENTEENR.[R] | 1xPhosph | A6ZI44 1x | 2.3E-06 | 0 | 1 | 11 | 2 | A6ZI44 | Fructose-bisphosphate aldolase OS=Mus musculus GN=Aldoa PE=1 SV=1 | A6ZI44 [98 | 0 | 1799.84 | 0 | 0.55 | 0.00318 | 4.66 | 81.1 | 118.9 | 2.99 | 7.56 | | High | High | High | High | High | High | High | High | High | High | High | 0 | 2.6E-07 | 3.64 |
| IMETSPTLLQISPGPPK | [R].IMETSPTLLQISPGPPK.[-] | 1xPhosph | Q6ZQ33 1 | 0.00052 | 0 | 1 | 4 | 1 | Q6ZQ33 | MKIAA0857 protein (Fragment) OS=Mus musculus GN=Rab11fip5 PE=2 SV=1 | Q6ZQ33 [1 | 0 | 2347.27 | 0 | 0.56 | 0.01014 | 17.43 | 80.9 | 119.1 | 7.35 | 7.1 | | High | High | High | High | High | High | High | High | High | High | High | 0 | 0.00012 | 2.63 |
| LSGINEIPRPFSPPVTSNTSPPPT | [K].LSGINEIPRPFSPPVTSNTSPPPTAPLAR.[A] | 2xPhosph | Q3UQN2 2 | 0.00035 | 0 | 1 | 3 | 2 | Q3UQN2 | F-BAR domain only protein 2 OS=Mus musculus GN=Fcho2 PE=1 SV=1 | Q3UQN2 [ | 0 | 3402.71 | 0 | 0.56 | 0.00518 | 14.01 | 79.4 | 120.6 | 8.59 | 12.19 | | High | High | High | High | High | High | High | High | High | High | High | 0 | 7.3E-05 | 2.77 |
| QEPVISESYNGSTPSILSLIK | [K].QEPVISESYNGSTPSILSLIK.[Q] | 1xPhosph | F8WGX5 1 | 1.5E-06 | 0 | 1 | 2 | 2 | F8WGX5 | DmX-like protein 1 OS=Mus musculus GN=Dmxl1 PE=1 SV=1 | F8WGX5 [ | 0 | 2800.48 | 0 | 0.57 | 0.0095 | 10.02 | 80.5 | 119.5 | 7.84 | 12.9 | | High | High | High | High | High | High | High | High | High | High | High | 0 | 1.7E-07 | 4.55 |
| VSLEEIYSGCTK | [RK].VSLEEIYSGCTK.[KR] | 1xCarbami | Q3TIT6 1x | 0.00942 | 0.00074 | 1 | 4 | 1 | Q3TIT6 | Putative uncharacterized protein OS=Mus musculus GN=Dnajb1 PE=2 SV=1 | Q3TIT6 [17 | 0 | 1923.95 | 0 | 0.58 | 0.09273 | 18.89 | 82.5 | 117.5 | 19.28 | 8.57 | | High | High | High | High | High | High | High | High | High | High | High | 0.00033 | 0.00294 | 1.85 |
| ELYEITPPEVSEEMIGSPIPEPR | [K].ELYEITPPEVSEEMIGSPIPEPR.[Q] | 1xPhosph | Q7TPS5 1x | 2.2E-07 | 0 | 2 | 6 | 2 | Q7TPS5; A | C2 domain-containing protein 5 OS=Mus musculus GN=C2cd5 PE=1 SV=2C2 domain-containing protein 5 OS=Mus musculus GN=C2cd5 PE=1 SV=1 | Q7TPS5 [6 | 0 | 2937.4 | 0 | 0.59 | 0.01973 | 24.81 | 84.1 | 115.9 | 9.07 | 13.21 | Shared | High | High | High | High | High | High | High | High | High | High | High | 0 | 1.8E-08 | 3.58 |
| LQSIGTENTEENRR | [R].LQSIGTENTEENRR.[F] | 1xPhosph | A6ZI44 1x | 0.02933 | 0.00186 | 1 | 11 | 1 | A6ZI44 | Fructose-bisphosphate aldolase OS=Mus musculus GN=Aldoa PE=1 SV=1 | A6ZI44 [98 | 1 | 1955.94 | 0 | 0.59 | 0.01159 | 21.66 | 80.1 | 119.9 | 7.39 | 10.98 | | High | High | High | High | High | High | High | High | High | High | High | 0.001 | 0.01047 | 0.91 |
| QFSPTMSPTLSSITQGVPLDTSK | [K].QFSPTMSPTLSSITQGVPLDTSK.[L] | 1xPhosph | Q3U182 1 | 0.00081 | 0 | 2 | 4 | 5 | Q3U182: C | CREB-regulated transcription coactivator 2 OS=Mus musculus GN=Crtc2 PE=1 SV=2Putative uncharacterized protein OS=Mus musculus GN=Crtc2 PE=2 SV=1 | Q3U182 [4 | 0 | 2976.5 | 0 | 0.6 | 0.00104 | 11.22 | 80.3 | 119.7 | 5.95 | 4.64 | Shared | High | High | High | High | High | High | High | High | High | High | High | 0.00072 | 0.00634 | 3.23 |
| AYLQSITPQGYSDSEER | [R].AYLQSITPQGYSDSEER.[E] | 1xPhosph | A0A0M3H | 0.00018 | 0 | 1 | 7 | 2 | A0A0M3H | Zinc finger E-box-binding homeobox 2 OS=Mus musculus GN=Zeb2 PE=1 SV=1 | A0A0M3H | 0 | 2253.03 | 0 | 0.63 | 0.1247 | 31.6 | 78.7 | 121.3 | 20.23 | 17.14 | | High | High | High | High | High | High | High | High | High | High | High | 0 | 3.5E-05 | 3.1 |
| RLSNSSLCSIEEEHR | [R].RLSNSSLCSIEEEHR.[T] | 1xCarbami | A0A0A0M | 0.06118 | 0.00465 | 1 | 5 | 2 | A0A0A0M | Ral GTPase-activating protein subunit alpha-2 OS=Mus musculus GN=Ralgapa2 PE=1 SV=1 | A0A0A0M | 1 | 2125.99 | 0 | 0.63 | 0.00438 | 4.45 | 78.6 | 121.4 | 7.55 | 6.52 | | High | High | High | High | High | High | High | High | High | High | High | 0.00237 | 0.02435 | 3.03 |
| QTLINIPALNDSDSEVEDISEIESN | [K].QTLINIPALNDSDSEVEDISEIESNR.[S] | 2xPhosph | Q9Z0U1 2 | 1.1E-05 | 0 | 1 | 1 | 5 | Q9Z0U1 | Tight junction protein ZO-2 OS=Mus musculus GN=Tjp2 PE=1 SV=2 | Q9Z0U1 [3 | 0 | 3290.49 | 0 | 0.64 | 0.03004 | 26.53 | 77.3 | 122.7 | 12.78 | 17.4 | | High | High | High | High | High | High | High | High | High | High | High | 0 | 1.5E-06 | 3.97 |
| SNDTLSGISSIEPSK | [K].SNDTLSGISSIEPSK.[H] | 1xPhosph | Q8BK03 1 | 0.11108 | 0.00899 | 1 | 5 | 1 | Q8BK03 | Mitoguardin 2 OS=Mus musculus GN=Miga2 PE=1 SV=1 | Q8BK03 [1 | 0 | 2073.05 | 0 | 0.64 | 0.01135 | 21.45 | 76.1 | 123.9 | 22.36 | 16.64 | | High | High | High | High | High | High | High | High | High | High | High | 0.00453 | 0.04909 | 2.37 |
| WVSELSYIEEK | [R].WVSELSYIEEK.[E] | 1xPhosph | Q9R0L6-2 | 4E-05 | 0 | 1 | 3 | 3 | Q9R0L6-2 | Isoform 2 of Pericentriolar material 1 protein OS=Mus musculus GN=Pcm1 | Q9R0L6-2 | 0 | 1920.98 | 0 | 0.66 | 0.12479 | 65.16 | 64.4 | 135.6 | 27.82 | 40.71 | | High | High | High | High | High | High | High | High | High | High | High | 0 | 6.5E-06 | 2.82 |

| | | | | | | | | | | | | | | | | | | | | | | | | | | | | | | | | | | | | |
|---|---|---|---|---|---|---|---|---|---|---|---|---|---|---|---|---|---|---|---|---|---|---|---|---|---|---|---|---|---|---|---|---|---|---|---|---|
| EASIDILHSIVK | [R].EASIDILHSIVK.[R] | 1xPhosphc | Q497E8 1x | 0.11648 | 0.00952 | 1 | 11 | 1 | Q497E8 | Proteasome (Prosome, macropain) 26S subunit, non-ATPase, 11 OS=Mus musculus GN=Psmd11 PE=2 SV=1 | Q497E8 [2 | 0 | 1863.04 | 0 | 0.67 | 0.00193 | 10.27 | 78 | 122 | 4.45 | 5.37 | | High | High | High | High | High | High | High | High | High | High | High | 0.00476 | 0.05182 | 2.19 |
| SLHGEFSASDSSLTDIQETR | [K].SLHGEFSASDSSLTDIQETR.[R] | 2xPhosphc | Q8C0T5 2x | 8.5E-05 | 0 | 1 | 3 | 1 | Q8C0T5 | Signal-induced proliferation-associated 1-like protein 1 OS=Mus musculus GN=Sipa1l1 PE=1 SV=2 | Q8C0T5 [1 | 0 | 2569.11 | 0 | 0.67 | 0.00544 | 13.44 | 78.1 | 121.9 | 9.22 | 8.36 | | High | High | High | High | High | High | High | High | High | High | High | 0 | 1.5E-05 | 3.82 |
| EALNIIGDISTSTVSTPVPPPVDD | [K].EALNIIGDISTSTVSTPVPPPVDDTWLQNTSG | 2xDeamid | F8WIV5 1x | 0.01493 | 0.00105 | 1 | 9 | 3 | F8WIV5 | Dynamin-2 OS=Mus musculus GN=Dnm2 PE=1 SV=2 | F8WIV5 [7 | 0 | 4439.15 | 0 | 0.68 | 0.00894 | 22.3 | 75.3 | 124.7 | 12.67 | 12.1 | | High | High | High | High | High | High | High | High | High | High | High | 0.00055 | 0.00492 | 3.42 |
| EVDSLGSIHSIPSTSVSTGSR | [R].EVDSLGSIHSIPSTSVSTGSR.[S] | 1xPhosphc | B2RY27 1x | 0.01506 | 0.00105 | 1 | 4 | 2 | B2RY27 | MCG125789, isoform CRA_a OS=Mus musculus GN=Taok3 PE=2 SV=1 | B2RY27 [3 | 0 | 2425.18 | 0 | 0.68 | 0.05609 | 8.6 | 76.9 | 123.1 | 13.37 | 19.8 | | High | High | High | High | High | High | High | High | High | High | High | 0.00057 | 0.00498 | 2.55 |
| ASLSDLTHLEDIEGLTVR | [R].ASLSDLTHLEDIEGLTVR.[Q] | 1xPhosphc | Q3UCG9 1 | 0.00072 | 0 | 1 | 6 | 2 | Q3UCG9 | E3 ubiquitin-protein ligase rififylin OS=Mus musculus GN=Rffl PE=1 SV=1 | Q3UCG9 [2 | 0 | 2278.15 | 0 | 0.71 | 0.00129 | 5.15 | 76.1 | 123.9 | 6.01 | 7.9 | | High | High | High | High | High | High | High | High | High | High | High | 0 | 0.00027 | 2.89 |
| SLTLGPSLSLGNISGVSMK | [R].SLTLGPSLSLGNISGVSMK.[S] | 1xPhosphc | B2RWU4 1 | 1.8E-06 | 0 | 2 | 11 | 2 | B2RWU4: ( | Cordon-bleu OS=Mus musculus GN=Cobl PE=2 SV=1Protein cordon-bleu OS=Mus musculus GN=Cobl PE=1 SV=1 | B2RWU4 [ | 0 | 2415.3 | 0 | 0.71 | 0.03904 | 7.55 | 75.7 | 124.3 | 7.29 | 26.98 | Shared | High | High | High | High | High | High | High | High | High | High | High | 0 | 2E-07 | 3.87 |
| GLEFPMTDLDMLSPIHTPQR | [K].GLEFPMTDLDMLSPIHTPQR.[T] | 1xPhosphc | Q3UDC3 1 | 0.0301 | 0.00186 | 1 | 3 | 1 | Q3UDC3 | Putative uncharacterized protein OS=Mus musculus GN=Tom1 PE=1 SV=1 | Q3UDC3 [1 | 0 | 2623.25 | 0 | 0.72 | 0.00374 | 3.14 | 77.1 | 122.9 | 8.46 | 8.73 | | High | High | High | High | High | High | High | High | High | High | High | 0.00101 | 0.01083 | 3.27 |
| NSGLTPPVTIVQPSSSAGQILAQI | [R].NSGLTPPVTIVQPSSSAGQILAQISR.[H] | 1xPhosphc | P53762 1x | 0.0034 | 0.00021 | 1 | 7 | 1 | P53762 | Aryl hydrocarbon receptor nuclear translocator OS=Mus musculus GN=Arnt PE=1 SV=3 | P53762 [6 | 0 | 2930.55 | 0 | 0.74 | 0.03353 | 15.4 | 74.8 | 125.2 | 6.86 | 18.12 | | High | High | High | High | High | High | High | High | High | High | High | 9.4E-05 | 0.00094 | 2.62 |
| SASANSAEAGGDTVTLDDILSLK | [R].SASANSAEAGGDTVTLDDILSLK.[S] | 1xPhosphc | Q9EPQ8 1 | 0.01602 | 0.00112 | 1 | 4 | 2 | Q9EPQ8 | Transcription factor 20 OS=Mus musculus GN=Tcf20 PE=1 SV=3 | Q9EPQ8 [1 | 0 | 2773.39 | 0 | 0.76 | 0.01921 | 27.72 | 73.6 | 126.4 | 15.23 | 14.32 | | High | High | High | High | High | High | High | High | High | High | High | 0.00058 | 0.00534 | 3.62 |
| LSGINEIPRPFSPPVTSNTSPPPT | [K].LSGINEIPRPFSPPVTSNTSPPPTAPLAR.[A] | 1xPhosphc | Q3UQN2 1 | 0.10846 | 0.00884 | 1 | 3 | 1 | Q3UQN2 | F-BAR domain only protein 2 OS=Mus musculus GN=Fcho2 PE=1 SV=1 | Q3UQN2 [ | 0 | 3322.74 | 0 | 0.77 | 0.00192 | 14.4 | 73.9 | 126.1 | 10.66 | 7.78 | | High | High | High | High | High | High | High | High | High | High | High | 0.00441 | 0.04729 | 3.01 |
| LYSLLGIDLTAPSNNSSPR | [K].LYSLLGIDLTAPSNNSSPR.[D] | 1xPhosphc | B2RRI2 1x | 0.00211 | 5.7E-05 | 1 | 8 | 2 | B2RRI2 | Protein strawberry notch homolog 1 OS=Mus musculus GN=Sbno1 PE=1 SV=1 | B2RRI2 [67 | 0 | 2327.18 | 0 | 0.77 | 0.00482 | 29.52 | 69.1 | 130.9 | 17.49 | 9.23 | | High | High | High | High | High | High | High | High | High | High | High | 4.9E-05 | 0.00063 | 3.46 |
| LPHISSHPLLQDLAAAR | [K].LPHISSHPLLQDLAAAR.[A] | 1xPhosphc | Q80U49 1 | 0.00715 | 0.00062 | 1 | 2 | 1 | Q80U49 | Centrosomal protein of 170 kDa protein B OS=Mus musculus GN=Cep170b PE=1 SV=2 | Q80U49 [8 | 0 | 2148.15 | 0 | 0.78 | 0.00101 | 10.78 | 73.2 | 126.8 | 3.99 | 6.12 | | High | High | High | High | High | High | High | High | High | High | High | 0.00027 | 0.00215 | 2.24 |
| TNILQYASTRPPTLSPIPHIPR | [K].TNILQYASTRPPTLSPIPHIPR.[S] | 1xPhosphc | P13405 1x | 0.00958 | 0.00074 | 1 | 4 | 2 | P13405 | Retinoblastoma-associated protein OS=Mus musculus GN=Rb1 PE=1 SV=2 | P13405 [7 | 0 | 2781.5 | 0 | 0.79 | 0.00121 | 6.72 | 73.5 | 126.5 | 5.42 | 8.45 | | High | High | High | High | High | High | High | High | High | High | High | 0.00081 | 0.00768 | 2.87 |
| CSDNSSYEEPLSPISASSSTSR | [R].CSDNSSYEEPLSPISASSSTSR.[R] | 1xCarbami | B2KFW6 1 | 0.00023 | 0 | 1 | 3 | 1 | B2KFW6 | Polyhomeotic-like 2 (Drosophila) OS=Mus musculus GN=Phc2 PE=4 SV=1 | B2KFW6 [7 | 0 | 2670.14 | 0 | 0.8 | 0.01581 | 17.79 | 76.7 | 123.3 | 10.48 | 16.53 | | High | High | High | High | High | High | High | High | High | High | High | 0 | 4.7E-05 | 3.64 |
| GSLTFEPLTLVPIQTK | [R].GSLTFEPLTLVPIQTK.[M] | 1xPhosphc | Q3UE92 1 | 0.00035 | 0 | 1 | 7 | 2 | Q3UE92 | Putative uncharacterized protein OS=Mus musculus GN=Xpnpep1 PE=1 SV=1 | Q3UE92 [5 | 0 | 2282.28 | 0 | 0.8 | 0.00019 | 1.59 | 71.3 | 128.7 | 6.83 | 4.14 | | High | High | High | High | High | High | High | High | High | High | High | 0 | 7.4E-05 | 2.87 |
| TSSLDTITGPYLTGQWPR | [R].TSSLDTITGPYLTGQWPR.[D] | 1xPhosphc | E9QKK4 1x | 3.7E-06 | 0 | 1 | 3 | 2 | E9QKK4 | Glucocorticoid-induced transcript 1 protein OS=Mus musculus GN=Glcci1 PE=1 SV=1 | E9QKK4 [1 | 0 | 2302.13 | 0 | 0.84 | 0.00054 | 5.56 | 70.9 | 129.1 | 5.66 | 3.25 | | High | High | High | High | High | High | High | High | High | High | High | 0 | 1.3E-06 | 4.18 |
| ALGVISNFQSSPK | [K].ALGVISNFQSSPK.[Y] | 1xPhosphc | F8WJ98 1x | 3.9E-07 | 0 | 1 | 6 | 2 | F8WJ98 | Transmembrane and coiled-coil domains protein 1 (Fragment) OS=Mus musculus GN=Tmcc1 PE=1 SV=2 | F8WJ98 [4 | 0 | 1886.02 | 0 | 0.85 | 0.00152 | 9.36 | 72.2 | 127.8 | 6.19 | 8.4 | | High | High | High | High | High | High | High | High | High | High | High | 0 | 3.5E-08 | 3.22 |
| SFSMQDLTTIR | [R].SFSMQDLTTIR.[G] | 1xPhosphc | A0A1D5RL | 0.0074 | 0.00062 | 1 | 2 | 2 | A0A1D5RL | Receptor expression-enhancing protein OS=Mus musculus GN=Reep1 PE=1 SV=1 | A0A1D5RL | 0 | 1607.77 | 0 | 0.86 | 0.00107 | 11.91 | 70.5 | 129.5 | 11.03 | 4.86 | | High | High | High | High | High | High | High | High | High | High | High | 0.00027 | 0.00223 | 1.91 |
| ADDDVDLEGISMDAPGYMSDG | [K].ADDDVDLEGISMDAPGYMSDGDVLSK.[N] | 1xPhosphc | E9Q842 1x | 0.00017 | 0 | 1 | 4 | 2 | E9Q842 | Neuron navigator 2 OS=Mus musculus GN=Nav2 PE=1 SV=1 | E9Q842 [8 | 0 | 3269.45 | 0 | 0.89 | 0.00537 | 12.53 | 70.2 | 129.8 | 14.82 | 19.46 | | High | High | High | High | High | High | High | High | High | High | High | 0 | 3.3E-05 | 3.5 |
| TPLENVPGNLSPIK | [K].TPLENVPGNLSPIK.[D] | 1xPhosphc | Q9EPN1 1 | 0.00909 | 0.00074 | 1 | 3 | 1 | Q9EPN1 | Neurobeachin OS=Mus musculus GN=Nbea PE=1 SV=1 | Q9EPN1 [1 | 0 | 2017.11 | 0 | 0.89 | 0.01041 | 26.18 | 69.1 | 130.9 | 13.3 | 12.34 | | High | High | High | High | High | High | High | High | High | High | High | 0.00033 | 0.00281 | 2.28 |
| VSLPPIPTVSNIK | [K].VSLPPIPTVSNIK.[S] | 1xPhosphc | Q8BH48 1 | 8.3E-05 | 0 | 1 | 3 | 1 | Q8BH48 | Ubiquitin-associated protein 1 OS=Mus musculus GN=Ubap1 PE=1 SV=1 | Q8BH48 [2 | 0 | 1903.11 | 0 | 0.89 | 0.00128 | 15.86 | 72 | 128 | 8.87 | 6.49 | | High | High | High | High | High | High | High | High | High | High | High | 0 | 1.5E-05 | 2.83 |
| NQYLSAIASDTEQEPK | [K].NQYLSAIASDTEQEPK.[I] | 1xPhosphc | Q5SWW4 | 3.4E-05 | 0 | 1 | 1 | 1 | Q5SWW4 | Mediator of RNA polymerase II transcription subunit 13 OS=Mus musculus GN=Med13 PE=1 SV=1 | Q5SWW4 | 0 | 2332.15 | 0 | 0.9 | 0.01456 | 27.9 | 70.8 | 129.2 | 15.83 | 16.4 | | High | High | High | High | High | High | High | High | High | High | High | 0 | 5.3E-06 | 2.34 |
| SPQLGVQAGLEPILSSATVDSTK | [K].SPQLGVQAGLEPILSSATVDSTK.[R] | 1xPhosphc | Q6P1H6 1 | 0.02105 | 0.00151 | 1 | 3 | 1 | Q6P1H6 | Ankyrin repeat and LEM domain-containing protein 2 OS=Mus musculus GN=Ankle2 PE=1 SV=2 | Q6P1H6 [8 | 0 | 2836.51 | 0 | 0.9 | 0.00096 | 5.08 | 68.5 | 131.5 | 12.79 | 10.66 | | High | High | High | High | High | High | High | High | High | High | High | 0.0008 | 0.0072 | 2.34 |
| DRSSPPPGYIPDELHQIAR | [R].DRSSPPPGYIPDELHQIAR.[N] | 1xPhosphc | G5E8L8 1x | 1E-06 | 0 | 1 | 1 | 4 | G5E8L8 | Mitogen activated protein kinase kinase kinase 2 OS=Mus musculus GN=Map3k2 PE=1 SV=1 | G5E8L8 [1 | 1 | 2457.21 | 0 | 0.91 | 0.0004 | 3.91 | 70.2 | 129.8 | 6.86 | 8.41 | | High | High | High | High | High | High | High | High | High | High | High | 0 | 3.4E-07 | 3.73 |
| GSSPDVHTLLEITEESDAVLVDK | [R].GSSPDVHTLLEITEESDAVLVDK.[S] | 1xPhosphc | O54724 1 | 2.8E-07 | 0 | 1 | 1 | 12 | O54724 | Caveolae-associated protein 1 OS=Mus musculus GN=Cavin1 PE=1 SV=1 | O54724 [3 | 0 | 2992.52 | 0 | 0.94 | 0.00521 | 14.42 | 68.7 | 131.3 | 11.66 | 9.43 | | High | High | High | High | High | High | High | High | High | High | High | 0 | 2.4E-08 | 4.3 |
| EVLESILSPGEPGDQK | [K].EVLESILSPGEPGDQK.[A] | 1xPhosphc | Q8BQ30-2 | 3.6E-06 | 0 | 1 | 4 | 3 | Q8BQ30-2 | Isoform 2 of Phostensin OS=Mus musculus GN=Ppp1r18 | Q8BQ30-2 | 0 | 2236.15 | 0 | 0.97 | 0.00043 | 7.89 | 68.9 | 131.1 | 4.78 | 4.31 | | High | High | High | High | High | High | High | High | High | High | High | 0 | 4.3E-07 | 3.2 |
| HSLECIQANQIFPR | [R].HSLECIQANQIFPR.[K] | 1xCarbami | Q8K296 1 | 1.5E-05 | 0 | 1 | 7 | 2 | Q8K296 | Myotubularin-related protein 3 OS=Mus musculus GN=Mtmr3 PE=1 SV=2 | Q8K296 [7 | 0 | 2021.98 | 0 | 0.99 | 0.00074 | 5.59 | 68.8 | 131.2 | 12.91 | 12.92 | | High | High | High | High | High | High | High | High | High | High | High | 0 | 2.1E-06 | 4.1 |
| LGSIAEIDLGVPPPIMK | [R].LGSIAEIDLGVPPPIMK.[S] | 1xPhosphc | Q99MR6 1 | 0.0002 | 0 | 1 | 5 | 1 | Q99MR6 | Serrate RNA effector molecule homolog OS=Mus musculus GN=Srrt PE=1 SV=1 | Q99MR6 [ | 0 | 2304.27 | 0 | 1.01 | 0.0006 | 4.3 | 71 | 129 | 12.54 | 9.58 | | High | High | High | High | High | High | High | High | High | High | High | 0 | 3.9E-05 | 2.27 |
| DEPQTVPDVPSFGDSPPLSPIDN | [K].DEPQTVPDVPSFGDSPPLSPIDMDTQER.[I] | 2xPhosphc | P15066 2x | 1.9E-06 | 0 | 1 | 2 | 4 | P15066 | Transcription factor jun-D OS=Mus musculus GN=Jund PE=1 SV=1 | P15066 [2 | 0 | 3474.49 | 0 | 1.03 | 0.00272 | 21.73 | 64 | 136 | 3.21 | 12.37 | | High | High | High | High | High | High | High | High | High | High | High | 0 | 2E-07 | 4.13 |
| NSSPSISPNTSFASDGSPSPLGGI | [R].NSSPSISPNTSFASDGSPSPLGGIK.[R] | 1xDeamid | Q9DBR1 1 | 0.004 | 0.00031 | 1 | 2 | 1 | Q9DBR1 | 5'-3' exoribonuclease 2 OS=Mus musculus GN=Xrn2 PE=1 SV=1 | Q9DBR1 [4 | 0 | 2943.44 | 0 | 1.06 | 0.01106 | 38.47 | 72.3 | 127.7 | 22.22 | 11.34 | | High | High | High | High | High | High | High | High | High | High | High | 0.00014 | 0.00112 | 3.15 |
| HQDGLPYIDDSPSSSPHLSSK | [R].HQDGLPYIDDSPSSSPHLSSK.[G] | 2xPhosphc | Q6PAJ1 2x | 0.00111 | 0 | 1 | 2 | 1 | Q6PAJ1 | Breakpoint cluster region protein OS=Mus musculus GN=Bcr PE=1 SV=3 | Q6PAJ1 [4 | 0 | 2656.15 | 0 | 1.07 | 0.00534 | 30.06 | 68.5 | 131.5 | 10.57 | 13.56 | | High | High | High | High | High | High | High | High | High | High | High | 0 | 0.00027 | 2.74 |
| DEPQTVPDVPSFGDSPPLSPIDN | [K].DEPQTVPDVPSFGDSPPLSPIDMDTQER.[I] | 2xPhosphc | P15066 2x | 6E-07 | 0 | 1 | 2 | 4 | P15066 | Transcription factor jun-D OS=Mus musculus GN=Jund PE=1 SV=1 | P15066 [2 | 0 | 3458.49 | 0 | 1.09 | 0.01818 | 21.22 | 65 | 135 | 4.73 | 22.4 | | High | High | High | High | High | High | High | High | High | High | High | 0 | 5.7E-08 | 5.25 |
| HSLSLDDIR | [R].HSLSLDDIR.[L] | 1xPhosphc | Q8C0V9 1 | 0.04003 | 0.00287 | 1 | 2 | 2 | Q8C0V9 | FERM domain-containing protein 6 OS=Mus musculus GN=Frmd6 PE=1 SV=2 | Q8C0V9 [5 | 0 | 1364.68 | 0 | 1.11 | 0.00045 | 2.45 | 61.5 | 138.5 | 6.36 | 8 | | High | High | High | High | High | High | High | High | High | High | High | 0.00148 | 0.0149 | 2.03 |
| GSLLQIPVK | [R].GSLLQIPVK.[T] | 1xPhosphc | I1VGT4 1x | 0.0005 | 0 | 2 | 19 | 1 | I1VGT4; I1 | Anion exchange protein OS=Mus musculus GN=Slc4a7 PE=2 SV=1Anion exchange protein OS=Mus musculus GN=Slc4a7 PE=2 SV=1 | I1VGT4 [1( | 0 | 1492.89 | 0 | 1.12 | 0.0029 | 6.5 | 66.7 | 133.3 | 7.48 | 12.63 | Shared | High | High | High | High | High | High | High | High | High | High | High | 0 | 0.00011 | 2.72 |
| LSGQNTMGSQDSLLEITFR | [K].LSGQNTMGSQDSLLEITFR.[S] | 1xPhosphc | Q9D3A8 1 | 0.00017 | 0 | 1 | 2 | 1 | Q9D3A8 | Carboxyl-terminal PDZ ligand of neuronal nitric oxide synthase protein OS=Mus musculus GN=Nos1ap PE=1 SV=3 | Q9D3A8 [3 | 0 | 2406.16 | 0 | 1.13 | 0.00373 | 21.08 | 65.1 | 134.9 | 10.5 | 13.26 | | High | High | High | High | High | High | High | High | High | High | High | 0 | 3.2E-05 | 2.92 |
| NSSPSISPNTSFASDGSPSPLGGI | [R].NSSPSISPNTSFASDGSPSPLGGIK.[R] | 2xPhosphc | Q9DBR1 2 | 8.6E-06 | 0 | 1 | 2 | 14 | Q9DBR1 | 5'-3' exoribonuclease 2 OS=Mus musculus GN=Xrn2 PE=1 SV=1 | Q9DBR1 [4 | 0 | 3022.42 | 0 | 1.15 | 9.8E-05 | 2.54 | 62.1 | 137.9 | 5.91 | 3.54 | | High | High | High | High | High | High | High | High | High | High | High | 0 | 0.00014 | 4.67 |
| DVVVEFPSPPSDSDFPPPPPEIEL | [K].DVVVEFPSPPSDSDFPPPPPEIELPLPPIEIPAV | 1xDeamid | F2Z3U3 1x | 1.8E-05 | 0 | 1 | 1 | 2 | F2Z3U3 | Ras association (RalGDS/AF-6) and pleckstrin homology domains 1 OS=Mus musculus GN=Raph1 PE=1 SV=2 | F2Z3U3 [1 | 0 | 4990.53 | 0 | 1.16 | 0.00922 | 24.06 | 64.1 | 135.9 | 17.9 | 12.64 | | High | High | High | High | High | High | High | High | High | High | High | 0 | 2.6E-06 | 4.51 |
| GQLTNIVSPTAATTPR | [K].GQLTNIVSPTAATTPR.[I] | 1xPhosphc | Q8CH77 1 | 3.3E-06 | 0 | 1 | 4 | 2 | Q8CH77 | Neuron navigator 1 OS=Mus musculus GN=Nav1 PE=1 SV=2 | Q8CH77 [9 | 0 | 1936.01 | 0 | 1.16 | 0.00017 | 2.48 | 61.7 | 138.3 | 5.97 | 4.8 | | High | High | High | High | High | High | High | High | High | High | High | 0 | 3.9E-07 | 3.75 |
| LSLSPISLAESWENSSPPPQAR | [K].LSLSPISLAESWENSSPPPQAR.[T] | 1xPhosphc | J3QNH8 1 | 7.2E-06 | 0 | 1 | 1 | 5 | J3QNH8 | Uncharacterized protein OS=Mus musculus GN=Gm7694 PE=1 SV=1 | J3QNH8 [2 | 0 | 2675.33 | 0 | 1.17 | 0.00295 | 25.74 | 58 | 142 | 11.23 | 14.96 | | High | High | High | High | High | High | High | High | High | High | High | 0 | 9.4E-07 | 3.26 |
| SLDLPSIGGSSVGK | [K].SLDLPSIGGSSVGK.[E] | 1xPhosphc | V9GX40 1 | 0.04669 | 0.00354 | 1 | 4 | 1 | V9GX40 | Synergin gamma OS=Mus musculus GN=Synrg PE=1 SV=1 | V9GX40 [9 | 0 | 1855 | 0 | 1.18 | 0.00016 | 4.78 | 61.2 | 138.8 | 5.93 | 3.23 | | High | High | High | High | High | High | High | High | High | High | High | 0.00181 | 0.0179 | 2.54 |
| RRESLSYIPK | [R].RRESLSYIPK.[G] | 1xPhosphc | Q4VAC9 1 | 0.09015 | 0.00698 | 1 | 3 | 1 | Q4VAC9 | Pleckstrin homology domain-containing family G member 3 OS=Mus musculus GN=Plekhg3 PE=1 SV=2 | Q4VAC9 [8 | 2 | 1787 | 0 | 1.21 | 0.01101 | 14.16 | 60.4 | 139.6 | 8.27 | 23.81 | | High | High | High | High | High | High | High | High | High | High | High | 0.00356 | 0.03803 | 1.47 |
| RTSLPAPGPNTSSVSLWSILR | [R].RTSLPAPGPNTSSVSLWSILR.[N] | 1xPhosphc | D3YTT6 1x | 0.01296 | 0.00089 | 1 | 6 | 2 | D3YTT6 | Oxysterol-binding protein OS=Mus musculus GN=Osbpl3 PE=1 SV=1 | D3YTT6 [5 | 1 | 2548.35 | 0 | 1.21 | 0.00043 | 10.17 | 61.1 | 138.9 | 12.29 | 6.41 | | High | High | High | High | High | High | High | High | High | High | High | 0.00045 | 0.0042 | 2.48 |
| MLQAISPK | [R].MLQAISPK.[Q] | 1xPhosphc | P49586 1x | 0.00457 | 0.00046 | 2 | 3 | 2 | P49586: Q | Choline-phosphate cytidylyltransferase A OS=Mus musculus GN=Pcyt1a PE=1 SV=1Choline-phosphate cytidylyltransferase B OS=Mus musculus GN=Pcyt1b PE=1 SV=2 | P49586 [3 | 0 | 1425.79 | 0 | 1.24 | 0.00124 | 12.6 | 58.6 | 141.4 | 5.91 | 9.75 | Shared | High | High | High | High | High | High | High | High | High | High | High | 0.0002 | 0.00131 | 1.35 |
| QEGGQISPNRPTSVPTAPALSSII | [R].QEGGQISPNRPTSVPTAPALSSIIAPK.[N] | 1xPhosphc | Q7TNI6 1x | 5.9E-05 | 0 | 1 | 1 | 3 | Q7TNI6 | Jerry protein OS=Mus musculus GN=Tom1l1 PE=2 SV=1 | Q7TNI6 [1 | 0 | 3254.76 | 0 | 1.26 | 0.00028 | 4.21 | 59.8 | 140.2 | 6.57 | 6.07 | | High | High | High | High | High | High | High | High | High | High | High | 0 | 9.9E-06 | 3.56 |
| SSDIISLLVSIYGSK | [R].SSDIISLLVSIYGSK.[D] | 1xPhosphc | Q6ZPR8 1x | 7.8E-05 | 0 | 1 | 3 | 2 | Q6ZPR8 | MKIAA1406 protein (Fragment) OS=Mus musculus GN=Anapc2 PE=2 SV=1 | Q6ZPR8 [5 | 0 | 2120.17 | 0 | 1.27 | 0.00052 | 19.02 | 55.1 | 144.9 | 17.6 | 5.33 | | High | High | High | High | High | High | High | High | High | High | High | 0.00145 | 0.01459 | 3.05 |
| QDMQPLSPISVHER | [R].QDMQPLSPISVHER.[Y] | 1xPhosphc | Q64701 1 | 0.00053 | 0 | 1 | 4 | 2 | Q64701 | Retinoblastoma-like protein 1 OS=Mus musculus GN=Rbl1 PE=1 SV=3 | Q64701 [6 | 0 | 1961.94 | 0 | 1.31 | 0.004 | 8.55 | 57.6 | 142.4 | 10.28 | 13.41 | | High | High | High | High | High | High | High | High | High | High | High | 0 | 0.00012 | 2.97 |
| QEGGQISPNRPTSVPTAPALSSII | [R].QEGGQISPNRPTSVPTAPALSSIIAPK.[N] | 1xDeamid | Q7TNI6 1x | 0.00183 | 5.7E-05 | 1 | 1 | 2 | Q7TNI6 | Jerry protein OS=Mus musculus GN=Tom1l1 PE=2 SV=1 | Q7TNI6 [1 | 0 | 3255.74 | 0 | 1.31 | 0.00049 | 4.06 | 59 | 141 | 11.23 | 6.92 | | High | High | High | High | High | High | High | High | High | High | High | 2.6E-05 | 0.00047 | 3.61 |
| HSSTGDLLSIELQQAK | [K].HSSTGDLLSIELQQAK.[SG] | 1xPhosphc | P0C090 1x | 0.00199 | 5.7E-05 | 1 | 4 | 3 | P0C090 | Roquin-2 OS=Mus musculus GN=Rc3h2 PE=1 SV=1 | P0C090 [9 | 0 | 2265.19 | 0 | 1.32 | 0.00917 | 14.92 | 61.3 | 138.7 | 17.46 | 20.12 | | High | High | High | High | High | High | High | High | High | High | High | 2.6E-05 | 0.00052 | 2.65 |
| SAQGTGFELGQLQSIR | [K].SAQGTGFELGQLQSIR.[S] | 1xPhosphc | Q8VDZ4 1 | 0.00024 | 0 | 1 | 2 | 1 | Q8VDZ4 | Palmitoyltransferase ZDHHC5 OS=Mus musculus GN=Zdhhc5 PE=1 SV=1 | Q8VDZ4 [4 | 0 | 2001 | 0 | 1.33 | 0.0048 | 13.61 | 63.8 | 136.2 | 13.44 | 13.07 | | High | High | High | High | High | High | High | High | High | High | High | 0 | 4.9E-05 | 3.82 |
| SLHGEFSASDSSLTDIQETR | [K].SLHGEFSASDSSLTDIQETR.[R] | 3xPhosphc | Q8C0T5 3x | 0.00642 | 0.00062 | 1 | 3 | 1 | Q8C0T5 | Signal-induced proliferation-associated 1-like protein 1 OS=Mus musculus GN=Sipa1l1 PE=1 SV=2 | Q8C0T5 [1 | 0 | 2649.07 | 0 | 1.33 | 0.00087 | 7.83 | 58.9 | 141.1 | 10.85 | 12.53 | | High | High | High | High | High | High | High | High | High | High | High | 0.00027 | 0.00192 | 3.31 |
| GIEFPMADLDALSPIHTPQR | [R].GIEFPMADLDALSPIHTPQR.[S] | 1xPhosphc | Q5SRX1 1x | 0.00047 | 0 | 1 | 6 | 1 | Q5SRX1 | TOM1-like protein 2 OS=Mus musculus GN=Tom1l2 PE=1 SV=1 | Q5SRX1 [1 | 0 | 2533.24 | 0 | 1.35 | 0.0038 | 18.41 | 57.2 | 142.8 | 26.14 | 5.87 | | High | High | High | High | High | High | High | High | High | High | High | 0 | 0.0001 | 2.58 |
| LTSIGSDEDEETETYQEK | [R].LTSIGSDEDEETETYQEK.[V] | 2xPhosphc | Q924N4 2 | 0.0002 | 0 | 2 | 4 | 2 | Q924N4; ( | Solute carrier family 12 member 6 OS=Mus musculus GN=Slc12a6 PE=1 SV=2Isoform 2 of Solute carrier family 12 member 6 OS=Mus musculus GN=Slc12a6 | Q924N4 [1 | 0 | 2692.16 | 0 | 1.37 | 0.00072 | 6.37 | 54.3 | 145.7 | 16.38 | 8.29 | Shared | High | High | High | High | High | High | High | High | High | High | High | 0 | 4E-05 | 3.66 |
| NSSPSISPNTSFASDGSPSPLGGI | [R].NSSPSISPNTSFASDGSPSPLGGIK.[R] | 1xPhosphc | Q9DBR1 1 | 4.6E-06 | 0 | 1 | 2 | 6 | Q9DBR1 | 5'-3' exoribonuclease 2 OS=Mus musculus GN=Xrn2 PE=1 SV=1 | Q9DBR1 [4 | 0 | 2942.45 | 0 | 1.37 | 0.00068 | 18.14 | 57.9 | 142.1 | 8.27 | 6.33 | | High | High | High | High | High | High | High | High | High | High | High | 0 | 4.6E-05 | 3.73 |
| THSGLTDGIPLQSISAPGSGIWSI | [R].THSGLTDGIPLQSISAPGSGIWSDGK.[D] | 1xPhosphc | Q6ZQ08-4 | 2.3E-05 | 0 | 1 | 7 | 1 | Q6ZQ08-4 | Isoform 4 of CCR4-NOT transcription complex subunit 1 OS=Mus musculus GN=Cnot1 | Q6ZQ08-4 | 0 | 3119.58 | 0 | 1.43 | 0.01236 | 44.59 | 54.2 | 145.8 | 45.39 | 28.51 | | High | High | High | High | High | High | High | High | High | High | High | 0 | 3.5E-06 | 4.23 |
| DSSGPVSPGTPSQEESEQICLYHI | [K].DSSGPVSPGTPSQEESEQICLYHIR.[K] | 1xCarbami | Q3U0S1 1 | 2.2E-06 | 0 | 1 | 2 | 1 | Q3U0S1 | Putative uncharacterized protein OS=Mus musculus GN=Parp12 PE=2 SV=1 | Q3U0S1 [2 | 0 | 3082.4 | 0 | 1.45 | 0.00179 | 20.55 | 53.3 | 146.7 | 11.39 | 11.24 | | High | High | High | High | High | High | High | High | High | High | High | 0 | 2.5E-07 | 4.14 |
| NVQTGLTESPVLLETLAEIFDTPA | [K].NVQTGLTESPVLLETLAEIFDTPASK.[V] | 1xPhosphc | S4R1P5 1x | 0.01262 | 0.00089 | 1 | 6 | 2 | S4R1P5 | Dystonin OS=Mus musculus GN=Dst PE=1 SV=1 | S4R1P5 [3 | 0 | 3311.74 | 0 | 1.49 | 0.00709 | 18.26 | 56.9 | 143.1 | 23.03 | 15.93 | | High | High | High | High | High | High | High | High | High | High | High | 0.00054 | 0.00455 | 2.98 |
| GLNGGITPLNSISPLK | [R].GLNGGITPLNSISPLK.[K] | 1xPhosphc | Q7TN98 1 | 8.3E-05 | 0 | 1 | 7 | 1 | Q7TN98 | Cytoplasmic polyadenylation element-binding protein 4 OS=Mus musculus GN=Cpeb4 PE=1 SV=1 | Q7TN98 [3 | 0 | 2119.19 | 0 | 1.53 | 0.00129 | 27.04 | 45.3 | 154.7 | 19.76 | 12.01 | | High | High | High | High | High | High | High | High | High | High | High | 0 | 1.4E-05 | 3.79 |
| LIASSPTLISGITSPPLLDSIK | [R].LIASSPTLISGITSPPLLDSIK.[T] | 1xPhosphc | B1ASA5 1x | 3E-05 | 0 | 1 | 2 | 3 | B1ASA5 | Zinc finger protein 362 OS=Mus musculus GN=Zfp362 PE=1 SV=1 | B1ASA5 [1 | 0 | 2761.58 | 0 | 1.53 | 0.00042 | 14.72 | 51.1 | 148.9 | 13.1 | 3.45 | | High | High | High | High | High | High | High | High | High | High | High | 0 | 4.7E-06 | 3.44 |
| QPSGAGLCDITEGTVVPEDR | [R].QPSGAGLCDITEGTVVPEDR.[C] | 1xCarbami | Q8K400 1x | 0.01734 | 0.0012 | 1 | 3 | 1 | Q8K400 | Syntaxin-binding protein 5 OS=Mus musculus GN=Stxbp5 PE=1 SV=3 | Q8K400 [6 | 0 | 2410.12 | 0 | 1.56 | 0.0049 | 30.19 | 52.9 | 147.1 | 23.26 | 10.5 | | High | High | High | High | High | High | High | High | High | High | High | 0.00064 | 0.00578 | 2.86 |
| DELSDISQGGSK | [K].DELSDISQGGSK.[A] | 1xPhosphc | B8X349 1x | 0.00169 | 5.7E-05 | 1 | 10 | 2 | B8X349 | JNK-interacting leucine zipper protein long form OS=Mus musculus GN=Spag9 PE=2 SV=1 | B8X349 [2 | 0 | 1773.87 | 0 | 1.67 | 0.00042 | 13.52 | 47.9 | 152.1 | 6.35 | 5.66 | | High | High | High | High | High | High | High | High | High | High | High | 2.6E-05 | 0.00043 | 3.69 |
| NSSPSISPNTSFASDGSPSPLGGI | [R].NSSPSISPNTSFASDGSPSPLGGIK.[R] | 2xPhosphc | Q9DBR1 2 | 0.00132 | 0 | 1 | 2 | 4 | Q9DBR1 | 5'-3' exoribonuclease 2 OS=Mus musculus GN=Xrn2 PE=1 SV=1 | Q9DBR1 [4 | 0 | 2793.26 | 0 | 1.68 | 0.15551 | 71.97 | 48.7 | 151.3 | 42.63 | 45.03 | | High | High | High | High | High | High | High | High | High | High | High | 0.00022 | 0.00145 | 3.75 |
| DKPVYDELFYTLSPINGK | [K].DKPVYDELFYTLSPINGK.[I] | 1xPhosphc | Q1MWP8 | 3.7E-05 | 0 | 1 | 5 | 1 | Q1MWP8 | EH-domain containing 4-KJR (Fragment) OS=Mus musculus GN=Ehd4 PE=2 SV=1 | Q1MWP8 | 0 | 2866.52 | 0 | 1.69 | 0.00024 | 10.1 | 50.9 | 149.1 | 10.58 | 5.04 | | High | High | High | High | High | High | High | High | High | High | High | 0 | 5.8E-06 | 3.97 |
| GSGFSLDVIDGPISQR | [K].GSGFSLDVIDGPISQR.[E] | 1xPhosphc | Q8C0T5 1x | 1.6E-05 | 0 | 1 | 4 | 8 | Q8C0T5 | Signal-induced proliferation-associated 1-like protein 1 OS=Mus musculus GN=Sipa1l1 PE=1 SV=2 | Q8C0T5 [2 | 0 | 1956.96 | 0 | 1.69 | 0.00037 | 10.2 | 48.7 | 151.3 | 7.44 | 5.78 | | High | High | High | High | High | High | High | High | High | High | High | 0 | 1.7E-05 | 4.33 |
| TSLPAPGPNTSSVSLWSILR | [R].TSLPAPGPNTSSVSLWSILR.[N] | 1xPhosphc | D3YTT6 1x | 2.6E-07 | 0 | 1 | 6 | 1 | D3YTT6 | Oxysterol-binding protein OS=Mus musculus GN=Osbpl3 PE=1 SV=1 | D3YTT6 [5 | 0 | 2392.25 | 0 | 1.73 | 0.00099 | 19.62 | 46.3 | 153.7 | 10.44 | 9.14 | | High | High | High | High | High | High | High | High | High | High | High | 0 | 2.2E-08 | 3.45 |
| SFSMHDLTAIQGDEPVGHRPYQ | [R].SFSMHDLTAIQGDEPVGHRPYQTLPEAK.[R] | 1xPhosphc | A0A1W2P | 0.01428 | 0.00101 | 1 | 2 | 1 | A0A1W2P | Receptor expression-enhancing protein OS=Mus musculus GN=Reep3 PE=1 SV=1 | A0A1W2P | 0 | 3678.8 | 0 | 1.77 | 0.00067 | 4.37 | 43.1 | 156.9 | 16.78 | 9.63 | | High | High | High | High | High | High | High | High | High | High | High | 0.00054 | 0.0047 | 2.98 |
| LLTSITSK | [R].LLTSITSK.[S] | 1xPhosphc | Q8K298 1x | 0.0047 | 0.00046 | 1 | 2 | 1 | Q8K298 | Anillin OS=Mus musculus GN=Anln PE=1 SV=2 | Q8K298 [9 | 0 | 1400.82 | 0 | 1.83 | 0.00649 | 38.77 | 44.1 | 155.9 | 23.72 | 17.74 | | High | High | High | High | High | High | High | High | High | High | High | 0.0002 | 0.00135 | 2.6 |
| DFPDSFQPGSPGHLGVIR | [R].DFPDSFQPGSPGHLGVIR.[D] | 1xPhosphc | Q8VCZ7 1x | 0.00022 | 0 | 1 | 1 | 1 | Q8VCZ7 | Zinc finger and BTB domain-containing protein 7C OS=Mus musculus GN=Zbtb7c PE=2 SV=1 | Q8VCZ7 [2 | 0 | 2235.08 | 0 | 1.85 | 0.00344 | 21.14 | 48.6 | 151.4 | 11.73 | 18.54 | | High | High | High | High | High | High | High | High | High | High | High | 0 | 4.3E-05 | 3.19 |
| LTSIGSDEDEETETYQEK | [R].LTSIGSDEDEETETYQEK.[V] | 1xPhosphc | Q924N4 1 | 0.00101 | 0 | 2 | 4 | 3 | Q924N4; ( | Solute carrier family 12 member 6 OS=Mus musculus GN=Slc12a6 PE=1 SV=2Isoform 2 of Solute carrier family 12 member 6 OS=Mus musculus GN=Slc12a6 | Q924N4 [1 | 0 | 2612.19 | 0 | 1.9 | 0.00223 | 25.55 | 40.7 | 159.3 | 20.26 | 18.35 | Shared | High | High | High | High | High | High | High | High | High | High | High | 0 | 0.00024 | 3.36 |
| WLGPILGYDTD | [K].WLGPILGYDTD.[-] | 1xPhosphc | A6H5Y3 1x | 0.00183 | 5.7E-05 | 1 | 2 | 2 | A6H5Y3 | Methionine synthase OS=Mus musculus GN=Mtr PE=1 SV=1 | A6H5Y3 [1 | 0 | 1558.74 | 0 | 1.97 | 0.00037 | 32.48 | 44.8 | 155.2 | 35.04 | 10.48 | | High | High | High | High | High | High | High | High | High | High | High | 2.6E-05 | 0.00047 | 2.81 |
| TPTGLVGAAAALQPVPAVPSAL( | [K].TPTGLVGAAAALQPVPAVPSALQNISLK.[H] | 1xPhosphc | B1AVH7 1 | 3.6E-07 | 0 | 1 | 2 | 1 | B1AVH7 | TBC1 domain family member 2A OS=Mus musculus GN=Tbc1d2 PE=2 SV=1 | B1AVH7 [1 | 0 | 3222.83 | 0 | 2 | 0.00483 | 49.42 | 39.7 | 160.3 | 20.89 | 19.14 | | High | High | High | High | High | High | High | High | High | High | High | 0 | 3.3E-08 | 4.52 |
| SFSMHDLTAIQGDEPVGHRPYQ | [R].SFSMHDLTAIQGDEPVGHRPYQTLPEAK.[R] | 1xPhosphc | A0A1W2P | 2.8E-06 | 0 | 1 | 2 | 1 | A0A1W2P | Receptor expression-enhancing protein OS=Mus musculus GN=Reep3 PE=1 SV=1 | A0A1W2P | 0 | 3662.81 | 0 | 2.06 | 0.00169 | 15.41 | 38.6 | 161.4 | 14.33 | 13.37 | | High | High | High | High | High | High | High | High | High | High | High | 0 | 3.3E-07 | 4.25 |
| GMVSLSEIEEALAK | [K].GMVSLSEIEEALAK.[N] | 1xPhosphc | P27546 1x | 0.02338 | 0.00158 | 1 | 3 | 2 | P27546 | Microtubule-associated protein 4 OS=Mus musculus GN=Map4 PE=1 SV=3 | P27546 [3 | 0 | 2031.05 | 0 | 2.11 | 0.00085 | 9.07 | 36.2 | 163.8 | 9.39 | 14.08 | | High | High | High | High | High | High | High | High | High | High | High | 0.00085 | 0.0081 | 3.17 |
| LSWGYGEHNGPIHWNELFPIAD | [R].LSWGYGEHNGPIHWNELFPIADGDQQSPIEI | 1xDeamid | Q9D6N1 1 | 0.00044 | 0 | 1 | 1 | 1 | Q9D6N1 | Carbonic anhydrase 13 OS=Mus musculus GN=Ca13 PE=1 SV=1 | Q9D6N1 [4 | 0 | 4187.03 | 0 | 2.12 | 0.00126 | 49.59 | 35.7 | 164.3 | 48.52 | 10.58 | | High | High | High | High | High | High | High | High | High | High | High | 0 | 9.4E-05 | 4.08 |
| GSLNEQIALVLIR | [R].GSLNEQIALVLIR.[L] | 1xPhosphc | Q5CZX6 1x | 0.10175 | 0.00819 | 1 | 5 | 1 | Q5CZX6 | Acbd5 protein (Fragment) OS=Mus musculus GN=Acbd5 PE=2 SV=1 | Q5CZX6 [4 | 0 | 1734.97 | 0 | 2.29 | 0.00043 | 5.72 | 38.1 | 161.9 | 9.78 | 9.85 | | High | High | High | High | High | High | High | High | High | High | High | 0.00413 | 0.0442 | 2.34 |
| TSLDEITSGK | [K].TSLDEITSGK.[G] | 1xPhosphc | Q8BT07 1x | 2.7E-05 | 0 | 1 | 3 | 2 | Q8BT07 | Centrosomal protein of 55 kDa OS=Mus musculus GN=Cep55 PE=1 SV=2 | Q8BT07 [3 | 0 | 1588.82 | 0 | 2.35 | 0.00385 | 33.39 | 30.9 | 169.1 | 14.8 | 21.3 | | High | High | High | High | High | High | High | High | High | High | High | 0 | 4.2E-06 | 2.8 |
| IGEEQSAEDAEDGPPELLFIHGGI | [K].IGEEQSAEDAEDGPPELLFIHGGHTAK.[I] | 1xPhosphc | Q3UJI2 1x | 1.9E-08 | 0 | 1 | 6 | 1 | Q3UJI2 | Putative uncharacterized protein OS=Mus musculus GN=Rbbp7 PE=2 SV=1 | Q3UJI2 [34 | 0 | 3385.64 | 0 | 2.39 | 0.00026 | 15.23 | 31.4 | 168.6 | 8.28 | 6.11 | | High | High | High | High | High | High | High | High | High | High | High | 0 | 1.2E-09 | 4.23 |
| LSWGYGEHNGPIHWNELFPIAD | [R].LSWGYGEHNGPIHWNELFPIADGDQQSPIEI | 1xPhosphc | Q9D6N1 1 | 0.01296 | 0.00089 | 1 | 1 | 2 | Q9D6N1 | Carbonic anhydrase 13 OS=Mus musculus GN=Ca13 PE=1 SV=1 | Q9D6N1 [4 | 0 | 4186.05 | 0 | 2.39 | 0.00031 | 11.32 | 29.2 | 170.8 | 18.89 | 6.57 | | High | High | High | High | High | High | High | High | High | High | High | 0.00058 | 0.00527 | 3.08 |
| RSSDIISLLVSIYGSK | [R].RSSDIISLLVSIYGSK.[D] | 1xPhosphc | Q6ZPR8 1x | 0.00124 | 0 | 1 | 3 | 3 | Q6ZPR8 | MKIAA1406 protein (Fragment) OS=Mus musculus GN=Anapc2 PE=2 SV=1 | Q6ZPR8 [5 | 1 | 2276.27 | 0 | 2.55 | 0.00021 | 4.68 | 29.2 | 170.8 | 11.27 | 4.43 | | High | High | High | High | High | High | High | High | High | High | High | 0 | 0.00031 | 2.75 |
| EGLLPISPATPPGSDEK | [K].EGLLPISPATPPGSDEK.[T] | 1xPhosphc | B9EHK8 1x | 0.00139 | 5.7E-05 | 1 | 4 | 2 | B9EHK8 | Homeo box A1 OS=Mus musculus GN=Hoxa1 PE=2 SV=1 | B9EHK8 [2 | 0 | 2246.17 | 0 | 2.56 | 0.00046 | 24.78 | 31.5 | 168.5 | 15.97 | 8.18 | | High | High | High | High | High | High | High | High | High | High | High | 2.6E-05 | 0.00034 | 3.02 |
| EEPVLTPIDEGHLPMINQDEFR | [K].EEPVLTPIDEGHLPMINQDEFR.[N] | 1xPhosphc | P23298 1x | 0.00044 | 0 | 1 | 3 | 1 | P23298 | Protein kinase C eta type OS=Mus musculus GN=Prkch PE=1 SV=2 | P23298 [6 | 0 | 2904.37 | 0 | 2.57 | 0.00115 | 14.76 | 26.9 | 173.1 | 5.29 | 14.86 | | High | High | High | High | High | High | High | High | High | High | High | 0 | 9.5E-05 | 2.8 |
| DLSPTAPCPAAATASLLASISR | [R].DLSPTAPCPAAATASLLASISR.[M] | 1xCarbami | Q6EBV9 1x | 0.00132 | 0 | 1 | 1 | 2 | Q6EBV9 | Autophagy-related protein 9B OS=Mus musculus GN=Atg9b PE=2 SV=2 | Q6EBV9 [7 | 0 | 2479.25 | 0 | 2.59 | 0.0004 | 8.3 | 32.7 | 167.3 | 36.91 | 11.35 | | High | High | High | High | High | High | High | High | High | High | High | 0 | 0.00033 | 2.83 |
| NAISLPTIGGSQALMLPLLSPVTF | [K].NAISLPTIGGSQALMLPLLSPVTFHSK.[Q] | 1xPhosphc | Q9CQC5 1 | 0.03488 | 0.00227 | 1 | 1 | 1 | Q9CQC5 | Cdc42 effector protein 3 OS=Mus musculus GN=Cdc42ep3 PE=1 SV=1 | Q9CQC5 [1 | 0 | 3346.83 | 0 | 2.68 | 0.00132 | 32.48 | 31.4 | 168.6 | 34.51 | 16.8 | | High | High | High | High | High | High | High | High | High | High | High | 0.00118 | 0.01272 | 2.57 |
| SPLSGIPVR | [R].SPLSGIPVR.[T] | 1xPhosphc | Q5SXY1 1x | 0.02833 | 0.00179 | 1 | 5 | 2 | Q5SXY1 | Cytospin-B OS=Mus musculus GN=Specc1 PE=1 SV=2 | Q5SXY1 [8 | 0 | 1234.68 | 0 | 2.76 | 0.00018 | 5.14 | 25.8 | 174.2 | 15.52 | 6.18 | | High | High | High | High | High | High | High | High | High | High | High | 0.00097 | 0.01007 | 2.06 |
| AAGHQELVQIPSRPLLSK | [R].AAGHQELVQIPSRPLLSK.[M] | 1xPhosphc | Q9D279 1 | 0.09933 | 0.008 | 1 | 1 | 1 | Q9D279 | Mitotic interactor and substrate of PLK1 OS=Mus musculus GN=Misp PE=1 SV=1 | Q9D279 [2 | 0 | 2482.39 | 0 | 2.92 | 0.00049 | 3.52 | 20.7 | 179.3 | 17.71 | 9.48 | | High | High | High | High | High | High | High | High | High | High | High | 0.00403 | 0.04276 | 2.39 |
| EEPVLTPIDEGHLPMINQDEFR | [K].EEPVLTPIDEGHLPMINQDEFR.[N] | 1xPhosphc | P23298 1x | 0.00527 | 0.0006 | 1 | 3 | 2 | P23298 | Protein kinase C eta type OS=Mus musculus GN=Prkch PE=1 SV=2 | P23298 [6 | 0 | 2888.37 | 0 | 4.11 | 0.00093 | 40.03 | 10.7 | 189.3 | 41.18 | 16.3 | | High | High | High | High | High | High | High | High | High | High | High | 0.00027 | 0.00153 | 3.52 |
| SSETLQSIIQSPK | [R].SSETLQSIIQSPK.[T] | 1xPhosphc | Q8BV49 1 | 0.00317 | 0.00016 | 1 | 2 | 2 | Q8BV49 | Pyrin and HIN domain-containing protein 1 OS=Mus musculus GN=Pyhin1 PE=1 SV=1 | Q8BV49 [1 | 0 | 1956.05 | 0 | 4.77 | 0.00021 | 27.54 | 7.1 | 192.9 | 24.44 | 7.04 | | High | High | High | High | High | High | High | High | High | High | High | 7.2E-05 | 0.00087 | 2.71 |
| SLDLDSIIAEVK | [R].SLDLDSIIAEVK.[A] | 1xPhosphc | P50446 1x | 0.0013 | 0 | 1 | 9 | 2 | P50446 | Keratin, type II cytoskeletal 6A OS=Mus musculus GN=Krt6a PE=1 SV=3 | P50446 [3 | 0 | 1841.01 | 1 | 6.64 | | 0 | 2.5 | 197.5 | 72.47 | 13.83 | | Not Found | High | Not Found | High | High | High | High | High | High | High | High | 0 | 0.00032 | 3.27 |
