## Supplementary_Table9 for "Aging-dependent dysregulation of EXOSC2 is maintained in cancer as a dependency"

| Checked | Confidence | Sequence | Annotated | Modificatio | Modificatio | Qvality PEP | Qvality q-v | # Protein G | # Proteins | # PSMs | Master Pro | Master Pro | Positions in | # Missed C | Theo. MH+ | # Razor Qu | Abundance | Abundance | Abundance | Abundance | Abundance | Abundance | Abundance | Quan Info | Found in S | Found in S | Found in S | Found in S | Found in S | Found in S | Found in S | Found in S | Found in S | Found in S | Confidence | Percolator | Percolator | XCorr (by Search Engine): Sequest HT |
|---|---|---|---|---|---|---|---|---|---|---|---|---|---|---|---|---|---|---|---|---|---|---|---|---|---|---|---|---|---|---|---|---|---|---|---|---|---|---|
| TRUE | High | DGSPVQPS | [K].DGSPV | 1xPhospho | D0VLQ4 1x | 0.001002 | 0 | 1 | 3 | 1 | D0VLQ4 | Tet oncoge | D0VLQ4 [8 | 0 | 2026.07 | 0 | -2.79 | 0.000809 | 0.62 | 174.7 | 25.3 | 8.38 | 33.21 | | High | High | High | High | High | High | High | High | High | High | High | 0 | 0.000238 | 3.14 |
| TRUE | High | IFYTTTPVK | [K].IFYTTTI | 1xPhospho | Q99KR3 1x | 0.023584 | 0.001578 | 1 | 1 | 1 | Q99KR3 | Endoribon | Q99KR3 [2 | 0 | 1607.885 | 0 | -2.34 | 0.000295 | 0.73 | 170.5 | 29.5 | 10.72 | 21.24 | | High | High | High | High | High | High | High | High | High | High | High | 0.000859 | 0.008243 | 2.03 |
| TRUE | High | LFVPSTIGA | [R].LFVPST | 1xPhospho | Q8K3F7 1x | 3.39E-06 | 0 | 1 | 4 | 2 | Q8K3F7 | L-threonin | Q8K3F7 [1 | 0 | 1956.019 | 0 | -3.99 | 0.000328 | 0.1 | 187.1 | 12.9 | 8.68 | 38.93 | | High | High | High | High | High | High | High | High | High | High | High | 0 | 4.01E-07 | 4.41 |
| TRUE | High | APSPARPFL | [R].APSPAF | 1xPhospho | Q3UHU5 1 | 0.002793 | 0.000109 | 1 | 2 | 1 | Q3UHU5 | Microtubu | Q3UHU5 [4 | 0 | 1621.923 | 0 | -2.94 | 0.000975 | 0.32 | 177.1 | 22.9 | 15.76 | 37.21 | | High | High | High | High | High | High | High | High | High | High | High | 4.94E-05 | 0.000755 | 1.57 |
| TRUE | High | ESSTNNSV | [R].ESSTNN | 1xPhospho | Q80TC1 1x | 0.000209 | 0 | 1 | 14 | 3 | Q80TC1 | MKIAA154 | Q80TC1 [3 | 0 | 1902.864 | 0 | -2.44 | 0.000154 | 0.6 | 169.4 | 30.6 | 3.77 | 14.09 | | High | High | High | High | High | High | High | High | High | High | High | 0 | 0.00023 | 2.68 |
| TRUE | High | GGEDAVT | [K].GGEDA | 1xPhospho | A2BDY3 1x | 2.18E-10 | 0 | 1 | 2 | 2 | A2BDY3 | Forkhead b | A2BDY3 [9 | 0 | 3134.609 | 0 | -2.75 | 0.000158 | 0.57 | 174.1 | 25.9 | 5.59 | 22.93 | | High | High | High | High | High | High | High | High | High | High | High | 0 | 9.71E-07 | 3.46 |
| TRUE | High | ELLSPAVSV | [K].ELLSPA | 1xPhospho | Q8BX22 1x | 9.08E-07 | 0 | 1 | 3 | 1 | Q8BX22 | Sal-like pro | Q8BX22 [9 | 0 | 3501.774 | 0 | -2.54 | 0.000793 | 0.01 | 170.8 | 29.2 | 12.06 | 19.47 | | High | High | High | High | High | High | High | High | High | High | High | 0 | 9.11E-08 | 4.67 |
| TRUE | High | ELLSPAVSV | [K].ELLSPA | 1xDeamida | Q8BX22 1x | 9.36E-05 | 0 | 1 | 3 | 1 | Q8BX22 | Sal-like pro | Q8BX22 [9 | 0 | 3486.763 | 0 | -2.63 | 0.000241 | 0.38 | 173.9 | 26.1 | 13.84 | 41 | | High | High | High | High | High | High | High | High | High | High | High | 0 | 1.67E-05 | 4.31 |
| TRUE | High | ELLSPAVSV | [K].ELLSPA | 1xPhospho | Q8BX22 1x | 7.53E-08 | 0 | 1 | 3 | 1 | Q8BX22 | Sal-like pro | Q8BX22 [9 | 0 | 3485.779 | 0 | -2.55 | 0.002229 | 0.47 | 170.3 | 29.7 | 19.41 | 13.98 | | High | High | High | High | High | High | High | High | High | High | High | 0 | 5.56E-09 | 5.04 |
| TRUE | High | SLSSSPVK | [K].SLSSSP | 1xPhospho | Q3TH99 1x | 0.002743 | 0.000109 | 1 | 3 | 2 | Q3TH99 | Putative ur | Q3TH99 [1 | 0 | 1342.738 | 0 | -2.03 | 0.00034 | 0.77 | 161.8 | 38.2 | 6.62 | 16.97 | | High | High | High | High | High | High | High | High | High | High | High | 0.000382 | 0.003276 | 1.8 |
| TRUE | High | VLMFCELP | [R].VLMFC | 1xCarbami | O70238 1x | 0.000302 | 0 | 1 | 2 | 2 | O70238 | Homeobox | O70238 [2 | 0 | 2155.005 | 0 | -2.46 | 0.000278 | 0.57 | 169.3 | 30.7 | 5.86 | 19.7 | | High | High | High | High | High | High | High | High | High | High | High | 0 | 6.24E-05 | 3.31 |
| TRUE | High | TVAISDAA | [R].TVAISD | 1xCarbami | P70445 1xl | 0.002671 | 0.000109 | 1 | 2 | 3 | P70445 | Eukaryotic | P70445 [21 | 0 | 3492.656 | 0 | -2.37 | 0.002146 | 1.98 | 160.9 | 39.1 | 18.36 | 38.23 | | High | High | High | High | High | High | High | High | High | High | High | 4.94E-05 | 0.000716 | 3.88 |
| TRUE | High | TVAISDAA | [R].TVAISD | 1xCarbami | P70445 2xl | 0.00129 | 0 | 1 | 2 | 2 | P70445 | Eukaryotic | P70445 [21 | 0 | 3572.622 | 0 | -2.2 | 0.000779 | 2.53 | 165.5 | 34.5 | 8.4 | 54.11 | | High | High | High | High | High | High | High | High | High | High | High | 0 | 0.000317 | 2.82 |
| TRUE | High | ESVGGSPQ | [R].ESVGG | 1xPhospho | Q9WV69-2 | 3.01E-05 | 0 | 2 | 5 | 3 | Q9WV69-2 | Isoform 2 o | Q9WV69-2 | 0 | 1513.766 | 0 | -3.94 | 0.000209 | 0.14 | 187.9 | 12.1 | 4.58 | 37.35 | Shared | High | High | High | High | High | High | High | High | High | High | High | 0 | 4.67E-06 | 2.59 |
| TRUE | High | SQGYPSSP | [K].SQGYP | 1xPhospho | D0VLQ4 1x | 6.35E-05 | 0 | 1 | 3 | 6 | D0VLQ4 | Tet oncoge | D0VLQ4 [9 | 0 | 1789.877 | 0 | -3.54 | 0.000222 | 0.14 | 184.1 | 15.9 | 3.04 | 36.11 | | High | High | High | High | High | High | High | High | High | High | High | 0 | 2.11E-05 | 2.83 |
| TRUE | High | SLSTCNK | [R].SLSTCN | 1xCarbami | E9PXD2 1x | 0.115566 | 0.009504 | 1 | 7 | 1 | E9PXD2 | Rho GTPas | E9PXD2 [7 | 0 | 1347.674 | 0 | -2.96 | 0.000313 | 0.7 | 178.1 | 21.9 | 4.36 | 32.52 | | High | High | High | High | High | High | High | High | High | High | High | 0.004733 | 0.05111 | 1.8 |
| TRUE | High | HSLSSLATA | [R].HSLSSL | 1xPhospho | Q80Y84 1x | 0.009931 | 0.000736 | 1 | 2 | 2 | Q80Y84 | Lysine-spe | Q80Y84 [9 | 0 | 1651.918 | 0 | -2.04 | 0.000333 | 0.51 | 159.3 | 40.7 | 6.97 | 7.88 | | High | High | High | High | High | High | High | High | High | High | High | 0.000344 | 0.00311 | 2.07 |
| TRUE | High | APSPSGLM | [R].APSPSG | 1xPhospho | U5LK19 1x | 0.001489 | 5.66E-05 | 2 | 16 | 3 | U5LK19; E | Marshalin- | U5LK19 [1 | 0 | 1511.713 | 0 | -2.33 | 0.000519 | 0.64 | 167.2 | 32.8 | 5.09 | 27.4 | Shared | High | High | High | High | High | High | High | High | High | High | High | 0.000814 | 0.007588 | 1.54 |
| TRUE | High | EIHPYLVMI | [K].EIHPYL | 1xPhospho | Q9EPB4 1x | 0.000967 | 0 | 1 | 2 | 1 | Q9EPB4 | Apoptosis- | Q9EPB4 [1 | 0 | 1882.886 | 0 | -3.35 | 0.000825 | 0.29 | 179.4 | 20.6 | 13.64 | 50.18 | | High | High | High | High | High | High | High | High | High | High | High | 0 | 0.000229 | 2.76 |
| TRUE | High | AEQSPVSSI | [K].AEQSP | 2xPhospho | Q80TS5 2x | 0.004955 | 0.000504 | 1 | 4 | 3 | Q80TS5 | Zinc finger | Q80TS5 [6 | 0 | 2264.05 | 1 | -2.04 | 0.00031 | 1.54 | 160 | 40 | 2.37 | 21.5 | | High | High | High | High | High | High | High | High | High | High | High | 0.000275 | 0.002117 | 2.55 |
| TRUE | High | DCVPLILNS | [K].DCVPLI | 3xCarbami | Q8C263 1x | 0.001158 | 0 | 1 | 1 | 1 | Q8C263 | Spindle an | Q8C263 [3 | 0 | 4668.127 | 0 | -2.14 | 0.003965 | 1.02 | 163.1 | 36.9 | 15.87 | 35.68 | | High | High | High | High | High | High | High | High | High | High | High | 0 | 0.000281 | 5.69 |
| TRUE | High | ELHPQLLSF | [R].ELHPQI | 1xPhospho | Q8VDV3 1x | 0.002921 | 0.00016 | 1 | 2 | 1 | Q8VDV3 | Guanine nu | Q8VDV3 [1 | 0 | 1801.003 | 0 | -2 | 0.000695 | 0.59 | 157.1 | 42.9 | 15.02 | 20.05 | | High | High | High | High | High | High | High | High | High | High | High | 7.19E-05 | 0.000795 | 1.7 |
| TRUE | High | EDSVDVVL | [K].EDSVD | 1xCarbami | Q9CWR8 1 | 2.47E-05 | 0 | 1 | 1 | 2 | Q9CWR8 | DNA (cytos | Q9CWR8 [4 | 0 | 1945.923 | 0 | -2.43 | 0.000918 | 1.23 | 166.8 | 33.2 | 10.56 | 19.92 | | High | High | High | High | High | High | High | High | High | High | High | 0 | 3.73E-06 | 3.63 |
| TRUE | High | EDSVDVVL | [K].EDSVD | 1xCarbami | Q9CWR8 1 | 2.21E-05 | 0 | 1 | 1 | 5 | Q9CWR8 | DNA (cytos | Q9CWR8 [4 | 1 | 3151.527 | 0 | -2.91 | 0.003382 | 0.65 | 178 | 22 | 24.38 | 23.39 | | High | High | High | High | High | High | High | High | High | High | High | 0 | 3.31E-06 | 3.7 |
| TRUE | High | LSADYEK | [R].LSADYE | 1xPhospho | B2RRE0 1x | 0.016596 | 0.001124 | 1 | 3 | 2 | B2RRE0 | A kinase (P | B2RRE0 [3 | 0 | 1134.528 | 0 | -2.05 | 0.000348 | 0.56 | 161.2 | 38.8 | 9.42 | 20 | | High | High | High | High | High | High | High | High | High | High | High | 0.002403 | 0.02577 | 2.18 |
| TRUE | High | EDSVDVVL | [K].EDSVD | 1xCarbami | Q9CWR8 2 | 1.64E-05 | 0 | 1 | 1 | 1 | Q9CWR8 | DNA (cytos | Q9CWR8 [4 | 1 | 3231.493 | 0 | -2.28 | 0.000669 | 0.87 | 163.1 | 36.9 | 12.02 | 27.24 | | High | High | High | High | High | High | High | High | High | High | High | 0 | 2.37E-06 | 4.36 |
| TRUE | High | ATAHSPDP | [K].ATAHSI | 1xCarbami | Q2WG77 1 | 0.037369 | 0.00249 | 1 | 1 | 1 | Q2WG77 | Protein rip | Q2WG77 [ | 0 | 1770.747 | 0 | -2.83 | 0.000466 | 1.31 | 172.1 | 27.9 | 8.11 | 46.36 | | High | High | High | High | High | High | High | High | High | High | High | 0.001308 | 0.01383 | 2.38 |
| TRUE | High | RTSLPDAP | [K].RTSLPD | 2xPhospho | A2TJV2 2xl | 2.01E-07 | 0 | 1 | 1 | 1 | A2TJV2 | Paralemmi | A2TJV2 [15 | 1 | 2501.098 | 0 | -2.37 | 0.000285 | 1.56 | 168.8 | 31.2 | 5.13 | 31.5 | | High | High | High | High | High | High | High | High | High | High | High | 0 | 1.68E-08 | 4.39 |
| TRUE | High | RTSLPDAP | [K].RTSLPD | 1xPhospho | A2TJV2 1xl | 1.58E-05 | 0 | 1 | 1 | 2 | A2TJV2 | Paralemmi | A2TJV2 [15 | 1 | 2421.132 | 0 | -3.74 | 0.00025 | 0.14 | 185.6 | 14.4 | 3.48 | 41.97 | | High | High | High | High | High | High | High | High | High | High | High | 0 | 2.27E-06 | 2.69 |
| TRUE | High | RTSLPDAP | [K].RTSLPD | 1xPhospho | A2TJV2 1xl | 0.001222 | 0 | 1 | 1 | 2 | A2TJV2 | Paralemmi | A2TJV2 [15 | 1 | 2405.137 | 0 | -3.29 | 0.00024 | 0.38 | 181.1 | 18.9 | 4.61 | 34.66 | | High | High | High | High | High | High | High | High | High | High | High | 2.58E-05 | 0.000386 | 3.29 |
| TRUE | High | RLPAFSPPS | [R].RLPAFS | 1xPhospho | Q6J1H4 1x | 7.08E-05 | 0 | 1 | 1 | 3 | Q6J1H4 | Undifferen | Q6J1H4 [7- | 1 | 2217.127 | 0 | -2.69 | 0.001001 | 0.73 | 173.2 | 26.8 | 13 | 27.11 | | High | High | High | High | High | High | High | High | High | High | High | 0 | 1.22E-05 | 3.48 |
| TRUE | High | IGAQMGHI | [K].IGAQM | 1xDeamida | E9PWE9 1x | 0.001011 | 0 | 1 | 1 | 3 | E9PWE9 | Tyrosine-p | E9PWE9 [2 | 0 | 4957.299 | 0 | -2.46 | 0.00195 | 1.38 | 166 | 34 | 15.1 | 17.51 | | High | High | High | High | High | High | High | High | High | High | High | 0 | 0.000243 | 4.48 |
| TRUE | High | GVQVPVSF | [K].GVQVP | 1xPhospho | Q6ZPI0 1xF | 7.56E-06 | 0 | 1 | 3 | 1 | Q6ZPI0 | Protein Jac | Q6ZPI0 [84 | 0 | 2011.094 | 0 | -2.6 | 0.00024 | 0.23 | 170.6 | 29.4 | 5.64 | 19.11 | | High | High | High | High | High | High | High | High | High | High | High | 0 | 9.9E-07 | 2.71 |
| TRUE | High | RVSGSGGH | [R].RVSGS | 1xPhospho | A0A0R4J1L | 0.010109 | 0.000736 | 1 | 2 | 1 | A0A0R4J1L | Microtubu | A0A0R4J1L | 1 | 1791.963 | 0 | -2.2 | 0.000405 | 0.4 | 168.5 | 31.5 | 9.8 | 15.24 | | High | High | High | High | High | High | High | High | High | High | High | 0.000344 | 0.00318 | 1.22 |
| TRUE | High | AGGGTTPV | [K].AGGGT | 1xPhospho | Q3UHU5 1 | 0.018774 | 0.001362 | 1 | 5 | 1 | Q3UHU5 | Microtubu | Q3UHU5 [ | 0 | 1482.715 | 0 | -2.28 | 0.00041 | 0.59 | 165.3 | 34.7 | 4.97 | 19.04 | | High | High | High | High | High | High | High | High | High | High | High | 0.000723 | 0.00633 | 1.29 |
| TRUE | High | ATQQSLGE | [R].ATQQS | 1xPhospho | B1AZP2 1xl | 0.005666 | 0.00062 | 1 | 3 | 1 | B1AZP2 | Disks large | B1AZP2 [4 | 0 | 1724.817 | 0 | -2.14 | 0.001282 | 0.62 | 165.7 | 34.3 | 11.18 | 44.69 | | High | High | High | High | High | High | High | High | High | High | High | 0.000275 | 0.001657 | 2.22 |
| TRUE | High | SVLTAGGV | [K].SVLTAC | 1xPhospho | A0A1B0GS | 2.43E-06 | 0 | 1 | 2 | 2 | A0A1B0GS | GRB2-asso | A0A1B0GS | 0 | 3138.465 | 0 | -2.17 | 0.000527 | 0.47 | 165.5 | 34.5 | 8.85 | 12.57 | | High | High | High | High | High | High | High | High | High | High | High | 0 | 2.75E-07 | 3.57 |
| TRUE | High | LGGLLPAG | [K].LGGLLF | 1xPhospho | G5DDB7 1x | 6.28E-11 | 0 | 1 | 3 | 3 | G5DDB7 | Phosphoin | G5DDB7 [4 | 0 | 3652.605 | 0 | -2.56 | 0.000261 | 0.04 | 171.7 | 28.3 | 10.25 | 32.19 | | High | High | High | High | High | High | High | High | High | High | High | 0 | 1.93E-12 | 6.8 |
| TRUE | High | EDMDISEP | [R].EDMDI | 1xPhospho | B1AU75 1x | 0.001411 | 5.66E-05 | 1 | 2 | 1 | B1AU75 | Nuclear au | B1AU75 [1 | 0 | 1859.839 | 0 | -2.55 | 0.000479 | 0.78 | 173 | 27 | 8.93 | 15.82 | | High | High | High | High | High | High | High | High | High | High | High | 2.58E-05 | 0.000349 | 2.1 |
| TRUE | High | EDMDISEP | [R].EDMDI | 1xPhospho | B1AU75 1x | 0.000386 | 0 | 1 | 2 | 2 | B1AU75 | Nuclear au | B1AU75 [1 | 0 | 1875.834 | 0 | -2.44 | 0.000225 | 0.75 | 169.6 | 30.4 | 5.64 | 15.46 | | High | High | High | High | High | High | High | High | High | High | High | 2.58E-05 | 0.000352 | 2.89 |
| TRUE | High | SVLDDFFT | [R].SVLDDI | 1xPhospho | E9Q696 1x | 1.44E-07 | 0 | 1 | 3 | 2 | E9Q696 | Tyrosine-p | E9Q696 [5 | 0 | 2352.063 | 0 | -5.11 | 0.000908 | 0.03 | 194 | 6 | 10.94 | 90.46 | | High | High | High | High | High | High | High | High | High | High | High | 0 | 7.98E-07 | 3.42 |
| TRUE | High | TTRSPEPVF | [R].TTRSPE | 1xPhospho | P97931 1xl | 0.007742 | 0.000664 | 1 | 1 | 2 | P97931 | Uracil-DNA | P97931 [2 | 1 | 3059.581 | 0 | -2.23 | 0.00068 | 2.02 | 166 | 34 | 7.54 | 34.98 | | High | High | High | High | High | High | High | High | High | High | High | 0.000327 | 0.002681 | 2.64 |
| TRUE | High | ATPMSSLLI | [R].ATPMS | 1xPhospho | B2RWY2 1 | 0.000651 | 0 | 1 | 7 | 1 | B2RWY2 | AT rich inte | B2RWY2 [1 | 0 | 2691.419 | 0 | -2.05 | 0.001448 | 0.88 | 160 | 40 | 9.94 | 17.49 | | High | High | High | High | High | High | High | High | High | High | High | 0 | 0.000147 | 2.97 |
| TRUE | High | VTTMSEND | [K].VTTMS | 1xPhospho | Q9JI59 1xP | 0.000611 | 0 | 1 | 1 | 1 | Q9JI59 | Junctional | Q9JI59 [28 | 0 | 1725.817 | 0 | -2.18 | 0.000275 | 0.84 | 163.9 | 36.1 | 8.21 | 15.39 | | High | High | High | High | High | High | High | High | High | High | High | 0 | 0.000137 | 2.09 |
| TRUE | High | SQSFGVAS | [R].SQSFG | 1xPhospho | Q8CI12 1xl | 1.78E-05 | 0 | 1 | 2 | 2 | Q8CI12 | Smoothelin | Q8CI12 [33 | 0 | 1806.94 | 0 | -2.73 | 0.00033 | 0.09 | 173.3 | 26.7 | 9.96 | 29.64 | | High | High | High | High | High | High | High | High | High | High | High | 0 | 5.29E-06 | 2.49 |
| TRUE | High | ATPMSSLLI | [R].ATPMS | 1xPhospho | B2RWY2 1 | 2.12E-06 | 0 | 1 | 7 | 1 | B2RWY2 | AT rich inte | B2RWY2 [1 | 0 | 2675.424 | 0 | -2.56 | 0.000313 | 0.39 | 173.6 | 26.4 | 11.04 | 22.12 | | High | High | High | High | High | High | High | High | High | High | High | 0 | 2.36E-07 | 3.59 |
| TRUE | High | ELPPELSHL | [K].ELPPEL | 1xPhospho | E9Q2Z1 1x | 7.06E-07 | 0 | 1 | 3 | 1 | E9Q2Z1 | CECR2, hist | E9Q2Z1 [3 | 0 | 2173.056 | 0 | -2.11 | 0.000517 | 1.29 | 162.4 | 37.6 | 6.83 | 23.3 | | High | High | High | High | High | High | High | High | High | High | High | 0 | 6.9E-08 | 1.98 |
| TRUE | High | NTPAVFSPI | [K].NTPAVI | 1xPhospho | Q9CPT0 1x | 0.002671 | 0.000109 | 1 | 2 | 2 | Q9CPT0 | Apoptosis | Q9CPT0 [3 | 0 | 1498.807 | 0 | -2.71 | 0.00022 | 0.67 | 173.7 | 26.3 | 4.41 | 21.98 | | High | High | High | High | High | High | High | High | High | High | High | 0.000327 | 0.002832 | 2.61 |
| TRUE | High | TEAEGLRP | [K].TEAEGI | 1xPhospho | Q0VGT2 1x | 0.00145 | 5.66E-05 | 1 | 7 | 3 | Q0VGT2 | Zinc finger | Q0VGT2 [3 | 0 | 2975.585 | 0 | -3.88 | 0.000358 | 0.19 | 186.5 | 13.5 | 5.79 | 48.43 | | High | High | High | High | High | High | High | High | High | High | High | 2.58E-05 | 0.00036 | 3.07 |
| TRUE | High | IVDGLSPLS | [R].IVDGLS | 1xPhospho | Q3UHU5 1 | 5.21E-05 | 0 | 1 | 5 | 1 | Q3UHU5 | Microtubu | Q3UHU5 [8 | 0 | 3028.544 | 0 | -2.65 | 0.014054 | 1.61 | 170.6 | 29.4 | 30.26 | 35.53 | | High | High | High | High | High | High | High | High | High | High | High | 0 | 8.66E-06 | 4 |
| TRUE | High | ESPTVSPR | [R].ESPTVS | 1xPhospho | D3YUT2 1x | 0.118334 | 0.009621 | 1 | 2 | 1 | D3YUT2 | Alpha-prot | D3YUT2 [1 | 0 | 1181.576 | 0 | -2.18 | 0.00024 | 1.2 | 165.7 | 34.3 | 8.04 | 28.86 | | High | High | High | High | High | High | High | High | High | High | High | 0.004826 | 0.05254 | 1.06 |
| TRUE | High | GGHGAASI | [K].GGHGA | 1xPhospho | Q8BMK4 1 | 0.007673 | 0.000664 | 1 | 1 | 4 | Q8BMK4 | Cytoskelet | Q8BMK4 [ | 0 | 1521.746 | 0 | -3.22 | 0.000202 | 0.21 | 181 | 19 | 6.87 | 23.64 | | High | High | High | High | High | High | High | High | High | High | High | 0.000294 | 0.002331 | 2.31 |
| TRUE | High | GVSPAAPP | [R].GVSPA | 1xPhospho | E9Q6W4 1 | 7.7E-06 | 0 | 1 | 1 | 2 | E9Q6W4 | Zinc finger | E9Q6W4 [2 | 0 | 3216.659 | 0 | -2.32 | 0.000158 | 0.45 | 166.8 | 33.2 | 4.7 | 16.17 | | High | High | High | High | High | High | High | High | High | High | High | 0 | 1.01E-06 | 4 |
| TRUE | High | TTSISPALA | [R].TTSISP | 1xPhospho | B2RY17 1xl | 0.045122 | 0.003373 | 1 | 11 | 2 | B2RY17 | Tnik protei | B2RY17 [6 | 0 | 1325.703 | 0 | -2.52 | 0.000151 | 0.52 | 169.4 | 30.6 | 5.23 | 22.03 | | High | High | High | High | High | High | High | High | High | High | High | 0.001727 | 0.01718 | 2.01 |
| TRUE | High | TPVLLGPP | [R].TPVLLG | 1xCarbami | Q7TNS8 1x | 1.5E-08 | 0 | 1 | 1 | 1 | Q7TNS8 | Elongin BC | Q7TNS8 [6 | 0 | 2703.398 | 0 | -2.53 | 0.000251 | 0.44 | 171.8 | 28.2 | 6.77 | 23.67 | | High | High | High | High | High | High | High | High | High | High | High | 0 | 9.09E-10 | 3.88 |
| TRUE | High | DAPSPPVLI | [K].DAPSPF | 2xCarbami | Q3KR45 1x | 1.86E-07 | 0 | 1 | 5 | 6 | Q3KR45 | DNA (cytos | Q3KR45 [3 | 0 | 2748.282 | 0 | -2.42 | 0.00046 | 0.58 | 168.5 | 31.5 | 5.91 | 23.03 | | High | High | High | High | High | High | High | High | High | High | High | 0 | 2.63E-08 | 6.02 |
| TRUE | High | DAPSPPVLI | [K].DAPSPF | 2xCarbami | Q3KR45 2x | 0.001848 | 5.66E-05 | 1 | 5 | 2 | Q3KR45 | DNA (cytos | Q3KR45 [3 | 0 | 2828.249 | 0 | -4.22 | 0.000191 | 0.05 | 190.2 | 9.8 | 4.28 | 50.08 | | High | High | High | High | High | High | High | High | High | High | High | 2.58E-05 | 0.000473 | 4.53 |
| TRUE | High | HSENDSVP | [K].HSEND | 1xPhospho | D0VLQ4 1x | 1.57E-05 | 0 | 1 | 3 | 2 | D0VLQ4 | Tet oncoge | D0VLQ4 [1 | 0 | 2986.313 | 0 | -2.48 | 0.002688 | 0.85 | 169.6 | 30.4 | 19.95 | 9.68 | | High | High | High | High | High | High | High | High | High | High | High | 0 | 2.25E-06 | 3.36 |
| TRUE | High | NTETVPSQ | [R].NTETVI | 1xPhospho | U5LK19 1x | 1.19E-06 | 0 | 2 | 17 | 2 | U5LK19; E | Marshalin- | U5LK19 [3 | 0 | 2578.177 | 0 | -2.05 | 0.000348 | 0.69 | 159.2 | 40.8 | 4.74 | 12.72 | Shared | High | High | High | High | High | High | High | High | High | High | High | 0 | 5.31E-07 | 3.89 |
| TRUE | High | DAPTSPAS | [K].DAPTSF | 1xPhospho | Q80TC1 1x | 1.43E-06 | 0 | 1 | 14 | 6 | Q80TC1 | MKIAA154 | Q80TC1 [4 | 0 | 2302.122 | 2 | -2.56 | 0.000213 | 0.64 | 172.6 | 27.4 | 8.84 | 23.49 | | High | High | High | High | High | High | High | High | High | High | High | 0 | 0.000148 | 3.79 |
| TRUE | High | HSENDSVP | [K].HSEND | 1xDeamida | D0VLQ4 1x | 0.002153 | 5.66E-05 | 1 | 3 | 2 | D0VLQ4 | Tet oncoge | D0VLQ4 [1 | 0 | 2987.297 | 0 | -2.34 | 0.003465 | 1.56 | 165 | 35 | 16.55 | 36.72 | | High | High | High | High | High | High | High | High | High | High | High | 2.58E-05 | 0.000562 | 3.63 |
| TRUE | High | RLDSENSP | [R].RLDSEN | 1xPhospho | G5E8P8 1x | 0.023379 | 0.001578 | 1 | 6 | 1 | G5E8P8 | Estrogen re | G5E8P8 [2 | 1 | 2649.405 | 0 | -3.02 | 0.000554 | 0.67 | 179.2 | 20.8 | 7.55 | 33.82 | | High | High | High | High | High | High | High | High | High | High | High | 0.000847 | 0.008166 | 1.62 |
| TRUE | High | GQDLGASV | [K].GQDLG | 1xPhospho | Q9D8C9 1x | 0.003372 | 0.00016 | 1 | 1 | 1 | Q9D8C9 | Purine nuc | Q9D8C9 [5 | 0 | 1927.978 | 0 | -2.42 | 0.000248 | 0.64 | 169.8 | 30.2 | 6.67 | 15 | | High | High | High | High | High | High | High | High | High | High | High | 9.42E-05 | 0.000935 | 2.21 |
| TRUE | High | LGEESKEPL | [K].LGEESK | 1xPhospho | Q923F1 1x | 0.003962 | 0.000311 | 1 | 1 | 2 | Q923F1 | Chloride ch | Q923F1 [9 | 1 | 3660.637 | 0 | -2.82 | 0.000627 | 0.69 | 175.5 | 24.5 | 13.5 | 27.65 | | High | High | High | High | High | High | High | High | High | High | High | 0.00554 | 0.06021 | 3.49 |
| TRUE | High | DSSVPGSP | [R].DSSVPC | 1xPhospho | Q9WV69-2 | 2.47E-06 | 0 | 2 | 2 | 3 | Q9WV69-2 | Isoform 2 o | Q9WV69-2 | 0 | 1868.977 | 0 | -5.03 | 0.000788 | 0.02 | 194.1 | 5.9 | 11.51 | 57.55 | Shared | High | High | High | High | High | High | High | High | High | High | High | 0 | 7.23E-05 | 4.12 |
| TRUE | High | SQEDEISPV | [R].SQEDEI | 1xPhospho | Q6PR54-3 | 0.000483 | 0 | 2 | 4 | 1 | Q6PR54-3; | Isoform 3 o | Q6PR54-3 | 0 | 1554.725 | 0 | -2.16 | 0.001144 | 0.86 | 161.3 | 38.7 | 7.86 | 39.27 | Shared | High | High | High | High | High | High | High | High | High | High | High | 0 | 0.000106 | 2.66 |
| TRUE | High | QLSPESLGT | [K].QLSPES | 1xPhospho | A2ARP8 1x | 9.67E-07 | 0 | 1 | 3 | 2 | A2ARP8 | Microtubu | A2ARP8 [1 | 0 | 2542.357 | 0 | -2.52 | 0.000971 | 0.49 | 168.1 | 31.9 | 12.97 | 22.53 | | High | High | High | High | High | High | High | High | High | High | High | 0 | 9.78E-08 | 3.45 |
| TRUE | High | ASYLESEK | [R].ASYLES | 1xPhospho | E9Q9N7 1x | 0.004572 | 0.000456 | 1 | 2 | 1 | E9Q9N7 | Immunoglo | E9Q9N7 [6 | 0 | 1464.739 | 0 | -2.44 | 0.000223 | 1.06 | 168.2 | 31.8 | 3.34 | 24.35 | | High | High | High | High | High | High | High | High | High | High | High | 0.000204 | 0.001306 | 2.09 |
| TRUE | High | DSDRDDFL | [K].DSDRD | 1xPhospho | Q9QY39 1x | 0.001986 | 5.66E-05 | 1 | 2 | 1 | Q9QY39 | PDZ domai | Q9QY39 [2 | 1 | 2626.078 | 0 | -2.66 | 0.00109 | 1.05 | 173.6 | 26.4 | 10.5 | 43.91 | | High | High | High | High | High | High | High | High | High | High | High | 2.58E-05 | 0.000515 | 2.51 |
| TRUE | High | MENGVLG | [R].MENG\ | 1xDeamida | B1AU75 1x | 1.03E-05 | 0 | 1 | 3 | 4 | B1AU75 | Nuclear au | B1AU75 [1 | 1 | 4967.23 | 0 | -3.56 | 0.002637 | 0.26 | 184.4 | 15.6 | 13.9 | 81.59 | | High | High | High | High | High | High | High | High | High | High | High | 0 | 1.76E-05 | 5.71 |
| TRUE | High | MERPGPGI | [K].MERPG | 1xCarbami | Q9JHU9 1x | 0.000191 | 0 | 1 | 1 | 1 | Q9JHU9 | Inositol-3-p | Q9JHU9 [5 | 0 | 3087.67 | 0 | -2.3 | 0.00149 | 0.59 | 167.1 | 32.9 | 17.35 | 10.57 | | High | High | High | High | High | High | High | High | High | High | High | 0 | 3.73E-05 | 1.63 |
| TRUE | High | RLSVENPM | [R].RLSVEN | 1xPhospho | Q62255 1x | 0.0095 | 0.000736 | 1 | 2 | 1 | Q62255 | Sal-like pro | Q62255 [1 | 1 | 2322.266 | 0 | -2.4 | 0.001351 | 0.81 | 168.2 | 31.8 | 12.8 | 20.56 | | High | High | High | High | High | High | High | High | High | High | High | 0.000327 | 0.002958 | 2.68 |
| TRUE | High | GTPEFSPLC | [R].GTPEFS | 1xCarbami | Q7TNS8 1x | 0.000396 | 0 | 1 | 1 | 1 | Q7TNS8 | Elongin BC | Q7TNS8 [3 | 0 | 1585.765 | 0 | -2.06 | 0.000697 | 0.83 | 161.5 | 38.5 | 7.69 | 14.43 | | High | High | High | High | High | High | High | High | High | High | High | 0 | 8.41E-05 | 2.29 |
| TRUE | High | AQDATAGV | [K].AQDAT | 1xPhospho | E9QN31 1x | 1.33E-08 | 0 | 1 | 2 | 7 | E9QN31 | Probable 2 | E9QN31 [1 | 0 | 3708.53 | 0 | -2.98 | 0.000291 | 0.93 | 177.6 | 22.4 | 4.28 | 33.9 | | High | High | High | High | High | High | High | High | High | High | High | 0 | 7.94E-10 | 5.51 |
| TRUE | High | GDVTAEEA | [R].GDVTA | 1xPhospho | P28667 1xl | 3.48E-09 | 0 | 1 | 1 | 7 | P28667 | MARCKS-re | P28667 [11 | 0 | 1911.947 | 1 | -2.19 | 0.000154 | 0.83 | 164.9 | 35.1 | 4.24 | 18.25 | | High | High | High | High | High | High | High | High | High | High | High | 0 | 1.76E-10 | 4.62 |
| TRUE | High | EVEVASPEV | [K].EVEVA | 1xPhospho | G3UYN0 1x | 0.000142 | 0 | 1 | 2 | 3 | G3UYN0 | L1TD1 OS= | G3UYN0 [3 | 0 | 1624.86 | 0 | -2.04 | 0.00031 | 0.78 | 163.1 | 36.9 | 6.67 | 9.01 | | High | High | High | High | High | High | High | High | High | High | High | 0 | 2.66E-05 | 2.47 |
| TRUE | High | AQDATAGV | [K].AQDAT | 2xPhospho | E9QN31 2x | 0.019618 | 0.001438 | 1 | 2 | 1 | E9QN31 | Probable 2 | E9QN31 [1 | 0 | 3788.496 | 0 | -2.35 | 0.000277 | 0.89 | 166.8 | 33.2 | 5.06 | 16.54 | | High | High | High | High | High | High | High | High | High | High | High | 0.000756 | 0.00666 | 3.79 |
| TRUE | High | TVSEPSLSG | [R].TVSEPS | 1xPhospho | D0VLQ5 1x | 1.95E-05 | 0 | 1 | 3 | 3 | D0VLQ5 | Tet oncoge | D0VLQ5 [9 | 0 | 2004.057 | 0 | -2.91 | 0.000402 | 0.3 | 174.8 | 25.2 | 9.15 | 24.49 | | High | High | High | High | High | High | High | High | High | High | High | 4.94E-05 | 0.000757 | 2.75 |
| TRUE | High | TEPACVSPI | [K].TEPAC\ | 1xCarbami | Q8BMQ3 1 | 6.62E-07 | 0 | 1 | 6 | 3 | Q8BMQ3 | Zinc finger | Q8BMQ3 [ | 0 | 2638.32 | 0 | -3.14 | 0.000518 | 0.34 | 179.8 | 20.2 | 9.67 | 22.24 | | High | High | High | High | High | High | High | High | High | High | High | 0 | 5.9E-06 | 3.89 |
| TRUE | High | NSGPAVPA | [R].NSGPA | 1xPhospho | Q3TAB2 1x | 0.00145 | 5.66E-05 | 1 | 1 | 1 | Q3TAB2 | Putative ur | Q3TAB2 [2 | 0 | 2573.338 | 0 | -2.42 | 0.000816 | 0.35 | 168.4 | 31.6 | 9.32 | 27.19 | | High | High | High | High | High | High | High | High | High | High | High | 2.58E-05 | 0.000361 | 3.15 |
| TRUE | High | VGTNYQAV | [R].VGTNY | 1xCarbami | Q8C796 1x | 6.2E-06 | 0 | 1 | 3 | 2 | Q8C796 | REST corep | Q8C796 [4 | 0 | 2654.341 | 0 | -2.03 | 0.000188 | 1.42 | 160.6 | 39.4 | 1.87 | 16.63 | | High | High | High | High | High | High | High | High | High | High | High | 0 | 0.000114 | 2.72 |
| TRUE | High | HTPGAPVN | [R].HTPGA | 1xCarbami | A2AUZ7 1x | 8E-07 | 0 | 1 | 1 | 2 | A2AUZ7 | Sal-like pro | A2AUZ7 [6 | 0 | 3016.412 | 0 | -2.11 | 0.000163 | 1.17 | 161.1 | 38.9 | 3.51 | 16.69 | | High | High | High | High | High | High | High | High | High | High | High | 0 | 7.95E-08 | 4.12 |
| TRUE | High | STSPPPSPE | [K].STSPPP | 1xPhospho | Q9WV69-2 | 2.93E-05 | 0 | 2 | 5 | 8 | Q9WV69-2 | Isoform 2 o | Q9WV69-2 | 0 | 1935.905 | 0 | -2.99 | 0.00014 | 0.27 | 177 | 23 | 2.28 | 20.99 | Shared | High | High | High | High | High | High | High | High | High | High | High | 0 | 1.55E-05 | 3.76 |
| TRUE | High | EEEEPNTD | [K].EEEEPN | 1xPhospho | B2RQ54 1x | 2.11E-05 | 0 | 1 | 1 | 2 | B2RQ54 | Developme | B2RQ54 [3 | 0 | 3284.443 | 0 | -3.64 | 0.001395 | 0.22 | 184.1 | 15.9 | 15.17 | 42.24 | | High | High | High | High | High | High | High | High | High | High | High | 0 | 3.14E-06 | 4.09 |
| TRUE | High | ASSSASTPV | [R].ASSSAS | 1xPhospho | Q3KR45 1x | 0.006653 | 0.00062 | 1 | 5 | 8 | Q3KR45 | DNA (cytos | Q3KR45 [1 | 0 | 3036.431 | 0 | -2.86 | 0.000236 | 0.53 | 178.1 | 21.9 | 6.51 | 33.49 | | High | High | High | High | High | High | High | High | High | High | High | 0.000275 | 0.001995 | 4.17 |
| TRUE | High | TPEDSVCEI | [K].TPEDS\ | 1xCarbami | Q9CZ91 1x | 0.000422 | 0 | 1 | 1 | 2 | Q9CZ91 | Serum resp | Q9CZ91 [2 | 0 | 3240.322 | 0 | -2.12 | 0.001937 | 1.53 | 162.6 | 37.4 | 9.11 | 38.99 | | High | High | High | High | High | High | High | High | High | High | High | 0 | 9.06E-05 | 4.38 |
| TRUE | High | ADSHGELD | [R].ADSHG | 1xPhospho | Q99L15 1xl | 0.025964 | 0.001645 | 1 | 9 | 2 | Q99L15 | Acot1 prot | Q99L15 [5 | 0 | 1492.699 | 0 | -2.1 | 0.001856 | 0.75 | 162.7 | 37.3 | 13.53 | 30.64 | | High | High | High | High | High | High | High | High | High | High | High | 0.00299 | 0.03159 | 2.23 |
| TRUE | High | EVRPSPSK | [R].EVRPSF | 1xPhospho | Q62315 1x | 0.014797 | 0.001049 | 1 | 5 | 2 | Q62315 | Protein Jur | Q62315 [3 | 0 | 1437.787 | 0 | -2.04 | 0.000201 | 0.03 | 160.3 | 39.7 | 2.45 | 17.9 | | High | High | High | High | High | High | High | High | High | High | High | 0.000555 | 0.004854 | 1.07 |
| TRUE | High | LEITSPSK | [RK].LEITSF | 1xPhospho | G3UY09 1x | 0.021797 | 0.001507 | 1 | 2 | 1 | G3UY09 | MCG61508 | G3UY09 [3 | 0 | 1412.78 | 0 | -2.12 | 0.000313 | 0.85 | 165.6 | 34.4 | 5.6 | 18.84 | | High | High | High | High | High | High | High | High | High | High | High | 0.000814 | 0.007491 | 1.15 |
| TRUE | High | EEEINIYQF | [K].EEEINI | 1xCarbami | P42209 1xl | 4.63E-08 | 0 | 1 | 3 | 7 | P42209 | Septin-1 O | P42209 [1 | 0 | 3203.369 | 0 | -2.57 | 0.00037 | 0.56 | 171.1 | 28.9 | 6.49 | 13.54 | | High | High | High | High | High | High | High | High | High | High | High | 0 | 3.21E-09 | 4.29 |
| TRUE | High | EEEINIYQF | [K].EEEINI | 1xCarbami | P42209 1xl | 3.94E-05 | 0 | 1 | 3 | 4 | P42209 | Septin-1 O | P42209 [1 | 1 | 3560.627 | 0 | -3.55 | 0.000338 | 0.28 | 184.2 | 15.8 | 8.11 | 39.29 | | High | High | High | High | High | High | High | High | High | High | High | 0.000275 | 0.001677 | 4.48 |
| TRUE | High | HTSLNDLSI | [R].HTSLNI | 1xPhospho | D3Z4U5 1x | 0.002334 | 0.000109 | 1 | 3 | 1 | D3Z4U5 | Janus kinas | D3Z4U5 [3 | 0 | 1565.789 | 0 | -3.25 | 0.000223 | 0.84 | 180.7 | 19.3 | 7.66 | 53.72 | | High | High | High | High | High | High | High | High | High | High | High | 4.94E-05 | 0.000616 | 1.79 |
| TRUE | High | QNSDPTSE | [R].QNSDP | 1xPhospho | B2RY17 1xl | 0.006478 | 0.00062 | 1 | 11 | 1 | B2RY17 | Tnik protei | B2RY17 [6 | 0 | 1961.917 | 0 | -2.06 | 0.000158 | 1.16 | 161.3 | 38.7 | 1.7 | 12.77 | | High | High | High | High | High | High | High | High | High | High | High | 0.000275 | 0.001933 | 2.22 |
| TRUE | High | ASSPQVAP | [R].ASSPQ\ | 1xPhospho | Q6J1H4 1x | 1.15E-06 | 0 | 1 | 2 | 5 | Q6J1H4 | Undifferen | Q6J1H4 [2 | 0 | 3707.993 | 0 | -3.64 | 0.002102 | 0.36 | 183.3 | 16.7 | 17.69 | 53.22 | | High | High | High | High | High | High | High | High | High | High | High | 0 | 1.18E-07 | 4.37 |
| TRUE | High | YDSDGDK | [R].YDSDG | 1xPhospho | Q80TC1 1x | 0.111965 | 0.009053 | 1 | 14 | 1 | Q80TC1 | MKIAA154 | Q80TC1 [3 | 0 | 1337.603 | 0 | -2.61 | 0.00014 | 0.59 | 171.8 | 28.2 | 5.35 | 19.24 | | High | High | High | High | High | High | High | High | High | High | High | 0.004543 | 0.0492 | 1.78 |
| TRUE | High | IQPSGADL | [R].IQPSGA | 1xPhospho | Q9WVL5 1 | 1.52E-06 | 0 | 1 | 1 | 1 | Q9WVL5 | MORC fam | Q9WVL5 [5 | 0 | 2210.147 | 0 | -2.54 | 0.000432 | 0.4 | 169.6 | 30.4 | 8.44 | 9.62 | | High | High | High | High | High | High | High | High | High | High | High | 0 | 1.63E-07 | 3.59 |
| TRUE | High | SPTDLTGV | [K].SPTDLT | 1xCarbami | Q7TSH4 1x | 0.000297 | 0 | 1 | 3 | 2 | Q7TSH4 | Centriolar | Q7TSH4 [4 | 0 | 2262.109 | 0 | -2.7 | 0.000158 | 0.82 | 173.6 | 26.4 | 3.67 | 31.73 | | High | High | High | High | High | High | High | High | High | High | High | 0 | 6.13E-05 | 2.17 |
| TRUE | High | SGATEASS | [R].SGATE/ | 1xPhospho | Q3TJR9 1xl | 0.000407 | 0 | 1 | 2 | 1 | Q3TJR9 | Achaete-sc | Q3TJR9 [1 | 0 | 1660.774 | 0 | -4.07 | 0.000191 | 0.14 | 189.8 | 10.2 | 5.54 | 53.81 | | High | High | High | High | High | High | High | High | High | High | High | 0 | 8.68E-05 | 1.76 |
| TRUE | High | ASSSASTPV | [R].ASSSAS | 1xPhospho | Q3KR45 1x | 0.004531 | 0.000456 | 1 | 5 | 7 | Q3KR45 | DNA (cytos | Q3KR45 [1 | 0 | 3052.426 | 0 | -2.2 | 0.001625 | 3.02 | 169.3 | 30.7 | 15.61 | 56.44 | | High | High | High | High | High | High | High | High | High | High | High | 0.000204 | 0.001299 | 3.49 |
| TRUE | High | NTTVCEISS | [K].NTTVCI | 1xCarbami | A0A1D9BZ | 9.62E-05 | 0 | 1 | 1 | 2 | A0A1D9BZ | GCNA OS= | A0A1D9BZ | 0 | 2775.234 | 0 | -3.47 | 0.038555 | 0.27 | 169.7 | 30.3 | 53.57 | 51.86 | | High | High | High | High | High | High | High | High | High | High | High | 0 | 2.27E-05 | 4.37 |
| TRUE | High | STSPPPSPE | [K].STSPPP | 2xPhospho | Q9WV69-2 | 0.008769 | 0.000736 | 2 | 5 | 3 | Q9WV69-2 | Isoform 2 o | Q9WV69-2 | 0 | 2015.871 | 0 | -2.69 | 0.001004 | 0.37 | 171.9 | 28.1 | 11.18 | 18.98 | Shared | High | High | High | High | High | High | High | High | High | High | High | 0.000706 | 0.006224 | 2.83 |
| TRUE | High | GRSSLSALA | [R].GRSSLS | 2xPhospho | Q6J1H4 2x | 0.000645 | 0 | 1 | 2 | 3 | Q6J1H4 | Undifferen | Q6J1H4 [1 | 1 | 3926.914 | 0 | -2.14 | 0.001888 | 2.26 | 157.9 | 42.1 | 12.31 | 28.07 | | High | High | High | High | High | High | High | High | High | High | High | 0 | 0.000145 | 4.3 |
| TRUE | High | DSISPTTVT | [R].DSISPT | 2xPhospho | D0VLQ5 2x | 0.075304 | 0.005727 | 1 | 3 | 1 | D0VLQ5 | Tet oncoge | D0VLQ5 [3 | 0 | 2356.14 | 0 | -2.75 | 0.005289 | 0.82 | 165.9 | 34.1 | 19.86 | 53.47 | | High | High | High | High | High | High | High | High | High | High | High | 0.002951 | 0.03083 | 2.33 |
| TRUE | High | DSISPTTVT | [K].DSISPT | 1xPhospho | D0VLQ5 1x | 2.23E-05 | 0 | 1 | 3 | 5 | D0VLQ5 | Tet oncoge | D0VLQ5 [3 | 0 | 2276.174 | 0 | -2.25 | 0.00022 | 0.41 | 166.3 | 33.7 | 7.8 | 19.08 | | High | High | High | High | High | High | High | High | High | High | High | 0 | 4.82E-05 | 3.69 |
| TRUE | High | NSSAITSPK | [K].NSSAIT | 1xPhospho | Q5SVI1 1xl | 0.002553 | 0.000109 | 1 | 9 | 2 | Q5SVI1 | Calcium/ca | Q5SVI1 [3 | 0 | 1442.766 | 0 | -2.72 | 0.000824 | 0.23 | 173 | 27 | 12.21 | 27.45 | | High | High | High | High | High | High | High | High | High | High | High | 0.000275 | 0.001675 | 1.13 |
| TRUE | High | GQTYATEV | [R].GQTYA | 1xPhospho | Q3TYF2 1x | 1.5E-05 | 0 | 1 | 2 | 4 | Q3TYF2 | Putative ur | Q3TYF2 [3 | 0 | 4085.735 | 0 | -3.29 | 0.000207 | 0.39 | 182.9 | 17.1 | 7.24 | 61.81 | | High | High | High | High | High | High | High | High | High | High | High | 0 | 2.14E-06 | 4.14 |
| TRUE | High | AELEEMEE | [K].AELEEN | 1xPhospho | A2ARP8 1x | 6.89E-05 | 0 | 1 | 3 | 2 | A2ARP8 | Microtubu | A2ARP8 [8 | 0 | 2943.274 | 0 | -2.15 | 0.002067 | 1.67 | 161.4 | 38.6 | 16.08 | 26.57 | | High | High | High | High | High | High | High | High | High | High | High | 0 | 1.18E-05 | 4.09 |
| TRUE | High | NSRPSSPVI | [K].NSRPSS | 1xPhospho | E9QLZ9 1xl | 4.8E-08 | 0 | 1 | 7 | 7 | E9QLZ9 | Protein ena | E9QLZ9 [3 | 0 | 2460.265 | 0 | -3.1 | 0.000771 | 0.54 | 179.1 | 20.9 | 9.17 | 27.96 | | High | High | High | High | High | High | High | High | High | High | High | 0 | 3.35E-09 | 3.65 |
| TRUE | High | NSRPSSPVI | [K].NSRPSS | 2xPhospho | E9QLZ9 2xl | 1.79E-06 | 0 | 1 | 7 | 1 | E9QLZ9 | Protein ena | E9QLZ9 [3 | 0 | 2311.068 | 0 | -2.54 | 0.007037 | 0.61 | 167.8 | 32.2 | 26.84 | 18.47 | | High | High | High | High | High | High | High | High | High | High | High | 0 | 1.95E-07 | 3.7 |

| | | | | | | | | | | | | | | | | | | | | | | | | | | | | | | | | | | | | | | |
|---|---|---|---|---|---|---|---|---|---|---|---|---|---|---|---|---|---|---|---|---|---|---|---|---|---|---|---|---|---|---|---|---|---|---|---|---|---|---|
| TRUE | High | APSTSTPEF | [K].APSTST | 1xPhosphc | E9QLZ9 1xl | 0.019618 | 0.001438 | 1 | 12 | 5 | E9QLZ9 | Protein ena | E9QLZ9 [7: | 0 | 1452.693 | 0 | -2.11 | 0.000468 | 0.68 | 162.3 | 37.7 | 1.91 | 28.24 | | High | High | High | High | High | High | High | High | High | High | High | 0.002675 | 0.02853 | 2.33 |
| TRUE | High | EDVTPPLSF | [K].EDVTPF | 2xPhosphc | Q3UHU5 2 | 0.000219 | 0 | 1 | 5 | 4 | Q3UHU5 | Microtubu | Q3UHU5 [: | 0 | 2043.969 | 0 | -2.01 | 0.000181 | 1.5 | 160.1 | 39.9 | 1.72 | 16.29 | | High | High | High | High | High | High | High | High | High | High | High | 0.000327 | 0.002964 | 2.6 |
| TRUE | High | VQGSPLK | [K].VQGSP | 1xPhosphc | B2RRE0 1x | 0.013545 | 0.000976 | 1 | 3 | 3 | B2RRE0 | A kinase (P | B2RRE0 [4! | 0 | 1266.722 | 0 | -2.88 | 0.000217 | 0.05 | 175.1 | 24.9 | 6.76 | 25.97 | | High | High | High | High | High | High | High | High | High | High | High | 0.000505 | 0.004404 | 1.93 |
| TRUE | High | ETPAPPPST | [K].ETPAPF | 2xPhosphc | Q6P9J5 2xl | 1.43E-06 | 0 | 1 | 1 | 2 | Q6P9J5 | KN motif a | Q6P9J5 [63 | 0 | 3330.672 | 0 | -2.74 | 0.000167 | 0.31 | 174 | 26 | 2.92 | 23.11 | | High | High | High | High | High | High | High | High | High | High | High | 0 | 1.51E-07 | 5.16 |
| TRUE | High | LETTAPALS | [R].LETTAP | 1xPhosphc | Q91XC0 1x | 0.000365 | 0 | 1 | 1 | 3 | Q91XC0 | LIM domai | Q91XC0 [1 | 0 | 1464.766 | 0 | -2.23 | 0.000362 | 1.27 | 166.2 | 33.8 | 4.76 | 19.78 | | High | High | High | High | High | High | High | High | High | High | High | 0 | 0.000134 | 2.26 |
| TRUE | High | NSPQSSPTS | [R].NSPQSS | 1xPhosphc | Q920Q8 1x | 0.000868 | 0 | 1 | 4 | 5 | Q920Q8 | Influenza v | Q920Q8 [3 | 0 | 1768.888 | 0 | -2.04 | 0.000348 | 0.92 | 160.7 | 39.3 | 6.75 | 18.46 | | High | High | High | High | High | High | High | High | High | High | High | 0.000204 | 0.001369 | 2.37 |
| TRUE | High | NSPQSSPTS | [R].NSPQSS | 1xPhosphc | Q920Q8 1x | 0.006895 | 0.00062 | 1 | 4 | 3 | Q920Q8 | Influenza v | Q920Q8 [3 | 0 | 1539.725 | 0 | -2.1 | 0.018632 | 3.13 | 162.3 | 37.7 | 29.81 | 32.41 | | High | High | High | High | High | High | High | High | High | High | High | 0.000585 | 0.005323 | 2.36 |
| TRUE | High | DSEGPFPAI | [K].DSEGPI | 1xPhosphc | A2TJV2 1xl | 2.51E-05 | 0 | 1 | 1 | 2 | A2TJV2 | Paralemmi | A2TJV2 [45 | 0 | 2273.063 | 0 | -2.21 | 0.002271 | 1.89 | 162.5 | 37.5 | 12.68 | 25.24 | | High | High | High | High | High | High | High | High | High | High | High | 0 | 3.78E-06 | 4.02 |
| TRUE | High | GESESEAK | [K].GESESE | 1xPhosphc | A0A1D9BZ | 0.000651 | 0 | 1 | 1 | 1 | A0A1D9BZ | GCNA OS= | A0A1D9BZ | 0 | 1374.655 | 0 | -2.93 | 0.000334 | 0.07 | 177.2 | 22.8 | 10.92 | 21.99 | | High | High | High | High | High | High | High | High | High | High | High | 0 | 0.000148 | 1.76 |
| TRUE | High | WLEETDSS | [K].WLEETI | 1xPhosphc | P21952 1xl | 0.000155 | 0 | 1 | 1 | 1 | P21952 | POU doma | P21952 [31 | 0 | 2404.132 | 0 | -2.67 | 0.000544 | 0.61 | 172.9 | 27.1 | 6.98 | 26.26 | | High | High | High | High | High | High | High | High | High | High | High | 0 | 2.93E-05 | 2.61 |
| TRUE | High | LYSILQGDS | [K].LYSILQ | 1xPhosphc | Q6NV83 1x | 0.000815 | 0 | 1 | 3 | 1 | Q6NV83 | U2 snRNP- | Q6NV83 [4 | 0 | 1859.992 | 0 | -2.19 | 0.000268 | 0.52 | 163.4 | 36.6 | 8.05 | 10.83 | | High | High | High | High | High | High | High | High | High | High | High | 0 | 0.000189 | 3.15 |
| TRUE | High | LRDSQGQF | [K].LRDSQ | 1xPhosphc | Q6J1H4 1xl | 0.000102 | 0 | 1 | 2 | 2 | Q6J1H4 | Undifferen | Q6J1H4 [1: | 1 | 2224.071 | 0 | -2.6 | 0.00024 | 0.81 | 170.3 | 29.7 | 5.34 | 19.32 | | High | High | High | High | High | High | High | High | High | High | High | 0.000449 | 0.003894 | 3.35 |
| TRUE | High | HSSSDEGS | [R].HSSSDE | 1xPhosphc | Q3U832 1x | 0.000102 | 0 | 1 | 3 | 2 | Q3U832 | Putative ur | Q3U832 [5 | 0 | 1528.704 | 0 | -2.13 | 0.000284 | 3.38 | 165 | 35 | 9.14 | 45.28 | | High | High | High | High | High | High | High | High | High | High | High | 0 | 1.82E-05 | 2.16 |
| TRUE | High | QLTESSEEA | [K].QLTESS | 1xPhosphc | B1AU75 1x | 0.000454 | 0 | 1 | 2 | 5 | B1AU75 | Nuclear au | B1AU75 [2 | 0 | 1659.824 | 0 | -2.51 | 0.000181 | 0.41 | 170.9 | 29.1 | 6.34 | 16.77 | | High | High | High | High | High | High | High | High | High | High | High | 0 | 9.82E-05 | 2.68 |
| TRUE | High | ETSPTRGEI | [R].ETSPTR | 1xPhosphc | A2ARP8 1x | 0.001049 | 0 | 1 | 3 | 3 | A2ARP8 | Microtubu | A2ARP8 [1: | 1 | 2279.147 | 0 | -2.19 | 0.002585 | 1.81 | 160.9 | 39.1 | 15.99 | 23.87 | | High | High | High | High | High | High | High | High | High | High | High | 0 | 0.000252 | 3.35 |
| TRUE | High | APVSPDSA | [K].APVSPI | 1xPhosphc | Q6J1H4 1x | 0.005978 | 0.00062 | 1 | 2 | 6 | Q6J1H4 | Undifferen | Q6J1H4 [5: | 0 | 1336.646 | 0 | -2.84 | 0.001013 | 0.2 | 174 | 26 | 14.15 | 20.82 | | High | High | High | High | High | High | High | High | High | High | High | 0.000275 | 0.00176 | 2.1 |
| TRUE | High | RSVSNSTQ | [K].RSVSNS | 1xPhosphc | E9PXD2 1x | 0.031713 | 0.002011 | 1 | 7 | 5 | E9PXD2 | Rho GTPas | E9PXD2 [7 | 1 | 3338.555 | 0 | -2.15 | 0.003581 | 3.12 | 170.2 | 29.8 | 18.09 | 46.24 | | High | High | High | High | High | High | High | High | High | High | High | 0.001308 | 0.01391 | 3.77 |
| TRUE | High | ETVESPTFF | [K].ETVESP | 1xPhosphc | B2RWW9 1 | 0.012731 | 0.000889 | 1 | 2 | 3 | B2RWW9 | Absent in r | B2RWW9 [ | 0 | 2148.139 | 0 | -2 | 0.003229 | 0.7 | 159.4 | 40.6 | 16.89 | 19.28 | | High | High | High | High | High | High | High | High | High | High | High | 0.000449 | 0.004117 | 3.42 |
| TRUE | High | GEGESPPV | [K].GEGESI | 1xPhosphc | P28667 1xl | 2.12E-06 | 0 | 1 | 1 | 1 | P28667 | MARCKS-re | P28667 [44 | 0 | 3787.759 | 0 | -2.28 | 0.001008 | 1.24 | 166 | 34 | 8.37 | 37.46 | | High | High | High | High | High | High | High | High | High | High | High | 0 | 2.36E-07 | 3.38 |
| TRUE | High | RLSSGPPA | [K].RLSSGP | 1xPhosphc | A6H5X4 1x | 0.000852 | 0 | 2 | 6 | 1 | A6H5X4; B | PHD finger | A6H5X4 [1 | 1 | 1675.93 | 0 | -2.16 | 0.000222 | 0.18 | 162.1 | 37.9 | 7.97 | 12.13 | Shared | High | High | High | High | High | High | High | High | High | High | High | 0 | 0.0002 | 1.4 |
| TRUE | High | LQSTEFSPS | [R].LQSTEF | 1xDeamida | Q9WV69 1 | 0.000137 | 0 | 1 | 3 | 5 | Q9WV69 | Dematin O | Q9WV69 [ | 0 | 2830.273 | 0 | -2.39 | 0.000696 | 0.55 | 166.1 | 33.9 | 8.9 | 19.92 | | High | High | High | High | High | High | High | High | High | High | High | 0 | 2.56E-05 | 3.61 |
| TRUE | High | LQSTEFSPS | [R].LQSTEF | 1xPhosphc | Q9WV69 1 | 7.55E-09 | 0 | 1 | 3 | 2 | Q9WV69 | Dematin O | Q9WV69 [ | 0 | 2829.289 | 0 | -2.93 | 0.000306 | 0.93 | 176.8 | 23.2 | 8.83 | 43.72 | | High | High | High | High | High | High | High | High | High | High | High | 0 | 4.22E-10 | 4.49 |
| TRUE | High | HSVTGYGD | [R].HSVTG | 1xCarbami | Q91XC0 1x | 0.00967 | 0.000736 | 1 | 1 | 2 | Q91XC0 | LIM domai | Q91XC0 [2 | 0 | 1730.752 | 0 | -2.74 | 0.002354 | 0.89 | 173.5 | 26.5 | 20.52 | 53.94 | | High | High | High | High | High | High | High | High | High | High | High | 0.000638 | 0.00563 | 2.17 |
| TRUE | High | NSPPGNSP | [R].NSPPGI | 1xPhosphc | Q7TN75-2 | 0.030367 | 0.001857 | 1 | 1 | 1 | Q7TN75-2 | Isoform 2 c | Q7TN75-2 | 0 | 1359.651 | 0 | -2.53 | 0.000417 | 0.34 | 173.5 | 26.5 | 9.96 | 14.99 | | High | High | High | High | High | High | High | High | High | High | High | 0.001011 | 0.01089 | 2.17 |
| TRUE | High | TCSPFAEEF | [R].TCSPFA | 1xCarbami | Q5FWH3 1 | 1.32E-05 | 0 | 1 | 1 | 1 | Q5FWH3 | Grainyhea | Q5FWH3 [ | 0 | 2277.091 | 0 | -2.02 | 0.000376 | 1.15 | 161.2 | 38.8 | 6.36 | 15.83 | | High | High | High | High | High | High | High | High | High | High | High | 0 | 1.85E-06 | 2.78 |
| TRUE | High | NTTVCEISS | [K].NTTVCI | 1xCarbami | A0A1D9BZ | 0.000524 | 0 | 1 | 1 | 2 | A0A1D9BZ | GCNA OS= | A0A1D9BZ | 0 | 2935.167 | 0 | -2.05 | 0.038087 | 0.48 | 161.7 | 38.3 | 42.28 | 33.63 | | High | High | High | High | High | High | High | High | High | High | High | 0.000117 | 0.001059 | 3.62 |
| TRUE | High | NTTVCEISS | [K].NTTVCI | 1xCarbami | A0A1D9BZ | 7.18E-07 | 0 | 1 | 1 | 5 | A0A1D9BZ | GCNA OS= | A0A1D9BZ | 0 | 3004.397 | 0 | -3.59 | 0.000343 | 0.01 | 184.7 | 15.3 | 12 | 35.4 | | High | High | High | High | High | High | High | High | High | High | High | 4.94E-05 | 0.000749 | 2.84 |
| TRUE | High | SLFWSGEE | [K].SLFWS | 1xPhosphc | Q5U4C1 1x | 0.001222 | 0 | 1 | 2 | 1 | Q5U4C1 | G-protein c | Q5U4C1 [3 | 0 | 1691.845 | 0 | -2.38 | 0.000266 | 0.78 | 167.7 | 32.3 | 5.25 | 14.24 | | High | High | High | High | High | High | High | High | High | High | High | 0 | 0.000298 | 3.23 |
| TRUE | High | QDPIPGSPI | [K].QDPIPC | 1xPhosphc | E9PXD2 1x | 0.000231 | 0 | 1 | 7 | 2 | E9PXD2 | Rho GTPas | E9PXD2 [5 | 0 | 1591.731 | 0 | -3.28 | 0.000237 | 0.28 | 181.9 | 18.1 | 4.8 | 27.06 | | High | High | High | High | High | High | High | High | High | High | High | 0 | 4.58E-05 | 2.4 |
| TRUE | High | ENNDRLST | [R].ENNDR | 1xPhosphc | H3BKF8 1x | 0.08727 | 0.006537 | 1 | 11 | 1 | H3BKF8 | Homeobox | H3BKF8 [1: | 1 | 1872.922 | 0 | -2.07 | 0.000162 | 0.44 | 161.6 | 38.4 | 2.14 | 11.09 | | High | High | High | High | High | High | High | High | High | High | High | 0.003375 | 0.03684 | 1.26 |
| TRUE | High | SATDGNTS | [R].SATDGI | 1xPhosphc | A2A432 1x | 0.026421 | 0.001676 | 1 | 4 | 2 | A2A432 | Cullin-4B C | A2A432 [9 | 1 | 2431.268 | 0 | -2.44 | 0.000932 | 0.2 | 168.6 | 31.4 | 15.14 | 10.61 | | High | High | High | High | High | High | High | High | High | High | High | 0.000903 | 0.009322 | 3.51 |
| TRUE | High | ENLSPTTIE | [R].ENLSPT | 1xPhosphc | P42228 1xl | 0.000993 | 0 | 1 | 3 | 2 | P42228 | Signal tran: | P42228 [73 | 0 | 2364.052 | 0 | -2.32 | 0.002834 | 2.36 | 166.6 | 33.4 | 12.97 | 39.25 | | High | High | High | High | High | High | High | High | High | High | High | 0 | 0.000238 | 2.61 |
| TRUE | High | EPSPHPHV | [R].EPSPHF | 1xPhosphc | B2RWW9 1 | 0.096181 | 0.007535 | 1 | 2 | 1 | B2RWW9 | Absent in r | B2RWW9 [ | 0 | 1879.901 | 0 | -2.05 | 0.002488 | 1 | 156.8 | 43.2 | 17.62 | 13.47 | | High | High | High | High | High | High | High | High | High | High | High | 0.003809 | 0.04103 | 2.51 |
| TRUE | High | ANAAFEPD | [R].ANAAFI | 1xPhosphc | Q3U5C8 1x | 0.006363 | 0.00062 | 1 | 3 | 4 | Q3U5C8 | Rho guanir | Q3U5C8 [4 | 0 | 4600.326 | 0 | -2.61 | 0.000806 | 0.74 | 171.9 | 28.1 | 14.39 | 38.55 | | High | High | High | High | High | High | High | High | High | High | High | 0.000275 | 0.001959 | 3.64 |
| TRUE | High | GLPPSVER | [R].GLPPSV | 1xPhosphc | Q9DAE2 1x | 0.027358 | 0.001676 | 1 | 1 | 4 | Q9DAE2 | Putative ur | Q9DAE2 [3 | 0 | 1163.602 | 0 | -3.52 | 0.000188 | 0.15 | 183.8 | 16.2 | 4.24 | 24.35 | | High | High | High | High | High | High | High | High | High | High | High | 0.004126 | 0.04422 | 1.12 |
| TRUE | High | TSISQGPST | [R].TSISQG | 1xPhosphc | A0A1L1SQ | 0.000529 | 0 | 1 | 3 | 1 | A0A1L1SQI | Dedicator c | A0A1L1SQI | 0 | 1484.731 | 0 | -2.08 | 0.000549 | 1.2 | 162.4 | 37.6 | 9.08 | 19.8 | | High | High | High | High | High | High | High | High | High | High | High | 0 | 0.000117 | 1.97 |
| TRUE | High | ANAAFEPD | [R].ANAAFI | 1xDeamida | Q3U5C8 1x | 0.045122 | 0.003373 | 1 | 3 | 4 | Q3U5C8 | Rho guanir | Q3U5C8 [4 | 0 | 4601.31 | 0 | -3.03 | 0.000191 | 0.37 | 178.1 | 21.9 | 3.25 | 32.95 | | High | High | High | High | High | High | High | High | High | High | High | 0.001727 | 0.01716 | 4.22 |
| TRUE | High | LHTGDK | [KR].LHTGI | 1xPhosphc | Q80TS5 1x | 0.071662 | 0.005278 | 2 | 10 | 2 | Q80TS5; Q | Zinc finger | Q80TS5 [1! | 0 | 1208.644 | 1 | -2.13 | 0.003357 | 2.21 | 162.8 | 37.2 | 16.2 | 27.99 | Shared | High | High | High | High | High | High | High | High | High | High | High | 0.002717 | 0.02922 | 1.7 |
| TRUE | High | TSLPDAPV | [R].TSLPDA | 1xPhosphc | A2TJV2 1xl | 1.49E-05 | 0 | 1 | 1 | 2 | A2TJV2 | Paralemmi | A2TJV2 [15 | 0 | 2249.036 | 0 | -2.16 | 0.00049 | 0.9 | 161.9 | 38.1 | 7.68 | 11.91 | | High | High | High | High | High | High | High | High | High | High | High | 0 | 2.11E-06 | 3.89 |
| TRUE | High | EPTKPLESP | [K].EPTKPL | 1xPhosphc | B2RRE0 1x | 7.18E-07 | 0 | 1 | 3 | 3 | B2RRE0 | A kinase (P | B2RRE0 [2 | 0 | 3130.616 | 0 | -2.3 | 0.007843 | 1.25 | 170.3 | 29.7 | 33.51 | 30.33 | | High | High | High | High | High | High | High | High | High | High | High | 0 | 3.1E-07 | 4.53 |
| TRUE | High | GLNTSHESI | [R].GLNTSI | 2xPhosphc | Q3U5C8 2x | 5.94E-07 | 0 | 1 | 1 | 1 | Q3U5C8 | Rho guanir | Q3U5C8 [2 | 0 | 2928.25 | 0 | -2.65 | 0.000154 | 0.62 | 173.6 | 26.4 | 6.21 | 22.81 | | High | High | High | High | High | High | High | High | High | High | High | 0 | 5.69E-08 | 3.92 |
| TRUE | High | HLLTHSSEF | [K].HLLTHS | 1xPhosphc | Q3URR7 1x | 0.016021 | 0.001124 | 1 | 7 | 1 | Q3URR7 | Zinc finger | Q3URR7 [4 | 0 | 1703.847 | 0 | -2.32 | 0.000956 | 0.48 | 166.2 | 33.8 | 11.56 | 7.87 | | High | High | High | High | High | High | High | High | High | High | High | 0.000585 | 0.00533 | 2.04 |
| TRUE | High | ALGSPTLK | [R].ALGSPT | 1xPhosphc | Q8C739 1x | 0.006536 | 0.00062 | 1 | 1 | 1 | Q8C739 | Protein FAI | Q8C739 [9 | 0 | 1324.764 | 0 | -2.08 | 0.000361 | 1.46 | 162.4 | 37.6 | 5.26 | 17.57 | | High | High | High | High | High | High | High | High | High | High | High | 0.000275 | 0.001959 | 1.83 |
| TRUE | High | QFSFGGDL | [R].QFSFG | 1xPhosphc | A0A0R4J0C | 0.001386 | 5.66E-05 | 1 | 4 | 1 | A0A0R4J0C | Disks large | A0A0R4J0C | 0 | 1849.934 | 0 | -2.84 | 0.000669 | 1.58 | 175 | 25 | 4.82 | 57.47 | | High | High | High | High | High | High | High | High | High | High | High | 2.58E-05 | 0.000345 | 2.39 |
| TRUE | High | RDSGVGAS | [R].RDSGV | 1xPhosphc | E9PXD2 1x | 0.110202 | 0.008961 | 1 | 7 | 1 | E9PXD2 | Rho GTPas | E9PXD2 [9 | 1 | 1427.721 | 0 | -2.46 | 0.000209 | 0.25 | 168.9 | 31.1 | 4.05 | 13.24 | | High | High | High | High | High | High | High | High | High | High | High | 0.00449 | 0.04859 | 2.43 |
| TRUE | High | DYNDLFDG | [K].DYNDLI | 1xDeamida | E9QPB3 1x | 0.000558 | 0 | 1 | 3 | 4 | E9QPB3 | WD repeat | E9QPB3 [3 | 0 | 3807.684 | 0 | -2.58 | 0.000842 | 0.25 | 173.5 | 26.5 | 12.2 | 36.14 | | High | High | High | High | High | High | High | High | High | High | High | 0 | 0.000171 | 3.78 |
| TRUE | High | SSATVGPK | [K].SSATV | 1xPhosphc | P27661 1xl | 0.013545 | 0.000976 | 1 | 1 | 3 | P27661 | Histone H2 | P27661 [12 | 0 | 1284.696 | 0 | -2.04 | 0.00036 | 0.46 | 159.1 | 40.9 | 9.03 | 11.36 | | High | High | High | High | High | High | High | High | High | High | High | 0.000505 | 0.004397 | 1.36 |
| TRUE | High | DYNDLFDG | [K].DYNDLI | 1xPhosphc | E9QPB3 1x | 0.023584 | 0.001578 | 1 | 3 | 3 | E9QPB3 | WD repeat | E9QPB3 [3 | 0 | 3806.7 | 0 | -2.83 | 0.000191 | 0.56 | 173.3 | 26.7 | 6.31 | 39.48 | | High | High | High | High | High | High | High | High | High | High | High | 0.001076 | 0.0114 | 3.66 |
| TRUE | High | VSPAHSPPI | [R].VSPAHS | 2xPhosphc | Q80TC1 2x | 0.003927 | 0.000311 | 1 | 14 | 1 | Q80TC1 | MKIAA154 | Q80TC1 [3 | 0 | 2065.976 | 0 | -2.79 | 0.00029 | 0.61 | 173.7 | 26.3 | 5.25 | 28.33 | | High | High | High | High | High | High | High | High | High | High | High | 0.00014 | 0.001103 | 2.26 |
| TRUE | High | ALGLEESPE | [R].ALGLEE | 1xPhosphc | A2ARP8 1x | 3.47E-05 | 0 | 1 | 3 | 2 | A2ARP8 | Microtubu | A2ARP8 [1 | 0 | 1925.951 | 0 | -2.88 | 0.000313 | 0.55 | 175.3 | 24.7 | 8.48 | 25.95 | | High | High | High | High | High | High | High | High | High | High | High | 0 | 5.47E-06 | 1.97 |
| TRUE | High | SHSLEAPGI | [K].SHSLEA | 1xPhosphc | Q6PFD5 1x | 0.065405 | 0.004734 | 1 | 2 | 1 | Q6PFD5 | Disks large | Q6PFD5 [1 | 0 | 1463.766 | 0 | -2.05 | 0.010715 | 2.85 | 161 | 39 | 19.85 | 34.67 | | High | High | High | High | High | High | High | High | High | High | High | 0.002446 | 0.02627 | 1.72 |
| TRUE | High | GPGGSPSG | [R].GPGGS | 1xPhosphc | Q91VC7 1x | 0.001881 | 5.66E-05 | 1 | 2 | 2 | Q91VC7 | Protein ph | Q91VC7 [2 | 0 | 1522.803 | 0 | -2.3 | 0.000181 | 0.89 | 167 | 33 | 9.6 | 29.59 | | High | High | High | High | High | High | High | High | High | High | High | 0.001308 | 0.01369 | 2 |
| TRUE | High | GLLPGTAP | [K].GLLPGT | 1xPhosphc | Q9JJA7 1xF | 1.69E-08 | 0 | 1 | 1 | 1 | Q9JJA7 | Cyclin-L2 C | Q9JJA7 [31 | 0 | 2462.31 | 0 | -2.91 | 0.000246 | 0.69 | 177.2 | 22.8 | 4.54 | 31.15 | | High | High | High | High | High | High | High | High | High | High | High | 0 | 1.03E-09 | 3.59 |
| TRUE | High | SSILDLSSG | [K].SSILDLS | 1xCarbami | Q7TN89 1x | 2.45E-06 | 0 | 1 | 1 | 3 | Q7TN89 | GTPase ER | Q7TN89 [7 | 0 | 1802.856 | 0 | -2.18 | 0.001414 | 1.69 | 164.6 | 35.4 | 13.11 | 14.74 | | High | High | High | High | High | High | High | High | High | High | High | 0 | 2.77E-07 | 3.06 |
| TRUE | High | ALETMAEC | [K].ALETM | 1xPhosphc | P27546 1xl | 0.028082 | 0.001791 | 1 | 3 | 3 | P27546 | Microtubu | P27546 [24 | 0 | 3838.835 | 0 | -2.11 | 0.003461 | 0.89 | 160 | 40 | 21.52 | 33.17 | | High | High | High | High | High | High | High | High | High | High | High | 0.000968 | 0.01002 | 2.52 |
| TRUE | High | ALEQSHSLI | [R].ALEQSI | 2xPhosphc | E9PXE2 2xl | 1.47E-05 | 0 | 1 | 19 | 1 | E9PXE2 | Guanine nu | E9PXE2 [9 | 0 | 2696.335 | 0 | -2.77 | 0.000729 | 0.28 | 173.7 | 26.3 | 7.17 | 27.36 | | High | High | High | High | High | High | High | High | High | High | High | 0 | 2.09E-06 | 3.15 |
| TRUE | High | EQESEVEK | [K].EQESEV | 1xPhosphc | B1AU75 1x | 0.002376 | 0.000109 | 1 | 2 | 1 | B1AU75 | Nuclear au | B1AU75 [1 | 0 | 1515.734 | 1 | -2.1 | 0.001336 | 1.08 | 163.9 | 36.1 | 12.61 | 12.47 | | High | High | High | High | High | High | High | High | High | High | High | 4.94E-05 | 0.000633 | 2.37 |
| TRUE | High | GLNTSHESI | [R].GLNTSI | 1xPhosphc | Q3U5C8 1x | 2.06E-08 | 0 | 1 | 1 | 5 | Q3U5C8 | Rho guanir | Q3U5C8 [2 | 0 | 2848.284 | 0 | -2.14 | 0.000434 | 0.62 | 163.2 | 36.8 | 9.02 | 18 | | High | High | High | High | High | High | High | High | High | High | High | 0 | 1.3E-09 | 5.43 |
| TRUE | High | QASPGSVS | [R].QASPG | 1xPhosphc | Q9CPN8 1x | 0.002355 | 0.000109 | 1 | 1 | 3 | Q9CPN8 | Insulin-like | Q9CPN8 [1 | 0 | 1398.739 | 0 | -2.33 | 0.000181 | 0.89 | 168.2 | 31.8 | 4.31 | 18.43 | | High | High | High | High | High | High | High | High | High | High | High | 0.000183 | 0.001283 | 1.79 |
| TRUE | High | LLGSGSPSS | [R].LLGSGS | 1xPhosphc | G3X981 1x | 0.005516 | 0.00062 | 1 | 5 | 2 | G3X981 | Peripherin | G3X981 [5 | 0 | 1427.709 | 0 | -2.16 | 0.000252 | 1.07 | 166.8 | 33.2 | 5.76 | 17.41 | | High | High | High | High | High | High | High | High | High | High | High | 0.000275 | 0.001616 | 2.22 |
| TRUE | High | GPAGVELN | [K].GPAGV | 1xPhosphc | P32037 1xl | 0.001398 | 5.66E-05 | 1 | 3 | 1 | P32037 | Solute carr | P32037 [47 | 0 | 1981.023 | 0 | -2.23 | 0.004585 | 1.35 | 166.9 | 33.1 | 23.92 | 27.81 | | High | High | High | High | High | High | High | High | High | High | High | 2.58E-05 | 0.000347 | 2.56 |
| TRUE | High | REETGIPVS | [R].REETGI | 1xPhosphc | Q8BRB7 1x | 0.102567 | 0.008249 | 1 | 3 | 1 | Q8BRB7 | Histone acc | Q8BRB7 [1 | 1 | 1888.009 | 0 | -2.72 | 0.000229 | 0.7 | 171.8 | 28.2 | 7.68 | 29.87 | | High | High | High | High | High | High | High | High | High | High | High | 0.004138 | 0.04428 | 2.01 |
| TRUE | High | EAFLVLSDE | [K].EAFLVL | 1xCarbami | Q7TT18 1x | 5.79E-07 | 0 | 1 | 4 | 5 | Q7TT18 | Activating | Q7TT18 [5 | 0 | 2350.129 | 0 | -2.93 | 0.000319 | 0.22 | 176.5 | 23.5 | 4.48 | 27.92 | | High | High | High | High | High | High | High | High | High | High | High | 0 | 9.66E-07 | 3.8 |
| TRUE | High | GNGLASPG | [R].GNGLA | 1xPhosphc | Q3UHU5 1 | 0.000165 | 0 | 1 | 5 | 1 | Q3UHU5 | Microtubu | Q3UHU5 [8 | 0 | 3309.606 | 0 | -2.15 | 0.000668 | 1.85 | 164.3 | 35.7 | 6.13 | 28.49 | | High | High | High | High | High | High | High | High | High | High | High | 0 | 3.15E-05 | 3.54 |
| TRUE | High | REGITPWA | [K].REGITP | 1xPhosphc | B2RRE0 1x | 0.004372 | 0.000409 | 1 | 3 | 1 | B2RRE0 | A kinase (P | B2RRE0 [5 | 1 | 1829.972 | 0 | -2.11 | 0.006858 | 1.68 | 162.4 | 37.6 | 20.72 | 18.51 | | High | High | High | High | High | High | High | High | High | High | High | 0.000183 | 0.001246 | 1.56 |
| TRUE | High | LSPSPLPAA | [R].LSPSPLI | 1xPhosphc | Q62066 1x | 0.001039 | 0 | 1 | 1 | 2 | Q62066 | Paired mes | Q62066 [2 | 0 | 2422.351 | 0 | -2.97 | 0.000324 | 0.28 | 177.1 | 22.9 | 6 | 38.31 | | High | High | High | High | High | High | High | High | High | High | High | 2.58E-05 | 0.000371 | 3.71 |
| TRUE | High | ISQDQSPTI | [K].ISQDQS | 1xPhosphc | I7HPW4 1x | 0.001233 | 0 | 1 | 1 | 1 | I7HPW4 | G protein-c | I7HPW4 [4 | 0 | 1655.841 | 0 | -2.23 | 0.000322 | 0.51 | 167.6 | 32.4 | 10.53 | 20.03 | | High | High | High | High | High | High | High | High | High | High | High | 0 | 0.000303 | 2.61 |
| TRUE | High | QEQAAVEI | [K].QEQAA | 1xPhosphc | B2RRE0 1x | 2.49E-05 | 0 | 1 | 2 | 2 | B2RRE0 | A kinase (P | B2RRE0 [8 | 0 | 4048.839 | 0 | -2.22 | 0.018897 | 3.8 | 162.5 | 37.5 | 13.49 | 62.57 | | High | High | High | High | High | High | High | High | High | High | High | 0 | 3.75E-06 | 4.41 |
| TRUE | High | QELGPGQE | [R].QELGP | 1xPhosphc | Q3UHU5 1 | 0.000136 | 0 | 1 | 5 | 4 | Q3UHU5 | Microtubu | Q3UHU5 [: | 0 | 1755.811 | 0 | -2.23 | 0.000384 | 0.32 | 165.5 | 34.5 | 8.42 | 12.63 | | High | High | High | High | High | High | High | High | High | High | High | 0 | 5.5E-05 | 2.95 |
| TRUE | High | EPDQSLLSF | [R].EPDQSI | 1xPhosphc | Q80VR2 1x | 2.23E-05 | 0 | 1 | 1 | 2 | Q80VR2 | RNA polym | Q80VR2 [1 | 0 | 1937.999 | 0 | -2.73 | 9.77E-05 | 0.22 | 173.9 | 26.1 | 4.07 | 16.32 | | High | High | High | High | High | High | High | High | High | High | High | 0 | 3.35E-06 | 3.2 |
| TRUE | High | EPELSHSGS | [R].EPELSH | 1xPhosphc | Q9R069 1x | 0.001916 | 5.66E-05 | 1 | 1 | 1 | Q9R069 | Basal cell a | Q9R069 [5 | 0 | 3056.409 | 0 | -2.2 | 0.002388 | 1.67 | 165.7 | 34.3 | 13.49 | 37.95 | | High | High | High | High | High | High | High | High | High | High | High | 2.58E-05 | 0.000495 | 2.1 |
| TRUE | High | SSFASSSAS | [R].SSFASS | 1xPhosphc | Q91XC0 1x | 6.02E-05 | 0 | 1 | 1 | 2 | Q91XC0 | LIM domai | Q91XC0 [1 | 0 | 2294.107 | 0 | -2.85 | 0.000483 | 0.25 | 176.2 | 23.8 | 10.52 | 27.95 | | High | High | High | High | High | High | High | High | High | High | High | 0 | 1.02E-05 | 1.27 |
| TRUE | High | GNSLPCVLI | [R].GNSLP | 1xCarbami | Q9WV69 1 | 0.00172 | 5.66E-05 | 1 | 3 | 2 | Q9WV69 | Dematin O | Q9WV69 [ | 0 | 1782.923 | 0 | -3.88 | 0.000154 | 0.15 | 187.8 | 12.2 | 5.24 | 35.4 | | High | High | High | High | High | High | High | High | High | High | High | 2.58E-05 | 0.000437 | 2.74 |
| TRUE | High | HFPGEDGD | [K].HFPGEI | 1xPhosphc | E9Q2Z1 1x | 2.98E-05 | 0 | 1 | 3 | 3 | E9Q2Z1 | CECR2, hist | E9Q2Z1 [5 | 0 | 2460.116 | 0 | -2.63 | 0.000669 | 0.13 | 169.8 | 30.2 | 9.05 | 17.67 | | High | High | High | High | High | High | High | High | High | High | High | 0 | 4.6E-06 | 3.11 |
| TRUE | High | RESASAWN | [R].RESASA | 1xPhosphc | Q3UYG1 1 | 0.005 | 0.000504 | 1 | 1 | 1 | Q3UYG1 | Coiled-coil | Q3UYG1 [1 | 1 | 1586.809 | 0 | -2.21 | 0.000313 | 0.69 | 162.5 | 37.5 | 8.05 | 15.67 | | High | High | High | High | High | High | High | High | High | High | High | 0.000225 | 0.00144 | 2.42 |
| TRUE | High | AVQEVHPS | [R].AVQEV | 1xPhosphc | O35738 1x | 0.000122 | 0 | 1 | 1 | 1 | O35738 | Krueppel-li | O35738 [2 | 0 | 1614.82 | 0 | -2.48 | 0.000434 | 0.34 | 168 | 32 | 6.43 | 18.91 | | High | High | High | High | High | High | High | High | High | High | High | 0 | 2.24E-05 | 2.99 |
| TRUE | High | TSPNSLVA | [R].TSPNSL | 1xPhosphc | Q0VGT2 1x | 1.31E-05 | 0 | 1 | 7 | 2 | Q0VGT2 | Zinc finger | Q0VGT2 [2 | 0 | 1844.911 | 0 | -2 | 0.000304 | 1.57 | 158.5 | 41.5 | 4.35 | 21.62 | | High | High | High | High | High | High | High | High | High | High | High | 0 | 1.83E-06 | 4.27 |
| TRUE | High | LLESGDPE | [K].LLESGD | 2xPhosphc | P28231 2xl | 0.000311 | 0 | 1 | 1 | 1 | P28231 | Gap junctic | P28231 [24 | 0 | 2175.986 | 0 | -2.5 | 0.016844 | 0.77 | 166.9 | 33.1 | 27.34 | 50.33 | | High | High | High | High | High | High | High | High | High | High | High | 0 | 6.39E-05 | 3.22 |
| TRUE | High | QEEPQLSPI | [K].QEEPQ | 1xDeamida | Q3UNW5 1 | 0.100129 | 0.008067 | 1 | 1 | 2 | Q3UNW5 | Transcripti | Q3UNW5 [ | 0 | 1764.8 | 0 | -2.61 | 0.000361 | 0.61 | 173.1 | 26.9 | 7.24 | 17.85 | | High | High | High | High | High | High | High | High | High | High | High | 0.004059 | 0.04328 | 1.95 |
| TRUE | High | RDSSTSTM | [R].RDSSTS | 1xPhosphc | Q0VGT2 1x | 0.000876 | 0 | 1 | 2 | 1 | Q0VGT2 | Zinc finger | Q0VGT2 [7 | 1 | 2060.916 | 0 | -2.13 | 0.002541 | 1.63 | 162.5 | 37.5 | 15.67 | 32.77 | | High | High | High | High | High | High | High | High | High | High | High | 0 | 0.000205 | 3.1 |
| TRUE | High | TSPGPALE | [R].TSPGPA | 1xPhosphc | Q3UNA4-2 | 2.26E-06 | 0 | 1 | 2 | 1 | Q3UNA4-2 | Isoform 2 c | Q3UNA4-2 | 0 | 2910.545 | 0 | -3.63 | 0.00022 | 0.07 | 185.3 | 14.7 | 6.48 | 61.99 | | High | High | High | High | High | High | High | High | High | High | High | 0 | 2.53E-07 | 3.74 |
| TRUE | High | SIDSLSVLK | [K].SIDSLS | 1xPhosphc | E9Q3I3 1xF | 0.00582 | 0.00062 | 1 | 2 | 1 | E9Q3I3 | Spermatog | E9Q3I3 [22 | 0 | 1499.849 | 0 | -2.15 | 0.000209 | 1.12 | 164.4 | 35.6 | 3.36 | 23.43 | | High | High | High | High | High | High | High | High | High | High | High | 0.000275 | 0.001713 | 1.44 |
| TRUE | High | HERSPAGE | [R].HERSPA | 1xCarbami | E9Q6S4 1x | 0.00359 | 0.00021 | 1 | 1 | 5 | E9Q6S4 | Uncharacte | E9Q6S4 [6 | 1 | 2711.192 | 0 | -2.86 | 0.000498 | 0.7 | 173 | 27 | 8.76 | 22.16 | | High | High | High | High | High | High | High | High | High | High | High | 0.000294 | 0.002346 | 3.88 |
| TRUE | High | EPLSPSSPP | [K].EPLSPS | 1xPhosphc | Q9CWR8 1 | 0.000231 | 0 | 1 | 1 | 5 | Q9CWR8 | DNA (cytos | Q9CWR8 [ | 0 | 1533.751 | 0 | -2.47 | 0.000207 | 0.84 | 169.4 | 30.6 | 4.09 | 21.79 | | High | High | High | High | High | High | High | High | High | High | High | 0.007335 | 0.08041 | 2.24 |
| TRUE | High | SYSHDVLPI | [K].SYSHD\ | 1xPhosphc | A0A1B0GS | 0.000239 | 0 | 1 | 2 | 5 | A0A1B0GS | GRB2-asso | A0A1B0GS | 0 | 1583.823 | 0 | -2.23 | 0.000779 | 1.61 | 167 | 33 | 6.97 | 35.85 | | High | High | High | High | High | High | High | High | High | High | High | 0.00168 | 0.0169 | 2.66 |
| TRUE | High | HLEVWDPS | [K].HLEVW | 1xPhosphc | Q99MV7 1 | 0.000488 | 0 | 1 | 3 | 1 | Q99MV7 | RING finge | Q99MV7 [ | 0 | 3484.802 | 0 | -3.46 | 0.000607 | 0.09 | 184.5 | 15.5 | 14.19 | 27.21 | | High | High | High | High | High | High | High | High | High | High | High | 0 | 0.000106 | 4.08 |
| TRUE | High | EAATQLSP | [R].EAATQI | 1xPhosphc | Q9CWU5 1 | 3.72E-07 | 0 | 1 | 2 | 1 | Q9CWU5 | KH domain | Q9CWU5 [ | 0 | 2725.406 | 0 | -3.31 | 0.000322 | 0.68 | 180.8 | 19.2 | 10.61 | 54.63 | | High | High | High | High | High | High | High | High | High | High | High | 0 | 3.35E-08 | 4.68 |
| TRUE | High | GPAGRPPP | [R].GPAGR | 1xPhosphc | Q3V293 1x | 0.024211 | 0.001612 | 1 | 3 | 1 | Q3V293 | Putative ur | Q3V293 [1 | 0 | 2497.28 | 0 | -2.09 | 0.00122 | 0.59 | 156.5 | 43.5 | 15.06 | 19.51 | | High | High | High | High | High | High | High | High | High | High | High | 0.000859 | 0.008496 | 2.6 |
| TRUE | High | LSPFLIAPP | [R].LSPFLIA | 2xPhosphc | D0VLQ5 2x | 0.001058 | 0 | 1 | 4 | 1 | D0VLQ5 | Tet oncoge | D0VLQ5 [1 | 0 | 2832.512 | 0 | -2.67 | 0.002238 | 0.35 | 169.7 | 30.3 | 14.8 | 27.03 | | High | High | High | High | High | High | High | High | High | High | High | 0 | 0.000254 | 2.86 |
| TRUE | High | EAAIPGLNE | [K].EAAIPG | 1xPhosphc | B1AU75 1x | 0.004033 | 0.000311 | 1 | 2 | 1 | B1AU75 | Nuclear au | B1AU75 [2 | 0 | 2138.078 | 0 | -3.12 | 0.000193 | 0.23 | 179.5 | 20.5 | 2.55 | 34.73 | | High | High | High | High | High | High | High | High | High | High | High | 0.000161 | 0.001136 | 2.15 |
| TRUE | High | LSPFLIAPP | [R].LSPFLIA | 1xPhosphc | D0VLQ5 1x | 1.03E-08 | 0 | 1 | 4 | 1 | D0VLQ5 | Tet oncoge | D0VLQ5 [1 | 0 | 2752.545 | 0 | -2.91 | 0.000771 | 0.32 | 177 | 23 | 13.39 | 28.48 | | High | High | High | High | High | High | High | High | High | High | High | 0 | 5.93E-10 | 3.09 |
| TRUE | High | LLESGDPE | [K].LLESGD | 1xPhosphc | P28231 1xl | 5.35E-08 | 0 | 1 | 1 | 1 | P28231 | Gap junctic | P28231 [24 | 0 | 2096.02 | 0 | -3.19 | 0.000859 | 0.31 | 183.4 | 16.6 | 14.06 | 28.65 | | High | High | High | High | High | High | High | High | High | High | High | 0 | 3.8E-09 | 3.65 |
| TRUE | High | EEGASRDS | [K].EEGASI | 1xPhosphc | A2TJV2 1xl | 0.000207 | 0 | 1 | 1 | 3 | A2TJV2 | Paralemmi | A2TJV2 [32 | 1 | 3342.548 | 0 | -2.07 | 0.000294 | 1.34 | 162 | 38 | 3.24 | 18.55 | | High | High | High | High | High | High | High | High | High | High | High | 0 | 5.17E-05 | 5.47 |
| TRUE | High | VPNSVDEH | [R].VPNSVI | 1xPhosphc | G3UYN0 1 | 0.051694 | 0.003745 | 1 | 2 | 1 | G3UYN0 | L1TD1 OS= | G3UYN0 [7 | 1 | 1828.927 | 0 | -2.21 | 0.001754 | 1.01 | 166 | 34 | 14.47 | 14.18 | | High | High | High | High | High | High | High | High | High | High | High | 0.00196 | 0.01999 | 1.87 |
| TRUE | High | AVEEQGDI | [K].AVEEQ | 1xPhosphc | Q00PI9 1xF | 2.67E-05 | 0 | 1 | 1 | 1 | Q00PI9 | Heterogen | Q00PI9 [17 | 0 | 1987.89 | 0 | -3.33 | 0.000264 | 0.49 | 180.8 | 19.2 | 4.72 | 46.51 | | High | High | High | High | High | High | High | High | High | High | High | 0 | 4.09E-06 | 3.45 |
| TRUE | High | LGSQTGPA | [R].LGSQT | 1xPhosphc | Q8BL80 1x | 1.48E-06 | 0 | 1 | 4 | 1 | Q8BL80 | Rho GTPas | Q8BL80 [4 | 0 | 1809.951 | 0 | -2.31 | 0.000363 | 1.77 | 169.4 | 30.6 | 5.34 | 30.48 | | High | High | High | High | High | High | High | High | High | High | High | 0 | 1.58E-07 | 3.5 |
| TRUE | High | RASSSASTF | [R].RASSSA | 1xPhosphc | Q3KR45 1x | 5.65E-05 | 0 | 1 | 5 | 5 | Q3KR45 | DNA (cytos | Q3KR45 [1 | 1 | 3192.532 | 0 | -2.87 | 0.000364 | 1.01 | 173.5 | 26.5 | 8.06 | 45.47 | | High | High | High | High | High | High | High | High | High | High | High | 0 | 1.03E-05 | 5.23 |
| TRUE | High | TASLPAIPN | [R].TASLPA | 1xPhosphc | Q60760 1x | 0.008691 | 0.000736 | 1 | 6 | 1 | Q60760 | Growth fac | Q60760 [7 | 0 | 4378.314 | 0 | -3.34 | 0.000777 | 0.24 | 181.8 | 18.2 | 12.77 | 33.1 | | High | High | High | High | High | High | High | High | High | High | High | 0.000327 | 0.002685 | 4.82 |
| TRUE | High | ATTDNSPS | [K].ATTDN | 1xPhosphc | P06837 1xl | 0.000458 | 0 | 1 | 1 | 1 | P06837 | Neuromod | P06837 [1 | 0 | 1545.756 | 0 | -2.3 | 0.000351 | 1.17 | 166.5 | 33.5 | 3.92 | 24.95 | | High | High | High | High | High | High | High | High | High | High | High | 0 | 9.93E-05 | 1.86 |
| TRUE | High | TTEGEESLE | [K].TTEGEE | 1xPhosphc | A2TJV2 1xl | 6.95E-05 | 0 | 1 | 1 | 1 | A2TJV2 | Paralemmi | A2TJV2 [47 | 0 | 1888.919 | 0 | -2.91 | 0.000368 | 0.35 | 176.7 | 23.3 | 5.51 | 24.53 | | High | High | High | High | High | High | High | High | High | High | High | 0 | 1.19E-05 | 2.54 |
| TRUE | High | TASLTSAAS | [R].TASLTS | 1xPhosphc | Q9QYG0 1 | 0.000151 | 0 | 1 | 3 | 2 | Q9QYG0 | Protein ND | Q9QYG0 [3 | 0 | 1645.8 | 0 | -2.8 | 0.000193 | 0.11 | 175.8 | 24.2 | 4.24 | 17.97 | | High | High | High | High | High | High | High | High | High | High | High | 0 | 2.87E-05 | 2.17 |
| TRUE | High | QGTSSPELI | [R].QGTSSI | 1xPhosphc | A2TJV2 1xl | 0.047894 | 0.00354 | 1 | 1 | 3 | A2TJV2 | Paralemmi | A2TJV2 [26 | 0 | 1968.024 | 0 | -3.51 | 0.000661 | 0.28 | 185.3 | 14.7 | 11.62 | 46.77 | | High | High | High | High | High | High | High | High | High | High | High | 0.001812 | 0.01836 | 2.65 |
| TRUE | High | SSPAPNPAI | [R].SSPAPN | 1xPhosphc | A0A068BIR | 7.56E-06 | 0 | 1 | 3 | 2 | A0A068BIR | KN motif a | A0A068BIR | 0 | 2380.259 | 0 | -2.07 | 0.000209 | 0.25 | 161.5 | 38.5 | 3.09 | 9.14 | | High | High | High | High | High | High | High | High | High | High | High | 0 | 9.83E-07 | 3.73 |
| TRUE | High | SSPAPNPAI | [R].SSPAPN | 2xPhosphc | A0A068BIR | 2.31E-05 | 0 | 1 | 3 | 1 | A0A068BIR | KN motif a | A0A068BIR | 0 | 2460.225 | 0 | -2.17 | 0.000687 | 1.76 | 163.6 | 36.4 | 5.03 | 32.34 | | High | High | High | High | High | High | High | High | High | High | High | 0 | 3.47E-06 | 3.29 |
| TRUE | High | LLPPLSPCY | [R].LLPPLSI | 1xCarbami | P47806 1xl | 0.000662 | 0 | 1 | 2 | 1 | P47806 | Zinc finger | P47806 [9 | 0 | 1993.1 | 0 | -3.5 | 0.00032 | 0.26 | 183.5 | 16.5 | 5.44 | 30.81 | | High | High | High | High | High | High | High | High | High | High | High | 0 | 0.00015 | 3.01 |
| TRUE | High | TTHAEGTR | [R].TTHAE | 2xPhosphc | D0VLQ5 2x | 0.00582 | 0.00062 | 1 | 4 | 1 | D0VLQ5 | Tet oncoge | D0VLQ5 [6 | 1 | 3685.916 | 0 | -2.47 | 0.00041 | 0.46 | 167.5 | 32.5 | 7.09 | 14.16 | | High | High | High | High | High | High | High | High | High | High | High | 0.000275 | 0.001718 | 2.94 |
| TRUE | High | TASPVNSP | [R].TASPVN | 1xPhosphc | Q7TPM6 1 | 0.003028 | 0.00016 | 1 | 2 | 1 | Q7TPM6 | Fibronectir | Q7TPM6 [ | 0 | 1308.651 | 0 | -2.11 | 0.001008 | 1.42 | 161.4 | 38.6 | 3.39 | 41.34 | | High | High | High | High | High | High | High | High | High | High | High | 7.19E-05 | 0.000828 | 1.78 |
| TRUE | High | DTPSPGDV | [K].DTPSPC | 1xPhosphc | S4R270 1xl | 0.019446 | 0.001438 | 1 | 4 | 2 | S4R270 | Bridging in | S4R270 [3 | 0 | 1464.73 | 0 | -2.47 | 0.00018 | 0.65 | 168.9 | 31.1 | 2.49 | 19.87 | | High | High | High | High | High | High | High | High | High | High | High | 0.000872 | 0.009026 | 2.83 |
| TRUE | High | VNGTTVSS | [K].VNGTT | 1xPhosphc | P11531 1xl | 0.005228 | 0.000551 | 1 | 12 | 2 | P11531 | Dystrophin | P11531 [3 | 0 | 1842.916 | 0 | -2.48 | 0.000192 | 0.83 | 169 | 31 | 2.59 | 26.92 | | High | High | High | High | High | High | High | High | High | High | High | 0.000267 | 0.001524 | 1.67 |
| TRUE | High | RASSSASTF | [R].RASSSA | 1xPhosphc | Q3KR45 1x | 0.001068 | 0 | 1 | 5 | 6 | Q3KR45 | DNA (cytos | Q3KR45 [1 | 1 | 3208.527 | 0 | -2.83 | 0.000206 | 0.41 | 175.8 | 24.2 | 5.38 | 23.67 | | High | High | High | High | High | High | High | High | High | High | High | 0 | 0.000257 | 3.29 |

| | | | | | | | | | | | | | | | | | | | | | | | | | | | | | | | | | | | | | | |
|---|---|---|---|---|---|---|---|---|---|---|---|---|---|---|---|---|---|---|---|---|---|---|---|---|---|---|---|---|---|---|---|---|---|---|---|---|---|---|
| TRUE | High | TTNDSAAS | [R].TTNDS/ | 1xPhosphc | Q3KR45 1x | 1.72E-05 | 0 | 1 | 5 | 3 | Q3KR45 | DNA (cytos | Q3KR45 [3 | 0 | 1939.941 | 0 | -2.33 | 0.000207 | 0.74 | 167.1 | 32.9 | 1.79 | 18.52 | | High | High | High | High | High | High | High | High | High | High | High | 0 | 2.47E-06 | 3.48 |
| TRUE | High | RYSWGTVE | [R].RYSWG | 1xCarbami | P42209 1xI | 0.000162 | 0 | 1 | 3 | 2 | P42209 | Septin-1 O | P42209 [24 | 1 | 2938.366 | 0 | -3.58 | 0.000191 | 0.21 | 184.9 | 15.1 | 1.97 | 50.28 | | High | High | High | High | High | High | High | High | High | High | High | 0 | 4.89E-05 | 3.49 |
| TRUE | High | ATSPSNNV | [R].ATSPSN | 2xPhosphc | H7BX94 2x | 2.05E-07 | 0 | 1 | 7 | 4 | H7BX94 | Lysine-rich | H7BX94 [1 | 0 | 2746.335 | 0 | -3.03 | 0.000466 | 0.67 | 178.2 | 21.8 | 7.12 | 46.27 | | High | High | High | High | High | High | High | High | High | High | High | 0 | 1.71E-08 | 3.86 |
| TRUE | High | ATSPSNNV | [R].ATSPSN | 1xDeamida | H7BX94 1x | 1.46E-05 | 0 | 1 | 7 | 25 | H7BX94 | Lysine-rich | H7BX94 [1 | 0 | 2667.353 | 0 | -2.78 | 0.000628 | 1.06 | 174.7 | 25.3 | 7.19 | 34.78 | | High | High | High | High | High | High | High | High | High | High | High | 2.58E-05 | 0.000398 | 3.73 |
| TRUE | High | ATSPSNNV | [R].ATSPSN | 1xPhosphc | H7BX94 1x | 1.08E-06 | 0 | 1 | 7 | 10 | H7BX94 | Lysine-rich | H7BX94 [1 | 0 | 2666.369 | 0 | -2.72 | 0.00024 | 0.34 | 173 | 27 | 3.11 | 25.79 | | High | High | High | High | High | High | High | High | High | High | High | 0 | 4.11E-07 | 4.44 |
| TRUE | High | TATSASAS | [R].TATSAS | 1xPhosphc | Q80TZ3-3 1 | 5.16E-05 | 0 | 1 | 3 | 1 | Q80TZ3-3 | Isoform 3 c | Q80TZ3-3 [ | 0 | 1471.736 | 0 | -3.29 | 0.000426 | 0.27 | 181.2 | 18.8 | 7.3 | 24.4 | | High | High | High | High | High | High | High | High | High | High | High | 0 | 8.59E-06 | 1.73 |
| TRUE | High | SSPGGAST | [R].SSPGG/ | 1xPhosphc | Q3UHU5 1 | 8.43E-06 | 0 | 1 | 2 | 2 | Q3UHU5 | Microtubu | Q3UHU5 [2 | 0 | 2252.998 | 0 | -2.27 | 0.000402 | 0.94 | 166.7 | 33.3 | 7.31 | 10.41 | | High | High | High | High | High | High | High | High | High | High | High | 0 | 1.11E-06 | 3.71 |
| TRUE | High | SSPGSMEC | [K].SSPGSN | 1xPhosphc | Q5DW34 1 | 9.91E-06 | 0 | 1 | 8 | 4 | Q5DW34 | Histone-lys | Q5DW34 [ | 0 | 3211.437 | 0 | -2.18 | 0.009131 | 3.42 | 168.8 | 31.2 | 25.85 | 39 | | High | High | High | High | High | High | High | High | High | High | High | 0 | 1.41E-06 | 4.31 |
| TRUE | High | SGSYIK | [R].SGSYIK | 1xPhosphc | B1AZP2 1xI | 0.052576 | 0.00384 | 2 | 21 | 1 | B1AZP2; Q | Disks large | B1AZP2 [36 | 0 | 1192.638 | 0 | -2.95 | 0.000309 | 0.2 | 177.1 | 22.9 | 4.29 | 26.93 | Shared | High | High | High | High | High | High | High | High | High | High | High | 0.001999 | 0.02035 | 1.57 |
| TRUE | High | QLAPGSPS | [R].QLAPG | 1xPhosphc | E9Q6W4 1 | 1.04E-05 | 0 | 1 | 1 | 3 | E9Q6W4 | Zinc finger | E9Q6W4 [2 | 0 | 1654.8 | 0 | -2.05 | 0.000187 | 0.87 | 162.3 | 37.7 | 4.29 | 10.97 | | High | High | High | High | High | High | High | High | High | High | High | 0 | 1.41E-06 | 2.29 |
| TRUE | High | SVPADEGN | [K].SVPADI | 1xPhosphc | F6VQ19 1x | 7.58E-07 | 0 | 1 | 4 | 2 | F6VQ19 | Leucine-ric | F6VQ19 [5 | 0 | 3015.523 | 0 | -2.38 | 0.000285 | 1.07 | 167.5 | 32.5 | 4.55 | 18.46 | | High | High | High | High | High | High | High | High | High | High | High | 0 | 7.42E-08 | 4.07 |
| TRUE | High | HQSFGAAV | [R].HQSFG | 1xPhosphc | Q3U5C8 1x | 0.002419 | 0.000109 | 1 | 3 | 1 | Q3U5C8 | Rho guanir | Q3U5C8 [1 | 0 | 1682.903 | 0 | -3.08 | 0.000209 | 0.16 | 178.9 | 21.1 | 9.05 | 30.31 | | High | High | High | High | High | High | High | High | High | High | High | 4.94E-05 | 0.000641 | 2.68 |
| TRUE | High | GGSADEG | [R].GGSAD | 1xPhosphc | Q9DAE2 1x | 0.000133 | 0 | 1 | 1 | 5 | Q9DAE2 | Putative ur | Q9DAE2 [1 | 0 | 2265.997 | 0 | -3.1 | 0.000393 | 0.16 | 179.2 | 20.8 | 6 | 25.66 | | High | High | High | High | High | High | High | High | High | High | High | 0 | 0.000121 | 2.17 |
| TRUE | High | NTTVCEISS | [K].NTTVCI | 1xCarbami | A0A1D9BZ | 0.000233 | 0 | 1 | 1 | 5 | A0A1D9BZ | GCNA OS= | A0A1D9BZ | 0 | 3084.363 | 0 | -2.19 | 0.000313 | 0.1 | 164.9 | 35.1 | 8.25 | 6.88 | | High | High | High | High | High | High | High | High | High | High | High | 0.000117 | 0.001088 | 2.93 |
| TRUE | High | NTTVCEISS | [K].NTTVCI | 1xCarbami | A0A1D9BZ | 0.019446 | 0.001438 | 1 | 1 | 2 | A0A1D9BZ | GCNA OS= | A0A1D9BZ | 0 | 3005.381 | 0 | -3.51 | 0.000158 | 0.24 | 184.5 | 15.5 | 6.69 | 41.7 | | High | High | High | High | High | High | High | High | High | High | High | 0.001151 | 0.01215 | 1.62 |
| TRUE | High | SCSAAGLH | [R].SCSAA | 1xCarbami | Q8BRK8 1x | 0.021989 | 0.001542 | 1 | 1 | 1 | Q8BRK8 | 5'-AMP-act | Q8BRK8 [4 | 0 | 1267.582 | 0 | -2.04 | 0.003994 | 0.56 | 161.6 | 38.4 | 11.46 | 39.95 | | High | High | High | High | High | High | High | High | High | High | High | 0.000814 | 0.007549 | 2.29 |
| TRUE | High | GSPVGME | [K].GSPVGI | 1xPhosphc | Q8BRB7 1x | 0.01894 | 0.001362 | 1 | 3 | 3 | Q8BRB7 | Histone ac | Q8BRB7 [1 | 0 | 1157.522 | 0 | -2.34 | 0.000294 | 0.26 | 169.2 | 30.8 | 6.55 | 8.72 | | High | High | High | High | High | High | High | High | High | High | High | 0.003646 | 0.03896 | 2.08 |
| TRUE | High | ECSLSPK | [R].ECSLSP | 1xCarbami | Q9WV69-2 | 0.050397 | 0.003641 | 2 | 5 | 1 | Q9WV69-2 | Isoform 2 c | Q9WV69-2 | 0 | 1438.645 | 0 | -4.17 | 0.000397 | 0.12 | 189.6 | 10.4 | 8.73 | 37.9 | Shared | High | High | High | High | High | High | High | High | High | High | High | 0.001916 | 0.01938 | 2.02 |
| TRUE | High | TSWGFLQS | [R].TSWGF | 1xPhosphc | Q99LI5 1xF | 0.008094 | 0.000736 | 1 | 1 | 1 | Q99LI5 | Zinc finger | Q99LI5 [11 | 0 | 2004.097 | 0 | -3.06 | 0.000748 | 0.65 | 179 | 21 | 6.35 | 50.8 | | High | High | High | High | High | High | High | High | High | High | High | 0.000327 | 0.002468 | 2.06 |
| TRUE | High | VSPVPSPS | [R].VSPVPS | 1xPhosphc | Q8VDP3 1x | 0.0009 | 0 | 1 | 4 | 2 | Q8VDP3 | [F-actin]-m | Q8VDP3 [7 | 0 | 1530.788 | 0 | -2.11 | 0.00034 | 0.49 | 162.3 | 37.7 | 4.39 | 18.46 | | High | High | High | High | High | High | High | High | High | High | High | 0 | 0.000211 | 2.29 |
| TRUE | High | VPITPASPV | [K].VPITPA | 1xPhosphc | Q6ZPI0 1xF | 0.003083 | 0.00016 | 1 | 1 | 2 | Q6ZPI0 | Protein Jad | Q6ZPI0 [72 | 0 | 1546.901 | 0 | -2.83 | 0.000114 | 0.27 | 175.3 | 24.7 | 6.26 | 27.85 | | High | High | High | High | High | High | High | High | High | High | High | 0.000275 | 0.001675 | 2.45 |
| TRUE | High | TAPNGGQI | [K].TAPNG | 1xDeamida | Q8C796 1x | 0.000143 | 0 | 1 | 2 | 5 | Q8C796 | REST corep | Q8C796 [2 | 0 | 3175.275 | 0 | -2.82 | 0.008837 | 0.65 | 165 | 35 | 27.67 | 40.12 | | High | High | High | High | High | High | High | High | High | High | High | 0 | 5.96E-05 | 3.48 |
| TRUE | High | TAPNGGQI | [K].TAPNG | 2xPhosphc | Q8C796 2x | 1.67E-05 | 0 | 1 | 2 | 9 | Q8C796 | REST corep | Q8C796 [2 | 0 | 3254.257 | 0 | -2.21 | 0.000846 | 1.4 | 161.7 | 38.3 | 8.97 | 21.59 | | High | High | High | High | High | High | High | High | High | High | High | 0 | 2.4E-06 | 3.19 |
| TRUE | High | ANSDIISLN | [R].ANSDII | 1xPhosphc | Q8C739 1x | 3.77E-05 | 0 | 1 | 1 | 2 | Q8C739 | Protein FAI | Q8C739 [2 | 0 | 1558.783 | 0 | -2.67 | 0.000154 | 0.31 | 173.1 | 26.9 | 3.25 | 14.15 | | High | High | High | High | High | High | High | High | High | High | High | 0 | 6.14E-06 | 2.35 |
| TRUE | High | CPAPLRPQ | [R].CPAPLR | 1xCarbami | Q7TNS8 1x | 0.067059 | 0.004796 | 1 | 1 | 1 | Q7TNS8 | Elongin BC | Q7TNS8 [1 | 0 | 2877.418 | 0 | -2.32 | 0.000486 | 0.08 | 164.7 | 35.3 | 6.96 | 13.22 | | High | High | High | High | High | High | High | High | High | High | High | 0.002455 | 0.02695 | 1.21 |
| TRUE | High | CPAPLRPQ | [R].CPAPLR | 1xCarbami | Q7TNS8 2x | 0.103392 | 0.008249 | 1 | 1 | 1 | Q7TNS8 | Elongin BC | Q7TNS8 [1 | 0 | 2957.384 | 0 | -2.19 | 0.000264 | 0.78 | 161.2 | 38.8 | 5.3 | 12.21 | | High | High | High | High | High | High | High | High | High | High | High | 0.004164 | 0.04493 | 1.17 |
| TRUE | High | EMVVVTSF | [K].EMVVV | 1xPhosphc | Q99MV7 1 | 0.028575 | 0.001829 | 1 | 3 | 2 | Q99MV7 | RING finge | Q99MV7 [3 | 0 | 1543.821 | 0 | -2.15 | 0.000444 | 0.37 | 163.2 | 36.8 | 7.39 | 8.95 | | High | High | High | High | High | High | High | High | High | High | High | 0.000984 | 0.0102 | 1.29 |
| TRUE | High | GIENPAFVI | [K].GIENPA | 1xPhosphc | Q8BGD4 1 | 5.55E-05 | 0 | 1 | 1 | 2 | Q8BGD4 | Solute carr | Q8BGD4 [6 | 0 | 1934.957 | 0 | -2.79 | 0.000285 | 0.46 | 174.7 | 25.3 | 6.8 | 26.17 | | High | High | High | High | High | High | High | High | High | High | High | 0 | 9.27E-06 | 3.38 |
| TRUE | High | RDAVQNSS | [K].RDAVQ | 1xPhosphc | Q6ZPI0 1xF | 7.41E-05 | 0 | 1 | 1 | 1 | Q6ZPI0 | Protein Jad | Q6ZPI0 [59 | 1 | 1886.937 | 0 | -2.63 | 0.000192 | 0.96 | 173.2 | 26.8 | 2.93 | 25.21 | | High | High | High | High | High | High | High | High | High | High | High | 0 | 1.28E-05 | 1.49 |
| TRUE | High | LHCLPFPSP | [K].LHCLPF | 1xCarbami | Q6NVG5 1 | 0.003655 | 0.00021 | 1 | 1 | 2 | Q6NVG5 | Melanoreg | Q6NVG5 [2 | 0 | 1376.664 | 0 | -2.12 | 0.000242 | 1 | 163.5 | 36.5 | 5.23 | 17.37 | | High | High | High | High | High | High | High | High | High | High | High | 9.42E-05 | 0.001019 | 1.94 |
| TRUE | High | SAGDVPVT | [R].SAGDV | 2xPhosphc | Q6J1H4 2x | 0.001398 | 5.66E-05 | 1 | 1 | 1 | Q6J1H4 | Undifferen | Q6J1H4 [2 | 0 | 3055.377 | 0 | -2.21 | 0.00041 | 0.55 | 165.5 | 34.5 | 11.22 | 23.3 | | High | High | High | High | High | High | High | High | High | High | High | 2.58E-05 | 0.000349 | 2.88 |
| TRUE | High | RGSETMAC | [R].RGSETN | 1xPhosphc | Q8K4Z3 1x | 0.004781 | 0.000456 | 1 | 1 | 2 | Q8K4Z3 | NAD(P)H-h | Q8K4Z3 [4 | 1 | 1715.892 | 0 | -2.08 | 0.000477 | 2.03 | 161.7 | 38.3 | 4.12 | 28.61 | | High | High | High | High | High | High | High | High | High | High | High | 0.00092 | 0.009574 | 2.25 |
| TRUE | High | SAGDVPVT | [R].SAGDV | 1xPhosphc | Q6J1H4 1x | 1.93E-08 | 0 | 1 | 1 | 27 | Q6J1H4 | Undifferen | Q6J1H4 [2 | 0 | 2991.406 | 0 | -2.3 | 0.000816 | 0.37 | 166.2 | 33.8 | 11.6 | 11.94 | | High | High | High | High | High | High | High | High | High | High | High | 0 | 3.33E-07 | 4.82 |
| TRUE | High | GPPPGSPEI | [R].GPPPG | 1xCarbami | Q3UHU5 1 | 6.18E-05 | 0 | 1 | 2 | 1 | Q3UHU5 | Microtubu | Q3UHU5 [2 | 0 | 2787.447 | 0 | -2.14 | 0.000207 | 0.43 | 161.7 | 38.3 | 3.93 | 20.26 | | High | High | High | High | High | High | High | High | High | High | High | 0 | 1.05E-05 | 3.2 |
| TRUE | High | SAGDVPVT | [R].SAGDV | 1xPhosphc | Q6J1H4 1x | 2.43E-07 | 0 | 1 | 1 | 2 | Q6J1H4 | Undifferen | Q6J1H4 [2 | 0 | 2975.411 | 0 | -2.88 | 0.000551 | 0.41 | 176.1 | 23.9 | 11.07 | 23.09 | | High | High | High | High | High | High | High | High | High | High | High | 0 | 7.11E-08 | 3.81 |
| TRUE | High | SAGDVPVT | [R].SAGDV | 1xPhosphc | Q6J1H4 1x | 1.37E-08 | 0 | 1 | 1 | 2 | Q6J1H4 | Undifferen | Q6J1H4 [2 | 0 | 2746.248 | 0 | -2.65 | 0.071566 | 1.63 | 175 | 25 | 65.96 | 44.08 | | High | High | High | High | High | High | High | High | High | High | High | 0 | 8.17E-10 | 5.47 |
| TRUE | High | SRSPEGHK | [K].SRSPEG | 1xPhosphc | Q8BX22 1x | 0.050826 | 0.003676 | 1 | 1 | 1 | Q8BX22 | Sal-like pro | Q8BX22 [8 | 1 | 1435.746 | 0 | -2.54 | 0.000181 | 0.42 | 170.4 | 29.6 | 3.05 | 18.01 | | High | High | High | High | High | High | High | High | High | High | High | 0.001916 | 0.0196 | 1.53 |
| TRUE | High | LSIYDNVPG | [R].LSIYDN | 1xPhosphc | E9PXD2 1x | 0.002192 | 0.000109 | 1 | 7 | 3 | E9PXD2 | Rho GTPas | E9PXD2 [8 | 0 | 4921.436 | 0 | -4.08 | 0.000831 | 0.14 | 188.8 | 11.2 | 4.7 | 95.33 | | High | High | High | High | High | High | High | High | High | High | High | 4.94E-05 | 0.000619 | 3.01 |
| TRUE | High | DVICIDNDS | [R].DVICID | 1xCarbami | E9PZ97 2x | 0.002293 | 0.000109 | 1 | 2 | 2 | E9PZ97 | Bloom syn | E9PZ97 [24 | 0 | 2556.123 | 0 | -2.31 | 0.000363 | 0.41 | 166.7 | 33.3 | 8.1 | 7.08 | | High | High | High | High | High | High | High | High | High | High | High | 4.94E-05 | 0.000608 | 3.04 |
| TRUE | High | LSLVSRSPS | [R].LSLVSR | 2xPhosphc | Q3U2W2 2 | 0.090888 | 0.007081 | 1 | 2 | 1 | Q3U2W2 | MYB bindir | Q3U2W2 [ | 1 | 2289.238 | 0 | -2.38 | 0.000233 | 0.95 | 168.8 | 31.2 | 5.75 | 17.55 | | High | High | High | High | High | High | High | High | High | High | High | 0.003633 | 0.03872 | 2.41 |
| TRUE | High | QAGSASAS | [K].QAGSA | 1xPhosphc | G3X8V5 1x | 0.000699 | 0 | 1 | 2 | 5 | G3X8V5 | RIKEN cDN | G3X8V5 [3 | 0 | 1909.897 | 0 | -2.27 | 0.001066 | 0.5 | 164.4 | 35.6 | 12.14 | 17.16 | | High | High | High | High | High | High | High | High | High | High | High | 0 | 0.00016 | 2.44 |
| TRUE | High | CPSPVPSDI | [R].CPSPVF | 2xCarbami | Q8CAQ6 1 | 0.017041 | 0.001204 | 1 | 1 | 1 | Q8CAQ6 | Putative ur | Q8CAQ6 [9 | 0 | 2023.882 | 0 | -2.74 | 0.000186 | 0.42 | 175.3 | 24.7 | 7.22 | 28.49 | | High | High | High | High | High | High | High | High | High | High | High | 0.000638 | 0.00572 | 2.48 |
| TRUE | High | LTADLSPGT | [R].LTADLS | 1xPhosphc | G3UYN0 1 | 1.11E-05 | 0 | 1 | 2 | 2 | G3UYN0 | L1TD1 OS= | G3UYN0 [7 | 0 | 1638.83 | 0 | -2.17 | 0.000659 | 1.08 | 163 | 37 | 10.08 | 9.99 | | High | High | High | High | High | High | High | High | High | High | High | 0 | 1.53E-06 | 3.1 |
| TRUE | High | CQETESNE | [R].CQETE | 1xCarbami | B2RRE0 1x | 3.26E-08 | 0 | 1 | 2 | 4 | B2RRE0 | A kinase (P | B2RRE0 [1 | 0 | 2433.089 | 0 | -2.39 | 0.000781 | 0.21 | 166.6 | 33.4 | 9.13 | 19.38 | | High | High | High | High | High | High | High | High | High | High | High | 0 | 1.26E-07 | 4.76 |
| TRUE | High | SSLSALAPA | [R].SSLSAL | 1xPhosphc | Q6J1H4 1x | 1.32E-05 | 0 | 1 | 2 | 6 | Q6J1H4 | Undifferen | Q6J1H4 [1 | 0 | 3633.825 | 0 | -2.65 | 0.00124 | 1.51 | 173.7 | 26.3 | 12.46 | 33.84 | | High | High | High | High | High | High | High | High | High | High | High | 0 | 1.84E-06 | 4.14 |
| TRUE | High | SLPGVQME | [K].SLPGVC | 1xPhosphc | Q9WVL5 1 | 0.100129 | 0.008067 | 1 | 1 | 1 | Q9WVL5 | MORC fam | Q9WVL5 [ | 0 | 1753.839 | 0 | -2.05 | 0.000382 | 1.46 | 162.6 | 37.4 | 5.58 | 16.01 | | High | High | High | High | High | High | High | High | High | High | High | 0.004032 | 0.04313 | 2.17 |
| TRUE | High | ITETEEDSD | [K].ITETEEI | 1xPhosphc | Q9CQS5 1x | 0.001437 | 5.66E-05 | 1 | 2 | 1 | Q9CQS5 | Serine/thre | Q9CQS5 [3 | 0 | 2933.294 | 0 | -2.11 | 0.002271 | 2.64 | 164.2 | 35.8 | 12.9 | 27.09 | | High | High | High | High | High | High | High | High | High | High | High | 2.58E-05 | 0.000359 | 3.52 |
| TRUE | High | IINDSDSESI | [R].IINDSD | 2xPhosphc | G3UWX1 2 | 6.77E-05 | 0 | 1 | 6 | 5 | G3UWX1 | Replication | G3UWX1 [ | 0 | 2411.103 | 0 | -2.77 | 0.000285 | 0.72 | 174.5 | 25.5 | 3.05 | 32.59 | | High | High | High | High | High | High | High | High | High | High | High | 0 | 1.16E-05 | 4.06 |
| TRUE | High | SVSELSLQ | [R].SVSELS | 1xPhosphc | B9EK92 1x | 0.00614 | 0.00062 | 1 | 2 | 2 | B9EK92 | Ras interac | B9EK92 [32 | 0 | 1384.703 | 0 | -2.44 | 0.000716 | 0.84 | 168.8 | 31.2 | 9.87 | 21.28 | | High | High | High | High | High | High | High | High | High | High | High | 0.000275 | 0.001855 | 2.27 |
| TRUE | High | ANSLGDLY | [R].ANSLGI | 1xPhosphc | G3UWG2 1 | 1.46E-06 | 0 | 1 | 4 | 1 | G3UWG2 | T-cell lymp | G3UWG2 [ | 0 | 1717.893 | 0 | -2.42 | 0.000191 | 1.14 | 170.5 | 29.5 | 3.81 | 22.89 | | High | High | High | High | High | High | High | High | High | High | High | 0 | 1.55E-07 | 3.86 |
| TRUE | High | TPAPAASPI | [K].TPAPA/ | 1xPhosphc | E9Q2Z1 1x | 0.000109 | 0 | 1 | 4 | 4 | E9Q2Z1 | CECR2, hist | E9Q2Z1 [1 | 0 | 2079.104 | 0 | -2.05 | 0.000285 | 1.28 | 161.1 | 38.9 | 4.33 | 13.74 | | High | High | High | High | High | High | High | High | High | High | High | 4.94E-05 | 0.000611 | 3.52 |
| TRUE | High | ETPAPPPST | [K].ETPAPF | 1xPhosphc | Q6P9J5 1x | 0.043236 | 0.003228 | 1 | 1 | 1 | Q6P9J5 | KN motif a | Q6P9J5 [63 | 0 | 3250.705 | 0 | -2.09 | 0.001068 | 1.76 | 162 | 38 | 13.59 | 24.01 | | High | High | High | High | High | High | High | High | High | High | High | 0.001651 | 0.01627 | 2.51 |
| TRUE | High | EETGIPVSP | [R].EETGIP | 1xPhosphc | Q8BRB7 1x | 0.001735 | 5.66E-05 | 1 | 3 | 1 | Q8BRB7 | Histone ac | Q8BRB7 [1 | 0 | 1731.908 | 0 | -2.87 | 0.000125 | 0.52 | 176.2 | 23.8 | 1.76 | 25.24 | | High | High | High | High | High | High | High | High | High | High | High | 2.58E-05 | 0.000444 | 2.09 |
| TRUE | High | FSPSQVPV | [K].FSPSQV | 1xPhosphc | U5LK19 1x | 0.001516 | 5.66E-05 | 2 | 9 | 2 | U5LK19; E | Marshalin- | U5LK19 [8 | 0 | 1554.788 | 0 | -2.67 | 0.000301 | 0.27 | 175.5 | 24.5 | 9.23 | 13.92 | Shared | High | High | High | High | High | High | High | High | High | High | High | 7.19E-05 | 0.000919 | 2.74 |
| TRUE | High | QPLTSPGS | [K].QPLTSF | 1xPhosphc | Q9WV69-2 | 0.001704 | 5.66E-05 | 2 | 2 | 3 | Q9WV69-2 | Isoform 2 c | Q9WV69-2 | 0 | 1611.794 | 0 | -3.08 | 0.000158 | 0.14 | 178.9 | 21.1 | 3.22 | 12.73 | Shared | High | High | High | High | High | High | High | High | High | High | High | 4.94E-05 | 0.000652 | 2.36 |
| TRUE | High | DLSIVAETG | [R].DLSIVA | 1xCarbami | E9Q2Z1 1x | 2.66E-06 | 0 | 1 | 4 | 1 | E9Q2Z1 | CECR2, hist | E9Q2Z1 [1 | 0 | 2793.333 | 0 | -2.98 | 0.00127 | 0.6 | 180 | 20 | 13.67 | 46.19 | | High | High | High | High | High | High | High | High | High | High | High | 0 | 3.06E-07 | 3.69 |
| TRUE | High | GSSDGGLC | [R].GSSDG | 1xPhosphc | A0A158SIT | 0.00614 | 0.00062 | 1 | 1 | 1 | A0A158SIT | Proline, glu | A0A158SIT | 0 | 2240.189 | 0 | -2.32 | 0.001035 | 0.31 | 166.9 | 33.1 | 18.24 | 38.02 | | High | High | High | High | High | High | High | High | High | High | High | 0.000275 | 0.001825 | 1.83 |
| TRUE | High | EGNSSPNH | [R].EGNSSI | 1xPhosphc | Q3UNW5 1 | 0.001179 | 0 | 1 | 1 | 2 | Q3UNW5 | Transcripti | Q3UNW5 [ | 0 | 4136.019 | 0 | -2.92 | 0.001944 | 0.38 | 173.6 | 26.4 | 19.2 | 26.01 | | High | High | High | High | High | High | High | High | High | High | High | 0 | 0.000286 | 3.99 |
| TRUE | High | STPLLVEPL | [R].STPLLV | 1xPhosphc | Q9QY39 1x | 0.002508 | 0.000109 | 1 | 2 | 1 | Q9QY39 | PDZ domai | Q9QY39 [4 | 0 | 2158.218 | 0 | -2.01 | 0.000836 | 1.24 | 160.2 | 39.8 | 10.4 | 11.96 | | High | High | High | High | High | High | High | High | High | High | High | 4.94E-05 | 0.000667 | 2.61 |
| TRUE | High | AASGGAAL | [R].AASGG | 1xPhosphc | B9EK92 1x | 3.78E-10 | 0 | 1 | 2 | 1 | B9EK92 | Ras interac | B9EK92 [28 | 0 | 2595.282 | 0 | -3.53 | 0.000432 | 0.57 | 184 | 16 | 10.02 | 41.53 | | High | High | High | High | High | High | High | High | High | High | High | 0 | 1.46E-11 | 5.43 |
| TRUE | High | SPEQPAESI | [R].SPEQP/ | 1xPhosphc | B2RRE0 1x | 1.3E-06 | 0 | 1 | 2 | 2 | B2RRE0 | A kinase (P | B2RRE0 [1 | 0 | 4369.003 | 0 | -2.68 | 0.000154 | 0.66 | 173.9 | 26.1 | 2.92 | 23.48 | | High | High | High | High | High | High | High | High | High | High | High | 0 | 1.36E-07 | 5.46 |
| TRUE | High | TLPPSSASI | [K].TLPPSS | 1xPhosphc | Q5DTZ0 1x | 5.21E-05 | 0 | 1 | 1 | 2 | Q5DTZ0 | Protein NY | Q5DTZ0 [9 | 0 | 3850.805 | 0 | -2.29 | 0.001483 | 0.55 | 167.9 | 32.1 | 12.28 | 46.5 | | High | High | High | High | High | High | High | High | High | High | High | 0 | 2.15E-05 | 4.04 |
| TRUE | High | DPQSPEGC | [K].DPQSPI | 1xPhosphc | A2TJV2 1xF | 0.083783 | 0.006345 | 1 | 1 | 1 | A2TJV2 | Paralemmi | A2TJV2 [84 | 0 | 1592.727 | 0 | -3.16 | 0.000158 | 0.55 | 181.1 | 18.9 | 8.57 | 53.19 | | High | High | High | High | High | High | High | High | High | High | High | 0.003265 | 0.03518 | 2.1 |
| TRUE | High | DNPDSPQK | [R].DNPDSI | 1xPhosphc | B9EIT8 1xP | 0.065952 | 0.004766 | 1 | 2 | 1 | B9EIT8 | Rg9mtd2 p | B9EIT8 [30 | 0 | 1438.698 | 0 | -2.4 | 0.00024 | 0.33 | 166.1 | 33.9 | 7.18 | 11.99 | | High | High | High | High | High | High | High | High | High | High | High | 0.002455 | 0.02661 | 2.31 |
| TRUE | High | SSTPSDINS | [R].SSTPSD | 1xPhosphc | Q69ZS8 1x | 0.001476 | 5.66E-05 | 1 | 9 | 2 | Q69ZS8 | Kazrin OS= | Q69ZS8 [3 | 0 | 1469.683 | 0 | -2.37 | 0.000181 | 0.86 | 168.5 | 31.5 | 3.14 | 18.63 | | High | High | High | High | High | High | High | High | High | High | High | 0.001916 | 0.01944 | 2.1 |
| TRUE | High | YLEAANSEI | [R].YLEAAN | 1xPhosphc | A2ADW8 1 | 4.15E-08 | 0 | 1 | 16 | 1 | A2ADW8 | Uncharacte | A2ADW8 [ | 0 | 3015.258 | 0 | -2.29 | 0.000404 | 0.54 | 166.3 | 33.7 | 9.13 | 5.82 | | High | High | High | High | High | High | High | High | High | High | High | 0 | 2.85E-09 | 4.58 |
| TRUE | High | LNSPTNSSI | [K].LNSPTN | 2xPhosphc | Q9DBX1 2x | 9.56E-06 | 0 | 1 | 1 | 4 | Q9DBX1 | Regulator c | Q9DBX1 [9 | 0 | 2311.13 | 0 | -2.55 | 0.000167 | 0.57 | 170.9 | 29.1 | 11.31 | 43.68 | | High | High | High | High | High | High | High | High | High | High | High | 0 | 8.9E-05 | 3.24 |
| TRUE | High | AATSPGVG | [R].AATSPC | 1xPhosphc | Q6ZPI0 1xF | 9.82E-06 | 0 | 1 | 1 | 4 | Q6ZPI0 | Protein Jad | Q6ZPI0 [70 | 0 | 1665.816 | 0 | -2.24 | 0.000965 | 1.36 | 165 | 35 | 6.98 | 24.96 | | High | High | High | High | High | High | High | High | High | High | High | 0 | 1.33E-06 | 2.41 |
| TRUE | High | YSSSGTPSS | [R].YSSSGT | 1xPhosphc | Q8K120 1x | 0.01861 | 0.001362 | 1 | 2 | 3 | Q8K120 | Nuclear fac | Q8K120 [2 | 0 | 1863.869 | 0 | -3.75 | 0.000598 | 0.06 | 187.1 | 12.9 | 15.11 | 68.2 | | High | High | High | High | High | High | High | High | High | High | High | 0.000872 | 0.008806 | 2.15 |
| TRUE | High | RPFTNDHS | [R].RPFTNI | 1xPhosphc | Q3UKE7 1x | 0.087983 | 0.006639 | 1 | 1 | 1 | Q3UKE7 | Hydroxyme | Q3UKE7 [4 | 0 | 3764.88 | 0 | -2.88 | 0.000264 | 0.12 | 175.7 | 24.3 | 7.79 | 23.82 | | High | High | High | High | High | High | High | High | High | High | High | 0.003402 | 0.03704 | 3.69 |
| TRUE | High | SPSLLQSG | [R].SPSLLQ | 1xPhosphc | Q3U2W2 1 | 2.23E-05 | 0 | 1 | 2 | 4 | Q3U2W2 | MYB bindir | Q3U2W2 [ | 0 | 1553.87 | 0 | -3.03 | 0.00034 | 0.26 | 179.3 | 20.7 | 11.31 | 27.95 | | High | High | High | High | High | High | High | High | High | High | High | 0 | 3.32E-06 | 2.86 |
| TRUE | High | YIESPDVEK | [K].YIESPD | 1xPhosphc | Q9R1C0 1x | 0.045122 | 0.003373 | 1 | 2 | 1 | Q9R1C0 | Transcripti | Q9R1C0 [1 | 0 | 1617.818 | 0 | -2.99 | 0.000333 | 0.12 | 179.6 | 20.4 | 9.49 | 24.15 | | High | High | High | High | High | High | High | High | High | High | High | 0.001727 | 0.01716 | 2.55 |
| TRUE | High | EGNNTDLP | [K].EGNNT | 1xDeamida | B1AV60 1x | 4.68E-05 | 0 | 1 | 4 | 2 | B1AV60 | NHS actin- | B1AV60 [1 | 0 | 3288.592 | 0 | -2.37 | 0.001547 | 1.35 | 164.1 | 35.9 | 15.98 | 35.43 | | High | High | High | High | High | High | High | High | High | High | High | 0 | 9.66E-06 | 3.92 |
| TRUE | High | SPEPVPGS | [R].SPEPVP | 2xPhosphc | P97931 2x | 1.32E-07 | 0 | 1 | 1 | 3 | P97931 | Uracil-DNA | P97931 [23 | 0 | 2781.351 | 0 | -2.8 | 0.000223 | 0.15 | 173.5 | 26.5 | 5.78 | 26.12 | | High | High | High | High | High | High | High | High | High | High | High | 0 | 1.04E-08 | 4.33 |
| TRUE | High | YPLSPPQE | [R].YPLSPP | 1xPhosphc | D0VLQ5 1x | 0.066503 | 0.004796 | 1 | 3 | 1 | D0VLQ5 | Tet oncoge | D0VLQ5 [9 | 0 | 1814.835 | 0 | -2.1 | 0.001032 | 1.28 | 160.8 | 39.2 | 9.95 | 14.52 | | High | High | High | High | High | High | High | High | High | High | High | 0.002455 | 0.02675 | 2.3 |
| TRUE | High | RSPGPGPS | [R].RSPGPC | 1xPhosphc | U5LK19 1x | 0.000297 | 0 | 2 | 9 | 1 | U5LK19; E | Marshalin- | U5LK19 [7 | 1 | 1514.768 | 0 | -2.22 | 0.000554 | 1.13 | 163.6 | 36.4 | 5.02 | 20.78 | Shared | High | High | High | High | High | High | High | High | High | High | High | 0 | 6.08E-05 | 3.71 |
| TRUE | High | GSGGPEST | [R].GSGGP | 1xPhosphc | Q7TNL7 1x | 5.47E-06 | 0 | 1 | 2 | 14 | Q7TNL7 | Dual specif | Q7TNL7 [4 | 0 | 3337.487 | 0 | -2.49 | 0.003648 | 0.62 | 170.1 | 29.9 | 13.53 | 56.25 | | High | High | High | High | High | High | High | High | High | High | High | 0 | 5.01E-06 | 4.92 |
| TRUE | High | LASNSPVLF | [R].LASNSF | 1xPhosphc | Q9CZZ4 1x | 0.002022 | 5.66E-05 | 1 | 4 | 1 | Q9CZZ4 | Putative ur | Q9CZZ4 [1 | 0 | 1779.936 | 0 | -2.24 | 0.001296 | 0.74 | 165.2 | 34.8 | 16.35 | 17.66 | | High | High | High | High | High | High | High | High | High | High | High | 2.58E-05 | 0.000528 | 2.81 |
| TRUE | High | GAVSSEDP | [R].GAVSSI | 1xPhosphc | E9Q8T1 1x | 0.000142 | 0 | 1 | 3 | 2 | E9Q8T1 | Transformi | E9Q8T1 [1 | 0 | 2202.084 | 0 | -2.34 | 0.00133 | 0.22 | 170.1 | 29.9 | 14.13 | 13.01 | | High | High | High | High | High | High | High | High | High | High | High | 0 | 6.92E-05 | 2.31 |
| TRUE | High | EGITPWAS | [R].EGITPW | 1xPhosphc | B2RRE0 1x | 0.001244 | 0 | 1 | 3 | 2 | B2RRE0 | A kinase (P | B2RRE0 [5 | 0 | 1444.708 | 0 | -2.24 | 0.003645 | 0.99 | 164.7 | 35.3 | 17.65 | 16.82 | | High | High | High | High | High | High | High | High | High | High | High | 0.003922 | 0.0417 | 2.1 |
| TRUE | High | TGSSPPIVS | [R].TGSSPF | 1xPhosphc | Q62255 1x | 1.66E-05 | 0 | 1 | 2 | 2 | Q62255 | Sal-like pro | Q62255 [1 | 0 | 1738.939 | 0 | -3.02 | 0.000805 | 0.53 | 178 | 22 | 9.96 | 23.6 | | High | High | High | High | High | High | High | High | High | High | High | 0 | 2.38E-06 | 2.9 |
| TRUE | High | TLIVSEASPI | [K].TLIVSE/ | 1xPhosphc | Q5U4C1 1x | 6.26E-06 | 0 | 1 | 3 | 1 | Q5U4C1 | G-protein c | Q5U4C1 [9 | 0 | 1882.993 | 0 | -2.37 | 0.000154 | 0.72 | 167.4 | 32.6 | 2.13 | 22.99 | | High | High | High | High | High | High | High | High | High | High | High | 0 | 7.97E-07 | 2.37 |
| TRUE | High | DGSSVEEV | [K].DGSSVI | 1xPhosphc | B1AU75 1x | 0.001087 | 0 | 1 | 2 | 2 | B1AU75 | Nuclear au | B1AU75 [3 | 0 | 1487.739 | 0 | -2.44 | 0.000197 | 0.72 | 167.9 | 32.1 | 3.56 | 18.21 | | High | High | High | High | High | High | High | High | High | High | High | 0 | 0.000261 | 2.92 |
| TRUE | High | TYNTNAQ\ | [K].TYNTN/ | 1xCarbami | B7XGA6 1x | 4.86E-06 | 0 | 1 | 8 | 1 | B7XGA6 | Alkaline ph | B7XGA6 [1 | 0 | 2940.421 | 0 | -2.02 | 0.000986 | 1.48 | 160.3 | 39.7 | 9.53 | 18.44 | | High | High | High | High | High | High | High | High | High | High | High | 0 | 6.01E-07 | 3.09 |
| TRUE | High | GADYTEPD | [R].GADYTI | 1xPhosphc | P97445 1x | 0.000396 | 0 | 1 | 6 | 1 | P97445 | Voltage-de | P97445 [22 | 1 | 2698.21 | 0 | -2.75 | 0.000459 | 0.35 | 174.2 | 25.8 | 8.32 | 13.92 | | High | High | High | High | High | High | High | High | High | High | High | 0 | 8.45E-05 | 1.8 |
| TRUE | High | EGITPWAS | [R].EGITPW | 1xPhosphc | B2RRE0 1x | 2.1E-05 | 0 | 1 | 3 | 1 | B2RRE0 | A kinase (P | B2RRE0 [5 | 0 | 1673.87 | 0 | -2.22 | 0.003173 | 1.97 | 162.7 | 37.3 | 17.17 | 16.6 | | High | High | High | High | High | High | High | High | High | High | High | 0 | 3.1E-06 | 3 |
| TRUE | High | ASSAGESN. | [R].ASSAGI | 1xCarbami | F6S200 1xF | 9.08E-07 | 0 | 1 | 8 | 2 | F6S200 | Pleckstrin I | F6S200 [66 | 0 | 1840.81 | 0 | -2.46 | 0.000372 | 0.19 | 168.5 | 31.5 | 6.39 | 27.07 | | High | High | High | High | High | High | High | High | High | High | High | 0 | 3.77E-06 | 3.43 |
| TRUE | High | SPFEIISPPA | [R].SPFEIIS | 2xPhosphc | A2ARP8 2x | 0.00011 | 0 | 1 | 3 | 5 | A2ARP8 | Microtubu | A2ARP8 [2 | 0 | 2446.096 | 0 | -2.14 | 0.001238 | 0.28 | 160.7 | 39.3 | 12.68 | 14.12 | | High | High | High | High | High | High | High | High | High | High | High | 0.001876 | 0.01876 | 3.02 |
| TRUE | High | QSSRENSSI | [R].QSSREI | 1xCarbami | Q8BX22 2x | 5.84E-07 | 0 | 1 | 1 | 11 | Q8BX22 | Sal-like pro | Q8BX22 [7 | 1 | 3769.585 | 0 | -2.85 | 0.001392 | 1.48 | 176.4 | 23.6 | 8.7 | 57.02 | | High | High | High | High | High | High | High | High | High | High | High | 0 | 5.55E-08 | 4.78 |
| TRUE | High | QSSRENSSI | [R].QSSREI | 1xCarbami | Q8BX22 1x | 4.63E-05 | 0 | 1 | 1 | 14 | Q8BX22 | Sal-like pro | Q8BX22 [7 | 1 | 3690.602 | 0 | -2.58 | 0.000209 | 0.76 | 170.3 | 29.7 | 3.36 | 20.95 | | High | High | High | High | High | High | High | High | High | High | High | 0 | 7.54E-06 | 5.2 |
| TRUE | High | QSSRENSSI | [R].QSSREI | 1xCarbami | Q8BX22 1x | 1.45E-06 | 0 | 1 | 1 | 10 | Q8BX22 | Sal-like pro | Q8BX22 [7 | 1 | 3689.618 | 0 | -2.6 | 0.000188 | 1.09 | 171.8 | 28.2 | 4.12 | 25.58 | | High | High | High | High | High | High | High | High | High | High | High | 0.000706 | 0.006205 | 4.37 |
| TRUE | High | VAYIPDETA | [K].VAYIPD | 1xPhosphc | Q9CPN8 1x | 4.29E-10 | 0 | 1 | 1 | 4 | Q9CPN8 | Insulin-like | Q9CPN8 [1 | 0 | 2407.186 | 0 | -2.21 | 0.00014 | 0.64 | 164.5 | 35.5 | 2.62 | 19.55 | | High | High | High | High | High | High | High | High | High | High | High | 0 | 1.67E-11 | 5.39 |
| TRUE | High | SPSPDSVAS | [R].SPSPDS | 1xPhosphc | F6RJ39 1xP | 0.000235 | 0 | 1 | 3 | 1 | F6RJ39 | Apoptotic | F6RJ39 [53 | 0 | 2309.154 | 0 | -2.59 | 0.000414 | 0.62 | 170.8 | 29.2 | 7.94 | 22.48 | | High | High | High | High | High | High | High | High | High | High | High | 0 | 4.69E-05 | 1.82 |
| TRUE | High | THTGERPN | [R].THTGEI | 2xCarbami | Q8BIF9 1xF | 0.015741 | 0.001124 | 1 | 1 | 1 | Q8BIF9 | Zinc finger | Q8BIF9 [87 | 0 | 2242.015 | 0 | -2.24 | 0.001802 | 1.55 | 164.7 | 35.3 | 8.86 | 32.42 | | High | High | High | High | High | High | High | High | High | High | High | 0.000585 | 0.005214 | 1.62 |
| TRUE | High | DGSPVQPS | [K].DGSPV | 1xPhosphc | D0VLQ4 1x | 0.000908 | 0 | 1 | 3 | 1 | D0VLQ4 | Tet oncoge | D0VLQ4 [8 | 0 | 2010.075 | 0 | -3.11 | 0.000163 | 0.17 | 179.2 | 20.8 | 2.29 | 25.87 | | High | High | High | High | High | High | High | High | High | High | High | 0 | 0.000214 | 1.84 |
| TRUE | High | YSWGTVE\ | [R].YSWGT | 1xCarbami | P42209 1x | 8.87E-05 | 0 | 1 | 3 | 1 | P42209 | Septin-1 O | P42209 [24 | 0 | 2782.265 | 0 | -3.1 | 0.000727 | 0.55 | 177.3 | 22.7 | 8 | 35.52 | | High | High | High | High | High | High | High | High | High | High | High | 0 | 1.57E-05 | 3.08 |
| TRUE | High | DIPSDTQSI | [K].DIPSDT | 2xPhosphc | Q3U2W2 2 | 0.001278 | 0 | 1 | 3 | 2 | Q3U2W2 | MYB bindir | Q3U2W2 [ | 0 | 1992.933 | 0 | -2.55 | 0.000471 | 0.74 | 170.8 | 29.2 | 4.13 | 28.73 | | High | High | High | High | High | High | High | High | High | High | High | 0 | 0.000314 | 3.33 |
| TRUE | High | SPQAQDTL | [R].SPQAQ | 1xCarbami | A2ARP8 1x | 4.35E-08 | 0 | 1 | 3 | 2 | A2ARP8 | Microtubu | A2ARP8 [1 | 0 | 3451.718 | 0 | -2.75 | 0.001298 | 0.42 | 173.9 | 26.1 | 13.59 | 23.82 | | High | High | High | High | High | High | High | High | High | High | High | 0 | 3.01E-09 | 3.96 |
| TRUE | High | EESGVLEIL | [K].EESGVL | 1xPhosphc | B2RQ54 1x | 0.001424 | 5.66E-05 | 1 | 1 | 2 | B2RQ54 | Developme | B2RQ54 [1 | 0 | 1940.039 | 0 | -2.58 | 0.000209 | 0.96 | 172.5 | 27.5 | 7.4 | 33.27 | | High | High | High | High | High | High | High | High | High | High | High | 2.58E-05 | 0.000355 | 2.94 |
| TRUE | High | LNTSDFQK | [R].LNTSDF | 1xPhosphc | E9QKI4 1xF | 0.000699 | 0 | 1 | 3 | 2 | E9QKI4 | Proline-rich | E9QKI4 [31 | 0 | 1490.766 | 0 | -2.19 | 0.001613 | 2.07 | 164.3 | 35.7 | 9.69 | 37.66 | | High | High | High | High | High | High | High | High | High | High | High | 0 | 0.000211 | 2.87 |
| TRUE | High | ASCVVIDSE | [K].ASCVVI | 1xCarbami | A0A1D9BZ | 6.43E-06 | 0 | 1 | 1 | 4 | A0A1D9BZ | GCNA OS= | A0A1D9BZ | 0 | 2676.104 | 0 | -2.05 | 0.000363 | 0.48 | 164.8 | 35.2 | 9.45 | 12.86 | | High | High | High | High | High | High | High | High | High | High | High | 0 | 4.97E-06 | 3.35 |
| TRUE | High | SETILSPPPI | [K].SETILSF | 1xPhosphc | Q8CCK0 1x | 0.103392 | 0.008249 | 1 | 3 | 1 | Q8CCK0 | Core histor | Q8CCK0 [1 | 0 | 1735.928 | 0 | -2.03 | 0.000492 | 0.93 | 156.6 | 43.4 | 7.96 | 11.9 | | High | High | High | High | High | High | High | High | High | High | High | 0.004164 | 0.04492 | 1.53 |
| TRUE | High | NDQQPTAS | [K].NDQQF | 1xPhosphc | E9PWS4 1x | 0.006834 | 0.00062 | 1 | 4 | 2 | E9PWS4 | Expressed | E9PWS4 [6 | 0 | 3159.468 | 0 | -5.03 | 0.000171 | 0.02 | 193.8 | 6.2 | 3.47 | 59.43 | | High | High | High | High | High | High | High | High | High | High | High | 0.000585 | 0.005408 | 2.22 |
| TRUE | High | RPSESDKEI | [R].RPSESD | 1xPhosphc | B2RRE0 1x | 0.008538 | 0.000736 | 1 | 3 | 4 | B2RRE0 | A kinase (P | B2RRE0 [6 | 1 | 2329.189 | 0 | -2.39 | 0.000213 | 0.77 | 167.8 | 32.2 | 5.64 | 23.93 | | High | High | High | High | High | High | High | High | High | High | High | 0.000327 | 0.002625 | 2.37 |
| TRUE | High | GAGPCSPG | [R].GAGPC | 1xCarbami | Q8R071 1x | 0.001077 | 0 | 1 | 1 | 1 | Q8R071 | Inositol-tri | Q8R071 [1 | 0 | 1409.645 | 0 | -3.21 | 0.000323 | 0.34 | 181 | 19 | 5.54 | 32.14 | | High | High | High | High | High | High | High | High | High | High | High | 0 | 0.000259 | 2.89 |
| TRUE | High | ASCVVIDSE | [K].ASCVVI | 1xCarbami | A0A1D9BZ | 1.48E-10 | 0 | 1 | 1 | 10 | A0A1D9BZ | GCNA OS= | A0A1D9BZ | 0 | 2596.137 | 0 | -2.44 | 0.000465 | 0.24 | 169.9 | 30.1 | 11.84 | 19.98 | | High | High | High | High | High | High | High | High | High | High | High | 0 | 5.05E-12 | 5.16 |
| TRUE | High | ASGSGSCN | [R].ASGSG | 2xCarbami | S4R270 2x | 0.0016 | 5.66E-05 | 1 | 4 | 2 | S4R270 | Bridging in | S4R270 [43 | 0 | 2639.047 | 0 | -2.41 | 0.000154 | 0.56 | 169.7 | 30.3 | 5.86 | 19.03 | | High | High | High | High | High | High | High | High | High | High | High | 0.000399 | 0.003471 | 3.21 |
| TRUE | High | SPEESTVQ\ | [K].SPEEST | 1xPhosphc | Q09XV5 1x | 3.27E-06 | 0 | 1 | 2 | 4 | Q09XV5 | Chromodo | Q09XV5 [2 | 0 | 2509.32 | 0 | -3.03 | 0.000181 | 0.38 | 179.5 | 20.5 | 4.29 | 22.78 | | High | High | High | High | High | High | High | High | High | High | High | 0 | 3.83E-07 | 3.5 |
| TRUE | High | SPEESHEA\ | [R].SPEESH | 1xCarbami | Q7TN89 1x | 4.39E-05 | 0 | 1 | 1 | 1 | Q7TN89 | GTPase ER | Q7TN89 [2 | 0 | 2096.951 | 0 | -3.69 | 0.000202 | 0.12 | 185.6 | 14.4 | 9.6 | 38.31 | | High | High | High | High | High | High | High | High | High | High | High | 0 | 7.12E-06 | 4.12 |
| TRUE | High | QPSSPGPQ | [R].QPSSPC | 1xPhosphc | Q61234 1x | 0.013909 | 0.001011 | 1 | 3 | 2 | Q61234 | Alpha-1-sy | Q61234 [1 | 0 | 1359.662 | 0 | -2.23 | 0.000285 | 0.7 | 166 | 34 | 4.71 | 19.11 | | High | High | High | High | High | High | High | High | High | High | High | 0.000555 | 0.004903 | 1.62 |

| | | | | | | | | | | | | | | | | | | | | | | | | | | | | | | | | | | | | | | |
|---|---|---|---|---|---|---|---|---|---|---|---|---|---|---|---|---|---|---|---|---|---|---|---|---|---|---|---|---|---|---|---|---|---|---|---|---|---|---|
| TRUE | High | ASGSGSCN | [R].ASGSG | 2xCarbami | S4R270 1x | 1.06E-07 | 0 | 1 | 4 | 4 | S4R270 | Bridging in | S4R270 [43 | 0 | 2559.08 | 0 | -2.6 | 0.000158 | 0.22 | 170.9 | 29.1 | 3.26 | 21.52 | | High | High | High | High | High | High | High | High | High | High | High | 0 | 8.21E-09 | 4.47 |
| TRUE | High | GSTGSLK | [R].GSTGSI | 1xPhospho | U5LK19 1x | 0.034878 | 0.002269 | 2 | 17 | 1 | U5LK19; E | Marshalin- | U5LK19 [3 | 0 | 1187.644 | 0 | -2.27 | 0.001232 | 1.89 | 168.7 | 31.3 | 11.9 | 30.8 | Shared | High | High | High | High | High | High | High | High | High | High | High | 0.001183 | 0.01273 | 0.94 |
| TRUE | High | LQASAPSLT | [K].LQASAI | 1xPhospho | P28231 1x | 7.68E-05 | 0 | 1 | 1 | 4 | P28231 | Gap junctic | P28231 [26 | 0 | 1406.749 | 0 | -2.71 | 0.000181 | 0.23 | 173.5 | 26.5 | 4.92 | 9.32 | | High | High | High | High | High | High | High | High | High | High | High | 0.000183 | 0.00126 | 3.07 |
| TRUE | High | LQASAPSLT | [K].LQASAI | 2xPhospho | P28231 2x | 0.002172 | 5.66E-05 | 1 | 1 | 3 | P28231 | Gap junctic | P28231 [26 | 0 | 1486.716 | 0 | -2.48 | 0.00025 | 0.2 | 170.6 | 29.4 | 7.16 | 17.71 | | High | High | High | High | High | High | High | High | High | High | High | 4.94E-05 | 0.000572 | 3.01 |
| TRUE | High | QSLGKPDC | [K].QSLGKI | 1xPhospho | E9PXD2 1x | 0.022182 | 0.001542 | 1 | 7 | 1 | E9PXD2 | Rho GTPas | E9PXD2 [1 | 0 | 1767.997 | 0 | -2.83 | 0.000571 | 0.32 | 175.7 | 24.3 | 8.34 | 21.63 | | High | High | High | High | High | High | High | High | High | High | High | 0.000814 | 0.007632 | 1.25 |
| TRUE | High | SSTLSSGSV | [R].SSTLSS | 1xPhospho | P70324 1x | 5.97E-05 | 0 | 1 | 4 | 1 | P70324 | T-box tran | P70324 [69 | 0 | 1874.988 | 0 | -2.45 | 0.000158 | 0.2 | 168.8 | 31.2 | 3.53 | 22.09 | | High | High | High | High | High | High | High | High | High | High | High | 0 | 1.01E-05 | 2.26 |
| TRUE | High | NDPNWAN | [K].NDPNW | 1xPhospho | Q02739 1x | 2.51E-05 | 0 | 1 | 2 | 2 | Q02739 | Gap junctic | Q02739 [2 | 0 | 1767.847 | 0 | -2.79 | 0.000158 | 0.2 | 174.7 | 25.3 | 3.05 | 13.07 | | High | High | High | High | High | High | High | High | High | High | High | 0 | 3.78E-06 | 4.1 |
| TRUE | High | SPDSFATN | [K].SPDSFA | 1xPhospho | D0VLQ4 1x | 0.000334 | 0 | 1 | 3 | 2 | D0VLQ4 | Tet oncoge | D0VLQ4 [9 | 0 | 1930.009 | 0 | -2.81 | 0.000202 | 0.55 | 175.1 | 24.9 | 4.33 | 20.64 | | High | High | High | High | High | High | High | High | High | High | High | 2.58E-05 | 0.000494 | 3.34 |
| TRUE | High | NDANTNSA | [K].NDANT | 1xPhospho | Q3UYG1 1 | 0.001077 | 0 | 1 | 1 | 1 | Q3UYG1 | Coiled-coil | Q3UYG1 [8 | 0 | 2773.287 | 0 | -2.2 | 0.001693 | 1.2 | 163.9 | 36.1 | 13.9 | 12.22 | | High | High | High | High | High | High | High | High | High | High | High | 0 | 0.000258 | 3.02 |
| TRUE | High | AALSHQLG | [R].AALSH | 1xPhospho | A2AUZ7 1x | 6.95E-05 | 0 | 2 | 4 | 1 | A2AUZ7; Q | Sal-like pro | A2AUZ7 [8 | 0 | 1847.978 | 0 | -2.41 | 0.000165 | 0.75 | 168.2 | 31.8 | 7.77 | 24 | Shared | High | High | High | High | High | High | High | High | High | High | High | 0 | 1.2E-05 | 1.69 |
| TRUE | High | SNTSSVQS | [R].SNTSSV | 1xPhospho | B2KFD2 1x | 1.13E-06 | 0 | 1 | 5 | 1 | B2KFD2 | 2810474O1 | B2KFD2 [1 | 0 | 1887.947 | 0 | -2.06 | 0.000191 | 0.4 | 160.7 | 39.3 | 2.76 | 10.57 | | High | High | High | High | High | High | High | High | High | High | High | 0 | 1.17E-07 | 1.9 |
| TRUE | High | GAPNALAE | [K].GAPNA | 1xPhospho | P70670 1x | 4.3E-07 | 0 | 1 | 1 | 1 | P70670 | Nascent pc | P70670 [12 | 0 | 1847.967 | 0 | -2.14 | 0.001893 | 0.98 | 158.3 | 41.7 | 16.83 | 30.24 | | High | High | High | High | High | High | High | High | High | High | High | 0 | 3.93E-08 | 3.91 |
| TRUE | High | NPSSQVSG | [K].NPSSQ | 1xPhospho | Q9D0D3 1 | 7.84E-06 | 0 | 1 | 2 | 2 | Q9D0D3 | Poly(A) RN | Q9D0D3 [1 | 0 | 2940.45 | 0 | -2.24 | 0.003092 | 1.79 | 162.6 | 37.4 | 8.69 | 49.92 | | High | High | High | High | High | High | High | High | High | High | High | 0 | 1.03E-06 | 3.43 |
| TRUE | High | QSHPTSAA | [K].QSHPT | 1xPhospho | A2AJ72 1x | 6.45E-07 | 0 | 1 | 6 | 1 | A2AJ72 | Far upstrea | A2AJ72 [52 | 0 | 3221.424 | 0 | -2.14 | 0.001562 | 0.39 | 166 | 34 | 16.09 | 11.37 | | High | High | High | High | High | High | High | High | High | High | High | 0 | 6.19E-08 | 4.87 |
| TRUE | High | GAQTSPVP | [K].GAQTS | 1xPhospho | Q80TS5 1x | 0.007881 | 0.000736 | 1 | 4 | 2 | Q80TS5 | Zinc finger | Q80TS5 [1 | 0 | 1221.619 | 0 | -2.2 | 0.000283 | 1.58 | 164.3 | 35.7 | 3.76 | 23.28 | | High | High | High | High | High | High | High | High | High | High | High | 0.000538 | 0.004423 | 1.38 |
| TRUE | High | RPSESDK | [R].RPSESD | 1xPhospho | B2RRE0 1x | 0.005717 | 0.00062 | 1 | 3 | 2 | B2RRE0 | A kinase (P | B2RRE0 [6 | 0 | 1356.692 | 0 | -2.23 | 0.00118 | 2.35 | 167.3 | 32.7 | 8.76 | 30.97 | | High | High | High | High | High | High | High | High | High | High | High | 0.002117 | 0.02196 | 1.54 |
| TRUE | High | GAQPGRHS | [R].GAQPG | 1xCarbami | Q91XC0 1x | 0.015741 | 0.001124 | 1 | 1 | 2 | Q91XC0 | LIM domai | Q91XC0 [2 | 1 | 2297.044 | 0 | -2.12 | 0.000821 | 1.19 | 161 | 39 | 10.49 | 20.42 | | High | High | High | High | High | High | High | High | High | High | High | 0.000585 | 0.005191 | 1.96 |
| TRUE | High | LPSPGLQG | [K].LPSPGL | 1xPhospho | Q0VGT2 1 | 0.030632 | 0.001857 | 1 | 3 | 2 | Q0VGT2 | Zinc finger | Q0VGT2 [1 | 0 | 1370.714 | 0 | -3.3 | 0.000499 | 0.37 | 181.2 | 18.8 | 9.55 | 26.66 | | High | High | High | High | High | High | High | High | High | High | High | 0.001011 | 0.01107 | 1.68 |
| TRUE | High | GSLPPAALE | [K].GSLPPA | 1xPhospho | Q5SVI1 1x | 6.31E-09 | 0 | 1 | 4 | 3 | Q5SVI1 | Calcium/ca | Q5SVI1 [34 | 0 | 2986.418 | 0 | -2.23 | 0.000334 | 1.15 | 169.9 | 30.1 | 10.5 | 26.19 | | High | High | High | High | High | High | High | High | High | High | High | 0 | 5.95E-10 | 4.76 |
| TRUE | High | TLYSFFSPT | [K].TLYSFF | 1xPhospho | P97931 1x | 0.000779 | 0 | 1 | 1 | 1 | P97931 | Uracil-DNA | P97931 [6- | 0 | 1984.023 | 0 | -2.66 | 0.000506 | 0.81 | 173.5 | 26.5 | 10.89 | 26.48 | | High | High | High | High | High | High | High | High | High | High | High | 0 | 0.000181 | 2.82 |
| TRUE | High | EGPRPLSQ | [R].EGPRPI | 1xPhospho | A2TJV2 1x | 8.2E-06 | 0 | 1 | 1 | 2 | A2TJV2 | Paralemmi | A2TJV2 [13 | 0 | 2906.437 | 0 | -3.62 | 0.000223 | 0.18 | 184.6 | 15.4 | 8 | 34.65 | | High | High | High | High | High | High | High | High | High | High | High | 0 | 1.08E-06 | 4.34 |
| TRUE | High | EGPRPLSQ | [R].EGPRPI | 1xPhospho | A2TJV2 1x | 7.68E-05 | 0 | 1 | 1 | 1 | A2TJV2 | Paralemmi | A2TJV2 [13 | 0 | 2922.432 | 0 | -2.26 | 0.000687 | 1.15 | 165.9 | 34.1 | 7.59 | 20.1 | | High | High | High | High | High | High | High | High | High | High | High | 0 | 1.34E-05 | 2.15 |
| TRUE | High | LPAFSPPSP | [R].LPAFSP | 2xPhospho | Q6J1H4 2x | 1.32E-07 | 0 | 1 | 1 | 20 | Q6J1H4 | Undifferen | Q6J1H4 [8- | 0 | 2140.992 | 0 | -2.28 | 0.002085 | 1.06 | 167 | 33 | 15.69 | 22.32 | | High | High | High | High | High | High | High | High | High | High | High | 0 | 0.00024 | 4.28 |
| TRUE | High | FTIDSDAIS | [R].FTIDSD | 1xPhospho | G3UWG2 1 | 1.8E-06 | 0 | 1 | 8 | 2 | G3UWG2 | T-cell lymp | G3UWG2 [ | 0 | 2106.041 | 0 | -2.45 | 0.000223 | 0.66 | 169.4 | 30.6 | 2.54 | 19.73 | | High | High | High | High | High | High | High | High | High | High | High | 0 | 1.98E-07 | 4.58 |
| TRUE | High | SPSDDGSG | [R].SPSDD | 1xPhospho | P70218 1x | 0.0006 | 0 | 1 | 4 | 3 | P70218 | Mitogen-a | P70218 [4C | 1 | 2593.246 | 0 | -2.32 | 0.000499 | 0.76 | 166.4 | 33.6 | 6.85 | 17.65 | | High | High | High | High | High | High | High | High | High | High | High | 0 | 0.000134 | 2.72 |
| TRUE | High | FSSSTTK | [R].FSSSTT | 1xPhospho | Q6J1H4 1x | 0.01319 | 0.000889 | 2 | 3 | 3 | Q6J1H4; Q | Undifferen | Q6J1H4 [2 | 0 | 1295.665 | 0 | -2.54 | 0.000495 | 0.91 | 170.5 | 29.5 | 9.48 | 17.12 | Shared | High | High | High | High | High | High | High | High | High | High | High | 0.000467 | 0.004285 | 1.57 |
| TRUE | High | LPAFSPPSP | [R].LPAFSP | 1xPhospho | Q6J1H4 1x | 0.000751 | 0 | 1 | 1 | 5 | Q6J1H4 | Undifferen | Q6J1H4 [8- | 0 | 2061.026 | 0 | -2.47 | 0.001663 | 1.32 | 169.5 | 30.5 | 14.3 | 28.32 | | High | High | High | High | High | High | High | High | High | High | High | 0 | 0.000174 | 3.31 |
| TRUE | High | QSIQKPPSF | [R].QSIQKF | 1xPhospho | O70238 1x | 7.16E-06 | 0 | 1 | 2 | 2 | O70238 | Homeobox | O70238 [9 | 0 | 2392.273 | 0 | -3.26 | 0.000605 | 0.46 | 183.1 | 16.9 | 9.82 | 35.41 | | High | High | High | High | High | High | High | High | High | High | High | 0 | 9.27E-07 | 2.57 |
| TRUE | High | FGLSSPLPS | [R].FGLSSP | 2xPhospho | Q8K120 2x | 0.064323 | 0.004664 | 1 | 2 | 1 | Q8K120 | Nuclear fa | Q8K120 [2 | 0 | 1546.727 | 0 | -2.65 | 0.003024 | 0.52 | 170.6 | 29.4 | 23.48 | 15.13 | | High | High | High | High | High | High | High | High | High | High | High | 0.002403 | 0.02586 | 1.91 |
| TRUE | High | EGPVGGES | [R].EGPVG | 1xPhospho | Q3UHU5 1 | 4.83E-07 | 0 | 1 | 5 | 2 | Q3UHU5 | Microtubu | Q3UHU5 [ | 0 | 2266.983 | 0 | -2.14 | 0.000914 | 1.66 | 162.6 | 37.4 | 7.35 | 25.75 | | High | High | High | High | High | High | High | High | High | High | High | 0 | 4.49E-08 | 3.07 |
| TRUE | High | LDHALSSPS | [R].LDHALS | 1xCarbami | Q8BGT8 2x | 0.000674 | 0 | 1 | 2 | 2 | Q8BGT8 | Phytanoyl- | Q8BGT8 [6 | 0 | 2388.096 | 0 | -2.45 | 0.000472 | 0.86 | 170.2 | 29.8 | 7.63 | 23.28 | | High | High | High | High | High | High | High | High | High | High | High | 0 | 0.000153 | 2.45 |
| TRUE | High | SSSSTGSTK | [R].SSSSTG | 1xPhospho | A0A140LJB | 0.014158 | 0.001011 | 1 | 12 | 1 | A0A140LJB | Espin OS=N | A0A140LJB | 0 | 1379.682 | 0 | -2.09 | 0.001617 | 1.23 | 165.8 | 34.2 | 17.26 | 28.5 | | High | High | High | High | High | High | High | High | High | High | High | 0.000538 | 0.00462 | 1.32 |
| TRUE | High | ASSEGAEQ | [R].ASSEGA | 1xPhospho | S4R270 1x | 1.6E-05 | 0 | 1 | 4 | 1 | S4R270 | Bridging in | S4R270 [4C | 0 | 1531.741 | 0 | -3.69 | 0.000188 | 0.18 | 185.5 | 14.5 | 7.59 | 41.08 | | High | High | High | High | High | High | High | High | High | High | High | 0 | 2.28E-06 | 2.63 |
| TRUE | High | ASPTQLNA | [R].ASPTQI | 1xPhospho | Q3UNW5 1 | 4.12E-05 | 0 | 1 | 1 | 2 | Q3UNW5 | Transcripti | Q3UNW5 [ | 0 | 2441.252 | 0 | -4.67 | 0.000233 | 0.01 | 192 | 8 | 4.69 | 62.9 | | High | High | High | High | High | High | High | High | High | High | High | 0 | 2.04E-05 | 3.77 |
| TRUE | High | FVPVVTSPV | [R].FVPVVT | 1xPhospho | F8WJ40 1x | 0.002334 | 0.000109 | 1 | 4 | 1 | F8WJ40 | Histone-lys | F8WJ40 [7 | 0 | 1610.932 | 0 | -3.79 | 0.000222 | 0.11 | 186.4 | 13.6 | 6.99 | 44.33 | | High | High | High | High | High | High | High | High | High | High | High | 4.94E-05 | 0.000615 | 2.89 |
| TRUE | High | TLETLDLET | [K].TLETLD | 1xPhospho | Q9CWR8 1 | 4.31E-08 | 0 | 1 | 1 | 8 | Q9CWR8 | DNA (cytos | Q9CWR8 [ | 0 | 3544.687 | 0 | -3.04 | 0.000928 | 0.84 | 172.8 | 27.2 | 16.28 | 43.5 | | High | High | High | High | High | High | High | High | High | High | High | 0 | 2.97E-09 | 5.02 |
| TRUE | High | QSATEIPLP | [R].QSATEI | 1xPhospho | P42209 1x | 0.000438 | 0 | 1 | 3 | 6 | P42209 | Septin-1 O | P42209 [31 | 0 | 2508.307 | 0 | -3.61 | 0.000427 | 0.31 | 184.9 | 15.1 | 6.9 | 38.5 | | High | High | High | High | High | High | High | High | High | High | High | 0 | 9.47E-05 | 2.94 |
| TRUE | High | QSATEIPLP | [R].QSATEI | 1xPhospho | P42209 1x | 0.000205 | 0 | 1 | 3 | 1 | P42209 | Septin-1 O | P42209 [31 | 0 | 2492.312 | 0 | -3.75 | 0.000297 | 0.26 | 184.9 | 15.1 | 5.32 | 42.66 | | High | High | High | High | High | High | High | High | High | High | High | 0 | 4.05E-05 | 3.69 |
| TRUE | High | SPSVEHDE | [R].SPSVEH | 1xPhospho | Q8CI12 1x | 0.000365 | 0 | 1 | 3 | 7 | Q8CI12 | Smoothelir | Q8CI12 [12 | 0 | 1978.896 | 0 | -2.15 | 0.000187 | 0.46 | 163.3 | 36.7 | 4.37 | 20.6 | | High | High | High | High | High | High | High | High | High | High | High | 4.94E-05 | 0.000607 | 2.8 |
| TRUE | High | LQISTPDIK | [R].LQISTP | 1xPhospho | P97431 1x | 0.051258 | 0.003745 | 1 | 2 | 1 | P97431 | Interferon | P97431 [42 | 0 | 1552.875 | 0 | -2.49 | 0.000676 | 0.22 | 169.4 | 30.6 | 8.49 | 23.97 | | High | High | High | High | High | High | High | High | High | High | High | 0.00196 | 0.01979 | 2.05 |
| TRUE | High | AQSSEVPLS | [K].AQSSE | 1xPhospho | V9GWU7 1 | 0.006195 | 0.00062 | 1 | 3 | 2 | V9GWU7 | Serine/thre | V9GWU7 [ | 0 | 1734.863 | 0 | -2.23 | 0.00022 | 1.45 | 166.1 | 33.9 | 3.66 | 23.3 | | High | High | High | High | High | High | High | High | High | High | High | 0.000275 | 0.001829 | 3.65 |
| TRUE | High | SSSNVNGG | [R].SSSNVN | 1xPhospho | Q6PGN3 1 | 4.8E-05 | 0 | 1 | 9 | 5 | Q6PGN3 | Serine/thre | Q6PGN3 [3 | 0 | 1640.748 | 0 | -2.22 | 0.000196 | 1.12 | 165 | 35 | 5.04 | 17.75 | | High | High | High | High | High | High | High | High | High | High | High | 0 | 7.89E-06 | 3.26 |
| TRUE | High | FVPSPPPN | [R].FVPSPP | 1xCarbami | Q8BMQ3 1 | 0.0008 | 0 | 1 | 5 | 1 | Q8BMQ3 | Zinc finger | Q8BMQ3 [ | 0 | 2044.038 | 0 | -2.27 | 0.001559 | 1.56 | 165.7 | 34.3 | 11.54 | 31.36 | | High | High | High | High | High | High | High | High | High | High | High | 0 | 0.000186 | 2.83 |
| TRUE | High | LDSENSPYL | [R].LDSENS | 1xPhospho | G5E8P8 1x | 1.44E-06 | 0 | 1 | 6 | 1 | G5E8P8 | Estrogen re | G5E8P8 [2 | 0 | 2493.304 | 0 | -3.74 | 0.000377 | 0.3 | 186 | 14 | 5.34 | 44.95 | | High | High | High | High | High | High | High | High | High | High | High | 0 | 1.53E-07 | 3.55 |
| TRUE | High | LDSENSPYL | [R].LDSENS | 1xPhospho | G5E8P8 1x | 0.003282 | 0.00016 | 1 | 6 | 1 | G5E8P8 | Estrogen re | G5E8P8 [2 | 0 | 2264.141 | 0 | -2.15 | 0.002951 | 0.63 | 163.2 | 36.8 | 12.72 | 28.02 | | High | High | High | High | High | High | High | High | High | High | High | 7.19E-05 | 0.000908 | 2.48 |
| TRUE | High | ADASQSIQ | [R].ADASQ | 1xCarbami | I3PQW4 1x | 6.93E-07 | 0 | 1 | 1 | 1 | I3PQW4 | Uncharacte | I3PQW4 [6 | 0 | 2599.259 | 0 | -2.77 | 0.000209 | 0.49 | 174.6 | 25.4 | 2.11 | 34.03 | | High | High | High | High | High | High | High | High | High | High | High | 0 | 6.71E-08 | 4.03 |
| TRUE | High | DIPSDTQSF | [K].DIPSDT | 1xPhospho | Q3U2W2 1 | 9.62E-05 | 0 | 1 | 3 | 6 | Q3U2W2 | MYB bindir | Q3U2W2 [ | 0 | 1912.967 | 0 | -2.59 | 0.000252 | 0.82 | 171.5 | 28.5 | 2.55 | 26.99 | | High | High | High | High | High | High | High | High | High | High | High | 9.42E-05 | 0.00098 | 3.34 |
| TRUE | High | NGVVTPDH | [K].NGVVT | 1xPhospho | Q6ZPI0 1x | 0.023584 | 0.001578 | 1 | 1 | 1 | Q6ZPI0 | Protein Jad | Q6ZPI0 [65 | 0 | 1561.814 | 0 | -2.18 | 0.00024 | 0.82 | 164.3 | 35.7 | 5.9 | 20.55 | | High | High | High | High | High | High | High | High | High | High | High | 0.000859 | 0.008201 | 1.71 |
| TRUE | High | LQNGSPLA | [K].LQNGS | 1xPhospho | D0VLQ5 1x | 0.024638 | 0.001612 | 1 | 3 | 1 | D0VLQ5 | Tet oncoge | D0VLQ5 [3 | 0 | 2697.376 | 0 | -2.69 | 0.000618 | 0.3 | 173.2 | 26.8 | 12.43 | 13.56 | | High | High | High | High | High | High | High | High | High | High | High | 0.000859 | 0.00861 | 1.98 |
| TRUE | High | LQNGSPLA | [K].LQNGS | 1xDeamida | D0VLQ5 1x | 0.00045 | 0 | 1 | 3 | 3 | D0VLQ5 | Tet oncoge | D0VLQ5 [3 | 0 | 2698.36 | 0 | -2.59 | 0.000154 | 0.29 | 171.4 | 28.6 | 1.81 | 16.04 | | High | High | High | High | High | High | High | High | High | High | High | 0 | 9.75E-05 | 2.8 |
| TRUE | High | SNGDLTPK | [R].SNGDL | 1xDeamida | P28667 1x | 0.009585 | 0.000736 | 1 | 1 | 5 | P28667 | MARCKS-re | P28667 [36 | 0 | 1370.697 | 0 | -2.07 | 0.000209 | 0.55 | 161.5 | 38.5 | 4.6 | 12.93 | | High | High | High | High | High | High | High | High | High | High | High | 0.000801 | 0.00747 | 1.67 |
| TRUE | High | FLEDDTSDI | [R].FLEDDT | 1xPhospho | P54763-2 1 | 5.07E-05 | 0 | 1 | 3 | 2 | P54763-2 | Isoform 2 c | P54763-2 [ | 0 | 2455.168 | 0 | -3.03 | 0.00099 | 0.48 | 179.7 | 20.3 | 14.35 | 36.87 | | High | High | High | High | High | High | High | High | High | High | High | 0 | 8.37E-06 | 2.47 |
| TRUE | High | TNSYGGK | [K].TNSYG | 1xPhospho | Q3KR45 1x | 0.005516 | 0.00062 | 1 | 6 | 1 | Q3KR45 | DNA (cytos | Q3KR45 [4 | 0 | 1264.634 | 0 | -3.41 | 0.000229 | 0.31 | 183.1 | 16.9 | 5.49 | 22.27 | | High | High | High | High | High | High | High | High | High | High | High | 0.000275 | 0.001612 | 1.49 |
| TRUE | High | FVQEHSSSI | [R].FVQEH | 1xPhospho | E9PZ97 1x | 0.001558 | 5.66E-05 | 1 | 2 | 1 | E9PZ97 | Bloom syn | E9PZ97 [10 | 0 | 1640.763 | 0 | -2.43 | 0.000393 | 0.45 | 167.6 | 32.4 | 7.66 | 19.93 | | High | High | High | High | High | High | High | High | High | High | High | 2.58E-05 | 0.000394 | 2.16 |
| TRUE | High | QNSVQNS | [R].QNSVQ | 1xPhospho | Q8BH82 1x | 5.67E-06 | 0 | 1 | 2 | 3 | Q8BH82 | N-acyl-pho | Q8BH82 [2 | 0 | 1715.791 | 0 | -2.23 | 0.000289 | 0.9 | 163.3 | 36.7 | 7.33 | 19.33 | | High | High | High | High | High | High | High | High | High | High | High | 0 | 7.72E-05 | 2.82 |
| TRUE | High | EEHQDISSF | [K].EEHQD | 2xPhospho | Q3U164 2x | 0.027121 | 0.001676 | 1 | 1 | 4 | Q3U164 | Putative ur | Q3U164 [6 | 1 | 3417.674 | 0 | -2.4 | 0.000199 | 1.03 | 170.4 | 29.6 | 4.63 | 29.74 | | High | High | High | High | High | High | High | High | High | High | High | 0.004769 | 0.05199 | 2.95 |
| TRUE | High | DFAPSPR | [R].DFAPSF | 1xPhospho | Q9DAE2 1x | 0.034579 | 0.002233 | 1 | 1 | 1 | Q9DAE2 | Putative ur | Q9DAE2 [2 | 0 | 1098.518 | 0 | -3.45 | 0.000275 | 0.35 | 183.3 | 16.7 | 4.67 | 31.97 | | High | High | High | High | High | High | High | High | High | High | High | 0.001183 | 0.01265 | 1.43 |
| TRUE | High | TISPESVYPI | [K].TISPES | 1xPhospho | Q8CIB9 1x | 3.54E-05 | 0 | 1 | 2 | 1 | Q8CIB9 | N-acetyltra | Q8CIB9 [31 | 0 | 2620.368 | 0 | -2.18 | 0.011068 | 1.49 | 163.9 | 36.1 | 33.48 | 17.69 | | High | High | High | High | High | High | High | High | High | High | High | 0 | 5.58E-06 | 3.76 |
| TRUE | High | RSSEASPLG | [R].RSSEAS | 1xPhospho | Q0VGT2 1 | 0.000442 | 0 | 1 | 2 | 1 | Q0VGT2 | Zinc finger | Q0VGT2 [8 | 1 | 3510.645 | 0 | -2.09 | 0.006722 | 2.84 | 170.2 | 29.8 | 27.25 | 34.18 | | High | High | High | High | High | High | High | High | High | High | High | 0 | 9.53E-05 | 3.15 |
| TRUE | High | DNHSQADI | [R].DNHSQ | 1xPhospho | A2A5V4 1x | 8.3E-07 | 0 | 1 | 5 | 2 | A2A5V4 | SH3 domai | A2A5V4 [2 | 0 | 2361.082 | 0 | -3.34 | 0.000598 | 0.29 | 182 | 18 | 12.93 | 31.47 | | High | High | High | High | High | High | High | High | High | High | High | 0 | 6.6E-07 | 3.4 |
| TRUE | High | VGGSDAED | [K].VGGSD | 2xPhospho | D3Z4R4 2x | 0.000584 | 0 | 1 | 1 | 1 | D3Z4R4 | NK6 home | D3Z4R4 [2 | 0 | 3270.34 | 0 | -2.86 | 0.000479 | 0.82 | 175.6 | 24.4 | 3.93 | 42.44 | | High | High | High | High | High | High | High | High | High | High | High | 0 | 0.00013 | 4.69 |
| TRUE | High | SPQNLSHA | [R].SPQNL | 1xPhospho | Q3U2B5 1 | 0.001373 | 5.66E-05 | 1 | 2 | 1 | Q3U2B5 | Putative ur | Q3U2B5 [1 | 0 | 2091.089 | 0 | -3.04 | 0.000329 | 0.73 | 178.8 | 21.2 | 5.65 | 31.56 | | High | High | High | High | High | High | High | High | High | High | High | 2.58E-05 | 0.000339 | 2.84 |
| TRUE | High | LAPSSNWP | [R].LAPSSN | 1xPhospho | Q3URR7 1 | 5.05E-07 | 0 | 1 | 3 | 1 | Q3URR7 | Zinc finger | Q3URR7 [1 | 0 | 2701.289 | 0 | -3.01 | 0.000217 | 0.14 | 177.1 | 22.9 | 6.33 | 31.17 | | High | High | High | High | High | High | High | High | High | High | High | 0 | 4.74E-08 | 5.37 |
| TRUE | High | DQVATSES | [K].DQVAT | 1xPhospho | B1AU75 1x | 1.96E-07 | 0 | 1 | 2 | 2 | B1AU75 | Nuclear au | B1AU75 [3 | 0 | 1801.899 | 0 | -2.21 | 0.000337 | 0.94 | 165 | 35 | 4.05 | 25.13 | | High | High | High | High | High | High | High | High | High | High | High | 0 | 1.63E-08 | 3.9 |
| TRUE | High | DFASPSSLK | [K].DFASPS | 1xPhospho | F6ZS36 1x | 0.08727 | 0.006537 | 1 | 2 | 1 | F6ZS36 | Uncharacte | F6ZS36 [25 | 0 | 1489.77 | 0 | -2.62 | 0.000366 | 0.9 | 173.3 | 26.7 | 4.66 | 35.04 | | High | High | High | High | High | High | High | High | High | High | High | 0.003346 | 0.03666 | 2.3 |
| TRUE | High | QRESVGGS | [K].QRESV | 1xPhospho | Q9WV69-2 | 0.008313 | 0.000736 | 2 | 5 | 1 | Q9WV69-2 | Isoform 2 c | Q9WV69-2 | 1 | 1797.926 | 0 | -2.08 | 0.000294 | 0.35 | 161.5 | 38.5 | 7.68 | 3.5 | Shared | High | High | High | High | High | High | High | High | High | High | High | 0.000327 | 0.002543 | 2.04 |
| TRUE | High | RSSEASQCS | [R].RSSEAS | 1xCarbami | Q0VGT2 1 | 0.028575 | 0.001829 | 1 | 2 | 2 | Q0VGT2 | Zinc finger | Q0VGT2 [8 | 1 | 3170.525 | 0 | -3.8 | 0.003713 | 0.25 | 185.5 | 14.5 | 25.6 | 60.36 | | High | High | High | High | High | High | High | High | High | High | High | 0.000984 | 0.01016 | 3.43 |
| TRUE | High | TLSGEEEAE | [R].TLSGEE | 1xPhospho | Q9D6W8 1 | 0.000118 | 0 | 1 | 1 | 1 | Q9D6W8 | BLOC-1-rel | Q9D6W8 [ | 0 | 1945.895 | 0 | -2.36 | 0.0006 | 0.91 | 168.9 | 31.1 | 5.86 | 47.28 | | High | High | High | High | High | High | High | High | High | High | High | 0 | 2.18E-05 | 2.9 |
| TRUE | High | AAPSSQSP | [K].AAPSSC | 1xCarbami | B2RWW9 1 | 2.43E-07 | 0 | 1 | 2 | 1 | B2RWW9 | Absent in r | B2RWW9 [ | 0 | 1993.982 | 0 | -2.5 | 0.00041 | 0.63 | 171.8 | 28.2 | 9.56 | 15.6 | | High | High | High | High | High | High | High | High | High | High | High | 0 | 2.07E-08 | 3.75 |
| TRUE | High | DERSPQNL | [K].DERSPC | 1xPhospho | Q3U2B5 1 | 0.000705 | 0 | 1 | 2 | 3 | Q3U2B5 | Putative ur | Q3U2B5 [1 | 1 | 2491.259 | 0 | -2.97 | 0.000162 | 0.36 | 177.4 | 22.6 | 3.69 | 22.01 | | High | High | High | High | High | High | High | High | High | High | High | 7.19E-05 | 0.00089 | 2.56 |
| TRUE | High | SSSLPSYGR | [K].SSSLPS | 1xPhospho | Q9WV69-2 | 0.072258 | 0.005346 | 2 | 5 | 2 | Q9WV69-2 | Isoform 2 c | Q9WV69-2 | 0 | 1262.598 | 0 | -4.19 | 0.000154 | 0.03 | 189.6 | 10.4 | 3.84 | 32.49 | Shared | High | High | High | High | High | High | High | High | High | High | High | 0.002855 | 0.02962 | 0.96 |
| TRUE | High | LPSALLPQT | [R].LPSALL | 1xPhospho | Q8BX22 1x | 0.000253 | 0 | 1 | 4 | 3 | Q8BX22 | Sal-like pro | Q8BX22 [3 | 0 | 3432.88 | 0 | -2.67 | 0.001475 | 0.43 | 172.8 | 27.2 | 14.67 | 13.51 | | High | High | High | High | High | High | High | High | High | High | High | 0 | 0.000113 | 3.15 |
| TRUE | High | LEEAQQVD | [R].LEEAQ | 1xPhospho | Q62315 1x | 3.23E-05 | 0 | 1 | 5 | 2 | Q62315 | Protein Jur | Q62315 [4 | 0 | 2461.331 | 0 | -2.05 | 0.001125 | 0.6 | 161.3 | 38.7 | 9.16 | 21.2 | | High | High | High | High | High | High | High | High | High | High | High | 0.000246 | 0.001489 | 3.57 |
| TRUE | High | AALTNSLSF | [RK].AALTN | 1xDeamida | Q3TJW2 1 | 0.000483 | 0 | 1 | 3 | 2 | Q3TJW2 | F-box only | Q3TJW2 [1 | 0 | 1639.907 | 0 | -2.37 | 0.000798 | 1.37 | 167.7 | 32.3 | 11.15 | 21.49 | | High | High | High | High | High | High | High | High | High | High | High | 0 | 0.000109 | 1.93 |
| TRUE | High | DRGSPPPSI | [K].DRGSPI | 1xPhospho | Q8BFY7 1x | 0.000573 | 0 | 2 | 3 | 1 | Q8BFY7; Q | Protein PIN | Q8BFY7 [1 | 1 | 1806.952 | 0 | -2.38 | 0.000153 | 0.17 | 168.7 | 31.3 | 4.06 | 12.77 | Shared | High | High | High | High | High | High | High | High | High | High | High | 0 | 0.000128 | 2.86 |
| TRUE | High | RQDEDYDE | [K].RQDED | 1xDeamida | Q8BKC5 1x | 3.44E-05 | 0 | 1 | 5 | 10 | Q8BKC5 | Importin-5 | Q8BKC5 [8 | 1 | 3842.706 | 0 | -2.43 | 0.001801 | 0.2 | 168.7 | 31.3 | 13.47 | 44.26 | | High | High | High | High | High | High | High | High | High | High | High | 0 | 5.41E-06 | 6.15 |
| TRUE | High | SPDATETM | [R].SPDATE | 1xCarbami | Q8BX22 2x | 0.054383 | 0.003951 | 1 | 1 | 1 | Q8BX22 | Sal-like pro | Q8BX22 [7 | 0 | 3117.371 | 0 | -2.18 | 0.000527 | 0.92 | 164.4 | 35.6 | 6.16 | 20.68 | | High | High | High | High | High | High | High | High | High | High | High | 0.002059 | 0.02128 | 2.75 |
| TRUE | High | DLPDGHAV | [K].DLPDGI | 1xPhospho | U5LK19 1x | 0.005228 | 0.000551 | 1 | 9 | 1 | U5LK19 | Marshalin- | U5LK19 [3 | 0 | 1472.71 | 0 | -2.5 | 0.000154 | 0.52 | 170.7 | 29.3 | 1.55 | 16.56 | | High | High | High | High | High | High | High | High | High | High | High | 0.000246 | 0.001521 | 2.67 |
| TRUE | High | DLPDGHVA | [K].DLPDGI | 1xPhospho | E9Q5B0 1x | 0.0095 | 0.000736 | 1 | 8 | 1 | E9Q5B0 | Calmodulir | E9Q5B0 [3 | 0 | 1571.778 | 0 | -2.37 | 0.00059 | 1.63 | 168.3 | 31.7 | 7.48 | 23.86 | | High | High | High | High | High | High | High | High | High | High | High | 0.000344 | 0.00297 | 2.52 |
| TRUE | High | SESPAVR | [R].SESPAV | 1xPhospho | Q3KR45 1x | 0.043236 | 0.003228 | 1 | 5 | 1 | Q3KR45 | DNA (cytos | Q3KR45 [1 | 0 | 1054.513 | 0 | -2.99 | 0.000779 | 0.55 | 179.8 | 20.2 | 12.02 | 33.6 | | High | High | High | High | High | High | High | High | High | High | High | 0.001651 | 0.01636 | 1.56 |
| TRUE | High | SSTTSSPK | [R].SSTTSS | 1xPhospho | A8CA96 1x | 0.008463 | 0.000736 | 1 | 3 | 2 | A8CA96 | Tissue-spe | A8CA96 [1 | 0 | 1332.681 | 0 | -2.83 | 0.000154 | 0.63 | 175.3 | 24.7 | 5.01 | 29.26 | | High | High | High | High | High | High | High | High | High | High | High | 0.000327 | 0.002593 | 1.34 |
| TRUE | High | KPEEESPRK | [R].KPEEES | 1xPhospho | B1AU76 1x | 0.081085 | 0.006219 | 2 | 4 | 4 | B1AU76; B | Nuclear au | B1AU76 [3 | 1 | 1637.866 | 0 | -2.08 | 0.000192 | 0.99 | 160.4 | 39.6 | 2.88 | 14.51 | Shared | High | High | High | High | High | High | High | High | High | High | High | 0.008399 | 0.09007 | 1.72 |
| TRUE | High | AALTNSLSF | [RK].AALTN | 1xPhospho | Q3TJW2 1 | 0.000808 | 0 | 1 | 3 | 1 | Q3TJW2 | F-box only | Q3TJW2 [1 | 0 | 1638.923 | 0 | -3.64 | 0.000154 | 0.21 | 185.3 | 14.7 | 3.54 | 32.46 | | High | High | High | High | High | High | High | High | High | High | High | 0 | 0.000187 | 2.69 |
| TRUE | High | GSQSSHES | [K].GSQSSI | 1xPhospho | D0VLQ5 1x | 0.002096 | 5.66E-05 | 1 | 3 | 1 | D0VLQ5 | Tet oncoge | D0VLQ5 [6 | 0 | 1761.739 | 0 | -2.39 | 0.000576 | 1.31 | 165.6 | 34.4 | 9 | 18.06 | | High | High | High | High | High | High | High | High | High | High | High | 2.58E-05 | 0.000548 | 2.57 |
| TRUE | High | RSPTCDVC | [R].RSPTCD | 2xCarbami | E9Q6W4 1 | 0.001158 | 0 | 1 | 1 | 2 | E9Q6W4 | Zinc finger | E9Q6W4 [2 | 1 | 1660.795 | 0 | -2.71 | 0.000211 | 0.22 | 174.4 | 25.6 | 6.64 | 13.2 | | High | High | High | High | High | High | High | High | High | High | High | 0 | 0.00028 | 2.89 |
| TRUE | High | LPLIVIDDDI | [K].LPLIVID | 1xPhospho | A0A1D9BZ | 2.9E-09 | 0 | 1 | 1 | 12 | A0A1D9BZ | GCNA OS= | A0A1D9BZ | 0 | 2378.214 | 0 | -2.46 | 0.000154 | 0.51 | 169.3 | 30.7 | 6.25 | 16.55 | | High | High | High | High | High | High | High | High | High | High | High | 0 | 1.44E-10 | 4.46 |
| TRUE | High | DHVPSTPT | [R].DHVPS | 1xPhospho | Q3UHU5 1 | 1.82E-06 | 0 | 1 | 5 | 1 | Q3UHU5 | Microtubu | Q3UHU5 [ | 0 | 2686.191 | 0 | -2.08 | 0.000191 | 0.79 | 160.4 | 39.6 | 4.7 | 20.72 | | High | High | High | High | High | High | High | High | High | High | High | 0 | 1.99E-07 | 3.34 |
| TRUE | High | GCSPDGYG | [R].GCSPD | 1xCarbami | Q9DAE2 1 | 0.000411 | 0 | 1 | 1 | 8 | Q9DAE2 | Putative ur | Q9DAE2 [3 | 0 | 1334.54 | 0 | -2.62 | 0.000143 | 0.36 | 172.2 | 27.8 | 1.86 | 14.78 | | High | High | High | High | High | High | High | High | High | High | High | 0 | 8.76E-05 | 2.12 |
| TRUE | High | SFSVEGR | [R].SFSVEG | 1xPhospho | Q571N6 1x | 0.072858 | 0.005599 | 1 | 6 | 1 | Q571N6 | MFLJ0027 | Q571N6 [6 | 0 | 1090.513 | 0 | -2.17 | 0.000427 | 1.84 | 164.6 | 35.4 | 3.44 | 28.39 | | High | High | High | High | High | High | High | High | High | High | High | 0.002867 | 0.02976 | 0.48 |
| TRUE | High | FLQSPEENT | [R].FLQSPE | 1xPhospho | Q7M6U3 1 | 2.58E-05 | 0 | 1 | 3 | 1 | Q7M6U3 | Inactive se | Q7M6U3 [ | 0 | 2341.209 | 0 | -2.03 | 0.000986 | 1.53 | 160.6 | 39.4 | 8.4 | 36.35 | | High | High | High | High | High | High | High | High | High | High | High | 0 | 3.93E-06 | 3.79 |
| TRUE | High | TWTPSGK | [K].TWTPS | 1xPhospho | Q71FD7 1x | 0.026421 | 0.001676 | 1 | 8 | 1 | Q71FD7 | Filamin-bin | Q71FD7 [6 | 0 | 1314.686 | 0 | -2.23 | 0.000391 | 0.68 | 167.5 | 32.5 | 8.1 | 9.84 | | High | High | High | High | High | High | High | High | High | High | High | 0.000903 | 0.009372 | 2.12 |
| TRUE | High | QSTESETPT | [R].QSTESE | 1xPhospho | A0A1B0GS | 0.011246 | 0.000772 | 1 | 2 | 2 | A0A1B0GS | GRB2-asso | A0A1B0GS | 0 | 1645.809 | 0 | -2.3 | 0.000938 | 0.89 | 165.9 | 34.1 | 11.33 | 19.97 | | High | High | High | High | High | High | High | High | High | High | High | 0.000418 | 0.00357 | 2.17 |
| TRUE | High | LCIDGYPSP | [R].LCIDGY | 1xCarbami | Q3TJB1 1x | 4.15E-08 | 0 | 1 | 5 | 1 | Q3TJB1 | Putative ur | Q3TJB1 [4 | 0 | 2323.217 | 0 | -2.2 | 0.001844 | 0.76 | 165.4 | 34.6 | 15.61 | 12.4 | | High | High | High | High | High | High | High | High | High | High | High | 0 | 2.84E-09 | 3.7 |
| TRUE | High | LPATSAESE | [R].LPATSA | 1xDeamida | Q3UKE7 1x | 7.33E-08 | 0 | 1 | 5 | 3 | Q3UKE7 | Hydroxym | Q3UKE7 [5 | 0 | 2108.959 | 3 | -2.28 | 0.000181 | 0.51 | 168.5 | 31.5 | 5.69 | 20.11 | | High | High | High | High | High | High | High | High | High | High | High | 0 | 0.000293 | 3.33 |
| TRUE | High | QQTNPSPT | [K].QQTNP | 1xPhospho | D0VLQ4 1x | 0.006195 | 0.00062 | 1 | 3 | 1 | D0VLQ4 | Tet oncoge | D0VLQ4 [7 | 0 | 1897.937 | 0 | -3.39 | 0.000413 | 0.3 | 182.6 | 17.4 | 6.96 | 28.34 | | High | High | High | High | High | High | High | High | High | High | High | 0.000275 | 0.001838 | 1.78 |
| TRUE | High | FSDEGDSD | [K].FSDEGI | 1xCarbami | E9PXD2 2x | 2.58E-05 | 0 | 1 | 7 | 4 | E9PXD2 | Rho GTPas | E9PXD2 [9 | 0 | 2902.214 | 0 | -2.3 | 0.002108 | 0.83 | 165.4 | 34.6 | 11.49 | 24.26 | | High | High | High | High | High | High | High | High | High | High | High | 0 | 3.9E-06 | 4.08 |
| TRUE | High | SPDATETM | [R].SPDATE | 1xCarbami | Q8BX22 1x | 9.31E-06 | 0 | 1 | 1 | 8 | Q8BX22 | Sal-like pro | Q8BX22 [7 | 0 | 3053.4 | 0 | -2.25 | 0.000266 | 0.78 | 165.2 | 34.8 | 3.18 | 27.4 | | High | High | High | High | High | High | High | High | High | High | High | 0 | 4.68E-06 | 4.2 |
| TRUE | High | RQDEDYDE | [K].RQDED | 1xPhospho | Q8BKC5 1x | 2.52E-07 | 0 | 1 | 5 | 4 | Q8BKC5 | Importin-5 | Q8BKC5 [8 | 1 | 3841.722 | 0 | -2.16 | 0.000408 | 0.87 | 163 | 37 | 6.94 | 16.67 | | High | High | High | High | High | High | High | High | High | High | High | 0 | 2.17E-08 | 6.13 |
| TRUE | High | TLSQSSESG | [R].TLSQSS | 1xCarbami | Q9QYG0 1 | 6.79E-06 | 0 | 1 | 3 | 3 | Q9QYG0 | Protein ND | Q9QYG0 [3 | 0 | 2771.21 | 0 | -2.16 | 0.001361 | 0.79 | 162.3 | 37.7 | 9.24 | 39.2 | | High | High | High | High | High | High | High | High | High | High | High | 0 | 3.33E-06 | 5.04 |
| TRUE | High | FLGGSQVS | [K].FLGGSC | 1xPhospho | P47806 1x | 0.000908 | 0 | 1 | 2 | 1 | P47806 | Zinc finger | P47806 [93 | 0 | 1843.972 | 0 | -2.94 | 0.000466 | 0.75 | 175.5 | 24.5 | 7.1 | 30.18 | | High | High | High | High | High | High | High | High | High | High | High | 0 | 0.000214 | 3.84 |
| TRUE | High | FSDEGDSD | [K].FSDEGI | 1xCarbami | E9PXD2 1x | 5.13E-09 | 0 | 1 | 7 | 1 | E9PXD2 | Rho GTPas | E9PXD2 [9 | 0 | 2822.248 | 0 | -2.99 | 0.00062 | 0.54 | 177.8 | 22.2 | 9.41 | 23.07 | | High | High | High | High | High | High | High | High | High | High | High | 0 | 2.73E-10 | 5.93 |
| FALSE | High | SEMTPEEL | [K].SEMTP | 1xPhospho | D3Z5N9 1x | 0.112856 | 0.009146 | 1 | 2 | 2 | D3Z5N9 | MCG4919 | D3Z5N9 [9 | 0 | 1745.843 | 0 | -0.77 | 0.001306 | 3.61 | 125.9 | 74.1 | 4.37 | 7.48 | | High | High | High | High | High | High | High | High | High | High | High | 0.0054 | 0.05933 | 2.68 |
| FALSE | High | SEHSLFQPI | [K].SEHSLF | 1xPhospho | Q8CH77 1x | 0.000656 | 0 | 1 | 3 | 2 | Q8CH77 | Neuron na | Q8CH77 [1 | 0 | 1610.834 | 0 | 0.22 | 0.050776 | 7.71 | 92.6 | 107.4 | 4.44 | 16.72 | | High | High | High | High | High | High | High | High | High | High | High | 0 | 0.000149 | 2.87 |
| FALSE | High | SCEEESDPE | [R].SCEEES | 1xCarbami | F8VPX2 1x | 1.26E-05 | 0 | 1 | 2 | 2 | F8VPX2 | F-box and | F8VPX2 [1 | 0 | 3016.224 | 0 | -0.37 | 0.105906 | 16.38 | 110.3 | 89.7 | 11.32 | 13.45 | | High | High | High | High | High | High | High | High | High | High | High | 0 | 4.73E-05 | 4.21 |
| FALSE | High | SCETSSQDI | [R].SCETSS | 1xCarbami | K4DI61 1x | 1.63E-05 | 0 | 1 | 4 | 3 | K4DI61 | PHD finger | K4DI61 [55 | 0 | 3269.5 | 0 | -0.02 | 0.895136 | 12.52 | 100.8 | 99.2 | 15.56 | 14.53 | | High | High | High | High | High | High | High | High | High | High | High | 0 | 2.35E-06 | 4.62 |
| FALSE | High | SCFSNHNE | [K].SCFSNH | 1xCarbami | P97929 1x | 0.001107 | 0 | 1 | 3 | 2 | P97929 | Breast can | P97929 [22 | 0 | 1972.899 | 0 | -1.25 | 0.000962 | 1.14 | 142.3 | 57.7 | 10.44 | 3.82 | | High | High | High | High | High | High | High | High | High | High | High | 0 | 0.000267 | 2.62 |
| FALSE | High | SAVGFSNG | [R].SAVGFS | 1xPhospho | D2DFA9 1x | 7.34E-05 | 0 | 1 | 3 | 1 | D2DFA9 | G protein-c | D2DFA9 [3 | 0 | 1660.789 | 0 | -0.13 | 0.267221 | 32.12 | 108.7 | 91.3 | 15.83 | 18.09 | | High | High | High | High | High | High | High | High | High | High | High | 0 | 1.27E-05 | 2.49 |
| FALSE | High | SCDYQFPS | [R].SCDYQI | 1xCarbami | Q69ZR9-2 1 | 1.35E-07 | 0 | 1 | 1 | 5 | Q69ZR9-2 | Isoform 2 c | Q69ZR9-2 | 0 | 2371.093 | 0 | -1.55 | 0.00024 | 0.8 | 146.8 | 53.2 | 3.81 | 7.2 | | High | High | High | High | High | High | High | High | High | High | High | 0.00168 | 0.01677 | 2.87 |
| FALSE | High | SCGGESPR | [R].SCGGE | 2xCarbami | Q3UHC1 2x | 0.058664 | 0.004294 | 1 | 5 | 1 | Q3UHC1 | Putative ur | Q3UHC1 [3 | 1 | 2385.038 | 0 | 1.08 | 0.000317 | 4.95 | 65.4 | 134.6 | 5.1 | 3.51 | | High | High | High | High | High | High | High | High | High | High | High | 0.002187 | 0.02321 | 2.8 |
| FALSE | High | SELVSEGRI | [R].SELVSE | 1xPhospho | B7ZCJ1 1x | 0.034878 | 0.002269 | 1 | 4 | 1 | B7ZCJ1 | Rho GTPas | B7ZCJ1 [16 | 0 | 3520.657 | 0 | -0.23 | 0.329375 | 23.44 | 108 | 92 | 7.23 | 35.74 | | High | High | High | High | High | High | High | High | High | High | High | 0.001183 | 0.01281 | 2.98 |

| | | | | | | | | | | | | | | | | | | | | | | | | | | | | | | | | | | | | | | |
|---|---|---|---|---|---|---|---|---|---|---|---|---|---|---|---|---|---|---|---|---|---|---|---|---|---|---|---|---|---|---|---|---|---|---|---|---|---|---|
| FALSE | High | SCDNEQGL | [K].SCDNE( | 1xCarbami | Q8CGN4 1: | 8.5E-06 | 0 | 1 | 4 | 2 | Q8CGN4 | BCL-6 core | Q8CGN4 [1 | 0 | 2470.176 | 0 | 0.55 | 0.012077 | 14.01 | 79.2 | 120.8 | 2.24 | 12.95 | | High | High | High | High | High | High | High | High | High | High | High | 0 | 1.13E-06 | 3.03 |
| FALSE | High | SELVLAPAF | [K].SELVLA | 1xPhospho | Q3U912 1x | 9.33E-07 | 0 | 1 | 2 | 1 | Q3U912 | Putative ur | Q3U912 [3 | 0 | 2344.238 | 0 | -0.02 | 0.49303 | 7.85 | 99.4 | 100.6 | 8.42 | 4.67 | | High | High | High | High | High | High | High | High | High | High | High | 0 | 9.38E-08 | 3.24 |
| FALSE | High | SAVRPSPSF | [K].SAVRPS | 2xPhospho | Q8BTI8 2xF | 0.024638 | 0.001612 | 1 | 3 | 3 | Q8BTI8 | Serine/argi | Q8BTI8 [34 | 0 | 1571.718 | 0 | -0.56 | 0.001493 | 3.74 | 121.1 | 78.9 | 4.54 | 6.85 | | High | High | High | High | High | High | High | High | High | High | High | 0.000859 | 0.008608 | 2.17 |
| FALSE | High | SEITPPPPQ | [K].SEITPPI | 1xPhospho | P53395 1xI | 0.112856 | 0.009146 | 1 | 2 | 1 | P53395 | Lipoamide | P53395 [22 | 0 | 1728.934 | 0 | 0.45 | 0.83735 | 32.83 | 88.1 | 111.9 | 28.62 | 40.25 | | High | High | High | High | High | High | High | High | High | High | High | 0.004569 | 0.04968 | 1.98 |
| FALSE | High | SAVSPDLHI | [R].SAVSPD | 1xPhospho | Q68FH0 1x | 0.000193 | 0 | 1 | 8 | 1 | Q68FH0 | Plakophilin | Q68FH0 [4 | 0 | 2064.036 | 0 | -1.13 | 0.004672 | 4.33 | 137.9 | 62.1 | 11.67 | 17.75 | | High | High | High | High | High | High | High | High | High | High | High | 0 | 3.75E-05 | 2.97 |
| FALSE | High | SAVRPSPSF | [K].SAVRPS | 1xPhospho | Q8BTI8 1xF | 0.030632 | 0.001857 | 1 | 3 | 2 | Q8BTI8 | Serine/argi | Q8BTI8 [34 | 0 | 1491.752 | 0 | -0.7 | 0.001799 | 3.42 | 123.9 | 76.1 | 6.45 | 5.63 | | High | High | High | High | High | High | High | High | High | High | High | 0.004757 | 0.05142 | 1.92 |
| FALSE | High | SAVSPDLHI | [R].SAVSPD | 2xPhospho | Q68FH0 2x | 0.005044 | 0.000504 | 1 | 8 | 2 | Q68FH0 | Plakophilin | Q68FH0 [4 | 0 | 2144.003 | 0 | -1.36 | 0.001949 | 2.93 | 145.3 | 54.7 | 11.51 | 8.32 | | High | High | High | High | High | High | High | High | High | High | High | 0.001916 | 0.0194 | 3.58 |
| FALSE | High | SELTDSASV | [K].SELTDS | 1xPhospho | O55106 1x | 0.000322 | 0 | 1 | 5 | 1 | O55106 | Striatin OS | O55106 [2 | 0 | 2064.03 | 0 | 1.56 | 0.006527 | 22.03 | 47 | 153 | 11.14 | 24.08 | | High | High | High | High | High | High | High | High | High | High | High | 0 | 6.66E-05 | 2.35 |
| FALSE | High | SAVLDSGL/ | [R].SAVLDS | 1xPhospho | Q3TN34 1x | 1.75E-05 | 0 | 1 | 4 | 4 | Q3TN34 | MICAL-like | Q3TN34 [8 | 0 | 2232.147 | 0 | -0.27 | 0.024471 | 7.33 | 109.5 | 90.5 | 5.01 | 15.36 | | High | High | High | High | High | High | High | High | High | High | High | 0 | 1.38E-05 | 2.83 |
| FALSE | High | SAVSQTTD | [R].SAVSQT | 1xCarbami | Q3UHC1 1: | 0.00824 | 0.000736 | 1 | 5 | 1 | Q3UHC1 | Putative ur | Q3UHC1 [4 | 0 | 1664.715 | 0 | -0.18 | 0.109213 | 9.08 | 106.7 | 93.3 | 9.57 | 21.29 | | High | High | High | High | High | High | High | High | High | High | High | 0.000327 | 0.002528 | 1.93 |
| FALSE | High | SAYFSDEEE | [K].SAYFSD | 1xPhospho | Q9CZ96 1x | 0.000379 | 0 | 1 | 2 | 1 | Q9CZ96 | Zinc finger | Q9CZ96 [2 | 0 | 1700.678 | 0 | 0.23 | 0.234832 | 15.06 | 92.3 | 107.7 | 6.34 | 14.36 | | High | High | High | High | High | High | High | High | High | High | High | 0 | 8E-05 | 3.32 |
| FALSE | High | SAYQDYDS | [K].SAYQDY | 1xPhospho | Q8CI71 1xF | 1.36E-06 | 0 | 1 | 3 | 7 | Q8CI71 | Syndetin O | Q8CI71 [55 | 0 | 2499.122 | 0 | 0.11 | 0.311257 | 3.57 | 96.5 | 103.5 | 2.46 | 6.88 | | High | High | High | High | High | High | High | High | High | High | High | 0 | 1.44E-07 | 3.91 |
| FALSE | High | SAVLDGILE | [R].SAVLDG | 1xPhospho | Q8R3C6 1x | 1.4E-05 | 0 | 1 | 1 | 5 | Q8R3C6 | Probable R | Q8R3C6 [9 | 0 | 3010.398 | 0 | -0.08 | 0.073697 | 2.2 | 103.4 | 96.6 | 8.7 | 9.54 | | High | High | High | High | High | High | High | High | High | High | High | 0 | 1.97E-06 | 4.17 |
| FALSE | High | SAYQDYDS | [K].SAYQDY | 2xPhospho | Q8CI71 2xF | 0.000142 | 0 | 1 | 3 | 2 | Q8CI71 | Syndetin O | Q8CI71 [55 | 0 | 2579.088 | 0 | 0.71 | 0.002682 | 8.68 | 75.4 | 124.6 | 9.93 | 7.64 | | High | High | High | High | High | High | High | High | High | High | High | 0 | 2.67E-05 | 3.02 |
| FALSE | High | SEISQSPTC | [R].SEISQSI | 1xCarbami | Q80U44 1x | 8.33E-05 | 0 | 1 | 1 | 2 | Q80U44 | Zinc finger | Q80U44 [9 | 0 | 2107.968 | 0 | 1.06 | 0.007243 | 19.22 | 62 | 138 | 8.27 | 18.28 | | High | High | High | High | High | High | High | High | High | High | High | 0 | 1.46E-05 | 2.03 |
| FALSE | High | SAYQDYDS | [K].SAYQDY | 1xPhospho | Q8CI71 1xF | 0.081085 | 0.006219 | 1 | 3 | 1 | Q8CI71 | Syndetin O | Q8CI71 [55 | 1 | 2655.223 | 0 | -0.1 | 0.903614 | 21.52 | 103.5 | 96.5 | 8.58 | 25.6 | | High | High | High | High | High | High | High | High | High | High | High | 0.003188 | 0.03361 | 2.35 |
| FALSE | High | SCDELSPVS | [R].SCDELS | 1xCarbami | A2A690-2 1 | 1.54E-05 | 0 | 1 | 3 | 1 | A2A690-2 | Isoform 2 c | A2A690-2 \| | 0 | 2573.143 | 0 | 2.37 | 0.000154 | 8.82 | 31.1 | 168.9 | 12.45 | 4.07 | | High | High | High | High | High | High | High | High | High | High | High | 0 | 2.2E-06 | 3.29 |
| FALSE | High | SAVHYDED | [K].SAVHYI | 2xPhospho | Q80XK6 2x | 5.75E-05 | 0 | 1 | 1 | 6 | Q80XK6 | Autophagy | Q80XK6 [1 | 0 | 3497.43 | 0 | 0.07 | 0.223682 | 15.66 | 93.6 | 106.4 | 8.42 | 14.92 | | High | High | High | High | High | High | High | High | High | High | High | 0.000246 | 0.001487 | 4.33 |
| FALSE | High | SEMAEHSD | [K].SEMAE | 1xPhospho | Q45KJ6-2 1 | 0.024001 | 0.001612 | 1 | 2 | 1 | Q45KJ6-2 | Isoform 2 c | Q45KJ6-2 [ | 1 | 2559.18 | 0 | -1.23 | 0.006559 | 2.38 | 140.1 | 59.9 | 11.12 | 22.76 | | High | High | High | High | High | High | High | High | High | High | High | 0.000859 | 0.008374 | 3.32 |
| FALSE | High | SEMTPEEL( | [K].SEMTPI | 1xPhospho | D3Z5N9 1x | 0.049549 | 0.00361 | 1 | 2 | 1 | D3Z5N9 | MCG49198 | D3Z5N9 [9 | 0 | 1729.848 | 0 | -0.13 | 0.277116 | 4.2 | 103.7 | 96.3 | 7.76 | 15.3 | | High | High | High | High | High | High | High | High | High | High | High | 0.001902 | 0.01915 | 2.19 |
| FALSE | High | SEISNIAR | [K].SEISNIA | 1xPhospho | Q9D7Z6 1x | 0.089425 | 0.006807 | 1 | 1 | 1 | Q9D7Z6 | Calcium-ac | Q9D7Z6 [8 | 0 | 1198.603 | 0 | 0.18 | 0.136949 | 5.12 | 93.2 | 106.8 | 3.3 | 9.36 | | High | High | High | High | High | High | High | High | High | High | High | 0.003564 | 0.03796 | 1.06 |
| FALSE | High | SELQAYIEC | [K].SELQAY | 1xCarbami | Q99J21 1xI | 1.84E-06 | 0 | 1 | 2 | 4 | Q99J21 | Mucolipin- | Q99J21 [53 | 0 | 2579.21 | 0 | 0.6 | 0.00675 | 12.78 | 79.5 | 120.5 | 13.66 | 8.25 | | High | High | High | High | High | High | High | High | High | High | High | 0 | 2.01E-07 | 4.45 |
| FALSE | High | SCDELSPVS | [R].SCDELS | 1xCarbami | A2A690-2 1 | 0.078467 | 0.006104 | 1 | 3 | 1 | A2A690-2 | Isoform 2 c | A2A690-2 \| | 0 | 2653.109 | 0 | 0.42 | 0.13309 | 19.49 | 87.8 | 112.2 | 17.19 | 5.63 | | High | High | High | High | High | High | High | High | High | High | High | 0.003148 | 0.03257 | 2.23 |
| FALSE | High | SCDNEQGL | [K].SCDNE( | 1xCarbami | Q8CGN4 1: | 2E-07 | 0 | 1 | 4 | 1 | Q8CGN4 | BCL-6 core | Q8CGN4 [1 | 0 | 2454.181 | 0 | 0.72 | 0.081909 | 31.1 | 73.5 | 126.5 | 9.73 | 28.82 | | High | High | High | High | High | High | High | High | High | High | High | 0 | 1.66E-08 | 3.69 |
| FALSE | High | SEIANVQS( | [K].SEIANV | 2xCarbami | Q99P72 1x | 7.77E-06 | 0 | 1 | 2 | 1 | Q99P72 | Reticulon-4 | Q99P72 [8 | 0 | 3261.557 | 0 | -0.34 | 0.713716 | 16.05 | 111.9 | 88.1 | 17.07 | 29.97 | | High | High | High | High | High | High | High | High | High | High | High | 0 | 1.01E-06 | 3.66 |
| FALSE | High | SDSSADCQ | [R].SDSSAD | 1xCarbami | Q9ESL4 1xI | 9.03E-05 | 0 | 1 | 1 | 1 | Q9ESL4 | Mitogen-a( | Q9ESL4 [56 | 0 | 1962.847 | 0 | 1.25 | 0.00024 | 8.12 | 62.2 | 137.8 | 8.45 | 5.21 | | High | High | High | High | High | High | High | High | High | High | High | 0 | 1.6E-05 | 3.35 |
| FALSE | High | SATLSSTES | [K].SATLSS | 2xPhospho | B2RRE0 2x | 0.001865 | 5.66E-05 | 1 | 3 | 3 | B2RRE0 | A kinase (P | B2RRE0 [62 | 0 | 2345.977 | 0 | -1.85 | 0.001211 | 0.73 | 151 | 49 | 9.45 | 20.85 | | High | High | High | High | High | High | High | High | High | High | High | 2.58E-05 | 0.000482 | 4.02 |
| FALSE | High | SAVFSGEG | [K].SAVFSG | 1xPhospho | Q7TQG1 1: | 0.081752 | 0.006219 | 1 | 6 | 1 | Q7TQG1 | Pleckstrin I | Q7TQG1 [9 | 0 | 1419.729 | 0 | 1.95 | 0.000266 | 4.2 | 42.1 | 157.9 | 6.05 | 5.76 | | High | High | High | High | High | High | High | High | High | High | High | 0.003229 | 0.03409 | 1.18 |
| FALSE | High | SEPTVVSLE | [K].SEPTVV | 1xPhospho | L8AZD2 1xI | 0.001615 | 5.66E-05 | 1 | 2 | 1 | L8AZD2 | Myotubula | L8AZD2 [63 | 0 | 1815.909 | 0 | 1.24 | 0.005931 | 32.31 | 66.6 | 133.4 | 13.03 | 16.88 | | High | High | High | High | High | High | High | High | High | High | High | 2.58E-05 | 0.00041 | 2.4 |
| FALSE | High | SASSSAAG! | [K].SASSSA | 1xPhospho | Q8CHS8 1x | 2.65E-08 | 0 | 1 | 4 | 2 | Q8CHS8 | Vacuolar p | Q8CHS8 [1 | 0 | 2414.233 | 0 | 0.42 | 0.01876 | 17.39 | 82.4 | 117.6 | 17.03 | 7.12 | | High | High | High | High | High | High | High | High | High | High | High | 0 | 1.72E-09 | 3.9 |
| FALSE | High | SEPTSPPPS | [K].SEPTSP | 1xPhospho | Q5PRE5 1x | 0.000837 | 0 | 1 | 2 | 4 | Q5PRE5 | Proline anc | Q5PRE5 [5 | 0 | 1782.908 | 0 | -1.47 | 0.000163 | 0.6 | 147.7 | 52.3 | 2.57 | 6.42 | | High | High | High | High | High | High | High | High | High | High | High | 0 | 0.000196 | 2.33 |
| FALSE | High | SASSSSDAE | [K].SASSSS | 1xPhospho | Q61687 1x | 8.22E-07 | 0 | 1 | 4 | 3 | Q61687 | Transcripti | Q61687 [1 | 0 | 2095.907 | 0 | 0.25 | 0.470544 | 21.24 | 94.4 | 105.6 | 12.41 | 11.75 | | High | High | High | High | High | High | High | High | High | High | High | 0 | 8.19E-08 | 4.84 |
| FALSE | High | SEPSSPPPA | [K].SEPSSP | 1xPhospho | A2AM37 1: | 0.000589 | 0 | 1 | 5 | 2 | A2AM37 | Transcripti | A2AM37 [3 | 0 | 1956.938 | 0 | -0.47 | 0.018232 | 7.03 | 116.2 | 83.8 | 11.1 | 11.61 | | High | High | High | High | High | High | High | High | High | High | High | 0 | 0.000131 | 2.62 |
| FALSE | High | SASSSSDAE | [K].SASSSS | 1xPhospho | Q61687 1x | 0.025514 | 0.001645 | 1 | 4 | 3 | Q61687 | Transcripti | Q61687 [1 | 1 | 2453.165 | 0 | 0.35 | 0.369408 | 16.21 | 91.2 | 108.8 | 18.21 | 16.62 | | High | High | High | High | High | High | High | High | High | High | High | 0.000872 | 0.009006 | 1.77 |
| FALSE | High | SASSTSSGS | [R].SASSTS | 1xPhospho | E9Q8I9-2 1 | 0.004256 | 0.00036 | 1 | 4 | 5 | E9Q8I9-2 | Isoform 2 c | E9Q8I9-2 [ | 0 | 2532.275 | 0 | -0.43 | 0.015087 | 10.22 | 114.8 | 85.2 | 9.26 | 8.46 | | High | High | High | High | High | High | High | High | High | High | High | 0.000275 | 0.00196 | 3 |
| FALSE | High | SASSYSDIE | [K].SASSYS | 1xPhospho | Q8CH77 1x | 0.004531 | 0.000456 | 1 | 2 | 3 | Q8CH77 | Neuron na | Q8CH77 [1 | 0 | 2635.186 | 0 | 0.67 | 0.364322 | 28.1 | 84 | 116 | 39.83 | 35.85 | | High | High | High | High | High | High | High | High | High | High | High | 0.000327 | 0.002413 | 2.91 |
| FALSE | High | SASSYSDIE | [K].SASSYS | 1xPhospho | Q8CH77 1x | 9.74E-06 | 0 | 1 | 2 | 4 | Q8CH77 | Neuron na | Q8CH77 [1 | 0 | 2864.349 | 0 | 0.83 | 0.001335 | 12.83 | 71.9 | 128.1 | 5.09 | 5.73 | | High | High | High | High | High | High | High | High | High | High | High | 0 | 2.36E-05 | 3.74 |
| FALSE | High | SASSYSDIE | [K].SASSYS | 2xPhospho | Q8CH77 2x | 0.000139 | 0 | 1 | 2 | 4 | Q8CH77 | Neuron na | Q8CH77 [1 | 0 | 2944.315 | 0 | 0.7 | 0.001936 | 9.48 | 76.1 | 123.9 | 7.62 | 4.92 | | High | High | High | High | High | High | High | High | High | High | High | 0 | 4.07E-05 | 3.4 |
| FALSE | High | SEPSLEPES | [R].SEPSLEI | 1xPhospho | Q8VDZ4 1x | 0.050397 | 0.003641 | 1 | 2 | 2 | Q8VDZ4 | Palmitoyltr | Q8VDZ4 [3 | 1 | 2433.21 | 0 | 0.58 | 0.000616 | 4.45 | 80 | 120 | 4.28 | 4.62 | | High | High | High | High | High | High | High | High | High | High | High | 0.003977 | 0.04245 | 2.26 |
| FALSE | High | SASTDAPPY | [K].SASTDA | 1xPhospho | Q91X43 1x | 0.035486 | 0.002304 | 1 | 1 | 2 | Q91X43 | SH3 domai | Q91X43 [1 | 0 | 2066.061 | 0 | 0.68 | 0.016213 | 8.64 | 77.5 | 122.5 | 10.1 | 12.54 | | High | High | High | High | High | High | High | High | High | High | High | 0.001214 | 0.01305 | 2.57 |
| FALSE | High | SEPSGSPPA | [K].SEPSGS | 1xPhospho | P53783 1xI | 0.002313 | 0.000109 | 1 | 1 | 1 | P53783 | Transcripti | P53783 [32 | 0 | 1685.785 | 0 | -0.2 | 0.077359 | 11.95 | 102.8 | 97.2 | 9.87 | 6.98 | | High | High | High | High | High | High | High | High | High | High | High | 4.94E-05 | 0.000612 | 1.58 |
| FALSE | High | SEPSDHST( | [R].SEPSDH | 1xDeamida | P39447 1xI | 0.000958 | 0 | 1 | 3 | 3 | P39447 | Tight junct | P39447 [32 | 0 | 2449.083 | 0 | 0.01 | 0.685941 | 18.56 | 100.8 | 99.2 | 17.81 | 13.65 | | High | High | High | High | High | High | High | High | High | High | High | 0 | 0.000228 | 3.13 |
| FALSE | High | SASTDLGT/ | [K].SASTDL | 1xPhospho | Q80TM6 1: | 4.65E-06 | 0 | 1 | 8 | 3 | Q80TM6 | R3H domai | Q80TM6 [9 | 0 | 1770.884 | 0 | 0.93 | 0.018207 | 19.37 | 69.7 | 130.3 | 5.73 | 25.15 | | High | High | High | High | High | High | High | High | High | High | High | 0 | 5.7E-07 | 3.93 |
| FALSE | High | SASTLCLPE | [K].SASTLC | 1xCarbami | Q497Z4 1x | 9.82E-06 | 0 | 1 | 2 | 1 | Q497Z4 | Rhpn2 pro | Q497Z4 [6 | 0 | 1782.902 | 0 | -1.02 | 0.005347 | 2.01 | 139.7 | 60.3 | 17.25 | 10.88 | | High | High | High | High | High | High | High | High | High | High | High | 0 | 1.32E-06 | 2.69 |
| FALSE | High | SASVNK | [R].SASVNI | 1xPhospho | Q6ZQ32 1x | 0.117405 | 0.009523 | 2 | 7 | 1 | Q6ZQ32; F | MKIAA085 | Q6ZQ32 [1 | 0 | 1143.618 | 0 | 1.69 | 0.000454 | 4.7 | 46.3 | 153.7 | 4.81 | 7.62 | Shared | High | High | High | High | High | High | High | High | High | High | High | 0.004769 | 0.05221 | 1.8 |
| FALSE | High | SEPSDHST( | [R].SEPSDH | 1xPhospho | P39447 1xI | 1.59E-06 | 0 | 1 | 3 | 2 | P39447 | Tight junct | P39447 [32 | 0 | 2448.099 | 0 | 0.15 | 0.770896 | 15.41 | 92.7 | 107.3 | 26.74 | 27.37 | | High | High | High | High | High | High | High | High | High | High | High | 0 | 1.71E-07 | 4.12 |
| FALSE | High | SASVSSISLS | [R].SASVSS | 1xPhospho | Q62245 1x | 0.00509 | 0.000551 | 1 | 3 | 1 | Q62245 | Son of seve | Q62245 [1 | 0 | 1603.871 | 0 | 0.51 | 0.02194 | 19.84 | 79.1 | 120.9 | 6.97 | 18.65 | | High | High | High | High | High | High | High | High | High | High | High | 0.000246 | 0.001476 | 1.73 |
| FALSE | High | SEPVSPSR | [K].SEPVSP | 1xPhospho | Q63943-2 1 | 0.098533 | 0.007902 | 2 | 6 | 1 | Q63943-2; | Isoform Mu | Q63943-2 \| | 0 | 1167.561 | 0 | 0.76 | 0.000763 | 9.12 | 74.8 | 125.2 | 4.63 | 5.26 | Shared | High | High | High | High | High | High | High | High | High | High | High | 0.003977 | 0.04245 | 0.87 |
| FALSE | High | SASWGSAD | [R].SASWG | 1xPhospho | E9QKK4 1x | 0.000725 | 0 | 1 | 7 | 3 | E9QKK4 | Glucocortic | E9QKK4 [2 | 0 | 1687.846 | 0 | 0.92 | 0.003297 | 11.19 | 72.4 | 127.6 | 6.43 | 10.04 | | High | High | High | High | High | High | High | High | High | High | High | 0 | 0.000167 | 3.24 |
| FALSE | High | SASSLHVCI | [R].SASSLH | 1xCarbami | F6RDI8 1xF | 0.017498 | 0.001245 | 1 | 1 | 1 | F6RDI8 | Serine/argi | F6RDI8 [34 | 0 | 1527.682 | 0 | 0.03 | 0.630328 | 20.92 | 99.1 | 100.9 | 11.26 | 6.3 | | High | High | High | High | High | High | High | High | High | High | High | 0.000674 | 0.005868 | 2.25 |
| FALSE | High | SASSLDDVI | [K].SASSLD | 1xPhospho | Q3U2K0 1x | 0.054844 | 0.003982 | 1 | 3 | 2 | Q3U2K0 | Protein FAI | Q3U2K0 [8 | 0 | 1816.95 | 0 | 0.32 | 0.054444 | 21.68 | 87.8 | 112.2 | 8.95 | 11.43 | | High | High | High | High | High | High | High | High | High | High | High | 0.002059 | 0.02134 | 2.56 |
| FALSE | High | SASPSQK | [R].SASPSC | 1xPhospho | O08856 1x | 0.089425 | 0.006807 | 1 | 1 | 1 | O08856 | RNA polym | O08856 [3 | 0 | 1242.65 | 0 | -0.32 | 0.015861 | 8 | 111.1 | 88.9 | 6.38 | 7.58 | | High | High | High | High | High | High | High | High | High | High | High | 0.003564 | 0.03792 | 1.56 |
| FALSE | High | SASPSTQPI | [R].SASPST | 1xPhospho | B1AR09 1x | 0.020679 | 0.001472 | 1 | 4 | 1 | B1AR09 | Myeloid/ly | B1AR09 [2 | 0 | 1569.793 | 0 | 0.06 | 0.122836 | 6.25 | 97.5 | 102.5 | 5.51 | 2.68 | | High | High | High | High | High | High | High | High | High | High | High | 0.000786 | 0.0071 | 1.43 |
| FALSE | High | SASPYHGF | [R].SASPYH | 1xPhospho | Q91YD3 1x | 0.115566 | 0.009504 | 1 | 2 | 1 | Q91YD3 | mRNA-dec | Q91YD3 [8 | 0 | 1757.857 | 0 | -1.08 | 0.000936 | 1.93 | 134.4 | 65.6 | 7.81 | 6.79 | | High | High | High | High | High | High | High | High | High | High | High | 0.004733 | 0.05111 | 1.33 |
| FALSE | High | SEQQAEAL | [K].SEQQA | 1xPhospho | Q3UZ39 1x | 0.011549 | 0.00086 | 1 | 2 | 1 | Q3UZ39 | Leucine-ric | Q3UZ39 [5 | 1 | 2326.226 | 0 | 0.45 | 0.024243 | 12.23 | 87.7 | 112.3 | 18.05 | 5.57 | | High | High | High | High | High | High | High | High | High | High | High | 0.000437 | 0.003686 | 1.2 |
| FALSE | High | SASQGALT | [R].SASQG/ | 1xPhospho | B7ZCJ1 1xF | 7.16E-06 | 0 | 1 | 6 | 1 | B7ZCJ1 | Rho GTPas | B7ZCJ1 [48 | 0 | 2179.049 | 0 | 0.14 | 0.902173 | 6.71 | 95.3 | 104.7 | 15.54 | 16.28 | | High | High | High | High | High | High | High | High | High | High | High | 0 | 9.26E-07 | 2.39 |
| FALSE | High | SASQLSQTI | [R].SASQLS | 1xPhospho | D9HP81 1x | 0.003224 | 0.00016 | 1 | 5 | 1 | D9HP81 | Regulating | D9HP81 [1 | 0 | 1787.894 | 0 | 0.37 | 0.013863 | 4.51 | 87.9 | 112.1 | 6.25 | 8.76 | | High | High | High | High | High | High | High | High | High | High | High | 7.19E-05 | 0.00089 | 3.41 |
| FALSE | High | SEQQAEAL | [K].SEQQA | 1xPhospho | Q3UZ39 1x | 3.65E-07 | 0 | 1 | 2 | 5 | Q3UZ39 | Leucine-ric | Q3UZ39 [5 | 0 | 1968.968 | 0 | 0.56 | 0.001667 | 7.4 | 81 | 119 | 6.12 | 3.33 | | High | High | High | High | High | High | High | High | High | High | High | 0 | 5.27E-08 | 4.41 |
| FALSE | High | SASQMEV/ | [R].SASQM | 1xPhospho | Q9JJ66 1xP | 0.065952 | 0.004766 | 1 | 1 | 1 | Q9JJ66 | Cell divisio | Q9JJ66 [84 | 0 | 2036.054 | 0 | 0.13 | 0.62649 | 10.11 | 95.6 | 104.4 | 8.02 | 14.1 | | High | High | High | High | High | High | High | High | High | High | High | 0.002446 | 0.02641 | 1.75 |
| FALSE | High | SASQSSLDI | [R].SASQSS | 1xPhospho | B8X349 1xI | 0.010758 | 0.000772 | 2 | 15 | 3 | B8X349; F6 | JNK-intera( | B8X349 [7 | 0 | 1460.74 | 0 | 2.1 | 0.000965 | 4.11 | 36 | 164 | 4.5 | 12.64 | Shared | High | High | High | High | High | High | High | High | High | High | High | 0.009773 | 0.1054 | 2.06 |
| FALSE | High | SASQSSLDI | [R].SASQSS | 2xPhospho | B8X349 2xI | 0.005924 | 0.00062 | 2 | 15 | 3 | B8X349; F6 | JNK-intera( | B8X349 [7 | 0 | 1540.706 | 0 | 2.05 | 0.000793 | 10.06 | 40.7 | 159.3 | 8.13 | 11.01 | Shared | High | High | High | High | High | High | High | High | High | High | High | 0.000275 | 0.001751 | 2.71 |
| FALSE | High | SASSASSLF | [R].SASSAS | 1xPhospho | Q3UR00 1x | 0.002376 | 0.000109 | 1 | 8 | 1 | Q3UR00 | Putative ur | Q3UR00 [7 | 0 | 2302.08 | 0 | -0.34 | 0.025456 | 8.64 | 106.6 | 93.4 | 8.54 | 15.38 | | High | High | High | High | High | High | High | High | High | High | High | 4.94E-05 | 0.000629 | 2.7 |
| FALSE | High | SEQPLSQVI | [K].SEQPLS | 2xPhospho | Q8BTI8 2xF | 0.086562 | 0.006435 | 1 | 3 | 1 | Q8BTI8 | Serine/argi | Q8BTI8 [12 | 0 | 2494.24 | 0 | -0.54 | 0.01476 | 3.73 | 129.6 | 70.4 | 18.6 | 14.58 | | High | High | High | High | High | High | High | High | High | High | High | 0.003303 | 0.03642 | 2.81 |
| FALSE | High | SASSCDVSS | [K].SASSCD | 1xCarbami | P61460-2 1 | 7.77E-06 | 0 | 1 | 5 | 3 | P61460-2 | Isoform 2 c | P61460-2 [ | 0 | 1932.857 | 0 | -0.07 | 0.61969 | 4.6 | 102.3 | 97.7 | 4.21 | 6.1 | | High | High | High | High | High | High | High | High | High | High | High | 0 | 1.02E-06 | 2.54 |
| FALSE | High | SASSDTSEE | [R].SASSDT | 1xPhospho | P70441 1xI | 2.9E-08 | 0 | 1 | 1 | 14 | P70441 | Na(+)/H(+) | P70441 [28 | 0 | 2320.059 | 0 | -0.33 | 0.006567 | 6.04 | 110.9 | 89.1 | 2.35 | 6.15 | | High | High | High | High | High | High | High | High | High | High | High | 0 | 1.89E-09 | 4.05 |
| FALSE | High | SASSDTSEE | [R].SASSDT | 2xPhospho | P70441 2xI | 3.29E-05 | 0 | 1 | 1 | 3 | P70441 | Na(+)/H(+) | P70441 [28 | 0 | 2400.026 | 0 | -0.28 | 0.187185 | 2.97 | 108.9 | 91.1 | 3.76 | 13.8 | | High | High | High | High | High | High | High | High | High | High | High | 0 | 5.13E-06 | 3.6 |
| FALSE | High | SEQPLSQVI | [K].SEQPLS | 1xPhospho | Q8BTI8 1xF | 2.4E-08 | 0 | 1 | 3 | 3 | Q8BTI8 | Serine/argi | Q8BTI8 [12 | 0 | 2414.273 | 0 | -0.56 | 0.003623 | 1.9 | 119.1 | 80.9 | 6.04 | 8.65 | | High | High | High | High | High | High | High | High | High | High | High | 0 | 1.54E-09 | 4.03 |
| FALSE | High | SASSFK | [K].SASSFK | 1xPhospho | Q7TT36 1x | 0.105058 | 0.008518 | 1 | 2 | 1 | Q7TT36 | Adhesion G | Q7TT36 [1 | 0 | 1164.607 | 0 | 0.33 | 0.009947 | 11.17 | 86.4 | 113.6 | 13.9 | 10.43 | | High | High | High | High | High | High | High | High | High | High | High | 0.004277 | 0.04568 | 1.09 |
| FALSE | High | SASSFVQTS | [K].SASSFV | 1xPhospho | Q9ERU9 1x | 0.007811 | 0.000664 | 1 | 2 | 2 | Q9ERU9 | E3 SUMO-p | Q9ERU9 [1 | 0 | 1726.882 | 0 | 0.4 | 0.106461 | 27.52 | 86.2 | 113.8 | 11.13 | 24.64 | | High | High | High | High | High | High | High | High | High | High | High | 0.000327 | 0.002379 | 2.04 |
| FALSE | High | SASSHSDIE | [K].SASSHS | 1xPhospho | E9Q842 1x | 0.000269 | 0 | 1 | 3 | 2 | E9Q842 | Neuron na | E9Q842 [1 | 0 | 2746.253 | 0 | -0.17 | 0.782726 | 6.74 | 106.6 | 93.4 | 14.97 | 4.97 | | High | High | High | High | High | High | High | High | High | High | High | 0 | 5.47E-05 | 2.52 |
| FALSE | High | SEQPLSQVI | [K].SEQPLS | 1xPhospho | Q8BTI8 1xF | 3.67E-05 | 0 | 1 | 3 | 4 | Q8BTI8 | Serine/argi | Q8BTI8 [12 | 0 | 2185.11 | 0 | -0.3 | 0.642015 | 1.89 | 111.4 | 88.6 | 68.63 | 54.38 | | High | High | High | High | High | High | High | High | High | High | High | 0 | 2.06E-05 | 3.7 |
| FALSE | High | SENACSCSI | [K].SENACS | 2xCarbami | E9Q3G8 1x | 0.000651 | 0 | 1 | 2 | 2 | E9Q3G8 | Nucleopor | E9Q3G8 [6 | 0 | 2726.106 | 0 | 0.51 | 0.006608 | 12.43 | 83.9 | 116.1 | 7.89 | 4.6 | | High | High | High | High | High | High | High | High | High | High | High | 0 | 0.000148 | 3.46 |
| FALSE | High | SASWGSTD | [R].SASWG | 1xPhospho | Q7TNF9 1x | 0.058173 | 0.004227 | 1 | 2 | 1 | Q7TNF9 | Protein FAI | Q7TNF9 [1 | 0 | 1412.616 | 0 | -1.17 | 0.001448 | 2.59 | 139.4 | 60.6 | 8.2 | 16.39 | | High | High | High | High | High | High | High | High | High | High | High | 0.002187 | 0.02296 | 1.77 |
| FALSE | High | SATDADMY | [R].SATDAI | 1xPhospho | Q6DFZ1 1x | 0.000267 | 0 | 1 | 4 | 1 | Q6DFZ1 | Golgi-speci | Q6DFZ1 [1 | 0 | 2304.135 | 0 | 1.24 | 0.005267 | 28.07 | 55.5 | 144.5 | 20.8 | 18.33 | | High | High | High | High | High | High | High | High | High | High | High | 0 | 5.39E-05 | 2.93 |
| FALSE | High | SATSGDIW | [K].SATSGI | 1xPhospho | Q8K019 1x | 1.46E-05 | 0 | 1 | 10 | 7 | Q8K019 | Bcl-2-assoc | Q8K019 [2 | 0 | 2303.054 | 0 | 0.12 | 0.990078 | 23.02 | 96 | 104 | 10.65 | 20.08 | | High | High | High | High | High | High | High | High | High | High | High | 0 | 0.000163 | 3.77 |
| FALSE | High | SATSGDIW | [K].SATSGI | 2xPhospho | Q8K019 2x | 4.12E-05 | 0 | 1 | 10 | 6 | Q8K019 | Bcl-2-assoc | Q8K019 [2 | 0 | 2383.021 | 0 | -0.76 | 0.001617 | 4.11 | 124.3 | 75.7 | 4.19 | 17.11 | | High | High | High | High | High | High | High | High | High | High | High | 0.000327 | 0.002928 | 3.98 |
| FALSE | High | SENVENTV | [K].SENVEI | 1xPhospho | E9PUJ6 1xF | 0.061176 | 0.004646 | 1 | 4 | 1 | E9PUJ6 | E3 ubiquiti | E9PUJ6 [12 | 0 | 2109.124 | 0 | 0.81 | 0.042976 | 13.13 | 73 | 127 | 39.87 | 11.3 | | High | High | High | High | High | High | High | High | High | High | High | 0.002354 | 0.02431 | 2.3 |
| FALSE | High | SENVEGTP | [K].SENVE( | 1xPhospho | Q01320 1x | 2.94E-07 | 0 | 2 | 3 | 2 | Q01320; Q | DNA topois | Q01320 [1 | 0 | 2223.997 | 0 | -0.52 | 0.011675 | 2.2 | 115.5 | 84.5 | 7.89 | 11.43 | Shared | High | High | High | High | High | High | High | High | High | High | High | 0 | 2.58E-08 | 3.85 |
| FALSE | High | SATSSSPGS | [K].SATSSS | 1xPhospho | G5E829 1x | 5.23E-06 | 0 | 1 | 9 | 6 | G5E829 | Plasma me | G5E829 [1 | 0 | 2066.984 | 0 | 2.31 | 0.004095 | 14.63 | 36.6 | 163.4 | 20.61 | 20.95 | | High | High | High | High | High | High | High | High | High | High | High | 0 | 6.5E-07 | 2.72 |
| FALSE | High | SATSSSPGS | [K].SATSSS | 2xPhospho | G5E829 2x | 0.008847 | 0.000736 | 1 | 9 | 4 | G5E829 | Plasma me | G5E829 [1 | 0 | 2146.951 | 0 | 1.87 | 0.000532 | 8.56 | 42.6 | 157.4 | 4.93 | 9.4 | | High | High | High | High | High | High | High | High | High | High | High | 0.000327 | 0.002742 | 2.67 |
| FALSE | High | SENQEALA | [K].SENQE/ | 1xPhospho | B2RUG8 1x | 1.5E-05 | 0 | 1 | 5 | 2 | B2RUG8 | Asxl2 prote | B2RUG8 [5 | 0 | 1911.983 | 0 | -0.1 | 0.321805 | 13.12 | 103.4 | 96.6 | 4.23 | 8.19 | | High | High | High | High | High | High | High | High | High | High | High | 0 | 2.13E-06 | 3.47 |
| FALSE | High | SATTTGSP( | [R].SATTTG | 1xPhospho | A0A1W2P7 | 2.06E-06 | 0 | 1 | 7 | 3 | A0A1W2P7 | Ral GTPase | A0A1W2P7 | 0 | 1674.835 | 0 | 0.59 | 0.000831 | 2.08 | 79.3 | 120.7 | 1.5 | 5.76 | | High | High | High | High | High | High | High | High | High | High | High | 0 | 2.3E-07 | 3.18 |
| FALSE | High | SENNSSNS | [K].SENNS! | 1xPhospho | P59017 1xI | 0.000558 | 0 | 1 | 3 | 7 | P59017 | Bcl-2-like p | P59017 [2 | 0 | 2197.054 | 0 | 0.82 | 0.004164 | 22.44 | 69.1 | 130.9 | 4.23 | 12.24 | | High | High | High | High | High | High | High | High | High | High | High | 0 | 0.000268 | 3.29 |
| FALSE | High | SATTTPSGS | [K].SATTTP | 1xPhospho | Q3UHJ0 1x | 0.002671 | 0.000109 | 1 | 2 | 2 | Q3UHJ0 | AP2-associ | Q3UHJ0 [6 | 0 | 1370.651 | 0 | 0.18 | 0.086215 | 12.14 | 96.6 | 103.4 | 8.4 | 5.64 | | High | High | High | High | High | High | High | High | High | High | High | 4.94E-05 | 0.000717 | 0.73 |
| FALSE | High | SATTTPSGS | [K].SATTTP | 2xPhospho | Q3UHJ0 2x | 0.079766 | 0.00613 | 1 | 2 | 1 | Q3UHJ0 | AP2-associ | Q3UHJ0 [6 | 0 | 1450.618 | 0 | -0.02 | 0.89462 | 7.41 | 103.4 | 96.6 | 8.13 | 15.4 | | High | High | High | High | High | High | High | High | High | High | High | 0.003148 | 0.03309 | 1.46 |
| FALSE | High | SENESLSPE | [R].SENESL | 1xPhospho | Q91WC0 1 | 2.04E-09 | 0 | 1 | 3 | 4 | Q91WC0 | Histone-lys | Q91WC0 [! | 0 | 3168.382 | 0 | 0.63 | 0.005132 | 13.12 | 78.5 | 121.5 | 7.2 | 7.5 | | High | High | High | High | High | High | High | High | High | High | High | 0 | 9.7E-11 | 4.99 |
| FALSE | High | SATVSPQP! | [R].SATVSF | 1xPhospho | A0A1L7NR( | 0.000194 | 0 | 1 | 1 | 1 | A0A1L7NR( | C-Cbl assoc | A0A1L7NR( | 0 | 3000.472 | 0 | 0.65 | 0.046156 | 21.21 | 77.7 | 122.3 | 16.87 | 17.32 | | High | High | High | High | High | High | High | High | High | High | High | 0 | 3.81E-05 | 2.3 |
| FALSE | High | SENESLSPE | [R].SENESL | 1xPhospho | Q91WC0 1 | 8.84E-07 | 0 | 1 | 3 | 2 | Q91WC0 | Histone-lys | Q91WC0 [! | 0 | 3152.387 | 0 | 0.41 | 0.046341 | 12.56 | 86.2 | 113.8 | 20.15 | 5.35 | | High | High | High | High | High | High | High | High | High | High | High | 0 | 8.82E-08 | 4.37 |
| FALSE | High | SATVSPQQ | [R].SATVSF | 1xPhospho | D3Z5J3 1xF | 0.000229 | 0 | 1 | 10 | 4 | D3Z5J3 | Sorbin and | D3Z5J3 [85 | 0 | 1834.901 | 0 | 1.23 | 0.000717 | 10.16 | 60.4 | 139.6 | 6.54 | 6.17 | | High | High | High | High | High | High | High | High | High | High | High | 0 | 4.54E-05 | 2.47 |
| FALSE | High | SAVAVYSY! | [R].SAVAVY | 1xCarbami | Q8BGW1 1 | 0.000738 | 0 | 1 | 4 | 1 | Q8BGW1 | Alpha-keto | Q8BGW1 [ | 0 | 3185.262 | 0 | -0.52 | 0.182182 | 11.79 | 113.3 | 86.7 | 4.98 | 29.88 | | High | High | High | High | High | High | High | High | High | High | High | 0 | 0.000169 | 4.11 |
| FALSE | High | SENAFDQA | [R].SENAFI | 1xPhospho | Q8BFW6 1: | 1.64E-05 | 0 | 1 | 4 | 3 | Q8BFW6 | Ectonuclec | Q8BFW6 [! | 0 | 1606.647 | 0 | 3.98 | 0.00041 | 10.62 | 11.9 | 188.1 | 15.01 | 10.48 | | High | High | High | High | High | High | High | High | High | High | High | 0 | 2.35E-06 | 3.31 |
| FALSE | High | SAVAVYSY! | [R].SAVAVY | 1xCarbami | Q8BGW1 2 | 0.000117 | 0 | 1 | 4 | 1 | Q8BGW1 | Alpha-keto | Q8BGW1 [ | 0 | 3265.228 | 0 | 0.05 | 0.837199 | 14.83 | 99.9 | 100.1 | 3.3 | 14.59 | | High | High | High | High | High | High | High | High | High | High | High | 0 | 2.15E-05 | 4.93 |
| FALSE | High | SAVDLTPIV | [K].SAVDLT | 1xPhospho | Q6PDQ2 1x | 3.23E-05 | 0 | 1 | 5 | 3 | Q6PDQ2 | Chromodo | Q6PDQ2 [1 | 0 | 1924.044 | 0 | -0.06 | 0.214524 | 5.51 | 102.2 | 97.8 | 2.26 | 8.46 | | High | High | High | High | High | High | High | High | High | High | High | 0 | 5.05E-06 | 3.06 |
| FALSE | High | SATPPPAEI | [K].SATPPP | 1xPhospho | Q542W3 1: | 1.39E-06 | 0 | 1 | 2 | 4 | Q542W3 | GTPase act | Q542W3 [2 | 0 | 2605.379 | 0 | -0.51 | 0.007794 | 4.63 | 119.2 | 80.8 | 9.93 | 2.75 | | High | High | High | High | High | High | High | High | High | High | High | 0 | 1.64E-07 | 2.55 |
| FALSE | High | SASYSYLED | [R].SASYSY | 1xPhospho | Q9DBR7 1x | 0.000235 | 0 | 1 | 4 | 4 | Q9DBR7 | Protein ph( | Q9DBR7 [9 | 0 | 1499.662 | 0 | 0.22 | 0.348614 | 38.37 | 92.4 | 107.6 | 5.82 | 21.19 | | High | High | High | High | High | High | High | High | High | High | High | 0 | 4.68E-05 | 2.99 |
| FALSE | High | SATPPPAEI | [K].SATPPP | 2xPhospho | Q3U6B1 2x | 3.5E-05 | 0 | 1 | 2 | 2 | Q3U6B1 | Putative ur | Q3U6B1 [2 | 0 | 2432.192 | 0 | 0.47 | 0.032791 | 8.39 | 86.2 | 113.8 | 6.84 | 14.68 | | High | High | High | High | High | High | High | High | High | High | High | 0 | 5.55E-06 | 2.37 |
| FALSE | High | SATPPPAEI | [K].SATPPP | 1xPhospho | Q3U6B1 1x | 0.047488 | 0.00354 | 1 | 2 | 3 | Q3U6B1 | Putative ur | Q3U6B1 [2 | 0 | 2123.062 | 0 | -0.5 | 0.509405 | 19.82 | 118.2 | 81.8 | 73.14 | 67.67 | | High | High | High | High | High | High | High | High | High | High | High | 0.001812 | 0.01818 | 2.56 |
| FALSE | High | SATDGNTS | [R].SATDGI | 1xPhospho | A2A432 1x | 2.51E-08 | 0 | 1 | 4 | 9 | A2A432 | Cullin-4B C | A2A432 [9 | 0 | 2074.011 | 0 | -1.14 | 0.000693 | 0.83 | 138.1 | 61.9 | 4.1 | 9.75 | | High | High | High | High | High | High | High | High | High | High | High | 0 | 1.61E-09 | 3.39 |
| FALSE | High | SEPSADSPI | [R].SEPSAD | 1xCarbami | Q8CAF4-3 | 8.34E-09 | 0 | 1 | 3 | 2 | Q8CAF4-3 | Isoform 3 c | Q8CAF4-3 | 0 | 2387.047 | 0 | 0.22 | 0.09332 | 7.89 | 95.2 | 104.8 | 10.26 | 7.26 | | High | High | High | High | High | High | High | High | High | High | High | 0 | 4.67E-10 | 4.28 |
| FALSE | High | SATDGNTS | [R].SATDGI | 1xDeamida | A2A432 1x | 0.032831 | 0.002195 | 1 | 4 | 1 | A2A432 | Cullin-4B C | A2A432 [9 | 0 | 2074.995 | 0 | -1.12 | 0.000661 | 1.28 | 137.6 | 62.4 | 2.5 | 12.42 | | High | High | High | High | High | High | High | High | High | High | High | 0.001151 | 0.01191 | 2.65 |
| FALSE | High | SEPQPEEG! | [R].SEPQPE | 1xPhospho | O88508 1x | 0.000779 | 0 | 1 | 6 | 1 | O88508 | DNA (cytos | O88508 [9 | 0 | 2049.989 | 0 | -0.17 | 0.019723 | 4.68 | 103.2 | 96.8 | 5.05 | 7.92 | | High | High | High | High | High | High | High | High | High | High | High | 0 | 0.00018 | 2.29 |
| FALSE | High | SATDGNTS | [R].SATDGI | 2xPhospho | A2A432 2x | 0.000279 | 0 | 1 | 4 | 1 | A2A432 | Cullin-4B C | A2A432 [9 | 0 | 2153.977 | 0 | -1.26 | 0.000466 | 1.77 | 138.8 | 61.2 | 7.34 | 7.36 | | High | High | High | High | High | High | High | High | High | High | High | 0 | 5.69E-05 | 2.55 |
| FALSE | High | SEPPGSGSI | [K].SEPPGS | 1xPhospho | A2A6A1 1x | 0.013666 | 0.001011 | 1 | 1 | 6 | A2A6A1 | G patch do | A2A6A1 [7 | 0 | 1544.731 | 0 | 0.12 | 0.020742 | 6.55 | 97.7 | 102.3 | 4.34 | 6.21 | | High | High | High | High | High | High | High | High | High | High | High | 0.002381 | 0.02461 | 2.39 |
| FALSE | High | SEPLPTFN( | [K].SEPLPT | 1xPhospho | Q8BHZ4 1x | 9.98E-05 | 0 | 1 | 4 | 1 | Q8BHZ4 | Zinc finger | Q8BHZ4 [1 | 0 | 3080.527 | 0 | -0.75 | 0.000589 | 2.14 | 123.7 | 76.3 | 2.7 | 11.88 | | High | High | High | High | High | High | High | High | High | High | High | 0 | 1.79E-05 | 3.77 |
| FALSE | High | SEPLPPSAT | [R].SEPLPP | 1xPhospho | Q8C2B3-3 | 0.003195 | 0.00016 | 1 | 11 | 1 | Q8C2B3-3 | Isoform 3 c | Q8C2B3-3 | 0 | 2351.257 | 0 | 1.07 | 0.000181 | 0.54 | 65.2 | 134.8 | 7.97 | 2.94 | | High | High | High | High | High | High | High | High | High | High | High | 7.19E-05 | 0.000873 | 2.42 |
| FALSE | High | SEPEQTGA | [K].SEPEQT | 1xCarbami | Q8BX09 1x | 1.01E-05 | 0 | 1 | 4 | 2 | Q8BX09 | Retinoblast | Q8BX09 [3 | 0 | 4610.006 | 0 | -0.85 | 0.001428 | 0.48 | 128.4 | 71.6 | 10.07 | 8.34 | | High | High | High | High | High | High | High | High | High | High | High | 0 | 1.89E-06 | 5.92 |
| FALSE | High | SATLLYDQI | [K].SATLLY | 1xPhospho | Q80TH2-2 | 0.042141 | 0.003162 | 1 | 6 | 2 | Q80TH2-2 | Isoform 2 c | Q80TH2-2 | 0 | 3352.733 | 0 | 0.91 | 0.006792 | 5.5 | 68.2 | 131.8 | 12.98 | 14.32 | | High | High | High | High | High | High | High | High | High | High | High | 0.002924 | 0.03035 | 3.76 |
| FALSE | High | SATISGSAT | [R].SATISG | 1xPhospho | E9Q4K7 1x | 0.033694 | 0.002195 | 1 | 2 | 1 | E9Q4K7 | Kinesin fan | E9Q4K7 [1 | 0 | 2386.299 | 0 | 0.63 | 0.000196 | 1.97 | 79.5 | 120.5 | 8.1 | 6.56 | | High | High | High | High | High | High | High | High | High | High | High | 0.001151 | 0.01229 | 2.65 |
| FALSE | High | SEPELTTVA | [K].SEPELT | 1xDeamida | E9Q6H8 1x | 8.7E-08 | 0 | 1 | 12 | 2 | E9Q6H8 | Pleckstrin I | E9Q6H8 [9 | 0 | 2602.206 | 0 | -0.08 | 0.24815 | 16.22 | 102.9 | 97.1 | 7.54 | 11.02 | | High | High | High | High | High | High | High | High | High | High | High | 0 | 6.55E-09 | 3.77 |
| FALSE | High | SEPELTTVA | [K].SEPELT | 1xPhospho | E9Q6H8 1x | 4.53E-07 | 0 | 1 | 12 | 2 | E9Q6H8 | Pleckstrin I | E9Q6H8 [9 | 0 | 2601.222 | 0 | -0.64 | 0.010058 | 4.73 | 122.7 | 77.3 | 6.49 | 20.67 | | High | High | High | High | High | High | High | High | High | High | High | 0 | 4.16E-08 | 4.87 |

| | | | | | | | | | | | | | | | | | | | | | | | | | | | | | | | | | | | | | | |
|---|---|---|---|---|---|---|---|---|---|---|---|---|---|---|---|---|---|---|---|---|---|---|---|---|---|---|---|---|---|---|---|---|---|---|---|---|---|---|
| FALSE | High | SATLSSTES | [K].SATLSS | 1xPhospho | B2RRE0 1x | 3.66E-08 | 0 | 1 | 3 | 7 | B2RRE0 | A kinase (P | B2RRE0 [62 | 0 | 2266.011 | 0 | -0.66 | 0.000738 | 2.95 | 120.6 | 79.4 | 5.02 | 6.15 | | High | High | High | High | High | High | High | High | High | High | High | 0 | 2.49E-09 | 4.07 |
| FALSE | High | SATLSSTES | [K].SATLSS | 1xPhospho | B2RRE0 1x | 6.36E-09 | 0 | 1 | 3 | 18 | B2RRE0 | A kinase (P | B2RRE0 [62 | 0 | 2282.006 | 0 | -1.15 | 0.003106 | 2.3 | 137.3 | 62.7 | 6.62 | 13.83 | | High | High | High | High | High | High | High | High | High | High | High | 0 | 2.21E-08 | 5.12 |
| FALSE | High | SEPASPQD | [K].SEPASP | 1xPhospho | Q8CBD1 1x | 0.0006 | 0 | 1 | 1 | 1 | Q8CBD1 | Nuclear re | Q8CBD1 [8 | 0 | 1964.941 | 0 | 1.52 | 0.000661 | 11.47 | 48.9 | 151.1 | 11.57 | 9.24 | | High | High | High | High | High | High | High | High | High | High | High | 0 | 0.000135 | 2.32 |
| FALSE | High | SATLSSTES | [K].SATLSS | 2xPhospho | B2RRE0 2x | 5.91E-05 | 0 | 1 | 3 | 5 | B2RRE0 | A kinase (P | B2RRE0 [62 | 0 | 2361.972 | 0 | -1.35 | 0.00061 | 2.26 | 143.5 | 56.5 | 3.72 | 19.86 | | High | High | High | High | High | High | High | High | High | High | High | 0.000344 | 0.003159 | 2.93 |
| FALSE | High | SATNLGQC | [K].SATNLC | 1xPhospho | Q9WVM1 1 | 0.014667 | 0.001049 | 1 | 1 | 1 | Q9WVM1 | Rac GTPase | Q9WVM1 [ | 0 | 1541.809 | 0 | -0.26 | 0.536529 | 21.66 | 108.8 | 91.2 | 7.73 | 27.41 | | High | High | High | High | High | High | High | High | High | High | High | 0.000555 | 0.004812 | 2.6 |
| FALSE | High | SATPANSD | [R].SATPAN | 1xPhospho | Q62388 1x | 0.000171 | 0 | 1 | 2 | 1 | Q62388 | Serine-pro | Q62388 [1 | 0 | 2033.901 | 0 | -0.37 | 0.00341 | 4.62 | 113.1 | 86.9 | 7.66 | 10.08 | | High | High | High | High | High | High | High | High | High | High | High | 0 | 3.3E-05 | 3.86 |
| FALSE | High | SATPPPAEF | [K].SATPPP | 1xPhospho | Q3U6B1 1x | 2.81E-07 | 0 | 1 | 2 | 7 | Q3U6B1 | Putative ur | Q3U6B1 [2 | 0 | 2352.225 | 0 | -0.54 | 0.001921 | 6.08 | 119.6 | 80.4 | 3.89 | 5.8 | | High | High | High | High | High | High | High | High | High | High | High | 0 | 2E-07 | 3.87 |
| FALSE | High | SCHDLDLL | [R].SCHDLD | 2xCarbami | Q921I6 1xP | 4.26E-07 | 0 | 1 | 1 | 1 | Q921I6 | SH3 domai | Q921I6 [29 | 0 | 3551.706 | 0 | -1.2 | 0.00184 | 1.62 | 138.6 | 61.4 | 8.35 | 16.7 | | High | High | High | High | High | High | High | High | High | High | High | 0 | 3.9E-08 | 5.67 |
| FALSE | High | SCSVTDTV | [R].SCSVTD | 1xCarbami | Q8CFI0 1xP | 0.000479 | 0 | 1 | 4 | 1 | Q8CFI0 | E3 ubiquiti | Q8CFI0 [36 | 0 | 2660.259 | 0 | -1.4 | 0.002764 | 4.25 | 144.5 | 55.5 | 11.14 | 17.99 | | High | High | High | High | High | High | High | High | High | High | High | 0 | 0.000105 | 4.35 |
| FALSE | High | SCLAEVSVS | [R].SCLAEV | 1xCarbami | P28359 2xP | 1.34E-06 | 0 | 1 | 1 | 1 | P28359 | Homeobox | P28359 [23 | 0 | 2367.096 | 0 | 4.22 | 0.000484 | 11.84 | 9.3 | 190.7 | 19.98 | 12.39 | | High | High | High | High | High | High | High | High | High | High | High | 0 | 1.41E-07 | 4.38 |
| FALSE | High | SEALEFVPI | [K].SEALEF | 1xPhospho | H7BX94 1x | 2.13E-05 | 0 | 1 | 2 | 3 | H7BX94 | Lysine-rich | H7BX94 [3 | 0 | 1970.029 | 0 | 0.12 | 0.135885 | 7.76 | 93.3 | 106.7 | 8.67 | 9.58 | | High | High | High | High | High | High | High | High | High | High | High | 0 | 3.17E-05 | 2.74 |
| FALSE | High | SEALEFVPI | [K].SEALEF | 1xPhospho | H7BX94 1x | 0.01861 | 0.001362 | 1 | 2 | 2 | H7BX94 | Lysine-rich | H7BX94 [3 | 0 | 1740.866 | 0 | 0.18 | 0.853855 | 35.1 | 96.5 | 103.5 | 31.73 | 36.47 | | High | High | High | High | High | High | High | High | High | High | High | 0.00111 | 0.01155 | 2.6 |
| FALSE | High | SDSALSAR | [K].SDSALS | 1xPhospho | Q01320 1x | 0.044358 | 0.003264 | 2 | 4 | 2 | Q01320; Q | DNA topois | Q01320 [1 | 0 | 1115.53 | 0 | -0.6 | 0.000549 | 0.93 | 120.7 | 79.3 | 2.14 | 5.86 | Shared | High | High | High | High | High | High | High | High | High | High | High | 0.002455 | 0.02668 | 1.94 |
| FALSE | High | SDSASDSQ | [K].SDSASD | 1xPhospho | O70311 1x | 0.001244 | 0 | 1 | 2 | 1 | O70311 | Glycylpept | O70311 [6 | 0 | 1704.809 | 0 | -0.56 | 0.004373 | 4.78 | 119.6 | 80.4 | 4.87 | 9.72 | | High | High | High | High | High | High | High | High | High | High | High | 0 | 0.000306 | 2.78 |
| FALSE | High | SDSDLLK | [R].SDSDLL | 1xPhospho | Q04692 1x | 0.027839 | 0.001755 | 1 | 1 | 1 | Q04692 | SWI/SNF-r | Q04692 [1 | 0 | 1315.691 | 0 | -0.55 | 0.001204 | 2.65 | 118.4 | 81.6 | 4.56 | 3.82 | | High | High | High | High | High | High | High | High | High | High | High | 0.000953 | 0.0099 | 1.75 |
| FALSE | High | SEAELQEA | [K].SEAELC | 1xPhospho | J7I1C3 1xP | 0.016596 | 0.001124 | 1 | 1 | 1 | J7I1C3 | Rif1 OS=M | J7I1C3 [178 | 0 | 2516.261 | 0 | -0.09 | 0.410179 | 7.27 | 101.7 | 98.3 | 26.19 | 15.4 | | High | High | High | High | High | High | High | High | High | High | High | 0.000585 | 0.00551 | 3.26 |
| FALSE | High | SDSDLLTCS | [K].SDSDLL | 1xCarbami | Q3UHP6 1x | 0.0004 | 0 | 1 | 3 | 1 | Q3UHP6 | Putative ur | Q3UHP6 [6 | 0 | 2351.993 | 0 | -0.19 | 0.056248 | 7.2 | 105 | 95 | 7.78 | 7.71 | | High | High | High | High | High | High | High | High | High | High | High | 0 | 8.52E-05 | 2.98 |
| FALSE | High | SDSDLLTCS | [K].SDSDLL | 1xCarbami | Q3UHP6 1x | 5.21E-05 | 0 | 1 | 3 | 1 | Q3UHP6 | Putative ur | Q3UHP6 [6 | 0 | 2367.988 | 0 | 0.53 | 0.055358 | 19.3 | 86.2 | 113.8 | 6.52 | 19.23 | | High | High | High | High | High | High | High | High | High | High | High | 0 | 8.59E-06 | 4.39 |
| FALSE | High | SDSDSSTLA | [R].SDSDSS | 1xPhospho | Q6NXJ0 1x | 2.44E-05 | 0 | 1 | 1 | 2 | Q6NXJ0 | Protein W | Q6NXJ0 [1 | 0 | 1548.756 | 0 | 0.14 | 0.164323 | 6.53 | 98.6 | 101.4 | 4.67 | 10.47 | | High | High | High | High | High | High | High | High | High | High | High | 0 | 2.55E-05 | 2.43 |
| FALSE | High | SDRGSGQ | [R].SDRGS | 1xPhospho | Q9CZX9 1x | 3.01E-06 | 0 | 1 | 1 | 2 | Q9CZX9 | ER membra | Q9CZX9 [3 | 1 | 2552.243 | 0 | 1.35 | 0.000909 | 8.5 | 55.5 | 144.5 | 7.91 | 8.65 | | High | High | High | High | High | High | High | High | High | High | High | 0 | 0.000325 | 3.75 |
| FALSE | High | SEAEDLAEF | [K].SEAEDL | 1xPhospho | Q6PIJ4 1xP | 0.004411 | 0.000409 | 1 | 1 | 4 | Q6PIJ4 | Nuclear fa | Q6PIJ4 [32 | 0 | 3762.934 | 0 | -0.1 | 0.104838 | 6.4 | 105 | 95 | 7.9 | 13.67 | | High | High | High | High | High | High | High | High | High | High | High | 0.000183 | 0.001262 | 2.64 |
| FALSE | High | SDSDSSTLS | [R].SDSDSS | 1xPhospho | Q5SXA9 1x | 0.01894 | 0.001362 | 1 | 2 | 1 | Q5SXA9 | Protein KIE | Q5SXA9 [9 | 0 | 1564.751 | 0 | -0.91 | 0.001341 | 2.08 | 129.1 | 70.9 | 3.85 | 9.11 | | High | High | High | High | High | High | High | High | High | High | High | 0.000723 | 0.006383 | 1.9 |
| FALSE | High | SDSDTLFET | [K].SDSDTL | 1xPhospho | Q3UHZ3 1x | 9.36E-08 | 0 | 1 | 3 | 5 | Q3UHZ3 | DNA (cytos | Q3UHZ3 [1 | 0 | 2108.959 | 0 | 0.26 | 0.024544 | 12.42 | 91 | 109 | 11.81 | 0.66 | | High | High | High | High | High | High | High | High | High | High | High | 4.94E-05 | 0.000687 | 3.57 |
| FALSE | High | SDSDTLFET | [K].SDSDTL | 2xPhospho | Q3UHZ3 2x | 0.031168 | 0.001935 | 1 | 3 | 1 | Q3UHZ3 | DNA (cytos | Q3UHZ3 [1 | 0 | 2188.925 | 0 | 0.5 | 0.020517 | 9.84 | 85 | 115 | 11.28 | 9.3 | | High | High | High | High | High | High | High | High | High | High | High | 0.001044 | 0.0112 | 2.18 |
| FALSE | High | SDSDTLSVE | [K].SDSDTL | 1xPhospho | P13864 1xP | 5.29E-07 | 0 | 1 | 3 | 9 | P13864 | DNA (cytos | P13864 [14 | 0 | 2147.991 | 0 | 0.1 | 0.428213 | 15.12 | 96.6 | 103.4 | 9.27 | 3.48 | | High | High | High | High | High | High | High | High | High | High | High | 2.58E-05 | 0.000384 | 3.83 |
| FALSE | High | SEAAAPHA | [K].SEAAAF | 2xPhospho | P51612 2xP | 9.73E-09 | 0 | 1 | 2 | 4 | P51612 | DNA repair | P51612 [86 | 0 | 3132.299 | 0 | 0.21 | 0.265453 | 15.44 | 95 | 105 | 7.72 | 14.7 | | High | High | High | High | High | High | High | High | High | High | High | 0 | 5.5E-07 | 4.48 |
| FALSE | High | SDSDTLSVE | [K].SDSDTL | 2xPhospho | P13864 2xP | 3.51E-06 | 0 | 1 | 3 | 1 | P13864 | DNA (cytos | P13864 [14 | 0 | 2227.957 | 0 | 0.44 | 0.019229 | 20.35 | 87 | 113 | 16.73 | 5.9 | | High | High | High | High | High | High | High | High | High | High | High | 0 | 4.19E-07 | 3.93 |
| FALSE | High | SDSDYDLSF | [R].SDSDYD | 1xPhospho | E9PVU7 1x | 0.004824 | 0.000456 | 1 | 14 | 1 | E9PVU7 | Phosphodi | E9PVU7 [1 | 0 | 1664.782 | 0 | -0.49 | 0.007146 | 2.87 | 114.2 | 85.8 | 8.85 | 5.7 | | High | High | High | High | High | High | High | High | High | High | High | 0.000204 | 0.001392 | 1.79 |
| FALSE | High | SDSDYDLSF | [R].SDSDYD | 2xPhospho | E9PVU7 2x | 0.016163 | 0.001124 | 1 | 14 | 2 | E9PVU7 | Phosphodi | E9PVU7 [1 | 0 | 1744.748 | 0 | 0.97 | 0.003082 | 23.18 | 67.6 | 132.4 | 10.39 | 11.75 | | High | High | High | High | High | High | High | High | High | High | High | 0.000585 | 0.00538 | 2.8 |
| FALSE | High | SDSEGSDY | [R].SDSEGS | 1xPhospho | Q6PDQ2 1x | 0.060159 | 0.004476 | 1 | 6 | 2 | Q6PDQ2 | Chromodo | Q6PDQ2 [9 | 0 | 1780.804 | 0 | -0.82 | 0.003295 | 6.4 | 127.9 | 72.1 | 7.97 | 7.74 | | High | High | High | High | High | High | High | High | High | High | High | 0.007957 | 0.0857 | 1.84 |
| FALSE | High | SEAAEPPKT | [K].SEAAEF | 1xCarbami | D3YXI6 1xP | 0.008313 | 0.000736 | 1 | 5 | 5 | D3YXI6 | Histone-lys | D3YXI6 [14 | 1 | 3120.589 | 0 | 0.45 | 0.002973 | 7.05 | 85.3 | 114.7 | 4.48 | 5.93 | | High | High | High | High | High | High | High | High | High | High | High | 0.000327 | 0.002544 | 3.14 |
| FALSE | High | SDSELVSK | [K].SDSELV | 1xPhospho | A0A0R4J0V | 0.008847 | 0.000736 | 1 | 8 | 1 | A0A0R4J0V | Phosphatic | A0A0R4J0V | 0 | 1402.723 | 0 | -0.63 | 0.003213 | 4.53 | 122.5 | 77.5 | 7.68 | 15.85 | | High | High | High | High | High | High | High | High | High | High | High | 0.000327 | 0.002724 | 1.38 |
| FALSE | High | SDQSPPSTI | [K].SDQSPI | 1xPhospho | Q7TT79 1x | 0.006031 | 0.00062 | 1 | 4 | 2 | Q7TT79 | Microceph | Q7TT79 [3 | 0 | 1396.667 | 0 | -0.98 | 0.000255 | 0.15 | 130.6 | 69.4 | 4.07 | 3.86 | | High | High | High | High | High | High | High | High | High | High | High | 0.000418 | 0.003609 | 1.9 |
| FALSE | High | SEALSSVLD | [R].SEALSS | 1xPhospho | P70399 1xP | 2.16E-07 | 0 | 1 | 4 | 2 | P70399 | TP53-bindi | P70399 [62 | 0 | 2407.135 | 0 | 0.06 | 0.193039 | 8.3 | 100.8 | 99.2 | 13.51 | 15.22 | | High | High | High | High | High | High | High | High | High | High | High | 0 | 1.83E-08 | 3.91 |
| FALSE | High | SDISPLTPR | [K].SDISPLT | 1xPhospho | P14873 1xP | 0.040028 | 0.002867 | 2 | 2 | 2 | P14873; B2 | Microtubu | P14873 [17 | 0 | 1294.661 | 0 | 0.68 | 0.00125 | 14.11 | 78.9 | 121.1 | 7.51 | 6.06 | Shared | High | High | High | High | High | High | High | High | High | High | High | 0.003291 | 0.03569 | 1.29 |
| FALSE | High | SEDDFDFA | [K].SEDDFD | 1xPhospho | A0A0R4J04 | 0.001881 | 5.66E-05 | 1 | 2 | 1 | A0A0R4J04 | CCAAT/enh | A0A0R4J04 | 0 | 2215.995 | 0 | -0.09 | 0.120557 | 8.22 | 104.9 | 95.1 | 7.35 | 8.21 | | High | High | High | High | High | High | High | High | High | High | High | 2.58E-05 | 0.000487 | 1.96 |
| FALSE | High | SEDCILSLD | [R].SEDCILS | 1xCarbami | B7ZP47 1xP | 2.82E-05 | 0 | 1 | 2 | 1 | B7ZP47 | Wapal prot | B7ZP47 [25 | 0 | 2550.212 | 0 | -0.35 | 0.239874 | 19.46 | 120.2 | 79.8 | 30.62 | 23.9 | | High | High | High | High | High | High | High | High | High | High | High | 0 | 4.36E-06 | 2.15 |
| FALSE | High | SDISPLTPR | [K].SDISPLT | 2xPhospho | P14873 2xP | 0.016892 | 0.001204 | 2 | 2 | 1 | P14873; B2 | Microtubu | P14873 [17 | 0 | 1374.627 | 0 | -0.02 | 0.623686 | 14.09 | 99.8 | 100.2 | 15.68 | 7.16 | Shared | High | High | High | High | High | High | High | High | High | High | High | 0.000638 | 0.005644 | 1.91 |
| FALSE | High | SDISSTVSE | [K].SDISSTV | 1xPhospho | Q5SWD9 1 | 0.000984 | 0 | 1 | 2 | 1 | Q5SWD9 | Pre-rRNA-p | Q5SWD9 [ | 0 | 1707.723 | 0 | -0.61 | 0.003751 | 5.14 | 125.9 | 74.1 | 12.68 | 17.26 | | High | High | High | High | High | High | High | High | High | High | High | 0 | 0.000234 | 2.53 |
| FALSE | High | SECSSDFG | [R].SECSSD | 1xCarbami | Q6PAR5 1x | 0.010382 | 0.000772 | 1 | 8 | 2 | Q6PAR5 | GTPase-act | Q6PAR5 [7 | 0 | 1611.713 | 0 | 0.03 | 0.90416 | 3.31 | 99 | 101 | 2.51 | 5.82 | | High | High | High | High | High | High | High | High | High | High | High | 0.000382 | 0.003277 | 1.59 |
| FALSE | High | SDISSTVSE | [K].SDISSTV | 1xPhospho | Q5SWD9 1 | 0.000772 | 0 | 1 | 2 | 6 | Q5SWD9 | Pre-rRNA-p | Q5SWD9 [ | 0 | 1723.718 | 0 | -0.56 | 0.002388 | 5.68 | 121.4 | 78.6 | 6.63 | 3.5 | | High | High | High | High | High | High | High | High | High | High | High | 0 | 0.000191 | 2.85 |
| FALSE | High | SDLTSELST | [K].SDLTSE | 1xPhospho | E9Q4Y4 1x | 0.086562 | 0.006435 | 1 | 1 | 1 | E9Q4Y4 | Centrosom | E9Q4Y4 [1 | 0 | 1730.877 | 0 | -0.99 | 0.005877 | 1.01 | 129.1 | 70.9 | 12.71 | 8.85 | | High | High | High | High | High | High | High | High | High | High | High | 0.003332 | 0.03652 | 1.88 |
| FALSE | High | SECSQASC | [K].SECSQA | 2xCarbami | F7BJB9 1xP | 0.0012 | 0 | 1 | 2 | 1 | F7BJB9 | MORC fam | F7BJB9 [79 | 0 | 1911.823 | 0 | 0.46 | 0.162937 | 31.97 | 89.1 | 110.9 | 11.81 | 18.83 | | High | High | High | High | High | High | High | High | High | High | High | 0 | 0.000293 | 1.91 |
| FALSE | High | SDQGVEGF | [K].SDQGV | 1xCarbami | A0A068BG | 3.47E-05 | 0 | 1 | 11 | 2 | A0A068BG | Retinoid X | A0A068BG | 0 | 3481.594 | 0 | -0.38 | 0.033554 | 7.97 | 107.3 | 92.7 | 10.92 | 9.61 | | High | High | High | High | High | High | High | High | High | High | High | 0 | 5.49E-06 | 4.79 |
| FALSE | High | SDNEETLG | [K].SDNEET | 1xPhospho | Q8C7S2 1x | 2.59E-06 | 0 | 1 | 5 | 5 | Q8C7S2 | Putative ur | Q8C7S2 [4 | 0 | 3805.843 | 0 | 0.7 | 0.005438 | 13.5 | 72.7 | 127.3 | 5.41 | 13.84 | | High | High | High | High | High | High | High | High | High | High | High | 0 | 2.96E-07 | 4.08 |
| FALSE | High | SDPEGGDS | [R].SDPEG | 1xCarbami | Q9D2D7 1x | 0.048715 | 0.00354 | 1 | 1 | 1 | Q9D2D7 | Zinc finger | Q9D2D7 [1 | 0 | 2548.204 | 0 | 0.61 | 0.009196 | 7.73 | 77 | 123 | 8.79 | 14.93 | | High | High | High | High | High | High | High | High | High | High | High | 0.001844 | 0.01867 | 3.15 |
| FALSE | High | SDPGPPTEI | [R].SDPGPI | 1xPhospho | P52734 1xP | 0.001629 | 5.66E-05 | 1 | 2 | 1 | P52734 | FYVE, Rho | P52734 [13 | 0 | 2380.268 | 0 | -1.25 | 0.004117 | 4.34 | 140.8 | 59.2 | 13.13 | 11.02 | | High | High | High | High | High | High | High | High | High | High | High | 2.58E-05 | 0.000412 | 2.1 |
| FALSE | High | SECDSSPEF | [R].SECDSS | 1xCarbami | Q8BTI8 1xP | 0.009416 | 0.000736 | 1 | 3 | 1 | Q8BTI8 | Serine/argi | Q8BTI8 [14 | 0 | 1673.749 | 0 | -0.87 | 0.003282 | 3.55 | 128 | 72 | 9.77 | 4.18 | | High | High | High | High | High | High | High | High | High | High | High | 0.000327 | 0.002926 | 2.44 |
| FALSE | High | SDPSLWNP | [R].SDPSLV | 1xPhospho | P70218 1xP | 0.000933 | 0 | 1 | 4 | 1 | P70218 | Mitogen-ac | P70218 [45 | 0 | 2631.317 | 0 | -2.43 | | 2.86 | 167.6 | 32.4 | 8.77 | 56.02 | | High | High | High | High | High | High | High | High | Not Found | High | High | 0 | 0.000221 | 3.09 |
| FALSE | High | SEASSSPPV | [K].SEASSS | 2xPhospho | P48432 2xP | 0.005135 | 0.000551 | 1 | 4 | 3 | P48432 | Transcripti | P48432 [24 | 0 | 2090.899 | 0 | 1.08 | 0.00682 | 12.38 | 60.8 | 139.2 | 7.78 | 14.83 | | High | High | High | High | High | High | High | High | High | High | High | 0.000294 | 0.002325 | 2.59 |
| FALSE | High | SDPSTPK | [K].SDPSTP | 1xPhospho | Q3UL54 1x | 0.015603 | 0.001124 | 1 | 8 | 2 | Q3UL54 | Putative ur | Q3UL54 [3 | 0 | 1269.649 | 0 | -0.55 | 0.003283 | 1.55 | 119.2 | 80.8 | 1.89 | 10.18 | | High | High | High | High | High | High | High | High | High | High | High | 0.000585 | 0.005151 | 1.5 |
| FALSE | High | SDPYHATT | [R].SDPYHA | 1xPhospho | Q7TMQ1 1 | 2.82E-08 | 0 | 1 | 2 | 4 | Q7TMQ1 | Gap juncti | Q7TMQ1 [ | 0 | 2096.047 | 0 | 3.65 | 0.00943 | 70.98 | 17.7 | 182.3 | 32.87 | 32 | | High | High | High | High | High | High | High | High | High | High | High | 0 | 3.9E-08 | 3.98 |
| FALSE | High | SEASSSPPV | [K].SEASSS | 1xPhospho | P48432 1xP | 0.000102 | 0 | 1 | 4 | 5 | P48432 | Transcripti | P48432 [24 | 0 | 2010.933 | 0 | 0.17 | 0.892286 | 21.55 | 94 | 106 | 13.25 | 14.65 | | High | High | High | High | High | High | High | High | High | High | High | 0 | 2.45E-05 | 2.67 |
| FALSE | High | SEASLLQLL | [K].SEASLL | 1xPhospho | Q66T02 1x | 0.001916 | 5.66E-05 | 1 | 4 | 1 | Q66T02 | Pleckstrin I | Q66T02 [9 | 0 | 1923.015 | 0 | 0.04 | 0.32731 | 3.89 | 99.3 | 100.7 | 8.43 | 3.87 | | High | High | High | High | High | High | High | High | High | High | High | 2.58E-05 | 0.000495 | 2.46 |
| FALSE | High | SDNEETLG | [K].SDNEET | 1xDeamida | Q8C7S2 1x | 0.000125 | 0 | 1 | 5 | 2 | Q8C7S2 | Putative ur | Q8C7S2 [4 | 0 | 3806.827 | 0 | 0.95 | 0.004439 | 14.53 | 66.2 | 133.8 | 6.4 | 11.49 | | High | High | High | High | High | High | High | High | High | High | High | 0 | 0.000023 | 4.08 |
| FALSE | High | SDSETEGL | [K].SDSETE | 1xPhospho | Q9Z2H5 1x | 0.002212 | 0.000109 | 1 | 2 | 2 | Q9Z2H5 | Band 4.1-li | Q9Z2H5 [6 | 0 | 2177.078 | 0 | -0.8 | 0.007755 | 2.85 | 126.9 | 73.1 | 10.85 | 8.19 | | High | High | High | High | High | High | High | High | High | High | High | 4.94E-05 | 0.000582 | 2.94 |
| FALSE | High | SDVYYFSPS | [K].SDVYYF | 1xPhospho | Q9Z2E1 1x | 0.00642 | 0.00062 | 1 | 2 | 1 | Q9Z2E1 | Methyl-Cp | Q9Z2E1 [1 | 0 | 1787.866 | 0 | 1.97 | 0.000324 | 16.67 | 38.8 | 161.2 | 14.32 | 6.1 | | High | High | High | High | High | High | High | High | High | High | High | 0.000275 | 0.001906 | 2.3 |
| FALSE | High | SDVSDQEE | [R].SDVSD | 1xCarbami | Q9R0L6-2 1 | 0.11108 | 0.008991 | 1 | 3 | 2 | Q9R0L6-2 | Isoform 2 c | Q9R0L6-2 [ | 1 | 3270.416 | 0 | 0.48 | 0.008708 | 11.47 | 84.7 | 115.3 | 8.57 | 14.74 | | High | High | High | High | High | High | High | High | High | High | High | 0.004517 | 0.049 | 3.36 |
| FALSE | High | SDSVDYGC | [R].SDSVDY | 1xPhospho | Q8CBY1 1x | 7.18E-07 | 0 | 1 | 3 | 2 | Q8CBY1 | Protein Sm | Q8CBY1 [1 | 0 | 2164.929 | 0 | 0.93 | 0.004241 | 13.36 | 75.4 | 124.6 | 12.23 | 15.25 | | High | High | High | High | High | High | High | High | High | High | High | 0 | 7.03E-08 | 3.57 |
| FALSE | High | SDSITEK | [R].SDSITEI | 1xPhospho | P70410 1xP | 0.013909 | 0.001011 | 1 | 4 | 3 | P70410 | Nucleolar ( | P70410 [50 | 0 | 1317.67 | 0 | 0.05 | 0.181807 | 5.99 | 98.7 | 101.3 | 5.27 | 4.11 | | High | High | High | High | High | High | High | High | High | High | High | 0.000538 | 0.00452 | 1.68 |
| FALSE | High | SDSSVITISF | [R].SDSSVI | 1xPhospho | Q80Z37 1x | 0.000154 | 0 | 1 | 2 | 2 | Q80Z37 | E3 ubiquiti | Q80Z37 [3 | 0 | 3120.413 | 0 | -0.08 | 0.337215 | 17.91 | 98.3 | 101.7 | 7.99 | 12.95 | | High | High | High | High | High | High | High | High | High | High | High | 0 | 2.91E-05 | 3.3 |
| FALSE | High | SDSNQYIR | [R].SDSNQ | 1xPhospho | F8VPT6 1xP | 0.021231 | 0.001507 | 1 | 3 | 2 | F8VPT6 | Ecotropic v | F8VPT6 [68 | 0 | 1291.588 | 0 | 0.63 | 0.001591 | 11.49 | 78.6 | 121.4 | 4.57 | 4.84 | | High | High | High | High | High | High | High | High | High | High | High | 0.001308 | 0.01355 | 2.16 |
| FALSE | High | SDSPAEEEF | [K].SDSPAE | 1xPhospho | B2RY61 1xP | 4.44E-06 | 0 | 1 | 3 | 1 | B2RY61 | AT rich inte | B2RY61 [93 | 0 | 2174.078 | 0 | 0.87 | 0.006124 | 5.22 | 73.2 | 126.8 | 4.55 | 12.96 | | High | High | High | High | High | High | High | High | High | High | High | 0 | 5.45E-07 | 3.14 |
| FALSE | High | SDSSQPML | [R].SDSSQF | 1xPhospho | P11531 1xP | 0.054844 | 0.003982 | 1 | 12 | 1 | P11531 | Dystrophin | P11531 [36 | 0 | 1458.686 | 0 | -0.58 | 0.037527 | 10.29 | 120.9 | 79.1 | 11.64 | 19.43 | | High | High | High | High | High | High | High | High | High | High | High | 0.002059 | 0.02151 | 1.62 |
| FALSE | High | SDSPAIQLR | [K].SDSPAI | 1xPhospho | Q60520-1 1 | 0.119269 | 0.00979 | 1 | 3 | 3 | Q60520-1 | Isoform 2 c | Q60520-1 [ | 0 | 1295.656 | 0 | -1.07 | 0.000241 | 0.98 | 134.8 | 65.2 | 3.77 | 2.08 | | High | High | High | High | High | High | High | High | High | High | High | 0.004866 | 0.05293 | 2.82 |
| FALSE | High | SDSPAWSL | [R].SDSPAV | 1xPhospho | A2A7B5 1x | 0.0009 | 0 | 1 | 2 | 1 | A2A7B5 | PR domain | A2A7B5 [7 | 0 | 1471.678 | 0 | -0.77 | 0.000411 | 1.87 | 127.5 | 72.5 | 4.58 | 6.38 | | High | High | High | High | High | High | High | High | High | High | High | 0 | 0.000212 | 2.16 |
| FALSE | High | SDSPAWSL | [R].SDSPAV | 2xPhospho | A2A7B5 2x | 0.08447 | 0.006345 | 1 | 2 | 1 | A2A7B5 | PR domain | A2A7B5 [7 | 0 | 1551.644 | 0 | -1.04 | 0.000605 | 2.63 | 128.5 | 71.5 | 8.07 | 13.51 | | High | High | High | High | High | High | High | High | High | High | High | 0.003265 | 0.03544 | 1.73 |
| FALSE | High | SDSLRPALN | [R].SDSLRP | 1xPhospho | Q3U007 1x | 0.087983 | 0.006639 | 1 | 2 | 1 | Q3U007 | Putative ur | Q3U007 [5 | 0 | 3574.722 | 0 | 1.54 | 0.009898 | 13.69 | 49.8 | 150.2 | 10.73 | 23.42 | | High | High | High | High | High | High | High | High | High | High | High | 0.003402 | 0.03702 | 1.74 |
| FALSE | High | SDSSPEEN | [R].SDSSPE | 1xDeamida | B2RWW2 1 | 3.87E-08 | 0 | 1 | 3 | 7 | B2RWW2 | Golgb1 pro | B2RWW2 [ | 0 | 2706.255 | 0 | 0.98 | 0.005468 | 8.45 | 69.8 | 130.2 | 6.51 | 15.01 | | High | High | High | High | High | High | High | High | High | High | High | 0 | 2.62E-09 | 4.08 |
| FALSE | High | SDSPEGLG | [K].SDSPEG | 1xCarbami | B8JJC1 1xP | 5.37E-06 | 0 | 1 | 3 | 1 | B8JJC1 | Tyrosyl-DN | B8JJC1 [79 | 0 | 3771.673 | 0 | -1.48 | 0.003751 | 3.01 | 145.1 | 54.9 | 18.17 | 10.77 | | High | High | High | High | High | High | High | High | High | High | High | 0 | 6.69E-07 | 5.41 |
| FALSE | High | SDSPEIPFQ | [R].SDSPEI | 1xPhospho | P70399 1xP | 9.19E-08 | 0 | 1 | 6 | 7 | P70399 | TP53-bindi | P70399 [14 | 0 | 3381.532 | 0 | 0.47 | 0.018174 | 22.72 | 84.7 | 115.3 | 6.54 | 11.76 | | High | High | High | High | High | High | High | High | High | High | High | 0 | 6.94E-09 | 6.23 |
| FALSE | High | SDSQASLT | [K].SDSQA | 1xPhospho | Q3UMB5 1 | 1.85E-06 | 0 | 1 | 2 | 2 | Q3UMB5 | Guanine nu | Q3UMB5 [ | 0 | 2102.084 | 0 | 0.94 | 0.002353 | 9.41 | 69.4 | 130.6 | 6.94 | 11.81 | | High | High | High | High | High | High | High | High | High | High | High | 0 | 2.03E-07 | 3.18 |
| FALSE | High | SDSQASLT | [K].SDSQA | 2xPhospho | Q3UMB5 2 | 0.000215 | 0 | 1 | 2 | 2 | Q3UMB5 | Guanine nu | Q3UMB5 [ | 0 | 2182.051 | 0 | 1.33 | 0.000311 | 8.88 | 57 | 143 | 5.48 | 6.62 | | High | High | High | High | High | High | High | High | High | High | High | 0 | 4.22E-05 | 3.76 |
| FALSE | High | SDSSGGYT | [R].SDSSG | 2xPhospho | Q6ZPR6 2x | 0.029844 | 0.001857 | 1 | 2 | 1 | Q6ZPR6 | Inhibitor o | Q6ZPR6 [9 | 0 | 2897.399 | 0 | -0.9 | 0.003856 | 3.25 | 131.2 | 68.8 | 13.61 | 16.88 | | High | High | High | High | High | High | High | High | High | High | High | 0.001011 | 0.01073 | 3.65 |
| FALSE | High | SDSSGGYT | [R].SDSSG | 1xPhospho | Q6ZPR6 1x | 0.00418 | 0.00036 | 1 | 2 | 3 | Q6ZPR6 | Inhibitor o | Q6ZPR6 [9 | 0 | 2817.432 | 0 | -0.93 | 0.006889 | 4.38 | 131.2 | 68.8 | 7.23 | 14.04 | | High | High | High | High | High | High | High | High | High | High | High | 0.000275 | 0.001638 | 4.26 |
| FALSE | High | SDSQASLT | [K].SDSQA | 3xPhospho | Q3UMB5 3 | 0.008769 | 0.000736 | 1 | 2 | 1 | Q3UMB5 | Guanine nu | Q3UMB5 [ | 0 | 2262.017 | 0 | 0.86 | 0.007613 | 8.47 | 74.7 | 125.3 | 8.01 | 15.2 | | High | High | High | High | High | High | High | High | High | High | High | 0.000327 | 0.002712 | 2.87 |
| FALSE | High | SDSSEVSAT | [K].SDSSEV | 1xPhospho | Q80U93 1x | 0.000132 | 0 | 1 | 3 | 2 | Q80U93 | Nuclear po | Q80U93 [1 | 0 | 3710.841 | 0 | -0.41 | 0.019346 | 9.51 | 114 | 86 | 17.59 | 19.93 | | High | High | High | High | High | High | High | High | High | High | High | 0 | 2.46E-05 | 3.83 |
| FALSE | High | SDSQQAVK | [R].SDSQQ | 1xPhospho | Q8BVL3 1x | 0.0006 | 0 | 1 | 1 | 2 | Q8BVL3 | Sorting ne | Q8BVL3 [4 | 0 | 1400.719 | 0 | 1.03 | 0.001852 | 5.91 | 66.2 | 133.8 | 2.66 | 12.14 | | High | High | High | High | High | High | High | High | High | High | High | 0 | 0.000135 | 2.56 |
| FALSE | High | SDSSPEEN | [R].SDSSPE | 1xPhospho | B2RWW2 1 | 1.69E-06 | 0 | 1 | 3 | 4 | B2RWW2 | Golgb1 pro | B2RWW2 [ | 0 | 2705.271 | 0 | 0.85 | 0.003802 | 17.19 | 67.8 | 132.2 | 9.45 | 10.3 | | High | High | High | High | High | High | High | High | High | High | High | 0 | 1.83E-07 | 4 |
| FALSE | High | SDSLPSFR | [K].SDSLPS | 1xPhospho | Q811L6 1xP | 0.096181 | 0.007535 | 1 | 4 | 1 | Q811L6 | Microtubu | Q811L6 [2 | 0 | 1217.576 | 0 | 1.35 | 0.013135 | 41.88 | 56.4 | 143.6 | 11.91 | 21.16 | | High | High | High | High | High | High | High | High | High | High | High | 0.003809 | 0.0411 | 1.92 |
| FALSE | High | SDSVTGHT | [R].SDSVT | 1xPhospho | A0A1S6GW | 0.003464 | 0.00021 | 1 | 6 | 3 | A0A1S6GW | Uncharacte | A0A1S6GW | 0 | 1684.831 | 0 | 0.74 | 0.028635 | 23.66 | 75 | 125 | 12.38 | 12.17 | | High | High | High | High | High | High | High | High | High | High | High | 9.42E-05 | 0.000961 | 2.17 |
| FALSE | High | SDSLPQVEI | [R].SDSLPC | 1xPhospho | Q76I79 1xP | 0.001168 | 0 | 1 | 3 | 1 | Q76I79 | Protein ph | Q76I79 [58 | 0 | 1911.972 | 0 | 0.38 | 0.371594 | 10.54 | 86.5 | 113.5 | 14.22 | 30.87 | | High | High | High | High | High | High | High | High | High | High | High | 0 | 0.000284 | 1.91 |
| FALSE | High | SDVSDQEE | [R].SDVSD | 1xCarbami | Q9R0L6-2 1 | 0.002313 | 0.000109 | 1 | 3 | 1 | Q9R0L6-2 | Isoform 2 c | Q9R0L6-2 [ | 1 | 3190.45 | 0 | 0.08 | 0.886261 | 24.35 | 100.6 | 99.4 | 4.87 | 25 | | High | High | High | High | High | High | High | High | High | High | High | 4.94E-05 | 0.000609 | 2.03 |
| FALSE | High | SDSFSVAAI | [R].SDSFSV | 1xPhospho | A7TU71 1x | 0.040028 | 0.002867 | 1 | 2 | 1 | A7TU71 | Protein Sh | A7TU71 [3 | 0 | 1248.582 | 0 | 1.95 | 0.001949 | 6.38 | 37.7 | 162.3 | 6.86 | 17.24 | | High | High | High | High | High | High | High | High | High | High | High | 0.001484 | 0.01488 | 2.23 |
| FALSE | High | SDSGAPDL | [K].SDSGA | 1xPhospho | Q6P5E3 1x | 0.003312 | 0.00016 | 1 | 3 | 1 | Q6P5E3 | Sal-like 1 ( | Q6P5E3 [5 | 0 | 1398.646 | 0 | -1.43 | 0.000261 | 0.12 | 145.7 | 54.3 | 8.29 | 8.67 | | High | High | High | High | High | High | High | High | High | High | High | 7.19E-05 | 0.00091 | 1.96 |
| FALSE | High | SDSGFAIQ | [K].SDSGFA | 1xPhospho | Q9DBN4 1x | 0.068752 | 0.005042 | 1 | 3 | 1 | Q9DBN4 | Putative m | Q9DBN4 [2 | 0 | 1724.866 | 0 | 0.43 | 0.087363 | 26.07 | 85.2 | 114.8 | 18.54 | 21.39 | | High | High | High | High | High | High | High | High | High | High | High | 0.002618 | 0.02793 | 2.46 |
| FALSE | High | SDVSDIGSI | [R].SDVSDI | 1xCarbami | Q3UGY8 1x | 8.25E-05 | 0 | 1 | 2 | 2 | Q3UGY8 | Brefeldin A | Q3UGY8 [6 | 0 | 2604.097 | 0 | 2.38 | 0.000333 | 5.57 | 31.8 | 168.2 | 9.43 | 8.12 | | High | High | High | High | High | High | High | High | High | High | High | 0 | 1.45E-05 | 4.51 |
| FALSE | High | SDVIHAPLF | [R].SDVIHA | 1xPhospho | Q3URD3 1x | 4.98E-05 | 0 | 1 | 6 | 1 | Q3URD3 | Sarcolemm | Q3URD3 [1 | 0 | 2013.082 | 0 | -0.3 | 0.006489 | 6.65 | 106.8 | 93.2 | 6.5 | 10.33 | | High | High | High | High | High | High | High | High | High | High | High | 0 | 8.18E-06 | 2.92 |
| FALSE | High | SDVGTSELI | [K].SDVGT | 1xPhospho | Q52KB6 1x | 7.72E-07 | 0 | 1 | 6 | 1 | Q52KB6 | C2 domain | Q52KB6 [4 | 0 | 2285.002 | 0 | -0.18 | 0.579778 | 1.13 | 105.2 | 94.8 | 6.78 | 6.22 | | High | High | High | High | High | High | High | High | High | High | High | 0 | 7.58E-08 | 3.87 |
| FALSE | High | SDSGGSSSI | [R].SDSGG | 1xPhospho | Q7TSC1 1x | 3.45E-06 | 0 | 1 | 3 | 2 | Q7TSC1 | Protein PRI | Q7TSC1 [7 | 0 | 1650.685 | 0 | 0.4 | 0.006276 | 7.57 | 81.1 | 118.9 | 7.94 | 7.97 | | High | High | High | High | High | High | High | High | High | High | High | 0 | 4.09E-07 | 2.73 |
| FALSE | High | SDSGSTSG | [K].SDSGST | 1xPhospho | Q9JJA7 1xP | 0.087983 | 0.006639 | 1 | 1 | 2 | Q9JJA7 | Cyclin-L2 C | Q9JJA7 [40 | 0 | 1507.704 | 0 | -0.69 | 0.007713 | 3.85 | 113.4 | 86.6 | 15.17 | 12.96 | | High | High | High | High | High | High | High | High | High | High | High | 0.003402 | 0.03703 | 1.62 |
| FALSE | High | SDSGVMLF | [K].SDSGVI | 1xPhospho | A2RSQ0 1x | 0.047488 | 0.00354 | 1 | 1 | 1 | A2RSQ0 | DENN dom | A2RSQ0 [8 | 0 | 1500.733 | 0 | 2.67 | 0.000209 | 5.76 | 27.4 | 172.6 | 15.66 | 3.57 | | High | High | High | High | High | High | High | High | High | High | High | 0.001812 | 0.01822 | 1.34 |
| FALSE | High | SDSHLGAY | [K].SDSHLG | 1xPhospho | E9QML5 1x | 6.31E-06 | 0 | 1 | 15 | 1 | E9QML5 | Zinc finger | E9QML5 [6 | 0 | 1810.889 | 0 | -0.42 | 0.0611 | 13.09 | 117.9 | 82.1 | 15.49 | 11.39 | | High | High | High | High | High | High | High | High | High | High | High | 0 | 8.08E-07 | 3.61 |
| FALSE | High | SDVEIPATV | [K].SDVEIP | 1xPhospho | Q9QYB5 1x | 0.000183 | 0 | 1 | 7 | 2 | Q9QYB5 | Gamma-ad | Q9QYB5 [4 | 0 | 3174.59 | 0 | 0.94 | 0.001949 | 6.14 | 68.7 | 131.3 | 7.41 | 9.2 | | High | High | High | High | High | High | High | High | High | High | High | 9.42E-05 | 0.000943 | 3.54 |
| FALSE | High | SDSLDSGVI | [K].SDSLDS | 1xPhospho | Q924A2 1x | 0.000908 | 0 | 1 | 1 | 2 | Q924A2 | Protein cap | Q924A2 [7 | 0 | 2726.251 | 0 | 0.14 | 0.105107 | 14.98 | 91.1 | 108.9 | 21.36 | 16.24 | | High | High | High | High | High | High | High | High | High | High | High | 0 | 0.000214 | 2.48 |
| FALSE | High | SDTPVEAQ | [R].SDTPVE | 1xCarbami | Q8CGF6 1x | 0.000253 | 0 | 1 | 2 | 1 | Q8CGF6 | WD repeat | Q8CGF6 [3 | 0 | 3073.426 | 0 | 0.42 | 0.026987 | 22.09 | 86.8 | 113.2 | 10.13 | 12.09 | | High | High | High | High | High | High | High | High | High | High | High | 0 | 5.1E-05 | 3.28 |
| FALSE | High | SDSYVELSE | [R].SDSYVE | 1xPhospho | A0A1S6GW | 0.000102 | 0 | 1 | 4 | 3 | A0A1S6GW | Nuclear ca | A0A1S6GW | 0 | 1656.736 | 0 | 1 | 0.004249 | 18.01 | 64.1 | 135.9 | 10.71 | 15.29 | | High | High | High | High | High | High | High | High | High | High | High | 0 | 1.83E-05 | 3.03 |
| FALSE | High | SDSYTLDPI | [K].SDSYTL | 1xPhospho | E9PUJ6 1xP | 0.039347 | 0.002793 | 1 | 5 | 2 | E9PUJ6 | E3 ubiquiti | E9PUJ6 [29 | 0 | 1691.773 | 0 | 0.27 | 0.092959 | 27.62 | 90.8 | 109.2 | 7.57 | 10.95 | | High | High | High | High | High | High | High | High | High | High | High | 0.001468 | 0.0147 | 2.85 |
| FALSE | High | SDSLDTDSS | [R].SDSLDT | 1xPhospho | Q6A065 1x | 0.000289 | 0 | 1 | 3 | 4 | Q6A065 | Centrosom | Q6A065 [8 | 0 | 2396.156 | 0 | 1.19 | 0.013622 | 17.45 | 57 | 143 | 12.83 | 27 | | High | High | High | High | High | High | High | High | High | High | High | 0 | 0.000239 | 2.59 |
| FALSE | High | SDSLGTQG | [R].SDSLGT | 1xPhospho | Q148V8 1x | 0.025073 | 0.001645 | 1 | 1 | 1 | Q148V8 | Protein FAI | Q148V8 [1 | 0 | 1229.572 | 0 | 1.05 | 0.000825 | 7.54 | 61.8 | 138.2 | 9.87 | 8.41 | | High | High | High | High | High | High | High | High | High | High | High | 0.000872 | 0.008828 | 1.87 |
| FALSE | High | SDSLILDHC | [R].SDSLILD | 1xPhospho | A0A087WC | 0.015061 | 0.001049 | 1 | 9 | 3 | A0A087WC | Kinesin-like | A0A087WC | 0 | 2251.141 | 0 | -0.74 | 0.006105 | 7.48 | 121.4 | 78.6 | 8.7 | 10.81 | | High | High | High | High | High | High | High | High | High | High | High | 0.000571 | 0.004962 | 3.17 |
| FALSE | High | SDSLLSFR | [R].SDSLLS | 1xPhospho | A0A0R4J0S | 0.026191 | 0.001676 | 1 | 2 | 2 | A0A0R4J0S | Cdc42 effe | A0A0R4J0S | 0 | 1233.608 | 0 | 1.43 | 0.000462 | 12.6 | 54.5 | 145.5 | 12.44 | 4.77 | | High | High | High | High | High | High | High | High | High | High | High | 0.000903 | 0.009275 | 1.38 |
| FALSE | High | SDSLPASR | [R].SDSLPA | 1xPhospho | Q1W617 1 | 0.031713 | 0.002011 | 1 | 2 | 1 | Q1W617 | Protein Sh | Q1W617 [2 | 0 | 1141.545 | 0 | 3.61 | 0.000582 | 30.06 | 15.8 | 184.2 | 26.13 | 12.66 | | High | High | High | High | High | High | High | High | High | High | High | 0.001093 | 0.01146 | 1.38 |
| FALSE | High | SEDDSAKFI | [K].SEDDSA | 1xPhospho | Q64511 1x | 0.003997 | 0.000311 | 1 | 2 | 2 | Q64511 | DNA topois | Q64511 [1 | 1 | 3560.692 | 0 | 0.51 | 0.105965 | 17.96 | 82.4 | 117.6 | 18.3 | 22.28 | | High | High | High | High | High | High | High | High | High | High | High | 0.00014 | 0.001127 | 3.3 |
| FALSE | High | SEDDSAVP | [R].SEDDSA | 1xPhospho | Q3U6Q1 1 | 0.00103 | 0 | 1 | 3 | 2 | Q3U6Q1 | Putative ur | Q3U6Q1 [6 | 0 | 1655.829 | 0 | 0.64 | 0.00813 | 21.76 | 72.1 | 127.9 | 14.74 | 10.39 | | High | High | High | High | High | High | High | High | High | High | High | 0 | 0.000247 | 2.43 |
| FALSE | High | SEDEESLAC | [R].SEDEES | 1xPhospho | Q3TYU7 1x | 0.01319 | 0.000889 | 1 | 3 | 1 | Q3TYU7 | Putative ur | Q3TYU7 [2 | 0 | 1730.825 | 0 | -0.8 | 0.002863 | 2.33 | 127.9 | 72.1 | 6.61 | 15.61 | | High | High | High | High | High | High | High | High | High | High | High | 0.000467 | 0.004295 | 2.02 |
| FALSE | High | SDISERPSV | [R].SDISERI | 1xPhospho | Q80X80 1x | 0.001689 | 5.66E-05 | 1 | 6 | 3 | Q80X80 | Phospholip | Q80X80 [6 | 0 | 2946.341 | 0 | 0.69 | 0.001821 | 10.44 | 78.9 | 121.1 | 5.54 | 7.16 | | High | High | High | High | High | High | High | High | High | High | High | 0.000786 | 0.006996 | 3.42 |
| FALSE | High | SCSLGALDI | [K].SCSLGA | 1xCarbami | Q7TSI1 1xP | 0.006834 | 0.00062 | 1 | 1 | 1 | Q7TSI1 | Pleckstrin I | Q7TSI1 [48 | 0 | 1488.753 | 0 | 0.48 | 0.003207 | 9.04 | 83.3 | 116.7 | 3.86 | 9.74 | | High | High | High | High | High | High | High | High | High | High | High | 0.000275 | 0.002058 | 2.16 |
| FALSE | High | SEGLSLER | [R].SEGLSL | 1xPhospho | Q3UMU9-3 | 0.078467 | 0.006104 | 1 | 4 | 1 | Q3UMU9-3 | Isoform 3 c | Q3UMU9-3 | 0 | 1199.587 | 0 | -0.58 | 0.020015 | 14.6 | 122.4 | 77.6 | 7.68 | 21.31 | | High | High | High | High | High | High | High | High | High | High | High | 0.003122 | 0.03243 | 1.52 |
| FALSE | High | SEGIEAELT | [K].SEGIEA | 1xPhospho | A2A5K6 1x | 4.43E-05 | 0 | 1 | 1 | 2 | A2A5K6 | Zinc finger | A2A5K6 [8 | 0 | 2668.219 | 0 | 0.19 | 0.196732 | 17.56 | 92.4 | 107.6 | 10.87 | 8.6 | | High | High | High | High | High | High | High | High | High | High | High | 0 | 7.18E-06 | 3.67 |

| | | | | | | | | | | | | | | | | | | | | | | | | | | | | | | | | | | | | | | |
|---|---|---|---|---|---|---|---|---|---|---|---|---|---|---|---|---|---|---|---|---|---|---|---|---|---|---|---|---|---|---|---|---|---|---|---|---|---|---|
| FALSE | High | SCSITGVDI | [R].SCSITG | 2xCarbami | A3KGI3 1xI | 0.004867 | 0.000504 | 1 | 2 | 2 | A3KGI3 | Kinetochor | A3KGI3 [14 | 0 | 3915.901 | 0 | -0.15 | 0.432534 | 14.09 | 105.5 | 94.5 | 6.07 | 9.3 | | High | High | High | High | High | High | High | High | High | High | High | 0.000225 | 0.001404 | 4.31 |
| FALSE | High | SCSMELHG | [K].SCSMEI | 1xCarbami | A6H8H2-2 | 1.77E-05 | 0 | 1 | 3 | 2 | A6H8H2-2 | Isoform 2 c | A6H8H2-2 | 0 | 3161.402 | 0 | 0.35 | 0.112424 | 10.4 | 88 | 112 | 8.13 | 22.93 | | High | High | High | High | High | High | High | High | High | High | High | 0 | 2.57E-06 | 3.6 |
| FALSE | High | SCSMHQLS | [K].SCSMH | 1xCarbami | B2RQE3 1x | 0.007145 | 0.00062 | 1 | 5 | 1 | B2RQE3 | Stim2 prot | B2RQE3 [7 | 0 | 2199.04 | 0 | 0.89 | 0.017128 | 4.54 | 68.9 | 131.1 | 12.96 | 13.64 | | High | High | High | High | High | High | High | High | High | High | High | 0.000275 | 0.002153 | 2.25 |
| FALSE | High | SCSMHQLS | [K].SCSMH | 1xCarbami | B2RQE3 1x | 0.002441 | 0.000109 | 1 | 5 | 1 | B2RQE3 | Stim2 prot | B2RQE3 [7 | 0 | 2215.035 | 0 | 1.09 | 0.004112 | 11.36 | 62.3 | 137.7 | 7.84 | 15.54 | | High | High | High | High | High | High | High | High | High | High | High | 4.94E-05 | 0.00065 | 2.97 |
| FALSE | High | SCSPEQPSI | [R].SCSPEC | 2xCarbami | A2ARL5 1xI | 1.8E-06 | 0 | 1 | 2 | 2 | A2ARL5 | Uncharacte | A2ARL5 [68 | 0 | 2278.029 | 0 | -1.47 | 0.000225 | 0.81 | 146.9 | 53.1 | 2.38 | 10.62 | | High | High | High | High | High | High | High | High | High | High | High | 0 | 1.97E-07 | 4.17 |
| FALSE | High | SCSPSSDSI | [R].SCSPSS | 1xCarbami | Q64355 1x | 0.019107 | 0.001402 | 1 | 2 | 1 | Q64355 | Embryonal | Q64355 [1 | 0 | 1768.823 | 0 | -1.95 | 0.000193 | 0.57 | 158.3 | 41.7 | 3.9 | 10.16 | | High | High | High | High | High | High | High | High | High | High | High | 0.00074 | 0.006477 | 1.82 |
| FALSE | High | SEGLSQEA | [R].SEGLSC | 1xCarbami | B1AQJ2 1xI | 0.000154 | 0 | 1 | 3 | 3 | B1AQJ2 | Ubiquitin c | B1AQJ2 [89 | 0 | 3600.611 | 0 | -0.54 | 0.001737 | 4.36 | 123 | 77 | 4.42 | 10.63 | | High | High | High | High | High | High | High | High | High | High | High | 0 | 2.9E-05 | 4.42 |
| FALSE | High | SCSQVLEH | [R].SCSQVI | 1xCarbami | O35465-2 1 | 0.003822 | 0.000261 | 1 | 3 | 1 | O35465-2 | Isoform 2 c | O35465-2 [ | 0 | 2193.078 | 0 | 1.02 | 0.003965 | 15.1 | 63.7 | 136.3 | 5.23 | 15.13 | | High | High | High | High | High | High | High | High | High | High | High | 0.000117 | 0.001073 | 3.62 |
| FALSE | High | SCTSSPSPL | [R].SCTSSP | 2xCarbami | A4QPD1 1x | 0.005717 | 0.00062 | 1 | 2 | 4 | A4QPD1 | Zinc finger | A4QPD1 [6 | 0 | 1945.871 | 0 | 0.89 | 0.001269 | 11.18 | 69.1 | 130.9 | 10.43 | 5.37 | | High | High | High | High | High | High | High | High | High | High | High | 0.000275 | 0.002195 | 2.61 |
| FALSE | High | SEGFELYNI | [K].SEGFEL | 1xPhospho | E9Q9C3 1x | 3.26E-05 | 0 | 1 | 5 | 6 | E9Q9C3 | Afadin OS= | E9Q9C3 [1 | 0 | 3957.757 | 0 | 0.29 | 0.118883 | 16.46 | 94.8 | 105.2 | 12.21 | 7.57 | | High | High | High | High | High | High | High | High | High | High | High | 0 | 5.12E-06 | 5.35 |
| FALSE | High | SCYNLTNP | [K].SCYNLT | 1xCarbami | E9Q6P5 2x | 0.072258 | 0.005346 | 1 | 1 | 2 | E9Q6P5 | Tetratricop | E9Q6P5 [6 | 1 | 2588.105 | 0 | -0.15 | 0.034287 | 8.88 | 106.6 | 93.4 | 5.54 | 3.34 | | High | High | High | High | High | High | High | High | High | High | High | 0.002855 | 0.02947 | 3.67 |
| FALSE | High | SDAEEDGV | [K].SDAEEI | 1xPhospho | Q3TXE5 1x | 3.4E-07 | 0 | 1 | 2 | 15 | Q3TXE5 | Putative ur | Q3TXE5 [5 | 0 | 2890.345 | 0 | 0.13 | 0.199406 | 25.93 | 91.1 | 108.9 | 8.7 | 9.85 | | High | High | High | High | High | High | High | High | High | High | High | 0 | 1.03E-07 | 3.47 |
| FALSE | High | SDAEEVDF | [R].SDAEEV | 1xCarbami | P31938 1xI | 0.000108 | 0 | 1 | 5 | 3 | P31938 | Dual specif | P31938 [36 | 0 | 3483.598 | 0 | -0.19 | 0.573183 | 30.23 | 107.2 | 92.8 | 28.13 | 12.47 | | High | High | High | High | High | High | High | High | High | High | High | 0 | 2.09E-05 | 4.64 |
| FALSE | High | SEGFELYNI | [K].SEGFEL | 1xDeamida | E9Q9C3 1x | 8.15E-07 | 0 | 1 | 5 | 9 | E9Q9C3 | Afadin OS= | E9Q9C3 [1 | 0 | 3942.746 | 0 | 0.21 | 0.065871 | 19.95 | 85.6 | 114.4 | 8.12 | 15.37 | | High | High | High | High | High | High | High | High | High | High | High | 0 | 5.44E-06 | 5.29 |
| FALSE | High | SEGAPSPVI | [K].SEGAPS | 1xPhospho | Q8CJF7 1xI | 0.001107 | 0 | 1 | 1 | 2 | Q8CJF7 | Protein EL | Q8CJF7 [13 | 0 | 1409.744 | 0 | -0.63 | 0.001797 | 2.63 | 121.4 | 78.6 | 6.32 | 4.41 | | High | High | High | High | High | High | High | High | High | High | High | 9.42E-05 | 0.001041 | 1.78 |
| FALSE | High | SDAICNFVI | [R].SDAICN | 2xCarbami | Q8BRH3 1x | 3.66E-08 | 0 | 1 | 3 | 2 | Q8BRH3 | Rho GTPas | Q8BRH3 [1 | 0 | 2189.992 | 0 | 0.01 | 0.638006 | 12.78 | 100.9 | 99.1 | 6.64 | 6.37 | | High | High | High | High | High | High | High | High | High | High | High | 0 | 2.48E-09 | 3.55 |
| FALSE | High | SDANTDLI | [K].SDANTI | 1xPhospho | P53986 1xI | 0.000731 | 0 | 1 | 5 | 1 | P53986 | Monocarb | P53986 [22 | 0 | 1812.914 | 0 | -0.4 | 0.000704 | 1.07 | 115.2 | 84.8 | 2.58 | 4.63 | | High | High | High | High | High | High | High | High | High | High | High | 0 | 0.000168 | 1.81 |
| FALSE | High | SCTSSDSTA | [K].SCTSSD | 1xCarbami | B2RSN1 1x | 0.068752 | 0.005042 | 1 | 6 | 2 | B2RSN1 | Myc proto- | B2RSN1 [2 | 0 | 2888.234 | 0 | -0.05 | 0.305417 | 12.37 | 105.7 | 94.3 | 20.83 | 4.97 | | High | High | High | High | High | High | High | High | High | High | High | 0.002924 | 0.03034 | 3.39 |
| FALSE | High | SEGRPETPI | [K].SEGRPE | 1xPhospho | Q6KCD5 1x | 0.104222 | 0.008348 | 1 | 3 | 2 | Q6KCD5 | Nipped-B-l | Q6KCD5 [7 | 0 | 1538.798 | 0 | 0.54 | 0.002475 | 1.9 | 81.5 | 118.5 | 6.34 | 4.4 | | High | High | High | High | High | High | High | High | High | High | High | 0.005428 | 0.05955 | 1.41 |
| FALSE | High | SCSIDTSPG | [R].SCSIDT | 1xCarbami | P53668 2xI | 0.001799 | 5.66E-05 | 1 | 6 | 1 | P53668 | LIM domai | P53668 [29 | 0 | 2398.02 | 0 | 1.17 | 0.005201 | 6.27 | 64.8 | 135.2 | 14.61 | 18.85 | | High | High | High | High | High | High | High | High | High | High | High | 2.58E-05 | 0.000461 | 2.44 |
| FALSE | High | SCSIDTSPG | [R].SCSIDT | 1xCarbami | P53668 1xI | 1.56E-05 | 0 | 1 | 6 | 3 | P53668 | LIM domai | P53668 [29 | 0 | 2318.053 | 0 | 0.58 | 0.003879 | 3.71 | 76.8 | 123.2 | 10.02 | 8.66 | | High | High | High | High | High | High | High | High | High | High | High | 0 | 2.23E-06 | 3.95 |
| FALSE | High | SCLLDREP | [R].SCLLDR | 1xCarbami | Q9DB28 1x | 0.011966 | 0.000889 | 1 | 1 | 1 | Q9DB28 | Ribonuclea | Q9DB28 [1 | 1 | 3074.342 | 0 | -0.38 | 0.08753 | 16.82 | 114.3 | 85.7 | 19.19 | 11.87 | | High | High | High | High | High | High | High | High | High | High | High | 0.000449 | 0.003834 | 2.59 |
| FALSE | High | SCLLDREP | [R].SCLLDR | 1xCarbami | Q9DB28 2x | 0.111965 | 0.009053 | 1 | 1 | 1 | Q9DB28 | Ribonuclea | Q9DB28 [1 | 1 | 3170.303 | 0 | -0.55 | 0.0435 | 11.3 | 118.3 | 81.7 | 14.04 | 10.72 | | High | High | High | High | High | High | High | High | High | High | High | 0.004543 | 0.04946 | 3.43 |
| FALSE | High | SEGVQDAC | [R].SEGVQ | 1xPhospho | F8VQN6 1x | 2.43E-06 | 0 | 1 | 1 | 2 | F8VQN6 | Rho guanir | F8VQN6 [3 | 0 | 2381.118 | 0 | 1.71 | 0.000181 | 2.13 | 48 | 152 | 11.74 | 3.9 | | High | High | High | High | High | High | High | High | High | High | High | 0 | 2.75E-07 | 3.76 |
| FALSE | High | SEGVQDAE | [R].SEGVQ | 1xPhospho | Q8R4H2 1x | 4.24E-09 | 0 | 1 | 1 | 2 | Q8R4H2 | Rho guanir | Q8R4H2 [3 | 0 | 2253.06 | 0 | 1.75 | 0.000209 | 13.59 | 48.9 | 151.1 | 11.08 | 5.83 | | High | High | High | High | High | High | High | High | High | High | High | 0 | 2.18E-10 | 3.52 |
| FALSE | High | SEGTTSTSY | [R].SEGTTS | 1xPhospho | Q8VDZ4 1x | 0.036731 | 0.00249 | 1 | 2 | 1 | Q8VDZ4 | Palmitoyltr | Q8VDZ4 [4 | 0 | 1598.772 | 0 | 0.29 | 0.034894 | 13.8 | 86.2 | 113.8 | 4.65 | 9.89 | | High | High | High | High | High | High | High | High | High | High | High | 0.001308 | 0.01357 | 1.71 |
| FALSE | High | SCLQCRPPI | [K].SCLQCF | 2xCarbami | P51944-2 1 | 0.00043 | 0 | 1 | 3 | 1 | P51944-2 | Isoform 2 c | P51944-2 [ | 0 | 2943.437 | 0 | 0.51 | 0.193946 | 33.08 | 82.6 | 117.4 | 11.63 | 21.09 | | High | High | High | High | High | High | High | High | High | High | High | 0 | 9.2E-05 | 2.08 |
| FALSE | High | SCLSPK | [R].SCLSPK | 1xCarbami | O09130 1x | 0.082424 | 0.006288 | 1 | 2 | 1 | O09130 | NFATC2-in | O09130 [2 | 0 | 1229.637 | 0 | -0.8 | 0.000728 | 3.01 | 129.4 | 70.6 | 7.67 | 7.89 | | High | High | High | High | High | High | High | High | High | High | High | 0.003265 | 0.03451 | 0.8 |
| FALSE | High | SCMLTGTP | [R].SCMLT | 1xCarbami | A0A0G2JG | 9.54E-05 | 0 | 1 | 10 | 4 | A0A0G2JG | Far upstre | A0A0G2JG | 0 | 2150.027 | 0 | -1.02 | 0.016645 | 5.44 | 132.7 | 67.3 | 10.88 | 28.45 | | High | High | High | High | High | High | High | High | High | High | High | 0.000689 | 0.005953 | 3.37 |
| FALSE | High | SCPPQDFPI | [K].SCPPQI | 2xCarbami | Q8BQ33 1x | 0.000137 | 0 | 1 | 1 | 4 | Q8BQ33 | Treslin OS= | Q8BQ33 [1 | 0 | 3630.656 | 0 | -0.83 | 0.001698 | 0.38 | 127.9 | 72.1 | 4.64 | 12.23 | | High | High | High | High | High | High | High | High | High | High | High | 0 | 2.56E-05 | 3.47 |
| FALSE | High | SCPSPEGSI | [R].SCPSPE | 1xCarbami | E9QKG8 1x | 0.054844 | 0.003982 | 1 | 8 | 1 | E9QKG8 | Holliday ju | E9QKG8 [4 | 0 | 1469.629 | 0 | -1.06 | 0.000787 | 2.87 | 134.2 | 65.8 | 6.66 | 7.18 | | High | High | High | High | High | High | High | High | High | High | High | 0.002059 | 0.02147 | 2.57 |
| FALSE | High | SCQHGDVS | [K].SCQHG | 1xCarbami | B2RSV7 1xI | 0.010382 | 0.000772 | 1 | 1 | 1 | B2RSV7 | Fgd6 prote | B2RSV7 [6 | 0 | 1930.879 | 0 | 1.76 | 0.000534 | 2.94 | 46.5 | 153.5 | 6.34 | 6.96 | | High | High | High | High | High | High | High | High | High | High | High | 0.000382 | 0.003276 | 1.75 |
| FALSE | High | SEGTTPEK | [K].SEGTTF | 1xPhospho | Q3U2W2 1 | 0.002172 | 5.66E-05 | 1 | 2 | 2 | Q3U2W2 | MYB bindir | Q3U2W2 [ | 0 | 1386.692 | 0 | 0.04 | 0.285001 | 11.65 | 97.8 | 102.2 | 0.94 | 8.07 | | High | High | High | High | High | High | High | High | High | High | High | 4.94E-05 | 0.000572 | 1.78 |
| FALSE | High | SEGTLIDLS | [K].SEGTLII | 1xPhospho | Q921I6 1xI | 9.65E-06 | 0 | 1 | 1 | 4 | Q921I6 | SH3 domai | Q921I6 [19 | 0 | 2813.353 | 0 | -0.55 | 0.02439 | 1.52 | 116.7 | 83.3 | 3.73 | 15.43 | | High | High | High | High | High | High | High | High | High | High | High | 0 | 1.3E-06 | 3.98 |
| FALSE | High | SEGSPNQG | [K].SEGSPN | 1xPhospho | Q99PL5 1xI | 0.077825 | 0.005963 | 2 | 9 | 1 | Q99PL5; Q | Ribosome- | Q99PL5 [76 | 1 | 1798.967 | 0 | -1.26 | 0.045941 | 2.79 | 140.4 | 59.6 | 26.56 | 15.79 | Shared | High | High | High | High | High | High | High | High | High | High | High | 0.003122 | 0.03231 | 1.49 |
| FALSE | High | SCSAEQPSI | [R].SCSAEC | 2xCarbami | A2ART0 1x | 2.28E-06 | 0 | 1 | 14 | 2 | A2ART0 | Uncharacte | A2ART0 [6 | 0 | 2266.029 | 0 | -1.47 | 0.00034 | 0.58 | 148.4 | 51.6 | 3.65 | 11.64 | | High | High | High | High | High | High | High | High | High | High | High | 0 | 6.84E-07 | 4.12 |
| FALSE | High | SEGSPNQG | [K].SEGSPN | 1xPhospho | Q99PL5 1xI | 0.002463 | 0.000109 | 2 | 9 | 3 | Q99PL5; Q | Ribosome- | Q99PL5 [76 | 0 | 1441.709 | 0 | -0.49 | 0.006653 | 8.33 | 116.7 | 83.3 | 4.9 | 7.1 | Shared | High | High | High | High | High | High | High | High | High | High | High | 4.94E-05 | 0.000655 | 2.76 |
| FALSE | High | SEGSEYEEI | [R].SEGSEY | 2xPhospho | Q7TSC1 2x | 0.032831 | 0.002195 | 1 | 3 | 1 | Q7TSC1 | Protein PRI | Q7TSC1 [1 | 0 | 1885.827 | 0 | 0.83 | 0.000949 | 2.25 | 72 | 128 | 5.18 | 4.79 | | High | High | High | High | High | High | High | High | High | High | High | 0.001151 | 0.0119 | 2.49 |
| FALSE | High | SCSASCMP | [R].SCSASC | 2xCarbami | Q3SYK5 1x | 0.022378 | 0.001542 | 1 | 4 | 2 | Q3SYK5 | Tyrosine-p | Q3SYK5 [7 | 0 | 1645.648 | 0 | 1.2 | 0.019959 | 22.07 | 70.8 | 129.2 | 17.11 | 25.89 | | High | High | High | High | High | High | High | High | High | High | High | 0.001011 | 0.01087 | 1.89 |
| FALSE | High | SCSHQLHFI | [K].SCSHQI | 1xCarbami | Q8CIB9 1xI | 1.2E-05 | 0 | 1 | 2 | 2 | Q8CIB9 | N-acetyltra | Q8CIB9 [58 | 0 | 2163.082 | 0 | -1.55 | 0.006842 | 1.65 | 150.3 | 49.7 | 22.12 | 5.81 | | High | High | High | High | High | High | High | High | High | High | High | 0 | 5.12E-05 | 2.25 |
| FALSE | High | SEGSEYEEI | [R].SEGSEY | 1xPhospho | Q7TSC1 1x | 0.003282 | 0.00016 | 1 | 3 | 3 | Q7TSC1 | Protein PRI | Q7TSC1 [1 | 0 | 1805.861 | 0 | 0.44 | 0.007824 | 5.59 | 86.2 | 113.8 | 8.56 | 6.14 | | High | High | High | High | High | High | High | High | High | High | High | 0.000786 | 0.007031 | 2.61 |
| FALSE | High | SCSHQTSA | [R].SCSHQ | 1xCarbami | Q99K70 1x | 0.002553 | 0.000109 | 1 | 1 | 2 | Q99K70 | Ras-relatec | Q99K70 [3 | 0 | 1840.903 | 0 | 1.76 | 0.017978 | 9.62 | 48.3 | 151.7 | 23.36 | 22.76 | | High | High | High | High | High | High | High | High | High | High | High | 0.000538 | 0.004571 | 2.76 |
| FALSE | High | SDAPDSTG | [R].SDAPD | 1xPhospho | Q64127 1x | 2.78E-06 | 0 | 1 | 5 | 2 | Q64127 | Transcripti | Q64127 [7 | 0 | 2779.257 | 0 | -0.98 | 0.001826 | 3.49 | 135.3 | 64.7 | 8.04 | 7.82 | | High | High | High | High | High | High | High | High | High | High | High | 0 | 3.21E-07 | 5.18 |
| FALSE | High | SCLAEVSVS | [R].SCLAEV | 1xCarbami | P28359 1xI | 3.74E-09 | 0 | 1 | 1 | 1 | P28359 | Homeobox | P28359 [23 | 0 | 2287.129 | 0 | 2.86 | 0.000432 | 2.18 | 26.8 | 173.2 | 9.38 | 10.89 | | High | High | High | High | High | High | High | High | High | High | High | 0 | 1.9E-10 | 3.77 |
| FALSE | High | SDAPDSTG | [R].SDAPD | 1xDeamida | Q64127 1x | 0.00025 | 0 | 1 | 5 | 9 | Q64127 | Transcripti | Q64127 [7 | 0 | 2780.241 | 0 | -0.66 | 0.001267 | 3.89 | 121.4 | 78.6 | 2.05 | 10.48 | | High | High | High | High | High | High | High | High | High | High | High | 0 | 5.03E-05 | 3.41 |
| FALSE | High | SDCHVSSP | [R].SDCHV | 1xCarbami | Q8BQ33 1x | 0.106748 | 0.008634 | 1 | 1 | 2 | Q8BQ33 | Treslin OS= | Q8BQ33 [1 | 0 | 2242.083 | 0 | -0.13 | 0.14179 | 8.05 | 103.9 | 96.1 | 5.49 | 6.73 | | High | High | High | High | High | High | High | High | High | High | High | 0.004327 | 0.04662 | 2.38 |
| FALSE | High | SDGSLDDG | [R].SDGSLI | 1xPhospho | A9CLV6 1xI | 1.44E-06 | 0 | 1 | 2 | 1 | A9CLV6 | Axotomy ir | A9CLV6 [3 | 0 | 1652.711 | 0 | 0.8 | 0.020252 | 21.91 | 76.6 | 123.4 | 16.6 | 16.27 | | High | High | High | High | High | High | High | High | High | High | High | 0 | 1.53E-07 | 2.79 |
| FALSE | High | SDGSNFAN | [K].SDGSNI | 1xPhospho | Q76N33 1x | 5.98E-09 | 0 | 1 | 2 | 1 | Q76N33 | AMSH-like | Q76N33 [2 | 0 | 1933.864 | 0 | 1.55 | 0.000689 | 4.28 | 46.9 | 153.1 | 9.12 | 8.5 | | High | High | High | High | High | High | High | High | High | High | High | 0 | 3.23E-10 | 4.41 |
| FALSE | High | SEDVLAQS | [K].SEDVLA | 1xPhospho | Q8BUH8 1 | 0.000984 | 0 | 2 | 4 | 1 | Q8BUH8; C | Sentrin-spe | Q8BUH8 [4 | 0 | 1811.956 | 0 | 1.96 | 0.000218 | 14.74 | 40.9 | 159.1 | 18.16 | 6.13 | Shared | High | High | High | High | High | High | High | High | High | High | High | 0 | 0.000235 | 3.7 |
| FALSE | High | SEDSSQPW | [K].SEDSSC | 2xCarbami | B2RS11 1xI | 0.006363 | 0.00062 | 1 | 3 | 2 | B2RS11 | Cyclin E1 C | B2RS11 [52 | 1 | 4767.043 | 0 | -0.25 | 0.095675 | 9.95 | 104.2 | 95.8 | 20.83 | 22.42 | | High | High | High | High | High | High | High | High | High | High | High | 0.000275 | 0.001889 | 4.47 |
| FALSE | High | SEDSSQPW | [K].SEDSSC | 2xCarbami | B2RS11 1xI | 0.101749 | 0.008194 | 1 | 3 | 1 | B2RS11 | Cyclin E1 C | B2RS11 [52 | 1 | 4766.059 | 0 | -0.06 | 0.32593 | 15.05 | 106.5 | 93.5 | 20.8 | 12.56 | | High | High | High | High | High | High | High | High | High | High | High | 0.004115 | 0.04385 | 3.09 |
| FALSE | High | SDHALTSP | [R].SDHALT | 1xPhospho | F8VQL9 1x | 9.14E-06 | 0 | 1 | 8 | 1 | F8VQL9 | Nuclear re | F8VQL9 [2 | 0 | 1721.862 | 0 | -0.15 | 0.138444 | 14.31 | 105.8 | 94.2 | 8.45 | 7.97 | | High | High | High | High | High | High | High | High | High | High | High | 0 | 1.22E-06 | 2.29 |
| FALSE | High | SDHGAYSC | [K].SDHGA | 1xPhospho | Q80TR8-2 | 0.000138 | 0 | 1 | 3 | 3 | Q80TR8-2 | Isoform 2 c | Q80TR8-2 | 0 | 1898.941 | 0 | 0.66 | 0.025855 | 9.03 | 79.3 | 120.7 | 11.49 | 8.29 | | High | High | High | High | High | High | High | High | High | High | High | 0.000505 | 0.004364 | 3.26 |
| FALSE | High | SDHSGTW | [K].SDHSG | 1xPhospho | E9Q6J5-3 1 | 2.51E-08 | 0 | 1 | 3 | 2 | E9Q6J5-3 | Isoform Ne | E9Q6J5-3 [ | 0 | 2142.975 | 0 | -0.62 | 0.006271 | 3.72 | 124.4 | 75.6 | 5.11 | 19.99 | | High | High | High | High | High | High | High | High | High | High | High | 0 | 1.62E-09 | 4 |
| FALSE | High | SDIASHFSN | [K].SDIASH | 1xPhospho | Q3UYC8 1x | 0.007209 | 0.00062 | 1 | 5 | 1 | Q3UYC8 | Putative ur | Q3UYC8 [7 | 0 | 1643.819 | 0 | -0.4 | 0.006816 | 7.4 | 115.9 | 84.1 | 5.42 | 10.59 | | High | High | High | High | High | High | High | High | High | High | High | 0.000275 | 0.002184 | 1.93 |
| FALSE | High | SEEAHAED | [K].SEEAH | 1xPhospho | Q9CY58 1x | 0.096181 | 0.007535 | 2 | 6 | 1 | Q9CY58; Q | Plasminoge | Q9CY58 [3 | 0 | 2205.922 | 0 | 0.69 | 0.026511 | 19.19 | 70 | 130 | 17.5 | 20.01 | Shared | High | High | High | High | High | High | High | High | High | High | High | 0.003809 | 0.04112 | 1.32 |
| FALSE | High | SEDSSQPW | [K].SEDSSC | 2xCarbami | B2RS11 1xI | 1.94E-06 | 0 | 1 | 3 | 3 | B2RS11 | Cyclin E1 C | B2RS11 [52 | 0 | 3491.517 | 0 | 0.61 | 0.067108 | 20.5 | 78.7 | 121.3 | 16.15 | 11.64 | | High | High | High | High | High | High | High | High | High | High | High | 0 | 4.38E-05 | 3.95 |
| FALSE | High | SEDLDNSID | [R].SEDLDN | 1xPhospho | Q1HL32 1x | 0.002463 | 0.000109 | 1 | 4 | 1 | Q1HL32 | Structural | Q1HL32 [8 | 0 | 1673.804 | 0 | -1.64 | 0.001736 | 1.57 | 151.3 | 48.7 | 7.46 | 39.54 | | High | High | High | High | High | High | High | High | High | High | High | 4.94E-05 | 0.000657 | 2.3 |
| FALSE | High | SDLDLGYEI | [R].SDLDLG | 1xPhospho | E9QMJ8 1x | 2.15E-07 | 0 | 1 | 4 | 2 | E9QMJ8 | Axin-1 OS= | E9QMJ8 [6 | 0 | 2573.201 | 0 | -0.05 | 0.96333 | 19.15 | 100.8 | 99.2 | 10.67 | 8.19 | | High | High | High | High | High | High | High | High | High | High | High | 0 | 1.79E-08 | 4.79 |
| FALSE | High | SDIDVNAA | [R].SDIDVN | 1xPhospho | B2RY22 1xI | 2.31E-05 | 0 | 2 | 16 | 1 | B2RY22; A( | Clasp1 pro | B2RY22 [60 | 0 | 1470.704 | 0 | -0.87 | 0.000614 | 0.98 | 128.3 | 71.7 | 12.44 | 12.38 | Shared | High | High | High | High | High | High | High | High | High | High | High | 0 | 3.48E-06 | 3.43 |
| FALSE | High | SDLIEDEEL | [R].SDLIED | 1xDeamida | P52479-2 1 | 0.008927 | 0.000736 | 1 | 2 | 2 | P52479-2 | Isoform 2 c | P52479-2 [ | 1 | 3473.577 | 0 | -0.48 | 0.520239 | 7.35 | 111.9 | 88.1 | 56.01 | 49.76 | | High | High | High | High | High | High | High | High | High | High | High | 0.000327 | 0.002759 | 4.26 |
| FALSE | High | SDLIEDEEL | [R].SDLIED | 1xPhospho | P52479-2 1 | 0.000668 | 0 | 1 | 2 | 7 | P52479-2 | Isoform 2 c | P52479-2 [ | 1 | 3701.756 | 0 | -0.81 | 0.001084 | 4.32 | 128 | 72 | 2.66 | 10.4 | | High | High | High | High | High | High | High | High | High | High | High | 7.19E-05 | 0.000885 | 3.24 |
| FALSE | High | SDLNGLPSI | [R].SDLNGI | 1xPhospho | Q5XJV7-2 1 | 1.12E-05 | 0 | 1 | 2 | 1 | Q5XJV7-2 | Isoform 2 c | Q5XJV7-2 [ | 0 | 1721.831 | 0 | -0.56 | 0.00191 | 3.14 | 120.4 | 79.6 | 4.7 | 6.65 | | High | High | High | High | High | High | High | High | High | High | High | 0 | 1.55E-06 | 1.84 |
| FALSE | High | SEDGYHSD | [R].SEDGYI | 1xPhospho | Q91YE7 1x | 1.21E-05 | 0 | 1 | 8 | 4 | Q91YE7 | RNA-bindir | Q91YE7 [7 | 0 | 2310.877 | 0 | 0.02 | 0.827367 | 7.6 | 99.4 | 100.6 | 11.38 | 14.75 | | High | High | High | High | High | High | High | High | High | High | High | 0 | 2.49E-06 | 4.52 |
| FALSE | High | SEDENTRP | [K].SEDENT | 1xCarbami | B1AY10 1xI | 0.000933 | 0 | 1 | 3 | 2 | B1AY10 | Transcripti | B1AY10 [1 | 0 | 2743.183 | 0 | 0.46 | 0.018668 | 11.17 | 83.8 | 116.2 | 4.73 | 11.84 | | High | High | High | High | High | High | High | High | High | High | High | 0.000859 | 0.008626 | 2.22 |
| FALSE | High | SDLSEEDLF | [R].SDLSEE | 1xPhospho | Q3UHQ6 1 | 0.013909 | 0.001011 | 1 | 3 | 1 | Q3UHQ6 | Protein do | Q3UHQ6 [ | 0 | 2335.058 | 0 | 0.67 | 0.00104 | 5.27 | 76.1 | 123.9 | 6.4 | 5.48 | | High | High | High | High | High | High | High | High | High | High | High | 0.000538 | 0.004538 | 2.9 |
| FALSE | High | SEDRPSSP | [R].SEDRPS | 1xPhospho | P70399 1xI | 2.4E-05 | 0 | 1 | 5 | 9 | P70399 | TP53-bindi | P70399 [26 | 0 | 2425.238 | 0 | 0.4 | 0.003225 | 8.12 | 85.1 | 114.9 | 4.75 | 7.37 | | High | High | High | High | High | High | High | High | High | High | High | 0 | 0.000306 | 3.2 |
| FALSE | High | SDGSENLC | [R].SDGSEI | 1xCarbami | Q9R0L6-2 1 | 0.000174 | 0 | 1 | 3 | 2 | Q9R0L6-2 | Isoform 2 c | Q9R0L6-2 [ | 0 | 1887.811 | 0 | 0 | 0.488412 | 1.1 | 102.8 | 97.2 | 3.99 | 5.3 | | High | High | High | High | High | High | High | High | High | High | High | 0.000327 | 0.002915 | 3.28 |
| FALSE | High | SDGGSDSG | [K].SDGGSI | 1xPhospho | E9Q842 1x | 2.98E-05 | 0 | 1 | 3 | 1 | E9Q842 | Neuron na | E9Q842 [9 | 0 | 1446.688 | 0 | 2.03 | 0.000193 | 10.5 | 39.5 | 160.5 | 13.36 | 5.62 | | High | High | High | High | High | High | High | High | High | High | High | 0 | 4.62E-06 | 2.33 |
| FALSE | High | SDGACDSP | [K].SDGACI | 1xCarbami | Q9Z277 1x | 1.02E-05 | 0 | 1 | 1 | 1 | Q9Z277 | Tyrosine-p | Q9Z277 [1 | 0 | 1763.756 | 0 | 0.28 | 0.133049 | 23.64 | 88.9 | 111.1 | 12.89 | 14.56 | | High | High | High | High | High | High | High | High | High | High | High | 0 | 1.37E-06 | 2.86 |
| FALSE | High | SDCQPNSP | [K].SDCQPI | 2xCarbami | P41183 1xI | 3.83E-05 | 0 | 1 | 2 | 1 | P41183 | B-cell lymp | P41183 [33 | 0 | 2221.951 | 0 | 2.62 | 0.001082 | 9.25 | 25.7 | 174.3 | 12.32 | 15.47 | | High | High | High | High | High | High | High | High | High | High | High | 0 | 6.11E-06 | 2.73 |
| FALSE | High | SEESPGWK | [K].SEESPG | 1xPhospho | Q9CYX7 1x | 0.002895 | 0.00016 | 1 | 1 | 2 | Q9CYX7 | RRP15-like | Q9CYX7 [2 | 0 | 1457.708 | 0 | -0.41 | 0.01309 | 9.63 | 109.2 | 90.8 | 7.99 | 5.76 | | High | High | High | High | High | High | High | High | High | High | High | 7.19E-05 | 0.000782 | 1.6 |
| FALSE | High | SEESLTSLH | [K].SEESLT | 1xPhospho | Q811P8 1x | 0.028575 | 0.001829 | 1 | 3 | 3 | Q811P8 | Rho GTPas | Q811P8 [7 | 0 | 2213.074 | 0 | -0.15 | 0.073257 | 9 | 106.8 | 93.2 | 7.96 | 4.99 | | High | High | High | High | High | High | High | High | High | High | High | 0.000997 | 0.0105 | 2.08 |
| FALSE | High | SEESAAAS | [R].SEESAA | 1xPhospho | O88413 1x | 0.069325 | 0.005112 | 1 | 1 | 1 | O88413 | Tubby-rela | O88413 [1 | 0 | 1545.756 | 0 | 0.01 | 0.228192 | 3.08 | 96.7 | 103.3 | 17.08 | 5.7 | | High | High | High | High | High | High | High | High | High | High | High | 0.002662 | 0.02815 | 1.63 |
| FALSE | High | SEEPSVSLP | [K].SEEPSV | 1xPhospho | Q6A065 1x | 0.009843 | 0.000736 | 1 | 2 | 1 | Q6A065 | Centrosom | Q6A065 [4 | 0 | 2196.151 | 0 | 0.89 | 0.004224 | 19.11 | 69.9 | 130.1 | 11.07 | 11.64 | | High | High | High | High | High | High | High | High | High | High | High | 0.000344 | 0.003077 | 2.92 |
| FALSE | High | SDCRSPTHI | [R].SDCRSF | 1xCarbami | G3UZM1 1 | 7.41E-05 | 0 | 1 | 3 | 2 | G3UZM1 | Probable J | G3UZM1 [ | 1 | 2311.106 | 0 | -0.38 | 0.155541 | 18.85 | 117.7 | 82.3 | 26.49 | 16.45 | | High | High | High | High | High | High | High | High | High | High | High | 0 | 1.86E-05 | 2.89 |
| FALSE | High | SEEEQSSAS | [K].SEEEQS | 1xPhospho | Q3U6P5 1x | 0.000389 | 0 | 2 | 6 | 5 | Q3U6P5; C | Putative ur | Q3U6P5 [2 | 0 | 1718.825 | 0 | -0.46 | 0.002458 | 2.6 | 115.1 | 84.9 | 5.78 | 6.77 | Shared | High | High | High | High | High | High | High | High | High | High | High | 0 | 8.25E-05 | 2.43 |
| FALSE | High | SDCSVDGG | [R].SDCSVI | 1xCarbami | E9Q3P4 1x | 0.040721 | 0.003058 | 1 | 5 | 1 | E9Q3P4 | Centromer | E9Q3P4 [8 | 1 | 2520.186 | 0 | -1.51 | 0.000847 | 2.37 | 147.9 | 52.1 | 9.06 | 8.62 | | High | High | High | High | High | High | High | High | High | High | High | 0.001581 | 0.01524 | 1.32 |
| FALSE | High | SDDDVSGA | [K].SDDDV | 1xPhospho | Q3V122 1x | 0.000386 | 0 | 1 | 4 | 2 | Q3V122 | DNA helica | Q3V122 [3 | 0 | 2276.058 | 0 | -0.47 | 0.00714 | 3.88 | 116.1 | 83.9 | 11.23 | 5.13 | | High | High | High | High | High | High | High | High | High | High | High | 0 | 8.17E-05 | 2.96 |
| FALSE | High | SDDGSHLQ | [K].SDDGSI | 1xPhospho | E9Q8V6 1x | 0.006653 | 0.00062 | 1 | 2 | 1 | E9Q8V6 | DENN/MA | E9Q8V6 [1 | 0 | 2153.927 | 0 | 0.28 | 0.062863 | 15.25 | 90.5 | 109.5 | 6.93 | 6.87 | | High | High | High | High | High | High | High | High | High | High | High | 0.000275 | 0.001989 | 2.41 |
| FALSE | High | SDDGTQDI | [K].SDDGT | 1xPhospho | Q3UTQ7 1 | 0.000639 | 0 | 1 | 4 | 2 | Q3UTQ7 | PR domain | Q3UTQ7 [4 | 0 | 2112.041 | 0 | -0.12 | 0.02977 | 5.67 | 106.8 | 93.2 | 6.15 | 7.34 | | High | High | High | High | High | High | High | High | High | High | High | 0 | 0.000144 | 2.88 |
| FALSE | High | SDDLVVDV | [K].SDDLVV | 1xPhospho | Q80TC1 1x | 2.19E-08 | 0 | 1 | 14 | 3 | Q80TC1 | MKIAA154 | Q80TC1 [3 | 0 | 2137.985 | 0 | -1.7 | 0.000443 | 0.33 | 152.6 | 47.4 | 7.62 | 7.64 | | High | High | High | High | High | High | High | High | High | High | High | 0 | 1.4E-09 | 4.71 |
| FALSE | High | SDDNLVVD | [K].SDDNL | 1xPhospho | Q3UY41 1x | 3.61E-06 | 0 | 1 | 4 | 2 | Q3UY41 | Putative ur | Q3UY41 [2 | 0 | 2254.007 | 0 | -0.87 | 0.004577 | 4.85 | 129.3 | 70.7 | 11.21 | 11.44 | | High | High | High | High | High | High | High | High | High | High | High | 0 | 6.23E-07 | 4.27 |
| FALSE | High | SDDNLVVD | [K].SDDNL | 3xPhospho | Q3UY41 3x | 0.081752 | 0.006219 | 1 | 4 | 1 | Q3UY41 | Putative ur | Q3UY41 [2 | 1 | 3203.328 | 0 | -0.6 | 0.008497 | 6.38 | 120.6 | 79.4 | 11.54 | 2.57 | | High | High | High | High | High | High | High | High | High | High | High | 0.003243 | 0.03421 | 3.27 |
| FALSE | High | SDDSNGIEI | [K].SDDSN | 1xDeamida | P21995 1xI | 0.007538 | 0.00062 | 1 | 4 | 1 | P21995 | Embigin OS | P21995 [30 | 0 | 1726.748 | 0 | -1.94 | 0.002988 | 2.41 | 156.5 | 43.5 | 9.95 | 30.43 | | High | High | High | High | High | High | High | High | High | High | High | 0.000275 | 0.002279 | 1.8 |
| FALSE | High | SDDTLDEEI | [R].SDDTLI | 1xPhospho | Q80YE7 1x | 0.011048 | 0.000772 | 1 | 3 | 1 | Q80YE7 | Death-assc | Q80YE7 [3 | 0 | 2184.966 | 0 | -0.45 | 0.008628 | 7.91 | 117.1 | 82.9 | 8.28 | 4.08 | | High | High | High | High | High | High | High | High | High | High | High | 0.000399 | 0.003504 | 2.26 |
| FALSE | High | SDEESEVIS | [K].SDEESE | 1xPhospho | Q8BHK9 1x | 0.019791 | 0.001438 | 1 | 3 | 2 | Q8BHK9 | DNA excisi | Q8BHK9 [1 | 0 | 1759.877 | 0 | 0.04 | 0.313098 | 10.45 | 98 | 102 | 5.28 | 9.6 | | High | High | High | High | High | High | High | High | High | High | High | 0.000773 | 0.006708 | 2.49 |
| FALSE | High | SEEEALETE | [K].SEEEAL | 1xPhospho | Q5SXI5 1xI | 0.0009 | 0 | 1 | 3 | 1 | Q5SXI5 | Zinc finger | Q5SXI5 [49 | 0 | 2546.216 | 0 | -0.15 | 0.18039 | 21.54 | 105.6 | 94.4 | 10.59 | 10.24 | | High | High | High | High | High | High | High | High | High | High | High | 0 | 0.000213 | 3 |
| FALSE | High | SDENVLDS | [R].SDENVI | 1xPhospho | A2AAE1-3 1 | 0.003372 | 0.00016 | 1 | 6 | 2 | A2AAE1-3 | Isoform 3 c | A2AAE1-3 | 0 | 1641.814 | 0 | -0.35 | 0.004969 | 2.65 | 111.9 | 88.1 | 3.44 | 6.77 | | High | High | High | High | High | High | High | High | High | High | High | 0.000275 | 0.001644 | 3.73 |
| FALSE | High | SEEAPAGD | [R].SEEAPA | 1xPhospho | Q91WE3 1 | 0.085162 | 0.006376 | 1 | 1 | 1 | Q91WE3 | Ribonuclea | Q91WE3 [9 | 0 | 3304.442 | 0 | -0.83 | 0.002309 | 4 | 127.9 | 72.1 | 6.92 | 12.27 | | High | High | High | High | High | High | High | High | High | High | High | 0.003291 | 0.03585 | 3.21 |
| FALSE | High | SDESLPDGI | [K].SDESLP | 1xPhospho | Q8BFU3-4 | 0.001735 | 5.66E-05 | 1 | 6 | 1 | Q8BFU3-4 | Isoform 4 c | Q8BFU3-4 | 0 | 1782.893 | 0 | -0.09 | 0.990729 | 13.19 | 103.3 | 96.7 | 10.48 | 10.72 | | High | High | High | High | High | High | High | High | High | High | High | 2.58E-05 | 0.000442 | 2.05 |
| FALSE | High | SDATASISL | [K].SDATAS | 1xPhospho | E9Q8A3 1x | 0.000112 | 0 | 1 | 3 | 3 | E9Q8A3 | Phosphatic | E9Q8A3 [2 | 0 | 1932.009 | 0 | -0.27 | 0.005149 | 2.48 | 109.1 | 90.9 | 3.69 | 4.8 | | High | High | High | High | High | High | High | High | High | High | High | 0 | 2.04E-05 | 2.46 |
| FALSE | High | SASPSAAG | [K].SASPSA | 1xPhospho | Q0VEE6 1x | 6.3E-05 | 0 | 1 | 2 | 1 | Q0VEE6 | Zinc finger | Q0VEE6 [4 | 0 | 1626.825 | 0 | 2.56 | 0.000154 | 3 | 29.8 | 170.2 | 4.85 | 3.85 | | High | High | High | High | High | High | High | High | High | High | High | 0 | 1.07E-05 | 2.22 |
| FALSE | High | SAINPVDT | [R].SAINPV | 1xPhospho | Q5NCY0 1x | 0.024855 | 0.001645 | 1 | 1 | 1 | Q5NCY0 | Lysine-spe | Q5NCY0 [9 | 0 | 3140.555 | 0 | -0.1 | 0.933236 | 18.23 | 97.8 | 102.2 | 16.89 | 12.13 | | High | High | High | High | High | High | High | High | High | High | High | 0.000872 | 0.008686 | 2.85 |
| FALSE | High | SASPGPAQ | [R].SASPGI | 2xPhospho | A2A9K7 2x | 0.000622 | 0 | 1 | 3 | 1 | A2A9K7 | Connector | A2A9K7 [5 | 0 | 2783.264 | 0 | 0.15 | 0.556631 | 52.23 | 94.7 | 105.3 | 26.65 | 14.18 | | High | High | High | High | High | High | High | High | High | High | High | 0 | 0.00014 | 2.9 |
| FALSE | High | RSSDPLGD | [R].RSSDPL | 1xPhospho | A2AFQ0 1x | 9.79E-08 | 0 | 1 | 10 | 2 | A2AFQ0 | E3 ubiquiti | A2AFQ0 [6 | 1 | 2600.222 | 0 | 0.45 | 0.005978 | 2.7 | 81.4 | 118.6 | 12.25 | 9.43 | | High | High | High | High | High | High | High | High | High | High | High | 0 | 7.49E-09 | 3.87 |
| FALSE | High | RSSDPLGD | [R].RSSDPL | 1xPhospho | A2AFQ0 1x | 6.95E-05 | 0 | 1 | 10 | 2 | A2AFQ0 | E3 ubiquiti | A2AFQ0 [6 | 1 | 2616.217 | 0 | 0.53 | 0.010481 | 16.95 | 85.6 | 114.4 | 9.11 | 6.57 | | High | High | High | High | High | High | High | High | High | High | High | 0 | 1.19E-05 | 3.97 |
| FALSE | High | RSSDPSLNI | [R].RSSDPS | 1xPhospho | Q8K296 1x | 0.016596 | 0.001124 | 1 | 7 | 3 | Q8K296 | Myotubula | Q8K296 [6 | 1 | 1670.851 | 0 | -0.13 | 0.193408 | 2.55 | 102 | 98 | 3.87 | 4.95 | | High | High | High | High | High | High | High | High | High | High | High | 0.008332 | 0.08972 | 1.89 |
| FALSE | High | RSSDPSLNI | [R].RSSDPS | 1xDeamida | Q8K296 1x | 0.012845 | 0.000889 | 1 | 7 | 2 | Q8K296 | Myotubula | Q8K296 [6 | 1 | 1671.836 | 0 | -0.06 | 0.998251 | 8.18 | 98.3 | 101.7 | 4.91 | 6.44 | | High | High | High | High | High | High | High | High | High | High | High | 0.000449 | 0.004151 | 1.88 |
| FALSE | High | RSSDPWSN | [R].RSSDPV | 1xPhospho | O70472 1x | 0.093123 | 0.007276 | 1 | 3 | 2 | O70472 | Transmem | O70472 [1 | 1 | 2105.903 | 0 | -0.18 | 0.117275 | 10.42 | 106.3 | 93.7 | 9.32 | 15.04 | | High | High | High | High | High | High | High | High | High | High | High | 0.003685 | 0.03961 | 1.94 |
| FALSE | High | RSSDQAVP | [R].RSSDQ | 1xPhospho | Q571J8 1xI | 1.13E-07 | 0 | 1 | 14 | 2 | Q571J8 | Non-specif | Q571J8 [41 | 1 | 2446.238 | 0 | 1.18 | 0.000384 | 9.73 | 62.5 | 137.5 | 5.79 | 5.55 | | High | High | High | High | High | High | High | High | High | High | High | 0 | 4.9E-08 | 3.14 |
| FALSE | High | RSSDSCPA | [R].RSSDSC | 1xCarbami | Q8CGF1 1x | 0.072858 | 0.005599 | 1 | 3 | 2 | Q8CGF1 | Rho GTPas | Q8CGF1 [1 | 1 | 1615.746 | 0 | 0.1 | 0.926859 | 9.04 | 98.9 | 101.1 | 21.49 | 7.69 | | High | High | High | High | High | High | High | High | High | High | High | 0.004543 | 0.0492 | 2.48 |
| FALSE | High | RSSDSWD | [R].RSSDSV | 1xPhospho | V9GXM1 1 | 2.03E-07 | 0 | 1 | 7 | 3 | V9GXM1 | ADP-ribosy | V9GXM1 [3 | 1 | 2478.145 | 0 | 0.92 | 0.021951 | 7.16 | 69.6 | 130.4 | 10.18 | 15.82 | | High | High | High | High | High | High | High | High | High | High | High | 0 | 1.73E-08 | 4.4 |
| FALSE | High | RSSEASQA | [R].RSSEAS | 1xPhospho | B7ZNV2 1x | 2.06E-06 | 0 | 1 | 2 | 3 | B7ZNV2 | GLI-Krupp | B7ZNV2 [8 | 1 | 3489.608 | 0 | 0.71 | 0.047733 | 32.32 | 81.9 | 118.1 | 16.65 | 22.21 | | High | High | High | High | High | High | High | High | High | High | High | 0 | 2.3E-07 | 4.53 |
| FALSE | High | RSSEPQLCI | [R].RSSEPC | 1xCarbami | Q9QZS8 1x | 0.085162 | 0.006376 | 1 | 2 | 1 | Q9QZS8 | SH2 domai | Q9QZS8 [4 | 1 | 2847.386 | 0 | -0.57 | 0.007367 | 6.75 | 119.8 | 80.2 | 5.86 | 12.77 | | High | High | High | High | High | High | High | High | High | High | High | 0.003279 | 0.03556 | 2.8 |
| FALSE | High | RSSEWDPV | [R].RSSEWI | 1xPhospho | Q8VED8 1x | 5.93E-06 | 0 | 1 | 3 | 2 | Q8VED8 | Mitochond | Q8VED8 [3 | 1 | 2340.236 | 0 | 0.32 | 0.056688 | 8.19 | 88.9 | 111.1 | 7.3 | 8.3 | | High | High | High | High | High | High | High | High | High | High | High | 0 | 1.11E-06 | 3.35 |
| FALSE | High | RSSFGLR | [R].RSSFGL | 1xPhospho | Q922M5 1 | 0.1059 | 0.008582 | 1 | 3 | 1 | Q922M5 | Cell divisio | Q922M5 [1 | 1 | 1131.587 | 0 | -0.72 | 0.000619 | 3.33 | 124.5 | 75.5 | 5.41 | 6.24 | | High | High | High | High | High | High | High | High | High | High | High | 0.004317 | 0.04625 | 0.75 |
| FALSE | High | RSSFLNAK | [R].RSSFLN | 1xPhospho | A0A0A6YW | 0.041072 | 0.003094 | 1 | 4 | 1 | A0A0A6YW | Rap guanin | A0A0A6YW | 1 | 1460.803 | 0 | -0.15 | 0.941926 | 3.14 | 102.8 | 97.2 | 3.89 | 12.27 | | High | High | High | High | High | High | High | High | High | High | High | 0.001609 | 0.01545 | 1.33 |
| FALSE | High | RSSFSEGQ | [R].RSSFSE | 1xPhospho | Q80U62 1x | 1.49E-06 | 0 | 1 | 5 | 4 | Q80U62 | Run domai | Q80U62 [3 | 1 | 2248.138 | 0 | 0.86 | 0.000192 | 6.71 | 71.2 | 128.8 | 4.53 | 1.81 | | High | High | High | High | High | High | High | High | High | High | High | 0 | 7.91E-06 | 4.16 |
| FALSE | High | RSSFSMEEI | [R].RSSFSN | 1xPhospho | A2AEW1 1 | 0.075927 | 0.005826 | 1 | 7 | 4 | A2AEW1 | Transcripti | A2AEW1 [5 | 1 | 1513.608 | 0 | -0.25 | 0.015687 | 2.23 | 108.8 | 91.2 | 2.62 | 9.1 | | High | High | High | High | High | High | High | High | High | High | High | 0.003265 | 0.03502 | 1.68 |

| | | | | | | | | | | | | | | | | | | | | | | | | | | | | | | | | | | | | | | |
|---|---|---|---|---|---|---|---|---|---|---|---|---|---|---|---|---|---|---|---|---|---|---|---|---|---|---|---|---|---|---|---|---|---|---|---|---|---|---|
| FALSE | High | RSSFSMEE( | [R].RSSFSM | 1xPhospho | Q3UKG7 1x | 0.028575 | 0.001829 | 1 | 6 | 3 | Q3UKG7 | Putative ur | Q3UKG7 [5 | 1 | 1665.739 | 0 | -1.27 | 0.002335 | 2.03 | 144.3 | 55.7 | 9.21 | 20.61 | | High | High | High | High | High | High | High | High | High | High | High | 0.001027 | 0.01109 | 2.38 |
| FALSE | High | RSSGAISNS | [R].RSSGAI | 1xCarbami | Q8C804 1x | 0.003788 | 0.00021 | 1 | 1 | 2 | Q8C804 | Spindle and | Q8C804 [8 | 1 | 2576.254 | 0 | -0.53 | 0.009744 | 3.79 | 129 | 71 | 18.11 | 23.96 | | High | High | High | High | High | High | High | High | High | High | High | 0.000117 | 0.001066 | 3.52 |
| FALSE | High | RSSGLISELI | [K].RSSGLI | 1xPhospho | Q6EDY6 1x | 0.007811 | 0.000664 | 1 | 4 | 2 | Q6EDY6 | F-actin-unc | Q6EDY6 [9 | 1 | 1925.953 | 0 | -1.14 | 0.000181 | 1.48 | 137.7 | 62.3 | 7.52 | 14.1 | | High | High | High | High | High | High | High | High | High | High | High | 0.000801 | 0.007293 | 1.74 |
| FALSE | High | RSSGLQPQ | [R].RSSGLC | 1xPhospho | E9Q4K7 1x | 4.82E-06 | 0 | 1 | 2 | 4 | E9Q4K7 | Kinesin fan | E9Q4K7 [1 | 1 | 1790.911 | 0 | -0.03 | 0.722935 | 23.83 | 102.1 | 97.9 | 7.84 | 12.92 | | High | High | High | High | High | High | High | High | High | High | High | 9.42E-05 | 0.000995 | 3.37 |
| FALSE | High | RSSGISPYF | [R].RSSGISI | 1xPhospho | Q0VGT2 1x | 0.007881 | 0.000736 | 1 | 2 | 1 | Q0VGT2 | Zinc finger | Q0VGT2 [8 | 1 | 1652.8 | 0 | -1.22 | 0.024839 | 2.38 | 139.9 | 60.1 | 13.29 | 58.77 | | High | High | High | High | High | High | High | High | High | High | High | 0.000327 | 0.002406 | 2.12 |
| FALSE | High | RSSGSADS | [R].RSSGSA | 1xPhospho | E9Q286 1x | 0.003558 | 0.00021 | 1 | 1 | 2 | E9Q286 | Little elong | E9Q286 [7 | 1 | 1742.781 | 0 | -0.65 | 0.022386 | 1.5 | 117.2 | 82.8 | 13.67 | 20.05 | | High | High | High | High | High | High | High | High | High | High | High | 9.42E-05 | 0.000986 | 2.39 |
| FALSE | High | RSSLAIEDG | [R].RSSLAII | 1xPhospho | Q4VAC9 1x | 0.001189 | 0 | 1 | 3 | 2 | Q4VAC9 | Pleckstrin I | Q4VAC9 [7 | 1 | 1613.866 | 0 | 1.39 | 0.000738 | 5.88 | 54.5 | 145.5 | 13.27 | 8.9 | | High | High | High | High | High | High | High | High | High | High | High | 0.000467 | 0.004291 | 2.43 |
| FALSE | High | RSSLAIEDG | [R].RSSLAII | 2xPhospho | Q4VAC9 2x | 0.027121 | 0.001676 | 1 | 3 | 1 | Q4VAC9 | Pleckstrin I | Q4VAC9 [7 | 1 | 1693.833 | 0 | 1.57 | 0.001591 | 31.45 | 48.5 | 151.5 | 12.16 | 10.11 | | High | High | High | High | High | High | High | High | High | High | High | 0.00092 | 0.009651 | 1.44 |
| FALSE | High | RSSLIETGQ | [K].RSSLIET | 1xPhospho | Q9DC04 1x | 0.000257 | 0 | 1 | 3 | 2 | Q9DC04 | Regulator c | Q9DC04 [6 | 1 | 2174.08 | 0 | 0.69 | 0.000592 | 4.93 | 78.3 | 121.7 | 5.36 | 6.76 | | High | High | High | High | High | High | High | High | High | High | High | 0 | 8.06E-05 | 2.8 |
| FALSE | High | RSSLPITEDI | [R].RSSLPIT | 1xPhospho | B2RY61 1x | 5.77E-06 | 0 | 1 | 6 | 4 | B2RY61 | AT rich inte | B2RY61 [14 | 1 | 1812.951 | 0 | 0.36 | 0.008089 | 12.35 | 88.6 | 111.4 | 6.56 | 7.7 | | High | High | High | High | High | High | High | High | High | High | High | 0 | 7.32E-07 | 3.15 |
| FALSE | High | RSSLPITEDI | [R].RSSLPIT | 2xPhospho | B2RY61 2x | 0.004033 | 0.000311 | 1 | 6 | 1 | B2RY61 | AT rich inte | B2RY61 [14 | 1 | 1892.917 | 0 | 0.47 | 0.005068 | 4.29 | 85.9 | 114.1 | 12.86 | 5.74 | | High | High | High | High | High | High | High | High | High | High | High | 0.000161 | 0.00114 | 2.59 |
| FALSE | High | RSSISGTEE | [RK].RSSIS( | 1xPhospho | G5E8W8 1x | 1.24E-08 | 0 | 2 | 7 | 7 | G5E8W8; A | Zinc finger | G5E8W8 [1 | 1 | 2803.344 | 0 | -1.83 | 0.000209 | 1.85 | 155.6 | 44.4 | 2.5 | 17.56 | Shared | High | High | High | High | High | High | High | High | High | High | High | 0 | 1.74E-06 | 3.59 |
| FALSE | High | RSSDPALT( | [R].RSSDPA | 1xDeamida | A5D6P2 1x | 0.000396 | 0 | 1 | 22 | 3 | A5D6P2 | Par-3 (Part | A5D6P2 [1 | 1 | 3050.379 | 0 | -0.52 | 0.095596 | 8.81 | 116.6 | 83.4 | 11.62 | 14.62 | | High | High | High | High | High | High | High | High | High | High | High | 0.000538 | 0.004475 | 3.83 |
| FALSE | High | RSSDPALT( | [R].RSSDPA | 1xPhospho | A5D6P2 1x | 3.31E-07 | 0 | 1 | 22 | 4 | A5D6P2 | Par-3 (Part | A5D6P2 [1 | 1 | 3049.395 | 0 | -0.6 | 0.005798 | 5.06 | 123 | 77 | 6.84 | 13.53 | | High | High | High | High | High | High | High | High | High | High | High | 0 | 2.92E-08 | 5.01 |
| FALSE | High | RSSDIISLLV | [R].RSSDIIS | 1xPhospho | Q6ZPR8 1x | 0.001244 | 0 | 1 | 3 | 3 | Q6ZPR8 | MKIAA140 | Q6ZPR8 [5 | 1 | 2276.267 | 0 | 2.55 | 0.000207 | 4.68 | 29.2 | 170.8 | 11.27 | 4.43 | | High | High | High | High | High | High | High | High | High | High | High | 0 | 0.000306 | 2.75 |
| FALSE | High | RSSDGSLSI | [K].RSSDGS | 2xPhospho | B8QI33 2x | 0.000411 | 0 | 1 | 3 | 3 | B8QI33 | Liprin-alph | B8QI33 [23 | 1 | 2119.983 | 0 | -0.04 | 0.506968 | 1.17 | 100.2 | 99.8 | 2.09 | 3.41 | | High | High | High | High | High | High | High | High | High | High | High | 0 | 8.81E-05 | 2.76 |
| FALSE | High | RSPLGGPP | [K].RSPLGG | 1xCarbami | B2RTL2 1x | 4.77E-06 | 0 | 1 | 4 | 1 | B2RTL2 | Phosphodi | B2RTL2 [20 | 1 | 1792.955 | 0 | 0.16 | 0.045143 | 6.89 | 93.6 | 106.4 | 8.75 | 4.96 | | High | High | High | High | High | High | High | High | High | High | High | 0 | 5.88E-07 | 3.32 |
| FALSE | High | RSPLSYPAV | [R].RSPLSY | 1xPhospho | P59764 1x | 0.00582 | 0.00062 | 1 | 3 | 1 | P59764 | Dedicator c | P59764 [16 | 1 | 1568.815 | 0 | 0.46 | 0.207191 | 41.1 | 88.4 | 111.6 | 19.62 | 12.54 | | High | High | High | High | High | High | High | High | High | High | High | 0.000275 | 0.001714 | 2.87 |
| FALSE | High | RSPPPETIK | [K].RSPPPE | 1xPhospho | G3UZM1 1 | 0.006306 | 0.00062 | 1 | 4 | 3 | G3UZM1 | Probable J | G3UZM1 [6 | 1 | 1562.871 | 0 | -0.72 | 0.000273 | 1.74 | 124.4 | 75.6 | 3.18 | 1.78 | | High | High | High | High | High | High | High | High | High | High | High | 0.000275 | 0.001881 | 3.07 |
| FALSE | High | RSPPPSLAK | [R].RSPPPS | 1xPhospho | P54285 1x | 0.006834 | 0.00062 | 1 | 3 | 1 | P54285 | Voltage-de | P54285 [15 | 1 | 1490.85 | 0 | 2.74 | 0.000188 | 6.28 | 27.6 | 172.4 | 11.15 | 5.02 | | High | High | High | High | High | High | High | High | High | High | High | 0.000275 | 0.002048 | 2.42 |
| FALSE | High | RSPPTSGL( | [R].RSPPTS | 1xPhospho | P47930 1x | 0.000852 | 0 | 1 | 2 | 2 | P47930 | Fos-relatec | P47930 [19 | 1 | 1607.847 | 0 | 1.37 | 0.000574 | 5.55 | 54 | 146 | 4.15 | 9.23 | | High | High | High | High | High | High | High | High | High | High | High | 0.000225 | 0.001399 | 3.26 |
| FALSE | High | RSPQMPEE | [K].RSPQM | 1xPhospho | A0A140T8I | 2.34E-05 | 0 | 1 | 2 | 1 | A0A140T8I | Protein Jad | A0A140T8I | 1 | 2309.144 | 0 | -0.27 | 0.015184 | 1.27 | 109.7 | 90.3 | 5.37 | 2.75 | | High | High | High | High | High | High | High | High | High | High | High | 0 | 3.51E-06 | 2.91 |
| FALSE | High | RSPQMPEE | [K].RSPQM | 1xPhospho | A0A140T8I | 0.000219 | 0 | 1 | 2 | 2 | A0A140T8I | Protein Jad | A0A140T8I | 1 | 2325.138 | 0 | -0.32 | 0.002393 | 3.33 | 110.1 | 89.9 | 7.32 | 6.31 | | High | High | High | High | High | High | High | High | High | High | High | 0 | 4.33E-05 | 3.09 |
| FALSE | High | RSPSAPVCI | [K].RSPSAP | 1xCarbami | Q3U3E2 1x | 0.000308 | 0 | 1 | 1 | 3 | Q3U3E2 | Protein FAI | Q3U3E2 [1 | 1 | 1539.812 | 0 | -0.66 | 0.001238 | 3.43 | 122.9 | 77.1 | 2.24 | 10.98 | | High | High | High | High | High | High | High | High | High | High | High | 0.000936 | 0.009702 | 1.96 |
| FALSE | High | RSPSDAGR | [K].RSPSDA | 2xPhospho | E9Q842 2x | 0.019276 | 0.001402 | 1 | 3 | 1 | E9Q842 | Neuron na | E9Q842 [1 | 2 | 2153.963 | 0 | 1.13 | 0.001714 | 23.41 | 62.9 | 137.1 | 14.33 | 8.85 | | High | High | High | High | High | High | High | High | High | High | High | 0.000756 | 0.006568 | 1.83 |
| FALSE | High | RSPSPAPPF | [R].RSPSPA | 2xPhospho | Q52KI8 2x | 6.18E-08 | 0 | 1 | 7 | 6 | Q52KI8 | Serine/argi | Q52KI8 [57 | 1 | 2130.05 | 0 | -0.23 | 0.002355 | 4.02 | 109.5 | 90.5 | 3.51 | 4.46 | | High | High | High | High | High | High | High | High | High | High | High | 0 | 4.44E-09 | 3.19 |
| FALSE | High | RSPSPSPTP | [K].RSPSPS | 1xPhospho | Q6PDG5 1x | 4.91E-06 | 0 | 1 | 3 | 1 | Q6PDG5 | SWI/SNF c | Q6PDG5 [3 | 1 | 1791.941 | 0 | 1.94 | 0.001542 | 21.76 | 37.4 | 162.6 | 19.99 | 13.44 | | High | High | High | High | High | High | High | High | High | High | High | 0 | 6.05E-07 | 3.03 |
| FALSE | High | RSPSPSPTP | [K].RSPSPS | 2xPhospho | Q6PDG5 2x | 0.014929 | 0.001049 | 1 | 3 | 1 | Q6PDG5 | SWI/SNF c | Q6PDG5 [3 | 1 | 1871.907 | 0 | 1.7 | 0.000377 | 2.12 | 43.2 | 156.8 | 11.48 | 7.15 | | High | High | High | High | High | High | High | High | High | High | High | 0.000555 | 0.004915 | 2.59 |
| FALSE | High | RSPSPYYSR | [R].RSPSPY | 2xPhospho | E9QP00 2x | 0.053023 | 0.00384 | 2 | 6 | 3 | E9QP00; P( | Transforme | E9QP00 [2 | 1 | 1501.644 | 0 | 0.23 | 0.040926 | 2.18 | 91.3 | 108.7 | 6.89 | 3.93 | Shared | High | High | High | High | High | High | High | High | High | High | High | 0.001999 | 0.02065 | 2.47 |
| FALSE | High | RSSITEPEG | [K].RSSITEI | 1xPhospho | E9QJU8 1x | 4.72E-05 | 0 | 1 | 2 | 2 | E9QJU8 | Transforma | E9QJU8 [7 | 1 | 2362.217 | 0 | 0.12 | 0.084197 | 9.89 | 92.3 | 107.7 | 6.91 | 6.34 | | High | High | High | High | High | High | High | High | High | High | High | 0 | 7.73E-06 | 4.06 |
| FALSE | High | RSPSVDSTI | [R].RSPSVD | 1xPhospho | P98203-3 1 | 0.096959 | 0.007777 | 1 | 5 | 1 | P98203-3 | Isoform 3 c | P98203-3 [ | 1 | 1313.641 | 0 | -0.47 | 0.013347 | 10.19 | 115.2 | 84.8 | 11.59 | 10.71 | | High | High | High | High | High | High | High | High | High | High | High | 0.003911 | 0.04145 | 1.02 |
| FALSE | High | RSPTALSEK | [R].RSPTAL | 1xPhospho | B0V2M3 1x | 0.025738 | 0.001645 | 1 | 3 | 1 | B0V2M3 | Zinc finger | B0V2M3 [1 | 1 | 1526.834 | 0 | -0.34 | 0.017762 | 5.95 | 112.2 | 87.8 | 6.48 | 7.42 | | High | High | High | High | High | High | High | High | High | High | High | 0.000872 | 0.009056 | 1.71 |
| FALSE | High | RSPTENVN | [K].RSPTEN | 1xPhospho | A2AGQ4 1x | 2.75E-05 | 0 | 1 | 28 | 2 | A2AGQ4 | MAP kinase | A2AGQ4 [1 | 1 | 1937.026 | 0 | 0.32 | 0.065269 | 15.09 | 93.6 | 106.4 | 12.39 | 11.5 | | High | High | High | High | High | High | High | High | High | High | High | 0 | 4.22E-06 | 4.03 |
| FALSE | High | RSPTSSAIPI | [K].RSPTSS | 1xPhospho | A0A0R4J1L | 0.005615 | 0.00062 | 1 | 8 | 1 | A0A0R4J1L | Microtubu | A0A0R4J1L | 1 | 1839.932 | 0 | -0.42 | 0.003951 | 1.79 | 114.2 | 85.8 | 5.01 | 4.08 | | High | High | High | High | High | High | High | High | High | High | High | 0.000275 | 0.001644 | 3.08 |
| FALSE | High | RSPVSPQL( | [R].RSPVSP | 1xPhospho | E9Q969 1x | 4E-07 | 0 | 1 | 4 | 1 | E9Q969 | Cytoplasmi | E9Q969 [5 | 1 | 2869.478 | 0 | 0.65 | 0.007264 | 3.2 | 77 | 123 | 5.99 | 12.29 | | High | High | High | High | High | High | High | High | High | High | High | 0 | 3.62E-08 | 4.18 |
| FALSE | High | RSPVVAVT( | [K].RSPVVA | 1xPhospho | Q3URY5 1x | 0.014538 | 0.001011 | 1 | 4 | 1 | Q3URY5 | Putative ur | Q3URY5 [1 | 1 | 1622.94 | 0 | -1.04 | 0.002425 | 2.29 | 134.7 | 65.3 | 8.06 | 10.9 | | High | High | High | High | High | High | High | High | High | High | High | 0.000555 | 0.004773 | 2.51 |
| FALSE | High | RSQSFNNY | [R].RSQSFN | 1xPhospho | E9Q842 1x | 0.070485 | 0.005214 | 1 | 7 | 1 | E9Q842 | Neuron na | E9Q842 [2 | 1 | 1796.873 | 0 | 1.05 | 0.002756 | 17.35 | 64.5 | 135.5 | 12.32 | 13.49 | | High | High | High | High | High | High | High | High | High | High | High | 0.00269 | 0.02872 | 2 |
| FALSE | High | RSRSHSDN | [R].RSRSHS | 1xCarbami | Q9R0U0-3 | 0.001615 | 5.66E-05 | 1 | 2 | 3 | Q9R0U0-3 | Isoform 3 c | Q9R0U0-3 | 2 | 3684.527 | 0 | -0.27 | 0.129833 | 24.98 | 122.5 | 77.5 | 28.11 | 15.53 | | High | High | High | High | High | High | High | High | High | High | High | 2.58E-05 | 0.000407 | 3.33 |
| FALSE | High | RSRSHSDN | [R].RSRSHS | 1xCarbami | Q9R0U0-3 | 0.104222 | 0.008348 | 1 | 2 | 3 | Q9R0U0-3 | Isoform 3 c | Q9R0U0-3 | 2 | 3764.494 | 0 | -0.54 | 0.021072 | 7.96 | 120.4 | 79.6 | 10.06 | 10.2 | | High | High | High | High | High | High | High | High | High | High | High | 0.004517 | 0.04891 | 4.09 |
| FALSE | High | RSRSVEDDI | [K].RSRSVE | 1xCarbami | P22518 2x | 2.47E-05 | 0 | 1 | 2 | 1 | P22518 | Dual specif | P22518 [13 | 2 | 3104.371 | 0 | -0.69 | 0.01263 | 5.11 | 119.7 | 80.3 | 11.3 | 7.43 | | High | High | High | High | High | High | High | High | High | High | High | 0 | 3.74E-06 | 3.13 |
| FALSE | High | RSSAEQHS | [R].RSSAEC | 1xCarbami | Q08BU3 1x | 0.000984 | 0 | 1 | 1 | 1 | Q08BU3 | OTTMUSG | Q08BU3 [6 | 1 | 2302.105 | 0 | 0.8 | 0.015375 | 19.11 | 73.8 | 126.2 | 10.02 | 14.17 | | High | High | High | High | High | High | High | High | High | High | High | 0 | 0.000235 | 3.15 |
| FALSE | High | RSSAIGIEN | [R].RSSAIG | 1xPhospho | A2ALS7 1x | 6.26E-06 | 0 | 2 | 6 | 4 | A2ALS7; A2 | Rap1 GTPa | A2ALS7 [24 | 1 | 2339.237 | 0 | 1.3 | 0.000928 | 11.87 | 56.8 | 143.2 | 9.28 | 7.56 | Shared | High | High | High | High | High | High | High | High | High | High | High | 0 | 7.96E-07 | 5.14 |
| FALSE | High | RSSCSSEPN | [R].RSSCSS | 1xCarbami | Q924A2 1x | 0.002974 | 0.00016 | 1 | 3 | 2 | Q924A2 | Protein cap | Q924A2 [2 | 1 | 1887.904 | 0 | -0.86 | 0.001626 | 3.49 | 124.9 | 75.1 | 9.69 | 8.18 | | High | High | High | High | High | High | High | High | High | High | High | 7.19E-05 | 0.000807 | 3.1 |
| FALSE | High | RSSDGSLSI | [K].RSSDGS | 1xPhospho | B8QI33 1x | 0.011966 | 0.000889 | 1 | 3 | 1 | B8QI33 | Liprin-alph | B8QI33 [23 | 1 | 2040.016 | 0 | 0.17 | 0.217157 | 28.02 | 98.9 | 101.1 | 9.49 | 15.41 | | High | High | High | High | High | High | High | High | High | High | High | 0.000449 | 0.003817 | 3.18 |
| FALSE | High | RSPSYPPPG | [R].RSPSYP | 1xCarbami | F8WIK2 1x | 0.01894 | 0.001362 | 1 | 2 | 1 | F8WIK2 | Poly [ADP- | F8WIK2 [3 | 1 | 1840.918 | 0 | 0.32 | 0.04273 | 13.37 | 89.1 | 110.9 | 9.31 | 14.79 | | High | High | High | High | High | High | High | High | High | High | High | 0.00074 | 0.006441 | 2.34 |
| FALSE | High | RSPGGTSPI | [R].RSPGG | 1xCarbami | A0A0U1RP | 0.007145 | 0.00062 | 1 | 4 | 2 | A0A0U1RP | Ataxin-2-lik | A0A0U1RP | 1 | 4417.173 | 0 | -0.26 | 0.035759 | 6.27 | 111.6 | 88.4 | 8.23 | 14.49 | | High | High | High | High | High | High | High | High | High | High | High | 0.000327 | 0.002609 | 3.88 |
| FALSE | High | RSSLTTAAL | [K].RSSLTT | 1xPhospho | A1A5B6 1x | 0.01441 | 0.001011 | 1 | 5 | 2 | A1A5B6 | TBC1 doma | A1A5B6 [2 | 1 | 2542.359 | 0 | 0.46 | 0.012611 | 4.49 | 85.3 | 114.7 | 14.05 | 10.56 | | High | High | High | High | High | High | High | High | High | High | High | 0.000538 | 0.004744 | 3.04 |
| FALSE | High | RSSLVDEAI | [R].RSSLVD | 2xPhospho | Q9R0L6-2 | 9.39E-06 | 0 | 1 | 3 | 2 | Q9R0L6-2 | Isoform 2 c | Q9R0L6-2 | 1 | 2626.173 | 0 | -0.09 | 0.328493 | 9.83 | 102.1 | 97.9 | 6.24 | 8.25 | | High | High | High | High | High | High | High | High | High | High | High | 0 | 1.25E-06 | 5.1 |
| FALSE | High | RSTQGVTL | [R].RSTQG | 1xPhospho | A6H644 1x | 3.88E-06 | 0 | 2 | 6 | 5 | A6H644; Q | MCG13049 | A6H644 [6 | 1 | 2314.206 | 0 | -0.2 | 0.177295 | 20.48 | 111.1 | 88.9 | 16.52 | 12.53 | Shared | High | High | High | High | High | High | High | High | High | High | High | 0 | 4.69E-07 | 3.88 |
| FALSE | High | RSTSLQPEI | [K].RSTSLQ | 1xPhospho | P48754 1x | 1.02E-06 | 0 | 1 | 5 | 3 | P48754 | Breast canc | P48754 [50 | 1 | 1959.035 | 0 | -0.52 | 0.007927 | 2.77 | 117.6 | 82.4 | 6.77 | 7.14 | | High | High | High | High | High | High | High | High | High | High | High | 0 | 1.04E-07 | 3.56 |
| FALSE | High | RSTSPHDVI | [R].RSTSPH | 2xCarbami | B2RWW1 2 | 0.021607 | 0.001507 | 1 | 3 | 1 | B2RWW1 | Microtubu | B2RWW1 [ | 1 | 2888.246 | 0 | -0.14 | 0.121294 | 13.38 | 104.8 | 95.2 | 4.26 | 16.06 | | High | High | High | High | High | High | High | High | High | High | High | 0.000801 | 0.007465 | 3.51 |
| FALSE | High | RSTSPIIGSF | [R].RSTSPII | 2xPhospho | Q3TC46 2x | 0.015329 | 0.001089 | 1 | 1 | 1 | Q3TC46 | Protein PA | Q3TC46 [1 | 1 | 1755.876 | 0 | -0.36 | 0.168833 | 11.18 | 110.1 | 89.9 | 9.2 | 15.41 | | High | High | High | High | High | High | High | High | High | High | High | 0.000571 | 0.005076 | 2.38 |
| FALSE | High | RSTVLGLP( | [R].RSTVLG | 1xPhospho | A2A7S8-2 | 5.91E-05 | 0 | 1 | 4 | 2 | A2A7S8-2 | Isoform 2 c | A2A7S8-2 [ | 1 | 2001.141 | 0 | 0.26 | 0.546209 | 10.01 | 94 | 106 | 19.01 | 7.66 | | High | High | High | High | High | High | High | High | High | High | High | 0 | 0.000103 | 2.26 |
| FALSE | High | RSVPVSSSS | [K].RSVPVS | 1xPhospho | P33215 1x | 7.81E-08 | 0 | 1 | 2 | 1 | P33215 | Protein NE | P33215 [32 | 1 | 2168.102 | 0 | 0.97 | 0.00545 | 30.61 | 76 | 124 | 27.37 | 16.51 | | High | High | High | High | High | High | High | High | High | High | High | 0 | 5.79E-09 | 5.33 |
| FALSE | High | RSVSASHE( | [R].RSVSAS | 1xPhospho | Q3U9G9 1x | 0.012959 | 0.000889 | 1 | 3 | 2 | Q3U9G9 | Lamin-B re | Q3U9G9 [1 | 1 | 1809.926 | 0 | -0.79 | 0.088603 | 3.67 | 125.7 | 74.3 | 17.85 | 24.42 | | High | High | High | High | High | High | High | High | High | High | High | 0.000449 | 0.004181 | 1.02 |
| FALSE | High | RSVSHGSN | [K].RSVSH( | 1xPhospho | Q3TLH4 1x | 2.06E-05 | 0 | 1 | 5 | 5 | Q3TLH4 | Protein PRI | Q3TLH4 [8 | 1 | 2216.016 | 0 | 0.14 | 0.116078 | 10.96 | 93.3 | 106.7 | 7.35 | 7.62 | | High | High | High | High | High | High | High | High | High | High | High | 0.000786 | 0.007068 | 3.81 |
| FALSE | High | RSWDQQII | [R].RSWDC | 1xPhospho | P97440 1x | 0.005467 | 0.00062 | 1 | 6 | 1 | P97440 | Histone RN | P97440 [18 | 1 | 1598.846 | 0 | -0.69 | 0.000406 | 2.16 | 121.6 | 78.4 | 4.8 | 4.77 | | High | High | High | High | High | High | High | High | High | High | High | 0.000275 | 0.001602 | 2.12 |
| FALSE | High | RSWGPAQ | [R].RSWGP | 1xPhospho | Q6EDY6 1x | 0.005717 | 0.00062 | 1 | 2 | 1 | Q6EDY6 | F-actin-unc | Q6EDY6 [1 | 1 | 2145.053 | 0 | -0.92 | 0.000401 | 2.96 | 129.1 | 70.9 | 6.97 | 15.85 | | High | High | High | High | High | High | High | High | High | High | High | 0.000275 | 0.001683 | 2.61 |
| FALSE | High | RSYDVPPPI | [R].RSYDVI | 1xPhospho | Q3U7Z6 1x | 0.047086 | 0.00354 | 1 | 2 | 1 | Q3U7Z6 | Phosphogl | Q3U7Z6 [1 | 1 | 3127.5 | 0 | 0.31 | 0.155622 | 27.13 | 89.2 | 110.8 | 7.06 | 18.06 | | High | High | High | High | High | High | High | High | High | High | High | 0.001812 | 0.01805 | 3.18 |
| FALSE | High | RSYEDDED | [K].RSYEDI | 1xPhospho | H9KV04 1x | 2.23E-05 | 0 | 1 | 2 | 2 | H9KV04 | Mini-chron | H9KV04 [1 | 1 | 2366.062 | 0 | -0.83 | 0.000668 | 3.48 | 124.4 | 75.6 | 6.46 | 11.47 | | High | High | High | High | High | High | High | High | High | High | High | 0 | 3.34E-06 | 2.64 |
| FALSE | High | RSYSLSELS | [K].RSYSLSI | 1xPhospho | Q921I6 1x | 7.1E-06 | 0 | 1 | 1 | 2 | Q921I6 | SH3 domai | Q921I6 [24 | 1 | 2119.156 | 0 | 0.46 | 0.00708 | 5.83 | 85.8 | 114.2 | 8.19 | 5.8 | | High | High | High | High | High | High | High | High | High | High | High | 0 | 9.17E-07 | 3.62 |
| FALSE | High | RSYSPDGK | [K].RSYSPD | 1xPhospho | Q6ZQ61 1x | 0.0008 | 0 | 1 | 8 | 1 | Q6ZQ61 | MCG12197 | Q6ZQ61 [6 | 1 | 1447.735 | 0 | -0.85 | 0.004142 | 4.79 | 128.6 | 71.4 | 3.77 | 15.01 | | High | High | High | High | High | High | High | High | High | High | High | 0 | 0.000185 | 3.11 |
| FALSE | High | RSYSPDGK | [K].RSYSPD | 2xPhospho | Q6ZQ61 2x | 0.020862 | 0.001472 | 1 | 8 | 3 | Q6ZQ61 | MCG12197 | Q6ZQ61 [6 | 1 | 1527.701 | 0 | 0.1 | 0.054719 | 4.64 | 96.5 | 103.5 | 4.65 | 6.85 | | High | High | High | High | High | High | High | High | High | High | High | 0.000801 | 0.007171 | 2.09 |
| FALSE | High | RSYSSATEE | [K].RSYSSA | 2xPhospho | Q4V9W2 2 | 1.97E-06 | 0 | 1 | 1 | 2 | Q4V9W2 | Protein SRI | Q4V9W2 [ | 1 | 2048.898 | 0 | 0.36 | 0.003474 | 6.71 | 86.1 | 113.9 | 8.67 | 2.16 | | High | High | High | High | High | High | High | High | High | High | High | 0 | 2.19E-07 | 3.82 |
| FALSE | High | RTASPPPPF | [R].RTASPP | 2xPhospho | Q52KI8 2x | 0.038017 | 0.002528 | 1 | 7 | 1 | Q52KI8 | Serine/argi | Q52KI8 [63 | 1 | 1665.853 | 0 | 0.12 | 0.221816 | 13.44 | 94.7 | 105.3 | 8.49 | 6.42 | | High | High | High | High | High | High | High | High | High | High | High | 0.001355 | 0.01409 | 2.22 |
| FALSE | High | RTEGYAAF( | [K].RTEGYA | 2xPhospho | A6H6B3 2x | 0.007404 | 0.00062 | 1 | 8 | 2 | A6H6B3 | RalA bindir | A6H6B3 [8 | 1 | 2979.27 | 0 | 0.1 | 0.217168 | 8.41 | 95.6 | 104.4 | 9.26 | 9.6 | | High | High | High | High | High | High | High | High | High | High | High | 0.000801 | 0.007347 | 2.96 |
| FALSE | High | RTESPSESC | [K].RTESPS | 1xCarbami | B7ZP47 1x | 6.37E-06 | 0 | 1 | 2 | 4 | B7ZP47 | Wapal prot | B7ZP47 [22 | 1 | 1914.94 | 0 | -0.98 | 0.000656 | 0.14 | 132.7 | 67.3 | 5.91 | 2.54 | | High | High | High | High | High | High | High | High | High | High | High | 0 | 5.55E-06 | 3.32 |
| FALSE | High | RTESVPSDI | [R].RTESVF | 1xPhospho | Q61097 1x | 7.74E-08 | 0 | 1 | 5 | 2 | Q61097 | Kinase sup | Q61097 [3 | 1 | 2007.97 | 0 | -0.65 | 0.006369 | 7.09 | 122.3 | 77.7 | 7.01 | 14.65 | | High | High | High | High | High | High | High | High | High | High | High | 0 | 5.74E-09 | 2.83 |
| FALSE | High | RTGSNISG | [K].RTGSNI | 2xPhospho | Q3B7Z2 2x | 0.000154 | 0 | 1 | 2 | 1 | Q3B7Z2 | Oxysterol-l | Q3B7Z2 [3 | 1 | 2732.258 | 0 | 0.49 | 0.002854 | 10.19 | 80.9 | 119.1 | 5.94 | 9.2 | | High | High | High | High | High | High | High | High | High | High | High | 0 | 2.92E-05 | 3.65 |
| FALSE | High | RTHSDASD | [R].RTHSDA | 1xPhospho | B9EKC3 1x | 3.6E-05 | 0 | 1 | 2 | 4 | B9EKC3 | Rho GTPas | B9EKC3 [1 | 1 | 2306.07 | 0 | -0.42 | 0.023421 | 9.76 | 114.9 | 85.1 | 11.67 | 9.52 | | High | High | High | High | High | High | High | High | High | High | High | 0 | 5.71E-06 | 3.32 |
| FALSE | High | RTHSDASD | [R].RTHSDA | 2xPhospho | B9EKC3 2x | 0.000322 | 0 | 1 | 2 | 2 | B9EKC3 | Rho GTPas | B9EKC3 [1 | 1 | 2386.037 | 0 | -0.21 | 0.042602 | 5.6 | 106.1 | 93.9 | 3.98 | 4.21 | | High | High | High | High | High | High | High | High | High | High | High | 0 | 6.66E-05 | 3.14 |
| FALSE | High | RTNGWVP | [R].RTNGW | 1xCarbami | Q497V6 1x | 0.031439 | 0.002011 | 1 | 3 | 1 | Q497V6 | Bromo adja | Q497V6 [5 | 1 | 2084.04 | 0 | 0.27 | 0.853666 | 34.77 | 92 | 108 | 30.71 | 30.05 | | High | High | High | High | High | High | High | High | High | High | High | 0.001076 | 0.01138 | 1.84 |
| FALSE | High | RTNSTGGS | [R].RTNST( | 1xPhospho | F8VPM7 1x | 3.04E-06 | 0 | 1 | 8 | 2 | F8VPM7 | ELKS/Rab6 | F8VPM7 [3 | 1 | 2205.066 | 0 | 0.41 | 0.073491 | 12.71 | 82 | 118 | 10.75 | 19.68 | | High | High | High | High | High | High | High | High | High | High | High | 0 | 3.53E-07 | 4.57 |
| FALSE | High | RTNSTGGS | [R].RTNST( | 1xDeamida | F8VPM7 1x | 2.24E-07 | 0 | 1 | 8 | 9 | F8VPM7 | ELKS/Rab6 | F8VPM7 [3 | 1 | 2206.05 | 0 | 0.52 | 0.030217 | 25.4 | 82.2 | 117.8 | 10.83 | 13.15 | | High | High | High | High | High | High | High | High | High | High | High | 0 | 0.000143 | 3.56 |
| FALSE | High | RTPDSTFSE | [R].RTPDST | 1xPhospho | Q921D9 1x | 0.000553 | 0 | 1 | 3 | 1 | Q921D9 | Grainyhead | Q921D9 [2 | 1 | 1953.972 | 0 | -0.31 | 0.001887 | 4.5 | 113.6 | 86.4 | 6.18 | 9.75 | | High | High | High | High | High | High | High | High | High | High | High | 0 | 0.000123 | 2.8 |
| FALSE | High | RSTQESLAI | [R].RSTQES | 1xPhospho | Q8K1C9 1x | 7.04E-06 | 0 | 1 | 2 | 4 | Q8K1C9 | Leucine-ric | Q8K1C9 [3 | 1 | 2088.074 | 0 | 2.27 | 0.008652 | 38.15 | 30.6 | 169.4 | 18.98 | 29.16 | | High | High | High | High | High | High | High | High | High | High | High | 0 | 9.14E-07 | 3.87 |
| FALSE | High | RSTPPLQLL | [R].RSTPPL | 1xPhospho | D3YYY8 1x | 0.003282 | 0.00016 | 1 | 1 | 2 | D3YYY8 | Rho guanir | D3YYY8 [2 | 1 | 1576.877 | 0 | 2.47 | 0.001125 | 8.63 | 31.6 | 168.4 | 25.61 | 10.84 | | High | High | High | High | High | High | High | High | High | High | High | 7.19E-05 | 0.000904 | 2.59 |
| FALSE | High | RSTELVLSP | [K].RSTELV | 1xPhospho | Q9ESK9 1x | 3.63E-05 | 0 | 1 | 6 | 3 | Q9ESK9 | RB1-induci | Q9ESK9 [2 | 1 | 1825.908 | 0 | -0.26 | 0.467578 | 24.3 | 107.9 | 92.1 | 15.7 | 19.17 | | High | High | High | High | High | High | High | High | High | High | High | 0 | 0.000188 | 3.31 |
| FALSE | High | RSTELVLSP | [K].RSTELV | 1xPhospho | Q9ESK9 1x | 0.000389 | 0 | 1 | 6 | 1 | Q9ESK9 | RB1-induci | Q9ESK9 [2 | 1 | 1809.913 | 0 | -0.18 | 0.425831 | 9.81 | 104.4 | 95.6 | 14.47 | 9.64 | | High | High | High | High | High | High | High | High | High | High | High | 0 | 8.25E-05 | 3 |
| FALSE | High | RSSMSSCG | [R].RSSMS | 2xCarbami | B2ZRS3 1x | 0.113753 | 0.009241 | 1 | 6 | 1 | B2ZRS3 | Depdc6-00 | B2ZRS3 [27 | 1 | 3689.705 | 0 | 2.13 | 0.000551 | 10.48 | 37.9 | 162.1 | 15.19 | 12.2 | | High | High | High | High | High | High | High | High | High | High | High | 0.00468 | 0.05043 | 3.27 |
| FALSE | High | RSSMSSCG | [R].RSSMS | 2xCarbami | B2ZRS3 1x | 0.011048 | 0.000772 | 1 | 6 | 2 | B2ZRS3 | Depdc6-00 | B2ZRS3 [27 | 1 | 3705.7 | 0 | 2.65 | 0.000527 | 1.76 | 26.4 | 173.6 | 14.54 | 10.58 | | High | High | High | High | High | High | High | High | High | High | High | 0.000399 | 0.003496 | 3.71 |
| FALSE | High | RSSNDSLN | [K].RSSNDS | 1xPhospho | B2RUG9 1x | 3.57E-05 | 0 | 1 | 8 | 2 | B2RUG9 | Adenomat | B2RUG9 [9 | 1 | 2412.144 | 0 | 0.09 | 0.747996 | 14.51 | 99.7 | 100.3 | 11.69 | 9.04 | | High | High | High | High | High | High | High | High | High | High | High | 0 | 5.64E-06 | 3 |
| FALSE | High | RSSPDDGN | [K].RSSPDI | 1xPhospho | Q3TQ38 1x | 0.000687 | 0 | 1 | 4 | 2 | Q3TQ38 | Putative ur | Q3TQ38 [1 | 1 | 2759.329 | 0 | 0.4 | 0.147059 | 35.18 | 91.8 | 108.2 | 21.2 | 5.25 | | High | High | High | High | High | High | High | High | High | High | High | 2.58E-05 | 0.000565 | 3.54 |
| FALSE | High | RSSPIFQTT | [K].RSSPIF( | 1xPhospho | Q9WV30-2 | 0.017966 | 0.001282 | 1 | 7 | 2 | Q9WV30-2 | Isoform 2 c | Q9WV30-2 | 1 | 1702.929 | 0 | 1.14 | 0.001726 | 11.25 | 62.3 | 137.7 | 9.35 | 6.55 | | High | High | High | High | High | High | High | High | High | High | High | 0.002403 | 0.02587 | 2.09 |
| FALSE | High | RSSPSPPLS | [R].RSSPSP | 2xPhospho | Q91WL6 2x | 0.021046 | 0.001507 | 1 | 2 | 2 | Q91WL6 | Sorting ne | Q91WL6 [1 | 1 | 1931.928 | 0 | -0.16 | 0.018556 | 1.56 | 106 | 94 | 2.9 | 5.39 | | High | High | High | High | High | High | High | High | High | High | High | 0.001468 | 0.01481 | 2.39 |
| FALSE | High | RSSPYHSEL | [K].RSSPYH | 1xPhospho | Q4V9W2 1 | 0.001168 | 0 | 1 | 1 | 4 | Q4V9W2 | Protein SRI | Q4V9W2 [ | 1 | 1842.952 | 0 | 0.24 | 0.295254 | 7.59 | 91.6 | 108.4 | 14.07 | 8.66 | | High | High | High | High | High | High | High | High | High | High | High | 0 | 0.000283 | 1.74 |
| FALSE | High | RSSQPSPT | [R].RSSQPS | 1xPhospho | A2AJI0 1xP | 1.39E-06 | 0 | 1 | 4 | 7 | A2AJI0 | MAP7 dom | A2AJI0 [11 | 1 | 2448.254 | 0 | 1 | 0.001414 | 11.56 | 66.8 | 133.2 | 13.72 | 5.79 | | High | High | High | High | High | High | High | High | High | High | High | 0 | 5.45E-06 | 3.39 |
| FALSE | High | RSSQPSPT | [R].RSSQPS | 2xPhospho | A2AJI0 2xP | 0.000534 | 0 | 1 | 4 | 2 | A2AJI0 | MAP7 dom | A2AJI0 [11 | 1 | 2528.22 | 0 | 1.43 | 0.000742 | 16.79 | 56.7 | 143.3 | 6.41 | 7.46 | | High | High | High | High | High | High | High | High | High | High | High | 7.19E-05 | 0.000924 | 3.58 |
| FALSE | High | RSSSDTAA | [R].RSSSDT | 1xCarbami | E9PYH6 1x | 0.00047 | 0 | 1 | 1 | 7 | E9PYH6 | SET domair | E9PYH6 [2 | 1 | 3800.666 | 0 | 0.31 | 0.051733 | 19.81 | 87.3 | 112.7 | 16.83 | 14.33 | | High | High | High | High | High | High | High | High | High | High | High | 2.58E-05 | 0.000436 | 4.47 |
| FALSE | High | RSSSDTAA | [R].RSSSDT | 1xCarbami | E9PYH6 1x | 3.32E-05 | 0 | 1 | 1 | 20 | E9PYH6 | SET domair | E9PYH6 [2 | 1 | 3801.65 | 0 | 0.2 | 0.178821 | 24.94 | 89.3 | 110.7 | 6.3 | 23.84 | | High | High | High | High | High | High | High | High | High | High | High | 0 | 8.23E-06 | 4.97 |
| FALSE | High | RSSSELSPE | [R].RSSSEL | 1xPhospho | Q8BTI8 1x | 4.05E-05 | 0 | 1 | 3 | 11 | Q8BTI8 | Serine/argi | Q8BTI8 [13 | 1 | 1985.036 | 0 | -0.44 | 0.007437 | 3.66 | 115.8 | 84.2 | 8.18 | 4.92 | | High | High | High | High | High | High | High | High | High | High | High | 0 | 6.52E-06 | 3.69 |
| FALSE | High | RSSSELSPE | [R].RSSSEL | 2xPhospho | Q8BTI8 2x | 3.28E-07 | 0 | 1 | 3 | 11 | Q8BTI8 | Serine/argi | Q8BTI8 [13 | 1 | 2065.002 | 0 | -0.85 | 0.000729 | 2.02 | 128.3 | 71.7 | 7.1 | 3.69 | | High | High | High | High | High | High | High | High | High | High | High | 0 | 2.9E-08 | 3.78 |
| FALSE | High | RSSLVDEAI | [R].RSSLVD | 1xPhospho | Q9R0L6-2 | 0.000146 | 0 | 1 | 3 | 2 | Q9R0L6-2 | Isoform 2 c | Q9R0L6-2 | 1 | 2546.206 | 0 | -0.37 | 0.013551 | 4.16 | 112.9 | 87.1 | 6.8 | 3.62 | | High | High | High | High | High | High | High | High | High | High | High | 0 | 2.73E-05 | 4.04 |
| FALSE | High | RSSSPAELE | [R].RSSSPA | 1xPhospho | A0A0A0M( | 3.54E-05 | 0 | 1 | 4 | 2 | A0A0A0M( | Ral GTPase | A0A0A0M( | 1 | 1754.945 | 0 | 1.14 | 0.00024 | 5.92 | 62.6 | 137.4 | 5.32 | 5.01 | | High | High | High | High | High | High | High | High | High | High | High | 0 | 5.6E-06 | 2.63 |
| FALSE | High | RSSSSSAA | [R].RSSSSS | 1xPhospho | Q6PHZ5 1x | 0.005322 | 0.000599 | 1 | 1 | 2 | Q6PHZ5 | Putative RI | Q6PHZ5 [2 | 1 | 4033.026 | 0 | 0.17 | 0.282209 | 22.72 | 89.8 | 110.2 | 11.34 | 15.46 | | High | High | High | High | High | High | High | High | High | High | High | 0.000267 | 0.001547 | 2.63 |
| FALSE | High | RSSTLAPA( | [R].RSSTLA | 1xPhospho | E9Q5G3 1x | 2.74E-08 | 0 | 1 | 1 | 14 | E9Q5G3 | Kinesin-like | E9Q5G3 [9 | 1 | 2842.309 | 0 | -0.65 | 0.00109 | 3.16 | 122.2 | 77.8 | 1.95 | 7.46 | | High | High | High | High | High | High | High | High | High | High | High | 0 | 1.79E-09 | 5.95 |
| FALSE | High | RSSTLSTLA | [R].RSSTLS | 1xPhospho | Q3U560 1x | 0.055308 | 0.003982 | 1 | 1 | 2 | Q3U560 | Putative ur | Q3U560 [4 | 1 | 1701.852 | 0 | 1.78 | 0.000487 | 1.51 | 41.9 | 158.1 | 10.47 | 8.5 | | High | High | High | High | High | High | High | High | High | High | High | 0.002074 | 0.0216 | 1.96 |
| FALSE | High | RSSTSQTVS | [K].RSSTSC | 1xPhospho | F6ZDS4 1x | 0.000114 | 0 | 1 | 7 | 3 | F6ZDS4 | Nucleopro | F6ZDS4 [71 | 1 | 3239.645 | 0 | -0.55 | 0.02005 | 8.21 | 114.9 | 85.1 | 10.93 | 19.52 | | High | High | High | High | High | High | High | High | High | High | High | 0 | 2.09E-05 | 4.13 |
| FALSE | High | RSSTSSEPT | [R].RSSTSS | 1xPhospho | A0A0R4J0J | 5.42E-06 | 0 | 1 | 3 | 8 | A0A0R4J0J | Cytospin-A | A0A0R4J0J | 1 | 1914.994 | 0 | 1.69 | 0.000223 | 6.22 | 47.2 | 152.8 | 7.35 | 5.12 | | High | High | High | High | High | High | High | High | High | High | High | 0 | 6.82E-07 | 4.46 |
| FALSE | High | RSSVFAAEI | [R].RSSVFA | 1xPhospho | O35425 1x | 0.00103 | 0 | 1 | 2 | 2 | O35425 | Bcl-2-relate | O35425 [6 | 1 | 2023.951 | 0 | 0.2 | 0.257456 | 23.97 | 96.6 | 103.4 | 8.43 | 12.2 | | High | High | High | High | High | High | High | High | High | High | High | 0 | 0.000264 | 3.37 |
| FALSE | High | RSSVNGEA | [R].RSSVN( | 1xPhospho | E9QP49 1x | 5.05E-07 | 0 | 1 | 7 | 4 | E9QP49 | EH domain | E9QP49 [1 | 1 | 1828.927 | 0 | 2.15 | 0.000181 | 9.54 | 37.4 | 162.6 | 6.04 | 4.05 | | High | High | High | High | High | High | High | High | High | High | High | 0 | 4.71E-08 | 3.83 |
| FALSE | High | RSSVNGEA | [R].RSSVN( | 1xDeamida | E9QP49 1x | 1.8E-06 | 0 | 1 | 7 | 4 | E9QP49 | EH domain | E9QP49 [1 | 1 | 1829.911 | 0 | 1.47 | 0.000536 | 5.92 | 52.3 | 147.7 | 9.42 | 6 | | High | High | High | High | High | High | High | High | High | High | High | 0 | 3.79E-05 | 3.67 |
| FALSE | High | RSSVSGISL | [R].RSSVSG | 1xPhospho | P70271 1x | 3.5E-05 | 0 | 1 | 3 | 4 | P70271 | PDZ and LI | P70271 [11 | 1 | 1728.848 | 0 | 0.12 | 0.053204 | 4.8 | 92.8 | 107.2 | 8.88 | 4.58 | | High | High | High | High | High | High | High | High | High | High | High | 0 | 5.54E-06 | 3.98 |
| FALSE | High | RSSVTFEDI | [R].RSSVTF | 1xPhospho | B2RPY5-2 | 0.000908 | 0 | 1 | 4 | 2 | B2RPY5-2 | Isoform 2 c | B2RPY5-2 [ | 1 | 2205.12 | 0 | 1.79 | 0.000455 | 8.5 | 45.7 | 154.3 | 5.83 | 8.37 | | High | High | High | High | High | High | High | High | High | High | High | 0 | 0.000213 | 3.3 |
| FALSE | High | RSSWSSDE | [R].RSSWS | 1xPhospho | Q7TSH8 1x | 2.5E-07 | 0 | 1 | 2 | 3 | Q7TSH8 | Transmem | Q7TSH8 [8 | 1 | 2317.148 | 0 | 0.61 | 0.026442 | 13.25 | 77.2 | 122.8 | 2.24 | 16.64 | | High | High | High | High | High | High | High | High | High | High | High | 0 | 3.47E-07 | 3.42 |
| FALSE | High | RSSYLLAIT | [R].RSSYLL | 1xPhospho | B1AQY2 1x | 0.037692 | 0.00249 | 1 | 4 | 2 | B1AQY2 | Rho GTPas | B1AQY2 [6 | 1 | 1718.904 | 0 | 0.24 | 0.092881 | 10.31 | 92.8 | 107.2 | 8.64 | 4.87 | | High | High | High | High | High | High | High | High | High | High | High | 0.002455 | 0.02655 | 1.99 |
| FALSE | High | RSTDSSSSV | [R].RSTDSS | 1xPhospho | Q8VDY9 1x | 0.003028 | 0.00016 | 1 | 1 | 2 | Q8VDY9 | Caspase ac | Q8VDY9 [6 | 1 | 2422.186 | 0 | -0.27 | 0.84902 | 17.54 | 109.5 | 90.5 | 7.29 | 62.43 | | High | High | High | High | High | High | High | High | High | High | High | 7.19E-05 | 0.000822 | 3.31 |
| FALSE | High | RSSSPAELE | [R].RSSSPA | 2xPhospho | A0A0A0M( | 0.000239 | 0 | 1 | 4 | 1 | A0A0A0M( | Ral GTPase | A0A0A0M( | 1 | 1834.912 | 0 | 1.15 | 0.000193 | 6.08 | 61.3 | 138.7 | 6.34 | 4.2 | | High | High | High | High | High | High | High | High | High | High | High | 0 | 4.77E-05 | 3.46 |
| FALSE | High | RSPGGTSPI | [R].RSPGG | 1xCarbami | A0A0U1RP | 5.5E-05 | 0 | 1 | 4 | 5 | A0A0U1RP | Ataxin-2-lik | A0A0U1RP | 1 | 4338.19 | 0 | 0.03 | 0.771815 | 12.49 | 98.1 | 101.9 | 6.86 | 4.84 | | High | High | High | High | High | High | High | High | High | High | High | 0 | 9.22E-06 | 4.16 |
| FALSE | High | RSPGGTSPI | [R].RSPGG | 1xCarbami | A0A0U1RP | 0.02933 | 0.001857 | 1 | 4 | 1 | A0A0U1RP | Ataxin-2-lik | A0A0U1RP | 1 | 4337.206 | 0 | -0.09 | 0.55053 | 12.89 | 103.2 | 96.8 | 7.05 | 6.55 | | High | High | High | High | High | High | High | High | High | High | High | 0.000997 | 0.01047 | 3.29 |

| | | | | | | | | | | | | | | | | | | | | | | | | | | | | | | | | | | | | | | |
|---|---|---|---|---|---|---|---|---|---|---|---|---|---|---|---|---|---|---|---|---|---|---|---|---|---|---|---|---|---|---|---|---|---|---|---|---|---|---|
| FALSE | High | RSPEHLGLI | [K].RSPEHL | 1xPhospho | B2RUA5 1x | 0.002058 | 5.66E-05 | 1 | 2 | 1 | B2RUA5 | SAPS doma | B2RUA5 [8 | 1 | 3528.663 | 0 | -0.61 | 0.786467 | 11.61 | 119.9 | 80.1 | 41.17 | 68.12 | | High | High | High | High | High | High | High | High | High | High | High | 2.58E-05 | 0.000534 | 3.85 |
| FALSE | High | RQSSDSCSI | [R].RQSSDS | 1xCarbami | Q9D2D7 1x | 0.002895 | 0.00016 | 1 | 1 | 2 | Q9D2D7 | Zinc finger | Q9D2D7 [1 | 1 | 2617.185 | 0 | -0.44 | 0.034656 | 13 | 117.8 | 82.2 | 13.55 | 8.65 | | High | High | High | High | High | High | High | High | High | High | High | 7.19E-05 | 0.000788 | 4.24 |
| FALSE | High | RQSSFPFNI | [R].RQSSFF | 1xPhospho | Q7TNL5 1x | 0.000221 | 0 | 1 | 4 | 1 | Q7TNL5 | Protein ph | Q7TNL5 [7 | 1 | 1875.988 | 0 | 0.23 | 0.099516 | 2.49 | 92 | 108 | 14.8 | 15.65 | | High | High | High | High | High | High | High | High | High | High | High | 0 | 4.35E-05 | 3.67 |
| FALSE | High | RQSSGDGV | [R].RQSSGI | 1xCarbami | B2RUP0 1x | 0.000184 | 0 | 1 | 4 | 2 | B2RUP0 | RNA bindir | B2RUP0 [1 | 1 | 2907.351 | 0 | 0.2 | 0.202968 | 20.32 | 94.9 | 105.1 | 15.68 | 9.82 | | High | High | High | High | High | High | High | High | High | High | High | 0 | 3.55E-05 | 4.89 |
| FALSE | High | RQSSGSTTI | [R].RQSSGS | 1xPhospho | B2RY22 1x | 3.19E-07 | 0 | 2 | 16 | 7 | B2RY22; A( | Clasp1 pro | B2RY22 [64 | 1 | 2146.993 | 0 | -0.78 | 0.000158 | 0.78 | 126.9 | 73.1 | 4.76 | 7.24 | Shared | High | High | High | High | High | High | High | High | High | High | High | 0 | 2.81E-08 | 4.03 |
| FALSE | High | RQSSGSTTI | [R].RQSSGS | 2xPhospho | B2RY22 2x | 0.001586 | 5.66E-05 | 2 | 16 | 5 | B2RY22; A( | Clasp1 pro | B2RY22 [64 | 1 | 2226.959 | 0 | 0.32 | 0.045431 | 12.12 | 90.8 | 109.2 | 8.23 | 11.81 | Shared | High | High | High | High | High | High | High | High | High | High | High | 0.000872 | 0.008701 | 3.48 |
| FALSE | High | RQSSHVQS | [K].RQSSH | 1xPhospho | P54116 1x | 0.01029 | 0.000772 | 1 | 3 | 2 | P54116 | Erythrocyt | P54116 [5- | 1 | 1521.748 | 0 | 6.2 | 0.000872 | 12.84 | 2.7 | 197.3 | 66.21 | 16.84 | | High | High | High | High | High | High | High | High | High | High | High | 0.001774 | 0.01753 | 2.44 |
| FALSE | High | RQSSPSCVI | [R].RQSSPS | 1xCarbami | E9PX30 1x | 0.01002 | 0.000736 | 1 | 5 | 1 | E9PX30 | Mitogen-a | E9PX30 [43 | 1 | 3058.492 | 0 | 0.57 | 0.00933 | 12.25 | 80.1 | 119.9 | 23.51 | 5.07 | | High | High | High | High | High | High | High | High | High | High | High | 0.000344 | 0.003139 | 2.6 |
| FALSE | High | RQSSPSCVI | [R].RQSSPS | 1xCarbami | E9PX30 1x | 0.000173 | 0 | 1 | 5 | 4 | E9PX30 | Mitogen-a | E9PX30 [43 | 1 | 3059.476 | 0 | 1.33 | 0.001312 | 22.97 | 57 | 143 | 13.87 | 7.63 | | High | High | High | High | High | High | High | High | High | High | High | 0 | 0.000145 | 3.93 |
| FALSE | High | RQSSPSCVI | [R].RQSSPS | 1xCarbami | E9PX30 2x | 0.000579 | 0 | 1 | 5 | 1 | E9PX30 | Mitogen-a | E9PX30 [43 | 1 | 3138.458 | 0 | 1.13 | 0.004544 | 24.82 | 59 | 141 | 38.66 | 6.43 | | High | High | High | High | High | High | High | High | High | High | High | 0 | 0.000129 | 4.75 |
| FALSE | High | RQSSSYEDI | [R].RQSSSY | 1xPhospho | O54916 1x | 0.000253 | 0 | 1 | 8 | 4 | O54916 | RalBP1-ass | O54916 [2 | 1 | 1920.926 | 0 | -0.67 | 0.015691 | 4.91 | 117.3 | 82.7 | 8.82 | 14.58 | | High | High | High | High | High | High | High | High | High | High | High | 0 | 5.1E-05 | 2.83 |
| FALSE | High | RQSSSYEDI | [R].RQSSSY | 2xPhospho | O54916 2x | 5.75E-05 | 0 | 1 | 8 | 2 | O54916 | RalBP1-ass | O54916 [2 | 1 | 2000.892 | 0 | 0.6 | 0.001414 | 0.32 | 79.6 | 120.4 | 2.56 | 7.99 | | High | High | High | High | High | High | High | High | High | High | High | 0 | 9.66E-06 | 4.48 |
| FALSE | High | RQSSTADA | [K].RQSSTA | 2xPhospho | Q7TSH2 2x | 0.000458 | 0 | 1 | 3 | 1 | Q7TSH2 | Phosphory | Q7TSH2 [6 | 1 | 3170.496 | 0 | 0.25 | 0.031277 | 9.14 | 87.5 | 112.5 | 5.6 | 11.39 | | High | High | High | High | High | High | High | High | High | High | High | 0 | 9.94E-05 | 2.78 |
| FALSE | High | RQSTDLPT( | [R].RQSTDI | 1xPhospho | E9Q6H8 1x | 1.2E-06 | 0 | 1 | 3 | 3 | E9Q6H8 | Pleckstrin | E9Q6H8 [5 | 1 | 2623.203 | 0 | 0.5 | 0.006898 | 19.87 | 82.3 | 117.7 | 16.21 | 2.74 | | High | High | High | High | High | High | High | High | High | High | High | 0 | 1.25E-07 | 4.31 |
| FALSE | High | RQSTESSPV | [K].RQSTES | 1xPhospho | G3UWZ0 1 | 0.000524 | 0 | 1 | 4 | 1 | G3UWZ0 | Bromodom | G3UWZ0 [1 | 1 | 1779.895 | 0 | 2.26 | 0.000622 | 5.07 | 33.6 | 166.4 | 5.54 | 11.01 | | High | High | High | High | High | High | High | High | High | High | High | 0 | 0.000116 | 2.09 |
| FALSE | High | RQSTESSPV | [K].RQSTES | 2xPhospho | G3UWZ0 2 | 0.001767 | 5.66E-05 | 1 | 4 | 2 | G3UWZ0 | Bromodom | G3UWZ0 [1 | 1 | 1859.862 | 0 | 2 | 0.000223 | 0.79 | 38.1 | 161.9 | 11.69 | 8.25 | | High | High | High | High | High | High | High | High | High | High | High | 2.58E-05 | 0.000452 | 2.64 |
| FALSE | High | RQSVGDFV | [R].RQSVG | 1xPhospho | Q80Y19 1x | 0.001179 | 0 | 1 | 3 | 1 | Q80Y19 | Rho GTPas | Q80Y19 [2 | 1 | 2043.079 | 0 | -0.33 | 0.059148 | 7.3 | 107 | 93 | 10.29 | 10.16 | | High | High | High | High | High | High | High | High | High | High | High | 0 | 0.000286 | 3.01 |
| FALSE | High | RQSVPTGE | [R].RQSVP | 1xPhospho | Q3TFZ4 1x | 0.022973 | 0.001578 | 1 | 3 | 1 | Q3TFZ4 | Putative ur | Q3TFZ4 [1 | 1 | 1452.716 | 0 | -0.56 | 0.026997 | 9.61 | 117.8 | 82.2 | 12.82 | 9.31 | | High | High | High | High | High | High | High | High | High | High | High | 0.000847 | 0.00796 | 1.89 |
| FALSE | High | RQTPPLAQ | [R].RQTPPI | 1xPhospho | Q80VY9 1x | 0.026885 | 0.001676 | 1 | 3 | 1 | Q80VY9 | Putative AT | Q80VY9 [4 | 1 | 1978.011 | 0 | -0.59 | 0.000446 | 2.84 | 121.7 | 78.3 | 4.19 | 4.18 | | High | High | High | High | High | High | High | High | High | High | High | 0.000903 | 0.009482 | 3.65 |
| FALSE | High | RQTPPLAQ | [R].RQTPPI | 2xPhospho | Q80VY9 2x | 0.013545 | 0.000976 | 1 | 3 | 1 | Q80VY9 | Putative AT | Q80VY9 [4 | 1 | 2057.977 | 0 | -0.1 | 0.289267 | 5.7 | 103.6 | 96.4 | 4.43 | 10.71 | | High | High | High | High | High | High | High | High | High | High | High | 0.000505 | 0.004397 | 2.26 |
| FALSE | High | RQVSASEL( | [R].RQVSAS | 1xCarbami | Q8R0W0 1 | 0.059159 | 0.004294 | 1 | 1 | 1 | Q8R0W0 | Epiplakin C | Q8R0W0 [2 | 1 | 2101.031 | 0 | 1.52 | 0.000293 | 0.48 | 55.6 | 144.4 | 7.21 | 6.61 | | High | High | High | High | High | High | High | High | High | High | High | 0.002202 | 0.0234 | 2.16 |
| FALSE | High | RQYEEEEEI | [K].RQYEEI | 1xPhospho | P49135 1x | 0.000415 | 0 | 1 | 2 | 1 | P49135 | TFIIH basal | P49135 [17 | 1 | 3659.627 | 0 | -0.58 | 0.046861 | 10.35 | 119.5 | 80.5 | 29.56 | 10.51 | | High | High | High | High | High | High | High | High | High | High | High | 0 | 8.84E-05 | 3.78 |
| FALSE | High | RRDSDGVD | [R].RRDSD( | 1xPhospho | O88532 1x | 3.14E-05 | 0 | 1 | 2 | 5 | O88532 | Zinc finger | O88532 [1( | 2 | 2176.044 | 0 | -0.09 | 0.091533 | 6.2 | 103.6 | 96.4 | 4.39 | 5.22 | | High | High | High | High | High | High | High | High | High | High | High | 0 | 0.000218 | 2.94 |
| FALSE | High | RRDSDIPVK | [K].RRDSDI | 1xPhospho | Q8K2W6 1 | 0.021607 | 0.001507 | 1 | 1 | 2 | Q8K2W6 | PHD finger | Q8K2W6 [1 | 2 | 1623.898 | 0 | -1.01 | 0.00014 | 1.28 | 133.7 | 66.3 | 2.39 | 5.32 | | High | High | High | High | High | High | High | High | High | High | High | 0.000801 | 0.007429 | 2.08 |
| FALSE | High | RRESASSSS | [K].RRESAS | 1xPhospho | Q9JI44 1xP | 0.035181 | 0.002269 | 1 | 1 | 3 | Q9JI44 | DNA methy | Q9JI44 [45 | 2 | 1818.948 | 0 | -0.57 | 0.004336 | 6.22 | 123.6 | 76.4 | 11.92 | 6.75 | | High | High | High | High | High | High | High | High | High | High | High | 0.004935 | 0.05361 | 2.36 |
| FALSE | High | RRESEEPTS | [K].RRESEE | 1xCarbami | Q6ZWQ0 1 | 0.000725 | 0 | 1 | 9 | 2 | Q6ZWQ0 | Nesprin-2 ( | Q6ZWQ0 [( | 2 | 3191.562 | 0 | 1.98 | 0.00302 | 26.52 | 41.7 | 158.3 | 19.57 | 18.02 | | High | High | High | High | High | High | High | High | High | High | High | 0.000275 | 0.002061 | 3.06 |
| FALSE | High | RRESLSYIPI | [R].RRESLS | 1xPhospho | Q4VAC9 1x | 0.090153 | 0.00698 | 1 | 3 | 1 | Q4VAC9 | Pleckstrin I | Q4VAC9 [8 | 2 | 1786.998 | 0 | 1.21 | 0.011006 | 14.16 | 60.4 | 139.6 | 8.27 | 23.81 | | High | High | High | High | High | High | High | High | High | High | High | 0.003564 | 0.03803 | 1.47 |
| FALSE | High | RRNGSDDD | [R].RRNGSI | 1xDeamida | Q3TYF6 1x | 0.017041 | 0.001204 | 1 | 3 | 2 | Q3TYF6 | DNA segme | Q3TYF6 [7 | 2 | 2373.135 | 0 | 0.18 | 0.035916 | 3.88 | 92.6 | 107.4 | 2.03 | 9.8 | | High | High | High | High | High | High | High | High | High | High | High | 0.000638 | 0.005688 | 2.08 |
| FALSE | High | RQSSADGS | [K].RQSSAI | 1xPhospho | Q8BX09 1x | 4.91E-06 | 0 | 1 | 4 | 2 | Q8BX09 | Retinoblast | Q8BX09 [4 | 1 | 1795.91 | 0 | -0.71 | 0.002654 | 4.44 | 125.7 | 74.3 | 9.33 | 4.9 | | High | High | High | High | High | High | High | High | High | High | High | 0 | 6.05E-07 | 4.15 |
| FALSE | High | RQSNVELG | [R].RQSNV | 1xPhospho | Q9QXN0 1 | 0.000356 | 0 | 1 | 7 | 1 | Q9QXN0 | Protein Shi | Q9QXN0 [6 | 1 | 2272.103 | 0 | -0.02 | 0.961846 | 7.75 | 103.3 | 96.7 | 14.74 | 10.63 | | High | High | High | High | High | High | High | High | High | High | High | 0 | 7.45E-05 | 2.74 |
| FALSE | High | RQSNVELG | [R].RQSNV | 1xPhospho | B7ZNA0 1x | 8.43E-06 | 0 | 1 | 1 | 1 | B7ZNA0 | Shroom3 p | B7ZNA0 [6 | 1 | 2241.097 | 0 | 0.2 | 0.17021 | 21.27 | 88.3 | 111.7 | 15.16 | 12.88 | | High | High | High | High | High | High | High | High | High | High | High | 0 | 1.11E-06 | 3.98 |
| FALSE | High | RQSNTFPFI | [K].RQSNTI | 1xPhospho | B1AVH7 1x | 0.048715 | 0.00354 | 1 | 1 | 1 | B1AVH7 | TBC1 doma | B1AVH7 [2 | 1 | 2051.99 | 0 | -0.38 | 0.025939 | 12.06 | 111.4 | 88.6 | 5.98 | 13.57 | | High | High | High | High | High | High | High | High | High | High | High | 0.00186 | 0.01873 | 2.37 |
| FALSE | High | RQELLPNA | [R].RQELLF | 1xPhospho | Q7TN02 1x | 0.004451 | 0.000409 | 1 | 1 | 1 | Q7TN02 | Mediator c | Q7TN02 [3 | 1 | 3002.447 | 0 | -0.13 | 0.732791 | 11.02 | 104.6 | 95.4 | 12.3 | 23.67 | | High | High | High | High | High | High | High | High | High | High | High | 0.000183 | 0.00127 | 3.09 |
| FALSE | High | RQEMESGI | [K].RQEME | 1xPhospho | Q04690 1x | 0.009843 | 0.000736 | 1 | 6 | 4 | Q04690 | Neurofibro | Q04690 [2 | 1 | 2028.024 | 0 | 0.88 | 0.000669 | 10.17 | 72.3 | 127.7 | 4.5 | 5.9 | | High | High | High | High | High | High | High | High | High | High | High | 0.000344 | 0.003074 | 3 |
| FALSE | High | RQESGSSLI | [R].RQESGS | 1xPhospho | Q3TLI0 1xP | 1.46E-06 | 0 | 1 | 3 | 3 | Q3TLI0 | Trafficking | Q3TLI0 [70 | 1 | 2814.398 | 0 | 1.35 | 0.000293 | 4.78 | 57.7 | 142.3 | 3.62 | 4.52 | | High | High | High | High | High | High | High | High | High | High | High | 0 | 1.56E-07 | 4.98 |
| FALSE | High | RQESPPNP | [K].RQESPI | 1xPhospho | Q7TT18 1x | 0.069903 | 0.005181 | 1 | 3 | 1 | Q7TT18 | Activating | Q7TT18 [6 | 1 | 3650.742 | 0 | -0.48 | 0.131857 | 15.8 | 117.1 | 82.9 | 15.6 | 24.57 | | High | High | High | High | High | High | High | High | High | High | High | 0.002662 | 0.02826 | 2.99 |
| FALSE | High | RQESQVQ( | [K].RQESQ | 1xPhospho | Q8R322 1x | 0.000386 | 0 | 1 | 1 | 2 | Q8R322 | Nucleopor | Q8R322 [3 | 1 | 4095.055 | 0 | -0.36 | 0.230325 | 3.74 | 112.4 | 87.6 | 13.68 | 14.82 | | High | High | High | High | High | High | High | High | High | High | High | 0 | 8.37E-05 | 4.1 |
| FALSE | High | RQESTETLV | [K].RQESTI | 1xPhospho | A0A087WC | 0.00051 | 0 | 1 | 2 | 2 | A0A087WC | TRAF3-inte | A0A087WC | 1 | 1741.868 | 0 | -0.49 | 0.006924 | 9.42 | 117 | 83 | 2.4 | 10.26 | | High | High | High | High | High | High | High | High | High | High | High | 0 | 0.000112 | 2.49 |
| FALSE | High | RQFSAPNLI | [R].RQFSAI | 1xPhospho | A2AH75 1x | 0.055308 | 0.003982 | 1 | 2 | 1 | A2AH75 | Kinesin fan | A2AH75 [1 | 1 | 1598.882 | 0 | 0.37 | 0.003106 | 5.1 | 89.6 | 110.4 | 5.11 | 8.25 | | High | High | High | High | High | High | High | High | High | High | High | 0.002105 | 0.02172 | 1.46 |
| FALSE | High | RQGSFSED | [K].RQGSF | 1xPhospho | B1ARU1 1x | 0.002232 | 0.000109 | 2 | 15 | 1 | B1ARU1; F | Microtubu | B1ARU1 [2 | 1 | 2028.032 | 0 | 0.82 | 0.000181 | 3.18 | 73.1 | 126.9 | 5.13 | 3.65 | Shared | High | High | High | High | High | High | High | High | High | High | High | 4.94E-05 | 0.000588 | 2.52 |
| FALSE | High | RQISEDVD( | [R].RQISED | 1xPhospho | B2RSC8 1x | 0.002272 | 0.000109 | 1 | 2 | 1 | B2RSC8 | E3 ubiquiti | B2RSC8 [3( | 1 | 1809.833 | 0 | 0.3 | 0.055092 | 23.87 | 89.7 | 110.3 | 5.5 | 15.65 | | High | High | High | High | High | High | High | High | High | High | High | 4.94E-05 | 0.000601 | 2.42 |
| FALSE | High | RQLSEEQP | [K].RQLSEE | 1xDeamida | Q542C3 1x | 0.003372 | 0.00016 | 1 | 2 | 1 | Q542C3 | Putative ur | Q542C3 [1 | 1 | 2068.048 | 0 | -0.26 | 0.061488 | 7.37 | 105.4 | 94.6 | 8.63 | 6.24 | | High | High | High | High | High | High | High | High | High | High | High | 9.42E-05 | 0.000929 | 2.96 |
| FALSE | High | RQNEYSVS | [K].RQNEY | 1xPhospho | Z4YKQ9 1x | 0.003962 | 0.000311 | 1 | 3 | 1 | Z4YKQ9 | ATPase fan | Z4YKQ9 [9 | 1 | 2135.024 | 0 | -0.56 | 0.004291 | 0.54 | 119.4 | 80.6 | 3.65 | 12.91 | | High | High | High | High | High | High | High | High | High | High | High | 0.00014 | 0.001113 | 2.06 |
| FALSE | High | RQNEYSVS | [K].RQNEY | 1xPhospho | Z4YKQ9 1x | 0.052133 | 0.00384 | 1 | 3 | 1 | Z4YKQ9 | ATPase fan | Z4YKQ9 [9 | 1 | 2151.019 | 0 | -0.52 | 0.011049 | 6.55 | 119.4 | 80.6 | 8.06 | 7.63 | | High | High | High | High | High | High | High | High | High | High | High | 0.001999 | 0.02032 | 1.99 |
| FALSE | High | RQNSSDSIS | [K].RQNSSI | 1xPhospho | Q8CH77 1x | 5.98E-06 | 0 | 1 | 5 | 1 | Q8CH77 | Neuron na | Q8CH77 [1 | 1 | 3017.494 | 0 | 0.38 | 0.269648 | 30.73 | 86.5 | 113.5 | 21.72 | 21.16 | | High | High | High | High | High | High | High | High | High | High | High | 0 | 7.6E-07 | 3.81 |
| FALSE | High | RRPGPHSP | [R].RRPGPI | 1xPhospho | Q9CR59 1x | 0.008614 | 0.000736 | 1 | 1 | 1 | Q9CR59 | Growth arr | Q9CR59 [3 | 1 | 2248.155 | 0 | -0.04 | 0.659045 | 27.89 | 108.4 | 91.6 | 20.07 | 24.97 | | High | High | High | High | High | High | High | High | High | High | High | 0.000327 | 0.00266 | 3.66 |
| FALSE | High | RQNYSESSI | [K].RQNYSI | 1xPhospho | Q61687 1x | 1.33E-05 | 0 | 1 | 4 | 4 | Q61687 | Transcripti | Q61687 [7 | 1 | 2285.987 | 0 | 0.04 | 0.872418 | 2.11 | 98.8 | 101.2 | 7.51 | 9.31 | | High | High | High | High | High | High | High | High | High | High | High | 0 | 0.000289 | 1.74 |
| FALSE | High | RQPSPQPS | [R].RQPSP( | 2xPhospho | Q8BTI8 2x | 0.037048 | 0.00249 | 1 | 3 | 2 | Q8BTI8 | Serine/argi | Q8BTI8 [26 | 1 | 1538.708 | 0 | -0.45 | 0.000377 | 2.5 | 115.3 | 84.7 | 2.06 | 5.24 | | High | High | High | High | High | High | High | High | High | High | High | 0.001308 | 0.01373 | 1.76 |
| FALSE | High | RQQDPSPG | [R].RQQDP | 1xPhospho | Q08024 1x | 0.000305 | 0 | 1 | 3 | 1 | Q08024 | Core-bindi | Q08024 [1 | 1 | 2379.171 | 0 | 1.48 | 0.000327 | 5.02 | 52.7 | 147.3 | 4.23 | 5.56 | | High | High | High | High | High | High | High | High | High | High | High | 0 | 6.27E-05 | 3.05 |
| FALSE | High | RQQYSDQI | [R].RQQYSI | 1xPhospho | Q9Z0U1 1x | 0.0016 | 5.66E-05 | 1 | 1 | 2 | Q9Z0U1 | Tight junct | Q9Z0U1 [4 | 1 | 2410.108 | 0 | -0.35 | 0.192856 | 6.66 | 111.9 | 88.1 | 8.51 | 21.68 | | High | High | High | High | High | High | High | High | High | High | High | 2.58E-05 | 0.000403 | 2.35 |
| FALSE | High | RQQYSDQI | [R].RQQYSI | 2xPhospho | Q9Z0U1 2x | 1.37E-05 | 0 | 1 | 1 | 2 | Q9Z0U1 | Tight junct | Q9Z0U1 [4 | 1 | 2490.074 | 0 | -0.17 | 0.034423 | 6.19 | 106 | 94 | 3.2 | 4.07 | | High | High | High | High | High | High | High | High | High | High | High | 0 | 1.92E-06 | 2.31 |
| FALSE | High | RQSAGDLL | [R].RQSAG | 1xPhospho | E9QQ68 1x | 0.001586 | 5.66E-05 | 1 | 3 | 2 | E9QQ68 | TBC1 doma | E9QQ68 [2 | 1 | 3139.573 | 0 | 1.83 | 0.001251 | 11.43 | 46.5 | 153.5 | 20.75 | 12.06 | | High | High | High | High | High | High | High | High | High | High | High | 7.19E-05 | 0.000819 | 4.69 |
| FALSE | High | RQSAILPQI | [K].RQSAIL | 1xPhospho | Q9EQU5 1x | 0.004491 | 0.000409 | 1 | 6 | 1 | Q9EQU5 | Protein SET | Q9EQU5 [5 | 1 | 1675.966 | 0 | -1.08 | 0.002076 | 3.66 | 136.2 | 63.8 | 11.46 | 11.25 | | High | High | High | High | High | High | High | High | High | High | High | 0.000204 | 0.001289 | 2.4 |
| FALSE | High | RQSCYLCD | [R].RQSCYI | 2xCarbami | E9Q1P8 1x | 0.000418 | 0 | 2 | 2 | 1 | E9Q1P8; Q | Interferon | E9Q1P8 [1 | 1 | 1776.812 | 0 | 1.01 | 0.001414 | 14.26 | 65.7 | 134.3 | 5.97 | 9.17 | Shared | High | High | High | High | High | High | High | High | High | High | High | 0 | 8.97E-05 | 2.08 |
| FALSE | High | RQSFASLAI | [R].RQSFAS | 1xPhospho | G5E8K3 1x | 0.049549 | 0.00361 | 2 | 19 | 1 | G5E8K3; A( | Ankyrin 3, | G5E8K3 [1 | 1 | 1457.783 | 0 | 1.11 | 0.000875 | 10.56 | 62.8 | 137.2 | 4.05 | 9.91 | Shared | High | High | High | High | High | High | High | High | High | High | High | 0.001902 | 0.01917 | 2.09 |
| FALSE | High | RQSGLYDG | [R].RQSGLY | 1xCarbami | Q80TY0-3 | 0.000837 | 0 | 1 | 12 | 1 | Q80TY0-3 | Isoform 3 ( | Q80TY0-3 | 1 | 2744.241 | 0 | 0.09 | 0.858253 | 28.43 | 98.4 | 101.6 | 14.01 | 14.19 | | High | High | High | High | High | High | High | High | High | High | High | 0 | 0.000196 | 2.36 |
| FALSE | High | RQSLELQK | [R].RQSLEL | 1xPhospho | Q9Z277 1x | 0.075927 | 0.005826 | 1 | 2 | 2 | Q9Z277 | Tyrosine-p | Q9Z277 [1 | 1 | 1539.866 | 0 | 1.36 | 0.000475 | 10.5 | 56.5 | 143.5 | 2.53 | 6.41 | | High | High | High | High | High | High | High | High | High | High | High | 0.003659 | 0.03903 | 2.02 |
| FALSE | High | RQSIPEEFK | [R].RQSIPE | 1xPhospho | Q925T6 1x | 0.001476 | 5.66E-05 | 1 | 9 | 1 | Q925T6 | Glutamate | Q925T6 [4 | 1 | 1671.887 | 0 | -1.75 | 0.000378 | 1.03 | 157 | 43 | 8.18 | 13.73 | | High | High | High | High | High | High | High | High | High | High | High | 2.58E-05 | 0.000371 | 2.15 |
| FALSE | High | RQSISPPYK | [R].RQSISP | 2xPhospho | Q14AX6-2 | 0.002172 | 5.66E-05 | 3 | 5 | 3 | Q14AX6-2; | Isoform 2 ( | Q14AX6-2 | 1 | 1693.848 | 0 | -0.24 | 0.003085 | 3.15 | 109.9 | 90.1 | 3.09 | 5.01 | Shared | High | High | High | High | High | High | High | High | High | High | High | 4.94E-05 | 0.000571 | 2.22 |
| FALSE | High | RQSLTSPDS | [R].RQSLTS | 2xPhospho | E9PZP8 2x | 0.012508 | 0.000889 | 1 | 3 | 1 | E9PZP8 | HECT and I | E9PZP8 [27 | 1 | 2290.054 | 0 | 0.52 | 0.001999 | 9.18 | 82.1 | 117.9 | 2.53 | 6.75 | | High | High | High | High | High | High | High | High | High | High | High | 0.000449 | 0.00404 | 1.96 |
| FALSE | High | RQPSPQPS | [R].RQPSP( | 1xPhospho | Q8BTI8 1x | 0.003688 | 0.00021 | 1 | 3 | 2 | Q8BTI8 | Serine/argi | Q8BTI8 [26 | 1 | 1458.742 | 0 | -0.58 | 0.000628 | 2.1 | 119.9 | 80.1 | 4.22 | 2.09 | | High | High | High | High | High | High | High | High | High | High | High | 0.001011 | 0.01072 | 2.4 |
| FALSE | High | RRPPSPDPI | [K].RRPPSP | 1xPhospho | Q62261 1x | 0.013074 | 0.000889 | 2 | 6 | 1 | Q62261; Q | Spectrin be | Q62261 [2( | 1 | 1573.805 | 0 | 0.38 | 0.299364 | 43.91 | 85.5 | 114.5 | 12.12 | 76.26 | Shared | High | High | High | High | High | High | High | High | High | High | High | 0.000467 | 0.004248 | 2.55 |
| FALSE | High | RRPSDENSI | [R].RRPSDE | 1xPhospho | Q8C8U0-3 | 3.09E-05 | 0 | 1 | 6 | 3 | Q8C8U0-3 | Isoform 3 ( | Q8C8U0-3 | 1 | 2846.353 | 0 | 1.08 | 0.00122 | 10.72 | 62 | 138 | 8.76 | 6.87 | | High | High | High | High | High | High | High | High | High | High | High | 0 | 5.94E-06 | 4.14 |
| FALSE | High | RRPSMALC | [K].RRPSM. | 1xPhospho | A2A7B5 1x | 0.093123 | 0.007276 | 1 | 2 | 1 | A2A7B5 | PR domain | A2A7B5 [4 | 1 | 2443.169 | 0 | -0.2 | 0.248233 | 24.15 | 110.4 | 89.6 | 7.19 | 21.92 | | High | High | High | High | High | High | High | High | High | High | High | 0.003685 | 0.03961 | 1.6 |
| FALSE | High | RSESCSEDK | [K].RSESCS | 1xCarbami | E9Q6J5-3 1 | 0.007538 | 0.00062 | 1 | 3 | 2 | E9Q6J5-3 | Isoform Ne | E9Q6J5-3 [ | 1 | 1635.745 | 0 | 0.09 | 0.32917 | 4.09 | 98.4 | 101.6 | 8.63 | 12.34 | | High | High | High | High | High | High | High | High | High | High | High | 0.000555 | 0.004921 | 1.61 |
| FALSE | High | RSESPFEGK | [R].RSESPF | 1xPhospho | P14873 1x | 0.00083 | 0 | 2 | 2 | 1 | P14873; B2 | Microtubu | P14873 [14 | 1 | 1574.798 | 0 | -0.34 | 0.073697 | 12.11 | 109.7 | 90.3 | 10.83 | 13.15 | Shared | High | High | High | High | High | High | High | High | High | High | High | 0 | 0.000193 | 1.55 |
| FALSE | High | RSESPFEGK | [R].RSESPF | 2xPhospho | P14873 2x | 0.01186 | 0.000889 | 2 | 2 | 2 | P14873; B2 | Microtubu | P14873 [14 | 1 | 1654.764 | 0 | -0.32 | 0.018729 | 8.65 | 110.4 | 89.6 | 3.79 | 9.67 | Shared | High | High | High | High | High | High | High | High | High | High | High | 0.000449 | 0.003779 | 2.07 |
| FALSE | High | RSESPPAEL | [K].RSESPP | 1xPhospho | G5E870 1x | 0.006536 | 0.00062 | 1 | 5 | 2 | G5E870 | E3 ubiquiti | G5E870 [3( | 1 | 1747.894 | 0 | -1.32 | 0.000358 | 1.42 | 144.6 | 55.4 | 6.4 | 5.37 | | High | High | High | High | High | High | High | High | High | High | High | 0.000275 | 0.001957 | 2.4 |
| FALSE | High | RSESSGNLI | [R].RSESSG | 1xPhospho | O08715-4 | 2.22E-06 | 0 | 1 | 6 | 5 | O08715-4 | Isoform 4 ( | O08715-4 | 1 | 1884.901 | 0 | 0.04 | 0.696977 | 23.38 | 96.7 | 103.3 | 7.98 | 11.27 | | High | High | High | High | High | High | High | High | High | High | High | 0 | 2.29E-06 | 3.88 |
| FALSE | High | RSETPPHW | [R].RSETPP | 2xPhospho | A2AR02 2x | 0.08727 | 0.006537 | 1 | 5 | 2 | A2AR02 | Peptidyl-pr | A2AR02 [3 | 1 | 1554.682 | 0 | -0.42 | 0.00795 | 6.78 | 114.5 | 85.5 | 4.34 | 9.5 | | High | High | High | High | High | High | High | High | High | High | High | 0.004059 | 0.04336 | 2.05 |
| FALSE | High | RSEVLADES | [R].RSEVLA | 1xCarbami | G3UWQ7 1 | 3.62E-07 | 0 | 1 | 5 | 2 | G3UWQ7 | Protein reg | G3UWQ7 [ | 1 | 2287.177 | 0 | 0.08 | 0.517846 | 16.57 | 99.6 | 100.4 | 12.12 | 9.71 | | High | High | High | High | High | High | High | High | High | High | High | 0 | 3.23E-08 | 4.09 |
| FALSE | High | RSFEVEEIE | [K].RSFEVE | 1xPhospho | Q80UG5 1x | 6.01E-08 | 0 | 1 | 3 | 4 | Q80UG5 | Septin-9 O | Q80UG5 [2 | 1 | 2293.106 | 0 | 0.86 | 0.049639 | 31.29 | 70.9 | 129.1 | 12.6 | 25.81 | | High | High | High | High | High | High | High | High | High | High | High | 0 | 2.52E-06 | 4.31 |
| FALSE | High | RSFEVEEIE | [K].RSFEVE | 2xPhospho | Q80UG5 2x | 0.041781 | 0.003126 | 1 | 3 | 1 | Q80UG5 | Septin-9 O | Q80UG5 [2 | 2 | 2529.174 | 0 | 0.81 | 0.009441 | 19.74 | 75.7 | 124.3 | 8.63 | 13.17 | | High | High | High | High | High | High | High | High | High | High | High | 0.001609 | 0.01566 | 2.41 |
| FALSE | High | RSFGNPFE( | [K].RSFGNI | 1xPhospho | Q8C078-5 | 0.007742 | 0.000664 | 1 | 5 | 1 | Q8C078-5 | Isoform 5 ( | Q8C078-5 | 1 | 1562.731 | 0 | 1.21 | 0.002304 | 24.4 | 63.2 | 136.8 | 13.56 | 9.1 | | High | High | High | High | High | High | High | High | High | High | High | 0.000294 | 0.002361 | 2.01 |
| FALSE | High | RSFGNPFEI | [K].RSFGNI | 1xPhospho | Q8VBY2 1x | 0.000352 | 0 | 1 | 2 | 1 | Q8VBY2 | Calcium/ca | Q8VBY2 [4 | 1 | 1714.826 | 0 | 0.32 | 0.002367 | 7.8 | 86 | 114 | 13.6 | 11.2 | | High | High | High | High | High | High | High | High | High | High | High | 0 | 7.42E-05 | 3 |
| FALSE | High | RSFVPEEEK | [K].RSFVPE | 1xPhospho | Q9D4H9-2 | 0.040373 | 0.002982 | 1 | 6 | 1 | Q9D4H9-2 | Isoform 2 ( | Q9D4H9-2 | 1 | 1658.856 | 0 | 0.86 | 0.000513 | 6.81 | 70.4 | 129.6 | 11.07 | 5.3 | | High | High | High | High | High | High | High | High | High | High | High | 0.00155 | 0.01505 | 2.04 |
| FALSE | High | RSFVYAK | [R].RSFVYA | 1xPhospho | E9PVX6 1x | 0.032831 | 0.002195 | 1 | 2 | 1 | E9PVX6 | Proliferatic | E9PVX6 [1 | 1 | 1408.775 | 0 | -0.24 | 0.001972 | 1.16 | 107.6 | 92.4 | 8.13 | 6.39 | | High | High | High | High | High | High | High | High | High | High | High | 0.001151 | 0.01197 | 2.36 |
| FALSE | High | RSEPQPEE( | [K].RSEPQI | 1xPhospho | O88508 1x | 1.77E-05 | 0 | 1 | 6 | 1 | O88508 | DNA (cytos | O88508 [9 | 1 | 2206.091 | 0 | -0.27 | 0.007167 | 1.79 | 108.6 | 91.4 | 7.18 | 7.06 | | High | High | High | High | High | High | High | High | High | High | High | 0 | 2.57E-06 | 1.89 |
| FALSE | High | RSGATGGL | [R].RSGAT( | 1xPhospho | Q9CXS4 1x | 0.000267 | 0 | 1 | 1 | 1 | Q9CXS4 | Centromer | Q9CXS4 [1 | 1 | 1600.764 | 0 | 0.23 | 0.459524 | 4.62 | 93.8 | 106.2 | 19.55 | 6.44 | | High | High | High | High | High | High | High | High | High | High | High | 0 | 5.39E-05 | 3.52 |
| FALSE | High | RSHDSDDS | [K].RSHDSI | 2xPhospho | A2A6A1 2x | 0.06169 | 0.004646 | 1 | 1 | 1 | A2A6A1 | G patch do | A2A6A1 [9 | 1 | 2217.91 | 0 | -0.44 | 0.020482 | 11.86 | 113.4 | 86.6 | 11.54 | 9.62 | | High | High | High | High | High | High | High | High | High | High | High | 0.002381 | 0.02466 | 2.67 |
| FALSE | High | RSLEGGGC | [R].RSLEG( | 1xCarbami | Q6P9Q4 1x | 0.009585 | 0.000736 | 1 | 1 | 1 | Q6P9Q4 | FH1/FH2 d | Q6P9Q4 [3 | 1 | 1496.724 | 0 | 0.41 | 0.447393 | 11.26 | 85.8 | 114.2 | 25.25 | 16.86 | | High | High | High | High | High | High | High | High | High | High | High | 0.000344 | 0.003006 | 2.85 |
| FALSE | High | RSLEQNVQ | [R].RSLEQN | 1xPhospho | Q3V122 1x | 0.006713 | 0.00062 | 1 | 3 | 1 | Q3V122 | DNA helica | Q3V122 [5 | 1 | 2104.148 | 0 | 0.07 | 0.38862 | 8.55 | 99.9 | 100.1 | 8.56 | 4.83 | | High | High | High | High | High | High | High | High | High | High | High | 0.000275 | 0.002017 | 2.74 |
| FALSE | High | RSLNLEDYI | [R].RSLNLE | 1xPhospho | A2AQ19 1x | 0.004867 | 0.000504 | 1 | 1 | 2 | A2AQ19 | RNA polym | A2AQ19 [7 | 1 | 1675.882 | 0 | -0.73 | 0.00725 | 6.11 | 126.2 | 73.8 | 8.9 | 11.7 | | High | High | High | High | High | High | High | High | High | High | High | 0.001484 | 0.0149 | 2.34 |
| FALSE | High | RSIQFVDW | [K].RSIQFV | 1xCarbami | A0A0A0M( | 3.94E-05 | 0 | 1 | 4 | 1 | A0A0A0M( | Tubulin alp | A0A0A0M( | 1 | 2279.145 | 0 | 0.43 | 0.101355 | 18.66 | 85.8 | 114.2 | 13.35 | 13.61 | | High | High | High | High | High | High | High | High | High | High | High | 0 | 6.28E-06 | 4.64 |
| FALSE | High | RSLSGSSP( | [R].RSLSGS | 1xCarbami | Q8BTI8 2x | 0.059657 | 0.004365 | 1 | 4 | 1 | Q8BTI8 | Serine/argi | Q8BTI8 [77 | 1 | 1793.842 | 0 | -1.06 | 0.002604 | 2.53 | 136.9 | 63.1 | 8.98 | 13.04 | | High | High | High | High | High | High | High | High | High | High | High | 0.002233 | 0.02357 | 2.37 |
| FALSE | High | RSLTNSHLE | [R].RSLTNS | 1xPhospho | Q8K2P7 1x | 0.019446 | 0.001438 | 1 | 3 | 1 | Q8K2P7 | Sodium-co | Q8K2P7 [5 | 1 | 1722.93 | 0 | -0.31 | 0.019785 | 6.07 | 114.1 | 85.9 | 11.43 | 8.28 | | High | High | High | High | High | High | High | High | High | High | High | 0.000756 | 0.006625 | 1.7 |
| FALSE | High | RSPAAFTSI | [R].RSPAAI | 1xPhospho | Q571G8 1x | 0.055308 | 0.003982 | 1 | 7 | 1 | Q571G8 | MKIAA164 | Q571G8 [1 | 1 | 2449.362 | 0 | 0.16 | 0.161287 | 12.13 | 92.3 | 107.7 | 11.73 | 5.91 | | High | High | High | High | High | High | High | High | High | High | High | 0.002105 | 0.02171 | 2.41 |
| FALSE | High | RSPAPCSDI | [K].RSPAP( | 1xCarbami | Q8BLH7 1x | 0.000184 | 0 | 1 | 2 | 4 | Q8BLH7 | HIRA-inter | Q8BLH7 [7 | 1 | 1652.823 | 0 | -1.23 | 0.000209 | 1.07 | 140.7 | 59.3 | 3.55 | 2.99 | | High | High | High | High | High | High | High | High | High | High | High | 0.000275 | 0.001977 | 2.88 |
| FALSE | High | RSPASSGSV | [R].RSPASS | 1xPhospho | A0A0B4J1E | 1.38E-05 | 0 | 1 | 3 | 6 | A0A0B4J1E | DNA-bindi | A0A0B4J1E | 1 | 2980.436 | 0 | 0.51 | 0.007505 | 10.43 | 85.6 | 114.4 | 8.35 | 8.6 | | High | High | High | High | High | High | High | High | High | High | High | 0 | 5.62E-06 | 4.67 |
| FALSE | High | RSPASSGSV | [R].RSPASS | 1xDeamida | A0A0B4J1E | 6.13E-05 | 0 | 1 | 3 | 2 | A0A0B4J1E | DNA-bindi | A0A0B4J1E | 1 | 2981.42 | 0 | 0.69 | 0.003908 | 7.95 | 72.8 | 127.2 | 15.22 | 10.82 | | High | High | High | High | High | High | High | High | High | High | High | 0 | 1.04E-05 | 4.62 |
| FALSE | High | RSPASSGSV | [R].RSPASS | 2xPhospho | A0A0B4J1E | 0.000109 | 0 | 1 | 3 | 2 | A0A0B4J1E | DNA-bindi | A0A0B4J1E | 1 | 3060.403 | 0 | 0.52 | 0.020162 | 19.99 | 80.4 | 119.6 | 9.4 | 14.17 | | High | High | High | High | High | High | High | High | High | High | High | 0 | 1.98E-05 | 4.26 |
| FALSE | High | RSPATNPVS | [R].RSPATN | 1xPhospho | Q8C205 1x | 5.4E-05 | 0 | 1 | 3 | 2 | Q8C205 | Putative ur | Q8C205 [6 | 1 | 2519.149 | 0 | -0.47 | 0.002972 | 3.9 | 115 | 85 | 5.32 | 8.59 | | High | High | High | High | High | High | High | High | High | High | High | 0 | 9.01E-06 | 3.41 |
| FALSE | High | RSGQEAAA | [K].RSGQE. | 3xCarbami | Q8VDY9 1x | 3.73E-05 | 0 | 1 | 1 | 2 | Q8VDY9 | Caspase ac | Q8VDY9 [1 | 1 | 4555.096 | 0 | -0.55 | 0.070636 | 7.86 | 119.2 | 80.8 | 7.93 | 23.41 | | High | High | High | High | High | High | High | High | High | High | High | 0 | 2.69E-05 | 4.79 |
| FALSE | High | RTPEEEFSA | [R].RTPEEE | 1xPhospho | E9Q4F8 1x | 0.027839 | 0.001755 | 1 | 3 | 1 | E9Q4F8 | Ankyrin rej | E9Q4F8 [1 | 1 | 1903.92 | 0 | -0.21 | 0.012886 | 6.22 | 109.2 | 90.8 | 5.82 | 7.74 | | High | High | High | High | High | High | High | High | High | High | High | 0.000953 | 0.009864 | 2.35 |
| FALSE | High | RSEPFQK | [K].RSEPFC | 1xPhospho | Q6NV99 1x | 0.112856 | 0.009146 | 1 | 6 | 1 | Q6NV99 | HAUS augn | Q6NV99 [5 | 1 | 1429.76 | 0 | -0.81 | 0.003692 | 2.5 | 127.5 | 72.5 | 15.48 | 6.35 | | High | High | High | High | High | High | High | High | High | High | High | 0.004569 | 0.04961 | 1.39 |
| FALSE | High | RSEGLSQE. | [K].RSEGLS | 1xCarbami | B1AQJ2 2x | 0.012845 | 0.000889 | 1 | 3 | 1 | B1AQJ2 | Ubiquitin c | B1AQJ2 [8 | 1 | 3836.679 | 0 | -0.36 | 0.002398 | 3.16 | 113.6 | 86.4 | 6.51 | 6.24 | | High | High | High | High | High | High | High | High | High | High | High | 0.000449 | 0.00414 | 3.28 |
| FALSE | High | RRSESSGNI | [R].RRSESS | 1xPhospho | O08715-4 | 0.001933 | 5.66E-05 | 1 | 6 | 1 | O08715-4 | Isoform 4 ( | O08715-4 | 2 | 2041.003 | 0 | -0.48 | 0.02854 | 6.19 | 112.5 | 87.5 | 13.98 | 8.93 | | High | High | High | High | High | High | High | High | High | High | High | 2.58E-05 | 0.000498 | 3.63 |
| FALSE | High | RRSESSGNI | [R].RRSESS | 2xPhospho | O08715-4 | 0.012508 | 0.000889 | 1 | 6 | 1 | O08715-4 | Isoform 4 ( | O08715-4 | 2 | 2120.969 | 0 | 0.75 | 0.061696 | 37.02 | 76.5 | 123.5 | 5.56 | 28.09 | | High | High | High | High | High | High | High | High | High | High | High | 0.000449 | 0.004018 | 2.47 |
| FALSE | High | RRSPSLSSK | [K].RRSPSL | 1xPhospho | Q52KI8 1x | 0.11108 | 0.008991 | 1 | 7 | 1 | Q52KI8 | Serine/argi | Q52KI8 [67 | 2 | 1326.709 | 0 | 0.01 | 0.598035 | 33.27 | 105.8 | 94.2 | 14.19 | 65.06 | | High | High | High | High | High | High | High | High | High | High | High | 0.004503 | 0.04867 | 1.89 |
| FALSE | High | RRSPSPAPI | [R].RRSPSP | 2xPhospho | Q52KI8 2x | 7.51E-07 | 0 | 1 | 7 | 11 | Q52KI8 | Serine/argi | Q52KI8 [57 | 2 | 2286.151 | 0 | -0.34 | 0.011956 | 4.07 | 111.7 | 88.3 | 7.78 | 3.08 | | High | High | High | High | High | High | High | High | High | High | High | 0.000538 | 0.004426 | 3.66 |
| FALSE | High | RRSPSPDFY | [R].RRSPSP | 2xPhospho | B1B0E8 2x | 0.093879 | 0.007306 | 1 | 2 | 1 | B1B0E8 | U2 small n | B1B0E8 [3 | 2 | 1870.902 | 0 | -0.41 | 0.005556 | 5.71 | 114.2 | 85.8 | 6.04 | 4.51 | | High | High | High | High | High | High | High | High | High | High | High | 0.003741 | 0.04021 | 1.35 |
| FALSE | High | RRSSDEDA | [R].RRSSDE | 2xPhospho | Q9D483 2x | 0.022973 | 0.001578 | 1 | 2 | 1 | Q9D483 | DNA-direct | Q9D483 [2 | 2 | 2065.936 | 0 | 0.41 | 0.006775 | 6.79 | 86.1 | 113.9 | 3.09 | 7.66 | | High | High | High | High | High | High | High | High | High | High | High | 0.000847 | 0.007991 | 1.14 |
| FALSE | High | RRSSGLQP( | [R].RRSSGL | 2xPhospho | E9Q4K7 2x | 0.003788 | 0.00021 | 1 | 2 | 2 | E9Q4K7 | Kinesin fan | E9Q4K7 [1 | 2 | 2026.979 | 0 | 0.61 | 0.001088 | 14.32 | 79.2 | 120.8 | 7.6 | 1.95 | | High | High | High | High | High | High | High | High | High | High | High | 0.000585 | 0.005497 | 3.61 |
| FALSE | High | RRSSTGLK | [R].RRSSTG | 2xPhospho | E9Q284 2x | 0.099328 | 0.008003 | 1 | 3 | 1 | E9Q284 | Coilin OS=I | E9Q284 [2 | 2 | 1522.791 | 0 | 0.02 | 0.517893 | 3.26 | 98.3 | 101.7 | 11.9 | 8.72 | | High | High | High | High | High | High | High | High | High | High | High | 0.004032 | 0.04277 | 0.89 |
| FALSE | High | RRTASPPPI | [K].RRTASF | 2xPhospho | Q52KI8 2x | 0.009843 | 0.000736 | 1 | 7 | 2 | Q52KI8 | Serine/argi | Q52KI8 [63 | 2 | 1821.954 | 0 | 0.06 | 0.684626 | 35.4 | 96.7 | 103.3 | 20.14 | 15.7 | | High | High | High | High | High | High | High | High | High | High | High | 0.001999 | 0.0209 | 2.02 |
| FALSE | High | RRTDEDAV | [R].RRTDEI | 1xPhospho | Q61584-3 | 0.000122 | 0 | 1 | 5 | 2 | Q61584-3 | Isoform B ( | Q61584-3 | 2 | 3531.704 | 0 | -0.95 | 0.000567 | 0.97 | 133.9 | 66.1 | 6.15 | 5.68 | | High | High | High | High | High | High | High | High | High | High | High | 0 | 2.23E-05 | 3.59 |
| FALSE | High | RRTEEGPTI | [R].RRTEE( | 1xPhospho | Q6ZQ61 1x | 0.00145 | 5.66E-05 | 1 | 6 | 1 | Q6ZQ61 | MCG12197 | Q6ZQ61 [1 | 2 | 1830.906 | 0 | 0.38 | 0.006456 | 4.49 | 85.3 | 114.7 | 5.66 | 8.55 | | High | High | High | High | High | High | High | High | High | High | High | 2.58E-05 | 0.000361 | 1.92 |
| FALSE | High | RRTENSLAS | [R].RRTENS | 1xPhospho | Q9JLI8 1xP | 0.008022 | 0.000736 | 1 | 1 | 1 | Q9JLI8 | Squamous | Q9JLI8 [64 | 2 | 2262.131 | 0 | -0.31 | 0.12665 | 2.73 | 110.7 | 89.3 | 9.48 | 13.69 | | High | High | High | High | High | High | High | High | High | High | High | 0.000327 | 0.002451 | 2.83 |
| FALSE | High | RSAPGGGN | [K].RSAPG( | 1xPhospho | Q5SQB7 1x | 0.000233 | 0 | 2 | 9 | 1 | Q5SQB7; C | MCG68069 | Q5SQB7 [1 | 1 | 1381.735 | 0 | -0.28 | 0.056819 | 6.65 | 110.6 | 89.4 | 5.73 | 11.02 | Shared | High | High | High | High | High | High | High | High | High | High | High | 0 | 4.66E-05 | 2.74 |
| FALSE | High | RSELNQAA | [K].RSELNC | 1xPhospho | Q3UYD0 1x | 0.001313 | 0 | 1 | 2 | 1 | Q3UYD0 | Putative ur | Q3UYD0 [4 | 1 | 2423.233 | 0 | -0.26 | 0.146469 | 7.85 | 109.1 | 90.9 | 13.03 | 11.48 | | High | High | High | High | High | High | High | High | High | High | High | 0 | 0.000323 | 3.6 |

| | | | | | | | | | | | | | | | | | | | | | | | | | | | | | | | | | | | | | | |
|---|---|---|---|---|---|---|---|---|---|---|---|---|---|---|---|---|---|---|---|---|---|---|---|---|---|---|---|---|---|---|---|---|---|---|---|---|---|---|
| FALSE | High | RSASPDDD | [K].RSASPD | 2xPhospho | Q9R0P4 2x | 2.45E-10 | 0 | 1 | 2 | 3 | Q9R0P4 | Small acidi | Q9R0P4 [1 | 1 | 2980.22 | 0 | 0.16 | 0.466265 | 11.94 | 99.8 | 100.2 | 9.06 | 10.69 | | High | High | High | High | High | High | High | High | High | High | High | 0 | 8.93E-12 | 6.9 |
| FALSE | High | RSCTLEGG | [R].RSCTLE | 1xCarbami | Q80TZ9 1x | 0.019446 | 0.001438 | 1 | 3 | 1 | Q80TZ9 | Arginine-gl | Q80TZ9 [3 | 1 | 1616.823 | 0 | 0.04 | 0.627483 | 26.87 | 98.7 | 101.3 | 7.06 | 13.69 | | High | High | High | High | High | High | High | High | High | High | High | 0.000756 | 0.006586 | 2.71 |
| FALSE | High | RSDAEEVD | [K].RSDAEE | 1xCarbami | P31938 1x | 0.000102 | 0 | 1 | 5 | 4 | P31938 | Dual specif | P31938 [36 | 1 | 3639.699 | 0 | 0.15 | 0.426849 | 15.89 | 96.5 | 103.5 | 14.23 | 13.13 | | High | High | High | High | High | High | High | High | High | High | High | 0 | 1.85E-05 | 3.95 |
| FALSE | High | RSDAEEVD | [K].RSDAEE | 1xCarbami | P31938 1x | 2.31E-05 | 0 | 1 | 5 | 1 | P31938 | Dual specif | P31938 [36 | 1 | 3641.667 | 0 | 0.2 | 0.129547 | 3.02 | 92.9 | 107.1 | 16.31 | 9.62 | | High | High | High | High | High | High | High | High | High | High | High | 0 | 3.45E-06 | 4.82 |
| FALSE | High | RSDLDLGYI | [R].RSDLDL | 1xPhospho | E9QMJ8 1x | 0.001865 | 5.66E-05 | 1 | 4 | 1 | E9QMJ8 | Axin-1 OS= | E9QMJ8 [6 | 1 | 2729.302 | 0 | -0.59 | 0.00853 | 2.22 | 118.8 | 81.2 | 11.29 | 6.3 | | High | High | High | High | High | High | High | High | High | High | High | 2.58E-05 | 0.000482 | 4.41 |
| FALSE | High | RSDQGGD | [R].RSDQG | 1xPhospho | Q546X9 1x | 2.9E-05 | 0 | 1 | 1 | 2 | Q546X9 | Ataxin 3 OS | Q546X9 [3 | 1 | 2089.906 | 0 | 0.22 | 0.532074 | 14.73 | 93.1 | 106.9 | 13.69 | 11.01 | | High | High | High | High | High | High | High | High | High | High | High | 7.19E-05 | 0.000878 | 3.07 |
| FALSE | High | RSDSEEK | [K].RSDSEE | 1xPhospho | A2AQ19 1x | 0.058173 | 0.004227 | 1 | 3 | 1 | A2AQ19 | RNA polym | A2AQ19 [8 | 1 | 1388.682 | 0 | -0.54 | 0.001931 | 0.76 | 117 | 83 | 9.83 | 6.51 | | High | High | High | High | High | High | High | High | High | High | High | 0.002187 | 0.023 | 1.85 |
| FALSE | High | RSDSMTFL | [R].RSDSM | 1xCarbami | E9Q2H1 1x | 0.114657 | 0.009411 | 1 | 2 | 2 | E9Q2H1 | E3 ubiquiti | E9Q2H1 [2 | 1 | 4537.261 | 0 | 0.14 | 0.537686 | 26.22 | 99 | 101 | 7.72 | 16.82 | | High | High | High | High | High | High | High | High | High | High | High | 0.004733 | 0.05097 | 3.22 |
| FALSE | High | RSDSPEIPF | [R].RSDSPE | 1xPhospho | P70399 1x | 2.96E-06 | 0 | 1 | 6 | 1 | P70399 | TP53-bindi | P70399 [14 | 1 | 3537.633 | 0 | 0.49 | 0.063379 | 24.48 | 85.6 | 114.4 | 10.34 | 12.75 | | High | High | High | High | High | High | High | High | High | High | High | 0 | 3.44E-07 | 5.64 |
| FALSE | High | RSDSQQAV | [R].RSDSQ | 1xPhospho | Q8BVL3 1x | 0.004696 | 0.000456 | 1 | 1 | 2 | Q8BVL3 | Sorting ne | Q8BVL3 [4 | 1 | 1556.82 | 0 | 1.09 | 0.001184 | 8.41 | 64.4 | 135.6 | 7.27 | 8.43 | | High | High | High | High | High | High | High | High | High | High | High | 0.000204 | 0.001355 | 2.39 |
| FALSE | High | RSDSSGGY | [K].RSDSSG | 1xPhospho | Q6ZPR6 1x | 0.003253 | 0.00016 | 1 | 2 | 5 | Q6ZPR6 | Inhibitor o | Q6ZPR6 [9 | 1 | 2973.533 | 0 | -0.14 | 0.122965 | 11.96 | 104.1 | 95.9 | 10.8 | 3.57 | | High | High | High | High | High | High | High | High | High | High | High | 7.19E-05 | 0.000894 | 4.04 |
| FALSE | High | RSEDSDED | [K].RSEDSD | 1xCarbami | E9PXD2 2x | 7.75E-05 | 0 | 1 | 8 | 1 | E9PXD2 | Rho GTPas | E9PXD2 [5 | 1 | 2413.003 | 0 | -1.4 | 0.002121 | 2.52 | 149.8 | 50.2 | 12.99 | 16.67 | | High | High | High | High | High | High | High | High | High | High | High | 0 | 1.36E-05 | 3.03 |
| FALSE | High | RSEGHEQT | [R].RSEGHI | 1xPhospho | Q7TSJ2 1x | 4.55E-08 | 0 | 1 | 5 | 4 | Q7TSJ2 | Microtubu | Q7TSJ2 [38 | 1 | 2740.32 | 0 | 0.41 | 0.024378 | 4.93 | 85.9 | 114.1 | 9.93 | 6.73 | | High | High | High | High | High | High | High | High | High | High | High | 0 | 3.17E-09 | 2.77 |
| FALSE | High | RSEGLSQE | [K].RSEGLS | 1xCarbami | B1AQJ2 1x | 0.020142 | 0.001472 | 1 | 3 | 2 | B1AQJ2 | Ubiquitin c | B1AQJ2 [8 | 1 | 3756.712 | 0 | -0.26 | 0.034995 | 1.79 | 108.5 | 91.5 | 7.93 | 16.3 | | High | High | High | High | High | High | High | High | High | High | High | 0.001076 | 0.01134 | 3.12 |
| FALSE | High | RSASPYPTH | [R].RSASPY | 1xCarbami | E9Q784 3x | 0.021418 | 0.001507 | 1 | 3 | 1 | E9Q784 | Zinc finger | E9Q784 [3 | 1 | 2312.949 | 0 | -0.95 | 0.000618 | 1.22 | 131.5 | 68.5 | 6.23 | 0.88 | | High | High | High | High | High | High | High | High | High | High | High | 0.000801 | 0.007331 | 2.08 |
| FALSE | High | SASPPCTG | [K].SASPPC | 1xCarbami | B2RX10 1x | 0.001411 | 5.66E-05 | 1 | 5 | 1 | B2RX10 | Fanconi an | B2RX10 [1 | 0 | 1556.698 | 0 | -0.78 | 0.002839 | 6.64 | 126.4 | 73.6 | 5.88 | 9.32 | | High | High | High | High | High | High | High | High | High | High | High | 2.58E-05 | 0.00035 | 2.05 |
| FALSE | High | RTPSPPPLP | [R].RTPSPP | 2xPhospho | A0A087WF | 0.015465 | 0.001089 | 1 | 2 | 2 | A0A087WF | Proline-ric | A0A087WF | 1 | 2277.114 | 0 | 2.75 | 0.000408 | 12.1 | 24.3 | 175.7 | 9.54 | 10.02 | | High | High | High | High | High | High | High | High | High | High | High | 0.000571 | 0.00513 | 3.31 |
| FALSE | High | RTPTMPQE | [R].RTPTM | 1xCarbami | Q14AX6 1x | 0.003526 | 0.00021 | 1 | 1 | 2 | Q14AX6 | Cyclin-dep | Q14AX6 [1 | 1 | 3024.509 | 0 | 0.27 | 0.026826 | 3.73 | 91 | 109 | 6.62 | 9.52 | | High | High | High | High | High | High | High | High | High | High | High | 9.42E-05 | 0.00098 | 2.9 |
| FALSE | High | SAGSSSPE | [R].SAGSSS | 1xCarbami | Q9CT10 3x | 0.056724 | 0.004124 | 1 | 1 | 1 | Q9CT10 | Ran-bindin | Q9CT10 [2 | 0 | 3061.105 | 0 | 1.08 | 0.025898 | 26.92 | 62.2 | 137.8 | 21.01 | 16.12 | | High | High | High | High | High | High | High | High | High | High | High | 0.002159 | 0.02239 | 2.18 |
| FALSE | High | SAGSSSPE | [R].SAGSSS | 1xCarbami | Q9CT10 1x | 2.8E-06 | 0 | 1 | 1 | 3 | Q9CT10 | Ran-bindin | Q9CT10 [2 | 0 | 3130.335 | 0 | 0.49 | 0.001298 | 3.59 | 83.8 | 116.2 | 5.05 | 4.23 | | High | High | High | High | High | High | High | High | High | High | High | 0 | 3.25E-07 | 3.46 |
| FALSE | High | SAGSSSPE | [R].SAGSSS | 1xCarbami | Q9CT10 2x | 2.1E-06 | 0 | 1 | 1 | 5 | Q9CT10 | Ran-bindin | Q9CT10 [2 | 0 | 3210.301 | 0 | -0.07 | 0.636656 | 2.84 | 103.3 | 96.7 | 3.78 | 4.72 | | High | High | High | High | High | High | High | High | High | High | High | 0 | 2.33E-07 | 3.98 |
| FALSE | High | SAGTETGS | [R].SAGTET | 1xPhospho | E9QPI5 1x | 1.71E-09 | 0 | 1 | 3 | 7 | E9QPI5 | Sister chro | E9QPI5 [11 | 0 | 2516.146 | 0 | 0.44 | 0.000936 | 6.95 | 83.3 | 116.7 | 3.79 | 3.42 | | High | High | High | High | High | High | High | High | High | High | High | 0 | 7.91E-11 | 4.61 |
| FALSE | High | SAGTETGS | [R].SAGTET | 1xDeamida | E9QPI5 1x | 1.07E-10 | 0 | 1 | 3 | 4 | E9QPI5 | Sister chro | E9QPI5 [11 | 0 | 2517.13 | 0 | 0.24 | 0.086473 | 23.31 | 91.6 | 108.4 | 14.64 | 9.63 | | High | High | High | High | High | High | High | High | High | High | High | 0 | 3.53E-12 | 4.96 |
| FALSE | High | SAIESDVDF | [K].SAIESD | 1xPhospho | O09012 1x | 0.005467 | 0.00062 | 1 | 3 | 2 | O09012 | Peroxisom | O09012 [2 | 0 | 1949.93 | 0 | 1.91 | 0.00278 | 7.87 | 39.7 | 160.3 | 14.67 | 18.41 | | High | High | High | High | High | High | High | High | High | High | High | 0.000275 | 0.001602 | 2.91 |
| FALSE | High | SALGEAPP | [R].SALGEA | 1xCarbami | H3BLD9 1x | 0.000168 | 0 | 1 | 4 | 1 | H3BLD9 | Zinc finger | H3BLD9 [1 | 0 | 1718.813 | 0 | -0.84 | 0.000366 | 2.55 | 129.2 | 70.8 | 1.03 | 7.82 | | High | High | High | High | High | High | High | High | High | High | High | 0 | 3.21E-05 | 2.85 |
| FALSE | High | SALLSPEEP | [K].SALLSP | 1xPhospho | Q9EPE9 1x | 0.031439 | 0.002011 | 1 | 2 | 1 | Q9EPE9 | Manganes | Q9EPE9 [9 | 0 | 1799.889 | 0 | 0.61 | 0.002417 | 3.88 | 82.2 | 117.8 | 5.59 | 7.05 | | High | High | High | High | High | High | High | High | High | High | High | 0.001076 | 0.01142 | 2.29 |
| FALSE | High | SALPPPSLC | [K].SALPPP | 1xPhospho | Q69ZS7 1x | 0.000233 | 0 | 1 | 3 | 2 | Q69ZS7 | HBS1-like p | Q69ZS7 [2 | 0 | 2573.306 | 0 | 0.43 | 0.030808 | 22.53 | 85.8 | 114.2 | 12.75 | 15.63 | | High | High | High | High | High | High | High | High | High | High | High | 0 | 0.000204 | 3.32 |
| FALSE | High | SALSESTQL | [K].SALSES | 1xPhospho | Q91WD7 1 | 5.38E-07 | 0 | 1 | 1 | 2 | Q91WD7 | Kinesin-like | Q91WD7 [ | 0 | 2152.058 | 0 | -0.52 | 0.007515 | 3.73 | 116.3 | 83.7 | 12.48 | 6.91 | | High | High | High | High | High | High | High | High | High | High | High | 0 | 5.06E-08 | 4.2 |
| FALSE | High | SALSPSK | [K].SALSPS | 1xPhospho | Q9ERU9 1x | 0.08447 | 0.006345 | 1 | 1 | 1 | Q9ERU9 | E3 SUMO- | Q9ERU9 [2 | 0 | 1227.675 | 0 | -0.55 | 0.000827 | 4.3 | 121.1 | 78.9 | 3.46 | 6.4 | | High | High | High | High | High | High | High | High | High | High | High | 0.003265 | 0.03525 | 1.31 |
| FALSE | High | SALSSSLRD | [R].SALSSS | 1xPhospho | E9PUA3 1x | 0.042868 | 0.003228 | 1 | 4 | 1 | E9PUA3 | IQ motif ar | E9PUA3 [9 | 1 | 2059.084 | 0 | 0.59 | 0.000592 | 3.66 | 79.7 | 120.3 | 5.17 | 4.54 | | High | High | High | High | High | High | High | High | High | High | High | 0.001651 | 0.01609 | 1.02 |
| FALSE | High | SALTLSLDS | [K].SALTLS | 1xPhospho | Q8BHK9 1x | 4.95E-06 | 0 | 1 | 3 | 1 | Q8BHK9 | DNA excisi | Q8BHK9 [9 | 0 | 1756.95 | 0 | -0.35 | 0.006148 | 3.61 | 110.2 | 89.8 | 6.06 | 5.59 | | High | High | High | High | High | High | High | High | High | High | High | 0 | 6.13E-07 | 2.55 |
| FALSE | High | SALYPALTP | [R].SALYPA | 1xPhospho | Q6YIY0 1x | 0.056248 | 0.004124 | 1 | 5 | 1 | Q6YIY0 | Gag protei | Q6YIY0 [12 | 0 | 1799.012 | 0 | 5.21 | 0.000261 | 31.42 | 4.4 | 195.6 | 41.3 | 8.62 | | High | High | High | High | High | High | High | High | High | High | High | 0.002117 | 0.022 | 3.45 |
| FALSE | High | SAMPFTAS | [R].SAMPF | 1xPhospho | O70400 1x | 0.043236 | 0.003228 | 1 | 3 | 2 | O70400 | PDZ and LI | O70400 [1 | 0 | 1729.818 | 0 | 0.08 | 0.871449 | 17.68 | 102.8 | 97.2 | 15.39 | 13.66 | | High | High | High | High | High | High | High | High | High | High | High | 0.001651 | 0.01637 | 2.56 |
| FALSE | High | SANDSTVH | [K].SANDS | 1xPhospho | Q80X50-5 | 0.000393 | 0 | 1 | 10 | 4 | Q80X50-5 | Isoform 5 | Q80X50-5 | 0 | 1928.952 | 0 | 0.18 | 0.584916 | 18.54 | 97.9 | 102.1 | 11.17 | 7.65 | | High | High | High | High | High | High | High | High | High | High | High | 0 | 8.71E-05 | 3.35 |
| FALSE | High | SANEMTTV | [R].SANEM | 1xCarbami | Q7TT79 1x | 0.090888 | 0.007081 | 1 | 5 | 2 | Q7TT79 | Microceph | Q7TT79 [2 | 0 | 2159.008 | 0 | -0.59 | 0.042399 | 8.71 | 117.7 | 82.3 | 14.35 | 8.52 | | High | High | High | High | High | High | High | High | High | High | High | 0.003633 | 0.03864 | 2.26 |
| FALSE | High | SANEMTTV | [R].SANEM | 1xCarbami | Q7TT79 1x | 5.85E-08 | 0 | 1 | 5 | 4 | Q7TT79 | Microceph | Q7TT79 [2 | 0 | 2175.002 | 0 | -0.55 | 0.044958 | 7.19 | 116.7 | 83.3 | 3.05 | 22.93 | | High | High | High | High | High | High | High | High | High | High | High | 0 | 1.29E-07 | 3.95 |
| FALSE | High | SANGGSES | [R].SANGG | 1xPhospho | Q9Z315 1x | 2.76E-07 | 0 | 1 | 1 | 7 | Q9Z315 | U4/U6.U5 | Q9Z315 [5 | 0 | 3162.415 | 0 | 0.1 | 0.249592 | 8.19 | 96 | 104 | 6.25 | 9.09 | | High | High | High | High | High | High | High | High | High | High | High | 0 | 1.71E-07 | 4.66 |
| FALSE | High | SANGGSES | [R].SANGG | 1xDeamida | Q9Z315 1x | 1.05E-07 | 0 | 1 | 1 | 6 | Q9Z315 | U4/U6.U5 | Q9Z315 [5 | 0 | 3163.399 | 0 | 0.4 | 0.059516 | 22.71 | 86.4 | 113.6 | 16.33 | 10.11 | | High | High | High | High | High | High | High | High | High | High | High | 0 | 8.13E-09 | 5.41 |
| FALSE | High | SANGGSES | [R].SANGG | 2xPhospho | Q9Z315 2x | 1.99E-06 | 0 | 1 | 1 | 3 | Q9Z315 | U4/U6.U5 | Q9Z315 [5 | 0 | 3242.382 | 0 | -0.27 | 0.076868 | 11.9 | 112.7 | 87.3 | 8.1 | 10.93 | | High | High | High | High | High | High | High | High | High | High | High | 0 | 2.2E-07 | 5.45 |
| FALSE | High | SANPTRPS | [R].SANPTF | 1xPhospho | Q99PU7 1x | 0.002172 | 5.66E-05 | 1 | 1 | 2 | Q99PU7 | Ubiquitin c | Q99PU7 [5 | 0 | 2304.211 | 0 | -0.47 | 0.036044 | 11.1 | 117.9 | 82.1 | 11.84 | 17.14 | | High | High | High | High | High | High | High | High | High | High | High | 2.58E-05 | 0.000569 | 2.01 |
| FALSE | High | SANQSPQS | [R].SANQS | 1xPhospho | A0A0R4J0V | 1.2E-10 | 0 | 1 | 8 | 2 | A0A0R4J0V | Phosphatic | A0A0R4J0V | 0 | 2831.289 | 0 | -0.47 | 0.021151 | 3.05 | 114.6 | 85.4 | 7.78 | 14.62 | | High | High | High | High | High | High | High | High | High | High | High | 0 | 3.99E-12 | 6.19 |
| FALSE | High | SANSPDGK | [K].SANSPD | 1xPhospho | P58252 1x | 0.012731 | 0.000889 | 1 | 14 | 2 | P58252 | Elongation | P58252 [27 | 0 | 1313.65 | 0 | -0.21 | 0.046206 | 7.02 | 109.5 | 90.5 | 5.74 | 12.95 | | High | High | High | High | High | High | High | High | High | High | High | 0.000585 | 0.005312 | 2.19 |
| FALSE | High | SAPASPNH | [R].SAPASF | 1xPhospho | Q3UCQ1 1 | 2.7E-06 | 0 | 1 | 2 | 1 | Q3UCQ1 | Forkhead b | Q3UCQ1 [3 | 0 | 2925.405 | 0 | 0.16 | 0.656137 | 6.82 | 96.5 | 103.5 | 12.23 | 20.22 | | High | High | High | High | High | High | High | High | High | High | High | 0 | 3.11E-07 | 2.63 |
| FALSE | High | SAPASPTHI | [R].SAPASF | 2xPhospho | P42128 2x | 0.027358 | 0.001676 | 1 | 1 | 2 | P42128 | Forkhead b | P42128 [40 | 0 | 1894.849 | 0 | 1.18 | 0.001966 | 21.28 | 59.8 | 140.2 | 11.25 | 13.71 | | High | High | High | High | High | High | High | High | High | High | High | 0.000936 | 0.009753 | 1.85 |
| FALSE | High | SAPASPTHI | [R].SAPASF | 2xPhospho | P42128 2x | 0.022773 | 0.001542 | 1 | 1 | 1 | P42128 | Forkhead b | P42128 [40 | 0 | 1910.843 | 0 | 1.24 | 0.005166 | 25.74 | 60.1 | 139.9 | 9.56 | 14.31 | | High | High | High | High | High | High | High | High | High | High | High | 0.000814 | 0.007881 | 1.77 |
| FALSE | High | SAGSPSSD | [K].SAGSPS | 1xPhospho | Q3TLH4 1x | 0.000173 | 0 | 1 | 6 | 4 | Q3TLH4 | Protein PRI | Q3TLH4 [1 | 0 | 1745.8 | 0 | -0.1 | 0.098199 | 5.1 | 108 | 92 | 11.85 | 9.97 | | High | High | High | High | High | High | High | High | High | High | High | 0 | 3.34E-05 | 2.67 |
| FALSE | High | SAGSIPLSP | [K].SAGSIP | 1xPhospho | Q811L6 1x | 0.113753 | 0.009241 | 1 | 4 | 1 | Q811L6 | Microtubu | Q811L6 [1 | 0 | 1477.798 | 0 | 0.26 | 0.204754 | 22.43 | 94.6 | 105.4 | 11.38 | 12.54 | | High | High | High | High | High | High | High | High | High | High | High | 0.00468 | 0.05049 | 1.37 |
| FALSE | High | SAGQDSPA | [R].SAGQD | 1xCarbami | Q6NZN1-2 | 7.28E-05 | 0 | 1 | 4 | 1 | Q6NZN1-2 | Isoform 2 | Q6NZN1-2 | 0 | 2441.131 | 0 | -0.24 | 0.368156 | 6.09 | 101.9 | 98.1 | 9.36 | 13.61 | | High | High | High | High | High | High | High | High | High | High | High | 0 | 1.25E-05 | 3.32 |
| FALSE | High | SAGGSAGP | [R].SAGGS | 1xPhospho | P26687 1x | 1.56E-10 | 0 | 1 | 1 | 2 | P26687 | Twist-relat | P26687 [4 | 0 | 2849.347 | 0 | 2.35 | 0.001021 | 13.03 | 36.4 | 163.6 | 14.53 | 12.65 | | High | High | High | High | High | High | High | High | High | High | High | 0 | 5.37E-12 | 5.58 |
| FALSE | High | SAEALGPG | [R].SAEALG | 1xPhospho | B1AQY2 1x | 0.01441 | 0.001011 | 1 | 3 | 1 | B1AQY2 | Rho GTPas | B1AQY2 [3 | 0 | 1633.851 | 0 | 0.41 | 0.025338 | 19.71 | 85.5 | 114.5 | 10.86 | 5.68 | | High | High | High | High | High | High | High | High | High | High | High | 0.000538 | 0.004707 | 2.26 |
| FALSE | High | SAEDLDLL | [R].SAEDLD | 1xPhospho | D3Z656 1x | 1.4E-09 | 0 | 1 | 4 | 2 | D3Z656 | Synaptojar | D3Z656 [8 | 0 | 2420.163 | 0 | 0.33 | 0.043269 | 16.05 | 84.7 | 115.3 | 17.42 | 9.38 | | High | High | High | High | High | High | High | High | High | High | High | 0 | 6.28E-11 | 4.71 |
| FALSE | High | SAEDLTDG | [R].SAEDLT | 1xPhospho | Q9D0L7 1x | 3.21E-06 | 0 | 1 | 2 | 1 | Q9D0L7 | Armadillo r | Q9D0L7 [4 | 0 | 2506.144 | 0 | -0.17 | 0.484141 | 25.83 | 103.5 | 96.5 | 52.98 | 29.04 | | High | High | High | High | High | High | High | High | High | High | High | 0 | 3.76E-07 | 3.48 |
| FALSE | High | SAEDLTDG | [R].SAEDLT | 1xPhospho | Q9D0L7 1x | 2.66E-07 | 0 | 1 | 2 | 7 | Q9D0L7 | Armadillo r | Q9D0L7 [4 | 0 | 2735.306 | 0 | -0.33 | 0.010148 | 5.04 | 109.7 | 90.3 | 1.84 | 9.27 | | High | High | High | High | High | High | High | High | High | High | High | 0 | 3.53E-08 | 4.98 |
| FALSE | High | SAEEIGFISL | [R].SAEEIG | 1xPhospho | G3UYN0 1x | 7.41E-05 | 0 | 1 | 1 | 3 | G3UYN0 | L1TD1 OS= | G3UYN0 [4 | 1 | 3104.544 | 0 | -0.8 | 0.000786 | 2.59 | 125.3 | 74.7 | 8.33 | 7.58 | | High | High | High | High | High | High | High | High | High | High | High | 0 | 1.29E-05 | 3.35 |
| FALSE | High | SAEESPSGT | [K].SAEESP | 1xPhospho | E9QML5 1x | 0.000382 | 0 | 1 | 11 | 3 | E9QML5 | Zinc finger | E9QML5 [8 | 0 | 2073.999 | 0 | -0.66 | 0.002666 | 4.78 | 123.8 | 76.2 | 6.62 | 5.52 | | High | High | High | High | High | High | High | High | High | High | High | 0 | 8.13E-05 | 2.48 |
| FALSE | High | SAELDASDS | [K].SAELDA | 1xPhospho | Q03141 1x | 0.000165 | 0 | 1 | 5 | 5 | Q03141 | MAP/micro | Q03141 [3 | 0 | 2407.164 | 0 | 0.35 | 0.005044 | 10.87 | 88.6 | 111.4 | 9.3 | 6.6 | | High | High | High | High | High | High | High | High | High | High | High | 0 | 0.000253 | 2.82 |
| FALSE | High | SAELDFPLE | [R].SAELDF | 1xPhospho | Q9EQM6 1 | 2.75E-06 | 0 | 1 | 1 | 2 | Q9EQM6 | Microproc | Q9EQM6 [ | 0 | 3081.336 | 0 | -0.37 | 0.028646 | 2.66 | 112.9 | 87.1 | 4.46 | 19.92 | | High | High | High | High | High | High | High | High | High | High | High | 0 | 3.17E-07 | 4.14 |
| FALSE | High | SAELDFPLE | [R].SAELDF | 1xPhospho | Q9EQM6 1 | 5.78E-10 | 0 | 1 | 1 | 10 | Q9EQM6 | Microproc | Q9EQM6 [ | 0 | 3097.331 | 0 | -0.62 | 0.003195 | 2.48 | 119.7 | 80.3 | 4.05 | 11.56 | | High | High | High | High | High | High | High | High | High | High | High | 0 | 2.33E-11 | 5.15 |
| FALSE | High | SAENPPSG | [R].SAENPF | 1xPhospho | Q3TXN7 1x | 5.91E-05 | 0 | 1 | 7 | 2 | Q3TXN7 | Putative ur | Q3TXN7 [6 | 0 | 1409.662 | 0 | 0.09 | 0.254531 | 21.99 | 91.8 | 108.2 | 18.48 | 15.46 | | High | High | High | High | High | High | High | High | High | High | High | 0 | 9.99E-06 | 1.61 |
| FALSE | High | SAENPPSG | [R].SAENPF | 1xPhospho | Q3TXN7 1x | 0.000274 | 0 | 1 | 7 | 1 | Q3TXN7 | Putative ur | Q3TXN7 [6 | 1 | 1766.92 | 0 | -0.04 | 0.443424 | 14.1 | 94.3 | 105.7 | 16.77 | 16.15 | | High | High | High | High | High | High | High | High | High | High | High | 0 | 5.58E-05 | 1.48 |
| FALSE | High | SAEPGSPG | [R].SAEPGS | 1xPhospho | Q8R4E9 1x | 0.091627 | 0.007146 | 1 | 1 | 2 | Q8R4E9 | DNA replic | Q8R4E9 [3 | 0 | 3390.76 | 0 | 0.17 | 0.106522 | 9.07 | 94.2 | 105.8 | 12.17 | 11.56 | | High | High | High | High | High | High | High | High | High | High | High | 0.007225 | 0.07852 | 2.31 |
| FALSE | High | SAEPLANT | [K].SAEPLA | 1xPhospho | H3BL37 1x | 5.15E-11 | 0 | 1 | 6 | 12 | H3BL37 | Treacle pro | H3BL37 [1 | 0 | 3336.661 | 0 | -0.53 | 0.000417 | 2.66 | 118.2 | 81.8 | 2.05 | 5.36 | | High | High | High | High | High | High | High | High | High | High | High | 0 | 1.54E-12 | 6.53 |
| FALSE | High | SAPASPTHI | [R].SAPASF | 3xPhospho | P42128 3x | 0.105058 | 0.008518 | 1 | 1 | 1 | P42128 | Forkhead b | P42128 [40 | 0 | 1974.815 | 0 | 0.82 | 0.004984 | 5.8 | 74.3 | 125.7 | 8.8 | 8.79 | | High | High | High | High | High | High | High | High | High | High | High | 0.004277 | 0.04555 | 1.55 |
| FALSE | High | SAEPLANT | [K].SAEPLA | 1xDeamida | H3BL37 1x | 3.23E-08 | 0 | 1 | 6 | 12 | H3BL37 | Treacle pro | H3BL37 [1 | 0 | 3337.645 | 0 | -0.46 | 0.002165 | 3.15 | 113.8 | 86.2 | 3.15 | 6.23 | | High | High | High | High | High | High | High | High | High | High | High | 0 | 2.14E-09 | 4.69 |
| FALSE | High | SAEPPRSPL | [K].SAEPPR | 1xPhospho | E9Q1Q1 1x | 0.045898 | 0.003408 | 2 | 12 | 1 | E9Q1Q1; C | Microtubu | E9Q1Q1 [1 | 1 | 1732.976 | 0 | -0.44 | 0.002755 | 5.24 | 116.7 | 83.3 | 6.57 | 12.31 | Shared | High | High | High | High | High | High | High | High | High | High | High | 0.001774 | 0.01752 | 1.33 |
| FALSE | High | SAEPPVVQ | [R].SAEPPV | 1xPhospho | Q5NCY0 1x | 0.000194 | 0 | 1 | 1 | 1 | Q5NCY0 | Lysine-spe | Q5NCY0 [2 | 0 | 3281.653 | 0 | -0.49 | 0.031993 | 7.23 | 115.6 | 84.4 | 10.98 | 15.91 | | High | High | High | High | High | High | High | High | High | High | High | 0 | 3.79E-05 | 4.13 |
| FALSE | High | SAEPSPTV | [K].SAEPSP | 1xPhospho | F8VQ28 1x | 5.67E-06 | 0 | 1 | 5 | 3 | F8VQ28 | Paxillin OS | F8VQ28 [1 | 0 | 2573.2 | 0 | 0.35 | 0.525199 | 63.16 | 90.1 | 109.9 | 39.55 | 24.21 | | High | High | High | High | High | High | High | High | High | High | High | 0 | 3.94E-06 | 4.51 |
| FALSE | High | SAEPSPTV | [K].SAEPSP | 1xPhospho | F8VQ28 1x | 9.42E-07 | 0 | 1 | 5 | 3 | F8VQ28 | Paxillin OS | F8VQ28 [1 | 0 | 2589.195 | 0 | 0.23 | 0.024353 | 10.04 | 93 | 107 | 11.74 | 8.07 | | High | High | High | High | High | High | High | High | High | High | High | 0 | 6.68E-07 | 3.63 |
| FALSE | High | SAEPSPTV | [K].SAEPSP | 2xPhospho | F8VQ28 2x | 1.99E-05 | 0 | 1 | 5 | 2 | F8VQ28 | Paxillin OS | F8VQ28 [1 | 0 | 2669.162 | 0 | 0.26 | 0.082414 | 12.64 | 91.1 | 108.9 | 5.24 | 21.62 | | High | High | High | High | High | High | High | High | High | High | High | 0 | 6.39E-05 | 3.72 |
| FALSE | High | SAEPTREP | [R].SAEPTR | 2xPhospho | B1AQD9 2x | 0.001644 | 5.66E-05 | 1 | 2 | 2 | B1AQD9 | Protein un | B1AQD9 [2 | 1 | 3096.303 | 0 | -0.38 | 0.016628 | 4.81 | 120.2 | 79.8 | 8.81 | 22.25 | | High | High | High | High | High | High | High | High | High | High | High | 0.000327 | 0.002868 | 2.64 |
| FALSE | High | SAEQQQP | [R].SAEQQ | 1xPhospho | F8WI37 1x | 6.02E-05 | 0 | 1 | 5 | 1 | F8WI37 | Histone-lys | F8WI37 [8 | 0 | 3173.556 | 0 | -0.05 | 0.228586 | 5.94 | 101.4 | 98.6 | 2.44 | 2.9 | | High | High | High | High | High | High | High | High | High | High | High | 0 | 1.01E-05 | 2.41 |
| FALSE | High | SAEVDSDD | [K].SAEVDS | 1xPhospho | Q61768 1x | 0.064323 | 0.004664 | 1 | 1 | 2 | Q61768 | Kinesin-1 h | Q61768 [8 | 0 | 2075.953 | 0 | -0.04 | 0.275184 | 4.97 | 103.7 | 96.3 | 8.44 | 12.94 | | High | High | High | High | High | High | High | High | High | High | High | 0.002403 | 0.02566 | 2.35 |
| FALSE | High | SAFSSLQNI | [R].SAFSSL | 1xPhospho | B7ZNA0 1x | 0.001021 | 0 | 2 | 8 | 4 | B7ZNA0; Q | Shroom3 p | B7ZNA0 [6 | 0 | 1857.931 | 0 | 0.08 | 0.025848 | 3.84 | 98.2 | 101.8 | 3.2 | 1.48 | Shared | High | High | High | High | High | High | High | High | High | High | High | 0.003911 | 0.04148 | 2.52 |
| FALSE | High | SAFTPATAT | [R].SAFTPA | 1xPhospho | E9Q0N0 1x | 2.09E-08 | 0 | 1 | 8 | 2 | E9Q0N0 | Intersectin | E9Q0N0 [8 | 0 | 2628.332 | 0 | -0.04 | 0.935394 | 19.95 | 101.4 | 98.6 | 4.37 | 12.17 | | High | High | High | High | High | High | High | High | High | High | High | 0 | 1.32E-09 | 4.7 |
| FALSE | High | SAFTPATAT | [R].SAFTPA | 2xPhospho | E9Q0N0 2x | 0.000276 | 0 | 1 | 8 | 1 | E9Q0N0 | Intersectin | E9Q0N0 [8 | 0 | 2708.299 | 0 | -0.94 | 0.011165 | 9.31 | 131.5 | 68.5 | 11.78 | 15.26 | | High | High | High | High | High | High | High | High | High | High | High | 0 | 5.63E-05 | 3.85 |
| FALSE | High | SAGDAGPS | [R].SAGDA | 1xPhospho | A0A1B0GR | 1.16E-07 | 0 | 1 | 1 | 2 | A0A1B0GR | RIKEN cDN | A0A1B0GR | 0 | 2518.139 | 0 | 1.59 | 0.00045 | 14.73 | 51 | 149 | 13.56 | 4.68 | | High | High | High | High | High | High | High | High | High | High | High | 0 | 1.02E-08 | 3.83 |
| FALSE | High | SAGDVPVT | [R].SAGDV | 1xPhospho | Q6J1H4 1x | 2.46E-08 | 0 | 1 | 1 | 9 | Q6J1H4 | Undifferen | Q6J1H4 [2 | 0 | 2762.243 | 0 | -1.5 | 0.103204 | 3.61 | 147.9 | 52.1 | 53.83 | 27.36 | | High | High | High | High | High | High | High | High | High | High | High | 0 | 1.59E-09 | 4.56 |
| FALSE | High | SAEPLANT | [K].SAEPLA | 2xPhospho | H3BL37 2x | 4.32E-06 | 0 | 1 | 6 | 7 | H3BL37 | Treacle pro | H3BL37 [1 | 0 | 3416.628 | 0 | 0.08 | 0.142637 | 10.62 | 89.7 | 110.3 | 9.12 | 14.51 | | High | High | High | High | High | High | High | High | High | High | High | 0 | 5.28E-07 | 4.05 |
| FALSE | High | SADWTEEL | [R].SADWT | 1xPhospho | Q9JL04 1x | 0.050397 | 0.003641 | 1 | 1 | 1 | Q9JL04 | Formin-2 C | Q9JL04 [4 | 0 | 1571.73 | 0 | 0.12 | 0.181095 | 15.68 | 92.5 | 107.5 | 6.76 | 10.61 | | High | High | High | High | High | High | High | High | High | High | High | 0.001916 | 0.01945 | 1.81 |
| FALSE | High | SAPCEDPF | [R].SAPCED | 1xCarbami | E9Q3G8 1x | 7.41E-05 | 0 | 1 | 3 | 1 | E9Q3G8 | Nucleopor | E9Q3G8 [5 | 0 | 1857.886 | 0 | -0.11 | 0.371503 | 5.14 | 100.2 | 99.8 | 11.15 | 14.05 | | High | High | High | High | High | High | High | High | High | High | High | 0 | 1.28E-05 | 2.68 |
| FALSE | High | SAPDGGDE | [R].SAPDG | 1xDeamida | G3UYW1 1 | 3.05E-07 | 0 | 1 | 3 | 10 | G3UYW1 | WD repeat | G3UYW1 [ | 0 | 3687.632 | 0 | 0.75 | 0.004041 | 5.07 | 80.6 | 119.4 | 13.05 | 7.01 | | High | High | High | High | High | High | High | High | High | High | High | 0 | 2.69E-08 | 6.44 |
| FALSE | High | SASANSAE | [R].SASANS | 1xPhospho | Q9EPQ8 1x | 0.016021 | 0.001124 | 1 | 4 | 2 | Q9EPQ8 | Transcripti | Q9EPQ8 [1 | 0 | 2773.391 | 0 | 0.76 | 0.019207 | 27.72 | 73.6 | 126.4 | 15.23 | 14.32 | | High | High | High | High | High | High | High | High | High | High | High | 0.000585 | 0.005338 | 3.62 |
| FALSE | High | SASAPTLAE | [R].SASAPT | 1xPhospho | F6YZ95 1x | 0.000118 | 0 | 1 | 1 | 3 | F6YZ95 | Liprin-beta | F6YZ95 [9 | 0 | 1742.898 | 0 | 1.64 | 0.000964 | 13.24 | 50 | 150 | 14.13 | 11.5 | | High | High | High | High | High | High | High | High | High | High | High | 0 | 2.17E-05 | 3.04 |
| FALSE | High | SASASPLTP | [R].SASASP | 1xCarbami | A2AJI0 1x | 7.08E-05 | 0 | 1 | 5 | 2 | A2AJI0 | MAP7 dom | A2AJI0 [36 | 0 | 1710.808 | 0 | 0.93 | 0.003545 | 13 | 70.7 | 129.3 | 3.96 | 12.99 | | High | High | High | High | High | High | High | High | High | High | High | 0 | 0.000174 | 2.3 |
| FALSE | High | SASATSLTL | [R].SASATS | 1xPhospho | Q6KAU8 1x | 0.008927 | 0.000736 | 1 | 2 | 1 | Q6KAU8 | Autophagy | Q6KAU8 [2 | 0 | 1402.714 | 0 | 2 | 0.000181 | 4.34 | 40.2 | 159.8 | 6.32 | 5.15 | | High | High | High | High | High | High | High | High | High | High | High | 0.000327 | 0.002776 | 1.45 |
| FALSE | High | SASCGSPLS | [R].SASCGS | 1xCarbami | A0A068BD | 3.83E-08 | 0 | 1 | 3 | 2 | A0A068BD | Ral guanine | A0A068BD | 0 | 2033.88 | 0 | -0.02 | 0.293624 | 8.22 | 101.4 | 98.6 | 4.84 | 11.1 | | High | High | High | High | High | High | High | High | High | High | High | 0 | 2.59E-09 | 2.18 |
| FALSE | High | SASDAGIR | [R].SASDA | 1xPhospho | O89032 1x | 0.020679 | 0.001472 | 1 | 3 | 1 | O89032 | SH3 and PX | O89032 [7 | 0 | 1085.519 | 0 | 1.11 | 0.113234 | 16.51 | 67.3 | 132.7 | 16.07 | 40.68 | | High | High | High | High | High | High | High | High | High | High | High | 0.000786 | 0.007045 | 1.99 |
| FALSE | High | SASDASISS | [K].SASDAS | 1xPhospho | F8WHD8 1 | 0.004654 | 0.000456 | 1 | 15 | 1 | F8WHD8 | Synaptojar | F8WHD8 [ | 0 | 2747.365 | 0 | 1.94 | 0.006061 | 43.07 | 47.7 | 152.3 | 21.49 | 20.56 | | High | High | High | High | High | High | High | High | High | High | High | 0.000204 | 0.001337 | 3.06 |
| FALSE | High | SASDSGCD | [K].SASDSG | 1xCarbami | B9EKS2 1x | 7.49E-06 | 0 | 1 | 3 | 3 | B9EKS2 | Jumonji do | B9EKS2 [2 | 0 | 1733.782 | 0 | 0.29 | 0.064472 | 1.75 | 93.2 | 106.8 | 4.24 | 7.28 | | High | High | High | High | High | High | High | High | High | High | High | 0 | 0.000103 | 1.81 |
| FALSE | High | SASESYTQ | [R].SASESY | 1xPhospho | Q2TBE6 1x | 9.62E-05 | 0 | 1 | 1 | 5 | Q2TBE6 | Phosphatic | Q2TBE6 [4 | 0 | 1786.785 | 0 | 0.56 | 0.013852 | 18.97 | 78.4 | 121.6 | 3.66 | 12.79 | | High | High | High | High | High | High | High | High | High | High | High | 0.00014 | 0.001109 | 2.96 |
| FALSE | High | SASFAFEFP | [K].SASFAF | 1xPhospho | E9QLH9 1x | 0.002096 | 5.66E-05 | 1 | 2 | 1 | E9QLH9 | PEAK1-rela | E9QLH9 [6 | 0 | 1668.844 | 0 | 1.5 | 0.000574 | 11.66 | 56.5 | 143.5 | 17.86 | 6.22 | | High | High | High | High | High | High | High | High | High | High | High | 2.58E-05 | 0.000547 | 2.86 |
| FALSE | High | SASFNTDP | [R].SASFNT | 1xPhospho | Q8CJG0 1x | 0.001233 | 0 | 1 | 4 | 2 | Q8CJG0 | Protein arg | Q8CJG0 [3 | 0 | 1565.72 | 0 | -0.66 | 0.00047 | 5.71 | 122.4 | 77.6 | 5.85 | 13.38 | | High | High | High | High | High | High | High | High | High | High | High | 7.19E-05 | 0.000763 | 1.96 |
| FALSE | High | SASGGDLN | [K].SASGG | 1xCarbami | B2RWU4 1 | 6.54E-09 | 0 | 1 | 9 | 4 | B2RWU4 | Cordon-ble | B2RWU4 [ | 0 | 2523.15 | 0 | -0.2 | 0.290019 | 20.02 | 107.2 | 92.8 | 13.32 | 11.79 | | High | High | High | High | High | High | High | High | High | High | High | 0 | 8.87E-08 | 5.1 |
| FALSE | High | SASGGDLN | [K].SASGG | 1xCarbami | B2RWU4 1 | 7.4E-08 | 0 | 1 | 9 | 5 | B2RWU4 | Cordon-ble | B2RWU4 [ | 0 | 2524.134 | 0 | -0.3 | 0.130583 | 9.9 | 107.2 | 92.8 | 5.35 | 12.81 | | High | High | High | High | High | High | High | High | High | High | High | 0 | 2.83E-07 | 5.29 |
| FALSE | High | SASLENK | [R].SASLEN | 1xPhospho | P39447 1x | 0.068752 | 0.005042 | 1 | 3 | 1 | P39447 | Tight junct | P39447 [1 | 0 | 1286.676 | 0 | 1.34 | 0.013046 | 42.25 | 56.7 | 143.3 | 12.76 | 24.27 | | High | High | High | High | High | High | High | High | High | High | High | 0.002589 | 0.02776 | 1.46 |
| FALSE | High | SASINHLLD | [R].SASINH | 1xPhospho | Q9QZM4 1 | 5.97E-05 | 0 | 1 | 1 | 3 | Q9QZM4 | Tumor nec | Q9QZM4 [ | 0 | 2176.085 | 0 | 1.59 | 0.001829 | 26.75 | 49.1 | 150.9 | 9.99 | 12.09 | | High | High | High | High | High | High | High | High | High | High | High | 0 | 1E-05 | 3.97 |
| FALSE | High | SASLSNLHS | [K].SASLSN | 1xPhospho | Q9D1C2 1x | 0.001011 | 0 | 1 | 1 | 2 | Q9D1C2 | Protein chi | Q9D1C2 [1 | 0 | 1608.794 | 0 | 0.55 | 0.000912 | 2.09 | 80.4 | 119.6 | 6.03 | 5.15 | | High | High | High | High | High | High | High | High | High | High | High | 0 | 0.000243 | 2.04 |
| FALSE | High | SASLSSLFIT | [R].SASLSS | 1xPhospho | Q68FD7 1x | 0.007811 | 0.000664 | 1 | 2 | 3 | Q68FD7 | Folliculin-ir | Q68FD7 [2 | 0 | 2518.279 | 0 | -0.18 | 0.15309 | 5.1 | 103.5 | 96.5 | 7.84 | 11.6 | | High | High | High | High | High | High | High | High | High | High | High | 0.000327 | 0.002378 | 3.48 |
| FALSE | High | SASLSYPDF | [R].SASLSY | 1xPhospho | Q69ZF7 1x | 0.002695 | 0.000109 | 1 | 2 | 2 | Q69ZF7 | Metal tran | Q69ZF7 [6 | 0 | 1304.609 | 0 | 0.42 | 0.002628 | 4.6 | 85.1 | 114.9 | 7.71 | 7.54 | | High | High | High | High | High | High | High | High | High | High | High | 0.003922 | 0.04164 | 1.48 |
| FALSE | High | SASITNLSLI | [R].SASITN | 1xPhospho | D3Z5N5 1x | 0.001058 | 0 | 2 | 4 | 1 | D3Z5N5; Q | 1-phospha | D3Z5N5 [3 | 0 | 1485.751 | 0 | -0.16 | 0.050393 | 2.98 | 104.9 | 95.1 | 3.53 | 4.69 | Shared | High | High | High | High | High | High | High | High | High | High | High | 0 | 0.000255 | 2.59 |
| FALSE | High | SASNLTELC | [K].SASNLT | 1xPhospho | E9Q3I3 1x | 0.000221 | 0 | 1 | 2 | 2 | E9Q3I3 | Spermatog | E9Q3I3 [24 | 0 | 1599.758 | 0 | -0.91 | 0.000756 | 3.17 | 130.2 | 69.8 | 4.63 | 8.58 | | High | High | High | High | High | High | High | High | High | High | High | 0.000872 | 0.009044 | 2.16 |
| FALSE | High | SASPDDDL | [R].SASPDD | 1xPhospho | Q9R0P4 1x | 2.05E-10 | 0 | 1 | 2 | 13 | Q9R0P4 | Small acidi | Q9R0P4 [1 | 0 | 2744.152 | 0 | -0.16 | 0.041591 | 9.94 | 106.9 | 93.1 | 3.14 | 6.67 | | High | High | High | High | High | High | High | High | High | High | High | 0 | 6E-09 | 6.11 |
| FALSE | High | SASPDDDL | [R].SASPDD | 1xDeamida | Q9R0P4 1x | 5.98E-09 | 0 | 1 | 2 | 3 | Q9R0P4 | Small acidi | Q9R0P4 [1 | 0 | 2745.136 | 0 | -0.35 | 0.047236 | 3.86 | 112.1 | 87.9 | 13.74 | 5.26 | | High | High | High | High | High | High | High | High | High | High | High | 0 | 3.21E-10 | 5.54 |
| FALSE | High | SASPDDDL | [R].SASPDD | 2xPhospho | Q9R0P4 2x | 0.022574 | 0.001542 | 1 | 2 | 1 | Q9R0P4 | Small acidi | Q9R0P4 [1 | 0 | 2824.119 | 0 | 0.07 | 0.986417 | 16.65 | 98 | 102 | 8.98 | 22.66 | | High | High | High | High | High | High | High | High | High | High | High | 0.000814 | 0.007834 | 3.26 |
| FALSE | High | SASPDDDL | [R].SASPDD | 1xPhospho | Q9R0P4 1x | 0.040721 | 0.003058 | 1 | 2 | 1 | Q9R0P4 | Small acidi | Q9R0P4 [1 | 1 | 3101.41 | 0 | -0.43 | 0.097367 | 17.8 | 119.6 | 80.4 | 21.94 | 24.15 | | High | High | High | High | High | High | High | High | High | High | High | 0.001597 | 0.01526 | 2.11 |
| FALSE | High | SASPDYNR | [R].SASPDY | 1xPhospho | G5E870 1x | 0.121159 | 0.009984 | 1 | 6 | 1 | G5E870 | E3 ubiquiti | G5E870 [7 | 0 | 1218.535 | 0 | -0.22 | 0.052543 | 7.43 | 110.3 | 89.7 | 6.99 | 7.42 | | High | High | High | High | High | High | High | High | High | High | High | 0.004961 | 0.05398 | 1.76 |
| FALSE | High | SASPGLPK | [K].SASPGL | 1xPhospho | Q9EQU5 1 | 0.069903 | 0.005181 | 1 | 5 | 1 | Q9EQU5 | Protein SET | Q9EQU5 [2 | 0 | 1065.554 | 0 | -0.78 | 0.685507 | 23.97 | 110.3 | 89.7 | 105.67 | 107.86 | | High | High | High | High | High | High | High | High | High | High | High | 0.002662 | 0.02839 | 1.9 |
| FALSE | High | SASPGLPK | [K].SASPGL | 1xPhospho | Q9EQU5 1 | 0.1059 | 0.008582 | 1 | 5 | 3 | Q9EQU5 | Protein SET | Q9EQU5 [2 | 0 | 1294.717 | 0 | -0.08 | 0.506714 | 15.07 | 102.6 | 97.4 | 6.17 | 9.01 | | High | High | High | High | High | High | High | High | High | High | High | 0.004291 | 0.04599 | 1.85 |
| FALSE | High | SASAISSDS | [K].SASAIS | 1xPhospho | Q9Z2D1 1x | 3.15E-09 | 0 | 1 | 3 | 1 | Q9Z2D1 | Myotubula | Q9Z2D1 [4 | 0 | 2537.161 | 0 | 0.11 | 0.144232 | 8.89 | 96 | 104 | 1.57 | 11.2 | | High | High | High | High | High | High | High | High | High | High | High | 0 | 1.57E-10 | 5.23 |
| FALSE | High | SASAENIPD | [R].SASAEN | 2xCarbami | Q920N2 1x | 0.030104 | 0.001857 | 1 | 1 | 1 | Q920N2 | Biotin--pro | Q920N2 [1 | 0 | 3247.423 | 0 | -1.27 | 0.000808 | 3.61 | 141.5 | 58.5 | 10.89 | 19.7 | | High | High | High | High | High | High | High | High | High | High | High | 0.001011 | 0.01083 | 2.68 |

| | | | | | | | | | | | | | | | | | | | | | | | | | | | | | | | | | | | | | |
|---|---|---|---|---|---|---|---|---|---|---|---|---|---|---|---|---|---|---|---|---|---|---|---|---|---|---|---|---|---|---|---|---|---|---|---|---|---|
| FALSE | High | SASADNLIL | [R].SASADN | 1xPhospho | Q8BGU5 1x | 0.000558 | 0 | 1 | 2 | 2 | Q8BGU5 | Cyclin-Y OS | Q8BGU5 [3 | 0 | 1465.761 | 0 | -0.01 | 0.416246 | 4.39 | 100.4 | 99.6 | 2.41 | 2.89 | | High | High | High | High | High | High | High | High | High | High | High | 0 | 0.000177 | 2.43 |
| FALSE | High | SASACQILII | [R].SASACC | 2xCarbami | Q7TSH4 2x | 4.04E-08 | 0 | 1 | 3 | 1 | Q7TSH4 | Centriolar | Q7TSH4 [3 | 0 | 2904.38 | 0 | 0.19 | 0.141917 | 19.64 | 88.8 | 111.2 | 14.85 | 12.76 | | High | High | High | High | High | High | High | High | High | High | High | 0 | 2.77E-09 | 5.16 |
| FALSE | High | SAPDISFTII | [R].SAPDIS | 1xPhospho | Q9D2N4 1x | 0.000639 | 0 | 1 | 10 | 1 | Q9D2N4 | Dystrobrev | Q9D2N4 [4 | 0 | 1916.977 | 0 | 1.52 | 0.001341 | 10.18 | 51.6 | 148.4 | 3.73 | 12.42 | | High | High | High | High | High | High | High | High | High | High | High | 0 | 0.000144 | 1.85 |
| FALSE | High | SAPEALSGA | [K].SAPEAL | 1xCarbami | Q8CH02 1x | 0.006773 | 0.00062 | 1 | 1 | 1 | Q8CH02 | SURP and C | Q8CH02 [3 | 0 | 4839.634 | 0 | 0.54 | 0.005406 | 10.5 | 81.6 | 118.4 | 8.24 | 6.34 | | High | High | High | High | High | High | High | High | High | High | High | 0.000275 | 0.002033 | 3.59 |
| FALSE | High | SAPETAAPI | [K].SAPETA | 2xPhospho | F8WI37 2x | 0.043981 | 0.003228 | 1 | 5 | 1 | F8WI37 | Histone-lys | F8WI37 [6 | 0 | 2252.02 | 0 | -0.05 | 0.274993 | 8.93 | 103.7 | 96.3 | 11.51 | 11.12 | | High | High | High | High | High | High | High | High | High | High | High | 0.001651 | 0.01656 | 2.46 |
| FALSE | High | SAPETLTLP | [K].SAPETL | 1xPhospho | Q8BTD8 1x | 0.001411 | 5.66E-05 | 1 | 4 | 3 | Q8BTD8 | Putative ur | Q8BTD8 [3 | 0 | 1936.008 | 0 | -1.49 | 0.00031 | 1.89 | 150.9 | 49.1 | 7.35 | 17.1 | | High | High | High | High | High | High | High | High | High | High | High | 0.007326 | 0.07976 | 2.31 |
| FALSE | High | SAPLPLTVE | [K].SAPLPL | 1xPhospho | F6RJ39 1xP | 0.022574 | 0.001542 | 1 | 3 | 1 | F6RJ39 | Apoptotic | F6RJ39 [47 | 0 | 2108.144 | 0 | -1.68 | 0.000302 | 0.73 | 152.3 | 47.7 | 7.31 | 7.42 | | High | High | High | High | High | High | High | High | High | High | High | 0.000814 | 0.007788 | 2.93 |
| FALSE | High | SAPLPSATA | [K].SAPLPS | 1xPhospho | G3X940 1x | 0.000229 | 0 | 1 | 3 | 1 | G3X940 | Histone acı | G3X940 [1 | 0 | 2134.064 | 0 | 1.34 | 0.000979 | 5.9 | 51.8 | 148.2 | 10.25 | 12.37 | | High | High | High | High | High | High | High | High | High | High | High | 0 | 4.57E-05 | 2.77 |
| FALSE | High | SAPIQEEQI | [K].SAPIQE | 1xPhospho | Q8K1N4 1x | 3.8E-08 | 0 | 1 | 2 | 2 | Q8K1N4 | Spermatog | Q8K1N4 [1 | 0 | 2039.01 | 0 | -0.05 | 0.896758 | 15.2 | 100.1 | 99.9 | 6.29 | 15.24 | | High | High | High | High | High | High | High | High | High | High | High | 0 | 2.57E-09 | 3.47 |
| FALSE | High | SAPLQSLTT | [K].SAPLQS | 1xPhospho | B1AQJ2 1x | 2.68E-09 | 0 | 1 | 2 | 5 | B1AQJ2 | Ubiquitin c | B1AQJ2 [5 | 0 | 2656.266 | 0 | 0.03 | 0.955863 | 9.78 | 101.3 | 98.7 | 3.9 | 6.9 | | High | High | High | High | High | High | High | High | High | High | High | 0 | 7.76E-09 | 4.4 |
| FALSE | High | SAPLQSLTT | [K].SAPLQS | 2xPhospho | B1AQJ2 2x | 0.000916 | 0 | 1 | 2 | 1 | B1AQJ2 | Ubiquitin c | B1AQJ2 [5 | 0 | 2736.233 | 0 | 0.66 | 0.002315 | 2.35 | 72.6 | 127.4 | 9.65 | 12.39 | | High | High | High | High | High | High | High | High | High | High | High | 0 | 0.000215 | 3.53 |
| FALSE | High | SAPNEPEA | [R].SAPNEI | 1xPhospho | Q5SS00 1x | 1.1E-06 | 0 | 1 | 1 | 2 | Q5SS00 | DBF4-type | Q5SS00 [6 | 0 | 3701.784 | 0 | -1.45 | 0.000426 | 1.55 | 147 | 53 | 5 | 9.42 | | High | High | High | High | High | High | High | High | High | High | High | 0 | 1.13E-07 | 4.29 |
| FALSE | High | SAPNEPEA | [R].SAPNEI | 2xPhospho | Q5SS00 2x | 1.72E-08 | 0 | 1 | 1 | 2 | Q5SS00 | DBF4-type | Q5SS00 [6 | 0 | 3781.75 | 0 | -0.96 | 0.003855 | 5.75 | 131.2 | 68.8 | 12.04 | 5.1 | | High | High | High | High | High | High | High | High | High | High | High | 0 | 1.05E-09 | 6.13 |
| FALSE | High | SAPPAGDE | [K].SAPPAC | 2xPhospho | Q6NS82 2x | 1.03E-07 | 0 | 1 | 4 | 3 | Q6NS82 | Reticuloph | Q6NS82 [2 | 0 | 3746.738 | 0 | 0.08 | 0.289419 | 5.93 | 96.6 | 103.4 | 3.62 | 7.66 | | High | High | High | High | High | High | High | High | High | High | High | 0 | 2.16E-05 | 4.42 |
| FALSE | High | SAPSSPTLD | [K].SAPSSP | 1xCarbami | Q9ERR1 1x | 0.011146 | 0.000772 | 1 | 2 | 3 | Q9ERR1 | Nuclear dis | Q9ERR1 [1 | 0 | 1829.876 | 0 | 1.35 | 0.000567 | 5.64 | 54.1 | 145.9 | 3.78 | 8.51 | | High | High | High | High | High | High | High | High | High | High | High | 0.000399 | 0.003554 | 2.83 |
| FALSE | High | SAPDGGDE | [R].SAPDG | 1xPhospho | G3UYW1 1 | 1.01E-05 | 0 | 1 | 3 | 3 | G3UYW1 | WD repeat | G3UYW1 [ | 0 | 3686.648 | 0 | 0.72 | 0.002065 | 14.78 | 75.4 | 124.6 | 9.95 | 6.54 | | High | High | High | High | High | High | High | High | High | High | High | 0 | 3.22E-06 | 5.8 |
| FALSE | High | SAPSSPTLD | [K].SAPSSP | 1xCarbami | Q9ERR1 2x | 0.00083 | 0 | 1 | 2 | 2 | Q9ERR1 | Nuclear dis | Q9ERR1 [1 | 0 | 1909.842 | 0 | 2.51 | 0.000246 | 19.87 | 31.9 | 168.1 | 14.75 | 4.99 | | High | High | High | High | High | High | High | High | High | High | High | 0.000275 | 0.002139 | 2.82 |
| FALSE | High | SAPTSPTAT | [K].SAPTSP | 2xPhospho | Q69Z38 2x | 0.042503 | 0.003197 | 1 | 1 | 1 | Q69Z38 | Pseudopoc | Q69Z38 [5 | 0 | 1979.949 | 0 | -0.54 | 0.006027 | 2.26 | 118.8 | 81.2 | 11.24 | 14.58 | | High | High | High | High | High | High | High | High | High | High | High | 0.001639 | 0.016 | 1.92 |
| FALSE | High | SAPTTLYLV | [R].SAPTTL | 1xPhospho | Q9ERQ3 1x | 0.00359 | 0.00021 | 1 | 2 | 2 | Q9ERQ3 | Zinc finger | Q9ERQ3 [4 | 0 | 3474.807 | 0 | -0.84 | 0.019891 | 7.71 | 128.3 | 71.7 | 8.89 | 26.67 | | High | High | High | High | High | High | High | High | High | High | High | 0.000327 | 0.002587 | 2.46 |
| FALSE | High | SAPVINNG | [K].SAPVIN | 1xPhospho | M1VMF6 1 | 0.017041 | 0.001204 | 1 | 8 | 1 | M1VMF6 | Metastasis | M1VMF6 [ | 0 | 2006.109 | 0 | -0.07 | 0.088682 | 1.36 | 102.2 | 97.8 | 6.56 | 4.71 | | High | High | High | High | High | High | High | High | High | High | High | 0.000638 | 0.005676 | 2.85 |
| FALSE | High | SAPVINNG | [K].SAPVIN | 1xDeamida | M1VMF6 1 | 5.31E-05 | 0 | 1 | 8 | 5 | M1VMF6 | Metastasis | M1VMF6 [ | 0 | 2007.093 | 0 | 0.07 | 0.318042 | 8.1 | 96.5 | 103.5 | 12.12 | 3.8 | | High | High | High | High | High | High | High | High | High | High | High | 0 | 8.82E-06 | 3.02 |
| FALSE | High | SAPVNGEA | [R].SAPVN | 1xPhospho | Q3UMU9-3 | 6.61E-06 | 0 | 1 | 4 | 2 | Q3UMU9-3 | Isoform 3 c | Q3UMU9-3 | 0 | 1726.878 | 0 | -0.57 | 0.005406 | 6.94 | 119.6 | 80.4 | 6.73 | 6.7 | | High | High | High | High | High | High | High | High | High | High | High | 0 | 8.51E-07 | 3.5 |
| FALSE | High | SAPVNGEA | [R].SAPVN | 1xDeamida | Q3UMU9-3 | 3.01E-05 | 0 | 1 | 4 | 3 | Q3UMU9-3 | Isoform 3 c | Q3UMU9-3 | 0 | 1727.862 | 0 | -0.57 | 0.002076 | 3.57 | 117.3 | 82.7 | 6.99 | 5.7 | | High | High | High | High | High | High | High | High | High | High | High | 0 | 1.21E-05 | 3.16 |
| FALSE | High | SAPYEFPEE | [R].SAPYEF | 1xPhospho | A2AE27 1x | 0.006363 | 0.00062 | 1 | 2 | 1 | A2AE27 | AMP deam | A2AE27 [8 | 0 | 2459.122 | 0 | 0.98 | 0.003738 | 26.29 | 67.5 | 132.5 | 14.58 | 12.75 | | High | High | High | High | High | High | High | High | High | High | High | 0.000275 | 0.001896 | 2.78 |
| FALSE | High | SAQDPPAP | [K].SAQDP | 1xPhospho | Q8VIG1 1x | 2.1E-06 | 0 | 1 | 1 | 1 | Q8VIG1 | RE1-silenci | Q8VIG1 [8 | 0 | 1913.977 | 0 | -1.22 | 0.000219 | 0.35 | 139.9 | 60.1 | 4.68 | 3.89 | | High | High | High | High | High | High | High | High | High | High | High | 0 | 2.34E-07 | 1.91 |
| FALSE | High | SAQETPESS | [K].SAQETI | 1xPhospho | Q9R0L6-2 | 7.29E-06 | 0 | 1 | 3 | 2 | Q9R0L6-2 | Isoform 2 c | Q9R0L6-2 | 0 | 4174.959 | 0 | -0.86 | 0.00582 | 1.86 | 127.1 | 72.9 | 15.97 | 8.17 | | High | High | High | High | High | High | High | High | High | High | High | 0 | 9.43E-07 | 3.49 |
| FALSE | High | SAQETPESS | [K].SAQETI | 1xDeamida | Q9R0L6-2 | 2.71E-07 | 0 | 1 | 3 | 7 | Q9R0L6-2 | Isoform 2 c | Q9R0L6-2 | 0 | 4175.943 | 0 | -0.17 | 0.262937 | 12.6 | 107.2 | 92.8 | 9.47 | 16.45 | | High | High | High | High | High | High | High | High | High | High | High | 0 | 3.49E-08 | 5.77 |
| FALSE | High | SAQETPESS | [K].SAQETI | 2xPhospho | Q9R0L6-2 | 0.010758 | 0.000772 | 1 | 3 | 2 | Q9R0L6-2 | Isoform 2 c | Q9R0L6-2 | 0 | 4254.926 | 0 | 0.15 | 0.1868 | 10.48 | 95.5 | 104.5 | 9.15 | 12.55 | | High | High | High | High | High | High | High | High | High | High | High | 0.000399 | 0.003387 | 3.23 |
| FALSE | High | SAQGTGFE | [K].SAQGT | 1xPhospho | Q8VDZ4 1x | 0.000244 | 0 | 1 | 2 | 1 | Q8VDZ4 | Palmitoyltr | Q8VDZ4 [4 | 0 | 2001 | 0 | 1.33 | 0.004795 | 13.61 | 63.8 | 136.2 | 13.44 | 13.07 | | High | High | High | High | High | High | High | High | High | High | High | 0 | 4.86E-05 | 3.82 |
| FALSE | High | SASACQILII | [R].SASACC | 2xCarbami | Q7TSH4 1x | 1.77E-09 | 0 | 1 | 3 | 1 | Q7TSH4 | Centriolar | Q7TSH4 [3 | 0 | 2824.414 | 0 | -0.12 | 0.137398 | 6.04 | 104.2 | 95.8 | 5.32 | 9.81 | | High | High | High | High | High | High | High | High | High | High | High | 0 | 8.18E-11 | 4.69 |
| FALSE | High | SAPTSPTAT | [K].SAPTSP | 1xPhospho | Q69Z38 1x | 9.89E-05 | 0 | 1 | 1 | 5 | Q69Z38 | Pseudopoc | Q69Z38 [5 | 0 | 1899.983 | 0 | 0.05 | 0.391627 | 5.02 | 97.4 | 102.6 | 7.89 | 6.5 | | High | High | High | High | High | High | High | High | High | High | High | 0 | 0.000307 | 3.14 |
| FALSE | High | SADTPSAPS | [R].SADTPS | 1xPhospho | Q3U2Z5 1x | 0.002671 | 0.000109 | 1 | 3 | 3 | Q3U2Z5 | Putative ur | Q3U2Z5 [6 | 0 | 3689.001 | 0 | 1.81 | 0.00034 | 5.71 | 44.3 | 155.7 | 15.75 | 4.23 | | High | High | High | High | High | High | High | High | High | High | High | 0.001011 | 0.01083 | 3.34 |
| FALSE | High | SADSVEEPI | [K].SADSVE | 1xPhospho | Q5DTX6 1x | 0.000426 | 0 | 1 | 1 | 2 | Q5DTX6 | Junctional | Q5DTX6 [1 | 0 | 1824.903 | 0 | 2.09 | 0.000177 | 11.15 | 38 | 162 | 9.79 | 3.44 | | High | High | High | High | High | High | High | High | High | High | High | 0 | 9.16E-05 | 3.1 |
| FALSE | High | SADSPSFR | [R].SADSPS | 1xPhospho | Q3UVM5 1 | 0.001799 | 5.66E-05 | 1 | 3 | 1 | Q3UVM5 | Putative ur | Q3UVM5 [ | 0 | 1175.53 | 0 | 0.73 | 0.006209 | 21.33 | 76.2 | 123.8 | 10.64 | 7.22 | | High | High | High | High | High | High | High | High | High | High | High | 2.58E-05 | 0.000462 | 1.66 |
| FALSE | High | RVESPVLPI | [K].RVESPV | 1xPhospho | P70340 1x | 0.056248 | 0.004124 | 1 | 6 | 1 | P70340 | Mothers ag | P70340 [12 | 1 | 1867.077 | 0 | 0.06 | 0.088931 | 5.26 | 98.9 | 101.1 | 4.68 | 2.64 | | High | High | High | High | High | High | High | High | High | High | High | 0.002117 | 0.02209 | 1.41 |
| FALSE | High | RVPGQPLA | [R].RVPGQ | 1xCarbami | Q542Q3 1x | 3.56E-08 | 0 | 1 | 1 | 5 | Q542Q3 | G two S ph | Q542Q3 [5 | 1 | 2944.537 | 0 | -0.11 | 0.973837 | 15.32 | 103.5 | 96.5 | 14.94 | 5.97 | | High | High | High | High | High | High | High | High | High | High | High | 0 | 2.4E-09 | 4.39 |
| FALSE | High | RVPLSPLSL | [R].RVPLSP | 1xDeamida | Q3TDX5 1x | 0.017192 | 0.001204 | 1 | 1 | 1 | Q3TDX5 | Putative ur | Q3TDX5 [2 | 1 | 2026.105 | 1 | 0.8 | 0.011293 | 36.23 | 70.2 | 129.8 | 14.93 | 11.67 | | High | High | High | High | High | High | High | High | High | High | High | 0.000638 | 0.005742 | 1.93 |
| FALSE | High | RVPSPLPK | [K].RVPSPL | 1xPhospho | Q8VDI7 1x | 0.097743 | 0.007804 | 1 | 2 | 1 | Q8VDI7 | Ubiquitin-a | Q8VDI7 [9 | 1 | 1431.849 | 0 | -1 | 0.001132 | 5.47 | 133.4 | 66.6 | 3.43 | 16.97 | | High | High | High | High | High | High | High | High | High | High | High | 0.00395 | 0.04197 | 1.63 |
| FALSE | High | RVPSPTPVI | [K].RVPSPT | 1xPhospho | Q8BTI8 1xI | 0.000837 | 0 | 1 | 2 | 4 | Q8BTI8 | Serine/argi | Q8BTI8 [25 | 1 | 1615.934 | 0 | -0.67 | 0.000334 | 2.33 | 121.4 | 78.6 | 5.08 | 6.51 | | High | High | High | High | High | High | High | High | High | High | High | 0 | 0.000195 | 2.08 |
| FALSE | High | RVPSYLDSK | [K].RVPSYL | 1xPhospho | A0A1D5RM | 0.041781 | 0.003126 | 1 | 2 | 1 | A0A1D5RM | Alstrom syı | A0A1D5RM | 1 | 1602.866 | 0 | 0.16 | 0.325127 | 23.54 | 95 | 105 | 11.15 | 15.33 | | High | High | High | High | High | High | High | High | High | High | High | 0.001609 | 0.01563 | 2.28 |
| FALSE | High | RVSADGQF | [R].RVSAD | 1xPhospho | Q8R516 1x | 0.002818 | 0.000109 | 1 | 5 | 1 | Q8R516 | E3 ubiquiti | Q8R516 [2 | 1 | 1569.774 | 0 | -0.84 | 0.000626 | 1.79 | 128.3 | 71.7 | 5.8 | 6.52 | | High | High | High | High | High | High | High | High | High | High | High | 4.94E-05 | 0.000759 | 2.44 |
| FALSE | High | RVSAILPGA | [R].RVSAIL | 1xPhospho | Q3UW53 1 | 3.16E-07 | 0 | 1 | 1 | 2 | Q3UW53 | Protein Nit | Q3UW53 [ | 1 | 3403.719 | 0 | 0.36 | 0.00227 | 9.09 | 87.8 | 112.2 | 9.45 | 2.54 | | High | High | High | High | High | High | High | High | High | High | High | 0 | 8.59E-06 | 3.69 |
| FALSE | High | RVSAILPGA | [R].RVSAIL | 1xDeamida | Q3UW53 1 | 9.03E-05 | 0 | 1 | 1 | 1 | Q3UW53 | Protein Nit | Q3UW53 [ | 1 | 3404.703 | 0 | 0.43 | 0.002563 | 1.57 | 87.2 | 112.8 | 13.5 | 10.99 | | High | High | High | High | High | High | High | High | High | High | High | 0 | 0.000016 | 3.21 |
| FALSE | High | RVSEVFDC | [R].RVSEVI | 1xCarbami | Q8BU30 1x | 0.043236 | 0.003228 | 1 | 2 | 1 | Q8BU30 | Isoleucine- | Q8BU30 [5 | 1 | 3820.758 | 0 | 1.63 | 0.000749 | 15.97 | 49.8 | 150.2 | 12.64 | 10.65 | | High | High | High | High | High | High | High | High | High | High | High | 0.001651 | 0.01638 | 3.25 |
| FALSE | High | RVSFADPIY | [R].RVSFAD | 1xPhospho | Q6PR54-3 | 2.33E-07 | 0 | 2 | 4 | 2 | Q6PR54-3; | Isoform 3 c | Q6PR54-3 | 1 | 2431.186 | 0 | -1.34 | 0.002019 | 1.62 | 143.3 | 56.7 | 13.84 | 7.59 | Shared | High | High | High | High | High | High | High | High | High | High | High | 0 | 1.97E-08 | 3.55 |
| FALSE | High | RVSFADTIK | [R].RVSFAD | 1xPhospho | A3KGI3 1xI | 0.002115 | 5.66E-05 | 1 | 4 | 1 | A3KGI3 | Kinetochor | A3KGI3 [58 | 1 | 1574.871 | 0 | -0.97 | 0.000191 | 1.01 | 132.3 | 67.7 | 4.44 | 4.07 | | High | High | High | High | High | High | High | High | High | High | High | 2.58E-05 | 0.000553 | 1.8 |
| FALSE | High | RVSFNEAA | [R].RVSFNI | 1xPhospho | E9PW07 1x | 0.057203 | 0.004227 | 1 | 2 | 1 | E9PW07 | Voltage-de | E9PW07 [1 | 1 | 1962.964 | 0 | 0.63 | 0.023917 | 19.04 | 80.3 | 119.7 | 12.43 | 13.38 | | High | High | High | High | High | High | High | High | High | High | High | 0.002159 | 0.02247 | 2.16 |
| FALSE | High | RVSGDGAC | [R].RVSGD | 1xPhospho | P81122 1xI | 0.105058 | 0.008518 | 1 | 2 | 1 | P81122 | Insulin rece | P81122 [5 | 1 | 1597.753 | 0 | -0.69 | 0.003844 | 6.37 | 125.5 | 74.5 | 6.57 | 21.4 | | High | High | High | High | High | High | High | High | High | High | High | 0.004291 | 0.04586 | 2.62 |
| FALSE | High | RVSGSATPI | [R].RVSGSA | 1xPhospho | Q6P9R1 1x | 3.26E-05 | 0 | 1 | 1 | 3 | Q6P9R1 | ATP-depen | Q6P9R1 [7 | 1 | 1737.848 | 0 | -0.56 | 0.001039 | 4.04 | 119.2 | 80.8 | 3.55 | 3.52 | | High | High | High | High | High | High | High | High | High | High | High | 0 | 5.09E-06 | 3.06 |
| FALSE | High | RVSGSATPI | [R].RVSGSA | 1xDeamida | Q6P9R1 1x | 0.053473 | 0.00384 | 1 | 1 | 1 | Q6P9R1 | ATP-depen | Q6P9R1 [7 | 1 | 1738.832 | 0 | -0.12 | 0.124873 | 8.96 | 103.3 | 96.7 | 10.54 | 7.45 | | High | High | High | High | High | High | High | High | High | High | High | 0.001999 | 0.0208 | 1.74 |
| FALSE | High | RVSHSPPPI | [R].RVSHSI | 2xPhospho | Q52KI8 2xI | 0.052576 | 0.00384 | 1 | 7 | 3 | Q52KI8 | Serine/argi | Q52KI8 [65 | 1 | 1622.822 | 0 | -0.38 | 0.00816 | 5.58 | 113.8 | 86.2 | 4.64 | 5.72 | | High | High | High | High | High | High | High | High | High | High | High | 0.003659 | 0.03914 | 2 |
| FALSE | High | RVSLGNEG | [R].RVSLGI | 1xPhospho | Q9D7Y9 1x | 0.004333 | 0.00036 | 1 | 2 | 1 | Q9D7Y9 | Protein SLX | Q9D7Y9 [3 | 2 | 2321.145 | 0 | 0.14 | 0.514795 | 10.96 | 95.1 | 104.9 | 9.91 | 12.11 | | High | High | High | High | High | High | High | High | High | High | High | 0.000161 | 0.001229 | 2.55 |
| FALSE | High | RVSLHCLLD | [R].RVSLHC | 1xCarbami | Q8BH65 1x | 0.01319 | 0.000889 | 1 | 2 | 1 | Q8BH65 | Protein DE | Q8BH65 [1 | 1 | 2170.113 | 0 | -0.14 | 0.053473 | 6.13 | 102.8 | 97.2 | 7.71 | 5.69 | | High | High | High | High | High | High | High | High | High | High | High | 0.000467 | 0.004256 | 1.7 |
| FALSE | High | RVSISEGDI | [K].RVSISE | 1xPhospho | P35550 1xI | 0.002355 | 0.000109 | 1 | 3 | 2 | P35550 | rRNA 2'-O- | P35550 [12 | 1 | 1643.841 | 0 | 1.4 | 0.000452 | 9.42 | 56 | 144 | 2.12 | 7.43 | | High | High | High | High | High | High | High | High | High | High | High | 0.001468 | 0.01478 | 2.37 |
| FALSE | High | RVSLSEIGF | [R].RVSLSE | 1xPhospho | Q3UPI3 1xI | 0.007605 | 0.000664 | 1 | 4 | 1 | Q3UPI3 | Putative ur | Q3UPI3 [1 | 1 | 1730.961 | 0 | -0.66 | 0.001894 | 3.26 | 124.9 | 75.1 | 6.86 | 7.92 | | High | High | High | High | High | High | High | High | High | High | High | 0.000294 | 0.002311 | 2.96 |
| FALSE | High | RVSLSSQLI | [K].RVSLSS | 1xPhospho | Z4YKQ9 1x | 0.0006 | 0 | 1 | 4 | 1 | Z4YKQ9 | ATPase fan | Z4YKQ9 [1 | 1 | 1898.051 | 0 | -0.54 | 0.001434 | 1.04 | 120 | 80 | 2.68 | 8.06 | | High | High | High | High | High | High | High | High | High | High | High | 0 | 0.000135 | 1.77 |
| FALSE | High | RVSLVGAD | [K].RVSLVC | 1xPhospho | Q80U72-4 | 0.000386 | 0 | 2 | 6 | 1 | Q80U72-4; | Isoform 4 c | Q80U72-4 | 1 | 1509.799 | 0 | 0.64 | 0.008201 | 22.59 | 76.3 | 123.7 | 5.08 | 14.78 | Shared | High | High | High | High | High | High | High | High | High | High | High | 0 | 8.22E-05 | 1.65 |
| FALSE | High | RVSLVTEEA | [R].RVSLVT | 1xCarbami | P59438 1xI | 1.1E-07 | 0 | 1 | 8 | 2 | P59438 | Hermansky | P59438 [6 | 1 | 3296.567 | 0 | 1.11 | 0.000784 | 9.48 | 61.1 | 138.9 | 9.01 | 6.27 | | High | High | High | High | High | High | High | High | High | High | High | 0 | 8.53E-09 | 5.2 |
| FALSE | High | RVSMLSEE | [R].RVSML | 1xPhospho | Q9Z206 1x | 6.3E-05 | 0 | 1 | 1 | 1 | Q9Z206 | Neuroepith | Q9Z206 [1 | 1 | 2845.375 | 0 | -0.77 | 0.001097 | 1.5 | 127.5 | 72.5 | 7.54 | 16.3 | | High | High | High | High | High | High | High | High | High | High | High | 0 | 1.07E-05 | 3.59 |
| FALSE | High | RVSPEEAIK | [R].RVSPEE | 1xPhospho | Q6ZWX6 1 | 0.004654 | 0.000456 | 1 | 1 | 1 | Q6ZWX6 | Eukaryotic | Q6ZWX6 [ | 1 | 1566.866 | 0 | -0.13 | 0.156273 | 6.73 | 104.7 | 95.3 | 3.79 | 7.78 | | High | High | High | High | High | High | High | High | High | High | High | 0.000204 | 0.001331 | 2.23 |
| FALSE | High | RVSPLNLSS | [R].RVSPLN | 2xPhospho | A0A0R4J1V | 0.063258 | 0.004664 | 1 | 4 | 1 | A0A0R4J1V | CDC23 (Cel | A0A0R4J1V | 1 | 1658.812 | 0 | -0.31 | 0.010126 | 7.98 | 109.6 | 90.4 | 5.43 | 5.65 | | High | High | High | High | High | High | High | High | High | High | High | 0.002403 | 0.02527 | 2.6 |
| FALSE | High | RVESEESGI | [R].RVESEE | 1xPhospho | Q3UM45 1 | 3.94E-05 | 0 | 1 | 2 | 1 | Q3UM45 | Protein pho | Q3UM45 [ | 2 | 2346.179 | 0 | -1.24 | 0.001221 | 1.26 | 142.3 | 57.7 | 10.35 | 15.47 | | High | High | High | High | High | High | High | High | High | High | High | 0 | 6.33E-06 | 1.67 |
| FALSE | High | RVEAATAY | [R].RVEAAT | 1xPhospho | Q8BMD7 1 | 7.51E-07 | 0 | 1 | 4 | 1 | Q8BMD7 | MORC fam | Q8BMD7 [ | 1 | 2372.154 | 0 | -0.24 | 0.136333 | 16.35 | 108.3 | 91.7 | 12.12 | 6.77 | | High | High | High | High | High | High | High | High | High | High | High | 0 | 7.34E-08 | 3.67 |
| FALSE | High | RVDQQSVE | [K].RVDQC | 2xPhospho | A0A140LI3 | 8.82E-06 | 0 | 1 | 1 | 1 | A0A140LI3 | Zinc finger | A0A140LI3 | 1 | 3577.598 | 0 | -0.01 | 0.510889 | 6.6 | 101.6 | 98.4 | 12.99 | 12.73 | | High | High | High | High | High | High | High | High | High | High | High | 0 | 1.18E-06 | 5.43 |
| FALSE | High | RVDQQSVE | [K].RVDQC | 1xPhospho | A0A140LI3 | 0.003342 | 0.00016 | 1 | 1 | 4 | A0A140LI3 | Zinc finger | A0A140LI3 | 1 | 3497.632 | 0 | -0.38 | 0.005771 | 3.44 | 115 | 85 | 6.85 | 5.14 | | High | High | High | High | High | High | High | High | High | High | High | 0.000161 | 0.001233 | 3.18 |
| FALSE | High | RTPTMPQE | [R].RTPTM | 1xPhospho | Q14AX6-3 | 6.07E-05 | 0 | 1 | 1 | 2 | Q14AX6-3 | Isoform 3 c | Q14AX6-3 | 1 | 2026.008 | 0 | -0.37 | 0.042323 | 0.64 | 114.2 | 85.8 | 7.17 | 20.59 | | High | High | High | High | High | High | High | High | High | High | High | 0 | 1.02E-05 | 3.37 |
| FALSE | High | RTPTMPQE | [R].RTPTM | 1xPhospho | Q14AX6-3 | 0.000174 | 0 | 1 | 1 | 4 | Q14AX6-3 | Isoform 3 c | Q14AX6-3 | 1 | 2042.003 | 0 | -0.42 | 0.123314 | 3.13 | 110.9 | 89.1 | 8.82 | 16.83 | | High | High | High | High | High | High | High | High | High | High | High | 0.000275 | 0.001653 | 2.83 |
| FALSE | High | RTPTMPQE | [R].RTPTM | 1xPhospho | Q14AX6-2 | 0.097743 | 0.007804 | 1 | 1 | 1 | Q14AX6-2 | Isoform 2 c | Q14AX6-2 | 1 | 1969.982 | 0 | -0.72 | 0.00552 | 6.21 | 123.2 | 76.8 | 10.96 | 9.71 | | High | High | High | High | High | High | High | High | High | High | High | 0.00395 | 0.04192 | 1.61 |
| FALSE | High | RTPVSYQN | [R].RTPVSY | 1xPhospho | Q5RIM6 1x | 0.042503 | 0.003197 | 1 | 10 | 1 | Q5RIM6 | Nuclear rec | Q5RIM6 [1 | 1 | 1730.879 | 0 | 0.38 | 0.029532 | 8.72 | 84.1 | 115.9 | 11.91 | 8.13 | | High | High | High | High | High | High | High | High | High | High | High | 0.001639 | 0.01599 | 2.56 |
| FALSE | High | RTSAETAD | [R].RTSAET | 1xPhospho | Q9D2E2 1x | 6.53E-05 | 0 | 1 | 1 | 7 | Q9D2E2 | Target of E | Q9D2E2 [4 | 1 | 3515.733 | 0 | 0.37 | 0.032234 | 13.19 | 88 | 112 | 6.62 | 11.1 | | High | High | High | High | High | High | High | High | High | High | High | 0 | 1.94E-05 | 4.85 |
| FALSE | High | RTSAQNCT | [R].RTSAQI | 1xCarbami | Q921S7 1x | 0.000248 | 0 | 1 | 1 | 4 | Q921S7 | 39S riboso | Q921S7 [2 | 1 | 2088.985 | 0 | 2.04 | 0.000388 | 6.54 | 43.4 | 156.6 | 11.61 | 8.45 | | High | High | High | High | High | High | High | High | High | High | High | 0 | 8.14E-05 | 3.2 |
| FALSE | High | RTSEDTSSC | [K].RTSEDT | 1xPhospho | Q3UYQ4 1 | 0.000442 | 0 | 1 | 2 | 3 | Q3UYQ4 | Putative ur | Q3UYQ4 [4 | 1 | 1886.926 | 0 | -0.3 | 0.09874 | 15.3 | 111.9 | 88.1 | 19.39 | 15.55 | | High | High | High | High | High | High | High | High | High | High | High | 0 | 9.49E-05 | 1.82 |
| FALSE | High | RTSEEDTPI | [R].RTSEED | 1xPhospho | Q9D8Y8 1x | 0.014797 | 0.001049 | 1 | 3 | 2 | Q9D8Y8 | Inhibitor of | Q9D8Y8 [1 | 1 | 1600.798 | 0 | -1.26 | 0.000158 | 0.29 | 140.8 | 59.2 | 4.63 | 7.25 | | High | High | High | High | High | High | High | High | High | High | High | 0.001151 | 0.01197 | 2.66 |
| FALSE | High | RTSLPAPGI | [R].RTSLPA | 1xPhospho | D3YTT6 1xI | 0.012959 | 0.000889 | 1 | 6 | 2 | D3YTT6 | Oxysterol-l | D3YTT6 [5 | 1 | 2548.349 | 0 | 1.21 | 0.000429 | 10.17 | 61.1 | 138.9 | 12.29 | 6.41 | | High | High | High | High | High | High | High | High | High | High | High | 0.000449 | 0.004199 | 2.48 |
| FALSE | High | RTSMGGT | [R].RTSMG | 1xPhospho | Q02248 1x | 1.33E-05 | 0 | 1 | 4 | 3 | Q02248 | Catenin be | Q02248 [5 | 1 | 2090.005 | 0 | -0.66 | 0.000397 | 2.65 | 122 | 78 | 2.19 | 3.66 | | High | High | High | High | High | High | High | High | High | High | High | 0 | 1.12E-05 | 4.46 |
| FALSE | High | RTSMGGT | [R].RTSMG | 1xPhospho | Q02248 1x | 1.86E-05 | 0 | 1 | 4 | 23 | Q02248 | Catenin be | Q02248 [5 | 1 | 2106 | 0 | -0.47 | 0.007032 | 4.13 | 116.3 | 83.7 | 4.97 | 6.21 | | High | High | High | High | High | High | High | High | High | High | High | 0 | 2.72E-06 | 4.23 |
| FALSE | High | RTSPAPDIL | [R].RTSPAF | 1xCarbami | S0DHL8 1xI | 0.010109 | 0.000736 | 1 | 4 | 2 | S0DHL8 | Capping pr | S0DHL8 [1 | 1 | 2500.247 | 0 | -1.89 | 0.000704 | 0.38 | 153.5 | 46.5 | 8.39 | 12.04 | | High | High | High | High | High | High | High | High | High | High | High | 0.000344 | 0.003186 | 2.81 |
| FALSE | High | RTSPTAYCI | [K].RTSPTA | 2xCarbami | E9Q2H1 1x | 0.001337 | 0 | 1 | 3 | 2 | E9Q2H1 | E3 ubiquiti | E9Q2H1 [1 | 1 | 2211.997 | 0 | -0.01 | 0.605505 | 9.39 | 98 | 102 | 9.54 | 6.25 | | High | High | High | High | High | High | High | High | High | High | High | 2.58E-05 | 0.00054 | 3.06 |
| FALSE | High | RVSQEANL | [R].RVSQE/ | 1xPhospho | E9Q2I4 1xF | 5.07E-08 | 0 | 1 | 3 | 1 | E9Q2I4 | ELM2 and | E9Q2I4 [45 | 1 | 2109.183 | 0 | 0.07 | 0.192683 | 8.16 | 95.9 | 104.1 | 7.99 | 6.53 | | High | High | High | High | High | High | High | High | High | High | High | 0 | 3.55E-09 | 3.74 |
| FALSE | High | RTSSAVWN | [K].RTSSAV | 1xPhospho | Q9WU62 1 | 0.012845 | 0.000889 | 1 | 2 | 1 | Q9WU62 | Inner centı | Q9WU62 [ | 1 | 1981.067 | 0 | -0.7 | 0.002809 | 6.45 | 119.4 | 80.6 | 6.4 | 15.5 | | High | High | High | High | High | High | High | High | High | High | High | 0.000449 | 0.004153 | 1.91 |
| FALSE | High | RTSSEDNLY | [K].RTSSED | 2xPhospho | Q9QXS1-3 | 0.106748 | 0.008634 | 1 | 2 | 2 | Q9QXS1-3 | Isoform PL | Q9QXS1-3 | 1 | 2025.961 | 0 | -0.11 | 0.556553 | 2.06 | 103.6 | 96.4 | 10.9 | 16.3 | | High | High | High | High | High | High | High | High | High | High | High | 0.004327 | 0.04659 | 1.79 |
| FALSE | High | RTSSEQAV | [R].RTSSEC | 1xPhospho | Q8K120 1x | 0.000558 | 0 | 1 | 2 | 2 | Q8K120 | Nuclear fac | Q8K120 [3 | 1 | 1623.842 | 0 | -1.17 | 0.002618 | 5.79 | 144.9 | 55.1 | 10.39 | 16.74 | | High | High | High | High | High | High | High | High | High | High | High | 0 | 0.000335 | 2.92 |
| FALSE | High | RTSSQYVA | [R].RTSSQY | 1xPhospho | B2RWU4 1 | 0.08447 | 0.006345 | 2 | 6 | 2 | B2RWU4; ( | Cordon-ble | B2RWU4 [ | 1 | 1920.036 | 0 | -0.9 | 0.000738 | 2.33 | 131.8 | 68.2 | 5.4 | 11.51 | Shared | High | High | High | High | High | High | High | High | High | High | High | 0.003265 | 0.03534 | 2.6 |
| FALSE | High | RTSTPVILE | [R].RTSTPV | 1xPhospho | E9Q6J5-3 1 | 1.64E-08 | 0 | 1 | 3 | 4 | E9Q6J5-3 | Isoform Ne | E9Q6J5-3 [ | 1 | 2312.133 | 0 | 0.43 | 0.001528 | 3.03 | 86.5 | 113.5 | 7.61 | 6.09 | | High | High | High | High | High | High | High | High | High | High | High | 0 | 1E-09 | 5.39 |
| FALSE | High | RTSTPVILE | [R].RTSTPV | 1xDeamida | E9Q6J5-3 1 | 4.43E-05 | 0 | 1 | 3 | 3 | E9Q6J5-3 | Isoform Ne | E9Q6J5-3 [ | 1 | 2313.117 | 0 | -0.15 | 0.047732 | 5.66 | 105.4 | 94.6 | 7.05 | 5.32 | | High | High | High | High | High | High | High | High | High | High | High | 0 | 1.11E-05 | 5.22 |
| FALSE | High | RTSTPVILE | [R].RTSTPV | 2xPhospho | E9Q6J5-3 2 | 0.000505 | 0 | 1 | 3 | 2 | E9Q6J5-3 | Isoform Ne | E9Q6J5-3 [ | 1 | 2392.1 | 0 | 0.92 | 0.002112 | 13.35 | 67.6 | 132.4 | 9.83 | 12.39 | | High | High | High | High | High | High | High | High | High | High | High | 0 | 0.000111 | 3.46 |
| FALSE | High | RTSTQENV | [R].RTSTQE | 1xPhospho | Q8R3I2 1xI | 0.000203 | 0 | 1 | 4 | 6 | Q8R3I2 | Lysophosp | Q8R3I2 [47 | 1 | 2137.117 | 0 | 3.09 | 0.000377 | 17.39 | 21.1 | 178.9 | 11.29 | 8.32 | | High | High | High | High | High | High | High | High | High | High | High | 0 | 4.4E-05 | 3.17 |
| FALSE | High | RTSTVLGTI | [K].RTSTVL | 1xPhospho | F8VPZ5 1xI | 9.36E-05 | 0 | 1 | 2 | 2 | F8VPZ5 | Excision re | F8VPZ5 [1 | 1 | 1712.935 | 0 | -1.25 | 0.002816 | 2.23 | 139.3 | 60.7 | 13.66 | 13.31 | | High | High | High | High | High | High | High | High | High | High | High | 0 | 1.68E-05 | 3.06 |
| FALSE | High | RTSVHGVS | [K].RTSVH | 1xPhospho | H7BX08 1x | 0.005666 | 0.00062 | 1 | 5 | 1 | H7BX08 | Calmodulir | H7BX08 [4 | 1 | 2233.142 | 0 | 0.23 | 0.005573 | 6.14 | 92.3 | 107.7 | 5.64 | 1.81 | | High | High | High | High | High | High | High | High | High | High | High | 0.000275 | 0.001659 | 3.13 |
| FALSE | High | RTSVSSPEC | [R].RTSVSS | 1xPhospho | Q8K382 1x | 0.001147 | 0 | 1 | 2 | 2 | Q8K382 | DENN dom | Q8K382 [5 | 1 | 1940.943 | 0 | 0.51 | 0.002235 | 15.06 | 81.6 | 118.4 | 10.56 | 1.39 | | High | High | High | High | High | High | High | High | High | High | High | 0 | 0.000278 | 2.46 |
| FALSE | High | RTSVSSPEC | [R].RTSVSS | 2xPhospho | Q8K382 2x | 0.022378 | 0.001542 | 1 | 2 | 1 | Q8K382 | DENN dom | Q8K382 [5 | 1 | 2020.909 | 0 | 0.49 | 0.00198 | 1.99 | 77.3 | 122.7 | 11.66 | 5.94 | | High | High | High | High | High | High | High | High | High | High | High | 0.000814 | 0.007719 | 2.31 |
| FALSE | High | RTYSDEAS | [K].RTYSDE | 1xPhospho | Q6ZQ33 1x | 0.000294 | 0 | 1 | 4 | 3 | Q6ZQ33 | MKIAA085 | Q6ZQ33 [3 | 1 | 1634.774 | 0 | 0.09 | 0.223462 | 7.21 | 97 | 103 | 5.67 | 7.14 | | High | High | High | High | High | High | High | High | High | High | High | 0.000859 | 0.008229 | 1.47 |
| FALSE | High | RVDLSDEE | [K].RVDLSD | 1xPhospho | Q9EQ28 1x | 0.000297 | 0 | 1 | 6 | 7 | Q9EQ28 | DNA polym | Q9EQ28 [3 | 1 | 1699.867 | 0 | -0.8 | 0.00021 | 1.58 | 127.7 | 72.3 | 3.61 | 5.75 | | High | High | High | High | High | High | High | High | High | High | High | 0.000275 | 0.001906 | 2.56 |
| FALSE | High | RTSSAVWN | [K].RTSSAV | 2xPhospho | Q9WU62 2 | 0.073462 | 0.005631 | 1 | 2 | 2 | Q9WU62 | Inner centı | Q9WU62 [ | 1 | 2061.034 | 0 | 0.23 | 0.100256 | 5.11 | 92.4 | 107.6 | 8.41 | 10.93 | | High | High | High | High | High | High | High | High | High | High | High | 0.005773 | 0.06343 | 2.09 |
| FALSE | High | RVSSDEEH | [R].RVSSDE | 1xCarbami | F6WJB7 1x | 0.000253 | 0 | 1 | 4 | 4 | F6WJB7 | Ubiquitin c | F6WJB7 [3 | 1 | 2585.232 | 0 | -0.09 | 0.195 | 18.69 | 103.4 | 96.6 | 6.8 | 6.3 | | High | High | High | High | High | High | High | High | High | High | High | 0 | 5.08E-05 | 3.57 |
| FALSE | High | RVSSDEEH | [R].RVSSDE | 1xCarbami | F6WJB7 2x | 7.89E-05 | 0 | 1 | 4 | 1 | F6WJB7 | Ubiquitin c | F6WJB7 [3 | 1 | 2665.198 | 0 | 0.64 | 0.002429 | 11.18 | 76.3 | 123.7 | 5.08 | 7.21 | | High | High | High | High | High | High | High | High | High | High | High | 0 | 1.38E-05 | 3.81 |
| FALSE | High | RVSSDLQH | [R].RVSSDL | 1xPhospho | A0A0R4J0I | 2.86E-06 | 0 | 1 | 2 | 2 | A0A0R4J0I | ETS domai | A0A0R4J0I | 1 | 2357.192 | 0 | 0.17 | 0.933992 | 38.66 | 103.4 | 96.6 | 22.64 | 22.27 | | High | High | High | High | High | High | High | High | High | High | High | 0 | 6.17E-07 | 3.13 |
| FALSE | High | RYSGNMEY | [K].RYSGNI | 1xPhospho | Q8BYR2 1x | 0.005182 | 0.000551 | 1 | 1 | 2 | Q8BYR2 | Serine/thre | Q8BYR2 [2 | 1 | 1799.835 | 0 | 1.2 | 0.003362 | 18.9 | 60.1 | 139.9 | 11.32 | 15.38 | | High | High | High | High | High | High | High | High | High | High | High | 0.000246 | 0.001511 | 2.49 |
| FALSE | High | RYSLIDPAS | [R].RYSLIDI | 1xPhospho | Q9Z0H8 1x | 1.47E-07 | 0 | 1 | 3 | 2 | Q9Z0H8 | CAP-Gly dc | Q9Z0H8 [9 | 1 | 2237.235 | 0 | 0.23 | 0.199134 | 16.58 | 94.7 | 105.3 | 15.73 | 11.28 | | High | High | High | High | High | High | High | High | High | High | High | 0 | 1.18E-08 | 4.15 |
| FALSE | High | RYSPSPPPK | [R].RYSPSP | 1xPhospho | Q52KI8 1xI | 0.000379 | 0 | 1 | 7 | 2 | Q52KI8 | Serine/argi | Q52KI8 [62 | 1 | 1566.845 | 0 | -0.51 | 0.009522 | 8.15 | 119.2 | 80.8 | 9.41 | 6.59 | | High | High | High | High | High | High | High | High | High | High | High | 0 | 8E-05 | 2.19 |
| FALSE | High | RYSPSPPPK | [R].RYSPSP | 2xPhospho | Q52KI8 2xI | 0.011346 | 0.000816 | 1 | 7 | 11 | Q52KI8 | Serine/argi | Q52KI8 [62 | 1 | 1646.811 | 0 | -0.38 | 0.004875 | 5.1 | 112.9 | 87.1 | 3.27 | 5.97 | | High | High | High | High | High | High | High | High | High | High | High | 0.002403 | 0.02583 | 3.34 |
| FALSE | High | RYSSSGTPS | [R].RYSSSG | 1xPhospho | Q8K120 1x | 0.001916 | 5.66E-05 | 1 | 2 | 4 | Q8K120 | Nuclear fac | Q8K120 [2 | 1 | 2019.97 | 0 | -1.86 | 0.000382 | 1.39 | 154.8 | 45.2 | 5.03 | 19.08 | | High | High | High | High | High | High | High | High | High | High | High | 0.000344 | 0.003095 | 3.18 |
| FALSE | High | RYSTSSSSG | [R].RYSTSS | 1xPhospho | Q8CAQ8 1 | 0.000563 | 0 | 1 | 15 | 4 | Q8CAQ8 | MICOS con | Q8CAQ8 [3 | 1 | 1939.989 | 0 | 2.22 | 0.001517 | 36.94 | 37.6 | 162.4 | 40.05 | 12.67 | | High | High | High | High | High | High | High | High | High | High | High | 0 | 0.000125 | 2.48 |
| FALSE | High | RYYSVDDN | [K].RYYSVD | 1xPhospho | B9EIW5 1x | 5.55E-05 | 0 | 1 | 5 | 2 | B9EIW5 | Ncoa7 prot | B9EIW5 [8 | 1 | 1939.932 | 0 | 1.29 | 0.007831 | 16.34 | 58.6 | 141.4 | 10.58 | 20.35 | | High | High | High | High | High | High | High | High | High | High | High | 0 | 9.23E-06 | 2.76 |
| FALSE | High | SAAAATSP/ | [K].SAAAA | 1xPhospho | Q924A2 1x | 5.6E-05 | 0 | 1 | 4 | 2 | Q924A2 | Protein cap | Q924A2 [1 | 0 | 2624.458 | 0 | 0.65 | 0.02781 | 32.68 | 81.1 | 118.9 | 17.02 | 13.36 | | High | High | High | High | High | High | High | High | High | High | High | 0 | 9.38E-06 | 3.75 |
| FALSE | High | SAAAGSPG | [R].SAAAG | 1xPhospho | F8VPV0 1xI | 0.000308 | 0 | 1 | 6 | 3 | F8VPV0 | Pericentrin | F8VPV0 [2 | 0 | 1283.676 | 0 | -0.27 | 0.001345 | 2.08 | 109.5 | 90.5 | 2.94 | 2.12 | | High | High | High | High | High | High | High | High | High | High | High | 0 | 6.38E-05 | 2.08 |
| FALSE | High | SAAATAALI | [R].SAAAT/ | 1xPhospho | Q8K2Z4-2 | 0.000196 | 0 | 1 | 3 | 1 | Q8K2Z4-2 | Isoform 2 c | Q8K2Z4-2 | 0 | 2912.404 | 0 | -0.78 | 0.001883 | 3.14 | 128.2 | 71.8 | 9.49 | 8.46 | | High | High | High | High | High | High | High | High | High | High | High | 0 | 3.84E-05 | 3.7 |
| FALSE | High | SAADDGVS | [K].SAADD | 1xPhospho | Q8JZS6 1xI | 0.010475 | 0.000772 | 1 | 3 | 1 | Q8JZS6 | NEDD4-bin | Q8JZS6 [28 | 0 | 1396.667 | 0 | -0.27 | 0.234212 | 5.49 | 107 | 93 | 7.34 | 11.73 | | High | High | High | High | High | High | High | High | High | High | High | 0.000382 | 0.003297 | 0.96 |
| FALSE | High | SAAEMYGS | [R].SAAEM | 1xPhospho | Q9Z204-2 | 0.001572 | 5.66E-05 | 1 | 3 | 9 | Q9Z204-2 | Isoform C1 | Q9Z204-2 | 0 | 2427.005 | 0 | -0.14 | 0.218463 | 5.83 | 103.7 | 96.3 | 10.34 | 9.8 | | High | High | High | High | High | High | High | High | High | High | High | 4.94E-05 | 0.000695 | 3.46 |
| FALSE | High | SAAEMYGS | [R].SAAEM | 1xPhospho | Q3U6P5 1x | 0.0004 | 0 | 1 | 3 | 4 | Q3U6P5 | Putative ur | Q3U6P5 [1 | 0 | 3738.687 | 0 | -1.25 | 0.001819 | 4.57 | 148.4 | 51.6 | 15.51 | 24.01 | | High | High | High | High | High | High | High | High | High | High | High | 0 | 8.52E-05 | 4.9 |

| | | | | | | | | | | | | | | | | | | | | | | | | | | | | | | | | | | | | | | |
|---|---|---|---|---|---|---|---|---|---|---|---|---|---|---|---|---|---|---|---|---|---|---|---|---|---|---|---|---|---|---|---|---|---|---|---|---|---|---|
| FALSE | High | RYSGDSDS | [R].RYSGDS | 1xPhosphc | Q8JZP9 1xf | 1.75E-06 | 0 | 1 | 1 | 3 | Q8JZP9 | GAS2-like p | Q8JZP9 [35 | 1 | 2399.013 | 0 | 0.34 | 0.184826 | 44.25 | 88.1 | 111.9 | 11.79 | 36.42 | | High | High | High | High | High | High | High | High | High | High | High | 0 | 1.91E-07 | 4.33 |
| FALSE | High | SAAEMYGS | [R].SAAEM | 1xPhosphc | Q3U6P5 1x | 4.62E-07 | 0 | 1 | 3 | 3 | Q3U6P5 | Putative ur | Q3U6P5 [1 | 0 | 3754.682 | 0 | -1.29 | 0.022031 | 4.97 | 137.7 | 62.3 | 16.95 | 24.97 | | High | High | High | High | High | High | High | High | High | High | High | 0 | 5.11E-07 | 5.06 |
| FALSE | High | SAAGSEK | [R].SAAGSI | 1xPhosphc | O55201 1x | 0.027839 | 0.001755 | 1 | 1 | 1 | O55201 | Transcripti | O55201 [3 | 0 | 1187.607 | 0 | 0.42 | 0.021216 | 20.34 | 87.4 | 112.6 | 13.62 | 15.29 | | High | High | High | High | High | High | High | High | High | High | High | 0.000953 | 0.009891 | 1.58 |
| FALSE | High | SAAPAPVS | [K].SAAPAI | 1xCarbami | A0A0U1RP | 0.004824 | 0.000456 | 1 | 7 | 3 | A0A0U1RP | Ataxin-2-lik | A0A0U1RP | 0 | 4753.405 | 0 | 0.09 | 0.76658 | 5.82 | 96.9 | 103.1 | 14.29 | 22.79 | | High | High | High | High | High | High | High | High | High | High | High | 0.000204 | 0.001393 | 4.4 |
| FALSE | High | SAAQAAAC | [K].SAAQA | 1xPhosphc | Q9CYS8 1x | 4.41E-07 | 0 | 2 | 11 | 1 | Q9CYS8; Q | Plasminoge | Q9CYS8 [5 | 0 | 1999.001 | 0 | 0.67 | 0.003084 | 7.29 | 81.1 | 118.9 | 11.38 | 5.99 | Shared | High | High | High | High | High | High | High | High | High | High | High | 0 | 4.04E-08 | 3.54 |
| FALSE | High | SAAQQENS | [R].SAAQQ | 1xPhosphc | Q8BI72 1xf | 0.070485 | 0.005214 | 1 | 1 | 2 | Q8BI72 | CDKN2A-in | Q8BI72 [16 | 0 | 1570.706 | 0 | 0.51 | 0.067331 | 23.12 | 80.4 | 119.6 | 15.77 | 10.9 | | High | High | High | High | High | High | High | High | High | High | High | 0.005374 | 0.05883 | 1.28 |
| FALSE | High | SAASPVVIS | [K].SAASPV | 1xPhosphc | H9KV00 1x | 0.000259 | 0 | 1 | 9 | 2 | H9KV00 | Protein SO | H9KV00 [1 | 0 | 1634.872 | 0 | 0.46 | 0.008602 | 9.11 | 86.5 | 113.5 | 5 | 10.62 | | High | High | High | High | High | High | High | High | High | High | High | 0 | 5.22E-05 | 2.69 |
| FALSE | High | SAASREDL | [R].SAASRE | 1xPhosphc | Q9ERD6 1x | 2.26E-07 | 0 | 1 | 7 | 2 | Q9ERD6 | Ras-specifi | Q9ERD6 [2 | 1 | 2321.108 | 0 | -0.93 | 0.01676 | 6.53 | 133.8 | 66.2 | 4.43 | 31.41 | | High | High | High | High | High | High | High | High | High | High | High | 0 | 1.92E-08 | 2.23 |
| FALSE | High | SAAVSASSI | [R].SAAVSA | 1xPhosphc | P54276 1x | 0.003927 | 0.000311 | 1 | 2 | 1 | P54276 | DNA misma | P54276 [56 | 0 | 1413.682 | 0 | -0.82 | 0.469799 | 5.22 | 126.9 | 73.1 | 98.16 | 85.49 | | High | High | High | High | High | High | High | High | High | High | High | 0.00014 | 0.001107 | 1.44 |
| FALSE | High | SAAVSASSI | [R].SAAVSA | 1xPhosphc | P54276 1xf | 7.65E-07 | 0 | 1 | 2 | 12 | P54276 | DNA misma | P54276 [56 | 0 | 1642.845 | 0 | -0.74 | 0.000349 | 2.15 | 125.3 | 74.7 | 2.91 | 7.07 | | High | High | High | High | High | High | High | High | High | High | High | 0 | 8.05E-08 | 3.61 |
| FALSE | High | SAAVSASSI | [R].SAAVSA | 2xPhosphc | P54276 2xf | 2.01E-06 | 0 | 1 | 2 | 1 | P54276 | DNA misma | P54276 [56 | 0 | 1722.812 | 0 | -0.9 | 0.001852 | 6.46 | 130.1 | 69.9 | 5.94 | 10.02 | | High | High | High | High | High | High | High | High | High | High | High | 0 | 2.23E-07 | 4.03 |
| FALSE | High | SADCSVPSI | [R].SADCSV | 1xCarbami | O70374 1x | 0.006957 | 0.00062 | 1 | 4 | 2 | O70374 | Protein CB | O70374 [5 | 0 | 1884.918 | 0 | -1.06 | 0.000533 | 3.58 | 136.8 | 63.2 | 3.41 | 8.76 | | High | High | High | High | High | High | High | High | High | High | High | 0.000382 | 0.003258 | 3.08 |
| FALSE | High | SADEESIED | [R].SADEES | 1xCarbami | A0A0R4J04 | 5.09E-06 | 0 | 1 | 2 | 6 | A0A0R4J04 | CCAAT/enh | A0A0R4J04 | 0 | 4290.782 | 0 | -0.14 | 0.14809 | 4.11 | 106.1 | 93.9 | 4.82 | 16.38 | | High | High | High | High | High | High | High | High | High | High | High | 0 | 6.3E-07 | 4.87 |
| FALSE | High | SADEESIED | [R].SADEES | 1xCarbami | A0A0R4J04 | 0.000181 | 0 | 1 | 2 | 7 | A0A0R4J04 | CCAAT/enh | A0A0R4J04 | 0 | 4306.777 | 0 | -0.06 | 0.144386 | 2.54 | 104.9 | 95.1 | 8.63 | 7.33 | | High | High | High | High | High | High | High | High | High | High | High | 0 | 0.000234 | 4.76 |
| FALSE | High | SADNSLEN | [K].SADNSI | 1xPhosphc | A0A0R4J1C | 0.020499 | 0.001472 | 1 | 11 | 1 | A0A0R4J1C | Disabled h | A0A0R4J1C | 0 | 1846.899 | 0 | 1.05 | 0.036575 | 33.16 | 65.4 | 134.6 | 12.17 | 28.72 | | High | High | High | High | High | High | High | High | High | High | High | 0.000786 | 0.007017 | 3.1 |
| FALSE | High | SAAGLCTSI | [R].SAAGL( | 1xPhosphc | Q7TSI1 1xF | 0.000244 | 0 | 1 | 1 | 2 | Q7TSI1 | Pleckstrin | Q7TSI1 [45 | 0 | 2084.952 | 0 | 0.69 | 0.000331 | 5.18 | 76.7 | 123.3 | 5.27 | 5.22 | | High | High | High | High | High | High | High | High | High | High | High | 0 | 6.16E-05 | 2.82 |
| FALSE | High | RTPTMPQE | [R].RTPTM | 1xCarbami | Q14AX6 1x | 0.003962 | 0.000311 | 1 | 1 | 1 | Q14AX6 | Cyclin-dep | Q14AX6 [1 | 0 | 3008.514 | 0 | 0.27 | 0.326172 | 9.08 | 92.8 | 107.2 | 7.73 | 15.65 | | High | High | High | High | High | High | High | High | High | High | High | 0.00014 | 0.00112 | 2.26 |
| FALSE | High | RYSGDSDS | [R].RYSGDS | 1xPhosphc | Q8JZP9 1xf | 0.000237 | 0 | 1 | 1 | 2 | Q8JZP9 | GAS2-like p | Q8JZP9 [35 | 1 | 2383.018 | 0 | 0.24 | 0.013164 | 7.75 | 91.2 | 108.8 | 9.44 | 4.09 | | High | High | High | High | High | High | High | High | High | High | High | 0 | 8.26E-05 | 3.53 |
| FALSE | High | RYSDLTTLV | [R].RYSDLT | 1xPhosphc | P25206 1xf | 9.89E-05 | 0 | 2 | 3 | 1 | P25206; Q | DNA replic | P25206 [15 | 1 | 2541.284 | 0 | -0.99 | 0.364288 | 8.3 | 122.6 | 77.4 | 17.74 | 57.82 | Shared | High | High | High | High | High | High | High | High | High | High | High | 0 | 1.77E-05 | 3.5 |
| FALSE | High | RVSSETHQ | [K].RVSSET | 1xPhosphc | E9Q7G0 1x | 0.000822 | 0 | 1 | 5 | 3 | E9Q7G0 | Nuclear mi | E9Q7G0 [1 | 1 | 2235.117 | 0 | -0.63 | 0.008195 | 4.99 | 121.3 | 78.7 | 7.43 | 14.73 | | High | High | High | High | High | High | High | High | High | High | High | 0.003461 | 0.03754 | 1.9 |
| FALSE | High | RVSSETHQ | [K].RVSSET | 2xPhosphc | E9Q7G0 2x | 0.009843 | 0.000736 | 1 | 5 | 3 | E9Q7G0 | Nuclear mi | E9Q7G0 [1 | 1 | 2315.083 | 0 | -0.3 | 0.167112 | 6.3 | 110.3 | 89.7 | 13.23 | 7.34 | | High | High | High | High | High | High | High | High | High | High | High | 0.001308 | 0.01379 | 3.16 |
| FALSE | High | RVSSGSCF | [R].RVSSGS | 1xCarbami | Q791T5 1x | 0.065405 | 0.004734 | 1 | 2 | 1 | Q791T5 | Mitochond | Q791T5 [3 | 1 | 1521.697 | 0 | 3.11 | 0.000468 | 6.38 | 18.8 | 181.2 | 7.59 | 11.12 | | High | High | High | High | High | High | High | High | High | High | High | 0.002432 | 0.02618 | 1.58 |
| FALSE | High | RVSSTPETF | [K].RVSSTP | 1xPhosphc | P33611 1xf | 1.21E-05 | 0 | 1 | 6 | 9 | P33611 | DNA polym | P33611 [12 | 1 | 1854.014 | 0 | 0.17 | 0.106919 | 4.41 | 97.1 | 102.9 | 6.52 | 7.68 | | High | High | High | High | High | High | High | High | High | High | High | 0 | 2.72E-05 | 3.11 |
| FALSE | High | RVSTDLPE | [R].RVSTDL | 1xCarbami | E9PZP8 1x | 0.000207 | 0 | 1 | 1 | 2 | E9PZP8 | HECT and F | E9PZP8 [14 | 1 | 2897.381 | 0 | 0.64 | 0.006121 | 9.65 | 77.9 | 122.1 | 6.38 | 7.23 | | High | High | High | High | High | High | High | High | High | High | High | 0 | 4.05E-05 | 3.19 |
| FALSE | High | RVSVCAET | [R].RVSVCA | 1xCarbami | Q8K1M3 1 | 3.47E-08 | 0 | 1 | 3 | 4 | Q8K1M3 | Protein kin | Q8K1M3 [5 | 1 | 2817.187 | 0 | 0.21 | 0.000656 | 2.46 | 92.5 | 107.5 | 5.42 | 4.14 | | High | High | High | High | High | High | High | High | High | High | High | 0 | 2.32E-09 | 4.41 |
| FALSE | High | RVSVELTN | [K].RVSVEL | 1xPhosphc | Q02257 1x | 0.001767 | 5.66E-05 | 1 | 1 | 1 | Q02257 | Junction pl | Q02257 [6 | 1 | 1931.077 | 0 | 0.1 | 0.056061 | 8.22 | 97.4 | 102.6 | 7.8 | 3.48 | | High | High | High | High | High | High | High | High | High | High | High | 2.58E-05 | 0.000453 | 2.16 |
| FALSE | High | RVSVTDRP | [K].RVSVTI | 1xPhosphc | E9Q842 1x | 0.072858 | 0.005599 | 1 | 6 | 1 | E9Q842 | Neuron na | E9Q842 [5 | 1 | 1824.014 | 0 | 1.98 | 0.006399 | 39.29 | 40.5 | 159.5 | 8.15 | 25.14 | | High | High | High | High | High | High | High | High | High | High | High | 0.002867 | 0.0299 | 0.94 |
| FALSE | High | RVVSEICET | [R].RVVSEI | 1xCarbami | E9QPB3 1x | 9.85E-07 | 0 | 1 | 5 | 2 | E9QPB3 | WD repeat | E9QPB3 [1 | 1 | 2588.268 | 0 | -0.94 | 0.001106 | 0.85 | 131 | 69 | 8.03 | 14.38 | | High | High | High | High | High | High | High | High | High | High | High | 0 | 9.95E-08 | 4.3 |
| FALSE | High | RWDQTAD | [R].RWDQT | 1xPhosphc | G5E866 1x | 0.042503 | 0.003197 | 1 | 5 | 2 | G5E866 | Splicing fac | G5E866 [1 | 1 | 2210.101 | 0 | -0.23 | 0.18949 | 5.55 | 105.5 | 94.5 | 8.58 | 5.71 | | High | High | High | High | High | High | High | High | High | High | High | 0.007161 | 0.0776 | 2.64 |
| FALSE | High | RWQSPVTI | [K].RWQSP | 1xPhosphc | Q52KI8 1xf | 0.006478 | 0.00062 | 1 | 5 | 2 | Q52KI8 | Serine/arg | Q52KI8 [54 | 1 | 1539.845 | 0 | -0.48 | 0.005782 | 7.19 | 118.1 | 81.9 | 6.24 | 7.74 | | High | High | High | High | High | High | High | High | High | High | High | 0.000275 | 0.00194 | 1.9 |
| FALSE | High | RWSAEVTS | [R].RWSAE | 1xPhosphc | Q3TJN8 1x | 1.75E-05 | 0 | 1 | 2 | 1 | Q3TJN8 | Putative ur | Q3TJN8 [2 | 1 | 2776.334 | 0 | -0.37 | 0.090371 | 17.15 | 112.7 | 87.3 | 10.59 | 17.42 | | High | High | High | High | High | High | High | High | High | High | High | 0 | 2.54E-06 | 2.5 |
| FALSE | High | RWSAPATA | [R].RWSAP | 1xPhosphc | A9JSM3 1x | 0.001865 | 5.66E-05 | 1 | 1 | 1 | A9JSM3 | Transmem | A9JSM3 [1 | 1 | 1508.757 | 0 | 0.61 | 0.043975 | 10.42 | 78.2 | 121.8 | 27.15 | 7.61 | | High | High | High | High | High | High | High | High | High | High | High | 2.58E-05 | 0.000479 | 1.19 |
| FALSE | High | RYSDLTTLV | [R].RYSDLT | 1xPhosphc | P25206 1xf | 7.04E-06 | 0 | 2 | 3 | 4 | P25206; Q | DNA replic | P25206 [15 | 1 | 2770.447 | 0 | -0.97 | 0.001104 | 0.88 | 131.2 | 68.8 | 6.18 | 9.08 | Shared | High | High | High | High | High | High | High | High | High | High | High | 0 | 9.11E-07 | 4.08 |
| FALSE | High | RWSDQLSL | [R].RWSDC | 1xPhosphc | G3UYW1 1 | 0.000845 | 0 | 1 | 5 | 1 | G3UYW1 | WD repeat | G3UYW1 [ | 1 | 1914.973 | 0 | 0.49 | 0.010461 | 16.2 | 82.3 | 117.7 | 9.33 | 10.23 | | High | High | High | High | High | High | High | High | High | High | High | 0 | 0.000197 | 2.51 |
| FALSE | High | RWSGNEPT | [K].RWSGN | 1xPhosphc | E9QAU9 1x | 0.000674 | 0 | 1 | 5 | 1 | E9QAU9 | Leucine-ric | E9QAU9 [3 | 1 | 2328.086 | 0 | 1.73 | 0.000716 | 14.93 | 44.5 | 155.5 | 15.51 | 10.83 | | High | High | High | High | High | High | High | High | High | High | High | 0 | 0.000153 | 3.28 |
| FALSE | High | RWSGNEST | [K].RWSGN | 1xPhosphc | Q921G6 1x | 3.98E-05 | 0 | 1 | 4 | 1 | Q921G6 | Leucine-ric | Q921G6 [3 | 1 | 2342.029 | 0 | -0.37 | 0.011097 | 3.61 | 110.9 | 89.1 | 7.83 | 9.85 | | High | High | High | High | High | High | High | High | High | High | High | 0 | 6.36E-06 | 2.89 |
| FALSE | High | RWSLASLP | [R].RWSLA | 1xCarbami | E9Q1Q1 1x | 0.000434 | 0 | 2 | 16 | 2 | E9Q1Q1; C | Microtubu | E9Q1Q1 [2 | 1 | 3686.741 | 0 | 0.32 | 0.163831 | 18.99 | 90.1 | 109.9 | 20.88 | 16 | Shared | High | High | High | High | High | High | High | High | High | High | High | 2.58E-05 | 0.000389 | 3.28 |
| FALSE | High | RWSLASLP | [R].RWSLA | 1xPhosphc | E9Q8S5 1x | 0.000132 | 0 | 1 | 5 | 1 | E9Q8S5 | Microtubu | E9Q8S5 [1 | 1 | 3155.484 | 0 | -0.86 | 0.000746 | 0.88 | 134.3 | 65.7 | 7.09 | 10.44 | | High | High | High | High | High | High | High | High | High | High | High | 0 | 2.45E-05 | 4.32 |
| FALSE | High | RYEDDGISI | [R].RYEDD( | 1xPhosphc | F6YRW4 1x | 0.000221 | 0 | 1 | 6 | 3 | F6YRW4 | F-box and | F6YRW4 [2 | 1 | 2279.048 | 0 | 0.87 | 0.000437 | 4.91 | 70.8 | 129.2 | 3.75 | 6.29 | | High | High | High | High | High | High | High | High | High | High | High | 4.94E-05 | 0.000602 | 2.13 |
| FALSE | High | RYEPYGMY | [R].RYEPY( | 1xPhosphc | B2RY22 1x | 4.31E-05 | 0 | 2 | 17 | 4 | B2RY22; A( | Clasp1 pro | B2RY22 [7 | 1 | 3099.218 | 0 | -0.91 | 0.000353 | 1.77 | 130.5 | 69.5 | 2.8 | 10.66 | Shared | High | High | High | High | High | High | High | High | High | High | High | 0 | 0.000157 | 3.12 |
| FALSE | High | RYEPYGMY | [R].RYEPY( | 1xCarbami | B2RY22 2x | 0.010109 | 0.000736 | 2 | 17 | 1 | B2RY22; A( | Clasp1 pro | B2RY22 [7 | 1 | 3179.185 | 0 | -0.31 | 0.208786 | 21.22 | 101.4 | 98.6 | 11.83 | 16.35 | Shared | High | High | High | High | High | High | High | High | High | High | High | 0.000344 | 0.003162 | 4.33 |
| FALSE | High | RYPSSISSSI | [R].RYPSSI | 1xPhosphc | Q80X50-5 | 3.39E-06 | 0 | 1 | 13 | 4 | Q80X50-5 | Isoform 5 c | Q80X50-5 | 1 | 1874.978 | 0 | -0.85 | 0.000931 | 4.25 | 128.3 | 71.7 | 3.72 | 8.07 | | High | High | High | High | High | High | High | High | High | High | High | 0 | 4E-07 | 2.98 |
| FALSE | High | RYPTPYPDI | [K].RYPTPY | 1xPhosphc | Q80TH2-2 | 0.000454 | 0 | 1 | 6 | 2 | Q80TH2-2 | Isoform 2 c | Q80TH2-2 | 1 | 1916.992 | 2 | 0.08 | 0.202493 | 14.22 | 95.5 | 104.5 | 4.69 | 12.16 | | High | High | High | High | High | High | High | High | High | High | High | 0 | 9.8E-05 | 2.12 |
| FALSE | High | RYSDFDLLI | [R].RYSDFC | 1xPhosphc | Q8BX57 1x | 0.001463 | 5.66E-05 | 1 | 1 | 4 | Q8BX57 | PX domain | Q8BX57 [5 | 1 | 3239.748 | 0 | 2.46 | 0.000261 | 4.69 | 30.8 | 169.2 | 4.8 | 7.58 | | High | High | High | High | High | High | High | High | High | High | High | 2.58E-05 | 0.000365 | 3.64 |
| FALSE | High | RYSDFEWL | [R].RYSDFE | 1xPhosphc | Q3V2H3 1x | 0.041781 | 0.003126 | 1 | 5 | 2 | Q3V2H3 | Putative ur | Q3V2H3 [7 | 1 | 1781.903 | 0 | 1.57 | 0.000515 | 6.8 | 50.2 | 149.8 | 3.23 | 8.82 | | High | High | High | High | High | High | High | High | High | High | High | 0.003265 | 0.03506 | 2.32 |
| FALSE | High | RYSDGWCI | [R].RYSDG\ | 2xCarbami | Q8BYZ1 1x | 0.009931 | 0.000736 | 1 | 2 | 6 | Q8BYZ1 | ABI gene fa | Q8BYZ1 [3 | 1 | 3353.375 | 0 | 2.81 | 0.00024 | 27.76 | 23.9 | 176.1 | 30.83 | 9.28 | | High | High | High | High | High | High | High | High | High | High | High | 0.000449 | 0.004165 | 3.9 |
| FALSE | High | RYSDHAGP | [R].RYSDH/ | 1xPhosphc | Q03141 1x | 3.63E-05 | 0 | 1 | 5 | 2 | Q03141 | MAP/micr | Q03141 [4 | 1 | 2466.295 | 0 | -0.73 | 0.005122 | 4.77 | 121.6 | 78.4 | 7.03 | 9.64 | | High | High | High | High | High | High | High | High | High | High | High | 0 | 5.78E-06 | 3.04 |
| FALSE | High | RWSEPSNC | [R].RWSEP | 1xPhosphc | Q571G1 1x | 0.009585 | 0.000736 | 1 | 5 | 2 | Q571G1 | MKIAA402 | Q571G1 [4 | 1 | 1669.846 | 0 | -1.08 | 0.000444 | 0.79 | 135.4 | 64.6 | 5.16 | 4.74 | | High | High | High | High | High | High | High | High | High | High | High | 0.002867 | 0.02993 | 1.76 |
| FALSE | High | SEQSSMSII | [R].SEQSS\ | 1xPhosphc | B2RQQ5 1 | 9.71E-05 | 0 | 1 | 1 | 2 | B2RQQ5 | Microtubu | B2RQQ5 [1 | 0 | 3191.317 | 0 | -0.43 | 0.947221 | 30.96 | 114.9 | 85.1 | 38.33 | 84.37 | | High | High | High | High | High | High | High | High | High | High | High | 0 | 1.74E-05 | 3.36 |
| FALSE | High | SFLSEPSSP | [R].SFLSEP | 1xPhosphc | E9PZY8 1x | 0.001615 | 5.66E-05 | 1 | 2 | 5 | E9PZY8 | RIKEN cDN | E9PZY8 [1 | 0 | 1472.698 | 0 | -0.54 | 0.00045 | 1.96 | 119.8 | 80.2 | 2.83 | 4.33 | | High | High | High | High | High | High | High | High | High | High | High | 0.000246 | 0.001493 | 1.98 |
| FALSE | High | SESAPSLHF | [R].SESAPS | 1xPhosphc | Q5I1X5 1x | 9.8E-05 | 0 | 1 | 1 | 3 | Q5I1X5 | RelA-assoc | Q5I1X5 [1 | 0 | 2235.146 | 0 | 2.21 | 0.005259 | 14.38 | 34.6 | 165.4 | 15.05 | 23.25 | | High | High | High | High | High | High | High | High | High | High | High | 7.19E-05 | 0.000887 | 2.85 |
| FALSE | High | SLSSPTVTL | [R].SLSSPT | 1xPhosphc | Q8CFI0 1xf | 5.12E-05 | 0 | 1 | 6 | 8 | Q8CFI0 | E3 ubiquiti | Q8CFI0 [47 | 0 | 2196.193 | 0 | 0.11 | 0.18982 | 6.3 | 97.7 | 102.3 | 3.35 | 8.24 | | High | High | High | High | High | High | High | High | High | High | High | 0 | 1.54E-05 | 3.01 |
| FALSE | High | SISSSNPDL | [K].SISSSNI | 2xPhosphc | A0A1L1SQ | 1.55E-06 | 0 | 1 | 5 | 4 | A0A1L1SQ | Dedicator | A0A1L1SQ | 0 | 2539.213 | 0 | -1.36 | 0.000981 | 1.33 | 143.7 | 56.3 | 5.08 | 12.61 | | High | High | High | High | High | High | High | High | High | High | High | 0 | 5.31E-07 | 3.51 |
| FALSE | High | SISSSNPDL | [K].SISSSNI | 2xPhosphc | A0A1L1SQ | 1.45E-05 | 0 | 1 | 5 | 4 | A0A1L1SQ | Dedicator | A0A1L1SQ | 0 | 2619.179 | 0 | -1.38 | 0.004083 | 6.31 | 144.5 | 55.5 | 8.81 | 34.11 | | High | High | High | High | High | High | High | High | High | High | High | 0 | 2.04E-06 | 3.55 |
| FALSE | High | SLSSSPEK | [K].SLSSSPI | 1xPhosphc | Q6PAR7 1x | 0.008769 | 0.000736 | 1 | 3 | 2 | Q6PAR7 | Phf20 prot | Q6PAR7 [1 | 0 | 1372.713 | 0 | -1.27 | 0.000361 | 0.81 | 140.5 | 59.5 | 5.6 | 5.12 | | High | High | High | High | High | High | High | High | High | High | High | 0.006821 | 0.07384 | 1.45 |
| FALSE | High | SLSSSPQAC | [R].SLSSSP | 1xPhosphc | A2AEW8 1x | 8.5E-06 | 0 | 1 | 3 | 2 | A2AEW8 | GRIP1-assc | A2AEW8 [6 | 0 | 3277.594 | 0 | 1.44 | 0.00356 | 25.77 | 51.5 | 148.5 | 20.76 | 15.34 | | High | High | High | High | High | High | High | High | High | High | High | 0 | 1.12E-06 | 5.14 |
| FALSE | High | SLSSSSQGF | [R].SLSSSS( | 1xPhosphc | Q9D312 1x | 2.98E-05 | 0 | 1 | 1 | 1 | Q9D312 | Keratin, typ | Q9D312 [1 | 0 | 2224.052 | 0 | 5.24 | 0.001634 | 24.64 | 4.9 | 195.1 | 66.88 | 20.28 | | High | High | High | High | High | High | High | High | High | High | High | 0 | 4.62E-06 | 2.87 |
| FALSE | High | SLSSSSQGF | [R].SLSSSS( | 1xPhosphc | Q9D312 1x | 1.83E-09 | 0 | 1 | 1 | 66 | Q9D312 | Keratin, typ | Q9D312 [1 | 0 | 2240.047 | 0 | 2.2 | 0.000756 | 14.32 | 35.8 | 164.2 | 10.98 | 13.46 | | High | High | High | High | High | High | High | High | High | High | High | 0 | 8.56E-11 | 5.25 |
| FALSE | High | SLSSSSQGF | [R].SLSSSS( | 2xPhosphc | Q9D312 2x | 6.02E-05 | 0 | 1 | 1 | 3 | Q9D312 | Keratin, typ | Q9D312 [1 | 0 | 2304.018 | 0 | 1.35 | 0.000975 | 14.36 | 54.3 | 145.7 | 15.27 | 11.82 | | High | High | High | High | High | High | High | High | High | High | High | 0 | 1.01E-05 | 3.72 |
| FALSE | High | SLSSSSQGF | [R].SLSSSS( | 2xPhosphc | Q9D312 2x | 0.000138 | 0 | 1 | 1 | 4 | Q9D312 | Keratin, typ | Q9D312 [1 | 0 | 2320.013 | 0 | 2.59 | 0.004053 | 12.75 | 27.3 | 172.7 | 13.07 | 23.35 | | High | High | High | High | High | High | High | High | High | High | High | 0 | 3.91E-05 | 3.64 |
| FALSE | High | SLSSTSSG | [R].SLSSTS | 1xPhosphc | E9PWI3 1x | 5.55E-05 | 0 | 1 | 3 | 1 | E9PWI3 | Mutated in | E9PWI3 [8 | 0 | 1478.75 | 0 | 2.56 | 0.000172 | 8.44 | 28.5 | 171.5 | 9.38 | 3.68 | | High | High | High | High | High | High | High | High | High | High | High | 0 | 9.26E-06 | 2.33 |
| FALSE | High | SISSTVQR | [R].SISSTV( | 1xPhosphc | Q80WQ6 1 | 0.017498 | 0.001245 | 1 | 2 | 1 | Q80WQ6 | Inactive rh | Q80WQ6 [ | 0 | 1186.603 | 0 | -0.06 | 0.516316 | 17.09 | 102 | 98 | 7.87 | 9.38 | | High | High | High | High | High | High | High | High | High | High | High | 0.000689 | 0.005893 | 1.35 |
| FALSE | High | SLSTDQFSS | [R].SLSTDC | 1xPhosphc | Q921D9 1x | 4.55E-05 | 0 | 1 | 3 | 2 | Q921D9 | Grainyhea | Q921D9 [1 | 0 | 2230.034 | 0 | 0.79 | 0.00375 | 16.39 | 67.8 | 132.2 | 18.21 | 10.33 | | High | High | High | High | High | High | High | High | High | High | High | 0 | 7.38E-06 | 3.99 |
| FALSE | High | SLSTEVEPK | [K].SLSTEV | 1xPhosphc | Q6ZPJ0 1x | 0.002272 | 0.000109 | 1 | 2 | 2 | Q6ZPJ0 | Testis-expr | Q6ZPJ0 [1 | 0 | 1527.807 | 0 | 0.55 | 0.044661 | 22.46 | 84 | 116 | 9.63 | 17.98 | | High | High | High | High | High | High | High | High | High | High | High | 0.000327 | 0.002538 | 2.34 |
| FALSE | High | SLSSPTDNL | [K].SLSSPT | 1xPhosphc | Q68FF6 1x | 7.77E-06 | 0 | 1 | 2 | 3 | Q68FF6 | ARF GTPas | Q68FF6 [3 | 0 | 1798.879 | 0 | 0.15 | 0.037818 | 3.31 | 96 | 104 | 2.71 | 4.75 | | High | High | High | High | High | High | High | High | High | High | High | 0 | 1.02E-06 | 3.26 |
| FALSE | High | SLSTGGSEV | [R].SLSTGC | 1xPhosphc | G3UVUS 1 | 0.006085 | 0.00062 | 1 | 10 | 3 | G3UVUS | TBC1 doma | G3UVUS [4 | 0 | 2408.045 | 0 | -0.33 | 0.004009 | 5.94 | 112.3 | 87.7 | 4.1 | 4.69 | | High | High | High | High | High | High | High | High | High | High | High | 0.000275 | 0.002288 | 3.22 |
| FALSE | High | SLSTPNVHI | [R].SLSTPN | 1xPhosphc | Q5DTN0 1x | 0.000173 | 0 | 1 | 5 | 2 | Q5DTN0 | MKIAA410 | Q5DTN0 [3 | 0 | 3241.541 | 0 | 0.39 | 0.027294 | 8.64 | 86.5 | 113.5 | 10.14 | 10.84 | | High | High | High | High | High | High | High | High | High | High | High | 0 | 3.31E-05 | 2.36 |
| FALSE | High | SLSTSGESL | [R].SLSTSG | 1xPhosphc | Q921J3 1xf | 0.000165 | 0 | 1 | 4 | 2 | Q921J3 | Dnajc5 pro | Q921J3 [8 | 0 | 2115.057 | 0 | 0.08 | 0.377892 | 31.38 | 80.5 | 119.5 | 32.53 | 31 | | High | High | High | High | High | High | High | High | High | High | High | 0 | 3.15E-05 | 2.66 |
| FALSE | High | SLSTSGESL | [R].SLSTSG | 1xPhosphc | Q921J3 1xf | 0.016163 | 0.001124 | 1 | 4 | 2 | Q921J3 | Dnajc5 pro | Q921J3 [8 | 0 | 2344.22 | 0 | 0.66 | 0.122983 | 20.44 | 78.6 | 121.4 | 17.1 | 18.77 | | High | High | High | High | High | High | High | High | High | High | High | 0.000585 | 0.005378 | 2.95 |
| FALSE | High | SISTVDLTE | [R].SISTVD | 1xPhosphc | Q9DBJ3 1x | 0.000148 | 0 | 1 | 1 | 3 | Q9DBJ3 | Brain-spec | Q9DBJ3 [4 | 0 | 1630.87 | 0 | 1.62 | 0.000238 | 3.57 | 48.2 | 151.8 | 5.92 | 5.99 | | High | High | High | High | High | High | High | High | High | High | High | 0 | 3.79E-05 | 2.99 |
| FALSE | High | SISVGENP( | [R].SISVGE | 1xPhosphc | Q921Q9 1x | 9.19E-08 | 0 | 1 | 1 | 2 | Q921Q9 | Mapkbp1 p | Q921Q9 [4 | 0 | 2413.233 | 0 | 0.88 | 0.001184 | 20.95 | 66.3 | 133.7 | 18.01 | 7.7 | | High | High | High | High | High | High | High | High | High | High | High | 0 | 1.12E-07 | 4.66 |
| FALSE | High | SISVGENP( | [R].SISVGE | 1xPhosphc | Q6NS57-2 | 0.000548 | 0 | 1 | 3 | 1 | Q6NS57-2 | Isoform 2 c | Q6NS57-2 | 0 | 2441.264 | 0 | -0.2 | 0.073646 | 6.86 | 107 | 93 | 4.95 | 7.56 | | High | High | High | High | High | High | High | High | High | High | High | 0 | 0.000121 | 3.37 |
| FALSE | High | SLSVLSPR | [R].SLSVLS | 1xPhosphc | D3Z3Q3 1x | 0.074685 | 0.005727 | 1 | 3 | 1 | D3Z3Q3 | Smoothelir | D3Z3Q3 [2 | 0 | 1167.634 | 0 | -0.78 | 0.002042 | 2.12 | 125.7 | 74.3 | 5.12 | 8.87 | | High | High | High | High | High | High | High | High | High | High | High | 0.002937 | 0.03078 | 1 |
| FALSE | High | SLSVPVDLS | [R].SLSVPV | 1xPhosphc | Q8BXQ8 1 | 0.036103 | 0.002342 | 1 | 1 | 2 | Q8BXQ8 | Protein FA | Q8BXQ8 [1 | 0 | 1381.729 | 0 | -0.25 | 0.028085 | 6.86 | 108.4 | 91.6 | 5.42 | 6.25 | | High | High | High | High | High | High | High | High | High | High | High | 0.001279 | 0.01334 | 1.59 |
| FALSE | High | SLSYSPVER | [R].SLSYSP | 1xPhosphc | Q8BTI8 1x | 0.002252 | 0.000109 | 1 | 3 | 1 | Q8BTI8 | Serine/arg | Q8BTI8 [26 | 0 | 1346.655 | 0 | -1.18 | 0.009974 | 1.93 | 136.2 | 63.8 | 16.26 | 8.23 | | High | High | High | High | High | High | High | High | High | High | High | 4.94E-05 | 0.000593 | 1.39 |
| FALSE | High | SLSYSPVER | [R].SLSYSP | 2xPhosphc | Q8BTI8 2x | 0.009756 | 0.000736 | 1 | 3 | 5 | Q8BTI8 | Serine/arg | Q8BTI8 [26 | 0 | 1426.622 | 0 | -0.37 | 0.000262 | 1.63 | 112.9 | 87.1 | 1.3 | 2.38 | | High | High | High | High | High | High | High | High | High | High | High | 0.000344 | 0.003059 | 2.26 |
| FALSE | High | SISYSSQK | [R].SISYSS( | 1xPhosphc | Q8VDZ4 1x | 0.058664 | 0.004294 | 1 | 1 | 1 | Q8VDZ4 | Palmitoyltr | Q8VDZ4 [6 | 0 | 1437.739 | 0 | 1.29 | 0.000778 | 15.83 | 59.4 | 140.6 | 13.58 | 6.37 | | High | High | High | High | High | High | High | High | High | High | High | 0.002187 | 0.02317 | 1.63 |
| FALSE | High | SITAYSPTT( | [K].SITAYS | 1xPhosphc | Q9CQ82 1 | 7.43E-06 | 0 | 1 | 2 | 2 | Q9CQ82 | Centromer | Q9CQ82 [2 | 0 | 3040.532 | 0 | -0.34 | 0.302821 | 31.51 | 110.6 | 89.4 | 16.76 | 20.34 | | High | High | High | High | High | High | High | High | High | High | High | 0 | 9.71E-07 | 4.94 |
| FALSE | High | SLTELVQQ | [K].SLTELV | 1xCarbami | H3BJQ9 1x | 1.15E-06 | 0 | 1 | 12 | 2 | H3BJQ9 | Homeobox | H3BJQ9 [1 | 0 | 2439.261 | 0 | 2.1 | 0.00196 | 21.16 | 37.8 | 162.2 | 8.04 | 16 | | High | High | High | High | High | High | High | High | High | High | High | 0 | 2.11E-07 | 4.52 |
| FALSE | High | SLTNGEN | [R].SLTNG | 1xPhosphc | Q3UVM5 1 | 0.016021 | 0.001124 | 1 | 3 | 2 | Q3UVM5 | Putative ur | Q3UVM5 [ | 0 | 2376.11 | 0 | -0.25 | 0.238112 | 12.03 | 109.7 | 90.3 | 9.05 | 15.82 | | High | High | High | High | High | High | High | High | High | High | High | 0.000585 | 0.005292 | 2.66 |
| FALSE | High | SLTESWEV | [R].SLTESW | 1xPhosphc | Q8VE91 1x | 9.11E-05 | 0 | 1 | 1 | 1 | Q8VE91 | Reticuloph | Q8VE91 [1 | 0 | 2428.216 | 0 | 2.98 | 0.00129 | 9.96 | 24.6 | 175.4 | 14.49 | 15.71 | | High | High | High | High | High | High | High | High | High | High | High | 0 | 1.61E-05 | 3.43 |
| FALSE | High | SISSPSMNI | [R].SISSPS | 1xPhosphc | E9Q4K7 1x | 0.00582 | 0.00062 | 1 | 2 | 2 | E9Q4K7 | Kinesin fan | E9Q4K7 [1 | 0 | 1303.591 | 0 | 0.2 | 0.046664 | 10.22 | 95.8 | 104.2 | 3.9 | 10.29 | | High | High | High | High | High | High | High | High | High | High | High | 0.006195 | 0.06694 | 0.7 |
| FALSE | High | SLSPPSSK | [R].SLSPP | 2xPhosphc | Q6GSP7 2x | 0.039347 | 0.002793 | 2 | 9 | 1 | Q6GSP7; A | Nuclear fac | Q6GSP7 [2 | 0 | 1507.721 | 0 | 3.34 | 0.000385 | 17.75 | 19.7 | 180.3 | 11.3 | 9.17 | Shared | High | High | High | High | High | High | High | High | High | High | High | 0.001468 | 0.01467 | 1.41 |
| FALSE | High | SLSPPWEQ | [R].SLSPPV | 1xPhosphc | Q91ZV0 1x | 0.011346 | 0.000816 | 1 | 12 | 2 | Q91ZV0 | Melanoma | Q91ZV0 [1 | 0 | 1523.709 | 0 | -0.21 | 0.14809 | 15.7 | 105.8 | 94.2 | 2.79 | 14.21 | | High | High | High | High | High | High | High | High | High | High | High | 0.003564 | 0.03805 | 2.26 |
| FALSE | High | SLSPQDR | [K].SLSPQD | 1xPhosphc | P25976 1xf | 0.020499 | 0.001472 | 1 | 6 | 1 | P25976 | Nucleolar t | P25976 [63 | 0 | 1111.535 | 0 | -1.15 | 0.001514 | 3.59 | 138 | 62 | 8.69 | 5.37 | | High | High | High | High | High | High | High | High | High | High | High | 0.000786 | 0.006988 | 1.62 |
| FALSE | High | SLSPQGR | [K].SLSPQC | 1xPhosphc | Q148V8 1x | 0.038017 | 0.002528 | 1 | 1 | 2 | Q148V8 | Protein FA | Q148V8 [8 | 0 | 1053.529 | 0 | 0.74 | 0.000605 | 4.32 | 73.9 | 126.1 | 5.61 | 8.36 | | High | High | High | High | High | High | High | High | High | High | High | 0.001324 | 0.01405 | 2.01 |
| FALSE | High | SLSPQGRD | [K].SLSPQC | 2xPhosphc | Q148V8 2x | 0.120211 | 0.009851 | 1 | 1 | 1 | Q148V8 | Protein FA | Q148V8 [8 | 1 | 3392.633 | 0 | 1.98 | 0.000346 | 4.9 | 41.1 | 158.9 | 6.92 | 6.23 | | High | High | High | High | High | High | High | High | High | High | High | 0.004922 | 0.05358 | 2.41 |
| FALSE | High | SISPSALQD | [R].SISPSAI | 1xPhosphc | F8VPR5 1x | 0.000568 | 0 | 1 | 4 | 1 | F8VPR5 | CREB-bindi | F8VPR5 [2 | 0 | 1608.856 | 0 | 1.62 | 0.000681 | 12.34 | 47.5 | 152.5 | 6.77 | 8.45 | | High | High | High | High | High | High | High | High | High | High | High | 0 | 0.000127 | 2.71 |
| FALSE | High | SLSPSHLTE | [R].SLSPSH | 1xPhosphc | E9Q784 1x | 0.000687 | 0 | 1 | 2 | 2 | E9Q784 | Zinc finger | E9Q784 [9 | 0 | 1550.741 | 0 | -0.76 | 0.005178 | 6.47 | 127.5 | 72.5 | 9.01 | 12.36 | | High | High | High | High | High | High | High | High | High | High | High | 0 | 0.000156 | 2.2 |
| FALSE | High | SLSPSHLTE | [R].SLSPSH | 2xPhosphc | E9Q784 2x | 0.010853 | 0.000772 | 1 | 2 | 2 | E9Q784 | Zinc finger | E9Q784 [9 | 0 | 1630.708 | 0 | -0.57 | 0.001214 | 5.66 | 119.8 | 80.2 | 2.12 | 9.4 | | High | High | High | High | High | High | High | High | High | High | High | 0.000399 | 0.003447 | 1.43 |
| FALSE | High | SLSPSPDSS | [R].SLSPSP | 1xPhosphc | E9Q6H8 1x | 0.000128 | 0 | 1 | 5 | 5 | E9Q6H8 | Pleckstrin | E9Q6H8 [1 | 0 | 2430.175 | 0 | 0.07 | 0.090449 | 4.29 | 97.8 | 102.2 | 3.09 | 6.85 | | High | High | High | High | High | High | High | High | High | High | High | 0 | 6.45E-05 | 3.56 |
| FALSE | High | SLSPSPDSS | [R].SLSPSP | 2xPhosphc | E9Q6H8 2x | 2.77E-05 | 0 | 1 | 5 | 1 | E9Q6H8 | Pleckstrin | E9Q6H8 [1 | 0 | 2510.141 | 0 | -0.5 | 0.090371 | 23.04 | 120.6 | 79.4 | 12.43 | 26.07 | | High | High | High | High | High | High | High | High | High | High | High | 0 | 4.24E-06 | 2.5 |
| FALSE | High | SISPSPSAIL | [R].SISPSP | 1xPhosphc | E9QAU4 1x | 0.001301 | 0 | 1 | 13 | 5 | E9QAU4 | Enhancer t | E9QAU4 [4 | 0 | 1565.814 | 0 | 0.92 | 0.000643 | 5.48 | 70 | 130 | 3.57 | 5.83 | | High | High | High | High | High | High | High | High | High | High | High | 0 | 0.000321 | 2.52 |
| FALSE | High | SISPSSLLDT | [R].SISPSSL | 1xPhosphc | Q60670 1x | 1.26E-05 | 0 | 1 | 3 | 3 | Q60670 | Serine/thr | Q60670 [4 | 0 | 2085.031 | 0 | -0.01 | 0.457593 | 15.26 | 98.9 | 101.1 | 6.23 | 17.95 | | High | High | High | High | High | High | High | High | High | High | High | 0 | 1.76E-06 | 3.42 |
| FALSE | High | SLSPSVR | [R].SLSPSV | 1xPhosphc | Q80U35 1x | 0.114657 | 0.009411 | 1 | 1 | 2 | Q80U35 | Rho guanir | Q80U35 [7 | 0 | 1054.55 | 0 | -0.66 | 0.001387 | 5.1 | 121.8 | 78.2 | 3.35 | 12 | | High | High | High | High | High | High | High | High | High | High | High | 0.004733 | 0.05088 | 1.03 |
| FALSE | High | SLSPVAAPI | [R].SLSPVA | 1xPhosphc | Q6PHZ5 1x | 0.004256 | 0.00036 | 1 | 1 | 2 | Q6PHZ5 | Putative RN | Q6PHZ5 [2 | 0 | 1416.781 | 0 | 0.81 | 0.001412 | 11.47 | 72.5 | 127.5 | 6.31 | 9.35 | | High | High | High | High | High | High | High | High | High | High | High | 0.002159 | 0.02264 | 2.04 |
| FALSE | High | SLSSPR | [R].SLSSPR | 1xPhosphc | Q8BTI8 1x | 0.114657 | 0.009411 | 1 | 4 | 1 | Q8BTI8 | Serine/arg | Q8BTI8 [7 | 0 | 955.4811 | 0 | 0.11 | 0.719986 | 9.78 | 96.6 | 103.4 | 5.46 | 9.68 | | High | High | High | High | High | High | High | High | High | High | High | 0.004707 | 0.0506 | 1.77 |
| FALSE | High | SLSPVLPAA | [R].SLSPVL | 1xCarbami | Q6P9R4 1x | 0.074071 | 0.0057 | 1 | 1 | 1 | Q6P9R4 | Rho guanir | Q6P9R4 [9 | 0 | 4049.028 | 0 | 1.48 | 0.001599 | 37.11 | 52.6 | 147.4 | 21.45 | 7.45 | | High | High | High | High | High | High | High | High | High | High | High | 0.002924 | 0.03026 | 2.62 |
| FALSE | High | SISQQK | [R].SISQQK | 1xPhosphc | Q9JIX8-3 1 | 0.073462 | 0.005631 | 3 | 11 | 1 | Q9JIX8-3; I | Isoform 3 c | Q9JIX8-3 [ | 0 | 1228.67 | 0 | 0.15 | 0.136694 | 12.16 | 96.6 | 103.4 | 5.23 | 9.5 | Shared | High | High | High | High | High | High | High | High | High | High | High | 0.002867 | 0.02997 | 1.49 |
| FALSE | High | SLSQSFENL | [K].SLSQSF | 1xPhosphc | Q4TVN0 1 | 3.04E-06 | 0 | 1 | 2 | 2 | Q4TVN0 | TBC1 doma | Q4TVN0 [2 | 0 | 2790.437 | 0 | -0.07 | 0.856717 | 42.99 | 114.7 | 85.3 | 10.57 | 39.33 | | High | High | High | High | High | High | High | High | High | High | High | 0 | 3.57E-07 | 4.82 |
| FALSE | High | SLSSDCILSI | [R].SLSSDC | 1xPhosphc | Q9D279 1x | 5.89E-07 | 0 | 1 | 1 | 5 | Q9D279 | Mitotic int | Q9D279 [3 | 0 | 1729.803 | 0 | 4.14 | 0.000159 | 28.42 | 10.4 | 189.6 | 28.91 | 5.28 | | High | High | High | High | High | High | High | High | High | High | High | 0 | 2.09E-06 | 3.21 |
| FALSE | High | SLSSDCILSI | [R].SLSSDC | 1xCarbami | Q9D279 2x | 0.001572 | 5.66E-05 | 1 | 1 | 2 | Q9D279 | Mitotic int | Q9D279 [3 | 0 | 1809.769 | 0 | 4.86 | 0.000193 | 46.48 | 6.7 | 193.3 | 49.44 | 4.98 | | High | High | High | High | High | High | High | High | High | High | High | 2.58E-05 | 0.000397 | 3.3 |
| FALSE | High | SLSSDEATN | [R].SLSSDE | 1xPhosphc | Q5F2E7-2 | 1.32E-05 | 0 | 1 | 2 | 3 | Q5F2E7-2 | Isoform 2 c | Q5F2E7-2 | 0 | 1685.794 | 0 | -0.07 | 0.985663 | 7.91 | 102 | 98 | 6.37 | 5.02 | | High | High | High | High | High | High | High | High | High | High | High | 0 | 2.58E-06 | 2.31 |
| FALSE | High | SLSSGESGF | [R].SLSSGE | 1xPhosphc | E9Q1Q1 1x | 0.002948 | 0.00016 | 1 | 6 | 1 | E9Q1Q1 | Microtubu | E9Q1Q1 [1 | 0 | 2373.103 | 0 | -0.88 | 0.018426 | 13.29 | 127.2 | 72.8 | 14.24 | 21.33 | | High | High | High | High | High | High | High | High | High | High | High | 7.19E-05 | 0.000805 | 2.04 |
| FALSE | High | SLSSGFSPE | [R].SLSSGF | 1xPhosphc | B9EKJ7 1xf | 6.07E-08 | 0 | 1 | 3 | 8 | B9EKJ7 | Tankyrase | B9EKJ7 [97 | 0 | 2810.281 | 0 | 0.67 | 0.00054 | 6.03 | 77.9 | 122.1 | 5.23 | 7.17 | | High | High | High | High | High | High | High | High | High | High | High | 0 | 1.77E-08 | 5.62 |
| FALSE | High | SLSSGFSPE | [R].SLSSGF | 2xPhosphc | B9EKJ7 2xf | 1.15E-07 | 0 | 1 | 3 | 1 | B9EKJ7 | Tankyrase | B9EKJ7 [97 | 0 | 2890.247 | 0 | 0.66 | 0.015102 | 29.18 | 77.7 | 122.3 | 9.78 | 9.79 | | High | High | High | High | High | High | High | High | High | High | High | 0 | 8.95E-09 | 4.36 |
| FALSE | High | SLSSPELDS | [K].SLSSPE | 1xPhosphc | Q6P1C6 1x | 0.000126 | 0 | 1 | 2 | 1 | Q6P1C6 | Leucine-ric | Q6P1C6 [1 | 0 | 2304.053 | 0 | -0.88 | 0.002303 | 2.47 | 123.6 | 76.4 | 11.15 | 8.53 | | High | High | High | High | High | High | High | High | High | High | High | 0 | 2.34E-05 | 3.09 |
| FALSE | High | SLSSPK | [KR].SLSSP | 1xPhosphc | Q7TQG1 1 | 0.069325 | 0.005112 | 2 | 13 | 4 | Q7TQG1; F | Pleckstrin | Q7TQG1 [5 | 0 | 1156.638 | 0 | -1.03 | 0.000449 | 0.87 | 132.4 | 67.6 | 5.78 | 7.56 | Shared | High | High | High | High | High | High | High | High | High | High | High | 0.00943 | 0.1019 | 1.61 |
| FALSE | High | SLSSPNCFC | [R].SLSSPN | 1xPhosphc | Q91Z58 1x | 0.000369 | 0 | 1 | 1 | 4 | Q91Z58 | Uncharacte | Q91Z58 [1 | 0 | 2226.987 | 0 | 0.93 | 0.025055 | 15.96 | 85.2 | 114.8 | 8.92 | 8.39 | | High | High | High | High | High | High | High | High | High | High | High | 0 | 0.000154 | 4.39 |
| FALSE | High | SLSSPNCFC | [R].SLSSPN | 1xCarbami | Q91Z58 1x | 3.73E-05 | 0 | 1 | 1 | 5 | Q91Z58 | Uncharacte | Q91Z58 [1 | 0 | 2242.982 | 0 | 0.93 | 0.021884 | 34.06 | 68.7 | 131.3 | 8.46 | 20.73 | | High | High | High | High | High | High | High | High | High | High | High | 0.000161 | 0.001186 | 4.16 |

| | | | | | | | | | | | | | | | | | | | | | | | | | | | | | | | | | | | | | | |
|---|---|---|---|---|---|---|---|---|---|---|---|---|---|---|---|---|---|---|---|---|---|---|---|---|---|---|---|---|---|---|---|---|---|---|---|---|---|---|
| FALSE | High | SLSSPPSSK | [R].SLSSPP | 1xPhospho | Q6GSP7 1x | 0.002232 | 0.000109 | 2 | 9 | 3 | Q6GSP7; A | Nuclear fac | Q6GSP7 [2 | 0 | 1427.755 | 0 | 2.99 | 0.000108 | 7.82 | 22.4 | 177.6 | 13.13 | 3.59 | Shared | High | High | High | High | High | High | High | High | High | High | High | 0.001296 | 0.01343 | 2.18 |
| FALSE | High | SLSQGSTN | [R].SLSQGS | 1xPhospho | B2RUJ6 1xl | 0.000772 | 0 | 1 | 6 | 2 | B2RUJ6 | Rap guanin | B2RUJ6 [10 | 0 | 2585.243 | 0 | 0.6 | 0.037048 | 23.67 | 82.3 | 117.7 | 4.8 | 20.38 | | High | High | High | High | High | High | High | High | High | High | High | 0.000872 | 0.009114 | 2.74 |
| FALSE | High | SLTGSGSQ | [R].SLTGSG | 1xPhospho | Q0VGR5 1x | 1.99E-06 | 0 | 1 | 2 | 6 | Q0VGR5 | CDK5 regul | Q0VGR5 [4 | 0 | 2114.115 | 0 | 0.39 | 0.003774 | 3.31 | 89.8 | 110.2 | 5.37 | 5.93 | | High | High | High | High | High | High | High | High | High | High | High | 0 | 2.21E-07 | 3.66 |
| FALSE | High | SLTLDGEEE | [K].SLTLDG | 1xPhospho | A0A0R4J29 | 0.016163 | 0.001124 | 1 | 5 | 1 | A0A0R4J29 | Eukaryotic | A0A0R4J29 | 0 | 2066 | 0 | 0.69 | 0.001779 | 10.04 | 74.1 | 125.9 | 4.77 | 6.51 | | High | High | High | High | High | High | High | High | High | High | High | 0.000585 | 0.005394 | 1.71 |
| FALSE | High | SLTLGPSLS | [R].SLTLGP | 1xPhospho | B2RWU4 1 | 0.000148 | 0 | 2 | 11 | 2 | B2RWU4; C | Cordon-ble | B2RWU4 [3 | 0 | 2399.302 | 0 | 0.36 | 0.02368 | 9.46 | 86 | 114 | 4.57 | 12.29 | Shared | High | High | High | High | High | High | High | High | High | High | High | 0 | 2.79E-05 | 2.29 |
| FALSE | High | SLYLEDEDE | [R].SLYLED | 1xPhospho | Q9D279 1x | 8.71E-05 | 0 | 1 | 3 | 1 | Q9D279 | Mitotic inte | Q9D279 [8 | 0 | 2020.977 | 0 | 2.91 | 0.00116 | 22.13 | 24.1 | 175.9 | 17.09 | 16.45 | | High | High | High | High | High | High | High | High | High | High | High | 0 | 1.54E-05 | 2.47 |
| FALSE | High | SLYSSSPGG | [R].SLYSSS | 1xPhospho | P20152 1xl | 5.84E-07 | 0 | 1 | 9 | 9 | P20152 | Vimentin C | P20152 [51 | 0 | 1753.836 | 0 | 2.98 | 0.000604 | 7.19 | 20.6 | 179.4 | 20.12 | 12.8 | | High | High | High | High | High | High | High | High | High | High | High | 0 | 5.56E-08 | 4.18 |
| FALSE | High | SLYSSSPGG | [R].SLYSSS | 2xPhospho | P20152 2xl | 0.005717 | 0.00062 | 1 | 9 | 1 | P20152 | Vimentin C | P20152 [51 | 0 | 1833.802 | 0 | 2.13 | 0.000945 | 26.1 | 33.3 | 166.7 | 16.69 | 13.47 | | High | High | High | High | High | High | High | High | High | High | High | 0.000275 | 0.001687 | 2.55 |
| FALSE | High | SLYTADSEN | [R].SLYTAD | 1xPhospho | Q3UWM4 | 1.65E-06 | 0 | 1 | 1 | 2 | Q3UWM4 | Lysine-spe | Q3UWM4 | 0 | 2038.926 | 0 | 0.06 | 0.310777 | 12.9 | 96.5 | 103.5 | 3.2 | 8.99 | | High | High | High | High | High | High | High | High | High | High | High | 0 | 1.78E-07 | 2.98 |
| FALSE | High | SIYYITGESK | [K].SIYYITG | 1xPhospho | P11499 1xl | 0.032267 | 0.002129 | 1 | 4 | 1 | P11499 | Heat shock | P11499 [48 | 0 | 1698.876 | 0 | -0.25 | 0.039808 | 5.86 | 106 | 94 | 11.35 | 6.07 | | High | High | High | High | High | High | High | High | High | High | High | 0.001126 | 0.01166 | 1.95 |
| FALSE | High | SMCENSPC | [R].SMCEN | 1xCarbami | Q546X9 1x | 2.31E-05 | 0 | 1 | 5 | 1 | Q546X9 | Ataxin 3 OS | Q546X9 [2 | 0 | 2563.089 | 0 | 0.09 | 0.401429 | 13.73 | 98.5 | 101.5 | 5.84 | 9.37 | | High | High | High | High | High | High | High | High | High | High | High | 0 | 3.46E-06 | 3.61 |
| FALSE | High | SMCENSPC | [R].SMCEN | 1xCarbami | Q546X9 1x | 1.5E-07 | 0 | 1 | 5 | 2 | Q546X9 | Ataxin 3 OS | Q546X9 [2 | 0 | 2579.084 | 0 | 0.12 | 0.288742 | 12.46 | 95.1 | 104.9 | 8.25 | 13.95 | | High | High | High | High | High | High | High | High | High | High | High | 0 | 1.21E-08 | 4.23 |
| FALSE | High | SMCENSPC | [R].SMCEN | 1xCarbami | Q546X9 2x | 0.016306 | 0.001124 | 1 | 5 | 2 | Q546X9 | Ataxin 3 OS | Q546X9 [2 | 0 | 2659.05 | 0 | 0.18 | 0.606234 | 19.13 | 96.3 | 103.7 | 14.3 | 9.72 | | High | High | High | High | High | High | High | High | High | High | High | 0.000585 | 0.005433 | 3.29 |
| FALSE | High | SMCENSPC | [R].SMCEN | 1xCarbami | Q546X9 1x | 0.002921 | 0.00016 | 1 | 5 | 1 | Q546X9 | Ataxin 3 OS | Q546X9 [2 | 1 | 2735.185 | 0 | -0.04 | 0.478988 | 11.73 | 101.3 | 98.7 | 9.13 | 39.76 | | High | High | High | High | High | High | High | High | High | High | High | 7.19E-05 | 0.000794 | 1.59 |
| FALSE | High | SMCENSPC | [R].SMCEN | 1xCarbami | Q546X9 2x | 0.079114 | 0.00613 | 1 | 5 | 1 | Q546X9 | Ataxin 3 OS | Q546X9 [2 | 1 | 2815.151 | 0 | 0.78 | 0.006548 | 12.51 | 77.7 | 122.3 | 14.86 | 17.29 | | High | High | High | High | High | High | High | High | High | High | High | 0.003148 | 0.0328 | 2.97 |
| FALSE | High | SMDDLLSA | [R].SMDDL | 1xCarbami | Q91XS1 1x | 0.111965 | 0.009053 | 1 | 3 | 1 | Q91XS1 | Myotubula | Q91XS1 [6 | 0 | 2177.966 | 0 | -0.44 | 0.029415 | 8.97 | 115.3 | 84.7 | 8.21 | 9.38 | | High | High | High | High | High | High | High | High | High | High | High | 0.004569 | 0.04951 | 2.96 |
| FALSE | High | SMDEFTAS | [R].SMDEF | 1xPhospho | B7ZP47 1xl | 0.043236 | 0.003228 | 1 | 2 | 2 | B7ZP47 | Wapal prot | B7ZP47 [38 | 0 | 2179.942 | 0 | -0.59 | 0.003246 | 3.85 | 121.2 | 78.8 | 3.73 | 11.96 | | High | High | High | High | High | High | High | High | High | High | High | 0.004782 | 0.0524 | 2.21 |
| FALSE | High | SMDQLDSS | [R].SMDQL | 1xPhospho | Q9CYC5 1x | 0.000112 | 0 | 1 | 1 | 3 | Q9CYC5 | Kinetochor | Q9CYC5 [3 | 0 | 1531.666 | 0 | -0.32 | 0.003295 | 2.34 | 110.7 | 89.3 | 4.51 | 6.46 | | High | High | High | High | High | High | High | High | High | High | High | 0.008359 | 0.08991 | 2 |
| FALSE | High | SLYGVGSSI | [R].SLYGVG | 1xPhospho | P50446 1xl | 0.002818 | 0.000109 | 1 | 5 | 3 | P50446 | Keratin, typ | P50446 [58 | 0 | 1435.76 | 0 | 6.45 | 0.005009 | 21.53 | 2.4 | 197.6 | 49.13 | 30.76 | | High | High | High | High | High | High | High | High | High | High | High | 0.000275 | 0.001604 | 1.42 |
| FALSE | High | SMDSSDLS | [R].SMDSS | 1xPhospho | Q68FF6 1xl | 3.63E-05 | 0 | 1 | 2 | 8 | Q68FF6 | ARF GTPas | Q68FF6 [4 | 0 | 2839.372 | 0 | -0.19 | 0.11222 | 14.92 | 106 | 94 | 8.94 | 18.5 | | High | High | High | High | High | High | High | High | High | High | High | 0 | 5.77E-06 | 4.87 |
| FALSE | High | SMDVDLNQ | [R].SMDVD | 1xPhospho | Q3TU15 1x | 4.68E-05 | 0 | 2 | 4 | 1 | Q3TU15; Q | Non-specif | Q3TU15 [4 | 0 | 2171.919 | 0 | -0.9 | 0.034869 | 11.24 | 130.1 | 69.9 | 9.34 | 30.82 | Shared | High | High | High | High | High | High | High | High | High | High | High | 0 | 7.67E-06 | 2.22 |
| FALSE | High | SMESDPED | [K].SMESD | 2xPhospho | O08648 2x | 6.65E-05 | 0 | 1 | 5 | 1 | O08648 | Mitogen-ac | O08648 [5 | 0 | 3495.391 | 0 | -0.91 | 0.026608 | 4.25 | 129.6 | 70.4 | 9.99 | 28.8 | | High | High | High | High | High | High | High | High | High | High | High | 0 | 1.13E-05 | 4.35 |
| FALSE | High | SMGSQEDI | [K].SMGSQ | 1xPhospho | O88543 1x | 0.000492 | 0 | 1 | 2 | 5 | O88543 | COP9 signa | O88543 [4 | 0 | 2313.995 | 0 | -1.18 | 0.000209 | 1.63 | 138.9 | 61.1 | 5.26 | 8.76 | | High | High | High | High | High | High | High | High | High | High | High | 0 | 0.000108 | 2.46 |
| FALSE | High | SMGSQEDI | [K].SMGSQ | 1xPhospho | O88543 1x | 0.000595 | 0 | 1 | 2 | 4 | O88543 | COP9 signa | O88543 [4 | 0 | 2329.99 | 0 | -1.05 | 0.00438 | 1.96 | 132.1 | 67.9 | 6.38 | 17.29 | | High | High | High | High | High | High | High | High | High | High | High | 0 | 0.000134 | 2.17 |
| FALSE | High | SMGTGDSA | [R].SMGTG | 1xPhospho | Q80YV2 1x | 2.49E-05 | 0 | 1 | 4 | 3 | Q80YV2 | Nuclear-int | Q80YV2 [3 | 0 | 2055.962 | 0 | -0.41 | 0.004517 | 3.67 | 112.9 | 87.1 | 4.24 | 13.56 | | High | High | High | High | High | High | High | High | High | High | High | 0 | 3.77E-06 | 2.79 |
| FALSE | High | SMGTGDSA | [R].SMGTG | 1xPhospho | Q80YV2 1x | 2.38E-06 | 0 | 1 | 4 | 11 | Q80YV2 | Nuclear-int | Q80YV2 [3 | 0 | 2071.957 | 0 | -0.48 | 0.002468 | 1.48 | 116.3 | 83.7 | 4.67 | 3.94 | | High | High | High | High | High | High | High | High | High | High | High | 0 | 2.87E-05 | 4.12 |
| FALSE | High | SMGTGDSA | [R].SMGTG | 1xPhospho | Q80YV2 1x | 2.26E-09 | 0 | 1 | 4 | 2 | Q80YV2 | Nuclear-int | Q80YV2 [3 | 1 | 2212.063 | 0 | -0.61 | 0.002487 | 5.47 | 119.8 | 80.2 | 5.39 | 7.21 | | High | High | High | High | High | High | High | High | High | High | High | 0 | 8.27E-05 | 3.61 |
| FALSE | High | SMGTGDSA | [R].SMGTG | 1xPhospho | Q80YV2 1x | 8.53E-07 | 0 | 1 | 4 | 37 | Q80YV2 | Nuclear-int | Q80YV2 [3 | 1 | 2228.058 | 0 | -0.5 | 0.000209 | 0.33 | 117.2 | 82.8 | 1.78 | 0.97 | | High | High | High | High | High | High | High | High | High | High | High | 0 | 9.49E-05 | 4.2 |
| FALSE | High | SMGYDDLD | [R].SMGYD | 1xPhospho | E9Q8Z8 1x | 0.000346 | 0 | 1 | 20 | 4 | E9Q8Z8 | Catenin de | E9Q8Z8 [2 | 0 | 2488.955 | 0 | -0.2 | 0.068753 | 1.89 | 108 | 92 | 6.88 | 8.82 | | High | High | High | High | High | High | High | High | High | High | High | 7.19E-05 | 0.000811 | 3.06 |
| FALSE | High | SMLQTPPD | [K].SMLQT | 1xPhospho | Q8BTI8 1xP | 2.58E-05 | 0 | 1 | 3 | 2 | Q8BTI8 | Serine/argi | Q8BTI8 [96 | 0 | 2141.071 | 0 | -0.51 | 0.039624 | 2.09 | 118.5 | 81.5 | 10.25 | 24.28 | | High | High | High | High | High | High | High | High | High | High | High | 0 | 0.000317 | 2.35 |
| FALSE | High | SMLQTPPD | [K].SMLQT | 1xPhospho | Q8BTI8 1xP | 6.26E-06 | 0 | 1 | 3 | 20 | Q8BTI8 | Serine/argi | Q8BTI8 [96 | 0 | 2157.066 | 0 | -0.51 | 0.006973 | 5.52 | 118.5 | 81.5 | 6.4 | 10.01 | | High | High | High | High | High | High | High | High | High | High | High | 2.58E-05 | 0.000376 | 3.22 |
| FALSE | High | SMLQTPPD | [K].SMLQT | 2xPhospho | Q8BTI8 2xP | 0.003282 | 0.00016 | 1 | 3 | 1 | Q8BTI8 | Serine/argi | Q8BTI8 [96 | 0 | 2221.038 | 0 | -0.76 | 0.003141 | 4.6 | 124.9 | 75.1 | 6.41 | 16.84 | | High | High | High | High | High | High | High | High | High | High | High | 7.19E-05 | 0.000904 | 2.42 |
| FALSE | High | SMPAGLTL | [R].SMPAG | 1xPhospho | E9Q6H8 1x | 6.67E-06 | 0 | 1 | 12 | 1 | E9Q6H8 | Pleckstrin h | E9Q6H8 [6 | 0 | 2250.151 | 0 | -0.17 | 0.090922 | 3.98 | 109.5 | 90.5 | 9.7 | 11.98 | | High | High | High | High | High | High | High | High | High | High | High | 0 | 8.58E-07 | 2.99 |
| FALSE | High | SMDSSDLS | [R].SMDSS | 1xPhospho | Q68FF6 1xl | 9.79E-08 | 0 | 1 | 2 | 13 | Q68FF6 | ARF GTPas | Q68FF6 [4 | 0 | 2855.367 | 0 | -0.51 | 0.007101 | 10.08 | 119.3 | 80.7 | 5.53 | 12.93 | | High | High | High | High | High | High | High | High | High | High | High | 0 | 7.49E-09 | 5.41 |
| FALSE | High | SLYGDVDS | [K].SLYGDV | 1xPhospho | Q3UHU5 1 | 2.27E-05 | 0 | 1 | 5 | 2 | Q3UHU5 | Microtubu | Q3UHU5 [5 | 0 | 2482.17 | 0 | -0.62 | 0.005735 | 5.64 | 121.8 | 78.2 | 10.48 | 5.64 | | High | High | High | High | High | High | High | High | High | High | High | 0 | 3.41E-06 | 3.59 |
| FALSE | High | SIYGDAEV | [R].SIYGDA | 1xPhospho | A0A087WC | 0.00045 | 0 | 1 | 2 | 1 | A0A087WC | MCG5930 | A0A087WC | 0 | 2326.117 | 0 | -0.23 | 0.263991 | 8.46 | 104.4 | 95.6 | 11.65 | 11.02 | | High | High | High | High | High | High | High | High | High | High | High | 0 | 9.76E-05 | 3.39 |
| FALSE | High | SIYDDISSP | [R].SIYDDIS | 2xPhospho | Q8R4R6 2x | 0.002192 | 0.000109 | 1 | 2 | 1 | Q8R4R6 | Nucleopor | Q8R4R6 [9 | 0 | 2355.072 | 0 | 0.25 | 0.244873 | 31.91 | 91.9 | 108.1 | 19 | 17.1 | | High | High | High | High | High | High | High | High | High | High | High | 4.94E-05 | 0.000577 | 3.47 |
| FALSE | High | SLTLGPSLS | [R].SLTLGP | 1xPhospho | B2RWU4 1 | 1.84E-06 | 0 | 2 | 11 | 2 | B2RWU4; C | Cordon-ble | B2RWU4 [3 | 0 | 2415.297 | 0 | 0.71 | 0.039035 | 7.55 | 75.7 | 124.3 | 7.29 | 26.98 | Shared | High | High | High | High | High | High | High | High | High | High | High | 0 | 2.01E-07 | 3.87 |
| FALSE | High | SLTNDWED | [K].SLTNDW | 1xPhospho | P11499 1xl | 0.00103 | 0 | 1 | 2 | 2 | P11499 | Heat shock | P11499 [30 | 0 | 2066.036 | 0 | 0.09 | 0.390634 | 7.98 | 95.9 | 104.1 | 7.1 | 9.63 | | High | High | High | High | High | High | High | High | High | High | High | 0 | 0.000246 | 3.17 |
| FALSE | High | SLTNDWEE | [K].SLTNDW | 1xPhospho | Q3UIF3 1xl | 0.000133 | 0 | 1 | 8 | 1 | Q3UIF3 | Putative ur | Q3UIF3 [31 | 0 | 2080.052 | 0 | -0.61 | 0.001585 | 3.87 | 121 | 79 | 3.74 | 6.67 | | High | High | High | High | High | High | High | High | High | High | High | 0 | 2.49E-05 | 3.7 |
| FALSE | High | SLTNLSFLT | [K].SLTNLS | 1xPhospho | Q8CH77 1x | 0.000111 | 0 | 1 | 3 | 1 | Q8CH77 | Neuron na | Q8CH77 [9 | 0 | 1993.03 | 0 | 0.63 | 0.025684 | 20.24 | 76.4 | 123.6 | 2.55 | 22.27 | | High | High | High | High | High | High | High | High | High | High | High | 0 | 2.02E-05 | 2.44 |
| FALSE | High | SLTNLVAAC | [K].SLTNLV | 1xPhospho | Q8CDD8 1x | 0.004824 | 0.000456 | 1 | 3 | 1 | Q8CDD8 | BTB/POZ d | Q8CDD8 [3 | 0 | 3808.83 | 0 | 0.25 | 0.27582 | 21.72 | 95.3 | 104.7 | 9.69 | 29.11 | | High | High | High | High | High | High | High | High | High | High | High | 0.000204 | 0.001394 | 2.51 |
| FALSE | High | SLTNLVAAC | [K].SLTNLV | 1xDeamida | Q8CDD8 1x | 0.039347 | 0.002793 | 1 | 3 | 1 | Q8CDD8 | BTB/POZ d | Q8CDD8 [3 | 0 | 3809.814 | 0 | 0.35 | 0.127946 | 21.8 | 90.4 | 109.6 | 11.61 | 8.48 | | High | High | High | High | High | High | High | High | High | High | High | 0.001453 | 0.01466 | 3.05 |
| FALSE | High | SLTPNPDSI | [R].SLTPNP | 1xPhospho | Q791V7 1x | 0.038346 | 0.002563 | 1 | 2 | 1 | Q791V7 | Uracil-DNA | Q791V7 [1 | 0 | 1497.678 | 0 | -0.77 | 0.009326 | 4.37 | 126.6 | 73.4 | 12.07 | 10.94 | | High | High | High | High | High | High | High | High | High | High | High | 0.001355 | 0.01425 | 2.81 |
| FALSE | High | SLTPPFR | [R].SLTPPF | 1xPhospho | Q8CJF7 1xl | 0.019107 | 0.001402 | 1 | 1 | 1 | Q8CJF7 | Protein ELY | Q8CJF7 [12 | 0 | 1126.586 | 0 | 0.37 | 0.006211 | 10.66 | 87.9 | 112.1 | 4.68 | 7.93 | | High | High | High | High | High | High | High | High | High | High | High | 0.00074 | 0.00648 | 1.32 |
| FALSE | High | SLTPPPSST | [R].SLTPPP | 1xPhospho | Q9EPQ8 1x | 0.001865 | 5.66E-05 | 1 | 5 | 3 | Q9EPQ8 | Transcripti | Q9EPQ8 [1 | 0 | 1768.913 | 0 | -0.45 | 0.005601 | 4.42 | 115.5 | 84.5 | 3.27 | 8.72 | | High | High | High | High | High | High | High | High | High | High | High | 2.58E-05 | 0.000478 | 1.64 |
| FALSE | High | SLTSCSSDF | [R].SLTSCS | 1xCarbami | Q6ZQ32 1x | 0.086562 | 0.006435 | 2 | 5 | 1 | Q6ZQ32; F | MKIAA085 | Q6ZQ32 [1 | 0 | 1682.766 | 0 | 0.29 | 0.023207 | 1.94 | 89.8 | 110.2 | 5.73 | 12.28 | Shared | High | High | High | High | High | High | High | High | High | High | High | 0.003346 | 0.03659 | 1.91 |
| FALSE | High | SLTSECHLL | [K].SLTSEC | 3xCarbami | A0A0R4J0C | 0.001831 | 5.66E-05 | 1 | 3 | 3 | A0A0R4J0C | Disks large | A0A0R4J0C | 0 | 2811.271 | 0 | -0.51 | 0.131737 | 12.88 | 120.1 | 79.9 | 14.25 | 28.75 | | High | High | High | High | High | High | High | High | High | High | High | 2.58E-05 | 0.000471 | 3.71 |
| FALSE | High | SITSPPSTSS | [R].SITSPPS | 1xPhospho | Q5CZY4 1x | 0.020319 | 0.001472 | 2 | 6 | 1 | Q5CZY4; P | Nuclear fac | Q5CZY4 [2 | 0 | 1730.898 | 0 | 1.32 | 0.000569 | 19.25 | 60.3 | 139.7 | 13 | 7.86 | Shared | High | High | High | High | High | High | High | High | High | High | High | 0.000786 | 0.006914 | 1.29 |
| FALSE | High | SLTSSDDAE | [R].SLTSSD | 1xPhospho | E9Q6J5-3 1 | 0.000656 | 0 | 1 | 3 | 2 | E9Q6J5-3 | Isoform Ne | E9Q6J5-3 [ | 0 | 1918.812 | 0 | 0.58 | 0.016245 | 18.74 | 80.3 | 119.7 | 13.78 | 7.95 | | High | High | High | High | High | High | High | High | High | High | High | 0.000382 | 0.003229 | 2.24 |
| FALSE | High | SLTSSDDAE | [R].SLTSSD | 1xPhospho | E9Q6J5-3 1 | 0.105058 | 0.008518 | 1 | 3 | 1 | E9Q6J5-3 | Isoform Ne | E9Q6J5-3 [ | 1 | 2276.07 | 0 | -0.25 | 0.027237 | 10.49 | 110.1 | 89.9 | 5.93 | 10.72 | | High | High | High | High | High | High | High | High | High | High | High | 0.004291 | 0.04587 | 1.73 |
| FALSE | High | SITSTPLSGI | [R].SITSTPL | 1xPhospho | Q9EP71 1x | 0.001986 | 5.66E-05 | 1 | 1 | 2 | Q9EP71 | Ankycorbir | Q9EP71 [2 | 0 | 1528.839 | 0 | 1.81 | 0.000763 | 11.43 | 42 | 158 | 12.63 | 8.56 | | High | High | High | High | High | High | High | High | High | High | High | 0.005054 | 0.05523 | 1.22 |
| FALSE | High | SITSTPLSGI | [R].SITSTPL | 2xPhospho | Q9EP71 2x | 0.027839 | 0.001755 | 1 | 1 | 1 | Q9EP71 | Ankycorbir | Q9EP71 [2 | 0 | 1608.805 | 0 | 1.65 | 0.001651 | 32.91 | 47.9 | 152.1 | 15.82 | 11.13 | | High | High | High | High | High | High | High | High | High | High | High | 0.000953 | 0.009907 | 2.58 |
| FALSE | High | SLVGSWLK | [R].SLVGSW | 1xPhospho | B1APX2 1x | 0.074071 | 0.0057 | 1 | 3 | 1 | B1APX2 | RIKEN cDN | B1APX2 [4 | 0 | 1427.806 | 0 | 0.55 | 0.01659 | 14.81 | 87 | 113 | 7.67 | 10.93 | | High | High | High | High | High | High | High | High | High | High | High | 0.002924 | 0.03034 | 1.74 |
| FALSE | High | SLVGTPYW | [K].SLVGTP | 1xPhospho | Q8BTW9 1 | 0.006085 | 0.00062 | 1 | 2 | 1 | Q8BTW9 | Serine/thre | Q8BTW9 [4 | 0 | 2129.07 | 0 | -0.59 | 0.00029 | 2.04 | 123.1 | 76.9 | 4.96 | 7.28 | | High | High | High | High | High | High | High | High | High | High | High | 0.000275 | 0.001792 | 1.26 |
| FALSE | High | SLVGTPYW | [K].SLVGTP | 1xPhospho | Q8BTW9 1 | 0.001751 | 5.66E-05 | 1 | 2 | 8 | Q8BTW9 | Serine/thre | Q8BTW9 [4 | 0 | 2145.065 | 0 | -0.39 | 0.002451 | 4.2 | 114.2 | 85.8 | 5.33 | 3.06 | | High | High | High | High | High | High | High | High | High | High | High | 7.19E-05 | 0.000858 | 3.09 |
| FALSE | High | SIVNTSPPC | [K].SIVNTS | 1xCarbami | Q9CXK9 1x | 0.002695 | 0.000109 | 1 | 2 | 2 | Q9CXK9 | RNA-bindir | Q9CXK9 [7 | 0 | 1439.692 | 0 | -0.27 | 0.004429 | 1.39 | 109.4 | 90.6 | 2.96 | 3.62 | | High | High | High | High | High | High | High | High | High | High | High | 4.94E-05 | 0.000723 | 1.47 |
| FALSE | High | SIVPSSPQS | [R].SIVPSS | 1xPhospho | B2RQE3 1x | 0.001127 | 0 | 1 | 6 | 2 | B2RQE3 | Stim2 prot | B2RQE3 [5 | 0 | 1494.751 | 0 | 1.97 | 0.000308 | 12.4 | 40.8 | 159.2 | 11.77 | 4.53 | | High | High | High | High | High | High | High | High | High | High | High | 0 | 0.000273 | 2.29 |
| FALSE | High | SLVPVGIPI | [K].SLVPVG | 1xPhospho | E9Q1M1 1 | 6.82E-08 | 0 | 1 | 3 | 1 | E9Q1M1 | PDZ domai | E9Q1M1 [2 | 0 | 2358.345 | 0 | 2.08 | 0.000357 | 16.99 | 36 | 164 | 11.98 | 6.01 | | High | High | High | High | High | High | High | High | High | High | High | 0 | 4.99E-09 | 4.34 |
| FALSE | High | SLVQDAPL | [R].SLVQDA | 1xCarbami | E9Q1K3 1x | 0.000233 | 0 | 1 | 2 | 1 | E9Q1K3 | Alpha-addu | E9Q1K3 [5 | 0 | 3445.685 | 0 | 0.81 | 0.000847 | 6.29 | 74.7 | 125.3 | 7.06 | 5.76 | | High | High | High | High | High | High | High | High | High | High | High | 0 | 4.66E-05 | 2.93 |
| FALSE | High | SLVQDAPL | [R].SLVQDA | 1xCarbami | E9Q1K3 1x | 0.000958 | 0 | 1 | 2 | 1 | E9Q1K3 | Alpha-addu | E9Q1K3 [5 | 0 | 3446.669 | 0 | 0.46 | 0.009603 | 3.92 | 86.9 | 113.1 | 11.31 | 13.14 | | High | High | High | High | High | High | High | High | High | High | High | 0 | 0.000228 | 5.53 |
| FALSE | High | SLVSPIPSPT | [R].SLVSPIP | 1xCarbami | P42128 1xl | 0.007673 | 0.000664 | 1 | 1 | 2 | P42128 | Forkhead b | P42128 [22 | 0 | 2730.374 | 0 | 1.06 | 0.042383 | 50.75 | 63.6 | 136.4 | 4.62 | 33.64 | | High | High | High | High | High | High | High | High | High | High | High | 0.000294 | 0.00233 | 2.49 |
| FALSE | High | SLVSPIPSPT | [R].SLVSPIP | 1xCarbami | P42128 2xl | 0.034281 | 0.002195 | 1 | 1 | 1 | P42128 | Forkhead b | P42128 [22 | 0 | 2810.34 | 0 | 0.15 | 0.360857 | 23.38 | 96.3 | 103.7 | 9.58 | 12.05 | | High | High | High | High | High | High | High | High | High | High | High | 0.001151 | 0.0125 | 2.76 |
| FALSE | High | SLVSPIPSPT | [R].SLVSPIP | 1xCarbami | P42128 3xl | 0.017809 | 0.001282 | 1 | 1 | 3 | P42128 | Forkhead b | P42128 [22 | 0 | 2890.306 | 0 | 1.18 | 0.011127 | 26.88 | 64.2 | 135.8 | 14.82 | 22.3 | | High | High | High | High | High | High | High | High | High | High | High | 0.000689 | 0.006001 | 2.83 |
| FALSE | High | SLVSQPTAI | [R].SLVSQP | 1xPhospho | F8VPU2 1x | 0.034878 | 0.002269 | 1 | 2 | 2 | F8VPU2 | FERM, Rho | F8VPU2 [3 | 0 | 2092.109 | 0 | 0.39 | 0.00988 | 3.39 | 85.3 | 114.7 | 12.37 | 7.26 | | High | High | High | High | High | High | High | High | High | High | High | 0.002403 | 0.02572 | 2.39 |
| FALSE | High | SIYDDISSP | [R].SIYDDIS | 1xPhospho | Q8R4R6 1x | 1.09E-06 | 0 | 1 | 2 | 5 | Q8R4R6 | Nucleopor | Q8R4R6 [9 | 0 | 2275.106 | 0 | -0.48 | 0.008377 | 1.62 | 120.5 | 79.5 | 12.28 | 6.19 | | High | High | High | High | High | High | High | High | High | High | High | 0 | 1.11E-07 | 3.64 |
| FALSE | High | SLSPPQSQS | [K].SLSPPQ | 1xPhospho | Q8BKX1 1x | 0.002212 | 0.000109 | 1 | 4 | 2 | Q8BKX1 | Brain-spec | Q8BKX1 [3 | 0 | 1596.84 | 0 | 1.82 | 0.00014 | 6.45 | 45.3 | 154.7 | 4.57 | 1.95 | | High | High | High | High | High | High | High | High | High | High | High | 4.94E-05 | 0.000583 | 1.81 |
| FALSE | High | SMPELLLLS | [K].SMPELL | 1xCarbami | B0V2M3 1x | 0.007274 | 0.00062 | 1 | 3 | 1 | B0V2M3 | Zinc finger | B0V2M3 [1 | 0 | 2424.305 | 0 | -0.25 | 0.216599 | 13.16 | 106.9 | 93.1 | 7.77 | 18.76 | | High | High | High | High | High | High | High | High | High | High | High | 0.000275 | 0.002187 | 3.33 |
| FALSE | High | SLSPNHNTI | [R].SLSPNH | 1xPhospho | E9PUJ6 1xP | 4.02E-06 | 0 | 1 | 5 | 1 | E9PUJ6 | E3 ubiquiti | E9PUJ6 [28 | 0 | 1991.073 | 0 | 0.3 | 0.492053 | 41.04 | 92.6 | 107.4 | 26.44 | 30.52 | | High | High | High | High | High | High | High | High | High | High | High | 0 | 4.87E-07 | 2.62 |
| FALSE | High | SLSPLSGTT | [R].SLSPLS | 1xPhospho | F6RJ39 1xP | 3.32E-05 | 0 | 2 | 4 | 6 | F6RJ39; Q8 | Apoptotic | F6RJ39 [42 | 0 | 1744.913 | 0 | 0.03 | 0.583092 | 8.54 | 101 | 99 | 4.18 | 6.12 | Shared | High | High | High | High | High | High | High | High | High | High | High | 4.94E-05 | 0.000751 | 2.68 |
| FALSE | High | SIPSLQSEY | [R].SIPSLQS | 1xPhospho | Q6GQT0 1 | 0.008927 | 0.000736 | 1 | 2 | 1 | Q6GQT0 | Dual specif | Q6GQT0 [5 | 0 | 1731.819 | 0 | 0.83 | 0.012786 | 13.07 | 70.7 | 129.3 | 20.35 | 17.98 | | High | High | High | High | High | High | High | High | High | High | High | 0.000327 | 0.002776 | 2.2 |
| FALSE | High | SLPSPQAAI | [K].SLPSPQ | 1xPhospho | E9Q1M1 1 | 0.08447 | 0.006345 | 1 | 3 | 1 | E9Q1M1 | PDZ domai | E9Q1M1 [1 | 0 | 1573.85 | 0 | 2.89 | 0.000209 | 19.16 | 24.2 | 175.8 | 13.97 | 4.82 | | High | High | High | High | High | High | High | High | High | High | High | 0.003265 | 0.03526 | 1.54 |
| FALSE | High | SLPSRSPPI | [K].SLPSRS | 3xPhospho | A2ALK8 3xl | 0.098533 | 0.007902 | 1 | 1 | 1 | A2ALK8 | Tyrosine-p | A2ALK8 [3 | 1 | 2076.927 | 0 | 0.49 | 0.004033 | 7.34 | 83.9 | 116.1 | 9.15 | 6.78 | | High | High | High | High | High | High | High | High | High | High | High | 0.003991 | 0.04254 | 1.82 |
| FALSE | High | SLPTSSSSS | [R].SLPTSS | 1xPhospho | Q8CHI8 1xl | 4E-07 | 0 | 1 | 5 | 5 | Q8CHI8 | E1A-bindin | Q8CHI8 [6 | 0 | 2607.286 | 0 | -0.15 | 0.217316 | 2.96 | 109.4 | 90.6 | 19.05 | 8.6 | | High | High | High | High | High | High | High | High | High | High | High | 0 | 3.94E-07 | 3.68 |
| FALSE | High | SLPTTVPES | [R].SLPTTV | 1xPhospho | Q6ZQ58 1x | 2.21E-05 | 0 | 1 | 3 | 8 | Q6ZQ58 | La-related | Q6ZQ58 [7 | 0 | 1769.867 | 0 | 0.79 | 0.000275 | 2.77 | 72.5 | 127.5 | 5.8 | 3.84 | | High | High | High | High | High | High | High | High | High | High | High | 0 | 3.29E-06 | 2.85 |
| FALSE | High | SLPTTVPES | [R].SLPTTV | 1xDeamida | Q6ZQ58 1x | 0.000352 | 0 | 1 | 3 | 2 | Q6ZQ58 | La-related | Q6ZQ58 [7 | 0 | 1770.851 | 0 | 0.79 | 0.001864 | 7.62 | 73.6 | 126.4 | 9.3 | 8.83 | | High | High | High | High | High | High | High | High | High | High | High | 0.000723 | 0.006342 | 2.79 |
| FALSE | High | SLPTTVPES | [R].SLPTTV | 2xPhospho | Q6ZQ58 2x | 0.016021 | 0.001124 | 1 | 3 | 2 | Q6ZQ58 | La-related | Q6ZQ58 [7 | 0 | 1849.834 | 0 | 0.29 | 0.094628 | 22.42 | 82.1 | 117.9 | 13.26 | 16.81 | | High | High | High | High | High | High | High | High | High | High | High | 0.000585 | 0.005303 | 2.53 |
| FALSE | High | SLPVSGTLE | [K].SLPVSG | 1xPhospho | F6ZBR8 1xl | 4.35E-05 | 0 | 2 | 2 | 1 | F6ZBR8; O8 | Protein Wi | F6ZBR8 [23 | 0 | 1752.909 | 0 | -0.46 | 0.017349 | 4.18 | 116.2 | 83.8 | 13.17 | 7.86 | Shared | High | High | High | High | High | High | High | High | High | High | High | 0 | 7.04E-06 | 3.12 |
| FALSE | High | SLQDDTEP | [K].SLQDD | 2xPhospho | G3X9K3 2x | 0.001659 | 5.66E-05 | 1 | 2 | 2 | G3X9K3 | Brefeldin A | G3X9K3 [2 | 0 | 4299.859 | 0 | 0.62 | 0.035932 | 28.74 | 78.7 | 121.3 | 18.43 | 14.4 | | High | High | High | High | High | High | High | High | High | High | High | 2.58E-05 | 0.00042 | 3.45 |
| FALSE | High | SLQDLSQP | [R].SLQDLS | 1xPhospho | Q6ZQE1 1x | 0.004572 | 0.000456 | 1 | 3 | 1 | Q6ZQE1 | MKIAA029 | Q6ZQE1 [8 | 0 | 2368.241 | 0 | 1.01 | 0.000442 | 11.48 | 69.3 | 130.7 | 8.7 | 4.91 | | High | High | High | High | High | High | High | High | High | High | High | 0.000204 | 0.001313 | 2.35 |
| FALSE | High | SIQDLTVTG | [R].SIQDLT | 1xPhospho | Q923A8 1x | 0.000118 | 0 | 1 | 5 | 2 | Q923A8 | Map3k7 pr | Q923A8 [4 | 0 | 2184.075 | 0 | -0.33 | 0.014939 | 7.04 | 113.3 | 86.7 | 2.75 | 9.04 | | High | High | High | High | High | High | High | High | High | High | High | 0 | 2.17E-05 | 3.76 |
| FALSE | High | SIQDLTVTG | [R].SIQDLT | 2xPhospho | Q923A8 2x | 0.000446 | 0 | 1 | 5 | 1 | Q923A8 | Map3k7 pr | Q923A8 [4 | 0 | 2264.041 | 0 | 0.6 | 0.007099 | 7.03 | 79.4 | 120.6 | 6.5 | 12.63 | | High | High | High | High | High | High | High | High | High | High | High | 0 | 9.66E-05 | 3.19 |
| FALSE | High | SIQFVDWC | [R].SIQFVD | 1xCarbami | A0A0A0MC | 0.000411 | 0 | 1 | 4 | 1 | A0A0A0MC | Tubulin alp | A0A0A0MC | 0 | 1893.881 | 0 | 0.14 | 0.262885 | 8.41 | 96.5 | 103.5 | 10 | 10.78 | | High | High | High | High | High | High | High | High | High | High | High | 0 | 0.000088 | 3.69 |
| FALSE | High | SLPPDPESE | [R].SLPPDP | 1xPhospho | Q80UW5 1 | 0.007339 | 0.00062 | 1 | 2 | 2 | Q80UW5 | Serine/thre | Q80UW5 [ | 0 | 1550.682 | 0 | 0.21 | 0.04392 | 10.43 | 89.3 | 110.7 | 12.8 | 9.99 | | High | High | High | High | High | High | High | High | High | High | High | 0.000275 | 0.002211 | 2.91 |
| FALSE | High | SLQQIAVEE | [K].SLQQIA | 2xCarbami | Q9CXH7 1x | 0.000126 | 0 | 1 | 1 | 1 | Q9CXH7 | Shugoshin | Q9CXH7 [1 | 0 | 4524.051 | 0 | 0.18 | 0.193007 | 12.45 | 94.7 | 105.3 | 7.72 | 15.34 | | High | High | High | High | High | High | High | High | High | High | High | 0 | 2.33E-05 | 4.64 |
| FALSE | High | SLQSPAAN | [K].SLQSPA | 1xPhospho | Q8BHK9 1x | 9.2E-05 | 0 | 1 | 3 | 1 | Q8BHK9 | DNA excisi | Q8BHK9 [9 | 0 | 1339.657 | 0 | 0.54 | 0.002151 | 8.47 | 80.3 | 119.7 | 11.14 | 8.84 | | High | High | High | High | High | High | High | High | High | High | High | 0 | 1.63E-05 | 2.18 |
| FALSE | High | SIQTSPTEE | [K].SIQTSP | 1xPhospho | Q9JL04 1xP | 0.051694 | 0.003745 | 1 | 1 | 2 | Q9JL04 | Formin-2 C | Q9JL04 [72 | 0 | 1513.71 | 0 | 0.21 | 0.073474 | 5 | 91.1 | 108.9 | 8.13 | 6.34 | | High | High | High | High | High | High | High | High | High | High | High | 0.001992 | 0.02008 | 1.8 |
| FALSE | High | SLRQSFR | [K].SLRQSF | 1xPhospho | A0A0R4J0S | 0.104222 | 0.008348 | 2 | 4 | 1 | A0A0R4J0S | Lethal(2) gi | A0A0R4J0S | 1 | 1202.624 | 0 | -1.09 | 0.00109 | 1.16 | 134.5 | 65.5 | 6.81 | 11.93 | Shared | High | High | High | High | High | High | High | High | High | High | High | 0.004264 | 0.04548 | 0.81 |
| FALSE | High | SIRTPEPVV | [K].SIRTPEI | 1xPhospho | E9QK89 1x | 3.36E-06 | 0 | 1 | 2 | 1 | E9QK89 | Mediator c | E9QK89 [1 | 1 | 3302.589 | 0 | -0.61 | 0.016168 | 6.49 | 120.8 | 79.2 | 11.37 | 8.51 | | High | High | High | High | High | High | High | High | High | High | High | 0 | 3.96E-07 | 2.89 |
| FALSE | High | SLSADEFEI | [R].SLSADE | 1xPhospho | B2KFD2 1x | 1.85E-06 | 0 | 1 | 3 | 3 | B2KFD2 | 28104740 | B2KFD2 [1 | 0 | 2228.162 | 0 | -1.58 | 0.000303 | 2.2 | 149.8 | 50.2 | 2.22 | 11.43 | | High | High | High | High | High | High | High | High | High | High | High | 0 | 2.03E-07 | 3.98 |
| FALSE | High | SLSADNFIG | [R].SLSADN | 1xPhospho | D3YUJ3 1xl | 3.01E-06 | 0 | 1 | 2 | 4 | D3YUJ3 | Cyclin Y-lik | D3YUJ3 [3 | 0 | 1629.82 | 0 | -0.05 | 0.204146 | 8.47 | 101.9 | 98.1 | 3.89 | 5.61 | | High | High | High | High | High | High | High | High | High | High | High | 0 | 0.000229 | 3.43 |
| FALSE | High | SLSADNFIG | [R].SLSADN | 1xDeamida | D3YUJ3 1xl | 0.000139 | 0 | 1 | 2 | 1 | D3YUJ3 | Cyclin Y-lik | D3YUJ3 [3 | 0 | 1630.804 | 0 | 0.02 | 0.633433 | 17.96 | 102 | 98 | 16.04 | 7.7 | | High | High | High | High | High | High | High | High | High | High | High | 0 | 2.59E-05 | 2.42 |
| FALSE | High | SISADSFDC | [R].SISADS | 1xPhospho | Q9D219 1x | 0.000458 | 0 | 1 | 1 | 1 | Q9D219 | B-cell CLL/l | Q9D219 [1 | 0 | 1434.646 | 0 | -0.21 | 0.004406 | 4.07 | 107.2 | 92.8 | 2.11 | 4.08 | | High | High | High | High | High | High | High | High | High | High | High | 0 | 9.93E-05 | 1.46 |
| FALSE | High | SISADSFDC | [R].SISADS | 2xPhospho | Q9D219 2x | 0.004867 | 0.000504 | 1 | 1 | 1 | Q9D219 | B-cell CLL/l | Q9D219 [1 | 0 | 1514.613 | 0 | 0.4 | 0.069654 | 8.37 | 91.6 | 108.4 | 11.1 | 7.88 | | High | High | High | High | High | High | High | High | High | High | High | 0.000225 | 0.001404 | 3.02 |
| FALSE | High | SISANQK | [R].SISANQ | 1xPhospho | C9K101 1xl | 0.065952 | 0.004766 | 1 | 3 | 1 | C9K101 | Non-specif | C9K101 [4 | 0 | 1285.692 | 0 | -0.26 | 0.065467 | 0.39 | 109 | 91 | 7.79 | 8.6 | | High | High | High | High | High | High | High | High | High | High | High | 0.002455 | 0.02646 | 1.19 |
| FALSE | High | SLSAPGNLI | [R].SLSAPG | 1xPhospho | Q8C078-5 | 0.009332 | 0.000736 | 1 | 5 | 1 | Q8C078-5 | Isoform 5 c | Q8C078-5 | 0 | 1638.923 | 0 | 0.85 | 0.000477 | 8.43 | 72 | 128 | 4.17 | 4.26 | | High | High | High | High | High | High | High | High | High | High | High | 0.000327 | 0.002912 | 2.8 |
| FALSE | High | SISAPVIFDI | [K].SISAPVI | 1xPhospho | Q3U5L3 1x | 0.003721 | 0.00021 | 1 | 3 | 2 | Q3U5L3 | Putative ur | Q3U5L3 [1 | 0 | 1413.734 | 0 | 1.82 | 0.005082 | 24.41 | 43.2 | 156.8 | 4.41 | 22.73 | | High | High | High | High | High | High | High | High | High | High | High | 9.42E-05 | 0.001035 | 2.17 |
| FALSE | High | SISASDLSFI | [R].SISASDI | 1xPhospho | Q640L5 1xl | 0.000462 | 0 | 1 | 1 | 1 | Q640L5 | Coiled-coil | Q640L5 [13 | 0 | 1592.834 | 0 | -0.93 | 0.000193 | 0.93 | 131.2 | 68.8 | 4.43 | 6.39 | | High | High | High | High | High | High | High | High | High | High | High | 0 | 0.000101 | 2.37 |
| FALSE | High | SLQQIAVEE | [K].SLQQIA | 2xCarbami | Q9CXH7 2x | 8.43E-06 | 0 | 1 | 1 | 1 | Q9CXH7 | Shugoshin | Q9CXH7 [1 | 0 | 4604.018 | 0 | 0.17 | 0.111148 | 12.21 | 92.7 | 107.3 | 10.14 | 4.93 | | High | High | High | High | High | High | High | High | High | High | High | 0 | 1.11E-06 | 5.52 |
| FALSE | High | SLSASPALG | [R].SLSASP | 1xPhospho | P58058 1xl | 0.003892 | 0.000261 | 1 | 5 | 3 | P58058 | NAD kinase | P58058 [46 | 0 | 1656.897 | 0 | 0.28 | 0.145775 | 9.49 | 92 | 108 | 6.24 | 8.78 | | High | High | High | High | High | High | High | High | High | High | High | 0.0032 | 0.03369 | 1.6 |
| FALSE | High | SIPNLEMPI | [K].SIPNLEI | 1xPhospho | E9QAU4 1x | 0.067619 | 0.004864 | 1 | 11 | 1 | E9QAU4 | Enhancer t | E9QAU4 [1 | 0 | 2192.144 | 0 | 0.23 | 0.037708 | 9.59 | 94 | 106 | 2.71 | 9.82 | | High | High | High | High | High | High | High | High | High | High | High | 0.002485 | 0.02719 | 3.37 |
| FALSE | High | SLPLSSPTM | [K].SLPLSSI | 1xPhospho | E9PUR0 1x | 0.090153 | 0.00698 | 1 | 4 | 1 | E9PUR0 | Ankyrin rep | E9PUR0 [1 | 0 | 1614.858 | 0 | -1.1 | 0.002273 | 3.27 | 135.8 | 64.2 | 5.85 | 12.68 | | High | High | High | High | High | High | High | High | High | High | High | 0.003606 | 0.03824 | 2.39 |
| FALSE | High | SLLVLDEVP | [R].SLLVLD | 1xPhospho | E9QAU4 1x | 0.004106 | 0.00036 | 1 | 3 | 1 | E9QAU4 | Enhancer t | E9QAU4 [1 | 0 | 2681.4 | 0 | -0.65 | 0.010742 | 4.62 | 122.1 | 77.9 | 10.05 | 9.42 | | High | High | High | High | High | High | High | High | High | High | High | 0.000161 | 0.001166 | 3.58 |
| FALSE | High | SILVSPTGL | [K].SILVSPT | 1xPhospho | Q91YK2 1x | 0.000876 | 0 | 1 | 2 | 2 | Q91YK2 | Ribosomal | Q91YK2 [6 | 0 | 1438.787 | 0 | 0.43 | 0.019501 | 16.68 | 87.6 | 112.4 | 4.21 | 10.18 | | High | High | High | High | High | High | High | High | High | High | High | 0 | 0.000206 | 1.68 |
| FALSE | High | SLMSSPED | [R].SLMSS | 2xPhospho | P14873 2xl | 0.001107 | 0 | 2 | 2 | 2 | P14873; B2 | Microtubu | P14873 [82 | 0 | 1825.846 | 0 | -0.72 | 0.015648 | 8.81 | 124.5 | 75.5 | 10.24 | 17.14 | Shared | High | High | High | High | High | High | High | High | High | High | High | 2.58E-05 | 0.00038 | 3.11 |
| FALSE | High | SLMSVSGII | [R].SLMSV | 1xPhospho | Q8K2J7 1xl | 6.31E-06 | 0 | 1 | 2 | 2 | Q8K2J7 | RELT-like p | Q8K2J7 [24 | 0 | 2641.356 | 0 | 0.61 | 0.019374 | 15.25 | 79 | 121 | 14.37 | 5.13 | | High | High | High | High | High | High | High | High | High | High | High | 0 | 8.05E-07 | 4.49 |
| FALSE | High | SLNEASMD | [K].SLNEAS | 1xPhospho | Q9D8C3 1x | 0.002355 | 0.000109 | 1 | 1 | 1 | Q9D8C3 | Putative bi | Q9D8C3 [2 | 0 | 2311.088 | 0 | -0.44 | 0.016788 | 10.7 | 117.2 | 82.8 | 10.29 | 7.58 | | High | High | High | High | High | High | High | High | High | High | High | 4.94E-05 | 0.000621 | 2.16 |
| FALSE | High | SLNFQEDD | [K].SLNFQE | 1xPhospho | Q9JJT9 1xP | 0.075304 | 0.005727 | 1 | 2 | 2 | Q9JJT9 | Phosphory | Q9JJT9 [33 | 0 | 1735.737 | 0 | -0.06 | 0.278934 | 5.4 | 103.4 | 96.6 | 2.91 | 7.91 | | High | High | High | High | High | High | High | High | High | High | High | 0.009441 | 0.1022 | 2.05 |
| FALSE | High | SLNFQEDD | [K].SLNFQE | 1xPhospho | Q9JJT9 1xP | 0.020862 | 0.001472 | 1 | 2 | 4 | Q9JJT9 | Phosphory | Q9JJT9 [33 | 1 | 4556.051 | 0 | 0.31 | 0.231211 | 8.92 | 88.7 | 111.3 | 13.48 | 10.41 | | High | High | High | High | High | High | High | High | High | High | High | 0.001011 | 0.01108 | 3.24 |
| FALSE | High | SLNFQEDD | [K].SLNFQE | 2xPhospho | Q9JJT9 2xP | 0.042141 | 0.003162 | 1 | 2 | 2 | Q9JJT9 | Phosphory | Q9JJT9 [33 | 1 | 4636.018 | 0 | 0.19 | 0.083238 | 7.4 | 92 | 108 | 12.82 | 12.36 | | High | High | High | High | High | High | High | High | High | High | High | 0.002924 | 0.03035 | 4.93 |

| | | | | | | | | | | | | | | | | | | | | | | | | | | | | | | | | | | | | | | |
|---|---|---|---|---|---|---|---|---|---|---|---|---|---|---|---|---|---|---|---|---|---|---|---|---|---|---|---|---|---|---|---|---|---|---|---|---|---|---|
| FALSE | High | SLNGGSDA | [K].SLNGG | 1xDeamida | Q99JF8 1x | 0.093879 | 0.007306 | 1 | 1 | 1 | Q99JF8 | PC4 and SF | Q99JF8 [47 | 0 | 3133.35 | 0 | -0.56 | 0.023421 | 7.9 | 118 | 82 | 5.16 | 17.49 | | High | High | High | High | High | High | High | High | High | High | High | 0.003727 | 0.04004 | 2.48 |
| FALSE | High | SLNLVDSP( | [K].SLNLVD | 1xPhospho | Q8BWJ3 1x | 0.101749 | 0.008194 | 1 | 2 | 1 | Q8BWJ3 | Phosphory | Q8BWJ3 [7 | 0 | 1962.108 | 0 | 0.33 | 0.206097 | 32.02 | 88.7 | 111.3 | 11.8 | 16.11 | | High | High | High | High | High | High | High | High | High | High | High | 0.004126 | 0.04404 | 1.78 |
| FALSE | High | SLNPALDG | [R].SLNPAL | 1xCarbami | Q8CGF6 1x | 0.001572 | 5.66E-05 | 1 | 2 | 1 | Q8CGF6 | WD repeat | Q8CGF6 [3 | 0 | 2437.22 | 0 | 1.26 | 0.001724 | 10.22 | 60.4 | 139.6 | 14.46 | 10.04 | | High | High | High | High | High | High | High | High | High | High | High | 2.58E-05 | 0.000395 | 2.9 |
| FALSE | High | SINPDEAV | [K].SINPDE | 1xPhospho | P63017 1x | 0.021607 | 0.001507 | 1 | 16 | 1 | P63017 | Heat shock | P63017 [36 | 0 | 2798.438 | 0 | 1.06 | 0.003621 | 16.69 | 63.5 | 136.5 | 11.59 | 17.18 | | High | High | High | High | High | High | High | High | High | High | High | 0.000801 | 0.007434 | 2.91 |
| FALSE | High | SLNQPTPPI | [K].SLNQPT | 1xPhospho | B1ARU1 1x | 0.00614 | 0.00062 | 2 | 16 | 1 | B1ARU1; F | Microtubu | B1ARU1 [5 | 0 | 2573.345 | 0 | -0.11 | 0.414699 | 15.71 | 101.8 | 98.2 | 10.94 | 10.26 | Shared | High | High | High | High | High | High | High | High | High | High | High | 0.000275 | 0.00181 | 2.98 |
| FALSE | High | SLPITIEMLI | [R].SLPITIE | 1xPhospho | Q501J7-3 1 | 0.019791 | 0.001438 | 1 | 3 | 1 | Q501J7-3 | Isoform 3 ( | Q501J7-3 [ | 0 | 1682.957 | 0 | 1.75 | 0.000654 | 6.98 | 45.1 | 154.9 | 7.63 | 9.02 | | High | High | High | High | High | High | High | High | High | High | High | 0.000773 | 0.00671 | 1.51 |
| FALSE | High | SLNQPTPPI | [K].SLNQPT | 1xPhospho | B1ARU1 1x | 0.062209 | 0.004646 | 2 | 16 | 1 | B1ARU1; F | Microtubu | B1ARU1 [5 | 0 | 2589.34 | 0 | 0.34 | 0.010447 | 9.46 | 88.2 | 111.8 | 6.05 | 9.79 | Shared | High | High | High | High | High | High | High | High | High | High | High | 0.002381 | 0.02479 | 2.84 |
| FALSE | High | SLNSDVSCI | [K].SLNSDV | 1xCarbami | G3X963 1x | 6.74E-07 | 0 | 1 | 3 | 25 | G3X963 | ATPase fan | G3X963 [7 | 0 | 4191.888 | 0 | 0.93 | 0.015227 | 3.96 | 74.2 | 125.8 | 12.42 | 16.97 | | High | High | High | High | High | High | High | High | High | High | High | 0 | 5.35E-07 | 4.87 |
| FALSE | High | SLNSVVTSI | [R].SLNSVV | 1xPhospho | A2AG68 1x | 0.007742 | 0.000664 | 1 | 3 | 1 | A2AG68 | Copper-tra | A2AG68 [1 | 0 | 1813.935 | 0 | -0.02 | 0.825725 | 8.32 | 101.7 | 98.3 | 5.5 | 5.38 | | High | High | High | High | High | High | High | High | High | High | High | 0.000294 | 0.002367 | 2.02 |
| FALSE | High | SLNVRSPEI | [K].SLNVRS | 1xPhospho | Q3UQS8 1x | 0.001898 | 5.66E-05 | 1 | 1 | 1 | Q3UQS8 | RNA-bindir | Q3UQS8 [1 | 1 | 2258.183 | 0 | 2.12 | 0.003177 | 14.63 | 37.5 | 162.5 | 12.59 | 20.82 | | High | High | High | High | High | High | High | High | High | High | High | 2.58E-05 | 0.000488 | 3.2 |
| FALSE | High | SINYSELDC | [K].SINYSEI | 1xPhospho | Q60848 1x | 1.13E-05 | 0 | 1 | 2 | 8 | Q60848 | Lymphocyt | Q60848 [4 | 0 | 2437.194 | 0 | -0.86 | 0.003537 | 7.92 | 128 | 72 | 7.95 | 8.82 | | High | High | High | High | High | High | High | High | High | High | High | 0 | 3.34E-05 | 3.95 |
| FALSE | High | SLPAGDAL | [R].SLPAGD | 1xPhospho | Q60875 1x | 0.002441 | 0.000109 | 1 | 8 | 3 | Q60875 | Rho guanir | Q60875 [8 | 0 | 2339.163 | 0 | -0.73 | 0.009193 | 2.08 | 123.8 | 76.2 | 12.38 | 9.23 | | High | High | High | High | High | High | High | High | High | High | High | 0.000275 | 0.0018 | 3.2 |
| FALSE | High | SLPASGTP( | [K].SLPASG | 1xPhospho | F2Z3U3 1x | 7.16E-06 | 0 | 1 | 1 | 4 | F2Z3U3 | Ras associa | F2Z3U3 [8 | 0 | 1975.067 | 0 | 0.63 | 0.009837 | 12.64 | 77.3 | 122.7 | 7.8 | 12.69 | | High | High | High | High | High | High | High | High | High | High | High | 0 | 9.31E-07 | 3.07 |
| FALSE | High | SLPASPSQS | [K].SLPASP | 1xPhospho | Q3U0X6 1x | 3.08E-07 | 0 | 1 | 2 | 3 | Q3U0X6 | Putative ur | Q3U0X6 [8 | 0 | 3084.529 | 0 | 0.15 | 0.434334 | 20.03 | 98 | 102 | 15.22 | 8.11 | | High | High | High | High | High | High | High | High | High | High | High | 0 | 3.95E-08 | 4.19 |
| FALSE | High | SLPASPSQS | [K].SLPASP | 1xDeamida | Q3U0X6 1x | 2.16E-07 | 0 | 1 | 2 | 9 | Q3U0X6 | Putative ur | Q3U0X6 [8 | 0 | 3085.513 | 0 | 0.1 | 0.351355 | 11.33 | 98.1 | 101.9 | 8.35 | 11.69 | | High | High | High | High | High | High | High | High | High | High | High | 0 | 1.81E-08 | 5.11 |
| FALSE | High | SIPDTPVPT | [R].SIPDTP | 1xPhospho | Q8K072 1x | 0.000337 | 0 | 1 | 1 | 1 | Q8K072 | Receptor e | Q8K072 [1 | 0 | 2852.396 | 0 | 0.6 | 0.018482 | 19.83 | 79.3 | 120.7 | 19.79 | 16.84 | | High | High | High | High | High | High | High | High | High | High | High | 0 | 7.02E-05 | 3 |
| FALSE | High | SIPENHPET | [K].SIPENH | 1xPhospho | Q6KCD5 1x | 0.000645 | 0 | 1 | 3 | 1 | Q6KCD5 | Nipped-B-l | Q6KCD5 [5 | 0 | 1786.914 | 0 | -0.08 | 0.165747 | 11.92 | 103.1 | 96.9 | 2.47 | 10.41 | | High | High | High | High | High | High | High | High | High | High | High | 0 | 0.000146 | 2.35 |
| FALSE | High | SLPGLASSV | [R].SLPGLA | 1xPhospho | Q9CWT6 1 | 0.033404 | 0.002195 | 1 | 1 | 2 | Q9CWT6 | Probable A | Q9CWT6 [ | 0 | 2104.146 | 0 | -0.09 | 0.84669 | 11.85 | 100.9 | 99.1 | 11.05 | 7.79 | | High | High | High | High | High | High | High | High | High | High | High | 0.001151 | 0.01213 | 2.88 |
| FALSE | High | SIPISPSASC | [R].SIPISPS | 1xPhospho | E9PU87 1x | 8.5E-06 | 0 | 1 | 4 | 1 | E9PU87 | Serine/thre | E9PU87 [8 | 0 | 2071.09 | 0 | 0.58 | 0.000474 | 1.67 | 80.6 | 119.4 | 2.69 | 5.51 | | High | High | High | High | High | High | High | High | High | High | High | 0 | 1.12E-06 | 3.15 |
| FALSE | High | SLPLSSPTM | [K].SLPLSSI | 1xPhospho | E9PUR0 1x | 0.023792 | 0.001612 | 1 | 4 | 1 | E9PUR0 | Ankyrin re | E9PUR0 [1 | 0 | 1598.863 | 0 | -1.11 | 0.000608 | 2.73 | 133.9 | 66.1 | 5.42 | 11.61 | | High | High | High | High | High | High | High | High | High | High | High | 0.000859 | 0.008299 | 1.42 |
| FALSE | High | SLNSDVSCI | [K].SLNSDV | 1xCarbami | G3X963 1x | 8.55E-08 | 0 | 1 | 3 | 5 | G3X963 | ATPase fan | G3X963 [7 | 0 | 4190.904 | 0 | 0.69 | 0.053913 | 34.57 | 76.4 | 123.6 | 8.36 | 18.83 | | High | High | High | High | High | High | High | High | High | High | High | 0 | 6.44E-09 | 5.22 |
| FALSE | High | SLSAVDAD | [K].SLSAVD | 1xPhospho | B2RSV7 1x | 0.004491 | 0.000409 | 1 | 2 | 2 | B2RSV7 | Fgd6 prote | B2RSV7 [5 | 0 | 1242.593 | 0 | 0.14 | 0.129185 | 9.02 | 95.7 | 104.3 | 2.85 | 8.97 | | High | High | High | High | High | High | High | High | High | High | High | 0.000204 | 0.001289 | 2.02 |
| FALSE | High | SLSCSDK | [R].SLSCSD | 1xCarbami | B7ZNA0 1x | 0.043236 | 0.003228 | 2 | 8 | 1 | B7ZNA0; Q | Shroom3 p | B7ZNA0 [1 | 0 | 1334.643 | 0 | 0.72 | 0.01021 | 23.31 | 75.8 | 124.2 | 14.2 | 11.97 | Shared | High | High | High | High | High | High | High | High | High | High | High | 0.001651 | 0.01628 | 1.24 |
| FALSE | High | SLSDATML | [K].SLSDAT | 2xPhospho | G5E8J4 2x | 1.06E-05 | 0 | 1 | 4 | 2 | G5E8J4 | Tyrosine-p | G5E8J4 [70 | 0 | 3100.243 | 0 | 0.62 | 0.01306 | 6.38 | 77.9 | 122.1 | 11.19 | 10.36 | | High | High | High | High | High | High | High | High | High | High | High | 0 | 1.43E-06 | 4.08 |
| FALSE | High | SLSGSSPCP | [R].SLSGSS | 1xCarbami | Q8BTI8 1x | 3.94E-05 | 0 | 1 | 4 | 7 | Q8BTI8 | Serine/argi | Q8BTI8 [77 | 0 | 1557.775 | 0 | -0.65 | 0.026353 | 6.8 | 122.1 | 77.9 | 16.47 | 10.52 | | High | High | High | High | High | High | High | High | High | High | High | 0 | 3.07E-05 | 2.35 |
| FALSE | High | SLSGSSPCP | [R].SLSGSS | 1xCarbami | Q8BTI8 2x | 0.012397 | 0.000889 | 1 | 4 | 4 | Q8BTI8 | Serine/argi | Q8BTI8 [77 | 0 | 1637.741 | 0 | 0.25 | 0.269248 | 9.93 | 91.7 | 108.3 | 9.55 | 8.14 | | High | High | High | High | High | High | High | High | High | High | High | 0.000449 | 0.003991 | 2.71 |
| FALSE | High | SLSGTLTEL | [R].SLSGTL | 1xPhospho | S4R1P5 1x | 0.017344 | 0.001204 | 1 | 8 | 2 | S4R1P5 | Dystonin O | S4R1P5 [48 | 0 | 1816.935 | 0 | 1.3 | 0.001537 | 4.46 | 57.7 | 142.3 | 11.13 | 9.1 | | High | High | High | High | High | High | High | High | High | High | High | 0.001308 | 0.0138 | 3.23 |
| FALSE | High | SISGTPAET | [R].SISGTP | 1xCarbami | E9Q2E4 1x | 4.76E-05 | 0 | 1 | 2 | 1 | E9Q2E4 | Uncharacte | E9Q2E4 [1 | 0 | 1856.923 | 0 | 1.58 | 0.001517 | 11.48 | 51.6 | 148.4 | 14.47 | 9.83 | | High | High | High | High | High | High | High | High | High | High | High | 0 | 7.78E-06 | 1.84 |
| FALSE | High | SLSHLAAP | [R].SLSHLA | 1xPhospho | E9PUQ5 1x | 0.000738 | 0 | 1 | 5 | 1 | E9PUQ5 | Golgin sub | E9PUQ5 [7 | 0 | 1775.982 | 0 | 0.89 | 0.011883 | 13.15 | 70.1 | 129.9 | 4.47 | 18.2 | | High | High | High | High | High | High | High | High | High | High | High | 0 | 0.00017 | 3.14 |
| FALSE | High | SLSLDPGQS | [K].SLSLDP | 1xPhospho | P52734 1x | 0.000616 | 0 | 1 | 2 | 2 | P52734 | FYVE, Rho( | P52734 [11 | 0 | 2353.139 | 0 | -1.18 | 0.003245 | 2.34 | 137.2 | 62.8 | 8.5 | 13.98 | | High | High | High | High | High | High | High | High | High | High | High | 0 | 0.000138 | 3.44 |
| FALSE | High | SLSIEIGHE | [K].SLSIEIG | 1xPhospho | O35623 1x | 0.012959 | 0.000889 | 1 | 2 | 2 | O35623 | BET1 homo | O35623 [4 | 0 | 1749.955 | 0 | -0.41 | 0.069853 | 7.62 | 117.2 | 82.8 | 21.36 | 10.95 | | High | High | High | High | High | High | High | High | High | High | High | 0.000449 | 0.004187 | 2.7 |
| FALSE | High | SLSLESTDR | [K].SLSLES | 1xPhospho | A0A1D5RL( | 0.079766 | 0.00613 | 1 | 9 | 1 | A0A1D5RL( | Serine/thre | A0A1D5RL( | 0 | 1316.63 | 0 | 1.38 | 0.002887 | 18.83 | 50.5 | 149.5 | 26.24 | 10.11 | | High | High | High | High | High | High | High | High | High | High | High | 0.003148 | 0.03303 | 1.3 |
| FALSE | High | SLSLGDK | [R].SLSLGD | 1xPhospho | V9GX40 1x | 0.051258 | 0.003745 | 1 | 5 | 2 | V9GX40 | Synergin g | V9GX40 [1 | 0 | 1257.686 | 0 | 0.61 | 0.00223 | 8.69 | 79 | 121 | 3.82 | 7.59 | | High | High | High | High | High | High | High | High | High | High | High | 0.00196 | 0.01978 | 1.51 |
| FALSE | High | SISLPVPVL | [R].SISLPVI | 1xPhospho | O35099 1x | 6.73E-06 | 0 | 1 | 3 | 3 | O35099 | Mitogen-a | O35099 [9 | 0 | 3560.791 | 0 | 1.73 | 0.002521 | 19.28 | 41.5 | 158.5 | 21.39 | 19.77 | | High | High | High | High | High | High | High | High | High | High | High | 0 | 4.69E-06 | 4.77 |
| FALSE | High | SLSMDSQ | [R].SLSMD | 1xPhospho | E9Q444 1x | 6.15E-06 | 0 | 1 | 8 | 3 | E9Q444 | Zinc finger | E9Q444 [3 | 0 | 2520.271 | 0 | 0.12 | 0.062176 | 1.85 | 95.3 | 104.7 | 3.97 | 9.32 | | High | High | High | High | High | High | High | High | High | High | High | 0 | 7.82E-07 | 4.03 |
| FALSE | High | SLSMVDLQ | [K].SLSMVI | 1xPhospho | Q3U2G7 1x | 0.003495 | 0.00021 | 1 | 11 | 1 | Q3U2G7 | Putative ur | Q3U2G7 [7 | 0 | 1543.739 | 0 | -0.76 | 0.000388 | 2.05 | 124.4 | 75.6 | 8.61 | 9.76 | | High | High | High | High | High | High | High | High | High | High | High | 9.42E-05 | 0.000974 | 2.62 |
| FALSE | High | SLSMVDLQ | [K].SLSMVI | 1xPhospho | Q3U2G7 1x | 0.019107 | 0.001402 | 1 | 11 | 1 | Q3U2G7 | Putative ur | Q3U2G7 [7 | 0 | 1559.734 | 0 | 0.04 | 0.573921 | 16.35 | 107.8 | 92.2 | 12.84 | 20.22 | | High | High | High | High | High | High | High | High | High | High | High | 0.00074 | 0.006507 | 1.86 |
| FALSE | High | SLSGSSPCP | [R].SLSGSS | 1xCarbami | Q8BTI8 2x | 0.000289 | 0 | 1 | 4 | 1 | Q8BTI8 | Serine/argi | Q8BTI8 [77 | 0 | 1408.578 | 0 | -1.14 | 0.102322 | 9.23 | 129.3 | 70.7 | 41.18 | 36.8 | | High | High | High | High | High | High | High | High | High | High | High | 0 | 5.89E-05 | 2.29 |
| FALSE | High | SLSNATIINI | [R].SLSNAT | 1xPhospho | E9PXC0 1x | 2.75E-05 | 0 | 1 | 6 | 1 | E9PXC0 | Signal reco | E9PXC0 [1 | 0 | 2498.334 | 0 | 1.68 | 0.000637 | 21.68 | 47.1 | 152.9 | 16.93 | 5.98 | | High | High | High | High | High | High | High | High | High | High | High | 0 | 4.21E-06 | 2.82 |
| FALSE | High | SISNPPGSN | [K].SISNPP | 1xPhospho | P83741-3 1 | 5.65E-05 | 0 | 1 | 7 | 2 | P83741-3 | Isoform 3 ( | P83741-3 [ | 0 | 1450.725 | 0 | 1.82 | 0.002517 | 32.04 | 44.1 | 155.9 | 13.03 | 14.28 | | High | High | High | High | High | High | High | High | High | High | High | 7.19E-05 | 0.000803 | 1.08 |
| FALSE | High | SLSNSNPDI | [R].SLSNSN | 1xPhospho | A2A9M4 1 | 4.53E-10 | 0 | 1 | 9 | 7 | A2A9M4 | Dedicator ( | A2A9M4 [8 | 0 | 2497.129 | 0 | 0.7 | 0.001496 | 5.99 | 74.8 | 125.2 | 8.34 | 5.4 | | High | High | High | High | High | High | High | High | High | High | High | 0 | 2.87E-10 | 4.81 |
| FALSE | High | SLSNSNPDI | [R].SLSNSN | 2xPhospho | A2A9M4 2 | 1.05E-05 | 0 | 1 | 9 | 4 | A2A9M4 | Dedicator ( | A2A9M4 [8 | 0 | 2577.096 | 0 | -0.71 | 0.00734 | 4.03 | 124.1 | 75.9 | 9.9 | 10.58 | | High | High | High | High | High | High | High | High | High | High | High | 0 | 1.42E-06 | 4.12 |
| FALSE | High | SLSNSSSQF | [R].SLSNSS | 1xPhospho | F8VQM8 1 | 0.002463 | 0.000109 | 1 | 10 | 1 | F8VQM8 | Xndc1-trar | F8VQM8 [2 | 0 | 1485.69 | 0 | -1.02 | 0.001003 | 1.64 | 133.3 | 66.7 | 7.3 | 4.57 | | High | High | High | High | High | High | High | High | High | High | High | 4.94E-05 | 0.000654 | 2.17 |
| FALSE | High | SISNSVSDL | [K].SISNSV | 1xPhospho | P48754 1x | 0.000123 | 0 | 1 | 5 | 1 | P48754 | Breast can | P48754 [58 | 0 | 2582.312 | 0 | -0.09 | 0.294588 | 16.32 | 103.8 | 96.2 | 7.17 | 8.95 | | High | High | High | High | High | High | High | High | High | High | High | 0 | 2.25E-05 | 2.67 |
| FALSE | High | SLSPAATVT | [K].SLSPAA | 1xPhospho | Q80X80 1x | 0.000311 | 0 | 1 | 6 | 1 | Q80X80 | Phospholip | Q80X80 [3 | 0 | 2423.181 | 0 | 1.07 | 0.004142 | 16.25 | 64.3 | 135.7 | 5.23 | 13.4 | | High | High | High | High | High | High | High | High | High | High | High | 0 | 6.42E-05 | 3.32 |
| FALSE | High | SISPCPK | [R].SISPCPI | 1xCarbami | Q8BTI8 2x | 0.06169 | 0.004646 | 1 | 3 | 1 | Q8BTI8 | Serine/argi | Q8BTI8 [92 | 0 | 1177.493 | 0 | -1.22 | 0.001025 | 7.07 | 138.5 | 61.5 | 12.13 | 38.93 | | High | High | High | High | High | High | High | High | High | High | High | 0.002381 | 0.02467 | 1.98 |
| FALSE | High | SISPCPK | [R].SISPCPI | 1xCarbami | Q8BTI8 1x | 0.074685 | 0.005727 | 1 | 3 | 1 | Q8BTI8 | Serine/argi | Q8BTI8 [92 | 0 | 1326.689 | 0 | -0.87 | 0.003827 | 2.8 | 130.7 | 69.3 | 11.67 | 7.42 | | High | High | High | High | High | High | High | High | High | High | High | 0.002937 | 0.03069 | 1.85 |
| FALSE | High | SLSPGAASS | [R].SLSPGA | 1xPhospho | Q9Z248 1x | 2.99E-07 | 0 | 1 | 3 | 2 | Q9Z248 | Zinc finger | Q9Z248 [1 | 0 | 2017.948 | 0 | 0.57 | 0.001694 | 3.92 | 80.7 | 119.3 | 6.67 | 4.87 | | High | High | High | High | High | High | High | High | High | High | High | 0 | 2.61E-08 | 3.26 |
| FALSE | High | SLSPGGAAI | [R].SLSPGG | 1xPhospho | Q0VBL3 1x | 1.78E-05 | 0 | 1 | 7 | 3 | Q0VBL3 | RNA bindir | Q0VBL3 [2 | 0 | 1457.735 | 0 | -0.23 | 0.00834 | 4.8 | 108.2 | 91.8 | 2.25 | 4.1 | | High | High | High | High | High | High | High | High | High | High | High | 0 | 2.57E-06 | 1.95 |
| FALSE | High | SLSPGK | [RK].SLSPG | 1xPhospho | Q3TUQ5 1 | 0.115566 | 0.009504 | 2 | 8 | 1 | Q3TUQ5; ( | Pinin OS=N | Q3TUQ5 [4 | 0 | 1126.627 | 0 | -0.2 | 0.052134 | 7.4 | 107 | 93 | 5.74 | 6.83 | Shared | High | High | High | High | High | High | High | High | High | High | High | 0.004733 | 0.05102 | 1.28 |
| FALSE | High | SLSPGVSR | [K].SLSPGV | 1xPhospho | F6RJ39 1xP | 0.051694 | 0.003745 | 2 | 4 | 2 | F6RJ39; Q8 | Apoptotic ( | F6RJ39 [60 | 0 | 1111.571 | 0 | 0.07 | 0.230275 | 11 | 94.9 | 105.1 | 5.18 | 8.54 | Shared | High | High | High | High | High | High | High | High | High | High | High | 0.00196 | 0.01995 | 1.31 |
| FALSE | High | SLSPIIGK | [R].SLSPIIG | 1xPhospho | A2AUK5 1x | 0.024424 | 0.001612 | 2 | 8 | 1 | A2AUK5; Q | Band 4.1-li | A2AUK5 [7 | 0 | 1352.795 | 0 | -0.69 | 0.000992 | 5.91 | 123.2 | 76.8 | 3.4 | 8.87 | Shared | High | High | High | High | High | High | High | High | High | High | High | 0.000859 | 0.008548 | 1.32 |
| FALSE | High | SLSNPDIAS | [K].SLSNPD | 1xPhospho | Q80U35 1x | 5.52E-06 | 0 | 1 | 1 | 1 | Q80U35 | Rho guanir | Q80U35 [4 | 0 | 2386.247 | 0 | -0.77 | 0.006247 | 5.9 | 121.8 | 78.2 | 11.41 | 22.75 | | High | High | High | High | High | High | High | High | High | High | High | 0 | 6.95E-07 | 3.7 |
| FALSE | High | SLSGSGYG | [R].SLSGSG | 1xPhospho | Q8CI12 1x | 0.007742 | 0.000664 | 1 | 2 | 2 | Q8CI12 | Smoothelir | Q8CI12 [24 | 0 | 1792.925 | 0 | -1.84 | 0.000241 | 0.43 | 155 | 45 | 3.53 | 12.4 | | High | High | High | High | High | High | High | High | High | High | High | 0.000294 | 0.002365 | 2.57 |
| FALSE | High | SLSGSCASF | [K].SLSGSC | 1xCarbami | F8VQ54 1x | 0.002096 | 5.66E-05 | 1 | 3 | 1 | F8VQ54 | Endoribon | F8VQ54 [1 | 0 | 1583.745 | 0 | -0.26 | 0.040179 | 8.95 | 109.2 | 90.8 | 9.66 | 6.9 | | High | High | High | High | High | High | High | High | High | High | High | 2.58E-05 | 0.000545 | 2.34 |
| FALSE | High | SLSFSEPQC | [R].SLSFSEI | 1xPhospho | E9Q5N9 1x | 0.000379 | 0 | 1 | 2 | 4 | E9Q5N9 | Zinc finger | E9Q5N9 [4 | 0 | 2180.14 | 0 | -1.31 | 0.001143 | 3.42 | 141.9 | 58.1 | 2.65 | 18.29 | | High | High | High | High | High | High | High | High | High | High | High | 0 | 8.05E-05 | 2.67 |
| FALSE | High | SLSDLTSQK | [K].SLSDLT | 1xPhospho | P70213 1x | 0.01441 | 0.001011 | 1 | 5 | 1 | P70213 | Friend viru | P70213 [24 | 0 | 1516.802 | 0 | 2.54 | 0.003589 | 13.6 | 30.8 | 169.2 | 9.41 | 22.89 | | High | High | High | High | High | High | High | High | High | High | High | 0.000538 | 0.004701 | 0.82 |
| FALSE | High | SISDQYVES | [K].SISDQY | 1xPhospho | Q6ZPR5-4 1 | 6.67E-06 | 0 | 1 | 9 | 3 | Q6ZPR5-4 | Isoform 4 ( | Q6ZPR5-4 | 0 | 1747.81 | 0 | -0.15 | 0.075887 | 2.82 | 105.5 | 94.5 | 3.1 | 7.61 | | High | High | High | High | High | High | High | High | High | High | High | 0.002403 | 0.02584 | 1.88 |
| FALSE | High | SLSDSESDD | [K].SLSDSE | 1xPhospho | Q32P00 1x | 3.35E-05 | 0 | 1 | 5 | 12 | Q32P00 | Chromobo | Q32P00 [9 | 0 | 1707.773 | 0 | -0.29 | 0.002838 | 4.67 | 110.1 | 89.9 | 3.05 | 4.34 | | High | High | High | High | High | High | High | High | High | High | High | 0 | 0.000262 | 3.83 |
| FALSE | High | SLSDSESDD | [K].SLSDSE | 2xPhospho | Q32P00 2x | 3.23E-05 | 0 | 1 | 5 | 9 | Q32P00 | Chromobo | Q32P00 [9 | 0 | 1787.739 | 0 | -0.32 | 0.000289 | 2.04 | 112.3 | 87.7 | 3.64 | 5.32 | | High | High | High | High | High | High | High | High | High | High | High | 0 | 1.23E-05 | 3.16 |
| FALSE | High | SLSDYTGPF | [R].SLSDYT | 1xPhospho | A0A1B0GS | 5.32E-06 | 0 | 1 | 4 | 2 | A0A1B0GS | Supervillin | A0A1B0GS | 0 | 1966.984 | 0 | -0.47 | 0.012032 | 4.76 | 116.3 | 83.7 | 10.04 | 8.02 | | High | High | High | High | High | High | High | High | High | High | High | 0 | 6.62E-07 | 3.69 |
| FALSE | High | SLSEAQDTS | [R].SLSEAC | 1xPhospho | Q8R2M2 1 | 8.12E-09 | 0 | 1 | 1 | 3 | Q8R2M2 | Deoxynucl | Q8R2M2 [3 | 0 | 3111.5 | 0 | -0.46 | 0.048274 | 8.56 | 115.9 | 84.1 | 15.47 | 14.63 | | High | High | High | High | High | High | High | High | High | High | High | 0 | 4.57E-10 | 4.16 |
| FALSE | High | SLSEAQDTS | [R].SLSEAC | 1xDeamida | Q8R2M2 1 | 2.01E-06 | 0 | 1 | 1 | 6 | Q8R2M2 | Deoxynucl | Q8R2M2 [3 | 0 | 3112.484 | 0 | -0.29 | 0.056554 | 6.32 | 108.9 | 91.1 | 7.22 | 10.43 | | High | High | High | High | High | High | High | High | High | High | High | 0 | 2.22E-07 | 3.44 |
| FALSE | High | SISEEK | [R].SISEEK. | 1xPhospho | E9Q1Q1 1x | 0.016596 | 0.001124 | 1 | 13 | 2 | E9Q1Q1 | Microtubu | E9Q1Q1 [8 | 0 | 1230.638 | 1 | -1.27 | 0.00037 | 1.24 | 141.4 | 58.6 | 2.21 | 15.94 | | High | High | High | High | High | High | High | High | High | High | High | 0.000622 | 0.005539 | 1.93 |
| FALSE | High | SISEGHPW | [K].SISEGH | 1xPhospho | Q6ZWQ0 1 | 8.18E-05 | 0 | 1 | 10 | 4 | Q6ZWQ0 | Nesprin-2 ( | Q6ZWQ0 [ | 0 | 2522.223 | 0 | 2.67 | 0.002241 | 12.5 | 25.6 | 174.4 | 6.84 | 19.52 | | High | High | High | High | High | High | High | High | High | High | High | 0.000449 | 0.003924 | 3.79 |
| FALSE | High | SISEGHPW | [K].SISEGH | 2xPhospho | Q6ZWQ0 2 | 6.02E-05 | 0 | 1 | 10 | 1 | Q6ZWQ0 | Nesprin-2 ( | Q6ZWQ0 [ | 0 | 2602.189 | 0 | 0.49 | 0.013303 | 20.56 | 79.8 | 120.2 | 3.72 | 14.97 | | High | High | High | High | High | High | High | High | High | High | High | 0 | 1.02E-05 | 2.47 |
| FALSE | High | SLSEIAIDLT | [R].SLSEIAI | 1xPhospho | Q8BIE6-2 1 | 0.092372 | 0.007211 | 1 | 7 | 1 | Q8BIE6-2 | Isoform 2 ( | Q8BIE6-2 [ | 0 | 2229.203 | 0 | 0.48 | 0.022099 | 10.15 | 83.5 | 116.5 | 7.86 | 16.87 | | High | High | High | High | High | High | High | High | High | High | High | 0.003672 | 0.03948 | 2.45 |
| FALSE | High | SLSELCLISV | [R].SLSELCI | 1xCarbami | Q66T02 1x | 0.002232 | 0.000109 | 1 | 4 | 1 | Q66T02 | Pleckstrin | Q66T02 [9 | 0 | 1910.002 | 0 | -0.03 | 0.794248 | 5.33 | 100.6 | 99.4 | 10.65 | 10.93 | | High | High | High | High | High | High | High | High | High | High | High | 4.94E-05 | 0.000588 | 2.65 |
| FALSE | High | SLSELESLK | [R].SLSELE | 1xPhospho | Q3B7Z2 1x | 0.000188 | 0 | 1 | 1 | 4 | Q3B7Z2 | Oxysterol-l | Q3B7Z2 [2 | 0 | 1543.838 | 0 | -0.05 | 0.1533 | 1.09 | 103 | 97 | 3.83 | 7.21 | | High | High | High | High | High | High | High | High | High | High | High | 0.000246 | 0.001498 | 2.13 |
| FALSE | High | SLSEPEELA | [R].SLSEPE | 1xPhospho | E9PV82 1x | 0.105058 | 0.008518 | 1 | 5 | 2 | E9PV82 | Protein FAI | E9PV82 [1 | 0 | 1439.698 | 0 | 0.56 | 0.001786 | 1.2 | 82.4 | 117.6 | 2.5 | 4.89 | | High | High | High | High | High | High | High | High | High | High | High | 0.005041 | 0.05515 | 2.75 |
| FALSE | High | SISEQELAE | [R].SISEQE | 1xPhospho | Q8BW72 1 | 1.81E-05 | 0 | 1 | 3 | 2 | Q8BW72 | Lysine-spe | Q8BW72 [ | 0 | 3065.468 | 0 | -1.33 | 0.000941 | 2.84 | 145 | 55 | 9.8 | 14.93 | | High | High | High | High | High | High | High | High | High | High | High | 0 | 2.65E-06 | 4.66 |
| FALSE | High | SLSEQPVVI | [R].SLSEQP | 1xPhospho | P19426 1x | 3.09E-05 | 0 | 1 | 3 | 3 | P19426 | Negative e | P19426 [4 | 0 | 2413.226 | 0 | 0.9 | 0.00295 | 18.27 | 69.6 | 130.4 | 11.23 | 7.03 | | High | High | High | High | High | High | High | High | High | High | High | 0 | 2.17E-05 | 2.59 |
| FALSE | High | SLSESAALR | [R].SLSESA | 1xPhospho | E9Q652 1x | 0.000359 | 0 | 1 | 4 | 4 | E9Q652 | Regulator ( | E9Q652 [1 | 0 | 1242.629 | 0 | 0.4 | 0.000486 | 3.4 | 85.2 | 114.8 | 3.2 | 3.1 | | High | High | High | High | High | High | High | High | High | High | High | 0 | 0.000262 | 1.7 |
| FALSE | High | SLSESELIGI | [K].SLSESEI | 1xPhospho | P51125-3 1 | 9.33E-07 | 0 | 1 | 12 | 11 | P51125-3 | Isoform 3 ( | P51125-3 [ | 0 | 2177.021 | 0 | 1.91 | 0.000659 | 2.81 | 42.5 | 157.5 | 9.19 | 8.23 | | High | High | High | High | High | High | High | High | High | High | High | 0 | 1.05E-06 | 4.85 |
| FALSE | High | SISESFLTVI | [R].SISESFL | 1xPhospho | Q5SW75 1 | 0.023175 | 0.001578 | 1 | 3 | 1 | Q5SW75 | Protein ph | Q5SW75 [3 | 0 | 1648.896 | 0 | 0.37 | 0.009323 | 6.56 | 85.9 | 114.1 | 11.78 | 12.06 | | High | High | High | High | High | High | High | High | High | High | High | 0.000847 | 0.008021 | 2.44 |
| FALSE | High | SLSESPFK | [K].SLSESPI | 1xPhospho | A0A1B0GS | 0.012508 | 0.000889 | 1 | 2 | 1 | A0A1B0GS | Nuclear po | A0A1B0GS | 0 | 1432.749 | 0 | -0.41 | 0.010429 | 8.26 | 113.5 | 86.5 | 4.03 | 10.37 | | High | High | High | High | High | High | High | High | High | High | High | 0.000449 | 0.004034 | 1.56 |
| FALSE | High | SLSESSVVN | [R].SLSESS | 1xPhospho | O88466 1x | 0.000196 | 0 | 2 | 5 | 2 | O88466; R | Zinc finger | O88466 [8 | 0 | 1518.707 | 0 | -1.39 | 0.000413 | 1.4 | 146 | 54 | 4.09 | 9.74 | Shared | High | High | High | High | High | High | High | High | High | High | High | 0 | 3.84E-05 | 2.86 |
| FALSE | High | SLSESSVVN | [R].SLSESS | 1xPhospho | O88466 1x | 0.003312 | 0.00016 | 2 | 5 | 3 | O88466; R | Zinc finger | O88466 [8 | 0 | 1534.702 | 0 | -1.34 | 0.001032 | 1.96 | 142.4 | 57.6 | 3.74 | 16.75 | Shared | High | High | High | High | High | High | High | High | High | High | High | 7.19E-05 | 0.000917 | 1.85 |
| FALSE | High | SLSESSVVN | [R].SLSESS | 2xPhospho | O88466 2x | 0.017041 | 0.001204 | 2 | 5 | 1 | O88466; R | Zinc finger | O88466 [8 | 0 | 1614.668 | 0 | -0.56 | 0.130641 | 8.74 | 119.2 | 80.8 | 23.99 | 14.82 | Shared | High | High | High | High | High | High | High | High | High | High | High | 0.000638 | 0.005678 | 1.56 |
| FALSE | High | SLSESYELS | [R].SLSESYI | 1xPhospho | E9PUA3 1x | 7.36E-06 | 0 | 1 | 5 | 8 | E9PUA3 | IQ motif ar | E9PUA3 [9 | 0 | 2239.078 | 0 | 0.81 | 0.000505 | 4.41 | 72.7 | 127.3 | 4.96 | 6.26 | | High | High | High | High | High | High | High | High | High | High | High | 0 | 0.000188 | 3.93 |
| FALSE | High | SLSEVGDEI | [R].SLSEVG | 1xPhospho | B2KF05 1x | 0.000157 | 0 | 1 | 1 | 4 | B2KF05 | Bromodom | B2KF05 [4 | 0 | 1770.856 | 0 | -0.49 | 0.001817 | 5.6 | 119.9 | 80.1 | 4.88 | 8.01 | | High | High | High | High | High | High | High | High | High | High | High | 0 | 8.85E-05 | 3.38 |
| FALSE | High | SLSFDPSLG | [R].SLSFDP | 1xCarbami | P23804 1x | 0.000147 | 0 | 1 | 4 | 1 | P23804 | E3 ubiquiti | P23804 [18 | 0 | 1902.923 | 0 | 1.55 | 0.001011 | 13.99 | 49 | 151 | 7.12 | 9.59 | | High | High | High | High | High | High | High | High | High | High | High | 0 | 2.77E-05 | 3.98 |
| FALSE | High | SISFPEVPR | [R].SISFPE | 1xPhospho | A0A1B0GS | 0.004256 | 0.00036 | 1 | 8 | 1 | A0A1B0GS | Supervillin | A0A1B0GS | 0 | 1340.681 | 0 | 2.26 | 0.000417 | 12.77 | 34.9 | 165.1 | 9.75 | 8.62 | | High | High | High | High | High | High | High | High | High | High | High | 0.000161 | 0.001208 | 1.66 |
| FALSE | High | SISFPEVPRS | [R].SISFPE | 1xPhospho | A0A1B0GS | 0.010475 | 0.000772 | 1 | 8 | 1 | A0A1B0GS | Supervillin | A0A1B0GS | 1 | 1882.024 | 0 | 0.27 | 0.033531 | 9.82 | 93.3 | 106.7 | 6.42 | 5.88 | | High | High | High | High | High | High | High | High | High | High | High | 0.000382 | 0.003318 | 1.13 |
| FALSE | High | SLSFSDEMS | [R].SLSFSD | 1xCarbami | Q8BGR5 1x | 0.019276 | 0.001402 | 1 | 3 | 1 | Q8BGR5 | Protein FAI | Q8BGR5 [1 | 0 | 1730.696 | 0 | 0.92 | 0.001169 | 12.45 | 70.1 | 129.9 | 6.89 | 5.06 | | High | High | High | High | High | High | High | High | High | High | High | 0.000756 | 0.006515 | 1.79 |
| FALSE | High | SLSPLSGTT | [R].SLSPLS( | 2xPhospho | F6RJ39 2xP | 0.001168 | 0 | 2 | 4 | 1 | F6RJ39; Q8 | Apoptotic ( | F6RJ39 [42 | 0 | 1824.88 | 0 | 0.11 | 0.111534 | 8.66 | 98.9 | 101.1 | 4.82 | 8.85 | Shared | High | High | High | High | High | High | High | High | High | High | High | 0 | 0.000285 | 2.36 |
| FALSE | High | SILTSLLLNS | [R].SILTSLL | 1xPhospho | Q64127 1x | 0.000434 | 0 | 1 | 5 | 1 | Q64127 | Transcripti | Q64127 [7 | 0 | 2644.364 | 0 | -0.26 | 0.042578 | 12.3 | 110.7 | 89.3 | 8.38 | 6.96 | | High | High | High | High | High | High | High | High | High | High | High | 0 | 9.3E-05 | 3.43 |
| FALSE | High | SMPELLLLS | [K].SMPELI | 1xCarbami | B0V2M3 1 | 1.42E-05 | 0 | 1 | 3 | 2 | B0V2M3 | Zinc finger | B0V2M3 [1 | 0 | 2440.3 | 0 | -0.27 | 0.113774 | 16.11 | 108.1 | 91.9 | 12.06 | 6.35 | | High | High | High | High | High | High | High | High | High | High | High | 0 | 2.01E-06 | 3.07 |
| FALSE | High | SMPPSLETS | [K].SMPPSI | 1xPhospho | E9PUJ6 1x | 0.001933 | 5.66E-05 | 1 | 5 | 2 | E9PUJ6 | E3 ubiquiti | E9PUJ6 [35 | 0 | 2457.224 | 0 | 0.12 | 0.210307 | 14 | 93.3 | 106.7 | 7.01 | 11.29 | | High | High | High | High | High | High | High | High | High | High | High | 7.19E-05 | 0.000877 | 2.69 |
| FALSE | High | SPASTSSVN | [K].SPASTS | 1xPhospho | Q3UGP0 1 | 0.000386 | 0 | 2 | 10 | 3 | Q3UGP0; ( | Putative ur | Q3UGP0 [3 | 0 | 2240.076 | 0 | -1.01 | 0.0029 | 0.38 | 129.7 | 70.3 | 15.59 | 16.19 | Shared | High | High | High | High | High | High | High | High | High | High | High | 0 | 9.02E-05 | 2.78 |
| FALSE | High | SPASTSSVN | [K].SPASTS | 1xDeamida | Q3UGP0 1 | 0.015061 | 0.001049 | 2 | 10 | 3 | Q3UGP0; ( | Putative ur | Q3UGP0 [3 | 0 | 2241.06 | 0 | -0.49 | 0.190515 | 13.41 | 116.8 | 83.2 | 8.87 | 26.82 | Shared | High | High | High | High | High | High | High | High | High | High | High | 0.000756 | 0.006675 | 3.36 |
| FALSE | High | SPASVNGT | [K].SPASVN | 1xDeamida | Q6PGN3 1 | 0.031168 | 0.001935 | 1 | 5 | 1 | Q6PGN3 | Serine/thre | Q6PGN3 [3 | 0 | 2197.115 | 0 | 0.13 | 0.163104 | 7.37 | 94.8 | 105.2 | 10.17 | 4.51 | | High | High | High | High | High | High | High | High | High | High | High | 0.001044 | 0.01122 | 1.48 |
| FALSE | High | SPATGGPQ | [K].SPATG( | 1xPhospho | Q8K2E5 1x | 2.38E-06 | 0 | 1 | 3 | 2 | Q8K2E5 | Budding ur | Q8K2E5 [5 | 0 | 1704.863 | 0 | -0.25 | 0.62027 | 10.58 | 103.4 | 96.6 | 12.08 | 17.38 | | High | High | High | High | High | High | High | High | High | High | High | 0 | 2.71E-07 | 3.6 |
| FALSE | High | SPATPQAK | [K].SPATPC | 1xCarbami | Q8C1T5 1x | 1.27E-06 | 0 | 1 | 2 | 1 | Q8C1T5 | Putative ur | Q8C1T5 [2 | 0 | 3476.528 | 0 | -0.17 | 0.303908 | 6.68 | 97.4 | 102.6 | 17.24 | 15.9 | | High | High | High | High | High | High | High | High | High | High | High | 0 | 1.33E-07 | 3.44 |
| FALSE | High | SPATPVCG | [R].SPATPV | 1xCarbami | Q14B71 1x | 1.83E-05 | 0 | 1 | 3 | 1 | Q14B71 | Cell divisio | Q14B71 [3 | 0 | 1984.982 | 0 | -1.02 | 0.000445 | 0.82 | 132.5 | 67.5 | 6.11 | 10.37 | | High | High | High | High | High | High | High | High | High | High | High | 0 | 2.67E-06 | 3.26 |
| FALSE | High | SPATSAGPI | [R].SPATSA | 1xPhospho | G3UYU0 1 | 7.68E-05 | 0 | 1 | 1 | 1 | G3UYU0 | Mex3 RNA | G3UYU0 [4 | 0 | 1890.952 | 0 | -0.3 | 0.081851 | 9.39 | 115.3 | 84.7 | 12.33 | 11.13 | | High | High | High | High | High | High | High | High | High | High | High | 0 | 1.34E-05 | 2.49 |
| FALSE | High | SPATSPISSI | [K].SPATSP | 1xPhospho | A0A1S6GW | 0.003622 | 0.00021 | 1 | 6 | 2 | A0A1S6GW | Uncharacte | A0A1S6GW | 0 | 1649.785 | 0 | 0.25 | 0.090157 | 4.91 | 92.9 | 107.1 | 7.61 | 9.76 | | High | High | High | High | High | High | High | High | High | High | High | 9.42E-05 | 0.001012 | 1.89 |
| FALSE | High | SPAYRPDYI | [R].SPAYRF | 1xPhospho | M0QWX4 | 4.89E-05 | 0 | 1 | 2 | 3 | M0QWX4 | Sentrin-spe | M0QWX4 | 0 | 3035.337 | 0 | -0.12 | 0.420159 | 11.67 | 103.8 | 96.2 | 10.32 | 6.5 | | High | High | High | High | High | High | High | High | High | High | High | 0 | 8.07E-06 | 2.9 |
| FALSE | High | SPAYTPQN | [K].SPAYTP | 1xPhospho | Q925J9 1x | 8.68E-07 | 0 | 1 | 2 | 5 | Q925J9 | Mediator c | Q925J9 [14 | 0 | 2620.161 | 0 | 0.63 | 0.162673 | 38.61 | 80 | 120 | 32.86 | 8.3 | | High | High | High | High | High | High | High | High | High | High | High | 0 | 3.06E-05 | 3.7 |
| FALSE | High | SPAYTPQN | [K].SPAYTP | 2xPhospho | Q925J9 2x | 0.058664 | 0.004294 | 1 | 2 | 2 | Q925J9 | Mediator c | Q925J9 [14 | 0 | 2700.128 | 0 | 1.17 | 0.000412 | 3.42 | 61.6 | 138.4 | 9.19 | 6.54 | | High | High | High | High | High | High | High | High | High | High | High | 0.002559 | 0.02748 | 3.05 |
| FALSE | High | SPCDSGYS | [K].SPCDS( | 1xCarbami | P14873 1x | 0.00824 | 0.000736 | 2 | 2 | 3 | P14873; B2 | Microtubu | P14873 [19 | 0 | 2173.976 | 0 | -0.86 | 0.006981 | 6.28 | 123 | 77 | 14.39 | 7.57 | Shared | High | High | High | High | High | High | High | High | High | High | High | 0.000327 | 0.002537 | 2.34 |
| FALSE | High | SPCETISSP | [K].SPCETI | 1xCarbami | Q0VGY8 1 | 9.98E-05 | 0 | 1 | 4 | 3 | Q0VGY8 | Protein TAI | Q0VGY8 [2 | 0 | 2248.082 | 0 | 0.53 | 0.00276 | 12.59 | 81.5 | 118.5 | 3.75 | 7.35 | | High | High | High | High | High | High | High | High | High | High | High | 0 | 1.79E-05 | 3.29 |
| FALSE | High | SPASSQRPS | [R].SPASSC | 1xDeamida | A2A863-3 | 0.001815 | 5.66E-05 | 1 | 5 | 3 | A2A863-3 | Isoform 3 ( | A2A863-3 | 0 | 2549.183 | 0 | 2.66 | 0.000533 | 21.78 | 31.8 | 168.2 | 14.76 | 10.17 | | High | High | High | High | High | High | High | High | High | High | High | 2.58E-05 | 0.000539 | 3.45 |
| FALSE | High | SPCGSPTG | [R].SPCGSF | 1xCarbami | Q3TTA7 1x | 0.002293 | 0.000109 | 1 | 3 | 3 | Q3TTA7 | E3 ubiquiti | Q3TTA7 [5 | 0 | 1612.781 | 0 | -1.04 | 0.002382 | 4.76 | 133.5 | 66.5 | 9.77 | 11.03 | | High | High | High | High | High | High | High | High | High | High | High | 4.94E-05 | 0.000608 | 2.3 |
| FALSE | High | SPCPGDPD | [K].SPCPGI | 1xCarbami | G3UVV3 1 | 0.027358 | 0.001676 | 1 | 1 | 1 | G3UVV3 | MCG1556, | G3UVV3 [1 | 0 | 2756.231 | 0 | -0.38 | 0.030093 | 5.06 | 117.6 | 82.4 | 12.4 | 14.74 | | High | High | High | High | High | High | High | High | High | High | High | 0.000936 | 0.0097 | 2.52 |
| FALSE | High | SPCSPLNC( | [R].SPCSPL | 2xCarbami | Q6NZK8-3 | 0.000515 | 0 | 1 | 3 | 1 | Q6NZK8-3 | Isoform 3 ( | Q6NZK8-3 | 0 | 1928.901 | 0 | 1.2 | 0.002273 | 22.88 | 60.3 | 139.7 | 17.37 | 9.06 | | High | High | High | High | High | High | High | High | High | High | High | 0 | 0.000114 | 3.07 |
| FALSE | High | SPDASIFDS | [K].SPDASI | 1xPhospho | Q8BJ05 1x | 0.001411 | 5.66E-05 | 1 | 3 | 2 | Q8BJ05 | Zinc finger | Q8BJ05 [8 | 0 | 2139.052 | 0 | -0.07 | 0.439806 | 26.62 | 105.6 | 94.4 | 13.7 | 10.03 | | High | High | High | High | High | High | High | High | High | High | High | 2.58E-05 | 0.000352 | 2.81 |
| FALSE | High | SPDATETM | [R].SPDATI | 1xCarbami | Q8BX22 1x | 7.95E-08 | 0 | 1 | 1 | 2 | Q8BX22 | Sal-like pro | Q8BX22 [7 | 0 | 3037.405 | 0 | -1.97 | 0.002227 | 2.44 | 159.2 | 40.8 | 14.1 | 12.26 | | High | High | High | High | High | High | High | High | High | High | High | 0 | 5.91E-09 | 4.35 |
| FALSE | High | SPDEALPG( | [R].SPDEAL | 1xPhospho | Q7TPM1 1 | 0.00172 | 5.66E-05 | 1 | 6 | 1 | Q7TPM1 | Protein PRI | Q7TPM1 [7 | 0 | 2362.128 | 0 | -0.15 | 0.710407 | 15.22 | 106.1 | 93.9 | 7.42 | 21.21 | | High | High | High | High | High | High | High | High | High | High | High | 2.58E-05 | 0.000437 | 3.52 |
| FALSE | High | SPDEALPG( | [R].SPDEAL | 2xPhospho | Q7TPM1 2 | 0.018125 | 0.001282 | 1 | 6 | 1 | Q7TPM1 | Protein PRI | Q7TPM1 [7 | 0 | 2442.094 | 0 | 0.23 | 0.087923 | 11.44 | 92.7 | 107.3 | 6.72 | 11.13 | | High | High | High | High | High | High | High | High | High | High | High | 0.000689 | 0.006111 | 1.96 |

| | | | | | | | | | | | | | | | | | | | | | | | | | | | | | | | | | | | | | | |
|---|---|---|---|---|---|---|---|---|---|---|---|---|---|---|---|---|---|---|---|---|---|---|---|---|---|---|---|---|---|---|---|---|---|---|---|---|---|---|
| FALSE | High | SPDEPSTP | [K].SPDEPS | 1xPhospho | Q69ZY3 1x | 0.057203 | 0.004227 | 1 | 5 | 2 | Q69ZY3 | MKIAA082 | Q69ZY3 [2 | 0 | 3370.744 | 0 | 0.01 | 0.954392 | 12.21 | 107.2 | 92.8 | 7.85 | 17.17 | | High | High | High | High | High | High | High | High | High | High | High | 0.002159 | 0.02251 | 3.22 |
| FALSE | High | SPDEPSTP | [K].SPDEPS | 1xDeamida | Q69ZY3 1x | 0.002948 | 0.00016 | 1 | 5 | 2 | Q69ZY3 | MKIAA082 | Q69ZY3 [2 | 0 | 3371.728 | 0 | -0.33 | 0.332609 | 30.08 | 102.6 | 97.4 | 25.67 | 12.38 | | High | High | High | High | High | High | High | High | High | High | High | 7.19E-05 | 0.0008 | 3.67 |
| FALSE | High | SPDGDGV | [R].SPDGD | 1xPhospho | Q8BK06 1x | 0.009931 | 0.000736 | 1 | 3 | 1 | Q8BK06 | F-box only | Q8BK06 [1 | 0 | 2564.144 | 0 | 0.95 | 0.002147 | 7.96 | 68.2 | 131.8 | 14.86 | 4.75 | | High | High | High | High | High | High | High | High | High | High | High | 0.000344 | 0.003101 | 3.22 |
| FALSE | High | SPDIGSAQ | [K].SPDIGS | 1xCarbami | Q14B71 1x | 0.000349 | 0 | 1 | 4 | 2 | Q14B71 | Cell divisio | Q14B71 [2 | 0 | 2392.124 | 0 | -0.77 | 0.002739 | 3.05 | 128.6 | 71.4 | 8.39 | 10.89 | | High | High | High | High | High | High | High | High | High | High | High | 0 | 7.33E-05 | 3.5 |
| FALSE | High | SPDPDPTG | [K].SPDPD | 1xPhospho | Q80WT5 1 | 0.000244 | 0 | 1 | 5 | 1 | Q80WT5 | Aftiphilin C | Q80WT5 [3 | 0 | 2555.174 | 0 | 0.9 | 0.112889 | 20.38 | 73.8 | 126.2 | 22.06 | 27.5 | | High | High | High | High | High | High | High | High | High | High | High | 0 | 0.000049 | 3.16 |
| FALSE | High | SPDPDPTG | [K].SPDPD | 1xPhospho | Q80WT5 1 | 2.89E-07 | 0 | 1 | 5 | 2 | Q80WT5 | Aftiphilin C | Q80WT5 [3 | 0 | 2571.169 | 0 | 0.95 | 0.011497 | 10.27 | 70.9 | 129.1 | 9.01 | 23.83 | | High | High | High | High | High | High | High | High | High | High | High | 0 | 1.51E-07 | 4.98 |
| FALSE | High | SPDSGHVA | [R].SPDSGH | 1xPhospho | E9QMJ8 1x | 0.077188 | 0.005893 | 1 | 5 | 2 | E9QMJ8 | Axin-1 OS= | E9QMJ8 [4 | 0 | 1435.735 | 0 | 0.48 | 0.02447 | 23.13 | 85.1 | 114.9 | 8.12 | 10.43 | | High | High | High | High | High | High | High | High | High | High | High | 0.003922 | 0.04181 | 2.55 |
| FALSE | High | SPCLSTFDE | [K].SPCLST | 1xCarbami | Q5DTT1 1x | 0.001968 | 5.66E-05 | 1 | 2 | 2 | Q5DTT1 | Protein KIA | Q5DTT1 [1 | 0 | 2604.242 | 0 | -0.02 | 0.381067 | 9.15 | 103.6 | 96.4 | 11.88 | 8.59 | | High | High | High | High | High | High | High | High | High | High | High | 7.19E-05 | 0.000779 | 2.23 |
| FALSE | High | SPDSPGPS | [R].SPDSPG | 1xPhospho | Q3TAF8 1x | 0.026652 | 0.001676 | 1 | 3 | 1 | Q3TAF8 | Putative ur | Q3TAF8 [4 | 0 | 1578.7 | 0 | 0.16 | 0.046274 | 3.06 | 94.4 | 105.6 | 5.97 | 9.66 | | High | High | High | High | High | High | High | High | High | High | High | 0.000903 | 0.009448 | 2.29 |
| FALSE | High | SPASSQRP | [R].SPASSC | 1xPhospho | A2A863-3 | 0.000662 | 0 | 1 | 5 | 2 | A2A863-3 | Isoform 3 c | A2A863-3 | 0 | 2548.199 | 0 | 0.56 | 0.00128 | 2.79 | 82.4 | 117.6 | 5.42 | 8.9 | | High | High | High | High | High | High | High | High | High | High | High | 0.000382 | 0.003331 | 1.91 |
| FALSE | High | SPASPLNN | [R].SPASPL | 1xPhospho | E9Q6T9 1x | 1.29E-05 | 0 | 1 | 3 | 1 | E9Q6T9 | BTB/POZ d | E9Q6T9 [6 | 0 | 2472.326 | 0 | 0.86 | 0.002337 | 16.68 | 68.3 | 131.7 | 7.37 | 9.04 | | High | High | High | High | High | High | High | High | High | High | High | 0 | 1.79E-06 | 3.22 |
| FALSE | High | SPAASGAP | [K].SPAASG | 2xPhospho | Q9JKB3 2x | 0.000369 | 0 | 1 | 5 | 3 | Q9JKB3 | Y-box-bind | Q9JKB3 [3 | 0 | 4381.079 | 0 | 0.63 | 0.033043 | 10.3 | 78.3 | 121.7 | 23.66 | 14.33 | | High | High | High | High | High | High | High | High | High | High | High | 0 | 7.75E-05 | 3.53 |
| FALSE | High | SPADSLSSA | [R].SPADSL | 1xPhospho | E9Q1P8 1x | 3.45E-06 | 0 | 1 | 1 | 5 | E9Q1P8 | Interferon | E9Q1P8 [2 | 0 | 2501.217 | 0 | 0.18 | 0.398985 | 16.19 | 91.1 | 108.9 | 14.3 | 5.14 | | High | High | High | High | High | High | High | High | High | High | High | 0 | 7.37E-07 | 4.24 |
| FALSE | High | SPAEEELTP | [K].SPAEEE | 1xPhospho | Q7TQG1 1 | 0.023792 | 0.001612 | 1 | 6 | 1 | Q7TQG1 | Pleckstrin | Q7TQG1 [9 | 0 | 1793.888 | 0 | 2.09 | 0.000313 | 11.33 | 37.4 | 162.6 | 12.77 | 9.33 | | High | High | High | High | High | High | High | High | High | High | High | 0.000859 | 0.008286 | 1.25 |
| FALSE | High | SPAELEPEP | [K].SPAELE | 1xPhospho | Q5I1X5 1x | 0.000845 | 0 | 1 | 1 | 2 | Q5I1X5 | RelA-assoc | Q5I1X5 [45 | 0 | 4223.085 | 0 | 1.47 | 0.000648 | 15.67 | 53 | 147 | 10.85 | 9.54 | | High | High | High | High | High | High | High | High | High | High | High | 4.94E-05 | 0.00074 | 3.28 |
| FALSE | High | SPAELEQTS | [K].SPAELE | 1xPhospho | Q99JG7 1x | 0.020862 | 0.001472 | 1 | 2 | 2 | Q99JG7 | TNFAIP3-in | Q99JG7 [1 | 0 | 2472.227 | 0 | 0.38 | 0.037567 | 16.48 | 86.9 | 113.1 | 12.88 | 11 | | High | High | High | High | High | High | High | High | High | High | High | 0.000786 | 0.00712 | 1.86 |
| FALSE | High | SPAEPVAA | [K].SPAEPV | 1xPhospho | Q7TT18 1x | 0.000404 | 0 | 1 | 4 | 1 | Q7TT18 | Activating | Q7TT18 [4 | 0 | 2379.137 | 0 | 0.16 | 0.042344 | 11.63 | 92.9 | 107.1 | 13.46 | 5.08 | | High | High | High | High | High | High | High | High | High | High | High | 0 | 8.6E-05 | 3 |
| FALSE | High | SPAFGGPG | [K].SPAFGC | 1xCarbami | Q924A2 2x | 0.047086 | 0.00354 | 1 | 1 | 1 | Q924A2 | Protein cap | Q924A2 [4 | 0 | 3003.265 | 0 | 0.36 | 0.035687 | 8.93 | 93.4 | 106.6 | 19.23 | 12.24 | | High | High | High | High | High | High | High | High | High | High | High | 0.001812 | 0.01799 | 1.56 |
| FALSE | High | SPAGFSLSD | [K].SPAGFS | 1xCarbami | E9QKD1 1x | 2.77E-05 | 0 | 1 | 2 | 2 | E9QKD1 | Nucleolar p | E9QKD1 [7 | 0 | 2364.131 | 0 | -0.31 | 0.010947 | 6.5 | 109 | 91 | 6.9 | 10.99 | | High | High | High | High | High | High | High | High | High | High | High | 0 | 4.24E-06 | 3.5 |
| FALSE | High | SPAGPASPI | [R].SPAGPA | 1xDeamida | Q2NL51 1x | 1.61E-05 | 0 | 1 | 1 | 2 | Q2NL51 | Glycogen s | Q2NL51 [4 | 0 | 4590.088 | 0 | 0.34 | 0.356566 | 38.27 | 85.4 | 114.6 | 23.54 | 21.27 | | High | High | High | High | High | High | High | High | High | High | High | 0 | 2.3E-06 | 6.36 |
| FALSE | High | SPAGSEDC | [R].SPAGSE | 1xCarbami | Q9CQS5 1x | 0.000198 | 0 | 1 | 2 | 2 | Q9CQS5 | Serine/thre | Q9CQS5 [4 | 0 | 2775.249 | 0 | -0.08 | 0.55545 | 13.64 | 104.1 | 95.9 | 14.62 | 51.24 | | High | High | High | High | High | High | High | High | High | High | High | 0 | 0.000208 | 3.78 |
| FALSE | High | SPALAFSLA | [K].SPALAF | 1xPhospho | Q9QYS9-7 | 0.000244 | 0 | 1 | 6 | 1 | Q9QYS9-7 | Isoform 7 c | Q9QYS9-7 | 0 | 1952.02 | 0 | 0.39 | 0.018597 | 10.5 | 89 | 111 | 8.08 | 9.44 | | High | High | High | High | High | High | High | High | High | High | High | 0 | 4.89E-05 | 2.39 |
| FALSE | High | SPALASVAS | [R].SPALAS | 1xCarbami | Q3URQ4 1 | 0.00017 | 0 | 1 | 2 | 1 | Q3URQ4 | Family with | Q3URQ4 [4 | 0 | 2507.205 | 0 | 0.88 | 0.00106 | 14.13 | 69.5 | 130.5 | 11.37 | 4.06 | | High | High | High | High | High | High | High | High | High | High | High | 0 | 3.27E-05 | 2.77 |
| FALSE | High | SPALLPSTV | [R].SPALLP | 1xPhospho | B9EKJ7 1xP | 0.007538 | 0.00062 | 1 | 2 | 1 | B9EKJ7 | Tankyrase | B9EKJ7 [56 | 0 | 2352.278 | 0 | 0.95 | 0.009898 | 11.32 | 67.7 | 132.3 | 21.94 | 9.18 | | High | High | High | High | High | High | High | High | High | High | High | 0.000294 | 0.002296 | 3.9 |
| FALSE | High | SPASSGSV | [R].SPASSG | 1xPhospho | A0A0B4J1E | 8.45E-07 | 0 | 1 | 3 | 4 | A0A0B4J1E | DNA-bindi | A0A0B4J1E | 0 | 2824.335 | 0 | 0.43 | 0.003862 | 6.09 | 86.5 | 113.5 | 2.77 | 6.54 | | High | High | High | High | High | High | High | High | High | High | High | 0 | 8.77E-08 | 3.13 |
| FALSE | High | SPALLPSTV | [R].SPALLP | 1xPhospho | B9EKJ7 1xP | 0.000181 | 0 | 1 | 2 | 2 | B9EKJ7 | Tankyrase | B9EKJ7 [56 | 0 | 2581.441 | 0 | 0.22 | 0.039674 | 11.93 | 92.7 | 107.3 | 8 | 6.57 | | High | High | High | High | High | High | High | High | High | High | High | 0 | 0.000035 | 2.96 |
| FALSE | High | SPALSSTTT | [K].SPALSS | 1xPhospho | B1AQJ2 1x | 7.28E-05 | 0 | 1 | 2 | 1 | B1AQJ2 | Ubiquitin c | B1AQJ2 [6 | 0 | 2638.345 | 0 | 0.6 | 0.024926 | 7.56 | 79.6 | 120.4 | 4.64 | 17.46 | | High | High | High | High | High | High | High | High | High | High | High | 0 | 1.25E-05 | 4.09 |
| FALSE | High | SPALSSTTT | [K].SPALSS | 1xPhospho | B1AQJ2 1x | 5.24E-07 | 0 | 1 | 2 | 2 | B1AQJ2 | Ubiquitin c | B1AQJ2 [6 | 0 | 2654.34 | 0 | -0.11 | 0.960255 | 12.42 | 103.7 | 96.3 | 9.73 | 21.5 | | High | High | High | High | High | High | High | High | High | High | High | 0 | 6.78E-06 | 3.41 |
| FALSE | High | SPAPGLAS | [K].SPAPGL | 1xCarbami | F6YAU7 1x | 0.001058 | 0 | 2 | 9 | 1 | F6YAU7; F | Nuclear re | F6YAU7 [2 | 0 | 2945.341 | 0 | 0.65 | 0.008667 | 14.07 | 82.2 | 117.8 | 12.81 | 10.64 | Shared | High | High | High | High | High | High | High | High | High | High | High | 0 | 0.000253 | 2.74 |
| FALSE | High | SPAPPPPA | [R].SPAPPP | 1xPhospho | Q63943-2 | 0.000337 | 0 | 2 | 6 | 5 | Q63943-2; | Isoform M | Q63943-2 | 0 | 3627.732 | 0 | 0.73 | 0.003679 | 11.42 | 73.1 | 126.9 | 5.47 | 10.23 | Shared | High | High | High | High | High | High | High | High | High | High | High | 0 | 7.03E-05 | 3.16 |
| FALSE | High | SPAPSNPT | [K].SPAPSN | 1xPhospho | Q3U2W2 1 | 1.92E-06 | 0 | 1 | 2 | 9 | Q3U2W2 | MYB bindi | Q3U2W2 [ | 0 | 2090.094 | 0 | 0.03 | 0.677211 | 11.36 | 98.9 | 101.1 | 4.24 | 8.77 | | High | High | High | High | High | High | High | High | High | High | High | 0 | 2.17E-06 | 3.39 |
| FALSE | High | SPAPSNPT | [K].SPAPSN | 2xPhospho | Q3U2W2 2 | 2.08E-06 | 0 | 1 | 2 | 7 | Q3U2W2 | MYB bindi | Q3U2W2 [ | 0 | 2170.06 | 0 | -0.18 | 0.031985 | 5.76 | 106.9 | 93.1 | 4.08 | 5.06 | | High | High | High | High | High | High | High | High | High | High | High | 0 | 2.31E-07 | 4.2 |
| FALSE | High | SPAPTYLDN | [K].SPAPTY | 1xPhospho | Q3U0X6 1x | 0.028327 | 0.001791 | 1 | 1 | 1 | Q3U0X6 | Putative ur | Q3U0X6 [6 | 0 | 1587.744 | 0 | -0.09 | 0.44008 | 9.79 | 102.8 | 97.2 | 6.19 | 11.27 | | High | High | High | High | High | High | High | High | High | High | High | 0.000968 | 0.01009 | 2.36 |
| FALSE | High | SPAPTYLDN | [K].SPAPTY | 1xPhospho | Q3U0X6 1x | 0.028575 | 0.001829 | 1 | 1 | 2 | Q3U0X6 | Putative ur | Q3U0X6 [6 | 0 | 1603.739 | 0 | 0.27 | 0.004683 | 6.4 | 89.9 | 110.1 | 4.27 | 4.57 | | High | High | High | High | High | High | High | High | High | High | High | 0.004769 | 0.05216 | 2.05 |
| FALSE | High | SPAPVQSP | [R].SPAPVC | 2xPhospho | F8VPR5 2x | 0.010758 | 0.000772 | 1 | 4 | 1 | F8VPR5 | CREB-bindi | F8VPR5 [2 | 0 | 2710.27 | 0 | 1.86 | 0.002439 | 23.82 | 43 | 157 | 13.37 | 15.46 | | High | High | High | High | High | High | High | High | High | High | High | 0.000399 | 0.003416 | 2.96 |
| FALSE | High | SPAPVQTA | [R].SPAPVC | 1xPhospho | A0A0N4SV | 0.034281 | 0.002195 | 1 | 8 | 1 | A0A0N4SV | Eukaryotic | A0A0N4SV | 0 | 1806.977 | 0 | 0.91 | 0.001187 | 6.61 | 69.5 | 130.5 | 9.47 | 4.56 | | High | High | High | High | High | High | High | High | High | High | High | 0.001151 | 0.01249 | 1.56 |
| FALSE | High | SPASDTYIV | [K].SPASDT | 1xPhospho | P70670 1x | 0.01186 | 0.000889 | 1 | 3 | 3 | P70670 | Nascent pc | P70670 [2 | 0 | 2023.019 | 0 | 0.7 | 0.001247 | 1.48 | 74.4 | 125.6 | 7.21 | 8.91 | | High | High | High | High | High | High | High | High | High | High | High | 0.000814 | 0.007735 | 3.38 |
| FALSE | High | SPASESGSF | [R].SPASES | 1xPhospho | A2RSJ4 1xP | 0.008094 | 0.000736 | 1 | 1 | 3 | A2RSJ4 | UHRF1-bin | A2RSJ4 [88 | 0 | 3142.575 | 0 | 0.79 | 0.006634 | 15.82 | 71.1 | 128.9 | 15.6 | 12.6 | | High | High | High | High | High | High | High | High | High | High | High | 0.000555 | 0.004798 | 2.38 |
| FALSE | High | SPASPHPPF | [R].SPASPH | 1xPhospho | Q7TN98 1x | 0.041425 | 0.003126 | 1 | 7 | 1 | Q7TN98 | Cytoplasmi | Q7TN98 [2 | 0 | 1736.847 | 0 | 1.66 | 0.002625 | 24.06 | 46.8 | 153.2 | 3.53 | 17.57 | | High | High | High | High | High | High | High | High | High | High | High | 0.001609 | 0.01553 | 1.38 |
| FALSE | High | SPALPTPLS | [R].SPALPT | 1xPhospho | Q69ZF7 1x | 0.008847 | 0.000736 | 1 | 2 | 1 | Q69ZF7 | Metal tran | Q69ZF7 [6 | 0 | 1347.723 | 0 | 0.59 | 0.016121 | 26.33 | 75.4 | 124.6 | 13.13 | 11.45 | | High | High | High | High | High | High | High | High | High | High | High | 0.000327 | 0.002734 | 1.61 |
| FALSE | High | SPDSTANV | [R].SPDSTA | 1xPhospho | Q91Z67 1x | 0.003433 | 0.00021 | 1 | 3 | 1 | Q91Z67 | SLIT-ROBO | Q91Z67 [2 | 0 | 1255.588 | 0 | 0.13 | 0.037953 | 5.5 | 94.3 | 105.7 | 6.06 | 7.05 | | High | High | High | High | High | High | High | High | High | High | High | 9.42E-05 | 0.000951 | 1.36 |
| FALSE | High | SPDSTSPSY | [K].SPDSTS | 1xPhospho | Q3UGE1 1 | 1.91E-05 | 0 | 1 | 5 | 10 | Q3UGE1 | Putative ur | Q3UGE1 [3 | 0 | 1593.7 | 0 | -0.03 | 0.947593 | 6.6 | 103.2 | 96.8 | 5.8 | 8.54 | | High | High | High | High | High | High | High | High | High | High | High | 0 | 3.62E-05 | 2.09 |
| FALSE | High | SPDTSAYC | [R].SPDTSA | 1xCarbami | P14873 1x | 0.018447 | 0.001324 | 2 | 2 | 2 | P14873; B | Microtubu | P14873 [2 | 0 | 2219.964 | 0 | -0.71 | 0.01865 | 6.26 | 124.3 | 75.7 | 9.78 | 20.8 | Shared | High | High | High | High | High | High | High | High | High | High | High | 0.000859 | 0.008628 | 2.33 |
| FALSE | High | SPGAGPGV | [R].SPGAG | 1xPhospho | P55200 1x | 0.046291 | 0.003523 | 1 | 2 | 1 | P55200 | Histone-lys | P55200 [25 | 0 | 3815.746 | 0 | 0.15 | 0.088116 | 3.62 | 90 | 110 | 12.14 | 16.53 | | High | High | High | High | High | High | High | High | High | High | High | 0.001789 | 0.01772 | 3.05 |
| FALSE | High | SPGASNFS | [K].SPGASN | 1xPhospho | Q8R5H1 1x | 0.000279 | 0 | 1 | 2 | 2 | Q8R5H1 | Ubiquitin c | Q8R5H1 [2 | 0 | 1743.908 | 0 | 0.5 | 0.001707 | 1.95 | 84.4 | 115.6 | 4.98 | 7.72 | | High | High | High | High | High | High | High | High | High | High | High | 0.000275 | 0.001632 | 2.6 |
| FALSE | High | SPGASVSS | [R].SPGAS | 1xCarbami | A2A7S8-2 | 3.75E-07 | 0 | 1 | 4 | 2 | A2A7S8-2 | Isoform 2 c | A2A7S8-2 | 0 | 2804.286 | 0 | -0.28 | 0.20357 | 21.53 | 104.1 | 95.9 | 18.23 | 14.72 | | High | High | High | High | High | High | High | High | High | High | High | 0 | 3.38E-08 | 4.45 |
| FALSE | High | SPGEATAG | [R].SPGEAT | 1xPhospho | Q91VY5 1x | 4.86E-06 | 0 | 1 | 3 | 4 | Q91VY5 | Lysine-spe | Q91VY5 [4 | 0 | 1869.879 | 0 | -0.77 | 0.00225 | 7.13 | 127 | 73 | 4.25 | 11.4 | | High | High | High | High | High | High | High | High | High | High | High | 0 | 5.88E-06 | 3.3 |
| FALSE | High | SPGEPSAA | [R].SPGEPS | 1xCarbami | Q53Z05 1x | 0.000213 | 0 | 1 | 4 | 2 | Q53Z05 | Sir2alpha C | Q53Z05 [4 | 0 | 2987.428 | 0 | -1.43 | 0.000456 | 1.63 | 146.2 | 53.8 | 6.74 | 8.59 | | High | High | High | High | High | High | High | High | High | High | High | 0 | 4.21E-05 | 3.42 |
| FALSE | High | SPGEVTGF | [R].SPGEVT | 2xPhospho | H3BKP8 2x | 0.00153 | 5.66E-05 | 1 | 3 | 1 | H3BKP8 | RIKEN cDN | H3BKP8 [5 | 0 | 2218.003 | 0 | -0.46 | 0.037364 | 17.41 | 118.8 | 81.2 | 15.34 | 18.74 | | High | High | High | High | High | High | High | High | High | High | High | 2.58E-05 | 0.000384 | 3.4 |
| FALSE | High | SPGEYINID | [K].SPGEYI | 1xPhospho | P81122 1x | 2.04E-05 | 0 | 1 | 3 | 1 | P81122 | Insulin rec | P81122 [9 | 0 | 2034.937 | 0 | -0.65 | 0.057648 | 15.26 | 126.1 | 73.9 | 23.84 | 32.05 | | High | High | High | High | High | High | High | High | High | High | High | 0 | 0.000003 | 4.33 |
| FALSE | High | SPGEYVNIE | [K].SPGEYV | 1xPhospho | P35569 1x | 4.69E-06 | 0 | 1 | 2 | 1 | P35569 | Insulin rec | P35569 [8 | 0 | 2863.35 | 0 | 0.32 | 0.403409 | 35.78 | 95.9 | 104.1 | 11.44 | 27.53 | | High | High | High | High | High | High | High | High | High | High | High | 0 | 5.79E-07 | 3.52 |
| FALSE | High | SPGGGGNI | [K].SPGGG | 1xPhospho | P32507-2 1 | 0.022973 | 0.001578 | 1 | 4 | 1 | P32507-2 | Isoform Al | P32507-2 | 1 | 2173.992 | 0 | -1.82 | 0.000324 | 0.24 | 155.2 | 44.8 | 6.05 | 16.04 | | High | High | High | High | High | High | High | High | High | High | High | 0.000847 | 0.00799 | 1.72 |
| FALSE | High | SPGLCSDS | [R].SPGLCS | 1xCarbami | B0V2M3 1 | 0.000553 | 0 | 1 | 2 | 2 | B0V2M3 | Zinc finger | B0V2M3 [1 | 0 | 1945.878 | 0 | -0.47 | 0.003114 | 1.43 | 117.1 | 82.9 | 4.44 | 6.13 | | High | High | High | High | High | High | High | High | High | High | High | 0.000327 | 0.002548 | 3.14 |
| FALSE | High | SPGLLSSDN | [K].SPGLLS | 1xPhospho | O70546-2 | 0.013787 | 0.001011 | 1 | 3 | 2 | O70546-2 | Isoform 2 c | O70546-2 | 0 | 2466.308 | 0 | 0.05 | 0.751506 | 12.92 | 98.1 | 101.9 | 9.97 | 24.22 | | High | High | High | High | High | High | High | High | High | High | High | 0.000538 | 0.004495 | 1.77 |
| FALSE | High | SPGLLSSDN | [K].SPGLLS | 1xPhospho | O70546-2 | 3.2E-05 | 0 | 1 | 3 | 2 | O70546-2 | Isoform 2 c | O70546-2 | 0 | 2482.303 | 0 | 0.29 | 0.095628 | 20.89 | 87.2 | 112.8 | 5.15 | 15.23 | | High | High | High | High | High | High | High | High | High | High | High | 0 | 1.76E-05 | 3.98 |
| FALSE | High | SPGLSQAT | [K].SPGLSC | 1xCarbami | Q5SPX8 1x | 6.74E-07 | 0 | 1 | 5 | 2 | Q5SPX8 | TBC1 doma | Q5SPX8 [4 | 0 | 2933.484 | 0 | 0.36 | 0.004696 | 0.58 | 88.4 | 111.6 | 5.19 | 5.09 | | High | High | High | High | High | High | High | High | High | High | High | 0 | 6.54E-08 | 3.98 |
| FALSE | High | SPGAAAGC | [K].SPGAA | 1xPhospho | A2A870 1x | 0.067059 | 0.004796 | 1 | 2 | 1 | A2A870 | Fas-bindin | A2A870 [2 | 1 | 2712.449 | 0 | 0.02 | 0.35146 | 11.73 | 94.2 | 105.8 | 9.02 | 13.87 | | High | High | High | High | High | High | High | High | High | High | High | 0.002485 | 0.02715 | 2.17 |
| FALSE | High | SPGIYALGE | [R].SPGIYA | 1xPhospho | Q80XC6 1x | 3.93E-07 | 0 | 1 | 1 | 1 | Q80XC6 | Protein NR | Q80XC6 [3 | 0 | 2016.009 | 0 | -0.45 | 0.016029 | 9.75 | 115 | 85 | 9.04 | 12.12 | | High | High | High | High | High | High | High | High | High | High | High | 0 | 3.54E-08 | 4.09 |
| FALSE | High | SPGNESAIS | [R].SPGNE | 1xPhospho | E9QLC6 1x | 0.104222 | 0.008348 | 1 | 5 | 1 | E9QLC6 | SUMO-spe | E9QLC6 [8 | 0 | 1581.747 | 0 | 0.4 | 0.039162 | 14.09 | 86.2 | 113.8 | 8.95 | 13.33 | | High | High | High | High | High | High | High | High | High | High | High | 0.00425 | 0.04543 | 1.63 |
| FALSE | High | SPGNSTLE | [K].SPGNS | 1xPhospho | Q80UF4 1x | 4.2E-05 | 0 | 1 | 7 | 1 | Q80UF4 | Serological | Q80UF4 [4 | 0 | 2060.949 | 0 | -0.35 | 0.1347 | 6.24 | 114.9 | 85.1 | 9.56 | 13.56 | | High | High | High | High | High | High | High | High | High | High | High | 0 | 6.77E-06 | 2.94 |
| FALSE | High | SPGNTSQP | [K].SPGNT | 1xPhospho | F6YAU7 1x | 0.038017 | 0.002528 | 2 | 9 | 1 | F6YAU7; F | Nuclear re | F6YAU7 [1 | 0 | 2003.004 | 0 | 0.81 | 0.002007 | 5.01 | 71 | 129 | 4.03 | 8.94 | Shared | High | High | High | High | High | High | High | High | High | High | High | 0.001355 | 0.01409 | 2.1 |
| FALSE | High | SPGQPSED | [R].SPGQP | 1xPhospho | Q5SXI5 1xP | 2.38E-05 | 0 | 1 | 3 | 2 | Q5SXI5 | Zinc finger | Q5SXI5 [18 | 0 | 2835.376 | 0 | 0.44 | 0.050085 | 29.6 | 84.9 | 115.1 | 8.99 | 12.51 | | High | High | High | High | High | High | High | High | High | High | High | 0 | 3.56E-06 | 4.15 |
| FALSE | High | SPGRPTQG | [R].SPGRPT | 1xPhospho | B7ZNV2 1x | 0.009756 | 0.000736 | 1 | 2 | 1 | B7ZNV2 | GLI-Krupp | B7ZNV2 [6 | 0 | 1998.021 | 0 | 1.01 | 0.000207 | 3.85 | 67.5 | 132.5 | 9.23 | 4.86 | | High | High | High | High | High | High | High | High | High | High | High | 0.000344 | 0.003044 | 1.47 |
| FALSE | High | SPGSLSLHD | [R].SPGSLS | 1xPhospho | P09450 1x | 0.003558 | 0.00021 | 1 | 4 | 2 | P09450 | Transcripti | P09450 [23 | 0 | 1741.893 | 0 | 2.75 | 0.000479 | 12.55 | 26 | 174 | 5.32 | 9.58 | | High | High | High | High | High | High | High | High | High | High | High | 9.42E-05 | 0.000989 | 1.75 |
| FALSE | High | SPGSISYLP | [R].SPGSIS | 1xPhospho | Q5RIM6 1x | 0.008022 | 0.000736 | 1 | 10 | 2 | Q5RIM6 | Nuclear re | Q5RIM6 [2 | 0 | 2069.076 | 0 | -0.1 | 0.445473 | 10.81 | 102.6 | 97.4 | 5.82 | 6.41 | | High | High | High | High | High | High | High | High | High | High | High | 0.000327 | 0.002451 | 2.61 |
| FALSE | High | SPGSNSK | [R].SPGSN | 1xPhospho | Q9ESZ8-5 | 0.026885 | 0.001676 | 2 | 8 | 1 | Q9ESZ8-5; | Isoform 5 c | Q9ESZ8-5 | 0 | 1214.618 | 0 | -0.07 | 0.430615 | 5.05 | 101.3 | 98.7 | 7.23 | 4.1 | Shared | High | High | High | High | High | High | High | High | High | High | High | 0.000903 | 0.009552 | 0.69 |
| FALSE | High | SPGSPAAP | [R].SPGSPA | 1xPhospho | A5D6P2 1x | 0.000352 | 0 | 1 | 23 | 4 | A5D6P2 | Par-3 (Part | A5D6P2 [6 | 0 | 2303.101 | 0 | -0.25 | 0.072188 | 3.71 | 107.5 | 92.5 | 10.58 | 5.75 | | High | High | High | High | High | High | High | High | High | High | High | 0 | 0.000103 | 2.78 |
| FALSE | High | SPGSPAAP | [R].SPGSPA | 1xPhospho | A5D6P2 1x | 0.002077 | 5.66E-05 | 1 | 23 | 1 | A5D6P2 | Par-3 (Part | A5D6P2 [6 | 1 | 2588.244 | 0 | -0.23 | 0.040557 | 14 | 108.6 | 91.4 | 4.63 | 11.91 | | High | High | High | High | High | High | High | High | High | High | High | 2.58E-05 | 0.00054 | 3.25 |
| FALSE | High | SPGSPLPPL | [R].SPGSPL | 2xPhospho | Q9D8N6 2x | 0.015195 | 0.001089 | 1 | 4 | 1 | Q9D8N6 | Protein lin- | Q9D8N6 [1 | 0 | 2595.183 | 0 | -0.59 | 0.003292 | 1.89 | 121.1 | 78.9 | 7.83 | 7.31 | | High | High | High | High | High | High | High | High | High | High | High | 0.000571 | 0.005009 | 2.82 |
| FALSE | High | SPGSPLPPL | [R].SPGSPL | 1xPhospho | Q9D8N6 1x | 1.16E-05 | 0 | 1 | 4 | 3 | Q9D8N6 | Protein lin- | Q9D8N6 [1 | 0 | 2744.38 | 0 | 0.07 | 0.163378 | 12.36 | 97.1 | 102.9 | 5.56 | 7.15 | | High | High | High | High | High | High | High | High | High | High | High | 0 | 1.59E-06 | 3.52 |
| FALSE | High | SPGSPLPPL | [R].SPGSPL | 2xPhospho | Q9D8N6 2x | 0.064862 | 0.004664 | 1 | 4 | 1 | Q9D8N6 | Protein lin- | Q9D8N6 [1 | 0 | 2824.346 | 0 | 0.17 | 0.025832 | 3.09 | 93.2 | 106.8 | 8.5 | 8.7 | | High | High | High | High | High | High | High | High | High | High | High | 0.002432 | 0.02597 | 2.57 |
| FALSE | High | SPGLYAIGY | [K].SPGLYA | 1xCarbami | Q6NZR2 1x | 0.00359 | 0.00021 | 1 | 3 | 4 | Q6NZR2 | Myb/SANT | Q6NZR2 [4 | 0 | 2959.411 | 0 | -1.38 | 0.00014 | 1.05 | 145 | 55 | 2.39 | 8.62 | | High | High | High | High | High | High | High | High | High | High | High | 9.42E-05 | 0.000998 | 2.55 |
| FALSE | High | SPFSVGVS | [K].SPFSVG | 1xPhospho | Q8BTM8 1 | 2.63E-05 | 0 | 1 | 6 | 3 | Q8BTM8 | Filamin-A C | Q8BTM8 [9 | 0 | 2058.092 | 0 | 1.7 | 0.000298 | 3.55 | 47 | 153 | 8.69 | 6.29 | | High | High | High | High | High | High | High | High | High | High | High | 0 | 4E-06 | 3.33 |
| FALSE | High | SPFQSNPS | [K].SPFQSN | 1xPhospho | A2CG63 1x | 3.44E-05 | 0 | 1 | 4 | 2 | A2CG63 | AT-rich int | A2CG63 [6 | 0 | 2089.008 | 0 | -0.28 | 0.300138 | 12.5 | 108.9 | 91.1 | 12.66 | 17.27 | | High | High | High | High | High | High | High | High | High | High | High | 0.000246 | 0.001514 | 1.96 |
| FALSE | High | SPFPGSPE | [R].SPFPGS | 1xPhospho | E9Q156 1x | 2.38E-05 | 0 | 1 | 2 | 4 | E9Q156 | tRNA (guar | E9Q156 [4 | 0 | 1712.866 | 0 | -0.09 | 0.716216 | 1.85 | 101.6 | 98.4 | 5.26 | 6.57 | | High | High | High | High | High | High | High | High | High | High | High | 0 | 3.57E-06 | 2.95 |
| FALSE | High | SPDTSAYC | [R].SPDTSA | 1xCarbami | P14873 1x | 1.25E-05 | 0 | 2 | 2 | 4 | P14873; B | Microtubu | P14873 [2 | 0 | 2235.959 | 0 | -0.73 | 0.031181 | 5.85 | 124.4 | 75.6 | 6.88 | 22.16 | Shared | High | High | High | High | High | High | High | High | High | High | High | 0 | 3.26E-06 | 4.32 |
| FALSE | High | SPEEDQQA | [R].SPEEDC | 1xPhospho | Q6P5H2-2 | 0.036103 | 0.002342 | 1 | 2 | 1 | Q6P5H2-2 | Isoform 2 c | Q6P5H2-2 | 0 | 1982.942 | 0 | 2.36 | 0.000328 | 6.75 | 32.7 | 167.3 | 11.91 | 9.79 | | High | High | High | High | High | High | High | High | High | High | High | 0.00123 | 0.01323 | 2.15 |
| FALSE | High | SPEFTEAEL | [R].SPEFTE | 1xPhospho | Q3UQS8 1 | 0.039011 | 0.002718 | 1 | 1 | 1 | Q3UQS8 | RNA-bindi | Q3UQS8 [1 | 0 | 1688.855 | 0 | 3.18 | 0.000444 | 15.23 | 19.9 | 180.1 | 17.93 | 11.56 | | High | High | High | High | High | High | High | High | High | High | High | 0.001453 | 0.01452 | 2.3 |
| FALSE | High | SPEGTTVS | [K].SPEGTT | 1xPhospho | P97434-2 1 | 0.000193 | 0 | 1 | 10 | 1 | P97434-2 | Isoform 2 c | P97434-2 | 0 | 2022.986 | 0 | 0.55 | 0.064535 | 13.89 | 85.8 | 114.2 | 14.79 | 17.66 | | High | High | High | High | High | High | High | High | High | High | High | 0 | 3.75E-05 | 2.49 |
| FALSE | High | SPEGTTVS | [K].SPEGTT | 1xPhospho | P97434-2 1 | 5.52E-06 | 0 | 1 | 10 | 4 | P97434-2 | Isoform 2 c | P97434-2 | 0 | 2038.981 | 0 | 0.63 | 0.019407 | 7.27 | 81 | 119 | 8.11 | 17.45 | | High | High | High | High | High | High | High | High | High | High | High | 0 | 6.92E-07 | 4.12 |
| FALSE | High | SPEKPPQS | [K].SPEKPP | 1xCarbami | Q8BTI8 1x | 0.000257 | 0 | 1 | 3 | 1 | Q8BTI8 | Serine/argi | Q8BTI8 [4 | 0 | 3120.553 | 0 | -0.65 | 0.008826 | 7.4 | 116.9 | 83.1 | 13.33 | 8.63 | | High | High | High | High | High | High | High | High | High | High | High | 0 | 5.19E-05 | 1.67 |
| FALSE | High | SPEIDDSVI | [K].SPEIDD | 1xPhospho | Z4YKQ9 1x | 0.003788 | 0.00021 | 1 | 3 | 2 | Z4YKQ9 | ATPase fan | Z4YKQ9 [7 | 1 | 3051.554 | 0 | -0.42 | 0.139253 | 18.92 | 110.3 | 89.7 | 14.07 | 15.57 | | High | High | High | High | High | High | High | High | High | High | High | 0.000117 | 0.001058 | 2.73 |
| FALSE | High | SPELSSPAY | [R].SPELSS | 1xPhospho | B7ZNA0 1x | 0.001039 | 0 | 1 | 1 | 1 | B7ZNA0 | Shroom3 p | B7ZNA0 [1 | 0 | 1915.982 | 0 | -0.18 | 0.243895 | 10.81 | 106.3 | 93.7 | 16.86 | 13.24 | | High | High | High | High | High | High | High | High | High | High | High | 0 | 0.00025 | 3.14 |
| FALSE | High | SPEMSMQ | [K].SPEMS | 1xCarbami | Q9EQQ9-3 | 0.003892 | 0.000261 | 1 | 3 | 1 | Q9EQQ9-3 | Isoform 3 c | Q9EQQ9-3 | 0 | 3433.507 | 0 | 0.16 | 0.927553 | 4.86 | 94.6 | 105.4 | 10.17 | 28.74 | | High | High | High | High | High | High | High | High | High | High | High | 0.00014 | 0.001098 | 3.24 |
| FALSE | High | SPEMSMQ | [K].SPEMS | 1xCarbami | Q9EQQ9-3 | 0.001898 | 5.66E-05 | 1 | 3 | 3 | Q9EQQ9-3 | Isoform 3 c | Q9EQQ9-3 | 0 | 3449.502 | 0 | -0.01 | 0.464576 | 14.83 | 100.8 | 99.2 | 5.1 | 12.18 | | High | High | High | High | High | High | High | High | High | High | High | 0.000399 | 0.003354 | 2.76 |
| FALSE | High | SPEPDTLPL | [R].SPEPDT | 1xPhospho | Q6PB97 1x | 2.34E-06 | 0 | 1 | 4 | 2 | Q6PB97 | F-box/LRR- | Q6PB97 [3 | 0 | 2279.127 | 0 | -1.43 | 0.000824 | 1.16 | 146 | 54 | 7.08 | 9.63 | | High | High | High | High | High | High | High | High | High | High | High | 0 | 4.77E-07 | 4.08 |
| FALSE | High | SPEPEATLT | [K].SPEPEA | 1xPhospho | D3YU22 1x | 0.002948 | 0.00016 | 1 | 7 | 1 | D3YU22 | LIM and ca | D3YU22 [7 | 0 | 2230.145 | 0 | 0.92 | 0.000197 | 4.47 | 69.3 | 130.7 | 4.31 | 3.28 | | High | High | High | High | High | High | High | High | High | High | High | 7.19E-05 | 0.000799 | 1.97 |
| FALSE | High | SPEPGQTW | [R].SPEPGC | 1xPhospho | E9Q616 1x | 0.017498 | 0.001245 | 1 | 5 | 1 | E9Q616 | AHNAK nu | E9Q616 [9 | 0 | 2153.985 | 0 | 2.1 | 0.001099 | 9.89 | 38.3 | 161.7 | 16.11 | 9.39 | | High | High | High | High | High | High | High | High | High | High | High | 0.000674 | 0.005847 | 2.24 |
| FALSE | High | SPEPVPGS | [R].SPEPVP | 1xPhospho | P97931 1x | 1.24E-07 | 0 | 1 | 1 | 4 | P97931 | Uracil-DNA | P97931 [2 | 0 | 2701.385 | 0 | -1.89 | 0.000172 | 0.52 | 158.4 | 41.6 | 3.17 | 12 | | High | High | High | High | High | High | High | High | High | High | High | 0 | 9.73E-09 | 4.08 |
| FALSE | High | SPEPVVTM | [K].SPEPVV | 1xPhospho | H9KV00 1x | 0.000136 | 0 | 1 | 7 | 3 | H9KV00 | Protein SO | H9KV00 [2 | 0 | 2132.075 | 0 | 0.57 | 0.139376 | 14.18 | 80.5 | 119.5 | 8.92 | 20.26 | | High | High | High | High | High | High | High | High | High | High | High | 0 | 2.54E-05 | 3.13 |
| FALSE | High | SPEPVVTM | [K].SPEPVV | 1xPhospho | H9KV00 1x | 6.83E-05 | 0 | 1 | 7 | 8 | H9KV00 | Protein SO | H9KV00 [2 | 0 | 2148.07 | 0 | 0.51 | 0.007795 | 10.46 | 82.4 | 117.6 | 4.98 | 13.71 | | High | High | High | High | High | High | High | High | High | High | High | 0.000225 | 0.001427 | 3.18 |
| FALSE | High | SPEQGRPS | [R].SPEQG | 1xPhospho | Q3U3E2 1x | 0.033116 | 0.002195 | 1 | 1 | 1 | Q3U3E2 | Protein FA | Q3U3E2 [1 | 0 | 1494.715 | 0 | -0.59 | 0.067579 | 8.94 | 114.3 | 85.7 | 10.7 | 22.12 | | High | High | High | High | High | High | High | High | High | High | High | 0.001151 | 0.01203 | 1.24 |
| FALSE | High | SPEQPAES | [R].SPEQPA | 1xDeamida | B2RRE0 1x | 6.55E-06 | 0 | 1 | 2 | 9 | B2RRE0 | A kinase (P | B2RRE0 [1 | 0 | 4369.987 | 0 | -0.72 | 0.100268 | 6.36 | 118.5 | 81.5 | 14.82 | 24.9 | | High | High | High | High | High | High | High | High | High | High | High | 0 | 1.14E-06 | 5.49 |
| FALSE | High | SPEQPAES | [R].SPEQPA | 2xPhospho | B2RRE0 2x | 0.000568 | 0 | 1 | 2 | 1 | B2RRE0 | A kinase (P | B2RRE0 [1 | 0 | 4448.97 | 0 | -0.28 | 0.044559 | 13.96 | 112.5 | 87.5 | 6.01 | 16.18 | | High | High | High | High | High | High | High | High | High | High | High | 0 | 0.000126 | 4.44 |
| FALSE | High | SPESQNLID | [K].SPESQN | 1xPhospho | S4R2I9 1xP | 0.000396 | 0 | 1 | 4 | 2 | S4R2I9 | Zinc finger | S4R2I9 [31 | 0 | 1826.93 | 0 | -1.35 | 0.000495 | 2.56 | 140.6 | 59.4 | 4.29 | 11.71 | | High | High | High | High | High | High | High | High | High | High | High | 0 | 8.4E-05 | 2.16 |
| FALSE | High | SPESVTLTK | [K].SPESVT | 1xPhospho | Q3U4X8 1x | 0.000316 | 0 | 1 | 6 | 2 | Q3U4X8 | DNA ligase | Q3U4X8 [2 | 0 | 1499.812 | 0 | -0.68 | 0.001119 | 3.66 | 123.7 | 76.3 | 4.35 | 8.15 | | High | High | High | High | High | High | High | High | High | High | High | 0 | 6.52E-05 | 1.92 |
| FALSE | High | SPETPMCG | [R].SPETPN | 1xCarbami | Q14B71 1x | 3.38E-08 | 0 | 1 | 4 | 3 | Q14B71 | Cell divisio | Q14B71 [2 | 0 | 2060.943 | 0 | -1.25 | 0.001536 | 4.32 | 144.6 | 55.4 | 8.61 | 13.89 | | High | High | High | High | High | High | High | High | High | High | High | 0 | 8.08E-08 | 4.78 |
| FALSE | High | SPETPMCG | [R].SPETPN | 1xCarbami | Q14B71 1x | 3.27E-06 | 0 | 1 | 4 | 7 | Q14B71 | Cell divisio | Q14B71 [2 | 0 | 2076.938 | 0 | -1.04 | 0.000481 | 0.65 | 133.2 | 66.8 | 2.88 | 8.47 | | High | High | High | High | High | High | High | High | High | High | High | 0 | 3.85E-07 | 3.64 |
| FALSE | High | SPFEGAVT | [R].SPFEGA | 1xPhospho | Q8BT14-3 | 1.38E-05 | 0 | 1 | 6 | 2 | Q8BT14-3 | Isoform 3 c | Q8BT14-3 | 0 | 2427.107 | 0 | -0.29 | 0.001989 | 3.81 | 110.1 | 89.9 | 2.85 | 4.7 | | High | High | High | High | High | High | High | High | High | High | High | 0 | 1.94E-06 | 3.72 |
| FALSE | High | SPFEIISPPA | [R].SPFEIIS | 2xPhospho | A2ARP8 2x | 5.91E-05 | 0 | 1 | 3 | 3 | A2ARP8 | Microtubu | A2ARP8 [2 | 0 | 2430.102 | 0 | -1.81 | 0.003529 | 1.19 | 157.6 | 42.4 | 16.92 | 17.87 | | High | High | High | High | High | High | High | High | High | High | High | 0 | 9.95E-06 | 3.14 |
| FALSE | High | SPFEVQVS | [R].SPFEVC | 1xPhospho | Q8VHX6 1x | 0.001301 | 0 | 1 | 2 | 1 | Q8VHX6 | Filamin-C C | Q8VHX6 [5 | 0 | 2112.078 | 0 | -0.73 | 0.003767 | 5.32 | 121.7 | 78.3 | 9.41 | 21.73 | | High | High | High | High | High | High | High | High | High | High | High | 0 | 0.000319 | 3.52 |
| FALSE | High | SPFNDRFP | [R].SPFND | 1xPhospho | F6UFZ5 1x | 0.080423 | 0.006219 | 2 | 6 | 1 | F6UFZ5; Q | PCF11 clea | F6UFZ5 [6 | 1 | 1529.772 | 0 | -0.34 | 0.005204 | 6.51 | 112 | 88 | 2.38 | 6.41 | Shared | High | High | High | High | High | High | High | High | High | High | High | 0.003188 | 0.03347 | 1.45 |
| FALSE | High | SPFNSPSP | [K].SPFNSP | 1xPhospho | P70255 1x | 2.25E-05 | 0 | 1 | 4 | 6 | P70255 | Nuclear fac | P70255 [3 | 0 | 1724.784 | 0 | 1.95 | 0.000485 | 11.4 | 41.4 | 158.6 | 7.25 | 9.92 | | High | High | High | High | High | High | High | High | High | High | High | 2.58E-05 | 0.00035 | 3.82 |
| FALSE | High | SPFNSPSP | [K].SPFNSP | 2xPhospho | P70255 2x | 0.000628 | 0 | 1 | 4 | 3 | P70255 | Nuclear fac | P70255 [3 | 0 | 1804.751 | 0 | 2.37 | 0.00016 | 3.35 | 31.9 | 168.1 | 7.6 | 5.33 | | High | High | High | High | High | High | High | High | High | High | High | 0 | 0.000142 | 2.78 |
| FALSE | High | SPAASGAP | [K].SPAASG | 1xDeamida | Q9JKB3 1x | 6.87E-07 | 0 | 1 | 5 | 3 | Q9JKB3 | Y-box-bind | Q9JKB3 [3 | 0 | 4302.097 | 0 | 1.29 | 0.000349 | 4.77 | 58.9 | 141.1 | 4.37 | 4.48 | | High | High | High | High | High | High | High | High | High | High | High | 0 | 6.67E-08 | 5.39 |
| FALSE | High | SMPLSTEG | [R].SMPLST | 1xPhospho | Q3UXJ3 1x | 0.005924 | 0.00062 | 2 | 5 | 1 | Q3UXJ3; Q | Putative ur | Q3UXJ3 [1 | 0 | 1978.946 | 0 | -0.33 | 0.308673 | 24.07 | 111.5 | 88.5 | 11.5 | 21.43 | Shared | High | High | High | High | High | High | High | High | High | High | High | 0.000275 | 0.001747 | 2.11 |
| FALSE | High | SPAASGAP | [K].SPAASG | 1xPhospho | Q9JKB3 1x | 2.71E-07 | 0 | 1 | 5 | 5 | Q9JKB3 | Y-box-bind | Q9JKB3 [3 | 0 | 4301.113 | 0 | 0.82 | 0.000933 | 6.34 | 73.3 | 126.7 | 2.8 | 5.51 | | High | High | High | High | High | High | High | High | High | High | High | 0.000382 | 0.003312 | 3.89 |
| FALSE | High | SPAAPGQS | [K].SPAAPC | 1xCarbami | Q3U1S6 1x | 0.037048 | 0.00249 | 1 | 1 | 2 | Q3U1S6 | Putative ur | Q3U1S6 [1 | 0 | 2072.931 | 0 | -0.27 | 0.088666 | 10.76 | 109.4 | 90.6 | 7.68 | 8.64 | | High | High | High | High | High | High | High | High | High | High | High | 0.002471 | 0.02708 | 2.14 |
| FALSE | High | SNATTPTK | [K].SNATTI | 1xPhospho | H9LAN6 1x | 0.001068 | 0 | 1 | 6 | 2 | H9LAN6 | Histone-lys | H9LAN6 [3 | 0 | 1357.713 | 0 | -1.11 | 0.000162 | 0.7 | 136.9 | 63.1 | 2.29 | 5.59 | | High | High | High | High | High | High | High | High | High | High | High | 0 | 0.000256 | 2.11 |
| FALSE | High | SNCLGTDE | [K].SNCLGT | 1xCarbami | Q8CHK4-3 | 0.010382 | 0.000772 | 1 | 5 | 1 | Q8CHK4-3 | Isoform 3 c | Q8CHK4-3 | 0 | 2617.092 | 0 | -0.16 | 0.415234 | 21.99 | 112.3 | 87.7 | 10.57 | 16.71 | | High | High | High | High | High | High | High | High | High | High | High | 0.000382 | 0.003286 | 1.95 |

| | | | | | | | | | | | | | | | | | | | | | | | | | | | | | | | | | | | | | | |
|---|---|---|---|---|---|---|---|---|---|---|---|---|---|---|---|---|---|---|---|---|---|---|---|---|---|---|---|---|---|---|---|---|---|---|---|---|---|---|
| FALSE | High | SNCLGTDE | [K].SNCLGT | 1xCarbami | Q8CHK4-3 | 0.061176 | 0.004646 | 1 | 5 | 3 | Q8CHK4-3 | Isoform 3 c | Q8CHK4-3 | 0 | 2697.059 | 0 | -0.14 | 0.075754 | 11.11 | 105.5 | 94.5 | 6.18 | 6.19 | | High | High | High | High | High | High | High | High | High | High | High | 0.002354 | 0.02432 | 2.23 |
| FALSE | High | SNDRNSMI | [K].SNDRN | 1xCarbami | E9Q5G3 1x | 0.000207 | 0 | 1 | 4 | 2 | E9Q5G3 | Kinesin-like | E9Q5G3 [1 | 1 | 2588.168 | 0 | -1.03 | 0.000555 | 0.7 | 134.5 | 65.5 | 6.58 | 7.53 | | High | High | High | High | High | High | High | High | High | High | High | 0 | 0.000295 | 3.57 |
| FALSE | High | SNDRNSMI | [K].SNDRN | 1xCarbami | E9Q5G3 1x | 1.82E-06 | 0 | 1 | 4 | 3 | E9Q5G3 | Kinesin-like | E9Q5G3 [1 | 1 | 2604.163 | 0 | -1.33 | 0.001316 | 3.78 | 144.3 | 55.7 | 6.63 | 14.89 | | High | High | High | High | High | High | High | High | High | High | High | 0 | 2E-07 | 3.92 |
| FALSE | High | SNDTLSGIS | [K].SNDTLS | 1xPhospho | Q8BK03 1x | 0.11108 | 0.008991 | 1 | 5 | 1 | Q8BK03 | Mitoguard | Q8BK03 [1 | 0 | 2073.052 | 0 | 0.64 | 0.011348 | 21.45 | 76.1 | 123.9 | 22.36 | 16.64 | | High | High | High | High | High | High | High | High | High | High | High | 0.004531 | 0.04909 | 2.37 |
| FALSE | High | SNDTSQTP | [K].SNDTSQ | 1xPhospho | P51125-3 1 | 5.37E-06 | 0 | 1 | 12 | 5 | P51125-3 | Isoform 3 c | P51125-3 [ | 0 | 1894.875 | 0 | 1.31 | 0.000805 | 10.1 | 56 | 144 | 3.1 | 10.11 | | High | High | High | High | High | High | High | High | High | High | High | 0.000267 | 0.001547 | 2.93 |
| FALSE | High | SNEDQSM | [K].SNEDQ | 1xPhospho | P48678 1x | 0.040028 | 0.002867 | 1 | 3 | 1 | P48678 | Prelamin-A | P48678 [45 | 0 | 1873.81 | 0 | 1.29 | 0.006377 | 9.71 | 58.1 | 141.9 | 20.07 | 16.93 | | High | High | High | High | High | High | High | High | High | High | High | 0.0015 | 0.01495 | 1.26 |
| FALSE | High | SNEDSLILE | [K].SNEDSL | 2xPhospho | Q9D4H9-2 | 0.025738 | 0.001645 | 1 | 4 | 1 | Q9D4H9-2 | Isoform 2 c | Q9D4H9-2 | 0 | 1536.68 | 0 | -0.21 | 0.415051 | 38.59 | 107.2 | 92.8 | 44.76 | 19.75 | | High | High | High | High | High | High | High | High | High | High | High | 0.000872 | 0.009071 | 2.65 |
| FALSE | High | SNEDSLILE | [K].SNEDSL | 1xPhospho | Q9D4H9-2 | 0.00625 | 0.00062 | 1 | 4 | 2 | Q9D4H9-2 | Isoform 2 c | Q9D4H9-2 | 0 | 1685.876 | 0 | -0.04 | 0.63505 | 12.45 | 99.3 | 100.7 | 5.11 | 9.18 | | High | High | High | High | High | High | High | High | High | High | High | 0.000275 | 0.001858 | 3.96 |
| FALSE | High | SNEHDHSK | [K].SNEHD | 1xPhospho | A2AR02 1x | 0.042141 | 0.003162 | 1 | 2 | 1 | A2AR02 | Peptidyl-pr | A2AR02 [5 | 0 | 1491.699 | 0 | -1.09 | 0.002031 | 0.86 | 143.3 | 56.7 | 12.08 | 12.14 | | High | High | High | High | High | High | High | High | High | High | High | 0.001625 | 0.0158 | 1.51 |
| FALSE | High | SNGDLTPK | [R].SNGDLT | 1xPhospho | P28667 1x | 0.004531 | 0.000456 | 1 | 1 | 2 | P28667 | MARCKS-re | P28667 [36 | 0 | 1369.713 | 0 | -1.79 | 0.000222 | 0.64 | 157.8 | 42.2 | 6.87 | 11.59 | | High | High | High | High | High | High | High | High | High | High | High | 0.000204 | 0.0013 | 1.23 |
| FALSE | High | SNGEVSESI | [K].SNGEVS | 1xPhospho | A2A654 1x | 0.003138 | 0.00016 | 1 | 5 | 3 | A2A654 | Bromodom | A2A654 [7 | 0 | 1756.852 | 0 | 0.04 | 0.398279 | 1.72 | 98.6 | 101.4 | 11.87 | 7.61 | | High | High | High | High | High | High | High | High | High | High | High | 0.001581 | 0.01523 | 2.36 |
| FALSE | High | SNATNSSYS | [R].SNATNS | 1xPhospho | G3UWG2 1 | 0.030898 | 0.001896 | 1 | 4 | 1 | G3UWG2 | T-cell lymp | G3UWG2 [ | 0 | 1747.785 | 0 | -0.52 | 0.033678 | 14.07 | 122.8 | 77.2 | 8.85 | 22.54 | | High | High | High | High | High | High | High | High | High | High | High | 0.001044 | 0.01117 | 1.07 |
| FALSE | High | SNGPTPELI | [K].SNGPTI | 1xPhospho | F8VPU2 1x | 1.02E-05 | 0 | 1 | 6 | 2 | F8VPU2 | FERM, Rho | F8VPU2 [8 | 0 | 2262.142 | 0 | -0.44 | 0.006088 | 3.87 | 115.2 | 84.8 | 2.33 | 9.38 | | High | High | High | High | High | High | High | High | High | High | High | 0 | 0.000162 | 3.71 |
| FALSE | High | SNHLNHVS | [K].SNHLNI | 1xPhospho | D3YUJ3 1x | 0.00204 | 5.66E-05 | 1 | 3 | 1 | D3YUJ3 | Cyclin Y-lik | D3YUJ3 [1 | 0 | 2070.09 | 0 | -0.12 | 0.243085 | 10.26 | 104.3 | 95.7 | 8.7 | 10.47 | | High | High | High | High | High | High | High | High | High | High | High | 2.58E-05 | 0.000532 | 2.42 |
| FALSE | High | SNICMPDP | [K].SNICMI | 1xCarbami | Q8K205 1x | 0.00086 | 0 | 1 | 2 | 1 | Q8K205 | Processing | Q8K205 [3 | 0 | 3161.566 | 0 | 0.61 | 0.041182 | 14.81 | 74.6 | 125.4 | 33.02 | 29.47 | | High | High | High | High | High | High | High | High | High | High | High | 0 | 0.000201 | 3.42 |
| FALSE | High | SNLELSPDN | [R].SNLELS | 1xPhospho | Q8CIG3 1x | 0.000584 | 0 | 1 | 3 | 1 | Q8CIG3 | Lysine-spe | Q8CIG3 [1 | 0 | 1776.909 | 0 | 0.8 | 0.00039 | 5.49 | 73.1 | 126.9 | 4.44 | 5.36 | | High | High | High | High | High | High | High | High | High | High | High | 0 | 0.000131 | 2.02 |
| FALSE | High | SNINGPGTI | [R].SNINGF | 1xDeamida | Q0VGB7 1x | 0.015061 | 0.001049 | 2 | 2 | 2 | Q0VGB7; A | Serine/thre | Q0VGB7 [1 | 0 | 2257.222 | 0 | 0.88 | 0.008246 | 17.78 | 70.7 | 129.3 | 12.08 | 12.46 | Shared | High | High | High | High | High | High | High | High | High | High | High | 0.000571 | 0.004964 | 1.95 |
| FALSE | High | SNLPNVGS | [K].SNLPNV | 1xPhospho | A0A0G2JG | 0.001516 | 5.66E-05 | 1 | 4 | 1 | A0A0G2JG | Crystallin b | A0A0G2JG | 0 | 2431.267 | 0 | 6.64 | | 0 | 4.5 | 195.5 | | 10.83 | | Not Found | Not Found | Not Found | High | Not Found | High | High | High | High | High | High | 2.58E-05 | 0.00038 | 2.8 |
| FALSE | High | SNISGWWS | [R].SNISGV | 1xPhospho | B1AZP2 1x | 0.001986 | 5.66E-05 | 1 | 3 | 2 | B1AZP2 | Disks large | B1AZP2 [1 | 0 | 2579.104 | 0 | -1.37 | 0.002064 | 4.22 | 150.4 | 49.6 | 13.93 | 18.59 | | High | High | High | High | High | High | High | High | High | High | High | 0.000161 | 0.001229 | 2.59 |
| FALSE | High | SNISPNFNF | [K].SNISPN | 1xPhospho | Q7TNL7 1x | 3.41E-08 | 0 | 1 | 12 | 2 | Q7TNL7 | Dual specif | Q7TNL7 [3 | 0 | 2438.141 | 0 | -1.54 | 0.001397 | 3.88 | 148.5 | 51.5 | 7.02 | 19.32 | | High | High | High | High | High | High | High | High | High | High | High | 0 | 4.07E-09 | 5.14 |
| FALSE | High | SNISPNFNF | [K].SNISPN | 1xPhospho | Q7TNL7 1x | 7.33E-08 | 0 | 1 | 12 | 27 | Q7TNL7 | Dual specif | Q7TNL7 [3 | 0 | 2454.136 | 0 | -0.94 | 0.001524 | 3.2 | 131.8 | 68.2 | 5.49 | 9.45 | | High | High | High | High | High | High | High | High | High | High | High | 0 | 3.65E-05 | 4.53 |
| FALSE | High | SNLSSPVR | [R].SNLSSP | 1xPhospho | Q9CQ82 1x | 0.007951 | 0.000736 | 1 | 2 | 1 | Q9CQ82 | Centromer | Q9CQ82 [6 | 0 | 1168.592 | 0 | -0.85 | 0.002295 | 5.33 | 128.6 | 71.4 | 6.49 | 7.02 | | High | High | High | High | High | High | High | High | High | High | High | 0.000327 | 0.002423 | 1.12 |
| FALSE | High | SNISSPVTP | [R].SNISSP | 1xPhospho | P70399 1x | 6.62E-07 | 0 | 1 | 6 | 8 | P70399 | TP53-bindi | P70399 [16 | 0 | 2358.139 | 0 | 0.29 | 0.006564 | 5.48 | 88.4 | 111.6 | 6.33 | 7.62 | | High | High | High | High | High | High | High | High | High | High | High | 0 | 5.2E-07 | 4.06 |
| FALSE | High | SNISSPVTP | [R].SNISSP | 1xPhospho | P70399 1x | 2.88E-06 | 0 | 1 | 6 | 3 | P70399 | TP53-bindi | P70399 [16 | 1 | 2715.397 | 0 | -0.03 | 0.811453 | 11.17 | 100.4 | 99.6 | 4.83 | 11.37 | | High | High | High | High | High | High | High | High | High | High | High | 0 | 3.35E-07 | 4 |
| FALSE | High | SNNHASAV | [K].SNNHA | 1xPhospho | Q99NH0 1x | 3.47E-05 | 0 | 1 | 6 | 2 | Q99NH0 | Ankyrin re | Q99NH0 [1 | 0 | 2173.072 | 0 | -0.36 | 0.548953 | 13.87 | 115 | 85 | 6.48 | 32.26 | | High | High | High | High | High | High | High | High | High | High | High | 0 | 5.5E-06 | 3.94 |
| FALSE | High | SNPLLSPK | [K].SNPLLS | 1xPhospho | E9PVX6 1x | 0.00311 | 0.00016 | 1 | 2 | 1 | E9PVX6 | Proliferatic | E9PVX6 [2 | 0 | 1393.786 | 0 | 0.04 | 0.745908 | 35.25 | 98.8 | 101.2 | 9.59 | 16.23 | | High | High | High | High | High | High | High | High | High | High | High | 7.19E-05 | 0.000851 | 1.5 |
| FALSE | High | SNGSLHTL | [R].SNGSLH | 1xCarbami | Q8C7U1 2x | 0.095408 | 0.007437 | 1 | 1 | 2 | Q8C7U1 | NEDD4-bin | Q8C7U1 [1 | 0 | 2387.093 | 0 | -0.42 | 0.000813 | 3.43 | 112.5 | 87.5 | 6.03 | 7.59 | | High | High | High | High | High | High | High | High | High | High | High | 0.004707 | 0.05055 | 3.16 |
| FALSE | High | SNPSIQATL | [R].SNPSIQ | 1xPhospho | Q91X84 1x | 0.002508 | 0.000109 | 1 | 2 | 1 | Q91X84 | CREB-regu | Q91X84 [3 | 0 | 1683.908 | 0 | 2.57 | 0.000815 | 30.47 | 30.7 | 169.3 | 21.37 | 10.31 | | High | High | High | High | High | High | High | High | High | High | High | 4.94E-05 | 0.000671 | 2.63 |
| FALSE | High | SNASDDPL | [K].SNASDI | 1xPhospho | O70445 1x | 6.7E-08 | 0 | 1 | 2 | 4 | O70445 | BRCA1-ass | O70445 [3 | 0 | 2896.404 | 0 | -1.28 | 0.000688 | 1.9 | 140.4 | 59.6 | 5.29 | 10.51 | | High | High | High | High | High | High | High | High | High | High | High | 0 | 4.89E-09 | 4.9 |
| FALSE | High | SMSSSSSA | [R].SMSSS | 1xPhospho | Q61425 1x | 0.000137 | 0 | 1 | 1 | 1 | Q61425 | Hydroxyac | Q61425 [1 | 0 | 1725.813 | 0 | -0.52 | 0.00257 | 5.8 | 116.6 | 83.4 | 7.67 | 17.18 | | High | High | High | High | High | High | High | High | High | High | High | 0 | 2.55E-05 | 1.76 |
| FALSE | High | SMPTSQSH | [K].SMPTS | 1xPhospho | Q3TLH4 1x | 0.001799 | 5.66E-05 | 1 | 7 | 3 | Q3TLH4 | Protein PRI | Q3TLH4 [1 | 0 | 2600.247 | 0 | 0.94 | 0.064071 | 37.2 | 72 | 128 | 10.3 | 30.91 | | High | High | High | High | High | High | High | High | High | High | High | 7.19E-05 | 0.000889 | 2.66 |
| FALSE | High | SMPTSQSH | [K].SMPTS | 1xPhospho | Q3TLH4 1x | 0.000279 | 0 | 1 | 7 | 5 | Q3TLH4 | Protein PRI | Q3TLH4 [1 | 0 | 2616.242 | 0 | 0.88 | 0.020918 | 27.7 | 75.6 | 124.4 | 11.52 | 17.76 | | High | High | High | High | High | High | High | High | High | High | High | 0 | 0.00017 | 2.99 |
| FALSE | High | SMSADEDL | [R].SMSAD | 1xPhospho | P19426 1x | 0.071662 | 0.005278 | 1 | 3 | 1 | P19426 | Negative e | P19426 [11 | 0 | 1773.756 | 0 | 0.01 | 0.269856 | 3.1 | 99.2 | 100.8 | 3.01 | 5.01 | | High | High | High | High | High | High | High | High | High | High | High | 0.002732 | 0.02931 | 2.27 |
| FALSE | High | SMSADEDL | [R].SMSAD | 1xPhospho | P19426 1x | 0.000215 | 0 | 1 | 3 | 8 | P19426 | Negative e | P19426 [11 | 0 | 1789.751 | 0 | 0.37 | 0.000652 | 1.35 | 87.9 | 112.1 | 6.98 | 3.73 | | High | High | High | High | High | High | High | High | High | High | High | 0 | 4.24E-05 | 2.89 |
| FALSE | High | SMSDVSAE | [K].SMSDV | 1xPhospho | Q6ZQ32 1x | 3.41E-05 | 0 | 2 | 8 | 1 | Q6ZQ32; F | MKIAA085 | Q6ZQ32 [5 | 0 | 1875.836 | 0 | 0.23 | 0.270812 | 26.81 | 94.2 | 105.8 | 13.39 | 44.47 | Shared | High | High | High | High | High | High | High | High | High | High | High | 0 | 5.39E-06 | 3.09 |
| FALSE | High | SMSENAVF | [R].SMSEN | 1xPhospho | Q6PCP5 1x | 0.005924 | 0.00062 | 3 | 11 | 1 | Q6PCP5; Q | Mitochond | Q6PCP5 [1 | 0 | 1202.544 | 0 | -0.48 | 0.00122 | 1.42 | 116.6 | 83.4 | 4.71 | 4.74 | Shared | High | High | High | High | High | High | High | High | High | High | High | 0.000275 | 0.001756 | 1.6 |
| FALSE | High | SMSENAVF | [R].SMSEN | 1xPhospho | Q6PCP5 1x | 0.001689 | 5.66E-05 | 3 | 11 | 13 | Q6PCP5; Q | Mitochond | Q6PCP5 [1 | 0 | 1218.539 | 0 | -0.07 | 0.335332 | 6.1 | 100.7 | 99.3 | 3.26 | 7.85 | Shared | High | High | High | High | High | High | High | High | High | High | High | 2.58E-05 | 0.000429 | 1.99 |
| FALSE | High | SMSETGTA | [R].SMSET | 1xPhospho | Q924A2 1x | 1.19E-05 | 0 | 1 | 4 | 4 | Q924A2 | Protein cap | Q924A2 [1 | 0 | 3563.78 | 0 | 0.39 | 0.01229 | 5.01 | 89.5 | 110.5 | 10.8 | 7.59 | | High | High | High | High | High | High | High | High | High | High | High | 0 | 1.64E-06 | 3.37 |
| FALSE | High | SMSFQGIR | [R].SMSFQ | 1xPhospho | E9Q557 1x | 0.088701 | 0.006669 | 1 | 2 | 4 | E9Q557 | Desmoplak | E9Q557 [2 | 0 | 1250.58 | 0 | 3.53 | 0.000211 | 28.6 | 14.6 | 185.4 | 20.07 | 6.61 | | High | High | High | High | High | High | High | High | High | High | High | 0.003432 | 0.03733 | 1.72 |
| FALSE | High | SMSHQAAI | [K].SMSHQ | 1xPhospho | Q8C0E2 1x | 0.007145 | 0.00062 | 1 | 1 | 2 | Q8C0E2 | Vacuolar p | Q8C0E2 [3 | 0 | 1595.756 | 0 | 0.21 | 0.127268 | 14.32 | 92.7 | 107.3 | 3.19 | 13.95 | | High | High | High | High | High | High | High | High | High | High | High | 0.000275 | 0.002147 | 2.19 |
| FALSE | High | SMSHQAAI | [K].SMSHQ | 1xPhospho | Q8C0E2 1x | 0.007811 | 0.000664 | 1 | 1 | 8 | Q8C0E2 | Vacuolar p | Q8C0E2 [3 | 0 | 1611.751 | 0 | 0.24 | 0.229022 | 10.61 | 94.4 | 105.6 | 7.49 | 9.37 | | High | High | High | High | High | High | High | High | High | High | High | 0.000953 | 0.009911 | 2.38 |
| FALSE | High | SMSIDDTPI | [R].SMSIDI | 1xPhospho | A2A9M4 1 | 0.092372 | 0.007211 | 1 | 6 | 1 | A2A9M4 | Dedicator c | A2A9M4 [1 | 0 | 1330.591 | 0 | 1.25 | 0.000192 | 5.35 | 59.5 | 140.5 | 12 | 2.99 | | High | High | High | High | High | High | High | High | High | High | High | 0.003659 | 0.03913 | 1.8 |
| FALSE | High | SMSIDDTPI | [R].SMSIDI | 1xPhospho | A2A9M4 1 | 0.045508 | 0.003373 | 1 | 6 | 1 | A2A9M4 | Dedicator c | A2A9M4 [1 | 0 | 1346.586 | 0 | 1.89 | 0.000223 | 11.49 | 43 | 157 | 12.09 | 4.16 | | High | High | High | High | High | High | High | High | High | High | High | 0.001727 | 0.01723 | 1.39 |
| FALSE | High | SMSVDLSH | [R].SMSVD | 1xPhospho | A2RSJ4 1x | 0.00017 | 0 | 1 | 1 | 2 | A2RSJ4 | UHRF1-bin | A2RSJ4 [93 | 0 | 1838.949 | 0 | 1.42 | 0.033411 | 33.39 | 52.2 | 147.8 | 12.71 | 38.8 | | High | High | High | High | High | High | High | High | High | High | High | 0 | 3.24E-05 | 2.49 |
| FALSE | High | SMSLDMG | [K].SMSLDI | 1xPhospho | Q04690 1x | 0.001968 | 5.66E-05 | 1 | 6 | 1 | Q04690 | Neurofibro | Q04690 [2 | 0 | 2149.007 | 0 | 0.65 | 0.072305 | 23.56 | 84.9 | 115.1 | 9.95 | 21.03 | | High | High | High | High | High | High | High | High | High | High | High | 2.58E-05 | 0.000511 | 2.4 |
| FALSE | High | SMSLIPTSP | [R].SMSLIP | 1xPhospho | G5E8A7 1x | 0.009756 | 0.000736 | 1 | 2 | 4 | G5E8A7 | Protein Sm | G5E8A7 [2 | 0 | 2862.358 | 0 | 0.96 | 0.001967 | 16.38 | 67.9 | 132.1 | 7.41 | 9.55 | | High | High | High | High | High | High | High | High | High | High | High | 0.002976 | 0.03151 | 3.4 |
| FALSE | High | SMSLIPTSP | [R].SMSLIP | 2xPhospho | G5E8A7 2x | 0.031168 | 0.001935 | 1 | 2 | 1 | G5E8A7 | Protein Sm | G5E8A7 [2 | 0 | 2926.33 | 0 | 1.45 | 0.006518 | 17.54 | 59.8 | 140.2 | 15.55 | 18.22 | | High | High | High | High | High | High | High | High | High | High | High | 0.001044 | 0.01122 | 2.09 |
| FALSE | High | SMSLIPTSP | [R].SMSLIP | 2xPhospho | G5E8A7 2x | 0.025964 | 0.001645 | 1 | 2 | 3 | G5E8A7 | Protein Sm | G5E8A7 [2 | 0 | 2942.325 | 0 | 1.12 | 0.0035 | 15.69 | 61.5 | 138.5 | 10.32 | 12.65 | | High | High | High | High | High | High | High | High | High | High | High | 0.003258 | 0.0344 | 2.94 |
| FALSE | High | SMSPNLAP | [R].SMSPN | 1xPhospho | F6UFZ5 1x | 0.009756 | 0.000736 | 2 | 7 | 2 | F6UFZ5; Q | PCF11 clea | F6UFZ5 [3 | 0 | 1498.774 | 0 | -0.09 | 0.203094 | 5.16 | 105.2 | 94.8 | 6.15 | 9.01 | Shared | High | High | High | High | High | High | High | High | High | High | High | 0.000344 | 0.003055 | 2.64 |
| FALSE | High | SMSPPPK | [R].SMSPPI | 1xPhospho | Q3TWW8 1 | 0.11108 | 0.008991 | 1 | 2 | 3 | Q3TWW8 | Serine/argi | Q3TWW8 [ | 0 | 1297.663 | 0 | -0.31 | 0.045958 | 4.15 | 115.3 | 84.7 | 4.33 | 24.85 | | High | High | High | High | High | High | High | High | High | High | High | 0.004503 | 0.04876 | 1.63 |
| FALSE | High | SMSQQESS | [R].SMSQC | 1xPhospho | Q80YR7 1x | 0.001158 | 0 | 1 | 1 | 1 | Q80YR7 | Claspin OS | Q80YR7 [4 | 0 | 1574.708 | 0 | -0.53 | 0.140004 | 3.4 | 111.5 | 88.5 | 14.39 | 12.62 | | High | High | High | High | High | High | High | High | High | High | High | 0 | 0.000282 | 1.82 |
| FALSE | High | SMSQVDVI | [K].SMSQV | 1xPhospho | Q80TU3 1x | 0.003224 | 0.00016 | 1 | 6 | 2 | Q80TU3 | MKIAA068 | Q80TU3 [1 | 1 | 2850.212 | 0 | -0.36 | 0.005912 | 4.48 | 113.2 | 86.8 | 7.31 | 4.2 | | High | High | High | High | High | High | High | High | High | High | High | 7.19E-05 | 0.000886 | 2.67 |
| FALSE | High | SMSQVDVI | [K].SMSQV | 1xPhospho | Q80TU3 1x | 0.00311 | 0.00016 | 1 | 6 | 1 | Q80TU3 | MKIAA068 | Q80TU3 [1 | 1 | 2866.207 | 0 | -0.37 | 0.171182 | 12.28 | 112.8 | 87.2 | 7.61 | 16.48 | | High | High | High | High | High | High | High | High | High | High | High | 7.19E-05 | 0.000849 | 1.62 |
| FALSE | High | SMSSAFCS | [R].SMSSA | 1xCarbami | Q921Q7 1x | 0.000107 | 0 | 1 | 1 | 1 | Q921Q7 | Ras and Ra | Q921Q7 [3 | 0 | 1864.854 | 0 | 2.77 | 0.002699 | 15.54 | 27.9 | 172.1 | 18.51 | 22.22 | | High | High | High | High | High | High | High | High | High | High | High | 0 | 1.94E-05 | 3.11 |
| FALSE | High | SMSSEGEN | [K].SMSSE | 1xPhospho | Q6A009 1x | 1.19E-05 | 0 | 1 | 2 | 2 | Q6A009 | E3 ubiquiti | Q6A009 [5 | 0 | 3491.532 | 0 | 0.29 | 0.979024 | 18.76 | 102 | 98 | 27.2 | 20.44 | | High | High | High | High | High | High | High | High | High | High | High | 0 | 1.63E-06 | 4.57 |
| FALSE | High | SMSSEGEN | [K].SMSSE | 1xPhospho | Q6A009 1x | 0.000144 | 0 | 1 | 2 | 7 | Q6A009 | E3 ubiquiti | Q6A009 [5 | 0 | 3507.527 | 0 | 0.09 | 0.249395 | 13.26 | 90.7 | 109.3 | 9.35 | 19.39 | | High | High | High | High | High | High | High | High | High | High | High | 0 | 2.73E-05 | 4.02 |
| FALSE | High | SMSSIPPYF | [R].SMSSIP | 1xPhospho | B8QI33 1x | 0.015195 | 0.001089 | 1 | 4 | 2 | B8QI33 | Liprin-alph | B8QI33 [7 | 0 | 2515.174 | 0 | -0.14 | 0.950507 | 17.68 | 104.1 | 95.9 | 19.12 | 6.66 | | High | High | High | High | High | High | High | High | High | High | High | 0.000571 | 0.005033 | 2.59 |
| FALSE | High | SMSSSSSA | [R].SMSSS | 1xPhospho | Q61425 1x | 0.110202 | 0.008961 | 1 | 1 | 1 | Q61425 | Hydroxyac | Q61425 [1 | 0 | 1709.818 | 0 | 0.18 | 0.179675 | 18.86 | 98.8 | 101.2 | 23.94 | 10.13 | | High | High | High | High | High | High | High | High | High | High | High | 0.004477 | 0.04842 | 1.19 |
| FALSE | High | SMSLIPTSP | [R].SMSLIP | 1xPhospho | G5E8A7 1x | 0.000141 | 0 | 1 | 2 | 3 | G5E8A7 | Protein Sm | G5E8A7 [2 | 0 | 2846.363 | 0 | 0.95 | 0.005119 | 13.66 | 69.8 | 130.2 | 4.76 | 12.22 | | High | High | High | High | High | High | High | High | High | High | High | 0 | 2.62E-05 | 3.44 |
| FALSE | High | SNPSPPSA | [R].SNPSPF | 1xPhospho | Q80U93 1x | 9.28E-05 | 0 | 1 | 2 | 1 | Q80U93 | Nuclear po | Q80U93 [1 | 0 | 2433.2 | 0 | -0.29 | 0.087458 | 15.79 | 108.4 | 91.6 | 9.22 | 18.11 | | High | High | High | High | High | High | High | High | High | High | High | 0 | 1.64E-05 | 3 |
| FALSE | High | SNPSPPSA | [R].SNPSPF | 1xPhospho | Q80U93 1x | 1.53E-05 | 0 | 1 | 2 | 5 | Q80U93 | Nuclear po | Q80U93 [1 | 0 | 2449.195 | 0 | 0.05 | 0.635051 | 24.44 | 98.4 | 101.6 | 8.15 | 30.76 | | High | High | High | High | High | High | High | High | High | High | High | 0 | 4.01E-06 | 3.38 |
| FALSE | High | SNRPVSAD | [K].SNRPVS | 1xPhospho | Q9R0L6-2 | 0.039011 | 0.002718 | 1 | 3 | 1 | Q9R0L6-2 | Isoform 2 c | Q9R0L6-2 [ | 0 | 2283.201 | 0 | -0.12 | 0.738423 | 5.7 | 102.7 | 97.3 | 8.27 | 12.72 | | High | High | High | High | High | High | High | High | High | High | High | 0.001421 | 0.01445 | 0.98 |
| FALSE | High | SNSSEVTPF | [R].SNSSEV | 1xPhospho | E9Q2I4 1x | 1.46E-05 | 0 | 1 | 3 | 8 | E9Q2I4 | ELM2 and | E9Q2I4 [69 | 0 | 2908.421 | 0 | -0.52 | 0.070718 | 12.76 | 118.3 | 81.7 | 15.98 | 14.65 | | High | High | High | High | High | High | High | High | High | High | High | 4.94E-05 | 0.000693 | 3.73 |
| FALSE | High | SNSSEVTPF | [R].SNSSEV | 2xPhospho | E9Q2I4 2x | 7.91E-06 | 0 | 1 | 3 | 4 | E9Q2I4 | ELM2 and | E9Q2I4 [69 | 0 | 2988.388 | 0 | 0.22 | 0.276869 | 22.33 | 87.6 | 112.4 | 12.19 | 39.64 | | High | High | High | High | High | High | High | High | High | High | High | 0 | 1.03E-06 | 4.84 |
| FALSE | High | SNSSGLSIG | [R].SNSSGI | 1xDeamida | Q69Z43 1x | 1.29E-05 | 0 | 1 | 2 | 9 | Q69Z43 | MKIAA199 | Q69Z43 [7 | 0 | 4990.157 | 0 | 1.89 | 0.000561 | 23.42 | 42.7 | 157.3 | 19.49 | 5.84 | | High | High | High | High | High | High | High | High | High | High | High | 0 | 1.8E-06 | 6.69 |
| FALSE | High | SNSSGNLH | [R].SNSSGI | 1xPhospho | Q8C0V0 1x | 0.036103 | 0.002342 | 1 | 2 | 2 | Q8C0V0 | Serine/thre | Q8C0V0 [7 | 0 | 2945.38 | 0 | -0.46 | 0.108988 | 16.77 | 115.7 | 84.3 | 23.24 | 16.57 | | High | High | High | High | High | High | High | High | High | High | High | 0.001263 | 0.01332 | 2.58 |
| FALSE | High | SNSSIDHLG | [K].SNSSID | 1xPhospho | A7TU71 1x | 0.000143 | 0 | 1 | 2 | 2 | A7TU71 | Protein Shr | A7TU71 [1 | 0 | 1876.932 | 0 | 1.47 | 0.017942 | 17.64 | 55.6 | 144.4 | 23.4 | 25.96 | | High | High | High | High | High | High | High | High | High | High | High | 0 | 2.69E-05 | 3.13 |
| FALSE | High | SNSSIDHLG | [K].SNSSID | 2xPhospho | A7TU71 2x | 0.103392 | 0.008249 | 1 | 2 | 1 | A7TU71 | Protein Shr | A7TU71 [1 | 0 | 1956.898 | 0 | 0.72 | 0.02342 | 6.47 | 75.6 | 124.4 | 11.98 | 26.37 | | High | High | High | High | High | High | High | High | High | High | High | 0.004164 | 0.045 | 1.97 |
| FALSE | High | SNSSSPVDI | [R].SNSSSP | 1xPhospho | Q6NXH3-2 | 0.012397 | 0.000889 | 1 | 3 | 1 | Q6NXH3-2 | Isoform 2 c | Q6NXH3-2 | 0 | 1458.724 | 0 | -0.63 | 0.009589 | 6.79 | 126.3 | 73.7 | 8.86 | 12.95 | | High | High | High | High | High | High | High | High | High | High | High | 0.000449 | 0.004 | 1.84 |
| FALSE | High | SNSSYLGSI | [R].SNSSYL | 1xCarbami | B2RXQ5 1x | 0.003822 | 0.000261 | 1 | 4 | 1 | B2RXQ5 | Phosphatic | B2RXQ5 [1 | 0 | 2732.072 | 0 | 0.62 | 0.057049 | 20 | 80.3 | 119.7 | 5.01 | 20.71 | | High | High | High | High | High | High | High | High | High | High | High | 0.000117 | 0.001067 | 3.18 |
| FALSE | High | SNSSYLGSI | [R].SNSSYL | 1xCarbami | B2RXQ5 1x | 0.000246 | 0 | 1 | 4 | 2 | B2RXQ5 | Phosphatic | B2RXQ5 [1 | 0 | 2748.067 | 0 | 0.82 | 0.007966 | 6.98 | 72.9 | 127.1 | 10.66 | 11.21 | | High | High | High | High | High | High | High | High | High | High | High | 0 | 8.34E-05 | 2.97 |
| FALSE | High | SNSTDQISF | [K].SNSTD | 1xPhospho | F8VPJ6 1x | 0.000573 | 0 | 1 | 3 | 6 | F8VPJ6 | Regulatory | F8VPJ6 [13 | 0 | 4009.87 | 0 | -0.36 | 0.034648 | 6.52 | 112.9 | 87.1 | 14.06 | 9.55 | | High | High | High | High | High | High | High | High | High | High | High | 9.42E-05 | 0.001012 | 3.38 |
| FALSE | High | SNSTEHLLI | [R].SNSTEH | 1xPhospho | Q9JID9 1x | 2.02E-05 | 0 | 1 | 1 | 2 | Q9JID9 | SH2B adap | Q9JID9 [58 | 0 | 2866.331 | 0 | 0.08 | 0.587192 | 40.69 | 97.4 | 102.6 | 12.07 | 13.14 | | High | High | High | High | High | High | High | High | High | High | High | 0 | 2.98E-06 | 4.68 |
| FALSE | High | SNSTSSMS | [R].SNSTSS | 1xPhospho | F8VQH0 1x | 4.2E-05 | 0 | 1 | 2 | 2 | F8VQH0 | Tyrosine-p | F8VQH0 [7 | 0 | 2007.853 | 0 | 0.01 | 0.987294 | 23.28 | 99.7 | 100.3 | 7.72 | 15.27 | | High | High | High | High | High | High | High | High | High | High | High | 0 | 0.00012 | 2.35 |
| FALSE | High | SNSVGIQD | [R].SNSVGI | 1xPhospho | E9QAQ7 1x | 4E-07 | 0 | 1 | 5 | 2 | E9QAQ7 | AT-rich inte | E9QAQ7 [1 | 0 | 2648.265 | 0 | 0.4 | 0.006834 | 11.77 | 88.9 | 111.1 | 15.32 | 9.84 | | High | High | High | High | High | High | High | High | High | High | High | 0 | 3.61E-08 | 5.46 |
| FALSE | High | SNSSEVTPF | [R].SNSSEV | 1xPhospho | E9Q2I4 1x | 0.000246 | 0 | 1 | 3 | 3 | E9Q2I4 | ELM2 and | E9Q2I4 [69 | 0 | 2892.426 | 0 | 0.52 | 0.175675 | 61.59 | 71.2 | 128.8 | 19.89 | 44.83 | | High | High | High | High | High | High | High | High | High | High | High | 0 | 4.93E-05 | 3.5 |
| FALSE | High | SNSVSLVPF | [K].SNSVSL | 1xPhospho | Q6QI06 1x | 0.000257 | 0 | 1 | 2 | 2 | Q6QI06 | Rapamycin | Q6QI06 [1 | 0 | 2044.043 | 0 | 1.12 | 0.002238 | 17.04 | 65.6 | 134.4 | 8.64 | 11.5 | | High | High | High | High | High | High | High | High | High | High | High | 0 | 5.19E-05 | 2.29 |
| FALSE | High | SNTEEALK | [R].SNTEEA | 1xPhospho | A0A140LJJ | 0.024001 | 0.001612 | 1 | 3 | 1 | A0A140LJJ | A-kinase ar | A0A140LJJ | 0 | 1429.734 | 0 | 0.15 | 0.054306 | 8.25 | 89 | 111 | 8.23 | 9 | | High | High | High | High | High | High | High | High | High | High | High | 0.000859 | 0.008393 | 1.53 |
| FALSE | High | SNTLDITDG | [R].SNTLDI | 1xPhospho | F8VQ05 1x | 0.000286 | 0 | 1 | 5 | 4 | F8VQ05 | FRY-like tra | F8VQ05 [1 | 0 | 1400.662 | 0 | 0.24 | 0.021167 | 9.14 | 91.7 | 108.3 | 2.84 | 8.7 | | High | High | High | High | High | High | High | High | High | High | High | 0.000382 | 0.00333 | 2.65 |
| FALSE | High | SNTPEVHT | [R].SNTPEV | 2xPhospho | P55200 2x | 0.001068 | 0 | 1 | 3 | 1 | P55200 | Histone-lys | P55200 [5 | 0 | 2937.287 | 0 | -0.23 | 0.400921 | 5.99 | 106.1 | 93.9 | 16.17 | 12.08 | | High | High | High | High | High | High | High | High | High | High | High | 0 | 0.000257 | 3.45 |
| FALSE | High | SNTPMGDI | [R].SNTPM | 1xPhospho | A2AN08-5 | 0.087983 | 0.006639 | 1 | 4 | 1 | A2AN08-5 | Isoform 5 c | A2AN08-5 | 0 | 1387.669 | 0 | 0.38 | 0.032018 | 17.6 | 91.5 | 108.5 | 7.47 | 9.53 | | High | High | High | High | High | High | High | High | High | High | High | 0.003402 | 0.0371 | 1.57 |
| FALSE | High | SNTPMGDI | [R].SNTPM | 1xPhospho | A2AN08-5 | 0.069325 | 0.005112 | 1 | 4 | 4 | A2AN08-5 | Isoform 5 c | A2AN08-5 | 0 | 1403.664 | 0 | 0.39 | 0.006474 | 10.9 | 87.2 | 112.8 | 3.47 | 7.78 | | High | High | High | High | High | High | High | High | High | High | High | 0.007374 | 0.08114 | 1.91 |
| FALSE | High | SNTPMGDI | [R].SNTPM | 1xPhospho | A2AN08-5 | 0.003622 | 0.00021 | 1 | 4 | 1 | A2AN08-5 | Isoform 5 c | A2AN08-5 | 1 | 2865.222 | 0 | 0.13 | 0.180139 | 8.99 | 86.2 | 113.8 | 18.42 | 14.99 | | High | High | High | High | High | High | High | High | High | High | High | 9.42E-05 | 0.001007 | 2.1 |
| FALSE | High | SNTQPSSC | [K].SNTQP | 1xCarbami | Q64337 1x | 0.006031 | 0.00062 | 1 | 4 | 3 | Q64337 | Sequestos | Q64337 [2 | 0 | 3961.876 | 0 | 1.62 | 0.0162 | 17.65 | 49.3 | 150.7 | 6.98 | 28.15 | | High | High | High | High | High | High | High | High | High | High | High | 0.000275 | 0.001787 | 3.34 |
| FALSE | High | SNVGGLLS | [K].SNVGG | 1xPhospho | Q6P4T3 1x | 0.000308 | 0 | 1 | 3 | 2 | Q6P4T3 | Eyes absen | Q6P4T3 [3 | 0 | 1436.746 | 0 | -0.41 | 0.000492 | 1.39 | 116.1 | 83.9 | 2.41 | 4.2 | | High | High | High | High | High | High | High | High | High | High | High | 0 | 6.38E-05 | 3.18 |
| FALSE | High | SNVLLSFDI | [R].SNVLLS | 1xPhospho | Q61037 1x | 0.000116 | 0 | 1 | 14 | 1 | Q61037 | Tuberin OS | Q61037 [9 | 0 | 2003.014 | 0 | -0.26 | 0.005203 | 5.24 | 108.7 | 91.3 | 5.83 | 9.83 | | High | High | High | High | High | High | High | High | High | High | High | 0 | 2.14E-05 | 3.19 |
| FALSE | High | SPAAFTSPF | [R].SPAAFT | 1xPhospho | Q571G8 1x | 4.16E-05 | 0 | 1 | 7 | 1 | Q571G8 | MKIAA164 | Q571G8 [1 | 0 | 2064.098 | 0 | -0.11 | 0.733724 | 2.82 | 97.8 | 102.2 | 14.82 | 5.77 | | High | High | High | High | High | High | High | High | High | High | High | 0 | 6.69E-06 | 3.28 |
| FALSE | High | SPAAFTSPF | [R].SPAAFT | 1xPhospho | Q571G8 1x | 0.000188 | 0 | 1 | 7 | 2 | Q571G8 | MKIAA164 | Q571G8 [1 | 0 | 2293.261 | 0 | 0.14 | 0.110113 | 14.81 | 95.6 | 104.4 | 10.01 | 9.67 | | High | High | High | High | High | High | High | High | High | High | High | 0 | 3.64E-05 | 2.69 |
| FALSE | High | SPAAGSSQ | [R].SPAAG | 1xPhospho | Q5CZY4 1x | 8.45E-07 | 0 | 1 | 5 | 2 | Q5CZY4 | Nuclear fac | Q5CZY4 [3 | 0 | 2824.355 | 0 | 1.96 | 0.000317 | 17.17 | 41.1 | 158.9 | 8.98 | 6.37 | | High | High | High | High | High | High | High | High | High | High | High | 0 | 8.4E-08 | 5.22 |
| FALSE | High | SPAAGSSQ | [R].SPAAG | 1xPhospho | P70257-2 1 | 0.115566 | 0.009504 | 1 | 2 | 1 | P70257-2 | Isoform NF | P70257-2 [ | 0 | 2439.078 | 0 | -0.09 | 0.265865 | 22.09 | 107.5 | 92.5 | 11.78 | 11.61 | | High | High | High | High | High | High | High | High | High | High | High | 0.004757 | 0.05132 | 2.56 |
| FALSE | High | SNSWVNT | [R].SNSWV | 1xPhospho | A2A9M4 1 | 0.000765 | 0 | 1 | 5 | 2 | A2A9M4 | Dedicator c | A2A9M4 [9 | 0 | 1627.825 | 0 | 1.25 | 0.00014 | 0.97 | 59.4 | 140.6 | 4.67 | 2.15 | | High | High | High | High | High | High | High | High | High | High | High | 7.19E-05 | 0.000865 | 3.53 |
| FALSE | High | SNSSEASS | [R].SNSSEA | 2xPhospho | P97825 2x | 0.014667 | 0.001049 | 1 | 1 | 2 | P97825 | Jupiter mic | P97825 [8 | 0 | 2174.966 | 0 | 0.96 | 0.002485 | 8.27 | 65.5 | 134.5 | 13 | 17.92 | | High | High | High | High | High | High | High | High | High | High | High | 0.000555 | 0.004807 | 2.77 |
| FALSE | High | SNSSEASS | [R].SNSSEA | 1xDeamida | P97825 1x | 0.002212 | 0.000109 | 1 | 1 | 4 | P97825 | Jupiter mic | P97825 [8 | 0 | 2095.984 | 0 | 1.74 | 0.00043 | 5.78 | 47 | 153 | 7.42 | 6.32 | | High | High | High | High | High | High | High | High | High | High | High | 4.94E-05 | 0.000595 | 2.95 |
| FALSE | High | SNSSEASS | [R].SNSSEA | 1xPhospho | P97825 1x | 5.19E-07 | 0 | 1 | 1 | 9 | P97825 | Jupiter mic | P97825 [8 | 0 | 2095 | 0 | 1.2 | 0.000275 | 7.81 | 62.6 | 137.4 | 5.59 | 4.68 | | High | High | High | High | High | High | High | High | High | High | High | 0 | 4.89E-08 | 3.55 |
| FALSE | High | SNSAPLIHG | [R].SNSAPL | 1xPhospho | Q9DB52 1x | 9.85E-07 | 0 | 1 | 1 | 2 | Q9DB52 | Protein FAI | Q9DB52 [3 | 0 | 2847.388 | 0 | 0.45 | 0.097966 | 2.32 | 85 | 115 | 9.89 | 12.01 | | High | High | High | High | High | High | High | High | High | High | High | 0 | 1E-07 | 4.81 |
| FALSE | High | SNSAPLIHV | [R].SNSAPL | 1xPhospho | Q6NZE7-2 | 2.85E-09 | 0 | 1 | 4 | 1 | Q6NZE7-2 | Isoform 2 c | Q6NZE7-2 | 0 | 2891.491 | 0 | -1.29 | 0.000779 | 1.5 | 148.5 | 51.5 | 11.04 | 15.25 | | High | High | High | High | High | High | High | High | High | High | High | 0 | 1.4E-10 | 5.17 |
| FALSE | High | SNSAWQIY | [R].SNSAW | 1xPhospho | Q9JM52-3 | 0.007019 | 0.00062 | 1 | 6 | 1 | Q9JM52-3 | Isoform 3 c | Q9JM52-3 | 0 | 1674.82 | 0 | 1.37 | 0.002684 | 27.31 | 57 | 143 | 13.33 | 13.64 | | High | High | High | High | High | High | High | High | High | High | High | 0.000275 | 0.002119 | 2.76 |
| FALSE | High | SNSCSSISV | [R].SNSCSS | 2xCarbami | E9Q5G3 1x | 9.66E-10 | 0 | 1 | 1 | 8 | E9Q5G3 | Kinesin-like | E9Q5G3 [7 | 0 | 2697.23 | 0 | -1.4 | 0.000422 | 1.42 | 142.7 | 57.3 | 5.78 | 7.58 | | High | High | High | High | High | High | High | High | High | High | High | 0 | 4.39E-11 | 6.15 |
| FALSE | High | SNSDITISDI | [R].SNSDIT | 1xPhospho | B9EK76 1x | 1.09E-07 | 0 | 1 | 4 | 3 | B9EK76 | Signal-indu | B9EK76 [1 | 0 | 3627.713 | 0 | -0.75 | 0.000961 | 2.24 | 123.8 | 76.2 | 8.07 | 3.86 | | High | High | High | High | High | High | High | High | High | High | High | 0 | 8.44E-09 | 6.96 |
| FALSE | High | SNSDITISEI | [R].SNSDIT | 1xCarbami | Q8C0T5 1x | 0.000158 | 0 | 1 | 4 | 3 | Q8C0T5 | Signal-indu | Q8C0T5 [1 | 0 | 3273.516 | 0 | 1.21 | 0.000877 | 10.27 | 58.8 | 141.2 | 12.04 | 7.69 | | High | High | High | High | High | High | High | High | High | High | High | 0 | 3E-05 | 3.14 |
| FALSE | High | SNSEETGEI | [K].SNSEET | 1xPhospho | Q9JJ89 1x | 0.001437 | 5.66E-05 | 1 | 1 | 1 | Q9JJ89 | Coiled-coil | Q9JJ89 [39 | 0 | 1631.793 | 0 | 1.37 | 0.003822 | 15.4 | 55.8 | 144.2 | 9.86 | 19.57 | | High | High | High | High | High | High | High | High | High | High | High | 2.58E-05 | 0.000357 | 2.06 |
| FALSE | High | SNSEFSPLC | [K].SNSEFS | 1xPhospho | Q8C804 1x | 0.0004 | 0 | 1 | 1 | 1 | Q8C804 | Spindle an | Q8C804 [5 | 0 | 1800.873 | 0 | -0.52 | 0.009225 | 1.13 | 117.9 | 82.1 | 10.31 | 4.27 | | High | High | High | High | High | High | High | High | High | High | High | 0 | 8.49E-05 | 2.17 |
| FALSE | High | SNSFNADR | [R].SNSFN | 1xPhospho | Q69ZR9-2 | 0.017498 | 0.001245 | 1 | 2 | 1 | Q69ZR9-2 | Isoform 2 c | Q69ZR9-2 [ | 0 | 1219.531 | 0 | -0.46 | 0.05301 | 14.4 | 115.8 | 84.2 | 12.14 | 9.94 | | High | High | High | High | High | High | High | High | High | High | High | 0.000674 | 0.005871 | 1.6 |
| FALSE | High | SNSFNNPL | [R].SNSFN | 1xPhospho | Q8BYR2 1x | 0.000334 | 0 | 1 | 1 | 1 | Q8BYR2 | Serine/thre | Q8BYR2 [4 | 0 | 1501.7 | 0 | 2.52 | 0.00014 | 4.61 | 29.7 | 170.3 | 6.35 | 3.04 | | High | High | High | High | High | High | High | High | High | High | High | 0 | 6.97E-05 | 2.6 |
| FALSE | High | SNSFSDER | [R].SNSFSD | 1xPhospho | E9Q616 1x | 0.005418 | 0.00062 | 1 | 4 | 4 | E9Q616 | AHNAK nuc | E9Q616 [5 | 0 | 1250.525 | 0 | 2.57 | 0.000236 | 10.63 | 27.1 | 172.9 | 9.35 | 6.55 | | High | High | High | High | High | High | High | High | High | High | High | 0.002455 | 0.02694 | 2.07 |
| FALSE | High | SNSFSDERI | [R].SNSFSD | 3xPhospho | E9Q616 3x | 0.001558 | 5.66E-05 | 1 | 4 | 1 | E9Q616 | AHNAK nuc | E9Q616 [5 | 1 | 3374.374 | 0 | 0.95 | 0.045 | 24.11 | 69.5 | 130.5 | 25.95 | 17.63 | | High | High | High | High | High | High | High | High | High | High | High | 2.58E-05 | 0.000394 | 4.03 |
| FALSE | High | SNSFSDERI | [R].SNSFSD | 1xPhospho | E9Q616 1x | 3.06E-05 | 0 | 1 | 4 | 2 | E9Q616 | AHNAK nuc | E9Q616 [5 | 1 | 3443.604 | 0 | 2.17 | 0.016022 | 40.67 | 37.8 | 162.2 | 14.67 | 34.35 | | High | High | High | High | High | High | High | High | High | High | High | 0 | 4.76E-06 | 4.26 |
| FALSE | High | SNSFSDERI | [R].SNSFSD | 2xPhospho | E9Q616 2x | 0.000133 | 0 | 1 | 4 | 3 | E9Q616 | AHNAK nuc | E9Q616 [5 | 1 | 3523.571 | 0 | 2.66 | 0.000154 | 10.02 | 27.3 | 172.7 | 7.95 | 3.91 | | High | High | High | High | High | High | High | High | High | High | High | 0 | 4.11E-05 | 4.45 |
| FALSE | High | SNSLDDSTI | [K].SNSLDI | 1xPhospho | A2AG50 1x | 6.53E-05 | 0 | 1 | 2 | 2 | A2AG50 | MAP7 dom | A2AG50 [6 | 0 | 2778.279 | 0 | -0.13 | 0.141276 | 10.45 | 108.6 | 91.4 | 12.58 | 16.95 | | High | High | High | High | High | High | High | High | High | High | High | 0 | 1.12E-05 | 3.46 |
| FALSE | High | SNSLDDSTI | [K].SNSLDI | 1xPhospho | A2AG50 1x | 1.99E-06 | 0 | 1 | 2 | 4 | A2AG50 | MAP7 dom | A2AG50 [6 | 0 | 2794.274 | 0 | -0.01 | 0.472282 | 7.88 | 95.2 | 104.8 | 8.36 | 6.43 | | High | High | High | High | High | High | High | High | High | High | High | 0 | 2.21E-07 | 3.82 |

| | | | | | | | | | | | | | | | | | | | | | | | | | | | | | | | | | | | | | | |
|---|---|---|---|---|---|---|---|---|---|---|---|---|---|---|---|---|---|---|---|---|---|---|---|---|---|---|---|---|---|---|---|---|---|---|---|---|---|---|
| FALSE | High | SNSLNNGC | [R].SNSLN | 1xPhospho | A2AH75 1x | 0.01218 | 0.000889 | 1 | 2 | 1 | A2AH75 | Kinesin fan | A2AH75 [1 | 0 | 1596.815 | 0 | 0.31 | 0.016364 | 12.35 | 89.1 | 110.9 | 6.2 | 4.74 | | High | High | High | High | High | High | High | High | High | High | High | 0.000449 | 0.003895 | 1.89 |
| FALSE | High | SNSLNNGC | [R].SNSLN | 1xDeamida | A2AH75 1x | 0.001179 | 0 | 1 | 2 | 3 | A2AH75 | Kinesin fan | A2AH75 [1 | 0 | 1597.799 | 0 | 0.35 | 0.116864 | 4.94 | 88 | 112 | 8.83 | 6.55 | | High | High | High | High | High | High | High | High | High | High | High | 0 | 0.000286 | 3.93 |
| FALSE | High | SNSLPHPA( | [R].SNSLPH | 1xPhospho | Q08274 1x | 0.003755 | 0.00021 | 1 | 4 | 1 | Q08274 | Dystrophia | Q08274 [4 | 0 | 1659.862 | 0 | 0.27 | 0.013338 | 9.6 | 90.8 | 109.2 | 8.59 | 8.34 | | High | High | High | High | High | High | High | High | High | High | High | 9.42E-05 | 0.001053 | 1.97 |
| FALSE | High | SNSLPHSA\ | [R].SNSLPH | 1xPhospho | Q3UWE6 1 | 0.000255 | 0 | 1 | 4 | 1 | Q3UWE6 | MCG14935 | Q3UWE6 [ | 0 | 2008.026 | 0 | 0.22 | 0.129738 | 26.97 | 91.6 | 108.4 | 12.42 | 7.83 | | High | High | High | High | High | High | High | High | High | High | High | 0 | 5.15E-05 | 2.69 |
| FALSE | High | SNSLSEQLT | [K].SNSLSE | 1xPhospho | Q6P9Q6-2 | 1.59E-07 | 0 | 1 | 6 | 5 | Q6P9Q6-2 | Isoform B ( | Q6P9Q6-2 | 0 | 2471.243 | 0 | -1.14 | 0.000813 | 0.9 | 138.2 | 61.8 | 7.72 | 6.95 | | High | High | High | High | High | High | High | High | High | High | High | 0 | 1.38E-06 | 4.28 |
| FALSE | High | SNSISQIPSI | [R].SNSISQ | 1xPhospho | Q99K90 1x | 0.0009 | 0 | 1 | 1 | 1 | Q99K90 | TGF-beta-a | Q99K90 [5 | 0 | 2285.105 | 0 | 0.76 | 0.002218 | 4.17 | 72.5 | 127.5 | 9.19 | 13.83 | | High | High | High | High | High | High | High | High | High | High | High | 0 | 0.000211 | 2.92 |
| FALSE | High | SNSNSSSVI | [R].SNSNSS | 1xPhospho | E0CYH7 1x | 9.2E-05 | 0 | 1 | 4 | 2 | E0CYH7 | Filamin A-ii | E0CYH7 [1( | 0 | 2121.011 | 0 | 0.56 | 0.067316 | 41 | 76.3 | 123.7 | 16.04 | 26.79 | | High | High | High | High | High | High | High | High | High | High | High | 0 | 1.63E-05 | 2.46 |
| FALSE | High | SNSPENK | [R].SNSPEN | 1xPhospho | Q924H7 1x | 0.08727 | 0.006537 | 1 | 10 | 1 | Q924H7 | WW doma | Q924H7 [6 | 0 | 1313.65 | 0 | 0.37 | 0.005009 | 9.09 | 87 | 113 | 3.67 | 4.72 | | High | High | High | High | High | High | High | High | High | High | High | 0.003375 | 0.03674 | 1.51 |
| FALSE | High | SNSPVSEKI | [K].SNSPVS | 1xPhospho | Q80Y82 1x | 0.027121 | 0.001676 | 1 | 12 | 1 | Q80Y82 | Protein kin | Q80Y82 [6 | 0 | 2321.236 | 0 | 0.22 | 0.058592 | 15.66 | 92.8 | 107.2 | 6.82 | 9.42 | | High | High | High | High | High | High | High | High | High | High | High | 0.00092 | 0.009653 | 1.47 |
| FALSE | High | SNSSDAPG | [K].SNSSD/ | 1xPhospho | Q3ULC7 1x | 9.69E-11 | 0 | 1 | 5 | 3 | Q3ULC7 | Putative ur | Q3ULC7 [2 | 0 | 2278.996 | 0 | -0.2 | 0.050598 | 6.37 | 106.7 | 93.3 | 1.93 | 11.55 | | High | High | High | High | High | High | High | High | High | High | High | 0 | 3.15E-12 | 4.84 |
| FALSE | High | SNSSDAPG | [K].SNSSD/ | 1xDeamida | Q3ULC7 1x | 0.039011 | 0.002718 | 1 | 5 | 2 | Q3ULC7 | Putative ur | Q3ULC7 [2 | 0 | 2279.98 | 0 | 0.2 | 0.18154 | 2.34 | 99.5 | 100.5 | 14.26 | 11.29 | | High | High | High | High | High | High | High | High | High | High | High | 0.001453 | 0.01452 | 2.1 |
| FALSE | High | SNSSDAPG | [K].SNSSD/ | 2xPhospho | Q3ULC7 2x | 0.000967 | 0 | 1 | 5 | 3 | Q3ULC7 | Putative ur | Q3ULC7 [2 | 0 | 2358.963 | 0 | -0.14 | 0.315855 | 25.31 | 104.7 | 95.3 | 13.72 | 21.99 | | High | High | High | High | High | High | High | High | High | High | High | 0 | 0.00023 | 3.78 |
| FALSE | High | SNSSEALLV | [R].SNSSEA | 1xPhospho | A2AEV7 1x | 3.41E-05 | 0 | 1 | 1 | 2 | A2AEV7 | Coiled-coil | A2AEV7 [3 | 0 | 1499.73 | 0 | 1.54 | 0.002337 | 33.83 | 50 | 150 | 9.46 | 13.23 | | High | High | High | High | High | High | High | High | High | High | High | 0 | 5.39E-06 | 2.83 |
| FALSE | High | SPAASAPSI | [R].SPAAS/ | 1xPhospho | Q8K3G7 1x | 0.000314 | 0 | 3 | 4 | 3 | Q8K3G7; C | Reticulon ( | Q8K3G7 [1 | 0 | 2270.256 | 0 | 1.88 | 0.017229 | 31.59 | 42.8 | 157.2 | 7.34 | 34.61 | Shared | High | High | High | High | High | High | High | High | High | High | High | 0 | 6.47E-05 | 4.21 |
| FALSE | High | SLLTAGATI | [R].SLLTAG | 1xPhospho | O55111 1x | 0.024855 | 0.001645 | 1 | 2 | 1 | O55111 | Desmoglei | O55111 [7 | 0 | 1571.826 | 0 | 0.72 | 0.077842 | 9.76 | 78.8 | 121.2 | 9.87 | 20.49 | | High | High | High | High | High | High | High | High | High | High | High | 0.000872 | 0.008749 | 2.62 |
| FALSE | High | SLISQDSQ\ | [R].SLISQD | 1xPhospho | Q9WU62 1 | 0.000442 | 0 | 1 | 2 | 3 | Q9WU62 | Inner centi | Q9WU62 [ | 0 | 2011.088 | 0 | -1.28 | 0.000218 | 0.34 | 141.9 | 58.1 | 6.65 | 7.81 | | High | High | High | High | High | High | High | High | High | High | High | 0.00014 | 0.001115 | 3.16 |
| FALSE | High | SLLQGSDLI | [K].SLLQGS | 1xPhospho | Q9WUF3 1 | 3.68E-06 | 0 | 1 | 1 | 4 | Q9WUF3 | CASP8-ass( | Q9WUF3 [ | 0 | 2202.131 | 0 | -0.88 | 0.000488 | 3.04 | 129.7 | 70.3 | 2.93 | 7.87 | | High | High | High | High | High | High | High | High | High | High | High | 0 | 6.37E-05 | 3.33 |
| FALSE | High | SGFGGMS! | [R].SGFGG | 1xPhospho | Q569X8 1x | 0.006773 | 0.00062 | 1 | 5 | 5 | Q569X8 | Nucleopor | Q569X8 [4 | 0 | 1519.718 | 0 | 0.14 | 0.184752 | 0.52 | 94.5 | 105.5 | 11.07 | 10.7 | | High | High | High | High | High | High | High | High | High | High | High | 0.000275 | 0.002037 | 1.72 |
| FALSE | High | SGGEDTDN | [K].SGGED | 1xPhospho | A2ANY6 1x | 1.35E-07 | 0 | 1 | 2 | 4 | A2ANY6 | Midasin OS | A2ANY6 [4 | 0 | 2743.224 | 0 | -1.53 | 0.000479 | 1.51 | 145.9 | 54.1 | 4.75 | 15.05 | | High | High | High | High | High | High | High | High | High | High | High | 0 | 1.08E-08 | 4.19 |
| FALSE | High | SGGELGAG | [R].SGGEL( | 1xPhospho | K3W4P2 1x | 0.001373 | 5.66E-05 | 1 | 4 | 3 | K3W4P2 | Integrator | K3W4P2 [3 | 0 | 3735.789 | 0 | -0.16 | 0.94156 | 4.62 | 100.9 | 99.1 | 15.51 | 19.87 | | High | High | High | High | High | High | High | High | High | High | High | 2.58E-05 | 0.000513 | 2.9 |
| FALSE | High | SGGGELTL( | [R].SGGGE | 1xPhospho | P70424 1x | 0.026885 | 0.001676 | 1 | 3 | 1 | P70424 | Receptor t | P70424 [10 | 0 | 2263.069 | 0 | -0.14 | 0.062297 | 8.76 | 107 | 93 | 5.68 | 11.09 | | High | High | High | High | High | High | High | High | High | High | High | 0.000903 | 0.009546 | 2.52 |
| FALSE | High | SGGGGGG | [K].SGGGG | 1xPhospho | Q9CX86 1x | 3.7E-05 | 0 | 1 | 1 | 1 | Q9CX86 | Heterogen | Q9CX86 [2 | 0 | 1414.606 | 0 | 0.03 | 0.566685 | 4.33 | 99 | 101 | 5.28 | 7.99 | | High | High | High | High | High | High | High | High | High | High | High | 0 | 5.86E-06 | 1.22 |
| FALSE | High | SGGGSGGI | [R].SGGGS( | 1xPhospho | Q9CW46 1 | 0.000334 | 0 | 1 | 2 | 3 | Q9CW46 | Ribonuclec | Q9CW46 [( | 0 | 3442.672 | 0 | 0.63 | 0.235322 | 50.36 | 78.6 | 121.4 | 25.16 | 24 | | High | High | High | High | High | High | High | High | High | High | High | 0 | 7.41E-05 | 3.55 |
| FALSE | High | SGGGSVGI | [R].SGGGS' | 1xPhospho | Q9QXS1-8 | 0.000114 | 0 | 1 | 6 | 1 | Q9QXS1-8 | Isoform PL | Q9QXS1-8 | 0 | 1867.875 | 0 | 1.37 | 0.000957 | 26.58 | 55.7 | 144.3 | 11.96 | 6.3 | | High | High | High | High | High | High | High | High | High | High | High | 0 | 2.08E-05 | 2.93 |
| FALSE | High | SGGSSTAS( | [R].SGGSST | 1xPhospho | Q5U4C3 1x | 0.003464 | 0.00021 | 1 | 1 | 3 | Q5U4C3 | Splicing fac | Q5U4C3 [8 | 0 | 1787.894 | 0 | -0.61 | 0.029924 | 8.63 | 120.9 | 79.1 | 8.5 | 19.67 | | High | High | High | High | High | High | High | High | High | High | High | 0.000246 | 0.001468 | 3.32 |
| FALSE | High | SGGSTTTD | [R].SGGST | 1xPhospho | Q925B0 1x | 0.037369 | 0.00249 | 1 | 2 | 1 | Q925B0 | PRKC apop | Q925B0 [8 | 0 | 2095.999 | 0 | -1.73 | 0.000236 | 0.36 | 155.7 | 44.3 | 10.44 | 31.39 | | High | High | High | High | High | High | High | High | High | High | High | 0.001308 | 0.0138 | 2.71 |
| FALSE | High | SGKNSQED | [R].SGKNS( | 1xPhospho | Q80XU3 1x | 0.003892 | 0.000261 | 1 | 2 | 2 | Q80XU3 | Nuclear ub | Q80XU3 [5 | 1 | 2436.138 | 0 | -0.91 | 0.008216 | 5.45 | 131.9 | 68.1 | 8.42 | 20.19 | | High | High | High | High | High | High | High | High | High | High | High | 0.000275 | 0.001968 | 2.21 |
| FALSE | High | SGKVTEEEI | [R].SGKVTI | 1xPhospho | Q8BLH7 1x | 0.035486 | 0.002304 | 1 | 2 | 1 | Q8BLH7 | HIRA-inter | Q8BLH7 [1 | 1 | 2149.07 | 0 | -0.96 | 0.000847 | 1.52 | 131.6 | 68.4 | 6.67 | 12.76 | | High | High | High | High | High | High | High | High | High | High | High | 0.001214 | 0.01299 | 1.35 |
| FALSE | High | SGLCSPSY\ | [R].SGLCSP | 1xCarbami | B2RSN1 1x | 0.000118 | 0 | 1 | 5 | 3 | B2RSN1 | Myc proto- | B2RSN1 [8 | 0 | 2195.041 | 0 | 1.15 | 0.002458 | 14.5 | 62.2 | 137.8 | 12.7 | 10.04 | | High | High | High | High | High | High | High | High | High | High | High | 0 | 2.17E-05 | 3.41 |
| FALSE | High | SGLCSPSY\ | [R].SGLCSP | 1xCarbami | B2RSN1 2x | 0.001644 | 5.66E-05 | 1 | 5 | 1 | B2RSN1 | Myc proto- | B2RSN1 [8 | 0 | 2275.007 | 0 | 0.27 | 0.015648 | 8.7 | 87.4 | 112.6 | 9.7 | 4.16 | | High | High | High | High | High | High | High | High | High | High | High | 2.58E-05 | 0.000417 | 4.16 |
| FALSE | High | SGFGGMS! | [R].SGFGG | 1xPhospho | Q569X8 1x | 0.068752 | 0.005042 | 1 | 5 | 1 | Q569X8 | Nucleopor | Q569X8 [4 | 0 | 1503.723 | 0 | 0.16 | 0.192573 | 11.57 | 94.4 | 105.6 | 6.53 | 12.38 | | High | High | High | High | High | High | High | High | High | High | High | 0.002589 | 0.02781 | 1.31 |
| FALSE | High | SGIDILVGT | [R].SGIDIL\ | 1xPhospho | Q3ULC7 1x | 0.012288 | 0.000889 | 1 | 5 | 1 | Q3ULC7 | Putative ur | Q3ULC7 [3 | 0 | 1493.793 | 0 | -0.65 | 0.001068 | 5.04 | 122.3 | 77.7 | 2.84 | 6.39 | | High | High | High | High | High | High | High | High | High | High | High | 0.000449 | 0.003948 | 1.53 |
| FALSE | High | SGLGSPYG | [R].SGLGSI | 1xPhospho | P70271 1x | 0.014158 | 0.001011 | 1 | 3 | 2 | P70271 | PDZ and LII | P70271 [13 | 0 | 1524.741 | 0 | 0.67 | 0.010399 | 20.29 | 75 | 125 | 7.79 | 12.66 | | High | High | High | High | High | High | High | High | High | High | High | 0.002924 | 0.03037 | 2.06 |
| FALSE | High | SGLLNSTSI | [R].SGLLNS | 1xPhospho | O35219 1x | 0.00967 | 0.000736 | 1 | 4 | 2 | O35219 | Potassium | O35219 [3 | 0 | 1772.863 | 0 | -1 | 0.001025 | 1.34 | 132.9 | 67.1 | 11.06 | 4.97 | | High | High | High | High | High | High | High | High | High | High | High | 0.000344 | 0.003008 | 2.97 |
| FALSE | High | SGIIPTSPTI | [K].SGIIPTS | 1xPhospho | O54992 1x | 0.005044 | 0.000504 | 1 | 6 | 2 | O54992 | MAP kinas( | O54992 [2( | 0 | 2177.13 | 0 | -0.09 | 0.705484 | 6.55 | 100.8 | 99.2 | 4.7 | 5.66 | | High | High | High | High | High | High | High | High | High | High | High | 0.000246 | 0.001466 | 2.51 |
| FALSE | High | SGLLSSFSL! | [K].SGLLSS | 1xPhospho | A0A1D5RM | 0.003342 | 0.00016 | 1 | 3 | 1 | A0A1D5RM | Kinesin-like | A0A1D5RM | 0 | 2211.138 | 0 | 1.01 | 0.017365 | 14.35 | 68.3 | 131.7 | 12.64 | 19.92 | | High | High | High | High | High | High | High | High | High | High | High | 7.19E-05 | 0.000925 | 2.21 |
| FALSE | High | SGLLSSFSL! | [K].SGLLSS | 1xPhospho | A0A1D5RM | 0.000822 | 0 | 1 | 3 | 1 | A0A1D5RM | Kinesin-like | A0A1D5RM | 0 | 2227.133 | 0 | 1.09 | 0.000188 | 4.62 | 62.4 | 137.6 | 10.03 | 2.86 | | High | High | High | High | High | High | High | High | High | High | High | 0 | 0.000191 | 2.03 |
| FALSE | High | SGLSPAND | [R].SGLSPA | 1xPhospho | O70310 1x | 1.6E-05 | 0 | 1 | 1 | 5 | O70310 | Glycylpept | O70310 [4 | 0 | 1655.841 | 0 | -0.44 | 0.020162 | 4.03 | 117.4 | 82.6 | 5.13 | 13.81 | | High | High | High | High | High | High | High | High | High | High | High | 0 | 2.29E-06 | 2.9 |
| FALSE | High | SGLSPAND | [R].SGLSPA | 1xDeamida | O70310 1x | 4.55E-05 | 0 | 1 | 1 | 3 | O70310 | Glycylpept | O70310 [4 | 0 | 1656.825 | 0 | -0.39 | 0.022986 | 5.6 | 115.4 | 84.6 | 5.22 | 10.47 | | High | High | High | High | High | High | High | High | High | High | High | 0 | 2.83E-05 | 2.44 |
| FALSE | High | SGLSPNLN\ | [R].SGLSPN | 1xPhospho | Q80UK7 1x | 0.000534 | 0 | 1 | 4 | 1 | Q80UK7 | Spindle ass | Q80UK7 [5 | 0 | 1579.804 | 0 | -0.07 | 0.809311 | 4 | 101.7 | 98.3 | 8.06 | 4.46 | | High | High | High | High | High | High | High | High | High | High | High | 0 | 0.000118 | 2.97 |
| FALSE | High | SGLSQSMS | [K].SGLSQS | 1xPhospho | Q3UMF0 1 | 0.003688 | 0.00021 | 1 | 6 | 1 | Q3UMF0 | Cordon-ble | Q3UMF0 [ | 0 | 1890.847 | 0 | 0.11 | 0.459154 | 13.66 | 98.7 | 101.3 | 8.79 | 17.76 | | High | High | High | High | High | High | High | High | High | High | High | 9.42E-05 | 0.00103 | 2.49 |
| FALSE | High | SGITSLLFG | [K].SGITSLI | 1xPhospho | A0A1W2P6 | 2.01E-07 | 0 | 1 | 3 | 2 | A0A1W2P6 | Small G prc | A0A1W2P6 | 0 | 2346.225 | 0 | 0.41 | 0.045439 | 19.12 | 91 | 109 | 14.12 | 8.67 | | High | High | High | High | High | High | High | High | High | High | High | 0 | 1.68E-08 | 3.04 |
| FALSE | High | SGMSPEQS | [K].SGMSP | 1xPhospho | Q8BTI8 1x | 0.002058 | 5.66E-05 | 1 | 3 | 1 | Q8BTI8 | Serine/argi | Q8BTI8 [10 | 0 | 1488.717 | 0 | -0.45 | 0.05799 | 8.45 | 117 | 83 | 10.72 | 32.14 | | High | High | High | High | High | High | High | High | High | High | High | 2.58E-05 | 0.000536 | 2 |
| FALSE | High | SGMSPEQS | [K].SGMSP | 1xPhospho | Q8BTI8 1x | 0.000584 | 0 | 1 | 3 | 46 | Q8BTI8 | Serine/argi | Q8BTI8 [10 | 0 | 1504.712 | 0 | -0.48 | 0.045351 | 5.36 | 116.7 | 83.3 | 10.19 | 24.76 | | High | High | High | High | High | High | High | High | High | High | High | 0.000449 | 0.004002 | 2.29 |
| FALSE | High | SGPAASPSI | [R].SGPAAS | 1xPhospho | Q3TJM4 1x | 3.26E-08 | 0 | 1 | 1 | 2 | Q3TJM4 | Centromer | Q3TJM4 [3 | 0 | 2438.195 | 0 | -0.06 | 0.139476 | 1.16 | 114 | 86 | 13 | 13.98 | | High | High | High | High | High | High | High | High | High | High | High | 0 | 2.17E-09 | 3.88 |
| FALSE | High | SGLGASPL( | [R].SGLGAS | 1xPhospho | A2AE27 1x | 5.07E-05 | 0 | 1 | 2 | 1 | A2AE27 | AMP deam | A2AE27 [3 | 0 | 1452.741 | 0 | 0.53 | 0.002065 | 11.38 | 81.7 | 118.3 | 6.48 | 4.21 | | High | High | High | High | High | High | High | High | High | High | High | 0 | 8.41E-06 | 2.76 |
| FALSE | High | SGPEGTLSI | [R].SGPEG | 1xPhospho | F8WJ40 1x | 0.000462 | 0 | 1 | 2 | 6 | F8WJ40 | Histone-lys | F8WJ40 [4 | 0 | 3488.745 | 0 | -0.5 | 0.003177 | 3.6 | 116.8 | 83.2 | 6.92 | 9.38 | | High | High | High | High | High | High | High | High | High | High | High | 0 | 0.000181 | 4.18 |
| FALSE | High | SGEWQETS | [K].SGEWC | 1xPhospho | B7ZP47 1x | 3.7E-05 | 0 | 1 | 4 | 2 | B7ZP47 | Wapal prot | B7ZP47 [10 | 0 | 2619.238 | 0 | -0.99 | 0.001173 | 2.78 | 131.2 | 68.8 | 7.85 | 3.95 | | High | High | High | High | High | High | High | High | High | High | High | 0 | 5.89E-06 | 3.33 |
| FALSE | High | SGEQITSSP | [R].SGEQIT | 1xPhospho | E9Q9J4 1x | 0.00537 | 0.000599 | 1 | 1 | 2 | E9Q9J4 | Inositol he: | E9Q9J4 [10 | 0 | 1854.961 | 0 | 0.37 | 0.021822 | 13.18 | 87.1 | 112.9 | 4.9 | 14.23 | | High | High | High | High | High | High | High | High | High | High | High | 0.000267 | 0.001568 | 2.81 |
| FALSE | High | SGAAEEDD | [K].SGAAEI | 1xPhospho | Q6NS46 1x | 6.08E-09 | 0 | 1 | 1 | 7 | Q6NS46 | Protein RRI | Q6NS46 [1 | 0 | 2055.875 | 0 | 0.42 | 0.037846 | 5.25 | 84.7 | 115.3 | 2.17 | 13.2 | | High | High | High | High | High | High | High | High | High | High | High | 0 | 3.3E-10 | 4.34 |
| FALSE | High | SGAASMPS | [R].SGAASI | 1xPhospho | Q8K1N2-2 | 0.00537 | 0.000599 | 1 | 3 | 1 | Q8K1N2-2 | Isoform 2 ( | Q8K1N2-2 | 0 | 1557.775 | 0 | 2.49 | 0.00099 | 21.98 | 30.5 | 169.5 | 16.42 | 12.42 | | High | High | High | High | High | High | High | High | High | High | High | 0.000267 | 0.001567 | 1.52 |
| FALSE | High | SGAASMPS | [R].SGAASI | 1xPhospho | Q8K1N2-2 | 4.68E-05 | 0 | 1 | 3 | 4 | Q8K1N2-2 | Isoform 2 ( | Q8K1N2-2 | 0 | 1573.77 | 0 | 2.92 | 0.007688 | 31.53 | 26.5 | 173.5 | 13.78 | 30.59 | | High | High | High | High | High | High | High | High | High | High | High | 0 | 7.61E-06 | 1.68 |
| FALSE | High | SGADSPPG | [R].SGADSI | 1xCarbami | Q2KHI9-2 1 | 0.001222 | 0 | 1 | 4 | 1 | Q2KHI9-2 | Isoform 2 ( | Q2KHI9-2 [ | 0 | 2764.231 | 0 | -1.12 | 0.001512 | 2.15 | 137 | 63 | 10.63 | 1.45 | | High | High | High | High | High | High | High | High | High | High | High | 0 | 0.000297 | 1.43 |
| FALSE | High | SGAGSPPG | [R].SGAGSI | 1xPhospho | Q8BTI8 1x | 0.000573 | 0 | 1 | 3 | 6 | Q8BTI8 | Serine/argi | Q8BTI8 [11 | 0 | 1295.676 | 0 | -0.49 | 0.003472 | 3.73 | 118.8 | 81.2 | 5.32 | 6.08 | | High | High | High | High | High | High | High | High | High | High | High | 0 | 0.000128 | 1.75 |
| FALSE | High | SGALASPTI | [R].SGALAS | 1xPhospho | Q8C0E3-2 | 0.000958 | 0 | 1 | 7 | 2 | Q8C0E3-2 | Isoform 2 ( | Q8C0E3-2 | 0 | 1742.831 | 0 | 0.64 | 0.007752 | 10.74 | 72.3 | 127.7 | 17.25 | 11.12 | | High | High | High | High | High | High | High | High | High | High | High | 0 | 0.000335 | 1.96 |
| FALSE | High | SGAQASST | [R].SGAQA | 1xPhospho | P48678 1x | 1.07E-06 | 0 | 1 | 3 | 11 | P48678 | Prelamin-A | P48678 [12 | 0 | 1668.816 | 0 | 1.54 | 0.001285 | 11.36 | 48.6 | 151.4 | 10.41 | 9.79 | | High | High | High | High | High | High | High | High | High | High | High | 0 | 1.04E-05 | 2.4 |
| FALSE | High | SGAQASST | [R].SGAQA | 2xPhospho | P48678 2x | 5.16E-05 | 0 | 1 | 3 | 4 | P48678 | Prelamin-A | P48678 [12 | 0 | 1748.782 | 0 | 1.7 | 0.000191 | 9.03 | 46.9 | 153.1 | 6.22 | 3.23 | | High | High | High | High | High | High | High | High | High | High | High | 0 | 1.97E-05 | 2.22 |
| FALSE | High | SGASCPSSI | [K].SGASCI | 1xCarbami | F8VPZ3 1x | 0.013787 | 0.001011 | 1 | 3 | 2 | F8VPZ3 | Ubiquitin-s | F8VPZ3 [13 | 0 | 1418.675 | 0 | 0.48 | 0.00771 | 9.68 | 84 | 116 | 5.56 | 7.26 | | High | High | High | High | High | High | High | High | High | High | High | 0.000555 | 0.004787 | 0.94 |
| FALSE | High | SGASEANL | [R].SGASE/ | 1xPhospho | E9PVX6 1x | 0.000255 | 0 | 1 | 2 | 2 | E9PVX6 | Proliferatic | E9PVX6 [5 | 0 | 1697.924 | 0 | -0.12 | 0.220772 | 10.1 | 104 | 96 | 5.73 | 5.45 | | High | High | High | High | High | High | High | High | High | High | High | 0.001902 | 0.01921 | 3.21 |
| FALSE | High | SGASSPFAI | [R].SGASSF | 1xCarbami | E9QLH9 1x | 2.1E-05 | 0 | 1 | 2 | 2 | E9QLH9 | PEAK1-rela | E9QLH9 [3 | 0 | 2717.305 | 0 | 1.11 | 0.001124 | 11.37 | 63.4 | 136.6 | 5.72 | 10.99 | | High | High | High | High | High | High | High | High | High | High | High | 0 | 3.09E-06 | 2.75 |
| FALSE | High | SGASSPFAI | [R].SGASSF | 1xCarbami | E9QLH9 2x | 4.59E-05 | 0 | 1 | 2 | 1 | E9QLH9 | PEAK1-rela | E9QLH9 [3 | 0 | 2797.271 | 0 | 1.34 | 0.000513 | 9.46 | 57.3 | 142.7 | 10.89 | 7.18 | | High | High | High | High | High | High | High | High | High | High | High | 0 | 7.51E-06 | 3.54 |
| FALSE | High | SGATGGLS | [R].SGATG( | 1xPhospho | Q9CXS4 1x | 0.000984 | 0 | 1 | 1 | 2 | Q9CXS4 | Centromer | Q9CXS4 [1 | 0 | 1444.663 | 0 | -0.17 | 0.143698 | 12.24 | 106 | 94 | 7.53 | 5.8 | | High | High | High | High | High | High | High | High | High | High | High | 0.000327 | 0.002378 | 1.75 |
| FALSE | High | SGEQITSSP | [R].SGEQIT | 2xPhospho | E9Q9J4 2x | 0.003167 | 0.00016 | 1 | 1 | 1 | E9Q9J4 | Inositol he: | E9Q9J4 [10 | 0 | 1934.928 | 0 | -0.6 | 0.001546 | 4.69 | 120 | 80 | 4.78 | 9.52 | | High | High | High | High | High | High | High | High | High | High | High | 7.19E-05 | 0.000865 | 2.7 |
| FALSE | High | SGAVFSSS( | [K].SGAVFS | 1xPhospho | E9QPB3 1x | 0.007742 | 0.000664 | 1 | 4 | 2 | E9QPB3 | WD repeat | E9QPB3 [8 | 0 | 2236.12 | 0 | -0.55 | 0.005934 | 4.15 | 119.7 | 80.3 | 9.7 | 8.94 | | High | High | High | High | High | High | High | High | High | High | High | 0.000294 | 0.002348 | 2.77 |
| FALSE | High | SGAYSAGY | [R].SGAYS/ | 1xPhospho | Q9Z2X1 1x | 1.85E-06 | 0 | 1 | 3 | 1 | Q9Z2X1 | Heterogen | Q9Z2X1 [2 | 0 | 3456.478 | 0 | 0.01 | 0.492548 | 19.74 | 97.9 | 102.1 | 15.24 | 12.35 | | High | High | High | High | High | High | High | High | High | High | High | 0 | 2.04E-07 | 5.14 |
| FALSE | High | SGDALTTV' | [R].SGDAL1 | 1xPhospho | Q8BIZ6 1x | 4.05E-05 | 0 | 1 | 1 | 3 | Q8BIZ6 | Smad nucl( | Q8BIZ6 [18 | 0 | 1398.744 | 0 | -0.51 | 0.009982 | 7.39 | 117.5 | 82.5 | 8.53 | 13.58 | | High | High | High | High | High | High | High | High | High | High | High | 0 | 1.93E-05 | 4.1 |
| FALSE | High | SGDALTTV' | [R].SGDAL1 | 1xPhospho | Q8BIZ6 1x | 0.000892 | 0 | 1 | 1 | 6 | Q8BIZ6 | Smad nucl( | Q8BIZ6 [18 | 0 | 1627.907 | 0 | -0.52 | 0.012821 | 4.66 | 117.4 | 82.6 | 6.26 | 15.32 | | High | High | High | High | High | High | High | High | High | High | High | 0.000275 | 0.00223 | 2.55 |
| FALSE | High | SGDETPGS | [R].SGDETI | 1xPhospho | Q00PI9 1x | 6.46E-08 | 0 | 1 | 3 | 4 | Q00PI9 | Heterogen | Q00PI9 [15 | 0 | 1884.863 | 0 | 0.33 | 0.006763 | 6.37 | 87.8 | 112.2 | 3.2 | 8.76 | | High | High | High | High | High | High | High | High | High | High | High | 0 | 4.7E-09 | 3.12 |
| FALSE | High | SGDGGSA( | [R].SGDGG | 1xPhospho | Q4VAA2 1x | 4.8E-05 | 0 | 1 | 4 | 1 | Q4VAA2 | Protein CD | Q4VAA2 [9 | 0 | 1498.73 | 0 | -0.68 | 0.003317 | 5.1 | 123.7 | 76.3 | 5.22 | 9.38 | | High | High | High | High | High | High | High | High | High | High | High | 0 | 7.87E-06 | 1.61 |
| FALSE | High | SGDGGSLG | [R].SGDGG | 1xPhospho | Q4VAA2 1x | 0.112856 | 0.009146 | 1 | 3 | 1 | Q4VAA2 | Protein CD | Q4VAA2 [5 | 0 | 1345.595 | 0 | -0.06 | 0.866377 | 18.09 | 102.2 | 97.8 | 8.65 | 8.73 | | High | High | High | High | High | High | High | High | High | High | High | 0.004569 | 0.04959 | 1.18 |
| FALSE | High | SGDPAPPD | [R].SGDPAI | 2xPhospho | Q5U4C3 2x | 0.000198 | 0 | 1 | 1 | 1 | Q5U4C3 | Splicing fac | Q5U4C3 [5 | 0 | 1929.787 | 0 | -0.33 | 0.156243 | 10.02 | 110.3 | 89.7 | 12.58 | 13.93 | | High | High | High | High | High | High | High | High | High | High | High | 0 | 3.88E-05 | 2.34 |
| FALSE | High | SGDPAPPD | [R].SGDPAI | 1xPhospho | Q5U4C3 1x | 0.000319 | 0 | 1 | 1 | 2 | Q5U4C3 | Splicing fac | Q5U4C3 [5 | 0 | 2078.984 | 0 | 0.53 | 0.0812 | 0.99 | 84.9 | 115.1 | 4.33 | 16.08 | | High | High | High | High | High | High | High | High | High | High | High | 0.000467 | 0.00427 | 2.92 |
| FALSE | High | SGDPSAGP | [K].SGDPS/ | 1xPhospho | Q9QY06-2 | 4.13E-09 | 0 | 2 | 10 | 2 | Q9QY06-2; | Isoform 2 ( | Q9QY06-2 | 0 | 2425.128 | 0 | -0.09 | 0.102262 | 3.79 | 103.5 | 96.5 | 3.18 | 4.66 | Shared | High | High | High | High | High | High | High | High | High | High | High | 0 | 2.13E-10 | 3.35 |
| FALSE | High | SGDPSAGP | [K].SGDPS/ | 2xPhospho | Q9QY06-2 | 0.010663 | 0.000772 | 2 | 10 | 1 | Q9QY06-2; | Isoform 2 ( | Q9QY06-2 | 0 | 2505.095 | 0 | 0.12 | 0.253851 | 8.85 | 95.8 | 104.2 | 10 | 6.08 | Shared | High | High | High | High | High | High | High | High | High | High | High | 0.000399 | 0.003376 | 2.15 |
| FALSE | High | SGDVYGYS | [R].SGDVY( | 1xCarbami | Q8VEG0 1x | 0.004218 | 0.00036 | 1 | 2 | 1 | Q8VEG0 | Coiled-coil | Q8VEG0 [6 | 0 | 2530.157 | 0 | -0.02 | 0.595924 | 20.84 | 98.1 | 101.9 | 23.68 | 6.72 | | High | High | High | High | High | High | High | High | High | High | High | 0.000161 | 0.001196 | 3.29 |
| FALSE | High | SGEGLGVF | [R].SGEGL( | 1xPhospho | Q91W98 1 | 0.000337 | 0 | 1 | 1 | 2 | Q91W98 | Solute carr | Q91W98 [2 | 0 | 1861.946 | 0 | 0.75 | 0.060785 | 8.9 | 74.4 | 125.6 | 18.27 | 30.86 | | High | High | High | High | High | High | High | High | High | High | High | 0 | 7.07E-05 | 3.52 |
| FALSE | High | SGELTALEF | [K].SGELTA | 1xPhospho | E9Q7D5 1x | 0.003857 | 0.000261 | 1 | 2 | 3 | E9Q7D5 | Rho guanir | E9Q7D5 [1 | 0 | 1971.999 | 0 | 2.16 | 0.000213 | 7.88 | 36.4 | 163.6 | 5.27 | 4.85 | | High | High | High | High | High | High | High | High | High | High | High | 0.000117 | 0.001083 | 3.1 |
| FALSE | High | SGAVQGA( | [R].SGAVQ | 1xPhospho | Q8CI08-2 1 | 0.000116 | 0 | 1 | 2 | 1 | Q8CI08-2 | Isoform 2 ( | Q8CI08-2 [ | 0 | 1803.932 | 0 | 0.72 | 0.001249 | 9.55 | 73.4 | 126.6 | 10.18 | 7.4 | | High | High | High | High | High | High | High | High | High | High | High | 0 | 2.12E-05 | 1.94 |
| FALSE | High | SGPEGTLSI | [R].SGPEG | 2xPhospho | F8WJ40 2x | 9.8E-05 | 0 | 1 | 2 | 1 | F8WJ40 | Histone-lys | F8WJ40 [4 | 0 | 3568.711 | 0 | -0.65 | 0.049597 | 4.85 | 125 | 75 | 23.18 | 3.73 | | High | High | High | High | High | High | High | High | High | High | High | 0 | 1.75E-05 | 4.64 |
| FALSE | High | SGPLFDSPI | [K].SGPLFD | 1xPhospho | O08696 1x | 0.045508 | 0.003373 | 1 | 6 | 1 | O08696 | Forkhead b | O08696 [6( | 0 | 1284.619 | 0 | 1.44 | 0.00022 | 6.29 | 53.5 | 146.5 | 10.8 | 7.2 | | High | High | High | High | High | High | High | High | High | High | High | 0.001727 | 0.0173 | 1.06 |
| FALSE | High | SGPLQPVN | [K].SGPLQI | 1xPhospho | Q9WVM1 | 0.009585 | 0.000736 | 1 | 1 | 1 | Q9WVM1 | Rac GTPase | Q9WVM1 | 0 | 2543.282 | 0 | -0.42 | 0.007102 | 6.12 | 113.5 | 86.5 | 5.05 | 6.67 | | High | High | High | High | High | High | High | High | High | High | High | 0.000344 | 0.002993 | 1.6 |
| FALSE | High | SGSGMSVI | [R].SGSGM | 1xPhospho | E9Q0N0 1x | 0.000302 | 0 | 1 | 10 | 3 | E9Q0N0 | Intersectin | E9Q0N0 [3 | 0 | 1892.862 | 0 | -0.6 | 0.009539 | 9.2 | 123.6 | 76.4 | 7.89 | 13.36 | | High | High | High | High | High | High | High | High | High | High | High | 0 | 6.26E-05 | 3.21 |
| FALSE | High | SGSGMSVI | [R].SGSGM | 1xPhospho | E9Q0N0 1x | 1.21E-06 | 0 | 1 | 10 | 3 | E9Q0N0 | Intersectin | E9Q0N0 [3 | 0 | 1908.857 | 0 | -0.68 | 0.088898 | 5.98 | 116.7 | 83.3 | 10.99 | 22.55 | | High | High | High | High | High | High | High | High | High | High | High | 0 | 1.26E-07 | 4.14 |
| FALSE | High | SGSGSGGA | [R].SGSGS( | 1xPhospho | P83741-3 1 | 0.002398 | 0.000109 | 1 | 8 | 2 | P83741-3 | Isoform 3 ( | P83741-3 [ | 0 | 1403.693 | 0 | 0.94 | 0.000987 | 17.09 | 69.9 | 130.1 | 12.23 | 8.29 | | High | High | High | High | High | High | High | High | High | High | High | 4.94E-05 | 0.000633 | 1.1 |
| FALSE | High | SGSGSGGA | [R].SGSGS( | 2xPhospho | P83741-3 2 | 0.002508 | 0.000109 | 1 | 8 | 1 | P83741-3 | Isoform 3 ( | P83741-3 [ | 0 | 1483.66 | 0 | 0.84 | 0.00148 | 13.86 | 72.4 | 127.6 | 6.97 | 5.74 | | High | High | High | High | High | High | High | High | High | High | High | 4.94E-05 | 0.000671 | 2.04 |
| FALSE | High | SGSHFSSPS | [R].SGSHFS | 1xPhospho | E9Q5F9 1x | 0.000837 | 0 | 1 | 3 | 1 | E9Q5F9 | Histone-lys | E9Q5F9 [1 | 0 | 1672.81 | 0 | 0 | 0.859666 | 24.28 | 98.3 | 101.7 | 21.11 | 18 | | High | High | High | High | High | High | High | High | High | High | High | 0 | 0.000196 | 2.12 |
| FALSE | High | SGSHGHVS | [R].SGSHGI | 1xPhospho | Q6PCQ0 1x | 0.056724 | 0.004124 | 1 | 2 | 1 | Q6PCQ0 | IQ domain- | Q6PCQ0 [1 | 0 | 1739.806 | 0 | 1.73 | 0.042296 | 29.12 | 46.3 | 153.7 | 24.89 | 43.95 | | High | High | High | High | High | High | High | High | High | High | High | 0.002117 | 0.02219 | 1.17 |
| FALSE | High | SGSLCPSVF | [R].SGSLCP | 1xCarbami | G5E8C6 1x | 1.11E-05 | 0 | 1 | 4 | 4 | G5E8C6 | Villin-like p | G5E8C6 [2 | 0 | 2413.22 | 0 | -0.04 | 0.86151 | 11.82 | 99.7 | 100.3 | 7.61 | 8.62 | | High | High | High | High | High | High | High | High | High | High | High | 0 | 1.52E-06 | 2.98 |
| FALSE | High | SGSIDDSFT | [K].SGSIDD | 1xPhospho | V9GX40 1x | 6.35E-05 | 0 | 1 | 4 | 2 | V9GX40 | Synergin g | V9GX40 [5 | 0 | 2603.162 | 0 | 0.21 | 0.221274 | 9.97 | 92.9 | 107.1 | 4.89 | 19.56 | | High | High | High | High | High | High | High | High | High | High | High | 0 | 1.08E-05 | 4.23 |
| FALSE | High | SGSLDSELS | [K].SGSLDS | 1xPhospho | A0A140LJJ! | 2.67E-05 | 0 | 1 | 3 | 3 | A0A140LJJ! | A-kinase ar | A0A140LJJ! | 0 | 1843.945 | 0 | 0.26 | 0.009672 | 5.82 | 90.2 | 109.8 | 3.06 | 5.03 | | High | High | High | High | High | High | High | High | High | High | High | 0 | 4.06E-06 | 3.65 |
| FALSE | High | SGSLGGSG | [R].SGSLG( | 1xPhospho | Q3TV65 1x | 0.001986 | 5.66E-05 | 1 | 4 | 1 | Q3TV65 | MPN doma | Q3TV65 [3 | 0 | 2314.106 | 0 | 0.12 | 0.334851 | 17.11 | 94.3 | 105.7 | 13.44 | 8.19 | | High | High | High | High | High | High | High | High | High | High | High | 2.58E-05 | 0.000515 | 2.58 |
| FALSE | High | SGSLGSSN( | [K].SGSLGS | 1xPhospho | Q8BLH7 1x | 5.09E-06 | 0 | 1 | 1 | 3 | Q8BLH7 | HIRA-inter | Q8BLH7 [3 | 0 | 1777.744 | 0 | -1.57 | 0.000209 | 0.81 | 150.9 | 49.1 | 5.73 | 15.16 | | High | High | High | High | High | High | High | High | High | High | High | 0.001597 | 0.01538 | 2.54 |
| FALSE | High | SGSLGSSN( | [K].SGSLGS | 1xDeamida | Q8BLH7 1x | 3.52E-07 | 0 | 1 | 1 | 2 | Q8BLH7 | HIRA-inter | Q8BLH7 [3 | 0 | 1778.728 | 0 | -1.58 | 0.000192 | 1.2 | 152 | 48 | 2.85 | 13.02 | | High | High | High | High | High | High | High | High | High | High | High | 0 | 7.35E-05 | 2.66 |
| FALSE | High | SGSLGSSN( | [K].SGSLGS | 1xPhospho | Q8BLH7 1x | 3.75E-07 | 0 | 1 | 1 | 4 | Q8BLH7 | HIRA-inter | Q8BLH7 [3 | 1 | 3251.47 | 0 | -1.7 | 0.000429 | 0.95 | 148.9 | 51.1 | 7.7 | 10.21 | | High | High | High | High | High | High | High | High | High | High | High | 0 | 3.37E-08 | 4.1 |
| FALSE | High | SGSGEMTS | [K].SGSGEI | 1xPhospho | A2RSJ4 1x | 0.022773 | 0.001542 | 1 | 1 | 2 | A2RSJ4 | UHRF1-bin | A2RSJ4 [98 | 0 | 1948.872 | 0 | 0.78 | 0.006258 | 13.05 | 73.6 | 126.4 | 13.29 | 13.68 | | High | High | High | High | High | High | High | High | High | High | High | 0.009209 | 0.1002 | 1.54 |
| FALSE | High | SGSLGSSN( | [K].SGSLGS | 1xDeamida | Q8BLH7 1x | 2.53E-05 | 0 | 1 | 1 | 2 | Q8BLH7 | HIRA-inter | Q8BLH7 [3 | 1 | 3252.454 | 0 | -1.86 | 0.000295 | 0.77 | 154 | 46 | 4.29 | 11.37 | | High | High | High | High | High | High | High | High | High | High | High | 0.001651 | 0.01645 | 2.43 |
| FALSE | High | SGSLSLTQI | [R].SGSLSL | 1xPhospho | E9Q557 1x | 0.003253 | 0.00016 | 1 | 3 | 2 | E9Q557 | Desmoplak | E9Q557 [2 | 0 | 2236.17 | 0 | 1.08 | 0.032252 | 33.94 | 63.6 | 136.4 | 29.96 | 11.96 | | High | High | High | High | High | High | High | High | High | High | High | 7.19E-05 | 0.000893 | 2.55 |
| FALSE | High | SGSLSLTQI | [R].SGSLSL | 1xPhospho | E9Q557 1x | 0.008538 | 0.000736 | 1 | 3 | 1 | E9Q557 | Desmoplak | E9Q557 [2 | 0 | 2252.165 | 0 | 0.88 | 0.00847 | 14.02 | 67.6 | 132.4 | 17.46 | 14.43 | | High | High | High | High | High | High | High | High | High | High | High | 0.000327 | 0.002621 | 2.62 |
| FALSE | High | SGSISNYR | [R].SGSISN | 1xPhospho | Q5EG47 1x | 0.005418 | 0.00062 | 1 | 3 | 2 | Q5EG47 | 5'-AMP-act | Q5EG47 [4 | 0 | 1192.556 | 0 | 0.83 | 0.000385 | 5.49 | 70.3 | 129.7 | 10.93 | 3.12 | | High | High | High | High | High | High | High | High | High | High | High | 0.000275 | 0.001583 | 1.89 |
| FALSE | High | SGSISSSPSI | [K].SGSISS | 1xPhospho | Q3U9G9 1 | 0.005615 | 0.00062 | 1 | 3 | 3 | Q3U9G9 | Lamin-B re | Q3U9G9 [6 | 0 | 1273.599 | 0 | -0.81 | 0.006782 | 1.14 | 126.6 | 73.4 | 14.48 | 4.15 | | High | High | High | High | High | High | High | High | High | High | High | 0.000449 | 0.004066 | 1.25 |
| FALSE | High | SGSISVK | [R].SGSISV | 1xPhospho | A2AJT5 1x | 0.042503 | 0.003197 | 1 | 4 | 1 | A2AJT5 | Arginine/se | A2AJT5 [72 | 0 | 1215.675 | 0 | 0.03 | 0.440676 | 36.5 | 89 | 111 | 15.68 | 20.6 | | High | High | High | High | High | High | High | High | High | High | High | 0.001651 | 0.01605 | 1.44 |
| FALSE | High | SGSIVELLA | [R].SGSIVE | 1xCarbami | G3X9H5 1x | 3.2E-11 | 0 | 1 | 3 | 1 | G3X9H5 | Huntingtin | G3X9H5 [3 | 0 | 2342.162 | 0 | 0.74 | 0.008737 | 8.5 | 74.9 | 125.1 | 6.65 | 17.69 | | High | High | High | High | High | High | High | High | High | High | High | 0 | 9.04E-13 | 5.38 |
| FALSE | High | SGSIVELLA | [R].SGSIVE | 1xCarbami | G3X9H5 2x | 6.56E-07 | 0 | 1 | 3 | 5 | G3X9H5 | Huntingtin | G3X9H5 [3 | 0 | 2422.129 | 0 | 1.21 | 0.000445 | 1.14 | 56.5 | 143.5 | 7.72 | 7.79 | | High | High | High | High | High | High | High | High | High | High | High | 0 | 1.6E-06 | 4.36 |
| FALSE | High | SGSMEED\ | [K].SGSME | 1xPhospho | P70255 1x | 3.04E-06 | 0 | 1 | 5 | 7 | P70255 | Nuclear fac | P70255 [28 | 0 | 3235.31 | 0 | 2.29 | 0.000244 | 28.66 | 36.6 | 163.4 | 33.61 | 6.79 | | High | High | High | High | High | High | High | High | High | High | High | 0 | 3.55E-07 | 6.11 |
| FALSE | High | SGSMEED\ | [K].SGSME | 2xPhospho | P70255 2x | 0.000181 | 0 | 1 | 5 | 2 | P70255 | Nuclear fac | P70255 [28 | 0 | 3299.281 | 0 | 2.4 | 0.004288 | 5.15 | 31.5 | 168.5 | 9.38 | 24.15 | | High | High | High | High | High | High | High | High | High | High | High | 2.58E-05 | 0.000392 | 4.22 |
| FALSE | High | SGSMEED\ | [K].SGSME | 2xPhospho | P70255 2x | 0.000219 | 0 | 1 | 5 | 6 | P70255 | Nuclear fac | P70255 [28 | 0 | 3315.276 | 0 | 2.58 | 0.000214 | 11.28 | 29.4 | 170.6 | 16.06 | 7.34 | | High | High | High | High | High | High | High | High | High | High | High | 0 | 4.32E-05 | 4.1 |
| FALSE | High | SGSNGSVV | [R].SGSNG! | 1xPhospho | B7ZNG0 1x | 2.05E-06 | 0 | 1 | 2 | 3 | B7ZNG0 | Kinesin-like | B7ZNG0 [8 | 0 | 2086.058 | 0 | -1.23 | 0.000442 | 2.33 | 140.7 | 59.3 | 3.95 | 9.4 | | High | High | High | High | High | High | High | High | High | High | High | 0 | 1.95E-06 | 4.35 |
| FALSE | High | SGSNGSVV | [R].SGSNG! | 1xDeamida | B7ZNG0 1x | 5.7E-05 | 0 | 1 | 2 | 3 | B7ZNG0 | Kinesin-like | B7ZNG0 [8 | 0 | 2087.042 | 0 | -0.9 | 0.001843 | 3.69 | 130.3 | 69.7 | 5.47 | 10.5 | | High | High | High | High | High | High | High | High | High | High | High | 0 | 9.51E-06 | 4.12 |
| FALSE | High | SGSNGSVV | [R].SGSNG! | 1xPhospho | B7ZNG0-2 | 3.36E-06 | 0 | 1 | 2 | 2 | B7ZNG0-2 | Isoform 2 ( | B7ZNG0-2 | 0 | 2585.286 | 0 | -0.91 | 0.001448 | 5.48 | 129.5 | 70.5 | 2.21 | 12.51 | | High | High | High | High | High | High | High | High | High | High | High | 0 | 3.96E-07 | 4.54 |
| FALSE | High | SGSLGSSN( | [K].SGSLGS | 2xPhospho | Q8BLH7 2x | 6.49E-06 | 0 | 1 | 1 | 5 | Q8BLH7 | HIRA-inter | Q8BLH7 [3 | 1 | 3331.436 | 0 | -1.23 | 0.000278 | 1.9 | 138.9 | 61.1 | 4.43 | 11.74 | | High | High | High | High | High | High | High | High | High | High | High | 0 | 8.32E-07 | 5.24 |

| | | | | | | | | | | | | | | | | | | | | | | | | | | | | | | | | | | | | | | |
|---|---|---|---|---|---|---|---|---|---|---|---|---|---|---|---|---|---|---|---|---|---|---|---|---|---|---|---|---|---|---|---|---|---|---|---|---|---|---|
| FALSE | High | SGSETPDG | [K].SGSETP | 2xPhospho | E9QLG3 2x | 0.001544 | 5.66E-05 | 1 | 5 | 6 | E9QLG3 | MAX gene- | E9QLG3 [1 | 0 | 1946.891 | 0 | -0.72 | 0.000209 | 1.18 | 123.4 | 76.6 | 4.36 | 6.4 | | High | High | High | High | High | High | High | High | High | High | High | 0.000418 | 0.003605 | 3.57 |
| FALSE | High | SGSETPDG | [K].SGSETP | 1xPhospho | E9QLG3 1x | 0.027839 | 0.001755 | 1 | 5 | 3 | E9QLG3 | MAX gene- | E9QLG3 [1 | 0 | 1866.925 | 0 | -0.44 | 0.00279 | 6.42 | 115.3 | 84.7 | 1.05 | 8.1 | | High | High | High | High | High | High | High | High | High | High | High | 0.001651 | 0.01645 | 2.58 |
| FALSE | High | SGSESSVE( | [R].SGSESS | 1xPhospho | Q8BTI8 1xF | 0.000369 | 0 | 1 | 3 | 3 | Q8BTI8 | Serine/argi | Q8BTI8 [14 | 0 | 1575.767 | 0 | -0.22 | 0.082608 | 5.24 | 106.3 | 93.7 | 14.14 | 6.09 | | High | High | High | High | High | High | High | High | High | High | High | 0.000467 | 0.00428 | 2.23 |
| FALSE | High | SGPSGNSC | [R].SGPSGI | 1xCarbami | P70182-3 1 | 4.39E-05 | 0 | 1 | 4 | 1 | P70182-3 | Isoform 3 c | P70182-3 [ | 0 | 2287.963 | 0 | 0.4 | 0.046634 | 13.87 | 85.4 | 114.6 | 13.35 | 16.12 | | High | High | High | High | High | High | High | High | High | High | High | 0 | 7.14E-06 | 4.7 |
| FALSE | High | SGPSHSGD | [R].SGPSHS | 1xPhospho | G3X959 1x | 0.000325 | 0 | 1 | 2 | 4 | G3X959 | Periphilin 1 | G3X959 [8 | 0 | 1644.685 | 0 | 0.32 | 0.037575 | 14.27 | 87.1 | 112.9 | 3.4 | 10.41 | | High | High | High | High | High | High | High | High | High | High | High | 2.58E-05 | 0.000524 | 2.99 |
| FALSE | High | SGPSPLSSP | [K].SGPSPL | 1xPhospho | Q99NH0 1x | 3.77E-05 | 0 | 1 | 5 | 5 | Q99NH0 | Ankyrin re | Q99NH0 [1 | 0 | 1665.861 | 0 | -0.36 | 0.009422 | 4.52 | 108.9 | 91.1 | 5.02 | 4.7 | | High | High | High | High | High | High | High | High | High | High | High | 0 | 3.93E-05 | 2.96 |
| FALSE | High | SGPSPLSSP | [K].SGPSPL | 1xDeamida | Q99NH0 1x | 0.001233 | 0 | 1 | 5 | 1 | Q99NH0 | Ankyrin re | Q99NH0 [1 | 0 | 1666.845 | 0 | 0.4 | 0.051581 | 16.88 | 86.7 | 113.3 | 9.41 | 7.07 | | High | High | High | High | High | High | High | High | High | High | High | 0 | 0.000302 | 2.39 |
| FALSE | High | SGPSPLSSP | [K].SGPSPL | 2xPhospho | Q99NH0 2x | 0.00017 | 0 | 1 | 5 | 1 | Q99NH0 | Ankyrin re | Q99NH0 [1 | 0 | 1745.828 | 0 | -0.39 | 0.002382 | 1.09 | 116.1 | 83.9 | 6.25 | 8.87 | | High | High | High | High | High | High | High | High | High | High | High | 0 | 3.27E-05 | 3.16 |
| FALSE | High | SGPTSDNE | [K].SGPTSD | 1xPhospho | Q3USH5 1x | 2.17E-05 | 0 | 1 | 1 | 6 | Q3USH5 | Splicing fac | Q3USH5 [2 | 0 | 3493.521 | 0 | -0.36 | 0.050965 | 3.39 | 111 | 89 | 12.68 | 9.82 | | High | High | High | High | High | High | High | High | High | High | High | 0 | 1.73E-05 | 5.03 |
| FALSE | High | SGPTSDNE | [K].SGPTSD | 1xDeamida | Q3USH5 1x | 0.02933 | 0.001857 | 1 | 1 | 2 | Q3USH5 | Splicing fac | Q3USH5 [2 | 0 | 3494.505 | 0 | -0.29 | 0.058725 | 15.34 | 107.6 | 92.4 | 11.66 | 9.44 | | High | High | High | High | High | High | High | High | High | High | High | 0.000997 | 0.01052 | 3.13 |
| FALSE | High | SGPTSPTPS | [R].SGPTSP | 1xCarbami | E9Q1Q1 1x | 4.35E-05 | 0 | 1 | 6 | 1 | E9Q1Q1 | Microtubu | E9Q1Q1 [1 | 0 | 2028.95 | 0 | -0.85 | 0.010549 | 5.74 | 125 | 75 | 14.41 | 15.04 | | High | High | High | High | High | High | High | High | High | High | High | 0 | 7.04E-06 | 2.86 |
| FALSE | High | SGPTSSPFS | [R].SGPTSS | 1xPhospho | Q8K3Z9 1x | 0.000815 | 0 | 1 | 2 | 7 | Q8K3Z9 | Nuclear en | Q8K3Z9 [4 | 0 | 1760.805 | 0 | 0.53 | 0.003341 | 14.47 | 79.9 | 120.1 | 8.96 | 7.16 | | High | High | High | High | High | High | High | High | High | High | High | 0 | 0.000231 | 3.05 |
| FALSE | High | SGPTSSPFS | [R].SGPTSS | 2xPhospho | Q8K3Z9 2x | 0.089425 | 0.006807 | 1 | 2 | 2 | Q8K3Z9 | Nuclear en | Q8K3Z9 [4 | 0 | 1840.772 | 0 | 0.68 | 0.000942 | 6.74 | 74.7 | 125.3 | 7.5 | 4.78 | | High | High | High | High | High | High | High | High | High | High | High | 0.003521 | 0.03774 | 2.62 |
| FALSE | High | SGPTSSPFS | [R].SGPTSS | 3xPhospho | Q8K3Z9 3x | 0.029076 | 0.001857 | 1 | 2 | 1 | Q8K3Z9 | Nuclear en | Q8K3Z9 [4 | 0 | 1920.738 | 0 | 0.57 | 0.018426 | 22.04 | 80.6 | 119.4 | 9.26 | 17.77 | | High | High | High | High | High | High | High | High | High | High | High | 0.000997 | 0.01036 | 2.36 |
| FALSE | High | SGQEVASP | [R].SGQEV. | 1xPhospho | I7H459 1xP | 9.82E-06 | 0 | 1 | 2 | 1 | I7H459 | Protein inc | I7H459 [20 | 0 | 3060.545 | 0 | 0.72 | 0.003906 | 7.19 | 74.4 | 125.6 | 12.23 | 8.69 | | High | High | High | High | High | High | High | High | High | High | High | 0 | 1.33E-06 | 3.57 |
| FALSE | High | SGQSFHGN | [R].SGQSFI | 1xPhospho | Q80U58 1x | 0.00625 | 0.00062 | 1 | 8 | 1 | Q80U58 | Pumilio ho | Q80U58 [6 | 0 | 2209.06 | 0 | 0.66 | 0.012915 | 6.51 | 76.2 | 123.8 | 16.24 | 11.97 | | High | High | High | High | High | High | High | High | High | High | High | 0.000275 | 0.001852 | 1.9 |
| FALSE | High | SGRSPEPDI | [R].SGRSPE | 1xPhospho | Q80U49 1x | 4.16E-05 | 0 | 1 | 2 | 2 | Q80U49 | Centrosom | Q80U49 [8 | 1 | 1872.962 | 0 | 0.73 | 0.001084 | 4.39 | 74.5 | 125.5 | 4.67 | 5.74 | | High | High | High | High | High | High | High | High | High | High | High | 0 | 6.72E-06 | 3.08 |
| FALSE | High | SGRSPTGN | [R].SGRSPT | 1xPhospho | B2RUG9 1x | 0.007404 | 0.00062 | 1 | 6 | 2 | B2RUG9 | Adenomat | B2RUG9 [2 | 1 | 2466.264 | 0 | 0.41 | 0.04696 | 18.15 | 87.8 | 112.2 | 12.06 | 8.65 | | High | High | High | High | High | High | High | High | High | High | High | 0.000275 | 0.00224 | 2.27 |
| FALSE | High | SGSAHGSG | [R].SGSAH( | 1xPhospho | P62996 1xP | 0.055308 | 0.003982 | 1 | 2 | 1 | P62996 | Transform | P62996 [20 | 0 | 1325.662 | 0 | 0.27 | 0.195452 | 8.29 | 91.4 | 108.6 | 7.15 | 8.08 | | High | High | High | High | High | High | High | High | High | High | High | 0.002105 | 0.02173 | 1.36 |
| FALSE | High | SGSAHGSG | [R].SGSAH( | 2xPhospho | P62996 2xP | 0.092372 | 0.007211 | 1 | 2 | 1 | P62996 | Transform | P62996 [20 | 0 | 1405.628 | 0 | -0.3 | 0.051504 | 13.15 | 110.4 | 89.6 | 8.94 | 13.1 | | High | High | High | High | High | High | High | High | High | High | High | 0.003672 | 0.03945 | 0.65 |
| FALSE | High | SGSALLQS( | [R].SGSALL | 1xPhospho | Q5DTM8 1 | 3.06E-08 | 0 | 1 | 3 | 5 | Q5DTM8 | E3 ubiquiti | Q5DTM8 [ | 0 | 2173.079 | 0 | 0.66 | 0.001043 | 12.77 | 74.5 | 125.5 | 7.93 | 5.45 | | High | High | High | High | High | High | High | High | High | High | High | 0 | 4.5E-07 | 4.1 |
| FALSE | High | SGSALLQS( | [R].SGSALL | 1xPhospho | Q5DTM8 1 | 0.000215 | 0 | 1 | 3 | 6 | Q5DTM8 | E3 ubiquiti | Q5DTM8 [ | 1 | 3214.634 | 0 | 0.46 | 0.022938 | 20.32 | 84.3 | 115.7 | 7.86 | 8.81 | | High | High | High | High | High | High | High | High | High | High | High | 0 | 4.23E-05 | 3.65 |
| FALSE | High | SGSASPAP( | [K].SGSASF | 1xPhospho | S4R1P5 1xP | 5.27E-06 | 0 | 1 | 5 | 2 | S4R1P5 | Dystonin O | S4R1P5 [16 | 0 | 2333.194 | 0 | 1.67 | 0.000558 | 13.03 | 49.8 | 150.2 | 7.85 | 9.49 | | High | High | High | High | High | High | High | High | High | High | High | 0 | 6.61E-07 | 3.54 |
| FALSE | High | SGSASPAP( | [K].SGSASF | 2xPhospho | S4R1P5 2xP | 5.5E-05 | 0 | 1 | 5 | 1 | S4R1P5 | Dystonin O | S4R1P5 [16 | 0 | 2413.161 | 0 | 1.46 | 0.000681 | 11.02 | 54.9 | 145.1 | 9.84 | 9.12 | | High | High | High | High | High | High | High | High | High | High | High | 0 | 9.2E-06 | 2.38 |
| FALSE | High | SGSCAWPA | [R].SGSCAV | 1xCarbami | Q6PAL0 1x | 0.100129 | 0.008067 | 1 | 3 | 1 | Q6PAL0 | BEN domai | Q6PAL0 [1 | 0 | 2921.332 | 0 | -0.19 | 0.185225 | 15.26 | 104.1 | 95.9 | 8.61 | 9.82 | | High | High | High | High | High | High | High | High | High | High | High | 0.004059 | 0.04329 | 2.18 |
| FALSE | High | SGSDGTPG | [R].SGSDGT | 1xPhospho | Q91XC0 1x | 0.017344 | 0.001204 | 1 | 1 | 1 | Q91XC0 | LIM domai | Q91XC0 [3 | 0 | 1471.719 | 0 | -0.96 | 0.005406 | 5.39 | 130.7 | 69.3 | 10.22 | 13.01 | | High | High | High | High | High | High | High | High | High | High | High | 0.000656 | 0.005795 | 2.11 |
| FALSE | High | SGSDLPNSI | [K].SGSDLF | 1xPhospho | Q9CPY3 1x | 0.000908 | 0 | 1 | 1 | 1 | Q9CPY3 | Sororin OS | Q9CPY3 [3 | 0 | 2000.932 | 0 | -0.57 | 0.007436 | 6.68 | 121.8 | 78.2 | 7.69 | 9.25 | | High | High | High | High | High | High | High | High | High | High | High | 0 | 0.000215 | 3 |
| FALSE | High | SGSDPQDV | [R].SGSDP( | 1xPhospho | B7ZNH9 1x | 0.002508 | 0.000109 | 1 | 3 | 1 | B7ZNH9 | Phosphoin | B7ZNH9 [1 | 0 | 3469.571 | 0 | -0.39 | 0.068055 | 9.09 | 117.7 | 82.3 | 12.52 | 38.86 | | High | High | High | High | High | High | High | High | High | High | High | 4.94E-05 | 0.000668 | 2.81 |
| FALSE | High | SGSEAEGL( | [R].SGSEAE | 1xPhospho | Q9EPU4 1x | 1.62E-06 | 0 | 1 | 1 | 5 | Q9EPU4 | Cleavage a | Q9EPU4 [7 | 0 | 4146.851 | 0 | -0.21 | 0.040129 | 8.04 | 106.6 | 93.4 | 5.32 | 16.58 | | High | High | High | High | High | High | High | High | High | High | High | 0 | 3.4E-06 | 5.47 |
| FALSE | High | SGSEAEGL( | [R].SGSEAE | 1xPhospho | Q9EPU4 1x | 8.84E-07 | 0 | 1 | 1 | 13 | Q9EPU4 | Cleavage a | Q9EPU4 [7 | 0 | 4162.846 | 0 | -0.27 | 0.00047 | 2.15 | 109.1 | 90.9 | 1.9 | 2.64 | | High | High | High | High | High | High | High | High | High | High | High | 0 | 8.89E-08 | 5.9 |
| FALSE | High | SGSEEIVTD | [R].SGSEEI' | 2xPhospho | E2JF22 2xP | 0.001058 | 0 | 1 | 8 | 1 | E2JF22 | Piezo-type | E2JF22 [16 | 0 | 3415.562 | 0 | 0.14 | 0.439695 | 21.16 | 94.1 | 105.9 | 19.64 | 16.87 | | High | High | High | High | High | High | High | High | High | High | High | 0 | 0.000255 | 3.74 |
| FALSE | High | SGSEGELS( | [K].SGSEGI | 1xCarbami | Z4YKQ9 1x | 0.005418 | 0.00062 | 1 | 4 | 1 | Z4YKQ9 | ATPase fan | Z4YKQ9 [2 | 0 | 2637.263 | 0 | -0.28 | 0.021812 | 4.66 | 109.6 | 90.4 | 5.99 | 6.25 | | High | High | High | High | High | High | High | High | High | High | High | 0.000275 | 0.001577 | 2.97 |
| FALSE | High | SFYSSHYSR | [K].SFYSSH | 1xPhospho | G3X959 1x | 0.072858 | 0.005599 | 1 | 7 | 1 | G3X959 | Periphilin 1 | G3X959 [1 | 0 | 1442.63 | 0 | 0.66 | 0.16905 | 43.33 | 80.1 | 119.9 | 19.37 | 22.49 | | High | High | High | High | High | High | High | High | High | High | High | 0.002867 | 0.02982 | 2.34 |
| FALSE | High | SGSPAPETI | [R].SGSPAF | 1xPhospho | E9PYH6 1xP | 3.52E-07 | 0 | 1 | 1 | 2 | E9PYH6 | SET domai | E9PYH6 [4 | 0 | 2810.283 | 0 | -0.02 | 0.9447 | 2.17 | 100.5 | 99.5 | 3.62 | 5.18 | | High | High | High | High | High | High | High | High | High | High | High | 0 | 3.16E-08 | 3.6 |
| FALSE | High | SFVKPPSLA | [K].SFVKPP | 1xPhospho | Q8CH77 1x | 0.007339 | 0.00062 | 1 | 4 | 1 | Q8CH77 | Neuron na | Q8CH77 [7 | 0 | 2183.244 | 0 | 0.75 | 0.033657 | 34.89 | 74.4 | 125.6 | 10.56 | 21.47 | | High | High | High | High | High | High | High | High | High | High | High | 0.000275 | 0.002221 | 1.16 |
| FALSE | High | SFTPDHVV' | [K].SFTPDH | 1xPhospho | O54724 1x | 0.003788 | 0.00021 | 1 | 1 | 2 | O54724 | Caveolae-a | O54724 [3 | 0 | 1600.772 | 0 | -0.52 | 0.02673 | 7.38 | 108.5 | 91.5 | 14.93 | 14.72 | | High | High | High | High | High | High | High | High | High | High | High | 0.001999 | 0.02033 | 2.27 |
| FALSE | High | SETDEYLAK | [K].SETDEY | 1xPhospho | Q80YR7 1x | 0.002212 | 0.000109 | 1 | 2 | 2 | Q80YR7 | Claspin OS | Q80YR7 [7 | 0 | 1593.781 | 0 | 0.36 | 0.005251 | 9.19 | 86.3 | 113.7 | 6.85 | 7.04 | | High | High | High | High | High | High | High | High | High | High | High | 0.000538 | 0.004466 | 2.36 |
| FALSE | High | SETGDSSIF | [K].SETGDS | 1xPhospho | Q8C0T5 1x | 0.061176 | 0.004646 | 1 | 4 | 1 | Q8C0T5 | Signal-indu | Q8C0T5 [2 | 0 | 1407.635 | 0 | 2.18 | 0.000534 | 12.49 | 35.5 | 164.5 | 4.15 | 9.59 | | High | High | High | High | High | High | High | High | High | High | High | 0.00237 | 0.02436 | 1.82 |
| FALSE | High | SETLVNAQ | [K].SETLVN | 1xPhospho | Q8CB16 1x | 2.44E-05 | 0 | 1 | 7 | 2 | Q8CB16 | Putative ur | Q8CB16 [1 | 0 | 2222.183 | 0 | 1.38 | 0.000463 | 2.25 | 54.9 | 145.1 | 5.43 | 5.7 | | High | High | High | High | High | High | High | High | High | High | High | 0 | 3.68E-06 | 2.68 |
| FALSE | High | SETLVNAQ | [K].SETLVN | 2xPhospho | Q8CB16 2x | 0.001255 | 0 | 1 | 7 | 2 | Q8CB16 | Putative ur | Q8CB16 [1 | 0 | 2302.15 | 0 | 1.56 | 0.000266 | 4.02 | 50.1 | 149.9 | 7.1 | 5.68 | | High | High | High | High | High | High | High | High | High | High | High | 4.94E-05 | 0.000613 | 2.84 |
| FALSE | High | SETNWESP | [K].SETNW | 1xPhospho | Q80WJ7 1x | 0.022973 | 0.001578 | 1 | 1 | 1 | Q80WJ7 | Protein LYI | Q80WJ7 [5 | 0 | 1386.614 | 0 | 1.38 | 0.081245 | 2.53 | 55.8 | 144.2 | 30.62 | 31.17 | | High | High | High | High | High | High | High | High | High | High | High | 0.000847 | 0.007995 | 2.4 |
| FALSE | High | SETNWESP | [K].SETNW | 1xPhospho | Q80WJ7 1x | 0.001049 | 0 | 1 | 1 | 2 | Q80WJ7 | Protein LYI | Q80WJ7 [5 | 0 | 1615.777 | 0 | 1.37 | 0.018158 | 23.81 | 58.2 | 141.8 | 12 | 22.57 | | High | High | High | High | High | High | High | High | High | High | High | 0 | 0.000252 | 3.04 |
| FALSE | High | SETPGNSS | [R].SETPGI | 1xPhospho | Q924W7 1 | 0.06169 | 0.004646 | 1 | 3 | 1 | Q924W7 | Suppressio | Q924W7 [3 | 0 | 2177.126 | 0 | -0.12 | 0.192757 | 3.36 | 103.5 | 96.5 | 6.64 | 6.9 | | High | High | High | High | High | High | High | High | High | High | High | 0.002381 | 0.02446 | 1.87 |
| FALSE | High | SETPPVPPI | [R].SETPPV | 1xPhospho | Q7TSC1 1x | 0.000524 | 0 | 1 | 3 | 1 | Q7TSC1 | Protein PRI | Q7TSC1 [8 | 0 | 3551.745 | 0 | 1.44 | 0.003418 | 26.02 | 54 | 146 | 13.64 | 15.54 | | High | High | High | High | High | High | High | High | High | High | High | 0 | 0.000115 | 3.03 |
| FALSE | High | SETPPVPPI | [R].SETPPV | 1xDeamida | Q7TSC1 1x | 0.011755 | 0.000889 | 1 | 3 | 2 | Q7TSC1 | Protein PRI | Q7TSC1 [8 | 0 | 3552.729 | 0 | 1.5 | 0.004624 | 34.21 | 53.9 | 146.1 | 18.45 | 12.14 | | High | High | High | High | High | High | High | High | High | High | High | 0.000449 | 0.003747 | 2.75 |
| FALSE | High | SETSPVAPI | [K].SETSPV | 1xPhospho | A0A0A6YW | 0.020679 | 0.001472 | 1 | 4 | 2 | A0A0A6YW | Rap guanin | A0A0A6YW | 0 | 1252.614 | 0 | 0.24 | 0.115047 | 17.96 | 92.7 | 107.3 | 8.3 | 8.65 | | High | High | High | High | High | High | High | High | High | High | High | 0.000786 | 0.007072 | 1.92 |
| FALSE | High | SETSSNPSS | [K].SETSSN | 1xCarbami | Q5SVL6 2x | 2.8E-05 | 0 | 1 | 3 | 3 | Q5SVL6 | Rap1 GTPa | Q5SVL6 [5 | 0 | 2352.023 | 0 | 2.69 | 0.000704 | 26.45 | 25.9 | 174.1 | 12.62 | 10.94 | | High | High | High | High | High | High | High | High | High | High | High | 0 | 8.8E-05 | 3.59 |
| FALSE | High | SEVDGGSG | [R].SEVDG( | 1xPhospho | A0A087WC | 3.03E-05 | 0 | 1 | 6 | 1 | A0A087WC | Snf2-relate | A0A087WC | 0 | 2490.253 | 0 | 0.54 | 0.003612 | 17.97 | 80.5 | 119.5 | 7.82 | 7.64 | | High | High | High | High | High | High | High | High | High | High | High | 0 | 4.69E-06 | 2.2 |
| FALSE | High | SEVGSLVAT | [K].SEVGSL | 1xCarbami | Q923W1 1 | 6.07E-05 | 0 | 1 | 1 | 4 | Q923W1 | Trimethylg | Q923W1 [ | 0 | 3990.93 | 0 | -0.83 | 0.000846 | 4.03 | 127.7 | 72.3 | 3.24 | 8.94 | | High | High | High | High | High | High | High | High | High | High | High | 0 | 1.02E-05 | 3.43 |
| FALSE | High | SETAPAETA | [-].SETAPA | 1xAcetyl [N | Q1WWK3 | 5.5E-05 | 0 | 1 | 1 | 3 | Q1WWK3 | Hist1h1b p | Q1WWK3 | 0 | 1919.92 | 0 | 0.16 | 0.012825 | 7.77 | 93.7 | 106.3 | 4.02 | 5.35 | | High | High | High | High | High | High | High | High | High | High | High | 0 | 9.16E-06 | 3.34 |
| FALSE | High | SEVGSLVAT | [K].SEVGSL | 1xCarbami | Q923W1 1 | 0.083101 | 0.006345 | 1 | 1 | 1 | Q923W1 | Trimethylg | Q923W1 [ | 0 | 3991.914 | 0 | -0.48 | 0.018703 | 4.31 | 112.1 | 87.9 | 12.11 | 17.65 | | High | High | High | High | High | High | High | High | High | High | High | 0.003265 | 0.03462 | 3.05 |
| FALSE | High | SEVQAIIES | [K].SEVQAI | 1xPhospho | B8X349 1xP | 0.000137 | 0 | 2 | 12 | 1 | B8X349; F6 | JNK-interac | B8X349 [3 | 0 | 2443.208 | 0 | 1.4 | 0.008145 | 24.7 | 54.8 | 145.2 | 28.26 | 13.35 | Shared | High | High | High | High | High | High | High | High | High | High | High | 0 | 2.55E-05 | 2.42 |
| FALSE | High | SEVQAIIES | [K].SEVQAI | 1xPhospho | B8X349 1xP | 5.82E-06 | 0 | 2 | 12 | 1 | B8X349; F6 | JNK-interac | B8X349 [3 | 0 | 2459.203 | 0 | 1.51 | 0.000812 | 7.03 | 49.4 | 150.6 | 13.69 | 14.36 | Shared | High | High | High | High | High | High | High | High | High | High | High | 0 | 7.38E-07 | 2.64 |
| FALSE | High | SEVQSEYLF | [R].SEVQSE | 1xPhospho | E9PYK3 1xP | 0.005717 | 0.00062 | 1 | 2 | 1 | E9PYK3 | Poly [ADP- | E9PYK3 [1 | 0 | 2361.126 | 0 | 0.35 | 0.002091 | 7.01 | 87.7 | 112.3 | 2.88 | 4.52 | | High | High | High | High | High | High | High | High | High | High | High | 0.000275 | 0.001674 | 2.46 |
| FALSE | High | SEVQSEYLF | [R].SEVQSE | 2xPhospho | E9PYK3 2xP | 0.010853 | 0.000772 | 1 | 2 | 1 | E9PYK3 | Poly [ADP- | E9PYK3 [1 | 0 | 2441.093 | 0 | 1.64 | 0.001266 | 6.1 | 48.8 | 151.2 | 4.86 | 13.36 | | High | High | High | High | High | High | High | High | High | High | High | 0.000399 | 0.003441 | 2.48 |
| FALSE | High | SEVSTSPFS | [K].SEVSTS | 1xPhospho | F8VQ93 1x | 0.00095 | 0 | 1 | 2 | 1 | F8VQ93 | PH-interac | F8VQ93 [1 | 0 | 1716.841 | 0 | 0.37 | 0.006147 | 7.98 | 87.1 | 112.9 | 5.33 | 11.09 | | High | High | High | High | High | High | High | High | High | High | High | 0 | 0.000225 | 3.07 |
| FALSE | High | SEVTSPIQR | [R].SEVTSP | 1xPhospho | E9Q6H8 1x | 0.035181 | 0.002269 | 1 | 12 | 2 | E9Q6H8 | Pleckstrin | E9Q6H8 [5 | 0 | 1325.666 | 0 | -0.53 | 0.014058 | 9.67 | 117.8 | 82.2 | 5.99 | 10.45 | | High | High | High | High | High | High | High | High | High | High | High | 0.001199 | 0.01289 | 1.95 |
| FALSE | High | SEVVEDEG | [R].SEVVEL | 1xPhospho | Q99LH9 1x | 7.15E-05 | 0 | 1 | 3 | 2 | Q99LH9 | SH3 domai | Q99LH9 [2 | 1 | 3355.558 | 0 | 0.19 | 0.059618 | 11.7 | 95.3 | 104.7 | 5.83 | 4.19 | | High | High | High | High | High | High | High | High | High | High | High | 0 | 1.23E-05 | 4.14 |
| FALSE | High | SFADENNV | [R].SFADEN | 1xPhospho | A0A1W2P8 | 0.001211 | 0 | 1 | 13 | 1 | A0A1W2P8 | Ankyrin-3 ( | A0A1W2P8 | 0 | 3810.687 | 0 | -0.34 | 0.015831 | 6.87 | 110.6 | 89.4 | 7.59 | 9.88 | | High | High | High | High | High | High | High | High | High | High | High | 0 | 0.000297 | 3.43 |
| FALSE | High | SFADIGILA | [R].SFADIG | 1xPhospho | I1VGT5 1xP | 3.26E-05 | 0 | 1 | 5 | 2 | I1VGT5 | Anion exch | I1VGT5 [23 | 0 | 2740.396 | 0 | 1.05 | 0.001262 | 19.12 | 65 | 135 | 9.57 | 4.98 | | High | High | High | High | High | High | High | High | High | High | High | 0 | 5.12E-06 | 2.97 |
| FALSE | High | SFADIGILA | [R].SFADIG | 2xPhospho | I1VGT5 2xP | 0.001644 | 5.66E-05 | 1 | 5 | 3 | I1VGT5 | Anion exch | I1VGT5 [23 | 0 | 2820.362 | 0 | 1.03 | 0.001339 | 4.26 | 66 | 134 | 8.94 | 8.79 | | High | High | High | High | High | High | High | High | High | High | High | 2.58E-05 | 0.000419 | 2.88 |
| FALSE | High | SFAGSVSP( | [R].SFAGS\ | 1xPhospho | Q52KC3 1x | 0.000219 | 0 | 1 | 3 | 1 | Q52KC3 | DNA helica | Q52KC3 [3 | 0 | 2007.89 | 0 | -0.05 | 0.857795 | 13.74 | 94 | 106 | 13.45 | 8.08 | | High | High | High | High | High | High | High | High | High | High | High | 0 | 4.34E-05 | 3.45 |
| FALSE | High | SFAPSTPLT | [R].SFAPST | 1xPhospho | Q64701 1x | 0.017966 | 0.001282 | 1 | 4 | 2 | Q64701 | Retinoblas | Q64701 [3 | 0 | 1442.724 | 0 | 2.65 | 0.000474 | 4.79 | 27.4 | 172.6 | 7.13 | 10.55 | | High | High | High | High | High | High | High | High | High | High | High | 0.000689 | 0.006028 | 2.75 |
| FALSE | High | SFDDEESVI | [K].SFDDEE | 1xPhospho | A0A1L1ST2 | 0.000331 | 0 | 2 | 12 | 4 | A0A1L1ST2 | CLIP-associ | A0A1L1ST2 | 0 | 2754.266 | 0 | -0.3 | 0.028615 | 10.33 | 108.7 | 91.3 | 3.91 | 7.56 | Shared | High | High | High | High | High | High | High | High | High | High | High | 0.000275 | 0.001757 | 3.7 |
| FALSE | High | SEVLADESI | [R].SEVLAD | 1xCarbami | G3UWQ7 1 | 0.000458 | 0 | 1 | 5 | 1 | G3UWQ7 | Protein reg | G3UWQ7 [ | 0 | 2131.076 | 0 | -0.04 | 0.68463 | 17.14 | 98.6 | 101.4 | 18.96 | 11.71 | | High | High | High | High | High | High | High | High | High | High | High | 0 | 9.89E-05 | 3.31 |
| FALSE | High | SFDFEGSLS | [R].SFDFEG | 1xPhospho | B7ZNX2 1x | 7.84E-06 | 0 | 1 | 3 | 2 | B7ZNX2 | Anaphase | B7ZNX2 [6 | 0 | 2132.108 | 0 | -0.04 | 0.459974 | 5.68 | 101.4 | 98.6 | 2.19 | 5.98 | | High | High | High | High | High | High | High | High | High | High | High | 0 | 5.95E-05 | 2.6 |
| FALSE | High | SESVVYADI | [K].SESVVY | 1xPhospho | Q3TEW6 1 | 0.001301 | 0 | 1 | 2 | 1 | Q3TEW6 | Myelin pro | Q3TEW6 [2 | 0 | 1447.703 | 0 | -0.47 | 0.011866 | 7.41 | 116 | 84 | 6.2 | 7.65 | | High | High | High | High | High | High | High | High | High | High | High | 0 | 0.000321 | 1.93 |
| FALSE | High | SESVEGFLS | [R].SESVEG | 1xPhospho | H7BX08 1x | 0.00013 | 0 | 1 | 6 | 2 | H7BX08 | Calmodulin | H7BX08 [1 | 0 | 1603.757 | 0 | 0.99 | 0.006267 | 20.93 | 66.9 | 133.1 | 6.63 | 14.51 | | High | High | High | High | High | High | High | High | High | High | High | 0 | 6.01E-05 | 2.43 |
| FALSE | High | SESDLEEAE | [K].SESDLE | 1xPhospho | Q64511 1x | 0.006031 | 0.00062 | 1 | 2 | 1 | Q64511 | DNA topois | Q64511 [1 | 0 | 1978.957 | 0 | 0.78 | 0.26588 | 36.24 | 79.2 | 120.8 | 8.39 | 45.46 | | High | High | High | High | High | High | High | High | High | High | High | 0.000275 | 0.001778 | 2.75 |
| FALSE | High | SESDQEDN | [K].SESDQI | 1xPhospho | A2AWT3-2 | 3.22E-07 | 0 | 1 | 4 | 4 | A2AWT3-2 | Isoform 2 c | A2AWT3-2 | 0 | 3100.258 | 0 | -0.68 | 0.001008 | 3.42 | 124.2 | 75.8 | 4.59 | 6.16 | | High | High | High | High | High | High | High | High | High | High | High | 0 | 2.84E-08 | 5.16 |
| FALSE | High | SESDSSPDS | [R].SESDSS | 1xPhospho | Q8BTI8 1xF | 0.031439 | 0.002011 | 1 | 3 | 1 | Q8BTI8 | Serine/argi | Q8BTI8 [15 | 0 | 2031.025 | 0 | -0.08 | 0.906626 | 27.4 | 102.9 | 97.1 | 21.29 | 15.75 | | High | High | High | High | High | High | High | High | High | High | High | 0.001076 | 0.01131 | 1.79 |
| FALSE | High | SESETSTM. | [R].SESETS | 1xPhospho | Q0VAW6 1 | 1.05E-05 | 0 | 1 | 5 | 4 | Q0VAW6 | RIKEN cDN | Q0VAW6 [ | 0 | 1466.628 | 0 | -0.98 | 0.007604 | 3.05 | 134.5 | 65.5 | 15.16 | 14.56 | | High | High | High | High | High | High | High | High | High | High | High | 7.19E-05 | 0.000862 | 1.83 |
| FALSE | High | SESGEEML | [K].SESGEE | 1xPhospho | Q9D7A8 1x | 0.007881 | 0.000736 | 1 | 1 | 1 | Q9D7A8 | Armadillo r | Q9D7A8 [2 | 0 | 4588.114 | 0 | -0.84 | 0.001018 | 2.6 | 129.4 | 70.6 | 4.59 | 11.31 | | High | High | High | High | High | High | High | High | High | High | High | 0.000327 | 0.002399 | 3.77 |
| FALSE | High | SESGEEML | [K].SESGEE | 1xPhospho | Q9D7A8 1x | 0.030104 | 0.001857 | 1 | 1 | 3 | Q9D7A8 | Armadillo r | Q9D7A8 [2 | 0 | 4604.109 | 0 | -0.26 | 0.201633 | 27.52 | 109 | 91 | 12.68 | 21.43 | | High | High | High | High | High | High | High | High | High | High | High | 0.004517 | 0.04891 | 3.56 |
| FALSE | High | SESGGLGV | [R].SESGGI | 1xPhospho | Q80U58 1x | 0.000639 | 0 | 1 | 8 | 4 | Q80U58 | Pumilio ho | Q80U58 [8 | 0 | 2737.341 | 0 | 0.76 | 0.029924 | 4.13 | 77.3 | 122.7 | 6.62 | 18.56 | | High | High | High | High | High | High | High | High | High | High | High | 0 | 0.000144 | 3.73 |
| FALSE | High | SESGGLGV | [R].SESGGI | 1xPhospho | Q80U58 1x | 0.000311 | 0 | 1 | 8 | 2 | Q80U58 | Pumilio ho | Q80U58 [8 | 0 | 2753.336 | 0 | 0.6 | 0.006439 | 21.97 | 78.7 | 121.3 | 12.95 | 4.93 | | High | High | High | High | High | High | High | High | High | High | High | 0 | 6.44E-05 | 4.01 |
| FALSE | High | SESGGLGV | [R].SESGGI | 1xCarbami | Q3UR00 1x | 9.56E-06 | 0 | 1 | 6 | 1 | Q3UR00 | Putative ur | Q3UR00 [2 | 0 | 2882.315 | 0 | -1.1 | 0.005383 | 7.99 | 140.4 | 59.6 | 12.67 | 16.88 | | High | High | High | High | High | High | High | High | High | High | High | 0 | 1.29E-06 | 4.19 |
| FALSE | High | SESGGLGV | [R].SESGGI | 1xCarbami | Q3UR00 1x | 1.92E-06 | 0 | 1 | 6 | 5 | Q3UR00 | Putative ur | Q3UR00 [2 | 0 | 2898.31 | 0 | -1.3 | 0.145383 | 4.06 | 138.3 | 61.7 | 21.86 | 47.34 | | High | High | High | High | High | High | High | High | High | High | High | 0 | 1.86E-06 | 3.33 |
| FALSE | High | SESGLSK | [R].SESGLS | 1xPhospho | Q6DID5 1x | 0.010853 | 0.000772 | 1 | 1 | 1 | Q6DID5 | PWWP dor | Q6DID5 [2 | 0 | 1016.486 | 0 | -0.54 | 0.00074 | 1.1 | 121.7 | 78.3 | 16.09 | 17.64 | | High | High | High | High | High | High | High | High | High | High | High | 0.000399 | 0.003442 | 2.35 |
| FALSE | High | SESLIDASEI | [R].SESLID/ | 1xPhospho | G5E8R8 1x | 1.05E-08 | 0 | 1 | 3 | 11 | G5E8R8 | UBX domai | G5E8R8 [2 | 0 | 2243.064 | 0 | 0.31 | 0.002141 | 6.26 | 86.8 | 113.2 | 4.39 | 5.17 | | High | High | High | High | High | High | High | High | High | High | High | 0 | 6.05E-10 | 4.95 |
| FALSE | High | SESLIDASEI | [R].SESLID/ | 2xPhospho | G5E8R8 2x | 0.000189 | 0 | 1 | 3 | 6 | G5E8R8 | UBX domai | G5E8R8 [2 | 0 | 2323.031 | 0 | 2.38 | 0.004106 | 3.91 | 31.8 | 168.2 | 20.46 | 23.46 | | High | High | High | High | High | High | High | High | High | High | High | 0 | 3.69E-05 | 5.14 |
| FALSE | High | SESVVSSLD | [R].SESVVS | 1xPhospho | P49025-5 1 | 2.07E-07 | 0 | 1 | 8 | 1 | P49025-5 | Isoform 5 c | P49025-5 [ | 0 | 1843.945 | 0 | 0.25 | 0.018944 | 0.43 | 91.6 | 108.4 | 4.01 | 3.28 | | High | High | High | High | High | High | High | High | High | High | High | 0 | 1.72E-08 | 3.18 |
| FALSE | High | SESLSNCSI( | [R].SESLSN | 1xCarbami | Q3UHP6 1x | 0.00013 | 0 | 1 | 4 | 3 | Q3UHP6 | Putative ur | Q3UHP6 [6 | 0 | 1719.839 | 0 | 0.1 | 0.200418 | 10.11 | 96.4 | 103.6 | 4.21 | 8.6 | | High | High | High | High | High | High | High | High | High | High | High | 0 | 0.000024 | 3.07 |
| FALSE | High | SESPCESQ\ | [K].SESPCE | 1xCarbami | B1AZI6 1xP | 4.91E-06 | 0 | 1 | 6 | 2 | B1AZI6 | THO comp | B1AZI6 [15 | 0 | 2092.93 | 0 | -0.49 | 0.001068 | 1.58 | 117.9 | 82.1 | 2.29 | 4.51 | | High | High | High | High | High | High | High | High | High | High | High | 0 | 6.07E-07 | 3.86 |
| FALSE | High | SESPEEPAL | [R].SESPEE | 1xPhospho | Q8BYU6 1x | 0.002313 | 0.000109 | 1 | 1 | 1 | Q8BYU6 | Torsin-1A-i | Q8BYU6 [8 | 0 | 2533.341 | 0 | 1.86 | 0.001221 | 10.37 | 43.8 | 156.2 | 16.82 | 11.38 | | High | High | High | High | High | High | High | High | High | High | High | 4.94E-05 | 0.000614 | 2.12 |
| FALSE | High | SESPEPGY\ | [R].SESPEP | 1xPhospho | Q920R0 1x | 0.027358 | 0.001676 | 1 | 2 | 2 | Q920R0 | Alsin OS=M | Q920R0 [1 | 0 | 2719.452 | 0 | -0.82 | 0.000986 | 1.74 | 126.3 | 73.7 | 5.91 | 4.95 | | High | High | High | High | High | High | High | High | High | High | High | 0.00092 | 0.009692 | 3.49 |
| FALSE | High | SESPFEGK | [R].SESPFE | 1xPhospho | P14873 1xP | 0.013074 | 0.000889 | 2 | 2 | 1 | P14873; B2 | Microtubu | P14873 [14 | 0 | 1189.534 | 0 | -0.2 | 0.063516 | 13.98 | 105.9 | 94.1 | 3.43 | 14.86 | Shared | High | High | High | High | High | High | High | High | High | High | High | 0.000467 | 0.004251 | 1.93 |
| FALSE | High | SESPLCSSK | [R].SESPLC | 1xCarbami | B9EKX8 1xP | 0.000267 | 0 | 1 | 5 | 1 | B9EKX8 | Ell2 protein | B9EKX8 [3 | 0 | 1532.743 | 0 | -1.42 | 0.000484 | 0.49 | 144.8 | 55.2 | 5.9 | 5 | | High | High | High | High | High | High | High | High | High | High | High | 0 | 5.38E-05 | 1.68 |
| FALSE | High | SESPIPVAD | [R].SESPIP\ | 1xPhospho | D2EAC2 1x | 1.45E-05 | 0 | 1 | 2 | 2 | D2EAC2 | Zinc finger | D2EAC2 [3 | 0 | 2398.124 | 0 | -0.06 | 0.114903 | 1.31 | 102.3 | 97.7 | 3.18 | 4.54 | | High | High | High | High | High | High | High | High | High | High | High | 0 | 2.05E-06 | 5.15 |
| FALSE | High | SESPPAELP | [R].SESPPA | 1xPhospho | G5E870 1x | 0.047894 | 0.00354 | 1 | 5 | 3 | G5E870 | E3 ubiquiti | G5E870 [3 | 0 | 1591.793 | 0 | -1.45 | 0.000231 | 0.7 | 145.2 | 54.8 | 4.49 | 5.08 | | High | High | High | High | High | High | High | High | High | High | High | 0.002485 | 0.02723 | 3.06 |
| FALSE | High | SESPPPLSE | [R].SESPPP | 1xPhospho | Q8K2F0-2 | 5.91E-05 | 0 | 1 | 6 | 5 | Q8K2F0-2 | Isoform 2 c | Q8K2F0-2 | 0 | 1705.881 | 0 | -0.67 | 0.000363 | 0.87 | 124.7 | 75.3 | 3.8 | 5.25 | | High | High | High | High | High | High | High | High | High | High | High | 0 | 9.95E-06 | 3.28 |
| FALSE | High | SESSAGICV | [R].SESSAG | 1xCarbami | B2RXR2 1x | 0.003622 | 0.00021 | 1 | 3 | 2 | B2RXR2 | Phosphatic | B2RXR2 [8 | 0 | 3177.59 | 0 | -0.19 | 0.159308 | 12.13 | 108.7 | 91.3 | 5.95 | 13.07 | | High | High | High | High | High | High | High | High | High | High | High | 9.42E-05 | 0.001007 | 2.55 |
| FALSE | High | SESSDFEV\ | [R].SESSDF | 1xPhospho | Q3TLH4 1x | 6.24E-05 | 0 | 1 | 5 | 8 | Q3TLH4 | Protein PRI | Q3TLH4 [1 | 0 | 1761.871 | 0 | -0.55 | 0.001273 | 3.74 | 117 | 83 | 3.99 | 4.97 | | High | High | High | High | High | High | High | High | High | High | High | 0.001665 | 0.01662 | 3.42 |
| FALSE | High | SESSDFEV\ | [R].SESSDF | 2xPhospho | Q3TLH4 2x | 0.023792 | 0.001612 | 1 | 5 | 2 | Q3TLH4 | Protein PRI | Q3TLH4 [1 | 0 | 1841.838 | 0 | -0.14 | 0.064652 | 7.9 | 103.7 | 96.3 | 6.56 | 3.11 | | High | High | High | High | High | High | High | High | High | High | High | 0.000859 | 0.008261 | 2.33 |
| FALSE | High | SESSGNLPS | [R].SESSGN | 1xPhospho | O08715-4 | 8.32E-08 | 0 | 1 | 6 | 4 | O08715-4 | Isoform 4 c | O08715-4 | 0 | 1728.8 | 0 | -0.04 | 0.872111 | 3.7 | 101.1 | 98.9 | 5.09 | 7.04 | | High | High | High | High | High | High | High | High | High | High | High | 0 | 6.23E-09 | 3.5 |
| FALSE | High | SESTEMVA | [R].SESTEM | 1xCarbami | B2RWU2 1 | 0.000229 | 0 | 1 | 2 | 4 | B2RWU2 | Immunode | B2RWU2 [ | 0 | 2035.912 | 0 | 0.74 | 0.010141 | 15.65 | 73.9 | 126.1 | 13.88 | 11.06 | | High | High | High | High | High | High | High | High | High | High | High | 0 | 4.57E-05 | 2.55 |
| FALSE | High | SESLTAESR | [R].SESLTA | 1xPhospho | E9PZP8 1xP | 0.108463 | 0.008839 | 1 | 1 | 1 | E9PZP8 | HECT and | E9PZP8 [14 | 0 | 1288.598 | 0 | 0.95 | 0.005936 | 16.81 | 69.5 | 130.5 | 36.98 | 8.01 | | High | High | High | High | High | High | High | High | High | High | High | 0.004411 | 0.04735 | 1.6 |
| FALSE | High | SFDNLTTT( | [R].SFDNLT | 1xCarbami | Q8K296 1x | 4.2E-05 | 0 | 1 | 7 | 1 | Q8K296 | Myotubula | Q8K296 [6 | 0 | 2365.077 | 0 | 0.3 | 0.229128 | 35.58 | 88.6 | 111.4 | 17.43 | 15.49 | | High | High | High | High | High | High | High | High | High | High | High | 0 | 6.78E-06 | 2.95 |
| FALSE | High | SFDNLTTT( | [R].SFDNLT | 1xCarbami | Q8K296 1x | 0.000681 | 0 | 1 | 7 | 2 | Q8K296 | Myotubula | Q8K296 [6 | 0 | 2381.072 | 0 | -0.57 | 0.007931 | 6.33 | 119.6 | 80.4 | 10.23 | 6.72 | | High | High | High | High | High | High | High | High | High | High | High | 7.19E-05 | 0.000874 | 3.02 |
| FALSE | High | SFDNSPGEI | [K].SFDNSF | 1xPhospho | O54826 1x | 1.48E-06 | 0 | 1 | 4 | 1 | O54826 | Protein AF- | O54826 [4 | 0 | 2552.232 | 0 | -0.12 | 0.196645 | 8.55 | 105.4 | 94.6 | 6.48 | 7.8 | | High | High | High | High | High | High | High | High | High | High | High | 0 | 1.58E-07 | 3.17 |
| FALSE | High | SFPSSPSK | [K].SFPSSP | 1xPhospho | A2CG76 1x | 0.002463 | 0.000109 | 1 | 5 | 4 | A2CG76 | Histone-lys | A2CG76 [1 | 0 | 1374.707 | 0 | -0.1 | 0.161613 | 4.44 | 103.3 | 96.7 | 4.73 | 5.14 | | High | High | High | High | High | High | High | High | High | High | High | 0.004317 | 0.04613 | 2.16 |
| FALSE | High | SFSAPYPTA | [R].SFSAPY | 1xCarbami | Q0PHV7 1x | 0.0004 | 0 | 1 | 2 | 2 | Q0PHV7 | Dapper ho | Q0PHV7 [1 | 0 | 2440.069 | 0 | -0.76 | 0.008036 | 5.4 | 130.2 | 69.8 | 12.24 | 11.12 | | High | High | High | High | High | High | High | High | High | High | High | 0 | 8.52E-05 | 4.59 |
| FALSE | High | SFSASQSTI | [R].SFSASC | 1xPhospho | E9Q444 1x | 0.04913 | 0.003576 | 1 | 9 | 1 | E9Q444 | Zinc finger | E9Q444 [4 | 0 | 1394.615 | 0 | 0.22 | 0.000429 | 0.97 | 93.3 | 106.7 | 3.34 | 3.66 | | High | High | High | High | High | High | High | High | High | High | High | 0.001889 | 0.01884 | 1.01 |
| FALSE | High | SFSEDVISH | [K].SFSEDV | 1xPhospho | S4R1P5 1xP | 0.001704 | 5.66E-05 | 1 | 7 | 2 | S4R1P5 | Dystonin O | S4R1P5 [4 | 0 | 1686.85 | 0 | -0.12 | 0.087574 | 5.73 | 105.4 | 94.6 | 5.08 | 4.27 | | High | High | High | High | High | High | High | High | High | High | High | 0.001151 | 0.0122 | 2.8 |
| FALSE | High | SFSENGER | [R].SFSENG | 1xPhospho | P23949 1xP | 0.090153 | 0.00698 | 1 | 1 | 1 | P23949 | mRNA dec | P23949 [12 | 0 | 1234.53 | 0 | 1.01 | 0.003186 | 19.22 | 66.8 | 133.2 | 7.86 | 13.77 | | High | High | High | High | High | High | High | High | High | High | High | 0.003606 | 0.03834 | 0.95 |
| FALSE | High | SFSLASSGN | [R].SFSLAS | 1xPhospho | G5E8A0 1x | 3.49E-07 | 0 | 1 | 2 | 4 | G5E8A0 | Oxysterol-l | G5E8A0 [1 | 0 | 1846.89 | 0 | -0.15 | 0.250696 | 2.83 | 102.4 | 97.6 | 4.71 | 10.64 | | High | High | High | High | High | High | High | High | High | High | High | 0 | 2.72E-06 | 4.11 |
| FALSE | High | SFSLASSGN | [R].SFSLAS | 2xPhospho | G5E8A0 2x | 0.068184 | 0.004935 | 1 | 2 | 3 | G5E8A0 | Oxysterol-l | G5E8A0 [1 | 0 | 1926.856 | 0 | 0.89 | 0.001342 | 9.61 | 69.3 | 130.7 | 9.9 | 9.35 | | High | High | High | High | High | High | High | High | High | High | High | 0.003796 | 0.04093 | 2.43 |
| FALSE | High | SFSLDEPPL | [R].SFSLDE | 1xPhospho | B2RQG2 1x | 2.56E-05 | 0 | 2 | 3 | 2 | B2RQG2; A | PHD finger | B2RQG2 [6 | 0 | 2641.393 | 0 | -0.76 | 0.00049 | 1.95 | 126.9 | 73.1 | 3.8 | 3.81 | Shared | High | High | High | High | High | High | High | High | High | High | High | 0 | 3.88E-06 | 3.81 |
| FALSE | High | SFSLDIK | [R].SFSLDII | 1xPhospho | Q99MG6 1 | 0.088701 | 0.006669 | 1 | 6 | 1 | Q99MG6 | Map kinase | Q99MG6 [ | 0 | 1347.733 | 1 | 0.01 | 0.756553 | 30.25 | 104.8 | 95.2 | 11.15 | 17.92 | | High | High | High | High | High | High | High | High | High | High | High | 0.003461 | 0.03751 | 1.39 |
| FALSE | High | SFSLEPDIT\ | [K].SFSLEPI | 1xPhospho | Q80Y19 1x | 0.000579 | 0 | 1 | 2 | 1 | Q80Y19 | Rho GTPas | Q80Y19 [5 | 0 | 1902.987 | 0 | -0.23 | 0.06458 | 14.72 | 109.7 | 90.3 | 12.06 | 17.34 | | High | High | High | High | High | High | High | High | High | High | High | 0 | 0.000129 | 2.54 |

| | | | | | | | | | | | | | | | | | | | | | | | | | | | | | | | | | | | | | | |
|---|---|---|---|---|---|---|---|---|---|---|---|---|---|---|---|---|---|---|---|---|---|---|---|---|---|---|---|---|---|---|---|---|---|---|---|---|---|---|
| FALSE | High | SFSLLSSNQ | [R].SFSLLS | 1xCarbami | P15261 1x | 4.95E-06 | 0 | 1 | 4 | 3 | P15261 | Interferon | P15261 [37 | 0 | 2608.206 | 0 | 0.69 | 0.021793 | 14.39 | 77.1 | 122.9 | 19.75 | 13.65 | | High | High | High | High | High | High | High | High | High | High | High | 0 | 6.16E-07 | 3.71 |
| FALSE | High | SFSLQSDK | [K].SFSLQS | 1xPhospho | E9PYP2 1x | 0.112856 | 0.009146 | 1 | 3 | 1 | E9PYP2 | Neurobeac | E9PYP2 [14 | 0 | 1449.739 | 0 | 0.29 | 0.36454 | 12.26 | 94.3 | 105.7 | 7.41 | 15.49 | | High | High | High | High | High | High | High | High | High | High | High | 0.004569 | 0.04957 | 1.32 |
| FALSE | High | SFSMHDLT | [R].SFSMH | 1xPhospho | A0A1W2P8 | 2.83E-06 | 0 | 1 | 2 | 1 | A0A1W2P8 | Receptor e | A0A1W2P8 | 0 | 3662.808 | 0 | 2.06 | 0.001689 | 15.41 | 38.6 | 161.4 | 14.33 | 13.37 | | High | High | High | High | High | High | High | High | High | High | High | 0 | 3.29E-07 | 4.25 |
| FALSE | High | SFPSRSPSC | [K].SFPSRS | 2xPhospho | Q9Z2C9 2x | 0.004572 | 0.000456 | 1 | 1 | 2 | Q9Z2C9 | Myotubula | Q9Z2C9 [5 | 1 | 3223.533 | 0 | -0.34 | 0.022774 | 3.19 | 111.2 | 88.8 | 10.32 | 4.24 | | High | High | High | High | High | High | High | High | High | High | High | 0.000204 | 0.00131 | 3.21 |
| FALSE | High | SFSMHDLT | [R].SFSMH | 1xPhospho | A0A1W2P8 | 0.014283 | 0.001011 | 1 | 2 | 1 | A0A1W2P8 | Receptor e | A0A1W2P8 | 0 | 3678.803 | 0 | 1.77 | 0.000675 | 4.37 | 43.1 | 156.9 | 16.78 | 9.63 | | High | High | High | High | High | High | High | High | High | High | High | 0.000538 | 0.004697 | 2.98 |
| FALSE | High | SFSMQDLR | [R].SFSMQ | 1xPhospho | Q8K072 1x | 0.015329 | 0.001089 | 1 | 1 | 3 | Q8K072 | Receptor e | Q8K072 [1 | 0 | 1308.586 | 0 | 0.58 | 0.041278 | 18.27 | 80 | 120 | 14.11 | 15.41 | | High | High | High | High | High | High | High | High | High | High | High | 0.000571 | 0.00505 | 1.67 |
| FALSE | High | SFSMQDLT | [R].SFSMQ | 1xPhospho | A0A1D5RL | 0.007404 | 0.00062 | 1 | 2 | 2 | A0A1D5RL | Receptor e | A0A1D5RL | 0 | 1607.77 | 0 | 0.86 | 0.001068 | 11.91 | 70.5 | 129.5 | 11.03 | 4.86 | | High | High | High | High | High | High | High | High | High | High | High | 0.000275 | 0.002233 | 1.91 |
| FALSE | High | SFSPEER | [R].SFSPEE | 1xPhospho | Q9Z0U1 1x | 0.072258 | 0.005346 | 1 | 1 | 3 | Q9Z0U1 | Tight junct | Q9Z0U1 [3 | 0 | 1160.519 | 0 | 0.7 | 0.001743 | 6.51 | 77 | 123 | 3.39 | 6.69 | | High | High | High | High | High | High | High | High | High | High | High | 0.003122 | 0.03234 | 1.96 |
| FALSE | High | SFSPGDFR | [R].SFSPGD | 1xPhospho | Q80WT5 1 | 0.005322 | 0.000599 | 1 | 5 | 2 | Q80WT5 | Aftiphilin C | Q80WT5 [1 | 0 | 1221.55 | 0 | 1.43 | 0.000641 | 13.11 | 54 | 146 | 5.77 | 7.26 | | High | High | High | High | High | High | High | High | High | High | High | 0.001076 | 0.01137 | 1.72 |
| FALSE | High | SFSPLQPGA | [R].SFSPLQ | 1xPhospho | A0A087WF | 0.053926 | 0.00384 | 1 | 4 | 1 | A0A087WF | Autophagy | A0A087WF | 0 | 1721.857 | 0 | 0.05 | 0.232831 | 12.08 | 93.9 | 106.1 | 8.73 | 12.05 | | High | High | High | High | High | High | High | High | High | High | High | 0.002031 | 0.02097 | 2.23 |
| FALSE | High | SFSQPGLR | [K].SFSQPG | 1xPhospho | Q60949 1x | 0.002272 | 0.000109 | 1 | 4 | 2 | Q60949 | TBC1 doma | Q60949 [2 | 0 | 1200.598 | 0 | -1.09 | 0.000358 | 2.75 | 137.4 | 62.6 | 2.58 | 13.7 | | High | High | High | High | High | High | High | High | High | High | High | 4.94E-05 | 0.000599 | 1.14 |
| FALSE | High | SFSSPEDYC | [R].SFSSPE | 1xPhospho | Q14A95 1x | 0.004372 | 0.000409 | 1 | 3 | 2 | Q14A95 | MCG9666, | Q14A95 [1 | 0 | 1524.657 | 0 | 0.29 | 0.201596 | 17.62 | 86.6 | 113.4 | 13.95 | 33.19 | | High | High | High | High | High | High | High | High | High | High | High | 0.000183 | 0.001248 | 1.88 |
| FALSE | High | SFSSPPSPS | [K].SFSSPP | 1xPhospho | Q91Z58 1x | 9.28E-05 | 0 | 1 | 1 | 2 | Q91Z58 | Uncharacte | Q91Z58 [9 | 0 | 1721.855 | 0 | 2.76 | 0.000125 | 9.18 | 25.6 | 174.4 | 10.07 | 2.56 | | High | High | High | High | High | High | High | High | High | High | High | 0.000847 | 0.007979 | 1.71 |
| FALSE | High | SFSSPPSPS | [K].SFSSPP | 2xPhospho | Q91Z58 2x | 0.064862 | 0.004664 | 1 | 1 | 1 | Q91Z58 | Uncharacte | Q91Z58 [9 | 0 | 1801.822 | 0 | 3.17 | 0.000223 | 25.38 | 20 | 180 | 17.8 | 6.59 | | High | High | High | High | High | High | High | High | High | High | High | 0.002432 | 0.02612 | 1.57 |
| FALSE | High | SFSVPK | [R].SFSVPK | 1xPhospho | E9Q8I9-2 1 | 0.109329 | 0.008865 | 1 | 2 | 2 | E9Q8I9-2 | Isoform 2 c | E9Q8I9-2 [ | 0 | 1202.659 | 0 | -1.07 | 0.000358 | 1.71 | 133.9 | 66.1 | 2.92 | 9.92 | | High | High | High | High | High | High | High | High | High | High | High | 0.004422 | 0.04784 | 0.88 |
| FALSE | High | SFTGGLGQ | [R].SFTGGL | 1xPhospho | Q3TCX6 1x | 8.56E-05 | 0 | 1 | 7 | 1 | Q3TCX6 | Putative ur | Q3TCX6 [1 | 0 | 1827.977 | 0 | 3.18 | 0.002161 | 27.17 | 20.9 | 179.1 | 22.5 | 18.94 | | High | High | High | High | High | High | High | High | High | High | High | 0 | 1.51E-05 | 3.41 |
| FALSE | High | SFTGMLGG | [K].SFTGM | 1xPhospho | E9Q4K7 1x | 8.45E-07 | 0 | 1 | 2 | 4 | E9Q4K7 | Kinesin fan | E9Q4K7 [1 | 0 | 2962.297 | 0 | 0.37 | 0.019917 | 10.52 | 81.5 | 118.5 | 11.93 | 8.16 | | High | High | High | High | High | High | High | High | High | High | High | 0 | 2.53E-06 | 3.76 |
| FALSE | High | SFTGMLGG | [K].SFTGM | 1xPhospho | E9Q4K7 1x | 9.91E-06 | 0 | 1 | 2 | 1 | E9Q4K7 | Kinesin fan | E9Q4K7 [1 | 0 | 2978.292 | 0 | 0.57 | 0.083211 | 23.11 | 86.3 | 113.7 | 18.74 | 15.02 | | High | High | High | High | High | High | High | High | High | High | High | 0 | 1.34E-06 | 4.22 |
| FALSE | High | SFSMQDLR | [R].SFSMQ | 1xPhospho | Q8K072 1x | 0.031439 | 0.002011 | 1 | 1 | 1 | Q8K072 | Receptor e | Q8K072 [1 | 0 | 1292.591 | 0 | 0.5 | 0.004653 | 9.23 | 83.4 | 116.6 | 9.79 | 4.14 | | High | High | High | High | High | High | High | High | High | High | High | 0.001076 | 0.01131 | 1.48 |
| FALSE | High | SFPSRSPSC | [K].SFPSRS | 1xPhospho | Q9Z2C9 1x | 2.41E-06 | 0 | 1 | 1 | 4 | Q9Z2C9 | Myotubula | Q9Z2C9 [5 | 1 | 3143.566 | 0 | -0.95 | 0.001524 | 3.71 | 131.7 | 68.3 | 3.98 | 16.88 | | High | High | High | High | High | High | High | High | High | High | High | 0 | 2.72E-07 | 4.16 |
| FALSE | High | SFPLSPAIP | [K].SFPLSP | 1xPhospho | Q8CE90-7 | 0.00129 | 0 | 1 | 3 | 1 | Q8CE90-7 | Isoform 7 c | Q8CE90-7 | 0 | 2350.179 | 0 | -1.98 | 0.004875 | 0.81 | 156.4 | 43.6 | 17.81 | 13.44 | | High | High | High | High | High | High | High | High | High | High | High | 0 | 0.000317 | 3.42 |
| FALSE | High | SFNYSPSSS | [K].SFNYSP | 1xPhospho | Q6NXI6 1x | 1.55E-06 | 0 | 1 | 3 | 6 | Q6NXI6 | Regulation | Q6NXI6 [6 | 0 | 2578.196 | 0 | -0.01 | 0.503589 | 10.24 | 100 | 100 | 5.44 | 11.52 | | High | High | High | High | High | High | High | High | High | High | High | 0 | 0.000217 | 3.24 |
| FALSE | High | SFDQLTPEE | [K].SFDQLT | 1xPhospho | O35887 1x | 0.019446 | 0.001438 | 1 | 8 | 1 | O35887 | Calumenin | O35887 [6 | 0 | 1818.893 | 0 | 1.08 | 0.010374 | 21.82 | 64.4 | 135.6 | 14.15 | 15.63 | | High | High | High | High | High | High | High | High | High | High | High | 0.000756 | 0.006588 | 2.17 |
| FALSE | High | SFDTVIGAC | [R].SFDTVI | 1xPhospho | Q8VI59 1x | 0.027597 | 0.001716 | 1 | 1 | 1 | Q8VI59 | Pecanex-lik | Q8VI59 [36 | 0 | 3017.648 | 0 | 0.83 | 0.000455 | 0.81 | 72.8 | 127.2 | 5.61 | 4.22 | | High | High | High | High | High | High | High | High | High | High | High | 0.000936 | 0.009756 | 1.92 |
| FALSE | High | SFDYGNLSI | [K].SFDYGN | 1xPhospho | B2RWU2 1 | 0.023584 | 0.001578 | 1 | 2 | 1 | B2RWU2 | Immunode | B2RWU2 [ | 0 | 2757.308 | 0 | 1.33 | 0.00199 | 11.44 | 58.7 | 141.3 | 6.08 | 11.79 | | High | High | High | High | High | High | High | High | High | High | High | 0.000859 | 0.008246 | 2.88 |
| FALSE | High | SFEDLTDHI | [K].SFEDLT | 1xPhospho | E9PXU2 1x | 0.004218 | 0.00036 | 1 | 8 | 2 | E9PXU2 | Disintegrin | E9PXU2 [8 | 0 | 1725.805 | 0 | 0.65 | 0.027643 | 40.68 | 77.6 | 122.4 | 10.94 | 15.9 | | High | High | High | High | High | High | High | High | High | High | High | 0.000246 | 0.001475 | 2.88 |
| FALSE | High | SFEDSGGG | [R].SFEDSG | 1xCarbami | Q6PAJ1 1x | 1.2E-08 | 0 | 1 | 2 | 5 | Q6PAJ1 | Breakpoint | Q6PAJ1 [32 | 0 | 3830.497 | 0 | 0.23 | 0.172917 | 31.39 | 87 | 113 | 12.88 | 30.88 | | High | High | High | High | High | High | High | High | High | High | High | 0 | 1.54E-09 | 5.54 |
| FALSE | High | SFEFEDASS | [K].SFEFED | 1xPhospho | Q924W7 1 | 0.108463 | 0.008839 | 1 | 5 | 1 | Q924W7 | Suppressio | Q924W7 [4 | 0 | 3600.667 | 0 | -0.04 | 0.933522 | 24.39 | 101.6 | 98.4 | 10.6 | 12.36 | | High | High | High | High | High | High | High | High | High | High | High | 0.004411 | 0.04735 | 2.46 |
| FALSE | High | SFEILRGSD | [K].SFEILR | 2xPhospho | A6H6E1 2x | 0.048715 | 0.00354 | 1 | 4 | 1 | A6H6E1 | Espl1 prote | A6H6E1 [1 | 1 | 2502.12 | 0 | -0.83 | 0.007035 | 6.35 | 126.6 | 73.4 | 9.24 | 10.3 | | High | High | High | High | High | High | High | High | High | High | High | 0.001876 | 0.01879 | 2.71 |
| FALSE | High | SFENISVDS | [K].SFENIS | 1xPhospho | Q8CGF1 1x | 0.00095 | 0 | 1 | 4 | 2 | Q8CGF1 | Rho GTPas | Q8CGF1 [1 | 0 | 2354.168 | 0 | -0.29 | 0.02853 | 10.23 | 112.6 | 87.4 | 10.1 | 5.38 | | High | High | High | High | High | High | High | High | High | High | High | 0 | 0.000226 | 3.61 |
| FALSE | High | SFENISVDS | [K].SFENIS | 2xPhospho | Q8CGF1 2x | 0.099328 | 0.008003 | 1 | 4 | 1 | Q8CGF1 | Rho GTPas | Q8CGF1 [1 | 0 | 2434.134 | 0 | 0.66 | 0.032026 | 22.71 | 78.1 | 121.9 | 17.09 | 14.78 | | High | High | High | High | High | High | High | High | High | High | High | 0.004032 | 0.04287 | 2.48 |
| FALSE | High | SFEPPPYTP | [R].SFEPPP | 1xPhospho | E9Q2I4 1x | 1.11E-06 | 0 | 1 | 3 | 2 | E9Q2I4 | ELM2 and | E9Q2I4 [64 | 0 | 2203.14 | 0 | 0.94 | 0.144621 | 37.62 | 74.3 | 125.7 | 7.66 | 38.65 | | High | High | High | High | High | High | High | High | High | High | High | 0.000467 | 0.004257 | 3.61 |
| FALSE | High | SFEPPPYTP | [R].SFEPPP | 2xPhospho | E9Q2I4 2x | 1.81E-05 | 0 | 1 | 3 | 7 | E9Q2I4 | ELM2 and | E9Q2I4 [64 | 0 | 2283.107 | 0 | 0.47 | 0.001587 | 5.03 | 83.7 | 116.3 | 7.75 | 5.21 | | High | High | High | High | High | High | High | High | High | High | High | 2.58E-05 | 0.000546 | 3.94 |
| FALSE | High | SFEQPSSEA | [K].SFEQPS | 1xPhospho | F8WGX5 1 | 0.009931 | 0.000736 | 1 | 2 | 1 | F8WGX5 | DmX-like p | F8WGX5 [4 | 1 | 4365.064 | 0 | 0.35 | 0.394901 | 20.63 | 87.1 | 112.9 | 21 | 21.1 | | High | High | High | High | High | High | High | High | High | High | High | 0.000344 | 0.003111 | 3.76 |
| FALSE | High | SFESLEEVE | [K].SFESLE | 1xPhospho | E9QK22 1x | 0.000993 | 0 | 1 | 2 | 1 | E9QK22 | Zinc finger | E9QK22 [9 | 0 | 1975.921 | 0 | -0.66 | 0.000236 | 1.97 | 120.8 | 79.2 | 2.43 | 5.13 | | High | High | High | High | High | High | High | High | High | High | High | 0 | 0.000237 | 2.11 |
| FALSE | High | SFEVEEIEP | [R].SFEVEE | 1xPhospho | Q80UG5 1 | 0.004531 | 0.000456 | 1 | 3 | 1 | Q80UG5 | Septin-9 O | Q80UG5 [3 | 0 | 2137.005 | 0 | 0.43 | 0.014593 | 4.5 | 86.5 | 113.5 | 15.56 | 4.59 | | High | High | High | High | High | High | High | High | High | High | High | 0.000204 | 0.001297 | 2.19 |
| FALSE | High | SFFGFEGPD | [R].SFFGFE | 1xCarbami | Q9CPY3 1x | 0.000705 | 0 | 1 | 2 | 2 | Q9CPY3 | Sororin OS | Q9CPY3 [1 | 0 | 2779.345 | 0 | -0.06 | 0.130543 | 6.86 | 102.1 | 97.9 | 8.16 | 6.55 | | High | High | High | High | High | High | High | High | High | High | High | 0 | 0.000162 | 2.46 |
| FALSE | High | SFGSGTELC | [R].SFGSGT | 1xPhospho | Q9CQS4 1x | 0.0009 | 0 | 1 | 1 | 2 | Q9CQS4 | Solute carr | Q9CQS4 [3 | 0 | 2607.244 | 0 | 2.62 | 0.000522 | 8.99 | 29.6 | 170.4 | 7.13 | 10.69 | | High | High | High | High | High | High | High | High | High | High | High | 0 | 0.000211 | 2.15 |
| FALSE | High | SFGSPNR | [K].SFGSPN | 1xPhospho | Q03147 1x | 0.021231 | 0.001507 | 1 | 2 | 1 | Q03147 | Cyclin-dep | Q03147 [1 | 0 | 1073.498 | 0 | -0.85 | 0.000536 | 3 | 128.5 | 71.5 | 4.89 | 4.64 | | High | High | High | High | High | High | High | High | High | High | High | 0.000801 | 0.007311 | 1.28 |
| FALSE | High | SFHFDPLSS | [K].SFHFDP | 1xPhospho | Q8VDZ4 1x | 0.003655 | 0.00021 | 1 | 2 | 2 | Q8VDZ4 | Palmitoyltr | Q8VDZ4 [4 | 0 | 1645.757 | 0 | 0.58 | 0.01047 | 15.13 | 81.7 | 118.3 | 4.07 | 13.06 | | High | High | High | High | High | High | High | High | High | High | High | 9.42E-05 | 0.001015 | 1.97 |
| FALSE | High | SFHVSPAR | [K].SFHVSP | 1xPhospho | Q9D279 1x | 0.015329 | 0.001089 | 1 | 2 | 1 | Q9D279 | Mitotic inte | Q9D279 [2 | 0 | 1209.598 | 0 | 1.78 | 0.002957 | 22.24 | 46.9 | 153.1 | 27.6 | 9.91 | | High | High | High | High | High | High | High | High | High | High | High | 0.000571 | 0.005037 | 2.26 |
| FALSE | High | SFLNYYADI | [K].SFLNYY | 1xPhospho | Q9DBR4-2 | 0.015061 | 0.001049 | 2 | 11 | 1 | Q9DBR4-2; | Isoform 2 c | Q9DBR4-2 | 0 | 1958.91 | 0 | 0.37 | 0.05866 | 10.62 | 87.3 | 112.7 | 13.17 | 16.66 | Shared | High | High | High | High | High | High | High | High | High | High | High | 0.000571 | 0.00497 | 2.15 |
| FALSE | High | RQEEAWVI | [R].RQEEA | 1xCarbami | Q3UJP8 1x | 3.09E-09 | 0 | 1 | 2 | 1 | Q3UJP8 | Trifunction | Q3UJP8 [7 | 1 | 2594.227 | 0 | -0.89 | 0.00046 | 1.7 | 130.4 | 69.6 | 5.14 | 8.2 | | High | High | High | High | High | High | High | High | High | High | High | 0 | 1.54E-10 | 5.54 |
| FALSE | High | SFISSSPSSF | [K].SFISSS | 1xPhospho | Q3UTJ2-2 | 0.002743 | 0.000109 | 1 | 16 | 2 | Q3UTJ2-2 | Isoform 2 c | Q3UTJ2-2 | 0 | 1547.73 | 0 | 0.42 | 0.010231 | 7.83 | 87.8 | 112.2 | 12.49 | 8.38 | | High | High | High | High | High | High | High | High | High | High | High | 4.94E-05 | 0.00074 | 1.89 |
| FALSE | High | SFLVASVLP | [R].SFLVAS | 1xPhospho | B9EIG6 1x | 0.001704 | 5.66E-05 | 1 | 5 | 2 | B9EIG6 | Pogo trans | B9EIG6 [13 | 0 | 2336.185 | 0 | -1.17 | 0.000896 | 4.11 | 137.2 | 62.8 | 7.94 | 14.63 | | High | High | High | High | High | High | High | High | High | High | High | 2.58E-05 | 0.000435 | 3.05 |
| FALSE | High | SFLVIVDGS | [K].SFLVIV | 1xPhospho | Q4VBD9 1x | 2.67E-05 | 0 | 1 | 2 | 1 | Q4VBD9 | GDNF-indu | Q4VBD9 [6 | 0 | 1699.944 | 0 | -0.15 | 0.007118 | 3.51 | 105.3 | 94.7 | 2.67 | 5.73 | | High | High | High | High | High | High | High | High | High | High | High | 0 | 4.1E-06 | 1.88 |
| FALSE | High | SFLVSYIAN | [R].SFLVSY | 1xPhospho | Q3TIN2 1x | 0.007019 | 0.00062 | 1 | 10 | 1 | Q3TIN2 | Putative ur | Q3TIN2 [7 | 0 | 1450.754 | 0 | 2.03 | 0.00046 | 3.24 | 41.7 | 158.3 | 16.58 | 8.93 | | High | High | High | High | High | High | High | High | High | High | High | 0.000275 | 0.002106 | 2.7 |
| FALSE | High | SFLVSYIAN | [R].SFLVSY | 1xPhospho | Q3TIN2 1x | 0.009087 | 0.000736 | 1 | 10 | 2 | Q3TIN2 | Putative ur | Q3TIN2 [7 | 0 | 1679.917 | 0 | 2.2 | 0.00014 | 1.46 | 35.9 | 164.1 | 7.26 | 3.49 | | High | High | High | High | High | High | High | High | High | High | High | 0.002354 | 0.02422 | 1.77 |
| FALSE | High | SFMESGGT | [K].SFMES | 1xPhospho | Q9CX34 1x | 3.47E-05 | 0 | 1 | 1 | 2 | Q9CX34 | Protein SG | Q9CX34 [3 | 0 | 2556.209 | 0 | -1.27 | 0.000904 | 2.51 | 139.1 | 60.9 | 5.97 | 16.8 | | High | High | High | High | High | High | High | High | High | High | High | 0 | 1.88E-05 | 2.73 |
| FALSE | High | SFNSDSPSI | [R].SFNSDS | 1xPhospho | Q8C4P0-2 | 2.53E-05 | 0 | 1 | 2 | 2 | Q8C4P0-2 | Isoform 2 c | Q8C4P0-2 | 0 | 2743.303 | 0 | -0.54 | 0.002542 | 5.54 | 118.6 | 81.4 | 5.03 | 4.52 | | High | High | High | High | High | High | High | High | High | High | High | 0 | 3.84E-06 | 3.65 |
| FALSE | High | SFNSDSPSI | [R].SFNSDS | 2xPhospho | Q8C4P0-2 | 0.04913 | 0.003576 | 1 | 2 | 1 | Q8C4P0-2 | Isoform 2 c | Q8C4P0-2 | 0 | 2823.269 | 0 | -0.09 | 0.079049 | 3.21 | 103.4 | 96.6 | 10.61 | 16.21 | | High | High | High | High | High | High | High | High | High | High | High | 0.001889 | 0.0189 | 2.87 |
| FALSE | High | SFTSGPGAI | [R].SFTSGP | 1xPhospho | P11679 1x | 0.007019 | 0.00062 | 1 | 1 | 2 | P11679 | Keratin, ty | P11679 [24 | 0 | 1188.561 | 0 | 4.2 | 0.00242 | 34.87 | 8.6 | 191.4 | 11.88 | 22.81 | | High | High | High | High | High | High | High | High | High | High | High | 0.000275 | 0.002107 | 1.94 |
| FALSE | High | SGSPAPETT | [R].SGSPAP | 2xPhospho | E9PYH6 2x | 0.011652 | 0.000889 | 1 | 1 | 1 | E9PYH6 | SET domai | E9PYH6 [4 | 0 | 2890.249 | 0 | -0.31 | 0.052919 | 14.08 | 111.4 | 88.6 | 8.73 | 13.11 | | High | High | High | High | High | High | High | High | High | High | High | 0.000449 | 0.003714 | 2.35 |
| FALSE | High | SGSPGTPA | [R].SGSPGT | 1xPhospho | Q9DBT3 1x | 3.15E-06 | 0 | 1 | 2 | 4 | Q9DBT3 | Coiled-coil | Q9DBT3 [2 | 0 | 2722.296 | 0 | 0.69 | 0.001819 | 9.06 | 78.1 | 121.9 | 4.22 | 7.16 | | High | High | High | High | High | High | High | High | High | High | High | 0 | 3.7E-07 | 4.62 |
| FALSE | High | SGSPLLSSH | [R].SGSPLL | 1xPhospho | Q9CR78 1x | 6.2E-06 | 0 | 1 | 1 | 1 | Q9CR78 | Myb/SANT | Q9CR78 [9 | 0 | 1878.014 | 0 | -0.86 | 0.004563 | 3.29 | 128.7 | 71.3 | 9.52 | 10.94 | | High | High | High | High | High | High | High | High | High | High | High | 0 | 7.88E-07 | 2.8 |
| FALSE | High | SLASQSSDI | [R].SLASQS | 1xPhospho | Q8K3Z9 1x | 0.000178 | 0 | 1 | 2 | 5 | Q8K3Z9 | Nuclear en | Q8K3Z9 [3 | 0 | 1939.953 | 0 | -0.49 | 0.044465 | 9.03 | 117.1 | 82.9 | 11.9 | 15.86 | | High | High | High | High | High | High | High | High | High | High | High | 0 | 4.89E-05 | 1.32 |
| FALSE | High | SLAVSDEEE | [R].SLAVSD | 1xPhospho | Q69ZX6 1x | 2.77E-05 | 0 | 2 | 2 | 1 | Q69ZX6; Q | MORC fam | Q69ZX6 [7 | 1 | 2459.159 | 0 | -1 | 0.001184 | 2.5 | 132.9 | 67.1 | 6.65 | 6.31 | Shared | High | High | High | High | High | High | High | High | High | High | High | 0 | 4.26E-06 | 2.81 |
| FALSE | High | SLCGSTLTD | [R].SLCGST | 2xCarbami | Q14B71 1x | 5.62E-06 | 0 | 1 | 1 | 2 | Q14B71 | Cell divisio | Q14B71 [9 | 0 | 2661.23 | 0 | -1.11 | 0.001745 | 3.26 | 136.7 | 63.3 | 8.19 | 6.7 | | High | High | High | High | High | High | High | High | High | High | High | 0 | 7.03E-07 | 4.18 |
| FALSE | High | SLCSEETDV | [R].SLCSEE | 1xCarbami | Q1W5W7 | 0.014158 | 0.001011 | 1 | 1 | 1 | Q1W5W7 | Putative ga | Q1W5W7 | 0 | 1921.865 | 0 | 1.35 | 0.002092 | 10.15 | 59.4 | 140.6 | 9.67 | 9.52 | | High | High | High | High | High | High | High | High | High | High | High | 0.000538 | 0.004624 | 1.92 |
| FALSE | High | SLCSLDYED | [K].SLCSLD | 1xCarbami | E9PV82 1x | 8.71E-05 | 0 | 1 | 3 | 2 | E9PV82 | Protein FAI | E9PV82 [3 | 0 | 2600.1 | 0 | -0.62 | 0.001436 | 2.36 | 121 | 79 | 4.53 | 7.47 | | High | High | High | High | High | High | High | High | High | High | High | 0 | 1.55E-05 | 2.95 |
| FALSE | High | SLCSPLQEV | [K].SLCSPL | 2xCarbami | Q6P9S1 1x | 1.9E-05 | 0 | 1 | 1 | 1 | Q6P9S1 | ATM intera | Q6P9S1 [3 | 0 | 2028.945 | 0 | -1.49 | 0.001887 | 2.34 | 148 | 52 | 15.17 | 6.98 | | High | High | High | High | High | High | High | High | High | High | High | 0 | 2.76E-06 | 3.9 |
| FALSE | High | SICSSPAIEG | [K].SICSSP | 1xCarbami | A3KGI3 1x | 0.000286 | 0 | 1 | 3 | 2 | A3KGI3 | Kinetochor | A3KGI3 [3 | 0 | 1849.917 | 0 | -0.17 | 0.040916 | 6.02 | 108.9 | 91.1 | 6.36 | 6.95 | | High | High | High | High | High | High | High | High | High | High | High | 0 | 0.000123 | 2.42 |
| FALSE | High | SLDAEPSVF | [K].SLDAEP | 1xPhospho | B9EIG6 1x | 0.096181 | 0.007535 | 1 | 5 | 1 | B9EIG6 | Pogo trans | B9EIG6 [4 | 0 | 2664.41 | 0 | -1.14 | 0.040272 | 9.25 | 138 | 62 | 30.25 | 14.79 | | High | High | High | High | High | High | High | High | High | High | High | 0.003823 | 0.04127 | 2.11 |
| FALSE | High | SLDDLESAT | [R].SLDDLE | 1xPhospho | E9Q652 1x | 2.02E-08 | 0 | 1 | 3 | 2 | E9Q652 | Regulator | E9Q652 [6 | 0 | 2674.311 | 0 | 0.9 | 0.003725 | 17.61 | 69.6 | 130.4 | 6.07 | 14.81 | | High | High | High | High | High | High | High | High | High | High | High | 0 | 1.26E-09 | 4.36 |
| FALSE | High | SLDDLSAPF | [R].SLDDLS | 1xPhospho | E9Q6B2 1x | 0.012845 | 0.000889 | 1 | 1 | 2 | E9Q6B2 | Coiled-coil | E9Q6B2 [2 | 0 | 1653.795 | 0 | -0.35 | 0.005149 | 5.93 | 114 | 86 | 5.83 | 8.09 | | High | High | High | High | High | High | High | High | High | High | High | 0.000449 | 0.004154 | 2.59 |
| FALSE | High | SIDDSEMES | [K].SIDDSE | 1xPhospho | Q5CZY4 1x | 0.009931 | 0.000736 | 2 | 7 | 1 | Q5CZY4; P | Nuclear fac | Q5CZY4 [2 | 0 | 2625.127 | 0 | 0.75 | 0.012601 | 6.23 | 76.7 | 123.3 | 10.86 | 16.47 | Shared | High | High | High | High | High | High | High | High | High | High | High | 0.000344 | 0.003116 | 2.41 |
| FALSE | High | SIDDSEMES | [K].SIDDSE | 1xPhospho | Q5CZY4 1x | 0.001586 | 5.66E-05 | 2 | 7 | 1 | Q5CZY4; P | Nuclear fac | Q5CZY4 [2 | 0 | 2641.121 | 0 | 0.75 | 0.040634 | 0.55 | 72 | 128 | 19.97 | 34.36 | Shared | High | High | High | High | High | High | High | High | High | High | High | 2.58E-05 | 0.000401 | 2.72 |
| FALSE | High | SLDGAAAA | [R].SLDGA | 1xPhospho | A0A068WA | 2.02E-05 | 0 | 1 | 10 | 2 | A0A068WA | Protein 4.1 | A0A068WA | 1 | 4143.112 | 0 | 1.14 | 0.001241 | 20.85 | 61.8 | 138.2 | 11.5 | 6.85 | | High | High | High | High | High | High | High | High | High | High | High | 0 | 2.99E-06 | 4.02 |
| FALSE | High | SLASFEAQ | [R].SLASFE | 1xPhospho | Q922B9 1x | 1.77E-07 | 0 | 1 | 5 | 1 | Q922B9 | Sperm-spe | Q922B9 [6 | 0 | 2154.019 | 0 | 0.45 | 0.051341 | 13.04 | 88.4 | 111.6 | 8.74 | 13.11 | | High | High | High | High | High | High | High | High | High | High | High | 0 | 1.46E-08 | 3.37 |
| FALSE | High | SLDGAAAA | [R].SLDGA | 1xDeamida | A0A068WA | 6.81E-07 | 0 | 1 | 10 | 1 | A0A068WA | Protein 4.1 | A0A068WA | 1 | 4144.096 | 0 | 1.17 | 0.001455 | 6.43 | 62.6 | 137.4 | 12.77 | 9.94 | | High | High | High | High | High | High | High | High | High | High | High | 0 | 6.6E-08 | 4.2 |
| FALSE | High | SLDGEVGT | [R].SLDGEV | 1xPhospho | Q9WV92-7 | 0.00086 | 0 | 1 | 3 | 2 | Q9WV92-7 | Isoform 7 c | Q9WV92-7 | 0 | 2065.025 | 0 | -1 | 0.009912 | 6.37 | 135.1 | 64.9 | 12.17 | 18.03 | | High | High | High | High | High | High | High | High | High | High | High | 0.002589 | 0.02777 | 2.23 |
| FALSE | High | SLDGLSGA | [R].SLDGLS | 1xCarbami | Q9D6W8 1 | 9.49E-10 | 0 | 1 | 1 | 2 | Q9D6W8 | BLOC-1-rel | Q9D6W8 [ | 0 | 2507.067 | 0 | 0.4 | 0.009796 | 3.05 | 86.3 | 113.7 | 7.83 | 7.42 | | High | High | High | High | High | High | High | High | High | High | High | 0 | 4.06E-11 | 4.73 |
| FALSE | High | SLDGSGSEI | [R].SLDGS | 2xCarbami | P59178 1x | 0.000751 | 0 | 2 | 6 | 1 | P59178; Q | Lethal(3)m | P59178 [7 | 0 | 2507.081 | 0 | -0.23 | 0.101015 | 13.51 | 112.9 | 87.1 | 9.61 | 12.33 | Shared | High | High | High | High | High | High | High | High | High | High | High | 0 | 0.000172 | 2.14 |
| FALSE | High | SLDLDSIIAE | [R].SLDLDS | 1xPhospho | P50446 1x | 0.001301 | 0 | 1 | 9 | 2 | P50446 | Keratin, ty | P50446 [31 | 0 | 1841.007 | 1 | 6.64 | | 0 | 2.5 | 197.5 | 72.47 | 13.83 | | Not Found | High | Not Found | High | High | High | High | High | High | High | High | 0 | 0.00032 | 3.27 |
| FALSE | High | SLDLLAALP | [K].SLDLLA | 1xPhospho | A0A0A0M | 0.000393 | 0 | 1 | 2 | 1 | A0A0A0M | Histone-lys | A0A0A0M | 0 | 2496.27 | 0 | 0.3 | 0.117026 | 21.64 | 89.7 | 110.3 | 8.46 | 12.21 | | High | High | High | High | High | High | High | High | High | High | High | 0 | 8.32E-05 | 3.4 |
| FALSE | High | SLDLPSIGG | [K].SLDLPS | 1xPhospho | V9GX40 1x | 0.046687 | 0.00354 | 1 | 4 | 1 | V9GX40 | Synergin g | V9GX40 [9 | 0 | 1854.998 | 0 | 1.18 | 0.00016 | 4.78 | 61.2 | 138.8 | 5.93 | 3.23 | | High | High | High | High | High | High | High | High | High | High | High | 0.001812 | 0.0179 | 2.54 |
| FALSE | High | SLDPDASLT | [R].SLDPDA | 1xPhospho | Q6ZWQ0 1 | 0.000164 | 0 | 1 | 10 | 1 | Q6ZWQ0 | Nesprin-2 | Q6ZWQ0 [ | 0 | 3248.565 | 0 | 2.93 | 0.001518 | 31.04 | 21.5 | 178.5 | 11.77 | 16.9 | | High | High | High | High | High | High | High | High | High | High | High | 0 | 3.12E-05 | 3.87 |
| FALSE | High | SLDQDSPSI | [R].SLDQDS | 1xPhospho | A2AAE1-3 | 0.000505 | 0 | 1 | 10 | 1 | A2AAE1-3 | Isoform 3 c | A2AAE1-3 | 0 | 1514.75 | 0 | -0.76 | 0.000234 | 2.79 | 124.4 | 75.6 | 2.37 | 5.89 | | High | High | High | High | High | High | High | High | High | High | High | 0 | 0.000111 | 1.46 |
| FALSE | High | SLDRLDSVI | [R].SLDRLD | 1xPhospho | B1AZP2 1x | 0.001313 | 0 | 1 | 3 | 1 | B1AZP2 | Disks large | B1AZP2 [4 | 1 | 2243.176 | 0 | -0.9 | 0.001251 | 5.51 | 130.4 | 69.6 | 6.56 | 12.32 | | High | High | High | High | High | High | High | High | High | High | High | 0 | 0.000324 | 2.01 |
| FALSE | High | SLDSDESED | [K].SLDSDE | 2xPhospho | B2RTB0 2x | 1.57E-09 | 0 | 1 | 1 | 14 | B2RTB0 | MCG17262 | B2RTB0 [5 | 0 | 2636.021 | 0 | 0.04 | 0.629281 | 11.64 | 96.6 | 103.4 | 6.73 | 3.8 | | High | High | High | High | High | High | High | High | High | High | High | 0 | 4.17E-10 | 5.8 |
| FALSE | High | SLDSDLSDC | [K].SLDSDL | 1xPhospho | E9PVA6 1x | 7.09E-09 | 0 | 1 | 3 | 2 | E9PVA6 | ARF GTPas | E9PVA6 [4 | 0 | 2750.325 | 0 | 0.41 | 0.023513 | 13.76 | 86.2 | 113.8 | 11.45 | 16.65 | | High | High | High | High | High | High | High | High | High | High | High | 0 | 3.91E-10 | 5.19 |
| FALSE | High | SLDSEGTR | [K].SLDSEG | 1xPhospho | Q6ZQ32 1x | 0.033116 | 0.002195 | 2 | 7 | 1 | Q6ZQ32; F | MKIAA085 | Q6ZQ32 [1 | 0 | 1173.535 | 0 | 2.23 | 0.000307 | 6.77 | 33.3 | 166.7 | 15.58 | 10.06 | Shared | High | High | High | High | High | High | High | High | High | High | High | 0.001151 | 0.01203 | 0.87 |
| FALSE | High | SLDSESISP | [R].SLDSES | 1xPhospho | Q8CGF1 1x | 0.100936 | 0.008097 | 1 | 3 | 1 | Q8CGF1 | Rho GTPas | Q8CGF1 [5 | 0 | 1855.842 | 0 | -0.06 | 0.76384 | 6.71 | 100.7 | 99.3 | 3.43 | 10.23 | | High | High | High | High | High | High | High | High | High | High | High | 0.004086 | 0.04367 | 2.75 |
| FALSE | High | SLDGAEFSI | [R].SLDGAE | 1xPhospho | A2AUK5 1x | 0.000227 | 0 | 1 | 4 | 3 | A2AUK5 | Band 4.1-li | A2AUK5 [4 | 0 | 3096.431 | 0 | -0.94 | 0.004895 | 2.29 | 130.2 | 69.8 | 10.61 | 8.82 | | High | High | High | High | High | High | High | High | High | High | High | 2.58E-05 | 0.000349 | 2.5 |
| FALSE | High | SLDSPSHISI | [R].SLDSPS | 1xPhospho | Q80WE4 1 | 0.001211 | 0 | 1 | 4 | 2 | Q80WE4 | Kinesin-like | Q80WE4 [9 | 0 | 1608.84 | 0 | -0.68 | 0.013655 | 8.56 | 123.1 | 76.9 | 11.74 | 11.48 | | High | High | High | High | High | High | High | High | High | High | High | 0.001597 | 0.01539 | 2.05 |
| FALSE | High | SLALSATSS | [K].SLALSA | 1xPhospho | Q8CJF7 1x | 6.09E-06 | 0 | 1 | 1 | 1 | Q8CJF7 | Protein ELY | Q8CJF7 [11 | 0 | 2461.258 | 0 | -0.63 | 0.000786 | 3 | 121.4 | 78.6 | 6.09 | 16.46 | | High | High | High | High | High | High | High | High | High | High | High | 0 | 7.77E-07 | 3.1 |
| FALSE | High | SLAFDSEHS | [K].SLAFDS | 1xPhospho | E9QAE4 1x | 0.010758 | 0.000772 | 1 | 2 | 1 | E9QAE4 | Histone-lys | E9QAE4 [4 | 0 | 1973.926 | 0 | -1.3 | 0.001904 | 3.97 | 142.7 | 57.3 | 9.99 | 8.59 | | High | High | High | High | High | High | High | High | High | High | High | 0.000399 | 0.003406 | 1.96 |
| FALSE | High | SKPEDLPSD | [K].SKPEDL | 1xPhospho | Q9ERU9 1x | 0.014797 | 0.001049 | 1 | 1 | 1 | Q9ERU9 | E3 SUMO- | Q9ERU9 [2 | 0 | 3608.896 | 0 | -0.44 | 0.003048 | 4.54 | 118.2 | 81.8 | 8.18 | 9.7 | | High | High | High | High | High | High | High | High | High | High | High | 0.000555 | 0.004872 | 3.27 |
| FALSE | High | SKPEESRSP | [K].SKPEES | 1xPhospho | Q91VY5 1x | 0.000668 | 0 | 1 | 3 | 1 | Q91VY5 | Lysine-spe | Q91VY5 [4 | 1 | 2912.44 | 0 | -1.09 | 0.000176 | 2.24 | 136.1 | 63.9 | 3.83 | 11.19 | | High | High | High | High | High | High | High | High | High | High | High | 0 | 0.000152 | 2.48 |
| FALSE | High | SKPELPPGL | [K].SKPELP | 1xPhospho | Q8CHP5 1x | 2.38E-08 | 0 | 1 | 2 | 1 | Q8CHP5 | Partner of | Q8CHP5 [5 | 0 | 3884.066 | 0 | 0.41 | 0.024185 | 17.92 | 86.4 | 113.6 | 9.4 | 5.53 | | High | High | High | High | High | High | High | High | High | High | High | 0 | 1.53E-09 | 4.12 |
| FALSE | High | SKPEPSTVS | [R].SKPEPS | 1xPhospho | O89091 1x | 0.034878 | 0.002269 | 1 | 5 | 1 | O89091 | Krueppel-li | O89091 [2 | 0 | 2243.141 | 0 | -0.75 | 0.002535 | 1.58 | 122.7 | 77.3 | 5.11 | 7 | | High | High | High | High | High | High | High | High | High | High | High | 0.001183 | 0.01277 | 2.77 |
| FALSE | High | SKPLDQSVI | [K].SKPLDQ | 1xPhospho | A0A1W2P6 | 0.000349 | 0 | 1 | 4 | 1 | A0A1W2P6 | Formin-like | A0A1W2P6 | 0 | 2213.203 | 0 | 0.07 | 0.949108 | 4.47 | 104.3 | 95.7 | 8.45 | 13.77 | | High | High | High | High | High | High | High | High | High | High | High | 0 | 7.31E-05 | 1.41 |
| FALSE | High | SKPIPIMPA | [K].SKPIPIM | 1xPhospho | Q8K1M6 1 | 1.17E-05 | 0 | 1 | 6 | 1 | Q8K1M6 | Dynamin-1 | Q8K1M6 [6 | 0 | 2161.242 | 0 | 0.44 | 0.005194 | 1.35 | 84.6 | 115.4 | 1.8 | 6.94 | | High | High | High | High | High | High | High | High | High | High | High | 0 | 1.6E-06 | 3.79 |
| FALSE | High | SKPLTPSTG | [K].SKPLTP | 1xPhospho | Q69ZA1 1x | 0.002004 | 5.66E-05 | 1 | 3 | 1 | Q69ZA1 | Cyclin-dep | Q69ZA1 [5 | 0 | 1854.07 | 0 | 0.21 | 0.146469 | 17.41 | 92.7 | 107.3 | 6.99 | 5.92 | | High | High | High | High | High | High | High | High | High | High | High | 2.58E-05 | 0.00052 | 1.63 |
| FALSE | High | SKPNSPILH | [K].SKPNSP | 1xPhospho | P59941 1x | 0.028824 | 0.001829 | 1 | 2 | 1 | P59941 | NAD-depe | P59941 [31 | 0 | 2238.309 | 0 | -0.36 | 0.028445 | 5.84 | 110.4 | 89.6 | 7.26 | 6.37 | | High | High | High | High | High | High | High | High | High | High | High | 0.000997 | 0.01029 | 1.43 |
| FALSE | High | SKPSPPPSE | [R].SKPSPP | 1xPhospho | A0A068F1 | 0.005565 | 0.00062 | 1 | 1 | 1 | A0A068F1 | Glyco-gag | A0A068F1 | 0 | 2326.261 | 0 | 3.26 | 0.001394 | 33.92 | 19.9 | 180.1 | 42.82 | 14.55 | | High | High | High | High | High | High | High | High | High | High | High | 0.000275 | 0.001625 | 2.25 |
| FALSE | High | SKSDATASI | [R].SKSDAT | 1xPhospho | E9Q8A3 1x | 5.03E-05 | 0 | 1 | 3 | 1 | E9Q8A3 | Phosphatic | E9Q8A3 [2 | 1 | 2376.299 | 0 | 0.74 | 0.001335 | 9.4 | 74.8 | 125.2 | 12.16 | 8.05 | | High | High | High | High | High | High | High | High | High | High | High | 0 | 8.33E-06 | 2.76 |
| FALSE | High | SKSDSDTLF | [R].SKSDSD | 1xPhospho | Q3UHZ3 1x | 9.08E-07 | 0 | 1 | 3 | 2 | Q3UHZ3 | DNA (cytos | Q3UHZ3 [1 | 1 | 2553.249 | 0 | -0.29 | 0.051468 | 13.12 | 112.7 | 87.3 | 13.84 | 6.89 | | High | High | High | High | High | High | High | High | High | High | High | 0 | 1.94E-05 | 3.72 |
| FALSE | High | SKSDSDTLS | [R].SKSDSD | 1xPhospho | P13864 1x | 0.000117 | 0 | 1 | 3 | 2 | P13864 | DNA (cytos | P13864 [13 | 1 | 2592.281 | 0 | -0.2 | 0.113982 | 20.54 | 108 | 92 | 14.03 | 9.39 | | High | High | High | High | High | High | High | High | High | High | High | 0 | 2.63E-05 | 3.81 |
| FALSE | High | SKSEASLLC | [K].SKSEAS | 1xPhospho | Q66T02 1x | 3.36E-06 | 0 | 1 | 4 | 1 | Q66T02 | Pleckstrin | Q66T02 [9 | 1 | 2367.305 | 0 | -0.1 | 0.472282 | 8.39 | 103.5 | 96.5 | 5.23 | 4.43 | | High | High | High | High | High | High | High | High | High | High | High | 0 | 3.96E-07 | 3.23 |
| FALSE | High | SLAFDSEHS | [K].SLAFDS | 2xPhospho | E9QAE4 2x | 0.028327 | 0.001791 | 1 | 2 | 2 | E9QAE4 | Histone-lys | E9QAE4 [4 | 0 | 2053.892 | 0 | -1.49 | 0.000158 | 0.86 | 149.3 | 50.7 | 3.33 | 7.32 | | High | High | High | High | High | High | High | High | High | High | High | 0.000968 | 0.01008 | 2.24 |
| FALSE | High | SKSEDMDS | [R].SKSEDM | 1xPhospho | Q7TT18 1x | 0.003253 | 0.00016 | 1 | 3 | 1 | Q7TT18 | Activating | Q7TT18 [5 | 1 | 2109.039 | 0 | -0.44 | 0.066235 | 8.7 | 113.8 | 86.2 | 8.28 | 13.73 | | High | High | High | High | High | High | High | High | High | High | High | 7.19E-05 | 0.000894 | 1.78 |
| FALSE | High | SKSEEAHAI | [K].SKSEEA | 1xPhospho | Q9CY58 1x | 0.002531 | 0.000109 | 2 | 6 | 2 | Q9CY58; Q | Plasminoge | Q9CY58 [3 | 1 | 2666.207 | 0 | 1.04 | 0.186812 | 40.64 | 77.1 | 122.9 | 19.37 | 75.05 | Shared | High | High | High | High | High | High | High | High | High | High | High | 4.94E-05 | 0.000677 | 3.12 |
| FALSE | High | SKSETSLPS | [R].SKSETS | 1xPhospho | Q9DBU1 1x | 0.014033 | 0.001011 | 1 | 5 | 1 | Q9DBU1 | Glycoprote | Q9DBU1 [1 | 1 | 1716.893 | 0 | 1.49 | 0.001538 | 3.47 | 47.5 | 152.5 | 8.64 | 14.06 | | High | High | High | High | High | High | High | High | High | High | High | 0.000538 | 0.004583 | 1.94 |
| FALSE | High | SKSEYELSIF | [K].SKSEYE | 1xPhospho | Q8BWW9 | 0.050397 | 0.003641 | 1 | 5 | 1 | Q8BWW9 | Serine/thre | Q8BWW9 | 1 | 2106.052 | 0 | -0.54 | 0.006742 | 3.48 | 115.7 | 84.3 | 4.72 | 9.8 | | High | High | High | High | High | High | High | High | High | High | High | 0.001916 | 0.01948 | 1.46 |
| FALSE | High | SKSNPDFLK | [K].SKSNPD | 1xPhospho | P97434-2 1 | 0.031989 | 0.00205 | 1 | 10 | 1 | P97434-2 | Isoform 2 c | P97434-2 [ | 1 | 1803.002 | 0 | -0.06 | 0.976313 | 13.4 | 105.6 | 94.4 | 18.31 | 5.69 | | High | High | High | High | High | High | High | High | High | High | High | 0.001126 | 0.01161 | 1.36 |
| FALSE | High | SKSPPKVPI | [K].SKSPPK | 1xPhospho | Q6NXN1 1 | 0.000941 | 0 | 1 | 3 | 1 | Q6NXN1 | SUZ domai | Q6NXN1 [3 | 2 | 3433.943 | 0 | -0.84 | 0.044773 | 8.98 | 128 | 72 | 21.58 | 11.58 | | High | High | High | High | High | High | High | High | High | High | High | 0 | 0.000224 | 2.76 |
| FALSE | High | SKSPPPPEE | [R].SKSPPP | 1xPhospho | Q00PI9 1x | 0.000442 | 0 | 1 | 1 | 2 | Q00PI9 | Heterogen | Q00PI9 [22 | 1 | 2063.103 | 0 | -0.18 | 0.924574 | 4.62 | 103.1 | 96.9 | 7.6 | 14.1 | | High | High | High | High | High | High | High | High | High | High | High | 0 | 9.53E-05 | 2.7 |
| FALSE | High | SKSTVTLAG | [K].SKSTVT | 1xCarbami | A7TU71 1x | 3.43E-07 | 0 | 1 | 2 | 2 | A7TU71 | Protein Shr | A7TU71 [6 | 1 | 2793.28 | 0 | 2.26 | 0.000361 | 8.3 | 36.7 | 163.3 | 9.95 | 7.01 | | High | High | High | High | High | High | High | High | High | High | High | 0 | 3.07E-08 | 3.11 |
| FALSE | High | SKTSPVASC | [K].SKTSPV | 1xPhospho | Q6A065 1x | 0.010568 | 0.000772 | 1 | 3 | 2 | Q6A065 | Centrosom | Q6A065 [1 | 1 | 2004.098 | 0 | 0.95 | 0.004525 | 27.08 | 68.8 | 131.2 | 17.48 | 4.75 | | High | High | High | High | High | High | High | High | High | High | High | 0.000382 | 0.003347 | 2.3 |

| | | | | | | | | | | | | | | | | | | | | | | | | | | | | | | | | | | | | | | |
|---|---|---|---|---|---|---|---|---|---|---|---|---|---|---|---|---|---|---|---|---|---|---|---|---|---|---|---|---|---|---|---|---|---|---|---|---|---|---|
| FALSE | High | SLAALDALI | [R].SLAALD | 1xPhospho | C0HKD9 1x | 9.64E-09 | 0 | 1 | 1 | 9 | C0HKD9 | Microfibril | C0HKD9 [2 | 0 | 3179.435 | 0 | -0.26 | 0.035885 | 12.4 | 108.7 | 91.3 | 3.22 | 14.88 | | High | High | High | High | High | High | High | High | High | High | High | 0 | 5.54E-10 | 4.2 |
| FALSE | High | SLAASGAQ | [K].SLAASG | 1xPhospho | E9QKB2 1x | 1.27E-06 | 0 | 1 | 2 | 1 | E9QKB2 | Intracellula | E9QKB2 [9 | 0 | 2613.307 | 0 | 1.4 | 0.001132 | 32.44 | 50.9 | 149.1 | 26.45 | 8.06 | | High | High | High | High | High | High | High | High | High | High | High | 0 | 1.33E-07 | 3.91 |
| FALSE | High | SIACSPK | [K].SIACSPI | 1xCarbami | Q8BKX6 1x | 0.022973 | 0.001578 | 1 | 1 | 1 | Q8BKX6 | Serine/thre | Q8BKX6 [3 | 0 | 1300.674 | 0 | -1.88 | 0.000266 | 0.28 | 155.3 | 44.7 | 3.89 | 11.78 | | High | High | High | High | High | High | High | High | High | High | High | 0.000847 | 0.007995 | 2.04 |
| FALSE | High | SLAENPSG | [K].SLAENF | 1xPhospho | A2APB8 1x | 0.000633 | 0 | 1 | 3 | 3 | A2APB8 | Targeting p | A2APB8 [6 | 0 | 2582.27 | 0 | -0.47 | 0.034866 | 7.58 | 116.3 | 83.7 | 11.45 | 9.15 | | High | High | High | High | High | High | High | High | High | High | High | 0 | 0.000212 | 3.7 |
| FALSE | High | SLAESGLN | [R].SLAESG | 1xPhospho | Q8CH77 1x | 6.64E-08 | 0 | 1 | 4 | 1 | Q8CH77 | Neuron na | Q8CH77 [4 | 0 | 2351.121 | 0 | 1.1 | 0.001981 | 15.93 | 64.7 | 135.3 | 3.1 | 10.91 | | High | High | High | High | High | High | High | High | High | High | High | 0 | 4.81E-09 | 5.08 |
| FALSE | High | SKSEEAHA | [K].SKSEEA | 1xPhospho | Q9CY58 1x | 1.41E-05 | 0 | 2 | 6 | 5 | Q9CY58; Q | Plasminoge | Q9CY58 [3 | 1 | 2650.212 | 0 | 1.03 | 0.013622 | 18.18 | 71.5 | 128.5 | 6.19 | 20.37 | Shared | High | High | High | High | High | High | High | High | High | High | High | 0 | 1.99E-06 | 2.25 |
| FALSE | High | SLDSPTSSP | [R].SLDSPT | 2xPhospho | Q6PGG2 2x | 5.91E-05 | 0 | 1 | 2 | 6 | Q6PGG2 | GEM-intera | Q6PGG2 [4 | 0 | 1691.724 | 0 | 2.68 | 0.000547 | 1.76 | 26.8 | 173.2 | 9.77 | 10.15 | | High | High | High | High | High | High | High | High | High | High | High | 0 | 9.92E-06 | 3.49 |
| FALSE | High | SLDSSGAG | [K].SLDSSG | 1xPhospho | P59017 1xP | 0.000117 | 0 | 1 | 3 | 3 | P59017 | Bcl-2-like p | P59017 [28 | 0 | 1488.735 | 0 | 1.31 | 0.001823 | 18.84 | 56.6 | 143.4 | 2.4 | 12.37 | | High | High | High | High | High | High | High | High | High | High | High | 0 | 0.000188 | 2.12 |
| FALSE | High | SIDTQTPG | [R].SIDTQT | 1xPhospho | Q3U3E2 1x | 0.003001 | 0.00016 | 1 | 1 | 1 | Q3U3E2 | Protein FAI | Q3U3E2 [3 | 0 | 1727.862 | 0 | -0.41 | 0.054321 | 5.48 | 114.2 | 85.8 | 8.18 | 17.16 | | High | High | High | High | High | High | High | High | High | High | High | 7.19E-05 | 0.000817 | 1.49 |
| FALSE | High | SIGSPESTP | [K].SIGSPE | 1xPhospho | Q8CH77 1x | 0.001767 | 5.66E-05 | 1 | 4 | 2 | Q8CH77 | Neuron na | Q8CH77 [7 | 0 | 1540.802 | 0 | 0.1 | 0.092959 | 6.98 | 98.5 | 101.5 | 7.08 | 6.15 | | High | High | High | High | High | High | High | High | High | High | High | 0.002264 | 0.0236 | 2.58 |
| FALSE | High | SLGSSADLE | [K].SLGSSA | 1xPhospho | Q3U226 1x | 7.89E-05 | 0 | 1 | 3 | 2 | Q3U226 | Putative ur | Q3U226 [3 | 0 | 1682.893 | 0 | 0.79 | 0.001716 | 3.46 | 73.3 | 126.7 | 8.79 | 7.74 | | High | High | High | High | High | High | High | High | High | High | High | 0 | 1.38E-05 | 2.44 |
| FALSE | High | SLGSSHSN | [K].SLGSSH | 1xPhospho | Q80XP8 1x | 0.000124 | 0 | 1 | 2 | 2 | Q80XP8 | Protein FAI | Q80XP8 [1 | 0 | 2233.075 | 0 | -1.09 | 0.043423 | 8.09 | 136 | 64 | 34.94 | 13.24 | | High | High | High | High | High | High | High | High | High | High | High | 0 | 8.16E-05 | 1.94 |
| FALSE | High | SLGTELQD | [K].SLGTEL | 1xPhospho | Q3UTB2 1x | 1.66E-08 | 0 | 1 | 1 | 1 | Q3UTB2 | Putative ur | Q3UTB2 [1 | 0 | 2061.031 | 0 | 3.02 | 0.000292 | 34.32 | 22.4 | 177.6 | 22.18 | 5.85 | | High | High | High | High | High | High | High | High | High | High | High | 0 | 1.02E-09 | 4.71 |
| FALSE | High | SLGTGAPV | [K].SLGTGA | 1xPhospho | B3V097 1xP | 6.05E-07 | 0 | 1 | 6 | 1 | B3V097 | BetaCstF-6 | B3V097 [10 | 0 | 3355.66 | 0 | 0.07 | 0.450734 | 8.49 | 97.5 | 102.5 | 6.68 | 10.68 | | High | High | High | High | High | High | High | High | High | High | High | 0 | 5.76E-08 | 3.75 |
| FALSE | High | SLGTHSPA | [K].SLGTHS | 1xPhospho | E9PVX6 1xP | 0.002293 | 0.000109 | 1 | 3 | 2 | E9PVX6 | Proliferatic | E9PVX6 [49 | 0 | 1647.924 | 0 | 0.24 | 0.217316 | 20.21 | 96.8 | 103.2 | 7.61 | 14.71 | | High | High | High | High | High | High | High | High | High | High | High | 0.002159 | 0.0226 | 1.9 |
| FALSE | High | SLGTHSPA | [K].SLGTHS | 2xPhospho | E9PVX6 2xP | 0.001767 | 5.66E-05 | 1 | 3 | 2 | E9PVX6 | Proliferatic | E9PVX6 [49 | 0 | 1727.89 | 0 | -0.03 | 0.466644 | 5.82 | 101.2 | 98.8 | 4.49 | 4.58 | | High | High | High | High | High | High | High | High | High | High | High | 0.000786 | 0.007018 | 2.45 |
| FALSE | High | SLHGEFSA | [K].SLHGEF | 2xPhospho | Q8C0T5 2x | 8.48E-05 | 0 | 1 | 3 | 1 | Q8C0T5 | Signal-indu | Q8C0T5 [1 | 0 | 2569.106 | 0 | 0.67 | 0.005443 | 13.44 | 78.1 | 121.9 | 9.22 | 8.36 | | High | High | High | High | High | High | High | High | High | High | High | 0 | 1.49E-05 | 3.82 |
| FALSE | High | SLHGEFSA | [K].SLHGEF | 3xPhospho | Q8C0T5 3x | 0.00642 | 0.00062 | 1 | 3 | 1 | Q8C0T5 | Signal-indu | Q8C0T5 [1 | 0 | 2649.072 | 0 | 1.33 | 0.000868 | 7.83 | 58.9 | 141.1 | 10.85 | 12.53 | | High | High | High | High | High | High | High | High | High | High | High | 0.000275 | 0.001919 | 3.31 |
| FALSE | High | SIHIVNDPG | [R].SIHIVNI | 1xPhospho | Q9D279 1x | 9.21E-09 | 0 | 1 | 2 | 4 | Q9D279 | Mitotic inte | Q9D279 [2 | 0 | 2773.381 | 0 | 5.06 | 0.000437 | 50.52 | 4.7 | 195.3 | 55.99 | 11.37 | | High | High | High | High | High | High | High | High | High | High | High | 0 | 5.25E-10 | 5.41 |
| FALSE | High | SIHIVNDPG | [R].SIHIVNI | 2xPhospho | Q9D279 2x | 0.007019 | 0.00062 | 1 | 1 | 1 | Q9D279 | Mitotic inte | Q9D279 [2 | 1 | 3578.718 | 0 | 2.42 | 0.002872 | 36.22 | 31.9 | 168.1 | 14.05 | 19.58 | | High | High | High | High | High | High | High | High | High | High | High | 0.000275 | 0.002109 | 3.58 |
| FALSE | High | SLHPWYGI | [K].SLHPW | 1xPhospho | Q8BGT6 1x | 0.006478 | 0.00062 | 1 | 2 | 2 | Q8BGT6 | MICAL-like | Q8BGT6 [4 | 0 | 2209.146 | 0 | 0.32 | 0.02398 | 3.6 | 89.6 | 110.4 | 4.79 | 6.05 | | High | High | High | High | High | High | High | High | High | High | High | 0.000275 | 0.001931 | 2.34 |
| FALSE | High | SLLDSDDSS | [R].SLLDSD | 1xPhospho | Q6PAL7 1x | 0.007811 | 0.000664 | 1 | 1 | 2 | Q6PAL7 | AT-hook D | Q6PAL7 [8 | 0 | 2905.367 | 0 | -1.18 | 0.002156 | 2.45 | 136.9 | 63.1 | 9.14 | 15.54 | | High | High | High | High | High | High | High | High | High | High | High | 0.000327 | 0.002383 | 3.65 |
| FALSE | High | SLGSLSQGS | [R].SLGSLS | 1xPhospho | A0A0A6YW | 2.52E-06 | 0 | 1 | 4 | 1 | A0A0A6YW | Rap guanin | A0A0A6YW | 0 | 2836.461 | 0 | -0.57 | 0.003186 | 6.16 | 119.6 | 80.4 | 6.76 | 9.61 | | High | High | High | High | High | High | High | High | High | High | High | 0 | 2.88E-07 | 4.32 |
| FALSE | High | SLLECIPAQ | [K].SLLECIF | 1xCarbami | A2CG63 1x | 2.45E-06 | 0 | 1 | 4 | 3 | A2CG63 | AT-rich inte | A2CG63 [4 | 0 | 2157.055 | 0 | 0.28 | 0.017574 | 7.51 | 92.9 | 107.1 | 4.31 | 6.13 | | High | High | High | High | High | High | High | High | High | High | High | 0 | 2.78E-07 | 3.09 |
| FALSE | High | SLLEGQEAI | [R].SLLEGC | 1xDeamida | P19001 1xP | 3.2E-05 | 0 | 1 | 1 | 6 | P19001 | Keratin, typ | P19001 [38 | 0 | 2450.237 | 0 | 4.12 | 0.001516 | 7.1 | 8.9 | 191.1 | 21.31 | 18.92 | | High | High | High | High | High | High | High | High | High | High | High | 0.000467 | 0.004239 | 2.94 |
| FALSE | High | SLLENDQLI | [K].SLLEND | 1xPhospho | Q3UGE9 1x | 1.97E-06 | 0 | 1 | 2 | 3 | Q3UGE9 | Tuftelin-int | Q3UGE9 [4 | 0 | 2908.368 | 0 | -0.13 | 0.215784 | 2.09 | 104.7 | 95.3 | 1.75 | 7.15 | | High | High | High | High | High | High | High | High | High | High | High | 0 | 2.17E-07 | 4.53 |
| FALSE | High | SLLGESAPA | [K].SLLGES | 1xPhospho | A0A087WF | 0.033404 | 0.002195 | 1 | 8 | 1 | A0A087WF | Arf-GAP dc | A0A087WF | 0 | 1988.098 | 0 | 1.43 | 0.005937 | 25.88 | 54 | 146 | 21.37 | 16.57 | | High | High | High | High | High | High | High | High | High | High | High | 0.001151 | 0.01218 | 2.68 |
| FALSE | High | SLLGLSNSE | [R].SLLGLS | 1xDeamida | Q9ERG2 1x | 0.000656 | 0 | 2 | 5 | 1 | Q9ERG2; B | Striatin-3 C | Q9ERG2 [2 | 0 | 2241.142 | 0 | 0.62 | 0.009386 | 9.47 | 77.1 | 122.9 | 11.44 | 9.35 | Shared | High | High | High | High | High | High | High | High | High | High | High | 0 | 0.000148 | 1.95 |
| FALSE | High | SLLGTDLES | [R].SLLGTD | 1xPhospho | E9Q3P4 1x | 0.034579 | 0.002233 | 1 | 4 | 1 | E9Q3P4 | Centromer | E9Q3P4 [1 | 0 | 1597.84 | 0 | -0.11 | 0.651801 | 24.44 | 102.1 | 97.9 | 4.2 | 19.13 | | High | High | High | High | High | High | High | High | High | High | High | 0.001183 | 0.01266 | 1.61 |
| FALSE | High | SLLLADSNG | [R].SLLLAD | 1xPhospho | Q9JM52-3 | 5.82E-06 | 0 | 1 | 9 | 2 | Q9JM52-3 | Isoform 3 c | Q9JM52-3 | 0 | 3521.757 | 0 | 0.81 | 0.002349 | 18.9 | 74.8 | 125.2 | 12.13 | 7.62 | | High | High | High | High | High | High | High | High | High | High | High | 0 | 1.06E-06 | 4.83 |
| FALSE | High | SLLLADSNG | [R].SLLLAD | 1xDeamida | Q9JM52-3 | 0.000308 | 0 | 1 | 9 | 9 | Q9JM52-3 | Isoform 3 c | Q9JM52-3 | 0 | 3522.741 | 0 | 1.45 | 0.00045 | 19.39 | 53.5 | 146.5 | 23.75 | 6.15 | | High | High | High | High | High | High | High | High | High | High | High | 0 | 6.38E-05 | 3.88 |
| FALSE | High | SLLLLPGLEI | [K].SLLLLP( | 1xPhospho | J7I1C3 1xP | 0.000144 | 0 | 1 | 1 | 6 | J7I1C3 | Rif1 OS=M | J7I1C3 [95 | 0 | 4237.022 | 0 | -0.83 | 0.010173 | 3.52 | 126.7 | 73.3 | 6.9 | 19.08 | | High | High | High | High | High | High | High | High | High | High | High | 0.000275 | 0.001984 | 3.44 |
| FALSE | High | SLINQTLSS | [R].SLINQT | 1xPhospho | A2A863-3 | 4.68E-05 | 0 | 1 | 6 | 3 | A2A863-3 | Isoform 3 c | A2A863-3 | 0 | 3188.651 | 0 | 2.45 | 0.000427 | 23.53 | 30.9 | 169.1 | 25.06 | 5.85 | | High | High | High | High | High | High | High | High | High | High | High | 2.58E-05 | 0.000433 | 2.87 |
| FALSE | High | SLLPNSSQI | [K].SLLPNS | 1xPhospho | Q8BTI8 1xP | 0.00625 | 0.00062 | 1 | 3 | 1 | Q8BTI8 | Serine/argi | Q8BTI8 [11 | 0 | 2357.171 | 0 | -0.7 | 0.038104 | 9.04 | 127 | 73 | 16.28 | 33.47 | | High | High | High | High | High | High | High | High | High | High | High | 0.000275 | 0.001852 | 2.55 |
| FALSE | High | SLLPNSSQI | [K].SLLPNS | 1xPhospho | Q8BTI8 1xP | 0.00013 | 0 | 1 | 3 | 1 | Q8BTI8 | Serine/argi | Q8BTI8 [11 | 0 | 2373.166 | 0 | -0.59 | 0.004904 | 5.02 | 122.6 | 77.4 | 10.9 | 9.19 | | High | High | High | High | High | High | High | High | High | High | High | 0 | 2.39E-05 | 2.94 |
| FALSE | High | SILPSEASP | [K].SILPSEA | 1xPhospho | B2RUG8 1x | 0.043981 | 0.003228 | 1 | 5 | 2 | B2RUG8 | Asxl2 prote | B2RUG8 [4 | 0 | 1464.766 | 0 | 0.11 | 0.00669 | 2.43 | 97.6 | 102.4 | 3.16 | 4.33 | | High | High | High | High | High | High | High | High | High | High | High | 0.001665 | 0.0167 | 2.28 |
| FALSE | High | SILPYPVSP | [K].SILPYPV | 1xPhospho | E9QLG3 1x | 9.62E-05 | 0 | 1 | 5 | 2 | E9QLG3 | MAX gene- | E9QLG3 [9 | 0 | 1638.927 | 0 | -0.82 | 0.000408 | 1.3 | 129.2 | 70.8 | 2.26 | 6.4 | | High | High | High | High | High | High | High | High | High | High | High | 0.000275 | 0.001878 | 1.78 |
| FALSE | High | SLLEGQEAI | [R].SLLEGC | 1xPhospho | P19001 1xP | 8.13E-06 | 0 | 1 | 1 | 1 | P19001 | Keratin, typ | P19001 [38 | 0 | 2449.253 | 0 | 6.64 | 0.00126 | 0 | 1.5 | 198.5 | 76.04 | 18.36 | | High | High | High | High | High | High | High | High | High | High | High | 0 | 1.08E-06 | 1.97 |
| FALSE | High | SLGSEEINP | [K].SLGSEE | 1xPhospho | O35188 1x | 0.025073 | 0.001645 | 1 | 2 | 1 | O35188 | Fractalkine | O35188 [2 | 0 | 2381.097 | 0 | 1.41 | 0.00106 | 19.44 | 52 | 148 | 15.5 | 6.71 | | High | High | High | High | High | High | High | High | High | High | High | 0.000872 | 0.008795 | 2.95 |
| FALSE | High | SIGSDSQGI | [R].SIGSDS | 1xPhospho | Q9R0L6-2 | 0.009931 | 0.000736 | 1 | 3 | 1 | Q9R0L6-2 | Isoform 2 c | Q9R0L6-2 | 0 | 1215.557 | 0 | 0.51 | 0.007509 | 6.94 | 75.7 | 124.3 | 7.34 | 10.73 | | High | High | High | High | High | High | High | High | High | High | High | 0.000344 | 0.003106 | 1.61 |
| FALSE | High | SIGSASPGP | [K].SIGSASI | 2xPhospho | Q8VDZ4 2x | 0.004143 | 0.00036 | 1 | 1 | 1 | Q8VDZ4 | Palmitoyltr | Q8VDZ4 [6 | 0 | 2182.014 | 0 | 0.65 | 0.007565 | 3.61 | 78.5 | 121.5 | 5.13 | 10.45 | | High | High | High | High | High | High | High | High | High | High | High | 0.000161 | 0.001174 | 2.36 |
| FALSE | High | SLDVTPGK | [R].SLDVTF | 1xPhospho | Q8CBR1 1x | 0.001373 | 5.66E-05 | 1 | 2 | 1 | Q8CBR1 | Immunoglo | Q8CBR1 [5 | 0 | 1354.738 | 0 | -1.51 | 0.000251 | 1.92 | 147.9 | 52.1 | 2.82 | 14.47 | | High | High | High | High | High | High | High | High | High | High | High | 2.58E-05 | 0.000341 | 1.78 |
| FALSE | High | SLDYLNLDI | [K].SLDYLN | 1xPhospho | Q8VEB3 1x | 0.007538 | 0.00062 | 1 | 1 | 2 | Q8VEB3 | UNC119-bi | Q8VEB3 [1 | 0 | 1618.849 | 0 | 0.63 | 0.015417 | 18.96 | 81.5 | 118.5 | 8.46 | 11.19 | | High | High | High | High | High | High | High | High | High | High | High | 0.000275 | 0.002286 | 1.83 |
| FALSE | High | SLEATDSAI | [R].SLEATD | 1xPhospho | Q61526 1x | 0.000573 | 0 | 1 | 1 | 1 | Q61526 | Receptor t | Q61526 [1 | 0 | 2420.039 | 0 | 2.01 | 0.002453 | 41.02 | 40.4 | 159.6 | 18.37 | 15.78 | | High | High | High | High | High | High | High | High | High | High | High | 0 | 0.000128 | 4.26 |
| FALSE | High | SLEDESQET | [K].SLEDES | 1xPhospho | Q6P5H2-2 | 5.26E-05 | 0 | 1 | 2 | 4 | Q6P5H2-2 | Isoform 2 c | Q6P5H2-2 | 0 | 2237.063 | 0 | 3.3 | 0.000554 | 15.18 | 18.3 | 181.7 | 11.97 | 11.74 | | High | High | High | High | High | High | High | High | High | High | High | 0 | 8.75E-06 | 3.58 |
| FALSE | High | SLEDESQET | [K].SLEDES | 2xPhospho | Q6P5H2-2 | 0.001516 | 5.66E-05 | 1 | 2 | 2 | Q6P5H2-2 | Isoform 2 c | Q6P5H2-2 | 0 | 2317.029 | 0 | 2.5 | 0.00102 | 25.74 | 28.7 | 171.3 | 7.88 | 13.88 | | High | High | High | High | High | High | High | High | High | High | High | 2.58E-05 | 0.00044 | 2.95 |
| FALSE | High | SLEEIYLFSL | [K].SLEEIYL | 1xPhospho | P25444 1xP | 0.013074 | 0.000889 | 1 | 14 | 1 | P25444 | 40S riboso | P25444 [77 | 0 | 2090.159 | 0 | -0.37 | 0.090072 | 11.61 | 112.7 | 87.3 | 13.09 | 12.04 | | High | High | High | High | High | High | High | High | High | High | High | 0.000467 | 0.004216 | 2.28 |
| FALSE | High | SIENETLNK | [K].SIENET | 1xPhospho | Q3TSY9 1x | 0.014158 | 0.001011 | 1 | 7 | 4 | Q3TSY9 | Putative ur | Q3TSY9 [4 | 0 | 1356.661 | 0 | -0.58 | 0.948458 | 30.33 | 91.7 | 108.3 | 56.28 | 64.89 | | High | High | High | High | High | High | High | High | High | High | High | 0.000538 | 0.004641 | 2.5 |
| FALSE | High | SLEPESQQS | [R].SLEPES | 1xPhospho | Q8R1F6 1x | 2.1E-05 | 0 | 1 | 3 | 3 | Q8R1F6 | Protein HID | Q8R1F6 [6 | 0 | 3032.296 | 0 | -1.27 | 0.002839 | 2.8 | 137.4 | 62.6 | 11.65 | 9.53 | | High | High | High | High | High | High | High | High | High | High | High | 0 | 3.1E-06 | 4.02 |
| FALSE | High | SLEQCSSPL | [R].SLEQCS | 1xCarbami | Q99LQ1 1x | 0.112856 | 0.009146 | 1 | 2 | 1 | Q99LQ1 | MAP3K12- | Q99LQ1 [1 | 0 | 1699.829 | 0 | 0.11 | 0.09877 | 11.71 | 92.6 | 107.4 | 5.58 | 9.06 | | High | High | High | High | High | High | High | High | High | High | High | 0.004596 | 0.04986 | 2.51 |
| FALSE | High | SLEQDLSK | [R].SLEQDL | 1xPhospho | Q8BQ33 1x | 0.003433 | 0.00021 | 1 | 1 | 1 | Q8BQ33 | Treslin OS= | Q8BQ33 [1 | 0 | 1228.602 | 0 | -0.98 | 0.000728 | 3.42 | 133.1 | 66.9 | 5.91 | 3.62 | | High | High | High | High | High | High | High | High | High | High | High | 9.42E-05 | 0.000949 | 2.36 |
| FALSE | High | SLEQYDQV | [R].SLEQYD | 1xPhospho | Q60902 1x | 1.03E-05 | 0 | 1 | 4 | 1 | Q60902 | Epidermal | Q60902 [5 | 0 | 3709.841 | 0 | 0.9 | 0.004341 | 23.86 | 69.6 | 130.4 | 11.21 | 12.86 | | High | High | High | High | High | High | High | High | High | High | High | 0 | 1.39E-06 | 4.51 |
| FALSE | High | SIFASPESV | [R].SIFASPI | 1xPhospho | Q3TGV3 1x | 2.65E-05 | 0 | 1 | 2 | 2 | Q3TGV3 | Putative ur | Q3TGV3 [1 | 0 | 1760.924 | 0 | -0.87 | 0.000965 | 3.16 | 128.5 | 71.5 | 6.32 | 8.78 | | High | High | High | High | High | High | High | High | High | High | High | 0 | 0.000256 | 3.49 |
| FALSE | High | SLFCGSPPL | [R].SLFCGS | 1xCarbami | Q9R0A1 1x | 1.53E-05 | 0 | 1 | 3 | 1 | Q9R0A1 | Chloride ch | Q9R0A1 [7 | 0 | 2403.192 | 0 | -0.07 | 0.559433 | 22.54 | 106.4 | 93.6 | 14.34 | 15.18 | | High | High | High | High | High | High | High | High | High | High | High | 0 | 2.18E-06 | 3 |
| FALSE | High | SLFFGSILAI | [R].SLFFGS | 1xDeamida | Q9Z0F6 1xI | 0.001313 | 0 | 1 | 1 | 2 | Q9Z0F6 | Cell cycle c | Q9Z0F6 [3 | 0 | 3247.54 | 0 | 0.09 | 0.937478 | 22.65 | 98.3 | 101.7 | 8.28 | 17.49 | | High | High | High | High | High | High | High | High | High | High | High | 0 | 0.000325 | 4.44 |
| FALSE | High | SIFREESPLF | [K].SIFREES | 1xPhospho | Q8K019 1x | 0.026652 | 0.001676 | 1 | 9 | 1 | Q8K019 | Bcl-2-assoc | Q8K019 [5 | 1 | 1542.788 | 0 | -0.36 | 0.014256 | 2.22 | 111.5 | 88.5 | 7.11 | 4.77 | | High | High | High | High | High | High | High | High | High | High | High | 0.000903 | 0.009383 | 1.54 |
| FALSE | High | SIFSSEK | [K].SIFSSEK | 1xPhospho | Q9ERU9 1x | 0.006085 | 0.00062 | 1 | 1 | 1 | Q9ERU9 | E3 SUMO-p | Q9ERU9 [2 | 0 | 1335.696 | 0 | -0.71 | 0.002198 | 1.98 | 124.1 | 75.9 | 5.25 | 8.12 | | High | High | High | High | High | High | High | High | High | High | High | 0.000275 | 0.001804 | 1.76 |
| FALSE | High | SLFTSQSSG | [K].SLFTSQ | 1xPhospho | Q69Z38 1x | 0.00967 | 0.000736 | 1 | 1 | 3 | Q69Z38 | Pseudopod | Q69Z38 [8 | 0 | 2918.382 | 0 | -0.2 | 0.523043 | 8.09 | 107 | 93 | 11.93 | 14.29 | | High | High | High | High | High | High | High | High | High | High | High | 0.000656 | 0.005824 | 3.07 |
| FALSE | High | SLGACHLLC | [R].SLGACH | 1xCarbami | B5B2N2 1x | 0.023379 | 0.001578 | 1 | 11 | 1 | B5B2N2 | Nuclear fac | B5B2N2 [2 | 0 | 1576.787 | 0 | -0.03 | 0.877081 | 4.93 | 100.6 | 99.4 | 6.46 | 9.89 | | High | High | High | High | High | High | High | High | High | High | High | 0.000847 | 0.008125 | 1.98 |
| FALSE | High | SLGDASPSA | [K].SLGDAS | 1xPhospho | Q0PHV7 1x | 0.002058 | 5.66E-05 | 1 | 2 | 1 | Q0PHV7 | Dapper ho | Q0PHV7 [1 | 0 | 1809.858 | 0 | -1.03 | 0.003667 | 3.36 | 134.2 | 65.8 | 10.43 | 15.32 | | High | High | High | High | High | High | High | High | High | High | High | 2.58E-05 | 0.000535 | 2.03 |
| FALSE | High | SLGDSLPST | [R].SLGDSL | 1xCarbami | E9Q7C4 1x | 0.021797 | 0.001507 | 1 | 2 | 1 | E9Q7C4 | Proline-rich | E9Q7C4 [4 | 0 | 2450.162 | 0 | 0.34 | 0.086038 | 34.51 | 86 | 114 | 20.63 | 19.7 | | High | High | High | High | High | High | High | High | High | High | High | 0.000814 | 0.007505 | 2.96 |
| FALSE | High | SLGEDPDSI | [K].SLGEDF | 1xPhospho | Q80UU6 1x | 5.25E-08 | 0 | 1 | 4 | 2 | Q80UU6 | Opioid gro | Q80UU6 [6 | 0 | 2723.144 | 0 | 0.54 | 0.04283 | 17.1 | 81.4 | 118.6 | 17.37 | 14.78 | | High | High | High | High | High | High | High | High | High | High | High | 0 | 3.7E-09 | 5.15 |
| FALSE | High | SLGEDPDSI | [K].SLGEDF | 1xPhospho | Q80UU6 1x | 5.84E-07 | 0 | 1 | 4 | 1 | Q80UU6 | Opioid gro | Q80UU6 [6 | 0 | 2739.139 | 0 | -0.4 | 0.090944 | 7.96 | 114.2 | 85.8 | 12.36 | 11.07 | | High | High | High | High | High | High | High | High | High | High | High | 0 | 5.57E-08 | 4.89 |
| FALSE | High | SLGEVEWE | [R].SLGEVE | 1xPhospho | Q6P5H2-2 | 0.004572 | 0.000456 | 1 | 2 | 2 | Q6P5H2-2 | Isoform 2 c | Q6P5H2-2 | 0 | 2168.022 | 0 | 1.09 | 0.030247 | 15.14 | 70.4 | 129.6 | 13.07 | 31.46 | | High | High | High | High | High | High | High | High | High | High | High | 0.000204 | 0.00131 | 3.28 |
| FALSE | High | SLGFLPQPT | [R].SLGFLP | 1xPhospho | A0A0R4J2D | 0.005615 | 0.00062 | 1 | 3 | 1 | A0A0R4J2D | Ubiquitin c | A0A0R4J2D | 0 | 2022.144 | 0 | -0.47 | 0.001251 | 1.96 | 116.8 | 83.2 | 6.11 | 5.89 | | High | High | High | High | High | High | High | High | High | High | High | 0.000275 | 0.001641 | 3.37 |
| FALSE | High | SLGFVGSP | [R].SLGFVG | 1xCarbami | Q8K2W9 1 | 0.028082 | 0.001791 | 1 | 2 | 1 | Q8K2W9 | Uncharacte | Q8K2W9 [2 | 0 | 1388.66 | 0 | -0.65 | 0.000569 | 1.74 | 121.6 | 78.4 | 5.11 | 6.43 | | High | High | High | High | High | High | High | High | High | High | High | 0.000968 | 0.009953 | 1.33 |
| FALSE | High | SLGGQQGS | [R].SLGGQ | 1xPhospho | E9PUA3 1x | 0.002313 | 0.000109 | 1 | 4 | 1 | E9PUA3 | IQ motif ar | E9PUA3 [3 | 0 | 1496.787 | 0 | 0.26 | 0.022477 | 2.74 | 89.6 | 110.4 | 3.6 | 7.57 | | High | High | High | High | High | High | High | High | High | High | High | 4.94E-05 | 0.000609 | 1.93 |
| FALSE | High | SLGHSLSIE | [R].SLGHSL | 1xCarbami | Q99MY8 1 | 0.010382 | 0.000772 | 1 | 1 | 1 | Q99MY8 | Histone-lys | Q99MY8 [6 | 0 | 1768.907 | 0 | 1.62 | 9.77E-05 | 11.79 | 49.2 | 150.8 | 13.42 | 3.55 | | High | High | High | High | High | High | High | High | High | High | High | 0.000382 | 0.003281 | 2.01 |
| FALSE | High | SLGLPSCIP | [K].SLGLPS | 2xCarbami | Q9DAP7 1x | 0.001503 | 5.66E-05 | 1 | 1 | 3 | Q9DAP7 | Histone ch | Q9DAP7 [1 | 0 | 2498.158 | 0 | 0.32 | 0.034729 | 22.32 | 85.1 | 114.9 | 12.41 | 12.51 | | High | High | High | High | High | High | High | High | High | High | High | 2.58E-05 | 0.000377 | 2.71 |
| FALSE | High | SIGSASPGP | [K].SIGSASI | 1xPhospho | Q8VDZ4 1x | 1.64E-07 | 0 | 1 | 1 | 2 | Q8VDZ4 | Palmitoyltr | Q8VDZ4 [6 | 0 | 2102.048 | 0 | 0.02 | 0.685034 | 6.25 | 99.9 | 100.1 | 6.99 | 4.07 | | High | High | High | High | High | High | High | High | High | High | High | 0 | 1.33E-08 | 3.53 |
| FALSE | High | SKPAGSDG | [K].SKPAGS | 1xPhospho | Q00PI9 1xP | 0.014158 | 0.001011 | 1 | 1 | 2 | Q00PI9 | Heterogen | Q00PI9 [18 | 1 | 1697.874 | 0 | 0.34 | 0.086759 | 23.53 | 88.4 | 111.6 | 12.03 | 9.81 | | High | High | High | High | High | High | High | High | High | High | High | 0.000538 | 0.004613 | 1.3 |
| FALSE | High | SKPAGSDG | [K].SKPAGS | 1xPhospho | Q00PI9 1xP | 0.001783 | 5.66E-05 | 1 | 1 | 1 | Q00PI9 | Heterogen | Q00PI9 [18 | 0 | 1541.773 | 0 | 0.48 | 0.00507 | 9.09 | 83.4 | 116.6 | 6.64 | 5.66 | | High | High | High | High | High | High | High | High | High | High | High | 2.58E-05 | 0.000458 | 1.1 |
| FALSE | High | SKPAAADS | [K].SKPAA | 1xPhospho | Q6P542 1x | 1.27E-07 | 0 | 1 | 1 | 16 | Q6P542 | ATP-bindin | Q6P542 [1 | 0 | 2760.307 | 0 | -0.41 | 0.000218 | 1.55 | 114 | 86 | 2.45 | 4.68 | | High | High | High | High | High | High | High | High | High | High | High | 0 | 1E-08 | 4.84 |
| FALSE | High | SKPAAADS | [K].SKPAA | 1xPhospho | Q6P542 1x | 2.98E-05 | 0 | 1 | 1 | 1 | Q6P542 | ATP-bindin | Q6P542 [1 | 0 | 2531.144 | 0 | -0.5 | 0.6552 | 36.4 | 132.6 | 67.4 | 113.87 | 121.14 | | High | High | High | High | High | High | High | High | High | High | High | 0 | 4.62E-06 | 3.41 |
| FALSE | High | SGTSTPTTF | [K].SGTSTF | 2xPhospho | P20357 2xP | 0.002192 | 0.000109 | 1 | 8 | 8 | P20357 | Microtubu | P20357 [15 | 0 | 2797.253 | 0 | 0.7 | 0.000402 | 4.88 | 77.9 | 122.1 | 6.93 | 5.28 | | High | High | High | High | High | High | High | High | High | High | High | 4.94E-05 | 0.000576 | 3.86 |
| FALSE | High | SGTTPKPV | [R].SGTTPK | 1xPhospho | A0A1W2P7 | 0.086562 | 0.006435 | 1 | 6 | 2 | A0A1W2P7 | Uncharacte | A0A1W2P7 | 0 | 2022.079 | 0 | -0.13 | 0.399286 | 11.49 | 103.9 | 96.1 | 8.15 | 8.46 | | High | High | High | High | High | High | High | High | High | High | High | 0.003317 | 0.0365 | 1.48 |
| FALSE | High | SGVAEAPG | [R].SGVAE | 1xPhospho | D3YZC8 1xP | 0.00642 | 0.00062 | 1 | 4 | 1 | D3YZC8 | Bromodom | D3YZC8 [4 | 0 | 1497.771 | 0 | -0.08 | 0.497645 | 19.12 | 102.8 | 97.2 | 13.49 | 16.46 | | High | High | High | High | High | High | High | High | High | High | High | 0.000275 | 0.001909 | 1.98 |
| FALSE | High | SGVAVPTS | [R].SGVAV | 1xPhospho | Q61823 1x | 0.001767 | 5.66E-05 | 1 | 1 | 3 | Q61823 | Programm | Q61823 [8 | 0 | 1480.818 | 0 | -1.58 | 0.000184 | 1.91 | 153 | 47 | 5.38 | 15.37 | | High | High | High | High | High | High | High | High | High | High | High | 2.58E-05 | 0.000453 | 2.68 |
| FALSE | High | SGVGCSLP | [K].SGVGC | 1xCarbami | Q8BGV4 1x | 0.000132 | 0 | 1 | 3 | 1 | Q8BGV4 | TELO2-inte | Q8BGV4 [1 | 0 | 2696.341 | 0 | -0.59 | 0.000492 | 1 | 120.3 | 79.7 | 8.19 | 7.73 | | High | High | High | High | High | High | High | High | High | High | High | 0 | 2.46E-05 | 4.45 |
| FALSE | High | SGVGTGPP | [R].SGVGT | 1xPhospho | Q5U4C9 1x | 0.052576 | 0.00384 | 1 | 1 | 1 | Q5U4C9 | Dual specif | Q5U4C9 [4 | 0 | 1925.046 | 0 | 0.81 | 0.001005 | 12.32 | 72 | 128 | 7.05 | 4.89 | | High | High | High | High | High | High | High | High | High | High | High | 0.001999 | 0.02033 | 1.67 |
| FALSE | High | SGVLGTQF | [K].SGVLG | 1xPhospho | A0A0N4SU | 5.88E-06 | 0 | 1 | 4 | 1 | A0A0N4SU | Peregrin O | A0A0N4SU | 0 | 2672.365 | 0 | 0.42 | 0.020402 | 8.67 | 85.5 | 114.5 | 2.68 | 9.72 | | High | High | High | High | High | High | High | High | High | High | High | 0 | 7.42E-07 | 3.8 |
| FALSE | High | SGVSITIDD | [K].SGVSIT | 1xPhospho | Q9JIX8-3 1 | 0.001127 | 0 | 3 | 11 | 1 | Q9JIX8-3; | Isoform 3 c | Q9JIX8-3 [ | 0 | 1567.793 | 0 | 0.65 | 0.006124 | 7.8 | 75.5 | 124.5 | 7.7 | 11.58 | Shared | High | High | High | High | High | High | High | High | High | High | High | 0 | 0.000273 | 2.52 |
| FALSE | High | SGVSPK | [K].SGVSPK | 1xPhospho | F8VQD1 1x | 0.108463 | 0.008839 | 1 | 13 | 1 | F8VQD1 | Protein pol | F8VQD1 [6 | 0 | 1112.612 | 0 | -0.2 | 0.034291 | 3.33 | 106.8 | 93.2 | 5.19 | 6.88 | | High | High | High | High | High | High | High | High | High | High | High | 0.004422 | 0.04768 | 1.29 |
| FALSE | High | SGVTNMSS | [R].SGVTN | 1xPhospho | Q8R3D1 1x | 0.010475 | 0.000772 | 1 | 1 | 1 | Q8R3D1 | TBC1 doma | Q8R3D1 [1 | 0 | 1682.834 | 0 | -1.4 | 0.000209 | 0.66 | 145.1 | 54.9 | 6.86 | 14.75 | | High | High | High | High | High | High | High | High | High | High | High | 0.000382 | 0.003297 | 1.8 |
| FALSE | High | SGVTNMSS | [R].SGVTN | 1xPhospho | Q8R3D1 1x | 0.029586 | 0.001857 | 1 | 1 | 4 | Q8R3D1 | TBC1 doma | Q8R3D1 [1 | 0 | 1698.829 | 0 | -1.15 | 0.00046 | 1.75 | 139.3 | 60.7 | 7.81 | 11.23 | | High | High | High | High | High | High | High | High | High | High | High | 0.009162 | 0.09911 | 2.32 |
| FALSE | High | SGYMSDSD | [K].SGYMS | 1xPhospho | Q8CH77 1x | 0.000738 | 0 | 1 | 3 | 2 | Q8CH77 | Neuron na | Q8CH77 [3 | 0 | 1844.821 | 0 | 0.77 | 0.046861 | 15.43 | 78 | 122 | 14.47 | 20.65 | | High | High | High | High | High | High | High | High | High | High | High | 0.000399 | 0.003548 | 1.83 |
| FALSE | High | SGYSSPGSI | [R].SGYSSP | 1xPhospho | A2A5Y6 1x | 8.86E-08 | 0 | 1 | 10 | 6 | A2A5Y6 | Microtubu | A2A5Y6 [5 | 0 | 1702.764 | 0 | -0.29 | 0.011559 | 4.84 | 110 | 90 | 3.68 | 6.16 | | High | High | High | High | High | High | High | High | High | High | High | 0.000689 | 0.00592 | 2.26 |
| FALSE | High | SGTSTPTTF | [K].SGTSTF | 1xPhospho | P20357 1xP | 9E-07 | 0 | 1 | 8 | 5 | P20357 | Microtubu | P20357 [15 | 0 | 2717.287 | 0 | 1.85 | 0.000675 | 9.73 | 44.8 | 155.2 | 11.08 | 11.16 | | High | High | High | High | High | High | High | High | High | High | High | 0 | 9.07E-08 | 4.24 |
| FALSE | High | SGYSSPGSI | [R].SGYSSP | 2xPhospho | A2A5Y6 2x | 0.002623 | 0.000109 | 1 | 10 | 5 | A2A5Y6 | Microtubu | A2A5Y6 [5 | 0 | 1782.73 | 0 | 0.4 | 0.003695 | 2.63 | 85.6 | 114.4 | 3.76 | 5.58 | | High | High | High | High | High | High | High | High | High | High | High | 0.000275 | 0.001588 | 2.54 |
| FALSE | High | SHAEGPQR | [R].SHAEGI | 2xPhospho | A0A0G2JG | 0.020499 | 0.001472 | 1 | 2 | 1 | A0A0G2JG | Crystallin b | A0A0G2JG | 0 | 2592.181 | 0 | 6.64 | | 0 | 4.2 | 195.8 | 3.65 | 13.6 | | Not Found | Not Found | High | Not Found | High | High | High | High | High | High | High | 0.000786 | 0.007041 | 1.61 |
| FALSE | High | SHAPFSPDI | [K].SHAPFS | 1xPhospho | Q571C7 1x | 1.6E-06 | 0 | 1 | 3 | 2 | Q571C7 | Transcripti | Q571C7 [1 | 0 | 1978.979 | 0 | -0.56 | 0.000783 | 4.21 | 118.7 | 81.3 | 3.19 | 7.56 | | High | High | High | High | High | High | High | High | High | High | High | 0 | 1.73E-07 | 3.48 |
| FALSE | High | SHAPWEEN | [K].SHAPW | 1xPhospho | P52963 1xP | 0.000454 | 0 | 1 | 2 | 2 | P52963 | Band 4.1-li | P52963 [41 | 0 | 2627.136 | 0 | 0.04 | 0.936352 | 4.64 | 100.1 | 99.9 | 13.11 | 6.25 | | High | High | High | High | High | High | High | High | High | High | High | 0 | 9.83E-05 | 3.12 |
| FALSE | High | SHATSPGSI | [R].SHATSF | 1xPhospho | Q8BMK0 1 | 7.72E-07 | 0 | 1 | 3 | 4 | Q8BMK0 | Centrosom | Q8BMK0 [ | 0 | 1863.973 | 0 | -0.97 | 0.011856 | 5.24 | 132.3 | 67.7 | 11.86 | 14.08 | | High | High | High | High | High | High | High | High | High | High | High | 0 | 1.3E-07 | 2.91 |
| FALSE | High | SHCHSPDIE | [R].SHCHSF | 1xCarbami | Q8CI03-2 1 | 2.56E-05 | 0 | 1 | 3 | 1 | Q8CI03-2 | Isoform 2 c | Q8CI03-2 [ | 0 | 1915.868 | 0 | -0.06 | 0.291612 | 19.75 | 104.3 | 95.7 | 10.06 | 10.12 | | High | High | High | High | High | High | High | High | High | High | High | 0 | 3.9E-06 | 2.52 |
| FALSE | High | SHDFYSHEI | [R].SHDFYS | 1xPhospho | P53349 1xP | 0.004033 | 0.000311 | 1 | 1 | 1 | P53349 | Mitogen-ac | P53349 [49 | 0 | 2554.181 | 0 | -0.86 | 0.002273 | 6.45 | 131.1 | 68.9 | 10.04 | 27.86 | | High | High | High | High | High | High | High | High | High | High | High | 0.000161 | 0.001139 | 3.03 |
| FALSE | High | SHDFYSHEI | [R].SHDFYS | 1xPhospho | F8VQ72 1x | 0.034579 | 0.002233 | 1 | 1 | 2 | F8VQ72 | Mitogen-ac | F8VQ72 [4 | 0 | 2584.192 | 0 | -0.1 | 0.943683 | 21.9 | 100.7 | 99.3 | 15.32 | 13.08 | | High | High | High | High | High | High | High | High | High | High | High | 0.003188 | 0.03354 | 2.9 |
| FALSE | High | SHEAEVLK | [K].SHEAEV | 1xPhospho | Q545B6 1x | 0.003892 | 0.000261 | 1 | 8 | 2 | Q545B6 | Stathmin C | Q545B6 [6 | 0 | 1221.608 | 0 | -1.25 | 0.013723 | 7.12 | 143 | 57 | 22.95 | 22.67 | | High | High | High | High | High | High | High | High | High | High | High | 0.000538 | 0.004619 | 2.42 |
| FALSE | High | SHEEVNTEI | [R].SHEEVN | 1xPhospho | Q6PCM2 1 | 0.072858 | 0.005599 | 1 | 2 | 1 | Q6PCM2 | Integrator | Q6PCM2 [8 | 0 | 1494.704 | 0 | -0.29 | 0.941841 | 33 | 110.1 | 89.9 | 21.6 | 45.45 | | High | High | High | High | High | High | High | High | High | High | High | 0.002867 | 0.02989 | 2.83 |
| FALSE | High | SHELDVDEI | [R].SHELDV | 1xPhospho | Q923W1 1 | 5.66E-09 | 0 | 1 | 1 | 3 | Q923W1 | Trimethylg | Q923W1 [4 | 0 | 3229.498 | 0 | -0.85 | 0.013136 | 8.67 | 130 | 70 | 8.46 | 24.1 | | High | High | High | High | High | High | High | High | High | High | High | 0 | 3.02E-10 | 5.42 |
| FALSE | High | SHESLLSPC | [R].SHESLL | 2xCarbami | E9PW37 1x | 8.22E-07 | 0 | 1 | 6 | 1 | E9PW37 | RAS protei | E9PW37 [2 | 0 | 2439.125 | 0 | 0.41 | 0.019358 | 4.58 | 80.9 | 119.1 | 15.78 | 13.85 | | High | High | High | High | High | High | High | High | High | High | High | 0 | 8.17E-08 | 4.28 |
| FALSE | High | SHETDGGS | [K].SHETD | 1xPhospho | Q9ERU9 1x | 0.003526 | 0.00021 | 1 | 1 | 3 | Q9ERU9 | E3 SUMO-p | Q9ERU9 [1 | 0 | 3608.622 | 0 | 0.35 | 0.114958 | 9.67 | 89.2 | 110.8 | 11.05 | 13.62 | | High | High | High | High | High | High | High | High | High | High | High | 9.42E-05 | 0.000982 | 3.22 |
| FALSE | High | SHETDGGS | [K].SHETD | 2xPhospho | Q9ERU9 2x | 0.004256 | 0.00036 | 1 | 1 | 1 | Q9ERU9 | E3 SUMO-p | Q9ERU9 [1 | 0 | 3688.588 | 0 | -0.01 | 0.457954 | 13.83 | 98 | 102 | 8.27 | 8.28 | | High | High | High | High | High | High | High | High | High | High | High | 0.000161 | 0.001209 | 3.83 |
| FALSE | High | SHAEGPQR | [R].SHAEGI | 1xPhospho | A0A0G2JG | 0.000221 | 0 | 1 | 2 | 5 | A0A0G2JG | Crystallin b | A0A0G2JG | 0 | 2512.214 | 0 | 2.57 | 0.000941 | 31.41 | 28 | 172 | 22.36 | 14.01 | | High | High | High | High | High | High | High | High | High | High | High | 0 | 4.36E-05 | 3.52 |
| FALSE | High | SGTSSPQSI | [R].SGTSSP | 1xPhospho | Q80YS6 1x | 0.000233 | 0 | 1 | 1 | 2 | Q80YS6 | Actin filam | Q80YS6 [6 | 0 | 1558.746 | 0 | 0.6 | 0.000627 | 9.44 | 77.5 | 122.5 | 7.44 | 2.18 | | High | High | High | High | High | High | High | High | High | High | High | 0 | 4.65E-05 | 1.85 |
| FALSE | High | SGTSQEELI | [R].SGTSQI | 1xPhospho | A0A087WC | 0.082424 | 0.006288 | 1 | 9 | 2 | A0A087WC | Kinesin-like | A0A087WC | 0 | 1315.609 | 0 | -0.58 | 0.022388 | 16.58 | 128.9 | 71.1 | 13.2 | 31.22 | | High | High | High | High | High | High | High | High | High | High | High | 0.003258 | 0.0344 | 1.62 |
| FALSE | High | SGTPVHCP | [R].SGTPVH | 1xCarbami | A6H663 2x | 0.003464 | 0.00021 | 1 | 3 | 1 | A6H663 | BCL2-assoc | A6H663 [2 | 0 | 1696.748 | 0 | 2.9 | 0.000598 | 14.84 | 23.6 | 176.4 | 21.62 | 11.01 | | High | High | High | High | High | High | High | High | High | High | High | 9.42E-05 | 0.000962 | 2.03 |
| FALSE | High | SGSPMVPS | [R].SGSPM | 1xPhospho | D3Z5N5 1x | 7.43E-06 | 0 | 2 | 4 | 1 | D3Z5N5; Q | 1-phospha | D3Z5N5 [3 | 0 | 2319.053 | 0 | 0.4 | 0.006273 | 12.83 | 90.8 | 109.2 | 11.02 | 6.24 | Shared | High | High | High | High | High | High | High | High | High | High | High | 0 | 9.68E-07 | 4.48 |
| FALSE | High | SGSPSLPQ | [K].SGSPSL | 1xPhospho | Q8C551 1x | 0.000616 | 0 | 1 | 3 | 4 | Q8C551 | RAD51-ass | Q8C551 [2 | 0 | 2063.037 | 0 | 0.07 | 0.23324 | 8.31 | 97.2 | 102.8 | 3.67 | 3.03 | | High | High | High | High | High | High | High | High | High | High | High | 0 | 0.000331 | 3.18 |
| FALSE | High | SGSPVATFI | [K].SGSPVA | 1xPhospho | Q810D6 1x | 0.083101 | 0.006345 | 1 | 4 | 1 | Q810D6 | Glutamate- | Q810D6 [3 | 0 | 1431.765 | 0 | -0.61 | 0.000489 | 3.06 | 122.7 | 77.3 | 4.86 | 4.13 | | High | High | High | High | High | High | High | High | High | High | High | 0.003265 | 0.03482 | 1.65 |
| FALSE | High | SGSQPACL | [K].SGSQPA | 1xCarbami | Q9QZK2 1x | 2.04E-05 | 0 | 1 | 2 | 1 | Q9QZK2 | Breast can | Q9QZK2 [3 | 0 | 1878.837 | 0 | -0.47 | 0.00272 | 3.59 | 118.4 | 81.6 | 3.05 | 9.03 | | High | High | High | High | High | High | High | High | High | High | High | 0 | 3.01E-06 | 3.2 |

| | | | | | | | | | | | | | | | | | | | | | | | | | | | | | | | | | | | | | | |
|---|---|---|---|---|---|---|---|---|---|---|---|---|---|---|---|---|---|---|---|---|---|---|---|---|---|---|---|---|---|---|---|---|---|---|---|---|---|---|
| FALSE | High | SGSSHASQ | [R].SGSSHA | 1xPhosphc | Q14A95 1x | 0.000269 | 0 | 1 | 3 | 2 | Q14A95 | MCG9666, | Q14A95 [8 | 0 | 3880.736 | 0 | 1.01 | 0.413154 | 20.13 | 67.7 | 132.3 | 73.27 | 41.6 | | High | High | High | High | High | High | High | High | High | High | High | 0 | 5.44E-05 | 4.04 |
| FALSE | High | SGSSPEMK | [R].SGSSPE | 1xPhosphc | Q8BTI8 1xP | 0.098533 | 0.007902 | 1 | 3 | 1 | Q8BTI8 | Serine/argi | Q8BTI8 [15 | 0 | 1360.658 | 0 | -0.64 | 0.012596 | 1.32 | 122 | 78 | 5.94 | 25.39 | | High | High | High | High | High | High | High | High | High | High | High | 0.003963 | 0.04233 | 2.45 |
| FALSE | High | SGSSPEMK | [R].SGSSPE | 1xPhosphc | Q8BTI8 1xP | 0.015741 | 0.001124 | 1 | 3 | 10 | Q8BTI8 | Serine/argi | Q8BTI8 [15 | 0 | 1376.653 | 0 | -0.63 | 0.01683 | 0.24 | 121.4 | 78.6 | 8.09 | 20.31 | | High | High | High | High | High | High | High | High | High | High | High | 0.003606 | 0.0383 | 1.95 |
| FALSE | High | SGSSPGLR | [R].SGSSPG | 1xPhosphc | Q8BTI8 1xP | 0.005924 | 0.00062 | 1 | 3 | 6 | Q8BTI8 | Serine/argi | Q8BTI8 [13 | 0 | 1069.524 | 0 | -0.3 | 0.030735 | 11.22 | 110.4 | 89.6 | 5.86 | 7.31 | | High | High | High | High | High | High | High | High | High | High | High | 0.003564 | 0.03805 | 2.08 |
| FALSE | High | SGSSPGLR | [R].SGSSPG | 2xPhosphc | Q8BTI8 2xP | 0.00967 | 0.000736 | 1 | 3 | 3 | Q8BTI8 | Serine/argi | Q8BTI8 [13 | 0 | 1149.49 | 0 | -0.56 | 0.002145 | 1.52 | 119.6 | 80.4 | 9.25 | 6.34 | | High | High | High | High | High | High | High | High | High | High | High | 0.002117 | 0.02212 | 2.16 |
| FALSE | High | SGSSPPK | [KR].SGSSF | 1xPhosphc | Q8BTI8 1xP | 0.034579 | 0.002233 | 1 | 8 | 2 | Q8BTI8 | Serine/argi | Q8BTI8 [79 | 0 | 1197.628 | 0 | 0 | 0.935345 | 14.32 | 99.6 | 100.4 | 5.45 | 4.39 | | High | High | High | High | High | High | High | High | High | High | High | 0.00269 | 0.0287 | 1.29 |
| FALSE | High | SGSSPPK | [KR].SGSSF | 2xPhosphc | Q8BTI8 2xP | 0.079766 | 0.00613 | 1 | 8 | 2 | Q8BTI8 | Serine/argi | Q8BTI8 [79 | 0 | 1277.594 | 0 | -0.45 | 0.006594 | 5.78 | 116.8 | 83.2 | 6.84 | 5.36 | | High | High | High | High | High | High | High | High | High | High | High | 0.003148 | 0.033 | 1.79 |
| FALSE | High | SGSSQELD( | [R].SGSSQE | 2xPhosphc | Q8BTI8 2xP | 0.117405 | 0.009523 | 1 | 3 | 1 | Q8BTI8 | Serine/argi | Q8BTI8 [14 | 0 | 2248.969 | 0 | -0.02 | 0.654252 | 10.32 | 103.5 | 96.5 | 10.06 | 16.26 | | High | High | High | High | High | High | High | High | High | High | High | 0.004769 | 0.0521 | 2.39 |
| FALSE | High | SGSSQELD( | [R].SGSSQE | 1xPhosphc | Q8BTI8 1xP | 0.000164 | 0 | 1 | 3 | 7 | Q8BTI8 | Serine/argi | Q8BTI8 [14 | 0 | 2398.165 | 0 | -0.37 | 0.054635 | 7.88 | 113.2 | 86.8 | 14.58 | 7.67 | | High | High | High | High | High | High | High | High | High | High | High | 4.94E-05 | 0.000696 | 2.16 |
| FALSE | High | SGSSSPDSE | [R].SGSSSP | 1xPhosphc | P70698 1xP | 3.79E-07 | 0 | 1 | 1 | 6 | P70698 | CTP syntha | P70698 [57 | 0 | 1974.967 | 0 | -1.18 | 0.001659 | 3.62 | 139.9 | 60.1 | 5.3 | 14.55 | | High | High | High | High | High | High | High | High | High | High | High | 0 | 3.41E-08 | 3.22 |
| FALSE | High | SGSSSPDSE | [R].SGSSSP | 2xPhosphc | P70698 2xP | 2.23E-05 | 0 | 1 | 1 | 6 | P70698 | CTP syntha | P70698 [57 | 0 | 2054.934 | 0 | -0.7 | 0.004646 | 7.8 | 123.9 | 76.1 | 6.53 | 10.84 | | High | High | High | High | High | High | High | High | High | High | High | 0 | 3.35E-05 | 3.19 |
| FALSE | High | SGSTDSLNI | [K].SGSTDS | 1xDeamida | E9QL65 1x | 0.039011 | 0.002718 | 1 | 3 | 2 | E9QL65 | Conserved | E9QL65 [53 | 0 | 3494.661 | 0 | -0.79 | 0.003062 | 3.3 | 126.5 | 73.5 | 13.5 | 10.08 | | High | High | High | High | High | High | High | High | High | High | High | 0.001421 | 0.01448 | 2.63 |
| FALSE | High | SGSTSPLGF | [R].SGSTSP | 1xPhosphc | Q3UFT4 1x | 0.001278 | 0 | 1 | 6 | 5 | Q3UFT4 | Putative ur | Q3UFT4 [4 | 0 | 1374.662 | 0 | -0.74 | 0.001363 | 1.81 | 125 | 75 | 3.97 | 7.43 | | High | High | High | High | High | High | High | High | High | High | High | 0.000382 | 0.003278 | 1.91 |
| FALSE | High | SGSTSPLGF | [R].SGSTSP | 2xPhosphc | Q3UFT4 2x | 0.003138 | 0.00016 | 1 | 6 | 2 | Q3UFT4 | Putative ur | Q3UFT4 [4 | 0 | 1454.628 | 0 | -1.33 | 0.000824 | 1.7 | 142.5 | 57.5 | 6.1 | 11.54 | | High | High | High | High | High | High | High | High | High | High | High | 0.000689 | 0.005938 | 2.86 |
| FALSE | High | SGSVLTSP | [K].SGSVLT | 1xPhosphc | Q80XP8 1x | 0.08727 | 0.006537 | 1 | 1 | 1 | Q80XP8 | Protein FAI | Q80XP8 [3 | 0 | 1056.518 | 0 | -0.56 | 0.001106 | 2.22 | 120.2 | 79.8 | 5.03 | 4.52 | | High | High | High | High | High | High | High | High | High | High | High | 0.003375 | 0.03684 | 1.99 |
| FALSE | High | SGSVTDPST | [R].SGSVTD | 1xPhosphc | Q60846 1x | 0.012508 | 0.000889 | 1 | 1 | 2 | Q60846 | Tumor nec | Q60846 [3 | 0 | 1710.846 | 0 | 2.97 | 0.000904 | 26.81 | 23.2 | 176.8 | 22.22 | 12.75 | | High | High | High | High | High | High | High | High | High | High | High | 0.000449 | 0.004032 | 2.06 |
| FALSE | High | SGSVVANH | [K].SGSVV/ | 1xPhosphc | O70263 1x | 3.63E-05 | 0 | 1 | 5 | 1 | O70263 | E3 ubiquiti | O70263 [2 | 0 | 1634.785 | 0 | -0.88 | 0.013889 | 5.23 | 124.9 | 75.1 | 13.45 | 18.93 | | High | High | High | High | High | High | High | High | High | High | High | 0 | 5.76E-06 | 2.83 |
| FALSE | High | SGSYSGRSI | [R].SGSYSG | 2xPhosphc | Q14AX6-2 | 0.075304 | 0.005727 | 3 | 5 | 1 | Q14AX6-2; | Isoform 2 c | Q14AX6-2 | 1 | 1846.772 | 0 | -0.26 | 0.015668 | 9.33 | 109.1 | 90.9 | 1.84 | 10.32 | Shared | High | High | High | High | High | High | High | High | High | High | High | 0.002976 | 0.03107 | 1.24 |
| FALSE | High | SGSYSSSST | [K].SGSYSS | 1xPhosphc | Q3TU81 1x | 3.01E-05 | 0 | 1 | 3 | 1 | Q3TU81 | Putative ur | Q3TU81 [3 | 0 | 1714.83 | 0 | -0.34 | 0.108002 | 20.4 | 113.2 | 86.8 | 15.59 | 15.77 | | High | High | High | High | High | High | High | High | High | High | High | 0 | 4.63E-06 | 2.55 |
| FALSE | High | SGTPPRPG | [K].SGTPPF | 1xCarbami | Q8BTI8 1xP | 0.003495 | 0.00021 | 1 | 3 | 3 | Q8BTI8 | Serine/argi | Q8BTI8 [82 | 0 | 2867.301 | 0 | -0.82 | 0.036932 | 3.24 | 130.9 | 69.1 | 19.24 | 29.26 | | High | High | High | High | High | High | High | High | High | High | High | 9.42E-05 | 0.000973 | 1.36 |
| FALSE | High | SGTPPRPG | [K].SGTPPF | 1xCarbami | Q8BTI8 1xP | 0.009416 | 0.000736 | 1 | 3 | 5 | Q8BTI8 | Serine/argi | Q8BTI8 [82 | 0 | 2883.296 | 0 | -0.81 | 0.007484 | 5.13 | 126 | 74 | 12.59 | 5.88 | | High | High | High | High | High | High | High | High | High | High | High | 0.000327 | 0.002931 | 1.96 |
| FALSE | High | SGTPPRPG | [K].SGTPPF | 1xCarbami | Q8BTI8 2xP | 2.34E-06 | 0 | 1 | 3 | 2 | Q8BTI8 | Serine/argi | Q8BTI8 [82 | 0 | 2947.268 | 0 | -1.07 | 0.009196 | 5.53 | 136.8 | 63.2 | 14.59 | 26.15 | | High | High | High | High | High | High | High | High | High | High | High | 0 | 3.36E-05 | 3 |
| FALSE | High | SGTPPRPG | [K].SGTPPF | 1xCarbami | Q8BTI8 2xP | 1.72E-05 | 0 | 1 | 3 | 8 | Q8BTI8 | Serine/argi | Q8BTI8 [82 | 0 | 2963.263 | 0 | -0.73 | 0.007603 | 5.02 | 124.2 | 75.8 | 11.28 | 4.73 | | High | High | High | High | High | High | High | High | High | High | High | 0 | 2.48E-06 | 2.77 |
| FALSE | High | SGTPTSPVE | [K].SGTPTS | 1xPhosphc | Q91YK2 1x | 5.97E-05 | 0 | 1 | 2 | 3 | Q91YK2 | Ribosomal | Q91YK2 [5 | 0 | 1597.824 | 0 | 0.39 | 0.0187 | 1.71 | 90.5 | 109.5 | 4.84 | 9.12 | | High | High | High | High | High | High | High | High | High | High | High | 0 | 1.01E-05 | 2.3 |
| FALSE | High | SGTPVHCP | [R].SGTPVH | 1xCarbami | A6H663 1x | 0.000146 | 0 | 1 | 3 | 3 | A6H663 | BCL2-assoc | A6H663 [2 | 0 | 1616.782 | 0 | 2.06 | 0.000374 | 14.02 | 39.4 | 160.6 | 7.53 | 6.39 | | High | High | High | High | High | High | High | High | High | High | High | 0 | 2.75E-05 | 2.53 |
| FALSE | High | SHGAEGG( | [K].SHGAE( | 1xPhosphc | Q76LS9 1xP | 8.5E-06 | 0 | 1 | 5 | 3 | Q76LS9 | Ubiquitin c | Q76LS9 [37 | 0 | 1707.81 | 0 | 0.7 | 0.000486 | 4.66 | 77.1 | 122.9 | 9.05 | 3.95 | | High | High | High | High | High | High | High | High | High | High | High | 0 | 1.13E-06 | 3.62 |
| FALSE | High | SESAENHS | [R].SESAEN | 1xPhosphc | Q80TH2-2 | 0.014538 | 0.001011 | 1 | 6 | 1 | Q80TH2-2 | Isoform 2 c | Q80TH2-2 | 0 | 1760.826 | 0 | 1.11 | 0.017023 | 42.7 | 63.4 | 136.6 | 10.8 | 25.39 | | High | High | High | High | High | High | High | High | High | High | High | 0.000538 | 0.004753 | 2.59 |
| FALSE | High | SHHAASTT | [R].SHHAA | 1xPhosphc | A2AFQ0 1x | 0.000274 | 0 | 1 | 8 | 1 | A2AFQ0 | E3 ubiquiti | A2AFQ0 [1 | 0 | 1885.912 | 0 | -0.58 | 0.014798 | 8.37 | 115.1 | 84.9 | 13.05 | 9.33 | | High | High | High | High | High | High | High | High | High | High | High | 0 | 5.54E-05 | 2.18 |
| FALSE | High | SHHAPMSF | [R].SHHAPI | 1xPhosphc | E9QAQ7 1x | 9.31E-06 | 0 | 1 | 3 | 2 | E9QAQ7 | AT-rich inte | E9QAQ7 [3 | 0 | 2197.017 | 0 | -0.76 | 0.006314 | 8.51 | 125.3 | 74.7 | 9.05 | 26.82 | | High | High | High | High | High | High | High | High | High | High | High | 0 | 1.46E-06 | 3.32 |
| FALSE | High | SHSMSQEF | [K].SHSMS | 1xPhosphc | Q91VX2 1x | 0.000269 | 0 | 1 | 2 | 6 | Q91VX2 | Ubiquitin-a | Q91VX2 [2 | 0 | 1768.834 | 0 | 0.37 | 0.010771 | 4.11 | 85 | 115 | 4.58 | 9.73 | | High | High | High | High | High | High | High | High | High | High | High | 0.000117 | 0.001082 | 3.49 |
| FALSE | High | SHSPLPAPF | [R].SHSPLP | 2xPhosphc | Q3TWW8 2 | 0.106748 | 0.008634 | 1 | 2 | 2 | Q3TWW8 | Serine/argi | Q3TWW8 [ | 0 | 1506.696 | 0 | -0.37 | 0.054177 | 13.23 | 113.8 | 86.2 | 13.73 | 12.28 | | High | High | High | High | High | High | High | High | High | High | High | 0.007598 | 0.08296 | 2.26 |
| FALSE | High | SHSPLPAPF | [R].SHSPLP | 1xPhosphc | Q3TWW8 1 | 0.000281 | 0 | 1 | 2 | 8 | Q3TWW8 | Serine/argi | Q3TWW8 [ | 0 | 1655.892 | 0 | -0.38 | 0.000986 | 2.26 | 111.9 | 88.1 | 2.48 | 4.4 | | High | High | High | High | High | High | High | High | High | High | High | 0 | 0.000116 | 2.62 |
| FALSE | High | SHSPPTDG | [R].SHSPPT | 1xPhosphc | Q6PCL9 1x | 0.000993 | 0 | 1 | 4 | 2 | Q6PCL9 | Poly(A) pol | Q6PCL9 [6 | 0 | 1651.809 | 0 | -0.51 | 0.001208 | 2.88 | 117.5 | 82.5 | 3.5 | 5.81 | | High | High | High | High | High | High | High | High | High | High | High | 0 | 0.000237 | 2.88 |
| FALSE | High | SHSPSMSQ | [R].SHSPSN | 1xPhosphc | Q6ZQ32 1x | 6.13E-05 | 0 | 2 | 7 | 1 | Q6ZQ32; F | MKIAA085 | Q6ZQ32 [1 | 0 | 1797.815 | 0 | 1.03 | 0.007243 | 14.12 | 68.9 | 131.1 | 9.08 | 15.52 | Shared | High | High | High | High | High | High | High | High | High | High | High | 0 | 1.04E-05 | 2.96 |
| FALSE | High | SHSPSMSQ | [R].SHSPSN | 1xPhosphc | Q6ZQ32 1x | 0.002419 | 0.000109 | 2 | 7 | 5 | Q6ZQ32; F | MKIAA085 | Q6ZQ32 [1 | 0 | 1813.81 | 0 | 1.06 | 0.039612 | 9.5 | 63 | 137 | 12.58 | 32.66 | Shared | High | High | High | High | High | High | High | High | High | High | High | 4.94E-05 | 0.000643 | 2.84 |
| FALSE | High | SHSPSPYTK | [K].SHSPSP | 1xPhosphc | Q3U6K5 1x | 0.002793 | 0.000109 | 1 | 3 | 2 | Q3U6K5 | Spermatog | Q3U6K5 [2 | 0 | 1541.777 | 0 | 1.62 | 0.001114 | 8.61 | 49 | 151 | 8.75 | 8.49 | | High | High | High | High | High | High | High | High | High | High | High | 0.000294 | 0.002322 | 1.85 |
| FALSE | High | SHSQDIPVI | [K].SHSQD | 1xPhosphc | Q3TY45 1x | 0.000157 | 0 | 1 | 2 | 1 | Q3TY45 | Putative ur | Q3TY45 [5 | 0 | 2359.246 | 0 | -0.76 | 0.001499 | 2.23 | 123.7 | 76.3 | 8.54 | 6.91 | | High | High | High | High | High | High | High | High | High | High | High | 0 | 2.98E-05 | 2.18 |
| FALSE | High | SHSSGTVS/ | [K].SHSSGT | 1xPhosphc | E9Q1A8 1x | 4.69E-06 | 0 | 1 | 6 | 1 | E9Q1A8 | HMG box-c | E9Q1A8 [4 | 0 | 1897.942 | 0 | -1.9 | 0.00128 | 1.63 | 156.3 | 43.7 | 10.74 | 15.04 | | High | High | High | High | High | High | High | High | High | High | High | 0 | 5.78E-07 | 4.73 |
| FALSE | High | SHSSIQFSF | [R].SHSSIQ | 1xPhosphc | H7BWZ9 1 | 0.037048 | 0.00249 | 1 | 5 | 1 | H7BWZ9 | Protein do | H7BWZ9 [1 | 0 | 1705.871 | 0 | 0.73 | 0.073671 | 6.76 | 74.2 | 125.8 | 15.26 | 16.5 | | High | High | High | High | High | High | High | High | High | High | High | 0.001308 | 0.01368 | 2.15 |
| FALSE | High | SHSSNSSPI | [R].SHSSNS | 1xPhosphc | Q6PR54-3 | 0.000284 | 0 | 2 | 4 | 1 | Q6PR54-3; | Isoform 3 c | Q6PR54-3 | 0 | 1694.888 | 0 | -1.55 | 0.003041 | 1.18 | 150.5 | 49.5 | 14.69 | 10.49 | Shared | High | High | High | High | High | High | High | High | High | High | High | 0 | 5.8E-05 | 2.57 |
| FALSE | High | SHSSPASLC | [R].SHSSPA | 1xPhosphc | Q9EPK5-2 | 0.075927 | 0.005826 | 1 | 2 | 1 | Q9EPK5-2 | Isoform 2 c | Q9EPK5-2 | 0 | 2973.464 | 0 | 0.63 | 0.051549 | 9.96 | 78.6 | 121.4 | 22.49 | 8.67 | | High | High | High | High | High | High | High | High | High | High | High | 0.002976 | 0.0312 | 2.75 |
| FALSE | High | SHSSPASLC | [R].SHSSPA | 2xPhosphc | Q9EPK5-2 | 0.020499 | 0.001472 | 1 | 2 | 1 | Q9EPK5-2 | Isoform 2 c | Q9EPK5-2 | 0 | 3053.431 | 0 | 0.67 | 0.026624 | 27.1 | 76.3 | 123.7 | 22.24 | 15.89 | | High | High | High | High | High | High | High | High | High | High | High | 0.000786 | 0.006979 | 3.11 |
| FALSE | High | SHSITNME | [R].SHSITN | 1xPhosphc | Q80TH2-2 | 7.47E-05 | 0 | 1 | 6 | 3 | Q80TH2-2 | Isoform 2 c | Q80TH2-2 | 0 | 1928.955 | 0 | 0.61 | 0.046053 | 18.63 | 77.5 | 122.5 | 7.99 | 26.76 | | High | High | High | High | High | High | High | High | High | High | High | 0 | 1.3E-05 | 3.38 |
| FALSE | High | SHSSPSLNF | [K].SHSSPS | 2xPhosphc | A0A1D5RN | 0.001313 | 0 | 1 | 9 | 1 | A0A1D5RN | PH and SEC | A0A1D5RN | 0 | 2313.093 | 0 | 1.05 | 0.000961 | 1.2 | 63.8 | 136.2 | 8.29 | 9.33 | | High | High | High | High | High | High | High | High | High | High | High | 0 | 0.000323 | 3.73 |
| FALSE | High | SHSSSASSA | [R].SHSSSA | 2xPhosphc | P97868 2xP | 0.000662 | 0 | 1 | 2 | 1 | P97868 | E3 ubiquiti | P97868 [17 | 0 | 2040.868 | 0 | 0.2 | 0.732517 | 9.53 | 94.1 | 105.9 | 21.27 | 10.45 | | High | High | High | High | High | High | High | High | High | High | High | 0 | 0.000151 | 2.5 |
| FALSE | High | SHSVSALE( | [R].SHSVSA | 1xCarbami | Q8BQ33 1x | 0.002576 | 0.000109 | 1 | 3 | 1 | Q8BQ33 | Treslin OS= | Q8BQ33 [9 | 0 | 2043.025 | 0 | -0.64 | 0.007099 | 6.99 | 123.8 | 76.2 | 10.7 | 5.2 | | High | High | High | High | High | High | High | High | High | High | High | 4.94E-05 | 0.00069 | 2.1 |
| FALSE | High | SHSWVNS/ | [R].SHSWV | 1xPhosphc | A0A1L1SQ | 7.29E-06 | 0 | 1 | 3 | 1 | A0A1L1SQ | Dedicator c | A0A1L1SQ | 0 | 2086.016 | 0 | -1.74 | 0.000727 | 1.67 | 157.6 | 42.4 | 10.5 | 21.73 | | High | High | High | High | High | High | High | High | High | High | High | 0 | 9.48E-07 | 3.72 |
| FALSE | High | SHSWVNS/ | [R].SHSWV | 1xDeamida | A0A1L1SQ | 0.025514 | 0.001645 | 1 | 3 | 1 | A0A1L1SQ | Dedicator c | A0A1L1SQ | 0 | 2087 | 0 | -0.6 | 0.002231 | 4.97 | 124.2 | 75.8 | 11.4 | 10.97 | | High | High | High | High | High | High | High | High | High | High | High | 0.000872 | 0.009006 | 2.27 |
| FALSE | High | SHTEEYTSK | [K].SHTEEY | 1xPhosphc | P49452 1xP | 0.007145 | 0.00062 | 1 | 1 | 1 | P49452 | Centromer | P49452 [61 | 0 | 1619.772 | 0 | -1.01 | 0.001448 | 4.19 | 132 | 68 | 7.76 | 11.67 | | High | High | High | High | High | High | High | High | High | High | High | 0.000275 | 0.002146 | 3.57 |
| FALSE | High | SHTGDLSPI | [R].SHTGD | 1xPhosphc | E9QN63 1x | 9.42E-07 | 0 | 1 | 13 | 2 | E9QN63 | Arf-GAP wi | E9QN63 [1 | 0 | 1706.806 | 0 | 0.45 | 0.00158 | 8.34 | 84.4 | 115.6 | 6.4 | 3.19 | | High | High | High | High | High | High | High | High | High | High | High | 0 | 9.54E-08 | 3.45 |
| FALSE | High | SHTGEAAA | [K].SHTGE/ | 1xPhosphc | P59017 1xP | 0.003055 | 0.00016 | 1 | 3 | 3 | P59017 | Bcl-2-like p | P59017 [38 | 0 | 1307.631 | 0 | 1.78 | 0.00251 | 35.98 | 42.5 | 157.5 | 8.15 | 15.91 | | High | High | High | High | High | High | High | High | High | High | High | 0.000327 | 0.002753 | 2.45 |
| FALSE | High | SHTSSNYD: | [R].SHTSSN | 1xPhosphc | Q14AX6-2 | 0.000815 | 0 | 3 | 5 | 4 | Q14AX6-2; | Isoform 2 c | Q14AX6-2 | 0 | 1826.836 | 0 | 0.16 | 0.288742 | 30.42 | 95.8 | 104.2 | 7.21 | 16.99 | Shared | High | High | High | High | High | High | High | High | High | High | High | 2.58E-05 | 0.000508 | 2.31 |
| FALSE | High | SHTSSNYD: | [R].SHTSSN | 2xPhosphc | Q14AX6-2 | 0.093879 | 0.007306 | 3 | 5 | 1 | Q14AX6-2; | Isoform 2 c | Q14AX6-2 | 0 | 1906.803 | 0 | 0.13 | 0.122859 | 11.43 | 95.6 | 104.4 | 12.23 | 14.29 | Shared | High | High | High | High | High | High | High | High | High | High | High | 0.003727 | 0.04016 | 2.19 |
| FALSE | High | SHVTSTPIP | [R].SHVTST | 1xPhosphc | E9Q238 1x | 3.6E-05 | 0 | 1 | 3 | 2 | E9Q238 | KN motif ar | E9Q238 [1 | 0 | 3197.619 | 0 | 1.01 | 0.00798 | 9.87 | 66.5 | 133.5 | 17.12 | 13.78 | | High | High | High | High | High | High | High | High | High | High | High | 0 | 8E-05 | 3.58 |
| FALSE | High | SHWDDGT | [R].SHWDD | 1xPhosphc | A2ARP8 1x | 0.000257 | 0 | 1 | 3 | 1 | A2ARP8 | Microtubu | A2ARP8 [2 | 0 | 2156.954 | 0 | -1.15 | 0.013083 | 7.2 | 143.9 | 56.1 | 19.74 | 19.69 | | High | High | High | High | High | High | High | High | High | High | High | 0 | 5.19E-05 | 1.94 |
| FALSE | High | SKESLQEA( | [K].SKESLQ | 1xPhosphc | P53986 1xP | 0.006031 | 0.00062 | 1 | 5 | 2 | P53986 | Monocarbo | P53986 [21 | 1 | 1844.013 | 0 | 0.57 | 0.001548 | 2.49 | 81.7 | 118.3 | 4 | 3.59 | | High | High | High | High | High | High | High | High | High | High | High | 0.000275 | 0.001779 | 2.01 |
| FALSE | High | SKPAAADS | [K].SKPAA/ | 1xPhosphc | Q5RL55 1x | 7.18E-07 | 0 | 1 | 1 | 2 | Q5RL55 | ATP-bindin | Q5RL55 [1 | 0 | 2631.264 | 0 | -0.44 | 0.10703 | 10.32 | 116.1 | 83.9 | 17.93 | 22.55 | | High | High | High | High | High | High | High | High | High | High | High | 0 | 6.97E-08 | 4.53 |
| FALSE | High | SHSSSASSA | [R].SHSSSA | 1xPhosphc | P97868 1xP | 0.000389 | 0 | 1 | 2 | 2 | P97868 | E3 ubiquiti | P97868 [17 | 0 | 1960.901 | 0 | 0.17 | 0.068617 | 8.19 | 91.5 | 108.5 | 10.35 | 7.91 | | High | High | High | High | High | High | High | High | High | High | High | 0 | 8.28E-05 | 2.1 |
| FALSE | High | SHSLSRPGF | [R].SHSLSR | 1xPhosphc | Q8VHK1 1x | 0.002376 | 0.000109 | 1 | 1 | 2 | Q8VHK1 | Caskin-2 O | Q8VHK1 [7 | 0 | 3649.733 | 0 | 0.67 | 0.011423 | 9.75 | 67.8 | 132.2 | 12.3 | 19.38 | | High | High | High | High | High | High | High | High | High | High | High | 0.000117 | 0.00107 | 2.79 |
| FALSE | High | SHSLSQHT/ | [R].SHSLSC | 1xPhosphc | E9Q3G8 1x | 0.007538 | 0.00062 | 1 | 2 | 2 | E9Q3G8 | Nucleopor | E9Q3G8 [2 | 0 | 1920.994 | 0 | 1.1 | 0.054959 | 5.29 | 63.5 | 136.5 | 10.86 | 30.75 | | High | High | High | High | High | High | High | High | High | High | High | 0.000275 | 0.002285 | 1.97 |
| FALSE | High | SHSLPNSLE | [R].SHSLPN | 2xPhosphc | Q3UJU9 2x | 0.001799 | 5.66E-05 | 1 | 4 | 2 | Q3UJU9 | Regulator c | Q3UJU9 [4 | 0 | 2163.931 | 0 | 1.04 | 0.000214 | 1.36 | 66 | 134 | 4.32 | 4.92 | | High | High | High | High | High | High | High | High | High | High | High | 2.58E-05 | 0.000459 | 2.74 |
| FALSE | High | SHHAPMSF | [R].SHHAPI | 1xPhosphc | E9QAQ7 1x | 1.2E-07 | 0 | 1 | 3 | 19 | E9QAQ7 | AT-rich inte | E9QAQ7 [3 | 0 | 2213.012 | 0 | -0.92 | 0.006792 | 5.24 | 127.4 | 72.6 | 10.25 | 10.92 | | High | High | High | High | High | High | High | High | High | High | High | 0 | 9.43E-09 | 3.59 |
| FALSE | High | SHHDGNLE | [K].SHHDG | 1xPhosphc | H9KV00 1x | 3.2E-05 | 0 | 1 | 7 | 2 | H9KV00 | Protein SO | H9KV00 [1 | 0 | 2124.016 | 0 | -0.89 | 0.00475 | 3.4 | 129.8 | 70.2 | 10.23 | 9.14 | | High | High | High | High | High | High | High | High | High | High | High | 0 | 5E-06 | 2.64 |
| FALSE | High | SHIAQWSP | [K].SHIAQV | 1xPhosphc | H6WCS3 1 | 0.028327 | 0.001791 | 1 | 5 | 2 | H6WCS3 | Interleukin | H6WCS3 [6 | 0 | 1822.895 | 0 | 0.01 | 0.8727 | 2.97 | 99.1 | 100.9 | 16.61 | 10.49 | | High | High | High | High | High | High | High | High | High | High | High | 0.001183 | 0.01265 | 2.57 |
| FALSE | High | SHLETMGS | [R].SHLETN | 1xPhosphc | Q3TAA7 1x | 0.032831 | 0.002195 | 1 | 1 | 2 | Q3TAA7 | Serine/thre | Q3TAA7 [4 | 0 | 2130.055 | 0 | -1.14 | 0.004626 | 3.66 | 144.7 | 55.3 | 19.5 | 17.63 | | High | High | High | High | High | High | High | High | High | High | High | 0.00302 | 0.03175 | 2.21 |
| FALSE | High | SHLIVTSPP | [R].SHLIVT | 1xPhosphc | E9Q5N9 1x | 0.000248 | 0 | 1 | 2 | 4 | E9Q5N9 | Zinc finger | E9Q5N9 [4 | 0 | 1415.761 | 0 | -0.83 | 0.000945 | 3.44 | 127.6 | 72.4 | 7.53 | 4.36 | | High | High | High | High | High | High | High | High | High | High | High | 0 | 5E-05 | 3 |
| FALSE | High | SHIPSEPYE | [R].SHIPSEI | 1xPhosphc | Q5RIM6 1x | 0.04913 | 0.003576 | 1 | 10 | 1 | Q5RIM6 | Nuclear rec | Q5RIM6 [2 | 0 | 2934.48 | 0 | -0.29 | 0.091709 | 8.86 | 109 | 91 | 9.64 | 8.03 | | High | High | High | High | High | High | High | High | High | High | High | 0.001889 | 0.01895 | 2.85 |
| FALSE | High | SHLSPSQTE | [R].SHLSPS | 1xPhosphc | B2RWU4 1 | 0.00086 | 0 | 2 | 6 | 2 | B2RWU4; ( | Cordon-ble | B2RWU4 [6 | 0 | 2146.028 | 0 | -0.21 | 0.106999 | 2.55 | 107.2 | 92.8 | 6.72 | 5.46 | Shared | High | High | High | High | High | High | High | High | High | High | High | 0.001651 | 0.0162 | 1.65 |
| FALSE | High | SHLVAQVS | [R].SHLVAC | 1xPhosphc | J3QNX5 1x | 0.00359 | 0.00021 | 1 | 1 | 1 | J3QNX5 | Cysteine-ki | J3QNX5 [3 | 0 | 1570.83 | 0 | -0.78 | 0.008059 | 2.18 | 124.3 | 75.7 | 12.36 | 15.94 | | High | High | High | High | High | High | High | High | High | High | High | 9.42E-05 | 0.001003 | 2.49 |
| FALSE | High | SHPAMSPG | [R].SHPAM | 1xPhosphc | E9Q4N7 1x | 0.033404 | 0.002195 | 1 | 2 | 2 | E9Q4N7 | AT-rich inte | E9Q4N7 [4 | 0 | 1921.85 | 0 | 0.41 | 0.022634 | 17.28 | 82.4 | 117.6 | 14.9 | 9.53 | | High | High | High | High | High | High | High | High | High | High | High | 0.001151 | 0.01218 | 2.45 |
| FALSE | High | SHPGGGPT | [R].SHPGG | 1xPhosphc | E9Q0S6 1x | 0.022773 | 0.001542 | 1 | 7 | 1 | E9Q0S6 | Tensin 1 O | E9Q0S6 [6 | 0 | 1611.784 | 0 | -0.25 | 0.02906 | 8.14 | 109.8 | 90.2 | 9.43 | 7.74 | | High | High | High | High | High | High | High | High | High | High | High | 0.000814 | 0.007889 | 2.15 |
| FALSE | High | SHPPANFF( | [R].SHPPAN | 1xPhosphc | F8VQ95 1x | 8.55E-08 | 0 | 1 | 2 | 1 | F8VQ95 | Transformi | F8VQ95 [3 | 0 | 2917.501 | 0 | -0.99 | 0.001564 | 3.5 | 132.4 | 67.6 | 7.97 | 5.5 | | High | High | High | High | High | High | High | High | High | High | High | 0 | 6.44E-09 | 4.24 |
| FALSE | High | SHQCGNEE | [R].SHQCG | 1xCarbami | P48754 1xP | 3.99E-10 | 0 | 1 | 3 | 3 | P48754 | Breast canc | P48754 [12 | 0 | 2159.983 | 0 | -0.63 | 0.001399 | 3.36 | 120.9 | 79.1 | 4.2 | 5.2 | | High | High | High | High | High | High | High | High | High | High | High | 0 | 1.55E-11 | 4.62 |
| FALSE | High | SHSAEGGK | [R].SHSAEC | 1xPhosphc | P24788 1xP | 0.050826 | 0.003676 | 1 | 3 | 2 | P24788 | Cyclin-dep | P24788 [11 | 0 | 1310.651 | 0 | -0.34 | 0.005594 | 6.57 | 114.1 | 85.9 | 4.84 | 7.33 | | High | High | High | High | High | High | High | High | High | High | High | 0.001916 | 0.01962 | 2.05 |
| FALSE | High | SHSAGGLQ | [R].SHSAG( | 1xPhosphc | E9Q842 1x | 2.45E-09 | 0 | 1 | 3 | 6 | E9Q842 | Neuron na | E9Q842 [1 | 0 | 3089.401 | 0 | 1.29 | 0.005127 | 23.15 | 54.5 | 145.5 | 5.4 | 17.92 | | High | High | High | High | High | High | High | High | High | High | High | 0 | 1.18E-10 | 4.82 |
| FALSE | High | SHSANDSE | [K].SHSANI | 1xPhosphc | B1APX2 1x | 0.005978 | 0.00062 | 1 | 3 | 1 | B1APX2 | RIKEN cDN | B1APX2 [3 | 0 | 1734.732 | 0 | 0.06 | 0.110322 | 2.87 | 92.5 | 107.5 | 8.48 | 8.47 | | High | High | High | High | High | High | High | High | High | High | High | 0.000275 | 0.001759 | 1.84 |
| FALSE | High | SHSDNDRP | [R].SHSDNI | 1xCarbami | Q9R0U0-3 | 0.000241 | 0 | 1 | 2 | 4 | Q9R0U0-3 | Isoform 3 c | Q9R0U0-3 | 0 | 3205.327 | 0 | -0.23 | 0.152487 | 8.93 | 107.8 | 92.2 | 11.14 | 9.8 | | High | High | High | High | High | High | High | High | High | High | High | 0 | 4.83E-05 | 3.51 |
| FALSE | High | SHSDNDRP | [R].SHSDNI | 1xCarbami | Q9R0U0-3 | 0.001615 | 5.66E-05 | 1 | 2 | 6 | Q9R0U0-3 | Isoform 3 c | Q9R0U0-3 | 0 | 3206.311 | 0 | 1.95 | 0.003874 | 20.72 | 37 | 163 | 15.45 | 23.37 | | High | High | High | High | High | High | High | High | High | High | High | 0.000773 | 0.006736 | 3.04 |
| FALSE | High | SHSDSPPK | [R].SHSDSF | 1xPhosphc | B2RXX0 1x | 0.008167 | 0.000736 | 1 | 6 | 1 | B2RXX0 | Nek1 prote | B2RXX0 [1 | 0 | 1392.692 | 0 | -1.18 | 0.001685 | 2.59 | 137.1 | 62.9 | 12.35 | 4.37 | | High | High | High | High | High | High | High | High | High | High | High | 0.000327 | 0.002511 | 1.28 |
| FALSE | High | SHSDTNIAS | [R].SHSDTN | 1xPhosphc | Q80U62 1x | 0.00359 | 0.00021 | 1 | 5 | 2 | Q80U62 | Run domai | Q80U62 [4 | 0 | 1396.642 | 0 | -0.22 | 0.010986 | 5.98 | 107.6 | 92.4 | 2.49 | 6.77 | | High | High | High | High | High | High | High | High | High | High | High | 0.000275 | 0.002047 | 2.14 |
| FALSE | High | SHSESASPS | [R].SHSESA | 1xPhosphc | Q99N57-2 | 1.21E-05 | 0 | 1 | 3 | 4 | Q99N57-2 | Isoform 2 c | Q99N57-2 | 0 | 3278.573 | 0 | -0.54 | 0.100553 | 19.64 | 118.4 | 81.6 | 19.05 | 25.65 | | High | High | High | High | High | High | High | High | High | High | High | 0 | 1.67E-06 | 4.93 |
| FALSE | High | SHSESASPS | [R].SHSESA | 2xPhosphc | Q99N57-2 | 1.54E-07 | 0 | 1 | 3 | 2 | Q99N57-2 | Isoform 2 c | Q99N57-2 | 0 | 3358.539 | 0 | 0.06 | 0.886764 | 2.85 | 97.1 | 102.9 | 5.37 | 11.77 | | High | High | High | High | High | High | High | High | High | High | High | 0 | 1.24E-08 | 5.27 |
| FALSE | High | SHSFADPA: | [R].SHSFAD | 1xPhosphc | Q5RIM6 1x | 0.002376 | 0.000109 | 1 | 11 | 1 | Q5RIM6 | Nuclear rec | Q5RIM6 [2 | 0 | 2251.096 | 0 | -0.17 | 0.200692 | 6.5 | 108.8 | 91.2 | 15.62 | 14.01 | | High | High | High | High | High | High | High | High | High | High | High | 4.94E-05 | 0.000631 | 2.45 |
| FALSE | High | SHSFSDPSF | [R].SHSFSD | 1xPhosphc | A0A0A6YW | 3.41E-05 | 0 | 1 | 8 | 4 | A0A0A6YW | Mitogen-ac | A0A0A6YW | 0 | 1626.793 | 0 | -1.46 | 0.000243 | 1.47 | 147.1 | 52.9 | 7.03 | 14.5 | | High | High | High | High | High | High | High | High | High | High | High | 0 | 5.37E-06 | 3.04 |
| FALSE | High | SHSFSDPSF | [R].SHSFSD | 2xPhosphc | A0A0A6YW | 0.107603 | 0.008634 | 1 | 8 | 1 | A0A0A6YW | Mitogen-ac | A0A0A6YW | 0 | 1706.759 | 0 | -0.54 | 0.065662 | 14.06 | 123.1 | 76.9 | 12.81 | 32.76 | | High | High | High | High | High | High | High | High | High | High | High | 0.004355 | 0.04697 | 1.82 |
| FALSE | High | SHSGSSPEV | [R].SHSGSS | 1xPhosphc | Q8BTI8 1xP | 5.1E-07 | 0 | 1 | 3 | 2 | Q8BTI8 | Serine/argi | Q8BTI8 [15 | 0 | 1754.836 | 0 | -0.54 | 0.006842 | 1.51 | 116.9 | 83.1 | 8.24 | 6.59 | | High | High | High | High | High | High | High | High | High | High | High | 0 | 4.76E-08 | 2.9 |
| FALSE | High | SHSGSSPEV | [R].SHSGSS | 2xPhosphc | Q8BTI8 2xP | 0.068752 | 0.005042 | 1 | 3 | 1 | Q8BTI8 | Serine/argi | Q8BTI8 [15 | 0 | 1834.803 | 0 | -0.49 | 0.036847 | 4.81 | 115.8 | 84.2 | 9.99 | 11.45 | | High | High | High | High | High | High | High | High | High | High | High | 0.002604 | 0.02787 | 2.25 |
| FALSE | High | SHSGTSPDI | [R].SHSGTS | 2xPhosphc | O54916 2x | 0.015061 | 0.001049 | 1 | 5 | 3 | O54916 | RalBP1-ass | O54916 [5 | 0 | 2990.351 | 0 | -0.13 | 0.368752 | 11.64 | 102.6 | 97.4 | 12.12 | 4.35 | | High | High | High | High | High | High | High | High | High | High | High | 0.000571 | 0.004946 | 2.43 |
| FALSE | High | SHSLDDLQ | [R].SHSLDD | 1xPhosphc | F8VQK5 1x | 0.000386 | 0 | 1 | 3 | 2 | F8VQK5 | SAM and S | F8VQK5 [8 | 0 | 2095.011 | 0 | 0.47 | 0.006531 | 8.28 | 83.9 | 116.1 | 5.9 | 9.93 | | High | High | High | High | High | High | High | High | High | High | High | 0 | 8.22E-05 | 2.95 |
| FALSE | High | SHSLPNSLE | [R].SHSLPN | 1xPhosphc | Q3UJU9 1x | 5.84E-07 | 0 | 1 | 4 | 4 | Q3UJU9 | Regulator c | Q3UJU9 [4 | 0 | 2083.965 | 0 | 0.6 | 0.017763 | 6.61 | 84.7 | 115.3 | 6.15 | 11.4 | | High | High | High | High | High | High | High | High | High | High | High | 0 | 1.18E-05 | 4.02 |
| FALSE | High | SHHANSPT | [K].SHHAN | 1xPhosphc | Q8R001 1x | 1.06E-05 | 0 | 1 | 4 | 2 | Q8R001 | Microtubu | Q8R001 [1 | 0 | 1786.9 | 0 | 0.09 | 0.696977 | 16.12 | 97.3 | 102.7 | 9.58 | 4.42 | | High | High | High | High | High | High | High | High | High | High | High | 0 | 9.33E-05 | 2.56 |
| FALSE | High | RQDSDLVC | [R].RQDSD | 1xPhosphc | Q9QXV1 1 | 3.76E-08 | 0 | 1 | 2 | 4 | Q9QXV1 | Chromobo | Q9QXV1 [2 | 1 | 2850.392 | 0 | 1.66 | 0.00039 | 22.48 | 49.5 | 150.5 | 12.42 | 5.81 | | High | High | High | High | High | High | High | High | High | High | High | 0 | 2.54E-09 | 5.16 |
| FALSE | High | AAAAAAAA | [K].AAAAA | 1xPhosphc | B2RUG8 1x | 0.002355 | 0.000109 | 1 | 4 | 1 | B2RUG8 | Asxl2 prote | B2RUG8 [6 | 0 | 2810.415 | 0 | 0.09 | 0.635787 | 7.54 | 96.6 | 103.4 | 6.02 | 8.16 | | High | High | High | High | High | High | High | High | High | High | High | 4.94E-05 | 0.000626 | 2.84 |
| FALSE | High | RQDPQVLS | [R].RQDPQ | 1xPhosphc | B1AY10 1x | 0.007145 | 0.00062 | 1 | 3 | 1 | B1AY10 | Transcripti | B1AY10 [3 | 1 | 1748.905 | 0 | 0.31 | 0.012889 | 7.09 | 85.5 | 114.5 | 10.33 | 6.31 | | High | High | High | High | High | High | High | High | High | High | High | 0.000275 | 0.002149 | 2.04 |
| FALSE | High | QELAVFCSI | [R].QELAVI | 1xCarbami | Q80TM2 1 | 2.06E-06 | 0 | 1 | 5 | 1 | Q80TM2 | MKIAA102 | Q80TM2 [2 | 0 | 2111.065 | 0 | 0.16 | 0.362814 | 9.53 | 95.1 | 104.9 | 10.26 | 3.81 | | High | High | High | High | High | High | High | High | High | High | High | 0 | 2.3E-07 | 2.63 |
| FALSE | High | QELLPNAE: | [R].QELLPN | 1xPhosphc | Q7TN02 1x | 0.006957 | 0.00062 | 1 | 1 | 1 | Q7TN02 | Mediator c | Q7TN02 [3 | 0 | 2846.346 | 0 | -0.15 | 0.183499 | 5.47 | 101.6 | 98.4 | 6.68 | 10.29 | | High | High | High | High | High | High | High | High | High | High | High | 0.000275 | 0.002084 | 2.63 |
| FALSE | High | QELSPSHSI | [R].QELSPS | 1xPhosphc | E9Q4K7 1x | 0.0026 | 0.000109 | 1 | 2 | 2 | E9Q4K7 | Kinesin fan | E9Q4K7 [1 | 0 | 1921.978 | 0 | 0.27 | 0.031764 | 10.24 | 88.5 | 111.5 | 8.18 | 6.96 | | High | High | High | High | High | High | High | High | High | High | High | 0.001609 | 0.01567 | 1.99 |
| FALSE | High | QELTDVNY | [R].QELTDV | 1xPhosphc | Q53Z05 1x | 0.000139 | 0 | 1 | 4 | 1 | Q53Z05 | Sir2alpha C | Q53Z05 [7 | 1 | 2033.983 | 0 | -1.44 | 0.000179 | 0.68 | 145.8 | 54.2 | 4.31 | 5.31 | | High | High | High | High | High | High | High | High | High | High | High | 0 | 2.59E-05 | 3.47 |
| FALSE | High | QEMEVKTE | [K].QEMEV | 1xPhosphc | Q3TUQ5 1 | 0.115566 | 0.009504 | 1 | 2 | 1 | Q3TUQ5 | Pinin OS=M | Q3TUQ5 [3 | 1 | 2261.077 | 0 | 0.16 | 0.854254 | 11.42 | 95.1 | 104.9 | 7.21 | 20.51 | | High | High | High | High | High | High | High | High | High | High | High | 0.004733 | 0.05114 | 2.89 |
| FALSE | High | QENSPNLD | [R].QENSPI | 1xPhosphc | Q8C351 1x | 0.002743 | 0.000109 | 1 | 2 | 1 | Q8C351 | Layilin OS= | Q8C351 [2 | 0 | 1969.958 | 0 | -1.04 | 0.000375 | 2.71 | 133.8 | 66.2 | 5.99 | 20.72 | | High | High | High | High | High | High | High | High | High | High | High | 4.94E-05 | 0.000742 | 2.41 |
| FALSE | High | QENSSNSS | [K].QENSSI | 1xPhosphc | Q9CXE2 1x | 2.15E-08 | 0 | 1 | 1 | 3 | Q9CXE2 | B-cell CLL/l | Q9CXE2 [1 | 0 | 3078.431 | 0 | 0.05 | 0.783171 | 4.6 | 99 | 101 | 3.65 | 9.46 | | High | High | High | High | High | High | High | High | High | High | High | 0 | 1.36E-09 | 4.09 |
| FALSE | High | QENSSNSS | [K].QENSSI | 1xDeamida | Q9CXE2 1x | 4.32E-06 | 0 | 1 | 1 | 3 | Q9CXE2 | B-cell CLL/l | Q9CXE2 [1 | 0 | 3079.415 | 0 | 0.06 | 0.698163 | 9.67 | 99.9 | 100.1 | 6.98 | 13.06 | | High | High | High | High | High | High | High | High | High | High | High | 0 | 5.25E-07 | 4.03 |
| FALSE | High | QEPAAPSL | [K].QEPAA | 1xPhosphc | A1A549 1x | 7.34E-05 | 0 | 1 | 3 | 1 | A1A549 | Transcripti | A1A549 [3 | 0 | 1991.061 | 0 | -0.8 | 0.001135 | 4.47 | 124.7 | 75.3 | 7.77 | 6.92 | | High | High | High | High | High | High | High | High | High | High | High | 0 | 1.27E-05 | 2.54 |
| FALSE | High | QEPEASQG | [K].QEPEA | 1xPhosphc | Q8C9B9 1x | 1.74E-07 | 0 | 1 | 3 | 4 | Q8C9B9 | Death-indu | Q8C9B9 [2 | 0 | 3253.515 | 0 | -0.4 | 0.004876 | 3.51 | 110.1 | 89.9 | 7.51 | 7.24 | | High | High | High | High | High | High | High | High | High | High | High | 0 | 1.43E-08 | 5.02 |
| FALSE | High | QEPEPGPN | [R].QEPEP( | 1xPhosphc | A2ARP8 1x | 0.011652 | 0.000889 | 1 | 3 | 2 | A2ARP8 | Microtubu | A2ARP8 [2 | 0 | 2551.243 | 0 | -1.13 | 0.001694 | 3.12 | 136.7 | 63.3 | 8.85 | 4.83 | | High | High | High | High | High | High | High | High | High | High | High | 0.000449 | 0.003706 | 2.89 |
| FALSE | High | QEPEVNG( | [K].QEPEVI | 1xPhosphc | B1AU76 1x | 2.12E-10 | 0 | 2 | 4 | 3 | B1AU76; B | Nuclear au | B1AU76 [3 | 0 | 2156.027 | 0 | -0.68 | 0.00041 | 0.7 | 122.7 | 77.3 | 5.93 | 3.87 | Shared | High | High | High | High | High | High | High | High | High | High | High | 0 | 4.46E-10 | 5.1 |
| FALSE | High | QEPEVNG( | [K].QEPEVI | 1xDeamida | B1AU76 1x | 0.002647 | 0.000109 | 2 | 4 | 1 | B1AU76; B | Nuclear au | B1AU76 [3 | 0 | 2157.011 | 0 | -0.47 | 0.000352 | 2.46 | 115.2 | 84.8 | 4.89 | 8.33 | Shared | High | High | High | High | High | High | High | High | High | High | High | 4.94E-05 | 0.000713 | 2.23 |
| FALSE | High | QELASDGE | [K].QELASI | 1xPhosphc | J3QQ27 1x | 3.2E-05 | 0 | 1 | 3 | 1 | J3QQ27 | Coiled-coil | J3QQ27 [7 | 0 | 3353.63 | 0 | -0.11 | 0.0877 | 5.22 | 104.7 | 95.3 | 4.83 | 5.63 | | High | High | High | High | High | High | High | High | High | High | High | 0 | 4.98E-06 | 2.68 |
| FALSE | High | QEPGSPPP. | [K].QEPGSI | 2xCarbami | Q3TDV8 1x | 9.06E-06 | 0 | 1 | 4 | 1 | Q3TDV8 | Putative ur | Q3TDV8 [1 | 0 | 3750.904 | 0 | -0.25 | 0.134627 | 22.42 | 112.5 | 87.5 | 18.64 | 24.31 | | High | High | High | High | High | High | High | High | High | High | High | 0 | 1.21E-06 | 4.55 |
| FALSE | High | QEPMSPEC | [K].QEPMS | 1xCarbami | O88466 1x | 0.000269 | 0 | 2 | 4 | 2 | O88466; R | Zinc finger | O88466 [1 | 0 | 3257.468 | 0 | -0.35 | 0.038215 | 3.03 | 116.4 | 83.6 | 11 | 15.67 | Shared | High | High | High | High | High | High | High | High | High | High | High | 0 | 5.46E-05 | 2.88 |

| | | | | | | | | | | | | | | | | | | | | | | | | | | | | | | | | | | | | | | |
|---|---|---|---|---|---|---|---|---|---|---|---|---|---|---|---|---|---|---|---|---|---|---|---|---|---|---|---|---|---|---|---|---|---|---|---|---|---|---|
| FALSE | High | QEPPEEDSI | [K].QEPPEE | 1xPhosphc | P47930 1xP | 1.55E-07 | 0 | 1 | 2 | 1 | P47930 | Fos-relatec | P47930 [22 | 0 | 2345.026 | 0 | 1.46 | 0.00505 | 9.63 | 52.4 | 147.6 | 6.8 | 20.28 | | High | High | High | High | High | High | High | High | High | High | High | 0 | 1.26E-08 | 3.3 |
| FALSE | High | QEPPSPPHS | [K].QEPPSF | 1xPhosphc | A0A075F6( | 0.077188 | 0.005893 | 1 | 5 | 1 | A0A075F6( | Heat shock | A0A075F6( | 0 | 1537.736 | 0 | -0.43 | 0.018 | 7.63 | 115.3 | 84.7 | 15.1 | 7.84 | | High | High | High | High | High | High | High | High | High | High | High | 0.003035 | 0.03182 | 1.51 |
| FALSE | High | QEPPSPPHS | [K].QEPPSF | 2xPhosphc | A0A075F6( | 0.014797 | 0.001049 | 1 | 5 | 3 | A0A075F6( | Heat shock | A0A075F6( | 0 | 1617.703 | 0 | 0.66 | 0.000158 | 1.64 | 77.8 | 122.2 | 4.91 | 3.89 | | High | High | High | High | High | High | High | High | High | High | High | 0.009441 | 0.102 | 1.8 |
| FALSE | High | QEPTCNNS | [K].QEPTCI | 1xCarbami | E9QAE4 1x | 0.001933 | 5.66E-05 | 1 | 3 | 1 | E9QAE4 | Histone-lys | E9QAE4 [1 | 0 | 1994.92 | 0 | -1.1 | 0.000918 | 2.26 | 138 | 62 | 6.91 | 8.78 | | High | High | High | High | High | High | High | High | High | High | High | 2.58E-05 | 0.000499 | 2.89 |
| FALSE | High | QEPVISESY | [K].QEPVIS | 1xPhosphc | F8WGX5 1: | 1.55E-06 | 0 | 1 | 2 | 2 | F8WGX5 | DmX-like p | F8WGX5 [1 | 0 | 2800.479 | 0 | 0.57 | 0.009495 | 10.02 | 80.5 | 119.5 | 7.84 | 12.9 | | High | High | High | High | High | High | High | High | High | High | High | 0 | 1.65E-07 | 4.55 |
| FALSE | High | QEPVISESY | [K].QEPVIS | 1xDeamida | F8WGX5 1: | 0.004218 | 0.00036 | 1 | 2 | 4 | F8WGX5 | DmX-like p | F8WGX5 [1 | 0 | 2801.463 | 0 | 0.27 | 0.038494 | 4.66 | 86 | 114 | 4.89 | 14.55 | | High | High | High | High | High | High | High | High | High | High | High | 0.000801 | 0.007166 | 3.09 |
| FALSE | High | QEQLSPEG | [R].QEQLSI | 1xPhosphc | E9QAU4 1x | 0.001489 | 5.66E-05 | 1 | 3 | 2 | E9QAU4 | Enhancer t | E9QAU4 [1 | 0 | 1821.895 | 0 | -0.22 | 0.017104 | 5.95 | 109.4 | 90.6 | 6.51 | 3.36 | | High | High | High | High | High | High | High | High | High | High | High | 0.000571 | 0.005046 | 2.44 |
| FALSE | High | QEQLSPK | [R].QEQLSI | 1xPhosphc | Q05BA5 1x | 0.022378 | 0.001542 | 1 | 6 | 1 | Q05BA5 | Nuclear re | Q05BA5 [7 | 0 | 1367.734 | 0 | -0.61 | 0.000371 | 1 | 118.7 | 81.3 | 3.77 | 4.44 | | High | High | High | High | High | High | High | High | High | High | High | 0.000814 | 0.007729 | 1.39 |
| FALSE | High | QEQQRASE | [R].QEQQR | 1xPhosphc | Q3TIZ8 1xP | 0.000967 | 0 | 1 | 4 | 1 | Q3TIZ8 | Putative ur | Q3TIZ8 [31 | 1 | 2070.002 | 0 | 0.6 | 0.000778 | 2.15 | 81.7 | 118.3 | 11.14 | 11.54 | | High | High | High | High | High | High | High | High | High | High | High | 0 | 0.000229 | 1.87 |
| FALSE | High | QESCSPHH | [K].QESCSF | 1xCarbami | Q9D2D7 1x | 0.002192 | 0.000109 | 1 | 3 | 1 | Q9D2D7 | Zinc finger | Q9D2D7 [2 | 0 | 2060.917 | 0 | -0.17 | 0.105461 | 11.58 | 110 | 90 | 10.37 | 11.11 | | High | High | High | High | High | High | High | High | High | High | High | 4.94E-05 | 0.000576 | 2.11 |
| FALSE | High | QESESEQE | [R].QESESE | 1xPhosphc | Q6NS46 1x | 1.53E-05 | 0 | 1 | 1 | 13 | Q6NS46 | Protein RRI | Q6NS46 [1 | 0 | 1957.952 | 0 | -0.65 | 0.001328 | 2.28 | 122.1 | 77.9 | 6.96 | 3.52 | | High | High | High | High | High | High | High | High | High | High | High | 0 | 2.18E-06 | 3.29 |
| FALSE | High | QESESEQE | [R].QESESE | 2xPhosphc | Q6NS46 2x | 0.000128 | 0 | 1 | 1 | 3 | Q6NS46 | Protein RRI | Q6NS46 [1 | 0 | 2037.918 | 0 | 0.92 | 0.009789 | 7.89 | 69 | 131 | 11.71 | 14.4 | | High | High | High | High | High | High | High | High | High | High | High | 0 | 2.38E-05 | 4.55 |
| FALSE | High | QEPLGSDS | [K].QEPLGS | 1xCarbami | Q7TQI3 1xI | 3.7E-08 | 0 | 1 | 1 | 6 | Q7TQI3 | Ubiquitin t | Q7TQI3 [1: | 0 | 3221.397 | 0 | 0.34 | 0.121134 | 25.54 | 81.7 | 118.3 | 17.16 | 17.77 | | High | High | High | High | High | High | High | High | High | High | High | 0 | 2.01E-08 | 6.38 |
| FALSE | High | QESFEAEK | [R].QESFEA | 1xPhosphc | Q8BHZ4 1x | 0.042141 | 0.003162 | 1 | 3 | 2 | Q8BHZ4 | Zinc finger | Q8BHZ4 [7 | 0 | 1505.729 | 0 | 0.17 | 0.14809 | 10.5 | 94.7 | 105.3 | 3.26 | 6.7 | | High | High | High | High | High | High | High | High | High | High | High | 0.001625 | 0.01578 | 2.9 |
| FALSE | High | QELANSSD | [K].QELAN: | 1xPhosphc | Q5SVL6 1x | 0.0002 | 0 | 1 | 4 | 1 | Q5SVL6 | Rap1 GTPa | Q5SVL6 [1( | 0 | 3429.774 | 0 | 3.09 | 0.001204 | 32.7 | 20.4 | 179.6 | 17 | 14.87 | | High | High | High | High | High | High | High | High | High | High | High | 0 | 3.89E-05 | 2.45 |
| FALSE | High | QEGTPEGL | [K].QEGTPI | 1xPhosphc | Q542I9 1xP | 0.018774 | 0.001362 | 1 | 1 | 1 | Q542I9 | MCG12265 | Q542I9 [43 | 0 | 1415.666 | 0 | -0.31 | 0.079643 | 3.82 | 104.9 | 95.1 | 15.9 | 17.03 | | High | High | High | High | High | High | High | High | High | High | High | 0.000723 | 0.006322 | 2.2 |
| FALSE | High | QEAVQEVS | [K].QEAVQ | 1xPhosphc | E9Q8S5 1x | 1.77E-06 | 0 | 1 | 3 | 1 | E9Q8S5 | Microtubu | E9Q8S5 [1: | 0 | 2223.017 | 0 | -0.06 | 0.522486 | 6.75 | 101.9 | 98.1 | 10.09 | 12.36 | | High | High | High | High | High | High | High | High | High | High | High | 0 | 1.94E-07 | 3.49 |
| FALSE | High | QEDENFNS | [R].QEDEN | 1xPhosphc | Q80TH2-2 | 0.006363 | 0.00062 | 1 | 6 | 1 | Q80TH2-2 | Isoform 2 c | Q80TH2-2 | 0 | 2941.341 | 0 | 0.34 | 0.002019 | 8.5 | 88.5 | 111.5 | 5.78 | 3.08 | | High | High | High | High | High | High | High | High | High | High | High | 0.000275 | 0.001887 | 3.24 |
| FALSE | High | QEDENFNS | [R].QEDEN | 1xPhosphc | Q80TH2-2 | 2.2E-07 | 0 | 1 | 6 | 2 | Q80TH2-2 | Isoform 2 c | Q80TH2-2 | 0 | 3170.504 | 0 | 0.27 | 0.067283 | 15.75 | 91.2 | 108.8 | 10.01 | 16.76 | | High | High | High | High | High | High | High | High | High | High | High | 0 | 4.83E-06 | 4.38 |
| FALSE | High | QEDENFNS | [R].QEDEN | 1xDeamida | Q80TH2-2 | 1.45E-09 | 0 | 1 | 6 | 2 | Q80TH2-2 | Isoform 2 c | Q80TH2-2 | 0 | 3171.488 | 0 | 0.4 | 0.021156 | 16.73 | 86.2 | 113.8 | 10.24 | 8.25 | | High | High | High | High | High | High | High | High | High | High | High | 0 | 6.56E-11 | 4.18 |
| FALSE | High | QEDLNSEA | [K].QEDLN: | 1xCarbami | Q7TT18 1x | 1.11E-08 | 0 | 1 | 3 | 4 | Q7TT18 | Activating t | Q7TT18 [1( | 0 | 2531.153 | 0 | 0.28 | 0.026682 | 5.85 | 91.7 | 108.3 | 2.65 | 9.75 | | High | High | High | High | High | High | High | High | High | High | High | 0 | 6.52E-10 | 4.43 |
| FALSE | High | QEDSSLSLE | [K].QEDSSI | 1xPhosphc | Q811S7 1x | 1.35E-05 | 0 | 1 | 3 | 4 | Q811S7 | Upstream- | Q811S7 [4! | 0 | 1988.947 | 0 | -0.06 | 0.445576 | 3.59 | 103.4 | 96.6 | 10.25 | 11.76 | | High | High | High | High | High | High | High | High | High | High | High | 0 | 2.28E-06 | 3.51 |
| FALSE | High | QEEATALEI | [K].QEEAT/ | 1xPhosphc | Q3U3B9 1x | 0.002313 | 0.000109 | 2 | 2 | 2 | Q3U3B9; C | Putative ur | Q3U3B9 [3 | 0 | 2419.095 | 0 | 0.54 | 0.029924 | 13.1 | 82.6 | 117.4 | 16.51 | 27.28 | Shared | High | High | High | High | High | High | High | High | High | High | High | 4.94E-05 | 0.000609 | 2.74 |
| FALSE | High | QEEEAAQC | [K].QEEEA/ | 1xPhosphc | Q7TNE3 1x | 3.65E-10 | 0 | 1 | 2 | 5 | Q7TNE3 | Sperm-ass | Q7TNE3 [1 | 0 | 2642.275 | 0 | -0.69 | 0.000554 | 3.02 | 121.8 | 78.2 | 4.89 | 5.19 | | High | High | High | High | High | High | High | High | High | High | High | 0 | 1.39E-11 | 5.06 |
| FALSE | High | QEEEEWTD | [K].QEEEE\ | 1xPhosphc | Q61458 1x | 0.000731 | 0 | 1 | 2 | 5 | Q61458 | Cyclin-H O: | Q61458 [3( | 0 | 2131.879 | 0 | -0.76 | 0.012953 | 12.3 | 130.1 | 69.9 | 17.05 | 35.78 | | High | High | High | High | High | High | High | High | High | High | High | 0.000275 | 0.002003 | 1.97 |
| FALSE | High | QEEGGSDC | [R].QEEGG | 1xPhosphc | S4R1P5 1xI | 0.0006 | 0 | 1 | 6 | 2 | S4R1P5 | Dystonin C | S4R1P5 [21 | 0 | 2731.235 | 0 | 0.6 | 0.018411 | 19.4 | 78.3 | 121.7 | 10.97 | 11.98 | | High | High | High | High | High | High | High | High | High | High | High | 0 | 0.000134 | 3.28 |
| FALSE | High | QEEPSTSQ | [R].QEEPST | 1xPhosphc | Q8BV49 1x | 0.020862 | 0.001472 | 1 | 7 | 2 | Q8BV49 | Pyrin and H | Q8BV49 [1 | 1 | 4569.077 | 0 | 6.64 | | 0 | 4 | 196 | | 19.95 | | Not Found | Not Found | Not Found | Not Found | High | High | High | High | High | High | High | 0.000953 | 0.009836 | 3.78 |
| FALSE | High | QEEPSTSQ | [R].QEEPST | 1xPhosphc | Q8BV49 1x | 0.002419 | 0.000109 | 1 | 7 | 4 | Q8BV49 | Pyrin and H | Q8BV49 [1 | 1 | 4585.072 | 0 | 4.91 | | 35.71 | 6.4 | 193.6 | 36.48 | 13.86 | | Not Found | High | High | High | High | High | High | High | High | High | High | 0.000204 | 0.001363 | 3.85 |
| FALSE | High | QEEPSTSQ | [R].QEEPST | 1xDeamida | Q8BV49 2x | 0.001767 | 5.66E-05 | 1 | 7 | 2 | Q8BV49 | Pyrin and H | Q8BV49 [1 | 1 | 4650.028 | 0 | 1.41 | 0.001823 | 21.12 | 49.8 | 150.2 | 12.16 | 12.84 | | High | High | High | High | High | High | High | High | High | High | High | 2.58E-05 | 0.00045 | 4.85 |
| FALSE | High | QELANSSD | [K].QELAN: | 1xPhosphc | Q5SVL6 1x | 0.002096 | 5.66E-05 | 1 | 4 | 2 | Q5SVL6 | Rap1 GTPa | Q5SVL6 [1( | 0 | 3413.779 | 0 | 2.62 | 0.000277 | 5.77 | 27.1 | 172.9 | 13.32 | 9.34 | | High | High | High | High | High | High | High | High | High | High | High | 0.000275 | 0.001681 | 2.19 |
| FALSE | High | QEEPSTSQ | [R].QEEPST | 2xPhosphc | Q8BV49 2x | 0.001313 | 0 | 1 | 7 | 3 | Q8BV49 | Pyrin and H | Q8BV49 [1 | 1 | 4665.039 | 0 | 5.48 | 0.001357 | 67.43 | 4.4 | 195.6 | 54.09 | 17.56 | | High | High | High | High | High | High | High | High | High | High | High | 2.58E-05 | 0.000411 | 4.51 |
| FALSE | High | QEESESPVI | [K].QEESES | 2xPhosphc | O70551 2x | 0.001267 | 0 | 1 | 3 | 2 | O70551 | SRSF prote | O70551 [3( | 0 | 2784.289 | 0 | -0.92 | 0.001634 | 2.26 | 132.4 | 67.6 | 7.57 | 7.59 | | High | High | High | High | High | High | High | High | High | High | High | 0 | 0.000312 | 3.47 |
| FALSE | High | QEESPPLVF | [K].QEESPF | 1xCarbami | F8WJ40 1x | 0.000106 | 0 | 1 | 2 | 1 | F8WJ40 | Histone-lys | F8WJ40 [4( | 0 | 1879.882 | 0 | -1.22 | 0.006523 | 1.34 | 136.8 | 63.2 | 12.9 | 11.51 | | High | High | High | High | High | High | High | High | High | High | High | 0 | 1.93E-05 | 1.96 |
| FALSE | High | QEESQQPL | [R].QEESQ( | 1xPhosphc | Q6P1B3 1x | 0.001986 | 5.66E-05 | 1 | 2 | 4 | Q6P1B3 | PILR alpha- | Q6P1B3 [2 | 0 | 4111.933 | 0 | -1.03 | 0.002564 | 1.63 | 127.4 | 72.6 | 13.18 | 13.08 | | High | High | High | High | High | High | High | High | High | High | High | 4.94E-05 | 0.00068 | 4.78 |
| FALSE | High | QEESVDPD | [K].QEESVI | 1xPhosphc | Q6PDQ2 1: | 0.0009 | 0 | 1 | 6 | 2 | Q6PDQ2 | Chromodo | Q6PDQ2 [1 | 0 | 2062.941 | 0 | 0.53 | 0.02911 | 19.43 | 76.8 | 123.2 | 8.13 | 18.05 | | High | High | High | High | High | High | High | High | High | High | High | 0.000344 | 0.002978 | 3.16 |
| FALSE | High | QEEVAFTQ | [K].QEEVAI | 1xPhosphc | Q3UMB7 1 | 0.000184 | 0 | 1 | 5 | 2 | Q3UMB7 | Putative ur | Q3UMB7 [ | 0 | 2351.075 | 0 | 0.65 | 0.004328 | 9.13 | 85 | 115 | 14.92 | 9.65 | | High | High | High | High | High | High | High | High | High | High | High | 0 | 0.000147 | 3.59 |
| FALSE | High | QEFQHFSP | [K].QEFQH | 1xPhosphc | Q1W617 1: | 0.000529 | 0 | 1 | 2 | 2 | Q1W617 | Protein Shi | Q1W617 [: | 0 | 1782.898 | 0 | 1.7 | 0.000498 | 11.11 | 47.2 | 152.8 | 6.39 | 7.97 | | High | High | High | High | High | High | High | High | High | High | High | 0 | 0.000117 | 3.02 |
| FALSE | High | QEGGQISP | [R].QEGGC | 1xPhosphc | Q7TNI6 1xI | 5.86E-05 | 0 | 1 | 1 | 3 | Q7TNI6 | Jerry prote | Q7TNI6 [1( | 0 | 3254.755 | 0 | 1.26 | 0.000281 | 4.21 | 59.8 | 140.2 | 6.57 | 6.07 | | High | High | High | High | High | High | High | High | High | High | High | 0 | 9.88E-06 | 3.56 |
| FALSE | High | QEGGQISP | [R].QEGGC | 1xDeamida | Q7TNI6 1xI | 0.001831 | 5.66E-05 | 1 | 1 | 2 | Q7TNI6 | Jerry prote | Q7TNI6 [1( | 0 | 3255.739 | 0 | 1.31 | 0.00049 | 4.06 | 59 | 141 | 11.23 | 6.92 | | High | High | High | High | High | High | High | High | High | High | High | 2.58E-05 | 0.000472 | 3.61 |
| FALSE | High | QEGLDLAS | [R].QEGLDI | 1xPhosphc | Q9DBY0 1x | 5.18E-06 | 0 | 1 | 6 | 1 | Q9DBY0 | Forkhead b | Q9DBY0 [2 | 0 | 2629.353 | 0 | -0.07 | 0.940036 | 8.41 | 98 | 102 | 6.15 | 12.6 | | High | High | High | High | High | High | High | High | High | High | High | 0 | 6.48E-07 | 3.66 |
| FALSE | High | QEGQDIVL | [R].QEGQD | 2xCarbami | Q9QY24-2 | 0.000958 | 0 | 1 | 1 | 1 | Q9QY24-2 | Isoform 2 c | Q9QY24-2 | 0 | 2121.966 | 0 | 2.94 | 0.000181 | 9.75 | 22.9 | 177.1 | 8.08 | 4.67 | | High | High | High | High | High | High | High | High | High | High | High | 0 | 0.000228 | 3.35 |
| FALSE | High | QEGQFQD( | [K].QEGQF | 1xPhosphc | B1AQJ2 1xI | 0.004613 | 0.000456 | 1 | 3 | 1 | B1AQJ2 | Ubiquitin c | B1AQJ2 [8! | 0 | 2365.086 | 0 | 0.39 | 0.102262 | 23.29 | 84.3 | 115.7 | 4.83 | 21.34 | | High | High | High | High | High | High | High | High | High | High | High | 0.000204 | 0.001328 | 2.49 |
| FALSE | High | QEGSSDDA | [K].QEGSSI | 1xPhosphc | P54276 1xI | 6.36E-09 | 0 | 1 | 2 | 4 | P54276 | DNA misma | P54276 [27 | 0 | 2985.289 | 0 | -0.88 | 0.003415 | 2.54 | 124.9 | 75.1 | 10.16 | 9.73 | | High | High | High | High | High | High | High | High | High | High | High | 0 | 3.48E-10 | 5.06 |
| FALSE | High | QEGSSPEP | [R].QEGSSI | 1xPhosphc | Q99ML1 1: | 0.067619 | 0.004864 | 1 | 3 | 1 | Q99ML1 | Bcl-2-bindi | Q99ML1 [6 | 0 | 1764.837 | 0 | -0.93 | 0.00062 | 4.7 | 132.7 | 67.3 | 6 | 15.57 | | High | High | High | High | High | High | High | High | High | High | High | 0.00253 | 0.0274 | 1.63 |
| FALSE | High | QEESESPVI | [K].QEESES | 1xPhosphc | O70551 1x | 2.29E-05 | 0 | 1 | 3 | 8 | O70551 | SRSF prote | O70551 [3( | 0 | 2704.323 | 0 | -0.67 | 0.000703 | 3.68 | 123.7 | 76.3 | 1.78 | 6.85 | | High | High | High | High | High | High | High | High | High | High | High | 0 | 3.45E-06 | 2.86 |
| FALSE | High | QESGSSLEI | [R].QESGS: | 1xPhosphc | Q3TLI0 1xP | 2.82E-08 | 0 | 1 | 3 | 2 | Q3TLI0 | Trafficking | Q3TLI0 [70 | 0 | 2658.297 | 0 | 1.3 | 0.000377 | 2.84 | 57.7 | 142.3 | 6.86 | 7.42 | | High | High | High | High | High | High | High | High | High | High | High | 0 | 1.19E-05 | 4.09 |
| FALSE | High | QESLKSPEE | [R].QESLKS | 1xPhosphc | Q6P5H2-2 | 0.002115 | 5.66E-05 | 1 | 2 | 1 | Q6P5H2-2 | Isoform 2 c | Q6P5H2-2 | 1 | 2230.023 | 0 | 0.8 | 0.003531 | 13.96 | 70.4 | 129.6 | 19.78 | 9.21 | | High | High | High | High | High | High | High | High | High | High | High | 2.58E-05 | 0.000551 | 3.42 |
| FALSE | High | QESLKSPEE | [R].QESLKS | 1xPhosphc | Q6P5H2-2 | 0.002232 | 0.000109 | 1 | 2 | 2 | Q6P5H2-2 | Isoform 2 c | Q6P5H2-2 | 1 | 2459.186 | 0 | 2.05 | 0.006054 | 21.99 | 42.6 | 157.4 | 25.76 | 17.57 | | High | High | High | High | High | High | High | High | High | High | High | 4.94E-05 | 0.000588 | 1.81 |
| FALSE | High | QFMAETQI | [R].QFMAE | 1xPhosphc | Q60865 1x | 9.98E-05 | 0 | 1 | 4 | 2 | Q60865 | Caprin-1 O | Q60865 [2! | 0 | 2027.955 | 0 | 0.46 | 0.093489 | 14.58 | 85.6 | 114.4 | 6.16 | 19.04 | | High | High | High | High | High | High | High | High | High | High | High | 0 | 1.8E-05 | 3.56 |
| FALSE | High | QFMAETQI | [R].QFMAE | 1xPhosphc | Q60865 1x | 3.45E-06 | 0 | 1 | 4 | 21 | Q60865 | Caprin-1 O | Q60865 [2! | 0 | 2043.95 | 0 | 0.32 | 0.010665 | 10.32 | 89.1 | 110.9 | 2.07 | 6.86 | | High | High | High | High | High | High | High | High | High | High | High | 0 | 4.08E-07 | 4.14 |
| FALSE | High | QFPPSPSAI | [R].QFPPSF | 1xPhosphc | Q9R0A0 1x | 0.000246 | 0 | 1 | 2 | 7 | Q9R0A0 | Peroxisom: | Q9R0A0 [2 | 0 | 1593.844 | 0 | 1.36 | 0.000628 | 9.87 | 59.9 | 140.1 | 5.79 | 9.46 | | High | High | High | High | High | High | High | High | High | High | High | 0.001142 | 0.01178 | 2.31 |
| FALSE | High | QFSAPNLK | [R].QFSAPI | 1xPhosphc | A2AH75 1x | 0.116482 | 0.009523 | 1 | 2 | 1 | A2AH75 | Kinesin fan | A2AH75 [1 | 0 | 1213.618 | 0 | 0.48 | 0.032974 | 15.25 | 83.1 | 116.9 | 5.85 | 12.36 | | High | High | High | High | High | High | High | High | High | High | High | 0.004757 | 0.05191 | 1.22 |
| FALSE | High | QFSAPNLK | [R].QFSAPI | 1xPhosphc | A2AH75 1x | 0.021989 | 0.001542 | 1 | 2 | 1 | A2AH75 | Kinesin fan | A2AH75 [1 | 0 | 1442.781 | 0 | 0.49 | 0.058403 | 25.95 | 84.4 | 115.6 | 7.21 | 16.97 | | High | High | High | High | High | High | High | High | High | High | High | 0.000814 | 0.007579 | 1.65 |
| FALSE | High | QFSLENVPI | [K].QFSLEN | 1xPhosphc | Q6A037 1x | 2.8E-05 | 0 | 1 | 2 | 3 | Q6A037 | NEDD4-bin | Q6A037 [2 | 0 | 2410.231 | 0 | -0.29 | 0.14348 | 8.07 | 106.4 | 93.6 | 6.6 | 11.1 | | High | High | High | High | High | High | High | High | High | High | High | 0.000183 | 0.001274 | 2.87 |
| FALSE | High | QFSPTMSP | [K].QFSPTN | 1xPhosphc | Q3U182 1x | 1.59E-07 | 0 | 2 | 4 | 2 | Q3U182; C | CREB-regu | Q3U182 [4 | 0 | 2960.509 | 0 | 0.19 | 0.893349 | 9.24 | 95.8 | 104.2 | 8.13 | 21.33 | Shared | High | High | High | High | High | High | High | High | High | High | High | 0 | 1.28E-08 | 5.65 |
| FALSE | High | QFSPTMSP | [K].QFSPTN | 1xPhosphc | Q3U182 1x | 0.000808 | 0 | 2 | 4 | 5 | Q3U182; C | CREB-regu | Q3U182 [4 | 0 | 2976.504 | 0 | 0.6 | 0.001041 | 11.22 | 80.3 | 119.7 | 5.95 | 4.64 | Shared | High | High | High | High | High | High | High | High | High | High | High | 0.000723 | 0.006337 | 3.23 |
| FALSE | High | QFSTSGSVI | [R].QFSTSC | 1xPhosphc | Q6IFX2 1xP | 0.064862 | 0.004664 | 1 | 1 | 1 | Q6IFX2 | Keratin, typ | Q6IFX2 [9- | 0 | 1478.766 | 0 | 0.72 | 0.077976 | 41.51 | 77.6 | 122.4 | 9.39 | 31.66 | | High | High | High | High | High | High | High | High | High | High | High | 0.002432 | 0.02607 | 2.12 |
| FALSE | High | QFSTSSDHI | [R].QFSTSS | 1xPhosphc | Q5SVR0 1x | 0.047488 | 0.00354 | 1 | 2 | 4 | Q5SVR0 | TBC1 doma | Q5SVR0 [1 | 0 | 1866.847 | 0 | 0.69 | 0.008985 | 11.16 | 73.4 | 126.6 | 5.81 | 13.75 | | High | High | High | High | High | High | High | High | High | High | High | 0.005812 | 0.06403 | 2.28 |
| FALSE | High | QFSVQAED | [R].QFSVQ, | 1xPhosphc | G5E870 1x | 0.001077 | 0 | 1 | 3 | 5 | G5E870 | E3 ubiquiti | G5E870 [1: | 1 | 2818.237 | 0 | -1.09 | 0.00016 | 0.87 | 136 | 64 | 1.84 | 4.79 | | High | High | High | High | High | High | High | High | High | High | High | 0 | 0.00026 | 2.67 |
| FALSE | High | QFSVQAED | [R].QFSVQ, | 2xPhosphc | G5E870 2x | 0.031989 | 0.00205 | 1 | 3 | 2 | G5E870 | E3 ubiquiti | G5E870 [1: | 1 | 2898.203 | 0 | -0.9 | 0.001972 | 0.35 | 126.7 | 73.3 | 7.08 | 6.69 | | High | High | High | High | High | High | High | High | High | High | High | 0.00111 | 0.01155 | 3.76 |
| FALSE | High | QFTSSSSM | [R].QFTSSS | 1xPhosphc | Q9EQD6 1: | 0.034878 | 0.002269 | 2 | 5 | 11 | Q9EQD6; C | Keratin inte | Q9EQD6 [7 | 0 | 1556.743 | 0 | 5.84 | 0.00149 | 17.52 | 2.6 | 197.4 | 47.16 | 19.81 | Shared | High | High | High | High | High | High | High | High | High | High | High | 0.005561 | 0.06032 | 2.28 |
| FALSE | High | QFLTPSLSC | [K].QFLTPS | 1xPhosphc | Q4QRL3-2 | 0.000668 | 0 | 1 | 3 | 1 | Q4QRL3-2 | Isoform 2 c | Q4QRL3-2 | 0 | 1329.665 | 0 | 0.91 | 0.004696 | 26 | 71.5 | 128.5 | 13.02 | 8.31 | | High | High | High | High | High | High | High | High | High | High | High | 0 | 0.000152 | 2.36 |
| FALSE | High | QFYHSIQDI | [R].QFYHSI | 1xPhosphc | P04627 1xI | 0.000181 | 0 | 1 | 6 | 2 | P04627 | Serine/thre | P04627 [1! | 0 | 1903.89 | 0 | 0.81 | 0.004893 | 9.91 | 71.7 | 128.3 | 12.97 | 6.81 | | High | High | High | High | High | High | High | High | High | High | High | 0 | 3.49E-05 | 2.54 |
| FALSE | High | QGADREES | [R].QGADR | 1xCarbami | A1L341 1xI | 0.002022 | 5.66E-05 | 1 | 5 | 4 | A1L341 | Dual-specit | A1L341 [73 | 1 | 2745.206 | 0 | -0.67 | 0.200743 | 8.03 | 122.8 | 77.2 | 6.62 | 36.72 | | High | High | High | High | High | High | High | High | High | High | High | 2.58E-05 | 0.000523 | 2.55 |
| FALSE | High | QGADREES | [R].QGADR | 1xCarbami | A1L341 2xI | 0.010109 | 0.000736 | 1 | 5 | 2 | A1L341 | Dual-specit | A1L341 [73 | 1 | 2825.172 | 0 | 0 | 0.634532 | 25.73 | 97.9 | 102.1 | 8.45 | 19.77 | | High | High | High | High | High | High | High | High | High | High | High | 0.000344 | 0.003182 | 3.08 |
| FALSE | High | QGAEDSLS | [R].QGAED | 1xCarbami | Q3UGP2 1: | 2.97E-07 | 0 | 1 | 3 | 1 | Q3UGP2 | Putative ur | Q3UGP2 [1 | 0 | 2254.001 | 0 | 1.71 | 0.000724 | 13.16 | 46.9 | 153.1 | 13.74 | 10.23 | | High | High | High | High | High | High | High | High | High | High | High | 0 | 2.6E-08 | 4.65 |
| FALSE | High | QGAGENSI | [K].QGAGE | 1xCarbami | A0A1L1SQI | 1.04E-05 | 0 | 1 | 7 | 1 | A0A1L1SQI | FAT atypica | A0A1L1SQI | 1 | 2833.26 | 0 | 0.6 | 0.001488 | 8.06 | 76 | 124 | 8.58 | 7.4 | | High | High | High | High | High | High | High | High | High | High | High | 0 | 1.41E-06 | 3.21 |
| FALSE | High | QGAIVAVT | [R].QGAIV/ | 1xPhosphc | Q3TXF9 1x | 8.37E-07 | 0 | 1 | 15 | 1 | Q3TXF9 | Sodium/pc | Q3TXF9 [7( | 0 | 2350.242 | 0 | 0.14 | 0.042543 | 8.29 | 91.7 | 108.3 | 9.08 | 12.07 | | High | High | High | High | High | High | High | High | High | High | High | 0 | 8.35E-08 | 3.96 |
| FALSE | High | QGAIYHGL | [K].QGAIYH | 1xPhosphc | E9Q9C3 1x | 5.8E-08 | 0 | 1 | 5 | 1 | E9Q9C3 | Afadin OS= | E9Q9C3 [1( | 0 | 2635.309 | 0 | -0.26 | 0.047189 | 5.62 | 107.8 | 92.2 | 2.21 | 14.51 | | High | High | High | High | High | High | High | High | High | High | High | 0 | 4.14E-09 | 3.91 |
| FALSE | High | QGAIYHGL | [K].QGAIYH | 1xPhosphc | E9Q9C3 1x | 1.21E-05 | 0 | 1 | 5 | 7 | E9Q9C3 | Afadin OS= | E9Q9C3 [1( | 0 | 2651.304 | 0 | -0.17 | 0.054306 | 9.67 | 107.5 | 92.5 | 5.71 | 7.31 | | High | High | High | High | High | High | High | High | High | High | High | 0 | 1.06E-05 | 4.08 |
| FALSE | High | QGAIYHGL | [K].QGAIYH | 1xDeamida | E9Q9C3 1x | 0.056248 | 0.004124 | 1 | 5 | 4 | E9Q9C3 | Afadin OS= | E9Q9C3 [1( | 0 | 2652.288 | 0 | -0.22 | 0.029506 | 4.59 | 108.5 | 91.5 | 10.45 | 11.02 | | High | High | High | High | High | High | High | High | High | High | High | 0.002675 | 0.02852 | 3.2 |
| FALSE | High | QGAIYHGL | [K].QGAIYH | 1xPhosphc | E9Q9C3 1x | 2.11E-08 | 0 | 1 | 5 | 4 | E9Q9C3 | Afadin OS= | E9Q9C3 [1( | 0 | 2667.299 | 0 | -0.22 | 0.111945 | 4.49 | 106.7 | 93.3 | 7.38 | 5.79 | | High | High | High | High | High | High | High | High | High | High | High | 0 | 1.34E-09 | 3.48 |
| FALSE | High | QGANSRPV | [K].QGANS | 1xPhosphc | Q8CFT3 1x | 0.021607 | 0.001507 | 1 | 3 | 3 | Q8CFT3 | Nerve grov | Q8CFT3 [2! | 0 | 2444.234 | 0 | 1.92 | 0.001075 | 12.75 | 44.2 | 155.8 | 7.23 | 13.23 | | High | High | High | High | High | High | High | High | High | High | High | 0.001651 | 0.01635 | 2.28 |
| FALSE | High | QGANSRPV | [K].QGANS | 1xDeamida | Q8CFT3 1x | 0.00153 | 5.66E-05 | 1 | 3 | 3 | Q8CFT3 | Nerve grov | Q8CFT3 [2! | 0 | 2445.218 | 0 | 1.95 | 0.000731 | 14.97 | 41 | 159 | 9.97 | 8.91 | | High | High | High | High | High | High | High | High | High | High | High | 0.001789 | 0.01764 | 1.59 |
| FALSE | High | QGDDEQSI | [R].QGDDE | 1xPhosphc | Q7TQC7 1x | 0.001767 | 5.66E-05 | 1 | 7 | 2 | Q7TQC7 | G patch do | Q7TQC7 [2 | 0 | 2184.953 | 0 | 0.6 | 0.028782 | 21.67 | 63.5 | 136.5 | 19.61 | 24.91 | | High | High | High | High | High | High | High | High | High | High | High | 2.58E-05 | 0.00045 | 2.44 |
| FALSE | High | QGDETPST | [R].QGDET | 1xPhosphc | A2AWA9 1 | 0.00045 | 0 | 1 | 3 | 2 | A2AWA9 | Rab GTPas | A2AWA9 [: | 0 | 2231.971 | 0 | 0.29 | 0.030831 | 14.3 | 83.2 | 116.8 | 9.8 | 13.45 | | High | High | High | High | High | High | High | High | High | High | High | 0 | 9.74E-05 | 2.68 |
| FALSE | High | QGADREES | [R].QGADR | 1xCarbami | A1L341 1xI | 0.006536 | 0.00062 | 1 | 5 | 2 | A1L341 | Dual-specit | A1L341 [73 | 1 | 2729.211 | 0 | 0.02 | 0.556809 | 6.07 | 97.3 | 102.7 | 5.72 | 10.66 | | High | High | High | High | High | High | High | High | High | High | High | 0.004193 | 0.04515 | 2.81 |
| FALSE | High | QFLISPPAS | [K].QFLISPI | 2xPhosphc | Q7TNY3 2x | 0.002172 | 5.66E-05 | 1 | 7 | 2 | Q7TNY3 | Calcipressi | Q7TNY3 [1 | 0 | 2242.148 | 0 | 0.36 | 0.008815 | 0.35 | 87.6 | 112.4 | 4.48 | 6.46 | | High | High | High | High | High | High | High | High | High | High | High | 0.000327 | 0.002504 | 1.99 |
| FALSE | High | QFLISPPAS | [K].QFLISPI | 1xPhosphc | Q7TNY3 1x | 0.000369 | 0 | 1 | 7 | 1 | Q7TNY3 | Calcipressi | Q7TNY3 [1 | 0 | 2162.182 | 0 | 0.21 | 0.191378 | 12.63 | 93.3 | 106.7 | 8.07 | 9.48 | | High | High | High | High | High | High | High | High | High | High | High | 0 | 7.78E-05 | 2.19 |
| FALSE | High | QFLLQQAS | [R].QFLLQ( | 1xPhosphc | Q9DBY0 1x | 4.26E-07 | 0 | 1 | 6 | 1 | Q9DBY0 | Forkhead b | Q9DBY0 [8 | 0 | 2586.297 | 0 | -0.51 | 0.013862 | 5 | 118.7 | 81.3 | 11.3 | 2.9 | | High | High | High | High | High | High | High | High | High | High | High | 0 | 3.9E-08 | 4.96 |
| FALSE | High | QESPEPPSI | [K].QESPEF | 1xPhosphc | P51944-2 1 | 0.000137 | 0 | 1 | 3 | 5 | P51944-2 | Isoform 2 c | P51944-2 [ | 0 | 2966.377 | 0 | 0.21 | 0.366032 | 17.14 | 91.5 | 108.5 | 7.23 | 12.42 | | High | High | High | High | High | High | High | High | High | High | High | 0 | 2.56E-05 | 3.83 |
| FALSE | High | QESPGITPI | [K].QESPGI | 1xPhosphc | E9PVX6 1xI | 0.001049 | 0 | 1 | 3 | 3 | E9PVX6 | Proliferatic | E9PVX6 [5: | 0 | 1390.693 | 0 | -0.11 | 0.980113 | 16.83 | 98.2 | 101.8 | 7.38 | 20.72 | | High | High | High | High | High | High | High | High | High | High | High | 2.58E-05 | 0.000381 | 1.91 |
| FALSE | High | QESPGITPI | [K].QESPGI | 2xPhosphc | E9PVX6 2xI | 0.011146 | 0.000772 | 1 | 3 | 1 | E9PVX6 | Proliferatic | E9PVX6 [5: | 0 | 1470.659 | 0 | 0.41 | 0.03056 | 10.98 | 84.3 | 115.7 | 7.41 | 17.4 | | High | High | High | High | High | High | High | High | High | High | High | 0.000399 | 0.003525 | 1.92 |
| FALSE | High | QESPPNPPI | [R].QESPPI | 1xPhosphc | Q7TT18 1x | 9.36E-05 | 0 | 1 | 3 | 1 | Q7TT18 | Activating t | Q7TT18 [6! | 0 | 3478.646 | 0 | 0.07 | 0.393776 | 10.69 | 97.4 | 102.6 | 11.66 | 7.03 | | High | High | High | High | High | High | High | High | High | High | High | 0 | 1.66E-05 | 4.31 |
| FALSE | High | QESPPNPPI | [R].QESPPI | 1xPhosphc | Q7TT18 1x | 0.112856 | 0.009146 | 1 | 3 | 1 | Q7TT18 | Activating t | Q7TT18 [6! | 0 | 3494.641 | 0 | -0.59 | 0.069446 | 11.17 | 122.9 | 77.1 | 16.43 | 13.48 | | High | High | High | High | High | High | High | High | High | High | High | 0.004583 | 0.04982 | 2.35 |
| FALSE | High | QESQQNSI | [R].QESQQ | 1xPhosphc | P40201 1xI | 0.012288 | 0.000889 | 1 | 1 | 1 | P40201 | Chromodo | P40201 [1! | 0 | 2565.189 | 0 | -0.12 | 0.786506 | 25.54 | 101.7 | 98.3 | 15.04 | 19.96 | | High | High | High | High | High | High | High | High | High | High | High | 0.000449 | 0.003935 | 2.89 |
| FALSE | High | QESQVQQ( | [R].QESQV | 1xPhosphc | Q8R322 1x | 1.43E-05 | 0 | 1 | 1 | 1 | Q8R322 | Nucleopor | Q8R322 [3! | 0 | 3938.954 | 0 | -0.14 | 0.031814 | 5.32 | 104 | 96 | 6.2 | 11.2 | | High | High | High | High | High | High | High | High | High | High | High | 0 | 2.02E-06 | 4.86 |
| FALSE | High | QESSENLPI | [K].QESSEN | 1xPhosphc | Q7TT18 1x | 0.000916 | 0 | 1 | 4 | 1 | Q7TT18 | Activating t | Q7TT18 [5: | 0 | 1569.793 | 0 | -0.58 | 0.000348 | 2.42 | 121.7 | 78.3 | 3.44 | 5.11 | | High | High | High | High | High | High | High | High | High | High | High | 0 | 0.000217 | 2.21 |
| FALSE | High | QESSRPGA | [K].QESSRF | 1xPhosphc | Q3UHL0 1x | 3.83E-05 | 0 | 1 | 6 | 1 | Q3UHL0 | Putative ur | Q3UHL0 [8 | 0 | 2873.299 | 0 | 0.24 | 0.178448 | 22.8 | 91.8 | 108.2 | 7.31 | 10.61 | | High | High | High | High | High | High | High | High | High | High | High | 0 | 6.1E-06 | 1.82 |
| FALSE | High | QESTETLV\ | [R].QESTET | 1xPhosphc | A0A087W( | 0.000501 | 0 | 1 | 2 | 1 | A0A087W( | TRAF3-inte | A0A087W( | 0 | 1585.767 | 0 | -0.43 | 0.026561 | 5.34 | 116.3 | 83.7 | 7.4 | 12.74 | | High | High | High | High | High | High | High | High | High | High | High | 0 | 0.00011 | 1.98 |
| FALSE | High | QESVELTV( | [R].QESVEI | 1xPhosphc | P52734 1xI | 1.45E-06 | 0 | 1 | 3 | 2 | P52734 | FYVE, Rho( | P52734 [3( | 0 | 1826.967 | 0 | -0.89 | 0.000707 | 3.98 | 130.1 | 69.9 | 1.74 | 10.46 | | High | High | High | High | High | High | High | High | High | High | High | 0 | 1.34E-06 | 2.91 |
| FALSE | High | QETAAVCG | [K].QETAA\ | 1xCarbami | E9PYL2 2xP | 2.11E-05 | 0 | 1 | 2 | 1 | E9PYL2 | Proline-ricl | E9PYL2 [15 | 0 | 2888.153 | 0 | -0.37 | 0.005223 | 1.42 | 113.7 | 86.3 | 6.32 | 11.59 | | High | High | High | High | High | High | High | High | High | High | High | 0 | 3.14E-06 | 3.7 |
| FALSE | High | QETESLME | [K].QETESL | 2xPhosphc | A0A1W2P6 | 0.012959 | 0.000889 | 1 | 6 | 1 | A0A1W2P6 | Bromodom | A0A1W2P6 | 0 | 2341.023 | 0 | 0.09 | 0.529998 | 21.18 | 93.9 | 106.1 | 11.97 | 15.86 | | High | High | High | High | High | High | High | High | High | High | High | 0.000449 | 0.004201 | 2.67 |
| FALSE | High | QETQALSA | [R].QETQA | 2xDeamida | Q4QRL3-2 | 0.001572 | 5.66E-05 | 1 | 3 | 2 | Q4QRL3-2 | Isoform 2 c | Q4QRL3-2 | 0 | 1485.704 | 0 | 1.47 | 0.852486 | 52.41 | 49.2 | 150.8 | 158.08 | 45.41 | | High | High | High | High | High | High | High | High | High | High | High | 2.58E-05 | 0.000395 | 1.94 |
| FALSE | High | QETVPATS | [R].QETVP/ | 1xPhosphc | A3KGF9 1x | 5.23E-06 | 0 | 1 | 1 | 2 | A3KGF9 | Uncharacte | A3KGF9 [3! | 0 | 2360.251 | 0 | 2.22 | 0.000154 | 6.32 | 35.3 | 164.7 | 7.66 | 3.88 | | High | High | High | High | High | High | High | High | High | High | High | 0 | 0.000064 | 3.25 |
| FALSE | High | QETVPATS | [R].QETVP/ | 2xPhosphc | A3KGF9 2x | 0.004955 | 0.000504 | 1 | 1 | 1 | A3KGF9 | Uncharacte | A3KGF9 [3! | 0 | 2440.218 | 0 | 2.67 | 0.000285 | 12.43 | 29.6 | 170.4 | 13.76 | 6.63 | | High | High | High | High | High | High | High | High | High | High | High | 0.000225 | 0.001439 | 2.28 |
| FALSE | High | QETVSDFT | [K].QETVSI | 1xPhosphc | Q9R1C7 1x | 1.46E-06 | 0 | 1 | 4 | 3 | Q9R1C7 | Pre-mRNA- | Q9R1C7 [3 | 0 | 1689.85 | 0 | -0.2 | 0.009687 | 5.71 | 108.3 | 91.7 | 4.78 | 4.01 | | High | High | High | High | High | High | High | High | High | High | High | 0 | 1.56E-07 | 4.06 |
| FALSE | High | QEVFAYSP: | [K].QEVFA\ | 1xPhosphc | Q5SVL6 1x | 0.002172 | 5.66E-05 | 1 | 2 | 1 | Q5SVL6 | Rap1 GTPa | Q5SVL6 [6! | 0 | 3205.533 | 0 | 4.73 | 0.002462 | 47.74 | 8 | 192 | 28.19 | 22.28 | | High | High | High | High | High | High | High | High | High | High | High | 4.94E-05 | 0.000572 | 3.46 |
| FALSE | High | QEVGPPLP | [R].QEVGP | 1xPhosphc | E9Q286 1x | 0.003526 | 0.00021 | 1 | 1 | 2 | E9Q286 | Little elong | E9Q286 [1! | 0 | 2411.366 | 0 | -0.84 | 0.000181 | 1.76 | 128.4 | 71.6 | 5.17 | 11.57 | | High | High | High | High | High | High | High | High | High | High | High | 0.00168 | 0.01679 | 2.36 |
| FALSE | High | QEVHCQN' | [K].QEVHC | 1xCarbami | Q8K298 1x | 1.08E-06 | 0 | 1 | 1 | 2 | Q8K298 | Anillin OS= | Q8K298 [4( | 0 | 1891.95 | 0 | 0.89 | 0.008712 | 9 | 70.2 | 129.8 | 23.11 | 1.85 | | High | High | High | High | High | High | High | High | High | High | High | 0 | 1.1E-07 | 3.71 |
| FALSE | High | QEVLPVLP/ | [K].QEVLP\ | 1xPhosphc | A2AJ15 1xP | 0.051694 | 0.003745 | 1 | 3 | 1 | A2AJ15 | Endoplasm | A2AJ15 [8! | 1 | 3718.991 | 0 | 1.38 | 0.001893 | 25.06 | 51 | 149 | 8.77 | 11.95 | | High | High | High | High | High | High | High | High | High | High | High | 0.001999 | 0.0201 | 3.18 |
| FALSE | High | QEVNGAVI | [K].QEVNG | 1xPhosphc | Q69Z61 1x | 0.003464 | 0.00021 | 1 | 2 | 1 | Q69Z61 | PWWP dor | Q69Z61 [2: | 0 | 2325.133 | 0 | -0.09 | 0.847124 | 4.96 | 102.9 | 97.1 | 4.27 | 11.54 | | High | High | High | High | High | High | High | High | High | High | High | 9.42E-05 | 0.000956 | 2.73 |
| FALSE | High | QEVNGAVI | [K].QEVNG | 1xDeamida | Q69Z61 1x | 0.093123 | 0.007276 | 1 | 2 | 1 | Q69Z61 | PWWP dor | Q69Z61 [2: | 0 | 2326.117 | 0 | 0.58 | 0.033963 | 1.69 | 80.9 | 119.1 | 14.52 | 22.58 | | High | High | High | High | High | High | High | High | High | High | High | 0.003698 | 0.03975 | 2.68 |
| FALSE | High | QEYDESGP | [K].QEYDE! | 1xPhosphc | P60710 1xI | 0.002376 | 0.000109 | 1 | 20 | 1 | P60710 | Actin, cyto | P60710 [3( | 0 | 1825.832 | 0 | -0.05 | 0.326498 | 4.35 | 98.4 | 101.6 | 9.72 | 6.81 | | High | High | High | High | High | High | High | High | High | High | High | 4.94E-05 | 0.000628 | 3.24 |
| FALSE | High | QFCQEGQI | [K].QFCQE( | 1xCarbami | Q9D0U6 1x | 2.38E-05 | 0 | 1 | 3 | 3 | Q9D0U6 | Repressor | Q9D0U6 [4 | 0 | 3159.513 | 0 | -0.75 | 0.000158 | 0.3 | 125.6 | 74.4 | 3.37 | 3.7 | | High | High | High | High | High | High | High | High | High | High | High | 0 | 3.56E-06 | 4.07 |
| FALSE | High | QFDASTPQ | [K].QFDAS | 1xPhosphc | Q8BHK9 1x | 2E-05 | 0 | 1 | 3 | 3 | Q8BHK9 | DNA excisi | Q8BHK9 [1 | 0 | 1887.844 | 0 | -0.04 | 0.940589 | 5.94 | 99.1 | 100.9 | 6.22 | 8.6 | | High | High | High | High | High | High | High | High | High | High | High | 0 | 2.95E-06 | 3.53 |
| FALSE | High | QFDASTPQ | [K].QFDAS | 2xPhosphc | Q8BHK9 2x | 0.092372 | 0.007211 | 1 | 3 | 2 | Q8BHK9 | DNA excisi | Q8BHK9 [1 | 0 | 1967.81 | 0 | -0.02 | 0.371087 | 3.64 | 96.9 | 103.1 | 8.76 | 12.6 | | High | High | High | High | High | High | High | High | High | High | High | 0.006741 | 0.07303 | 1.82 |
| FALSE | High | QFDEESLES | [R].QFDEES | 1xPhosphc | Q9R0L6-2 : | 1.73E-05 | 0 | 1 | 3 | 1 | Q9R0L6-2 | Isoform 2 c | Q9R0L6-2 [ | 0 | 3238.515 | 0 | 0.21 | 0.175506 | 7.61 | 90.7 | 109.3 | 8.03 | 11.5 | | High | High | High | High | High | High | High | High | High | High | High | 0 | 2.52E-06 | 4.33 |
| FALSE | High | QFDEESLES | [R].QFDEES | 1xPhosphc | Q9R0L6-2 : | 1.17E-06 | 0 | 1 | 3 | 2 | Q9R0L6-2 | Isoform 2 c | Q9R0L6-2 [ | 0 | 3254.51 | 0 | 0.42 | 0.131208 | 6.77 | 87.3 | 112.7 | 9.35 | 9.7 | | High | High | High | High | High | High | High | High | High | High | High | 0 | 0.000248 | 3.66 |
| FALSE | High | QEAPLHSPI | [R].QEAPLH | 1xPhosphc | B9EKF4 1xI | 0.001087 | 0 | 1 | 3 | 1 | B9EKF4 | Zinc finger | B9EKF4 [1( | 0 | 1999.03 | 0 | 0.11 | 0.079885 | 6.69 | 96.1 | 103.9 | 1.06 | 5.78 | | High | High | High | High | High | High | High | High | High | High | High | 0 | 0.000262 | 2.24 |

| | | | | | | | | | | | | | | | | | | | | | | | | | | | | | | | | | | | | | | |
|---|---|---|---|---|---|---|---|---|---|---|---|---|---|---|---|---|---|---|---|---|---|---|---|---|---|---|---|---|---|---|---|---|---|---|---|---|---|---|
| FALSE | High | QEAGDSPP | [K].QEAGD | 2xPhosphc | Q62311 2x | 0.000211 | 0 | 1 | 1 | 3 | Q62311 | Transcripti | Q62311 [6 | 0 | 2066.96 | 0 | -0.22 | 0.002647 | 3.48 | 107.7 | 92.3 | 5.26 | 7.4 | | High | High | High | High | High | High | High | High | High | High | High | 0 | 0.000112 | 4.03 |
| FALSE | High | QEAGDSPP | [K].QEAGD | 1xPhosphc | Q62311 1x | 1.19E-06 | 0 | 1 | 1 | 3 | Q62311 | Transcripti | Q62311 [6 | 0 | 1986.994 | 0 | -0.42 | 0.006006 | 4.98 | 113.9 | 86.1 | 2.42 | 6.86 | | High | High | High | High | High | High | High | High | High | High | High | 0 | 1.61E-06 | 3.52 |
| FALSE | High | QEAEDLSL | [R].QEAED | 2xPhosphc | E9PV45 2x | 4.01E-05 | 0 | 1 | 2 | 5 | E9PV45 | Ubiquitin c | E9PV45 [20 | 0 | 3787.728 | 0 | 0.6 | 0.001984 | 13.38 | 78.9 | 121.1 | 8.96 | 8.23 | | High | High | High | High | High | High | High | High | High | High | High | 0 | 0.000271 | 3.92 |
| FALSE | High | QALMPVTF | [R].QALMP | 1xCarbami | Q5FWX6 1 | 7.53E-08 | 0 | 1 | 3 | 1 | Q5FWX6 | Serine/thre | Q5FWX6 [5 | 0 | 2611.303 | 0 | 1.25 | 0.000361 | 7.8 | 58.8 | 141.2 | 5.57 | 6.83 | | High | High | High | High | High | High | High | High | High | High | High | 0 | 5.57E-09 | 4.13 |
| FALSE | High | QALNGLSC | [K].QALNG | 1xPhosphc | A0A0R4J0S | 6.68E-07 | 0 | 1 | 7 | 1 | A0A0R4J0S | Uveal auto | A0A0R4J0S | 0 | 2390.237 | 0 | -0.49 | 0.000311 | 1.62 | 118.5 | 81.5 | 4.44 | 8.06 | | High | High | High | High | High | High | High | High | High | High | High | 0 | 6.44E-08 | 3.02 |
| FALSE | High | QALNGLSC | [K].QALNG | 1xDeamida | A0A0R4J0S | 6.89E-05 | 0 | 1 | 7 | 2 | A0A0R4J0S | Uveal auto | A0A0R4J0S | 0 | 2391.221 | 0 | -1.12 | 0.001839 | 2.74 | 133.4 | 66.6 | 6.49 | 12.03 | | High | High | High | High | High | High | High | High | High | High | High | 4.94E-05 | 0.000618 | 2.73 |
| FALSE | High | QAIPESEAE | [R].QAIPES | 1xPhosphc | Q6ZQB7 1x | 0.081085 | 0.006219 | 1 | 4 | 1 | Q6ZQB7 | MKIAA042 | Q6ZQB7 [7 | 1 | 4093.908 | 0 | -0.21 | 0.180179 | 14.41 | 107.3 | 92.7 | 11.11 | 5.67 | | High | High | High | High | High | High | High | High | High | High | High | 0.0032 | 0.0337 | 3.07 |
| FALSE | High | QALQDLVD | [R].QALQD | 1xPhosphc | Q6P1D7 1x | 4.49E-07 | 0 | 1 | 2 | 4 | Q6P1D7 | Structure-s | Q6P1D7 [4 | 0 | 2676.344 | 0 | -0.49 | 0.007381 | 6.36 | 115 | 85 | 10.37 | 11.59 | | High | High | High | High | High | High | High | High | High | High | High | 0 | 4.13E-08 | 3.92 |
| FALSE | High | QAISEPDEE | [K].QAISEP | 1xCarbami | Q9WU62 1 | 1.96E-07 | 0 | 1 | 2 | 1 | Q9WU62 | Inner centr | Q9WU62 [ | 0 | 3167.449 | 0 | -1.33 | 0.000278 | 1.71 | 143 | 57 | 3.72 | 5.08 | | High | High | High | High | High | High | High | High | High | High | High | 0 | 1.63E-08 | 5.21 |
| FALSE | High | QANAASVC | [K].QANAA | 1xPhosphc | E9Q4Y4 1x | 0.003083 | 0.00016 | 1 | 2 | 1 | E9Q4Y4 | Centrosom | E9Q4Y4 [1 | 0 | 3813.994 | 0 | -0.49 | 0.017776 | 10.01 | 114.3 | 85.7 | 13.37 | 18.74 | | High | High | High | High | High | High | High | High | High | High | High | 7.19E-05 | 0.000841 | 3.74 |
| FALSE | High | QAPFRSPT | [K].QAPFR | 2xPhosphc | Q8BG30 2x | 0.087983 | 0.006639 | 1 | 1 | 1 | Q8BG30 | Negative e | Q8BG30 [2 | 1 | 2422.116 | 0 | -0.24 | 0.023867 | 8.96 | 109.9 | 90.1 | 6.96 | 5.84 | | High | High | High | High | High | High | High | High | High | High | High | 0.003402 | 0.03714 | 2.46 |
| FALSE | High | QAPGVGAV | [R].QAPGV | 2xPhosphc | A2AES4 2x | 3.63E-05 | 0 | 1 | 3 | 2 | A2AES4 | OTU doma | A2AES4 [1 | 1 | 3753.627 | 0 | -0.43 | 0.006871 | 9.95 | 114.1 | 85.9 | 2.41 | 15.25 | | High | High | High | High | High | High | High | High | High | High | High | 0 | 5.75E-06 | 4.51 |
| FALSE | High | QAPQVLQS | [K].QAPQV | 1xPhosphc | B2RWU2 1 | 0.014929 | 0.001049 | 1 | 2 | 2 | B2RWU2 | Immunode | B2RWU2 [ | 0 | 2232.122 | 0 | 0.17 | 0.074335 | 13.49 | 92 | 108 | 2.76 | 9.2 | | High | High | High | High | High | High | High | High | High | High | High | 0.000555 | 0.004912 | 2.01 |
| FALSE | High | QAPSISDSE | [K].QAPSIS | 1xPhosphc | E9PVA2 1x | 0.000369 | 0 | 1 | 3 | 2 | E9PVA2 | Ribosomal | E9PVA2 [1 | 0 | 1656.825 | 0 | 0.62 | 0.000539 | 7.19 | 80.3 | 119.7 | 5.98 | 4.65 | | High | High | High | High | High | High | High | High | High | High | High | 0 | 7.75E-05 | 2.75 |
| FALSE | High | QAPSPETD | [K].QAPSP | 1xPhosphc | Q8BRM2 1 | 0.108463 | 0.008839 | 1 | 2 | 1 | Q8BRM2 | RAB6-inter | Q8BRM2 [ | 0 | 2022.088 | 0 | 0.84 | 0.000905 | 11.57 | 70.1 | 129.9 | 9.08 | 3.98 | | High | High | High | High | High | High | High | High | High | High | High | 0.004411 | 0.04741 | 1.78 |
| FALSE | High | QAQSPAGV | [R].QAQSP | 1xPhosphc | B8QI33 1x | 0.000483 | 0 | 1 | 3 | 1 | B8QI33 | Liprin-alph | B8QI33 [14 | 0 | 2167.141 | 0 | -0.22 | 0.054417 | 18 | 110.3 | 89.7 | 9.5 | 8.1 | | High | High | High | High | High | High | High | High | High | High | High | 0 | 0.000105 | 3.32 |
| FALSE | High | QALMPVTF | [R].QALMP | 1xCarbami | Q5FWX6 1 | 2.54E-06 | 0 | 1 | 3 | 1 | Q5FWX6 | Serine/thre | Q5FWX6 [5 | 0 | 2595.308 | 0 | 1.15 | 0.000405 | 10.4 | 61.9 | 138.1 | 9.83 | 8.44 | | High | High | High | High | High | High | High | High | High | High | High | 0 | 2.89E-07 | 3.76 |
| FALSE | High | QAQSSTEIF | [K].QAQSS | 1xPhosphc | B2RRE0 1x | 3.69E-07 | 0 | 1 | 3 | 9 | B2RRE0 | A kinase (P | B2RRE0 [2 | 0 | 3027.456 | 0 | -1.36 | 0.001144 | 2.45 | 143.9 | 56.1 | 6.52 | 11.66 | | High | High | High | High | High | High | High | High | High | High | High | 0 | 1.11E-07 | 5.07 |
| FALSE | High | QAQVVGD | [K].QAQVV | 1xPhosphc | Q9QY06-2 | 0.000311 | 0 | 2 | 9 | 2 | Q9QY06-2; | Isoform 2 c | Q9QY06-2 | 0 | 2982.525 | 0 | 0.57 | 0.002152 | 9.37 | 77.7 | 122.3 | 6.8 | 6.04 | Shared | High | High | High | High | High | High | High | High | High | High | High | 0 | 6.4E-05 | 2.49 |
| FALSE | High | QASASYDS | [K].QASAS | 1xPhosphc | Q8K2F0-2 | 4.37E-07 | 0 | 1 | 4 | 16 | Q8K2F0-2 | Isoform 2 c | Q8K2F0-2 | 0 | 3032.301 | 0 | -0.76 | 0.000353 | 2.05 | 125.6 | 74.4 | 2.31 | 3.43 | | High | High | High | High | High | High | High | High | High | High | High | 0 | 2.09E-07 | 4.03 |
| FALSE | High | QASASYDS | [K].QASAS | 1xPhosphc | Q8K2F0-2 | 1.53E-06 | 0 | 1 | 4 | 34 | Q8K2F0-2 | Isoform 2 c | Q8K2F0-2 | 0 | 3048.296 | 0 | -0.74 | 0.000466 | 0.95 | 124.5 | 75.5 | 3.89 | 3.7 | | High | High | High | High | High | High | High | High | High | High | High | 0 | 8.07E-07 | 4.69 |
| FALSE | High | QASDTEED | [R].QASDT | 1xPhosphc | Q80WC1-4 | 5.35E-08 | 0 | 1 | 5 | 3 | Q80WC1-4 | Isoform 4 c | Q80WC1-4 | 0 | 2280.991 | 0 | 0.11 | 0.079492 | 5.25 | 95.8 | 104.2 | 5.07 | 7.32 | | High | High | High | High | High | High | High | High | High | High | High | 0 | 3.78E-09 | 3.89 |
| FALSE | High | QASEDSNE | [K].QASED | 1xPhosphc | E9QN63 1x | 3.81E-06 | 0 | 1 | 13 | 3 | E9QN63 | Arf-GAP wi | E9QN63 [1 | 0 | 2686.317 | 0 | 0.48 | 0.025092 | 8.55 | 78.4 | 121.6 | 8.38 | 15.19 | | High | High | High | High | High | High | High | High | High | High | High | 0 | 4.56E-07 | 3.56 |
| FALSE | High | QASEGANS | [R].QASEG | 1xPhosphc | A2ADB0 1x | 2.15E-12 | 0 | 1 | 5 | 4 | A2ADB0 | Msx2-inter | A2ADB0 [1 | 0 | 2754.293 | 0 | 1.16 | 0.002849 | 7.17 | 60.6 | 139.4 | 13.69 | 15.54 | | High | High | High | High | High | High | High | High | High | High | High | 0 | 4.37E-14 | 5.53 |
| FALSE | High | QASEQNW | [K].QASEQ | 1xPhosphc | Q7TMQ1 1 | 2.94E-07 | 0 | 1 | 3 | 5 | Q7TMQ1 | Gap junctic | Q7TMQ1 [ | 0 | 2204.956 | 0 | 1.48 | 0.001492 | 5.43 | 55.3 | 144.7 | 7.69 | 13.91 | | High | High | High | High | High | High | High | High | High | High | High | 0 | 2.29E-06 | 4.31 |
| FALSE | High | QASEQNW | [K].QASEQ | 2xPhosphc | Q7TMQ1 2 | 0.000126 | 0 | 1 | 3 | 1 | Q7TMQ1 | Gap junctic | Q7TMQ1 [ | 0 | 2284.922 | 0 | -0.01 | 0.748423 | 8.15 | 97.7 | 102.3 | 9.52 | 14.02 | | High | High | High | High | High | High | High | High | High | High | High | 0 | 2.33E-05 | 3.89 |
| FALSE | High | QASESEDD | [R].QASESE | 1xPhosphc | Q4G0F8-3 | 2.9E-05 | 0 | 1 | 3 | 1 | Q4G0F8-3 | Isoform 3 c | Q4G0F8-3 | 0 | 1806.856 | 0 | 0.27 | 0.017537 | 8.56 | 91.1 | 108.9 | 3.89 | 6.24 | | High | High | High | High | High | High | High | High | High | High | High | 0 | 4.48E-06 | 2.85 |
| FALSE | High | QASESEDD | [R].QASESE | 2xPhosphc | Q4G0F8-3 | 7.47E-05 | 0 | 1 | 3 | 1 | Q4G0F8-3 | Isoform 3 c | Q4G0F8-3 | 0 | 1886.823 | 0 | 1.27 | 0.001745 | 7.46 | 60.1 | 139.9 | 5.96 | 9.08 | | High | High | High | High | High | High | High | High | High | High | High | 0 | 1.29E-05 | 3.69 |
| FALSE | High | QASFSGSP | [R].QASFS | 1xPhosphc | B2RWU2 1 | 0.003721 | 0.00021 | 1 | 2 | 2 | B2RWU2 | Immunode | B2RWU2 [ | 0 | 2699.333 | 0 | -0.19 | 0.091669 | 1.51 | 106.6 | 93.4 | 12.51 | 10.6 | | High | High | High | High | High | High | High | High | High | High | High | 9.42E-05 | 0.001039 | 3.12 |
| FALSE | High | QASGSPEQ | [K].QASGS | 1xPhosphc | Q3UZ39 1x | 0.002077 | 5.66E-05 | 1 | 2 | 3 | Q3UZ39 | Leucine-ric | Q3UZ39 [6 | 0 | 1469.74 | 0 | 1.28 | 0.000255 | 5.57 | 56.5 | 143.5 | 3.35 | 5.46 | | High | High | High | High | High | High | High | High | High | High | High | 2.58E-05 | 0.000541 | 3.44 |
| FALSE | High | QASLDGLQ | [R].QASLD | 1xPhosphc | A1A5B6 1x | 0.000758 | 0 | 1 | 5 | 2 | A1A5B6 | TBC1 doma | A1A5B6 [5 | 0 | 1537.794 | 0 | 0.47 | 0.009986 | 0.49 | 80.6 | 119.4 | 6.19 | 10.57 | | High | High | High | High | High | High | High | High | High | High | High | 0 | 0.000175 | 2.05 |
| FALSE | High | QAQTQTSF | [R].QAQTC | 1xDeamida | Q3TXU6 1x | 1.31E-05 | 0 | 1 | 3 | 4 | Q3TXU6 | Putative ur | Q3TXU6 [3 | 0 | 4441.12 | 0 | -0.56 | 0.017648 | 4.55 | 116.1 | 83.9 | 9.53 | 14.45 | | High | High | High | High | High | High | High | High | High | High | High | 0 | 2.02E-06 | 4.11 |
| FALSE | High | QAIEEEGG | [K].QAIEEE | 1xPhosphc | Q8CH25-2 | 9.61E-08 | 0 | 1 | 3 | 2 | Q8CH25-2 | Isoform 2 c | Q8CH25-2 | 0 | 3110.482 | 0 | -0.33 | 0.001578 | 0.4 | 109.2 | 90.8 | 7.91 | 6.3 | | High | High | High | High | High | High | High | High | High | High | High | 0 | 7.35E-09 | 4.16 |
| FALSE | High | QALAVGYC | [R].QALAV | 1xPhosphc | Q3TUF7 1x | 8.45E-07 | 0 | 1 | 3 | 2 | Q3TUF7 | YEATS dom | Q3TUF7 [1 | 0 | 1784.889 | 0 | -0.77 | 0.000376 | 1.77 | 126.7 | 73.3 | 3.49 | 3.89 | | High | High | High | High | High | High | High | High | High | High | High | 0 | 8.42E-08 | 2.67 |
| FALSE | High | QAGSFGSP | [K].QAGSF | 1xPhosphc | E9QL43 1x | 0.001158 | 0 | 1 | 2 | 1 | E9QL43 | Nucleopor | E9QL43 [2 | 0 | 3216.54 | 0 | -0.29 | 0.020103 | 12.79 | 115.1 | 84.9 | 6.11 | 17.17 | | High | High | High | High | High | High | High | High | High | High | High | 0 | 0.000282 | 2.98 |
| FALSE | High | NYQQNYQ | [R].NYQQN | 1xDeamida | Q3UBT1 1x | 1.46E-05 | 0 | 1 | 6 | 23 | Q3UBT1 | Putative ur | Q3UBT1 [1 | 0 | 2227.991 | 0 | -0.23 | 0.126432 | 10.2 | 107.9 | 92.1 | 9.1 | 7.55 | | High | High | High | High | High | High | High | High | High | High | High | 0 | 7.41E-06 | 3.31 |
| FALSE | High | NYQQNYQ | [R].NYQQN | 2xPhosphc | Q3UBT1 2x | 0.020679 | 0.001472 | 1 | 6 | 1 | Q3UBT1 | Putative ur | Q3UBT1 [1 | 0 | 2306.973 | 0 | 0.1 | 0.261903 | 11.97 | 95.4 | 104.6 | 8.67 | 6.04 | | High | High | High | High | High | High | High | High | High | High | High | 0.000786 | 0.007065 | 2.96 |
| FALSE | High | NYQQNYQ | [R].NYQQN | 1xDeamida | Q3UBT1 1x | 1.18E-07 | 0 | 1 | 5 | 3 | Q3UBT1 | Putative ur | Q3UBT1 [1 | 1 | 3796.672 | 0 | 0.33 | 0.070447 | 18.13 | 88.6 | 111.4 | 10.29 | 9.12 | | High | High | High | High | High | High | High | High | High | High | High | 0 | 9.23E-09 | 5.83 |
| FALSE | High | NYSETIEVD | [K].NYSETI | 1xPhosphc | Q3URU4 1 | 0.000779 | 0 | 1 | 4 | 6 | Q3URU4 | Meiotic rec | Q3URU4 [6 | 0 | 2870.209 | 0 | -0.4 | 0.004099 | 6.52 | 114 | 86 | 5.24 | 5.78 | | High | High | High | High | High | High | High | High | High | High | High | 0 | 0.00018 | 3.6 |
| FALSE | High | NYSSSPPCI | [R].NYSSSP | 1xCarbami | B1AY10 1x | 0.000976 | 0 | 1 | 3 | 4 | B1AY10 | Transcripti | B1AY10 [4 | 0 | 1723.782 | 0 | 0.62 | 0.001132 | 4.28 | 78.9 | 121.1 | 8.53 | 1.77 | | High | High | High | High | High | High | High | High | High | High | High | 0 | 0.000233 | 1.3 |
| FALSE | High | NYVPSYDP | [K].NYVPSY | 1xPhosphc | A2BG77 1x | 1.09E-06 | 0 | 1 | 4 | 2 | A2BG77 | Nuclear fac | A2BG77 [4 | 0 | 3161.519 | 0 | 3.82 | | 52.39 | 15.6 | 184.4 | 38.58 | 11.16 | | High | Not Found | High | High | High | High | High | High | High | High | High | 0 | 1.9E-07 | 4.25 |
| FALSE | High | NYVPSYDP | [K].NYVPSY | 1xPhosphc | Q6GSP7 1x | 0.078467 | 0.006104 | 1 | 2 | 1 | Q6GSP7 | Nuclear fac | Q6GSP7 [4 | 0 | 2585.143 | 0 | 3.51 | | 23.14 | 17.5 | 182.5 | 17.11 | 9.39 | | High | High | High | Not Found | High | High | High | High | High | High | High | 0.003148 | 0.03261 | 2.44 |
| FALSE | High | QAAAVALT | [K].QAAAV | 1xPhosphc | Q9JI13 1xP | 4.16E-05 | 0 | 1 | 4 | 1 | Q9JI13 | Something | Q9JI13 [34 | 0 | 2441.237 | 0 | -0.42 | 0.036047 | 2.24 | 118.4 | 81.6 | 22.99 | 19.75 | | High | High | High | High | High | High | High | High | High | High | High | 0 | 6.7E-06 | 3.74 |
| FALSE | High | QAALSPALI | [R].QAALSI | 1xPhosphc | Q8K382 1x | 0.000164 | 0 | 1 | 2 | 1 | Q8K382 | DENN dom | Q8K382 [7 | 0 | 1816.03 | 0 | 0.87 | 0.051592 | 19.45 | 73.4 | 126.6 | 36.91 | 11.11 | | High | High | High | High | High | High | High | High | High | High | High | 0 | 3.11E-05 | 2.52 |
| FALSE | High | QAALSSLQ | [R].QAALS | 1xPhosphc | Q8BI71 1xP | 5.98E-06 | 0 | 1 | 2 | 3 | Q8BI71 | Exocyst co | Q8BI71 [69 | 0 | 2131.075 | 0 | -1.36 | 0.002989 | 1.46 | 140.9 | 59.1 | 14.19 | 7.09 | | High | High | High | High | High | High | High | High | High | High | High | 0 | 1.02E-06 | 3.09 |
| FALSE | High | QAAPAAPS | [R].QAAPA | 1xPhosphc | Q149H7 1x | 0.038677 | 0.002563 | 1 | 2 | 2 | Q149H7 | Myeloid le | Q149H7 [1 | 0 | 2348.196 | 0 | -0.34 | 0.012715 | 1.03 | 112.5 | 87.5 | 9.17 | 3.94 | | High | High | High | High | High | High | High | High | High | High | High | 0.003303 | 0.03626 | 1.42 |
| FALSE | High | QAAPAAPS | [R].QAAPA | 2xPhosphc | Q149H7 2x | 0.007742 | 0.000664 | 1 | 2 | 1 | Q149H7 | Myeloid le | Q149H7 [1 | 0 | 2428.162 | 0 | -1.01 | 0.000158 | 0.43 | 133.8 | 66.2 | 3.37 | 8.9 | | High | High | High | High | High | High | High | High | High | High | High | 0.000294 | 0.00235 | 1.32 |
| FALSE | High | QAASSQEP | [K].QAASS | 1xPhosphc | E9Q7G0 1x | 0.047086 | 0.00354 | 1 | 4 | 2 | E9Q7G0 | Nuclear mi | E9Q7G0 [2 | 0 | 1982.927 | 0 | -0.59 | 0.040115 | 4.89 | 116.4 | 83.6 | 12.24 | 12.07 | | High | High | High | High | High | High | High | High | High | High | High | 0.001812 | 0.01804 | 2.26 |
| FALSE | High | QACSPCSS | [K].QACSP | 3xCarbami | A2AFQ0 1x | 6.2E-09 | 0 | 1 | 5 | 2 | A2AFQ0 | E3 ubiquiti | A2AFQ0 [3 | 0 | 3358.52 | 0 | -0.13 | 0.273646 | 10.49 | 103.7 | 96.3 | 10.5 | 6.42 | | High | High | High | High | High | High | High | High | High | High | High | 0 | 3.35E-10 | 4 |
| FALSE | High | QADEEMT | [R].QADEE | 1xPhosphc | Q6PG57 1x | 0.000892 | 0 | 1 | 4 | 2 | Q6PG57 | Mitogen-ac | Q6PG57 [1 | 0 | 1795.777 | 0 | 3.06 | 0.001068 | 16.92 | 22.6 | 177.4 | 25.82 | 13.83 | | High | High | High | High | High | High | High | High | High | High | High | 0 | 0.00021 | 2.75 |
| FALSE | High | QADLSFSS | [K].QADLSI | 1xPhosphc | Q3U6P5 1x | 1.21E-05 | 0 | 2 | 3 | 5 | Q3U6P5; C | Putative ur | Q3U6P5 [2 | 0 | 1976.98 | 0 | 0.19 | 0.574895 | 13.45 | 91.6 | 108.4 | 5.81 | 18.4 | Shared | High | High | High | High | High | High | High | High | High | High | High | 0 | 1.67E-06 | 3.19 |
| FALSE | High | QADLSFSS | [K].QADLSI | 1xPhosphc | Q3U6P5 1x | 2.67E-08 | 0 | 2 | 3 | 41 | Q3U6P5; C | Putative ur | Q3U6P5 [2 | 0 | 1992.975 | 0 | 0.07 | 0.419613 | 11.05 | 94.1 | 105.9 | 5.01 | 9.86 | Shared | High | High | High | High | High | High | High | High | High | High | High | 0 | 1.55E-05 | 3.67 |
| FALSE | High | QADLSFSS | [K].QADLSI | 2xPhosphc | Q3U6P5 2x | 0.034878 | 0.002269 | 2 | 3 | 3 | Q3U6P5; C | Putative ur | Q3U6P5 [2 | 0 | 2056.947 | 0 | -0.54 | 0.00743 | 4.53 | 120.9 | 79.1 | 6.14 | 16.75 | Shared | High | High | High | High | High | High | High | High | High | High | High | 0.001183 | 0.01273 | 2.31 |
| FALSE | High | QADSCPYG | [R].QADSC | 1xCarbami | Q68ED7 1x | 7.2E-08 | 0 | 1 | 1 | 1 | Q68ED7 | CREB-regu | Q68ED7 [1 | 0 | 2680.195 | 0 | 0.31 | 0.007798 | 4.54 | 89.6 | 110.4 | 7.05 | 4.4 | | High | High | High | High | High | High | High | High | High | High | High | 0 | 5.32E-09 | 3.35 |
| FALSE | High | QADSETK | [K].QADSE | 1xPhosphc | Q3ULC7 1x | 0.026652 | 0.001676 | 1 | 4 | 2 | Q3ULC7 | Putative ur | Q3ULC7 [1 | 0 | 1316.65 | 0 | -0.22 | 0.066212 | 3.46 | 107.7 | 92.3 | 5.19 | 9.62 | | High | High | High | High | High | High | High | High | High | High | High | 0.000903 | 0.009459 | 1.36 |
| FALSE | High | QADVADQ | [K].QADVA | 1xPhosphc | P18608 1x | 5.85E-08 | 0 | 1 | 3 | 9 | P18608 | Non-histor | P18608 [62 | 0 | 3783.712 | 0 | 0.13 | 0.043249 | 4 | 97 | 103 | 9.91 | 8.43 | | High | High | High | High | High | High | High | High | High | High | High | 0 | 1.67E-08 | 5.21 |
| FALSE | High | QADVADQ | [K].QADVA | 1xDeamida | P18608 1x | 6.9E-09 | 0 | 1 | 3 | 19 | P18608 | Non-histor | P18608 [62 | 0 | 3784.696 | 0 | 0.01 | 0.926034 | 13.32 | 103.4 | 96.6 | 10.86 | 4.27 | | High | High | High | High | High | High | High | High | High | High | High | 0 | 1.52E-09 | 5.84 |
| FALSE | High | QADVADQ | [K].QADVA | 2xPhosphc | P18608 2x | 4.03E-07 | 0 | 1 | 3 | 9 | P18608 | Non-histor | P18608 [62 | 0 | 3863.679 | 0 | 0.25 | 0.005811 | 1.4 | 91.3 | 108.7 | 8.82 | 4.5 | | High | High | High | High | High | High | High | High | High | High | High | 0 | 1.32E-06 | 6.12 |
| FALSE | High | QADVPAAV | [K].QADVP | 1xPhosphc | P06837 1x | 0.000231 | 0 | 1 | 1 | 1 | P06837 | Neuromod | P06837 [1 | 0 | 2794.391 | 0 | -0.36 | 0.071317 | 10.77 | 109.8 | 90.2 | 7.59 | 12.92 | | High | High | High | High | High | High | High | High | High | High | High | 0 | 4.58E-05 | 3.32 |
| FALSE | High | QAEEESHR | [R].QAEEE | 1xPhosphc | Q99LB0 1x | 0.003138 | 0.00016 | 1 | 2 | 1 | Q99LB0 | Deoxynucl | Q99LB0 [1 | 1 | 2103.064 | 0 | 0.91 | 0.006281 | 16.8 | 69.5 | 130.5 | 14.08 | 8.04 | | High | High | High | High | High | High | High | High | High | High | High | 7.19E-05 | 0.000858 | 2.26 |
| FALSE | High | QAEEGSPS | [R].QAEEG | 1xPhosphc | Q14CI5 1x | 0.00582 | 0.00062 | 1 | 5 | 3 | Q14CI5 | Zinc finger | Q14CI5 [47 | 0 | 3162.395 | 0 | 0.89 | 0.001124 | 8.98 | 69.9 | 130.1 | 8.28 | 10.21 | | High | High | High | High | High | High | High | High | High | High | High | 0.000786 | 0.006896 | 2.46 |
| FALSE | High | QAEMLQN | [R].QAEML | 1xPhosphc | Q9Z0Z7 1x | 0.000386 | 0 | 1 | 2 | 1 | Q9Z0Z7 | Krueppel-li | Q9Z0Z7 [3 | 0 | 2769.43 | 0 | -1.46 | 0.004921 | 2.36 | 143.4 | 56.6 | 18.52 | 11.27 | | High | High | High | High | High | High | High | High | High | High | High | 0 | 8.22E-05 | 2.53 |
| FALSE | High | QAEMLQN | [R].QAEML | 1xPhosphc | Q9Z0Z7 1x | 8.2E-06 | 0 | 1 | 2 | 1 | Q9Z0Z7 | Krueppel-li | Q9Z0Z7 [3 | 0 | 2785.425 | 0 | -1.38 | 0.004825 | 0.81 | 143.2 | 56.8 | 10.91 | 17.54 | | High | High | High | High | High | High | High | High | High | High | High | 0 | 1.08E-06 | 3.22 |
| FALSE | High | QAGPSPEA | [R].QAGPS | 1xPhosphc | Q80U76 1x | 0.000589 | 0 | 1 | 2 | 2 | Q80U76 | MKIAA011 | Q80U76 [6 | 0 | 1463.709 | 0 | -0.94 | 0.000748 | 0.84 | 131.4 | 68.6 | 4.44 | 5.65 | | High | High | High | High | High | High | High | High | High | High | High | 0 | 0.000132 | 3.86 |
| FALSE | High | QASLPGSP | [R].QASLP | 1xPhosphc | E9Q1M1 1 | 0.060159 | 0.004476 | 1 | 1 | 1 | E9Q1M1 | PDZ domai | E9Q1M1 [ | 0 | 1561.83 | 0 | 2.5 | 0.000193 | 5.06 | 29.7 | 170.3 | 8.86 | 6.31 | | High | High | High | High | High | High | High | High | High | High | High | 0.002293 | 0.02389 | 1.66 |
| FALSE | High | QGFPDRES | [K].QGFPD | 1xPhosphc | P14873 1x | 3.28E-07 | 0 | 2 | 2 | 3 | P14873; B2 | Microtubu | P14873 [14 | 1 | 2978.455 | 0 | -0.34 | 0.028896 | 12.05 | 111.9 | 88.1 | 8.32 | 9.8 | Shared | High | High | High | High | High | High | High | High | High | High | High | 0 | 2.9E-08 | 4.12 |
| FALSE | High | QASLSQSIF | [R].QASLS | 1xPhosphc | Q80WQ6 1 | 0.056248 | 0.004124 | 1 | 1 | 1 | Q80WQ6 | Inactive rh | Q80WQ6 [ | 0 | 1298.667 | 0 | 0.04 | 0.811324 | 20.93 | 100.2 | 99.8 | 6.71 | 8.23 | | High | High | High | High | High | High | High | High | High | High | High | 0.002117 | 0.0221 | 1.87 |
| FALSE | High | QASPAWK | [R].QASPA | 1xPhosphc | Q64327 1x | 0.095408 | 0.007437 | 1 | 3 | 1 | Q64327 | Male-enha | Q64327 [1 | 0 | 1325.702 | 0 | 0.85 | 0.037459 | 39.08 | 71.2 | 128.8 | 12.29 | 26.47 | | High | High | High | High | High | High | High | High | High | High | High | 0.003796 | 0.04095 | 1.56 |
| FALSE | High | QDGTSNSF | [K].QDGTS | 1xPhosphc | Q0VEE6 1x | 0.001986 | 5.66E-05 | 1 | 1 | 2 | Q0VEE6 | Zinc finger | Q0VEE6 [5 | 0 | 1558.751 | 0 | 2.08 | 0.000468 | 5.52 | 39.6 | 160.4 | 4.95 | 9.92 | | High | High | High | High | High | High | High | High | High | High | High | 2.58E-05 | 0.000517 | 1.87 |
| FALSE | High | QDHADSCV | [R].QDHAD | 1xCarbami | Q3ULB8 2x | 1.62E-06 | 0 | 1 | 3 | 1 | Q3ULB8 | Putative ur | Q3ULB8 [8 | 0 | 2437.942 | 0 | 0.12 | 0.371154 | 7.93 | 103.4 | 96.6 | 16.33 | 15.32 | | High | High | High | High | High | High | High | High | High | High | High | 0 | 1.75E-07 | 3.75 |
| FALSE | High | QDHDYCSF | [K].QDHDY | 1xCarbami | E9QLC6 1x | 0.022182 | 0.001542 | 1 | 5 | 1 | E9QLC6 | SUMO-spe | E9QLC6 [1 | 0 | 1816.798 | 0 | -0.4 | 0.000669 | 0.38 | 114.6 | 85.4 | 3.06 | 5.2 | | High | High | High | High | High | High | High | High | High | High | High | 0.000814 | 0.007694 | 2 |
| FALSE | High | QDHQSSQS | [R].QDHQS | 1xPhosphc | Q80SY5 1x | 2.22E-07 | 0 | 1 | 1 | 6 | Q80SY5 | Pre-mRNA- | Q80SY5 [5 | 0 | 2339.055 | 0 | 0.1 | 0.41815 | 13.2 | 97.4 | 102.6 | 6.17 | 9.45 | | High | High | High | High | High | High | High | High | High | High | High | 0 | 1.88E-08 | 3.91 |
| FALSE | High | QDLQNEPF | [R].QDLQN | 1xPhosphc | Q3TYD6 1x | 0.001222 | 0 | 1 | 1 | 2 | Q3TYD6 | Serine/thre | Q3TYD6 [8 | 0 | 2800.369 | 0 | -0.19 | 0.344543 | 26.33 | 108.7 | 91.3 | 14.19 | 19.26 | | High | High | High | High | High | High | High | High | High | High | High | 0 | 0.000298 | 2.57 |
| FALSE | High | QDLSSSLTD | [R].QDLSS | 1xPhosphc | A0A1S6GW | 8.79E-05 | 0 | 1 | 6 | 3 | A0A1S6GW | Uncharacte | A0A1S6GW | 0 | 1833.888 | 0 | 0.64 | 0.014376 | 11.39 | 76.6 | 123.4 | 24.23 | 10.72 | | High | High | High | High | High | High | High | High | High | High | High | 0 | 1.55E-05 | 3.89 |
| FALSE | High | QDITQLEES | [R].QDITQL | 2xPhosphc | O70551 2x | 0.031439 | 0.002011 | 1 | 2 | 1 | O70551 | SRSF prote | O70551 [4 | 1 | 3761.72 | 0 | -0.44 | 0.014443 | 7.81 | 115.3 | 84.7 | 7.99 | 7.98 | | High | High | High | High | High | High | High | High | High | High | High | 0.001076 | 0.01138 | 2.55 |
| FALSE | High | QDMQPLSI | [R].QDMQ | 1xPhosphc | Q64701 1x | 0.000534 | 0 | 1 | 4 | 2 | Q64701 | Retinoblast | Q64701 [6 | 0 | 1961.935 | 0 | 1.31 | 0.003995 | 8.55 | 57.6 | 142.4 | 10.28 | 13.41 | | High | High | High | High | High | High | High | High | High | High | High | 0 | 0.000118 | 2.97 |
| FALSE | High | QDNDSNVI | [K].QDNDS | 1xPhosphc | Q6PR54-3 | 0.063788 | 0.004664 | 1 | 3 | 1 | Q6PR54-3 | Isoform 3 c | Q6PR54-3 | 1 | 2515.208 | 0 | -0.47 | 0.003994 | 2.89 | 121 | 79 | 10.59 | 8.47 | | High | High | High | High | High | High | High | High | High | High | High | 0.002403 | 0.02548 | 2.34 |
| FALSE | High | QDPVVHLS | [R].QDPVV | 1xPhosphc | Q8C0T5 1x | 2.1E-05 | 0 | 1 | 3 | 2 | Q8C0T5 | Signal-indu | Q8C0T5 [1 | 0 | 1771.951 | 0 | 1.47 | 0.000295 | 7.45 | 52 | 148 | 5.03 | 5.82 | | High | High | High | High | High | High | High | High | High | High | High | 0 | 4.63E-06 | 2.89 |
| FALSE | High | QDSAGPILI | [R].QDSAG | 1xPhosphc | B2RY19 1x | 0.003001 | 0.00016 | 1 | 7 | 1 | B2RY19 | Cgnl1 prote | B2RY19 [2 | 0 | 1508.731 | 0 | 1.93 | 0.000361 | 0.71 | 41.4 | 158.6 | 9.19 | 9.38 | | High | High | High | High | High | High | High | High | High | High | High | 7.19E-05 | 0.000816 | 2.92 |
| FALSE | High | QDSDSDVV | [R].QDSDS | 1xPhosphc | A0A0A0M0 | 0.000605 | 0 | 1 | 4 | 2 | A0A0A0M0 | Calmodulin | A0A0A0M0 | 0 | 3464.558 | 0 | -0.32 | 0.016452 | 2.73 | 109.3 | 90.7 | 10.32 | 7.59 | | High | High | High | High | High | High | High | High | High | High | High | 0 | 0.000136 | 4.8 |
| FALSE | High | QDSDSDVV | [R].QDSDS | 1xDeamida | A0A0A0M0 | 6.83E-05 | 0 | 1 | 4 | 4 | A0A0A0M0 | Calmodulin | A0A0A0M0 | 0 | 3465.542 | 0 | -0.24 | 0.203903 | 28.4 | 108.8 | 91.2 | 12.24 | 17.05 | | High | High | High | High | High | High | High | High | High | High | High | 0 | 2.04E-05 | 4.45 |
| FALSE | High | QDGGTAPV | [K].QDGGT | 1xPhosphc | Q0VBL3 1x | 2.63E-06 | 0 | 1 | 8 | 2 | Q0VBL3 | RNA bindin | Q0VBL3 [7 | 0 | 1823.93 | 0 | 0.04 | 0.713576 | 19.25 | 104.4 | 95.6 | 7.3 | 12.13 | | High | High | High | High | High | High | High | High | High | High | High | 0 | 3.03E-07 | 4.32 |
| FALSE | High | QDSDSDVV | [R].QDSDS | 2xPhosphc | A0A0A0M0 | 0.005182 | 0.000551 | 1 | 4 | 1 | A0A0A0M0 | Calmodulin | A0A0A0M0 | 0 | 3544.524 | 0 | 0.01 | 0.881115 | 31.43 | 99.7 | 100.3 | 12.64 | 10.22 | | High | High | High | High | High | High | High | High | High | High | High | 0.000246 | 0.0015 | 4.03 |
| FALSE | High | QDSLVINLI | [K].QDSLVI | 1xPhosphc | G3X956 1x | 0.089425 | 0.006807 | 1 | 2 | 1 | G3X956 | Suppressor | G3X956 [6 | 0 | 1480.772 | 0 | -0.32 | 0.089203 | 16.34 | 107.3 | 92.7 | 16.8 | 9.4 | | High | High | High | High | High | High | High | High | High | High | High | 0.003535 | 0.03784 | 1.93 |
| FALSE | High | QDSNVNEE | [K].QDSNV | 1xPhosphc | B2RXR2 1x | 0.015741 | 0.001124 | 1 | 3 | 1 | B2RXR2 | Phosphatid | B2RXR2 [8 | 0 | 1786.841 | 0 | 0.37 | 0.058774 | 32.83 | 87.7 | 112.3 | 14.95 | 14.87 | | High | High | High | High | High | High | High | High | High | High | High | 0.000585 | 0.005193 | 2.38 |
| FALSE | High | QDSQTENS | [R].QDSQT | 1xPhosphc | B2RY51 1x | 1.03E-05 | 0 | 1 | 3 | 2 | B2RY51 | Thyroid ho | B2RY51 [9 | 0 | 2308.075 | 0 | 0.78 | 0.000548 | 4.27 | 72.8 | 127.2 | 8.74 | 6.42 | | High | High | High | High | High | High | High | High | High | High | High | 0 | 1.39E-06 | 3.85 |
| FALSE | High | QDSSTSVT | [K].QDSST | 1xPhosphc | P30415 1x | 2.1E-06 | 0 | 1 | 1 | 2 | P30415 | NK-tumor r | P30415 [11 | 0 | 2148.047 | 0 | -0.16 | 0.096029 | 5.08 | 105.5 | 94.5 | 4.67 | 8.9 | | High | High | High | High | High | High | High | High | High | High | High | 0 | 2.33E-07 | 2.97 |
| FALSE | High | QDSVESQL | [K].QDSVE | 1xPhosphc | E9Q4R1 1x | 0.031713 | 0.002011 | 1 | 2 | 1 | E9Q4R1 | Protein FA | E9Q4R1 [2 | 0 | 1571.808 | 0 | 2.04 | 0.00034 | 16 | 38.2 | 161.8 | 13.31 | 7.21 | | High | High | High | High | High | High | High | High | High | High | High | 0.001076 | 0.01145 | 1.98 |
| FALSE | High | QDSWEVV | [R].QDSWE | 1xPhosphc | D3YTT6 1x | 0.000213 | 0 | 1 | 8 | 2 | D3YTT6 | Oxysterol-l | D3YTT6 [3 | 0 | 1626.773 | 0 | 2.13 | 0.000377 | 11.39 | 36.6 | 163.4 | 6.31 | 7.79 | | High | High | High | High | High | High | High | High | High | High | High | 0.000161 | 0.001201 | 2.88 |
| FALSE | High | QDTSSEGV | [R].QDTSS | 1xPhosphc | A4QPD1 1x | 1.66E-05 | 0 | 1 | 2 | 3 | A4QPD1 | Zinc finger | A4QPD1 [8 | 0 | 2961.325 | 0 | 1.47 | 0.000793 | 15.05 | 49.1 | 150.9 | 10.54 | 10.63 | | High | High | High | High | High | High | High | High | High | High | High | 0 | 2.24E-05 | 2.93 |
| FALSE | High | QDVETCRP | [R].QDVET | 1xCarbami | E9QKG8 1x | 0.106748 | 0.008634 | 1 | 8 | 2 | E9QKG8 | Holliday ju | E9QKG8 [4 | 0 | 1944.884 | 0 | -0.72 | 0.000442 | 1.28 | 122.9 | 77.1 | 1.99 | 5.8 | | High | High | High | High | High | High | High | High | High | High | High | 0.007212 | 0.07829 | 1.86 |
| FALSE | High | QDVFGEGS | [R].QDVFG | 1xPhosphc | J3QNE8 1x | 0.027358 | 0.001676 | 1 | 8 | 1 | J3QNE8 | Putative so | J3QNE8 [8 | 0 | 1561.673 | 0 | 0.76 | 0.001281 | 6.05 | 74.1 | 125.9 | 13.07 | 7.19 | | High | High | High | High | High | High | High | High | High | High | High | 0.000936 | 0.00973 | 2.46 |
| FALSE | High | QDVPPSPS | [R].QDVPP | 1xPhosphc | A5D6P2 1x | 0.001313 | 0 | 1 | 8 | 1 | A5D6P2 | Par-3 (Part | A5D6P2 [1 | 0 | 1589.789 | 0 | -0.44 | 0.001925 | 5.09 | 115.3 | 84.7 | 6.56 | 3.8 | | High | High | High | High | High | High | High | High | High | High | High | 0 | 0.000325 | 1.85 |
| FALSE | High | QDYIDYSD | [K].QDYIDY | 1xPhosphc | Q3TI79 1xP | 9.14E-06 | 0 | 1 | 3 | 3 | Q3TI79 | Putative ur | Q3TI79 [41 | 0 | 1828.841 | 0 | 0.49 | 0.023633 | 18.77 | 82.6 | 117.4 | 5.66 | 13.96 | | High | High | High | High | High | High | High | High | High | High | High | 0 | 6.86E-05 | 2.76 |
| FALSE | High | QEADELSG | [K].QEADE | 1xPhosphc | Q8CH25-2 | 3.1E-07 | 0 | 1 | 4 | 2 | Q8CH25-2 | Isoform 2 c | Q8CH25-2 | 0 | 2579.18 | 0 | -0.44 | 0.006061 | 6.1 | 115.5 | 84.5 | 9.23 | 10.48 | | High | High | High | High | High | High | High | High | High | High | High | 0 | 4.82E-05 | 4.4 |
| FALSE | High | QEAEDLSL | [R].QEAED | 1xPhosphc | E9PV45 1x | 3.04E-06 | 0 | 1 | 2 | 3 | E9PV45 | Ubiquitin c | E9PV45 [2 | 0 | 3707.761 | 0 | 0.21 | 0.276835 | 11.53 | 94.6 | 105.4 | 13.3 | 12.6 | | High | High | High | High | High | High | High | High | High | High | High | 0 | 3.55E-07 | 3.23 |
| FALSE | High | QDSGSAPS | [R].QDSGS | 1xPhosphc | O35739 1x | 0.001325 | 0 | 1 | 2 | 2 | O35739 | Krueppel-li | O35739 [1 | 0 | 2476.285 | 0 | -1.81 | 0.002077 | 1.69 | 152.6 | 47.4 | 11.21 | 16.32 | | High | High | High | High | High | High | High | High | High | High | High | 0 | 0.000327 | 2.18 |
| FALSE | High | QDFSPERP | [R].QDFSP | 1xPhosphc | B1AV60 1x | 0.004867 | 0.000504 | 1 | 4 | 1 | B1AV60 | NHS actin-i | B1AV60 [7 | 0 | 1641.84 | 0 | -0.82 | 0.002019 | 3.74 | 124.9 | 75.1 | 7.99 | 5.31 | | High | High | High | High | High | High | High | High | High | High | High | 0.000225 | 0.001407 | 1.37 |
| FALSE | High | QDEQPGYN | [R].QDEQP | 1xPhosphc | E9Q411 1x | 0.013909 | 0.001011 | 1 | 1 | 1 | E9Q411 | Neuroblast | E9Q411 [3 | 1 | 2953.413 | 0 | -0.28 | 0.066982 | 14.31 | 111.9 | 88.1 | 12.26 | 6.53 | | High | High | High | High | High | High | High | High | High | High | High | 0.000538 | 0.004531 | 2.41 |
| FALSE | High | QDEMDESI | [R].QDEMI | 1xPhosphc | E9QN63 1x | 0.001189 | 0 | 1 | 12 | 1 | E9QN63 | Arf-GAP wi | E9QN63 [7 | 0 | 2959.373 | 0 | 0.54 | 0.007241 | 5.53 | 80.1 | 119.9 | 5.96 | 9.32 | | High | High | High | High | High | High | High | High | High | High | High | 0 | 0.000291 | 3.17 |
| FALSE | High | QASPETSA | [R].QASPE | 1xPhosphc | B2RWW2 1 | 0.000146 | 0 | 1 | 8 | 2 | B2RWW2 | Golgb1 prc | B2RWW2 [ | 0 | 2044 | 0 | 0.38 | 0.128754 | 13.91 | 87.3 | 112.7 | 22.75 | 7.77 | | High | High | High | High | High | High | High | High | High | High | High | 0 | 2.74E-05 | 3.9 |
| FALSE | High | QASPLGTP | [K].QASPL | 1xPhosphc | E9Q4D5 1x | 0.005467 | 0.00062 | 1 | 3 | 2 | E9Q4D5 | GTP-bindin | E9Q4D5 [2 | 0 | 2378.262 | 0 | -0.47 | 0.073243 | 18.95 | 116.9 | 83.1 | 15.19 | 24.75 | | High | High | High | High | High | High | High | High | High | High | High | 0.000275 | 0.001593 | 2.34 |
| FALSE | High | QASPLISPL | [K].QASPLI | 1xCarbami | F8VPU2 1x | 8.9E-06 | 0 | 1 | 2 | 1 | F8VPU2 | FERM, Rho | F8VPU2 [5 | 0 | 2189.099 | 0 | 0.42 | 0.050883 | 22.03 | 84.5 | 115.5 | 10.09 | 16.76 | | High | High | High | High | High | High | High | High | High | High | High | 0 | 1.19E-06 | 3.95 |
| FALSE | High | QASPNIVIA | [R].QASPN | 1xPhosphc | Q8C266 1x | 0.017966 | 0.001282 | 1 | 3 | 1 | Q8C266 | Putative ur | Q8C266 [1 | 0 | 1934.088 | 0 | 0.28 | 0.00428 | 4.38 | 90.2 | 109.8 | 10.9 | 6.02 | | High | High | High | High | High | High | High | High | High | High | High | 0.000689 | 0.006016 | 1.98 |
| FALSE | High | QASPQLQF | [K].QASPQ | 1xPhosphc | Z4YKQ9 1x | 0.00034 | 0 | 1 | 3 | 2 | Z4YKQ9 | ATPase fan | Z4YKQ9 [9 | 0 | 1964.025 | 0 | -0.53 | 0.002281 | 5.64 | 122.1 | 77.9 | 4.96 | 7.84 | | High | High | High | High | High | High | High | High | High | High | High | 0 | 7.08E-05 | 3.01 |
| FALSE | High | QASPTEVV | [R].QASPTI | 1xPhosphc | Q80U58 1x | 0.000375 | 0 | 1 | 7 | 4 | Q80U58 | Pumilio ho | Q80U58 [1 | 0 | 1424.698 | 0 | 0.49 | 0.000275 | 3.73 | 82.8 | 117.2 | 2.89 | 3.3 | | High | High | High | High | High | High | High | High | High | High | High | 0 | 7.93E-05 | 3.28 |
| FALSE | High | QASPVQTH | [R].QASPV | 1xPhosphc | E9PWH2 1 | 0.000117 | 0 | 1 | 2 | 2 | E9PWH2 | Caspase re | E9PWH2 [8 | 0 | 1630.872 | 0 | 2.05 | 0.000192 | 14.3 | 38.3 | 161.7 | 8.21 | 4.03 | | High | High | High | High | High | High | High | High | High | High | High | 0 | 2.15E-05 | 2.86 |
| FALSE | High | QASPYYCS | [R].QASPY | 1xCarbami | Q3UPW2 1 | 1.42E-05 | 0 | 1 | 2 | 6 | Q3UPW2 | ETS-relatec | Q3UPW2 [ | 0 | 2972.297 | 0 | -0.48 | 0.035679 | 11.92 | 116.9 | 83.1 | 12.89 | 8.98 | | High | High | High | High | High | High | High | High | High | High | High | 0 | 2.65E-06 | 4.72 |
| FALSE | High | QASSPVFTI | [K].QASSP | 1xPhosphc | Q8R4U7 1x | 5.7E-05 | 0 | 2 | 4 | 3 | Q8R4U7; B | Leucine zip | Q8R4U7 [4 | 0 | 1616.845 | 0 | -0.12 | 0.118855 | 7.98 | 103.9 | 96.1 | 8.03 | 6.06 | Shared | High | High | High | High | High | High | High | High | High | High | High | 0 | 9.59E-06 | 2.67 |
| FALSE | High | QASTDAGT | [R].QASTD | 1xPhosphc | P46938 1x | 3.14E-11 | 0 | 1 | 6 | 9 | P46938 | Transcripti | P46938 [92 | 0 | 2090.023 | 0 | 0.39 | 0.00919 | 2.63 | 91 | 109 | 4.93 | 6.91 | | High | High | High | High | High | High | High | High | High | High | High | 0 | 8.87E-13 | 5.6 |

| | | | | | | | | | | | | | | | | | | | | | | | | | | | | | | | | | | | | | | |
|---|---|---|---|---|---|---|---|---|---|---|---|---|---|---|---|---|---|---|---|---|---|---|---|---|---|---|---|---|---|---|---|---|---|---|---|---|---|---|
| FALSE | High | QASTDAGT | [R].QASTDA | 2xPhospho | P46938 2x | 8.4E-05 | 0 | 1 | 6 | 1 | P46938 | Transcripti | P46938 [92 | 0 | 2169.989 | 0 | 1 | 0.001615 | 10.77 | 65.5 | 134.5 | 8.56 | 7.45 | | High | High | High | High | High | High | High | High | High | High | High | 0 | 1.48E-05 | 3.79 |
| FALSE | High | QASTVEYLI | [R].QASTVI | 1xCarbami | Q8VE18 1x | 3.81E-09 | 0 | 1 | 1 | 2 | Q8VE18 | Protein SM | Q8VE18 [7 | 0 | 2570.255 | 0 | 1.29 | 0.006456 | 9.85 | 51.1 | 148.9 | 13.68 | 23.65 | | High | High | High | High | High | High | High | High | High | High | High | 0 | 1.95E-10 | 3.98 |
| FALSE | High | QASTVEYLI | [R].QASTVI | 1xCarbami | Q8VE18 1x | 1.77E-07 | 0 | 1 | 1 | 1 | Q8VE18 | Protein SM | Q8VE18 [7 | 0 | 2586.25 | 0 | 1.29 | 0.003279 | 12.99 | 54.9 | 145.1 | 9.36 | 14.78 | | High | High | High | High | High | High | High | High | High | High | High | 0 | 1.46E-08 | 3.71 |
| FALSE | High | QATGASW | [K].QATGA | 1xPhospho | P59729 1x | 0.035793 | 0.002304 | 1 | 5 | 1 | P59729 | Ras and Ra | P59729 [49 | 0 | 1767.863 | 0 | 2.75 | 0.000295 | 16.87 | 25.7 | 174.3 | 12.5 | 6.84 | | High | High | High | High | High | High | High | High | High | High | High | 0.00123 | 0.01317 | 1.83 |
| FALSE | High | QATPGVPA | [R].QATPG | 1xPhospho | Q6ZQ88 1x | 2.72E-05 | 0 | 1 | 3 | 1 | Q6ZQ88 | Lysine-spe | Q6ZQ88 [8 | 0 | 1707.797 | 0 | -0.35 | 0.016125 | 1.14 | 118.8 | 81.2 | 8.88 | 13.61 | | High | High | High | High | High | High | High | High | High | High | High | 0 | 4.17E-06 | 2.38 |
| FALSE | High | QATPGVPA | [R].QATPG | 1xPhospho | Q6ZQ88 1x | 2.01E-06 | 0 | 1 | 3 | 2 | Q6ZQ88 | Lysine-spe | Q6ZQ88 [8 | 0 | 1723.792 | 0 | -0.72 | 0.004241 | 3.75 | 126.1 | 73.9 | 4.24 | 15.09 | | High | High | High | High | High | High | High | High | High | High | High | 0 | 2.22E-07 | 3.66 |
| FALSE | High | QATVNVSY | [R].QATVN | 1xPhospho | P40201 1x | 0.035486 | 0.002304 | 1 | 5 | 1 | P40201 | Chromodo | P40201 [23 | 0 | 1547.823 | 0 | -0.07 | 0.577525 | 16.61 | 101.8 | 98.2 | 17.86 | 7.68 | | High | High | High | High | High | High | High | High | High | High | High | 0.001214 | 0.01301 | 0.93 |
| FALSE | High | QAVDVSPL | [R].QAVDV | 1xPhospho | P97461 1x | 0.051694 | 0.003745 | 1 | 4 | 1 | P97461 | 40S ribosor | P97461 [13 | 0 | 1293.677 | 0 | -0.36 | 0.016155 | 5.92 | 112.2 | 87.8 | 9.84 | 4.7 | | High | High | High | High | High | High | High | High | High | High | High | 0.00196 | 0.02004 | 1.67 |
| FALSE | High | QAVEGSSS | [R].QAVEG | 1xPhospho | Q7TQ95 1x | 1.83E-05 | 0 | 1 | 1 | 5 | Q7TQ95 | Protein lun | Q7TQ95 [3 | 0 | 4784.236 | 0 | 1.76 | 0.007798 | 52.23 | 51.4 | 148.6 | 39 | 21.1 | | High | High | High | High | High | High | High | High | High | High | High | 0 | 2.66E-06 | 5.07 |
| FALSE | High | QAVVQTTF | [K].QAVVQ | 1xPhospho | Q8BRG8 1x | 0.041072 | 0.003094 | 1 | 4 | 2 | Q8BRG8 | Transmem | Q8BRG8 [1 | 0 | 1405.74 | 0 | -0.16 | 0.004532 | 3.29 | 104.4 | 95.6 | 2.87 | 4.69 | | High | High | High | High | High | High | High | High | High | High | High | 0.004517 | 0.04902 | 1.58 |
| FALSE | High | QCEGITSPE | [K].QCEGIT | 1xCarbami | Q8BVQ5 1x | 5.21E-05 | 0 | 1 | 2 | 3 | Q8BVQ5 | Protein phi | Q8BVQ5 [2 | 0 | 1830.871 | 0 | -1.26 | 0.013546 | 1.32 | 142.3 | 57.7 | 28.33 | 7.53 | | High | High | High | High | High | High | High | High | High | High | High | 0 | 8.62E-06 | 2.35 |
| FALSE | High | QDAIGKPP | [K].QDAIGI | 1xPhospho | Q80Y82 1x | 0.000284 | 0 | 1 | 12 | 4 | Q80Y82 | Protein kin | Q80Y82 [7 | 0 | 3270.677 | 0 | 0.15 | 0.165829 | 18.4 | 94.1 | 105.9 | 9.81 | 13.98 | | High | High | High | High | High | High | High | High | High | High | High | 0 | 5.8E-05 | 2.94 |
| FALSE | High | QDAPGPTS | [K].QDAPG | 1xPhospho | A0A0R4J0S | 2.19E-08 | 0 | 1 | 5 | 3 | A0A0R4J0S | Uveal auto | A0A0R4J0S | 0 | 3052.467 | 0 | 0.78 | 0.000496 | 6.04 | 71.8 | 128.2 | 13.98 | 9.77 | | High | High | High | High | High | High | High | High | High | High | High | 0 | 1.39E-09 | 5.65 |
| FALSE | High | QDDSPPRP | [K].QDDSP | 1xPhospho | Q69ZC8 1x | 0.000639 | 0 | 1 | 1 | 1 | Q69ZC8 | GPALPP m | Q69ZC8 [1 | 0 | 2653.452 | 0 | -0.24 | 0.030712 | 3.01 | 106.6 | 93.4 | 6.2 | 1.58 | | High | High | High | High | High | High | High | High | High | High | High | 0 | 0.000144 | 2.96 |
| FALSE | High | QDDSPSGA | [K].QDDSP | 1xPhospho | Q14AX6-2 | 4.23E-08 | 0 | 3 | 5 | 1 | Q14AX6-2; | Isoform 2 c | Q14AX6-2 | 0 | 2464.05 | 0 | -0.06 | 0.580473 | 4.71 | 101.4 | 98.6 | 11.43 | 2.11 | Shared | High | High | High | High | High | High | High | High | High | High | High | 0 | 2.91E-09 | 3.62 |
| FALSE | High | QDDSPSGA | [K].QDDSP | 2xPhospho | Q14AX6-2 | 0.000837 | 0 | 3 | 5 | 4 | Q14AX6-2; | Isoform 2 c | Q14AX6-2 | 0 | 2544.017 | 0 | -0.25 | 0.773049 | 8.28 | 98.5 | 101.5 | 22.83 | 19.6 | Shared | High | High | High | High | High | High | High | High | High | High | High | 0 | 0.000196 | 2.71 |
| FALSE | High | QDDSPSGA | [K].QDDSP | 3xPhospho | Q14AX6-2 | 0.003892 | 0.000261 | 3 | 5 | 1 | Q14AX6-2; | Isoform 2 c | Q14AX6-2 | 0 | 2623.983 | 0 | -0.08 | 0.626176 | 13.8 | 102.9 | 97.1 | 20.46 | 10.28 | Shared | High | High | High | High | High | High | High | High | High | High | High | 0.00014 | 0.001094 | 3.84 |
| FALSE | High | QDEDSDSE | [R].QDEDS | 1xPhospho | Q9CZH7 1x | 2.21E-05 | 0 | 1 | 2 | 4 | Q9CZH7 | Matrix-rem | Q9CZH7 [9 | 0 | 3511.466 | 0 | -1.42 | 0.002672 | 7.22 | 140.7 | 59.3 | 7.74 | 34.51 | | High | High | High | High | High | High | High | High | High | High | High | 0 | 3.28E-06 | 3.74 |
| FALSE | High | QDEDSDSE | [R].QDEDS | 1xPhospho | Q9CZH7 1x | 1.58E-08 | 0 | 1 | 2 | 8 | Q9CZH7 | Matrix-rem | Q9CZH7 [9 | 0 | 3527.461 | 0 | -1.13 | 0.002031 | 3.95 | 137 | 63 | 8.35 | 8.81 | | High | High | High | High | High | High | High | High | High | High | High | 0 | 9.61E-10 | 5.78 |
| FALSE | High | QASPAK | [R].QASPAI | 1xPhospho | Q791V7 1x | 0.081752 | 0.006219 | 1 | 2 | 1 | Q791V7 | Uracil-DNA | Q791V7 [3 | 0 | 1139.623 | 0 | -1.84 | 0.000192 | 1.24 | 157.8 | 42.2 | 3.25 | 16.38 | | High | High | High | High | High | High | High | High | High | High | High | 0.0032 | 0.03403 | 1.28 |
| FALSE | High | NYQQNYQ | [R].NYQQN | 1xPhospho | Q3UBT1 1x | 5.72E-06 | 0 | 1 | 6 | 8 | Q3UBT1 | Putative ur | Q3UBT1 [1 | 0 | 2227.007 | 0 | -0.36 | 0.021037 | 4.53 | 111.6 | 88.4 | 4.57 | 7.3 | | High | High | High | High | High | High | High | High | High | High | High | 0 | 3.54E-06 | 3.86 |
| FALSE | High | QGGAQPLT | [K].QGGAC | 1xPhospho | A0A0R4J0C | 0.000279 | 0 | 2 | 2 | 2 | A0A0R4J0C | E3 ubiquiti | A0A0R4J0C | 0 | 1534.839 | 0 | 1.52 | 0.000545 | 14.39 | 52 | 148 | 6.68 | 7.16 | Shared | High | High | High | High | High | High | High | High | High | High | High | 0 | 5.71E-05 | 3.68 |
| FALSE | High | QGGGQPS | [K].QGGGC | 1xPhospho | Q8C7E9 1x | 2.74E-08 | 0 | 1 | 1 | 5 | Q8C7E9 | Cleavage st | Q8C7E9 [5 | 0 | 2912.383 | 0 | -0.69 | 0.000176 | 1.34 | 122.7 | 77.3 | 3.69 | 4.49 | | High | High | High | High | High | High | High | High | High | High | High | 0 | 1.78E-09 | 4.82 |
| FALSE | High | QITIIDSPCL | [K].QITIIDS | 2xCarbami | Q3TK27 1x | 0.003721 | 0.00021 | 1 | 3 | 2 | Q3TK27 | Putative ur | Q3TK27 [2 | 0 | 2962.534 | 0 | -1.23 | 0.004273 | 5.49 | 140.3 | 59.7 | 10.1 | 17.91 | | High | High | High | High | High | High | High | High | High | High | High | 9.42E-05 | 0.00104 | 3.97 |
| FALSE | High | QLTNGSLP | [R].QLTNG | 1xPhospho | P70174 1x | 0.003558 | 0.00021 | 1 | 1 | 2 | P70174 | Histamine | P70174 [22 | 0 | 2099.155 | 0 | 0.16 | 0.192025 | 6.77 | 96 | 104 | 9.61 | 2.93 | | High | High | High | High | High | High | High | High | High | High | High | 0.000183 | 0.001243 | 3 |
| FALSE | High | QITQEDDD | [K].QITQED | 1xPhospho | Q9EQC8 1x | 9.22E-06 | 0 | 1 | 2 | 9 | Q9EQC8 | Papillary re | Q9EQC8 [2 | 0 | 3306.498 | 0 | -0.48 | 0.000797 | 2.11 | 116.3 | 83.7 | 4.19 | 3.47 | | High | High | High | High | High | High | High | High | High | High | High | 0 | 1.23E-06 | 3.87 |
| FALSE | High | QLTQPETS | [K].QLTQPI | 1xPhospho | Q62418 1x | 0.000976 | 0 | 1 | 3 | 2 | Q62418 | Drebrin-lik | Q62418 [2 | 0 | 1588.757 | 0 | 1.88 | 0.016213 | 54.24 | 42.6 | 157.4 | 18.11 | 33.89 | | High | High | High | High | High | High | High | High | High | High | High | 0 | 0.000231 | 1.5 |
| FALSE | High | QITTSPITVI | [K].QITTSP | 1xPhospho | Q8BYR2 1x | 0.085859 | 0.006406 | 1 | 1 | 1 | Q8BYR2 | Serine/thre | Q8BYR2 [6 | 0 | 1424.771 | 0 | 2.56 | 0.000394 | 58.84 | 29 | 171 | 43.18 | 5.75 | | High | High | High | High | High | High | High | High | High | High | High | 0.003303 | 0.03611 | 2.14 |
| FALSE | High | QITVEELVF | [K].QITVEE | 1xPhospho | Q9QXS1-3 | 0.002647 | 0.000109 | 3 | 23 | 4 | Q9QXS1-3; | Isoform PL | Q9QXS1-3 | 0 | 1395.745 | 0 | 1.82 | 0.006027 | 22.68 | 41.6 | 158.4 | 13.34 | 22.57 | Shared | High | High | High | High | High | High | High | High | High | High | High | 4.94E-05 | 0.000707 | 2.59 |
| FALSE | High | QIVIASTSK | [R].QIVIAS | 1xPhospho | Q8R2M2 1 | 0.000718 | 0 | 1 | 1 | 2 | Q8R2M2 | Deoxynucl | Q8R2M2 [1 | 0 | 1484.849 | 0 | -0.25 | 0.696499 | 33 | 88.8 | 111.2 | 29.47 | 26.77 | | High | High | High | High | High | High | High | High | High | High | High | 0 | 0.000165 | 2.8 |
| FALSE | High | QLVTSPSPI | [R].QLVTSI | 1xPhospho | F8VPJ6 1xP | 3.88E-06 | 0 | 1 | 1 | 4 | F8VPJ6 | Regulatory | F8VPJ6 [37 | 0 | 2112.045 | 0 | -0.55 | 0.001003 | 2.95 | 118.2 | 81.8 | 2.25 | 6.09 | | High | High | High | High | High | High | High | High | High | High | High | 0.000571 | 0.00497 | 2.88 |
| FALSE | High | QLVYVEQA | [R].QLVYVI | 1xPhospho | G3X932 1x | 1.1E-07 | 0 | 1 | 5 | 2 | G3X932 | DNA segme | G3X932 [3 | 0 | 1944.024 | 0 | -0.37 | 0.00417 | 5.78 | 110.4 | 89.6 | 9.38 | 9.52 | | High | High | High | High | High | High | High | High | High | High | High | 0 | 8.54E-09 | 3.68 |
| FALSE | High | QLWNPPQ | [K].QLWNI | 1xPhospho | Q5RJH6-2 | 6.07E-08 | 0 | 1 | 4 | 1 | Q5RJH6-2 | Isoform 2 c | Q5RJH6-2 | 0 | 2130.151 | 0 | -0.69 | 0.005298 | 6.23 | 123.5 | 76.5 | 6.06 | 11.27 | | High | High | High | High | High | High | High | High | High | High | High | 0 | 4.37E-09 | 3.24 |
| FALSE | High | QMAELMP | [R].QMAEL | 2xPhospho | Q8VIG1 2x | 0.007538 | 0.00062 | 1 | 3 | 2 | Q8VIG1 | RE1-silenci | Q8VIG1 [7 | 0 | 3667.599 | 0 | -0.91 | 0.002041 | 5.21 | 126.6 | 73.4 | 5.93 | 10.46 | | High | High | High | High | High | High | High | High | High | High | High | 0.000294 | 0.002296 | 3.17 |
| FALSE | High | QMGVTEW | [R].QMGV | 1xPhospho | F6ZBR8 1x | 0.0009 | 0 | 2 | 5 | 1 | F6ZBR8; O | Protein Wi | F6ZBR8 [57 | 0 | 2315.094 | 0 | 0.21 | 0.411068 | 14.88 | 94.5 | 105.5 | 7.39 | 11.95 | Shared | High | High | High | High | High | High | High | High | High | High | High | 0 | 0.000211 | 3.84 |
| FALSE | High | QMLTPTSP | [R].QMLTP | 1xPhospho | F8VPN7 1x | 0.000454 | 0 | 1 | 6 | 1 | F8VPN7 | Dedicator | F8VPN7 [4 | 0 | 3982.931 | 0 | 1.73 | 0.003621 | 40.31 | 46.4 | 153.6 | 29.02 | 10.91 | | High | High | High | High | High | High | High | High | High | High | High | 0 | 9.83E-05 | 3.7 |
| FALSE | High | QLTGNTVC | [K].QLTGN | 1xPhospho | Q9WV30-2 | 0.11108 | 0.008991 | 1 | 9 | 1 | Q9WV30-2 | Isoform 2 c | Q9WV30-2 | 0 | 2174.137 | 0 | 0.52 | 0.026817 | 11.13 | 82.3 | 117.7 | 17.77 | 12.47 | | High | High | High | High | High | High | High | High | High | High | High | 0.004503 | 0.04881 | 2.28 |
| FALSE | High | QMNLGFG | [R].QMNL | 1xPhospho | B1AQY2 1x | 0.001211 | 0 | 1 | 3 | 1 | B1AQY2 | Rho GTPas | B1AQY2 [5 | 0 | 2101.91 | 0 | 0.35 | 0.414579 | 53.51 | 100 | 100 | 13.69 | 34.05 | | High | High | High | High | High | High | High | High | High | High | High | 0 | 0.000297 | 1.53 |
| FALSE | High | QMSEPIVP | [R].QMSEP | 1xPhospho | Q8BJR1 1x | 4.31E-05 | 0 | 1 | 2 | 1 | Q8BJR1 | ETS translo | Q8BJR1 [26 | 0 | 2445.302 | 0 | 1.22 | 0.007584 | 23.85 | 60.3 | 139.7 | 11.63 | 16.43 | | High | High | High | High | High | High | High | High | High | High | High | 0 | 6.97E-06 | 3.07 |
| FALSE | High | QMSEPIVP | [R].QMSEP | 1xPhospho | Q8BJR1 1x | 0.00043 | 0 | 1 | 2 | 1 | Q8BJR1 | ETS translo | Q8BJR1 [26 | 0 | 2461.297 | 0 | 1.38 | 0.004445 | 14.47 | 58.2 | 141.8 | 12.43 | 14.8 | | High | High | High | High | High | High | High | High | High | High | High | 0 | 9.23E-05 | 2.69 |
| FALSE | High | QMSFDLTK | [R].QMSFD | 1xPhospho | E9Q9B7 1x | 0.025073 | 0.001645 | 1 | 5 | 1 | E9Q9B7 | Kinase D-ir | E9Q9B7 [9 | 0 | 1507.763 | 0 | 0.41 | 0.003443 | 5.84 | 86.4 | 113.6 | 6.63 | 2.35 | | High | High | High | High | High | High | High | High | High | High | High | 0.000872 | 0.008837 | 1.72 |
| FALSE | High | QMSFDLTK | [R].QMSFD | 1xPhospho | E9Q9B7 1x | 0.005135 | 0.000551 | 1 | 5 | 2 | E9Q9B7 | Kinase D-ir | E9Q9B7 [9 | 0 | 1523.758 | 0 | 0.4 | 0.005134 | 1.88 | 87.2 | 112.8 | 3.42 | 9.21 | | High | High | High | High | High | High | High | High | High | High | High | 0.000246 | 0.001496 | 1.72 |
| FALSE | High | QMSLCGTF | [R].QMSLC | 1xCarbami | E9PV45 1x | 0.000109 | 0 | 1 | 2 | 2 | E9PV45 | Ubiquitin c | E9PV45 [1 | 0 | 1704.81 | 0 | 1.42 | 0.00222 | 4.83 | 50.5 | 149.5 | 14.15 | 14.39 | | High | High | High | High | High | High | High | High | High | High | High | 0 | 1.98E-05 | 1.93 |
| FALSE | High | QMSSGLAS | [R].QMSSG | 1xPhospho | Q3UUL4 1x | 2.05E-10 | 0 | 1 | 1 | 1 | Q3UUL4 | Putative ur | Q3UUL4 [8 | 0 | 2320.084 | 0 | -0.15 | 0.125694 | 11.66 | 107.4 | 92.6 | 10.27 | 12.51 | | High | High | High | High | High | High | High | High | High | High | High | 0 | 7.28E-12 | 3.85 |
| FALSE | High | QMSSGLAS | [R].QMSSG | 2xPhospho | Q3UUL4 2x | 2.45E-06 | 0 | 1 | 1 | 1 | Q3UUL4 | Putative ur | Q3UUL4 [8 | 0 | 2400.051 | 0 | 0.66 | 0.001066 | 9.16 | 74.7 | 125.3 | 11.4 | 8.77 | | High | High | High | High | High | High | High | High | High | High | High | 0 | 2.79E-07 | 4.09 |
| FALSE | High | QMSYGPEI | [R].QMSYG | 1xPhospho | A2AJ88 1x | 0.011447 | 0.000816 | 1 | 4 | 1 | A2AJ88 | Patatin-like | A2AJ88 [33 | 0 | 1791.782 | 0 | 0.88 | 0.002647 | 18.19 | 70.8 | 129.2 | 6.9 | 10.57 | | High | High | High | High | High | High | High | High | High | High | High | 0.000437 | 0.003647 | 1.88 |
| FALSE | High | QNETADEA | [K].QNETA | 1xPhospho | E9Q5C9 1x | 5.4E-05 | 0 | 1 | 2 | 5 | E9Q5C9 | Nucleolar a | E9Q5C9 [5 | 0 | 2041.984 | 0 | 0.29 | 0.099145 | 3.8 | 92.3 | 107.7 | 3.67 | 11.36 | | High | High | High | High | High | High | High | High | High | High | High | 0 | 5.59E-05 | 3.54 |
| FALSE | High | QNGSNDSI | [R].QNGSN | 1xPhospho | Q3U955 1x | 0.002232 | 0.000109 | 1 | 9 | 8 | Q3U955 | Putative ur | Q3U955 [1 | 1 | 2584.084 | 0 | -0.28 | 0.031572 | 1.1 | 108.7 | 91.3 | 3.73 | 8.56 | | High | High | High | High | High | High | High | High | High | High | High | 0.007398 | 0.08162 | 3.15 |
| FALSE | High | QNGSNDSI | [R].QNGSN | 1xDeamida | Q3U955 1x | 0.005615 | 0.00062 | 1 | 9 | 6 | Q3U955 | Putative ur | Q3U955 [1 | 1 | 2585.068 | 0 | -0.1 | 0.402462 | 6.01 | 104.6 | 95.4 | 6.73 | 11.29 | | High | High | High | High | High | High | High | High | High | High | High | 0.003672 | 0.0394 | 2.62 |
| FALSE | High | QNGSNDSI | [R].QNGSN | 1xPhospho | Q61033 1x | 0.015061 | 0.001049 | 1 | 1 | 1 | Q61033 | Lamina-ass | Q61033 [1 | 1 | 2439.046 | 0 | 0.8 | 0.017776 | 18.47 | 75.1 | 124.9 | 9.96 | 17.55 | | High | High | High | High | High | High | High | High | High | High | High | 0.000571 | 0.004958 | 1.35 |
| FALSE | High | QNGSNDSI | [R].QNGSN | 1xDeamida | Q61033 1x | 0.001189 | 0 | 1 | 1 | 5 | Q61033 | Lamina-ass | Q61033 [1 | 1 | 2440.03 | 0 | 0.59 | 0.004648 | 11.04 | 82.7 | 117.3 | 10.36 | 16.42 | | High | High | High | High | High | High | High | High | High | High | High | 0 | 0.000289 | 2.28 |
| FALSE | High | QMNMSPF | [K].QMNM | 1xPhospho | Q8CCS6 1x | 2.74E-08 | 0 | 1 | 7 | 13 | Q8CCS6 | Polyadenyl | Q8CCS6 [1 | 0 | 2781.343 | 0 | -0.11 | 0.242789 | 5.8 | 103.7 | 96.3 | 5.19 | 14.45 | | High | High | High | High | High | High | High | High | High | High | High | 0 | 1.79E-09 | 5.4 |
| FALSE | High | QNLLAPQN | [K].QNLLAI | 1xPhospho | F8WIP8 1x | 3.33E-06 | 0 | 1 | 9 | 5 | F8WIP8 | Osteopont | F8WIP8 [5 | 0 | 2166.121 | 0 | -0.89 | 0.034974 | 14.3 | 127.3 | 72.7 | 14.94 | 21.2 | | High | High | High | High | High | High | High | High | High | High | High | 0 | 3.92E-07 | 3.43 |
| FALSE | High | QLTEMLPS | [K].QLTEM | 1xPhospho | Q64152 1x | 2.59E-07 | 0 | 1 | 2 | 1 | Q64152 | Transcripti | Q64152 [1 | 0 | 2725.404 | 0 | 0 | 0.62858 | 8.92 | 95.1 | 104.9 | 18.27 | 19.51 | | High | High | High | High | High | High | High | High | High | High | High | 0 | 2.24E-08 | 3.92 |
| FALSE | High | QLTANTGH | [R].QLTAN | 1xPhospho | A2AFQ0 1x | 0.005044 | 0.000504 | 1 | 6 | 1 | A2AFQ0 | E3 ubiquiti | A2AFQ0 [2 | 0 | 2325.145 | 0 | 0.29 | 0.071181 | 10.51 | 87.7 | 112.3 | 15.19 | 21.29 | | High | High | High | High | High | High | High | High | High | High | High | 0.000225 | 0.001456 | 2.52 |
| FALSE | High | QISGQYSG | [K].QISGQ | 1xPhospho | A2A9M4 1 | 1.24E-05 | 0 | 1 | 6 | 1 | A2A9M4 | Dedicator | A2A9M4 [2 | 0 | 2044.088 | 0 | 0.79 | 0.000934 | 2.19 | 70.4 | 129.6 | 8.66 | 5.81 | | High | High | High | High | High | High | High | High | High | High | High | 0 | 1.72E-06 | 2.73 |
| FALSE | High | QISGSSTG | [K].QISGSS | 1xCarbami | Q920Q8 1x | 1.99E-06 | 0 | 1 | 3 | 1 | Q920Q8 | Influenza v | Q920Q8 [2 | 0 | 2662.262 | 0 | -0.4 | 0.05059 | 11.55 | 112 | 88 | 11.06 | 10.93 | | High | High | High | High | High | High | High | High | High | High | High | 0 | 2.21E-07 | 3.61 |
| FALSE | High | QISGSSTG | [K].QISGSS | 1xCarbami | Q920Q8 1x | 5.51E-09 | 0 | 1 | 3 | 10 | Q920Q8 | Influenza v | Q920Q8 [2 | 0 | 2678.257 | 0 | -0.41 | 0.035102 | 12.27 | 115.7 | 84.3 | 3.3 | 17.87 | | High | High | High | High | High | High | High | High | High | High | High | 0 | 6.35E-07 | 3.61 |
| FALSE | High | QLSHDQES | [R].QLSHD | 1xPhospho | B7ZCJ1 1xP | 0.000111 | 0 | 1 | 6 | 1 | B7ZCJ1 | Rho GTPas | B7ZCJ1 [85 | 0 | 2771.34 | 0 | 0.29 | 0.314182 | 28.02 | 94.6 | 105.4 | 12.73 | 12.35 | | High | High | High | High | High | High | High | High | High | High | High | 0 | 2.02E-05 | 2.13 |
| FALSE | High | QISLASETL | [K].QISLAS | 1xPhospho | S4R189 1x | 3.8E-05 | 0 | 1 | 3 | 1 | S4R189 | Rho guanir | S4R189 [14 | 0 | 2325.117 | 0 | -0.38 | 0.041812 | 10.23 | 114.2 | 85.8 | 7.87 | 10.93 | | High | High | High | High | High | High | High | High | High | High | High | 0 | 6.04E-06 | 3.04 |
| FALSE | High | QLSLEGAG | [K].QLSLEG | 1xPhospho | V9GX40 1x | 4.71E-08 | 0 | 1 | 4 | 1 | V9GX40 | Synergin g | V9GX40 [8 | 0 | 2161.102 | 0 | 0.35 | 0.228113 | 11.76 | 86.6 | 113.4 | 7.19 | 18.24 | | High | High | High | High | High | High | High | High | High | High | High | 0 | 3.3E-09 | 5.2 |
| FALSE | High | QLSLEGAG | [K].QLSLEG | 1xPhospho | V9GX40 1x | 4.09E-09 | 0 | 1 | 4 | 3 | V9GX40 | Synergin g | V9GX40 [8 | 0 | 2177.097 | 0 | 0.33 | 0.148582 | 13.06 | 84.5 | 115.5 | 12.82 | 30.34 | | High | High | High | High | High | High | High | High | High | High | High | 0 | 2.12E-10 | 5.59 |
| FALSE | High | QLSLPLTQS | [R].QLSLPL | 1xPhospho | Q80TA6 1x | 0.001361 | 0 | 1 | 2 | 1 | Q80TA6 | Myotubula | Q80TA6 [5 | 0 | 1652.939 | 0 | 0.52 | 0.021743 | 13.63 | 82.2 | 117.8 | 4.26 | 18.5 | | High | High | High | High | High | High | High | High | High | High | High | 2.58E-05 | 0.000338 | 2.69 |
| FALSE | High | QLSNANIG | [K].QLSNAI | 1xPhospho | O88622 1x | 0.072858 | 0.005599 | 1 | 5 | 1 | O88622 | Poly(ADP-r | O88622 [1 | 1 | 4541.101 | 0 | -1.87 | 0.003083 | 0.49 | 150.9 | 49.1 | 17.14 | 24.94 | | High | High | High | High | High | High | High | High | High | High | High | 0.002867 | 0.02974 | 3.36 |
| FALSE | High | QLSNDAAS | [K].QLSND | 1xPhospho | Q8BMA5 1 | 4.22E-07 | 0 | 1 | 1 | 1 | Q8BMA5 | Protein NP | Q8BMA5 [ | 0 | 2327.153 | 0 | -0.25 | 0.013532 | 6.19 | 108.8 | 91.2 | 8.61 | 6.62 | | High | High | High | High | High | High | High | High | High | High | High | 0 | 3.84E-08 | 3.61 |
| FALSE | High | QLSPATQL | [K].QLSPAT | 1xPhospho | Q8K2W9 1 | 0.000718 | 0 | 1 | 2 | 1 | Q8K2W9 | Uncharacte | Q8K2W9 [ | 0 | 1634.883 | 0 | -0.56 | 0.003625 | 4.45 | 122.1 | 77.9 | 5.4 | 10.91 | | High | High | High | High | High | High | High | High | High | High | High | 0 | 0.000165 | 2.89 |
| FALSE | High | QLSPEAQA | [K].QLSPEA | 1xPhospho | Q9D4H9-2 | 0.035486 | 0.002304 | 1 | 4 | 1 | Q9D4H9-2 | Isoform 2 c | Q9D4H9-2 | 0 | 1308.651 | 0 | -0.41 | 0.034083 | 4.87 | 110.8 | 89.2 | 7.04 | 9.07 | | High | High | High | High | High | High | High | High | High | High | High | 0.001214 | 0.01298 | 2.21 |
| FALSE | High | QLSPPLSPS | [R].QLSPPL | 1xCarbami | E9QLG3 2x | 0.000148 | 0 | 1 | 5 | 1 | E9QLG3 | MAX gene- | E9QLG3 [1 | 0 | 3187.364 | 0 | -0.37 | 0.03925 | 13.14 | 110.5 | 89.5 | 9.54 | 7.84 | | High | High | High | High | High | High | High | High | High | High | High | 0 | 2.81E-05 | 4.7 |
| FALSE | High | QLTEMLPS | [K].QLTEM | 1xPhospho | Q64152 1x | 6.78E-09 | 0 | 1 | 2 | 1 | Q64152 | Transcripti | Q64152 [1 | 0 | 2709.41 | 0 | -0.12 | 0.93842 | 17.51 | 98.7 | 101.3 | 7.37 | 19.71 | | High | High | High | High | High | High | High | High | High | High | High | 0 | 3.7E-10 | 3.81 |
| FALSE | High | QLSPTPGYI | [R].QLSPTF | 1xPhospho | Q5RIM6 1x | 2.31E-05 | 0 | 1 | 10 | 1 | Q5RIM6 | Nuclear re | Q5RIM6 [1 | 0 | 2769.279 | 0 | 0.64 | 0.014165 | 25.95 | 74.6 | 125.4 | 15.58 | 13.98 | | High | High | High | High | High | High | High | High | High | High | High | 0 | 3.48E-06 | 4.11 |
| FALSE | High | QISQDVK | [R].QISQDV | 1xPhospho | A2AE27 1x | 0.005182 | 0.000551 | 1 | 2 | 3 | A2AE27 | AMP deam | A2AE27 [1 | 0 | 1355.734 | 0 | 0.4 | 0.023913 | 10.97 | 86.3 | 113.7 | 4.85 | 15 | | High | High | High | High | High | High | High | High | High | High | High | 0.000399 | 0.003555 | 2.27 |
| FALSE | High | QLSQQNLT | [R].QLSQQ | 1xPhospho | B2RUG9 1x | 0.001516 | 5.66E-05 | 1 | 6 | 1 | B2RUG9 | Adenomat | B2RUG9 [2 | 0 | 1597.871 | 0 | 0.63 | 0.05311 | 41.09 | 72.7 | 127.3 | 17.62 | 22.4 | | High | High | High | High | High | High | High | High | High | High | High | 2.58E-05 | 0.00038 | 2.04 |
| FALSE | High | QLSSGVSEI | [R].QLSSG | 1xPhospho | Q545F4 1x | 0.000731 | 0 | 1 | 4 | 7 | Q545F4 | Hspb1 prot | Q545F4 [8 | 0 | 1384.703 | 0 | 0.03 | 0.737981 | 12.77 | 98.1 | 101.9 | 7.17 | 9.51 | | High | High | High | High | High | High | High | High | High | High | High | 0 | 0.000167 | 2.43 |
| FALSE | High | QLSSSQGS | [R].QLSSSC | 1xPhospho | Q3UFK8 1x | 1.4E-06 | 0 | 1 | 2 | 3 | Q3UFK8 | FERM dom | Q3UFK8 [4 | 0 | 2665.208 | 0 | 0.8 | 0.004215 | 11.19 | 71.8 | 128.2 | 9.32 | 7.42 | | High | High | High | High | High | High | High | High | High | High | High | 0 | 0.000141 | 4.34 |
| FALSE | High | QLSSSVTGI | [K].QLSSSV | 1xCarbami | Q6ZQK2 1x | 9E-07 | 0 | 1 | 7 | 4 | Q6ZQK2 | MKIAA005 | Q6ZQK2 [5 | 0 | 2474.143 | 0 | 0.12 | 0.380094 | 16.08 | 94.5 | 105.5 | 3.59 | 14.47 | | High | High | High | High | High | High | High | High | High | High | High | 0 | 9.02E-08 | 3.62 |
| FALSE | High | QLSTEQAA | [R].QLSTEC | 1xPhospho | A0A0R4J1L | 0.004411 | 0.000409 | 1 | 8 | 2 | A0A0R4J1L | Microtubu | A0A0R4J1L | 0 | 2213.147 | 0 | -0.24 | 0.023243 | 2.82 | 109.3 | 90.7 | 7.97 | 3.95 | | High | High | High | High | High | High | High | High | High | High | High | 0.000183 | 0.001256 | 2.66 |
| FALSE | High | QLSVPASD | [K].QLSVPA | 1xPhospho | Q6P542 1x | 2.73E-06 | 0 | 2 | 4 | 7 | Q6P542; Q | ATP-bindin | Q6P542 [1 | 0 | 2358.143 | 0 | -0.4 | 0.008673 | 7.84 | 113.7 | 86.3 | 4.01 | 7.4 | Shared | High | High | High | High | High | High | High | High | High | High | High | 0 | 2E-05 | 3.33 |
| FALSE | High | QLSVPASD | [K].QLSVPA | 2xPhospho | Q6P542 2x | 0.000176 | 0 | 2 | 4 | 4 | Q6P542; Q | ATP-bindin | Q6P542 [1 | 0 | 2438.109 | 0 | -0.06 | 0.381083 | 2.52 | 102.1 | 97.9 | 5.14 | 4.03 | Shared | High | High | High | High | High | High | High | High | High | High | High | 0 | 4.88E-05 | 3.41 |
| FALSE | High | QLSVSDR | [R].QLSVSD | 1xPhospho | E9PV45 1x | 0.026191 | 0.001676 | 1 | 2 | 2 | E9PV45 | Ubiquitin c | E9PV45 [1 | 0 | 1113.55 | 0 | 1.93 | 0.001861 | 12.16 | 41.4 | 158.6 | 5.98 | 15.31 | | High | High | High | High | High | High | High | High | High | High | High | 0.001372 | 0.01436 | 1.47 |
| FALSE | High | QISYNCSDI | [K].QISYNC | 1xCarbami | Q9CY58 1x | 7.06E-07 | 0 | 1 | 2 | 18 | Q9CY58 | Plasminoge | Q9CY58 [2 | 0 | 4173.848 | 0 | 0.04 | 0.468097 | 8.42 | 100.8 | 99.2 | 4.85 | 8.59 | | High | High | High | High | High | High | High | High | High | High | High | 0 | 1.17E-06 | 4.79 |
| FALSE | High | QISYNCSDI | [K].QISYNC | 1xCarbami | Q9CY58 1x | 3.9E-05 | 0 | 1 | 2 | 23 | Q9CY58 | Plasminoge | Q9CY58 [2 | 0 | 4174.832 | 0 | 0.34 | 0.008916 | 13.4 | 88.1 | 111.9 | 4.83 | 5.32 | | High | High | High | High | High | High | High | High | High | High | High | 0 | 1.29E-05 | 5.75 |
| FALSE | High | QISYNCSDI | [K].QISYNC | 1xCarbami | Q9CY58 2x | 7.95E-08 | 0 | 1 | 2 | 5 | Q9CY58 | Plasminoge | Q9CY58 [2 | 0 | 4253.815 | 0 | 0.41 | 0.009124 | 2.8 | 88.6 | 111.4 | 5.36 | 7.13 | | High | High | High | High | High | High | High | High | High | High | High | 0 | 5.91E-09 | 4.82 |
| FALSE | High | QLSYSDSDI | [R].QLSYSD | 1xPhospho | Q6NZK8-3 | 0.027839 | 0.001755 | 1 | 3 | 3 | Q6NZK8-3 | Isoform 3 c | Q6NZK8-3 | 0 | 1693.845 | 0 | 1.02 | 0.001101 | 7.34 | 67.5 | 132.5 | 10.36 | 6.27 | | High | High | High | High | High | High | High | High | High | High | High | 0.001609 | 0.01571 | 2.6 |
| FALSE | High | QISPVK | [R].QISPVK | 1xPhospho | B8JJC1 1xP | 0.034579 | 0.002233 | 1 | 3 | 2 | B8JJC1 | Tyrosyl-DN | B8JJC1 [59 | 0 | 1209.701 | 0 | -0.92 | 0.000181 | 1.55 | 131.3 | 68.7 | 6.11 | 10.07 | | High | High | High | High | High | High | High | High | High | High | High | 0.001183 | 0.01269 | 1.73 |
| FALSE | High | QNLLAPQN | [K].QNLLAI | 2xPhospho | F8WIP8 2x | 2.3E-06 | 0 | 1 | 9 | 4 | F8WIP8 | Osteopont | F8WIP8 [5 | 0 | 2246.087 | 0 | 0.3 | 0.437904 | 52.57 | 89.5 | 110.5 | 25.19 | 20.72 | | High | High | High | High | High | High | High | High | High | High | High | 0 | 2.58E-07 | 3.56 |
| FALSE | High | QNLLSVGD | [K].QNLLSV | 1xPhospho | Q91ZW3 1 | 0.00051 | 0 | 1 | 3 | 1 | Q91ZW3 | SWI/SNF-r | Q91ZW3 [1 | 0 | 1473.73 | 0 | 0.23 | 0.028782 | 16.51 | 90 | 110 | 11.05 | 6.77 | | High | High | High | High | High | High | High | High | High | High | High | 0 | 0.000112 | 1.35 |
| FALSE | High | QNLWFAN | [R].QNLWF | 1xPhospho | G5E8A7 1x | 2.27E-08 | 0 | 1 | 3 | 2 | G5E8A7 | Protein Sm | G5E8A7 [6 | 0 | 2387.08 | 0 | 0.86 | 0.005117 | 18.93 | 73.4 | 126.6 | 7.65 | 14.39 | | High | High | High | High | High | High | High | High | High | High | High | 0 | 1.44E-09 | 2.47 |
| FALSE | High | QNYSESSN | [R].QNYSE | 1xPhospho | Q61687 1x | 1.64E-07 | 0 | 1 | 4 | 6 | Q61687 | Transcripti | Q61687 [7 | 0 | 2129.886 | 0 | -0.18 | 0.105204 | 11.44 | 105.8 | 94.2 | 2.09 | 10.73 | | High | High | High | High | High | High | High | High | High | High | High | 0 | 3.24E-08 | 5.27 |
| FALSE | High | QPAEPGTS | [K].QPAEP | 1xCarbami | P41234 1x | 1.17E-05 | 0 | 1 | 3 | 3 | P41234 | ATP-bindin | P41234 [12 | 0 | 2519.144 | 0 | 0.66 | 0.002654 | 10.89 | 76.6 | 123.4 | 8.77 | 7.56 | | High | High | High | High | High | High | High | High | High | High | High | 0 | 1.61E-06 | 3.9 |
| FALSE | High | QPAPTTHS | [R].QPAPT | 1xPhospho | Q8C547 1x | 0.00359 | 0.00021 | 1 | 2 | 1 | Q8C547 | HEAT repe | Q8C547 [2 | 0 | 1917.025 | 0 | 0.89 | 0.001684 | 5.75 | 65.8 | 134.2 | 10.81 | 8.17 | | High | High | High | High | High | High | High | High | High | High | High | 9.42E-05 | 0.001002 | 2.9 |
| FALSE | High | QPASLHGS | [R].QPASLI | 3xPhospho | Q3TZJ2 3x | 0.002623 | 0.000109 | 1 | 4 | 1 | Q3TZJ2 | Putative ur | Q3TZJ2 [17 | 0 | 2174.985 | 0 | 0.02 | 0.880956 | 9.35 | 99.4 | 100.6 | 11.58 | 11.51 | | High | High | High | High | High | High | High | High | High | High | High | 4.94E-05 | 0.000702 | 2.31 |
| FALSE | High | QPASSCVQ | [K].QPASS | 1xCarbami | Q923A2 1x | 5.73E-07 | 0 | 1 | 1 | 2 | Q923A2 | Protein Spi | Q923A2 [5 | 0 | 2232.125 | 0 | -0.44 | 0.042707 | 15.98 | 114.3 | 85.7 | 10.78 | 9.38 | | High | High | High | High | High | High | High | High | High | High | High | 0 | 5.45E-08 | 3.49 |
| FALSE | High | QPCPSDGD | [K].QPCPSI | 1xCarbami | Q99JF8 1x | 2.82E-05 | 0 | 1 | 4 | 1 | Q99JF8 | PC4 and SF | Q99JF8 [2 | 0 | 2259.991 | 0 | -0.55 | 0.000815 | 1.54 | 118.8 | 81.2 | 4.72 | 6.77 | | High | High | High | High | High | High | High | High | High | High | High | 0 | 4.35E-06 | 3.44 |
| FALSE | High | QPEDESHC | [K].QPEDE | 1xPhospho | G5E8T6 1x | 0.000279 | 0 | 1 | 3 | 2 | G5E8T6 | Membrane | G5E8T6 [8 | 0 | 2222.967 | 0 | -0.31 | 0.022767 | 3.51 | 110.9 | 89.1 | 5.11 | 5.69 | | High | High | High | High | High | High | High | High | High | High | High | 4.94E-05 | 0.000594 | 2.36 |
| FALSE | High | QPELENSLS | [K].QPELEN | 1xPhospho | Z4YKQ9 1x | 6.23E-08 | 0 | 1 | 4 | 1 | Z4YKQ9 | ATPase fan | Z4YKQ9 [3 | 0 | 2719.323 | 0 | -0.02 | 0.277045 | 6.71 | 102.7 | 97.3 | 8.73 | 7.39 | | High | High | High | High | High | High | High | High | High | High | High | 0 | 4.52E-09 | 4.89 |
| FALSE | High | QPELENSLS | [K].QPELEN | 2xPhospho | Z4YKQ9 2x | 1.67E-05 | 0 | 1 | 4 | 1 | Z4YKQ9 | ATPase fan | Z4YKQ9 [3 | 0 | 2799.289 | 0 | -0.29 | 0.016943 | 4.68 | 108.6 | 91.4 | 2.99 | 5.62 | | High | High | High | High | High | High | High | High | High | High | High | 0 | 2.4E-06 | 4.45 |
| FALSE | High | QPEPHPGS | [R].QPEPHI | 1xPhospho | Q9JJ89 1xP | 0.000122 | 0 | 1 | 1 | 2 | Q9JJ89 | Coiled-coil | Q9JJ89 [16 | 0 | 3826.787 | 0 | 0.42 | 0.045925 | 5.44 | 86.5 | 113.5 | 6.25 | 12.3 | | High | High | High | High | High | High | High | High | High | High | High | 0 | 2.24E-05 | 3.7 |
| FALSE | High | QPETVLTET | [R].QPETVI | 1xPhospho | A2A6Q5-3 | 8.4E-08 | 0 | 1 | 6 | 1 | A2A6Q5-3 | Isoform 3 c | A2A6Q5-3 | 0 | 2393.18 | 0 | -0.41 | 0.008268 | 3.81 | 114.3 | 85.7 | 5.75 | 6.73 | | High | High | High | High | High | High | High | High | High | High | High | 0 | 6.27E-09 | 3.74 |
| FALSE | High | QPETYHTY | [R].QPETYI | 1xPhospho | Q61191 1x | 0.001615 | 5.66E-05 | 1 | 2 | 2 | Q61191 | Host cell fa | Q61191 [1 | 0 | 2321.065 | 0 | -0.17 | 0.602353 | 5.63 | 105.5 | 94.5 | 13.28 | 16.38 | | High | High | High | High | High | High | High | High | High | High | High | 2.58E-05 | 0.000408 | 2.93 |
| FALSE | High | QPGAEDSP | [R].QPGAE | 1xPhospho | Q9JI99 1xP | 0.004143 | 0.00036 | 1 | 2 | 2 | Q9JI99 | Sphingosin | Q9JI99 [59 | 0 | 1537.766 | 0 | 1.25 | 0.000248 | 6.93 | 57.7 | 142.3 | 8.87 | 5.46 | | High | High | High | High | High | High | High | High | High | High | High | 0.002455 | 0.02666 | 2.25 |
| FALSE | High | QNVPTGGS | [K].QNVPT | 1xPhospho | Q7TPM1 1 | 1.49E-05 | 0 | 1 | 1 | 1 | Q7TPM1 | Protein PRI | Q7TPM1 [1 | 0 | 1938.927 | 0 | 0.59 | 0.00607 | 16.55 | 86.9 | 113.1 | 17.11 | 14.99 | | High | High | High | High | High | High | High | High | High | High | High | 0 | 2.12E-06 | 3.14 |
| FALSE | High | QPGDSPQA | [R].QPGDS | 1xPhospho | Q99K30 1x | 0.022973 | 0.001578 | 1 | 2 | 2 | Q99K30 | Epidermal | Q99K30 [2 | 0 | 1465.745 | 0 | 1.07 | 0.000738 | 11.9 | 64.3 | 135.7 | 4.37 | 8.33 | | High | High | High | High | High | High | High | High | High | High | High | 0.007016 | 0.0763 | 1.94 |
| FALSE | High | QPGFPQPS | [K].QPGFP | 2xPhospho | Q91VC7 2x | 0.005516 | 0.00062 | 1 | 1 | 1 | Q91VC7 | Protein phi | Q91VC7 [1 | 0 | 2298.004 | 0 | -1.29 | 0.005655 | 4.17 | 142 | 58 | 9.17 | 24.26 | | High | High | High | High | High | High | High | High | High | High | High | 0.000275 | 0.001611 | 3.63 |
| FALSE | High | QPGSLEVN | [R].QPGSLI | 1xPhospho | Q3TWC8 1 | 0.006773 | 0.00062 | 1 | 7 | 1 | Q3TWC8 | Putative ur | Q3TWC8 [1 | 0 | 1680.872 | 0 | 1.19 | 0.000276 | 9 | 59.6 | 140.4 | 9.64 | 5.01 | | High | High | High | High | High | High | High | High | High | High | High | 0.000275 | 0.002024 | 3.43 |
| FALSE | High | QPGSPSPPS | [K].QPGSPS | 1xPhospho | Q8VD12 1x | 7.75E-05 | 0 | 1 | 2 | 3 | Q8VD12 | Zinc finger | Q8VD12 [1 | 0 | 2374.206 | 0 | 0.91 | 0.002537 | 12.14 | 70 | 130 | 7.37 | 7.1 | | High | High | High | High | High | High | High | High | High | High | High | 0.000275 | 0.002163 | 2.76 |
| FALSE | High | QPGTCSPD | [R].QPGTC | 1xCarbami | Q3U0X6 1x | 6.97E-06 | 0 | 1 | 1 | 4 | Q3U0X6 | Putative ur | Q3U0X6 [4 | 0 | 2473.075 | 0 | -0.08 | 0.348765 | 3.15 | 101.7 | 98.3 | 3.26 | 8.45 | | High | High | High | High | High | High | High | High | High | High | High | 0 | 9.04E-07 | 4.28 |
| FALSE | High | QPGTCSPD | [R].QPGTC | 1xCarbami | Q3U0X6 2x | 0.019276 | 0.001402 | 1 | 1 | 1 | Q3U0X6 | Putative ur | Q3U0X6 [4 | 1 | 4790.07 | 0 | -0.85 | 0.001251 | 1.68 | 128 | 72 | 8.16 | 10.09 | | High | High | High | High | High | High | High | High | High | High | High | 0.000756 | 0.006561 | 4.77 |
| FALSE | High | QPGTSLVD | [R].QPGTSI | 1xPhospho | A2AM95 1 | 9.71E-05 | 0 | 1 | 3 | 4 | A2AM95 | Emerin OS | A2AM95 [1 | 0 | 2217.065 | 0 | -0.24 | 0.023026 | 6.31 | 108.5 | 91.5 | 6.73 | 3.78 | | High | High | High | High | High | High | High | High | High | High | High | 0 | 1.89E-05 | 3.07 |
| FALSE | High | QPLASPSTC | [K].QPLASF | 1xPhospho | Q8BM75 1 | 0.002486 | 0.000109 | 1 | 2 | 1 | Q8BM75 | AT-rich int | Q8BM75 [5 | 0 | 1927.96 | 0 | 0.97 | 0.036583 | 21.85 | 73.2 | 126.8 | 14.13 | 25.28 | | High | High | High | High | High | High | High | High | High | High | High | 4.94E-05 | 0.000664 | 2.18 |
| FALSE | High | QPLASPSTC | [K].QPLASF | 1xPhospho | Q8BM75 1 | 0.012397 | 0.000889 | 1 | 2 | 2 | Q8BM75 | AT-rich int | Q8BM75 [5 | 0 | 1943.955 | 0 | 1.03 | 0.002949 | 12.92 | 65.2 | 134.8 | 4.03 | 13.9 | | High | High | High | High | High | High | High | High | High | High | High | 0.000689 | 0.005934 | 3.01 |

| | | | | | | | | | | | | | | | | | | | | | | | | | | | | | | | | | | | | | | |
|---|---|---|---|---|---|---|---|---|---|---|---|---|---|---|---|---|---|---|---|---|---|---|---|---|---|---|---|---|---|---|---|---|---|---|---|---|---|---|
| FALSE | High | QPLEQSQT | [K].QPLEQ | 1xPhospho | Q3UJ69 1x | 2.43E-07 | 0 | 1 | 4 | 6 | Q3UJ69 | Putative ur | Q3UJ69 [4 | 0 | 2675.322 | 0 | 1.22 | 0.001124 | 14.49 | 60.4 | 139.6 | 6.21 | 10.93 | | High | High | High | High | High | High | High | High | High | High | High | 0 | 1.15E-05 | 4.01 |
| FALSE | High | QPLEVGSP | [R].QPLEV | 1xCarbami | E9Q9T0 1x | 0.004491 | 0.000409 | 1 | 3 | 1 | E9Q9T0 | Fibrosin-lik | E9Q9T0 [1 | 1 | 2895.36 | 0 | -0.8 | 0.007702 | 2.15 | 128.6 | 71.4 | 10.25 | 9.79 | | High | High | High | High | High | High | High | High | High | High | High | 0.000183 | 0.001278 | 2.36 |
| FALSE | High | QPLFVPAN | [K].QPLFVF | 1xPhospho | A2ADB0 1x | 0.000372 | 0 | 1 | 5 | 2 | A2ADB0 | Msx2-inter | A2ADB0 [3 | 0 | 3455.849 | 0 | 0.46 | 0.035851 | 2.22 | 87.1 | 112.9 | 5.48 | 9.43 | | High | High | High | High | High | High | High | High | High | High | High | 0 | 7.83E-05 | 3.8 |
| FALSE | High | QPLLLSEDE | [K].QPLLLS | 1xPhospho | Q8R1B4 1x | 2.47E-05 | 0 | 1 | 7 | 2 | Q8R1B4 | Eukaryotic | Q8R1B4 [3 | 0 | 1825.867 | 0 | -0.27 | 0.733771 | 25.04 | 123 | 77 | 107.11 | 105.7 | | High | High | High | High | High | High | High | High | High | High | High | 0 | 3.74E-06 | 3.75 |
| FALSE | High | QPLLLSEDE | [K].QPLLLS | 1xPhospho | Q8R1B4 1x | 3.26E-08 | 0 | 1 | 7 | 11 | Q8R1B4 | Eukaryotic | Q8R1B4 [3 | 0 | 2055.03 | 0 | -0.26 | 0.004952 | 5.2 | 109.5 | 90.5 | 2.36 | 6.62 | | High | High | High | High | High | High | High | High | High | High | High | 0 | 2.16E-09 | 3.82 |
| FALSE | High | QPGFPQPS | [K].QPGFP | 1xPhospho | Q91VC7 1x | 0.000157 | 0 | 1 | 1 | 6 | Q91VC7 | Protein ph | Q91VC7 [1 | 0 | 2218.038 | 0 | -1.97 | 0.000204 | 1.73 | 161.1 | 38.9 | 6.62 | 25.11 | | High | High | High | High | High | High | High | High | High | High | High | 0 | 8.12E-05 | 3.17 |
| FALSE | High | QNTEEAGL | [K].QNTEE | 1xPhospho | Q3TI84 1xP | 6.16E-07 | 0 | 1 | 2 | 1 | Q3TI84 | CDC16 cell | Q3TI84 [57 | 0 | 2239.137 | 0 | 0.23 | 0.002162 | 1.7 | 92.1 | 107.9 | 1.99 | 4.82 | | High | High | High | High | High | High | High | High | High | High | High | 0 | 5.88E-08 | 3.3 |
| FALSE | High | QNTAWEFI | [K].QNTAW | 1xPhospho | B1ATC3 1x | 0.000462 | 0 | 1 | 3 | 1 | B1ATC3 | Trinucleoti | B1ATC3 [6 | 0 | 1802.795 | 0 | -1.79 | 0.001216 | 2.36 | 159.8 | 40.2 | 10.95 | 18.48 | | High | High | High | High | High | High | High | High | High | High | High | 0 | 0.0001 | 1.56 |
| FALSE | High | QNTAWDT | [K].QNTAW | 1xPhospho | F6ZMJ4 1x | 9.2E-05 | 0 | 1 | 5 | 3 | F6ZMJ4 | Trinucleoti | F6ZMJ4 [7 | 0 | 1714.764 | 0 | 0.37 | 0.245938 | 5.73 | 87.2 | 112.8 | 4.2 | 23.71 | | High | High | High | High | High | High | High | High | High | High | High | 0 | 1.63E-05 | 2.83 |
| FALSE | High | QNPEQSAD | [K].QNPEQ | 1xPhospho | Q8R1B4 1x | 7.1E-06 | 0 | 1 | 5 | 3 | Q8R1B4 | Eukaryotic | Q8R1B4 [1 | 0 | 1998.906 | 0 | -0.28 | 0.011187 | 0.23 | 107.7 | 92.3 | 3.01 | 6.9 | | High | High | High | High | High | High | High | High | High | High | High | 0 | 9.19E-07 | 3.97 |
| FALSE | High | QNPTVGSH | [K].QNPTV | 1xCarbami | E9Q9J4 1x | 5.13E-06 | 0 | 1 | 1 | 4 | E9Q9J4 | Inositol he | E9Q9J4 [10 | 0 | 3588.663 | 0 | 0.39 | 0.25837 | 21.89 | 94.3 | 105.7 | 21 | 15.4 | | High | High | High | High | High | High | High | High | High | High | High | 0 | 6.39E-07 | 4.54 |
| FALSE | High | QNQPTSCV | [K].QNQPT | 1xCarbami | Q9ERU9 1x | 8.32E-08 | 0 | 1 | 2 | 2 | Q9ERU9 | E3 SUMO-p | Q9ERU9 [1 | 0 | 2215.985 | 0 | -0.16 | 0.064786 | 11.83 | 112.1 | 87.9 | 14.05 | 12.2 | | High | High | High | High | High | High | High | High | High | High | High | 0 | 6.25E-09 | 3.58 |
| FALSE | High | QNQQSSSD | [R].QNQQS | 1xPhospho | Q52KI8 1xP | 3.32E-05 | 0 | 1 | 6 | 4 | Q52KI8 | Serine/argi | Q52KI8 [48 | 0 | 2779.242 | 0 | 0.07 | 0.844621 | 31.18 | 97.5 | 102.5 | 20.97 | 3.47 | | High | High | High | High | High | High | High | High | High | High | High | 0 | 5.22E-06 | 2.75 |
| FALSE | High | QNSATESA | [R].QNSAT | 1xPhospho | B1AZP2 1x | 6.76E-08 | 0 | 1 | 6 | 6 | B1AZP2 | Disks large | B1AZP2 [9 | 0 | 2618.218 | 0 | 0.9 | 0.00207 | 18.52 | 72.1 | 127.9 | 5.86 | 7.61 | | High | High | High | High | High | High | High | High | High | High | High | 0 | 5.72E-09 | 4.9 |
| FALSE | High | QNSCHAQI | [K].QNSCH | 2xCarbami | A0A0R4IZY | 4.25E-10 | 0 | 1 | 3 | 4 | A0A0R4IZY | tRNA-dihy | A0A0R4IZY | 0 | 3973.826 | 0 | -0.43 | 0.144737 | 14.27 | 119.5 | 80.5 | 17.46 | 10.94 | | High | High | High | High | High | High | High | High | High | High | High | 0 | 1.65E-11 | 5.69 |
| FALSE | High | QNSCHAQI | [K].QNSCH | 2xCarbami | A0A0R4IZY | 0.048715 | 0.00354 | 1 | 3 | 1 | A0A0R4IZY | tRNA-dihy | A0A0R4IZY | 0 | 4053.792 | 0 | 0.1 | 0.120513 | 3.99 | 87.9 | 112.1 | 15.68 | 13.3 | | High | High | High | High | High | High | High | High | High | High | High | 0.001876 | 0.01878 | 3.18 |
| FALSE | High | QNSDDK | [R].QNSDD | 1xPhospho | Q8CBY1 1x | 0.095408 | 0.007437 | 1 | 3 | 1 | Q8CBY1 | Protein Sm | Q8CBY1 [1 | 0 | 1244.592 | 0 | 1.93 | 0.000158 | 9.49 | 43.8 | 156.2 | 14.67 | 4.78 | | High | High | High | High | High | High | High | High | High | High | High | 0.003809 | 0.04101 | 1.45 |
| FALSE | High | QNSDPTSE | [R].QNSDP | 1xPhospho | Q9JM52-3 | 2.65E-05 | 0 | 1 | 8 | 2 | Q9JM52-3 | Isoform 3 c | Q9JM52-3 | 0 | 3592.711 | 0 | 1.76 | 0.001103 | 5.65 | 47.3 | 152.7 | 3.66 | 11.25 | | High | High | High | High | High | High | High | High | High | High | High | 0 | 4.04E-06 | 3.29 |
| FALSE | High | QNSFQLQK | [K].QNSFQ | 1xPhospho | A2AKX3 1x | 0.014033 | 0.001011 | 1 | 2 | 1 | A2AKX3 | Probable h | A2AKX3 [9 | 0 | 1530.808 | 0 | 0.88 | 0.011006 | 17.11 | 69.1 | 130.9 | 6.18 | 18.98 | | High | High | High | High | High | High | High | High | High | High | High | 0.000538 | 0.004603 | 2.53 |
| FALSE | High | QNSGIIASE | [K].QNSGII | 1xPhospho | E9QLG3 1x | 9E-07 | 0 | 1 | 5 | 3 | E9QLG3 | MAX gene- | E9QLG3 [2 | 0 | 3386.543 | 0 | -0.87 | 0.004311 | 0.93 | 123.3 | 76.7 | 10.7 | 7.33 | | High | High | High | High | High | High | High | High | High | High | High | 0 | 1.65E-06 | 4.69 |
| FALSE | High | QNSGIIASE | [K].QNSGII | 1xDeamida | E9QLG3 1x | 2.75E-05 | 0 | 1 | 5 | 3 | E9QLG3 | MAX gene- | E9QLG3 [2 | 0 | 3387.527 | 0 | -0.59 | 0.021918 | 6.77 | 119.5 | 80.5 | 12.08 | 21.31 | | High | High | High | High | High | High | High | High | High | High | High | 0 | 4.19E-06 | 5.62 |
| FALSE | High | QNSGIIASE | [K].QNSGII | 2xPhospho | E9QLG3 2x | 6.15E-06 | 0 | 1 | 5 | 1 | E9QLG3 | MAX gene- | E9QLG3 [2 | 0 | 3466.51 | 0 | -1.33 | 0.002607 | 2.4 | 144.3 | 55.7 | 7.27 | 18.62 | | High | High | High | High | High | High | High | High | High | High | High | 0 | 7.81E-07 | 4.17 |
| FALSE | High | QNSLGCIG | [R].QNSLG | 2xCarbami | G3UYU0 1x | 1.46E-08 | 0 | 1 | 1 | 2 | G3UYU0 | Mex3 RNA- | G3UYU0 [3 | 0 | 2476.084 | 0 | 0.27 | 0.001431 | 1.57 | 88.8 | 111.2 | 2.86 | 4.94 | | High | High | High | High | High | High | High | High | High | High | High | 0 | 8.84E-10 | 5.79 |
| FALSE | High | QNSPAGSK | [R].QNSPA | 1xPhospho | E9QM77 1: | 0.006478 | 0.00062 | 1 | 9 | 3 | E9QM77 | Ataxin-2 O | E9QM77 [7 | 0 | 1326.682 | 0 | 1.3 | 0.000434 | 6.29 | 58.8 | 141.2 | 8.16 | 4.56 | | High | High | High | High | High | High | High | High | High | High | High | 0.000275 | 0.001924 | 2.37 |
| FALSE | High | QNSPVTSP | [R].QNSPV | 1xPhospho | Q3TTA7 1x | 0.013666 | 0.001011 | 1 | 3 | 3 | Q3TTA7 | E3 ubiquiti | Q3TTA7 [4 | 0 | 1934.953 | 0 | 0.97 | 0.013532 | 26.4 | 65.4 | 134.6 | 8.65 | 25.27 | | High | High | High | High | High | High | High | High | High | High | High | 0.00074 | 0.00646 | 1.97 |
| FALSE | High | QNSPVTSP | [R].QNSPV | 2xPhospho | Q3TTA7 2x | 8.3E-07 | 0 | 1 | 3 | 2 | Q3TTA7 | E3 ubiquiti | Q3TTA7 [4 | 0 | 2014.92 | 0 | -0.25 | 0.448961 | 6.54 | 103.7 | 96.3 | 10.79 | 9.59 | | High | High | High | High | High | High | High | High | High | High | High | 0 | 8.26E-08 | 3.26 |
| FALSE | High | QNSPVTSP | [R].QNSPV | 3xPhospho | Q3TTA7 3x | 0.002172 | 5.66E-05 | 1 | 3 | 1 | Q3TTA7 | E3 ubiquiti | Q3TTA7 [4 | 0 | 2094.886 | 0 | 0.56 | 0.055609 | 23.8 | 80.8 | 119.2 | 11.32 | 12.37 | | High | High | High | High | High | High | High | High | High | High | High | 4.94E-05 | 0.000572 | 3.6 |
| FALSE | High | QNSQLPAC | [R].QNSQL | 1xPhospho | E9QLZ9 1x | 1E-06 | 0 | 1 | 11 | 2 | E9QLZ9 | Protein en | E9QLZ9 [1 | 0 | 2802.362 | 0 | -0.47 | 0.026561 | 10.67 | 116 | 84 | 6.63 | 11.68 | | High | High | High | High | High | High | High | High | High | High | High | 0 | 1.02E-07 | 5.42 |
| FALSE | High | QNSQLPAC | [R].QNSQL | 1xDeamida | E9QLZ9 1x | 8.63E-05 | 0 | 1 | 11 | 4 | E9QLZ9 | Protein en | E9QLZ9 [1 | 0 | 2803.346 | 0 | -0.37 | 0.024655 | 8.77 | 117.9 | 82.1 | 10.62 | 12.1 | | High | High | High | High | High | High | High | High | High | High | High | 0 | 1.53E-05 | 4.65 |
| FALSE | High | QNSSCTVD | [K].QNSSC | 1xCarbami | Q6A037 1x | 1.3E-05 | 0 | 1 | 4 | 1 | Q6A037 | NEDD4-bin | Q6A037 [4 | 0 | 2958.355 | 0 | -0.66 | 0.000442 | 0.32 | 124.1 | 75.9 | 9.15 | 8.3 | | High | High | High | High | High | High | High | High | High | High | High | 0 | 1.81E-06 | 3.6 |
| FALSE | High | QNSSDSISS | [R].QNSSD | 1xPhospho | Q8CH77 1x | 5.73E-07 | 0 | 1 | 5 | 3 | Q8CH77 | Neuron na | Q8CH77 [1 | 0 | 2861.393 | 0 | 0.41 | 0.303324 | 17.77 | 84.9 | 115.1 | 23.12 | 24.98 | | High | High | High | High | High | High | High | High | High | High | High | 0 | 1.2E-07 | 4.49 |
| FALSE | High | QNSSSSDS | [R].QNSSS | 1xPhospho | Q3U2Z5 1x | 2.49E-05 | 0 | 1 | 3 | 8 | Q3U2Z5 | Putative ur | Q3U2Z5 [6 | 0 | 1689.764 | 0 | 1.21 | 0.000385 | 2.4 | 60.3 | 139.7 | 7.39 | 3.52 | | High | High | High | High | High | High | High | High | High | High | High | 0.000204 | 0.001316 | 2.24 |
| FALSE | High | QNSSSSDS | [R].QNSSS | 2xPhospho | Q3U2Z5 2x | 0.000144 | 0 | 1 | 3 | 4 | Q3U2Z5 | Putative ur | Q3U2Z5 [6 | 0 | 1769.731 | 0 | 1.93 | 0.000158 | 11.16 | 40.8 | 159.2 | 9.52 | 2.69 | | High | High | High | High | High | High | High | High | High | High | High | 0 | 2.71E-05 | 2.87 |
| FALSE | High | QNSTGQSQ | [R].QNSTG | 1xPhospho | E9PW37 1x | 0.002355 | 0.000109 | 1 | 3 | 2 | E9PW37 | RAS protei | E9PW37 [1 | 0 | 1427.684 | 0 | 1.12 | 0.000532 | 10.35 | 62.1 | 137.9 | 8.94 | 3.55 | | High | High | High | High | High | High | High | High | High | High | High | 9.42E-05 | 0.000959 | 2.46 |
| FALSE | High | QNSVESDL | [R].QNSVE | 1xPhospho | A2A654 1x | 9.03E-05 | 0 | 1 | 4 | 2 | A2A654 | Bromodom | A2A654 [1 | 0 | 1542.7 | 0 | 0.09 | 0.035942 | 2.27 | 94.1 | 105.9 | 7.16 | 7.7 | | High | High | High | High | High | High | High | High | High | High | High | 0 | 1.6E-05 | 3.14 |
| FALSE | High | QNSVNSGI | [R].QNSVN | 1xPhospho | J3QM82 1x | 0.001558 | 5.66E-05 | 1 | 2 | 1 | J3QM82 | MCG16947 | J3QM82 [1 | 0 | 1791.932 | 0 | 1.02 | 0.000417 | 14.97 | 64.7 | 135.3 | 12.85 | 3.75 | | High | High | High | High | High | High | High | High | High | High | High | 2.58E-05 | 0.000391 | 2.86 |
| FALSE | High | QNSVSSNT | [R].QNSVS | 1xPhospho | Q3UDC3 1x | 0.002719 | 0.000109 | 1 | 4 | 1 | Q3UDC3 | Putative ur | Q3UDC3 [1 | 0 | 1516.695 | 0 | 0.54 | 0.000252 | 2.65 | 80.6 | 119.4 | 6.01 | 5.62 | | High | High | High | High | High | High | High | High | High | High | High | 4.94E-05 | 0.00073 | 1.87 |
| FALSE | High | QNTAGSPK | [K].QNTAG | 1xPhospho | A5D6P2 1x | 0.028082 | 0.001791 | 1 | 22 | 2 | A5D6P2 | Par-3 (Part | A5D6P2 [1 | 0 | 1340.698 | 0 | -0.23 | 0.064192 | 4.52 | 106 | 94 | 3.91 | 7.78 | | High | High | High | High | High | High | High | High | High | High | High | 0.000968 | 0.01 | 1.64 |
| FALSE | High | QLSGQSTS | [R].QLSGQ | 2xPhospho | Q9EPQ8 2x | 0.003927 | 0.000311 | 1 | 3 | 3 | Q9EPQ8 | Transcripti | Q9EPQ8 [5 | 0 | 2120.955 | 0 | -0.52 | 0.003859 | 5.07 | 112.2 | 87.8 | 11.5 | 14.66 | | High | High | High | High | High | High | High | High | High | High | High | 0.00014 | 0.001103 | 2.29 |
| FALSE | High | QLSGQSTS | [R].QLSGQ | 1xPhospho | Q9EPQ8 1x | 1.78E-05 | 0 | 1 | 3 | 3 | Q9EPQ8 | Transcripti | Q9EPQ8 [5 | 0 | 2040.989 | 0 | -0.47 | 0.003417 | 0.89 | 118.3 | 81.7 | 7.06 | 7.77 | | High | High | High | High | High | High | High | High | High | High | High | 0 | 2.58E-06 | 2.9 |
| FALSE | High | QLSFISPPA | [K].QLSFISI | 1xPhospho | Q3UT19 1x | 0.002376 | 0.000109 | 1 | 2 | 1 | Q3UT19 | Putative ur | Q3UT19 [1 | 0 | 1948.071 | 0 | -0.86 | 0.001042 | 2.62 | 129.1 | 70.9 | 4.49 | 6.42 | | High | High | High | High | High | High | High | High | High | High | High | 4.94E-05 | 0.000631 | 1.43 |
| FALSE | High | QLSESESSL | [R].QLSESE | 1xPhospho | D3YZP9 1x | 4.61E-06 | 0 | 1 | 3 | 7 | D3YZP9 | Coiled-coil | D3YZP9 [3 | 0 | 2079.863 | 0 | 0.84 | 0.001759 | 8.26 | 68.5 | 131.5 | 9.33 | 7.21 | | High | High | High | High | High | High | High | High | High | High | High | 0 | 2.43E-05 | 4.23 |
| FALSE | High | QGTAIGSEI | [R].QGTAI | 1xPhospho | Q3TFK5 1x | 4.64E-09 | 0 | 1 | 1 | 3 | Q3TFK5 | G patch do | Q3TFK5 [2 | 0 | 2183.966 | 0 | -0.12 | 0.212928 | 7.6 | 103.9 | 96.1 | 1.94 | 9.74 | | High | High | High | High | High | High | High | High | High | High | High | 0 | 7.68E-06 | 3.55 |
| FALSE | High | QGTDDSK | [R].QGTDD | 1xPhospho | Q6ZWQ0 1 | 0.059159 | 0.004294 | 1 | 10 | 2 | Q6ZWQ0 | Nesprin-2 | Q6ZWQ0 [ | 0 | 1288.619 | 0 | 1.51 | 0.001183 | 12.47 | 50 | 150 | 12.62 | 10.7 | | High | High | High | High | High | High | High | High | High | High | High | 0.002187 | 0.02337 | 1.13 |
| FALSE | High | QGTLDGSF | [R].QGTLD | 1xCarbami | M0QWX4 | 0.008388 | 0.000736 | 1 | 3 | 1 | M0QWX4 | Sentrin-spe | M0QWX4 | 0 | 1738.767 | 0 | -0.31 | 0.254504 | 9.86 | 112.3 | 87.7 | 8.64 | 22.83 | | High | High | High | High | High | High | High | High | High | High | High | 0.000327 | 0.002569 | 1.59 |
| FALSE | High | QGYACPYY | [R].QGYAC | 1xCarbami | Q8BL48 1x | 0.000176 | 0 | 1 | 2 | 2 | Q8BL48 | RING finge | Q8BL48 [2 | 0 | 2025.929 | 0 | 0.43 | 0.003812 | 3.07 | 83 | 117 | 7.62 | 4.31 | | High | High | High | High | High | High | High | High | High | High | High | 0 | 9.95E-05 | 2.51 |
| FALSE | High | QHAHTALS | [R].QHAHT | 1xPhospho | Q9ERQ3 1x | 0.017809 | 0.001282 | 1 | 2 | 4 | Q9ERQ3 | Zinc finger | Q9ERQ3 [5 | 0 | 1610.8 | 0 | -0.24 | 0.04404 | 2.76 | 107.8 | 92.2 | 5.93 | 9.37 | | High | High | High | High | High | High | High | High | High | High | High | 0.000689 | 0.006005 | 2.21 |
| FALSE | High | QHALSLNV | [R].QHALSI | 1xPhospho | Q8CJF7 1x | 1.61E-10 | 0 | 1 | 1 | 2 | Q8CJF7 | Protein ELY | Q8CJF7 [17 | 0 | 2745.422 | 0 | -0.33 | 0.019587 | 9.96 | 106.8 | 93.2 | 11.41 | 9.42 | | High | High | High | High | High | High | High | High | High | High | High | 0 | 5.54E-12 | 4.69 |
| FALSE | High | QHAQPNQ | [K].QHAQP | 1xPhospho | Q544H6 1x | 1.12E-07 | 0 | 1 | 3 | 2 | Q544H6 | Cyclin E2, i | Q544H6 [1 | 0 | 2796.408 | 0 | 1.19 | 0.002176 | 13.81 | 62 | 138 | 5.53 | 10.63 | | High | High | High | High | High | High | High | High | High | High | High | 0 | 8.67E-09 | 4.23 |
| FALSE | High | QHEVEIPSF | [K].QHEVE | 1xPhospho | B2RTL2 1x | 0.004781 | 0.000456 | 1 | 5 | 1 | B2RTL2 | Phosphodi | B2RTL2 [2 | 0 | 1698.841 | 0 | -0.12 | 0.404409 | 2.95 | 101.7 | 98.3 | 6.58 | 6.75 | | High | High | High | High | High | High | High | High | High | High | High | 0.000204 | 0.001382 | 2.84 |
| FALSE | High | QHEVEIPSF | [K].QHEVE | 1xPhospho | E9PVU7 1x | 0.001751 | 5.66E-05 | 1 | 11 | 2 | E9PVU7 | Phosphodi | E9PVU7 [3 | 0 | 1931.004 | 0 | -0.54 | 0.002393 | 6.65 | 117.7 | 82.3 | 6.52 | 4.72 | | High | High | High | High | High | High | High | High | High | High | High | 2.58E-05 | 0.000448 | 2.24 |
| FALSE | High | QHGEQLG | [R].QHGEC | 1xPhospho | Q3UR00 1x | 5.53E-07 | 0 | 1 | 10 | 2 | Q3UR00 | Putative ur | Q3UR00 [8 | 0 | 2386.119 | 0 | -0.73 | 0.001041 | 4.05 | 126.6 | 73.4 | 5.13 | 7.48 | | High | High | High | High | High | High | High | High | High | High | High | 0 | 5.24E-08 | 4.08 |
| FALSE | High | QHGIPIPVT | [K].QHGIPI | 1xPhospho | P16460 1x | 0.000244 | 0 | 1 | 2 | 1 | P16460 | Argininosu | P16460 [16 | 0 | 1724.986 | 0 | -1.65 | 0.001035 | 1.48 | 147.6 | 52.4 | 9.97 | 19.39 | | High | High | High | High | High | High | High | High | High | High | High | 0 | 4.87E-05 | 2.02 |
| FALSE | High | QHLASGGD | [K].QHLAS | 1xPhospho | Q91ZJ0 1x | 3.02E-07 | 0 | 1 | 3 | 3 | Q91ZJ0 | Crossover | Q91ZJ0 [83 | 0 | 2171.065 | 0 | -0.4 | 0.020442 | 6.35 | 114.9 | 85.1 | 9.69 | 12.69 | | High | High | High | High | High | High | High | High | High | High | High | 0 | 2.64E-08 | 3.54 |
| FALSE | High | QHLENDPG | [K].QHLENI | 1xPhospho | Q3UNH3 1: | 1.84E-07 | 0 | 1 | 4 | 2 | Q3UNH3 | Putative ur | Q3UNH3 [1 | 0 | 2417.139 | 0 | 0.45 | 0.00809 | 6.48 | 84.6 | 115.4 | 5.56 | 9.62 | | High | High | High | High | High | High | High | High | High | High | High | 0 | 1.51E-08 | 2.95 |
| FALSE | High | QGSVVCSR | [R].QGSVV | 1xCarbami | Q3UFK8 1x | 0.030898 | 0.001896 | 1 | 1 | 1 | Q3UFK8 | FERM dom | Q3UFK8 [4 | 0 | 1201.56 | 0 | 0.77 | 0.00066 | 7.76 | 75.4 | 124.6 | 8.87 | 5.88 | | High | High | High | High | High | High | High | High | High | High | High | 0.001027 | 0.01116 | 1.37 |
| FALSE | High | QHILGLTAS | [K].QHILGL | 1xPhospho | A2AQ14 1x | 4.91E-06 | 0 | 1 | 1 | 1 | A2AQ14 | Protein Mi | A2AQ14 [2 | 0 | 1702.966 | 0 | -0.95 | 0.000444 | 1.36 | 132.5 | 67.5 | 6.05 | 6.24 | | High | High | High | High | High | High | High | High | High | High | High | 0 | 6.07E-07 | 3.84 |
| FALSE | High | QHRPLPSTI | [R].QHRPLI | 1xPhospho | E9Q7D5 1x | 0.001558 | 5.66E-05 | 1 | 2 | 3 | E9Q7D5 | Rho guanir | E9Q7D5 [7 | 0 | 2776.373 | 0 | 3.27 | 0.001309 | 28.35 | 19.9 | 180.1 | 9.21 | 16.02 | | High | High | High | High | High | High | High | High | High | High | High | 2.58E-05 | 0.000393 | 2.4 |
| FALSE | High | QHRPLPSTI | [R].QHRPLI | 2xPhospho | E9Q7D5 2x | 0.008022 | 0.000736 | 1 | 2 | 2 | E9Q7D5 | Rho guanir | E9Q7D5 [7 | 0 | 2856.339 | 0 | 1.86 | 0.000275 | 5.51 | 41.3 | 158.7 | 14.05 | 4.25 | | High | High | High | High | High | High | High | High | High | High | High | 0.000327 | 0.002445 | 2.26 |
| FALSE | High | QHSLPSSEI | [R].QHSLPS | 1xPhospho | Q9ESJ4 1x | 0.000284 | 0 | 1 | 1 | 3 | Q9ESJ4 | NCK-intera | Q9ESJ4 [14 | 0 | 2923.43 | 0 | 0 | 0.337172 | 7.24 | 92.3 | 107.7 | 13.98 | 15.55 | | High | High | High | High | High | High | High | High | High | High | High | 0 | 5.83E-05 | 3.02 |
| FALSE | High | QHSPGSTD | [R].QHSPG | 1xPhospho | O70374 1x | 3.65E-07 | 0 | 1 | 4 | 2 | O70374 | Protein CB | O70374 [3 | 0 | 2024.887 | 0 | -1.46 | 0.000768 | 2.3 | 149.6 | 50.4 | 6.98 | 10.4 | | High | High | High | High | High | High | High | High | High | High | High | 0 | 4.65E-08 | 4.16 |
| FALSE | High | QHSPIQVPI | [K].QHSPIC | 1xCarbami | Q8VBU8 1x | 0.000209 | 0 | 1 | 7 | 3 | Q8VBU8 | Protein BA | Q8VBU8 [9 | 0 | 2975.488 | 0 | -0.62 | 0.063532 | 12.58 | 120 | 80 | 17.3 | 18.41 | | High | High | High | High | High | High | High | High | High | High | High | 0.001651 | 0.01657 | 3.81 |
| FALSE | High | QHSSGTLLI | [R].QHSSG | 1xPhospho | Q3U1G4 1x | 0.000681 | 0 | 1 | 4 | 2 | Q3U1G4 | Putative ur | Q3U1G4 [5 | 0 | 2008.052 | 0 | 2.08 | 0.00061 | 25.29 | 34.4 | 165.6 | 10.93 | 9.22 | | High | High | High | High | High | High | High | High | High | High | High | 0 | 0.000315 | 3.06 |
| FALSE | High | QHSYIEQG | [K].QHSYIE | 1xPhospho | Q61164 1x | 0.007019 | 0.00062 | 1 | 3 | 1 | Q61164 | Transcripti | Q61164 [4 | 0 | 1627.825 | 0 | -0.54 | 0.002872 | 3.67 | 119.7 | 80.3 | 9.3 | 5.56 | | High | High | High | High | High | High | High | High | High | High | High | 0.000275 | 0.002112 | 2.83 |
| FALSE | High | QKPCSPLSE | [K].QKPCSI | 1xCarbami | B0V2M3 1x | 0.001255 | 0 | 1 | 3 | 1 | B0V2M3 | Zinc finger | B0V2M3 [2 | 0 | 2484.318 | 0 | -0.29 | 0.039214 | 8.04 | 112.7 | 87.3 | 10.55 | 7.37 | | High | High | High | High | High | High | High | High | High | High | High | 0 | 0.000308 | 2.21 |
| FALSE | High | QKPETSSLE | [K].QKPETS | 1xPhospho | Q8C804 1x | 0.105058 | 0.008518 | 1 | 2 | 1 | Q8C804 | Spindle an | Q8C804 [7 | 0 | 1914.055 | 0 | -0.77 | 0.001507 | 1.69 | 125.5 | 74.5 | 6.99 | 5.03 | | High | High | High | High | High | High | High | High | High | High | High | 0.004277 | 0.04578 | 1.49 |
| FALSE | High | QKPSLEEQ | [K].QKPSLE | 1xPhospho | F8VQC7 1x | 0.003526 | 0.00021 | 1 | 35 | 1 | F8VQC7 | Kinectin O | F8VQC7 [1 | 0 | 2066.187 | 0 | 1.94 | 0.000471 | 6.9 | 41.3 | 158.7 | 9.75 | 8.01 | | High | High | High | High | High | High | High | High | High | High | High | 9.42E-05 | 0.000979 | 2.2 |
| FALSE | High | QIADTPPH | [R].QIADTF | 1xPhospho | Q8JZX4 1x | 1.66E-05 | 0 | 1 | 2 | 2 | Q8JZX4 | Splicing fac | Q8JZX4 [67 | 0 | 1956.072 | 0 | -0.06 | 0.460584 | 3.51 | 101.1 | 98.9 | 2.73 | 4.45 | | High | High | High | High | High | High | High | High | High | High | High | 0 | 2.38E-06 | 2.85 |
| FALSE | High | QIAGPDSE | [R].QIAGPI | 1xPhospho | E9PWG6 1: | 0.000524 | 0 | 1 | 2 | 3 | E9PWG6 | Non-SMC c | E9PWG6 [9 | 0 | 2492.142 | 0 | -0.59 | 0.000695 | 2.89 | 119.2 | 80.8 | 5.95 | 5.14 | | High | High | High | High | High | High | High | High | High | High | High | 0 | 0.000116 | 2.63 |
| FALSE | High | QIAGPDSE | [R].QIAGPI | 1xPhospho | E9PWG6 1: | 0.000382 | 0 | 1 | 2 | 6 | E9PWG6 | Non-SMC c | E9PWG6 [9 | 0 | 2508.137 | 0 | -0.47 | 0.019844 | 2.61 | 117.9 | 82.1 | 5.73 | 13.4 | | High | High | High | High | High | High | High | High | High | High | High | 0 | 0.00028 | 3.32 |
| FALSE | High | QHQQPFQ | [R].QHQQF | 1xPhospho | B2RUG8 1x | 0.000314 | 0 | 1 | 5 | 1 | B2RUG8 | Asxl2 prote | B2RUG8 [5 | 0 | 2260.123 | 0 | 0.37 | 0.09335 | 21.67 | 87.3 | 112.7 | 7.82 | 12.71 | | High | High | High | High | High | High | High | High | High | High | High | 0 | 0.000065 | 3.85 |
| FALSE | High | QGSSSVSG | [R].QGSSS | 1xPhospho | P97445 1x | 1.3E-05 | 0 | 1 | 6 | 1 | P97445 | Voltage-de | P97445 [22 | 0 | 2244.034 | 0 | -1.86 | 0.002443 | 1.73 | 159.4 | 40.6 | 14.34 | 15.47 | | High | High | High | High | High | High | High | High | High | High | High | 0 | 1.8E-06 | 2.59 |
| FALSE | High | QGSRQDSV | [K].QGSRQ | 2xPhospho | D3YTT6 2x | 0.066503 | 0.004796 | 1 | 8 | 1 | D3YTT6 | Oxysterol-l | D3YTT6 [2 | 1 | 2134.952 | 0 | 1.06 | 0.001125 | 8.88 | 66.8 | 133.2 | 3.25 | 9.56 | | High | High | High | High | High | High | High | High | High | High | High | 0.002455 | 0.02676 | 2.26 |
| FALSE | High | QGSRPGSP | [R].QGSRP | 2xPhospho | H7BX49 2x | 0.022973 | 0.001578 | 1 | 2 | 1 | H7BX49 | WD repeat | H7BX49 [1 | 0 | 2383.101 | 0 | -0.43 | 0.008212 | 8.21 | 114.8 | 85.2 | 9.89 | 2.35 | | High | High | High | High | High | High | High | High | High | High | High | 0.000847 | 0.008005 | 2.3 |
| FALSE | High | QGGQGNA | [R].QGGQ | 1xPhospho | Q3UZ18 1x | 5.87E-09 | 0 | 1 | 1 | 1 | Q3UZ18 | Little elong | Q3UZ18 [5 | 0 | 2898.327 | 0 | -0.94 | 0.045016 | 6.24 | 129.2 | 70.8 | 14.28 | 18.52 | | High | High | High | High | High | High | High | High | High | High | High | 0 | 3.16E-10 | 4.58 |
| FALSE | High | QGGSCSIA | [R].QGGSC | 1xCarbami | Q61687 1x | 0.000244 | 0 | 1 | 3 | 2 | Q61687 | Transcripti | Q61687 [8 | 0 | 1888.924 | 0 | -0.36 | 0.000964 | 1.96 | 112.5 | 87.5 | 3.1 | 2.64 | | High | High | High | High | High | High | High | High | High | High | High | 0 | 4.93E-05 | 2.48 |
| FALSE | High | QGHDSLEH | [R].QGHDS | 1xPhospho | Q9R1C0 1x | 0.002058 | 5.66E-05 | 1 | 2 | 2 | Q9R1C0 | Transcripti | Q9R1C0 [2 | 0 | 1744.785 | 0 | -1.33 | 0.000406 | 1.93 | 144 | 56 | 4.47 | 8.71 | | High | High | High | High | High | High | High | High | High | High | High | 0.000204 | 0.001337 | 2.21 |
| FALSE | High | QGLAETSSI | [R].QGLAE | 1xPhospho | Q9R0L6-2 | 7.34E-05 | 0 | 1 | 3 | 7 | Q9R0L6-2 | Isoform 2 c | Q9R0L6-2 | 0 | 1837.962 | 0 | 0.96 | 0.002274 | 3.35 | 71.3 | 128.7 | 10.27 | 5.87 | | High | High | High | High | High | High | High | High | High | High | High | 0 | 1.27E-05 | 3.11 |
| FALSE | High | QGLAETSSI | [R].QGLAE | 2xPhospho | Q9R0L6-2 | 5.55E-05 | 0 | 1 | 3 | 1 | Q9R0L6-2 | Isoform 2 c | Q9R0L6-2 | 0 | 1917.929 | 0 | 0.36 | 0.003989 | 4.9 | 86.1 | 113.9 | 10.55 | 10.53 | | High | High | High | High | High | High | High | High | High | High | High | 0 | 9.3E-06 | 3.83 |
| FALSE | High | QGLAETSSI | [R].QGLAE | 1xCarbami | Q9R0L6-2 | 0.036731 | 0.00249 | 1 | 3 | 2 | Q9R0L6-2 | Isoform 2 c | Q9R0L6-2 | 1 | 3557.558 | 0 | 0.33 | 0.025895 | 18.93 | 88.7 | 111.3 | 9.05 | 7.64 | | High | High | High | High | High | High | High | High | High | High | High | 0.001308 | 0.01352 | 4.44 |
| FALSE | High | QGLGNHSF | [K].QGLGN | 1xPhospho | B1AYC9 1x | 0.001002 | 0 | 1 | 1 | 2 | B1AYC9 | Amyloid be | B1AYC9 [5 | 0 | 1503.727 | 0 | 1.96 | 0.00049 | 13.5 | 37.2 | 162.8 | 13.56 | 8.52 | | High | High | High | High | High | High | High | High | High | High | High | 0.000117 | 0.001076 | 2.98 |
| FALSE | High | QGLGNHSF | [K].QGLGN | 1xDeamida | B1AYC9 1x | 0.003312 | 0.00016 | 1 | 1 | 1 | B1AYC9 | Amyloid be | B1AYC9 [5 | 0 | 1504.711 | 0 | 1.76 | 0.000261 | 18.48 | 45.5 | 154.5 | 11.76 | 2.64 | | High | High | High | High | High | High | High | High | High | High | High | 7.19E-05 | 0.000913 | 2.97 |
| FALSE | High | QGLPSPPG | [R].QGLPS | 1xPhospho | Q2TGE7 1x | 0.001629 | 5.66E-05 | 1 | 3 | 1 | Q2TGE7 | Palmitoyltr | Q2TGE7 [6 | 0 | 1415.725 | 0 | 0.34 | 0.011382 | 11.66 | 88.1 | 111.9 | 2.52 | 7.98 | | High | High | High | High | High | High | High | High | High | High | High | 2.58E-05 | 0.000411 | 2.15 |
| FALSE | High | QGLSYSSLK | [K].QGLSYS | 1xPhospho | Q61730 1x | 0.073462 | 0.005631 | 1 | 1 | 1 | Q61730 | Interleukin | Q61730 [5 | 0 | 1520.813 | 0 | 1.56 | 0.005266 | 36.91 | 49.7 | 150.3 | 12.04 | 20.87 | | High | High | High | High | High | High | High | High | High | High | High | 0.002911 | 0.03021 | 1.5 |
| FALSE | High | QGPDSPGC | [K].QGPDS | 1xPhospho | Q3UXZ9 1x | 0.00418 | 0.00036 | 1 | 1 | 1 | Q3UXZ9 | Lysine-spe | Q3UXZ9 [1 | 0 | 2568.261 | 0 | 0.28 | 0.046938 | 4.76 | 92.6 | 107.4 | 8.84 | 9.57 | | High | High | High | High | High | High | High | High | High | High | High | 0.000161 | 0.001179 | 2.15 |
| FALSE | High | QGPELGSC | [R].QGPEL | 1xPhospho | Q3UJB9 1x | 2.08E-06 | 0 | 1 | 6 | 1 | Q3UJB9 | Enhancer c | Q3UJB9 [7 | 0 | 2234.029 | 0 | -0.21 | 0.063056 | 4.17 | 107.1 | 92.9 | 7.61 | 5.98 | | High | High | High | High | High | High | High | High | High | High | High | 0 | 2.32E-07 | 4.19 |
| FALSE | High | QGPSPVSP | [K].QGPSP | 1xPhospho | Q3U2G7 1x | 2.75E-05 | 0 | 1 | 8 | 1 | Q3U2G7 | Putative ur | Q3U2G7 [1 | 0 | 1646.81 | 0 | -1.26 | 0.00024 | 1.35 | 143.3 | 56.7 | 4.74 | 6.57 | | High | High | High | High | High | High | High | High | High | High | High | 0 | 4.19E-06 | 2.19 |
| FALSE | High | QGPSQPLS | [R].QGPSQ | 1xPhospho | Q3UIL6-2 1 | 7.36E-06 | 0 | 1 | 8 | 1 | Q3UIL6-2 | Isoform 2 c | Q3UIL6-2 [ | 0 | 2555.295 | 0 | -1.32 | 0.00476 | 4.52 | 143.4 | 56.6 | 3.87 | 30.45 | | High | High | High | High | High | High | High | High | High | High | High | 0 | 9.56E-07 | 3.23 |
| FALSE | High | QGPSTPPP | [K].QGPSTI | 1xPhospho | Q6ZQ89 1x | 0.000194 | 0 | 1 | 3 | 2 | Q6ZQ89 | E3 ubiquiti | Q6ZQ89 [8 | 0 | 1619.752 | 0 | -0.08 | 0.311637 | 6.49 | 100.3 | 99.7 | 13.54 | 13.23 | | High | High | High | High | High | High | High | High | High | High | High | 0 | 3.78E-05 | 2.85 |
| FALSE | High | QGSDEWG | [R].QGSDE | 1xPhospho | G5E8A7 1x | 2.2E-06 | 0 | 1 | 3 | 2 | G5E8A7 | Protein Sm | G5E8A7 [1 | 0 | 3327.584 | 0 | 0.95 | 0.003621 | 21.53 | 70.6 | 129.4 | 10.24 | 9.55 | | High | High | High | High | High | High | High | High | High | High | High | 0 | 2.47E-07 | 4.24 |
| FALSE | High | QGSFSEDV | [R].QGSFSI | 1xPhospho | B1ARU1 1x | 0.001349 | 0 | 2 | 15 | 1 | B1ARU1; F | Microtubu | B1ARU1 [2 | 0 | 1642.768 | 0 | 1.12 | 0.011923 | 11.39 | 62.9 | 137.1 | 18.45 | 11.29 | Shared | High | High | High | High | High | High | High | High | High | High | High | 0 | 0.000332 | 2.24 |
| FALSE | High | QGSFSEDV | [R].QGSFSI | 1xPhospho | B1ARU1 1x | 8.9E-06 | 0 | 2 | 15 | 1 | B1ARU1; F | Microtubu | B1ARU1 [2 | 0 | 1871.93 | 0 | 1.07 | 0.00046 | 9.8 | 65.9 | 134.1 | 4.65 | 6.29 | Shared | High | High | High | High | High | High | High | High | High | High | High | 0 | 1.19E-06 | 3.16 |
| FALSE | High | QGSGAESE | [R].QGSGA | 1xPhospho | Q8K1C9 1x | 0.01029 | 0.000772 | 1 | 2 | 1 | Q8K1C9 | Leucine-ric | Q8K1C9 [3 | 0 | 2799.353 | 0 | 1.57 | 0.000631 | 24.02 | 50.4 | 149.6 | 32.17 | 9.17 | | High | High | High | High | High | High | High | High | High | High | High | 0.000382 | 0.003241 | 3.18 |
| FALSE | High | QGSGLFSN | [R].QGSGL | 2xPhospho | Q8BXJ2-4 | 0.000279 | 0 | 2 | 3 | 2 | Q8BXJ2-4; | Isoform 4 c | Q8BXJ2-4 [ | 0 | 3663.865 | 0 | 0.46 | 0.021288 | 10.77 | 84.2 | 115.8 | 7.74 | 10.97 | Shared | High | High | High | High | High | High | High | High | High | High | High | 0 | 0.000209 | 2.46 |
| FALSE | High | QGSGSSQF | [K].QGSGS | 1xPhospho | Q62318 1x | 0.002793 | 0.000109 | 1 | 3 | 7 | Q62318 | Transcripti | Q62318 [4 | 0 | 4086.777 | 0 | -1.76 | 0.000457 | 0.67 | 150.3 | 49.7 | 6.59 | 14.43 | | High | High | High | High | High | High | High | High | High | High | High | 4.94E-05 | 0.000751 | 3.76 |
| FALSE | High | QGSIEDLLH | [R].QGSIED | 1xPhospho | Q9D6X5 1x | 0.034281 | 0.002195 | 1 | 1 | 1 | Q9D6X5 | Solute carr | Q9D6X5 [2 | 0 | 2595.361 | 0 | 0.33 | 0.703928 | 14.32 | 90.7 | 109.3 | 18.06 | 12.32 | | High | High | High | High | High | High | High | High | High | High | High | 0.001151 | 0.01247 | 2.06 |
| FALSE | High | QGSLEPDW | [R].QGSLEI | 1xPhospho | Q8C708 1x | 0.0026 | 0.000109 | 1 | 1 | 2 | Q8C708 | Transmem | Q8C708 [1 | 0 | 1791.863 | 0 | 0.93 | 0.006035 | 20.45 | 69.5 | 130.5 | 9.27 | 13.84 | | High | High | High | High | High | High | High | High | High | High | High | 4.94E-05 | 0.000694 | 2.46 |
| FALSE | High | QGSLHVGD | [R].QGSLH | 1xDeamida | Q542P4 1x | 1.2E-05 | 0 | 1 | 6 | 3 | Q542P4 | Membrane | Q542P4 [1 | 0 | 3584.8 | 0 | 1.68 | 0.000644 | 0.64 | 45.5 | 154.5 | 13.34 | 13.47 | | High | High | High | High | High | High | High | High | High | High | High | 0 | 1.65E-06 | 3.66 |
| FALSE | High | QGSLVLEEI | [R].QGSLVI | 1xPhospho | Q544C5 1x | 0.005275 | 0.000599 | 1 | 2 | 1 | Q544C5 | MCG14526 | Q544C5 [3 | 0 | 2633.488 | 0 | -0.63 | 0.025252 | 7.33 | 121 | 79 | 10.88 | 13.75 | | High | High | High | High | High | High | High | High | High | High | High | 0.000267 | 0.001536 | 3.2 |
| FALSE | High | QGSPTPAL | [R].QGSPTI | 1xPhospho | E9Q0S6 1x | 0.025073 | 0.001645 | 1 | 10 | 1 | E9Q0S6 | Tensin 1 O | E9Q0S6 [1 | 0 | 1662.887 | 0 | -0.05 | 0.896752 | 12.85 | 100.8 | 99.2 | 8.78 | 9.3 | | High | High | High | High | High | High | High | High | High | High | High | 0.000872 | 0.008759 | 2.23 |
| FALSE | High | QGSPVAAG | [R].QGSPV | 1xPhospho | O88477 1x | 0.005666 | 0.00062 | 1 | 2 | 2 | O88477 | Insulin-like | O88477 [1 | 0 | 1362.698 | 0 | -1.46 | 0.20496 | 4.69 | 147.9 | 52.1 | 64.35 | 72.05 | | High | High | High | High | High | High | High | High | High | High | High | 0.000275 | 0.00167 | 2.53 |
| FALSE | High | QGSPVAAG | [R].QGSPV | 1xPhospho | O88477 1x | 5.82E-06 | 0 | 1 | 2 | 9 | O88477 | Insulin-like | O88477 [1 | 0 | 1591.861 | 0 | -1.38 | 0.00021 | 1.05 | 144.9 | 55.1 | 3.74 | 9.6 | | High | High | High | High | High | High | High | High | High | High | High | 2.58E-05 | 0.000555 | 3.23 |
| FALSE | High | QGSRPGSP | [R].QGSRP | 1xPhospho | H7BX49 1x | 0.000215 | 0 | 1 | 2 | 3 | H7BX49 | WD repeat | H7BX49 [1 | 0 | 2303.134 | 0 | 0.07 | 0.254531 | 9.08 | 100.5 | 99.5 | 6.62 | 6.25 | | High | High | High | High | High | High | High | High | High | High | High | 0.000344 | 0.003177 | 2.71 |
| FALSE | High | QIAPEPPKF | [K].QIAPEP | 1xPhospho | Q8BTI8 1x | 0.110202 | 0.008961 | 1 | 6 | 2 | Q8BTI8 | Serine/argi | Q8BTI8 [16 | 0 | 2133.187 | 0 | -0.66 | 0.000313 | 0.72 | 124.3 | 75.7 | 2.41 | 3.55 | | High | High | High | High | High | High | High | High | High | High | High | 0.00449 | 0.04854 | 1.13 |
| FALSE | High | QGGASQSI | [R].QGGAS | 1xPhospho | P17809 1x | 0.000616 | 0 | 1 | 3 | 3 | P17809 | Solute carr | P17809 [46 | 0 | 1415.693 | 0 | 0.03 | 0.271421 | 5.77 | 95.8 | 104.2 | 14.05 | 16.15 | | High | High | High | High | High | High | High | High | High | High | High | 0 | 0.000138 | 1.96 |
| FALSE | High | QLASEEDD | [R].QLASEE | 1xPhospho | Q3TR89 1x | 0.025293 | 0.001645 | 1 | 6 | 1 | Q3TR89 | Putative ur | Q3TR89 [9 | 0 | 2272.115 | 0 | 0.4 | 0.044308 | 18.67 | 85.3 | 114.7 | 13.85 | 8.94 | | High | High | High | High | High | High | High | High | High | High | High | 0.000872 | 0.008849 | 3.03 |
| FALSE | High | QIASSSPSL | [R].QIASSS | 1xPhospho | Q0VGY8 1x | 0.000198 | 0 | 1 | 4 | 1 | Q0VGY8 | Protein TA | Q0VGY8 [4 | 0 | 1739.934 | 0 | -0.42 | 0.003183 | 2.31 | 113.9 | 86.1 | 3.44 | 6.57 | | High | High | High | High | High | High | High | High | High | High | High | 0 | 3.88E-05 | 1.05 |
| FALSE | High | QIPDQDEN | [K].QIPDQI | 1xPhospho | Q5DTS2 1x | 0.068184 | 0.004935 | 1 | 4 | 1 | Q5DTS2 | MKIAA402 | Q5DTS2 [4 | 0 | 2135.042 | 0 | -0.08 | 0.513037 | 10.55 | 102 | 98 | 9.65 | 8.65 | | High | High | High | High | High | High | High | High | High | High | High | 0.002559 | 0.02748 | 3.18 |
| FALSE | High | QIPGGGSG | [K].QIPGG | 1xPhospho | Q8BMA6 1 | 2.54E-09 | 0 | 1 | 2 | 2 | Q8BMA6 | Signal reco | Q8BMA6 [ | 0 | 2139.937 | 0 | 0.01 | 0.822451 | 5.22 | 100 | 100 | 9.22 | 6.39 | | High | High | High | High | High | High | High | High | High | High | High | 0 | 1.25E-06 | 4.27 |
| FALSE | High | QLPHVQLP | [K].QLPHV | 1xDeamida | Q6P5E3 1x | 0.000958 | 0 | 1 | 3 | 3 | Q6P5E3 | Sal-like 1 ( | Q6P5E3 [3 | 0 | 4610.347 | 0 | -1.35 | 0.000357 | 0.94 | 143.4 | 56.6 | 6.75 | 17.6 | | High | High | High | High | High | High | High | High | High | High | High | 2.58E-05 | 0.000516 | 3.44 |

| | | | | | | | | | | | | | | | | | | | | | | | | | | | | | | | | | | | | | | |
|---|---|---|---|---|---|---|---|---|---|---|---|---|---|---|---|---|---|---|---|---|---|---|---|---|---|---|---|---|---|---|---|---|---|---|---|---|---|---|
| FALSE | High | QLPLEPGN | [K].QLPLEP | 1xCarbami | A2AJI0 1xP | 0.000139 | 0 | 1 | 4 | 2 | A2AJI0 | MAP7 dom | A2AJI0 [65 | 0 | 3721.818 | 0 | 0.96 | 0.000602 | 2.21 | 65.4 | 134.6 | 9.33 | 3.92 | | High | High | High | High | High | High | High | High | High | High | High | 0 | 2.61E-05 | 4.72 |
| FALSE | High | QIPSAEDLE | [R].QIPSAE | 1xPhospho | Q8C2B3-3 | 7.51E-07 | 0 | 1 | 11 | 1 | Q8C2B3-3 | Isoform 3 c | Q8C2B3-3 | 0 | 2974.345 | 0 | 2.04 | 0.000877 | 8.22 | 39.1 | 160.9 | 8.92 | 12.39 | | High | High | High | High | High | High | High | High | High | High | High | 0 | 7.37E-08 | 5.66 |
| FALSE | High | QIPSAEDLE | [R].QIPSAE | 1xPhospho | Q8C2B3-3 | 1.1E-07 | 0 | 1 | 11 | 1 | Q8C2B3-3 | Isoform 3 c | Q8C2B3-3 | 0 | 2990.34 | 0 | 2.2 | 0.00338 | 10.85 | 35.8 | 164.2 | 10.3 | 22.53 | | High | High | High | High | High | High | High | High | High | High | High | 0 | 8.49E-09 | 5.34 |
| FALSE | High | QIPSSPLQC | [K].QIPSSP | 1xPhospho | A0A1B0GS | 2.29E-05 | 0 | 1 | 8 | 2 | A0A1B0GS | Supervillin | A0A1B0GS | 0 | 3030.488 | 0 | -0.02 | 0.954392 | 10.51 | 104.4 | 95.6 | 7.01 | 7.88 | | High | High | High | High | High | High | High | High | High | High | High | 0 | 0.000125 | 4.26 |
| FALSE | High | QIPSSPLQC | [K].QIPSSP | 2xPhospho | A0A1B0GS | 0.000334 | 0 | 1 | 8 | 2 | A0A1B0GS | Supervillin | A0A1B0GS | 0 | 3110.455 | 0 | 0.87 | 0.002019 | 18.22 | 68.4 | 131.6 | 5.52 | 10.11 | | High | High | High | High | High | High | High | High | High | High | High | 0.000327 | 0.002676 | 3.46 |
| FALSE | High | QIPSSPLQC | [K].QIPSSP | 3xPhospho | A0A1B0GS | 0.020679 | 0.001472 | 1 | 8 | 1 | A0A1B0GS | Supervillin | A0A1B0GS | 0 | 3190.421 | 0 | 1.21 | 0.000435 | 4.02 | 56.9 | 143.1 | 5.73 | 7.27 | | High | High | High | High | High | High | High | High | High | High | High | 0.000786 | 0.00709 | 4.35 |
| FALSE | High | QLPTPVGA | [K].QLPTP | 1xPhospho | Q8C4X2 1x | 1.16E-06 | 0 | 1 | 1 | 2 | Q8C4X2 | Casein kina | Q8C4X2 [3 | 0 | 2287.164 | 0 | -0.21 | 0.321327 | 6.55 | 103.1 | 96.9 | 8.78 | 9.28 | | High | High | High | High | High | High | High | High | High | High | High | 0 | 1.2E-07 | 3.95 |
| FALSE | High | QIPVSPGSF | [K].QIPVSP | 1xPhospho | Q9D0N8 1x | 0.058173 | 0.004227 | 1 | 1 | 1 | Q9D0N8 | Transmem | Q9D0N8 [1 | 0 | 1249.65 | 0 | 1.24 | 0.000463 | 8.83 | 58.5 | 141.5 | 5.91 | 6.73 | | High | High | High | High | High | High | High | High | High | High | High | 0.002187 | 0.02296 | 1.09 |
| FALSE | High | QIPVVGSVI | [R].QIPVVG | 1xPhospho | Q99K01-4 | 1.66E-05 | 0 | 1 | 5 | 1 | Q99K01-4 | Isoform 4 c | Q99K01-4 | 0 | 2722.473 | 0 | 0.68 | 0.004135 | 10.97 | 80 | 120 | 13.12 | 3.91 | | High | High | High | High | High | High | High | High | High | High | High | 0 | 2.37E-06 | 4.19 |
| FALSE | High | QIQALEDA | [K].QIQALE | 1xPhospho | Q5NC05 1x | 0.004033 | 0.000311 | 1 | 1 | 2 | Q5NC05 | Transcripti | Q5NC05 [4 | 0 | 2721.448 | 0 | -0.51 | 0.001825 | 4.2 | 117.6 | 82.4 | 7.66 | 4.04 | | High | High | High | High | High | High | High | High | High | High | High | 0.000161 | 0.001135 | 2.86 |
| FALSE | High | QINVPALD | [K].QINVPA | 1xPhospho | B2RT41 1x | 0.000681 | 0 | 1 | 3 | 1 | B2RT41 | Zinc finger, | B2RT41 [13 | 0 | 2161.11 | 0 | -0.53 | 0.027074 | 4.3 | 122.6 | 77.4 | 14.16 | 20.38 | | High | High | High | High | High | High | High | High | High | High | High | 0 | 0.000155 | 2.93 |
| FALSE | High | QLQDSLHD | [R].QLQDS | 1xPhospho | Q9EP71 1x | 0.066503 | 0.004796 | 1 | 1 | 1 | Q9EP71 | Ankycorbir | Q9EP71 [4 | 0 | 1862.978 | 0 | 0.86 | 0.000902 | 5.72 | 70.9 | 129.1 | 6.66 | 10.86 | | High | High | High | High | High | High | High | High | High | High | High | 0.002455 | 0.02679 | 2.28 |
| FALSE | High | QLQLSSPLK | [R].QLQLS | 1xPhospho | Q9Z2E2 1x | 0.000958 | 0 | 1 | 6 | 2 | Q9Z2E2 | Methyl-Cp | Q9Z2E2 [4 | 0 | 1551.891 | 0 | 0.09 | 0.560503 | 2.97 | 97.7 | 102.3 | 8.16 | 3.34 | | High | High | High | High | High | High | High | High | High | High | High | 0 | 0.000227 | 2.25 |
| FALSE | High | QLQPASEL | [R].QLQPA | 1xPhospho | B7ZNX2 1x | 3.99E-10 | 0 | 1 | 6 | 3 | B7ZNX2 | Anaphase | B7ZNX2 [4 | 0 | 3588.835 | 0 | -0.56 | 0.001896 | 4.45 | 118 | 82 | 5.05 | 8.68 | | High | High | High | High | High | High | High | High | High | High | High | 0 | 1.54E-11 | 5.07 |
| FALSE | High | QLQQQLSN | [R].QLQQC | 1xPhospho | A2AGH6 1x | 0.000107 | 0 | 1 | 6 | 1 | A2AGH6 | Mediator c | A2AGH6 [2 | 0 | 2594.293 | 0 | 1.1 | 0.006057 | 10.8 | 62.6 | 137.4 | 13.92 | 14.18 | | High | High | High | High | High | High | High | High | High | High | High | 0 | 1.95E-05 | 3.34 |
| FALSE | High | QLSADSAE | [K].QLSADS | 1xPhospho | E9PU87 1x | 5.62E-06 | 0 | 1 | 3 | 1 | E9PU87 | Serine/thre | E9PU87 [9 | 0 | 2052.937 | 0 | 0.19 | 0.45525 | 17.19 | 94.9 | 105.1 | 8.29 | 9.99 | | High | High | High | High | High | High | High | High | High | High | High | 0 | 7.09E-07 | 2.92 |
| FALSE | High | QLSADSAE | [K].QLSADS | 1xPhospho | E9PU87 1x | 0.00083 | 0 | 1 | 3 | 2 | E9PU87 | Serine/thre | E9PU87 [9 | 0 | 2068.932 | 0 | 0.03 | 0.426934 | 7.79 | 99.4 | 100.6 | 5.51 | 27.02 | | High | High | High | High | High | High | High | High | High | High | High | 0 | 0.000194 | 2.57 |
| FALSE | High | QLSDSFK | [R].QLSDSF | 1xPhospho | Q08943-2 | 0.005615 | 0.00062 | 1 | 2 | 5 | Q08943-2 | Isoform 2 c | Q08943-2 | 0 | 1362.707 | 0 | -1.23 | 0.000936 | 1.39 | 140.2 | 59.8 | 4.93 | 12.12 | | High | High | High | High | High | High | High | High | High | High | High | 0.000275 | 0.001639 | 2.38 |
| FALSE | High | QISEDVDG | [R].QISEDV | 1xPhospho | B2RSC8 1x | 0.001503 | 5.66E-05 | 1 | 2 | 1 | B2RSC8 | E3 ubiquiti | B2RSC8 [30 | 0 | 1653.732 | 0 | 0.69 | 0.00171 | 7.86 | 74.1 | 125.9 | 17.35 | 7.47 | | High | High | High | High | High | High | High | High | High | High | High | 2.58E-05 | 0.000379 | 1.95 |
| FALSE | High | QISEDVDG | [R].QISEDV | 1xPhospho | B2RSC8 1x | 0.001011 | 0 | 1 | 2 | 6 | B2RSC8 | E3 ubiquiti | B2RSC8 [30 | 1 | 2893.32 | 0 | -0.92 | 0.001356 | 3.04 | 127.9 | 72.1 | 3.49 | 12.04 | | High | High | High | High | High | High | High | High | High | High | High | 0.000638 | 0.005721 | 3.31 |
| FALSE | High | QISEDVDG | [R].QISEDV | 2xPhospho | B2RSC8 2x | 0.054383 | 0.003951 | 1 | 2 | 1 | B2RSC8 | E3 ubiquiti | B2RSC8 [30 | 1 | 2973.287 | 0 | 0.18 | 0.070751 | 9.28 | 93.4 | 106.6 | 4.38 | 12.27 | | High | High | High | High | High | High | High | High | High | High | High | 0.002059 | 0.02126 | 3.51 |
| FALSE | High | QISEDVDG | [R].QISEDV | 1xDeamida | B2RSC8 2x | 0.038677 | 0.002563 | 1 | 2 | 1 | B2RSC8 | E3 ubiquiti | B2RSC8 [30 | 1 | 2974.271 | 0 | 0.17 | 0.031654 | 2.17 | 91.5 | 108.5 | 5.88 | 5.38 | | High | High | High | High | High | High | High | High | High | High | High | 0.001388 | 0.01441 | 3.8 |
| FALSE | High | QLSEEQPS | [R].QLSEEC | 1xPhospho | Q542C3 1x | 2.49E-05 | 0 | 1 | 2 | 2 | Q542C3 | Putative ur | Q542C3 [1 | 0 | 1910.962 | 0 | -0.28 | 0.000897 | 2.65 | 109.8 | 90.2 | 3.07 | 4.99 | | High | High | High | High | High | High | High | High | High | High | High | 0 | 3.74E-06 | 2.76 |
| FALSE | High | QLSEEQPS | [R].QLSEEC | 1xDeamida | Q542C3 1x | 4.55E-05 | 0 | 1 | 2 | 3 | Q542C3 | Putative ur | Q542C3 [1 | 0 | 1911.947 | 0 | -0.22 | 0.010297 | 4.99 | 109.5 | 90.5 | 3.76 | 6.94 | | High | High | High | High | High | High | High | High | High | High | High | 0 | 2.3E-05 | 2.67 |
| FALSE | High | QLSENGEL | [K].QLSEN | 1xPhospho | Q6PR54-3 | 0.013666 | 0.001011 | 2 | 4 | 1 | Q6PR54-3; | Isoform 3 c | Q6PR54-3 | 0 | 2142.073 | 0 | -0.26 | 0.035985 | 8.34 | 109.8 | 90.2 | 13.03 | 6.32 | Shared | High | High | High | High | High | High | High | High | High | High | High | 0.000538 | 0.004439 | 2.48 |
| FALSE | High | QLQHGEDE | [K].QLQHG | 1xPhospho | Q3U2W2 1 | 5.58E-07 | 0 | 1 | 2 | 9 | Q3U2W2 | MYB bindir | Q3U2W2 [ | 0 | 2411.04 | 0 | 0.15 | 0.418161 | 6.14 | 94.2 | 105.8 | 3.99 | 13.78 | | High | High | High | High | High | High | High | High | High | High | High | 0 | 5.3E-08 | 3.62 |
| FALSE | High | QLMSQSQ | [R].QLMSC | 1xPhospho | Q9CZ62 1x | 0.000337 | 0 | 1 | 3 | 4 | Q9CZ62 | Centrosom | Q9CZ62 [3 | 0 | 2557.205 | 0 | -0.42 | 0.005921 | 2.18 | 113.3 | 86.7 | 10.55 | 5.26 | | High | High | High | High | High | High | High | High | High | High | High | 0 | 7.03E-05 | 3.14 |
| FALSE | High | QLLTLSNEL | [R].QLLTLS | 1xPhospho | Q3TCP5 1x | 0.000237 | 0 | 1 | 5 | 1 | Q3TCP5 | Putative ur | Q3TCP5 [5 | 0 | 1781.936 | 0 | -0.33 | 0.039432 | 9.88 | 115.4 | 84.6 | 5.71 | 15.04 | | High | High | High | High | High | High | High | High | High | High | High | 0 | 4.72E-05 | 2.8 |
| FALSE | High | QLLSSGSSE | [R].QLLSSG | 1xCarbami | Q60I26 1xP | 0.00625 | 0.00062 | 1 | 2 | 2 | Q60I26 | ALS2 C-ter | Q60I26 [13 | 0 | 2478.157 | 0 | 0.07 | 0.573882 | 18.23 | 92.9 | 107.1 | 15.22 | 16.31 | | High | High | High | High | High | High | High | High | High | High | High | 0.000275 | 0.001852 | 2.14 |
| FALSE | High | QIASSSPSL | [R].QIASSS | 2xPhospho | Q0VGY8 2x | 0.006594 | 0.00062 | 1 | 4 | 2 | Q0VGY8 | Protein TA | Q0VGY8 [4 | 0 | 1819.901 | 0 | 0.65 | 0.001857 | 10.33 | 78.1 | 121.9 | 4.78 | 5.69 | | High | High | High | High | High | High | High | High | High | High | High | 0.000275 | 0.001979 | 2.36 |
| FALSE | High | QICLVMLE | [K].QICLVN | 1xCarbami | P60335 1x | 8.11E-05 | 0 | 1 | 1 | 2 | P60335 | Poly(rC)-bi | P60335 [16 | 0 | 2269.128 | 0 | 0.11 | 0.334688 | 11.16 | 97.2 | 102.8 | 3.64 | 6.11 | | High | High | High | High | High | High | High | High | High | High | High | 0 | 1.42E-05 | 3.75 |
| FALSE | High | QICLVMLE | [K].QICLVN | 1xCarbami | P60335 1x | 6.88E-08 | 0 | 1 | 1 | 22 | P60335 | Poly(rC)-bi | P60335 [16 | 0 | 2285.123 | 0 | 0.07 | 0.008376 | 0.38 | 97.9 | 102.1 | 0.68 | 1.19 | | High | High | High | High | High | High | High | High | High | High | High | 0 | 4.99E-08 | 4.74 |
| FALSE | High | QICVVMLE | [K].QICVVN | 1xCarbami | Q3TT81 1x | 0.000418 | 0 | 1 | 1 | 1 | Q3TT81 | Putative ur | Q3TT81 [1 | 0 | 2139.079 | 0 | -0.65 | 0.402414 | 5.48 | 126 | 74 | 56.63 | 54.21 | | High | High | High | High | High | High | High | High | High | High | High | 0 | 8.99E-05 | 3.86 |
| FALSE | High | QICVVMLE | [K].QICVVN | 1xCarbami | Q3TT81 1x | 6.35E-08 | 0 | 1 | 1 | 4 | Q3TT81 | Putative ur | Q3TT81 [1 | 0 | 2368.242 | 0 | -1.17 | 0.001158 | 0.45 | 138.9 | 61.1 | 5.63 | 8.85 | | High | High | High | High | High | High | High | High | High | High | High | 0 | 4.57E-09 | 4.17 |
| FALSE | High | QICVVMLE | [K].QICVVN | 1xCarbami | Q3TT81 1x | 4.79E-07 | 0 | 1 | 1 | 20 | Q3TT81 | Putative ur | Q3TT81 [1 | 0 | 2384.237 | 0 | -0.88 | 0.001613 | 3.38 | 129.8 | 70.2 | 4.39 | 9.4 | | High | High | High | High | High | High | High | High | High | High | High | 0 | 1.08E-07 | 3.76 |
| FALSE | High | QLDDTPPQ | [R].QLDDT | 1xCarbami | Q91VW9 1 | 1.05E-07 | 0 | 2 | 6 | 1 | Q91VW9; | Zinc finger | Q91VW9 [ | 0 | 3023.396 | 0 | -0.62 | 0.001209 | 4.12 | 118.4 | 81.6 | 3.89 | 7.08 | Shared | High | High | High | High | High | High | High | High | High | High | High | 0 | 8.05E-09 | 3.59 |
| FALSE | High | QLEEALVP | [K].QLEEAL | 1xPhospho | V9GX48 1x | 3.52E-07 | 0 | 1 | 3 | 1 | V9GX48 | M-phase p | V9GX48 [8 | 0 | 2332.199 | 0 | 0.73 | 0.001371 | 10.97 | 75.2 | 124.8 | 4.14 | 5.13 | | High | High | High | High | High | High | High | High | High | High | High | 0 | 3.16E-08 | 3.07 |
| FALSE | High | QLEEHGEG | [R].QLEEH | 1xPhospho | Q8K4F6 1x | 0.00025 | 0 | 1 | 1 | 2 | Q8K4F6 | Probable 2 | Q8K4F6 [3 | 0 | 1849.91 | 0 | -0.02 | 0.791477 | 1.81 | 100 | 100 | 8.08 | 6.62 | | High | High | High | High | High | High | High | High | High | High | High | 0 | 5.02E-05 | 3.21 |
| FALSE | High | QLEHVMG | [R].QLEHV | 1xPhospho | Q6P9L6 1x | 0.006653 | 0.00062 | 1 | 1 | 1 | Q6P9L6 | Kinesin-like | Q6P9L6 [1 | 0 | 2456.193 | 0 | 0 | 0.910294 | 10.6 | 100.2 | 99.8 | 14.71 | 8.21 | | High | High | High | High | High | High | High | High | High | High | High | 0.000275 | 0.001989 | 2.85 |
| FALSE | High | QLENTEHN | [R].QLENT | 1xPhospho | Q6P1D7 1x | 0.052133 | 0.00384 | 1 | 2 | 1 | Q6P1D7 | Structure-s | Q6P1D7 [7 | 0 | 2286.073 | 0 | -0.46 | 0.007948 | 2.91 | 117.3 | 82.7 | 9.56 | 4.27 | | High | High | High | High | High | High | High | High | High | High | High | 0.001999 | 0.02027 | 2.29 |
| FALSE | High | QLESGGFE | [R].QLESG | 1xPhospho | Q6PGF7 1x | 0.003001 | 0.00016 | 1 | 1 | 1 | Q6PGF7 | Exocyst co | Q6PGF7 [1 | 0 | 1402.657 | 0 | 0.5 | 0.009645 | 9.01 | 83.4 | 116.6 | 10.17 | 3.47 | | High | High | High | High | High | High | High | High | High | High | High | 7.19E-05 | 0.000818 | 1.93 |
| FALSE | High | QIESSDYDS | [K].QIESSD | 1xPhospho | Q8C0D7 1x | 2.13E-07 | 0 | 1 | 5 | 1 | Q8C0D7 | Inhibitor o | Q8C0D7 [1 | 0 | 1970.9 | 0 | -0.42 | 0.005905 | 5.79 | 112.7 | 87.3 | 6.5 | 2.41 | | High | High | High | High | High | High | High | High | High | High | High | 0 | 1.78E-08 | 2.58 |
| FALSE | High | QLGDTASE | [K].QLGDT | 1xPhospho | E9QLG3 1x | 0.010758 | 0.000772 | 1 | 4 | 2 | E9QLG3 | MAX gene- | E9QLG3 [2 | 0 | 1910.988 | 0 | -0.85 | 0.000523 | 4.15 | 127.4 | 72.6 | 3.59 | 6.56 | | High | High | High | High | High | High | High | High | High | High | High | 0.001916 | 0.01956 | 2.15 |
| FALSE | High | QLGTAQAC | [K].QLGTA | 1xPhospho | Q7TSI1 1xP | 3.54E-05 | 0 | 1 | 1 | 2 | Q7TSI1 | Pleckstrin | Q7TSI1 [38 | 0 | 3746.818 | 0 | 0.4 | 0.369124 | 12.74 | 90 | 110 | 15.31 | 13.13 | | High | High | High | High | High | High | High | High | High | High | High | 0 | 5.61E-06 | 4 |
| FALSE | High | QLGYQPPN | [R].QLGYQ | 1xPhospho | Q91WE3 1 | 6.67E-06 | 0 | 1 | 1 | 1 | Q91WE3 | Ribonuclea | Q91WE3 [ | 0 | 2877.48 | 0 | -0.96 | 0.000423 | 2.4 | 131.3 | 68.7 | 3.82 | 7.86 | | High | High | High | High | High | High | High | High | High | High | High | 0 | 8.55E-07 | 3.11 |
| FALSE | High | QLHFDSPS | [R].QLHFD | 1xPhospho | Q6NWW5 | 0.000267 | 0 | 1 | 3 | 1 | Q6NWW5 | Kinesin-like | Q6NWW5 | 0 | 1754.888 | 0 | -1.08 | 0.000537 | 0.47 | 137.6 | 62.4 | 7.87 | 9.82 | | High | High | High | High | High | High | High | High | High | High | High | 0 | 5.42E-05 | 2.33 |
| FALSE | High | QLHIEGASI | [R].QLHIEG | 1xPhospho | Q3UH59 1x | 5.79E-11 | 0 | 1 | 6 | 5 | Q3UH59 | Myosin-10 | Q3UH59 [1 | 0 | 2625.27 | 0 | -1.78 | 0.000246 | 1.69 | 154.9 | 45.1 | 4.02 | 13 | | High | High | High | High | High | High | High | High | High | High | High | 0 | 2.34E-10 | 5.59 |
| FALSE | High | QLHIEGASI | [R].QLHIEG | 2xPhospho | Q3UH59 2x | 3.65E-07 | 0 | 1 | 6 | 3 | Q3UH59 | Myosin-10 | Q3UH59 [1 | 0 | 2705.236 | 0 | -0.8 | 0.020648 | 15.71 | 124.9 | 75.1 | 6.72 | 31.54 | | High | High | High | High | High | High | High | High | High | High | High | 0 | 6.09E-07 | 4.39 |
| FALSE | High | QLLDHLPG | [R].QLLDH | 1xPhospho | Q80WQ8 1 | 3.93E-07 | 0 | 1 | 3 | 3 | Q80WQ8 | Mis18-binc | Q80WQ8 [ | 0 | 3161.544 | 0 | -0.82 | 0.000289 | 0.26 | 128.8 | 71.2 | 5.15 | 8.99 | | High | High | High | High | High | High | High | High | High | High | High | 0 | 3.57E-08 | 4.43 |
| FALSE | High | QLLDSDEE | [R].QLLDSD | 1xPhospho | Q9R1X4 1x | 7.29E-06 | 0 | 1 | 7 | 1 | Q9R1X4 | Protein tim | Q9R1X4 [1 | 0 | 1958.807 | 0 | -1.53 | 0.001084 | 3.29 | 148.6 | 51.4 | 9.87 | 20.33 | | High | High | High | High | High | High | High | High | High | High | High | 0 | 9.5E-07 | 3.48 |
| FALSE | High | QLLDSDEE | [R].QLLDSD | 1xPhospho | Q9R1X4 1x | 0.006653 | 0.00062 | 1 | 7 | 1 | Q9R1X4 | Protein tim | Q9R1X4 [1 | 1 | 2114.908 | 0 | -1.91 | 0.000364 | 0.89 | 157.2 | 42.8 | 6.05 | 17.58 | | High | High | High | High | High | High | High | High | High | High | High | 0.000275 | 0.001982 | 2.2 |
| FALSE | High | QIIDSGTGD | [R].QIIDSG | 1xPhospho | Q3TTL0 1x | 0.008847 | 0.000736 | 1 | 1 | 1 | Q3TTL0 | Uncharacte | Q3TTL0 [3 | 0 | 2848.449 | 0 | -0.08 | 0.993723 | 16.49 | 102.4 | 97.6 | 13.66 | 8.11 | | High | High | High | High | High | High | High | High | High | High | High | 0.000327 | 0.002746 | 3.03 |
| FALSE | High | QLLPPPSPC | [R].QLLPPP | 1xPhospho | E9Q238 1x | 4.72E-05 | 0 | 1 | 3 | 2 | E9Q238 | KN motif a | E9Q238 [1 | 0 | 1651.913 | 0 | 1.42 | 0.00101 | 7.67 | 50.5 | 149.5 | 11.13 | 11.09 | | High | High | High | High | High | High | High | High | High | High | High | 0 | 7.69E-06 | 2.97 |
| FALSE | High | QILPSPVLQ | [K].QILPSP | 1xPhospho | B2RRF6 1x | 1.6E-05 | 0 | 1 | 1 | 1 | B2RRF6 | Zinc finger | B2RRF6 [4 | 0 | 2092.146 | 0 | -1.31 | 0.000495 | 1.98 | 144.1 | 55.9 | 4.01 | 15.48 | | High | High | High | High | High | High | High | High | High | High | High | 0 | 2.3E-06 | 3.44 |
| FALSE | High | QLLSPK | [RK].QLLSP | 1xPhospho | Q148V8 1x | 0.052576 | 0.00384 | 1 | 2 | 2 | Q148V8 | Protein FA | Q148V8 [1 | 0 | 1223.717 | 0 | 1.3 | 0.000457 | 9.44 | 57.8 | 142.2 | 5.14 | 5.58 | | High | High | High | High | High | High | High | High | High | High | High | 0.001999 | 0.02047 | 1.3 |
| FALSE | High | QLLSPSSFS | [R].QLLSPS | 1xPhospho | P33611 1x | 1.26E-06 | 0 | 1 | 6 | 4 | P33611 | DNA polym | P33611 [14 | 0 | 2300.194 | 0 | -0.11 | 0.347028 | 8.49 | 103.9 | 96.1 | 6.35 | 3.97 | | High | High | High | High | High | High | High | High | High | High | High | 0 | 6.86E-06 | 3.03 |
| FALSE | High | QLLSPSSFS | [R].QLLSPS | 2xPhospho | P33611 2x | 0.001933 | 5.66E-05 | 1 | 6 | 3 | P33611 | DNA polym | P33611 [14 | 0 | 2380.16 | 0 | 0.1 | 0.533305 | 13.24 | 96.6 | 103.4 | 12.94 | 3.29 | | High | High | High | High | High | High | High | High | High | High | High | 2.58E-05 | 0.000501 | 2.39 |
| FALSE | High | QILSQSTDS | [R].QILSQS | 1xPhospho | Q60875 1x | 1.78E-05 | 0 | 1 | 14 | 3 | Q60875 | Rho guanir | Q60875 [1 | 0 | 1817.867 | 0 | -0.42 | 0.041827 | 4.43 | 115.7 | 84.3 | 16.51 | 15.02 | | High | High | High | High | High | High | High | High | High | High | High | 0 | 2.59E-06 | 2.89 |
| FALSE | High | QIASQFPPF | [K].QIASQF | 1xPhospho | F2Z3U3 1x | 5.86E-05 | 0 | 1 | 1 | 1 | F2Z3U3 | Ras associa | F2Z3U3 [8 | 0 | 2779.484 | 0 | 1.46 | 0.005523 | 24.58 | 53 | 147 | 10.62 | 19.69 | | High | High | High | High | High | High | High | High | High | High | High | 0 | 9.9E-06 | 2.77 |
| FALSE | High | NYQQNYQ | [R].NYQQN | 1xPhospho | Q3UBT1 1x | 1.11E-08 | 0 | 1 | 6 | 12 | Q3UBT1 | Putative ur | Q3UBT1 [1 | 0 | 1997.844 | 0 | -0.32 | 0.677884 | 7.58 | 109.4 | 90.6 | 53.9 | 55.75 | | High | High | High | High | High | High | High | High | High | High | High | 0 | 6.48E-10 | 3.9 |
| FALSE | High | NYPNSPPR | [K].NYPNSI | 1xPhospho | Q8CIB9 1x | 0.043236 | 0.003228 | 1 | 2 | 3 | Q8CIB9 | N-acetyltra | Q8CIB9 [1 | 0 | 1253.588 | 0 | -1.66 | 0.000376 | 0.57 | 152.5 | 47.5 | 6.48 | 2.76 | | High | High | High | High | High | High | High | High | High | High | High | 0.001651 | 0.0164 | 1.38 |
| FALSE | High | NYPLPSPA | [K].NYPLPS | 1xPhospho | A2ACM0 1 | 0.001244 | 0 | 1 | 4 | 3 | A2ACM0 | Putative ur | A2ACM0 [6 | 0 | 2236.121 | 0 | 1.74 | 0.0011 | 25.32 | 46.2 | 153.8 | 21.19 | 8.11 | | High | High | High | High | High | High | High | High | High | High | High | 0.000344 | 0.003079 | 2.81 |
| FALSE | High | NPQMPQY | [R].NPQMP | 1xDeamida | E9QAQ7 1x | 0.045898 | 0.003408 | 1 | 5 | 2 | E9QAQ7 | AT-rich int | E9QAQ7 [7 | 0 | 2382.1 | 0 | 0.95 | 0.058826 | 21.95 | 75.1 | 124.9 | 13.73 | 24.57 | | High | High | High | High | High | High | High | High | High | High | High | 0.001741 | 0.01744 | 2.88 |
| FALSE | High | NPQMPQY | [R].NPQMP | 2xPhospho | E9QAQ7 2x | 0.000136 | 0 | 1 | 5 | 3 | E9QAQ7 | AT-rich int | E9QAQ7 [7 | 0 | 2461.082 | 0 | 0.08 | 0.652365 | 15.38 | 96.2 | 103.8 | 14.93 | 10.71 | | High | High | High | High | High | High | High | High | High | High | High | 0.004999 | 0.05465 | 3.75 |
| FALSE | High | NPQNSSQS | [R].NPQNS | 1xPhospho | Q14C24 1x | 0.002623 | 0.000109 | 1 | 1 | 5 | Q14C24 | MCG14259 | Q14C24 [5 | 0 | 1682.77 | 0 | -0.75 | 0.000609 | 0.85 | 127.7 | 72.3 | 6.55 | 8.15 | | High | High | High | High | High | High | High | High | High | High | High | 0.000689 | 0.006174 | 2.09 |
| FALSE | High | NPSLLGQD | [R].NPSLLG | 1xCarbami | Q924W7 2 | 0.014667 | 0.001049 | 1 | 5 | 2 | Q924W7 | Suppressio | Q924W7 [ | 0 | 2869.386 | 0 | 0.14 | 0.57227 | 19.34 | 92.8 | 107.2 | 17.22 | 13.34 | | High | High | High | High | High | High | High | High | High | High | High | 0.000555 | 0.004814 | 3.04 |
| FALSE | High | NPSISQTLL | [K].NPSISQ | 1xPhospho | Q5SSW2 1 | 7.41E-05 | 0 | 1 | 1 | 2 | Q5SSW2 | Proteasom | Q5SSW2 [1 | 0 | 1952.051 | 0 | -1.29 | 0.000512 | 1.29 | 140.5 | 59.5 | 3.83 | 8.67 | | High | High | High | High | High | High | High | High | High | High | High | 0 | 3.2E-05 | 2.94 |
| FALSE | High | NPSPGMSA | [R].NPSPGI | 1xPhospho | B7ZNH9 1x | 0.001039 | 0 | 1 | 3 | 1 | B7ZNH9 | Phosphoin | B7ZNH9 [1 | 0 | 1442.711 | 0 | 0.74 | 0.001425 | 0.99 | 73.9 | 126.1 | 10.46 | 8.22 | | High | High | High | High | High | High | High | High | High | High | High | 0 | 0.000248 | 1.92 |
| FALSE | High | NPSPPPPTC | [R].NPSPPI | 1xCarbami | Q9D2C2 1x | 1.04E-08 | 0 | 1 | 2 | 1 | Q9D2C2 | Protein SA | Q9D2C2 [4 | 0 | 2592.076 | 0 | -0.25 | 0.009204 | 6.6 | 108.7 | 91.3 | 8.03 | 5.85 | | High | High | High | High | High | High | High | High | High | High | High | 0 | 5.98E-10 | 4.6 |
| FALSE | High | NPSPPPPTC | [R].NPSPPI | 1xCarbami | Q9D2C2 2x | 3.55E-07 | 0 | 1 | 2 | 2 | Q9D2C2 | Protein SA | Q9D2C2 [4 | 0 | 2672.042 | 0 | -0.15 | 0.093278 | 5.21 | 101 | 99 | 9.53 | 19.92 | | High | High | High | High | High | High | High | High | High | High | High | 0 | 3.19E-08 | 5.42 |
| FALSE | High | NPSPPTCAI | [R].NPSPPT | 2xCarbami | P48754 1x | 5.03E-09 | 0 | 1 | 5 | 4 | P48754 | Breast can | P48754 [62 | 0 | 2945.331 | 0 | -0.79 | 0.00286 | 6.09 | 125.9 | 74.1 | 8.24 | 10.48 | | High | High | High | High | High | High | High | High | High | High | High | 0 | 3.14E-10 | 6.48 |
| FALSE | High | NPSPSSGA | [R].NPSPSS | 1xPhospho | B9EKK8 1x | 2.56E-06 | 0 | 1 | 4 | 1 | B9EKK8 | Uncharacte | B9EKK8 [9 | 0 | 2018.975 | 0 | -0.33 | 0.135247 | 13.77 | 113.2 | 86.8 | 14.55 | 12.83 | | High | High | High | High | High | High | High | High | High | High | High | 0 | 2.93E-07 | 3.36 |
| FALSE | High | NPSSTLGSK | [K].NPSSTL | 1xPhospho | Q8K0V2 1x | 0.042868 | 0.003228 | 1 | 3 | 1 | Q8K0V2 | DCN1-like | Q8K0V2 [1 | 0 | 1428.75 | 0 | 0.58 | 0.015133 | 18.02 | 76.8 | 123.2 | 13.44 | 13.24 | | High | High | High | High | High | High | High | High | High | High | High | 0.001651 | 0.01611 | 1.63 |
| FALSE | High | NPVLSQTG | [K].NPVLSC | 1xPhospho | E9Q842 1x | 0.028824 | 0.001829 | 1 | 3 | 1 | E9Q842 | Neuron na | E9Q842 [1 | 0 | 1553.767 | 0 | 1.64 | 0.00163 | 19.26 | 48.4 | 151.6 | 16.01 | 10.55 | | High | High | High | High | High | High | High | High | High | High | High | 0.000984 | 0.01025 | 1.6 |
| FALSE | High | NPYAHISIP | [K].NPYAH | 1xPhospho | Q9D1E4 1x | 0.049549 | 0.00361 | 1 | 2 | 1 | Q9D1E4 | Cysteine-ri | Q9D1E4 [1 | 0 | 1476.756 | 0 | 1.44 | 0.001484 | 23.82 | 55.1 | 144.9 | 17.4 | 7.67 | | High | High | High | High | High | High | High | High | High | High | High | 0.001889 | 0.01908 | 2.03 |
| FALSE | High | NPQMPQY | [R].NPQMP | 1xPhospho | E9QAQ7 1x | 5.01E-07 | 0 | 1 | 5 | 27 | E9QAQ7 | AT-rich int | E9QAQ7 [7 | 0 | 2381.116 | 0 | 0.29 | 0.014137 | 9.31 | 89.7 | 110.3 | 7.02 | 7.29 | | High | High | High | High | High | High | High | High | High | High | High | 0 | 1.44E-05 | 5.02 |
| FALSE | High | NPYLLSEEE | [R].NPYLLS | 1xPhospho | Q8CCP0 1x | 0.000407 | 0 | 1 | 3 | 3 | Q8CCP0 | Nuclear ex | Q8CCP0 [4 | 0 | 3074.32 | 0 | -1.21 | 0.00062 | 2.1 | 141.1 | 58.9 | 9.07 | 8.25 | | High | High | High | High | High | High | High | High | High | High | High | 0 | 8.7E-05 | 3.11 |
| FALSE | High | NQAAVEAC | [R].NQAAV | 1xPhospho | E9PVX6 1x | 6.72E-09 | 0 | 1 | 2 | 1 | E9PVX6 | Proliferatic | E9PVX6 [3 | 0 | 2806.33 | 0 | -0.13 | 0.266028 | 8.98 | 106.7 | 93.3 | 9.01 | 15.05 | | High | High | High | High | High | High | High | High | High | High | High | 0 | 3.66E-10 | 6.16 |
| FALSE | High | NQAAVEAC | [R].NQAAV | 2xPhospho | E9PVX6 2x | 2.75E-05 | 0 | 1 | 2 | 1 | E9PVX6 | Proliferatic | E9PVX6 [3 | 0 | 2886.296 | 0 | 0.55 | 0.01229 | 12.87 | 83 | 117 | 12.74 | 6.19 | | High | High | High | High | High | High | High | High | High | High | High | 0 | 4.2E-06 | 3.01 |
| FALSE | High | NQAIQISPT | [R].NQAIQ | 1xPhospho | Q8BV49 1x | 7.62E-09 | 0 | 1 | 7 | 1 | Q8BV49 | Pyrin and H | Q8BV49 [1 | 0 | 2342.155 | 0 | 6.64 | 0.00036 | 0 | 1.9 | 198.1 | 82.55 | 10.97 | | High | High | High | High | High | High | High | High | High | High | High | 0 | 4.22E-10 | 4.09 |
| FALSE | High | NQAIYAAV | [R].NQAIY | 1xPhospho | Q923B1 1x | 1.13E-05 | 0 | 1 | 2 | 2 | Q923B1 | Lariat debr | Q923B1 [5 | 0 | 1918.827 | 0 | -0.28 | 0.088524 | 4.17 | 107.6 | 92.4 | 5.42 | 9.55 | | High | High | High | High | High | High | High | High | High | High | High | 0 | 1.55E-06 | 4.02 |
| FALSE | High | NQAQTSSE | [R].NQAQT | 1xPhospho | G3X963 1x | 7.79E-07 | 0 | 1 | 2 | 2 | G3X963 | ATPase fan | G3X963 [7 | 0 | 1966.916 | 0 | 0.2 | 0.025522 | 5.02 | 93 | 107 | 4.22 | 6.92 | | High | High | High | High | High | High | High | High | High | High | High | 0 | 7.66E-08 | 3.13 |
| FALSE | High | NQATSASS | [K].NQATS | 1xPhospho | Q6A065 1x | 5.81E-05 | 0 | 1 | 2 | 5 | Q6A065 | Centrosom | Q6A065 [4 | 0 | 1560.767 | 0 | 1.11 | 0.00062 | 10.84 | 62.2 | 137.8 | 9.1 | 4.49 | | High | High | High | High | High | High | High | High | High | High | High | 4.94E-05 | 0.000722 | 1.81 |
| FALSE | High | NQDEESQE | [K].NQDEE | 1xPhospho | Q6P542 1x | 0.003822 | 0.000261 | 3 | 3 | 1 | Q6P542; Q | ATP-bindin | Q6P542 [5 | 0 | 2194.068 | 0 | -0.89 | 0.010828 | 9.41 | 131.2 | 68.8 | 10.08 | 21.6 | Shared | High | High | High | High | High | High | High | High | High | High | High | 0.000117 | 0.001071 | 2.3 |
| FALSE | High | NQDHSPSP | [R].NQDHS | 2xPhospho | O08905 2x | 0.038677 | 0.002563 | 1 | 6 | 1 | O08905 | RNP particl | O08905 [2 | 0 | 4000.825 | 0 | -0.71 | 0.012167 | 9.78 | 123.6 | 76.4 | 7.48 | 22.05 | | High | High | High | High | High | High | High | High | High | High | High | 0.001388 | 0.01439 | 3.86 |
| FALSE | High | NQDNYVSV | [R].NQDNY | 2xPhospho | F8VQ54 2x | 2.91E-06 | 0 | 1 | 3 | 5 | F8VQ54 | Endoribon | F8VQ54 [3 | 0 | 3394.309 | 0 | -0.66 | 0.003227 | 4.41 | 121.8 | 78.2 | 7.59 | 6.21 | | High | High | High | High | High | High | High | High | High | High | High | 0 | 5.96E-06 | 4.49 |
| FALSE | High | NQDSSAPP | [K].NQDSS | 1xPhospho | Q571C7 1x | 0.000382 | 0 | 1 | 2 | 1 | Q571C7 | Transcripti | Q571C7 [2 | 0 | 4301.123 | 0 | 0.49 | 0.035144 | 8.62 | 84.5 | 115.5 | 17.61 | 16.73 | | High | High | High | High | High | High | High | High | High | High | High | 0 | 8.13E-05 | 3.84 |
| FALSE | High | NQEIQQGE | [R].NQEIQ | 1xPhospho | Q9ESC8 1x | 0.020499 | 0.001472 | 1 | 3 | 2 | Q9ESC8 | AF4/FMR2 | Q9ESC8 [1 | 0 | 3130.517 | 0 | 0.05 | 0.627359 | 6.1 | 101.5 | 98.5 | 4.2 | 11.95 | | High | High | High | High | High | High | High | High | High | High | High | 0.000786 | 0.007013 | 1.99 |
| FALSE | High | NQGGYGG | [R].NQGGY | 1xPhospho | Q3UK83 1x | 1.72E-08 | 0 | 3 | 6 | 14 | Q3UK83; P | Putative ur | Q3UK83 [3 | 0 | 2003.829 | 0 | 0.43 | 0.567008 | 55.56 | 87.2 | 112.8 | 20.8 | 47.97 | Shared | High | High | High | High | High | High | High | High | High | High | High | 0 | 1.95E-09 | 2.72 |
| FALSE | High | NQHFNESF | [K].NQHFN | 1xPhospho | Q8CJ27 1x | 0.013545 | 0.000976 | 1 | 2 | 1 | Q8CJ27 | Abnormal | Q8CJ27 [1 | 0 | 2118.042 | 0 | -0.07 | 0.235973 | 7.3 | 105.2 | 94.8 | 12.89 | 18.16 | | High | High | High | High | High | High | High | High | High | High | High | 0.000538 | 0.004414 | 1.68 |
| FALSE | High | NPYLLSEEE | [R].NPYLLS | 1xPhospho | Q8CCP0 1x | 2.25E-08 | 0 | 1 | 3 | 7 | Q8CCP0 | Nuclear ex | Q8CCP0 [4 | 0 | 3303.483 | 0 | 0.09 | 0.408696 | 16.17 | 97 | 103 | 10.43 | 8.89 | | High | High | High | High | High | High | High | High | High | High | High | 0 | 1.43E-09 | 5.29 |
| FALSE | High | NQHSLYTA | [R].NQHSL | 1xPhospho | Q571K4 1x | 0.000101 | 0 | 1 | 1 | 4 | Q571K4 | TGF-beta-a | Q571K4 [3 | 0 | 2240.055 | 0 | 0.2 | 0.017537 | 7.57 | 90.7 | 109.3 | 9.58 | 7.11 | | High | High | High | High | High | High | High | High | High | High | High | 0 | 1.8E-05 | 3.24 |
| FALSE | High | NPQMPQY | [R].NPQMP | 1xPhospho | E9QAQ7 1x | 3.27E-06 | 0 | 1 | 5 | 3 | E9QAQ7 | AT-rich int | E9QAQ7 [7 | 0 | 2365.121 | 0 | -0.25 | 0.014045 | 4.01 | 108.5 | 91.5 | 2.07 | 9.57 | | High | High | High | High | High | High | High | High | High | High | High | 0 | 3.85E-07 | 4.49 |
| FALSE | High | NPPSGDLL | [K].NPPSGI | 1xPhospho | Q6PAL0 1x | 0.002793 | 0.000109 | 1 | 2 | 1 | Q6PAL0 | BEN domai | Q6PAL0 [1 | 0 | 2344.188 | 0 | -0.07 | 0.076186 | 1.53 | 101.6 | 98.4 | 8.75 | 11.37 | | High | High | High | High | High | High | High | High | High | High | High | 4.94E-05 | 0.000755 | 2.84 |
| FALSE | High | NNSPAPPQ | [K].NNSPAI | 1xDeamida | Q14AX6-2 | 4.08E-05 | 0 | 3 | 3 | 2 | Q14AX6-2; | Isoform 2 c | Q14AX6-2 | 0 | 1855.972 | 0 | 0.04 | 0.870187 | 8.05 | 99.4 | 100.6 | 7.13 | 12.09 | Shared | High | High | High | High | High | High | High | High | High | High | High | 0.000327 | 0.002723 | 2.59 |
| FALSE | High | NNSSSNVE | [K].NNSSSN | 1xPhospho | A0A1B0GS | 0.000155 | 0 | 1 | 9 | 1 | A0A1B0GS | Nuclear po | A0A1B0GS | 0 | 2173.016 | 0 | 0.03 | 0.695725 | 17.74 | 100.3 | 99.7 | 12.95 | 8.5 | | High | High | High | High | High | High | High | High | High | High | High | 0 | 2.95E-05 | 3.95 |
| FALSE | High | NNSSSNVE | [K].NNSSSN | 1xPhospho | A0A1B0GS | 0.000515 | 0 | 1 | 9 | 3 | A0A1B0GS | Nuclear po | A0A1B0GS | 0 | 2189.011 | 0 | -0.14 | 0.127379 | 9.98 | 104.7 | 95.3 | 4.42 | 6.98 | | High | High | High | High | High | High | High | High | High | High | High | 2.58E-05 | 0.000371 | 2.37 |
| FALSE | High | NNSSSSNIE | [K].NNSSSS | 1xPhospho | G3X963 1x | 0.000133 | 0 | 1 | 4 | 6 | G3X963 | ATPase fan | G3X963 [1 | 0 | 2329.096 | 0 | 0.43 | 0.049303 | 4.92 | 85.2 | 114.8 | 5.39 | 12.19 | | High | High | High | High | High | High | High | High | High | High | High | 0 | 0.000135 | 3.75 |
| FALSE | High | NNSSSSNIE | [K].NNSSSS | 2xPhospho | G3X963 2x | 0.00043 | 0 | 1 | 4 | 4 | G3X963 | ATPase fan | G3X963 [1 | 0 | 2409.062 | 0 | -0.35 | 0.00602 | 1.52 | 111.9 | 88.1 | 4.06 | 5.15 | | High | High | High | High | High | High | High | High | High | High | High | 0.003279 | 0.03549 | 2.4 |
| FALSE | High | NNSYTCYT. | [R].NNSYT | 2xCarbami | Q80UP8 1x | 2.11E-05 | 0 | 1 | 1 | 3 | Q80UP8 | Sodium-de | Q80UP8 [3 | 0 | 2773.214 | 0 | 0.08 | 0.674458 | 6.65 | 103 | 97 | 11.39 | 8.67 | | High | High | High | High | High | High | High | High | High | High | High | 0 | 3.12E-06 | 4.25 |
| FALSE | High | NNSYTCYT. | [R].NNSYT | 2xCarbami | Q80UP8 1x | 0.001137 | 0 | 1 | 1 | 1 | Q80UP8 | Sodium-de | Q80UP8 [3 | 0 | 2789.208 | 0 | 1.04 | 0.001912 | 13.81 | 67.1 | 132.9 | 15.27 | 8.81 | | High | High | High | High | High | High | High | High | High | High | High | 0 | 0.000275 | 3.65 |
| FALSE | High | NNTITTTSS | [R].NNTITT | 1xPhospho | Q99NH0 1x | 0.005228 | 0.000551 | 1 | 8 | 2 | Q99NH0 | Ankyrin re | Q99NH0 [1 | 0 | 1375.667 | 0 | -0.22 | 0.066088 | 12.94 | 106.2 | 93.8 | 5.32 | 13.81 | | High | High | High | High | High | High | High | High | High | High | High | 0.000418 | 0.003597 | 2.36 |
| FALSE | High | NNVMTSPI | [R].NNVMT | 1xPhospho | P49452 1x | 0.000292 | 0 | 1 | 1 | 2 | P49452 | Centromer | P49452 [6 | 0 | 1907.981 | 0 | -1.06 | 0.005079 | 4.79 | 131.3 | 68.7 | 10.86 | 23.94 | | High | High | High | High | High | High | High | High | High | High | High | 0 | 5.96E-05 | 1.5 |
| FALSE | High | NNVSPVSR | [K].NNVSP | 1xPhospho | A0A0A6YW | 0.107603 | 0.008634 | 1 | 8 | 1 | A0A0A6YW | Mitogen-a | A0A0A6YW | 0 | 1181.588 | 0 | -1.41 | 0.000933 | 2 | 145.4 | 54.6 | 5.79 | 11.61 | | High | High | High | High | High | High | High | High | High | High | High | 0.004341 | 0.0469 | 1.06 |
| FALSE | High | NNVTNYNT | [R].NNVTN | 2xPhospho | A4QPD1 2x | 5.03E-05 | 0 | 1 | 5 | 1 | A4QPD1 | Zinc finger | A4QPD1 [1 | 0 | 2732.081 | 0 | 1.17 | 0.003725 | 33.18 | 60.9 | 139.1 | 9.81 | 13.53 | | High | High | High | High | High | High | High | High | High | High | High | 0 | 8.25E-06 | 4.2 |
| FALSE | High | NNVTNYNT | [R].NNVTN | 2xPhospho | A4QPD1 2x | 3.7E-08 | 0 | 1 | 5 | 3 | A4QPD1 | Zinc finger | A4QPD1 [1 | 0 | 2961.244 | 0 | 0.6 | 0.006898 | 10.88 | 78.4 | 121.6 | 10.52 | 8.06 | | High | High | High | High | High | High | High | High | High | High | High | 0 | 2.49E-09 | 4.62 |
| FALSE | High | NNWSFDH | [K].NNWSF | 1xPhospho | B7ZWN9 1 | 0.003464 | 0.00021 | 2 | 4 | 1 | B7ZWN9; F | SUMO/sen | B7ZWN9 [3 | 0 | 2547.071 | 0 | 0.51 | 0.030986 | 18.3 | 83.8 | 116.2 | 10.75 | 10.33 | Shared | High | High | High | High | High | High | High | High | High | High | High | 9.42E-05 | 0.000957 | 2.82 |
| FALSE | High | NPQEQLPP | [R].NPQEQ | 2xCarbami | Q9CWP9 1 | 0.005516 | 0.00062 | 1 | 1 | 1 | Q9CWP9 | Proline-ric | Q9CWP9 [ | 0 | 2085.945 | 0 | -1.12 | 0.001068 | 3.13 | 140.8 | 59.2 | 6.29 | 11.77 | | High | High | High | High | High | High | High | High | High | High | High | 0.000275 | 0.001614 | 2.95 |
| FALSE | High | NNYSPGRF | [K].NNYSP | 1xCarbami | A5D6P2 1x | 9.48E-06 | 0 | 1 | 23 | 1 | A5D6P2 | Par-3 (Part | A5D6P2 [3 | 1 | 2387.055 | 0 | -0.5 | 0.017882 | 9.27 | 117.3 | 82.7 | 7.83 | 12.47 | | High | High | High | High | High | High | High | High | High | High | High | 0 | 1.27E-06 | 1.14 |
| FALSE | High | NNYSPGRF | [K].NNYSP | 1xCarbami | A5D6P2 2x | 0.008388 | 0.000736 | 1 | 23 | 2 | A5D6P2 | Par-3 (Part | A5D6P2 [3 | 1 | 2467.021 | 0 | -0.65 | 0.009796 | 4.42 | 121.4 | 78.6 | 10.12 | 10.74 | | High | High | High | High | High | High | High | High | High | High | High | 0.005336 | 0.05805 | 2.08 |

| | | | | | | | | | | | | | | | | | | | | | | | | | | | | | | | | | | | | | | |
|---|---|---|---|---|---|---|---|---|---|---|---|---|---|---|---|---|---|---|---|---|---|---|---|---|---|---|---|---|---|---|---|---|---|---|---|---|---|---|
| FALSE | High | NPAPPVGS | [R].NPAPPV | 1xPhospho | Q8BGT6 1x | 3.01E-06 | 0 | 1 | 2 | 1 | Q8BGT6 | MICAL-like | Q8BGT6 [6 | 0 | 2142.172 | 0 | 2.18 | 0.000608 | 24.52 | 36.2 | 163.8 | 18.2 | 7.72 | | High | High | High | High | High | High | High | High | High | High | High | 0 | 3.51E-07 | 2.74 |
| FALSE | High | NPATTFLTV | [K].NPATTF | 1xPhospho | Q5DTI8 1xl | 0.006773 | 0.00062 | 1 | 2 | 1 | Q5DTI8 | Extended s | Q5DTI8 [68 | 0 | 2485.362 | 0 | 0.98 | 0.007276 | 14.57 | 63.2 | 136.8 | 4.81 | 18.91 | | High | High | High | High | High | High | High | High | High | High | High | 0.000275 | 0.002027 | 2.2 |
| FALSE | High | NPEPDSDE | [K].NPEPDS | 1xPhospho | Q9D0F4 1x | 0.00025 | 0 | 1 | 1 | 2 | Q9D0F4 | NF-kappa-I | Q9D0F4 [1 | 1 | 2959.418 | 0 | -0.65 | 0.001125 | 3.29 | 120.5 | 79.5 | 2.65 | 7.17 | | High | High | High | High | High | High | High | High | High | High | High | 0.00014 | 0.001092 | 1.97 |
| FALSE | High | NPGASPAE | [R].NPGAS | 1xCarbami | Q7M6Z9 1: | 0.062731 | 0.004664 | 1 | 4 | 2 | Q7M6Z9 | Kinesin ligh | Q7M6Z9 [5 | 0 | 2870.224 | 0 | 1.3 | 0.001911 | 14.83 | 52.1 | 147.9 | 10.98 | 13.22 | | High | High | High | High | High | High | High | High | High | High | High | 0.005149 | 0.05617 | 2.79 |
| FALSE | High | NPGTAVSA | [R].NPGTA | 1xPhospho | O54826 1x | 8.25E-05 | 0 | 1 | 4 | 1 | O54826 | Protein AF- | O54826 [3 | 0 | 2902.439 | 0 | 0.56 | 0.006898 | 13.56 | 81 | 119 | 7.58 | 11.85 | | High | High | High | High | High | High | High | High | High | High | High | 0 | 1.45E-05 | 3.45 |
| FALSE | High | NPICDSPSD | [R].NPICDS | 2xCarbami | E9Q286 1x | 4.3E-07 | 0 | 1 | 1 | 5 | E9Q286 | Little elong | E9Q286 [9 | 0 | 2794.19 | 0 | -1.01 | 0.000954 | 2.24 | 134.4 | 65.6 | 7.08 | 3.42 | | High | High | High | High | High | High | High | High | High | High | High | 0 | 5.95E-08 | 4.72 |
| FALSE | High | NPIFQDSW | [R].NPIFQD | 1xPhospho | Q3TYF6 1x | 3.24E-09 | 0 | 1 | 4 | 1 | Q3TYF6 | DNA segme | Q3TYF6 [9 | 0 | 3529.471 | 0 | 0.09 | 0.794264 | 22 | 97.1 | 102.9 | 21.58 | 15.4 | | High | High | High | High | High | High | High | High | High | High | High | 0 | 1.62E-10 | 5.75 |
| FALSE | High | NPLGPPGL | [K].NPLGPI | 1xPhospho | Q3V1Z0 1x | 0.006031 | 0.00062 | 1 | 5 | 2 | Q3V1Z0 | Iporin OS= | Q3V1Z0 [9 | 0 | 1688.857 | 0 | 1.85 | 0.012602 | 21.79 | 41.7 | 158.3 | 14.69 | 30.76 | | High | High | High | High | High | High | High | High | High | High | High | 0.000449 | 0.004078 | 1.84 |
| FALSE | High | NPNNSSLSI | [K].NPNNS | 1xPhospho | Q91YN0 1x | 3.35E-05 | 0 | 1 | 1 | 1 | Q91YN0 | Protein C1 | Q91YN0 [2 | 0 | 1869.936 | 0 | 1.17 | 0.001374 | 13.13 | 59.7 | 140.3 | 10.38 | 12.83 | | High | High | High | High | High | High | High | High | High | High | High | 0 | 5.28E-06 | 1.84 |
| FALSE | High | NPPEAATA | [K].NPPEA | 1xPhospho | P13595 1xl | 0.00095 | 0 | 1 | 2 | 2 | P13595 | Neural cell | P13595 [9 | 0 | 1788.93 | 0 | -0.33 | 0.259411 | 19.43 | 112.4 | 87.6 | 12.57 | 14.97 | | High | High | High | High | High | High | High | High | High | High | High | 0 | 0.000226 | 2.53 |
| FALSE | High | NPPHCLEIT | [K].NPPHCI | 1xCarbami | A3KGV1-2 | 0.001411 | 5.66E-05 | 1 | 5 | 2 | A3KGV1-2 | Isoform 2 c | A3KGV1-2 | 0 | 2244.114 | 0 | -0.33 | 0.022323 | 1.5 | 111.4 | 88.6 | 7.11 | 7.11 | | High | High | High | High | High | High | High | High | High | High | High | 0.000161 | 0.001148 | 3.23 |
| FALSE | High | NPPLSPQS | [K].NPPLSP | 1xPhospho | Q8CI08-2 1 | 1.75E-08 | 0 | 1 | 3 | 4 | Q8CI08-2 | Isoform 2 c | Q8CI08-2 [ | 0 | 3617.715 | 0 | -1.19 | 0.000312 | 1.33 | 139.4 | 60.6 | 5.28 | 15.97 | | High | High | High | High | High | High | High | High | High | High | High | 0 | 1.08E-09 | 5.22 |
| FALSE | High | NNYSPGRF | [K].NNYSP | 1xCarbami | A5D6P2 1x | 0.00204 | 5.66E-05 | 1 | 23 | 1 | A5D6P2 | Par-3 (Part | A5D6P2 [3 | 1 | 2388.039 | 0 | -0.44 | 0.002458 | 3.24 | 116 | 84 | 9.58 | 6.31 | | High | High | High | High | High | High | High | High | High | High | High | 2.58E-05 | 0.000533 | 2.09 |
| FALSE | High | NQLSDPDII | [K].NQLSD | 1xDeamida | B2RS91 1xl | 0.027839 | 0.001755 | 1 | 1 | 1 | B2RS91 | RNA polym | B2RS91 [74 | 0 | 1568.797 | 0 | -0.07 | 0.823892 | 2.25 | 102.5 | 97.5 | 7.21 | 9.65 | | High | High | High | High | High | High | High | High | High | High | High | 0.000953 | 0.009878 | 3.49 |
| FALSE | High | NQLVLIVNI | [K].NQLVLI | 1xPhospho | Q6P5E3 1x | 9.73E-09 | 0 | 1 | 3 | 1 | Q6P5E3 | Sal-like 1 (I | Q6P5E3 [6 | 0 | 1989.062 | 0 | -1.35 | 0.109803 | 15.9 | 143.8 | 56.2 | 35.88 | 41.77 | | High | High | High | High | High | High | High | High | High | High | High | 0 | 5.59E-10 | 4.82 |
| FALSE | High | NQLVLIVNI | [K].NQLVLI | 1xPhospho | Q6P5E3 1x | 1.47E-07 | 0 | 1 | 3 | 1 | Q6P5E3 | Sal-like 1 (I | Q6P5E3 [6 | 0 | 2218.225 | 0 | -1.27 | 0.000743 | 2.43 | 141.2 | 58.8 | 5.2 | 13.81 | | High | High | High | High | High | High | High | High | High | High | High | 0 | 1.17E-08 | 4.19 |
| FALSE | High | NRHSPDHP | [K].NRHSPI | 1xPhospho | Q8C1S0 1x | 3.09E-05 | 0 | 1 | 1 | 2 | Q8C1S0 | Mediator c | Q8C1S0 [2 | 1 | 2591.162 | 0 | -0.9 | 0.014244 | 8.92 | 130.1 | 69.9 | 21.39 | 20.35 | | High | High | High | High | High | High | High | High | High | High | High | 2.58E-05 | 0.000504 | 2.58 |
| FALSE | High | NRHSPDHP | [K].NRHSPI | 1xPhospho | Q8C1S0 1x | 0.000158 | 0 | 1 | 1 | 6 | Q8C1S0 | Mediator c | Q8C1S0 [2 | 1 | 2607.157 | 0 | -0.8 | 0.089153 | 3.84 | 128.5 | 71.5 | 12.25 | 34.37 | | High | High | High | High | High | High | High | High | High | High | High | 0 | 2.99E-05 | 3 |
| FALSE | High | NRNASTSF | [R].NRNAS | 1xDeamida | P54103 1xl | 0.007274 | 0.00062 | 1 | 6 | 2 | P54103 | DnaJ homc | P54103 [43 | 1 | 2178.048 | 0 | 0.06 | 0.450292 | 18.86 | 98 | 102 | 10.41 | 3.19 | | High | High | High | High | High | High | High | High | High | High | High | 0.000275 | 0.002196 | 2.59 |
| FALSE | High | NRPETSLN | [R].NRPETS | 1xPhospho | Q542P4 1x | 3.85E-10 | 0 | 1 | 6 | 1 | Q542P4 | Membrane | Q542P4 [3 | 0 | 3909.848 | 0 | 0.36 | 0.046652 | 5.51 | 82.8 | 117.2 | 15.61 | 16.98 | | High | High | High | High | High | High | High | High | High | High | High | 0 | 1.48E-11 | 4.51 |
| FALSE | High | NRPETSLN | [R].NRPETS | 1xDeamida | Q542P4 1x | 1.41E-05 | 0 | 1 | 6 | 3 | Q542P4 | Membrane | Q542P4 [3 | 0 | 3910.832 | 0 | 0.66 | 0.017781 | 37.83 | 75.7 | 124.3 | 19.71 | 8.26 | | High | High | High | High | High | High | High | High | High | High | High | 0 | 6.21E-06 | 3.48 |
| FALSE | High | NRPGYVSE | [K].NRPGY | 1xPhospho | Q9ERU9 1x | 0.053926 | 0.00384 | 1 | 1 | 1 | Q9ERU9 | E3 SUMO- | Q9ERU9 [2 | 0 | 2813.17 | 0 | -0.91 | 0.000544 | 2.46 | 130.4 | 69.6 | 9.82 | 16.23 | | High | High | High | High | High | High | High | High | High | High | High | 0.002015 | 0.02093 | 2.73 |
| FALSE | High | NRPGYVSE | [K].NRPGY | 1xPhospho | Q9ERU9 1x | 1.37E-07 | 0 | 1 | 1 | 7 | Q9ERU9 | E3 SUMO- | Q9ERU9 [2 | 0 | 2829.165 | 0 | 0.02 | 0.972674 | 10.75 | 98.6 | 101.4 | 16.32 | 19.56 | | High | High | High | High | High | High | High | High | High | High | High | 0 | 1.08E-08 | 4.42 |
| FALSE | High | NRPGYVSE | [K].NRPGY | 1xPhospho | Q9ERU9 1x | 1.19E-09 | 0 | 1 | 1 | 12 | Q9ERU9 | E3 SUMO- | Q9ERU9 [2 | 0 | 3042.333 | 0 | 0.02 | 0.541837 | 3.45 | 100.1 | 99.9 | 6.04 | 18.04 | | High | High | High | High | High | High | High | High | High | High | High | 0 | 5.25E-11 | 3.85 |
| FALSE | High | NRPGYVSE | [K].NRPGY | 1xPhospho | Q9ERU9 1x | 3.04E-09 | 0 | 1 | 1 | 24 | Q9ERU9 | E3 SUMO- | Q9ERU9 [2 | 0 | 3058.328 | 0 | -0.17 | 0.031113 | 4.05 | 107.4 | 92.6 | 6.64 | 2.21 | | High | High | High | High | High | High | High | High | High | High | High | 0 | 4.13E-10 | 3.82 |
| FALSE | High | NRPLSDEEI | [K].NRPLSD | 1xPhospho | G5E866 1x | 1.13E-07 | 0 | 1 | 5 | 2 | G5E866 | Splicing fac | G5E866 [3 | 0 | 2649.267 | 0 | -0.23 | 0.11256 | 1.69 | 111.4 | 88.6 | 16.53 | 13.63 | | High | High | High | High | High | High | High | High | High | High | High | 0 | 8.79E-09 | 3.75 |
| FALSE | High | NRPLSDEEI | [K].NRPLSD | 1xPhospho | G5E866 1x | 2E-07 | 0 | 1 | 5 | 1 | G5E866 | Splicing fac | G5E866 [3 | 0 | 2665.262 | 0 | -0.45 | 0.0747 | 16.07 | 115.5 | 84.5 | 11.9 | 14.49 | | High | High | High | High | High | High | High | High | High | High | High | 0 | 1.65E-08 | 3.12 |
| FALSE | High | NRSAEEGE | [R].NRSAEI | 1xPhospho | B9EKJ7 1xF | 3.88E-06 | 0 | 1 | 3 | 2 | B9EKJ7 | Tankyrase | B9EKJ7 [16 | 1 | 1973.958 | 0 | -0.45 | 0.016163 | 6.43 | 114.9 | 85.1 | 4.9 | 13.36 | | High | High | High | High | High | High | High | High | High | High | High | 0 | 4.68E-07 | 3.77 |
| FALSE | High | NRSDTEQE | [R].NRSDTI | 1xPhospho | A0A0R4J0J | 0.008388 | 0.000736 | 1 | 2 | 1 | A0A0R4J0J | THO comp | A0A0R4J0J | 1 | 1701.821 | 0 | 0 | 0.845725 | 5.71 | 101.4 | 98.6 | 4.24 | 6.08 | | High | High | High | High | High | High | High | High | High | High | High | 0.000327 | 0.002566 | 2.22 |
| FALSE | High | NRHSAQTE | [R].NRHSA | 1xPhospho | Q9R0L6-2 | 0.000164 | 0 | 1 | 3 | 1 | Q9R0L6-2 | Isoform 2 c | Q9R0L6-2 [ | 1 | 2133.085 | 0 | -0.33 | 0.012675 | 7.66 | 112 | 88 | 5.13 | 13.65 | | High | High | High | High | High | High | High | High | High | High | High | 0 | 3.12E-05 | 2.57 |
| FALSE | High | NSAAELGA | [K].NSAAEI | 1xPhospho | Q8BSD5 1x | 2.1E-05 | 0 | 1 | 1 | 1 | Q8BSD5 | SH2 domai | Q8BSD5 [7 | 0 | 1485.771 | 0 | 1.32 | 0.001214 | 18.21 | 53.7 | 146.3 | 12.89 | 10.64 | | High | High | High | High | High | High | High | High | High | High | High | 0 | 3.11E-06 | 2.51 |
| FALSE | High | NSAAVPGE | [R].NSAAV | 2xPhospho | G3UWZ0 2 | 0.00359 | 0.00021 | 1 | 2 | 2 | G3UWZ0 | Bromodom | G3UWZ0 [ | 1 | 2943.27 | 0 | 1.38 | 0.004221 | 9.36 | 57.1 | 142.9 | 16.1 | 19.43 | | High | High | High | High | High | High | High | High | High | High | High | 9.42E-05 | 0.001003 | 4.02 |
| FALSE | High | NSADDEEL | [R].NSADD | 1xPhospho | Q9D4H9-2 | 2.12E-09 | 0 | 1 | 4 | 6 | Q9D4H9-2 | Isoform 2 c | Q9D4H9-2 | 0 | 2605.228 | 0 | 0.2 | 0.028781 | 8.34 | 96.1 | 103.9 | 3.09 | 6.94 | | High | High | High | High | High | High | High | High | High | High | High | 0 | 1.02E-06 | 4.22 |
| FALSE | High | NSADDEEL | [R].NSADD | 2xPhospho | Q9D4H9-2 | 5.4E-05 | 0 | 1 | 4 | 4 | Q9D4H9-2 | Isoform 2 c | Q9D4H9-2 | 0 | 2685.195 | 0 | 0.72 | 0.002546 | 11.2 | 73 | 127 | 5.96 | 10.65 | | High | High | High | High | High | High | High | High | High | High | High | 0 | 8.95E-06 | 4.23 |
| FALSE | High | NSAEIAPPL | [R].NSAEIA | 1xPhospho | B2RY04 1xl | 6.02E-05 | 0 | 1 | 2 | 1 | B2RY04 | Dedicator c | B2RY04 [1 | 0 | 1572.835 | 0 | -0.81 | 0.004113 | 4.51 | 127.3 | 72.7 | 7.35 | 16.58 | | High | High | High | High | High | High | High | High | High | High | High | 0 | 1.02E-05 | 2.85 |
| FALSE | High | NSAGGTSV | [K].NSAGG | 1xPhospho | Q61234 1x | 0.000651 | 0 | 1 | 3 | 2 | Q61234 | Alpha-1-sy | Q61234 [1 | 0 | 2293.081 | 0 | -1.19 | 0.000751 | 1.91 | 139.3 | 60.7 | 7.89 | 3.4 | | High | High | High | High | High | High | High | High | High | High | High | 0 | 0.000148 | 3.34 |
| FALSE | High | NSAGGTSV | [K].NSAGG | 2xPhospho | Q61234 2x | 5.76E-09 | 0 | 1 | 3 | 1 | Q61234 | Alpha-1-sy | Q61234 [1 | 0 | 2373.047 | 0 | -1.42 | 0.002359 | 3.7 | 145.7 | 54.3 | 11.58 | 8.77 | | High | High | High | High | High | High | High | High | High | High | High | 0 | 3.08E-10 | 4.91 |
| FALSE | High | NSASATTPI | [R].NSASAT | 1xPhospho | A0A1L1SSC | 4.49E-07 | 0 | 1 | 10 | 4 | A0A1L1SSC | Leucine-ric | A0A1L1SSC | 0 | 1758.822 | 0 | 1.18 | 0.001466 | 6.3 | 61.6 | 138.4 | 9.81 | 9.39 | | High | High | High | High | High | High | High | High | High | High | High | 0.001889 | 0.01907 | 1.91 |
| FALSE | High | NSASATTPI | [R].NSASAT | 2xPhospho | A0A1L1SSC | 2.51E-05 | 0 | 1 | 10 | 2 | A0A1L1SSC | Leucine-ric | A0A1L1SSC | 0 | 1838.788 | 0 | 1 | 0.001025 | 11.26 | 64.2 | 135.8 | 13.53 | 4.36 | | High | High | High | High | High | High | High | High | High | High | High | 0 | 3.8E-06 | 2.66 |
| FALSE | High | NSASVGIIC | [K].NSASV | 1xPhospho | Q8C0N2 1x | 0.000466 | 0 | 1 | 1 | 2 | Q8C0N2 | Glycerol-3- | Q8C0N2 [6 | 0 | 1353.709 | 0 | 3.2 | 0.000917 | 8.3 | 20 | 180 | 16.06 | 15.54 | | High | High | High | High | High | High | High | High | High | High | High | 0 | 0.000101 | 2.09 |
| FALSE | High | NSATCHSE | [R].NSATCI | 1xCarbami | Q3UZN6 1x | 3.94E-05 | 0 | 1 | 3 | 3 | Q3UZN6 | Putative ur | Q3UZN6 [2 | 0 | 2001.795 | 0 | -0.23 | 0.380242 | 6.47 | 111.2 | 88.8 | 10.52 | 12.43 | | High | High | High | High | High | High | High | High | High | High | High | 0 | 6.34E-06 | 3.66 |
| FALSE | High | NSATSADE | [R].NSATSA | 1xPhospho | Q571J8 1xl | 0.000725 | 0 | 1 | 17 | 1 | Q571J8 | Non-specif | Q571J8 [43 | 0 | 2068.929 | 0 | 0.11 | 0.069538 | 6.5 | 98.3 | 101.7 | 7.56 | 7.72 | | High | High | High | High | High | High | High | High | High | High | High | 0 | 0.000166 | 2.86 |
| FALSE | High | NSCEGFFL | [R].NSCEGI | 1xCarbami | Q9CRA9 1x | 0.002022 | 5.66E-05 | 1 | 2 | 1 | Q9CRA9 | FGFR1 onc | Q9CRA9 [1 | 0 | 1711.735 | 0 | 0.73 | 0.011333 | 17.24 | 77.2 | 122.8 | 8.46 | 11.18 | | High | High | High | High | High | High | High | High | High | High | High | 2.58E-05 | 0.000526 | 2.74 |
| FALSE | High | NSCNVGG | [R].NSCNV | 1xCarbami | Q8K3A9 1x | 0.048715 | 0.00354 | 1 | 1 | 2 | Q8K3A9 | 7SK snRNA | Q8K3A9 [1 | 0 | 1636.699 | 0 | -0.04 | 0.597723 | 31.21 | 111.2 | 88.8 | 12.91 | 24.61 | | High | High | High | High | High | High | High | High | High | High | High | 0.002403 | 0.02587 | 1.93 |
| FALSE | High | NSAAVPGE | [R].NSAAV | 1xPhospho | G3UWZ0 1 | 0.000151 | 0 | 1 | 2 | 5 | G3UWZ0 | Bromodom | G3UWZ0 [ | 1 | 2863.304 | 0 | 0.9 | 0.002051 | 10.18 | 69.6 | 130.4 | 8.15 | 8.58 | | High | High | High | High | High | High | High | High | High | High | High | 0.000689 | 0.005916 | 2.96 |
| FALSE | High | NRESNSLV | [K].NRESN | 1xPhospho | Q80WE4 1 | 0.012731 | 0.000889 | 1 | 4 | 1 | Q80WE4 | Kinesin-like | Q80WE4 [7 | 1 | 1897.031 | 0 | 0.53 | 0.001921 | 2.36 | 82.6 | 117.4 | 5.83 | 9.13 | | High | High | High | High | High | High | High | High | High | High | High | 0.000449 | 0.004129 | 2.09 |
| FALSE | High | NQYLSAIAS | [K].NQYLS | 1xPhospho | Q5SWW4 | 3.38E-05 | 0 | 1 | 1 | 1 | Q5SWW4 | Mediator c | Q5SWW4 [ | 0 | 2332.147 | 0 | 0.9 | 0.014562 | 27.9 | 70.8 | 129.2 | 15.83 | 16.4 | | High | High | High | High | High | High | High | High | High | High | High | 0 | 5.29E-06 | 2.34 |
| FALSE | High | NQYDNDV | [R].NQYDN | 1xPhospho | Q9R1P4 1x | 2.41E-06 | 0 | 1 | 2 | 1 | Q9R1P4 | Proteasom | Q9R1P4 [4 | 0 | 2087.939 | 0 | -0.27 | 0.025096 | 7.55 | 113.4 | 86.6 | 6.87 | 8.36 | | High | High | High | High | High | High | High | High | High | High | High | 0 | 2.72E-07 | 4.07 |
| FALSE | High | NQLVLIVNI | [K].NQLVLI | 1xDeamida | Q6P5E3 1x | 1.48E-06 | 0 | 1 | 3 | 1 | Q6P5E3 | Sal-like 1 (I | Q6P5E3 [6 | 0 | 2219.209 | 0 | -1 | 0.000518 | 2 | 134.9 | 65.1 | 12.9 | 15.1 | | High | High | High | High | High | High | High | High | High | High | High | 0 | 1.58E-07 | 3.66 |
| FALSE | High | NQMGGSN | [R].NQMG | 1xPhospho | Q91YD3 1x | 0.096181 | 0.007535 | 1 | 2 | 1 | Q91YD3 | mRNA-dec | Q91YD3 [1 | 0 | 4201.928 | 0 | 0.25 | 0.227467 | 4.05 | 89 | 111 | 12.76 | 23.58 | | High | High | High | High | High | High | High | High | High | High | High | 0.003809 | 0.04115 | 2.79 |
| FALSE | High | NQMGGSN | [R].NQMG | 1xDeamida | Q91YD3 1x | 1.51E-07 | 0 | 1 | 2 | 6 | Q91YD3 | mRNA-dec | Q91YD3 [1 | 0 | 4416.08 | 0 | -0.55 | 0.049949 | 2.73 | 122.8 | 77.2 | 18.54 | 23.02 | | High | High | High | High | High | High | High | High | High | High | High | 0 | 1.22E-08 | 5.43 |
| FALSE | High | NQMGGSN | [R].NQMG | 1xPhospho | Q91YD3 1x | 4.82E-06 | 0 | 1 | 2 | 3 | Q91YD3 | mRNA-dec | Q91YD3 [1 | 0 | 4431.091 | 0 | -1.12 | 0.005121 | 5.05 | 136.8 | 63.2 | 8.43 | 15.55 | | High | High | High | High | High | High | High | High | High | High | High | 0 | 2.18E-05 | 4.12 |
| FALSE | High | NQNCAAEI | [R].NQNCA | 1xCarbami | O54928 1x | 0.076555 | 0.005826 | 1 | 2 | 1 | O54928 | Suppressor | O54928 [7 | 0 | 2351.152 | 0 | -0.47 | 0.002741 | 1.21 | 116 | 84 | 6.26 | 5.85 | | High | High | High | High | High | High | High | High | High | High | High | 0.00299 | 0.03158 | 1.58 |
| FALSE | High | NQPSPDTA | [R].NQPSPI | 1xPhospho | D3Z183 1x | 1.78E-05 | 0 | 1 | 1 | 2 | D3Z183 | Major facil | D3Z183 [7 | 0 | 3667.703 | 0 | 6.64 | | 0 | 9.5 | 190.5 | 4.4 | 21.09 | | High | High | Not Found | Not Found | Not Found | High | High | High | High | High | High | 0 | 3.63E-06 | 3.99 |
| FALSE | High | NQPTVYIA | [R].NQPTV | 1xPhospho | Q99K90 1x | 0.000267 | 0 | 1 | 3 | 1 | Q99K90 | TGF-beta-a | Q99K90 [3 | 0 | 2529.189 | 0 | -0.24 | 0.007305 | 4.51 | 109.2 | 90.8 | 5.48 | 10.29 | | High | High | High | High | High | High | High | High | High | High | High | 0 | 5.4E-05 | 2.62 |
| FALSE | High | NQQQGLT | [K].NQQQ | 1xPhospho | P23949 1xl | 8.25E-08 | 0 | 1 | 1 | 1 | P23949 | mRNA deca | P23949 [3 | 0 | 3637.852 | 0 | 0.64 | 0.011073 | 28.8 | 78.1 | 121.9 | 19.75 | 5.3 | | High | High | High | High | High | High | High | High | High | High | High | 0 | 6.18E-09 | 4.96 |
| FALSE | High | NQQSQPS | [K].NQQSC | 1xPhospho | B2RWS6 1 | 0.004143 | 0.00036 | 1 | 3 | 1 | B2RWS6 | Histone ac | B2RWS6 [4 | 0 | 1938.869 | 0 | 0.1 | 0.359485 | 24.64 | 96.5 | 103.5 | 12.02 | 11.64 | | High | High | High | High | High | High | High | High | High | High | High | 0.000161 | 0.001171 | 1.64 |
| FALSE | High | NQRESQQI | [K].NQRES | 1xPhospho | Q8VDR2 1x | 5.91E-05 | 0 | 1 | 6 | 1 | Q8VDR2 | Hmmr prot | Q8VDR2 [4 | 1 | 2125.08 | 0 | -0.68 | 0.001052 | 5.41 | 124.2 | 75.8 | 5.49 | 17.62 | | High | High | High | High | High | High | High | High | High | High | High | 0 | 1E-05 | 3.31 |
| FALSE | High | NQSFCPTV | [R].NQSFCI | 1xCarbami | P14115 1xl | 0.060159 | 0.004476 | 1 | 1 | 2 | P14115 | 60S riboso | P14115 [6 | 0 | 1960.96 | 0 | 1 | 0.026814 | 2.13 | 66.8 | 133.2 | 16.39 | 29.13 | | High | High | High | High | High | High | High | High | High | High | High | 0.0032 | 0.03379 | 2.37 |
| FALSE | High | NQSLGQSA | [R].NQSLG | 1xPhospho | E9Q1Q1 1x | 0.000458 | 0 | 1 | 5 | 1 | E9Q1Q1 | Microtubu | E9Q1Q1 [1 | 0 | 1880.968 | 0 | -0.67 | 0.067331 | 11.62 | 122.6 | 77.4 | 23.86 | 12.13 | | High | High | High | High | High | High | High | High | High | High | High | 0 | 9.94E-05 | 3.07 |
| FALSE | High | NQSLGQSA | [R].NQSLG | 1xPhospho | E9Q1Q1 1x | 0.009585 | 0.000736 | 1 | 5 | 1 | E9Q1Q1 | Microtubu | E9Q1Q1 [1 | 0 | 2110.131 | 0 | -1.21 | 0.004786 | 5.63 | 135.9 | 64.1 | 8.69 | 20.52 | | High | High | High | High | High | High | High | High | High | High | High | 0.000344 | 0.002996 | 2.6 |
| FALSE | High | NQSPGSHC | [R].NQSPG | 1xPhospho | Q8C7E7 1x | 0.045898 | 0.003408 | 1 | 2 | 1 | Q8C7E7 | Starch-bin | Q8C7E7 [1 | 0 | 1620.712 | 0 | 2.43 | 0.000945 | 2.89 | 29.4 | 170.6 | 14.9 | 16.08 | | High | High | High | High | High | High | High | High | High | High | High | 0.001774 | 0.01747 | 1.45 |
| FALSE | High | NQSPVLEP | [R].NQSPV | 1xPhospho | A7E215 1xl | 0.001586 | 5.66E-05 | 1 | 3 | 2 | A7E215 | Rps6ka3 pr | A7E215 [7 | 0 | 1504.772 | 0 | 1.77 | 0.000436 | 17.53 | 45.7 | 154.3 | 7.94 | 6.2 | | High | High | High | High | High | High | High | High | High | High | High | 0.000814 | 0.007496 | 2.16 |
| FALSE | High | NQSSEDILF | [R].NQSSEI | 1xPhospho | E9QJU8 1x | 0.010758 | 0.000772 | 1 | 2 | 1 | E9QJU8 | Transform | E9QJU8 [4 | 0 | 1370.651 | 0 | 0.81 | 0.058263 | 33.92 | 73.8 | 126.2 | 19.3 | 24.36 | | High | High | High | High | High | High | High | High | High | High | High | 0.000399 | 0.003406 | 2.09 |
| FALSE | High | NQSSPPPA | [R].NQSSPI | 1xCarbami | E9QK36 1x | 0.004867 | 0.000504 | 1 | 5 | 4 | E9QK36 | WD repeat | E9QK36 [3 | 0 | 1842.913 | 0 | -0.35 | 0.012717 | 5.81 | 109.7 | 90.3 | 2.48 | 8.38 | | High | High | High | High | High | High | High | High | High | High | High | 0.000585 | 0.005354 | 2.32 |
| FALSE | High | NQSYLQLL | [R].NQSYL | 1xCarbami | Q9CQU5 1 | 3.8E-05 | 0 | 1 | 1 | 2 | Q9CQU5 | ZW10 inter | Q9CQU5 [2 | 0 | 2247.161 | 0 | 0.23 | 0.049161 | 13.42 | 90 | 110 | 9.5 | 7.78 | | High | High | High | High | High | High | High | High | High | High | High | 0 | 6.08E-06 | 2.95 |
| FALSE | High | NQTHQGS | [K].NQTHC | 1xPhospho | Q9D0N7 1x | 7.28E-05 | 0 | 1 | 1 | 2 | Q9D0N7 | Chromatin | Q9D0N7 [4 | 0 | 1564.707 | 0 | 0.27 | 0.282756 | 16.6 | 96.3 | 103.7 | 13.34 | 17.64 | | High | High | High | High | High | High | High | High | High | High | High | 0.000327 | 0.002573 | 2.21 |
| FALSE | High | NQTQLNTS | [K].NQTQL | 1xPhospho | Q3UQN6 1 | 0.00418 | 0.00036 | 1 | 4 | 2 | Q3UQN6 | Putative ur | Q3UQN6 [ | 0 | 1669.826 | 0 | 0.04 | 0.83221 | 4.13 | 99.2 | 100.8 | 11.35 | 8.95 | | High | High | High | High | High | High | High | High | High | High | High | 0.000161 | 0.001188 | 1.9 |
| FALSE | High | NQTYSFSP | [K].NQTYSI | 1xPhospho | Q7TPS5 1x | 0.005228 | 0.000551 | 1 | 4 | 2 | Q7TPS5 | C2 domain | Q7TPS5 [2 | 0 | 1696.835 | 0 | -0.55 | 0.011524 | 2.93 | 118.5 | 81.5 | 4.58 | 10.95 | | High | High | High | High | High | High | High | High | High | High | High | 0.001999 | 0.02009 | 1.99 |
| FALSE | High | NQVPSQPS | [R].NQVPS | 1xPhospho | Q5RIM6 1x | 1.61E-05 | 0 | 1 | 8 | 1 | Q5RIM6 | Nuclear re | Q5RIM6 [2 | 0 | 2913.419 | 0 | 0.03 | 0.588045 | 4.59 | 99 | 101 | 6.29 | 13.65 | | High | High | High | High | High | High | High | High | High | High | High | 0 | 2.32E-06 | 3.89 |
| FALSE | High | NQVVPESD | [R].NQVVP | 1xPhospho | Q3U4X8 1x | 0.000389 | 0 | 1 | 7 | 3 | Q3U4X8 | DNA ligase | Q3U4X8 [5 | 0 | 1607.788 | 0 | -0.93 | 0.155143 | 10.24 | 131.1 | 68.9 | 42.24 | 40.08 | | High | High | High | High | High | High | High | High | High | High | High | 2.58E-05 | 0.000342 | 2.95 |
| FALSE | High | NQVVPESD | [R].NQVVP | 1xPhospho | Q3U4X8 1x | 1.55E-06 | 0 | 1 | 7 | 5 | Q3U4X8 | DNA ligase | Q3U4X8 [5 | 0 | 1836.951 | 0 | -1.23 | 0.001082 | 0.77 | 139 | 61 | 4.05 | 12.38 | | High | High | High | High | High | High | High | High | High | High | High | 0 | 3.92E-06 | 3.31 |
| FALSE | High | NQVVPESD | [R].NQVVP | 1xDeamida | Q3U4X8 1x | 2.9E-05 | 0 | 1 | 7 | 3 | Q3U4X8 | DNA ligase | Q3U4X8 [5 | 0 | 1837.935 | 0 | -0.95 | 0.000997 | 1.49 | 130.1 | 69.9 | 6.73 | 6.29 | | High | High | High | High | High | High | High | High | High | High | High | 0 | 4.49E-06 | 2.68 |
| FALSE | High | NQVVPESD | [R].NQVVP | 1xPhospho | Q3U4X8 1x | 0.00642 | 0.00062 | 1 | 7 | 2 | Q3U4X8 | DNA ligase | Q3U4X8 [5 | 1 | 1993.052 | 0 | -1.27 | 0.000818 | 0.95 | 139.1 | 60.9 | 14.56 | 20.99 | | High | High | High | High | High | High | High | High | High | High | High | 0.000275 | 0.001917 | 2.04 |
| FALSE | High | NQVVSSTN | [K].NQVVS | 1xPhospho | Q8C4X2 1x | 4.52E-09 | 0 | 1 | 1 | 5 | Q8C4X2 | Casein kina | Q8C4X2 [3 | 0 | 2384.093 | 0 | 0.2 | 0.066934 | 8.11 | 92.4 | 107.6 | 2.88 | 7.87 | | High | High | High | High | High | High | High | High | High | High | High | 0 | 2.45E-08 | 4.49 |
| FALSE | High | NQVVSSTN | [K].NQVVS | 1xDeamida | Q8C4X2 1x | 1.39E-09 | 0 | 1 | 1 | 2 | Q8C4X2 | Casein kina | Q8C4X2 [3 | 0 | 2385.077 | 0 | 0.13 | 0.088628 | 10 | 93.2 | 106.8 | 4.88 | 7.75 | | High | High | High | High | High | High | High | High | High | High | High | 0 | 8.2E-11 | 5.84 |
| FALSE | High | NQWQLSA | [K].NQWQ | 1xPhospho | Q3TYV5 1x | 0.003055 | 0.00016 | 1 | 3 | 1 | Q3TYV5 | 2',3'-cyclic- | Q3TYV5 [1 | 0 | 1855.936 | 0 | -0.68 | 0.003058 | 6.2 | 123.8 | 76.2 | 5.16 | 12.63 | | High | High | High | High | High | High | High | High | High | High | High | 7.19E-05 | 0.000837 | 2.58 |
| FALSE | High | NNSPAPPQ | [K].NNSPA | 1xPhospho | Q14AX6-2 | 0.000194 | 0 | 3 | 3 | 3 | Q14AX6-2; | Isoform 2 c | Q14AX6-2 | 0 | 1854.988 | 0 | 0.02 | 0.8126 | 6.97 | 97.9 | 102.1 | 8.87 | 14.02 | Shared | High | High | High | High | High | High | High | High | High | High | High | 0.000585 | 0.005186 | 2.81 |
| FALSE | High | NNSPAPPQ | [K].NNSPA | 1xPhospho | Q14AX6-2 | 0.111965 | 0.009053 | 3 | 3 | 1 | Q14AX6-2; | Isoform 2 c | Q14AX6-2 | 0 | 1625.825 | 0 | 0.05 | 0.810965 | 22.26 | 99 | 101 | 31.64 | 28.25 | Shared | High | High | High | High | High | High | High | High | High | High | High | 0.004543 | 0.04942 | 1.83 |
| FALSE | High | NNSISEAK | [R].NNSISE | 1xPhospho | P51432 1xl | 0.008614 | 0.000736 | 1 | 4 | 2 | P51432 | 1-phospha | P51432 [11 | 0 | 1400.719 | 0 | 0.71 | 0.005447 | 7.84 | 74.1 | 125.9 | 9.37 | 9.15 | | High | High | High | High | High | High | High | High | High | High | High | 0.000571 | 0.004992 | 2.08 |
| FALSE | High | NNSISEAK | [R].NNSISE | 1xPhospho | P51432 1xl | 0.08447 | 0.006345 | 1 | 4 | 2 | P51432 | 1-phospha | P51432 [11 | 0 | 1171.556 | 0 | 0.47 | 0.000486 | 1.33 | 82.9 | 117.1 | 3.39 | 2.49 | | High | High | High | High | High | High | High | High | High | High | High | 0.003265 | 0.03534 | 1.7 |
| FALSE | High | NLLPAYRPI | [K].NLLPAY | 1xPhospho | Q9D2D7 1x | 0.027839 | 0.001755 | 1 | 3 | 1 | Q9D2D7 | Zinc finger | Q9D2D7 [5 | 0 | 3057.612 | 0 | -0.64 | 0.00303 | 7.85 | 121.7 | 78.3 | 6.23 | 12.55 | | High | High | High | High | High | High | High | High | High | High | High | 0.000953 | 0.009919 | 2.51 |
| FALSE | High | NLLQHNNS | [R].NLLQHI | 1xPhospho | E9Q9R9 1x | 0.000845 | 0 | 1 | 5 | 1 | E9Q9R9 | Disks large | E9Q9R9 [8 | 0 | 2489.166 | 0 | 0.48 | 0.019715 | 6.54 | 85.6 | 114.4 | 7.86 | 10.11 | | High | High | High | High | High | High | High | High | High | High | High | 0 | 0.000197 | 2.77 |
| FALSE | High | NLLQQSWI | [R].NLLQQ | 1xPhospho | E9Q9R9 1x | 0.001799 | 5.66E-05 | 1 | 4 | 1 | E9Q9R9 | Disks large | E9Q9R9 [2 | 0 | 1929.955 | 0 | -0.49 | 0.003077 | 4.99 | 116.3 | 83.7 | 8.4 | 10.15 | | High | High | High | High | High | High | High | High | High | High | High | 2.58E-05 | 0.000463 | 2.86 |
| FALSE | High | NLLQQSWI | [R].NLLQQ | 1xPhospho | E9Q9R9 1x | 0.008691 | 0.000736 | 1 | 4 | 1 | E9Q9R9 | Disks large | E9Q9R9 [2 | 0 | 1945.949 | 0 | -0.35 | 0.020381 | 1.78 | 108.6 | 91.4 | 4.09 | 7.02 | | High | High | High | High | High | High | High | High | High | High | High | 0.000327 | 0.002681 | 3.09 |
| FALSE | High | NIISAASLS | [R].NIISAAS | 1xPhospho | Q7TPM1 1 | 2.92E-08 | 0 | 1 | 3 | 1 | Q7TPM1 | Protein PRI | Q7TPM1 [2 | 0 | 2083.063 | 0 | 1.02 | 0.000223 | 1.44 | 65.8 | 134.2 | 5.09 | 2.41 | | High | High | High | High | High | High | High | High | High | High | High | 0 | 1.92E-09 | 3.88 |
| FALSE | High | NIISPPWDF | [R].NIISPPV | 1xPhospho | Q3TI84 1xF | 0.009416 | 0.000736 | 1 | 2 | 1 | Q3TI84 | CDC16 cell | Q3TI84 [55 | 0 | 1553.772 | 0 | -0.58 | 0.000614 | 1.01 | 118.8 | 81.2 | 4.04 | 1.71 | | High | High | High | High | High | High | High | High | High | High | High | 0.000327 | 0.002932 | 2.67 |
| FALSE | High | NLNLVSST | [R].NLNLV | 1xPhospho | B2RTN5 1x | 7.04E-06 | 0 | 1 | 2 | 3 | B2RTN5 | Glucocortic | B2RTN5 [7 | 0 | 1784.992 | 0 | 1.22 | 0.003473 | 13.55 | 59.7 | 140.3 | 12.39 | 15.9 | | High | High | High | High | High | High | High | High | High | High | High | 0 | 9.08E-07 | 3.11 |
| FALSE | High | NLNNSNLF | [K].NLNNS | 1xPhospho | A0A1B0GS | 0.00043 | 0 | 1 | 9 | 2 | A0A1B0GS | Nuclear po | A0A1B0GS | 0 | 3654.618 | 0 | -0.25 | 0.026933 | 2.51 | 109.4 | 90.6 | 12.18 | 10.91 | | High | High | High | High | High | High | High | High | High | High | High | 0 | 9.26E-05 | 3.47 |
| FALSE | High | NLNNSNLF | [K].NLNNS | 1xDeamida | A0A1B0GS | 6.37E-06 | 0 | 1 | 9 | 10 | A0A1B0GS | Nuclear po | A0A1B0GS | 0 | 3655.602 | 0 | -0.34 | 0.010084 | 6.71 | 108.4 | 91.6 | 6.7 | 10.25 | | High | High | High | High | High | High | High | High | High | High | High | 0 | 8.17E-07 | 4.57 |
| FALSE | High | NLNNSNLF | [K].NLNNS | 2xDeamida | A0A1B0GS | 0.000328 | 0 | 1 | 9 | 5 | A0A1B0GS | Nuclear po | A0A1B0GS | 0 | 3656.586 | 0 | -0.31 | 0.176182 | 3.14 | 109.4 | 90.6 | 9.41 | 9.22 | | High | High | High | High | High | High | High | High | High | High | High | 0 | 6.82E-05 | 4.6 |
| FALSE | High | NLNNSNLF | [K].NLNNS | 2xPhospho | A0A1B0GS | 0.00172 | 5.66E-05 | 1 | 9 | 3 | A0A1B0GS | Nuclear po | A0A1B0GS | 0 | 3734.585 | 0 | -0.18 | 0.005109 | 0.89 | 106 | 94 | 4.18 | 8.42 | | High | High | High | High | High | High | High | High | High | High | High | 0.000161 | 0.001151 | 3.6 |
| FALSE | High | NLNNSNLF | [K].NLNNS | 1xDeamida | A0A1B0GS | 0.001117 | 0 | 1 | 9 | 5 | A0A1B0GS | Nuclear po | A0A1B0GS | 0 | 3735.569 | 0 | -0.37 | 0.016618 | 6.35 | 110.5 | 89.5 | 8.38 | 9.18 | | High | High | High | High | High | High | High | High | High | High | High | 0 | 0.000269 | 3.64 |
| FALSE | High | NLPAHGG | [R].NLPAH | 1xPhospho | B9EKC1 1x | 0.017344 | 0.001204 | 1 | 2 | 2 | B9EKC1 | PAS domai | B9EKC1 [5 | 0 | 2869.522 | 0 | -0.52 | 0.085215 | 8.91 | 109.6 | 90.4 | 14.5 | 19.35 | | High | High | High | High | High | High | High | High | High | High | High | 0.001279 | 0.01338 | 2.16 |
| FALSE | High | NLIELSLLP | [K].NLIELSI | 1xPhospho | Q6NS46 1x | 2.69E-08 | 0 | 1 | 1 | 1 | Q6NS46 | Protein RR | Q6NS46 [1 | 0 | 3230.711 | 0 | -0.28 | 0.010863 | 7.05 | 105.6 | 94.4 | 8.2 | 4.69 | | High | High | High | High | High | High | High | High | High | High | High | 0 | 1.75E-09 | 2.5 |
| FALSE | High | NIPCAAVSI | [K].NIPCAA | 1xCarbami | O89091 1x | 0.014283 | 0.001011 | 1 | 5 | 1 | O89091 | Krueppel-li | O89091 [1 | 0 | 1507.729 | 0 | -0.45 | 0.003054 | 1.91 | 115.4 | 84.6 | 6.22 | 4.47 | | High | High | High | High | High | High | High | High | High | High | High | 0.000538 | 0.004668 | 1.7 |
| FALSE | High | NLPETIPW | [K].NLPETI | 1xPhospho | P70268-2 1 | 0.000393 | 0 | 1 | 3 | 2 | P70268-2 | Isoform 2 c | P70268-2 [ | 0 | 2685.312 | 0 | 0.71 | 0.000339 | 5.8 | 76.5 | 123.5 | 2.67 | 3.81 | | High | High | High | High | High | High | High | High | High | High | High | 0 | 8.31E-05 | 4.15 |
| FALSE | High | NLPPSGAV | [K].NLPPSG | 1xPhospho | Q571F9 1x | 8.78E-08 | 0 | 1 | 3 | 3 | Q571F9 | MKIAA411 | Q571F9 [2 | 0 | 2405.336 | 0 | -0.05 | 0.670142 | 7.27 | 101.8 | 98.2 | 6.69 | 7.33 | | High | High | High | High | High | High | High | High | High | High | High | 0 | 2.41E-06 | 4.87 |
| FALSE | High | NLPSFGTEI | [R].NLPSFG | 1xPhospho | Q5SVQ0 1x | 0.040373 | 0.002982 | 1 | 5 | 1 | Q5SVQ0 | Histone ac | Q5SVQ0 [6 | 0 | 1977.916 | 0 | -1.3 | 0.000779 | 2.39 | 142.3 | 57.7 | 5.99 | 12.59 | | High | High | High | High | High | High | High | High | High | High | High | 0.001566 | 0.01512 | 1.94 |
| FALSE | High | NLPSSMFV | [R].NLPSSN | 1xPhospho | Q923B1 1x | 0.004491 | 0.000409 | 1 | 2 | 2 | Q923B1 | Lariat debr | Q923B1 [4 | 1 | 2864.354 | 0 | -1.18 | 0.011023 | 8.69 | 140.2 | 59.8 | 16.53 | 19.64 | | High | High | High | High | High | High | High | High | High | High | High | 0.000449 | 0.00388 | 3.51 |
| FALSE | High | NLPSSMFV | [R].NLPSSN | 1xPhospho | Q923B1 1x | 0.032267 | 0.002129 | 1 | 2 | 4 | Q923B1 | Lariat debr | Q923B1 [4 | 1 | 2880.349 | 0 | -1.66 | 0.045808 | 1.43 | 146.2 | 53.8 | 17.28 | 48.81 | | High | High | High | High | High | High | High | High | High | High | High | 0.001665 | 0.01675 | 3.07 |
| FALSE | High | NLPSSMFV | [R].NLPSSN | 2xPhospho | Q923B1 2x | 0.014667 | 0.001049 | 1 | 2 | 1 | Q923B1 | Lariat debr | Q923B1 [4 | 1 | 2944.32 | 0 | -0.03 | 0.587089 | 22.44 | 107.2 | 92.8 | 16.37 | 16.18 | | High | High | High | High | High | High | High | High | High | High | High | 0.000555 | 0.004792 | 4.29 |
| FALSE | High | NIQQDNSE | [K].NIQQD | 1xPhospho | Q61699 1x | 0.001039 | 0 | 1 | 3 | 5 | Q61699 | Heat shock | Q61699 [5 | 0 | 4279.931 | 0 | -0.36 | 0.55865 | 34.23 | 112.5 | 87.5 | 54.75 | 63.18 | | High | High | High | High | High | High | High | High | High | High | High | 2.58E-05 | 0.000486 | 4.55 |
| FALSE | High | NIQQDNSE | [K].NIQQD | 1xDeamida | Q61699 1x | 8.09E-11 | 0 | 1 | 3 | 4 | Q61699 | Heat shock | Q61699 [5 | 0 | 4280.915 | 0 | -0.66 | 0.619408 | 5.54 | 118.1 | 81.9 | 52.96 | 78.87 | | High | High | High | High | High | High | High | High | High | High | High | 0 | 3.65E-12 | 5.48 |
| FALSE | High | NIQQDNSE | [K].NIQQD | 1xPhospho | Q61699 1x | 5.01E-07 | 0 | 1 | 3 | 11 | Q61699 | Heat shock | Q61699 [5 | 0 | 4509.094 | 0 | -0.6 | 0.003991 | 1.86 | 120.8 | 79.2 | 6.18 | 5.62 | | High | High | High | High | High | High | High | High | High | High | High | 0 | 4.66E-08 | 5.72 |
| FALSE | High | NIQQDNSE | [K].NIQQD | 1xDeamida | Q61699 1x | 1.18E-08 | 0 | 1 | 3 | 23 | Q61699 | Heat shock | Q61699 [5 | 0 | 4510.078 | 0 | -0.32 | 0.042071 | 14.68 | 113 | 87 | 11.54 | 6.61 | | High | High | High | High | High | High | High | High | High | High | High | 0 | 6.89E-10 | 8.04 |
| FALSE | High | NIQQDNSE | [K].NIQQD | 2xPhospho | Q61699 2x | 1.79E-07 | 0 | 1 | 3 | 3 | Q61699 | Heat shock | Q61699 [5 | 0 | 4589.061 | 0 | -0.4 | 0.0077 | 6.48 | 114 | 86 | 4.72 | 6.92 | | High | High | High | High | High | High | High | High | High | High | High | 0 | 1.48E-08 | 7 |
| FALSE | High | NLRPLYGD | [K].NLRPLY | 2xPhospho | E9Q8M1 2 | 0.025293 | 0.001645 | 1 | 2 | 1 | E9Q8M1 | Zinc finger | E9Q8M1 [1 | 0 | 2986.4 | 0 | 0.22 | 0.010915 | 8.04 | 93.2 | 106.8 | 9.07 | 6.74 | | High | High | High | High | High | High | High | High | High | High | High | 0.000872 | 0.008853 | 2.23 |
| FALSE | High | NISEESPLT | [K].NISEES | 1xPhospho | B9EKC1 1x | 0.065405 | 0.004734 | 1 | 2 | 1 | B9EKC1 | PAS domai | B9EKC1 [5 | 0 | 1591.768 | 0 | -0.41 | 0.003906 | 5.77 | 116.7 | 83.3 | 10.18 | 16.83 | | High | High | High | High | High | High | High | High | High | High | High | 0.002432 | 0.02615 | 2.3 |
| FALSE | High | NLPDFLGD | [K].NLPDFL | 1xPhospho | Q9EP53 1x | 7.38E-07 | 0 | 1 | 4 | 1 | Q9EP53 | Hamartin C | Q9EP53 [4 | 0 | 2530.237 | 0 | -0.15 | 0.055132 | 2.43 | 105.1 | 94.9 | 16.24 | 10.69 | | High | High | High | High | High | High | High | High | High | High | High | 0 | 7.21E-08 | 4.35 |

| | | | | | | | | | | | | | | | | | | | | | | | | | | | | | | | | | | | | | | |
|---|---|---|---|---|---|---|---|---|---|---|---|---|---|---|---|---|---|---|---|---|---|---|---|---|---|---|---|---|---|---|---|---|---|---|---|---|---|---|
| FALSE | High | NLIEGFNSF | [K].NLIEGF | 1xCarbami | A2AG50 1x | 0.000297 | 0 | 1 | 2 | 2 | A2AG50 | MAP7 dom | A2AG50 [7 | 0 | 2451.11 | 0 | -0.01 | 0.933379 | 24.48 | 99.9 | 100.1 | 5.85 | 17.29 | | High | High | High | High | High | High | High | High | High | High | High | 0 | 6.1E-05 | 4.75 |
| FALSE | High | NLLEDDSD | [R].NLLEDD | 1xPhospho | O70480 1x | 1.36E-08 | 0 | 1 | 3 | 9 | O70480 | Vesicle-ass | O70480 [2 | 0 | 2294.99 | 0 | 0.91 | 0.001194 | 11.94 | 69.3 | 130.7 | 10.6 | 6.53 | | High | High | High | High | High | High | High | High | High | High | High | 0 | 1.5E-08 | 4.73 |
| FALSE | High | NILDSDNYS | [R].NILDSD | 1xPhospho | A6H5Z3 1x | 0.003001 | 0.00016 | 1 | 4 | 1 | A6H5Z3 | Exocyst co | A6H5Z3 [4 | 0 | 2823.374 | 0 | -0.28 | 0.095925 | 18.11 | 106.6 | 93.4 | 5.34 | 14.03 | | High | High | High | High | High | High | High | High | High | High | High | 7.19E-05 | 0.000815 | 2.91 |
| FALSE | High | NHQSPSFS | [R].NHQSP | 1xPhospho | A2AG58 1x | 0.009087 | 0.000736 | 1 | 1 | 1 | A2AG58 | Uncharacte | A2AG58 [5 | 0 | 1569.783 | 0 | -1.17 | 0.000886 | 2.39 | 136.7 | 63.3 | 4.18 | 11.41 | | High | High | High | High | High | High | High | High | High | High | High | 0.000327 | 0.002809 | 1.69 |
| FALSE | High | NHSGSRTP | [R].NHSGS | 2xPhospho | Q8BTI8 2x | 0.035181 | 0.002269 | 1 | 3 | 2 | Q8BTI8 | Serine/argi | Q8BTI8 [20 | 1 | 2041.942 | 0 | -0.31 | 0.081712 | 12.52 | 108.8 | 91.2 | 8.18 | 9.78 | | High | High | High | High | High | High | High | High | High | High | High | 0.001999 | 0.0206 | 1.61 |
| FALSE | High | NHSNAQFI | [R].NHSNA | 1xCarbami | P41230 1x | 0.000693 | 0 | 1 | 3 | 2 | P41230 | Lysine-spe | P41230 [31 | 0 | 2033.91 | 0 | 0.12 | 0.329454 | 45.09 | 93 | 107 | 15.36 | 18.39 | | High | High | High | High | High | High | High | High | High | High | High | 0 | 0.000158 | 3.15 |
| FALSE | High | NHSPSPPV | [R].NHSPSF | 2xPhospho | Q8CAF4-3 | 6.3E-05 | 0 | 1 | 3 | 1 | Q8CAF4-3 | Isoform 3 c | Q8CAF4-3 | 0 | 2688.32 | 0 | 0.73 | 0.017287 | 29.8 | 75.1 | 124.9 | 12.97 | 13.57 | | High | High | High | High | High | High | High | High | High | High | High | 0 | 1.06E-05 | 3.49 |
| FALSE | High | NKLEGDSD | [K].NKLEGI | 1xPhospho | Q9JMH9-6 | 0.031168 | 0.001935 | 2 | 4 | 2 | Q9JMH9-6; | Isoform 6 c | Q9JMH9-6 | 1 | 2359.107 | 0 | -0.78 | 0.046124 | 10.09 | 127.1 | 72.9 | 22.33 | 10.63 | Shared | High | High | High | High | High | High | High | High | High | High | High | 0.006317 | 0.06749 | 2.62 |
| FALSE | High | NKPLSPIK | [K].NKPLSF | 1xPhospho | G3UWX1 1 | 0.092372 | 0.007211 | 2 | 9 | 3 | G3UWX1; I | Replication | G3UWX1 [ | 0 | 1434.849 | 0 | -0.14 | 0.739565 | 32.37 | 104.7 | 95.3 | 48.17 | 49.51 | Shared | High | High | High | High | High | High | High | High | High | High | High | 0.005527 | 0.05995 | 2.48 |
| FALSE | High | NKPLSPIK | [K].NKPLSF | 1xPhospho | G3UWX1 1 | 0.005 | 0.000504 | 2 | 9 | 2 | G3UWX1; I | Replication | G3UWX1 [ | 0 | 1664.012 | 0 | -0.34 | 0.008365 | 7.11 | 111.9 | 88.1 | 3.74 | 7.66 | Shared | High | High | High | High | High | High | High | High | High | High | High | 0.00269 | 0.02878 | 2.11 |
| FALSE | High | NKPQASAP | [K].NKPQA | 1xPhospho | Q9CZX5 1x | 0.005768 | 0.00062 | 1 | 1 | 1 | Q9CZX5 | PIN2/TERF | Q9CZX5 [1 | 0 | 2521.27 | 0 | 0.33 | 0.14425 | 23.39 | 88.6 | 111.4 | 27.45 | 6.22 | | High | High | High | High | High | High | High | High | High | High | High | 0.000275 | 0.00169 | 2.88 |
| FALSE | High | NKPRSPVV | [K].NKPRSF | 1xPhospho | Q3U1V3 1x | 0.002974 | 0.00016 | 1 | 3 | 1 | Q3U1V3 | Protein-ser | Q3U1V3 [6 | 1 | 2121.24 | 0 | 1.12 | 0.001812 | 14.1 | 63.2 | 136.8 | 11.12 | 7.71 | | High | High | High | High | High | High | High | High | High | High | High | 7.19E-05 | 0.000811 | 2.48 |
| FALSE | High | NIAPTAEHI | [R].NIAPTA | 1xPhospho | G3X963 1x | 0.003138 | 0.00016 | 1 | 2 | 1 | G3X963 | ATPase fan | G3X963 [1 | 0 | 2411.155 | 0 | 0.35 | 0.054927 | 3.14 | 88.3 | 111.7 | 7.76 | 15.55 | | High | High | High | High | High | High | High | High | High | High | High | 7.19E-05 | 0.000864 | 2.74 |
| FALSE | High | NLASPEGTI | [R].NLASPE | 1xPhospho | B0V2P5 1x | 0.009006 | 0.000736 | 1 | 4 | 2 | B0V2P5 | DmX-like p | B0V2P5 [18 | 0 | 2023.124 | 0 | 0.16 | 0.038074 | 9.48 | 89 | 111 | 9.05 | 7.19 | | High | High | High | High | High | High | High | High | High | High | High | 0.000327 | 0.002794 | 2.16 |
| FALSE | High | NLATSADTI | [K].NLATSA | 1xPhospho | Q8BKX6 1x | 8.92E-07 | 0 | 1 | 1 | 5 | Q8BKX6 | Serine/thre | Q8BKX6 [3 | 0 | 2266.173 | 0 | 0.25 | 0.039211 | 1.41 | 89.4 | 110.6 | 9.83 | 8.99 | | High | High | High | High | High | High | High | High | High | High | High | 0 | 4.95E-06 | 2.91 |
| FALSE | High | NLATSADTI | [K].NLATSA | 2xPhospho | Q8BKX6 2x | 0.005275 | 0.000599 | 1 | 1 | 4 | Q8BKX6 | Serine/thre | Q8BKX6 [3 | 0 | 2346.14 | 0 | 0.54 | 0.010209 | 6.4 | 82.7 | 117.3 | 6.28 | 8.41 | | High | High | High | High | High | High | High | High | High | High | High | 0.000275 | 0.001577 | 2.36 |
| FALSE | High | NIAVAHRD | [K].NIAVAH | 1xPhospho | Q91VE6 1x | 1.84E-06 | 0 | 1 | 2 | 2 | Q91VE6 | MKI67 FHA | Q91VE6 [1 | 1 | 2629.35 | 0 | 0.12 | 0.534273 | 24.94 | 95.7 | 104.3 | 10.94 | 9.52 | | High | High | High | High | High | High | High | High | High | High | High | 0 | 2.01E-07 | 3.44 |
| FALSE | High | NLDLASPEI | [K].NLDLAS | 1xPhospho | B2RRY4 1x | 0.117405 | 0.009523 | 1 | 2 | 2 | B2RRY4 | Membrane | B2RRY4 [2 | 0 | 1913.966 | 0 | 0.75 | 0.004815 | 2.06 | 76.8 | 123.2 | 12.39 | 3.95 | | High | High | High | High | High | High | High | High | High | High | High | 0.005599 | 0.06146 | 2.29 |
| FALSE | High | NLDPSEDE | [R].NLDPSE | 1xPhospho | Q149H7 1x | 3.76E-08 | 0 | 1 | 2 | 3 | Q149H7 | Myeloid le | Q149H7 [1 | 0 | 2243.048 | 0 | -0.67 | 0.000333 | 1.49 | 125.4 | 74.6 | 5.86 | 5.19 | | High | High | High | High | High | High | High | High | High | High | High | 0 | 2.54E-09 | 4.27 |
| FALSE | High | NLDPSEDE | [R].NLDPSE | 2xPhospho | Q149H7 2x | 8.15E-07 | 0 | 1 | 2 | 3 | Q149H7 | Myeloid le | Q149H7 [1 | 0 | 2323.014 | 0 | -1.01 | 0.000669 | 2.99 | 133.5 | 66.5 | 6.89 | 6.76 | | High | High | High | High | High | High | High | High | High | High | High | 0 | 1.42E-07 | 4.33 |
| FALSE | High | NLEAPYIPD | [R].NLEAPY | 1xPhospho | B2RQQ7 1x | 0.000407 | 0 | 1 | 2 | 4 | B2RQQ7 | CDC42 bin | B2RQQ7 [3 | 0 | 3337.466 | 0 | 0.94 | 0.002863 | 7.88 | 71.6 | 128.4 | 9.08 | 13.37 | | High | High | High | High | High | High | High | High | High | High | High | 2.58E-05 | 0.000343 | 3.28 |
| FALSE | High | NLEEAAAE | [R].NLEEA | 2xCarbami | Q9WU40-2 | 4.12E-05 | 0 | 1 | 4 | 1 | Q9WU40-2 | Isoform 2 c | Q9WU40-2 | 0 | 2908.244 | 0 | 0.03 | 0.742743 | 9.36 | 103.1 | 96.9 | 10.65 | 17.52 | | High | High | High | High | High | High | High | High | High | High | High | 0 | 6.62E-06 | 4.83 |
| FALSE | High | NLEQILNG | [K].NLEQIL | 1xPhospho | Q9ERG2 1x | 1.26E-05 | 0 | 2 | 5 | 2 | Q9ERG2; B | Striatin-3 C | Q9ERG2 [2 | 0 | 1707.852 | 0 | 0.4 | 0.699095 | 10.74 | 89.9 | 110.1 | 36.59 | 42.07 | Shared | High | High | High | High | High | High | High | High | High | High | High | 0 | 1.75E-06 | 3.76 |
| FALSE | High | NLEQILNG | [K].NLEQIL | 1xPhospho | Q9ERG2 1x | 1.4E-08 | 0 | 2 | 5 | 2 | Q9ERG2; B | Striatin-3 C | Q9ERG2 [2 | 0 | 1937.015 | 0 | 0.23 | 0.072719 | 10.77 | 91.6 | 108.4 | 7.17 | 7.27 | Shared | High | High | High | High | High | High | High | High | High | High | High | 0 | 8.4E-10 | 3.69 |
| FALSE | High | NLEQILNG | [K].NLEQIL | 1xDeamida | Q9ERG2 1x | 7.03E-09 | 0 | 2 | 5 | 9 | Q9ERG2; B | Striatin-3 C | Q9ERG2 [2 | 0 | 1937.999 | 0 | 0.22 | 0.101242 | 2.89 | 92.2 | 107.8 | 8.36 | 9.54 | Shared | High | High | High | High | High | High | High | High | High | High | High | 0 | 4.2E-06 | 4.17 |
| FALSE | High | NLETLPSFS | [R].NLETLP | 2xPhospho | E9PYL2 2x | 0.000852 | 0 | 1 | 2 | 4 | E9PYL2 | Proline-ricl | E9PYL2 [13 | 0 | 2586.167 | 0 | -0.31 | 0.041658 | 11.51 | 110.8 | 89.2 | 2.12 | 13.27 | | High | High | High | High | High | High | High | High | High | High | High | 0 | 0.000199 | 3.19 |
| FALSE | High | NLFNQEMI | [R].NLFNQ | 1xPhospho | Q8BQ33 1x | 9.48E-06 | 0 | 1 | 3 | 1 | Q8BQ33 | Treslin OS= | Q8BQ33 [9 | 0 | 1961.981 | 0 | -0.97 | 0.001286 | 3.48 | 134.7 | 65.3 | 5.9 | 8.23 | | High | High | High | High | High | High | High | High | High | High | High | 0 | 1.28E-06 | 4.58 |
| FALSE | High | NIGRDTPTS | [K].NIGRDT | 1xPhospho | F8WI88 1x | 3.73E-05 | 0 | 1 | 4 | 1 | F8WI88 | PEST prote | F8WI88 [1 | 1 | 2314.159 | 0 | 0.89 | 0.009854 | 10.25 | 62.9 | 137.1 | 18.12 | 21.67 | | High | High | High | High | High | High | High | High | High | High | High | 0 | 5.92E-06 | 2.5 |
| FALSE | High | NLGSINTEL | [R].NLGSIN | 1xPhospho | O08547 1x | 3.18E-06 | 0 | 1 | 3 | 3 | O08547 | Vesicle-tra | O08547 [1 | 0 | 1895.943 | 0 | 0.12 | 0.963715 | 14.8 | 98.1 | 101.9 | 12.4 | 12.11 | | High | High | High | High | High | High | High | High | High | High | High | 0 | 3.72E-07 | 4.11 |
| FALSE | High | NIGYESASV | [K].NIGYES | 2xCarbami | Q9R0L6-2 | 1.11E-05 | 0 | 1 | 3 | 3 | Q9R0L6-2 | Isoform 2 c | Q9R0L6-2 | 0 | 2427.097 | 0 | 0.31 | 0.089289 | 3.19 | 90.6 | 109.4 | 10.71 | 11.87 | | High | High | High | High | High | High | High | High | High | High | High | 0 | 6.18E-06 | 3.98 |
| FALSE | High | NIGYESASV | [K].NIGYES | 2xCarbami | Q9R0L6-2 | 0.005717 | 0.00062 | 1 | 3 | 1 | Q9R0L6-2 | Isoform 2 c | Q9R0L6-2 | 0 | 2507.064 | 0 | 0.22 | 0.178794 | 6.08 | 90.2 | 109.8 | 7.48 | 26.39 | | High | High | High | High | High | High | High | High | High | High | High | 0.000275 | 0.001679 | 2.74 |
| FALSE | High | NLLDLFSPC | [R].NLLDLF | 1xPhospho | Q3TVV9 1x | 0.000548 | 0 | 1 | 4 | 1 | Q3TVV9 | Putative ur | Q3TVV9 [4 | 0 | 2765.468 | 0 | -0.21 | 0.78074 | 20.98 | 110.9 | 89.1 | 11.97 | 23.67 | | High | High | High | High | High | High | High | High | High | High | High | 0 | 0.000122 | 3.98 |
| FALSE | High | NLSFDELYF | [K].NLSFDE | 1xPhospho | Q8CJF7 1x | 9.62E-05 | 0 | 1 | 1 | 1 | Q8CJF7 | Protein ELY | Q8CJF7 [15 | 0 | 1904.924 | 0 | -0.7 | 0.4506 | 7.02 | 124.8 | 75.2 | 73.73 | 64.99 | | High | High | High | High | High | High | High | High | High | High | High | 0 | 1.73E-05 | 3.33 |
| FALSE | High | NSCNVGG | [R].NSCNV | 1xCarbami | Q8K3A9 1x | 1.46E-07 | 0 | 1 | 1 | 2 | Q8K3A9 | 7SK snRNA | Q8K3A9 [1 | 0 | 1865.862 | 0 | 0.2 | 0.162014 | 13.54 | 93.2 | 106.8 | 12.28 | 7.52 | | High | High | High | High | High | High | High | High | High | High | High | 0 | 7.7E-06 | 3.59 |
| FALSE | High | NLSFDELYF | [K].NLSFDE | 1xPhospho | Q8CJF7 1x | 0.010109 | 0.000736 | 1 | 1 | 3 | Q8CJF7 | Protein ELY | Q8CJF7 [15 | 0 | 2134.087 | 0 | -0.86 | 0.000447 | 1.65 | 127.3 | 72.7 | 4.15 | 5.11 | | High | High | High | High | High | High | High | High | High | High | High | 0.001214 | 0.01307 | 3.25 |
| FALSE | High | NLSFNITED | [K].NLSFNI | 1xPhospho | P09405 1x | 0.002398 | 0.000109 | 1 | 6 | 1 | P09405 | Nucleolin ( | P09405 [40 | 0 | 1731.84 | 0 | -0.86 | 0.245125 | 13.2 | 129.4 | 70.6 | 65.82 | 50.52 | | High | High | High | High | High | High | High | High | High | High | High | 4.94E-05 | 0.000639 | 3.75 |
| FALSE | High | NMAPGAG | [R].NMAPG | 1xCarbami | B9EKJ7 1x | 0.002153 | 5.66E-05 | 1 | 3 | 1 | B9EKJ7 | Tankyrase | B9EKJ7 [12 | 0 | 1709.734 | 0 | 0.73 | 0.045613 | 18.07 | 77.3 | 122.7 | 16.87 | 16.12 | | High | High | High | High | High | High | High | High | High | High | High | 2.58E-05 | 0.000565 | 2.47 |
| FALSE | High | NMAPGAG | [R].NMAPG | 1xCarbami | B9EKJ7 1x | 0.000462 | 0 | 1 | 3 | 3 | B9EKJ7 | Tankyrase | B9EKJ7 [12 | 0 | 1725.729 | 0 | 0.42 | 0.008737 | 5.09 | 83.8 | 116.2 | 11.1 | 5.2 | | High | High | High | High | High | High | High | High | High | High | High | 0 | 0.0001 | 2.79 |
| FALSE | High | NMAPSQQ | [R].NMAPS | 1xPhospho | Q6AXD2 1x | 0.065952 | 0.004766 | 1 | 5 | 2 | Q6AXD2 | Abi2 prote | Q6AXD2 [2 | 0 | 1539.719 | 0 | -1.26 | 0.013448 | 5.71 | 140.7 | 59.3 | 11.99 | 25.4 | | High | High | High | High | High | High | High | High | High | High | High | 0.002455 | 0.02662 | 1.99 |
| FALSE | High | NMDQGG | [R].NMDQ | 1xPhospho | E9PV45 1x | 0.004654 | 0.000456 | 1 | 2 | 1 | E9PV45 | Ubiquitin c | E9PV45 [1 | 0 | 1400.583 | 0 | -0.1 | 0.138464 | 12.67 | 107.7 | 92.3 | 7.44 | 10.17 | | High | High | High | High | High | High | High | High | High | High | High | 0.000204 | 0.001332 | 0.92 |
| FALSE | High | NMGGPYG | [R].NMGGI | 1xPhospho | O88569-3 | 1.36E-09 | 0 | 2 | 4 | 24 | O88569-3; | Isoform 3 c | O88569-3 | 0 | 2515.03 | 0 | 0.06 | 0.169251 | 7.47 | 97.8 | 102.2 | 4.43 | 6.7 | Shared | High | High | High | High | High | High | High | High | High | High | High | 0 | 3.31E-06 | 5.16 |
| FALSE | High | NMPNVSD | [R].NMPN | 1xPhospho | Q8R2M2 1 | 1.45E-08 | 0 | 1 | 1 | 6 | Q8R2M2 | Deoxynucl | Q8R2M2 [2 | 0 | 2686.081 | 0 | -0.18 | 0.073671 | 3.97 | 103.9 | 96.1 | 6.36 | 9.88 | | High | High | High | High | High | High | High | High | High | High | High | 0 | 2.06E-09 | 5.34 |
| FALSE | High | NMPNVSD | [R].NMPN | 1xPhospho | Q8R2M2 1 | 3.39E-09 | 0 | 1 | 1 | 14 | Q8R2M2 | Deoxynucl | Q8R2M2 [2 | 0 | 2702.076 | 0 | 0.12 | 0.103628 | 4.11 | 97.1 | 102.9 | 6.09 | 4.78 | | High | High | High | High | High | High | High | High | High | High | High | 0 | 1.7E-10 | 6.48 |
| FALSE | High | NMPNVSD | [R].NMPN | 2xPhospho | Q8R2M2 2 | 2.58E-08 | 0 | 1 | 1 | 6 | Q8R2M2 | Deoxynucl | Q8R2M2 [2 | 0 | 2766.048 | 0 | -0.39 | 0.00194 | 4.13 | 114.5 | 85.5 | 4.8 | 6.08 | | High | High | High | High | High | High | High | High | High | High | High | 0 | 1.67E-09 | 5.2 |
| FALSE | High | NMPNVSD | [R].NMPN | 1xDeamida | Q8R2M2 2 | 3.27E-06 | 0 | 1 | 1 | 1 | Q8R2M2 | Deoxynucl | Q8R2M2 [2 | 0 | 2767.032 | 0 | -0.67 | 0.000675 | 3.03 | 119.8 | 80.2 | 7.09 | 14.37 | | High | High | High | High | High | High | High | High | High | High | High | 0 | 3.86E-07 | 4.37 |
| FALSE | High | NMPNVSD | [R].NMPN | 2xPhospho | Q8R2M2 2 | 1.26E-07 | 0 | 1 | 1 | 19 | Q8R2M2 | Deoxynucl | Q8R2M2 [2 | 0 | 2782.043 | 0 | -0.06 | 0.046805 | 4.79 | 102.5 | 97.5 | 1.49 | 4.44 | | High | High | High | High | High | High | High | High | High | High | High | 0 | 9.91E-09 | 5.68 |
| FALSE | High | NMPNVSD | [R].NMPN | 3xPhospho | Q8R2M2 3 | 0.010568 | 0.000772 | 1 | 1 | 2 | Q8R2M2 | Deoxynucl | Q8R2M2 [2 | 0 | 2846.014 | 0 | -0.19 | 0.151142 | 6.89 | 106.2 | 93.8 | 17.75 | 9.38 | | High | High | High | High | High | High | High | High | High | High | High | 0.000382 | 0.003341 | 3.53 |
| FALSE | High | NMPSLSAA | [R].NMPSL | 1xPhospho | Q3UZ39 1x | 0.000207 | 0 | 1 | 10 | 1 | Q3UZ39 | Leucine-ric | Q3UZ39 [6 | 0 | 2130.046 | 0 | 1.86 | 0.018724 | 8.67 | 46.1 | 153.9 | 15.81 | 29.63 | | High | High | High | High | High | High | High | High | High | High | High | 0 | 4.07E-05 | 3.4 |
| FALSE | High | NMPSLSAA | [R].NMPSL | 1xPhospho | Q3UZ39 1x | 0.000111 | 0 | 1 | 10 | 3 | Q3UZ39 | Leucine-ric | Q3UZ39 [6 | 0 | 2146.041 | 0 | 1.28 | 0.001537 | 11.32 | 58.4 | 141.6 | 11.59 | 14.75 | | High | High | High | High | High | High | High | High | High | High | High | 0 | 7.07E-05 | 3.66 |
| FALSE | High | NMALLSQL | [K].NMALL | 1xPhospho | P70372 1x | 1.59E-06 | 0 | 1 | 7 | 2 | P70372 | ELAV-like p | P70372 [1 | 0 | 1925.951 | 0 | 0.31 | 0.015616 | 5.26 | 88.6 | 111.4 | 6.14 | 6.85 | | High | High | High | High | High | High | High | High | High | High | High | 0 | 1.71E-07 | 4.29 |
| FALSE | High | NMQLSNTI | [K].NMQLS | 1xPhospho | B2RR82 1x | 0.053473 | 0.00384 | 1 | 9 | 1 | B2RR82 | Intersectin | B2RR82 [5 | 0 | 2409.225 | 0 | 0.27 | 0.066272 | 11.88 | 90.8 | 109.2 | 7.72 | 7.5 | | High | High | High | High | High | High | High | High | High | High | High | 0.001999 | 0.02073 | 2.33 |
| FALSE | High | NMSPDLR | [R].NMSPD | 1xPhospho | Q9QYB5 1x | 0.102567 | 0.008249 | 1 | 6 | 1 | Q9QYB5 | Gamma-ad | Q9QYB5 [4 | 0 | 1157.522 | 0 | 1.55 | 0.001214 | 12.71 | 51.2 | 148.8 | 6.96 | 11.51 | | High | High | High | High | High | High | High | High | High | High | High | 0.004138 | 0.0443 | 2.02 |
| FALSE | High | NMTSPSFL | [R].NMTSP | 1xPhospho | Q8BLM0 1 | 3.6E-08 | 0 | 1 | 1 | 3 | Q8BLM0 | Krueppel-li | Q8BLM0 [4 | 0 | 3033.487 | 0 | -3.88 | | 0.12 | 183.6 | 16.4 | 17.44 | 21.07 | | High | High | High | High | High | Not Found | High | High | High | High | High | 0 | 7.08E-09 | 5.16 |
| FALSE | High | NMTSPSFL | [R].NMTSP | 1xPhospho | Q8BLM0 1 | 6.3E-05 | 0 | 1 | 1 | 5 | Q8BLM0 | Krueppel-li | Q8BLM0 [4 | 0 | 3049.482 | 0 | -0.14 | 0.078996 | 11.08 | 106.3 | 93.7 | 10.4 | 14.33 | | High | High | High | High | High | High | High | High | High | High | High | 0 | 1.07E-05 | 4.01 |
| FALSE | High | NMTVEQLI | [K].NMTVE | 1xPhospho | Q6ZPJ3 1x | 4.43E-05 | 0 | 1 | 1 | 2 | Q6ZPJ3 | (E3-indepe | Q6ZPJ3 [82 | 0 | 3150.616 | 0 | 0.19 | 0.51159 | 5.29 | 96.8 | 103.2 | 14 | 8.96 | | High | High | High | High | High | High | High | High | High | High | High | 0 | 7.16E-06 | 3.53 |
| FALSE | High | NMTVEQLI | [K].NMTVE | 1xPhospho | Q6ZPJ3 1x | 0.000244 | 0 | 1 | 1 | 1 | Q6ZPJ3 | (E3-indepe | Q6ZPJ3 [82 | 0 | 3166.611 | 0 | 0.29 | 0.025645 | 10.78 | 88.2 | 111.8 | 12.44 | 5.69 | | High | High | High | High | High | High | High | High | High | High | High | 0 | 4.89E-05 | 3.7 |
| FALSE | High | NNAYLAQS | [K].NNAYL | 1xPhospho | Q3TF87 1x | 4.23E-05 | 0 | 1 | 5 | 1 | Q3TF87 | Putative ur | Q3TF87 [2 | 0 | 2048.062 | 0 | -0.68 | 0.007666 | 5.75 | 123.1 | 76.9 | 5.86 | 10.3 | | High | High | High | High | High | High | High | High | High | High | High | 0 | 6.83E-06 | 3.16 |
| FALSE | High | NNAYPHK | [R].NNAYP | 1xPhospho | Q9CW64 1 | 0.097743 | 0.007804 | 1 | 3 | 1 | Q9CW64 | Putative ur | Q9CW64 [ | 0 | 1381.703 | 0 | -1.06 | 0.001931 | 3.82 | 134 | 66 | 1.67 | 18.13 | | High | High | High | High | High | High | High | High | High | High | High | 0.00395 | 0.04202 | 1.98 |
| FALSE | High | NNEAEPSG | [K].NNEAE | 1xPhospho | D3Z482 1x | 5.75E-08 | 0 | 1 | 2 | 1 | D3Z482 | Ankyrin re | D3Z482 [5 | 0 | 4098.868 | 0 | 0.6 | 0.008516 | 10.63 | 79.1 | 120.9 | 5.36 | 11.38 | | High | High | High | High | High | High | High | High | High | High | High | 0 | 4.11E-09 | 5.93 |
| FALSE | High | NNLSLGDA | [R].NNLSL | 1xPhospho | Q8BJ56 1x | 0.001373 | 5.66E-05 | 1 | 3 | 2 | Q8BJ56 | Patatin-like | Q8BJ56 [42 | 0 | 1424.735 | 0 | 0.61 | 0.179147 | 12.53 | 79.3 | 120.7 | 24.16 | 28.75 | | High | High | High | High | High | High | High | High | High | High | High | 0.000161 | 0.001235 | 2.7 |
| FALSE | High | NNLSLGDA | [R].NNLSL | 1xPhospho | Q8BJ56 1x | 0.000362 | 0 | 1 | 3 | 1 | Q8BJ56 | Patatin-like | Q8BJ56 [42 | 0 | 1653.898 | 0 | 1.48 | 0.010551 | 14.17 | 52.4 | 147.6 | 12.01 | 22.71 | | High | High | High | High | High | High | High | High | High | High | High | 0 | 7.63E-05 | 3 |
| FALSE | High | NNPDEEAN | [K].NNPDE | 1xPhospho | P13405 1x | 5.94E-10 | 0 | 1 | 4 | 3 | P13405 | Retinoblast | P13405 [35 | 0 | 2198.044 | 0 | 0.96 | 0.000507 | 2.24 | 65.4 | 134.6 | 4.12 | 5.15 | | High | High | High | High | High | High | High | High | High | High | High | 0 | 2.42E-11 | 5.91 |
| FALSE | High | NNPDEEAN | [K].NNPDE | 2xPhospho | P13405 2x | 9.42E-07 | 0 | 1 | 4 | 2 | P13405 | Retinoblast | P13405 [35 | 0 | 2278.01 | 0 | 1.22 | 0.018843 | 21.07 | 61.1 | 138.9 | 10.2 | 31.23 | | High | High | High | High | High | High | High | High | High | High | High | 0 | 1.18E-06 | 4.51 |
| FALSE | High | NNRPAFFS | [R].NNRPA | 1xPhospho | B1APX2 1x | 0.106748 | 0.008634 | 1 | 3 | 1 | B1APX2 | RIKEN cDN | B1APX2 [3 | 0 | 1686.857 | 0 | 0.28 | 0.25596 | 42.38 | 88.2 | 111.8 | 18.76 | 34.21 | | High | High | High | High | High | High | High | High | High | High | High | 0.004327 | 0.0468 | 2.16 |
| FALSE | High | NMSPDEIK | [R].NMSPD | 1xPhospho | Q02614 1x | 0.067619 | 0.004864 | 1 | 1 | 2 | Q02614 | SAP30-bin | Q02614 [1 | 0 | 1487.722 | 0 | -0.1 | 0.668253 | 4.85 | 102.3 | 97.7 | 3.62 | 10.53 | | High | High | High | High | High | High | High | High | High | High | High | 0.00253 | 0.02732 | 2.04 |
| FALSE | High | NMALLSQL | [K].NMALL | 1xPhospho | P70372 1x | 1.28E-06 | 0 | 1 | 7 | 1 | P70372 | ELAV-like p | P70372 [1 | 0 | 1909.956 | 0 | 0.15 | 0.075616 | 7.27 | 94.4 | 105.6 | 8.89 | 7.69 | | High | High | High | High | High | High | High | High | High | High | High | 0 | 1.34E-07 | 4.81 |
| FALSE | High | NIYCLENCS | [K].NIYCLE | 2xCarbami | A2AKX3 1x | 0.009416 | 0.000736 | 1 | 2 | 1 | A2AKX3 | Probable h | A2AKX3 [6 | 0 | 2209.043 | 0 | 0.24 | 0.218983 | 12.16 | 93.7 | 106.3 | 8.26 | 9.97 | | High | High | High | High | High | High | High | High | High | High | High | 0.000327 | 0.002945 | 2.84 |
| FALSE | High | NIVVVTSDS | [K].NIVVVT | 1xCarbami | Q8BJW7 2x | 0.000241 | 0 | 1 | 1 | 1 | Q8BJW7 | Crossover | Q8BJW7 [4 | 0 | 2675.28 | 0 | -0.5 | 0.025744 | 15.61 | 117.2 | 82.8 | 14.72 | 19.41 | | High | High | High | High | High | High | High | High | High | High | High | 0 | 4.83E-05 | 4 |
| FALSE | High | NLSFNITED | [K].NLSFNI | 1xPhospho | P09405 1x | 5.19E-07 | 0 | 1 | 6 | 6 | P09405 | Nucleolin ( | P09405 [40 | 0 | 1961.003 | 0 | -1.29 | 0.000177 | 0.69 | 141 | 59 | 4.84 | 6.37 | | High | High | High | High | High | High | High | High | High | High | High | 0 | 5.85E-07 | 3.38 |
| FALSE | High | NLSGSTLYF | [R].NLSGST | 1xPhospho | F8VPJ6 1x | 0.004333 | 0.00036 | 1 | 3 | 1 | F8VPJ6 | Regulatory | F8VPJ6 [11 | 0 | 1926.989 | 0 | -0.94 | 0.001542 | 4.28 | 130.3 | 69.7 | 6.52 | 8.09 | | High | High | High | High | High | High | High | High | High | High | High | 0.000183 | 0.001238 | 1.98 |
| FALSE | High | NLSLEDMS | [K].NLSLED | 2xCarbami | Q3TU15 1x | 2.87E-08 | 0 | 2 | 4 | 5 | Q3TU15; C | Non-specif | Q3TU15 [3 | 0 | 3644.61 | 0 | -1.08 | 0.001371 | 4.29 | 131.4 | 68.6 | 11.53 | 29.64 | Shared | High | High | High | High | High | High | High | High | High | High | High | 0 | 1.89E-09 | 5.25 |
| FALSE | High | NLSLEDMS | [K].NLSLED | 2xCarbami | Q3TU15 1x | 2.03E-07 | 0 | 2 | 4 | 7 | Q3TU15; C | Non-specif | Q3TU15 [3 | 0 | 3660.604 | 0 | -1.11 | 0.005732 | 9.02 | 139 | 61 | 7.64 | 23.93 | Shared | High | High | High | High | High | High | High | High | High | High | High | 0 | 5.52E-07 | 4.78 |
| FALSE | High | NISLSSEEE | [K].NISLSS | 2xPhospho | Q5U3K5 2x | 7.98E-06 | 0 | 1 | 1 | 5 | Q5U3K5 | Rab-like pr | Q5U3K5 [4 | 0 | 2285.005 | 0 | -0.25 | 0.015342 | 2.82 | 108.3 | 91.7 | 8.05 | 3.95 | | High | High | High | High | High | High | High | High | High | High | High | 0 | 1.05E-06 | 3.66 |
| FALSE | High | NISLSSEEE | [K].NISLSS | 3xPhospho | Q5U3K5 3x | 0.001087 | 0 | 1 | 1 | 2 | Q5U3K5 | Rab-like pr | Q5U3K5 [4 | 0 | 2364.971 | 0 | 0.56 | 0.004058 | 13.41 | 82.2 | 117.8 | 5.85 | 6.22 | | High | High | High | High | High | High | High | High | High | High | High | 0 | 0.000261 | 3.65 |
| FALSE | High | NLSNSPNSI | [R].NLSNSF | 1xPhospho | Q3UL97 1x | 0.001799 | 5.66E-05 | 1 | 3 | 2 | Q3UL97 | Activating t | Q3UL97 [2 | 0 | 1758.886 | 0 | -1.9 | 0.000267 | 1.29 | 161.3 | 38.7 | 5.38 | 17.54 | | High | High | High | High | High | High | High | High | High | High | High | 2.58E-05 | 0.000461 | 2.61 |
| FALSE | High | NLSPAAVIN | [K].NLSPAA | 1xPhospho | Q9DBR4-2 | 3.74E-06 | 0 | 2 | 11 | 1 | Q9DBR4-2; | Isoform 2 c | Q9DBR4-2 | 0 | 1995.093 | 0 | -0.3 | 0.044563 | 2.08 | 106.2 | 93.8 | 8.67 | 5.07 | Shared | High | High | High | High | High | High | High | High | High | High | High | 0 | 4.47E-07 | 3.52 |
| FALSE | High | NLSPAPASS | [R].NLSPAF | 1xPhospho | Q7TQ95 1x | 3.64E-09 | 0 | 1 | 3 | 1 | Q7TQ95 | Protein lun | Q7TQ95 [1 | 0 | 2863.476 | 0 | 1.73 | 0.002224 | 13.84 | 54 | 146 | 27.18 | 19.95 | | High | High | High | High | High | High | High | High | High | High | High | 0 | 1.85E-10 | 5.32 |
| FALSE | High | NLSPGAVE | [K].NLSPGA | 1xPhospho | F8WHL2 1x | 0.005044 | 0.000504 | 1 | 4 | 3 | F8WHL2 | Coatomer s | F8WHL2 [1 | 0 | 1552.757 | 0 | -0.23 | 0.016125 | 2.42 | 108 | 92 | 3.07 | 6.9 | | High | High | High | High | High | High | High | High | High | High | High | 0.000246 | 0.001458 | 2.21 |
| FALSE | High | NISPGLHN | [R].NISPGL | 1xDeamida | D3Z5N5 1x | 0.004955 | 0.000504 | 2 | 3 | 1 | D3Z5N5; Q | 1-phospha | D3Z5N5 [1 | 0 | 1704.872 | 0 | 0.16 | 0.071154 | 6.27 | 94.5 | 105.5 | 9.4 | 5.74 | Shared | High | High | High | High | High | High | High | High | High | High | High | 0.000225 | 0.001437 | 1.76 |
| FALSE | High | NLSPPSVSS | [K].NLSPPS | 1xPhospho | Q8BFU3-4 | 0.001916 | 5.66E-05 | 1 | 4 | 1 | Q8BFU3-4 | Isoform 4 c | Q8BFU3-4 | 0 | 2027.065 | 0 | -0.4 | 0.044834 | 7.91 | 114.8 | 85.2 | 7.83 | 19.05 | | High | High | High | High | High | High | High | High | High | High | High | 2.58E-05 | 0.000493 | 2.98 |
| FALSE | High | NLSPPSVSS | [K].NLSPPS | 1xPhospho | Q8BFU3-4 | 1.91E-05 | 0 | 1 | 4 | 2 | Q8BFU3-4 | Isoform 4 c | Q8BFU3-4 | 0 | 2043.06 | 0 | -0.29 | 0.589842 | 14.35 | 103.7 | 96.3 | 15.61 | 17.94 | | High | High | High | High | High | High | High | High | High | High | High | 0 | 2.8E-06 | 3.36 |
| FALSE | High | NISPSTNSV | [K].NISPSTI | 1xPhospho | Q6NXH3-2 | 0.001916 | 5.66E-05 | 1 | 3 | 1 | Q6NXH3-2 | Isoform 2 c | Q6NXH3-2 | 0 | 1584.84 | 0 | -0.66 | 0.005132 | 4.38 | 122.9 | 77.1 | 6.93 | 8.14 | | High | High | High | High | High | High | High | High | High | High | High | 2.58E-05 | 0.000494 | 2.39 |
| FALSE | High | NISSDDSED | [R].NISSDD | 1xPhospho | Q6ZPV2 1x | 0.074685 | 0.005727 | 1 | 3 | 3 | Q6ZPV2 | DNA helica | Q6ZPV2 [4 | 0 | 3634.741 | 0 | -0.49 | 0.006462 | 7.54 | 119.5 | 80.5 | 7.77 | 16.87 | | High | High | High | High | High | High | High | High | High | High | High | 0.002937 | 0.03062 | 3.27 |
| FALSE | High | NLSSTTDDI | [R].NLSSTT | 1xPhospho | P57016 1x | 0.067059 | 0.004796 | 1 | 2 | 3 | P57016 | Ladinin-1 C | P57016 [36 | 0 | 1831.873 | 0 | -1.03 | 0.000396 | 1.95 | 131.7 | 68.3 | 4.52 | 9.24 | | High | High | High | High | High | High | High | High | High | High | High | 0.002455 | 0.02691 | 2.3 |
| FALSE | High | NLSTDASTI | [R].NLSTDA | 1xPhospho | Q6A0A2 1x | 2.42E-05 | 0 | 1 | 2 | 2 | Q6A0A2 | La-related | Q6A0A2 [5 | 0 | 2037.022 | 0 | 1.17 | 0.000255 | 3.96 | 61.7 | 138.3 | 13.34 | 5.39 | | High | High | High | High | High | High | High | High | High | High | High | 0 | 3.64E-06 | 3.87 |
| FALSE | High | NLSTEALK | [R].NLSTEA | 1xPhospho | Q5SYL3 1x | 0.029844 | 0.001857 | 1 | 1 | 1 | Q5SYL3 | Protein KIA | Q5SYL3 [1 | 0 | 1413.775 | 0 | 0.58 | 0.004418 | 20.62 | 80.2 | 119.8 | 8.16 | 7.34 | | High | High | High | High | High | High | High | High | High | High | High | 0.001011 | 0.01076 | 1.4 |
| FALSE | High | NLSYGDNS | [R].NLSYGI | 1xPhospho | H3BKJ0 1x | 1.06E-06 | 0 | 1 | 1 | 3 | H3BKJ0 | Disks large | H3BKJ0 [5 | 0 | 1861.805 | 0 | 0.67 | 0.047157 | 11.7 | 79.6 | 120.4 | 25.57 | 9.71 | | High | High | High | High | High | High | High | High | High | High | High | 0 | 1.52E-07 | 3.26 |
| FALSE | High | NLTANDPS | [R].NLTANI | 1xCarbami | Q9DA19 1x | 2.9E-09 | 0 | 1 | 1 | 3 | Q9DA19 | Corepresso | Q9DA19 [1 | 0 | 3523.586 | 0 | -0.39 | 0.002085 | 2.41 | 115.5 | 84.5 | 4.1 | 5.28 | | High | High | High | High | High | High | High | High | High | High | High | 0 | 1.44E-10 | 6.45 |
| FALSE | High | NLTEQNSY | [K].NLTEQI | 1xPhospho | Q6A0A9 1x | 0.057203 | 0.004227 | 1 | 1 | 1 | Q6A0A9 | Constitutiv | Q6A0A9 [4 | 1 | 3265.604 | 0 | 2.29 | 0.004376 | 28.25 | 34 | 166 | 28.64 | 18.06 | | High | High | High | High | High | High | High | High | High | High | High | 0.002159 | 0.02241 | 2.5 |
| FALSE | High | NLTGLAED | [R].NLTGLA | 1xPhospho | Q9ERV1 1x | 0.008538 | 0.000736 | 1 | 2 | 1 | Q9ERV1 | Probable E | Q9ERV1 [1 | 1 | 2238.235 | 0 | 0.32 | 0.007704 | 8.28 | 85.6 | 114.4 | 7.4 | 6.23 | | High | High | High | High | High | High | High | High | High | High | High | 0.000327 | 0.002625 | 2.22 |
| FALSE | High | NITGSSPVA | [K].NITGSS | 1xPhospho | H3BK65 1x | 0.003372 | 0.00016 | 1 | 4 | 2 | H3BK65 | Epidermal | H3BK65 [3 | 0 | 2045.072 | 0 | 0.28 | 0.824026 | 7.88 | 102.9 | 97.1 | 23.45 | 37.38 | | High | High | High | High | High | High | High | High | High | High | High | 7.19E-05 | 0.000929 | 3.09 |
| FALSE | High | NLTMSPLH | [R].NLTMS | 1xPhospho | A0A0A0MC | 0.036416 | 0.002419 | 1 | 2 | 1 | A0A0A0MC | Histone-lys | A0A0A0MC | 0 | 1594.843 | 0 | 0.17 | 0.122821 | 6.06 | 93.6 | 106.4 | 6.33 | 10.93 | | High | High | High | High | High | High | High | High | High | High | High | 0.001308 | 0.01344 | 1.68 |
| FALSE | High | NLTSSPAM | [R].NLTSSP | 1xPhospho | B9EKJ7 1x | 0.000227 | 0 | 1 | 2 | 5 | B9EKJ7 | Tankyrase | B9EKJ7 [24 | 0 | 1972.982 | 0 | -0.06 | 0.684126 | 13.19 | 100.2 | 99.8 | 10.28 | 17.16 | | High | High | High | High | High | High | High | High | High | High | High | 2.58E-05 | 0.000542 | 3.57 |
| FALSE | High | NIVEPTSGI | [R].NIVEPT | 1xPhospho | E9PVX6 1x | 0.120211 | 0.009851 | 1 | 2 | 1 | E9PVX6 | Proliferatic | E9PVX6 [1 | 0 | 1857.976 | 0 | -0.04 | 0.867796 | 9.87 | 101.9 | 98.1 | 7.73 | 7.13 | | High | High | High | High | High | High | High | High | High | High | High | 0.004961 | 0.05385 | 2.46 |
| FALSE | High | NLVGALDE | [K].NLVGA | 1xPhospho | Q8C0J6 1x | 1.35E-06 | 0 | 1 | 1 | 2 | Q8C0J6 | Ankyrin re | Q8C0J6 [3 | 0 | 2023.917 | 0 | 1.7 | 0.000513 | 17.7 | 47.1 | 152.9 | 14.99 | 6.28 | | High | High | High | High | High | High | High | High | High | High | High | 0 | 1.42E-07 | 4.9 |
| FALSE | High | NLVSGGHL | [K].NLVSG | 1xPhospho | Q8K1G2 1x | 0.064862 | 0.004664 | 1 | 3 | 1 | Q8K1G2 | Limbin OS= | Q8K1G2 [3 | 0 | 1803.895 | 0 | 0.53 | 0.001718 | 6.94 | 79.7 | 120.3 | 12.15 | 10.48 | | High | High | High | High | High | High | High | High | High | High | High | 0.002432 | 0.02597 | 2.2 |
| FALSE | High | NLVSWESC | [K].NLVSW | 1xPhospho | O70439 1x | 3.01E-05 | 0 | 1 | 4 | 1 | O70439 | Syntaxin-7 | O70439 [1 | 0 | 3395.584 | 0 | -0.23 | 0.53261 | 19.53 | 105.7 | 94.3 | 11.38 | 14.21 | | High | High | High | High | High | High | High | High | High | High | High | 0 | 4.67E-06 | 3.72 |
| FALSE | High | NLSFEIK | [R].NLSFEI | 1xPhospho | P35831 1x | 0.010758 | 0.000772 | 1 | 1 | 2 | P35831 | Tyrosine-p | P35831 [43 | 0 | 1388.759 | 0 | 1.33 | 0.005618 | 29.27 | 56.5 | 143.5 | 2.36 | 19.14 | | High | High | High | High | High | High | High | High | High | High | High | 0.000585 | 0.005168 | 1.73 |
| FALSE | High | NSCSSPLR | [R].NSCSSF | 1xCarbami | O88845 1x | 0.048303 | 0.00354 | 1 | 4 | 2 | O88845 | A-kinase ar | O88845 [2 | 0 | 1229.555 | 0 | 0.09 | 0.231107 | 13.15 | 97.9 | 102.1 | 6.44 | 5.68 | | High | High | High | High | High | High | High | High | High | High | High | 0.005553 | 0.06027 | 1.23 |
| FALSE | High | NSDDAPW | [K].NSDDA | 1xPhospho | Q80TJ7 1x | 0.001783 | 5.66E-05 | 1 | 2 | 2 | Q80TJ7 | Histone lys | Q80TJ7 [8 | 0 | 1654.788 | 0 | -0.17 | 0.183446 | 7.17 | 104.3 | 95.7 | 9.79 | 11.25 | | High | High | High | High | High | High | High | High | High | High | High | 2.58E-05 | 0.000455 | 2.78 |
| FALSE | High | NSDEEAPK | [K].NSDEE | 1xPhospho | Q8C0D7 1x | 0.043981 | 0.003228 | 1 | 6 | 1 | Q8C0D7 | Inhibitor o | Q8C0D7 [1 | 0 | 1427.682 | 0 | -0.49 | 0.007617 | 3.64 | 113.7 | 86.3 | 10.53 | 8.85 | | High | High | High | High | High | High | High | High | High | High | High | 0.001665 | 0.01662 | 1.7 |
| FALSE | High | NSVQSGES | [K].NSVQS | 2xPhospho | E9PUJ6 2x | 2.13E-05 | 0 | 1 | 4 | 1 | E9PUJ6 | E3 ubiquiti | E9PUJ6 [17 | 1 | 3066.323 | 0 | -0.43 | 0.010081 | 4.72 | 111.9 | 88.1 | 9.38 | 10.79 | | High | High | High | High | High | High | High | High | High | High | High | 0 | 3.16E-06 | 2.66 |
| FALSE | High | NSVSINRP | [K].NSVSIN | 1xPhospho | Q80Y98 1x | 0.011146 | 0.000772 | 1 | 2 | 2 | Q80Y98 | Phospholip | Q80Y98 [4 | 0 | 2329.142 | 0 | 0.47 | 0.178786 | 12.7 | 94.4 | 105.6 | 22.18 | 10.48 | | High | High | High | High | High | High | High | High | High | High | High | 0.000399 | 0.003538 | 1.93 |
| FALSE | High | NSVSINRP | [K].NSVSIN | 1xPhospho | Q80Y98 1x | 0.011246 | 0.000772 | 1 | 2 | 1 | Q80Y98 | Phospholip | Q80Y98 [4 | 0 | 2558.305 | 0 | 0.68 | 0.000552 | 3.96 | 77 | 123 | 11.96 | 6.07 | | High | High | High | High | High | High | High | High | High | High | High | 0.000399 | 0.003565 | 2.83 |
| FALSE | High | NSVSPVGV | [R].NSVSP | 1xPhospho | Q6ZQ32 1x | 0.0004 | 0 | 1 | 2 | 2 | Q6ZQ32 | MKIAA085 | Q6ZQ32 [1 | 0 | 2393.273 | 0 | -1.51 | 0.00016 | 0.38 | 149.1 | 50.9 | 3.34 | 5.99 | | High | High | High | High | High | High | High | High | High | High | High | 0 | 8.48E-05 | 3.3 |
| FALSE | High | NSWDSPAI | [R].NSWDS | 1xPhospho | E9PUA3 1x | 3.1E-07 | 0 | 1 | 4 | 2 | E9PUA3 | IQ motif ar | E9PUA3 [4 | 0 | 1916.874 | 0 | 0.77 | 0.005559 | 4.15 | 74.2 | 125.8 | 7.46 | 10.63 | | High | High | High | High | High | High | High | High | High | High | High | 0 | 2.74E-08 | 3.36 |
| FALSE | High | NSYSNSQA | [K].NSYSNS | 1xPhospho | Q9ESC8 1x | 0.004451 | 0.000409 | 1 | 2 | 1 | Q9ESC8 | AF4/FMR2 | Q9ESC8 [1 | 0 | 1902.88 | 0 | 0.1 | 0.970584 | 59.71 | 96.7 | 103.3 | 30.1 | 22.54 | | High | High | High | High | High | High | High | High | High | High | High | 0.000183 | 0.001268 | 2.45 |
| FALSE | High | NSYSNSQA | [K].NSYSNS | 1xPhospho | Q9ESC8 1x | 7.06E-07 | 0 | 1 | 2 | 1 | Q9ESC8 | AF4/FMR2 | Q9ESC8 [1 | 0 | 2132.043 | 0 | -0.45 | 0.000961 | 2.6 | 114.3 | 85.7 | 2.38 | 4.3 | | High | High | High | High | High | High | High | High | High | High | High | 0 | 6.86E-08 | 3.17 |
| FALSE | High | NSYSNSQA | [K].NSYSNS | 1xDeamida | Q9ESC8 1x | 0.046291 | 0.003523 | 1 | 2 | 1 | Q9ESC8 | AF4/FMR2 | Q9ESC8 [1 | 0 | 2133.027 | 0 | -0.18 | 0.230684 | 14.79 | 105.5 | 94.5 | 19.12 | 7.36 | | High | High | High | High | High | High | High | High | High | High | High | 0.001789 | 0.0177 | 1.87 |
| FALSE | High | NTDNSDDI | [K].NTDNS | 1xPhospho | F8VQD1 1x | 0.013666 | 0.001011 | 1 | 9 | 1 | F8VQD1 | Protein pol | F8VQD1 [1 | 1 | 2649.183 | 0 | -0.04 | 0.829239 | 15.25 | 101.5 | 98.5 | 6.57 | 9.63 | | High | High | High | High | High | High | High | High | High | High | High | 0.000538 | 0.004444 | 3.32 |

| | | | | | | | | | | | | | | | | | | | | | | | | | | | | | | | | | | | | | | |
|---|---|---|---|---|---|---|---|---|---|---|---|---|---|---|---|---|---|---|---|---|---|---|---|---|---|---|---|---|---|---|---|---|---|---|---|---|---|---|
| FALSE | High | NTDNSDDI | [K].NTDNS | 2xPhospho | F8VQD1 2x | 0.047488 | 0.00354 | 1 | 9 | 1 | F8VQD1 | Protein pol | F8VQD1 [1 | 1 | 2729.149 | 0 | -0.06 | 0.316234 | 6.66 | 101.4 | 98.6 | 11.95 | 5.89 | | High | High | High | High | High | High | High | High | High | High | High | 0.001812 | 0.01825 | 2.6 |
| FALSE | High | NTDSHSAS | [R].NTDSH | 1xPhospho | Q6ZQF0 1x | 0.021797 | 0.001507 | 1 | 2 | 1 | Q6ZQF0 | DNA topois | Q6ZQF0 [8 | 0 | 1593.747 | 0 | 0.56 | 0.222593 | 26.68 | 86.4 | 113.6 | 28.54 | 13.44 | | High | High | High | High | High | High | High | High | High | High | High | 0.000814 | 0.007532 | 2.51 |
| FALSE | High | NTDSHSAS | [R].NTDSH | 1xPhospho | Q6ZQF0 1x | 0.000579 | 0 | 1 | 2 | 2 | Q6ZQF0 | DNA topois | Q6ZQF0 [8 | 0 | 1822.91 | 0 | 0.24 | 0.029107 | 7.13 | 93 | 107 | 5.64 | 4.07 | | High | High | High | High | High | High | High | High | High | High | High | 0 | 0.000129 | 2.68 |
| FALSE | High | NTETQPTS | [K].NTETQ | 1xPhospho | E9PZ26 1xP | 0.000181 | 0 | 1 | 3 | 2 | E9PZ26 | Bromodom | E9PZ26 [89 | 0 | 2040.041 | 0 | 0.44 | 0.056608 | 6.13 | 84.9 | 115.1 | 4.56 | 13.7 | | High | High | High | High | High | High | High | High | High | High | High | 0 | 3.5E-05 | 2.79 |
| FALSE | High | NSVPADEG | [K].NSVPAI | 1xPhospho | Q3UMG5-2 | 0.00015 | 0 | 1 | 2 | 1 | Q3UMG5-2 | Isoform 2 o | Q3UMG5-2 | 0 | 3129.566 | 0 | 0.57 | 0.035991 | 38.94 | 74.8 | 125.2 | 18.78 | 12.17 | | High | High | High | High | High | High | High | High | High | High | High | 0 | 2.82E-05 | 3.3 |
| FALSE | High | NTETVPSQ | [R].NTETVF | 2xPhospho | U5LK19 2xP | 2.56E-06 | 0 | 2 | 17 | 4 | U5LK19; E9 | Marshalin- | U5LK19 [33 | 0 | 2658.144 | 0 | -1.11 | 0.00118 | 4.08 | 136.8 | 63.2 | 6.63 | 10.05 | Shared | High | High | High | High | High | High | High | High | High | High | High | 0 | 2.94E-07 | 3.2 |
| FALSE | High | NTFTAWSE | [K].NTFTA\ | 1xPhospho | Q6ZQ58 1x | 1.51E-05 | 0 | 1 | 2 | 12 | Q6ZQ58 | La-related | Q6ZQ58 [5 | 0 | 2502.018 | 0 | 0.06 | 0.320776 | 6.1 | 97 | 103 | 5.95 | 8.76 | | High | High | High | High | High | High | High | High | High | High | High | 0 | 5.26E-06 | 3.69 |
| FALSE | High | NTFTAWSE | [K].NTFTA\ | 2xPhospho | Q6ZQ58 2x | 0.000178 | 0 | 1 | 2 | 8 | Q6ZQ58 | La-related | Q6ZQ58 [5 | 0 | 2581.985 | 0 | 0.7 | 0.007339 | 7.22 | 75.1 | 124.9 | 8.03 | 12.95 | | High | High | High | High | High | High | High | High | High | High | High | 0 | 3.43E-05 | 4.21 |
| FALSE | High | NTFTAWSE | [K].NTFTA\ | 1xPhospho | Q6ZQ58 1x | 1.69E-05 | 0 | 1 | 2 | 4 | Q6ZQ58 | La-related | Q6ZQ58 [5 | 1 | 3187.415 | 0 | -0.01 | 0.872418 | 9.41 | 102.1 | 97.9 | 6.43 | 3.78 | | High | High | High | High | High | High | High | High | High | High | High | 0 | 2.43E-06 | 4.15 |
| FALSE | High | NTFTAWSE | [K].NTFTA\ | 2xPhospho | Q6ZQ58 2x | 0.000941 | 0 | 1 | 2 | 2 | Q6ZQ58 | La-related | Q6ZQ58 [5 | 1 | 3267.381 | 0 | 0.32 | 0.006293 | 6.68 | 88.9 | 111.1 | 4.95 | 6.74 | | High | High | High | High | High | High | High | High | High | High | High | 0.00014 | 0.001119 | 3.39 |
| FALSE | High | NTGNGTQS | [R].NTGNG | 1xPhospho | B9EJA4 1xP | 2.53E-05 | 0 | 1 | 7 | 3 | B9EJA4 | Clasp2 pro | B9EJA4 [89 | 0 | 2384.123 | 0 | 0.01 | 0.720608 | 36.17 | 98 | 102 | 11.24 | 15.37 | | High | High | High | High | High | High | High | High | High | High | High | 0 | 3.85E-06 | 2.44 |
| FALSE | High | NTGNGTQS | [R].NTGNG | 1xDeamida | B9EJA4 1xP | 0.057686 | 0.004227 | 1 | 7 | 3 | B9EJA4 | Clasp2 pro | B9EJA4 [89 | 0 | 2385.107 | 0 | -0.32 | 0.450292 | 4.72 | 106.5 | 93.5 | 13.49 | 13.42 | | High | High | High | High | High | High | High | High | High | High | High | 0.002159 | 0.02275 | 2.7 |
| FALSE | High | NTGSPK | [R].NTGSPI | 1xPhospho | Q924K8 1x | 0.112856 | 0.009146 | 1 | 10 | 3 | Q924K8 | Metastasis | Q924K8 [4 | 0 | 1141.602 | 0 | -0.53 | 0.001649 | 3.38 | 117.4 | 82.6 | 3.54 | 4.99 | | High | High | High | High | High | High | High | High | High | High | High | 0.004999 | 0.05469 | 1.76 |
| FALSE | High | NTGVSSAS | [K].NTGVS | 2xPhospho | Q6NXI6 2xP | 0.00083 | 0 | 1 | 4 | 1 | Q6NXI6 | Regulation | Q6NXI6 [48 | 0 | 2887.38 | 0 | 0.36 | 0.44594 | 39.64 | 90.2 | 109.8 | 23.78 | 19.46 | | High | High | High | High | High | High | High | High | High | High | High | 0 | 0.000193 | 2.31 |
| FALSE | High | NTGVSSAS | [K].NTGVS | 1xPhospho | Q6NXI6 1xP | 3.86E-07 | 0 | 1 | 4 | 2 | Q6NXI6 | Regulation | Q6NXI6 [48 | 0 | 3036.577 | 0 | -0.36 | 0.10295 | 2.56 | 110.8 | 89.2 | 11.48 | 7.02 | | High | High | High | High | High | High | High | High | High | High | High | 0 | 3.47E-08 | 2.59 |
| FALSE | High | NTPESPPYS | [K].NTPESF | 1xPhospho | Q3UH85 1x | 4.99E-09 | 0 | 1 | 5 | 3 | Q3UH85 | Nucleolar p | Q3UH85 [3 | 0 | 2367.116 | 0 | -1.42 | 0.000385 | 1.6 | 144.9 | 55.1 | 6.53 | 6.67 | | High | High | High | High | High | High | High | High | High | High | High | 0 | 2.63E-10 | 3.07 |
| FALSE | High | NTPSQHSH | [K].NTPSQI | 1xPhospho | Q8K019 1x | 3.21E-06 | 0 | 1 | 10 | 3 | Q8K019 | Bcl-2-assoc | Q8K019 [2 | 0 | 2150.993 | 0 | -0.38 | 0.108756 | 30.74 | 114.9 | 85.1 | 16.18 | 37.25 | | High | High | High | High | High | High | High | High | High | High | High | 0 | 3.78E-07 | 3.02 |
| FALSE | High | NTPSQHSH | [K].NTPSQI | 2xPhospho | Q8K019 2x | 0.002623 | 0.000109 | 1 | 10 | 2 | Q8K019 | Bcl-2-assoc | Q8K019 [2 | 0 | 2230.959 | 0 | 0.01 | 0.972789 | 19.43 | 100.2 | 99.8 | 9.33 | 13.11 | | High | High | High | High | High | High | High | High | High | High | High | 0.001151 | 0.01201 | 2.35 |
| FALSE | High | NTQGVTTI | [R].NTQGV | 1xPhospho | A0A1S6GW | 0.0008 | 0 | 1 | 3 | 1 | A0A1S6GW | Uncharacte | A0A1S6GW | 0 | 1871.988 | 0 | -0.53 | 0.017482 | 5.4 | 118.5 | 81.5 | 9.87 | 8.04 | | High | High | High | High | High | High | High | High | High | High | High | 0 | 0.000185 | 2.53 |
| FALSE | High | NTFSFGSK | [R].NTFSFG | 1xPhospho | Q9ERU9 1x | 0.032267 | 0.002129 | 1 | 1 | 1 | Q9ERU9 | E3 SUMO-p | Q9ERU9 [1 | 0 | 1425.718 | 0 | 0.61 | 0.014958 | 21.85 | 78 | 122 | 9.31 | 15.75 | | High | High | High | High | High | High | High | High | High | High | High | 0.001126 | 0.01168 | 1.38 |
| FALSE | High | NSVLSDPG | [R].NSVLSD | 2xPhospho | Q68FM7 2x | 0.007742 | 0.000664 | 1 | 4 | 1 | Q68FM7 | Rho guanir | Q68FM7 [2 | 0 | 2555.236 | 0 | -0.21 | 0.146949 | 7.17 | 107.6 | 92.4 | 13.43 | 13.85 | | High | High | High | High | High | High | High | High | High | High | High | 0.000294 | 0.00235 | 2.61 |
| FALSE | High | NSVEEWTT | [K].NSVEE\ | 1xPhospho | Q91VX2 1x | 0.000122 | 0 | 1 | 2 | 1 | Q91VX2 | Ubiquitin-a | Q91VX2 [2 | 0 | 2837.281 | 0 | 1.02 | 0.048489 | 32.49 | 76.2 | 123.8 | 17.49 | 31.26 | | High | High | High | High | High | High | High | High | High | High | High | 0 | 2.23E-05 | 2.96 |
| FALSE | High | NSVCLERD | [K].NSVCLE | 1xCarbami | Q8BMI4 2x | 9.36E-05 | 0 | 1 | 2 | 3 | Q8BMI4 | Flap endor | Q8BMI4 [7 | 1 | 2884.226 | 0 | 0.64 | 0.007309 | 19.67 | 74.4 | 125.6 | 9.05 | 8.72 | | High | High | High | High | High | High | High | High | High | High | High | 0 | 0.000129 | 3.12 |
| FALSE | High | NSSSPPSPS | [R].NSSSPP | 2xPhospho | E9Q1P8 2x | 1.09E-05 | 0 | 1 | 1 | 1 | E9Q1P8 | Interferon | E9Q1P8 [4 | 0 | 1864.75 | 0 | -0.07 | 0.613442 | 10.16 | 102 | 98 | 7.85 | 8.77 | | High | High | High | High | High | High | High | High | High | High | High | 0 | 1.5E-06 | 3.36 |
| FALSE | High | NSSSPPSPS | [R].NSSSPP | 2xPhospho | E9Q1P8 2x | 3.9E-05 | 0 | 1 | 1 | 2 | E9Q1P8 | Interferon | E9Q1P8 [4 | 0 | 1880.745 | 0 | -0.18 | 0.428903 | 24.91 | 99.6 | 100.4 | 15.67 | 28.67 | | High | High | High | High | High | High | High | High | High | High | High | 0 | 6.25E-06 | 2.17 |
| FALSE | High | NSSSPVAAT | [K].NSSSPV | 1xPhospho | Q3UZZ2 1x | 0.000219 | 0 | 1 | 1 | 1 | Q3UZZ2 | Putative ur | Q3UZZ2 [9 | 0 | 2607.343 | 0 | -0.8 | 0.191077 | 6.65 | 125.9 | 74.1 | 30.72 | 30.3 | | High | High | High | High | High | High | High | High | High | High | High | 0 | 4.3E-05 | 3.08 |
| FALSE | High | NSSSPVAAT | [K].NSSSPV | 1xPhospho | Q3UZZ2 1x | 0.115566 | 0.009504 | 1 | 1 | 2 | Q3UZZ2 | Putative ur | Q3UZZ2 [9 | 0 | 2836.506 | 0 | -0.99 | 0.044541 | 14.16 | 133.1 | 66.9 | 20.81 | 22.5 | | High | High | High | High | High | High | High | High | High | High | High | 0.004747 | 0.05116 | 2.32 |
| FALSE | High | NSSSPVSLD | [R].NSSSPV | 1xPhospho | Q8BL06 1x | 5.21E-05 | 0 | 1 | 1 | 4 | Q8BL06 | Inactive ub | Q8BL06 [6 | 0 | 2384.133 | 0 | -0.39 | 0.011192 | 3.35 | 116.4 | 83.6 | 4.63 | 11.94 | | High | High | High | High | High | High | High | High | High | High | High | 0 | 1.79E-05 | 2.52 |
| FALSE | High | NSSSPVSPA | [R].NSSSPV | 1xPhospho | Q8K3X4 1x | 2.22E-07 | 0 | 1 | 1 | 6 | Q8K3X4 | Interferon | Q8K3X4 [6 | 0 | 1778.864 | 0 | -1.17 | 0.000746 | 2.86 | 136.4 | 63.6 | 3.25 | 12.45 | | High | High | High | High | High | High | High | High | High | High | High | 0 | 9.15E-07 | 2.62 |
| FALSE | High | NSSSPVSPA | [R].NSSSPV | 2xPhospho | Q8K3X4 2x | 6.37E-06 | 0 | 1 | 1 | 1 | Q8K3X4 | Interferon | Q8K3X4 [6 | 0 | 1858.83 | 0 | -1.01 | 0.00252 | 3.86 | 132 | 68 | 5.21 | 15.55 | | High | High | High | High | High | High | High | High | High | High | High | 0 | 8.17E-07 | 3.07 |
| FALSE | High | NSSSSTSPV | [R].NSSSST | 1xPhospho | B2RUG9 1x | 0.000407 | 0 | 1 | 6 | 2 | B2RUG9 | Adenomat | B2RUG9 [2 | 0 | 1618.809 | 0 | 0.23 | 0.030208 | 3.84 | 91.2 | 108.8 | 5.41 | 10.14 | | High | High | High | High | High | High | High | High | High | High | High | 0 | 8.69E-05 | 1.93 |
| FALSE | High | NSSSSTSPV | [R].NSSSST | 2xPhospho | B2RUG9 2x | 0.001898 | 5.66E-05 | 1 | 6 | 1 | B2RUG9 | Adenomat | B2RUG9 [2 | 0 | 1698.775 | 0 | 0.06 | 0.382657 | 13.85 | 95.6 | 104.4 | 12.82 | 5.83 | | High | High | High | High | High | High | High | High | High | High | High | 2.58E-05 | 0.000489 | 2.04 |
| FALSE | High | NSSSTVSCL | [R].NSSSTV | 1xCarbami | P41233 1xP | 0.002376 | 0.000109 | 1 | 2 | 1 | P41233 | ATP-bindin | P41233 [11 | 0 | 1620.807 | 0 | 2.81 | 0.003932 | 17.94 | 24.9 | 175.1 | 14.01 | 23.36 | | High | High | High | High | High | High | High | High | High | High | High | 4.94E-05 | 0.000627 | 2.33 |
| FALSE | High | NSSWYSSG | [R].NSSWY | 1xPhospho | Q8K1N4 1x | 0.002115 | 5.66E-05 | 1 | 1 | 3 | Q8K1N4 | Spermatog | Q8K1N4 [4 | 0 | 1449.636 | 0 | -0.57 | 0.024572 | 4.16 | 119.4 | 80.6 | 10.09 | 6.15 | | High | High | High | High | High | High | High | High | High | High | High | 2.58E-05 | 0.000554 | 1.8 |
| FALSE | High | NSTANLSP | [K].NSTANI | 1xPhospho | B2RRF6 1xP | 0.00195 | 5.66E-05 | 1 | 1 | 1 | B2RRF6 | Zinc finger | B2RRF6 [47 | 0 | 2802.444 | 0 | -0.02 | 0.920908 | 11.17 | 96.5 | 103.5 | 7.82 | 6.39 | | High | High | High | High | High | High | High | High | High | High | High | 2.58E-05 | 0.000504 | 2.99 |
| FALSE | High | NSTDLDSA | [K].NSTDLD | 1xPhospho | Q924A2 1x | 5.31E-05 | 0 | 1 | 3 | 2 | Q924A2 | Protein cap | Q924A2 [2 | 0 | 2212.042 | 0 | 0.3 | 0.603791 | 20.6 | 86.9 | 113.1 | 17 | 23.01 | | High | High | High | High | High | High | High | High | High | High | High | 0 | 2.13E-05 | 2.61 |
| FALSE | High | NSTDLDSA | [K].NSTDLD | 2xPhospho | Q924A2 2x | 0.011246 | 0.000772 | 1 | 3 | 2 | Q924A2 | Protein cap | Q924A2 [2 | 0 | 2292.009 | 0 | -0.06 | 0.584635 | 12.72 | 103.3 | 96.7 | 13.72 | 7.41 | | High | High | High | High | High | High | High | High | High | High | High | 0.000418 | 0.00358 | 2.84 |
| FALSE | High | NSTDSRPV | [R].NSTDSF | 1xPhospho | Q3UWE6 1 | 0.117405 | 0.009523 | 1 | 2 | 1 | Q3UWE6 | MCG14935 | Q3UWE6 [ | 0 | 1790.864 | 0 | -0.08 | 0.602134 | 14.82 | 104.5 | 95.5 | 10.96 | 10.23 | | High | High | High | High | High | High | High | High | High | High | High | 0.004782 | 0.05238 | 0.93 |
| FALSE | High | NSTDYSSES | [R].NSTDYS | 1xPhospho | Q3UY41 1x | 0.000262 | 0 | 1 | 4 | 2 | Q3UY41 | Putative ur | Q3UY41 [2 | 0 | 1655.757 | 0 | 0.48 | 0.480197 | 71.77 | 94.4 | 105.6 | 48.29 | 103.35 | | High | High | High | High | High | High | High | High | High | High | High | 0 | 0.000137 | 2.63 |
| FALSE | High | NSTLSDSGI | [R].NSTLSD | 1xPhospho | Q921I6 1xP | 0.043607 | 0.003228 | 1 | 1 | 1 | Q921I6 | SH3 domai | Q921I6 [11 | 0 | 2628.195 | 0 | -0.09 | 0.276212 | 19.66 | 103.7 | 96.3 | 13.78 | 7.72 | | High | High | High | High | High | High | High | High | High | High | High | 0.001651 | 0.01654 | 2.73 |
| FALSE | High | NSTLSDSGI | [R].NSTLSD | 1xPhospho | Q921I6 1xP | 0.000892 | 0 | 1 | 1 | 4 | Q921I6 | SH3 domai | Q921I6 [11 | 0 | 2857.358 | 0 | -0.31 | 0.011979 | 0.75 | 110.3 | 89.7 | 4.48 | 6.49 | | High | High | High | High | High | High | High | High | High | High | High | 2.58E-05 | 0.000451 | 4 |
| FALSE | High | NSTLSDSGI | [R].NSTLSD | 1xPhospho | Q921I6 1xP | 6.51E-07 | 0 | 1 | 1 | 3 | Q921I6 | SH3 domai | Q921I6 [11 | 0 | 2873.353 | 0 | -0.21 | 0.060204 | 6.41 | 107.6 | 92.4 | 7.23 | 13.33 | | High | High | High | High | High | High | High | High | High | High | High | 0 | 8.37E-08 | 4.79 |
| FALSE | High | NSTLSEEDY | [K].NSTLSE | 1xPhospho | O55003 1x | 0.000229 | 0 | 1 | 2 | 6 | O55003 | BCL2/aden | O55003 [8 | 0 | 1764.789 | 0 | -0.79 | 0.000667 | 1.31 | 130.9 | 69.1 | 5.98 | 9.3 | | High | High | High | High | High | High | High | High | High | High | High | 2.58E-05 | 0.00055 | 2.47 |
| FALSE | High | NSTLSEEDY | [K].NSTLSE | 2xPhospho | O55003 2x | 0.000176 | 0 | 1 | 2 | 1 | O55003 | BCL2/aden | O55003 [8 | 0 | 1844.755 | 0 | -0.51 | 0.044483 | 15.41 | 126.1 | 73.9 | 20.93 | 21.61 | | High | High | High | High | High | High | High | High | High | High | High | 0 | 3.39E-05 | 4.18 |
| FALSE | High | NSTPNSSG | [R].NSTPNS | 1xPhospho | Q8VC70 1x | 8.49E-12 | 0 | 1 | 4 | 4 | Q8VC70 | RNA-bindir | Q8VC70 [3 | 0 | 2473.136 | 0 | 1.39 | 0.00049 | 4.56 | 54.6 | 145.4 | 6.49 | 8.17 | | High | High | High | High | High | High | High | High | High | High | High | 0 | 2.04E-13 | 4.52 |
| FALSE | High | NSTPNSSG | [R].NSTPNS | 2xPhospho | Q8VC70 2x | 1.46E-05 | 0 | 1 | 4 | 1 | Q8VC70 | RNA-bindir | Q8VC70 [3 | 0 | 2553.102 | 0 | 0.66 | 0.00348 | 4.22 | 76.7 | 123.3 | 3.65 | 8.53 | | High | High | High | High | High | High | High | High | High | High | High | 0 | 2.07E-06 | 3.18 |
| FALSE | High | NSTPSEPDS | [K].NSTPSE | 1xPhospho | P51859 1xP | 0.000382 | 0 | 1 | 2 | 2 | P51859 | Hepatoma- | P51859 [19 | 0 | 3209.405 | 0 | -0.3 | 0.409287 | 5.24 | 110.5 | 89.5 | 17.9 | 26.24 | | High | High | High | High | High | High | High | High | High | High | High | 0 | 8.08E-05 | 3.64 |
| FALSE | High | NSTSGSDFI | [K].NSTSGS | 1xPhospho | Q61687 1x | 0.00204 | 5.66E-05 | 1 | 4 | 2 | Q61687 | Transcripti | Q61687 [7 | 0 | 1467.62 | 0 | -0.24 | 0.866727 | 43.14 | 106.9 | 93.1 | 57.08 | 52.41 | | High | High | High | High | High | High | High | High | High | High | High | 2.58E-05 | 0.000531 | 2.66 |
| FALSE | High | NSTSGSDFI | [K].NSTSGS | 1xPhospho | Q61687 1x | 7.21E-05 | 0 | 1 | 4 | 3 | Q61687 | Transcripti | Q61687 [7 | 0 | 1696.783 | 0 | -0.24 | 0.027524 | 3.56 | 107.7 | 92.3 | 4.9 | 5.57 | | High | High | High | High | High | High | High | High | High | High | High | 0.00014 | 0.001102 | 2.82 |
| FALSE | High | NSTSGSDFI | [K].NSTSGS | 2xPhospho | Q61687 2x | 0.001767 | 5.66E-05 | 1 | 4 | 3 | Q61687 | Transcripti | Q61687 [7 | 0 | 1776.749 | 0 | -0.18 | 0.099459 | 10.75 | 106.2 | 93.8 | 11.05 | 6.57 | | High | High | High | High | High | High | High | High | High | High | High | 0.000275 | 0.001787 | 2.5 |
| FALSE | High | NSTSSIEPR | [R].NSTSSII | 1xPhospho | A0A0A6YW | 0.017652 | 0.001282 | 2 | 11 | 2 | A0A0A6YW | Mitogen-a | A0A0A6YW | 0 | 1299.614 | 0 | -0.89 | 0.00471 | 3.73 | 130.4 | 69.6 | 5.97 | 14.65 | Shared | High | High | High | High | High | High | High | High | High | High | High | 0.001789 | 0.01758 | 2.23 |
| FALSE | High | NSVASPTSI | [R].NSVASI | 1xPhospho | A2AEV7 1x | 0.085859 | 0.006406 | 1 | 1 | 1 | A2AEV7 | Coiled-coil | A2AEV7 [2 | 0 | 1425.694 | 0 | 1.26 | 0.001899 | 7.1 | 58.9 | 141.1 | 13.67 | 5.5 | | High | High | High | High | High | High | High | High | High | High | High | 0.003291 | 0.03599 | 1.09 |
| FALSE | High | NTSLPPLW | [K].NTSLPP | 1xPhospho | E9Q3G8 1x | 7.84E-06 | 0 | 1 | 2 | 7 | E9Q3G8 | Nucleopor | E9Q3G8 [2 | 0 | 1905.931 | 0 | -0.48 | 0.007624 | 6.31 | 116.3 | 83.7 | 5.13 | 12.79 | | High | High | High | High | High | High | High | High | High | High | High | 0 | 1.03E-06 | 3.7 |
| FALSE | High | NSSSPPSPS | [R].NSSSPP | 1xPhospho | E9Q1P8 1x | 1.03E-05 | 0 | 1 | 1 | 4 | E9Q1P8 | Interferon | E9Q1P8 [4 | 0 | 1800.779 | 0 | -1.1 | 0.000541 | 1.42 | 139.2 | 60.8 | 5.19 | 11.85 | | High | High | High | High | High | High | High | High | High | High | High | 0 | 1.39E-06 | 3.06 |
| FALSE | High | NTSLPPLW | [K].NTSLPP | 2xPhospho | E9Q3G8 2x | 0.001629 | 5.66E-05 | 1 | 2 | 2 | E9Q3G8 | Nucleopor | E9Q3G8 [2 | 0 | 1985.897 | 0 | -0.94 | 0.001986 | 2.4 | 129 | 71 | 8.72 | 12.79 | | High | High | High | High | High | High | High | High | High | High | High | 2.58E-05 | 0.000411 | 2.83 |
| FALSE | High | NTSPHSEP | [R].NTSPHS | 2xCarbami | Q5SPW0 1 | 0.000186 | 0 | 1 | 5 | 2 | Q5SPW0 | Vacuolar p | Q5SPW0 [5 | 0 | 3437.44 | 0 | 0.65 | 0.001933 | 2.48 | 78 | 122 | 8.78 | 4.01 | | High | High | High | High | High | High | High | High | High | High | High | 0 | 3.61E-05 | 3.87 |
| FALSE | High | NVPQEDIC | [R].NVPQE | 1xCarbami | A0A0B4J1E | 5.4E-05 | 0 | 1 | 4 | 2 | A0A0B4J1E | Importin su | A0A0B4J1E | 0 | 2449.006 | 0 | 0.54 | 0.00508 | 4.6 | 77.7 | 122.3 | 7.87 | 9.43 | | High | High | High | High | High | High | High | High | High | High | High | 0 | 8.99E-06 | 3.57 |
| FALSE | High | NVPQEESLI | [R].NVPQE | 1xPhospho | O35344 1x | 1.16E-07 | 0 | 1 | 2 | 3 | O35344 | Importin su | O35344 [5 | 0 | 2346.022 | 0 | 0.44 | 0.467453 | 3.66 | 85 | 115 | 36.68 | 12.47 | | High | High | High | High | High | High | High | High | High | High | High | 0 | 9.02E-09 | 2.86 |
| FALSE | High | NVPQEESLI | [R].NVPQE | 1xPhospho | O35344 1x | 1.6E-09 | 0 | 1 | 2 | 7 | O35344 | Importin su | O35344 [5 | 0 | 2575.185 | 0 | -0.52 | 0.002358 | 1.43 | 117.6 | 82.4 | 2.83 | 8.85 | | High | High | High | High | High | High | High | High | High | High | High | 0 | 7.35E-11 | 4.33 |
| FALSE | High | NVPQEESLI | [R].NVPQE | 2xPhospho | O35344 2x | 1.95E-05 | 0 | 1 | 2 | 1 | O35344 | Importin su | O35344 [5 | 0 | 2655.152 | 0 | 0.17 | 0.254975 | 15.92 | 96.7 | 103.3 | 10.81 | 17.92 | | High | High | High | High | High | High | High | High | High | High | High | 0 | 2.85E-06 | 3.7 |
| FALSE | High | NVPVNPTS | [R].NVPVN | 1xPhospho | Q05BA5 1x | 1.22E-05 | 0 | 1 | 6 | 1 | Q05BA5 | Nuclear re | Q05BA5 [8 | 0 | 2277.086 | 0 | 0.39 | 0.019091 | 13.34 | 88.7 | 111.3 | 6.8 | 15.59 | | High | High | High | High | High | High | High | High | High | High | High | 0 | 1.68E-06 | 3.01 |
| FALSE | High | NVQTGLTE | [K].NVQTG | 1xPhospho | S4R1P5 1xP | 0.012619 | 0.000889 | 1 | 6 | 2 | S4R1P5 | Dystonin O | S4R1P5 [30 | 0 | 3311.743 | 0 | 1.49 | 0.007094 | 18.26 | 56.9 | 143.1 | 23.03 | 15.93 | | High | High | High | High | High | High | High | High | High | High | High | 0.000538 | 0.004547 | 2.98 |
| FALSE | High | NVSASCISK | [K].NVSAS | 1xCarbami | Q80WQ8 1 | 0.002895 | 0.00016 | 1 | 4 | 1 | Q80WQ8 | Mis18-binc | Q80WQ8 [ | 0 | 1503.764 | 0 | -1.44 | 0.000197 | 1.2 | 146.5 | 53.5 | 3.2 | 7.61 | | High | High | High | High | High | High | High | High | High | High | High | 7.19E-05 | 0.000783 | 2.72 |
| FALSE | High | NVSEELDR | [K].NVSEEL | 1xPhospho | B2RSV4 1xP | 0.014158 | 0.001011 | 1 | 2 | 1 | B2RSV4 | Splicing fac | B2RSV4 [1 | 1 | 2238.142 | 0 | -0.65 | 0.001373 | 1.1 | 122.4 | 77.6 | 5.41 | 3.56 | | High | High | High | High | High | High | High | High | High | High | High | 0.000538 | 0.004635 | 2.03 |
| FALSE | High | NVSGSLK | [K].NVSGSI | 1xPhospho | Q9JI11 1xP | 0.044739 | 0.003302 | 1 | 3 | 2 | Q9JI11 | Serine/thre | Q9JI11 [41 | 0 | 1242.686 | 0 | 0.17 | 0.040262 | 8.53 | 94.1 | 105.9 | 2.4 | 6.44 | | High | High | High | High | High | High | High | High | High | High | High | 0.001712 | 0.01703 | 1.15 |
| FALSE | High | NVSSFPDD | [R].NVSSFF | 1xPhospho | Q52L97 1xP | 4.77E-09 | 0 | 1 | 6 | 6 | Q52L97 | Importin su | Q52L97 [5 | 0 | 2185.996 | 0 | 0.22 | 0.010575 | 3.03 | 92.5 | 107.5 | 4.71 | 4.16 | | High | High | High | High | High | High | High | High | High | High | High | 0 | 2.51E-10 | 4.88 |
| FALSE | High | NVSSFPDD | [R].NVSSFF | 2xPhospho | Q52L97 2xP | 2.47E-05 | 0 | 1 | 6 | 1 | Q52L97 | Importin su | Q52L97 [5 | 0 | 2265.963 | 0 | -0.05 | 0.721048 | 11.32 | 100.9 | 99.1 | 8 | 8.91 | | High | High | High | High | High | High | High | High | High | High | High | 0 | 3.73E-06 | 3.62 |
| FALSE | High | NVSTCEVT | [R].NVSTCI | 1xCarbami | Q9QZR5 1x | 0.000181 | 0 | 1 | 5 | 1 | Q9QZR5 | Homeodor | Q9QZR5 [8 | 0 | 2260.048 | 0 | 0.35 | 0.019189 | 11.51 | 85.9 | 114.1 | 13.56 | 2.56 | | High | High | High | High | High | High | High | High | High | High | High | 0 | 3.48E-05 | 3.16 |
| FALSE | High | NVSTESGE | [R].NVSTES | 1xPhospho | B8X349 1xP | 0.003224 | 0.00016 | 2 | 12 | 1 | B8X349; F6 | JNK-intera | B8X349 [3 | 0 | 1860.863 | 0 | 2.37 | 0.000623 | 26.47 | 32.4 | 167.6 | 16.68 | 8.56 | Shared | High | High | High | High | High | High | High | High | High | High | High | 7.19E-05 | 0.000883 | 3.25 |
| FALSE | High | NVPPPSIEP | [K].NVPPPS | 1xPhospho | Q99LG1 1x | 4.31E-05 | 0 | 1 | 1 | 2 | Q99LG1 | Transmem | Q99LG1 [2 | 0 | 2939.557 | 0 | -0.08 | 0.068094 | 5.78 | 103.3 | 96.7 | 4.75 | 2.56 | | High | High | High | High | High | High | High | High | High | High | High | 0 | 6.98E-06 | 2.2 |
| FALSE | High | NVVNYDN\ | [K].NVVNY | 2xPhospho | A0A0M3HE | 0.000219 | 0 | 1 | 15 | 1 | A0A0M3HE | Zinc finger | A0A0M3HE | 0 | 2830.211 | 0 | 0.43 | 0.662723 | 28.65 | 85.3 | 114.7 | 24.66 | 25.42 | | High | High | High | High | High | High | High | High | High | High | High | 0 | 4.3E-05 | 4.54 |
| FALSE | High | NVYYELND | [R].NVYYEI | 1xPhospho | E9QAU4 1x | 0.003433 | 0.00021 | 1 | 9 | 2 | E9QAU4 | Enhancer t | E9QAU4 [2 | 0 | 1593.751 | 0 | 0.22 | 0.028529 | 1.9 | 92.5 | 107.5 | 4.02 | 6.72 | | High | High | High | High | High | High | High | High | High | High | High | 9.42E-05 | 0.000946 | 3.45 |
| FALSE | High | NWEDEDF\ | [K].NWEDE | 1xPhospho | E9PX68 1xP | 4.58E-07 | 0 | 1 | 7 | 9 | E9PX68 | Solute carr | E9PX68 [40 | 0 | 2606.008 | 0 | -0.11 | 0.072787 | 6.44 | 102.6 | 97.4 | 4 | 6.91 | | High | High | High | High | High | High | High | High | High | High | High | 0 | 3.39E-07 | 3.85 |
| FALSE | High | NWSGCGE | [R].NWSGC | 1xCarbami | P61406 1xP | 0.000543 | 0 | 1 | 1 | 2 | P61406 | Telomerase | P61406 [32 | 0 | 1789.798 | 0 | 0.56 | 0.007447 | 6.29 | 80.5 | 119.5 | 9.48 | 7.22 | | High | High | High | High | High | High | High | High | High | High | High | 0 | 0.000121 | 3.99 |
| FALSE | High | NWTEDIEG | [R].NWTED | 1xPhospho | P70255 1xP | 0.002077 | 5.66E-05 | 1 | 4 | 1 | P70255 | Nuclear fac | P70255 [31 | 0 | 1940.92 | 0 | 0.15 | 0.133491 | 7.64 | 93.9 | 106.1 | 2.18 | 11.88 | | High | High | High | High | High | High | High | High | High | High | High | 2.58E-05 | 0.000539 | 2.92 |
| FALSE | High | NWTEDIEG | [R].NWTED | 1xPhospho | P70255 1xP | 9.13E-09 | 0 | 1 | 4 | 9 | P70255 | Nuclear fac | P70255 [31 | 0 | 2170.083 | 0 | 2.27 | 0.000672 | 4.05 | 33.7 | 166.3 | 7.55 | 11.72 | | High | High | High | High | High | High | High | High | High | High | High | 0 | 6.56E-09 | 4.64 |
| FALSE | High | NYAGEEEE | [R].NYAGE | 1xPhospho | Q9JKB3 1xP | 9.7E-08 | 0 | 1 | 1 | 7 | Q9JKB3 | Y-box-bind | Q9JKB3 [18 | 0 | 3370.386 | 0 | 0.09 | 0.856319 | 11.84 | 108.6 | 91.4 | 17.49 | 16.9 | | High | High | High | High | High | High | High | High | High | High | High | 0 | 1.24E-08 | 4.48 |
| FALSE | High | NYAGEEEE | [R].NYAGE | 2xPhospho | Q9JKB3 2xP | 0.000271 | 0 | 1 | 1 | 6 | Q9JKB3 | Y-box-bind | Q9JKB3 [18 | 0 | 3450.352 | 0 | 0.11 | 0.346779 | 13.28 | 97.6 | 102.4 | 16.21 | 8.15 | | High | High | High | High | High | High | High | High | High | High | High | 0 | 5.54E-05 | 4.45 |
| FALSE | High | NYGSPLISG | [K].NYGSPI | 1xPhospho | Q925J9 1xP | 1.46E-06 | 0 | 1 | 2 | 1 | Q925J9 | Mediator c | Q925J9 [14 | 0 | 1858.972 | 0 | 0.93 | 0.000222 | 8.35 | 68.8 | 131.2 | 12.9 | 5.7 | | High | High | High | High | High | High | High | High | High | High | High | 0 | 1.56E-07 | 3.45 |
| FALSE | High | NYGSPLISG | [K].NYGSPI | 2xPhospho | Q925J9 2xP | 4.68E-05 | 0 | 1 | 2 | 1 | Q925J9 | Mediator c | Q925J9 [14 | 0 | 1938.938 | 0 | 1.26 | 0.000563 | 18.16 | 58.1 | 141.9 | 10.05 | 4.63 | | High | High | High | High | High | High | High | High | High | High | High | 0 | 7.68E-06 | 4.63 |
| FALSE | High | NYGSYSTQ | [R].NYGSYS | 1xPhospho | B2M0S2 1x | 0.000758 | 0 | 1 | 7 | 5 | B2M0S2 | Secretory c | B2M0S2 [5 | 0 | 2555.279 | 0 | 0.52 | 0.181367 | 26.99 | 89.9 | 110.1 | 13.29 | 24.62 | | High | High | High | High | High | High | High | High | High | High | High | 0 | 0.000217 | 2.99 |
| FALSE | High | NYLGSHSPI | [R].NYLGSH | 1xPhospho | Q6A050 1x | 0.017966 | 0.001282 | 1 | 2 | 1 | Q6A050 | MKIAA054 | Q6A050 [6 | 0 | 1540.793 | 0 | -1.02 | 0.000348 | 2.98 | 136.7 | 63.3 | 3.6 | 12.9 | | High | High | High | High | High | High | High | High | High | High | High | 0.000689 | 0.006059 | 1.85 |
| FALSE | High | NYILDQTN | [R].NYILDC | 1xPhospho | Q8VDM6 1 | 7.04E-06 | 0 | 1 | 2 | 1 | Q8VDM6 | Heterogen | Q8VDM6 [ | 0 | 2050.98 | 0 | -0.48 | 0.036141 | 5.17 | 119.1 | 80.9 | 11.25 | 11.18 | | High | High | High | High | High | High | High | High | High | High | High | 0 | 9.06E-07 | 3.72 |
| FALSE | High | NYILSASAS | [R].NYILSA | 1xPhospho | A2AWI7 1x | 4.68E-05 | 0 | 1 | 4 | 3 | A2AWI7 | Endophilin | A2AWI7 [2 | 0 | 2534.258 | 0 | 0.17 | 0.144593 | 28.64 | 89.7 | 110.3 | 13.4 | 11.92 | | High | High | High | High | High | High | High | High | High | High | High | 0 | 7.63E-06 | 3.65 |
| FALSE | High | NVYSSSR | [R].NVYSSS | 1xPhospho | Q9R0U0-3 | 0.057686 | 0.004227 | 1 | 7 | 1 | Q9R0U0-3 | Isoform 3 o | Q9R0U0-3 | 0 | 1121.519 | 0 | -0.81 | 0.000236 | 1.43 | 126.3 | 73.7 | 3.52 | 3.27 | | High | High | High | High | High | High | High | High | High | High | High | 0.002159 | 0.02262 | 0.35 |
| FALSE | High | NVPPPSIEP | [K].NVPPPS | 1xPhospho | Q99LG1 1x | 0.012845 | 0.000889 | 1 | 1 | 1 | Q99LG1 | Transmem | Q99LG1 [2 | 0 | 2710.394 | 0 | -0.34 | 0.789833 | 45.17 | 109.9 | 90.1 | 44.95 | 56.4 | | High | High | High | High | High | High | High | High | High | High | High | 0.000449 | 0.004133 | 2.64 |
| FALSE | High | NVPGLAAP | [R-].NVPGL | 1xPhospho | G3X9K4 1x | 6.79E-06 | 0 | 1 | 5 | 1 | G3X9K4 | SAPS doma | G3X9K4 [6 | 0 | 1905.025 | 0 | 1.09 | 0.000677 | 4.12 | 66.4 | 133.6 | 5.77 | 6.78 | | High | High | High | High | High | High | High | High | High | High | High | 0 | 8.75E-07 | 2.24 |
| FALSE | High | NVMSAFGI | [K].NVMSA | 2xPhospho | Q9Z1A1 2x | 0.015465 | 0.001089 | 1 | 5 | 1 | Q9Z1A1 | Putative ur | Q9Z1A1 [1 | 0 | 2896.231 | 0 | 0.13 | 0.342212 | 25.32 | 87.9 | 112.1 | 17.26 | 49.17 | | High | High | High | High | High | High | High | High | High | High | High | 0.000571 | 0.005125 | 3.53 |
| FALSE | High | NTSSDNSD | [R].NTSSDI | 2xPhospho | Q09XV5 2x | 1.25E-05 | 0 | 1 | 3 | 1 | Q09XV5 | Chromodo | Q09XV5 [5 | 0 | 2338.946 | 0 | -0.46 | 0.034869 | 9.5 | 114.7 | 85.3 | 15.41 | 16.13 | | High | High | High | High | High | High | High | High | High | High | High | 0 | 1.74E-06 | 5.09 |
| FALSE | High | NTSYSYSEI | [K].NTSYSY | 1xPhospho | Q8BUH8 1x | 0.083783 | 0.006345 | 1 | 2 | 1 | Q8BUH8 | Sentrin-spe | Q8BUH8 [2 | 0 | 2198.926 | 0 | 0.67 | 0.001008 | 13.64 | 77.2 | 122.8 | 15.57 | 10.58 | | High | High | High | High | High | High | High | High | High | High | High | 0.003265 | 0.03519 | 2.73 |
| FALSE | High | NTTEEVGS | [K].NTTEE\ | 1xPhospho | Q5BLK4 1x | 0.000207 | 0 | 1 | 2 | 2 | Q5BLK4 | Terminal u | Q5BLK4 [6 | 0 | 1670.84 | 0 | 0.75 | 0.00145 | 7.39 | 74.3 | 125.7 | 4.75 | 5.47 | | High | High | High | High | High | High | High | High | High | High | High | 0 | 4.05E-05 | 2.74 |
| FALSE | High | NTTKPETIP | [K].NTTKPE | 2xPhospho | Q8C9B9 2x | 0.045898 | 0.003408 | 1 | 3 | 4 | Q8C9B9 | Death-indu | Q8C9B9 [7 | 0 | 3534.507 | 0 | 0.11 | 0.959451 | 41.37 | 96.2 | 103.8 | 39.64 | 27.69 | | High | High | High | High | High | High | High | High | High | High | High | 0.002403 | 0.02552 | 4.01 |
| FALSE | High | NTTKPETIP | [K].NTTKPE | 2xPhospho | Q8C9B9 2x | 2.49E-05 | 0 | 1 | 3 | 2 | Q8C9B9 | Death-indu | Q8C9B9 [7 | 0 | 3550.502 | 0 | -0.01 | 0.888174 | 39.5 | 101.3 | 98.7 | 38.45 | 46.04 | | High | High | High | High | High | High | High | High | High | High | High | 0 | 3.77E-06 | 5.29 |
| FALSE | High | NTTKPETIP | [K].NTTKPE | 1xPhospho | Q8C9B9 1x | 1.22E-05 | 0 | 1 | 3 | 1 | Q8C9B9 | Death-indu | Q8C9B9 [7 | 0 | 3699.698 | 0 | -0.75 | 0.011572 | 9.15 | 125.6 | 74.4 | 14.56 | 23.43 | | High | High | High | High | High | High | High | High | High | High | High | 0 | 1.69E-06 | 3.13 |
| FALSE | High | NTTKPETIP | [K].NTTKPE | 2xPhospho | Q8C9B9 2x | 0.000259 | 0 | 1 | 3 | 1 | Q8C9B9 | Death-indu | Q8C9B9 [7 | 0 | 3763.67 | 0 | -0.07 | 0.293528 | 6.7 | 102.5 | 97.5 | 11.19 | 25.97 | | High | High | High | High | High | High | High | High | High | High | High | 0 | 5.22E-05 | 2.08 |
| FALSE | High | NTTPPSLAF | [R].NTTPPS | 1xPhospho | Q8BJH1 1x | 0.056724 | 0.004124 | 1 | 1 | 2 | Q8BJH1 | Zinc finger | Q8BJH1 [2 | 0 | 1265.645 | 0 | 0.12 | 0.538074 | 15.16 | 96.4 | 103.6 | 6.33 | 5.39 | | High | High | High | High | High | High | High | High | High | High | High | 0.002117 | 0.02218 | 1.75 |
| FALSE | High | NTTPTGAA | [R].NTTPTC | 1xPhospho | P27546 1xP | 0.0026 | 0.000109 | 1 | 10 | 4 | P27546 | Microtubu | P27546 [84 | 0 | 1939.915 | 0 | 0.55 | 0.116751 | 14.12 | 77.4 | 122.6 | 11.69 | 20.25 | | High | High | High | High | High | High | High | High | High | High | High | 0.002717 | 0.02912 | 2.75 |
| FALSE | High | NTTPTGAA | [R].NTTPTC | 1xPhospho | P27546 1xP | 3.32E-05 | 0 | 1 | 10 | 5 | P27546 | Microtubu | P27546 [84 | 0 | 1955.91 | 0 | 0.47 | 0.031814 | 13.13 | 84.1 | 115.9 | 7.5 | 19.82 | | High | High | High | High | High | High | High | High | High | High | High | 0.006563 | 0.0705 | 2.14 |
| FALSE | High | NTTVCEISS | [K].NTTVCI | 1xCarbami | A0A1D9BZ | 0.004143 | 0.00036 | 1 | 1 | 3 | A0A1D9BZ | GCNA OS=I | A0A1D9BZ | 0 | 2855.2 | 0 | -1.2 | 0.025094 | 6.63 | 138.7 | 61.3 | 29.18 | 12.29 | | High | High | High | High | High | High | High | High | High | High | High | 0.000467 | 0.004247 | 3.24 |
| FALSE | High | NTTVCEISS | [K].NTTVCI | 1xCarbami | A0A1D9BZ | 9.36E-05 | 0 | 1 | 1 | 2 | A0A1D9BZ | GCNA OS=I | A0A1D9BZ | 0 | 2856.185 | 0 | -1.41 | 0.023766 | 2.42 | 144.2 | 55.8 | 33.14 | 8.29 | | High | High | High | High | High | High | High | High | High | High | High | 0 | 0.000239 | 4.03 |
| FALSE | High | NTVCPEQS | [K].NTVCPI | 1xCarbami | Q9D2D7 1x | 8.58E-06 | 0 | 1 | 5 | 5 | Q9D2D7 | Zinc finger | Q9D2D7 [9 | 0 | 2835.264 | 0 | -0.17 | 0.082599 | 12.52 | 112.3 | 87.7 | 7.51 | 11.65 | | High | High | High | High | High | High | High | High | High | High | High | 0 | 1.14E-06 | 4.98 |
| FALSE | High | NTVCPEQS | [K].NTVCPI | 1xCarbami | Q9D2D7 1x | 0.104222 | 0.008348 | 1 | 5 | 1 | Q9D2D7 | Zinc finger | Q9D2D7 [9 | 0 | 2836.248 | 0 | 0.17 | 0.593673 | 13.97 | 102.6 | 97.4 | 18.07 | 12.43 | | High | High | High | High | High | High | High | High | High | High | High | 0.004206 | 0.04529 | 2.68 |
| FALSE | High | NTVCPEQS | [K].NTVCPI | 1xCarbami | Q9D2D7 2x | 0.001933 | 5.66E-05 | 1 | 5 | 1 | Q9D2D7 | Zinc finger | Q9D2D7 [9 | 0 | 2915.231 | 0 | 0.5 | 0.197732 | 12.84 | 91.6 | 108.4 | 18.17 | 19.65 | | High | High | High | High | High | High | High | High | High | High | High | 2.58E-05 | 0.000499 | 3.86 |
| FALSE | High | NTVDIVDKI | [K].NTVDIV | 1xDeamida | B2RQG2 1x | 0.016743 | 0.001166 | 1 | 1 | 8 | B2RQG2 | PHD finger | B2RQG2 [3 | 1 | 4244.987 | 0 | -0.56 | 0.028428 | 4.09 | 119.2 | 80.8 | 20.16 | 17.07 | | High | High | High | High | High | High | High | High | High | High | High | 0.003122 | 0.03229 | 3.88 |
| FALSE | High | NTVDLVNT | [K].NTVDL\ | 1xPhospho | B2RUA5 1x | 1.71E-06 | 0 | 1 | 2 | 5 | B2RUA5 | SAPS doma | B2RUA5 [5 | 0 | 2715.185 | 0 | -0.91 | 0.002219 | 6.88 | 130.6 | 69.4 | 3.91 | 13.05 | | High | High | High | High | High | High | High | High | High | High | High | 0 | 1.85E-07 | 3.32 |
| FALSE | High | NTVDLVTT | [R].NTVDL\ | 1xCarbami | G5E8R4 1x | 0.008167 | 0.000736 | 1 | 2 | 4 | G5E8R4 | SAPS doma | G5E8R4 [5 | 0 | 3071.443 | 0 | -0.36 | 0.015255 | 3.51 | 112.7 | 87.3 | 5.35 | 10.42 | | High | High | High | High | High | High | High | High | High | High | High | 0.002455 | 0.02656 | 2.67 |
| FALSE | High | NTVMCAV\ | [K].NTVMC | 1xCarbami | Q3U3X7 1x | 0.000916 | 0 | 1 | 5 | 1 | Q3U3X7 | Putative ur | Q3U3X7 [6 | 1 | 3638.733 | 1 | 1.09 | 0.00077 | 11.78 | 63.9 | 136.1 | 10.08 | 8.33 | | High | High | High | High | High | High | High | High | High | High | High | 0 | 0.000215 | 3.58 |
| FALSE | High | NVAEALGH | [K].NVAEA | 1xPhospho | Q8R3Y8 1x | 6.02E-05 | 0 | 1 | 2 | 3 | Q8R3Y8 | Interferon | Q8R3Y8 [4 | 0 | 1660.882 | 0 | 1.13 | 0.000209 | 3.45 | 62.8 | 137.2 | 5.52 | 3.53 | | High | High | High | High | High | High | High | High | High | High | High | 0 | 1.01E-05 | 2.51 |
| FALSE | High | NVESVSPES | [R].NVESVS | 1xPhospho | Q7TMI6 1x | 0.014033 | 0.001011 | 1 | 4 | 3 | Q7TMI6 | Zinc finger | Q7TMI6 [3 | 0 | 2523.281 | 0 | 0.31 | 0.17114 | 7.73 | 92.4 | 107.6 | 7.74 | 13.24 | | High | High | High | High | High | High | High | High | High | High | High | 0.000801 | 0.007338 | 2.2 |
| FALSE | High | NVEVEVLSI | [K].NVEVE\ | 1xPhospho | J3QPN0 1x | 0.01029 | 0.000772 | 1 | 2 | 1 | J3QPN0 | MCG14131 | J3QPN0 [3 | 0 | 1738.939 | 0 | -1.97 | 0.00035 | 0.95 | 159 | 41 | 7.71 | 15.23 | | High | High | High | High | High | High | High | High | High | High | High | 0.000382 | 0.003233 | 2.82 |

| | | | | | | | | | | | | | | | | | | | | | | | | | | | | | | | | | | | | | | |
|---|---|---|---|---|---|---|---|---|---|---|---|---|---|---|---|---|---|---|---|---|---|---|---|---|---|---|---|---|---|---|---|---|---|---|---|---|---|---|
| FALSE | High | NVFEVGAG | [R].NVFEV | 1xPhospho | Q80XI6 1xP | 1.2E-06 | 0 | 1 | 1 | 3 | Q80XI6 | Mitogen-a | Q80XI6 [51 | 0 | 1901.9 | 0 | 0.8 | 0.004316 | 21.75 | 74.9 | 125.1 | 7.72 | 9.87 | | High | High | High | High | High | High | High | High | High | High | High | 0 | 1.25E-07 | 4.09 |
| FALSE | High | NVHSEDFE | [R].NVHSE | 1xPhospho | Q3UYC8 1x | 0.000699 | 0 | 1 | 3 | 1 | Q3UYC8 | Putative ur | Q3UYC8 [9 | 0 | 1555.674 | 0 | 0.33 | 0.006902 | 7.33 | 87.3 | 112.7 | 5.58 | 9.1 | | High | High | High | High | High | High | High | High | High | High | High | 0 | 0.000159 | 1.94 |
| FALSE | High | NVLITPGK | [K].NVLITP | 1xPhospho | Q8BHE0 1x | 0.067619 | 0.004864 | 1 | 1 | 1 | Q8BHE0 | Proline-ricl | Q8BHE0 [2 | 0 | 1379.806 | 0 | 1.51 | 0.000505 | 17.75 | 52.9 | 147.1 | 20.34 | 4.51 | | High | High | High | High | High | High | High | High | High | High | High | 0.00253 | 0.02739 | 1.94 |
| FALSE | High | NVIPDTPPS | [R].NVIPDT | 1xPhospho | E9Q777 1x | 1.24E-05 | 0 | 1 | 3 | 1 | E9Q777 | A kinase (P | E9Q777 [1 | 0 | 3146.606 | 0 | 0.18 | 0.075845 | 12.39 | 90.7 | 109.3 | 17.38 | 7.61 | | High | High | High | High | High | High | High | High | High | High | High | 0 | 1.72E-06 | 3.31 |
| FALSE | High | NVIPDTPPS | [R].NVIPDT | 2xPhospho | E9Q777 2x | 0.003755 | 0.00021 | 1 | 3 | 1 | E9Q777 | A kinase (P | E9Q777 [1 | 0 | 3226.573 | 0 | 1.23 | 0.011545 | 15.4 | 59.8 | 140.2 | 11.48 | 18.05 | | High | High | High | High | High | High | High | High | High | High | High | 9.42E-05 | 0.001052 | 3.46 |
| FALSE | High | NVMSAFGI | [K].NVMSA | 1xPhospho | Q9Z1A1 1x | 2.19E-05 | 0 | 1 | 5 | 2 | Q9Z1A1 | Putative ur | Q9Z1A1 [1 | 0 | 2800.27 | 0 | 1.06 | 0.00098 | 9.71 | 62 | 138 | 17.29 | 14.05 | | High | High | High | High | High | High | High | High | High | High | High | 0 | 3.27E-06 | 3.81 |
| FALSE | High | NVMSAFGI | [K].NVMSA | 1xPhospho | Q9Z1A1 1x | 3.84E-09 | 0 | 1 | 5 | 2 | Q9Z1A1 | Putative ur | Q9Z1A1 [1 | 0 | 2816.265 | 0 | 1.34 | 0.036029 | 4.56 | 56.6 | 143.4 | 11.88 | 38 | | High | High | High | High | High | High | High | High | High | High | High | 0 | 3.86E-09 | 5.54 |
| FALSE | High | NTSMSPR | [K].NTSMS | 1xPhospho | Q3TV38 1x | 0.019966 | 0.001438 | 1 | 4 | 1 | Q3TV38 | Putative ur | Q3TV38 [2 | 0 | 1117.491 | 0 | 1.93 | 0.002319 | 12.87 | 42.4 | 157.6 | 7.07 | 16.48 | | High | High | High | High | High | High | High | High | High | High | High | 0.000786 | 0.006804 | 1.03 |
| FALSE | High | QPLPDEFLS | [K].QPLPDI | 1xCarbami | P35123 1xI | 6.74E-07 | 0 | 1 | 9 | 2 | P35123 | Ubiquitin c | P35123 [63 | 0 | 2580.2 | 0 | 0.29 | 0.14841 | 28.78 | 98.9 | 101.1 | 17.29 | 10.85 | | High | High | High | High | High | High | High | High | High | High | High | 0 | 3.09E-06 | 4.15 |
| FALSE | High | NSSSPPSPS | [R].NSSSPP | 1xPhospho | E9Q1P8 1x | 1.32E-05 | 0 | 1 | 1 | 1 | E9Q1P8 | Interferon | E9Q1P8 [4 | 0 | 1784.784 | 0 | -1.17 | 0.001693 | 1.71 | 143.2 | 56.8 | 7.82 | 14.91 | | High | High | High | High | High | High | High | High | High | High | High | 0 | 1.85E-06 | 1.31 |
| FALSE | High | NSSQTGGK | [K].NSSQT | 1xPhospho | Q925J9 1xI | 8.87E-05 | 0 | 1 | 2 | 3 | Q925J9 | Mediator c | Q925J9 [11 | 0 | 2084.079 | 0 | 1.51 | 0.147754 | 62.84 | 56.7 | 143.3 | 40.69 | 40.51 | | High | High | High | High | High | High | High | High | High | High | High | 0 | 1.57E-05 | 2.92 |
| FALSE | High | NSGTPGLQ | [R].NSGTP | 1xPhospho | P55200 1xI | 6.91E-06 | 0 | 1 | 3 | 1 | P55200 | Histone-lys | P55200 [30 | 0 | 2685.438 | 0 | 0.14 | 0.402147 | 5.61 | 98 | 102 | 12 | 9.95 | | High | High | High | High | High | High | High | High | High | High | High | 0 | 8.91E-07 | 4.55 |
| FALSE | High | NSGTPGLQ | [R].NSGTP | 2xPhospho | P55200 2xI | 0.030367 | 0.001857 | 1 | 3 | 2 | P55200 | Histone-lys | P55200 [30 | 0 | 2765.404 | 0 | 0.97 | 0.001919 | 8.07 | 57.4 | 142.6 | 19.32 | 13.6 | | High | High | High | High | High | High | High | High | High | High | High | 0.001011 | 0.01094 | 3.18 |
| FALSE | High | NSLEGYSR | [R].NSLEGY | 1xPhospho | Q6PHZ5 1x | 0.049971 | 0.003641 | 1 | 2 | 1 | Q6PHZ5 | Putative RI | Q6PHZ5 [5 | 0 | 1234.567 | 0 | 0.52 | 0.000436 | 1.92 | 83.5 | 116.5 | 5.94 | 6.74 | | High | High | High | High | High | High | High | High | High | High | High | 0.001902 | 0.01935 | 1.37 |
| FALSE | High | NSIEIAGLTI | [R].NSIEIA | 1xPhospho | G5E8J4 1xI | 0.001831 | 5.66E-05 | 1 | 5 | 1 | G5E8J4 | Tyrosine-p | G5E8J4 [63 | 0 | 2023.021 | 0 | 0.57 | 0.002898 | 9.12 | 80.6 | 119.4 | 7.88 | 3.95 | | High | High | High | High | High | High | High | High | High | High | High | 2.58E-05 | 0.000469 | 2.66 |
| FALSE | High | NSLESISSID | [R].NSLESIS | 1xPhospho | Q80U72-4 | 0.002463 | 0.000109 | 2 | 5 | 1 | Q80U72-4; | Isoform 4 c | Q80U72-4 | 0 | 1529.741 | 0 | 0.22 | 0.056061 | 12.63 | 88.2 | 111.8 | 6.85 | 12.46 | Shared | High | High | High | High | High | High | High | High | High | High | High | 4.94E-05 | 0.000655 | 2.85 |
| FALSE | High | NSLESISSID | [R].NSLESIS | 2xPhospho | Q80U72-4 | 0.004911 | 0.000504 | 2 | 5 | 2 | Q80U72-4; | Isoform 4 c | Q80U72-4 | 0 | 1609.707 | 0 | 0.19 | 0.043291 | 13.04 | 94.3 | 105.7 | 2.58 | 8.31 | Shared | High | High | High | High | High | High | High | High | High | High | High | 0.000275 | 0.00196 | 3.21 |
| FALSE | High | NSLGEGHL | [R].NSLGE | 1xPhospho | Q5SUQ9 1x | 0.008927 | 0.000736 | 1 | 2 | 2 | Q5SUQ9 | CST compl | Q5SUQ9 [1 | 0 | 4501.459 | 0 | 0.43 | 0.056347 | 22.07 | 92.5 | 107.5 | 7.66 | 15.15 | | High | High | High | High | High | High | High | High | High | High | High | 0.000327 | 0.002775 | 3.31 |
| FALSE | High | NSLGGDVL | [R].NSLGG | 1xPhospho | Q9DBR1 1x | 0.018774 | 0.001362 | 1 | 2 | 1 | Q9DBR1 | 5'-3' exorib | Q9DBR1 [6 | 0 | 1514.782 | 0 | 1.34 | 0.007175 | 12.95 | 53.7 | 146.3 | 19.29 | 21.98 | | High | High | High | High | High | High | High | High | High | High | High | 0.000723 | 0.006335 | 3.06 |
| FALSE | High | NSLGGDVL | [R].NSLGG | 1xPhospho | Q9DBR1 1x | 1.32E-05 | 0 | 1 | 2 | 1 | Q9DBR1 | 5'-3' exorib | Q9DBR1 [6 | 0 | 1743.945 | 0 | 2.62 | 0.000779 | 12.88 | 28 | 172 | 13.35 | 13.49 | | High | High | High | High | High | High | High | High | High | High | High | 0 | 1.85E-06 | 2.87 |
| FALSE | High | NSLGQSR | [R].NSLGQ | 1xPhospho | Q8C9G3 1x | 0.038346 | 0.002563 | 1 | 6 | 1 | Q8C9G3 | Putative ur | Q8C9G3 [4 | 0 | 1070.519 | 0 | -0.67 | 0.005999 | 5.87 | 125.8 | 74.2 | 6.74 | 16.11 | | High | High | High | High | High | High | High | High | High | High | High | 0.001355 | 0.01426 | 0.86 |
| FALSE | High | NSLIELPED | [R].NSLIELI | 1xCarbami | O70481 1x | 0.000411 | 0 | 1 | 2 | 1 | O70481 | E3 ubiquiti | O70481 [1 | 0 | 2815.332 | 0 | 0.97 | 0.002756 | 9.93 | 70.6 | 129.4 | 6.39 | 9.96 | | High | High | High | High | High | High | High | High | High | High | High | 0 | 8.75E-05 | 1.69 |
| FALSE | High | NSLLGLEEE | [K].NSLLGL | 1xPhospho | Q62273 1x | 0.002232 | 0.000109 | 1 | 2 | 1 | Q62273 | Sulfate trai | Q62273 [5 | 0 | 2814.374 | 0 | 2.09 | 0.0028 | 53.02 | 32.1 | 167.9 | 45.91 | 18.21 | | High | High | High | High | High | High | High | High | High | High | High | 4.94E-05 | 0.000586 | 2.75 |
| FALSE | High | NSLMSLSEI | [R].NSLMS | 1xPhospho | A0A140LJJ | 0.028824 | 0.001829 | 1 | 7 | 2 | A0A140LJJ | A-kinase ar | A0A140LJJ | 0 | 2194.005 | 0 | 0.13 | 0.807811 | 10.16 | 96.1 | 103.9 | 2.6 | 15.39 | | High | High | High | High | High | High | High | High | High | High | High | 0.001889 | 0.01908 | 2.44 |
| FALSE | High | NSGSFSSPS | [R].NSGSFS | 1xPhospho | A0A0R4J1L | 0.000325 | 0 | 1 | 8 | 1 | A0A0R4J1L | Microtubu | A0A0R4J1L | 0 | 1631.763 | 0 | -1.28 | 0.009302 | 4.23 | 147.8 | 52.2 | 17.78 | 18.12 | | High | High | High | High | High | High | High | High | High | High | High | 0 | 6.76E-05 | 1.95 |
| FALSE | High | NSIQFTDG | [R].NSIQFT | 1xPhospho | A7E215 1xI | 1.14E-06 | 0 | 1 | 3 | 4 | A7E215 | Rps6ka3 pr | A7E215 [4 | 0 | 1938.961 | 0 | 2.92 | 0.000181 | 21.3 | 22.8 | 177.2 | 18.7 | 4.91 | | High | High | High | High | High | High | High | High | High | High | High | 0 | 1.17E-07 | 3.3 |
| FALSE | High | NSLSASPPF | [R].NSLSAS | 1xPhospho | Q8BQ33 1x | 6.47E-05 | 0 | 1 | 1 | 2 | Q8BQ33 | Treslin OS= | Q8BQ33 [1 | 0 | 2135.094 | 0 | -0.59 | 0.030855 | 4.04 | 121.4 | 78.6 | 15.97 | 17.32 | | High | High | High | High | High | High | High | High | High | High | High | 0 | 1.1E-05 | 3.38 |
| FALSE | High | NSLSASPPF | [R].NSLSAS | 2xPhospho | Q8BQ33 2x | 6.13E-05 | 0 | 1 | 1 | 2 | Q8BQ33 | Treslin OS= | Q8BQ33 [1 | 0 | 2215.06 | 0 | -0.91 | 0.006988 | 2.16 | 131.7 | 68.3 | 12.85 | 17.72 | | High | High | High | High | High | High | High | High | High | High | High | 0.001999 | 0.02047 | 2.7 |
| FALSE | High | NSLSILVPH | [R].NSLSIL | 1xPhospho | Q68FD7 1x | 0.040373 | 0.002982 | 1 | 1 | 1 | Q68FD7 | Folliculin-ir | Q68FD7 [9 | 0 | 1817.993 | 0 | 0.1 | 0.344317 | 18.75 | 97.9 | 102.1 | 8.35 | 7.76 | | High | High | High | High | High | High | High | High | High | High | High | 0.001566 | 0.01507 | 1.33 |
| FALSE | High | NSLSPATQI | [K].NSLSPA | 1xPhospho | Q3V0B5 1x | 0.006536 | 0.00062 | 1 | 4 | 2 | Q3V0B5 | Putative ur | Q3V0B5 [5 | 0 | 1282.635 | 0 | -1.16 | 0.006323 | 4 | 135.2 | 64.8 | 12.78 | 35.93 | | High | High | High | High | High | High | High | High | High | High | High | 0.000538 | 0.004489 | 1.76 |
| FALSE | High | NSISPGRN | [R].NSISPG | 1xDeamida | B2RUG9 1x | 0.034579 | 0.002233 | 1 | 6 | 1 | B2RUG9 | Adenomat | B2RUG9 [2 | 1 | 2077.084 | 0 | 0.52 | 0.01478 | 5.89 | 84.3 | 115.7 | 10.15 | 8.65 | | High | High | High | High | High | High | High | High | High | High | High | 0.001183 | 0.01269 | 2.43 |
| FALSE | High | NSLSQVED | [R].NSLSQ | 1xPhospho | A2A654 1x | 7.04E-06 | 0 | 1 | 5 | 1 | A2A654 | Bromodom | A2A654 [1 | 0 | 2460.162 | 0 | -0.24 | 0.159943 | 15.17 | 109.4 | 90.6 | 9.34 | 17.7 | | High | High | High | High | High | High | High | High | High | High | High | 0 | 9.15E-07 | 2.91 |
| FALSE | High | NSLSQVED | [R].NSLSQ | 1xPhospho | A2A654 1x | 9.89E-05 | 0 | 1 | 5 | 2 | A2A654 | Bromodom | A2A654 [1 | 0 | 2476.157 | 0 | -0.03 | 0.568317 | 7.65 | 100.8 | 99.2 | 12.95 | 7.94 | | High | High | High | High | High | High | High | High | High | High | High | 2.58E-05 | 0.000513 | 3.08 |
| FALSE | High | NSITEISDN | [K].NSITEIS | 1xPhospho | D3Z0X5 1xI | 1.87E-06 | 0 | 1 | 7 | 7 | D3Z0X5 | Pleckstrin I | D3Z0X5 [5 | 0 | 2486.129 | 0 | 0.33 | 0.02999 | 6.05 | 90.1 | 109.9 | 8.48 | 6.59 | | High | High | High | High | High | High | High | High | High | High | High | 0 | 1.34E-06 | 4.92 |
| FALSE | High | NSITEISDN | [K].NSITEIS | 2xPhospho | D3Z0X5 2xI | 0.000639 | 0 | 1 | 7 | 2 | D3Z0X5 | Pleckstrin I | D3Z0X5 [5 | 0 | 2566.095 | 0 | 1.22 | 0.007798 | 3.64 | 58.7 | 141.3 | 10.32 | 18.43 | | High | High | High | High | High | High | High | High | High | High | High | 0 | 0.00017 | 3.19 |
| FALSE | High | NSITEISDN | [K].NSITEIS | 1xDeamida | D3Z0X5 2xI | 0.065952 | 0.004766 | 1 | 7 | 1 | D3Z0X5 | Pleckstrin I | D3Z0X5 [5 | 0 | 2567.079 | 0 | 1.05 | 0.012778 | 21.29 | 69.6 | 130.4 | 23 | 13.54 | | High | High | High | High | High | High | High | High | High | High | High | 0.002455 | 0.02661 | 2.85 |
| FALSE | High | NSLTGEEG | [R].NSLTGI | 1xPhospho | Q9JI99 1xP | 0.000188 | 0 | 1 | 2 | 2 | Q9JI99 | Sphingosin | Q9JI99 [10 | 0 | 1813.935 | 0 | 0.53 | 0.005945 | 4.6 | 80.6 | 119.4 | 6.55 | 9.97 | | High | High | High | High | High | High | High | High | High | High | High | 0 | 3.65E-05 | 2.45 |
| FALSE | High | NSLWSGDI | [R].NSLWS | 1xPhospho | E9Q842 1x | 0.034281 | 0.002195 | 1 | 3 | 1 | E9Q842 | Neuron na | E9Q842 [9 | 0 | 1672.835 | 0 | 1.28 | 0.003151 | 10.54 | 52.4 | 147.6 | 17.43 | 15.67 | | High | High | High | High | High | High | High | High | High | High | High | 0.001168 | 0.01256 | 1.75 |
| FALSE | High | NSLYSTDFI | [K].NSLYST | 1xPhospho | Q9Z2U2 1x | 4.52E-09 | 0 | 1 | 2 | 5 | Q9Z2U2 | Zinc finger | Q9Z2U2 [6 | 0 | 3277.41 | 0 | -0.66 | 0.008707 | 8.52 | 122.4 | 77.6 | 7.32 | 12.79 | | High | High | High | High | High | High | High | High | High | High | High | 0 | 2.37E-10 | 5.83 |
| FALSE | High | NSIQNQES | [K].NSIQN | 1xCarbami | E9PWG2 1: | 6.35E-08 | 0 | 1 | 3 | 2 | E9PWG2 | Trafficking | E9PWG2 [2 | 0 | 2830.231 | 0 | 0.55 | 0.003938 | 13.87 | 77.2 | 122.8 | 8.96 | 5.28 | | High | High | High | High | High | High | High | High | High | High | High | 0 | 4.58E-09 | 4.8 |
| FALSE | High | NSGSDGSP | [R].NSGSD | 1xPhospho | Q6GQX2 1: | 0.039011 | 0.002718 | 1 | 1 | 2 | Q6GQX2 | Nck-associ | Q6GQX2 [4 | 0 | 1666.8 | 0 | -0.04 | 0.734154 | 14.2 | 104.5 | 95.5 | 5.17 | 9.77 | | High | High | High | High | High | High | High | High | High | High | High | 0.001421 | 0.01448 | 2.22 |
| FALSE | High | NSGSAAAE | [R].NSGSA | 1xPhospho | Q3U0X6 1x | 0.003372 | 0.00016 | 1 | 2 | 2 | Q3U0X6 | Putative ur | Q3U0X6 [8 | 0 | 2082.99 | 0 | -0.39 | 0.043046 | 6.76 | 112.8 | 87.2 | 10.03 | 8.44 | | High | High | High | High | High | High | High | High | High | High | High | 9.42E-05 | 0.000979 | 2.74 |
| FALSE | High | NSGPVSEV | [K].NSGPV | 1xDeamida | Q8BTI8 1xI | 8.83E-10 | 0 | 1 | 3 | 7 | Q8BTI8 | Serine/argi | Q8BTI8 [12 | 0 | 2287.126 | 0 | -0.42 | 0.005769 | 4.15 | 115.1 | 84.9 | 7.79 | 3.78 | | High | High | High | High | High | High | High | High | High | High | High | 0 | 3.78E-11 | 5.16 |
| FALSE | High | NSDLFTVLS | [K].NSDLFT | 1xPhospho | E9Q8I9-2 1 | 0.014929 | 0.001049 | 1 | 2 | 1 | E9Q8I9-2 | Isoform 2 c | E9Q8I9-2 [ | 0 | 1460.735 | 0 | -0.7 | 0.000466 | 2.78 | 125.5 | 74.5 | 4.78 | 4.74 | | High | High | High | High | High | High | High | High | High | High | High | 0.000555 | 0.004891 | 1.23 |
| FALSE | High | NSDLLTSPD | [K].NSDLLT | 1xPhospho | P05627 1xI | 0.00129 | 0 | 1 | 3 | 1 | P05627 | Transcripti | P05627 [57 | 0 | 1780.93 | 0 | 0.59 | 0.376572 | 40.53 | 83.7 | 116.3 | 38.35 | 31.58 | | High | High | High | High | High | High | High | High | High | High | High | 0 | 0.000317 | 3.42 |
| FALSE | High | NSDLLTSPD | [K].NSDLLT | 1xPhospho | P05627 1xI | 8.58E-06 | 0 | 1 | 3 | 4 | P05627 | Transcripti | P05627 [57 | 0 | 2010.092 | 0 | 0.63 | 0.022269 | 16.31 | 78.5 | 121.5 | 11.88 | 9.65 | | High | High | High | High | High | High | High | High | High | High | High | 0 | 0.000142 | 2.77 |
| FALSE | High | NSDPWAAS | [K].NSDPW | 1xPhospho | Q8CHU3 1x | 0.000146 | 0 | 1 | 6 | 1 | Q8CHU3 | Epsin-2 OS | Q8CHU3 [3 | 0 | 2267.086 | 0 | 0.79 | 0.02701 | 31.35 | 68.2 | 131.8 | 11.15 | 28.41 | | High | High | High | High | High | High | High | High | High | High | High | 0 | 2.74E-05 | 4.05 |
| FALSE | High | NSDQEGG | [R].NSDQE | 1xPhospho | Q52KI8 1xI | 0.000257 | 0 | 1 | 7 | 2 | Q52KI8 | Serine/argi | Q52KI8 [84 | 0 | 1429.672 | 0 | -0.21 | 0.109391 | 6.88 | 110.4 | 89.6 | 11 | 18.31 | | High | High | High | High | High | High | High | High | High | High | High | 2.58E-05 | 0.000537 | 2.07 |
| FALSE | High | NSDSIVSLP | [K].NSDSIV | 2xPhospho | B2RY22 2xI | 0.095408 | 0.007437 | 2 | 17 | 2 | B2RY22; A | Clasp1 pro | B2RY22 [54 | 1 | 3694.773 | 0 | 0.43 | 0.0251 | 12.75 | 85.2 | 114.8 | 8.33 | 9.12 | Shared | High | High | High | High | High | High | High | High | High | High | High | 0.003768 | 0.04068 | 3.07 |
| FALSE | High | NSDSNLLSI | [K].NSDSN | 1xPhospho | E9PWG2 1: | 0.026191 | 0.001676 | 1 | 3 | 1 | E9PWG2 | Trafficking | E9PWG2 [2 | 0 | 2142.016 | 0 | 0.1 | 0.694681 | 14.18 | 99.5 | 100.5 | 22.64 | 19.26 | | High | High | High | High | High | High | High | High | High | High | High | 0.000903 | 0.009203 | 2.64 |
| FALSE | High | NSDSNLLSI | [K].NSDSN | 1xPhospho | E9PWG2 1: | 4.35E-08 | 0 | 1 | 3 | 4 | E9PWG2 | Trafficking | E9PWG2 [2 | 0 | 2371.179 | 0 | -0.13 | 0.145869 | 1.94 | 103.9 | 96.1 | 4.31 | 10.88 | | High | High | High | High | High | High | High | High | High | High | High | 0 | 2.98E-09 | 4.22 |
| FALSE | High | NSDSNLLSI | [K].NSDSN | 1xDeamida | E9PWG2 1: | 0.000106 | 0 | 1 | 3 | 3 | E9PWG2 | Trafficking | E9PWG2 [2 | 0 | 2372.163 | 0 | 0.2 | 0.493296 | 18.4 | 97.9 | 102.1 | 19.24 | 7.04 | | High | High | High | High | High | High | High | High | High | High | High | 0 | 1.92E-05 | 3.51 |
| FALSE | High | NSDSPASA | [R].NSDSP | 1xPhospho | Q5PRE5 1x | 0.000492 | 0 | 1 | 2 | 1 | Q5PRE5 | Proline anc | Q5PRE5 [4 | 0 | 1629.789 | 0 | -0.68 | 0.002128 | 4.73 | 123.3 | 76.7 | 4.69 | 7.68 | | High | High | High | High | High | High | High | High | High | High | High | 0 | 0.000108 | 1.97 |
| FALSE | High | NSEDDELTI | [K].NSEDDI | 1xPhospho | E9PWW9 1 | 8.03E-05 | 0 | 1 | 1 | 2 | E9PWW9 | Remodelin | E9PWW9 [ | 0 | 1588.751 | 0 | -0.76 | 0.00101 | 2.04 | 126.8 | 73.2 | 5.85 | 3.48 | | High | High | High | High | High | High | High | High | High | High | High | 0 | 1.4E-05 | 2.89 |
| FALSE | High | NSEIDELK | [R].NSEIDE | 1xPhospho | F8VPV0 1xI | 0.001147 | 0 | 1 | 7 | 1 | F8VPV0 | Pericentrin | F8VPV0 [1 | 0 | 1485.76 | 0 | 0.31 | 0.059264 | 29.57 | 87.7 | 112.3 | 8.63 | 11.41 | | High | High | High | High | High | High | High | High | High | High | High | 0 | 0.000277 | 2.74 |
| FALSE | High | NSEPAEGLI | [R].NSEPAI | 1xPhospho | Q3UIV3 1x | 0.002334 | 0.000109 | 1 | 11 | 1 | Q3UIV3 | Putative ur | Q3UIV3 [5 | 1 | 2049.98 | 0 | -1.13 | 0.067694 | 3.03 | 136.3 | 63.7 | 34.15 | 24.38 | | High | High | High | High | High | High | High | High | High | High | High | 4.94E-05 | 0.000617 | 1.77 |
| FALSE | High | NSEPAEGLI | [R].NSEPAI | 1xPhospho | Q3UIV3 1x | 0.000233 | 0 | 1 | 11 | 2 | Q3UIV3 | Putative ur | Q3UIV3 [5 | 1 | 2279.143 | 0 | -1.66 | 0.000459 | 0.65 | 152.5 | 47.5 | 5.55 | 11.61 | | High | High | High | High | High | High | High | High | High | High | High | 0 | 4.66E-05 | 2.29 |
| FALSE | High | NSEPAEGLI | [R].NSEPAI | 2xPhospho | Q3UIV3 2x | 0.069325 | 0.005112 | 1 | 11 | 2 | Q3UIV3 | Putative ur | Q3UIV3 [5 | 1 | 2359.11 | 0 | -0.55 | 0.009204 | 2.95 | 119.3 | 80.7 | 9.8 | 4.03 | | High | High | High | High | High | High | High | High | High | High | High | 0.002618 | 0.028 | 1.94 |
| FALSE | High | NSESCIPED | [R].NSESCI | 2xCarbami | E9Q309 1x | 0.000712 | 0 | 1 | 1 | 1 | E9Q309 | Centrosom | E9Q309 [1 | 0 | 2828.232 | 0 | 0.02 | 0.996213 | 16.45 | 100.7 | 99.3 | 17.67 | 8.9 | | High | High | High | High | High | High | High | High | High | High | High | 0 | 0.000163 | 3.34 |
| FALSE | High | NSFGYR | [K].NSFGYI | 1xPhospho | Q6A0A2 1x | 0.091627 | 0.007146 | 1 | 3 | 1 | Q6A0A2 | La-related | Q6A0A2 [4 | 0 | 1052.476 | 0 | 0.36 | 0.03865 | 17.82 | 89 | 111 | 7.88 | 5.8 | | High | High | High | High | High | High | High | High | High | High | High | 0.003646 | 0.03895 | 1.96 |
| FALSE | High | NSFSPSK | [K].NSFSPS | 1xPhospho | Q9CQ82 1x | 0.045122 | 0.003373 | 1 | 1 | 1 | Q9CQ82 | Centromer | Q9CQ82 [1 | 0 | 1304.665 | 0 | -0.86 | 0.001561 | 1.79 | 129.7 | 70.3 | 9.64 | 5.44 | | High | High | High | High | High | High | High | High | High | High | High | 0.001712 | 0.01706 | 1.53 |
| FALSE | High | NSFYTQLPI | [R].NSFYT | 1xPhospho | Q3UM95 1 | 0.006653 | 0.00062 | 1 | 3 | 4 | Q3UM95 | Putative ur | Q3UM95 [ | 0 | 2776.359 | 0 | 0.56 | 0.191659 | 18.87 | 80.4 | 119.6 | 25.88 | 33.98 | | High | High | High | High | High | High | High | High | High | High | High | 0.000275 | 0.001992 | 3.29 |
| FALSE | High | NSGAGAVI | [K].NSGAG | 1xPhospho | Q8CAF4-3 | 0.000297 | 0 | 1 | 3 | 1 | Q8CAF4-3 | Isoform 3 c | Q8CAF4-3 | 0 | 2157.017 | 0 | 0.59 | 0.023663 | 18.22 | 79.7 | 120.3 | 15.24 | 10.27 | | High | High | High | High | High | High | High | High | High | High | High | 0 | 6.08E-05 | 3.04 |
| FALSE | High | NSGATADA | [R].NSGAT | 1xPhospho | B2RUC3 1x | 1.69E-06 | 0 | 1 | 8 | 1 | B2RUC3 | Excision re | B2RUC3 [3 | 0 | 1612.753 | 0 | -0.46 | 0.030043 | 2.96 | 116 | 84 | 11.42 | 7.4 | | High | High | High | High | High | High | High | High | High | High | High | 0 | 1.83E-07 | 1.2 |
| FALSE | High | NSGEETSS | [K].NSGEE | 1xPhospho | B2RQG2 1x | 4.81E-09 | 0 | 1 | 2 | 3 | B2RQG2 | PHD finger | B2RQG2 [1 | 0 | 2728.324 | 0 | -1.21 | 0.000427 | 1.18 | 140.9 | 59.1 | 7.1 | 4.54 | | High | High | High | High | High | High | High | High | High | High | High | 0 | 7.84E-07 | 4.41 |
| FALSE | High | NSGGGYSII | [K].NSGGG | 1xPhospho | Q5U464 1x | 0.006834 | 0.00062 | 1 | 1 | 1 | Q5U464 | Arf-GAP wi | Q5U464 [2 | 0 | 2122.023 | 0 | -1.36 | 0.002793 | 0.5 | 147.7 | 52.3 | 10.02 | 30.05 | | High | High | High | High | High | High | High | High | High | High | High | 0.000275 | 0.002051 | 1.7 |
| FALSE | High | NSGLGSPSI | [R].NSGLG | 1xCarbami | Q8BL48 2x | 0.00642 | 0.00062 | 1 | 3 | 2 | Q8BL48 | RING finge | Q8BL48 [3 | 0 | 2282.983 | 0 | 0.8 | 0.000892 | 2.49 | 70.9 | 129.1 | 6.75 | 4.19 | | High | High | High | High | High | High | High | High | High | High | High | 0.000275 | 0.001916 | 2.3 |
| FALSE | High | NSGLTPPV | [R].NSGLTI | 1xPhospho | P53762 1xI | 0.003402 | 0.00021 | 1 | 7 | 1 | P53762 | Aryl hydro | P53762 [60 | 0 | 2930.555 | 0 | 0.74 | 0.033526 | 15.4 | 74.8 | 125.2 | 6.86 | 18.12 | | High | High | High | High | High | High | High | High | High | High | High | 9.42E-05 | 0.000939 | 2.62 |
| FALSE | High | NSGIWDSP | [R].NSGIW | 1xPhospho | Q68FM7 1: | 0.050826 | 0.003676 | 1 | 5 | 1 | Q68FM7 | Rho guanir | Q68FM7 [1 | 1 | 3302.469 | 0 | -0.25 | 0.05713 | 8.97 | 109 | 91 | 10.3 | 8.96 | | High | High | High | High | High | High | High | High | High | High | High | 0.001932 | 0.01966 | 2.36 |
| FALSE | High | NSGPASPA | [R].NSGPA | 1xPhospho | H9LAN6 1x | 7.02E-05 | 0 | 1 | 7 | 1 | H9LAN6 | Histone-lys | H9LAN6 [9 | 0 | 1899.928 | 0 | -1.02 | 0.001394 | 0.98 | 134.2 | 65.8 | 7.66 | 5.99 | | High | High | High | High | High | High | High | High | High | High | High | 0 | 1.21E-05 | 1.59 |
| FALSE | High | NSGPVSEV | [K].NSGPV | 1xPhospho | Q8BTI8 1xI | 2.6E-05 | 0 | 1 | 3 | 3 | Q8BTI8 | Serine/argi | Q8BTI8 [12 | 0 | 2056.979 | 0 | -0.29 | 0.293993 | 16.34 | 112.7 | 87.3 | 27.45 | 26.67 | | High | High | High | High | High | High | High | High | High | High | High | 0.000246 | 0.001513 | 3.84 |
| FALSE | High | NSGPVSEV | [K].NSGPV | 1xPhospho | Q8BTI8 1xI | 1.07E-08 | 0 | 1 | 3 | 6 | Q8BTI8 | Serine/argi | Q8BTI8 [12 | 0 | 2286.142 | 0 | -0.47 | 0.016319 | 5.61 | 116.1 | 83.9 | 8.74 | 8.39 | | High | High | High | High | High | High | High | High | High | High | High | 0 | 6.23E-10 | 4.69 |
| FALSE | High | NSNPIIAEL | [R].NSNPII | 1xPhospho | Q501J7-3 1 | 0.009756 | 0.000736 | 1 | 4 | 1 | Q501J7-3 | Isoform 3 c | Q501J7-3 [ | 0 | 3755.042 | 0 | 0.87 | 0.047532 | 26.41 | 73.7 | 126.3 | 12.59 | 22.93 | | High | High | High | High | High | High | High | High | High | High | High | 0.000344 | 0.003064 | 3.24 |
| FALSE | High | NSSQTGGK | [K].NSSQT | 1xPhospho | Q925J9 1xI | 0.000616 | 0 | 1 | 2 | 2 | Q925J9 | Mediator c | Q925J9 [11 | 0 | 2313.242 | 0 | 1.07 | 0.000813 | 9.15 | 63.8 | 136.2 | 4.52 | 6.54 | | High | High | High | High | High | High | High | High | High | High | High | 0 | 0.000139 | 2.66 |
| FALSE | High | NSNSDGW | [K].NSNSD | 1xPhospho | V9GXM1 1 | 0.000186 | 0 | 1 | 7 | 3 | V9GXM1 | ADP-ribosy | V9GXM1 [3 | 0 | 2233.899 | 0 | 1.16 | 0.000857 | 4.55 | 59.9 | 140.1 | 13.3 | 13.5 | | High | High | High | High | High | High | High | High | High | High | High | 0 | 3.59E-05 | 3.94 |
| FALSE | High | NSPGAVG | [K].NSPGA | 1xPhospho | A0A1B0GR | 2.42E-08 | 0 | 1 | 3 | 4 | A0A1B0GR | Single strai | A0A1B0GR | 0 | 1860.917 | 0 | 1.34 | 0.000405 | 17 | 55.8 | 144.2 | 6.59 | 4.94 | | High | High | High | High | High | High | High | High | High | High | High | 0 | 1.79E-06 | 4.57 |
| FALSE | High | NSSGPQSG | [R].NSSGP | 1xPhospho | E9PZM7 1x | 0.005275 | 0.000599 | 1 | 3 | 1 | E9PZM7 | SR-related | E9PZM7 [1 | 0 | 1531.656 | 0 | -0.61 | 0.039615 | 9.84 | 120.9 | 79.1 | 14.05 | 21.54 | | High | High | High | High | High | High | High | High | High | High | High | 0.000267 | 0.001538 | 1.89 |
| FALSE | High | NSSGTGLC | [R].NSSGT | 1xCarbami | Q8CFU0 1x | 0.008927 | 0.000736 | 1 | 1 | 1 | Q8CFU0 | Transmem | Q8CFU0 [2 | 0 | 1624.699 | 0 | 1.32 | 0.001069 | 7 | 53.3 | 146.7 | 11.36 | 11.34 | | High | High | High | High | High | High | High | High | High | High | High | 0.000327 | 0.00275 | 1.38 |
| FALSE | High | NSSGVSPL | [R].NSSGV | 1xCarbami | G5E8J3 1xI | 3.06E-05 | 0 | 1 | 3 | 2 | G5E8J3 | Bromodom | G5E8J3 [3 | 0 | 3102.574 | 0 | 1.24 | 0.000458 | 11.83 | 59.4 | 140.6 | 5.22 | 5.42 | | High | High | High | High | High | High | High | High | High | High | High | 0 | 4.74E-06 | 3.94 |
| FALSE | High | NSSLLSFDS | [K].NSSLLS | 1xPhospho | Q8K039 1x | 5.43E-07 | 0 | 1 | 2 | 8 | Q8K039 | Uncharacte | Q8K039 [1 | 0 | 1894.779 | 0 | -0.4 | 0.005556 | 4.99 | 111.3 | 88.7 | 5.01 | 8.64 | | High | High | High | High | High | High | High | High | High | High | High | 0 | 5.14E-08 | 4.33 |
| FALSE | High | NSSLLSFDS | [K].NSSLLS | 2xPhospho | Q8K039 2x | 1.89E-06 | 0 | 1 | 2 | 3 | Q8K039 | Uncharacte | Q8K039 [1 | 0 | 1974.746 | 0 | 1.33 | 0.013067 | 5.96 | 52.9 | 147.1 | 15.27 | 30.65 | | High | High | High | High | High | High | High | High | High | High | High | 0 | 2.07E-07 | 4.05 |
| FALSE | High | NSSIQLK | [R].NSSIQL | 1xPhospho | Q5PRF0 1x | 0.049549 | 0.00361 | 1 | 1 | 1 | Q5PRF0 | HEAT repe | Q5PRF0 [2 | 0 | 1327.739 | 0 | 1.14 | 0.003539 | 23.9 | 62.5 | 137.5 | 10.86 | 13.62 | | High | High | High | High | High | High | High | High | High | High | High | 0.001889 | 0.01909 | 1.77 |
| FALSE | High | NSSLVGR | [R].NSSLV | 1xPhospho | A2A9M4 1: | 0.036103 | 0.002342 | 1 | 6 | 3 | A2A9M4 | Dedicator c | A2A9M4 [4 | 0 | 1041.529 | 0 | 1.08 | 0.000517 | 10.19 | 61.4 | 138.6 | 9.64 | 4.23 | | High | High | High | High | High | High | High | High | High | High | High | 0.006155 | 0.0664 | 1.19 |
| FALSE | High | NSSMSLDE | [R].NSSMS | 1xPhospho | E9Q842 1x | 0.002719 | 0.000109 | 1 | 3 | 1 | E9Q842 | Neuron na | E9Q842 [1 | 0 | 1548.738 | 0 | 0.88 | 0.006898 | 13.57 | 67.7 | 132.3 | 10.27 | 14.72 | | High | High | High | High | High | High | High | High | High | High | High | 4.94E-05 | 0.000729 | 1.76 |
| FALSE | High | NSSMSLDE | [R].NSSMS | 1xPhospho | E9Q842 1x | 0.073462 | 0.005631 | 1 | 3 | 1 | E9Q842 | Neuron na | E9Q842 [1 | 0 | 1564.733 | 0 | 1.26 | 0.001516 | 15.32 | 59.5 | 140.5 | 13.79 | 8.87 | | High | High | High | High | High | High | High | High | High | High | High | 0.002911 | 0.03021 | 2.38 |
| FALSE | High | NSSNEELTI | [R].NSSNEI | 1xPhospho | Q3UQN2 1 | 0.005924 | 0.00062 | 1 | 2 | 3 | Q3UQN2 | F-BAR dom | Q3UQN2 [4 | 0 | 1559.772 | 0 | 0.94 | 0.001564 | 5.54 | 69.4 | 130.6 | 4.5 | 10.47 | | High | High | High | High | High | High | High | High | High | High | High | 0.000585 | 0.005189 | 1.9 |
| FALSE | High | NSSNFK | [K].NSSNFI | 1xPhospho | D3YYI8 1xF | 0.112856 | 0.009146 | 1 | 2 | 1 | D3YYI8 | Histone de | D3YYI8 [43 | 0 | 1234.623 | 0 | -0.56 | 0.015875 | 3.59 | 120.6 | 79.4 | 11.16 | 9.54 | | High | High | High | High | High | High | High | High | High | High | High | 0.004596 | 0.05 | 1.24 |
| FALSE | High | NSSNTGVG | [K].NSSNT | 1xPhospho | B2RY22 1xI | 2.2E-06 | 0 | 2 | 18 | 3 | B2RY22; A | Clasp1 pro | B2RY22 [1 | 0 | 1856.87 | 0 | 0.05 | 0.261238 | 12.86 | 97.5 | 102.5 | 5.12 | 9.61 | Shared | High | High | High | High | High | High | High | High | High | High | High | 0 | 3.2E-07 | 2.25 |
| FALSE | High | NSSNTGVG | [K].NSSNT | 1xDeamida | B2RY22 1xI | 3.94E-05 | 0 | 2 | 18 | 3 | B2RY22; A | Clasp1 pro | B2RY22 [1 | 0 | 1857.854 | 0 | -0.05 | 0.735812 | 8.71 | 102.3 | 97.7 | 7.02 | 10.67 | Shared | High | High | High | High | High | High | High | High | High | High | High | 0 | 5.21E-05 | 2.9 |
| FALSE | High | NSSFTQEA | [R].NSSFT | 1xPhospho | Q9WU22 1 | 0.010568 | 0.000772 | 1 | 3 | 1 | Q9WU22 | Tyrosine-p | Q9WU22 [ | 0 | 1449.657 | 0 | -0.48 | 0.004081 | 2.7 | 113.6 | 86.4 | 9.52 | 5.41 | | High | High | High | High | High | High | High | High | High | High | High | 0.000382 | 0.003332 | 2.75 |
| FALSE | High | NSSPAPES | [K].NSSPAI | 1xPhospho | Q8BRH3 1x | 0.022773 | 0.001542 | 1 | 4 | 2 | Q8BRH3 | Rho GTPas | Q8BRH3 [4 | 0 | 2071.018 | 0 | -0.3 | 0.041794 | 13.54 | 111.3 | 88.7 | 4.12 | 13.8 | | High | High | High | High | High | High | High | High | High | High | High | 0.000814 | 0.007889 | 2.35 |
| FALSE | High | NSSPAVPA | [R].NSSPA | 1xPhospho | H3BL37 1xI | 2.33E-07 | 0 | 1 | 7 | 10 | H3BL37 | Treacle prc | H3BL37 [8 | 0 | 2603.348 | 0 | 0.59 | 0.008662 | 18.58 | 79.8 | 120.2 | 8.33 | 10.89 | | High | High | High | High | High | High | High | High | High | High | High | 0 | 1.97E-08 | 3.69 |
| FALSE | High | NSSPAVPA | [R].NSSPA | 1xDeamida | H3BL37 1xI | 4.94E-05 | 0 | 1 | 7 | 1 | H3BL37 | Treacle prc | H3BL37 [8 | 0 | 2604.332 | 0 | 0.24 | 0.108576 | 16.17 | 95.4 | 104.6 | 10.96 | 4.13 | | High | High | High | High | High | High | High | High | High | High | High | 0 | 8.16E-06 | 3.69 |
| FALSE | High | NSSPAVPA | [R].NSSPA | 2xPhospho | H3BL37 2xI | 0.000125 | 0 | 1 | 7 | 1 | H3BL37 | Treacle prc | H3BL37 [8 | 0 | 2683.315 | 0 | 1.25 | 0.001573 | 2.8 | 60.9 | 139.1 | 7.69 | 10.32 | | High | High | High | High | High | High | High | High | High | High | High | 0 | 2.32E-05 | 3.35 |
| FALSE | High | NSSPLNCSI | [K].NSSPLN | 1xCarbami | Q9JHC9 1x | 0.01218 | 0.000889 | 1 | 7 | 2 | Q9JHC9 | ETS-relatec | Q9JHC9 [3 | 0 | 1343.598 | 0 | -0.21 | 0.020384 | 4.85 | 107.9 | 92.1 | 7.31 | 10.32 | | High | High | High | High | High | High | High | High | High | High | High | 0.000449 | 0.003922 | 1.78 |
| FALSE | High | NSSPPPSPN | [R].NSSPPF | 1xPhospho | G5E829 1x | 0.042868 | 0.003228 | 1 | 5 | 3 | G5E829 | Plasma me | G5E829 [1 | 0 | 1562.798 | 0 | 1.01 | 0.004997 | 5.48 | 66.8 | 133.2 | 5.33 | 11.39 | | High | High | High | High | High | High | High | High | High | High | High | 0.001651 | 0.01613 | 2.18 |
| FALSE | High | NSSPPPSPN | [R].NSSPPF | 2xPhospho | G5E829 2x | 0.017498 | 0.001245 | 1 | 5 | 1 | G5E829 | Plasma me | G5E829 [1 | 0 | 1642.764 | 0 | 1.02 | 0.002328 | 13.17 | 66.7 | 133.3 | 3.77 | 10.87 | | High | High | High | High | High | High | High | High | High | High | High | 0.000674 | 0.00584 | 2.09 |
| FALSE | High | NSSPRSPTN | [R].NSSPRS | 2xPhospho | Q3UGS4 2x | 0.000892 | 0 | 1 | 1 | 1 | Q3UGS4 | Mapk-regu | Q3UGS4 [1 | 1 | 2945.303 | 0 | 0.87 | 0.026169 | 21.13 | 76.5 | 123.5 | 11.2 | 17.9 | | High | High | High | High | High | High | High | High | High | High | High | 0 | 0.000209 | 3.09 |
| FALSE | High | NSSPSISPN | [R].NSSPSI | 1xPhospho | Q9DBR1 1x | 1.95E-05 | 0 | 1 | 2 | 8 | Q9DBR1 | 5'-3' exorib | Q9DBR1 [4 | 0 | 2713.292 | 0 | 0.15 | 0.250964 | 24.39 | 94.7 | 105.3 | 16.39 | 25.53 | | High | High | High | High | High | High | High | High | High | High | High | 0 | 2.86E-05 | 4.1 |
| FALSE | High | NSSPSISPN | [R].NSSPSI | 2xPhospho | Q9DBR1 2x | 0.001325 | 0 | 1 | 2 | 4 | Q9DBR1 | 5'-3' exorib | Q9DBR1 [4 | 0 | 2793.258 | 0 | 1.68 | 0.155508 | 71.97 | 48.7 | 151.3 | 42.63 | 45.03 | | High | High | High | High | High | High | High | High | High | High | High | 0.000225 | 0.001448 | 3.75 |
| FALSE | High | NSSPSISPN | [R].NSSPSI | 1xPhospho | Q9DBR1 1x | 4.61E-06 | 0 | 1 | 2 | 6 | Q9DBR1 | 5'-3' exorib | Q9DBR1 [4 | 0 | 2942.455 | 0 | 1.37 | 0.000677 | 18.14 | 57.9 | 142.1 | 8.27 | 6.33 | | High | High | High | High | High | High | High | High | High | High | High | 0 | 4.61E-05 | 3.73 |
| FALSE | High | NSSPSISPN | [R].NSSPSI | 1xDeamida | Q9DBR1 1x | 0.003997 | 0.000311 | 1 | 2 | 1 | Q9DBR1 | 5'-3' exorib | Q9DBR1 [4 | 0 | 2943.439 | 0 | 1.06 | 0.011056 | 38.47 | 72.3 | 127.7 | 22.22 | 11.34 | | High | High | High | High | High | High | High | High | High | High | High | 0.00014 | 0.001123 | 3.15 |
| FALSE | High | NSSPSISPN | [R].NSSPSI | 2xPhospho | Q9DBR1 2x | 8.58E-06 | 0 | 1 | 2 | 14 | Q9DBR1 | 5'-3' exorib | Q9DBR1 [4 | 0 | 3022.421 | 0 | 1.15 | 9.77E-05 | 2.54 | 62.1 | 137.9 | 5.91 | 3.54 | | High | High | High | High | High | High | High | High | High | High | High | 0 | 0.000135 | 4.67 |
| FALSE | High | NSSQQEDS | [K].NSSQQ | 1xPhospho | E9PWW9 1 | 0.117405 | 0.009523 | 1 | 4 | 2 | E9PWW9 | Remodelin | E9PWW9 [ | 1 | 2934.325 | 0 | -1.32 | 0.001035 | 1.26 | 144 | 56 | 8.5 | 13.2 | | High | High | High | High | High | High | High | High | High | High | High | 0.004782 | 0.05237 | 2.47 |
| FALSE | High | NSSPAVPA | [R].NSSPA | 1xPhospho | H3BL37 1xI | 0.103392 | 0.008249 | 1 | 7 | 3 | H3BL37 | Treacle prc | H3BL37 [8 | 0 | 2374.185 | 0 | 0.52 | 0.72643 | 58.09 | 85.4 | 114.6 | 40.54 | 43.16 | | High | High | High | High | High | High | High | High | High | High | High | 0.005054 | 0.05524 | 2.08 |
| FALSE | High | NSSDTCLR | [R].NSSDT | 1xCarbami | Q80VE5 1x | 0.007274 | 0.00062 | 1 | 1 | 1 | Q80VE5 | TBC1 doma | Q80VE5 [1 | 0 | 1261.545 | 0 | 0.27 | 0.009277 | 4.64 | 88.1 | 111.9 | 4.19 | 7.59 | | High | High | High | High | High | High | High | High | High | High | High | 0.000275 | 0.002195 | 2.1 |
| FALSE | High | NSSDQEGN | [R].NSSDQ | 2xPhospho | E9QLG3 2x | 0.022973 | 0.001578 | 1 | 7 | 2 | E9QLG3 | MAX gene- | E9QLG3 [2 | 0 | 2147.859 | 0 | -0.62 | 0.014042 | 3.86 | 124.3 | 75.7 | 9.85 | 27.37 | | High | High | High | High | High | High | High | High | High | High | High | 0.000847 | 0.007971 | 3.58 |
| FALSE | High | NSSDQEGN | [R].NSSDQ | 1xPhospho | E9QLG3 1x | 1.91E-07 | 0 | 1 | 7 | 5 | E9QLG3 | MAX gene- | E9QLG3 [2 | 0 | 2067.893 | 0 | -0.71 | 0.001 | 4.08 | 123.1 | 76.9 | 2.28 | 8.79 | | High | High | High | High | High | High | High | High | High | High | High | 0 | 3.59E-08 | 4.78 |
| FALSE | High | NSPGAVG | [K].NSPGA | 2xPhospho | A0A1B0GR | 1.66E-05 | 0 | 1 | 3 | 2 | A0A1B0GR | Single strai | A0A1B0GR | 0 | 1940.883 | 0 | 1.81 | 0.000209 | 7.96 | 44 | 156 | 7.08 | 5.92 | | High | High | High | High | High | High | High | High | High | High | High | 0 | 2.38E-06 | 3.88 |
| FALSE | High | NSPGCQVA | [R].NSPGC | 1xCarbami | Q9DBR1 1x | 4.98E-05 | 0 | 1 | 2 | 3 | Q9DBR1 | 5'-3' exorib | Q9DBR1 [4 | 0 | 1595.72 | 0 | 0.78 | 0.000445 | 1.71 | 76.1 | 123.9 | 6.83 | 4.69 | | High | High | High | High | High | High | High | High | High | High | High | 0 | 1.56E-05 | 2.84 |

| | | | | | | | | | | | | | | | | | | | | | | | | | | | | | | | | | | | | | | |
|---|---|---|---|---|---|---|---|---|---|---|---|---|---|---|---|---|---|---|---|---|---|---|---|---|---|---|---|---|---|---|---|---|---|---|---|---|---|---|
| FALSE | High | NSPGVPAG | [K].NSPGVI | 1xPhospho | E9PUQ5 1x | 0.037692 | 0.00249 | 1 | 9 | 1 | E9PUQ5 | Golgin sub | E9PUQ5 [3 | 0 | 1435.771 | 0 | 0.47 | 0.002044 | 8.56 | 84.3 | 115.7 | 4.71 | 6.04 | | High | High | High | High | High | High | High | High | High | High | High | 0.001324 | 0.01402 | 1.66 |
| FALSE | High | NSPGVSHS | [K].NSPGV! | 1xPhospho | Q9WUF3 1 | 0.003433 | 0.00021 | 1 | 1 | 1 | Q9WUF3 | CASP8-asso | Q9WUF3 [ | 0 | 1665.836 | 0 | -0.13 | 0.249292 | 7.65 | 101.7 | 98.3 | 6.78 | 9.17 | | High | High | High | High | High | High | High | High | High | High | High | 9.42E-05 | 0.000946 | 1.75 |
| FALSE | High | NSPLDCGS. | [R].NSPLDC | 1xCarbami | Q99NH0 1x | 0.000984 | 0 | 1 | 4 | 4 | Q99NH0 | Ankyrin rej | Q99NH0 [2 | 0 | 1655.73 | 0 | 0.06 | 0.87209 | 28.44 | 97.8 | 102.2 | 30.4 | 36.14 | | High | High | High | High | High | High | High | High | High | High | High | 0.000275 | 0.001718 | 3.19 |
| FALSE | High | NSPLDCGS. | [R].NSPLDC | 1xCarbami | Q99NH0 2x | 0.063258 | 0.004664 | 1 | 4 | 1 | Q99NH0 | Ankyrin rej | Q99NH0 [2 | 0 | 1735.696 | 0 | -0.39 | 0.262738 | 17.47 | 113.3 | 86.7 | 8.83 | 28.73 | | High | High | High | High | High | High | High | High | High | High | High | 0.002403 | 0.02521 | 2.73 |
| FALSE | High | NSPLDCGS. | [R].NSPLDC | 1xCarbami | Q99NH0 1x | 1.29E-06 | 0 | 1 | 4 | 3 | Q99NH0 | Ankyrin rej | Q99NH0 [2 | 0 | 1884.893 | 0 | -0.08 | 0.96979 | 10.92 | 102.6 | 97.4 | 10.38 | 9.33 | | High | High | High | High | High | High | High | High | High | High | High | 0 | 1.35E-07 | 3.68 |
| FALSE | High | NSPLDCGS. | [R].NSPLDC | 1xCarbami | Q99NH0 2x | 0.000121 | 0 | 1 | 4 | 1 | Q99NH0 | Ankyrin rej | Q99NH0 [2 | 0 | 1964.859 | 0 | -0.27 | 0.125408 | 9.24 | 108.9 | 91.1 | 8.85 | 6.92 | | High | High | High | High | High | High | High | High | High | High | High | 0 | 2.21E-05 | 3.23 |
| FALSE | High | NSPLPNCT | [R].NSPLPN | 1xCarbami | I7H459 1xF | 0.069903 | 0.005181 | 1 | 2 | 1 | I7H459 | Protein inc | I7H459 [33 | 0 | 1702.782 | 0 | 1.15 | 0.002635 | 19.96 | 61.8 | 138.2 | 12.02 | 10.15 | | High | High | High | High | High | High | High | High | High | High | High | 0.002662 | 0.02827 | 1.4 |
| FALSE | High | NSPMAVDI | [K].NSPMA | 1xPhospho | P97929 1xI | 6.95E-08 | 0 | 1 | 3 | 1 | P97929 | Breast can | P97929 [90 | 0 | 2877.411 | 0 | -1.18 | 0.001964 | 5.58 | 138.4 | 61.6 | 2.02 | 24.61 | | High | High | High | High | High | High | High | High | High | High | High | 0 | 5.07E-09 | 5.85 |
| FALSE | High | NSPMAVDI | [K].NSPMA | 1xPhospho | P97929 1xI | 1.24E-08 | 0 | 1 | 3 | 2 | P97929 | Breast can | P97929 [90 | 0 | 2893.405 | 0 | -1.03 | 0.008264 | 4.97 | 132.5 | 67.5 | 11.87 | 17.1 | | High | High | High | High | High | High | High | High | High | High | High | 0 | 7.36E-10 | 4.02 |
| FALSE | High | NSPMTISR | [R].NSPMT | 1xPhospho | O35144 1x | 0.01861 | 0.001362 | 1 | 4 | 3 | O35144 | Telomeric | O35144 [3 | 0 | 1230.575 | 0 | -0.88 | 0.000448 | 1.14 | 129.5 | 70.5 | 5.22 | 3.74 | | High | High | High | High | High | High | High | High | High | High | High | 0.000723 | 0.006292 | 1.9 |
| FALSE | High | NSPNNISGI | [K].NSPNN | 1xPhospho | Q9D032 1x | 7.29E-09 | 0 | 1 | 7 | 3 | Q9D032 | Single-stra | Q9D032 [3 | 0 | 2030.986 | 0 | -0.03 | 0.943532 | 8.86 | 101.9 | 98.1 | 3.02 | 6.85 | | High | High | High | High | High | High | High | High | High | High | High | 0 | 7.95E-10 | 3.65 |
| FALSE | High | NSPNNMSI | [K].NSPNN | 1xPhospho | Q9CYZ8 1x | 3.01E-05 | 0 | 1 | 2 | 1 | Q9CYZ8 | Single-stra | Q9CYZ8 [3 | 0 | 2022.927 | 0 | 0.67 | 0.026517 | 24.04 | 79.5 | 120.5 | 7.94 | 13.83 | | High | High | High | High | High | High | High | High | High | High | High | 0 | 4.64E-06 | 4.34 |
| FALSE | High | NSPNNMSI | [K].NSPNN | 1xPhospho | Q9CYZ8 1x | 1.83E-08 | 0 | 1 | 2 | 4 | Q9CYZ8 | Single-stra | Q9CYZ8 [3 | 0 | 2038.921 | 0 | 2.66 | 0.000514 | 9.18 | 28.4 | 171.6 | 19.72 | 8.07 | | High | High | High | High | High | High | High | High | High | High | High | 0 | 2.32E-09 | 4.68 |
| FALSE | High | NSPNNMSI | [K].NSPNN | 2xPhospho | Q9CYZ8 2x | 0.023584 | 0.001578 | 1 | 2 | 1 | Q9CYZ8 | Single-stra | Q9CYZ8 [3 | 0 | 2118.888 | 0 | 0.68 | 0.009645 | 18.17 | 80.3 | 119.7 | 17.13 | 9.08 | | High | High | High | High | High | High | High | High | High | High | High | 0.000859 | 0.008219 | 2.82 |
| FALSE | High | NSPPFLK | [K].NSPPFL | 1xPhospho | B0V2M3 1x | 0.072258 | 0.005346 | 1 | 5 | 1 | B0V2M3 | Zinc finger | B0V2M3 [7 | 0 | 1340.738 | 0 | -0.71 | 0.00268 | 6.12 | 120.1 | 79.9 | 5.41 | 9.81 | | High | High | High | High | High | High | High | High | High | High | High | 0.002855 | 0.02951 | 1.91 |
| FALSE | High | NSPQALR | [R].NSPQA | 1xPhospho | B2RX66 1x | 0.044739 | 0.003302 | 1 | 1 | 1 | B2RX66 | MCG12481 | B2RX66 [9 | 0 | 1094.556 | 0 | 0.28 | 0.04504 | 15.83 | 85.7 | 114.3 | 7.83 | 12.34 | | High | High | High | High | High | High | High | High | High | High | High | 0.00168 | 0.01692 | 2.8 |
| FALSE | High | NSPQSSPT | [R].NSPQS | 2xPhospho | Q920Q8 2x | 0.000589 | 0 | 1 | 4 | 7 | Q920Q8 | Influenza v | Q920Q8 [3 | 0 | 1848.855 | 0 | -1.27 | 0.002567 | 3.12 | 141.3 | 58.7 | 8.3 | 13.65 | | High | High | High | High | High | High | High | High | High | High | High | 2.58E-05 | 0.000559 | 2.76 |
| FALSE | High | NSPSAAST | [R].NSPSA | 1xPhospho | Q80TZ9 1x | 1.29E-07 | 0 | 1 | 2 | 2 | Q80TZ9 | Arginine-gl | Q80TZ9 [6 | 0 | 1890.885 | 0 | 0.39 | 0.01642 | 15.77 | 87.3 | 112.7 | 5.99 | 6.87 | | High | High | High | High | High | High | High | High | High | High | High | 0 | 1.02E-08 | 3.35 |
| FALSE | High | NSPSLQPP | [R].NSPSLC | 1xPhospho | S4R189 1xI | 0.000103 | 0 | 1 | 8 | 1 | S4R189 | Rho guanir | S4R189 [14 | 0 | 2887.476 | 0 | -0.27 | 0.173381 | 11.65 | 112.3 | 87.7 | 14.62 | 11.46 | | High | High | High | High | High | High | High | High | High | High | High | 0 | 1.86E-05 | 3.17 |
| FALSE | High | NSPSNLSS | [R].NSPSNI | 1xPhospho | Q8C0T5 1x | 6.3E-05 | 0 | 1 | 3 | 2 | Q8C0T5 | Signal-indu | Q8C0T5 [1 | 0 | 2253.982 | 0 | 2.67 | 0.001068 | 34.71 | 27.2 | 172.8 | 20.91 | 11.48 | | High | High | High | High | High | High | High | High | High | High | High | 0 | 5.17E-05 | 3.36 |
| FALSE | High | NSPSQEELI | [R].NSPSQI | 1xCarbami | P48754 1xI | 0.001767 | 5.66E-05 | 1 | 3 | 1 | P48754 | Breast can | P48754 [14 | 0 | 3603.622 | 0 | 0.32 | 0.507776 | 2.7 | 99 | 101 | 22.29 | 18.56 | | High | High | High | High | High | High | High | High | High | High | High | 2.58E-05 | 0.000452 | 2.81 |
| FALSE | High | NSPTGLAPI | [R].NSPTGI | 2xPhospho | M0QWZ1 2 | 6.91E-06 | 0 | 1 | 2 | 1 | M0QWZ1 | Protein FAI | M0QWZ1 [ | 0 | 4037.095 | 0 | -0.06 | 0.342563 | 23.2 | 103.7 | 96.3 | 12.8 | 10.66 | | High | High | High | High | High | High | High | High | High | High | High | 0 | 8.89E-07 | 4.89 |
| FALSE | High | NSQEQLLD | [K].NSQEQ | 1xPhospho | A2AAE1-3 | 2.89E-07 | 0 | 1 | 8 | 2 | A2AAE1-3 | Isoform 3 c | A2AAE1-3 | 0 | 1827.87 | 0 | -0.45 | 0.01315 | 2.47 | 114.4 | 85.6 | 7.92 | 5.81 | | High | High | High | High | High | High | High | High | High | High | High | 0 | 2.51E-08 | 4.61 |
| FALSE | High | NSQLGIFSS | [R].NSQLGI | 1xPhospho | Q9D0F1 1x | 0.000259 | 0 | 1 | 1 | 2 | Q9D0F1 | Kinetochor | Q9D0F1 [6 | 0 | 1834.935 | 0 | -1.02 | 0.00388 | 7.14 | 136.6 | 63.4 | 12.18 | 12.72 | | High | High | High | High | High | High | High | High | High | High | High | 7.19E-05 | 0.000816 | 3.2 |
| FALSE | High | NSRPSSPVI | [K].NSRPSS | 1xPhospho | E9QLZ9 1xI | 1.2E-05 | 0 | 1 | 7 | 2 | E9QLZ9 | Protein ena | E9QLZ9 [3 | 0 | 2231.102 | 0 | -1.81 | 0.004809 | 1.49 | 155.6 | 44.4 | 19.92 | 8.89 | | High | High | High | High | High | High | High | High | High | High | High | 0 | 1.65E-06 | 3.06 |
| FALSE | High | NSRPSSPVI | [K].NSRPSS | 1xDeamida | E9QLZ9 1xI | 0.002948 | 0.00016 | 1 | 7 | 4 | E9QLZ9 | Protein ena | E9QLZ9 [3 | 0 | 2461.249 | 0 | -1.98 | 0.003082 | 2.17 | 159.5 | 40.5 | 13.02 | 32.74 | | High | High | High | High | High | High | High | High | High | High | High | 7.19E-05 | 0.000799 | 2.52 |
| FALSE | High | NSRPSSPVI | [K].NSRPSS | 2xPhospho | E9QLZ9 2xI | 0.000573 | 0 | 1 | 7 | 2 | E9QLZ9 | Protein ena | E9QLZ9 [3 | 0 | 2540.231 | 0 | -1.92 | 0.001713 | 0.21 | 157.7 | 42.3 | 11.38 | 13.93 | | High | High | High | High | High | High | High | High | High | High | High | 0 | 0.000128 | 3.88 |
| FALSE | High | NSNSLLTLD | [K].NSNSLI | 1xCarbami | E9QML5 1x | 5.55E-05 | 0 | 1 | 5 | 2 | E9QML5 | Zinc finger | E9QML5 [1 | 0 | 3219.564 | 0 | -0.65 | 0.000626 | 2.06 | 123.7 | 76.3 | 6.45 | 9.72 | | High | High | High | High | High | High | High | High | High | High | High | 0 | 9.31E-06 | 4.82 |
| FALSE | High | QPLSGTPK | [K].QPLSGT | 1xPhospho | Q8BP27 1x | 0.001244 | 0 | 1 | 3 | 2 | Q8BP27 | Swi5-depe | Q8BP27 [1 | 0 | 1365.754 | 0 | -1.06 | 0.000667 | 1.12 | 136.9 | 63.1 | 9.8 | 6.89 | | High | High | High | High | High | High | High | High | High | High | High | 0 | 0.000305 | 1.82 |
| FALSE | High | QPPGVPNC | [R].QPPGV | 1xPhospho | Q6PEV3 1x | 0.000108 | 0 | 1 | 1 | 10 | Q6PEV3 | WAS/WASI | Q6PEV3 [2 | 0 | 2665.282 | 0 | 0.45 | 0.002708 | 11.78 | 84.1 | 115.9 | 3.36 | 6.85 | | High | High | High | High | High | High | High | High | High | High | High | 0 | 4.96E-05 | 2.77 |
| FALSE | High | QPPGVPNC | [R].QPPGV | 1xDeamida | Q6PEV3 1x | 0.002486 | 0.000109 | 1 | 1 | 3 | Q6PEV3 | WAS/WASI | Q6PEV3 [2 | 0 | 2666.266 | 0 | 0.54 | 0.00952 | 8.59 | 80 | 120 | 6.73 | 8.18 | | High | High | High | High | High | High | High | High | High | High | High | 4.94E-05 | 0.000662 | 2.58 |
| FALSE | High | RKSGEVEEI | [K].RKSGEV | 1xPhospho | Q80WE4 1 | 0.112856 | 0.009146 | 1 | 4 | 1 | Q80WE4 | Kinesin-like | Q80WE4 [1 | 2 | 2156.193 | 0 | 0.02 | 0.806471 | 7.05 | 99 | 101 | 4.17 | 5.21 | | High | High | High | High | High | High | High | High | High | High | High | 0.004583 | 0.04981 | 1.83 |
| FALSE | High | RKSPENTEC | [K].RKSPEN | 1xPhospho | A0A087WS | 0.047086 | 0.00354 | 1 | 4 | 1 | A0A087WS | High mobil | A0A087WS | 2 | 1913.046 | 0 | 1.42 | 0.001429 | 14.9 | 51.1 | 148.9 | 14.49 | 9.69 | | High | High | High | High | High | High | High | High | High | High | High | 0.001812 | 0.01805 | 1.28 |
| FALSE | High | RLDDQESP | [R].RLDDQ | 1xPhospho | Q60520-1 | 0.01029 | 0.000772 | 1 | 4 | 2 | Q60520-1 | Isoform 2 c | Q60520-1 | 1 | 2084.996 | 0 | 0.17 | 0.770896 | 29.1 | 98.2 | 101.8 | 8.87 | 22.82 | | High | High | High | High | High | High | High | High | High | High | High | 0.000872 | 0.008649 | 2.33 |
| FALSE | High | RIDFIPVSP | [K].RIDFIPV | 2xPhospho | Q9DB52 2x | 0.00359 | 0.00021 | 1 | 1 | 2 | Q9DB52 | Protein FAI | Q9DB52 [1 | 1 | 2042.007 | 0 | 0.71 | 0.003337 | 19.36 | 77 | 123 | 10.44 | 7 | | High | High | High | High | High | High | High | High | High | High | High | 0.001999 | 0.02017 | 2.56 |
| FALSE | High | RIDFTPVSP | [K].RIDFTP | 2xPhospho | Q6NZE7-2 | 0.000269 | 0 | 1 | 4 | 2 | Q6NZE7-2 | Isoform 2 c | Q6NZE7-2 | 1 | 2029.971 | 0 | -0.63 | 0.000426 | 3.31 | 120 | 80 | 2.8 | 7.46 | | High | High | High | High | High | High | High | High | High | High | High | 0 | 5.44E-05 | 3.15 |
| FALSE | High | RLDGLPNSI | [R].RLDGLF | 1xPhospho | Q8BMG1 1 | 0.000316 | 0 | 1 | 4 | 1 | Q8BMG1 | ATR-intera | Q8BMG1 [ | 1 | 1546.83 | 0 | -0.08 | 0.227944 | 6.5 | 101.4 | 98.6 | 5.65 | 5.32 | | High | High | High | High | High | High | High | High | High | High | High | 0 | 6.52E-05 | 1.62 |
| FALSE | High | RIDISPSALF | [R].RIDISPS | 1xPhospho | Q8K019 1x | 0.01095 | 0.000772 | 1 | 9 | 2 | Q8K019 | Bcl-2-assoc | Q8K019 [6 | 1 | 1436.782 | 0 | -0.63 | 0.006617 | 4.7 | 124 | 76 | 10.22 | 5.57 | | High | High | High | High | High | High | High | High | High | High | High | 0.001876 | 0.01875 | 2.19 |
| FALSE | High | RIDISPSTFF | [R-].RIDISP | 1xPhospho | Q569Z6 1x | 0.081085 | 0.006219 | 1 | 4 | 1 | Q569Z6 | Thyroid ho | Q569Z6 [6 | 1 | 1500.777 | 0 | -0.31 | 0.028886 | 8.27 | 112.4 | 87.6 | 9.09 | 6.61 | | High | High | High | High | High | High | High | High | High | High | High | 0.0032 | 0.03375 | 1.47 |
| FALSE | High | RLDSENSP | [R].RLDSEN | 2xPhospho | G5E8P8 2x | 9.71E-05 | 0 | 1 | 6 | 2 | G5E8P8 | Estrogen re | G5E8P8 [2 | 1 | 2729.372 | 0 | -1.49 | 0.001491 | 1.74 | 145.4 | 54.6 | 11.36 | 14.53 | | High | High | High | High | High | High | High | High | High | High | High | 0 | 2.61E-05 | 3.28 |
| FALSE | High | RLDSSTCLH | [K].RLDSST | 1xCarbami | Q9Z2Z9 1xI | 0.001489 | 5.66E-05 | 1 | 2 | 9 | Q9Z2Z9 | Glutamine- | Q9Z2Z9 [2 | 1 | 2097.056 | 0 | -0.62 | 0.0039 | 6.34 | 121.8 | 78.2 | 4.61 | 10.73 | | High | High | High | High | High | High | High | High | High | High | High | 4.94E-05 | 0.00068 | 2.43 |
| FALSE | High | RIDTPSLEE | [K].RIDTPS | 1xPhospho | A0A0R4J1Z | 0.003167 | 0.00016 | 1 | 18 | 3 | A0A0R4J1Z | POU doma | A0A0R4J1Z | 1 | 2984.491 | 0 | 0.72 | 0.009302 | 22.62 | 77.1 | 122.9 | 18.25 | 10.43 | | High | High | High | High | High | High | High | High | High | High | High | 0.000275 | 0.001878 | 2.88 |
| FALSE | High | RLEDQAAA | [R].RLEDQ | 1xPhospho | Q8BX02 1x | 0.000375 | 0 | 1 | 2 | 1 | Q8BX02 | KN motif a | Q8BX02 [1 | 1 | 3956.033 | 0 | -0.14 | 0.870347 | 12.38 | 99.1 | 100.9 | 10.16 | 15.07 | | High | High | High | High | High | High | High | High | High | High | High | 0 | 7.95E-05 | 2.68 |
| FALSE | High | RLEEAQQV | [R].RLEEAC | 1xPhospho | Q62315 1x | 0.060666 | 0.00451 | 1 | 5 | 3 | Q62315 | Protein Jur | Q62315 [4 | 1 | 2617.432 | 0 | -1.94 | 0.000275 | 0.73 | 157.2 | 42.8 | 3.34 | 16.87 | | High | High | High | High | High | High | High | High | High | High | High | 0.004032 | 0.0427 | 2.17 |
| FALSE | High | RKSELGAEI | [R].RKSELG | 1xCarbami | Q9QY06-2 | 1.26E-07 | 0 | 2 | 9 | 1 | Q9QY06-2; | Isoform 2 c | Q9QY06-2 | 2 | 3256.664 | 0 | -0.48 | 0.008616 | 6.51 | 115 | 85 | 10.68 | 9.73 | Shared | High | High | High | High | High | High | High | High | High | High | High | 0 | 9.91E-09 | 3.34 |
| FALSE | High | RLEISPDSSI | [R].RLEISPI | 1xPhospho | Q3TIY5 1xF | 0.01095 | 0.000772 | 1 | 4 | 1 | Q3TIY5 | Putative ur | Q3TIY5 [14 | 1 | 1694.831 | 0 | -0.02 | 0.892149 | 10.86 | 99.7 | 100.3 | 8.28 | 6.99 | | High | High | High | High | High | High | High | High | High | High | High | 0.000399 | 0.003486 | 1.9 |
| FALSE | High | RLEQTPVRI | [R].RLEQTF | 1xPhospho | Q8K3W3 1 | 0.000286 | 0 | 1 | 1 | 2 | Q8K3W3 | Protein CA | Q8K3W3 [3 | 2 | 3166.655 | 0 | 0.17 | 0.024185 | 6.97 | 94.8 | 105.2 | 6.63 | 2.37 | | High | High | High | High | High | High | High | High | High | High | High | 2.58E-05 | 0.000395 | 2.96 |
| FALSE | High | RLESPEPEC | [K].RLESPE | 1xPhospho | Q8BFY7-2 | 0.011346 | 0.000816 | 1 | 2 | 1 | Q8BFY7-2 | Isoform 2 c | Q8BFY7-2 | 1 | 1989.996 | 0 | 0.35 | 0.053804 | 7.7 | 87.9 | 112.1 | 8.3 | 8.18 | | High | High | High | High | High | High | High | High | High | High | High | 0.000418 | 0.003603 | 2.61 |
| FALSE | High | RLESPEPEC | [K].RLESPE | 1xPhospho | Q8BFY7 1x | 3.12E-05 | 0 | 1 | 2 | 1 | Q8BFY7 | Protein PIN | Q8BFY7 [6 | 1 | 2189.091 | 0 | -1.82 | 0.000209 | 1 | 153.4 | 46.6 | 6.56 | 12.29 | | High | High | High | High | High | High | High | High | High | High | High | 0 | 4.83E-06 | 3.3 |
| FALSE | High | RLGSLADEI | [K].RLGSLA | 1xPhospho | A0A0R4J18 | 0.020862 | 0.001472 | 1 | 2 | 1 | A0A0R4J18 | X-ray repai | A0A0R4J18 | 1 | 1673.903 | 0 | 0.74 | 0.010058 | 6.74 | 78.9 | 121.1 | 10.63 | 8.58 | | High | High | High | High | High | High | High | High | High | High | High | 0.000786 | 0.007154 | 1.99 |
| FALSE | High | RLGSTGSQ | [K].RLGSTC | 2xPhospho | Q501J7-3 2 | 0.000488 | 0 | 1 | 5 | 1 | Q501J7-3 | Isoform 3 c | Q501J7-3 [ | 1 | 2864.338 | 0 | 0.93 | 0.009066 | 12.8 | 65.7 | 134.3 | 9.02 | 17.45 | | High | High | High | High | High | High | High | High | High | High | High | 0 | 0.000106 | 3.41 |
| FALSE | High | RLNHSPPQ | [R].RLNHSF | 1xDeamida | Q6NZN0 1x | 0.082424 | 0.006288 | 1 | 8 | 1 | Q6NZN0 | RNA-bindir | Q6NZN0 [1 | 1 | 1675.811 | 0 | -0.77 | 0.003058 | 6.39 | 124.4 | 75.6 | 4.85 | 16.06 | | High | High | High | High | High | High | High | High | High | High | High | 0.003243 | 0.0343 | 1.51 |
| FALSE | High | RLNPDSPQ | [K].RLNPDS | 1xPhospho | E9QKG8 1x | 0.010853 | 0.000772 | 1 | 9 | 1 | E9QKG8 | Holliday ju | E9QKG8 [5 | 1 | 2023.037 | 0 | -0.87 | 0.002004 | 1.4 | 129.9 | 70.1 | 10.16 | 6.54 | | High | High | High | High | High | High | High | High | High | High | High | 0.000399 | 0.003442 | 2.7 |
| FALSE | High | RINPPSSGC | [K].RINPPS | 1xPhospho | A0A0R4J1Z | 0.012959 | 0.000889 | 1 | 19 | 1 | A0A0R4J1Z | POU doma | A0A0R4J1Z | 1 | 2123.126 | 0 | -0.86 | 0.002128 | 3.41 | 127.1 | 72.9 | 9.62 | 6.31 | | High | High | High | High | High | High | High | High | High | High | High | 0.000449 | 0.004205 | 2.47 |
| FALSE | High | RLPAFSPPS | [R].RLPAFS | 2xPhospho | Q6J1H4 2xI | 2.5E-07 | 0 | 1 | 1 | 14 | Q6J1H4 | Undifferen | Q6J1H4 [7- | 1 | 2297.093 | 0 | -1.4 | 0.002201 | 1.32 | 145.1 | 54.9 | 13.14 | 8.9 | | High | High | High | High | High | High | High | High | High | High | High | 0 | 2.14E-08 | 3.81 |
| FALSE | High | RLPALSHSE | [R].RLPALS | 2xPhospho | A0A0N4SU | 2.87E-05 | 0 | 1 | 10 | 3 | A0A0N4SU | Peregrin O | A0A0N4SU | 1 | 3132.334 | 0 | -0.2 | 0.017684 | 3.38 | 106.8 | 93.2 | 2.86 | 7.99 | | High | High | High | High | High | High | High | High | High | High | High | 0 | 4.44E-06 | 3.56 |
| FALSE | High | RLPASPVIS | [R].RLPASP | 1xPhospho | A2A5V4 1x | 5.33E-07 | 0 | 1 | 2 | 1 | A2A5V4 | SH3 domai | A2A5V4 [6 | 1 | 2860.423 | 0 | 1.05 | 0.000756 | 5.13 | 66.1 | 133.9 | 9.52 | 10.04 | | High | High | High | High | High | High | High | High | High | High | High | 0 | 5.02E-08 | 4.33 |
| FALSE | High | RLPASPVIS | [R].RLPASP | 1xPhospho | A2A5V4 1x | 1.09E-07 | 0 | 1 | 2 | 3 | A2A5V4 | SH3 domai | A2A5V4 [6 | 1 | 2876.417 | 0 | 0.96 | 0.031006 | 10.05 | 71.1 | 128.9 | 9.25 | 27.87 | | High | High | High | High | High | High | High | High | High | High | High | 0 | 5.88E-07 | 4.66 |
| FALSE | High | RIPSAVSSP | [K].RIPSAV | 1xPhospho | E9Q3G8 1x | 1.9E-05 | 0 | 1 | 2 | 2 | E9Q3G8 | Nucleopor | E9Q3G8 [3 | 1 | 2018.063 | 0 | -0.77 | 0.002896 | 2.34 | 125.9 | 74.1 | 9.12 | 9.74 | | High | High | High | High | High | High | High | High | High | High | High | 0 | 0.000123 | 2.79 |
| FALSE | High | RLEISPDSSI | [R].RLEISPI | 3xPhospho | Q3TIY5 3xF | 0.001361 | 0 | 1 | 4 | 1 | Q3TIY5 | Putative ur | Q3TIY5 [14 | 1 | 1854.764 | 0 | 0.81 | 0.001251 | 8.27 | 71 | 129 | 8.24 | 9.69 | | High | High | High | High | High | High | High | High | High | High | High | 0 | 0.000338 | 3.1 |
| FALSE | High | RIPSAVSSP | [K].RIPSAV | 2xPhospho | E9Q3G8 2x | 0.026421 | 0.001676 | 1 | 2 | 2 | E9Q3G8 | Nucleopor | E9Q3G8 [3 | 1 | 2098.03 | 0 | -0.38 | 0.006795 | 4.84 | 113.4 | 86.6 | 3.46 | 11.25 | | High | High | High | High | High | High | High | High | High | High | High | 0.003389 | 0.0369 | 2.69 |
| FALSE | High | RKSEEEEDC | [K].RKSEEE | 1xPhospho | Q04750 1x | 0.099328 | 0.008003 | 1 | 3 | 1 | Q04750 | DNA topois | Q04750 [1 | 2 | 1974.015 | 0 | -0.08 | 0.416891 | 8.69 | 103.3 | 96.7 | 12.55 | 8.41 | | High | High | High | High | High | High | High | High | High | High | High | 0.004032 | 0.04274 | 1.23 |
| FALSE | High | RKPSSSDSS | [K].RKPSSS | 1xPhospho | Q01320 1x | 0.009843 | 0.000736 | 2 | 4 | 3 | Q01320; Q | DNA topois | Q01320 [1 | 1 | 2238.044 | 0 | -0.78 | 0.003728 | 0.13 | 130.1 | 69.9 | 8.5 | 10.81 | Shared | High | High | High | High | High | High | High | High | High | High | High | 0.002703 | 0.02909 | 1.1 |
| FALSE | High | RHSPSPRPF | [K].RHSPSF | 2xPhospho | Q52KI8 2xF | 0.086562 | 0.006435 | 1 | 7 | 1 | Q52KI8 | Serine/argi | Q52KI8 [70 | 1 | 1478.698 | 0 | -0.21 | 0.155811 | 13.16 | 105.7 | 94.3 | 26.94 | 8.84 | | High | High | High | High | High | High | High | High | High | High | High | 0.003346 | 0.03656 | 2.88 |
| FALSE | High | RHSRPQGA | [R].RHSRPC | 1xPhospho | O35134 1x | 0.013308 | 0.000889 | 1 | 4 | 1 | O35134 | DNA-direct | O35134 [1 | 1 | 1888.032 | 0 | 0.57 | 0.008345 | 2.71 | 82.1 | 117.9 | 6.23 | 11.99 | | High | High | High | High | High | High | High | High | High | High | High | 0.000486 | 0.004323 | 2.12 |
| FALSE | High | RHSSDINHI | [R].RHSSDI | 1xPhospho | E9PUI5 1xF | 0.007742 | 0.000664 | 1 | 5 | 1 | E9PUI5 | Formin-bin | E9PUI5 [48 | 1 | 1928.965 | 0 | -0.47 | 0.008414 | 4.26 | 114.4 | 85.6 | 9.96 | 6.6 | | High | High | High | High | High | High | High | High | High | High | High | 0.000294 | 0.002354 | 2.37 |
| FALSE | High | RHSSETFSA | [R].RHSSET | 2xPhospho | P35569 2xI | 0.032267 | 0.002129 | 1 | 2 | 1 | P35569 | Insulin rece | P35569 [10 | 1 | 1764.767 | 0 | 0.08 | 0.610243 | 30.61 | 99.6 | 100.4 | 12.11 | 22.17 | | High | High | High | High | High | High | High | High | High | High | High | 0.001142 | 0.01169 | 2.13 |
| FALSE | High | RHSSLPTES | [R].RHSSLP | 2xPhospho | O54774 2x | 0.035486 | 0.002304 | 1 | 2 | 1 | O54774 | AP-3 comp | O54774 [7 | 1 | 2398.064 | 0 | -0.25 | 0.066843 | 6.92 | 108.5 | 91.5 | 7.35 | 10.32 | | High | High | High | High | High | High | High | High | High | High | High | 0.001214 | 0.01299 | 2.74 |
| FALSE | High | RHSSLPTES | [R].RHSSLP | 3xPhospho | O54774 3x | 0.013666 | 0.001011 | 1 | 2 | 1 | O54774 | AP-3 comp | O54774 [7 | 1 | 2478.03 | 0 | -0.44 | 0.185454 | 9.69 | 115.2 | 84.8 | 14.08 | 20.47 | | High | High | High | High | High | High | High | High | High | High | High | 0.000538 | 0.004458 | 3.97 |
| FALSE | High | RHSTEGPEI | [R].RHSTEC | 1xPhospho | B2RW11 1x | 0.001002 | 0 | 1 | 1 | 1 | B2RW11 | Ankyrin rej | B2RW11 [2 | 1 | 2371.148 | 0 | -0.04 | 0.342336 | 11.29 | 100.6 | 99.4 | 4.56 | 11.96 | | High | High | High | High | High | High | High | High | High | High | High | 0 | 0.000238 | 3.04 |
| FALSE | High | RHSTSDLSI | [K].RHSTSD | 1xPhospho | B1AQY2 1x | 6E-07 | 0 | 1 | 4 | 2 | B1AQY2 | Rho GTPas | B1AQY2 [6 | 1 | 2116.986 | 0 | 0.08 | 0.664877 | 9.76 | 95.6 | 104.4 | 12.85 | 14.17 | | High | High | High | High | High | High | High | High | High | High | High | 0 | 1.59E-05 | 3.28 |
| FALSE | High | RHSVNLDE | [R].RHSVNI | 1xPhospho | E9PVX6 1xI | 3.51E-06 | 0 | 1 | 2 | 3 | E9PVX6 | Proliferatic | E9PVX6 [3 | 1 | 2415.218 | 0 | -0.04 | 0.411037 | 9.49 | 101.4 | 98.6 | 10.35 | 4.16 | | High | High | High | High | High | High | High | High | High | High | High | 0.000267 | 0.00156 | 2.64 |
| FALSE | High | RHSVSEMT | [R].RHSVSE | 1xCarbami | Q812A5 1x | 1.6E-06 | 0 | 1 | 2 | 1 | Q812A5 | Proline-ricl | Q812A5 [2 | 1 | 3473.545 | 0 | -1.09 | 0.002895 | 5.46 | 134.9 | 65.1 | 10.99 | 19.12 | | High | High | High | High | High | High | High | High | High | High | High | 0 | 1.73E-07 | 4.45 |
| FALSE | High | RHSYENDG | [R].RHSYEN | 1xPhospho | Q3UYV9 1x | 0.002077 | 5.66E-05 | 1 | 2 | 4 | Q3UYV9 | Nuclear ca | Q3UYV9 [5 | 1 | 1833.823 | 0 | -0.29 | 0.707822 | 24.71 | 104.3 | 95.7 | 18.1 | 24.2 | | High | High | High | High | High | High | High | High | High | High | High | 0.000814 | 0.007733 | 2.79 |
| FALSE | High | RHSYENDG | [R].RHSYEN | 1xPhospho | Q3UYV9 1x | 0.000308 | 0 | 1 | 2 | 7 | Q3UYV9 | Nuclear ca | Q3UYV9 [5 | 1 | 2062.986 | 0 | -0.04 | 0.978662 | 1.78 | 101.4 | 98.6 | 4.21 | 3.66 | | High | High | High | High | High | High | High | High | High | High | High | 0.000872 | 0.009135 | 2.47 |
| FALSE | High | RHSYENDG | [R].RHSYEN | 1xDeamida | Q3UYV9 1x | 0.001516 | 5.66E-05 | 1 | 2 | 2 | Q3UYV9 | Nuclear ca | Q3UYV9 [5 | 1 | 2063.97 | 0 | -0.27 | 0.00933 | 7.5 | 110.8 | 89.2 | 5.47 | 3.36 | | High | High | High | High | High | High | High | High | High | High | High | 7.19E-05 | 0.000791 | 2.29 |
| FALSE | High | RKPSVPDT | [K].RKPSVF | 1xPhospho | Q99M51 1 | 6.97E-06 | 0 | 1 | 4 | 1 | Q99M51 | Cytoplasmi | Q99M51 [8 | 1 | 2881.413 | 0 | 0.23 | 0.058048 | 6.97 | 89.8 | 110.2 | 7.04 | 4.97 | | High | High | High | High | High | High | High | High | High | High | High | 0 | 9.03E-07 | 3.18 |
| FALSE | High | RHTITNGVI | [R].RHTITN | 1xCarbami | Q9D818-3 | 1.57E-06 | 0 | 1 | 3 | 1 | Q9D818-3 | Isoform 3 c | Q9D818-3 | 1 | 2152.135 | 0 | 0.65 | 0.010089 | 17.35 | 77.9 | 122.1 | 10.14 | 8.96 | | High | High | High | High | High | High | High | High | High | High | High | 0 | 1.7E-07 | 3.3 |
| FALSE | High | RHTSLEEEE | [R].RHTSLE | 1xPhospho | D3Z357 1xI | 0.004781 | 0.000456 | 1 | 6 | 2 | D3Z357 | Histone-lys | D3Z357 [4 | 1 | 2186.126 | 0 | 0.19 | 0.793978 | 30.97 | 112.6 | 87.4 | 61.09 | 20.66 | | High | High | High | High | High | High | High | High | High | High | High | 0.000204 | 0.001381 | 2.86 |
| FALSE | High | RHTVDEYS | [R].RHTVDI | 1xPhospho | Q8C0Q9 1x | 0.006773 | 0.00062 | 1 | 3 | 1 | Q8C0Q9 | Rap guanin | Q8C0Q9 [4 | 1 | 1897.957 | 0 | -0.01 | 0.508423 | 8.57 | 100.5 | 99.5 | 9.07 | 5.22 | | High | High | High | High | High | High | High | High | High | High | High | 0.000275 | 0.002039 | 2.88 |
| FALSE | High | RHTVGCGE | [R].RHTVG | 1xCarbami | Q7TPN9 1x | 0.003655 | 0.00021 | 1 | 1 | 1 | Q7TPN9 | Proline-ricl | Q7TPN9 [3 | 1 | 1582.718 | 0 | -0.35 | 0.058048 | 3.29 | 110.1 | 89.9 | 13.3 | 14.41 | | High | High | High | High | High | High | High | High | High | High | High | 9.42E-05 | 0.001018 | 2.04 |
| FALSE | High | RHVSPEPVI | [K].RHVSPE | 1xPhospho | E9PVX6 1xI | 0.006653 | 0.00062 | 1 | 2 | 1 | E9PVX6 | Proliferatic | E9PVX6 [2 | 1 | 1586.882 | 0 | 0.39 | 0.186253 | 1.75 | 87.2 | 112.8 | 11.86 | 23.95 | | High | High | High | High | High | High | High | High | High | High | High | 0.000275 | 0.001986 | 1.92 |
| FALSE | High | RKGSDDDC | [K].RKGSDI | 1xPhospho | O88574 1x | 0.000984 | 0 | 1 | 1 | 2 | O88574 | Histone de | O88574 [1 | 2 | 4083.991 | 0 | -0.8 | 0.004336 | 4.34 | 123.6 | 76.4 | 12.09 | 15.51 | | High | High | High | High | High | High | High | High | High | High | High | 0 | 0.000235 | 3.7 |
| FALSE | High | RKGSDDDC | [K].RKGSDI | 1xDeamida | O88574 1x | 0.000884 | 0 | 1 | 1 | 2 | O88574 | Histone de | O88574 [1 | 2 | 4084.975 | 0 | -0.9 | 0.001307 | 4.72 | 130.1 | 69.9 | 6.79 | 10.21 | | High | High | High | High | High | High | High | High | High | High | High | 0 | 0.000208 | 4.01 |
| FALSE | High | RKPDLENV | [R].RKPDLE | 1xPhospho | Q6PR54-3 | 0.000751 | 0 | 2 | 4 | 1 | Q6PR54-3; | Isoform 3 c | Q6PR54-3 | 1 | 3068.587 | 0 | -0.76 | 0.002319 | 4.19 | 124.7 | 75.3 | 8.33 | 6.82 | Shared | High | High | High | High | High | High | High | High | High | High | High | 0 | 0.000173 | 2.05 |
| FALSE | High | RKPDQTLD | [R].RKPDQ | 1xPhospho | P18572 1xI | 0.000984 | 0 | 1 | 5 | 2 | P18572 | Basigin OS= | P18572 [35 | 1 | 2733.443 | 0 | 4.17 | 0.000444 | 13.15 | 11.9 | 188.1 | 21.95 | 10.2 | | High | High | High | High | High | High | High | High | High | High | High | 0 | 0.000234 | 3.14 |
| FALSE | High | RKPGAGGS | [R].RKPGA | 1xPhospho | A2AJI0 1xP | 0.01894 | 0.001362 | 1 | 5 | 1 | A2AJI0 | MAP7 dom | A2AJI0 [39 | 1 | 1776.005 | 0 | 1.58 | 0.000528 | 4.45 | 52.6 | 147.4 | 7.84 | 9.38 | | High | High | High | High | High | High | High | High | High | High | High | 0.00074 | 0.006445 | 1.5 |
| FALSE | High | RKPSCSEGI | [R].RKPSCS | 1xCarbami | P51612 1xI | 0.007274 | 0.00062 | 1 | 3 | 2 | P51612 | DNA repair | P51612 [39 | 1 | 2402.199 | 0 | 0.38 | 0.192513 | 24.17 | 88.5 | 111.5 | 13.92 | 14.94 | | High | High | High | High | High | High | High | High | High | High | High | 0.00074 | 0.006455 | 2.2 |
| FALSE | High | RKPSPEPEC | [K].RKPSPE | 1xPhospho | E9Q1P8 1x | 2.4E-05 | 0 | 1 | 1 | 2 | E9Q1P8 | Interferon | E9Q1P8 [3 | 1 | 2371.299 | 0 | -0.32 | 0.008832 | 2.11 | 111.7 | 88.3 | 5.47 | 4.34 | | High | High | High | High | High | High | High | High | High | High | High | 0 | 3.61E-06 | 3.42 |
| FALSE | High | RKPSPSPPF | [K].RKPSPS | 1xPhospho | P97496 1xI | 8.35E-06 | 0 | 1 | 6 | 3 | P97496 | SWI/SNF c | P97496 [32 | 1 | 2243.195 | 0 | -1.3 | 0.000344 | 1.13 | 141.2 | 58.8 | 6.16 | 4.31 | | High | High | High | High | High | High | High | High | High | High | High | 0 | 1.11E-06 | 3.16 |
| FALSE | High | RKPSPSPPF | [K].RKPSPS | 2xPhospho | P97496 2xI | 2.56E-06 | 0 | 1 | 6 | 7 | P97496 | SWI/SNF c | P97496 [32 | 1 | 2323.161 | 0 | -1.06 | 0.000266 | 2.11 | 134.5 | 65.5 | 2.7 | 5.33 | | High | High | High | High | High | High | High | High | High | High | High | 0 | 2.93E-07 | 3.44 |
| FALSE | High | RHTITNGVI | [R].RHTITN | 1xCarbami | Q9D818-3 | 0.091627 | 0.007146 | 1 | 3 | 1 | Q9D818-3 | Isoform 3 c | Q9D818-3 | 1 | 2153.119 | 0 | 0.89 | 0.011866 | 14.17 | 70 | 130 | 13.76 | 11.98 | | High | High | High | High | High | High | High | High | High | High | High | 0.003633 | 0.03878 | 3.34 |
| FALSE | High | RLPSSPASP | [R].RLPSSP | 1xPhospho | A2AUK5 1x | 0.008538 | 0.000736 | 2 | 10 | 1 | A2AUK5; C | Band 4.1-li | A2AUK5 [5 | 1 | 1761.966 | 0 | -0.48 | 0.002353 | 0.98 | 114.8 | 85.2 | 8.08 | 5.61 | Shared | High | High | High | High | High | High | High | High | High | High | High | 0.000327 | 0.002632 | 2.48 |
| FALSE | High | RIPVSPEQT | [R].RIPVSPI | 1xPhospho | Q8R2S9 1x | 0.00359 | 0.00021 | 1 | 1 | 2 | Q8R2S9 | Actin-relat | Q8R2S9 [1 | 1 | 1491.788 | 0 | -0.92 | 0.000405 | 0.87 | 130.8 | 69.2 | 4.99 | 6.38 | | High | High | High | High | High | High | High | High | High | High | High | 9.42E-05 | 0.001002 | 1.2 |
| FALSE | High | RIPYTPGEII | [K].RIPYTP | 1xPhospho | Q3TJE8 1xI | 0.001097 | 0 | 1 | 5 | 1 | Q3TJE8 | UDP-gluco | Q3TJE8 [47 | 1 | 1809.008 | 0 | -0.89 | 0.000266 | 1.16 | 130.4 | 69.6 | 6.22 | 9.73 | | High | High | High | High | High | High | High | High | High | High | High | 0 | 0.000264 | 1.34 |
| FALSE | High | RISHSLYSG | [R].RISHSL | 1xPhospho | A5D6P2 1x | 1.91E-05 | 0 | 1 | 23 | 1 | A5D6P2 | Par-3 (Part | A5D6P2 [7 | 1 | 2426.191 | 0 | -0.33 | 0.233921 | 16.03 | 111.4 | 88.6 | 9.09 | 15.8 | | High | High | High | High | High | High | High | High | High | High | High | 0 | 2.81E-06 | 3.66 |
| FALSE | High | RISICSSDK | [K].RISICSS | 1xCarbami | D3YYI8 1xF | 0.002077 | 5.66E-05 | 1 | 2 | 3 | D3YYI8 | Histone de | D3YYI8 [40 | 1 | 1603.828 | 0 | -0.25 | 0.06019 | 8.12 | 110.2 | 89.8 | 6.79 | 7.65 | | High | High | High | High | High | High | High | High | High | High | High | 2.58E-05 | 0.00054 | 1.88 |
| FALSE | High | RISICSSDK | [K].RISICSS | 1xCarbami | D3YYI8 2xF | 0.039347 | 0.002793 | 1 | 2 | 1 | D3YYI8 | Histone de | D3YYI8 [40 | 1 | 1683.794 | 0 | -1.27 | 0.000191 | 0.68 | 141.2 | 58.8 | 2.18 | 6.53 | | High | High | High | High | High | High | High | High | High | High | High | 0.001453 | 0.0146 | 1.86 |
| FALSE | High | RLSLDIQSP | [R].RLSLDI | 1xPhospho | Q921C3 1x | 1.13E-05 | 0 | 1 | 5 | 1 | Q921C3 | Bromodom | Q921C3 [6 | 1 | 1988.089 | 0 | -0.82 | 0.000401 | 2.52 | 128.4 | 71.6 | 3.58 | 2.67 | | High | High | High | High | High | High | High | High | High | High | High | 0 | 1.55E-06 | 3.03 |
| FALSE | High | RISIGGGSC | [K].RISIGG | 1xCarbami | P50446 1xI | 8.57E-09 | 0 | 1 | 5 | 1 | P50446 | Keratin, ty | P50446 [67 | 1 | 2019.963 | 0 | 2.78 | 0.000673 | 4.82 | 29.5 | 170.5 | 20.8 | 14.44 | | High | High | High | High | High | High | High | High | High | High | High | 0 | 4.82E-10 | 3.04 |
| FALSE | High | RLSLGSYPD | [R].RLSLGS | 1xPhospho | Q8R123-2 | 0.000418 | 0 | 1 | 2 | 1 | Q8R123-2 | Isoform 2 c | Q8R123-2 | 1 | 2685.348 | 0 | 1.56 | 0.000381 | 16.62 | 50.9 | 149.1 | 11.69 | 4.28 | | High | High | High | High | High | High | High | High | High | High | High | 0 | 8.97E-05 | 3.5 |
| FALSE | High | RISIIINASAI | [R].RISIIINA | 1xPhospho | Q8BP60 1x | 0.015465 | 0.001089 | 1 | 4 | 1 | Q8BP60 | Putative ur | Q8BP60 [1 | 1 | 3960.218 | 0 | -0.07 | 0.09874 | 4.9 | 102.9 | 97.1 | 4.28 | 7.08 | | High | High | High | High | High | High | High | High | High | High | High | 0.000571 | 0.005098 | 3.33 |
| FALSE | High | RLSLPGLLS | [R].RLSLPG | 1xPhospho | Q920R0 1x | 0.0004 | 0 | 1 | 2 | 1 | Q920R0 | Alsin OS=M | Q920R0 [4 | 1 | 1832.036 | 0 | -0.07 | 0.341199 | 4.84 | 104.1 | 95.9 | 4.78 | 9.57 | | High | High | High | High | High | High | High | High | High | High | High | 0 | 8.48E-05 | 2.19 |
| FALSE | High | RLSLPGLLS | [R].RLSLPG | 2xPhospho | Q920R0 2x | 0.043607 | 0.003228 | 1 | 2 | 1 | Q920R0 | Alsin OS=M | Q920R0 [4 | 1 | 1912.002 | 0 | -0.2 | 0.04326 | 11.61 | 107.1 | 92.9 | 5.2 | 7.92 | | High | High | High | High | High | High | High | High | High | High | High | 0.001651 | 0.01641 | 2.8 |
| FALSE | High | RLSLTPDPE | [R].RLSLTP | 1xPhospho | Q6P9Q6-2 | 3.35E-05 | 0 | 1 | 3 | 2 | Q6P9Q6-2 | Isoform B c | Q6P9Q6-2 | 1 | 1693.929 | 0 | 0.45 | 0.00142 | 1.89 | 84 | 116 | 4.41 | 6.12 | | High | High | High | High | High | High | High | High | High | High | High | 0 | 5.24E-06 | 3.02 |
| FALSE | High | RLSLTPDPE | [R].RLSLTP | 2xPhospho | Q6P9Q6-2 | 0.013426 | 0.000976 | 1 | 3 | 1 | Q6P9Q6-2 | Isoform B c | Q6P9Q6-2 | 1 | 1773.895 | 0 | -0.03 | 0.432383 | 8.7 | 101 | 99 | 8.57 | 5.7 | | High | High | High | High | High | High | High | High | High | High | High | 0.000505 | 0.004361 | 2.1 |
| FALSE | High | RISIVESCFC | [R].RISIVES | 1xCarbami | Q91WB4 1 | 7.74E-08 | 0 | 1 | 2 | 1 | Q91WB4 | Pleckstrin I | Q91WB4 [1 | 1 | 2538.31 | 0 | -0.79 | 0.00104 | 2.16 | 127.4 | 72.6 | 7.64 | 5.28 | | High | High | High | High | High | High | High | High | High | High | High | 0 | 5.71E-09 | 3.66 |
| FALSE | High | RISMEDLNI | [R].RISMED | 1xPhospho | Q8K0D0 1x | 0.015465 | 0.001089 | 1 | 4 | 3 | Q8K0D0 | Cyclin-depe | Q8K0D0 [1 | 1 | 1659.854 | 0 | -0.11 | 0.50957 | 13.05 | 102 | 98 | 7.87 | 29.86 | | High | High | High | High | High | High | High | High | High | High | High | 0.000903 | 0.009447 | 2.49 |
| FALSE | High | RISHLFFHK | [R].RISHLFI | 1xPhospho | F8VPX1 1xI | 0.029586 | 0.001857 | 1 | 2 | 1 | F8VPX1 | Ubiquitin c | F8VPX1 [1 | 1 | 1722.961 | 0 | 1.6 | 0.001132 | 23.84 | 49.5 | 150.5 | 7.5 | 9.44 | | High | High | High | High | High | High | High | High | High | High | High | 0.000997 | 0.01062 | 1.76 |
| FALSE | High | RLSMSEVK | [R].RLSMSI | 1xPhospho | E9Q9R9 1x | 0.013308 | 0.000889 | 1 | 8 | 1 | E9Q9R9 | Disks large | E9Q9R9 [1 | 1 | 1503.801 | 0 | -0.53 | 0.024677 | 13.79 | 118 | 82 | 6.44 | 16.48 | | High | High | High | High | High | High | High | High | High | High | High | 0.000486 | 0.004316 | 1.21 |

| | | | | | | | | | | | | | | | | | | | | | | | | | | | | | | | | | | | | | | |
|---|---|---|---|---|---|---|---|---|---|---|---|---|---|---|---|---|---|---|---|---|---|---|---|---|---|---|---|---|---|---|---|---|---|---|---|---|---|---|
| FALSE | High | RISMVEELE | [R].RISMVE | 1xPhospho | Q8R2P8 1x | 0.005467 | 0.00062 | 1 | 5 | 1 | Q8R2P8 | Lysine--tRN | Q8R2P8 [4 | 1 | 1787.938 | 0 | 0.44 | 0.052587 | 8.36 | 86.7 | 113.3 | 6.9 | 12.61 | | High | High | High | High | High | High | High | High | High | High | High | 0.000275 | 0.001598 | 1.66 |
| FALSE | High | RLSNASAS | [R].RLSNAS | 1xPhospho | Q8R1F6 1x | 0.011048 | 0.000772 | 1 | 3 | 1 | Q8R1F6 | Protein HID | Q8R1F6 [6 | 1 | 3015.513 | 0 | -1.29 | 0.001054 | 0.59 | 139.4 | 60.6 | 8.27 | 9.71 | | High | High | High | High | High | High | High | High | High | High | High | 0.000399 | 0.003508 | 3.08 |
| FALSE | High | RLSNLPTVD | [R].RLSNLP | 1xCarbami | F8VQD1 1x | 3.85E-06 | 0 | 1 | 14 | 2 | F8VQD1 | Protein pol | F8VQD1 [3 | 1 | 2890.485 | 0 | 0.55 | 0.00237 | 9.13 | 78.3 | 121.7 | 9.88 | 10.16 | | High | High | High | High | High | High | High | High | High | High | High | 0 | 4.63E-07 | 3.21 |
| FALSE | High | RLSNPCIQF | [R].RLSNPC | 1xCarbami | Q8BUL6 1x | 0.021231 | 0.001507 | 1 | 3 | 2 | Q8BUL6 | Pleckstrin h | Q8BUL6 [3 | 1 | 1452.734 | 0 | 0.25 | 0.021822 | 9.98 | 90.9 | 109.1 | 6.51 | 7.13 | | High | High | High | High | High | High | High | High | High | High | High | 0.001308 | 0.01352 | 1.97 |
| FALSE | High | RLSNPPVEI | [R].RLSNPP | 1xPhospho | F7BJB9 1xP | 2.09E-07 | 0 | 1 | 2 | 3 | F7BJB9 | MORC fam | F7BJB9 [56 | 1 | 2029.052 | 0 | -0.68 | 0.001015 | 3.43 | 123.2 | 76.8 | 4.43 | 5.87 | | High | High | High | High | High | High | High | High | High | High | High | 0 | 1.74E-08 | 3.16 |
| FALSE | High | RLSNPPVEI | [R].RLSNPP | 1xDeamida | F7BJB9 1xP | 1.46E-05 | 0 | 1 | 2 | 3 | F7BJB9 | MORC fam | F7BJB9 [56 | 1 | 2030.036 | 0 | -0.53 | 0.012692 | 6.7 | 117.5 | 82.5 | 11.2 | 2.37 | | High | High | High | High | High | High | High | High | High | High | High | 0 | 0.000176 | 3.41 |
| FALSE | High | RLSNSSLCS | [R].RLSNSS | 1xCarbami | A0A0A0MC | 0.061176 | 0.004646 | 1 | 5 | 2 | A0A0A0MC | Ral GTPase | A0A0A0MC | 1 | 2125.99 | 0 | 0.63 | 0.004385 | 4.45 | 78.6 | 121.4 | 7.55 | 6.52 | | High | High | High | High | High | High | High | High | High | High | High | 0.00237 | 0.02435 | 3.03 |
| FALSE | High | RLSNVSSSC | [R].RLSNVS | 2xPhospho | P20357 2xP | 0.061176 | 0.004646 | 1 | 5 | 2 | P20357 | Microtubu | P20357 [17 | 1 | 3874.002 | 0 | 2.03 | 0.000875 | 7.62 | 41.9 | 158.1 | 13.09 | 10.45 | | High | High | High | High | High | High | High | High | High | High | High | 0.008888 | 0.09537 | 3.14 |
| FALSE | High | RLSPGSDIY | [R].RLSPGS | 1xPhospho | Q8C0T5 1x | 0.000446 | 0 | 1 | 3 | 1 | Q8C0T5 | Signal-indu | Q8C0T5 [1 | 1 | 2348.188 | 0 | 1.69 | 0.002019 | 31.84 | 47.1 | 152.9 | 18.54 | 10.08 | | High | High | High | High | High | High | High | High | High | High | High | 0 | 9.65E-05 | 3.21 |
| FALSE | High | RLSPIMLGH | [R].RLSPIM | 1xPhospho | Q9CQT7 1x | 0.057686 | 0.004227 | 1 | 3 | 1 | Q9CQT7 | Desumoyla | Q9CQT7 [2 | 1 | 1568.9 | 0 | -0.33 | 0.058793 | 12.56 | 111.3 | 88.7 | 9.1 | 6.32 | | High | High | High | High | High | High | High | High | High | High | High | 0.002159 | 0.02265 | 1.24 |
| FALSE | High | RISPPPPSD | [R].RISPPPI | 1xPhospho | Q569L8 1xP | 0.039347 | 0.002793 | 1 | 1 | 1 | Q569L8 | Centromer | Q569L8 [84 | 1 | 1975.085 | 0 | -0.95 | 0.002165 | 2.28 | 131.4 | 68.6 | 9.16 | 8.76 | | High | High | High | High | High | High | High | High | High | High | High | 0.001453 | 0.01464 | 2.58 |
| FALSE | High | RLSPSASPP | [R].RLSPSA | 2xPhospho | Q52KI8 2xP | 0.117405 | 0.009523 | 1 | 6 | 1 | Q52KI8 | Serine/argi | Q52KI8 [38 | 1 | 1456.691 | 0 | 0.2 | 0.042435 | 3.75 | 94.5 | 105.5 | 5.69 | 5.36 | | High | High | High | High | High | High | High | High | High | High | High | 0.004769 | 0.05197 | 2.52 |
| FALSE | High | RLSPSTPSN | [K].RLSPST | 2xPhospho | O54839 2x | 0.000458 | 0 | 1 | 2 | 1 | O54839 | Eomesode | O54839 [6 | 1 | 2270.135 | 0 | -1.18 | 0.001191 | 3.73 | 138.5 | 61.5 | 9.18 | 19.99 | | High | High | High | High | High | High | High | High | High | High | High | 0 | 9.92E-05 | 2.67 |
| FALSE | High | RISMVEELE | [R].RISMVE | 1xPhospho | Q8R2P8 1x | 4.43E-05 | 0 | 1 | 5 | 2 | Q8R2P8 | Lysine--tRN | Q8R2P8 [4 | 1 | 1771.943 | 0 | 2.5 | 0.000158 | 7.5 | 29.6 | 170.4 | 10.52 | 3.91 | | High | High | High | High | High | High | High | High | High | High | High | 0 | 7.2E-06 | 2.87 |
| FALSE | High | RLSGVEEAY | [R].RLSGVE | 1xPhospho | D3YUT2 1x | 0.020862 | 0.001472 | 1 | 2 | 1 | D3YUT2 | Alpha-prot | D3YUT2 [2 | 1 | 1922.031 | 0 | -1.97 | 0.000856 | 2.02 | 160.5 | 39.5 | 6.31 | 22.69 | | High | High | High | High | High | High | High | High | High | High | High | 0.000801 | 0.007171 | 2.83 |
| FALSE | High | RLSGSNLCS | [R].RLSGSN | 1xCarbami | Q9R1C7 1x | 2.14E-06 | 0 | 1 | 4 | 2 | Q9R1C7 | Pre-mRNA- | Q9R1C7 [3 | 1 | 2508.19 | 0 | 0.24 | 0.241344 | 8.01 | 98.1 | 101.9 | 8.08 | 13.62 | | High | High | High | High | High | High | High | High | High | High | High | 0 | 2.4E-07 | 4.44 |
| FALSE | High | RISGSECAT | [K].RISGSE | 1xCarbami | A2ASY1 2xP | 0.024424 | 0.001612 | 1 | 5 | 1 | A2ASY1 | TAF3 RNA p | A2ASY1 [42 | 1 | 1823.853 | 0 | -0.13 | 0.347571 | 6.55 | 99.7 | 100.3 | 8.13 | 8.19 | | High | High | High | High | High | High | High | High | High | High | High | 0.000859 | 0.008544 | 1.81 |
| FALSE | High | RLSAEGGD | [R].RLSAEG | 1xPhospho | Q3UHH1 1: | 0.027358 | 0.001676 | 1 | 5 | 1 | Q3UHH1 | Zinc finger | Q3UHH1 [5 | 1 | 1470.772 | 0 | -0.16 | 0.097437 | 7.75 | 106.5 | 93.5 | 5 | 8.42 | | High | High | High | High | High | High | High | High | High | High | High | 0.000936 | 0.009742 | 1.43 |
| FALSE | High | RLSAGTTVA | [R].RLSAGT | 1xPhospho | Q8K1N2-2 | 6.26E-06 | 0 | 1 | 3 | 2 | Q8K1N2-2 | Isoform 2 c | Q8K1N2-2 | 1 | 1884.036 | 0 | 1.12 | 0.001595 | 11.97 | 60.7 | 139.3 | 9.83 | 12.05 | | High | High | High | High | High | High | High | High | High | High | High | 0 | 7.97E-07 | 4.13 |
| FALSE | High | RLSASDIVS | [R].RLSASD | 1xPhospho | B2RT41 1xP | 1.49E-06 | 0 | 1 | 1 | 3 | B2RT41 | Zinc finger, | B2RT41 [35 | 1 | 1742.945 | 0 | -0.23 | 0.032599 | 1.71 | 108 | 92 | 2.71 | 6.11 | | High | High | High | High | High | High | High | High | High | High | High | 0 | 1.59E-07 | 3.2 |
| FALSE | High | RLSASSTGS | [K].RLSASS | 1xPhospho | Q3TIT7 1xP | 0.033116 | 0.002195 | 1 | 7 | 1 | Q3TIT7 | Putative ur | Q3TIT7 [43 | 1 | 1431.715 | 0 | -0.61 | 0.00175 | 5.33 | 120.2 | 79.8 | 8.46 | 9.34 | | High | High | High | High | High | High | High | High | High | High | High | 0.001151 | 0.01207 | 1.66 |
| FALSE | High | RISDDDTAT | [R].RISDDD | 1xCarbami | Q08775 1x | 0.070485 | 0.005214 | 1 | 8 | 2 | Q08775 | Runt-relate | Q08775 [4 | 1 | 2939.39 | 0 | 1.77 | 0.001031 | 30.73 | 46.4 | 153.6 | 15.35 | 9.27 | | High | High | High | High | High | High | High | High | High | High | High | 0.004667 | 0.05018 | 2.48 |
| FALSE | High | RLSDEHEPE | [K].RLSDEH | 1xPhospho | Q923B1 1x | 0.069903 | 0.005181 | 1 | 2 | 1 | Q923B1 | Lariat debr | Q923B1 [5 | 1 | 1704.79 | 0 | -0.71 | 0.001658 | 3.47 | 122.4 | 77.6 | 6.37 | 10.2 | | High | High | High | High | High | High | High | High | High | High | High | 0.002675 | 0.02852 | 1.63 |
| FALSE | High | RISDPLTSSI | [R].RISDPL | 1xPhospho | Q3UJN1 1x | 0.018447 | 0.001324 | 1 | 3 | 2 | Q3UJN1 | DNA helica | Q3UJN1 [1 | 1 | 1594.815 | 0 | 0.35 | 0.001972 | 3.81 | 87.8 | 112.2 | 5.7 | 9 | | High | High | High | High | High | High | High | High | High | High | High | 0.001126 | 0.01163 | 2.45 |
| FALSE | High | RISDPLTSSI | [R].RISDPL | 3xPhospho | Q3UJN1 3x | 0.001002 | 0 | 1 | 3 | 4 | Q3UJN1 | DNA helica | Q3UJN1 [1 | 1 | 1754.748 | 0 | -0.39 | 0.01028 | 3.58 | 111.6 | 88.4 | 9.1 | 3.26 | | High | High | High | High | High | High | High | High | High | High | High | 0 | 0.000239 | 3.89 |
| FALSE | High | RISDPQVFS | [R].RISDPQ | 1xPhospho | E0CYH7 1x | 2.53E-05 | 0 | 1 | 4 | 1 | E0CYH7 | Filamin A-in | E0CYH7 [78 | 1 | 1714.929 | 0 | 0.26 | 0.232485 | 42.32 | 91.4 | 108.6 | 19.98 | 22.26 | | High | High | High | High | High | High | High | High | High | High | High | 0 | 3.83E-06 | 2.36 |
| FALSE | High | RLSEEACPC | [R].RLSEEA | 1xCarbami | O08715-4 1 | 9.82E-06 | 0 | 1 | 6 | 4 | O08715-4 | Isoform 4 c | O08715-4 | 1 | 2730.385 | 0 | 0.07 | 0.641123 | 5.64 | 97.2 | 102.8 | 4.57 | 9.6 | | High | High | High | High | High | High | High | High | High | High | High | 0 | 2.22E-06 | 4.42 |
| FALSE | High | RISEMEEEL | [R].RISEME | 1xPhospho | Q9DBR7 1x | 0.012731 | 0.000889 | 1 | 2 | 1 | Q9DBR7 | Protein ph | Q9DBR7 [9 | 1 | 1801.917 | 0 | 0.26 | 0.227497 | 17.48 | 92.6 | 107.4 | 10.16 | 10.94 | | High | High | High | High | High | High | High | High | High | High | High | 0.000449 | 0.004119 | 2.73 |
| FALSE | High | RLSENNCSI | [R].RLSENN | 1xCarbami | Q8C7S2 1x | 0.00025 | 0 | 1 | 5 | 1 | Q8C7S2 | Putative ur | Q8C7S2 [2 | 1 | 3661.699 | 0 | 0.76 | 0.020384 | 13.94 | 77.2 | 122.8 | 6.92 | 15.71 | | High | High | High | High | High | High | High | High | High | High | High | 0 | 5.03E-05 | 3.02 |
| FALSE | High | RLSENNCSI | [R].RLSENN | 1xCarbami | Q8C7S2 1x | 0.002793 | 0.000109 | 1 | 5 | 5 | Q8C7S2 | Putative ur | Q8C7S2 [2 | 1 | 3662.683 | 0 | 0.32 | 0.002536 | 6.07 | 88.7 | 111.3 | 5.82 | 8.02 | | High | High | High | High | High | High | High | High | High | High | High | 4.94E-05 | 0.000757 | 3.97 |
| FALSE | High | RLSEQLAHT | [R].RLSEQL | 1xPhospho | E9Q8A3 1x | 0.085859 | 0.006406 | 1 | 4 | 1 | E9Q8A3 | Phosphatic | E9Q8A3 [5 | 1 | 2137.157 | 0 | -0.16 | 0.029919 | 1.01 | 105.2 | 94.8 | 3.4 | 7.28 | | High | High | High | High | High | High | High | High | High | High | High | 0.003303 | 0.03611 | 2.64 |
| FALSE | High | RLSEQLAHT | [R].RLSEQL | 2xPhospho | E9Q8A3 2x | 0.007538 | 0.00062 | 1 | 4 | 1 | E9Q8A3 | Phosphatic | E9Q8A3 [5 | 1 | 2217.123 | 0 | 0.14 | 0.831576 | 3.12 | 96.6 | 103.4 | 11.48 | 6.19 | | High | High | High | High | High | High | High | High | High | High | High | 0.000275 | 0.002287 | 1.9 |
| FALSE | High | RLSESLHSA | [R].RLSESL | 1xPhospho | E9Q6J5-3 1 | 5.75E-05 | 0 | 1 | 3 | 2 | E9Q6J5-3 | Isoform Ne | E9Q6J5-3 [ | 1 | 2024.021 | 0 | 0.52 | 0.085068 | 24.22 | 84.4 | 115.6 | 11.28 | 15.96 | | High | High | High | High | High | High | High | High | High | High | High | 0 | 9.67E-06 | 2.11 |
| FALSE | High | RLSESPVFD | [R].RLSESP | 1xPhospho | P23949 1xP | 0.00034 | 0 | 1 | 1 | 2 | P23949 | mRNA deca | P23949 [42 | 1 | 2578.239 | 0 | -0.02 | 0.789782 | 15.75 | 101.2 | 98.8 | 11.26 | 8.67 | | High | High | High | High | High | High | High | High | High | High | High | 0 | 0.000124 | 3.64 |
| FALSE | High | RLSESPVFD | [R].RLSESP | 2xPhospho | P23949 2xP | 0.000772 | 0 | 1 | 1 | 2 | P23949 | mRNA deca | P23949 [42 | 1 | 2658.205 | 0 | 0.44 | 0.117381 | 33.01 | 82.8 | 117.2 | 18.49 | 24.86 | | High | High | High | High | High | High | High | High | High | High | High | 0 | 0.000179 | 3.77 |
| FALSE | High | RLSESQLSF | [R].RLSESQ | 1xPhospho | Q8C2Q3 1x | 0.046687 | 0.00354 | 1 | 2 | 1 | Q8C2Q3 | RNA-bindir | Q8C2Q3 [6 | 1 | 1531.783 | 0 | 0.53 | 0.007729 | 9.16 | 80 | 120 | 3.78 | 11.97 | | High | High | High | High | High | High | High | High | High | High | High | 0.001812 | 0.01786 | 1.37 |
| FALSE | High | RLSESSALK | [R].RLSESS. | 1xPhospho | Q91XU0 1x | 0.015603 | 0.001124 | 1 | 2 | 3 | Q91XU0 | ATPase WF | Q91XU0 [7 | 1 | 1528.85 | 0 | -0.62 | 0.0027 | 2.59 | 119.5 | 80.5 | 9.21 | 7.38 | | High | High | High | High | High | High | High | High | High | High | High | 0.000585 | 0.00514 | 1.73 |
| FALSE | High | RLSESSALK | [R].RLSESS. | 2xPhospho | Q91XU0 2x | 0.094641 | 0.007408 | 1 | 2 | 1 | Q91XU0 | ATPase WF | Q91XU0 [7 | 1 | 1608.816 | 0 | -0.9 | 0.001717 | 2.9 | 132.1 | 67.9 | 8.19 | 6.16 | | High | High | High | High | High | High | High | High | High | High | High | 0.003754 | 0.04033 | 2.85 |
| FALSE | High | RISETSEDQ | [R].RISETSE | 1xPhospho | P70174 1xP | 0.009087 | 0.000736 | 1 | 1 | 2 | P70174 | Histamine I | P70174 [34 | 1 | 1957.943 | 0 | 1.86 | 0.000448 | 11.2 | 43.1 | 156.9 | 9.63 | 8.73 | | High | High | High | High | High | High | High | High | High | High | High | 0.000327 | 0.00282 | 3.02 |
| FALSE | High | RISFGQEDS | [K].RISFGQ | 1xPhospho | G3UWZ0 1 | 1.43E-06 | 0 | 1 | 2 | 1 | G3UWZ0 | Bromodom | G3UWZ0 [ | 1 | 1976.025 | 0 | 1.62 | 0.000248 | 2.59 | 50.9 | 149.1 | 4.3 | 4.53 | | High | High | High | High | High | High | High | High | High | High | High | 0 | 1.52E-07 | 3.35 |
| FALSE | High | RISFSTSDV | [R].RISFSTS | 1xPhospho | E9PZW8 1x | 0.005615 | 0.00062 | 1 | 6 | 1 | E9PZW8 | Unconvent | E9PZW8 [1 | 1 | 1764.93 | 0 | 0.81 | 0.000438 | 7.01 | 71 | 129 | 7.16 | 2.38 | | High | High | High | High | High | High | High | High | High | High | High | 0.000275 | 0.001653 | 2.3 |
| FALSE | High | RISGASELG | [R].RISGAS | 1xPhospho | E9PUK7 1x | 0.000662 | 0 | 1 | 2 | 1 | E9PUK7 | Runt-relate | E9PUK7 [3 | 1 | 1897.937 | 0 | 3.25 | 0.000209 | 15.14 | 19.1 | 180.9 | 19.44 | 6.31 | | High | High | High | High | High | High | High | High | High | High | High | 0 | 0.000151 | 3.42 |
| FALSE | High | RLSGDSEEI | [K].RLSGDS | 1xPhospho | A0A1W2P6 | 0.000176 | 0 | 1 | 8 | 2 | A0A1W2P6 | Bromodom | A0A1W2P6 | 1 | 2658.285 | 0 | -1.31 | 0.000173 | 1.3 | 143 | 57 | 2.09 | 6.72 | | High | High | High | High | High | High | High | High | High | High | High | 0 | 4.38E-05 | 3.7 |
| FALSE | High | RISGLIYEET | [K].RISGLIY | 1xPhospho | B2RTM0 1x | 0.000124 | 0 | 1 | 2 | 2 | B2RTM0 | Histone H4 | B2RTM0 [4 | 1 | 1645.851 | 0 | 0.25 | 0.0054 | 2.92 | 91.6 | 108.4 | 2.32 | 4.17 | | High | High | High | High | High | High | High | High | High | High | High | 0 | 2.27E-05 | 3.04 |
| FALSE | High | RLSGLSAQ | [K].RLSGLS | 1xPhospho | G3X8Q1 1x | 0.053473 | 0.00384 | 1 | 3 | 1 | G3X8Q1 | Calcineurir | G3X8Q1 [1 | 1 | 1867.02 | 0 | -0.04 | 0.449366 | 15.94 | 101.5 | 98.5 | 1.52 | 13.49 | | High | High | High | High | High | High | High | High | High | High | High | 0.001999 | 0.02083 | 1.79 |
| FALSE | High | RISGSECAT | [K].RISGSE | 1xCarbami | A2ASY1 1xP | 7.21E-05 | 0 | 1 | 5 | 1 | A2ASY1 | TAF3 RNA p | A2ASY1 [42 | 1 | 1743.886 | 0 | -0.32 | 0.053703 | 8.53 | 111.2 | 88.8 | 7.29 | 7.75 | | High | High | High | High | High | High | High | High | High | High | High | 0 | 1.24E-05 | 2.62 |
| FALSE | High | RHSLSCTK | [K].RHSLSC | 1xCarbami | G3X9H5 1x | 0.005467 | 0.00062 | 1 | 2 | 2 | G3X9H5 | Huntingtin | G3X9H5 [1 | 1 | 1526.791 | 0 | 1.51 | 0.000278 | 6.3 | 51.2 | 148.8 | 4.45 | 4.36 | | High | High | High | High | High | High | High | High | High | High | High | 0.000275 | 0.001592 | 1.59 |
| FALSE | High | RHSIAIGEV | [K].RHSIAIC | 1xCarbami | A7MCV2 1: | 0.001644 | 5.66E-05 | 1 | 3 | 1 | A7MCV2 | DEP domai | A7MCV2 [1 | 1 | 1774.899 | 0 | -0.62 | 0.006815 | 4.65 | 123.3 | 76.7 | 9.07 | 6.95 | | High | High | High | High | High | High | High | High | High | High | High | 2.58E-05 | 0.000418 | 1.92 |
| FALSE | High | RHSHSHSPI | [R].RHSHSH | 3xPhospho | P62996 3xP | 0.104222 | 0.008348 | 1 | 2 | 1 | P62996 | Transform | P62996 [93 | 1 | 1888.728 | 0 | -0.1 | 0.214398 | 5.55 | 104.5 | 95.5 | 5.12 | 22.31 | | High | High | High | High | High | High | High | High | High | High | High | 0.004236 | 0.04538 | 2.3 |
| FALSE | High | RHSGTGAT | [R].RHSGTC | 2xPhospho | Q60520-1 2 | 0.01894 | 0.001362 | 1 | 2 | 1 | Q60520-1 | Isoform 2 c | Q60520-1 | 1 | 1825.913 | 0 | -1.02 | 0.00598 | 1.55 | 132.6 | 67.4 | 12.28 | 7.91 | | High | High | High | High | High | High | High | High | High | High | High | 0.000723 | 0.006415 | 2.11 |
| FALSE | High | RGLSVDSA | [K].RGLSVD | 1xPhospho | A0A1D5RLI | 0.003312 | 0.00016 | 1 | 7 | 1 | A0A1D5RLI | Transforma | A0A1D5RLI | 1 | 1826.978 | 0 | 0.76 | 0.000729 | 2.42 | 75.3 | 124.7 | 10.19 | 2.78 | | High | High | High | High | High | High | High | High | High | High | High | 7.19E-05 | 0.000909 | 2.53 |
| FALSE | High | RGNSESVS | [R].RGNSE | 1xPhospho | Q9DBY8 1x | 0.000967 | 0 | 1 | 2 | 2 | Q9DBY8 | Nuclear va | Q9DBY8 [1 | 1 | 1800.926 | 0 | 0.53 | 0.003778 | 2.85 | 85.6 | 114.4 | 5 | 5.43 | | High | High | High | High | High | High | High | High | High | High | High | 0 | 0.000231 | 2.51 |
| FALSE | High | RGPAEGEE | [R].RGPAE | 1xCarbami | Q3UMT1 1 | 0.000868 | 0 | 1 | 3 | 1 | Q3UMT1 | Protein ph | Q3UMT1 [ | 2 | 3194.437 | 0 | -0.14 | 0.112723 | 10.38 | 106.4 | 93.6 | 7.73 | 10.96 | | High | High | High | High | High | High | High | High | High | High | High | 0 | 0.000203 | 3.78 |
| FALSE | High | RGPEVEGS | [R].RGPEVI | 1xPhospho | Q921C3 1x | 3.42E-06 | 0 | 1 | 5 | 1 | Q921C3 | Bromodom | Q921C3 [1 | 1 | 1891.948 | 0 | -1.05 | 0.000223 | 0.58 | 134.7 | 65.3 | 3.86 | 4.26 | | High | High | High | High | High | High | High | High | High | High | High | 0 | 4.04E-07 | 1.87 |
| FALSE | High | RGPEVTSQ | [K].RGPEV | 1xCarbami | E9QM77 1: | 1.01E-06 | 0 | 1 | 8 | 4 | E9QM77 | Ataxin-2 O | E9QM77 [8 | 1 | 2427.21 | 0 | 0.86 | 0.002267 | 12.36 | 69.9 | 130.1 | 8.17 | 6.85 | | High | High | High | High | High | High | High | High | High | High | High | 0 | 1.03E-07 | 4.05 |
| FALSE | High | RGPGYTSG | [R].RGPGY | 1xDeamida | E9QAS8 2x | 0.000164 | 0 | 1 | 6 | 5 | E9QAS8 | Synaptic fu | E9QAS8 [4 | 2 | 4657.931 | 0 | 0.09 | 0.626777 | 7.5 | 96.9 | 103.1 | 9.65 | 1.3 | | High | High | High | High | High | High | High | High | High | High | High | 0 | 3.13E-05 | 4.51 |
| FALSE | High | RGPNYTSG | [R].RGPNY | 1xPhospho | Q61584-3 1 | 0.003055 | 0.00016 | 2 | 9 | 6 | Q61584-3; | Isoform B c | Q61584-3 | 1 | 3041.332 | 0 | -0.86 | 0.001766 | 4.64 | 128.6 | 71.4 | 6.8 | 10.33 | Shared | High | High | High | High | High | High | High | High | High | High | High | 0.001151 | 0.01247 | 2.75 |
| FALSE | High | RGPNYTSG | [R].RGPNY | 2xPhospho | Q61584-3 2 | 0.000529 | 0 | 2 | 9 | 3 | Q61584-3; | Isoform B c | Q61584-3 | 1 | 3121.299 | 0 | -0.94 | 9.77E-05 | 0.74 | 131.4 | 68.6 | 2.31 | 4.14 | Shared | High | High | High | High | High | High | High | High | High | High | High | 0 | 0.000183 | 2.2 |
| FALSE | High | RGPSNSTSI | [K].RGPSN | 1xPhospho | Q9R0A1 1x | 0.013666 | 0.001011 | 1 | 3 | 3 | Q9R0A1 | Chloride ch | Q9R0A1 [7 | 1 | 2843.356 | 0 | -0.55 | 0.023867 | 5.96 | 118.8 | 81.2 | 13.23 | 8.18 | | High | High | High | High | High | High | High | High | High | High | High | 0.000814 | 0.007791 | 3.77 |
| FALSE | High | RGPSNSTSI | [K].RGPSN | 1xPhospho | Q9R0A1 1x | 1.59E-06 | 0 | 1 | 3 | 5 | Q9R0A1 | Chloride ch | Q9R0A1 [7 | 1 | 2859.351 | 0 | -0.91 | 0.01021 | 2.42 | 130.6 | 69.4 | 10.78 | 14.12 | | High | High | High | High | High | High | High | High | High | High | High | 0 | 4.98E-07 | 4.38 |
| FALSE | High | RGSADIIPL | [R].RGSADI | 1xCarbami | O89032 1x | 1.53E-06 | 0 | 1 | 3 | 1 | O89032 | SH3 and PX | O89032 [8 | 1 | 2532.366 | 0 | 1.14 | 0.003573 | 9.5 | 58.2 | 141.8 | 11.23 | 12.95 | | High | High | High | High | High | High | High | High | High | High | High | 0 | 1.64E-07 | 3.1 |
| FALSE | High | RGSADIIPL | [R].RGSADI | 1xCarbami | O89032 2x | 0.006536 | 0.00062 | 1 | 3 | 1 | O89032 | SH3 and PX | O89032 [8 | 1 | 2612.332 | 0 | 0.97 | 0.001567 | 11.2 | 67.7 | 132.3 | 6.79 | 7.16 | | High | High | High | High | High | High | High | High | High | High | High | 0.000275 | 0.001958 | 2.79 |
| FALSE | High | RGSALGIG | [R].RGSAL | 1xPhospho | A2ALS7 1xP | 0.000779 | 0 | 1 | 1 | 1 | A2ALS7 | Rap1 GTPa | A2ALS7 [21 | 1 | 2391.341 | 0 | 1.93 | 0.001353 | 7.36 | 38.3 | 161.7 | 10.72 | 13.05 | | High | High | High | High | High | High | High | High | High | High | High | 0 | 0.00018 | 2.02 |
| FALSE | High | RGHHGSLG | [R].RGHHG | 1xPhospho | Q8R149 1x | 0.035181 | 0.002269 | 1 | 2 | 2 | Q8R149 | BUD13 hor | Q8R149 [2 | 1 | 1657.812 | 0 | -0.19 | 0.316282 | 14.57 | 106.8 | 93.2 | 11.65 | 18.9 | | High | High | High | High | High | High | High | High | High | High | High | 0.001199 | 0.01285 | 2.99 |
| FALSE | High | RGSASFVSS | [K].RGSASF | 1xPhospho | Q8BI72 1xP | 0.0016 | 5.66E-05 | 1 | 1 | 1 | Q8BI72 | CDKN2A-in | Q8BI72 [23 | 1 | 1789.998 | 0 | -0.41 | 0.013067 | 5.88 | 114 | 86 | 8.75 | 10.33 | | High | High | High | High | High | High | High | High | High | High | High | 2.58E-05 | 0.000405 | 2.87 |
| FALSE | High | RGSDASDF | [R].RGSDA | 1xCarbami | B1ARU1 1x | 6.7E-08 | 0 | 2 | 19 | 4 | B1ARU1; F | Microtubu | B1ARU1 [5 | 1 | 3646.529 | 0 | 0.11 | 0.140193 | 4.48 | 95.5 | 104.5 | 9.1 | 3.76 | Shared | High | High | High | High | High | High | High | High | High | High | High | 0 | 7.68E-08 | 6.61 |
| FALSE | High | RGSDASDF | [R].RGSDA | 1xCarbami | B1ARU1 2x | 6.73E-06 | 0 | 2 | 19 | 4 | B1ARU1; F | Microtubu | B1ARU1 [5 | 1 | 3725.511 | 0 | 1.19 | 0.003575 | 8.21 | 65 | 135 | 26.11 | 13.42 | Shared | High | High | High | High | High | High | High | High | High | High | High | 0 | 8.64E-07 | 4.79 |
| FALSE | High | RGSDASDF | [R].RGSDA | 1xCarbami | S4R1P5 1xP | 6.11E-07 | 0 | 1 | 10 | 3 | S4R1P5 | Dystonin O | S4R1P5 [76 | 1 | 3894.817 | 0 | 0.23 | 0.106799 | 10.43 | 96.6 | 103.4 | 18.12 | 16.81 | | High | High | High | High | High | High | High | High | High | High | High | 0 | 5.82E-08 | 5.22 |
| FALSE | High | RGSDGEW | [R].RGSDG | 1xPhospho | Q6RT24 1x | 7.23E-06 | 0 | 1 | 2 | 2 | Q6RT24 | Centromer | Q6RT24 [8 | 1 | 2629.236 | 0 | -0.37 | 0.000945 | 2.08 | 112.1 | 87.9 | 5.2 | 2.9 | | High | High | High | High | High | High | High | High | High | High | High | 0 | 9.35E-07 | 3.47 |
| FALSE | High | RGSDIDNP | [R].RGSDID | 1xPhospho | G5E8L8 1xP | 3.9E-05 | 0 | 1 | 2 | 1 | G5E8L8 | Mitogen ac | G5E8L8 [32 | 1 | 2450.213 | 0 | 1.56 | 0.000333 | 9.22 | 50.6 | 149.4 | 7.86 | 5.51 | | High | High | High | High | High | High | High | High | High | High | High | 0 | 6.25E-06 | 4.33 |
| FALSE | High | RGSDIDNP | [R].RGSDID | 2xPhospho | G5E8L8 2xP | 0.112856 | 0.009146 | 1 | 2 | 1 | G5E8L8 | Mitogen ac | G5E8L8 [32 | 1 | 2530.179 | 0 | 0.87 | 0.003671 | 21.15 | 70.6 | 129.4 | 12.09 | 10.14 | | High | High | High | High | High | High | High | High | High | High | High | 0.004596 | 0.04991 | 2.71 |
| FALSE | High | RGSDIDNP | [R].RGSDID | 3xPhospho | G5E8L8 3xP | 0.100129 | 0.008067 | 1 | 2 | 1 | G5E8L8 | Mitogen ac | G5E8L8 [32 | 2 | 2950.331 | 0 | 0.57 | 0.00645 | 9.92 | 78.5 | 121.5 | 13.79 | 4.91 | | High | High | High | High | High | High | High | High | High | High | High | 0.004046 | 0.04315 | 3.55 |
| FALSE | High | RGSDIGVR | [R].RGSDIG | 1xPhospho | Q99LH9 1x | 0.030367 | 0.001857 | 1 | 2 | 2 | Q99LH9 | SH3 domai | Q99LH9 [3 | 1 | 1168.604 | 0 | -0.01 | 0.799831 | 18.39 | 100.8 | 99.2 | 1.7 | 12.04 | | High | High | High | High | High | High | High | High | High | High | High | 0.006167 | 0.06662 | 1.43 |
| FALSE | High | RGSETMAC | [R].RGSETM | 1xPhospho | Q8K4Z3 1x | 5.5E-05 | 0 | 1 | 1 | 5 | Q8K4Z3 | NAD(P)H-h | Q8K4Z3 [4 | 1 | 1731.886 | 0 | -1.68 | 0.000922 | 1.62 | 152 | 48 | 3.22 | 23.1 | | High | High | High | High | High | High | High | High | High | High | High | 2.58E-05 | 0.000411 | 3.53 |
| FALSE | High | RGSFDATG | [R].RGSFD/ | 1xPhospho | Q9QXS1-3 | 0.002376 | 0.000109 | 3 | 23 | 1 | Q9QXS1-3; | Isoform PL | Q9QXS1-3 | 1 | 3082.309 | 0 | 2.42 | 0.00041 | 27.81 | 31.5 | 168.5 | 32.46 | 5.75 | Shared | High | High | High | High | High | High | High | High | High | High | High | 4.94E-05 | 0.00063 | 3.55 |
| FALSE | High | RGSFDATG | [R].RGSFD/ | 1xPhospho | Q9QXS1-3 | 0.002719 | 0.000109 | 3 | 23 | 2 | Q9QXS1-3; | Isoform PL | Q9QXS1-3 | 1 | 3098.304 | 0 | 1.71 | 0.005932 | 15.58 | 47.7 | 152.3 | 16.99 | 26.49 | Shared | High | High | High | High | High | High | High | High | High | High | High | 4.94E-05 | 0.000731 | 3.14 |
| FALSE | High | RGSFPLAAS | [R].RGSFPL | 1xPhospho | Q9D4F2 1x | 3.46E-07 | 0 | 1 | 1 | 1 | Q9D4F2 | Phospholip | Q9D4F2 [6 | 1 | 2640.313 | 0 | 1.05 | 0.001495 | 7.04 | 67.3 | 132.7 | 8.49 | 12.46 | | High | High | High | High | High | High | High | High | High | High | High | 0 | 3.09E-08 | 4.34 |
| FALSE | High | RGSFSSENY | [R].RGSFSS | 1xPhospho | Q3TSG4 1x | 0.042503 | 0.003197 | 1 | 1 | 1 | Q3TSG4 | RNA deme | Q3TSG4 [3 | 1 | 1697.763 | 0 | -0.09 | 0.195802 | 6.43 | 101.2 | 98.8 | 5.78 | 8.1 | | High | High | High | High | High | High | High | High | High | High | High | 0.001639 | 0.01596 | 1.41 |
| FALSE | High | RGSDASDF | [R].RGSDA | 1xCarbami | B1ARU1 1x | 4.53E-10 | 0 | 2 | 19 | 11 | B1ARU1; F | Microtubu | B1ARU1 [5 | 1 | 3645.545 | 0 | 0.06 | 0.218612 | 8.66 | 95.1 | 104.9 | 7.88 | 10.65 | Shared | High | High | High | High | High | High | High | High | High | High | High | 0 | 1.78E-11 | 7.25 |
| FALSE | High | RGGGGGD | [K].RGGGG | 1xPhospho | B2RRE0 1xP | 0.000127 | 0 | 1 | 3 | 6 | B2RRE0 | A kinase (P | B2RRE0 [52 | 1 | 3525.581 | 0 | -1.21 | 0.003578 | 3.76 | 138.7 | 61.3 | 8.33 | 14.5 | | High | High | High | High | High | High | High | High | High | High | High | 0 | 4.92E-05 | 5.44 |
| FALSE | High | RGFSDSGG | [R].RGFSDS | 1xPhospho | Q3TUQ5 1: | 7.65E-07 | 0 | 1 | 3 | 2 | Q3TUQ5 | Pinin OS=M | Q3TUQ5 [6 | 1 | 1770.894 | 0 | 1.31 | 0.005153 | 24.28 | 58.2 | 141.8 | 13.57 | 11.65 | | High | High | High | High | High | High | High | High | High | High | High | 0 | 7.53E-08 | 3.31 |
| FALSE | High | RGEPQTPT | [R].RGEPQ | 1xPhospho | Q62187 1x | 0.000183 | 0 | 1 | 1 | 1 | Q62187 | Transcripti | Q62187 [1 | 1 | 3145.618 | 0 | 0.48 | 0.007151 | 8.82 | 82.8 | 117.2 | 11.89 | 5.62 | | High | High | High | High | High | High | High | High | High | High | High | 0 | 3.53E-05 | 3.99 |
| FALSE | High | REYQEFSSF | [K].REYQEF | 2xPhospho | P48972 2xP | 0.003001 | 0.00016 | 1 | 1 | 1 | P48972 | Myb-relate | P48972 [27 | 1 | 2579.187 | 0 | -0.95 | 0.003064 | 2.85 | 131.7 | 68.3 | 10.62 | 8.42 | | High | High | High | High | High | High | High | High | High | High | High | 7.19E-05 | 0.00082 | 2.41 |
| FALSE | High | RFEIINAIYE | [K].RFEIINA | 1xCarbami | E9PW66 1x | 0.0004 | 0 | 1 | 5 | 1 | E9PW66 | Nucleosom | E9PW66 [1 | 1 | 4623.198 | 0 | -0.99 | 0.004509 | 4.45 | 138.5 | 61.5 | 12.33 | 15.37 | | High | High | High | High | High | High | High | High | High | High | High | 0 | 8.51E-05 | 3.82 |
| FALSE | High | RFGEYNSN | [K].RFGEYN | 1xPhospho | Q8R3N1 1x | 5.88E-06 | 0 | 1 | 4 | 1 | Q8R3N1 | Nucleolar p | Q8R3N1 [8 | 1 | 2208.074 | 0 | -0.27 | 0.015526 | 7.97 | 109.3 | 90.7 | 4.2 | 8.73 | | High | High | High | High | High | High | High | High | High | High | High | 0 | 7.44E-07 | 1.92 |
| FALSE | High | RFPGYDSES | [K].RFPGYD | 1xPhospho | P47962 1xP | 0.039011 | 0.002718 | 1 | 5 | 2 | P47962 | 60S riboso | P47962 [17 | 1 | 1723.846 | 0 | 0.01 | 0.414008 | 2.92 | 100.4 | 99.6 | 4.74 | 2.67 | | High | High | High | High | High | High | High | High | High | High | High | 0.001421 | 0.01446 | 1.46 |
| FALSE | High | RFPSTGSCT | [K].RFPSTG | 1xCarbami | Q8BIE6-2 1 | 0.002793 | 0.000109 | 1 | 4 | 3 | Q8BIE6-2 | Isoform 2 c | Q8BIE6-2 [ | 1 | 2747.302 | 0 | 0.25 | 0.073697 | 9.68 | 88.6 | 111.4 | 7.57 | 12.5 | | High | High | High | High | High | High | High | High | High | High | High | 0.000449 | 0.003809 | 4.07 |
| FALSE | High | RFPTDYAST | [R].RFPTDY | 1xPhospho | Q6A065 1x | 0.02933 | 0.001857 | 1 | 3 | 1 | Q6A065 | Centrosom | Q6A065 [1 | 1 | 2388.034 | 0 | 0.8 | 0.00301 | 3.99 | 71.9 | 128.1 | 9.35 | 9.32 | | High | High | High | High | High | High | High | High | High | High | High | 0.000997 | 0.01054 | 2.3 |
| FALSE | High | RFPTDYAST | [R].RFPTDY | 2xPhospho | Q6A065 2x | 0.024638 | 0.001612 | 1 | 3 | 1 | Q6A065 | Centrosom | Q6A065 [1 | 1 | 2468.001 | 0 | 0.84 | 0.002367 | 17.85 | 71.8 | 128.2 | 13.09 | 6.17 | | High | High | High | High | High | High | High | High | High | High | High | 0.000872 | 0.008656 | 3.46 |
| FALSE | High | RFSDFLGLH | [R].RFSDFL | 1xPhospho | Q3U5L3 1x | 9.14E-06 | 0 | 1 | 3 | 1 | Q3U5L3 | Putative ur | Q3U5L3 [1 | 1 | 1844.982 | 0 | 1.71 | 0.000487 | 6.81 | 46.8 | 153.2 | 3.97 | 8.45 | | High | High | High | High | High | High | High | High | High | High | High | 0 | 1.22E-06 | 2.67 |
| FALSE | High | RFSDFLGLY | [R].RFSDFL | 1xPhospho | D3YWH1 1 | 9.65E-06 | 0 | 1 | 6 | 1 | D3YWH1 | Sorting nex | D3YWH1 [ | 1 | 1912.997 | 0 | 1.92 | 0.000363 | 2.65 | 40 | 160 | 4.41 | 8.2 | | High | High | High | High | High | High | High | High | High | High | High | 0 | 1.3E-06 | 3.23 |
| FALSE | High | RFSDGAAS | [R].RFSDG/ | 1xPhospho | E9PU87 1x | 3.57E-09 | 0 | 1 | 4 | 1 | E9PU87 | Serine/thre | E9PU87 [6 | 1 | 1936.009 | 0 | 0.17 | 0.194194 | 17.87 | 88.3 | 111.7 | 11.39 | 15.82 | | High | High | High | High | High | High | High | High | High | High | High | 0 | 1.81E-10 | 4.51 |
| FALSE | High | RFSDLGFEV | [R].RFSDLG | 1xPhospho | O08738 1x | 0.025073 | 0.001645 | 1 | 3 | 1 | O08738 | Caspase-6 | O08738 [6 | 1 | 1735.918 | 0 | 2.14 | 0.000364 | 13.67 | 37.9 | 162.1 | 18.71 | 10.48 | | High | High | High | High | High | High | High | High | High | High | High | 0.000872 | 0.008823 | 2.3 |
| FALSE | High | RFSEGVLQ | [R].RFSEGV | 1xPhospho | B9EKJ7 1xP | 3.45E-06 | 0 | 1 | 2 | 3 | B9EKJ7 | Tankyrase | B9EKJ7 [42 | 1 | 2383.206 | 0 | -0.39 | 0.103629 | 12.61 | 112.9 | 87.1 | 13.24 | 12.97 | | High | High | High | High | High | High | High | High | High | High | High | 0 | 0.000324 | 3.33 |
| FALSE | High | RFSFCFSPE | [R].RFSFCF | 2xCarbami | Q64337 1x | 8.79E-05 | 0 | 1 | 3 | 7 | Q64337 | Sequestos | Q64337 [2 | 1 | 3077.348 | 0 | 2.63 | 0.000352 | 12.15 | 28.4 | 171.6 | 7.16 | 8.81 | | High | High | High | High | High | High | High | High | High | High | High | 0.000538 | 0.004648 | 3.83 |
| FALSE | High | RFSFCFSPE | [R].RFSFCF | 2xCarbami | Q64337 1x | 0.000584 | 0 | 1 | 3 | 1 | Q64337 | Sequestos | Q64337 [2 | 1 | 3078.332 | 0 | 3.73 | | 64.08 | 19.3 | 180.7 | 54.35 | 10.76 | | Not Found | High | High | High | High | High | High | High | High | High | High | 0 | 0.00013 | 4.74 |
| FALSE | High | RFSFPLSLF | [R].RFSFPL | 1xPhospho | Q8CHW4 1 | 0.001558 | 5.66E-05 | 1 | 1 | 1 | Q8CHW4 | Translatior | Q8CHW4 [ | 1 | 2421.217 | 0 | 0.57 | 0.000899 | 6.46 | 78.3 | 121.7 | 7.54 | 7.02 | | High | High | High | High | High | High | High | High | High | High | High | 2.58E-05 | 0.000392 | 4.01 |
| FALSE | High | RFSFSPSAV | [R].RFSFSP | 1xPhospho | Q4VAC9 1x | 0.022574 | 0.001542 | 1 | 3 | 1 | Q4VAC9 | Pleckstrin h | Q4VAC9 [1 | 1 | 1646.825 | 0 | 0.55 | 0.059677 | 20.51 | 83.2 | 116.8 | 12.58 | 15.54 | | High | High | High | High | High | High | High | High | High | High | High | 0.000814 | 0.007821 | 1.52 |
| FALSE | High | RFSHDGEE | [K].RFSHD | 1xPhospho | Q8BJ05 1xP | 0.0002 | 0 | 1 | 3 | 3 | Q8BJ05 | Zinc finger | Q8BJ05 [30 | 1 | 2508.999 | 0 | 0.8 | 0.026679 | 23.97 | 69.1 | 130.9 | 14.02 | 16.37 | | High | High | High | High | High | High | High | High | High | High | High | 0.000538 | 0.004419 | 2.74 |
| FALSE | High | RFSLSPSLG | [R].RFSLSP | 2xPhospho | Q8BXQ8 2x | 0.037692 | 0.00249 | 1 | 1 | 2 | Q8BXQ8 | Protein FA | Q8BXQ8 [2 | 1 | 1891.903 | 0 | -1.82 | 0.000561 | 1.02 | 155.9 | 44.1 | 8.53 | 12.42 | | High | High | High | High | High | High | High | High | High | High | High | 0.008583 | 0.0915 | 2.36 |
| FALSE | High | RFSMEDLN | [R].RFSME | 1xPhospho | Q0VDL6 1x | 0.001437 | 5.66E-05 | 1 | 1 | 2 | Q0VDL6 | PCTAIRE-m | Q0VDL6 [6 | 1 | 1677.844 | 0 | -0.3 | 0.064991 | 11.42 | 111.6 | 88.4 | 5.76 | 11.43 | | High | High | High | High | High | High | High | High | High | High | High | 2.58E-05 | 0.000359 | 2.03 |
| FALSE | High | RFSMEDLN | [R].RFSME | 1xPhospho | Q0VDL6 1x | 5.03E-05 | 0 | 1 | 1 | 3 | Q0VDL6 | PCTAIRE-m | Q0VDL6 [6 | 1 | 1693.838 | 1 | -0.27 | 0.071901 | 5.44 | 109.2 | 90.8 | 2.45 | 12.63 | | High | High | High | High | High | High | High | High | High | High | High | 0 | 8.28E-06 | 2.41 |
| FALSE | High | RFSPPSSSL | [R].RFSPPS | 1xPhospho | Q08775 1x | 0.002895 | 0.00016 | 2 | 14 | 1 | Q08775; E | Runt-relate | Q08775 [1 | 1 | 1926.025 | 0 | 2.85 | 0.000317 | 4.67 | 25.6 | 174.4 | 9.86 | 9.4 | Shared | High | High | High | High | High | High | High | High | High | High | High | 7.19E-05 | 0.000788 | 2.87 |
| FALSE | High | RFSPSQVPI | [R].RFSPSC | 1xPhospho | H7BX08 1x | 0.032548 | 0.002167 | 1 | 6 | 1 | H7BX08 | Calmodulir | H7BX08 [1 | 1 | 1724.905 | 0 | 0.19 | 0.312903 | 9.72 | 93.9 | 106.1 | 9.5 | 9.32 | | High | High | High | High | High | High | High | High | High | High | High | 0.001142 | 0.01182 | 2.51 |
| FALSE | High | RFSVQEQD | [R].RFSVQE | 1xPhospho | Q542P8 1x | 3.18E-06 | 0 | 1 | 2 | 8 | Q542P8 | EMG1 nucl | Q542P8 [1 | 1 | 2386.184 | 0 | 0.42 | 0.001285 | 2.02 | 84.9 | 115.1 | 5.45 | 4.06 | | High | High | High | High | High | High | High | High | High | High | High | 0 | 0.00018 | 4.13 |
| FALSE | High | RFSVQEQD | [R].RFSVQE | 2xPhospho | Q542P8 2x | 0.007339 | 0.00062 | 1 | 2 | 1 | Q542P8 | EMG1 nucl | Q542P8 [1 | 1 | 2466.151 | 0 | -0.58 | 0.030006 | 2.46 | 115.9 | 84.1 | 11.2 | 12.72 | | High | High | High | High | High | High | High | High | High | High | High | 0.000275 | 0.002224 | 2.7 |
| FALSE | High | RFSVQEQD | [R].RFSVQE | 1xPhospho | Q542P8 1x | 0.001168 | 0 | 1 | 2 | 1 | Q542P8 | EMG1 nucl | Q542P8 [1 | 2 | 2743.442 | 0 | -0.02 | 0.759401 | 42.64 | 104.2 | 95.8 | 20.02 | 21.69 | | High | High | High | High | High | High | High | High | High | High | High | 0 | 0.000284 | 2.02 |
| FALSE | High | RGEDEGSF | [R].RGEDE | 1xPhospho | Q148V8 1x | 7.06E-07 | 0 | 1 | 1 | 2 | Q148V8 | Protein FA | Q148V8 [1 | 1 | 2623.162 | 0 | 1.39 | 0.000192 | 1.53 | 55.4 | 144.6 | 5.98 | 6.06 | | High | High | High | High | High | High | High | High | High | High | High | 0 | 6.84E-08 | 3.97 |
| FALSE | High | RGEDEGSF | [R].RGEDE | 1xDeamida | Q148V8 1x | 4.69E-06 | 0 | 1 | 1 | 3 | Q148V8 | Protein FA | Q148V8 [1 | 1 | 2624.146 | 0 | 0.85 | 0.005452 | 10.96 | 73.6 | 126.4 | 8.73 | 11.61 | | High | High | High | High | High | High | High | High | High | High | High | 0 | 5.78E-07 | 4.76 |
| FALSE | High | RGEDEGSF | [R].RGEDE | 1xDeamida | Q148V8 1x | 0.035486 | 0.002304 | 1 | 1 | 3 | Q148V8 | Protein FA | Q148V8 [1 | 2 | 3320.549 | 0 | 0.64 | 0.003929 | 9.37 | 83.8 | 116.2 | 14.09 | 8.11 | | High | High | High | High | High | High | High | High | High | High | High | 0.001214 | 0.01302 | 3.26 |

| | | | | | | | | | | | | | | | | | | | | | | | | | | | | | | | | | | | | | | |
|---|---|---|---|---|---|---|---|---|---|---|---|---|---|---|---|---|---|---|---|---|---|---|---|---|---|---|---|---|---|---|---|---|---|---|---|---|---|---|
| FALSE | High | RGEDIDCD | [K].RGEDID | 1xCarbami | A2A654 1x | 0.008094 | 0.000736 | 1 | 5 | 1 | A2A654 | Bromodom | A2A654 [5 | 1 | 2428.074 | 0 | -0.15 | 0.30058 | 18.19 | 106.8 | 93.2 | 13.25 | 6.85 | | High | High | High | High | High | High | High | High | High | High | High | 0.000327 | 0.002472 | 2.17 |
| FALSE | High | RGSFSSENY | [R].RGSFSS | 1xDeamida | Q3TSG4 1x | 0.116482 | 0.009523 | 1 | 1 | 1 | Q3TSG4 | RNA deme | Q3TSG4 [3 | 1 | 1698.747 | 0 | -0.15 | 0.350459 | 4.62 | 106.9 | 93.1 | 10.23 | 7.38 | | High | High | High | High | High | High | High | High | High | High | High | 0.004757 | 0.05169 | 2.25 |
| FALSE | High | RLSPVEK | [R].RLSPVE | 1xPhospho | P97360 1xl | 0.114657 | 0.009411 | 1 | 6 | 1 | P97360 | Transcripti | P97360 [21 | 1 | 1366.786 | 0 | 0.18 | 0.003177 | 1.33 | 92.6 | 107.4 | 2.52 | 2.91 | | High | High | High | High | High | High | High | High | High | High | High | 0.004707 | 0.05077 | 1.73 |
| FALSE | High | RGSGDPMI | [R].RGSGD | 1xPhospho | Q8VI59 1xl | 0.062731 | 0.004664 | 1 | 1 | 1 | Q8VI59 | Pecanex-lik | Q8VI59 [28 | 1 | 1626.808 | 0 | 0.42 | 0.153571 | 11.65 | 84.1 | 115.9 | 13.54 | 33.86 | | High | High | High | High | High | High | High | High | High | High | High | 0.002403 | 0.02507 | 2.18 |
| FALSE | High | RGSGDTSIS | [R].RGSGD | 1xPhospho | Q3UZ39 1x | 2E-07 | 0 | 1 | 10 | 18 | Q3UZ39 | Leucine-ric | Q3UZ39 [8 | 1 | 2107.953 | 0 | 0.61 | 0.012623 | 17.42 | 77.4 | 122.6 | 7.44 | 14.01 | | High | High | High | High | High | High | High | High | High | High | High | 0 | 1.67E-08 | 4.43 |
| FALSE | High | RGSQNSSE | [R].RGSQN | 1xPhospho | Q8BZ47 1x | 0.094641 | 0.007408 | 1 | 2 | 2 | Q8BZ47 | Zinc finger | Q8BZ47 [4 | 1 | 2794.352 | 0 | -0.19 | 0.445384 | 16.27 | 99.8 | 100.2 | 15.72 | 23.49 | | High | High | High | High | High | High | High | High | High | High | High | 0.003754 | 0.04035 | 2.38 |
| FALSE | High | RGSQNSSE | [R].RGSQN | 1xDeamida | Q8BZ47 1x | 0.022574 | 0.001542 | 1 | 2 | 1 | Q8BZ47 | Zinc finger | Q8BZ47 [4 | 1 | 2795.336 | 0 | -0.49 | 0.365123 | 34.17 | 116.8 | 83.2 | 26.89 | 27.19 | | High | High | High | High | High | High | High | High | High | High | High | 0.000814 | 0.007794 | 2.52 |
| FALSE | High | RGSQPSTP | [R].RGSQP | 3xPhospho | Q05A36 3x | 0.00824 | 0.000736 | 1 | 4 | 1 | Q05A36 | RNA-bindir | Q05A36 [5 | 2 | 3129.452 | 0 | 0.42 | 0.018207 | 5.67 | 88.3 | 111.7 | 8.42 | 5.45 | | High | High | High | High | High | High | High | High | High | High | High | 0.000327 | 0.002528 | 3.68 |
| FALSE | High | RGSSPGSLI | [R].RGSSPC | 1xPhospho | A0A1W2P7 | 0.000165 | 0 | 1 | 7 | 3 | A0A1W2P7 | Ral GTPase | A0A1W2P7 | 1 | 1765.961 | 0 | 1.13 | 0.001035 | 13.37 | 66.2 | 133.8 | 7.42 | 7.15 | | High | High | High | High | High | High | High | High | High | High | High | 0 | 3.15E-05 | 3.99 |
| FALSE | High | RGSSQPES | [R].RGSSQI | 1xCarbami | Q8BMI0 1x | 5.37E-06 | 0 | 1 | 2 | 4 | Q8BMI0 | F-box only | Q8BMI0 [8 | 1 | 2520.066 | 0 | 0.28 | 0.17158 | 29.8 | 92.1 | 107.9 | 19.32 | 10.85 | | High | High | High | High | High | High | High | High | High | High | High | 0 | 6.72E-07 | 4.14 |
| FALSE | High | RGSTSQEN | [R].RGSTSC | 1xPhospho | E9Q6J5-3 1 | 0.000426 | 0 | 1 | 4 | 6 | E9Q6J5-3 | Isoform Ne | E9Q6J5-3 [ | 1 | 1648.813 | 0 | 0.26 | 0.027288 | 15.13 | 89.3 | 110.7 | 9.34 | 8.81 | | High | High | High | High | High | High | High | High | High | High | High | 0 | 0.000184 | 3.04 |
| FALSE | High | RGSTTSIPS | [R].RGSTTS | 1xPhospho | D3Z742 1xl | 7.88E-08 | 0 | 1 | 2 | 6 | D3Z742 | MCG12536 | D3Z742 [43 | 1 | 2693.2 | 0 | 0.62 | 0.003324 | 5.56 | 76.6 | 123.4 | 3.71 | 9.8 | | High | High | High | High | High | High | High | High | High | High | High | 0 | 1.23E-08 | 4.53 |
| FALSE | High | RGSTTSIPS | [R].RGSTTS | 2xPhospho | D3Z742 2xl | 0.001674 | 5.66E-05 | 1 | 2 | 3 | D3Z742 | MCG12536 | D3Z742 [43 | 1 | 2773.166 | 0 | 1.38 | 0.001537 | 13.92 | 59.1 | 140.9 | 8.78 | 10.83 | | High | High | High | High | High | High | High | High | High | High | High | 2.58E-05 | 0.000426 | 4.28 |
| FALSE | High | RGSYEYSQ | [R].RGSYEY | 1xPhospho | A2ADB0 1x | 0.000693 | 0 | 1 | 5 | 2 | A2ADB0 | Msx2-inter | A2ADB0 [6 | 1 | 1583.705 | 0 | 0.62 | 0.006288 | 8.46 | 79.6 | 120.4 | 7.72 | 10.19 | | High | High | High | High | High | High | High | High | High | High | High | 0.007094 | 0.07695 | 2.36 |
| FALSE | High | RGTGQSDI | [R].RGTGQ | 2xPhospho | P97801 2xl | 3.17E-05 | 0 | 1 | 3 | 4 | P97801 | Survival m | P97801 [20 | 1 | 2711.237 | 0 | -0.57 | 0.001244 | 1.26 | 120.2 | 79.8 | 4.77 | 4.91 | | High | High | High | High | High | High | High | High | High | High | High | 0 | 0.00022 | 3.88 |
| FALSE | High | RGTPEFSPI | [R].RGTPEF | 1xCarbami | Q7TNS8 1x | 0.035181 | 0.002269 | 1 | 1 | 1 | Q7TNS8 | Elongin BC | Q7TNS8 [3 | 1 | 1741.866 | 0 | -1.79 | 0.006558 | 1.48 | 156.4 | 43.6 | 23.71 | 16.46 | | High | High | High | High | High | High | High | High | High | High | High | 0.001183 | 0.01284 | 2.61 |
| FALSE | High | RGTSPRPPI | [K].RGTSPF | 1xPhospho | P24788 1xl | 0.000454 | 0 | 1 | 4 | 4 | P24788 | Cyclin-dep | P24788 [73 | 1 | 2953.482 | 0 | -0.46 | 0.00118 | 1.99 | 118.4 | 81.6 | 3.62 | 4.56 | | High | High | High | High | High | High | High | High | High | High | High | 0 | 9.78E-05 | 3.09 |
| FALSE | High | RGTSPRPPI | [K].RGTSPF | 2xPhospho | P24788 2xl | 0.00418 | 0.00036 | 1 | 4 | 3 | P24788 | Cyclin-dep | P24788 [73 | 1 | 3033.448 | 0 | -0.41 | 0.00252 | 2.56 | 113.3 | 86.7 | 5 | 4.9 | | High | High | High | High | High | High | High | High | High | High | High | 0.000847 | 0.007959 | 2.59 |
| FALSE | High | RGSQGDIK | [R].RGSQG | 1xPhospho | Q8CI51 1x | 0.017344 | 0.001204 | 2 | 11 | 2 | Q8CI51; Q | PDZ and LI | Q8CI51 [22 | 1 | 1398.751 | 0 | 0.28 | 0.068941 | 5.14 | 89.5 | 110.5 | 10.41 | 13.31 | Shared | High | High | High | High | High | High | High | High | High | High | High | 0.000674 | 0.005839 | 2.07 |
| FALSE | High | RGYSYDDS | [R].RGYSYI | 1xPhospho | Q8C5N3 1x | 0.000148 | 0 | 1 | 14 | 5 | Q8C5N3 | Pre-mRNA- | Q8C5N3 [5 | 1 | 1790.725 | 0 | -0.55 | 0.00411 | 3.38 | 120.7 | 79.3 | 3.45 | 9.86 | | High | High | High | High | High | High | High | High | High | High | High | 0.001597 | 0.01538 | 2.18 |
| FALSE | High | RHEDLVAP | [K].RHEDLV | 2xPhospho | Q8CFT2 2x | 0.013545 | 0.000976 | 1 | 2 | 1 | Q8CFT2 | Histone-lys | Q8CFT2 [1 | 1 | 3069.491 | 0 | -0.43 | 0.006105 | 7.52 | 113.1 | 86.9 | 5.21 | 10.42 | | High | High | High | High | High | High | High | High | High | High | High | 0.000505 | 0.004407 | 2.44 |
| FALSE | High | RHESDEDS | [R].RHESDI | 1xPhospho | O08796 1x | 0.008614 | 0.000736 | 1 | 6 | 1 | O08796 | Eukaryotic | O08796 [4 | 1 | 1840.802 | 0 | 1.04 | 0.003855 | 31.34 | 61.7 | 138.3 | 9.03 | 12.82 | | High | High | High | High | High | High | High | High | High | High | High | 0.000327 | 0.002645 | 2.02 |
| FALSE | High | RHPPLYQA | [K].RHPPLY | 1xPhospho | E9PYL2 1xF | 0.067619 | 0.004864 | 1 | 2 | 1 | E9PYL2 | Proline-ricl | E9PYL2 [12 | 1 | 2507.394 | 0 | -0.48 | 0.007167 | 7.75 | 116.3 | 83.7 | 6.15 | 18.26 | | High | High | High | High | High | High | High | High | High | High | High | 0.002485 | 0.02722 | 2.17 |
| FALSE | High | RHQEDQDI | [K].RHQED | 1xPhospho | B2RT77 1xl | 3.37E-07 | 0 | 1 | 1 | 4 | B2RT77 | TWIST neig | B2RT77 [28 | 1 | 3327.507 | 0 | -0.14 | 0.014859 | 2.2 | 105.5 | 94.5 | 2.97 | 3.54 | | High | High | High | High | High | High | High | High | High | High | High | 0 | 2.98E-08 | 4.29 |
| FALSE | High | RHSCAEAL | [R].RHSCAI | 1xCarbami | Q60591 1x | 3.92E-06 | 0 | 1 | 14 | 1 | Q60591 | Nuclear fac | Q60591 [2 | 1 | 2340.185 | 0 | 1.46 | 0.004361 | 1.21 | 51.3 | 148.7 | 10.05 | 17.77 | | High | High | High | High | High | High | High | High | High | High | High | 0 | 4.73E-07 | 3.93 |
| FALSE | High | RHSCAEAL | [R].RHSCAI | 1xCarbami | Q60591 2x | 0.000389 | 0 | 1 | 14 | 2 | Q60591 | Nuclear fac | Q60591 [2 | 1 | 2420.151 | 0 | 2.7 | 0.000377 | 32.22 | 26.7 | 173.3 | 16.46 | 6.23 | | High | High | High | High | High | High | High | High | High | High | High | 2.58E-05 | 0.000516 | 3.88 |
| FALSE | High | RHSDFCGP | [R].RHSDF( | 1xCarbami | Q8CIP4 1xF | 0.081085 | 0.006219 | 1 | 2 | 1 | Q8CIP4 | MAP/micro | Q8CIP4 [42 | 1 | 2341.168 | 0 | -0.5 | 0.034876 | 1.98 | 115.5 | 84.5 | 10.09 | 13.19 | | High | High | High | High | High | High | High | High | High | High | High | 0.0032 | 0.03376 | 2.09 |
| FALSE | High | RHSDVSLA | [R].RHSDVS | 1xPhospho | D9HP81 1x | 0.010758 | 0.000772 | 1 | 5 | 1 | D9HP81 | Regulating | D9HP81 [3 | 1 | 2107.997 | 0 | 0.4 | 0.000172 | 2.67 | 85.2 | 114.8 | 3.54 | 2.32 | | High | High | High | High | High | High | High | High | High | High | High | 0.000399 | 0.003417 | 2.4 |
| FALSE | High | RHSFTSLTN | [K].RHSFTS | 1xPhospho | Q69ZI1 1xF | 0.000188 | 0 | 1 | 6 | 1 | Q69ZI1 | E3 ubiquiti | Q69ZI1 [30 | 1 | 1946.992 | 0 | 0.07 | 0.32068 | 11 | 99.9 | 100.1 | 9.1 | 11.6 | | High | High | High | High | High | High | High | High | High | High | High | 0 | 3.63E-05 | 2.61 |
| FALSE | High | RHSGQDVI | [R].RHSGQ | 1xPhospho | Q3UVN5 1 | 0.000164 | 0 | 1 | 4 | 5 | Q3UVN5 | Putative ur | Q3UVN5 [1 | 1 | 1683.889 | 0 | 0.18 | 0.274225 | 11.59 | 89.7 | 110.3 | 9.48 | 41.46 | | High | High | High | High | High | High | High | High | High | High | High | 0.000859 | 0.008642 | 4.07 |
| FALSE | High | RHSGQDVI | [R].RHSGQ | 1xDeamida | Q3UVN5 1 | 0.085859 | 0.006406 | 1 | 4 | 1 | Q3UVN5 | Putative ur | Q3UVN5 [1 | 1 | 1684.873 | 0 | 0.21 | 0.566558 | 44.31 | 96.2 | 103.8 | 18.49 | 49.52 | | High | High | High | High | High | High | High | High | High | High | High | 0.003303 | 0.03607 | 2.7 |
| FALSE | High | RHSGQDVI | [R].RHSGQ | 1xPhospho | Q3UVN5 1 | 0.000622 | 0 | 1 | 4 | 5 | Q3UVN5 | Putative ur | Q3UVN5 [1 | 1 | 1913.052 | 0 | 0.41 | 0.005436 | 6.93 | 84.1 | 115.9 | 5.36 | 5.67 | | High | High | High | High | High | High | High | High | High | High | High | 0.004569 | 0.04949 | 2.42 |
| FALSE | High | RHSGTGAT | [R].RHSGT( | 1xPhospho | Q60520-1 | 9.14E-06 | 0 | 1 | 2 | 3 | Q60520-1 | Isoform 2 c | Q60520-1 | | 1 | 1745.946 | 0 | -0.58 | 0.003361 | 1.36 | 117.4 | 82.6 | 4.65 | 9.65 | | High | High | High | High | High | High | High | High | High | High | High | 0 | 1.22E-06 | 2.98 |
| FALSE | High | RHDSLPDP | [R].RHDSLF | 1xPhospho | Q3UHP6 1x | 0.000418 | 0 | 1 | 5 | 2 | Q3UHP6 | Putative ur | Q3UHP6 [9 | 1 | 1717.822 | 0 | 0.02 | 0.834185 | 4.99 | 99.8 | 100.2 | 7.07 | 8.65 | | High | High | High | High | High | High | High | High | High | High | High | 0.000538 | 0.004712 | 2.43 |
| FALSE | High | RGSPTTGLI | [R].RGSPTT | 1xPhospho | Q148V8 1x | 0.000223 | 0 | 1 | 1 | 6 | Q148V8 | Protein FAI | Q148V8 [9 | 1 | 1858.95 | 0 | 1.12 | 0.005725 | 4.67 | 63.7 | 136.3 | 5.14 | 16.44 | | High | High | High | High | High | High | High | High | High | High | High | 0 | 4.42E-05 | 2.97 |
| FALSE | High | RGSPTTGLI | [R].RGSPTT | 1xPhospho | Q148V8 1x | 0.013074 | 0.000889 | 1 | 1 | 1 | Q148V8 | Protein FAI | Q148V8 [9 | 1 | 1842.955 | 0 | 0.82 | 0.002198 | 17.98 | 70.4 | 129.6 | 14.71 | 13.69 | | High | High | High | High | High | High | High | High | High | High | High | 0.000467 | 0.004216 | 2.09 |
| FALSE | High | RGSPPAAG | [K].RGSPPA | 2xPhospho | Q545J5 2xl | 0.001881 | 5.66E-05 | 1 | 2 | 1 | Q545J5 | Basic Krup | Q545J5 [69 | 1 | 1986.982 | 0 | 1.08 | 0.001894 | 6.37 | 64.1 | 135.9 | 7.1 | 12.85 | | High | High | High | High | High | High | High | High | High | High | High | 2.58E-05 | 0.000483 | 3.01 |
| FALSE | High | RGSGDTSS | [R].RGSGD | 1xPhospho | A0A1L1SSC | 5.67E-06 | 0 | 1 | 12 | 3 | A0A1L1SSC | Leucine-ric | A0A1L1SSC | 1 | 2415.16 | 0 | 0.15 | 0.044223 | 4.66 | 92 | 108 | 3.79 | 8.78 | | High | High | High | High | High | High | High | High | High | High | High | 0.00074 | 0.006504 | 3.74 |
| FALSE | High | RGSGDTSS | [R].RGSGD | 2xPhospho | A0A1L1SSC | 0.005615 | 0.00062 | 1 | 12 | 5 | A0A1L1SSC | Leucine-ric | A0A1L1SSC | 1 | 2495.127 | 0 | 0.06 | 0.804756 | 30.33 | 95.1 | 104.9 | 22.42 | 11.24 | | High | High | High | High | High | High | High | High | High | High | High | 0.000275 | 0.001639 | 3.56 |
| FALSE | High | RGSGHPAY | [R].RGSGH | 1xPhospho | Q9DC11 1x | 8.35E-06 | 0 | 1 | 3 | 2 | Q9DC11 | Plexin dom | Q9DC11 [5 | 1 | 2321.169 | 0 | 0.21 | 0.158932 | 22.67 | 96.8 | 103.2 | 9.2 | 14.83 | | High | High | High | High | High | High | High | High | High | High | High | 0 | 1.11E-06 | 3.75 |
| FALSE | High | RGSGPEIFT | [R].RGSGPI | 1xPhospho | Q4FJY1 1xF | 0.0003 | 0 | 1 | 3 | 1 | Q4FJY1 | Ier3 protei | Q4FJY1 [29 | 1 | 2127.047 | 0 | 0.41 | 0.007002 | 6.56 | 85.2 | 114.8 | 4.15 | 10.91 | | High | High | High | High | High | High | High | High | High | High | High | 0 | 6.14E-05 | 3.29 |
| FALSE | High | RGSGPVDL | [R].RGSGP' | 2xCarbami | Q923B1 1x | 4.3E-07 | 0 | 1 | 2 | 1 | Q923B1 | Lariat debr | Q923B1 [2 | 1 | 2429.167 | 0 | 1.94 | 0.00024 | 13.54 | 40.6 | 159.4 | 10.39 | 4.3 | | High | High | High | High | High | High | High | High | High | High | High | 0 | 3.91E-08 | 4.33 |
| FALSE | High | RGSGSACS | [R].RGSGS/ | 4xCarbami | Q99J21 1xl | 0.000967 | 0 | 1 | 2 | 1 | Q99J21 | Mucolipin- | Q99J21 [55 | 1 | 2043.822 | 0 | 0.57 | 0.014017 | 18.44 | 79.8 | 120.2 | 16.35 | 4.17 | | High | High | High | High | High | High | High | High | High | High | High | 0 | 0.000229 | 2.96 |
| FALSE | High | RGSGSVDE | [R].RGSGS\ | 1xPhospho | Q99P72 1x | 1.41E-06 | 0 | 1 | 1 | 2 | Q99P72 | Reticulon-4 | Q99P72 [1 | 1 | 3024.513 | 0 | -0.33 | 0.076231 | 9.49 | 110.5 | 89.5 | 15.19 | 19.86 | | High | High | High | High | High | High | High | High | High | High | High | 0 | 1.49E-07 | 3.9 |
| FALSE | High | RGSGSVDE | [R].RGSGS\ | 1xPhospho | Q99P72 1x | 3.37E-07 | 0 | 1 | 1 | 5 | Q99P72 | Reticulon-4 | Q99P72 [1 | 1 | 3253.676 | 0 | -0.33 | 0.031941 | 6.82 | 116.1 | 83.9 | 10.56 | 11.63 | | High | High | High | High | High | High | High | High | High | High | High | 0 | 2.98E-08 | 5 |
| FALSE | High | RGSGSVDE | [R].RGSGS\ | 2xPhospho | Q99P72 2x | 1.29E-05 | 0 | 1 | 1 | 2 | Q99P72 | Reticulon-4 | Q99P72 [1 | 1 | 3333.642 | 0 | -0.51 | 0.008289 | 4.64 | 123.6 | 76.4 | 14.67 | 13.87 | | High | High | High | High | High | High | High | High | High | High | High | 0 | 1.79E-06 | 4.72 |
| FALSE | High | RGSGSVDE | [R].RGSGS\ | 1xPhospho | Q8BHF5 1x | 0.000681 | 0 | 1 | 1 | 1 | Q8BHF5 | Reticulon C | Q8BHF5 [1 | 1 | 3811.951 | 0 | 0.37 | 0.164195 | 9.25 | 91 | 109 | 17.21 | 17.25 | | High | High | High | High | High | High | High | High | High | High | High | 0 | 0.000154 | 3.67 |
| FALSE | High | RGSGSVVV | [R].RGSGS\ | 1xPhospho | Q8K3G7 1x | 0.001799 | 5.66E-05 | 1 | 2 | 2 | Q8K3G7 | Reticulon C | Q8K3G7 [1 | 1 | 1915.999 | 0 | 2.39 | 0.000913 | 6.91 | 32.1 | 167.9 | 10.68 | 12.68 | | High | High | High | High | High | High | High | High | High | High | High | 0.000585 | 0.00517 | 2.6 |
| FALSE | High | RGSLELGN | [R].RGSLEL | 1xPhospho | A0A1B0GS | 8.82E-06 | 0 | 1 | 6 | 2 | A0A1B0GS | Supervillin | A0A1B0GS | 1 | 2502.312 | 0 | 0.83 | 0.003366 | 10.2 | 71.7 | 128.3 | 4.23 | 9.49 | | High | High | High | High | High | High | High | High | High | High | High | 0 | 1.17E-06 | 4 |
| FALSE | High | RGSLEMSS | [R].RGSLEN | 1xPhospho | Q9Z248 1x | 1.38E-06 | 0 | 1 | 3 | 2 | Q9Z248 | Zinc finger | Q9Z248 [1 | 1 | 1929.894 | 0 | 0.11 | 0.545358 | 1.69 | 95.2 | 104.8 | 5.17 | 12.32 | | High | High | High | High | High | High | High | High | High | High | High | 0 | 1.45E-07 | 2.88 |
| FALSE | High | RGSLEMSS | [R].RGSLEN | 1xPhospho | Q9Z248 1x | 0.003927 | 0.000311 | 1 | 3 | 3 | Q9Z248 | Zinc finger | Q9Z248 [1 | 1 | 1945.889 | 0 | -0.13 | 0.942036 | 14.51 | 102.2 | 97.8 | 15.02 | 22.43 | | High | High | High | High | High | High | High | High | High | High | High | 0.000183 | 0.001257 | 3.17 |
| FALSE | High | RGSLEMSS | [R].RGSLEN | 2xPhospho | Q9Z248 2x | 0.045898 | 0.003408 | 1 | 3 | 1 | Q9Z248 | Zinc finger | Q9Z248 [1 | 1 | 2009.86 | 0 | -0.07 | 0.759017 | 20.41 | 102.5 | 97.5 | 7 | 13.11 | | High | High | High | High | High | High | High | High | High | High | High | 0.001741 | 0.01743 | 2.26 |
| FALSE | High | RGSLGEEG | [R].RGSLGI | 1xPhospho | Q8K120 1x | 1.55E-07 | 0 | 1 | 2 | 2 | Q8K120 | Nuclear fac | Q8K120 [2 | 1 | 2500.316 | 0 | -1.97 | 0.000323 | 1.16 | 160.4 | 39.6 | 5.19 | 18.19 | | High | High | High | High | High | High | High | High | High | High | High | 0 | 1.26E-08 | 4.08 |
| FALSE | High | RGSLLSEPA | [K].RGSLLS | 1xPhospho | E9Q4K7 1x | 0.041425 | 0.003126 | 1 | 1 | 1 | E9Q4K7 | Kinesin fan | E9Q4K7 [7 | 1 | 1734.947 | 0 | 1.23 | 0.000609 | 12.2 | 58.1 | 141.9 | 14.32 | 8.28 | | High | High | High | High | High | High | High | High | High | High | High | 0.001609 | 0.01559 | 1.64 |
| FALSE | High | RGSLPDQK | [R].RGSLPE | 1xPhospho | Q3TDH0 1x | 0.008463 | 0.000736 | 1 | 6 | 2 | Q3TDH0 | Putative ur | Q3TDH0 [4 | 1 | 1438.782 | 0 | -0.22 | 0.150763 | 5.63 | 108.6 | 91.4 | 4.65 | 9.98 | | High | High | High | High | High | High | High | High | High | High | High | 0.001011 | 0.01101 | 1.95 |
| FALSE | High | RGSLPDTG | [R].RGSLPE | 1xPhospho | Q8VI56 1xl | 0.028575 | 0.001829 | 1 | 2 | 1 | Q8VI56 | Low-densit | Q8VI56 [18 | 1 | 1654.872 | 0 | -0.92 | 0.001142 | 2.83 | 130 | 70 | 6.28 | 11.08 | | High | High | High | High | High | High | High | High | High | High | High | 0.000984 | 0.0102 | 2.31 |
| FALSE | High | RGSLPTEAS | [K].RGSLPT | 1xCarbami | E9Q3E2 1x | 0.039347 | 0.002793 | 2 | 4 | 1 | E9Q3E2; Q | Synaptopo | E9Q3E2 [1 | 1 | 1588.724 | 0 | -0.64 | 0.035273 | 4.28 | 116.1 | 83.9 | 11.49 | 20.86 | Shared | High | High | High | High | High | High | High | High | High | High | High | 0.001453 | 0.01462 | 2.6 |
| FALSE | High | RGSLSPPSS | [R].RGSLSP | 2xPhospho | E9Q5F9 2x | 0.010382 | 0.000772 | 1 | 3 | 3 | E9Q5F9 | Histone-lys | E9Q5F9 [2 | 1 | 1795.798 | 0 | -0.15 | 0.007294 | 3.04 | 104.3 | 95.7 | 5.11 | 3.14 | | High | High | High | High | High | High | High | High | High | High | High | 0.000382 | 0.003268 | 2.75 |
| FALSE | High | RGSISSMSS | [R].RGSISS | 1xPhospho | Q80Y56 1x | 2.9E-05 | 0 | 1 | 1 | 3 | Q80Y56 | Rabenosyn | Q80Y56 [2 | 1 | 2323.162 | 0 | 0.09 | 0.50724 | 14.81 | 101.4 | 98.6 | 7.78 | 29.71 | | High | High | High | High | High | High | High | High | High | High | High | 0 | 4.48E-06 | 3.51 |
| FALSE | High | RGSISSMSS | [R].RGSISS | 2xPhospho | Q80Y56 2x | 0.002355 | 0.000109 | 1 | 1 | 2 | Q80Y56 | Rabenosyn | Q80Y56 [2 | 1 | 2387.133 | 0 | 0.12 | 0.155644 | 8.74 | 96.3 | 103.7 | 1.49 | 7.07 | | High | High | High | High | High | High | High | High | High | High | High | 4.94E-05 | 0.000621 | 2.6 |
| FALSE | High | RGSLTFAGI | [R].RGSLTF | 1xPhospho | Q148V8 1x | 2.47E-05 | 0 | 1 | 1 | 4 | Q148V8 | Protein FAI | Q148V8 [9 | 1 | 1777.925 | 0 | 1.57 | 0.000176 | 4.23 | 49.1 | 150.9 | 9.99 | 4.05 | | High | High | High | High | High | High | High | High | High | High | High | 0 | 3.74E-06 | 3.2 |
| FALSE | High | RGSLTLEEK | [K].RGSLTL | 1xPhospho | Q8CFE4 1x | 0.050826 | 0.003676 | 1 | 3 | 2 | Q8CFE4 | SCY1-like p | Q8CFE4 [6 | 1 | 1570.861 | 0 | 0.11 | 0.144555 | 10.93 | 94.2 | 105.8 | 6.31 | 4.89 | | High | High | High | High | High | High | High | High | High | High | High | 0.001916 | 0.01963 | 1.81 |
| FALSE | High | RGSMFYDE | [R].RGSMF | 1xPhospho | Q9WVF8 1 | 0.058173 | 0.004227 | 1 | 2 | 1 | Q9WVF8 | Tumor sup | Q9WVF8 [4 | 1 | 3390.564 | 0 | 0.21 | 0.763955 | 7.27 | 95.9 | 104.1 | 12.13 | 12.8 | | High | High | High | High | High | High | High | High | High | High | High | 0.002187 | 0.02297 | 2.45 |
| FALSE | High | RGSMFYDE | [R].RGSMF | 1xPhospho | Q9WVF8 1 | 0.000772 | 0 | 1 | 2 | 2 | Q9WVF8 | Tumor sup | Q9WVF8 [4 | 1 | 3406.559 | 0 | -0.19 | 0.332969 | 9.58 | 107.8 | 92.2 | 12.44 | 13.98 | | High | High | High | High | High | High | High | High | High | High | High | 0 | 0.000179 | 3.07 |
| FALSE | High | RGSNIWLE | [R].RGSNIV | 1xPhospho | Q6ZQ33 1x | 0.048715 | 0.00354 | 1 | 2 | 1 | Q6ZQ33 | MKIAA085 | Q6ZQ33 [7 | 1 | 1737.945 | 0 | -0.57 | 0.03349 | 11.29 | 125.7 | 74.3 | 17.67 | 21.09 | | High | High | High | High | High | High | High | High | High | High | High | 0.001876 | 0.01877 | 2.84 |
| FALSE | High | RGSPPAAG | [K].RGSPPA | 1xPhospho | Q545J5 1xl | 2.98E-05 | 0 | 1 | 2 | 1 | Q545J5 | Basic Krup | Q545J5 [69 | 1 | 1907.015 | 0 | 0.57 | 0.011149 | 10.22 | 80.5 | 119.5 | 7.95 | 11.01 | | High | High | High | High | High | High | High | High | High | High | High | 0 | 4.61E-06 | 3.89 |
| FALSE | High | RGSGDTSIS | [R].RGSGD | 1xPhospho | Q3UZ39 1x | 4.62E-07 | 0 | 1 | 10 | 12 | Q3UZ39 | Leucine-ric | Q3UZ39 [8 | 1 | 2091.958 | 0 | 0.79 | 0.002281 | 9.56 | 73.5 | 126.5 | 5.61 | 10.22 | | High | High | High | High | High | High | High | High | High | High | High | 0 | 4.28E-08 | 4.12 |
| FALSE | High | RISQGDK | [K].RISQGD | 1xPhospho | Q80Y20 1x | 0.113753 | 0.009241 | 1 | 4 | 1 | Q80Y20 | Alkylated D | Q80Y20 [5 | 1 | 1341.729 | 0 | -0.24 | 0.063936 | 4.23 | 109.5 | 90.5 | 8.06 | 9.17 | | High | High | High | High | High | High | High | High | High | High | High | 0.004667 | 0.05031 | 1.64 |
| FALSE | High | RLSQISAGE | [K].RLSQIS | 1xCarbami | E9PYP2 1xl | 0.000645 | 0 | 1 | 3 | 1 | E9PYP2 | Neurobeac | E9PYP2 [26 | 1 | 2638.258 | 0 | 0.54 | 0.05315 | 21.61 | 82.9 | 117.1 | 14.39 | 19.34 | | High | High | High | High | High | High | High | High | High | High | High | 0 | 0.000145 | 4.58 |
| FALSE | High | RLSQQSAS | [R].RLSQQ | 1xPhospho | Q9DCM0 1 | 7.15E-05 | 0 | 1 | 1 | 1 | Q9DCM0 | Persulfide | Q9DCM0 [ | 1 | 1892.032 | 0 | -0.69 | 0.018764 | 4.08 | 116.2 | 83.8 | 18.22 | 16.02 | | High | High | High | High | High | High | High | High | High | High | High | 0 | 1.23E-05 | 1.99 |
| FALSE | High | RPPQSPQV | [K].RPPQSF | 1xPhospho | A0EQY3 1x | 2.67E-05 | 0 | 1 | 4 | 1 | A0EQY3 | Jacobsen s | A0EQY3 [1 | 0 | 1875.004 | 0 | 0.06 | 0.472015 | 7.26 | 96.5 | 103.5 | 9.17 | 6.7 | | High | High | High | High | High | High | High | High | High | High | High | 0 | 4.09E-06 | 1.87 |
| FALSE | High | SPGSVAVSI | [K].SPGSVA | 1xPhospho | G3X928 1x | 0.000231 | 0 | 1 | 4 | 2 | G3X928 | SEC23-inte | G3X928 [6 | 0 | 1752.966 | 0 | -0.43 | 0.011007 | 10 | 116.8 | 83.2 | 7.23 | 8.15 | | High | High | High | High | High | High | High | High | High | High | High | 0.000689 | 0.005931 | 2.9 |
| FALSE | High | RPPSIGDVF | [R].RPPSIG | 1xPhospho | M0QWZ1 | 0.000628 | 0 | 1 | 2 | 1 | M0QWZ1 | Protein FAI | M0QWZ1 [ | 0 | 2075.12 | 0 | -0.59 | 0.015263 | 1.41 | 117.4 | 82.6 | 10.62 | 9.18 | | High | High | High | High | High | High | High | High | High | High | High | 0 | 0.000141 | 1.94 |
| FALSE | High | RPPSPDPN | [R].RPPSPD | 1xPhospho | Q62261 1x | 0.000434 | 0 | 2 | 6 | 1 | Q62261; Q | Spectrin be | Q62261 [2 | 0 | 1417.704 | 0 | 1.25 | 0.004047 | 28.66 | 59.3 | 140.7 | 16.11 | 11.56 | Shared | High | High | High | High | High | High | High | High | High | High | High | 0 | 9.38E-05 | 2.78 |
| FALSE | High | RPPSPDPN | [R].RPPSPD | 1xPhospho | Q62261 1x | 0.000116 | 0 | 2 | 6 | 6 | Q62261; Q | Spectrin be | Q62261 [2 | 0 | 1646.867 | 0 | 1.37 | 0.000263 | 9.74 | 56.9 | 143.1 | 4.01 | 4.8 | Shared | High | High | High | High | High | High | High | High | High | High | High | 0.000689 | 0.005935 | 2.24 |
| FALSE | High | RPPSPDPN | [R].RPPSPD | 1xDeamida | Q62261 1x | 0.000976 | 0 | 2 | 6 | 1 | Q62261; Q | Spectrin be | Q62261 [2 | 0 | 1647.851 | 0 | 1.37 | 0.000193 | 8.84 | 55 | 145 | 4.35 | 3.23 | Shared | High | High | High | High | High | High | High | High | High | High | High | 0 | 0.000231 | 2.1 |
| FALSE | High | RPPSPDVIV | [K].RPPSPD | 1xPhospho | E9QMN5 1 | 3.02E-07 | 0 | 1 | 4 | 3 | E9QMN5 | Transcripti | E9QMN5 [ | 0 | 2472.233 | 0 | -0.54 | 0.000414 | 2.87 | 117.7 | 82.3 | 2.8 | 4.86 | | High | High | High | High | High | High | High | High | High | High | High | 0 | 2.64E-08 | 3.73 |
| FALSE | High | RPPSPDVIV | [K].RPPSPD | 2xPhospho | E9QMN5 2 | 6.33E-07 | 0 | 1 | 4 | 5 | E9QMN5 | Transcripti | E9QMN5 [ | 0 | 2552.2 | 0 | -0.65 | 0.000295 | 1.18 | 122.9 | 77.1 | 1.34 | 5.55 | | High | High | High | High | High | High | High | High | High | High | High | 0 | 6.11E-08 | 4.25 |
| FALSE | High | RPPSPDVIV | [K].RPPSPD | 3xPhospho | E9QMN5 3 | 7.91E-06 | 0 | 1 | 4 | 5 | E9QMN5 | Transcripti | E9QMN5 [ | 0 | 2632.166 | 0 | -0.3 | 0.000209 | 0.15 | 109.2 | 90.8 | 4.66 | 4.81 | | High | High | High | High | High | High | High | High | High | High | High | 0 | 1.04E-06 | 5.2 |
| FALSE | High | RPPSQSLTS | [R].RPPSQS | 1xPhospho | Q8R550-3 | 2.07E-07 | 0 | 1 | 7 | 7 | Q8R550-3 | Isoform 3 c | Q8R550-3 | 0 | 3428.651 | 0 | 2.27 | 0.000442 | 14.45 | 32.5 | 167.5 | 20.04 | 8.15 | | High | High | High | High | High | High | High | High | High | High | High | 0 | 1.73E-08 | 4.98 |
| FALSE | High | RPQANSVE | [K].RPQAN | 1xPhospho | C9K101 1x | 0.01218 | 0.000889 | 1 | 3 | 1 | C9K101 | Non-specif | C9K101 [43 | 0 | 1910.974 | 0 | -0.74 | 0.015337 | 8.48 | 124.1 | 75.9 | 9.33 | 18.87 | | High | High | High | High | High | High | High | High | High | High | High | 0.000449 | 0.003917 | 2.48 |
| FALSE | High | RPQFSPIES | [K].RPQFSF | 1xCarbami | Q3USC0 1x | 0.000248 | 0 | 1 | 2 | 5 | Q3USC0 | Putative ur | Q3USC0 [1 | 0 | 4298.98 | 0 | -0.43 | 0.003908 | 1.9 | 116 | 84 | 2.87 | 7.41 | | High | High | High | High | High | High | High | High | High | High | High | 0 | 0.000129 | 3.89 |
| FALSE | High | RPQFSPIES | [K].RPQFSF | 1xCarbami | Q3USC0 1x | 0.011549 | 0.00086 | 1 | 2 | 10 | Q3USC0 | Putative ur | Q3USC0 [1 | 0 | 4299.964 | 0 | -0.66 | 0.009668 | 8 | 122.6 | 77.4 | 7.66 | 15.71 | | High | High | High | High | High | High | High | High | High | High | High | 0.000449 | 0.004027 | 4.18 |
| FALSE | High | RPPPSAALL | [R].RPPPSA | 1xPhospho | Q8VCE9 1x | 3.57E-05 | 0 | 1 | 6 | 1 | Q8VCE9 | Pleckstrin l | Q8VCE9 [5 | 0 | 2511.425 | 0 | 1.54 | 0.000291 | 2.98 | 50.5 | 149.5 | 16.44 | 8.96 | | High | High | High | High | High | High | High | High | High | High | High | 0 | 5.64E-06 | 3.39 |
| FALSE | High | RPQFSPIES | [K].RPQFSF | 1xCarbami | Q3USC0 2x | 0.021231 | 0.001507 | 1 | 2 | 1 | Q3USC0 | Putative ur | Q3USC0 [1 | 0 | 4378.946 | 0 | -0.66 | 0.049844 | 3.6 | 119.8 | 80.2 | 20.74 | 10.87 | | High | High | High | High | High | High | High | High | High | High | High | 0.000801 | 0.007281 | 2.87 |
| FALSE | High | RPQSPGVS | [R].RPQSP( | 2xPhospho | A2ADB0 2x | 1.31E-05 | 0 | 1 | 5 | 3 | A2ADB0 | Msx2-inter | A2ADB0 [7 | 0 | 1893.857 | 0 | -0.05 | 0.684834 | 1.7 | 99.4 | 100.6 | 4.05 | 5.54 | | High | High | High | High | High | High | High | High | High | High | High | 2.58E-05 | 0.000563 | 2.9 |
| FALSE | High | RPQSWDEV | [R].RPQSW | 1xPhospho | Q5U4C1 1x | 0.000466 | 0 | 1 | 2 | 1 | Q5U4C1 | G-protein c | Q5U4C1 [1 | 0 | 2158.11 | 0 | -0.85 | 0.000507 | 2.44 | 125.5 | 74.5 | 5.68 | 9.95 | | High | High | High | High | High | High | High | High | High | High | High | 0 | 0.000102 | 3.04 |
| FALSE | High | RPQYSNPP | [R].RPQYSI | 1xPhospho | Q3UBT1 1x | 0.034878 | 0.002269 | 1 | 5 | 2 | Q3UBT1 | Putative ur | Q3UBT1 [2 | 0 | 3532.659 | 0 | 0.72 | 0.011207 | 13.39 | 73.1 | 126.9 | 12.98 | 12.36 | | High | High | High | High | High | High | High | High | High | High | High | 0.001183 | 0.01273 | 2.9 |
| FALSE | High | RPQYSNPP | [R].RPQYSI | 1xDeamida | Q3UBT1 1x | 0.000779 | 0 | 1 | 5 | 5 | Q3UBT1 | Putative ur | Q3UBT1 [2 | 0 | 3533.643 | 0 | 0.58 | 0.031316 | 39.98 | 80.3 | 119.7 | 9.3 | 16.46 | | High | High | High | High | High | High | High | High | High | High | High | 7.19E-05 | 0.000789 | 3.51 |
| FALSE | High | RPQYSNPP | [R].RPQYSI | 1xPhospho | Q3UBT1 1x | 0.001002 | 0 | 1 | 5 | 1 | Q3UBT1 | Putative ur | Q3UBT1 [2 | 0 | 3548.654 | 0 | 0.69 | 0.038658 | 25.04 | 78.1 | 121.9 | 7.18 | 21.44 | | High | High | High | High | High | High | High | High | High | High | High | 0 | 0.00024 | 2.94 |
| FALSE | High | RPRPGTED | [K].RPRPG1 | 1xPhospho | Q91XA5 1x | 6.41E-05 | 0 | 1 | 2 | 1 | Q91XA5 | snRNA-acti | Q91XA5 [2 | 0 | 3711.709 | 0 | -0.42 | 0.034617 | 6.71 | 113.1 | 86.9 | 13.48 | 16.02 | | High | High | High | High | High | High | High | High | High | High | High | 0 | 1.09E-05 | 2.54 |
| FALSE | High | RPRPSTPAI | [K].RPRPST | 1xPhospho | E9PYH6 1xl | 0.000102 | 0 | 1 | 3 | 3 | E9PYH6 | SET domai | E9PYH6 [9 | 1 | 2791.33 | 0 | -0.5 | 0.000948 | 3.4 | 115.1 | 84.9 | 3.33 | 5.61 | | High | High | High | High | High | High | High | High | High | High | High | 0.004327 | 0.04639 | 1.84 |
| FALSE | High | RPSALSEN | [K].RPSALS | 1xPhospho | Q99M54 1 | 0.001489 | 5.66E-05 | 1 | 1 | 1 | Q99M54 | Cell divisio | Q99M54 [2 | 0 | 1638.898 | 0 | -0.74 | 0.021418 | 1.74 | 126 | 74 | 21.61 | 5.27 | | High | High | High | High | High | High | High | High | High | High | High | 2.58E-05 | 0.000373 | 2.66 |
| FALSE | High | RPSANCDP | [R].RPSAN( | 1xCarbami | P15105 1xl | 0.000104 | 0 | 1 | 2 | 1 | P15105 | Glutamine | P15105 [34 | 0 | 2228.073 | 0 | 1.8 | 0.000795 | 29.1 | 49.6 | 150.4 | 14.54 | 7.87 | | High | High | High | High | High | High | High | High | High | High | High | 0 | 1.88E-05 | 3.33 |
| FALSE | High | RPSASSPNI | [R].RPSASS | 1xPhospho | Q9WTU0 1 | 0.000515 | 0 | 1 | 2 | 2 | Q9WTU0 | Lysine-spe | Q9WTU0 [ | 0 | 1838.953 | 0 | 1.46 | 0.00034 | 3.93 | 50.7 | 149.3 | 5.18 | 7.79 | | High | High | High | High | High | High | High | High | High | High | High | 0.003633 | 0.03862 | 1.96 |
| FALSE | High | RPSASSPNI | [R].RPSASS | 2xPhospho | Q9WTU0 2 | 0.035793 | 0.002304 | 1 | 2 | 2 | Q9WTU0 | Lysine-spe | Q9WTU0 [ | 0 | 1918.919 | 0 | 1.9 | 0.000196 | 12 | 43 | 157 | 10.75 | 3.05 | | High | High | High | High | High | High | High | High | High | High | High | 0.001214 | 0.01315 | 2.43 |
| FALSE | High | RPSDENSIT | [R].RPSDEN | 1xPhospho | Q8C8U0-3 | 2.34E-05 | 0 | 1 | 6 | 2 | Q8C8U0-3 | Isoform 3 c | Q8C8U0-3 | 0 | 2690.252 | 0 | 1.34 | 0.000865 | 21.7 | 58.9 | 141.1 | 14.26 | 7.32 | | High | High | High | High | High | High | High | High | High | High | High | 0 | 6.27E-06 | 3.48 |
| FALSE | High | RPSDGNSG | [R].RPSDGI | 1xPhospho | B2RY61 1x | 8.55E-08 | 0 | 1 | 3 | 2 | B2RY61 | AT rich inte | B2RY61 [1 | 0 | 1752.941 | 0 | -0.51 | 0.011023 | 2.18 | 115.2 | 84.8 | 9.1 | 5.4 | | High | High | High | High | High | High | High | High | High | High | High | 0 | 6.39E-09 | 3.22 |
| FALSE | High | RPQSPGVS | [R].RPQSP( | 1xPhospho | A2ADB0 1x | 0.000311 | 0 | 1 | 5 | 1 | A2ADB0 | Msx2-inter | A2ADB0 [7 | 0 | 1813.891 | 0 | 0.68 | 0.025 | 19.07 | 73.3 | 126.7 | 13.48 | 14.87 | | High | High | High | High | High | High | High | High | High | High | High | 0 | 6.45E-05 | 2.06 |
| FALSE | High | RPPGDVLE | [K].RPPGD' | 1xPhospho | Q9ERG2 1x | 0.000284 | 0 | 2 | 4 | 2 | Q9ERG2; B | Striatin-3 C | Q9ERG2 [2 | 0 | 4375.984 | 0 | 0.51 | 0.122821 | 35.5 | 82.5 | 117.5 | 10.57 | 33.1 | Shared | High | High | High | High | High | High | High | High | High | High | High | 0 | 5.78E-05 | 4 |
| FALSE | High | RPPGDVLE | [K].RPPGD' | 1xPhospho | Q9ERG2 1x | 0.029586 | 0.001857 | 2 | 4 | 2 | Q9ERG2; B | Striatin-3 C | Q9ERG2 [2 | 0 | 4359.989 | 0 | 0.38 | 0.079561 | 19.28 | 88.4 | 111.6 | 9.7 | 13.3 | Shared | High | High | High | High | High | High | High | High | High | High | High | 0.000997 | 0.01056 | 3.14 |
| FALSE | High | RPPESPPIV | [K].RPPESP | 1xPhospho | Q9CQY2 1x | 0.003688 | 0.00021 | 1 | 2 | 2 | Q9CQY2 | RNMT-acti | Q9CQY2 [3 | 0 | 2102.027 | 0 | 0.91 | 0.001765 | 18.28 | 69.4 | 130.6 | 4.16 | 8.76 | | High | High | High | High | High | High | High | High | High | High | High | 0.000275 | 0.00182 | 2.29 |
| FALSE | High | RPGGPSPLI | [K].RPGGP! | 1xPhospho | Q7TSC1 1x | 8.45E-07 | 0 | 1 | 3 | 2 | Q7TSC1 | Protein PRI | Q7TSC1 [1 | 0 | 2838.362 | 0 | 0.73 | 0.000992 | 8.17 | 76.4 | 123.6 | 10.25 | 6.26 | | High | High | High | High | High | High | High | High | High | High | High | 0 | 8.45E-08 | 4.49 |
| FALSE | High | RPGNEELPI | [R].RPGNEI | 1xPhospho | Q6PAR5 1x | 6.23E-08 | 0 | 1 | 9 | 6 | Q6PAR5 | GTPase-act | Q6PAR5 [9 | 0 | 3512.779 | 0 | 0.14 | 0.520645 | 9.25 | 94.5 | 105.5 | 11.54 | 8.43 | | High | High | High | High | High | High | High | High | High | High | High | 0 | 4.47E-09 | 4.91 |
| FALSE | High | RPGPAAPP | [R].RPGPA/ | 1xPhospho | Q5Y5T2 1x | 2.21E-05 | 0 | 1 | 1 | 1 | Q5Y5T2 | Palmitoyltr | Q5Y5T2 [2 | 0 | 2127.106 | 0 | 0.22 | 0.007486 | 4.29 | 94.4 | 105.6 | 10.35 | 7.7 | | High | High | High | High | High | High | High | High | High | High | High | 0 | 3.28E-06 | 3.34 |
| FALSE | High | RPGPHSPD | [R].RPGPH! | 1xPhospho | Q9CR59 1x | 0.014033 | 0.001011 | 1 | 1 | 1 | Q9CR59 | Growth arr | Q9CR59 [3 | 0 | 2092.054 | 0 | 0.15 | 0.218227 | 3.27 | 94.1 | 105.9 | 6.47 | 11.14 | | High | High | High | High | High | High | High | High | High | High | High | 0.000538 | 0.004567 | 1.73 |
| FALSE | High | RPGPLPASI | [R].RPGPLF | 1xPhospho | H3BJQ9 1x | 0.001107 | 0 | 1 | 6 | 1 | H3BJQ9 | Homeobox | H3BJQ9 [4 | 0 | 1889.036 | 0 | 1.74 | 0.000406 | 6.18 | 46.5 | 153.5 | 9.16 | 8.44 | | High | High | High | High | High | High | High | High | High | High | High | 0 | 0.000266 | 1.93 |
| FALSE | High | RPGPPLSPE | [R].RPGPPL | 1xPhospho | A0A140T8 | 0.029076 | 0.001857 | 1 | 1 | 1 | A0A140T8 | Chromosol | A0A140T8 | 0 | 1527.825 | 0 | -0.36 | 0.002383 | 3.93 | 114.1 | 85.9 | 5.71 | 3.81 | | High | High | High | High | High | High | High | High | High | High | High | 0.000997 | 0.01042 | 1.7 |
| FALSE | High | RPGPPLSPE | [R].RPGPPL | 2xPhospho | A0A140T8 | 0.025073 | 0.001645 | 1 | 1 | 1 | A0A140T8 | Chromosol | A0A140T8 | 1 | 2703.404 | 0 | 0.02 | 0.725145 | 14.87 | 98.2 | 101.8 | 6.78 | 11.37 | | High | High | High | High | High | High | High | High | High | High | High | 0.000872 | 0.008796 | 1.73 |
| FALSE | High | RPGQSFHV | [R].RPGQS | 1xPhospho | Q3UR00 1x | 5.58E-07 | 0 | 1 | 6 | 1 | Q3UR00 | Putative ur | Q3UR00 [1 | 0 | 2419.208 | 0 | -0.9 | 0.019221 | 5.12 | 129 | 71 | 15.03 | 13.81 | | High | High | High | High | High | High | High | High | High | High | High | 0 | 5.26E-08 | 3.65 |

| | | | | | | | | | | | | | | | | | | | | | | | | | | | | | | | | | | | | | | |
|---|---|---|---|---|---|---|---|---|---|---|---|---|---|---|---|---|---|---|---|---|---|---|---|---|---|---|---|---|---|---|---|---|---|---|---|---|---|---|
| FALSE | High | RPGSVSSTI | [K].RPGSVS | 1xPhospho | Q8K3X4 1x | 0.030632 | 0.001857 | 1 | 1 | 4 | Q8K3X4 | Interferon | Q8K3X4 [3 | 0 | 1627.764 | 0 | -1.03 | 0.000432 | 0.79 | 133.6 | 66.4 | 5.65 | 5.05 | | High | High | High | High | High | High | High | High | High | High | High | 0.007094 | 0.07698 | 2.56 |
| FALSE | High | RPHSPDPG | [R].RPHSPD | 1xPhospho | D3Z2R5 1x | 0.00095 | 0 | 1 | 1 | 1 | D3Z2R5 | Selenoprot | D3Z2R5 [10 | 0 | 1888.938 | 0 | 1.89 | 0.001017 | 19.74 | 40.7 | 159.3 | 14.45 | 12.15 | | High | High | High | High | High | High | High | High | High | High | High | 0 | 0.000225 | 1.77 |
| FALSE | High | RPKPEEDEI | [K].RPKPEE | 1xPhospho | Q8VCD7 1x | 1.51E-06 | 0 | 1 | 4 | 1 | Q8VCD7 | Lysine-spe | Q8VCD7 [3 | 0 | 3414.571 | 0 | 0.09 | 0.632544 | 14.85 | 97 | 103 | 5.69 | 9.62 | | High | High | High | High | High | High | High | High | High | High | High | 0 | 1.61E-07 | 6.41 |
| FALSE | High | RPKPEEDEI | [K].RPKPEE | 1xPhospho | Q8VCD7 1x | 0.000328 | 0 | 1 | 4 | 2 | Q8VCD7 | Lysine-spe | Q8VCD7 [3 | 0 | 3430.566 | 0 | -0.73 | 0.003176 | 1.51 | 124.3 | 75.7 | 2.54 | 14.43 | | High | High | High | High | High | High | High | High | High | High | High | 0 | 6.82E-05 | 3.64 |
| FALSE | High | RPLDSPEVE | [K].RPLDSP | 1xPhospho | A2ASY1 1x | 3.92E-06 | 0 | 1 | 5 | 3 | A2ASY1 | TAF3 RNA | A2ASY1 [17 | 0 | 2299.112 | 0 | 0.01 | 0.711713 | 1.63 | 99.6 | 100.4 | 5.95 | 4.93 | | High | High | High | High | High | High | High | High | High | High | High | 0 | 4.69E-07 | 3.36 |
| FALSE | High | RPLETTQDS | [R].RPLETT | 1xPhospho | Q3TC93 1x | 0.003892 | 0.000261 | 1 | 2 | 1 | Q3TC93 | HCLS1-binc | Q3TC93 [2 | 0 | 1825.983 | 0 | 1.83 | 0.000222 | 13.14 | 45.3 | 154.7 | 15.72 | 6.64 | | High | High | High | High | High | High | High | High | High | High | High | 0.00014 | 0.001091 | 1.66 |
| FALSE | High | RPLIASPSQ | [R].RPLIASI | 1xPhospho | Q8VDF2 1x | 8.4E-05 | 0 | 1 | 1 | 1 | Q8VDF2 | E3 ubiquiti | Q8VDF2 [2 | 0 | 1909.062 | 0 | 0.24 | 0.121001 | 19.21 | 91.4 | 108.6 | 9.82 | 9.99 | | High | High | High | High | High | High | High | High | High | High | High | 0 | 1.48E-05 | 1.24 |
| FALSE | High | RPILQLSPP | [R].RPILQL | 1xPhospho | A0A0R4IZZ | 0.000322 | 0 | 1 | 4 | 2 | A0A0R4IZZ | Formin bin | A0A0R4IZZ | 0 | 1639.925 | 0 | -0.35 | 0.000858 | 0.81 | 113.7 | 86.3 | 3.98 | 2.93 | | High | High | High | High | High | High | High | High | High | High | High | 0 | 6.66E-05 | 0.94 |
| FALSE | High | RPLPVESPD | [K].RPLPVE | 1xPhospho | Q545J5 1x | 6.71E-05 | 0 | 1 | 3 | 2 | Q545J5 | Basic Krup | Q545J5 [24 | 0 | 1703.868 | 0 | 1.02 | 0.000444 | 4.69 | 65.7 | 134.3 | 4.78 | 6.11 | | High | High | High | High | High | High | High | High | High | High | High | 0.000204 | 0.001324 | 1.91 |
| FALSE | High | RPLRPASPE | [K].RPLRPA | 1xPhospho | Q9Z2F6 1x | 0.120211 | 0.009851 | 1 | 2 | 1 | Q9Z2F6 | B-cell lymp | Q9Z2F6 [33 | 0 | 1857.99 | 0 | 0.54 | 0.005967 | 13.43 | 81.3 | 118.7 | 6.73 | 8.77 | | High | High | High | High | High | High | High | High | High | High | High | 0.004949 | 0.05382 | 1.03 |
| FALSE | High | RPLSASLGC | [R].RPLSAS | 1xPhospho | Q3UT88 1x | 1.1E-05 | 0 | 1 | 3 | 1 | Q3UT88 | Putative ur | Q3UT88 [4 | 0 | 3614.863 | 0 | 0.99 | 0.002607 | 15.57 | 68.9 | 131.1 | 10.56 | 6.74 | | High | High | High | High | High | High | High | High | High | High | High | 0 | 1.52E-06 | 5 |
| FALSE | High | RPLSGPDVS | [K].RPLSGF | 1xPhospho | Q8BTW9 1 | 2.09E-08 | 0 | 1 | 1 | 1 | Q8BTW9 | Serine/thre | Q8BTW9 [1 | 0 | 2620.375 | 0 | 0.26 | 0.030427 | 11.54 | 90.9 | 109.1 | 3.36 | 10.78 | | High | High | High | High | High | High | High | High | High | High | High | 0 | 1.32E-09 | 4.13 |
| FALSE | High | RPLSGPDVS | [K].RPLSGF | 2xPhospho | Q8BTW9 2 | 0.010758 | 0.000772 | 1 | 1 | 1 | Q8BTW9 | Serine/thre | Q8BTW9 [1 | 0 | 2700.341 | 0 | 1.36 | 0.000635 | 4.95 | 51.6 | 148.4 | 13.93 | 6.76 | | High | High | High | High | High | High | High | High | High | High | High | 0.000399 | 0.003393 | 3.34 |
| FALSE | High | RPLSPQSK | [K].RPLSPC | 1xPhospho | G3X8T2 1x | 0.005978 | 0.00062 | 1 | 3 | 2 | G3X8T2 | RIKEN cDN | G3X8T2 [90 | 0 | 1450.818 | 0 | -0.44 | 0.000364 | 2.92 | 115.2 | 84.8 | 2.76 | 4.02 | | High | High | High | High | High | High | High | High | High | High | High | 0.000555 | 0.004934 | 1.34 |
| FALSE | High | RPLSSSCEA | [K].RPLSSS | 1xCarbami | Q501J7-3 1 | 0.001301 | 0 | 1 | 4 | 2 | Q501J7-3 | Isoform 3 c | Q501J7-3 [ | 0 | 1672.849 | 0 | 0.72 | 0.002563 | 7.33 | 73.9 | 126.1 | 3.15 | 9.23 | | High | High | High | High | High | High | High | High | High | High | High | 0 | 0.000322 | 2.12 |
| FALSE | High | RPLTTSPSL | [R].RPLTTS | 1xPhospho | Q9CQB2 1x | 5.91E-08 | 0 | 1 | 1 | 3 | Q9CQB2 | MAPK regu | Q9CQB2 [7 | 0 | 3124.515 | 0 | 0.72 | 0.003483 | 11.24 | 75.7 | 124.3 | 9.68 | 6.26 | | High | High | High | High | High | High | High | High | High | High | High | 0 | 4.22E-09 | 4.26 |
| FALSE | High | RPMEFEAA | [K].RPMEF | 1xPhospho | Q99LF1 1x | 0.000375 | 0 | 1 | 1 | 1 | Q99LF1 | Akirin-1 OS | Q99LF1 [9- | 0 | 2284.181 | 0 | -0.07 | 0.349939 | 15.27 | 102.7 | 97.3 | 6.73 | 7.23 | | High | High | High | High | High | High | High | High | High | High | High | 0 | 7.9E-05 | 2.15 |
| FALSE | High | RPMSDPSV | [R].RPMSD | 1xPhospho | Q6PAR5 1x | 0.012845 | 0.000889 | 1 | 9 | 3 | Q6PAR5 | GTPase-act | Q6PAR5 [9 | 0 | 1570.703 | 0 | 1.13 | 0.002085 | 12.07 | 61.1 | 138.9 | 5.71 | 11.84 | | High | High | High | High | High | High | High | High | High | High | High | 0.007957 | 0.0859 | 1.65 |
| FALSE | High | RPMSSPEM | [K].RPMSSI | 1xPhospho | Q61191 1x | 0.013308 | 0.000889 | 1 | 3 | 7 | Q61191 | Host cell fa | Q61191 [20 | 0 | 1616.794 | 0 | -0.63 | 0.000547 | 1.75 | 121.4 | 78.6 | 6.63 | 8.21 | | High | High | High | High | High | High | High | High | High | High | High | 0.000814 | 0.00753 | 2.18 |
| FALSE | High | RPNEDSDE | [K].RPNEDS | 1xPhospho | Q99LI7 1xP | 0.000196 | 0 | 1 | 3 | 2 | Q99LI7 | Cleavage st | Q99LI7 [68 | 0 | 2000.885 | 0 | -0.15 | 0.050198 | 0.63 | 105.1 | 94.9 | 3.2 | 8.09 | | High | High | High | High | High | High | High | High | High | High | High | 0 | 3.83E-05 | 2.63 |
| FALSE | High | RPPASQSSI | [R].RPPASC | 1xPhospho | Q6P1D7 1x | 0.018285 | 0.001282 | 1 | 2 | 1 | Q6P1D7 | Structure-s | Q6P1D7 [4 | 0 | 2213.184 | 0 | -0.39 | 0.060132 | 9.55 | 113.6 | 86.4 | 8.11 | 8.96 | | High | High | High | High | High | High | High | High | High | High | High | 0.000689 | 0.006145 | 2.41 |
| FALSE | High | RPSDSVQP | [R].RPSDSV | 1xCarbami | Q8C6B2 1x | 1.74E-07 | 0 | 1 | 5 | 3 | Q8C6B2 | Rhotekin C | Q8C6B2 [1 | 0 | 2030.943 | 0 | 0.45 | 0.012334 | 7.19 | 83.3 | 116.7 | 7.54 | 13.84 | | High | High | High | High | High | High | High | High | High | High | High | 2.58E-05 | 0.000375 | 4.04 |
| FALSE | High | RPGGHSQS | [K].RPGGH | 1xCarbami | Q6NV66 2x | 0.000718 | 0 | 1 | 3 | 4 | Q6NV66 | Zinc finger | Q6NV66 [1 | 0 | 3202.439 | 0 | -0.74 | 0.018748 | 11.45 | 122.8 | 77.2 | 9.91 | 22.21 | | High | High | High | High | High | High | High | High | High | High | High | 0 | 0.000165 | 3.3 |
| FALSE | High | RPSEGNYQ | [K].RPSEGN | 1xPhospho | Q5SRY7-2 1 | 2.85E-05 | 0 | 1 | 5 | 3 | Q5SRY7-2 | Isoform 2 c | Q5SRY7-2 | 0 | 1616.82 | 0 | 0.37 | 0.009354 | 11.32 | 88.8 | 111.2 | 1.99 | 10.13 | | High | High | High | High | High | High | High | High | High | High | High | 0 | 4.38E-06 | 2.51 |
| FALSE | High | RPSEPLEPR | [R].RPSEPL | 1xPhospho | A0A0G2JFT | 0.074071 | 0.0057 | 1 | 3 | 1 | A0A0G2JFT | Protein RU | A0A0G2JFT | 0 | 1389.709 | 0 | 0.92 | 0.004969 | 18.68 | 70.8 | 129.2 | 11.49 | 15.54 | | High | High | High | High | High | High | High | High | High | High | High | 0.002924 | 0.03029 | 1.49 |
| FALSE | High | RPSTIAEQT | [R].RPSTIAI | 1xPhospho | F8VPT7 1x | 2.22E-06 | 0 | 1 | 4 | 2 | F8VPT7 | Serine/thre | F8VPT7 [35 | 0 | 1839.014 | 0 | 0.94 | 0.000793 | 2.56 | 66.3 | 133.7 | 8.03 | 8.89 | | High | High | High | High | High | High | High | High | High | High | High | 0 | 1.44E-06 | 3.14 |
| FALSE | High | RPSTSPDVI | [R].RPSTSP | 1xPhospho | Q3TV38 1x | 1.94E-06 | 0 | 1 | 3 | 6 | Q3TV38 | Putative ur | Q3TV38 [2 | 0 | 1943.99 | 0 | 0.89 | 0.000374 | 3.41 | 72.4 | 127.6 | 7.12 | 4.83 | | High | High | High | High | High | High | High | High | High | High | High | 0 | 3.54E-07 | 3.36 |
| FALSE | High | RPSVDSLVS | [K].RPSVDS | 1xPhospho | F2Z3U3 1x | 0.000639 | 0 | 1 | 1 | 1 | F2Z3U3 | Ras associa | F2Z3U3 [10 | 0 | 1625.903 | 0 | 0.39 | 0.002304 | 5.81 | 86.4 | 113.6 | 7.81 | 6.32 | | High | High | High | High | High | High | High | High | High | High | High | 0 | 0.000144 | 1.64 |
| FALSE | High | RPSVGSQS | [R].RPSVGS | 1xPhospho | Q80TJ7 1x | 1.43E-05 | 0 | 1 | 2 | 3 | Q80TJ7 | Histone lys | Q80TJ7 [98 | 0 | 2012.033 | 0 | 0.17 | 0.147275 | 8.24 | 99.5 | 100.5 | 10.17 | 7.18 | | High | High | High | High | High | High | High | High | High | High | High | 0 | 1.36E-05 | 3.35 |
| FALSE | High | RPSVPPSPC | [K].RPSVPF | 1xPhospho | D3Z5N5 1x | 0.113753 | 0.009241 | 2 | 3 | 1 | D3Z5N5; Q | 1-phospha | D3Z5N5 [1 | 0 | 1358.714 | 0 | 0.35 | 0.006364 | 9.74 | 89.7 | 110.3 | 8.98 | 9.32 | Shared | High | High | High | High | High | High | High | High | High | High | High | 0.00468 | 0.05046 | 1.47 |
| FALSE | High | RPSVSFAEC | [K].RPSVSF | 1xPhospho | Q5ND28 1x | 0.024638 | 0.001612 | 1 | 1 | 1 | Q5ND28 | Scavenger | Q5ND28 [5 | 0 | 1716.909 | 0 | -1.08 | 0.004407 | 4.79 | 141.8 | 58.2 | 17.64 | 29.65 | | High | High | High | High | High | High | High | High | High | High | High | 0.000872 | 0.008656 | 1.71 |
| FALSE | High | RPSVTQGT | [R].RPSVTC | 1xCarbami | Q69ZF3 1x | 6.78E-09 | 0 | 1 | 1 | 2 | Q69ZF3 | Non-lysosc | Q69ZF3 [89 | 0 | 2538.279 | 0 | -0.11 | 0.045748 | 6.85 | 104 | 96 | 3.16 | 4.17 | | High | High | High | High | High | High | High | High | High | High | High | 0 | 2.3E-08 | 3.98 |
| FALSE | High | RPSVYLPTR | [R].RPSVYL | 1xPhospho | Q9D9Z5 1x | 2.11E-05 | 0 | 1 | 2 | 1 | Q9D9Z5 | DET1- and | Q9D9Z5 [3 | 0 | 1397.75 | 0 | 1.3 | 0.000328 | 10.14 | 58.8 | 141.2 | 4.37 | 5.35 | | High | High | High | High | High | High | High | High | High | High | High | 0 | 3.14E-06 | 2.07 |
| FALSE | High | RPSYTLGM | [R].RPSYTL | 1xPhospho | Q8C0T5 1x | 0.003892 | 0.000261 | 1 | 3 | 1 | Q8C0T5 | Signal-indu | Q8C0T5 [1 | 0 | 1590.848 | 0 | 1.22 | 0.006026 | 12.83 | 62.7 | 137.3 | 11.5 | 15.86 | | High | High | High | High | High | High | High | High | High | High | High | 0.00014 | 0.001094 | 1.54 |
| FALSE | High | RPSYTLGM | [R].RPSYTL | 1xPhospho | Q8C0T5 1x | 0.000119 | 0 | 1 | 3 | 1 | Q8C0T5 | Signal-indu | Q8C0T5 [1 | 0 | 1606.843 | 0 | 1.65 | 0.003537 | 6.68 | 49.2 | 150.8 | 6.11 | 17.96 | | High | High | High | High | High | High | High | High | High | High | High | 0 | 2.19E-05 | 1.07 |
| FALSE | High | RPTAPSTPD | [R].RPTAPS | 1xPhospho | Q8CIV2-2 1 | 0.026191 | 0.001676 | 1 | 4 | 2 | Q8CIV2-2 | Isoform 2 c | Q8CIV2-2 [ | 0 | 3051.462 | 0 | 1.44 | 0.000785 | 5.33 | 48.3 | 151.7 | 16.4 | 11.82 | | High | High | High | High | High | High | High | High | High | High | High | 0.000903 | 0.00924 | 3.22 |
| FALSE | High | RPTDEKPPA | [R].RPTDEK | 1xPhospho | Q3TXW9 1 | 2.8E-06 | 0 | 1 | 3 | 1 | Q3TXW9 | Putative ur | Q3TXW9 [6 | 0 | 2599.374 | 0 | 0.59 | 0.000955 | 8.73 | 80.9 | 119.1 | 5.94 | 6.62 | | High | High | High | High | High | High | High | High | High | High | High | 0 | 3.25E-07 | 2.06 |
| FALSE | High | RPTEAVSPI | [K].RPTEAV | 1xPhospho | F6RDI8 1xP | 0.007673 | 0.000664 | 1 | 7 | 1 | F6RDI8 | Serine/argi | F6RDI8 [46 | 0 | 1293.677 | 0 | -0.46 | 0.612451 | 13.09 | 117.2 | 82.8 | 62.76 | 72.25 | | High | High | High | High | High | High | High | High | High | High | High | 0.000294 | 0.002324 | 1.81 |
| FALSE | High | RPSTEPGA | [R].RPSTEP | 1xCarbami | E9PUV9 1x | 0.002376 | 0.000109 | 1 | 1 | 1 | E9PUV9 | Caspase re | E9PUV9 [2 | 0 | 1869.848 | 0 | 0.74 | 0.000347 | 5.77 | 75.2 | 124.8 | 11.98 | 10.28 | | High | High | High | High | High | High | High | High | High | High | High | 4.94E-05 | 0.000631 | 1.96 |
| FALSE | High | RPTEAVSPI | [K].RPTEAV | 1xPhospho | F6RDI8 1xP | 0.016451 | 0.001124 | 1 | 7 | 3 | F6RDI8 | Serine/argi | F6RDI8 [46 | 0 | 1522.839 | 0 | -0.55 | 0.002069 | 2.4 | 117.7 | 82.3 | 7.25 | 3.78 | | High | High | High | High | High | High | High | High | High | High | High | 0.000832 | 0.007903 | 1.22 |
| FALSE | High | RPTLGVQLI | [R].RPTLGV | 1xPhospho | A0A0R4J0J | 0.001815 | 5.66E-05 | 1 | 4 | 1 | A0A0R4J0J | THO comp | A0A0R4J0J | 0 | 1550.814 | 0 | 0.87 | 0.032315 | 20.73 | 67.7 | 132.3 | 14.76 | 26.31 | | High | High | High | High | High | High | High | High | High | High | High | 2.58E-05 | 0.000465 | 3.15 |
| FALSE | High | RPTLGVQLI | [R].RPTLGV | 1xPhospho | A0A0R4J0J | 1.68E-07 | 0 | 1 | 4 | 2 | A0A0R4J0J | THO comp | A0A0R4J0J | 0 | 1779.977 | 0 | 1.08 | 0.024042 | 35.51 | 59.7 | 140.3 | 21.37 | 24.34 | | High | High | High | High | High | High | High | High | High | High | High | 0 | 3.3E-08 | 3.46 |
| FALSE | High | RPTSFADEI | [K].RPTSFA | 1xPhospho | Q6PGL7 1x | 0.028824 | 0.001829 | 1 | 4 | 1 | Q6PGL7 | WASH com | Q6PGL7 [2 | 0 | 1642.815 | 0 | -0.23 | 0.024323 | 7 | 112.1 | 87.9 | 12.68 | 9.25 | | High | High | High | High | High | High | High | High | High | High | High | 0.000997 | 0.01027 | 1.8 |
| FALSE | High | RPTTPAPTI | [R].RPTTPA | 1xPhospho | Q8K1L5 1x | 0.003997 | 0.000311 | 1 | 1 | 3 | Q8K1L5 | Protein ph | Q8K1L5 [10 | 0 | 3249.56 | 0 | 1.16 | 0.04505 | 32.6 | 63.3 | 136.7 | 21.08 | 37.2 | | High | High | High | High | High | High | High | High | High | High | High | 0.000161 | 0.001152 | 3.58 |
| FALSE | High | RPVSAMFII | [K].RPVSAN | 1xPhospho | P25976 1x | 0.002212 | 0.000109 | 1 | 6 | 1 | P25976 | Nucleolar t | P25976 [40 | 0 | 2095.07 | 0 | -0.44 | 0.013773 | 8.09 | 115 | 85 | 6.07 | 11.23 | | High | High | High | High | High | High | High | High | High | High | High | 4.94E-05 | 0.000581 | 2.56 |
| FALSE | High | RPVSFPETF | [K].RPVSFP | 1xPhospho | G3X9J0 1x | 2E-05 | 0 | 1 | 3 | 2 | G3X9J0 | Signal-indu | G3X9J0 [11 | 0 | 2282.117 | 0 | 0.26 | 0.011092 | 5.73 | 90.9 | 109.1 | 2.21 | 6.65 | | High | High | High | High | High | High | High | High | High | High | High | 4.94E-05 | 0.000737 | 3.14 |
| FALSE | High | RPVTPPSQT | [R].RPVTPF | 1xPhospho | Q9JHE7 1x | 0.001107 | 0 | 1 | 4 | 2 | Q9JHE7 | Protein TSS | Q9JHE7 [12 | 0 | 1615.852 | 0 | -0.09 | 0.092268 | 1.88 | 103.4 | 96.6 | 3.09 | 4.27 | | High | High | High | High | High | High | High | High | High | High | High | 0 | 0.000268 | 1.43 |
| FALSE | High | RPWDLISQ | [R].RPWDL | 1xPhospho | Q5DTX6 1x | 0.002948 | 0.00016 | 1 | 1 | 1 | Q5DTX6 | Junctional | Q5DTX6 [8 | 0 | 2271.194 | 0 | 1.73 | 0.001064 | 16.92 | 46.3 | 153.7 | 14.3 | 11.62 | | High | High | High | High | High | High | High | High | High | High | High | 7.19E-05 | 0.000798 | 2.61 |
| FALSE | High | RPYGSSLSV | [R].RPYGSS | 1xPhospho | Q80XC3-2 | 4.01E-05 | 0 | 1 | 2 | 4 | Q80XC3-2 | Isoform 2 c | Q80XC3-2 | 0 | 2161.094 | 0 | 0.78 | 0.000251 | 2.09 | 73.3 | 126.7 | 4.93 | 5.73 | | High | High | High | High | High | High | High | High | High | High | High | 0 | 3.04E-05 | 3.98 |
| FALSE | High | RPYSPEK | [K].RPYSPE | 1xPhospho | Q8BZR9 1x | 0.099328 | 0.008003 | 1 | 3 | 1 | Q8BZR9 | Nuclear ca | Q8BZR9 [4 | 0 | 1414.75 | 0 | -0.45 | 0.001079 | 3.03 | 113.8 | 86.2 | 5.18 | 3.95 | | High | High | High | High | High | High | High | High | High | High | High | 0.004032 | 0.04302 | 1.24 |
| FALSE | High | RPYTGNPQ | [R].RPYTGI | 1xPhospho | P70398 1x | 0.001586 | 5.66E-05 | 1 | 5 | 1 | P70398 | Probable u | P70398 [24 | 0 | 3988.813 | 0 | -0.47 | 0.014375 | 4.82 | 115 | 85 | 10.2 | 11.15 | | High | High | High | High | High | High | High | High | High | High | High | 2.58E-05 | 0.000398 | 3.75 |
| FALSE | High | RPYTGNPQ | [R].RPYTGI | 1xDeamida | P70398 1x | 0.000127 | 0 | 1 | 5 | 4 | P70398 | Probable u | P70398 [24 | 0 | 3989.797 | 0 | -0.48 | 0.013974 | 7.12 | 113.4 | 86.6 | 6.2 | 9.17 | | High | High | High | High | High | High | High | High | High | High | High | 0.000294 | 0.002293 | 3.5 |
| FALSE | High | RPYTGNPQ | [R].RPYTGI | 2xDeamida | P70398 1x | 0.007881 | 0.000736 | 1 | 5 | 4 | P70398 | Probable u | P70398 [24 | 0 | 3990.781 | 0 | -0.19 | 0.023039 | 5.87 | 107.8 | 92.2 | 10.58 | 10.78 | | High | High | High | High | High | High | High | High | High | High | High | 0.000327 | 0.002395 | 5.06 |
| FALSE | High | RPTLAWLE | [R].RPTLAV | 1xCarbami | A0A1S6GW | 5.24E-07 | 0 | 1 | 6 | 2 | A0A1S6GW | Uncharacte | A0A1S6GW | 0 | 2600.283 | 0 | 1.4 | 0.000748 | 9.26 | 56.8 | 143.2 | 9.8 | 8.16 | | High | High | High | High | High | High | High | High | High | High | High | 0 | 4.91E-08 | 3.53 |
| FALSE | High | RPSTAALTF | [R].RPSTAA | 1xPhospho | B2KF25 1x | 0.056248 | 0.004124 | 1 | 3 | 2 | B2KF25 | T-complex | B2KF25 [46 | 0 | 1778.911 | 0 | -0.75 | 0.02648 | 4.12 | 125.3 | 74.7 | 13.42 | 22.04 | | High | High | High | High | High | High | High | High | High | High | High | 0.002455 | 0.02652 | 2.27 |
| FALSE | High | RPSSPPPPS | [R].RPSSPP | 1xCarbami | B0V2M3 1x | 0.001337 | 0 | 1 | 2 | 3 | B0V2M3 | Zinc finger | B0V2M3 [3 | 0 | 1833.888 | 0 | -0.52 | 0.023778 | 9.55 | 116.4 | 83.6 | 9.02 | 13.18 | | High | High | High | High | High | High | High | High | High | High | High | 0 | 0.00033 | 2.96 |
| FALSE | High | RPSSPAPGI | [R].RPSSPA | 1xPhospho | O35216 1x | 0.003655 | 0.00021 | 1 | 6 | 3 | O35216 | Histone H3 | O35216 [1 | 0 | 1417.715 | 0 | -1.26 | 9.77E-05 | 1.32 | 141.6 | 58.4 | 4.49 | 8.76 | | High | High | High | High | High | High | High | High | High | High | High | 9.42E-05 | 0.00102 | 2.46 |
| FALSE | High | RPSESDKEE | [R].RPSESD | 1xPhospho | B2RRE0 1x | 0.000407 | 0 | 1 | 3 | 2 | B2RRE0 | A kinase (P | B2RRE0 [6 | 1 | 2100.026 | 0 | -1.88 | 0.152965 | 2.11 | 163.9 | 36.1 | 73.88 | 63.94 | | High | High | High | High | High | High | High | High | High | High | High | 0 | 9.06E-05 | 2.66 |
| FALSE | High | RPSETK | [R].RPSETK | 1xPhospho | Q9CU65 1x | 0.085859 | 0.006406 | 1 | 2 | 1 | Q9CU65 | Zinc finger | Q9CU65 [1 | 0 | 1255.681 | 0 | -0.63 | 0.013103 | 2.78 | 122 | 78 | 9.48 | 10.19 | | High | High | High | High | High | High | High | High | High | High | High | 0.003291 | 0.03592 | 0.76 |
| FALSE | High | RPSFSSAPV | [K].RPSFSS. | 2xPhospho | B2RRI2 2xP | 1.22E-05 | 0 | 1 | 4 | 1 | B2RRI2 | Protein stra | B2RRI2 [81 | 0 | 4221.929 | 0 | 0.01 | 0.503538 | 19.17 | 96.4 | 103.6 | 14.22 | 8.98 | | High | High | High | High | High | High | High | High | High | High | High | 0 | 1.7E-06 | 5.44 |
| FALSE | High | RPSFSSAPV | [K].RPSFSS. | 1xDeamida | B2RRI2 2xP | 1.05E-05 | 0 | 1 | 4 | 1 | B2RRI2 | Protein stra | B2RRI2 [81 | 0 | 4222.913 | 0 | 0.13 | 0.314325 | 13.56 | 94.7 | 105.3 | 12.1 | 19.63 | | High | High | High | High | High | High | High | High | High | High | High | 0 | 1.42E-06 | 5.7 |
| FALSE | High | RPSGPEGA | [R].RPSGPE | 1xPhospho | Q8CHP6 1x | 0.025293 | 0.001645 | 1 | 3 | 1 | Q8CHP6 | Polyhomec | Q8CHP6 [8 | 0 | 1306.647 | 0 | 0.34 | 0.06021 | 7.52 | 87.8 | 112.2 | 5.22 | 14.45 | | High | High | High | High | High | High | High | High | High | High | High | 0.000872 | 0.00891 | 2.02 |
| FALSE | High | RPSGQQGA | [R].RPSGQ | 1xPhospho | Q6P1B3 1x | 0.102567 | 0.008249 | 1 | 2 | 1 | Q6P1B3 | PILR alpha- | Q6P1B3 [2 | 0 | 1378.715 | 0 | -0.76 | 0.00611 | 6.66 | 125.8 | 74.2 | 8.14 | 11.34 | | High | High | High | High | High | High | High | High | High | High | High | 0.004138 | 0.04465 | 1.48 |
| FALSE | High | RPSGSDISS | [K].RPSGSD | 1xPhospho | Q68FE8 1x | 0.000119 | 0 | 1 | 6 | 1 | Q68FE8 | Zinc finger | Q68FE8 [1 | 0 | 1881.984 | 0 | -1.27 | 0.000384 | 2.29 | 142.7 | 57.3 | 6.02 | 7.38 | | High | High | High | High | High | High | High | High | High | High | High | 0 | 2.2E-05 | 2.3 |
| FALSE | High | RPSLGELLL | [R].RPSLGE | 1xPhospho | A0A1C7CY | 0.011246 | 0.000772 | 1 | 6 | 1 | A0A1C7CY | FTS and Hc | A0A1C7CY | 0 | 1462.834 | 0 | -0.43 | 0.034595 | 13.56 | 112 | 88 | 8.37 | 14.55 | | High | High | High | High | High | High | High | High | High | High | High | 0.000418 | 0.003592 | 1.83 |
| FALSE | High | RPSILPESS | [R].RPSILPE | 1xPhospho | E9Q7D5 1x | 0.000628 | 0 | 1 | 2 | 2 | E9Q7D5 | Rho guanir | E9Q7D5 [1 | 0 | 1765.905 | 0 | 2.31 | 0.000885 | 13.01 | 34.4 | 165.6 | 16.43 | 13.69 | | High | High | High | High | High | High | High | High | High | High | High | 0 | 0.000141 | 2.2 |
| FALSE | High | RPSLLSEFH | [R].RPSLLSI | 1xPhospho | Q5RIM6 1x | 0.103392 | 0.008249 | 1 | 11 | 1 | Q5RIM6 | Nuclear re | Q5RIM6 [6 | 0 | 2417.192 | 0 | -0.25 | 0.445384 | 5.55 | 107.3 | 92.7 | 8.52 | 14.36 | | High | High | High | High | High | High | High | High | High | High | High | 0.004164 | 0.04504 | 1.89 |
| FALSE | High | RPSLLSEFQ | [R].RPSLLSI | 1xPhospho | F8VQL9 1x | 0.002793 | 0.000109 | 1 | 4 | 2 | F8VQL9 | Nuclear re | F8VQL9 [6 | 0 | 1911.953 | 0 | 0.51 | 0.014167 | 6.87 | 79.1 | 120.9 | 13.25 | 10.68 | | High | High | High | High | High | High | High | High | High | High | High | 4.94E-05 | 0.000757 | 2.39 |
| FALSE | High | RPSLPALHF | [R].RPSLPA | 1xPhospho | Q3SYK5 1x | 0.065952 | 0.004766 | 1 | 4 | 1 | Q3SYK5 | Tyrosine-p | Q3SYK5 [1 | 0 | 1817.06 | 0 | -0.48 | 0.163635 | 15.78 | 115 | 85 | 12.64 | 25.1 | | High | High | High | High | High | High | High | High | High | High | High | 0.002455 | 0.02653 | 2.03 |
| FALSE | High | RPSLPCISR | [R].RPSLPC | 1xCarbami | Q61409 1x | 0.019618 | 0.001438 | 1 | 2 | 1 | Q61409 | cGMP-inhil | Q61409 [2 | 0 | 1394.718 | 0 | 0.54 | 0.005191 | 7.84 | 76.9 | 123.1 | 10.13 | 12.45 | | High | High | High | High | High | High | High | High | High | High | High | 0.000756 | 0.006685 | 1.67 |
| FALSE | High | RPSLPPSK | [R].RPSLPP | 1xPhospho | Q8BJ05 1x | 0.020679 | 0.001472 | 1 | 3 | 4 | Q8BJ05 | Zinc finger | Q8BJ05 [34 | 0 | 1419.813 | 0 | 1 | 0.00045 | 11.8 | 68.2 | 131.8 | 9.29 | 7.62 | | High | High | High | High | High | High | High | High | High | High | High | 0.004138 | 0.04453 | 1.66 |
| FALSE | High | RPSMDSEA | [R].RPSMD | 1xPhospho | Q80U35 1x | 0.008094 | 0.000736 | 1 | 1 | 1 | Q80U35 | Rho guanir | Q80U35 [1 | 0 | 2030.905 | 0 | -0.16 | 0.386069 | 10.16 | 104.3 | 95.7 | 11.83 | 9.1 | | High | High | High | High | High | High | High | High | High | High | High | 0.000327 | 0.002478 | 3.54 |
| FALSE | High | RPSNRSPTS | [R].RPSNRS | 1xPhospho | Q4VAC9 1x | 0.080423 | 0.006219 | 1 | 2 | 1 | Q4VAC9 | Pleckstrin I | Q4VAC9 [6 | 1 | 1867.979 | 0 | 1.57 | 0.000236 | 6.97 | 48.6 | 151.4 | 6.89 | 5.09 | | High | High | High | High | High | High | High | High | High | High | High | 0.003188 | 0.03327 | 2.5 |
| FALSE | High | RPSNSTPPF | [K].RPSNST | 1xPhospho | Q7TNL5 1x | 0.004218 | 0.00036 | 1 | 4 | 2 | Q7TNL5 | Protein ph | Q7TNL5 [5 | 0 | 2048.094 | 0 | 0.82 | 0.007262 | 14.65 | 73.5 | 126.5 | 13.92 | 5.13 | | High | High | High | High | High | High | High | High | High | High | High | 0.000161 | 0.001194 | 1.71 |
| FALSE | High | RPSNSTPPF | [K].RPSNST | 1xDeamida | Q7TNL5 1x | 0.004256 | 0.00036 | 1 | 4 | 1 | Q7TNL5 | Protein ph | Q7TNL5 [5 | 0 | 2049.078 | 0 | 0.98 | 0.000198 | 1.07 | 67.9 | 132.1 | 10.13 | 3.47 | | High | High | High | High | High | High | High | High | High | High | High | 0.000161 | 0.001209 | 2.56 |
| FALSE | High | RPSPDDDL | [K].RPSPDI | 1xPhospho | B2RSC8 1x | 1.3E-09 | 0 | 1 | 2 | 8 | B2RSC8 | E3 ubiquiti | B2RSC8 [2 | 0 | 2883.23 | 0 | 0.12 | 0.105461 | 11.83 | 92.3 | 107.7 | 4.7 | 9.39 | | High | High | High | High | High | High | High | High | High | High | High | 0 | 6.19E-10 | 5.23 |
| FALSE | High | RPSPDDDL | [K].RPSPDI | 1xDeamida | B2RSC8 1x | 0.016163 | 0.001124 | 1 | 2 | 2 | B2RSC8 | E3 ubiquiti | B2RSC8 [2 | 0 | 2884.214 | 0 | 0.04 | 0.55538 | 17.25 | 98.8 | 101.2 | 4.5 | 16.54 | | High | High | High | High | High | High | High | High | High | High | High | 0.000859 | 0.008332 | 3.65 |
| FALSE | High | RPSPDDDL | [K].RPSPDI | 1xPhospho | B2RSC8 1x | 1.45E-10 | 0 | 1 | 2 | 33 | B2RSC8 | E3 ubiquiti | B2RSC8 [2 | 0 | 2899.225 | 0 | 0.2 | 0.025771 | 7.3 | 89.6 | 110.4 | 5.14 | 6.5 | | High | High | High | High | High | High | High | High | High | High | High | 0 | 1.26E-10 | 5.69 |
| FALSE | High | RPSPDDDL | [K].RPSPDI | 1xDeamida | B2RSC8 1x | 0.00103 | 0 | 1 | 2 | 5 | B2RSC8 | E3 ubiquiti | B2RSC8 [2 | 0 | 2900.209 | 0 | 0.07 | 0.209026 | 9.59 | 99.5 | 100.5 | 6.41 | 10.49 | | High | High | High | High | High | High | High | High | High | High | High | 0.005324 | 0.05792 | 3.11 |
| FALSE | High | RPSPGEVSI | [R].RPSPGE | 1xPhospho | H3BJD6 1x | 0.015741 | 0.001124 | 1 | 9 | 1 | H3BJD6 | Protein ph | H3BJD6 [8 | 0 | 1494.808 | 0 | 0.72 | 0.006443 | 5.38 | 76.8 | 123.2 | 9.45 | 5.78 | | High | High | High | High | High | High | High | High | High | High | High | 0.000585 | 0.005233 | 2.07 |
| FALSE | High | RPSQEEDA | [K].RPSQEE | 1xPhospho | P97784 1x | 4.85E-05 | 0 | 1 | 1 | 1 | P97784 | Cryptochrc | P97784 [58 | 0 | 2066.032 | 0 | 0.01 | 0.946387 | 13.61 | 97 | 103 | 6.34 | 10.19 | | High | High | High | High | High | High | High | High | High | High | High | 0 | 7.97E-06 | 2.92 |
| FALSE | High | RPSSAGELI | [K].RPSSAG | 1xPhospho | Q8C9B9 1x | 0.000213 | 0 | 1 | 1 | 1 | Q8C9B9 | Death-indu | Q8C9B9 [1 | 0 | 1597.835 | 0 | -0.67 | 0.004251 | 6.81 | 122.9 | 77.1 | 5.88 | 8.62 | | High | High | High | High | High | High | High | High | High | High | High | 0 | 4.2E-05 | 2.1 |
| FALSE | High | RPSSDSTEE | [K].RPSSDS | 1xPhospho | Q3TN34 1x | 0.105058 | 0.008518 | 1 | 4 | 2 | Q3TN34 | MICAL-like | Q3TN34 [1 | 0 | 1930.952 | 0 | 0.11 | 0.704509 | 18.26 | 96 | 104 | 11.56 | 14.52 | | High | High | High | High | High | High | High | High | High | High | High | 0.004277 | 0.0456 | 2.86 |
| FALSE | High | RPSSGGEEI | [K].RPSSGC | 1xPhospho | Q8VDP4 1x | 0.000184 | 0 | 1 | 1 | 7 | Q8VDP4 | Cell cycle a | Q8VDP4 [6 | 0 | 1742.836 | 0 | -0.05 | 0.695403 | 9.03 | 101.7 | 98.3 | 3.38 | 5.79 | | High | High | High | High | High | High | High | High | High | High | High | 0 | 3.57E-05 | 2.81 |
| FALSE | High | RPSSGNAST | [R].RPSSGN | 1xPhospho | S4R1P5 1x | 0.023175 | 0.001578 | 1 | 2 | 1 | S4R1P5 | Dystonin O | S4R1P5 [13 | 0 | 1403.663 | 0 | 0.65 | 0.001147 | 3.27 | 78.5 | 121.5 | 3.34 | 7.17 | | High | High | High | High | High | High | High | High | High | High | High | 0.000847 | 0.008048 | 1.93 |
| FALSE | High | RPSSLQSLF | [R].RPSSLQ | 1xPhospho | H3BJQ9 1x | 8.17E-08 | 0 | 1 | 12 | 2 | H3BJQ9 | Homeobox | H3BJQ9 [1 | 0 | 2166.127 | 0 | 1.98 | 0.000331 | 2.73 | 40.2 | 159.8 | 9.59 | 7.03 | | High | High | High | High | High | High | High | High | High | High | High | 0 | 6.11E-09 | 4.42 |
| FALSE | High | RPSELGCFS | [R].RPSELG | 1xCarbami | A0A0R4J28 | 0.012959 | 0.000889 | 1 | 5 | 1 | A0A0R4J28 | DOM-3 hoi | A0A0R4J28 | 0 | 1944.92 | 0 | 0.73 | 0.004552 | 25.08 | 74.8 | 125.2 | 7.34 | 9.8 | | High | High | High | High | High | High | High | High | High | High | High | 0.000449 | 0.004176 | 2.29 |
| FALSE | High | REWEPADI | [K].REWEP | 1xPhospho | E9QK89 1x | 0.004531 | 0.000456 | 1 | 2 | 1 | E9QK89 | Mediator c | E9QK89 [8 | 1 | 2021.953 | 0 | -0.74 | 0.010413 | 1.9 | 124.6 | 75.4 | 7.04 | 18.11 | | High | High | High | High | High | High | High | High | High | High | High | 0.000204 | 0.0013 | 1.16 |
| FALSE | High | RPGGHSQS | [K].RPGGH | 1xCarbami | Q6NV66 1x | 0.000246 | 0 | 1 | 3 | 14 | Q6NV66 | Zinc finger | Q6NV66 [1 | 0 | 3122.473 | 0 | -0.4 | 0.08316 | 16.32 | 112.1 | 87.9 | 12.29 | 14.39 | | High | High | High | High | High | High | High | High | High | High | High | 2.58E-05 | 0.000368 | 3.37 |
| FALSE | High | RPEVTEPEE | [K].RPEVTE | 1xPhospho | Q3TMA0 1 | 1.27E-07 | 0 | 1 | 2 | 7 | Q3TMA0 | Putative ur | Q3TMA0 [4 | 0 | 2267.121 | 0 | 2.56 | 0.00023 | 9.37 | 28.6 | 171.4 | 8.3 | 7.12 | | High | High | High | High | High | High | High | High | High | High | High | 0 | 1.18E-06 | 2.79 |
| FALSE | High | RMSPKPEL | [K].RMSPK | 1xPhospho | Q9R1K9 1x | 0.002719 | 0.000109 | 1 | 3 | 1 | Q9R1K9 | Centrin-2 C | Q9R1K9 [1 | 1 | 2326.244 | 0 | 0.1 | 0.494587 | 23.63 | 89.7 | 110.3 | 11.62 | 19.3 | | High | High | High | High | High | High | High | High | High | High | High | 4.94E-05 | 0.000733 | 2.45 |
| FALSE | High | RNCEDPAS | [K].RNCEDI | 1xCarbami | Q9JHI2 1xP | 0.018285 | 0.001282 | 1 | 4 | 1 | Q9JHI2 | tRNA-speci | Q9JHI2 [18 | 1 | 1881.929 | 0 | -1.07 | 0.000563 | 1.16 | 136.3 | 63.7 | 3.61 | 9.63 | | High | High | High | High | High | High | High | High | High | High | High | 0.000689 | 0.006166 | 2.6 |
| FALSE | High | RNDSSELEI | [R].RNDSSI | 1xPhospho | Q04692 1x | 0.000102 | 0 | 1 | 1 | 8 | Q04692 | SWI/SNF-r | Q04692 [1 | 1 | 2530.233 | 0 | -1.04 | 0.000263 | 1.91 | 135.2 | 64.8 | 3.19 | 3.77 | | High | High | High | High | High | High | High | High | High | High | High | 0 | 9.45E-05 | 4.56 |
| FALSE | High | RNDSSELEI | [R].RNDSSI | 2xPhospho | Q04692 2x | 2.18E-07 | 0 | 1 | 1 | 8 | Q04692 | SWI/SNF-r | Q04692 [1 | 1 | 2610.199 | 0 | -0.95 | 0.000264 | 0.93 | 132.5 | 67.5 | 3.97 | 2.38 | | High | High | High | High | High | High | High | High | High | High | High | 0 | 1.09E-07 | 5.41 |
| FALSE | High | RNEAFVCK | [R].RNEAFV | 1xCarbami | Q80U72-4 | 2.47E-06 | 0 | 2 | 5 | 5 | Q80U72-4; | Isoform 4 c | Q80U72-4 | 1 | 3167.569 | 0 | -0.13 | 0.31963 | 8.94 | 104.4 | 95.6 | 6.49 | 9.29 | Shared | High | High | High | High | High | High | High | High | High | High | High | 0 | 2.81E-07 | 4.19 |
| FALSE | High | RNEDEDSP | [K].RNEDEI | 1xPhospho | A0A0A6YX | 0.033694 | 0.002195 | 1 | 3 | 2 | A0A0A6YX | 60S riboso | A0A0A6YX | 1 | 1741.816 | 0 | -1.57 | 0.000246 | 1.29 | 151.2 | 48.8 | 7.01 | 13.18 | | High | High | High | High | High | High | High | High | High | High | High | 0.004032 | 0.04291 | 2.09 |
| FALSE | High | RNESLTATI | [R].RNESLT | 1xPhospho | O89032 1x | 0.011048 | 0.000772 | 1 | 3 | 1 | O89032 | SH3 and PX | O89032 [9 | 1 | 1671.826 | 0 | 1.64 | 0.000675 | 13.71 | 51.3 | 148.7 | 11.08 | 9.35 | | High | High | High | High | High | High | High | High | High | High | High | 0.000399 | 0.003515 | 1.27 |
| FALSE | High | RNGSDDDI | [R].RNGSD | 1xDeamida | Q3TYF6 1x | 0.007019 | 0.00062 | 1 | 3 | 2 | Q3TYF6 | DNA segme | Q3TYF6 [8 | 1 | 2217.034 | 0 | 0.09 | 0.281337 | 6.85 | 96.4 | 103.6 | 8.04 | 11.25 | | High | High | High | High | High | High | High | High | High | High | High | 0.000275 | 0.002103 | 2.83 |
| FALSE | High | RNGSPTPA | [R].RNGSP | 1xDeamida | Q3U3E2 1x | 2.62E-08 | 0 | 1 | 1 | 7 | Q3U3E2 | Protein FAI | Q3U3E2 [7 | 1 | 2468.224 | 0 | 0.07 | 0.335804 | 17.02 | 96.5 | 103.5 | 7.42 | 8.56 | | High | High | High | High | High | High | High | High | High | High | High | 0 | 1.7E-09 | 5.5 |
| FALSE | High | RNLLEDDSI | [R].RNLLED | 1xPhospho | O70480 1x | 0.000314 | 0 | 1 | 3 | 1 | O70480 | Vesicle-ass | O70480 [2 | 1 | 2451.091 | 0 | 0.07 | 0.355892 | 16.82 | 97.9 | 102.1 | 13.91 | 9.78 | | High | High | High | High | High | High | High | High | High | High | High | 0 | 6.47E-05 | 3.55 |
| FALSE | High | RNNSISEAK | [K].RNNSIS | 1xPhospho | P51432 1x | 0.011048 | 0.000772 | 1 | 4 | 2 | P51432 | 1-phospha | P51432 [11 | 1 | 1556.82 | 0 | 0.79 | 0.001921 | 4.45 | 73.4 | 126.6 | 6.22 | 6.71 | | High | High | High | High | High | High | High | High | High | High | High | 0.000585 | 0.005516 | 1.38 |
| FALSE | High | RNNVTNYN | [R].RNNVT | 2xPhospho | A4QPD1 2x | 0.000933 | 0 | 1 | 5 | 1 | A4QPD1 | Zinc finger | A4QPD1 [1 | 1 | 3117.345 | 0 | 1.33 | 0.000826 | 10.65 | 53.3 | 146.7 | 8.74 | 8.96 | | High | High | High | High | High | High | High | High | High | High | High | 0 | 0.00022 | 3.43 |
| FALSE | High | RNPWSDDI | [K].RNPWS | 1xPhospho | Q64511 1x | 0.060159 | 0.004476 | 1 | 2 | 1 | Q64511 | DNA topois | Q64511 [1 | 1 | 1771.842 | 0 | -0.02 | 0.698055 | 31.9 | 100.5 | 99.5 | 9.3 | 28.38 | | High | High | High | High | High | High | High | High | High | High | High | 0.002279 | 0.02379 | 0.89 |
| FALSE | High | RMSDEFEG | [R].RMSDE | 1xPhospho | Q61337 1x | 0.000158 | 0 | 1 | 6 | 2 | Q61337 | Bcl2-associ | Q61337 [1 | 1 | 1886.876 | 0 | -0.16 | 0.676132 | 25.75 | 107.5 | 92.5 | 14.47 | 46.76 | | High | High | High | High | High | High | High | High | High | High | High | 0.000399 | 0.003427 | 3.13 |
| FALSE | High | RNSAEETW | [K].RNSAEI | 1xPhospho | Q3TH64 1x | 2.9E-05 | 0 | 1 | 3 | 1 | Q3TH64 | Putative ur | Q3TH64 [3 | 1 | 2010.942 | 0 | 1.35 | 0.000324 | 4.93 | 59 | 141 | 11.73 | 8.11 | | High | High | High | High | High | High | High | High | High | High | High | 0 | 4.47E-06 | 2.98 |
| FALSE | High | RNSAPVSV | [R].RNSAP | 1xPhospho | B2RSI0 1xP | 0.009843 | 0.000736 | 1 | 1 | 1 | B2RSI0 | CDC42 GTF | B2RSI0 [11 | 1 | 1551.821 | 0 | -0.22 | 0.986015 | 5.5 | 102.9 | 97.1 | 12.78 | 17.91 | | High | High | High | High | High | High | High | High | High | High | High | 0.000344 | 0.003071 | 2.97 |
| FALSE | High | RNSCNVGC | [R].RNSCN | 1xCarbami | Q8K3A9 1x | 4.67E-08 | 0 | 1 | 1 | 3 | Q8K3A9 | 7SK snRNA | Q8K3A9 [1 | 1 | 2021.963 | 0 | 0.42 | 0.00727 | 2.07 | 89.5 | 110.5 | 6.38 | 8.17 | | High | High | High | High | High | High | High | High | High | High | High | 0 | 3.24E-09 | 2.93 |
| FALSE | High | RNSCNVGC | [R].RNSCN | 1xCarbami | Q8K3A9 1x | 2.2E-06 | 0 | 1 | 1 | 1 | Q8K3A9 | 7SK snRNA | Q8K3A9 [1 | 1 | 2022.947 | 0 | 0.49 | 0.018191 | 18.59 | 83.3 | 116.7 | 11.21 | 17.71 | | High | High | High | High | High | High | High | High | High | High | High | 0 | 2.46E-07 | 3.69 |
| FALSE | High | RNSDSAGP | [R].RNSDS/ | 1xPhospho | G5E8T2 1x | 1.21E-09 | 0 | 1 | 12 | 2 | G5E8T2 | E3 ubiquiti | G5E8T2 [1 | 1 | 2047.991 | 0 | -1.05 | 0.000694 | 1.29 | 134.8 | 65.2 | 6.1 | 4.36 | | High | High | High | High | High | High | High | High | High | High | High | 0 | 5.34E-11 | 5.35 |
| FALSE | High | RNSEGSEA | [R].RNSEG | 1xCarbami | B7ZCJ1 1xF | 6.68E-07 | 0 | 1 | 5 | 2 | B7ZCJ1 | Rho GTPas | B7ZCJ1 [16 | 1 | 2649.166 | 0 | 0.3 | 0.040644 | 5.88 | 90.3 | 109.7 | 8.13 | 9.46 | | High | High | High | High | High | High | High | High | High | High | High | 0 | 6.46E-08 | 4.32 |

| | | | | | | | | | | | | | | | | | | | | | | | | | | | | | | | | | | | | | | |
|---|---|---|---|---|---|---|---|---|---|---|---|---|---|---|---|---|---|---|---|---|---|---|---|---|---|---|---|---|---|---|---|---|---|---|---|---|---|---|
| FALSE | High | RNSEIDELK | [R].RNSEID | 1xPhospho | F8VPV0 1x | 0.001361 | 0 | 1 | 7 | 2 | F8VPV0 | Pericentrin | F8VPV0 [14 | 1 | 1641.861 | 0 | 0.51 | 0.009773 | 7.3 | 84.3 | 115.7 | 3.55 | 9.6 | | High | High | High | High | High | High | High | High | High | High | High | 0 | 0.000336 | 2.77 |
| FALSE | High | RNSEPVGL | [K].RNSEPV | 1xPhospho | Q3ULZ2 1x | 0.004867 | 0.000504 | 1 | 1 | 1 | Q3ULZ2 | FH2 domai | Q3ULZ2 [6 | 1 | 2821.517 | 0 | 0.86 | 0.005082 | 9.67 | 73.9 | 126.1 | 19.35 | 13.85 | | High | High | High | High | High | High | High | High | High | High | High | 0.000225 | 0.001404 | 2.48 |
| FALSE | High | RNSFTPLSS | [R].RNSFTF | 1xPhospho | A0A087WF | 0.013074 | 0.000889 | 1 | 6 | 2 | A0A087WF | 6-phospho | A0A087WF | 1 | 1888.948 | 0 | -0.22 | 0.016927 | 8.16 | 108.9 | 91.1 | 8.47 | 4.35 | | High | High | High | High | High | High | High | High | High | High | High | 0.000467 | 0.00422 | 2.1 |
| FALSE | High | RNSGEPEP | [K].RNSGEI | 1xPhospho | B2RQG2 1x | 0.002921 | 0.00016 | 1 | 2 | 1 | B2RQG2 | PHD finger | B2RQG2 [5 | 1 | 2459.244 | 0 | -1.08 | 0.000759 | 3.67 | 135.1 | 64.9 | 4.98 | 10.03 | | High | High | High | High | High | High | High | High | High | High | High | 7.19E-05 | 0.00079 | 2.03 |
| FALSE | High | RNSGYHSF | [R].RNSGYI | 1xPhospho | Q8CBX9 1x | 0.030367 | 0.001857 | 1 | 1 | 1 | Q8CBX9 | Basic immu | Q8CBX9 [4 | 1 | 1432.668 | 0 | 0.48 | 0.003488 | 14.13 | 85.3 | 114.7 | 10.91 | 7.08 | | High | High | High | High | High | High | High | High | High | High | High | 0.001011 | 0.01091 | 1.7 |
| FALSE | High | RNSLDVSQ | [R].RNSLDV | 1xPhospho | A2AJK6 1x | 6.56E-07 | 0 | 1 | 2 | 2 | A2AJK6 | Chromodo | A2AJK6 [62 | 1 | 1842.936 | 0 | 0.05 | 0.686038 | 12.3 | 102.4 | 97.6 | 10.65 | 4.92 | | High | High | High | High | High | High | High | High | High | High | High | 0 | 6.34E-08 | 3.61 |
| FALSE | High | RNSIEIAGL | [K].RNSIEIA | 1xPhospho | G5E8J4 1x | 8.74E-06 | 0 | 1 | 5 | 1 | G5E8J4 | Tyrosine-p | G5E8J4 [63 | 1 | 2179.122 | 0 | 0.77 | 0.000887 | 0.8 | 75.1 | 124.9 | 3.23 | 5.87 | | High | High | High | High | High | High | High | High | High | High | High | 0 | 1.16E-06 | 3.95 |
| FALSE | High | RNSLGGDV | [R].RNSLG | 1xPhospho | Q9DBR1 1x | 0.000731 | 0 | 1 | 2 | 1 | Q9DBR1 | 5'-3' exorib | Q9DBR1 [6 | 1 | 1900.046 | 0 | 2.43 | 0.000554 | 3.79 | 28.7 | 171.3 | 11.43 | 11.69 | | High | High | High | High | High | High | High | High | High | High | High | 0 | 0.000168 | 3.4 |
| FALSE | High | RNSAPESIL | [R].RNSAPI | 1xPhospho | Q60591 1x | 0.001968 | 5.66E-05 | 1 | 14 | 2 | Q60591 | Nuclear fac | Q60591 [3 | 1 | 2443.351 | 0 | 2.87 | 0.000234 | 3.73 | 23.6 | 176.4 | 4.4 | 6.94 | | High | High | High | High | High | High | High | High | High | High | High | 4.94E-05 | 0.000639 | 2.58 |
| FALSE | High | RMSADMS | [K].RMSAD | 1xPhospho | Q923A8 1x | 0.009416 | 0.000736 | 1 | 5 | 1 | Q923A8 | Map3k7 pr | Q923A8 [3 | 1 | 1736.755 | 0 | -0.54 | 0.001377 | 6.84 | 117.7 | 82.3 | 3.3 | 9.7 | | High | High | High | High | High | High | High | High | High | High | High | 0.000327 | 0.00293 | 1.97 |
| FALSE | High | RMSADMS | [K].RMSAD | 1xPhospho | Q923A8 1x | 0.000892 | 0 | 1 | 5 | 1 | Q923A8 | Map3k7 pr | Q923A8 [3 | 1 | 1704.765 | 0 | -0.11 | 0.853237 | 28.65 | 105.3 | 94.7 | 6.09 | 21.32 | | High | High | High | High | High | High | High | High | High | High | High | 0 | 0.000209 | 3.22 |
| FALSE | High | RLTVSSLQE | [R].RLTVSS | 1xPhospho | Q8BTM8 1 | 0.025514 | 0.001645 | 1 | 5 | 1 | Q8BTM8 | Filamin-A C | Q8BTM8 [2 | 1 | 1956.093 | 0 | 0.94 | 0.028373 | 13.61 | 69 | 131 | 13.8 | 27.21 | | High | High | High | High | High | High | High | High | High | High | High | 0.000872 | 0.00893 | 2.76 |
| FALSE | High | RLSQSDED | [K].RLSQSD | 1xPhospho | Q8C6G8 1x | 0.000276 | 0 | 1 | 3 | 3 | Q8C6G8 | WD repeat | Q8C6G8 [9 | 1 | 1626.805 | 0 | 0.72 | 9.77E-05 | 2.13 | 74.8 | 125.2 | 3.99 | 2.7 | | High | High | High | High | High | High | High | High | High | High | High | 0 | 5.61E-05 | 2.89 |
| FALSE | High | RISQSQPVI | [R].RISQSQ | 1xPhospho | E9Q2H1 1x | 0.047086 | 0.00354 | 1 | 2 | 2 | E9Q2H1 | E3 ubiquiti | E9Q2H1 [1 | 1 | 1379.736 | 0 | 0.24 | 0.00302 | 4.19 | 92.2 | 107.8 | 3.92 | 1.73 | | High | High | High | High | High | High | High | High | High | High | High | 0.001812 | 0.01815 | 1.63 |
| FALSE | High | RLSQSSYQI | [K].RLSQSS | 1xPhospho | Q61033 1x | 0.005322 | 0.000599 | 1 | 1 | 2 | Q61033 | Lamina-ass | Q61033 [4 | 1 | 2246.065 | 0 | 0.52 | 0.023424 | 17.15 | 79.2 | 120.8 | 13.38 | 11.65 | | High | High | High | High | High | High | High | High | High | High | High | 0.000267 | 0.001549 | 2.87 |
| FALSE | High | RLSQSTESN | [K].RLSQST | 1xPhospho | Q8C4S8 1x | 1.3E-06 | 0 | 1 | 2 | 2 | Q8C4S8 | DENN dom | Q8C4S8 [4 | 1 | 1831.932 | 0 | -1.6 | 0.00039 | 0.4 | 150.2 | 49.8 | 7.79 | 9.43 | | High | High | High | High | High | High | High | High | High | High | High | 0 | 1.36E-07 | 2.18 |
| FALSE | High | RISQVSSGE | [R].RISQVS | 1xPhospho | B2RXS1 1x | 0.000475 | 0 | 1 | 8 | 1 | B2RXS1 | Nbeal2 pro | B2RXS1 [27 | 1 | 2289.071 | 0 | 2.25 | 0.000846 | 5.97 | 35.7 | 164.3 | 12.72 | 11.55 | | High | High | High | High | High | High | High | High | High | High | High | 0 | 0.000103 | 3.33 |
| FALSE | High | RLSSASTGK | [R].RLSSAS | 1xPhospho | Q3TIV2 1x | 6.53E-05 | 0 | 1 | 3 | 1 | Q3TIV2 | Putative ur | Q3TIV2 [73 | 1 | 2930.548 | 0 | -0.4 | 0.167621 | 6.99 | 113.7 | 86.3 | 18.02 | 39.1 | | High | High | High | High | High | High | High | High | High | High | High | 0 | 1.12E-05 | 2.99 |
| FALSE | High | RLSSLGEVD | [R].RLSSLG | 1xPhospho | Q3UPD0 1x | 0.006653 | 0.00062 | 1 | 5 | 1 | Q3UPD0 | Putative ur | Q3UPD0 [1 | 1 | 2739.49 | 0 | 0.35 | 0.03022 | 8.61 | 90.4 | 109.6 | 15.72 | 8.09 | | High | High | High | High | High | High | High | High | High | High | High | 0.000275 | 0.001982 | 2.93 |
| FALSE | High | RISSSSAQP | [R].RISSSSA | 1xCarbami | A0A0R4J0Y | 1.06E-05 | 0 | 1 | 4 | 4 | A0A0R4J0Y | Probable h | A0A0R4J0Y | 1 | 3141.529 | 0 | 0.15 | 0.10581 | 10.28 | 95.2 | 104.8 | 4.79 | 5.25 | | High | High | High | High | High | High | High | High | High | High | High | 0 | 1.43E-06 | 6.39 |
| FALSE | High | RISSSSAQP | [R].RISSSSA | 1xCarbami | A0A0R4J0Y | 0.000924 | 0 | 1 | 4 | 3 | A0A0R4J0Y | Probable h | A0A0R4J0Y | 1 | 3221.495 | 0 | 0.53 | 0.012549 | 17.31 | 82.3 | 117.7 | 15.93 | 16.22 | | High | High | High | High | High | High | High | High | High | High | High | 7.19E-05 | 0.000788 | 4.14 |
| FALSE | High | RLSSSSSAA | [R].RLSSSS | 1xPhospho | B7ZNA0 1x | 0.018285 | 0.001282 | 2 | 8 | 2 | B7ZNA0; Q | Shroom3 p | B7ZNA0 [7 | 1 | 1718.842 | 0 | 0.22 | 0.03826 | 11.1 | 90.4 | 109.6 | 12.57 | 6.22 | Shared | High | High | High | High | High | High | High | High | High | High | High | 0.000689 | 0.006177 | 2.76 |
| FALSE | High | RLSSTGGQ | [R].RLSSTG | 1xPhospho | Q8C1D8 1x | 0.001267 | 0 | 1 | 3 | 2 | Q8C1D8 | Protein IW | Q8C1D8 [6 | 1 | 1468.747 | 0 | -0.45 | 0.015459 | 13.16 | 120.7 | 79.3 | 7.56 | 11.84 | | High | High | High | High | High | High | High | High | High | High | High | 0.003922 | 0.0418 | 1.74 |
| FALSE | High | RLSSTGGQ | [R].RLSSTG | 2xPhospho | Q8C1D8 2x | 0.100129 | 0.008067 | 1 | 3 | 2 | Q8C1D8 | Protein IW | Q8C1D8 [6 | 1 | 1548.713 | 0 | 0.26 | 0.042275 | 6.8 | 89.1 | 110.9 | 7.17 | 9.81 | | High | High | High | High | High | High | High | High | High | High | High | 0.00488 | 0.05317 | 2.45 |
| FALSE | High | RLSSTSLAS | [R].RLSSTS | 1xPhospho | Q8BZ03 1x | 0.000543 | 0 | 1 | 2 | 1 | Q8BZ03 | Serine/thre | Q8BZ03 [1 | 1 | 1766.911 | 0 | 0.4 | 0.582997 | 25.38 | 93.8 | 106.2 | 20.78 | 22.57 | | High | High | High | High | High | High | High | High | High | High | High | 0 | 0.00012 | 2.02 |
| FALSE | High | RLSTIFEECI | [K].RLSTIFE | 1xCarbami | E9QAU4 1x | 1.15E-06 | 0 | 1 | 3 | 3 | E9QAU4 | Enhancer t | E9QAU4 [1 | 1 | 2235.02 | 0 | -1.28 | 0.000761 | 1.44 | 141.7 | 58.3 | 6.18 | 5.65 | | High | High | High | High | High | High | High | High | High | High | High | 0 | 6.43E-07 | 4.66 |
| FALSE | High | RLSTLILHG | [R].RLSTLIL | 1xCarbami | Q91VL8 1x | 0.003195 | 0.00016 | 1 | 2 | 2 | Q91VL8 | Telomeric | Q91VL8 [4 | 1 | 1949.035 | 0 | -1.45 | 0.000158 | 1.13 | 145 | 55 | 3.43 | 10.22 | | High | High | High | High | High | High | High | High | High | High | High | 0.000847 | 0.008014 | 2.53 |
| FALSE | High | RLSVDGPM | [R].RLSVDC | 1xPhospho | Q6P5E3 1x | 2.72E-05 | 0 | 1 | 3 | 1 | Q6P5E3 | Sal-like 1 (I | Q6P5E3 [1 | 1 | 2326.239 | 0 | -1.3 | 0.000256 | 0.76 | 143.7 | 56.3 | 5.19 | 4.4 | | High | High | High | High | High | High | High | High | High | High | High | 0 | 4.18E-06 | 4.39 |
| FALSE | High | RLSVDGPM | [R].RLSVDC | 1xPhospho | Q6P5E3 1x | 2.34E-09 | 0 | 1 | 3 | 2 | Q6P5E3 | Sal-like 1 (I | Q6P5E3 [1 | 1 | 2342.234 | 0 | -1.36 | 0.001044 | 2.11 | 141.9 | 58.1 | 5.29 | 14.46 | | High | High | High | High | High | High | High | High | High | High | High | 0 | 1.13E-10 | 5.54 |
| FALSE | High | RLSVEEAV | [R].RLSVEE | 1xPhospho | Q8R0W0 1 | 5.24E-07 | 0 | 1 | 1 | 1 | Q8R0W0 | Epiplakin C | Q8R0W0 [2 | 1 | 2579.385 | 0 | 3.09 | 0.000209 | 23.57 | 19.5 | 180.5 | 19.92 | 6.9 | | High | High | High | High | High | High | High | High | High | High | High | 0 | 4.94E-08 | 4.85 |
| FALSE | High | RLSVELTSS | [K].RLSVEL | 1xPhospho | Q02248 1x | 0.00824 | 0.000736 | 1 | 4 | 1 | Q02248 | Catenin be | Q02248 [6 | 1 | 1716.925 | 0 | -0.4 | 0.006343 | 4.1 | 111.7 | 88.3 | 7.53 | 7.84 | | High | High | High | High | High | High | High | High | High | High | High | 0.000327 | 0.002523 | 2.17 |
| FALSE | High | RLSVENME | [R].RLSVEN | 1xPhospho | Q9WTV7 1 | 0.000202 | 0 | 1 | 1 | 2 | Q9WTV7 | E3 ubiquiti | Q9WTV7 [ | 1 | 1831.857 | 0 | 1.36 | 0.002492 | 11.13 | 56.8 | 143.2 | 2.47 | 14.97 | | High | High | High | High | High | High | High | High | High | High | High | 0.000449 | 0.004192 | 2.65 |
| FALSE | High | RLSVENME | [R].RLSVEN | 1xPhospho | Q9WTV7 1 | 0.002441 | 0.000109 | 1 | 1 | 1 | Q9WTV7 | E3 ubiquiti | Q9WTV7 [ | 1 | 1847.852 | 0 | 1.21 | 0.001375 | 10.39 | 60.1 | 139.9 | 11.59 | 11.07 | | High | High | High | High | High | High | High | High | High | High | High | 4.94E-05 | 0.00065 | 2.98 |
| FALSE | High | RLSVQGQA | [R].RLSVQ | 1xPhospho | F8WJD4 1x | 5.88E-06 | 0 | 2 | 6 | 1 | F8WJD4; Q | Symplekin | F8WJD4 [4 | 1 | 3593.865 | 0 | -0.27 | 0.352099 | 34.63 | 109.4 | 90.6 | 14.09 | 21.9 | Shared | High | High | High | High | High | High | High | High | High | High | High | 0 | 7.43E-07 | 4.15 |
| FALSE | High | RLSVQGQA | [R].RLSVQ | 1xPhospho | F8WJD4 1x | 0.096181 | 0.007535 | 2 | 6 | 2 | F8WJD4; Q | Symplekin | F8WJD4 [4 | 1 | 3609.86 | 0 | 0.04 | 0.985916 | 25.98 | 99.7 | 100.3 | 27.78 | 7.78 | Shared | High | High | High | High | High | High | High | High | High | High | High | 0.004086 | 0.04376 | 3.11 |
| FALSE | High | RLSVVGPPI | [K].RLSVVG | 1xPhospho | G5E8L8 1x | 0.000269 | 0 | 1 | 2 | 2 | G5E8L8 | Mitogen ac | G5E8L8 [1 | 1 | 1403.772 | 0 | 0.74 | 0.006182 | 23.01 | 76.4 | 123.6 | 11.16 | 9 | | High | High | High | High | High | High | High | High | High | High | High | 0 | 5.46E-05 | 2.38 |
| FALSE | High | RISWPENSI | [R].RISWPE | 1xPhospho | Q3U3G8 1x | 0.000292 | 0 | 1 | 2 | 1 | Q3U3G8 | Putative ur | Q3U3G8 [2 | 1 | 2250.136 | 0 | 1.65 | 0.00315 | 21.44 | 48.8 | 151.2 | 5.44 | 17.16 | | High | High | High | High | High | High | High | High | High | High | High | 0 | 5.98E-05 | 3.02 |
| FALSE | High | RLSYNTASI | [R].RLSYNT | 1xPhospho | Q9D1R9 1x | 9.48E-06 | 0 | 1 | 3 | 2 | Q9D1R9 | 60S riboso | Q9D1R9 [1 | 1 | 1691.888 | 0 | 0.28 | 0.415168 | 34.06 | 91.5 | 108.5 | 9.09 | 19.7 | | High | High | High | High | High | High | High | High | High | High | High | 0 | 1.27E-06 | 2.46 |
| FALSE | High | RLTGGLLDI | [R].RLTGGI | 1xPhospho | G5E8K3 1x | 0.106748 | 0.008634 | 2 | 26 | 1 | G5E8K3; A | Ankyrin 3, | G5E8K3 [1 | 1 | 1309.719 | 0 | 0.86 | 0.001285 | 6.32 | 66 | 134 | 9.22 | 9.84 | Shared | High | High | High | High | High | High | High | High | High | High | High | 0.004327 | 0.04652 | 2.02 |
| FALSE | High | RLTQEDLH | [R].RLTQED | 1xPhospho | Q8R4D1 1x | 0.020862 | 0.001472 | 1 | 5 | 1 | Q8R4D1 | Sodium/hy | Q8R4D1 [5 | 1 | 1670.833 | 0 | 3.54 | 0.000547 | 13.49 | 14.8 | 185.2 | 17.27 | 12.78 | | High | High | High | High | High | High | High | High | High | High | High | 0.000786 | 0.007133 | 1.7 |
| FALSE | High | RLTVNEAV | [R].RLTVNE | 1xPhospho | Q9QXS1-3 | 0.0016 | 5.66E-05 | 3 | 24 | 1 | Q9QXS1-3; | Isoform PL | Q9QXS1-3 | 1 | 1567.897 | 0 | 1.93 | 0.003428 | 16.59 | 40.3 | 159.7 | 7.31 | 19.26 | Shared | High | High | High | High | High | High | High | High | High | High | High | 2.58E-05 | 0.000406 | 2.4 |
| FALSE | High | RNSLIELPEI | [K].RNSLIEI | 1xCarbami | O70481 1x | 1.21E-05 | 0 | 1 | 2 | 1 | O70481 | E3 ubiquiti | O70481 [1 | 1 | 2971.433 | 0 | 1.79 | 0.000348 | 13.38 | 46.8 | 153.2 | 9.47 | 7.63 | | High | High | High | High | High | High | High | High | High | High | High | 0 | 1.68E-06 | 4.9 |
| FALSE | High | RPGASPTG | [R].RPGASI | 1xPhospho | A0A0R4J1C | 9.74E-06 | 0 | 1 | 9 | 2 | A0A0R4J1C | Anion exch | A0A0R4J1C | 0 | 3515.517 | 0 | 2.89 | 0.003574 | 65.54 | 23.8 | 176.2 | 35.13 | 20.25 | | High | High | High | High | High | High | High | High | High | High | High | 0 | 1.31E-06 | 4.98 |
| FALSE | High | RNSLPQIPT | [K].RNSLPC | 1xPhospho | Q9QVP9 1x | 6.55E-06 | 0 | 1 | 6 | 1 | Q9QVP9 | Protein-tyr | Q9QVP9 [3 | 1 | 2031.095 | 0 | -0.48 | 0.006945 | 3.57 | 117.4 | 82.6 | 10.63 | 4.4 | | High | High | High | High | High | High | High | High | High | High | High | 0 | 8.41E-07 | 3.76 |
| FALSE | High | RNSLTGEE | [R].RNSLTC | 1xPhospho | Q9JI99 1xP | 5.84E-07 | 0 | 1 | 2 | 2 | Q9JI99 | Sphingosin | Q9JI99 [99 | 1 | 1970.036 | 0 | 1.05 | 0.001497 | 11.17 | 65 | 135 | 8.36 | 6.67 | | High | High | High | High | High | High | High | High | High | High | High | 0 | 5.57E-08 | 2.62 |
| FALSE | High | RPASDPEL | [K].RPASDI | 1xPhospho | Q3UIZ4 1x | 0.005467 | 0.00062 | 1 | 2 | 2 | Q3UIZ4 | Putative ur | Q3UIZ4 [11 | 0 | 1477.736 | 0 | -0.3 | 0.013785 | 5.42 | 110.2 | 89.8 | 5.16 | 6.73 | | High | High | High | High | High | High | High | High | High | High | High | 0.005561 | 0.06074 | 1.94 |
| FALSE | High | RPASGDDL | [R].RPASGI | 1xPhospho | Q8C5X2 1x | 2.02E-05 | 0 | 1 | 9 | 2 | Q8C5X2 | DCN1-like | Q8C5X2 [8 | 0 | 1654.857 | 0 | -1.72 | 0.000271 | 0.42 | 153.4 | 46.6 | 5.05 | 19.74 | | High | High | High | High | High | High | High | High | High | High | High | 0.000538 | 0.004553 | 2.03 |
| FALSE | High | RPASPSHN | [K].RPASPS | 1xPhospho | F8WIA0 1x | 0.003083 | 0.00016 | 1 | 3 | 4 | F8WIA0 | INO80 com | F8WIA0 [2 | 0 | 2255.097 | 0 | -0.06 | 0.756973 | 24.2 | 100.8 | 99.2 | 13.13 | 11.34 | | High | High | High | High | High | High | High | High | High | High | High | 0.000814 | 0.007575 | 2.53 |
| FALSE | High | RPASPSHN | [K].RPASPS | 1xDeamida | F8WIA0 1x | 0.013545 | 0.000976 | 1 | 3 | 3 | F8WIA0 | INO80 com | F8WIA0 [2 | 0 | 2256.081 | 0 | -0.05 | 0.902819 | 15.67 | 101.3 | 98.7 | 11.14 | 16.46 | | High | High | High | High | High | High | High | High | High | High | High | 0.000638 | 0.005603 | 2.04 |
| FALSE | High | RPASPSSPE | [K].RPASPS | 1xPhospho | A0A0R4J24 | 4.03E-07 | 0 | 1 | 2 | 6 | A0A0R4J24 | Sin3 histon | A0A0R4J24 | 0 | 2592.277 | 0 | -0.54 | 0.056339 | 16.56 | 117.2 | 82.8 | 12.61 | 12.88 | | High | High | High | High | High | High | High | High | High | High | High | 0 | 3.65E-08 | 4.24 |
| FALSE | High | RPASPSSPE | [K].RPASPS | 2xPhospho | A0A0R4J24 | 0.000229 | 0 | 1 | 2 | 8 | A0A0R4J24 | Sin3 histon | A0A0R4J24 | 0 | 2672.243 | 0 | -0.35 | 0.009497 | 4.64 | 113.7 | 86.3 | 3.71 | 8.23 | | High | High | High | High | High | High | High | High | High | High | High | 0 | 4.53E-05 | 4.24 |
| FALSE | High | RPASPTPLL | [K].RPASPT | 1xCarbami | A0A0R4J0C | 0.000292 | 0 | 1 | 1 | 1 | A0A0R4J0C | BRF2, subu | A0A0R4J0C | 0 | 2767.574 | 0 | -0.78 | 0.004552 | 2.9 | 122.4 | 77.6 | 10.7 | 4.89 | | High | High | High | High | High | High | High | High | High | High | High | 0 | 5.95E-05 | 2.01 |
| FALSE | High | RPASTAGLI | [K].RPASTA | 1xPhospho | Q6ZQB7 1x | 8.5E-06 | 0 | 1 | 4 | 2 | Q6ZQB7 | MKIAA042 | Q6ZQB7 [5 | 0 | 2760.468 | 0 | -0.24 | 0.249889 | 12.16 | 111.5 | 88.5 | 12.98 | 15.84 | | High | High | High | High | High | High | High | High | High | High | High | 0 | 1.8E-06 | 4.33 |
| FALSE | High | RPASVSSA | [K].RPASVS | 1xPhospho | E9Q1P8 1x | 5.48E-07 | 0 | 1 | 1 | 2 | E9Q1P8 | Interferon | E9Q1P8 [2 | 0 | 1847.896 | 0 | 0.74 | 0.021512 | 25.55 | 75 | 125 | 11.57 | 14.86 | | High | High | High | High | High | High | High | High | High | High | High | 0 | 5.19E-08 | 2.61 |
| FALSE | High | RPDAPAAS | [R].RPDAP | 1xPhospho | Q8BIZ6 1x | 2.78E-06 | 0 | 1 | 1 | 6 | Q8BIZ6 | Smad nucl | Q8BIZ6 [41 | 0 | 2487.209 | 0 | -0.43 | 0.014955 | 3.88 | 116.4 | 83.6 | 9.75 | 3.8 | | High | High | High | High | High | High | High | High | High | High | High | 0 | 0.000319 | 3.39 |
| FALSE | High | RPDAPAAS | [R].RPDAP | 2xPhospho | Q8BIZ6 2x | 0.000693 | 0 | 1 | 1 | 1 | Q8BIZ6 | Smad nucl | Q8BIZ6 [41 | 0 | 2567.175 | 0 | -0.38 | 0.056712 | 17.1 | 114.7 | 85.3 | 12.59 | 10.66 | | High | High | High | High | High | High | High | High | High | High | High | 0 | 0.000159 | 2.24 |
| FALSE | High | RPDAPAAS | [R].RPDAP | 2xPhospho | Q8BIZ6 2x | 0.000198 | 0 | 1 | 1 | 1 | Q8BIZ6 | Smad nucl | Q8BIZ6 [41 | 1 | 2867.33 | 0 | -0.19 | 0.266002 | 8.64 | 106.7 | 93.3 | 9.96 | 15.83 | | High | High | High | High | High | High | High | High | High | High | High | 0 | 3.88E-05 | 2.66 |
| FALSE | High | RPDAPAAS | [R].RPDAP | 3xPhospho | Q8BIZ6 3x | 0.001831 | 5.66E-05 | 1 | 1 | 1 | Q8BIZ6 | Smad nucl | Q8BIZ6 [41 | 1 | 2947.296 | 0 | -0.12 | 0.197444 | 12.43 | 106.7 | 93.3 | 4.53 | 11.12 | | High | High | High | High | High | High | High | High | High | High | High | 2.58E-05 | 0.000471 | 2.83 |
| FALSE | High | RPASDDEC | [K].RPASDI | 1xCarbami | Q7TMI3 1x | 2.79E-07 | 0 | 1 | 2 | 4 | Q7TMI3 | E3 ubiquiti | Q7TMI3 [6 | 0 | 1971.888 | 0 | 0.22 | 0.016758 | 9.09 | 92.6 | 107.4 | 4.37 | 6.98 | | High | High | High | High | High | High | High | High | High | High | High | 0 | 2.42E-08 | 3.85 |
| FALSE | High | RPDGEEED | [K].RPDGEI | 1xPhospho | Q810V0 1x | 0.001865 | 5.66E-05 | 1 | 3 | 3 | Q810V0 | U3 small nu | Q810V0 [2 | 0 | 3723.554 | 0 | -0.54 | 0.063582 | 21.64 | 118.1 | 81.9 | 21.22 | 19.2 | | High | High | High | High | High | High | High | High | High | High | High | 0.00749 | 0.08236 | 4 |
| FALSE | High | RPDSPASFI | [K].RPDSPA | 1xPhospho | A0A0R4J1A | 0.02933 | 0.001857 | 1 | 2 | 2 | A0A0R4J1A | GAS2-like | A0A0R4J1A | 0 | 1873.973 | 0 | 0.33 | 0.004757 | 0.86 | 87.5 | 112.5 | 3.17 | 5.48 | | High | High | High | High | High | High | High | High | High | High | High | 0.000997 | 0.01051 | 2.02 |
| FALSE | High | RPDYAPME | [K].RPDYAI | 1xPhospho | C0HKD9 1x | 0.024855 | 0.001645 | 1 | 1 | 2 | C0HKD9 | Microfibril | C0HKD9 [4 | 0 | 3116.385 | 0 | -0.1 | 0.587331 | 23.01 | 102.4 | 97.6 | 7.95 | 22.75 | | High | High | High | High | High | High | High | High | High | High | High | 0.001308 | 0.01351 | 2.75 |
| FALSE | High | RPDYAPME | [K].RPDYAI | 2xPhospho | C0HKD9 2x | 4.44E-06 | 0 | 1 | 1 | 11 | C0HKD9 | Microfibril | C0HKD9 [4 | 0 | 3180.356 | 0 | -0.27 | 0.059148 | 10.76 | 109 | 91 | 5.62 | 13.23 | | High | High | High | High | High | High | High | High | High | High | High | 0 | 1.41E-05 | 4.54 |
| FALSE | High | RPECSSSDT | [K].RPECSS | 1xCarbami | Q61687 1x | 2.56E-05 | 0 | 1 | 4 | 3 | Q61687 | Transcripti | Q61687 [1 | 0 | 1833.845 | 0 | -0.3 | 0.013554 | 3.43 | 110.1 | 89.9 | 6.38 | 4.55 | | High | High | High | High | High | High | High | High | High | High | High | 0 | 3.86E-06 | 1.9 |
| FALSE | High | RPECSSSDT | [K].RPECSS | 1xCarbami | Q61687 2x | 0.048715 | 0.00354 | 1 | 4 | 1 | Q61687 | Transcripti | Q61687 [1 | 0 | 1913.812 | 0 | -0.15 | 0.011483 | 1.86 | 108.5 | 91.5 | 4.71 | 2.91 | | High | High | High | High | High | High | High | High | High | High | High | 0.001844 | 0.01864 | 1.81 |
| FALSE | High | RPEFFTFG | [K].RPEFFT | 1xCarbami | E9PWG2 1 | 0.001147 | 0 | 1 | 4 | 1 | E9PWG2 | Trafficking | E9PWG2 [9 | 0 | 3688.776 | 0 | 0.1 | 0.163232 | 15.28 | 93.7 | 106.3 | 16.59 | 5.7 | | High | High | High | High | High | High | High | High | High | High | High | 0 | 0.000278 | 2.84 |
| FALSE | High | RPEPPSPYF | [R].RPEPPS | 1xPhospho | Q9D2X8 1x | 0.063788 | 0.004664 | 1 | 3 | 1 | Q9D2X8 | Uncharacte | Q9D2X8 [2 | 0 | 1407.698 | 0 | -0.41 | 0.002085 | 4.25 | 114.1 | 85.9 | 5.73 | 5.96 | | High | High | High | High | High | High | High | High | High | High | High | 0.002403 | 0.02547 | 1.73 |
| FALSE | High | RPEPSSDYI | [K].RPEPSS | 1xPhospho | Q8CGN4 1 | 0.000319 | 0 | 1 | 4 | 1 | Q8CGN4 | BCL-6 core | Q8CGN4 [1 | 0 | 2197.094 | 0 | 0.69 | 0.008134 | 7.4 | 74.7 | 125.3 | 7.28 | 14.27 | | High | High | High | High | High | High | High | High | High | High | High | 0 | 6.64E-05 | 3.34 |
| FALSE | High | RPEPSSDYI | [K].RPEPSS | 2xPhospho | Q8CGN4 2 | 0.002695 | 0.000109 | 1 | 4 | 2 | Q8CGN4 | BCL-6 core | Q8CGN4 [1 | 0 | 2277.061 | 0 | 0.35 | 0.005924 | 7.77 | 88.2 | 111.8 | 2.08 | 6.35 | | High | High | High | High | High | High | High | High | High | High | High | 0.000275 | 0.001599 | 3.15 |
| FALSE | High | RPESAPAES | [R].RPESAF | 1xPhospho | Q62245 1x | 0.000294 | 0 | 1 | 3 | 1 | Q62245 | Son of seve | Q62245 [1 | 0 | 1880.952 | 0 | -0.4 | 0.003679 | 5.7 | 113.8 | 86.2 | 3.47 | 7.03 | | High | High | High | High | High | High | High | High | High | High | High | 0 | 6.02E-05 | 2.19 |
| FALSE | High | RPESLEK | [K].RPESLE | 1xPhospho | Q80YR9 1x | 0.1059 | 0.008582 | 1 | 4 | 1 | Q80YR9 | RNA-bindir | Q80YR9 [3 | 0 | 1396.76 | 0 | -0.36 | 0.00295 | 3.12 | 109.3 | 90.7 | 3.68 | 4.84 | | High | High | High | High | High | High | High | High | High | High | High | 0.004317 | 0.04606 | 1.13 |
| FALSE | High | RPESPPSVL | [K].RPESPP | 1xPhospho | B1AR09 1x | 4.44E-06 | 0 | 1 | 3 | 1 | B1AR09 | Myeloid/ly | B1AR09 [2 | 0 | 2467.336 | 0 | 0.2 | 0.137602 | 13.92 | 92.9 | 107.1 | 11.16 | 4.77 | | High | High | High | High | High | High | High | High | High | High | High | 0 | 5.41E-07 | 3.19 |
| FALSE | High | RPESRGSG | [R].RPESRC | 2xPhospho | Q9JLF6 2x | 0.0003 | 0 | 2 | 2 | 3 | Q9JLF6; A0 | Protein-glu | Q9JLF6 [88 | 1 | 2189.006 | 0 | 0.37 | 0.023162 | 13.9 | 87.4 | 112.6 | 7.14 | 5.56 | Shared | High | High | High | High | High | High | High | High | High | High | High | 0.009099 | 0.098 | 2.35 |
| FALSE | High | RPDPDSDE | [R].RPDPD | 1xPhospho | Q8JZX4 1x | 0.000334 | 0 | 1 | 1 | 1 | Q8JZX4 | Splicing fac | Q8JZX4 [1 | 0 | 2046.813 | 0 | -0.57 | 0.003323 | 7.34 | 122 | 78 | 10.21 | 10.19 | | High | High | High | High | High | High | High | High | High | High | High | 0 | 6.93E-05 | 2.66 |
| FALSE | High | RPAGGSLG | [R].RPAGG | 2xPhospho | A2CG35 2x | 0.000633 | 0 | 1 | 2 | 1 | A2CG35 | Ras-relatec | A2CG35 [5 | 0 | 2198.068 | 0 | 1.72 | 0.000648 | 8.11 | 45.1 | 154.9 | 16.33 | 8.64 | | High | High | High | High | High | High | High | High | High | High | High | 0 | 0.000143 | 3 |
| FALSE | High | RPAGGSLG | [R].RPAGG | 1xPhospho | A2CG35 1x | 3E-08 | 0 | 1 | 2 | 3 | A2CG35 | Ras-relatec | A2CG35 [5 | 0 | 2118.102 | 0 | 1.12 | 0.000406 | 7.81 | 61.8 | 138.2 | 4.5 | 4.53 | | High | High | High | High | High | High | High | High | High | High | High | 0.000161 | 0.001186 | 2.64 |
| FALSE | High | RPADVSPA | [K].RPADV | 1xPhospho | A2AI52 1x | 0.022182 | 0.001542 | 1 | 4 | 1 | A2AI52 | Ubiquitin c | A2AI52 [58 | 0 | 2293.195 | 0 | -0.42 | 0.012759 | 4.54 | 111.9 | 88.1 | 6.5 | 12.91 | | High | High | High | High | High | High | High | High | High | High | High | 0.000814 | 0.007652 | 2.67 |
| FALSE | High | RNSLTGEE | [R].RNSLTC | 1xDeamida | Q9JI99 1xP | 0.039686 | 0.00283 | 1 | 2 | 1 | Q9JI99 | Sphingosin | Q9JI99 [99 | 1 | 1971.02 | 0 | 0.43 | 0.016653 | 5.83 | 83.9 | 116.1 | 5.22 | 11.11 | | High | High | High | High | High | High | High | High | High | High | High | 0.001468 | 0.01476 | 2.61 |
| FALSE | High | RNSPLVNQ | [K].RNSPLV | 1xPhospho | A0A1B0GR | 0.00047 | 0 | 1 | 1 | 1 | A0A1B0GR | RIKEN cDN | A0A1B0GR | 1 | 1706.972 | 0 | 1.12 | 0.000396 | 3.39 | 62.4 | 137.6 | 6.81 | 8.55 | | High | High | High | High | High | High | High | High | High | High | High | 0 | 0.000103 | 3.12 |
| FALSE | High | RNSPMAQ | [R].RNSPM | 1xCarbami | P70445 1x | 0.096959 | 0.007777 | 1 | 2 | 1 | P70445 | Eukaryotic | P70445 [63 | 1 | 3723.864 | 0 | -1.07 | 0.005725 | 6.37 | 135 | 65 | 10.97 | 15.49 | | High | High | High | High | High | High | High | High | High | High | High | 0.003922 | 0.0418 | 2.71 |
| FALSE | High | RNSSDSFSI | [R].RNSSDS | 1xPhospho | G3X9T5 1x | 0.1059 | 0.008582 | 1 | 6 | 1 | G3X9T5 | MCG12342 | G3X9T5 [6 | 1 | 1364.616 | 0 | 0.57 | 0.00264 | 5.81 | 80.8 | 119.2 | 9.71 | 7.52 | | High | High | High | High | High | High | High | High | High | High | High | 0.004317 | 0.04615 | 1.12 |
| FALSE | High | RNSSLVGR | [R].RNSSLV | 1xPhospho | A2A9M4 1 | 0.049549 | 0.00361 | 1 | 6 | 1 | A2A9M4 | Dedicator | A2A9M4 [4 | 1 | 1197.63 | 0 | 0.75 | 0.010455 | 12.53 | 82.2 | 117.8 | 13.79 | 10.94 | | High | High | High | High | High | High | High | High | High | High | High | 0.001889 | 0.019 | 1.13 |
| FALSE | High | RNSSPPPSI | [K].RNSSPF | 2xPhospho | G5E829 2x | 0.000687 | 0 | 1 | 5 | 2 | G5E829 | Plasma me | G5E829 [1 | 1 | 1798.865 | 0 | 1.33 | 0.000737 | 1.21 | 55.4 | 144.6 | 4.65 | 8.89 | | High | High | High | High | High | High | High | High | High | High | High | 0.00092 | 0.009632 | 2.62 |
| FALSE | High | RNSSSPPSI | [R].RNSSSF | 2xPhospho | E9Q1P8 2x | 0.075927 | 0.005826 | 1 | 1 | 3 | E9Q1P8 | Interferon | E9Q1P8 [4 | 1 | 2036.846 | 0 | 0.41 | 0.023494 | 11.77 | 86.1 | 113.9 | 7.63 | 8.09 | | High | High | High | High | High | High | High | High | High | High | High | 0.002976 | 0.03121 | 2.75 |
| FALSE | High | RNSSSPVSI | [R].RNSSSF | 1xPhospho | Q8K3X4 1x | 0.001476 | 5.66E-05 | 1 | 1 | 2 | Q8K3X4 | Interferon | Q8K3X4 [6 | 1 | 1934.965 | 0 | -0.62 | 0.02017 | 3.78 | 120.9 | 79.1 | 14.72 | 7.11 | | High | High | High | High | High | High | High | High | High | High | High | 2.58E-05 | 0.000369 | 3.26 |
| FALSE | High | RNSSSPVSI | [R].RNSSSF | 1xDeamida | Q8K3X4 1x | 0.008094 | 0.000736 | 1 | 1 | 2 | Q8K3X4 | Interferon | Q8K3X4 [6 | 1 | 1935.949 | 0 | 0.11 | 0.165638 | 9.37 | 94.7 | 105.3 | 7.37 | 6.7 | | High | High | High | High | High | High | High | High | High | High | High | 0.000327 | 0.002489 | 2.52 |
| FALSE | High | RNSSSPVSI | [R].RNSSSF | 2xPhospho | Q8K3X4 2x | 0.000483 | 0 | 1 | 1 | 4 | Q8K3X4 | Interferon | Q8K3X4 [6 | 1 | 2014.931 | 0 | -0.02 | 0.92981 | 11.11 | 100.2 | 99.8 | 4.2 | 7.87 | | High | High | High | High | High | High | High | High | High | High | High | 0.002455 | 0.02688 | 3.09 |
| FALSE | High | RNSSSPVSI | [R].RNSSSF | 3xPhospho | Q8K3X4 3x | 0.005467 | 0.00062 | 1 | 1 | 2 | Q8K3X4 | Interferon | Q8K3X4 [6 | 1 | 2094.897 | 0 | -0.21 | 0.324369 | 4.73 | 107.3 | 92.7 | 2.93 | 11.63 | | High | High | High | High | High | High | High | High | High | High | High | 0.000814 | 0.007857 | 3.41 |
| FALSE | High | RNSTATAM | [R].RNSTAT | 1xPhospho | Q6PAL5 1x | 2.85E-05 | 0 | 1 | 5 | 2 | Q6PAL5 | Uncharacte | Q6PAL5 [6 | 1 | 1975.008 | 0 | 0.97 | 0.016927 | 12.99 | 69.2 | 130.8 | 5.79 | 18.28 | | High | High | High | High | High | High | High | High | High | High | High | 0 | 4.38E-06 | 3.79 |
| FALSE | High | RNSTATAM | [R].RNSTAT | 1xPhospho | Q6PAL5 1x | 0.000331 | 0 | 1 | 5 | 3 | Q6PAL5 | Uncharacte | Q6PAL5 [6 | 1 | 1991.003 | 0 | 0.84 | 0.004353 | 12.96 | 72.3 | 127.7 | 6.3 | 11.78 | | High | High | High | High | High | High | High | High | High | High | High | 0 | 6.87E-05 | 3.77 |
| FALSE | High | RNSTMGE | [K].RNSTM | 1xPhospho | Q8VI59 1x | 1.95E-05 | 0 | 1 | 2 | 2 | Q8VI59 | Pecanex-lik | Q8VI59 [94 | 1 | 2631.156 | 0 | -0.01 | 0.863206 | 21.05 | 107.2 | 92.8 | 9.36 | 12.74 | | High | High | High | High | High | High | High | High | High | High | High | 0 | 2.87E-06 | 4.87 |
| FALSE | High | RNSTSSTN | [R].RNSTSS | 1xCarbami | P61460-2 1 | 9.8E-05 | 0 | 1 | 3 | 2 | P61460-2 | Isoform 2 c | P61460-2 [ | 1 | 2269.953 | 0 | -0.31 | 0.053968 | 8.33 | 111.3 | 88.7 | 9.01 | 14.84 | | High | High | High | High | High | High | High | High | High | High | High | 0 | 1.76E-05 | 3.93 |
| FALSE | High | RNSTSSTN | [R].RNSTSS | 1xCarbami | P61460-2 1 | 6.53E-05 | 0 | 1 | 3 | 3 | P61460-2 | Isoform 2 c | P61460-2 [ | 1 | 2285.948 | 0 | -0.26 | 0.206097 | 14.1 | 112.3 | 87.7 | 9.26 | 18.15 | | High | High | High | High | High | High | High | High | High | High | High | 0.001183 | 0.01284 | 3.46 |
| FALSE | High | RNSTTFPSI | [R].RNSTTF | 1xPhospho | Q9DB52 1x | 0.067059 | 0.004796 | 1 | 1 | 1 | Q9DB52 | Protein FAI | Q9DB52 [5 | 1 | 1374.673 | 0 | 0.74 | 0.00131 | 7.36 | 73.1 | 126.9 | 6.48 | 8.01 | | High | High | High | High | High | High | High | High | High | High | High | 0.002455 | 0.02704 | 1.6 |
| FALSE | High | RNSVTPLAS | [R].RNSVTI | 1xPhospho | A7UAK5 1x | 4.92E-07 | 0 | 1 | 11 | 2 | A7UAK5 | 6-phospho | A7UAK5 [4 | 1 | 2035.099 | 0 | -0.64 | 0.000992 | 2.73 | 120 | 80 | 4.15 | 8.41 | | High | High | High | High | High | High | High | High | High | High | High | 0 | 4.57E-08 | 4.03 |
| FALSE | High | RNSVTPLAS | [R].RNSVTI | 2xPhospho | A7UAK5 2x | 0.002974 | 0.00016 | 1 | 11 | 1 | A7UAK5 | 6-phospho | A7UAK5 [4 | 1 | 2115.065 | 0 | -0.77 | 0.003738 | 4.62 | 125.1 | 74.9 | 3.93 | 11.76 | | High | High | High | High | High | High | High | High | High | High | High | 7.19E-05 | 0.000807 | 2.98 |
| FALSE | High | RNSWATEI | [K].RNSWA | 1xPhospho | Q8VBT6 1x | 3.54E-05 | 0 | 1 | 1 | 3 | Q8VBT6 | Apolipopro | Q8VBT6 [3 | 1 | 2469.279 | 0 | 2.55 | 0.00459 | 18.38 | 26 | 174 | 7.89 | 25.1 | | High | High | High | High | High | High | High | High | High | High | High | 0 | 5.6E-06 | 3.91 |
| FALSE | High | RNSYANCL | [K].RNSYAI | 1xCarbami | Q3UH06-3 | 0.112856 | 0.009146 | 1 | 4 | 1 | Q3UH06-3 | Isoform 3 c | Q3UH06-3 | 1 | 1791.898 | 0 | -0.33 | 0.096543 | 6.98 | 111.4 | 88.6 | 12.72 | 13.51 | | High | High | High | High | High | High | High | High | High | High | High | 0.004569 | 0.04974 | 1.72 |
| FALSE | High | RNTAPEAQ | [R].RNTAPI | 1xPhospho | B2RW11 1x | 0.116482 | 0.009523 | 1 | 1 | 1 | B2RW11 | Ankyrin rep | B2RW11 [3 | 1 | 2092.039 | 0 | -0.35 | 0.05678 | 7.61 | 110.4 | 89.6 | 6.22 | 10.35 | | High | High | High | High | High | High | High | High | High | High | High | 0.004757 | 0.05147 | 3.39 |
| FALSE | High | RNTIDSTAS | [R].RNTIDS | 1xPhospho | Q3TYS6 1x | 0.009931 | 0.000736 | 1 | 4 | 1 | Q3TYS6 | Putative ur | Q3TYS6 [3 | 1 | 1938.927 | 0 | -0.72 | 0.004419 | 7.5 | 128.6 | 71.4 | 8.19 | 14.38 | | High | High | High | High | High | High | High | High | High | High | High | 0.000344 | 0.003127 | 2.79 |
| FALSE | High | RNTNDFDE | [K].RNTND | 1xPhospho | Q5DTM7 1 | 0.042141 | 0.003162 | 1 | 8 | 1 | Q5DTM7 | MKIAA411 | Q5DTM7 [ | 1 | 1676.805 | 0 | 0.7 | 0.001043 | 1.82 | 75.7 | 124.3 | 5.74 | 9.09 | | High | High | High | High | High | High | High | High | High | High | High | 0.001639 | 0.01585 | 1.66 |
| FALSE | High | RNYSSSPP | [R].RNYSSS | 1xCarbami | B1AY10 2x | 0.071662 | 0.005278 | 1 | 3 | 3 | B1AY10 | Transcripti | B1AY10 [4 | 1 | 1959.85 | 0 | -0.47 | 0.013797 | 5.07 | 113.5 | 86.5 | 10.55 | 7.51 | | High | High | High | High | High | High | High | High | High | High | High | 0.003672 | 0.03945 | 1.93 |
| FALSE | High | RPAAAAAA | [K].RPAAA | 1xPhospho | Q91XU0 1x | 7.47E-05 | 0 | 1 | 2 | 5 | Q91XU0 | ATPase WF | Q91XU0 [1 | 0 | 1562.8 | 0 | -0.75 | 0.000413 | 1.76 | 124.6 | 75.4 | 4.55 | 3.68 | | High | High | High | High | High | High | High | High | High | High | High | 0 | 1.29E-05 | 1.56 |
| FALSE | High | RPAASSAA | [K].RPAASS | 2xCarbami | Q62318 1x | 4.32E-09 | 0 | 1 | 2 | 18 | Q62318 | Transcripti | Q62318 [3 | 0 | 3633.674 | 0 | -1.19 | 0.000727 | 2.7 | 139.1 | 60.9 | 5.5 | 6.69 | | High | High | High | High | High | High | High | High | High | High | High | 0 | 8.84E-09 | 4.62 |
| FALSE | High | RPAASSAA | [K].RPAASS | 2xCarbami | Q62318 1x | 4.87E-07 | 0 | 1 | 2 | 6 | Q62318 | Transcripti | Q62318 [3 | 0 | 3634.658 | 0 | 0.15 | 0.032345 | 7.75 | 92.7 | 107.3 | 7.74 | 7.84 | | High | High | High | High | High | High | High | High | High | High | High | 7.19E-05 | 0.000901 | 3.26 |
| FALSE | High | RPAATDSPI | [K].RPAATI | 1xPhospho | Q8CJ76 1x | 0.005872 | 0.00062 | 1 | 3 | 2 | Q8CJ76 | Leukemia-a | Q8CJ76 [27 | 0 | 1480.793 | 0 | 0.97 | 0.000222 | 7.08 | 67.5 | 132.5 | 5.48 | 4.05 | | High | High | High | High | High | High | High | High | High | High | High | 0.000275 | 0.001727 | 1.1 |
| FALSE | High | RNSLSGSST | [R].RNSLSC | 1xPhospho | Q80UE6 1x | 0.012508 | 0.000889 | 1 | 4 | 2 | Q80UE6 | Serine/thre | Q80UE6 [1 | 1 | 1902.887 | 0 | 1.11 | 0.000336 | 6.22 | 63.1 | 136.9 | 9.72 | 5.93 | | High | High | High | High | High | High | High | High | High | High | High | 0.000449 | 0.004034 | 2.73 |
| FALSE | High | RQDSAGPII | [R].RQDSA | 1xPhospho | B2RY19 1x | 1.36E-05 | 0 | 1 | 7 | 2 | B2RY19 | Cgnl1 prot | B2RY19 [28 | 1 | 1664.832 | 0 | 1.9 | 0.000173 | 5.79 | 42.2 | 157.8 | 6.42 | 3.75 | | High | High | High | High | High | High | High | High | High | High | High | 0 | 1.9E-06 | 2.44 |
| FALSE | High | REVSPAPV | [R].REVSPA | 1xPhospho | G3X9J0 1x | 5.05E-07 | 0 | 2 | 10 | 1 | G3X9J0; Q | Signal-indu | G3X9J0 [13 | 1 | 1990.089 | 0 | 1.39 | 0.00014 | 2.34 | 55.2 | 144.8 | 5.28 | 2.18 | Shared | High | High | High | High | High | High | High | High | High | High | High | 0 | 4.72E-08 | 3.98 |
| FALSE | High | RETVVESQ | [K].RETVVE | 2xPhospho | E9PZM7 2x | 1.55E-07 | 0 | 1 | 3 | 9 | E9PZM7 | SR-related | E9PZM7 [8 | 1 | 2364.125 | 0 | 0.2 | 0.147162 | 2.56 | 93 | 107 | 6.97 | 8.49 | | High | High | High | High | High | High | High | High | High | High | High | 0.000275 | 0.002137 | 3.47 |

| | | | | | | | | | | | | | | | | | | | | | | | | | | | | | | | | | | | | | | |
|---|---|---|---|---|---|---|---|---|---|---|---|---|---|---|---|---|---|---|---|---|---|---|---|---|---|---|---|---|---|---|---|---|---|---|---|---|---|---|
| FALSE | High | QSPPHGEA | [K].QSPPH( | 1xPhospho | G3X9K3 1x | 0.003655 | 0.00021 | 1 | 2 | 2 | G3X9K3 | Brefeldin A | G3X9K3 [5 | 0 | 1488.761 | 0 | 0.1 | 0.272144 | 8.59 | 97.4 | 102.6 | 3.07 | 6.2 | | High | High | High | High | High | High | High | High | High | High | High | 0.005216 | 0.05667 | 1.75 |
| FALSE | High | QSPQSASL | [K].QSPQS/ | 1xCarbami | F8VPU2 1x | 1.69E-05 | 0 | 1 | 2 | 1 | F8VPU2 | FERM, Rho | F8VPU2 [3 | 0 | 3205.487 | 0 | 1.31 | 0.00135 | 28 | 59.6 | 140.4 | 15.75 | 11.03 | | High | High | High | High | High | High | High | High | High | High | High | 0 | 2.44E-06 | 3.86 |
| FALSE | High | QSPQSASL | [K].QSPQS/ | 1xCarbami | F8VPU2 1x | 0.000993 | 0 | 1 | 2 | 1 | F8VPU2 | FERM, Rho | F8VPU2 [3 | 0 | 3206.471 | 0 | -0.19 | 0.588905 | 1.74 | 104.2 | 95.8 | 9.62 | 8.05 | | High | High | High | High | High | High | High | High | High | High | High | 0 | 0.000236 | 3.67 |
| FALSE | High | QSPQTLK | [R].QSPQT | 1xPhospho | Q8K3G5 1x | 0.070485 | 0.005214 | 1 | 3 | 1 | Q8K3G5 | Inactive se | Q8K3G5 [1 | 0 | 1339.739 | 0 | 0.7 | 0.003503 | 14.9 | 76.4 | 123.6 | 8.79 | 6 | | High | High | High | High | High | High | High | High | High | High | High | 0.00269 | 0.02877 | 1.91 |
| FALSE | High | QSPSPSEVS | [K].QSPSPS | 1xPhospho | A0A0N4SU | 6.24E-05 | 0 | 1 | 10 | 2 | A0A0N4SU | Peregrin O | A0A0N4SU | 0 | 1751.816 | 0 | 0.32 | 0.007167 | 10.01 | 86.5 | 113.5 | 7.72 | 10.37 | | High | High | High | High | High | High | High | High | High | High | High | 0.002059 | 0.0213 | 2.36 |
| FALSE | High | QSPSPTLEE | [R].QSPSPT | 1xPhospho | B2RTL2 1x | 6.98E-10 | 0 | 1 | 4 | 7 | B2RTL2 | Phosphodi | B2RTL2 [67 | 0 | 2844.432 | 0 | -0.15 | 0.053388 | 6.14 | 106.9 | 93.1 | 4.79 | 9.19 | | High | High | High | High | High | High | High | High | High | High | High | 0 | 2.91E-11 | 4.06 |
| FALSE | High | QSPSQANG | [K].QSPSQ/ | 1xCarbami | Q91YD3 1x | 0.073462 | 0.005631 | 1 | 2 | 1 | Q91YD3 | mRNA-dec | Q91YD3 [1 | 0 | 2911.364 | 0 | -0.86 | 0.001627 | 2.08 | 129 | 71 | 9.63 | 2.58 | | High | High | High | High | High | High | High | High | High | High | High | 0.002896 | 0.03001 | 2.36 |
| FALSE | High | QSPSSSPTH | [K].QSPSSS | 3xPhospho | P49586 3x | 0.057686 | 0.004227 | 1 | 1 | 2 | P49586 | Choline-ph | P49586 [31 | 1 | 2439.993 | 0 | 1.37 | 0.003054 | 8.62 | 57.9 | 142.1 | 7.46 | 12.37 | | High | High | High | High | High | High | High | High | High | High | High | 0.002159 | 0.02277 | 2 |
| FALSE | High | QSPTLNLLF | [R].QSPTLN | 1xPhospho | Q80WC1-4 | 0.004696 | 0.000456 | 1 | 6 | 1 | Q80WC1-4 | Isoform 4 c | Q80WC1-4 | 0 | 1648.862 | 0 | 0 | 0.714392 | 23.73 | 93.1 | 106.9 | 14.66 | 9.79 | | High | High | High | High | High | High | High | High | High | High | High | 0.000204 | 0.001345 | 1.46 |
| FALSE | High | QSPVSSPTF | [K].QSPVSS | 1xPhospho | Q811Q9 1x | 0.060666 | 0.00451 | 1 | 2 | 1 | Q811Q9 | Choline-ph | Q811Q9 [3 | 0 | 1267.625 | 0 | 0.13 | 0.224055 | 5.42 | 99.6 | 100.4 | 11.77 | 7.34 | | High | High | High | High | High | High | High | High | High | High | High | 0.002354 | 0.02414 | 1.02 |
| FALSE | High | QSQAQSKF | [K].QSQAQ | 1xPhospho | A2A5R2 1x | 0.000155 | 0 | 1 | 3 | 4 | A2A5R2 | Brefeldin A | A2A5R2 [2 | 0 | 2197.183 | 0 | 0.28 | 0.014235 | 9 | 90.6 | 109.4 | 5.75 | 4.62 | | High | High | High | High | High | High | High | High | High | High | High | 0 | 2.93E-05 | 2.1 |
| FALSE | High | QSQIQNQ( | [K].QSQIQI | 1xPhospho | Q3U4X8 1x | 5.11E-08 | 0 | 1 | 4 | 9 | Q3U4X8 | DNA ligase | Q3U4X8 [9 | 0 | 2508.061 | 0 | -1.07 | 0.000887 | 2.15 | 134.2 | 65.8 | 5.86 | 5.49 | | High | High | High | High | High | High | High | High | High | High | High | 0 | 3.6E-09 | 5.4 |
| FALSE | High | QSQIQNQ( | [K].QSQIQI | 2xPhospho | Q3U4X8 2x | 1.11E-05 | 0 | 1 | 4 | 18 | Q3U4X8 | DNA ligase | Q3U4X8 [9 | 0 | 2588.028 | 0 | -0.78 | 0.002092 | 5.59 | 126.3 | 73.7 | 9.32 | 11.06 | | High | High | High | High | High | High | High | High | High | High | High | 0 | 1.53E-06 | 3.96 |
| FALSE | High | QSPNSSSA | [R].QSPNS | 1xPhospho | Q8K3X4 1x | 9.08E-07 | 0 | 1 | 1 | 3 | Q8K3X4 | Interferon | Q8K3X4 [1 | 0 | 1758.822 | 0 | -0.11 | 0.035723 | 1.88 | 104.6 | 95.4 | 3.05 | 3.59 | | High | High | High | High | High | High | High | High | High | High | High | 0 | 9.11E-08 | 2.47 |
| FALSE | High | QSQTSSSD | [R].QSQTS | 1xPhospho | Q99K46 1x | 0.033986 | 0.002195 | 1 | 4 | 1 | Q99K46 | Ubiquitin c | Q99K46 [8 | 0 | 3062.335 | 0 | -1.11 | 0.001517 | 1.17 | 135.8 | 64.2 | 9.82 | 5.02 | | High | High | High | High | High | High | High | High | High | High | High | 0.001151 | 0.01239 | 3.1 |
| FALSE | High | QSQVTEEE | [K].QSQVT | 1xPhospho | B2RRE0 1x | 3.7E-08 | 0 | 1 | 2 | 10 | B2RRE0 | A kinase (P | B2RRE0 [1 | 0 | 3508.57 | 0 | -1.24 | 0.005447 | 3.61 | 137 | 63 | 12.05 | 14.66 | | High | High | High | High | High | High | High | High | High | High | High | 0 | 2.21E-07 | 4.74 |
| FALSE | High | QSSADGSC | [R].QSSAD | 1xPhospho | Q8BX09 1x | 1.41E-06 | 0 | 1 | 4 | 1 | Q8BX09 | Retinoblast | Q8BX09 [4 | 0 | 1639.809 | 0 | -0.62 | 0.003243 | 3.44 | 121.2 | 78.8 | 9.59 | 3.9 | | High | High | High | High | High | High | High | High | High | High | High | 0 | 1.49E-07 | 2.41 |
| FALSE | High | QSSASGYQ | [R].QSSAS( | 1xPhospho | E9Q0S6 1x | 0.007274 | 0.00062 | 1 | 10 | 2 | E9Q0S6 | Tensin 1 O | E9Q0S6 [1 | 0 | 4327.007 | 0 | -0.63 | 0.04261 | 18.03 | 124.6 | 75.4 | 14.46 | 38.24 | | High | High | High | High | High | High | High | High | High | High | High | 0.000275 | 0.002192 | 4.18 |
| FALSE | High | QSSDFDTQ | [R].QSSDFI | 1xPhospho | A2AFQ0 1x | 0.000176 | 0 | 1 | 2 | 13 | A2AFQ0 | E3 ubiquiti | A2AFQ0 [1 | 0 | 4863.123 | 0 | 0.65 | 0.012008 | 24.33 | 76.6 | 123.4 | 13.06 | 10.6 | | High | High | High | High | High | High | High | High | High | High | High | 0.000689 | 0.006021 | 4.67 |
| FALSE | High | QSSEGSAN | [R].QSSEG | 1xPhospho | Q3TJN8 1x | 0.00086 | 0 | 1 | 2 | 2 | Q3TJN8 | Putative ur | Q3TJN8 [3 | 0 | 1445.704 | 0 | 0.14 | 0.120561 | 11.3 | 94.9 | 105.1 | 6.84 | 16.96 | | High | High | High | High | High | High | High | High | High | High | High | 0 | 0.000202 | 2.93 |
| FALSE | High | QSSFASLLN | [K].QSSFAS | 1xPhospho | F6WJB7 1x | 3.08E-07 | 0 | 1 | 5 | 3 | F6WJB7 | Ubiquitin c | F6WJB7 [5 | 0 | 2376.203 | 0 | 0.67 | 0.021771 | 26.47 | 78.6 | 121.4 | 6.65 | 13.11 | | High | High | High | High | High | High | High | High | High | High | High | 0 | 2.71E-08 | 3.52 |
| FALSE | High | QSSFEIPDD | [R].QSSFEI | 1xPhospho | P46938 1x | 1.56E-07 | 0 | 1 | 7 | 2 | P46938 | Transcripti | P46938 [14 | 0 | 2755.345 | 0 | 0.53 | 0.042841 | 23.26 | 80.3 | 119.7 | 11.64 | 24.05 | | High | High | High | High | High | High | High | High | High | High | High | 0 | 1.26E-08 | 3.59 |
| FALSE | High | QSSFEIPDD | [R].QSSFEI | 1xPhospho | P46938 1x | 4.56E-06 | 0 | 1 | 7 | 13 | P46938 | Transcripti | P46938 [14 | 0 | 2771.34 | 0 | 0.68 | 0.000626 | 5.85 | 77.8 | 122.2 | 9.3 | 2.61 | | High | High | High | High | High | High | High | High | High | High | High | 0 | 5.61E-07 | 3.33 |
| FALSE | High | QSSFPFNLI | [R].QSSFPF | 1xPhospho | Q7TNL5 1x | 0.004491 | 0.000409 | 1 | 4 | 2 | Q7TNL5 | Protein ph | Q7TNL5 [8 | 0 | 1719.887 | 0 | 1.32 | 0.000875 | 10.01 | 57.3 | 142.7 | 7.86 | 6.97 | | High | High | High | High | High | High | High | High | High | High | High | 0.000449 | 0.003817 | 2.85 |
| FALSE | High | QSSGASSS | [R].QSSGA | 1xPhospho | Q3TQX5 1x | 9.11E-05 | 0 | 1 | 3 | 12 | Q3TQX5 | Putative ur | Q3TQX5 [6 | 0 | 1670.722 | 0 | 0.82 | 0.004946 | 21.1 | 67.9 | 132.1 | 10.23 | 11.85 | | High | High | High | High | High | High | High | High | High | High | High | 0.000449 | 0.004034 | 2.6 |
| FALSE | High | QSSGEQSP | [R].QSSGE( | 1xPhospho | Q61122 1x | 0.009756 | 0.000736 | 1 | 3 | 2 | Q61122 | NGFI-A-bin | Q61122 [3 | 0 | 3179.444 | 0 | 1.28 | 0.000434 | 2.7 | 56.3 | 143.7 | 7.72 | 6.6 | | High | High | High | High | High | High | High | High | High | High | High | 0.000344 | 0.003049 | 2.22 |
| FALSE | High | QSSGEQSP | [R].QSSGE( | 1xDeamida | Q61122 1x | 0.000941 | 0 | 1 | 3 | 4 | Q61122 | NGFI-A-bin | Q61122 [3 | 0 | 3180.428 | 0 | 2.17 | 0.000209 | 7.06 | 35 | 165 | 19.6 | 6.44 | | High | High | High | High | High | High | High | High | High | High | High | 2.58E-05 | 0.000392 | 2.68 |
| FALSE | High | QSSGPGAS | [R].QSSGP( | 1xPhospho | F6ZFU0 1x | 2.76E-07 | 0 | 1 | 4 | 2 | F6ZFU0 | Elongation | F6ZFU0 [3 | 0 | 2304.082 | 0 | 0.11 | 0.303249 | 8.25 | 95.8 | 104.2 | 10.76 | 7.38 | | High | High | High | High | High | High | High | High | High | High | High | 0 | 2.4E-08 | 3.93 |
| FALSE | High | QSQTSSSD | [R].QSQTS | 1xPhospho | Q99K46 1x | 0.011755 | 0.000889 | 1 | 4 | 1 | Q99K46 | Ubiquitin c | Q99K46 [8 | 0 | 3078.33 | 0 | -1.1 | 0.003413 | 4.89 | 139.7 | 60.3 | 10.99 | 15.91 | | High | High | High | High | High | High | High | High | High | High | High | 0.000449 | 0.003769 | 2.96 |
| FALSE | High | QSSGSTTN | [R].QSSGS | 1xPhospho | B2RY22 1x | 6E-07 | 0 | 2 | 16 | 6 | B2RY22; A( | Clasp1 pro | B2RY22 [6 | 0 | 1990.892 | 0 | -0.6 | 0.016499 | 13.7 | 122.7 | 77.3 | 8.35 | 25.06 | Shared | High | High | High | High | High | High | High | High | High | High | High | 0 | 5.73E-08 | 2.58 |
| FALSE | High | QSPGPALA | [R].QSPGP/ | 1xPhospho | P42128 1x | 0.001255 | 0 | 1 | 2 | 2 | P42128 | Forkhead b | P42128 [8 | 0 | 1205.624 | 0 | 1.05 | 0.000628 | 3.7 | 65 | 135 | 6.03 | 4.63 | | High | High | High | High | High | High | High | High | High | High | High | 0.000161 | 0.00114 | 2.35 |
| FALSE | High | QSPGAQVF | [K].QSPGA( | 1xPhospho | Q3TSB2 1x | 2.1E-05 | 0 | 1 | 3 | 1 | Q3TSB2 | Putative ur | Q3TSB2 [1 | 0 | 2490.377 | 0 | 2.24 | 0.001622 | 31.27 | 33.2 | 166.8 | 9.47 | 16.38 | | High | High | High | High | High | High | High | High | High | High | High | 0 | 3.1E-06 | 3.02 |
| FALSE | High | QSEQPVKP | [R].QSEQP | 1xPhospho | P70399 1x | 0.000186 | 0 | 1 | 3 | 2 | P70399 | TP53-bindi | P70399 [1 | 0 | 3603.74 | 0 | 0.37 | 0.410233 | 47.03 | 88.4 | 111.6 | 12.34 | 22 | | High | High | High | High | High | High | High | High | High | High | High | 0 | 3.6E-05 | 3.74 |
| FALSE | High | QSESPDTK | [K].QSESPI | 1xPhospho | B7ZCJ1 1x | 0.001127 | 0 | 1 | 5 | 1 | B7ZCJ1 | Rho GTPas | B7ZCJ1 [19 | 0 | 1429.698 | 0 | 0 | 0.709695 | 1.8 | 99.6 | 100.4 | 3.96 | 4.79 | | High | High | High | High | High | High | High | High | High | High | High | 0 | 0.000272 | 1.8 |
| FALSE | High | QSFTMVAE | [K].QSFTM | 1xPhospho | Q543N3 1x | 0.001127 | 0 | 1 | 9 | 1 | Q543N3 | LIM and SH | Q543N3 [6 | 0 | 1917.898 | 0 | 0.01 | 0.997341 | 32.83 | 99.7 | 100.3 | 10.67 | 19.06 | | High | High | High | High | High | High | High | High | High | High | High | 0 | 0.000273 | 3.55 |
| FALSE | High | QSGEAFVE | [R].QSGEA | 1xPhospho | Q9Z2X1 1x | 1.79E-06 | 0 | 1 | 5 | 1 | Q9Z2X1 | Heterogen | Q9Z2X1 [5 | 0 | 2320.1 | 0 | 1.08 | 0.000486 | 2.84 | 64.4 | 135.6 | 4.1 | 7.89 | | High | High | High | High | High | High | High | High | High | High | High | 0 | 1.94E-07 | 2.73 |
| FALSE | High | QSGGLEPF | [K].QSGGL | 1xPhospho | Q8BMG1 1 | 1.16E-05 | 0 | 1 | 5 | 1 | Q8BMG1 | ATR-intera | Q8BMG1 [ | 0 | 1653.82 | 0 | -0.13 | 0.214175 | 11.57 | 104.2 | 95.8 | 8.01 | 4.17 | | High | High | High | High | High | High | High | High | High | High | High | 0 | 1.59E-06 | 2.26 |
| FALSE | High | QSGGSYLD | [R].QSGGS | 1xPhospho | Q9D361 1x | 0.000219 | 0 | 1 | 3 | 1 | Q9D361 | U11/U12 s | Q9D361 [2 | 0 | 1693.763 | 0 | -0.76 | 0.005275 | 2.25 | 125.6 | 74.4 | 7.38 | 12.07 | | High | High | High | High | High | High | High | High | High | High | High | 0 | 4.31E-05 | 2.79 |
| FALSE | High | QSGLLLEEE | [K].QSGLLL | 1xPhospho | O88271 1x | 3.35E-05 | 0 | 1 | 1 | 2 | O88271 | Craniofacia | O88271 [6 | 0 | 1984.988 | 0 | -0.43 | 0.046881 | 13.13 | 117 | 83 | 17.07 | 11.77 | | High | High | High | High | High | High | High | High | High | High | High | 0 | 5.27E-06 | 3.51 |
| FALSE | High | QSGTPGFL | [K].QSGTP | 1xPhospho | P83741-3 1 | 2.73E-06 | 0 | 1 | 7 | 1 | P83741-3 | Isoform 3 c | P83741-3 [ | 0 | 2132.156 | 0 | 1.17 | 0.000769 | 9.42 | 61.6 | 138.4 | 4.56 | 6.85 | | High | High | High | High | High | High | High | High | High | High | High | 0 | 3.15E-07 | 2.41 |
| FALSE | High | QSGTPPGN | [R].QSGTPI | 1xPhospho | E9QM90 1 | 5.12E-05 | 0 | 1 | 8 | 2 | E9QM90 | RIKEN cDN | E9QM90 [ | 0 | 2458.211 | 0 | 0.13 | 0.286408 | 29.29 | 98.8 | 101.2 | 4.26 | 21.32 | | High | High | High | High | High | High | High | High | High | High | High | 0 | 8.48E-06 | 2.6 |
| FALSE | High | QSGTPPGN | [R].QSGTPI | 1xPhospho | E9QM90 1 | 0.003997 | 0.000311 | 1 | 8 | 2 | E9QM90 | RIKEN cDN | E9QM90 [ | 0 | 2474.206 | 0 | 0.06 | 0.816636 | 27.01 | 96.2 | 103.8 | 21.94 | 10.78 | | High | High | High | High | High | High | High | High | High | High | High | 0.000246 | 0.001502 | 3.16 |
| FALSE | High | QSHPTSAA | [K].QSHPT | 1xPhospho | A2AJ72 1x | 8.74E-06 | 0 | 1 | 6 | 1 | A2AJ72 | Far upstre | A2AJ72 [52 | 0 | 3237.419 | 0 | -1.58 | 0.016766 | 5.27 | 149.8 | 50.2 | 25.04 | 41.25 | | High | High | High | High | High | High | High | High | High | High | High | 0 | 1.17E-06 | 3.79 |
| FALSE | High | QSHSESSP | [R].QSHSE | 1xPhospho | Q8BTI8 1x | 0.000215 | 0 | 1 | 3 | 12 | Q8BTI8 | Serine/arg | Q8BTI8 [8 | 0 | 1924.905 | 0 | -1.09 | 0.000718 | 1.34 | 136.1 | 63.9 | 6.69 | 4.51 | | High | High | High | High | High | High | High | High | High | High | High | 7.19E-05 | 0.000774 | 3.34 |
| FALSE | High | QSHSESSP | [R].QSHSE | 2xPhospho | Q8BTI8 2x | 0.001544 | 5.66E-05 | 1 | 3 | 1 | Q8BTI8 | Serine/arg | Q8BTI8 [8 | 0 | 2004.872 | 0 | -0.94 | 0.001651 | 2.38 | 131 | 69 | 7.57 | 4.55 | | High | High | High | High | High | High | High | High | High | High | High | 2.58E-05 | 0.000388 | 3.51 |
| FALSE | High | QSPGHQSF | [K].QSPGH | 2xPhospho | Q9D2D7 2x | 0.001898 | 5.66E-05 | 1 | 3 | 1 | Q9D2D7 | Zinc finger | Q9D2D7 [2 | 0 | 2034.982 | 0 | -0.42 | 0.055816 | 8.45 | 115.4 | 84.6 | 8.37 | 14.46 | | High | High | High | High | High | High | High | High | High | High | High | 2.58E-05 | 0.000489 | 2.86 |
| FALSE | High | QSHSPQPS | [K].QSHSP( | 1xPhospho | Q9ET80 1x | 0.005275 | 0.000599 | 1 | 1 | 1 | Q9ET80 | Junctophili | Q9ET80 [4 | 0 | 2110.961 | 0 | -1.29 | 0.002539 | 2.77 | 139.5 | 60.5 | 12.99 | 26.35 | | High | High | High | High | High | High | High | High | High | High | High | 0.000267 | 0.001528 | 2.61 |
| FALSE | High | QSLEGASN | [R].QSLEG/ | 1xPhospho | O08715-4 | 0.000193 | 0 | 1 | 6 | 1 | O08715-4 | Isoform 4 c | O08715-4 | 0 | 1367.652 | 0 | 0.95 | 0.001715 | 12.53 | 69.4 | 130.6 | 5.25 | 9.92 | | High | High | High | High | High | High | High | High | High | High | High | 0 | 3.76E-05 | 2.74 |
| FALSE | High | QSLEPDK | [K].QSLEPL | 1xPhospho | Q3TRR0 1x | 0.049971 | 0.003641 | 1 | 5 | 1 | Q3TRR0 | Microtubu | Q3TRR0 [4 | 0 | 1354.702 | 0 | 0.15 | 0.083731 | 13.96 | 92.9 | 107.1 | 9.64 | 4.46 | | High | High | High | High | High | High | High | High | High | High | High | 0.001902 | 0.01932 | 1.78 |
| FALSE | High | QSILPPQS | [R].QSILPP | 1xPhospho | Q99K30 1x | 0.000183 | 0 | 1 | 2 | 1 | Q99K30 | Epidermal | Q99K30 [1 | 0 | 2182.162 | 0 | 2.75 | 0.001763 | 7.88 | 28.5 | 171.5 | 11.07 | 16.16 | | High | High | High | High | High | High | High | High | High | High | High | 0 | 3.53E-05 | 2.83 |
| FALSE | High | QSIPSNDCS | [K].QSIPSN | 1xCarbami | A2APB8 1x | 0.013426 | 0.000976 | 1 | 3 | 2 | A2APB8 | Targeting p | A2APB8 [9 | 0 | 2059.977 | 0 | -0.31 | 0.000434 | 1.88 | 110.8 | 89.2 | 6.97 | 9.3 | | High | High | High | High | High | High | High | High | High | High | High | 0.005703 | 0.06265 | 2.29 |
| FALSE | High | QSIQKPPSF | [R].QSIQKF | 1xPhospho | Q9EQM5 1 | 0.003788 | 0.00021 | 1 | 2 | 1 | Q9EQM5 | Homeobox | Q9EQM5 [ | 0 | 2378.257 | 0 | -1.72 | 0.000222 | 1.6 | 156.7 | 43.3 | 5.74 | 14.12 | | High | High | High | High | High | High | High | High | High | High | High | 0.000117 | 0.001059 | 2.3 |
| FALSE | High | QSISFSGLF | [R].QSISFS | 1xPhospho | P22893 1x | 0.026885 | 0.001676 | 1 | 3 | 1 | P22893 | mRNA deca | P22893 [17 | 0 | 1544.767 | 0 | 0.22 | 0.009165 | 8.63 | 92.1 | 107.9 | 4.14 | 4.65 | | High | High | High | High | High | High | High | High | High | High | High | 0.000903 | 0.009502 | 1.88 |
| FALSE | High | QSLSSADN | [R].QSLSS/ | 1xPhospho | A2A5R2 1x | 1.96E-07 | 0 | 1 | 3 | 3 | A2A5R2 | Brefeldin A | A2A5R2 [3 | 0 | 2531.209 | 0 | -0.2 | 0.162043 | 14.73 | 109.6 | 90.4 | 6.2 | 16.69 | | High | High | High | High | High | High | High | High | High | High | High | 0 | 1.63E-08 | 4.58 |
| FALSE | High | QSLSSADN | [R].QSLSS/ | 2xPhospho | A2A5R2 2x | 0.000705 | 0 | 1 | 3 | 6 | A2A5R2 | Brefeldin A | A2A5R2 [3 | 0 | 2611.175 | 0 | -0.05 | 0.361792 | 8.34 | 101.5 | 98.5 | 6.65 | 7.65 | | High | High | High | High | High | High | High | High | High | High | High | 0 | 0.000162 | 5.34 |
| FALSE | High | QSLTSPDS( | [R].QSLTSF | 1xPhospho | E9PZP8 1x | 0.061176 | 0.004646 | 1 | 3 | 1 | E9PZP8 | HECT and F | E9PZP8 [27 | 0 | 2053.987 | 0 | -0.34 | 0.045143 | 11.39 | 110.7 | 89.3 | 5.45 | 9.51 | | High | High | High | High | High | High | High | High | High | High | High | 0.002354 | 0.02429 | 1.58 |
| FALSE | High | QSMAFSILI | [R].QSMAF | 1xPhospho | E9Q7G0 1x | 0.009416 | 0.000736 | 1 | 5 | 1 | E9Q7G0 | Nuclear mi | E9Q7G0 [2 | 0 | 1890.98 | 0 | 0.24 | 0.092683 | 8.34 | 86.8 | 113.2 | 14.27 | 16.37 | | High | High | High | High | High | High | High | High | High | High | High | 0.000327 | 0.002941 | 2.48 |
| FALSE | High | QSNASSDV | [K].QSNAS | 1xPhospho | Q99JF8 1x | 5.71E-09 | 0 | 1 | 4 | 9 | Q99JF8 | PC4 and SF | Q99JF8 [1 | 0 | 1959.931 | 0 | -0.29 | 0.004914 | 1.96 | 111 | 89 | 3.31 | 5.84 | | High | High | High | High | High | High | High | High | High | High | High | 0 | 3.08E-10 | 4.96 |
| FALSE | High | QSPAGISLC | [R].QSPAG | 1xPhospho | Q80V94 1x | 9.58E-10 | 0 | 1 | 1 | 1 | Q80V94 | AP-4 comp | Q80V94 [6 | 0 | 2627.333 | 0 | -0.25 | 0.123818 | 11.54 | 109.8 | 90.2 | 7.55 | 10.83 | | High | High | High | High | High | High | High | High | High | High | High | 0 | 4.13E-11 | 4.6 |
| FALSE | High | QSPDEFCH | [K].QSPDE | 1xCarbami | Q5DTY9 1x | 1.75E-05 | 0 | 1 | 1 | 1 | Q5DTY9 | BTB/POZ d | Q5DTY9 [1 | 0 | 2724.072 | 0 | 1.54 | 0.00151 | 8.7 | 51.1 | 148.9 | 25.57 | 4.83 | | High | High | High | High | High | High | High | High | High | High | High | 0 | 2.54E-06 | 4.32 |
| FALSE | High | QSLDISSEV | [R].QSLDIS | 1xPhospho | Q547N0 1x | 0.001222 | 0 | 1 | 2 | 4 | Q547N0 | BRMS1 OS | Q547N0 [1 | 0 | 2147.01 | 0 | -1.13 | 0.000623 | 1.65 | 137 | 63 | 8.22 | 7.16 | | High | High | High | High | High | High | High | High | High | High | High | 0.001011 | 0.01069 | 3.04 |
| FALSE | High | QSSLAEPVS | [K].QSSLAE | 1xPhospho | O88845 1x | 4.69E-06 | 0 | 1 | 4 | 2 | O88845 | A-kinase ar | O88845 [1 | 0 | 1767.929 | 0 | 0.42 | 0.000293 | 3.21 | 85 | 115 | 2.52 | 1.42 | | High | High | High | High | High | High | High | High | High | High | High | 0 | 3.41E-05 | 2.19 |
| FALSE | High | QSSIGNSPS | [R].QSSIGN | 1xPhospho | E9QM77 1 | 0.003558 | 0.00021 | 1 | 9 | 4 | E9QM77 | Ataxin-2 O | E9QM77 [ | 0 | 4475.248 | 0 | 0.81 | 0.021553 | 14.1 | 72.1 | 127.9 | 8.03 | 22.64 | | High | High | High | High | High | High | High | High | High | High | High | 0.000638 | 0.005782 | 4.97 |
| FALSE | High | QSSIGNSPS | [R].QSSIGN | 1xDeamida | E9QM77 1 | 1.2E-06 | 0 | 1 | 9 | 6 | E9QM77 | Ataxin-2 O | E9QM77 [ | 0 | 4492.227 | 0 | 2.55 | 0.003099 | 43.91 | 32.9 | 167.1 | 56.34 | 10.69 | | High | High | High | High | High | High | High | High | High | High | High | 0 | 3.44E-05 | 6.01 |
| FALSE | High | QTEEVTAL | [R].QTEEV | 1xPhospho | Q80XC6 1x | 0.000314 | 0 | 1 | 1 | 5 | Q80XC6 | Protein NR | Q80XC6 [4 | 0 | 2078 | 0 | -0.4 | 0.000527 | 2.24 | 113.6 | 86.4 | 1.64 | 3.55 | | High | High | High | High | High | High | High | High | High | High | High | 0 | 0.000185 | 3.48 |
| FALSE | High | QTEEVTAL | [R].QTEEV | 2xPhospho | Q80XC6 2x | 0.008167 | 0.000736 | 1 | 1 | 5 | Q80XC6 | Protein NR | Q80XC6 [4 | 0 | 2157.967 | 0 | -0.5 | 0.013301 | 7.93 | 117.2 | 82.8 | 4.85 | 12.48 | | High | High | High | High | High | High | High | High | High | High | High | 0.000327 | 0.002507 | 3.21 |
| FALSE | High | QTENAFSP | [K].QTENA | 1xPhospho | E9QA16 1x | 0.012397 | 0.000889 | 1 | 8 | 1 | E9QA16 | Caldesmon | E9QA16 [2 | 0 | 1445.662 | 0 | 0.1 | 0.094315 | 7.76 | 95.1 | 104.9 | 3.35 | 5.85 | | High | High | High | High | High | High | High | High | High | High | High | 0.000449 | 0.003973 | 2.08 |
| FALSE | High | QTEPSVNT | [R].QTEPS\ | 1xPhospho | V9GX48 1x | 0.101749 | 0.008194 | 1 | 3 | 1 | V9GX48 | M-phase p | V9GX48 [8 | 1 | 2068.048 | 0 | -0.1 | 0.583167 | 0.73 | 101.6 | 98.4 | 6.81 | 2.45 | | High | High | High | High | High | High | High | High | High | High | High | 0.004115 | 0.04391 | 1.99 |
| FALSE | High | QTESNPAN | [R].QTESN | 1xPhospho | Q61687 1x | 0.000941 | 0 | 1 | 4 | 2 | Q61687 | Transcripti | Q61687 [6 | 0 | 2695.163 | 0 | -0.31 | 0.044786 | 11.13 | 111.2 | 88.8 | 17.21 | 11.16 | | High | High | High | High | High | High | High | High | High | High | High | 0 | 0.000223 | 2.95 |
| FALSE | High | QTESNPAN | [R].QTESN | 1xDeamida | Q61687 1x | 0.000343 | 0 | 1 | 4 | 2 | Q61687 | Transcripti | Q61687 [6 | 0 | 2696.147 | 0 | -0.07 | 0.42419 | 25.64 | 103.2 | 96.8 | 22.55 | 13.97 | | High | High | High | High | High | High | High | High | High | High | High | 0 | 7.14E-05 | 2.99 |
| FALSE | High | QTESNPAN | [R].QTESN | 1xPhospho | Q61687 1x | 4.36E-09 | 0 | 1 | 4 | 7 | Q61687 | Transcripti | Q61687 [6 | 0 | 2711.158 | 0 | -0.27 | 0.041369 | 10.37 | 108 | 92 | 4.88 | 10.94 | | High | High | High | High | High | High | High | High | High | High | High | 0 | 2.27E-10 | 5.46 |
| FALSE | High | QTGYGWI( | [K].QTGYG | 2xCarbami | Q9D4H9-2 | 2.62E-08 | 0 | 1 | 2 | 2 | Q9D4H9-2 | Isoform 2 c | Q9D4H9-2 | 0 | 2531.098 | 0 | 0.14 | 0.737482 | 54.54 | 86 | 114 | 25.17 | 30 | | High | High | High | High | High | High | High | High | High | High | High | 0 | 1.7E-09 | 4.13 |
| FALSE | High | QTHNHSPT | [R].QTHNH | 1xPhospho | Q8R149 1x | 0.056248 | 0.004124 | 1 | 2 | 1 | Q8R149 | BUD13 hor | Q8R149 [2 | 0 | 1793.839 | 0 | 0.31 | 0.883446 | 6.03 | 93.7 | 106.3 | 54.02 | 26.11 | | High | High | High | High | High | High | High | High | High | High | High | 0.003265 | 0.03444 | 1.91 |
| FALSE | High | QTHTAVAS | [R].QTHTA | 3xPhospho | G3UYY4 3x | 0.074071 | 0.0057 | 1 | 1 | 1 | G3UYY4 | Tetratricop | G3UYY4 [8 | 0 | 2905.287 | 0 | 0.01 | 0.445825 | 25.77 | 99.2 | 100.8 | 23.82 | 87.06 | | High | High | High | High | High | High | High | High | High | High | High | 0.002924 | 0.03027 | 2.09 |
| FALSE | High | QTHTGERF | [K].QTHTG | 1xCarbami | A2ART0 1x | 0.009931 | 0.000736 | 2 | 11 | 1 | A2ART0; A | Uncharacte | A2ART0 [5 | 0 | 1800.757 | 0 | -1.57 | 0.000222 | 0.31 | 149.3 | 50.7 | 7.24 | 8.43 | Shared | High | High | High | High | High | High | High | High | High | High | High | 0.000344 | 0.003123 | 2.89 |
| FALSE | High | QTLINIPAL | [K].QTLINI | 1xPhospho | Q9Z0U1 1x | 1.53E-06 | 0 | 1 | 1 | 7 | Q9Z0U1 | Tight junct | Q9Z0U1 [3 | 0 | 3210.525 | 0 | 0.29 | 0.172635 | 16.21 | 97.2 | 102.8 | 8.71 | 14.4 | | High | High | High | High | High | High | High | High | High | High | High | 0 | 1.65E-07 | 4.16 |
| FALSE | High | QTLINIPAL | [K].QTLINI | 2xPhospho | Q9Z0U1 2x | 1.1E-05 | 0 | 1 | 1 | 5 | Q9Z0U1 | Tight junct | Q9Z0U1 [3 | 0 | 3290.492 | 0 | 0.64 | 0.030039 | 26.53 | 77.3 | 122.7 | 12.78 | 17.4 | | High | High | High | High | High | High | High | High | High | High | High | 0 | 1.51E-06 | 3.97 |
| FALSE | High | QTEEGVRS | [R].QTEEG | 1xPhospho | G3UW85 1x | 0.017966 | 0.001282 | 1 | 1 | 1 | G3UW85 | Enhancer c | G3UW85 [ | 1 | 3497.884 | 0 | -1.15 | 0.003323 | 3.13 | 137.7 | 62.3 | 13.6 | 9.55 | | High | High | High | High | High | High | High | High | High | High | High | 0.000689 | 0.006062 | 3.03 |
| FALSE | High | QTLPSSPT | [K].QTLPSS | 1xPhospho | Q9JKN6 1x | 0.009087 | 0.000736 | 1 | 2 | 5 | Q9JKN6 | RNA-bindir | Q9JKN6 [1 | 0 | 1698.908 | 0 | 0.2 | 0.003313 | 3.75 | 93.1 | 106.9 | 8.31 | 6.81 | | High | High | High | High | High | High | High | High | High | High | High | 0.001844 | 0.0187 | 2.34 |
| FALSE | High | QTNVQDQ | [K].QTNVQ | 1xCarbami | Q8C080 1x | 3.22E-07 | 0 | 1 | 3 | 2 | Q8C080 | Sorting nex | Q8C080 [5 | 0 | 2666.146 | 0 | 0.17 | 0.105224 | 14.89 | 98.5 | 101.5 | 21.24 | 10.35 | | High | High | High | High | High | High | High | High | High | High | High | 0 | 2.84E-08 | 3.9 |
| FALSE | High | QTPPASPSI | [K].QTPPA | 1xPhospho | Q60598 1x | 0.004955 | 0.000504 | 1 | 3 | 4 | Q60598 | Src substra | Q60598 [4 | 0 | 3657.824 | 0 | 0.17 | 0.031252 | 6.27 | 93.5 | 106.5 | 5.93 | 6.04 | | High | High | High | High | High | High | High | High | High | High | High | 0.000225 | 0.001435 | 2.61 |
| FALSE | High | QTPPASPSI | [K].QTPPA | 2xPhospho | Q60598 2x | 0.001767 | 5.66E-05 | 1 | 3 | 4 | Q60598 | Src substra | Q60598 [4 | 0 | 3737.791 | 0 | 0.38 | 0.044299 | 21.27 | 84.8 | 115.2 | 13.18 | 11.69 | | High | High | High | High | High | High | High | High | High | High | High | 2.58E-05 | 0.000452 | 4.02 |
| FALSE | High | QTPPLAQP | [R].QTPPL/ | 1xPhospho | Q80VY9 1x | 0.002058 | 5.66E-05 | 1 | 3 | 5 | Q80VY9 | Putative AT | Q80VY9 [4 | 0 | 1821.91 | 0 | -0.07 | 0.762626 | 25.43 | 102.3 | 97.7 | 5.59 | 15.03 | | High | High | High | High | High | High | High | High | High | High | High | 0.000399 | 0.003478 | 2.38 |
| FALSE | High | QTPQAPQS | [R].QTPQA | 1xPhospho | F6ZDS4 1x | 0.000656 | 0 | 1 | 5 | 7 | F6ZDS4 | Nucleoproi | F6ZDS4 [2 | 0 | 1418.699 | 0 | -0.54 | 0.000492 | 1.41 | 119.6 | 80.4 | 3.15 | 3.51 | | High | High | High | High | High | High | High | High | High | High | High | 4.94E-05 | 0.00066 | 1.82 |
| FALSE | High | QTPQVNIN | [R].QTPQV | 1xPhospho | P47810 1x | 4.39E-05 | 0 | 1 | 1 | 2 | P47810 | Wee1-like | P47810 [22 | 0 | 2726.386 | 0 | -0.25 | 0.23733 | 18.81 | 109.8 | 90.2 | 23.02 | 18.85 | | High | High | High | High | High | High | High | High | High | High | High | 0 | 7.11E-06 | 4.06 |
| FALSE | High | QTPTPPTH | [R].QTPTPI | 1xPhospho | B2RWS6 1 | 0.002058 | 5.66E-05 | 1 | 1 | 3 | B2RWS6 | Histone ac | B2RWS6 [8 | 0 | 3608.806 | 0 | 0.26 | 0.054644 | 23.66 | 92.8 | 107.2 | 9.7 | 11.62 | | High | High | High | High | High | High | High | High | High | High | High | 7.19E-05 | 0.00086 | 3.08 |
| FALSE | High | QTPTPPTH | [R].QTPTPI | 2xPhospho | B2RWS6 2 | 5.93E-06 | 0 | 1 | 1 | 1 | B2RWS6 | Histone ac | B2RWS6 [8 | 0 | 3688.772 | 0 | 0.27 | 0.219919 | 26.07 | 90.8 | 109.2 | 14.74 | 8.28 | | High | High | High | High | High | High | High | High | High | High | High | 0 | 7.52E-07 | 3.79 |
| FALSE | High | QTPTSPCQ | [K].QTPTSF | 1xCarbami | G3X932 1x | 0.020862 | 0.001472 | 1 | 4 | 1 | G3X932 | DNA segm | G3X932 [3 | 0 | 1771.807 | 0 | 0.08 | 0.183045 | 6.85 | 96 | 104 | 1.65 | 8.83 | | High | High | High | High | High | High | High | High | High | High | High | 0.000786 | 0.007118 | 1.77 |
| FALSE | High | QTQSASST | [R].QTQSA | 1xPhospho | A0A0A6YW | 0.004824 | 0.000456 | 1 | 7 | 1 | A0A0A6YW | Mitogen-a | A0A0A6YW | 0 | 1716.893 | 0 | -0.82 | 0.022823 | 8.72 | 127.8 | 72.2 | 9.2 | 28.82 | | High | High | High | High | High | High | High | High | High | High | High | 0.000204 | 0.001395 | 2.87 |
| FALSE | High | QTRSSGSE | [R].QTRSS( | 1xPhospho | Q55VQ0 3x | 2.08E-06 | 0 | 1 | 5 | 3 | Q55VQ0 | Histone ac | Q55VQ0 [9 | 1 | 2496.988 | 0 | -1.39 | 0.000236 | 1.02 | 144.9 | 55.1 | 2.99 | 6.71 | | High | High | High | High | High | High | High | High | High | High | High | 0 | 2.31E-07 | 5.49 |
| FALSE | High | QTSDSSVG | [R].QTSDS | 1xPhospho | Q6PA06 1x | 9.98E-05 | 0 | 1 | 2 | 3 | Q6PA06 | Atlastin-2 | Q6PA06 [2 | 0 | 4313.917 | 0 | 0.01 | 0.715609 | 30.56 | 99.7 | 100.3 | 6.59 | 28.95 | | High | High | High | High | High | High | High | High | High | High | High | 0 | 1.93E-05 | 4.56 |
| FALSE | High | QTSLETSIL | [R].QTSLET | 1xCarbami | Q9EP53 1x | 1.26E-05 | 0 | 1 | 4 | 2 | Q9EP53 | Hamartin C | Q9EP53 [5 | 0 | 2200.134 | 0 | -0.9 | 0.001907 | 3.67 | 128.5 | 71.5 | 6.45 | 9.54 | | High | High | High | High | High | High | High | High | High | High | High | 0 | 1.9E-05 | 3.18 |
| FALSE | High | QTNGYLNG | [K].QTNGY | 2xDeamida | Q543C2 1x | 0.025738 | 0.001645 | 1 | 3 | 2 | Q543C2 | Uridine-cy | Q543C2 [2 | 0 | 1241.794 | 0 | -0.49 | 0.125104 | 10.14 | 112.7 | 87.3 | 13.63 | 17.14 | | High | High | High | High | High | High | High | High | High | High | High | 0.000903 | 0.009268 | 2.74 |
| FALSE | High | QTDPSTPQ | [K].QTDPS | 1xPhospho | Q69ZA1 1x | 3.9E-05 | 0 | 1 | 2 | 4 | Q69ZA1 | Cyclin-dep | Q69ZA1 [1 | 0 | 1970.947 | 0 | 0.03 | 0.660287 | 7.84 | 99.1 | 100.9 | 3.29 | 5.67 | | High | High | High | High | High | High | High | High | High | High | High | 0 | 6.25E-06 | 3.69 |
| FALSE | High | QTASPSDG | [K].QTASP | 1xPhospho | A3KFM7 1 | 9.71E-05 | 0 | 1 | 2 | 3 | A3KFM7 | Chromodo | A3KFM7 [1 | 0 | 2134.011 | 0 | -0.11 | 0.074641 | 3.83 | 103.8 | 96.2 | 4.09 | 2.81 | | High | High | High | High | High | High | High | High | High | High | High | 0 | 1.74E-05 | 3.68 |
| FALSE | High | QSWPSPAL | [R].QSWPS | 1xPhospho | Q6P1H6 1x | 0.011549 | 0.00086 | 1 | 3 | 1 | Q6P1H6 | Ankyrin rep | Q6P1H6 [8 | 0 | 1551.834 | 0 | 0.21 | 0.253366 | 9.79 | 92.7 | 107.3 | 3.6 | 11.38 | | High | High | High | High | High | High | High | High | High | High | High | 0.000437 | 0.003699 | 2.56 |
| FALSE | High | QSSLPGTG | [R].QSSLP( | 1xPhospho | E9QM73 1 | 0.002004 | 5.66E-05 | 1 | 16 | 3 | E9QM73 | Serine/thre | E9QM73 [ | 0 | 2088.078 | 0 | 0.37 | 0.022006 | 9.44 | 87.3 | 112.7 | 13.06 | 14.11 | | High | High | High | High | High | High | High | High | High | High | High | 7.19E-05 | 0.000898 | 2.67 |
| FALSE | High | QSSLSEK | [R].QSSLSE | 1xPhospho | E9Q0J5 1x | 0.077188 | 0.005893 | 1 | 3 | 1 | E9Q0J5 | Kinesin-like | E9Q0J5 [1 | 0 | 1316.686 | 0 | 2.82 | 0.000605 | 16.29 | 25.3 | 174.7 | 11.93 | 10.5 | | High | High | High | High | High | High | High | High | High | High | High | 0.003064 | 0.03185 | 1.9 |
| FALSE | High | QSSLTFQS | [K].QSSLTF | 1xPhospho | Q3UMF0 1 | 4.1E-06 | 0 | 1 | 6 | 4 | Q3UMF0 | Cordon-ble | Q3UMF0 [ | 0 | 2026.943 | 0 | 0.17 | 0.212831 | 22.14 | 91.6 | 108.4 | 7.41 | 12.68 | | High | High | High | High | High | High | High | High | High | High | High | 0 | 4.95E-07 | 3.93 |
| FALSE | High | QSSPAMPH | [R].QSSPAI | 1xPhospho | B2RY17 1x | 0.047086 | 0.00354 | 1 | 7 | 1 | B2RY17 | Tnik protei | B2RY17 [5 | 0 | 1536.765 | 0 | -1.75 | 0.000285 | 1.29 | 156.5 | 43.5 | 10.76 | 21.02 | | High | High | High | High | High | High | High | High | High | High | High | 0.001812 | 0.01803 | 2.39 |
| FALSE | High | QSSPESPVS | [K].QSSPES | 1xPhospho | B1AUW6 1 | 0.001097 | 0 | 1 | 5 | 1 | B1AUW6 | Interleukin | B1AUW6 [ | 0 | 1680.816 | 0 | 1.13 | 0.003981 | 19.81 | 63.6 | 136.4 | 7.07 | 11.4 | | High | High | High | High | High | High | High | High | High | High | High | 0 | 0.000266 | 2.64 |
| FALSE | High | QSSPFLDD | [R].QSSPFL | 1xCarbami | Q3UQS8 1 | 0.000529 | 0 | 1 | 3 | 3 | Q3UQS8 | RNA-bindir | Q3UQS8 [1 | 0 | 1734.817 | 0 | 3.59 | 0.00051 | 8.64 | 15 | 185 | 12.37 | 11.87 | | High | High | High | High | High | High | High | High | High | High | High | 0 | 0.000116 | 3.09 |
| FALSE | High | QSSPSCVP | [R].QSSPS( | 1xCarbami | E9PX30 1x | 5.7E-08 | 0 | 1 | 5 | 3 | E9PX30 | Mitogen-a | E9PX30 [4 | 0 | 2902.391 | 0 | 1.68 | 0.000347 | 15.6 | 48.8 | 151.2 | 12.99 | 4.19 | | High | High | High | High | High | High | High | High | High | High | High | 0 | 7.77E-09 | 4.25 |
| FALSE | High | QSSPSCVP | [R].QSSPS( | 1xCarbami | E9PX30 1x | 2.34E-06 | 0 | 1 | 5 | 6 | E9PX30 | Mitogen-a | E9PX30 [4 | 0 | 2903.375 | 0 | 1.49 | 0.000325 | 5.64 | 52.4 | 147.6 | 10.85 | 4.45 | | High | High | High | High | High | High | High | High | High | High | High | 0 | 2.64E-07 | 4.15 |
| FALSE | High | QSSQDNEV | [K].QSSQD | 1xPhospho | A2AR02 1x | 0.001659 | 5.66E-05 | 1 | 2 | 1 | A2AR02 | Peptidyl-pr | A2AR02 [6 | 0 | 1572.767 | 0 | -0.44 | 0.000968 | 4.21 | 115 | 85 | 0.95 | 5.01 | | High | High | High | High | High | High | High | High | High | High | High | 2.58E-05 | 0.000422 | 2.18 |
| FALSE | High | QSSSEISLA | [K].QSSSEI | 1xPhospho | Q7TSC1 1x | 0.000674 | 0 | 1 | 6 | 3 | Q7TSC1 | Protein PR | Q7TSC1 [4 | 0 | 1614.794 | 0 | 0.44 | 0.027488 | 10.68 | 85.4 | 114.6 | 8.69 | 12.42 | | High | High | High | High | High | High | High | High | High | High | High | 0 | 0.000308 | 2.34 |
| FALSE | High | QSSSGPAS( | [K].QSSSG | 1xPhospho | P19426 1x | 2.47E-05 | 0 | 1 | 3 | 2 | P19426 | Negative e | P19426 [3 | 0 | 1727.873 | 0 | 0.95 | 0.004171 | 9.66 | 68.4 | 131.6 | 11.75 | 6.76 | | High | High | High | High | High | High | High | High | High | High | High | 0 | 9.35E-06 | 3.41 |
| FALSE | High | QSSSPYEDI | [K].QSSPY | 1xPhospho | Q8BTI8 1x | 0.008388 | 0.000736 | 1 | 3 | 8 | Q8BTI8 | Serine/arg | Q8BTI8 [3 | 0 | 1578.745 | 0 | -0.59 | 0.00146 | 2.33 | 120.4 | 79.6 | 3.9 | 5.48 | | High | High | High | High | High | High | High | High | High | High | High | 0.000756 | 0.006611 | 3.21 |
| FALSE | High | QSSSSDTDI | [R].QSSSD | 1xPhospho | Q7TPS5 1x | 0.000116 | 0 | 1 | 4 | 2 | Q7TPS5 | C2 domain | Q7TPS5 [3 | 0 | 2003.994 | 0 | 0.62 | 0.013448 | 19.5 | 78.8 | 121.2 | 15.56 | 9.55 | | High | High | High | High | High | High | High | High | High | High | High | 0 | 2.12E-05 | 2.95 |
| FALSE | High | QSSSSDTDI | [R].QSSSD | 2xPhospho | Q7TPS5 2x | 0.001087 | 0 | 1 | 4 | 1 | Q7TPS5 | C2 domain | Q7TPS5 [3 | 0 | 2083.96 | 0 | -0.96 | 0.009227 | 9.06 | 131.2 | 68.8 | 14.71 | 10.51 | | High | High | High | High | High | High | High | High | High | High | High | 0 | 0.000261 | 3.25 |

| | | | | | | | | | | | | | | | | | | | | | | | | | | | | | | | | | | | | | | |
|---|---|---|---|---|---|---|---|---|---|---|---|---|---|---|---|---|---|---|---|---|---|---|---|---|---|---|---|---|---|---|---|---|---|---|---|---|---|---|
| FALSE | High | QSSSYEDP | [R].QSSSYE | 1xPhospho | O54916 1x | 3.7E-05 | 0 | 1 | 8 | 1 | O54916 | RalBP1-ass | O54916 [2 | 0 | 1764.825 | 0 | 0.63 | 0.005078 | 16.25 | 80.9 | 119.1 | 10.82 | 5.64 | | High | High | High | High | High | High | High | High | High | High | High | 0 | 5.88E-06 | 2.47 |
| FALSE | High | QSSTADAP | [R].QSSTAI | 1xPhospho | Q7TSH2 1x | 0.008538 | 0.000736 | 1 | 3 | 5 | Q7TSH2 | Phosphory | Q7TSH2 [6 | 0 | 2934.429 | 0 | -0.45 | 0.003023 | 6.09 | 115.8 | 84.2 | 2.15 | 8.51 | | High | High | High | High | High | High | High | High | High | High | High | 0.000327 | 0.00262 | 2.72 |
| FALSE | High | QSSTDSEL | [R].QSSTD | 1xPhospho | Q80TM6 1 | 0.000793 | 0 | 1 | 7 | 2 | Q80TM6 | R3H domai | Q80TM6 [3 | 0 | 1532.761 | 0 | 1.01 | 0.004081 | 17.59 | 66.4 | 133.6 | 5.49 | 16.09 | | High | High | High | High | High | High | High | High | High | High | High | 0 | 0.000185 | 3.04 |
| FALSE | High | QSSTFGGA | [R].QSSTF | 1xPhospho | Q8BXW8 1 | 5.29E-07 | 0 | 1 | 2 | 2 | Q8BXW8 | Putative ur | Q8BXW8 [ | 0 | 2521.201 | 0 | 0.76 | 0.003277 | 6.33 | 72.7 | 127.3 | 7.19 | 8.15 | | High | High | High | High | High | High | High | High | High | High | High | 0 | 4.97E-08 | 4.03 |
| FALSE | High | QSSVSSTA | [R].QSSVS | 1xPhospho | Q80TH2-2 | 7.01E-08 | 0 | 1 | 6 | 2 | Q80TH2-2 | Isoform 2 c | Q80TH2-2 | 0 | 2014.964 | 0 | 0.82 | 0.003624 | 1.58 | 77.8 | 122.2 | 12.66 | 9.32 | | High | High | High | High | High | High | High | High | High | High | High | 0 | 9.47E-09 | 4.21 |
| FALSE | High | QSSVTQVT | [K].QSSVT | 1xPhospho | E9QML5 1 | 0.01319 | 0.000889 | 1 | 16 | 1 | E9QML5 | Zinc finger | E9QML5 [1 | 0 | 1727.841 | 0 | 0.15 | 0.982656 | 60.73 | 97.9 | 102.1 | 46.51 | 55.26 | | High | High | High | High | High | High | High | High | High | High | High | 0.000467 | 0.004277 | 2.62 |
| FALSE | High | QSSVTQVT | [K].QSSVT | 1xPhospho | E9QML5 1 | 3.1E-06 | 0 | 1 | 16 | 2 | E9QML5 | Zinc finger | E9QML5 [1 | 0 | 1957.004 | 0 | -0.11 | 0.066034 | 6.38 | 103.1 | 96.9 | 5.68 | 6.39 | | High | High | High | High | High | High | High | High | High | High | High | 0 | 3.61E-07 | 4.03 |
| FALSE | High | QSTESSPV | [R].QSTESS | 1xPhospho | G3UWZ0 1 | 0.004654 | 0.000456 | 1 | 4 | 4 | G3UWZ0 | Bromodom | G3UWZ0 [ | 0 | 1623.794 | 0 | 1.44 | 0.000609 | 12.08 | 54 | 146 | 7.68 | 9.11 | | High | High | High | High | High | High | High | High | High | High | High | 0.000204 | 0.001339 | 2.64 |
| FALSE | High | QSTSFLVL | [K].QSTSFL | 1xPhospho | O70400 1x | 0.000331 | 0 | 1 | 2 | 3 | O70400 | PDZ and LI | O70400 [2 | 0 | 2432.273 | 0 | 0.52 | 0.032971 | 30.19 | 78.6 | 121.4 | 4.33 | 17.73 | | High | High | High | High | High | High | High | High | High | High | High | 0 | 0.00011 | 2.81 |
| FALSE | High | QSVDGK | [K].QSVDG | 1xPhospho | Q64152 1x | 0.063788 | 0.004664 | 1 | 2 | 1 | Q64152 | Transcripti | Q64152 [1 | 0 | 1171.612 | 0 | 0.01 | 0.810372 | 33.71 | 100.3 | 99.7 | 7.97 | 17.06 | | High | High | High | High | High | High | High | High | High | High | High | 0.002403 | 0.02548 | 1.28 |
| FALSE | High | QSVFGASS | [K].QSVFG | 1xPhospho | P70227 1x | 1.55E-08 | 0 | 1 | 1 | 1 | P70227 | Inositol 1,4 | P70227 [93 | 0 | 2423.217 | 0 | 0.99 | 0.000791 | 8.2 | 67.1 | 132.9 | 8.72 | 10.16 | | High | High | High | High | High | High | High | High | High | High | High | 0 | 9.42E-10 | 3.76 |
| FALSE | High | QSVGSGSP | [K].QSVGS | 1xPhospho | B2RUG9 1x | 0.000239 | 0 | 1 | 6 | 1 | B2RUG9 | Adenomat | B2RUG9 [2 | 0 | 2011.006 | 0 | -0.03 | 0.774718 | 7.97 | 100.9 | 99.1 | 4.04 | 7.29 | | High | High | High | High | High | High | High | High | High | High | High | 0 | 4.78E-05 | 3.89 |
| FALSE | High | QSVIHFSP | [R].QSVIHF | 1xPhospho | Q7TNB8 1x | 0.01894 | 0.001362 | 1 | 2 | 1 | Q7TNB8 | Protein str | Q7TNB8 [1 | 0 | 1765.851 | 0 | 0.62 | 0.002809 | 14.03 | 78.1 | 121.9 | 7.39 | 5.21 | | High | High | High | High | High | High | High | High | High | High | High | 0.00074 | 0.006439 | 2 |
| FALSE | High | QSVNHQM | [R].QSVNH | 1xPhospho | Q3UYL7 1x | 0.000976 | 0 | 1 | 5 | 2 | Q3UYL7 | Annexin O | Q3UYL7 [3 | 0 | 1966.864 | 0 | 1.61 | 0.001184 | 25.47 | 49.4 | 150.6 | 14.82 | 7.91 | | High | High | High | High | High | High | High | High | High | High | High | 0 | 0.000231 | 3.5 |
| FALSE | High | QSWASSSS | [R].QSWAS | 1xCarbami | B2RUJ6 1x | 0.000123 | 0 | 1 | 8 | 2 | B2RUJ6 | Rap guanin | B2RUJ6 [14 | 0 | 2842.301 | 0 | 1.54 | 0.011821 | 10.06 | 62.3 | 137.7 | 21.58 | 19.05 | | High | High | High | High | High | High | High | High | High | High | High | 0 | 7.83E-05 | 3.59 |
| FALSE | High | QSELSAEE | [K].QSELSA | 1xPhospho | Q7TQK5 1x | 1.75E-05 | 0 | 1 | 5 | 3 | Q7TQK5 | Coiled-coil | Q7TQK5 [2 | 0 | 1871.904 | 0 | -0.39 | 0.002045 | 3.24 | 113.4 | 86.6 | 2.9 | 5.18 | | High | High | High | High | High | High | High | High | High | High | High | 0 | 2.53E-06 | 2.77 |
| FALSE | High | QSEKPPSP | [K].QSEKP | 1xCarbami | A2ADB0 1x | 0.000876 | 0 | 1 | 2 | 3 | A2ADB0 | Msx2-inter | A2ADB0 [2 | 0 | 2769.377 | 0 | 0.49 | 0.002816 | 1.8 | 82.7 | 117.3 | 6.23 | 10.24 | | High | High | High | High | High | High | High | High | High | High | High | 0 | 0.000206 | 2.89 |
| FALSE | High | QSDSVK | [K].QSDSV | 1xPhospho | A2ARQ4 1x | 0.01894 | 0.001362 | 1 | 5 | 2 | A2ARQ4 | Small EDRK | A2ARQ4 [1 | 0 | 1201.623 | 0 | 0.39 | 0.082811 | 27.82 | 84.5 | 115.5 | 21.16 | 17.88 | | High | High | High | High | High | High | High | High | High | High | High | 0.000723 | 0.006411 | 1.58 |
| FALSE | High | QSDGQSSS | [K].QSDGQ | 1xPhospho | B2RUG8 1x | 2.55E-08 | 0 | 1 | 3 | 4 | B2RUG8 | Asxl2 prote | B2RUG8 [2 | 0 | 2683.225 | 0 | 0.43 | 0.062556 | 12.94 | 87.5 | 112.5 | 7.24 | 10.29 | | High | High | High | High | High | High | High | High | High | High | High | 0 | 5.62E-07 | 4.52 |
| FALSE | High | QQEAPSNL | [R].QQEAP | 1xPhospho | P04627 1x | 6.3E-05 | 0 | 1 | 4 | 2 | P04627 | Serine/thre | P04627 [16 | 0 | 3175.562 | 0 | 2.26 | 0.001663 | 21.8 | 31.6 | 168.4 | 13.4 | 17.45 | | High | High | High | High | High | High | High | High | High | High | High | 0.000786 | 0.007013 | 3.66 |
| FALSE | High | QQEEEAAE | [R].QQEEE | 1xPhospho | Q8BQ30-2 | 0.005467 | 0.00062 | 1 | 5 | 1 | Q8BQ30-2 | Isoform 2 c | Q8BQ30-2 | 0 | 4139.995 | 0 | 1.17 | 0.0119 | 2.95 | 67.9 | 132.1 | 9.95 | 19.43 | | High | High | High | High | High | High | High | High | High | High | High | 0.000275 | 0.0016 | 2.62 |
| FALSE | High | QQEEEAAE | [R].QQEEE | 1xPhospho | Q8BQ30-2 | 0.012288 | 0.000889 | 1 | 5 | 4 | Q8BQ30-2 | Isoform 2 c | Q8BQ30-2 | 0 | 4155.99 | 0 | 0.92 | 0.004052 | 15.91 | 71.8 | 128.2 | 7.39 | 9.4 | | High | High | High | High | High | High | High | High | High | High | High | 0.003461 | 0.03758 | 3.54 |
| FALSE | High | QQEHSAAE | [K].QQEHS | 2xCarbami | Q9QWF0 1 | 1.12E-05 | 0 | 1 | 2 | 3 | Q9QWF0 | Chromatin | Q9QWF0 [ | 0 | 4230.907 | 0 | 0.09 | 0.997926 | 21.96 | 95.6 | 104.4 | 8.86 | 21.42 | | High | High | High | High | High | High | High | High | High | High | High | 0 | 1.54E-06 | 2.87 |
| FALSE | High | QQEHSAAE | [K].QQEHS | 2xCarbami | Q9QWF0 1 | 0.000645 | 0 | 1 | 2 | 4 | Q9QWF0 | Chromatin | Q9QWF0 [ | 0 | 4231.891 | 0 | -0.05 | 0.714594 | 24.61 | 100.7 | 99.3 | 9.39 | 16.63 | | High | High | High | High | High | High | High | High | High | High | High | 0 | 0.000146 | 4.44 |
| FALSE | High | QQEHSAAE | [K].QQEHS | 2xCarbami | Q9QWF0 1 | 8.56E-05 | 0 | 1 | 2 | 3 | Q9QWF0 | Chromatin | Q9QWF0 [ | 0 | 4246.902 | 0 | -0.06 | 0.836943 | 2.56 | 99.2 | 100.8 | 8.61 | 7.26 | | High | High | High | High | High | High | High | High | High | High | High | 0 | 1.5E-05 | 3.61 |
| FALSE | High | QQEVLVSL | [R].QQEVL | 1xCarbami | S4R189 1x | 0.000142 | 0 | 1 | 6 | 2 | S4R189 | Rho guanir | S4R189 [40 | 0 | 3092.563 | 0 | -0.57 | 0.050745 | 3.84 | 120.6 | 79.4 | 25.23 | 3.2 | | High | High | High | High | High | High | High | High | High | High | High | 0 | 2.67E-05 | 3.89 |
| FALSE | High | QQFSPLSD | [R].QQFSP | 1xPhospho | Q69ZR9-2 | 3.45E-06 | 0 | 1 | 2 | 3 | Q69ZR9-2 | Isoform 2 c | Q69ZR9-2 | 0 | 2823.313 | 0 | -0.85 | 0.00202 | 0.83 | 129.3 | 70.7 | 7.63 | 4.66 | | High | High | High | High | High | High | High | High | High | High | High | 0 | 2.08E-06 | 4.44 |
| FALSE | High | QQFSPLSD | [R].QQFSP | 1xDeamida | Q69ZR9-2 | 5.65E-05 | 0 | 1 | 2 | 3 | Q69ZR9-2 | Isoform 2 c | Q69ZR9-2 | 0 | 2824.297 | 0 | -0.78 | 0.002895 | 1.35 | 125.9 | 74.1 | 14.07 | 5.68 | | High | High | High | High | High | High | High | High | High | High | High | 0 | 2.01E-05 | 4.29 |
| FALSE | High | QQFSPLSD | [R].QQFSP | 2xPhospho | Q69ZR9-2 | 2.08E-05 | 0 | 1 | 2 | 3 | Q69ZR9-2 | Isoform 2 c | Q69ZR9-2 | 0 | 2903.279 | 0 | -0.13 | 0.09961 | 12.3 | 103.4 | 96.6 | 3.23 | 10.06 | | High | High | High | High | High | High | High | High | High | High | High | 0 | 6.98E-06 | 4.62 |
| FALSE | High | QQGLDDAI | [K].QQGLD | 1xPhospho | Q9QYB5 1x | 5.03E-05 | 0 | 1 | 2 | 4 | Q9QYB5 | Gamma-ad | Q9QYB5 [5 | 0 | 3995.83 | 0 | -0.12 | 0.265397 | 14.06 | 104.9 | 95.1 | 12.18 | 7.7 | | High | High | High | High | High | High | High | High | High | High | High | 0 | 8.28E-06 | 4.03 |
| FALSE | High | QQGPPPAF | [K].QQGPP | 1xPhospho | Q0VGY8 1x | 1.36E-05 | 0 | 1 | 4 | 4 | Q0VGY8 | Protein TA | Q0VGY8 [1 | 0 | 2973.388 | 0 | -0.32 | 0.000833 | 1.86 | 111.2 | 88.8 | 1.65 | 2.56 | | High | High | High | High | High | High | High | High | High | High | High | 0 | 1.46E-05 | 4.02 |
| FALSE | High | QQGSPLSR | [R].QQGSP | 1xPhospho | E9PVA6 1x | 0.007209 | 0.00062 | 1 | 9 | 2 | E9PVA6 | ARF GTPas | E9PVA6 [3 | 0 | 1181.588 | 0 | -0.1 | 0.269248 | 7.53 | 104.4 | 95.6 | 5.64 | 5.19 | | High | High | High | High | High | High | High | High | High | High | High | 0.000275 | 0.002182 | 1.41 |
| FALSE | High | QQDSALTP | [R].QQDSA | 1xDeamida | Q60953 1x | 0.000454 | 0 | 2 | 5 | 11 | Q60953; Q | Protein PM | Q60953 [2 | 0 | 4995.239 | 0 | 0.29 | 0.831269 | 17.18 | 92.5 | 107.5 | 11.5 | 27.56 | Shared | High | High | High | High | High | High | High | High | High | High | High | 0.000275 | 0.001762 | 3.95 |
| FALSE | High | QQGVDSLS | [K].QQGVD | 1xPhospho | Q9CU65 1x | 3.3E-06 | 0 | 1 | 3 | 2 | Q9CU65 | Zinc finger | Q9CU65 [2 | 0 | 2064.114 | 0 | -0.9 | 0.001414 | 2.45 | 132.8 | 67.2 | 9.8 | 3.02 | | High | High | High | High | High | High | High | High | High | High | High | 0 | 3.89E-07 | 2.73 |
| FALSE | High | QQHEQLNI | [R].QQHEC | 1xPhospho | B1ARU1 1x | 0.000123 | 0 | 2 | 13 | 1 | B1ARU1; F | Microtubu | B1ARU1 [3 | 0 | 3612.752 | 0 | 0.56 | 0.015986 | 5.16 | 80.1 | 119.9 | 3.51 | 16.05 | Shared | High | High | High | High | High | High | High | High | High | High | High | 0 | 2.26E-05 | 4.18 |
| FALSE | High | QQHVISTE | [K].QQHVI | 2xPhospho | J3QNK5 2x | 3.94E-05 | 0 | 1 | 4 | 1 | J3QNK5 | Histone de | J3QNK5 [8 | 0 | 3142.304 | 0 | -0.32 | 0.083761 | 5.11 | 111.5 | 88.5 | 6.16 | 26.67 | | High | High | High | High | High | High | High | High | High | High | High | 0 | 6.29E-06 | 4.8 |
| FALSE | High | QQIENGSG | [K].QQIEN | 1xCarbami | Q9CQJ4 1x | 0.002793 | 0.000109 | 1 | 3 | 5 | Q9CQJ4 | E3 ubiquiti | Q9CQJ4 [1 | 0 | 4053.715 | 0 | 0.87 | 0.05217 | 20.07 | 70.2 | 129.8 | 20.37 | 14.5 | | High | High | High | High | High | High | High | High | High | High | High | 4.94E-05 | 0.000755 | 4.2 |
| FALSE | High | QQIENGSG | [K].QQIEN | 1xCarbami | Q9CQJ4 2x | 0.000538 | 0 | 1 | 3 | 2 | Q9CQJ4 | E3 ubiquiti | Q9CQJ4 [1 | 0 | 4132.698 | 0 | -0.54 | 0.022234 | 8.57 | 118 | 82 | 14.2 | 6.96 | | High | High | High | High | High | High | High | High | High | High | High | 0.001011 | 0.01087 | 2.95 |
| FALSE | High | QQIIHSGH | [R].QQIIHS | 1xPhospho | G5E8D8 1x | 0.010853 | 0.000772 | 1 | 4 | 2 | G5E8D8 | MLX intera | G5E8D8 [7 | 0 | 2186.053 | 0 | 0.42 | 0.039465 | 15.21 | 87.6 | 112.4 | 9.13 | 15.64 | | High | High | High | High | High | High | High | High | High | High | High | 0.000399 | 0.003436 | 3.2 |
| FALSE | High | QQLPDCIV | [K].QQLPD | 1xCarbami | Q6ZPR5-4 | 8.15E-07 | 0 | 1 | 9 | 2 | Q6ZPR5-4 | Isoform 4 c | Q6ZPR5-4 | 0 | 2517.299 | 0 | -0.13 | 0.105965 | 7.81 | 104.8 | 95.2 | 7.16 | 7.97 | | High | High | High | High | High | High | High | High | High | High | High | 0 | 8.84E-08 | 2.93 |
| FALSE | High | QQLQELGA | [R].QQLQE | 1xPhospho | B2RPS6 1x | 1.22E-07 | 0 | 1 | 3 | 2 | B2RPS6 | Putative ur | B2RPS6 [3 | 0 | 2262.19 | 0 | -0.11 | 0.127974 | 2.57 | 103.2 | 96.8 | 1.11 | 4.82 | | High | High | High | High | High | High | High | High | High | High | High | 0 | 9.01E-08 | 3.49 |
| FALSE | High | QQLSAEEL | [K].QQLSA | 1xPhospho | O08583 1x | 3.54E-05 | 0 | 1 | 3 | 1 | O08583 | THO comp | O08583 [2 | 0 | 2344.102 | 0 | 0.14 | 0.950133 | 8.8 | 99.3 | 100.7 | 17.79 | 4.76 | | High | High | High | High | High | High | High | High | High | High | High | 0 | 5.59E-06 | 4.42 |
| FALSE | High | QQLSEEEK | [R].QQLSE | 1xPhospho | Q5XJE5 1x | 0.027597 | 0.001716 | 1 | 2 | 1 | Q5XJE5 | RNA polym | Q5XJE5 [2 | 0 | 1528.766 | 0 | -0.54 | 0.00557 | 7.72 | 121 | 79 | 10.71 | 10.03 | | High | High | High | High | High | High | High | High | High | High | High | 0.000953 | 0.009828 | 1.76 |
| FALSE | High | QQISGPELI | [R].QQISG | 1xPhospho | Q9WTV7 1 | 0.016021 | 0.001124 | 1 | 1 | 2 | Q9WTV7 | E3 ubiquiti | Q9WTV7 [ | 0 | 1506.788 | 0 | 1.66 | 0.000181 | 4.62 | 47.2 | 152.8 | 5.59 | 3.78 | | High | High | High | High | High | High | High | High | High | High | High | 0.000585 | 0.005313 | 2.02 |
| FALSE | High | QQLSPSPG | [K].QQLSP | 1xPhospho | Q7TN98 1x | 6.41E-08 | 0 | 1 | 8 | 3 | Q7TN98 | Cytoplasmi | Q7TN98 [9 | 0 | 2547.311 | 0 | 2.22 | 0.000125 | 10.48 | 35.3 | 164.7 | 11.15 | 3.99 | | High | High | High | High | High | High | High | High | High | High | High | 0 | 5.61E-08 | 5.29 |
| FALSE | High | QQITEDPE | [K].QQITE | 1xPhospho | Q0PD32 1x | 0.012397 | 0.000889 | 1 | 2 | 2 | Q0PD32 | Rab23 OS= | Q0PD32 [1 | 0 | 2367.123 | 0 | -0.42 | 0.002838 | 4.3 | 115.1 | 84.9 | 7.41 | 9.69 | | High | High | High | High | High | High | High | High | High | High | High | 0.000638 | 0.005663 | 2.36 |
| FALSE | High | QQMDAYE | [K].QQMD | 1xPhospho | Q921I6 1x | 0.000297 | 0 | 1 | 1 | 1 | Q921I6 | SH3 domai | Q921I6 [86 | 0 | 1686.714 | 0 | -0.82 | 0.001138 | 5.99 | 126.7 | 73.3 | 6.07 | 17.21 | | High | High | High | High | High | High | High | High | High | High | High | 0 | 6.12E-05 | 2.9 |
| FALSE | High | QQHEQLNI | [R].QQHEC | 1xPhospho | B1ARU1 1x | 7.47E-08 | 0 | 2 | 13 | 2 | B1ARU1; F | Microtubu | B1ARU1 [3 | 0 | 3596.757 | 0 | 1.17 | 0.006573 | 13.02 | 64.1 | 135.9 | 21.74 | 17.36 | Shared | High | High | High | High | High | High | High | High | High | High | High | 0 | 5.52E-09 | 4.97 |
| FALSE | High | QQDSALTP | [R].QQDSA | 1xPhospho | Q60953 1x | 0.007274 | 0.00062 | 2 | 5 | 8 | Q60953; Q | Protein PM | Q60953 [2 | 0 | 4994.255 | 0 | 0.09 | 0.859807 | 21.78 | 98.8 | 101.2 | 9.11 | 17.91 | Shared | High | High | High | High | High | High | High | High | High | High | High | 0.000275 | 0.002204 | 4.52 |
| FALSE | High | QQDPSPGS | [R].QQDPS | 1xPhospho | Q08024 1x | 2.03E-06 | 0 | 1 | 3 | 2 | Q08024 | Core-bindi | Q08024 [1 | 0 | 2223.069 | 0 | 1.69 | 0.00014 | 2 | 47.7 | 152.3 | 4.48 | 2.94 | | High | High | High | High | High | High | High | High | High | High | High | 0 | 4.62E-06 | 3.4 |
| FALSE | High | QQATQGW | [R].QQATC | 1xPhospho | A0A1S6GW | 0.000628 | 0 | 1 | 2 | 1 | A0A1S6GW | Uncharacte | A0A1S6GW | 0 | 2952.458 | 0 | 1 | 0.000789 | 13.68 | 64.4 | 135.6 | 12.91 | 10.33 | | High | High | High | High | High | High | High | High | High | High | High | 0 | 0.000142 | 2.79 |
| FALSE | High | QPPMFSPL | [K].QPPMF | 1xPhospho | B2RR82 1x | 0.015061 | 0.001049 | 1 | 7 | 1 | B2RR82 | Intersectin | B2RR82 [1 | 0 | 1668.838 | 0 | 1.04 | 0.004627 | 16.74 | 63.1 | 136.9 | 15.11 | 12.79 | | High | High | High | High | High | High | High | High | High | High | High | 0.000571 | 0.004971 | 1.21 |
| FALSE | High | QPPPLAPQ | [K].QPPPL | 1xPhospho | P46938 1x | 0.000188 | 0 | 1 | 6 | 1 | P46938 | Transcripti | P46938 [26 | 0 | 4251.163 | 0 | 0.34 | 0.09948 | 6.23 | 88.4 | 111.6 | 6.19 | 17.26 | | High | High | High | High | High | High | High | High | High | High | High | 0 | 3.64E-05 | 3.83 |
| FALSE | High | QPPTAAGR | [K].QPPTA | 1xPhospho | Q3TWG5 1 | 9.28E-05 | 0 | 1 | 2 | 1 | Q3TWG5 | Dynein cyt | Q3TWG5 [ | 0 | 1828.927 | 0 | 0.22 | 0.001562 | 4.49 | 92.8 | 107.2 | 5.24 | 2.31 | | High | High | High | High | High | High | High | High | High | High | High | 0 | 1.65E-05 | 1.23 |
| FALSE | High | QPPTASSG | [R].QPPTA | 1xPhospho | Q2TBA3 1x | 2.9E-05 | 0 | 1 | 2 | 2 | Q2TBA3 | Mucosa-as | Q2TBA3 [1 | 0 | 2303.123 | 0 | 0.12 | 0.277103 | 17.26 | 92.9 | 107.1 | 12.13 | 7.78 | | High | High | High | High | High | High | High | High | High | High | High | 0 | 4.48E-06 | 4.18 |
| FALSE | High | QPPVSPGT | [K].QPPVS | 1xPhospho | Q3TE85 1x | 2.47E-05 | 0 | 1 | 2 | 3 | Q3TE85 | Putative ur | Q3TE85 [3 | 0 | 2004.093 | 0 | -0.32 | 0.063249 | 16.18 | 111.5 | 88.5 | 6.54 | 12.56 | | High | High | High | High | High | High | High | High | High | High | High | 0 | 3.73E-06 | 3.89 |
| FALSE | High | QPPVSPLT | [R].QPPVS | 2xPhospho | Q91X84 2x | 1.18E-06 | 0 | 1 | 2 | 2 | Q91X84 | CREB-regu | Q91X84 [3 | 0 | 2591.226 | 0 | 2.11 | 0.000464 | 21.62 | 40.1 | 159.9 | 17.02 | 7.02 | | High | High | High | High | High | High | High | High | High | High | High | 0 | 1.23E-07 | 2.93 |
| FALSE | High | QPSDSSIDK | [R].QPSDS | 1xPhospho | G3X9H5 1x | 0.000967 | 0 | 1 | 3 | 1 | G3X9H5 | Huntingtin | G3X9H5 [6 | 0 | 1514.75 | 0 | 1.91 | 0.000191 | 4.74 | 41.7 | 158.3 | 2.93 | 4.17 | | High | High | High | High | High | High | High | High | High | High | High | 0 | 0.00023 | 1.76 |
| FALSE | High | QPSDTSET | [R].QPSDT | 1xPhospho | Q68FM7 1 | 9.42E-07 | 0 | 1 | 5 | 4 | Q68FM7 | Rho guanir | Q68FM7 [3 | 0 | 1827.869 | 0 | 0.17 | 0.047284 | 8.03 | 96.4 | 103.6 | 5.17 | 3.66 | | High | High | High | High | High | High | High | High | High | High | High | 0 | 9.5E-08 | 3.98 |
| FALSE | High | QPSEEEIIK | [R].QPSEEE | 1xPhospho | Q62048 1x | 0.001463 | 5.66E-05 | 1 | 2 | 2 | Q62048 | Astrocytic | Q62048 [1 | 0 | 1610.844 | 0 | -0.33 | 0.039105 | 9.88 | 116.1 | 83.9 | 12.68 | 23.44 | | High | High | High | High | High | High | High | High | High | High | High | 2.58E-05 | 0.000511 | 2.41 |
| FALSE | High | QPSGAGLC | [R].QPSGA | 1xCarbami | Q8K400 1x | 0.017344 | 0.001204 | 1 | 3 | 1 | Q8K400 | Syntaxin-b | Q8K400 [6 | 0 | 2410.116 | 0 | 1.56 | 0.004895 | 30.19 | 52.9 | 147.1 | 23.26 | 10.5 | | High | High | High | High | High | High | High | High | High | High | High | 0.000638 | 0.005784 | 2.86 |
| FALSE | High | QPSIELPSN | [R].QPSIEL | 1xPhospho | P19973 1x | 0.075304 | 0.005727 | 1 | 4 | 1 | P19973 | Lymphocyt | P19973 [24 | 0 | 2113.102 | 0 | 0.95 | 0.060516 | 33.06 | 78.2 | 121.8 | 16.07 | 41.49 | | High | High | High | High | High | High | High | High | High | High | High | 0.002976 | 0.031 | 2.05 |
| FALSE | High | QPSLSAACI | [R].QPSLSA | 1xCarbami | Q8R146 1x | 0.001516 | 5.66E-05 | 1 | 4 | 1 | Q8R146 | Acylamino- | Q8R146 [2 | 0 | 2677.289 | 0 | 0.18 | 0.151404 | 10.02 | 94.5 | 105.5 | 12.88 | 13.93 | | High | High | High | High | High | High | High | High | High | High | High | 2.58E-05 | 0.000382 | 2.66 |
| FALSE | High | QPSMSEAN | [R].QPSMS | 1xCarbami | Q3U0S4 1x | 5.12E-05 | 0 | 1 | 4 | 2 | Q3U0S4 | Putative ur | Q3U0S4 [8 | 0 | 2294.104 | 0 | -0.48 | 0.020033 | 1.52 | 117.6 | 82.4 | 5.81 | 20.41 | | High | High | High | High | High | High | High | High | High | High | High | 0 | 1.24E-05 | 4.32 |
| FALSE | High | QPSMSEAN | [R].QPSMS | 1xCarbami | Q3U0S4 1x | 2.22E-06 | 0 | 1 | 4 | 2 | Q3U0S4 | Putative ur | Q3U0S4 [8 | 0 | 2310.099 | 0 | -0.55 | 0.005934 | 5.33 | 115.6 | 84.4 | 10.55 | 7.16 | | High | High | High | High | High | High | High | High | High | High | High | 0 | 2.48E-07 | 2.87 |
| FALSE | High | QPSPLLLLP | [R].QPSPLL | 1xPhospho | Q924A2 1x | 0.005135 | 0.000551 | 1 | 1 | 1 | Q924A2 | Protein cap | Q924A2 [7 | 0 | 2618.416 | 0 | 0.6 | 0.01235 | 12.74 | 87.9 | 112.1 | 9.38 | 11.74 | | High | High | High | High | High | High | High | High | High | High | High | 0.000246 | 0.001491 | 3.28 |
| FALSE | High | QPSPVVPA | [R].QPSPV | 1xPhospho | B2RX88 1x | 0.001267 | 0 | 1 | 5 | 1 | B2RX88 | Centrosom | B2RX88 [8 | 0 | 1816.013 | 0 | -0.45 | 0.00943 | 4.96 | 117.7 | 82.3 | 7.59 | 5.49 | | High | High | High | High | High | High | High | High | High | High | High | 0 | 0.000312 | 2.61 |
| FALSE | High | QPSQADTC | [R].QPSQA | 1xPhospho | E9QJU7 1x | 5.73E-10 | 0 | 1 | 7 | 1 | E9QJU7 | SH3 domai | E9QJU7 [4 | 0 | 2581.087 | 0 | 2.73 | 0.000637 | 16.48 | 24.2 | 175.8 | 13.41 | 11.47 | | High | High | High | High | High | High | High | High | High | High | High | 0 | 2.32E-11 | 4.88 |
| FALSE | High | QPSQASPL | [R].QPSQA | 1xPhospho | Q5ND43 1 | 0.025514 | 0.001645 | 1 | 2 | 2 | Q5ND43 | Putative ph | Q5ND43 [2 | 0 | 2873.547 | 0 | 0.28 | 0.083937 | 22.86 | 90.2 | 109.8 | 10.19 | 12.97 | | High | High | High | High | High | High | High | High | High | High | High | 0.000903 | 0.009333 | 3.22 |
| FALSE | High | QPSSPSHN | [R].QPSSPS | 1xPhospho | A0A1W2P6 | 0.000164 | 0 | 1 | 11 | 4 | A0A1W2P6 | Bromodom | A0A1W2P6 | 0 | 1646.785 | 0 | -0.99 | 0.028488 | 7.31 | 136.3 | 63.7 | 19.83 | 24.62 | | High | High | High | High | High | High | High | High | High | High | High | 0.000204 | 0.001367 | 1.26 |
| FALSE | High | QPTPPFFG | [R].QPTPP | 1xPhospho | Q8C2Q3 1 | 0.003028 | 0.00016 | 1 | 5 | 3 | Q8C2Q3 | RNA-bindir | Q8C2Q3 [2 | 0 | 1355.671 | 0 | -0.07 | 0.400577 | 10.3 | 101.8 | 98.2 | 2.78 | 6 | | High | High | High | High | High | High | High | High | High | High | High | 0.008786 | 0.09401 | 1.8 |
| FALSE | High | QPVIGDQN | [K].QPVIG | 1xPhospho | Q61687 1x | 0.000147 | 0 | 1 | 4 | 3 | Q61687 | Transcripti | Q61687 [7 | 0 | 2597.293 | 0 | -0.3 | 0.080109 | 17.38 | 111.8 | 88.2 | 10.13 | 15.26 | | High | High | High | High | High | High | High | High | High | High | High | 0.000449 | 0.003864 | 3.28 |
| FALSE | High | QPVIGDQN | [K].QPVIG | 1xPhospho | Q61687 1x | 1.91E-05 | 0 | 1 | 4 | 21 | Q61687 | Transcripti | Q61687 [7 | 0 | 2613.288 | 0 | -0.33 | 0.133173 | 6.86 | 113.3 | 86.7 | 4.66 | 16.56 | | High | High | High | High | High | High | High | High | High | High | High | 0 | 0.000134 | 3.58 |
| FALSE | High | QPVTDPLL | [K].QPVTD | 1xPhospho | Q3UYD0 1 | 0.000237 | 0 | 1 | 3 | 2 | Q3UYD0 | Putative ur | Q3UYD0 [1 | 0 | 1975.092 | 0 | -0.3 | 0.044092 | 11.83 | 116.1 | 83.9 | 18.7 | 25.2 | | High | High | High | High | High | High | High | High | High | High | High | 0 | 4.74E-05 | 3.65 |
| FALSE | High | QPYNAVFS | [R].QPYNA | 1xPhospho | Q80YR6 1x | 0.004106 | 0.00036 | 1 | 1 | 1 | Q80YR6 | DNA endor | Q80YR6 [8 | 0 | 1688.881 | 0 | -0.42 | 0.02512 | 9.97 | 111.8 | 88.2 | 9.28 | 20.27 | | High | High | High | High | High | High | High | High | High | High | High | 0.000161 | 0.00116 | 2.16 |
| FALSE | High | QQAAFYG | [R].QQAAF | 1xPhospho | A2AJ72 1x | 3.3E-06 | 0 | 1 | 3 | 1 | A2AJ72 | Far upstrea | A2AJ72 [55 | 0 | 2500.146 | 0 | -0.91 | 0.000473 | 1.23 | 130.6 | 69.4 | 4.71 | 3.8 | | High | High | High | High | High | High | High | High | High | High | High | 0 | 3.87E-07 | 3.7 |
| FALSE | High | QQAAYYAC | [R].QQAAY | 1xPhospho | A0A0G2JG | 0.000375 | 0 | 1 | 4 | 4 | A0A0G2JG | Far upstrea | A0A0G2JG | 0 | 2890.318 | 0 | -1.72 | 0.001836 | 1.78 | 156.7 | 43.3 | 13.76 | 22.23 | | High | High | High | High | High | High | High | High | High | High | High | 0.001916 | 0.01938 | 3.37 |
| FALSE | High | QQAAYYAC | [R].QQAAY | 1xPhospho | A0A0G2JG | 1.43E-07 | 0 | 1 | 4 | 25 | A0A0G2JG | Far upstrea | A0A0G2JG | 0 | 2906.313 | 0 | -1.56 | 0.000238 | 0.48 | 149.4 | 50.6 | 6.96 | 6.3 | | High | High | High | High | High | High | High | High | High | High | High | 0 | 3.45E-05 | 3.8 |
| FALSE | High | QQASRPNS | [R].QQASR | 1xPhospho | Q6VGS5-2 | 0.017652 | 0.001282 | 1 | 3 | 1 | Q6VGS5-2 | Isoform 2 c | Q6VGS5-2 | 0 | 1768.854 | 0 | -0.93 | 0.004952 | 7.07 | 131.1 | 68.9 | 8.73 | 14.84 | | High | High | High | High | High | High | High | High | High | High | High | 0.000689 | 0.005922 | 2.37 |
| FALSE | High | QQASSDSP | [K].QQASS | 1xCarbami | P97412 1x | 1.78E-05 | 0 | 1 | 4 | 2 | P97412 | Lysosomal- | P97412 [8 | 0 | 1644.725 | 0 | 1.09 | 0.000878 | 8.95 | 65 | 135 | 9.13 | 5.25 | | High | High | High | High | High | High | High | High | High | High | High | 0 | 2.58E-06 | 1.92 |
| FALSE | High | QQNSNPSL | [K].QQNSN | 1xPhospho | B2RUP0 1x | 8.4E-05 | 0 | 1 | 3 | 2 | B2RUP0 | RNA bindir | B2RUP0 [3 | 0 | 1553.809 | 0 | 0.1 | 0.176618 | 12.35 | 89.9 | 110.1 | 10.5 | 11.24 | | High | High | High | High | High | High | High | High | High | High | High | 0 | 1.48E-05 | 3.06 |
| FALSE | High | QTSITQTIR | [R].QTSITQ | 1xPhospho | Q6PIX5 1x | 0.021989 | 0.001542 | 1 | 4 | 1 | Q6PIX5 | Inactive rh | Q6PIX5 [74 | 0 | 1356.709 | 0 | -0.05 | 0.629898 | 10.38 | 100.9 | 99.1 | 4.77 | 9.22 | | High | High | High | High | High | High | High | High | High | High | High | 0.000814 | 0.007593 | 1.53 |
| FALSE | High | QQNSPLAK | [K].QQNSP | 1xPhospho | Q9JM52-3 | 0.030898 | 0.001896 | 1 | 9 | 1 | Q9JM52-3 | Isoform 3 c | Q9JM52-3 | 0 | 1423.771 | 0 | 1.67 | 0.00034 | 11.69 | 48.4 | 151.6 | 4.82 | 6.06 | | High | High | High | High | High | High | High | High | High | High | High | 0.001044 | 0.01118 | 1.61 |
| FALSE | High | QQPAAQQ | [K].QQPAA | 1xPhospho | P97329 1x | 0.005666 | 0.00062 | 1 | 1 | 1 | P97329 | Kinesin-like | P97329 [6 | 0 | 1862.969 | 0 | -0.49 | 0.006889 | 5.36 | 118.5 | 81.5 | 4.59 | 10.27 | | High | High | High | High | High | High | High | High | High | High | High | 0.000275 | 0.001667 | 2.3 |
| FALSE | High | QQTQQVQ | [R].QQTQC | 1xPhospho | E9PW37 1 | 0.004106 | 0.00036 | 1 | 3 | 1 | E9PW37 | RAS protei | E9PW37 [1 | 0 | 2525.19 | 0 | 0.11 | 0.155644 | 18.46 | 96.6 | 103.4 | 5.71 | 12.04 | | High | High | High | High | High | High | High | High | High | High | High | 0.000161 | 0.001166 | 3.55 |
| FALSE | High | QQTQQVQ | [R].QQTQC | 1xPhospho | E9PW37 1 | 1.87E-07 | 0 | 1 | 3 | 1 | E9PW37 | RAS protei | E9PW37 [1 | 0 | 2541.185 | 0 | 0.98 | 0.008759 | 14.35 | 67.3 | 132.7 | 16.1 | 13.2 | | High | High | High | High | High | High | High | High | High | High | High | 0 | 1.55E-08 | 5.1 |
| FALSE | High | QQTQTDN | [K].QQTQT | 2xPhospho | A2A8R7 2x | 9.11E-08 | 0 | 1 | 6 | 3 | A2A8R7 | Terminal u | A2A8R7 [2 | 0 | 3458.477 | 0 | -1.24 | 0.002388 | 2.16 | 139.1 | 60.9 | 6.07 | 17.47 | | High | High | High | High | High | High | High | High | High | High | High | 0 | 6.91E-09 | 5.2 |
| FALSE | High | QQVPITPV | [K].QQVPI | 1xPhospho | E4NKG5 1x | 8.25E-05 | 0 | 1 | 1 | 1 | E4NKG5 | MCG11736 | E4NKG5 [3 | 0 | 2716.504 | 0 | -1.37 | 0.000931 | 2.05 | 145.2 | 54.8 | 6.22 | 12.81 | | High | High | High | High | High | High | High | High | High | High | High | 0 | 1.44E-05 | 2.07 |
| FALSE | High | QQVSWEQ | [K].QQVSW | 1xPhospho | Q05AH6 1x | 9.16E-07 | 0 | 1 | 2 | 4 | Q05AH6 | Uncharacte | Q05AH6 [7 | 0 | 2471.155 | 0 | -0.9 | 0.000561 | 2.37 | 130.1 | 69.9 | 4.21 | 11.32 | | High | High | High | High | High | High | High | High | High | High | High | 0 | 9.23E-08 | 3.43 |
| FALSE | High | QQYSDQD | [R].QQYSD | 1xPhospho | Q9Z0U1 1x | 4.17E-06 | 0 | 1 | 1 | 8 | Q9Z0U1 | Tight junct | Q9Z0U1 [4 | 0 | 2254.007 | 0 | 0.16 | 0.013256 | 5.75 | 93.6 | 106.4 | 2.24 | 3.43 | | High | High | High | High | High | High | High | High | High | High | High | 0 | 5.04E-07 | 2.89 |
| FALSE | High | QQYSDQD | [R].QQYSD | 2xPhospho | Q9Z0U1 2x | 0.000442 | 0 | 1 | 1 | 5 | Q9Z0U1 | Tight junct | Q9Z0U1 [4 | 0 | 2333.973 | 0 | 0 | 0.921401 | 9.05 | 98.8 | 101.2 | 4.64 | 6.7 | | High | High | High | High | High | High | High | High | High | High | High | 0 | 9.5E-05 | 2.78 |
| FALSE | High | QRESSNQI | [K].QRESSN | 1xPhospho | Q3V300 1x | 0.011549 | 0.00086 | 1 | 1 | 2 | Q3V300 | Kinesin-like | Q3V300 [5 | 1 | 1982.084 | 0 | -0.36 | 0.223927 | 13.44 | 110.4 | 89.6 | 10.66 | 14.6 | | High | High | High | High | High | High | High | High | High | High | High | 0.000814 | 0.007812 | 2.4 |
| FALSE | High | QRGSETDT | [R].QRGSE | 1xPhospho | Q3TLH4 1x | 0.001386 | 5.66E-05 | 1 | 5 | 3 | Q3TLH4 | Protein PR | Q3TLH4 [1 | 1 | 2616.183 | 0 | -1.65 | 0.005741 | 0.92 | 141.1 | 58.9 | 19.32 | 12.9 | | High | High | High | High | High | High | High | High | High | High | High | 2.58E-05 | 0.000344 | 2.91 |
| FALSE | High | QRGSETGS | [R].QRGSE | 2xPhospho | Q7TSC1 2x | 0.0026 | 0.000109 | 1 | 3 | 1 | Q7TSC1 | Protein PR | Q7TSC1 [1 | 1 | 2749.212 | 0 | -0.33 | 0.012893 | 6.4 | 109.8 | 90.2 | 6.83 | 3.58 | | High | High | High | High | High | High | High | High | High | High | High | 4.94E-05 | 0.000696 | 1.41 |
| FALSE | High | QRLSFEGA | [R].QRLSFE | 1xPhospho | Q9CXS4 1x | 0.026421 | 0.001676 | 1 | 2 | 1 | Q9CXS4 | Centromer | Q9CXS4 [1 | 1 | 1644.887 | 0 | 0.2 | 0.102027 | 16.63 | 90.3 | 109.7 | 4.6 | 10.36 | | High | High | High | High | High | High | High | High | High | High | High | 0.000903 | 0.009364 | 1.01 |
| FALSE | High | QRLSPTSN | [R].QRLSPT | 1xPhospho | Q9Z0U1 1x | 0.051694 | 0.003745 | 1 | 1 | 1 | Q9Z0U1 | Tight junct | Q9Z0U1 [8 | 1 | 1568.856 | 0 | 0.27 | 0.007243 | 6.41 | 89.8 | 110.2 | 2.69 | 5.98 | | High | High | High | High | High | High | High | High | High | High | High | 0.00196 | 0.01994 | 1.27 |
| FALSE | High | QRPASLVQ | [R].QRPASI | 1xPhospho | P14901 1x | 8.28E-06 | 0 | 1 | 2 | 4 | P14901 | Heme oxyg | P14901 [23 | 0 | 2146.086 | 0 | -0.53 | 0.003287 | 4.19 | 115.9 | 84.1 | 11.15 | 20.58 | | High | High | High | High | High | High | High | High | High | High | High | 0.000275 | 0.001837 | 2.62 |
| FALSE | High | QQTPVHSC | [R].QQTPV | 1xPhospho | Q3U0N4 1 | 5.91E-05 | 0 | 1 | 2 | 1 | Q3U0N4 | Putative ur | Q3U0N4 [8 | 0 | 2057.058 | 0 | -0.55 | 0.070285 | 16.32 | 118.9 | 81.1 | 25.84 | 4.91 | | High | High | High | High | High | High | High | High | High | High | High | 0 | 9.94E-06 | 3.1 |
| FALSE | High | QRPLTASL | [K].QRPLT | 1xCarbami | Q8BHG9 1 | 0.000104 | 0 | 1 | 1 | 2 | Q8BHG9 | CGG triplet | Q8BHG9 [8 | 0 | 2567.305 | 0 | 0.05 | 0.940091 | 13.2 | 98.3 | 101.7 | 14.41 | 2.9 | | High | High | High | High | High | High | High | High | High | High | High | 2.58E-05 | 0.000441 | 3.32 |
| FALSE | High | QRPSYDVF | [K].QRPSY | 1xPhospho | F8WJ40 1x | 1.05E-06 | 0 | 1 | 5 | 1 | F8WJ40 | Histone-lys | F8WJ40 [1 | 0 | 2827.241 | 0 | -0.88 | 0.001634 | 5.14 | 129.7 | 70.3 | 7.54 | 8.84 | | High | High | High | High | High | High | High | High | High | High | High | 0 | 1.08E-07 | 3.2 |
| FALSE | High | QRPSYDVF | [K].QRPSY | 2xPhospho | F8WJ40 2x | 2.73E-06 | 0 | 1 | 5 | 2 | F8WJ40 | Histone-lys | F8WJ40 [1 | 0 | 2907.207 | 0 | -0.88 | 0.000477 | 2.83 | 129.8 | 70.2 | 5.21 | 6.21 | | High | High | High | High | High | High | High | High | High | High | High | 0 | 3.14E-07 | 3.21 |
| FALSE | High | QRPSYDVF | [K].QRPSY | 3xPhospho | F8WJ40 3x | 0.0026 | 0.000109 | 1 | 5 | 1 | F8WJ40 | Histone-lys | F8WJ40 [1 | 0 | 2987.174 | 0 | 0.18 | 0.449366 | 29.33 | 93.6 | 106.4 | 13.96 | 12.28 | | High | High | High | High | High | High | High | High | High | High | High | 4.94E-05 | 0.000697 | 2.84 |
| FALSE | High | QRSDEDNK | [K].QRSDE | 1xPhospho | B2RRE0 1x | 7.6E-08 | 0 | 1 | 2 | 1 | B2RRE0 | A kinase (P | B2RRE0 [1 | 1 | 2638.308 | 0 | -1.28 | 0.00315 | 2.96 | 141.6 | 58.4 | 10.36 | 11.7 | | High | High | High | High | High | High | High | High | High | High | High | 0 | 5.59E-09 | 3.82 |
| FALSE | High | QRSPAAPE | [R].QRSPA | 1xPhospho | Q8BX80 1x | 0.117405 | 0.009523 | 1 | 4 | 1 | Q8BX80 | Cytosolic e | Q8BX80 [1 | 1 | 1521.819 | 0 | -0.16 | 0.280301 | 16.05 | 106.3 | 93.7 | 6.31 | 12.01 | | High | High | High | High | High | High | High | High | High | High | High | 0.004782 | 0.05239 | 2.19 |
| FALSE | High | QRSPFPGS | [R].QRSPF | 1xPhospho | E9Q156 1x | 0.020142 | 0.001472 | 1 | 2 | 2 | E9Q156 | tRNA (guar | E9Q156 [4 | 1 | 1997.026 | 0 | 1.04 | 0.004516 | 11.45 | 68 | 132 | 11.77 | 8.59 | | High | High | High | High | High | High | High | High | High | High | High | 0.000786 | 0.00685 | 3.45 |
| FALSE | High | QRSPFPGS | [R].QRSPF | 2xPhospho | E9Q156 2x | 0.036731 | 0.00249 | 1 | 2 | 1 | E9Q156 | tRNA (guar | E9Q156 [4 | 1 | 2076.992 | 0 | 0.55 | 0.035518 | 20 | 83.9 | 116.1 | 10.89 | 9.34 | | High | High | High | High | High | High | High | High | High | High | High | 0.001308 | 0.01356 | 1.84 |
| FALSE | High | QRSPIALPV | [R].QRSPIA | 1xPhospho | A2AJT5 1x | 0.008388 | 0.000736 | 1 | 5 | 1 | A2AJT5 | Arginine/se | A2AJT5 [2 | 1 | 1646.976 | 0 | -0.82 | 0.001216 | 2.49 | 126.4 | 73.6 | 8.77 | 4.12 | | High | High | High | High | High | High | High | High | High | High | High | 0.000327 | 0.002573 | 3.1 |
| FALSE | High | QSANGSFS | [K].QSANG | 1xCarbami | Q5NBY9 1x | 0.00129 | 0 | 1 | 4 | 3 | Q5NBY9 | POZ (BTB) | Q5NBY9 [5 | 0 | 2105.014 | 0 | -0.94 | 0.000651 | 3.69 | 129.5 | 70.5 | 6.3 | 11.16 | | High | High | High | High | High | High | High | High | High | High | High | 0.000161 | 0.001174 | 3.55 |
| FALSE | High | QSARPSTP | [R].QSARP | 1xCarbami | Q60949 1x | 0.000281 | 0 | 1 | 4 | 2 | Q60949 | TBC1 doma | Q60949 [1 | 0 | 2495.096 | 0 | -1.29 | 0.000236 | 1.62 | 142.6 | 57.4 | 4.93 | 11.21 | | High | High | High | High | High | High | High | High | High | High | High | 0 | 5.73E-05 | 1.87 |
| FALSE | High | QSCELSPSK | [K].QSCELS | 1xCarbami | E9PYG6 1x | 0.059159 | 0.004294 | 1 | 4 | 1 | E9PYG6 | RAS p21 pr | E9PYG6 [8 | 0 | 1573.77 | 0 | -0.76 | 0.000285 | 1.37 | 125.1 | 74.9 | 2.5 | 5.99 | | High | High | High | High | High | High | High | High | High | High | High | 0.002202 | 0.02342 | 1.09 |

| | | | | | | | | | | | | | | | | | | | | | | | | | | | | | | | | | | | | | | |
|---|---|---|---|---|---|---|---|---|---|---|---|---|---|---|---|---|---|---|---|---|---|---|---|---|---|---|---|---|---|---|---|---|---|---|---|---|---|---|
| FALSE | High | QSCLHPFTI | [K].QSCLHI | 1xCarbami | P41233 1xI | 3.37E-07 | 0 | 1 | 2 | 2 | P41233 | ATP-bindin | P41233 [12 | 0 | 3168.33 | 0 | 1.89 | 0.000282 | 1.31 | 39.8 | 160.2 | 9.97 | 8.74 | | High | High | High | High | High | High | High | High | High | High | High | 0 | 3.01E-08 | 4.32 |
| FALSE | High | QSDDEVYA | [K].QSDDE | 1xPhospho | Q3TVM1 1 | 2.47E-05 | 0 | 1 | 2 | 1 | Q3TVM1 | Putative ur | Q3TVM1 [4 | 0 | 2504.221 | 0 | -0.38 | 0.004123 | 5.78 | 112.8 | 87.2 | 4.5 | 10.41 | | High | High | High | High | High | High | High | High | High | High | High | 0 | 3.71E-06 | 2.75 |
| FALSE | High | QRPQSSSD | [R].QRPQS | 1xPhospho | P06803 1xI | 0.000235 | 0 | 1 | 1 | 2 | P06803 | Serine/thre | P06803 [28 | 0 | 1824.88 | 0 | -1.22 | 0.000403 | 1.2 | 139.9 | 60.1 | 3.42 | 10.47 | | High | High | High | High | High | High | High | High | High | High | High | 0.000872 | 0.009161 | 2.2 |
| FALSE | High | QQSYQASE | [K].QQSYQ | 1xPhospho | Q80TY5 1x | 2.64E-07 | 0 | 1 | 3 | 1 | Q80TY5 | Vacuolar p | Q80TY5 [1 | 0 | 2225.089 | 0 | -0.02 | 0.647754 | 11.01 | 100.7 | 99.3 | 6.2 | 10.09 | | High | High | High | High | High | High | High | High | High | High | High | 0 | 2.29E-08 | 3.66 |
| FALSE | High | QQSVPAPS | [R].QQSVP | 2xPhospho | E9Q4F8 2x | 0.004033 | 0.000311 | 1 | 3 | 2 | E9Q4F8 | Ankyrin re | E9Q4F8 [1 | 0 | 2941.415 | 0 | 0.76 | 0.001469 | 10.58 | 75.6 | 124.4 | 5.77 | 5.3 | | High | High | High | High | High | High | High | High | High | High | High | 0.000161 | 0.001142 | 3.02 |
| FALSE | High | QQSVPAPS | [R].QQSVP | 1xPhospho | E9Q4F8 1x | 0.00086 | 0 | 1 | 3 | 2 | E9Q4F8 | Ankyrin re | E9Q4F8 [1 | 0 | 2861.449 | 0 | -0.08 | 0.482817 | 13.19 | 101.7 | 98.3 | 3.86 | 11.94 | | High | High | High | High | High | High | High | High | High | High | High | 0 | 0.000201 | 3.99 |
| FALSE | High | QQPALEQF | [K].QQPAL | 1xPhospho | Q9WV92-7 | 9.14E-06 | 0 | 1 | 13 | 2 | Q9WV92-7 | Isoform 7 c | Q9WV92-7 | 0 | 2587.332 | 0 | -1.8 | 0.001876 | 2.47 | 154.7 | 45.3 | 10.78 | 19.99 | | High | High | High | High | High | High | High | High | High | High | High | 0 | 1.49E-06 | 3.71 |
| FALSE | High | QQPPEPEV | [K].QQPPE | 1xPhospho | Q6P542 1x | 1.22E-09 | 0 | 2 | 3 | 6 | Q6P542; Q | ATP-bindin | Q6P542 [7 | 0 | 2676.259 | 0 | 0.1 | 0.230179 | 6.45 | 96.9 | 103.1 | 6.79 | 6.52 | Shared | High | High | High | High | High | High | High | High | High | High | High | 0 | 5.42E-11 | 4.87 |
| FALSE | High | QQPPLGPS | [R].QQPPL | 1xPhospho | Q8BHL3 1x | 7.75E-05 | 0 | 1 | 1 | 2 | Q8BHL3 | TBC1 doma | Q8BHL3 [6 | 0 | 2387.335 | 0 | 0.31 | 0.047805 | 12.22 | 89.4 | 110.6 | 10.46 | 14.15 | | High | High | High | High | High | High | High | High | High | High | High | 0 | 1.34E-05 | 3.77 |
| FALSE | High | QQPQLPRF | [K].QQPQL | 1xCarbami | A2A7S8-2 | 2.2E-06 | 0 | 1 | 4 | 1 | A2A7S8-2 | Isoform 2 c | A2A7S8-2 [ | 0 | 4098.965 | 0 | -0.21 | 0.09291 | 5.55 | 113.9 | 86.1 | 17.42 | 18.44 | | High | High | High | High | High | High | High | High | High | High | High | 0 | 2.45E-07 | 3.86 |
| FALSE | High | QQPSQSPR | [K].QQPSQ | 1xPhospho | Q3U0M8 1 | 0.1059 | 0.008582 | 1 | 3 | 1 | Q3U0M8 | Putative ur | Q3U0M8 [ | 0 | 1236.594 | 0 | -0.7 | 0.011165 | 6.01 | 123 | 77 | 5.12 | 19.9 | | High | High | High | High | High | High | High | High | High | High | High | 0.004317 | 0.04634 | 0.94 |
| FALSE | High | QQPVAAVI | [R].QQPVA | 1xPhospho | E9Q6J5-3 1 | 1.5E-05 | 0 | 1 | 3 | 1 | E9Q6J5-3 | Isoform Ne | E9Q6J5-3 [ | 0 | 4824.347 | 0 | 0.49 | 0.506736 | 24.95 | 84.9 | 115.1 | 5.39 | 43.13 | | High | High | High | High | High | High | High | High | High | High | High | 0 | 2.14E-06 | 3.89 |
| FALSE | High | QQQEPTCE | [K].QQQEP | 1xCarbami | Q6NSP9 1x | 8.66E-06 | 0 | 1 | 2 | 4 | Q6NSP9 | High mobil | Q6NSP9 [3 | 0 | 1966.935 | 0 | 2.66 | 0.000317 | 15.03 | 28 | 172 | 12.75 | 6.67 | | High | High | High | High | High | High | High | High | High | High | High | 0 | 0.000034 | 2.66 |
| FALSE | High | QQQMSDT | [K].QQQM | 1xPhospho | E9PVX6 1x | 0.000779 | 0 | 1 | 2 | 1 | E9PVX6 | Proliferatic | E9PVX6 [7 | 0 | 2477.118 | 0 | -0.14 | 0.90023 | 5.95 | 104.3 | 95.7 | 8.61 | 19.84 | | High | High | High | High | High | High | High | High | High | High | High | 0 | 0.00018 | 3.03 |
| FALSE | High | QQQQQQ( | [K].QQQQ( | 1xPhospho | A0A0A0M( | 1.12E-05 | 0 | 1 | 2 | 2 | A0A0A0M( | Histone-lys | A0A0A0M( | 0 | 3692.784 | 0 | 1.5 | 0.004391 | 22.76 | 51.2 | 148.8 | 12.19 | 16.29 | | High | High | High | High | High | High | High | High | High | High | High | 0 | 1.55E-06 | 4.19 |
| FALSE | High | QQQQSGG | [K].QQQQS | 1xPhospho | Q921G6 1x | 0.000227 | 0 | 1 | 5 | 2 | Q921G6 | Leucine-ric | Q921G6 [4 | 0 | 1624.743 | 0 | -0.44 | 0.013392 | 10.51 | 116.6 | 83.4 | 5.87 | 13.22 | | High | High | High | High | High | High | High | High | High | High | High | 0.000275 | 0.001833 | 1.55 |
| FALSE | High | QQQSFTNF | [K].QQQSF | 1xPhospho | Q9D0K1 1x | 0.019107 | 0.001402 | 1 | 1 | 2 | Q9D0K1 | Peroxisoma | Q9D0K1 [3 | 0 | 1943.04 | 0 | 1.93 | 0.008123 | 21.36 | 44.5 | 155.5 | 9.88 | 25.91 | | High | High | High | High | High | High | High | High | High | High | High | 0.00074 | 0.006458 | 3.12 |
| FALSE | High | QQQSPPK | [R].QQQSP | 1xPhospho | A0A0R4J0Y | 0.008847 | 0.000736 | 1 | 6 | 2 | A0A0R4J0Y | Probable h | A0A0R4J0Y | 0 | 1350.718 | 0 | -0.35 | 0.008577 | 9.22 | 112.2 | 87.8 | 4.36 | 6.31 | | High | High | High | High | High | High | High | High | High | High | High | 0.003754 | 0.04038 | 1.66 |
| FALSE | High | QQQSQTA | [R].QQQSC | 1xPhospho | Q8R1B4 1x | 0.00359 | 0.00021 | 1 | 2 | 2 | Q8R1B4 | Eukaryotic | Q8R1B4 [9 | 0 | 1262.562 | 0 | 1.3 | 0.002042 | 0.07 | 60.1 | 139.9 | 11.57 | 10.51 | | High | High | High | High | High | High | High | High | High | High | High | 9.42E-05 | 0.001004 | 1.44 |
| FALSE | High | QQRPSLTE | [R].QQRPS | 1xPhospho | Q8BP00 1x | 0.025738 | 0.001645 | 1 | 2 | 1 | Q8BP00 | IQ calmodu | Q8BP00 [4 | 0 | 1787.946 | 0 | -0.81 | 0.005002 | 5.43 | 131.1 | 68.9 | 12.9 | 8.38 | | High | High | High | High | High | High | High | High | High | High | High | 0.000872 | 0.009016 | 1.99 |
| FALSE | High | QQRPSSPV | [R].QQRPS | 1xPhospho | Q9DBU5 1x | 0.029844 | 0.001857 | 1 | 2 | 1 | Q9DBU5 | E3 ubiquiti | Q9DBU5 [1 | 0 | 1434.742 | 0 | 1.27 | 0.001829 | 17.35 | 58.2 | 141.8 | 12.44 | 8.83 | | High | High | High | High | High | High | High | High | High | High | High | 0.001011 | 0.01076 | 1.96 |
| FALSE | High | QQSEEDLL | [K].QQSEE | 1xPhospho | Q4FK79 1x | 8.68E-07 | 0 | 1 | 2 | 1 | Q4FK79 | Putative ur | Q4FK79 [9 | 0 | 2016.948 | 0 | 0.34 | 0.146275 | 28.3 | 86.3 | 113.7 | 9.43 | 15.75 | | High | High | High | High | High | High | High | High | High | High | High | 0 | 8.68E-08 | 2.92 |
| FALSE | High | QQSEISTHS | [R].QQSEIS | 1xPhospho | Q3UGJ5 1x | 0.002134 | 5.66E-05 | 1 | 3 | 1 | Q3UGJ5 | Putative ur | Q3UGJ5 [8 | 0 | 1438.678 | 0 | 0.82 | 0.013622 | 7.11 | 74.2 | 125.8 | 8.01 | 13.29 | | High | High | High | High | High | High | High | High | High | High | High | 2.58E-05 | 0.000559 | 1.92 |
| FALSE | High | QQSFGVKF | [K].QQSFG | 1xPhospho | F2Z3U3 1xI | 5.32E-06 | 0 | 1 | 1 | 2 | F2Z3U3 | Ras associa | F2Z3U3 [84 | 0 | 2942.672 | 0 | 1.75 | 0.000191 | 3.85 | 45.6 | 154.4 | 3.51 | 4.11 | | High | High | High | High | High | High | High | High | High | High | High | 0 | 6.62E-07 | 3.27 |
| FALSE | High | QQSFSGSL | [R].QQSFS | 2xPhospho | Q4VAC9 2x | 0.000538 | 0 | 1 | 2 | 1 | Q4VAC9 | Pleckstrin I | Q4VAC9 [5 | 0 | 2552.147 | 0 | 1.69 | 0.000313 | 7.66 | 47 | 153 | 6.62 | 5.09 | | High | High | High | High | High | High | High | High | High | High | High | 0 | 0.000119 | 2.29 |
| FALSE | High | QQSHDPVS | [K].QQSHD | 1xPhospho | Q924H7 1x | 0.001544 | 5.66E-05 | 1 | 8 | 1 | Q924H7 | WW doma | Q924H7 [5 | 0 | 1459.689 | 0 | 0.17 | 0.021682 | 10.85 | 91.5 | 108.5 | 3.16 | 6.63 | | High | High | High | High | High | High | High | High | High | High | High | 2.58E-05 | 0.000387 | 2.3 |
| FALSE | High | QQSHFAM | [R].QQSHF | 1xPhospho | P60335 1xI | 0.000106 | 0 | 1 | 1 | 1 | P60335 | Poly(rC)-bi | P60335 [24 | 0 | 3144.468 | 0 | -0.03 | 0.606416 | 12.5 | 105.4 | 94.6 | 10.69 | 7.78 | | High | High | High | High | High | High | High | High | High | High | High | 0 | 1.93E-05 | 2.63 |
| FALSE | High | QQSHFAM | [R].QQSHF | 1xPhospho | P60335 1xI | 0.013426 | 0.000976 | 1 | 1 | 4 | P60335 | Poly(rC)-bi | P60335 [24 | 0 | 3160.463 | 0 | 0 | 0.873903 | 25.88 | 101 | 99 | 10.33 | 14.49 | | High | High | High | High | High | High | High | High | High | High | High | 0.000505 | 0.004374 | 2.6 |
| FALSE | High | QQSLGGSE | [R].QQSLG | 1xPhospho | A0A1C7CY | 1.13E-05 | 0 | 1 | 6 | 2 | A0A1C7CY | FTS and Hc | A0A1C7CY | 0 | 1804.879 | 0 | -0.31 | 0.075159 | 7.17 | 109.9 | 90.1 | 6.32 | 10.25 | | High | High | High | High | High | High | High | High | High | High | High | 0 | 1.55E-06 | 3.57 |
| FALSE | High | QQSLVEK | [R].QQSLV | 1xPhospho | Q80Y19 1x | 0.011447 | 0.000816 | 1 | 2 | 1 | Q80Y19 | Rho GTPas | Q80Y19 [7 | 0 | 1369.749 | 0 | -0.09 | 0.038321 | 5.84 | 105.9 | 94.1 | 5.01 | 2.81 | | High | High | High | High | High | High | High | High | High | High | High | 0.000437 | 0.003644 | 1.89 |
| FALSE | High | QQSPPQGI | [R].QQSPP | 1xDeamida | Q9D9U4 1x | 0.044739 | 0.003302 | 1 | 1 | 1 | Q9D9U4 | Testis-expr | Q9D9U4 [2 | 0 | 2378.2 | 0 | 0.5 | 0.031814 | 12.96 | 82 | 118 | 4.75 | 13.57 | | High | High | High | High | High | High | High | High | High | High | High | 0.001712 | 0.01702 | 2.15 |
| FALSE | High | QQSQPTK | [R].QQSQP | 1xPhospho | Q546B3 1x | 0.045898 | 0.003408 | 1 | 3 | 1 | Q546B3 | DNA-bindi | Q546B3 [5 | 0 | 1354.713 | 0 | 1.87 | 0.001084 | 25.96 | 43 | 157 | 7.02 | 10.9 | | High | High | High | High | High | High | High | High | High | High | High | 0.001741 | 0.01743 | 2.06 |
| FALSE | High | QQSSQAQ | [R].QQSSQ | 1xPhospho | A0A0B4J1E | 0.000662 | 0 | 1 | 3 | 2 | A0A0B4J1E | DNA-bindi | A0A0B4J1E | 0 | 1614.825 | 0 | 2.65 | 0.001764 | 12.67 | 27.3 | 172.7 | 12.86 | 16.35 | | High | High | High | High | High | High | High | High | High | High | High | 0 | 0.00015 | 2.53 |
| FALSE | High | QQSSWTSF | [K].QQSSW | 1xPhospho | Q8K3Z9 1x | 6.59E-05 | 0 | 1 | 2 | 3 | Q8K3Z9 | Nuclear en | Q8K3Z9 [4 | 0 | 2096.96 | 0 | -0.02 | 0.526037 | 22.5 | 90.4 | 109.6 | 12.86 | 19.35 | | High | High | High | High | High | High | High | High | High | High | High | 0 | 1.13E-05 | 2.52 |
| FALSE | High | QQSTGAIP | [R].QQSTG | 1xPhospho | Q3UDC3 1x | 0.001831 | 5.66E-05 | 1 | 3 | 1 | Q3UDC3 | Putative ur | Q3UDC3 [4 | 0 | 1637.821 | 0 | 0.95 | 0.007479 | 12.72 | 69.3 | 130.7 | 13.1 | 16.82 | | High | High | High | High | High | High | High | High | High | High | High | 2.58E-05 | 0.000472 | 2.14 |
| FALSE | High | QQNTPQA | [R].QQNTP | 1xPhospho | Q6PCM1 1 | 6.62E-07 | 0 | 1 | 4 | 2 | Q6PCM1 | Lysine-spe | Q6PCM1 [3 | 0 | 2274.164 | 0 | -0.84 | 0.000406 | 1.18 | 125.7 | 74.3 | 3.76 | 5.74 | | High | High | High | High | High | High | High | High | High | High | High | 0 | 6.38E-08 | 3.53 |
| FALSE | High | QTSLTSSQI | [R].QTSLTS | 1xPhospho | Q9D4H4 1x | 2.7E-05 | 0 | 1 | 2 | 1 | Q9D4H4 | Angiomoti | Q9D4H4 [8 | 0 | 1990.015 | 0 | 2.74 | 0.000163 | 15.51 | 24.7 | 175.3 | 15.34 | 5.69 | | High | High | High | High | High | High | High | High | High | High | High | 0 | 4.12E-06 | 3.81 |
| FALSE | High | QTSPDISTE | [R].QTSPDI | 1xPhospho | Q8C3S2 1x | 0.000127 | 0 | 1 | 3 | 4 | Q8C3S2 | Transport a | Q8C3S2 [7 | 0 | 2250.153 | 0 | -0.99 | 0.000442 | 2.3 | 132.2 | 67.8 | 3.01 | 6.4 | | High | High | High | High | High | High | High | High | High | High | High | 0 | 7.82E-05 | 2.17 |
| FALSE | High | QTSPEPVA | [K].QTSPEF | 1xPhospho | Q8K2X7 1x | 1.53E-10 | 0 | 1 | 3 | 4 | Q8K2X7 | DNA repair | Q8K2X7 [8 | 0 | 2356.175 | 0 | -0.03 | 0.985795 | 8.09 | 100.4 | 99.6 | 7.16 | 6.26 | | High | High | High | High | High | High | High | High | High | High | High | 0 | 6.25E-10 | 5.18 |
| FALSE | High | RDSFSGSK | [R].RDSFSG | 1xPhospho | Q8BM75 1 | 0.091627 | 0.007146 | 1 | 3 | 1 | Q8BM75 | AT-rich inte | Q8BM75 [2 | 1 | 1421.719 | 0 | 1.09 | 0.000385 | 8.65 | 63.6 | 136.4 | 3.66 | 5.47 | | High | High | High | High | High | High | High | High | High | High | High | 0.003659 | 0.03908 | 1.78 |
| FALSE | High | RDSLASLFP | [R].RDSLAS | 1xPhospho | Q91W40 1 | 0.001087 | 0 | 1 | 2 | 2 | Q91W40 | Kinesin ligh | Q91W40 [ | 1 | 2275.126 | 0 | -0.45 | 0.019192 | 8.18 | 113.3 | 86.7 | 10.58 | 6.04 | | High | High | High | High | High | High | High | High | High | High | High | 0 | 0.000263 | 3.54 |
| FALSE | High | RDSICSSVS | [R].RDSICS | 1xCarbami | E9QP99 1x | 0.095408 | 0.007437 | 1 | 5 | 1 | E9QP99 | Golgin sub | E9QP99 [3 | 1 | 3489.726 | 0 | 0.08 | 0.190481 | 5.94 | 94.4 | 105.6 | 6.58 | 17.81 | | High | High | High | High | High | High | High | High | High | High | High | 0.003768 | 0.04071 | 4.65 |
| FALSE | High | RDSLEEGEI | [K].RDSLEE | 1xPhospho | P24788 1xI | 0.007209 | 0.00062 | 1 | 4 | 1 | P24788 | Cyclin-dep | P24788 [45 | 1 | 1512.726 | 0 | -0.14 | 0.543478 | 10.39 | 105.8 | 94.2 | 8.17 | 9.42 | | High | High | High | High | High | High | High | High | High | High | High | 0.000275 | 0.002167 | 3.35 |
| FALSE | High | RDSLGSFST | [K].RDSLGS | 1xPhospho | B9EKJ7 1xF | 0.00129 | 0 | 1 | 3 | 2 | B9EKJ7 | Tankyrase | B9EKJ7 [86 | 1 | 1434.694 | 0 | 0.54 | 0.068094 | 3.92 | 78.5 | 121.5 | 5.38 | 33.59 | | High | High | High | High | High | High | High | High | High | High | High | 0 | 0.000317 | 2.11 |
| FALSE | High | RDSLPGK | [R].RDSLPG | 1xPhospho | P70188 1xI | 0.068184 | 0.004935 | 1 | 3 | 1 | P70188 | Kinesin-ass | P70188 [10 | 1 | 1310.723 | 0 | 0.73 | 0.00279 | 2.27 | 72.1 | 127.9 | 9.3 | 8.8 | | High | High | High | High | High | High | High | High | High | High | High | 0.002559 | 0.02748 | 1.99 |
| FALSE | High | RDSLTGSSI | [R].RDSLTG | 1xPhospho | Q3UR00 1x | 4.94E-05 | 0 | 1 | 8 | 4 | Q3UR00 | Putative ur | Q3UR00 [7 | 1 | 1879.957 | 0 | -0.42 | 0.003957 | 5.99 | 116.3 | 83.7 | 6.04 | 3.31 | | High | High | High | High | High | High | High | High | High | High | High | 4.94E-05 | 0.00064 | 2.78 |
| FALSE | High | RDSNPEIK | [R].RDSNPI | 1xPhospho | Z4YKQ9 1x | 0.017041 | 0.001204 | 1 | 3 | 2 | Z4YKQ9 | ATPase fan | Z4YKQ9 [1 | 1 | 1496.787 | 0 | 0.6 | 0.005158 | 14.45 | 79.4 | 120.6 | 5.74 | 7.31 | | High | High | High | High | High | High | High | High | High | High | High | 0.001355 | 0.01424 | 2.46 |
| FALSE | High | RDSNTSTIS | [R].RDSNTS | 1xPhospho | B7ZNV2 1x | 0.043981 | 0.003228 | 1 | 2 | 1 | B7ZNV2 | GLI-Kruppe | B7ZNV2 [8 | 1 | 2053.975 | 0 | -1.08 | 0.000384 | 0.45 | 138.2 | 61.8 | 2.95 | 10.23 | | High | High | High | High | High | High | High | High | High | High | High | 0.001665 | 0.01668 | 2.42 |
| FALSE | High | RDSPLQGG | [R].RDSPLC | 1xPhospho | A0A0A6YW | 4.08E-11 | 0 | 2 | 13 | 3 | A0A0A6YW | Mitogen-ac | A0A0A6YW | 1 | 2193.036 | 0 | -0.58 | 0.002883 | 1.23 | 117.4 | 82.6 | 3.2 | 9.08 | Shared | High | High | High | High | High | High | High | High | High | High | High | 0 | 1.39E-10 | 4.75 |
| FALSE | High | RDSPTYDP | [R].RDSPTY | 1xPhospho | Q8CFC7 1x | 0.00418 | 0.00036 | 1 | 4 | 3 | Q8CFC7 | CLK4-assoc | Q8CFC7 [2 | 1 | 1779.872 | 0 | -0.55 | 0.00058 | 1.83 | 117.9 | 82.1 | 3.19 | 1.93 | | High | High | High | High | High | High | High | High | High | High | High | 0.000161 | 0.001179 | 2.24 |
| FALSE | High | RDSSDDWI | [R].RDSSDI | 1xPhospho | P28028 1xI | 2.8E-06 | 0 | 2 | 3 | 7 | P28028; F6 | Serine/thre | P28028 [48 | 1 | 2483.14 | 0 | -0.18 | 0.278953 | 2.26 | 104.8 | 95.2 | 3.06 | 7.45 | Shared | High | High | High | High | High | High | High | High | High | High | High | 0 | 5.51E-07 | 4.07 |
| FALSE | High | RDSSEHGA | [R].RDSSEH | 1xPhospho | I6L9H1 1xP | 0.013545 | 0.000976 | 1 | 3 | 3 | I6L9H1 | Dishevellec | I6L9H1 [16 | 1 | 2125.984 | 0 | -0.6 | 0.001079 | 2.81 | 120.5 | 79.5 | 3.99 | 3.68 | | High | High | High | High | High | High | High | High | High | High | High | 0.004138 | 0.04462 | 2.85 |
| FALSE | High | RDSFSENEI | [R].RDSFSE | 1xDeamida | Q8C5N3 1x | 0.01319 | 0.000889 | 1 | 4 | 1 | Q8C5N3 | Pre-mRNA- | Q8C5N3 [8 | 1 | 1650.778 | 0 | -0.2 | 0.007985 | 2.1 | 109.3 | 90.7 | 6.41 | 8.34 | | High | High | High | High | High | High | High | High | High | High | High | 0.000467 | 0.004274 | 1.61 |
| FALSE | High | RDSSESQL/ | [R].RDSSES | 1xPhospho | Q0VAW6 1 | 0.008769 | 0.000736 | 1 | 3 | 1 | Q0VAW6 | RIKEN cDN | Q0VAW6 [ | 1 | 2461.141 | 0 | -0.96 | 0.17675 | 11.55 | 134.6 | 65.4 | 28.45 | 58.61 | | High | High | High | High | High | High | High | High | High | High | High | 0.000327 | 0.002714 | 2.29 |
| FALSE | High | RDSSESQL/ | [R].RDSSES | 2xPhospho | Q0VAW6 2 | 0.002463 | 0.000109 | 1 | 3 | 3 | Q0VAW6 | RIKEN cDN | Q0VAW6 [ | 1 | 2770.27 | 0 | -0.64 | 0.046499 | 12.01 | 121.9 | 78.1 | 15.25 | 17.26 | | High | High | High | High | High | High | High | High | High | High | High | 4.94E-05 | 0.000657 | 2.61 |
| FALSE | High | RDSSGGAK | [R].RDSSG( | 1xPhospho | Q3U5I9 1xI | 0.072858 | 0.005599 | 1 | 3 | 1 | Q3U5I9 | Putative ur | Q3U5I9 [22 | 1 | 1315.677 | 0 | 1.19 | 0.000278 | 7.52 | 61.5 | 138.5 | 5.12 | 4.59 | | High | High | High | High | High | High | High | High | High | High | High | 0.002867 | 0.02987 | 1.62 |
| FALSE | High | RDSSSLSSP | [K].RDSSSL | 1xPhospho | Q9Z277 1x | 2.37E-07 | 0 | 1 | 2 | 1 | Q9Z277 | Tyrosine-p | Q9Z277 [3 | 1 | 1926.01 | 0 | -0.27 | 0.025677 | 13.18 | 108.9 | 91.1 | 5.06 | 7.92 | | High | High | High | High | High | High | High | High | High | High | High | 0 | 2.02E-08 | 3.23 |
| FALSE | High | RDSSSLSSP | [K].RDSSSL | 2xPhospho | Q9Z277 2x | 0.003622 | 0.00021 | 1 | 2 | 3 | Q9Z277 | Tyrosine-p | Q9Z277 [3 | 1 | 2005.976 | 0 | 0.15 | 0.057587 | 4.23 | 95.4 | 104.6 | 5.79 | 8.29 | | High | High | High | High | High | High | High | High | High | High | High | 9.42E-05 | 0.001013 | 2.76 |
| FALSE | High | RDSSVIK | [R].RDSSVI | 1xPhospho | H3BKF8 1x | 0.080423 | 0.006219 | 1 | 11 | 1 | H3BKF8 | Homeobox | H3BKF8 [1 | 1 | 1342.75 | 0 | -1.05 | 0.000263 | 2.98 | 134.9 | 65.1 | 0.95 | 8.87 | | High | High | High | High | High | High | High | High | High | High | High | 0.003188 | 0.03327 | 1.77 |
| FALSE | High | RDSTEAPKI | [R].RDSTEA | 1xPhospho | K3W4P2 1x | 0.000331 | 0 | 1 | 4 | 1 | K3W4P2 | Integrator | K3W4P2 [1 | 1 | 2757.361 | 0 | 0.77 | 0.001151 | 7.17 | 74.1 | 125.9 | 2.6 | 7.07 | | High | High | High | High | High | High | High | High | High | High | High | 0 | 6.88E-05 | 2.96 |
| FALSE | High | RDSTETLHS | [K].RDSTET | 1xPhospho | E9PV87 1xI | 0.031168 | 0.001935 | 1 | 4 | 2 | E9PV87 | Protein TAI | E9PV87 [4 | 1 | 2428.134 | 0 | 0.36 | 0.076099 | 26.54 | 87.7 | 112.3 | 10.78 | 12.49 | | High | High | High | High | High | High | High | High | High | High | High | 0.001061 | 0.01123 | 2.4 |
| FALSE | High | RDSTPSLK | [R].RDSTPS | 1xPhospho | Q8C142 1x | 0.043607 | 0.003228 | 1 | 2 | 2 | Q8C142 | Low densit | Q8C142 [1 | 1 | 1441.782 | 0 | 1.53 | 0.000166 | 7.97 | 51.7 | 148.3 | 4.28 | 3.41 | | High | High | High | High | High | High | High | High | High | High | High | 0.005122 | 0.05575 | 2.04 |
| FALSE | High | RDSVLAASI | [R].RDSVLA | 1xPhospho | P11881 1xI | 0.026885 | 0.001676 | 1 | 8 | 1 | P11881 | Inositol 1,4 | P11881 [15 | 1 | 1283.667 | 0 | 0.92 | 0.005912 | 12.44 | 61.4 | 138.6 | 11.09 | 14.44 | | High | High | High | High | High | High | High | High | High | High | High | 0.000903 | 0.009477 | 1.95 |
| FALSE | High | RDSVLTSK | [K].RDSVLT | 1xPhospho | Q6NZB0 1x | 0.039011 | 0.002718 | 1 | 6 | 1 | Q6NZB0 | DnaJ homc | Q6NZB0 [3 | 1 | 1443.797 | 0 | -0.59 | 0.007436 | 4.08 | 123.4 | 76.6 | 10.68 | 7.49 | | High | High | High | High | High | High | High | High | High | High | High | 0.001453 | 0.01453 | 1.64 |
| FALSE | High | RDSWDFA( | [R].RDSWD | 1xCarbami | Q8BGY4 1x | 0.00095 | 0 | 1 | 4 | 1 | Q8BGY4 | Kelch-like p | Q8BGY4 [4 | 1 | 1932.944 | 0 | 1.27 | 0.000748 | 11.01 | 58.5 | 141.5 | 12.1 | 4.17 | | High | High | High | High | High | High | High | High | High | High | High | 0 | 0.000226 | 4.31 |
| FALSE | High | RDSYLSCK | [R].RDSYLS | 1xCarbami | Q542Q3 1x | 0.008614 | 0.000736 | 1 | 1 | 1 | Q542Q3 | G two S ph | Q542Q3 [4 | 1 | 1566.775 | 0 | -0.11 | 0.42146 | 5.92 | 102.2 | 97.8 | 8.12 | 12.59 | | High | High | High | High | High | High | High | High | High | High | High | 0.000327 | 0.002644 | 1.62 |
| FALSE | High | RDTDAGEG | [R].RDTDA | 1xPhospho | Q3ULT8 1x | 8.06E-06 | 0 | 1 | 6 | 2 | Q3ULT8 | Putative ur | Q3ULT8 [4 | 2 | 3640.708 | 0 | 0.61 | 0.002411 | 8.62 | 79.2 | 120.8 | 7.02 | 4.58 | | High | High | High | High | High | High | High | High | High | High | High | 0.000786 | 0.007105 | 3.7 |
| FALSE | High | RDSSESQL/ | [R].RDSSES | 1xPhospho | Q0VAW6 1 | 0.000379 | 0 | 1 | 3 | 5 | Q0VAW6 | RIKEN cDN | Q0VAW6 [ | 1 | 2690.303 | 0 | -0.72 | 0.000621 | 2.59 | 123.3 | 76.7 | 4.19 | 4.12 | | High | High | High | High | High | High | High | High | High | High | High | 0 | 0.000176 | 2.87 |
| FALSE | High | RDSFSENEI | [R].RDSFSE | 1xPhospho | Q8C5N3 1x | 0.000155 | 0 | 1 | 4 | 6 | Q8C5N3 | Pre-mRNA- | Q8C5N3 [8 | 1 | 1649.794 | 0 | -0.45 | 0.002565 | 3.93 | 114.6 | 85.4 | 3.47 | 4.21 | | High | High | High | High | High | High | High | High | High | High | High | 0 | 2.94E-05 | 2.86 |
| FALSE | High | RDSFIGTPY | [R].RDSFIG | 1xCarbami | O54988 1x | 1.13E-05 | 0 | 1 | 2 | 2 | O54988 | STE20-like | O54988 [1 | 1 | 3158.48 | 0 | 0.39 | 0.024316 | 7.91 | 89.4 | 110.6 | 8.17 | 11.67 | | High | High | High | High | High | High | High | High | High | High | High | 0 | 2.06E-06 | 5.41 |
| FALSE | High | RDSFIGTPY | [R].RDSFIG | 1xCarbami | O54988 1x | 3.66E-08 | 0 | 1 | 2 | 1 | O54988 | STE20-like | O54988 [1 | 1 | 3142.485 | 0 | 0.5 | 0.026379 | 15.44 | 81.1 | 118.9 | 13.74 | 7.73 | | High | High | High | High | High | High | High | High | High | High | High | 0 | 2.48E-09 | 5.64 |
| FALSE | High | RASYNETKI | [R].RASYNI | 1xPhospho | Q6ZPJ0 1xI | 0.000822 | 0 | 1 | 4 | 1 | Q6ZPJ0 | Testis-expr | Q6ZPJ0 [58 | 1 | 2781.479 | 0 | 0.33 | 0.034866 | 4.57 | 88.7 | 111.3 | 7.93 | 17.84 | | High | High | High | High | High | High | High | High | High | High | High | 0 | 0.000191 | 3.24 |
| FALSE | High | RATGDGSS | [R].RATGD | 1xPhospho | O88466 1x | 0.002004 | 5.66E-05 | 2 | 4 | 1 | O88466; R | Zinc finger | O88466 [1 | 1 | 1980.959 | 0 | -0.17 | 0.008493 | 2.08 | 105.7 | 94.3 | 3.17 | 5.67 | Shared | High | High | High | High | High | High | High | High | High | High | High | 2.58E-05 | 0.000518 | 3.13 |
| FALSE | High | RATISSPLEI | [R].RATISSI | 1xPhospho | Q9D6W8 1 | 2.59E-06 | 0 | 1 | 1 | 1 | Q9D6W8 | BLOC-1-rel | Q9D6W8 [ | 1 | 2025.058 | 0 | 0.71 | 0.012021 | 18.3 | 75.3 | 124.7 | 14.7 | 13.45 | | High | High | High | High | High | High | High | High | High | High | High | 0 | 2.95E-07 | 2.63 |
| FALSE | High | RATPEPSG | [R].RATPEF | 1xPhospho | Q3T9L1 1xI | 6.71E-05 | 0 | 1 | 4 | 1 | Q3T9L1 | Coronin O | Q3T9L1 [4 | 1 | 2053.975 | 0 | -1.04 | 0.005916 | 8.47 | 136.6 | 63.4 | 5.58 | 23.1 | | High | High | High | High | High | High | High | High | High | High | High | 0 | 1.15E-05 | 3.31 |
| FALSE | High | RATPILSETI | [K].RATPIL | 1xPhospho | P54276 1xI | 0.000164 | 0 | 1 | 2 | 2 | P54276 | DNA mism | P54276 [32 | 1 | 1653.934 | 0 | -0.9 | 0.003785 | 1.54 | 129.7 | 70.3 | 5.75 | 13.19 | | High | High | High | High | High | High | High | High | High | High | High | 0 | 3.14E-05 | 3.4 |
| FALSE | High | RATPINLAS | [K].RATPIN | 1xPhospho | O08856 1x | 0.010382 | 0.000772 | 1 | 1 | 1 | O08856 | RNA polym | O08856 [1 | 1 | 1591.888 | 0 | -0.22 | 0.138879 | 17.01 | 107.5 | 92.5 | 10.7 | 9.62 | | High | High | High | High | High | High | High | High | High | High | High | 0.000382 | 0.003281 | 2.62 |
| FALSE | High | RCSDDFSP | [K].RCSDDI | 1xCarbami | Q80U44 1x | 0.000161 | 0 | 1 | 1 | 1 | Q80U44 | Zinc finger | Q80U44 [8 | 1 | 2729.344 | 0 | 0.87 | 0.015966 | 8.84 | 71.6 | 128.4 | 9.76 | 16 | | High | High | High | High | High | High | High | High | High | High | High | 0 | 3.05E-05 | 4.08 |
| FALSE | High | RCSDDFSP | [K].RCSDDI | 1xCarbami | Q80U44 1x | 0.000292 | 0 | 1 | 1 | 1 | Q80U44 | Zinc finger | Q80U44 [8 | 1 | 2745.339 | 0 | 0.96 | 0.024747 | 29.06 | 74.4 | 125.6 | 12.09 | 25.37 | | High | High | High | High | High | High | High | High | High | High | High | 0 | 5.98E-05 | 4.82 |
| FALSE | High | RCSEGEGA | [K].RCSEGI | 1xCarbami | B1AQJ2 1xI | 6.39E-07 | 0 | 1 | 3 | 1 | B1AQJ2 | Ubiquitin c | B1AQJ2 [8 | 1 | 1800.872 | 0 | -1.4 | 0.00617 | 5.39 | 142.7 | 57.3 | 17.29 | 13.22 | | High | High | High | High | High | High | High | High | High | High | High | 0 | 6.14E-08 | 2.89 |
| FALSE | High | RCSETSSK | [R].RCSETS | 1xCarbami | Q9EP71 1x | 0.057686 | 0.004227 | 1 | 1 | 1 | Q9EP71 | Ankycorbir | Q9EP71 [8 | 1 | 1492.723 | 0 | 0.74 | 0.013423 | 5.64 | 75.9 | 124.1 | 12.11 | 9.97 | | High | High | High | High | High | High | High | High | High | High | High | 0.002159 | 0.02263 | 1.49 |
| FALSE | High | RCSSSLPPI( | [R].RCSSSL | 1xCarbami | Q3UIA2 1x | 0.003138 | 0.00016 | 1 | 5 | 1 | Q3UIA2 | Rho GTPas | Q3UIA2 [6 | 1 | 2941.471 | 0 | -0.25 | 0.301766 | 12.38 | 108.8 | 91.2 | 6.88 | 19.29 | | High | High | High | High | High | High | High | High | High | High | High | 7.19E-05 | 0.000858 | 2.39 |
| FALSE | High | RCSWDKPE | [R].RCSWD | 1xCarbami | G3X9N1 1x | 0.045122 | 0.003373 | 1 | 5 | 1 | G3X9N1 | Rho GTPas | G3X9N1 [3 | 1 | 2084.015 | 0 | -0.13 | 0.434219 | 23.05 | 108.2 | 91.8 | 10.52 | 19.57 | | High | High | High | High | High | High | High | High | High | High | High | 0.001712 | 0.01707 | 1.92 |
| FALSE | High | RDASEEELI | [R].RDASEI | 1xPhospho | E9Q8D0 1x | 0.047086 | 0.00354 | 1 | 1 | 1 | E9Q8D0 | DnaJ homc | E9Q8D0 [1 | 1 | 1614.814 | 0 | -1.46 | 0.000706 | 2.16 | 146.8 | 53.2 | 6.14 | 9.07 | | High | High | High | High | High | High | High | High | High | High | High | 0.001812 | 0.01793 | 1.09 |
| FALSE | High | RDDIEDGD | [R].RDDIED | 1xPhospho | F8VQD1 1x | 2.24E-10 | 0 | 1 | 13 | 7 | F8VQD1 | Protein pol | F8VQD1 [4 | 1 | 2796.265 | 0 | -0.14 | 0.030272 | 5.73 | 106.1 | 93.9 | 2.17 | 8.08 | | High | High | High | High | High | High | High | High | High | High | High | 0 | 8.18E-11 | 5.91 |
| FALSE | High | RDDIEDGD | [R].RDDIED | 1xPhospho | F8VQD1 1x | 7.56E-06 | 0 | 1 | 13 | 9 | F8VQD1 | Protein pol | F8VQD1 [4 | 1 | 2812.26 | 0 | -0.13 | 0.587192 | 13.57 | 104.9 | 95.1 | 8.87 | 16.11 | | High | High | High | High | High | High | High | High | High | High | High | 0 | 9.91E-07 | 3.91 |
| FALSE | High | RDEDMLYS | [R].RDEDM | 1xPhospho | Q3U6Q1 1 | 0.001424 | 5.66E-05 | 1 | 4 | 1 | Q3U6Q1 | Putative ur | Q3U6Q1 [2 | 1 | 2047.936 | 0 | 0.1 | 0.196312 | 11.66 | 96.5 | 103.5 | 7.71 | 6.15 | | High | High | High | High | High | High | High | High | High | High | High | 2.58E-05 | 0.000356 | 2.61 |
| FALSE | High | RDFASEAPT | [K].RDFASI | 1xCarbami | P97434-2 1 | 2.96E-10 | 0 | 1 | 6 | 2 | P97434-2 | Isoform 2 c | P97434-2 [ | 1 | 2704.275 | 0 | 0.74 | 0.001455 | 2.05 | 74.9 | 125.1 | 12.06 | 6 | | High | High | High | High | High | High | High | High | High | High | High | 0 | 1.1E-11 | 3.65 |
| FALSE | High | RDFASEAPT | [K].RDFASI | 1xCarbami | P97434-2 1 | 2.41E-06 | 0 | 1 | 6 | 1 | P97434-2 | Isoform 2 c | P97434-2 [ | 2 | 2860.376 | 0 | 0.82 | 0.000439 | 4.53 | 72.2 | 127.8 | 5.36 | 7.2 | | High | High | High | High | High | High | High | High | High | High | High | 0 | 2.71E-07 | 1.76 |
| FALSE | High | RDGSEEDE | [K].RDGSEI | 1xPhospho | Q05D44 1x | 0.012072 | 0.000889 | 1 | 1 | 3 | Q05D44 | Eukaryotic | Q05D44 [1 | 1 | 1918.843 | 0 | -0.42 | 0.01331 | 6.73 | 110.3 | 89.7 | 11.04 | 15.15 | | High | High | High | High | High | High | High | High | High | High | High | 0.000773 | 0.00671 | 2.86 |
| FALSE | High | RDLIDNSFN | [K].RDLIDN | 1xPhospho | Q9DBE9 1x | 0.016596 | 0.001124 | 1 | 1 | 1 | Q9DBE9 | pre-rRNA p | Q9DBE9 [6 | 1 | 1558.758 | 0 | -0.17 | 0.031091 | 7.06 | 104.9 | 95.1 | 10.03 | 8.14 | | High | High | High | High | High | High | High | High | High | High | High | 0.000585 | 0.005516 | 1.11 |
| FALSE | High | RDLNSSFET | [R].RDLNSS | 1xPhospho | Q8K3G5 1x | 0.000359 | 0 | 1 | 5 | 2 | Q8K3G5 | Inactive se | Q8K3G5 [5 | 1 | 1918.968 | 0 | 0.13 | 0.046053 | 8.36 | 92.4 | 107.6 | 5.43 | 4.1 | | High | High | High | High | High | High | High | High | High | High | High | 0 | 0.000261 | 1.39 |
| FALSE | High | RDSAPVPQ | [R].RDSAP\ | 1xPhospho | E9Q6J5-3 1 | 0.011146 | 0.000772 | 1 | 3 | 1 | E9Q6J5-3 | Isoform Ne | E9Q6J5-3 [ | 1 | 1462.737 | 0 | 0.76 | 0.002234 | 3.67 | 70 | 130 | 14.58 | 11.65 | | High | High | High | High | High | High | High | High | High | High | High | 0.000399 | 0.003555 | 2.2 |
| FALSE | High | RDSCGGG/ | [R].RDSCG( | 1xCarbami | Q6GQT6 1 | 1.85E-06 | 0 | 1 | 1 | 2 | Q6GQT6 | Sterol regu | Q6GQT6 [8 | 1 | 2307.965 | 0 | 0.23 | 0.026952 | 2.16 | 91.9 | 108.1 | 3.37 | 4.88 | | High | High | High | High | High | High | High | High | High | High | High | 0 | 2.04E-07 | 5.43 |
| FALSE | High | RDSDGVD( | [R].RDSDG | 1xPhospho | O88532 1x | 1.13E-08 | 0 | 1 | 2 | 5 | O88532 | Zinc finger | O88532 [1 | 1 | 2019.942 | 0 | 0.12 | 0.020287 | 4.38 | 95.7 | 104.3 | 2.54 | 3.34 | | High | High | High | High | High | High | High | High | High | High | High | 0 | 6.63E-10 | 4.12 |
| FALSE | High | RDSEAGSS | [K].RDSEA( | 1xPhospho | A0A0R4J17 | 2.47E-05 | 0 | 1 | 8 | 5 | A0A0R4J17 | Transcripti | A0A0R4J17 | 1 | 1861.849 | 0 | -0.43 | 0.002357 | 2.94 | 112.9 | 87.1 | 7.05 | 4.17 | | High | High | High | High | High | High | High | High | High | High | High | 0 | 3.74E-06 | 3.54 |
| FALSE | High | RDSESESDI | [K].RDSESE | 1xPhospho | Q3TI79 1xF | 0.010568 | 0.000772 | 1 | 2 | 1 | Q3TI79 | Putative ur | Q3TI79 [44 | 1 | 2456.196 | 0 | -1.19 | 0.000429 | 1.41 | 138.3 | 61.7 | 3.57 | 7.1 | | High | High | High | High | High | High | High | High | High | High | High | 0.000382 | 0.003327 | 1.91 |
| FALSE | High | RDSESESDI | [K].RDSESE | 2xPhospho | Q3TI79 2xF | 4.17E-06 | 0 | 1 | 2 | 8 | Q3TI79 | Putative ur | Q3TI79 [44 | 1 | 2536.162 | 0 | -0.22 | 0.056901 | 11.58 | 109.1 | 90.9 | 6.98 | 16.45 | | High | High | High | High | High | High | High | High | High | High | High | 0 | 5.04E-07 | 4.94 |
| FALSE | High | RDSETIEDV | [K].RDSETI | 1xPhospho | Q8CIL4 1xF | 9.71E-05 | 0 | 1 | 3 | 2 | Q8CIL4 | Uncharacte | Q8CIL4 [49 | 1 | 2685.37 | 0 | -0.61 | 0.004532 | 5.37 | 121 | 79 | 7.78 | 7.73 | | High | High | High | High | High | High | High | High | High | High | High | 0 | 1.74E-05 | 4.92 |
| FALSE | High | RDSFDDRG | [R].RDSFDI | 1xPhospho | Q6ZQ61 1x | 2.86E-07 | 0 | 1 | 6 | 7 | Q6ZQ61 | MCG12197 | Q6ZQ61 [1 | 2 | 2827.3 | 0 | 0.17 | 0.552364 | 5.86 | 95.3 | 104.7 | 8.03 | 12.64 | | High | High | High | High | High | High | High | High | High | High | High | 2.58E-05 | 0.000348 | 3.51 |
| FALSE | High | RDTDSDTQ | [R].RDTDSI | 1xCarbami | B1AZP2 1xI | 3.61E-06 | 0 | 2 | 11 | 5 | B1AZP2; H | Disks large | B1AZP2 [7 | 1 | 2353.002 | 0 | 0.54 | 0.011102 | 2.25 | 82.1 | 117.9 | 8.47 | 18.68 | Shared | High | High | High | High | High | High | High | High | High | High | High | 0 | 0.000101 | 3.08 |
| FALSE | High | RASVSTNQ | [K].RASVST | 1xPhospho | B9EKJ7 1xF | 2.67E-05 | 0 | 1 | 3 | 2 | B9EKJ7 | Tankyrase | B9EKJ7 [88 | 1 | 2792.281 | 0 | 0.38 | 0.050233 | 21.01 | 91.9 | 108.1 | 8.55 | 15.81 | | High | High | High | High | High | High | High | High | High | High | High | 0 | 4.07E-06 | 4.23 |
| FALSE | High | RDTSSNDII | [R].RDTSSN | 1xPhospho | P17426 1xI | 1.01E-05 | 0 | 1 | 2 | 2 | P17426 | AP-2 comp | P17426 [63 | 1 | 3549.727 | 0 | -0.42 | 0.053072 | 15.86 | 107.8 | 92.2 | 15.14 | 8.87 | | High | High | High | High | High | High | High | High | High | High | High | 0 | 2.26E-05 | 3.76 |
| FALSE | High | RDYDDMSI | [R].RDYDD | 1xPhospho | Q5FWJ5 1x | 0.043236 | 0.003228 | 2 | 13 | 6 | Q5FWJ5; C | Hnrpk prot | Q5FWJ5 [2 | 1 | 1479.614 | 0 | -0.16 | 0.002599 | 1.53 | 105.2 | 94.8 | 4.95 | 3.15 | Shared | High | High | High | High | High | High | High | High | High | High | High | 0.008188 | 0.08856 | 1.9 |
| FALSE | High | REPPPPQTI | [R].REPPPP | 1xPhospho | Q5U4C3 1x | 1.65E-07 | 0 | 1 | 1 | 3 | Q5U4C3 | Splicing fac | Q5U4C3 [3 | 1 | 2579.282 | 0 | 0.31 | 0.024952 | 17.37 | 87.4 | 112.6 | 2.8 | 10.82 | | High | High | High | High | High | High | High | High | High | High | High | 0 | 1.34E-08 | 3.96 |
| FALSE | High | REPSYFEIP | [R].REPSYF | 1xPhospho | Q80U49 1x | 0.000165 | 0 | 1 | 3 | 2 | Q80U49 | Centrosom | Q80U49 [2 | 1 | 1904.992 | 0 | 0.94 | 0.087513 | 14.85 | 75.1 | 124.9 | 16.38 | 32.05 | | High | High | High | High | High | High | High | High | High | High | High | 0 | 3.15E-05 | 3.17 |
| FALSE | High | REQEWDSS | [K-].REQEV | 1xPhospho | Q61687 1x | 0.001476 | 5.66E-05 | 1 | 6 | 1 | Q61687 | Transcripti | Q61687 [9 | 2 | 3575.658 | 0 | -0.2 | 0.149362 | 11.19 | 112.1 | 87.9 | 6.05 | 20.59 | | High | High | High | High | High | High | High | High | High | High | High | 2.58E-05 | 0.000368 | 2.76 |
| FALSE | High | REQGSSGL | [R].REQGS | 1xPhospho | Q3UHJ0 1x | 0.000404 | 0 | 1 | 3 | 5 | Q3UHJ0 | AP2-associ | Q3UHJ0 [9 | 1 | 2867.312 | 0 | 1.23 | 0.001678 | 25.8 | 60.8 | 139.2 | 24.09 | 4.93 | | High | High | High | High | High | High | High | High | High | High | High | 0 | 0.000237 | 2.49 |
| FALSE | High | REREESEDI | [K].REREES | 1xPhospho | Q8CFE3 1x | 8.4E-05 | 0 | 1 | 2 | 1 | Q8CFE3 | REST corep | Q8CFE3 [2 | 2 | 3681.717 | 0 | 0.72 | 0.007526 | 13.13 | 77.8 | 122.2 | 13.31 | 10.93 | | High | High | High | High | High | High | High | High | High | High | High | 0 | 1.47E-05 | 2.78 |
| FALSE | High | REREESEDI | [K].REREES | 1xDeamida | Q8CFE3 1x | 0.000738 | 0 | 1 | 2 | 7 | Q8CFE3 | REST corep | Q8CFE3 [2 | 2 | 3682.701 | 0 | 0.29 | 0.058666 | 6.38 | 90.8 | 109.2 | 8.38 | 11.57 | | High | High | High | High | High | High | High | High | High | High | High | 7.19E-05 | 0.000862 | 3.5 |

| | | | | | | | | | | | | | | | | | | | | | | | | | | | | | | | | | | | | | | |
|---|---|---|---|---|---|---|---|---|---|---|---|---|---|---|---|---|---|---|---|---|---|---|---|---|---|---|---|---|---|---|---|---|---|---|---|---|---|---|
| FALSE | High | RESASSSSS | [R].RESASS | 1xPhospho | Q9JI44 1xP | 0.005467 | 0.00062 | 1 | 1 | 2 | Q9JI44 | DNA methy | Q9JI44 [45 | 1 | 1662.846 | 0 | -0.63 | 0.000531 | 1.2 | 121.1 | 78.9 | 5.37 | 3.35 | | High | High | High | High | High | High | High | High | High | High | High | 0.000275 | 0.001597 | 2.56 |
| FALSE | High | RESASSSSS | [R].RESASS | 2xPhospho | Q9JI44 2xP | 0.017652 | 0.001282 | 1 | 1 | 1 | Q9JI44 | DNA methy | Q9JI44 [45 | 1 | 1742.813 | 0 | -0.47 | 0.014763 | 3.57 | 117.1 | 82.9 | 9.65 | 5.62 | | High | High | High | High | High | High | High | High | High | High | High | 0.000689 | 0.005915 | 2.06 |
| FALSE | High | RESDDSDV | [R].RESDDS | 1xPhospho | Q62187 1x | 7.04E-06 | 0 | 1 | 2 | 1 | Q62187 | Transcripti | Q62187 [4 | 1 | 1929.875 | 0 | -0.78 | 0.000985 | 3.04 | 126 | 74 | 4.69 | 7.8 | | High | High | High | High | High | High | High | High | High | High | High | 0 | 9.1E-07 | 3.26 |
| FALSE | High | RESDDSDV | [R].RESDDS | 2xPhospho | Q62187 2x | 4.17E-06 | 0 | 1 | 2 | 2 | Q62187 | Transcripti | Q62187 [4 | 1 | 2009.842 | 0 | -0.14 | 0.033005 | 5.39 | 106 | 94 | 5.67 | 6.83 | | High | High | High | High | High | High | High | High | High | High | High | 0 | 0.000101 | 4.03 |
| FALSE | High | RESDGTPG | [K].RESDGT | 1xPhospho | A0A1L7NR | 0.000346 | 0 | 2 | 12 | 2 | A0A1L7NR | C-Cbl asso | A0A1L7NR | 1 | 2096.981 | 0 | 0.15 | 0.031253 | 4.36 | 95.6 | 104.4 | 3.85 | 1.83 | Shared | High | High | High | High | High | High | High | High | High | High | High | 0 | 7.23E-05 | 2.64 |
| FALSE | High | RESEEPTSP | [R].RESEEP | 1xCarbami | Q6ZWQ0 1 | 0.001544 | 5.66E-05 | 1 | 9 | 1 | Q6ZWQ0 | Nesprin-2 | Q6ZWQ0 [ | 1 | 3035.461 | 0 | 1.9 | 0.001482 | 5.87 | 40.1 | 159.9 | 7.33 | 15.86 | | High | High | High | High | High | High | High | High | High | High | High | 2.58E-05 | 0.000389 | 3.16 |
| FALSE | High | RESFDLNV | [K].RESFDL | 1xPhospho | Q9QZD4 1x | 4.61E-06 | 0 | 1 | 2 | 1 | Q9QZD4 | DNA repair | Q9QZD4 [5 | 1 | 2523.29 | 0 | 0.82 | 0.001269 | 11.95 | 70.5 | 129.5 | 6.94 | 6.59 | | High | High | High | High | High | High | High | High | High | High | High | 0 | 5.67E-07 | 2.84 |
| FALSE | High | REPAVNET | [K].REPAVI | 1xPhospho | Q9WUF3 1 | 0.000656 | 0 | 1 | 1 | 1 | Q9WUF3 | CASP8-ass | Q9WUF3 [ | 1 | 2568.296 | 0 | -0.28 | 0.001008 | 3.66 | 110.2 | 89.8 | 7 | 10.35 | | High | High | High | High | High | High | High | High | High | High | High | 0 | 0.000149 | 2.19 |
| FALSE | High | RESFSSHCA | [K].RESFSS | 1xCarbami | Q91X43 1x | 0.001933 | 5.66E-05 | 1 | 2 | 1 | Q91X43 | SH3 domai | Q91X43 [5 | 1 | 1746.84 | 0 | -0.28 | 0.052878 | 6.71 | 109.6 | 90.4 | 6.32 | 5.49 | | High | High | High | High | High | High | High | High | High | High | High | 2.58E-05 | 0.0005 | 1.68 |
| FALSE | High | RESGSPYS | [K].RESGSF | 1xPhospho | Q5DU09 1x | 0.000379 | 0 | 1 | 4 | 2 | Q5DU09 | Zinc finger | Q5DU09 [5 | 1 | 2048.047 | 0 | 0.43 | 0.006788 | 3.92 | 84.7 | 115.3 | 7.54 | 7.69 | | High | High | High | High | High | High | High | High | High | High | High | 0.001453 | 0.01461 | 2.5 |
| FALSE | High | RESKPEEPF | [R].RESKPE | 1xPhospho | A0A0R4J1 | 0.001489 | 5.66E-05 | 1 | 5 | 2 | A0A0R4J1 | Menin OS= | A0A0R4J1 | 1 | 2158.188 | 0 | 0.07 | 0.928824 | 10.1 | 103.1 | 96.9 | 9.42 | 8.5 | | High | High | High | High | High | High | High | High | High | High | High | 2.58E-05 | 0.000373 | 2.34 |
| FALSE | High | RESLSYIPK | [R].RESLSY | 1xPhospho | Q4VAC9 1x | 0.004696 | 0.000456 | 1 | 3 | 1 | Q4VAC9 | Pleckstrin | Q4VAC9 [8 | 1 | 1630.897 | 0 | 0 | 0.868709 | 23.99 | 99.9 | 100.1 | 5.91 | 13.21 | | High | High | High | High | High | High | High | High | High | High | High | 0.000204 | 0.001354 | 1.74 |
| FALSE | High | RESPSPAPK | [K].RESPSP | 1xPhospho | Q52KI8 1xF | 0.019618 | 0.001438 | 1 | 6 | 1 | Q52KI8 | Serine/arg | Q52KI8 [44 | 1 | 1759.962 | 0 | -1.25 | 0.268159 | 15.89 | 135.7 | 64.3 | 83.15 | 113.65 | | High | High | High | High | High | High | High | High | High | High | High | 0.000756 | 0.006645 | 1.46 |
| FALSE | High | RESSTVGG | [K].RESSTV | 1xPhospho | Q3TW94 1 | 2.1E-05 | 0 | 1 | 3 | 1 | Q3TW94 | Putative ur | Q3TW94 [ | 1 | 2565.153 | 0 | 0.13 | 0.021512 | 2.87 | 97.4 | 102.6 | 9.33 | 4.97 | | High | High | High | High | High | High | High | High | High | High | High | 0 | 3.09E-06 | 2.66 |
| FALSE | High | RESTGSYS | [R].RESTGS | 2xPhospho | F6WJB7 2x | 0.00172 | 5.66E-05 | 1 | 5 | 1 | F6WJB7 | Ubiquitin c | F6WJB7 [1 | 1 | 3528.462 | 0 | -0.18 | 0.82376 | 28.31 | 106.3 | 93.7 | 19.27 | 18.22 | | High | High | High | High | High | High | High | High | High | High | High | 2.58E-05 | 0.000437 | 3.46 |
| FALSE | High | RESVSPEDS | [K].RESVSP | 1xPhospho | G3UWX1 1 | 0.004256 | 0.00036 | 2 | 8 | 3 | G3UWX1; | Replication | G3UWX1 [ | 1 | 1800.878 | 0 | -0.29 | 0.001701 | 2.66 | 110.6 | 89.4 | 3.46 | 1.99 | Shared | High | High | High | High | High | High | High | High | High | High | High | 0.000161 | 0.001206 | 1.59 |
| FALSE | High | RESVSPEDS | [K].RESVSP | 2xPhospho | G3UWX1 2 | 0.057686 | 0.004227 | 2 | 8 | 1 | G3UWX1; | Replication | G3UWX1 [ | 1 | 1880.844 | 0 | 0.02 | 0.782986 | 11.58 | 102.8 | 97.2 | 8.3 | 16.73 | Shared | High | High | High | High | High | High | High | High | High | High | High | 0.002159 | 0.0227 | 2.55 |
| FALSE | High | RESVVSGD | [K].RESVVS | 1xPhospho | P14873 1xI | 0.002058 | 5.66E-05 | 2 | 2 | 1 | P14873; B2 | Microtubu | P14873 [98 | 2 | 2948.36 | 0 | -0.7 | 0.201234 | 18.97 | 121.2 | 78.8 | 21.35 | 42.42 | Shared | High | High | High | High | High | High | High | High | High | High | High | 2.58E-05 | 0.000539 | 3.17 |
| FALSE | High | RETDGSETI | [R].RETDGS | 1xPhospho | Q3TRW3 1 | 1.02E-06 | 0 | 1 | 5 | 1 | Q3TRW3 | Putative ur | Q3TRW3 [ | 1 | 2373.138 | 0 | 1.35 | 0.003024 | 22.47 | 51.5 | 148.5 | 9.37 | 16.19 | | High | High | High | High | High | High | High | High | High | High | High | 0 | 1.04E-07 | 4.75 |
| FALSE | High | RETTVASDS | [K].RETTVA | 2xPhospho | Q8C1D8 2x | 0.001898 | 5.66E-05 | 1 | 4 | 1 | Q8C1D8 | Protein IW | Q8C1D8 [3 | 1 | 2226.993 | 0 | -0.4 | 0.00241 | 2.25 | 113.6 | 86.4 | 3.54 | 6.02 | | High | High | High | High | High | High | High | High | High | High | High | 2.58E-05 | 0.000492 | 1.99 |
| FALSE | High | RETVVESQ | [K].RETVVE | 2xPhospho | E9PZM7 2x | 0.014929 | 0.001049 | 1 | 3 | 2 | E9PZM7 | SR-related | E9PZM7 [8 | 1 | 2134.962 | 0 | 0.05 | 0.989326 | 27.03 | 100.8 | 99.2 | 38.29 | 44.65 | | High | High | High | High | High | High | High | High | High | High | High | 0.000555 | 0.004905 | 2.4 |
| FALSE | High | RETVVESQ | [K].RETVVE | 1xPhospho | E9PZM7 1x | 1.14E-08 | 0 | 1 | 3 | 3 | E9PZM7 | SR-related | E9PZM7 [8 | 1 | 2284.159 | 0 | 0.56 | 0.004527 | 5.8 | 81.5 | 118.5 | 4.1 | 7.96 | | High | High | High | High | High | High | High | High | High | High | High | 0 | 6.71E-10 | 4.31 |
| FALSE | High | RESGENPP | [K].RESGEN | 1xPhospho | P33174 1xI | 8.56E-05 | 0 | 1 | 1 | 2 | P33174 | Chromoso | P33174 [78 | 1 | 1664.877 | 0 | -0.23 | 0.009595 | 3.49 | 108 | 92 | 4.61 | 3.36 | | High | High | High | High | High | High | High | High | High | High | High | 0.00554 | 0.06007 | 3.01 |
| FALSE | High | RENPVTPD | [K].RENPVT | 1xPhospho | E9PVX6 1xI | 0.000138 | 0 | 1 | 2 | 1 | E9PVX6 | Proliferatic | E9PVX6 [3 | 1 | 1721.817 | 0 | 0.23 | 0.327895 | 41.94 | 92.5 | 107.5 | 12.97 | 17.99 | | High | High | High | High | High | High | High | High | High | High | High | 0 | 2.59E-05 | 1.72 |
| FALSE | High | RELTGSNA | [K].RELTGS | 1xPhospho | A0A1W2P6 | 0.053473 | 0.00384 | 1 | 6 | 2 | A0A1W2P6 | Bromodom | A0A1W2P6 | 1 | 1755.859 | 0 | -0.73 | 0.000553 | 0.82 | 124.6 | 75.4 | 3.74 | 5.68 | | High | High | High | High | High | High | High | High | High | High | High | 0.006084 | 0.0662 | 1.3 |
| FALSE | High | REISSSPTS | [R].REISSSF | 2xPhospho | Q8BTI8 2xF | 0.00967 | 0.000736 | 1 | 3 | 2 | Q8BTI8 | Serine/arg | Q8BTI8 [44 | 1 | 1709.828 | 0 | -0.8 | 0.000727 | 1.4 | 123.2 | 76.8 | 6.31 | 4.53 | | High | High | High | High | High | High | High | High | High | High | High | 0.000344 | 0.003035 | 2.18 |
| FALSE | High | REATLPPVS | [K].REATLP | 1xPhospho | Q6NZJ6 1xI | 1.08E-05 | 0 | 1 | 9 | 5 | Q6NZJ6 | Eukaryotic | Q6NZJ6 [1 | 1 | 1830.029 | 0 | 0.1 | 0.243848 | 9 | 96.2 | 103.8 | 3.4 | 5.39 | | High | High | High | High | High | High | High | High | High | High | High | 0 | 4.31E-06 | 3.18 |
| FALSE | High | REATLPPVS | [K].REATLP | 2xPhospho | Q6NZJ6 2xI | 0.000253 | 0 | 1 | 9 | 2 | Q6NZJ6 | Eukaryotic | Q6NZJ6 [1 | 1 | 1909.995 | 0 | 1.16 | 0.00537 | 19.02 | 65.7 | 134.3 | 8.21 | 18.46 | | High | High | High | High | High | High | High | High | High | High | High | 0.002455 | 0.02699 | 2.64 |
| FALSE | High | REATPPGTI | [R].REATPF | 1xPhospho | P58334 1xI | 0.000173 | 0 | 1 | 2 | 1 | P58334 | Krueppel-li | P58334 [48 | 1 | 4044.127 | 0 | 0.4 | 0.302498 | 8.58 | 85.7 | 114.3 | 16.22 | 20.62 | | High | High | High | High | High | High | High | High | High | High | High | 0 | 3.33E-05 | 3.05 |
| FALSE | High | REDEEESLI | [K].REDEEE | 1xCarbami | Q01853 1x | 1.21E-05 | 0 | 1 | 2 | 4 | Q01853 | Transitiona | Q01853 [1 | 1 | 2651.113 | 0 | 0.11 | 0.664693 | 2.83 | 97.8 | 102.2 | 5.29 | 5.04 | | High | High | High | High | High | High | High | High | High | High | High | 0 | 1.67E-06 | 4.03 |
| FALSE | High | REDPGDNV | [K].REDPGI | 1xPhospho | Q4VAA2 1x | 4.05E-05 | 0 | 1 | 6 | 3 | Q4VAA2 | Protein CD | Q4VAA2 [1 | 1 | 2485.103 | 0 | -0.2 | 0.071622 | 3.21 | 109.2 | 90.8 | 6.87 | 16.26 | | High | High | High | High | High | High | High | High | High | High | High | 0 | 6.49E-06 | 2.77 |
| FALSE | High | REDSFESLD | [R].REDSFE | 1xPhospho | Q6ZQ32 1x | 0.059159 | 0.004294 | 2 | 5 | 2 | Q6ZQ32; F | MKIAA085 | Q6ZQ32 [1 | 1 | 1906.875 | 0 | 0.53 | 0.000181 | 2.52 | 83 | 117 | 4.14 | 3.21 | Shared | High | High | High | High | High | High | High | High | High | High | High | 0.002217 | 0.02349 | 3.27 |
| FALSE | High | REDSPGPE | [K].REDSPC | 1xPhospho | D3YUY7 1x | 0.000386 | 0 | 1 | 3 | 1 | D3YUY7 | Tripartite r | D3YUY7 [4 | 1 | 2123.024 | 0 | 0.02 | 0.297017 | 4.56 | 99.4 | 100.6 | 7.83 | 7.71 | | High | High | High | High | High | High | High | High | High | High | High | 0 | 8.14E-05 | 3.68 |
| FALSE | High | REDSPGPE | [K].REDSPC | 1xPhospho | D3YUY7 1x | 5.32E-06 | 0 | 1 | 3 | 1 | D3YUY7 | Tripartite r | D3YUY7 [4 | 1 | 2139.019 | 0 | 0.11 | 0.091675 | 4.14 | 96.3 | 103.7 | 3.88 | 5.92 | | High | High | High | High | High | High | High | High | High | High | High | 0 | 6.63E-07 | 4.31 |
| FALSE | High | REEAAPPTI | [K].REEAAF | 1xPhospho | Q80WJ7 1x | 6.88E-08 | 0 | 1 | 3 | 3 | Q80WJ7 | Protein LYF | Q80WJ7 [8 | 1 | 2457.279 | 0 | 1.95 | 0.000909 | 8.54 | 42.2 | 157.8 | 5.03 | 12.29 | | High | High | High | High | High | High | High | High | High | High | High | 0 | 5E-09 | 3.04 |
| FALSE | High | REEQTDTS | [R].REEQTI | 1xPhospho | P36916 1xI | 0.000158 | 0 | 1 | 1 | 4 | P36916 | Guanine nu | P36916 [44 | 1 | 2406.088 | 0 | -1.28 | 0.004726 | 0.9 | 141.7 | 58.3 | 13.59 | 14.81 | | High | High | High | High | High | High | High | High | High | High | High | 0 | 5.83E-05 | 2.02 |
| FALSE | High | REEQTDTS | [R].REEQTI | 2xPhospho | P36916 2xI | 0.002553 | 0.000109 | 1 | 1 | 2 | P36916 | Guanine nu | P36916 [44 | 1 | 2486.055 | 0 | -1.2 | 0.003914 | 0.49 | 138.5 | 61.5 | 6.35 | 19 | | High | High | High | High | High | High | High | High | High | High | High | 4.94E-05 | 0.000684 | 2.67 |
| FALSE | High | REEQTDTS | [R].REEQTI | 3xPhospho | P36916 3xI | 0.000302 | 0 | 1 | 1 | 1 | P36916 | Guanine nu | P36916 [44 | 1 | 2566.021 | 0 | -0.53 | 0.019505 | 14.38 | 119 | 81 | 12.59 | 17.62 | | High | High | High | High | High | High | High | High | High | High | High | 0 | 6.24E-05 | 3.44 |
| FALSE | High | REESPMDV | [R].REESPN | 1xPhospho | A2AFQ0 1x | 0.002376 | 0.000109 | 1 | 4 | 4 | A2AFQ0 | E3 ubiquiti | A2AFQ0 [3 | 1 | 3966.868 | 0 | 0.04 | 0.502965 | 16.67 | 99.3 | 100.7 | 8.34 | 6.9 | | High | High | High | High | High | High | High | High | High | High | High | 4.94E-05 | 0.000629 | 3.99 |
| FALSE | High | REESPMDV | [R].REESPN | 1xDeamida | A2AFQ0 1x | 0.097743 | 0.007804 | 1 | 4 | 3 | A2AFQ0 | E3 ubiquiti | A2AFQ0 [3 | 1 | 3967.852 | 0 | -0.12 | 0.362814 | 1.07 | 104.3 | 95.7 | 12.84 | 7.91 | | High | High | High | High | High | High | High | High | High | High | High | 0.005122 | 0.05568 | 4.21 |
| FALSE | High | REESPMDV | [R].REESPN | 1xPhospho | A2AFQ0 1x | 0.000924 | 0 | 1 | 4 | 4 | A2AFQ0 | E3 ubiquiti | A2AFQ0 [3 | 1 | 3982.863 | 0 | -0.04 | 0.888174 | 10.76 | 101.3 | 98.7 | 2.66 | 14.8 | | High | High | High | High | High | High | High | High | High | High | High | 0 | 0.000218 | 3.45 |
| FALSE | High | REESPMDV | [R].REESPN | 2xPhospho | A2AFQ0 2x | 0.040028 | 0.002867 | 1 | 4 | 1 | A2AFQ0 | E3 ubiquiti | A2AFQ0 [3 | 1 | 4046.834 | 0 | -0.17 | 0.39686 | 19.14 | 109.6 | 90.4 | 18.14 | 16.38 | | High | High | High | High | High | High | High | High | High | High | High | 0.001484 | 0.0149 | 3.91 |
| FALSE | High | REFITGDVE | [R].REFITG | 1xPhospho | Q8C1W9 1 | 0.002895 | 0.00016 | 1 | 4 | 1 | Q8C1W9 | Putative ur | Q8C1W9 [ | 1 | 3558.595 | 0 | 0.01 | 0.601267 | 30.68 | 107.7 | 92.3 | 13.4 | 21.18 | | High | High | High | High | High | High | High | High | High | High | High | 7.19E-05 | 0.000787 | 3.44 |
| FALSE | High | REFSGGSYI | [K].REFSGC | 1xPhospho | A0A1D5RL | 7.96E-05 | 0 | 1 | 9 | 2 | A0A1D5RL | Serine/thre | A0A1D5RL | 1 | 1783.878 | 0 | 2.29 | 0.000244 | 10.89 | 33.9 | 166.1 | 11.79 | 4.67 | | High | High | High | High | High | High | High | High | High | High | High | 0.003265 | 0.03527 | 1.99 |
| FALSE | High | REFVEDEE | [K].REFVED | 2xPhospho | Q8BGS0 2x | 0.00537 | 0.000599 | 1 | 2 | 1 | Q8BGS0 | Protein MA | Q8BGS0 [2 | 1 | 3310.368 | 0 | -0.25 | 0.031814 | 7.24 | 107.3 | 92.7 | 7.55 | 6.5 | | High | High | High | High | High | High | High | High | High | High | High | 0.000267 | 0.001562 | 2.94 |
| FALSE | High | REGAGDTE | [K].REGAG | 1xPhospho | Q8C9B9 1x | 0.06169 | 0.004646 | 1 | 3 | 1 | Q8C9B9 | Death-indu | Q8C9B9 [3 | 1 | 3000.401 | 0 | -0.55 | 0.043598 | 11.1 | 119.5 | 80.5 | 12.47 | 17.1 | | High | High | High | High | High | High | High | High | High | High | High | 0.002381 | 0.02459 | 2.56 |
| FALSE | High | REGDSPTA | [K].REGDSI | 1xPhospho | Q6PDX6 1x | 0.022773 | 0.001542 | 1 | 2 | 1 | Q6PDX6 | E3 ubiquiti | Q6PDX6 [2 | 1 | 2798.233 | 0 | -0.43 | 0.243534 | 12.51 | 120.4 | 79.6 | 24.04 | 45.08 | | High | High | High | High | High | High | High | High | High | High | High | 0.000832 | 0.007929 | 2.57 |
| FALSE | High | REGPVGGE | [K].REGPV | 1xPhospho | Q3UHU5 1 | 1.58E-07 | 0 | 1 | 5 | 2 | Q3UHU5 | Microtubu | Q3UHU5 [ | 1 | 2407.089 | 0 | -1.82 | 0.00036 | 0.37 | 154.1 | 45.9 | 8.68 | 11.04 | | High | High | High | High | High | High | High | High | High | High | High | 0 | 1.27E-08 | 2.49 |
| FALSE | High | REGPVGGE | [K].REGPV | 1xPhospho | Q3UHU5 1 | 0.01002 | 0.000736 | 1 | 5 | 1 | Q3UHU5 | Microtubu | Q3UHU5 [ | 1 | 2423.084 | 0 | -1.16 | 0.001532 | 5.14 | 140.3 | 59.7 | 8.5 | 10.43 | | High | High | High | High | High | High | High | High | High | High | High | 0.000344 | 0.00315 | 1.7 |
| FALSE | High | REGTPPPPI | [R].REGTPF | 1xPhospho | F8VQL9 1x | 0.002948 | 0.00016 | 1 | 8 | 2 | F8VQL9 | Nuclear re | F8VQL9 [1 | 1 | 1606.83 | 0 | 0.75 | 6.71E-06 | 1.3 | 74.7 | 125.3 | 3.76 | 2.07 | | High | High | High | High | High | High | High | High | High | High | High | 7.19E-05 | 0.0008 | 2.4 |
| FALSE | High | REHANIDA | [R].REHAN | 1xPhospho | Q8VDQ8 1 | 0.00967 | 0.000736 | 1 | 5 | 1 | Q8VDQ8 | NAD-depe | Q8VDQ8 [ | 1 | 3101.542 | 0 | 0.29 | 0.261024 | 5.96 | 93 | 107 | 16.24 | 7.35 | | High | High | High | High | High | High | High | High | High | High | High | 0.000344 | 0.003025 | 2.76 |
| FALSE | High | REHQGYSP | [K].REHQG | 1xCarbami | A2A7F4 1xI | 5.13E-06 | 0 | 1 | 2 | 2 | A2A7F4 | MCG12287 | A2A7F4 [1 | 1 | 2384.147 | 0 | -0.63 | 0.010015 | 3.82 | 123.6 | 76.4 | 7.93 | 9.21 | | High | High | High | High | High | High | High | High | High | High | High | 0 | 6.4E-07 | 2.05 |
| FALSE | High | RELPTLSPA | [K].RELPTL | 1xPhospho | E9Q8A3 1x | 1.41E-05 | 0 | 1 | 3 | 2 | E9Q8A3 | Phosphatic | E9Q8A3 [2 | 1 | 2404.289 | 0 | 0.05 | 0.446844 | 20.49 | 98.3 | 101.7 | 7 | 7.07 | | High | High | High | High | High | High | High | High | High | High | High | 0 | 1.98E-06 | 3.79 |
| FALSE | High | REISETEAK | [K].REISETI | 1xPhospho | A2AEW1 1 | 0.036103 | 0.002342 | 1 | 7 | 1 | A2AEW1 | Transcripti | A2AEW1 [ | 1 | 1600.835 | 0 | 1.34 | 0.006375 | 24.24 | 64.2 | 135.8 | 14.22 | 17.91 | | High | High | High | High | High | High | High | High | High | High | High | 0.001279 | 0.01336 | 1.16 |
| FALSE | High | REISSSPTS | [K].REISSSF | 1xPhospho | Q8BTI8 1xF | 0.068752 | 0.005042 | 1 | 3 | 1 | Q8BTI8 | Serine/arg | Q8BTI8 [44 | 1 | 1629.861 | 0 | -0.71 | 0.008797 | 6.92 | 123.5 | 76.5 | 5.56 | 16.04 | | High | High | High | High | High | High | High | High | High | High | High | 0.002589 | 0.02777 | 1.54 |
| FALSE | High | RDVSSPDE | [K].RDVSSF | 1xPhospho | Q544H8 1x | 0.017498 | 0.001245 | 1 | 1 | 2 | Q544H8 | Putative ur | Q544H8 [1 | 1 | 2043.975 | 0 | -0.72 | 0.005936 | 6.74 | 124.1 | 75.9 | 4.87 | 11.14 | | High | High | High | High | High | High | High | High | High | High | High | 0.002187 | 0.02335 | 2.3 |
| FALSE | High | REVLYDSE | [K].REVLYD | 2xPhospho | Q5U4C3 2x | 0.004143 | 0.00036 | 1 | 1 | 4 | Q5U4C3 | Splicing fac | Q5U4C3 [6 | 1 | 2127.92 | 0 | -0.15 | 0.005475 | 3.37 | 105.5 | 94.5 | 5.19 | 3.36 | | High | High | High | High | High | High | High | High | High | High | High | 0.000344 | 0.00311 | 3.19 |
| FALSE | High | RASVSTNQ | [K].RASVST | 1xPhospho | B9EKJ7 1xF | 1.25E-05 | 0 | 1 | 3 | 2 | B9EKJ7 | Tankyrase | B9EKJ7 [88 | 1 | 2776.286 | 0 | 0.34 | 0.605906 | 16.84 | 90.5 | 109.5 | 16.54 | 30.45 | | High | High | High | High | High | High | High | High | High | High | High | 0 | 1.74E-06 | 4.24 |
| FALSE | High | RASTEDSPI | [R].RASTED | 1xPhospho | B9EK83 1xI | 3.3E-06 | 0 | 2 | 3 | 3 | B9EK83; A | Transient r | B9EK83 [1 | 1 | 1958.947 | 0 | 1.72 | 0.000239 | 10.48 | 47 | 153 | 6.82 | 4.96 | Shared | High | High | High | High | High | High | High | High | High | High | High | 0 | 1.25E-05 | 3.34 |
| FALSE | High | QVSNPLPK | [K].QVSNP | 1xPhospho | Q8VD75 1x | 0.013308 | 0.000889 | 1 | 2 | 1 | Q8VD75 | Huntingtin | Q8VD75 [1 | 0 | 1420.797 | 0 | 1.23 | 0.00241 | 12.76 | 58.2 | 141.8 | 8.84 | 12.41 | | High | High | High | High | High | High | High | High | High | High | High | 0.000486 | 0.004315 | 2.11 |
| FALSE | High | QVSPVASV | [R].QVSPV | 1xPhospho | A0A1B0GR | 0.056724 | 0.004124 | 1 | 1 | 1 | A0A1B0GR | RIKEN cDN | A0A1B0GR | 0 | 1348.719 | 0 | 1.75 | 0.000158 | 8.51 | 45.9 | 154.1 | 7.35 | 4.33 | | High | High | High | High | High | High | High | High | High | High | High | 0.002148 | 0.0223 | 2.49 |
| FALSE | High | QVSTENDS | [K].QVSTEI | 1xPhospho | E9PUJ6 1xI | 4.33E-07 | 0 | 1 | 4 | 4 | E9PUJ6 | E3 ubiquiti | E9PUJ6 [1 | 0 | 1794.858 | 0 | 1.17 | 0.00148 | 14.67 | 61.4 | 138.6 | 6.21 | 9.2 | | High | High | High | High | High | High | High | High | High | High | High | 0 | 3.97E-08 | 3.23 |
| FALSE | High | QVSTEPSS | [R].QVSTEI | 1xPhospho | P70441 1xI | 9.91E-06 | 0 | 1 | 1 | 4 | P70441 | Na(+)/H(+) | P70441 [3 | 0 | 3371.724 | 0 | 2.17 | 0.003876 | 33.61 | 34.5 | 165.5 | 27.98 | 18.33 | | High | High | High | High | High | High | High | High | High | High | High | 0 | 1.34E-06 | 4.18 |
| FALSE | High | QVTASPSS | [R].QVTASI | 1xPhospho | Q3SYK5 1x | 0.003312 | 0.00016 | 1 | 4 | 2 | Q3SYK5 | Tyrosine-p | Q3SYK5 [8 | 0 | 1836.962 | 0 | -0.07 | 0.911229 | 28.07 | 101.6 | 98.4 | 11.47 | 8.88 | | High | High | High | High | High | High | High | High | High | High | High | 7.19E-05 | 0.000912 | 1.75 |
| FALSE | High | QVTPGPAT | [R].QVTPG | 1xCarbami | B2RXU3 1x | 0.000115 | 0 | 1 | 2 | 4 | B2RXU3 | ADP-ribosy | B2RXU3 [1 | 0 | 4295.793 | 0 | 0.62 | 0.003855 | 4 | 78.5 | 121.5 | 8.59 | 11.5 | | High | High | High | High | High | High | High | High | High | High | High | 0 | 0.000021 | 4.2 |
| FALSE | High | QVTPGPAT | [R].QVTPG | 1xCarbami | B2RXU3 1x | 2.43E-06 | 0 | 1 | 2 | 11 | B2RXU3 | ADP-ribosy | B2RXU3 [1 | 0 | 4296.777 | 0 | 1.49 | 0.001689 | 23.41 | 54.1 | 145.9 | 11.23 | 11 | | High | High | High | High | High | High | High | High | High | High | High | 0 | 3.61E-06 | 4.37 |
| FALSE | High | QVVDSAYE | [K].QVVDS | 1xPhospho | A0A1B0GS | 1.41E-06 | 0 | 1 | 9 | 1 | A0A1B0GS | L-lactate d | A0A1B0GS | 0 | 1788.955 | 0 | 0.22 | 0.050007 | 13.96 | 87.3 | 112.7 | 16.56 | 19.65 | | High | High | High | High | High | High | High | High | High | High | High | 0 | 1.5E-07 | 3.09 |
| FALSE | High | QVVQTPNT | [R].QVVQT | 1xPhospho | A0A1W2P7 | 0.099328 | 0.008003 | 1 | 4 | 1 | A0A1W2P7 | Uncharacte | A0A1W2P7 | 0 | 1996.047 | 0 | 0.02 | 0.336066 | 9.36 | 94.4 | 105.6 | 11.44 | 10.12 | | High | High | High | High | High | High | High | High | High | High | High | 0.004032 | 0.04286 | 2.39 |
| FALSE | High | QWLCAPLC | [K].QWLCA | 2xCarbami | P54276 1xI | 2.08E-05 | 0 | 1 | 2 | 1 | P54276 | DNA mism | P54276 [7 | 0 | 2170.002 | 0 | 0.19 | 0.617067 | 21.99 | 94.3 | 105.7 | 15.01 | 13.15 | | High | High | High | High | High | High | High | High | High | High | High | 0 | 3.06E-06 | 3.98 |
| FALSE | High | QWNLTSPS | [R].QWNLT | 1xPhospho | H7BX08 1x | 2.28E-06 | 0 | 1 | 6 | 6 | H7BX08 | Calmodulir | H7BX08 [8 | 0 | 3119.523 | 0 | 0.14 | 0.315265 | 8.47 | 95 | 105 | 14.37 | 7.7 | | High | High | High | High | High | High | High | High | High | High | High | 0 | 2.78E-07 | 3.88 |
| FALSE | High | QWSVQSG | [R].QWSV( | 1xPhospho | B0QZP6 1x | 0.04913 | 0.003576 | 1 | 4 | 1 | B0QZP6 | Transmem | B0QZP6 [4 | 0 | 1521.741 | 0 | 0.65 | 0.006167 | 8.64 | 81.5 | 118.5 | 8.35 | 14.01 | | High | High | High | High | High | High | High | High | High | High | High | 0.001889 | 0.01893 | 2.5 |
| FALSE | High | QWVESSPK | [R].QWVES | 1xPhospho | Q5SSZ5 1xI | 0.001783 | 5.66E-05 | 1 | 2 | 2 | Q5SSZ5 | Tensin-3 O | Q5SSZ5 [9 | 0 | 1498.771 | 0 | -0.92 | 0.005735 | 5.37 | 129.7 | 70.3 | 7.87 | 12.06 | | High | High | High | High | High | High | High | High | High | High | High | 2.58E-05 | 0.000457 | 2.6 |
| FALSE | High | QVSITGFFC | [R].QVSITG | 1xPhospho | Q9EQ28 1x | 0.002869 | 0.00016 | 1 | 5 | 1 | Q9EQ28 | DNA polym | Q9EQ28 [4 | 0 | 1692.913 | 0 | 1.15 | 0.004783 | 18.94 | 57.3 | 142.7 | 7.05 | 17.61 | | High | High | High | High | High | High | High | High | High | High | High | 7.19E-05 | 0.000778 | 3.39 |
| FALSE | High | QYEEEEED | [R].QYEEEE | 1xPhospho | P49135 1xI | 6.77E-05 | 0 | 1 | 2 | 1 | P49135 | TFIIH basal | P49135 [1 | 0 | 3503.526 | 0 | -1.08 | 0.001238 | 0.41 | 133.3 | 66.7 | 8.42 | 20.95 | | High | High | High | High | High | High | High | High | High | High | High | 0 | 1.16E-05 | 2.72 |
| FALSE | High | QYQDSPNC | [K].QYQDS | 1xPhospho | Q3U671 1x | 0.003433 | 0.00021 | 1 | 2 | 1 | Q3U671 | Putative ur | Q3U671 [3 | 0 | 1645.799 | 0 | -1.13 | 0.003469 | 6.47 | 138.2 | 61.8 | 12.54 | 16.42 | | High | High | High | High | High | High | High | High | High | High | High | 9.42E-05 | 0.000948 | 1.58 |
| FALSE | High | QYSETEPHI | [R].QYSETE | 1xPhospho | E9Q6J5-3 1 | 0.030632 | 0.001857 | 1 | 3 | 2 | E9Q6J5-3 | Isoform Ne | E9Q6J5-3 [ | 0 | 1872.878 | 0 | 1.17 | 0.047913 | 13.94 | 69.5 | 130.5 | 16.38 | 25.26 | | High | High | High | High | High | High | High | High | High | High | High | 0.001011 | 0.01098 | 1.56 |
| FALSE | High | QYTSPEEID | [R].QYTSPE | 1xPhospho | B2RTB0 1xI | 9.2E-05 | 0 | 1 | 1 | 1 | B2RTB0 | MCG17262 | B2RTB0 [1 | 0 | 2388.174 | 0 | -0.21 | 0.239178 | 4.19 | 109.1 | 90.9 | 6.49 | 15.88 | | High | High | High | High | High | High | High | High | High | High | High | 0 | 1.64E-05 | 3.6 |
| FALSE | High | RADALTSSI | [R].RADALT | 1xPhospho | Q3UJN1 1x | 0.018774 | 0.001362 | 1 | 3 | 2 | Q3UJN1 | DNA helica | Q3UJN1 [3 | 1 | 1439.721 | 0 | -0.4 | 0.006552 | 1.71 | 110.8 | 89.2 | 4.32 | 4.93 | | High | High | High | High | High | High | High | High | High | High | High | 0.000723 | 0.006364 | 1.92 |
| FALSE | High | RADANLLT | [K].RADAN | 1xPhospho | Q3U2K4 1x | 3.4E-07 | 0 | 1 | 4 | 2 | Q3U2K4 | Putative ur | Q3U2K4 [6 | 1 | 2113.012 | 0 | 0.02 | 0.530147 | 3.47 | 99.3 | 100.7 | 4.56 | 3.71 | | High | High | High | High | High | High | High | High | High | High | High | 0 | 3.01E-08 | 2.76 |
| FALSE | High | RADLNQGI | [R].RADLN | 1xPhospho | Q8C845 1x | 0.00418 | 0.00036 | 1 | 2 | 1 | Q8C845 | EF-hand do | Q8C845 [6 | 1 | 2033.997 | 0 | 1.31 | 0.00046 | 12.91 | 56.9 | 143.1 | 4.73 | 7.06 | | High | High | High | High | High | High | High | High | High | High | High | 0.000161 | 0.001184 | 2.15 |
| FALSE | High | RADLNQGI | [R].RADLN | 1xPhospho | Q8C845 1x | 0.039686 | 0.00283 | 1 | 2 | 1 | Q8C845 | EF-hand do | Q8C845 [6 | 2 | 2190.098 | 0 | 1.42 | 0.000377 | 6.73 | 53.7 | 146.3 | 9.31 | 8.38 | | High | High | High | High | High | High | High | High | High | High | High | 0.001468 | 0.01478 | 0.98 |
| FALSE | High | RADNCSPV | [R].RADNC | 1xCarbami | Q0VGY8 1x | 0.000281 | 0 | 1 | 4 | 3 | Q0VGY8 | Protein TA | Q0VGY8 [2 | 1 | 2985.428 | 0 | 0.22 | 0.067495 | 6.29 | 92.5 | 107.5 | 9.35 | 6.48 | | High | High | High | High | High | High | High | High | High | High | High | 0.000801 | 0.007303 | 3.79 |
| FALSE | High | RAEAQALL | [R].RAEAQ | 1xPhospho | A2A4P4 1x | 3.8E-05 | 0 | 1 | 3 | 9 | A2A4P4 | Uncharacte | A2A4P4 [5 | 1 | 2418.139 | 0 | 0.3 | 0.039038 | 9.67 | 89.6 | 110.4 | 5.39 | 11.69 | | High | High | High | High | High | High | High | High | High | High | High | 2.58E-05 | 0.000367 | 2.85 |
| FALSE | High | RAELSGNC | [R].RAELSG | 1xCarbami | G3X932 1x | 0.039011 | 0.002718 | 1 | 4 | 2 | G3X932 | DNA segm | G3X932 [2 | 1 | 2120.088 | 0 | 0.03 | 0.573407 | 25.19 | 103.2 | 96.8 | 10.91 | 12.99 | | High | High | High | High | High | High | High | High | High | High | High | 0.001453 | 0.01452 | 1.61 |
| FALSE | High | RAETFGGY | [R].RAETF( | 1xPhospho | Q6P9R4 1x | 3.33E-06 | 0 | 1 | 2 | 2 | Q6P9R4 | Rho guanir | Q6P9R4 [5 | 1 | 2196.074 | 0 | 0.9 | 0.001479 | 9.47 | 69.7 | 130.3 | 6.28 | 6.51 | | High | High | High | High | High | High | High | High | High | High | High | 0 | 7.1E-06 | 1.94 |
| FALSE | High | RAPAAQPP | [R].RAPAA( | 1xPhospho | B2RPU8 1x | 1.22E-09 | 0 | 1 | 4 | 3 | B2RPU8 | MCG13067 | B2RPU8 [2 | 1 | 2462.287 | 0 | -0.86 | 0.022568 | 4.04 | 126.7 | 73.3 | 8.5 | 20.67 | | High | High | High | High | High | High | High | High | High | High | High | 0 | 5.45E-11 | 4.79 |
| FALSE | High | RAPAPPPP( | [R].RAPAPF | 1xPhospho | B2RWU4 1 | 0.000505 | 0 | 2 | 10 | 2 | B2RWU4; ( | Cordon-ble | B2RWU4 [3 | 1 | 2408.28 | 0 | -0.06 | 0.900261 | 4.4 | 100.2 | 99.8 | 8.37 | 3.53 | Shared | High | High | High | High | High | High | High | High | High | High | High | 0 | 0.000111 | 3.51 |
| FALSE | High | QYNIPHGP | [R].QYNIPH | 1xPhospho | A2AM95 1 | 0.012619 | 0.000889 | 1 | 3 | 1 | A2AM95 | Emerin OS | A2AM95 [1 | 0 | 1847.937 | 0 | 0.25 | 0.362826 | 42.45 | 82.3 | 117.7 | 12.79 | 33.12 | | High | High | High | High | High | High | High | High | High | High | High | 0.000449 | 0.004069 | 1.18 |
| FALSE | High | QVSASSSPI | [R].QVSAS | 2xPhospho | B2RQB7 2x | 0.004451 | 0.000409 | 1 | 2 | 1 | B2RQB7 | Atxn7 prot | B2RQB7 [6 | 0 | 2788.3 | 0 | 0.6 | 0.033416 | 22.04 | 80.4 | 119.6 | 14.4 | 8.18 | | High | High | High | High | High | High | High | High | High | High | High | 0.000183 | 0.001271 | 2.72 |
| FALSE | High | QVSASELC | [R].QVSASI | 1xCarbami | Q8R0W0 1 | 0.000184 | 0 | 1 | 1 | 6 | Q8R0W0 | Epiplakin C | Q8R0W0 [ | 0 | 1944.93 | 0 | 1.61 | 0.00044 | 5.53 | 48.3 | 151.7 | 5.53 | 6.73 | | High | High | High | High | High | High | High | High | High | High | High | 0.000275 | 0.002046 | 3.26 |
| FALSE | High | QVSAGDLF | [K].QVSAG | 1xPhospho | Q8R0W0 1 | 0.046687 | 0.00354 | 1 | 1 | 2 | Q8R0W0 | Epiplakin C | Q8R0W0 [ | 0 | 1301.645 | 0 | 3.44 | 0.000353 | 16.4 | 17.4 | 182.6 | 11.06 | 9.09 | | High | High | High | High | High | High | High | High | High | High | High | 0.001805 | 0.01779 | 1.93 |
| FALSE | High | QTSSSPEG | [R].QTSSSF | 1xCarbami | Q9QZD4 1x | 1.71E-07 | 0 | 1 | 2 | 1 | Q9QZD4 | DNA repair | Q9QZD4 [5 | 0 | 2016.971 | 0 | 0.48 | 0.001604 | 1.66 | 84 | 116 | 2.88 | 7.06 | | High | High | High | High | High | High | High | High | High | High | High | 0 | 1.4E-08 | 3.68 |
| FALSE | High | QTSSSPEG | [R].QTSSSF | 1xCarbami | Q9QZD4 2x | 2.34E-06 | 0 | 1 | 2 | 5 | Q9QZD4 | DNA repair | Q9QZD4 [5 | 0 | 2096.938 | 0 | 0.66 | 0.000384 | 2.69 | 79.2 | 120.8 | 3.12 | 3.72 | | High | High | High | High | High | High | High | High | High | High | High | 0 | 4.03E-05 | 3.47 |
| FALSE | High | QTSSTTND | [R].QTSSTT | 1xPhospho | Q9QZM4 1 | 0.000186 | 0 | 1 | 1 | 11 | Q9QZM4 | Tumor nec | Q9QZM4 [ | 0 | 2356.082 | 0 | 1.9 | 0.009329 | 22.88 | 41.5 | 158.5 | 13.99 | 24.42 | | High | High | High | High | High | High | High | High | High | High | High | 0 | 0.000316 | 3.31 |
| FALSE | High | QTSVDSGI | [K].QTSVDS | 1xPhospho | Q8BUR4 1x | 0.000712 | 0 | 1 | 1 | 3 | Q8BUR4 | Dedicator | Q8BUR4 [1 | 0 | 1342.645 | 0 | 0.69 | 0.01643 | 29.33 | 79 | 121 | 7.85 | 17.57 | | High | High | High | High | High | High | High | High | High | High | High | 4.94E-05 | 0.000724 | 3.26 |
| FALSE | High | QTTSPASK | [K].QTTSPA | 1xPhospho | P25976 1xI | 0.001815 | 5.66E-05 | 1 | 6 | 5 | P25976 | Nucleolar t | P25976 [3 | 0 | 1357.713 | 0 | -0.84 | 0.000234 | 1.06 | 126.5 | 73.5 | 3.71 | 6.93 | | High | High | High | High | High | High | High | High | High | High | High | 0.000449 | 0.003766 | 2.01 |
| FALSE | High | QVAVRPQS | [R].QVAVR | 1xPhospho | O89032 1x | 0.002419 | 0.000109 | 1 | 3 | 1 | O89032 | SH3 and PX | O89032 [9 | 0 | 2274.278 | 0 | 1.04 | 0.0022 | 32.74 | 65.7 | 134.3 | 18.4 | 4.74 | | High | High | High | High | High | High | High | High | High | High | High | 4.94E-05 | 0.000642 | 2.45 |
| FALSE | High | QVCEDSLE | [K].QVCED | 1xCarbami | Q9D2E2 1x | 0.000933 | 0 | 1 | 1 | 1 | Q9D2E2 | Target of E | Q9D2E2 [3 | 0 | 1875.881 | 0 | 0.27 | 0.050766 | 11.24 | 86.3 | 113.7 | 8.77 | 14.07 | | High | High | High | High | High | High | High | High | High | High | High | 0 | 0.000221 | 3.17 |
| FALSE | High | QVDLESNP | [K].QVDLE | 1xPhospho | A0A0R4J0Y | 0.079766 | 0.00613 | 1 | 6 | 1 | A0A0R4J0Y | Probable h | A0A0R4J0Y | 1 | 3339.673 | 0 | -0.34 | 0.064318 | 10.96 | 111.8 | 88.2 | 13.52 | 5.74 | | High | High | High | High | High | High | High | High | High | High | High | 0.003148 | 0.03309 | 2.37 |
| FALSE | High | QVDITSSSS | [K].QVDITS | 2xPhospho | Q61687 2x | 7.22E-09 | 0 | 1 | 3 | 2 | Q61687 | Transcripti | Q61687 [1 | 0 | 3561.468 | 0 | 0.66 | 0.04239 | 37.84 | 72.6 | 127.4 | 12.64 | 20.5 | | High | High | High | High | High | High | High | High | High | High | High | 0 | 3.98E-10 | 5.73 |
| FALSE | High | QVDSPSAT | [K].QVDSP | 1xPhospho | D3Z357 1xI | 2.19E-08 | 0 | 1 | 3 | 1 | D3Z357 | Histone-lys | D3Z357 [6 | 0 | 2789.231 | 0 | -0.06 | 0.800956 | 28.58 | 99.7 | 100.3 | 9.39 | 15.67 | | High | High | High | High | High | High | High | High | High | High | High | 0 | 1.38E-09 | 4.66 |
| FALSE | High | QVEALGLA | [R].QVEAL( | 2xPhospho | Q8CI12 2xF | 6.35E-05 | 0 | 1 | 3 | 5 | Q8CI12 | Smoothelir | Q8CI12 [8 | 0 | 3146.537 | 0 | -1.68 | 0.000328 | 1.34 | 152.5 | 47.5 | 5.37 | 4.63 | | High | High | High | High | High | High | High | High | High | High | High | 0 | 1.28E-05 | 3.94 |
| FALSE | High | QVEDSLPFI | [R].QVEDSI | 1xPhospho | Q8BW72 1 | 0.001503 | 5.66E-05 | 1 | 3 | 2 | Q8BW72 | Lysine-spe | Q8BW72 [ | 0 | 2604.252 | 0 | -1.17 | 0.003981 | 7.04 | 136.1 | 63.9 | 10.43 | 15.56 | | High | High | High | High | High | High | High | High | High | High | High | 2.58E-05 | 0.000376 | 3.15 |
| FALSE | High | QVHPDTGI | [K].QVHPD | 1xPhospho | Q8CGP2-2 | 3.33E-06 | 0 | 1 | 18 | 4 | Q8CGP2-2 | Isoform 2 | Q8CGP2-2 | 0 | 1706.888 | 0 | -0.56 | 0.044844 | 10.64 | 114.6 | 85.4 | 15.66 | 11.98 | | High | High | High | High | High | High | High | High | High | High | High | 0 | 3.93E-07 | 2.29 |
| FALSE | High | QVLLGPYN | [K].QVLLGI | 1xPhospho | A0A0R4J07 | 5.26E-05 | 0 | 1 | 4 | 1 | A0A0R4J07 | Acyl-Coenz | A0A0R4J07 | 0 | 3058.476 | 0 | 0.44 | 0.039602 | 12.93 | 86.3 | 113.7 | 10.86 | 11.08 | | High | High | High | High | High | High | High | High | High | High | High | 0 | 8.72E-06 | 3.7 |
| FALSE | High | QVISAGEG | [K].QVISAG | 1xPhospho | Q3TUF7 1x | 2.7E-06 | 0 | 1 | 4 | 1 | Q3TUF7 | YEATS dom | Q3TUF7 [6 | 0 | 1941.009 | 0 | -0.59 | 0.001425 | 1.41 | 116.8 | 83.2 | 7.13 | 4.74 | | High | High | High | High | High | High | High | High | High | High | High | 0 | 3.12E-07 | 2.49 |
| FALSE | High | QVLTPSGL | [K].QVLTPS | 1xPhospho | G3X961 1x | 6.27E-07 | 0 | 1 | 5 | 3 | G3X961 | Kelch dom | G3X961 [4 | 0 | 2046.14 | 0 | -0.54 | 0.004778 | 5.48 | 117.6 | 82.4 | 2.99 | 9.12 | | High | High | High | High | High | High | High | High | High | High | High | 0 | 5.26E-06 | 3.9 |
| FALSE | High | QVLTPSGL | [K].QVLTPS | 2xPhospho | G3X961 2x | 5.67E-06 | 0 | 1 | 5 | 1 | G3X961 | Kelch dom | G3X961 [4 | 0 | 2126.106 | 0 | -1.05 | 0.003173 | 2.43 | 132.9 | 67.1 | 10.76 | 8.86 | | High | High | High | High | High | High | High | High | High | High | High | 0 | 7.14E-07 | 3.51 |
| FALSE | High | QVMGPSG | [R].QVMGI | 1xPhospho | E9Q0S6 1x | 7.82E-05 | 0 | 1 | 10 | 1 | E9Q0S6 | Tensin 1 O | E9Q0S6 [1 | 0 | 3439.642 | 0 | -0.24 | 0.997997 | 21.72 | 107.5 | 92.5 | 22.93 | 22.36 | | High | High | High | High | High | High | High | High | High | High | High | 0 | 1.37E-05 | 2.88 |
| FALSE | High | QVMSVVQ | [K].QVMSV | 1xPhospho | Q8R1F1 1x | 5.41E-09 | 0 | 1 | 1 | 26 | Q8R1F1 | Niban-like | Q8R1F1 [6 | 0 | 3908.835 | 0 | 1.46 | 0.057117 | 35.51 | 56.4 | 143.6 | 19.59 | 53.39 | | High | High | High | High | High | High | High | High | High | High | High | 0 | 5.84E-09 | 5.2 |

| | | | | | | | | | | | | | | | | | | | | | | | | | | | | | | | | | | | | | | |
|---|---|---|---|---|---|---|---|---|---|---|---|---|---|---|---|---|---|---|---|---|---|---|---|---|---|---|---|---|---|---|---|---|---|---|---|---|---|---|
| FALSE | High | QVNSTSSP | [R].QVNST | 1xPhosphc | B1AVZ0 1x | 0.000916 | 0 | 1 | 1 | 2 | B1AVZ0 | Uracil phos | B1AVZ0 [1 | 0 | 2276.076 | 0 | 2.03 | 0.000555 | 6.61 | 39.9 | 160.1 | 8 | 8.71 | | High | High | High | High | High | High | High | High | High | High | High | 0.000246 | 0.001466 | 2.8 |
| FALSE | High | QVNSTSSP | [R].QVNST | 2xPhosphc | B1AVZ0 2x | 0.000852 | 0 | 1 | 1 | 2 | B1AVZ0 | Uracil phos | B1AVZ0 [1 | 0 | 2356.042 | 0 | 1.67 | 0.000275 | 8.46 | 48.6 | 151.4 | 6.05 | 5.96 | | High | High | High | High | High | High | High | High | High | High | High | 0 | 0.000199 | 3.12 |
| FALSE | High | QVNYNDG | [K].QVNYN | 1xPhosphc | Q6PDQ2 1x | 0.00103 | 0 | 1 | 6 | 2 | Q6PDQ2 | Chromodo | Q6PDQ2 [1 | 0 | 1733.733 | 0 | -0.33 | 0.0183 | 3.81 | 110.2 | 89.8 | 7.24 | 8.84 | | High | High | High | High | High | High | High | High | High | High | High | 0 | 0.000246 | 1.83 |
| FALSE | High | QVNYNDG | [K].QVNYN | 1xDeamida | Q6PDQ2 1x | 3.73E-05 | 0 | 1 | 5 | 15 | Q6PDQ2 | Chromodo | Q6PDQ2 [1 | 1 | 4896.889 | 0 | 0.14 | 0.030649 | 7.07 | 94.1 | 105.9 | 10.22 | 9.25 | | High | High | High | High | High | High | High | High | High | High | High | 0 | 5.95E-06 | 4.69 |
| FALSE | High | QVPESGDC | [K].QVPESC | 1xCarbami | Q149F1 1x | 0.000314 | 0 | 1 | 2 | 2 | Q149F1 | RNA pseud | Q149F1 [8 | 0 | 3674.722 | 0 | -0.35 | 0.02984 | 7.77 | 112.3 | 87.7 | 10.18 | 5.57 | | High | High | High | High | High | High | High | High | High | High | High | 0 | 6.54E-05 | 4.44 |
| FALSE | High | QVPIAPLHL | [K].QVPIAF | 1xPhosphc | P97434-2 1 | 0.000369 | 0 | 1 | 11 | 1 | P97434-2 | Isoform 2 c | P97434-2 [ | 0 | 1897.015 | 0 | 0.91 | 0.002756 | 21.28 | 64.9 | 135.1 | 19.87 | 11.51 | | High | High | High | High | High | High | High | High | High | High | High | 0 | 7.78E-05 | 2.72 |
| FALSE | High | QVPNESFF | [K].QVPNE | 1xPhosphc | Q8C1W9 1 | 7.36E-06 | 0 | 1 | 5 | 1 | Q8C1W9 | Putative ur | Q8C1W9 [2 | 0 | 2339.188 | 0 | -0.41 | 0.02187 | 9.59 | 115.4 | 84.6 | 7.47 | 11.78 | | High | High | High | High | High | High | High | High | High | High | High | 0 | 9.61E-07 | 3.58 |
| FALSE | High | QVQELQSF | [R].QVQEL | 1xPhosphc | E9QQ10 1x | 2.62E-08 | 0 | 1 | 2 | 2 | E9QQ10 | A-kinase ar | E9QQ10 [1 | 0 | 2956.473 | 0 | 1.09 | 0.000357 | 1.2 | 63.6 | 136.4 | 13.7 | 9.85 | | High | High | High | High | High | High | High | High | High | High | High | 0 | 1.69E-09 | 5.11 |
| FALSE | High | QVQQNSLI | [K].QVQQN | 1xPhosphc | E9Q0N0 1x | 0.001117 | 0 | 1 | 10 | 2 | E9Q0N0 | Intersectin | E9Q0N0 [5 | 0 | 1418.71 | 0 | 0.09 | 0.276131 | 7.5 | 94.3 | 105.7 | 5.51 | 10.16 | | High | High | High | High | High | High | High | High | High | High | High | 0 | 0.00027 | 2.49 |
| FALSE | High | QVQSLTCE | [R].QVQSL | 1xCarbami | P20152 1xl | 0.0003 | 0 | 1 | 7 | 2 | P20152 | Vimentin C | P20152 [32 | 0 | 2029.044 | 0 | 1.48 | 0.001811 | 17.8 | 49.6 | 150.4 | 15.41 | 14.76 | | High | High | High | High | High | High | High | High | High | High | High | 0 | 6.18E-05 | 2.85 |
| FALSE | High | RAPAPPPP | [R].RAPAP | 1xDeamida | B2RWU4 1 | 0.077825 | 0.005963 | 2 | 10 | 2 | B2RWU4; C | Cordon-ble | B2RWU4 [ | 1 | 2409.264 | 0 | -0.12 | 0.191166 | 8.57 | 101 | 99 | 8.08 | 9.23 | Shared | High | High | High | High | High | High | High | High | High | High | High | 0.003064 | 0.03201 | 3.7 |
| FALSE | High | RASVCAEA | [R].RASVCA | 1xCarbami | P31324 1xl | 2.2E-10 | 0 | 1 | 1 | 6 | P31324 | cAMP-depe | P31324 [11 | 1 | 2722.114 | 0 | 0.47 | 0.028861 | 12.65 | 86 | 114 | 9.19 | 10.62 | | High | High | High | High | High | High | High | High | High | High | High | 0 | 9.93E-10 | 5.78 |
| FALSE | High | RAPSPDGF | [R].RAPSPD | 1xPhosphc | P53349 1x | 0.012731 | 0.000889 | 2 | 3 | 1 | P53349; F8 | Mitogen-ac | P53349 [28 | 1 | 2289.039 | 0 | -0.76 | 0.059259 | 5.83 | 124 | 76 | 15.36 | 27.36 | Shared | High | High | High | High | High | High | High | High | High | High | High | 0.000449 | 0.004115 | 3.03 |
| FALSE | High | RAPSPVVSI | [R].RAPSPV | 2xPhosphc | D3Z5K8 2x | 0.002844 | 0.00016 | 1 | 5 | 1 | D3Z5K8 | SH3 and m | D3Z5K8 [16 | 1 | 2086.075 | 0 | -0.37 | 0.793278 | 7.94 | 112.7 | 87.3 | 15.02 | 79.39 | | High | High | High | High | High | High | High | High | High | High | High | 7.19E-05 | 0.000772 | 2.71 |
| FALSE | High | RASMQPAI | [R].RASMC | 1xPhosphc | E9Q7G0 1x | 2.53E-05 | 0 | 1 | 5 | 1 | E9Q7G0 | Nuclear mi | E9Q7G0 [1 | 1 | 2195.121 | 0 | -0.16 | 0.577064 | 7.88 | 102 | 98 | 19.71 | 14.18 | | High | High | High | High | High | High | High | High | High | High | High | 0 | 3.84E-06 | 3.6 |
| FALSE | High | RASMQPAI | [R].RASMC | 1xPhosphc | E9Q7G0 1x | 1.86E-05 | 0 | 1 | 5 | 2 | E9Q7G0 | Nuclear mi | E9Q7G0 [1 | 1 | 2211.115 | 0 | 0.11 | 0.471952 | 43.06 | 96.6 | 103.4 | 14.83 | 21.69 | | High | High | High | High | High | High | High | High | High | High | High | 0 | 2.71E-06 | 3.87 |
| FALSE | High | RASPGLSM | [R].RASPGI | 1xPhosphc | Q545J5 1xl | 0.000563 | 0 | 1 | 3 | 1 | Q545J5 | Basic Krupp | Q545J5 [89 | 1 | 2166.139 | 0 | 0.76 | 0.004126 | 10.49 | 74.2 | 125.8 | 8.64 | 7.65 | | High | High | High | High | High | High | High | High | High | High | High | 0 | 0.000125 | 3.84 |
| FALSE | High | RASPGLSM | [R].RASPGI | 2xPhosphc | Q545J5 2xl | 0.007538 | 0.00062 | 1 | 3 | 2 | Q545J5 | Basic Krupp | Q545J5 [89 | 1 | 2230.111 | 0 | 0.79 | 0.003014 | 7.22 | 71.9 | 128.1 | 2.3 | 9.64 | | High | High | High | High | High | High | High | High | High | High | High | 0.000275 | 0.002287 | 2.59 |
| FALSE | High | RASPGSEEI | [R].RASPGS | 1xPhosphc | A0A068BFI | 0.000122 | 0 | 1 | 7 | 2 | A0A068BFI | Kinesin-like | A0A068BFI | 1 | 1896.938 | 0 | -0.58 | 0.000209 | 1.6 | 119.5 | 80.5 | 2.86 | 3.18 | | High | High | High | High | High | High | High | High | High | High | High | 0 | 2.93E-05 | 3.44 |
| FALSE | High | RASPLCETS | [R].RASPLC | 1xCarbami | Q0VBL3 1x | 4.2E-05 | 0 | 1 | 6 | 1 | Q0VBL3 | RNA bindir | Q0VBL3 [1 | 1 | 1799.846 | 0 | 0.52 | 0.004581 | 7.42 | 83 | 117 | 6.98 | 4.32 | | High | High | High | High | High | High | High | High | High | High | High | 0 | 6.77E-06 | 2.93 |
| FALSE | High | RASPNSDD | [R].RASPNS | 1xPhosphc | Q8BHS6 1x | 1.38E-05 | 0 | 1 | 1 | 2 | Q8BHS6 | Armadillo r | Q8BHS6 [1 | 1 | 2523.286 | 0 | -1.51 | 0.000824 | 1.72 | 149.1 | 50.9 | 4.59 | 19.37 | | High | High | High | High | High | High | High | High | High | High | High | 0 | 1.94E-06 | 2.93 |
| FALSE | High | RASPPAAG | [R].RASPPA | 1xPhosphc | Q6A0A2 1x | 0.013545 | 0.000976 | 1 | 3 | 2 | Q6A0A2 | La-related | Q6A0A2 [7 | 1 | 1392.776 | 0 | 0.4 | 0.012277 | 8.53 | 87.5 | 112.5 | 7.39 | 4.52 | | High | High | High | High | High | High | High | High | High | High | High | 0.002159 | 0.02244 | 2.58 |
| FALSE | High | RASPSPPP | [R].RASPSP | 1xPhosphc | Q52KI8 1x | 0.019276 | 0.001402 | 1 | 7 | 2 | Q52KI8 | Serine/argi | Q52KI8 [64 | 1 | 1474.818 | 0 | -0.55 | 0.015918 | 8.93 | 116.8 | 83.2 | 9.21 | 9.34 | | High | High | High | High | High | High | High | High | High | High | High | 0.000756 | 0.006536 | 2.3 |
| FALSE | High | RASPSPPP | [R].RASPSP | 2xPhosphc | Q52KI8 2x | 0.036103 | 0.002342 | 1 | 7 | 3 | Q52KI8 | Serine/argi | Q52KI8 [64 | 1 | 1554.785 | 0 | -0.4 | 0.001293 | 3.36 | 113.7 | 86.3 | 1.18 | 4.4 | | High | High | High | High | High | High | High | High | High | High | High | 0.001279 | 0.01334 | 2.49 |
| FALSE | High | RASQGLISA | [R].RASQG | 1xPhosphc | Q6PR54-3 | 2.72E-08 | 0 | 2 | 4 | 5 | Q6PR54-3; | Isoform 3 c | Q6PR54-3 | 1 | 2703.324 | 0 | -1.08 | 0.009122 | 2.86 | 138.3 | 61.7 | 18.69 | 12.21 | Shared | High | High | High | High | High | High | High | High | High | High | High | 0 | 1.77E-09 | 5.55 |
| FALSE | High | RASQGLISA | [R].RASQG | 1xDeamida | Q6PR54-3 | 0.103392 | 0.008249 | 2 | 4 | 1 | Q6PR54-3; | Isoform 3 c | Q6PR54-3 | 1 | 2704.308 | 0 | -0.57 | 0.005645 | 6.09 | 119.3 | 80.7 | 8.16 | 11.34 | Shared | High | High | High | High | High | High | High | High | High | High | High | 0.004138 | 0.04469 | 4.21 |
| FALSE | High | RASQGLISA | [R].RASQG | 2xPhosphc | Q6PR54-3 | 0.096959 | 0.007777 | 2 | 4 | 1 | Q6PR54-3; | Isoform 3 c | Q6PR54-3 | 1 | 2783.29 | 0 | -1.07 | 0.015977 | 6.41 | 130.3 | 69.7 | 19.03 | 18.21 | Shared | High | High | High | High | High | High | High | High | High | High | High | 0.003922 | 0.04159 | 2.94 |
| FALSE | High | RASLVAQP | [K].RASLVA | 1xPhosphc | Q8K190 1x | 9.89E-05 | 0 | 1 | 1 | 1 | Q8K190 | SAYSvFN d | Q8K190 [1 | 1 | 2628.376 | 0 | -0.82 | 0.02284 | 7.26 | 128.5 | 71.5 | 8.99 | 23.74 | | High | High | High | High | High | High | High | High | High | High | High | 0 | 1.78E-05 | 3.45 |
| FALSE | High | RASQSSLES | [R].RASQSS | 1xPhosphc | D9HP81 1x | 0.000154 | 0 | 1 | 6 | 2 | D9HP81 | Regulating | D9HP81 [1 | 1 | 2108.981 | 0 | 0.79 | 0.002675 | 9.74 | 74.4 | 125.6 | 11.78 | 5.55 | | High | High | High | High | High | High | High | High | High | High | High | 0 | 2.91E-05 | 3.4 |
| FALSE | High | RASSAAPS | [R].RASSAA | 1xPhosphc | Q7M753 1 | 5.15E-07 | 0 | 1 | 4 | 4 | Q7M753 | Pantothen | Q7M753 [5 | 1 | 1927.907 | 0 | -0.3 | 0.055339 | 8.81 | 106.7 | 93.3 | 14.09 | 7.18 | | High | High | High | High | High | High | High | High | High | High | High | 0 | 4.83E-08 | 3.99 |
| FALSE | High | RASSCSLA | [R].RASSCS | 1xCarbami | P58801 1xl | 2.23E-05 | 0 | 1 | 1 | 2 | P58801 | Receptor-i | P58801 [40 | 1 | 2402.292 | 0 | -0.49 | 0.009225 | 8.3 | 118.2 | 81.8 | 14.4 | 13.04 | | High | High | High | High | High | High | High | High | High | High | High | 0 | 5.93E-06 | 4.78 |
| FALSE | High | RASSDLDQ | [R].RASSDL | 1xPhosphc | Q3UMU9-3 | 6.12E-08 | 0 | 1 | 4 | 2 | Q3UMU9-3 | Isoform 3 c | Q3UMU9-3 | 1 | 3291.479 | 0 | 0.35 | 0.028301 | 6.53 | 89.9 | 110.1 | 13.8 | 12.65 | | High | High | High | High | High | High | High | High | High | High | High | 0 | 4.43E-09 | 5.76 |
| FALSE | High | RASSDLSIA | [K].RASSDL | 1xPhosphc | O54988 1x | 8.4E-05 | 0 | 1 | 2 | 2 | O54988 | STE20-like | O54988 [3 | 1 | 2133.048 | 0 | 0.14 | 0.219975 | 16.01 | 94.7 | 105.3 | 1.31 | 9.6 | | High | High | High | High | High | High | High | High | High | High | High | 0 | 1.48E-05 | 3.24 |
| FALSE | High | RASSDLSIA | [K].RASSDL | 2xPhosphc | O54988 2x | 0.021797 | 0.001507 | 1 | 2 | 2 | O54988 | STE20-like | O54988 [3 | 1 | 2213.014 | 0 | -0.48 | 0.011967 | 3.73 | 116.4 | 83.6 | 7.65 | 7.08 | | High | High | High | High | High | High | High | High | High | High | High | 0.001011 | 0.01098 | 2.27 |
| FALSE | High | RASSEIVTE | [R].RASSEI | 1xPhosphc | Q8BUH8 1 | 0.001039 | 0 | 2 | 7 | 1 | Q8BUH8; C | Sentrin-spe | Q8BUH8 [1 | 1 | 1714.914 | 0 | 1.94 | 0.000486 | 15.94 | 43.3 | 156.7 | 10.71 | 7.42 | Shared | High | High | High | High | High | High | High | High | High | High | High | 0 | 0.00025 | 1.99 |
| FALSE | High | RASSLDLEE | [R].RASSLD | 2xPhosphc | G3X9F6 2x | 0.000595 | 0 | 1 | 6 | 1 | G3X9F6 | SH3 domai | G3X9F6 [4 | 1 | 1661.75 | 0 | 3.48 | 0.000275 | 11.83 | 16.5 | 183.5 | 16.3 | 7.35 | | High | High | High | High | High | High | High | High | High | High | High | 0 | 0.000133 | 2.96 |
| FALSE | High | RASSPFR | [R].RASSPF | 2xPhosphc | E9Q5C9 2x | 0.093123 | 0.007276 | 1 | 2 | 1 | E9Q5C9 | Nucleolar a | E9Q5C9 [6 | 1 | 1209.538 | 0 | 0.21 | 0.035034 | 7.99 | 92.1 | 107.9 | 6.52 | 5.5 | | High | High | High | High | High | High | High | High | High | High | High | 0.003685 | 0.03957 | 1.28 |
| FALSE | High | RASSQDGL | [R].RASSQI | 1xPhosphc | B7ZMS8 1x | 0.007145 | 0.00062 | 1 | 9 | 1 | B7ZMS8 | Mdm1 pro | B7ZMS8 [5 | 1 | 1842.973 | 0 | -0.05 | 0.488083 | 19.13 | 101.1 | 98.9 | 8.96 | 12.22 | | High | High | High | High | High | High | High | High | High | High | High | 0.000275 | 0.002151 | 2.35 |
| FALSE | High | RASSQDGL | [R].RASSQI | 2xPhosphc | B7ZMS8 2x | 0.017192 | 0.001204 | 1 | 9 | 1 | B7ZMS8 | Mdm1 pro | B7ZMS8 [5 | 1 | 1922.939 | 0 | 0.51 | 0.001546 | 5.76 | 79.4 | 120.6 | 6.71 | 9.07 | | High | High | High | High | High | High | High | High | High | High | High | 0.000638 | 0.005738 | 2.64 |
| FALSE | High | RASSSAQA | [R].RASSSA | 1xCarbami | P54276 1xl | 1.21E-05 | 0 | 1 | 2 | 6 | P54276 | DNA misma | P54276 [75 | 1 | 3158.538 | 0 | 0.22 | 0.040956 | 9.6 | 96.1 | 103.9 | 9.24 | 7.47 | | High | High | High | High | High | High | High | High | High | High | High | 0 | 1.67E-06 | 4.18 |
| FALSE | High | RASTASCPI | [R].RASTAS | 1xCarbami | O35954 1x | 1.81E-07 | 0 | 1 | 1 | 3 | O35954 | Membrane | O35954 [6 | 1 | 2580.171 | 0 | 1.58 | 0.000426 | 9.3 | 47.9 | 152.1 | 15.58 | 5.98 | | High | High | High | High | High | High | High | High | High | High | High | 0 | 3.43E-08 | 4.6 |
| FALSE | High | RASTASCPI | [R].RASTAS | 1xCarbami | O35954 2x | 0.000668 | 0 | 1 | 1 | 2 | O35954 | Membrane | O35954 [6 | 1 | 2660.137 | 0 | 1.34 | 0.00034 | 2.54 | 57.2 | 142.8 | 13.34 | 9.43 | | High | High | High | High | High | High | High | High | High | High | High | 0 | 0.000152 | 3.34 |
| FALSE | High | RASQTPAQ | [R].RASQTI | 1xPhosphc | Q62187 1x | 0.000213 | 0 | 1 | 2 | 1 | Q62187 | Transcripti | Q62187 [5 | 1 | 2624.312 | 0 | 0.24 | 0.053724 | 8.52 | 91.7 | 108.3 | 10.83 | 6.95 | | High | High | High | High | High | High | High | High | High | High | High | 0 | 4.22E-05 | 2.47 |
| FALSE | High | RASISEPSD | [R].RASISE | 1xPhosphc | Q3TAA7 1x | 1.77E-05 | 0 | 1 | 1 | 2 | Q3TAA7 | Serine/thre | Q3TAA7 [3 | 1 | 1981.943 | 0 | -1.13 | 0.000213 | 1.33 | 136.7 | 63.3 | 5.43 | 8.94 | | High | High | High | High | High | High | High | High | High | High | High | 0 | 2.57E-06 | 3.79 |
| FALSE | High | RASLSEIGF | [R].RASLSE | 1xPhosphc | Q8K0D0 1x | 4.13E-06 | 0 | 1 | 4 | 1 | Q8K0D0 | Cyclin-dep | Q8K0D0 [1 | 1 | 1702.929 | 0 | -0.57 | 0.019276 | 9.05 | 119.4 | 80.6 | 4.97 | 17.48 | | High | High | High | High | High | High | High | High | High | High | High | 0 | 4.99E-07 | 3.34 |
| FALSE | High | RASLSDLTH | [R].RASLSD | 1xPhosphc | Q3UCG9 1 | 0.006594 | 0.00062 | 1 | 6 | 4 | Q3UCG9 | E3 ubiquiti | Q3UCG9 [2 | 1 | 2434.254 | 0 | -0.22 | 0.005696 | 2.44 | 107.9 | 92.1 | 3.28 | 2.87 | | High | High | High | High | High | High | High | High | High | High | High | 0.000585 | 0.00549 | 2.64 |
| FALSE | High | RAPSVANI( | [K].RAPSVA | 1xPhosphc | Q8BTM8 1 | 0.114657 | 0.009411 | 1 | 5 | 1 | Q8BTM8 | Filamin-A C | Q8BTM8 [2 | 1 | 2134.068 | 0 | -0.38 | 0.27281 | 12.23 | 108.5 | 91.5 | 12.3 | 28.58 | | High | High | High | High | High | High | High | High | High | High | High | 0.004707 | 0.05053 | 2.5 |
| FALSE | High | RAPSVANI( | [K].RAPSVA | 1xCarbami | Q8BTM8 1 | 3.29E-05 | 0 | 1 | 5 | 2 | Q8BTM8 | Filamin-A C | Q8BTM8 [2 | 1 | 2363.231 | 0 | 0.49 | 0.017947 | 6.25 | 85.1 | 114.9 | 8.41 | 10.29 | | High | High | High | High | High | High | High | High | High | High | High | 0 | 5.17E-06 | 2.41 |
| FALSE | High | RAPTRGTP | [K].RAPTRC | 1xPhosphc | Q5U3K5 1x | 0.004738 | 0.000456 | 1 | 1 | 1 | Q5U3K5 | Rab-like pr | Q5U3K5 [5 | 2 | 3392.664 | 0 | -0.51 | 0.02439 | 6.35 | 113.8 | 86.2 | 13.18 | 8.11 | | High | High | High | High | High | High | High | High | High | High | High | 0.000204 | 0.001361 | 3.04 |
| FALSE | High | RASAANGF | [R].RASAAI | 1xPhosphc | Q8BH49 1x | 0.004294 | 0.00036 | 1 | 1 | 1 | Q8BH49 | Sesquipedi | Q8BH49 [1 | 1 | 2147.107 | 0 | 0.17 | 0.109422 | 15.35 | 87.9 | 112.1 | 11.05 | 11.69 | | High | High | High | High | High | High | High | High | High | High | High | 0.000161 | 0.001223 | 2.52 |
| FALSE | High | RASAGPVP | [R].RASAGI | 2xPhosphc | Q8BHL3 2x | 3.44E-05 | 0 | 1 | 1 | 1 | Q8BHL3 | TBC1 doma | Q8BHL3 [6 | 1 | 3941.034 | 0 | -0.09 | 0.735531 | 4.42 | 102.8 | 97.2 | 12.16 | 17.98 | | High | High | High | High | High | High | High | High | High | High | High | 0 | 5.44E-06 | 4.33 |
| FALSE | High | RASAGTPSI | [R].RASAG | 2xPhosphc | H9LAN6 2x | 0.030898 | 0.001896 | 1 | 1 | 7 | H9LAN6 | Histone-lys | H9LAN6 [1 | 1 | 2134.071 | 0 | -1.49 | 0.001528 | 0.85 | 145 | 55 | 8.23 | 13.98 | | High | High | High | High | High | High | High | High | High | High | High | 0.001044 | 0.01117 | 3.09 |
| FALSE | High | RASAPDLV | [K].RASAPI | 1xPhosphc | Q7TSH3 1x | 4.72E-05 | 0 | 1 | 1 | 1 | Q7TSH3 | Zinc finger | Q7TSH3 [7 | 1 | 1950.008 | 0 | -0.03 | 0.824421 | 28.44 | 97.7 | 102.3 | 10.47 | 17.28 | | High | High | High | High | High | High | High | High | High | High | High | 0 | 7.73E-06 | 3.11 |
| FALSE | High | RASAPDLV | [K].RASAPI | 1xPhosphc | Q7TSH3 1x | 0.041781 | 0.003126 | 1 | 1 | 1 | Q7TSH3 | Zinc finger | Q7TSH3 [7 | 1 | 1966.003 | 0 | 0.13 | 0.178668 | 11.29 | 95.6 | 104.4 | 9.21 | 13.63 | | High | High | High | High | High | High | High | High | High | High | High | 0.003303 | 0.03643 | 3 |
| FALSE | High | RASAPLPGI | [R].RASAPI | 1xPhosphc | Q5EBG6 1x | 0.004867 | 0.000504 | 1 | 1 | 1 | Q5EBG6 | Heat shock | Q5EBG6 [1 | 1 | 1692.878 | 0 | -0.42 | 0.087433 | 16.75 | 119.2 | 80.8 | 8.43 | 33.25 | | High | High | High | High | High | High | High | High | High | High | High | 0.000225 | 0.001407 | 1.88 |
| FALSE | High | RASASPSQ' | [R].RASASF | 1xPhosphc | Q8BGD4 1 | 9.82E-06 | 0 | 1 | 1 | 1 | Q8BGD4 | Solute carr | Q8BGD4 [2 | 1 | 2011.017 | 0 | 0.54 | 0.001319 | 8.41 | 81 | 119 | 3.25 | 4.95 | | High | High | High | High | High | High | High | High | High | High | High | 0 | 1.33E-06 | 1.88 |
| FALSE | High | RASATTGQ | [R].RASATT | 1xPhosphc | Q7TSH3 1x | 0.037692 | 0.00249 | 1 | 1 | 1 | Q7TSH3 | Zinc finger | Q7TSH3 [4 | 1 | 1514.809 | 0 | 0.3 | 0.036392 | 4.28 | 86.7 | 113.3 | 5.98 | 15.48 | | High | High | High | High | High | High | High | High | High | High | High | 0.001324 | 0.01395 | 1.98 |
| FALSE | High | RASCSSSTC | [R].RASCSS | 1xCarbami | Q9Z1E4 1x | 0.000209 | 0 | 1 | 2 | 2 | Q9Z1E4 | Glycogen [ | Q9Z1E4 [6 | 1 | 1722.825 | 0 | -0.72 | 0.005429 | 4.24 | 124.5 | 75.5 | 13.44 | 11.79 | | High | High | High | High | High | High | High | High | High | High | High | 0 | 4.12E-05 | 3.01 |
| FALSE | High | RASDAFPEI | [R].RASDAI | 1xPhosphc | P48754 1x | 0.001189 | 0 | 1 | 4 | 2 | P48754 | Breast canc | P48754 [68 | 1 | 1687.846 | 0 | -0.74 | 0.000352 | 1.63 | 124.1 | 75.9 | 4.66 | 2.21 | | High | High | High | High | High | High | High | High | High | High | High | 0 | 0.00029 | 2.93 |
| FALSE | High | RASDDGK | [R].RASDD | 1xPhosphc | Q99N57-2 | 0.074071 | 0.0057 | 1 | 3 | 1 | Q99N57-2 | Isoform 2 c | Q99N57-2 | 1 | 1286.651 | 0 | 0.08 | 0.160966 | 5.25 | 96.7 | 103.3 | 3.75 | 8.45 | | High | High | High | High | High | High | High | High | High | High | High | 0.002911 | 0.03026 | 1.56 |
| FALSE | High | RASDGGAN | [R].RASDG | 1xPhosphc | E9PU87 1x | 3.24E-06 | 0 | 1 | 6 | 1 | E9PU87 | Serine/thre | E9PU87 [5 | 1 | 2458.333 | 0 | 0.42 | 0.018987 | 8.99 | 85.7 | 114.3 | 5.99 | 14.61 | | High | High | High | High | High | High | High | High | High | High | High | 0 | 3.8E-07 | 4.04 |
| FALSE | High | RASDSNTH | [R].RASDSN | 1xPhosphc | Q9QZN4 1x | 0.000279 | 0 | 1 | 1 | 1 | Q9QZN4 | F-box only | Q9QZN4 [2 | 1 | 2631.209 | 0 | 0.33 | 0.057757 | 10.45 | 91.8 | 108.2 | 9.05 | 6.67 | | High | High | High | High | High | High | High | High | High | High | High | 0 | 5.69E-05 | 2.42 |
| FALSE | High | RASDTSLT( | [R].RASDTS | 1xPhosphc | Q60670 1x | 2.43E-07 | 0 | 1 | 3 | 2 | Q60670 | Serine/thre | Q60670 [5 | 1 | 1814.978 | 0 | -0.15 | 0.387746 | 8.67 | 109 | 91 | 8.33 | 13.48 | | High | High | High | High | High | High | High | High | High | High | High | 0 | 2.94E-06 | 3.93 |
| FALSE | High | RASDTSLT( | [R].RASDTS | 1xPhosphc | F8VPT7 1xl | 1.03E-06 | 0 | 1 | 4 | 1 | F8VPT7 | Serine/thre | F8VPT7 [58 | 1 | 1930.995 | 0 | 0.38 | 0.060878 | 9.8 | 82 | 118 | 16.2 | 15.79 | | High | High | High | High | High | High | High | High | High | High | High | 0 | 1.05E-07 | 3.83 |
| FALSE | High | RASFAEK | [K].RASFAE | 1xPhosphc | Q9QXS1-3 | 0.116482 | 0.009523 | 3 | 23 | 1 | Q9QXS1-3; | Isoform PL | Q9QXS1-3 | 1 | 1346.723 | 0 | 1.06 | 0.003992 | 13.45 | 64.7 | 135.3 | 8.67 | 11.93 | Shared | High | High | High | High | High | High | High | High | High | High | High | 0.004757 | 0.05186 | 1.9 |
| FALSE | High | RASGDTAK | [K].RASGD | 1xPhosphc | Q3TDI5 1xl | 0.019618 | 0.001438 | 1 | 1 | 1 | Q3TDI5 | Peroxisom | Q3TDI5 [53 | 1 | 1343.708 | 0 | 0.2 | 0.252552 | 36.11 | 88.7 | 111.3 | 21.52 | 17.75 | | High | High | High | High | High | High | High | High | High | High | High | 0.000756 | 0.006653 | 1.5 |
| FALSE | High | RASGPGIPI | [K].RASGP | 1xPhosphc | Q61526 1x | 5.75E-08 | 0 | 1 | 1 | 1 | Q61526 | Receptor t | Q61526 [9 | 1 | 2345.263 | 0 | 2.63 | 0.000614 | 20.47 | 28.2 | 171.8 | 11.91 | 10.97 | | High | High | High | High | High | High | High | High | High | High | High | 0 | 4.09E-09 | 3.66 |
| FALSE | High | RASGQAFE | [K].RASGQ | 1xPhosphc | Q545B6 1x | 5.65E-08 | 0 | 1 | 8 | 1 | Q545B6 | Stathmin C | Q545B6 [1 | 1 | 1853.984 | 0 | -1.16 | 0.000565 | 1.67 | 138.3 | 61.7 | 9.56 | 11.67 | | High | High | High | High | High | High | High | High | High | High | High | 0 | 4.04E-09 | 3.48 |
| FALSE | High | RASGQAFE | [K].RASGQ | 2xPhosphc | Q545B6 2x | 4.31E-05 | 0 | 1 | 8 | 3 | Q545B6 | Stathmin C | Q545B6 [1 | 1 | 1933.95 | 0 | -1.45 | 0.000405 | 2.17 | 146.1 | 53.9 | 4.82 | 8.14 | | High | High | High | High | High | High | High | High | High | High | High | 0.001581 | 0.01516 | 2.37 |
| FALSE | High | RASGSAGA | [R].RASGSA | 1xPhosphc | B2RQE3 1x | 1.05E-06 | 0 | 1 | 4 | 2 | B2RQE3 | Stim2 prot | B2RQE3 [9 | 1 | 2024.971 | 0 | 0.37 | 0.02848 | 17.84 | 91.6 | 108.4 | 9.69 | 13.65 | | High | High | High | High | High | High | High | High | High | High | High | 0 | 1.08E-07 | 3.98 |
| FALSE | High | RASHALFPI | [R].RASHAI | 1xPhosphc | Q80TU6 1x | 0.001437 | 5.66E-05 | 1 | 1 | 1 | Q80TU6 | MKIAA067 | Q80TU6 [4 | 1 | 2001.036 | 0 | 1.91 | 0.001245 | 13.75 | 44.1 | 155.9 | 16.95 | 9.98 | | High | High | High | High | High | High | High | High | High | High | High | 2.58E-05 | 0.000358 | 2.6 |
| FALSE | High | RASLCTQP | [K].RASLCT | 1xCarbami | Q7TS72 1x | 0.01002 | 0.000736 | 1 | 1 | 1 | Q7TS72 | Inositol-tri | Q7TS72 [1 | 1 | 2578.148 | 0 | -0.54 | 0.02736 | 8.97 | 123.9 | 76.1 | 15 | 13.03 | | High | High | High | High | High | High | High | High | High | High | High | 0.000344 | 0.003146 | 3.53 |
| FALSE | High | RASLEIGES | [R].RASLEI | 1xPhosphc | Q9QY06-2 | 5.24E-07 | 0 | 2 | 8 | 3 | Q9QY06-2; | Isoform 2 c | Q9QY06-2 | 1 | 2159.115 | 0 | 0.04 | 0.190625 | 5.51 | 97.5 | 102.5 | 4.13 | 9.86 | Shared | High | High | High | High | High | High | High | High | High | High | High | 0 | 4.22E-06 | 4.23 |
| FALSE | High | RASLSCGGI | [R].RASLSC | 1xCarbami | E9Q1A8 1x | 0.000331 | 0 | 1 | 7 | 1 | E9Q1A8 | HMG box-c | E9Q1A8 [3 | 1 | 2841.328 | 0 | -0.5 | 0.005915 | 6.76 | 116 | 84 | 10.05 | 6.92 | | High | High | High | High | High | High | High | High | High | High | High | 0 | 6.86E-05 | 3.16 |
| FALSE | High | RASLSDIGF | [R].RASLSD | 1xPhosphc | Q0VDL6 1x | 0.000668 | 0 | 1 | 1 | 1 | Q0VDL6 | PCTAIRE-m | Q0VDL6 [1 | 1 | 1688.914 | 0 | -0.24 | 0.183376 | 16.39 | 106.2 | 93.8 | 6.25 | 14.83 | | High | High | High | High | High | High | High | High | High | High | High | 0 | 0.000151 | 2.03 |
| FALSE | High | RAPSPGSYI | [K].RAPSPG | 1xPhosphc | O35126 1x | 0.007082 | 0.00062 | 1 | 1 | 1 | O35126 | Atrophin-1 | O35126 [6 | 1 | 1500.798 | 0 | 0.72 | 0.000181 | 2.46 | 75.7 | 124.3 | 1.74 | 2.02 | | High | High | High | High | High | High | High | High | High | High | High | 0.000275 | 0.002144 | 1.47 |
| FALSE | High | SPGSVAVSI | [K].SPGSVA | 1xDeamida | G3X928 1x | 0.000297 | 0 | 1 | 4 | 1 | G3X928 | SEC23-inte | G3X928 [6 | 0 | 1753.95 | 0 | -0.38 | 0.047049 | 6.25 | 113 | 87 | 12.13 | 8.08 | | High | High | High | High | High | High | High | High | High | High | High | 0 | 6.08E-05 | 2.86 |
| FALSE | High | SPQLSLSPR | [K].SPQLSL | 2xPhosphc | F6ZBR8 2xl | 0.008094 | 0.000736 | 2 | 5 | 2 | F6ZBR8; O | Protein Wi | F6ZBR8 [19 | 0 | 2113.086 | 0 | 0.17 | 0.003603 | 1.89 | 93.5 | 106.5 | 5.49 | 7.18 | Shared | High | High | High | High | High | High | High | High | High | High | High | 0.009151 | 0.09877 | 1.88 |
| FALSE | High | SPGTPAGE | [R].SPGTPA | 1xPhosphc | Q8K232 1x | 0.024855 | 0.001645 | 2 | 7 | 2 | Q8K232; E | Adducin 1 | Q8K232 [3 | 0 | 1634.762 | 0 | 0.23 | 0.030425 | 13.66 | 88.3 | 111.7 | 12.94 | 9.92 | Shared | High | High | High | High | High | High | High | High | High | High | High | 0.001916 | 0.0194 | 2.06 |
| FALSE | High | TVPSTPTLV | [R].TVPSTP | 1xPhosphc | F6ZDS4 1xl | 7.16E-06 | 0 | 1 | 5 | 4 | F6ZDS4 | Nucleopro | F6ZDS4 [22 | 0 | 1712.93 | 0 | 1.48 | 0.010732 | 14.64 | 54.1 | 145.9 | 10.11 | 23.78 | | High | High | High | High | High | High | High | High | High | High | High | 0 | 8.66E-06 | 3.86 |
| FALSE | High | TVQLTDLG | [R].TVQLTI | 1xPhosphc | Q810A4 1x | 6.29E-08 | 0 | 1 | 3 | 5 | Q810A4 | RNA helica | Q810A4 [5 | 0 | 2029.942 | 0 | -0.26 | 0.016976 | 1 | 107.5 | 92.5 | 3.71 | 7.29 | | High | High | High | High | High | High | High | High | High | High | High | 0 | 4.55E-09 | 3.75 |
| FALSE | High | TVQNSSVS | [K].TVQNS | 1xPhosphc | Q5F2E7-2 1 | 4.37E-07 | 0 | 1 | 2 | 4 | Q5F2E7-2 | Isoform 2 c | Q5F2E7-2 | 0 | 3086.36 | 0 | 0.37 | 0.000572 | 3.17 | 86.2 | 113.8 | 3.4 | 4.08 | | High | High | High | High | High | High | High | High | High | High | High | 0 | 8.63E-08 | 6.18 |
| FALSE | High | TVQSPNSS | [R].TVQSPI | 1xPhosphc | Q64127 1x | 0.00103 | 0 | 1 | 5 | 1 | Q64127 | Transcripti | Q64127 [6 | 0 | 3122.591 | 0 | -1.13 | 0.004492 | 4.61 | 137.3 | 62.7 | 9.24 | 25.04 | | High | High | High | High | High | High | High | High | High | High | High | 0 | 0.000247 | 4.11 |
| FALSE | High | TVQSPNSS | [R].TVQSPI | 1xDeamida | Q64127 1x | 0.005135 | 0.000551 | 1 | 5 | 1 | Q64127 | Transcripti | Q64127 [6 | 0 | 3123.575 | 0 | -0.82 | 0.008577 | 6.27 | 127.7 | 72.3 | 13.93 | 13.27 | | High | High | High | High | High | High | High | High | High | High | High | 0.000246 | 0.001488 | 4.91 |
| FALSE | High | TVQSPNSS | [R].TVQSPI | 1xPhosphc | Q64127 1x | 0.00642 | 0.00062 | 1 | 5 | 1 | Q64127 | Transcripti | Q64127 [6 | 0 | 3138.586 | 0 | -1.15 | 0.025632 | 3.72 | 142 | 58 | 19.37 | 32.67 | | High | High | High | High | High | High | High | High | High | High | High | 0.000275 | 0.001907 | 3.6 |
| FALSE | High | TVQSQIGP | [K].TVQSQ | 1xPhosphc | Q3U2C9 1x | 0.001751 | 5.66E-05 | 1 | 8 | 1 | Q3U2C9 | PHD finger | Q3U2C9 [6 | 0 | 2776.383 | 0 | -0.44 | 0.007515 | 8.58 | 110.8 | 89.2 | 7.77 | 11.56 | | High | High | High | High | High | High | High | High | High | High | High | 2.58E-05 | 0.000448 | 1.78 |
| FALSE | High | TVRPEVNA | [K].TVRPEV | 1xPhosphc | Q9WTU0 1 | 0.003622 | 0.00021 | 1 | 1 | 1 | Q9WTU0 | Lysine-spe | Q9WTU0 [ | 0 | 2457.091 | 0 | 0.82 | 0.004661 | 24.15 | 70.2 | 129.8 | 16.66 | 13.18 | | High | High | High | High | High | High | High | High | High | High | High | 9.42E-05 | 0.001009 | 2.54 |
| FALSE | High | TVRPEVNA | [K].TVRPEV | 1xCarbami | Q9WTU0 2 | 6.09E-06 | 0 | 1 | 1 | 1 | Q9WTU0 | Lysine-spe | Q9WTU0 [ | 0 | 2537.058 | 0 | 1.86 | 0.002894 | 14.25 | 46.3 | 153.7 | 22.25 | 12.21 | | High | High | High | High | High | High | High | High | High | High | High | 0 | 7.72E-07 | 3.32 |
| FALSE | High | TVRPTSAPI | [K].TVRPTS | 1xPhosphc | E9QMQ3 1 | 0.000815 | 0 | 2 | 7 | 1 | E9QMQ3; | TRIO and F | E9QMQ3 [ | 0 | 1809.988 | 0 | 1.84 | 0.000791 | 7.72 | 44.5 | 155.5 | 12.4 | 12.96 | Shared | High | High | High | High | High | High | High | High | High | High | High | 0 | 0.000189 | 1.77 |
| FALSE | High | TVSASAAD | [R].TVSASA | 1xPhosphc | G5E8J4 1xl | 3.24E-06 | 0 | 1 | 4 | 2 | G5E8J4 | Tyrosine-p | G5E8J4 [65 | 0 | 1453.725 | 0 | 1.55 | 0.000241 | 10.02 | 59.7 | 140.3 | 9.13 | 5.21 | | High | High | High | High | High | High | High | High | High | High | High | 0 | 3.83E-07 | 2.28 |
| FALSE | High | TVSASSLPS | [R].TVSASS | 1xPhosphc | A0A0R4J0Y | 2.49E-05 | 0 | 1 | 4 | 2 | A0A0R4J0Y | Probable h | A0A0R4J0Y | 0 | 2073.994 | 0 | 0.46 | 0.004993 | 11.85 | 82.4 | 117.6 | 7.19 | 7.79 | | High | High | High | High | High | High | High | High | High | High | High | 0 | 3.78E-06 | 3.33 |
| FALSE | High | TVSASSTGI | [R].TVSASS | 1xPhosphc | A0A0R4J12 | 1E-06 | 0 | 2 | 3 | 2 | A0A0R4J12 | SRSF prote | A0A0R4J12 | 0 | 1700.887 | 0 | -0.09 | 0.1347 | 1.05 | 102.1 | 97.9 | 3.92 | 5.56 | Shared | High | High | High | High | High | High | High | High | High | High | High | 0 | 1.02E-07 | 2.53 |
| FALSE | High | TVSPPISPE | [R].TVSPPI | 2xPhosphc | A2AN08-5 | 0.005467 | 0.00062 | 1 | 3 | 1 | A2AN08-5 | Isoform 5 c | A2AN08-5 | 0 | 3615.783 | 0 | -0.01 | 0.490373 | 13.88 | 98.7 | 101.3 | 8.75 | 12.85 | | High | High | High | High | High | High | High | High | High | High | High | 0.000275 | 0.001606 | 3.05 |
| FALSE | High | TVSASSTGI | [R].TVSASS | 2xPhosphc | A0A0R4J12 | 0.039686 | 0.00283 | 2 | 3 | 3 | A0A0R4J12 | SRSF prote | A0A0R4J12 | 0 | 1780.854 | 0 | -0.56 | 0.005268 | 7.48 | 118.6 | 81.4 | 3.78 | 10.63 | Shared | High | High | High | High | High | High | High | High | High | High | High | 0.00196 | 0.0199 | 2.89 |
| FALSE | High | TVSDNSLSS | [R].TVSDN | 1xPhosphc | Q80TY0-3 | 0.001916 | 5.66E-05 | 1 | 11 | 1 | Q80TY0-3 | Isoform 3 c | Q80TY0-3 | 0 | 1433.672 | 0 | -0.23 | 0.80724 | 4.41 | 110 | 90 | 21.29 | 71.45 | | High | High | High | High | High | High | High | High | High | High | High | 2.58E-05 | 0.000494 | 2.91 |
| FALSE | High | TVSDNSLSS | [R].TVSDN | 1xPhosphc | Q80TY0-3 | 4.8E-05 | 0 | 1 | 11 | 3 | Q80TY0-3 | Isoform 3 c | Q80TY0-3 | 0 | 1662.835 | 0 | 0.19 | 0.109243 | 16.34 | 97.4 | 102.6 | 5.5 | 14.24 | | High | High | High | High | High | High | High | High | High | High | High | 0 | 7.92E-06 | 2.44 |
| FALSE | High | TVSDNSLSS | [R].TVSDN | 1xDeamida | Q80TY0-3 | 0.014158 | 0.001011 | 1 | 11 | 1 | Q80TY0-3 | Isoform 3 c | Q80TY0-3 | 0 | 1663.819 | 0 | 0.31 | 0.043879 | 15.32 | 92.6 | 107.4 | 5.66 | 12.13 | | High | High | High | High | High | High | High | High | High | High | High | 0.000538 | 0.004651 | 1.55 |
| FALSE | High | TVSDNSLSS | [R].TVSDN | 2xPhosphc | Q80TY0-3 | 0.002398 | 0.000109 | 1 | 11 | 1 | Q80TY0-3 | Isoform 3 c | Q80TY0-3 | 0 | 1742.802 | 0 | 0.71 | 0.004935 | 13.7 | 75.9 | 124.1 | 7.91 | 9.43 | | High | High | High | High | High | High | High | High | High | High | High | 4.94E-05 | 0.000638 | 3.03 |
| FALSE | High | TVSEPNLK | [K].TVSEPN | 1xPhosphc | Q8C2B3-3 | 0.085162 | 0.006376 | 1 | 11 | 1 | Q8C2B3-3 | Isoform 3 c | Q8C2B3-3 | 0 | 1425.775 | 0 | 2.28 | 0.000361 | 9.87 | 34.7 | 165.3 | 3.65 | 8.25 | | High | High | High | High | High | High | High | High | High | High | High | 0.003291 | 0.03569 | 2.43 |
| FALSE | High | TVSETPAVI | [K].TVSETF | 1xPhosphc | G5E884 1x | 0.002441 | 0.000109 | 1 | 1 | 1 | G5E884 | Serine/thre | G5E884 [1 | 0 | 4262.043 | 0 | -0.78 | 0.010252 | 5.99 | 129 | 71 | 11.61 | 9.5 | | High | High | High | High | High | High | High | High | High | High | High | 4.94E-05 | 0.000646 | 4.26 |
| FALSE | High | TVSFEALPI | [K].TVSFEA | 1xPhosphc | Q505D1 1x | 0.017192 | 0.001204 | 1 | 2 | 1 | Q505D1 | Serine/thre | Q505D1 [1 | 0 | 1588.801 | 0 | 0.12 | 0.533456 | 19.4 | 100.4 | 99.6 | 6.68 | 13.4 | | High | High | High | High | High | High | High | High | High | High | High | 0.000638 | 0.00577 | 1.95 |
| FALSE | High | TVSFSAAEI | [K].TVSFSA | 1xPhosphc | E9PYH6 1x | 2.7E-05 | 0 | 1 | 3 | 5 | E9PYH6 | SET domai | E9PYH6 [1 | 0 | 2936.506 | 0 | -1.24 | 0.002225 | 1.58 | 139.4 | 60.6 | 13.76 | 8.42 | | High | High | High | High | High | High | High | High | High | High | High | 0 | 4.17E-06 | 3.92 |
| FALSE | High | TVSFSAAEI | [K].TVSFSA | 2xPhosphc | E9PYH6 2x | 2.52E-06 | 0 | 1 | 3 | 5 | E9PYH6 | SET domai | E9PYH6 [1 | 0 | 3016.472 | 0 | -0.92 | 0.001803 | 2.2 | 134.3 | 65.7 | 14.88 | 10.83 | | High | High | High | High | High | High | High | High | High | High | High | 0 | 2.88E-07 | 4.68 |
| FALSE | High | TVSFSSMP | [K].TVSFSS | 1xPhosphc | Q3USB7 1x | 0.086562 | 0.006435 | 1 | 1 | 1 | Q3USB7 | Inactive ph | Q3USB7 [9 | 0 | 1753.848 | 0 | 2.48 | 0.00355 | 4.77 | 31.1 | 168.9 | 7.49 | 23.51 | | High | High | High | High | High | High | High | High | High | High | High | 0.003303 | 0.0364 | 2.5 |
| FALSE | High | TVSLPLPDL | [R].TVSLPL | 1xPhosphc | Q80XC2 1x | 1.42E-07 | 0 | 1 | 1 | 1 | Q80XC2 | tRNA (ader | Q80XC2 [2 | 0 | 3041.449 | 0 | 0.08 | 0.669433 | 8.21 | 95.4 | 104.6 | 13.08 | 9.16 | | High | High | High | High | High | High | High | High | High | High | High | 0 | 1.13E-08 | 4.46 |
| FALSE | High | TVSISEIPN | [R].TVSISEI | 1xPhosphc | Q6KAU8 1 | 0.01588 | 0.001124 | 1 | 3 | 2 | Q6KAU8 | Autophagy | Q6KAU8 [2 | 0 | 2242.162 | 0 | 2.34 | 0.000118 | 5.1 | 32.9 | 167.1 | 18.74 | 4.2 | | High | High | High | High | High | High | High | High | High | High | High | 0.000903 | 0.00954 | 2.93 |
| FALSE | High | TVSISESEEI | [K].TVSISE | 1xPhosphc | Q640L3-2 1 | 0.002355 | 0.000109 | 1 | 4 | 5 | Q640L3-2 | Isoform 2 c | Q640L3-2 [ | 0 | 2808.348 | 0 | -0.27 | 0.127586 | 15 | 108.7 | 91.3 | 8.71 | 8.71 | | High | High | High | High | High | High | High | High | High | High | High | 4.94E-05 | 0.000625 | 3.48 |
| FALSE | High | TVSDLALQI | [R].TVSDLA | 1xPhosphc | A7YQ68 1x | 0.036103 | 0.002342 | 1 | 2 | 1 | A7YQ68 | Prickle-like | A7YQ68 [8 | 0 | 1829.9 | 0 | -0.55 | 0.002612 | 4.33 | 119.8 | 80.2 | 2.51 | 8.6 | | High | High | High | High | High | High | High | High | High | High | High | 0.00123 | 0.01325 | 2.16 |
| FALSE | High | TVSISESEEI | [K].TVSISE | 2xPhosphc | Q640L3-2 2 | 0.007019 | 0.00062 | 1 | 4 | 3 | Q640L3-2 | Isoform 2 c | Q640L3-2 [ | 0 | 2888.314 | 0 | -0.27 | 0.033354 | 6.94 | 110.1 | 89.9 | 6 | 5.99 | | High | High | High | High | High | High | High | High | High | High | High | 0.000275 | 0.002103 | 2.57 |
| FALSE | High | TVPSPESSE | [K].TVPSPE | 1xPhosphc | B2KFD2 1x | 0.000808 | 0 | 1 | 4 | 2 | B2KFD2 | 2810474O | B2KFD2 [1 | 0 | 1681.845 | 0 | -1.68 | 0.000158 | 0.32 | 153.1 | 46.9 | 5.32 | 7.69 | | High | High | High | High | High | High | High | High | High | High | High | 0 | 0.000188 | 3.16 |

| | | | | | | | | | | | | | | | | | | | | | | | | | | | | | | | | | | | | | | |
|---|---|---|---|---|---|---|---|---|---|---|---|---|---|---|---|---|---|---|---|---|---|---|---|---|---|---|---|---|---|---|---|---|---|---|---|---|---|---|
| FALSE | High | TVPCSSPR | [K].TVPCSS | 1xCarbami | E9QLK3 1x | 0.053473 | 0.00384 | 1 | 3 | 1 | E9QLK3 | BRCA1-ass | E9QLK3 [7 | 0 | 1212.565 | 0 | -0.54 | 0.009877 | 3.28 | 118.7 | 81.3 | 7.52 | 7.66 | | High | High | High | High | High | High | High | High | High | High | High | 0.001999 | 0.0209 | 1.56 |
| FALSE | High | TVDETDNL | [K].TVDETI | 1xDeamida | Q6NVE8 1x | 1.37E-05 | 0 | 1 | 1 | 1 | Q6NVE8 | WD repeat | Q6NVE8 [1 | 0 | 3604.704 | 0 | 0.76 | 0.022938 | 22.02 | 78.6 | 121.4 | 14.74 | 12.61 | | High | High | High | High | High | High | High | High | High | High | High | 0 | 1.92E-06 | 4.75 |
| FALSE | High | TVDLGLTG | [R].TVDLGI | 1xPhospho | Q8JZN5 1x | 4.72E-05 | 0 | 1 | 2 | 1 | Q8JZN5 | Acyl-CoA d | Q8JZN5 [48 | 0 | 2803.464 | 0 | 0.62 | 0.045748 | 24.23 | 73 | 127 | 16.45 | 15.46 | | High | High | High | High | High | High | High | High | High | High | High | 0 | 7.71E-06 | 4.2 |
| FALSE | High | TVDMEETD | [K].TVDME | 1xPhospho | A2ANY6 1x | 0.014667 | 0.001049 | 1 | 2 | 2 | A2ANY6 | Midasin OS | A2ANY6 [4 | 0 | 4389.839 | 0 | -0.54 | 0.031911 | 13.29 | 118.4 | 81.6 | 14.12 | 15 | | High | High | High | High | High | High | High | High | High | High | High | 0.000555 | 0.00481 | 3.94 |
| FALSE | High | TVDQGAP( | [R].TVDQG | 1xPhospho | Q3UHR0 1x | 4.66E-07 | 0 | 1 | 2 | 1 | Q3UHR0 | BAH and co | Q3UHR0 [1 | 0 | 3495.777 | 0 | -0.35 | 0.16805 | 14.81 | 112 | 88 | 13.7 | 16.62 | | High | High | High | High | High | High | High | High | High | High | High | 0 | 4.31E-08 | 4.91 |
| FALSE | High | TVDSDSTM | [K].TVDSDS | 1xPhospho | Q61482 1x | 0.004824 | 0.000456 | 1 | 8 | 1 | Q61482 | CDE1-bindi | Q61482 [20 | 0 | 1624.754 | 0 | 0.51 | 0.014028 | 2.51 | 81.6 | 118.4 | 15.18 | 7.42 | | High | High | High | High | High | High | High | High | High | High | High | 0.000204 | 0.001393 | 2.15 |
| FALSE | High | TVELQEKPI | [K].TVELQE | 1xPhospho | D3U0D6 1x | 0.011246 | 0.000772 | 1 | 5 | 3 | D3U0D6 | Kidney higl | D3U0D6 [8 | 0 | 2778.478 | 0 | 0.46 | 0.021086 | 21.32 | 84.8 | 115.2 | 3.87 | 12.24 | | High | High | High | High | High | High | High | High | High | High | High | 0.000418 | 0.003588 | 2.2 |
| FALSE | High | TVFAGAVP | [K].TVFAG/ | 1xPhospho | Q60902 1x | 0.001077 | 0 | 1 | 5 | 1 | Q60902 | Epidermal | Q60902 [2 | 0 | 1957.076 | 0 | -0.2 | 0.554307 | 23.88 | 98.7 | 101.3 | 39.9 | 130.9 | | High | High | High | High | High | High | High | High | High | High | High | 0 | 0.000259 | 3.51 |
| FALSE | High | TVFAGAVP | [K].TVFAG/ | 1xPhospho | Q60902 1x | 1.17E-07 | 0 | 1 | 5 | 2 | Q60902 | Epidermal | Q60902 [2 | 0 | 2186.239 | 0 | 0.29 | 0.019069 | 5.38 | 92.7 | 107.3 | 4.94 | 5.03 | | High | High | High | High | High | High | High | High | High | High | High | 0 | 1.41E-06 | 3.35 |
| FALSE | High | TVFNSETPS | [R].TVFNSE | 1xPhospho | Q3UDC3 1x | 0.039347 | 0.002793 | 1 | 5 | 1 | Q3UDC3 | Putative ur | Q3UDC3 [1 | 0 | 1446.683 | 0 | 0.4 | 0.017316 | 9.37 | 86.1 | 113.9 | 6.59 | 5.05 | | High | High | High | High | High | High | High | High | High | High | High | 0.001453 | 0.01464 | 1.26 |
| FALSE | High | TVGHSVVS | [R].TVGHS\ | 1xPhospho | O35621 1x | 0.001097 | 0 | 1 | 2 | 1 | O35621 | Phosphom | O35621 [2 | 0 | 1918.959 | 0 | 0.41 | 0.019849 | 12.41 | 85.9 | 114.1 | 9.33 | 8.85 | | High | High | High | High | High | High | High | High | High | High | High | 0 | 0.000264 | 3.31 |
| FALSE | High | TVGNVSPT | [R].TVGNV | 1xPhospho | Q14A95 1x | 0.012072 | 0.000889 | 1 | 3 | 1 | Q14A95 | MCG9666, | Q14A95 [1 | 0 | 1796.893 | 0 | 0.38 | 0.003089 | 6.14 | 86.4 | 113.6 | 8.46 | 7.25 | | High | High | High | High | High | High | High | High | High | High | High | 0.000449 | 0.003878 | 2.96 |
| FALSE | High | TVGNVSPT | [R].TVGNV | 1xPhospho | Q14A95 1x | 0.000815 | 0 | 1 | 3 | 3 | Q14A95 | MCG9666, | Q14A95 [1 | 0 | 1812.888 | 0 | 0.58 | 0.005613 | 7.95 | 78.1 | 121.9 | 9.38 | 9.27 | | High | High | High | High | High | High | High | High | High | High | High | 0 | 0.00019 | 2.51 |
| FALSE | High | TVGPGETP | [R].TVGPG | 1xPhospho | D3YWD3 1 | 0.048715 | 0.00354 | 1 | 3 | 1 | D3YWD3 | Transmem | D3YWD3 [2 | 0 | 1222.603 | 0 | 0.7 | 0.004762 | 6.09 | 76.3 | 123.7 | 7.02 | 9.07 | | High | High | High | High | High | High | High | High | High | High | High | 0.001876 | 0.01873 | 1.43 |
| FALSE | High | TVPPPVPQ | [K].TVPPP\ | 1xPhospho | P97465 1x | 1.62E-06 | 0 | 1 | 2 | 2 | P97465 | Docking pr | P97465 [27 | 0 | 3035.569 | 0 | 2.76 | 0.000391 | 40.55 | 23.9 | 176.1 | 26.37 | 8.35 | | High | High | High | High | High | High | High | High | High | High | High | 0 | 1.74E-07 | 5.25 |
| FALSE | High | TVGTNTPP | [R].TVGTN | 1xPhospho | E9Q0S6 1x | 0.000343 | 0 | 1 | 11 | 1 | E9Q0S6 | Tensin 1 O | E9Q0S6 [1 | 0 | 1696.826 | 0 | -0.05 | 0.319412 | 24.41 | 103.4 | 96.6 | 9.95 | 13.05 | | High | High | High | High | High | High | High | High | High | High | High | 0 | 7.19E-05 | 1.43 |
| FALSE | High | TVGTPIASV | [R].TVGTPI | 1xDeamida | Q3TXS7 1x | 1.51E-06 | 0 | 1 | 6 | 2 | Q3TXS7 | 26S protea | Q3TXS7 [2 | 0 | 2696.38 | 0 | 0.03 | 0.761475 | 13.25 | 101.6 | 98.4 | 10.81 | 6.82 | | High | High | High | High | High | High | High | High | High | High | High | 0 | 1.61E-07 | 4.1 |
| FALSE | High | TVLESGGE | [K].TVLESG | 1xPhospho | Q3TI79 1x | 2.25E-08 | 0 | 1 | 3 | 6 | Q3TI79 | Putative ur | Q3TI79 [47 | 0 | 2334.075 | 0 | -0.1 | 0.145829 | 8.54 | 103.5 | 96.5 | 3.96 | 5.6 | | High | High | High | High | High | High | High | High | High | High | High | 0 | 1.26E-07 | 4.11 |
| FALSE | High | TVLESGGE | [K].TVLESG | 1xDeamida | Q3TI79 1x | 4.74E-07 | 0 | 1 | 3 | 5 | Q3TI79 | Putative ur | Q3TI79 [47 | 0 | 2335.059 | 0 | -0.03 | 0.257281 | 3.97 | 102.1 | 97.9 | 6.88 | 3.67 | | High | High | High | High | High | High | High | High | High | High | High | 0 | 4.41E-08 | 3.47 |
| FALSE | High | TVIIEQSW( | [R].TVIIEQ | 1xPhospho | P63038 1x | 0.031439 | 0.002011 | 1 | 3 | 1 | P63038 | 60 kDa hea | P63038 [61 | 0 | 1653.845 | 0 | -0.39 | 0.207379 | 11.61 | 117.7 | 82.3 | 39.77 | 11.39 | | High | High | High | High | High | High | High | High | High | High | High | 0.001076 | 0.01138 | 2.08 |
| FALSE | High | TVIIEQSW( | [R].TVIIEQ | 1xPhospho | P63038 1x | 2.74E-07 | 0 | 1 | 3 | 1 | P63038 | 60 kDa hea | P63038 [61 | 0 | 1883.008 | 0 | -1.56 | 0.000223 | 1.43 | 149.5 | 50.5 | 3.47 | 6.97 | | High | High | High | High | High | High | High | High | High | High | High | 0 | 2.38E-08 | 3.04 |
| FALSE | High | TVLPNNLSI | [K].TVLPNI | 1xPhospho | B2RWS6 1x | 0.001137 | 0 | 1 | 2 | 1 | B2RWS6 | Histone ac | B2RWS6 [2 | 0 | 2101.081 | 0 | -0.22 | 0.10112 | 6.94 | 108.8 | 91.2 | 10.54 | 9.52 | | High | High | High | High | High | High | High | High | High | High | High | 0 | 0.000277 | 1.97 |
| FALSE | High | TVLSLFDED | [R].TVLSLF | 1xPhospho | Q6PGL7 1x | 4.14E-07 | 0 | 1 | 5 | 2 | Q6PGL7 | WASH com | Q6PGL7 [7 | 0 | 1948.956 | 0 | -1.32 | 0.011434 | 3.8 | 135.3 | 64.7 | 19.09 | 18.27 | | High | High | High | High | High | High | High | High | High | High | High | 0 | 3.79E-08 | 4 |
| FALSE | High | TVLSLFDED | [R].TVLSLF | 1xCarbami | Q6PGL7 1x | 0.111965 | 0.009053 | 1 | 5 | 1 | Q6PGL7 | WASH com | Q6PGL7 [7 | 1 | 3481.561 | 0 | -1.34 | 0.002855 | 2.57 | 146.1 | 53.9 | 8.88 | 18.24 | | High | High | High | High | High | High | High | High | High | High | High | 0.004569 | 0.04951 | 4.09 |
| FALSE | High | TVLYNDDS | [R].TVLYNI | 1xPhospho | A2AHJ4 1x | 4.21E-06 | 0 | 1 | 1 | 4 | A2AHJ4 | Bromodom | A2AHJ4 [17 | 0 | 3210.431 | 0 | -0.13 | 0.384556 | 8.22 | 107.8 | 92.2 | 8.6 | 12.82 | | High | High | High | High | High | High | High | High | High | High | High | 0 | 5.1E-07 | 4.19 |
| FALSE | High | TVMDGGLI | [R].TVMDC | 1xPhospho | Q6P5B5 1x | 3.29E-05 | 0 | 1 | 1 | 2 | Q6P5B5 | Fragile X m | Q6P5B5 [5 | 0 | 3243.413 | 0 | 0.03 | 0.533714 | 10.99 | 98.5 | 101.5 | 9.76 | 19.71 | | High | High | High | High | High | High | High | High | High | High | High | 0.008346 | 0.08981 | 2.06 |
| FALSE | High | TVMDGGLI | [R].TVMDC | 1xPhospho | Q6P5B5 1x | 4.82E-06 | 0 | 1 | 1 | 3 | Q6P5B5 | Fragile X m | Q6P5B5 [5 | 0 | 3259.408 | 0 | 0.1 | 0.825015 | 5.59 | 96.6 | 103.4 | 6.64 | 13.92 | | High | High | High | High | High | High | High | High | High | High | High | 0 | 5.92E-07 | 3.06 |
| FALSE | High | TVMDGGLI | [R].TVMDC | 1xDeamida | Q6P5B5 1x | 0.109329 | 0.008865 | 1 | 1 | 1 | Q6P5B5 | Fragile X m | Q6P5B5 [5 | 0 | 3260.392 | 0 | 0.12 | 0.959841 | 14.43 | 101 | 99 | 14.51 | 17.67 | | High | High | High | High | High | High | High | High | High | High | High | 0.004422 | 0.04778 | 2.32 |
| FALSE | High | TVMDGGLI | [R].TVMDC | 1xPhospho | Q6P5B5 1x | 0.000389 | 0 | 1 | 1 | 3 | Q6P5B5 | Fragile X m | Q6P5B5 [5 | 0 | 3275.403 | 0 | 0.01 | 0.401699 | 8.88 | 99.6 | 100.4 | 5.61 | 40.23 | | High | High | High | High | High | High | High | High | High | High | High | 0 | 8.24E-05 | 3.04 |
| FALSE | High | TVGTPIASV | [R].TVGTPI | 1xPhospho | Q3TXS7 1x | 8.78E-08 | 0 | 1 | 6 | 3 | Q3TXS7 | 26S protea | Q3TXS7 [2 | 0 | 2695.396 | 0 | -0.28 | 0.000918 | 3.39 | 109.8 | 90.2 | 2.51 | 5.61 | | High | High | High | High | High | High | High | High | High | High | High | 0 | 5.48E-07 | 3.75 |
| FALSE | High | TVSPDRLEL | [K].TVSPDF | 1xPhospho | P70168 1x | 0.001629 | 5.66E-05 | 1 | 4 | 1 | P70168 | Importin su | P70168 [10 | 1 | 2095.12 | 0 | -0.98 | 0.000408 | 1.36 | 134.1 | 65.9 | 3.05 | 9.35 | | High | High | High | High | High | High | High | High | High | High | High | 2.58E-05 | 0.000414 | 1.67 |
| FALSE | High | TVSPEGYSI | [R].TVSPEG | 1xPhospho | P30415 1x | 0.012845 | 0.000889 | 1 | 2 | 1 | P30415 | NK-tumor | P30415 [25 | 0 | 1433.651 | 0 | 0.48 | 0.021991 | 22.84 | 83.5 | 116.5 | 10.89 | 7.32 | | High | High | High | High | High | High | High | High | High | High | High | 0.000449 | 0.004149 | 2.08 |
| FALSE | High | TVSPGSVSI | [R].TVSPGS | 1xPhospho | B2RR82 1x | 0.109329 | 0.008865 | 1 | 6 | 1 | B2RR82 | Intersectin | B2RR82 [8 | 0 | 2414.228 | 0 | -0.13 | 0.538166 | 33.48 | 101 | 99 | 14.16 | 59.51 | | High | High | High | High | High | High | High | High | High | High | High | 0.004422 | 0.04777 | 2.12 |
| FALSE | High | TWTLCGTP | [R].TWTLC | 1xCarbami | P68181-4 1 | 0.000184 | 0 | 1 | 6 | 1 | P68181-4 | Isoform 4 c | P68181-4 [ | 0 | 2730.423 | 0 | 1.42 | 0.030094 | 58.23 | 37.6 | 162.4 | 27.74 | 37.35 | | High | High | High | High | High | High | High | High | High | High | High | 0 | 3.58E-05 | 3.49 |
| FALSE | High | TYAENLSPI | [R].TYAENI | 1xPhospho | F6UFZ5 1x | 0.000239 | 0 | 1 | 1 | 1 | F6UFZ5 | PCF11 clea | F6UFZ5 [59 | 0 | 1682.774 | 0 | -0.42 | 0.000894 | 1.19 | 115 | 85 | 5.61 | 3.29 | | High | High | High | High | High | High | High | High | High | High | High | 0 | 4.78E-05 | 3.17 |
| FALSE | High | TYANAEQD | [R].TYANAI | 1xPhospho | O35379 1x | 1.01E-07 | 0 | 1 | 2 | 2 | O35379 | Multidrug | O35379 [8 | 0 | 2682.218 | 0 | 1.77 | 0.001112 | 10.38 | 41.1 | 158.9 | 10.09 | 14.01 | | High | High | High | High | High | High | High | High | High | High | High | 0 | 2.8E-08 | 3.7 |
| FALSE | High | TYANAEQD | [R].TYANAI | 2xPhospho | O35379 2x | 3.14E-08 | 0 | 1 | 2 | 2 | O35379 | Multidrug | O35379 [8 | 0 | 2762.185 | 0 | 0.7 | 0.004346 | 14.45 | 72.1 | 127.9 | 6.31 | 14.15 | | High | High | High | High | High | High | High | High | High | High | High | 0 | 2.09E-09 | 4.34 |
| FALSE | High | TYDTDASE | [K].TYDTD/ | 2xCarbami | Q6PR54-3 | 0.001751 | 5.66E-05 | 2 | 4 | 1 | Q6PR54-3; | Isoform 3 c | Q6PR54-3 | 0 | 4118.788 | 0 | -1.24 | 0.00125 | 3.03 | 141.3 | 58.7 | 9.21 | 19.59 | Shared | High | High | High | High | High | High | High | High | High | High | High | 2.58E-05 | 0.000446 | 2.63 |
| FALSE | High | TYDTDASE | [K].TYDTD/ | 2xCarbami | Q6PR54-3 | 0.000352 | 0 | 2 | 4 | 2 | Q6PR54-3; | Isoform 3 c | Q6PR54-3 | 0 | 4119.772 | 0 | -0.26 | 0.036916 | 8.46 | 109.9 | 90.1 | 11.02 | 10.8 | Shared | High | High | High | High | High | High | High | High | High | High | High | 0 | 0.000115 | 4.08 |
| FALSE | High | TYEDMTLE | [K].TYEDM | 1xPhospho | Q8BVF2 1x | 0.000221 | 0 | 1 | 1 | 2 | Q8BVF2 | Phosducin- | Q8BVF2 [4 | 0 | 3190.266 | 0 | -0.41 | 0.003186 | 4.95 | 113.7 | 86.3 | 8.49 | 11.29 | | High | High | High | High | High | High | High | High | High | High | High | 0 | 4.53E-05 | 3.34 |
| FALSE | High | TYEDMTLE | [K].TYEDM | 1xPhospho | Q8BVF2 1x | 2.09E-08 | 0 | 1 | 1 | 3 | Q8BVF2 | Phosducin- | Q8BVF2 [4 | 0 | 3206.261 | 0 | -0.33 | 0.05531 | 3.69 | 111.3 | 88.7 | 7.03 | 10.61 | | High | High | High | High | High | High | High | High | High | High | High | 0 | 1.33E-09 | 4.92 |
| FALSE | High | TYPQSPK | [K].TYPQSF | 1xPhospho | A2AG50 1x | 0.113753 | 0.009241 | 1 | 5 | 1 | A2AG50 | MAP7 dom | A2AG50 [3 | 0 | 1358.712 | 0 | 0.74 | 0.001225 | 1.28 | 72.9 | 127.1 | 4.28 | 7.59 | | High | High | High | High | High | High | High | High | High | High | High | 0.004625 | 0.05008 | 1.38 |
| FALSE | High | TYQVPPVQ | [K].TYQVPI | 1xPhospho | Q920I9 1x | 4.94E-05 | 0 | 1 | 2 | 1 | Q920I9 | WD repeat | Q920I9 [14 | 0 | 2529.327 | 0 | 0.4 | 0.063671 | 1.73 | 85.8 | 114.2 | 7.36 | 9.51 | | High | High | High | High | High | High | High | High | High | High | High | 0 | 8.14E-06 | 2.88 |
| FALSE | High | TYSAPAINA | [R].TYSAPA | 1xPhospho | D3YTT6 1x | 2.54E-06 | 0 | 1 | 3 | 2 | D3YTT6 | Oxysterol-l | D3YTT6 [24 | 0 | 2460.258 | 0 | 1.52 | 0.000192 | 6.04 | 51.6 | 148.4 | 4.62 | 3.72 | | High | High | High | High | High | High | High | High | High | High | High | 0 | 2.9E-07 | 2.9 |
| FALSE | High | TYSAPAINA | [R].TYSAPA | 2xPhospho | D3YTT6 2x | 6.67E-06 | 0 | 1 | 3 | 1 | D3YTT6 | Oxysterol-l | D3YTT6 [24 | 0 | 2540.224 | 0 | -0.12 | 0.510785 | 3.55 | 95.3 | 104.7 | 13.01 | 19.25 | | High | High | High | High | High | High | High | High | High | High | High | 0 | 8.57E-07 | 3.92 |
| FALSE | High | TYSAPAINA | [R].TYSAPA | 1xPhospho | D3YTT6 1x | 1.77E-06 | 0 | 1 | 3 | 1 | D3YTT6 | Oxysterol-l | D3YTT6 [24 | 1 | 2817.516 | 0 | 0.15 | 0.166446 | 3.12 | 94.7 | 105.3 | 6.54 | 12.87 | | High | High | High | High | High | High | High | High | High | High | High | 0 | 1.94E-07 | 3.96 |
| FALSE | High | TWSTPEVT | [R].TWSTPI | 1xPhospho | Q8VCH5 1x | 0.01095 | 0.000772 | 1 | 4 | 1 | Q8VCH5 | Rab9 effec | Q8VCH5 [1 | 0 | 1907.91 | 0 | -0.27 | 0.074084 | 17.07 | 108.9 | 91.1 | 10.55 | 14.55 | | High | High | High | High | High | High | High | High | High | High | High | 0.000399 | 0.003484 | 3.14 |
| FALSE | High | TYSDEASQ | [R].TYSDEA | 1xPhospho | Q6ZQ33 1x | 0.000314 | 0 | 1 | 4 | 2 | Q6ZQ33 | MKIAA085 | Q6ZQ33 [3 | 0 | 1478.673 | 0 | 0.41 | 0.00419 | 1.56 | 85.9 | 114.1 | 6.85 | 3.28 | | High | High | High | High | High | High | High | High | High | High | High | 0 | 6.5E-05 | 2.25 |
| FALSE | High | TYSLGSALF | [R].TYSLGS | 1xPhospho | P20152 1x | 0.069325 | 0.005112 | 1 | 9 | 1 | P20152 | Vimentin C | P20152 [37 | 0 | 1804.916 | 0 | 1.46 | 0.004478 | 27.77 | 48.3 | 151.7 | 11.41 | 19.93 | | High | High | High | High | High | High | High | High | High | High | High | 0.002618 | 0.02804 | 1.14 |
| FALSE | High | TYSQDCSFI | [R].TYSQD( | 1xCarbami | F8VQD1 1x | 0.002948 | 0.00016 | 1 | 10 | 2 | F8VQD1 | Protein pol | F8VQD1 [9 | 0 | 1673.765 | 0 | 1.47 | 0.00104 | 18.48 | 56.4 | 143.6 | 8 | 11 | | High | High | High | High | High | High | High | High | High | High | High | 7.19E-05 | 0.000803 | 2.05 |
| FALSE | High | TYSSSGSSG | [R].TYSSSG | 1xPhospho | Q6AXD2 1x | 4.98E-08 | 0 | 1 | 5 | 3 | Q6AXD2 | Abi2 prote | Q6AXD2 [2 | 0 | 1849.792 | 0 | -0.06 | 0.74733 | 25.83 | 104.5 | 95.5 | 25.95 | 29.95 | | High | High | High | High | High | High | High | High | High | High | High | 0 | 3.5E-09 | 3.16 |
| FALSE | High | TYTHEVVTI | [R].TYTHE\ | 1xPhospho | P97377 1x | 0.013787 | 0.001011 | 1 | 4 | 2 | P97377 | Cyclin-dep | P97377 [15 | 0 | 1876.92 | 0 | -0.12 | 0.595472 | 23.92 | 100 | 100 | 17.39 | 11.07 | | High | High | High | High | High | High | High | High | High | High | High | 0.000538 | 0.004488 | 1.76 |
| FALSE | High | VAAAAGSG | [R].VAAAA( | 1xCarbami | Q2TBE6 2x | 1.54E-07 | 0 | 1 | 1 | 1 | Q2TBE6 | Phosphatic | Q2TBE6 [3 | 0 | 2179.919 | 0 | 1.29 | 0.000434 | 5.17 | 59.5 | 140.5 | 4.42 | 8.16 | | High | High | High | High | High | High | High | High | High | High | High | 0 | 1.24E-08 | 2.55 |
| FALSE | High | VAAEPESPI | [R].VAAEPE | 1xPhospho | Q9DBN9 1x | 2.53E-05 | 0 | 1 | 2 | 2 | Q9DBN9 | Probable A | Q9DBN9 [1 | 0 | 2337.182 | 0 | 0.19 | 0.024255 | 5.59 | 91.2 | 108.8 | 6.68 | 7.02 | | High | High | High | High | High | High | High | High | High | High | High | 0 | 1.12E-05 | 3.27 |
| FALSE | High | VAAGHELC | [K].VAAGH | 2xPhospho | Q7TMQ1 2 | 0.00145 | 5.66E-05 | 1 | 3 | 2 | Q7TMQ1 | Gap juncti | Q7TMQ1 [ | 1 | 2934.455 | 0 | 1.68 | 0.000154 | 9.13 | 46.4 | 153.6 | 6.49 | 2.1 | | High | High | High | High | High | High | High | High | High | High | High | 2.58E-05 | 0.00036 | 1.93 |
| FALSE | High | VAAGHELC | [K].VAAGH | 3xPhospho | Q7TMQ1 3 | 0.022182 | 0.001542 | 1 | 3 | 2 | Q7TMQ1 | Gap juncti | Q7TMQ1 [ | 1 | 3014.421 | 0 | 0.4 | 0.00171 | 3.01 | 85.1 | 114.9 | 6.04 | 2.57 | | High | High | High | High | High | High | High | High | High | High | High | 0.001712 | 0.01705 | 2.56 |
| FALSE | High | VAAGLGLS | [K].VAAGL( | 1xDeamida | Q8R205 1x | 0.065952 | 0.004766 | 1 | 2 | 1 | Q8R205 | Zinc finger | Q8R205 [1 | 0 | 2019.093 | 0 | -0.11 | 0.08724 | 7.79 | 105.3 | 94.7 | 7.4 | 6.71 | | High | High | High | High | High | High | High | High | High | High | High | 0.002455 | 0.02655 | 1.91 |
| FALSE | High | VAASGSPG | [R].VAASG | 1xPhospho | D3Z5I1 1xP | 0.063788 | 0.004664 | 1 | 3 | 1 | D3Z5I1 | Zinc finger | D3Z5I1 [54 | 0 | 1311.707 | 0 | 0.84 | 0.001064 | 12.02 | 71.8 | 128.2 | 5.74 | 8.01 | | High | High | High | High | High | High | High | High | High | High | High | 0.002403 | 0.02542 | 2.18 |
| FALSE | High | VAASMPLS | [K].VAASM | 1xPhospho | F8WJ40 1x | 1.71E-06 | 0 | 1 | 4 | 2 | F8WJ40 | Histone-lys | F8WJ40 [8 | 0 | 2639.377 | 0 | -0.37 | 0.715722 | 17.89 | 108.3 | 91.7 | 10.54 | 30.79 | | High | High | High | High | High | High | High | High | High | High | High | 0 | 1.86E-07 | 3.76 |
| FALSE | High | VAASPRPG | [R].VAASPI | 1xPhospho | Q3UHL0 1x | 0.000103 | 0 | 1 | 5 | 3 | Q3UHL0 | Putative ur | Q3UHL0 [1 | 0 | 3507.67 | 0 | 0.25 | 0.025009 | 12.06 | 93.8 | 106.2 | 8.37 | 8.09 | | High | High | High | High | High | High | High | High | High | High | High | 2.58E-05 | 0.000434 | 3.56 |
| FALSE | High | VADDSVPG | [K].VADDS\ | 1xPhospho | P61406 1x | 0.053023 | 0.00384 | 1 | 1 | 1 | P61406 | Telomeras | P61406 [22 | 0 | 2123.172 | 0 | -0.14 | 0.061435 | 10.88 | 105.1 | 94.9 | 6.38 | 2.86 | | High | High | High | High | High | High | High | High | High | High | High | 0.001999 | 0.0206 | 1.38 |
| FALSE | High | TYSECEDG | [K].TYSECE | 1xCarbami | Q6A0E5 1x | 4.94E-05 | 0 | 1 | 2 | 4 | Q6A0E5 | MKIAA002 | Q6A0E5 [4 | 0 | 2921.285 | 0 | 0.15 | 0.281051 | 3.46 | 97.4 | 102.6 | 4.34 | 9.2 | | High | High | High | High | High | High | High | High | High | High | High | 0 | 8.14E-06 | 4.38 |
| FALSE | High | TWSRDEVI | [R].TWSRD | 1xDeamida | Q6A065 1x | 0.012619 | 0.000889 | 1 | 6 | 1 | Q6A065 | Centrosom | Q6A065 [1 | 1 | 2913.369 | 0 | 1.25 | 0.004117 | 38.06 | 58.1 | 141.9 | 21.67 | 15.41 | | High | High | High | High | High | High | High | High | High | High | High | 0.000449 | 0.004057 | 2.13 |
| FALSE | High | TWSPTTCG | [R].TWSPT | 1xCarbami | Q6PD21 1x | 0.100936 | 0.008097 | 1 | 1 | 1 | Q6PD21 | SH2 domai | Q6PD21 [2 | 0 | 1632.786 | 0 | 2.03 | 0.000382 | 11.94 | 40.3 | 159.7 | 8.17 | 7.35 | | High | High | High | High | High | High | High | High | High | High | High | 0.004072 | 0.04357 | 1.51 |
| FALSE | High | TWPGSGSI | [R].TWPGS | 1xPhospho | Q6P5E8 1x | 0.025964 | 0.001645 | 1 | 2 | 1 | Q6P5E8 | Diacylglyce | Q6P5E8 [1 | 0 | 1253.588 | 0 | 0.62 | 0.002704 | 7.76 | 82.3 | 117.7 | 4.48 | 8.03 | | High | High | High | High | High | High | High | High | High | High | High | 0.000872 | 0.009132 | 1.09 |
| FALSE | High | TVSPGSVSI | [R].TVSPGS | 1xPhospho | B2RR82 1x | 0.000334 | 0 | 1 | 6 | 2 | B2RR82 | Intersectin | B2RR82 [8 | 0 | 2643.391 | 0 | 0.91 | 0.000221 | 2.21 | 69 | 131 | 6.13 | 4.54 | | High | High | High | High | High | High | High | High | High | High | High | 0 | 6.96E-05 | 3.85 |
| FALSE | High | TVSPGSVSI | [R].TVSPGS | 2xPhospho | B2RR82 2x | 1.04E-06 | 0 | 1 | 6 | 2 | B2RR82 | Intersectin | B2RR82 [8 | 0 | 2723.357 | 0 | 0.44 | 0.006728 | 6.41 | 84.8 | 115.2 | 5.18 | 5.1 | | High | High | High | High | High | High | High | High | High | High | High | 0 | 1.07E-07 | 3.58 |
| FALSE | High | TVSPPPGIS | [R].TVSPPP | 1xPhospho | Q80XI6 1x | 0.090888 | 0.007081 | 1 | 1 | 1 | Q80XI6 | Mitogen-a | Q80XI6 [74 | 0 | 1319.692 | 0 | 0.82 | 0.002101 | 8.63 | 72.4 | 127.6 | 3.39 | 8.71 | | High | High | High | High | High | High | High | High | High | High | High | 0.003606 | 0.03845 | 2.43 |
| FALSE | High | TVSQESLTI | [K].TVSQES | 1xPhospho | A0A0G2JE | 0.069325 | 0.005112 | 1 | 9 | 2 | A0A0G2JE | Nexilin OS | A0A0G2JE | 0 | 1684.892 | 0 | -0.97 | 0.002443 | 2.74 | 132.3 | 67.7 | 6.77 | 16.79 | | High | High | High | High | High | High | High | High | High | High | High | 0.003122 | 0.03242 | 2.82 |
| FALSE | High | TVSQQSFD | [R].TVSQQ | 1xPhospho | Q570Z6 1x | 3.94E-05 | 0 | 1 | 2 | 3 | Q570Z6 | MKIAA412 | Q570Z6 [1 | 0 | 2376.055 | 0 | -0.81 | 0.002026 | 5.32 | 128.2 | 71.8 | 7.59 | 11.64 | | High | High | High | High | High | High | High | High | High | High | High | 2.58E-05 | 0.000412 | 2.81 |
| FALSE | High | TVSQQSFD | [R].TVSQQ | 2xPhospho | Q570Z6 2x | 6.41E-05 | 0 | 1 | 2 | 3 | Q570Z6 | MKIAA412 | Q570Z6 [1 | 0 | 2456.022 | 0 | 0.14 | 0.194246 | 19.41 | 94.9 | 105.1 | 6.29 | 15.37 | | High | High | High | High | High | High | High | High | High | High | High | 0 | 9.26E-05 | 5.25 |
| FALSE | High | TVSQSSSLI | [R].TVSQSS | 1xPhospho | Q9ESC8 1x | 0.004491 | 0.000409 | 1 | 2 | 2 | Q9ESC8 | AF4/FMR2 | Q9ESC8 [8 | 0 | 1474.792 | 0 | -0.72 | 0.000357 | 1.22 | 124.8 | 75.2 | 6.85 | 4.96 | | High | High | High | High | High | High | High | High | High | High | High | 0.000183 | 0.001284 | 1.34 |
| FALSE | High | TVSSPIPYTI | [R].TVSSPII | 1xPhospho | D3YZP9 1x | 0.000115 | 0 | 1 | 3 | 1 | D3YZP9 | Coiled-coil | D3YZP9 [34 | 0 | 4339.185 | 0 | 0.44 | 0.312709 | 35.63 | 85.8 | 114.2 | 20.53 | 24.8 | | High | High | High | High | High | High | High | High | High | High | High | 0 | 0.000021 | 4.78 |
| FALSE | High | TVSSTPNST | [K].TVSSTP | 1xPhospho | P35831 1x | 2.44E-05 | 0 | 1 | 1 | 2 | P35831 | Tyrosine-p | P35831 [56 | 0 | 3569.716 | 0 | 1.14 | 0.001203 | 11.95 | 61.3 | 138.7 | 13.82 | 8.57 | | High | High | High | High | High | High | High | High | High | High | High | 0 | 3.83E-06 | 3.93 |
| FALSE | High | TVSTQHST | [R].TVSTQI | 1xDeamida | E9PVA6 2x | 0.015195 | 0.001089 | 1 | 8 | 1 | E9PVA6 | ARF GTPas | E9PVA6 [3 | 0 | 3961.549 | 0 | -0.02 | 0.965775 | 38.48 | 100.8 | 99.2 | 13.78 | 25.96 | | High | High | High | High | High | High | High | High | High | High | High | 0.000571 | 0.004994 | 5.44 |
| FALSE | High | TVSTSSQPI | [R].TVSTSS | 1xPhospho | F8VQH0 1x | 1.43E-06 | 0 | 1 | 4 | 2 | F8VQH0 | Tyrosine-p | F8VQH0 [8 | 0 | 1857.843 | 0 | -0.36 | 0.020027 | 8.26 | 112.5 | 87.5 | 5.91 | 6.31 | | High | High | High | High | High | High | High | High | High | High | High | 0 | 1.52E-07 | 4 |
| FALSE | High | TVSVEGDA | [R].TVSVE( | 1xPhospho | E9QAD8 1x | 0.000281 | 0 | 1 | 3 | 1 | E9QAD8 | IQ motif ar | E9QAD8 [2 | 0 | 2998.388 | 0 | 1.1 | 0.020072 | 27.35 | 65 | 135 | 13.57 | 21.98 | | High | High | High | High | High | High | High | High | High | High | High | 0 | 5.75E-05 | 4.35 |
| FALSE | High | TVSVEGDA | [R].TVSVE( | 1xDeamida | E9QAD8 1x | 0.061176 | 0.004646 | 1 | 3 | 1 | E9QAD8 | IQ motif ar | E9QAD8 [2 | 0 | 2999.372 | 0 | 0.75 | 0.022975 | 23.45 | 74.5 | 125.5 | 12.22 | 13.78 | | High | High | High | High | High | High | High | High | High | High | High | 0.00237 | 0.02437 | 2.98 |
| FALSE | High | TVSVPSEG( | [R].TVSVPS | 1xPhospho | E9PW37 1x | 1.12E-07 | 0 | 1 | 5 | 2 | E9PW37 | RAS protei | E9PW37 [1 | 0 | 2719.327 | 0 | 0.24 | 0.077052 | 14.92 | 91 | 109 | 6.9 | 8.53 | | High | High | High | High | High | High | High | High | High | High | High | 0 | 8.73E-09 | 3.47 |
| FALSE | High | TVTASHSA | [K].TVTASH | 1xPhospho | E9QLG3 1x | 3.08E-07 | 0 | 1 | 6 | 5 | E9QLG3 | MAX gene- | E9QLG3 [5 | 0 | 1993.016 | 0 | -0.66 | 0.004623 | 7.84 | 122.6 | 77.4 | 8.37 | 6.41 | | High | High | High | High | High | High | High | High | High | High | High | 0 | 2.7E-08 | 4.11 |
| FALSE | High | TVTCVTVEI | [R].TVTCV1 | 1xCarbami | A6H619 1x | 4.4E-09 | 0 | 1 | 3 | 1 | A6H619 | PHD and RI | A6H619 [9 | 0 | 3013.49 | 0 | -0.06 | 0.352588 | 12.4 | 106.4 | 93.6 | 15.33 | 14.7 | | High | High | High | High | High | High | High | High | High | High | High | 0 | 2.28E-10 | 4.9 |
| FALSE | High | TVTDVSTID | [K].TVTDVS | 1xPhospho | Q8BR92 1x | 0.000379 | 0 | 1 | 2 | 1 | Q8BR92 | Paralemmi | Q8BR92 [3 | 0 | 3868.911 | 0 | -1.41 | 0.005406 | 6.69 | 144 | 56 | 9.84 | 22.3 | | High | High | High | High | High | High | High | High | High | High | High | 0 | 8.06E-05 | 3.13 |
| FALSE | High | TVTLTHEAI | [R].TVTLTH | 1xPhospho | E9PV58 1x | 1.17E-05 | 0 | 1 | 4 | 2 | E9PV58 | Transporti | E9PV58 [34 | 0 | 3941.667 | 0 | 0.62 | 0.017063 | 17.7 | 76.9 | 123.1 | 9.96 | 16.04 | | High | High | High | High | High | High | High | High | High | High | High | 0 | 1.61E-06 | 3.58 |
| FALSE | High | TVTNAVVT | [K].TVTNA\ | 1xPhospho | P63017 1x | 0.015329 | 0.001089 | 1 | 14 | 2 | P63017 | Heat shock | P63017 [13 | 0 | 2291.127 | 0 | 0.1 | 0.947588 | 8.15 | 98 | 102 | 15.78 | 7.68 | | High | High | High | High | High | High | High | High | High | High | High | 0.000571 | 0.005046 | 2.09 |
| FALSE | High | TVTPASSAI | [K].TVTPAS | 1xPhospho | O08553 1x | 0.00537 | 0.000599 | 1 | 1 | 4 | O08553 | Dihydropyi | O08553 [5 | 0 | 1399.76 | 0 | 0.5 | 0.005747 | 5.28 | 84.7 | 115.3 | 7.17 | 13.04 | | High | High | High | High | High | High | High | High | High | High | High | 0.000267 | 0.001559 | 1.96 |
| FALSE | High | TVTSEEVNI | [K].TVTSEE | 1xDeamida | Q571C7 1x | 0.020679 | 0.001472 | 1 | 2 | 2 | Q571C7 | Transcripti | Q571C7 [4 | 0 | 2958.257 | 0 | -0.01 | 0.798264 | 6.82 | 98.6 | 101.4 | 9.51 | 20.96 | | High | High | High | High | High | High | High | High | High | High | High | 0.000786 | 0.007085 | 3.71 |
| FALSE | High | TVTSSGSSE | [R].TVTSSG | 1xPhospho | Q8BMI0 1x | 1.06E-05 | 0 | 1 | 2 | 2 | Q8BMI0 | F-box only | Q8BMI0 [7 | 0 | 2227.033 | 0 | 0.96 | 0.002028 | 18.96 | 67.9 | 132.1 | 12.11 | 8.43 | | High | High | High | High | High | High | High | High | High | High | High | 0 | 1.43E-06 | 3.33 |
| FALSE | High | TVTVLAQV | [K].TVTVLA | 1xPhospho | Z4YKQ9 1x | 0.007951 | 0.000736 | 1 | 4 | 1 | Z4YKQ9 | ATPase fan | Z4YKQ9 [3 | 0 | 2026.15 | 0 | -0.21 | 0.037539 | 6.17 | 107 | 93 | 10.64 | 4.92 | | High | High | High | High | High | High | High | High | High | High | High | 0.000327 | 0.002439 | 2.13 |
| FALSE | High | TVVAASTPI | [R].TVVAAS | 1xCarbami | Q3UH06-3 | 3.7E-05 | 0 | 1 | 4 | 1 | Q3UH06-3 | Isoform 3 c | Q3UH06-3 | 0 | 3765.006 | 0 | 0.23 | 0.177 | 6.46 | 99.2 | 100.8 | 9.52 | 10.2 | | High | High | High | High | High | High | High | High | High | High | High | 0 | 5.87E-06 | 3.63 |
| FALSE | High | TVVGPSVA | [K].TVVGPS | 1xPhospho | Q9QZS3 1x | 0.012731 | 0.000889 | 1 | 5 | 2 | Q9QZS3 | Protein nu | Q9QZS3 [2 | 0 | 4207.98 | 0 | 0.71 | 0.061133 | 34.24 | 87.7 | 112.3 | 14.02 | 30.78 | | High | High | High | High | High | High | High | High | High | High | High | 0.000449 | 0.004111 | 2.8 |
| FALSE | High | TVVHLLSGI | [K].TVVHLI | 1xPhospho | H3BK88 1x | 0.029586 | 0.001857 | 2 | 8 | 1 | H3BK88; H | Treacle pro | H3BK88 [1 | 0 | 1491.87 | 0 | 1.39 | 0.004952 | 18.4 | 51.7 | 148.3 | 20.67 | 14.58 | Shared | High | High | High | High | High | High | High | High | High | High | High | 0.000997 | 0.01058 | 2.42 |
| FALSE | High | TVYLQSALS | [K].TVYLQS | 1xPhospho | A0A0R4J25 | 1.23E-07 | 0 | 1 | 3 | 1 | A0A0R4J25 | Etoposide | A0A0R4J25 | 0 | 2196.12 | 0 | 0.85 | 0.023826 | 34.91 | 70.8 | 129.2 | 11.59 | 19.08 | | High | High | High | High | High | High | High | High | High | High | High | 0 | 9.62E-09 | 3.64 |
| FALSE | High | TWAVLYPA | [K].TWAVL | 1xPhospho | P97465 1x | 0.000155 | 0 | 1 | 2 | 1 | P97465 | Docking pr | P97465 [25 | 0 | 1933.989 | 0 | 1.54 | 0.000791 | 19.62 | 54 | 146 | 8.13 | 8.29 | | High | High | High | High | High | High | High | High | High | High | High | 0 | 2.96E-05 | 2.98 |
| FALSE | High | TWCGSPPY | [K].TWCGS | 1xCarbami | E9PU87 1x | 0.007274 | 0.00062 | 1 | 6 | 1 | E9PU87 | Serine/thre | E9PU87 [2 | 0 | 2448.171 | 0 | 1.58 | 0.049724 | 96.67 | 38.1 | 161.9 | 46.4 | 42.18 | | High | High | High | High | High | High | High | High | High | High | High | 0.000275 | 0.002195 | 4.16 |
| FALSE | High | TVASDLQK | [R].TVASDI | 1xPhospho | Q99LH9 1x | 0.000933 | 0 | 1 | 2 | 2 | Q99LH9 | SH3 domai | Q99LH9 [3 | 0 | 1399.76 | 0 | -0.52 | 0.037846 | 3.1 | 118.1 | 81.9 | 2.2 | 18.78 | | High | High | High | High | High | High | High | High | High | High | High | 0 | 0.000221 | 2.66 |
| FALSE | High | TTYLEDLPP | [K].TTYLED | 1xPhospho | Q8C5Q4 1x | 3.54E-09 | 0 | 1 | 2 | 1 | Q8C5Q4 | G-rich sequ | Q8C5Q4 [1 | 0 | 2730.393 | 0 | 0.3 | 0.012749 | 4.99 | 90.4 | 109.6 | 2.53 | 5.51 | | High | High | High | High | High | High | High | High | High | High | High | 0 | 1.79E-10 | 3.1 |
| FALSE | High | TTVTVPSD( | [K].TTVTVF | 1xPhospho | Q9WVM1 | 0.018125 | 0.001282 | 1 | 1 | 2 | Q9WVM1 | Rac GTPase | Q9WVM1 | 0 | 3023.518 | 0 | -1.66 | 0.002186 | 1.77 | 148.6 | 51.4 | 13.52 | 8.99 | | High | High | High | High | High | High | High | High | High | High | High | 0.000689 | 0.006124 | 2.74 |
| FALSE | High | TTVSGPSH | [K].TTVSGI | 1xPhospho | Q5SS80 1x | 1.93E-07 | 0 | 1 | 3 | 3 | Q5SS80 | Dehydroge | Q5SS80 [34 | 0 | 2417.175 | 0 | 1.01 | 0.013936 | 23.76 | 63.7 | 136.3 | 18.83 | 24.34 | | High | High | High | High | High | High | High | High | High | High | High | 0 | 1.59E-08 | 4.4 |
| FALSE | High | TSVDGFQG | [R].TSVDGI | 1xPhospho | Q9WTK2 1 | 1.26E-05 | 0 | 1 | 2 | 2 | Q9WTK2 | Chromodo | Q9WTK2 [2 | 0 | 1918.92 | 0 | 0.47 | 0.029547 | 14.33 | 86.7 | 113.3 | 5.23 | 14.62 | | High | High | High | High | High | High | High | High | High | High | High | 0 | 1.75E-06 | 3.32 |
| FALSE | High | TSVDGFQG | [R].TSVDGI | 2xPhospho | Q9WTK2 2 | 0.01095 | 0.000772 | 1 | 2 | 2 | Q9WTK2 | Chromodo | Q9WTK2 [2 | 0 | 1998.886 | 0 | 1.44 | 0.001041 | 15.92 | 49.5 | 150.5 | 17.63 | 12.47 | | High | High | High | High | High | High | High | High | High | High | High | 0.000399 | 0.003467 | 2.71 |
| FALSE | High | TSVEQADP | [R].TSVEQ/ | 2xCarbami | Q6NZN1-2 | 0.002441 | 0.000109 | 1 | 4 | 1 | Q6NZN1-2 | Isoform 2 c | Q6NZN1-2 | 0 | 3184.453 | 0 | -0.47 | 0.065254 | 8 | 111.8 | 88.2 | 11.23 | 19.66 | | High | High | High | High | High | High | High | High | High | High | High | 4.94E-05 | 0.00065 | 2.88 |
| FALSE | High | TSVGNIAVI | [K].TSVGNI | 1xPhospho | E9PVX6 1x | 0.000193 | 0 | 1 | 2 | 2 | E9PVX6 | Proliferatic | E9PVX6 [1 | 0 | 1225.65 | 0 | -0.27 | 0.048894 | 7.19 | 106.6 | 93.4 | 8.65 | 7.05 | | High | High | High | High | High | High | High | High | High | High | High | 0 | 3.76E-05 | 1.88 |
| FALSE | High | TSVLDGIVF | [K].TSVLDC | 1xPhospho | Q8CJF7 1x | 0.040721 | 0.003058 | 1 | 1 | 1 | Q8CJF7 | Protein ELY | Q8CJF7 [14 | 0 | 1694.893 | 0 | -0.04 | 0.952418 | 26.25 | 101.5 | 98.5 | 12.48 | 29.73 | | High | High | High | High | High | High | High | High | High | High | High | 0.001581 | 0.01524 | 1.73 |
| FALSE | High | TSVLGSSEI | [K].TSVLGS | 1xPhospho | F6YAU7 1x | 0.000389 | 0 | 2 | 9 | 2 | F6YAU7; F8 | Nuclear re | F6YAU7 [1 | 0 | 3059.423 | 0 | 0.55 | 0.106464 | 16.88 | 75.9 | 124.1 | 17 | 25.82 | Shared | High | High | High | High | High | High | High | High | High | High | High | 0 | 0.000135 | 3.45 |
| FALSE | High | TSVLGSSEI | [K].TSVLGS | 1xPhospho | F6YAU7 1x | 0.002508 | 0.000109 | 2 | 9 | 3 | F6YAU7; F8 | Nuclear re | F6YAU7 [1 | 0 | 3075.418 | 0 | 0.78 | 0.005906 | 6.07 | 74.1 | 125.9 | 4.49 | 13.15 | Shared | High | High | High | High | High | High | High | High | High | High | High | 4.94E-05 | 0.000669 | 2.47 |
| FALSE | High | TSVISQSEF | [K].TSVISQ | 1xPhospho | Q8K298 1x | 2.96E-06 | 0 | 1 | 1 | 4 | Q8K298 | Anillin OS= | Q8K298 [7 | 0 | 1918.993 | 0 | 0.46 | 0.018597 | 4.89 | 80 | 120 | 8.54 | 15.62 | | High | High | High | High | High | High | High | High | High | High | High | 0 | 3.45E-07 | 3.4 |
| FALSE | High | TSVPESIAS | [R].TSVPES | 1xPhospho | D3YU22 1x | 6.79E-06 | 0 | 1 | 7 | 1 | D3YU22 | LIM and ca | D3YU22 [4 | 0 | 2114.078 | 0 | 0.81 | 0.000831 | 1.48 | 72.6 | 127.4 | 9.87 | 3.53 | | High | High | High | High | High | High | High | High | High | High | High | 0 | 8.72E-07 | 4.63 |
| FALSE | High | TSVQTEDD | [R].TSVQTI | 1xPhospho | P26231 1x | 6.37E-06 | 0 | 1 | 7 | 5 | P26231 | Catenin alp | P26231 [65 | 0 | 2128.012 | 0 | -0.71 | 0.002135 | 2.03 | 125.6 | 74.4 | 6.81 | 7.07 | | High | High | High | High | High | High | High | High | High | High | High | 0 | 8.13E-07 | 4.76 |
| FALSE | High | TSVSLPESC | [K].TSVSLP | 1xPhospho | G3UZM1 1 | 1.36E-06 | 0 | 1 | 3 | 4 | G3UZM1 | Probable J | G3UZM1 [ | 0 | 2296.159 | 0 | -0.87 | 0.000241 | 1.52 | 129.3 | 70.7 | 2.01 | 5.92 | | High | High | High | High | High | High | High | High | High | High | High | 0 | 1.45E-07 | 4.68 |
| FALSE | High | TSVSLPESC | [K].TSVSLP | 1xDeamida | G3UZM1 1 | 2.67E-05 | 0 | 1 | 3 | 1 | G3UZM1 | Probable J | G3UZM1 [ | 0 | 2297.143 | 0 | -0.32 | 0.050053 | 7.65 | 112.2 | 87.8 | 12.12 | 16.57 | | High | High | High | High | High | High | High | High | High | High | High | 0 | 4.08E-06 | 3.27 |

| | | | | | | | | | | | | | | | | | | | | | | | | | | | | | | | | | | | | | | |
|---|---|---|---|---|---|---|---|---|---|---|---|---|---|---|---|---|---|---|---|---|---|---|---|---|---|---|---|---|---|---|---|---|---|---|---|---|---|---|
| FALSE | High | TSVSSPEQI | [R].TSVSSP | 1xPhospho | Q8K382 1x | 0.008094 | 0.000736 | 1 | 2 | 1 | Q8K382 | DENN dom | Q8K382 [5 | 0 | 1784.842 | 0 | 0.12 | 0.324085 | 16.22 | 95.6 | 104.4 | 9.65 | 5.62 | | High | High | High | High | High | High | High | High | High | High | High | 0.000327 | 0.002466 | 2.77 |
| FALSE | High | TSVDGFQG | [R].TSVDGI | 1xPhospho | Q9WTK2 1 | 7.47E-05 | 0 | 1 | 2 | 2 | Q9WTK2 | Chromodo | Q9WTK2 [2 | 0 | 1689.757 | 0 | 1.34 | 0.001732 | 4.47 | 58.5 | 141.5 | 9.84 | 14.84 | | High | High | High | High | High | High | High | High | High | High | High | 0 | 1.3E-05 | 2.61 |
| FALSE | High | TSVSSVSAS | [R].TSVSSV | 1xPhospho | P19246 1xI | 0.00407 | 0.00036 | 1 | 2 | 4 | P19246 | Neurofilam | P19246 [53 | 0 | 1473.715 | 0 | -1.76 | 0.001178 | 2.21 | 151 | 49 | 9.19 | 17.57 | | High | High | High | High | High | High | High | High | High | High | High | 0.000585 | 0.005515 | 0.74 |
| FALSE | High | TSVTTPAAI | [R].TSVTTP | 1xPhospho | Q3TAB2 1x | 0.001799 | 5.66E-05 | 2 | 8 | 4 | Q3TAB2; H | Putative ur | Q3TAB2 [2 | 0 | 1412.735 | 0 | -0.49 | 0.000354 | 0.72 | 117.1 | 82.9 | 3.68 | 4.92 | Shared | High | High | High | High | High | High | High | High | High | High | High | 0.000275 | 0.002046 | 3.24 |
| FALSE | High | TSWPPATG | [R].TSWPP. | 1xPhospho | E9Q7D5 1x | 0.014797 | 0.001049 | 1 | 2 | 1 | E9Q7D5 | Rho guanir | E9Q7D5 [8 | 0 | 1281.619 | 0 | 0.67 | 0.001881 | 7.78 | 78.3 | 121.7 | 4.07 | 6.78 | | High | High | High | High | High | High | High | High | High | High | High | 0.000555 | 0.004841 | 1.2 |
| FALSE | High | TTAAHSLV( | [K].TTAAH! | 1xPhospho | Q9ES74 1x | 0.119269 | 0.00979 | 1 | 1 | 1 | Q9ES74 | Serine/thr | Q9ES74 [1 | 0 | 2306.073 | 0 | 0.48 | 0.430351 | 49.56 | 89.4 | 110.6 | 12.42 | 49.59 | | High | High | High | High | High | High | High | High | High | High | High | 0.004907 | 0.05341 | 2.42 |
| FALSE | High | TTASPSVR | [K].TTASPS | 1xPhospho | A2AWA9 1 | 0.092372 | 0.007211 | 1 | 2 | 2 | A2AWA9 | Rab GTPas | A2AWA9 [4 | 0 | 1127.566 | 0 | 0.77 | 0.000364 | 7.98 | 75.8 | 124.2 | 6.06 | 2.39 | | High | High | High | High | High | High | High | High | High | High | High | 0.003672 | 0.03937 | 1.68 |
| FALSE | High | TTASSDFTK | [K].TTASSD | 1xPhospho | Q6NS59 1x | 0.017498 | 0.001245 | 1 | 2 | 1 | Q6NS59 | Protein FAI | Q6NS59 [5 | 0 | 1495.745 | 0 | 1.68 | 0.003589 | 37.33 | 41.8 | 158.2 | 16.09 | 17.33 | | High | High | High | High | High | High | High | High | High | High | High | 0.000689 | 0.005895 | 2.34 |
| FALSE | High | TTATLDPTS | [R].TTATLD | 1xPhospho | Q3U1J1-2 | 6.22E-07 | 0 | 1 | 4 | 3 | Q3U1J1-2 | Isoform 2 c | Q3U1J1-2 | 0 | 2121.006 | 0 | -0.01 | 0.726041 | 3.75 | 99.8 | 100.2 | 2.05 | 4.23 | | High | High | High | High | High | High | High | High | High | High | High | 0 | 5.98E-08 | 3.45 |
| FALSE | High | TTATPLK | [R].TTATPL | 1xPhospho | Q8BMA5 1 | 0.063788 | 0.004664 | 1 | 1 | 1 | Q8BMA5 | Protein NP | Q8BMA5 [ | 0 | 1269.722 | 0 | -1.31 | 0.002118 | 2.6 | 152.6 | 47.4 | 16.15 | 25.21 | | High | High | High | High | High | High | High | High | High | High | High | 0.002403 | 0.02554 | 1.25 |
| FALSE | High | TTCSSGGA | [R].TTCSSG | 1xCarbami | Q3THH0 1x | 0.011447 | 0.000816 | 1 | 4 | 2 | Q3THH0 | Putative ur | Q3THH0 [1 | 0 | 2369.101 | 0 | 0.46 | 0.054756 | 22.95 | 80.3 | 119.7 | 6.89 | 21.63 | | High | High | High | High | High | High | High | High | High | High | High | 0.000571 | 0.005021 | 3.27 |
| FALSE | High | TTCSSSEED | [K].TTCSSS | 2xCarbami | Q9CWE0 1 | 0.004256 | 0.00036 | 1 | 2 | 2 | Q9CWE0 | Mitochond | Q9CWE0 [2 | 0 | 2357.029 | 0 | -0.24 | 0.006602 | 3.08 | 108.6 | 91.4 | 4.99 | 1.07 | | High | High | High | High | High | High | High | High | High | High | High | 0.000161 | 0.001206 | 2.17 |
| FALSE | High | TTEAPCSP( | [K].TTEAPC | 1xCarbami | Q80U72 1x | 3.08E-07 | 0 | 1 | 3 | 3 | Q80U72 | Protein scr | Q80U72 [1 | 0 | 3045.464 | 0 | -0.33 | 0.020418 | 3.55 | 112.1 | 87.9 | 5.05 | 8.96 | | High | High | High | High | High | High | High | High | High | High | High | 0 | 2.7E-08 | 4.14 |
| FALSE | High | TTEAPCSP( | [K].TTEAPC | 1xCarbami | Q80U72 1x | 0.001689 | 5.66E-05 | 1 | 3 | 2 | Q80U72 | Protein scr | Q80U72 [1 | 0 | 3046.448 | 0 | -0.15 | 0.288946 | 21.1 | 105.1 | 94.9 | 9.24 | 10.59 | | High | High | High | High | High | High | High | High | High | High | High | 2.58E-05 | 0.000516 | 4.5 |
| FALSE | High | TTEAPCSP( | [K].TTEAPC | 1xCarbami | Q80U72 2x | 0.000543 | 0 | 1 | 3 | 2 | Q80U72 | Protein scr | Q80U72 [1 | 0 | 3125.43 | 0 | 0.24 | 0.112425 | 19.64 | 95 | 105 | 9.16 | 10.2 | | High | High | High | High | High | High | High | High | High | High | High | 0 | 0.00012 | 3.4 |
| FALSE | High | TTEHSPNQ | [K].TTEHSF | 1xPhospho | Q91Z96 1x | 0.004911 | 0.000504 | 1 | 1 | 1 | Q91Z96 | BMP-2-ind | Q91Z96 [6 | 0 | 1579.788 | 0 | -1.01 | 0.000806 | 1.19 | 134.2 | 65.8 | 8.12 | 6.82 | | High | High | High | High | High | High | High | High | High | High | High | 0.000225 | 0.001418 | 2.38 |
| FALSE | High | TSVTSLR | [K].TSVTSL | 1xPhospho | Q6RT24 1x | 0.081752 | 0.006219 | 1 | 2 | 1 | Q6RT24 | Centromer | Q6RT24 [4 | 0 | 1072.56 | 0 | -0.41 | 0.002182 | 5.24 | 114.4 | 85.6 | 2.25 | 7.56 | | High | High | High | High | High | High | High | High | High | High | High | 0.003229 | 0.0341 | 0.95 |
| FALSE | High | TSVATQCD | [K].TSVAT( | 1xCarbami | Q3UIV3 2x | 2.63E-05 | 0 | 1 | 12 | 1 | Q3UIV3 | Putative ur | Q3UIV3 [7 | 0 | 2468.039 | 0 | -0.38 | 0.003259 | 3.81 | 114.6 | 85.4 | 4.46 | 8.21 | | High | High | High | High | High | High | High | High | High | High | High | 0 | 4.01E-06 | 4.24 |
| FALSE | High | TSVATQCD | [K].TSVAT( | 1xCarbami | Q3UIV3 1x | 6.79E-06 | 0 | 1 | 12 | 4 | Q3UIV3 | Putative ur | Q3UIV3 [7 | 0 | 2388.073 | 0 | -1.27 | 0.002885 | 5.07 | 139.8 | 60.2 | 11.48 | 38.12 | | High | High | High | High | High | High | High | High | High | High | High | 0 | 8.74E-07 | 4.41 |
| FALSE | High | TSTWSGES | [K].TSTWS( | 1xPhospho | P09405 1xI | 0.039347 | 0.002793 | 1 | 6 | 1 | P09405 | Nucleolin ( | P09405 [47 | 0 | 1520.74 | 0 | -0.51 | 0.002408 | 1.86 | 118.8 | 81.2 | 4.36 | 9.39 | | High | High | High | High | High | High | High | High | High | High | High | 0.001468 | 0.01472 | 1.77 |
| FALSE | High | TSSSSPANS | [K].TSSSSP. | 2xPhospho | B2RT41 2xI | 0.027839 | 0.001755 | 1 | 2 | 1 | B2RT41 | Zinc finger, | B2RT41 [80 | 0 | 2214.882 | 0 | 0.35 | 0.004015 | 6.16 | 86.8 | 113.2 | 8.34 | 10.06 | | High | High | High | High | High | High | High | High | High | High | High | 0.000953 | 0.009878 | 1.94 |
| FALSE | High | TSSSSPLSSI | [K].TSSSSPI | 1xPhospho | Q6ZPJ0 1xI | 0.00359 | 0.00021 | 1 | 3 | 1 | Q6ZPJ0 | Testis-expr | Q6ZPJ0 [15 | 0 | 1702.866 | 0 | -1.09 | 0.000401 | 2.02 | 136 | 64 | 7.2 | 6.72 | | High | High | High | High | High | High | High | High | High | High | High | 9.42E-05 | 0.001002 | 2.55 |
| FALSE | High | TSSTALGEK | [K].TSSTAL | 1xPhospho | Q8CB77 1x | 0.0016 | 5.66E-05 | 1 | 1 | 3 | Q8CB77 | Elongin-A ( | Q8CB77 [4 | 0 | 1431.75 | 0 | 0.78 | 0.00722 | 10.12 | 71.1 | 128.9 | 9.06 | 13.85 | | High | High | High | High | High | High | High | High | High | High | High | 2.58E-05 | 0.000404 | 2.91 |
| FALSE | High | TSSTCSNES | [R].TSSTCS | 1xCarbami | Q8BYJ6 1xI | 0.001463 | 5.66E-05 | 1 | 9 | 3 | Q8BYJ6 | TBC1 doma | Q8BYJ6 [75 | 0 | 2345.064 | 0 | 0.06 | 0.189012 | 9.25 | 97.7 | 102.3 | 4.86 | 6.88 | | High | High | High | High | High | High | High | High | High | High | High | 2.58E-05 | 0.000364 | 3.05 |
| FALSE | High | TSSTCSNES | [R].TSSTCS | 1xCarbami | Q8BYJ6 2xI | 0.001933 | 5.66E-05 | 1 | 9 | 2 | Q8BYJ6 | TBC1 doma | Q8BYJ6 [75 | 0 | 2425.031 | 0 | 0.44 | 0.040271 | 19.97 | 84.9 | 115.1 | 7.58 | 18.88 | | High | High | High | High | High | High | High | High | High | High | High | 2.58E-05 | 0.000498 | 3.79 |
| FALSE | High | TSSTDEVLS | [R].TSSTDE | 1xPhospho | E9PVV1 1x | 3.38E-05 | 0 | 2 | 6 | 3 | E9PVV1; E | Transcripti | E9PVV1 [5 | 0 | 1746.825 | 0 | -0.16 | 0.90416 | 31.48 | 102.9 | 97.1 | 50.78 | 75.72 | Shared | High | High | High | High | High | High | High | High | High | High | High | 0 | 6.28E-06 | 1.98 |
| FALSE | High | TSSTDEVLS | [R].TSSTDE | 1xPhospho | E9PVV1 1x | 0.000103 | 0 | 2 | 6 | 11 | E9PVV1; E | Transcripti | E9PVV1 [5 | 0 | 1975.988 | 0 | 0.02 | 0.573634 | 5.21 | 99.2 | 100.8 | 3.52 | 2.86 | Shared | High | High | High | High | High | High | High | High | High | High | High | 0 | 7.39E-05 | 3.82 |
| FALSE | High | TSSTDLSDI | [K].TSSTDL | 1xPhospho | Q3V141 1x | 9.76E-07 | 0 | 1 | 7 | 7 | Q3V141 | Putative ur | Q3V141 [7 | 0 | 2687.467 | 0 | 0.26 | 0.083045 | 8.46 | 92.5 | 107.5 | 4.88 | 8.44 | | High | High | High | High | High | High | High | High | High | High | High | 0 | 2.1E-07 | 4.13 |
| FALSE | High | TSSTDQQP | [R].TSSTDC | 1xPhospho | E9Q8L9 1xI | 2.38E-06 | 0 | 1 | 3 | 4 | E9Q8L9 | Rab11 fam | E9Q8L9 [2 | 0 | 2188.012 | 0 | -0.03 | 0.90033 | 27.55 | 102.2 | 97.8 | 8.88 | 16.98 | | High | High | High | High | High | High | High | High | High | High | High | 0 | 2.69E-07 | 3.12 |
| FALSE | High | TSSTDQQP | [R].TSSTDC | 1xPhospho | E9Q8L9 1xI | 0.001674 | 5.66E-05 | 1 | 3 | 2 | E9Q8L9 | Rab11 fam | E9Q8L9 [2 | 0 | 2417.175 | 0 | 0.21 | 0.140193 | 2.93 | 96.8 | 103.2 | 6.38 | 5.38 | | High | High | High | High | High | High | High | High | High | High | High | 2.58E-05 | 0.00046 | 1.53 |
| FALSE | High | TSSTNEDEI | [R].TSSTNE | 1xPhospho | Q61286-2 | 0.000267 | 0 | 2 | 7 | 6 | Q61286-2; | Isoform AL | Q61286-2 | 0 | 2245.027 | 0 | -0.38 | 0.013706 | 1.31 | 113.2 | 86.8 | 8.82 | 4.19 | Shared | High | High | High | High | High | High | High | High | High | High | High | 0 | 5.4E-05 | 3.42 |
| FALSE | High | TSSTVSTPS | [K].TSSTVS | 1xPhospho | Q5SFM8 1x | 0.001933 | 5.66E-05 | 1 | 3 | 4 | Q5SFM8 | RNA-bindir | Q5SFM8 [8 | 0 | 1532.797 | 0 | -0.57 | 0.002264 | 4.98 | 118.6 | 81.4 | 4.58 | 8.63 | | High | High | High | High | High | High | High | High | High | High | High | 0.003685 | 0.03963 | 1.53 |
| FALSE | High | TSSVPEYVY | [R].TSSVPE | 1xPhospho | Q3TIY5 1xI | 0.000343 | 0 | 1 | 4 | 3 | Q3TIY5 | Putative ur | Q3TIY5 [80 | 0 | 1710.876 | 0 | 0.5 | 0.004259 | 7.69 | 84.9 | 115.1 | 4.31 | 10.43 | | High | High | High | High | High | High | High | High | High | High | High | 0 | 7.17E-05 | 2.5 |
| FALSE | High | TSSVSSLAS | [R].TSSVSS | 1xCarbami | Q8K3Z9 1x | 1.14E-06 | 0 | 1 | 2 | 4 | Q8K3Z9 | Nuclear en | Q8K3Z9 [3 | 0 | 2221.037 | 0 | 0.16 | 0.077242 | 8.44 | 94.7 | 105.3 | 2.9 | 6.12 | | High | High | High | High | High | High | High | High | High | High | High | 0 | 9.56E-06 | 2.95 |
| FALSE | High | TSSVSSLAS | [R].TSSVSS | 1xCarbami | Q8K3Z9 2x | 0.000393 | 0 | 1 | 2 | 2 | Q8K3Z9 | Nuclear en | Q8K3Z9 [3 | 0 | 2301.003 | 0 | 0.13 | 0.232377 | 20.18 | 94.1 | 105.9 | 17.56 | 22.62 | | High | High | High | High | High | High | High | High | High | High | High | 0 | 0.000149 | 2.68 |
| FALSE | High | TSSVSSISQ | [K].TSSVSS | 1xPhospho | A0A1S6GW | 8.13E-06 | 0 | 1 | 2 | 3 | A0A1S6GW | Uncharacte | A0A1S6GW | 0 | 1772.863 | 0 | 0.53 | 0.000692 | 5.58 | 82.9 | 117.1 | 4.68 | 3.03 | | High | High | High | High | High | High | High | High | High | High | High | 0 | 1.07E-06 | 3.78 |
| FALSE | High | TSSVVEEHI | [R].TSSVVE | 1xPhospho | A0A140LJ0 | 1.81E-08 | 0 | 1 | 4 | 3 | A0A140LJ0 | Zinc finger | A0A140LJ0 | 0 | 2911.331 | 0 | 1.21 | 0.001333 | 7.39 | 63.2 | 136.8 | 6.76 | 7.99 | | High | High | High | High | High | High | High | High | High | High | High | 0 | 1.12E-09 | 3.95 |
| FALSE | High | TSSVVEEHI | [R].TSSVVE | 2xPhospho | A0A140LJ0 | 0.003526 | 0.00021 | 1 | 4 | 1 | A0A140LJ0 | Zinc finger | A0A140LJ0 | 0 | 2991.297 | 0 | 0.72 | 0.002121 | 13.07 | 75.4 | 124.6 | 10.99 | 10.55 | | High | High | High | High | High | High | High | High | High | High | High | 9.42E-05 | 0.00098 | 3.25 |
| FALSE | High | TSSVVTLEV | [R].TSSVVT | 1xPhospho | E9Q9C3 1x | 5.23E-06 | 0 | 1 | 5 | 1 | E9Q9C3 | Afadin OS= | E9Q9C3 [1 | 0 | 1671.933 | 0 | 1.05 | 0.004179 | 14.2 | 62.2 | 137.8 | 11.23 | 13.15 | | High | High | High | High | High | High | High | High | High | High | High | 0 | 6.52E-07 | 3.18 |
| FALSE | High | TSSYSGSNA | [R].TSSYSG | 1xPhospho | B2RXS2 1xI | 0.002058 | 5.66E-05 | 1 | 5 | 1 | B2RXS2 | Fam179b p | B2RXS2 [80 | 0 | 2852.371 | 0 | 0.66 | 0.003365 | 18.1 | 82 | 118 | 14.12 | 8.25 | | High | High | High | High | High | High | High | High | High | High | High | 2.58E-05 | 0.000535 | 2.91 |
| FALSE | High | TSTGSTST\ | [R].TSTGST | 1xPhospho | A6H8H2-2 | 0.000822 | 0 | 1 | 3 | 6 | A6H8H2-2 | Isoform 2 c | A6H8H2-2 | 0 | 3588.776 | 0 | -0.02 | 0.695652 | 23.08 | 93.6 | 106.4 | 23.68 | 9.18 | | High | High | High | High | High | High | High | High | High | High | High | 2.58E-05 | 0.00036 | 3.52 |
| FALSE | High | TSTLPPLPLI | [K].TSTLPP | 1xPhospho | Q14AX6-2 | 1.29E-05 | 0 | 3 | 4 | 4 | Q14AX6-2; | Isoform 2 c | Q14AX6-2 | 0 | 2970.519 | 0 | 0.29 | 0.25277 | 8.75 | 89.9 | 110.1 | 10.99 | 12.94 | Shared | High | High | High | High | High | High | High | High | High | High | High | 0.001061 | 0.01124 | 3.17 |
| FALSE | High | TSTPVILEG | [R].TSTPVII | 1xPhospho | E9Q6J5-3 1 | 6.58E-08 | 0 | 1 | 3 | 6 | E9Q6J5-3 | Isoform Ne | E9Q6J5-3 [ | 0 | 2156.032 | 0 | 0.65 | 0.001272 | 7.84 | 81.1 | 118.9 | 12.16 | 9.62 | | High | High | High | High | High | High | High | High | High | High | High | 0 | 4.76E-09 | 4.93 |
| FALSE | High | TSTQENVH | [R].TSTQEI | 1xPhospho | Q8R3I2 1xI | 0.000892 | 0 | 1 | 4 | 3 | Q8R3I2 | Lysophosp | Q8R3I2 [47 | 0 | 1981.016 | 0 | 2.57 | 0.000434 | 8.43 | 28.4 | 171.6 | 12.05 | 9.66 | | High | High | High | High | High | High | High | High | High | High | High | 0 | 0.00021 | 3.31 |
| FALSE | High | TSTSAVPNI | [R].TSTSAV | 1xPhospho | Q8K232 1x | 0.003372 | 0.00016 | 2 | 8 | 2 | Q8K232; E | Adducin 1 | Q8K232 [4 | 0 | 2438.31 | 0 | 1.5 | 0.001774 | 3.73 | 53 | 147 | 13.71 | 11.95 | Shared | High | High | High | High | High | High | High | High | High | High | High | 9.42E-05 | 0.000934 | 3.13 |
| FALSE | High | TSTSLHSV( | [K].TSTSLH | 1xCarbami | B9EK83 1xI | 0.001373 | 5.66E-05 | 1 | 2 | 2 | B9EK83 | Transient r | B9EK83 [1 | 0 | 1958.884 | 0 | 2.52 | 0.000209 | 4.65 | 28.2 | 171.8 | 8.6 | 5.25 | | High | High | High | High | High | High | High | High | High | High | High | 0.002045 | 0.0211 | 2.3 |
| FALSE | High | TSTSLHSV( | [K].TSTSLH | 1xCarbami | A2AI57 1xI | 0.050826 | 0.003676 | 1 | 1 | 2 | A2AI57 | Transient r | A2AI57 [14 | 0 | 1887.847 | 0 | 0.85 | 0.001703 | 10.09 | 71.4 | 128.6 | 4.69 | 10.62 | | High | High | High | High | High | High | High | High | High | High | High | 0.001948 | 0.01967 | 1.93 |
| FALSE | High | TSTSPQNS( | [K].TSTSPC | 1xPhospho | Q80TX5 1x | 1.41E-06 | 0 | 1 | 4 | 3 | Q80TX5 | MKIAA057 | Q80TX5 [1 | 0 | 1702.841 | 0 | -0.24 | 0.007488 | 1.89 | 109.8 | 90.2 | 3.23 | 5.33 | | High | High | High | High | High | High | High | High | High | High | High | 0 | 1.5E-07 | 2.74 |
| FALSE | High | TSTSPQNS( | [K].TSTSPC | 1xDeamida | Q80TX5 1x | 0.00095 | 0 | 1 | 4 | 4 | Q80TX5 | MKIAA057 | Q80TX5 [1 | 0 | 1703.825 | 0 | -0.26 | 0.022152 | 7.05 | 112.2 | 87.8 | 12.78 | 11.35 | | High | High | High | High | High | High | High | High | High | High | High | 0.000275 | 0.001972 | 2.47 |
| FALSE | High | TTESWAPP | [K].TTESW. | 2xPhospho | Q5HZJ0 2xI | 0.041072 | 0.003094 | 1 | 2 | 1 | Q5HZJ0 | Ribonuclea | Q5HZJ0 [3 | 2 | 3037.433 | 0 | -0.9 | 0.001562 | 1.78 | 131.4 | 68.6 | 5.49 | 10.45 | | High | High | High | High | High | High | High | High | High | High | High | 0.001609 | 0.01541 | 2.6 |
| FALSE | High | VADNSFDA | [R].VADNS | 1xPhospho | E9Q5C9 1x | 0.000356 | 0 | 1 | 2 | 3 | E9Q5C9 | Nucleolar a | E9Q5C9 [6 | 0 | 1504.745 | 0 | -0.68 | 0.002273 | 1.31 | 121.7 | 78.3 | 7.05 | 5.16 | | High | High | High | High | High | High | High | High | High | High | High | 4.94E-05 | 0.000593 | 2.93 |
| FALSE | High | TTFDEAMA | [K].TTFDEA | 1xPhospho | O70456 1x | 0.065952 | 0.004766 | 1 | 2 | 1 | O70456 | 14-3-3 pro | O70456 [1 | 0 | 2712.252 | 0 | 0.64 | 0.004615 | 5.88 | 78.1 | 121.9 | 2.82 | 13.21 | | High | High | High | High | High | High | High | High | High | High | High | 0.002455 | 0.02662 | 2.34 |
| FALSE | High | TTGASQLQ | [K].TTGAS( | 1xPhospho | Q9D842 1x | 0.000454 | 0 | 1 | 3 | 1 | Q9D842 | Aprataxin a | Q9D842 [1 | 0 | 1826.91 | 0 | 0.68 | 0.411753 | 51.48 | 77 | 123 | 39.16 | 55.6 | | High | High | High | High | High | High | High | High | High | High | High | 0 | 9.8E-05 | 2.37 |
| FALSE | High | TTRTPEEG( | [K].TTRTPE | 1xPhospho | P14873 1xI | 0.006957 | 0.00062 | 2 | 2 | 2 | P14873; B2 | Microtubu | P14873 [1 | 1 | 2485.19 | 0 | -0.2 | 0.34831 | 28.38 | 106.9 | 93.1 | 9.95 | 16.02 | Shared | High | High | High | High | High | High | High | High | High | High | High | 0.000756 | 0.006576 | 3.34 |
| FALSE | High | TTRTPEVS( | [K].TTRTPE | 1xPhospho | P14873 1xI | 3.88E-06 | 0 | 2 | 2 | 2 | P14873; B2 | Microtubu | P14873 [1 | 1 | 2170.083 | 0 | -0.76 | 0.011165 | 6.64 | 125.3 | 74.7 | 11.29 | 9.82 | Shared | High | High | High | High | High | High | High | High | High | High | High | 0 | 1.44E-06 | 3.26 |
| FALSE | High | TTSAGTGG | [R].TTSAGT | 1xPhospho | E9PW43 1x | 0.016451 | 0.001124 | 1 | 3 | 4 | E9PW43 | Protein tra | E9PW43 [6 | 0 | 1449.64 | 0 | -0.43 | 0.028659 | 9.13 | 120.7 | 79.3 | 19.05 | 12.95 | | High | High | High | High | High | High | High | High | High | High | High | 0.000585 | 0.005477 | 1.7 |
| FALSE | High | TTSASSVK | [K].TTSASS | 1xPhospho | P27546 1xI | 0.023792 | 0.001612 | 1 | 10 | 2 | P27546 | Microtubu | P27546 [83 | 0 | 1318.702 | 0 | 0.59 | 0.003407 | 4.75 | 79.8 | 120.2 | 3.41 | 8.74 | | High | High | High | High | High | High | High | High | High | High | High | 0.002117 | 0.02191 | 1.44 |
| FALSE | High | TTSGATPA | [K].TTSGAT | 1xPhospho | Q3TNN6 1x | 0.004781 | 0.000456 | 1 | 6 | 2 | Q3TNN6 | Putative ur | Q3TNN6 [1 | 0 | 1354.693 | 0 | 2.91 | 0.000618 | 7.82 | 23.9 | 176.1 | 13.11 | 12.56 | | High | High | High | High | High | High | High | High | High | High | High | 0.000204 | 0.001369 | 2.59 |
| FALSE | High | TTSGDDAC | [R].TTSGDI | 1xCarbami | A2A9M4 1 | 0.049549 | 0.00361 | 1 | 6 | 2 | A2A9M4 | Dedicator c | A2A9M4 [4 | 0 | 3272.606 | 0 | 0.62 | 0.041946 | 38.28 | 76.4 | 123.6 | 20.51 | 12.57 | | High | High | High | High | High | High | High | High | High | High | High | 0.001889 | 0.01901 | 2.53 |
| FALSE | High | TTSGYSGG | [K].TTSGYS | 1xPhospho | P11679 1xI | 7.14E-08 | 0 | 1 | 1 | 5 | P11679 | Keratin, typ | P11679 [42 | 0 | 4501.976 | 0 | 3.48 | 0.004534 | 8.3 | 12.5 | 187.5 | 26.76 | 27.62 | | High | High | High | High | High | High | High | High | High | High | High | 0 | 5.23E-09 | 5.91 |
| FALSE | High | TTSGYSGG | [K].TTSGYS | 1xPhospho | P11679 1xI | 7.1E-06 | 0 | 1 | 1 | 15 | P11679 | Keratin, typ | P11679 [42 | 0 | 4517.971 | 0 | 2.62 | 0.002989 | 11.97 | 28.6 | 171.4 | 30.05 | 20.61 | | High | High | High | High | High | High | High | High | High | High | High | 0 | 9.2E-07 | 5.7 |
| FALSE | High | TTSPATDA( | [K].TTSPAT | 1xPhospho | Q8R010 1x | 0.000837 | 0 | 1 | 1 | 1 | Q8R010 | Aminoacyl | Q8R010 [3 | 0 | 2921.373 | 0 | 0.2 | 0.008299 | 1.79 | 92.7 | 107.3 | 6.43 | 5.37 | | High | High | High | High | High | High | High | High | High | High | High | 0 | 0.000196 | 2.35 |
| FALSE | High | TTSPLQSP\ | [K].TTSPLQ | 1xPhospho | Q14B71 1x | 8.68E-07 | 0 | 1 | 2 | 1 | Q14B71 | Cell divisio | Q14B71 [4 | 0 | 2966.47 | 0 | -0.91 | 0.003105 | 4.54 | 133.9 | 66.1 | 13.43 | 9.75 | | High | High | High | High | High | High | High | High | High | High | High | 0 | 8.66E-08 | 4.85 |
| FALSE | High | TTSPLQSP\ | [K].TTSPLQ | 2xPhospho | Q14B71 2x | 1.91E-05 | 0 | 1 | 2 | 2 | Q14B71 | Cell divisio | Q14B71 [4 | 0 | 3046.436 | 0 | -0.64 | 0.002335 | 4.34 | 120.2 | 79.8 | 7.99 | 9.88 | | High | High | High | High | High | High | High | High | High | High | High | 0 | 2.82E-06 | 4.33 |
| FALSE | High | TTSPSSDTD | [R].TTSPSS | 1xPhospho | Q3U2C9 1x | 0.004613 | 0.000456 | 1 | 6 | 4 | Q3U2C9 | PHD finger | Q3U2C9 [1 | 0 | 2329.169 | 0 | -0.74 | 0.000154 | 0.67 | 125 | 75 | 3.54 | 5.89 | | High | High | High | High | High | High | High | High | High | High | High | 0.000204 | 0.001326 | 3.04 |
| FALSE | High | TTSPTQEL( | [K].TTSPTC | 1xPhospho | A2BI37 1xI | 9.02E-08 | 0 | 1 | 2 | 3 | A2BI37 | Zinc finger | A2BI37 [19 | 0 | 2186.151 | 0 | -1.84 | 0.000209 | 1.12 | 155 | 45 | 6.21 | 13.47 | | High | High | High | High | High | High | High | High | High | High | High | 0 | 1.54E-08 | 3.48 |
| FALSE | High | TTRSPDTSA | [K].TTRSPD | 1xCarbami | P14873 1xI | 1.45E-06 | 0 | 2 | 2 | 2 | P14873; B2 | Microtubu | P14873 [2 | 1 | 2594.156 | 0 | -0.7 | 0.020163 | 13.3 | 119.5 | 80.5 | 10.09 | 14.99 | Shared | High | High | High | High | High | High | High | High | High | High | High | 0 | 1.54E-07 | 3.37 |
| FALSE | High | TTSQCK | [R].TTSQCI | 1xCarbami | Q91YL3 1xI | 0.093879 | 0.007306 | 1 | 4 | 2 | Q91YL3 | Uridine-cyt | Q91YL3 [6 | 0 | 1262.622 | 0 | 1.43 | 0.000987 | 6.76 | 54.3 | 145.7 | 11.08 | 9.16 | | High | High | High | High | High | High | High | High | High | High | High | 0.003727 | 0.04017 | 1.54 |
| FALSE | High | TTSSANNPI | [R].TTSSAN | 1xCarbami | Q3U741 1x | 0.063258 | 0.004664 | 1 | 5 | 2 | Q3U741 | DEAD (Asp | Q3U741 [4 | 1 | 2582.085 | 0 | 0.69 | 0.009227 | 6.32 | 74.6 | 125.4 | 11.91 | 14.99 | | High | High | High | High | High | High | High | High | High | High | High | 0.005216 | 0.0567 | 3.91 |
| FALSE | High | TTSSETNTF | [R].TTSSET | 1xPhospho | Q8CBQ5 1x | 3.7E-05 | 0 | 1 | 1 | 4 | Q8CBQ5 | Phosphatic | Q8CBQ5 [8 | 0 | 2895.432 | 0 | -0.61 | 0.016369 | 8.69 | 122.9 | 77.1 | 15.03 | 15.42 | | High | High | High | High | High | High | High | High | High | High | High | 0 | 5.86E-06 | 3.27 |
| FALSE | High | TTSSGAAG | [K].TTSSGA | 1xPhospho | Q5DTZ0 1x | 1.9E-05 | 0 | 1 | 1 | 1 | Q5DTZ0 | Protein NY | Q5DTZ0 [2 | 0 | 2953.41 | 0 | -1.48 | 0.000651 | 1.85 | 148.4 | 51.6 | 8.42 | 36.52 | | High | High | High | High | High | High | High | High | High | High | High | 0 | 2.79E-06 | 4.96 |
| FALSE | High | TTSSPTQEL | [K].TTSSPT | 1xPhospho | G5E8W8 1 | 0.000246 | 0 | 1 | 5 | 6 | G5E8W8 | Zinc finger | G5E8W8 [1 | 0 | 2273.183 | 0 | -0.64 | 0.002142 | 3.01 | 120.8 | 79.2 | 3.73 | 8.2 | | High | High | High | High | High | High | High | High | High | High | High | 0 | 0.000166 | 4.04 |
| FALSE | High | TTSSPTQEL | [K].TTSSPT | 2xPhospho | G5E8W8 2 | 0.000176 | 0 | 1 | 5 | 1 | G5E8W8 | Zinc finger | G5E8W8 [1 | 0 | 2353.149 | 0 | -0.91 | 0.030839 | 21.62 | 128 | 72 | 14.83 | 49.61 | | High | High | High | High | High | High | High | High | High | High | High | 0 | 3.4E-05 | 3.2 |
| FALSE | High | TTSSSDLTT | [R].TTSSSD | 1xPhospho | A2AKB9 1x | 8.48E-05 | 0 | 1 | 2 | 1 | A2AKB9 | DDB1- and | A2AKB9 [3 | 0 | 1970.875 | 0 | 1.56 | 0.001422 | 13.39 | 50.7 | 149.3 | 4.16 | 13.32 | | High | High | High | High | High | High | High | High | High | High | High | 0 | 1.5E-05 | 2.53 |
| FALSE | High | TTSTSNPSS | [K].TTSTSN | 1xPhospho | Q61687 1x | 0.017192 | 0.001204 | 1 | 3 | 1 | Q61687 | Transcripti | Q61687 [1 | 0 | 2047.921 | 0 | -0.72 | 0.024236 | 1.59 | 128 | 72 | 22.56 | 22.3 | | High | High | High | High | High | High | High | High | High | High | High | 0.000638 | 0.005777 | 2.48 |
| FALSE | High | TTSTSNPSS | [K].TTSTSN | 2xPhospho | Q61687 2x | 0.018285 | 0.001282 | 1 | 3 | 1 | Q61687 | Transcripti | Q61687 [1 | 0 | 2127.887 | 0 | 0.15 | 0.570118 | 109.82 | 81 | 119 | 63.74 | 99.36 | | High | High | High | High | High | High | High | High | High | High | High | 0.000689 | 0.006146 | 3.19 |
| FALSE | High | TTSTSNPSS | [K].TTSTSN | 1xPhospho | Q61687 1x | 2.16E-06 | 0 | 1 | 3 | 8 | Q61687 | Transcripti | Q61687 [1 | 0 | 2277.084 | 0 | -0.18 | 0.036041 | 6.97 | 104.8 | 95.2 | 1.67 | 7.04 | | High | High | High | High | High | High | High | High | High | High | High | 0 | 2.4E-07 | 4.3 |
| FALSE | High | TTSTSNPSS | [K].TTSTSN | 2xPhospho | Q61687 2x | 0.000611 | 0 | 1 | 3 | 3 | Q61687 | Transcripti | Q61687 [1 | 0 | 2357.05 | 0 | 0.64 | 0.003376 | 12.38 | 79.3 | 120.7 | 2.47 | 9.44 | | High | High | High | High | High | High | High | High | High | High | High | 0 | 0.000137 | 2.58 |
| FALSE | High | TTTPPDSG\ | [R].TTTPPD | 1xPhospho | Q80Z37 1x | 0.000933 | 0 | 1 | 2 | 2 | Q80Z37 | E3 ubiquiti | Q80Z37 [2 | 0 | 2510.285 | 0 | 0.1 | 0.308043 | 22.15 | 92.5 | 107.5 | 9.18 | 8.84 | | High | High | High | High | High | High | High | High | High | High | High | 2.58E-05 | 0.000347 | 1.83 |
| FALSE | High | TTTSPFFSE | [R].TTTSPF | 1xPhospho | Q505G8 1x | 0.076555 | 0.005826 | 1 | 4 | 1 | Q505G8 | Zinc finger | Q505G8 [7 | 0 | 1844.831 | 0 | 2.58 | 0.000286 | 6.61 | 31.2 | 168.8 | 14.13 | 9.22 | | High | High | High | High | High | High | High | High | High | High | High | 0.002976 | 0.03142 | 2.48 |
| FALSE | High | TTTTPEVK | [K].TTTTPE | 1xPhospho | P14873 1xI | 0.087983 | 0.006639 | 2 | 3 | 2 | P14873; B2 | Microtubu | P14873 [2 | 0 | 1414.759 | 0 | -0.5 | 0.014394 | 1.48 | 117.6 | 82.4 | 8.28 | 8.22 | Shared | High | High | High | High | High | High | High | High | High | High | High | 0.003402 | 0.03714 | 2.03 |
| FALSE | High | TTSSADAK | [K].TTSSAD | 1xPhospho | Q3TGQ4 1 | 0.11108 | 0.008991 | 1 | 3 | 1 | Q3TGQ4 | Putative ur | Q3TGQ4 [3 | 0 | 1318.666 | 0 | 0.92 | 0.004689 | 6.59 | 67.9 | 132.1 | 6.96 | 17.72 | | High | High | High | High | High | High | High | High | High | High | High | 0.004517 | 0.049 | 1.6 |
| FALSE | High | TTQSLQDF | [R].TTQSL( | 1xPhospho | Q3UGE9 1 | 1.57E-08 | 0 | 1 | 3 | 5 | Q3UGE9 | Tuftelin-in | Q3UGE9 [1 | 0 | 3195.466 | 0 | 0.01 | 0.474437 | 3.85 | 100.4 | 99.6 | 2.81 | 4.32 | | High | High | High | High | High | High | High | High | High | High | High | 0 | 9.55E-10 | 5 |
| FALSE | High | TTQSLQDF | [R].TTQSL( | 1xPhospho | Q3UGE9 1 | 3.33E-06 | 0 | 1 | 3 | 2 | Q3UGE9 | Tuftelin-in | Q3UGE9 [1 | 0 | 2966.303 | 0 | 0.16 | 0.781613 | 55.7 | 94.4 | 105.6 | 56.24 | 78.27 | | High | High | High | High | High | High | High | High | High | High | High | 0 | 3.93E-07 | 4.7 |
| FALSE | High | TTQSGQM | [K].TTQSG | 1xPhospho | H3BJW3 1x | 0.00095 | 0 | 1 | 3 | 1 | H3BJW3 | Cleavage a | H3BJW3 [1 | 0 | 1764.824 | 0 | -0.24 | 0.002281 | 3.46 | 107.8 | 92.2 | 4.63 | 8.95 | | High | High | High | High | High | High | High | High | High | High | High | 0 | 0.000224 | 2.09 |
| FALSE | High | TTGASQLQ | [K].TTGAS( | 1xPhospho | Q9D842 1x | 4.04E-08 | 0 | 1 | 3 | 2 | Q9D842 | Aprataxin a | Q9D842 [1 | 0 | 2056.073 | 0 | 0.22 | 0.177381 | 7.52 | 92.5 | 107.5 | 3.87 | 9.66 | | High | High | High | High | High | High | High | High | High | High | High | 0 | 2.76E-09 | 3.24 |
| FALSE | High | TTGYPSSPA | [R].TTGYPS | 1xPhospho | Q09XV5 1x | 0.000958 | 0 | 1 | 2 | 3 | Q09XV5 | Chromodo | Q09XV5 [2 | 0 | 2078 | 0 | 0.78 | 0.004118 | 21.03 | 75.8 | 124.2 | 12.31 | 7.73 | | High | High | High | High | High | High | High | High | High | High | High | 0 | 0.000226 | 2.16 |
| FALSE | High | TTHASSDG | [R].TTHASS | 1xPhospho | Q7TPM1 1 | 0.000209 | 0 | 1 | 6 | 1 | Q7TPM1 | Protein PRI | Q7TPM1 [8 | 0 | 1952.937 | 0 | -0.41 | 0.002995 | 4.52 | 115 | 85 | 6.02 | 5.78 | | High | High | High | High | High | High | High | High | High | High | High | 0 | 4.09E-05 | 3.17 |
| FALSE | High | TTHSPYEEF | [R].TTHSPY | 1xPhospho | Q6PHZ5 1x | 0.01588 | 0.001124 | 1 | 1 | 1 | Q6PHZ5 | Putative RI | Q6PHZ5 [6 | 0 | 1428.636 | 0 | 0.38 | 0.007032 | 8.77 | 87.7 | 112.3 | 4.07 | 6.75 | | High | High | High | High | High | High | High | High | High | High | High | 0.000585 | 0.005239 | 2.72 |
| FALSE | High | TTIESHTKP | [K].TTIESH | 1xPhospho | Q80U93 1x | 0.051694 | 0.003745 | 1 | 2 | 1 | Q80U93 | Nuclear po | Q80U93 [9 | 0 | 1892.004 | 0 | -0.74 | 0.005591 | 6.25 | 124.9 | 75.1 | 7.9 | 10.05 | | High | High | High | High | High | High | High | High | High | High | High | 0.00196 | 0.02003 | 1.18 |
| FALSE | High | TTLLPDSSP | [R].TTLLPD | 1xPhospho | Q8CAF4-3 | 0.018447 | 0.001324 | 1 | 3 | 2 | Q8CAF4-3 | Isoform 3 c | Q8CAF4-3 | 0 | 1482.74 | 0 | 0.48 | 0.004658 | 3.67 | 83.4 | 116.6 | 11.83 | 5.68 | | High | High | High | High | High | High | High | High | High | High | High | 0.000706 | 0.006227 | 1.8 |
| FALSE | High | TTITTTTSS | [R].TTITTT | 1xPhospho | Q0VE46 1x | 0.000235 | 0 | 1 | 2 | 9 | Q0VE46 | Myadm pr | Q0VE46 [8 | 0 | 1980.969 | 0 | 0.14 | 0.243923 | 11.26 | 95.8 | 104.2 | 3.45 | 12.78 | | High | High | High | High | High | High | High | High | High | High | High | 0 | 4.69E-05 | 2.69 |
| FALSE | High | TTNPFYNTI | [R].TTNPFY | 1xPhospho | Q5SW75 1 | 0.004781 | 0.000456 | 1 | 4 | 2 | Q5SW75 | Protein ph | Q5SW75 [1 | 0 | 1413.596 | 0 | 1.39 | 0.000282 | 10.1 | 56.8 | 143.2 | 16.06 | 7.39 | | High | High | High | High | High | High | High | High | High | High | High | 0.000538 | 0.004523 | 2.42 |
| FALSE | High | TTNTPTVV | [R].TTNTPT | 1xPhospho | Q61191 1x | 0.011755 | 0.000889 | 1 | 2 | 2 | Q61191 | Host cell fa | Q61191 [1 | 0 | 1297.671 | 0 | 0.58 | 0.003376 | 12.51 | 81.7 | 118.3 | 4.87 | 7.49 | | High | High | High | High | High | High | High | High | High | High | High | 0.002662 | 0.02827 | 1.91 |
| FALSE | High | TTNTSLVTS | [R].TTNTSL | 1xPhospho | Q9ESK9 1x | 2.34E-05 | 0 | 1 | 4 | 1 | Q9ESK9 | RB1-induci | Q9ESK9 [2 | 0 | 1873.983 | 0 | -0.1 | 0.064002 | 7.48 | 105.6 | 94.4 | 6.1 | 6.65 | | High | High | High | High | High | High | High | High | High | High | High | 0 | 3.51E-06 | 2.61 |
| FALSE | High | TTNTSLVTS | [R].TTNTSL | 2xPhospho | Q9ESK9 2x | 0.068752 | 0.005042 | 1 | 4 | 1 | Q9ESK9 | RB1-induci | Q9ESK9 [2 | 0 | 1953.949 | 0 | 0.68 | 0.000916 | 7.48 | 76.9 | 123.1 | 6.74 | 6.22 | | High | High | High | High | High | High | High | High | High | High | High | 0.002604 | 0.02786 | 2.31 |
| FALSE | High | TTPAPSPGS | [R].TTPAPS | 1xPhospho | Q9CQ20 1x | 0.083783 | 0.006345 | 1 | 1 | 1 | Q9CQ20 | Mid1-inter | Q9CQ20 [6 | 0 | 2230.038 | 0 | 0.62 | 0.028325 | 21.76 | 79.5 | 120.5 | 13.24 | 14.34 | | High | High | High | High | High | High | High | High | High | High | High | 0.003265 | 0.03507 | 2.39 |
| FALSE | High | TTPAPSPGS | [R].TTPAPS | 2xPhospho | Q9CQ20 2x | 0.002743 | 0.000109 | 1 | 1 | 1 | Q9CQ20 | Mid1-inter | Q9CQ20 [6 | 0 | 2310.004 | 0 | 1.14 | 0.005337 | 16.47 | 65.3 | 134.7 | 17.33 | 11.61 | | High | High | High | High | High | High | High | High | High | High | High | 4.94E-05 | 0.00074 | 3.55 |
| FALSE | High | TTPAPSTNS | [K].TTPAPS | 1xPhospho | Q571J8 1xI | 5.18E-06 | 0 | 1 | 16 | 3 | Q571J8 | Non-specif | Q571J8 [46 | 0 | 2042.996 | 0 | 0.58 | 0.007059 | 13.85 | 82.3 | 117.7 | 6.7 | 10.29 | | High | High | High | High | High | High | High | High | High | High | High | 0 | 6.46E-07 | 4.81 |
| FALSE | High | TTPDVHTEI | [R].TTPDVI | 1xPhospho | Q8BH05 1x | 0.002313 | 0.000109 | 1 | 1 | 1 | Q8BH05 | Zinc finger | Q8BH05 [5 | 0 | 2227.017 | 0 | 1.16 | 0.000442 | 7.61 | 61.7 | 138.3 | 14.12 | 5.15 | | High | High | High | High | High | High | High | High | High | High | High | 4.94E-05 | 0.000611 | 3 |
| FALSE | High | TTPECDESF | [K].TTPECD | 1xCarbami | Q8BP78 1x | 0.077188 | 0.005893 | 1 | 1 | 1 | Q8BP78 | Protein FRA | Q8BP78 [2 | 1 | 1857.882 | 0 | -0.14 | 0.040724 | 1.95 | 107.2 | 92.8 | 5.14 | 6.19 | | High | High | High | High | High | High | High | High | High | High | High | 0.003035 | 0.03178 | 1.86 |
| FALSE | High | TTPGVQPT | [R].TTPGV( | 1xPhospho | Q3U0S4 1x | 3.11E-08 | 0 | 1 | 2 | 10 | Q3U0S4 | Putative ur | Q3U0S4 [4 | 0 | 3724.618 | 0 | -1.05 | 0.000213 | 0.86 | 134 | 66 | 3.08 | 9.24 | | High | High | High | High | High | High | High | High | High | High | High | 0 | 2.06E-09 | 5.11 |
| FALSE | High | TTPGVQPT | [R].TTPGV( | 1xDeamida | Q3U0S4 1x | 0.063788 | 0.004664 | 1 | 2 | 3 | Q3U0S4 | Putative ur | Q3U0S4 [4 | 0 | 3725.602 | 0 | 0.31 | 0.413047 | 41.31 | 90.2 | 109.8 | 20.04 | 32.86 | | High | High | High | High | High | High | High | High | High | High | High | 0.002967 | 0.03096 | 2.53 |
| FALSE | High | TTPLASPSL | [R].TTPLAS | 1xPhospho | Q8CJF7 1xI | 5.97E-05 | 0 | 1 | 1 | 4 | Q8CJF7 | Protein ELY | Q8CJF7 [12 | 0 | 1592.825 | 0 | -0.1 | 0.055209 | 2.83 | 104.5 | 95.5 | 1.89 | 4.95 | | High | High | High | High | High | High | High | High | High | High | High | 0 | 1.01E-05 | 1.86 |
| FALSE | High | TTPLASPSL | [R].TTPLAS | 2xPhospho | Q8CJF7 2xI | 0.003402 | 0.00021 | 1 | 1 | 2 | Q8CJF7 | Protein ELY | Q8CJF7 [12 | 0 | 1672.791 | 0 | -0.12 | 0.636656 | 17.59 | 107.4 | 92.6 | 14.57 | 11.33 | | High | High | High | High | High | High | High | High | High | High | High | 9.42E-05 | 0.000936 | 2.77 |
| FALSE | High | TTPPESAP( | [R].TTPPES | 1xPhospho | Q8K3X4 1x | 1.1E-05 | 0 | 1 | 1 | 1 | Q8K3X4 | Interferon | Q8K3X4 [5 | 0 | 3699.816 | 0 | -0.23 | 0.137922 | 27.13 | 112 | 88 | 18.36 | 17.07 | | High | High | High | High | High | High | High | High | High | High | High | 0 | 1.51E-06 | 3.18 |
| FALSE | High | TTPPTQKPI | [R].TTPPTC | 1xPhospho | Q6AXD2 1x | 0.005 | 0.000504 | 1 | 10 | 1 | Q6AXD2 | Abi2 prote | Q6AXD2 [1 | 0 | 2418.307 | 0 | -0.4 | 0.02854 | 6.56 | 115.9 | 84.1 | 8.04 | 15.4 | | High | High | High | High | High | High | High | High | High | High | High | 0.000225 | 0.001453 | 2 |
| FALSE | High | TTPSATSLP | [R].TTPSAT | 1xPhospho | P81269 1xI | 2.47E-06 | 0 | 1 | 3 | 2 | P81269 | Cyclic AMP | P81269 [18 | 0 | 3201.673 | 0 | 0.04 | 0.189472 | 5.74 | 97 | 103 | 6.85 | 8.06 | | High | High | High | High | High | High | High | High | High | High | High | 0 | 1.08E-06 | 4.27 |
| FALSE | High | TTPTGGLPI | [K].TTPTG( | 1xPhospho | Q8BU00 1x | 0.007339 | 0.00062 | 1 | 2 | 2 | Q8BU00 | Zinc finger | Q8BU00 [2 | 0 | 1208.624 | 0 | 0.56 | 0.038321 | 23.12 | 83.7 | 116.3 | 8.8 | 12.44 | | High | High | High | High | High | High | High | High | High | High | High | 0.000275 | 0.002227 | 1.68 |
| FALSE | High | TTPTLSPEE | [R].TTPTLS | 1xPhospho | Q60793 1x | 0.001848 | 5.66E-05 | 1 | 1 | 4 | Q60793 | Krueppel-li | Q60793 [3 | 0 | 1866.941 | 0 | 1.01 | 0.003799 | 13.54 | 66.5 | 133.5 | 7.54 | 9.23 | | High | High | High | High | High | High | High | High | High | High | High | 2.58E-05 | 0.000476 | 2.26 |

| | | | | | | | | | | | | | | | | | | | | | | | | | | | | | | | | | | | | | | |
|---|---|---|---|---|---|---|---|---|---|---|---|---|---|---|---|---|---|---|---|---|---|---|---|---|---|---|---|---|---|---|---|---|---|---|---|---|---|---|
| FALSE | High | TTPTLSPEE | [R].TTPTLS | 2xPhospho | Q60793 2x | 0.034281 | 0.002195 | 1 | 1 | 2 | Q60793 | Krueppel-li | Q60793 [3 | 0 | 1946.907 | 0 | 0.85 | 0.007764 | 4.43 | 71.9 | 128.1 | 7.09 | 12.74 | | High | High | High | High | High | High | High | High | High | High | High | 0.001151 | 0.01252 | 1.92 |
| FALSE | High | TTQDVNTG | [K].TTQDV | 1xDeamida | G3UWG2 1 | 0.000141 | 0 | 1 | 4 | 2 | G3UWG2 | T-cell lymp | G3UWG2 [ | 0 | 3397.501 | 0 | -0.98 | 0.010909 | 6.64 | 132.2 | 67.8 | 14.45 | 10.73 | | High | High | High | High | High | High | High | High | High | High | High | 2.58E-05 | 0.000382 | 4.18 |
| FALSE | High | TTQENGHI | [K].TTQEN | 1xPhospho | Q8C156 1x | 0.000568 | 0 | 1 | 2 | 4 | Q8C156 | Condensin | Q8C156 [5 | 0 | 3991.969 | 0 | -1.01 | 0.000876 | 3.44 | 133.5 | 66.5 | 4.07 | 9.49 | | High | High | High | High | High | High | High | High | High | High | High | 0 | 0.000126 | 3.63 |
| FALSE | High | TTQENGHI | [K].TTQEN | 1xDeamida | Q8C156 1x | 3.44E-05 | 0 | 1 | 2 | 19 | Q8C156 | Condensin | Q8C156 [5 | 0 | 3992.953 | 0 | -0.78 | 0.000324 | 1.1 | 127.1 | 72.9 | 2.8 | 5.03 | | High | High | High | High | High | High | High | High | High | High | High | 0 | 1.13E-05 | 4.47 |
| FALSE | High | TTFSPPVA | [K].TTFSPP | 1xPhospho | Q5SVL6 1x | 0.047488 | 0.00354 | 1 | 3 | 2 | Q5SVL6 | Rap1 GTPa | Q5SVL6 [5 | 0 | 1799.971 | 0 | 3.08 | 0.000269 | 12.77 | 19.7 | 180.3 | 16.53 | 7.79 | | High | High | High | High | High | High | High | High | High | High | High | 0.001812 | 0.01818 | 2.14 |
| FALSE | High | TSSSSLTSS | [R].TSSSSL | 1xPhospho | P51141 1x | 0.014929 | 0.001049 | 1 | 3 | 2 | P51141 | Segment p | P51141 [38 | 0 | 2924.527 | 0 | 1.59 | 0.002652 | 26.59 | 46.1 | 153.9 | 17.4 | 16.41 | | High | High | High | High | High | High | High | High | High | High | High | 0.000555 | 0.00492 | 4.51 |
| FALSE | High | VADNSFDA | [R].VADNS | 1xDeamida | E9Q5C9 1x | 0.006536 | 0.00062 | 1 | 2 | 1 | E9Q5C9 | Nucleolar a | E9Q5C9 [6 | 0 | 1505.729 | 0 | -0.52 | 0.003419 | 1.17 | 117.4 | 82.6 | 7.53 | 2.47 | | High | High | High | High | High | High | High | High | High | High | High | 0.000275 | 0.00196 | 1.98 |
| FALSE | High | VADPGPSI | [K].VADPG | 1xPhospho | Q3UH06-3 | 9.39E-06 | 0 | 1 | 4 | 1 | Q3UH06-3 | Isoform 3 c | Q3UH06-3 | 0 | 2923.47 | 1 | 0.19 | 0.98784 | 20.77 | 97.8 | 102.2 | 30.86 | 17.16 | | High | High | High | High | High | High | High | High | High | High | High | 0 | 1.26E-06 | 3.44 |
| FALSE | High | VEPGQEVH | [K].VEPGQ | 1xPhospho | E9Q8T1 1x | 0.001349 | 0 | 1 | 4 | 4 | E9Q8T1 | Transformi | E9Q8T1 [7 | 0 | 1887.88 | 0 | -1.83 | 0.001886 | 2.88 | 151.4 | 48.6 | 13.15 | 11.33 | | High | High | High | High | High | High | High | High | High | High | High | 7.19E-05 | 0.000865 | 2.55 |
| FALSE | High | VEPQEGSD | [K].VEPQE | 1xPhospho | E9QLG3 1x | 1.82E-06 | 0 | 1 | 5 | 2 | E9QLG3 | MAX gene- | E9QLG3 [2 | 0 | 2867.375 | 0 | -0.18 | 0.028084 | 7.12 | 107.7 | 92.3 | 3.36 | 6.05 | | High | High | High | High | High | High | High | High | High | High | High | 0 | 1.99E-07 | 4.57 |
| FALSE | High | VEPSSQSP | [K].VEPSSC | 1xPhospho | F8VPM7 1x | 0.006363 | 0.00062 | 1 | 8 | 2 | F8VPM7 | ELKS/Rab6 | F8VPM7 [1 | 1 | 1692.827 | 0 | 0.21 | 0.038634 | 10.11 | 94.1 | 105.9 | 2.76 | 9.74 | | High | High | High | High | High | High | High | High | High | High | High | 0.001142 | 0.01181 | 1.42 |
| FALSE | High | VEPVPAISP | [K].VEPVPA | 1xPhospho | Q3UCQ1 1 | 2.57E-07 | 0 | 1 | 1 | 1 | Q3UCQ1 | Forkhead b | Q3UCQ1 [5 | 0 | 1945.036 | 0 | 0.66 | 0.00459 | 13.85 | 77.6 | 122.4 | 10.57 | 10.13 | | High | High | High | High | High | High | High | High | High | High | High | 0 | 2.22E-08 | 3.42 |
| FALSE | High | VEQALSEV | [R].VEQALS | 1xPhospho | E9Q6P5 1x | 9.11E-05 | 0 | 1 | 1 | 2 | E9Q6P5 | Tetratricop | E9Q6P5 [6 | 0 | 2369.237 | 0 | 0.29 | 0.030735 | 1.42 | 84.9 | 115.1 | 6.91 | 11.5 | | High | High | High | High | High | High | High | High | High | High | High | 0 | 1.62E-05 | 2.62 |
| FALSE | High | VEQALSEV | [R].VEQALS | 2xPhospho | E9Q6P5 2x | 0.004451 | 0.000409 | 1 | 1 | 1 | E9Q6P5 | Tetratricop | E9Q6P5 [6 | 0 | 2449.203 | 0 | 0.73 | 0.008785 | 23.72 | 74.8 | 125.2 | 16.59 | 8.54 | | High | High | High | High | High | High | High | High | High | High | High | 0.000183 | 0.001267 | 2.79 |
| FALSE | High | VEQEEASP | [K].VEQEE | 1xPhospho | A0A0R4J1L | 7.72E-07 | 0 | 1 | 5 | 3 | A0A0R4J1L | Zinc finger | A0A0R4J1L | 0 | 2793.282 | 0 | 0.06 | 0.504923 | 13.05 | 98.2 | 101.8 | 8.36 | 4.39 | | High | High | High | High | High | High | High | High | High | High | High | 0 | 7.57E-08 | 4.89 |
| FALSE | High | VEQEESSPI | [K].VEQEE | 1xPhospho | Q91VW9 1 | 0.016451 | 0.001124 | 1 | 2 | 2 | Q91VW9 | Zinc finger | Q91VW9 [ | 0 | 2809.277 | 0 | -0.63 | 0.001858 | 4.79 | 124.3 | 75.7 | 6.41 | 4.77 | | High | High | High | High | High | High | High | High | High | High | High | 0.000814 | 0.007568 | 3.05 |
| FALSE | High | VEQLSSGLI | [K].VEQLSS | 1xPhospho | Q99K01-4 | 0.000219 | 0 | 1 | 4 | 1 | Q99K01-4 | Isoform 4 c | Q99K01-4 | 0 | 2878.357 | 0 | -0.17 | 0.901691 | 3.53 | 100.3 | 99.7 | 10.71 | 17.98 | | High | High | High | High | High | High | High | High | High | High | High | 0 | 4.31E-05 | 3.97 |
| FALSE | High | VEQNEQG | [R].VEQNE | 1xPhospho | P97931 1x | 8.28E-06 | 0 | 2 | 3 | 1 | P97931; Q | Uracil-DNA | P97931 [50 | 0 | 2193.075 | 0 | -1.89 | 0.002296 | 2.48 | 157.6 | 42.4 | 12.48 | 17.61 | Shared | High | High | High | High | High | High | High | High | High | High | High | 0 | 1.1E-06 | 3.42 |
| FALSE | High | VERPSSPFS | [K].VERPSS | 1xPhospho | P22682 1x | 3.26E-05 | 0 | 1 | 4 | 4 | P22682 | E3 ubiquiti | P22682 [47 | 0 | 2656.352 | 0 | -0.05 | 0.780744 | 6.2 | 101.4 | 98.6 | 5.6 | 5.06 | | High | High | High | High | High | High | High | High | High | High | High | 0 | 5.11E-06 | 3.06 |
| FALSE | High | VERPSSPFS | [K].VERPSS | 1xPhospho | P22682 1x | 0.002743 | 0.000109 | 1 | 4 | 9 | P22682 | E3 ubiquiti | P22682 [47 | 0 | 2672.347 | 0 | -0.11 | 0.189806 | 11.33 | 103.9 | 96.1 | 4.23 | 7.64 | | High | High | High | High | High | High | High | High | High | High | High | 0.000786 | 0.007073 | 2.32 |
| FALSE | High | VESAGPGG | [K].VESAGI | 1xPhospho | F8WJ40 1x | 4.82E-14 | 0 | 1 | 4 | 4 | F8WJ40 | Histone-lys | F8WJ40 [9 | 0 | 2560.188 | 0 | -0.87 | 0.000713 | 0.71 | 129.8 | 70.2 | 4.24 | 5.55 | | High | High | High | High | High | High | High | High | High | High | High | 0 | 7.77E-13 | 5.07 |
| FALSE | High | VEPAWHD | [K].VEPAW | 2xPhospho | Q6ZQ58 2x | 0.079766 | 0.00613 | 1 | 3 | 2 | Q6ZQ58 | La-related | Q6ZQ58 [2 | 1 | 3017.344 | 0 | 0.17 | 0.28526 | 13.58 | 95.9 | 104.1 | 7.02 | 16.11 | | High | High | High | High | High | High | High | High | High | High | High | 0.009441 | 0.102 | 3.7 |
| FALSE | High | VESAGPGG | [K].VESAGI | 2xPhospho | F8WJ40 2x | 0.000286 | 0 | 1 | 4 | 2 | F8WJ40 | Histone-lys | F8WJ40 [9 | 0 | 2640.154 | 0 | -1.16 | 0.000748 | 0.81 | 137.5 | 62.5 | 8.35 | 4.2 | | High | High | High | High | High | High | High | High | High | High | High | 2.58E-05 | 0.00038 | 2.38 |
| FALSE | High | VESEESGDI | [R].VESEES | 1xPhospho | Q3UM45 1 | 0.071662 | 0.005278 | 1 | 2 | 1 | Q3UM45 | Protein ph | Q3UM45 [ | 0 | 1832.82 | 0 | -1.13 | 0.053236 | 1.65 | 132.5 | 67.5 | 15.78 | 24.91 | | High | High | High | High | High | High | High | High | High | High | High | 0.002703 | 0.0291 | 2.04 |
| FALSE | High | VESGYFSLE | [R].VESGYF | 1xPhospho | P97434-2 1 | 0.000267 | 0 | 1 | 9 | 1 | P97434-2 | Isoform 2 c | P97434-2 [ | 0 | 1696.86 | 0 | 1.54 | 0.000445 | 5 | 49.7 | 150.3 | 5.15 | 6.73 | | High | High | High | High | High | High | High | High | High | High | High | 0 | 5.41E-05 | 2.56 |
| FALSE | High | VESLESLYF | [K].VESLES | 1xPhospho | E9Q7G0 1x | 7.16E-06 | 0 | 1 | 4 | 4 | E9Q7G0 | Nuclear mi | E9Q7G0 [1 | 0 | 2019.004 | 0 | 0 | 0.782726 | 19.52 | 97.6 | 102.4 | 9.92 | 13.17 | | High | High | High | High | High | High | High | High | High | High | High | 0 | 9.28E-07 | 4.29 |
| FALSE | High | VESNSEGT | [R].VESNSE | 1xCarbami | P20263 1x | 2.82E-05 | 0 | 1 | 2 | 10 | P20263 | POU doma | P20263 [98 | 0 | 2772.291 | 0 | 0.12 | 0.094418 | 3.08 | 95.9 | 104.1 | 6.05 | 2.36 | | High | High | High | High | High | High | High | High | High | High | High | 0 | 2.98E-05 | 2.8 |
| FALSE | High | VESNSEGT | [R].VESNSE | 1xCarbami | P20263 2x | 0.015603 | 0.001124 | 1 | 2 | 4 | P20263 | POU doma | P20263 [98 | 0 | 2852.258 | 0 | -0.23 | 0.010173 | 4.26 | 108.7 | 91.3 | 5.42 | 5.57 | | High | High | High | High | High | High | High | High | High | High | High | 0.000903 | 0.009213 | 2.84 |
| FALSE | High | VESPPAAR | [K].VESPPA | 1xPhospho | O55176 1x | 0.000426 | 0 | 1 | 4 | 2 | O55176 | E3 ubiquiti | O55176 [1 | 0 | 1135.571 | 0 | 1.43 | 0.00121 | 15.88 | 55.7 | 144.3 | 6.42 | 9.7 | | High | High | High | High | High | High | High | High | High | High | High | 0 | 9.16E-05 | 2.74 |
| FALSE | High | VESPPAPLF | [R].VESPPA | 1xPhospho | Q3UHT3 1x | 0.011146 | 0.000772 | 1 | 2 | 2 | Q3UHT3 | A disintegr | Q3UHT3 [8 | 0 | 1806.947 | 0 | -1.74 | 0.002233 | 2.6 | 151.5 | 48.5 | 10.73 | 30.98 | | High | High | High | High | High | High | High | High | High | High | High | 0.001812 | 0.01812 | 2.15 |
| FALSE | High | VESPPLEEV | [K].VESPPL | 1xCarbami | O08696 1x | 4.13E-06 | 0 | 1 | 7 | 2 | O08696 | Forkhead b | O08696 [4 | 0 | 2464.224 | 0 | 1.34 | 0.003867 | 10.44 | 56.1 | 143.9 | 5.98 | 15.66 | | High | High | High | High | High | High | High | High | High | High | High | 0 | 6.46E-06 | 4.04 |
| FALSE | High | VESSENVP | [K].VESSEN | 1xPhospho | Q3TIV2 1x | 1.62E-07 | 0 | 1 | 5 | 3 | Q3TIV2 | Putative ur | Q3TIV2 [14 | 0 | 4653.02 | 0 | -0.34 | 0.020358 | 10.16 | 109.3 | 90.7 | 6.27 | 9.42 | | High | High | High | High | High | High | High | High | High | High | High | 0 | 1.32E-08 | 4.58 |
| FALSE | High | VESSENVP | [K].VESSEN | 1xDeamida | Q3TIV2 1x | 2.11E-08 | 0 | 1 | 5 | 12 | Q3TIV2 | Putative ur | Q3TIV2 [14 | 0 | 4654.004 | 0 | 0.13 | 0.031208 | 6.99 | 97.9 | 102.1 | 4.96 | 8.82 | | High | High | High | High | High | High | High | High | High | High | High | 0 | 5.23E-09 | 5.69 |
| FALSE | High | VESSENVP | [K].VESSEN | 2xPhospho | Q3TIV2 2x | 0.000276 | 0 | 1 | 5 | 2 | Q3TIV2 | Putative ur | Q3TIV2 [14 | 0 | 4732.986 | 0 | 0.06 | 0.672231 | 18.47 | 96.6 | 103.4 | 10.4 | 8.48 | | High | High | High | High | High | High | High | High | High | High | High | 0 | 5.61E-05 | 4.67 |
| FALSE | High | VESTSVGSI | [R].VESTSV | 1xPhospho | Q9WV92-7 | 2.91E-07 | 0 | 1 | 15 | 2 | Q9WV92-7 | Isoform 7 c | Q9WV92-7 | 0 | 1913.999 | 0 | -1.59 | 0.005383 | 2.92 | 143.4 | 56.6 | 12.61 | 26.04 | | High | High | High | High | High | High | High | High | High | High | High | 0 | 9.18E-06 | 3.31 |
| FALSE | High | VESTTTPD | [K].VESTTT | 1xCarbami | Q6P6J0 1x | 1.77E-06 | 0 | 1 | 1 | 4 | Q6P6J0 | tRNA-speci | Q6P6J0 [5- | 0 | 2701.304 | 0 | -0.69 | 0.01994 | 3.9 | 124.9 | 75.1 | 23.01 | 4.75 | | High | High | High | High | High | High | High | High | High | High | High | 0 | 1.93E-07 | 4.42 |
| FALSE | High | VESASAAG | [K].VESASA | 1xCarbami | Q6PD21 1x | 0.000143 | 0 | 1 | 1 | 1 | Q6PD21 | SH2 domai | Q6PD21 [1 | 0 | 2039.906 | 0 | 2.45 | 0.000237 | 8.83 | 32.5 | 167.5 | 18.22 | 9.52 | | High | High | High | High | High | High | High | High | High | High | High | 0 | 2.68E-05 | 2.95 |
| FALSE | High | VETLSPEGA | [K].VETLSP | 1xPhospho | Q61850 1x | 1.25E-05 | 0 | 1 | 1 | 1 | Q61850 | Forkhead b | Q61850 [2 | 0 | 1863.941 | 0 | -0.32 | 0.765498 | 4.26 | 107.9 | 92.1 | 20.94 | 18.1 | | High | High | High | High | High | High | High | High | High | High | High | 0 | 1.75E-06 | 3.8 |
| FALSE | High | VEPAWHD | [K].VEPAW | 2xPhospho | Q6ZQ58 2x | 0.01894 | 0.001362 | 1 | 3 | 1 | Q6ZQ58 | La-related | Q6ZQ58 [2 | 1 | 2788.181 | 0 | 0.16 | 0.963397 | 30.84 | 103.7 | 96.3 | 46.22 | 48.06 | | High | High | High | High | High | High | High | High | High | High | High | 0.000723 | 0.006396 | 2.88 |
| FALSE | High | VENSNTVS | [R].VENSN | 1xPhospho | Q0VBD2 1x | 0.0009 | 0 | 1 | 1 | 1 | Q0VBD2 | Protein MC | Q0VBD2 [7 | 0 | 2209.022 | 0 | -1.02 | 0.002833 | 4.25 | 138.1 | 61.9 | 11.58 | 8.99 | | High | High | High | High | High | High | High | High | High | High | High | 0 | 0.000211 | 2.94 |
| FALSE | High | VEGDTCSLI | [K].VEGDT | 1xCarbami | Q5DTI2 1x | 4.05E-05 | 0 | 1 | 4 | 4 | Q5DTI2 | ATPase, Ca | Q5DTI2 [38 | 0 | 3340.606 | 0 | 0.1 | 0.192513 | 12.31 | 93.5 | 106.5 | 7.97 | 11.83 | | High | High | High | High | High | High | High | High | High | High | High | 0 | 1.66E-05 | 4.09 |
| FALSE | High | VEGNFNPF | [K].VEGNF | 1xPhospho | B2KF52 1x | 7.98E-06 | 0 | 1 | 2 | 1 | B2KF52 | Uncharacte | B2KF52 [2 | 0 | 1972.993 | 0 | -0.85 | 0.001125 | 6.52 | 132.3 | 67.7 | 5.44 | 12.9 | | High | High | High | High | High | High | High | High | High | High | High | 0 | 1.05E-06 | 3.63 |
| FALSE | High | VEGSEPTTI | [K].VEGSEF | 1xPhospho | P09405 1x | 5.57E-06 | 0 | 1 | 7 | 3 | P09405 | Nucleolin ( | P09405 [30 | 0 | 2826.448 | 0 | -0.71 | 0.002304 | 5.17 | 128.6 | 71.4 | 4.79 | 15.76 | | High | High | High | High | High | High | High | High | High | High | High | 0 | 7.01E-07 | 4.54 |
| FALSE | High | VEGSESPY | [R].VEGSES | 1xPhospho | Q3UFM6 1 | 0.000279 | 0 | 1 | 3 | 3 | Q3UFM6 | Expressed | Q3UFM6 [ | 0 | 1546.699 | 0 | -0.45 | 0.005714 | 7.85 | 118.4 | 81.6 | 6.88 | 8.31 | | High | High | High | High | High | High | High | High | High | High | High | 0 | 5.68E-05 | 3.24 |
| FALSE | High | VEGSPTEE | [K].VEGSPT | 1xPhospho | E9Q3Y4 1x | 7.06E-07 | 0 | 1 | 4 | 2 | E9Q3Y4 | Lipopolysa | E9Q3Y4 [1 | 0 | 3079.467 | 0 | 1.07 | 0.011769 | 29.77 | 68 | 132 | 22.14 | 15.15 | | High | High | High | High | High | High | High | High | High | High | High | 0 | 2.2E-07 | 3.82 |
| FALSE | High | VEGSTGAN | [K].VEGST | 1xCarbami | Q8BLH7 2x | 3.48E-06 | 0 | 1 | 1 | 6 | Q8BLH7 | HIRA-intera | Q8BLH7 [2 | 0 | 3029.237 | 0 | -1.24 | 0.000447 | 2.02 | 142.4 | 57.6 | 4.01 | 12.16 | | High | High | High | High | High | High | High | High | High | High | High | 0 | 4.14E-07 | 4.66 |
| FALSE | High | VEGTEIVKF | [K].VEGTEI | 1xPhospho | P97868 1x | 0.080423 | 0.006219 | 1 | 3 | 1 | P97868 | E3 ubiquiti | P97868 [11 | 0 | 1822.013 | 0 | 0.29 | 0.971553 | 48.05 | 99.4 | 100.6 | 66.08 | 62.13 | | High | High | High | High | High | High | High | High | High | High | High | 0.003188 | 0.03342 | 1.75 |
| FALSE | High | VEHGGSSG | [K].VEHGG | 1xPhospho | Q9CWC4 1 | 0.002058 | 5.66E-05 | 1 | 5 | 2 | Q9CWC4 | Transcripti | Q9CWC4 [ | 0 | 1482.735 | 0 | -0.95 | 0.000975 | 1.29 | 131.3 | 68.7 | 5.12 | 7.61 | | High | High | High | High | High | High | High | High | High | High | High | 2.58E-05 | 0.000537 | 1.88 |
| FALSE | High | VEHHDLSV | [K].VEHHD | 1xPhospho | Q8R4H2 1x | 9.59E-07 | 0 | 2 | 2 | 2 | Q8R4H2; F | Rho guanir | Q8R4H2 [1 | 0 | 2069.001 | 0 | 1.66 | 0.000154 | 6.42 | 47.5 | 152.5 | 5.77 | 2.81 | Shared | High | High | High | High | High | High | High | High | High | High | High | 0 | 9.71E-08 | 4.22 |
| FALSE | High | VELADSVE | [K].VELADS | 1xPhospho | Q0VEE6 1x | 1.03E-06 | 0 | 1 | 2 | 3 | Q0VEE6 | Zinc finger | Q0VEE6 [4 | 0 | 3002.512 | 0 | 2.22 | 0.000497 | 8.5 | 35.3 | 164.7 | 18.29 | 10.45 | | High | High | High | High | High | High | High | High | High | High | High | 0 | 1.05E-07 | 4.16 |
| FALSE | High | VELADSVE | [K].VELADS | 2xPhospho | Q0VEE6 2x | 2.98E-05 | 0 | 1 | 2 | 2 | Q0VEE6 | Zinc finger | Q0VEE6 [4 | 0 | 3082.479 | 0 | 1.61 | 0.000731 | 10.49 | 49.2 | 150.8 | 9.69 | 9.69 | | High | High | High | High | High | High | High | High | High | High | High | 0 | 4.62E-06 | 3.95 |
| FALSE | High | VELESSSPA | [K].VELESS | 1xPhospho | Q8VDH2 1x | 0.01029 | 0.000772 | 1 | 4 | 1 | Q8VDH2 | 2810452K2 | Q8VDH2 [6 | 0 | 2335.066 | 0 | -0.41 | 0.026319 | 10.04 | 113.1 | 86.9 | 5.08 | 10.05 | | High | High | High | High | High | High | High | High | High | High | High | 0.000382 | 0.003246 | 2.7 |
| FALSE | High | VELPLEDQ | [K].VELPLE | 1xPhospho | B2RRE0 1x | 5.96E-08 | 0 | 1 | 3 | 3 | B2RRE0 | A kinase (P | B2RRE0 [3 | 0 | 2522.304 | 0 | -1.83 | 0.001134 | 1.95 | 155.9 | 44.1 | 10.61 | 12.54 | | High | High | High | High | High | High | High | High | High | High | High | 0 | 2.47E-08 | 4.36 |
| FALSE | High | VEPAWHD | [K].VEPAW | 1xPhospho | Q6ZQ58 1x | 2.14E-06 | 0 | 1 | 3 | 2 | Q6ZQ58 | La-related | Q6ZQ58 [2 | 0 | 2266.079 | 0 | 0.31 | 0.019384 | 8.89 | 87.6 | 112.4 | 4.99 | 9.24 | | High | High | High | High | High | High | High | High | High | High | High | 0 | 0.000107 | 3.42 |
| FALSE | High | VELSESEED | [K].VELSES | 1xPhospho | Q52KI8 1x | 0.001615 | 5.66E-05 | 1 | 6 | 5 | Q52KI8 | Serine/argi | Q52KI8 [45 | 0 | 1702.819 | 0 | -0.49 | 0.00108 | 3 | 117.1 | 82.9 | 1.9 | 5.26 | | High | High | High | High | High | High | High | High | High | High | High | 0.000275 | 0.001802 | 2.51 |
| FALSE | High | VELSPAPSG | [R].VELSPA | 1xPhospho | G3X9X7 1x | 2.77E-05 | 0 | 1 | 8 | 1 | G3X9X7 | Activated ( | G3X9X7 [7 | 0 | 1896.879 | 0 | 0.49 | 0.001134 | 3.43 | 82.7 | 117.3 | 4.13 | 5.49 | | High | High | High | High | High | High | High | High | High | High | High | 0 | 4.26E-06 | 3.07 |
| FALSE | High | VELSPVAK | [K].VELSPV | 1xPhospho | Q6NV99 1x | 0.003055 | 0.00016 | 1 | 5 | 2 | Q6NV99 | HAUS augn | Q6NV99 [8 | 0 | 1380.79 | 0 | 0.92 | 0.000441 | 6.83 | 69.8 | 130.2 | 5.63 | 4.25 | | High | High | High | High | High | High | High | High | High | High | High | 0.000161 | 0.001135 | 2.84 |
| FALSE | High | VEITPESILS | [K].VEITPE | 1xPhospho | Q6NXI6 1x | 0.000267 | 0 | 1 | 4 | 1 | Q6NXI6 | Regulation | Q6NXI6 [5 | 0 | 2025.129 | 0 | -0.73 | 0.002715 | 5.96 | 123.3 | 76.7 | 8.87 | 6.43 | | High | High | High | High | High | High | High | High | High | High | High | 0 | 5.39E-05 | 2.57 |
| FALSE | High | VENDENNE | [K].VENDE | 1xPhospho | Q8BP27 1x | 9.97E-08 | 0 | 1 | 2 | 4 | Q8BP27 | Swi5-depe | Q8BP27 [1 | 0 | 2516.144 | 0 | -0.24 | 0.019583 | 7.86 | 108 | 92 | 2.06 | 8.26 | | High | High | High | High | High | High | High | High | High | High | High | 0 | 5.22E-07 | 4.64 |
| FALSE | High | VENDENNE | [K].VENDE | 1xDeamida | Q8BP27 1x | 4.8E-05 | 0 | 1 | 2 | 3 | Q8BP27 | Swi5-depe | Q8BP27 [1 | 0 | 2517.128 | 0 | -0.21 | 0.167585 | 6.98 | 104 | 96 | 6.79 | 8.17 | | High | High | High | High | High | High | High | High | High | High | High | 0 | 1.07E-05 | 3.92 |
| FALSE | High | VENDENNE | [K].VENDE | 1xCarbami | Q8BP27 1x | 0.000645 | 0 | 1 | 2 | 2 | Q8BP27 | Swi5-depe | Q8BP27 [1 | 1 | 3492.593 | 0 | -0.27 | 0.040634 | 4.5 | 111.6 | 88.4 | 5.09 | 7.84 | | High | High | High | High | High | High | High | High | High | High | High | 0 | 0.000146 | 4.04 |
| FALSE | High | VENESDEG | [K].VENESI | 1xPhospho | P97496 1x | 0.000119 | 0 | 1 | 5 | 1 | P97496 | SWI/SNF c | P97496 [7 | 0 | 1659.752 | 0 | -1.15 | 0.001065 | 3 | 137.6 | 62.4 | 7.42 | 6.31 | | High | High | High | High | High | High | High | High | High | High | High | 0 | 2.2E-05 | 1.78 |
| FALSE | High | VENGSLCD | [K].VENGSI | 2xCarbami | A0A024QY | 2.7E-06 | 0 | 1 | 6 | 2 | A0A024QY | Mitochond | A0A024QY | 0 | 2534.099 | 0 | 0.35 | 0.051671 | 20.16 | 88.7 | 111.3 | 3.93 | 15.23 | | High | High | High | High | High | High | High | High | High | High | High | 0 | 3.11E-07 | 4.76 |
| FALSE | High | VENMDISD | [R].VENMD | 1xPhospho | Q9Z315 1x | 0.018125 | 0.001282 | 1 | 1 | 1 | Q9Z315 | U4/U6.U5 | Q9Z315 [4 | 0 | 4179.909 | 0 | 0.39 | 0.159349 | 32.7 | 86.4 | 113.6 | 10.02 | 21.13 | | High | High | High | High | High | High | High | High | High | High | High | 0.000689 | 0.006129 | 2.64 |
| FALSE | High | VENMDISD | [R].VENMD | 2xPhospho | Q9Z315 2x | 1.07E-05 | 0 | 1 | 1 | 5 | Q9Z315 | U4/U6.U5 | Q9Z315 [4 | 0 | 4259.876 | 0 | -0.31 | 0.013258 | 3.61 | 112.1 | 87.9 | 4.01 | 13.83 | | High | High | High | High | High | High | High | High | High | High | High | 0 | 0.000102 | 3.46 |
| FALSE | High | VENMSSN | [K].VENMS | 1xPhospho | Q78PG9 1x | 9.91E-06 | 0 | 1 | 1 | 1 | Q78PG9 | Coiled-coil | Q78PG9 [1 | 0 | 2227.836 | 0 | -1.32 | 0.012404 | 7.02 | 142.5 | 57.5 | 11.73 | 33.96 | | High | High | High | High | High | High | High | High | High | High | High | 0 | 1.33E-06 | 4.19 |
| FALSE | High | VENMSSN | [K].VENMS | 1xPhospho | Q78PG9 1x | 3.82E-07 | 0 | 1 | 1 | 20 | Q78PG9 | Coiled-coil | Q78PG9 [1 | 0 | 2243.831 | 0 | -1.21 | 0.000905 | 2.55 | 140.4 | 59.6 | 5.03 | 19.62 | | High | High | High | High | High | High | High | High | High | High | High | 0 | 3.04E-07 | 4.46 |
| FALSE | High | VENMSSN | [K].VENMS | 1xPhospho | Q78PG9 1x | 9.11E-08 | 0 | 1 | 1 | 46 | Q78PG9 | Coiled-coil | Q78PG9 [1 | 0 | 2259.826 | 0 | -0.96 | 0.000485 | 1.81 | 132.1 | 67.9 | 4.69 | 9.02 | | High | High | High | High | High | High | High | High | High | High | High | 0 | 1.2E-07 | 5.24 |
| FALSE | High | VELSESEED | [K].VELSES | 2xPhospho | Q52KI8 2x | 7.23E-06 | 0 | 1 | 6 | 5 | Q52KI8 | Serine/argi | Q52KI8 [45 | 0 | 1782.785 | 0 | -0.38 | 0.001345 | 2.09 | 113.7 | 86.3 | 3.61 | 2.2 | | High | High | High | High | High | High | High | High | High | High | High | 0.000267 | 0.001562 | 3.49 |
| FALSE | High | VETLSPEGA | [K].VETLSP | 2xPhospho | Q61850 2x | 9.54E-05 | 0 | 1 | 1 | 1 | Q61850 | Forkhead b | Q61850 [2 | 0 | 1943.908 | 0 | -0.13 | 0.26863 | 20.78 | 104.6 | 95.4 | 15.6 | 6.52 | | High | High | High | High | High | High | High | High | High | High | High | 0 | 1.7E-05 | 3.04 |
| FALSE | High | VETPDVNL | [R].VETPD | 1xPhospho | Q3UC10 1x | 2.21E-08 | 0 | 1 | 4 | 8 | Q3UC10 | DNA helica | Q3UC10 [6 | 0 | 4285.903 | 0 | -0.41 | 0.006609 | 8.56 | 114 | 86 | 4.06 | 8.46 | | High | High | High | High | High | High | High | High | High | High | High | 0 | 1.4E-09 | 6.06 |
| FALSE | High | VETPDVNL | [R].VETPD | 1xDeamida | Q3UC10 1x | 1.43E-08 | 0 | 1 | 4 | 43 | Q3UC10 | DNA helica | Q3UC10 [6 | 0 | 4286.887 | 0 | -0.65 | 0.011665 | 1.66 | 121.8 | 78.2 | 14.09 | 3.89 | | High | High | High | High | High | High | High | High | High | High | High | 0 | 8.58E-10 | 6.59 |
| FALSE | High | VGAHSPVG | [R].VGAHS | 1xPhospho | H3BJX1 1x | 0.000365 | 0 | 1 | 1 | 1 | H3BJX1 | Ecotropic v | H3BJX1 [39 | 0 | 1710.865 | 0 | 0.42 | 0.244205 | 31.14 | 81.8 | 118.2 | 17.06 | 25.03 | | High | High | High | High | High | High | High | High | High | High | High | 0 | 7.67E-05 | 2.62 |
| FALSE | High | VGASLTPEI | [R].VGASL | 1xPhospho | Q3U1J1-2 | 0.00614 | 0.00062 | 1 | 2 | 1 | Q3U1J1-2 | Isoform 2 c | Q3U1J1-2 | 0 | 2061.176 | 0 | -0.62 | 0.000442 | 1.78 | 120.8 | 79.2 | 4.09 | 4.51 | | High | High | High | High | High | High | High | High | High | High | High | 0.000275 | 0.001815 | 2.27 |
| FALSE | High | VGCDESPS | [K].VGCDE | 1xCarbami | O08734 1x | 1.97E-05 | 0 | 1 | 2 | 6 | O08734 | Bcl-2 homo | O08734 [1 | 0 | 2903.26 | 0 | 0.71 | 0.002838 | 11.75 | 76 | 124 | 5.36 | 8.01 | | High | High | High | High | High | High | High | High | High | High | High | 0 | 2.9E-06 | 4.32 |
| FALSE | High | VGDGSPVL | [R].VGDGS | 1xPhospho | B2RXU3 1x | 3.38E-05 | 0 | 1 | 2 | 2 | B2RXU3 | ADP-ribosy | B2RXU3 [7 | 0 | 1621.86 | 0 | 0.42 | 0.007945 | 4.86 | 86.3 | 113.7 | 5.52 | 5.25 | | High | High | High | High | High | High | High | High | High | High | High | 0 | 5.31E-06 | 2.68 |
| FALSE | High | VGDSQNPI | [R].VGDSQ | 1xPhospho | Q3ULN2 1x | 8.87E-05 | 0 | 1 | 2 | 3 | Q3ULN2 | General tra | Q3ULN2 [6 | 0 | 2601.288 | 0 | -0.44 | 0.050706 | 15.25 | 112 | 88 | 7.5 | 15.99 | | High | High | High | High | High | High | High | High | High | High | High | 0 | 1.58E-05 | 3.05 |
| FALSE | High | VGECSGPA | [R].VGECS | 1xCarbami | F8VQL9 1x | 1.94E-07 | 0 | 1 | 8 | 4 | F8VQL9 | Nuclear re | F8VQL9 [7 | 0 | 2626.165 | 0 | 0.19 | 0.053395 | 11.66 | 93.8 | 106.2 | 10 | 5.01 | | High | High | High | High | High | High | High | High | High | High | High | 0 | 1.21E-07 | 4.47 |
| FALSE | High | VGECSGPA | [R].VGECS | 1xCarbami | F8VQL9 2x | 0.000302 | 0 | 1 | 8 | 1 | F8VQL9 | Nuclear re | F8VQL9 [7 | 0 | 2706.132 | 0 | 0.9 | 0.000824 | 3.56 | 71.7 | 128.3 | 4.83 | 7.21 | | High | High | High | High | High | High | High | High | High | High | High | 0 | 6.23E-05 | 3.25 |
| FALSE | High | VGECSGPA | [R].VGECS | 1xCarbami | F8VQL9 3x | 6.53E-05 | 0 | 1 | 8 | 4 | F8VQL9 | Nuclear re | F8VQL9 [7 | 0 | 2786.098 | 0 | 0.39 | 0.006026 | 7.28 | 85.4 | 114.6 | 8.04 | 6.38 | | High | High | High | High | High | High | High | High | High | High | High | 0 | 8.55E-05 | 3.94 |
| FALSE | High | VGESDCSD | [R].VGESD | 1xCarbami | Q9ERQ3 2x | 0.000993 | 0 | 1 | 2 | 3 | Q9ERQ3 | Zinc finger | Q9ERQ3 [3 | 0 | 2861.155 | 0 | -0.15 | 0.467124 | 14.84 | 103 | 97 | 11.51 | 13.89 | | High | High | High | High | High | High | High | High | High | High | High | 2.58E-05 | 0.000444 | 2.94 |
| FALSE | High | VGFEGNTK | [R].VGFEG | 1xPhospho | Q80WQ8 1 | 0.002058 | 5.66E-05 | 1 | 2 | 1 | Q80WQ8 | Mis18-binc | Q80WQ8 [ | 0 | 1389.718 | 0 | -1.33 | 0.000209 | 0.61 | 142.5 | 57.5 | 5.72 | 5.81 | | High | High | High | High | High | High | High | High | High | High | High | 2.58E-05 | 0.000538 | 1.83 |
| FALSE | High | VGGAEHSF | [K].VGGAE | 1xPhospho | Q3U3R4 1x | 0.004781 | 0.000456 | 1 | 8 | 1 | Q3U3R4 | Lipase mat | Q3U3R4 [2 | 0 | 1872.917 | 0 | -0.09 | 0.934639 | 26.71 | 97.6 | 102.4 | 12.44 | 15.7 | | High | High | High | High | High | High | High | High | High | High | High | 0.000204 | 0.001371 | 1.58 |
| FALSE | High | VGGAGPPF | [K].VGGAG | 1xPhospho | Q80V76 1x | 4.44E-06 | 0 | 1 | 2 | 1 | Q80V76 | Pygopus 2 | Q80V76 [8 | 0 | 2125.083 | 0 | 0.62 | 0.008577 | 15.61 | 74.4 | 125.6 | 10.98 | 11.71 | | High | High | High | High | High | High | High | High | High | High | High | 0 | 5.41E-07 | 4.33 |
| FALSE | High | VGGASLGA | [K].VGGAS | 1xDeamida | A2AJ72 1x | 0.015603 | 0.001124 | 1 | 4 | 2 | A2AJ72 | Far upstrea | A2AJ72 [42 | 0 | 3469.689 | 0 | -0.19 | 0.197967 | 12.86 | 105.7 | 94.3 | 7.66 | 12.99 | | High | High | High | High | High | High | High | High | High | High | High | 0.000585 | 0.005136 | 3.19 |
| FALSE | High | VGAHGSEE | [K].VGAHG | 1xPhospho | Q8BMK4 1 | 1.73E-05 | 0 | 1 | 1 | 1 | Q8BMK4 | Cytoskelet | Q8BMK4 [ | 0 | 1825.925 | 0 | -0.11 | 0.186696 | 2.26 | 104.7 | 95.3 | 6.61 | 13.4 | | High | High | High | High | High | High | High | High | High | High | High | 0 | 2.51E-06 | 2.79 |
| FALSE | High | VGGLDFSP | [K].VGGLD | 1xPhospho | O08696 1x | 0.000639 | 0 | 1 | 6 | 1 | O08696 | Forkhead b | O08696 [6 | 0 | 1355.692 | 0 | 1.63 | 0.00059 | 7.41 | 49.2 | 150.8 | 3.23 | 9.03 | | High | High | High | High | High | High | High | High | High | High | High | 0 | 0.000145 | 0.99 |
| FALSE | High | VGGSSVDL | [R-].VGGSS | 1xPhospho | E9Q8Z8 1x | 0.023792 | 0.001612 | 1 | 20 | 2 | E9Q8Z8 | Catenin de | E9Q8Z8 [2 | 0 | 1335.662 | 0 | -0.04 | 0.429949 | 15.9 | 101.3 | 98.7 | 6.52 | 12.71 | | High | High | High | High | High | High | High | High | High | High | High | 0.001151 | 0.01218 | 1.9 |
| FALSE | High | VGGTSDVE | [K].VGGTS | 1xPhospho | P63038 1x | 0.019276 | 0.001402 | 1 | 2 | 2 | P63038 | 60 kDa hea | P63038 [40 | 0 | 1771.888 | 0 | -0.24 | 0.009986 | 5.12 | 108.9 | 91.1 | 2.05 | 6 | | High | High | High | High | High | High | High | High | High | High | High | 0.000756 | 0.006556 | 3.18 |
| FALSE | High | VGGYATSP | [R].VGGYA | 1xPhospho | E9QK48 1x | 0.000141 | 0 | 1 | 1 | 1 | E9QK48 | Echinoderr | E9QK48 [1 | 0 | 1688.866 | 0 | -0.68 | 0.039674 | 7.81 | 123 | 77 | 11.85 | 22.96 | | High | High | High | High | High | High | High | High | High | High | High | 0 | 2.63E-05 | 2.51 |
| FALSE | High | VGLPSPSSL | [K].VGLPSF | 1xPhospho | Q9D168 1x | 0.001002 | 0 | 1 | 1 | 1 | Q9D168 | Integrator | Q9D168 [3 | 0 | 3711.848 | 0 | 0.11 | 0.532074 | 15.98 | 98.4 | 101.6 | 14.63 | 9.83 | | High | High | High | High | High | High | High | High | High | High | High | 0 | 0.00024 | 4.35 |
| FALSE | High | VGLPSPSSL | [K].VGLPSF | 1xDeamida | Q9D168 1x | 1.69E-06 | 0 | 1 | 1 | 12 | Q9D168 | Integrator | Q9D168 [3 | 0 | 3712.832 | 0 | 0.14 | 0.99512 | 13.96 | 97.5 | 102.5 | 10.29 | 13.64 | | High | High | High | High | High | High | High | High | High | High | High | 0 | 1.77E-05 | 4.48 |
| FALSE | High | VGIQLPEVI | [K].VGIQLF | 1xPhospho | E9Q616 1x | 0.000189 | 0 | 1 | 4 | 1 | E9Q616 | AHNAK nu | E9Q616 [5 | 0 | 2137.192 | 0 | 1.69 | 0.001471 | 4.84 | 47.6 | 152.4 | 12.73 | 17.15 | | High | High | High | High | High | High | High | High | High | High | High | 0 | 3.69E-05 | 2.3 |
| FALSE | High | VGLSPDSP | [R].VGLSPD | 1xPhospho | Q8VHK1 1x | 0.007145 | 0.00062 | 1 | 1 | 1 | Q8VHK1 | Caskin-2 O | Q8VHK1 [3 | 0 | 1479.704 | 0 | 1.14 | 0.000216 | 5.95 | 63.8 | 136.2 | 6.9 | 3.67 | | High | High | High | High | High | High | High | High | High | High | High | 0.000275 | 0.002165 | 3.2 |
| FALSE | High | VGLSPDSP | [R].VGLSPD | 1xPhospho | Q8VHK1 1x | 0.016743 | 0.001166 | 1 | 1 | 1 | Q8VHK1 | Caskin-2 O | Q8VHK1 [3 | 1 | 2709.304 | 0 | 1.06 | 0.000899 | 10.72 | 64.4 | 135.6 | 7.85 | 4.66 | | High | High | High | High | High | High | High | High | High | High | High | 0.000622 | 0.005579 | 2.78 |
| FALSE | High | VGITSPVEK | [R].VGITSP | 1xPhospho | A0A0A0M | 0.004613 | 0.000456 | 1 | 2 | 1 | A0A0A0M | Histone-lys | A0A0A0M | 0 | 1467.822 | 0 | 0.38 | 0.0159 | 3.42 | 88.2 | 111.8 | 6.19 | 7.91 | | High | High | High | High | High | High | High | High | High | High | High | 0.000204 | 0.001321 | 2.91 |
| FALSE | High | VGLVEDSP | [K].VGLVEI | 1xPhospho | Q8K212 1x | 2.36E-05 | 0 | 1 | 3 | 21 | Q8K212 | Phosphofu | Q8K212 [7 | 0 | 3865.843 | 0 | 6.64 | | 0 | | 200 | | 15.57 | | Not Found | Not Found | Not Found | High | Not Found | High | High | High | High | High | High | 7.19E-05 | 0.000793 | 4.01 |
| FALSE | High | VGNSGDAI | [R].VGNSG | 1xPhospho | Q8R4U7 1x | 0.005666 | 0.00062 | 1 | 1 | 1 | Q8R4U7 | Leucine zip | Q8R4U7 [9 | 0 | 1607.763 | 0 | -0.23 | 0.112425 | 15.38 | 108.1 | 91.9 | 7.72 | 10.04 | | High | High | High | High | High | High | High | High | High | High | High | 0.000275 | 0.001659 | 1.63 |
| FALSE | High | VGPGAGAS | [R].VGPGA | 1xPhospho | B2RQ83 1x | 0.010475 | 0.000772 | 2 | 5 | 1 | B2RQ83; B | Gem (Nucl | B2RQ83 [4 | 0 | 1646.825 | 0 | -0.64 | 0.001008 | 3.27 | 123.1 | 76.9 | 3.63 | 10.72 | Shared | High | High | High | High | High | High | High | High | High | High | High | 0.000382 | 0.003294 | 2.23 |
| FALSE | High | VGPNADSL | [K].VGPNA | 1xPhospho | P48754 1x | 0.025073 | 0.001645 | 1 | 6 | 1 | P48754 | Breast can | P48754 [33 | 0 | 1439.673 | 0 | -0.6 | 0.000454 | 2.15 | 121.6 | 78.4 | 6.08 | 5.47 | | High | High | High | High | High | High | High | High | High | High | High | 0.000872 | 0.008754 | 1.67 |
| FALSE | High | VGGSEVTE | [K].VGGSE | 1xPhospho | A0A0R4J17 | 3.19E-07 | 0 | 1 | 3 | 1 | A0A0R4J17 | Bromodom | A0A0R4J17 | 1 | 3250.376 | 0 | -1.26 | 0.006573 | 3.29 | 145.1 | 54.9 | 17.13 | 17.56 | | High | High | High | High | High | High | High | High | High | High | High | 0 | 2.81E-08 | 3.42 |
| FALSE | High | VFVGGLSP | [K].VFVGG | 1xPhospho | Q9Z130 1x | 4.85E-05 | 0 | 1 | 3 | 2 | Q9Z130 | Heterogen | Q9Z130 [1 | 0 | 2244.157 | 0 | -0.79 | 0.000211 | 0.53 | 124.7 | 75.3 | 4.67 | 5 | | High | High | High | High | High | High | High | High | High | High | High | 0 | 0.00019 | 2.73 |
| FALSE | High | VFTSDISDP | [K].VFTSDI | 1xPhospho | P83741-3 1 | 0.002768 | 0.000109 | 1 | 5 | 2 | P83741-3 | Isoform 3 c | P83741-3 [ | 0 | 4591.268 | 0 | 0.39 | 0.21659 | 24.81 | 89.2 | 110.8 | 12.68 | 34.94 | | High | High | High | High | High | High | High | High | High | High | High | 4.94E-05 | 0.000747 | 3.35 |
| FALSE | High | VFTPPPNS | [K].VFTPPP | 1xPhospho | Q9DB07 1x | 0.000822 | 0 | 1 | 1 | 3 | Q9DB07 | Intraflagell | Q9DB07 [2 | 0 | 2472.19 | 0 | -0.86 | 0.032911 | 14.89 | 128.8 | 71.2 | 14.21 | 22.3 | | High | High | High | High | High | High | High | High | High | High | High | 0 | 0.000191 | 3.25 |
| FALSE | High | VETPDVNL | [R].VETPD | 2xDeamida | Q3UC10 1x | 3.31E-07 | 0 | 1 | 4 | 9 | Q3UC10 | DNA helica | Q3UC10 [6 | 0 | 4287.871 | 0 | -0.46 | 0.01052 | 10.43 | 115.9 | 84.1 | 5.98 | 16.43 | | High | High | High | High | High | High | High | High | High | High | High | 0 | 4.79E-07 | 5.45 |
| FALSE | High | VETPDVNL | [R].VETPD | 1xPhospho | Q3UC10 1x | 1.2E-06 | 0 | 1 | 4 | 13 | Q3UC10 | DNA helica | Q3UC10 [6 | 0 | 4301.898 | 0 | -0.34 | 0.096769 | 7.91 | 112 | 88 | 7.42 | 13.48 | | High | High | High | High | High | High | High | High | High | High | High | 0 | 1.66E-06 | 5.88 |
| FALSE | High | VETPDVNL | [R].VETPD | 1xDeamida | Q3UC10 1x | 6.74E-07 | 0 | 1 | 4 | 33 | Q3UC10 | DNA helica | Q3UC10 [6 | 0 | 4302.882 | 0 | -0.28 | 0.188616 | 25.35 | 111 | 89 | 23.47 | 8.28 | | High | High | High | High | High | High | High | High | High | High | High | 0 | 1.16E-07 | 5.64 |
| FALSE | High | VETQNPSL | [K].VETQN | 1xPhospho | A0A140LJJ | 0.000152 | 0 | 1 | 6 | 1 | A0A140LJJ | A-kinase ar | A0A140LJJ | 0 | 2055.041 | 0 | 0.13 | 0.190601 | 15.06 | 96.4 | 103.6 | 6.56 | 4 | | High | High | High | High | High | High | High | High | High | High | High | 0 | 2.9E-05 | 3.22 |
| FALSE | High | VEVAAPAH | [R].VEVAAI | 1xPhospho | P0C7T6 1x | 0.000815 | 0 | 1 | 1 | 1 | P0C7T6 | Ataxin-1-lik | P0C7T6 [31 | 1 | 2470.229 | 0 | 0.69 | 0.00359 | 14.72 | 75 | 125 | 16.43 | 8.2 | | High | High | High | High | High | High | High | High | High | High | High | 0 | 0.000189 | 3.76 |
| FALSE | High | VEVESGGS | [R].VEVES | 1xPhospho | Q3TCX6 1x | 2.74E-08 | 0 | 1 | 7 | 2 | Q3TCX6 | Putative ur | Q3TCX6 [2 | 0 | 1936.958 | 0 | 1.11 | 0.003024 | 19.67 | 63 | 137 | 9.33 | 14.61 | | High | High | High | High | High | High | High | High | High | High | High | 0 | 1.78E-09 | 3.73 |

| | | | | | | | | | | | | | | | | | | | | | | | | | | | | | | | | | | | | | | |
|---|---|---|---|---|---|---|---|---|---|---|---|---|---|---|---|---|---|---|---|---|---|---|---|---|---|---|---|---|---|---|---|---|---|---|---|---|---|---|
| FALSE | High | VEVSHVTLI | [K].VEVSH\ | 1xPhospho | E9Q784 1x | 0.000852 | 0 | 1 | 1 | 1 | E9Q784 | Zinc finger | E9Q784 [1: | 1 | 2285.206 | 0 | -0.26 | 0.083538 | 13.23 | 113.7 | 86.3 | 13.75 | 15.7 | | High | High | High | High | High | High | High | High | High | High | High | 0 | 0.0002 | 2.47 |
| FALSE | High | VEVTPTVPI | [K].VEVTPT | 1xPhospho | Q8C729-2 : | 0.016892 | 0.001204 | 1 | 2 | 2 | Q8C729-2 | Isoform 2 c | Q8C729-2 | 0 | 1306.697 | 0 | -1.25 | 0.000426 | 1.76 | 140.9 | 59.1 | 2.56 | 8.46 | | High | High | High | High | High | High | High | High | High | High | High | 0.000638 | 0.005627 | 2.18 |
| FALSE | High | VEVVSPATI | [K].VEVVSF | 1xDeamida | Q8CHK4-3 | 0.099328 | 0.008003 | 1 | 5 | 2 | Q8CHK4-3 | Isoform 3 c | Q8CHK4-3 | 0 | 3004.487 | 0 | -0.7 | 0.015108 | 6.6 | 121 | 79 | 15.71 | 9.63 | | High | High | High | High | High | High | High | High | High | High | High | 0.004032 | 0.04289 | 2.82 |
| FALSE | High | VFAEDQDV | [K].VFAED( | 1xCarbami | Q04692 1x | 5.98E-06 | 0 | 1 | 2 | 1 | Q04692 | SWI/SNF-r | Q04692 [2 | 0 | 2750.274 | 0 | -1.37 | 0.001216 | 2.94 | 143.6 | 56.4 | 8.06 | 11.43 | | High | High | High | High | High | High | High | High | High | High | High | 0 | 7.61E-07 | 4.14 |
| FALSE | High | VFDDDSDD | [R].VFDDD: | 1xPhospho | Q8BGC0 1x | 9.5E-07 | 0 | 1 | 1 | 6 | Q8BGC0 | HIV Tat-spe | Q8BGC0 [6 | 0 | 2667.112 | 0 | -0.41 | 0.005 | 3.67 | 115.5 | 84.5 | 4.79 | 5.52 | | High | High | High | High | High | High | High | High | High | High | High | 0 | 0.000269 | 4.19 |
| FALSE | High | VFDLQFSTI | [R].VFDLQI | 1xPhospho | Q8C156 1x | 4.01E-05 | 0 | 1 | 1 | 1 | Q8C156 | Condensin | Q8C156 [6 | 0 | 2356.154 | 0 | 0.19 | 0.262866 | 4.65 | 95 | 105 | 11.48 | 8.07 | | High | High | High | High | High | High | High | High | High | High | High | 0 | 6.44E-06 | 2.89 |
| FALSE | High | VFEDNSNE | [K].VFEDN! | 1xPhospho | Q8BGC0 1x | 0.024001 | 0.001612 | 1 | 1 | 1 | Q8BGC0 | HIV Tat-spe | Q8BGC0 [7 | 0 | 1390.609 | 0 | 0.16 | 0.804799 | 33.14 | 107.1 | 92.9 | 102.08 | 57.01 | | High | High | High | High | High | High | High | High | High | High | High | 0.000859 | 0.008351 | 1.94 |
| FALSE | High | VFEDNSNE | [K].VFEDN! | 1xPhospho | Q8BGC0 1x | 0.000297 | 0 | 1 | 1 | 5 | Q8BGC0 | HIV Tat-spe | Q8BGC0 [7 | 0 | 1619.772 | 0 | -0.28 | 0.010155 | 1.94 | 108.7 | 91.3 | 5.88 | 2.36 | | High | High | High | High | High | High | High | High | High | High | High | 0 | 8.61E-05 | 3.27 |
| FALSE | High | VFEDNSNE | [K].VFEDN! | 1xDeamida | Q8BGC0 1x | 0.000286 | 0 | 1 | 1 | 5 | Q8BGC0 | HIV Tat-spe | Q8BGC0 [7 | 0 | 1620.756 | 0 | -0.11 | 0.013452 | 3.05 | 105 | 95 | 4.8 | 3.47 | | High | High | High | High | High | High | High | High | High | High | High | 0.000622 | 0.005562 | 2.84 |
| FALSE | High | VFEEATSP( | [R].VFEEAT | 1xCarbami | Q7TNF9 1x | 3.63E-05 | 0 | 1 | 2 | 2 | Q7TNF9 | Protein FAI | Q7TNF9 [3 | 0 | 2819.361 | 0 | -0.77 | 0.00521 | 8.42 | 127.9 | 72.1 | 11.82 | 11.82 | | High | High | High | High | High | High | High | High | High | High | High | 0 | 1.1E-05 | 3.88 |
| FALSE | High | VFEECSPK | [K].VFEECS | 1xCarbami | Q3U3E2 1x | 0.079114 | 0.00613 | 1 | 1 | 1 | Q3U3E2 | Protein FAI | Q3U3E2 [4 | 0 | 1533.742 | 0 | -0.02 | 0.645217 | 7.67 | 100.6 | 99.4 | 4.96 | 2.71 | | High | High | High | High | High | High | High | High | High | High | High | 0.003148 | 0.03289 | 1.62 |
| FALSE | High | VFFDTDDD | [K].VFFDTD | 1xPhospho | O55176 1x | 3.38E-08 | 0 | 1 | 3 | 2 | O55176 | E3 ubiquiti | O55176 [2 | 0 | 2276.955 | 0 | 0.03 | 0.612252 | 13.65 | 100.2 | 99.8 | 15.89 | 17.12 | | High | High | High | High | High | High | High | High | High | High | High | 0 | 2.25E-09 | 5.17 |
| FALSE | High | VFGALGPI( | [R].VFGAL( | 1xPhospho | Q91VU8 1x | 0.000758 | 0 | 1 | 2 | 2 | Q91VU8 | Prostate tu | Q91VU8 [6 | 0 | 2825.491 | 0 | -0.03 | 0.918851 | 8.78 | 101.7 | 98.3 | 19.93 | 6.12 | | High | High | High | High | High | High | High | High | High | High | High | 0 | 0.000175 | 4.13 |
| FALSE | High | VFGTDSSL( | [K].VFGTD! | 1xPhospho | Q8BX22 1x | 0.001021 | 0 | 1 | 2 | 1 | Q8BX22 | Sal-like pro | Q8BX22 [3 | 0 | 1781.915 | 0 | -1.7 | 0.000199 | 0.75 | 152.6 | 47.4 | 5.11 | 14.75 | | High | High | High | High | High | High | High | High | High | High | High | 0 | 0.000245 | 2.02 |
| FALSE | High | VFIDQNLSI | [K].VFIDQN | 1xPhospho | Q3V3X3 1x | 2.75E-06 | 0 | 1 | 4 | 2 | Q3V3X3 | MCG3705, | Q3V3X3 [1 | 0 | 1755.945 | 0 | -0.66 | 0.000815 | 2.47 | 121.5 | 78.5 | 2.61 | 8.56 | | High | High | High | High | High | High | High | High | High | High | High | 0.000571 | 0.004996 | 2.1 |
| FALSE | High | VFLGQQSP | [R].VFLGQ( | 1xPhospho | Q9D279 1x | 0.091627 | 0.007146 | 1 | 3 | 2 | Q9D279 | Mitotic inte | Q9D279 [7 | 0 | 1340.693 | 0 | 4.84 | 0.000159 | 40.7 | 6.5 | 193.5 | 35.8 | 4.2 | | High | High | High | High | High | High | High | High | High | High | High | 0.007994 | 0.08653 | 1.12 |
| FALSE | High | VFLQDNPT | [K].VFLQDI | 1xPhospho | Q6ZPR8 1x | 0.0008 | 0 | 1 | 4 | 4 | Q6ZPR8 | MKIAA140 | Q6ZPR8 [3 | 0 | 2522.26 | 0 | -0.05 | 0.886246 | 10.43 | 100.8 | 99.2 | 3.63 | 8.03 | | High | High | High | High | High | High | High | High | High | High | High | 2.58E-05 | 0.000352 | 2.52 |
| FALSE | High | VFLRPSPED | [R].VFLRPS | 1xPhospho | Q9Z0U1 1x | 8.97E-09 | 0 | 1 | 1 | 1 | Q9Z0U1 | Tight junct | Q9Z0U1 [4 | 0 | 2571.326 | 0 | -0.04 | 0.209682 | 3.41 | 101.5 | 98.5 | 3.84 | 5.85 | | High | High | High | High | High | High | High | High | High | High | High | 0 | 5.09E-10 | 3.13 |
| FALSE | High | VFNNIGAD | [K].VFNNI( | 1xPhospho | Q8CFE4 1x | 5.85E-08 | 0 | 1 | 3 | 2 | Q8CFE4 | SCY1-like p | Q8CFE4 [6 | 0 | 2231.054 | 0 | 0.44 | 0.008493 | 8.67 | 86.2 | 113.8 | 4.67 | 9.31 | | High | High | High | High | High | High | High | High | High | High | High | 0 | 4.18E-09 | 3.15 |
| FALSE | High | VFPQSSSW | [K].VFPQS! | 1xPhospho | E9Q9R9 1x | 7.28E-05 | 0 | 1 | 5 | 1 | E9Q9R9 | Disks large | E9Q9R9 [7 | 0 | 2606.306 | 0 | -1.08 | 0.000537 | 1.75 | 135.7 | 64.3 | 6.78 | 4.89 | | High | High | High | High | High | High | High | High | High | High | High | 0 | 1.26E-05 | 2.85 |
| FALSE | High | VFSGGEDL | [R].VFSGGI | 1xPhospho | E9QMX4 1: | 5.94E-07 | 0 | 1 | 4 | 1 | E9QMX4 | Ribosomal | E9QMX4 [5 | 0 | 2560.202 | 0 | 0.21 | 0.374814 | 19.46 | 102.1 | 97.9 | 11.04 | 23.65 | | High | High | High | High | High | High | High | High | High | High | High | 0 | 5.68E-08 | 3.19 |
| FALSE | High | VFSPVIR | [K].VFSPVII | 1xPhospho | Q924A2 1x | 0.014158 | 0.001011 | 1 | 4 | 1 | Q924A2 | Protein cap | Q924A2 [1 | 0 | 1126.622 | 0 | 0.18 | 0.01868 | 4.94 | 91.6 | 108.4 | 8.51 | 4.29 | | High | High | High | High | High | High | High | High | High | High | High | 0.000538 | 0.004623 | 1.55 |
| FALSE | High | VFSSSQGV | [R].VFSSSC | 1xPhospho | A0A0N4SU | 0.006834 | 0.00062 | 1 | 2 | 3 | A0A0N4SU | U6 snRNA- | A0A0N4SU | 0 | 2389.273 | 0 | 0.96 | 0.002833 | 10.95 | 63.5 | 136.5 | 14.41 | 9.79 | | High | High | High | High | High | High | High | High | High | High | High | 0.000294 | 0.002332 | 3 |
| FALSE | High | VEGAHSLS | [R].VEGAH | 1xPhospho | B2RRE7 1x | 0.000924 | 0 | 1 | 2 | 1 | B2RRE7 | OTU doma | B2RRE7 [9 | 0 | 1968.02 | 0 | 0.17 | 0.024864 | 3.16 | 93.1 | 106.9 | 11.61 | 9.12 | | High | High | High | High | High | High | High | High | High | High | High | 0 | 0.000218 | 2.81 |
| FALSE | High | VEFDEEFS( | [K].VEFDEE | 2xPhospho | Q64511 2x | 0.00537 | 0.000599 | 1 | 2 | 1 | Q64511 | DNA topois | Q64511 [1 | 0 | 3556.606 | 0 | 0.04 | 0.222742 | 9.61 | 96.3 | 103.7 | 7.94 | 9.74 | | High | High | High | High | High | High | High | High | High | High | High | 0.000267 | 0.001571 | 4.28 |
| FALSE | High | VEFDEEFS( | [K].VEFDEE | 1xDeamida | Q64511 1x | 0.016451 | 0.001124 | 1 | 2 | 1 | Q64511 | DNA topois | Q64511 [1 | 0 | 3477.624 | 0 | 0.28 | 0.036957 | 15.53 | 95.7 | 104.3 | 12.52 | 7.2 | | High | High | High | High | High | High | High | High | High | High | High | 0.000585 | 0.005468 | 2.8 |
| FALSE | High | VEFDEEFS( | [K].VEFDEE | 1xPhospho | Q64511 1x | 8.28E-06 | 0 | 1 | 2 | 9 | Q64511 | DNA topois | Q64511 [1 | 0 | 3476.64 | 0 | 0.49 | 0.002936 | 8.86 | 84 | 116 | 8.51 | 12.02 | | High | High | High | High | High | High | High | High | High | High | High | 0 | 1.09E-06 | 4.79 |
| FALSE | High | VAQVTGET | [R].VAQVT | 1xPhospho | P48381 1x | 2.46E-08 | 0 | 1 | 3 | 1 | P48381 | Transcripti | P48381 [64 | 0 | 3467.753 | 0 | 0.51 | 0.018689 | 12.8 | 83.8 | 116.2 | 16.54 | 5.39 | | High | High | High | High | High | High | High | High | High | High | High | 0 | 1.59E-09 | 4.51 |
| FALSE | High | VASAESCH | [R].VASAES | 1xCarbami | Q8R2R3 1x | 3.38E-05 | 0 | 1 | 1 | 2 | Q8R2R3 | Alpha- and | Q8R2R3 [1 | 0 | 2509.086 | 0 | -0.52 | 0.007574 | 4.65 | 118.7 | 81.3 | 3.5 | 13.79 | | High | High | High | High | High | High | High | High | High | High | High | 0 | 8.52E-06 | 3.68 |
| FALSE | High | VASDLEVE( | [K].VASDLE | 1xPhospho | Q8K298 1x | 0.000147 | 0 | 1 | 1 | 2 | Q8K298 | Anillin OS= | Q8K298 [5 | 0 | 1968.948 | 0 | 0.01 | 0.51811 | 12.23 | 96.5 | 103.5 | 9.06 | 6.33 | | High | High | High | High | High | High | High | High | High | High | High | 0 | 2.76E-05 | 4.39 |
| FALSE | High | VASEEEEVI | [K].VASEEE | 1xPhospho | E9Q3E2 1x | 0.000765 | 0 | 2 | 7 | 2 | E9Q3E2; Q | Synaptopo | E9Q3E2 [2 | 0 | 2242.202 | 0 | -0.5 | 0.075317 | 6.98 | 114.3 | 85.7 | 16.26 | 23.58 | Shared | High | High | High | High | High | High | High | High | High | High | High | 0 | 0.000177 | 3.67 |
| FALSE | High | VASGEQPL | [K].VASGE( | 1xPhospho | B7ZMS8 1x | 0.016306 | 0.001124 | 1 | 9 | 1 | B7ZMS8 | Mdm1 pro | B7ZMS8 [4 | 0 | 1466.802 | 0 | -1.08 | 0.000237 | 1.72 | 135.4 | 64.6 | 2.08 | 5.65 | | High | High | High | High | High | High | High | High | High | High | High | 0.000585 | 0.00542 | 2.62 |
| FALSE | High | VASGSDLH | [K].VASGSI | 1xPhospho | Q9JMH9-6 | 2.21E-05 | 0 | 1 | 8 | 1 | Q9JMH9-6 | Isoform 6 c | Q9JMH9-6 | 0 | 2211.013 | 0 | 0.28 | 0.490936 | 25.74 | 90.3 | 109.7 | 18.11 | 18.4 | | High | High | High | High | High | High | High | High | High | High | High | 0 | 3.29E-06 | 2.47 |
| FALSE | High | VASIFDDPL | [K].VASIFD | 1xPhospho | Q6PGL7 1x | 2.26E-07 | 0 | 1 | 2 | 2 | Q6PGL7 | WASH com | Q6PGL7 [1 | 0 | 1889.888 | 0 | 0.12 | 0.245142 | 20.05 | 91.8 | 108.2 | 15.83 | 24.81 | | High | High | High | High | High | High | High | High | High | High | High | 0 | 7.49E-05 | 3.41 |
| FALSE | High | VASLGSLPL | [K].VASLGS | 1xPhospho | F8WJ40 1x | 3.77E-05 | 0 | 1 | 4 | 1 | F8WJ40 | Histone-lys | F8WJ40 [7 | 0 | 2024.108 | 0 | -0.43 | 0.004623 | 1.3 | 111.7 | 88.3 | 6.19 | 7.45 | | High | High | High | High | High | High | High | High | High | High | High | 0 | 6E-06 | 4.05 |
| FALSE | High | VASMASFC | [R].VASMA | 1xPhospho | Q7TME2 1: | 9.25E-07 | 0 | 1 | 1 | 1 | Q7TME2 | Sperm-ass | Q7TME2 [9 | 0 | 2405.171 | 0 | 1.13 | 0.017349 | 38.91 | 63.1 | 136.9 | 37.3 | 22.16 | | High | High | High | High | High | High | High | High | High | High | High | 0 | 9.29E-08 | 3.45 |
| FALSE | High | VASPLSPTS | [K].VASPLS | 1xPhospho | Q8BSN5 1x | 1.61E-05 | 0 | 1 | 4 | 5 | Q8BSN5 | PHD finger | Q8BSN5 [1 | 0 | 3757.984 | 0 | -0.71 | 0.106397 | 8.82 | 126.4 | 73.6 | 25.7 | 20.17 | | High | High | High | High | High | High | High | High | High | High | High | 0 | 3.23E-06 | 3.61 |
| FALSE | High | VASPSQGC | [R].VASPSC | 1xPhospho | Q68FH0 1x | 3.05E-07 | 0 | 1 | 6 | 3 | Q68FH0 | Plakophilin | Q68FH0 [3 | 0 | 1954.005 | 0 | -1.03 | 0.00033 | 1.76 | 135.1 | 64.9 | 4.54 | 4.2 | | High | High | High | High | High | High | High | High | High | High | High | 0 | 2.67E-08 | 4.21 |
| FALSE | High | VASPSQGC | [R].VASPSC | 2xPhospho | Q68FH0 2x | 1.6E-05 | 0 | 1 | 6 | 2 | Q68FH0 | Plakophilin | Q68FH0 [3 | 0 | 2033.971 | 0 | -0.73 | 0.002846 | 0.91 | 124.1 | 75.9 | 6.05 | 6.37 | | High | High | High | High | High | High | High | High | High | High | High | 0 | 2.28E-06 | 4.25 |
| FALSE | High | VASPTSGLI | [R].VASPTS | 1xPhospho | P81122 1x | 0.019966 | 0.001438 | 1 | 3 | 1 | P81122 | Insulin rece | P81122 [10 | 0 | 1397.781 | 0 | -0.49 | 0.007151 | 7.43 | 118.9 | 81.1 | 2.96 | 13.36 | | High | High | High | High | High | High | High | High | High | High | High | 0.000786 | 0.006801 | 1.65 |
| FALSE | High | VAQVLSCE | [K].VAQVL! | 1xCarbami | Q3U4X8 2x | 4.3E-07 | 0 | 1 | 7 | 3 | Q3U4X8 | DNA ligase | Q3U4X8 [7 | 0 | 2749.208 | 0 | 0.09 | 0.160238 | 10.19 | 99 | 101 | 2.08 | 9.89 | | High | High | High | High | High | High | High | High | High | High | High | 0 | 8.78E-08 | 5.24 |
| FALSE | High | VASPVNK | [R].VASPVI | 1xPhospho | Q8BYJ6 1x | 0.019791 | 0.001438 | 1 | 9 | 1 | Q8BYJ6 | TBC1 doma | Q8BYJ6 [7 | 0 | 1252.707 | 0 | 0.56 | 0.007386 | 17.35 | 80.1 | 119.9 | 8.6 | 7.21 | | High | High | High | High | High | High | High | High | High | High | High | 0.000773 | 0.006734 | 1.98 |
| FALSE | High | VASSPVMV | [K].VASSP\ | 1xPhospho | Q61191 1x | 0.000286 | 0 | 1 | 2 | 1 | Q61191 | Host cell fa | Q61191 [5 | 0 | 1724.86 | 0 | -0.61 | 0.005193 | 7.52 | 121.5 | 78.5 | 6.51 | 8.13 | | High | High | High | High | High | High | High | High | High | High | High | 0 | 5.86E-05 | 3.65 |
| FALSE | High | VASSPVMV | [K].VASSP\ | 1xPhospho | Q61191 1x | 7.77E-06 | 0 | 1 | 2 | 6 | Q61191 | Host cell fa | Q61191 [5 | 0 | 1740.855 | 0 | -0.45 | 0.012334 | 8.16 | 115.5 | 84.5 | 6.89 | 7.55 | | High | High | High | High | High | High | High | High | High | High | High | 0 | 2.43E-06 | 3.12 |
| FALSE | High | VASSVEQLI | [R].VASSVE | 1xPhospho | Q6ZPQ6 1x | 0.027597 | 0.001716 | 1 | 3 | 2 | Q6ZPQ6 | Membrane | Q6ZPQ6 [3 | 0 | 3159.659 | 0 | -0.6 | 0.043692 | 13.02 | 123.2 | 76.8 | 19.87 | 16.33 | | High | High | High | High | High | High | High | High | High | High | High | 0.000997 | 0.01042 | 2.72 |
| FALSE | High | VATLNSEEI | [K].VATLN! | 1xPhospho | Q3V1M8 1 | 6.91E-06 | 0 | 1 | 7 | 3 | Q3V1M8 | Putative ur | Q3V1M8 [2 | 0 | 2345.131 | 0 | 0.61 | 0.003603 | 5.28 | 79.1 | 120.9 | 6.51 | 5.7 | | High | High | High | High | High | High | High | High | High | High | High | 0 | 8.93E-07 | 3.32 |
| FALSE | High | VATSAASG | [R].VATSA/ | 1xPhospho | A2AG50 1x | 0.000837 | 0 | 1 | 5 | 1 | A2AG50 | MAP7 dom | A2AG50 [3 | 0 | 1676.832 | 0 | -0.26 | 0.513124 | 7.61 | 109.8 | 90.2 | 10.47 | 21.24 | | High | High | High | High | High | High | High | High | High | High | High | 0 | 0.000195 | 2.58 |
| FALSE | High | VAVCPSSP\ | [K].VAVCP! | 1xCarbami | A0A1L1SS/ | 0.000257 | 0 | 1 | 2 | 2 | A0A1L1SS/ | Centrosom | A0A1L1SS/ | 0 | 2887.468 | 0 | -0.69 | 0.0051 | 3.62 | 123.4 | 76.6 | 12.45 | 8.73 | | High | High | High | High | High | High | High | High | High | High | High | 0 | 5.2E-05 | 3.04 |
| FALSE | High | VAVDPVVT | [R].VAVDP' | 1xPhospho | F8VQ93 1x | 0.000114 | 0 | 1 | 2 | 2 | F8VQ93 | PH-interac | F8VQ93 [1 | 0 | 2354.226 | 0 | 0.14 | 0.731168 | 8.39 | 98.5 | 101.5 | 16.68 | 6.13 | | High | High | High | High | High | High | High | High | High | High | High | 0 | 2.08E-05 | 3.09 |
| FALSE | High | VAVEELSP( | [R].VAVEEI | 1xPhospho | Q80SX8 1x | 0.003526 | 0.00021 | 1 | 4 | 1 | Q80SX8 | ATP-depen | Q80SX8 [3 | 0 | 1647.83 | 0 | -0.01 | 0.929488 | 8.56 | 100.2 | 99.8 | 4.43 | 7.66 | | High | High | High | High | High | High | High | High | High | High | High | 9.42E-05 | 0.000981 | 2.72 |
| FALSE | High | VAVEYLDP! | [R].VAVEYI | 1xPhospho | Q9WVA3 1 | 2.87E-05 | 0 | 1 | 3 | 1 | Q9WVA3 | Mitotic che | Q9WVA3 [ | 0 | 2112.103 | 0 | -0.55 | 0.004083 | 5.06 | 118.8 | 81.2 | 4.8 | 6 | | High | High | High | High | High | High | High | High | High | High | High | 0 | 4.44E-06 | 1.68 |
| FALSE | High | VAVPTGPT | [K].VAVPT( | 1xPhospho | Q3UJD6 1x | 0.000725 | 0 | 1 | 6 | 3 | Q3UJD6 | Ubiquitin c | Q3UJD6 [4 | 0 | 3144.556 | 0 | -0.07 | 0.733724 | 7.74 | 99.3 | 100.7 | 12.41 | 7.76 | | High | High | High | High | High | High | High | High | High | High | High | 2.58E-05 | 0.00036 | 3.26 |
| FALSE | High | VAVSADPN | [R].VAVSAI | 1xPhospho | Q99PT1 1x | 0.019791 | 0.001438 | 1 | 1 | 1 | Q99PT1 | Rho GDP-d | Q99PT1 [5 | 0 | 1960.047 | 0 | 1.87 | 0.005549 | 17.27 | 43 | 157 | 10.72 | 23.73 | | High | High | High | High | High | High | High | High | High | High | High | 0.000773 | 0.006763 | 1.88 |
| FALSE | High | VCDSLSEDI | [R].VCDSLS | 1xCarbami | Q8BQ33 1x | 0.003688 | 0.00021 | 1 | 1 | 2 | Q8BQ33 | Treslin OS= | Q8BQ33 [1 | 0 | 1973.929 | 0 | -0.44 | 0.003272 | 4.61 | 120.2 | 79.8 | 11.41 | 11.32 | | High | High | High | High | High | High | High | High | High | High | High | 9.42E-05 | 0.001027 | 3.4 |
| FALSE | High | VCLEIPQEV | [R].VCLEIP( | 1xCarbami | Q8BW72 1 | 0.000568 | 0 | 1 | 3 | 1 | Q8BW72 | Lysine-spe | Q8BW72 [4 | 0 | 2541.308 | 0 | -0.06 | 0.84249 | 9.36 | 98.7 | 101.3 | 12.28 | 12.5 | | High | High | High | High | High | High | High | High | High | High | High | 0 | 0.000127 | 2.2 |
| FALSE | High | VASSERHS! | [K].VASSEF | 2xPhospho | Q3U832 2x | 0.000595 | 0 | 1 | 3 | 1 | Q3U832 | Putative ur | Q3U832 [5 | 1 | 2237.984 | 0 | -1.43 | 0.004129 | 1.61 | 146.1 | 53.9 | 15.76 | 8.19 | | High | High | High | High | High | High | High | High | High | High | High | 0 | 0.000133 | 2.11 |
| FALSE | High | VAQVLSCE | [K].VAQVL! | 1xCarbami | Q3U4X8 1x | 1.4E-07 | 0 | 1 | 7 | 5 | Q3U4X8 | DNA ligase | Q3U4X8 [7 | 0 | 2669.242 | 0 | -0.47 | 0.001247 | 3.09 | 115.2 | 84.8 | 2.56 | 5.01 | | High | High | High | High | High | High | High | High | High | High | High | 0 | 1.12E-08 | 4.67 |
| FALSE | High | VAQVLSCE | [K].VAQVL! | 1xCarbami | Q3U4X8 2x | 5.23E-06 | 0 | 1 | 7 | 1 | Q3U4X8 | DNA ligase | Q3U4X8 [7 | 0 | 2520.045 | 0 | -0.08 | 0.904388 | 44.93 | 111 | 89 | 45.51 | 56.31 | | High | High | High | High | High | High | High | High | High | High | High | 0 | 6.5E-07 | 3.94 |
| FALSE | High | VAQVLSCE | [K].VAQVL! | 1xCarbami | Q3U4X8 1x | 2.46E-07 | 0 | 1 | 7 | 2 | Q3U4X8 | DNA ligase | Q3U4X8 [7 | 0 | 2440.079 | 0 | -0.33 | 0.647586 | 15.47 | 119.1 | 80.9 | 61.85 | 61.86 | | High | High | High | High | High | High | High | High | High | High | High | 0 | 9.11E-07 | 4.64 |
| FALSE | High | VAEFNNVT | [K].VAEFNI | 1xPhospho | P27546 1x | 5.65E-08 | 0 | 1 | 3 | 6 | P27546 | Microtubu | P27546 [4 | 0 | 2630.337 | 0 | 1.2 | 0.000545 | 13.8 | 61 | 139 | 12.65 | 2.41 | | High | High | High | High | High | High | High | High | High | High | High | 0 | 1.34E-06 | 4.14 |
| FALSE | High | VAEFNNVT | [K].VAEFNI | 1xDeamida | P27546 1x | 0.001659 | 5.66E-05 | 1 | 3 | 9 | P27546 | Microtubu | P27546 [4 | 0 | 2631.321 | 0 | 0.85 | 0.000662 | 6.14 | 71.2 | 128.8 | 5.04 | 7.77 | | High | High | High | High | High | High | High | High | High | High | High | 7.19E-05 | 0.000907 | 2.85 |
| FALSE | High | VAEFNNVT | [K].VAEFNI | 2xPhospho | P27546 2x | 1.85E-06 | 0 | 1 | 3 | 1 | P27546 | Microtubu | P27546 [4 | 0 | 2710.303 | 0 | 1.48 | 0.005379 | 10.3 | 55.7 | 144.3 | 14 | 13.17 | | High | High | High | High | High | High | High | High | High | High | High | 0 | 2.03E-07 | 4.23 |
| FALSE | High | VAELEEEQ! | [K].VAELEE | 1xPhospho | A2ARP8 1x | 2.58E-05 | 0 | 1 | 3 | 1 | A2ARP8 | Microtubu | A2ARP8 [3 | 0 | 2796.302 | 0 | -1.28 | 0.005188 | 3.84 | 144 | 56 | 15.38 | 29.83 | | High | High | High | High | High | High | High | High | High | High | High | 0 | 3.93E-06 | 3.52 |
| FALSE | High | VAGEESALI | [R].VAGEE! | 1xCarbami | Q8CCE9 1x | 0.0012 | 0 | 1 | 5 | 1 | Q8CCE9 | Transcripti | Q8CCE9 [3 | 0 | 3727.854 | 0 | -0.64 | 0.001876 | 3.31 | 128.3 | 71.7 | 9.99 | 13.96 | | High | High | High | High | High | High | High | High | High | High | High | 0 | 0.000292 | 3.19 |
| FALSE | High | VAGEESALI | [R].VAGEE! | 1xCarbami | Q8CCE9 1x | 0.000529 | 0 | 1 | 5 | 2 | Q8CCE9 | Transcripti | Q8CCE9 [3 | 0 | 3728.838 | 0 | -0.51 | 0.005827 | 8.83 | 118.4 | 81.6 | 10.4 | 14.9 | | High | High | High | High | High | High | High | High | High | High | High | 0 | 0.000117 | 4.23 |
| FALSE | High | VAGGSGSE | [R].VAGGS | 1xPhospho | Q9Z1X2 1x | 1.9E-05 | 0 | 1 | 3 | 9 | Q9Z1X2 | Phosphatic | Q9Z1X2 [8 | 0 | 1739.934 | 0 | -0.01 | 0.61969 | 8.68 | 98.5 | 101.5 | 6.56 | 5.19 | | High | High | High | High | High | High | High | High | High | High | High | 0 | 2.78E-06 | 3.53 |
| FALSE | High | VAGSEGSS | [K].VAGSE( | 1xPhospho | P97868 1x | 4.92E-07 | 0 | 1 | 3 | 4 | P97868 | E3 ubiquiti | P97868 [12 | 0 | 2798.293 | 0 | -0.35 | 0.010054 | 4.52 | 110.5 | 89.5 | 5.69 | 4.77 | | High | High | High | High | High | High | High | High | High | High | High | 0 | 4.59E-08 | 5.15 |
| FALSE | High | VAGSEGSS | [K].VAGSE( | 1xPhospho | P97868 1x | 9.22E-06 | 0 | 1 | 3 | 6 | P97868 | E3 ubiquiti | P97868 [12 | 1 | 3155.551 | 0 | -0.16 | 0.037669 | 4.46 | 105.4 | 94.6 | 4.85 | 8.08 | | High | High | High | High | High | High | High | High | High | High | High | 0 | 3.89E-05 | 3.47 |
| FALSE | High | VALENDDR | [R].VALENI | 1xPhospho | E9QM77 1: | 0.006536 | 0.00062 | 1 | 7 | 2 | E9QM77 | Ataxin-2 O | E9QM77 [4 | 1 | 2071.995 | 0 | 0.6 | 0.001426 | 4.93 | 79.7 | 120.3 | 6.34 | 4.76 | | High | High | High | High | High | High | High | High | High | High | High | 0.000449 | 0.003782 | 2.15 |
| FALSE | High | VALGDGVC | [R].VALGD( | 2xPhospho | Q3TII9 2xP | 0.001179 | 0 | 1 | 2 | 8 | Q3TII9 | Putative ur | Q3TII9 [20 | 0 | 3408.596 | 0 | -1.78 | 0.001219 | 1.82 | 155.5 | 44.5 | 6.51 | 30.3 | | High | High | High | High | High | High | High | High | High | High | High | 0.004596 | 0.04989 | 3.9 |
| FALSE | High | VAILTDDEE | [K].VAILTD | 1xPhospho | D3Z183 1x | 9.14E-06 | 0 | 1 | 3 | 2 | D3Z183 | Major facil | D3Z183 [7- | 0 | 1927.967 | 0 | 4.63 | 0.00014 | 49.44 | 7.5 | 192.5 | 41.9 | 4.36 | | High | High | High | High | High | High | High | High | High | High | High | 0 | 1.22E-06 | 3.93 |
| FALSE | High | VALSPVTEI | [R].VALSPV | 1xCarbami | P48972 1x | 2.47E-05 | 0 | 1 | 1 | 2 | P48972 | Myb-relate | P48972 [42 | 0 | 3175.6 | 0 | -1.36 | 0.00018 | 1.26 | 144.8 | 55.2 | 3.05 | 5.74 | | High | High | High | High | High | High | High | High | High | High | High | 0 | 3.74E-06 | 4.79 |
| FALSE | High | VANFSNMI | [K].VANFSI | 2xDeamida | P40201 1x | 0.035793 | 0.002304 | 1 | 4 | 1 | P40201 | Chromodo | P40201 [10 | 0 | 2434.021 | 0 | -0.79 | 0.105053 | 4.15 | 125.5 | 74.5 | 25.27 | 14 | | High | High | High | High | High | High | High | High | High | High | High | 0.001214 | 0.01311 | 3.12 |
| FALSE | High | VANFSNMI | [K].VANFSI | 1xPhospho | P40201 1x | 0.000674 | 0 | 1 | 4 | 1 | P40201 | Chromodo | P40201 [10 | 1 | 2990.385 | 0 | -0.47 | 0.003121 | 2.02 | 117.7 | 82.3 | 2.52 | 13.42 | | High | High | High | High | High | High | High | High | High | High | High | 0 | 0.000153 | 2.67 |
| FALSE | High | VANFSNMI | [K].VANFSI | 1xPhospho | P40201 1x | 3.27E-06 | 0 | 1 | 4 | 1 | P40201 | Chromodo | P40201 [10 | 1 | 3006.38 | 0 | -0.35 | 0.12248 | 0.49 | 107.6 | 92.4 | 13.77 | 15.9 | | High | High | High | High | High | High | High | High | High | High | High | 0 | 3.84E-07 | 3.11 |
| FALSE | High | VANSPTMV | [K].VANSPT | 1xPhospho | E9Q4F8 1x | 0.003526 | 0.00021 | 1 | 4 | 2 | E9Q4F8 | Ankyrin rep | E9Q4F8 [2 | 0 | 1995.111 | 0 | -0.66 | 0.003279 | 1.3 | 120.4 | 79.6 | 4.33 | 10.26 | | High | High | High | High | High | High | High | High | High | High | High | 9.42E-05 | 0.000982 | 2.95 |
| FALSE | High | VAPENQLT | [K].VAPEN( | 1xPhospho | Q8CJF7 1x | 3.39E-06 | 0 | 1 | 1 | 2 | Q8CJF7 | Protein ELY | Q8CJF7 [17 | 0 | 1985.005 | 0 | -0.73 | 0.000863 | 3.29 | 126.9 | 73.1 | 4.89 | 4.91 | | High | High | High | High | High | High | High | High | High | High | High | 0 | 4.02E-07 | 4.41 |
| FALSE | High | VAPGPSSG | [K].VAPGP! | 1xCarbami | Q8CH77 1x | 4.17E-06 | 0 | 1 | 3 | 3 | Q8CH77 | Neuron na | Q8CH77 [1 | 0 | 2562.222 | 0 | 0.66 | 0.000429 | 7.4 | 78.8 | 121.2 | 8.5 | 5.22 | | High | High | High | High | High | High | High | High | High | High | High | 0 | 5.07E-07 | 3.37 |
| FALSE | High | VAPGPSSG | [K].VAPGP! | 1xCarbami | Q8CH77 2x | 0.054383 | 0.003951 | 1 | 3 | 1 | Q8CH77 | Neuron na | Q8CH77 [1 | 0 | 2642.188 | 0 | 0.26 | 0.085663 | 11.18 | 93.6 | 106.4 | 11.39 | 6.55 | | High | High | High | High | High | High | High | High | High | High | High | 0.002059 | 0.02128 | 1.99 |
| FALSE | High | VAPGPSSG | [K].VAPGP! | 1xCarbami | Q8CH77 1x | 0.002508 | 0.000109 | 1 | 3 | 2 | Q8CH77 | Neuron na | Q8CH77 [1 | 1 | 2718.323 | 0 | 0.54 | 0.033081 | 24.98 | 83.4 | 116.6 | 17.03 | 7.29 | | High | High | High | High | High | High | High | High | High | High | High | 0.000344 | 0.003012 | 3.16 |
| FALSE | High | VAPLSSSLD | [K].VAPLSS | 1xPhospho | Q8BXW8 1 | 0.000322 | 0 | 1 | 2 | 2 | Q8BXW8 | Putative ur | Q8BXW8 [ | 0 | 2205.146 | 0 | 0.02 | 0.688932 | 15.47 | 99.4 | 100.6 | 5.56 | 13.02 | | High | High | High | High | High | High | High | High | High | High | High | 0 | 7.37E-05 | 3.37 |
| FALSE | High | VAPLWHSS | [K].VAPLW | 1xPhospho | Q8C0T5 1x | 0.0102 | 0.000736 | 1 | 3 | 1 | Q8C0T5 | Signal-indu | Q8C0T5 [1 | 0 | 2176.1 | 0 | 1.62 | 0.002593 | 6.43 | 48.1 | 151.9 | 10.58 | 13.79 | | High | High | High | High | High | High | High | High | High | High | High | 0.000382 | 0.003217 | 1.54 |
| FALSE | High | VAPPPPALT | [K].VAPPPF | 1xCarbami | Q5U4C3 1x | 9.05E-09 | 0 | 1 | 1 | 4 | Q5U4C3 | Splicing fac | Q5U4C3 [9 | 0 | 2605.298 | 0 | 0.03 | 0.850474 | 5.4 | 100.5 | 99.5 | 4.72 | 5.06 | | High | High | High | High | High | High | High | High | High | High | High | 0 | 2.95E-07 | 4.36 |
| FALSE | High | VAPPPPALT | [K].VAPPPF | 1xCarbami | Q5U4C3 2x | 2.52E-06 | 0 | 1 | 1 | 4 | Q5U4C3 | Splicing fac | Q5U4C3 [9 | 0 | 2685.265 | 0 | -0.22 | 0.004088 | 3.95 | 107 | 93 | 6.37 | 6.28 | | High | High | High | High | High | High | High | High | High | High | High | 0 | 2.87E-07 | 2.95 |
| FALSE | High | VAPQQCSE | [R].VAPQQ | 1xCarbami | Q5U3K5 1x | 0.003721 | 0.00021 | 1 | 1 | 2 | Q5U3K5 | Rab-like pr | Q5U3K5 [4 | 0 | 1911.929 | 0 | 0.55 | 0.000972 | 3.19 | 81.3 | 118.7 | 2.09 | 5.07 | | High | High | High | High | High | High | High | High | High | High | High | 0.000275 | 0.00204 | 3.42 |
| FALSE | High | VAPWEME | [R].VAPWE | 1xPhospho | Q8C6P8 1x | 0.000267 | 0 | 1 | 2 | 1 | Q8C6P8 | Zinc finger | Q8C6P8 [4 | 0 | 2688.111 | 0 | 0.09 | 0.679835 | 21.76 | 93.2 | 106.8 | 14.21 | 23 | | High | High | High | High | High | High | High | High | High | High | High | 0 | 5.4E-05 | 2.89 |
| FALSE | High | VAQAPSPC | [K].VAQAP | 1xCarbami | E9Q921 1x | 0.06169 | 0.004646 | 1 | 5 | 1 | E9Q921 | AF4/FMR2 | E9Q921 [1 | 0 | 1466.702 | 0 | -0.33 | 0.007099 | 2.29 | 111.3 | 88.7 | 11.38 | 4.73 | | High | High | High | High | High | High | High | High | High | High | High | 0.002381 | 0.02463 | 1.78 |
| FALSE | High | VAQLLSEE/ | [R].VAQLL! | 1xCarbami | Q6P1D7 1x | 1.19E-07 | 0 | 1 | 2 | 1 | Q6P1D7 | Structure-s | Q6P1D7 [3 | 0 | 2794.471 | 0 | -0.44 | 0.008166 | 6.37 | 113.3 | 86.7 | 13.42 | 14.01 | | High | High | High | High | High | High | High | High | High | High | High | 0 | 9.35E-09 | 4.19 |
| FALSE | High | VCSDEGGK | [R].VCSDE( | 1xCarbami | Q3U280 1x | 0.014929 | 0.001049 | 1 | 4 | 1 | Q3U280 | Putative ur | Q3U280 [1 | 0 | 1389.649 | 0 | 0.26 | 0.033478 | 7.6 | 91.6 | 108.4 | 6.78 | 7.82 | | High | High | High | High | High | High | High | High | High | High | High | 0.000555 | 0.004904 | 1.53 |
| FALSE | High | VADPDHDH | [R].VADPD | 1xPhospho | P63085 1x | 9.36E-05 | 0 | 1 | 3 | 4 | P63085 | Mitogen-ac | P63085 [17 | 0 | 2453.134 | 0 | 2.49 | 0.053501 | 43.33 | 29.5 | 170.5 | 22.38 | 51.59 | | High | High | High | High | High | High | High | High | High | High | High | 0 | 1.67E-05 | 3.41 |
| FALSE | High | VCSIDLEIDS | [K].VCSIDL | 1xCarbami | Q3TCR9 1x | 6.68E-07 | 0 | 1 | 5 | 1 | Q3TCR9 | Putative ur | Q3TCR9 [1 | 0 | 2792.375 | 0 | 0.06 | 0.631152 | 14.59 | 103 | 97 | 22.57 | 35.36 | | High | High | High | High | High | High | High | High | High | High | High | 0 | 6.46E-08 | 4.52 |
| FALSE | High | VCSVEYIN( | [K].VCSVEY | 2xCarbami | Q7TN31 1x | 7.38E-07 | 0 | 1 | 1 | 2 | Q7TN31 | Angiogenic | Q7TN31 [2 | 0 | 2764.236 | 0 | -0.37 | 0.007902 | 4.56 | 113.2 | 86.8 | 4.39 | 6.12 | | High | High | High | High | High | High | High | High | High | High | High | 0 | 7.21E-08 | 4.94 |
| FALSE | High | VDSSTNSSI | [K].VDSSTI | 1xPhospho | Q921K9 1x | 1.95E-05 | 0 | 1 | 2 | 1 | Q921K9 | B-cell CLL/I | Q921K9 [1 | 0 | 3057.364 | 0 | 0.53 | 0.029477 | 4.89 | 82.2 | 117.8 | 11.07 | 11.27 | | High | High | High | High | High | High | High | High | High | High | High | 0 | 2.86E-06 | 4.36 |
| FALSE | High | VDSSTNSSI | [K].VDSSTI | 2xPhospho | Q921K9 2x | 7.41E-05 | 0 | 1 | 2 | 6 | Q921K9 | B-cell CLL/I | Q921K9 [1 | 0 | 3137.33 | 0 | 0.53 | 0.011822 | 19.24 | 81.9 | 118.1 | 6.33 | 13.24 | | High | High | High | High | High | High | High | High | High | High | High | 0 | 1.28E-05 | 3.64 |
| FALSE | High | VDSSTNSSI | [K].VDSSTI | 3xPhospho | Q921K9 3x | 0.001168 | 0 | 1 | 2 | 2 | Q921K9 | B-cell CLL/I | Q921K9 [1 | 0 | 3217.296 | 0 | 0.65 | 0.002841 | 10.69 | 74.3 | 125.7 | 9.36 | 14.76 | | High | High | High | High | High | High | High | High | High | High | High | 0.000275 | 0.001864 | 3.64 |
| FALSE | High | VDSTEEINE | [R].VDSTEE | 1xPhospho | Q640N3 1x | 6.3E-05 | 0 | 1 | 3 | 2 | Q640N3 | Rho GTPas | Q640N3 [8 | 0 | 2294.051 | 0 | 0.02 | 0.618723 | 34.77 | 98 | 102 | 11.2 | 34.1 | | High | High | High | High | High | High | High | High | High | High | High | 0 | 1.07E-05 | 3.09 |
| FALSE | High | VDSTTCLFF | [R].VDSTT( | 1xCarbami | P47856 1x | 2.87E-05 | 0 | 1 | 2 | 10 | P47856 | Glutamine- | P47856 [2 | 0 | 2063.017 | 0 | 0.67 | 0.001219 | 11.86 | 76.8 | 123.2 | 4.64 | 5.13 | | High | High | High | High | High | High | High | High | High | High | High | 0 | 4.41E-06 | 3.37 |
| FALSE | High | VDTAASSS\ | [R].VDTAA! | 1xCarbami | Q6ZPR6 1x | 2.04E-05 | 0 | 1 | 2 | 1 | Q6ZPR6 | Inhibitor of | Q6ZPR6 [1 | 0 | 2867.385 | 0 | -0.06 | 0.77313 | 3.19 | 102.2 | 97.8 | 4.08 | 7.75 | | High | High | High | High | High | High | High | High | High | High | High | 0 | 3E-06 | 5.1 |
| FALSE | High | VDVEALEN | [K].VDVEAI | 1xPhospho | F8VQC1 1x | 0.000573 | 0 | 1 | 5 | 1 | F8VQC1 | Signal reco | F8VQC1 [5 | 0 | 2043 | 0 | -0.09 | 0.193851 | 7.48 | 103 | 97 | 7.68 | 4.71 | | High | High | High | High | High | High | High | High | High | High | High | 0 | 0.000127 | 3.11 |
| FALSE | High | VDVSPTSQ | [R].VDVSPT | 1xPhospho | Q99KI0 1x | 0.011246 | 0.000772 | 1 | 1 | 1 | Q99KI0 | Aconitate h | Q99KI0 [5 | 0 | 1297.635 | 0 | -1.06 | 0.000807 | 2.32 | 136 | 64 | 8.41 | 8.83 | | High | High | High | High | High | High | High | High | High | High | High | 0.000418 | 0.00359 | 2.45 |
| FALSE | High | VDVVGATC | [K].VDVVG | 1xCarbami | Q3TSY9 1x | 0.000302 | 0 | 1 | 7 | 2 | Q3TSY9 | Putative ur | Q3TSY9 [5 | 0 | 1900.879 | 0 | 0.17 | 0.033008 | 9.76 | 93 | 107 | 7.78 | 8.74 | | High | High | High | High | High | High | High | High | High | High | High | 0 | 6.24E-05 | 2.7 |
| FALSE | High | VDYESQST | [K].VDYES( | 1xPhospho | P47878 1x | 1.4E-08 | 0 | 1 | 3 | 8 | P47878 | Insulin-like | P47878 [18 | 0 | 2590.16 | 0 | 5.05 | 0.000365 | 11.94 | 5.9 | 194.1 | 30.21 | 9.95 | | High | High | High | High | High | High | High | High | High | High | High | 0 | 9.06E-08 | 5.43 |
| FALSE | High | VEAEVLSTI | [K].VEAEVI | 1xPhospho | Q3UXZ9 1x | 1.88E-08 | 0 | 1 | 5 | 2 | Q3UXZ9 | Lysine-spe | Q3UXZ9 [1 | 0 | 2079.021 | 0 | -0.08 | 0.009093 | 0.23 | 102.7 | 97.3 | 4.2 | 5.88 | | High | High | High | High | High | High | High | High | High | High | High | 0 | 1.51E-09 | 4.23 |
| FALSE | High | VEAPDSEA | [K].VEAPD! | 1xCarbami | Q8BHZ4 1x | 2.2E-07 | 0 | 1 | 3 | 2 | Q8BHZ4 | Zinc finger | Q8BHZ4 [1 | 0 | 2593.145 | 0 | -0.06 | 0.824565 | 14.08 | 103.1 | 96.9 | 8 | 11.04 | | High | High | High | High | High | High | High | High | High | High | High | 0 | 1.85E-08 | 4.43 |
| FALSE | High | VEASIVRPP | [K].VEASIV | 2xPhospho | F8WJ40 2x | 0.08727 | 0.006537 | 1 | 2 | 2 | F8WJ40 | Histone-lys | F8WJ40 [5 | 0 | 3250.648 | 0 | -0.29 | 0.052636 | 5.37 | 107.5 | 92.5 | 9.76 | 11.74 | | High | High | High | High | High | High | High | High | High | High | High | 0.003375 | 0.0368 | 2.78 |
| FALSE | High | VDSSSEDG | [K].VDSSSE | 1xPhospho | P97864 1x | 0.000292 | 0 | 1 | 1 | 4 | P97864 | Caspase-7 | P97864 [14 | 0 | 2115.001 | 0 | 0.73 | 0.006074 | 14.21 | 75.5 | 124.5 | 4.23 | 13.65 | | High | High | High | High | High | High | High | High | High | High | High | 0 | 5.98E-05 | 2.74 |
| FALSE | High | VEDGTSAV | [K].VEDGT! | 1xPhospho | Q5SWW4 : | 1.29E-05 | 0 | 1 | 1 | 2 | Q5SWW4 | Mediator c | Q5SWW4 [ | 0 | 3190.527 | 0 | 0.69 | 0.057277 | 20.49 | 76.6 | 123.4 | 9.7 | 22.34 | | High | High | High | High | High | High | High | High | High | High | High | 0 | 1.79E-06 | 2.98 |
| FALSE | High | VEDSGTPP' | [R].VEDSG1 | 1xPhospho | Q99KC3 1x | 0.004333 | 0.00036 | 1 | 1 | 1 | Q99KC3 | Nop2 prote | Q99KC3 [3 | 0 | 1989.973 | 0 | 0.66 | 0.001329 | 10.88 | 76 | 124 | 3.82 | 6.39 | | High | High | High | High | High | High | High | High | High | High | High | 0.000161 | 0.001234 | 2.69 |
| FALSE | High | VEEASPPA\ | [K].VEEASF | 1xPhospho | O88508 1x | 3.94E-05 | 0 | 1 | 7 | 4 | O88508 | DNA (cytos | O88508 [2 | 0 | 3853.878 | 0 | -0.9 | 0.010173 | 7.01 | 130.3 | 69.7 | 14.3 | 21.62 | | High | High | High | High | High | High | High | High | High | High | High | 0 | 6.33E-06 | 4.66 |
| FALSE | High | VEEASPPA\ | [K].VEEASF | 2xPhospho | O88508 2x | 1.16E-05 | 0 | 1 | 7 | 6 | O88508 | DNA (cytos | O88508 [2 | 0 | 3933.845 | 0 | -0.69 | 0.013429 | 9.67 | 123.5 | 76.5 | 11.31 | 15.81 | | High | High | High | High | High | High | High | High | High | High | High | 0 | 1.59E-06 | 6.08 |

| | | | | | | | | | | | | | | | | | | | | | | | | | | | | | | | | | | | | | | |
|---|---|---|---|---|---|---|---|---|---|---|---|---|---|---|---|---|---|---|---|---|---|---|---|---|---|---|---|---|---|---|---|---|---|---|---|---|---|---|
| FALSE | High | VEEDGPTE | [K].VEEDG | 1xCarbami | Q3UZ39 1x | 6.72E-09 | 0 | 1 | 2 | 3 | Q3UZ39 | Leucine-ric | Q3UZ39 [6 | 0 | 3010.282 | 0 | 0.95 | 0.00031 | 1.88 | 67.5 | 132.5 | 5.43 | 2.23 | | High | High | High | High | High | High | High | High | High | High | High | 0 | 5.55E-09 | 5.88 |
| FALSE | High | VEEEDTGD | [K].VEEEDT | 1xPhospho | B7ZP47 1xP | 6.54E-09 | 0 | 1 | 2 | 11 | B7ZP47 | Wapal prot | B7ZP47 [64 | 0 | 3139.392 | 0 | -0.8 | 0.000188 | 1.04 | 127.5 | 72.5 | 2.07 | 4.31 | | High | High | High | High | High | High | High | High | High | High | High | 0 | 3.57E-10 | 4.1 |
| FALSE | High | VEEEDTGD | [K].VEEEDT | 2xPhospho | B7ZP47 2xP | 0.002768 | 0.000109 | 1 | 2 | 1 | B7ZP47 | Wapal prot | B7ZP47 [64 | 0 | 3219.358 | 0 | -0.56 | 0.009949 | 5.24 | 120.1 | 79.9 | 17.45 | 11.53 | | High | High | High | High | High | High | High | High | High | High | High | 4.94E-05 | 0.000748 | 2.58 |
| FALSE | High | VEEGAEGD | [K].VEEGAI | 1xCarbami | Q80UU6 1x | 2.85E-09 | 0 | 1 | 5 | 3 | Q80UU6 | Opioid gro | Q80UU6 [4 | 0 | 3897.81 | 0 | 0.36 | 0.031508 | 14.88 | 88.4 | 111.6 | 10.87 | 7.32 | | High | High | High | High | High | High | High | High | High | High | High | 0 | 1.41E-10 | 6.4 |
| FALSE | High | VEELSSGTD | [K].VEELSS | 2xPhospho | G3UYN0 2x | 0.002232 | 0.000109 | 1 | 3 | 6 | G3UYN0 | L1TD1 OS= | G3UYN0 [1 | 0 | 2383.015 | 0 | -1.02 | 0.001778 | 3.65 | 135.9 | 64.1 | 10.8 | 4.8 | | High | High | High | High | High | High | High | High | High | High | High | 0.000183 | 0.001248 | 4.18 |
| FALSE | High | VEELSSGTD | [K].VEELSS | 3xPhospho | G3UYN0 3x | 8.66E-06 | 0 | 1 | 3 | 7 | G3UYN0 | L1TD1 OS= | G3UYN0 [1 | 0 | 2462.982 | 0 | -1.2 | 0.000181 | 0.19 | 140.9 | 59.1 | 3.26 | 4.62 | | High | High | High | High | High | High | High | High | High | High | High | 0 | 1.77E-06 | 3.67 |
| FALSE | High | VEEQVEDS | [R].VEEQVI | 2xPhospho | Q8CGC6 2x | 0.003253 | 0.00016 | 1 | 1 | 2 | Q8CGC6 | RNA-bindir | Q8CGC6 [2 | 1 | 4618.903 | 0 | -0.6 | 0.027415 | 4.04 | 119.6 | 80.4 | 10.79 | 12.46 | | High | High | High | High | High | High | High | High | High | High | High | 0.003521 | 0.03772 | 4.62 |
| FALSE | High | VEESSEISPI | [K].VEESSE | 1xPhospho | Q8BJQ2 1x | 0.000281 | 0 | 1 | 1 | 2 | Q8BJQ2 | Ubiquitin c | Q8BJQ2 [4 | 0 | 1868.929 | 0 | -0.33 | 0.000945 | 2.82 | 112.1 | 87.9 | 3.24 | 3.61 | | High | High | High | High | High | High | High | High | High | High | High | 0 | 5.76E-05 | 2.49 |
| FALSE | High | VEESTGAA | [R].VEESTG | 2xPhospho | Q3UH06-3 | 6.95E-05 | 0 | 1 | 4 | 1 | Q3UH06-3 | Isoform 3 c | Q3UH06-3 | 0 | 3529.555 | 0 | -0.02 | 0.573726 | 27.34 | 105.4 | 94.6 | 16.09 | 11.88 | | High | High | High | High | High | High | High | High | High | High | High | 0 | 1.2E-05 | 4.28 |
| FALSE | High | VEEVEEDSI | [K].VEEVEE | 1xPhospho | B2RRE0 1x | 0.000379 | 0 | 1 | 2 | 1 | B2RRE0 | A kinase (P | B2RRE0 [1( | 0 | 2244.094 | 0 | -1.05 | 0.001152 | 2.86 | 136.7 | 63.3 | 9.64 | 25.6 | | High | High | High | High | High | High | High | High | High | High | High | 0 | 8.02E-05 | 3.48 |
| FALSE | High | VEDPEEGA | [K].VEDPEE | 1xPhospho | Q924N4-2 | 1.76E-08 | 0 | 1 | 1 | 1 | Q924N4-2 | Isoform 2 c | Q924N4-2 | 0 | 2733.327 | 0 | 2.29 | 0.000701 | 14.84 | 32 | 168 | 9.24 | 11.43 | | High | High | High | High | High | High | High | High | High | High | High | 0 | 1.09E-09 | 4.64 |
| FALSE | High | VDSPTVTT | [K].VDSPTV | 1xPhospho | Q7M6Z9 1: | 0.000155 | 0 | 1 | 13 | 2 | Q7M6Z9 | Kinesin ligh | Q7M6Z9 [4 | 0 | 1699.928 | 0 | 1.28 | 0.0012 | 16.6 | 61.2 | 138.8 | 4.96 | 10.29 | | High | High | High | High | High | High | High | High | High | High | High | 0 | 2.94E-05 | 2.64 |
| FALSE | High | VDSPTVNT | [K].VDSPTV | 1xPhospho | Q91YS4 1x | 0.001021 | 0 | 1 | 3 | 1 | Q91YS4 | Kinesin ligh | Q91YS4 [4 | 0 | 1511.767 | 0 | 0.89 | 0.000297 | 2.27 | 69.3 | 130.7 | 6.89 | 4.05 | | High | High | High | High | High | High | High | High | High | High | High | 0 | 0.000244 | 1.6 |
| FALSE | High | VDSPMAYS | [R].VDSPM | 1xPhospho | Q61146 1x | 0.016021 | 0.001124 | 1 | 4 | 1 | Q61146 | Occludin O | Q61146 [3 | 0 | 1809.849 | 0 | 0.55 | 0.050373 | 25.27 | 78.7 | 121.3 | 10.3 | 18.36 | | High | High | High | High | High | High | High | High | High | High | High | 0.000585 | 0.005301 | 1.8 |
| FALSE | High | VCTISPAPR | [R].VCTISP. | 1xCarbami | O70374 1x | 0.013545 | 0.000976 | 1 | 5 | 1 | O70374 | Protein CBI | O70374 [2 | 0 | 1309.654 | 0 | -0.98 | 0.000377 | 2.7 | 132.5 | 67.5 | 3.25 | 7.35 | | High | High | High | High | High | High | High | High | High | High | High | 0.000538 | 0.004415 | 1.31 |
| FALSE | High | VDDDSLGE | [R].VDDDS | 1xPhospho | D3Z5N6 1x | 2.62E-07 | 0 | 1 | 4 | 3 | D3Z5N6 | Zinc finger | D3Z5N6 [1 | 0 | 1945.874 | 0 | 0.19 | 0.025753 | 6.92 | 89.7 | 110.3 | 7.52 | 6.89 | | High | High | High | High | High | High | High | High | High | High | High | 0 | 2.25E-08 | 3.88 |
| FALSE | High | VDDVAYTV | [K].VDDVA | 1xPhospho | Q925P3 1x | 9.27E-08 | 0 | 1 | 8 | 2 | Q925P3 | CEA-relate | Q925P3 [4 | 0 | 3269.579 | 0 | 2.8 | 0.00278 | 21.86 | 25.7 | 174.3 | 31.38 | 20.02 | | High | High | High | High | High | High | High | High | High | High | High | 0 | 7.01E-09 | 3.35 |
| FALSE | High | VDEETSQA | [K].VDEETS | 2xCarbami | Q497V5 1x | 1.8E-05 | 0 | 1 | 2 | 1 | Q497V5 | S1 RNA-bir | Q497V5 [1 | 0 | 3705.666 | 0 | 1 | 0.000413 | 10.21 | 65.6 | 134.4 | 14.87 | 6.12 | | High | High | High | High | High | High | High | High | High | High | High | 0 | 2.62E-06 | 3.27 |
| FALSE | High | VDENENCS | [K].VDENEI | 1xCarbami | Q8VDS4 1x | 9.28E-05 | 0 | 1 | 1 | 3 | Q8VDS4 | Regulation | Q8VDS4 [1 | 0 | 2952.35 | 0 | -0.19 | 0.397589 | 17 | 103.3 | 96.7 | 7.04 | 11.24 | | High | High | High | High | High | High | High | High | High | High | High | 0 | 1.64E-05 | 3.67 |
| FALSE | High | VDGPRSPS | [K].VDGPRI | 3xPhospho | H7BX95 3x | 0.005 | 0.000504 | 1 | 2 | 1 | H7BX95 | Serine/argi | H7BX95 [1 | 2 | 1902.786 | 0 | -0.27 | 0.011079 | 5.76 | 108.1 | 91.9 | 3.24 | 7.5 | | High | High | High | High | High | High | High | High | High | High | High | 0.000225 | 0.001448 | 2.53 |
| FALSE | High | VDHGAEIIT | [R].VDHGA | 1xPhospho | P20357 1xP | 4.96E-07 | 0 | 1 | 5 | 4 | P20357 | Microtubu | P20357 [17 | 0 | 1818.895 | 0 | 1.57 | 0.000213 | 9.08 | 49 | 151 | 7.53 | 5.02 | | High | High | High | High | High | High | High | High | High | High | High | 0 | 4.6E-08 | 4.42 |
| FALSE | High | VDHGAEIIT | [R].VDHGA | 3xPhospho | P20357 3xP | 0.000297 | 0 | 1 | 5 | 5 | P20357 | Microtubu | P20357 [17 | 1 | 2663.183 | 0 | 1.62 | 0.000609 | 10.89 | 50.1 | 149.9 | 6.34 | 9 | | High | High | High | High | High | High | High | High | High | High | High | 0 | 0.00007 | 4 |
| FALSE | High | VDIEAPDVS | [K].VDIEAP | 1xPhospho | E9Q616 1x | 1.04E-05 | 0 | 1 | 3 | 1 | E9Q616 | AHNAK nuc | E9Q616 [1 | 0 | 2193.109 | 0 | 3.56 | 0.000405 | 7.35 | 15 | 185 | 39.38 | 12.86 | | High | High | High | High | High | High | High | High | High | High | High | 0 | 1.4E-06 | 3.41 |
| FALSE | High | VDLETPSLD | [K].VDLETF | 1xPhospho | E9Q616 1x | 1.1E-07 | 0 | 1 | 5 | 2 | E9Q616 | AHNAK nuc | E9Q616 [4 | 0 | 3287.616 | 0 | 3.41 | 0.00185 | 17.84 | 18.6 | 181.4 | 24.7 | 21.51 | | High | High | High | High | High | High | High | High | High | High | High | 0 | 1.37E-06 | 4.84 |
| FALSE | High | VDLETPSLD | [K].VDLETF | 1xPhospho | E9Q616 1x | 1.56E-05 | 0 | 1 | 5 | 2 | E9Q616 | AHNAK nuc | E9Q616 [4 | 0 | 3303.611 | 0 | 4.18 | 0.010133 | 65.87 | 11.3 | 188.7 | 41.56 | 35.61 | | High | High | High | High | High | High | High | High | High | High | High | 0 | 7.68E-06 | 4.48 |
| FALSE | High | VDIITEEMF | [R].VDIITEE | 1xPhospho | O54774 1x | 2.93E-05 | 0 | 1 | 2 | 2 | O54774 | AP-3 comp | O54774 [7 | 0 | 2798.31 | 0 | 0.2 | 0.077437 | 9.71 | 94.5 | 105.5 | 1.89 | 6.98 | | High | High | High | High | High | High | High | High | High | High | High | 0 | 4.53E-06 | 3.11 |
| FALSE | High | VDIITEEMF | [R].VDIITEE | 1xPhospho | O54774 1x | 8.3E-07 | 0 | 1 | 2 | 4 | O54774 | AP-3 comp | O54774 [7 | 0 | 2814.304 | 0 | 0.05 | 0.384448 | 26.82 | 91.9 | 108.1 | 18.1 | 47.48 | | High | High | High | High | High | High | High | High | High | High | High | 0 | 2.8E-07 | 3.81 |
| FALSE | High | VDIITEEMF | [R].VDIITEE | 1xPhospho | O54774 1x | 0.002334 | 0.000109 | 1 | 2 | 1 | O54774 | AP-3 comp | O54774 [7 | 1 | 3655.676 | 0 | 0.03 | 0.868734 | 20.53 | 99 | 101 | 9.14 | 15.75 | | High | High | High | High | High | High | High | High | High | High | High | 4.94E-05 | 0.000617 | 3.37 |
| FALSE | High | VDIITEEMF | [R].VDIITEE | 1xPhospho | O54774 1x | 5.81E-05 | 0 | 1 | 2 | 4 | O54774 | AP-3 comp | O54774 [7 | 1 | 3671.671 | 0 | -0.04 | 0.801776 | 10.69 | 99.8 | 100.2 | 4 | 13.29 | | High | High | High | High | High | High | High | High | High | High | High | 0 | 6.38E-05 | 3.55 |
| FALSE | High | VDLSDEEAI | [R].VDLSDE | 1xPhospho | Q9EQ28 1x | 0.000126 | 0 | 1 | 6 | 2 | Q9EQ28 | DNA polym | Q9EQ28 [3 | 0 | 1543.766 | 0 | -0.73 | 0.000154 | 1.94 | 125.9 | 74.1 | 4.01 | 8.73 | | High | High | High | High | High | High | High | High | High | High | High | 0 | 2.34E-05 | 3.49 |
| FALSE | High | VDLSDGNT | [K].VDLSD( | 1xCarbami | Q3U285-3 | 0.000308 | 0 | 1 | 3 | 1 | Q3U285-3 | Isoform 3 c | Q3U285-3 | 0 | 2312.161 | 0 | 0.49 | 0.01263 | 9.7 | 83 | 117 | 2.75 | 10.58 | | High | High | High | High | High | High | High | High | High | High | High | 0 | 6.36E-05 | 2.94 |
| FALSE | High | VDLTLDPD | [R].VDLTLD | 1xPhospho | Q5NCC3 1x | 0.005 | 0.000504 | 1 | 2 | 1 | Q5NCC3 | E3 ubiquiti | Q5NCC3 [4 | 1 | 2624.365 | 0 | -0.44 | 0.001797 | 3.78 | 114.4 | 85.6 | 9.97 | 8.07 | | High | High | High | High | High | High | High | High | High | High | High | 0.000225 | 0.001441 | 3.13 |
| FALSE | High | VDNARVSP | [K].VDNAR | 1xPhospho | Q6PGL7 1x | 2.7E-06 | 0 | 1 | 5 | 1 | Q6PGL7 | WASH com | Q6PGL7 [7 | 1 | 2651.381 | 0 | -0.26 | 0.038398 | 1.91 | 107.8 | 92.2 | 9.27 | 3.04 | | High | High | High | High | High | High | High | High | High | High | High | 0 | 3.1E-07 | 4.04 |
| FALSE | High | VDNARVSP | [K].VDNAR | 1xDeamida | Q6PGL7 1x | 0.000147 | 0 | 1 | 5 | 1 | Q6PGL7 | WASH com | Q6PGL7 [7 | 1 | 2652.365 | 0 | -0.08 | 0.347322 | 28.54 | 102.1 | 97.9 | 6.65 | 15.44 | | High | High | High | High | High | High | High | High | High | High | High | 0 | 2.76E-05 | 3.42 |
| FALSE | High | VDNDENEI | [K].VDNDE | 1xPhospho | Q5SQB7 1x | 0.038677 | 0.002563 | 2 | 9 | 1 | Q5SQB7; C | MCG68069 | Q5SQB7 [3 | 0 | 1877.859 | 0 | -0.19 | 0.328989 | 4.79 | 108.1 | 91.9 | 5.66 | 9.35 | Shared | High | High | High | High | High | High | High | High | High | High | High | 0.001388 | 0.01441 | 2.1 |
| FALSE | High | VDPSLMED | [K].VDPSLN | 1xPhospho | Q9CQU3 1: | 0.000553 | 0 | 1 | 1 | 3 | Q9CQU3 | Protein REI | Q9CQU3 [8 | 0 | 2457.151 | 0 | 0.4 | 0.004901 | 7.19 | 84.4 | 115.6 | 5.04 | 5.68 | | High | High | High | High | High | High | High | High | High | High | High | 0 | 0.000123 | 3.03 |
| FALSE | High | VDPTYGLE | [K].VDPTY( | 1xCarbami | P97496 1xP | 0.010758 | 0.000772 | 1 | 5 | 1 | P97496 | SWI/SNF c | P97496 [7 | 1 | 3990.902 | 0 | -1.31 | 0.009717 | 0.3 | 143.1 | 56.9 | 22.25 | 9.29 | | High | High | High | High | High | High | High | High | High | High | High | 0.000399 | 0.0034 | 2.97 |
| FALSE | High | VDPVVDVA | [K].VDPVVI | 1xPhospho | P97929 1xP | 0.000328 | 0 | 1 | 3 | 2 | P97929 | Breast canc | P97929 [11 | 0 | 1874.044 | 0 | -0.87 | 0.000952 | 0.87 | 128.3 | 71.7 | 4.38 | 7.49 | | High | High | High | High | High | High | High | High | High | High | High | 0 | 6.8E-05 | 3.67 |
| FALSE | High | VDSDGDA( | [K].VDSDG | 1xCarbami | E9QKD1 1x | 0.000286 | 0 | 1 | 2 | 1 | E9QKD1 | Nucleolar p | E9QKD1 [6 | 0 | 1939.784 | 0 | 0.18 | 0.019054 | 4.49 | 95.7 | 104.3 | 12.27 | 8 | | High | High | High | High | High | High | High | High | High | High | High | 0 | 5.87E-05 | 2.88 |
| FALSE | High | VDSDSK | [R].VDSDSI | 1xPhospho | A2CG76 1x | 0.065952 | 0.004766 | 1 | 5 | 2 | A2CG76 | Histone-lys | A2CG76 [3 | 0 | 1188.591 | 0 | 0.1 | 0.018586 | 2.04 | 95.2 | 104.8 | 4.97 | 4.83 | | High | High | High | High | High | High | High | High | High | High | High | 0.002455 | 0.02652 | 1.53 |
| FALSE | High | VDSDTSNP | [R].VDSDTS | 1xCarbami | Q3TLH4 2x | 0.000458 | 0 | 1 | 5 | 1 | Q3TLH4 | Protein PRI | Q3TLH4 [2 | 0 | 2864.235 | 0 | 0.19 | 0.206961 | 21.82 | 100.4 | 99.6 | 10.59 | 14.78 | | High | High | High | High | High | High | High | High | High | High | High | 0 | 9.97E-05 | 4.02 |
| FALSE | High | VDSEGISPN | [K].VDSEGI | 1xPhospho | P51660 1xP | 0.039011 | 0.002718 | 1 | 3 | 2 | P51660 | Peroxisoma | P51660 [3( | 0 | 1382.651 | 0 | -0.79 | 0.000522 | 3.5 | 125.9 | 74.1 | 3.43 | 4.75 | | High | High | High | High | High | High | High | High | High | High | High | 0.009004 | 0.09614 | 1.12 |
| FALSE | High | VDSEGPST. | [R].VDSEGI | 1xPhospho | Q99LX5 1x | 0.002212 | 0.000109 | 1 | 1 | 1 | Q99LX5 | Multiple m | Q99LX5 [1 | 0 | 1693.8 | 0 | 0 | 0.54051 | 24.66 | 96.5 | 103.5 | 11.86 | 16.86 | | High | High | High | High | High | High | High | High | High | High | High | 4.94E-05 | 0.000582 | 2.24 |
| FALSE | High | VCSIDLEID | [K].VCSIDL | 1xCarbami | Q3TCR9 1x | 4.71E-08 | 0 | 1 | 5 | 5 | Q3TCR9 | Putative ur | Q3TCR9 [1 | 0 | 2808.37 | 0 | -0.31 | 0.700633 | 29 | 110.6 | 89.4 | 13.79 | 32.21 | | High | High | High | High | High | High | High | High | High | High | High | 0 | 3.3E-09 | 5.43 |
| FALSE | High | VGQFGQH | [R].VGQFG | 1xDeamida | Q9EPQ8 1x | 0.005467 | 0.00062 | 1 | 2 | 2 | Q9EPQ8 | Transcripti | Q9EPQ8 [2 | 0 | 2940.3 | 0 | -0.38 | 0.16964 | 18.4 | 113.1 | 86.9 | 22.86 | 23.19 | | High | High | High | High | High | High | High | High | High | High | High | 0.000275 | 0.001603 | 3.94 |
| FALSE | High | TSSSLDDVF | [R].TSSSLD | 1xPhospho | M0QWZ1 1 | 0.052133 | 0.00384 | 1 | 3 | 2 | M0QWZ1 | Protein FAI | M0QWZ1 [ | 0 | 1846.96 | 0 | 0.12 | 0.613391 | 11.13 | 97.3 | 102.7 | 8.44 | 8.65 | | High | High | High | High | High | High | High | High | High | High | High | 0.001999 | 0.02016 | 2.74 |
| FALSE | High | TSSSETEEK | [R].TSSSET | 1xPhospho | B2RRE8 1x | 0.001463 | 5.66E-05 | 1 | 2 | 4 | B2RRE8 | CD2-associ | B2RRE8 [2 | 0 | 1535.724 | 0 | 0.22 | 0.22127 | 26.96 | 92.2 | 107.8 | 4.25 | 23.95 | | High | High | High | High | High | High | High | High | High | High | High | 0.007174 | 0.07763 | 2.02 |
| FALSE | High | TPISTPVSG | [K].TPISTP\ | 2xPhospho | Q9DBJ3 2x | 0.001865 | 5.66E-05 | 1 | 1 | 1 | Q9DBJ3 | Brain-spec | Q9DBJ3 [2 | 0 | 2384.117 | 0 | 0.63 | 0.003474 | 14.09 | 72.3 | 127.7 | 11.77 | 8.41 | | High | High | High | High | High | High | High | High | High | High | High | 2.58E-05 | 0.000482 | 3.09 |
| FALSE | High | TPISTPVSG | [K].TPISTP\ | 2xPhospho | Q9DBJ3 2x | 0.00086 | 0 | 1 | 1 | 1 | Q9DBJ3 | Brain-spec | Q9DBJ3 [2 | 0 | 2400.112 | 0 | 1.32 | 0.0011 | 16.05 | 54.8 | 145.2 | 17.18 | 13.33 | | High | High | High | High | High | High | High | High | High | High | High | 0 | 0.000202 | 2.74 |
| FALSE | High | TPLTQDPA: | [R].TPLTQD | 1xPhospho | Q3UFM6 1 | 6.07E-05 | 0 | 1 | 3 | 1 | Q3UFM6 | Expressed | Q3UFM6 [4 | 0 | 2473.29 | 0 | 0.01 | 0.804012 | 7.63 | 103.7 | 96.3 | 13.2 | 10.05 | | High | High | High | High | High | High | High | High | High | High | High | 0 | 1.02E-05 | 3.3 |
| FALSE | High | TPLTSADEI | [R].TPLTSA | 1xPhospho | Q6A065 1x | 0.004372 | 0.000409 | 1 | 3 | 2 | Q6A065 | Centrosom | Q6A065 [1 | 0 | 2088.053 | 0 | 0.05 | 0.314475 | 12.46 | 97 | 103 | 6.22 | 9.27 | | High | High | High | High | High | High | High | High | High | High | High | 0.000183 | 0.001248 | 2.1 |
| FALSE | High | TPITSVPLG | [K].TPITSVI | 1xPhospho | E9QMD3 1 | 3.41E-05 | 0 | 1 | 2 | 2 | E9QMD3 | Zinc finger | E9QMD3 [4 | 0 | 2204.234 | 0 | 2.5 | 0.000431 | 11 | 30.2 | 169.8 | 5.88 | 8.47 | | High | High | High | High | High | High | High | High | High | High | High | 0 | 2.82E-05 | 4.09 |
| FALSE | High | TPMYGSQT | [R].TPMYG | 1xPhospho | O55201 1x | 0.001021 | 0 | 1 | 2 | 1 | O55201 | Transcripti | O55201 [7 | 0 | 2060.866 | 0 | -0.04 | 0.923376 | 17.23 | 98 | 102 | 24.61 | 12.72 | | High | High | High | High | High | High | High | High | High | High | High | 0 | 0.000243 | 2.74 |
| FALSE | High | TPNITPNTK | [K].TPNITP | 1xPhospho | Q8K298 1x | 0.003464 | 0.00021 | 1 | 2 | 1 | Q8K298 | Anillin OS= | Q8K298 [3 | 0 | 1523.823 | 0 | 0.38 | 0.077039 | 14.58 | 87.1 | 112.9 | 12.16 | 5.43 | | High | High | High | High | High | High | High | High | High | High | High | 9.42E-05 | 0.00096 | 1.56 |
| FALSE | High | TPPAESLSC | [K].TPPAES | 1xPhospho | A2ADB0 1x | 0.001489 | 5.66E-05 | 1 | 5 | 2 | A2ADB0 | Msx2-inter | A2ADB0 [2 | 0 | 2197.079 | 0 | -0.08 | 0.360851 | 10.25 | 102.8 | 97.2 | 5.11 | 6.84 | | High | High | High | High | High | High | High | High | High | High | High | 2.58E-05 | 0.000372 | 2.83 |
| FALSE | High | TPPCSQESI | [K].TPPCSC | 1xCarbami | Q80YR6 1x | 4.47E-05 | 0 | 1 | 3 | 1 | Q80YR6 | DNA endor | Q80YR6 [5 | 0 | 2689.283 | 0 | -1.8 | 0.001128 | 1.84 | 156.3 | 43.7 | 10.36 | 12.96 | | High | High | High | High | High | High | High | High | High | High | High | 0 | 7.27E-06 | 2.41 |
| FALSE | High | TPPDTELH( | [K].TPPDTE | 1xPhospho | Q8VDC1 1x | 7.82E-05 | 0 | 1 | 2 | 2 | Q8VDC1 | FYVE and c | Q8VDC1 [3 | 0 | 2637.394 | 0 | 1.68 | 0.00073 | 16.08 | 48.3 | 151.7 | 13.1 | 9.47 | | High | High | High | High | High | High | High | High | High | High | High | 0 | 1.36E-05 | 2.73 |
| FALSE | High | TPPGSPSSE | [R].TPPGSF | 1xPhospho | Q8C804 1x | 5.67E-06 | 0 | 1 | 2 | 2 | Q8C804 | Spindle anc | Q8C804 [2 | 0 | 2168.052 | 0 | -0.71 | 0.001304 | 2.28 | 127.6 | 72.4 | 6.57 | 9.15 | | High | High | High | High | High | High | High | High | High | High | High | 0 | 7.15E-07 | 3.94 |
| FALSE | High | TPPGSPSSE | [R].TPPGSF | 2xPhospho | Q8C804 2x | 6.79E-06 | 0 | 1 | 2 | 2 | Q8C804 | Spindle anc | Q8C804 [2 | 0 | 2248.019 | 1 | -0.28 | 0.010382 | 1.12 | 109.5 | 90.5 | 4.3 | 6.94 | | High | High | High | High | High | High | High | High | High | High | High | 0 | 8.75E-07 | 4.67 |
| FALSE | High | TPPLAIK | [R].TPPLAII | 1xPhospho | E9Q784 1x | 0.048715 | 0.00354 | 1 | 1 | 2 | E9Q784 | Zinc finger | E9Q784 [1 | 0 | 1048.601 | 0 | -0.65 | 0.001912 | 3.37 | 124.8 | 75.2 | 7.49 | 6.77 | | High | High | High | High | High | High | High | High | High | High | High | 0.001844 | 0.01866 | 1.54 |
| FALSE | High | TPISTPVSG | [K].TPISTP\ | 1xPhospho | Q9DBJ3 1x | 1.93E-08 | 0 | 1 | 1 | 5 | Q9DBJ3 | Brain-spec | Q9DBJ3 [2 | 0 | 2320.146 | 0 | 0.13 | 0.044442 | 3.74 | 95.5 | 104.5 | 1.24 | 6.74 | | High | High | High | High | High | High | High | High | High | High | High | 0 | 1.2E-09 | 4.03 |
| FALSE | High | TPPPEEPG\ | [R].TPPPEE | 1xPhospho | Q8K2W9 1 | 0.105058 | 0.008518 | 1 | 2 | 1 | Q8K2W9 | Uncharacte | Q8K2W9 [ | 0 | 1472.76 | 0 | 0.04 | 0.39518 | 12.15 | 97.8 | 102.2 | 6.9 | 15.74 | | High | High | High | High | High | High | High | High | High | High | High | 0.004291 | 0.04592 | 2.14 |
| FALSE | High | TPPSPGSPL | [R].TPPSPG | 1xPhospho | A0A0G2JFT | 0.000267 | 0 | 1 | 3 | 2 | A0A0G2JFT | Protein RU | A0A0G2JFT | 0 | 2965.548 | 0 | 0.61 | 0.029171 | 3.58 | 80.1 | 119.9 | 17.02 | 10.09 | | High | High | High | High | High | High | High | High | High | High | High | 0 | 0.000336 | 3.16 |
| FALSE | High | TPPSSPPEV | [K].TPPSSP | 1xPhospho | Q8R0F5 1x | 0.000217 | 0 | 1 | 2 | 1 | Q8R0F5 | RNA-bindir | Q8R0F5 [1 | 0 | 2223.047 | 0 | 0.22 | 0.013915 | 7.34 | 91.2 | 108.8 | 6.54 | 10.34 | | High | High | High | High | High | High | High | High | High | High | High | 0 | 4.27E-05 | 3.28 |
| FALSE | High | TPPSTPPAT | [R].TPPSTP | 1xPhospho | F8WJE0 1x | 1.53E-08 | 0 | 1 | 9 | 6 | F8WJE0 | Deoxynucl | F8WJE0 [5 | 0 | 2853.314 | 0 | 0.92 | 0.00057 | 4.52 | 69.5 | 130.5 | 3.61 | 7.3 | | High | High | High | High | High | High | High | High | High | High | High | 0 | 9.26E-10 | 5.34 |
| FALSE | High | TPPSTPPAT | [R].TPPSTP | 2xPhospho | F8WJE0 2x | 5.58E-07 | 0 | 1 | 9 | 4 | F8WJE0 | Deoxynucl | F8WJE0 [5 | 0 | 2933.28 | 0 | 0.5 | 0.007584 | 9.55 | 86.1 | 113.9 | 6.36 | 9.21 | | High | High | High | High | High | High | High | High | High | High | High | 0 | 5.28E-08 | 4.35 |
| FALSE | High | TPPTPPSSI\ | [K].TPPTPP | 1xPhospho | Q3U2C9 1x | 0.019966 | 0.001438 | 1 | 1 | 1 | Q3U2C9 | PHD finger | Q3U2C9 [8 | 0 | 1732.965 | 0 | -0.48 | 0.00093 | 2.82 | 116.5 | 83.5 | 6.61 | 4.13 | | High | High | High | High | High | High | High | High | High | High | High | 0.000786 | 0.006834 | 2.57 |
| FALSE | High | TPPVAVTSI | [K].TPPVA\ | 1xPhospho | Q8CH77 1x | 6.27E-07 | 0 | 1 | 4 | 4 | Q8CH77 | Neuron na | Q8CH77 [5 | 0 | 2457.352 | 0 | 0.47 | 0.021491 | 22.27 | 85.2 | 114.8 | 9.59 | 10.2 | | High | High | High | High | High | High | High | High | High | High | High | 0 | 8.69E-07 | 2.36 |
| FALSE | High | TPPVLIMNI | [K].TPPVLII | 1xPhospho | A2AUZ7 1x | 0.002974 | 0.00016 | 2 | 3 | 2 | A2AUZ7; Q | Sal-like pro | A2AUZ7 [6 | 0 | 2596.22 | 0 | -1.6 | 0.000436 | 1.84 | 150.9 | 49.1 | 6.02 | 11.37 | Shared | High | High | High | High | High | High | High | High | High | High | High | 7.19E-05 | 0.000808 | 4.15 |
| FALSE | High | TPPVLIMNI | [K].TPPVLII | 1xPhospho | A2AUZ7 1x | 5.81E-05 | 0 | 2 | 3 | 1 | A2AUZ7; Q | Sal-like pro | A2AUZ7 [6 | 0 | 2612.215 | 0 | -1.16 | 0.003596 | 6.58 | 138.1 | 61.9 | 5.01 | 24.23 | Shared | High | High | High | High | High | High | High | High | High | High | High | 0 | 9.77E-06 | 3.27 |
| FALSE | High | TPPVLPAPL | [K].TPPVLP | 1xPhospho | Q80U49 1x | 0.104222 | 0.008348 | 1 | 2 | 1 | Q80U49 | Centrosom | Q80U49 [5 | 0 | 1567.881 | 0 | 1.94 | 0.001212 | 12.25 | 45.3 | 154.7 | 14.82 | 9.52 | | High | High | High | High | High | High | High | High | High | High | High | 0.004264 | 0.04553 | 2.27 |
| FALSE | High | TPPVLSPVF | [K].TPPVLS | 1xPhospho | A2ASY1 1x | 0.074685 | 0.005727 | 1 | 5 | 1 | A2ASY1 | TAF3 RNA | A2ASY1 [2 | 0 | 1274.707 | 0 | -0.46 | 0.008095 | 3.45 | 116.3 | 83.7 | 7.97 | 5.79 | | High | High | High | High | High | High | High | High | High | High | High | 0.002937 | 0.0307 | 1.34 |
| FALSE | High | TPQAPTPA | [R].TPQAP1 | 1xPhospho | Q8BTI8 1xP | 3.41E-05 | 0 | 1 | 3 | 2 | Q8BTI8 | Serine/argi | Q8BTI8 [22 | 0 | 1826.973 | 0 | -0.81 | 0.002492 | 4.06 | 127.5 | 72.5 | 7.31 | 5.05 | | High | High | High | High | High | High | High | High | High | High | High | 0 | 5.39E-06 | 3.06 |
| FALSE | High | TPQASTYS\ | [K].TPQAST | 1xPhospho | P14873 1xP | 0.083101 | 0.006345 | 2 | 2 | 1 | P14873; B2 | Microtubu | P14873 [2( | 0 | 1914.832 | 0 | -0.25 | 0.039129 | 5.95 | 110.7 | 89.3 | 11.5 | 14.86 | Shared | High | High | High | High | High | High | High | High | High | High | High | 0.003265 | 0.03485 | 2.15 |
| FALSE | High | TPQEAIMD | [R].TPQEAI | 1xPhospho | A0A068BF | 1.16E-06 | 0 | 1 | 3 | 1 | A0A068BF | Ring finger | A0A068BF | 0 | 2124.025 | 0 | 0.2 | 0.235563 | 8.09 | 95.7 | 104.3 | 8.55 | 13.93 | | High | High | High | High | High | High | High | High | High | High | High | 0 | 1.2E-07 | 3.44 |
| FALSE | High | TPPPSYNS/ | [K].TPPPSY | 1xPhospho | Q8BXW8 1 | 0.000681 | 0 | 1 | 2 | 6 | Q8BXW8 | Putative ur | Q8BXW8 [ | 0 | 2885.323 | 0 | 0.14 | 0.7185 | 21.83 | 95 | 105 | 5.01 | 16.47 | | High | High | High | High | High | High | High | High | High | High | High | 0.000538 | 0.004629 | 3.8 |
| FALSE | High | TPQEAIMD | [R].TPQEAI | 1xPhospho | A0A068BF | 0.001313 | 0 | 1 | 3 | 1 | A0A068BF | Ring finger | A0A068BF | 0 | 2140.019 | 0 | -0.68 | 0.000945 | 2.97 | 122.3 | 77.7 | 6.52 | 10.4 | | High | High | High | High | High | High | High | High | High | High | High | 0 | 0.000323 | 2.14 |
| FALSE | High | TPISTPVSG | [K].TPISTP\ | 1xPhospho | Q9DBJ3 1x | 6.55E-06 | 0 | 1 | 1 | 1 | Q9DBJ3 | Brain-spec | Q9DBJ3 [2 | 0 | 2304.151 | 0 | 1.08 | 0.002397 | 18.25 | 65.4 | 134.6 | 11.37 | 8.27 | | High | High | High | High | High | High | High | High | High | High | High | 0 | 8.38E-07 | 3.03 |
| FALSE | High | TPLSPSPMI | [R].TPLSPS | 1xPhospho | A0A0R4J0J | 0.020679 | 0.001472 | 1 | 3 | 2 | A0A0R4J0J | Cytospin-A | A0A0R4J0J | 0 | 1511.794 | 0 | 1.54 | 0.007618 | 27.02 | 51.2 | 148.8 | 13.73 | 21.6 | | High | High | High | High | High | High | High | High | High | High | High | 0.000786 | 0.007071 | 2.11 |
| FALSE | High | TPGNCEPP | [R].TPGNCI | 1xCarbami | B2RWU4 1 | 0.013545 | 0.000976 | 2 | 6 | 1 | B2RWU4; ( | Cordon-ble | B2RWU4 [ | 0 | 1629.729 | 0 | -0.44 | 0.675404 | 8.86 | 116.3 | 83.7 | 29.96 | 46.53 | Shared | High | High | High | High | High | High | High | High | High | High | High | 0.000538 | 0.004423 | 3 |
| FALSE | High | TPGNCEPP | [R].TPGNCI | 1xCarbami | B2RWU4 1 | 2.08E-05 | 0 | 2 | 6 | 2 | B2RWU4; ( | Cordon-ble | B2RWU4 [ | 0 | 1858.892 | 0 | -0.03 | 0.930463 | 6.34 | 100.6 | 99.4 | 5.35 | 5.93 | Shared | High | High | High | High | High | High | High | High | High | High | High | 0 | 3.09E-06 | 2.85 |
| FALSE | High | TPGNCEPP | [R].TPGNCI | 1xCarbami | B2RWU4 1 | 1.78E-05 | 0 | 2 | 6 | 2 | B2RWU4; ( | Cordon-ble | B2RWU4 [ | 0 | 1859.876 | 0 | 0.08 | 0.249842 | 6.87 | 99.3 | 100.7 | 15.43 | 18.4 | Shared | High | High | High | High | High | High | High | High | High | High | High | 0 | 2.58E-06 | 3.02 |
| FALSE | High | TPGNSAGA | [R].TPGNS/ | 1xPhospho | Q3TU15 1x | 0.0002 | 0 | 2 | 4 | 1 | Q3TU15; C | Non-specif | Q3TU15 [4 | 0 | 2253.096 | 0 | -0.28 | 0.030715 | 9.78 | 111.6 | 88.4 | 5.82 | 9.49 | Shared | High | High | High | High | High | High | High | High | High | High | High | 0 | 3.9E-05 | 2.86 |
| FALSE | High | TPGNSAGA | [R].TPGNS/ | 1xPhospho | Q3TU15 1x | 7.21E-05 | 0 | 2 | 4 | 1 | Q3TU15; C | Non-specif | Q3TU15 [4 | 1 | 2409.197 | 0 | -0.79 | 0.001117 | 2.7 | 125.7 | 74.3 | 6.55 | 5.44 | Shared | High | High | High | High | High | High | High | High | High | High | High | 0 | 1.24E-05 | 1.97 |
| FALSE | High | TPGPPSSQ | [R].TPGPPS | 1xPhospho | D3Z656 1x | 2.58E-05 | 0 | 1 | 4 | 3 | D3Z656 | Synaptojar | D3Z656 [1 | 0 | 2488.249 | 0 | -0.07 | 0.7364 | 4.99 | 103.5 | 96.5 | 5.11 | 16.87 | | High | High | High | High | High | High | High | High | High | High | High | 0 | 3.93E-06 | 3.79 |
| FALSE | High | TPGRPLSSY | [R].TPGRPL | 1xPhospho | P31938 1xP | 0.000892 | 0 | 1 | 4 | 4 | P31938 | Dual specif | P31938 [2 | 0 | 4226.147 | 0 | -0.23 | 0.036462 | 5.17 | 103.3 | 96.7 | 9.21 | 16.03 | | High | High | High | High | High | High | High | High | High | High | High | 4.94E-05 | 0.000736 | 3.39 |
| FALSE | High | TPGTPGTP: | [R].TPGTP( | 1xPhospho | P20357 1xP | 0.004613 | 0.000456 | 1 | 8 | 3 | P20357 | Microtubu | P20357 [16 | 0 | 1539.741 | 0 | 1.55 | 0.00024 | 6.28 | 49.3 | 150.7 | 5.18 | 4.42 | | High | High | High | High | High | High | High | High | High | High | High | 0.002748 | 0.02939 | 2.65 |
| FALSE | High | TPGTPGTP: | [R].TPGTP( | 2xPhospho | P20357 2xP | 0.01894 | 0.001362 | 1 | 8 | 1 | P20357 | Microtubu | P20357 [16 | 0 | 1619.707 | 0 | 0.95 | 0.005561 | 21.56 | 71.5 | 128.5 | 10.34 | 14.49 | | High | High | High | High | High | High | High | High | High | High | High | 0.00074 | 0.006444 | 2.88 |
| FALSE | High | TPGVLLPG\ | [R].TPGVLL | 1xPhospho | Q8CHI8 1xP | 9.48E-06 | 0 | 1 | 7 | 3 | Q8CHI8 | E1A-bindin | Q8CHI8 [2 | 0 | 2986.495 | 0 | -0.65 | 0.005434 | 1.39 | 122.7 | 77.3 | 4.55 | 12.08 | | High | High | High | High | High | High | High | High | High | High | High | 0 | 1.27E-06 | 4.99 |
| FALSE | High | TPGVLLPG\ | [R].TPGVLL | 2xPhospho | Q8CHI8 2xP | 0.029586 | 0.001857 | 1 | 7 | 2 | Q8CHI8 | E1A-bindin | Q8CHI8 [2 | 0 | 3050.466 | 0 | -0.25 | 0.332063 | 29.81 | 106.3 | 93.7 | 12.67 | 27.88 | | High | High | High | High | High | High | High | High | High | High | High | 0.000997 | 0.01064 | 2.99 |
| FALSE | High | TPGVLLPG\ | [R].TPGVLL | 2xPhospho | Q8CHI8 2xP | 0.037692 | 0.00249 | 1 | 7 | 1 | Q8CHI8 | E1A-bindin | Q8CHI8 [2 | 0 | 3066.461 | 0 | -0.09 | 0.332076 | 9.77 | 102 | 98 | 4.2 | 17.95 | | High | High | High | High | High | High | High | High | High | High | High | 0.001308 | 0.01392 | 3.13 |
| FALSE | High | TPHAVLTP\ | [K].TPHAVI | 1xPhospho | Q80U93 1x | 8.03E-05 | 0 | 1 | 2 | 1 | Q80U93 | Nuclear po | Q80U93 [1 | 0 | 2072.131 | 0 | -0.54 | 0.038101 | 4.27 | 112.5 | 87.5 | 11.36 | 8.43 | | High | High | High | High | High | High | High | High | High | High | High | 0 | 1.41E-05 | 2.65 |
| FALSE | High | TPLSTGGTI | [K].TPLSTG | 1xPhospho | E9QMN5 1 | 7.86E-07 | 0 | 1 | 3 | 1 | E9QMN5 | Transcripti | E9QMN5 [ | 0 | 2521.383 | 0 | -0.97 | 0.008196 | 5.35 | 132 | 68 | 11.04 | 13.45 | | High | High | High | High | High | High | High | High | High | High | High | 0 | 7.78E-08 | 4.76 |
| FALSE | High | TPHYGSQT | [R].TPHYGS | 1xPhospho | O55201 1x | 0.000422 | 0 | 1 | 2 | 3 | O55201 | Transcripti | O55201 [7 | 0 | 1961.907 | 0 | -0.83 | 0.043276 | 7.08 | 128.1 | 71.9 | 11.36 | 23.59 | | High | High | High | High | High | High | High | High | High | High | High | 0 | 0.000148 | 3.04 |
| FALSE | High | TPLAALGG | [R].TPLAAL | 1xPhospho | B1AS46 1xP | 2.1E-05 | 0 | 1 | 3 | 1 | B1AS46 | Zinc finger | B1AS46 [1 | 0 | 1854.956 | 0 | -1.04 | 0.001307 | 4.63 | 135.2 | 64.8 | 8.88 | 17.2 | | High | High | High | High | High | High | High | High | High | High | High | 0 | 3.11E-06 | 5.06 |
| FALSE | High | TPLATIADC | [R].TPLATI/ | 1xPhospho | P11157 1xP | 1.39E-06 | 0 | 1 | 2 | 2 | P11157 | Ribonuclec | P11157 [6- | 0 | 2274.231 | 0 | 0.08 | 0.848784 | 31.09 | 106 | 94 | 71.78 | 82.52 | | High | High | High | High | High | High | High | High | High | High | High | 0 | 1.47E-07 | 4.26 |
| FALSE | High | TPLATIADC | [R].TPLATI/ | 1xPhospho | P11157 1xP | 6.87E-07 | 0 | 1 | 2 | 4 | P11157 | Ribonuclec | P11157 [6- | 0 | 2503.394 | 0 | -0.34 | 0.020886 | 4.87 | 111.4 | 88.6 | 3.6 | 7.69 | | High | High | High | High | High | High | High | High | High | High | High | 0 | 6.68E-08 | 3.51 |
| FALSE | High | TPLATIADC | [R].TPLATI/ | 1xDeamida | P11157 1xP | 0.006085 | 0.00062 | 1 | 2 | 2 | P11157 | Ribonuclec | P11157 [6- | 0 | 2504.378 | 0 | -0.14 | 0.575369 | 18.11 | 106 | 94 | 12.93 | 11.87 | | High | High | High | High | High | High | High | High | High | High | High | 0.000275 | 0.001793 | 2.42 |
| FALSE | High | TPLENVPG | [K].TPLENV | 1xPhospho | Q9EPN1 1x | 0.009087 | 0.000736 | 1 | 3 | 1 | Q9EPN1 | Neurobeac | Q9EPN1 [1 | 0 | 2017.114 | 0 | 0.89 | 0.010409 | 26.18 | 69.1 | 130.9 | 13.3 | 12.34 | | High | High | High | High | High | High | High | High | High | High | High | 0.000327 | 0.002813 | 2.28 |
| FALSE | High | TPLGASLDI | [R].TPLGAS | 1xPhospho | Q922B9 1x | 0.007673 | 0.000664 | 1 | 4 | 1 | Q922B9 | Sperm-spe | Q922B9 [8 | 0 | 2126.078 | 0 | 0.6 | 0.004077 | 2.68 | 79.9 | 120.1 | 5.85 | 6.29 | | High | High | High | High | High | High | High | High | High | High | High | 0.000294 | 0.002332 | 2.99 |
| FALSE | High | TPLLSSLSA | [R].TPLLSSI | 1xCarbami | Q01279 1x | 0.004613 | 0.000456 | 1 | 3 | 3 | Q01279 | Epidermal | Q01279 [1( | 0 | 2515.242 | 0 | 0.79 | 0.002936 | 10.46 | 73.7 | 126.3 | 14.65 | 4.49 | | High | High | High | High | High | High | High | High | High | High | High | 0.000204 | 0.001326 | 3.13 |
| FALSE | High | TPIQAGGY( | [K].TPIQAG | 1xPhospho | P28028 1xP | 0.011549 | 0.00086 | 1 | 1 | 1 | P28028 | Serine/thre | P28028 [7 | 0 | 2095.067 | 0 | -0.17 | 0.024952 | 8.55 | 108.2 | 91.8 | 5.16 | 8.66 | | High | High | High | High | High | High | High | High | High | High | High | 0.000437 | 0.003675 | 2.1 |
| FALSE | High | TPIQVESSP | [R].TPIQVE | 1xPhospho | Q99M54 1: | 0.002077 | 5.66E-05 | 1 | 5 | 1 | Q99M54 | Cell divisio | Q99M54 [3 | 0 | 2871.527 | 0 | -0.49 | 0.004563 | 5.7 | 116.9 | 83.1 | 9.02 | 6.7 | | High | High | High | High | High | High | High | High | High | High | High | 2.58E-05 | 0.000541 | 3.08 |
| FALSE | High | TPIQVESSP | [R].TPIQVE | 1xDeamida | Q99M54 1: | 9.36E-05 | 0 | 1 | 5 | 3 | Q99M54 | Cell divisio | Q99M54 [3 | 0 | 2872.511 | 0 | -0.36 | 0.010246 | 4.41 | 119.1 | 80.9 | 18.95 | 17.57 | | High | High | High | High | High | High | High | High | High | High | High | 0 | 1.67E-05 | 2.91 |
| FALSE | High | TPIQVESSP | [R].TPIQVE | 2xPhospho | Q99M54 2: | 0.000178 | 0 | 1 | 5 | 1 | Q99M54 | Cell divisio | Q99M54 [3 | 0 | 2951.493 | 0 | -0.25 | 0.219315 | 27.2 | 119.2 | 80.8 | 16.51 | 25.8 | | High | High | High | High | High | High | High | High | High | High | High | 0 | 3.44E-05 | 2.89 |
| FALSE | High | TPLSEAEFE | [R].TPLSEA | 1xPhospho | H3BJW3 1x | 0.023175 | 0.001578 | 1 | 4 | 1 | H3BJW3 | Cleavage a | H3BJW3 [4 | 0 | 1974.908 | 0 | -1.72 | 0.000447 | 2.02 | 154.6 | 45.4 | 4.25 | 29.2 | | High | High | High | High | High | High | High | High | High | High | High | 0.000847 | 0.008054 | 2.18 |
| FALSE | High | TPISLSPQID | [K].TPISLSF | 2xCarbami | Q5SVT3 1x | 0.006594 | 0.00062 | 1 | 3 | 3 | Q5SVT3 | Ewing's tur | Q5SVT3 [3 | 0 | 3179.65 | 0 | -1.31 | 0.00409 | 8.67 | 143 | 57 | 12.72 | 33.68 | | High | High | High | High | High | High | High | High | High | High | High | 0.000275 | 0.002033 | 3.55 |
| FALSE | High | TPKPPTDDI | [K].TPKPPT | 1xPhospho | Q6P1G2 1x | 0.002252 | 0.000109 | 1 | 7 | 4 | Q6P1G2 | Lysine-spe | Q6P1G2 [4 | 0 | 3363.645 | 0 | -0.06 | 0.361792 | 0.36 | 102.2 | 97.8 | 7.05 | 7.4 | | High | High | High | High | High | High | High | High | High | High | High | 0.009466 | 0.1024 | 3.27 |
| FALSE | High | TPQEAITD( | [R].TPQEAI | 1xPhospho | Q9CQJ4 1x | 2.15E-08 | 0 | 1 | 3 | 5 | Q9CQJ4 | E3 ubiquiti | Q9CQJ4 [2 | 0 | 2134.099 | 0 | -0.24 | 0.0162 | 6.23 | 107 | 93 | 2.93 | 4.88 | | High | High | High | High | High | High | High | High | High | High | High | 0 | 1.36E-09 | 4.95 |
| FALSE | High | TPQNVPAS | [K].TPQNV | 1xPhospho | F8VPJ6 1xP | 0.065405 | 0.004734 | 1 | 1 | 1 | F8VPJ6 | Regulatory | F8VPJ6 [41 | 0 | 1604.763 | 0 | -0.36 | 0.018819 | 4.42 | 112.3 | 87.7 | 4.13 | 7.78 | | High | High | High | High | High | High | High | High | High | High | High | 0.002446 | 0.02628 | 1.75 |

| | | | | | | | | | | | | | | | | | | | | | | | | | | | | | | | | | | | | | | |
|---|---|---|---|---|---|---|---|---|---|---|---|---|---|---|---|---|---|---|---|---|---|---|---|---|---|---|---|---|---|---|---|---|---|---|---|---|---|---|
| FALSE | High | TPQPCSRP | [R].TPQPCS | 1xCarbami | A0A087WC | 0.008769 | 0.000736 | 1 | 9 | 1 | A0A087WC | Kinesin-like | A0A087WC | 0 | 3066.466 | 0 | -1.01 | 0.005259 | 4.21 | 132.9 | 67.1 | 17.31 | 14.35 | | High | High | High | High | High | High | High | High | High | High | High | 0.000327 | 0.002708 | 2.99 |
| FALSE | High | TPTCQSSTI | [R].TPTCQS | 1xCarbami | P97329 1xI | 0.001058 | 0 | 1 | 1 | 3 | P97329 | Kinesin-like | P97329 [85 | 0 | 1966.842 | 0 | 0.21 | 0.287599 | 8.02 | 93.9 | 106.1 | 16.41 | 8.65 | | High | High | High | High | High | High | High | High | High | High | High | 0 | 0.000253 | 1.66 |
| FALSE | High | TPTDLLQSS | [K].TPTDLL | 1xPhospho | Q8BT14-3 | 0.069903 | 0.005181 | 1 | 4 | 2 | Q8BT14-3 | Isoform 3 o | Q8BT14-3 | 0 | 1755.873 | 0 | 0.17 | 0.099156 | 12.67 | 95.4 | 104.6 | 3.26 | 9.69 | | High | High | High | High | High | High | High | High | High | High | High | 0.002675 | 0.0285 | 1.94 |
| FALSE | High | TPTGLVGA | [K].TPTGLV | 1xPhospho | B1AVH7 1x | 3.62E-07 | 0 | 1 | 2 | 1 | B1AVH7 | TBC1 doma | B1AVH7 [1 | 0 | 3222.827 | 0 | 2 | 0.004833 | 49.42 | 39.7 | 160.3 | 20.89 | 19.14 | | High | High | High | High | High | High | High | High | High | High | High | 0 | 3.26E-08 | 4.52 |
| FALSE | High | TPTGNAPS | [R].TPTGNA | 1xPhospho | Q5SVQ0 1x | 9.06E-06 | 0 | 1 | 6 | 3 | Q5SVQ0 | Histone ac | Q5SVQ0 [1 | 0 | 3393.61 | 0 | -0.67 | 0.012398 | 6.61 | 119.7 | 80.3 | 4.68 | 20.7 | | High | High | High | High | High | High | High | High | High | High | High | 0 | 1.21E-06 | 4.97 |
| FALSE | High | TPTGPDLD | [K].TPTGPD | 1xPhospho | A0A1L1STC | 0.061176 | 0.004646 | 1 | 7 | 1 | A0A1L1STC | Nesprin-1 ( | A0A1L1STC | 0 | 1832.908 | 0 | 2.33 | 0.000439 | 19.43 | 32.3 | 167.7 | 24.09 | 10.57 | | High | High | High | High | High | High | High | High | High | High | High | 0.002381 | 0.0244 | 2.38 |
| FALSE | High | TPTGSPATI | [K].TPTGSP | 1xPhospho | Q6P1G2 1x | 0.00195 | 5.66E-05 | 1 | 7 | 1 | Q6P1G2 | Lysine-spe | Q6P1G2 [4 | 0 | 1813.935 | 0 | 0.16 | 0.208941 | 4.31 | 96.6 | 103.4 | 7.18 | 4.36 | | High | High | High | High | High | High | High | High | High | High | High | 2.58E-05 | 0.000504 | 2.24 |
| FALSE | High | TPTISPVITA | [K].TPTISPV | 1xPhospho | B2KG46 1x | 0.002623 | 0.000109 | 1 | 6 | 2 | B2KG46 | Protein bic | B2KG46 [5 | 0 | 2673.395 | 0 | 0.9 | 0.005136 | 10.37 | 74.1 | 125.9 | 20.19 | 6.46 | | High | High | High | High | High | High | High | High | High | High | High | 0.000161 | 0.001202 | 2.86 |
| FALSE | High | TPTPEPAEV | [K].TPTPEP | 1xPhospho | D3YUV1 1x | 0.005978 | 0.00062 | 1 | 3 | 2 | D3YUV1 | Nuclear re | D3YUV1 [4 | 0 | 1635.783 | 0 | 0.16 | 0.060133 | 9.21 | 96.1 | 103.9 | 2.51 | 5.73 | | High | High | High | High | High | High | High | High | High | High | High | 0.0015 | 0.01493 | 2.02 |
| FALSE | High | TPTPSTPGI | [K].TPTPST | 1xPhospho | Q6PDQ2 1x | 4.65E-06 | 0 | 1 | 5 | 13 | Q6PDQ2 | Chromodo | Q6PDQ2 [1 | 0 | 3123.565 | 0 | 0.02 | 0.815753 | 10.5 | 102.4 | 97.6 | 6.27 | 7.97 | | High | High | High | High | High | High | High | High | High | High | High | 0 | 5.73E-07 | 4.21 |
| FALSE | High | TPTPSTPGI | [K].TPTPST | 2xPhospho | Q6PDQ2 2x | 1.27E-07 | 0 | 1 | 5 | 8 | Q6PDQ2 | Chromodo | Q6PDQ2 [1 | 0 | 3203.531 | 0 | 0.35 | 0.022785 | 21.18 | 87.1 | 112.9 | 4.38 | 11.49 | | High | High | High | High | High | High | High | High | High | High | High | 0 | 9.97E-09 | 4.08 |
| FALSE | High | TPTPSVETL | [K].TPTPSV | 1xPhospho | A0A0A0MC | 0.024855 | 0.001645 | 1 | 2 | 1 | A0A0A0MC | Calmodulin | A0A0A0MC | 0 | 1721.867 | 0 | -0.76 | 0.002142 | 5.11 | 123.7 | 76.3 | 10.95 | 13.95 | | High | High | High | High | High | High | High | High | High | High | High | 0.000872 | 0.008708 | 1.47 |
| FALSE | High | TPTSASPG | [K].TPTSAS | 1xPhospho | Q91VE6 1x | 0.000203 | 0 | 1 | 2 | 1 | Q91VE6 | MKI67 FHA | Q91VE6 [2 | 0 | 1440.75 | 0 | -0.91 | 0.000328 | 2.44 | 129.2 | 70.8 | 3.54 | 11.14 | | High | High | High | High | High | High | High | High | High | High | High | 0 | 3.99E-05 | 1.93 |
| FALSE | High | TPTSGDFSI | [K].TPTSGD | 1xPhospho | Q8CAF4-3 | 0.002553 | 0.000109 | 1 | 4 | 6 | Q8CAF4-3 | Isoform 3 o | Q8CAF4-3 | 0 | 2574.228 | 0 | 1.53 | 0.001041 | 7.04 | 52.3 | 147.7 | 5.56 | 10.69 | | High | High | High | High | High | High | High | High | High | High | High | 0.000859 | 0.008607 | 2.78 |
| FALSE | High | TPSYQLSPN | [K].TPSYQL | 1xPhospho | Q9R207 1x | 0.00083 | 0 | 1 | 2 | 2 | Q9R207 | Nibrin OS= | Q9R207 [4 | 0 | 1705.864 | 0 | -0.36 | 0.037399 | 0.63 | 109.6 | 90.4 | 5.99 | 11.18 | | High | High | High | High | High | High | High | High | High | High | High | 0 | 0.000194 | 1.99 |
| FALSE | High | TPTSPLK | [K].TPTSPL | 1xPhospho | Q91ZW3 1 | 0.081085 | 0.006219 | 1 | 1 | 4 | Q91ZW3 | SWI/SNF-r | Q91ZW3 [1 | 0 | 1281.722 | 0 | -0.55 | 0.000648 | 1.68 | 119.2 | 80.8 | 5.99 | 6.29 | | High | High | High | High | High | High | High | High | High | High | High | 0.003188 | 0.03361 | 1.79 |
| FALSE | High | TPTSTPVHI | [R].TPTSTP | 1xPhospho | Q3TUF7 1x | 0.004572 | 0.000456 | 1 | 4 | 2 | Q3TUF7 | YEATS dom | Q3TUF7 [4 | 0 | 1618.897 | 0 | -0.61 | 0.009564 | 8.21 | 117.1 | 82.9 | 9.89 | 9.89 | | High | High | High | High | High | High | High | High | High | High | High | 0.000204 | 0.001312 | 1.57 |
| FALSE | High | TPTTPGLG | [R].TPTTPG | 1xPhospho | Q8C180 1x | 9.27E-08 | 0 | 1 | 1 | 5 | Q8C180 | Fibroblast | Q8C180 [1 | 0 | 2163.08 | 0 | 0.46 | 0.008233 | 13.09 | 82.8 | 117.2 | 6.32 | 9.34 | | High | High | High | High | High | High | High | High | High | High | High | 0 | 1.23E-08 | 4.99 |
| FALSE | High | TPTTPGLG | [R].TPTTPG | 2xPhospho | Q8C180 2x | 3.01E-06 | 0 | 1 | 1 | 1 | Q8C180 | Fibroblast | Q8C180 [1 | 0 | 2243.046 | 0 | 0.41 | 0.027544 | 12.73 | 83.6 | 116.4 | 16.07 | 9.12 | | High | High | High | High | High | High | High | High | High | High | High | 0 | 3.5E-07 | 3.79 |
| FALSE | High | TPTVDVTV | [K].TPTVD\ | 1xPhospho | E9Q616 1x | 0.000191 | 0 | 1 | 5 | 2 | E9Q616 | AHNAK nu | E9Q616 [4 | 0 | 3132.612 | 0 | 3.15 | 0.000308 | 0.76 | 19.4 | 180.6 | 7.25 | 8.86 | | High | High | High | High | High | High | High | High | High | High | High | 2.58E-05 | 0.000417 | 3.07 |
| FALSE | High | TPVAPGPP | [K].TPVAP( | 1xPhospho | Q3U3X7 1x | 9.89E-05 | 0 | 1 | 2 | 1 | Q3U3X7 | Putative ur | Q3U3X7 [3 | 0 | 2071.078 | 0 | -0.69 | 0.002153 | 2.04 | 120.9 | 79.1 | 7.68 | 10.05 | | High | High | High | High | High | High | High | High | High | High | High | 0 | 1.77E-05 | 2.97 |
| FALSE | High | TPVASECT | [R].TPVASE | 2xCarbami | Q99PU7 1x | 4.79E-07 | 0 | 1 | 1 | 6 | Q99PU7 | Ubiquitin c | Q99PU7 [3 | 0 | 3768.713 | 0 | -0.48 | 0.001437 | 0.99 | 119.6 | 80.4 | 5.16 | 10.79 | | High | High | High | High | High | High | High | High | High | High | High | 0 | 4.72E-08 | 4.74 |
| FALSE | High | TPVASTHSI | [R].TPVAST | 1xPhospho | Q03141 1x | 0.005135 | 0.000551 | 1 | 5 | 4 | Q03141 | MAP/micro | Q03141 [5 | 0 | 2036.985 | 0 | -0.77 | 0.011552 | 4.37 | 121.3 | 78.7 | 14.16 | 13.28 | | High | High | High | High | High | High | High | High | High | High | High | 0.000246 | 0.001483 | 2.56 |
| FALSE | High | TPVATPPP | [R].TPVATF | 1xPhospho | E9PV87 1xI | 2.03E-07 | 0 | 1 | 5 | 5 | E9PV87 | Protein TA | E9PV87 [1 | 0 | 2816.51 | 0 | 0.39 | 0.004721 | 11.85 | 83.5 | 116.5 | 11.26 | 3.69 | | High | High | High | High | High | High | High | High | High | High | High | 0 | 9.49E-07 | 4.1 |
| FALSE | High | TPVDYIDLP | [K].TPVDYI | 1xPhospho | A0A1L7NR | 0.003224 | 0.00016 | 2 | 12 | 5 | A0A1L7NR | C-Cbl asso | A0A1L7NR | 0 | 2105.999 | 0 | 1.4 | 0.000693 | 17.31 | 53.3 | 146.7 | 11.75 | 5.92 | Shared | High | High | High | High | High | High | High | High | High | High | High | 0.000786 | 0.006903 | 3.78 |
| FALSE | High | TPVGAEQT | [R].TPVGAI | 1xPhospho | E9QAE4 1x | 4.72E-05 | 0 | 1 | 2 | 1 | E9QAE4 | Histone-lys | E9QAE4 [2 | 0 | 2438.31 | 0 | -0.15 | 0.061125 | 7.89 | 107.2 | 92.8 | 8.66 | 10.34 | | High | High | High | High | High | High | High | High | High | High | High | 0 | 7.76E-06 | 2.84 |
| FALSE | High | TPVQSQQF | [K].TPVQS( | 1xPhospho | Q3TI61 1xF | 1.94E-07 | 0 | 1 | 8 | 6 | Q3TI61 | Putative ur | Q3TI61 [9- | 0 | 2467.212 | 0 | 0.06 | 0.851184 | 8.66 | 99.8 | 100.2 | 6.87 | 5.33 | | High | High | High | High | High | High | High | High | High | High | High | 0 | 6.22E-06 | 3.83 |
| FALSE | High | TPVSSTLPV | [K].TPVSST | 1xPhospho | P0C090 1xI | 0.000738 | 0 | 1 | 4 | 2 | P0C090 | Roquin-2 C | P0C090 [78 | 0 | 3037.548 | 0 | -0.23 | 0.093088 | 14.59 | 106.6 | 93.4 | 9.08 | 12.18 | | High | High | High | High | High | High | High | High | High | High | High | 0 | 0.000169 | 3.22 |
| FALSE | High | TQAAGPSS | [K].TQAAG | 1xPhospho | Q3TN28 1x | 0.007145 | 0.00062 | 1 | 4 | 2 | Q3TN28 | Putative ur | Q3TN28 [4 | 0 | 2127.136 | 0 | -0.47 | 0.010811 | 7.83 | 116 | 84 | 4.13 | 10.32 | | High | High | High | High | High | High | High | High | High | High | High | 0.000275 | 0.002147 | 3.41 |
| FALSE | High | TPTSPLSPS | [K].TPTSPL | 2xPhospho | E9Q309 2x | 0.001476 | 5.66E-05 | 1 | 1 | 1 | E9Q309 | Centrosom | E9Q309 [1 | 0 | 1847.896 | 0 | -0.06 | 0.106574 | 1.02 | 103.5 | 96.5 | 6.77 | 4.3 | | High | High | High | High | High | High | High | High | High | High | High | 2.58E-05 | 0.00037 | 2.95 |
| FALSE | High | TPSYQLSPN | [K].TPSYQL | 1xPhospho | Q9R207 1x | 0.023175 | 0.001578 | 1 | 2 | 1 | Q9R207 | Nibrin OS= | Q9R207 [4 | 0 | 1689.869 | 0 | 0.04 | 0.866377 | 8.36 | 98.9 | 101.1 | 5.04 | 9.77 | | High | High | High | High | High | High | High | High | High | High | High | 0.000847 | 0.008079 | 2.52 |
| FALSE | High | TPSVTSLAL | [R].TPSVTS | 1xPhospho | Q3UYD0 1 | 4.74E-07 | 0 | 1 | 2 | 5 | Q3UYD0 | Putative ur | Q3UYD0 [5 | 0 | 2624.291 | 0 | 0.04 | 0.418196 | 7.45 | 95.9 | 104.1 | 8.13 | 10.77 | | High | High | High | High | High | High | High | High | High | High | High | 0 | 8.95E-08 | 4.31 |
| FALSE | High | TPSTVTLNI | [R].TPSTVT | 1xPhospho | E9Q9B7 1x | 4.58E-07 | 0 | 1 | 12 | 2 | E9Q9B7 | Kinase D-ir | E9Q9B7 [1 | 0 | 2009.985 | 0 | 0.67 | 0.002549 | 9.68 | 77.6 | 122.4 | 13.6 | 8.85 | | High | High | High | High | High | High | High | High | High | High | High | 0 | 1.11E-07 | 3.52 |
| FALSE | High | TPQSPVSR | [R].TPQSP\ | 1xPhospho | Q6P9Q4 1x | 0.003167 | 0.00016 | 1 | 1 | 2 | Q6P9Q4 | FH1/FH2 d | Q6P9Q4 [4 | 0 | 1180.592 | 0 | 0.1 | 0.123449 | 9.98 | 91.7 | 108.3 | 5.32 | 9.82 | | High | High | High | High | High | High | High | High | High | High | High | 7.19E-05 | 0.000867 | 1.37 |
| FALSE | High | TPRDSPGIF | [R].TPRDSF | 1xPhospho | Q810V8 1x | 0.001558 | 5.66E-05 | 1 | 8 | 2 | Q810V8 | Rps6ka1 pr | Q810V8 [3 | 1 | 2313.17 | 0 | -0.8 | 0.01645 | 2.78 | 129.6 | 70.4 | 13.33 | 16.06 | | High | High | High | High | High | High | High | High | High | High | High | 2.58E-05 | 0.000393 | 3.25 |
| FALSE | High | TPSASHEE | [R].TPSASH | 1xPhospho | Q8BJU0 1x | 6.95E-05 | 0 | 1 | 2 | 3 | Q8BJU0 | Small gluta | Q8BJU0 [3 | 0 | 1551.653 | 0 | 0.16 | 0.326449 | 35.95 | 96.5 | 103.5 | 9.02 | 37.54 | | High | High | High | High | High | High | High | High | High | High | High | 0 | 1.2E-05 | 3.23 |
| FALSE | High | TPSEIQFHC | [R].TPSEIQ | 1xPhospho | Q61161 1x | 0.002463 | 0.000109 | 1 | 5 | 1 | Q61161 | Mitogen-a | Q61161 [3 | 0 | 1851.977 | 0 | 0.6 | 0.002928 | 13.36 | 77.5 | 122.5 | 9.44 | 8.28 | | High | High | High | High | High | High | High | High | High | High | High | 4.94E-05 | 0.000658 | 2.71 |
| FALSE | High | TPSESNSPS | [K].TPSESN | 1xPhospho | Q505G8 1x | 4.47E-05 | 0 | 1 | 6 | 2 | Q505G8 | Zinc finger | Q505G8 [9 | 0 | 2778.308 | 0 | 1.58 | 0.00499 | 53.99 | 51 | 149 | 42.75 | 12.79 | | High | High | High | High | High | High | High | High | High | High | High | 0 | 1.29E-05 | 3.38 |
| FALSE | High | TPSFDAPN | [R].TPSFDA | 1xPhospho | Q3TZW5 1 | 0.014538 | 0.001011 | 1 | 5 | 1 | Q3TZW5 | Putative ur | Q3TZW5 [2 | 0 | 1600.757 | 0 | -0.67 | 0.030484 | 2.96 | 123.5 | 76.5 | 10.27 | 14.1 | | High | High | High | High | High | High | High | High | High | High | High | 0.000538 | 0.004752 | 2.46 |
| FALSE | High | TPSIDLQR | [R].TPSIDL | 1xPhospho | E9Q8V6 1x | 0.001349 | 0 | 1 | 3 | 1 | E9Q8V6 | DENN/MAI | E9Q8V6 [1 | 0 | 1238.634 | 0 | 0.12 | 0.082787 | 3.01 | 92.7 | 107.3 | 4.24 | 11.21 | | High | High | High | High | High | High | High | High | High | High | High | 0 | 0.000333 | 1.62 |
| FALSE | High | TPSLPAVDT | [R].TPSLPA | 1xPhospho | Q2M4G9 1 | 4.89E-08 | 0 | 1 | 2 | 1 | Q2M4G9 | Protein inh | Q2M4G9 [ | 0 | 2663.317 | 0 | 0.85 | 0.002447 | 10.97 | 72.4 | 127.6 | 14.86 | 7.86 | | High | High | High | High | High | High | High | High | High | High | High | 0 | 3.43E-09 | 4.09 |
| FALSE | High | TPSLSPASS | [R].TPSLSP | 1xPhospho | Q8VC03 1x | 0.000579 | 0 | 1 | 1 | 3 | Q8VC03 | Echinoderr | Q8VC03 [8 | 0 | 1482.729 | 0 | 1.51 | 0.001772 | 4.71 | 49.6 | 150.4 | 7.24 | 15.28 | | High | High | High | High | High | High | High | High | High | High | High | 0 | 0.000128 | 3.35 |
| FALSE | High | TPSLSSLNS | [R].TPSLSS | 1xPhospho | E9Q9B7 1x | 2.25E-05 | 0 | 1 | 11 | 3 | E9Q9B7 | Kinase D-ir | E9Q9B7 [1 | 0 | 2431.237 | 0 | 0.84 | 0.028312 | 34.89 | 75.4 | 124.6 | 15.94 | 16.36 | | High | High | High | High | High | High | High | High | High | High | High | 0 | 3.38E-06 | 3.66 |
| FALSE | High | TPSNDGK | [R].TPSND( | 1xPhospho | Q925J9 1xI | 0.029076 | 0.001857 | 1 | 2 | 1 | Q925J9 | Mediator c | Q925J9 [9 | 0 | 1256.629 | 0 | 1.32 | 0.000428 | 9.71 | 55.7 | 144.3 | 10.48 | 7.36 | | High | High | High | High | High | High | High | High | High | High | High | 0.000997 | 0.01036 | 1.65 |
| FALSE | High | TPSNGAEG | [R].TPSNG | 1xPhospho | A0A1W2P7 | 0.017498 | 0.001245 | 1 | 6 | 1 | A0A1W2P7 | Uncharacte | A0A1W2P7 | 0 | 1508.731 | 0 | -0.16 | 0.047898 | 5.97 | 105.6 | 94.4 | 5.57 | 5.88 | | High | High | High | High | High | High | High | High | High | High | High | 0.000674 | 0.005863 | 1.25 |
| FALSE | High | TPSNGAEG | [R].TPSNG | 1xDeamida | A0A1W2P7 | 0.089425 | 0.006807 | 1 | 6 | 1 | A0A1W2P7 | Uncharacte | A0A1W2P7 | 0 | 1509.715 | 0 | 0.04 | 0.96365 | 14.36 | 99 | 101 | 5.14 | 7.35 | | High | High | High | High | High | High | High | High | High | High | High | 0.003476 | 0.03769 | 2.17 |
| FALSE | High | TPSPEPVDI | [R].TPSPEP | 1xPhospho | Q80TM9 1 | 0.022182 | 0.001542 | 1 | 6 | 2 | Q80TM9 | Nischarin C | Q80TM9 [1 | 0 | 1507.781 | 0 | -0.41 | 0.015276 | 8.18 | 114.3 | 85.7 | 3.59 | 15.49 | | High | High | High | High | High | High | High | High | High | High | High | 0.007994 | 0.08657 | 2.02 |
| FALSE | High | TPSPGEQQ | [R].TPSPGE | 1xPhospho | Q9CPS8 1x | 2.74E-07 | 0 | 1 | 1 | 1 | Q9CPS8 | Small mem | Q9CPS8 [3 | 0 | 2093.068 | 0 | 0.79 | 0.013816 | 22.13 | 73 | 127 | 9.88 | 17.04 | | High | High | High | High | High | High | High | High | High | High | High | 0 | 2.37E-08 | 4.72 |
| FALSE | High | TPSPKEEDI | [K].TPSPKE | 1xPhospho | Q80XU3 1x | 0.000496 | 0 | 1 | 2 | 2 | Q80XU3 | Nuclear ub | Q80XU3 [2 | 1 | 2666.305 | 0 | -0.98 | 0.011274 | 4.82 | 132.9 | 67.1 | 17.91 | 12.93 | | High | High | High | High | High | High | High | High | High | High | High | 0.000117 | 0.001061 | 2.29 |
| FALSE | High | TPSPPCPAN | [K].TPSPPC | 1xCarbami | Q9WU62 1 | 0.00095 | 0 | 1 | 2 | 2 | Q9WU62 | Inner centr | Q9WU62 [ | 0 | 1606.806 | 0 | -1.36 | 0.000264 | 0.43 | 142.3 | 57.7 | 4.18 | 4.74 | | High | High | High | High | High | High | High | High | High | High | High | 2.58E-05 | 0.000366 | 2.36 |
| FALSE | High | TPSPPPPILI | [R].TPSPPP | 1xPhospho | E9Q784 1x | 0.064323 | 0.004664 | 1 | 3 | 1 | E9Q784 | Zinc finger | E9Q784 [2 | 0 | 2225.26 | 0 | -1.3 | 0.005995 | 0.9 | 140.4 | 59.6 | 11.1 | 16.38 | | High | High | High | High | High | High | High | High | High | High | High | 0.002403 | 0.02575 | 1.72 |
| FALSE | High | TPSPSASTT | [R].TPSPSA | 1xPhospho | Q9QZ11 1x | 2.56E-05 | 0 | 1 | 1 | 1 | Q9QZ11 | Exonuclea | Q9QZ11 [6 | 0 | 1829.9 | 0 | 0 | 0.562369 | 5.48 | 99.9 | 100.1 | 9.54 | 11.9 | | High | High | High | High | High | High | High | High | High | High | High | 0 | 3.86E-06 | 4.22 |
| FALSE | High | TPSPSSPPL | [K].TPSPSS | 1xPhospho | Q91VY5 1x | 0.012397 | 0.000889 | 1 | 3 | 3 | Q91VY5 | Lysine-spe | Q91VY5 [4 | 0 | 3686.001 | 0 | -0.85 | 0.000603 | 0.88 | 128.8 | 71.2 | 3.68 | 5.59 | | High | High | High | High | High | High | High | High | High | High | High | 0.000449 | 0.004004 | 2.75 |
| FALSE | High | TPSQDSDY | [K].TPSQDS | 1xPhospho | P35831 1xI | 0.00195 | 5.66E-05 | 1 | 1 | 1 | P35831 | Tyrosine-p | P35831 [81 | 0 | 2251.105 | 0 | 0.89 | 0.042346 | 37.54 | 65.8 | 134.2 | 40.02 | 22.67 | | High | High | High | High | High | High | High | High | High | High | High | 2.58E-05 | 0.000506 | 1.93 |
| FALSE | High | TPSQEGCQ | [K].TPSQE( | 1xCarbami | Q5BL07 1x | 4.45E-07 | 0 | 1 | 2 | 2 | Q5BL07 | Peroxisom | Q5BL07 [1 | 0 | 2085.911 | 0 | 0.46 | 0.003612 | 11.96 | 83.4 | 116.6 | 13.4 | 10.73 | | High | High | High | High | High | High | High | High | High | High | High | 0 | 4.11E-08 | 5.55 |
| FALSE | High | TPSSCSSSL | [K].TPSSCS | 1xCarbami | A3KGI3 1xI | 0.006594 | 0.00062 | 1 | 1 | 4 | A3KGI3 | Kinetochor | A3KGI3 [16 | 0 | 1906.923 | 0 | -0.3 | 0.010146 | 6.62 | 108.7 | 91.3 | 5.62 | 3.65 | | High | High | High | High | High | High | High | High | High | High | High | 0.000275 | 0.001964 | 3.21 |
| FALSE | High | TPSSDVLVF | [K].TPSSDV | 1xPhospho | Q3UJI2 1xF | 0.025964 | 0.001645 | 2 | 8 | 1 | Q3UJI2; Q6 | Putative ur | Q3UJI2 [14 | 0 | 2010.024 | 0 | -0.08 | 0.094077 | 5.15 | 104.8 | 95.2 | 3.97 | 7.79 | Shared | High | High | High | High | High | High | High | High | High | High | High | 0.000872 | 0.009112 | 1.94 |
| FALSE | High | TPSSEEISPT | [R].TPSSEE | 1xPhospho | A6H6B3 1x | 0.088701 | 0.006669 | 1 | 9 | 2 | A6H6B3 | RalA bindir | A6H6B3 [2 | 0 | 1713.871 | 0 | 0.16 | 0.063971 | 10.26 | 95.2 | 104.8 | 2.37 | 8.99 | | High | High | High | High | High | High | High | High | High | High | High | 0.003432 | 0.03733 | 3.51 |
| FALSE | High | TPSSEEISPT | [R].TPSSEE | 2xPhospho | A6H6B3 2x | 0.007274 | 0.00062 | 1 | 9 | 4 | A6H6B3 | RalA bindir | A6H6B3 [2 | 0 | 1793.838 | 0 | -0.86 | 0.000399 | 2.34 | 129.9 | 70.1 | 1.94 | 7.39 | | High | High | High | High | High | High | High | High | High | High | High | 0.001151 | 0.01237 | 3.08 |
| FALSE | High | TPSSHGYN | [K].TPSSHG | 2xPhospho | Q8BQ33 2x | 0.007742 | 0.000664 | 1 | 1 | 1 | Q8BQ33 | Treslin OS= | Q8BQ33 [1 | 0 | 2097.945 | 0 | -0.8 | 0.002438 | 4.51 | 128.4 | 71.6 | 11.08 | 4.37 | | High | High | High | High | High | High | High | High | High | High | High | 0.000294 | 0.002347 | 3.36 |
| FALSE | High | TPSSQNLLT | [K].TPSSQN | 1xPhospho | Q3UFT4 1x | 1.23E-06 | 0 | 1 | 7 | 1 | Q3UFT4 | Putative ur | Q3UFT4 [6 | 0 | 1722.935 | 0 | -1.68 | 0.000104 | 0.32 | 152.5 | 47.5 | 1.88 | 5.81 | | High | High | High | High | High | High | High | High | High | High | High | 0 | 1.29E-07 | 2.33 |
| FALSE | High | TPSSSSTLA | [R].TPSSSS | 2xPhospho | F8WJF7 2x | 0.027839 | 0.001755 | 1 | 7 | 3 | F8WJF7 | E3 ubiquiti | F8WJF7 [5 | 0 | 1742.76 | 0 | 0.27 | 0.012502 | 14.91 | 88.6 | 111.4 | 14.1 | 6.22 | | High | High | High | High | High | High | High | High | High | High | High | 0.000968 | 0.009943 | 3.43 |
| FALSE | High | TPGFASPK | [K].TPGFAS | 1xPhospho | E9Q3G8 1x | 0.019791 | 0.001438 | 1 | 3 | 2 | E9Q3G8 | Nucleopor | E9Q3G8 [6 | 0 | 1342.717 | 0 | 0.26 | 0.042998 | 18.36 | 91.1 | 108.9 | 13.16 | 5.59 | | High | High | High | High | High | High | High | High | High | High | High | 0.000773 | 0.006713 | 1.57 |
| FALSE | High | TPGEVNTS | [K].TPGEVI | 1xPhospho | Q8BHK9 1x | 0.002058 | 5.66E-05 | 1 | 3 | 1 | Q8BHK9 | DNA excisi | Q8BHK9 [1 | 0 | 1606.779 | 0 | 0.08 | 0.306113 | 18.05 | 91.2 | 108.8 | 16.8 | 11.66 | | High | High | High | High | High | High | High | High | High | High | High | 2.58E-05 | 0.000538 | 2.35 |
| FALSE | High | TPGDFNYA | [K].TPGDFI | 1xPhospho | P14873 1xI | 2.28E-06 | 0 | 2 | 2 | 1 | P14873; B2 | Microtubu | P14873 [18 | 0 | 4324.933 | 0 | -0.83 | 0.00535 | 2.12 | 134.7 | 65.3 | 16.61 | 15.09 | Shared | High | High | High | High | High | High | High | High | High | High | High | 0 | 2.58E-07 | 4.52 |
| FALSE | High | TPGCQSPG | [R].TPGCQ | 1xCarbami | E9QMJ8 2x | 0.019966 | 0.001438 | 1 | 5 | 2 | E9QMJ8 | Axin-1 OS= | E9QMJ8 [4 | 1 | 2747.253 | 0 | 0.45 | 0.001977 | 3.84 | 89.1 | 110.9 | 12.17 | 11.07 | | High | High | High | High | High | High | High | High | High | High | High | 0.004327 | 0.04687 | 2.93 |
| FALSE | High | TNHSSPEA | [K].TNHSSI | 1xPhospho | A0A0A6YW | 8.71E-05 | 0 | 2 | 6 | 2 | A0A0A6YW | Mitogen-a | A0A0A6YW | 0 | 1707.847 | 0 | -0.23 | 0.503884 | 15.38 | 109.2 | 90.8 | 3.79 | 27.01 | Shared | High | High | High | High | High | High | High | High | High | High | High | 0 | 1.55E-05 | 2.82 |
| FALSE | High | TNKPSTPTT | [R].TNKPST | 1xPhospho | Q9QYY8 1x | 0.093879 | 0.007306 | 1 | 2 | 2 | Q9QYY8 | Spastin OS | Q9QYY8 [2 | 0 | 1810.983 | 0 | 2.01 | 0.000278 | 15.6 | 37.8 | 162.2 | 10.3 | 5.73 | | High | High | High | High | High | High | High | High | High | High | High | 0.006781 | 0.07362 | 1.5 |
| FALSE | High | TNILQYAST | [K].TNILQY | 1xPhospho | P13405 1xI | 0.009585 | 0.000736 | 1 | 4 | 2 | P13405 | Retinoblas | P13405 [75 | 0 | 2781.501 | 0 | 0.79 | 0.001212 | 6.72 | 73.5 | 126.5 | 5.42 | 8.45 | | High | High | High | High | High | High | High | High | High | High | High | 0.000814 | 0.007681 | 2.87 |
| FALSE | High | TNIPLSPTL | [R].TNIPLSI | 1xPhospho | Q69ZJ2 1xF | 0.009756 | 0.000736 | 1 | 5 | 1 | Q69ZJ2 | MKIAA146 | Q69ZJ2 [32 | 0 | 1824.028 | 0 | 0.8 | 0.00035 | 6.21 | 75 | 125 | 7.85 | 7.7 | | High | High | High | High | High | High | High | High | High | High | High | 0.000344 | 0.003039 | 1.87 |
| FALSE | High | TNLQPSEST | [R].TNLQP | 1xPhospho | Q9D600 1x | 0.003312 | 0.00016 | 1 | 1 | 1 | Q9D600 | DNA replic | Q9D600 [1 | 0 | 1890.832 | 0 | -0.94 | 0.001184 | 3.67 | 127.9 | 72.1 | 12.06 | 14.22 | | High | High | High | High | High | High | High | High | High | High | High | 7.19E-05 | 0.000914 | 3.24 |
| FALSE | High | TNLTSQSS | [K].TNLTS( | 1xPhospho | E9Q481 1x | 5.82E-06 | 0 | 1 | 3 | 3 | E9Q481 | Serine/thre | E9Q481 [7 | 0 | 3518.742 | 0 | -1.29 | 0.003545 | 3.13 | 142.7 | 57.3 | 10.7 | 19.52 | | High | High | High | High | High | High | High | High | High | High | High | 0 | 5.35E-06 | 3.29 |
| FALSE | High | TNLTSQSS | [K].TNLTS( | 2xPhospho | E9Q481 2x | 0.000933 | 0 | 1 | 3 | 3 | E9Q481 | Serine/thre | E9Q481 [7 | 0 | 3598.708 | 0 | -0.39 | 0.0462 | 4.76 | 109.4 | 90.6 | 8.81 | 11.16 | | High | High | High | High | High | High | High | High | High | High | High | 0.002855 | 0.02946 | 3.7 |
| FALSE | High | TNPFLNGN | [R].TNPFLN | 1xDeamida | Q921I6 1xF | 0.113753 | 0.009241 | 1 | 1 | 1 | Q921I6 | SH3 domai | Q921I6 [16 | 0 | 2596.27 | 0 | -0.74 | 0.003768 | 8.45 | 118.6 | 81.4 | 8.24 | 17.04 | | High | High | High | High | High | High | High | High | High | High | High | 0.004667 | 0.05026 | 2.73 |
| FALSE | High | TNPPTQKP | [R].TNPPT( | 1xPhospho | J3QNK8 1x | 0.089425 | 0.006807 | 1 | 15 | 1 | J3QNK8 | Abl interac | J3QNK8 [1 | 0 | 1969.011 | 0 | -0.4 | 0.691237 | 50.2 | 107.9 | 92.1 | 47.63 | 91.48 | | High | High | High | High | High | High | High | High | High | High | High | 0.003564 | 0.03801 | 1.84 |
| FALSE | High | TNPPTQKP | [R].TNPPT( | 1xPhospho | J3QNK8 1x | 0.002844 | 0.00016 | 1 | 15 | 3 | J3QNK8 | Abl interac | J3QNK8 [1 | 0 | 2198.173 | 0 | 0.08 | 0.116031 | 6.8 | 95.6 | 104.4 | 4.41 | 6.98 | | High | High | High | High | High | High | High | High | High | High | High | 7.19E-05 | 0.000766 | 2.47 |
| FALSE | High | TNPSPTNPI | [R].TNPSPT | 1xPhospho | Q9QZS3 1x | 8.63E-05 | 0 | 1 | 4 | 4 | Q9QZS3 | Protein nu | Q9QZS3 [6 | 0 | 2171.079 | 0 | -1.52 | 0.000333 | 0.5 | 146.7 | 53.3 | 4.38 | 10.44 | | High | High | High | High | High | High | High | High | High | High | High | 0 | 2.43E-05 | 4.22 |
| FALSE | High | TNQAILSQI | [K].TNQAIL | 1xPhospho | B1AXZ5 1xI | 0.001659 | 5.66E-05 | 1 | 10 | 2 | B1AXZ5 | ELAV-like p | B1AXZ5 [2 | 0 | 2042.027 | 0 | -1.88 | 0.000236 | 0.56 | 157.6 | 42.4 | 7.78 | 10.96 | | High | High | High | High | High | High | High | High | High | High | High | 2.58E-05 | 0.00042 | 2.82 |
| FALSE | High | TNQPESGL | [K].TNQPE | 1xPhospho | B9EKC1 1xI | 0.000151 | 0 | 1 | 2 | 2 | B9EKC1 | PAS domai | B9EKC1 [1 | 0 | 2239.117 | 0 | 0.1 | 0.337131 | 21.34 | 96.4 | 103.6 | 11.97 | 15.1 | | High | High | High | High | High | High | High | High | High | High | High | 0.000204 | 0.00132 | 3.32 |
| FALSE | High | TNHRLSLPT | [K].TNHRLS | 1xCarbami | Q03160 1x | 0.003282 | 0.00016 | 1 | 2 | 3 | Q03160 | Growth fac | Q03160 [4 | 1 | 2675.329 | 0 | -0.75 | 0.006123 | 2.58 | 123.1 | 76.9 | 16.5 | 17.12 | | High | High | High | High | High | High | High | High | High | High | High | 0.000344 | 0.003159 | 2.93 |
| FALSE | High | TNSDSALH | [R].TNSDS/ | 1xPhospho | Q68ED7 1x | 0.01095 | 0.000772 | 1 | 1 | 1 | Q68ED7 | CREB-regu | Q68ED7 [1 | 0 | 3686.679 | 0 | -0.51 | 0.002017 | 0.32 | 117 | 83 | 10.89 | 10.77 | | High | High | High | High | High | High | High | High | High | High | High | 0.000399 | 0.003474 | 3.75 |
| FALSE | High | TNSEQVDE | [R].TNSEQ | 1xPhospho | Q8C6M1 1 | 0.000772 | 0 | 1 | 1 | 1 | Q8C6M1 | Ubiquitin c | Q8C6M1 [3 | 0 | 2935.235 | 0 | 0.02 | 0.853883 | 13.54 | 100.3 | 99.7 | 6.54 | 20.83 | | High | High | High | High | High | High | High | High | High | High | High | 0 | 0.000179 | 3.27 |
| FALSE | High | TNSFDVPR | [R].TNSFD\ | 1xPhospho | V9GXJ4 1xI | 0.009585 | 0.000736 | 1 | 6 | 2 | V9GXJ4 | Homeobox | V9GXJ4 [1 | 0 | 1244.587 | 0 | -1.8 | 0.000842 | 0.22 | 154.6 | 45.4 | 10.68 | 6.62 | | High | High | High | High | High | High | High | High | High | High | High | 0.003606 | 0.03835 | 1.32 |
| FALSE | High | TNSGISSAS | [R].TNSGIS | 1xPhospho | B2RRF1 1xI | 0.000141 | 0 | 1 | 6 | 1 | B2RRF1 | B230339M | B2RRF1 [7 | 0 | 3196.532 | 0 | 1.83 | 0.000281 | 11.82 | 44.3 | 155.7 | 11.21 | 3.49 | | High | High | High | High | High | High | High | High | High | High | High | 0 | 2.62E-05 | 3.01 |
| FALSE | High | TNSGSFYS | [R].TNSGSI | 1xPhospho | Q8BQ33 1x | 0.00051 | 0 | 1 | 1 | 1 | Q8BQ33 | Treslin OS= | Q8BQ33 [1 | 0 | 1939.957 | 0 | -0.53 | 0.013724 | 2.43 | 116.5 | 83.5 | 3.21 | 14.52 | | High | High | High | High | High | High | High | High | High | High | High | 0 | 0.000112 | 2.54 |
| FALSE | High | TNSLNRPG | [K].TNSLNI | 1xPhospho | Q8R550-3 | 0.069325 | 0.005112 | 1 | 7 | 1 | Q8R550-3 | Isoform 3 o | Q8R550-3 | 0 | 1701.9 | 0 | 1.48 | 0.00109 | 16.1 | 57.4 | 142.6 | 15.92 | 8.95 | | High | High | High | High | High | High | High | High | High | High | High | 0.002633 | 0.02809 | 0.91 |
| FALSE | High | TNSLPFSQI | [K].TNSLPF | 1xPhospho | L7N209 1xI | 0.007145 | 0.00062 | 1 | 2 | 1 | L7N209 | HBS1-like p | L7N209 [3 | 0 | 1358.667 | 0 | 0.54 | 0.013714 | 11.55 | 81.9 | 118.1 | 7 | 11.13 | | High | High | High | High | High | High | High | High | High | High | High | 0.000275 | 0.002154 | 2.26 |
| FALSE | High | TNSMGSAT | [R].TNSMG | 1xPhospho | Q8BZ47 1x | 0.000868 | 0 | 1 | 2 | 1 | Q8BZ47 | Zinc finger | Q8BZ47 [4 | 0 | 1956.987 | 0 | -0.18 | 0.307956 | 17.1 | 101.9 | 98.1 | 9.13 | 17.56 | | High | High | High | High | High | High | High | High | High | High | High | 0 | 0.000203 | 2.83 |
| FALSE | High | TNSMGSAT | [R].TNSMG | 1xPhospho | Q8BZ47 1x | 3.79E-07 | 0 | 1 | 2 | 9 | Q8BZ47 | Zinc finger | Q8BZ47 [4 | 0 | 1972.982 | 0 | -0.18 | 0.175952 | 8.58 | 107.6 | 92.4 | 6.79 | 11.23 | | High | High | High | High | High | High | High | High | High | High | High | 0 | 1.82E-05 | 3.8 |
| FALSE | High | TNSMQQLI | [R].TNSMC | 1xPhospho | E9Q6H8 1x | 0.000136 | 0 | 1 | 9 | 1 | E9Q6H8 | Pleckstrin | E9Q6H8 [4 | 0 | 2044.034 | 0 | 0.03 | 0.686435 | 4.13 | 97.7 | 102.3 | 17.23 | 11.83 | | High | High | High | High | High | High | High | High | High | High | High | 0 | 2.53E-05 | 3.12 |
| FALSE | High | TNSMSFLP | [R].TNSMS | 1xPhospho | Q6ZWQ0 1 | 0.116482 | 0.009523 | 1 | 2 | 1 | Q6ZWQ0 | Nesprin-2 ( | Q6ZWQ0 [ | 0 | 1547.774 | 0 | 1.62 | 0.023713 | 18.3 | 49.2 | 150.8 | 6.21 | 37.28 | | High | High | High | High | High | High | High | High | High | High | High | 0.004757 | 0.05171 | 2.41 |
| FALSE | High | TNSMSFLP | [R].TNSMS | 1xPhospho | Q6ZWQ0 1 | 0.000426 | 0 | 1 | 2 | 1 | Q6ZWQ0 | Nesprin-2 ( | Q6ZWQ0 [ | 0 | 1760.942 | 0 | 4.42 | 0.002511 | 23.91 | 7.5 | 192.5 | 24.15 | 23.49 | | High | High | High | High | High | High | High | High | High | High | High | 0 | 9.16E-05 | 3.81 |
| FALSE | High | TNSMSFLP | [R].TNSMS | 1xPhospho | Q6ZWQ0 1 | 3.17E-05 | 0 | 1 | 2 | 6 | Q6ZWQ0 | Nesprin-2 ( | Q6ZWQ0 [ | 0 | 1776.937 | 0 | 3.62 | 0.000379 | 3.2 | 13.3 | 186.7 | 11.91 | 10.26 | | High | High | High | High | High | High | High | High | High | High | High | 0 | 4.94E-06 | 3.82 |
| FALSE | High | TNSMSSSG | [R].TNSMS | 1xPhospho | A0A1S6GW | 0.011755 | 0.000889 | 1 | 8 | 2 | A0A1S6GW | Uncharacte | A0A1S6GW | 0 | 1703.762 | 0 | 0.15 | 0.146059 | 16.73 | 94.9 | 105.1 | 3.37 | 13.74 | | High | High | High | High | High | High | High | High | High | High | High | 0.000859 | 0.008324 | 1.52 |
| FALSE | High | TNSDSALH | [R].TNSDS/ | 1xPhospho | Q68ED7 1x | 5.31E-05 | 0 | 1 | 1 | 1 | Q68ED7 | CREB-regu | Q68ED7 [1 | 0 | 3702.674 | 0 | -0.48 | 0.647734 | 3.97 | 118.9 | 81.1 | 13.01 | 48.41 | | High | High | High | High | High | High | High | High | High | High | High | 0 | 8.8E-06 | 4.35 |
| FALSE | High | TNGSSPGP | [K].TNGSSI | 1xPhospho | Q80YR7 1x | 0.000538 | 0 | 1 | 2 | 3 | Q80YR7 | Claspin OS | Q80YR7 [1 | 0 | 1382.708 | 0 | -0.19 | 0.065671 | 4.7 | 106.4 | 93.6 | 3.88 | 13.57 | | High | High | High | High | High | High | High | High | High | High | High | 0 | 0.000139 | 1.8 |
| FALSE | High | TNGISDVQ | [R].TNGISD | 1xPhospho | G5E852 1x | 7.95E-08 | 0 | 1 | 2 | 3 | G5E852 | Tyrosine-p | G5E852 [5 | 0 | 1937.989 | 0 | 1.5 | 0.001191 | 21.77 | 53.1 | 146.9 | 10.14 | 8.95 | | High | High | High | High | High | High | High | High | High | High | High | 0 | 5.93E-09 | 4.16 |
| FALSE | High | TNGDNGIN | [K].TNGDN | 2xPhospho | E9Q4Y4 2x | 3.88E-06 | 0 | 1 | 1 | 1 | E9Q4Y4 | Centrosom | E9Q4Y4 [4 | 0 | 2550.075 | 1 | -0.17 | 0.052281 | 2.22 | 110.7 | 89.3 | 8.65 | 13.09 | | High | High | High | High | High | High | High | High | High | High | High | 0 | 4.67E-07 | 4.68 |
| FALSE | High | TLSVTSAGS | [R].TLSVTS | 1xCarbami | Q9CQS5 1x | 0.000346 | 0 | 1 | 1 | 6 | Q9CQS5 | Serine/thre | Q9CQS5 [4 | 0 | 2756.456 | 0 | -0.56 | 0.004077 | 4.52 | 122.7 | 77.3 | 6.42 | 9.56 | | High | High | High | High | High | High | High | High | High | High | High | 0.000449 | 0.004204 | 3.06 |
| FALSE | High | TLSVTSAGS | [R].TLSVTS | 1xCarbami | Q9CQS5 2x | 0.007145 | 0.00062 | 1 | 1 | 2 | Q9CQS5 | Serine/thre | Q9CQS5 [4 | 0 | 2836.422 | 0 | -0.73 | 0.017184 | 1.99 | 124.6 | 75.4 | 6.35 | 20.16 | | High | High | High | High | High | High | High | High | High | High | High | 0.000275 | 0.002148 | 3.79 |
| FALSE | High | TLTAEEAEE | [R].TLTAEE | 1xPhospho | Q3THK3 1x | 0.009756 | 0.000736 | 1 | 3 | 1 | Q3THK3 | General tra | Q3THK3 [1 | 0 | 1901.837 | 0 | 0.17 | 0.754748 | 21.64 | 90.8 | 109.2 | 24.35 | 21.78 | | High | High | High | High | High | High | High | High | High | High | High | 0.000344 | 0.003057 | 2.6 |
| FALSE | High | TLTDEVNSI | [K].TLTDEV | 1xPhospho | Q6PDG5 1x | 9.65E-06 | 0 | 1 | 3 | 1 | Q6PDG5 | SWI/SNF c | Q6PDG5 [2 | 0 | 1757.779 | 0 | 2.03 | 0.000313 | 3.83 | 38.2 | 161.8 | 6.62 | 8.19 | | High | High | High | High | High | High | High | High | High | High | High | 0 | 1.3E-06 | 3.35 |
| FALSE | High | TLTDEVNSI | [K].TLTDEV | 1xPhospho | Q6PDG5 1x | 2.15E-05 | 0 | 1 | 3 | 4 | Q6PDG5 | SWI/SNF c | Q6PDG5 [2 | 1 | 1913.88 | 0 | 2.1 | 0.000428 | 11.58 | 38.1 | 161.9 | 6.37 | 9.02 | | High | High | High | High | High | High | High | High | High | High | High | 0 | 3.77E-05 | 2.21 |
| FALSE | High | TITLEVEPSI | [K].TITLEVI | 1xPhospho | E9QNP0 1x | 6E-07 | 0 | 1 | 17 | 2 | E9QNP0 | KxDL motif | E9QNP0 [1 | 0 | 2326.22 | 0 | -0.47 | 0.015098 | 0.86 | 111.4 | 88.6 | 10.66 | 5.23 | | High | High | High | High | High | High | High | High | High | High | High | 0 | 5.7E-08 | 3.29 |
| FALSE | High | TLTPANSP\ | [R].TLTPAN | 1xPhospho | Q3TQ38 1x | 2.63E-05 | 0 | 1 | 5 | 3 | Q3TQ38 | Putative ur | Q3TQ38 [3 | 0 | 1924.019 | 0 | -0.12 | 0.028481 | 3.58 | 102.8 | 97.2 | 10.08 | 12.64 | | High | High | High | High | High | High | High | High | High | High | High | 0.000723 | 0.006346 | 3.25 |
| FALSE | High | TLTPPLNK | [R].TLTPPL | 1xPhospho | B2RUE3 1x | 0.025964 | 0.001645 | 1 | 3 | 1 | B2RUE3 | UTP14, U3 | B2RUE3 [1 | 0 | 1421.817 | 0 | -1.2 | 0.000502 | 0.99 | 143 | 57 | 8.82 | 17.35 | | High | High | High | High | High | High | High | High | High | High | High | 0.000872 | 0.009112 | 2.15 |
| FALSE | High | TLTPSLR | [R].TLTPSL | 1xPhospho | E9Q784 1x | 0.024001 | 0.001612 | 1 | 3 | 2 | E9Q784 | Zinc finger | E9Q784 [3 | 0 | 1096.596 | 0 | -0.63 | 0.000313 | 2.56 | 121.5 | 78.5 | 2.39 | 2.45 | | High | High | High | High | High | High | High | High | High | High | High | 0.000859 | 0.00837 | 1.41 |
| FALSE | High | TLTPTPDGI | [R].TLTPTP | 1xPhospho | Q61210-5 | 0.037692 | 0.00249 | 1 | 8 | 1 | Q61210-5 | Isoform 5 o | Q61210-5 | 0 | 1467.786 | 0 | 0.41 | 0.001229 | 5.97 | 84.8 | 115.2 | 5.42 | 6.05 | | High | High | High | High | High | High | High | High | High | High | High | 0.001324 | 0.01399 | 1.64 |
| FALSE | High | TLTSPADA | [K].TLTSPA | 1xPhospho | Q3UFT4 1x | 2.02E-05 | 0 | 1 | 7 | 2 | Q3UFT4 | Putative ur | Q3UFT4 [3 | 0 | 1900.937 | 0 | -1.1 | 0.002705 | 3.82 | 134.5 | 65.5 | 7.58 | 13.56 | | High | High | High | High | High | High | High | High | High | High | High | 0 | 2.97E-06 | 3.57 |
| FALSE | High | TITVPVSGS | [K].TITVPV | 1xPhospho | Q8BMB0 1 | 0.000751 | 0 | 2 | 7 | 1 | Q8BMB0; A | BRCA2-inte | Q8BMB0 [ | 0 | 1623.912 | 0 | -1.04 | 0.000361 | 0.9 | 136 | 64 | 3.95 | 4.56 | Shared | High | High | High | High | High | High | High | High | High | High | High | 0 | 0.000172 | 3.26 |

| | | | | | | | | | | | | | | | | | | | | | | | | | | | | | | | | | | | | | | |
|---|---|---|---|---|---|---|---|---|---|---|---|---|---|---|---|---|---|---|---|---|---|---|---|---|---|---|---|---|---|---|---|---|---|---|---|---|---|---|
| FALSE | High | TLVHSSSDC | [R].TLVHSS | 1xPhospho | Q571K4 1x | 0.002647 | 0.000109 | 1 | 1 | 1 | Q571K4 | TGF-beta-a | Q571K4 [9 | 0 | 2525.255 | 0 | -0.17 | 0.311536 | 9.07 | 102.6 | 97.4 | 10.07 | 6.85 | | High | High | High | High | High | High | High | High | High | High | High | 4.94E-05 | 0.00071 | 1.93 |
| FALSE | High | TLVLDCHYI | [K].TLVLDC | 1xCarbami | Q8R5L1 1x | 1E-05 | 0 | 1 | 2 | 3 | Q8R5L1 | Compleme | Q8R5L1 [1 | 0 | 3699.718 | 0 | 0.49 | 0.101544 | 4.24 | 82.4 | 117.6 | 16.67 | 25.3 | | High | High | High | High | High | High | High | High | High | High | High | 0 | 1.36E-06 | 3.64 |
| FALSE | High | TLVLSNLSY | [K].TLVLSN | 1xPhospho | P09405 1xl | 4.27E-05 | 0 | 1 | 6 | 1 | P09405 | Nucleolin ( | P09405 [48 | 0 | 1935.06 | 0 | -1.53 | 0.000565 | 1.49 | 147.3 | 52.7 | 5.83 | 13.36 | | High | High | High | High | High | High | High | High | High | High | High | 0 | 6.88E-06 | 2.13 |
| FALSE | High | TLVLSSSPT | [K].TLVLSS | 1xPhospho | A0A1W2P7 | 0.001799 | 5.66E-05 | 1 | 2 | 4 | A0A1W2P7 | RAB3A inte | A0A1W2P7 | 0 | 2536.368 | 0 | -0.01 | 0.521741 | 4.65 | 99.8 | 100.2 | 8.64 | 8.58 | | High | High | High | High | High | High | High | High | High | High | High | 0.000327 | 0.002638 | 3.58 |
| FALSE | High | TLYESVFGS | [R].TLYESV | 1xCarbami | E9Q6T9 1x | 1.19E-06 | 0 | 1 | 1 | 2 | E9Q6T9 | BTB/POZ d | E9Q6T9 [5 | 0 | 2553.235 | 0 | 0.76 | 0.001003 | 0.37 | 74.3 | 125.7 | 10.64 | 10.62 | | High | High | High | High | High | High | High | High | High | High | High | 0 | 8.39E-07 | 3.95 |
| FALSE | High | TIYLSSPDS/ | [R].TIYLSSF | 1xPhospho | E9QLH9 1x | 5.93E-06 | 0 | 1 | 2 | 2 | E9QLH9 | PEAK1-rela | E9QLH9 [4 | 0 | 2185.089 | 0 | 0.73 | 0.016634 | 18.65 | 74 | 126 | 8.54 | 14.82 | | High | High | High | High | High | High | High | High | High | High | High | 0 | 7.48E-07 | 3.07 |
| FALSE | High | TMATWGC | [K].TMATW | 1xCarbami | Q9R0L6-2 : | 1.81E-05 | 0 | 1 | 3 | 1 | Q9R0L6-2 | Isoform 2 c | Q9R0L6-2 [ | 0 | 3472.476 | 0 | -0.52 | 0.014167 | 10.1 | 117.3 | 82.7 | 12.47 | 2.57 | | High | High | High | High | High | High | High | High | High | High | High | 0 | 2.65E-06 | 3.97 |
| FALSE | High | TMATWGC | [K].TMATW | 1xCarbami | Q9R0L6-2 : | 0.028327 | 0.001791 | 1 | 3 | 1 | Q9R0L6-2 | Isoform 2 c | Q9R0L6-2 [ | 0 | 3552.442 | 0 | 0.26 | 0.054961 | 18.64 | 85 | 115 | 11.19 | 17.39 | | High | High | High | High | High | High | High | High | High | High | High | 0.000984 | 0.01013 | 3.48 |
| FALSE | High | TMELSPR | [K].TMELSI | 1xPhospho | E9QMD3 1 | 0.08727 | 0.006537 | 1 | 5 | 1 | E9QMD3 | Zinc finger | E9QMD3 [: | 0 | 1158.543 | 0 | 1.84 | 0.000614 | 16.82 | 46.8 | 153.2 | 13.19 | 8.78 | | High | High | High | High | High | High | High | High | High | High | High | 0.003375 | 0.0368 | 1.4 |
| FALSE | High | TMFAQVES | [K].TMFAQ | 1xPhospho | G3X934 1x | 0.000271 | 0 | 1 | 2 | 2 | G3X934 | MCG11596 | G3X934 [6 | 0 | 2153.971 | 0 | -0.56 | 0.000746 | 3.21 | 118.5 | 81.5 | 6.81 | 10.7 | | High | High | High | High | High | High | High | High | High | High | High | 0.000327 | 0.002436 | 3.52 |
| FALSE | High | TMFAQVES | [K].TMFAQ | 1xPhospho | G3X934 1x | 8.5E-06 | 0 | 1 | 2 | 6 | G3X934 | MCG11596 | G3X934 [6 | 0 | 2169.966 | 0 | -0.29 | 0.013785 | 6.95 | 113.2 | 86.8 | 5.11 | 6.48 | | High | High | High | High | High | High | High | High | High | High | High | 0 | 1.12E-06 | 3.91 |
| FALSE | High | TMSSDDEE | [K].TMSSD | 1xCarbami | A0A0R4J0\ | 5.4E-05 | 0 | 1 | 4 | 4 | A0A0R4J0\ | Integrator | A0A0R4J0\ | 0 | 1913.791 | 0 | 0.04 | 0.919871 | 13.7 | 98.7 | 101.3 | 2.04 | 11.05 | | High | High | High | High | High | High | High | High | High | High | High | 0 | 8.98E-06 | 3.51 |
| FALSE | High | TMTTNSSD | [R].TMTTN | 1xPhospho | P46938 1xl | 5.09E-06 | 0 | 1 | 3 | 19 | P46938 | Transcripti | P46938 [36 | 0 | 2441.064 | 0 | 1.09 | 0.00573 | 29.88 | 63.8 | 136.2 | 9.53 | 13.53 | | High | High | High | High | High | High | High | High | High | High | High | 0.001665 | 0.01672 | 3.61 |
| FALSE | High | TNATATTP | [R].TNATA | 1xPhospho | Q80ZQ9 1x | 0.045122 | 0.003373 | 1 | 1 | 2 | Q80ZQ9 | Protein Sin | Q80ZQ9 [1 | 0 | 1085.508 | 0 | 0.44 | 0.003862 | 8.09 | 84.5 | 115.5 | 5.19 | 6.69 | | High | High | High | High | High | High | High | High | High | High | High | 0.001712 | 0.0171 | 1.71 |
| FALSE | High | TNDGIGAS | [K].TNDGI( | 1xPhospho | O54917 1x | 0.006195 | 0.00062 | 1 | 3 | 2 | O54917 | Transcripti | O54917 [2 | 0 | 1584.803 | 0 | -0.07 | 0.33997 | 6.74 | 102.5 | 97.5 | 2.74 | 18.42 | | High | High | High | High | High | High | High | High | High | High | High | 0.000275 | 0.001835 | 2.84 |
| FALSE | High | TNDSTEQN | [K].TNDSTI | 1xPhospho | A2AFQ0 1x | 5.26E-05 | 0 | 1 | 6 | 3 | A2AFQ0 | E3 ubiquiti | A2AFQ0 [2 | 0 | 4082.795 | 0 | 0.54 | 0.639846 | 8.12 | 86 | 114 | 20.79 | 40.61 | | High | High | High | High | High | High | High | High | High | High | High | 0 | 8.73E-06 | 3.34 |
| FALSE | High | TNDSTEQN | [K].TNDSTI | 1xPhospho | A2AFQ0 1x | 1.07E-05 | 0 | 1 | 6 | 6 | A2AFQ0 | E3 ubiquiti | A2AFQ0 [2 | 0 | 4098.79 | 0 | 0.03 | 0.378474 | 14.07 | 92.2 | 107.8 | 14.26 | 13.63 | | High | High | High | High | High | High | High | High | High | High | High | 0 | 1.46E-06 | 4.08 |
| FALSE | High | TNSMSSSG | [R].TNSMS | 1xPhospho | A0A1S6GW | 0.003312 | 0.00016 | 1 | 8 | 4 | A0A1S6GW | Uncharacte | A0A1S6GW | 0 | 1719.757 | 0 | 0.06 | 0.364447 | 15.59 | 94.5 | 105.5 | 11.52 | 35.54 | | High | High | High | High | High | High | High | High | High | High | High | 0.000756 | 0.006615 | 2.05 |
| FALSE | High | TQAPPTPD | [R].TQAPP | 1xPhospho | Q9WVL3 1 | 0.009332 | 0.000736 | 1 | 2 | 1 | Q9WVL3 | Solute carr | Q9WVL3 [ | 0 | 1492.781 | 0 | 0.09 | 0.021714 | 4.22 | 95.6 | 104.4 | 4.59 | 3.92 | | High | High | High | High | High | High | High | High | High | High | High | 0.000327 | 0.002917 | 2.26 |
| FALSE | High | TNSMSSSG | [R].TNSMS | 1xDeamida | A0A1S6GW | 0.067059 | 0.004796 | 1 | 8 | 1 | A0A1S6GW | Uncharacte | A0A1S6GW | 0 | 1720.741 | 0 | 0.23 | 0.646828 | 39.46 | 93.6 | 106.4 | 20.62 | 23.39 | | High | High | High | High | High | High | High | High | High | High | High | 0.002485 | 0.02711 | 1.61 |
| FALSE | High | TNSPAYSDI | [K].TNSPAY | 2xPhospho | Q8BZ47 2x | 1.34E-06 | 0 | 1 | 2 | 2 | Q8BZ47 | Zinc finger | Q8BZ47 [8 | 0 | 2531.063 | 0 | -0.07 | 0.321188 | 9.26 | 102.8 | 97.2 | 8.04 | 7.01 | | High | High | High | High | High | High | High | High | High | High | High | 0 | 8.19E-05 | 3.72 |
| FALSE | High | TPATTASSF | [R].TPATTA | 1xCarbami | P25799 1xl | 0.000446 | 0 | 1 | 2 | 3 | P25799 | Nuclear fac | P25799 [89 | 0 | 2966.449 | 0 | -0.06 | 0.59159 | 34.25 | 102 | 98 | 13.24 | 21.21 | | High | High | High | High | High | High | High | High | High | High | High | 0 | 9.62E-05 | 3.99 |
| FALSE | High | TPCAVLSPT | [K].TPCAVL | 1xCarbami | P29477 1xl | 1.51E-06 | 0 | 1 | 1 | 1 | P29477 | Nitric oxide | P29477 [31 | 0 | 2180.107 | 0 | 4.75 | 0.001839 | 33.45 | 8.7 | 191.3 | 31.74 | 19.88 | | High | High | High | High | High | High | High | High | High | High | High | 0 | 1.61E-07 | 2.73 |
| FALSE | High | TPDLSNQN | [K].TPDLSN | 1xPhospho | Q3TLH4 1x | 0.000109 | 0 | 1 | 7 | 5 | Q3TLH4 | Protein PRI | Q3TLH4 [1 | 0 | 3596.477 | 0 | -0.85 | 0.012889 | 5.61 | 126 | 74 | 14.69 | 14.58 | | High | High | High | High | High | High | High | High | High | High | High | 0 | 1.99E-05 | 4.33 |
| FALSE | High | TPDLSNQN | [K].TPDLSN | 1xDeamida | Q3TLH4 2x | 0.033986 | 0.002195 | 1 | 7 | 4 | Q3TLH4 | Protein PRI | Q3TLH4 [1 | 0 | 3677.428 | 0 | 0.85 | 0.009421 | 28.61 | 73.2 | 126.8 | 32.01 | 13.71 | | High | High | High | High | High | High | High | High | High | High | High | 0.003148 | 0.03268 | 3.96 |
| FALSE | High | TPDLSNQN | [K].TPDLSN | 3xPhospho | Q3TLH4 3x | 1.3E-05 | 0 | 1 | 7 | 8 | Q3TLH4 | Protein PRI | Q3TLH4 [1 | 0 | 3756.41 | 0 | -1.02 | 0.015376 | 15.47 | 133.1 | 66.9 | 8.39 | 43.64 | | High | High | High | High | High | High | High | High | High | High | High | 0 | 1.81E-06 | 5.32 |
| FALSE | High | TPDLSVQLI | [K].TPDLSV | 1xPhospho | E9PYB0 1xl | 9.45E-05 | 0 | 1 | 1 | 1 | E9PYB0 | AHNAK nu | E9PYB0 [12 | 0 | 2250.203 | 0 | 2.19 | 0.000623 | 10.72 | 37.6 | 162.4 | 32.59 | 13.8 | | High | High | High | High | High | High | High | High | High | High | High | 0 | 1.69E-05 | 3.64 |
| FALSE | High | TPDLSVQLI | [K].TPDLSV | 1xPhospho | E9PYB0 1xl | 1.51E-06 | 0 | 2 | 3 | 2 | E9PYB0; F7 | AHNAK nu | E9PYB0 [54 | 0 | 2264.219 | 0 | 2.07 | 0.000682 | 14.69 | 35.5 | 164.5 | 16.48 | 11.13 | Shared | High | High | High | High | High | High | High | High | High | High | High | 0 | 1.6E-07 | 3.71 |
| FALSE | High | TPDSGHVS | [K].TPDSGI | 1xPhospho | A2ALS7 1xl | 0.045508 | 0.003373 | 2 | 6 | 1 | A2ALS7; A2 | Rap1 GTPa | A2ALS7 [26 | 0 | 1819.899 | 0 | 0.84 | 0.005028 | 9.19 | 70.6 | 129.4 | 7.48 | 11.73 | Shared | High | High | High | High | High | High | High | High | High | High | High | 0.001741 | 0.01736 | 1.88 |
| FALSE | High | TPDSTFSET | [R].TPDSTF | 1xPhospho | Q921D9 1x | 0.002232 | 0.000109 | 1 | 3 | 1 | Q921D9 | Grainyhead | Q921D9 [2 | 0 | 1797.871 | 0 | 0.18 | 0.223489 | 11.64 | 97 | 103 | 13.23 | 14.36 | | High | High | High | High | High | High | High | High | High | High | High | 4.94E-05 | 0.000586 | 1.35 |
| FALSE | High | TPDVQESP | [R].TPDVQ | 1xPhospho | Q99LG0 1x | 0.008691 | 0.000736 | 1 | 2 | 1 | Q99LG0 | Ubiquitin c | Q99LG0 [5 | 0 | 2052.03 | 0 | -0.33 | 0.023619 | 4.76 | 111.8 | 88.2 | 6.05 | 9.54 | | High | High | High | High | High | High | High | High | High | High | High | 0.000327 | 0.002685 | 2.77 |
| FALSE | High | TPEASVPT | [R].TPEASV | 1xPhospho | E9QK89 1x | 0.005516 | 0.00062 | 1 | 2 | 2 | E9QK89 | Mediator c | E9QK89 [1 | 0 | 2468.273 | 0 | -0.46 | 0.026118 | 12.78 | 115.8 | 84.2 | 9.52 | 10.93 | | High | High | High | High | High | High | High | High | High | High | High | 0.000275 | 0.002135 | 4.02 |
| FALSE | High | TPEAVEEP/ | [K].TPEAVE | 1xPhospho | Q14A90 1x | 0.077825 | 0.005963 | 1 | 1 | 1 | Q14A90 | Dihydrouri | Q14A90 [3 | 0 | 2324.167 | 0 | -0.71 | 0.000452 | 1.46 | 127.7 | 72.3 | 8.61 | 11.55 | | High | High | High | High | High | High | High | High | High | High | High | 0.003109 | 0.03219 | 2.02 |
| FALSE | High | TPEDGGYT | [K].TPEDG( | 1xCarbami | P14873 1xl | 0.001615 | 5.66E-05 | 2 | 2 | 1 | P14873; B2 | Microtubu | P14873 [19 | 0 | 2137.976 | 0 | -0.6 | 0.016022 | 5.21 | 120.5 | 79.5 | 14.66 | 9.48 | Shared | High | High | High | High | High | High | High | High | High | High | High | 2.58E-05 | 0.000407 | 2.83 |
| FALSE | High | TPASSSETA | [K].TPASSS | 1xPhospho | F8WJ93 1x | 0.000699 | 0 | 1 | 9 | 8 | F8WJ93 | Echinoderr | F8WJ93 [8 | 0 | 3572.732 | 0 | -0.64 | 0.000598 | 2.13 | 120.8 | 79.2 | 7.65 | 5.89 | | High | High | High | High | High | High | High | High | High | High | High | 0 | 0.00016 | 3.31 |
| FALSE | High | TPEEGGYS | [R].TPEEG( | 1xPhospho | P14873 1xl | 0.017966 | 0.001282 | 2 | 2 | 1 | P14873; B2 | Microtubu | P14873 [19 | 0 | 2126.994 | 0 | -0.54 | 0.007171 | 8.26 | 118.3 | 81.7 | 11.52 | 6.31 | Shared | High | High | High | High | High | High | High | High | High | High | High | 0.000689 | 0.006052 | 2.52 |
| FALSE | High | TPEEHGLYI | [K].TPEEH( | 1xDeamida | A0A0R4J02 | 0.000411 | 0 | 1 | 4 | 2 | A0A0R4J02 | DNA repair | A0A0R4J02 | 0 | 3507.62 | 0 | -0.2 | 0.094944 | 6.23 | 106 | 94 | 10.15 | 4.2 | | High | High | High | High | High | High | High | High | High | High | High | 0 | 8.79E-05 | 4.28 |
| FALSE | High | TPEEHGLYI | [K].TPEEH( | 2xPhospho | A0A0R4J02 | 7.75E-05 | 0 | 1 | 4 | 2 | A0A0R4J02 | DNA repair | A0A0R4J02 | 0 | 3586.603 | 0 | 0.04 | 0.055133 | 1.9 | 97.1 | 102.9 | 7.33 | 5.71 | | High | High | High | High | High | High | High | High | High | High | High | 0 | 1.35E-05 | 4.89 |
| FALSE | High | TPEELDDSI | [R].TPEELD | 1xPhospho | P26231 1xl | 5.51E-09 | 0 | 1 | 3 | 14 | P26231 | Catenin alp | P26231 [63 | 0 | 2468.023 | 0 | -0.85 | 0.000302 | 2.03 | 129.6 | 70.4 | 1.7 | 6.25 | | High | High | High | High | High | High | High | High | High | High | High | 0 | 2.93E-10 | 4.71 |
| FALSE | High | TPEELDDSI | [R].TPEELD | 2xPhospho | P26231 2xl | 6.85E-06 | 0 | 1 | 3 | 2 | P26231 | Catenin alp | P26231 [63 | 1 | 2791.122 | 0 | -1.26 | 0.000678 | 2.63 | 142.1 | 57.9 | 5.72 | 9.65 | | High | High | High | High | High | High | High | High | High | High | High | 0 | 8.82E-07 | 3.94 |
| FALSE | High | TPEGSQGE | [K].TPEGS( | 1xPhospho | A0A0R4J2D | 1.11E-05 | 0 | 1 | 3 | 1 | A0A0R4J2D | Ubiquitin c | A0A0R4J2D | 0 | 2531.129 | 0 | -0.55 | 0.010242 | 2.15 | 121 | 79 | 6.97 | 17.74 | | High | High | High | High | High | High | High | High | High | High | High | 0 | 1.53E-06 | 4.43 |
| FALSE | High | TPELIVPVD | [R].TPELIVI | 1xPhospho | E9QK89 1x | 0.007951 | 0.000736 | 1 | 2 | 2 | E9QK89 | Mediator c | E9QK89 [1 | 0 | 3023.579 | 0 | -0.54 | 0.025442 | 11.07 | 120.4 | 79.6 | 11.57 | 18.02 | | High | High | High | High | High | High | High | High | High | High | High | 0.000327 | 0.002438 | 4.01 |
| FALSE | High | TPELVELSS | [R].TPELVE | 1xPhospho | Q80Z37 1x | 9E-07 | 0 | 1 | 2 | 3 | Q80Z37 | E3 ubiquiti | Q80Z37 [4 | 0 | 2660.3 | 0 | -0.1 | 0.141563 | 6.73 | 103 | 97 | 9.9 | 7.34 | | High | High | High | High | High | High | High | High | High | High | High | 0 | 9.05E-08 | 4.09 |
| FALSE | High | TPESFLGPN | [K].TPESFL | 1xPhospho | Q80VP1-2 | 2.52E-06 | 0 | 1 | 2 | 3 | Q80VP1-2 | Isoform 2 c | Q80VP1-2 | 0 | 3557.866 | 0 | -0.41 | 0.00687 | 6.96 | 117.6 | 82.4 | 5.46 | 10.41 | | High | High | High | High | High | High | High | High | High | High | High | 0 | 2.86E-07 | 4.38 |
| FALSE | High | TPESTNVPS | [K].TPESTN | 1xPhospho | Q9R0L6-2 : | 1.04E-06 | 0 | 1 | 3 | 6 | Q9R0L6-2 | Isoform 2 c | Q9R0L6-2 [ | 0 | 2255.084 | 0 | 0.42 | 0.001807 | 2.79 | 84.9 | 115.1 | 5.18 | 6.38 | | High | High | High | High | High | High | High | High | High | High | High | 0 | 1.65E-07 | 3.12 |
| FALSE | High | TPESVVPTC | [K].TPESVV | 1xPhospho | E9QK89 1x | 0.00625 | 0.00062 | 1 | 2 | 1 | E9QK89 | Mediator c | E9QK89 [1 | 0 | 2489.33 | 0 | -0.49 | 0.024064 | 7.17 | 112.4 | 87.6 | 12.91 | 14.05 | | High | High | High | High | High | High | High | High | High | High | High | 0.000275 | 0.001856 | 2.83 |
| FALSE | High | TPEVSFLPE | [K].TPEVSF | 1xPhospho | Q5U4C3 1x | 1.7E-07 | 0 | 1 | 1 | 3 | Q5U4C3 | Splicing fac | Q5U4C3 [9 | 0 | 2272.058 | 0 | -0.11 | 0.426435 | 6.95 | 101.9 | 98.1 | 4.93 | 7.17 | | High | High | High | High | High | High | High | High | High | High | High | 0 | 2.41E-05 | 3.58 |
| FALSE | High | TPFAEDQLI | [K].TPFAED | 1xPhospho | B7ZNA0 1x | 2.82E-05 | 0 | 2 | 8 | 2 | B7ZNA0; Q | Shroom3 p | B7ZNA0 [4 | 1 | 3113.571 | 0 | 0.17 | 0.002097 | 1.69 | 95 | 105 | 3.86 | 3.97 | Shared | High | High | High | High | High | High | High | High | High | High | High | 0 | 4.33E-06 | 3.79 |
| FALSE | High | TPFQVEHV | [K].TPFQVI | 1xPhospho | Q9DAR7 1x | 0.011048 | 0.000772 | 1 | 2 | 1 | Q9DAR7 | m7GpppX | Q9DAR7 [8 | 0 | 2632.415 | 0 | -0.95 | 0.000712 | 1.73 | 130.5 | 69.5 | 4.45 | 10.53 | | High | High | High | High | High | High | High | High | High | High | High | 0.000399 | 0.00349 | 2.97 |
| FALSE | High | TPEEHGLYI | [K].TPEEH( | 1xPhospho | A0A0R4J02 | 1.03E-06 | 0 | 1 | 4 | 3 | A0A0R4J02 | DNA repair | A0A0R4J02 | 0 | 3506.636 | 0 | -0.41 | 0.00397 | 1.17 | 114.6 | 85.4 | 5.31 | 7.21 | | High | High | High | High | High | High | High | High | High | High | High | 0 | 1.05E-07 | 3.82 |
| FALSE | High | TPASPVVH | [K].TPASPV | 1xPhospho | G5E924 1x | 0.000308 | 0 | 1 | 4 | 2 | G5E924 | Heterogen | G5E924 [8 | 0 | 1385.75 | 0 | -0.85 | 0.001518 | 5.57 | 128.7 | 71.3 | 5.24 | 7.33 | | High | High | High | High | High | High | High | High | High | High | High | 0 | 6.34E-05 | 1.81 |
| FALSE | High | TPARESEG( | [K].TPARES | 1xPhospho | Q9DBY8 1x | 2.47E-05 | 0 | 1 | 2 | 1 | Q9DBY8 | Nuclear va | Q9DBY8 [1 | 1 | 2131.063 | 0 | -0.25 | 0.070719 | 10.76 | 109.8 | 90.2 | 10.53 | 19.86 | | High | High | High | High | High | High | High | High | High | High | High | 0 | 3.71E-06 | 2.28 |
| FALSE | High | TPAQSGAV | [R].TPAQS( | 1xPhospho | O55201 1x | 0.00537 | 0.000599 | 1 | 2 | 1 | O55201 | Transcripti | O55201 [8 | 0 | 2219.008 | 0 | -0.54 | 0.015085 | 5.37 | 125.7 | 74.3 | 12.15 | 13.25 | | High | High | High | High | High | High | High | High | High | High | High | 0.000267 | 0.00157 | 3.83 |
| FALSE | High | TNSPGFQK | [R].TNSPGI | 1xPhospho | Q6NZN0 1x | 0.092372 | 0.007211 | 1 | 8 | 2 | Q6NZN0 | RNA-bindir | Q6NZN0 [5 | 0 | 1416.729 | 0 | -0.66 | 0.000477 | 0.53 | 122.3 | 77.7 | 3.33 | 2.26 | | High | High | High | High | High | High | High | High | High | High | High | 0.003659 | 0.03924 | 2.22 |
| FALSE | High | TNSPSISPSI | [K].TNSPSI | 1xPhospho | E9QM77 1: | 0.000255 | 0 | 1 | 8 | 2 | E9QM77 | Ataxin-2 O | E9QM77 [8 | 0 | 2338.151 | 0 | -0.68 | 0.014963 | 9.09 | 123.4 | 76.6 | 8.97 | 24.29 | | High | High | High | High | High | High | High | High | High | High | High | 0 | 5.12E-05 | 2.64 |
| FALSE | High | TNSPSISPSI | [K].TNSPSI | 1xPhospho | E9QM77 1: | 0.000305 | 0 | 1 | 8 | 9 | E9QM77 | Ataxin-2 O | E9QM77 [8 | 0 | 2354.146 | 0 | -0.5 | 0.067834 | 4.85 | 114.5 | 85.5 | 5.66 | 18.46 | | High | High | High | High | High | High | High | High | High | High | High | 0 | 6.26E-05 | 2.79 |
| FALSE | High | TNSPSSAK | [R].TNSPSS | 1xPhospho | G5E870 1x | 0.001799 | 5.66E-05 | 1 | 6 | 3 | G5E870 | E3 ubiquiti | G5E870 [8 | 0 | 1329.682 | 0 | -1.5 | 0.001698 | 1.68 | 145.7 | 54.3 | 9.5 | 12.36 | | High | High | High | High | High | High | High | High | High | High | High | 0.000382 | 0.003211 | 1.71 |
| FALSE | High | TNSSDSERS | [R].TNSSDS | 2xPhospho | Q8K212 2x | 0.000114 | 0 | 1 | 3 | 2 | Q8K212 | Phosphofu | Q8K212 [5 | 1 | 2573.123 | 0 | 2.5 | 0.001912 | 25.45 | 29.8 | 170.2 | 19.99 | 15.28 | | High | High | High | High | High | High | High | High | High | High | High | 0 | 2.09E-05 | 1.78 |
| FALSE | High | TNSSPLFK | [K].TNSSPL | 1xPhospho | B2RQG2 1x | 0.013426 | 0.000976 | 2 | 2 | 3 | B2RQG2; A | PHD finger | B2RQG2 [2 | 0 | 1431.765 | 0 | -0.98 | 0.000608 | 2.09 | 134 | 66 | 4.1 | 7.23 | Shared | High | High | High | High | High | High | High | High | High | High | High | 0.000505 | 0.00435 | 2.85 |
| FALSE | High | TNSSSSSPV | [R].TNSSSS | 1xPhospho | Q8BMB0 1 | 0.002486 | 0.000109 | 2 | 7 | 4 | Q8BMB0; / | BRCA2-inte | Q8BMB0 [: | 0 | 1743.929 | 0 | -0.17 | 0.095979 | 10.78 | 105.8 | 94.2 | 7.24 | 5.85 | Shared | High | High | High | High | High | High | High | High | High | High | High | 4.94E-05 | 0.000659 | 3 |
| FALSE | High | TNSSSSSPV | [R].TNSSSS | 2xPhospho | Q8BMB0 2 | 0.000501 | 0 | 2 | 7 | 1 | Q8BMB0; / | BRCA2-inte | Q8BMB0 [: | 0 | 1823.896 | 0 | -0.75 | 0.000613 | 4.45 | 123.1 | 76.9 | 2.88 | 7.41 | Shared | High | High | High | High | High | High | High | High | High | High | High | 0 | 0.000109 | 2.94 |
| FALSE | High | TNSSTGLEI | [K].TNSSTC | 1xPhospho | E9Q6J5-3 1 | 7.1E-06 | 0 | 1 | 3 | 3 | E9Q6J5-3 | Isoform Ne | E9Q6J5-3 [ | 1 | 2871.293 | 0 | 0.31 | 0.126947 | 9.96 | 89.2 | 110.8 | 3.94 | 14.4 | | High | High | High | High | High | High | High | High | High | High | High | 0 | 5.08E-05 | 4.49 |
| FALSE | High | TNSSTGLEI | [K].TNSSTC | 2xPhospho | E9Q6J5-3 2 | 0.015741 | 0.001124 | 1 | 3 | 4 | E9Q6J5-3 | Isoform Ne | E9Q6J5-3 [ | 1 | 2951.26 | 0 | 0.28 | 0.040582 | 8.37 | 90.9 | 109.1 | 2.78 | 9.85 | | High | High | High | High | High | High | High | High | High | High | High | 0.00186 | 0.01873 | 3.38 |
| FALSE | High | TNSTGGSS | [R].TNSTG( | 1xPhospho | F8VPM7 1x | 4.22E-07 | 0 | 1 | 8 | 5 | F8VPM7 | ELKS/Rab6 | F8VPM7 [3 | 0 | 1819.802 | 0 | 0.47 | 0.417383 | 21.48 | 86.3 | 113.7 | 37.84 | 56.08 | | High | High | High | High | High | High | High | High | High | High | High | 0 | 1.89E-07 | 4.11 |
| FALSE | High | TNSTGGSS | [R].TNSTG( | 1xPhospho | F8VPM7 1x | 5.48E-07 | 0 | 1 | 8 | 7 | F8VPM7 | ELKS/Rab6 | F8VPM7 [3 | 0 | 2048.965 | 0 | 0.49 | 0.015507 | 17.85 | 86.1 | 113.9 | 4.89 | 10.58 | | High | High | High | High | High | High | High | High | High | High | High | 0 | 5.2E-08 | 4.54 |
| FALSE | High | TNSTGGSS | [R].TNSTG( | 1xDeamida | F8VPM7 1x | 5.48E-07 | 0 | 1 | 8 | 9 | F8VPM7 | ELKS/Rab6 | F8VPM7 [3 | 0 | 2049.949 | 0 | 0.52 | 0.102275 | 21.56 | 86.6 | 113.4 | 8.88 | 31.62 | | High | High | High | High | High | High | High | High | High | High | High | 0 | 5.62E-08 | 4.51 |
| FALSE | High | TNTMNGSI | [R].TNTMN | 1xPhospho | E9QLZ9 1xl | 0.011549 | 0.00086 | 1 | 12 | 1 | E9QLZ9 | Protein en: | E9QLZ9 [7 | 0 | 1406.675 | 0 | -0.57 | 0.169831 | 23.35 | 110 | 90 | 31.3 | 13.85 | | High | High | High | High | High | High | High | High | High | High | High | 0.000437 | 0.003689 | 1.88 |
| FALSE | High | TNTPVK | [R].TNTPVI | 1xPhospho | Q9ESZ8-5 : | 0.052576 | 0.00384 | 2 | 8 | 1 | Q9ESZ8-5; | Isoform 5 c | Q9ESZ8-5 [ | 0 | 1197.664 | 0 | -0.53 | 0.011564 | 2.4 | 118.3 | 81.7 | 6.01 | 9.41 | Shared | High | High | High | High | High | High | High | High | High | High | High | 0.001999 | 0.02033 | 1.61 |
| FALSE | High | TNVLPTQQ | [K].TNVLPT | 1xCarbami | Q6P6I6 1xP | 2.66E-06 | 0 | 1 | 2 | 1 | Q6P6I6 | DNA-direct | Q6P6I6 [24 | 0 | 2264.033 | 0 | -0.36 | 0.002665 | 2.19 | 115.3 | 84.7 | 6.87 | 4.14 | | High | High | High | High | High | High | High | High | High | High | High | 0 | 3.06E-07 | 4.97 |
| FALSE | High | TNVQTTTI | [R].TNVQC | 1xCarbami | Q69ZR2 1x | 0.017041 | 0.001204 | 1 | 4 | 3 | Q69ZR2 | E3 ubiquiti | Q69ZR2 [1 | 0 | 3880.851 | 0 | 0.15 | 0.178934 | 3.25 | 94.9 | 105.1 | 9.94 | 10.15 | | High | High | High | High | High | High | High | High | High | High | High | 0.000638 | 0.00572 | 2.75 |
| FALSE | High | TPAAAAAM | [R].TPAAA/ | 1xPhospho | Q8BTI8 1xF | 0.000121 | 0 | 1 | 3 | 2 | Q8BTI8 | Serine/argi | Q8BTI8 [22 | 0 | 1650.824 | 0 | -0.59 | 0.021468 | 10.78 | 123.9 | 76.1 | 15.5 | 20.62 | | High | High | High | High | High | High | High | High | High | High | High | 0 | 2.21E-05 | 3.1 |
| FALSE | High | TPAAAAAM | [R].TPAAA/ | 1xPhospho | Q8BTI8 1xF | 6.81E-07 | 0 | 1 | 3 | 14 | Q8BTI8 | Serine/argi | Q8BTI8 [22 | 0 | 1666.819 | 0 | -0.52 | 0.042483 | 0.41 | 118 | 82 | 7.86 | 12.19 | | High | High | High | High | High | High | High | High | High | High | High | 0 | 4.81E-06 | 3.13 |
| FALSE | High | TPAALAALS | [R].TPAALA | 1xPhospho | Q8BTI8 1xF | 2.83E-06 | 0 | 1 | 3 | 1 | Q8BTI8 | Serine/argi | Q8BTI8 [22 | 0 | 2868.434 | 0 | 0.11 | 0.032688 | 6.53 | 96.2 | 103.8 | 6.29 | 9.69 | | High | High | High | High | High | High | High | High | High | High | High | 0 | 3.28E-07 | 2.9 |
| FALSE | High | TPAASAVN | [R].TPAASA | 1xPhospho | Q8BTI8 1xF | 0.003655 | 0.00021 | 1 | 3 | 2 | Q8BTI8 | Serine/argi | Q8BTI8 [22 | 0 | 1507.783 | 0 | -0.33 | 0.013644 | 9.06 | 111.6 | 88.4 | 4.89 | 8.92 | | High | High | High | High | High | High | High | High | High | High | High | 0.000204 | 0.001357 | 1.39 |
| FALSE | High | TPADTGFA | [K].TPADT( | 1xPhospho | A0A0R4J0I | 0.000772 | 0 | 1 | 3 | 3 | A0A0R4J0I | ETS domair | A0A0R4J0I | 0 | 3022.439 | 0 | -0.65 | 0.020243 | 7.52 | 126.7 | 73.3 | 13.73 | 15.73 | | High | High | High | High | High | High | High | High | High | High | High | 0 | 0.000285 | 2.2 |
| FALSE | High | TPAGGPQT | [K].TPAGG | 1xPhospho | Q3UFM6 1 | 1.37E-07 | 0 | 1 | 3 | 2 | Q3UFM6 | Expressed | Q3UFM6 [ | 0 | 2161.049 | 0 | 0.15 | 0.163368 | 8.81 | 96.1 | 103.9 | 4.86 | 10.57 | | High | High | High | High | High | High | High | High | High | High | High | 0 | 1.09E-08 | 5.69 |
| FALSE | High | TPAGGPQT | [K].TPAGG | 2xPhospho | Q3UFM6 2 | 0.069325 | 0.005112 | 1 | 3 | 1 | Q3UFM6 | Expressed | Q3UFM6 [ | 0 | 2241.015 | 0 | -0.67 | 0.003156 | 4.46 | 117.1 | 82.9 | 6.68 | 8.02 | | High | High | High | High | High | High | High | High | High | High | High | 0.002662 | 0.02817 | 2.01 |
| FALSE | High | TPALQPSPI | [K].TPALQI | 1xPhospho | Q3TIV2 1xl | 0.005516 | 0.00062 | 1 | 4 | 2 | Q3TIV2 | Putative ur | Q3TIV2 [17 | 0 | 1910.969 | 0 | -0.62 | 0.003771 | 5.03 | 121.5 | 78.5 | 7.19 | 6.49 | | High | High | High | High | High | High | High | High | High | High | High | 0.000275 | 0.001618 | 2.29 |
| FALSE | High | TPALQPSPI | [K].TPALQI | 1xDeamida | Q3TIV2 1xl | 0.007673 | 0.000664 | 1 | 4 | 2 | Q3TIV2 | Putative ur | Q3TIV2 [17 | 0 | 1911.953 | 0 | -0.65 | 0.00406 | 4.72 | 123.9 | 76.1 | 9.97 | 6.47 | | High | High | High | High | High | High | High | High | High | High | High | 0.000294 | 0.002324 | 2.62 |
| FALSE | High | TPALSTPPL | [K].TPALST | 1xCarbami | Q3TAA7 1x | 3.67E-05 | 0 | 1 | 1 | 2 | Q3TAA7 | Serine/thre | Q3TAA7 [4 | 0 | 3546.769 | 0 | -0.28 | 0.087732 | 5.01 | 105.1 | 94.9 | 17.12 | 16.75 | | High | High | High | High | High | High | High | High | High | High | High | 0 | 6.26E-06 | 3.75 |
| FALSE | High | TPAPATTPS | [K].TPAPAT | 1xPhospho | Q6NXI6 1xl | 0.000359 | 0 | 1 | 4 | 2 | Q6NXI6 | Regulation | Q6NXI6 [5 | 0 | 2468.331 | 0 | -0.09 | 0.249326 | 17.43 | 113.8 | 86.2 | 14.01 | 16.09 | | High | High | High | High | High | High | High | High | High | High | High | 0 | 7.58E-05 | 2.65 |
| FALSE | High | TPAPSPQTS | [R].TPAPSP | 1xPhospho | Q9DCS9 1x | 0.002022 | 5.66E-05 | 1 | 2 | 3 | Q9DCS9 | NADH dehy | Q9DCS9 [1 | 0 | 2564.378 | 0 | 0.43 | 0.14685 | 12.44 | 93.3 | 106.7 | 16.3 | 9.41 | | High | High | High | High | High | High | High | High | High | High | High | 2.58E-05 | 0.000524 | 3.01 |
| FALSE | High | TNSPAYSDI | [K].TNSPAY | 1xPhospho | Q8BZ47 1x | 1.19E-06 | 0 | 1 | 2 | 2 | Q8BZ47 | Zinc finger | Q8BZ47 [8 | 0 | 2451.096 | 0 | 0.01 | 0.634017 | 12.45 | 98.2 | 101.8 | 7.32 | 16.71 | | High | High | High | High | High | High | High | High | High | High | High | 0 | 1.23E-07 | 3.45 |
| FALSE | High | TSSSGFADI | [K].TSSSGF | 1xDeamida | Q60902 1x | 0.113753 | 0.009241 | 1 | 2 | 5 | Q60902 | Epidermal | Q60902 [8 | 0 | 3047.408 | 0 | 0.91 | 0.003445 | 2.42 | 67.5 | 132.5 | 17.36 | 18.89 | | High | High | High | High | High | High | High | High | High | High | High | 0.008759 | 0.09385 | 3.37 |
| FALSE | High | TQAQHSSP | [R].TQAQH | 1xPhospho | A2ADY9 1x | 2.11E-07 | 0 | 1 | 1 | 2 | A2ADY9 | Protein DD | A2ADY9 [1 | 0 | 2901.376 | 0 | -0.99 | 0.002547 | 4.4 | 134 | 66 | 10.78 | 11.07 | | High | High | High | High | High | High | High | High | High | High | High | 0 | 1.77E-08 | 4.05 |
| FALSE | High | TQAQVSSP | [R].TQAQV | 1xPhospho | Q811P8 1x | 2.81E-07 | 0 | 1 | 3 | 4 | Q811P8 | Rho GTPas | Q811P8 [6 | 0 | 2040.078 | 0 | -1.1 | 0.001495 | 2.59 | 136.1 | 63.9 | 9.43 | 7.84 | | High | High | High | High | High | High | High | High | High | High | High | 0 | 2.18E-07 | 3.98 |
| FALSE | High | TSMGGTQ | [R].TSMGG | 1xPhospho | Q02248 1x | 6.16E-07 | 0 | 1 | 4 | 3 | Q02248 | Catenin be | Q02248 [5 | 0 | 1933.904 | 0 | -0.67 | 0.003287 | 5.83 | 122.1 | 77.9 | 3.62 | 9.67 | | High | High | High | High | High | High | High | High | High | High | High | 0 | 5.93E-08 | 3.96 |
| FALSE | High | TSMGGTQ | [R].TSMGG | 1xPhospho | Q02248 1x | 9.79E-08 | 0 | 1 | 4 | 11 | Q02248 | Catenin be | Q02248 [5 | 0 | 1949.899 | 0 | -0.5 | 0.003612 | 6.7 | 117 | 83 | 5.87 | 4.47 | | High | High | High | High | High | High | High | High | High | High | High | 0 | 7.49E-09 | 4 |
| FALSE | High | TSPAAQAP | [K].TSPAA( | 1xPhospho | A2BIE1 1xF | 9.82E-06 | 0 | 1 | 2 | 1 | A2BIE1 | Glutamine | A2BIE1 [11 | 0 | 2355.185 | 0 | -0.52 | 0.002162 | 5.31 | 116.6 | 83.4 | 8.65 | 7.13 | | High | High | High | High | High | High | High | High | High | High | High | 0 | 1.33E-06 | 3.5 |
| FALSE | High | TSPADHGG | [R].TSPADH | 1xPhospho | A2AN08-5 | 2.15E-07 | 0 | 1 | 4 | 1 | A2AN08-5 | Isoform 5 c | A2AN08-5 | 0 | 3149.397 | 0 | -0.29 | 0.166374 | 11.8 | 110.7 | 89.3 | 9.56 | 12.5 | | High | High | High | High | High | High | High | High | High | High | High | 0 | 1.8E-08 | 4 |
| FALSE | High | TSPAGGTW | [R].TSPAG( | 1xPhospho | E9QM77 1: | 6.47E-05 | 0 | 1 | 8 | 1 | E9QM77 | Ataxin-2 O | E9QM77 [6 | 0 | 1953.963 | 0 | 0.34 | 0.064315 | 32.73 | 85 | 115 | 11.79 | 16.06 | | High | High | High | High | High | High | High | High | High | High | High | 0 | 1.1E-05 | 2.46 |
| FALSE | High | TSPAPSDS/ | [R].TSPAPS | 1xPhospho | Q5SXY1 1x | 5.4E-05 | 0 | 1 | 5 | 2 | Q5SXY1 | Cytospin-B | Q5SXY1 [8 | 0 | 1799.919 | 0 | 1.71 | 0.001222 | 6.37 | 47.5 | 152.5 | 13.47 | 8.18 | | High | High | High | High | High | High | High | High | High | High | High | 0 | 9.04E-06 | 1.85 |
| FALSE | High | TSPAPWK | [R].TSPAPW | 1xPhospho | Q8BTI8 1xF | 0.077825 | 0.005963 | 1 | 3 | 1 | Q8BTI8 | Serine/argi | Q8BTI8 [18 | 0 | 1324.707 | 0 | -0.69 | 0.006921 | 5.25 | 123.4 | 76.6 | 7.7 | 12.06 | | High | High | High | High | High | High | High | High | High | High | High | 0.003079 | 0.03205 | 1.29 |
| FALSE | High | TSPASLDFF | [K].TSPASL | 1xPhospho | A0A140T8 | 0.053926 | 0.00384 | 1 | 1 | 2 | A0A140T8 | Chromoso | A0A140T8 | 0 | 1725.83 | 0 | -0.78 | 0.448022 | 21.32 | 124.1 | 75.9 | 52.55 | 64.28 | | High | High | High | High | High | High | High | High | High | High | High | 0.002031 | 0.02095 | 2.21 |
| FALSE | High | TSPASLDFF | [K].TSPASL | 1xPhospho | A0A140T8 | 1.56E-05 | 0 | 1 | 1 | 6 | A0A140T8 | Chromoso | A0A140T8 | 0 | 1954.993 | 0 | -0.85 | 0.000477 | 3.09 | 128.7 | 71.3 | 4.88 | 3.57 | | High | High | High | High | High | High | High | High | High | High | High | 0 | 2.21E-06 | 2.96 |
| FALSE | High | TSPDHPK | [R].TSPDHF | 1xPhospho | Q8CDM8 1 | 0.075927 | 0.005826 | 1 | 1 | 1 | Q8CDM8 | Protein FAI | Q8CDM8 [ | 0 | 1319.676 | 0 | -0.45 | 0.002708 | 7.26 | 118.9 | 81.1 | 3.86 | 10.86 | | High | High | High | High | High | High | High | High | High | High | High | 0.002976 | 0.03128 | 1.97 |
| FALSE | High | TSPDPTPVS | [K].TSPDPT | 1xPhospho | Q60591 1x | 0.002695 | 0.000109 | 1 | 14 | 5 | Q60591 | Nuclear fac | Q60591 [3 | 0 | 1922.988 | 0 | 1.93 | 0.000725 | 3.74 | 41.6 | 158.4 | 2.23 | 10.26 | | High | High | High | High | High | High | High | High | High | High | High | 4.94E-05 | 0.000726 | 3.01 |
| FALSE | High | TSPDSTSSV | [K].TSPDST | 1xPhospho | Q7TSY8 1x | 0.000731 | 0 | 1 | 1 | 1 | Q7TSY8 | Shugoshin | Q7TSY8 [1 | 0 | 1432.652 | 0 | -0.91 | 0.00043 | 1 | 133 | 67 | 6.63 | 9.46 | | High | High | High | High | High | High | High | High | High | High | High | 0 | 0.000168 | 1.56 |
| FALSE | High | TSPDTDSEI | [K].TSPDTI | 1xPhospho | Q5XG69 1x | 1.21E-05 | 0 | 1 | 1 | 4 | Q5XG69 | Soluble lan | Q5XG69 [4 | 0 | 2619.178 | 0 | -0.96 | 0.003141 | 2.87 | 136 | 64 | 6.99 | 19.46 | | High | High | High | High | High | High | High | High | High | High | High | 0 | 1.67E-06 | 3.46 |
| FALSE | High | TSMDGAPI | [K].TSMDG | 1xPhospho | E9Q842 1x | 4.27E-08 | 0 | 1 | 3 | 2 | E9Q842 | Neuron na | E9Q842 [1 | 0 | 2339.006 | 0 | 0.98 | 0.002958 | 24.21 | 65.7 | 134.3 | 8.61 | 10.44 | | High | High | High | High | High | High | High | High | High | High | High | 0 | 2.94E-09 | 4.15 |
| FALSE | High | TSPDTDSEI | [K].TSPDTI | 1xPhospho | Q5XG69 1x | 2.31E-08 | 0 | 1 | 1 | 11 | Q5XG69 | Soluble lan | Q5XG69 [4 | 0 | 2635.173 | 0 | -0.83 | 0.009734 | 8.85 | 126.9 | 73.1 | 12.08 | 14.59 | | High | High | High | High | High | High | High | High | High | High | High | 0 | 1.48E-09 | 5.22 |
| FALSE | High | TSPEFSQPI | [K].TSPEFS | 1xPhospho | Q8VBT0 1x | 0.041781 | 0.003126 | 1 | 1 | 1 | Q8VBT0 | Thioredoxi | Q8VBT0 [2 | 0 | 1671.876 | 0 | 0.56 | 0.011725 | 25.47 | 79.6 | 120.4 | 7.76 | 10.03 | | High | High | High | High | High | High | High | High | High | High | High | 0.001609 | 0.01574 | 2.92 |
| FALSE | High | TSPEPQREI | [R].TSPEPC | 2xPhospho | Q9Z1R2 2x | 0.017809 | 0.001282 | 1 | 7 | 1 | Q9Z1R2 | Large proli | Q9Z1R2 [9 | 1 | 2919.243 | 0 | -0.39 | 0.025724 | 8.01 | 111.7 | 88.3 | 6.27 | 19.79 | | High | High | High | High | High | High | High | High | High | High | High | 0.000689 | 0.005995 | 2.94 |
| FALSE | High | TSPGIPQK | [R].TSPGIP | 1xPhospho | Q3UHU5 1 | 0.107603 | 0.008634 | 1 | 5 | 1 | Q3UHU5 | Microtubu | Q3UHU5 [ | 0 | 1365.754 | 0 | -1.42 | 0.000872 | 1.05 | 147.2 | 52.8 | 8.39 | 9.42 | | High | High | High | High | High | High | High | High | High | High | High | 0.004399 | 0.0472 | 1.43 |
| FALSE | High | TSPGPAGR | [R].TSPGP/ | 1xPhospho | J3QK52 1xl | 0.002974 | 0.00016 | 1 | 2 | 1 | J3QK52 | Nucleolar c | J3QK52 [5 | 0 | 1304.667 | 0 | -0.24 | 0.036075 | 9.49 | 110.2 | 89.8 | 3.28 | 10.69 | | High | High | High | High | High | High | High | High | High | High | High | 7.19E-05 | 0.000811 | 1.92 |
| FALSE | High | TSPGRPLG | [K].TSPGRF | 1xPhospho | Q80X32 1x | 0.039686 | 0.00283 | 1 | 1 | 2 | Q80X32 | UPF0461 p | Q80X32 [1 | 0 | 1652.914 | 0 | 0.42 | 0.007781 | 8.94 | 82.8 | 117.2 | 7.64 | 6.68 | | High | High | High | High | High | High | High | High | High | High | High | 0.002662 | 0.02844 | 1.2 |
| FALSE | High | TSPGSPSPF | [K].TSPGSF | 1xPhospho | Q6ZQ32 1x | 0.016892 | 0.001204 | 2 | 7 | 4 | Q6ZQ32; F | MKIAA085 | Q6ZQ32 [1 | 0 | 1194.572 | 0 | 1.38 | 0.000209 | 2.88 | 56.5 | 143.5 | 2.86 | 4.56 | Shared | High | High | High | High | High | High | High | High | High | High | High | 0.000638 | 0.005647 | 2.27 |
| FALSE | High | TSPGTSSAY | [R].TSPGTS | 2xPhospho | Q7TSZ8 2x | 1.01E-07 | 0 | 1 | 1 | 6 | Q7TSZ8 | Nucleus ac | Q7TSZ8 [2 | 0 | 4445.673 | 0 | -0.77 | 0.003906 | 4.41 | 127.4 | 72.6 | 8.25 | 16.08 | | High | High | High | High | High | High | High | High | High | High | High | 0 | 7.73E-09 | 7.17 |
| FALSE | High | TSPLMLDR | [R].TSPLMI | 1xPhospho | Q8BTI8 1xF | 0.039686 | 0.00283 | 1 | 3 | 2 | Q8BTI8 | Serine/argi | Q8BTI8 [23 | 0 | 1241.616 | 0 | -0.54 | 0.015001 | 5 | 121.7 | 78.3 | 8.48 | 14.55 | | High | High | High | High | High | High | High | High | High | High | High | 0.001468 | 0.01481 | 2.51 |
| FALSE | High | TSPLMLDR | [R].TSPLMI | 1xPhospho | Q8BTI8 1xF | 0.009843 | 0.000736 | 1 | 3 | 6 | Q8BTI8 | Serine/argi | Q8BTI8 [23 | 0 | 1257.611 | 0 | -0.43 | 0.014908 | 2.92 | 114.7 | 85.3 | 5.23 | 13.54 | | High | High | High | High | High | High | High | High | High | High | High | 0.000344 | 0.003071 | 1.86 |

| | | | | | | | | | | | | | | | | | | | | | | | | | | | | | | | | | | | | | | |
|---|---|---|---|---|---|---|---|---|---|---|---|---|---|---|---|---|---|---|---|---|---|---|---|---|---|---|---|---|---|---|---|---|---|---|---|---|---|---|
| FALSE | High | TSPINLGLT | [R].TSPINL | 1xPhospho | Q3UQ91 1x | 4.4E-06 | 0 | 1 | 2 | 1 | Q3UQ91 | Putative ur | Q3UQ91 [2 | 0 | 2349.319 | 0 | 0.9 | 0.00297 | 17.16 | 68.8 | 131.2 | 10.78 | 10.6 | | High | High | High | High | High | High | High | High | High | High | High | 0 | 5.38E-07 | 3.58 |
| FALSE | High | TSPLNSSGS | [K].TSPLNS | 1xPhospho | Q3V141 1x | 4.36E-06 | 0 | 1 | 8 | 6 | Q3V141 | Putative ur | Q3V141 [5 | 0 | 1586.737 | 0 | -0.5 | 0.001003 | 3.52 | 116.2 | 83.8 | 4.92 | 0.87 | | High | High | High | High | High | High | High | High | High | High | High | 0 | 2.04E-06 | 2.88 |
| FALSE | High | TSPLNSSGS | [K].TSPLNS | 1xDeamida | Q3V141 1x | 0.000112 | 0 | 1 | 8 | 2 | Q3V141 | Putative ur | Q3V141 [5 | 0 | 1587.721 | 0 | -0.34 | 0.018135 | 4.43 | 112 | 88 | 11.92 | 4.25 | | High | High | High | High | High | High | High | High | High | High | High | 0 | 2.05E-05 | 1.6 |
| FALSE | High | TSPLNSSGS | [K].TSPLNS | 2xPhospho | Q3V141 2x | 0.026421 | 0.001676 | 1 | 8 | 1 | Q3V141 | Putative ur | Q3V141 [5 | 0 | 1666.704 | 0 | -0.96 | 0.000641 | 2.12 | 130.7 | 69.3 | 4.67 | 7.72 | | High | High | High | High | High | High | High | High | High | High | High | 0.000903 | 0.009325 | 2.15 |
| FALSE | High | TSPDTTFAE | [R].TSPDTT | 1xCarbami | Q6Y5D8 1x | 9.02E-08 | 0 | 1 | 4 | 2 | Q6Y5D8 | Rho GTPas | Q6Y5D8 [5 | 0 | 2723.259 | 0 | -1.58 | 0.000253 | 0.85 | 148 | 52 | 4.08 | 10.41 | | High | High | High | High | High | High | High | High | High | High | High | 0 | 6.83E-09 | 4.52 |
| FALSE | High | TSPNRLSPK | [R].TSPNRL | 2xPhospho | Q3U0M8 2 | 0.071662 | 0.005278 | 1 | 3 | 2 | Q3U0M8 | Putative ur | Q3U0M8 [ | 1 | 1617.817 | 0 | -0.84 | 0.000642 | 0.92 | 125.3 | 74.7 | 4.81 | 11.12 | | High | High | High | High | High | High | High | High | High | High | High | 0.002703 | 0.02908 | 1.64 |
| FALSE | High | TSMDGAPI | [K].TSMDG | 1xPhospho | E9Q842 1x | 2.27E-05 | 0 | 1 | 3 | 1 | E9Q842 | Neuron na | E9Q842 [1 | 0 | 2323.011 | 0 | 1.55 | 0.002691 | 30.59 | 49.4 | 150.6 | 26.44 | 9.89 | | High | High | High | High | High | High | High | High | High | High | High | 0 | 3.4E-06 | 4.15 |
| FALSE | High | TSLVASQD | [R].TSLVAS | 1xCarbami | A0A1B0GS | 0.086562 | 0.006435 | 1 | 8 | 2 | A0A1B0GS | Supervillin | A0A1B0GS | 0 | 3560.608 | 0 | -0.62 | 0.013639 | 6.17 | 125.3 | 74.7 | 20.74 | 43.66 | | High | High | High | High | High | High | High | High | High | High | High | 0.003317 | 0.03649 | 3.15 |
| FALSE | High | TSFSQEQS | [K].TSFSQE | 1xCarbami | S4R221 1x | 0.000308 | 0 | 1 | 9 | 2 | S4R221 | Rho GTPas | S4R221 [18 | 0 | 2270.907 | 0 | 0.31 | 0.036862 | 2.8 | 97 | 103 | 13.61 | 13.15 | | High | High | High | High | High | High | High | High | High | High | High | 0.000585 | 0.005379 | 3.14 |
| FALSE | High | TSFSVGSDI | [R].TSFSVG | 1xPhospho | B2RTN5 1x | 8.82E-06 | 0 | 1 | 4 | 2 | B2RTN5 | Glucocorti | B2RTN5 [1 | 0 | 1888.889 | 0 | -0.21 | 0.557247 | 0.48 | 103.7 | 96.3 | 6.09 | 11.17 | | High | High | High | High | High | High | High | High | High | High | High | 0 | 1.17E-06 | 3.13 |
| FALSE | High | TSGPPVSEL | [K].TSGPPV | 1xPhospho | P43274 1x | 0.000158 | 0 | 1 | 1 | 1 | P43274 | Histone H1 | P43274 [35 | 0 | 1766.971 | 0 | 0.34 | 0.034311 | 4 | 88.3 | 111.7 | 14.45 | 5.38 | | High | High | High | High | High | High | High | High | High | High | High | 0 | 3.01E-05 | 3.07 |
| FALSE | High | TSGQTFAN | [R].TSGQTI | 1xPhospho | Q8BU22 1x | 1.22E-05 | 0 | 1 | 4 | 1 | Q8BU22 | Kinesin-like | Q8BU22 [4 | 0 | 1665.743 | 0 | -0.75 | 0.003279 | 3.34 | 127.1 | 72.9 | 7.09 | 12.64 | | High | High | High | High | High | High | High | High | High | High | High | 0 | 1.7E-06 | 3.06 |
| FALSE | High | TSGSDDPG | [R].TSGSDI | 1xPhospho | Q69ZW3 1x | 0.001799 | 5.66E-05 | 1 | 2 | 4 | Q69ZW3 | EH domain | Q69ZW3 [6 | 0 | 2790.345 | 0 | 0.24 | 0.083679 | 21.54 | 91.6 | 108.4 | 13.45 | 5.27 | | High | High | High | High | High | High | High | High | High | High | High | 2.58E-05 | 0.000462 | 2.82 |
| FALSE | High | TSGSGFHR | [R].TSGSGI | 1xPhospho | E0CYH0 1x | 0.001189 | 0 | 1 | 2 | 2 | E0CYH0 | MCG16685 | E0CYH0 [2 | 0 | 1157.53 | 0 | -0.87 | 0.000255 | 2.39 | 129.4 | 70.6 | 1.87 | 6.57 | | High | High | High | High | High | High | High | High | High | High | High | 0.003782 | 0.04083 | 1.89 |
| FALSE | High | TSGSSSSD | [R].TSGSSS | 1xPhospho | Q5SXY1 1x | 0.000118 | 0 | 1 | 8 | 6 | Q5SXY1 | Cytospin-B | Q5SXY1 [3 | 0 | 1593.777 | 0 | 1.97 | 0.001395 | 8.09 | 41.1 | 158.9 | 5.55 | 13.27 | | High | High | High | High | High | High | High | High | High | High | High | 0 | 0.000115 | 2.6 |
| FALSE | High | TSHAAEQS | [K].TSHAAI | 1xPhospho | B2RXS2 1x | 0.003195 | 0.00016 | 1 | 4 | 1 | B2RXS2 | Fam179b p | B2RXS2 [10 | 0 | 1855.854 | 0 | 0.55 | 0.273592 | 17 | 81.3 | 118.7 | 21.93 | 96.39 | | High | High | High | High | High | High | High | High | High | High | High | 7.19E-05 | 0.000877 | 1.99 |
| FALSE | High | TSHLDGSSI | [K].TSHLD | 1xPhospho | P55200 1x | 0.000154 | 0 | 1 | 2 | 1 | P55200 | Histone-lys | P55200 [22 | 0 | 1881.936 | 0 | 0.5 | 0.001999 | 7.9 | 83 | 117 | 2.72 | 8.56 | | High | High | High | High | High | High | High | High | High | High | High | 0 | 2.91E-05 | 2.93 |
| FALSE | High | TSLAGDTSI | [R].TSLAGI | 1xPhospho | B1ARU1 1x | 5.77E-06 | 0 | 2 | 19 | 1 | B1ARU1; F | Microtubu | B1ARU1 [5 | 0 | 2277.101 | 0 | 0.4 | 0.034683 | 20.67 | 86.3 | 113.7 | 9.01 | 8.7 | Shared | High | High | High | High | High | High | High | High | High | High | High | 0 | 7.3E-07 | 3.36 |
| FALSE | High | TSLDEDSAT | [K].TSLDED | 1xPhospho | Q6P1D7 1x | 0.015741 | 0.001124 | 1 | 2 | 2 | Q6P1D7 | Structure-s | Q6P1D7 [1 | 0 | 2714.276 | 0 | 0.26 | 0.012888 | 9.47 | 89.8 | 110.2 | 12.92 | 12.74 | | High | High | High | High | High | High | High | High | High | High | High | 0.002202 | 0.0234 | 2.61 |
| FALSE | High | TSLDEITSG | [K].TSLDEI | 1xPhospho | Q8BT07 1x | 2.72E-05 | 0 | 1 | 3 | 2 | Q8BT07 | Centrosom | Q8BT07 [3 | 0 | 1588.824 | 0 | 2.35 | 0.003854 | 33.39 | 30.9 | 169.1 | 14.8 | 21.3 | | High | High | High | High | High | High | High | High | High | High | High | 0 | 4.18E-06 | 2.8 |
| FALSE | High | TSLDVSNS | [K].TSLDVS | 1xPhospho | Q8CH77 1x | 0.000102 | 0 | 1 | 4 | 4 | Q8CH77 | Neuron na | Q8CH77 [6 | 0 | 2226.1 | 0 | -0.13 | 0.302438 | 19.64 | 107.7 | 92.3 | 9.44 | 10.07 | | High | High | High | High | High | High | High | High | High | High | High | 0 | 1.82E-05 | 3.06 |
| FALSE | High | TSLVIVESTI | [K].TSLVIVI | 1xPhospho | Q8C8R3-4 | 8.66E-06 | 0 | 1 | 7 | 1 | Q8C8R3-4 | Isoform 4 c | Q8C8R3-4 | 0 | 2362.138 | 0 | 0.81 | 0.005899 | 18.64 | 78 | 122 | 13.7 | 9.25 | | High | High | High | High | High | High | High | High | High | High | High | 0 | 1.15E-06 | 4.24 |
| FALSE | High | TSLEGSDV | [R].TSLEGS | 1xPhospho | Q8BYH3 1x | 0.052576 | 0.00384 | 1 | 1 | 1 | Q8BYH3 | tRNA:m(4) | Q8BYH3 [3 | 0 | 1599.746 | 0 | -1.45 | 0.000255 | 0.49 | 147.9 | 52.1 | 4.93 | 4.64 | | High | High | High | High | High | High | High | High | High | High | High | 0.001999 | 0.02041 | 1.86 |
| FALSE | High | TSLEQCPAI | [K].TSLEQC | 1xCarbami | E9Q7F2 1x | 0.006085 | 0.00062 | 1 | 1 | 3 | E9Q7F2 | E3 ubiquiti | E9Q7F2 [2 | 1 | 2614.165 | 0 | -0.27 | 0.03868 | 4.4 | 109.1 | 90.9 | 7.57 | 10.46 | | High | High | High | High | High | High | High | High | High | High | High | 0.000801 | 0.007184 | 1.89 |
| FALSE | High | TSLHALTVD | [R].TSLHAL | 1xPhospho | Q91VS8 1x | 0.004218 | 0.00036 | 1 | 2 | 2 | Q91VS8 | FERM, Rho | Q91VS8 [3 | 0 | 1603.866 | 0 | 0.68 | 0.004928 | 10.84 | 73.5 | 126.5 | 9.21 | 10.22 | | High | High | High | High | High | High | High | High | High | High | High | 0.000275 | 0.001888 | 3.42 |
| FALSE | High | TSLHALTVD | [R].TSLHAL | 1xPhospho | Q91VS8 1x | 6.37E-06 | 0 | 1 | 2 | 1 | Q91VS8 | FERM, Rho | Q91VS8 [3 | 0 | 1833.029 | 0 | 0.8 | 0.005736 | 2.19 | 72 | 128 | 8.02 | 13.33 | | High | High | High | High | High | High | High | High | High | High | High | 0 | 8.14E-07 | 3.21 |
| FALSE | High | TSLLFEDDA | [R].TSLLFEI | 1xPhospho | Q6PGL7 1x | 0.00642 | 0.00062 | 1 | 5 | 2 | Q6PGL7 | WASH com | Q6PGL7 [6 | 0 | 3492.744 | 0 | -0.77 | 0.000574 | 4.02 | 120.5 | 79.5 | 9.38 | 13.37 | | High | High | High | High | High | High | High | High | High | High | High | 0.002432 | 0.02603 | 2.78 |
| FALSE | High | TSLPAPGPI | [R].TSLPAP | 1xPhospho | D3YTT6 1x | 2.59E-07 | 0 | 1 | 6 | 1 | D3YTT6 | Oxysterol-l | D3YTT6 [5 | 0 | 2392.247 | 0 | 1.73 | 0.000994 | 19.62 | 46.3 | 153.7 | 10.44 | 9.14 | | High | High | High | High | High | High | High | High | High | High | High | 0 | 2.23E-08 | 3.45 |
| FALSE | High | TSLSASPFE | [K].TSLSAS | 1xPhospho | A2ADL9 1x | 0.004824 | 0.000456 | 1 | 7 | 2 | A2ADL9 | InaD-like (I | A2ADL9 [4 | 0 | 1801.832 | 0 | 0.73 | 0.009577 | 11.76 | 75.4 | 124.6 | 17.05 | 10.01 | | High | High | High | High | High | High | High | High | High | High | High | 0.000327 | 0.002473 | 2.71 |
| FALSE | High | TSISDPPESI | [R].TSISDPI | 1xPhospho | Q62245 1x | 0.012845 | 0.000889 | 1 | 2 | 1 | Q62245 | Son of seve | Q62245 [1 | 0 | 2012.03 | 0 | -0.03 | 0.748557 | 7.95 | 97.2 | 102.8 | 7.94 | 11.15 | | High | High | High | High | High | High | High | High | High | High | High | 0.000449 | 0.004137 | 2.18 |
| FALSE | High | TSISDPPESI | [R].TSISDPI | 1xPhospho | Q62245 1x | 0.091627 | 0.007146 | 1 | 2 | 1 | Q62245 | Son of seve | Q62245 [1 | 1 | 2493.295 | 0 | -0.18 | 0.05315 | 7.9 | 102.8 | 97.2 | 5.5 | 9.07 | | High | High | High | High | High | High | High | High | High | High | High | 0.003659 | 0.03905 | 2.28 |
| FALSE | High | TSLSPESHP | [R].TSLSPE | 1xPhospho | Q6DID5 1x | 0.000454 | 0 | 1 | 2 | 5 | Q6DID5 | PWWP dor | Q6DID5 [1 | 0 | 2577.321 | 0 | -0.93 | 0.000489 | 1.89 | 131.1 | 68.9 | 4.54 | 2.33 | | High | High | High | High | High | High | High | High | High | High | High | 0.000267 | 0.001526 | 3.5 |
| FALSE | High | TSLSSVSPG | [R].TSLSSV | 1xCarbami | Q80Z37 1x | 0.012731 | 0.000889 | 1 | 1 | 2 | Q80Z37 | E3 ubiquiti | Q80Z37 [1 | 1 | 1875.836 | 0 | 0.2 | 0.061855 | 11.05 | 91.4 | 108.6 | 2.6 | 8.67 | | High | High | High | High | High | High | High | High | High | High | High | 0.000449 | 0.00412 | 2.11 |
| FALSE | High | TSISTASAT | [K].TSISTAS | 1xCarbami | P35831 1x | 4.21E-06 | 0 | 1 | 1 | 3 | P35831 | Tyrosine-p | P35831 [62 | 0 | 2451.127 | 0 | 0.91 | 0.00073 | 10.89 | 69.2 | 130.8 | 6.28 | 7.28 | | High | High | High | High | High | High | High | High | High | High | High | 0 | 0.000145 | 4.29 |
| FALSE | High | TSLSYSR | [R].TSLSYSI | 1xPhospho | E9Q5F9 1x | 0.067619 | 0.004864 | 1 | 3 | 1 | E9Q5F9 | Histone-lys | E9Q5F9 [4 | 0 | 1122.539 | 0 | -0.08 | 0.378851 | 10.8 | 101.4 | 98.6 | 10.52 | 8.66 | | High | High | High | High | High | High | High | High | High | High | High | 0.002485 | 0.02722 | 1.37 |
| FALSE | High | TSLTPSSQD | [K].TSLTPS | 1xPhospho | E9QMX4 1x | 4.43E-05 | 0 | 1 | 4 | 1 | E9QMX4 | Ribosomal | E9QMX4 [4 | 0 | 2958.325 | 0 | -0.31 | 0.031697 | 13.8 | 111.1 | 88.9 | 5.86 | 9.22 | | High | High | High | High | High | High | High | High | High | High | High | 0 | 7.22E-06 | 3.45 |
| FALSE | High | TSLEGSDV | [R].TSLEGS | 1xPhospho | Q8BYH3 1x | 0.057686 | 0.004227 | 1 | 1 | 1 | Q8BYH3 | tRNA:m(4) | Q8BYH3 [3 | 1 | 1957.004 | 0 | -1.5 | 0.000693 | 2.19 | 147.7 | 52.3 | 6.69 | 13.06 | | High | High | High | High | High | High | High | High | High | High | High | 0.002159 | 0.02272 | 2.19 |
| FALSE | High | TSPPSSPK | [K].TSPPSS | 1xPhospho | Q811Q9 1x | 0.090153 | 0.00698 | 1 | 2 | 1 | Q811Q9 | Choline-ph | Q811Q9 [3 | 0 | 1338.707 | 0 | -0.45 | 0.0389 | 10.66 | 115.7 | 84.3 | 13.44 | 8.23 | | High | High | High | High | High | High | High | High | High | High | High | 0.003606 | 0.0383 | 1.16 |
| FALSE | High | TSPQAAAS | [R].TSPQA | 1xPhospho | O70472 1x | 0.000217 | 0 | 1 | 1 | 2 | O70472 | Transmem | O70472 [1 | 0 | 1801.91 | 0 | 0.28 | 0.083111 | 6.88 | 90.7 | 109.3 | 6.2 | 10.87 | | High | High | High | High | High | High | High | High | High | High | High | 0 | 4.3E-05 | 2.39 |
| FALSE | High | TSPQAPGL | [R].TSPQAI | 1xPhospho | A0A0A0M | 0.00153 | 5.66E-05 | 1 | 4 | 2 | A0A0A0M | Calmodulir | A0A0A0M | 0 | 1605.856 | 0 | 0.12 | 0.09045 | 1.79 | 95.7 | 104.3 | 3.68 | 7.81 | | High | High | High | High | High | High | High | High | High | High | High | 0.000449 | 0.003963 | 2.29 |
| FALSE | High | TSSETLECK | [K].TSSETL | 1xCarbami | Q6P5H2-2 | 0.002921 | 0.00016 | 1 | 2 | 1 | Q6P5H2-2 | Isoform 2 c | Q6P5H2-2 | 0 | 1592.764 | 0 | 2.28 | 0.004647 | 18.52 | 32.7 | 167.3 | 8.07 | 23.12 | | High | High | High | High | High | High | High | High | High | High | High | 7.19E-05 | 0.000797 | 2.45 |
| FALSE | High | TSSFAEPG | [R].TSSFAE | 1xPhospho | Q2NL51 1x | 3.23E-05 | 0 | 1 | 2 | 3 | Q2NL51 | Glycogen s | Q2NL51 [1 | 0 | 2742.195 | 0 | -1.33 | 0.405137 | 10.84 | 139.8 | 60.2 | 61.68 | 88.21 | | High | High | High | High | High | High | High | High | High | High | High | 0 | 5.07E-06 | 2.93 |
| FALSE | High | TSSFAEPG | [R].TSSFAE | 1xPhospho | Q2NL51 1x | 3.63E-08 | 0 | 1 | 2 | 8 | Q2NL51 | Glycogen s | Q2NL51 [1 | 0 | 2971.358 | 0 | -1.28 | 0.000824 | 1.75 | 141.3 | 58.7 | 4.5 | 11.27 | | High | High | High | High | High | High | High | High | High | High | High | 0 | 5.02E-07 | 5.33 |
| FALSE | High | TSSFTDQLI | [R].TSSFTD | 1xPhospho | Q91VW5 1 | 0.006306 | 0.00062 | 1 | 7 | 2 | Q91VW5 | Golgin sub | Q91VW5 [ | 0 | 2004.911 | 0 | 0.73 | 0.00155 | 11.19 | 75 | 125 | 7.22 | 4.7 | | High | High | High | High | High | High | High | High | High | High | High | 0.000399 | 0.003506 | 2.79 |
| FALSE | High | TSSIADEGT | [R].TSSIADI | 1xPhospho | D3Z4J3 1xP | 1.01E-07 | 0 | 1 | 4 | 8 | D3Z4J3 | Unconvent | D3Z4J3 [16 | 0 | 2151.042 | 0 | 0.51 | 0.008783 | 19.88 | 82.4 | 117.6 | 9.78 | 4.43 | | High | High | High | High | High | High | High | High | High | High | High | 0 | 1.81E-08 | 4.27 |
| FALSE | High | TSSLDNEGI | [R].TSSLDN | 1xPhospho | J3QNH8 1x | 0.000442 | 0 | 1 | 1 | 3 | J3QNH8 | Uncharacte | J3QNH8 [2 | 0 | 2183.011 | 0 | 5.2 | | 52.22 | 5.1 | 194.9 | 46.97 | 10.5 | | Not Found | High | High | High | High | High | High | High | High | High | High | 0 | 9.55E-05 | 2.69 |
| FALSE | High | TSSLDSAVI | [K].TSSLDS | 1xPhospho | Q69ZI1 1xP | 0.002508 | 0.000109 | 1 | 4 | 2 | Q69ZI1 | E3 ubiquiti | Q69ZI1 [80 | 0 | 1816.941 | 0 | 1.27 | 0.006288 | 29.79 | 59.1 | 140.9 | 20.04 | 10.19 | | High | High | High | High | High | High | High | High | High | High | High | 4.94E-05 | 0.000668 | 2.36 |
| FALSE | High | TSSLDTLAA | [R].TSSLDT | 1xPhospho | Q3U3E2 1x | 4.21E-06 | 0 | 1 | 1 | 3 | Q3U3E2 | Protein FAI | Q3U3E2 [2 | 0 | 2265.127 | 0 | -0.42 | 0.001876 | 2.59 | 116.1 | 83.9 | 6.59 | 7.38 | | High | High | High | High | High | High | High | High | High | High | High | 0 | 5.09E-07 | 4.17 |
| FALSE | High | TSSLDTITG | [R].TSSLDT | 1xPhospho | E9QKK4 1x | 3.74E-06 | 0 | 1 | 3 | 2 | E9QKK4 | Glucocorti | E9QKK4 [1 | 0 | 2302.132 | 0 | 0.84 | 0.000544 | 5.56 | 70.9 | 129.1 | 5.66 | 3.25 | | High | High | High | High | High | High | High | High | High | High | High | 0 | 1.31E-06 | 4.18 |
| FALSE | High | TSSLNLSED | [K].TSSLNL | 1xPhospho | Q3UT88 1x | 8.75E-10 | 0 | 1 | 3 | 3 | Q3UT88 | Putative ur | Q3UT88 [1 | 0 | 2491.042 | 0 | 0.23 | 0.045749 | 17.97 | 88.5 | 111.5 | 10.32 | 6.84 | | High | High | High | High | High | High | High | High | High | High | High | 0 | 3.73E-11 | 4.83 |
| FALSE | High | TSSLNLSED | [K].TSSLNL | 2xPhospho | Q3UT88 2x | 0.000144 | 0 | 1 | 3 | 1 | Q3UT88 | Putative ur | Q3UT88 [1 | 0 | 2571.008 | 0 | 0.22 | 0.096351 | 8.46 | 96.8 | 103.2 | 14.08 | 7.51 | | High | High | High | High | High | High | High | High | High | High | High | 0 | 2.72E-05 | 2.15 |
| FALSE | High | TSSLPESSP | [R].TSSLPE | 1xPhospho | Q8C7S2 1x | 0.003622 | 0.00021 | 1 | 4 | 1 | Q8C7S2 | Putative ur | Q8C7S2 [3 | 0 | 1428.682 | 0 | -0.14 | 0.842129 | 38.32 | 105.3 | 94.7 | 52.61 | 70.05 | | High | High | High | High | High | High | High | High | High | High | High | 9.42E-05 | 0.001005 | 2.24 |
| FALSE | High | TSSLPESSP | [R].TSSLPE | 1xPhospho | Q8C7S2 1x | 6.18E-05 | 0 | 1 | 4 | 5 | Q8C7S2 | Putative ur | Q8C7S2 [3 | 0 | 1657.845 | 0 | 0.41 | 0.000886 | 3.4 | 83.3 | 116.7 | 6.03 | 7.13 | | High | High | High | High | High | High | High | High | High | High | High | 0.000689 | 0.006174 | 2.69 |
| FALSE | High | TSSESVGTI | [K].TSSESV | 1xPhospho | Q9JI10 1xP | 0.067059 | 0.004796 | 1 | 3 | 1 | Q9JI10 | Serine/thre | Q9JI10 [32 | 0 | 1379.608 | 0 | 0.69 | 0.026755 | 23.89 | 77.6 | 122.4 | 16.71 | 19.61 | | High | High | High | High | High | High | High | High | High | High | High | 0.002455 | 0.02694 | 1.76 |
| FALSE | High | TSSIPVQEA | [R].TSSIPV | 1xPhospho | Q91Z58 1x | 5.5E-05 | 0 | 1 | 1 | 5 | Q91Z58 | Uncharacte | Q91Z58 [3 | 0 | 2823.37 | 0 | 2.07 | 0.000689 | 7.56 | 36.1 | 163.9 | 13.69 | 11.96 | | High | High | High | High | High | High | High | High | High | High | High | 0 | 9.19E-06 | 3.37 |
| FALSE | High | TSSISGPLSI | [R].TSSISGI | 1xPhospho | Q62261-2 | 2.21E-08 | 0 | 1 | 4 | 7 | Q62261-2 | Isoform 2 c | Q62261-2 | 0 | 3108.488 | 0 | 1.83 | 0.001039 | 12.75 | 42.3 | 157.7 | 11.51 | 13.18 | | High | High | High | High | High | High | High | High | High | High | High | 0 | 1.4E-09 | 4.83 |
| FALSE | High | TSSISGPLSI | [R].TSSISGI | 1xDeamida | Q62261-2 | 0.000538 | 0 | 1 | 4 | 8 | Q62261-2 | Isoform 2 c | Q62261-2 | 0 | 3109.472 | 0 | 1.63 | 0.000849 | 28.81 | 52.7 | 147.3 | 30.18 | 11.53 | | High | High | High | High | High | High | High | High | High | High | High | 0 | 0.000119 | 3.33 |
| FALSE | High | TSSISGPLSI | [R].TSSISGI | 2xPhospho | Q62261-2 | 6.26E-06 | 0 | 1 | 4 | 3 | Q62261-2 | Isoform 2 c | Q62261-2 | 0 | 3188.454 | 0 | 2.67 | 0.000979 | 42.1 | 27.7 | 172.3 | 32.69 | 14.25 | | High | High | High | High | High | High | High | High | High | High | High | 0 | 8.01E-07 | 3.67 |
| FALSE | High | TSSLSSDSS | [R].TSSLSSI | 1xPhospho | Q330P7 1x | 0.003997 | 0.000311 | 1 | 1 | 2 | Q330P7 | SCF apopto | Q330P7 [1 | 0 | 1562.726 | 0 | 0.69 | 0.002249 | 8.54 | 75 | 125 | 9.56 | 8.11 | | High | High | High | High | High | High | High | High | High | High | High | 0.00014 | 0.001127 | 3.73 |
| FALSE | High | TSSLTHSEE | [R].TSSLTH | 1xPhospho | Q9CT10 1x | 0.010568 | 0.000772 | 1 | 1 | 3 | Q9CT10 | Ran-bindin | Q9CT10 [5 | 0 | 1427.662 | 0 | 0.51 | 0.456015 | 94.66 | 81.2 | 118.8 | 43.45 | 82.58 | | High | High | High | High | High | High | High | High | High | High | High | 0.000382 | 0.003337 | 2.3 |
| FALSE | High | TSSLTHSEE | [R].TSSLTH | 1xPhospho | Q9CT10 1x | 0.003167 | 0.00016 | 1 | 1 | 6 | Q9CT10 | Ran-bindin | Q9CT10 [5 | 0 | 1656.825 | 0 | 0.61 | 0.003589 | 6.81 | 80.3 | 119.7 | 5.1 | 7.95 | | High | High | High | High | High | High | High | High | High | High | High | 7.19E-05 | 0.000871 | 2.36 |
| FALSE | High | TSSLTHSEE | [R].TSSLTH | 2xPhospho | Q9CT10 2x | 0.003495 | 0.00021 | 1 | 1 | 1 | Q9CT10 | Ran-bindin | Q9CT10 [5 | 0 | 1736.791 | 0 | 0.6 | 0.010457 | 5.64 | 81.2 | 118.8 | 7.09 | 11.38 | | High | High | High | High | High | High | High | High | High | High | High | 9.42E-05 | 0.000969 | 2.29 |
| FALSE | High | TSSNASTIS | [R].TSSNAS | 1xPhospho | Q9R1E0 1x | 0.01894 | 0.001362 | 1 | 3 | 2 | Q9R1E0 | Forkhead b | Q9R1E0 [3 | 0 | 1389.657 | 0 | -1.26 | 0.000901 | 1.29 | 139.5 | 60.5 | 3.65 | 12.08 | | High | High | High | High | High | High | High | High | High | High | High | 0.00074 | 0.006445 | 1.87 |
| FALSE | High | TSSPEVQE | [K].TSSPEV | 1xPhospho | Q9QZD4 1x | 0.000202 | 0 | 1 | 4 | 2 | Q9QZD4 | DNA repair | Q9QZD4 [3 | 0 | 1643.829 | 0 | 0.3 | 0.000926 | 2.4 | 90.4 | 109.6 | 1.53 | 2.79 | | High | High | High | High | High | High | High | High | High | High | High | 0 | 3.95E-05 | 2.95 |
| FALSE | High | TSSPGNTLS | [R].TSSPGN | 1xCarbami | Q8K296 1x | 0.000231 | 0 | 1 | 9 | 2 | Q8K296 | Myotubula | Q8K296 [8 | 0 | 2721.43 | 0 | 0.65 | 0.012886 | 4.97 | 78.6 | 121.4 | 3.89 | 10.85 | | High | High | High | High | High | High | High | High | High | High | High | 0 | 4.61E-05 | 3.14 |
| FALSE | High | TSSPLHAAI | [R].TSSPLH | 1xPhospho | Q99KW3-2 | 0.103392 | 0.008249 | 1 | 5 | 2 | Q99KW3-2 | Isoform 2 c | Q99KW3-2 | 0 | 1345.683 | 0 | 0.15 | 0.408877 | 22.34 | 94.9 | 105.1 | 13.66 | 11.56 | | High | High | High | High | High | High | High | High | High | High | High | 0.004152 | 0.04485 | 1.89 |
| FALSE | High | TSSPTSLPP | [R].TSSPTS | 1xPhospho | A0A0N4SV | 0.00311 | 0.00016 | 1 | 9 | 4 | A0A0N4SV | Eukaryotic | A0A0N4SV | 0 | 1535.803 | 0 | 0.67 | 0.001801 | 7.24 | 74.8 | 125.2 | 9.07 | 5.4 | | High | High | High | High | High | High | High | High | High | High | High | 7.19E-05 | 0.000855 | 2.57 |
| FALSE | High | TSSQYVAS | [R].TSSQY\ | 1xPhospho | B2RWU4 1 | 0.003083 | 0.00016 | 2 | 6 | 2 | B2RWU4; C | Cordon-ble | B2RWU4 [ | 0 | 1763.934 | 0 | -0.46 | 0.039397 | 2.33 | 117.3 | 82.7 | 13.98 | 17.25 | Shared | High | High | High | High | High | High | High | High | High | High | High | 7.19E-05 | 0.000843 | 2.9 |
| FALSE | High | TSSIPVQEA | [R].TSSIPV | 2xPhospho | Q91Z58 2x | 0.000167 | 0 | 1 | 1 | 1 | Q91Z58 | Uncharacte | Q91Z58 [3 | 0 | 2903.336 | 0 | 1.56 | 0.002706 | 13.82 | 47.1 | 152.9 | 16.96 | 17.99 | | High | High | High | High | High | High | High | High | High | High | High | 0 | 3.19E-05 | 4.48 |
| FALSE | High | TSSEQAVA | [R].TSSEQA | 1xPhospho | Q8K120 1x | 6.71E-05 | 0 | 1 | 2 | 2 | Q8K120 | Nuclear fac | Q8K120 [3 | 0 | 1467.741 | 0 | -1.41 | 0.000656 | 1.61 | 142.7 | 57.3 | 5.72 | 14.17 | | High | High | High | High | High | High | High | High | High | High | High | 0 | 1.81E-05 | 3.57 |
| FALSE | High | TSSEDNLYL | [R].TSSEDN | 1xPhospho | Q9QXS1-3 | 6.09E-06 | 0 | 1 | 2 | 10 | Q9QXS1-3 | Isoform PL | Q9QXS1-3 | 0 | 1789.893 | 0 | 1.57 | 0.000405 | 7.63 | 49.2 | 150.8 | 5.27 | 6.79 | | High | High | High | High | High | High | High | High | High | High | High | 0 | 7.75E-07 | 3.53 |
| FALSE | High | TSSDSPCW | [R].TSSDSP | 1xCarbami | B2RQG2 1x | 0.003464 | 0.00021 | 1 | 2 | 1 | B2RQG2 | PHD finger | B2RQG2 [1 | 0 | 1917.882 | 0 | -0.09 | 0.05015 | 3.23 | 103.4 | 96.6 | 7.15 | 12.29 | | High | High | High | High | High | High | High | High | High | High | High | 9.42E-05 | 0.00096 | 2.39 |
| FALSE | High | TSPQELSEE | [R].TSPQEL | 1xPhospho | Q8CFI0 1x | 0.000235 | 0 | 1 | 8 | 1 | Q8CFI0 | E3 ubiquiti | Q8CFI0 [33 | 0 | 1670.784 | 0 | -0.58 | 0.002375 | 5.68 | 123.9 | 76.1 | 6.24 | 6.38 | | High | High | High | High | High | High | High | High | High | High | High | 0 | 4.69E-05 | 2.7 |
| FALSE | High | TSPSLCPES | [K].TSPSLC | 2xCarbami | A0A140T8 | 0.000108 | 0 | 1 | 1 | 2 | A0A140T8 | Chromosor | A0A140T8 | 0 | 4690.319 | 0 | -1.45 | 0.000391 | 0.14 | 146.5 | 53.5 | 4.65 | 6.17 | | High | High | High | High | High | High | High | High | High | High | High | 0 | 1.96E-05 | 3.96 |
| FALSE | High | TSPSPPTQ | [R].TSPSPP | 1xPhospho | A0A0R4J1C | 1.69E-06 | 0 | 1 | 9 | 3 | A0A0R4J1C | Anion exch | A0A0R4J1C | 0 | 2111.012 | 0 | 1.62 | 0.001414 | 5.51 | 50.3 | 149.7 | 4.79 | 13.83 | | High | High | High | High | High | High | High | High | High | High | High | 0 | 1.83E-07 | 3.76 |
| FALSE | High | TSPSSFPSP | [R].TSPSSFI | 1xPhospho | B2RUA5 1x | 0.038017 | 0.002528 | 1 | 2 | 1 | B2RUA5 | SAPS doma | B2RUA5 [6 | 0 | 4295.066 | 0 | -0.34 | 0.149544 | 11.77 | 108.7 | 91.3 | 17.47 | 29.05 | | High | High | High | High | High | High | High | High | High | High | High | 0.001355 | 0.01415 | 2.57 |
| FALSE | High | TSPSSSPAS | [K].TSPSSS | 1xPhospho | P49586 1x | 0.000337 | 0 | 1 | 1 | 2 | P49586 | Choline-ph | P49586 [34 | 0 | 1485.715 | 0 | 0.85 | 0.001184 | 15.17 | 71.9 | 128.1 | 6.75 | 4.08 | | High | High | High | High | High | High | High | High | High | High | High | 0.00074 | 0.006502 | 2.3 |
| FALSE | High | TSPSSSPAS | [K].TSPSSS | 2xPhospho | P49586 2x | 0.013909 | 0.001011 | 1 | 1 | 1 | P49586 | Choline-ph | P49586 [34 | 0 | 1565.681 | 0 | 1.56 | 0.000181 | 3.3 | 51.7 | 148.3 | 7.17 | 3.63 | | High | High | High | High | High | High | High | High | High | High | High | 0.000538 | 0.004541 | 2.12 |
| FALSE | High | TSPTASVNI | [K].TSPTAS | 1xPhospho | Q5XG69 1x | 0.003962 | 0.000311 | 1 | 1 | 3 | Q5XG69 | Soluble lan | Q5XG69 [3 | 0 | 1442.766 | 0 | -1.29 | 0.00092 | 3.58 | 142 | 58 | 8.6 | 14.46 | | High | High | High | High | High | High | High | High | High | High | High | 0.00014 | 0.00112 | 2.05 |
| FALSE | High | TSPTVATQ | [R].TSPTVA | 1xPhospho | Q3U3E2 1x | 2.86E-07 | 0 | 1 | 1 | 6 | Q3U3E2 | Protein FAI | Q3U3E2 [1 | 0 | 1973.974 | 0 | -0.12 | 0.324621 | 9.27 | 101.6 | 98.4 | 6.24 | 9.84 | | High | High | High | High | High | High | High | High | High | High | High | 0 | 3.01E-07 | 3.55 |
| FALSE | High | TSPVASGS | [K].TSPVAS | 1xPhospho | Q6A065 1x | 1.12E-05 | 0 | 1 | 3 | 2 | Q6A065 | Centrosom | Q6A065 [1 | 0 | 1559.808 | 0 | 1.45 | 0.000751 | 13.58 | 55.9 | 144.1 | 13.05 | 6.16 | | High | High | High | High | High | High | High | High | High | High | High | 0 | 1.54E-06 | 1.54 |
| FALSE | High | TSPVQSK | [K].TSPVQS | 1xPhospho | P11862 1x | 0.079114 | 0.00613 | 1 | 2 | 1 | P11862 | Growth arr | P11862 [28 | 0 | 1284.696 | 0 | -1.56 | 0.001486 | 3.13 | 150.5 | 49.5 | 12.3 | 18.07 | | High | High | High | High | High | High | High | High | High | High | High | 0.003148 | 0.03277 | 1.45 |
| FALSE | High | TSPVVAPTS | [K].TSPVVA | 2xPhospho | Q91Z67 2x | 0.000305 | 0 | 1 | 1 | 4 | Q91Z67 | SLIT-ROBO | Q91Z67 [9 | 0 | 2807.44 | 0 | 0.16 | 0.053395 | 9.87 | 93.8 | 106.2 | 8.32 | 4.19 | | High | High | High | High | High | High | High | High | High | High | High | 0 | 6.29E-05 | 4.06 |
| FALSE | High | TSPYILR | [K].TSPYILI | 1xPhospho | E9Q3P4 1x | 0.041425 | 0.003126 | 1 | 5 | 1 | E9Q3P4 | Centromer | E9Q3P4 [2 | 0 | 1158.612 | 0 | -0.01 | 0.844161 | 4.16 | 101.7 | 98.3 | 3.93 | 7.94 | | High | High | High | High | High | High | High | High | High | High | High | 0.001609 | 0.01548 | 2.34 |
| FALSE | High | TSPYPGSK | [R].TSPYPG | 1xPhospho | Q8BVU5 1x | 0.024424 | 0.001612 | 1 | 5 | 2 | Q8BVU5 | ADP-ribose | Q8BVU5 [6 | 0 | 1374.707 | 0 | -1.41 | 0.003294 | 2.29 | 148 | 52 | 11.51 | 18.96 | | High | High | High | High | High | High | High | High | High | High | High | 0.005784 | 0.06382 | 1.51 |
| FALSE | High | TSQAACLSI | [R].TSQAA | 1xCarbami | P97929 1x | 0.00418 | 0.00036 | 1 | 3 | 1 | P97929 | Breast can | P97929 [48 | 0 | 1599.776 | 0 | -1.15 | 0.001732 | 2.31 | 138.3 | 61.7 | 9.68 | 8.32 | | High | High | High | High | High | High | High | High | High | High | High | 0.000161 | 0.001189 | 0.85 |
| FALSE | High | TSQCSSPSL | [K].TSQCS | 1xCarbami | A4QPD1 1x | 0.000276 | 0 | 1 | 4 | 4 | A4QPD1 | Zinc finger | A4QPD1 [2 | 0 | 2640.265 | 0 | 1.21 | 0.007151 | 7.72 | 61.3 | 138.7 | 14.06 | 13.72 | | High | High | High | High | High | High | High | High | High | High | High | 0.000689 | 0.005915 | 2.49 |
| FALSE | High | TSQCSSPSL | [K].TSQCS | 1xCarbami | A4QPD1 2x | 0.032548 | 0.002167 | 1 | 4 | 2 | A4QPD1 | Zinc finger | A4QPD1 [2 | 0 | 2720.231 | 0 | 1.72 | 0.001732 | 19.44 | 43.8 | 156.2 | 12.67 | 12.24 | | High | High | High | High | High | High | High | High | High | High | High | 0.001142 | 0.01181 | 3.27 |
| FALSE | High | TSQEILNSS | [K].TSQEILI | 1xPhospho | A2A7B5 1x | 2.82E-08 | 0 | 1 | 2 | 2 | A2A7B5 | PR domain | A2A7B5 [4 | 0 | 2548.258 | 0 | -1.02 | 0.000628 | 1.54 | 133.4 | 66.6 | 4.81 | 7.65 | | High | High | High | High | High | High | High | High | High | High | High | 0 | 1.84E-09 | 4.13 |
| FALSE | High | TSQEILNSS | [K].TSQEILI | 1xDeamida | A2A7B5 1x | 0.001158 | 0 | 1 | 2 | 7 | A2A7B5 | PR domain | A2A7B5 [4 | 0 | 2549.242 | 0 | -0.25 | 0.056328 | 4.27 | 110.4 | 89.6 | 4.89 | 7.78 | | High | High | High | High | High | High | High | High | High | High | High | 0.000638 | 0.005662 | 2.54 |
| FALSE | High | TSQETADV | [R].TSQETA | 1xPhospho | Q8BJ71 1x | 0.003055 | 0.00016 | 1 | 1 | 3 | Q8BJ71 | Nuclear po | Q8BJ71 [5 | 0 | 1516.766 | 0 | -0.93 | 0.000285 | 1.95 | 131 | 69 | 3.41 | 3.23 | | High | High | High | High | High | High | High | High | High | High | High | 0.005041 | 0.05519 | 1.94 |
| FALSE | High | TSQLGDSP | [K].TSQLGI | 1xPhospho | E9Q3G8 1x | 0.015329 | 0.001089 | 1 | 2 | 1 | E9Q3G8 | Nucleopor | E9Q3G8 [2 | 0 | 1934.967 | 0 | -0.2 | 0.243075 | 5.41 | 106.5 | 93.5 | 17.33 | 5.84 | | High | High | High | High | High | High | High | High | High | High | High | 0.000571 | 0.005038 | 2.25 |
| FALSE | High | TSQQNVSN | [R].TSQQN | 1xPhospho | Q3UHJ0 1x | 8.45E-07 | 0 | 1 | 2 | 2 | Q3UHJ0 | AP2-associ | Q3UHJ0 [6 | 0 | 3313.469 | 0 | 0.05 | 0.88369 | 35.81 | 97.4 | 102.6 | 11.58 | 25.44 | | High | High | High | High | High | High | High | High | High | High | High | 0 | 8.4E-08 | 4.88 |
| FALSE | High | TSRDASPSS | [K].TSRDAS | 2xPhospho | G3X959 2x | 8.28E-06 | 0 | 1 | 7 | 2 | G3X959 | Periphilin 1 | G3X959 [2 | 1 | 2243.036 | 0 | 0.79 | 0.004776 | 5.65 | 72.5 | 127.5 | 2.12 | 12.88 | | High | High | High | High | High | High | High | High | High | High | High | 0 | 1.1E-06 | 3.44 |
| FALSE | High | TSSALAVTV | [R].TSSALA | 1xPhospho | P70399 1x | 0.002272 | 0.000109 | 1 | 5 | 1 | P70399 | TP53-bindi | P70399 [12 | 0 | 2470.273 | 0 | 0.84 | 0.002833 | 9.99 | 68.3 | 131.7 | 13.91 | 10.35 | | High | High | High | High | High | High | High | High | High | High | High | 4.94E-05 | 0.000599 | 1.99 |
| FALSE | High | TSSASDQE | [R].TSSASD | 1xPhospho | B2RQQ7 1x | 1.96E-07 | 0 | 1 | 2 | 5 | B2RQQ7 | CDC42 bin | B2RQQ7 [9 | 0 | 1877.889 | 0 | 1.26 | 0.000377 | 14.26 | 58.9 | 141.1 | 13.12 | 3.75 | | High | High | High | High | High | High | High | High | High | High | High | 0 | 1.62E-08 | 3.02 |
| FALSE | High | TSSASTSDF | [K].TSSAST | 1xPhospho | Q3TN28 1x | 3.26E-08 | 0 | 1 | 4 | 2 | Q3TN28 | Putative ur | Q3TN28 [2 | 0 | 3561.736 | 0 | -0.19 | 0.158764 | 10.87 | 106.4 | 93.6 | 14.57 | 10.95 | | High | High | High | High | High | High | High | High | High | High | High | 0 | 2.16E-09 | 4.01 |
| FALSE | High | TSSAVWNS | [R].TSSAVV | 1xPhospho | Q9WU62 1 | 0.000286 | 0 | 1 | 2 | 3 | Q9WU62 | Inner centr | Q9WU62 [ | 0 | 1824.966 | 0 | -0.82 | 0.000665 | 2.15 | 127.7 | 72.3 | 5.76 | 7.76 | | High | High | High | High | High | High | High | High | High | High | High | 0 | 5.85E-05 | 2.95 |
| FALSE | High | TSSAVWNS | [R].TSSAVV | 2xPhospho | Q9WU62 2 | 0.002463 | 0.000109 | 1 | 2 | 2 | Q9WU62 | Inner centr | Q9WU62 [ | 0 | 1904.932 | 0 | -0.59 | 0.000612 | 2.65 | 120 | 80 | 4.99 | 6.64 | | High | High | High | High | High | High | High | High | High | High | High | 4.94E-05 | 0.000657 | 3.22 |
| FALSE | High | TSSDHTNP | [R].TSSDHT | 1xPhospho | O54916 1x | 0.000407 | 0 | 1 | 7 | 2 | O54916 | RalBP1-ass | O54916 [4 | 0 | 3363.729 | 0 | 0.05 | 0.847223 | 17.54 | 100.2 | 99.8 | 20.22 | 14.25 | | High | High | High | High | High | High | High | High | High | High | High | 0.000275 | 0.001726 | 2.85 |
| FALSE | High | TSSDPNLNI | [R].TSSDPN | 1xPhospho | Q91XS1 1x | 0.000198 | 0 | 1 | 3 | 2 | Q91XS1 | Myotubula | Q91XS1 [6 | 0 | 2006.913 | 0 | 0.23 | 0.079294 | 7.63 | 93 | 107 | 8.32 | 2.15 | | High | High | High | High | High | High | High | High | High | High | High | 0 | 4.17E-05 | 1.47 |
| FALSE | High | TSFLEGTLR | [R].TSFLEG | 1xPhospho | F8VQB6 1x | 0.005044 | 0.000504 | 1 | 4 | 1 | F8VQB6 | Unconvent | F8VQB6 [1 | 0 | 1332.676 | 0 | -1.75 | 0.010552 | 1.13 | 147 | 53 | 21.42 | 18.96 | | High | High | High | High | High | High | High | High | High | High | High | 0.000246 | 0.001463 | 2.01 |
| FALSE | High | TSFDENDSI | [K].TSFDEN | 1xPhospho | Q05D44 1x | 8.89E-09 | 0 | 1 | 1 | 6 | Q05D44 | Eukaryotic | Q05D44 [1 | 0 | 2195.963 | 0 | -0.66 | 0.018217 | 5.77 | 121.5 | 78.5 | 10.64 | 9.44 | | High | High | High | High | High | High | High | High | High | High | High | 0 | 5.04E-10 | 3.4 |
| FALSE | High | TSEVLVPSN | [K].TSEVLV | 1xDeamida | Q3TLH4 1x | 7.81E-08 | 0 | 1 | 6 | 5 | Q3TLH4 | Protein PRI | Q3TLH4 [1 | 0 | 3586.804 | 0 | 0.1 | 0.321582 | 13.99 | 96.5 | 103.5 | 6.58 | 5.55 | | High | High | High | High | High | High | High | High | High | High | High | 0 | 5.8E-09 | 5.47 |
| FALSE | High | TSEVLVPSN | [K].TSEVLV | 1xPhospho | Q3TLH4 1x | 5.88E-06 | 0 | 1 | 6 | 1 | Q3TLH4 | Protein PRI | Q3TLH4 [1 | 0 | 3585.82 | 0 | 0.2 | 0.003092 | 3.33 | 92.2 | 107.8 | 3.66 | 6.15 | | High | High | High | High | High | High | High | High | High | High | High | 0 | 7.41E-07 | 3.65 |
| FALSE | High | TQPLWDDI | [R].TQPLW | 1xPhospho | Q9D0B1 1x | 2.1E-05 | 0 | 1 | 1 | 1 | Q9D0B1 | Zinc finger | Q9D0B1 [2 | 0 | 3320.561 | 0 | 0.73 | 0.004786 | 14.53 | 72.6 | 127.4 | 11.27 | 10.33 | | High | High | High | High | High | High | High | High | High | High | High | 0 | 3.12E-06 | 4.12 |
| FALSE | High | TQQQQDS | [K].TQQQC | 1xPhospho | A0A1B0GR | 2.78E-06 | 0 | 1 | 1 | 3 | A0A1B0GR | RIKEN cDN | A0A1B0GR | 0 | 2406.077 | 0 | 0.62 | 0.023554 | 22.42 | 82.1 | 117.9 | 13.83 | 11.83 | | High | High | High | High | High | High | High | High | High | High | High | 0 | 1.31E-06 | 3.43 |
| FALSE | High | TQQSFEK | [K].TQQSFI | 1xPhospho | Q80X50-5 | 0.049549 | 0.00361 | 1 | 9 | 1 | Q80X50-5 | Isoform 5 c | Q80X50-5 | 0 | 1405.713 | 0 | 0.46 | 0.006161 | 9.56 | 82.6 | 117.4 | 6.99 | 10.48 | | High | High | High | High | High | High | High | High | High | High | High | 0.001902 | 0.01915 | 1.42 |
| FALSE | High | TQQSPVSN | [K].TQQSP | 1xPhospho | Q9JHC9 1x | 1.93E-07 | 0 | 1 | 6 | 3 | Q9JHC9 | ETS-relate | Q9JHC9 [1 | 0 | 2180.136 | 0 | -0.63 | 0.000364 | 2.31 | 121.4 | 78.6 | 1.55 | 6.26 | | High | High | High | High | High | High | High | High | High | High | High | 0 | 2.77E-06 | 3.51 |
| FALSE | High | TQQSPVSN | [K].TQQSP | 1xDeamida | Q9JHC9 1x | 5.35E-05 | 0 | 1 | 6 | 4 | Q9JHC9 | ETS-relate | Q9JHC9 [1 | 0 | 2181.12 | 0 | -0.68 | 0.00207 | 0.84 | 121.5 | 78.5 | 10.61 | 5.17 | | High | High | High | High | High | High | High | High | High | High | High | 0 | 8.92E-06 | 3.9 |

| | | | | | | | | | | | | | | | | | | | | | | | | | | | | | | | | | | | | | | |
|---|---|---|---|---|---|---|---|---|---|---|---|---|---|---|---|---|---|---|---|---|---|---|---|---|---|---|---|---|---|---|---|---|---|---|---|---|---|---|
| FALSE | High | TQQSPVSN | [K].TQQSP | 2xPhospho | Q9JHC9 2x | 0.000492 | 0 | 1 | 6 | 2 | Q9JHC9 | ETS-related | Q9JHC9 [1 | 0 | 2260.103 | 0 | -0.59 | 0.000195 | 1.23 | 120 | 80 | 2.26 | 1.48 | | High | High | High | High | High | High | High | High | High | High | High | 0 | 0.000107 | 2.61 |
| FALSE | High | TQSAPLPQ | [R].TQSAPL | 1xPhospho | Q6NZM9 1 | 0.000286 | 0 | 1 | 2 | 1 | Q6NZM9 | Histone de | Q6NZM9 [ | 0 | 3761.989 | 0 | 0.34 | 0.017229 | 9.3 | 88.4 | 111.6 | 8.07 | 6.03 | | High | High | High | High | High | High | High | High | High | High | High | 0 | 5.88E-05 | 5.02 |
| FALSE | High | TQSGNFNT | [R].TQSGN | 1xPhospho | O35711-4 | 0.000116 | 0 | 1 | 4 | 1 | O35711-4 | Isoform 4 c | O35711-4 | 0 | 2151.937 | 0 | -0.21 | 0.087869 | 5.51 | 107.1 | 92.9 | 5.93 | 7.88 | | High | High | High | High | High | High | High | High | High | High | High | 0 | 2.12E-05 | 4.42 |
| FALSE | High | TQSGNFNT | [R].TQSGN | 1xPhospho | O35711-4 | 8.56E-05 | 0 | 1 | 4 | 2 | O35711-4 | Isoform 4 c | O35711-4 | 0 | 2167.932 | 0 | -0.63 | 0.007781 | 2.87 | 121.4 | 78.6 | 3.72 | 18.57 | | High | High | High | High | High | High | High | High | High | High | High | 0 | 1.51E-05 | 3.78 |
| FALSE | High | TQSLSALPK | [R].TQSLSA | 1xPhospho | Q924A2 1x | 0.085162 | 0.006376 | 1 | 4 | 1 | Q924A2 | Protein cap | Q924A2 [1 | 0 | 1482.833 | 0 | 0.07 | 0.304236 | 10.42 | 96.6 | 103.4 | 4.96 | 4.06 | | High | High | High | High | High | High | High | High | High | High | High | 0.003291 | 0.03562 | 1.96 |
| FALSE | High | TQSPATSH | [K].TQSPAT | 1xPhospho | B1AX39 1x | 0.007019 | 0.00062 | 1 | 2 | 1 | B1AX39 | Zinc finger | B1AX39 [1 | 0 | 1695.847 | 0 | -0.39 | 0.021735 | 9.55 | 118.8 | 81.2 | 13.19 | 12.68 | | High | High | High | High | High | High | High | High | High | High | High | 0.000275 | 0.00211 | 1.91 |
| FALSE | High | TQSPGGCS | [R].TQSPG | 1xCarbami | Q3TVI4 1x | 5.52E-06 | 0 | 1 | 3 | 1 | Q3TVI4 | Protein HE | Q3TVI4 [73 | 0 | 1840.883 | 0 | -0.79 | 0.019096 | 2.99 | 125.2 | 74.8 | 9.39 | 15.97 | | High | High | High | High | High | High | High | High | High | High | High | 0 | 6.89E-07 | 2.97 |
| FALSE | High | TQSPGGCS | [R].TQSPG | 1xCarbami | Q3TVI4 2x | 9.82E-06 | 0 | 1 | 3 | 1 | Q3TVI4 | Protein HE | Q3TVI4 [73 | 0 | 1920.849 | 0 | -0.4 | 0.014898 | 10.32 | 112 | 88 | 6.05 | 9.33 | | High | High | High | High | High | High | High | High | High | High | High | 0 | 1.33E-06 | 3.1 |
| FALSE | High | TQPLALPPF | [R].TQPLAL | 1xDeamida | O54804 1x | 0.012959 | 0.000889 | 1 | 5 | 1 | O54804 | Choline kin | O54804 [5 | 0 | 3659.933 | 0 | -1.48 | 0.000222 | 1.14 | 147.1 | 52.9 | 5.78 | 11.91 | | High | High | High | High | High | High | High | High | High | High | High | 0.000449 | 0.004182 | 3.81 |
| FALSE | High | TQSPHSPK | [K].TQSPHS | 1xPhospho | B7ZCJ1 1xF | 0.108463 | 0.008839 | 1 | 7 | 1 | B7ZCJ1 | Rho GTPas | B7ZCJ1 [10 | 0 | 1419.74 | 0 | 0.15 | 0.432606 | 1.01 | 95.9 | 104.1 | 4.6 | 4.79 | | High | High | High | High | High | High | High | High | High | High | High | 0.004422 | 0.04758 | 1.88 |
| FALSE | High | TQSSASLA | [R].TQSSAS | 1xPhospho | Q8C2Q3 1x | 3.85E-06 | 0 | 1 | 4 | 5 | Q8C2Q3 | RNA-bindir | Q8C2Q3 [5 | 0 | 2774.31 | 0 | 1.66 | 0.005756 | 20 | 48.4 | 151.6 | 9.97 | 19.85 | | High | High | High | High | High | High | High | High | High | High | High | 0 | 4.63E-07 | 4.78 |
| FALSE | High | TQSSGQDC | [R].TQSSG | 1xPhospho | F8WHW6 1 | 0.054383 | 0.003951 | 1 | 5 | 1 | F8WHW6 | Phosphatic | F8WHW6 [ | 0 | 2875.362 | 0 | -0.22 | 0.040646 | 10.04 | 111.4 | 88.6 | 4.7 | 7.93 | | High | High | High | High | High | High | High | High | High | High | High | 0.002059 | 0.02124 | 1.62 |
| FALSE | High | TQSSGQDC | [R].TQSSG | 2xPhospho | F8WHW6 2 | 0.004372 | 0.000409 | 1 | 5 | 1 | F8WHW6 | Phosphatic | F8WHW6 [ | 0 | 2955.329 | 0 | -0.82 | 0.001951 | 3.43 | 129.9 | 70.1 | 4.26 | 14.03 | | High | High | High | High | High | High | High | High | High | High | High | 0.000183 | 0.001239 | 2.36 |
| FALSE | High | TQSSLTASE | [R].TQSSLT | 1xPhospho | Q8R4U7 1x | 0.001179 | 0 | 2 | 4 | 2 | Q8R4U7; B | Leucine zip | Q8R4U7 [9 | 0 | 1701.862 | 0 | -1.11 | 0.00183 | 2.79 | 136.2 | 63.8 | 9.92 | 7.48 | Shared | High | High | High | High | High | High | High | High | High | High | High | 0.002419 | 0.02595 | 2.48 |
| FALSE | High | TQSSPAAP | [R].TQSSPA | 1xPhospho | B7ZDF5 1x | 0.000123 | 0 | 1 | 8 | 3 | B7ZDF5 | Histone de | B7ZDF5 [66 | 0 | 1715.844 | 0 | 0.95 | 0.004552 | 21.31 | 70.6 | 129.4 | 9.85 | 15.02 | | High | High | High | High | High | High | High | High | High | High | High | 0 | 2.26E-05 | 2.33 |
| FALSE | High | TQSSPLPQ | [R].TQSSPL | 1xPhospho | B7ZDF5 1x | 0.000751 | 0 | 1 | 8 | 1 | B7ZDF5 | Histone de | B7ZDF5 [50 | 0 | 3785.945 | 0 | 0.87 | 0.003281 | 4.05 | 70.4 | 129.6 | 9.93 | 13.77 | | High | High | High | High | High | High | High | High | High | High | High | 0 | 0.000172 | 4.14 |
| FALSE | High | TQSSPLPQ | [R].TQSSPL | 1xPhospho | B7ZDF5 1x | 2.65E-05 | 0 | 1 | 8 | 2 | B7ZDF5 | Histone de | B7ZDF5 [50 | 0 | 3801.94 | 0 | 0.95 | 0.007779 | 22.1 | 68.3 | 131.7 | 10.02 | 15.01 | | High | High | High | High | High | High | High | High | High | High | High | 0 | 4.06E-06 | 3.9 |
| FALSE | High | TQSSSCEDI | [R].TQSSSC | 1xCarbami | O70589-3 | 0.004106 | 0.00036 | 1 | 2 | 2 | O70589-3 | Isoform 3 c | O70589-3 | 0 | 2304.083 | 0 | -0.97 | 0.002625 | 4.3 | 129.1 | 70.9 | 8.42 | 17.36 | | High | High | High | High | High | High | High | High | High | High | High | 0.000327 | 0.002718 | 2.23 |
| FALSE | High | TQSSVPAPI | [K].TQSSVF | 1xPhospho | Q9CQJ7 1x | 0.006773 | 0.00062 | 1 | 4 | 1 | Q9CQJ7 | Securin OS | Q9CQJ7 [9 | 0 | 2385.163 | 0 | -0.01 | 0.971941 | 19.33 | 99.9 | 100.1 | 17.87 | 7.29 | | High | High | High | High | High | High | High | High | High | High | High | 0.000275 | 0.002037 | 2.53 |
| FALSE | High | TQTCTGSS | [R].TQTCT | 1xCarbami | Q99PJ2 1x | 0.000501 | 0 | 1 | 1 | 1 | Q99PJ2 | Probable E | Q99PJ2 [31 | 0 | 1901.944 | 0 | -0.41 | 0.003589 | 4.65 | 116.5 | 83.5 | 7.3 | 6.38 | | High | High | High | High | High | High | High | High | High | High | High | 0 | 0.00011 | 2.4 |
| FALSE | High | TQTGDSSS | [R].TQTGD | 1xPhospho | A2AEW8 1 | 0.02933 | 0.001857 | 1 | 3 | 3 | A2AEW8 | GRIP1-assc | A2AEW8 [6 | 0 | 1917.843 | 0 | 1.45 | 0.002225 | 8.84 | 53.6 | 146.4 | 12.77 | 11.3 | | High | High | High | High | High | High | High | High | High | High | High | 0.001468 | 0.01469 | 1.48 |
| FALSE | High | TQTGHVLG | [R].TQTGH | 1xPhospho | E9Q555 1x | 1.23E-05 | 0 | 1 | 3 | 1 | E9Q555 | E3 ubiquiti | E9Q555 [4 | 0 | 2421.161 | 0 | 1.32 | 0.001448 | 16.27 | 59.1 | 140.9 | 10.63 | 10.15 | | High | High | High | High | High | High | High | High | High | High | High | 0 | 1.7E-06 | 3.63 |
| FALSE | High | TQTPPLGQ | [R].TQTPPL | 1xPhospho | A0JNY7 1x | 8.79E-05 | 0 | 1 | 7 | 3 | A0JNY7 | Eukaryotic | A0JNY7 [50 | 0 | 1888.014 | 0 | 0.35 | 0.504099 | 43.95 | 88.1 | 111.9 | 37.39 | 62.26 | | High | High | High | High | High | High | High | High | High | High | High | 0 | 9.25E-05 | 3.73 |
| FALSE | High | TQSQSIYNI | [K].TQSQSI | 1xCarbami | Q8BKX6 1x | 3.94E-05 | 0 | 1 | 1 | 4 | Q8BKX6 | Serine/thre | Q8BKX6 [3 | 0 | 4847.271 | 0 | -0.13 | 0.306125 | 26.29 | 113.4 | 86.6 | 20.65 | 13.64 | | High | High | High | High | High | High | High | High | High | High | High | 0 | 6.3E-06 | 5.07 |
| FALSE | High | TQPDGTSV | [R].TQPDG | 1xPhospho | E9Q7G0 1x | 4.92E-07 | 0 | 1 | 4 | 8 | E9Q7G0 | Nuclear mi | E9Q7G0 [1 | 0 | 2233.07 | 0 | 0.12 | 0.020579 | 1.88 | 95.7 | 104.3 | 4.47 | 6.8 | | High | High | High | High | High | High | High | High | High | High | High | 0 | 2.03E-06 | 3.14 |
| FALSE | High | TQLWASEF | [R].TQLWA | 1xPhospho | Q99MR6 1 | 0.000628 | 0 | 1 | 5 | 1 | Q99MR6 | Serrate RN | Q99MR6 [ | 0 | 3196.706 | 0 | -0.45 | 0.00145 | 2.82 | 117.7 | 82.3 | 4.01 | 5.47 | | High | High | High | High | High | High | High | High | High | High | High | 0 | 0.000141 | 2.95 |
| FALSE | High | TQLWASEF | [R].TQLWA | 1xPhospho | Q99MR6 1 | 0.00051 | 0 | 1 | 5 | 1 | Q99MR6 | Serrate RN | Q99MR6 [ | 0 | 2967.543 | 0 | -0.52 | 0.859395 | 34.14 | 117.8 | 82.2 | 62.63 | 100.25 | | High | High | High | High | High | High | High | High | High | High | High | 0 | 0.000112 | 2.74 |
| FALSE | High | TQAYSNPG | [R].TQAYSI | 1xCarbami | Q3UCG9 1 | 0.000908 | 0 | 1 | 8 | 1 | Q3UCG9 | E3 ubiquiti | Q3UCG9 [6 | 0 | 2987.353 | 0 | -0.75 | 0.068699 | 11.46 | 120.6 | 79.4 | 26.18 | 17.58 | | High | High | High | High | High | High | High | High | High | High | High | 0 | 0.000214 | 2.35 |
| FALSE | High | TQCAPSPIC | [K].TQCAP | 1xCarbami | Q9D4V4 1x | 0.000375 | 0 | 1 | 5 | 2 | Q9D4V4 | TATA box-l | Q9D4V4 [3 | 0 | 1667.859 | 0 | -1.26 | 0.000177 | 0.6 | 142 | 58 | 1.91 | 5.34 | | High | High | High | High | High | High | High | High | High | High | High | 0 | 7.93E-05 | 2.26 |
| FALSE | High | TQCPAFAE | [K].TQCPAI | 3xCarbami | E9Q2E4 1x | 2.89E-07 | 0 | 1 | 3 | 2 | E9Q2E4 | Uncharacte | E9Q2E4 [1 | 0 | 2816.18 | 0 | 1.07 | 0.002844 | 16.43 | 61.5 | 138.5 | 17.31 | 9.73 | | High | High | High | High | High | High | High | High | High | High | High | 0 | 2.52E-08 | 3.97 |
| FALSE | High | TQDDFCSP | [K].TQDDF | 1xCarbami | Q3TLR7 1x | 0.012072 | 0.000889 | 1 | 1 | 1 | Q3TLR7 | Denticleles | Q3TLR7 [7 | 0 | 1974.799 | 0 | -0.15 | 0.678099 | 22.02 | 103.3 | 96.7 | 10.54 | 14.68 | | High | High | High | High | High | High | High | High | High | High | High | 0.000449 | 0.003864 | 2.41 |
| FALSE | High | TQDPSSPC | [R].TQDPS | 1xCarbami | Q9D0N7 1x | 1.19E-05 | 0 | 1 | 1 | 6 | Q9D0N7 | Chromatin | Q9D0N7 [4 | 0 | 2994.453 | 0 | -0.05 | 0.381051 | 9.19 | 100.9 | 99.1 | 5.82 | 10.95 | | High | High | High | High | High | High | High | High | High | High | High | 0 | 1.07E-05 | 4.18 |
| FALSE | High | TQDPSSPC | [R].TQDPS | 1xCarbami | Q9D0N7 2x | 8.5E-06 | 0 | 1 | 1 | 2 | Q9D0N7 | Chromatin | Q9D0N7 [4 | 0 | 3074.419 | 0 | 0.8 | 0.003443 | 13.19 | 71.4 | 128.6 | 10.39 | 10.67 | | High | High | High | High | High | High | High | High | High | High | High | 0 | 3.06E-05 | 2.89 |
| FALSE | High | TQDPSSPC | [R].TQDPS | 1xCarbami | Q9D0N7 1x | 0.000758 | 0 | 1 | 1 | 3 | Q9D0N7 | Chromatin | Q9D0N7 [4 | 1 | 4147.087 | 0 | -0.22 | 0.228078 | 22.72 | 105.6 | 94.4 | 12.96 | 16.68 | | High | High | High | High | High | High | High | High | High | High | High | 0 | 0.000175 | 3.18 |
| FALSE | High | TQDPVPPE | [R].TQDPV | 1xPhospho | Q8C1S0 1x | 4.98E-08 | 0 | 1 | 1 | 5 | Q8C1S0 | Mediator c | Q8C1S0 [1 | 0 | 2288.085 | 0 | -0.46 | 0.010252 | 4.01 | 115.9 | 84.1 | 6.7 | 7.23 | | High | High | High | High | High | High | High | High | High | High | High | 0 | 3.5E-09 | 3.17 |
| FALSE | High | TQDPVPPE | [R].TQDPV | 2xPhospho | Q8C1S0 2x | 0.008691 | 0.000736 | 1 | 1 | 1 | Q8C1S0 | Mediator c | Q8C1S0 [1 | 0 | 2368.051 | 0 | 0.02 | 0.88167 | 10.27 | 100.9 | 99.1 | 10.98 | 8.81 | | High | High | High | High | High | High | High | High | High | High | High | 0.000327 | 0.002693 | 2.77 |
| FALSE | High | TQEIESLAS | [R].TQEIES | 1xPhospho | E9QQ10 1x | 0.000687 | 0 | 1 | 4 | 3 | E9QQ10 | A-kinase ar | E9QQ10 [2 | 0 | 1958.025 | 0 | -0.4 | 0.010528 | 9.11 | 112.9 | 87.1 | 11.98 | 10.51 | | High | High | High | High | High | High | High | High | High | High | High | 0.000872 | 0.009103 | 2.79 |
| FALSE | High | TQEISRPNS | [K].TQEISR | 1xPhospho | Q80TZ9 1x | 0.010663 | 0.000772 | 1 | 2 | 1 | Q80TZ9 | Arginine-gl | Q80TZ9 [6 | 0 | 2558.121 | 0 | 0.02 | 0.711732 | 5.46 | 99.2 | 100.8 | 4.26 | 7.65 | | High | High | High | High | High | High | High | High | High | High | High | 0.000399 | 0.00337 | 1.4 |
| FALSE | High | TQEISRPNS | [K].TQEISR | 2xPhospho | Q80TZ9 2x | 0.014667 | 0.001049 | 1 | 2 | 2 | Q80TZ9 | Arginine-gl | Q80TZ9 [6 | 0 | 2638.087 | 0 | -0.27 | 0.072019 | 11.46 | 110.4 | 89.6 | 12.07 | 5.45 | | High | High | High | High | High | High | High | High | High | High | High | 0.002589 | 0.02766 | 1.49 |
| FALSE | High | TQESCGIAI | [R].TQESC | 1xCarbami | Q9CZJ2 1x | 0.100129 | 0.008067 | 1 | 2 | 2 | Q9CZJ2 | Heat shock | Q9CZJ2 [32 | 0 | 2792.405 | 0 | -0.35 | 0.04035 | 10.07 | 113.6 | 86.4 | 22.1 | 14.22 | | High | High | High | High | High | High | High | High | High | High | High | 0.004032 | 0.04307 | 2.82 |
| FALSE | High | TQESGQGS | [K].TQESG | 1xPhospho | Q924A2 1x | 0.000369 | 0 | 1 | 4 | 2 | Q924A2 | Protein cap | Q924A2 [1 | 0 | 3180.645 | 0 | -0.26 | 0.218684 | 9.36 | 108.8 | 91.2 | 12.53 | 8.41 | | High | High | High | High | High | High | High | High | High | High | High | 0 | 7.77E-05 | 2.28 |
| FALSE | High | TQESPTAEI | [R].TQESPT | 1xPhospho | E9PUP1 1x | 1.93E-05 | 0 | 1 | 4 | 1 | E9PUP1 | Synapse de | E9PUP1 [5 | 0 | 2040.03 | 0 | -1.15 | 0.000264 | 0.23 | 138.3 | 61.7 | 9.66 | 14 | | High | High | High | High | High | High | High | High | High | High | High | 0 | 2.82E-06 | 3.1 |
| FALSE | High | TQGESISEC | [R].TQGESI | 1xCarbami | A0A1S6GW | 6.25E-09 | 0 | 1 | 6 | 2 | A0A1S6GW | Uncharacte | A0A1S6GW | 0 | 3348.474 | 0 | 1.49 | 0.000395 | 4.71 | 53.6 | 146.4 | 10.72 | 3.13 | | High | High | High | High | High | High | High | High | High | High | High | 0 | 3.4E-10 | 4.76 |
| FALSE | High | TQGPTDSL | [R].TQGPT | 1xPhospho | A2AJG0 1x | 5.09E-06 | 0 | 1 | 3 | 6 | A2AJG0 | Ribonuclea | A2AJG0 [2 | 0 | 3850.768 | 0 | -0.37 | 0.011548 | 8.33 | 114.5 | 85.5 | 11.26 | 7.03 | | High | High | High | High | High | High | High | High | High | High | High | 0 | 2.8E-05 | 4.72 |
| FALSE | High | TQGPTDSL | [R].TQGPT | 1xDeamida | A2AJG0 1x | 3.21E-06 | 0 | 1 | 3 | 8 | A2AJG0 | Ribonuclea | A2AJG0 [2 | 0 | 3851.752 | 0 | -0.64 | 0.005466 | 12.26 | 121.8 | 78.2 | 7.38 | 27.26 | | High | High | High | High | High | High | High | High | High | High | High | 0 | 3.79E-07 | 5.04 |
| FALSE | High | TQGSSMSV | [K].TQGSSI | 1xPhospho | Q8CCE5 1x | 8.66E-06 | 0 | 1 | 2 | 6 | Q8CCE5 | DNA topois | Q8CCE5 [8 | 0 | 2121.019 | 0 | -0.68 | 0.004842 | 4.23 | 121.8 | 78.2 | 7.2 | 9.66 | | High | High | High | High | High | High | High | High | High | High | High | 0 | 1.15E-06 | 2.91 |
| FALSE | High | TQGSSMSV | [K].TQGSSI | 1xPhospho | Q8CCE5 1x | 2.36E-05 | 0 | 1 | 2 | 5 | Q8CCE5 | DNA topois | Q8CCE5 [8 | 0 | 2137.014 | 0 | -0.39 | 0.125968 | 10.2 | 117.5 | 82.5 | 8.36 | 17.99 | | High | High | High | High | High | High | High | High | High | High | High | 0 | 3.54E-06 | 3.04 |
| FALSE | High | TQGSSMSV | [K].TQGSSI | 2xPhospho | Q8CCE5 2x | 0.028824 | 0.001829 | 1 | 2 | 1 | Q8CCE5 | DNA topois | Q8CCE5 [8 | 0 | 2200.985 | 0 | -0.27 | 0.112344 | 7.18 | 112.9 | 87.1 | 5.61 | 19.46 | | High | High | High | High | High | High | High | High | High | High | High | 0.000997 | 0.01029 | 2.72 |
| FALSE | High | TQGSSMSV | [K].TQGSSI | 1xPhospho | Q01320 1x | 5.5E-05 | 0 | 1 | 2 | 6 | Q01320 | DNA topois | Q01320 [1 | 0 | 2220.087 | 0 | -0.6 | 0.001262 | 3.14 | 123.8 | 76.2 | 5.53 | 5.51 | | High | High | High | High | High | High | High | High | High | High | High | 0 | 5.53E-05 | 3.19 |
| FALSE | High | TQGSSMSV | [K].TQGSSI | 1xPhospho | Q01320 1x | 1.69E-06 | 0 | 1 | 2 | 4 | Q01320 | DNA topois | Q01320 [1 | 0 | 2236.082 | 0 | -0.21 | 0.043199 | 9.46 | 110.7 | 89.3 | 12.56 | 4.25 | | High | High | High | High | High | High | High | High | High | High | High | 0 | 9.15E-07 | 4.36 |
| FALSE | High | TQGSVSAA | [R].TQGSV | 1xPhospho | Q3TA75 1x | 0.0009 | 0 | 1 | 1 | 1 | Q3TA75 | Putative ur | Q3TA75 [1 | 0 | 3546.71 | 0 | 1.23 | 0.037284 | 55.88 | 46.7 | 153.3 | 34.79 | 38.46 | | High | High | High | High | High | High | High | High | High | High | High | 0 | 0.000212 | 3.46 |
| FALSE | High | TQGYLESPI | [K].TQGYLI | 1xPhospho | Q3TZR9 1x | 0.003372 | 0.00016 | 1 | 1 | 1 | Q3TZR9 | Activating t | Q3TZR9 [4 | 0 | 1560.808 | 0 | 0.44 | 0.238737 | 13.92 | 87.1 | 112.9 | 15.96 | 13.41 | | High | High | High | High | High | High | High | High | High | High | High | 9.42E-05 | 0.000931 | 2.56 |
| FALSE | High | TQHSSLDQ | [K].TQHSSI | 1xPhospho | Q6NZM9 1 | 0.006773 | 0.00062 | 1 | 2 | 2 | Q6NZM9 | Histone de | Q6NZM9 [ | 0 | 3842.81 | 0 | 0.88 | 0.066324 | 37.09 | 69.5 | 130.5 | 22.6 | 32.24 | | High | High | High | High | High | High | High | High | High | High | High | 0.000275 | 0.002034 | 3.47 |
| FALSE | High | TQLASWSD | [R].TQLAS | 1xPhospho | Q9QXS1-3 | 3.35E-09 | 0 | 3 | 23 | 7 | Q9QXS1-3; | Isoform PL | Q9QXS1-3 | 0 | 3426.697 | 0 | 2.87 | 0.000909 | 20.7 | 21.4 | 178.6 | 14.07 | 14.07 | Shared | High | High | High | High | High | High | High | High | High | High | High | 0 | 2.83E-07 | 5.46 |
| FALSE | High | TQLSPEAK | [R].TQLSPE | 1xPhospho | F8VPY0 1x | 0.006957 | 0.00062 | 1 | 2 | 1 | F8VPY0 | Zinc finger | F8VPY0 [3 | 0 | 1411.76 | 0 | 3.46 | 0.000616 | 11.29 | 16.3 | 183.7 | 11.5 | 12.48 | | High | High | High | High | High | High | High | High | High | High | High | 0.000275 | 0.002101 | 1.6 |
| FALSE | High | TQLSTSAEF | [K].TQLSTS | 1xPhospho | Q9D1F5 1x | 0.004781 | 0.000456 | 1 | 1 | 5 | Q9D1F5 | MRN comp | Q9D1F5 [1 | 0 | 2122.049 | 0 | -1.28 | 0.000503 | 3.13 | 142.8 | 57.2 | 4.91 | 16.65 | | High | High | High | High | High | High | High | High | High | High | High | 0.001774 | 0.01751 | 1.73 |
| FALSE | High | TQTPPLGQ | [R].TQTPPL | 1xPhospho | A0JNY7 1x | 0.000103 | 0 | 1 | 7 | 1 | A0JNY7 | Eukaryotic | A0JNY7 [50 | 0 | 2117.177 | 0 | 0.68 | 0.006614 | 14.15 | 75.2 | 124.8 | 5.56 | 11.01 | | High | High | High | High | High | High | High | High | High | High | High | 0 | 1.87E-05 | 3.42 |
| FALSE | High | TQAQHSSP | [R].TQAQH | 1xPhospho | A2ADY9 1x | 5.35E-08 | 0 | 1 | 1 | 4 | A2ADY9 | Protein DD | A2ADY9 [1 | 0 | 2917.371 | 0 | -0.89 | 0.001475 | 2.27 | 130.1 | 69.9 | 2.85 | 17.12 | | High | High | High | High | High | High | High | High | High | High | High | 0 | 3.77E-09 | 4.96 |
| FALSE | High | TQTSLMSP | [K].TQTSLN | 1xPhospho | Q8BMA5 1 | 0.029076 | 0.001857 | 1 | 1 | 4 | Q8BMA5 | Protein NP | Q8BMA5 [ | 0 | 1402.66 | 0 | -0.5 | 0.003577 | 5.34 | 116.4 | 83.6 | 5.72 | 6.41 | | High | High | High | High | High | High | High | High | High | High | High | 0.009099 | 0.09835 | 1.32 |
| FALSE | High | TQVLSPDSI | [K].TQVLSF | 1xPhospho | E9Q9C3 1x | 9.56E-06 | 0 | 1 | 5 | 1 | E9Q9C3 | Afadin OS= | E9Q9C3 [1 | 0 | 1945.045 | 0 | 0.44 | 0.016053 | 15.36 | 86 | 114 | 7.26 | 10.45 | | High | High | High | High | High | High | High | High | High | High | High | 0 | 1.28E-06 | 2.93 |
| FALSE | High | TSAPEPR | [R].TSAPEP | 1xPhospho | Q0VGB7 1 | 0.062209 | 0.004646 | 2 | 3 | 1 | Q0VGB7; A | Serine/thre | Q0VGB7 [6 | 0 | 1066.513 | 0 | -0.34 | 0.024585 | 9.13 | 118.5 | 81.5 | 11.37 | 17.32 | Shared | High | High | High | High | High | High | High | High | High | High | High | 0.002381 | 0.02476 | 1.73 |
| FALSE | High | TSAPLLQQ | [R].TSAPLL | 1xPhospho | Q8BYH8 1x | 5.84E-07 | 0 | 1 | 3 | 2 | Q8BYH8 | Chromodo | Q8BYH8 [2 | 0 | 2750.469 | 0 | 0.56 | 0.047387 | 8.91 | 80.5 | 119.5 | 13.89 | 13.32 | | High | High | High | High | High | High | High | High | High | High | High | 0 | 5.55E-08 | 4.18 |
| FALSE | High | TSAQCGDV | [R].TSAQC | 1xCarbami | Q14AM7 1 | 6.52E-08 | 0 | 1 | 1 | 1 | Q14AM7 | UPF0472 p | Q14AM7 [ | 0 | 2255.078 | 0 | 0.05 | 0.620587 | 19.45 | 98.1 | 101.9 | 13.56 | 10.01 | | High | High | High | High | High | High | High | High | High | High | High | 0 | 4.74E-09 | 5.04 |
| FALSE | High | TSAQCGDV | [R].TSAQC | 1xCarbami | Q14AM7 1 | 0.058664 | 0.004294 | 1 | 1 | 1 | Q14AM7 | UPF0472 p | Q14AM7 [ | 1 | 2411.179 | 0 | 0.13 | 0.593856 | 8.7 | 94 | 106 | 24.82 | 2.81 | | High | High | High | High | High | High | High | High | High | High | High | 0.002187 | 0.0232 | 2.19 |
| FALSE | High | TSAQNCTL | [R].TSAQN | 1xCarbami | Q921S7 1x | 3.67E-05 | 0 | 1 | 1 | 1 | Q921S7 | 39S riboso | Q921S7 [2 | 0 | 1932.884 | 0 | 0.92 | 0.003534 | 10.78 | 66 | 134 | 19.15 | 12.85 | | High | High | High | High | High | High | High | High | High | High | High | 0 | 5.83E-06 | 3.59 |
| FALSE | High | TSASGPLTF | [K].TSASGF | 1xPhospho | E9QK36 1x | 0.01441 | 0.001011 | 1 | 5 | 1 | E9QK36 | WD repeat | E9QK36 [1 | 0 | 2298.206 | 0 | -0.38 | 0.00977 | 5.4 | 114.2 | 85.8 | 3.58 | 12.99 | | High | High | High | High | High | High | High | High | High | High | High | 0.000538 | 0.004727 | 2.43 |
| FALSE | High | TSATTPNFI | [R].TSATTP | 1xPhospho | Q9ERH4 1x | 0.047894 | 0.00354 | 1 | 1 | 2 | Q9ERH4 | Nucleolar a | Q9ERH4 [1 | 0 | 1504.781 | 0 | -1.57 | 0.00028 | 0.09 | 148.5 | 51.5 | 6.78 | 8.8 | | High | High | High | High | High | High | High | High | High | High | High | 0.001812 | 0.01826 | 2.22 |
| FALSE | High | TSAVPNHS | [K].TSAVPN | 1xPhospho | E9PZY8 1x | 1.95E-06 | 0 | 1 | 4 | 1 | E9PZY8 | RIKEN cDN | E9PZY8 [57 | 0 | 2490.135 | 0 | -0.36 | 0.000689 | 2.8 | 114.4 | 85.6 | 6.61 | 9.38 | | High | High | High | High | High | High | High | High | High | High | High | 0 | 2.16E-07 | 3.41 |
| FALSE | High | TSCPGENT | [R].TSCPGE | 1xCarbami | J3QM81 1x | 0.000322 | 0 | 1 | 5 | 1 | J3QM81 | Ankyrin rep | J3QM81 [7 | 0 | 1819.83 | 0 | 0.2 | 0.108387 | 6.19 | 96.2 | 103.8 | 11.23 | 7.88 | | High | High | High | High | High | High | High | High | High | High | High | 0 | 6.69E-05 | 2.02 |
| FALSE | High | TSCPGENT | [R].TSCPGE | 1xCarbami | J3QM81 1x | 0.018285 | 0.001282 | 1 | 5 | 2 | J3QM81 | Ankyrin rep | J3QM81 [7 | 1 | 3281.622 | 0 | -0.38 | 0.004096 | 7.61 | 115 | 85 | 4.68 | 12.37 | | High | High | High | High | High | High | High | High | High | High | High | 0.00074 | 0.006428 | 2.37 |
| FALSE | High | TSCSPSVR | [R].TSCSPS | 1xCarbami | A2ADB0 1x | 0.003402 | 0.00021 | 1 | 5 | 2 | A2ADB0 | Msx2-inter | A2ADB0 [1 | 0 | 1202.544 | 0 | 0.85 | 0.003747 | 10.65 | 74 | 126 | 8.25 | 7.96 | | High | High | High | High | High | High | High | High | High | High | High | 0.001712 | 0.01711 | 1.57 |
| FALSE | High | TSDANETE | [K].TSDANI | 1xCarbami | Q3UYV9 1x | 0.003433 | 0.00021 | 1 | 2 | 6 | Q3UYV9 | Nuclear ca | Q3UYV9 [2 | 0 | 2271.005 | 0 | -0.13 | 0.842879 | 26.69 | 104.6 | 95.4 | 40.54 | 42.73 | | High | High | High | High | High | High | High | High | High | High | High | 9.42E-05 | 0.00095 | 2.93 |
| FALSE | High | TSDANETE | [K].TSDANI | 1xCarbami | Q3UYV9 1x | 8.84E-07 | 0 | 1 | 2 | 7 | Q3UYV9 | Nuclear ca | Q3UYV9 [2 | 0 | 2500.168 | 0 | -0.17 | 0.017316 | 4.63 | 106.8 | 93.2 | 3.54 | 6.25 | | High | High | High | High | High | High | High | High | High | High | High | 0 | 3.53E-05 | 3.72 |
| FALSE | High | TSALLFSSD | [K].TSALLF | 2xPhospho | Q6PGL7 2x | 0.013909 | 0.001011 | 1 | 5 | 3 | Q6PGL7 | WASH com | Q6PGL7 [6 | 0 | 3112.395 | 0 | 0.37 | 0.186557 | 15.04 | 85.8 | 114.2 | 14.03 | 13.88 | | High | High | High | High | High | High | High | High | High | High | High | 0.000538 | 0.004532 | 2.62 |
| FALSE | High | TSDANETE | [K].TSDANI | 1xCarbami | Q3UYV9 1x | 1.18E-05 | 0 | 1 | 2 | 3 | Q3UYV9 | Nuclear ca | Q3UYV9 [2 | 0 | 2501.152 | 0 | -0.3 | 0.002793 | 1.07 | 114.1 | 85.9 | 5.4 | 5.26 | | High | High | High | High | High | High | High | High | High | High | High | 0 | 1.63E-06 | 3.6 |
| FALSE | High | TSDIFGSPV | [K].TSDIFG | 1xPhospho | Q3U7P4 1x | 4.05E-05 | 0 | 1 | 3 | 1 | Q3U7P4 | Putative ur | Q3U7P4 [9 | 0 | 3016.648 | 0 | -0.67 | 0.144988 | 4.59 | 120.4 | 79.6 | 17.82 | 13.56 | | High | High | High | High | High | High | High | High | High | High | High | 0 | 6.48E-06 | 2.62 |
| FALSE | High | TSDLSPEGS | [K].TSDLSP | 1xCarbami | Q9DBQ7 1 | 0.000524 | 0 | 1 | 2 | 1 | Q9DBQ7 | Protein-ass | Q9DBQ7 [4 | 0 | 2527.116 | 0 | 1.01 | 0.007032 | 36.41 | 66.4 | 133.6 | 22.53 | 7.1 | | High | High | High | High | High | High | High | High | High | High | High | 0 | 0.000116 | 3.3 |
| FALSE | High | TSDPDVILC | [K].TSDPDV | 1xCarbami | Q7TPE5 1x | 0.000908 | 0 | 1 | 1 | 2 | Q7TPE5 | Probable R | Q7TPE5 [1 | 0 | 2140.056 | 0 | 0.43 | 0.031764 | 19.95 | 86.4 | 113.6 | 6 | 13.23 | | High | High | High | High | High | High | High | High | High | High | High | 2.58E-05 | 0.000536 | 3.47 |
| FALSE | High | TSDQDFTP | [K].TSDQD | 1xPhospho | Q3UMU9-3 | 0.014158 | 0.001011 | 1 | 4 | 1 | Q3UMU9-3 | Isoform 3 c | Q3UMU9-3 | 0 | 1705.809 | 0 | 0.42 | 0.014489 | 6.65 | 85.6 | 114.4 | 11.14 | 8.57 | | High | High | High | High | High | High | High | High | High | High | High | 0.000538 | 0.004633 | 2.19 |
| FALSE | High | TSDSDQQT | [R].TSDSD | 1xPhospho | Q8K3K8 1x | 0.000118 | 0 | 1 | 1 | 4 | Q8K3K8 | Optineurin | Q8K3K8 [5 | 0 | 1897.853 | 0 | 2.69 | 0.000235 | 7.21 | 27 | 173 | 7.8 | 5.32 | | High | High | High | High | High | High | High | High | High | High | High | 0 | 2.22E-05 | 3.61 |
| FALSE | High | TSDVPLEPL | [R].TSDVPL | 1xPhospho | A2ATK9 1x | 0.003788 | 0.00021 | 1 | 4 | 1 | A2ATK9 | Family with | A2ATK9 [8 | 0 | 1932.963 | 0 | 1.95 | 0.000857 | 16.94 | 36.6 | 163.4 | 11.16 | 12.05 | | High | High | High | High | High | High | High | High | High | High | High | 0.000117 | 0.001061 | 2.44 |
| FALSE | High | TSEDQAAII | [K].TSEDQ | 1xPhospho | Q3UZ64 1x | 7.84E-06 | 0 | 1 | 10 | 2 | Q3UZ64 | Nuclear fac | Q3UZ64 [3 | 0 | 1767.929 | 0 | -0.02 | 0.711713 | 11.75 | 100.2 | 99.8 | 10.41 | 9.11 | | High | High | High | High | High | High | High | High | High | High | High | 0 | 1.03E-06 | 3.12 |
| FALSE | High | TSEDTSSGS | [R].TSEDTS | 1xPhospho | Q3UYQ4 1 | 2.86E-06 | 0 | 1 | 2 | 7 | Q3UYQ4 | Putative ur | Q3UYQ4 [4 | 0 | 1730.825 | 0 | -0.68 | 0.000313 | 1.46 | 123.6 | 76.4 | 2.65 | 5.75 | | High | High | High | High | High | High | High | High | High | High | High | 0 | 1.13E-06 | 1.85 |
| FALSE | High | TSEDTSSGS | [R].TSEDTS | 1xPhospho | Q3UYQ4 1 | 0.015741 | 0.001124 | 1 | 2 | 3 | Q3UYQ4 | Putative ur | Q3UYQ4 [4 | 1 | 1858.92 | 0 | -0.65 | 0.021016 | 12.14 | 122.4 | 77.6 | 15.13 | 8.28 | | High | High | High | High | High | High | High | High | High | High | High | 0.006155 | 0.06637 | 2.52 |
| FALSE | High | TSEFPTPLF | [K].TSEFPT | 1xCarbami | A0A0R4J12 | 0.022973 | 0.001578 | 1 | 2 | 2 | A0A0R4J12 | SRSF prote | A0A0R4J12 | 0 | 4567.106 | 0 | -0.11 | 0.479222 | 22.81 | 105.8 | 94.2 | 12.47 | 13.15 | | High | High | High | High | High | High | High | High | High | High | High | 0.000847 | 0.007981 | 3.6 |
| FALSE | High | TSESSEEFL | [R].TSESSE | 1xPhospho | Q5XG69 1x | 1.33E-08 | 0 | 1 | 1 | 21 | Q5XG69 | Soluble lan | Q5XG69 [3 | 0 | 3599.573 | 0 | -1.31 | 0.000574 | 0.48 | 143.4 | 56.6 | 4.31 | 13.74 | | High | High | High | High | High | High | High | High | High | High | High | 0 | 7.97E-10 | 5.18 |
| FALSE | High | TSESSEEFL | [R].TSESSE | 2xPhospho | Q5XG69 2x | 4.37E-07 | 0 | 1 | 1 | 24 | Q5XG69 | Soluble lan | Q5XG69 [3 | 0 | 3679.539 | 0 | -1.02 | 0.000878 | 1.98 | 135.5 | 64.5 | 7.99 | 12.97 | | High | High | High | High | High | High | High | High | High | High | High | 0 | 2.36E-07 | 4.83 |
| FALSE | High | TSETSAPFP | [R].TSETSA | 1xPhospho | Q8BGS1 1x | 1.38E-05 | 0 | 1 | 2 | 2 | Q8BGS1 | Band 4.1-li | Q8BGS1 [4 | 0 | 2284.106 | 0 | -0.64 | 0.019689 | 14.39 | 117.3 | 82.7 | 5.28 | 19.01 | | High | High | High | High | High | High | High | High | High | High | High | 0 | 2.8E-06 | 2.89 |
| FALSE | High | TSDFNTFL | [K].TSDFNT | 1xCarbami | Q9D1P4 1x | 4.08E-05 | 0 | 1 | 2 | 2 | Q9D1P4 | Cysteine ar | Q9D1P4 [1 | 0 | 2055.904 | 0 | 2.06 | 0.001592 | 22.83 | 35.8 | 164.2 | 7.04 | 16.03 | | High | High | High | High | High | High | High | High | High | High | High | 0 | 6.54E-06 | 4.08 |
| FALSE | High | TSALLFSSD | [K].TSALLF | 1xPhospho | Q6PGL7 1x | 1.77E-06 | 0 | 1 | 5 | 2 | Q6PGL7 | WASH com | Q6PGL7 [6 | 0 | 3032.429 | 0 | -0.69 | 0.009968 | 7.92 | 121.2 | 78.8 | 7.7 | 20.27 | | High | High | High | High | High | High | High | High | High | High | High | 0 | 1.93E-07 | 4.27 |
| FALSE | High | TSALLFSSD | [K].TSALLF | 2xPhospho | Q6PGL7 2x | 0.002869 | 0.00016 | 1 | 5 | 1 | Q6PGL7 | WASH com | Q6PGL7 [6 | 0 | 2883.232 | 0 | 0.06 | 0.577961 | 47.16 | 95.4 | 104.6 | 25.36 | 24.7 | | High | High | High | High | High | High | High | High | High | High | High | 7.19E-05 | 0.000774 | 2.65 |
| FALSE | High | TSAFSPPGS | [R].TSAFSP | 2xCarbami | E9Q777 1x | 1.71E-06 | 0 | 1 | 2 | 3 | E9Q777 | A kinase (P | E9Q777 [4 | 0 | 3142.312 | 0 | 0.16 | 0.631041 | 24.19 | 95.1 | 104.9 | 9.21 | 27.2 | | High | High | High | High | High | High | High | High | High | High | High | 0 | 1.61E-05 | 4.44 |
| FALSE | High | TQVTETPC | [K].TQVTE | 1xCarbami | G3X961 1x | 0.000124 | 0 | 1 | 3 | 1 | G3X961 | Kelch dom | G3X961 [5 | 0 | 1875.887 | 0 | 0.62 | 0.000218 | 5.14 | 76.3 | 123.7 | 9.5 | 5.82 | | High | High | High | High | High | High | High | High | High | High | High | 0 | 2.29E-05 | 2.26 |
| FALSE | High | TQYNQVPS | [K].TQYNQ | 1xPhospho | P97792 1x | 0.007811 | 0.000664 | 1 | 3 | 1 | P97792 | Coxsackiev | P97792 [31 | 0 | 1921.853 | 0 | -0.99 | 0.001994 | 2 | 134.4 | 65.6 | 13.92 | 9.79 | | High | High | High | High | High | High | High | High | High | High | High | 0.000327 | 0.002379 | 1.47 |
| FALSE | High | TRASLPVVF | [R].TRASLP | 1xPhospho | E9QAU4 1x | 0.076555 | 0.005826 | 1 | 9 | 2 | E9QAU4 | Enhancer t | E9QAU4 [2 | 1 | 1307.74 | 0 | 0.24 | 0.018621 | 6.2 | 91.8 | 108.2 | 4.78 | 4.69 | | High | High | High | High | High | High | High | High | High | High | High | 0.002976 | 0.03147 | 1.84 |
| FALSE | High | TREEEPCQ | [K].TREEEP | 1xCarbami | Q8K3Z9 1x | 2.11E-07 | 0 | 1 | 2 | 6 | Q8K3Z9 | Nuclear en | Q8K3Z9 [4 | 1 | 2813.343 | 0 | 0.39 | 0.023227 | 1.91 | 87.7 | 112.3 | 6.09 | 9.65 | | High | High | High | High | High | High | High | High | High | High | High | 0 | 1.77E-08 | 3.7 |
| FALSE | High | TREEEPCQ | [K].TREEEP | 1xCarbami | Q8K3Z9 1x | 0.014033 | 0.001011 | 1 | 2 | 2 | Q8K3Z9 | Nuclear en | Q8K3Z9 [4 | 2 | 3669.792 | 0 | 0.9 | 0.014034 | 7.72 | 72.1 | 127.9 | 9.06 | 15.62 | | High | High | High | High | High | High | High | High | High | High | High | 0.000538 | 0.004597 | 3.04 |
| FALSE | High | TRHSPTPQ | [K].TRHSPT | 1xPhospho | Q52KI8 1x | 0.090888 | 0.007081 | 1 | 6 | 1 | Q52KI8 | Serine/argi | Q52KI8 [40 | 1 | 1717.833 | 0 | -0.41 | 0.17885 | 28.52 | 120.1 | 79.9 | 25.5 | 15.65 | | High | High | High | High | High | High | High | High | High | High | High | 0.003606 | 0.03841 | 2.57 |
| FALSE | High | TRITPEEMT | [R].TRITPE | 1xPhospho | Q3UHI0-2 | 0.110202 | 0.008961 | 1 | 2 | 1 | Q3UHI0-2 | Isoform 2 c | Q3UHI0-2 | 1 | 1872.991 | 0 | -0.22 | 0.024478 | 8.15 | 108 | 92 | 5.7 | 6.12 | | High | High | High | High | High | High | High | High | High | High | High | 0.004462 | 0.04838 | 2.28 |
| FALSE | High | TRNSGIWD | [R].TRNSGI | 1xPhospho | Q68FM7 1 | 0.023792 | 0.001612 | 1 | 6 | 1 | Q68FM7 | Rho guanir | Q68FM7 [1 | 1 | 1954.922 | 0 | -0.16 | 0.10492 | 8.72 | 105.4 | 94.6 | 9.06 | 5.14 | | High | High | High | High | High | High | High | High | High | High | High | 0.000859 | 0.008261 | 2.58 |
| FALSE | High | TRNSGIWD | [R].TRNSGI | 1xPhospho | Q68FM7 1 | 0.001463 | 5.66E-05 | 1 | 5 | 2 | Q68FM7 | Rho guanir | Q68FM7 [1 | 2 | 3788.781 | 0 | -0.2 | 0.119967 | 8.75 | 107.1 | 92.9 | 5.54 | 11.42 | | High | High | High | High | High | High | High | High | High | High | High | 2.58E-05 | 0.000383 | 3.14 |
| FALSE | High | TRPGAVSP | [K].TRPGA | 1xPhospho | Q0KL02-4 | 3.12E-06 | 0 | 1 | 3 | 1 | Q0KL02-4 | Isoform 4 c | Q0KL02-4 | 0 | 2896.537 | 0 | 0.61 | 0.017931 | 12.13 | 74.7 | 125.3 | 8.62 | 15.82 | | High | High | High | High | High | High | High | High | High | High | High | 0 | 3.67E-07 | 3.96 |
| FALSE | High | TRSEPLPPS | [R].TRSEPL | 1xPhospho | Q8C2B3-3 | 8.98E-06 | 0 | 1 | 11 | 1 | Q8C2B3-3 | Isoform 3 c | Q8C2B3-3 | 1 | 2608.406 | 0 | 1.66 | 0.00031 | 16.11 | 46.9 | 153.1 | 13.08 | 3.15 | | High | High | High | High | High | High | High | High | High | High | High | 0 | 1.2E-06 | 4.12 |
| FALSE | High | TRSFDNLT | [K].TRSFDN | 1xCarbami | Q8K296 1x | 0.008388 | 0.000736 | 1 | 7 | 1 | Q8K296 | Myotubula | Q8K296 [6 | 1 | 2638.22 | 0 | -0.13 | 0.898547 | 13.57 | 97.5 | 102.5 | 11.58 | 16.6 | | High | High | High | High | High | High | High | High | High | High | High | 0.000327 | 0.002587 | 3.04 |
| FALSE | High | TRSPDVISS | [R].TRSPDV | 1xPhospho | Q3UJB9 1x | 0.000132 | 0 | 1 | 6 | 1 | Q3UJB9 | Enhancer c | Q3UJB9 [7 | 1 | 3210.609 | 0 | -0.87 | 0.00288 | 2.14 | 127.2 | 72.8 | 5.86 | 9.63 | | High | High | High | High | High | High | High | High | High | High | High | 0 | 2.45E-05 | 4.55 |
| FALSE | High | TRSPQQPV | [R].TRSPQ | 1xCarbami | Q3UFY0 1x | 3.33E-06 | 0 | 1 | 2 | 2 | Q3UFY0 | Ribosomal | Q3UFY0 [6 | 1 | 2024.04 | 0 | 0.17 | 0.162673 | 9.37 | 96.1 | 103.9 | 7.73 | 4.88 | | High | High | High | High | High | High | High | High | High | High | High | 0 | 3.92E-07 | 3.42 |
| FALSE | High | TRSPSPTLG | [R].TRSPSP | 2xPhospho | D3Z0X5 2x | 6.71E-05 | 0 | 1 | 7 | 2 | D3Z0X5 | Pleckstrin | D3Z0X5 [5 | 1 | 1957.935 | 0 | 0.74 | 0.003863 | 22.17 | 72 | 128 | 6.79 | 9.87 | | High | High | High | High | High | High | High | High | High | High | High | 7.19E-05 | 0.000918 | 3.16 |
| FALSE | High | TRSWESSS | [R].TRSWE | 1xPhospho | Q80YV2 1x | 0.012731 | 0.000889 | 1 | 3 | 1 | Q80YV2 | Nuclear-int | Q80YV2 [3 | 1 | 2868.373 | 0 | 0.08 | 0.249889 | 6.48 | 97.3 | 102.7 | 6.09 | 4.8 | | High | High | High | High | High | High | High | High | High | High | High | 0.000449 | 0.004128 | 2.13 |
| FALSE | High | TRSWESSS | [R].TRSWE | 2xPhospho | Q80YV2 2x | 0.006031 | 0.00062 | 1 | 3 | 1 | Q80YV2 | Nuclear-int | Q80YV2 [3 | 1 | 2948.339 | 0 | 0.22 | 0.039817 | 9.4 | 92.6 | 107.4 | 6.12 | 6.78 | | High | High | High | High | High | High | High | High | High | High | High | 0.000275 | 0.001777 | 2.48 |
| FALSE | High | TRSWESSS | [R].TRSWE | 3xPhospho | Q80YV2 3x | 0.004491 | 0.000409 | 1 | 3 | 1 | Q80YV2 | Nuclear-int | Q80YV2 [3 | 1 | 3028.305 | 0 | 0.14 | 0.035445 | 1.96 | 97.9 | 102.1 | 5.09 | 7.82 | | High | High | High | High | High | High | High | High | High | High | High | 0.000183 | 0.001277 | 2.84 |

| | | | | | | | | | | | | | | | | | | | | | | | | | | | | | | | | | | | | | | |
|---|---|---|---|---|---|---|---|---|---|---|---|---|---|---|---|---|---|---|---|---|---|---|---|---|---|---|---|---|---|---|---|---|---|---|---|---|---|---|
| FALSE | High | TRTEDSEQ | [R].TRTEDS | 1xPhospho | Q5BLK4 1x | 0.060159 | 0.004476 | 1 | 2 | 1 | Q5BLK4 | Terminal u | Q5BLK4 [1 | 1 | 3244.551 | 0 | 0.75 | 0.035095 | 19.85 | 74.5 | 125.5 | 20.29 | 31.15 | | High | High | High | High | High | High | High | High | High | High | High | 0.002279 | 0.02379 | 3.61 |
| FALSE | High | TRTLPGTP( | [R].TRTLPG | 1xPhospho | Q08274 1x | 3.94E-05 | 0 | 1 | 4 | 1 | Q08274 | Dystrophia | Q08274 [4 | 1 | 2120.07 | 0 | 0.07 | 0.83735 | 17.81 | 97.5 | 102.5 | 10.47 | 3.82 | | High | High | High | High | High | High | High | High | High | High | High | 0 | 6.32E-06 | 3.84 |
| FALSE | High | TRTPASINA | [R].TRTPAS | 1xPhospho | Q62383 1x | 0.00311 | 0.00016 | 1 | 2 | 1 | Q62383 | Transcripti | Q62383 [1 | 1 | 2519.318 | 0 | 0.03 | 0.49733 | 13.96 | 97.5 | 102.5 | 9.44 | 5.6 | | High | High | High | High | High | High | High | High | High | High | High | 7.19E-05 | 0.000849 | 2.99 |
| FALSE | High | TRTPASINA | [R].TRTPAS | 2xPhospho | Q62383 2x | 0.030632 | 0.001857 | 1 | 2 | 1 | Q62383 | Transcripti | Q62383 [1 | 1 | 2599.284 | 0 | 0.37 | 0.077709 | 20.49 | 87.7 | 112.3 | 7.27 | 17.08 | | High | High | High | High | High | High | High | High | High | High | High | 0.001011 | 0.01104 | 2.65 |
| FALSE | High | TRTSQEELI | [R].TRTSQE | 1xPhospho | Q8BJ05 1xI | 1.58E-05 | 0 | 1 | 3 | 1 | Q8BJ05 | Zinc finger | Q8BJ05 [38 | 1 | 2399.159 | 0 | 0.61 | 0.066915 | 33.81 | 90.8 | 109.2 | 12.1 | 21.08 | | High | High | High | High | High | High | High | High | High | High | High | 0 | 2.26E-06 | 2.88 |
| FALSE | High | TRTSQEELI | [R].TRTSQE | 1xPhospho | Q8BJ05 1xI | 0.030104 | 0.001857 | 1 | 3 | 3 | Q8BJ05 | Zinc finger | Q8BJ05 [38 | 1 | 2415.154 | 0 | 0.52 | 0.008816 | 10.37 | 82 | 118 | 10.11 | 12.22 | | High | High | High | High | High | High | High | High | High | High | High | 0.001011 | 0.01076 | 3.05 |
| FALSE | High | TSAAGFPL\ | [R].TSAAGI | 1xPhospho | D3Z5I1 1xP | 9.76E-07 | 0 | 1 | 5 | 1 | D3Z5I1 | Zinc finger | D3Z5I1 [34 | 0 | 1597.83 | 0 | -0.5 | 0.057854 | 3.1 | 116.3 | 83.7 | 8.9 | 15.83 | | High | High | High | High | High | High | High | High | High | High | High | 0 | 9.9E-08 | 4.17 |
| FALSE | High | TSAASPPV\ | [K].TSAASF | 1xPhospho | Q9DCL8 1x | 0.005044 | 0.000504 | 1 | 2 | 2 | Q9DCL8 | Protein ph | Q9DCL8 [1 | 0 | 3156.659 | 0 | 1.02 | 0.00024 | 7.76 | 64.3 | 135.7 | 7.72 | 5.38 | | High | High | High | High | High | High | High | High | High | High | High | 0.000246 | 0.001458 | 2.55 |
| FALSE | High | TSADVEIR | [R].TSADVI | 1xPhospho | A0A087WC | 0.007881 | 0.000736 | 1 | 5 | 2 | A0A087WC | Snf2-relate | A0A087WC | 0 | 1199.587 | 0 | 0.2 | 0.025812 | 7.65 | 94.2 | 105.8 | 5.08 | 5.46 | | High | High | High | High | High | High | High | High | High | High | High | 0.000327 | 0.002405 | 2.65 |
| FALSE | High | TSAEDSDA. | [K].TSAEDS | 1xPhospho | P70410 1xI | 5.15E-07 | 0 | 1 | 3 | 2 | P70410 | Nucleolar ( | P70410 [64 | 0 | 1730.825 | 0 | -0.29 | 0.013936 | 5.44 | 108.5 | 91.5 | 7.84 | 6.94 | | High | High | High | High | High | High | High | High | High | High | High | 0 | 4.84E-08 | 3.26 |
| FALSE | High | TSAETADV. | [R].TSAETA | 1xPhospho | Q9D2E2 1x | 2.83E-06 | 0 | 1 | 1 | 2 | Q9D2E2 | Target of E | Q9D2E2 [4 | 0 | 3359.632 | 0 | 0.54 | 0.143048 | 6.85 | 84.7 | 115.3 | 8.23 | 20.91 | | High | High | High | High | High | High | High | High | High | High | High | 0 | 3.27E-07 | 4.51 |
| FALSE | High | TQTYPTDW | [K].TQTYPT | 1xPhospho | Q3TDA7 1x | 0.001087 | 0 | 1 | 2 | 6 | Q3TDA7 | Protein kin | Q3TDA7 [3 | 0 | 4708.907 | 0 | 0.03 | 0.444105 | 11.05 | 103 | 97 | 9.55 | 18.11 | | High | High | High | High | High | High | High | High | High | High | High | 0.003303 | 0.03628 | 4.26 |
| FALSE | High | VGQGSQS( | [K].VGQGS | 1xPhospho | A3KGJ7 1xI | 1.12E-05 | 0 | 1 | 3 | 1 | A3KGJ7 | Centrosom | A3KGJ7 [22 | 0 | 2668.276 | 0 | -0.81 | 0.004243 | 5.43 | 124.1 | 75.9 | 5.67 | 12.79 | | High | High | High | High | High | High | High | High | High | High | High | 0 | 1.54E-06 | 3.84 |
| FALSE | High | VGSAGAAA | [R].VGSAG. | 1xPhospho | A2BE91 1xI | 9.91E-06 | 0 | 1 | 2 | 1 | A2BE91 | WD repeat | A2BE91 [13 | 0 | 1822.901 | 0 | 0.7 | 0.004141 | 2.18 | 80.4 | 119.6 | 7.99 | 11.12 | | High | High | High | High | High | High | High | High | High | High | High | 0 | 1.34E-06 | 3.26 |
| FALSE | High | VGSEHSLLI | [R].VGSEH | 1xPhospho | Q80TH2-2 | 0.003962 | 0.000311 | 1 | 6 | 2 | Q80TH2-2 | Isoform 2 c | Q80TH2-2 | 0 | 1873.983 | 0 | 1.01 | 0.040547 | 37.6 | 68.6 | 131.4 | 13.36 | 27.84 | | High | High | High | High | High | High | High | High | High | High | High | 0.00014 | 0.001116 | 3.13 |
| FALSE | High | WLDESDAE | [R].WLDES | 1xPhospho | Q8K2C9 1x | 1.75E-06 | 0 | 1 | 1 | 5 | Q8K2C9 | Very-long- | Q8K2C9 [1 | 0 | 1802.787 | 0 | 1.19 | 0.003247 | 5.79 | 61.6 | 138.4 | 2.54 | 12.66 | | High | High | High | High | High | High | High | High | High | High | High | 0 | 1.91E-07 | 3.53 |
| FALSE | High | WLDESDAE | [R].WLDES | 1xPhospho | Q8K2C9 1x | 7.18E-07 | 0 | 1 | 1 | 17 | Q8K2C9 | Very-long- | Q8K2C9 [1 | 0 | 1818.782 | 0 | 1.13 | 0.000877 | 7.96 | 61.7 | 138.3 | 3.3 | 8.84 | | High | High | High | High | High | High | High | High | High | High | High | 0 | 1.33E-07 | 4.44 |
| FALSE | High | WLDGASFI | [R].WLDGA | 1xPhospho | E9Q2H1 1x | 0.045122 | 0.003373 | 1 | 3 | 1 | E9Q2H1 | E3 ubiquiti | E9Q2H1 [3 | 0 | 1618.71 | 0 | 1.03 | 0.004527 | 15.75 | 64.7 | 135.3 | 12.07 | 13.28 | | High | High | High | High | High | High | High | High | High | High | High | 0.001727 | 0.01713 | 1.5 |
| FALSE | High | WLGPILGY | [K].WLGPII | 1xPhospho | A6H5Y3 1x | 0.001831 | 5.66E-05 | 1 | 2 | 2 | A6H5Y3 | Methionin | A6H5Y3 [1 | 0 | 1558.739 | 0 | 1.97 | 0.000373 | 32.48 | 44.8 | 155.2 | 35.04 | 10.48 | | High | High | High | High | High | High | High | High | High | High | High | 2.58E-05 | 0.000469 | 2.81 |
| FALSE | High | WLHSPLVK | [R].WLHSP | 1xPhospho | Q6PGG6 1x | 0.102567 | 0.008249 | 1 | 1 | 1 | Q6PGG6 | Guanine nu | Q6PGG6 [4 | 0 | 1517.865 | 0 | -0.33 | 0.011981 | 7.04 | 107.1 | 92.9 | 10.29 | 13.55 | | High | High | High | High | High | High | High | High | High | High | High | 0.004138 | 0.0443 | 2.46 |
| FALSE | High | WLLHSGPE | [K].WLLHS | 1xCarbami | Q3U829 1x | 0.020319 | 0.001472 | 1 | 5 | 1 | Q3U829 | AP-5 comp | Q3U829 [2 | 0 | 2577.173 | 0 | 0.94 | 0.01432 | 2.6 | 71.8 | 128.2 | 22.23 | 9.52 | | High | High | High | High | High | High | High | High | High | High | High | 0.000786 | 0.006911 | 3.08 |
| FALSE | High | WLNDAENI | [K].WLNDA | 1xPhospho | A0A0R4J1Z | 2.31E-11 | 0 | 1 | 19 | 6 | A0A0R4J1Z | POU doma | A0A0R4J1Z | 0 | 3539.649 | 0 | -0.09 | 0.285992 | 3.51 | 103.5 | 96.5 | 3.28 | 11.41 | | High | High | High | High | High | High | High | High | High | High | High | 0 | 1.71E-10 | 5.97 |
| FALSE | High | WLNDAENI | [K].WLNDA | 1xDeamida | A0A0R4J1Z | 8.94E-08 | 0 | 1 | 19 | 2 | A0A0R4J1Z | POU doma | A0A0R4J1Z | 0 | 3540.633 | 0 | 0 | 0.662525 | 17.25 | 101.7 | 98.3 | 11.69 | 15.86 | | High | High | High | High | High | High | High | High | High | High | High | 0 | 6.74E-09 | 4.95 |
| FALSE | High | WLNDAENI | [K].WLNDA | 2xPhospho | A0A0R4J1Z | 0.00025 | 0 | 1 | 19 | 5 | A0A0R4J1Z | POU doma | A0A0R4J1Z | 0 | 3619.615 | 0 | -0.58 | 0.027464 | 9.5 | 120.3 | 79.7 | 17.07 | 12.24 | | High | High | High | High | High | High | High | High | High | High | High | 0 | 6.92E-05 | 3.93 |
| FALSE | High | WIPLSSDS( | [K].WIPLSS | 1xPhospho | B2RUR8 1x | 2.02E-08 | 0 | 1 | 1 | 8 | B2RUR8 | OTU doma | B2RUR8 [4 | 0 | 3116.478 | 0 | 0.79 | 0.002784 | 9.68 | 73.2 | 126.8 | 19.99 | 7.08 | | High | High | High | High | High | High | High | High | High | High | High | 0 | 6.47E-09 | 5.01 |
| FALSE | High | WLPNSPG\ | [K].WLPNS | 1xPhospho | E9QLG3 1x | 0.000219 | 0 | 1 | 7 | 1 | E9QLG3 | MAX gene- | E9QLG3 [4 | 0 | 1606.876 | 0 | -0.57 | 0.003414 | 0.81 | 120.5 | 79.5 | 3.53 | 13.64 | | High | High | High | High | High | High | High | High | High | High | High | 0 | 4.34E-05 | 2.58 |
| FALSE | High | WLSASVNI | [R].WLSAS | 1xPhospho | Q6NVF4 1x | 1.62E-09 | 0 | 1 | 1 | 7 | Q6NVF4 | DNA helica | Q6NVF4 [9 | 0 | 2559.145 | 0 | -0.19 | 0.003058 | 2.56 | 104.7 | 95.3 | 2.86 | 3.83 | | High | High | High | High | High | High | High | High | High | High | High | 0 | 3.6E-10 | 5.1 |
| FALSE | High | WNPETVES | [K].WNPET | 1xCarbami | Q6P5H2-2 | 4.4E-06 | 0 | 1 | 2 | 1 | Q6P5H2-2 | Isoform 2 c | Q6P5H2-2 | 0 | 3366.467 | 0 | 2.1 | 0.005406 | 19.07 | 37.9 | 162.1 | 10.22 | 24.04 | | High | High | High | High | High | High | High | High | High | High | High | 0 | 5.38E-07 | 4.38 |
| FALSE | High | WLDDLLAS | [R].WLDDL | 1xDeamida | B9EKJ7 1xF | 5.65E-05 | 0 | 1 | 3 | 2 | B9EKJ7 | Tankyrase | B9EKJ7 [68 | 0 | 2106.011 | 0 | -0.53 | 0.228044 | 18.4 | 109.8 | 90.2 | 17.33 | 21.74 | | High | High | High | High | High | High | High | High | High | High | High | 0 | 9.45E-06 | 2.67 |
| FALSE | High | WNSNDFID | [K].WNSNI | 1xPhospho | O35954 1x | 0.107603 | 0.008634 | 1 | 1 | 1 | O35954 | Membrane | O35954 [3 | 0 | 3576.701 | 0 | 2.84 | 0.003856 | 9.79 | 26.5 | 173.5 | 13.25 | 23.59 | | High | High | High | High | High | High | High | High | High | High | High | 0.00437 | 0.04705 | 2.4 |
| FALSE | High | WNSPNGLI | [K].WNSPN | 1xPhospho | O35144 1x | 0.008094 | 0.000736 | 1 | 3 | 1 | O35144 | Telomeric | O35144 [4 | 0 | 1711.846 | 0 | -0.63 | 0.007702 | 6.86 | 121.5 | 78.5 | 5.15 | 15.91 | | High | High | High | High | High | High | High | High | High | High | High | 0.000327 | 0.002478 | 2.28 |
| FALSE | High | WPPGPTTS | [R].WPPGP | 1xPhospho | P51432 1xI | 0.001349 | 0 | 1 | 5 | 2 | P51432 | 1-phospha | P51432 [92 | 0 | 2326.091 | 0 | 0.7 | 0.00728 | 16.62 | 77.1 | 122.9 | 6.63 | 10.83 | | High | High | High | High | High | High | High | High | High | High | High | 2.58E-05 | 0.000433 | 1.85 |
| FALSE | High | WPQGVVP | [K].WPQG\ | 1xPhospho | B2RQC6 1x | 1.44E-07 | 0 | 1 | 7 | 6 | B2RQC6 | CAD protei | B2RQC6 [1 | 0 | 3249.651 | 0 | 0.49 | 0.014196 | 17.37 | 79.4 | 120.6 | 13.19 | 9.29 | | High | High | High | High | High | High | High | High | High | High | High | 0 | 1.15E-08 | 4.34 |
| FALSE | High | WPSEASGK | [R].WPSEA | 1xPhospho | Q8BI84 1xF | 0.018285 | 0.001282 | 1 | 4 | 1 | Q8BI84 | Melanoma | Q8BI84 [17 | 0 | 1399.702 | 0 | -0.41 | 0.087309 | 6.68 | 118 | 82 | 16.78 | 15.63 | | High | High | High | High | High | High | High | High | High | High | High | 0.000689 | 0.006175 | 1.02 |
| FALSE | High | WPSEEEK | [K].WPSEEI | 1xPhospho | Q8K387 1x | 0.025964 | 0.001645 | 1 | 2 | 3 | Q8K387 | Ubiquitin c | Q8K387 [4 | 0 | 1442.697 | 0 | 0.4 | 0.008185 | 12.17 | 85.8 | 114.2 | 5.89 | 5.63 | | High | High | High | High | High | High | High | High | High | High | High | 0.000872 | 0.009166 | 1.56 |
| FALSE | High | WPTGDNIH | [R].WPTGD | 1xPhospho | Q3UR00 1x | 0.000183 | 0 | 1 | 11 | 2 | Q3UR00 | Putative ur | Q3UR00 [1 | 0 | 1968.928 | 0 | -1.9 | 0.00436 | 2.88 | 159.1 | 40.9 | 18.94 | 25.89 | | High | High | High | High | High | High | High | High | High | High | High | 0 | 3.53E-05 | 2.25 |
| FALSE | High | WQDPSAD | [R].WQDPS | 1xPhospho | A0A1L1STC | 0.005 | 0.000504 | 1 | 7 | 1 | A0A1L1STC | Nesprin-1 ( | A0A1L1STC | 0 | 3198.513 | 0 | 0.88 | 0.01072 | 18.67 | 68 | 132 | 15.99 | 16.82 | | High | High | High | High | High | High | High | High | High | High | High | 0.000225 | 0.001439 | 2.01 |
| FALSE | High | WQELSPET | [K].WQELS | 1xPhospho | P58283-2 1 | 0.000446 | 0 | 1 | 3 | 1 | P58283-2 | Isoform 2 c | P58283-2 [ | 0 | 1799.898 | 0 | 0.4 | 0.006671 | 12.06 | 86.6 | 113.4 | 4.61 | 5.92 | | High | High | High | High | High | High | High | High | High | High | High | 0 | 9.6E-05 | 2.43 |
| FALSE | High | WQHDLFD: | [K].WQHDI | 1xPhospho | O08583 1x | 4.27E-08 | 0 | 1 | 1 | 1 | O08583 | THO comp | O08583 [8 | 0 | 2660.255 | 0 | -1.16 | 0.006642 | 7.74 | 135.3 | 64.7 | 8.28 | 24.22 | | High | High | High | High | High | High | High | High | High | High | High | 0 | 2.93E-09 | 4.31 |
| FALSE | High | WQRPASPI | [R].WQRPA | 1xDeamida | A2AI08 1xF | 0.039686 | 0.00283 | 1 | 1 | 1 | A2AI08 | Taperin OS | A2AI08 [45 | 0 | 3593.881 | 0 | 1.35 | 0.01278 | 23.3 | 65.9 | 134.1 | 14.45 | 24.88 | | High | High | High | High | High | High | High | High | High | High | High | 0.001468 | 0.01485 | 3.37 |
| FALSE | High | WRLSPGET | [K].WRLSP | 1xPhospho | Q8BQ30-2 | 0.070485 | 0.005214 | 1 | 4 | 1 | Q8BQ30-2 | Isoform 2 c | Q8BQ30-2 | 1 | 1965.963 | 0 | 0.77 | 0.018239 | 13.53 | 76 | 124 | 9.62 | 12.3 | | High | High | High | High | High | High | High | High | High | High | High | 0.00269 | 0.02878 | 2.59 |
| FALSE | High | WRSTAEID. | [R].WRSTA | 1xPhospho | Q0PHV7 1x | 0.005717 | 0.00062 | 1 | 1 | 1 | Q0PHV7 | Dapper ho | Q0PHV7 [4 | 1 | 1782.837 | 0 | -1.27 | 0.000647 | 5.01 | 138.7 | 61.3 | 4.15 | 25.93 | | High | High | High | High | High | High | High | High | High | High | High | 0.000275 | 0.001671 | 2.64 |
| FALSE | High | WSDQLSLD | [R].WSDQI | 1xPhospho | G3UYW1 1 | 0.001349 | 0 | 1 | 5 | 1 | G3UYW1 | WD repeat | G3UYW1 [ | 0 | 1758.872 | 0 | 0.9 | 0.002621 | 14.51 | 68.2 | 131.8 | 6.4 | 10.13 | | High | High | High | High | High | High | High | High | High | High | High | 0 | 0.000335 | 2.44 |
| FALSE | High | WNSPAEE( | [R].WNSPA | 1xCarbami | Q571M2 1: | 5.13E-06 | 0 | 1 | 4 | 2 | Q571M2 | MKIAA402 | Q571M2 [4 | 0 | 2503.162 | 0 | -0.33 | 0.03022 | 9.77 | 112.2 | 87.8 | 10.58 | 11.92 | | High | High | High | High | High | High | High | High | High | High | High | 0 | 6.39E-07 | 2.63 |
| FALSE | High | WSDSSK | [K].WSDSS | 1xPhospho | Q14AX6-2 | 0.093123 | 0.007276 | 3 | 5 | 1 | Q14AX6-2; | Isoform 2 c | Q14AX6-2 | 0 | 1247.607 | 0 | -0.15 | 0.505963 | 29.04 | 108.4 | 91.6 | 5.8 | 24.8 | Shared | High | High | High | High | High | High | High | High | High | High | High | 0.003698 | 0.03981 | 1.19 |
| FALSE | High | WLDDLLAS | [R].WLDDL | 1xPhospho | B9EKJ7 1xF | 3.3E-06 | 0 | 1 | 3 | 3 | B9EKJ7 | Tankyrase | B9EKJ7 [68 | 0 | 2105.027 | 0 | -0.41 | 0.157743 | 16.62 | 106.7 | 93.3 | 12.88 | 25.08 | | High | High | High | High | High | High | High | High | High | High | High | 0 | 0.000277 | 2.77 |
| FALSE | High | WKESPENE | [R].WKESP | 1xPhospho | Q05AH6 1x | 0.047488 | 0.00354 | 1 | 2 | 1 | Q05AH6 | Uncharacte | Q05AH6 [2 | 1 | 1880.931 | 0 | -0.75 | 0.000751 | 2.24 | 124.2 | 75.8 | 7.93 | 8.02 | | High | High | High | High | High | High | High | High | High | High | High | 0.001812 | 0.01824 | 0.76 |
| FALSE | High | WDETPAS( | [R].WDETP | 1xPhospho | G5E866 1x | 1.97E-05 | 0 | 1 | 5 | 1 | G5E866 | Splicing fac | G5E866 [3 | 0 | 2597.272 | 0 | 0.51 | 0.208994 | 1.12 | 82.5 | 117.5 | 13.12 | 27.74 | | High | High | High | High | High | High | High | High | High | High | High | 0 | 2.89E-06 | 3.1 |
| FALSE | High | WDETPAS( | [R].WDETP | 1xPhospho | G5E866 1x | 2.9E-09 | 0 | 1 | 5 | 7 | G5E866 | Splicing fac | G5E866 [3 | 0 | 2613.267 | 0 | 0.03 | 0.26863 | 3.26 | 99.2 | 100.8 | 8.15 | 8.31 | | High | High | High | High | High | High | High | High | High | High | High | 0 | 1.44E-10 | 4.5 |
| FALSE | High | WDETPAS( | [R].WDETP | 2xPhospho | G5E866 2x | 5.97E-05 | 0 | 1 | 5 | 1 | G5E866 | Splicing fac | G5E866 [3 | 0 | 2677.239 | 0 | 0.26 | 0.6847 | 7.65 | 92.7 | 107.3 | 11.44 | 24.08 | | High | High | High | High | High | High | High | High | High | High | High | 0 | 1.01E-05 | 2.62 |
| FALSE | High | WDETPGR | [R].WDETP | 1xPhospho | G5E866 1x | 0.085859 | 0.006406 | 1 | 5 | 2 | G5E866 | Splicing fac | G5E866 [2 | 0 | 1169.519 | 0 | 0.34 | 0.368515 | 11.66 | 88.2 | 111.8 | 10.69 | 21.79 | | High | High | High | High | High | High | High | High | High | High | High | 0.007304 | 0.07936 | 1.7 |
| FALSE | High | WDQTADC | [R].WDQTA | 1xPhospho | G5E866 1x | 6.41E-08 | 0 | 1 | 5 | 3 | G5E866 | Splicing fac | G5E866 [2 | 0 | 2054 | 0 | 0.04 | 0.562149 | 6.26 | 99.7 | 100.3 | 12.37 | 15.68 | | High | High | High | High | High | High | High | High | High | High | High | 0 | 1.03E-05 | 3.13 |
| FALSE | High | WDQTADC | [R].WDQTA | 2xPhospho | G5E866 2x | 2.35E-08 | 0 | 1 | 5 | 3 | G5E866 | Splicing fac | G5E866 [2 | 0 | 2133.966 | 0 | 0.42 | 0.489182 | 8.01 | 85.5 | 114.5 | 20.14 | 21.46 | | High | High | High | High | High | High | High | High | High | High | High | 0 | 3E-08 | 3.12 |
| FALSE | High | WDSDNDE | [K].WDSDN | 1xPhospho | F8WGX5 1: | 0.000415 | 0 | 1 | 2 | 2 | F8WGX5 | DmX-like p | F8WGX5 [1 | 0 | 2559.1 | 0 | 0.02 | 0.509238 | 5.12 | 99.1 | 100.9 | 1.13 | 4.4 | | High | High | High | High | High | High | High | High | High | High | High | 0 | 8.84E-05 | 3.06 |
| FALSE | High | WDSDNDE | [K].WDSDN | 1xPhospho | F8WGX5 1: | 6.93E-07 | 0 | 1 | 2 | 2 | F8WGX5 | DmX-like p | F8WGX5 [1 | 0 | 2575.095 | 0 | 0.14 | 0.149798 | 21.34 | 94.8 | 105.2 | 6.38 | 7.07 | | High | High | High | High | High | High | High | High | High | High | High | 0 | 6.74E-08 | 3.92 |
| FALSE | High | WDSYDNF! | [R].WDSYD | 1xPhospho | E9Q0S6 1x | 9.28E-05 | 0 | 1 | 6 | 2 | E9Q0S6 | Tensin 1 O | E9Q0S6 [5 | 0 | 1692.701 | 0 | -0.19 | 0.026635 | 7.64 | 106.2 | 93.8 | 4.51 | 5.75 | | High | High | High | High | High | High | High | High | High | High | High | 0 | 0.000249 | 3.27 |
| FALSE | High | WDSYENFI | [R].WDSYE | 1xPhospho | Q8CGB6-3 | 7.43E-06 | 0 | 1 | 4 | 2 | Q8CGB6-3 | Isoform 3 c | Q8CGB6-3 | 0 | 3037.306 | 0 | -0.96 | 0.000222 | 1.65 | 132.3 | 67.7 | 3.95 | 7.32 | | High | High | High | High | High | High | High | High | High | High | High | 0 | 9.67E-07 | 3.45 |
| FALSE | High | WDSYENM | [R].WDSYE | 1xPhospho | Q5SSZ5 1xI | 0.000147 | 0 | 1 | 1 | 3 | Q5SSZ5 | Tensin-3 O | Q5SSZ5 [3 | 0 | 3507.618 | 0 | -1.01 | 0.002308 | 6.05 | 133.7 | 66.3 | 7.74 | 10.17 | | High | High | High | High | High | High | High | High | High | High | High | 0 | 2.77E-05 | 4.85 |
| FALSE | High | WDSYENM | [R].WDSYE | 1xDeamida | Q5SSZ5 1xI | 0.000279 | 0 | 1 | 1 | 2 | Q5SSZ5 | Tensin-3 O | Q5SSZ5 [3 | 0 | 3508.602 | 0 | -1.83 | 0.004608 | 3.15 | 158.4 | 41.6 | 4.1 | 48.63 | | High | High | High | High | High | High | High | High | High | High | High | 0.00395 | 0.04213 | 3.96 |
| FALSE | High | WDSYENM | [R].WDSYE | 1xPhospho | Q5SSZ5 1xI | 3.23E-08 | 0 | 1 | 1 | 10 | Q5SSZ5 | Tensin-3 O | Q5SSZ5 [3 | 0 | 3523.613 | 0 | -1.02 | 0.00417 | 6.06 | 133.9 | 66.1 | 7.99 | 16.67 | | High | High | High | High | High | High | High | High | High | High | High | 0 | 2.16E-09 | 4.46 |
| FALSE | High | WLAESPVG | [R].WLAES | 1xPhospho | A2ARP8 1x | 0.035181 | 0.002269 | 1 | 3 | 1 | A2ARP8 | Microtubu | A2ARP8 [1 | 0 | 2334.167 | 0 | -1.39 | 0.022466 | 8.19 | 148.1 | 51.9 | 22.91 | 30.47 | | High | High | High | High | High | High | High | High | High | High | High | 0.001199 | 0.01285 | 1.61 |
| FALSE | High | WEAHIYAS | [R].WEAHI | 1xPhospho | Q9D279 1x | 0.000751 | 0 | 1 | 1 | 3 | Q9D279 | Mitotic int | Q9D279 [6 | 0 | 1772.737 | 0 | 3.43 | 0.00024 | 9.86 | 17.6 | 182.4 | 15.11 | 8.3 | | High | High | High | High | High | High | High | High | High | High | High | 0 | 0.000173 | 2.46 |
| FALSE | High | WEGGDSN | [R].WEGGI | 1xPhospho | A0A0U1RP | 2.42E-05 | 0 | 1 | 8 | 7 | A0A0U1RP | Ataxin-2-lik | A0A0U1RP | 0 | 3778.524 | 0 | 0.02 | 0.6062 | 16.08 | 100.1 | 99.9 | 13.37 | 41.7 | | High | High | High | High | High | High | High | High | High | High | High | 0 | 3.66E-06 | 4.67 |
| FALSE | High | WEGGDSN | [R].WEGGI | 1xPhospho | A0A0U1RP | 2.66E-06 | 0 | 1 | 8 | 26 | A0A0U1RP | Ataxin-2-lik | A0A0U1RP | 0 | 3794.519 | 0 | -0.29 | 0.271421 | 11.02 | 107.7 | 92.3 | 17.69 | 9.62 | | High | High | High | High | High | High | High | High | High | High | High | 0 | 0.000189 | 5.44 |
| FALSE | High | WELFDQHI | [K].WELFD | 2xPhospho | Q6NV83 2x | 0.000426 | 0 | 1 | 3 | 2 | Q6NV83 | U2 snRNP- | Q6NV83 [7 | 0 | 4560.793 | 0 | -0.24 | 0.153471 | 15 | 110.7 | 89.3 | 10.36 | 22.88 | | High | High | High | High | High | High | High | High | High | High | High | 0 | 9.14E-05 | 4.88 |
| FALSE | High | WEPALGEF | [R].WEPAL | 1xPhospho | Q9EP53 1x | 1.31E-05 | 0 | 1 | 4 | 1 | Q9EP53 | Hamartin ( | Q9EP53 [1 | 0 | 2865.469 | 0 | -0.44 | 0.123549 | 17.52 | 116.5 | 83.5 | 16.13 | 11.47 | | High | High | High | High | High | High | High | High | High | High | High | 0 | 1.83E-06 | 4.62 |
| FALSE | High | WESQDSGI | [K].WESQD | 1xPhospho | D3Z357 1xI | 0.004411 | 0.000409 | 1 | 4 | 1 | D3Z357 | Histone-lys | D3Z357 [1 | 1 | 1960.894 | 0 | 1.99 | 0.003461 | 3.94 | 41.9 | 158.1 | 5.2 | 18.98 | | High | High | High | High | High | High | High | High | High | High | High | 0.000183 | 0.001255 | 3.79 |
| FALSE | High | WESQQDV | [R].WESQC | 1xPhospho | E9Q4K7 1x | 1.08E-05 | 0 | 1 | 2 | 2 | E9Q4K7 | Kinesin fan | E9Q4K7 [1 | 0 | 1971.937 | 0 | 0.35 | 0.008588 | 7.87 | 84.8 | 115.2 | 8.19 | 5.07 | | High | High | High | High | High | High | High | High | High | High | High | 0 | 1.6E-06 | 4.03 |
| FALSE | High | WESSASFD | [R].WESSA | 1xPhospho | Q6P9R4 1x | 0.0003 | 0 | 1 | 1 | 1 | Q6P9R4 | Rho guanir | Q6P9R4 [9 | 0 | 1707.84 | 0 | 2.45 | 0.000656 | 11.79 | 29.7 | 170.3 | 17.83 | 10 | | High | High | High | High | High | High | High | High | High | High | High | 0 | 6.16E-05 | 1.94 |
| FALSE | High | WFSNPSR | [R].WFSNP | 2xPhospho | Q9EST3 2x | 0.104222 | 0.008348 | 1 | 6 | 1 | Q9EST3 | Eukaryotic | Q9EST3 [3 | 0 | 1282.522 | 0 | 2.04 | 0.000845 | 12.54 | 39 | 161 | 9.88 | 10.69 | | High | High | High | High | High | High | High | High | High | High | High | 0.00425 | 0.04545 | 0.83 |
| FALSE | High | WGQLGAS | [R].WGQL( | 1xPhospho | Q9EP97 1x | 0.052576 | 0.00384 | 1 | 1 | 1 | Q9EP97 | Sentrin-spe | Q9EP97 [9 | 0 | 1311.641 | 0 | -1.45 | 0.003537 | 4.29 | 146 | 54 | 14.45 | 15.04 | | High | High | High | High | High | High | High | High | High | High | High | 0.001999 | 0.02044 | 0.79 |
| FALSE | High | WGSQGNR | [R].WGSQ( | 1xPhospho | A0A0R4IZX | 0.045122 | 0.003373 | 1 | 3 | 2 | A0A0R4IZX | Differentia | A0A0R4IZX | 0 | 1113.504 | 0 | 1.56 | 0.000324 | 10.52 | 48.6 | 151.4 | 10.41 | 7.81 | | High | High | High | High | High | High | High | High | High | High | High | 0.001712 | 0.01712 | 0.73 |
| FALSE | High | WHQPPPSI | [R].WHQPI | 1xPhospho | Q6PDM1 1 | 0.000311 | 0 | 1 | 7 | 1 | Q6PDM1 | Male-speci | Q6PDM1 [4 | 0 | 1733.909 | 0 | 0.24 | 0.049795 | 2.99 | 92.1 | 107.9 | 2.56 | 9.39 | | High | High | High | High | High | High | High | High | High | High | High | 0 | 6.4E-05 | 2.43 |
| FALSE | High | WHSQNQS | [R].WHSQN | 1xCarbami | Q9CS00 1x | 0.00824 | 0.000736 | 1 | 1 | 1 | Q9CS00 | Cactin OS= | Q9CS00 [1 | 0 | 2202.871 | 0 | 0.14 | 0.87045 | 12.66 | 94.7 | 105.3 | 13.62 | 8.91 | | High | High | High | High | High | High | High | High | High | High | High | 0.000327 | 0.002522 | 3.16 |
| FALSE | High | WHSQNQS | [R].WHSQN | 1xCarbami | Q9CS00 2x | 0.00642 | 0.00062 | 1 | 1 | 1 | Q9CS00 | Cactin OS= | Q9CS00 [1 | 0 | 2282.837 | 0 | 0.27 | 0.071115 | 14.87 | 91.8 | 108.2 | 4.08 | 17.47 | | High | High | High | High | High | High | High | High | High | High | High | 0.000275 | 0.001908 | 3.94 |
| FALSE | High | WEELSTSG | [R].WEELS | 1xPhospho | G5E8N7 1x | 0.0003 | 0 | 1 | 8 | 7 | G5E8N7 | Bifunction | G5E8N7 [1 | 0 | 3863.706 | 0 | 0.65 | 0.002085 | 5.87 | 77.2 | 122.8 | 2.36 | 8.61 | | High | High | High | High | High | High | High | High | High | High | High | 0.000246 | 0.001488 | 3.36 |
| FALSE | High | WSDYVER | [R].WSDYV | 1xPhospho | Q60520-1 | 0.052576 | 0.00384 | 1 | 3 | 1 | Q60520-1 | Isoform 2 c | Q60520-1 | 0 | 1263.561 | 0 | -0.85 | 0.000826 | 3.38 | 126.2 | 73.8 | 5.85 | 6.19 | | High | High | High | High | High | High | High | High | High | High | High | 0.001999 | 0.02049 | 2.17 |
| FALSE | High | WSGNESTI | [R].WSGNE | 1xPhospho | Q921G6 1x | 0.000294 | 0 | 1 | 4 | 3 | Q921G6 | Leucine-ric | Q921G6 [3 | 0 | 2185.928 | 0 | -0.07 | 0.850499 | 22.92 | 98.8 | 101.2 | 19.62 | 15.87 | | High | High | High | High | High | High | High | High | High | High | High | 0 | 0.000165 | 2.67 |
| FALSE | High | WSGSGSG! | [R].WSGSG | 1xPhospho | Q5Y5T2 1x | 0.020499 | 0.001472 | 1 | 1 | 1 | Q5Y5T2 | Palmitoyltr | Q5Y5T2 [4 | 0 | 1503.679 | 0 | -0.08 | 0.688602 | 8.67 | 103.5 | 96.5 | 7.49 | 6.06 | | High | High | High | High | High | High | High | High | High | High | High | 0.000786 | 0.007039 | 1 |
| FALSE | High | YCRPESQEI | [K].YCRPES | 1xCarbami | P42227-3 1 | 0.000765 | 0 | 1 | 1 | 1 | P42227-3 | Isoform De | P42227-3 [ | 0 | 2953.396 | 0 | 0.71 | 0.012124 | 6.65 | 75.9 | 124.1 | 9.65 | 11.08 | | High | High | High | High | High | High | High | High | High | High | High | 0 | 0.000177 | 1.93 |
| FALSE | High | YCRPESQEI | [K].YCRPES | 1xCarbami | P42227 1xI | 0.000738 | 0 | 1 | 7 | 1 | P42227 | Signal tran | P42227 [68 | 0 | 3040.428 | 0 | 0.99 | 0.002786 | 10.86 | 67.3 | 132.7 | 9.82 | 8.79 | | High | High | High | High | High | High | High | High | High | High | High | 0 | 0.00017 | 2.37 |
| FALSE | High | YCRPESQEI | [K].YCRPES | 1xCarbami | P42227 1xI | 0.003721 | 0.00021 | 1 | 7 | 1 | P42227 | Signal tran | P42227 [68 | 0 | 3041.412 | 0 | 1.19 | 0.009742 | 39.8 | 66.8 | 133.2 | 16.42 | 17.69 | | High | High | High | High | High | High | High | High | High | High | High | 9.42E-05 | 0.001037 | 3.38 |
| FALSE | High | YCSADECL( | [R].YCSADE | 2xCarbami | A2CG63 1x | 1.41E-06 | 0 | 1 | 4 | 2 | A2CG63 | AT-rich inte | A2CG63 [8 | 0 | 2210.986 | 0 | -0.29 | 0.056489 | 2.14 | 109.5 | 90.5 | 5.37 | 10.86 | | High | High | High | High | High | High | High | High | High | High | High | 0 | 1.5E-07 | 3.2 |
| FALSE | High | YCSADECL( | [R].YCSADE | 2xCarbami | A2CG63 2x | 4.37E-07 | 0 | 1 | 4 | 2 | A2CG63 | AT-rich inte | A2CG63 [8 | 0 | 2290.953 | 0 | -0.35 | 0.004809 | 2.61 | 113.7 | 86.3 | 7.2 | 5.68 | | High | High | High | High | High | High | High | High | High | High | High | 0 | 4.01E-08 | 3.66 |
| FALSE | High | YDCFLHPFI | [K].YDCFLH | 1xCarbami | Q571L5 1xI | 0.032267 | 0.002129 | 1 | 8 | 1 | Q571L5 | MKIAA406 | Q571L5 [3 | 0 | 2549.209 | 0 | -0.73 | 0.018066 | 11.76 | 129.1 | 70.9 | 16.47 | 14.98 | | High | High | High | High | High | High | High | High | High | High | High | 0.001142 | 0.0117 | 2.61 |
| FALSE | High | YDDSDGIP\ | [K].YDDSD | 1xPhospho | Q62315 1x | 0.015465 | 0.001089 | 1 | 4 | 1 | Q62315 | Protein Jur | Q62315 [1 | 0 | 1877.779 | 0 | -1.2 | 0.002517 | 0.9 | 141.6 | 58.4 | 11.41 | 13.81 | | High | High | High | High | High | High | High | High | High | High | High | 0.000571 | 0.005094 | 1.58 |
| FALSE | High | YDDYSSSR | [R].YDDYS | 1xPhospho | Q91VM5 1 | 0.093879 | 0.007306 | 1 | 3 | 2 | Q91VM5 | RNA bindir | Q91VM5 [ | 0 | 1301.525 | 0 | 0.93 | 0.001086 | 6.46 | 65.9 | 134.1 | 6.39 | 9.09 | | High | High | High | High | High | High | High | High | High | High | High | 0.003698 | 0.03995 | 0.85 |
| FALSE | High | YDEGSGGS | [R].YDEGS | 1xPhospho | Q8VID5 1xI | 0.01218 | 0.000889 | 1 | 1 | 2 | Q8VID5 | ATP-depen | Q8VID5 [4 | 1 | 2404.045 | 0 | 0.05 | 0.534317 | 9.92 | 97.5 | 102.5 | 9.18 | 22.6 | | High | High | High | High | High | High | High | High | High | High | High | 0.000449 | 0.003908 | 1.29 |
| FALSE | High | YDENEDLSI | [R].YDENEI | 1xPhospho | Q6P1G2 1x | 2.85E-05 | 0 | 1 | 5 | 3 | Q6P1G2 | Lysine-spe | Q6P1G2 [1 | 0 | 2320.043 | 0 | -0.51 | 0.002153 | 2.26 | 115.4 | 84.6 | 3.27 | 6.52 | | High | High | High | High | High | High | High | High | High | High | High | 0 | 4.4E-06 | 3.22 |
| FALSE | High | YDERPGPSI | [R].YDERP( | 1xPhospho | A2RS68 1xI | 0.094641 | 0.007408 | 1 | 2 | 3 | A2RS68 | Small nucle | A2RS68 [2 | 0 | 1829.89 | 0 | -0.34 | 0.007768 | 6.02 | 110.6 | 89.4 | 4.24 | 7.56 | | High | High | High | High | High | High | High | High | High | High | High | 0.003741 | 0.04029 | 2.08 |
| FALSE | High | YDFEPYVSI | [R].YDFEPY | 1xPhospho | D3Z7R7 1xI | 0.000164 | 0 | 1 | 6 | 1 | D3Z7R7 | Rho GTPas | D3Z7R7 [5 | 0 | 2430.067 | 0 | 0.85 | 0.000826 | 19.79 | 68.5 | 131.5 | 13.76 | 3.91 | | High | High | High | High | High | High | High | High | High | High | High | 0 | 3.11E-05 | 2.79 |
| FALSE | High | YDFWDQQ | [K].YDFWD | 1xPhospho | E9Q4K7 1x | 0.000108 | 0 | 1 | 1 | 5 | E9Q4K7 | Kinesin fan | E9Q4K7 [8 | 0 | 3351.571 | 0 | 0.21 | 0.017369 | 6.02 | 91.3 | 108.7 | 8.75 | 4.44 | | High | High | High | High | High | High | High | High | High | High | High | 0 | 1.97E-05 | 4.11 |
| FALSE | High | YCNSSVPD | [R].YCNSS\ | 2xCarbami | Q8BLU0 1x | 0.000538 | 0 | 1 | 1 | 2 | Q8BLU0 | Leucine-ric | Q8BLU0 [6 | 0 | 2027.819 | 0 | 2.95 | 0.003229 | 25.24 | 22.2 | 177.8 | 11.52 | 22.28 | | High | High | High | High | High | High | High | High | High | High | High | 0.000723 | 0.00639 | 1.53 |
| FALSE | High | YDLILDEQ( | [K].YDLILDI | 1xPhospho | Q9CXG3 1x | 4.94E-05 | 0 | 1 | 4 | 1 | Q9CXG3 | Peptidyl-pr | Q9CXG3 [3 | 0 | 2062.999 | 0 | 1.65 | 0.002241 | 30.46 | 43.1 | 156.9 | 20 | 14.36 | | High | High | High | High | High | High | High | High | High | High | High | 0 | 8.11E-06 | 2.75 |
| FALSE | High | YDIPATAAT | [K].YDIPAT | 1xPhospho | Q61191 1x | 1.17E-05 | 0 | 1 | 2 | 1 | Q61191 | Host cell fa | Q61191 [4 | 0 | 3212.664 | 0 | -0.41 | 0.006784 | 2.62 | 113.6 | 86.4 | 6.02 | 5 | | High | High | High | High | High | High | High | High | High | High | High | 0 | 1.6E-06 | 4.12 |
| FALSE | High | YDLSSPIQT | [K].YDLSSP | 1xPhospho | Q8BM55 1: | 0.000115 | 0 | 1 | 2 | 1 | Q8BM55 | Transmem | Q8BM55 [4 | 0 | 2183.047 | 0 | 0.51 | 0.028738 | 17.67 | 87.2 | 112.8 | 8.03 | 13.16 | | High | High | High | High | High | High | High | High | High | High | High | 0 | 2.1E-05 | 2.87 |
| FALSE | High | YDSDGDKS | [R].YDSDGI | 2xPhospho | Q80TC1 2x | 0.000176 | 0 | 1 | 14 | 1 | Q80TC1 | MKIAA154 | Q80TC1 [3 | 1 | 2998.244 | 0 | -0.39 | 0.10581 | 19.57 | 117.2 | 82.8 | 21.04 | 17.53 | | High | High | High | High | High | High | High | High | High | High | High | 0 | 3.39E-05 | 4.09 |
| FALSE | High | YDSLLGR | [R].YDSLLG | 1xPhospho | Q9DBR7 1x | 0.012619 | 0.000889 | 1 | 3 | 1 | Q9DBR7 | Protein ph | Q9DBR7 [9 | 0 | 1132.56 | 0 | 1.36 | 0.004529 | 6.52 | 59.6 | 140.4 | 11.67 | 17.4 | | High | High | High | High | High | High | High | High | High | High | High | 0.000449 | 0.004083 | 1.8 |
| FALSE | High | YDSQLSSNI | [K].YDSQLS | 1xPhospho | Q8C145 1x | 0.000325 | 0 | 1 | 2 | 2 | Q8C145 | Zinc transp | Q8C145 [4 | 0 | 1837.862 | 0 | 1.95 | 0.001007 | 5.62 | 41.1 | 158.9 | 4.34 | 13.6 | | High | High | High | High | High | High | High | High | High | High | High | 0 | 0.000178 | 2.54 |
| FALSE | High | YDSSNQVK | [R].YDSSN( | 1xPhospho | D3YTR4 1x | 0.045508 | 0.003373 | 2 | 4 | 2 | D3YTR4; Q | Methyl-Cp | D3YTR4 [5 | 0 | 1249.566 | 0 | -0.98 | 0.00134 | 2.37 | 131.6 | 68.4 | 5.16 | 9.27 | Shared | High | High | High | High | High | High | High | High | High | High | High | 0.001741 | 0.01736 | 1.68 |
| FALSE | High | YDSSNQVK | [R].YDSSN( | 1xPhospho | D3YTR4 1x | 0.006895 | 0.00062 | 2 | 4 | 2 | D3YTR4; Q | Methyl-Cp | D3YTR4 [5 | 0 | 1478.729 | 0 | -0.72 | 0.001897 | 2.61 | 122.7 | 77.3 | 7.65 | 4.82 | Shared | High | High | High | High | High | High | High | High | High | High | High | 0.000585 | 0.005266 | 2.93 |
| FALSE | High | YDSVNEK | [R].YDSVNI | 1xPhospho | A0A0A6YY | 0.083101 | 0.006345 | 1 | 12 | 1 | A0A0A6YY | Protocadh | A0A0A6YY | 0 | 1392.681 | 0 | -0.86 | 0.050373 | 14.03 | 125.3 | 74.7 | 16.71 | 28.45 | | High | High | High | High | High | High | High | High | High | High | High | 0.003265 | 0.03484 | 2.13 |
| FALSE | High | YDTPTSK | [R].YDTPTS | 1xPhospho | E9Q5F9 1x | 0.023379 | 0.001578 | 1 | 3 | 1 | E9Q5F9 | Histone-lys | E9Q5F9 [2 | 0 | 1349.675 | 0 | -0.07 | 0.055698 | 3.91 | 104.1 | 95.9 | 4.58 | 5.04 | | High | High | High | High | High | High | High | High | High | High | High | 0.000847 | 0.008099 | 1.82 |
| FALSE | High | YDYGNQY( | [K].YDYGN | 1xPhospho | B0V2W5 1: | 0.000108 | 0 | 1 | 4 | 2 | B0V2W5 | Zinc finger | B0V2W5 [7 | 0 | 3132.41 | 0 | 0.01 | 0.846243 | 2.18 | 99.8 | 100.2 | 4.04 | 5.31 | | High | High | High | High | High | High | High | High | High | High | High | 0 | 1.97E-05 | 3.88 |
| FALSE | High | YEDDGISDI | [R].YEDDG | 1xPhospho | F6YRW4 1x | 0.030104 | 0.001857 | 1 | 6 | 2 | F6YRW4 | F-box and l | F6YRW4 [2 | 0 | 2122.947 | 0 | 0.62 | 0.001112 | 3.2 | 77.4 | 122.6 | 3.86 | 7.93 | | High | High | High | High | High | High | High | High | High | High | High | 0.001011 | 0.01086 | 2.71 |
| FALSE | High | YEDLSSSDE | [K].YEDLSS | 2xPhospho | Q9CZK6 2x | 0.100936 | 0.008097 | 1 | 3 | 1 | Q9CZK6 | Ankyrin rep | Q9CZK6 [2 | 0 | 2283.941 | 0 | -0.08 | 0.871098 | 28.78 | 99.8 | 100.2 | 9.35 | 24.79 | | High | High | High | High | High | High | High | High | High | High | High | 0.004086 | 0.0437 | 2.05 |
| FALSE | High | YEEGESEAI | [R].YEEGES | 1xPhospho | F8VQD1 1x | 9.59E-07 | 0 | 1 | 13 | 1 | F8VQD1 | Protein pol | F8VQD1 [3 | 0 | 3689.619 | 0 | -0.05 | 0.925633 | 7.68 | 99 | 101 | 5.51 | 13.5 | | High | High | High | High | High | High | High | High | High | High | High | 0 | 9.69E-08 | 6.32 |
| FALSE | High | YDLIQNSES | [R].YDLIQN | 1xPhospho | B1ATC3 1x | 0.000221 | 0 | 1 | 3 | 1 | B1ATC3 | Trinucleoti | B1ATC3 [1 | 0 | 2846.371 | 0 | -0.37 | 0.06162 | 15.72 | 112.5 | 87.5 | 13.64 | 16.25 | | High | High | High | High | High | High | High | High | High | High | High | 0 | 4.36E-05 | 4.04 |
| FALSE | High | YAYFNGCS | [K].YAYFN( | 1xCarbami | Q7TMQ1 2 | 0.003001 | 0.00016 | 1 | 2 | 3 | Q7TMQ1 | Gap juncti | Q7TMQ1 [ | 0 | 3110.385 | 0 | 2.54 | 0.001195 | 14.96 | 27.7 | 172.3 | 6.29 | 15.88 | | High | High | High | High | High | High | High | High | High | High | High | 7.19E-05 | 0.000818 | 2.71 |
| FALSE | High | YAYFNGCS | [K].YAYFN( | 1xCarbami | Q7TMQ1 1 | 3.89E-07 | 0 | 1 | 2 | 6 | Q7TMQ1 | Gap juncti | Q7TMQ1 [ | 0 | 3046.413 | 0 | 2.64 | 0.000541 | 11.07 | 29 | 171 | 14.19 | 9.09 | | High | High | High | High | High | High | High | High | High | High | High | 0.000859 | 0.008348 | 3.47 |

| | | | | | | | | | | | | | | | | | | | | | | | | | | | | | | | | | | | | | | |
|---|---|---|---|---|---|---|---|---|---|---|---|---|---|---|---|---|---|---|---|---|---|---|---|---|---|---|---|---|---|---|---|---|---|---|---|---|---|---|
| FALSE | High | YAYFNGCS | [K].YAYFN( | 1xCarbami | Q7TMQ1 1 | 0.000184 | 0 | 1 | 2 | 4 | Q7TMQ1 | Gap junctio | Q7TMQ1 [ | 0 | 3031.402 | 0 | 0.5 | 0.002704 | 5.47 | 82.5 | 117.5 | 4.37 | 8.16 | | High | High | High | High | High | High | High | High | High | High | High | 0 | 3.56E-05 | 4.12 |
| FALSE | High | WSHMFLLI | [K].WSHMI | 2xPhospho | Q3TXC4 2x | 0.040028 | 0.002867 | 1 | 9 | 1 | Q3TXC4 | Putative ur | Q3TXC4 [4 | 0 | 3177.496 | 0 | 2.26 | 0.011202 | 23.68 | 39.5 | 160.5 | 20.65 | 30.09 | | High | High | High | High | High | High | High | High | High | High | High | 0.0015 | 0.01492 | 3.03 |
| FALSE | High | WSLNTYK | [K].WSLNT | 1xPhospho | G5E8V9 1x | 0.005228 | 0.000551 | 1 | 4 | 1 | G5E8V9 | ADP-ribosy | G5E8V9 [1 | 0 | 1449.754 | 0 | 1.32 | 0.007067 | 26.85 | 60.1 | 139.9 | 17.54 | 14.65 | | High | High | High | High | High | High | High | High | High | High | High | 0.000246 | 0.001521 | 1.88 |
| FALSE | High | WSPEHNA( | [R].WSPEH | 1xPhospho | Q1W617 1 | 9.39E-06 | 0 | 1 | 2 | 2 | Q1W617 | Protein Shi | Q1W617 [7 | 0 | 3243.536 | 0 | 2.75 | 0.002186 | 27.57 | 26.3 | 173.7 | 10.59 | 19.05 | | High | High | High | High | High | High | High | High | High | High | High | 0 | 1.26E-06 | 4.4 |
| FALSE | High | WSPVASTL | [R].WSPVA | 1xPhospho | Q3TPZ5 1x | 0.000884 | 0 | 1 | 1 | 1 | Q3TPZ5 | Dynactin 2 | Q3TPZ5 [3 | 0 | 1891.988 | 0 | -0.02 | 0.373163 | 9.66 | 101.8 | 98.2 | 8.24 | 5.24 | | High | High | High | High | High | High | High | High | High | High | High | 0 | 0.000207 | 2.38 |
| FALSE | High | WSSGVGG | [R].WSSGV | 1xPhospho | E9Q2H1 1x | 0.000958 | 0 | 1 | 2 | 1 | E9Q2H1 | E3 ubiquiti | E9Q2H1 [1 | 0 | 1603.706 | 0 | -0.41 | 0.002708 | 4.65 | 114.5 | 85.5 | 6 | 5.77 | | High | High | High | High | High | High | High | High | High | High | High | 0 | 0.000228 | 0.9 |
| FALSE | High | WSTDQED( | [K].WSTDC | 1xPhospho | B2RWU4 1 | 3.72E-07 | 0 | 2 | 6 | 10 | B2RWU4; ( | Cordon-ble | B2RWU4 [ | 0 | 3012.388 | 0 | -0.39 | 0.00933 | 6.14 | 112.7 | 87.3 | 2.24 | 17.04 | Shared | High | High | High | High | High | High | High | High | High | High | High | 0 | 3.36E-08 | 4.47 |
| FALSE | High | WSTDQED( | [K].WSTDC | 1xDeamida | B2RWU4 1 | 8.63E-05 | 0 | 2 | 6 | 5 | B2RWU4; ( | Cordon-ble | B2RWU4 [ | 0 | 3013.372 | 0 | -0.77 | 0.063578 | 1.32 | 118.4 | 81.6 | 20.29 | 15.02 | Shared | High | High | High | High | High | High | High | High | High | High | High | 0.000275 | 0.001815 | 4.28 |
| FALSE | High | WSTDQED( | [K].WSTDC | 2xPhospho | B2RWU4 2 | 0.000248 | 0 | 2 | 6 | 3 | B2RWU4; ( | Cordon-ble | B2RWU4 [ | 0 | 3092.354 | 0 | -0.06 | 0.28535 | 12.33 | 101.7 | 98.3 | 15.63 | 19.72 | Shared | High | High | High | High | High | High | High | High | High | High | High | 0 | 4.99E-05 | 3.84 |
| FALSE | High | WSTTAGFL | [R].WSTTA | 1xPhospho | A5D6P2 1x | 0.092372 | 0.007211 | 1 | 22 | 1 | A5D6P2 | Par-3 (Part | A5D6P2 [1 | 0 | 1548.823 | 0 | -0.26 | 0.159259 | 7.84 | 107.4 | 92.6 | 7.98 | 9.14 | | High | High | High | High | High | High | High | High | High | High | High | 0.003659 | 0.03931 | 2.29 |
| FALSE | High | WTKPSSFS | [R].WTKPS | 1xPhospho | Q6DVA0 1x | 0.003892 | 0.000261 | 1 | 1 | 1 | Q6DVA0 | LEM doma | Q6DVA0 [5 | 0 | 1735.789 | 0 | 0.82 | 0.012843 | 21.71 | 75.9 | 124.1 | 13.35 | 16.79 | | High | High | High | High | High | High | High | High | High | High | High | 0.000117 | 0.001089 | 2.33 |
| FALSE | High | WTPVIVSN | [R].WTPVI | 1xPhospho | G3X940 1x | 3.19E-07 | 0 | 1 | 5 | 2 | G3X940 | Histone ac | G3X940 [7 | 0 | 3201.476 | 0 | 0.56 | 0.012515 | 14.15 | 79.7 | 120.3 | 4.65 | 13.66 | | High | High | High | High | High | High | High | High | High | High | High | 0 | 2.81E-08 | 6.13 |
| FALSE | High | WVAQSVSI | [R].WVAQS | 1xPhospho | A0A068F1 | 0.004696 | 0.000456 | 1 | 1 | 4 | A0A068F1 | Glyco-gag | A0A068F1 | 0 | 1637.789 | 0 | 4.81 | 0.000295 | 31.29 | 6.9 | 193.1 | 33.08 | 8.99 | | High | High | High | High | High | High | High | High | High | High | High | 0.000204 | 0.001351 | 2.11 |
| FALSE | High | WVEENIPS | [K].WVEEN | 1xPhospho | Q8BKU8-3 | 0.002153 | 5.66E-05 | 1 | 4 | 2 | Q8BKU8-3 | Isoform 3 ( | Q8BKU8-3 | 0 | 3516.669 | 0 | 2.13 | 0.000209 | 12.37 | 37.2 | 162.8 | 10.49 | 6.13 | | High | High | High | High | High | High | High | High | High | High | High | 2.58E-05 | 0.000563 | 3.11 |
| FALSE | High | WVEENVPS | [K].WVEEN | 1xPhospho | A2AQJ6 1x | 1.33E-09 | 0 | 1 | 4 | 8 | A2AQJ6 | Transmem | A2AQJ6 [5 | 0 | 3079.471 | 0 | 0.81 | 0.004357 | 8.24 | 72.2 | 127.8 | 2.01 | 11.26 | | High | High | High | High | High | High | High | High | High | High | High | 0 | 7.23E-09 | 4.59 |
| FALSE | High | WVSELSYIE | [R].WVSEL | 1xPhospho | Q9R0L6-2 | 4.05E-05 | 0 | 1 | 3 | 3 | Q9R0L6-2 | Isoform 2 ( | Q9R0L6-2 | 0 | 1920.976 | 0 | 0.66 | 0.124793 | 65.16 | 64.4 | 135.6 | 27.82 | 40.71 | | High | High | High | High | High | High | High | High | High | High | High | 0 | 6.5E-06 | 2.82 |
| FALSE | High | YAALSVDG | [K].YAALSV | 1xCarbami | Q3TSY9 1x | 3.43E-10 | 0 | 1 | 6 | 22 | Q3TSY9 | Putative ur | Q3TSY9 [5 | 0 | 2398.896 | 0 | 0.29 | 0.031329 | 5.41 | 90 | 110 | 4.66 | 8.11 | | High | High | High | High | High | High | High | High | High | High | High | 0 | 5.72E-10 | 6.25 |
| FALSE | High | YACGLWGI | [K].YACGL | 1xCarbami | D3YZC9 1x | 5.03E-05 | 0 | 1 | 2 | 1 | D3YZC9 | Splicing fac | D3YZC9 [4 | 0 | 1746.824 | 0 | -0.44 | 0.000997 | 4.05 | 114.7 | 85.3 | 3.01 | 3.73 | | High | High | High | High | High | High | High | High | High | High | High | 0 | 8.27E-06 | 1.95 |
| FALSE | High | YACSPLPSY | [K].YACSPL | 3xCarbami | A0A1B0GS | 0.000111 | 0 | 1 | 2 | 2 | A0A1B0GS | Nuclear po | A0A1B0GS | 0 | 3017.321 | 0 | -0.49 | 0.00837 | 10.42 | 119.9 | 80.1 | 6.85 | 11.67 | | High | High | High | High | High | High | High | High | High | High | High | 0 | 2.02E-05 | 4.23 |
| FALSE | High | YADEEIPRS | [K].YADEEI | 1xPhospho | Q80X90 1x | 0.001267 | 0 | 1 | 1 | 1 | Q80X90 | Filamin-B ( | Q80X90 [1 | 1 | 1990.009 | 0 | 0.61 | 0.011162 | 12.31 | 81.7 | 118.3 | 11.83 | 9.94 | | High | High | High | High | High | High | High | High | High | High | High | 0 | 0.00031 | 2.03 |
| FALSE | High | YADQEVPR | [K].YADQE | 1xPhospho | Q8VHX6 1x | 0.0095 | 0.000736 | 1 | 2 | 1 | Q8VHX6 | Filamin-C ( | Q8VHX6 [1 | 1 | 1975.009 | 0 | -1.09 | 0.005468 | 4.93 | 133.8 | 66.2 | 12.07 | 17.6 | | High | High | High | High | High | High | High | High | High | High | High | 0.000327 | 0.002951 | 1.5 |
| FALSE | High | YADVASPT | [K].YADVA | 1xPhospho | E9Q842 1x | 0.016743 | 0.001166 | 1 | 4 | 2 | E9Q842 | Neuron na | E9Q842 [1 | 0 | 1401.698 | 0 | 0.73 | 0.001404 | 8.81 | 74.6 | 125.4 | 2.3 | 7.04 | | High | High | High | High | High | High | High | High | High | High | High | 0.000638 | 0.005603 | 1.34 |
| FALSE | High | YAGLDDTD | [R].YAGLDI | 2xPhospho | Q91WL8 2 | 4.06E-06 | 0 | 1 | 4 | 4 | Q91WL8 | WW doma | Q91WL8 [6 | 0 | 2683.069 | 0 | 0.05 | 0.471042 | 10.51 | 98.2 | 101.8 | 10.77 | 20.95 | | High | High | High | High | High | High | High | High | High | High | High | 0 | 2.78E-05 | 3.66 |
| FALSE | High | YAGSGEED | [R].YAGSGI | 1xPhospho | Q5BLK4 1x | 0.002623 | 0.000109 | 1 | 2 | 1 | Q5BLK4 | Terminal u | Q5BLK4 [8 | 0 | 2734.238 | 0 | 0.26 | 0.107707 | 24.15 | 89.1 | 110.9 | 9.84 | 11.98 | | High | High | High | High | High | High | High | High | High | High | High | 4.94E-05 | 0.000701 | 2.35 |
| FALSE | High | YAHQQPPS | [R].YAHQQ | 1xPhospho | F8VQ28 1x | 1.01E-07 | 0 | 1 | 5 | 1 | F8VQ28 | Paxillin OS= | F8VQ28 [7 | 0 | 2495.274 | 0 | 1.35 | 0.00024 | 0.95 | 56.6 | 143.4 | 2.56 | 5.19 | | High | High | High | High | High | High | High | High | High | High | High | 0 | 7.72E-09 | 3.24 |
| FALSE | High | YAPNPGG( | [K].YAPNP( | 1xCarbami | G3X932 1x | 1.45E-06 | 0 | 1 | 5 | 1 | G3X932 | DNA segme | G3X932 [4 | 0 | 2457.184 | 0 | -0.56 | 0.001968 | 4.06 | 119.1 | 80.9 | 10.22 | 7.2 | | High | High | High | High | High | High | High | High | High | High | High | 0 | 1.55E-07 | 3.6 |
| FALSE | High | YASGLLK | [K].YASGLL | 1xPhospho | I6L9H1 1xP | 0.054383 | 0.003951 | 1 | 2 | 2 | I6L9H1 | Dishevelle | I6L9H1 [47 | 0 | 1060.564 | 0 | -0.99 | 0.011867 | 2.54 | 129.6 | 70.4 | 10.18 | 21.9 | | High | High | High | High | High | High | High | High | High | High | High | 0.003768 | 0.04059 | 2.22 |
| FALSE | High | YAWTDEEE | [R].YAWTD | 1xPhospho | J3QMP9 1x | 9.32E-10 | 0 | 1 | 3 | 6 | J3QMP9 | Centrosom | J3QMP9 [4 | 0 | 2852.292 | 0 | -0.2 | 0.04882 | 7.92 | 106.9 | 93.1 | 6.09 | 9.26 | | High | High | High | High | High | High | High | High | High | High | High | 0 | 3.99E-11 | 4.39 |
| FALSE | High | YAWTDEEE | [R].YAWTD | 2xPhospho | J3QMP9 2x | 1.85E-05 | 0 | 1 | 3 | 3 | J3QMP9 | Centrosom | J3QMP9 [4 | 0 | 2932.258 | 0 | 0.26 | 0.0803 | 14.4 | 90.6 | 109.4 | 10.05 | 17.22 | | High | High | High | High | High | High | High | High | High | High | High | 0 | 2.7E-06 | 4.08 |
| FALSE | High | YAYFNGCS | [K].YAYFN( | 1xCarbami | Q7TMQ1 1 | 3.23E-05 | 0 | 1 | 2 | 3 | Q7TMQ1 | Gap junctio | Q7TMQ1 [ | 0 | 3030.418 | 0 | 2.94 | 0.002128 | 19.41 | 21.8 | 178.2 | 17.17 | 20.49 | | High | High | High | High | High | High | High | High | High | High | High | 0 | 5.03E-06 | 3.42 |
| FALSE | High | WDDESHIT | [R].WDDES | 1xPhospho | E9QML5 1x | 0.001147 | 0 | 1 | 16 | 1 | E9QML5 | Zinc finger | E9QML5 [1 | 0 | 2190.088 | 0 | -0.05 | 0.781514 | 13.07 | 103.3 | 96.7 | 10.95 | 9.94 | | High | High | High | High | High | High | High | High | High | High | High | 0 | 0.000277 | 2.29 |
| FALSE | High | WCNVQST | [R].WCNV( | 1xCarbami | Q545E8 1x | 2.49E-09 | 0 | 1 | 2 | 8 | Q545E8 | Deoxycytid | Q545E8 [5 | 0 | 2882.332 | 0 | -0.05 | 0.123449 | 3.41 | 103.2 | 96.8 | 5.94 | 3.45 | | High | High | High | High | High | High | High | High | High | High | High | 0 | 0.000137 | 4 |
| FALSE | High | WCLVSQTI | [R].WCLVS | 1xCarbami | Q8BGX7 1x | 0.003997 | 0.000311 | 1 | 3 | 3 | Q8BGX7 | DNA repair | Q8BGX7 [2 | 0 | 3000.443 | 0 | -0.53 | 0.00216 | 5.57 | 117 | 83 | 2.32 | 11.19 | | High | High | High | High | High | High | High | High | High | High | High | 0.000161 | 0.001177 | 2.94 |
| FALSE | High | WASPSPPR | [R].WASPS | 2xPhospho | B0V2M3 2x | 0.050397 | 0.003641 | 1 | 2 | 1 | B0V2M3 | Zinc finger | B0V2M3 [6 | 0 | 1286.553 | 0 | -0.37 | 0.013652 | 2.98 | 115.1 | 84.9 | 3.33 | 14.99 | | High | High | High | High | High | High | High | High | High | High | High | 0.001916 | 0.01954 | 1.32 |
| FALSE | High | VTQVDGN | [K].VTQVD | 1xPhospho | Q3U4S9 1x | 8.63E-05 | 0 | 1 | 4 | 1 | Q3U4S9 | Coatomer | Q3U4S9 [4 | 0 | 1480.736 | 0 | -0.28 | 0.004801 | 7 | 109.7 | 90.3 | 2.93 | 5.35 | | High | High | High | High | High | High | High | High | High | High | High | 0 | 1.53E-05 | 1.73 |
| FALSE | High | VTRPNTSPI | [K].VTRPN | 1xPhospho | I1VGT4 1x | 9.08E-07 | 0 | 2 | 13 | 2 | I1VGT4; I1 | Anion exch | I1VGT4 [11 | 0 | 3438.812 | 0 | -0.46 | 0.181347 | 17.25 | 115.9 | 84.1 | 16.79 | 14.06 | Shared | High | High | High | High | High | High | High | High | High | High | High | 0 | 9.09E-08 | 3.08 |
| FALSE | High | VTSDYHSG | [K].VTSDYH | 2xPhospho | P50615 2x | 8.79E-05 | 0 | 1 | 1 | 1 | P50615 | Protein BT | P50615 [13 | 0 | 2507.026 | 0 | 0.29 | 0.082649 | 10.23 | 92.2 | 107.8 | 10.64 | 7.75 | | High | High | High | High | High | High | High | High | High | High | High | 0 | 1.55E-05 | 2.92 |
| FALSE | High | VTSENGAT | [R].VTSEN( | 1xPhospho | Q80U04 1x | 0.098533 | 0.007902 | 1 | 3 | 1 | Q80U04 | E3 ubiquiti | Q80U04 [3 | 0 | 1458.679 | 0 | 1.59 | 0.001811 | 9.42 | 42.9 | 157.1 | 13.31 | 14 | | High | High | High | High | High | High | High | High | High | High | High | 0.003991 | 0.04255 | 1.9 |
| FALSE | High | VTSFSTPPT | [R].VTSFST | 2xPhospho | B1APX2 2x | 0.112856 | 0.009146 | 1 | 2 | 1 | B1APX2 | RIKEN cDN | B1APX2 [3 | 0 | 1707.759 | 0 | 0.6 | 0.000486 | 9.07 | 79.6 | 120.4 | 15.1 | 7.97 | | High | High | High | High | High | High | High | High | High | High | High | 0.004583 | 0.04978 | 1.64 |
| FALSE | High | VTSLANLIS | [R].VTSLAN | 1xPhospho | Q9Z206 1x | 0.016596 | 0.001124 | 1 | 2 | 1 | Q9Z206 | Neuroepith | Q9Z206 [9 | 0 | 1578.882 | 0 | -1.12 | 0.001034 | 0.75 | 136.7 | 63.3 | 6.13 | 6.25 | | High | High | High | High | High | High | High | High | High | High | High | 0.000622 | 0.005529 | 2.51 |
| FALSE | High | VTSPSLAAL | [R].VTSPSL | 1xPhospho | E9Q5N9 1x | 0.002334 | 0.000109 | 1 | 2 | 1 | E9Q5N9 | Zinc finger | E9Q5N9 [3 | 0 | 1990.139 | 0 | -0.55 | 0.005219 | 7.87 | 118.8 | 81.2 | 4.73 | 7.23 | | High | High | High | High | High | High | High | High | High | High | High | 4.94E-05 | 0.000619 | 2.34 |
| FALSE | High | VTSPSSGYS | [R].VTSPSS | 1xPhospho | Q8CAF4-3 | 3.77E-05 | 0 | 1 | 4 | 5 | Q8CAF4-3 | Isoform 3 ( | Q8CAF4-3 | 0 | 3203.664 | 0 | 0.85 | 0.002566 | 11.48 | 72.9 | 127.1 | 8.07 | 9.56 | | High | High | High | High | High | High | High | High | High | High | High | 0 | 6.01E-06 | 4.48 |
| FALSE | High | VTSPTVFS( | [K].VTSPTV | 1xPhospho | P70174 1x | 0.003055 | 0.00016 | 1 | 1 | 2 | P70174 | Histamine | P70174 [28 | 0 | 1745.831 | 0 | 2.97 | 0.000793 | 28.38 | 21.7 | 178.3 | 29.07 | 13.64 | | High | High | High | High | High | High | High | High | High | High | High | 7.19E-05 | 0.000834 | 2.57 |
| FALSE | High | VTSSMSSP | [R].VTSSM | 1xPhospho | B8QI33 1x | 0.039347 | 0.002793 | 1 | 4 | 1 | B8QI33 | Liprin-alph | B8QI33 [12 | 0 | 1920.921 | 0 | -0.22 | 0.238858 | 12.93 | 104.1 | 95.9 | 14.33 | 21.43 | | High | High | High | High | High | High | High | High | High | High | High | 0.001453 | 0.01464 | 1.73 |
| FALSE | High | VTSSVPLPS | [R].VTSSVP | 1xPhospho | Q8BVL3 1x | 0.000113 | 0 | 1 | 1 | 4 | Q8BVL3 | Sorting nex | Q8BVL3 [3 | 0 | 1924.958 | 0 | 0.09 | 0.120676 | 8.94 | 96 | 104 | 3.98 | 4.61 | | High | High | High | High | High | High | High | High | High | High | High | 0 | 2.05E-05 | 3.23 |
| FALSE | High | VTSTPQKP( | [K].VTSTPC | 1xPhospho | B7ZNX2 1x | 0.003655 | 0.00021 | 1 | 3 | 3 | B7ZNX2 | Anaphase | B7ZNX2 [8 | 0 | 2380.191 | 0 | -0.7 | 0.000227 | 0.86 | 123.6 | 76.4 | 4.49 | 5.26 | | High | High | High | High | High | High | High | High | High | High | High | 0.004999 | 0.05469 | 2.08 |
| FALSE | High | VTTGGNQ | [K].VTTGG | 1xPhospho | Q8VCF0 1x | 0.009585 | 0.000736 | 1 | 2 | 1 | Q8VCF0 | Mitochond | Q8VCF0 [3 | 0 | 1797.869 | 0 | 0.71 | 0.001252 | 18.29 | 74.7 | 125.3 | 13.11 | 2.16 | | High | High | High | High | High | High | High | High | High | High | High | 0.000344 | 0.002996 | 2.02 |
| FALSE | High | VTQQVNPI | [R].VTQQV | 1xPhospho | Q99L27 1x | 0.000274 | 0 | 1 | 1 | 1 | Q99L27 | GMP reduc | Q99L27 [3 | 0 | 1770.863 | 0 | 0.27 | 0.130106 | 10.97 | 90.8 | 109.2 | 6.73 | 8.43 | | High | High | High | High | High | High | High | High | High | High | High | 0 | 5.58E-05 | 2.35 |
| FALSE | High | VTTPWSPA | [R].VTTPW | 1xPhospho | Q6PIX9-2 1 | 0.017192 | 0.001204 | 1 | 2 | 2 | Q6PIX9-2 | Isoform 2 ( | Q6PIX9-2 [ | 0 | 2083.051 | 0 | -0.63 | 0.028488 | 14.95 | 119.3 | 80.7 | 13.05 | 14.22 | | High | High | High | High | High | High | High | High | High | High | High | 0.000638 | 0.005762 | 2.47 |
| FALSE | High | VTVLPAEP | [K].VTVLPA | 1xPhospho | Q8CJ96 1x | 2.84E-08 | 0 | 1 | 1 | 2 | Q8CJ96 | Ras associa | Q8CJ96 [3 | 0 | 3695.871 | 0 | 0.66 | 0.007624 | 19.8 | 77.5 | 122.5 | 7.8 | 15.36 | | High | High | High | High | High | High | High | High | High | High | High | 0 | 1.85E-09 | 5.37 |
| FALSE | High | VVAACELA | [R].VVAAC | 2xCarbami | Q7TT18 1x | 3.37E-07 | 0 | 1 | 3 | 3 | Q7TT18 | Activating | Q7TT18 [2 | 0 | 4124.779 | 0 | -0.22 | 0.069547 | 8.1 | 109.4 | 90.6 | 11.41 | 13.29 | | High | High | High | High | High | High | High | High | High | High | High | 0 | 7.92E-08 | 4.37 |
| FALSE | High | VVAACELA | [R].VVAAC | 2xCarbami | Q7TT18 1x | 1.9E-06 | 0 | 1 | 3 | 3 | Q7TT18 | Activating | Q7TT18 [2 | 0 | 4125.763 | 0 | -0.01 | 0.412547 | 17.63 | 102.1 | 97.9 | 4.56 | 15.9 | | High | High | High | High | High | High | High | High | High | High | High | 0 | 2.09E-07 | 4.82 |
| FALSE | High | VVAGEVNF | [K].VVAGE | 1xPhospho | Q7TS75 1x | 0.00103 | 0 | 1 | 1 | 2 | Q7TS75 | APC memb | Q7TS75 [2 | 0 | 3360.786 | 0 | 0.3 | 0.029791 | 2.71 | 89.1 | 110.9 | 11.44 | 9.93 | | High | High | High | High | High | High | High | High | High | High | High | 0 | 0.000247 | 2.61 |
| FALSE | High | VVANVSEP | [R].VVANV | 1xPhospho | H7BX94 1x | 2.14E-06 | 0 | 1 | 7 | 4 | H7BX94 | Lysine-rich | H7BX94 [7 | 0 | 3420.62 | 0 | -0.04 | 0.590778 | 4.21 | 100.1 | 99.9 | 4.07 | 3.53 | | High | High | High | High | High | High | High | High | High | High | High | 0 | 5.01E-07 | 4.63 |
| FALSE | High | VVANVSEP | [R].VVANV | 1xDeamida | H7BX94 1x | 3.52E-07 | 0 | 1 | 7 | 7 | H7BX94 | Lysine-rich | H7BX94 [7 | 0 | 3421.604 | 0 | -0.07 | 0.038043 | 2.51 | 102.5 | 97.5 | 3.12 | 4.13 | | High | High | High | High | High | High | High | High | High | High | High | 0 | 6E-08 | 4.74 |
| FALSE | High | VVAPTISSP | [K].VVAPTI | 1xCarbami | Q80TM2 1 | 1.16E-05 | 0 | 1 | 3 | 3 | Q80TM2 | MKIAA102 | Q80TM2 [7 | 0 | 2449.236 | 0 | 0.25 | 0.052081 | 5.32 | 85.6 | 114.4 | 16.94 | 11.47 | | High | High | High | High | High | High | High | High | High | High | High | 0 | 1.59E-06 | 3.55 |
| FALSE | High | VVCDADDS | [R].VVCDA | 1xCarbami | Q8C1D8 1x | 1.01E-08 | 0 | 1 | 4 | 2 | Q8C1D8 | Protein IW | Q8C1D8 [3 | 0 | 2264.004 | 0 | -0.39 | 0.003571 | 7.6 | 118.4 | 81.6 | 4.46 | 9.82 | | High | High | High | High | High | High | High | High | High | High | High | 0 | 5.8E-10 | 3.7 |
| FALSE | High | VVCDADDS | [R].VVCDA | 1xCarbami | Q8C1D8 2x | 6.47E-05 | 0 | 1 | 4 | 3 | Q8C1D8 | Protein IW | Q8C1D8 [3 | 0 | 2343.97 | 0 | -0.24 | 0.070942 | 9.3 | 104.5 | 95.5 | 5.33 | 9.18 | | High | High | High | High | High | High | High | High | High | High | High | 0 | 0.000011 | 3.34 |
| FALSE | High | VVDFGSAT | [K].VVDFG | 1xPhospho | P22518 1x | 0.003464 | 0.00021 | 1 | 4 | 1 | P22518 | Dual specif | P22518 [32 | 0 | 2645.208 | 0 | -0.23 | 0.365773 | 28.79 | 104.3 | 95.7 | 17.2 | 22.67 | | High | High | High | High | High | High | High | High | High | High | High | 9.42E-05 | 0.00096 | 4.47 |
| FALSE | High | VVDNSPPA | [R].VVDNS | 1xPhospho | Q3UHC1 1 | 0.000852 | 0 | 1 | 5 | 2 | Q3UHC1 | Putative ur | Q3UHC1 [2 | 0 | 1771.976 | 0 | -0.26 | 0.003707 | 2.99 | 109 | 91 | 3.95 | 2.75 | | High | High | High | High | High | High | High | High | High | High | High | 0 | 0.000216 | 3.16 |
| FALSE | High | VVDYSQFC | [K].VVDYS( | 1xPhospho | Q80XU3 1x | 6.9E-09 | 0 | 1 | 2 | 31 | Q80XU3 | Nuclear ub | Q80XU3 [1 | 0 | 2547.04 | 0 | -0.16 | 0.15087 | 7.64 | 107.6 | 92.4 | 3.18 | 10.76 | | High | High | High | High | High | High | High | High | High | High | High | 0 | 5.28E-08 | 5 |
| FALSE | High | VVDYSQFC | [K].VVDYS( | 1xPhospho | Q80XU3 1x | 0.000188 | 0 | 1 | 2 | 1 | Q80XU3 | Nuclear ub | Q80XU3 [1 | 1 | 3428.521 | 0 | -0.71 | 0.018765 | 16.04 | 126.3 | 73.7 | 15.76 | 37.51 | | High | High | High | High | High | High | High | High | High | High | High | 0 | 3.63E-05 | 3.14 |
| FALSE | High | VTTTNAVA | [R].VTTTN | 1xCarbami | Q3UV57 1x | 2.27E-08 | 0 | 1 | 1 | 1 | Q3UV57 | Putative ur | Q3UV57 [4 | 0 | 2546.164 | 0 | 1.27 | 0.001216 | 3.77 | 57.6 | 142.4 | 11.22 | 8.26 | | High | High | High | High | High | High | High | High | High | High | High | 0 | 1.45E-09 | 4.6 |
| FALSE | High | VTQEEIK | [R].VTQEEI | 1xPhospho | E9Q317 1x | 0.025964 | 0.001645 | 1 | 3 | 1 | E9Q317 | Histone de | E9Q317 [2 | 0 | 1384.749 | 0 | -0.02 | 0.958831 | 7.83 | 100.3 | 99.7 | 10.71 | 8.02 | | High | High | High | High | High | High | High | High | High | High | High | 0.000872 | 0.00914 | 2.19 |
| FALSE | High | VTPTQSLRS | [R].VTPTQ | 1xPhospho | Q3UA65 1x | 0.000121 | 0 | 1 | 6 | 1 | Q3UA65 | DNA helica | Q3UA65 [1 | 1 | 1798.89 | 0 | -0.69 | 0.001617 | 5.68 | 124.3 | 75.7 | 3.95 | 8.55 | | High | High | High | High | High | High | High | High | High | High | High | 0 | 2.22E-05 | 1.2 |
| FALSE | High | VTPTQSLR | [R].VTPTQ | 1xPhospho | Q3UA65 1x | 0.111965 | 0.009053 | 1 | 6 | 3 | Q3UA65 | DNA helica | Q3UA65 [1 | 0 | 1210.639 | 0 | -0.65 | 0.000791 | 1.85 | 122.7 | 77.3 | 3.55 | 8.19 | | High | High | High | High | High | High | High | High | High | High | High | 0.008242 | 0.0889 | 1.98 |
| FALSE | High | VTFPSDEDI | [R].VTFPSD | 1xPhospho | Q8BKR5 1x | 0.000276 | 0 | 1 | 1 | 3 | Q8BKR5 | Protein ph | Q8BKR5 [5 | 0 | 2328.178 | 0 | 0.5 | 0.005877 | 9.52 | 80 | 120 | 7.1 | 9.58 | | High | High | High | High | High | High | High | High | High | High | High | 0 | 5.65E-05 | 3.55 |
| FALSE | High | VTGDNQPI | [K].VTGDN | 1xPhospho | F8VQC1 1x | 0.000808 | 0 | 1 | 5 | 1 | F8VQC1 | Signal reco | F8VQC1 [5 | 0 | 1396.724 | 0 | 0.1 | 0.233978 | 17.58 | 93.3 | 106.7 | 7.54 | 11.11 | | High | High | High | High | High | High | High | High | High | High | High | 0 | 0.000188 | 1.85 |
| FALSE | High | VTGGSSSS | [R].VTGGS | 1xPhospho | Q9EP53 1x | 0.01186 | 0.000889 | 1 | 5 | 3 | Q9EP53 | Hamartin ( | Q9EP53 [1 | 0 | 2743.355 | 0 | -0.98 | 0.001595 | 1.48 | 132.8 | 67.2 | 5.46 | 13.77 | | High | High | High | High | High | High | High | High | High | High | High | 0.000832 | 0.007896 | 2.5 |
| FALSE | High | VTGSPAEE | [K].VTGSP | 1xPhospho | E9Q284 1x | 4.25E-06 | 0 | 1 | 3 | 3 | E9Q284 | Coilin OS= | E9Q284 [1 | 0 | 2058.952 | 0 | -1.56 | 0.000418 | 2.37 | 150.4 | 49.6 | 4.62 | 10.48 | | High | High | High | High | High | High | High | High | High | High | High | 0 | 5.14E-07 | 3.42 |
| FALSE | High | VTGVYELSI | [R].VTGVYI | 1xCarbami | A0A087W( | 0.013308 | 0.000889 | 1 | 9 | 1 | A0A087W( | Kinesin-like | A0A087W( | 0 | 2672.241 | 0 | -0.51 | 0.005616 | 7.48 | 118.6 | 81.4 | 9.56 | 7.18 | | High | High | High | High | High | High | High | High | High | High | High | 0.000486 | 0.004333 | 2.64 |
| FALSE | High | VTHEVAGE | [R].VTHEV | 1xPhospho | Q69Z38 1x | 1.58E-05 | 0 | 1 | 2 | 2 | Q69Z38 | Pseudopod | Q69Z38 [1 | 0 | 1603.804 | 0 | -0.51 | 0.004429 | 4.02 | 116.3 | 83.7 | 5.35 | 5.61 | | High | High | High | High | High | High | High | High | High | High | High | 0 | 2.27E-06 | 2.59 |
| FALSE | High | VTIAQGGV | [KR].VTIAC | 1xPhospho | C0HKE6 1x | 0.090153 | 0.00698 | 2 | 13 | 1 | C0HKE6; B | Histone H2 | C0HKE6 [1 | 0 | 2469.461 | 0 | -0.21 | 0.473611 | 10.47 | 98.3 | 101.7 | 18.44 | 9.5 | Shared | High | High | High | High | High | High | High | High | High | High | High | 0.003606 | 0.03831 | 2.08 |
| FALSE | High | VTIASLPR | [R].VTIASL | 1xPhospho | G5E880 1x | 0.054844 | 0.003982 | 2 | 7 | 1 | G5E880; Q | Lymphoid- | G5E880 [3 | 0 | 1165.654 | 0 | 3.52 | 0.000263 | 24.81 | 14.4 | 185.6 | 19.52 | 7.84 | Shared | High | High | High | High | High | High | High | High | High | High | High | 0.002059 | 0.02145 | 1.02 |
| FALSE | High | VTIPPPSSS | [K].VTIPPP | 1xPhospho | Q571F3 1x | 0.000237 | 0 | 1 | 2 | 1 | Q571F3 | MFLJ00061 | Q571F3 [1 | 0 | 2873.594 | 0 | 0.35 | 0.087054 | 10.42 | 88.3 | 111.7 | 14.54 | 10.21 | | High | High | High | High | High | High | High | High | High | High | High | 0 | 4.73E-05 | 4.02 |
| FALSE | High | VTLQDYHL | [R].VTLQD | 1xPhospho | Q9DAZ9 1x | 1.03E-07 | 0 | 1 | 2 | 4 | Q9DAZ9 | Abscission, | Q9DAZ9 [2 | 0 | 2783.261 | 0 | 0.55 | 0.007192 | 13.24 | 81.9 | 118.1 | 2.66 | 10.3 | | High | High | High | High | High | High | High | High | High | High | High | 0 | 7.94E-09 | 3.86 |
| FALSE | High | VTLSTAETE | [K].VTLSTA | 1xPhospho | P49452 1x | 0.009931 | 0.000736 | 1 | 4 | 2 | P49452 | Centromer | P49452 [18 | 0 | 2815.438 | 0 | -0.18 | 0.092928 | 12.62 | 110.6 | 89.4 | 13.09 | 7.8 | | High | High | High | High | High | High | High | High | High | High | High | 0.000344 | 0.003109 | 3.14 |
| FALSE | High | VTLTPEEEA | [K].VTLTPE | 1xPhospho | A0A1B0GS | 0.008691 | 0.000736 | 1 | 9 | 2 | A0A1B0GS | L-lactate d | A0A1B0GS | 0 | 1453.714 | 0 | 0.11 | 0.204454 | 10.85 | 96.1 | 103.9 | 4.42 | 9.63 | | High | High | High | High | High | High | High | High | High | High | High | 0.003215 | 0.03404 | 3.37 |
| FALSE | High | VTLYNADQ | [RK].VTLYN | 1xPhospho | Q8BUR3 1x | 5.77E-06 | 0 | 1 | 2 | 1 | Q8BUR3 | Forkhead | Q8BUR3 [2 | 0 | 1946.869 | 0 | 0.79 | 0.005611 | 20.24 | 69.7 | 130.3 | 16.88 | 8.05 | | High | High | High | High | High | High | High | High | High | High | High | 0 | 7.27E-07 | 3.08 |
| FALSE | High | VTNAISPES | [R].VTNAIS | 1xPhospho | Q9R0L6-2 | 2.08E-06 | 0 | 1 | 4 | 4 | Q9R0L6-2 | Isoform 2 ( | Q9R0L6-2 | 0 | 1779.884 | 0 | -0.1 | 0.075797 | 7.25 | 102.7 | 97.3 | 6.35 | 2.25 | | High | High | High | High | High | High | High | High | High | High | High | 0 | 2.33E-07 | 3.2 |
| FALSE | High | VTNAISPES | [R].VTNAIS | 2xPhospho | Q9R0L6-2 | 0.000209 | 0 | 1 | 4 | 5 | Q9R0L6-2 | Isoform 2 ( | Q9R0L6-2 | 0 | 1859.85 | 0 | 0.17 | 0.000578 | 1.5 | 94 | 106 | 3.09 | 3.32 | | High | High | High | High | High | High | High | High | High | High | High | 7.19E-05 | 0.000789 | 2.4 |
| FALSE | High | VTNAISPES | [R].VTNAIS | 3xPhospho | Q9R0L6-2 | 0.012845 | 0.000889 | 1 | 4 | 1 | Q9R0L6-2 | Isoform 2 ( | Q9R0L6-2 | 0 | 1939.817 | 0 | -0.29 | 0.005608 | 2.63 | 109.2 | 90.8 | 8 | 8.36 | | High | High | High | High | High | High | High | High | High | High | High | 0.000449 | 0.004158 | 2.62 |
| FALSE | High | VTNGAFTG | [K].VTNGA | 1xPhospho | P17751 1x | 0.000446 | 0 | 1 | 1 | 1 | P17751 | Triosephos | P17751 [12 | 0 | 2160.118 | 0 | 1.37 | 0.00022 | 5.27 | 55 | 145 | 4.15 | 3.48 | | High | High | High | High | High | High | High | High | High | High | High | 0 | 9.67E-05 | 2.5 |
| FALSE | High | VTNGAFTG | [K].VTNGA | 1xDeamida | P17751 1x | 0.015465 | 0.001089 | 1 | 1 | 1 | P17751 | Triosephos | P17751 [12 | 0 | 2161.102 | 0 | 1.42 | 0.000425 | 10.62 | 54.3 | 145.7 | 3.48 | 5.85 | | High | High | High | High | High | High | High | High | High | High | High | 0.000571 | 0.005099 | 2.4 |
| FALSE | High | VTNGAFTG | [K].VTNGA | 1xPhospho | P17751 1x | 0.000628 | 0 | 1 | 1 | 1 | P17751 | Triosephos | P17751 [12 | 0 | 2176.113 | 0 | 1.24 | 0.000386 | 9.62 | 61.4 | 138.6 | 6.66 | 6.63 | | High | High | High | High | High | High | High | High | High | High | High | 0 | 0.000142 | 3.68 |
| FALSE | High | VTNGVHQI | [R].VTNGV | 1xPhospho | Q5SSZ5 1x | 9.06E-06 | 0 | 1 | 1 | 3 | Q5SSZ5 | Tensin-3 O | Q5SSZ5 [6 | 0 | 3599.718 | 0 | -1.18 | 0.002936 | 2.71 | 137.5 | 62.5 | 12.41 | 10.2 | | High | High | High | High | High | High | High | High | High | High | High | 0 | 1.22E-06 | 5.73 |
| FALSE | High | VTNGVHQI | [R].VTNGV | 1xDeamida | Q5SSZ5 1x | 1.98E-07 | 0 | 1 | 1 | 14 | Q5SSZ5 | Tensin-3 O | Q5SSZ5 [6 | 0 | 3600.702 | 0 | -1.05 | 0.000539 | 1.63 | 136 | 64 | 5.44 | 4.95 | | High | High | High | High | High | High | High | High | High | High | High | 0 | 1.64E-08 | 5 |
| FALSE | High | VTNGVHQI | [R].VTNGV | 2xDeamida | Q5SSZ5 1x | 0.001386 | 5.66E-05 | 1 | 1 | 7 | Q5SSZ5 | Tensin-3 O | Q5SSZ5 [6 | 0 | 3601.686 | 0 | -0.77 | 0.019288 | 8.62 | 126.9 | 73.1 | 12.36 | 18.99 | | High | High | High | High | High | High | High | High | High | High | High | 4.94E-05 | 0.000645 | 4.01 |
| FALSE | High | VTNGVHQI | [R].VTNGV | 1xDeamida | Q5SSZ5 2x | 0.027358 | 0.001676 | 1 | 1 | 6 | Q5SSZ5 | Tensin-3 O | Q5SSZ5 [6 | 0 | 3680.668 | 0 | -0.51 | 0.009684 | 4.49 | 117.6 | 82.4 | 8.61 | 9.66 | | High | High | High | High | High | High | High | High | High | High | High | 0.000984 | 0.01017 | 4.07 |
| FALSE | High | VTNIVSFDI | [K].VTNIVS | 1xPhospho | Q9D3S3 1x | 1.8E-05 | 0 | 1 | 2 | 1 | Q9D3S3 | Sorting nex | Q9D3S3 [2 | 0 | 2850.333 | 0 | 0.57 | 0.014535 | 22.01 | 82.1 | 117.9 | 7.79 | 11.94 | | High | High | High | High | High | High | High | High | High | High | High | 0 | 2.6E-06 | 3.42 |
| FALSE | High | VTPAPLSPF | [K].VTPAPL | 1xPhospho | Q9WTN3 1 | 0.021046 | 0.001507 | 1 | 4 | 1 | Q9WTN3 | Sterol regu | Q9WTN3 [ | 0 | 2152.218 | 0 | 1.26 | 0.000693 | 8.75 | 60.6 | 139.4 | 5.06 | 10.06 | | High | High | High | High | High | High | High | High | High | High | High | 0.000801 | 0.007221 | 2.16 |
| FALSE | High | VTPEEEEEI | [R].VTPEEE | 1xPhospho | Q3U3A6 1x | 9.42E-07 | 0 | 1 | 5 | 1 | Q3U3A6 | Sodium/hy | Q3U3A6 [7 | 0 | 2329.036 | 0 | -0.04 | 0.302249 | 16.87 | 105.8 | 94.2 | 12.92 | 20.36 | | High | High | High | High | High | High | High | High | High | High | High | 0 | 9.5E-08 | 4.03 |
| FALSE | High | VTPLPFQP( | [R].VTPLPF | 1xPhospho | B6ZHD0 1x | 0.034579 | 0.002233 | 1 | 4 | 2 | B6ZHD0 | Erythrocyt | B6ZHD0 [6 | 0 | 2988.548 | 0 | 1.67 | 0.003434 | 19.4 | 47.1 | 152.9 | 13.1 | 15.72 | | High | High | High | High | High | High | High | High | High | High | High | 0.001812 | 0.01811 | 3.52 |
| FALSE | High | VTPNLSR | [R].VTPNLS | 1xPhospho | Q8VEF1 1x | 0.083101 | 0.006345 | 1 | 5 | 1 | Q8VEF1 | GRAM don | Q8VEF1 [2 | 0 | 1095.576 | 0 | 1.61 | 0.003166 | 20.6 | 51.5 | 148.5 | 15.98 | 14.66 | | High | High | High | High | High | High | High | High | High | High | High | 0.003265 | 0.03479 | 1.8 |
| FALSE | High | VTPQSDGS | [K].VTPQSI | 1xPhospho | Q571M2 1 | 0.041781 | 0.003126 | 1 | 4 | 1 | Q571M2 | MKIAA402 | Q571M2 [5 | 0 | 1630.809 | 0 | -0.82 | 0.00471 | 4.11 | 126.5 | 73.5 | 8.27 | 9.3 | | High | High | High | High | High | High | High | High | High | High | High | 0.001609 | 0.01568 | 2.07 |
| FALSE | High | VVDYSQFC | [K].VVDYS( | 1xDeamida | Q80XU3 1x | 0.057203 | 0.004227 | 1 | 2 | 2 | Q80XU3 | Nuclear ub | Q80XU3 [1 | 1 | 3429.505 | 0 | -0.7 | 0.040247 | 15.25 | 134.8 | 65.2 | 18.69 | 26.25 | | High | High | High | High | High | High | High | High | High | High | High | 0.002159 | 0.02243 | 3.29 |
| FALSE | High | YEEPEYDVI | [K].YEEPEY | 1xPhospho | O89032 1x | 2.86E-07 | 0 | 1 | 3 | 4 | O89032 | SH3 and PX | O89032 [5 | 0 | 3460.393 | 0 | 0.22 | 0.20057 | 5.15 | 92.5 | 107.5 | 9.73 | 15.14 | | High | High | High | High | High | High | High | High | High | High | High | 0 | 2.5E-08 | 5.03 |
| FALSE | High | VVEEMSE | [K].VVEEM | 1xPhospho | Q3TI79 1x | 0.009585 | 0.000736 | 1 | 3 | 13 | Q3TI79 | Putative ur | Q3TI79 [50 | 0 | 1147.479 | 0 | -0.2 | 0.108185 | 10.42 | 105.7 | 94.3 | 3.68 | 14.4 | | High | High | High | High | High | High | High | High | High | High | High | 0.000801 | 0.00719 | 1.93 |
| FALSE | High | VVEPELMG | [K].VVEPEL | 1xCarbami | B2RQC6 1x | 8.25E-05 | 0 | 1 | 6 | 1 | B2RQC6 | CAD protei | B2RQC6 [1 | 0 | 2687.281 | 0 | -0.2 | 0.647754 | 24.65 | 107 | 93 | 9.01 | 17.63 | | High | High | High | High | High | High | High | High | High | High | High | 0 | 1.45E-05 | 3.68 |
| FALSE | High | VVSSTSEEE | [R].VVSSTS | 2xPhospho | E9Q0A7 2x | 0.000267 | 0 | 2 | 6 | 1 | E9Q0A7; Q | Oxidation | E9Q0A7 [1 | 0 | 2290.018 | 0 | 0.12 | 0.39518 | 7.48 | 95.9 | 104.1 | 7.24 | 21.25 | Shared | High | High | High | High | High | High | High | High | High | High | High | 0 | 5.42E-05 | 3.5 |
| FALSE | High | VVVSQTK | [K].VVVSQ | 1xPhospho | P09405 1x | 0.049549 | 0.00361 | 1 | 8 | 2 | P09405 | Nucleolin ( | P09405 [64 | 0 | 1298.749 | 0 | -1.01 | 0.000234 | 1.74 | 133.6 | 66.4 | 3.71 | 8.45 | | High | High | High | High | High | High | High | High | High | High | High | 0.002976 | 0.03109 | 1.45 |
| FALSE | High | VVYAFSPK | [K].VVYAF | 1xPhospho | Q811P8 1x | 0.003622 | 0.00021 | 1 | 3 | 1 | Q811P8 | Rho GTPas | Q811P8 [8 | 0 | 1448.795 | 0 | -1.53 | 0.000438 | 0.91 | 150.6 | 49.4 | 8.8 | 14.03 | | High | High | High | High | High | High | High | High | High | High | High | 9.42E-05 | 0.001008 | 1.8 |
| FALSE | High | VVYMNSLL | [K].VVYMN | 1xDeamida | Q5U467 1x | 0.077825 | 0.005963 | 1 | 3 | 1 | Q5U467 | F-box and | Q5U467 [2 | 1 | 2765.44 | 0 | -0.25 | 0.801758 | 21.37 | 107.6 | 92.4 | 16.63 | 17.04 | | High | High | High | High | High | High | High | High | High | High | High | 0.003122 | 0.03224 | 2.42 |
| FALSE | High | VWVLRDSE | [R].VWVLR | 1xPhospho | E9Q7D5 1x | 1.03E-05 | 0 | 1 | 1 | 2 | E9Q7D5 | Rho guanir | E9Q7D5 [3 | 1 | 2371.076 | 0 | 2.28 | 0.000474 | 9.29 | 37.8 | 162.2 | 22.98 | 9.32 | | High | High | High | High | High | High | High | High | High | High | High | 0 | 4.23E-06 | 3.29 |
| FALSE | High | VYEDSGIPL | [K].VYEDS( | 1xPhospho | Q9DCT6-3 | 0.0002 | 0 | 2 | 10 | 1 | Q9DCT6-3; | Isoform 3 ( | Q9DCT6-3 | 0 | 1910.935 | 0 | -0.39 | 0.388395 | 18.09 | 110.5 | 89.5 | 36.12 | 28.52 | Shared | High | High | High | High | High | High | High | High | High | High | High | 0 | 3.9E-05 | 3.7 |
| FALSE | High | VYEDSGIPL | [K].VYEDS( | 1xPhospho | Q9DCT6-3 | 2.18E-07 | 0 | 2 | 10 | 3 | Q9DCT6-3; | Isoform 3 ( | Q9DCT6-3 | 0 | 2140.098 | 0 | -0.43 | 0.018679 | 2.55 | 113.6 | 86.4 | 6.49 | 10.71 | Shared | High | High | High | High | High | High | High | High | High | High | High | 0 | 6.72E-06 | 3.03 |
| FALSE | High | VYEDSGIPL | [K].VYEDS( | 1xPhospho | Q9DCT6-3 | 8.06E-06 | 0 | 2 | 10 | 4 | Q9DCT6-3; | Isoform 3 ( | Q9DCT6-3 | 1 | 2268.193 | 0 | -0.2 | 0.41131 | 34.24 | 116.4 | 83.6 | 57.66 | 42.76 | Shared | High | High | High | High | High | High | High | High | High | High | High | 0 | 1.27E-05 | 4.07 |
| FALSE | High | VYEDSGIPL | [K].VYEDS( | 1xPhospho | Q9DCT6-3 | 0.009756 | 0.000736 | 2 | 10 | 2 | Q9DCT6-3; | Isoform 3 ( | Q9DCT6-3 | 1 | 2497.356 | 0 | -0.56 | 0.01956 | 4.45 | 119.3 | 80.7 | 6.96 | 15.33 | Shared | High | High | High | High | High | High | High | High | High | High | High | 0.002187 | 0.02323 | 2.65 |
| FALSE | High | VYELLGGEI | [K].VYELLG | 1xPhospho | P52734 1x | 0.08447 | 0.006345 | 1 | 4 | 1 | P52734 | FYVE, Rho( | P52734 [57 | 0 | 2287.187 | 0 | -1.18 | 0.020349 | 2.9 | 136.7 | 63.3 | 15.19 | 24.31 | | High | High | High | High | High | High | High | High | High | High | High | 0.003265 | 0.03523 | 2.55 |
| FALSE | High | VYFQSPPG | [R].VYFQSI | 1xPhospho | Q62084 1x | 2.51E-05 | 0 | 1 | 1 | 4 | Q62084 | Protein ph | Q62084 [2 | 0 | 2625.182 | 0 | 0.09 | 0.049852 | 5.68 | 96.6 | 103.4 | 7.18 | 2.86 | | High | High | High | High | High | High | High | High | High | High | High | 0 | 3.79E-06 | 4 |

| | | | | | | | | | | | | | | | | | | | | | | | | | | | | | | | | | | | | | | |
|---|---|---|---|---|---|---|---|---|---|---|---|---|---|---|---|---|---|---|---|---|---|---|---|---|---|---|---|---|---|---|---|---|---|---|---|---|---|---|
| FALSE | High | VYLSDSDN | [R].VYLSDS | 2xPhospho | Q921C3 2x | 3.27E-06 | 0 | 1 | 3 | 2 | Q921C3 | Bromodom | Q921C3 [1 | 0 | 2660.178 | 0 | -1.01 | 0.001933 | 5.13 | 133.5 | 66.5 | 8.02 | 8.77 | | High | High | High | High | High | High | High | High | High | High | High | 0 | 3.84E-07 | 5.02 |
| FALSE | High | VYLSPGSR | [K].VYLSPG | 1xPhospho | P39689 1xl | 0.071071 | 0.005247 | 1 | 3 | 2 | P39689 | Cyclin-dep | P39689 [75 | 0 | 1187.602 | 0 | 3.25 | 0.000222 | 10.12 | 19.1 | 180.9 | 12.26 | 6.43 | | High | High | High | High | High | High | High | High | High | High | High | 0.002703 | 0.02902 | 1.35 |
| FALSE | High | VVSSTSEEE | [R].VVSSTS | 1xPhospho | E9Q0A7 1x | 4.39E-05 | 0 | 2 | 6 | 11 | E9Q0A7; Q | Oxidation r | E9Q0A7 [1 | 0 | 2210.052 | 0 | 0.45 | 0.007752 | 5.89 | 84.3 | 115.7 | 7.56 | 7.35 | Shared | High | High | High | High | High | High | High | High | High | High | High | 2.58E-05 | 0.000423 | 3.82 |
| FALSE | High | VYSLPDGTI | [R].VYSLPD | 1xPhospho | O70194 1x | 0.000656 | 0 | 1 | 1 | 5 | O70194 | Eukaryotic | O70194 [5 | 0 | 3877.459 | 0 | -1.4 | 0.005326 | 1.66 | 145.4 | 54.6 | 7.8 | 29.12 | | High | High | High | High | High | High | High | High | High | High | High | 0 | 0.000148 | 5.31 |
| FALSE | High | VYSPAHTEI | [K].VYSPAH | 2xPhospho | E9QAQ7 2x | 9.36E-08 | 0 | 1 | 5 | 3 | E9QAQ7 | AT-rich inte | E9QAQ7 [1 | 0 | 2757.21 | 0 | -0.81 | 0.000445 | 3.36 | 127.6 | 72.4 | 3.83 | 5.04 | | High | High | High | High | High | High | High | High | High | High | High | 0 | 2.81E-05 | 4.1 |
| FALSE | High | VYSSSEFLA | [K].VYSSSE | 1xPhospho | E9Q8S5 1x | 0.00015 | 0 | 1 | 2 | 8 | E9Q8S5 | Microtubu | E9Q8S5 [7 | 0 | 2438.184 | 0 | -0.45 | 0.008655 | 5.8 | 116.9 | 83.1 | 6.07 | 6.96 | | High | High | High | High | High | High | High | High | High | High | High | 0 | 6.83E-05 | 3.82 |
| FALSE | High | VYTHEVVT | [R].VYTHEV | 1xPhospho | P11440 1xl | 0.025073 | 0.001645 | 1 | 4 | 4 | P11440 | Cyclin-dep | P11440 [15 | 0 | 1874.94 | 0 | -0.15 | 0.274898 | 28.51 | 103.6 | 96.4 | 31.07 | 23.23 | | High | High | High | High | High | High | High | High | High | High | High | 0.000872 | 0.0088 | 2.31 |
| FALSE | High | VYTYIQSR | [KR].VYTYI | 1xPhospho | Q5U4C9 1x | 0.004696 | 0.000456 | 1 | 3 | 1 | Q5U4C9 | Dual specif | Q5U4C9 [3 | 0 | 1338.666 | 0 | 1.34 | 0.001189 | 9.5 | 55.3 | 144.7 | 12.63 | 9.25 | | High | High | High | High | High | High | High | High | High | High | High | 0.000204 | 0.001345 | 2.31 |
| FALSE | High | VYVPDTLS | [R].VYVPD | 1xPhospho | Q80WE4 1 | 0.079766 | 0.00613 | 1 | 4 | 2 | Q80WE4 | Kinesin-like | Q80WE4 [4 | 0 | 1991.014 | 0 | -0.24 | 0.09082 | 2.13 | 108.2 | 91.8 | 20.1 | 8.65 | | High | High | High | High | High | High | High | High | High | High | High | 0.004769 | 0.052 | 2.59 |
| FALSE | High | VYYSPPVAI | [R].VYYSPP | 1xPhospho | A2ACV6 1x | 0.001386 | 5.66E-05 | 1 | 4 | 3 | A2ACV6 | Protein SO | A2ACV6 [1 | 0 | 1360.686 | 0 | 1.68 | 0.000167 | 4.2 | 47.3 | 152.7 | 3.89 | 3.56 | | High | High | High | High | High | High | High | High | High | High | High | 2.58E-05 | 0.000344 | 2.69 |
| FALSE | High | WAAHGTSI | [R].WAAH | 1xPhospho | D3Z0X5 1xl | 3.7E-05 | 0 | 1 | 6 | 2 | D3Z0X5 | Pleckstrin | D3Z0X5 [4 | 0 | 2225.059 | 0 | 1.04 | 0.002367 | 11.6 | 61.8 | 138.2 | 5.73 | 12.65 | | High | High | High | High | High | High | High | High | High | High | High | 0 | 0.000124 | 2.75 |
| FALSE | High | WAALVVPS | [K].WAALV | 1xPhospho | Q62321 1x | 0.085162 | 0.006376 | 1 | 2 | 2 | Q62321 | TL antigen | Q62321 [2 | 0 | 1952.029 | 0 | 1.1 | 0.241298 | 11.62 | 74.7 | 125.3 | 25.1 | 40.69 | | High | High | High | High | High | High | High | High | High | High | High | 0.003291 | 0.03587 | 1.97 |
| FALSE | High | WAGKPPEI | [K].WAGKP | 1xCarbami | Q80YV2 1x | 0.025964 | 0.001645 | 1 | 5 | 1 | Q80YV2 | Nuclear-int | Q80YV2 [9 | 0 | 2434.354 | 0 | -0.18 | 0.26249 | 5.2 | 105.9 | 94.1 | 16.12 | 9.82 | | High | High | High | High | High | High | High | High | High | High | High | 0.000872 | 0.009126 | 2.59 |
| FALSE | High | WAHDKFS( | [K].WAHDK | 1xPhospho | Q569Z6 1x | 0.000248 | 0 | 1 | 3 | 5 | Q569Z6 | Thyroid ho | Q569Z6 [9 | 1 | 3175.389 | 0 | -0.11 | 0.122712 | 6.53 | 106.6 | 93.4 | 5 | 15.06 | | High | High | High | High | High | High | High | High | High | High | High | 0 | 4.97E-05 | 4.39 |
| FALSE | High | WAHDKFS( | [K].WAHDK | 2xPhospho | Q569Z6 2x | 0.000322 | 0 | 1 | 3 | 4 | Q569Z6 | Thyroid ho | Q569Z6 [9 | 1 | 3255.356 | 0 | -0.03 | 0.419928 | 13.17 | 101.2 | 98.8 | 3.81 | 20 | | High | High | High | High | High | High | High | High | High | High | High | 0 | 6.7E-05 | 5.25 |
| FALSE | High | WALAWSF | [R].WALAV | 1xPhospho | Q5SW15 1 | 0.000352 | 0 | 1 | 2 | 1 | Q5SW15 | Methyltrar | Q5SW15 [3 | 0 | 2704.346 | 0 | -0.12 | 0.043953 | 2.85 | 105.2 | 94.8 | 11.22 | 5.72 | | High | High | High | High | High | High | High | High | High | High | High | 0 | 7.37E-05 | 3.35 |
| FALSE | High | WANETLEK | [K].WANET | 1xPhospho | G3X959 1x | 0.085859 | 0.006406 | 1 | 6 | 1 | G3X959 | Periphilin 1 | G3X959 [2 | 0 | 1528.781 | 0 | -0.37 | 0.027211 | 9.51 | 112.2 | 87.8 | 7.31 | 21.31 | | High | High | High | High | High | High | High | High | High | High | High | 0.003303 | 0.03615 | 1.55 |
| FALSE | High | VYSPAHTEI | [K].VYSPAH | 1xPhospho | E9QAQ7 1x | 7.11E-10 | 0 | 1 | 5 | 12 | E9QAQ7 | AT-rich inte | E9QAQ7 [1 | 0 | 2677.243 | 0 | -0.93 | 0.000391 | 2.44 | 130.1 | 69.9 | 4.7 | 4.41 | | High | High | High | High | High | High | High | High | High | High | High | 0 | 2.94E-11 | 3.56 |
| FALSE | High | VVSQSQPG | [K].VVSQS( | 2xPhospho | B2RY22 2xl | 0.003788 | 0.00021 | 2 | 7 | 1 | B2RY22; A( | Clasp1 pro | B2RY22 [66 | 0 | 1433.639 | 0 | -0.59 | 0.004825 | 3.48 | 114.1 | 85.9 | 12.02 | 11.9 | Shared | High | High | High | High | High | High | High | High | High | High | High | 0.000117 | 0.001063 | 2.34 |
| FALSE | High | VVSQSQPG | [K].VVSQS( | 1xPhospho | B2RY22 1xl | 0.000548 | 0 | 2 | 7 | 3 | B2RY22; A( | Clasp1 pro | B2RY22 [66 | 0 | 1353.673 | 0 | -0.65 | 0.000636 | 4.53 | 125 | 75 | 7.61 | 16.01 | Shared | High | High | High | High | High | High | High | High | High | High | High | 7.19E-05 | 0.00091 | 2.04 |
| FALSE | High | VVSPPEPEK | [K].VVSPPE | 1xPhospho | Q8VDP4 1x | 0.046291 | 0.003523 | 1 | 1 | 1 | Q8VDP4 | Cell cycle a | Q8VDP4 [5 | 0 | 1519.817 | 0 | 0.42 | 0.003754 | 6.72 | 84.6 | 115.4 | 1.56 | 7.24 | | High | High | High | High | High | High | High | High | High | High | High | 0.001789 | 0.01761 | 1.88 |
| FALSE | High | VVEPELMG | [K].VVEPEL | 1xCarbami | B2RQC6 1x | 5.98E-06 | 0 | 1 | 6 | 1 | B2RQC6 | CAD protei | B2RQC6 [1 | 0 | 2703.276 | 0 | -0.34 | 0.033452 | 4.63 | 109.5 | 90.5 | 10.39 | 5.71 | | High | High | High | High | High | High | High | High | High | High | High | 0 | 7.57E-07 | 3.76 |
| FALSE | High | VVEPSETLT | [K].VVEPSE | 1xPhospho | H9KV00 1x | 1.29E-05 | 0 | 1 | 7 | 4 | H9KV00 | Protein SO | H9KV00 [2 | 0 | 4565.137 | 0 | 0.67 | 0.243569 | 40.88 | 77.2 | 122.8 | 14.25 | 32.74 | | High | High | High | High | High | High | High | High | High | High | High | 0.000204 | 0.001369 | 5.08 |
| FALSE | High | VVEPSETLT | [K].VVEPSE | 1xPhospho | H9KV00 1x | 0.00195 | 5.66E-05 | 1 | 7 | 3 | H9KV00 | Protein SO | H9KV00 [2 | 0 | 4581.132 | 0 | 0.72 | 0.269767 | 77.64 | 83.7 | 116.3 | 36.28 | 53.32 | | High | High | High | High | High | High | High | High | High | High | High | 2.58E-05 | 0.000555 | 4.07 |
| FALSE | High | VVEPSETLT | [K].VVEPSE | 1xDeamida | H9KV00 1x | 0.008614 | 0.000736 | 1 | 7 | 6 | H9KV00 | Protein SO | H9KV00 [2 | 0 | 4582.116 | 0 | -0.2 | 0.308239 | 8.42 | 110.8 | 89.2 | 16.38 | 24.22 | | High | High | High | High | High | High | High | High | High | High | High | 0.002675 | 0.02864 | 3.98 |
| FALSE | High | VVFENEQD | [R].VVFENI | 1xPhospho | A2AAE1-3 | 1.37E-05 | 0 | 1 | 6 | 2 | A2AAE1-3 | Isoform 3 c | A2AAE1-3 | 0 | 2175.074 | 0 | -0.66 | 0.000268 | 2.07 | 122.4 | 77.6 | 1.69 | 5.37 | | High | High | High | High | High | High | High | High | High | High | High | 0 | 1.93E-06 | 3.39 |
| FALSE | High | VVHGLQET | [R].VVHGL | 1xPhospho | Q80X32 1x | 8.06E-06 | 0 | 1 | 1 | 1 | Q80X32 | UPF0461 p | Q80X32 [1 | 0 | 2331.2 | 0 | 0.01 | 0.587938 | 5.07 | 98.5 | 101.5 | 3.3 | 12.12 | | High | High | High | High | High | High | High | High | High | High | High | 0 | 1.06E-06 | 2.3 |
| FALSE | High | VVLASASPI | [R].VVLASA | 1xPhospho | Q9D927 1x | 0.004033 | 0.000311 | 1 | 1 | 1 | Q9D927 | Putative ur | Q9D927 [1 | 0 | 1208.66 | 0 | 1.36 | 0.000827 | 13.58 | 56.2 | 143.8 | 8.48 | 8.61 | | High | High | High | High | High | High | High | High | High | High | High | 0.000161 | 0.001139 | 1.4 |
| FALSE | High | VVLGDSVD | [R].VVLGD | 1xPhospho | P49070 1xl | 1.15E-08 | 0 | 1 | 1 | 2 | P49070 | Calcium sig | P49070 [83 | 0 | 2379.175 | 0 | 0.67 | 0.056676 | 30.36 | 83.6 | 116.4 | 13.58 | 17.34 | | High | High | High | High | High | High | High | High | High | High | High | 0 | 6.78E-10 | 5.11 |
| FALSE | High | VVIIQPHED | [K].VVIIQPI | 1xPhospho | B9EK76 1xl | 0.002793 | 0.000109 | 1 | 4 | 1 | B9EK76 | Signal-indu | B9EK76 [1 | 0 | 1755.899 | 0 | -0.42 | 0.01642 | 7.61 | 116.7 | 83.3 | 8.56 | 8.98 | | High | High | High | High | High | High | High | High | High | High | High | 4.94E-05 | 0.000758 | 2.34 |
| FALSE | High | VVLSDSPQ | [R].VVLSDS | 1xPhospho | Q6NV99 1x | 1.24E-05 | 0 | 1 | 6 | 1 | Q6NV99 | HAUS augn | Q6NV99 [5 | 0 | 1638.83 | 0 | 0.35 | 0.001684 | 4.92 | 85 | 115 | 3.45 | 5.31 | | High | High | High | High | High | High | High | High | High | High | High | 0 | 1.73E-06 | 4.11 |
| FALSE | High | VVLSLQHA | [R].VVLSLC | 1xPhospho | A0A1B0GS | 0.0003 | 0 | 1 | 2 | 2 | A0A1B0GS | Nuclear po | A0A1B0GS | 0 | 2542.25 | 0 | -0.16 | 0.202671 | 10.55 | 106.6 | 93.4 | 14.5 | 8.98 | | High | High | High | High | High | High | High | High | High | High | High | 0 | 6.17E-05 | 2.74 |
| FALSE | High | VVNGAAAS | [K].VVNGA | 1xPhospho | G5E866 1x | 4.2E-05 | 0 | 1 | 5 | 1 | G5E866 | Splicing fac | G5E866 [1 | 0 | 1763.946 | 0 | 0.15 | 0.859666 | 13.25 | 97.7 | 102.3 | 21.08 | 8.16 | | High | High | High | High | High | High | High | High | High | High | High | 0 | 6.79E-06 | 1.64 |
| FALSE | High | VVNGAAAS | [K].VVNGA | 1xDeamida | G5E866 1x | 0.000194 | 0 | 1 | 5 | 2 | G5E866 | Splicing fac | G5E866 [1 | 0 | 1764.93 | 0 | 0.15 | 0.621829 | 12.82 | 99.1 | 100.9 | 14.73 | 5.33 | | High | High | High | High | High | High | High | High | High | High | High | 0.000467 | 0.004301 | 2.19 |
| FALSE | High | VVNGGVPV | [R].VVNGG | 1xCarbami | E9QM77 1 | 5.12E-05 | 0 | 1 | 3 | 1 | E9QM77 | Ataxin-2 O | E9QM77 [5 | 0 | 2626.28 | 0 | 0.31 | 0.030461 | 8.62 | 95.8 | 104.2 | 15.6 | 12.11 | | High | High | High | High | High | High | High | High | High | High | High | 0 | 8.48E-06 | 3.49 |
| FALSE | High | VVNGGVPV | [R].VVNGG | 1xCarbami | E9QM77 1 | 0.020862 | 0.001472 | 1 | 3 | 2 | E9QM77 | Ataxin-2 O | E9QM77 [5 | 0 | 2627.264 | 0 | 0.2 | 0.090148 | 5 | 94.6 | 105.4 | 8.2 | 11.56 | | High | High | High | High | High | High | High | High | High | High | High | 0.000801 | 0.00718 | 2.33 |
| FALSE | High | VVNGGVPV | [R].VVNGG | 1xCarbami | E9QM77 2 | 0.002398 | 0.000109 | 1 | 3 | 1 | E9QM77 | Ataxin-2 O | E9QM77 [5 | 0 | 2707.23 | 0 | 0.51 | 0.095634 | 4.82 | 87.4 | 112.6 | 23.15 | 12.46 | | High | High | High | High | High | High | High | High | High | High | High | 4.94E-05 | 0.000635 | 2.98 |
| FALSE | High | VVNLNGEL | [K].VVNLN | 1xPhospho | Q9R0M4 1 | 1.67E-05 | 0 | 1 | 2 | 1 | Q9R0M4 | Podocalyxi | Q9R0M4 [4 | 0 | 2834.511 | 0 | 3.53 | 0.006 | 31.94 | 14.5 | 185.5 | 8.96 | 30.12 | | High | High | High | High | High | High | High | High | High | High | High | 0 | 2.41E-06 | 2.95 |
| FALSE | High | VVNLNGEL | [K].VVNLN | 1xDeamida | Q9R0M4 1 | 0.000815 | 0 | 1 | 2 | 1 | Q9R0M4 | Podocalyxi | Q9R0M4 [4 | 0 | 2835.495 | 0 | 3.34 | 0.00419 | 39.36 | 10 | 190 | 39.47 | 26.33 | | High | High | High | High | High | High | High | High | High | High | High | 0 | 0.00019 | 3.04 |
| FALSE | High | VVNITSPTH | [R].VVNITS | 1xPhospho | Q9JHC9 1x | 0.000993 | 0 | 1 | 7 | 1 | Q9JHC9 | ETS-relatec | Q9JHC9 [3 | 0 | 1778.864 | 0 | -0.29 | 0.010221 | 6.18 | 110.8 | 89.2 | 4.31 | 4.69 | | High | High | High | High | High | High | High | High | High | High | High | 0 | 0.000237 | 1.76 |
| FALSE | High | VVPGPATS | [K].VVPGP | 1xPhospho | Q91Z58 1x | 0.021046 | 0.001507 | 1 | 1 | 2 | Q91Z58 | Uncharacte | Q91Z58 [6 | 0 | 1490.838 | 0 | 2.87 | 0.000154 | 14.49 | 25 | 175 | 14.68 | 3.14 | | High | High | High | High | High | High | High | High | High | High | High | 0.002159 | 0.02245 | 1.89 |
| FALSE | High | VVPIASLTP | [K].VVPIAS | 1xPhospho | Q3U8B3 1x | 0.032548 | 0.002167 | 1 | 3 | 1 | Q3U8B3 | Putative ur | Q3U8B3 [2 | 0 | 1941.086 | 0 | -1.11 | 0.002225 | 5.57 | 133 | 67 | 5.49 | 17.09 | | High | High | High | High | High | High | High | High | High | High | High | 0.001142 | 0.01185 | 2.36 |
| FALSE | High | VVPPDGHN | [K].VVPPD( | 1xPhospho | E9Q2Z1 1x | 0.000308 | 0 | 1 | 4 | 2 | E9Q2Z1 | CECR2, hist | E9Q2Z1 [1 | 0 | 2129.032 | 0 | -0.95 | 0.001753 | 3.22 | 131.7 | 68.3 | 5.93 | 14.01 | | High | High | High | High | High | High | High | High | High | High | High | 0 | 6.32E-05 | 2.64 |
| FALSE | High | VVPQQITH | [K].VVPQQ | 1xPhospho | J3QNK5 1x | 1.46E-05 | 0 | 1 | 4 | 4 | J3QNK5 | Histone de | J3QNK5 [4 | 0 | 1671.878 | 0 | -0.79 | 0.001907 | 5 | 125.4 | 74.6 | 6.73 | 4.78 | | High | High | High | High | High | High | High | High | High | High | High | 0.000505 | 0.004348 | 3.08 |
| FALSE | High | VVPQRSEI( | [K].VVPQR | 1xPhospho | Q6A065 1x | 0.008094 | 0.000736 | 1 | 2 | 1 | Q6A065 | Centrosom | Q6A065 [6 | 1 | 1779.977 | 0 | 1.64 | 0.009968 | 32.37 | 46.9 | 153.1 | 12.07 | 25.84 | | High | High | High | High | High | High | High | High | High | High | High | 0.000327 | 0.002465 | 2.19 |
| FALSE | High | VVQEYIDAI | [K].VVQEY | 1xPhospho | P18052 1xl | 3.66E-08 | 0 | 1 | 4 | 1 | P18052 | Receptor-t | P18052 [81 | 0 | 2447.194 | 0 | 1.17 | 0.001788 | 17.04 | 63 | 137 | 9.49 | 10.7 | | High | High | High | High | High | High | High | High | High | High | High | 0 | 2.47E-09 | 4.1 |
| FALSE | High | VVSEICETE | [R].VVSEIC | 1xCarbami | E9QPB3 1x | 3.49E-07 | 0 | 1 | 5 | 2 | E9QPB3 | WD repeat | E9QPB3 [1 | 0 | 2432.167 | 0 | -1.06 | 0.000269 | 0.97 | 133.8 | 66.2 | 5.17 | 7.88 | | High | High | High | High | High | High | High | High | High | High | High | 0 | 3.11E-08 | 3.57 |
| FALSE | High | VVSHDDSL | [R].VVSHD | 1xPhospho | G3X8R8 1x | 0.006085 | 0.00062 | 1 | 4 | 5 | G3X8R8 | MCG12388 | G3X8R8 [5 | 0 | 3126.55 | 0 | 0.41 | 0.012362 | 0.53 | 81 | 119 | 8.95 | 10.83 | | High | High | High | High | High | High | High | High | High | High | High | 0.000275 | 0.001808 | 2.42 |
| FALSE | High | VVSISSEHL | [K].VVSISS | 1xPhospho | O55201 1x | 0.004572 | 0.000456 | 1 | 2 | 1 | O55201 | Transcripti | O55201 [1 | 0 | 2275.235 | 0 | -1.23 | 0.01511 | 4.06 | 143.5 | 56.5 | 18.71 | 20.1 | | High | High | High | High | High | High | High | High | High | High | High | 0.000204 | 0.001315 | 2.64 |
| FALSE | High | VVSISSSDF | [R].VVSISS | 1xPhospho | Q69ZR9-2 | 0.002293 | 0.000109 | 1 | 2 | 2 | Q69ZR9-2 | Isoform 2 c | Q69ZR9-2 | 0 | 1764.919 | 0 | 0.84 | 0.015782 | 23.74 | 76.5 | 123.5 | 16.58 | 12.57 | | High | High | High | High | High | High | High | High | High | High | High | 4.94E-05 | 0.000739 | 2.31 |
| FALSE | High | VVELSPLK | [R].VVELSP | 1xPhospho | H9LAN6 1x | 0.028327 | 0.001791 | 1 | 6 | 1 | H9LAN6 | Histone-lys | H9LAN6 [2 | 0 | 1422.837 | 0 | -1.36 | 0.000233 | 0.73 | 143.1 | 56.9 | 2.92 | 8.54 | | High | High | High | High | High | High | High | High | High | High | High | 0.000968 | 0.01008 | 1.63 |
| FALSE | High | VTFGEDLSI | [R].VTFGEI | 1xCarbami | Q14B71 1x | 2.24E-06 | 0 | 1 | 2 | 1 | Q14B71 | Cell divisio | Q14B71 [3 | 0 | 3203.562 | 0 | -0.72 | 0.00959 | 0.4 | 126.1 | 73.9 | 14.21 | 8.46 | | High | High | High | High | High | High | High | High | High | High | High | 0 | 2.52E-07 | 3.68 |
| FALSE | High | YEESLADSI | [K].YEESLA | 1xPhospho | Q6KCD5 1x | 0.064862 | 0.004664 | 1 | 2 | 1 | Q6KCD5 | Nipped-B-l | Q6KCD5 [1 | 0 | 1579.673 | 0 | -0.46 | 0.209016 | 18.48 | 113.9 | 86.1 | 25.56 | 39.23 | | High | High | High | High | High | High | High | High | High | High | High | 0.002432 | 0.02613 | 2.31 |
| FALSE | High | YEHDSDLSI | [K].YEHDSI | 2xPhospho | Q8R149 2x | 0.003224 | 0.00016 | 1 | 3 | 1 | Q8R149 | BUD13 hor | Q8R149 [3 | 0 | 1704.687 | 0 | 0.16 | 0.164073 | 5.99 | 96.4 | 103.6 | 4.75 | 6.52 | | High | High | High | High | High | High | High | High | High | High | High | 7.19E-05 | 0.000883 | 2.34 |
| FALSE | High | YSDFEWLR | [R].YSDFEV | 1xPhospho | O70492 1x | 0.002719 | 0.000109 | 1 | 5 | 1 | O70492 | Sorting ne | O70492 [7 | 0 | 1424.645 | 0 | -0.27 | 0.092571 | 11.36 | 105.4 | 94.6 | 7.75 | 11.45 | | High | High | High | High | High | High | High | High | High | High | High | 4.94E-05 | 0.00073 | 2.61 |
| FALSE | High | YSDHAGPA | [R].YSDHA( | 1xPhospho | Q03141 1x | 0.00013 | 0 | 1 | 5 | 1 | Q03141 | MAP/micro | Q03141 [4 | 0 | 2310.194 | 0 | -0.69 | 0.006228 | 5 | 123.3 | 76.7 | 8.77 | 11.12 | | High | High | High | High | High | High | High | High | High | High | High | 0 | 2.41E-05 | 1.89 |
| FALSE | High | YSDLTTLVA | [R].YSDLTT | 1xPhospho | P25206 1xl | 3.01E-05 | 0 | 2 | 3 | 3 | P25206; Q | DNA replic | P25206 [15 | 0 | 2614.346 | 0 | -0.97 | 0.007117 | 4.35 | 132.4 | 67.6 | 10.61 | 14.29 | Shared | High | High | High | High | High | High | High | High | High | High | High | 0 | 5.38E-06 | 3.43 |
| FALSE | High | YSDSSLPPS | [K].YSDSSL | 1xPhospho | E9Q7E2 1x | 0.002313 | 0.000109 | 1 | 5 | 2 | E9Q7E2 | AT-rich inte | E9Q7E2 [1 | 0 | 1876.909 | 0 | 0.23 | 0.058835 | 6.34 | 95.2 | 104.8 | 10.11 | 12.55 | | High | High | High | High | High | High | High | High | High | High | High | 4.94E-05 | 0.000613 | 2.63 |
| FALSE | High | YSDVEVPA | [K].YSDVE\ | 1xCarbami | Q8K232 1x | 5.32E-06 | 0 | 2 | 7 | 2 | Q8K232; E | Adducin 1 | Q8K232 [4 | 0 | 3207.414 | 0 | 1.21 | 0.008648 | 28.76 | 61.3 | 138.7 | 14.74 | 15.28 | Shared | High | High | High | High | High | High | High | High | High | High | High | 0 | 1.1E-05 | 4.58 |
| FALSE | High | YSEPERPSS | [R].YSEPER | 3xPhospho | A2AIX1 3xl | 0.002844 | 0.00016 | 1 | 2 | 1 | A2AIX1 | SEC16 hom | A2AIX1 [12 | 1 | 2902.216 | 0 | -0.01 | 0.800141 | 29.88 | 97.1 | 102.9 | 19.1 | 13.28 | | High | High | High | High | High | High | High | High | High | High | High | 7.19E-05 | 0.000771 | 2.93 |
| FALSE | High | YSEWTVPA | [K].YSEWT | 1xPhospho | E9PZ97 1xl | 0.008388 | 0.000736 | 1 | 2 | 1 | E9PZ97 | Bloom syn | E9PZ97 [12 | 0 | 2030.906 | 0 | -0.83 | 0.001229 | 3.07 | 124.6 | 75.4 | 5.52 | 7.28 | | High | High | High | High | High | High | High | High | High | High | High | 0.000327 | 0.002565 | 2.16 |
| FALSE | High | YSGSSLSN | [K].YSGSSL | 1xPhospho | Q8K1N2-2 | 0.086562 | 0.006435 | 1 | 3 | 1 | Q8K1N2-2 | Isoform 2 c | Q8K1N2-2 | 0 | 1788.783 | 0 | 1.1 | 0.004707 | 18.14 | 63.7 | 136.3 | 20.77 | 20.86 | | High | High | High | High | High | High | High | High | High | High | High | 0.003303 | 0.03627 | 1.84 |
| FALSE | High | YSGSSLSN | [K].YSGSSL | 1xPhospho | Q8K1N2-2 | 0.001586 | 5.66E-05 | 1 | 3 | 2 | Q8K1N2-2 | Isoform 2 c | Q8K1N2-2 | 0 | 1804.777 | 0 | 2.31 | 0.000963 | 17.55 | 37.3 | 162.7 | 17.33 | 11.66 | | High | High | High | High | High | High | High | High | High | High | High | 2.58E-05 | 0.000399 | 2.15 |
| FALSE | High | YSIDPSVVN | [K].YSIDPS | 1xPhospho | I1VGT4 1xl | 0.012619 | 0.000889 | 2 | 13 | 1 | I1VGT4; I1 | Anion exch | I1VGT4 [10 | 0 | 2322.134 | 0 | 0.54 | 0.021365 | 22.18 | 71.9 | 128.1 | 11.08 | 17.28 | Shared | High | High | High | High | High | High | High | High | High | High | High | 0.000449 | 0.00407 | 2.38 |
| FALSE | High | YSLGSVTTS | [K].YSLGSV | 1xPhospho | Q8BS90 1x | 2.31E-07 | 0 | 1 | 2 | 5 | Q8BS90 | Protein au | Q8BS90 [2 | 0 | 3191.664 | 0 | -1.06 | 0.002419 | 4.84 | 127.9 | 72.1 | 9.78 | 12.12 | | High | High | High | High | High | High | High | High | High | High | High | 0 | 3.62E-08 | 5.18 |
| FALSE | High | YSIPDLAVD | [R].YSIPDL | 1xPhospho | A0A0A6YW | 0.001767 | 5.66E-05 | 1 | 4 | 1 | A0A0A6YW | Rap guanin | A0A0A6YW | 0 | 2526.351 | 0 | 2.66 | 0.005211 | 4.84 | 34.1 | 165.9 | 36.93 | 20.13 | | High | High | High | High | High | High | High | High | High | High | High | 2.58E-05 | 0.00045 | 2.3 |
| FALSE | High | YSLPSK | [K].YSLPSK | 1xPhospho | Q9Z277 1x | 0.104222 | 0.008348 | 1 | 1 | 1 | Q9Z277 | Tyrosine-p | Q9Z277 [2 | 0 | 1232.669 | 0 | 0.12 | 0.100569 | 6.02 | 93.4 | 106.6 | 8.21 | 9.14 | | High | High | High | High | High | High | High | High | High | High | High | 0.004206 | 0.0452 | 1.19 |
| FALSE | High | YSDEQLNS | [K].YSDEQ | 1xPhospho | B2RUG9 1x | 0.005768 | 0.00062 | 1 | 8 | 1 | B2RUG9 | Adenomat | B2RUG9 [1 | 1 | 2404.073 | 0 | -0.33 | 0.02869 | 3.17 | 107.5 | 92.5 | 8.11 | 14.01 | | High | High | High | High | High | High | High | High | High | High | High | 0.000275 | 0.001694 | 2.56 |
| FALSE | High | YSLSPSK | [K].YSLSPS | 1xPhospho | Q9ERU9 1x | 0.010382 | 0.000772 | 1 | 1 | 3 | Q9ERU9 | E3 SUMO-p | Q9ERU9 [7 | 0 | 1319.701 | 0 | -1.35 | 0.000207 | 1.03 | 144.3 | 55.7 | 5.61 | 6.28 | | High | High | High | High | High | High | High | High | High | High | High | 0.000382 | 0.00327 | 1.63 |
| FALSE | High | YSPDEMNN | [R].YSPDEN | 1xPhospho | Q3U3C9-2 | 0.000415 | 0 | 1 | 4 | 1 | Q3U3C9-2 | Isoform 2 c | Q3U3C9-2 | 0 | 2355.025 | 0 | -1.2 | 0.000567 | 1.39 | 138.3 | 61.7 | 4.4 | 9.34 | | High | High | High | High | High | High | High | High | High | High | High | 0 | 8.9E-05 | 2.37 |
| FALSE | High | YSPDQLCV | [K].YSPDQ | 1xCarbami | Q3USC0 1x | 7.75E-05 | 0 | 1 | 6 | 2 | Q3USC0 | Putative ur | Q3USC0 [6 | 0 | 2089.044 | 0 | -1.44 | 0.001349 | 0.58 | 146.1 | 53.9 | 6.92 | 14.66 | | High | High | High | High | High | High | High | High | High | High | High | 0 | 1.35E-05 | 2.83 |
| FALSE | High | YSPGQLR | [K].YSPGQ | 1xPhospho | Q9CZH7 1x | 0.057686 | 0.004227 | 1 | 2 | 1 | Q9CZH7 | Matrix-rem | Q9CZH7 [1 | 0 | 1129.56 | 0 | -1.17 | 0.001561 | 5.6 | 142 | 58 | 4.64 | 21.51 | | High | High | High | High | High | High | High | High | High | High | High | 0.002159 | 0.02267 | 2.12 |
| FALSE | High | YSPGYCSSI | [R].YSPGY( | 1xCarbami | Q91WC9 1 | 9.59E-07 | 0 | 1 | 1 | 5 | Q91WC9 | Sn1-specifi | Q91WC9 [5 | 0 | 2399.088 | 0 | 1.42 | 0.000405 | 3.02 | 53.6 | 146.4 | 9.07 | 6.64 | | High | High | High | High | High | High | High | High | High | High | High | 0 | 9.68E-08 | 4.15 |
| FALSE | High | YSPNTQVE | [R].YSPNT( | 1xPhospho | Q3TNN6 1x | 0.013308 | 0.000889 | 1 | 4 | 1 | Q3TNN6 | Putative ur | Q3TNN6 [3 | 1 | 2841.495 | 0 | 2.29 | 0.000805 | 28.42 | 33.9 | 166.1 | 11.43 | 10.13 | | High | High | High | High | High | High | High | High | High | High | High | 0.000486 | 0.00433 | 2.89 |
| FALSE | High | YSPSPPPK | [R].YSPSPP | 2xPhospho | Q52KI8 2xl | 0.008094 | 0.000736 | 1 | 7 | 2 | Q52KI8 | Serine/argi | Q52KI8 [62 | 0 | 1490.71 | 0 | -0.29 | 0.001168 | 2.65 | 109.1 | 90.9 | 3.76 | 7.08 | | High | High | High | High | High | High | High | High | High | High | High | 0.000327 | 0.002482 | 1.97 |
| FALSE | High | YSPSQNSPI | [R].YSPSQN | 1xPhospho | Q8K019 1x | 1.25E-06 | 0 | 1 | 10 | 6 | Q8K019 | Bcl-2-assoc | Q8K019 [2 | 0 | 2028.985 | 0 | -0.33 | 0.070774 | 10.15 | 115.4 | 84.6 | 13.03 | 10.13 | | High | High | High | High | High | High | High | High | High | High | High | 0 | 1.3E-07 | 3.96 |
| FALSE | High | YSPSQNSPI | [R].YSPSQN | 2xPhospho | Q8K019 2x | 0.005768 | 0.00062 | 1 | 10 | 5 | Q8K019 | Bcl-2-assoc | Q8K019 [2 | 0 | 2108.952 | 0 | -0.15 | 0.017178 | 3.54 | 104.1 | 95.9 | 5.04 | 6.25 | | High | High | High | High | High | High | High | High | High | High | High | 0.000275 | 0.001702 | 2.47 |
| FALSE | High | YSPSQNSPI | [R].YSPSQN | 1xDeamida | Q8K019 2x | 0.040721 | 0.003058 | 1 | 10 | 1 | Q8K019 | Bcl-2-assoc | Q8K019 [2 | 0 | 2109.936 | 0 | -0.92 | 0.000222 | 2.8 | 131 | 69 | 2.36 | 10.09 | | High | High | High | High | High | High | High | High | High | High | High | 0.001581 | 0.01522 | 1.93 |
| FALSE | High | YSPSQNVP | [R].YSPSQN | 1xPhospho | Q64127 1x | 0.024424 | 0.001612 | 1 | 5 | 1 | Q64127 | Transcripti | Q64127 [5 | 0 | 1356.651 | 0 | -0.05 | 0.812704 | 20.64 | 99.2 | 100.8 | 11.02 | 8.78 | | High | High | High | High | High | High | High | High | High | High | High | 0.000859 | 0.008509 | 2.26 |
| FALSE | High | YSPTSPK | [SKT].YSPT | 1xPhospho | P08775 1xl | 0.017344 | 0.001204 | 1 | 3 | 3 | P08775 | DNA-direct | P08775 [18 | 0 | 1317.686 | 0 | -0.56 | 0.112813 | 18.96 | 117.1 | 82.9 | 22.56 | 10.55 | | High | High | High | High | High | High | High | High | High | High | High | 0.009556 | 0.1032 | 1.44 |
| FALSE | High | YSPTSPTYS | [K].YSPTSP | 1xPhospho | P08775 1xl | 1.87E-07 | 0 | 1 | 3 | 9 | P08775 | DNA-direct | P08775 [19 | 0 | 2051.014 | 0 | -0.11 | 0.910555 | 35.95 | 102.9 | 97.1 | 17.37 | 13.5 | | High | High | High | High | High | High | High | High | High | High | High | 0 | 4.93E-07 | 3.13 |
| FALSE | High | YSPTSPTYS | [K].YSPTSP | 2xPhospho | P08775 2xl | 0.000681 | 0 | 1 | 3 | 4 | P08775 | DNA-direct | P08775 [19 | 0 | 2130.98 | 0 | -0.52 | 0.077851 | 7.4 | 118.4 | 81.6 | 17.68 | 4.8 | | High | High | High | High | High | High | High | High | High | High | High | 0.000161 | 0.001148 | 3.38 |
| FALSE | High | YSISSDNSD | [K].YSISSD | 3xPhospho | Q6ZQF7 3x | 0.000396 | 0 | 1 | 3 | 2 | Q6ZQF7 | E3 ubiquiti | Q6ZQF7 [8 | 0 | 2757.053 | 0 | 1.21 | 0.006818 | 30.46 | 60.9 | 139.1 | 19.03 | 13.68 | | High | High | High | High | High | High | High | High | High | High | High | 0 | 0.000199 | 3.91 |
| FALSE | High | YSPTSPTYS | [K].YSPTSP | 1xPhospho | P08775 1xl | 0.002022 | 5.66E-05 | 1 | 3 | 3 | P08775 | DNA-direct | P08775 [18 | 0 | 2796.379 | 0 | 0.23 | 0.201035 | 7.5 | 91.9 | 108.1 | 8.23 | 12.24 | | High | High | High | High | High | High | High | High | High | High | High | 0.003265 | 0.03454 | 2.8 |
| FALSE | High | YSASPTLR | [R].YSASPT | 1xPhospho | E9PVX6 1xl | 0.021231 | 0.001507 | 1 | 2 | 1 | E9PVX6 | Proliferatic | E9PVX6 [7 | 0 | 1203.597 | 0 | 0.02 | 0.426403 | 16.27 | 99.2 | 100.8 | 3.98 | 13.01 | | High | High | High | High | High | High | High | High | High | High | High | 0.000801 | 0.007314 | 2.18 |
| FALSE | High | YSAENDPSI | [K].YSAENI | 1xPhospho | F6RGI3 1xl | 2.08E-05 | 0 | 1 | 1 | 2 | F6RGI3 | Uncharacte | F6RGI3 [48 | 0 | 2861.24 | 0 | -0.16 | 0.062326 | 5.34 | 105.7 | 94.3 | 3.83 | 11 | | High | High | High | High | High | High | High | High | High | High | High | 0 | 3.06E-06 | 3.65 |
| FALSE | High | YPSGSEIPV | [R].YPSGSE | 1xPhospho | A0A1B0GS | 1.61E-05 | 0 | 1 | 5 | 6 | A0A1B0GS | Supervillin | A0A1B0GS | 0 | 2216.078 | 0 | 0.95 | 0.000442 | 7.85 | 71.7 | 128.3 | 4.97 | 5.15 | | High | High | High | High | High | High | High | High | High | High | High | 0 | 2.31E-06 | 3.75 |
| FALSE | High | YPSPPVGS\ | [R].YPSPPV | 1xPhospho | P47941 1xl | 5.92E-09 | 0 | 1 | 1 | 2 | P47941 | Crk-like pro | P47941 [1 | 0 | 2995.465 | 0 | -0.07 | 0.868661 | 27.5 | 102.4 | 97.6 | 16.67 | 11.15 | | High | High | High | High | High | High | High | High | High | High | High | 0 | 4.05E-09 | 4.76 |
| FALSE | High | YPSSISSSP( | [R].YPSSISS | 1xPhospho | Q80X50-5 | 0.001021 | 0 | 1 | 13 | 3 | Q80X50-5 | Isoform 5 c | Q80X50-5 | 0 | 1718.877 | 0 | -0.71 | 0.001286 | 3.37 | 123.2 | 76.8 | 4.97 | 8.51 | | High | High | High | High | High | High | High | High | High | High | High | 0 | 0.000243 | 3.17 |
| FALSE | High | YPVSPGSPI | [R].YPVSPG | 1xPhospho | A0A1B0GR | 0.000184 | 0 | 1 | 1 | 2 | A0A1B0GR | RIKEN cDN | A0A1B0GR | 0 | 1567.736 | 0 | 1.11 | 0.000717 | 9 | 65.4 | 134.6 | 7.12 | 7.86 | | High | High | High | High | High | High | High | High | High | High | High | 0 | 3.55E-05 | 1.95 |
| FALSE | High | YPVSPGSPI | [R].YPVSPG | 2xPhospho | A0A1B0GR | 0.000908 | 0 | 1 | 1 | 1 | A0A1B0GR | RIKEN cDN | A0A1B0GR | 0 | 1647.702 | 0 | 0.81 | 0.000873 | 13.15 | 72.5 | 127.5 | 6.53 | 4.84 | | High | High | High | High | High | High | High | High | High | High | High | 0 | 0.000213 | 2.33 |
| FALSE | High | YPVSPGSPI | [R].YPVSPG | 2xPhospho | A0A1B0GR | 0.012731 | 0.000889 | 1 | 1 | 1 | A0A1B0GR | RIKEN cDN | A0A1B0GR | 1 | 2004.96 | 0 | 0.66 | 0.001435 | 6.13 | 80.7 | 119.3 | 5.72 | 4.96 | | High | High | High | High | High | High | High | High | High | High | High | 0.000449 | 0.004092 | 2.15 |
| FALSE | High | YPWPSAAF | [R].YPWPS | 1xPhospho | Q8CI61 1xl | 0.003622 | 0.00021 | 1 | 1 | 1 | Q8CI61 | BAG family | Q8CI61 [25 | 0 | 4983.374 | 0 | -0.64 | 0.059124 | 1.8 | 121.9 | 78.1 | 9.68 | 28.42 | | High | High | High | High | High | High | High | High | High | High | High | 9.42E-05 | 0.001012 | 3.73 |
| FALSE | High | YQDEVFGG | [R].YQDEV | 1xPhospho | Q571F9 1x | 1.55E-07 | 0 | 1 | 3 | 27 | Q571F9 | MKIAA411 | Q571F9 [1 | 0 | 3525.494 | 0 | -0.2 | 0.023621 | 7.57 | 110.2 | 89.8 | 5.35 | 5.72 | | High | High | High | High | High | High | High | High | High | High | High | 0 | 1.87E-07 | 5.02 |
| FALSE | High | YQDEVFGG | [R].YQDEV | 1xDeamida | Q571F9 1x | 2.79E-08 | 0 | 1 | 3 | 7 | Q571F9 | MKIAA411 | Q571F9 [1 | 0 | 3526.478 | 0 | -0.22 | 0.086647 | 3.76 | 109.2 | 90.8 | 14.57 | 15.96 | | High | High | High | High | High | High | High | High | High | High | High | 0 | 1.82E-09 | 5.26 |
| FALSE | High | YQEADSEE | [R].YQEAD | 1xPhospho | Q6A046 1x | 0.032548 | 0.002167 | 1 | 3 | 1 | Q6A046 | Phospholip | Q6A046 [4 | 0 | 2180.005 | 0 | 1.45 | 0.000229 | 4.68 | 52.7 | 147.3 | 8.3 | 5.29 | | High | High | High | High | High | High | High | High | High | High | High | 0.001142 | 0.01185 | 2.96 |
| FALSE | High | YQEDSDPE | [K].YQEDSI | 1xPhospho | Q3TSG4 1x | 0.098533 | 0.007902 | 1 | 1 | 1 | Q3TSG4 | RNA deme | Q3TSG4 [6 | 1 | 2934.32 | 0 | -0.1 | 0.248744 | 9.43 | 100.4 | 99.6 | 5.52 | 5.57 | | High | High | High | High | High | High | High | High | High | High | High | 0.003963 | 0.04232 | 2.27 |
| FALSE | High | YQEDSDPE | [K].YQEDSI | 2xPhospho | Q3TSG4 2x | 0.001021 | 0 | 1 | 1 | 3 | Q3TSG4 | RNA deme | Q3TSG4 [6 | 1 | 3014.286 | 0 | -0.06 | 0.040676 | 3.2 | 102.2 | 97.8 | 4.14 | 2.95 | | High | High | High | High | High | High | High | High | High | High | High | 0 | 0.000245 | 2.76 |
| FALSE | High | YQEQELPP | [K].YQEQE | 1xPhospho | P10833 1xl | 0.001398 | 5.66E-05 | 1 | 1 | 2 | P10833 | Ras-relatec | P10833 [1 | 0 | 2004.963 | 0 | 0.77 | 0.000781 | 6.79 | 74.7 | 125.3 | 4.94 | 8.07 | | High | High | High | High | High | High | High | High | High | High | High | 2.58E-05 | 0.000347 | 2.5 |
| FALSE | High | YSALLSPSE | [K].YSALLS | 1xPhospho | D4AFX6 1x | 0.025073 | 0.001645 | 1 | 3 | 1 | D4AFX6 | Centaurin | D4AFX6 [5 | 0 | 1889.966 | 0 | 0.47 | 0.019401 | 7.64 | 81.5 | 118.5 | 22.79 | 5.88 | | High | High | High | High | High | High | High | High | High | High | High | 0.000872 | 0.008789 | 1.7 |
| FALSE | High | YQEVWTSS | [R].YQEVW | 1xPhospho | Q69Z38 1x | 0.041072 | 0.003094 | 1 | 1 | 2 | Q69Z38 | Pseudopod | Q69Z38 [5 | 0 | 1749.805 | 0 | -0.34 | 0.190706 | 22.63 | 109.9 | 90.1 | 12.79 | 14.97 | | High | High | High | High | High | High | High | High | High | High | High | 0.004707 | 0.05076 | 2.64 |
| FALSE | High | YQPSYETTI | [K].YQPSYE | 1xCarbami | Q3UNW5 1 | 0.000656 | 0 | 1 | 1 | 2 | Q3UNW5 | Transcripti | Q3UNW5 [ | 0 | 4944.234 | 0 | -6.64 | | 0 | 185.9 | 14.1 | 12.08 | 35.75 | | High | High | High | High | High | High | High | Not Found | Not Found | Not Found | High | 0 | 0.000296 | 4.03 |
| FALSE | High | YQQPFEDF | [R].YQQPF | 1xPhospho | Q62077 1x | 0.012845 | 0.000889 | 1 | 5 | 1 | Q62077 | 1-phospha | Q62077 [1 | 1 | 3074.4 | 0 | -0.54 | 0.175632 | 4.73 | 117.6 | 82.4 | 7.61 | 23.93 | | High | High | High | High | High | High | High | High | High | High | High | 0.000449 | 0.00417 | 2.14 |
| FALSE | High | YRDYSPPY | [K].YRDYSF | 1xPhospho | Q6ZPZ3 1xl | 0.005467 | 0.00062 | 1 | 6 | 2 | Q6ZPZ3 | Zinc finger | Q6ZPZ3 [1 | 1 | 3428.647 | 0 | 0.09 | 0.44594 | 8.34 | 106 | 94 | 20.95 | 13.63 | | High | High | High | High | High | High | High | High | High | High | High | 0.000449 | 0.003784 | 2.65 |
| FALSE | High | YRHTPTGN | [R].YRHTPT | 1xPhospho | Q8BN21 1x | 0.000628 | 0 | 1 | 2 | 1 | Q8BN21 | Serine/thre | Q8BN21 [4 | 1 | 2409.176 | 0 | 1.4 | 0.003573 | 20.58 | 55.4 | 144.6 | 15.71 | 10.6 | | High | High | High | High | High | High | High | High | High | High | High | 0 | 0.000142 | 3.49 |
| FALSE | High | YRSDIHTEA | [R].YRSDIH | 1xPhospho | Q3UH60 1x | 0.004572 | 0.000456 | 1 | 1 | 1 | Q3UH60 | Disco-inter | Q3UH60 [9 | 1 | 2311.221 | 0 | 0.65 | 0.033217 | 30.92 | 79 | 121 | 21.3 | 9.66 | | High | High | High | High | High | High | High | High | High | High | High | 0.000204 | 0.001312 | 2.34 |
| FALSE | High | YRSDVHTE. | [R].YRSDVH | 1xPhospho | E9PWR4 1x | 0.002419 | 0.000109 | 1 | 3 | 1 | E9PWR4 | Disco-inter | E9PWR4 [8 | 1 | 2297.206 | 0 | -0.73 | 0.007115 | 7.21 | 124 | 76 | 15.49 | 16.59 | | High | High | High | High | High | High | High | High | High | High | High | 4.94E-05 | 0.000641 | 2.98 |
| FALSE | High | YRSPEPDPY | [R].YRSPEP | 1xPhospho | O35492 1x | 0.000211 | 0 | 1 | 5 | 3 | O35492 | Dual specif | O35492 [1 | 1 | 1951.915 | 0 | -0.23 | 0.034083 | 3.57 | 107.9 | 92.1 | 1.18 | 6.98 | | High | High | High | High | High | High | High | High | High | High | High | 0 | 4.15E-05 | 2.6 |
| FALSE | High | YRSPPEGY | [K].YRSPPE | 1xPhospho | O88466 1x | 0.087983 | 0.006639 | 2 | 7 | 1 | O88466; R | Zinc finger | O88466 [3 | 1 | 3345.53 | 0 | -0.95 | 0.012974 | 8.81 | 131.7 | 68.3 | 10.87 | 27.68 | Shared | High | High | High | High | High | High | High | High | High | High | High | 0.003389 | 0.03698 | 2.83 |
| FALSE | High | YRSPPTVYI | [R].YRSPPT | 1xPhospho | Q8BRG8 1x | 0.004143 | 0.00036 | 1 | 5 | 1 | Q8BRG8 | Transmem | Q8BRG8 [2 | 1 | 2163.089 | 0 | -0.09 | 0.551074 | 20.53 | 102.6 | 97.4 | 13.53 | 10.84 | | High | High | High | High | High | High | High | High | High | High | High | 0.000161 | 0.001177 | 2.65 |

| | | | | | | | | | | | | | | | | | | | | | | | | | | | | | | | | | | | | | | |
|---|---|---|---|---|---|---|---|---|---|---|---|---|---|---|---|---|---|---|---|---|---|---|---|---|---|---|---|---|---|---|---|---|---|---|---|---|---|---|
| FALSE | High | YRSPPTVYI | [R].YRSPPT | 2xPhosphc | Q8BRG8 2x | 0.000174 | 0 | 1 | 5 | 1 | Q8BRG8 | Transmem | Q8BRG8 [2 | 1 | 2243.055 | 0 | -0.2 | 0.045195 | 7.21 | 107 | 93 | 10.21 | 4.56 | | High | High | High | High | High | High | High | High | High | High | High | 0 | 3.36E-05 | 2.98 |
| FALSE | High | YRSPYSGPI | [R].YRSPYS | 1xPhosphc | Q8VH51 1x | 0.088701 | 0.006669 | 1 | 5 | 3 | Q8VH51 | RNA-bindir | Q8VH51 [9 | 1 | 1592.824 | 0 | -0.38 | 0.002855 | 2.95 | 113.1 | 86.9 | 3.98 | 3.07 | | High | High | High | High | High | High | High | High | High | High | High | 0.004769 | 0.05217 | 2.34 |
| FALSE | High | YRSQSGED | [R].YRSQS( | 1xPhosphc | Q5BL07 1x | 2.41E-06 | 0 | 1 | 2 | 1 | Q5BL07 | Peroxisom | Q5BL07 [1 | 1 | 2543.254 | 0 | 1.31 | 0.003373 | 18.14 | 57.5 | 142.5 | 6.57 | 15.99 | | High | High | High | High | High | High | High | High | High | High | High | 0 | 2.74E-07 | 4.03 |
| FALSE | High | YRSQSGED | [R].YRSQS( | 2xPhosphc | Q5BL07 2x | 0.036103 | 0.002342 | 1 | 2 | 1 | Q5BL07 | Peroxisom | Q5BL07 [1 | 1 | 2623.221 | 0 | 0.64 | 0.002539 | 15.27 | 81.8 | 118.2 | 8.78 | 6.28 | | High | High | High | High | High | High | High | High | High | High | High | 0.00123 | 0.01326 | 2.6 |
| FALSE | High | YQLQSQEE | [R].YQLQS( | 1xPhosphc | Q8CH77 1x | 0.004696 | 0.000456 | 1 | 4 | 1 | Q8CH77 | Neuron na | Q8CH77 [9 | 0 | 1791.893 | 0 | 1.84 | 0.0107 | 19.62 | 42.9 | 157.1 | 18.45 | 24.85 | | High | High | High | High | High | High | High | High | High | High | High | 0.000204 | 0.001353 | 2.53 |
| FALSE | High | YSPTSPTYS | [K].YSPTSP | 2xPhosphc | P08775 2xI | 2.7E-05 | 0 | 1 | 3 | 4 | P08775 | DNA-direct | P08775 [18 | 0 | 2876.345 | 0 | -0.34 | 0.386383 | 14.13 | 110.3 | 89.7 | 15.5 | 10.38 | | High | High | High | High | High | High | High | High | High | High | High | 0 | 4.12E-06 | 4.2 |
| FALSE | High | YSPTSPTYS | [K].YSPTSP | 1xPhosphc | P08775 1xI | 0.00017 | 0 | 1 | 3 | 6 | P08775 | DNA-direct | P08775 [18 | 0 | 2065.03 | 0 | -0.22 | 0.65506 | 19.44 | 104.7 | 95.3 | 16.16 | 10.09 | | High | High | High | High | High | High | High | High | High | High | High | 0 | 4.11E-05 | 3.89 |
| FALSE | High | YSPTSPTYS | [K].YSPTSP | 2xPhosphc | P08775 2xI | 0.002895 | 0.00016 | 1 | 3 | 4 | P08775 | DNA-direct | P08775 [18 | 0 | 2144.996 | 0 | -0.59 | 0.062857 | 10.79 | 118.4 | 81.6 | 17.09 | 7.91 | | High | High | High | High | High | High | High | High | High | High | High | 7.19E-05 | 0.000784 | 2.82 |
| FALSE | High | YTPTSPSYS | [K].YTPTSP | 1xPhosphc | P08775 1xI | 2.8E-09 | 0 | 1 | 3 | 9 | P08775 | DNA-direct | P08775 [18 | 0 | 2782.326 | 0 | -0.13 | 0.525199 | 30.03 | 108.4 | 91.6 | 17.95 | 13.19 | | High | High | High | High | High | High | High | High | High | High | High | 0 | 1.38E-10 | 4.72 |
| FALSE | High | YTPTSPSYS | [K].YTPTSP | 2xPhosphc | P08775 2xI | 0.00407 | 0.00036 | 1 | 3 | 2 | P08775 | DNA-direct | P08775 [18 | 0 | 2862.293 | 0 | -0.39 | 0.042087 | 9.41 | 114.3 | 85.7 | 13.6 | 9.04 | | High | High | High | High | High | High | High | High | High | High | High | 0.000275 | 0.002023 | 3.16 |
| FALSE | High | YTVAPTSL\ | [K].YTVAPT | 1xPhosphc | Q8BRG8 1x | 4.53E-07 | 0 | 1 | 4 | 1 | Q8BRG8 | Transmem | Q8BRG8 [8 | 0 | 2680.509 | 0 | -0.41 | 0.045516 | 8.5 | 115 | 85 | 12.83 | 15.18 | | High | High | High | High | High | High | High | High | High | High | High | 0 | 4.19E-08 | 3.54 |
| FALSE | High | YTVPLPSPV | [K].YTVPLP | 1xPhosphc | P20357 1xI | 0.004143 | 0.00036 | 1 | 2 | 1 | P20357 | Microtubu | P20357 [87 | 0 | 3656.693 | 0 | -0.17 | 0.710222 | 14.09 | 99.6 | 100.4 | 29.37 | 23.04 | | High | High | High | High | High | High | High | High | High | High | High | 0.000161 | 0.001168 | 2.58 |
| FALSE | High | YVASYLLA/ | [R].YVASYL | 1xPhosphc | P99027 1xI | 2.61E-06 | 0 | 1 | 1 | 4 | P99027 | 60S acidic r | P99027 [3- | 0 | 2407.267 | 0 | -0.93 | 0.013065 | 2.15 | 124.3 | 75.7 | 16.31 | 8.9 | | High | High | High | High | High | High | High | High | High | High | High | 0 | 2.99E-07 | 3.78 |
| FALSE | High | YVASYLLA/ | [R].YVASYL | 1xDeamida | P99027 1xI | 3.29E-05 | 0 | 1 | 1 | 2 | P99027 | 60S acidic r | P99027 [3- | 0 | 2408.251 | 0 | -0.55 | 0.084678 | 3.64 | 118.2 | 81.8 | 14.66 | 20.87 | | High | High | High | High | High | High | High | High | High | High | High | 0 | 5.12E-06 | 3.43 |
| FALSE | High | YVDEDNSD | [K].YVDEDI | 1xPhosphc | F6WIU1 1x | 2.85E-05 | 0 | 1 | 3 | 4 | F6WIU1 | Protein KRI | F6WIU1 [1 | 0 | 2059.844 | 0 | 0.09 | 0.040584 | 3.37 | 94.2 | 105.8 | 8.02 | 8.51 | | High | High | High | High | High | High | High | High | High | High | High | 0 | 4.37E-06 | 3.64 |
| FALSE | High | YVESDDEKI | [K].YVESDI | 1xPhosphc | Q61687 1x | 0.001011 | 0 | 1 | 13 | 4 | Q61687 | Transcripti | Q61687 [8 | 0 | 2549.17 | 0 | -0.3 | 0.974451 | 24.4 | 110.3 | 89.7 | 29.82 | 64.21 | | High | High | High | High | High | High | High | High | High | High | High | 0.000689 | 0.006122 | 2.96 |
| FALSE | High | YVESDDEKI | [K].YVESDI | 1xPhosphc | Q61687 1x | 3.07E-06 | 0 | 1 | 13 | 6 | Q61687 | Transcripti | Q61687 [8 | 0 | 2778.333 | 0 | 0.03 | 0.621677 | 7.52 | 100.8 | 99.2 | 5.39 | 5.87 | | High | High | High | High | High | High | High | High | High | High | High | 0 | 3.58E-07 | 3.45 |
| FALSE | High | YVHYEIDA( | [K].YVHYEI | 1xCarbami | A0A0R4J04 | 1.07E-09 | 0 | 1 | 5 | 3 | A0A0R4J04 | N6-adenos | A0A0R4J04 | 0 | 2594.189 | 0 | -0.83 | 0.002405 | 2.64 | 130.3 | 69.7 | 5.27 | 14.94 | | High | High | High | High | High | High | High | High | High | High | High | 0 | 8.33E-09 | 5.23 |
| FALSE | High | YVLGQLVG | [K].YVLGQI | 1xPhosphc | P35761 1xI | 3.87E-05 | 0 | 1 | 5 | 2 | P35761 | Dual specif | P35761 [81 | 0 | 2353.33 | 0 | -0.52 | 0.007752 | 1.45 | 116.4 | 83.6 | 8.41 | 6.96 | | High | High | High | High | High | High | High | High | High | High | High | 0 | 6.18E-06 | 3.19 |
| FALSE | High | YVLLNDQA | [K].YVLLNI | 1xPhosphc | Q3TGG0 1x | 3.74E-06 | 0 | 1 | 2 | 3 | Q3TGG0 | Putative ur | Q3TGG0 [6 | 0 | 2360.06 | 0 | 0.1 | 0.490074 | 14.71 | 97.1 | 102.9 | 9.38 | 3.85 | | High | High | High | High | High | High | High | High | High | High | High | 0 | 4.5E-07 | 4.24 |
| FALSE | High | YVLSSQHRI | [R].YVLSSC | 1xPhosphc | E9Q9C3 1x | 0.121159 | 0.009984 | 1 | 5 | 1 | E9Q9C3 | Afadin OS= | E9Q9C3 [6 | 0 | 2194.086 | 0 | 0.25 | 0.393129 | 23.56 | 94.3 | 105.7 | 13.03 | 20.87 | | High | High | High | High | High | High | High | High | High | High | High | 0.004961 | 0.05409 | 0.99 |
| FALSE | High | YTPSGQSG | [K].YTPSG( | 1xPhosphc | A6ZI44 1xP | 0.001516 | 5.66E-05 | 1 | 3 | 2 | A6ZI44 | Fructose-b | A6ZI44 [39 | 0 | 2567.165 | 0 | 0.43 | 0.279147 | 28.12 | 86 | 114 | 31.3 | 19.23 | | High | High | High | High | High | High | High | High | High | High | High | 7.19E-05 | 0.000912 | 2.38 |
| FALSE | High | YVSGSSPDI | [K].YVSGSS | 1xPhosphc | Q62130 1x | 1.46E-06 | 0 | 1 | 4 | 8 | Q62130 | Tyrosine-p | Q62130 [5 | 0 | 1589.777 | 0 | 0.69 | 0.000462 | 6.05 | 74.9 | 125.1 | 5.76 | 5.78 | | High | High | High | High | High | High | High | High | High | High | High | 0 | 1.85E-07 | 4.16 |
| FALSE | High | YVSQNCVT | [K].YVSQN | 1xCarbami | Q5DW34 1 | 0.116482 | 0.009523 | 1 | 10 | 1 | Q5DW34 | Histone-lys | Q5DW34 [ | 0 | 2108.934 | 0 | 0.11 | 0.785969 | 15.92 | 97.9 | 102.1 | 7.86 | 9.15 | | High | High | High | High | High | High | High | High | High | High | High | 0.004757 | 0.0516 | 2.48 |
| FALSE | High | YVSSDGTEI | [R].YVSSD( | 1xPhosphc | Q8C8R3-4 | 0.006195 | 0.00062 | 1 | 7 | 1 | Q8C8R3-4 | Isoform 4 ( | Q8C8R3-4 | 0 | 1523.739 | 0 | 0.9 | 0.000462 | 1.48 | 70.3 | 129.7 | 6.43 | 5.5 | | High | High | High | High | High | High | High | High | High | High | High | 0.000275 | 0.001846 | 2.34 |
| FALSE | High | YVWLVYE( | [R].YVWLV | 1xCarbami | Q3TGC5 1x | 3.87E-05 | 0 | 1 | 3 | 2 | Q3TGC5 | Putative ur | Q3TGC5 [1 | 0 | 3248.599 | 0 | 0.39 | 0.030889 | 16.45 | 86.5 | 113.5 | 19.52 | 8.58 | | High | High | High | High | High | High | High | High | High | High | High | 0 | 6.2E-06 | 3.52 |
| FALSE | High | YWEEETTP | [R].YWEEE | 1xPhosphc | A0A068BF: | 6.65E-05 | 0 | 1 | 3 | 1 | A0A068BF: | WD repeat | A0A068BF: | 0 | 2338.095 | 0 | -0.01 | 0.438854 | 2.11 | 98.6 | 101.4 | 2.18 | 13.1 | | High | High | High | High | High | High | High | High | High | High | High | 0 | 1.13E-05 | 4.42 |
| FALSE | High | YWEEETTP | [R].YWEEE | 1xPhosphc | A0A068BF: | 0.002441 | 0.000109 | 1 | 3 | 1 | A0A068BF: | WD repeat | A0A068BF: | 1 | 2695.353 | 0 | 0.04 | 0.607353 | 7.15 | 98.1 | 101.9 | 7.02 | 5.02 | | High | High | High | High | High | High | High | High | High | High | High | 4.94E-05 | 0.000652 | 2.68 |
| FALSE | High | YWEEETTP | [R].YWEEE | 1xPhosphc | Q9Z0H1 1x | 0.00311 | 0.00016 | 1 | 1 | 2 | Q9Z0H1 | WD repeat | Q9Z0H1 [2 | 0 | 2368.106 | 0 | -0.58 | 0.004406 | 3.3 | 117.5 | 82.5 | 5.39 | 7.93 | | High | High | High | High | High | High | High | High | High | High | High | 7.19E-05 | 0.00085 | 3.01 |
| FALSE | High | YWGPASPT | [K].YWGPA | 1xPhosphc | Q99K30 1x | 0.003892 | 0.000261 | 1 | 1 | 1 | Q99K30 | Epidermal | Q99K30 [5 | 0 | 1681.85 | 0 | 1.26 | 0.000225 | 8.35 | 58.7 | 141.3 | 6.23 | 4.48 | | High | High | High | High | High | High | High | High | High | High | High | 0.00014 | 0.001097 | 2.12 |
| FALSE | High | YYEEDSDSI | [R].YYEEDS | 1xPhosphc | O55176 1x | 9.82E-06 | 0 | 1 | 6 | 1 | O55176 | E3 ubiquiti | O55176 [3 | 0 | 1788.762 | 0 | 0.74 | 0.000783 | 7.19 | 75.3 | 124.7 | 5.28 | 3.81 | | High | High | High | High | High | High | High | High | High | High | High | 0 | 1.32E-06 | 2.69 |
| FALSE | High | YYEEDSDSI | [R].YYEEDS | 2xPhosphc | O55176 2x | 0.045122 | 0.003373 | 1 | 6 | 1 | O55176 | E3 ubiquiti | O55176 [3 | 0 | 1868.728 | 0 | 1.17 | 0.003671 | 31.16 | 61.8 | 138.2 | 24.78 | 7.89 | | High | High | High | High | High | High | High | High | High | High | High | 0.001727 | 0.01712 | 2.17 |
| FALSE | High | YYSDSDDEI | [K].YYSDSD | 1xPhosphc | E9Q6J5-3 1 | 1.32E-05 | 0 | 1 | 3 | 8 | E9Q6J5-3 | Isoform Ne | E9Q6J5-3 [ | 0 | 2028.864 | 0 | -0.49 | 0.013974 | 8.48 | 114.9 | 85.1 | 7.15 | 14.79 | | High | High | High | High | High | High | High | High | High | High | High | 0 | 7.82E-06 | 3.96 |
| FALSE | High | YYSDSDDEI | [K].YYSDSD | 2xPhosphc | E9Q6J5-3 2 | 9.94E-07 | 0 | 1 | 3 | 8 | E9Q6J5-3 | Isoform Ne | E9Q6J5-3 [ | 0 | 2108.83 | 0 | 0.24 | 0.019956 | 6.82 | 90.5 | 109.5 | 2.54 | 7.62 | | High | High | High | High | High | High | High | High | High | High | High | 0 | 1.01E-07 | 3.99 |
| FALSE | High | YYSEGESA( | [R].YYSEGE | 1xPhosphc | F8WIJ5 1xI | 1.61E-07 | 0 | 1 | 7 | 3 | F8WIJ5 | Phospholip | F8WIJ5 [93 | 0 | 3700.635 | 0 | 0.18 | 0.771516 | 24.01 | 92.5 | 107.5 | 21.52 | 13.21 | | High | High | High | High | High | High | High | High | High | High | High | 0 | 1.3E-08 | 4.39 |
| FALSE | High | YYSEGESA( | [R].YYSEGE | 1xPhosphc | F8WIJ5 1xI | 0.004143 | 0.00036 | 1 | 7 | 1 | F8WIJ5 | Phospholip | F8WIJ5 [93 | 1 | 4057.893 | 0 | 0.87 | 0.004705 | 15.27 | 71.7 | 128.3 | 15.27 | 10.52 | | High | High | High | High | High | High | High | High | High | High | High | 0.000161 | 0.001173 | 3.15 |
| FALSE | High | YVSGSSPDI | [K].YVSGSS | 2xPhosphc | Q62130 2x | 0.023792 | 0.001612 | 1 | 4 | 1 | Q62130 | Tyrosine-p | Q62130 [5 | 0 | 1669.744 | 0 | 0.52 | 0.019212 | 8.1 | 82.2 | 117.8 | 4.18 | 12.26 | | High | High | High | High | High | High | High | High | High | High | High | 0.000859 | 0.008324 | 4.23 |
| FALSE | High | YTPQTLPV( | [R].YTPQTL | 1xPhosphc | Q8CFT2 1x | 0.000211 | 0 | 1 | 3 | 2 | Q8CFT2 | Histone-lys | Q8CFT2 [1 | 0 | 3636.943 | 0 | 0 | 0.678099 | 4.67 | 100.1 | 99.9 | 10.29 | 7.01 | | High | High | High | High | High | High | High | High | High | High | High | 0 | 4.14E-05 | 3.73 |
| FALSE | High | YTPQSPTYT | [R].YTPQSI | 1xPhosphc | P08775 1xI | 0.00034 | 0 | 1 | 3 | 5 | P08775 | DNA-direct | P08775 [18 | 0 | 3516.65 | 0 | 0.11 | 0.697988 | 10.97 | 94.1 | 105.9 | 14.41 | 19.79 | | High | High | High | High | High | High | High | High | High | High | High | 0 | 0.000261 | 4.26 |
| FALSE | High | YTPENLTEI | [K].YTPENL | 1xPhosphc | Q04736 1x | 8.03E-05 | 0 | 1 | 4 | 2 | Q04736 | Tyrosine-p | Q04736 [1 | 0 | 3795.852 | 0 | 0.2 | 0.992746 | 36.83 | 98.8 | 101.2 | 29.67 | 31.83 | | High | High | High | High | High | High | High | High | High | High | High | 0 | 1.41E-05 | 4.21 |
| FALSE | High | YSPTVQTR | [K].YSPTVC | 1xPhosphc | Q9R269 1x | 0.080423 | 0.006219 | 1 | 4 | 1 | Q9R269 | Periplakin ( | Q9R269 [1 | 0 | 1260.619 | 0 | 2.68 | 0.000158 | 6.55 | 27.1 | 172.9 | 12.72 | 5.48 | | High | High | High | High | High | High | High | High | High | High | High | 0.003188 | 0.03351 | 1.64 |
| FALSE | High | YSQDDND( | [R].YSQDDI | 1xPhosphc | Q8BJS4 1xI | 0.008167 | 0.000736 | 1 | 7 | 4 | Q8BJS4 | SUN doma | Q8BJS4 [11 | 0 | 3133.4 | 0 | -0.81 | 0.000965 | 1.26 | 125.7 | 74.3 | 10.31 | 17.07 | | High | High | High | High | High | High | High | High | High | High | High | 0.00074 | 0.00646 | 3.85 |
| FALSE | High | YSQEGFSG | [K].YSQEGI | 1xPhosphc | Q80YR9 1x | 1.2E-05 | 0 | 1 | 4 | 1 | Q80YR9 | RNA-bindir | Q80YR9 [3 | 0 | 1812.857 | 0 | -0.57 | 0.005904 | 3.95 | 121.1 | 78.9 | 11.09 | 4.31 | | High | High | High | High | High | High | High | High | High | High | High | 0 | 1.65E-06 | 3.78 |
| FALSE | High | YSQSVQYG | [K].YSQSV( | 1xPhosphc | Q6PCN7 1x | 1.91E-05 | 0 | 1 | 4 | 3 | Q6PCN7 | Helicase-lik | Q6PCN7 [1 | 0 | 2073.959 | 0 | 0.34 | 0.035613 | 16.21 | 88.1 | 111.9 | 9 | 5.32 | | High | High | High | High | High | High | High | High | High | High | High | 0 | 2.82E-06 | 2.5 |
| FALSE | High | YSRSSHSGI | [R].YSRSSH | 2xPhosphc | Q14AX9 2x | 0.002576 | 0.000109 | 1 | 2 | 1 | Q14AX9 | Mannose r | Q14AX9 [1 | 1 | 2225.004 | 0 | 0.23 | 0.216823 | 22.1 | 94 | 106 | 12.84 | 18.53 | | High | High | High | High | High | High | High | High | High | High | High | 4.94E-05 | 0.000691 | 3.2 |
| FALSE | High | YSSCSTIFLI | [K].YSSCST | 1xCarbami | D3YUJ3 1xI | 1.11E-05 | 0 | 1 | 3 | 11 | D3YUJ3 | Cyclin Y-lik | D3YUJ3 [12 | 0 | 2599.195 | 0 | -0.11 | 0.012083 | 2.73 | 105.9 | 94.1 | 2.74 | 6.02 | | High | High | High | High | High | High | High | High | High | High | High | 0 | 2.96E-06 | 3.15 |
| FALSE | High | YSSNGNLE | [R].YSSNGI | 1xPhosphc | Q61146 1x | 0.000822 | 0 | 1 | 3 | 1 | Q61146 | Occludin O | Q61146 [3 | 0 | 1834.899 | 0 | -0.3 | 0.018931 | 9.56 | 110.7 | 89.3 | 4.07 | 15.11 | | High | High | High | High | High | High | High | High | High | High | High | 0 | 0.000192 | 2.98 |
| FALSE | High | YSSPAAGS/ | [R].YSSPAA | 1xPhosphc | Q64701 1x | 0.007471 | 0.00062 | 1 | 4 | 3 | Q64701 | Retinoblast | Q64701 [6 | 0 | 1476.75 | 0 | 2.62 | 0.000154 | 8.35 | 26.3 | 173.7 | 10.18 | 4.51 | | High | High | High | High | High | High | High | High | High | High | High | 0.000275 | 0.002258 | 1.64 |
| FALSE | High | YSSSSGGS\ | [R].YSSSSG | 1xPhosphc | F8VQ05 1x | 3.54E-05 | 0 | 1 | 3 | 2 | F8VQ05 | FRY-like tra | F8VQ05 [1 | 0 | 1947.863 | 0 | 2.24 | 0.000177 | 1.34 | 35.2 | 164.8 | 4.12 | 4.83 | | High | High | High | High | High | High | High | High | High | High | High | 0 | 1.76E-05 | 3.21 |
| FALSE | High | YSSTETLK | [R].YSSTET | 1xPhosphc | Q80U35 1x | 0.020679 | 0.001472 | 1 | 1 | 1 | Q80U35 | Rho guanir | Q80U35 [3 | 0 | 1466.754 | 0 | -0.6 | 0.015134 | 12.76 | 116.7 | 83.3 | 5.78 | 17.14 | | High | High | High | High | High | High | High | High | High | High | High | 0.000786 | 0.007056 | 2.29 |
| FALSE | High | YSSVYGAS\ | [R].YSSVYG | 1xPhosphc | Q9CRB9 1x | 3.17E-08 | 0 | 1 | 5 | 7 | Q9CRB9 | MICOS con | Q9CRB9 [4 | 0 | 2158.036 | 0 | -0.43 | 0.045143 | 14.8 | 114.6 | 85.4 | 19.14 | 8.52 | | High | High | High | High | High | High | High | High | High | High | High | 0 | 5.25E-08 | 3.47 |
| FALSE | High | YSTPHAFTI | [R].YSTPHA | 1xPhosphc | Q99N57-2 | 0.001986 | 5.66E-05 | 1 | 3 | 1 | Q99N57-2 | Isoform 2 ( | Q99N57-2 | 0 | 2797.267 | 0 | -0.02 | 0.591138 | 8.27 | 102 | 98 | 10.1 | 9.26 | | High | High | High | High | High | High | High | High | High | High | High | 2.58E-05 | 0.000515 | 3.35 |
| FALSE | High | YSTSPASSD | [K].YSTSPA | 1xPhosphc | G3X9T5 1x | 0.000815 | 0 | 1 | 7 | 5 | G3X9T5 | MCG12342 | G3X9T5 [6 | 0 | 2016.936 | 0 | 0.51 | 0.015448 | 14.57 | 81.2 | 118.8 | 3.24 | 11.63 | | High | High | High | High | High | High | High | High | High | High | High | 0.000786 | 0.007052 | 2.61 |
| FALSE | High | YSTSPVPEI | [R].YSTSPV | 1xPhosphc | Q5I1X5 1xI | 0.016306 | 0.001124 | 1 | 4 | 3 | Q5I1X5 | RelA-assoc | Q5I1X5 [70 | 0 | 1772.821 | 0 | 2.18 | 0.002063 | 2.62 | 34.2 | 165.8 | 8.88 | 16.75 | | High | High | High | High | High | High | High | High | High | High | High | 0.000585 | 0.005448 | 1.71 |
| FALSE | High | YSTSPVPEI | [R].YSTSPV | 2xPhosphc | Q5I1X5 2xI | 0.003892 | 0.000261 | 1 | 4 | 2 | Q5I1X5 | RelA-assoc | Q5I1X5 [70 | 0 | 1852.787 | 0 | 0.82 | 0.000909 | 8.28 | 75.5 | 124.5 | 6.91 | 5.09 | | High | High | High | High | High | High | High | High | High | High | High | 0.00014 | 0.001093 | 2.48 |
| FALSE | High | YSVGTYNS | [R].YSVGT\ | 1xPhosphc | F8VQB6 1x | 0.003822 | 0.000261 | 1 | 4 | 1 | F8VQB6 | Unconvent | F8VQB6 [1 | 0 | 1733.773 | 0 | -1.62 | 0.000245 | 0.14 | 150.9 | 49.1 | 8.06 | 14.67 | | High | High | High | High | High | High | High | High | High | High | High | 0.000117 | 0.001067 | 1.09 |
| FALSE | High | YSVSSPNSI | [R].YSVSSP | 1xPhosphc | Q61572 1x | 0.000876 | 0 | 1 | 1 | 1 | Q61572 | Forkhead b | Q61572 [5 | 0 | 2831.349 | 0 | 0.75 | 0.006043 | 25.5 | 72.1 | 127.9 | 12.36 | 8.46 | | High | High | High | High | High | High | High | High | High | High | High | 0 | 0.000206 | 3.28 |
| FALSE | High | YSYPPLVDI | [K].YSYPPL | 1xCarbami | B1AU74 1x | 8.32E-08 | 0 | 1 | 4 | 4 | B1AU74 | Motile spe | B1AU74 [2 | 0 | 3912.781 | 0 | -0.16 | 0.250677 | 14.58 | 103.4 | 96.6 | 8.6 | 9.64 | | High | High | High | High | High | High | High | High | High | High | High | 0 | 8.91E-09 | 5.54 |
| FALSE | High | YSYQLLQCI | [R].YSYQLL | 1xCarbami | Q5XJV7-2 1 | 0.005 | 0.000504 | 1 | 3 | 1 | Q5XJV7-2 | Isoform 2 ( | Q5XJV7-2 [ | 0 | 1838.835 | 0 | -0.11 | 0.571932 | 20.16 | 102.6 | 97.4 | 10.34 | 15.35 | | High | High | High | High | High | High | High | High | High | High | High | 0.000225 | 0.001441 | 2.63 |
| FALSE | High | YTAGSASPT | [R].YTAGS/ | 1xPhosphc | A2AAE1-3 1 | 0.087983 | 0.006639 | 1 | 6 | 1 | A2AAE1-3 | Isoform 3 ( | A2AAE1-3 | 0 | 1865.945 | 0 | -0.38 | 0.008944 | 6.22 | 113.9 | 86.1 | 3.33 | 8.61 | | High | High | High | High | High | High | High | High | High | High | High | 0.003402 | 0.03724 | 2.34 |
| FALSE | High | YTDQSGEE | [R].YTDQS( | 1xPhosphc | Q8K1Z0 1x | 2.86E-07 | 0 | 1 | 2 | 2 | Q8K1Z0 | Ubiquinon | Q8K1Z0 [7 | 0 | 2910.179 | 0 | 0.32 | 0.01854 | 7.76 | 90.9 | 109.1 | 4.03 | 8.18 | | High | High | High | High | High | High | High | High | High | High | High | 0 | 4.45E-06 | 5.09 |
| FALSE | High | YTEFNSSTT | [K].YTEFNS | 1xPhosphc | E9QJS7 1xI | 0.000411 | 0 | 1 | 8 | 2 | E9QJS7 | Adhesion ( | E9QJS7 [79 | 0 | 2146.949 | 0 | 0.82 | 0.022329 | 46.4 | 72.3 | 127.7 | 15.6 | 15.93 | | High | High | High | High | High | High | High | High | High | High | High | 0 | 8.79E-05 | 2.22 |
| FALSE | High | YTFAPSASL | [R].YTFAPS | 1xPhosphc | Q3TJD7 1xI | 0.000674 | 0 | 1 | 5 | 1 | Q3TJD7 | PDZ and LII | Q3TJD7 [1 | 0 | 1736.902 | 0 | 0.75 | 0.002538 | 16.28 | 75.3 | 124.7 | 16.42 | 14.23 | | High | High | High | High | High | High | High | High | High | High | High | 0 | 0.000153 | 1.78 |
| FALSE | High | YTFDFSEEE | [K].YTFDFS | 1xPhosphc | Q64511 1x | 1.71E-10 | 0 | 1 | 2 | 13 | Q64511 | DNA topois | Q64511 [1 | 0 | 3492.478 | 0 | 0.81 | 0.002995 | 7.04 | 74.6 | 125.4 | 5.83 | 12.83 | | High | High | High | High | High | High | High | High | High | High | High | 0 | 9.24E-12 | 6.7 |
| FALSE | High | YTFDFSEEE | [K].YTFDFS | 2xPhosphc | Q64511 2x | 5.98E-09 | 0 | 1 | 2 | 8 | Q64511 | DNA topois | Q64511 [1 | 0 | 3572.444 | 0 | -0.75 | 0.037235 | 2.8 | 125.2 | 74.8 | 17.4 | 19.99 | | High | High | High | High | High | High | High | High | High | High | High | 0 | 3.21E-10 | 6.82 |
| FALSE | High | YTIGDIFVH | [K].YTIGDII | 1xPhosphc | B9EK76 1xI | 2.04E-05 | 0 | 1 | 4 | 1 | B9EK76 | Signal-indu | B9EK76 [1 | 0 | 1841.915 | 0 | -1.34 | 0.00137 | 0.13 | 140.4 | 59.6 | 6.67 | 11.59 | | High | High | High | High | High | High | High | High | High | High | High | 0 | 3.01E-06 | 2.65 |
| FALSE | High | YTLPPGVDI | [K].YTLPPG | 1xPhosphc | Q545F4 1x | 0.011048 | 0.000772 | 1 | 3 | 2 | Q545F4 | Hspb1 prot | Q545F4 [1 | 0 | 3603.921 | 0 | -0.85 | 0.002245 | 2.2 | 126.4 | 73.6 | 10.81 | 9.78 | | High | High | High | High | High | High | High | High | High | High | High | 0.000399 | 0.003505 | 3.37 |
| FALSE | High | YTPASPSSP | [R].YTPASP | 1xCarbami | A0A1D5RL: | 0.0095 | 0.000736 | 1 | 5 | 3 | A0A1D5RL: | Calpain-15 | A0A1D5RL: | 0 | 2340.021 | 0 | -0.15 | 0.183793 | 11.34 | 104.7 | 95.3 | 11.23 | 12.57 | | High | High | High | High | High | High | High | High | High | High | High | 0.000585 | 0.005373 | 2.29 |
| FALSE | High | YTPDDHR | [R].YTPDDI | 1xPhosphc | A2AG58 1x | 0.005768 | 0.00062 | 1 | 2 | 2 | A2AG58 | Uncharacte | A2AG58 [1 | 0 | 1212.525 | 0 | -1.67 | 0.00037 | 0.46 | 152.8 | 47.2 | 5.36 | 8.36 | | High | High | High | High | High | High | High | High | High | High | High | 0.000275 | 0.001695 | 2.33 |
| FALSE | High | YPQQYQPS | [R].YPQQY( | 1xPhosphc | Q6PHZ5 1x | 0.085859 | 0.006406 | 1 | 2 | 1 | Q6PHZ5 | Putative RI | Q6PHZ5 [4 | 0 | 3074.486 | 0 | 0.8 | 0.032561 | 39.43 | 73.2 | 126.8 | 20.4 | 18.78 | | High | High | High | High | High | High | High | High | High | High | High | 0.003303 | 0.03609 | 2.43 |
| FALSE | High | YPPYTTPPI | [R].YPPYTT | 1xPhosphc | O35071 1x | 0.024424 | 0.001612 | 1 | 4 | 2 | O35071 | Kinesin-like | O35071 [1 | 0 | 1400.681 | 0 | 0.08 | 0.222239 | 16.34 | 97.2 | 102.8 | 9.04 | 9.05 | | High | High | High | High | High | High | High | High | High | High | High | 0.001609 | 0.01552 | 2.4 |
| FALSE | High | YPPSPAK | [R].YPPSPA | 1xPhosphc | B7ZN13 1x | 0.031713 | 0.002011 | 1 | 3 | 2 | B7ZN13 | Aldehyde ( | B7ZN13 [4 | 0 | 1297.696 | 0 | 4.65 | 0.000154 | 48.38 | 7.6 | 192.4 | 40.95 | 2.96 | | High | High | High | High | High | High | High | High | High | High | High | 0.001812 | 0.01843 | 1.32 |
| FALSE | High | YPPFSPPEP | [R].YPPFSP | 1xPhosphc | Q5SSZ5 1xI | 0.005717 | 0.00062 | 1 | 1 | 1 | Q5SSZ5 | Tensin-3 O | Q5SSZ5 [8 | 0 | 2617.347 | 0 | -0.9 | 0.010232 | 4.32 | 127.1 | 72.9 | 18.63 | 17.67 | | High | High | High | High | High | High | High | High | High | High | High | 0.000275 | 0.001684 | 2.7 |
| FALSE | High | YGGISTAS\ | [K].YGGIST | 1xPhosphc | Q99KG3 1x | 0.000611 | 0 | 2 | 3 | 1 | Q99KG3; ( | RNA-bindir | Q99KG3 [8 | 0 | 2036.953 | 0 | 0.42 | 0.02113 | 6.45 | 84.6 | 115.4 | 10.71 | 12.64 | Shared | High | High | High | High | High | High | High | High | High | High | High | 0 | 0.000138 | 2.61 |
| FALSE | High | YGGPPPDS | [K].YGGPPI | 1xPhosphc | Q7TMK9 1 | 0.006363 | 0.00062 | 1 | 8 | 1 | Q7TMK9 | Heterogen | Q7TMK9 [1 | 0 | 3104.538 | 0 | 0.24 | 0.158046 | 26.9 | 91 | 109 | 18.12 | 15.3 | | High | High | High | High | High | High | High | High | High | High | High | 0.000275 | 0.0019 | 3.25 |
| FALSE | High | YGGPPPGV | [K].YGGPPI | 1xPhosphc | Q6VY05 1x | 8.35E-06 | 0 | 1 | 1 | 1 | Q6VY05 | Dead end p | Q6VY05 [4 | 0 | 2622.259 | 0 | -1.71 | 0.000108 | 0.38 | 152.6 | 47.4 | 2.66 | 8.97 | | High | High | High | High | High | High | High | High | High | High | High | 0 | 1.11E-06 | 3.55 |
| FALSE | High | YGLGYQLC | [K].YGLGY( | 1xCarbami | Q3TPZ2 1x | 1.31E-06 | 0 | 1 | 2 | 1 | Q3TPZ2 | Serine/thre | Q3TPZ2 [4 | 0 | 2687.237 | 0 | -0.42 | 0.069915 | 10.79 | 116.4 | 83.6 | 15.59 | 18.86 | | High | High | High | High | High | High | High | High | High | High | High | 0 | 1.39E-07 | 4.17 |
| FALSE | High | YGLLTNTEI | [R].YGLLTN | 2xPhosphc | Q9DAZ5 2x | 0.000119 | 0 | 1 | 1 | 4 | Q9DAZ5 | Uncharacte | Q9DAZ5 [7 | 0 | 3563.465 | 0 | 1.61 | 0.002784 | 10.97 | 47.3 | 152.7 | 5.53 | 18.04 | | High | High | High | High | High | High | High | High | High | High | High | 0 | 4.46E-05 | 4.97 |
| FALSE | High | YGLLTNTEI | [R].YGLLTN | 1xDeamida | Q9DAZ5 2x | 0.000651 | 0 | 1 | 1 | 2 | Q9DAZ5 | Uncharacte | Q9DAZ5 [7 | 0 | 3564.449 | 0 | 1.31 | 0.00473 | 8.31 | 57.9 | 142.1 | 16.09 | 20.37 | | High | High | High | High | High | High | High | High | High | High | High | 0.000786 | 0.006926 | 3.97 |
| FALSE | High | YGLLTNTEI | [R].YGLLTN | 2xPhosphc | Q9DAZ5 2x | 0.000237 | 0 | 1 | 1 | 4 | Q9DAZ5 | Uncharacte | Q9DAZ5 [7 | 0 | 3579.46 | 0 | 1.62 | 0.000255 | 2.82 | 49.3 | 150.7 | 3.79 | 5.97 | | High | High | High | High | High | High | High | High | High | High | High | 0 | 4.73E-05 | 4.06 |
| FALSE | High | YGLIYHASL | [K].YGLIYH | 1xPhosphc | Q6DFW4 1 | 1.98E-07 | 0 | 1 | 4 | 2 | Q6DFW4 | Nucleolar p | Q6DFW4 [ | 0 | 2258.199 | 0 | -0.54 | 0.000181 | 0.81 | 118.6 | 81.4 | 2.66 | 6.62 | | High | High | High | High | High | High | High | High | High | High | High | 0 | 4.95E-08 | 3.18 |
| FALSE | High | YGLQDSDE | [K].YGLQD: | 1xPhosphc | A0A1B0GS: | 0.0006 | 0 | 1 | 9 | 1 | A0A1B0GS: | Nuclear po | A0A1B0GS: | 0 | 2210.933 | 0 | -0.26 | 0.551596 | 47.06 | 110.1 | 89.9 | 63.08 | 54.26 | | High | High | High | High | High | High | High | High | High | High | High | 0 | 0.000135 | 4.06 |
| FALSE | High | YGLQDSDE | [K].YGLQD: | 1xPhosphc | A0A1B0GS: | 2.61E-06 | 0 | 1 | 9 | 10 | A0A1B0GS: | Nuclear po | A0A1B0GS: | 0 | 2440.096 | 0 | -0.33 | 0.000406 | 2.48 | 112.2 | 87.8 | 2.02 | 4.22 | | High | High | High | High | High | High | High | High | High | High | High | 0 | 6.58E-05 | 4.42 |
| FALSE | High | YGLSPGPV | [R].YGLSPG | 1xPhosphc | Q9WU40-2 | 0.016451 | 0.001124 | 1 | 4 | 1 | Q9WU40-2 | Isoform 2 ( | Q9WU40-2 | 0 | 2502.341 | 0 | 0.23 | 0.011042 | 6.53 | 90 | 110 | 7.5 | 11.11 | | High | High | High | High | High | High | High | High | High | High | High | 0.000585 | 0.005475 | 1.6 |
| FALSE | High | YGMGTSVI | [R].YGMGT | 1xPhosphc | P35486 1xI | 0.014667 | 0.001049 | 1 | 1 | 2 | P35486 | Pyruvate d | P35486 [22 | 0 | 1308.586 | 0 | -0.88 | 0.000405 | 2.26 | 129.4 | 70.6 | 5.06 | 8.42 | | High | High | High | High | High | High | High | High | High | High | High | 0.002105 | 0.02163 | 1.86 |
| FALSE | High | YGMGTSVI | [R].YGMGT | 1xPhosphc | P35486 1xI | 0.041781 | 0.003126 | 1 | 1 | 15 | P35486 | Pyruvate d | P35486 [22 | 0 | 1324.581 | 0 | -0.82 | 0.000378 | 2.75 | 127 | 73 | 3.48 | 7.14 | | High | High | High | High | High | High | High | High | High | High | High | 0.004046 | 0.0432 | 1.35 |
| FALSE | High | YGGDSDHI | [K].YGGDS | 2xPhosphc | Q9EQM6 2 | 2.29E-05 | 0 | 1 | 1 | 1 | Q9EQM6 | Microproc | Q9EQM6 [ | 0 | 2857.186 | 0 | -0.64 | 0.032184 | 14.36 | 119.8 | 80.2 | 7.51 | 22.45 | | High | High | High | High | High | High | High | High | High | High | High | 0 | 3.42E-06 | 3.96 |
| FALSE | High | YGPLSEPG: | [R].YGPLSE | 1xPhosphc | Q8K412 1x | 0.090153 | 0.00698 | 1 | 3 | 1 | Q8K412 | Memory-re | Q8K412 [6 | 0 | 1442.688 | 0 | 1.64 | 0.000333 | 8.48 | 47.4 | 152.6 | 9.94 | 6.52 | | High | High | High | High | High | High | High | High | High | High | High | 0.003564 | 0.03806 | 1.38 |
| FALSE | High | YGPSSVEDT | [K].YGPSSV | 1xPhosphc | O70251 1x | 4.72E-05 | 0 | 1 | 2 | 7 | O70251 | Elongation | O70251 [7 | 1 | 4734.072 | 0 | -0.68 | 0.030889 | 6.76 | 120.1 | 79.9 | 17.29 | 7.16 | | High | High | High | High | High | High | High | High | High | High | High | 0 | 7.76E-06 | 5.52 |
| FALSE | High | YGQDGVVI | [R].YGQDG | 1xPhosphc | Q80YR5 1x | 0.042141 | 0.003162 | 1 | 5 | 1 | Q80YR5 | Scaffold at | Q80YR5 [1 | 0 | 1973.953 | 0 | -0.39 | 0.060949 | 13.16 | 112 | 88 | 7.57 | 9.45 | | High | High | High | High | High | High | High | High | High | High | High | 0.001639 | 0.01584 | 2.59 |
| FALSE | High | YGQFSGLN | [K].YGQFS( | 1xPhosphc | P62137 1xI | 0.000126 | 0 | 1 | 3 | 1 | P62137 | Serine/thre | P62137 [3 | 0 | 2223.127 | 0 | -0.12 | 0.430997 | 17.9 | 105.4 | 94.6 | 6.07 | 11.61 | | High | High | High | High | High | High | High | High | High | High | High | 0 | 2.32E-05 | 1.15 |
| FALSE | High | YGSDDECS. | [K].YGSDDI | 1xCarbami | Q8C440 1x | 0.000264 | 0 | 1 | 4 | 1 | Q8C440 | Putative ur | Q8C440 [2 | 0 | 4026.682 | 0 | 1.96 | 0.001086 | 6.18 | 40.7 | 159.3 | 5.94 | 13.73 | | High | High | High | High | High | High | High | High | High | High | High | 0 | 5.35E-05 | 4.57 |
| FALSE | High | YGSDHTAN | [K].YGSDH | 1xPhosphc | Q8BJQ2 1x | 1.1E-08 | 0 | 1 | 1 | 1 | Q8BJQ2 | Ubiquitin c | Q8BJQ2 [3 | 0 | 3171.452 | 0 | -0.26 | 0.100553 | 13.68 | 110.8 | 89.2 | 12.5 | 5.65 | | High | High | High | High | High | High | High | High | High | High | High | 0 | 6.4E-10 | 4.44 |
| FALSE | High | YGSDHTAN | [K].YGSDH | 1xDeamida | Q8BJQ2 1x | 9.67E-07 | 0 | 1 | 1 | 2 | Q8BJQ2 | Ubiquitin c | Q8BJQ2 [3 | 0 | 3172.436 | 0 | -0.16 | 0.034434 | 4.93 | 105.7 | 94.3 | 4.02 | 3.87 | | High | High | High | High | High | High | High | High | High | High | High | 0 | 9.78E-08 | 4.28 |
| FALSE | High | YGSIVDDEI | [R].YGSIVD | 1xPhosphc | B9EKK3 1xI | 0.003342 | 0.00016 | 1 | 4 | 3 | B9EKK3 | IQ motif co | B9EKK3 [1 | 0 | 1362.614 | 0 | 2.75 | 0.000158 | 15.85 | 26.7 | 173.3 | 17.55 | 5.13 | | High | High | High | High | High | High | High | High | High | High | High | 0.000814 | 0.007807 | 2.61 |
| FALSE | High | YGSQEHPI( | [R].YGSQEI | 1xPhosphc | Q3UGJ5 1x | 0.002172 | 5.66E-05 | 1 | 3 | 2 | Q3UGJ5 | Putative ur | Q3UGJ5 [8 | 0 | 1539.704 | 0 | 0.5 | 0.125238 | 15.87 | 79.8 | 120.2 | 14.88 | 27.73 | | High | High | High | High | High | High | High | High | High | High | High | 0.000382 | 0.003291 | 1.92 |
| FALSE | High | YGTDLSR | [K].YGTDLS | 1xPhosphc | Q3TXF9 1x | 0.036731 | 0.00249 | 1 | 2 | 1 | Q3TXF9 | Sodium/pc | Q3TXF9 [5 | 0 | 1120.524 | 0 | -1.19 | 0.006375 | 4.37 | 141.1 | 58.9 | 18.61 | 8.26 | | High | High | High | High | High | High | High | High | High | High | High | 0.001308 | 0.01357 | 2.36 |
| FALSE | High | YGVYEDEN | [R].YGVYEI | 1xPhosphc | Q9D7A3 1x | 2.98E-05 | 0 | 1 | 2 | 6 | Q9D7A3 | Putative ur | Q9D7A3 [2 | 0 | 3107.37 | 0 | 0.24 | 0.030676 | 6.33 | 93.7 | 106.3 | 4.24 | 4.67 | | High | High | High | High | High | High | High | High | High | High | High | 2.58E-05 | 0.000349 | 3.18 |
| FALSE | High | YGVYEDEN | [R].YGVYEI | 1xPhosphc | Q3UJW4 1 | 3.8E-05 | 0 | 1 | 2 | 2 | Q3UJW4 | Putative ur | Q3UJW4 [2 | 0 | 3035.349 | 0 | -0.3 | 0.002394 | 4.02 | 111.9 | 88.1 | 5.24 | 1.69 | | High | High | High | High | High | High | High | High | High | High | High | 0 | 6.04E-06 | 3.57 |
| FALSE | High | YGYTHLST( | [K].YGYTHI | 1xPhosphc | Q9R0Y5-2 | 0.021046 | 0.001507 | 1 | 4 | 2 | Q9R0Y5-2 | Isoform 2 ( | Q9R0Y5-2 | 0 | 1804.883 | 0 | 0.36 | 0.005771 | 5.16 | 89 | 111 | 5.63 | 7.97 | | High | High | High | High | High | High | High | High | High | High | High | 0.000801 | 0.007233 | 2.28 |
| FALSE | High | YHGHSMSI | [R].YHGHSI | 1xPhosphc | P35486 1xI | 4.35E-05 | 0 | 1 | 1 | 5 | P35486 | Pyruvate d | P35486 [28 | 0 | 1901.82 | 0 | 0.31 | 0.077209 | 20.54 | 89 | 111 | 8.55 | 10.76 | | High | High | High | High | High | High | High | High | High | High | High | 0 | 3.18E-05 | 3.11 |
| FALSE | High | YGPSSVEDT | [K].YGPSSV | 1xPhosphc | O70251 1x | 1.33E-08 | 0 | 1 | 2 | 7 | O70251 | Elongation | O70251 [7 | 0 | 2251.053 | 0 | -0.35 | 0.003427 | 5.52 | 114.1 | 85.9 | 2.65 | 7 | | High | High | High | High | High | High | High | High | High | High | High | 0 | 7.94E-10 | 3.37 |
| FALSE | High | YGGDEIPFS | [K].YGGDE | 1xPhosphc | Q8BTM8 1 | 0.000573 | 0 | 1 | 5 | 1 | Q8BTM8 | Filamin-A ( | Q8BTM8 [1 | 0 | 1709.777 | 0 | 0.33 | 0.082035 | 18.38 | 91 | 109 | 6.25 | 13.87 | | High | High | High | High | High | High | High | High | High | High | High | 0 | 0.000128 | 1.41 |
| FALSE | High | YFSSENMD | [R].YFSSEN | 1xPhosphc | G5E870 1x | 0.015061 | 0.001049 | 1 | 2 | 1 | G5E870 | E3 ubiquiti | G5E870 [1 | 0 | 2467.116 | 0 | -0.06 | 0.393541 | 31.17 | 106.3 | 93.7 | 11.6 | 10.71 | | High | High | High | High | High | High | High | High | High | High | High | 0.000571 | 0.004981 | 2.59 |
| FALSE | High | YFSPPPPAI | [K].YFSPPP | 1xPhosphc | Q3TYD6 1x | 0.003927 | 0.000311 | 1 | 1 | 1 | Q3TYD6 | Serine/thre | Q3TYD6 [1 | 0 | 1340.66 | 0 | -0.6 | 0.00252 | 1.07 | 119.7 | 80.3 | 4.27 | 5.53 | | High | High | High | High | High | High | High | High | High | High | High | 0.00014 | 0.0011 | 1.92 |
| FALSE | High | YEISPEAYEI | [K].YEISPE/ | 1xPhosphc | Q9D1E6 1x | 0.000193 | 0 | 1 | 1 | 5 | Q9D1E6 | Tubulin-fol | Q9D1E6 [1 | 0 | 1565.709 | 0 | -1.01 | 0.00043 | 4.03 | 134.5 | 65.5 | 2.67 | 11.79 | | High | High | High | High | High | High | High | High | High | High | High | 0 | 3.74E-05 | 2.86 |
| FALSE | High | YENHSATA | [R].YENHS/ | 1xPhosphc | B2RRF0 1xI | 0.071071 | 0.005247 | 1 | 1 | 1 | B2RRF0 | Ptprk prote | B2RRF0 [8 | 0 | 1660.717 | 0 | 0.78 | 0.003242 | 12.03 | 70.9 | 129.1 | 16.59 | 7.95 | | High | High | High | High | High | High | High | High | High | High | High | 0.002703 | 0.02892 | 2.84 |
| FALSE | High | YEPAAVSEI | [K].YEPAA\ | 1xPhosphc | Q3TXF9 1x | 0.0004 | 0 | 1 | 2 | 1 | Q3TXF9 | Sodium/pc | Q3TXF9 [1 | 0 | 1840.888 | 0 | 0.81 | 0.031694 | 6.45 | 72.2 | 127.8 | 18.29 | 24.07 | | High | High | High | High | High | High | High | High | High | High | High | 0 | 8.54E-05 | 2.04 |
| FALSE | High | YEPAAVSEI | [K].YEPAA\ | 1xPhosphc | Q3TXF9 1x | 0.025964 | 0.001645 | 1 | 2 | 1 | Q3TXF9 | Sodium/pc | Q3TXF9 [1 | 1 | 2198.146 | 0 | 0.67 | 0.042372 | 8.14 | 77.1 | 122.9 | 11.42 | 32.59 | | High | High | High | High | High | High | High | High | High | High | High | 0.000872 | 0.00914 | 1.68 |
| FALSE | High | YEPYGMYS | [R].YEPYGI | 1xCarbami | B2RY22 1xI | 0.002153 | 5.66E-05 | 2 | 17 | 2 | B2RY22; A( | Clasp1 pro | B2RY22 [7 | 0 | 2927.122 | 0 | -0.48 | 0.072411 | 19.11 | 114.5 | 85.5 | 7.01 | 27.36 | Shared | High | High | High | High | High | High | High | High | High | High | High | 2.58E-05 | 0.000566 | 3.59 |

| | | | | | | | | | | | | | | | | | | | | | | | | | | | | | | | | | | | | | | |
|---|---|---|---|---|---|---|---|---|---|---|---|---|---|---|---|---|---|---|---|---|---|---|---|---|---|---|---|---|---|---|---|---|---|---|---|---|---|---|
| FALSE | High | YEPYGMYS | [R].YEPYGM | 1xCarbami | B2RY22 1xI | 4.4E-06 | 0 | 2 | 17 | 2 | B2RY22; A( | Clasp1 pro | B2RY22 [76 | 0 | 2943.117 | 0 | -1 | 0.002042 | 3.94 | 136 | 64 | 11.02 | 10.52 | Shared | High | High | High | High | High | High | High | High | High | High | High | 0 | 5.36E-07 | 4.31 |
| FALSE | High | YEQGFITDI | [K].YEQGFI | 1xPhospho | P24547 1xI | 5.82E-06 | 0 | 1 | 4 | 1 | P24547 | Inosine-5'- | P24547 [11 | 0 | 2231.177 | 0 | -1.07 | 0.004833 | 4.23 | 133.4 | 66.6 | 10.68 | 10.35 | | High | High | High | High | High | High | High | High | High | High | High | 0 | 7.36E-07 | 2.68 |
| FALSE | High | YESGPDGG | [K].YESGPD | 1xPhospho | Q7M6Z9 1: | 0.002974 | 0.00016 | 1 | 1 | 1 | Q7M6Z9 | Kinesin ligh | Q7M6Z9 [5 | 0 | 2236.006 | 0 | 1.55 | 0.001183 | 15.04 | 50.1 | 149.9 | 23.82 | 12.84 | | High | High | High | High | High | High | High | High | High | High | High | 7.19E-05 | 0.00081 | 2.95 |
| FALSE | High | YESYGMHS | [R].YESYGM | 1xCarbami | B9EJA4 1xF | 1.02E-05 | 0 | 1 | 7 | 1 | B9EJA4 | Clasp2 pro | B9EJA4 [62 | 0 | 2879.061 | 0 | -1.2 | 0.005635 | 2.13 | 134.5 | 65.5 | 15.54 | 15.69 | | High | High | High | High | High | High | High | High | High | High | High | 0 | 1.38E-06 | 4.96 |
| FALSE | High | YESYGMHS | [R].YESYGM | 1xCarbami | B9EJA4 1xF | 0.000492 | 0 | 1 | 7 | 1 | B9EJA4 | Clasp2 pro | B9EJA4 [62 | 0 | 2880.045 | 0 | -1.17 | 0.006329 | 3.37 | 139.9 | 60.1 | 20.74 | 13.37 | | High | High | High | High | High | High | High | High | High | High | High | 0 | 0.000108 | 3.84 |
| FALSE | High | YEVAAQHE | [K].YEVAA( | 1xPhospho | Q9Z1K8 1x | 0.001233 | 0 | 1 | 3 | 1 | Q9Z1K8 | Y+L amino | Q9Z1K8 [7 | 0 | 3252.546 | 0 | -1.88 | 0.000399 | 1.38 | 155.8 | 44.2 | 3.18 | 23.66 | | High | High | High | High | High | High | High | High | High | High | High | 0 | 0.000303 | 3.68 |
| FALSE | High | YFDDEFTA( | [R].YFDDEF | 1xPhospho | Q3TY95 1x | 5.7E-05 | 0 | 1 | 3 | 3 | Q3TY95 | MCG22777 | Q3TY95 [4 | 0 | 2425.116 | 0 | 0.26 | 0.011865 | 4.54 | 92.1 | 107.9 | 2.13 | 4.5 | | High | High | High | High | High | High | High | High | High | High | High | 0 | 1.81E-05 | 3.6 |
| FALSE | High | YFDEEFTA( | [R].YFDEEF | 1xPhospho | Q6NXW0 1 | 4.52E-06 | 0 | 1 | 4 | 3 | Q6NXW0 | Thymoma | Q6NXW0 [ | 0 | 2668.32 | 0 | 0.62 | 0.007872 | 15.07 | 75.2 | 124.8 | 8.46 | 9.5 | | High | High | High | High | High | High | High | High | High | High | High | 0 | 5.52E-07 | 3.95 |
| FALSE | High | YFDSGDYN | [K].YFDSGI | 1xPhospho | P60840 1xI | 4.59E-05 | 0 | 1 | 6 | 2 | P60840 | Alpha-endo | P60840 [64 | 0 | 1848.828 | 0 | -1.66 | 0.000947 | 1.31 | 153.1 | 46.9 | 14.79 | 32.33 | | High | High | High | High | High | High | High | High | High | High | High | 0 | 7.5E-06 | 3.02 |
| FALSE | High | YFDSGDYN | [K].YFDSGI | 1xPhospho | P60840 1xI | 9.65E-06 | 0 | 1 | 6 | 7 | P60840 | Alpha-endo | P60840 [64 | 0 | 1864.823 | 0 | -1.43 | 0.00505 | 6.03 | 154.6 | 45.4 | 15.92 | 28.27 | | High | High | High | High | High | High | High | High | High | High | High | 0.000225 | 0.001451 | 3.43 |
| FALSE | High | YFDTNSEIE | [K].YFDTNS | 1xPhospho | Q5UAK0-1 | 1.74E-06 | 0 | 1 | 4 | 3 | Q5UAK0-1 | Isoform 1 c | Q5UAK0-1 | 0 | 3636.535 | 0 | 0.2 | 0.321204 | 4.64 | 93.7 | 106.3 | 4.46 | 15.38 | | High | High | High | High | High | High | High | High | High | High | High | 0 | 1.88E-07 | 4.94 |
| FALSE | High | YFDTNSEIE | [K].YFDTNS | 2xPhospho | Q5UAK0-1 | 0.000322 | 0 | 1 | 4 | 3 | Q5UAK0-1 | Isoform 1 c | Q5UAK0-1 | 0 | 3716.502 | 0 | -0.17 | 0.213393 | 12.32 | 104.1 | 95.9 | 9.44 | 10.44 | | High | High | High | High | High | High | High | High | High | High | High | 0 | 6.66E-05 | 4.22 |
| FALSE | High | YFDVHPEP | [K].YFDVHI | 1xPhospho | A7TU71 1x | 0.001572 | 5.66E-05 | 1 | 2 | 3 | A7TU71 | Protein Sh | A7TU71 [1 | 0 | 3088.522 | 0 | 2.15 | 0.000426 | 14.42 | 36.7 | 163.3 | 21.81 | 11.38 | | High | High | High | High | High | High | High | High | High | High | High | 0.00014 | 0.001096 | 4.5 |
| FALSE | High | YFGFDDLSI | [K].YFGFDI | 1xCarbami | B7ZP47 1xI | 9.97E-08 | 0 | 1 | 2 | 3 | B7ZP47 | Wapal prot | B7ZP47 [46 | 0 | 2980.119 | 0 | 0.04 | 0.483921 | 13.42 | 103.9 | 96.1 | 9.87 | 27.05 | | High | High | High | High | High | High | High | High | High | High | High | 0 | 7.58E-09 | 4.58 |
| FALSE | High | YFGFDDLSI | [K].YFGFDI | 1xCarbami | B7ZP47 2xI | 1.19E-07 | 0 | 1 | 2 | 11 | B7ZP47 | Wapal prot | B7ZP47 [46 | 0 | 3060.085 | 0 | -0.55 | 0.000149 | 0.19 | 119 | 81 | 1.54 | 2.2 | | High | High | High | High | High | High | High | High | High | High | High | 0 | 1.12E-07 | 4.47 |
| FALSE | High | YFGTDAVP | [R].YFGTD/ | 1xPhospho | Q91YL3 1xI | 0.005872 | 0.00062 | 1 | 3 | 2 | Q91YL3 | Uridine-cyt | Q91YL3 [5 | 0 | 2310.949 | 0 | 2.46 | 0.000401 | 4.75 | 30.8 | 169.2 | 5.52 | 9.6 | | High | High | High | High | High | High | High | High | High | High | High | 0.006234 | 0.06725 | 1.93 |
| FALSE | High | YFIDDTESD | [K].YFIDDT | 1xPhospho | Q80WQ8 1 | 1.22E-05 | 0 | 1 | 3 | 2 | Q80WQ8 | Mis18-binc | Q80WQ8 [ | 0 | 2157.952 | 0 | -0.46 | 0.002719 | 6.12 | 113.9 | 86.1 | 4.05 | 8.81 | | High | High | High | High | High | High | High | High | High | High | High | 0 | 1.69E-06 | 3.75 |
| FALSE | High | YFIDDTESD | [K].YFIDDT | 2xPhospho | Q80WQ8 2 | 5.91E-05 | 0 | 1 | 3 | 3 | Q80WQ8 | Mis18-binc | Q80WQ8 [ | 0 | 2237.918 | 0 | -0.93 | 0.000308 | 1.29 | 130.8 | 69.2 | 3.33 | 5.27 | | High | High | High | High | High | High | High | High | High | High | High | 0 | 9.93E-06 | 4.07 |
| FALSE | High | YFLERPPVS | [K].YFLERP | 1xPhospho | Q8CGN4 1: | 6.61E-06 | 0 | 1 | 4 | 3 | Q8CGN4 | BCL-6 core | Q8CGN4 [1 | 0 | 3329.64 | 0 | 0.85 | 0.002487 | 10.88 | 72.2 | 127.8 | 10.33 | 13.61 | | High | High | High | High | High | High | High | High | High | High | High | 0 | 8.48E-07 | 3.13 |
| FALSE | High | YFPSGIPLL | [K].YFPSGII | 1xPhospho | A0A1L1SSA | 3.09E-05 | 0 | 1 | 6 | 1 | A0A1L1SSA | Centrosom | A0A1L1SSA | 0 | 2535.33 | 0 | -0.26 | 0.016475 | 3.54 | 110.8 | 89.2 | 5.6 | 5.58 | | High | High | High | High | High | High | High | High | High | High | High | 0 | 4.8E-06 | 2.38 |
| FALSE | High | YFQISQDEI | [R].YFQISQ | 1xPhospho | Q9D7S7 1x | 3.32E-08 | 0 | 1 | 2 | 39 | Q9D7S7 | 60S ribosoi | Q9D7S7 [1 | 0 | 2057.806 | 0 | 0.37 | 0.016178 | 5.93 | 87.1 | 112.9 | 8.51 | 3.83 | | High | High | High | High | High | High | High | High | High | High | High | 0 | 5.22E-09 | 4.22 |
| FALSE | High | YFQISQDEI | [R].YFQISQ | 1xDeamida | Q9D7S7 1x | 0.052133 | 0.00384 | 1 | 2 | 1 | Q9D7S7 | 60S ribosoi | Q9D7S7 [1 | 0 | 2058.79 | 0 | -0.44 | 0.20623 | 1.55 | 115 | 85 | 6.67 | 25.69 | | High | High | High | High | High | High | High | High | High | High | High | 0.001999 | 0.02029 | 4.34 |
| FALSE | High | YFQISQDEI | [R].YFQISQ | 2xPhospho | Q9D7S7 2x | 7.4E-08 | 0 | 1 | 2 | 40 | Q9D7S7 | 60S ribosoi | Q9D7S7 [1 | 0 | 2137.773 | 0 | 0.15 | 0.210232 | 11.74 | 94.6 | 105.4 | 5.6 | 6.46 | | High | High | High | High | High | High | High | High | High | High | High | 0 | 5.5E-07 | 3.81 |
| FALSE | High | YFSLDLTHD | [K].YFSLDL | 1xPhospho | Q8C0V9 1x | 0.025964 | 0.001645 | 1 | 2 | 1 | Q8C0V9 | FERM dom | Q8C0V9 [6 | 0 | 2219.051 | 0 | 0.68 | 0.043238 | 33.44 | 74.7 | 125.3 | 19.17 | 23.33 | | High | High | High | High | High | High | High | High | High | High | High | 0.000903 | 0.009182 | 2.5 |
| FALSE | High | YHGHSMSI | [R].YHGHSI | 1xPhospho | P35486 1xI | 0.000386 | 0 | 1 | 1 | 26 | P35486 | Pyruvate d | P35486 [28 | 0 | 1917.815 | 0 | 0.36 | 0.066655 | 6.8 | 90.3 | 109.7 | 2.72 | 12.99 | | High | High | High | High | High | High | High | High | High | High | High | 0 | 0.000211 | 3.43 |
| FALSE | High | YEFWGPLY | [K].YEFWG | 1xPhospho | Q07231 1x | 0.054383 | 0.003951 | 1 | 2 | 1 | Q07231 | Zinc finger | Q07231 [2 | 0 | 3142.428 | 0 | 0.5 | 0.118387 | 39.78 | 82.9 | 117.1 | 18.32 | 15.11 | | High | High | High | High | High | High | High | High | High | High | High | 0.002059 | 0.02127 | 2.87 |
| FALSE | High | YHGHSMSI | [R].YHGHSI | 2xPhospho | P35486 2xI | 0.003622 | 0.00021 | 1 | 1 | 1 | P35486 | Pyruvate d | P35486 [28 | 0 | 1981.787 | 0 | 1.33 | 0.009968 | 24.18 | 41.8 | 158.2 | 28.85 | 26.01 | | High | High | High | High | High | High | High | High | High | High | High | 9.42E-05 | 0.001013 | 2.78 |
| FALSE | High | YHSADDILI | [R].YHSADI | 1xPhospho | A7TU71 1x | 8.56E-05 | 0 | 1 | 2 | 1 | A7TU71 | Protein Sh | A7TU71 [9 | 0 | 3177.495 | 0 | 1.42 | 0.015335 | 58.48 | 62.2 | 137.8 | 44.51 | 18.23 | | High | High | High | High | High | High | High | High | High | High | High | 0 | 1.5E-05 | 2.86 |
| FALSE | High | YMAENPTA | [R].YMAEN | 1xPhospho | Q810A7 1x | 8.22E-07 | 0 | 1 | 2 | 5 | Q810A7 | ATP-depen | Q810A7 [1 | 0 | 4049.825 | 0 | 0.02 | 0.831605 | 3.56 | 99.4 | 100.6 | 3.34 | 11.8 | | High | High | High | High | High | High | High | High | High | High | High | 0 | 8.12E-08 | 4.71 |
| FALSE | High | YMAENPTA | [R].YMAEN | 1xPhospho | Q810A7 1x | 4.17E-09 | 0 | 1 | 2 | 20 | Q810A7 | ATP-depen | Q810A7 [1 | 0 | 4065.82 | 0 | 0.06 | 0.541478 | 6.36 | 98.4 | 101.6 | 2.81 | 6.35 | | High | High | High | High | High | High | High | High | High | High | High | 0 | 2.15E-10 | 5.44 |
| FALSE | High | YMDAETSL | [K].YMDAE | 1xPhospho | I1VGT4 1xF | 0.006306 | 0.00062 | 2 | 19 | 1 | I1VGT4; I1 | Anion exch | I1VGT4 [11 | 0 | 1238.521 | 0 | 1.39 | 0.006271 | 28.36 | 51.1 | 148.9 | 17.38 | 18.04 | Shared | High | High | High | High | High | High | High | High | High | High | High | 0.000275 | 0.001874 | 1.95 |
| FALSE | High | YMDAETSL | [K].YMDAE | 1xPhospho | I1VGT4 1xF | 0.013074 | 0.000889 | 2 | 19 | 7 | I1VGT4; I1 | Anion exch | I1VGT4 [11 | 0 | 1254.516 | 0 | 1.93 | 0.000691 | 9.66 | 41.6 | 158.4 | 4.94 | 10.82 | Shared | High | High | High | High | High | High | High | High | High | High | High | 0.000814 | 0.007671 | 2 |
| FALSE | High | YMDDDGS | [K].YMDDD | 1xCarbami | Q9D4V4 1x | 0.000297 | 0 | 1 | 5 | 3 | Q9D4V4 | TATA box-I | Q9D4V4 [1 | 0 | 3936.733 | 0 | -1.25 | 0.002357 | 3.42 | 138.6 | 61.4 | 7.65 | 13.46 | | High | High | High | High | High | High | High | High | High | High | High | 0 | 6.13E-05 | 4.22 |
| FALSE | High | YMLTHQEL | [K].YMLTH | 1xPhospho | E9Q5G3 1x | 9.22E-06 | 0 | 1 | 1 | 1 | E9Q5G3 | Kinesin-like | E9Q5G3 [8 | 0 | 2502.235 | 0 | -1.73 | 0.000425 | 1.33 | 153.6 | 46.4 | 5.73 | 15.14 | | High | High | High | High | High | High | High | High | High | High | High | 0 | 1.24E-06 | 2.49 |
| FALSE | High | YMLTHQEL | [K].YMLTH | 1xPhospho | E9Q5G3 1x | 8.5E-06 | 0 | 1 | 1 | 5 | E9Q5G3 | Kinesin-like | E9Q5G3 [8 | 0 | 2518.23 | 0 | -1.56 | 0.000707 | 2.94 | 151.4 | 48.6 | 5.32 | 17.16 | | High | High | High | High | High | High | High | High | High | High | High | 0 | 1.13E-06 | 3.43 |
| FALSE | High | YMSFTDK | [K].YMSFTI | 1xPhospho | G5E884 1x | 0.025073 | 0.001645 | 1 | 3 | 2 | G5E884 | Serine/thre | G5E884 [1 | 0 | 1445.679 | 0 | -0.8 | 0.001455 | 2.96 | 128 | 72 | 5.66 | 11 | | High | High | High | High | High | High | High | High | High | High | High | 0.002202 | 0.0234 | 2.33 |
| FALSE | High | YMSGQTSI | [R].YMSGC | 1xPhospho | Q9JM13 1x | 0.02933 | 0.001857 | 1 | 1 | 2 | Q9JM13 | Rab5 GDP/ | Q9JM13 [3 | 0 | 1351.591 | 0 | -0.1 | 0.692116 | 18.55 | 103.6 | 96.4 | 6.87 | 9.61 | | High | High | High | High | High | High | High | High | High | High | High | 0.001812 | 0.01837 | 1.39 |
| FALSE | High | YMSSDTTS | [R].YMSSD | 1xPhospho | Q60520-1 | 0.000651 | 0 | 1 | 3 | 4 | Q60520-1 | Isoform 2 c | Q60520-1 | 0 | 1695.75 | 0 | -1.13 | 0.001045 | 2.88 | 137.7 | 62.3 | 6.73 | 6.87 | | High | High | High | High | High | High | High | High | High | High | High | 0.000847 | 0.007979 | 2.3 |
| FALSE | High | YMSSDTTS | [R].YMSSD | 1xPhospho | Q60520-1 | 3.54E-05 | 0 | 1 | 3 | 4 | Q60520-1 | Isoform 2 c | Q60520-1 | 0 | 1711.745 | 0 | -0.84 | 0.000812 | 2.69 | 130.1 | 69.9 | 4.06 | 7.52 | | High | High | High | High | High | High | High | High | High | High | High | 0 | 5.61E-06 | 2.79 |
| FALSE | High | YMTQMSV | [R].YMTQM | 1xPhospho | Q9R0L6-2 | 0.007881 | 0.000736 | 1 | 4 | 1 | Q9R0L6-2 | Isoform 2 c | Q9R0L6-2 | 0 | 2322.116 | 0 | 0.91 | 0.028525 | 26.17 | 62.3 | 137.7 | 11.79 | 26.96 | | High | High | High | High | High | High | High | High | High | High | High | 0.000327 | 0.002404 | 2.36 |
| FALSE | High | YMTQMSV | [R].YMTQM | 1xPhospho | Q9R0L6-2 | 3.67E-05 | 0 | 1 | 4 | 14 | Q9R0L6-2 | Isoform 2 c | Q9R0L6-2 | 0 | 2338.111 | 0 | 1 | 0.002254 | 8.3 | 66.3 | 133.7 | 12.17 | 11.57 | | High | High | High | High | High | High | High | High | High | High | High | 0.000246 | 0.001485 | 2.9 |
| FALSE | High | YLSTTPETT | [R].YLSTTP | 1xCarbami | Q8K0L9 1x | 0.01861 | 0.001362 | 1 | 3 | 2 | Q8K0L9 | Zinc finger | Q8K0L9 [3 | 0 | 1774.803 | 0 | 4.16 | 0.000241 | 41.87 | 9.4 | 190.6 | 30.27 | 6.69 | | High | High | High | High | High | High | High | High | High | High | High | 0.000723 | 0.006266 | 2.03 |
| FALSE | High | YNAPTSHV | [K].YNAPTS | 1xPhospho | B8X349 1xI | 0.100129 | 0.008067 | 2 | 13 | 1 | B8X349; F6 | JNK-interac | B8X349 [5 | 0 | 1939.009 | 0 | 2.36 | 0.001116 | 17.38 | 32.2 | 167.8 | 8.53 | 13.82 | Shared | High | High | High | High | High | High | High | High | High | High | High | 0.004059 | 0.04336 | 1.93 |
| FALSE | High | YNDWSDD | [R].YNDWS | 1xPhospho | Q8BHS6 1x | 4.39E-05 | 0 | 1 | 1 | 2 | Q8BHS6 | Armadillo r | Q8BHS6 [5 | 0 | 2342.934 | 0 | -1.04 | 0.005159 | 2.51 | 134.9 | 65.1 | 8.96 | 13.62 | | High | High | High | High | High | High | High | High | High | High | High | 0 | 7.15E-06 | 4.13 |
| FALSE | High | YNDWSDD | [R].YNDWS | 2xPhospho | Q8BHS6 2x | 0.024424 | 0.001612 | 1 | 1 | 1 | Q8BHS6 | Armadillo r | Q8BHS6 [5 | 0 | 2422.9 | 0 | -0.72 | 0.000255 | 0.61 | 123.8 | 76.2 | 7.74 | 10.74 | | High | High | High | High | High | High | High | High | High | High | High | 0.000859 | 0.008555 | 1.87 |
| FALSE | High | YNLDASEEI | [K].YNLDAS | 1xPhospho | Q9R020 1x | 0.014033 | 0.001011 | 1 | 5 | 3 | Q9R020 | Zinc finger | Q9R020 [1 | 0 | 1822.758 | 0 | -0.4 | 0.758158 | 26.52 | 122.1 | 77.9 | 110.55 | 106.76 | | High | High | High | High | High | High | High | High | High | High | High | 0.000538 | 0.004567 | 1.99 |
| FALSE | High | YNLDASEEI | [K].YNLDAS | 1xPhospho | Q9R020 1x | 1.82E-06 | 0 | 1 | 5 | 8 | Q9R020 | Zinc finger | Q9R020 [1 | 0 | 2051.921 | 0 | -0.32 | 0.000209 | 0.19 | 112.3 | 87.7 | 2.5 | 3.41 | | High | High | High | High | High | High | High | High | High | High | High | 0 | 9.57E-07 | 3.58 |
| FALSE | High | YNLDASEEI | [K].YNLDAS | 2xPhospho | Q9R020 2x | 0.065952 | 0.004766 | 1 | 5 | 1 | Q9R020 | Zinc finger | Q9R020 [1 | 0 | 2131.887 | 0 | -0.65 | 0.005647 | 6.61 | 123.7 | 76.3 | 8.46 | 9.5 | | High | High | High | High | High | High | High | High | High | High | High | 0.002455 | 0.02656 | 2.28 |
| FALSE | High | YPAPSSSSS | [K].YPAPSS | 1xPhospho | A0A0R4J0( | 5.09E-10 | 0 | 1 | 2 | 5 | A0A0R4J0( | Ran-bindin | A0A0R4J0( | 0 | 3887.687 | 0 | 0.01 | 0.680992 | 21.34 | 105 | 95 | 8.62 | 18.18 | | High | High | High | High | High | High | High | High | High | High | High | 0 | 2.02E-11 | 5.27 |
| FALSE | High | YPCATSDT/ | [R].YPCATS | 1xCarbami | E9Q7D5 1x | 1.28E-10 | 0 | 1 | 1 | 5 | E9Q7D5 | Rho guanir | E9Q7D5 [5 | 0 | 3169.461 | 0 | 0.96 | 0.001024 | 10.83 | 68.5 | 131.5 | 18.66 | 11.25 | | High | High | High | High | High | High | High | High | High | High | High | 0 | 7.36E-11 | 4.55 |
| FALSE | High | YPDIASPTF | [K].YPDIAS | 1xPhospho | E9QMF5 1: | 0.072858 | 0.005599 | 1 | 2 | 1 | E9QMF5 | Neuron na | E9QMF5 [1 | 0 | 1475.713 | 0 | 3.13 | 0.00039 | 26.2 | 18.9 | 181.1 | 17.47 | 8.31 | | High | High | High | High | High | High | High | High | High | High | High | 0.002867 | 0.02972 | 1.75 |
| FALSE | High | YPEDGLSPI | [R].YPEDGI | 1xPhospho | A0A1W2P6 | 0.048303 | 0.00354 | 1 | 10 | 1 | A0A1W2P6 | Bromodom | A0A1W2P6 | 0 | 1411.671 | 0 | -0.74 | 0.311886 | 10.18 | 127.4 | 72.6 | 55.46 | 49.67 | | High | High | High | High | High | High | High | High | High | High | High | 0.001844 | 0.01861 | 1.86 |
| FALSE | High | YPEDGLSPI | [R].YPEDGI | 1xPhospho | A0A1W2P6 | 0.000563 | 0 | 1 | 10 | 3 | A0A1W2P6 | Bromodom | A0A1W2P6 | 0 | 1640.834 | 0 | -0.89 | 0.001229 | 1.79 | 127.1 | 72.9 | 7.49 | 5.08 | | High | High | High | High | High | High | High | High | High | High | High | 0 | 0.000126 | 2.14 |
| FALSE | High | YPESNRTP\ | [K].YPESNF | 1xPhospho | Q9ESC8 1x | 0.06169 | 0.004646 | 1 | 2 | 1 | Q9ESC8 | AF4/FMR2 | Q9ESC8 [6 | 1 | 3138.519 | 0 | -0.59 | 0.00063 | 3.65 | 122.6 | 77.4 | 5.28 | 3 | | High | High | High | High | High | High | High | High | High | High | High | 0.002381 | 0.02454 | 1.46 |
| FALSE | High | YPLSPPQEI | [R].YPLSPP | 1xPhospho | D0VLQ5 1x | 0.000553 | 0 | 1 | 3 | 2 | D0VLQ5 | Tet oncoge | D0VLQ5 [9 | 0 | 1830.829 | 0 | -1.92 | 0.000158 | 1.25 | 159.5 | 40.5 | 5.5 | 17.16 | | High | High | High | High | High | High | High | High | High | High | High | 0.004086 | 0.04366 | 1.93 |
| FALSE | High | YPNQDHLT | [R].YPNQD | 1xDeamida | D0VLQ5 1x | 0.009843 | 0.000736 | 1 | 6 | 4 | D0VLQ5 | Tet oncoge | D0VLQ5 [1 | 0 | 3866.89 | 0 | -1.3 | 0.018753 | 10.27 | 146.1 | 53.9 | 24.16 | 43.86 | | High | High | High | High | High | High | High | High | High | High | High | 0.000449 | 0.004066 | 3.77 |
| FALSE | High | YNAVLGFG | [K].YNAVL( | 2xPhospho | A2AHG2 2x | 4.59E-05 | 0 | 1 | 8 | 1 | A2AHG2 | PHD finger | A2AHG2 [4 | 0 | 3430.601 | 0 | 0.4 | 0.051212 | 2.41 | 87.1 | 112.9 | 9.26 | 8.22 | | High | High | High | High | High | High | High | High | High | High | High | 0 | 7.5E-06 | 2.54 |
| FALSE | High | YLSFTPPEK | [K].YLSFTPI | 1xPhospho | Q8CIN4 1xI | 0.002895 | 0.00016 | 1 | 1 | 5 | Q8CIN4 | Serine/thre | Q8CIN4 [1 | 0 | 1619.849 | 0 | 0.51 | 0.002153 | 7.27 | 78.1 | 121.9 | 6.31 | 9.25 | | High | High | High | High | High | High | High | High | High | High | High | 7.19E-05 | 0.000789 | 2 |
| FALSE | High | YLSESDGIP | [R].YLSESD | 1xPhospho | E9Q4E4 1x | 0.000418 | 0 | 1 | 3 | 1 | E9Q4E4 | DENN dom | E9Q4E4 [2 | 0 | 1809.908 | 0 | -0.66 | 0.001627 | 0.77 | 119.8 | 80.2 | 9.68 | 6.67 | | High | High | High | High | High | High | High | High | High | High | High | 0 | 9E-05 | 1.87 |
| FALSE | High | YLSDSYSPR | [K].YLSDSY | 1xPhospho | A0A0R4J0( | 0.071662 | 0.005278 | 1 | 3 | 1 | A0A0R4J0( | Conserved | A0A0R4J0( | 0 | 1396.635 | 0 | -0.23 | 0.001414 | 1.7 | 109.2 | 90.8 | 1.84 | 3.53 | | High | High | High | High | High | High | High | High | High | High | High | 0.002717 | 0.02916 | 1.88 |
| FALSE | High | YHSLGNISF | [R].YHSLGN | 1xPhospho | Q08274 1x | 0.004781 | 0.000456 | 1 | 4 | 2 | Q08274 | Dystrophia | Q08274 [5 | 0 | 1355.667 | 0 | 0.36 | 0.003027 | 7.21 | 87.8 | 112.2 | 3.22 | 5.22 | | High | High | High | High | High | High | High | High | High | High | High | 0.000275 | 0.002137 | 1.8 |
| FALSE | High | YHSVSTETF | [R].YHSVST | 1xPhospho | Q60949 1x | 8.63E-08 | 0 | 1 | 5 | 1 | Q60949 | TBC1 doma | Q60949 [6 | 0 | 1751.795 | 0 | -0.33 | 0.040557 | 4.19 | 113.2 | 86.8 | 10.66 | 23 | | High | High | High | High | High | High | High | High | High | High | High | 0 | 6.51E-09 | 2.35 |
| FALSE | High | YLAEDSNIS | [K].YLAEDS | 1xPhospho | C7G3P2 1x | 7.82E-05 | 0 | 1 | 2 | 3 | C7G3P2 | MKIAA053 | C7G3P2 [5 | 0 | 2660.229 | 0 | -0.07 | 0.564022 | 12.49 | 100.6 | 99.4 | 12.01 | 5.52 | | High | High | High | High | High | High | High | High | High | High | High | 0 | 1.93E-05 | 3.1 |
| FALSE | High | YIASVQGS/ | [R].YIASVQ | 1xPhospho | Q9ERU9 1x | 0.000442 | 0 | 1 | 1 | 1 | Q9ERU9 | E3 SUMO-p | Q9ERU9 [1 | 0 | 1641.82 | 0 | -0.16 | 0.699701 | 18.13 | 102.6 | 97.4 | 7.68 | 14.31 | | High | High | High | High | High | High | High | High | High | High | High | 0 | 9.54E-05 | 3.14 |
| FALSE | High | YLCEGTESF | [R].YLCEGT | 1xCarbami | B2RRF0 1xI | 0.001799 | 5.66E-05 | 1 | 3 | 2 | B2RRF0 | Ptprk prote | B2RRF0 [8 | 0 | 2629.232 | 0 | 1.27 | 0.000855 | 7.73 | 61.4 | 138.6 | 9.49 | 10.94 | | High | High | High | High | High | High | High | High | High | High | High | 2.58E-05 | 0.000461 | 2.2 |
| FALSE | High | YIDLEPDTP | [R].YIDLEPI | 1xPhospho | Q7TQG1 1: | 0.000108 | 0 | 1 | 6 | 1 | Q7TQG1 | Pleckstrin | Q7TQG1 [1 | 0 | 2397.224 | 0 | 2.96 | 0.000311 | 6.31 | 23.7 | 176.3 | 8.54 | 8.69 | | High | High | High | High | High | High | High | High | High | High | High | 0 | 1.97E-05 | 3.27 |
| FALSE | High | YIEEFHSNI | [K].YIEEFH | 1xPhospho | Q3UHU5 1: | 0.1059 | 0.008582 | 1 | 5 | 1 | Q3UHU5 | Microtubu | Q3UHU5 [1 | 0 | 2338.002 | 0 | -1.94 | 0.000866 | 0.65 | 155.2 | 44.8 | 8.84 | 13.27 | | High | High | High | High | High | High | High | High | High | High | High | 0.004317 | 0.04634 | 1.5 |
| FALSE | High | YLEESDDDI | [K].YLEESD | 1xPhospho | Q01320 1x | 2.86E-07 | 0 | 2 | 4 | 104 | Q01320; Q | DNA topois | Q01320 [1 | 0 | 1784.699 | 0 | -1.1 | 0.000154 | 0.72 | 135.7 | 64.3 | 1.5 | 2.82 | Shared | High | High | High | High | High | High | High | High | High | High | High | 0 | 1.44E-07 | 3.5 |
| FALSE | High | YLEIDSDEE | [K].YLEIDSI | 1xPhospho | A0A0R4J0E | 6.24E-05 | 0 | 1 | 4 | 4 | A0A0R4J0E | Protein SD. | A0A0R4J0E | 0 | 1664.725 | 0 | -0.8 | 0.001887 | 3.81 | 125.7 | 74.3 | 5.85 | 6.64 | | High | High | High | High | High | High | High | High | High | High | High | 4.94E-05 | 0.00068 | 2.74 |
| FALSE | High | YLEIDSDEE | [K].YLEIDSI | 1xPhospho | A0A0R4J0E | 0.005615 | 0.00062 | 1 | 4 | 1 | A0A0R4J0E | Protein SD. | A0A0R4J0E | 1 | 2433.175 | 0 | -0.8 | 0.000436 | 2.45 | 127.1 | 72.9 | 4.11 | 3.11 | | High | High | High | High | High | High | High | High | High | High | High | 0.000275 | 0.00165 | 1.33 |
| FALSE | High | YIESPLESES | [R].YIESPLE | 1xPhospho | Q3TYA6 1x | 4.37E-07 | 0 | 1 | 3 | 4 | Q3TYA6 | M-phase p | Q3TYA6 [2 | 0 | 3637.616 | 0 | -1.48 | 0.000497 | 1.73 | 150.1 | 49.9 | 6.72 | 8.34 | | High | High | High | High | High | High | High | High | High | High | High | 0 | 4.03E-08 | 5.54 |
| FALSE | High | YIESPLESES | [R].YIESPLE | 1xDeamida | Q3TYA6 1x | 6.61E-06 | 0 | 1 | 3 | 3 | Q3TYA6 | M-phase p | Q3TYA6 [2 | 0 | 3638.6 | 0 | -1.5 | 0.000443 | 0.93 | 150.1 | 49.9 | 7.01 | 16.58 | | High | High | High | High | High | High | High | High | High | High | High | 0 | 8.48E-07 | 5.08 |
| FALSE | High | YIESPLESES | [R].YIESPLE | 1xPhospho | Q3TYA6 1x | 0.002398 | 0.000109 | 1 | 3 | 3 | Q3TYA6 | M-phase p | Q3TYA6 [2 | 1 | 4507.015 | 0 | -1.23 | 0.002264 | 3.62 | 146.8 | 53.2 | 12.55 | 22.17 | | High | High | High | High | High | High | High | High | High | High | High | 4.94E-05 | 0.000636 | 3.98 |
| FALSE | High | YIESPLESES | [R].YIESPLE | 2xPhospho | Q3TYA6 2x | 0.002647 | 0.000109 | 1 | 3 | 1 | Q3TYA6 | M-phase p | Q3TYA6 [2 | 1 | 4586.981 | 0 | -1.44 | 0.002578 | 3.16 | 145 | 55 | 12.1 | 11.42 | | High | High | High | High | High | High | High | High | High | High | High | 4.94E-05 | 0.000707 | 3.25 |
| FALSE | High | YLFNQLFGI | [K].YLFNQI | 1xPhospho | Q9Z1F9 1xI | 2.02E-05 | 0 | 1 | 3 | 1 | Q9Z1F9 | SUMO-acti | Q9Z1F9 [1 | 0 | 2681.197 | 0 | -0.7 | 0.003039 | 3.49 | 124.7 | 75.3 | 9.98 | 9.09 | | High | High | High | High | High | High | High | High | High | High | High | 0 | 2.99E-06 | 3.89 |
| FALSE | High | YLGDAFGS | [K].YLGDAI | 1xPhospho | E9Q5N9 1x | 0.000573 | 0 | 1 | 2 | 4 | E9Q5N9 | Zinc finger | E9Q5N9 [2 | 0 | 3613.636 | 0 | -1.54 | 0.002031 | 2.26 | 151.5 | 48.5 | 9.03 | 16.49 | | High | High | High | High | High | High | High | High | High | High | High | 0 | 0.000128 | 3.73 |
| FALSE | High | YLGGSMDI | [R].YLGGSI | 1xPhospho | D3YTR4 1x | 0.000343 | 0 | 2 | 4 | 1 | D3YTR4; Q | Methyl-Cp | D3YTR4 [2 | 0 | 1917.866 | 0 | 0.81 | 0.005952 | 4.78 | 71.1 | 128.9 | 13.08 | 11.67 | Shared | High | High | High | High | High | High | High | High | High | High | High | 0 | 0.000072 | 2.09 |
| FALSE | High | YLGGSMDI | [R].YLGGSI | 1xPhospho | D3YTR4 1x | 7.04E-06 | 0 | 2 | 4 | 1 | D3YTR4; Q | Methyl-Cp | D3YTR4 [2 | 0 | 1933.86 | 0 | 0.8 | 0.051987 | 27.09 | 72.9 | 127.1 | 10.92 | 31.19 | Shared | High | High | High | High | High | High | High | High | High | High | High | 0 | 9.09E-07 | 2.22 |
| FALSE | High | YLGVQLPL/ | [K].YLGVQI | 1xPhospho | E9QLG3 1x | 4.48E-06 | 0 | 1 | 4 | 1 | E9QLG3 | MAX gene- | E9QLG3 [7 | 0 | 4097.078 | 0 | -0.53 | 0.003291 | 7.73 | 115.8 | 84.2 | 4.42 | 15.02 | | High | High | High | High | High | High | High | High | High | High | High | 0 | 5.46E-07 | 4.69 |
| FALSE | High | YLLGNAPV | [K].YLLGN/ | 1xPhospho | Q91VL8 1x | 1.8E-06 | 0 | 1 | 2 | 1 | Q91VL8 | Telomeric | Q91VL8 [1 | 0 | 1999.067 | 0 | -1.08 | 0.00022 | 1.77 | 136 | 64 | 1.39 | 6.19 | | High | High | High | High | High | High | High | High | High | High | High | 0 | 1.97E-07 | 1.91 |
| FALSE | High | YILNVTPNI | [R].YILNVT | 1xPhospho | Q7TNL7 1x | 0.005565 | 0.00062 | 1 | 2 | 1 | Q7TNL7 | Dual specif | Q7TNL7 [3 | 0 | 2313.266 | 0 | -1.11 | 0.000509 | 4.31 | 137.2 | 62.8 | 7.13 | 22.47 | | High | High | High | High | High | High | High | High | High | High | High | 0.000275 | 0.001631 | 2.36 |
| FALSE | High | YLLSQSSPA | [K].YLLSQS | 1xPhospho | Q91YR1 1x | 4.12E-08 | 0 | 1 | 1 | 5 | Q91YR1 | Twinfilin-1 | Q91YR1 [1 | 0 | 2384.195 | 0 | 0.79 | 0.002355 | 8.4 | 68.8 | 131.2 | 10.68 | 7.09 | | High | High | High | High | High | High | High | High | High | High | High | 0 | 2.33E-08 | 3.7 |
| FALSE | High | YLMLTAPA | [R].YLMLT/ | 1xPhospho | Q64127 1x | 0.040721 | 0.003058 | 1 | 2 | 1 | Q64127 | Transcripti | Q64127 [8 | 0 | 4389.256 | 0 | -0.68 | 0.001858 | 7.11 | 123.1 | 76.9 | 7.7 | 12.67 | | High | High | High | High | High | High | High | High | High | High | High | 0.001597 | 0.0153 | 3.8 |
| FALSE | High | YLMLTAPA | [R].YLMLT/ | 1xPhospho | Q64127 1x | 0.000738 | 0 | 1 | 2 | 2 | Q64127 | Transcripti | Q64127 [8 | 0 | 4405.25 | 0 | -1.03 | 0.01382 | 2.93 | 128.8 | 71.2 | 22.56 | 11.28 | | High | High | High | High | High | High | High | High | High | High | High | 0 | 0.00017 | 4 |
| FALSE | High | YLPAITSTP | [K].YLPAITS | 1xPhospho | Q9R0L6-2 | 9.79E-08 | 0 | 1 | 3 | 12 | Q9R0L6-2 | Isoform 2 c | Q9R0L6-2 | 0 | 2663.322 | 0 | 0.11 | 0.250489 | 2.24 | 96.6 | 103.4 | 4.62 | 10.97 | | High | High | High | High | High | High | High | High | High | High | High | 0 | 7.26E-08 | 3.99 |
| FALSE | High | YLPHSAGR | [K].YLPHSA | 1xPhospho | P97461 1xI | 0.109329 | 0.008865 | 1 | 4 | 2 | P97461 | 40S ribosoi | P97461 [48 | 0 | 1209.598 | 0 | -0.3 | 0.010155 | 4.18 | 110.7 | 89.3 | 5.17 | 4.22 | | High | High | High | High | High | High | High | High | High | High | High | 0.004462 | 0.04809 | 2.61 |
| FALSE | High | YIPTEDVSG | [K].YIPTED | 1xPhospho | Q62210 1x | 1.07E-06 | 0 | 1 | 1 | 1 | Q62210 | Baculoviral | Q62210 [5 | 0 | 2258.115 | 0 | -0.53 | 0.102053 | 22.48 | 125.3 | 74.7 | 38.18 | 15.64 | | High | High | High | High | High | High | High | High | High | High | High | 0 | 1.09E-07 | 4.63 |
| FALSE | High | YLQAHTQS | [R].YLQAH | 1xPhospho | A0A0R4J0I | 0.000171 | 0 | 1 | 2 | 1 | A0A0R4J0I | ETS domai | A0A0R4J0I | 0 | 2386.154 | 0 | -0.21 | 0.120528 | 3.21 | 114.7 | 85.3 | 22.17 | 16.4 | | High | High | High | High | High | High | High | High | High | High | High | 0 | 3.3E-05 | 2.39 |
| FALSE | High | YISDGVECS | [R].YISDGV | 1xCarbami | A1BQX3 2x | 5.31E-05 | 0 | 1 | 1 | 1 | A1BQX3 | Tricellulin i | A1BQX3 [1 | 0 | 2094.881 | 0 | -0.62 | 0.006327 | 1.89 | 121.2 | 78.8 | 11.54 | 9.03 | | High | High | High | High | High | High | High | High | High | High | High | 0 | 8.8E-06 | 3.11 |
| FALSE | High | YHHCTDEP | [K].YHHCTI | 1xCarbami | Q3TIA9 1xI | 0.001516 | 5.66E-05 | 1 | 4 | 1 | Q3TIA9 | Putative ur | Q3TIA9 [2 | 0 | 2752.284 | 0 | -1.64 | 0.002035 | 0.22 | 148 | 52 | 12.83 | 15.16 | | High | High | High | High | High | High | High | High | High | High | High | 2.58E-05 | 0.000381 | 2.46 |
| FALSE | High | VTFFEPSP( | [K].VTFFEP | 1xDeamida | Q3UKX3 1x | 0.090153 | 0.00698 | 1 | 15 | 1 | Q3UKX3 | Putative ur | Q3UKX3 [3 | 0 | 2365.1 | 0 | 0.59 | 0.002238 | 4.83 | 79.8 | 120.2 | 13.32 | 3.84 | | High | High | High | High | High | High | High | High | High | High | High | 0.003606 | 0.03836 | 2.56 |
| FALSE | High | VTFFEPSP( | [K].VTFFEP | 1xPhospho | Q3UKX3 1x | 3.73E-05 | 0 | 1 | 15 | 2 | Q3UKX3 | Putative ur | Q3UKX3 [3 | 0 | 2364.116 | 0 | 0.57 | 0.005601 | 10.54 | 81.5 | 118.5 | 11.05 | 7.08 | | High | High | High | High | High | High | High | High | High | High | High | 0 | 5.93E-06 | 3.05 |
| FALSE | High | VTFALPDDI | [R].VTFALP | 1xPhospho | Q810V0 1x | 2.48E-07 | 0 | 1 | 2 | 3 | Q810V0 | U3 small nu | Q810V0 [3 | 0 | 2556.289 | 0 | -0.2 | 0.002919 | 1.99 | 107.3 | 92.7 | 2.57 | 5.05 | | High | High | High | High | High | High | High | High | High | High | High | 0 | 2.11E-08 | 4.81 |
| FALSE | High | VNASASSLI | [K].VNASA | 1xPhospho | Q61553 1x | 0.112856 | 0.009146 | 1 | 2 | 2 | Q61553 | Fascin OS= | Q61553 [3 | 0 | 1414.771 | 0 | -1.59 | 0.000209 | 0.51 | 150.7 | 49.3 | 4.77 | 13.96 | | High | High | High | High | High | High | High | High | High | High | High | 0.004596 | 0.04995 | 1.59 |
| FALSE | High | VNFPENGF | [K].VNFPEI | 1xPhospho | P63037 1xI | 6.04E-06 | 0 | 1 | 2 | 1 | P63037 | DnaJ homo | P63037 [32 | 0 | 2002.009 | 0 | -1.26 | 0.000235 | 0.47 | 141.4 | 58.6 | 6.08 | 6.93 | | High | High | High | High | High | High | High | High | High | High | High | 0 | 7.68E-07 | 3.32 |
| FALSE | High | VNFPENGF | [K].VNFPEI | 1xDeamida | P63037 1xI | 1.12E-05 | 0 | 1 | 2 | 4 | P63037 | DnaJ homo | P63037 [32 | 0 | 2002.993 | 0 | -1.14 | 0.00143 | 5.1 | 138.1 | 61.9 | 4.33 | 16.17 | | High | High | High | High | High | High | High | High | High | High | High | 0 | 1.55E-06 | 3.91 |
| FALSE | High | VNFTEEEW | [R].VNFTEI | 1xPhospho | Q91VL6 1x | 7.41E-05 | 0 | 1 | 2 | 1 | Q91VL6 | Zinc finger | Q91VL6 [1 | 0 | 2488.205 | 0 | -1.05 | 0.004416 | 0.39 | 141.5 | 58.5 | 17.18 | 9.79 | | High | High | High | High | High | High | High | High | High | High | High | 0 | 1.28E-05 | 3.35 |
| FALSE | High | VNGDASPA | [K].VNGDA | 1xPhospho | P26645 1xI | 1.08E-06 | 0 | 1 | 3 | 2 | P26645 | Myristoyla | P26645 [41 | 0 | 1892.952 | 0 | 2.01 | 0.00043 | 4.11 | 40.3 | 159.7 | 2.48 | 8.01 | | High | High | High | High | High | High | High | High | High | High | High | 0 | 2.77E-07 | 4.3 |
| FALSE | High | VNGDASPA | [K].VNGDA | 1xDeamida | P26645 1xI | 1.17E-06 | 0 | 1 | 3 | 4 | P26645 | Myristoyla | P26645 [41 | 0 | 1893.936 | 0 | 1.85 | 0.000201 | 4.91 | 44.3 | 155.7 | 7.93 | 4.58 | | High | High | High | High | High | High | High | High | High | High | High | 0 | 1.21E-07 | 4.44 |
| FALSE | High | VNGTTVSS | [K].VNGTT | 1xDeamida | P11531 1xI | 0.013426 | 0.000976 | 1 | 12 | 1 | P11531 | Dystrophin | P11531 [35 | 0 | 1843.9 | 0 | -1.81 | 0.000438 | 1.55 | 153.9 | 46.1 | 4.3 | 15.95 | | High | High | High | High | High | High | High | High | High | High | High | 0.000505 | 0.004352 | 2.66 |
| FALSE | High | VNHEPEPA | [R].VNHEPI | 1xPhospho | O08539 1x | 1.13E-05 | 0 | 1 | 3 | 3 | O08539 | Myc box-d | O08539 [3 | 0 | 2397.222 | 0 | 1.27 | 0.000267 | 3.05 | 55.4 | 144.6 | 4.24 | 6.06 | | High | High | High | High | High | High | High | High | High | High | High | 0 | 1.55E-06 | 2.75 |
| FALSE | High | VNLEESTG | [R].VNLEES | 1xPhospho | Q8BRF7 1x | 4.1E-06 | 0 | 1 | 3 | 3 | Q8BRF7 | Sec1 family | Q8BRF7 [2 | 0 | 2493.275 | 0 | 0.46 | 0.001627 | 5.09 | 82.8 | 117.2 | 3.72 | 7.75 | | High | High | High | High | High | High | High | High | High | High | High | 0 | 4.95E-07 | 2.7 |
| FALSE | High | VNLFPGLSI | [K].VNLFP( | 1xPhospho | B9EKJ7 1xF | 0.008614 | 0.000736 | 1 | 3 | 1 | B9EKJ7 | Tankyrase | B9EKJ7 [16 | 0 | 1881.065 | 0 | -0.2 | 0.007246 | 2.22 | 107 | 93 | 6.86 | 7.46 | | High | High | High | High | High | High | High | High | High | High | High | 0.000327 | 0.002652 | 2.73 |
| FALSE | High | VNISLLPK | [R].VNISLLI | 1xDeamida | Q69ZN6 1x | 0.120211 | 0.009851 | 1 | 3 | 1 | Q69ZN6 | N-acetylglu | Q69ZN6 [6 | 0 | 1422.837 | 0 | -1.23 | 0.000905 | 3.29 | 136.1 | 63.9 | 7.3 | 13.94 | | High | High | High | High | High | High | High | High | High | High | High | 0.004907 | 0.05354 | 1.8 |
| FALSE | High | VNNSTPEL | [R].VNNSTI | 1xCarbami | Q8C263 1x | 3.29E-05 | 0 | 1 | 1 | 4 | Q8C263 | Spindle and | Q8C263 [1 | 0 | 1869.955 | 0 | -1.34 | 0.000357 | 1.1 | 145.7 | 54.3 | 5.33 | 9.7 | | High | High | High | High | High | High | High | High | High | High | High | 0 | 0.000187 | 3.3 |
| FALSE | High | VNQQPNA | [K].VNQQF | 1xPhospho | Q9JMA1 1x | 0.000382 | 0 | 1 | 4 | 1 | Q9JMA1 | Ubiquitin c | Q9JMA1 [3 | 1 | 2418.275 | 0 | -0.71 | 0.141276 | 16.95 | 128.3 | 71.7 | 36.06 | 16.15 | | High | High | High | High | High | High | High | High | High | High | High | 0 | 8.14E-05 | 3.98 |
| FALSE | High | VMVAPLSG | [K].VMVAF | 1xPhospho | A2A654 1x | 0.000289 | 0 | 1 | 8 | 1 | A2A654 | Bromodom | A2A654 [2 | 0 | 1998.075 | 0 | 0.12 | 0.126591 | 3.69 | 94.7 | 105.3 | 9.72 | 6.63 | | High | High | High | High | High | High | High | High | High | High | High | 0 | 5.91E-05 | 3.4 |
| FALSE | High | VNQSALEA | [R].VNQSA | 1xPhospho | Q8K3H0 1x | 9.76E-07 | 0 | 1 | 1 | 3 | Q8K3H0 | DCC-intera | Q8K3H0 [3 | 0 | 2268.122 | 0 | 0.45 | 0.008175 | 11.27 | 86 | 114 | 5.11 | 9.94 | | High | High | High | High | High | High | High | High | High | High | High | 0 | 2.76E-07 | 3.75 |
| FALSE | High | VNRPGWN | [R].VNRPG | 1xDeamida | E9PXF8 1xI | 0.031439 | 0.002011 | 1 | 5 | 2 | E9PXF8 | SET-bindin | E9PXF8 [1 | 0 | 3543.678 | 0 | 0.49 | 0.03442 | 6.43 | 83.1 | 116.9 | 18.73 | 17.76 | | High | High | High | High | High | High | High | High | High | High | High | 0.006366 | 0.06853 | 2.72 |
| FALSE | High | VNSATKPEI | [R].VNSATI | 1xPhospho | Q80TR8-2 | 0.053023 | 0.00384 | 1 | 4 | 1 | Q80TR8-2 | Isoform 2 c | Q80TR8-2 | 0 | 1867.993 | 0 | -0.7 | 0.430997 | 35.55 | 121.9 | 78.1 | 48.98 | 40.2 | | High | High | High | High | High | High | High | High | High | High | High | 0.001999 | 0.02058 | 2.74 |

| | | | | | | | | | | | | | | | | | | | | | | | | | | | | | | | | | | | | | | |
|---|---|---|---|---|---|---|---|---|---|---|---|---|---|---|---|---|---|---|---|---|---|---|---|---|---|---|---|---|---|---|---|---|---|---|---|---|---|---|
| FALSE | High | VNSATKPE | [R].VNSATI | 1xPhospho | Q80TR8-2 1 | 0.000241 | 0 | 1 | 4 | 1 | Q80TR8-2 | Isoform 2 c | Q80TR8-2 | 0 | 2097.156 | 0 | -0.68 | 0.000958 | 1.03 | 122.9 | 77.1 | 5.33 | 2.48 | | High | High | High | High | High | High | High | High | High | High | High | 0 | 4.84E-05 | 1.46 |
| FALSE | High | VNSGDTVG | [R].VNSGD | 1xPhospho | E9PUJ6 1xP | 0.000142 | 0 | 1 | 5 | 2 | E9PUJ6 | E3 ubiquiti | E9PUJ6 [35 | 0 | 1613.81 | 0 | 0.7 | 9.77E-05 | 2.31 | 76.2 | 123.8 | 4.95 | 2.99 | | High | High | High | High | High | High | High | High | High | High | High | 0 | 2.65E-05 | 2.07 |
| FALSE | High | VNSNSLDLI | [K].VNSNSI | 1xPhospho | Q8CH77 1x | 5.05E-07 | 0 | 1 | 4 | 2 | Q8CH77 | Neuron na | Q8CH77 [8 | 0 | 2397.17 | 0 | 0.5 | 0.146427 | 4.73 | 82.7 | 117.3 | 18.51 | 21.79 | | High | High | High | High | High | High | High | High | High | High | High | 0 | 4.74E-08 | 3.97 |
| FALSE | High | VNSQPGPC | [R].VNSQP | 1xPhospho | Q8BGG7 1 | 0.047894 | 0.00354 | 1 | 2 | 1 | Q8BGG7 | Ubiquitin-a | Q8BGG7 [3 | 0 | 1492.793 | 0 | 2.43 | 0.001088 | 7.14 | 31.3 | 168.7 | 11.16 | 13.3 | | High | High | High | High | High | High | High | High | High | High | High | 0.001812 | 0.0184 | 2.2 |
| FALSE | High | VNSSGESG | [R].VNSSGI | 1xPhospho | Q05D44 1x | 4.89E-08 | 0 | 1 | 1 | 5 | Q05D44 | Eukaryotic | Q05D44 [1 | 0 | 2193.95 | 0 | 0 | 0.859135 | 12.89 | 97.5 | 102.5 | 13.69 | 11.71 | | High | High | High | High | High | High | High | High | High | High | High | 0 | 3.42E-09 | 3.95 |
| FALSE | High | VNSSGESG | [R].VNSSGI | 2xPhospho | Q05D44 2x | 0.0008 | 0 | 1 | 1 | 2 | Q05D44 | Eukaryotic | Q05D44 [1 | 0 | 2273.916 | 0 | -1.29 | 0.007617 | 6.49 | 141.9 | 58.1 | 16.99 | 15.51 | | High | High | High | High | High | High | High | High | High | High | High | 0 | 0.000186 | 3.36 |
| FALSE | High | VNSSGESG | [R].VNSSGI | 3xPhospho | Q05D44 3x | 4.44E-06 | 0 | 1 | 1 | 4 | Q05D44 | Eukaryotic | Q05D44 [1 | 0 | 2353.883 | 0 | -0.52 | 0.002587 | 5.46 | 116.7 | 83.3 | 4.7 | 5.46 | | High | High | High | High | High | High | High | High | High | High | High | 0 | 5.41E-07 | 4.53 |
| FALSE | High | VNSVGSSA | [R].VNSVG | 1xPhospho | P05532 1xI | 0.077188 | 0.005893 | 1 | 4 | 1 | P05532 | Mast/stem | P05532 [96 | 0 | 2307.107 | 0 | -0.86 | 0.130641 | 1.16 | 125.9 | 74.1 | 24.67 | 23.07 | | High | High | High | High | High | High | High | High | High | High | High | 0.003064 | 0.03197 | 3.34 |
| FALSE | High | VNTAGASD | [K].VNTAG | 1xPhospho | A0A0R4J07 | 3.47E-05 | 0 | 1 | 3 | 1 | A0A0R4J07 | Acyl-Coenz | A0A0R4J07 | 0 | 2171.111 | 0 | 0.61 | 0.013136 | 8.02 | 73.4 | 126.6 | 8.43 | 17.39 | | High | High | High | High | High | High | High | High | High | High | High | 0 | 5.45E-06 | 2.9 |
| FALSE | High | VNTAGASD | [K].VNTAG | 1xDeamida | A0A0R4J07 | 4.22E-07 | 0 | 1 | 3 | 1 | A0A0R4J07 | Acyl-Coenz | A0A0R4J07 | 0 | 2172.095 | 0 | 0.65 | 0.001715 | 11.75 | 77.7 | 122.3 | 6.68 | 7.7 | | High | High | High | High | High | High | High | High | High | High | High | 0 | 3.86E-08 | 2.42 |
| FALSE | High | VNTYPEDS | [R].VNTYPI | 1xPhospho | E9PX30 1xI | 0.000393 | 0 | 1 | 5 | 2 | E9PX30 | Mitogen-a | E9PX30 [39 | 0 | 2174.031 | 0 | 1.8 | 0.071846 | 53.71 | 42.9 | 157.1 | 24.17 | 49.27 | | High | High | High | High | High | High | High | High | High | High | High | 0 | 8.39E-05 | 2.84 |
| FALSE | High | VNRGSVSS | [R].VNRGS | 1xPhospho | F8VQ93 1x | 0.002419 | 0.000109 | 1 | 5 | 1 | F8VQ93 | PH-interac | F8VQ93 [6 | 1 | 2502.266 | 0 | 0.71 | 0.16003 | 53.27 | 76 | 124 | 39.13 | 21.63 | | High | High | High | High | High | High | High | High | High | High | High | 4.94E-05 | 0.000641 | 2.93 |
| FALSE | High | VNVTVDYII | [K].VNVTVI | 1xPhospho | Q3TRW3 1 | 5.16E-05 | 0 | 1 | 5 | 1 | Q3TRW3 | Putative ur | Q3TRW3 [4 | 0 | 2928.471 | 0 | 0.6 | 0.004041 | 6.25 | 80.8 | 119.2 | 6.77 | 5.34 | | High | High | High | High | High | High | High | High | High | High | High | 0 | 8.58E-06 | 1.96 |
| FALSE | High | VMTVTAVT | [R].VMTVT | 1xPhospho | Q3UMU9-3 | 7.15E-05 | 0 | 1 | 4 | 7 | Q3UMU9-3 | Isoform 3 c | Q3UMU9-3 | 0 | 1878.908 | 0 | -0.08 | 0.432379 | 7.14 | 104.5 | 95.5 | 6.05 | 7.31 | | High | High | High | High | High | High | High | High | High | High | High | 9.42E-05 | 0.000965 | 3.65 |
| FALSE | High | VMTLQWL | [R].VMTLQ | 1xPhospho | E9QKG8 1x | 0.000142 | 0 | 1 | 7 | 4 | E9QKG8 | Holliday ju | E9QKG8 [3 | 0 | 1969.1 | 0 | -1.23 | 0.000503 | 0.84 | 138 | 62 | 5.53 | 8.89 | | High | High | High | High | High | High | High | High | High | High | High | 0 | 5.05E-05 | 3.25 |
| FALSE | High | VLSSVNPVI | [K].VLSSVN | 1xPhospho | S4R2I9 1xP | 1.69E-05 | 0 | 1 | 4 | 1 | S4R2I9 | Zinc finger | S4R2I9 [58 | 0 | 3213.694 | 0 | -1.14 | 0.003619 | 2.83 | 137.8 | 62.2 | 11.46 | 11.37 | | High | High | High | High | High | High | High | High | High | High | High | 0 | 2.44E-06 | 4.85 |
| FALSE | High | VLSSVNPVI | [K].VLSSVN | 1xPhospho | S4R2I9 1xP | 6.18E-05 | 0 | 1 | 4 | 1 | S4R2I9 | Zinc finger | S4R2I9 [58 | 0 | 3229.689 | 0 | -0.93 | 0.001414 | 1.21 | 131.6 | 68.4 | 12.69 | 9.72 | | High | High | High | High | High | High | High | High | High | High | High | 0 | 1.05E-05 | 4.26 |
| FALSE | High | VLSTGGPG | [K].VLSTGC | 1xPhospho | F6ZBR8 1xI | 0.041781 | 0.003126 | 2 | 5 | 1 | F6ZBR8; O | Protein Wi | F6ZBR8 [43 | 0 | 1639.825 | 0 | 1.03 | 0.000285 | 7.76 | 64.2 | 135.8 | 9.37 | 5.83 | Shared | High | High | High | High | High | High | High | High | High | High | High | 0.001609 | 0.0157 | 2.71 |
| FALSE | High | VLSTSTDLE | [R].VLSTST | 1xPhospho | B2RY22 1xI | 0.000662 | 0 | 1 | 5 | 12 | B2RY22 | Clasp1 pro | B2RY22 [74 | 0 | 2242.198 | 0 | -0.13 | 0.30209 | 7.87 | 107.7 | 92.3 | 6.79 | 13.48 | | High | High | High | High | High | High | High | High | High | High | High | 0.000538 | 0.004462 | 3.51 |
| FALSE | High | VLTANSNP | [K].VLTANS | 1xPhospho | Q9DAT2 1x | 6.76E-08 | 0 | 1 | 2 | 4 | Q9DAT2 | MRG/MOR | Q9DAT2 [1 | 0 | 1982.036 | 0 | 0.74 | 0.001987 | 12.65 | 75.6 | 124.4 | 8.24 | 5.45 | | High | High | High | High | High | High | High | High | High | High | High | 0 | 4.94E-09 | 2.46 |
| FALSE | High | VLTANSNP | [K].VLTANS | 2xPhospho | Q9DAT2 2x | 0.000751 | 0 | 1 | 2 | 3 | Q9DAT2 | MRG/MOR | Q9DAT2 [1 | 0 | 2062.002 | 0 | 1.29 | 0.000746 | 6.79 | 56 | 144 | 11.96 | 8.51 | | High | High | High | High | High | High | High | High | High | High | High | 0.000538 | 0.004699 | 2.56 |
| FALSE | High | VITNQYNSI | [R].VITNQY | 1xPhospho | O70400 1x | 0.000133 | 0 | 1 | 3 | 2 | O70400 | PDZ and LII | O70400 [1 | 0 | 3628.757 | 0 | 0.12 | 0.276415 | 10.75 | 94.3 | 105.7 | 6.34 | 12.27 | | High | High | High | High | High | High | High | High | High | High | High | 0 | 2.5E-05 | 4.12 |
| FALSE | High | VLTPEEQL | [K].VLTPEE | 1xPhospho | Q66JS6 1xP | 0.003001 | 0.00016 | 1 | 3 | 1 | Q66JS6 | Eukaryotic | Q66JS6 [11 | 0 | 1780.95 | 0 | 0.65 | 0.024452 | 10.52 | 77.9 | 122.1 | 12 | 8.17 | | High | High | High | High | High | High | High | High | High | High | High | 7.19E-05 | 0.000814 | 2.75 |
| FALSE | High | VLTPIK | [R].VLTPIK. | 1xPhospho | Q7TSG1 1x | 0.057686 | 0.004227 | 1 | 2 | 1 | Q7TSG1 | Centrosom | Q7TSG1 [3 | 0 | 1208.742 | 0 | -1.45 | 0.000406 | 0.22 | 144.4 | 55.6 | 5.91 | 5.28 | | High | High | High | High | High | High | High | High | High | High | High | 0.002159 | 0.02268 | 1.98 |
| FALSE | High | VLTPPHDV | [R].VLTPPH | 1xPhospho | U5LK19 1xI | 0.000993 | 0 | 2 | 9 | 2 | U5LK19; E | Marshalin- | U5LK19 [79 | 0 | 2005.047 | 0 | -1.4 | 0.00085 | 2.17 | 148.6 | 51.4 | 5.92 | 15.49 | Shared | High | High | High | High | High | High | High | High | High | High | High | 0 | 0.000237 | 3.47 |
| FALSE | High | VLTPPQAA | [R].VLTPPC | 1xPhospho | Q3U2C9 1x | 2.38E-09 | 0 | 1 | 10 | 2 | Q3U2C9 | PHD finger | Q3U2C9 [6 | 0 | 2289.18 | 0 | -0.24 | 0.00727 | 5.4 | 108.8 | 91.2 | 5.2 | 3.35 | | High | High | High | High | High | High | High | High | High | High | High | 0 | 1.15E-10 | 4.64 |
| FALSE | High | VLTPTQVM | [K].VLTPTC | 1xPhospho | Q3TGC5 1x | 0.100936 | 0.008097 | 1 | 4 | 2 | Q3TGC5 | Putative ur | Q3TGC5 [4 | 0 | 3036.563 | 0 | 1.33 | 0.003498 | 4.07 | 56.7 | 143.3 | 10.33 | 15.09 | | High | High | High | High | High | High | High | High | High | High | High | 0.00607 | 0.06596 | 1.88 |
| FALSE | High | VITSDRSSE | [K].VITSDR | 1xPhospho | O54988 1x | 1.08E-05 | 0 | 1 | 2 | 4 | O54988 | STE20-like | O54988 [5 | 1 | 2804.36 | 0 | 0.32 | 0.000946 | 3.74 | 87.2 | 112.8 | 3.25 | 5.09 | | High | High | High | High | High | High | High | High | High | High | High | 0 | 1.48E-06 | 4.16 |
| FALSE | High | VMTVTAVT | [R].VMTVT | 1xPhospho | Q3UMU9-3 | 0.049971 | 0.003641 | 1 | 4 | 2 | Q3UMU9-3 | Isoform 3 c | Q3UMU9-3 | 0 | 1862.913 | 0 | -0.25 | 0.207076 | 14.68 | 106.1 | 93.9 | 19.41 | 12.3 | | High | High | High | High | High | High | High | High | High | High | High | 0.001902 | 0.01925 | 2.94 |
| FALSE | High | VLVGEGGS | [R].VLVGEC | 1xPhospho | A6H6E1 1x | 0.000181 | 0 | 1 | 4 | 2 | A6H6E1 | Espl1 prote | A6H6E1 [2 | 0 | 1945.995 | 1 | 0.13 | 0.127355 | 12.37 | 91.8 | 108.2 | 6.7 | 11.02 | | High | High | High | High | High | High | High | High | High | High | High | 4.94E-05 | 0.000696 | 2.89 |
| FALSE | High | VLVQNAAC | [K].VLVQN. | 1xPhospho | Q80TM2 1 | 1.77E-07 | 0 | 1 | 5 | 2 | Q80TM2 | MKIAA102 | Q80TM2 [2 | 0 | 1781.956 | 0 | 0.12 | 0.114801 | 9.34 | 93.3 | 106.7 | 13.14 | 11 | | High | High | High | High | High | High | High | High | High | High | High | 0 | 1.46E-08 | 3.52 |
| FALSE | High | VLVSYGSPI | [K].VLVSYC | 1xPhospho | E9Q777 1x | 0.022574 | 0.001542 | 1 | 2 | 2 | E9Q777 | A kinase (P | E9Q777 [2 | 0 | 1487.828 | 0 | -0.33 | 0.006027 | 4.77 | 109.4 | 90.6 | 2.12 | 7.37 | | High | High | High | High | High | High | High | High | High | High | High | 0.005613 | 0.06168 | 1.81 |
| FALSE | High | VIYDFIEK | [K].VIYDFIE | 1xPhospho | Q7TPN5 1x | 0.000645 | 0 | 1 | 4 | 2 | Q7TPN5 | Wiskott-Al | Q7TPN5 [2 | 0 | 1564.843 | 0 | -0.93 | 0.003483 | 7.22 | 133.8 | 66.2 | 7.76 | 18.45 | | High | High | High | High | High | High | High | High | High | High | High | 0 | 0.00026 | 2.32 |
| FALSE | High | VIYSQPSTR | [K].VIYSQP | 1xPhospho | O88792 1x | 0.069325 | 0.005112 | 1 | 1 | 1 | O88792 | Junctional | O88792 [2 | 0 | 1359.687 | 0 | 0.92 | 0.00104 | 19.79 | 71.7 | 128.3 | 16.6 | 4.79 | | High | High | High | High | High | High | High | High | High | High | High | 0.002618 | 0.02798 | 1.18 |
| FALSE | High | VMSPENFF | [R].VMSPE | 1xPhospho | Q8BYH8 1x | 1.52E-06 | 0 | 1 | 6 | 1 | Q8BYH8 | Chromodo | Q8BYH8 [5 | 0 | 2117.039 | 0 | 0.32 | 0.047922 | 18.98 | 87.2 | 112.8 | 10.6 | 11.39 | | High | High | High | High | High | High | High | High | High | High | High | 0 | 1.62E-07 | 3.05 |
| FALSE | High | VMSPENFF | [R].VMSPE | 1xPhospho | Q8BYH8 1x | 0.000815 | 0 | 1 | 6 | 1 | Q8BYH8 | Chromodo | Q8BYH8 [5 | 0 | 2133.034 | 0 | 0.26 | 0.164128 | 5.55 | 91.1 | 108.9 | 6.34 | 26.54 | | High | High | High | High | High | High | High | High | High | High | High | 0 | 0.00019 | 2.86 |
| FALSE | High | VMSPLQSP | [K].VMSPL | 1xPhospho | Q3UIL6-2 1 | 0.020499 | 0.001472 | 1 | 7 | 2 | Q3UIL6-2 | Isoform 2 c | Q3UIL6-2 [ | 0 | 1641.869 | 0 | -0.83 | 0.018035 | 7.76 | 123.4 | 76.6 | 12.66 | 13.73 | | High | High | High | High | High | High | High | High | High | High | High | 0.001151 | 0.01225 | 2.34 |
| FALSE | High | VMSPLQSP | [K].VMSPL | 2xPhospho | Q3UIL6-2 2 | 0.004867 | 0.000504 | 1 | 7 | 2 | Q3UIL6-2 | Isoform 2 c | Q3UIL6-2 [ | 0 | 1705.84 | 0 | -1.22 | 0.001282 | 2.8 | 139.2 | 60.8 | 8.47 | 9.66 | | High | High | High | High | High | High | High | High | High | High | High | 0.000225 | 0.00141 | 2.85 |
| FALSE | High | VMTIPYQP | [R].VMTIPY | 1xCarbami | P60335 1xI | 9.45E-05 | 0 | 1 | 1 | 2 | P60335 | Poly(rC)-bi | P60335 [17 | 0 | 2784.312 | 0 | 0.46 | 0.021406 | 12.13 | 83.9 | 116.1 | 8.16 | 10.91 | | High | High | High | High | High | High | High | High | High | High | High | 0 | 1.69E-05 | 3.1 |
| FALSE | High | VMTIPYQP | [R].VMTIPY | 1xCarbami | P60335 1xI | 3.53E-08 | 0 | 1 | 1 | 9 | P60335 | Poly(rC)-bi | P60335 [17 | 0 | 2800.307 | 0 | 0.38 | 0.00542 | 9.5 | 87.5 | 112.5 | 6.22 | 3.5 | | High | High | High | High | High | High | High | High | High | High | High | 0 | 5.19E-05 | 3.69 |
| FALSE | High | VMTIPYQP | [R].VMTIPY | 1xCarbami | P60335 1xI | 0.00407 | 0.00036 | 1 | 1 | 4 | P60335 | Poly(rC)-bi | P60335 [17 | 0 | 2801.291 | 0 | 0.35 | 0.117639 | 25.22 | 87.9 | 112.1 | 16.94 | 7.83 | | High | High | High | High | High | High | High | High | High | High | High | 0.000275 | 0.002034 | 3.06 |
| FALSE | High | VMTIPYQP | [R].VMTIPY | 1xCarbami | P60335 1xI | 8.37E-07 | 0 | 1 | 1 | 22 | P60335 | Poly(rC)-bi | P60335 [17 | 0 | 2816.302 | 0 | 0.47 | 0.004488 | 4.42 | 84 | 116 | 6.66 | 4.61 | | High | High | High | High | High | High | High | High | High | High | High | 0 | 0.000197 | 3.82 |
| FALSE | High | VMTLQWL | [R].VMTLQ | 1xPhospho | E9QKG8 1x | 9.42E-07 | 0 | 1 | 7 | 1 | E9QKG8 | Holliday ju | E9QKG8 [3 | 0 | 1953.105 | 0 | -1.45 | 0.000378 | 0.18 | 144.2 | 55.8 | 7.86 | 5.39 | | High | High | High | High | High | High | High | High | High | High | High | 0 | 9.5E-08 | 3.38 |
| FALSE | High | VLVNSASA | [R].VLVNS/ | 1xPhospho | B9EJA4 1xP | 0.000396 | 0 | 1 | 7 | 2 | B9EJA4 | Clasp2 pro | B9EJA4 [48 | 0 | 1554.866 | 0 | -0.26 | 0.010295 | 3.24 | 108.4 | 91.6 | 7.63 | 3.16 | | High | High | High | High | High | High | High | High | High | High | High | 0 | 8.47E-05 | 2.76 |
| FALSE | High | VPAASNST | [R].VPAASI | 1xPhospho | Q3U8K7 1x | 9.28E-05 | 0 | 1 | 3 | 1 | Q3U8K7 | Histone-lys | Q3U8K7 [4 | 0 | 1596.84 | 0 | -0.55 | 0.01935 | 7.28 | 122 | 78 | 10.91 | 12.05 | | High | High | High | High | High | High | High | High | High | High | High | 0 | 1.66E-05 | 3.17 |
| FALSE | High | VPALASPA | [R].VPALAS | 1xPhospho | Q3UHZ3 1x | 6.71E-05 | 0 | 3 | 4 | 2 | Q3UHZ3; C | DNA (cytos | Q3UHZ3 [1 | 0 | 1895.994 | 0 | 0.05 | 0.43661 | 9.68 | 98.7 | 101.3 | 5.18 | 6.93 | Shared | High | High | High | High | High | High | High | High | High | High | High | 0 | 1.14E-05 | 2.53 |
| FALSE | High | VPANSVFLI | [K].VPANS\ | 1xPhospho | Q80WC9 1 | 0.067059 | 0.004796 | 1 | 2 | 1 | Q80WC9 | Acyl-CoA s | Q80WC9 [ | 1 | 2289.115 | 0 | -0.5 | 0.435264 | 39.92 | 117.2 | 82.8 | 41.92 | 69.8 | | High | High | High | High | High | High | High | High | High | High | High | 0.002455 | 0.02696 | 2.37 |
| FALSE | High | VPNGRGSP | [K].VPNGR | 1xDeamida | Q0P5V9-2 | 0.014667 | 0.001049 | 1 | 4 | 1 | Q0P5V9-2 | Isoform 2 c | Q0P5V9-2 | 1 | 1860.953 | 0 | -0.85 | 0.000642 | 1.51 | 128 | 72 | 5.71 | 7.99 | | High | High | High | High | High | High | High | High | High | High | High | 0.000555 | 0.004814 | 2.26 |
| FALSE | High | VPPFNSSGI | [K].VPPFNS | 1xPhospho | Q5XG69 1x | 0.005275 | 0.000599 | 1 | 1 | 1 | Q5XG69 | Soluble lan | Q5XG69 [4 | 0 | 2747.406 | 0 | -0.64 | 0.022873 | 5.14 | 118.6 | 81.4 | 9.54 | 12.05 | | High | High | High | High | High | High | High | High | High | High | High | 0.000267 | 0.001536 | 2.5 |
| FALSE | High | VPPFVDLN | [K].VPPFVD | 1xPhospho | B2RUM8 1 | 6.15E-06 | 0 | 1 | 1 | 2 | B2RUM8 | RNA helica | B2RUM8 [6 | 0 | 2149.109 | 0 | -2 | 0.002046 | 0.94 | 156.5 | 43.5 | 13.65 | 16.6 | | High | High | High | High | High | High | High | High | High | High | High | 0 | 7.85E-07 | 3 |
| FALSE | High | VPPIDAAAS | [K].VPPIDA | 1xPhospho | E9Q555 1x | 0.000255 | 0 | 1 | 2 | 2 | E9Q555 | E3 ubiquiti | E9Q555 [2 | 0 | 1702.954 | 0 | 2.36 | 0.000207 | 19 | 34.6 | 165.4 | 16.34 | 3.81 | | High | High | High | High | High | High | High | High | High | High | High | 0 | 5.14E-05 | 2.96 |
| FALSE | High | VPPSPDELF | [R].VPPSPD | 1xPhospho | Q6KAP5 1x | 0.102567 | 0.008249 | 1 | 4 | 1 | Q6KAP5 | MFLJ00227 | Q6KAP5 [4 | 0 | 1502.745 | 0 | 1.28 | 0.001166 | 6.09 | 59.2 | 140.8 | 6.7 | 8.71 | | High | High | High | High | High | High | High | High | High | High | High | 0.004138 | 0.04438 | 2.54 |
| FALSE | High | VPPTDSRPI | [K].VPPTDS | 2xPhospho | Q3TCX6 2x | 0.004294 | 0.00036 | 1 | 7 | 2 | Q3TCX6 | Putative ur | Q3TCX6 [2 | 0 | 3029.494 | 0 | 1.33 | 0.005584 | 6.57 | 59.2 | 140.8 | 4.31 | 20.21 | | High | High | High | High | High | High | High | High | High | High | High | 0.000161 | 0.001215 | 3.7 |
| FALSE | High | VPQFSVGE | [K].VPQFS\ | 1xPhospho | Q542Q3 1x | 5.53E-07 | 0 | 1 | 1 | 1 | Q542Q3 | G two S ph | Q542Q3 [4 | 0 | 2124.114 | 0 | 0.13 | 0.403687 | 17.73 | 95.8 | 104.2 | 10.69 | 11.04 | | High | High | High | High | High | High | High | High | High | High | High | 0 | 5.23E-08 | 4.27 |
| FALSE | High | VPQFSVGE | [K].VPQFS\ | 2xPhospho | Q542Q3 2x | 2.25E-05 | 0 | 1 | 1 | 1 | Q542Q3 | G two S ph | Q542Q3 [4 | 0 | 2204.081 | 0 | 0.27 | 0.059798 | 15.97 | 85.7 | 114.3 | 13.5 | 14.18 | | High | High | High | High | High | High | High | High | High | High | High | 0 | 3.38E-06 | 2.57 |
| FALSE | High | VPRSPDLPI | [K].VPRSPD | 1xPhospho | Q91ZW3 1 | 1.46E-05 | 0 | 1 | 2 | 2 | Q91ZW3 | SWI/SNF-r | Q91ZW3 [7 | 1 | 2130.147 | 0 | -0.06 | 0.848879 | 5.79 | 100 | 100 | 3.9 | 9.19 | | High | High | High | High | High | High | High | High | High | High | High | 0 | 2.07E-06 | 3.09 |
| FALSE | High | VPSAGDVE | [R].VPSAGI | 1xPhospho | Q9CYG7 1x | 0.002334 | 0.000109 | 1 | 2 | 2 | Q9CYG7 | Mitochond | Q9CYG7 [1 | 0 | 1238.598 | 0 | 0.64 | 0.003074 | 9.15 | 77.3 | 122.7 | 4.89 | 9.15 | | High | High | High | High | High | High | High | High | High | High | High | 4.94E-05 | 0.000617 | 1.67 |
| FALSE | High | VPSAPGQE | [R].VPSAPC | 1xPhospho | A2ARP8 1x | 0.000479 | 0 | 1 | 3 | 2 | A2ARP8 | Microtubu | A2ARP8 [2 | 0 | 2047.051 | 0 | -1.82 | 0.00198 | 0.9 | 155.4 | 44.6 | 12.61 | 20.18 | | High | High | High | High | High | High | High | High | High | High | High | 0 | 0.000105 | 2.78 |
| FALSE | High | VPSASDSD | [K].VPSASD | 1xPhospho | Q3UMU9-3 | 0.001147 | 0 | 1 | 4 | 4 | Q3UMU9-3 | Isoform 3 c | Q3UMU9-3 | 0 | 1530.745 | 0 | 0.14 | 0.0149 | 4.15 | 95.3 | 104.7 | 7.88 | 5.74 | | High | High | High | High | High | High | High | High | High | High | High | 0.000161 | 0.001164 | 2.19 |
| FALSE | High | VPSASDSD | [K].VPSASD | 2xPhospho | Q3UMU9-3 | 0.001211 | 0 | 1 | 4 | 1 | Q3UMU9-3 | Isoform 3 c | Q3UMU9-3 | 0 | 1610.712 | 0 | 0.1 | 0.229932 | 8.6 | 97.9 | 102.1 | 3.47 | 9.63 | | High | High | High | High | High | High | High | High | High | High | High | 0 | 0.000297 | 2.58 |
| FALSE | High | VPMAGVG | [K].VPMAG | 1xCarbami | A0A0U1RP | 5.31E-05 | 0 | 1 | 7 | 1 | A0A0U1RP | Ataxin-2-lik | A0A0U1RP | 0 | 3682.863 | 0 | 0.15 | 0.137377 | 15.19 | 95.6 | 104.4 | 9.78 | 10.23 | | High | High | High | High | High | High | High | High | High | High | High | 0 | 8.8E-06 | 3.44 |
| FALSE | High | VPSDSSLGT | [R].VPSDSS | 1xPhospho | Q8CJ53 1xI | 0.002508 | 0.000109 | 1 | 4 | 5 | Q8CJ53 | Cdc42-inte | Q8CJ53 [29 | 0 | 2092.027 | 0 | -0.03 | 0.539728 | 1.15 | 100.2 | 99.8 | 2.88 | 7.08 | | High | High | High | High | High | High | High | High | High | High | High | 4.94E-05 | 0.000671 | 2.38 |
| FALSE | High | VPSEPSALV | [K].VPSEPS | 1xPhospho | Q5SWP3 1 | 0.00509 | 0.000551 | 1 | 1 | 1 | Q5SWP3 | NAC-alpha | Q5SWP3 [1 | 0 | 2041.057 | 0 | -0.58 | 0.030592 | 4.38 | 116.8 | 83.2 | 6.83 | 14.5 | | High | High | High | High | High | High | High | High | High | High | High | 0.000246 | 0.001478 | 3.03 |
| FALSE | High | VPSFSIVEQ | [R].VPSFSI\ | 1xCarbami | E9Q4Y4 1x | 2.02E-05 | 0 | 1 | 1 | 1 | E9Q4Y4 | Centrosom | E9Q4Y4 [9 | 0 | 2359.084 | 0 | 0.1 | 0.89589 | 9.44 | 96.4 | 103.6 | 27.17 | 9.65 | | High | High | High | High | High | High | High | High | High | High | High | 0 | 2.97E-06 | 3.87 |
| FALSE | High | VPSFTFTPT | [K].VPSFTF | 1xPhospho | G5E843 1x | 0.096959 | 0.007777 | 1 | 2 | 1 | G5E843 | Roundabou | G5E843 [8 | 0 | 1952.972 | 0 | 0.65 | 0.000897 | 11.32 | 77.8 | 122.2 | 12.2 | 7.76 | | High | High | High | High | High | High | High | High | High | High | High | 0.003922 | 0.04162 | 1.48 |
| FALSE | High | VPSGLFDTI | [K].VPSGLF | 1xPhospho | A6H8H2-2 | 0.048303 | 0.00354 | 1 | 3 | 2 | A6H8H2-2 | Isoform 2 c | A6H8H2-2 | 0 | 1528.736 | 0 | 1.48 | 0.000408 | 7.42 | 53.8 | 146.2 | 6.8 | 8.82 | | High | High | High | High | High | High | High | High | High | High | High | 0.00302 | 0.03166 | 2.21 |
| FALSE | High | VPSGLFDTI | [K].VPSGLF | 1xDeamida | A6H8H2-2 | 0.042868 | 0.003228 | 1 | 3 | 2 | A6H8H2-2 | Isoform 2 c | A6H8H2-2 | 0 | 1529.72 | 0 | 1.38 | 0.00032 | 1.42 | 55.3 | 144.7 | 4.11 | 5.79 | | High | High | High | High | High | High | High | High | High | High | High | 0.003389 | 0.03686 | 2.75 |
| FALSE | High | VPSLDHAM | [R].VPSLDH | 1xPhospho | Q99LW0 1 | 0.003892 | 0.000261 | 1 | 3 | 1 | Q99LW0 | Ankyrin re | Q99LW0 [2 | 0 | 2532.128 | 0 | 0.03 | 0.793068 | 19.77 | 99.8 | 100.2 | 22.09 | 7.12 | | High | High | High | High | High | High | High | High | High | High | High | 0.00014 | 0.001091 | 3.46 |
| FALSE | High | VPSLDHAM | [R].VPSLDH | 1xDeamida | Q99LW0 1 | 2.34E-05 | 0 | 1 | 3 | 2 | Q99LW0 | Ankyrin re | Q99LW0 [2 | 0 | 2533.112 | 0 | 0.03 | 0.287097 | 7.25 | 95.2 | 104.8 | 8.73 | 12.42 | | High | High | High | High | High | High | High | High | High | High | High | 0 | 3.51E-06 | 4.9 |
| FALSE | High | VPSLDHAM | [R].VPSLDH | 1xPhospho | Q99LW0 1 | 9.67E-07 | 0 | 1 | 3 | 2 | Q99LW0 | Ankyrin re | Q99LW0 [2 | 0 | 2548.122 | 0 | -0.03 | 0.457915 | 27.52 | 99.5 | 100.5 | 22.26 | 29.11 | | High | High | High | High | High | High | High | High | High | High | High | 0 | 9.82E-08 | 4.1 |
| FALSE | High | VPSLDHAM | [R].VPSLDH | 1xDeamida | Q99LW0 1 | 2.44E-05 | 0 | 1 | 3 | 1 | Q99LW0 | Ankyrin re | Q99LW0 [2 | 0 | 2549.106 | 0 | 0.05 | 0.595827 | 48.64 | 98.1 | 101.9 | 23.99 | 26.54 | | High | High | High | High | High | High | High | High | High | High | High | 0 | 3.68E-06 | 4.6 |
| FALSE | High | VPSISTETEI | [K].VPSISTI | 1xPhospho | Q5SFM8 1x | 0.000837 | 0 | 1 | 3 | 2 | Q5SFM8 | RNA-bindir | Q5SFM8 [1 | 0 | 2115.051 | 0 | 0.09 | 0.617672 | 8.49 | 97.9 | 102.1 | 19.14 | 8.31 | | High | High | High | High | High | High | High | High | High | High | High | 0 | 0.000196 | 2.93 |
| FALSE | High | VPSMESLFI | [R].VPSME. | 1xPhospho | Q91VW5 1 | 0.015329 | 0.001089 | 1 | 7 | 1 | Q91VW5 | Golgin sub | Q91VW5 [ | 0 | 1390.664 | 0 | 0.84 | 0.000376 | 3.45 | 71.4 | 128.6 | 6.92 | 6.65 | | High | High | High | High | High | High | High | High | High | High | High | 0.000571 | 0.005047 | 1.36 |
| FALSE | High | VPSPGGPQ | [K].VPSPGC | 1xPhospho | Q6GQX2 1 | 9.48E-06 | 0 | 1 | 1 | 1 | Q6GQX2 | Nck-associ | Q6GQX2 [4 | 0 | 1838.973 | 0 | 0.2 | 0.454995 | 12.06 | 91.6 | 108.4 | 4.82 | 14.85 | | High | High | High | High | High | High | High | High | High | High | High | 0 | 1.28E-06 | 3.16 |
| FALSE | High | VPSPGNAA | [K].VPSPGI | 1xPhospho | Q6ZWQ0 1 | 8.03E-05 | 0 | 1 | 3 | 2 | Q6ZWQ0 | Nesprin-2 | Q6ZWQ0 [ | 0 | 2678.344 | 0 | 1.11 | 0.003874 | 21.26 | 67.1 | 132.9 | 14.41 | 12.26 | | High | High | High | High | High | High | High | High | High | High | High | 0 | 9.56E-05 | 2.48 |
| FALSE | High | VPSDSSLGT | [R].VPSDSS | 2xPhospho | Q8CJ53 2xI | 0.000122 | 0 | 1 | 4 | 2 | Q8CJ53 | Cdc42-inte | Q8CJ53 [29 | 0 | 2171.994 | 0 | -0.37 | 0.004274 | 2.02 | 112 | 88 | 6.67 | 7.89 | | High | High | High | High | High | High | High | High | High | High | High | 0 | 2.24E-05 | 4.02 |
| FALSE | High | VPMAGVG | [K].VPMAG | 1xCarbami | A0A0U1RP | 3.63E-05 | 0 | 1 | 7 | 2 | A0A0U1RP | Ataxin-2-lik | A0A0U1RP | 0 | 3666.868 | 0 | -0.02 | 0.763465 | 31.83 | 97.1 | 102.9 | 13.77 | 28.28 | | High | High | High | High | High | High | High | High | High | High | High | 0 | 5.76E-06 | 3.63 |
| FALSE | High | VPLSPLSLP | [R].VPLSPL | 1xDeamida | Q3TDX5 1x | 5.97E-05 | 0 | 1 | 1 | 1 | Q3TDX5 | Putative ur | Q3TDX5 [2 | 0 | 1870.004 | 1 | 0.86 | 0.000569 | 2.11 | 70.8 | 129.2 | 3.88 | 7.32 | | High | High | High | High | High | High | High | High | High | High | High | 0 | 1E-05 | 3.26 |
| FALSE | High | VPLPSLSPT | [K].VPLPSL | 1xPhospho | Q8BMF4 1 | 1.19E-05 | 0 | 1 | 1 | 1 | Q8BMF4 | Dihydrolipo | Q8BMF4 [ | 0 | 2064.076 | 0 | -0.42 | 0.002739 | 5.15 | 114.4 | 85.6 | 2.77 | 9.62 | | High | High | High | High | High | High | High | High | High | High | High | 0 | 1.65E-06 | 4.13 |
| FALSE | High | VPAPFSPPF | [K].VPAPFS | 1xPhospho | A2A7S8-2 1 | 0.001411 | 5.66E-05 | 1 | 4 | 1 | A2A7S8-2 | Isoform 2 c | A2A7S8-2 [ | 0 | 1661.907 | 0 | -0.24 | 0.013837 | 7.02 | 112.3 | 87.7 | 10.89 | 11.65 | | High | High | High | High | High | High | High | High | High | High | High | 2.58E-05 | 0.000351 | 0.95 |
| FALSE | High | VPAPPGSP | [K-].VPAPP | 1xCarbami | Q60953 1x | 0.00407 | 0.00036 | 2 | 7 | 4 | Q60953; Q | Protein PM | Q60953 [1 | 0 | 1346.649 | 0 | -0.15 | 0.010895 | 4.05 | 105 | 95 | 3.58 | 5.28 | Shared | High | High | High | High | High | High | High | High | High | High | High | 0.005981 | 0.06466 | 1.45 |
| FALSE | High | VPASPLPGI | [K].VPASPL | 1xPhospho | Q571J8 1xI | 0.020499 | 0.001472 | 1 | 16 | 1 | Q571J8 | Non-specif | Q571J8 [45 | 0 | 1430.761 | 0 | 0.21 | 0.03162 | 6.52 | 93.7 | 106.3 | 4.33 | 13.64 | | High | High | High | High | High | High | High | High | High | High | High | 0.000786 | 0.007037 | 1.44 |
| FALSE | High | VPCNVEGIS | [R].VPCNVI | 1xCarbami | E9QKK4 1x | 6.47E-05 | 0 | 1 | 6 | 1 | E9QKK4 | Glucocortic | E9QKK4 [2 | 0 | 2109.07 | 0 | 0.21 | 0.033286 | 7.21 | 94.5 | 105.5 | 4.83 | 6.61 | | High | High | High | High | High | High | High | High | High | High | High | 0 | 1.1E-05 | 3.16 |
| FALSE | High | VPCTPVAC | [K].VPCTP\ | 2xCarbami | G3X963 1x | 2.22E-07 | 0 | 1 | 3 | 2 | G3X963 | ATPase fan | G3X963 [1 | 0 | 2167.106 | 0 | 0.7 | 0.011215 | 12.74 | 73.3 | 126.7 | 7.74 | 15.93 | | High | High | High | High | High | High | High | High | High | High | High | 0 | 1.87E-08 | 4.54 |
| FALSE | High | VPCTPVAC | [K].VPCTP\ | 2xCarbami | G3X963 2x | 0.000933 | 0 | 1 | 3 | 1 | G3X963 | ATPase fan | G3X963 [1 | 0 | 2247.072 | 0 | 0.33 | 0.009361 | 9.8 | 88.7 | 111.3 | 18.21 | 6.21 | | High | High | High | High | High | High | High | High | High | High | High | 0 | 0.00022 | 3.73 |
| FALSE | High | VPDSESEPI | [K].VPDSES | 1xPhospho | Q91YK2 1x | 0.001158 | 0 | 1 | 2 | 3 | Q91YK2 | Ribosomal | Q91YK2 [4 | 0 | 1739.769 | 0 | -0.79 | 0.000181 | 1.34 | 126.8 | 73.2 | 4.34 | 6.44 | | High | High | High | High | High | High | High | High | High | High | High | 0.000399 | 0.003397 | 3.5 |
| FALSE | High | VPDSPATK | [K].VPDSP/ | 1xPhospho | Q61493 1x | 0.016743 | 0.001166 | 1 | 2 | 1 | Q61493 | DNA polym | Q61493 [1 | 0 | 1352.723 | 0 | -1.25 | 0.000774 | 2.53 | 140.7 | 59.3 | 6.24 | 5.91 | | High | High | High | High | High | High | High | High | High | High | High | 0.000622 | 0.005557 | 2 |
| FALSE | High | VPDSPVFTI | [R].VPDSP\ | 1xCarbami | Q8BUM9-2 | 0.014797 | 0.001049 | 1 | 3 | 1 | Q8BUM9-2 | Isoform 2 c | Q8BUM9-2 | 0 | 2339.114 | 0 | 0.36 | 0.014729 | 13.48 | 86.6 | 113.4 | 8.22 | 8.19 | | High | High | High | High | High | High | High | High | High | High | High | 0.000555 | 0.004852 | 2.09 |
| FALSE | High | VPDVDISSF | [K].VPDVD | 1xPhospho | E9Q616 1x | 1.03E-09 | 0 | 1 | 5 | 2 | E9Q616 | AHNAK nu | E9Q616 [4 | 0 | 2771.409 | 0 | 5.83 | | 36.89 | 3.4 | 196.6 | 25.14 | 23.43 | | High | Not Found | High | High | High | High | High | High | High | High | High | 0 | 1.04E-10 | 6.39 |
| FALSE | High | VPDVDISSF | [K].VPDVD | 1xPhospho | E9Q616 1x | 4.77E-06 | 0 | 1 | 5 | 7 | E9Q616 | AHNAK nu | E9Q616 [4 | 0 | 2787.404 | 0 | 3.14 | 0.002102 | 20.74 | 21.6 | 178.4 | 12.95 | 19.04 | | High | High | High | High | High | High | High | High | High | High | High | 0 | 2.06E-06 | 5.32 |
| FALSE | High | VPEIEVTVE | [K].VPEIEV | 1xPhospho | Q9ESZ8-5 | 2.33E-07 | 0 | 2 | 8 | 3 | Q9ESZ8-5; | Isoform 5 c | Q9ESZ8-5 [ | 0 | 2619.286 | 0 | 0.07 | 0.258049 | 8.14 | 94.9 | 105.1 | 17.06 | 13.63 | Shared | High | High | High | High | High | High | High | High | High | High | High | 0 | 1.98E-08 | 4.58 |
| FALSE | High | VPELLGPEI | [K].VPELLG | 1xDeamida | Q91VJ2 1xI | 0.001107 | 0 | 1 | 1 | 2 | Q91VJ2 | Caveolae-a | Q91VJ2 [14 | 0 | 4013.848 | 0 | 2.62 | 0.001084 | 55.82 | 27.9 | 172.1 | 43.7 | 11.79 | | High | High | High | High | High | High | High | High | High | High | High | 2.58E-05 | 0.000401 | 3.28 |
| FALSE | High | VPELLGPEI | [K].VPELLG | 2xPhospho | Q91VJ2 2xI | 0.015603 | 0.001124 | 1 | 1 | 5 | Q91VJ2 | Caveolae-a | Q91VJ2 [14 | 0 | 4092.83 | 0 | 1.76 | 0.000534 | 22.46 | 45.4 | 154.6 | 18.17 | 4.54 | | High | High | High | High | High | High | High | High | High | High | High | 0.003672 | 0.03947 | 3.24 |
| FALSE | High | VPELPNTSS | [K].VPELPN | 1xPhospho | Q04891 1x | 1.72E-05 | 0 | 1 | 2 | 1 | Q04891 | Transcripti | Q04891 [2 | 0 | 1994.061 | 0 | 1.49 | 0.000334 | 7.87 | 49.8 | 150.2 | 10.52 | 6.85 | | High | High | High | High | High | High | High | High | High | High | High | 0 | 2.48E-06 | 3.71 |
| FALSE | High | VPEPESQTS | [K].VPEPES | 1xPhospho | Q8BGF7 1x | 6.18E-05 | 0 | 1 | 6 | 2 | Q8BGF7 | PAB-deper | Q8BGF7 [1 | 0 | 1736.887 | 0 | -0.46 | 0.00418 | 6.4 | 115.9 | 84.1 | 3.21 | 9.56 | | High | High | High | High | High | High | High | High | High | High | High | 0 | 1.05E-05 | 3.89 |
| FALSE | High | VPFAPGPSI | [R].VPFAPC | 1xPhospho | Q99MR1 1 | 9.22E-06 | 0 | 1 | 1 | 1 | Q99MR1 | GRB10-inte | Q99MR1 [5 | 0 | 2444.188 | 0 | 0.37 | 0.083129 | 16.69 | 87.3 | 112.7 | 10.32 | 15.14 | | High | High | High | High | High | High | High | High | High | High | High | 0 | 1.23E-06 | 3.39 |
| FALSE | High | VPFSPGPAI | [R].VPFSPG | 1xPhospho | Q6Y7W8 1 | 0.000118 | 0 | 1 | 8 | 1 | Q6Y7W8 | GRB10-inte | Q6Y7W8 [5 | 0 | 2467.168 | 0 | -0.02 | 0.724765 | 10.54 | 99.8 | 100.2 | 6.21 | 12.91 | | High | High | High | High | High | High | High | High | High | High | High | 0 | 2.18E-05 | 3.93 |
| FALSE | High | VPGAPDSS | [R].VPGAPI | 1xPhospho | Q8BJM7 1x | 0.000223 | 0 | 1 | 4 | 1 | Q8BJM7 | S-adenosyl | Q8BJM7 [5 | 0 | 1820.956 | 0 | 0.47 | 0.00594 | 12.57 | 79.9 | 120.1 | 14.98 | 11.2 | | High | High | High | High | High | High | High | High | High | High | High | 0 | 4.41E-05 | 3.28 |
| FALSE | High | VPGPELPG | [K].VPGPEI | 1xPhospho | Q4VAC9 1x | 0.000884 | 0 | 1 | 1 | 2 | Q4VAC9 | Pleckstrin | Q4VAC9 [5 | 0 | 3306.555 | 0 | 2.07 | 0.000593 | 10.54 | 37.2 | 162.8 | 17.24 | 9.31 | | High | High | High | High | High | High | High | High | High | High | High | 0 | 0.000208 | 3.81 |
| FALSE | High | VPGPSSSEI | [K].VPGPSS | 1xPhospho | Q4FJY3 1xP | 0.018447 | 0.001324 | 1 | 1 | 3 | Q4FJY3 | MCG1292S | Q4FJY3 [26 | 0 | 2502.184 | 0 | 0.64 | 0.007529 | 21.26 | 78 | 122 | 3.93 | 13.49 | | High | High | High | High | High | High | High | High | High | High | High | 0.000706 | 0.006234 | 2.53 |
| FALSE | High | VPGPSSSEI | [K].VPGPSS | 1xPhospho | Q4FJY3 1xP | 5.63E-07 | 0 | 1 | 1 | 6 | Q4FJY3 | MCG1292S | Q4FJY3 [26 | 0 | 2518.178 | 0 | 1.28 | 0.000158 | 4.89 | 58.4 | 141.6 | 12.03 | 6.34 | | High | High | High | High | High | High | High | High | High | High | High | 0 | 5.31E-08 | 4.16 |
| FALSE | High | VPGQPLAP | [R].VPGQP | 1xCarbami | Q542Q3 1x | 4.49E-07 | 0 | 1 | 1 | 2 | Q542Q3 | G two S ph | Q542Q3 [5 | 0 | 2788.436 | 0 | 0.18 | 0.089653 | 12.98 | 91.3 | 108.7 | 14.34 | 7.87 | | High | High | High | High | High | High | High | High | High | High | High | 0 | 4.14E-08 | 4.42 |
| FALSE | High | VPGQPLAP | [R].VPGQP | 1xCarbami | Q542Q3 2x | 8.63E-05 | 0 | 1 | 1 | 1 | Q542Q3 | G two S ph | Q542Q3 [5 | 0 | 2868.402 | 0 | 0.54 | 0.004187 | 12.19 | 81.5 | 118.5 | 9.06 | 5.58 | | High | High | High | High | High | High | High | High | High | High | High | 0 | 1.52E-05 | 3.43 |
| FALSE | High | VPGSPENE | [K].VPGSPE | 1xPhospho | A2ADL9 1x | 5.12E-05 | 0 | 1 | 7 | 2 | A2ADL9 | InaD-like (I | A2ADL9 [5 | 0 | 1607.845 | 0 | 0.61 | 0.000438 | 7.45 | 78.9 | 121.1 | 6.24 | 4.06 | | High | High | High | High | High | High | High | High | High | High | High | 0.00092 | 0.009638 | 2.64 |
| FALSE | High | VPHANSGS | [K].VPHAN. | 1xPhospho | E9Q1M1 1 | 4.89E-05 | 0 | 1 | 3 | 2 | E9Q1M1 | PDZ domai | E9Q1M1 [1 | 0 | 1761.848 | 0 | 0.2 | 0.628926 | 7.61 | 93.8 | 106.2 | 13.3 | 8.44 | | High | High | High | High | High | High | High | High | High | High | High | 0 | 8.03E-06 | 3.73 |
| FALSE | High | VPHNTPVD | [R].VPHNTI | 1xPhospho | A2A4N8 1x | 0.001476 | 5.66E-05 | 1 | 2 | 1 | A2A4N8 | Next to BR | A2A4N8 [5 | 0 | 1990.041 | 0 | 0.67 | 0.0232 | 25.03 | 79.3 | 120.7 | 13.44 | 9.46 | | High | High | High | High | High | High | High | High | High | High | High | 2.58E-05 | 0.00037 | 2.7 |
| FALSE | High | VPLFSPFSP | [R].VPLFSP | 1xPhospho | A0A1D5RL | 0.022773 | 0.001542 | 1 | 5 | 1 | A0A1D5RL | Calpain-15 | A0A1D5RL | 0 | 2319.21 | 0 | 0.62 | 0.002895 | 14.4 | 78.8 | 121.2 | 9.21 | 5.85 | | High | High | High | High | High | High | High | High | High | High | High | 0.000814 | 0.007885 | 2.65 |
| FALSE | High | VPLNQFDF | [K].VPLNQI | 1xPhospho | B2RUM8 1 | 0.000314 | 0 | 1 | 1 | 1 | B2RUM8 | RNA helica | B2RUM8 [5 | 0 | 2006.019 | 0 | -1.58 | 0.000333 | 1.6 | 149.8 | 50.2 | 4.46 | 6.24 | | High | High | High | High | High | High | High | High | High | High | High | 0 | 6.49E-05 | 2.31 |
| FALSE | High | VLSSTSEED | [R].VLSSTS | 2xPhospho | B9EIW5 2x | 0.000674 | 0 | 1 | 4 | 1 | B9EIW5 | Ncoa7 prot | B9EIW5 [2 | 0 | 2194.033 | 0 | 2.99 | 0.000702 | 28.56 | 22.8 | 177.2 | 25.46 | 11.76 | | High | High | High | High | High | High | High | High | High | High | High | 0 | 0.000153 | 3.08 |
| FALSE | High | VLSSTSEED | [R].VLSSTS | 1xPhospho | B9EIW5 1x | 0.000209 | 0 | 1 | 4 | 2 | B9EIW5 | Ncoa7 prot | B9EIW5 [2 | 0 | 2114.067 | 0 | 1.66 | 0.001286 | 22.59 | 48 | 152 | 15.36 | 12.68 | | High | High | High | High | High | High | High | High | High | High | High | 0 | 4.1E-05 | 2.81 |

| | | | | | | | | | | | | | | | | | | | | | | | | | | | | | | | | | | | | | | |
|---|---|---|---|---|---|---|---|---|---|---|---|---|---|---|---|---|---|---|---|---|---|---|---|---|---|---|---|---|---|---|---|---|---|---|---|---|---|---|
| FALSE | High | VLSSTINNC | [K].VLSSTIN | 1xPhospho | S4R221 1xP | 5.31E-09 | 0 | 1 | 7 | 2 | S4R221 | Rho GTPas | S4R221 [56 | 0 | 3567.699 | 0 | -0.55 | 0.001445 | 6.82 | 118.7 | 81.3 | 3.4 | 12.43 | | High | High | High | High | High | High | High | High | High | High | High | 0 | 2.82E-10 | 5.14 |
| FALSE | High | VLSSSEDE | [R].VLSSSS | 1xPhospho | Q8BJW7 1x | 0.001158 | 0 | 1 | 1 | 3 | Q8BJW7 | Crossover | Q8BJW7 [8 | 0 | 2288.09 | 0 | -0.93 | 0.011342 | 2.22 | 133.7 | 66.3 | 17.18 | 13.64 | | High | High | High | High | High | High | High | High | High | High | High | 0 | 0.000282 | 3.94 |
| FALSE | High | VHGLPTTSI | [R].VHGLPT | 1xPhospho | Q3UKG7 1x | 1.05E-08 | 0 | 1 | 5 | 2 | Q3UKG7 | Putative ur | Q3UKG7 [3 | 0 | 2901.531 | 0 | -1.37 | 0.00242 | 1.76 | 140.1 | 59.9 | 12.24 | 7.56 | | High | High | High | High | High | High | High | High | High | High | High | 0 | 6.04E-10 | 5.06 |
| FALSE | High | VHGLPTTSI | [R].VHGLPT | 1xPhospho | Q3UKG7 1x | 4.39E-05 | 0 | 1 | 5 | 4 | Q3UKG7 | Putative ur | Q3UKG7 [3 | 0 | 2917.526 | 0 | -0.93 | 0.006462 | 6.63 | 132 | 68 | 5.97 | 20.71 | | High | High | High | High | High | High | High | High | High | High | High | 0 | 1.64E-05 | 2.68 |
| FALSE | High | VHGPGAES | [K].VHGPG | 1xPhospho | Q571J5 1xP | 1.64E-05 | 0 | 1 | 1 | 1 | Q571J5 | Zinc finger | Q571J5 [14 | 0 | 1516.793 | 0 | 0.15 | 0.072571 | 11.22 | 93.7 | 106.3 | 12.16 | 10.25 | | High | High | High | High | High | High | High | High | High | High | High | 0 | 2.36E-06 | 2.18 |
| FALSE | High | VHIPNDDA | [R].VHIPND | 1xCarbami | Q3UJI2 1xP | 4.74E-07 | 0 | 1 | 7 | 4 | Q3UJI2 | Putative ur | Q3UJI2 [83 | 0 | 2709.202 | 0 | -0.37 | 0.015758 | 7.95 | 112.2 | 87.8 | 11.92 | 6.56 | | High | High | High | High | High | High | High | High | High | High | High | 0 | 4.39E-08 | 5.66 |
| FALSE | High | VHIPNDDA | [R].VHIPND | 1xCarbami | Q3UJI2 1xP | 0.056248 | 0.004124 | 1 | 6 | 1 | Q3UJI2 | Putative ur | Q3UJI2 [83 | 1 | 4061.894 | 0 | 0.25 | 0.359436 | 31.46 | 95.2 | 104.8 | 16.43 | 7.97 | | High | High | High | High | High | High | High | High | High | High | High | 0.002117 | 0.02206 | 3.87 |
| FALSE | High | VHNFMLGI | [K].VHNFM | 1xPhospho | P47713 1xP | 4.66E-07 | 0 | 1 | 2 | 1 | P47713 | Cytosolic p | P47713 [48 | 0 | 2595.336 | 0 | -0.33 | 0.147409 | 21.51 | 104.3 | 95.7 | 12.72 | 23.94 | | High | High | High | High | High | High | High | High | High | High | High | 0 | 4.3E-08 | 4.58 |
| FALSE | High | VHNFMLGI | [K].VHNFM | 1xPhospho | P47713 1xP | 0.000372 | 0 | 1 | 2 | 1 | P47713 | Cytosolic p | P47713 [48 | 0 | 2611.331 | 0 | -1.02 | 0.059124 | 5.22 | 128.5 | 71.5 | 19.95 | 35.76 | | High | High | High | High | High | High | High | High | High | High | High | 0 | 7.89E-05 | 3.11 |
| FALSE | High | VHPFEDSD | [K].VHPFED | 2xPhospho | Q3TC66 2x | 0.003558 | 0.00021 | 1 | 5 | 1 | Q3TC66 | Putative ur | Q3TC66 [1 | 0 | 2782.076 | 0 | -0.48 | 0.013136 | 10.24 | 117 | 83 | 1.9 | 18.94 | | High | High | High | High | High | High | High | High | High | High | High | 9.42E-05 | 0.000989 | 3.59 |
| FALSE | High | VHPFEDSD | [K].VHPFED | 3xPhospho | Q3TC66 3x | 8.76E-07 | 0 | 1 | 5 | 11 | Q3TC66 | Putative ur | Q3TC66 [1 | 0 | 2862.042 | 0 | -0.39 | 0.016053 | 5.35 | 113.9 | 86.1 | 6.26 | 9.56 | | High | High | High | High | High | High | High | High | High | High | High | 0 | 8.78E-08 | 6.35 |
| FALSE | High | VHSPCSTSI | [R].VHSPCS | 1xCarbami | Q3UZ01 1x | 0.001615 | 5.66E-05 | 1 | 3 | 1 | Q3UZ01 | RNA-bindir | Q3UZ01 [1 | 0 | 1884.893 | 0 | -0.62 | 0.011192 | 10.17 | 123.4 | 76.6 | 6.93 | 15.59 | | High | High | High | High | High | High | High | High | High | High | High | 2.58E-05 | 0.00041 | 2.68 |
| FALSE | High | VHSPPASLV | [R].VHSPPA | 1xPhospho | H9KV04 1x | 0.009843 | 0.000736 | 1 | 2 | 3 | H9KV04 | Mini-chron | H9KV04 [2 | 0 | 1468.787 | 0 | -0.85 | 0.001303 | 2.59 | 127.9 | 72.1 | 5.27 | 7.14 | | High | High | High | High | High | High | High | High | High | High | High | 0.000936 | 0.00979 | 1.99 |
| FALSE | High | VHSSDSYK | [R].VHSSDS | 1xPhospho | Q9D0R9 1x | 0.055777 | 0.004093 | 1 | 1 | 1 | Q9D0R9 | WD repeat | Q9D0R9 [3 | 0 | 1460.719 | 0 | 0.69 | 0.037924 | 27.14 | 75.8 | 124.2 | 20.14 | 19.26 | | High | High | High | High | High | High | High | High | High | High | High | 0.002117 | 0.02181 | 1.7 |
| FALSE | High | VHSSLGDTI | [R].VHSSLG | 1xPhospho | A2CG76 1x | 0.010382 | 0.000772 | 1 | 5 | 1 | A2CG76 | Histone-lys | A2CG76 [9 | 0 | 2363.19 | 0 | -0.36 | 0.345648 | 10.12 | 107.2 | 92.8 | 23.51 | 17.03 | | High | High | High | High | High | High | High | High | High | High | High | 0.000382 | 0.003274 | 2.02 |
| FALSE | High | VHGLPSTN | [R].VHGLPS | 1xPhospho | Q8CB62 1x | 2E-08 | 0 | 1 | 2 | 2 | Q8CB62 | Centrobin | Q8CB62 [7 | 0 | 2604.343 | 0 | -0.72 | 0.005447 | 2.5 | 124.3 | 75.7 | 8.7 | 6.22 | | High | High | High | High | High | High | High | High | High | High | High | 0 | 1.25E-09 | 4.99 |
| FALSE | High | VHTGEK | [KRS].VHTG | 1xPhospho | F6ZS36 1xP | 0.075927 | 0.005826 | 1 | 24 | 2 | F6ZS36 | Uncharacte | F6ZS36 [15 | 0 | 1208.644 | 0 | -0.78 | 0.004574 | 4.96 | 127.1 | 72.9 | 6.69 | 11.47 | | High | High | High | High | High | High | High | High | High | High | High | 0.009053 | 0.09705 | 2.28 |
| FALSE | High | VKASPITNI | [K].VKASPI | 1xPhospho | Q64511 1x | 0.009087 | 0.000736 | 1 | 2 | 1 | Q64511 | DNA topois | Q64511 [1 | 1 | 3100.569 | 0 | -0.03 | 0.914347 | 56.89 | 101 | 99 | 26.96 | 27.06 | | High | High | High | High | High | High | High | High | High | High | High | 0.000327 | 0.002827 | 3.11 |
| FALSE | High | VKESSIIAPV | [R].VKESSII | 1xPhospho | Q571L5 1xP | 0.001087 | 0 | 1 | 6 | 2 | Q571L5 | MKIAA406 | Q571L5 [50 | 1 | 2688.426 | 0 | -0.14 | 0.237224 | 23.91 | 114.6 | 85.4 | 15.4 | 28.33 | | High | High | High | High | High | High | High | High | High | High | High | 0 | 0.000261 | 2.6 |
| FALSE | High | VKPESPIGK | [R].VKPESP | 1xPhospho | Q5SXJ3 1xP | 0.000681 | 0 | 1 | 3 | 1 | Q5SXJ3 | Fanconi an | Q5SXJ3 [19 | 0 | 1722.017 | 0 | 0.3 | 0.014294 | 12.79 | 88.6 | 111.4 | 9.32 | 4.27 | | High | High | High | High | High | High | High | High | High | High | High | 0 | 0.000155 | 1.24 |
| FALSE | High | VKPGSPGL | [K].VKPGSP | 1xPhospho | Q3TXU6 1x | 0.090888 | 0.007081 | 2 | 6 | 1 | Q3TXU6; C | Putative ur | Q3TXU6 [7 | 0 | 1853.081 | 0 | -0.32 | 0.005891 | 6.44 | 110.5 | 89.5 | 4.71 | 7.1 | Shared | High | High | High | High | High | High | High | High | High | High | High | 0.003633 | 0.03854 | 1.56 |
| FALSE | High | VKTPAASA | [K].VKTPAA | 1xPhospho | Q8R2U2 1x | 0.076555 | 0.005826 | 1 | 3 | 1 | Q8R2U2 | Nucleolus a | Q8R2U2 [2 | 1 | 1741.023 | 0 | -0.41 | 0.038256 | 3.52 | 109.6 | 90.4 | 5.33 | 13.03 | | High | High | High | High | High | High | High | High | High | High | High | 0.00302 | 0.03168 | 1.57 |
| FALSE | High | VLAGETLSV | [K].VLAGET | 1xPhospho | Q9Z1X4-3 | 1.3E-05 | 0 | 1 | 6 | 1 | Q9Z1X4-3 | Isoform 3 c | Q9Z1X4-3 | | 0 | 2219.116 | 0 | -0.49 | 0.03291 | 3.97 | 121.7 | 78.3 | 17.01 | 11.59 | | High | High | High | High | High | High | High | High | High | High | High | 0 | 1.81E-06 | 3.72 |
| FALSE | High | VIAGSSPEG | [K].VIAGSS | 1xPhospho | Q6PR54-3 | 2.54E-06 | 0 | 2 | 4 | 4 | Q6PR54-3; | Isoform 3 c | Q6PR54-3 | 0 | 2197.077 | 0 | -0.97 | 0.007478 | 6.61 | 132.2 | 67.8 | 9.06 | 16.78 | Shared | High | High | High | High | High | High | High | High | High | High | High | 0 | 6.35E-06 | 3.16 |
| FALSE | High | VIAGSSPEG | [K].VIAGSS | 1xPhospho | Q6PR54-3 | 9.07E-10 | 0 | 2 | 4 | 12 | Q6PR54-3; | Isoform 3 c | Q6PR54-3 | 0 | 2213.072 | 0 | -0.85 | 0.008623 | 5.35 | 128.1 | 71.9 | 7.17 | 17.16 | Shared | High | High | High | High | High | High | High | High | High | High | High | 0 | 3.89E-11 | 4.71 |
| FALSE | High | VIAISGDAD | [R].VIAISGI | 1xPhospho | Q64701 1x | 0.000668 | 0 | 1 | 3 | 4 | Q64701 | Retinoblast | Q64701 [10 | 0 | 1781.945 | 0 | 1.08 | 0.000394 | 9.98 | 64.1 | 135.9 | 6.11 | 5.06 | | High | High | High | High | High | High | High | High | High | High | High | 4.94E-05 | 0.000652 | 3.52 |
| FALSE | High | VLDEFEQN | [K].VLDEFE | 1xPhospho | A2A8R0 1x | 0.00011 | 0 | 1 | 2 | 1 | A2A8R0 | MCG17975 | A2A8R0 [1 | 0 | 2960.443 | 0 | 0.93 | 0.007336 | 34.27 | 64.1 | 135.9 | 20.14 | 13.96 | | High | High | High | High | High | High | High | High | High | High | High | 0 | 2E-05 | 3.43 |
| FALSE | High | VLDNYLTSI | [K].VLDNYI | 1xPhospho | Q9Z1Q5 1x | 1.69E-05 | 0 | 1 | 2 | 4 | Q9Z1Q5 | Chloride in | Q9Z1Q5 [1 | 0 | 3315.535 | 0 | 0.79 | 0.008517 | 28.23 | 71.1 | 128.9 | 11.16 | 12.48 | | High | High | High | High | High | High | High | High | High | High | High | 0 | 2.44E-06 | 3.32 |
| FALSE | High | VLDSPSR | [R].VLDSPS | 1xPhospho | O70585 1x | 0.08727 | 0.006537 | 1 | 8 | 2 | O70585 | Dystrobrev | O70585 [3 | 0 | 1082.544 | 0 | -0.6 | 0.00065 | 1.5 | 123.2 | 76.8 | 8.03 | 7.04 | | High | High | High | High | High | High | High | High | High | High | High | 0.008059 | 0.08726 | 1.21 |
| FALSE | High | VLEAADGS | [R].VLEAAD | 1xPhospho | Q546X9 1x | 5.25E-08 | 0 | 1 | 5 | 2 | Q546X9 | Ataxin 3 OS | Q546X9 [2 | 0 | 2503.107 | 0 | -0.24 | 0.142831 | 19.99 | 113 | 87 | 10.75 | 18.44 | | High | High | High | High | High | High | High | High | High | High | High | 0 | 3.7E-09 | 3.77 |
| FALSE | High | VHVQFFDD | [R].VHVQF | 1xPhospho | P54276 1xP | 2.32E-06 | 0 | 1 | 2 | 8 | P54276 | DNA misma | P54276 [12 | 0 | 1756.826 | 0 | -0.99 | 0.000158 | 1.09 | 134.5 | 65.5 | 2.81 | 4.61 | | High | High | High | High | High | High | High | High | High | High | High | 0 | 0.000178 | 4.14 |
| FALSE | High | VHGHGGA | [K].VHGHG | 1xPhospho | Q6PD29 1x | 1.9E-05 | 0 | 1 | 2 | 2 | Q6PD29 | Zinc finger | Q6PD29 [4 | 0 | 3278.606 | 0 | 0.03 | 0.324761 | 8.09 | 90.2 | 109.8 | 20.35 | 16.65 | | High | High | High | High | High | High | High | High | High | High | High | 0 | 2.77E-06 | 3 |
| FALSE | High | VHGEPASD | [R].VHGEP | 2xPhospho | E9PZ26 2xP | 0.01029 | 0.000772 | 1 | 5 | 3 | E9PZ26 | Bromodom | E9PZ26 [11 | 0 | 1743.708 | 0 | 0.27 | 0.02101 | 6.91 | 87.7 | 112.3 | 5.49 | 7.72 | | High | High | High | High | High | High | High | High | High | High | High | 0.000382 | 0.003233 | 2.3 |
| FALSE | High | VHGEPASD | [R].VHGEP | 1xPhospho | E9PZ26 1xP | 0.000446 | 0 | 1 | 5 | 3 | E9PZ26 | Bromodom | E9PZ26 [11 | 0 | 1663.741 | 0 | -0.14 | 0.144948 | 9.12 | 106.1 | 93.9 | 7.21 | 8.44 | | High | High | High | High | High | High | High | High | High | High | High | 0 | 9.64E-05 | 3.06 |
| FALSE | High | VGSGAEPG | [K].VGSGA | 1xPhospho | Q3U2K0 1x | 0.001222 | 0 | 1 | 3 | 3 | Q3U2K0 | Protein FAI | Q3U2K0 [7 | 0 | 2872.424 | 0 | -0.29 | 0.020342 | 3.97 | 110.1 | 89.9 | 5.51 | 5.03 | | High | High | High | High | High | High | High | High | High | High | High | 7.19E-05 | 0.000776 | 2.65 |
| FALSE | High | VGSLDNVG | [K].VGSLDI | 1xPhospho | P27546 1xP | 3.48E-06 | 0 | 1 | 13 | 2 | P27546 | Microtubu | P27546 [10 | 0 | 1899.989 | 0 | 1.16 | 0.087432 | 28.98 | 58.4 | 141.6 | 24.29 | 43.09 | | High | High | High | High | High | High | High | High | High | High | High | 0 | 4.13E-07 | 4.99 |
| FALSE | High | VGSLDNVG | [K].VGSLDI | 1xPhospho | P27546 1xP | 8.87E-05 | 0 | 1 | 13 | 2 | P27546 | Microtubu | P27546 [10 | 0 | 2129.152 | 0 | 1.07 | 0.00868 | 14.81 | 59 | 141 | 13.63 | 17.86 | | High | High | High | High | High | High | High | High | High | High | High | 0 | 1.57E-05 | 2.67 |
| FALSE | High | VGSLDVNV | [K].VGSLDV | 1xPhospho | E9Q616 1x | 0.000573 | 0 | 1 | 3 | 1 | E9Q616 | AHNAK nuc | E9Q616 [2 | 0 | 1468.818 | 0 | 2.33 | 0.000255 | 16.28 | 34.5 | 165.5 | 17.69 | 6.24 | | High | High | High | High | High | High | High | High | High | High | High | 0 | 0.000127 | 2.52 |
| FALSE | High | VGSLTPPSS | [K].VGSLTP | 2xPhospho | Q3UHJ0 2x | 5.03E-05 | 0 | 1 | 2 | 5 | Q3UHJ0 | AP2-associ | Q3UHJ0 [6 | 0 | 1687.847 | 0 | 0.55 | 0.010953 | 11.43 | 82 | 118 | 9.03 | 8.91 | | High | High | High | High | High | High | High | High | High | High | High | 0.005442 | 0.05962 | 2.33 |
| FALSE | High | VGSPATVT | [K].VGSPAT | 1xPhospho | A2A654 1x | 6.43E-06 | 0 | 1 | 8 | 1 | A2A654 | Bromodom | A2A654 [20 | 0 | 1915.009 | 0 | -0.28 | 0.110147 | 5.37 | 107.6 | 92.4 | 7 | 9.21 | | High | High | High | High | High | High | High | High | High | High | High | 0 | 8.25E-07 | 3.56 |
| FALSE | High | VGSPERPLS | [K].VGSPEF | 1xPhospho | Q5SVQ0 1x | 0.002441 | 0.000109 | 1 | 8 | 1 | Q5SVQ0 | Histone ac | Q5SVQ0 [5 | 0 | 2168.131 | 0 | -1.03 | 0.004758 | 2.72 | 131 | 69 | 8.32 | 14.46 | | High | High | High | High | High | High | High | High | High | High | High | 4.94E-05 | 0.000651 | 1.49 |
| FALSE | High | VGSPLDYSI | [K].VGSPLD | 2xPhospho | E9PWW9 2 | 0.083101 | 0.006345 | 1 | 1 | 1 | E9PWW9 | Remodelin | E9PWW9 [ | 0 | 2620.215 | 0 | -0.83 | 0.086162 | 8.96 | 123.9 | 76.1 | 31.55 | 16.04 | | High | High | High | High | High | High | High | High | High | High | High | 0.003265 | 0.03463 | 2.42 |
| FALSE | High | VGSPLDYSI | [K].VGSPLD | 1xDeamida | E9PWW9 2 | 0.004411 | 0.000409 | 1 | 1 | 2 | E9PWW9 | Remodelin | E9PWW9 [ | 0 | 2621.199 | 0 | -0.94 | | 4.77 | 129.4 | 70.6 | 22.83 | 24.59 | | High | High | High | High | High | High | Not Found | High | High | High | High | 0.000183 | 0.001252 | 2.88 |
| FALSE | High | VGSPLDYSI | [K].VGSPLD | 1xPhospho | E9PWW9 1 | 4.74E-10 | 0 | 1 | 1 | 1 | E9PWW9 | Remodelin | E9PWW9 [ | 0 | 2769.411 | 0 | -1.75 | 0.000158 | 0.87 | 153.4 | 46.6 | 4.29 | 7.92 | | High | High | High | High | High | High | High | High | High | High | High | 0 | 1.88E-11 | 5.2 |
| FALSE | High | VGSPLDYSI | [K].VGSPLD | 1xDeamida | E9PWW9 1 | 8.94E-08 | 0 | 1 | 1 | 4 | E9PWW9 | Remodelin | E9PWW9 [ | 0 | 2770.395 | 0 | -1.58 | 0.000254 | 1.32 | 150.2 | 49.8 | 4.5 | 4.47 | | High | High | High | High | High | High | High | High | High | High | High | 0 | 6.72E-09 | 4.08 |
| FALSE | High | VGSPLDYSI | [K].VGSPLD | 2xPhospho | E9PWW9 2 | 0.001002 | 0 | 1 | 1 | 5 | E9PWW9 | Remodelin | E9PWW9 [ | 0 | 2849.378 | 0 | -0.86 | 0.005233 | 4.94 | 128.8 | 71.2 | 9.54 | 10.56 | | High | High | High | High | High | High | High | High | High | High | High | 0 | 0.000239 | 3.44 |
| FALSE | High | VGSPLTLTI | [R].VGSPLT | 1xPhospho | Q68FH0 1x | 0.002272 | 0.000109 | 1 | 6 | 1 | Q68FH0 | Plakophilin | Q68FH0 [3 | 0 | 1667.857 | 0 | -0.64 | 0.009573 | 3.44 | 123.1 | 76.9 | 10.26 | 4.8 | | High | High | High | High | High | High | High | High | High | High | High | 4.94E-05 | 0.0006 | 2.68 |
| FALSE | High | VGSQDVSL | [R].VGSQD | 1xPhospho | E9QJT6 1xP | 0.000908 | 0 | 1 | 5 | 2 | E9QJT6 | Non-canor | E9QJT6 [59 | 0 | 2141.126 | 0 | 0.72 | 0.009873 | 18.11 | 75.4 | 124.6 | 6.38 | 12.25 | | High | High | High | High | High | High | High | High | High | High | High | 0 | 0.000214 | 2.72 |
| FALSE | High | VGSSQHLR | [R].VGSSQ | 1xPhospho | J3QNH8 1x | 0.001077 | 0 | 1 | 1 | 1 | J3QNH8 | Uncharacte | J3QNH8 [4 | 0 | 1192.604 | 0 | 3.13 | 0.000406 | 20.82 | 20.4 | 179.6 | 15.12 | 10.39 | | High | High | High | High | High | High | High | High | High | High | High | 0 | 0.000259 | 2.06 |
| FALSE | High | VGSSSSSFF | [R].VGSSSS | 1xPhospho | P11679 1xP | 0.005418 | 0.00062 | 1 | 1 | 3 | P11679 | Keratin, ty | P11679 [41 | 0 | 1222.567 | 0 | 4.85 | 0.004235 | 14.3 | 6.1 | 193.9 | 20 | 28.77 | | High | High | High | High | High | High | High | High | High | High | High | 0.000585 | 0.00543 | 1.33 |
| FALSE | High | VGSTENIK | [K].VGSTEN | 1xPhospho | P27546 1xP | 0.005135 | 0.000551 | 1 | 11 | 3 | P27546 | Microtubu | P27546 [91 | 0 | 1385.744 | 0 | 1.09 | 0.008336 | 17.65 | 57.1 | 142.9 | 15.87 | 18.2 | | High | High | High | High | High | High | High | High | High | High | High | 0.000327 | 0.002694 | 2.55 |
| FALSE | High | VGTEGLVD | [R].VGTEG | 1xPhospho | G3X9K4 1x | 0.000984 | 0 | 1 | 3 | 1 | G3X9K4 | SAPS doma | G3X9K4 [2 | 0 | 2002.968 | 0 | 0.24 | 0.020405 | 4.2 | 86.9 | 113.1 | 13.57 | 8.96 | | High | High | High | High | High | High | High | High | High | High | High | 0 | 0.000235 | 3.43 |
| FALSE | High | VGTNYQAV | [R].VGTNY | 1xCarbami | Q8C796 1x | 0.007082 | 0.00062 | 1 | 3 | 1 | Q8C796 | REST corep | Q8C796 [4 | 0 | 2425.178 | 0 | 0.25 | 0.71652 | 6.77 | 93.2 | 106.8 | 68.83 | 18.05 | | High | High | High | High | High | High | High | High | High | High | High | 0.000275 | 0.002137 | 2.39 |
| FALSE | High | VGTNYQAV | [R].VGTNY | 1xCarbami | Q8C796 1x | 0.005666 | 0.00062 | 1 | 3 | 2 | Q8C796 | REST corep | Q8C796 [4 | 0 | 2655.325 | 0 | -0.33 | 0.032319 | 6.1 | 111.3 | 88.7 | 14.71 | 3.1 | | High | High | High | High | High | High | High | High | High | High | High | 0.000275 | 0.001666 | 2.92 |
| FALSE | High | VGTPTRPC | [K].VGTPTF | 1xCarbami | H3BK65 1x | 0.001437 | 5.66E-05 | 1 | 5 | 1 | H3BK65 | Epidermal | H3BK65 [8 | 0 | 1769.897 | 0 | 1.02 | 0.019425 | 13.49 | 61.8 | 138.2 | 11.81 | 25.8 | | High | High | High | High | High | High | High | High | High | High | High | 2.58E-05 | 0.00036 | 3.13 |
| FALSE | High | VGTPTRPC | [K].VGTPTF | 1xCarbami | H3BK65 1x | 0.003372 | 0.00016 | 1 | 5 | 2 | H3BK65 | Epidermal | H3BK65 [8 | 0 | 1999.06 | 0 | 1.08 | 0.015257 | 7.8 | 62.7 | 137.3 | 17.22 | 18.85 | | High | High | High | High | High | High | High | High | High | High | High | 0.000538 | 0.00456 | 1.57 |
| FALSE | High | VGVSQQPE | [K].VGVSQ | 1xPhospho | B2RXS4 1xP | 0.000181 | 0 | 1 | 3 | 1 | B2RXS4 | Plexin-B2 C | B2RXS4 [15 | 0 | 2278.055 | 0 | 1.55 | 0.000335 | 7.34 | 51.5 | 148.5 | 8.71 | 7.57 | | High | High | High | High | High | High | High | High | High | High | High | 0 | 3.48E-05 | 3.04 |
| FALSE | High | VGVSTPSS | [R].VGVSTI | 1xPhospho | Q8CI08-2 1 | 8.56E-05 | 0 | 1 | 2 | 2 | Q8CI08-2 | Isoform 2 c | Q8CI08-2 [ | 0 | 1581.784 | 0 | 1.17 | 0.003268 | 3.28 | 61 | 139 | 16.58 | 10.61 | | High | High | High | High | High | High | High | High | High | High | High | 0 | 1.51E-05 | 2.38 |
| FALSE | High | VHAPSATP | [R].VHAPSA | 1xPhospho | Q9QZS8 1x | 1.1E-06 | 0 | 1 | 2 | 1 | Q9QZS8 | SH2 domai | Q9QZS8 [4 | 0 | 2241.123 | 0 | -0.25 | 0.126438 | 8.89 | 114.1 | 85.9 | 25.77 | 32.04 | | High | High | High | High | High | High | High | High | High | High | High | 0 | 1.12E-07 | 3.58 |
| FALSE | High | VHAYFAPV | [K].VHAYFA | 1xPhospho | Q9CZV8-3 | 1.17E-05 | 0 | 1 | 3 | 1 | Q9CZV8-3 | Isoform 3 c | Q9CZV8-3 | 0 | 2148.084 | 0 | 0.65 | 0.005877 | 25.15 | 77.9 | 122.1 | 5.36 | 9.26 | | High | High | High | High | High | High | High | High | High | High | High | 0 | 1.61E-06 | 2.85 |
| FALSE | High | VHAYFAPV | [K].VHAYFA | 2xPhospho | Q9CZV8-3 | 0.000645 | 0 | 1 | 3 | 2 | Q9CZV8-3 | Isoform 3 c | Q9CZV8-3 | 0 | 2228.05 | 0 | 0.43 | 0.035266 | 20.69 | 85 | 115 | 6.12 | 12.27 | | High | High | High | High | High | High | High | High | High | High | High | 0 | 0.000146 | 3.37 |
| FALSE | High | VHDFPSGN | [K].VHDFPS | 1xPhospho | G5E870 1x | 8.19E-09 | 0 | 1 | 2 | 1 | G5E870 | E3 ubiquiti | G5E870 [1 | 0 | 2127.981 | 0 | -0.69 | 0.003874 | 1.77 | 124.5 | 75.5 | 6.96 | 13.02 | | High | High | High | High | High | High | High | High | High | High | High | 0 | 4.61E-10 | 4.94 |
| FALSE | High | VHDFPSGN | [K].VHDFPS | 1xDeamida | G5E870 1x | 8.29E-10 | 0 | 1 | 2 | 4 | G5E870 | E3 ubiquiti | G5E870 [1 | 0 | 2128.965 | 0 | -0.3 | 0.050207 | 5.8 | 110.3 | 89.7 | 6.35 | 7.55 | | High | High | High | High | High | High | High | High | High | High | High | 0 | 0.000116 | 3.91 |
| FALSE | High | VLEASPGRI | [R].VLEASP | 1xPhospho | A0A075F6C | 0.014538 | 0.001011 | 1 | 5 | 2 | A0A075F6C | Heat shock | A0A075F6C | 0 | 3166.606 | 0 | 0.6 | 0.070104 | 23.27 | 78.8 | 121.2 | 12.8 | 16.03 | | High | High | High | High | High | High | High | High | High | High | High | 0.007957 | 0.08583 | 2.19 |
| FALSE | High | VPSPPPAPE | [R].VPSPPP | 1xCarbami | Q80U38 1x | 1.69E-06 | 0 | 1 | 1 | 2 | Q80U38 | Protein KH | Q80U38 [3 | 0 | 2165.009 | 0 | -0.9 | 0.000546 | 2.51 | 130.8 | 69.2 | 3.01 | 9.8 | | High | High | High | High | High | High | High | High | High | High | High | 0 | 1.84E-07 | 3.88 |
| FALSE | High | VLEDDPEA | [R].VLEDDI | 1xPhospho | Q03145 1x | 5.75E-05 | 0 | 1 | 1 | 1 | Q03145 | Ephrin type | Q03145 [7 | 0 | 2221.068 | 0 | 0.61 | 0.001073 | 14.46 | 79 | 121 | 12.03 | 3.81 | | High | High | High | High | High | High | High | High | High | High | High | 0 | 9.62E-06 | 2.67 |
| FALSE | High | VIEDEDGA | [R].VIEDED | 1xCarbami | Q9R0L6-2 | 1.99E-06 | 0 | 1 | 3 | 4 | Q9R0L6-2 | Isoform 2 c | Q9R0L6-2 | | 0 | 4697.176 | 0 | 0.09 | 0.129034 | 6.18 | 98.8 | 101.2 | 6.51 | 11.94 | | High | High | High | High | High | High | High | High | High | High | High | 0 | 2.19E-07 | 5.61 |
| FALSE | High | VLPQGPSTI | [R].VLPQGI | 1xPhospho | Q8CC12 1x | 0.004256 | 0.00036 | 1 | 3 | 3 | Q8CC12 | Codanin-1 | Q8CC12 [6 | 0 | 1632.913 | 0 | -0.75 | 0.000857 | 0.66 | 125.2 | 74.8 | 2.63 | 9.88 | | High | High | High | High | High | High | High | High | High | High | High | 0.000161 | 0.001206 | 2.39 |
| FALSE | High | VIPTPGK | [K].VIPTPG | 1xPhospho | P09405 1xP | 0.061176 | 0.004646 | 1 | 7 | 3 | P09405 | Nucleolin C | P09405 [10 | 0 | 1249.732 | 0 | -0.7 | 0.003215 | 0.22 | 125.2 | 74.8 | 10.57 | 5.62 | | High | High | High | High | High | High | High | High | High | High | High | 0.003064 | 0.03185 | 1.66 |
| FALSE | High | VLPVSPGAI | [R].VLPVSP | 1xPhospho | Q8BIV1 1xP | 0.003558 | 0.00021 | 1 | 2 | 1 | Q8BIV1 | Putative ur | Q8BIV1 [14 | 0 | 1923.051 | 0 | 0.12 | 0.484608 | 12 | 96 | 104 | 4.4 | 9.19 | | High | High | High | High | High | High | High | High | High | High | High | 9.42E-05 | 0.000987 | 2.69 |
| FALSE | High | VLQAMGYI | [K].VLQAM | 1xCarbami | Q91WM1 1 | 3.21E-09 | 0 | 1 | 2 | 2 | Q91WM1 | Spermatid | Q91WM1 [ | 0 | 3084.435 | 0 | -0.88 | 0.000609 | 3.84 | 130.9 | 69.1 | 4.91 | 12.6 | | High | High | High | High | High | High | High | High | High | High | High | 0 | 1.6E-10 | 4.81 |
| FALSE | High | VLQAMGYI | [K].VLQAM | 1xCarbami | Q91WM1 1 | 0.020862 | 0.001472 | 1 | 2 | 2 | Q91WM1 | Spermatid | Q91WM1 [ | 0 | 3100.43 | 0 | -0.5 | 0.012188 | 9.67 | 116.7 | 83.3 | 9.7 | 12.16 | | High | High | High | High | High | High | High | High | High | High | High | 0.000786 | 0.007132 | 3.21 |
| FALSE | High | VLQATVVA | [K].VLQATV | 1xPhospho | Q64433 1x | 0.000128 | 0 | 1 | 2 | 1 | Q64433 | 10 kDa hea | Q64433 [4 | 0 | 1824.04 | 0 | -1.68 | 0.000193 | 0.98 | 151.6 | 48.4 | 4.52 | 7.23 | | High | High | High | High | High | High | High | High | High | High | High | 0 | 2.38E-05 | 2.61 |
| FALSE | High | VLQDMGLI | [K].VLQDM | 1xPhospho | Q9Z1X4-3 | 0.001127 | 0 | 1 | 6 | 1 | Q9Z1X4-3 | Isoform 3 c | Q9Z1X4-3 | | 1 | 2415.199 | 0 | -0.06 | 0.531866 | 17.46 | 102.2 | 97.8 | 13.52 | 10.62 | | High | High | High | High | High | High | High | High | High | High | High | 0 | 0.000273 | 2.23 |
| FALSE | High | VLQPSPVSI | [K].VLQPSF | 1xPhospho | Q3URX8 1x | 0.015603 | 0.001124 | 1 | 3 | 2 | Q3URX8 | Putative ur | Q3URX8 [7 | 0 | 1907.931 | 0 | -0.28 | 0.004345 | 4.84 | 110 | 90 | 3.35 | 3.97 | | High | High | High | High | High | High | High | High | High | High | High | 0.000585 | 0.005149 | 2.92 |
| FALSE | High | VLSAPPHFI | [R].VLSAPP | 1xPhospho | Q9D8N0 1x | 0.086562 | 0.006435 | 1 | 1 | 1 | Q9D8N0 | Elongation | Q9D8N0 [3 | 1 | 2760.397 | 0 | 0.41 | 0.082424 | 19.74 | 89.9 | 110.1 | 14.88 | 14.72 | | High | High | High | High | High | High | High | High | High | High | High | 0.003303 | 0.0363 | 1.93 |
| FALSE | High | VLSCQSSLK | [R].VLSCQS | 1xCarbami | Q8BX22 1x | 0.011966 | 0.000889 | 1 | 2 | 1 | Q8BX22 | Sal-like pro | Q8BX22 [5 | 0 | 1559.827 | 0 | -1.81 | 0.004055 | 1.7 | 151.9 | 48.1 | 20.57 | 11.71 | | High | High | High | High | High | High | High | High | High | High | High | 0.000449 | 0.00385 | 2.8 |
| FALSE | High | VLSDDEED | [R].VLSDDE | 1xPhospho | Q80X90 1x | 5.31E-05 | 0 | 1 | 1 | 2 | Q80X90 | Filamin-B C | Q80X90 [7 | 0 | 3289.555 | 0 | 1.6 | 0.002231 | 19.86 | 50.8 | 149.2 | 12.8 | 10.92 | | High | High | High | High | High | High | High | High | High | High | High | 0 | 8.81E-06 | 4.05 |
| FALSE | High | VISDDNWS | [K].VISDDN | 1xPhospho | Q9WUF3 1 | 1.15E-05 | 0 | 1 | 1 | 2 | Q9WUF3 | CASP8-asso | Q9WUF3 [ | 0 | 2131.072 | 0 | 0.17 | 0.301064 | 7.72 | 96.8 | 103.2 | 6.01 | 14.64 | | High | High | High | High | High | High | High | High | High | High | High | 0.000161 | 0.001213 | 3.55 |
| FALSE | High | VISDDNWS | [K].VISDDN | 2xPhospho | Q9WUF3 2 | 0.014929 | 0.001049 | 1 | 1 | 1 | Q9WUF3 | CASP8-asso | Q9WUF3 [ | 0 | 2211.039 | 0 | 0.64 | 0.381785 | 39.39 | 86.6 | 113.4 | 37.33 | 34.37 | | High | High | High | High | High | High | High | High | High | High | High | 0.000555 | 0.004926 | 3.02 |
| FALSE | High | VLPCQSPN | [K].VLPCQS | 1xCarbami | Q6A060 1x | 0.065405 | 0.004734 | 1 | 7 | 1 | Q6A060 | MKIAA048 | Q6A060 [6 | 1 | 4019.06 | 0 | 1.11 | 0.000669 | 1.35 | 63.6 | 136.4 | 6.3 | 8.26 | | High | High | High | High | High | High | High | High | High | High | High | 0.002446 | 0.02627 | 3.48 |
| FALSE | High | VLSDSEEEE | [K].VLSDSE | 2xPhospho | F8VQ93 2x | 0.0008 | 0 | 1 | 2 | 3 | F8VQ93 | PH-interact | F8VQ93 [1 | 0 | 1782.785 | 0 | 0.22 | 0.173214 | 11.77 | 94.9 | 105.1 | 2.55 | 13.01 | | High | High | High | High | High | High | High | High | High | High | High | 0 | 0.000185 | 3.62 |
| FALSE | High | VLSGPGSA | [R].VLSGPG | 1xPhospho | E9PUV9 1x | 0.015195 | 0.001089 | 1 | 2 | 1 | E9PUV9 | Caspase re | E9PUV9 [6 | 0 | 1152.598 | 0 | 1.57 | 0.000447 | 19.2 | 51.6 | 148.4 | 12.63 | 5.14 | | High | High | High | High | High | High | High | High | High | High | High | 0.000571 | 0.004999 | 0.74 |
| FALSE | High | VLSGQDTE | [K].VLSGQI | 1xPhospho | Q8VD57 1x | 0.012397 | 0.000889 | 1 | 3 | 1 | Q8VD57 | Vesicle tra | Q8VD57 [7 | 0 | 1428.657 | 0 | 1.04 | 0.007327 | 2.23 | 60.4 | 139.6 | 23.22 | 13.11 | | High | High | High | High | High | High | High | High | High | High | High | 0.000449 | 0.004004 | 3.15 |
| FALSE | High | VISHDRDSI | [R].VISHDR | 1xPhospho | E9PZY8 1xP | 2.2E-06 | 0 | 1 | 4 | 4 | E9PZY8 | RIKEN cDN | E9PZY8 [13 | 1 | 3326.77 | 0 | -0.54 | 0.003186 | 2.72 | 118.4 | 81.6 | 4.51 | 6.81 | | High | High | High | High | High | High | High | High | High | High | High | 0 | 2.46E-07 | 3.46 |
| FALSE | High | VLSIGDGIA | [R].VLSIGD | 1xPhospho | Q03265 1x | 0.001244 | 0 | 1 | 2 | 1 | Q03265 | ATP syntha | Q03265 [7 | 0 | 1309.708 | 0 | 1.15 | 0.000281 | 3.49 | 61.9 | 138.1 | 13.71 | 9.71 | | High | High | High | High | High | High | High | High | High | High | High | 0 | 0.000306 | 2.25 |
| FALSE | High | VLSMDGA( | [R].VLSMD | 1xPhospho | Q8VI59 1xP | 0.053473 | 0.00384 | 1 | 1 | 1 | Q8VI59 | Pecanex-lik | Q8VI59 [50 | 0 | 1614.776 | 0 | 0.73 | 0.090739 | 19.89 | 74.9 | 125.1 | 8.77 | 43.93 | | High | High | High | High | High | High | High | High | High | High | High | 0.001999 | 0.02077 | 2.27 |
| FALSE | High | VLSPLIIK | [R].VLSPLII | 1xPhospho | E9Q7F2 1x | 0.004411 | 0.000409 | 1 | 1 | 1 | E9Q7F2 | E3 ubiquiti | E9Q7F2 [3 | 0 | 1420.894 | 0 | -0.57 | 0.001148 | 3.93 | 120.6 | 79.4 | 6.96 | 4.54 | | High | High | High | High | High | High | High | High | High | High | High | 0.000183 | 0.001256 | 2.37 |
| FALSE | High | VLSPILLNN | [R].VLSPILL | 1xPhospho | Q9WU62 1 | 0.000815 | 0 | 1 | 2 | 1 | Q9WU62 | Inner centr | Q9WU62 [ | 0 | 2245.334 | 0 | -0.21 | 0.022633 | 9.13 | 107.2 | 92.8 | 6.03 | 7.65 | | High | High | High | High | High | High | High | High | High | High | High | 0 | 0.000189 | 2.71 |
| FALSE | High | VLSPILLNN | [R].VLSPILL | 1xDeamida | Q9WU62 1 | 0.000662 | 0 | 1 | 2 | 3 | Q9WU62 | Inner centr | Q9WU62 [ | 0 | 2246.318 | 0 | -0.1 | 0.308043 | 13.62 | 102.2 | 97.8 | 10.25 | 14.8 | | High | High | High | High | High | High | High | High | High | High | High | 0.000275 | 0.001604 | 3.74 |
| FALSE | High | VLSPLRSPP | [K].VLSPLR | 2xPhospho | P14873 2xP | 0.005228 | 0.000551 | 2 | 2 | 3 | P14873; B2 | Microtubu | P14873 [13 | 1 | 3422.694 | 0 | -0.81 | 0.01092 | 5.46 | 127.3 | 72.7 | 14.99 | 6.85 | Shared | High | High | High | High | High | High | High | High | High | High | High | 0.000538 | 0.00459 | 4.52 |
| FALSE | High | VISPPPFAP | [K].VISPPPI | 2xPhospho | Q3UT88 2x | 0.065405 | 0.004734 | 1 | 3 | 1 | Q3UT88 | Putative ur | Q3UT88 [7 | 0 | 3705.83 | 0 | 1.04 | 0.000492 | 13.67 | 69.1 | 130.9 | 9.39 | 4.61 | | High | High | High | High | High | High | High | High | High | High | High | 0.002432 | 0.02618 | 2.9 |
| FALSE | High | VLSQTVPSS | [K].VLSQTV | 1xCarbami | Q9CT10 1x | 0.000331 | 0 | 1 | 1 | 18 | Q9CT10 | Ran-bindin | Q9CT10 [1 | 0 | 3855.798 | 0 | 0.8 | 0.037985 | 10.33 | 66.8 | 133.2 | 11.92 | 28.56 | | High | High | High | High | High | High | High | High | High | High | High | 2.58E-05 | 0.000404 | 4.14 |
| FALSE | High | VLSQTVPSS | [K].VLSQTV | 1xCarbami | Q9CT10 1x | 1.3E-05 | 0 | 1 | 1 | 2 | Q9CT10 | Ran-bindin | Q9CT10 [1 | 0 | 3871.793 | 0 | 0.34 | 0.209016 | 25.33 | 84.9 | 115.1 | 23.85 | 42.61 | | High | High | High | High | High | High | High | High | High | High | High | 0 | 1.82E-06 | 3.95 |
| FALSE | High | VISSSQVDO | [K].VISSSQ | 1xPhospho | Q80U04 1x | 5.5E-05 | 0 | 1 | 2 | 3 | Q80U04 | E3 ubiquiti | Q80U04 [3 | 0 | 1987.969 | 0 | 1.43 | 0.008457 | 30.39 | 58.6 | 141.4 | 19.92 | 20.21 | | High | High | High | High | High | High | High | High | High | High | High | 0 | 9.16E-06 | 4.23 |
| FALSE | High | VISFIENSST | [K].VISFIEN | 1xPhospho | Q8K0F1 1x | 0.008691 | 0.000736 | 1 | 1 | 1 | Q8K0F1 | TBC1 doma | Q8K0F1 [5 | 0 | 1872.931 | 0 | 3.69 | 0.007605 | 17.86 | 13.3 | 186.7 | 67.8 | 27.27 | | High | High | High | High | High | High | High | High | High | High | High | 0.000327 | 0.002676 | 1.96 |
| FALSE | High | VLPCGSPP | [R].VLPCGS | 1xCarbami | E9Q9R9 1x | 0.003433 | 0.00021 | 1 | 4 | 1 | E9Q9R9 | Disks large | E9Q9R9 [1 | 0 | 1487.764 | 0 | 0.49 | 0.006718 | 9.13 | 86.5 | 113.5 | 7.66 | 10.09 | | High | High | High | High | High | High | High | High | High | High | High | 9.42E-05 | 0.00095 | 2.16 |
| FALSE | High | VLPCGPSP( | [R].VLPCGF | 1xCarbami | P52734 1xP | 0.001168 | 0 | 1 | 2 | 3 | P52734 | FYVE, RhoC | P52734 [76 | 0 | 1693.82 | 0 | -0.93 | 0.000832 | 1.72 | 131.1 | 68.9 | 3.63 | 9.78 | | High | High | High | High | High | High | High | High | High | High | High | 0 | 0.000283 | 2.21 |
| FALSE | High | VLPALELSD | [R].VLPALE | 1xPhospho | Q9D7Y9 1x | 0.007145 | 0.00062 | 1 | 2 | 1 | Q9D7Y9 | Protein SLX | Q9D7Y9 [3 | 0 | 2116.243 | 0 | -0.81 | 0.012404 | 5.02 | 127.2 | 72.8 | 5.33 | 31.73 | | High | High | High | High | High | High | High | High | High | High | High | 0.000275 | 0.00216 | 3.15 |
| FALSE | High | VIEDEDGA | [R].VIEDED | 1xCarbami | Q9R0L6-2 | 3.1E-10 | 0 | 1 | 3 | 3 | Q9R0L6-2 | Isoform 2 c | Q9R0L6-2 | | 0 | 4698.16 | 0 | -0.08 | 0.879479 | 9.11 | 101 | 99 | 10.36 | 10.23 | | High | High | High | High | High | High | High | High | High | High | High | 0 | 1.17E-11 | 7.59 |
| FALSE | High | VIEDEDGA | [R].VIEDED | 1xCarbami | Q9R0L6-2 | 2.69E-07 | 0 | 1 | 3 | 2 | Q9R0L6-2 | Isoform 2 c | Q9R0L6-2 | | 0 | 4777.143 | 0 | -0.02 | 0.56994 | 21.43 | 92.5 | 107.5 | 24.48 | 11.47 | | High | High | High | High | High | High | High | High | High | High | High | 0 | 3.46E-08 | 5.43 |
| FALSE | High | VLEGSQAG | [R].VLEGS( | 1xPhospho | G3X960 1x | 0.002768 | 0.000109 | 1 | 2 | 5 | G3X960 | MCG5982 | G3X960 [4 | 1 | 2671.329 | 0 | 0.34 | 0.004615 | 2.84 | 88.4 | 111.6 | 7.09 | 9.96 | | High | High | High | High | High | High | High | High | High | High | High | 0.00014 | 0.001111 | 2.9 |
| FALSE | High | VIEPPAASC | [R].VIEPPA | 1xCarbami | Q3UYI5 1xP | 0.001516 | 5.66E-05 | 1 | 1 | 2 | Q3UYI5 | Ral guanine | Q3UYI5 [4 | 0 | 1776.855 | 0 | 1.33 | 0.004802 | 20.48 | 53.5 | 146.5 | 16.52 | 16.76 | | High | High | High | High | High | High | High | High | High | High | High | 2.58E-05 | 0.000382 | 2.81 |
| FALSE | High | VLEQLTGQ | [K].VLEQLT | 1xPhospho | Q9CXW4 1 | 0.014929 | 0.001049 | 1 | 5 | 1 | Q9CXW4 | 60S riboso | Q9CXW4 [ | 0 | 2085.14 | 0 | 0 | 0.567856 | 12.33 | 98.7 | 101.3 | 7.6 | 14.78 | | High | High | High | High | High | High | High | High | High | High | High | 0.000555 | 0.00489 | 1.81 |
| FALSE | High | VLGAGSSG | [R].VLGAG | 1xPhospho | Q8VC03 1x | 1.93E-07 | 0 | 1 | 1 | 3 | Q8VC03 | Echinoderr | Q8VC03 [8 | 0 | 1733.878 | 0 | 1.96 | 0.000587 | 18.69 | 40.6 | 159.4 | 15.49 | 8.19 | | High | High | High | High | High | High | High | High | High | High | High | 0 | 1.59E-08 | 4.16 |
| FALSE | High | VLGGNSAT | [R].VLGGN | 1xPhospho | Q99K90 1x | 0.009585 | 0.000736 | 1 | 3 | 2 | Q99K90 | TGF-beta-a | Q99K90 [4 | 0 | 1367.688 | 0 | 1.61 | 0.000769 | 14.66 | 51.9 | 148.1 | 11.39 | 8.48 | | High | High | High | High | High | High | High | High | High | High | High | 0.002381 | 0.02449 | 1.22 |
| FALSE | High | VLGGTGSG | [K].VLGGT | 1xPhospho | Q3UHH1 1 | 0.002058 | 5.66E-05 | 1 | 5 | 2 | Q3UHH1 | Zinc finger | Q3UHH1 [5 | 0 | 1370.745 | 0 | 0.22 | 0.029613 | 7.96 | 93.8 | 106.2 | 2.99 | 9.91 | | High | High | High | High | High | High | High | High | High | High | High | 2.58E-05 | 0.000539 | 1.78 |
| FALSE | High | VLGSLPSSC | [K].VLGSLP | 1xCarbami | Q8K368 1x | 1.32E-08 | 0 | 1 | 3 | 3 | Q8K368 | Fanconi an | Q8K368 [5 | 0 | 2426.231 | 0 | -1.25 | 0.000479 | 4.07 | 142.7 | 57.3 | 2.93 | 18.79 | | High | High | High | High | High | High | High | High | High | High | High | 0 | 7.89E-10 | 3.72 |
| FALSE | High | VLGSPAYAI | [K].VLGSPA | 1xPhospho | J3QK54 1xP | 0.004654 | 0.000456 | 1 | 1 | 2 | J3QK54 | Fidgetin-lik | J3QK54 [32 | 0 | 2270.187 | 0 | -0.43 | 0.012826 | 9.12 | 119.6 | 80.4 | 6.85 | 11.62 | | High | High | High | High | High | High | High | High | High | High | High | 0.00092 | 0.009637 | 2.91 |
| FALSE | High | VLGSPPVEI | [R].VLGSPF | 1xCarbami | Q6ZPG3 1x | 0.000148 | 0 | 1 | 1 | 1 | Q6ZPG3 | MKIAA189 | Q6ZPG3 [7 | 0 | 2780.28 | 0 | -0.98 | 0.007474 | 6.94 | 131.5 | 68.5 | 12.68 | 24.55 | | High | High | High | High | High | High | High | High | High | High | High | 0 | 2.8E-05 | 2.44 |

| | | | | | | | | | | | | | | | | | | | | | | | | | | | | | | | | | | | | | | |
|---|---|---|---|---|---|---|---|---|---|---|---|---|---|---|---|---|---|---|---|---|---|---|---|---|---|---|---|---|---|---|---|---|---|---|---|---|---|---|
| FALSE | High | VLGSQEAL | [K].VLGSQI | 1xPhospho | Q3U2X5 1x | 0.000479 | 0 | 1 | 4 | 1 | Q3U2X5 | Amine oxic | Q3U2X5 [3 | 0 | 2324.194 | 0 | 0.57 | 0.002468 | 2.19 | 81.1 | 118.9 | 4.16 | 9.79 | | High | High | High | High | High | High | High | High | High | High | High | 0 | 0.000104 | 2.22 |
| FALSE | High | VLGTPVLA | [K].VLGTPV | 1xPhospho | Q9WVH3 1 | 2.12E-06 | 0 | 1 | 2 | 1 | Q9WVH3 | Forkhead b | Q9WVH3 [ | 0 | 2190.064 | 0 | -1.05 | 0.002756 | 4.65 | 135.5 | 64.5 | 17.05 | 14.3 | | High | High | High | High | High | High | High | High | High | High | High | 0 | 2.36E-07 | 3.22 |
| FALSE | High | VLGTSPEAI | [R].VLGTSP | 1xPhospho | B2RQC6 1x | 4.35E-05 | 0 | 1 | 5 | 2 | B2RQC6 | CAD protei | B2RQC6 [1 | 0 | 1867.9 | 0 | 0.41 | 0.01469 | 6.64 | 85.6 | 114.4 | 5.81 | 9.65 | | High | High | High | High | High | High | High | High | High | High | High | 0 | 7.03E-06 | 3.88 |
| FALSE | High | VLHSSNPVI | [K].VLHSSN | 1xPhospho | Q8BHZ4 1x | 1.75E-05 | 0 | 1 | 4 | 1 | Q8BHZ4 | Zinc finger | Q8BHZ4 [5 | 0 | 2640.338 | 0 | -0.2 | 0.062604 | 9.01 | 106.3 | 93.7 | 9.09 | 11.48 | | High | High | High | High | High | High | High | High | High | High | High | 0 | 2.54E-06 | 3.96 |
| FALSE | High | VLLAADSEI | [R].VLLAAD | 1xPhospho | E9QK89 1x | 1.1E-06 | 0 | 1 | 2 | 6 | E9QK89 | Mediator c | E9QK89 [1 | 0 | 2100.969 | 0 | -0.84 | 0.000236 | 1.46 | 128.8 | 71.2 | 2.94 | 4.33 | | High | High | High | High | High | High | High | High | High | High | High | 0 | 6.92E-07 | 3.88 |
| FALSE | High | VLLAADSEI | [R].VLLAAD | 2xPhospho | E9QK89 2x | 0.000124 | 0 | 1 | 2 | 2 | E9QK89 | Mediator c | E9QK89 [1 | 0 | 2180.935 | 0 | 0.13 | 0.029993 | 6.91 | 95.4 | 104.6 | 9.41 | 6.27 | | High | High | High | High | High | High | High | High | High | High | High | 9.42E-05 | 0.001004 | 3.86 |
| FALSE | High | VILASPETS | [K].VILASPI | 1xPhospho | G3X9W4 1 | 0.000137 | 0 | 1 | 4 | 1 | G3X9W4 | Nuclear re | G3X9W4 [9 | 0 | 1740.955 | 0 | 0.88 | 0.001453 | 8.21 | 71.8 | 128.2 | 5.97 | 8.83 | | High | High | High | High | High | High | High | High | High | High | High | 0 | 2.54E-05 | 3.29 |
| FALSE | High | VLIEDTDDE | [K].VLIEDT | 1xPhospho | E3VX19 1x | 0.037369 | 0.00249 | 1 | 5 | 2 | E3VX19 | Xenotropic | E3VX19 [6 | 0 | 1643.725 | 0 | 0.06 | 0.996173 | 27.12 | 97.9 | 102.1 | 14.93 | 29.69 | | High | High | High | High | High | High | High | High | High | High | High | 0.002159 | 0.02293 | 2.29 |
| FALSE | High | VLLGFSSDE | [K].VLLGFS | 1xPhospho | Q9WU40-2 | 6.35E-05 | 0 | 1 | 4 | 5 | Q9WU40-2 | Isoform 2 c | Q9WU40-2 | 0 | 2117 | 0 | 0.55 | 0.007596 | 10.41 | 83.1 | 116.9 | 8.3 | 12.02 | | High | High | High | High | High | High | High | High | High | High | High | 0 | 0.000214 | 3.55 |
| FALSE | High | VLLHSPGRI | [R].VLLHSP | 2xPhospho | Q8R1G6 2x | 0.040373 | 0.002982 | 1 | 2 | 1 | Q8R1G6 | PDZ and LI | Q8R1G6 [1 | 0 | 1791.887 | 0 | 0.11 | 0.125323 | 9.17 | 97.6 | 102.4 | 12.76 | 11.6 | | High | High | High | High | High | High | High | High | High | High | High | 0.00155 | 0.01503 | 1.34 |
| FALSE | High | VLINGSAY | [R].VLINGS | 1xPhospho | H9KV00 1x | 1.87E-06 | 0 | 1 | 1 | 1 | H9KV00 | Protein SO | H9KV00 [2 | 0 | 2331.215 | 0 | 0.51 | 0.015691 | 24.41 | 82.9 | 117.1 | 10.69 | 5.84 | | High | High | High | High | High | High | High | High | High | High | High | 0 | 2.06E-07 | 3.63 |
| FALSE | High | VIIPSPPKDI | [K].VIIPSPP | 1xPhospho | A2SW42 1x | 0.002153 | 5.66E-05 | 1 | 4 | 1 | A2SW42 | Zinc finger | A2SW42 [1 | 1 | 2658.416 | 0 | -1.77 | 0.001621 | 1.38 | 155 | 45 | 9.55 | 16.14 | | High | High | High | High | High | High | High | High | High | High | High | 2.58E-05 | 0.000567 | 2.38 |
| FALSE | High | VLLPTPPST | [R].VLLPTP | 1xPhospho | Q8C9B9 1x | 3.5E-05 | 0 | 1 | 1 | 3 | Q8C9B9 | Death-indu | Q8C9B9 [1 | 0 | 2702.497 | 0 | -0.54 | 0.005285 | 6.23 | 121 | 79 | 10.68 | 11.39 | | High | High | High | High | High | High | High | High | High | High | High | 0 | 5.52E-06 | 3.1 |
| FALSE | High | VLIQANQE | [R].VLIQAN | 1xPhospho | Q569X8 1x | 2.2E-06 | 0 | 1 | 4 | 1 | Q569X8 | Nucleopor | Q569X8 [3 | 0 | 2183.07 | 0 | 0.32 | 0.248435 | 35.11 | 91 | 109 | 10.72 | 26.25 | | High | High | High | High | High | High | High | High | High | High | High | 0 | 2.45E-07 | 3.38 |
| FALSE | High | VLISEEFELE | [K].VLISEEI | 1xCarbami | Q8BQ33 1x | 6.81E-07 | 0 | 1 | 1 | 1 | Q8BQ33 | Treslin OS= | Q8BQ33 [1 | 0 | 3052.535 | 0 | -0.85 | 0.003719 | 5.57 | 131.4 | 68.6 | 11.43 | 12.54 | | High | High | High | High | High | High | High | High | High | High | High | 0 | 6.58E-08 | 3.29 |
| FALSE | High | VLISVGSYK | [K].VLISVG | 1xPhospho | Q3UMB5 1 | 0.051258 | 0.003745 | 1 | 2 | 1 | Q3UMB5 | Guanine nu | Q3UMB5 [ | 0 | 1503.859 | 0 | 1.72 | 0.00378 | 20.67 | 44.4 | 155.6 | 9.44 | 21.9 | | High | High | High | High | High | High | High | High | High | High | High | 0.00196 | 0.0198 | 2.42 |
| FALSE | High | VLLTSENDI | [R].VLLTSE | 1xPhospho | Q7TSY8 1x | 9.76E-07 | 0 | 1 | 1 | 2 | Q7TSY8 | Shugoshin | Q7TSY8 [1 | 0 | 2260.027 | 0 | -0.61 | 0.117285 | 10.32 | 123.4 | 76.6 | 18.19 | 32.6 | | High | High | High | High | High | High | High | High | High | High | High | 0 | 4.26E-07 | 2.86 |
| FALSE | High | VINAAHSFY | [K].VINAAH | 1xPhospho | Q3THK7 1x | 0.021797 | 0.001507 | 1 | 1 | 1 | Q3THK7 | GMP synth | Q3THK7 [2 | 1 | 3084.499 | 0 | -0.44 | 0.027373 | 11.75 | 115.2 | 84.8 | 13.32 | 15.05 | | High | High | High | High | High | High | High | High | High | High | High | 0.000801 | 0.007479 | 2.35 |
| FALSE | High | VIEDEDGA | [R].VIEDED | 1xCarbami | Q9R0L6-2 | 3.79E-07 | 0 | 1 | 3 | 8 | Q9R0L6-2 | Isoform 2 c | Q9R0L6-2 | 0 | 4696.192 | 0 | -0.04 | 0.954392 | 19.8 | 98.4 | 101.6 | 8.13 | 11.45 | | High | High | High | High | High | High | High | High | High | High | High | 0 | 9.06E-08 | 6.42 |
| FALSE | High | VPSPPPAPI | [R].VPSPPP | 1xCarbami | Q80U38 1x | 0.046291 | 0.003523 | 1 | 1 | 1 | Q80U38 | Protein KH | Q80U38 [3 | 1 | 2436.137 | 0 | 0 | 0.455983 | 8.98 | 96.3 | 103.7 | 9.56 | 10.96 | | High | High | High | High | High | High | High | High | High | High | High | 0.001789 | 0.01772 | 3.5 |
| FALSE | High | VPSPPPGH | [K].VPSPPP | 1xPhospho | Q04750 1x | 0.050826 | 0.003676 | 1 | 1 | 1 | Q04750 | DNA topois | Q04750 [3 | 0 | 1453.797 | 0 | -0.05 | 0.127767 | 1.62 | 105.2 | 94.8 | 7.37 | 3.84 | | High | High | High | High | High | High | High | High | High | High | High | 0.001932 | 0.01966 | 1.94 |
| FALSE | High | VPSPQGAD | [R].VPSPQ | 1xPhospho | Q3UFT4 1x | 0.042503 | 0.003197 | 1 | 6 | 1 | Q3UFT4 | Putative ur | Q3UFT4 [5 | 0 | 1334.667 | 0 | 0.01 | 0.607892 | 4.73 | 100.7 | 99.3 | 3.6 | 7.15 | | High | High | High | High | High | High | High | High | High | High | High | 0.001639 | 0.01597 | 2.18 |
| FALSE | High | VSPSPSQES | [R].VSPSPS | 1xPhospho | Q6NVE8 1x | 0.002022 | 5.66E-05 | 1 | 2 | 4 | Q6NVE8 | WD repeat | Q6NVE8 [5 | 0 | 1957.988 | 0 | 0.62 | 0.004993 | 4.68 | 75.2 | 124.8 | 5.78 | 12.26 | | High | High | High | High | High | High | High | High | High | High | High | 0.000859 | 0.008367 | 2.95 |
| FALSE | High | VSPSPSQES | [R].VSPSPS | 2xPhospho | Q6NVE8 2x | 0.000241 | 0 | 1 | 2 | 1 | Q6NVE8 | WD repeat | Q6NVE8 [5 | 0 | 2037.955 | 0 | 0.78 | 0.011531 | 13.57 | 73.5 | 126.5 | 5.08 | 19.72 | | High | High | High | High | High | High | High | High | High | High | High | 0 | 4.84E-05 | 2.67 |
| FALSE | High | VSPSTSYTP | [R].VSPSTS | 1xPhospho | H9KV04 1x | 0.014667 | 0.001049 | 1 | 2 | 2 | H9KV04 | Mini-chron | H9KV04 [1 | 0 | 1490.709 | 0 | -0.93 | 0.000477 | 2.28 | 133.4 | 66.6 | 6.85 | 6.86 | | High | High | High | High | High | High | High | High | High | High | High | 0.000555 | 0.004799 | 1.73 |
| FALSE | High | VSPTGAAG | [K].VSPTGA | 1xPhospho | Q80U72-4 | 0.08727 | 0.006537 | 2 | 5 | 1 | Q80U72-4; | Isoform 4 c | Q80U72-4 | 0 | 1124.566 | 0 | -0.51 | 0.000829 | 3.22 | 116.7 | 83.3 | 5.51 | 2.85 | Shared | High | High | High | High | High | High | High | High | High | High | High | 0.003361 | 0.03672 | 1.97 |
| FALSE | High | VSPVPPSGT | [R].VSPVPF | 1xPhospho | A2SW42 1x | 3.78E-06 | 0 | 1 | 7 | 2 | A2SW42 | Zinc finger | A2SW42 [2 | 0 | 1834.915 | 0 | -1.49 | 0.001478 | 2.63 | 147.7 | 52.3 | 8.26 | 17 | | High | High | High | High | High | High | High | High | High | High | High | 0 | 1.54E-05 | 1.52 |
| FALSE | High | VSPVPPSGT | [R].VSPVPF | 1xPhospho | A2SW42 1x | 4.45E-07 | 0 | 1 | 7 | 2 | A2SW42 | Zinc finger | A2SW42 [2 | 0 | 2064.078 | 0 | -1.16 | 0.000461 | 2.22 | 136 | 64 | 5.29 | 12.2 | | High | High | High | High | High | High | High | High | High | High | High | 0 | 4.11E-08 | 3.66 |
| FALSE | High | VSPVPSPSC | [R].VSPVPS | 2xPhospho | Q8VDP3 2x | 0.000908 | 0 | 1 | 4 | 2 | Q8VDP3 | [F-actin]-m | Q8VDP3 [7 | 0 | 1610.754 | 0 | -1.66 | 0.001117 | 2.65 | 152 | 48 | 6.12 | 17.28 | | High | High | High | High | High | High | High | High | High | High | High | 0.002455 | 0.02677 | 2.89 |
| FALSE | High | VSPYSLEISI | [K].VSPYSL | 1xPhospho | J3QM81 1x | 0.000237 | 0 | 1 | 5 | 2 | J3QM81 | Ankyrin re | J3QM81 [8 | 0 | 1789.939 | 0 | -0.5 | 0.019045 | 1.27 | 114.9 | 85.1 | 5.01 | 9.31 | | High | High | High | High | High | High | High | High | High | High | High | 0 | 4.72E-05 | 2.76 |
| FALSE | High | VSQEANLL | [R].VSQEAI | 1xPhospho | E9Q2I4 1xI | 0.008691 | 0.000736 | 1 | 3 | 1 | E9Q2I4 | ELM2 and | E9Q2I4 [45 | 0 | 1953.082 | 0 | 0.15 | 0.100697 | 4.98 | 93 | 107 | 5.75 | 10.03 | | High | High | High | High | High | High | High | High | High | High | High | 0.000327 | 0.002688 | 2.26 |
| FALSE | High | VSQGVEDC | [K].VSQGV | 1xPhospho | Q9R1T2 1x | 0.000244 | 0 | 1 | 2 | 1 | Q9R1T2 | SUMO-acti | Q9R1T2 [1 | 0 | 1753.877 | 0 | -0.71 | 0.000297 | 1.18 | 125.2 | 74.8 | 3.73 | 8 | | High | High | High | High | High | High | High | High | High | High | High | 0 | 4.9E-05 | 3.25 |
| FALSE | High | VSQSSAESI | [K].VSQSSA | 1xPhospho | E9QLH9 1x | 1.65E-07 | 0 | 1 | 2 | 2 | E9QLH9 | PEAK1-rela | E9QLH9 [7 | 0 | 1790.852 | 0 | 0.96 | 0.000382 | 9.23 | 67.3 | 132.7 | 14.21 | 3.02 | | High | High | High | High | High | High | High | High | High | High | High | 0 | 1.35E-08 | 4.33 |
| FALSE | High | VSQVESPE | [K].VSQVE | 1xPhospho | P28359 1xI | 0.002895 | 0.00016 | 1 | 1 | 2 | P28359 | Homeobox | P28359 [21 | 0 | 1611.84 | 0 | 4.09 | 0.000291 | 14.87 | 11.6 | 188.4 | 18.3 | 8.8 | | High | High | High | High | High | High | High | High | High | High | High | 0.000204 | 0.001351 | 3.36 |
| FALSE | High | VSQVSLESI | [K].VSQVSI | 1xPhospho | Q6P5H2-2 | 0.043981 | 0.003228 | 1 | 2 | 3 | Q6P5H2-2 | Isoform 2 c | Q6P5H2-2 | 0 | 1756.95 | 0 | 3 | 0.003088 | 21.78 | 18.4 | 181.6 | 11.14 | 23.7 | | High | High | High | High | High | High | High | High | High | High | High | 0.001889 | 0.01906 | 2.16 |
| FALSE | High | VSPSPAGP | [K].VSPSPA | 1xPhospho | Q3U0S1 1x | 0.001087 | 0 | 1 | 2 | 2 | Q3U0S1 | Putative ur | Q3U0S1 [2 | 0 | 1664.784 | 0 | 1.4 | 0.001386 | 7.36 | 54.8 | 145.2 | 4.54 | 11.86 | | High | High | High | High | High | High | High | High | High | High | High | 0.000449 | 0.004185 | 2.29 |
| FALSE | High | VSQVSLESI | [K].VSQVSI | 2xPhospho | Q6P5H2-2 | 0.043607 | 0.003228 | 1 | 2 | 1 | Q6P5H2-2 | Isoform 2 c | Q6P5H2-2 | 0 | 1836.916 | 0 | 2.11 | 0.000911 | 8.55 | 35.6 | 164.4 | 19.66 | 14.49 | | High | High | High | High | High | High | High | High | High | High | High | 0.001651 | 0.01651 | 2.65 |
| FALSE | High | VSSAPIPCP | [K].VSSAPI | 1xCarbami | A6H663 1x | 0.000225 | 0 | 1 | 3 | 2 | A6H663 | BCL2-assoc | A6H663 [3 | 0 | 2765.435 | 0 | 1.72 | 0.001811 | 4.97 | 46.6 | 153.4 | 12.45 | 12.25 | | High | High | High | High | High | High | High | High | High | High | High | 0.001639 | 0.01582 | 3.11 |
| FALSE | High | VSSAPIPCP | [K].VSSAPI | 1xCarbami | A6H663 2x | 0.033116 | 0.002195 | 1 | 3 | 1 | A6H663 | BCL2-assoc | A6H663 [3 | 0 | 2845.401 | 0 | 2.36 | 0.001931 | 23.09 | 34.9 | 165.1 | 18.77 | 15.63 | | High | High | High | High | High | High | High | High | High | High | High | 0.001151 | 0.01202 | 2.06 |
| FALSE | High | VSSDEEHT | [R].VSSDEE | 1xCarbami | F6WJB7 1x | 1.56E-06 | 0 | 1 | 4 | 2 | F6WJB7 | Ubiquitin c | F6WJB7 [3 | 0 | 2429.131 | 0 | 0.06 | 0.737353 | 4.39 | 98.1 | 101.9 | 5.6 | 9.7 | | High | High | High | High | High | High | High | High | High | High | High | 0 | 1.68E-07 | 3.83 |
| FALSE | High | VSSEDPGP | [K].VSSEDF | 1xPhospho | E9QAU4 1x | 0.000876 | 0 | 1 | 11 | 1 | E9QAU4 | Enhancer t | E9QAU4 [1 | 0 | 1679.784 | 0 | 0.93 | 0.033286 | 37.14 | 62.1 | 137.9 | 21.75 | 20.16 | | High | High | High | High | High | High | High | High | High | High | High | 0 | 0.000206 | 2.4 |
| FALSE | High | VSSETHQG | [R].VSSETH | 1xPhospho | E9Q7G0 1x | 9.8E-05 | 0 | 1 | 5 | 4 | E9Q7G0 | Nuclear mi | E9Q7G0 [1 | 0 | 1849.853 | 0 | -0.51 | 0.002154 | 2.84 | 115.6 | 84.4 | 5.72 | 7.9 | | High | High | High | High | High | High | High | High | High | High | High | 0 | 1.75E-05 | 2.86 |
| FALSE | High | VSSETHQG | [R].VSSETH | 1xPhospho | E9Q7G0 1x | 1.22E-05 | 0 | 1 | 5 | 11 | E9Q7G0 | Nuclear mi | E9Q7G0 [1 | 0 | 2079.016 | 0 | 0.14 | 0.462526 | 8.44 | 95.7 | 104.3 | 8.69 | 10.86 | | High | High | High | High | High | High | High | High | High | High | High | 0 | 4.57E-06 | 3.66 |
| FALSE | High | VSSETHQG | [R].VSSETH | 2xPhospho | E9Q7G0 2x | 0.001586 | 5.66E-05 | 1 | 5 | 1 | E9Q7G0 | Nuclear mi | E9Q7G0 [1 | 0 | 2158.982 | 0 | -0.41 | 0.029117 | 2.46 | 113.9 | 86.1 | 5.17 | 13.7 | | High | High | High | High | High | High | High | High | High | High | High | 2.58E-05 | 0.0004 | 2.97 |
| FALSE | High | VSSFSMPS | [R].VSSFSM | 1xPhospho | P70227 1xI | 0.000534 | 0 | 1 | 2 | 1 | P70227 | Inositol 1,4 | P70227 [18 | 0 | 1480.67 | 0 | 1.76 | 0.002198 | 13.6 | 45.6 | 154.4 | 5.42 | 15.39 | | High | High | High | High | High | High | High | High | High | High | High | 0 | 0.000118 | 2.37 |
| FALSE | High | VSSFSMPS | [R].VSSFSM | 1xPhospho | P70227 1xI | 0.002793 | 0.000109 | 1 | 2 | 5 | P70227 | Inositol 1,4 | P70227 [18 | 0 | 1496.665 | 0 | 1.61 | 0.000918 | 12.35 | 49.4 | 150.6 | 6.58 | 11.6 | | High | High | High | High | High | High | High | High | High | High | High | 0.001484 | 0.01486 | 2.67 |
| FALSE | High | VSSGFGLSI | [R].VSSGFC | 1xPhospho | G3UYW1 1 | 0.003055 | 0.00016 | 1 | 3 | 1 | G3UYW1 | WD repeat | G3UYW1 [ | 0 | 1419.765 | 0 | 1.35 | 0.005735 | 35.87 | 56.3 | 143.7 | 11.82 | 16.67 | | High | High | High | High | High | High | High | High | High | High | High | 7.19E-05 | 0.000833 | 2.44 |
| FALSE | High | VSSGGLTE | [R].VSSGGI | 1xPhospho | Q3U1T9 1x | 0.000731 | 0 | 1 | 4 | 3 | Q3U1T9 | DENN dom | Q3U1T9 [6 | 0 | 1989.107 | 0 | 0.79 | 0.000406 | 4.22 | 72.3 | 127.7 | 5.72 | 4.66 | | High | High | High | High | High | High | High | High | High | High | High | 0 | 0.000168 | 2.55 |
| FALSE | High | VSSHTWR | [R].VSSHTV | 1xPhospho | Q3UIV3 1x | 0.082424 | 0.006288 | 1 | 12 | 1 | Q3UIV3 | Putative ur | Q3UIV3 [6 | 0 | 1181.567 | 0 | -1.41 | 0.000438 | 1.42 | 142.8 | 57.2 | 6.92 | 9.82 | | High | High | High | High | High | High | High | High | High | High | High | 0.003265 | 0.03444 | 1.15 |
| FALSE | High | VSSLASEEL | [K].VSSLAS | 1xPhospho | B2RWU4 1 | 0.001386 | 5.66E-05 | 2 | 6 | 3 | B2RWU4; ( | Cordon-ble | B2RWU4 [ | 0 | 1761.862 | 0 | 1.15 | 0.012952 | 32.27 | 60.8 | 139.2 | 15.33 | 17.3 | Shared | High | High | High | High | High | High | High | High | High | High | High | 2.58E-05 | 0.000345 | 3.17 |
| FALSE | High | VSRVEDGS | [K].VSRVED | 1xPhospho | E9QLZ9 1xI | 1.79E-06 | 0 | 1 | 12 | 1 | E9QLZ9 | Protein en | E9QLZ9 [6 | 1 | 2833.413 | 0 | -0.61 | 0.01789 | 8 | 123.1 | 76.9 | 14.61 | 9.15 | | High | High | High | High | High | High | High | High | High | High | High | 0 | 1.96E-07 | 3.96 |
| FALSE | High | VSPSIQPQI | [R].VSPSIQ | 1xPhospho | E9Q0S6 1x | 5.4E-05 | 0 | 1 | 7 | 3 | E9Q0S6 | Tensin 1 O | E9Q0S6 [1 | 0 | 2246.138 | 0 | -0.45 | 0.001923 | 1.39 | 115.6 | 84.4 | 3.07 | 5.32 | | High | High | High | High | High | High | High | High | High | High | High | 0 | 9E-06 | 3.71 |
| FALSE | High | VSPQDANS | [R].VSPQD | 1xPhospho | E9Q8Z8 1x | 6.61E-06 | 0 | 1 | 8 | 1 | E9Q8Z8 | Catenin de | E9Q8Z8 [4 | 0 | 2084.004 | 0 | -0.05 | 0.877688 | 5.03 | 97 | 103 | 9.35 | 10.02 | | High | High | High | High | High | High | High | High | High | High | High | 0 | 8.47E-07 | 2.29 |
| FALSE | High | VSPPENGP | [K].VSPPEN | 1xPhospho | Q7TPM1 1 | 0.004033 | 0.000311 | 1 | 3 | 1 | Q7TPM1 | Protein PR | Q7TPM1 [5 | 0 | 1459.751 | 0 | -0.66 | 0.000887 | 3.73 | 122.4 | 77.6 | 3.25 | 6.46 | | High | High | High | High | High | High | High | High | High | High | High | 0.000161 | 0.001139 | 2.15 |
| FALSE | High | VSLLGPVTT | [K].VSLLGP | 1xPhospho | Q9WVM1 | 6.12E-08 | 0 | 1 | 1 | 1 | Q9WVM1 | Rac GTPase | Q9WVM1 | 0 | 2266.286 | 0 | -0.28 | 0.014303 | 6.09 | 110.8 | 89.2 | 5.75 | 9.32 | | High | High | High | High | High | High | High | High | High | High | High | 0 | 4.39E-09 | 3.11 |
| FALSE | High | VSLPDVDLI | [K].VSLPDV | 1xPhospho | E9Q616 1x | 0.000106 | 0 | 1 | 3 | 1 | E9Q616 | AHNAK nu | E9Q616 [1 | 0 | 1751.96 | 0 | 1.95 | 0.001124 | 21.83 | 41.2 | 158.8 | 26.24 | 8.86 | | High | High | High | High | High | High | High | High | High | High | High | 0 | 1.93E-05 | 2.64 |
| FALSE | High | VSLPPIPTV | [K].VSLPPII | 1xPhospho | Q8BH48 1x | 8.33E-05 | 0 | 1 | 3 | 1 | Q8BH48 | Ubiquitin-a | Q8BH48 [2 | 0 | 1903.107 | 0 | 0.89 | 0.001276 | 15.86 | 72 | 128 | 8.87 | 6.49 | | High | High | High | High | High | High | High | High | High | High | High | 0 | 1.46E-05 | 2.83 |
| FALSE | High | VSLSDCIQE | [K].VSLSDC | 1xCarbami | Q3TCJ8 1xI | 0.002272 | 0.000109 | 1 | 2 | 1 | Q3TCJ8 | Coiled-coil | Q3TCJ8 [5 | 0 | 1787.901 | 0 | 3.8 | 0.000417 | 12.52 | 12.8 | 187.2 | 20.24 | 11.73 | | High | High | High | High | High | High | High | High | High | High | High | 4.94E-05 | 0.000598 | 2.49 |
| FALSE | High | VSISEGDD | [R].VSISEG | 1xPhospho | P35550 1xI | 0.092372 | 0.007211 | 1 | 3 | 1 | P35550 | rRNA 2'-O- | P35550 [12 | 0 | 1487.739 | 0 | 1.23 | 0.006776 | 25.68 | 60.9 | 139.1 | 13.59 | 17.31 | | High | High | High | High | High | High | High | High | High | High | High | 0.003672 | 0.03943 | 1.95 |
| FALSE | High | VSLTLADLE | [R].VSLTLA | 1xPhospho | E9QKD1 1x | 0.000131 | 0 | 1 | 2 | 2 | E9QKD1 | Nucleolar | E9QKD1 [4 | 0 | 2019.047 | 0 | -0.3 | 0.0236 | 9.66 | 108.7 | 91.3 | 7.96 | 4.94 | | High | High | High | High | High | High | High | High | High | High | High | 0 | 4.18E-05 | 4.11 |
| FALSE | High | VSNAESNT | [R].VSNAES | 1xPhospho | Q8C180 1x | 0.037048 | 0.00249 | 1 | 1 | 1 | Q8C180 | Fibroblast | Q8C180 [2 | 0 | 1584.803 | 0 | 1.45 | 0.000295 | 11.36 | 57.3 | 142.7 | 13.89 | 6.9 | | High | High | High | High | High | High | High | High | High | High | High | 0.001308 | 0.01375 | 2.02 |
| FALSE | High | VSNCSVAS | [R].VSNCSV | 1xCarbami | Q8C551 1x | 0.000231 | 0 | 1 | 1 | 3 | Q8C551 | RAD51-ass | Q8C551 [1 | 0 | 2291.051 | 0 | 0.04 | 0.778054 | 2.34 | 100 | 100 | 3.33 | 3.89 | | High | High | High | High | High | High | High | High | High | High | High | 0 | 4.6E-05 | 2.18 |
| FALSE | High | VSNGSPSLI | [R].VSNGSI | 1xPhospho | P46414 1xI | 0.008022 | 0.000736 | 1 | 2 | 1 | P46414 | Cyclin-dep | P46414 [6- | 0 | 1354.657 | 0 | 2.55 | 0.000317 | 12.81 | 30.4 | 169.6 | 7.7 | 7.41 | | High | High | High | High | High | High | High | High | High | High | High | 0.000327 | 0.00246 | 1.9 |
| FALSE | High | VSNGSPSLI | [R].VSNGSI | 1xDeamida | P46414 1xI | 0.028327 | 0.001791 | 1 | 2 | 1 | P46414 | Cyclin-dep | P46414 [6- | 0 | 1355.641 | 0 | 2.52 | 0.000201 | 6.23 | 30.4 | 169.6 | 6.38 | 6.02 | | High | High | High | High | High | High | High | High | High | High | High | 0.000968 | 0.01009 | 3.03 |
| FALSE | High | VSPAAAQL | [R].VSPAAA | 1xPhospho | A0A0A0MC | 0.001898 | 5.66E-05 | 1 | 2 | 1 | A0A0A0MC | Histone-lys | A0A0A0MC | 0 | 2061.082 | 0 | 0.41 | 0.021661 | 5.54 | 84.1 | 115.9 | 6.57 | 13.6 | | High | High | High | High | High | High | High | High | High | High | High | 2.58E-05 | 0.00049 | 2.37 |
| FALSE | High | VSPAHSPPI | [R].VSPAHS | 2xPhospho | Q80TC1 2x | 0.063788 | 0.004664 | 1 | 14 | 1 | Q80TC1 | MKIAA154 | Q80TC1 [3 | 0 | 1836.813 | 0 | -0.29 | 0.132164 | 21.06 | 105.9 | 94.1 | 18.18 | 16.14 | | High | High | High | High | High | High | High | High | High | High | High | 0.002403 | 0.02557 | 1.78 |
| FALSE | High | VSPAHSPPI | [R].VSPAHS | 1xPhospho | Q80TC1 1x | 0.00407 | 0.00036 | 1 | 14 | 1 | Q80TC1 | MKIAA154 | Q80TC1 [3 | 0 | 1986.01 | 0 | -0.65 | 0.000511 | 2.77 | 123.4 | 76.6 | 5.18 | 6.79 | | High | High | High | High | High | High | High | High | High | High | High | 0.000161 | 0.001155 | 2.59 |
| FALSE | High | VSPAHSPPI | [R].VSPAHS | 1xDeamida | Q80TC1 1x | 0.001058 | 0 | 1 | 14 | 1 | Q80TC1 | MKIAA154 | Q80TC1 [3 | 0 | 1986.994 | 0 | -1.67 | 0.00023 | 1.05 | 153.9 | 46.1 | 4.49 | 10.2 | | High | High | High | High | High | High | High | High | High | High | High | 0 | 0.000253 | 2.8 |
| FALSE | High | VSPANAAC | [R].VSPANA | 1xPhospho | E0CZ26 1xI | 0.000349 | 0 | 1 | 3 | 1 | E0CZ26 | Potassium | E0CZ26 [3 | 0 | 1994.986 | 0 | 0.77 | 0.004699 | 11.64 | 75.2 | 124.8 | 18.59 | 19.07 | | High | High | High | High | High | High | High | High | High | High | High | 0 | 7.32E-05 | 3.8 |
| FALSE | High | VSPASSVDS | [K].VSPASS | 1xPhospho | Q9ESC8 1x | 1.93E-05 | 0 | 1 | 2 | 4 | Q9ESC8 | AF4/FMR2 | Q9ESC8 [4 | 0 | 2375.19 | 0 | -0.46 | 0.001066 | 0.26 | 118.8 | 81.2 | 5.24 | 7.7 | | High | High | High | High | High | High | High | High | High | High | High | 0 | 1.65E-05 | 3.04 |
| FALSE | High | VSPDDVQF | [K].VSPDD | 1xPhospho | Q03963 1x | 0.0102 | 0.000736 | 1 | 2 | 2 | Q03963 | Interferon- | Q03963 [2 | 0 | 1224.582 | 0 | 0.17 | 0.013928 | 2.7 | 92.3 | 107.7 | 3.22 | 6.07 | | High | High | High | High | High | High | High | High | High | High | High | 0.007292 | 0.07917 | 1.96 |
| FALSE | High | VSPEAEECI | [K].VSPEAE | 1xCarbami | Q5SXA9 1x | 0.001674 | 5.66E-05 | 1 | 2 | 2 | Q5SXA9 | Protein KIE | Q5SXA9 [8 | 0 | 1867.928 | 0 | -0.47 | 0.005802 | 6.82 | 117.3 | 82.7 | 6.66 | 7.42 | | High | High | High | High | High | High | High | High | High | High | High | 2.58E-05 | 0.000424 | 2.89 |
| FALSE | High | VSPELNSTI | [K].VSPELN | 1xPhospho | Q6PFD6 1x | 0.078467 | 0.006104 | 1 | 1 | 2 | Q6PFD6 | Kinesin-like | Q6PFD6 [7 | 0 | 3072.61 | 0 | -0.27 | 0.089289 | 7.12 | 110.9 | 89.1 | 21.52 | 6.57 | | High | High | High | High | High | High | High | High | High | High | High | 0.003265 | 0.0351 | 3.37 |
| FALSE | High | VSPENLVDI | [R].VSPENI | 1xPhospho | Q5RIM6 1x | 0.077188 | 0.005893 | 1 | 10 | 1 | Q5RIM6 | Nuclear re | Q5RIM6 [2 | 0 | 1309.66 | 0 | -0.4 | 0.309108 | 19.71 | 113.9 | 86.1 | 29.59 | 26.39 | | High | High | High | High | High | High | High | High | High | High | High | 0.00302 | 0.03176 | 2.22 |
| FALSE | High | VSPENLVDI | [R].VSPENI | 1xPhospho | Q5RIM6 1x | 0.002508 | 0.000109 | 1 | 10 | 3 | Q5RIM6 | Nuclear re | Q5RIM6 [2 | 0 | 1538.823 | 0 | -0.3 | 0.008579 | 1.89 | 110.8 | 89.2 | 5.92 | 4.56 | | High | High | High | High | High | High | High | High | High | High | High | 4.94E-05 | 0.00067 | 1.86 |
| FALSE | High | VSPEVGSAI | [R].VSPEVC | 1xPhospho | Q6PGL7 1x | 0.000538 | 0 | 1 | 5 | 1 | Q6PGL7 | WASH com | Q6PGL7 [7 | 0 | 1866.941 | 0 | -0.28 | 0.073115 | 11.63 | 111.5 | 88.5 | 14.6 | 7.65 | | High | High | High | High | High | High | High | High | High | High | High | 0 | 0.000119 | 2.74 |
| FALSE | High | VSPEVGSAI | [R].VSPEVC | 1xPhospho | Q6PGL7 1x | 9.71E-05 | 0 | 1 | 5 | 3 | Q6PGL7 | WASH com | Q6PGL7 [7 | 0 | 2096.104 | 0 | -0.21 | 0.044977 | 2.07 | 107.1 | 92.9 | 8.21 | 7.12 | | High | High | High | High | High | High | High | High | High | High | High | 0 | 0.000136 | 2.78 |
| FALSE | High | VSPGEILK | [R].VSPGEI | 1xPhospho | Q9CYX7 1x | 0.00537 | 0.000599 | 1 | 1 | 2 | Q9CYX7 | RRP15-like | Q9CYX7 [1 | 0 | 1380.79 | 0 | -0.6 | 0.012699 | 3.45 | 119.6 | 80.4 | 11.12 | 13.02 | | High | High | High | High | High | High | High | High | High | High | High | 0.000267 | 0.001565 | 1.94 |
| FALSE | High | VSPHHPAP | [R].VSPHHI | 1xPhospho | Q03347-4 | 0.008463 | 0.000736 | 1 | 5 | 2 | Q03347-4 | Isoform 4 c | Q03347-4 | 0 | 1715.858 | 0 | 1.85 | 0.001378 | 10.99 | 43.4 | 156.6 | 7.48 | 15.82 | | High | High | High | High | High | High | High | High | High | High | High | 0.003809 | 0.04112 | 2.12 |
| FALSE | High | VSPLNLSSV | [R].VSPLNL | 1xPhospho | A0A0R4J1V | 0.0016 | 5.66E-05 | 1 | 4 | 3 | A0A0R4J1V | CDC23 (Cel | A0A0R4J1V | 0 | 1422.744 | 0 | -0.22 | 0.023111 | 6.78 | 107.5 | 92.5 | 3.9 | 5.28 | | High | High | High | High | High | High | High | High | High | High | High | 0.000467 | 0.004221 | 2.27 |
| FALSE | High | VSPMLPSS | [R].VSPMLI | 1xPhospho | Q9QY06-2 | 4.61E-06 | 0 | 1 | 4 | 2 | Q9QY06-2 | Isoform 2 c | Q9QY06-2 | 0 | 2013.038 | 0 | -1.48 | 0.002085 | 4.02 | 147.4 | 52.6 | 7.16 | 23.05 | | High | High | High | High | High | High | High | High | High | High | High | 0 | 2.42E-06 | 4.4 |
| FALSE | High | VSPMLPSS | [R].VSPMLI | 1xPhospho | E9PZW8 1x | 0.002115 | 5.66E-05 | 1 | 3 | 2 | E9PZW8 | Unconvent | E9PZW8 [1 | 0 | 3108.619 | 0 | 1.35 | 0.008264 | 42.72 | 57.2 | 142.8 | 19.72 | 18.42 | | High | High | High | High | High | High | High | High | High | High | High | 2.58E-05 | 0.000555 | 3.95 |
| FALSE | High | VSPPASPPI | [K].VSPPAS | 2xPhospho | F8WI37 2xI | 0.000223 | 0 | 1 | 7 | 2 | F8WI37 | Histone-lys | F8WI37 [4 | 0 | 2528.264 | 0 | -0.42 | 0.004822 | 5.4 | 111.8 | 88.2 | 6.23 | 11.69 | | High | High | High | High | High | High | High | High | High | High | High | 0 | 4.43E-05 | 2.76 |
| FALSE | High | VSSIPHDLC | [R].VSSIPH | 1xCarbami | B2RQE3 1x | 0.000411 | 0 | 1 | 6 | 2 | B2RQE3 | Stim2 prot | B2RQE3 [7 | 0 | 2131.041 | 0 | 1.48 | 0.000805 | 9.29 | 53.2 | 146.8 | 10.02 | 7.83 | | High | High | High | High | High | High | High | High | High | High | High | 0 | 8.8E-05 | 2.57 |
| FALSE | High | VSIGNITLSI | [K].VSIGNI | 1xPhospho | Q3UQN2 1 | 2.52E-06 | 0 | 1 | 3 | 2 | Q3UQN2 | F-BAR dom | Q3UQN2 [ | 0 | 1722.935 | 0 | 0.81 | 0.006889 | 6.54 | 69.7 | 130.3 | 14.83 | 15.44 | | High | High | High | High | High | High | High | High | High | High | High | 0 | 2.85E-07 | 2.73 |
| FALSE | High | VSSLSPEQE | [K].VSSLSP | 1xPhospho | Q80XP8 1x | 1.33E-06 | 0 | 1 | 2 | 5 | Q80XP8 | Protein FAI | Q80XP8 [1 | 0 | 2126.094 | 0 | -0.74 | 0.000285 | 0.83 | 125.6 | 74.4 | 3.76 | 3.04 | | High | High | High | High | High | High | High | High | High | High | High | 0 | 0.000119 | 3.27 |
| FALSE | High | VSSPIANGL | [K].VSSPIAI | 1xPhospho | D3YYY8 1xI | 4.05E-05 | 0 | 1 | 1 | 2 | D3YYY8 | Rho guanir | D3YYY8 [1 | 0 | 1731.956 | 0 | 2.39 | 0.000487 | 1.1 | 36.1 | 163.9 | 13.25 | 10.84 | | High | High | High | High | High | High | High | High | High | High | High | 0 | 6.52E-06 | 3.38 |
| FALSE | High | VSVATPDV | [K].VSVATI | 1xPhospho | E9Q616 1x | 0.000362 | 0 | 1 | 6 | 2 | E9Q616 | AHNAK nu | E9Q616 [5 | 0 | 2297.204 | 0 | 2.41 | 0.000273 | 21.72 | 32.8 | 167.2 | 37.69 | 4.28 | | High | High | High | High | High | High | High | High | High | High | High | 0 | 0.00017 | 3.26 |
| FALSE | High | VSVATPDV | [K].VSVATI | 2xPhospho | E9Q616 2x | 0.000434 | 0 | 1 | 6 | 1 | E9Q616 | AHNAK nu | E9Q616 [5 | 0 | 2377.171 | 0 | 0.95 | 0.001764 | 11.54 | 65.1 | 134.9 | 9.36 | 8.31 | | High | High | High | High | High | High | High | High | High | High | High | 0 | 9.31E-05 | 3.12 |
| FALSE | High | VSVCAETFI | [R].VSVCAI | 1xCarbami | Q8K1M3 1 | 0.001189 | 0 | 1 | 3 | 1 | Q8K1M3 | Protein kin | Q8K1M3 [9 | 0 | 2661.086 | 0 | 0.2 | 0.677168 | 33.43 | 99.9 | 100.1 | 15.96 | 16.34 | | High | High | High | High | High | High | High | High | High | High | High | 0 | 0.00029 | 2.69 |
| FALSE | High | VSVDSNLF | [R].VSVDSI | 1xPhospho | B0V2P5 1xI | 0.109329 | 0.008865 | 1 | 5 | 2 | B0V2P5 | DmX-like p | B0V2P5 [1 | 0 | 1895.992 | 0 | 0.95 | 0.000546 | 2.7 | 68.1 | 131.9 | 4.74 | 5.88 | | High | High | High | High | High | High | High | High | High | High | High | 0.005839 | 0.06408 | 1.7 |
| FALSE | High | VSVENSLK | [R].VSVEN | 1xPhospho | H7BX94 1x | 0.003195 | 0.00016 | 1 | 2 | 1 | H7BX94 | Lysine-rich | H7BX94 [2 | 0 | 1413.775 | 0 | 0.42 | 0.057683 | 22.56 | 83 | 117 | 13.59 | 17.47 | | High | High | High | High | High | High | High | High | High | High | High | 7.19E-05 | 0.000878 | 2.73 |
| FALSE | High | VSVGRLSP | [K].VSVGRI | 1xPhospho | F8VPM9 1x | 0.119269 | 0.00979 | 1 | 2 | 1 | F8VPM9 | Zinc finger | F8VPM9 [1 | 1 | 2285.19 | 0 | 0.39 | 0.068415 | 24.64 | 86.7 | 113.3 | 6.27 | 17.19 | | High | High | High | High | High | High | High | High | High | High | High | 0.004866 | 0.0531 | 2.01 |
| FALSE | High | VSVQGGS | [R].VSVQG | 1xPhospho | Q9Z2A7 1x | 0.00204 | 5.66E-05 | 1 | 1 | 1 | Q9Z2A7 | Diacylglyce | Q9Z2A7 [1 | 0 | 1453.782 | 0 | -0.18 | 0.220299 | 26.24 | 109.6 | 90.4 | 15.93 | 14.96 | | High | High | High | High | High | High | High | High | High | High | High | 2.58E-05 | 0.000532 | 3.07 |
| FALSE | High | VSVSESNVI | [K].VSVSES | 1xPhospho | Q04690 1x | 0.000616 | 0 | 1 | 6 | 4 | Q04690 | Neurofibrc | Q04690 [2 | 0 | 2611.352 | 0 | 1.35 | 0.00184 | 13.94 | 50.4 | 149.6 | 14.68 | 16.54 | | High | High | High | High | High | High | High | High | High | High | High | 0 | 0.000138 | 2.56 |
| FALSE | High | VSVSESNVI | [K].VSVSES | 1xDeamida | Q04690 1x | 0.007881 | 0.000736 | 1 | 6 | 2 | Q04690 | Neurofibrc | Q04690 [2 | 0 | 2612.336 | 0 | 1.79 | 0.001558 | 44.68 | 49.8 | 150.2 | 43.74 | 10.79 | | High | High | High | High | High | High | High | High | High | High | High | 0.000327 | 0.002414 | 4.12 |
| FALSE | High | VSVSPGR | [R].VSVSPC | 2xPhospho | Q52KI8 2xI | 0.021607 | 0.001507 | 1 | 6 | 4 | Q52KI8 | Serine/argi | Q52KI8 [42 | 0 | 1090.49 | 0 | -0.18 | 0.053147 | 4.39 | 113 | 87 | 9.85 | 10.89 | | High | High | High | High | High | High | High | High | High | High | High | 0.001151 | 0.01201 | 2 |
| FALSE | High | VSVTTNTTI | [R].VSVTTN | 1xPhospho | A0A1S6GW | 8.53E-07 | 0 | 1 | 6 | 2 | A0A1S6GW | Uncharact | A0A1S6GW | 0 | 2250.079 | 0 | 1.29 | 0.000202 | 3.95 | 58.9 | 141.1 | 3.44 | 4.54 | | High | High | High | High | High | High | High | High | High | High | High | 0 | 8.52E-08 | 3.88 |
| FALSE | High | VSWENPSF | [K].VSWEN | 1xPhospho | Q8K124 1x | 0.00145 | 5.66E-05 | 1 | 1 | 1 | Q8K124 | Pleckstrin | Q8K124 [2 | 0 | 3733.8 | 0 | -0.36 | 0.076546 | 14.36 | 114.9 | 85.1 | 7.83 | 21.96 | | High | High | High | High | High | High | High | High | High | High | High | 2.58E-05 | 0.000362 | 3.59 |
| FALSE | High | VSWENPSF | [K].VSWEN | 1xDeamida | Q8K124 1x | 0.000411 | 0 | 1 | 1 | 2 | Q8K124 | Pleckstrin | Q8K124 [2 | 0 | 3734.784 | 0 | -0.45 | 0.084302 | 21.02 | 115.4 | 84.6 | 12.84 | 18.78 | | High | High | High | High | High | High | High | High | High | High | High | 0 | 8.78E-05 | 4.54 |
| FALSE | High | VSVASGDH | [K].VSVAS | 1xPhospho | Q99MV7 1 | 0.008463 | 0.000736 | 1 | 3 | 1 | Q99MV7 | RING finge | Q99MV7 [ | 0 | 1574.809 | 0 | -0.89 | 0.001307 | 5.36 | 135.9 | 64.1 | 8.01 | 14.39 | | High | High | High | High | High | High | High | High | High | High | High | 0.000327 | 0.0026 | 1.91 |
| FALSE | High | VSWQDLK | [R].VSWQD | 1xPhospho | Q9D8S5 1x | 0.037692 | 0.00249 | 1 | 2 | 1 | Q9D8S5 | MCG7614, | Q9D8S5 [1 | 0 | 1413.754 | 0 | 0.33 | 0.780638 | 16.98 | 91.3 | 108.7 | 9.96 | 27.82 | | High | High | High | High | High | High | High | High | High | High | High | 0.001324 | 0.01397 | 2.27 |
| FALSE | High | VTAPFPSSI | [R].VTAPFF | 1xPhospho | Q05BA5 1x | 0.015061 | 0.001049 | 1 | 7 | 2 | Q05BA5 | Nuclear re | Q05BA5 [2 | 0 | 1944.967 | 0 | -1.04 | 0.004252 | 1.84 | 133.4 | 66.6 | 10.03 | 9.2 | | High | High | High | High | High | High | High | High | High | High | High | 0.000571 | 0.00494 | 2.23 |
| FALSE | High | VTDALPEPI | [K].VTDALI | 1xDeamida | Q9CQ10 1x | 0.000501 | 0 | 1 | 1 | 1 | Q9CQ10 | Charged m | Q9CQ10 [1 | 0 | 4071.693 | 0 | 1.27 | 0.034086 | 81.86 | 60 | 140 | 35.51 | 26.07 | | High | High | High | High | High | High | High | High | High | High | High | 0 | 0.000109 | 4.75 |
| FALSE | High | VTDALPEPI | [K].VTDALI | 1xPhospho | Q9CQ10 1x | 0.000651 | 0 | 1 | 1 | 5 | Q9CQ10 | Charged m | Q9CQ10 [1 | 0 | 4086.704 | 0 | 0.82 | 0.009059 | 32.47 | 70.4 | 129.6 | 11.34 | 12.41 | | High | High | High | High | High | High | High | High | High | High | High | 0 | 0.000148 | 3.47 |
| FALSE | High | VTDALPEPI | [K].VTDALI | 1xDeamida | Q9CQ10 1x | 1.43E-05 | 0 | 1 | 1 | 6 | Q9CQ10 | Charged m | Q9CQ10 [1 | 0 | 4087.688 | 0 | 1.12 | 0.001798 | 9.02 | 64.5 | 135.5 | 11.85 | 5.46 | | High | High | High | High | High | High | High | High | High | High | High | 0 | 2.02E-06 | 4.28 |
| FALSE | High | VTDALPEPI | [K].VTDALI | 1xPhospho | Q9CQ10 1x | 8.53E-07 | 0 | 1 | 1 | 3 | Q9CQ10 | Charged m | Q9CQ10 [1 | 0 | 4102.699 | 0 | 0.98 | 0.010452 | 21.68 | 68.1 | 131.9 | 6.19 | 20.55 | | High | High | High | High | High | High | High | High | High | High | High | 0 | 8.53E-08 | 4.23 |
| FALSE | High | VTDEENAS | [K].VTDEEI | 1xPhospho | Q9ERU9 1x | 1.47E-05 | 0 | 1 | 1 | 6 | Q9ERU9 | E3 SUMO- | Q9ERU9 [2 | 0 | 2662.213 | 0 | 1.23 | 0.000222 | 14.9 | 60.6 | 139.4 | 10.35 | 1.94 | | High | High | High | High | High | High | High | High | High | High | High | 0 | 2.1E-06 | 3.71 |
| FALSE | High | VTDSGSSEI | [K].VTDSG | 1xPhospho | F8WIP8 1xI | 7.82E-05 | 0 | 1 | 10 | 2 | F8WIP8 | Osteopont | F8WIP8 [2 | 0 | 1576.751 | 0 | 0.13 | 0.961355 | 41.78 | 95.6 | 104.4 | 26.82 | 20.26 | | High | High | High | High | High | High | High | High | High | High | High | 0.000689 | 0.006155 | 2.1 |
| FALSE | High | VTEECEEAI | [K].VTEECE | 1xCarbami | Q3UL54 1x | 1.21E-06 | 0 | 1 | 8 | 3 | Q3UL54 | Putative ur | Q3UL54 [3 | 0 | 2892.435 | 0 | 0.14 | 0.121418 | 11.32 | 95.6 | 104.4 | 3.84 | 7.17 | | High | High | High | High | High | High | High | High | High | High | High | 0 | 1.26E-07 | 3.46 |

| | | | | | | | | | | | | | | | | | | | | | | | | | | | | | | | | | | | | | | |
|---|---|---|---|---|---|---|---|---|---|---|---|---|---|---|---|---|---|---|---|---|---|---|---|---|---|---|---|---|---|---|---|---|---|---|---|---|---|---|
| FALSE | High | VTEETEEPI | [K].VTEETE | 2xCarbami | P70399 1xl | 1.36E-06 | 0 | 1 | 4 | 6 | P70399 | TP53-bindi | P70399 [12 | 0 | 4830.142 | 0 | 0.36 | 0.063045 | 7.24 | 85.8 | 114.2 | 4.92 | 16.55 | | High | High | High | High | High | High | High | High | High | High | High | 0 | 1.44E-07 | 6.63 |
| FALSE | High | VTEETEEPI | [K].VTEETE | 2xCarbami | P70399 1xl | 1.99E-06 | 0 | 1 | 4 | 5 | P70399 | TP53-bindi | P70399 [12 | 0 | 4831.126 | 0 | 0.3 | 0.082159 | 9.95 | 93.8 | 106.2 | 9.42 | 12.3 | | High | High | High | High | High | High | High | High | High | High | High | 0 | 2.2E-07 | 5.15 |
| FALSE | High | VTEETEEPI | [K].VTEETE | 2xCarbami | P70399 2xl | 6.24E-05 | 0 | 1 | 4 | 2 | P70399 | TP53-bindi | P70399 [12 | 0 | 4910.109 | 0 | 0.34 | 0.708802 | 19.6 | 96.3 | 103.7 | 12.62 | 29.99 | | High | High | High | High | High | High | High | High | High | High | High | 0 | 1.06E-05 | 4.76 |
| FALSE | High | VTENLASLT | [R].VTENLA | 1xPhospho | Q3UFM6 1 | 0.000105 | 0 | 1 | 3 | 1 | Q3UFM6 | Expressed | Q3UFM6 [3 | 0 | 1710.944 | 0 | 0.01 | 0.918519 | 4.88 | 98.3 | 101.7 | 3.25 | 7.28 | | High | High | High | High | High | High | High | High | High | High | High | 0 | 1.9E-05 | 3.1 |
| FALSE | High | VTEQGGQ | [R].VTEQG | 1xPhospho | A0A023UE | 9.7E-08 | 0 | 1 | 2 | 3 | A0A023UE | MCG5110C | A0A023UE | 0 | 2580.13 | 0 | -1.4 | 0.002161 | 5.55 | 145.1 | 54.9 | 8.73 | 21.99 | | High | High | High | High | High | High | High | High | High | High | High | 0 | 7.37E-09 | 3.91 |
| FALSE | High | VTAASGHC | [R].VTAASC | 1xCarbami | Q6A0A9 1x | 0.001361 | 0 | 1 | 2 | 1 | Q6A0A9 | Constitutiv | Q6A0A9 [9 | 0 | 2062.885 | 0 | 0.8 | 0.049456 | 20.6 | 77.3 | 122.7 | 7.9 | 21.45 | | High | High | High | High | High | High | High | High | High | High | High | 2.58E-05 | 0.000338 | 2.9 |
| FALSE | High | VSTTTDSPV | [K].VSTTTD | 2xPhospho | Q9DB52 2x | 5.03E-05 | 0 | 1 | 1 | 1 | Q9DB52 | Protein FAI | Q9DB52 [2 | 0 | 3349.626 | 0 | 2.54 | 0.000204 | 29.93 | 29.3 | 170.7 | 49.48 | 10.55 | | High | High | High | High | High | High | High | High | High | High | High | 0 | 8.29E-06 | 4.23 |
| FALSE | High | VSTTTDSPV | [K].VSTTTD | 1xPhospho | Q9DB52 1x | 0.000229 | 0 | 1 | 1 | 1 | Q9DB52 | Protein FAI | Q9DB52 [2 | 0 | 3269.659 | 0 | 0.79 | 0.001093 | 8.67 | 72.5 | 127.5 | 8.21 | 7.17 | | High | High | High | High | High | High | High | High | High | High | High | 0 | 4.53E-05 | 3.41 |
| FALSE | High | VSTSSQEQ | [R].VSTSSC | 1xPhospho | Q8BJ05 1xl | 0.035181 | 0.002269 | 1 | 3 | 1 | Q8BJ05 | Zinc finger | Q8BJ05 [13 | 0 | 1531.777 | 0 | 0.79 | 0.005735 | 6.8 | 68.6 | 131.4 | 12.4 | 13.71 | | High | High | High | High | High | High | High | High | High | High | High | 0.001199 | 0.01286 | 2.11 |
| FALSE | High | VSSPIANGL | [K].VSSPIA | 1xDeamida | D3YYY8 1xl | 0.021989 | 0.001542 | 1 | 1 | 2 | D3YYY8 | Rho guanir | D3YYY8 [1( | 0 | 1732.94 | 0 | 0.76 | 0.001792 | 18.54 | 70.2 | 129.8 | 16.84 | 10.52 | | High | High | High | High | High | High | High | High | High | High | High | 0.000814 | 0.007609 | 2.81 |
| FALSE | High | VSSPLPSPS | [R].VSSPLP | 1xPhospho | Q8VHR5 1x | 4.17E-06 | 0 | 1 | 5 | 2 | Q8VHR5 | Transcripti | Q8VHR5 [3 | 0 | 2625.3 | 0 | -0.38 | 0.026517 | 9.36 | 114.4 | 85.6 | 12.15 | 15.57 | | High | High | High | High | High | High | High | High | High | High | High | 0 | 5.05E-07 | 3.3 |
| FALSE | High | VSSPLPSPS | [R].VSSPLP | 1xPhospho | Q8VHR5 1x | 1.46E-05 | 0 | 1 | 5 | 3 | Q8VHR5 | Transcripti | Q8VHR5 [3 | 0 | 2641.294 | 0 | -0.18 | 0.86789 | 21.83 | 106.5 | 93.5 | 21.44 | 36.83 | | High | High | High | High | High | High | High | High | High | High | High | 0 | 2.06E-06 | 2.9 |
| FALSE | High | VSSPLPSPS | [R].VSSPLP | 2xPhospho | Q8VHR5 2x | 1.08E-05 | 0 | 1 | 5 | 2 | Q8VHR5 | Transcripti | Q8VHR5 [3 | 0 | 2705.266 | 0 | -0.31 | 0.042504 | 10.23 | 111.7 | 88.3 | 10.54 | 8.79 | | High | High | High | High | High | High | High | High | High | High | High | 0 | 1.48E-06 | 3.94 |
| FALSE | High | VSSPLSPLS | [K].VSSPLS | 1xPhospho | Q99LJ0 1xP | 3.21E-06 | 0 | 1 | 1 | 4 | Q99LJ0 | CTTNBP2 N | Q99LJ0 [55 | 0 | 1820.033 | 0 | 0.57 | 0.002642 | 10.2 | 81.4 | 118.6 | 6.97 | 5.27 | | High | High | High | High | High | High | High | High | High | High | High | 0 | 3.78E-07 | 2.96 |
| FALSE | High | VSSPPPTIT | [K].VSSPPP | 1xPhospho | E9Q0S6 1x | 0.005924 | 0.00062 | 1 | 10 | 2 | E9Q0S6 | Tensin 1 O | E9Q0S6 [1( | 0 | 1878.014 | 0 | -0.16 | 0.261903 | 10.6 | 107.1 | 92.9 | 6.82 | 9.8 | | High | High | High | High | High | High | High | High | High | High | High | 0.000344 | 0.002973 | 2.44 |
| FALSE | High | VSSPSCDS( | [K].VSSPSC | 2xCarbami | Q7TSZ8 1xl | 1.42E-05 | 0 | 1 | 1 | 3 | Q7TSZ8 | Nucleus ac | Q7TSZ8 [12 | 0 | 4852.28 | 0 | -1.06 | 0.003629 | 1.84 | 135.3 | 64.7 | 8.09 | 12.49 | | High | High | High | High | High | High | High | High | High | High | High | 0 | 2.01E-06 | 4.04 |
| FALSE | High | VSSPSCDS( | [K].VSSPSC | 2xCarbami | Q7TSZ8 1xl | 3.92E-06 | 0 | 1 | 1 | 1 | Q7TSZ8 | Nucleus ac | Q7TSZ8 [12 | 0 | 4853.264 | 0 | -1.15 | 0.001359 | 3.07 | 138 | 62 | 10.25 | 12.63 | | High | High | High | High | High | High | High | High | High | High | High | 0 | 4.72E-07 | 4.13 |
| FALSE | High | VSSPTLLEA | [K].VSSPTL | 1xCarbami | Q80VI1 1xl | 2.1E-05 | 0 | 1 | 1 | 3 | Q80VI1 | E3 ubiquiti | Q80VI1 [5- | 0 | 2476.281 | 0 | -1.95 | 0.001219 | 0.92 | 157.7 | 42.3 | 16.32 | 33.81 | | High | High | High | High | High | High | High | High | High | High | High | 0 | 3.12E-06 | 3.22 |
| FALSE | High | VSSPTVNT | [R].VSSPTV | 1xPhospho | Q9DBS5 1x | 0.005516 | 0.00062 | 1 | 1 | 1 | Q9DBS5 | Kinesin ligh | Q9DBS5 [4 | 0 | 1684.929 | 0 | 0.52 | 0.006165 | 10.22 | 84 | 116 | 8.72 | 6.82 | | High | High | High | High | High | High | High | High | High | High | High | 0.000275 | 0.001618 | 2.73 |
| FALSE | High | VSSPVLETV | [K].VSSPVL | 1xPhospho | Q8BTI8 1xP | 1.22E-06 | 0 | 1 | 3 | 16 | Q8BTI8 | Serine/argi | Q8BTI8 [13 | 0 | 1651.862 | 0 | -0.58 | 0.001072 | 4.38 | 119.1 | 80.9 | 4.93 | 3.41 | | High | High | High | High | High | High | High | High | High | High | High | 0 | 1.27E-07 | 4.57 |
| FALSE | High | VSSQGCLQ | [R].VSSQG | 1xCarbami | Z4YKQ9 1x | 0.002921 | 0.00016 | 1 | 4 | 3 | Z4YKQ9 | ATPase fan | Z4YKQ9 [4 | 0 | 1753.789 | 0 | 0.02 | 0.311841 | 3.75 | 97.1 | 102.9 | 5.52 | 15.12 | | High | High | High | High | High | High | High | High | High | High | High | 7.19E-05 | 0.000791 | 2.63 |
| FALSE | High | VSSSCLDLP | [R].VSSSCL | 1xCarbami | Q6NXI6 1xl | 1.48E-06 | 0 | 1 | 3 | 2 | Q6NXI6 | Regulation | Q6NXI6 [1( | 0 | 2205.04 | 0 | 0.68 | 0.004514 | 9.11 | 73.1 | 126.9 | 11.06 | 8.39 | | High | High | High | High | High | High | High | High | High | High | High | 0 | 1.57E-07 | 3.18 |
| FALSE | High | VSSSPGIK | [R].VSSSPG | 1xPhospho | Q3V1H4 1x | 0.002695 | 0.000109 | 1 | 3 | 2 | Q3V1H4 | E74-like fa | Q3V1H4 [3 | 0 | 1312.728 | 0 | 0.66 | 0.00148 | 7.7 | 77 | 123 | 3.99 | 5.44 | | High | High | High | High | High | High | High | High | High | High | High | 4.94E-05 | 0.000724 | 2.65 |
| FALSE | High | VSSSSGSPA | [K].VSSSSG | 1xPhospho | B2RY22 1xl | 0.000158 | 0 | 2 | 12 | 1 | B2RY22; A( | Clasp1 pro | B2RY22 [61 | 0 | 2720.313 | 0 | -0.5 | 0.120025 | 14.43 | 113.9 | 86.1 | 22.53 | 13.26 | Shared | High | High | High | High | High | High | High | High | High | High | High | 0 | 3.01E-05 | 2.69 |
| FALSE | High | VSSSSSK | [K].VSSSSS | 1xPhospho | Q542V3 1x | 0.018447 | 0.001324 | 1 | 1 | 3 | Q542V3 | Putative ur | Q542V3 [3 | 0 | 1219.634 | 0 | -0.45 | 0.002209 | 3.59 | 115.3 | 84.7 | 3.81 | 5.47 | | High | High | High | High | High | High | High | High | High | High | High | 0.000706 | 0.006222 | 1.65 |
| FALSE | High | VSSTPETPL | [R].VSSTPE | 1xPhospho | P33611 1xl | 0.028082 | 0.001791 | 1 | 6 | 3 | P33611 | DNA polym | P33611 [12 | 0 | 1468.75 | 0 | 0.22 | 0.033081 | 12.62 | 93.3 | 106.7 | 6.2 | 7.1 | | High | High | High | High | High | High | High | High | High | High | High | 0.000968 | 0.009992 | 2.13 |
| FALSE | High | VSSTPETPL | [R].VSSTPE | 1xPhospho | P33611 1xl | 4.43E-05 | 0 | 1 | 6 | 9 | P33611 | DNA polym | P33611 [12 | 0 | 1697.913 | 0 | 0.19 | 0.010543 | 2.42 | 93.1 | 106.9 | 3.36 | 4.87 | | High | High | High | High | High | High | High | High | High | High | High | 7.19E-05 | 0.000847 | 2.75 |
| FALSE | High | VSSTPETPL | [R].VSSTPE | 2xPhospho | P33611 2xl | 0.009332 | 0.000736 | 1 | 6 | 2 | P33611 | DNA polym | P33611 [12 | 0 | 1777.879 | 0 | -0.09 | 0.161791 | 3.94 | 101.9 | 98.1 | 11.2 | 2.84 | | High | High | High | High | High | High | High | High | High | High | High | 0.000327 | 0.002913 | 2.74 |
| FALSE | High | VSSVCLELY | [R].VSSVCL | 1xCarbami | Q99M31-2 | 0.000334 | 0 | 1 | 2 | 1 | Q99M31-2 | Isoform 2 c | Q99M31-2 | 0 | 2138.049 | 0 | -0.26 | 0.087709 | 3.95 | 99 | 101 | 22.1 | 19.19 | | High | High | High | High | High | High | High | High | High | High | High | 0 | 6.96E-05 | 2.84 |
| FALSE | High | VSTEGSDQ | [R].VSTEGS | 1xPhospho | P97467 1xl | 0.001158 | 0 | 1 | 5 | 2 | P97467 | Peptidyl-gl | P97467 [94 | 0 | 1617.777 | 0 | 3.91 | 0.000192 | 20.05 | 12.5 | 187.5 | 23.99 | 5.73 | | High | High | High | High | High | High | High | High | High | High | High | 0 | 0.000281 | 2.57 |
| FALSE | High | VSTGDTSP( | [R].VSTGDT | 1xCarbami | B1APX2 2x | 0.004143 | 0.00036 | 1 | 3 | 2 | B1APX2 | RIKEN cDN | B1APX2 [3( | 0 | 2810.089 | 0 | 0.67 | 0.023227 | 20.95 | 76.5 | 123.5 | 13.49 | 15.67 | | High | High | High | High | High | High | High | High | High | High | High | 0.000161 | 0.001174 | 3.37 |
| FALSE | High | VSTLVQSP( | [K].VSTLVC | 1xPhospho | Q6NZN1-2 | 0.057203 | 0.004227 | 1 | 5 | 1 | Q6NZN1-2 | Isoform 2 c | Q6NZN1-2 | 0 | 1737.992 | 0 | -0.4 | 0.022212 | 6.8 | 113.3 | 86.7 | 3.97 | 13.15 | | High | High | High | High | High | High | High | High | High | High | High | 0.002159 | 0.02258 | 1.32 |
| FALSE | High | VSTPASSSC | [K].VSTPAS | 1xPhospho | Q62481 1x | 0.077825 | 0.005963 | 1 | 1 | 1 | Q62481 | Vacuolar p | Q62481 [1( | 0 | 1529.798 | 0 | -0.11 | 0.49303 | 13.44 | 95.8 | 104.2 | 17.47 | 15.85 | | High | High | High | High | High | High | High | High | High | High | High | 0.003109 | 0.03221 | 1.55 |
| FALSE | High | VSTPHGVS | [R].VSTPHC | 1xPhospho | Q7TMX5 1: | 4.94E-05 | 0 | 1 | 2 | 1 | Q7TMX5 | Protein SH | Q7TMX5 [5 | 0 | 2417.093 | 0 | 0.12 | 0.708764 | 27.27 | 97.1 | 102.9 | 15.69 | 18.18 | | High | High | High | High | High | High | High | High | High | High | High | 0 | 8.11E-06 | 4.37 |
| FALSE | High | VSTPSDYTL | [K].VSTPSD | 1xPhospho | A0A0R4J1I | 0.117405 | 0.009523 | 1 | 6 | 1 | A0A0R4J1I | Menin OS= | A0A0R4J1I | 0 | 1996.044 | 0 | -0.09 | 0.329907 | 15.5 | 112 | 88 | 8.55 | 18.41 | | High | High | High | High | High | High | High | High | High | High | High | 0.004769 | 0.05204 | 2.06 |
| FALSE | High | VSTSGQIR | [K].VSTSGC | 1xPhospho | E9PUQ8 1x | 0.01319 | 0.000889 | 1 | 1 | 1 | E9PUQ8 | Diacylglyce | E9PUQ8 [4 | 0 | 1156.592 | 0 | 0.16 | 0.044465 | 2.45 | 95.7 | 104.3 | 2.85 | 4.28 | | High | High | High | High | High | High | High | High | High | High | High | 0.000467 | 0.004274 | 2.01 |
| FALSE | High | VSTSPVGLI | [R].VSTSPV | 1xPhospho | E9PUR0 1x | 6.2E-06 | 0 | 1 | 7 | 5 | E9PUR0 | Ankyrin rep | E9PUR0 [2 | 0 | 3533.773 | 0 | -0.45 | 0.025662 | 4.6 | 110 | 90 | 10.44 | 5.84 | | High | High | High | High | High | High | High | High | High | High | High | 0 | 1.49E-06 | 5.4 |
| FALSE | High | VSTSQASP( | [K].VSTSQA | 1xPhospho | Q9ESK9 1x | 3.67E-05 | 0 | 1 | 6 | 3 | Q9ESK9 | RB1-induci | Q9ESK9 [6 | 0 | 1782.858 | 0 | -0.59 | 0.000729 | 1.98 | 121.2 | 78.8 | 3.52 | 6.61 | | High | High | High | High | High | High | High | High | High | High | High | 0 | 1.17E-05 | 3.5 |
| FALSE | High | VSSLSPEQE | [K].VSSLSP | 2xPhospho | Q80XP8 2x | 0.000924 | 0 | 1 | 2 | 5 | Q80XP8 | Protein FAI | Q80XP8 [1 | 0 | 2206.06 | 0 | -0.85 | 0.005126 | 2.88 | 128.7 | 71.3 | 9.8 | 7.3 | | High | High | High | High | High | High | High | High | High | High | High | 0 | 0.000302 | 3.55 |
| FALSE | High | TLSVSTPAV | [K].TLSVST | 1xPhospho | Q6NZN0 1x | 8.56E-05 | 0 | 1 | 8 | 3 | Q6NZN0 | RNA-bindir | Q6NZN0 [7 | 0 | 2212.126 | 0 | 0.1 | 0.782368 | 19.9 | 100.5 | 99.5 | 11.87 | 18.99 | | High | High | High | High | High | High | High | High | High | High | High | 0 | 3.43E-05 | 3.04 |
| FALSE | High | VSLESQNV | [R].VSLESC | 1xPhospho | P59729 1xl | 1.6E-05 | 0 | 1 | 4 | 1 | P59729 | Ras and Ra | P59729 [38 | 0 | 1801.853 | 0 | 3.08 | 0.000663 | 32.49 | 18.8 | 181.2 | 22.06 | 11.45 | | High | High | High | High | High | High | High | High | High | High | High | 0 | 2.29E-06 | 3.08 |
| FALSE | High | VSLEITTFSI | [R].VSLEITT | 1xPhospho | Q3UHH1 1: | 0.006031 | 0.00062 | 1 | 4 | 1 | Q3UHH1 | Zinc finger | Q3UHH1 [1 | 0 | 1402.707 | 0 | 0.76 | 0.006493 | 17.78 | 71.6 | 128.4 | 14.79 | 11.91 | | High | High | High | High | High | High | High | High | High | High | High | 0.000275 | 0.001778 | 2.12 |
| FALSE | High | VQEAESPG | [K].VQEAE | 1xPhospho | Q80UP8 1x | 0.008022 | 0.000736 | 1 | 1 | 1 | Q80UP8 | Sodium-de | Q80UP8 [2 | 0 | 1629.829 | 0 | 1.5 | 0.000218 | 13.67 | 49.3 | 150.7 | 13.28 | 4.37 | | High | High | High | High | High | High | High | High | High | High | High | 0.000327 | 0.002449 | 2.7 |
| FALSE | High | VQEAQDSI | [K].VQEAQ | 2xPhospho | Q8VDQ8 2: | 1.44E-09 | 0 | 1 | 2 | 5 | Q8VDQ8 | NAD-deper | Q8VDQ8 [1 | 0 | 3104.228 | 0 | -0.52 | 0.189306 | 5.6 | 114.1 | 85.9 | 20.88 | 24.19 | | High | High | High | High | High | High | High | High | High | High | High | 0 | 6.49E-11 | 6.45 |
| FALSE | High | VQEAQDSI | [K].VQEAQ | 2xPhospho | Q8VDQ8 2: | 2.28E-06 | 0 | 1 | 2 | 1 | Q8VDQ8 | NAD-deper | Q8VDQ8 [1 | 0 | 3120.223 | 0 | -0.63 | 0.013837 | 5.95 | 121.1 | 78.9 | 7.87 | 21.48 | | High | High | High | High | High | High | High | High | High | High | High | 0 | 2.56E-07 | 4.73 |
| FALSE | High | VQEDSSSEI | [K].VQEDS | 1xPhospho | Q61687 1x | 0.001659 | 5.66E-05 | 1 | 3 | 4 | Q61687 | Transcripti | Q61687 [1 | 0 | 1660.783 | 0 | -0.25 | 0.046427 | 7.99 | 110.9 | 89.1 | 8.54 | 8.73 | | High | High | High | High | High | High | High | High | High | High | High | 0.001665 | 0.01663 | 2.88 |
| FALSE | High | VQEESYPQ | [R].VQEESY | 1xPhospho | A2AG58 1x | 4.39E-05 | 0 | 1 | 2 | 3 | A2AG58 | Uncharacte | A2AG58 [1 | 0 | 1628.752 | 0 | -1.32 | 0.000163 | 0.3 | 142.1 | 57.9 | 3.29 | 2.84 | | High | High | High | High | High | High | High | High | High | High | High | 0 | 4.38E-05 | 2.3 |
| FALSE | High | VQENAVEF | [R].VQENA | 1xPhospho | Q8R4E9 1x | 8.28E-06 | 0 | 1 | 1 | 1 | Q8R4E9 | DNA replic | Q8R4E9 [1 | 0 | 2023.015 | 0 | -0.46 | 0.004342 | 4.76 | 115.9 | 84.1 | 7.43 | 2.17 | | High | High | High | High | High | High | High | High | High | High | High | 0 | 1.09E-06 | 2.68 |
| FALSE | High | VQGEAVSN | [K].VQGEA | 1xPhospho | P70670 1xl | 0.039686 | 0.00283 | 1 | 3 | 2 | P70670 | Nascent pc | P70670 [21 | 0 | 4169.897 | 0 | 0.19 | 0.41116 | 14.82 | 96.4 | 103.6 | 10.41 | 17.38 | | High | High | High | High | High | High | High | High | High | High | High | 0.001468 | 0.01473 | 3.51 |
| FALSE | High | VQGEAVSN | [K].VQGEA | 1xPhospho | P70670 1xl | 2.41E-06 | 0 | 1 | 3 | 83 | P70670 | Nascent pc | P70670 [21 | 0 | 4399.06 | 0 | -0.48 | 0.001065 | 2.93 | 115.3 | 84.7 | 1.32 | 7.14 | | High | High | High | High | High | High | High | High | High | High | High | 0 | 2.6E-06 | 5.17 |
| FALSE | High | VQGEAVSN | [K].VQGEA | 1xDeamida | P70670 1xl | 3.04E-09 | 0 | 1 | 3 | 214 | P70670 | Nascent pc | P70670 [21 | 0 | 4400.044 | 0 | -0.47 | 0.001789 | 5.16 | 114.5 | 85.5 | 3.28 | 7.06 | | High | High | High | High | High | High | High | High | High | High | High | 2.58E-05 | 0.000388 | 5.61 |
| FALSE | High | VQGEAVSN | [K].VQGEA | 2xPhospho | P70670 2xl | 0.000442 | 0 | 1 | 3 | 1 | P70670 | Nascent pc | P70670 [21 | 0 | 4479.026 | 0 | 0.35 | 0.027796 | 8.28 | 87.9 | 112.1 | 4.23 | 12.93 | | High | High | High | High | High | High | High | High | High | High | High | 0 | 9.57E-05 | 3.61 |
| FALSE | High | VQGLLENG | [K].VQGLLI | 1xPhospho | P41230 1xl | 3.26E-08 | 0 | 1 | 3 | 3 | P41230 | Lysine-spe | P41230 [13 | 0 | 2211.131 | 0 | -0.66 | 0.000406 | 2 | 122.4 | 77.6 | 1.73 | 3.7 | | High | High | High | High | High | High | High | High | High | High | High | 0 | 2.16E-09 | 4.07 |
| FALSE | High | VQGLLENG | [K].VQGLLI | 1xDeamida | P41230 1xl | 2.66E-07 | 0 | 1 | 3 | 8 | P41230 | Lysine-spe | P41230 [13 | 0 | 2212.115 | 0 | -0.56 | 0.000669 | 0.61 | 118.9 | 81.1 | 5.64 | 4.32 | | High | High | High | High | High | High | High | High | High | High | High | 0 | 6.24E-06 | 3.87 |
| FALSE | High | VQGSDSDE | [K].VQGSD | 1xPhospho | G3UY93 1x | 0.040721 | 0.003058 | 1 | 5 | 2 | G3UY93 | Valine--tRN | G3UY93 [5 | 0 | 2000.938 | 0 | 0.69 | 0.024269 | 22.17 | 75.7 | 124.3 | 9.62 | 16.41 | | High | High | High | High | High | High | High | High | High | High | High | 0.001581 | 0.01521 | 3.18 |
| FALSE | High | VQDHAGRI | [R].VQDHA | 1xPhospho | Q80W49-2 | 0.117405 | 0.009523 | 1 | 2 | 1 | Q80W49-2 | Isoform 2 c | Q80W49-2 | 1 | 2465.259 | 0 | 1.97 | 0.000401 | 15.49 | 40.9 | 159.1 | 8.11 | 8.62 | | High | High | High | High | High | High | High | High | High | High | High | 0.004782 | 0.05229 | 1.87 |
| FALSE | High | VQGTGVTI | [K].VQGTG | 1xPhospho | Q99K01-4 | 4.7E-07 | 0 | 1 | 5 | 6 | Q99K01-4 | Isoform 4 c | Q99K01-4 | 0 | 1886.994 | 0 | 0.56 | 0.000892 | 4.66 | 81.2 | 118.8 | 4.75 | 4.15 | | High | High | High | High | High | High | High | High | High | High | High | 0 | 4.35E-08 | 2.63 |
| FALSE | High | VQHNGNC | [K].VQHNG | 2xCarbami | Q8BZX4-2 | 9.97E-08 | 0 | 1 | 2 | 4 | Q8BZX4-2 | Isoform 2 c | Q8BZX4-2 | 0 | 2523.116 | 0 | -0.23 | 0.041244 | 5.27 | 107.9 | 92.1 | 4.61 | 6.38 | | High | High | High | High | High | High | High | High | High | High | High | 0 | 7.59E-09 | 5.05 |
| FALSE | High | VQHNGNC | [K].VQHNG | 2xCarbami | Q8BZX4-2 | 7.12E-07 | 0 | 1 | 2 | 26 | Q8BZX4-2 | Isoform 2 c | Q8BZX4-2 | 0 | 2524.1 | 0 | -0.14 | 0.011249 | 3.68 | 104.9 | 95.1 | 0.64 | 4.33 | | High | High | High | High | High | High | High | High | High | High | High | 0 | 2.44E-05 | 3.99 |
| FALSE | High | VQIPVSHPI | [K].VQIPVS | 1xPhospho | P39447 1xl | 0.001799 | 5.66E-05 | 1 | 3 | 4 | P39447 | Tight junct | P39447 [11 | 0 | 3625.581 | 0 | -0.23 | 0.039133 | 4.44 | 105.3 | 94.7 | 3.67 | 9.07 | | High | High | High | High | High | High | High | High | High | High | High | 0.000327 | 0.002708 | 3.96 |
| FALSE | High | VQIPVSHPI | [K].VQIPVS | 2xPhospho | P39447 2xl | 4.01E-05 | 0 | 1 | 3 | 6 | P39447 | Tight junct | P39447 [11 | 0 | 3705.547 | 0 | -0.16 | 0.038684 | 5.64 | 104.4 | 95.6 | 4.87 | 3.26 | | High | High | High | High | High | High | High | High | High | High | High | 0.000773 | 0.006717 | 4.12 |
| FALSE | High | VQISPDSG( | [K].VQISPD | 1xPhospho | Q3U0V1 1x | 0.000241 | 0 | 1 | 1 | 3 | Q3U0V1 | Far upstrea | Q3U0V1 [1 | 0 | 1663.825 | 0 | 0.16 | 0.028719 | 8.56 | 94.8 | 105.2 | 6.89 | 2.4 | | High | High | High | High | High | High | High | High | High | High | High | 0.000382 | 0.003235 | 2.4 |
| FALSE | High | VQISPDSG( | [K].VQISPD | 2xPhospho | Q3U0V1 2x | 0.000662 | 0 | 1 | 1 | 1 | Q3U0V1 | Far upstrea | Q3U0V1 [1 | 0 | 1743.792 | 0 | 0.33 | 0.036716 | 5.2 | 88.9 | 111.1 | 5.07 | 7.19 | | High | High | High | High | High | High | High | High | High | High | High | 0 | 0.000151 | 3.35 |
| FALSE | High | VQMPEPLL | [R].VQMPE | 1xPhospho | E9QK89 1x | 2.7E-05 | 0 | 1 | 2 | 1 | E9QK89 | Mediator c | E9QK89 [1( | 0 | 3262.575 | 0 | -0.69 | 0.022683 | 10.33 | 130 | 70 | 21.09 | 17.26 | | High | High | High | High | High | High | High | High | High | High | High | 0 | 4.12E-06 | 3.33 |
| FALSE | High | VQMTSLGS | [K].VQMTS | 1xPhospho | E9Q3G8 1x | 0.035486 | 0.002304 | 1 | 2 | 1 | E9Q3G8 | Nucleopor | E9Q3G8 [5 | 0 | 2808.466 | 0 | -0.27 | 0.189671 | 20.97 | 102.2 | 97.8 | 8.39 | 18.8 | | High | High | High | High | High | High | High | High | High | High | High | 0.001214 | 0.01299 | 2.66 |
| FALSE | High | VQMTSLGS | [K].VQMTS | 1xPhospho | E9Q3G8 1x | 1.7E-07 | 0 | 1 | 2 | 2 | E9Q3G8 | Nucleopor | E9Q3G8 [5 | 0 | 2824.461 | 0 | -0.26 | 0.141052 | 16.27 | 111.3 | 88.7 | 7.95 | 15.11 | | High | High | High | High | High | High | High | High | High | High | High | 0 | 1.39E-08 | 5.35 |
| FALSE | High | VQNHLPAS | [R].VQNHL | 1xPhospho | Q62219 1x | 6.68E-07 | 0 | 1 | 9 | 4 | Q62219 | Transformi | Q62219 [1 | 0 | 2331.181 | 0 | -0.46 | 0.067494 | 9.77 | 114.9 | 85.1 | 11.26 | 17.97 | | High | High | High | High | High | High | High | High | High | High | High | 0 | 2.27E-06 | 3.94 |
| FALSE | High | VQNHLPAS | [R].VQNHL | 1xDeamida | Q62219 1x | 2.34E-05 | 0 | 1 | 9 | 4 | Q62219 | Transformi | Q62219 [1 | 0 | 2332.165 | 0 | -0.18 | 0.335824 | 14.67 | 100.9 | 99.1 | 8.28 | 10.36 | | High | High | High | High | High | High | High | High | High | High | High | 0 | 1E-05 | 3.93 |
| FALSE | High | VQNSPPVG | [R].VQNSP | 1xPhospho | E9Q8T1 1x | 0.002212 | 0.000109 | 2 | 10 | 1 | E9Q8T1; Q | Transformi | E9Q8T1 [2 | 0 | 1262.646 | 0 | -1.39 | 0.000275 | 1.75 | 142.9 | 57.1 | 5.82 | 11.35 | Shared | High | High | High | High | High | High | High | High | High | High | High | 4.94E-05 | 0.000579 | 1.75 |
| FALSE | High | VQNTLDSS | [R].VQNTLI | 1xPhospho | Q7TSG3 1x | 0.007811 | 0.000664 | 1 | 1 | 2 | Q7TSG3 | F-box only | Q7TSG3 [1 | 0 | 2331.055 | 0 | -0.15 | 0.119702 | 12.19 | 107.9 | 92.1 | 16.14 | 4.5 | | High | High | High | High | High | High | High | High | High | High | High | 0.000449 | 0.003756 | 2.94 |
| FALSE | High | VQGTGVTI | [K].VQGTG | 2xPhospho | Q99K01-4 | 0.003622 | 0.00021 | 1 | 5 | 1 | Q99K01-4 | Isoform 4 c | Q99K01-4 | 0 | 1966.96 | 0 | -0.51 | 0.004077 | 2.68 | 114.2 | 85.8 | 10.22 | 7.61 | | High | High | High | High | High | High | High | High | High | High | High | 9.42E-05 | 0.001011 | 2.92 |
| FALSE | High | VQASPGNV | [R].VQASP | 1xPhospho | Q3U2C9 1x | 1.77E-05 | 0 | 1 | 1 | 1 | Q3U2C9 | PHD finger | Q3U2C9 [7 | 0 | 2741.347 | 0 | -0.96 | 0.013281 | 5.27 | 132 | 68 | 12.13 | 14.9 | | High | High | High | High | High | High | High | High | High | High | High | 0 | 2.56E-06 | 3.56 |
| FALSE | High | VQASPAAP | [R].VQASP | 1xPhospho | A2AIV6 1xl | 0.002743 | 0.000109 | 1 | 3 | 1 | A2AIV6 | snRNA-acti | A2AIV6 [12 | 0 | 1644.831 | 0 | 0.42 | 0.025039 | 15.39 | 85.9 | 114.1 | 12.17 | 8.25 | | High | High | High | High | High | High | High | High | High | High | High | 4.94E-05 | 0.00074 | 2.11 |
| FALSE | High | VQAPPPKP | [K].VQAPP | 1xPhospho | E9PYL2 1xl | 0.019276 | 0.001402 | 1 | 2 | 2 | E9PYL2 | Proline-rich | E9PYL2 [16 | 0 | 2185.224 | 0 | -0.54 | 0.00056 | 2.76 | 116.7 | 83.3 | 2.81 | 6.01 | | High | High | High | High | High | High | High | High | High | High | High | 0.001566 | 0.01511 | 2 |
| FALSE | High | VPSPSALLV | [K].VPSPSA | 1xPhospho | Q921I6 1xl | 3.52E-07 | 0 | 1 | 1 | 2 | Q921I6 | SH3 domai | Q921I6 [40 | 0 | 2462.31 | 0 | -0.68 | 0.005543 | 7.97 | 121.5 | 78.5 | 9.77 | 7.91 | | High | High | High | High | High | High | High | High | High | High | High | 0 | 3.14E-08 | 4.65 |
| FALSE | High | VPSPTGEP | [R].VPSPTG | 1xPhospho | Q921Y4 1x | 0.000179 | 0 | 1 | 1 | 1 | Q921Y4 | Molybdate | Q921Y4 [4 | 0 | 1665.797 | 0 | 0.38 | 0.012966 | 14.59 | 87.1 | 112.9 | 5.76 | 6.84 | | High | High | High | High | High | High | High | High | High | High | High | 0 | 3.44E-05 | 1.92 |
| FALSE | High | VPSPTPVPI | [R].VPSPTP | 1xPhospho | Q8BTI8 1xl | 0.053473 | 0.00384 | 1 | 2 | 1 | Q8BTI8 | Serine/argi | Q8BTI8 [25 | 0 | 1230.67 | 0 | -0.64 | 0.005155 | 5.97 | 119.4 | 80.6 | 10.36 | 9.52 | | High | High | High | High | High | High | High | High | High | High | High | 0.001999 | 0.02077 | 1.48 |
| FALSE | High | VPSPTPVPI | [R].VPSPTP | 1xPhospho | Q8BTI8 1xl | 0.003755 | 0.00021 | 1 | 2 | 3 | Q8BTI8 | Serine/argi | Q8BTI8 [25 | 0 | 1459.833 | 0 | -0.54 | 0.000426 | 0.65 | 117.5 | 82.5 | 5.25 | 8.56 | | High | High | High | High | High | High | High | High | High | High | High | 0.002924 | 0.0304 | 2.25 |
| FALSE | High | VPSSDEEV | [R].VPSSDE | 1xPhospho | B9EKJ7 1xl | 0.000941 | 0 | 1 | 3 | 3 | B9EKJ7 | Tankyrase | B9EKJ7 [16 | 0 | 1995.911 | 0 | -0.26 | 0.050487 | 9.67 | 109.1 | 90.9 | 10.3 | 6.15 | | High | High | High | High | High | High | High | High | High | High | High | 0 | 0.000223 | 3.02 |
| FALSE | High | VPSSDEEV | [R].VPSSDE | 2xPhospho | B9EKJ7 2xl | 1.6E-05 | 0 | 1 | 3 | 6 | B9EKJ7 | Tankyrase | B9EKJ7 [16 | 0 | 2075.877 | 0 | 0.17 | 0.193445 | 13.75 | 94.1 | 105.9 | 7.35 | 6.47 | | High | High | High | High | High | High | High | High | High | High | High | 0 | 2.29E-06 | 3.99 |
| FALSE | High | VPSSGSSTL | [K].VPSSGS | 1xPhospho | E9PV87 1xl | 1.69E-06 | 0 | 1 | 6 | 9 | E9PV87 | Protein TAI | E9PV87 [12 | 0 | 2833.355 | 0 | -0.03 | 0.317084 | 9.96 | 101 | 99 | 8.93 | 6.01 | | High | High | High | High | High | High | High | High | High | High | High | 0 | 1.84E-07 | 4.1 |
| FALSE | High | VPSSLEGSE | [K].VPSSLE | 1xPhospho | H7BX59 1x | 1.95E-08 | 0 | 1 | 6 | 5 | H7BX59 | tRNA pseu | H7BX59 [4 | 0 | 1773.727 | 0 | 0.68 | 0.002118 | 13.03 | 76.9 | 123.1 | 9.34 | 4.48 | | High | High | High | High | High | High | High | High | High | High | High | 0 | 2.7E-09 | 4.21 |
| FALSE | High | VPSSLEGSE | [K].VPSSLE | 2xPhospho | H7BX59 2x | 1.46E-07 | 0 | 1 | 6 | 14 | H7BX59 | tRNA pseu | H7BX59 [4 | 0 | 1853.693 | 0 | 0.7 | 0.000646 | 3.41 | 78.8 | 121.2 | 4.87 | 5.54 | | High | High | High | High | High | High | High | High | High | High | High | 0 | 5.24E-08 | 4.51 |
| FALSE | High | VPSWEDN | [R].VPSWE | 1xCarbami | Q3UKC1 1x | 0.000157 | 0 | 1 | 1 | 1 | Q3UKC1 | Tax1-bindi | Q3UKC1 [6 | 0 | 2052.941 | 0 | 0.52 | 0.063467 | 30.79 | 79.6 | 120.4 | 14.9 | 23.39 | | High | High | High | High | High | High | High | High | High | High | High | 0 | 2.99E-05 | 3.29 |
| FALSE | High | VPSYLDSK | [R].VPSYLD | 1xPhospho | A0A1D5RM | 0.049549 | 0.00361 | 1 | 2 | 1 | A0A1D5RM | Alstrom sy | A0A1D5RM | 0 | 1446.765 | 0 | -0.09 | 0.537686 | 9.65 | 98.6 | 101.4 | 12.53 | 14.69 | | High | High | High | High | High | High | High | High | High | High | High | 0.001902 | 0.01913 | 1.91 |
| FALSE | High | VPTAPSPK | [K].VPTAPS | 1xPhospho | Q5HZJ2 1xl | 0.098533 | 0.007902 | 1 | 5 | 1 | Q5HZJ2 | Atxn7l2 pro | Q5HZJ2 [15 | 0 | 1334.749 | 0 | -1.03 | 0.000381 | 1.66 | 132.5 | 67.5 | 4.68 | 6.38 | | High | High | High | High | High | High | High | High | High | High | High | 0.003963 | 0.04238 | 1.26 |
| FALSE | High | VPTDESEA( | [R].VPTDES | 1xPhospho | A0A0R4J1C | 1.15E-08 | 0 | 1 | 11 | 2 | A0A0R4J1C | Anion exch | A0A0R4J1C | 0 | 2702.325 | 0 | 1.43 | 0.012135 | 22.95 | 61 | 139 | 11.12 | 23.26 | | High | High | High | High | High | High | High | High | High | High | High | 0 | 6.79E-10 | 5.42 |
| FALSE | High | VPTEEEDG | [K].VPTEEE | 1xCarbami | Q61699 1x | 1.05E-05 | 0 | 1 | 2 | 4 | Q61699 | Heat shock | Q61699 [5( | 0 | 3889.703 | 0 | 0.16 | 0.547037 | 6.62 | 92.4 | 107.6 | 8.72 | 12.77 | | High | High | High | High | High | High | High | High | High | High | High | 0 | 1.42E-06 | 4.62 |
| FALSE | High | VPTEEEDG | [K].VPTEEE | 1xCarbami | Q61699 1x | 0.000334 | 0 | 1 | 2 | 2 | Q61699 | Heat shock | Q61699 [5( | 0 | 3890.687 | 0 | -0.01 | 0.724717 | 8.12 | 100.8 | 99.2 | 9.95 | 11.66 | | High | High | High | High | High | High | High | High | High | High | High | 0 | 6.97E-05 | 5.3 |
| FALSE | High | VPTLSPLTC | [K].VPTLSP | 1xPhospho | Q542Q3 1x | 0.002921 | 0.00016 | 1 | 1 | 1 | Q542Q3 | G two S ph | Q542Q3 [3 | 0 | 2229.148 | 0 | 0.86 | 0.002397 | 7.99 | 69.1 | 130.9 | 5.62 | 11.39 | | High | High | High | High | High | High | High | High | High | High | High | 7.19E-05 | 0.000794 | 4.08 |
| FALSE | High | VPTLSPLTC | [K].VPTLSP | 2xPhospho | Q542Q3 2x | 0.021797 | 0.001507 | 1 | 1 | 2 | Q542Q3 | G two S ph | Q542Q3 [3 | 0 | 2309.114 | 0 | 0.54 | 0.000748 | 1.19 | 80.9 | 119.1 | 5.86 | 7.49 | | High | High | High | High | High | High | High | High | High | High | High | 0.000814 | 0.00749 | 3.02 |
| FALSE | High | VPTPPGVP | [K].VPTPPG | 1xPhospho | P47242 1xl | 0.046687 | 0.00354 | 1 | 2 | 1 | P47242 | Paired box | P47242 [17 | 0 | 2034.135 | 0 | 3.82 | | 59.55 | 16.2 | 183.8 | 27.21 | 7.49 | | High | Not Found | Not Found | High | High | High | High | High | High | High | High | 0.001812 | 0.01782 | 3.21 |
| FALSE | High | VPTPSISPIF | [K].VPTPSI | 2xPhospho | Q8CJ27 2xl | 0.000892 | 0 | 1 | 2 | 1 | Q8CJ27 | Abnormal s | Q8CJ27 [2( | 0 | 1455.721 | 0 | 0.47 | 0.012792 | 21.14 | 84.6 | 115.4 | 8.56 | 10.56 | | High | High | High | High | High | High | High | High | High | High | High | 0 | 0.000209 | 2.9 |
| FALSE | High | VPTRHSPPI | [R].VPTRHS | 1xPhospho | Q8CFK6 1x | 0.010853 | 0.000772 | 1 | 1 | 1 | Q8CFK6 | DENN dom | Q8CFK6 [6 | 1 | 3150.578 | 0 | -1.38 | 0.001247 | 2.66 | 144.1 | 55.9 | 11.97 | 14.23 | | High | High | High | High | High | High | High | High | High | High | High | 0.000399 | 0.003437 | 2.26 |
| FALSE | High | VPTTPVK | [K].VPTTPV | 1xPhospho | Q8R5A6-2 | 0.033986 | 0.002195 | 1 | 2 | 1 | Q8R5A6-2 | Isoform 2 c | Q8R5A6-2 | 0 | 1279.743 | 0 | -0.63 | 0.002424 | 5.92 | 123.9 | 76.1 | 6.24 | 16.54 | | High | High | High | High | High | High | High | High | High | High | High | 0.001151 | 0.0124 | 1.43 |
| FALSE | High | VPVASPSAI | [R].VPVASI | 1xPhospho | Q571J8 1xl | 1.93E-05 | 0 | 1 | 18 | 5 | Q571J8 | Non-specif | Q571J8 [56 | 0 | 2245.081 | 0 | 0.34 | 0.158541 | 2.8 | 88.3 | 111.7 | 9.33 | 16.82 | | High | High | High | High | High | High | High | High | High | High | High | 0.00092 | 0.009676 | 2.2 |
| FALSE | High | VPVGDSFLI | [K].VPVGD | 1xPhospho | A0A1B0GS | 9.11E-05 | 0 | 1 | 7 | 2 | A0A1B0GS | Supervillin | A0A1B0GS | 0 | 1885.971 | 0 | 0.95 | 0.001438 | 8.03 | 68.1 | 131.9 | 6.92 | 10.82 | | High | High | High | High | High | High | High | High | High | High | High | 2.58E-05 | 0.000402 | 3.41 |
| FALSE | High | VPVLISPTLI | [K].VPVLISI | 1xPhospho | F8VQH0 1x | 0.108463 | 0.008839 | 1 | 5 | 1 | F8VQH0 | Tyrosine-p | F8VQH0 [9 | 0 | 1604.979 | 0 | -0.69 | 0.001348 | 5.95 | 122 | 78 | 3.19 | 10.72 | | High | High | High | High | High | High | High | High | High | High | High | 0.004411 | 0.0473 | 1.83 |
| FALSE | High | VPVNLLNSI | [RK].VPVNI | 1xPhospho | P35761 1xl | 7.61E-05 | 0 | 1 | 6 | 1 | P35761 | Dual specif | P35761 [27 | 0 | 2143.16 | 0 | -0.36 | 0.003866 | 5.36 | 113.4 | 86.6 | 3.57 | 11.46 | | High | High | High | High | High | High | High | High | High | High | High | 0 | 1.32E-05 | 4.05 |
| FALSE | High | VPYLASSPS | [K].VPYLAS | 1xPhospho | Q8BFX3 1x | 1.38E-05 | 0 | 1 | 1 | 3 | Q8BFX3 | BTB/POZ d | Q8BFX3 [7 | 0 | 3102.492 | 0 | 0.25 | 0.238754 | 11.01 | 92.3 | 107.7 | 10.07 | 8.47 | | High | High | High | High | High | High | High | High | High | High | High | 0 | 2.33E-05 | 2.87 |
| FALSE | High | VQAAQAQ | [R].VQAAC | 1xPhospho | Q9Z0H0 1x | 0.00967 | 0.000736 | 1 | 3 | 5 | Q9Z0H0 | Cell divisio | Q9Z0H0 [4 | 0 | 2213.1 | 0 | -1.38 | 0.000191 | 1.1 | 144.4 | 55.6 | 2.69 | 7.62 | | High | High | High | High | High | High | High | High | High | High | High | 0.000344 | 0.00303 | 2.49 |
| FALSE | High | VQAMQISE | [R].VQAM( | 1xPhospho | Q4VAA2 1x | 0.048303 | 0.00354 | 1 | 4 | 1 | Q4VAA2 | Protein CD | Q4VAA2 [1 | 0 | 1587.822 | 0 | -0.2 | 0.070945 | 2.83 | 110.1 | 89.9 | 4.89 | 22.14 | | High | High | High | High | High | High | High | High | High | High | High | 0.001812 | 0.01847 | 2.43 |
| FALSE | High | VQANLDTF | [K].VQANL | 1xPhospho | E9Q616 1x | 2.06E-08 | 0 | 1 | 5 | 2 | E9Q616 | AHNAK nuc | E9Q616 [4 | 0 | 2462.258 | 0 | 4.52 | 0.001001 | 27.02 | 11.3 | 188.7 | 35.9 | 16.34 | | High | High | High | High | High | High | High | High | High | High | High | 0 | 1.3E-09 | 4.76 |
| FALSE | High | VQNVEDFV | [R].VQNVE | 1xPhospho | Q5U3K5 1x | 0.004143 | 0.00036 | 1 | 1 | 2 | Q5U3K5 | Rab-like pr | Q5U3K5 [3 | 1 | 3373.661 | 0 | -0.3 | 0.004126 | 4.44 | 110.4 | 89.6 | 6.44 | 5.28 | | High | High | High | High | High | High | High | High | High | High | High | 0.000161 | 0.001173 | 3.11 |
| FALSE | High | VSLENGEA | [K].VSLENC | 1xPhospho | A2A5R2 1x | 0.000173 | 0 | 1 | 3 | 1 | A2A5R2 | Brefeldin A | A2A5R2 [2( | 0 | 1380.672 | 0 | -0.88 | 0.011434 | 1.14 | 123.5 | 76.5 | 10.87 | 11.44 | | High | High | High | High | High | High | High | High | High | High | High | 0 | 3.32E-05 | 3.37 |
| FALSE | High | VQPQWSPI | [R].VQPQW | 1xCarbami | Q3UXN3 1: | 0.000334 | 0 | 1 | 3 | 2 | Q3UXN3 | Putative ur | Q3UXN3 [9 | 0 | 2018.936 | 0 | -0.73 | 0.006847 | 2.5 | 122.3 | 77.7 | 6.11 | 12.05 | | High | High | High | High | High | High | High | High | High | High | High | 0 | 6.97E-05 | 2.39 |
| FALSE | High | VQSEPQEE | [R].VQSEP( | 1xCarbami | Q8R4Z6 1x | 3.69E-07 | 0 | 1 | 3 | 1 | Q8R4Z6 | DEAD-box | Q8R4Z6 [7 | 0 | 2258.938 | 0 | -0.37 | 0.002101 | 3.9 | 112 | 88 | 4.45 | 6.97 | | High | High | High | High | High | High | High | High | High | High | High | 0 | 3.31E-08 | 3.44 |
| FALSE | High | VSEGCLTSS | [K].VSEGCI | 1xCarbami | Q3UQN6 1 | 6.45E-07 | 0 | 1 | 6 | 9 | Q3UQN6 | Putative ur | Q3UQN6 [9 | 0 | 2469.091 | 0 | -0.16 | 0.095376 | 6.14 | 105.5 | 94.5 | 6.18 | 5.16 | | High | High | High | High | High | High | High | High | High | High | High | 0 | 1.09E-06 | 4.78 |
| FALSE | High | VSEGCLTSS | [K].VSEGCI | 1xCarbami | Q3UQN6 1 | 0.00103 | 0 | 1 | 6 | 1 | Q3UQN6 | Putative ur | Q3UQN6 [9 | 1 | 4249.96 | 0 | -0.05 | 0.720622 | 19.15 | 99.2 | 100.8 | 3.78 | 21.32 | | High | High | High | High | High | High | High | High | High | High | High | 0 | 0.000246 | 3.15 |
| FALSE | High | VSEGGPAE | [R].VSEGGI | 1xPhospho | Q9DBG9 1x | 0.000111 | 0 | 1 | 1 | 2 | Q9DBG9 | Tax1-bindi | Q9DBG9 [6 | 0 | 2179.141 | 0 | 1.61 | 0.000329 | 4.74 | 49.3 | 150.7 | 8.9 | 6.71 | | High | High | High | High | High | High | High | High | High | High | High | 0 | 2.03E-05 | 2.85 |

| | | | | | | | | | | | | | | | | | | | | | | | | | | | | | | | | | | | | | | |
|---|---|---|---|---|---|---|---|---|---|---|---|---|---|---|---|---|---|---|---|---|---|---|---|---|---|---|---|---|---|---|---|---|---|---|---|---|---|---|
| FALSE | High | VSEGLDGC | [R].VSEGLI | 1xCarbami | Q7TSH3 1x | 1.77E-06 | 0 | 1 | 1 | 2 | Q7TSH3 | Zinc finger | Q7TSH3 [1 | 0 | 1858.902 | 0 | -0.3 | 0.017568 | 4.45 | 110.4 | 89.6 | 7.79 | 7.69 | | High | High | High | High | High | High | High | High | High | High | High | 0 | 1.94E-07 | 3.86 |
| FALSE | High | VSEQEDAV | [R].VSEQEI | 1xPhosphc | G3UYN0 1x | 0.002921 | 0.00016 | 1 | 2 | 2 | G3UYN0 | L1TD1 OS= | G3UYN0 [6 | 0 | 1942.977 | 0 | -1.05 | 0.001052 | 3.97 | 134.8 | 65.2 | 5.16 | 11.38 | | High | High | High | High | High | High | High | High | High | High | High | 0.000814 | 0.007649 | 3.25 |
| FALSE | High | VSESVADD | [R].VSESVA | 1xCarbami | B9EKS2 1xl | 0.001544 | 5.66E-05 | 1 | 3 | 2 | B9EKS2 | Jumonji do | B9EKS2 [54 | 1 | 3530.661 | 0 | -0.11 | 0.945574 | 6.04 | 103.6 | 96.4 | 15.46 | 20.15 | | High | High | High | High | High | High | High | High | High | High | High | 2.58E-05 | 0.000388 | 2.73 |
| FALSE | High | VSETEYSAL | [R].VSETEY | 1xPhosphc | P56485 1xl | 0.002948 | 0.00016 | 1 | 1 | 2 | P56485 | Atypical ch | P56485 [34 | 0 | 2137.047 | 0 | 2.87 | 0.000388 | 7.38 | 23.9 | 176.1 | 12.29 | 9.72 | | High | High | High | High | High | High | High | High | High | High | High | 7.19E-05 | 0.000798 | 3.06 |
| FALSE | High | VSFADTISH | [K].VSFADT | 2xPhosphc | Q8CG73 2x | 0.002623 | 0.000109 | 1 | 2 | 1 | Q8CG73 | Protein fan | Q8CG73 [9 | 0 | 2612.245 | 0 | 0.63 | 0.000598 | 9.02 | 81.1 | 118.9 | 6 | 5.07 | | High | High | High | High | High | High | High | High | High | High | High | 4.94E-05 | 0.000706 | 2.82 |
| FALSE | High | VSFTPDGC | [R].VSFTPD | 1xCarbami | A0A0R4J0H | 0.000134 | 0 | 1 | 2 | 1 | A0A0R4J0H | Fibronectir | A0A0R4J0H | 0 | 3205.475 | 0 | 0.43 | 0.050053 | 15.02 | 85.2 | 114.8 | 1.98 | 15.86 | | High | High | High | High | High | High | High | High | High | High | High | 0 | 2.51E-05 | 4.23 |
| FALSE | High | VSGADGYE | [R].VSGAD | 1xPhosphc | Q3UC10 1x | 3.74E-06 | 0 | 1 | 3 | 2 | Q3UC10 | DNA helica | Q3UC10 [2 | 0 | 1662.757 | 0 | -1.76 | 0.00034 | 0.25 | 153.3 | 46.7 | 2.2 | 15.61 | | High | High | High | High | High | High | High | High | High | High | High | 0 | 4.49E-07 | 2.92 |
| FALSE | High | VSGAGLSP | [R].VSGAG | 1xPhosphc | B9EKJ7 1xP | 0.000496 | 0 | 1 | 3 | 1 | B9EKJ7 | Tankyrase | B9EKJ7 [11 | 0 | 1239.63 | 0 | -0.37 | 0.00532 | 7.24 | 112.9 | 87.1 | 4.14 | 8.89 | | High | High | High | High | High | High | High | High | High | High | High | 0 | 0.000109 | 1.35 |
| FALSE | High | VSGGFPED | [R].VSGGFI | 1xPhosphc | O70439 1x | 0.001039 | 0 | 1 | 4 | 1 | O70439 | Syntaxin-7 | O70439 [1 | 0 | 1418.64 | 0 | 0.11 | 0.273125 | 18.56 | 95.4 | 104.6 | 12.95 | 5.94 | | High | High | High | High | High | High | High | High | High | High | High | 0 | 0.00025 | 1.58 |
| FALSE | High | VSGGFPED | [R].VSGGFI | 1xPhosphc | O70439 1x | 0.004333 | 0.00036 | 1 | 4 | 2 | O70439 | Syntaxin-7 | O70439 [1 | 0 | 1647.803 | 0 | -0.2 | 0.210946 | 8.76 | 105.4 | 94.6 | 9.16 | 9.12 | | High | High | High | High | High | High | High | High | High | High | High | 0.000204 | 0.001348 | 2.79 |
| FALSE | High | VSEEDQSLI | [K].VSEEDC | 1xPhosphc | P41234 1xl | 0.091627 | 0.007146 | 1 | 3 | 1 | P41234 | ATP-bindin | P41234 [13 | 0 | 2317.085 | 0 | -0.1 | 0.63606 | 15.06 | 104.7 | 95.3 | 8.34 | 11.69 | | High | High | High | High | High | High | High | High | High | High | High | 0.003659 | 0.03906 | 2.49 |
| FALSE | High | VSGGSPLV. | [R].VSGGSI | 1xCarbami | Q9CWY3 1 | 0.00025 | 0 | 1 | 1 | 2 | Q9CWY3 | N-lysine m | Q9CWY3 [ | 0 | 1789.887 | 0 | -0.23 | 0.00338 | 1.17 | 107.7 | 92.3 | 5.04 | 2.2 | | High | High | High | High | High | High | High | High | High | High | High | 0 | 5.05E-05 | 3.46 |
| FALSE | High | VSGPSSSEI | [K].VSGPSS | 1xPhosphc | Q3ULZ6 1x | 2.88E-06 | 0 | 1 | 2 | 3 | Q3ULZ6 | Putative ur | Q3ULZ6 [2 | 0 | 2492.163 | 0 | 2.05 | 0.009338 | 39.68 | 34.9 | 165.1 | 13.95 | 29.54 | | High | High | High | High | High | High | High | High | High | High | High | 0 | 4.01E-06 | 3.32 |
| FALSE | High | VSGPSSSEI | [K].VSGPSS | 1xPhosphc | Q3ULZ6 1x | 1.7E-07 | 0 | 1 | 2 | 12 | Q3ULZ6 | Putative ur | Q3ULZ6 [2 | 0 | 2508.158 | 1 | 1.11 | 0.000546 | 2.84 | 62.6 | 137.4 | 10.25 | 3.2 | | High | High | High | High | High | High | High | High | High | High | High | 0 | 2.48E-08 | 5.02 |
| FALSE | High | VSGSATPN | [R].VSGSAT | 1xPhosphc | Q6P9R1 1x | 0.005924 | 0.00062 | 1 | 1 | 1 | Q6P9R1 | ATP-depen | Q6P9R1 [7 | 0 | 1581.747 | 0 | -0.4 | 0.001473 | 2.96 | 115.3 | 84.7 | 5.67 | 5.23 | | High | High | High | High | High | High | High | High | High | High | High | 0.000275 | 0.001739 | 1.76 |
| FALSE | High | VSGSPCHH | [R].VSGSPC | 1xCarbami | A2AKB9 1x | 0.000118 | 0 | 1 | 2 | 1 | A2AKB9 | DDB1- and | A2AKB9 [3 | 0 | 2294.027 | 0 | 1.8 | 0.030383 | 34.8 | 53.1 | 146.9 | 33.43 | 27.39 | | High | High | High | High | High | High | High | High | High | High | High | 0 | 2.17E-05 | 2.17 |
| FALSE | High | VSGSPEQD | [K].VSGSPE | 1xPhosphc | Q8BHN5 1x | 7.15E-05 | 0 | 1 | 2 | 3 | Q8BHN5 | RNA-bindir | Q8BHN5 [1 | 0 | 1792.759 | 0 | -1.06 | 0.000319 | 1.62 | 134.9 | 65.1 | 6.13 | 5.21 | | High | High | High | High | High | High | High | High | High | High | High | 0 | 1.23E-05 | 3.2 |
| FALSE | High | VSGSSDDL | [K].VSGSSD | 1xPhosphc | B2RUC3 1x | 0.001463 | 5.66E-05 | 1 | 8 | 2 | B2RUC3 | Excision re | B2RUC3 [4 | 0 | 1645.809 | 0 | -0.65 | 0.000523 | 0.55 | 122.6 | 77.4 | 9.64 | 8.32 | | High | High | High | High | High | High | High | High | High | High | High | 2.58E-05 | 0.000364 | 3.25 |
| FALSE | High | VSGSSSDSS | [R].VSGSSS | 1xPhosphc | Q9D809 1x | 4.52E-06 | 0 | 1 | 2 | 1 | Q9D809 | MCG22941 | Q9D809 [4 | 0 | 2589.109 | 0 | 0.53 | 0.011864 | 12.2 | 83.5 | 116.5 | 17.82 | 6.43 | | High | High | High | High | High | High | High | High | High | High | High | 0 | 5.54E-07 | 3.98 |
| FALSE | High | VSGTLETK | [K].VSGTLE | 1xPhosphc | Q60930 1x | 0.069903 | 0.005181 | 1 | 4 | 1 | Q60930 | Voltage-de | Q60930 [6 | 0 | 1372.749 | 0 | 0.32 | 0.189562 | 11.74 | 90.4 | 109.6 | 10.22 | 14.53 | | High | High | High | High | High | High | High | High | High | High | High | 0.002662 | 0.02833 | 1.63 |
| FALSE | High | VSHSPEPLK | [K].VSHSPE | 1xPhosphc | F6SH61 1xl | 0.06169 | 0.004646 | 1 | 1 | 1 | F6SH61 | C-Jun-amir | F6SH61 [84 | 0 | 1531.829 | 0 | 1.95 | 0.000313 | 15.43 | 44.1 | 155.9 | 13.25 | 5.24 | | High | High | High | High | High | High | High | High | High | High | High | 0.002381 | 0.02459 | 2.22 |
| FALSE | High | VSHSPPPK | [R].VSHSPF | 1xPhosphc | Q52KI8 1xP | 0.115566 | 0.009504 | 1 | 7 | 1 | Q52KI8 | Serine/argi | Q52KI8 [65 | 0 | 1386.755 | 0 | -0.19 | 0.081316 | 7.83 | 107 | 93 | 7.49 | 6.59 | | High | High | High | High | High | High | High | High | High | High | High | 0.004733 | 0.0511 | 1.75 |
| FALSE | High | VSIASEVES | [R].VSIASE | 1xPhosphc | A2AG68 1x | 1.74E-06 | 0 | 1 | 4 | 4 | A2AG68 | Copper-tra | A2AG68 [3 | 0 | 2619.317 | 0 | 0.72 | 0.002581 | 6.6 | 74.1 | 125.9 | 11.69 | 6.88 | | High | High | High | High | High | High | High | High | High | High | High | 0 | 1.9E-07 | 4.46 |
| FALSE | High | VSLEDLYN | [K].VSLEDL | 1xPhosphc | Q9QYJ0 1x | 2.25E-05 | 0 | 1 | 2 | 1 | Q9QYJ0 | DnaJ homc | Q9QYJ0 [1 | 0 | 1675.871 | 0 | -0.22 | 0.029037 | 8.75 | 111.5 | 88.5 | 8.48 | 7.21 | | High | High | High | High | High | High | High | High | High | High | High | 0 | 3.37E-06 | 2.72 |
| FALSE | High | VSLEEIYSG | [RK].VSLEE | 1xCarbami | Q3TIT6 1xP | 0.009416 | 0.000736 | 1 | 4 | 1 | Q3TIT6 | Putative ur | Q3TIT6 [17 | 0 | 1923.954 | 0 | 0.58 | 0.092726 | 18.89 | 82.5 | 117.5 | 19.28 | 8.57 | | High | High | High | High | High | High | High | High | High | High | High | 0.000327 | 0.002943 | 1.85 |
| FALSE | High | VSGGSPLV. | [R].VSGGSI | 1xCarbami | Q9CWY3 2 | 1.52E-06 | 0 | 1 | 1 | 1 | Q9CWY3 | N-lysine m | Q9CWY3 [ | 0 | 1869.853 | 0 | -0.07 | 0.795258 | 12.11 | 102.5 | 97.5 | 10.96 | 9.81 | | High | High | High | High | High | High | High | High | High | High | High | 0 | 1.63E-07 | 3.88 |
| FALSE | High | VSDSSLSPE | [R].VSDSSL | 1xPhosphc | Q6PR54-3 | 6.02E-05 | 0 | 2 | 4 | 5 | Q6PR54-3; | Isoform 3 c | Q6PR54-3 | 0 | 1586.808 | 0 | -1.37 | 0.000343 | 1.07 | 145.9 | 54.1 | 7.53 | 4.87 | Shared | High | High | High | High | High | High | High | High | High | High | High | 0.000872 | 0.008881 | 1.77 |
| FALSE | High | VSDQNSPV | [R].VSDQN | 1xPhosphc | E9PV45 1xl | 0.001021 | 0 | 1 | 2 | 5 | E9PV45 | Ubiquitin c | E9PV45 [2 | 0 | 1721.924 | 0 | 0.55 | 0.005023 | 10.39 | 80.5 | 119.5 | 2.22 | 9.31 | | High | High | High | High | High | High | High | High | High | High | High | 0 | 0.000245 | 2.55 |
| FALSE | High | VSDQNSPV | [R].VSDQN | 1xPhosphc | E9PV45 1xl | 0.009416 | 0.000736 | 1 | 2 | 2 | E9PV45 | Ubiquitin c | E9PV45 [2 | 0 | 1492.761 | 0 | 0.65 | 0.04696 | 23.73 | 77.8 | 122.2 | 9.56 | 17.76 | | High | High | High | High | High | High | High | High | High | High | High | 0.000327 | 0.002941 | 2.13 |
| FALSE | High | VQSGMGL | [K].VQSGM | 2xPhosphc | C7G3P2 2x | 0.050397 | 0.003641 | 1 | 3 | 1 | C7G3P2 | MKIAA053 | C7G3P2 [1 | 0 | 3324.515 | 0 | 0.05 | 0.907358 | 6.86 | 98.5 | 101.5 | 18.63 | 16.42 | | High | High | High | High | High | High | High | High | High | High | High | 0.001916 | 0.01952 | 2.42 |
| FALSE | High | VQSGSEGK | [R].VQSGSI | 1xPhosphc | Q9ERU9 1x | 0.030632 | 0.001857 | 1 | 1 | 1 | Q9ERU9 | E3 SUMO-p | Q9ERU9 [2 | 0 | 1329.682 | 0 | -0.14 | 0.073447 | 4.27 | 106.1 | 93.9 | 7.69 | 4 | | High | High | High | High | High | High | High | High | High | High | High | 0.001011 | 0.01104 | 2.32 |
| FALSE | High | VQSHQQG | [R].VQSHQ | 1xCarbami | Q8K2E5 1x | 0.003892 | 0.000261 | 1 | 3 | 1 | Q8K2E5 | Budding ur | Q8K2E5 [3 | 0 | 1867.877 | 0 | -1.54 | 0.004404 | 3.87 | 152.9 | 47.1 | 18.72 | 12.18 | | High | High | High | High | High | High | High | High | High | High | High | 0.00014 | 0.001098 | 2.11 |
| FALSE | High | VQSLEGEK | [K].VQSLEC | 1xPhosphc | P14873 1xl | 0.000573 | 0 | 2 | 2 | 1 | P14873; B2 | Microtubu | P14873 [17 | 1 | 2082.181 | 0 | -1.2 | 0.000458 | 1.23 | 139 | 61 | 8.2 | 7.39 | Shared | High | High | High | High | High | High | High | High | High | High | High | 0 | 0.000128 | 3.57 |
| FALSE | High | VQSPPLTR | [R].VQSPPI | 1xPhosphc | A2A654 1x | 0.018125 | 0.001282 | 1 | 8 | 1 | A2A654 | Bromodom | A2A654 [2 | 0 | 1206.645 | 0 | 0.18 | 0.114175 | 5.88 | 96.1 | 103.9 | 3.43 | 9.21 | | High | High | High | High | High | High | High | High | High | High | High | 0.000689 | 0.006116 | 1.58 |
| FALSE | High | VQSSLCLGI | [R].VQSSLC | 1xCarbami | Q4QRL3-2 | 0.030104 | 0.001857 | 1 | 3 | 2 | Q4QRL3-2 | Isoform 2 c | Q4QRL3-2 | 0 | 2099.999 | 0 | 0.82 | 0.016022 | 22.37 | 66.7 | 133.3 | 13.62 | 15.84 | | High | High | High | High | High | High | High | High | High | High | High | 0.002381 | 0.02466 | 3.11 |
| FALSE | High | VQSSPNLL | [R].VQSSPI | 1xPhosphc | Q3UTJ2-2 | 0.093879 | 0.007306 | 1 | 15 | 1 | Q3UTJ2-2 | Isoform 2 c | Q3UTJ2-2 | 0 | 1521.799 | 0 | 1.3 | 0.002007 | 2.95 | 57.4 | 142.6 | 11.28 | 10.97 | | High | High | High | High | High | High | High | High | High | High | High | 0.003698 | 0.03989 | 2.44 |
| FALSE | High | VQTTPPPTI | [K].VQTTPI | 1xPhosphc | Q3UHJ0 1x | 9.67E-07 | 0 | 1 | 2 | 4 | Q3UHJ0 | AP2-associ | Q3UHJ0 [6 | 0 | 1933.056 | 0 | -0.14 | 0.344345 | 7.77 | 103.2 | 96.8 | 5.58 | 5.4 | | High | High | High | High | High | High | High | High | High | High | High | 0 | 9.76E-08 | 3.88 |
| FALSE | High | VQVAPLQC | [K].VQVAP | 1xPhosphc | Q9Z0U1 1x | 2.49E-06 | 0 | 1 | 1 | 9 | Q9Z0U1 | Tight junct | Q9Z0U1 [9 | 0 | 2125.064 | 0 | 0.22 | 0.002441 | 3.89 | 92.6 | 107.4 | 2.7 | 1.75 | | High | High | High | High | High | High | High | High | High | High | High | 0.000859 | 0.008475 | 2.93 |
| FALSE | High | VQVAPLQC | [K].VQVAP | 2xPhosphc | Q9Z0U1 2x | 3.06E-05 | 0 | 1 | 1 | 2 | Q9Z0U1 | Tight junct | Q9Z0U1 [9 | 0 | 2205.03 | 0 | -1.12 | 0.000726 | 3.91 | 139.5 | 60.5 | 4.51 | 9.98 | | High | High | High | High | High | High | High | High | High | High | High | 0 | 0.000119 | 1.81 |
| FALSE | High | VREDDEDS | [R].VREDD | 2xPhosphc | G5E870 2x | 9.48E-06 | 0 | 1 | 3 | 28 | G5E870 | E3 ubiquiti | G5E870 [1 | 1 | 3930.591 | 0 | -1.1 | 0.005756 | 4.7 | 137.6 | 62.4 | 14.97 | 12.35 | | High | High | High | High | High | High | High | High | High | High | High | 0 | 6.98E-06 | 6.29 |
| FALSE | High | VRHDTPDL | [R].VRHDT | 2xPhosphc | Q8R149 2x | 0.006957 | 0.00062 | 1 | 2 | 2 | Q8R149 | BUD13 hor | Q8R149 [1 | 1 | 1778.819 | 0 | -0.13 | 0.004738 | 2.69 | 104.8 | 95.2 | 0.82 | 3.95 | | High | High | High | High | High | High | High | High | High | High | High | 0.000903 | 0.009496 | 1.62 |
| FALSE | High | VRHDTPDP | [R].VRHDT | 2xPhosphc | Q8R149 2x | 0.083101 | 0.006345 | 1 | 2 | 1 | Q8R149 | BUD13 hor | Q8R149 [2 | 2 | 1918.889 | 0 | -0.16 | 0.929891 | 15.58 | 103.5 | 96.5 | 10.83 | 19.67 | | High | High | High | High | High | High | High | High | High | High | High | 0.003265 | 0.03471 | 3.19 |
| FALSE | High | VRLSTPLQT | [R].VRLSTP | 1xCarbami | Q3UZG4 1x | 0.005275 | 0.000599 | 1 | 3 | 2 | Q3UZG4 | Aminoacyl | Q3UZG4 [8 | 1 | 3826.966 | 0 | -0.05 | 0.894122 | 26.7 | 96.6 | 103.4 | 12.49 | 12.46 | | High | High | High | High | High | High | High | High | High | High | High | 0.000267 | 0.001538 | 2.17 |
| FALSE | High | VRVSPQDA | [R].VRVSP( | 1xPhosphc | E9Q8Z8 1x | 2.81E-07 | 0 | 1 | 8 | 1 | E9Q8Z8 | Catenin de | E9Q8Z8 [4 | 1 | 2339.174 | 0 | -0.23 | 0.002561 | 3.2 | 107.8 | 92.2 | 7.01 | 7.79 | | High | High | High | High | High | High | High | High | High | High | High | 0 | 2.44E-08 | 3.91 |
| FALSE | High | VRVSPQDA | [R].VRVSP( | 1xPhosphc | E9Q8Z8 1x | 1.32E-05 | 0 | 1 | 8 | 2 | E9Q8Z8 | Catenin de | E9Q8Z8 [4 | 1 | 2355.169 | 0 | -0.34 | 0.405048 | 2.56 | 108.7 | 91.3 | 5.1 | 19.67 | | High | High | High | High | High | High | High | High | High | High | High | 0 | 1.84E-05 | 2.94 |
| FALSE | High | VRVSPQDA | [R].VRVSP( | 1xDeamida | E9Q8Z8 1x | 1.87E-09 | 0 | 1 | 8 | 7 | E9Q8Z8 | Catenin de | E9Q8Z8 [4 | 1 | 2356.153 | 0 | -0.24 | 0.172585 | 4.92 | 108.6 | 91.4 | 7.08 | 11.53 | | High | High | High | High | High | High | High | High | High | High | High | 0.002059 | 0.02129 | 4.1 |
| FALSE | High | VRYDSSNQ | [R].VRYDSS | 1xPhosphc | D3YTR4 1x | 0.014283 | 0.001011 | 2 | 4 | 2 | D3YTR4; Q | Methyl-Cp | D3YTR4 [4 | 1 | 1733.899 | 0 | -0.89 | 0.000552 | 2.1 | 130.5 | 69.5 | 5.38 | 15.96 | Shared | High | High | High | High | High | High | High | High | High | High | High | 0.000538 | 0.004661 | 1.98 |
| FALSE | High | VSAASPCEI | [R].VSAASF | 1xCarbami | G8DM25 1 | 0.002272 | 0.000109 | 1 | 1 | 3 | G8DM25 | DNA polym | G8DM25 [ | 0 | 1981.925 | 0 | -1.84 | 0.000466 | 0.98 | 156.2 | 43.8 | 2.53 | 19.96 | | High | High | High | High | High | High | High | High | High | High | High | 4.94E-05 | 0.000601 | 2.92 |
| FALSE | High | VSADGAAK | [K].VSADG. | 1xPhosphc | P18608 1xl | 0.001516 | 5.66E-05 | 1 | 3 | 1 | P18608 | Non-histor | P18608 [6- | 0 | 1256.665 | 0 | 1.15 | 0.029673 | 33.81 | 66.3 | 133.7 | 14.67 | 35.81 | | High | High | High | High | High | High | High | High | High | High | High | 2.58E-05 | 0.00038 | 2.21 |
| FALSE | High | VSAGNAVF | [R].VSAGN. | 1xPhosphc | Q3TVV9 1x | 7.41E-05 | 0 | 1 | 4 | 1 | Q3TVV9 | Putative ur | Q3TVV9 [3 | 0 | 1927.036 | 0 | 1.2 | 0.000278 | 4.6 | 63.8 | 136.2 | 5.51 | 5.86 | | High | High | High | High | High | High | High | High | High | High | High | 0 | 1.28E-05 | 2.15 |
| FALSE | High | VSALAGPS | [K].VSALAC | 1xPhosphc | Q69ZR2 1x | 8.06E-06 | 0 | 1 | 3 | 3 | Q69ZR2 | E3 ubiquiti | Q69ZR2 [6 | 0 | 2985.454 | 0 | 0.53 | 0.001218 | 4.7 | 81.8 | 118.2 | 0.57 | 5.83 | | High | High | High | High | High | High | High | High | High | High | High | 0 | 1.06E-06 | 3.27 |
| FALSE | High | VSAQVAAF | [R].VSAQV. | 1xPhosphc | Q8K2J0 1xl | 0.072258 | 0.005346 | 1 | 1 | 1 | Q8K2J0 | 1-phospha | Q8K2J0 [1 | 0 | 2753.416 | 0 | 2.88 | 0.001012 | 19.8 | 23.6 | 176.4 | 33.87 | 15.32 | | High | High | High | High | High | High | High | High | High | High | High | 0.002855 | 0.02955 | 3.11 |
| FALSE | High | VSARESPVS | [K].VSARES | 1xPhosphc | F8WGX5 1 | 0.000415 | 0 | 1 | 2 | 2 | F8WGX5 | DmX-like p | F8WGX5 [2 | 1 | 2651.344 | 0 | 0.54 | 0.006724 | 10.11 | 82.1 | 117.9 | 8.44 | 5.61 | | High | High | High | High | High | High | High | High | High | High | High | 0 | 8.83E-05 | 3.32 |
| FALSE | High | VSASPPPV | [K].VSASPF | 1xPhosphc | G3X8R8 1x | 0.000496 | 0 | 1 | 4 | 2 | G3X8R8 | MCG12388 | G3X8R8 [2 | 0 | 3350.769 | 0 | 0.36 | 0.008049 | 10.16 | 84 | 116 | 8.01 | 9.4 | | High | High | High | High | High | High | High | High | High | High | High | 0 | 0.000108 | 4.24 |
| FALSE | High | VSASSPTAI | [R].VSASSP | 1xPhosphc | Q6PAR7 1x | 0.002004 | 5.66E-05 | 1 | 3 | 4 | Q6PAR7 | Phf20 prot | Q6PAR7 [5 | 0 | 1385.744 | 0 | -1.61 | 9.77E-05 | 0.29 | 150.6 | 49.4 | 2.59 | 6.3 | | High | High | High | High | High | High | High | High | High | High | High | 2.58E-05 | 0.000518 | 2.36 |
| FALSE | High | VSDESVLPI | [R].VSDES\ | 1xPhosphc | F6RJ39 1xP | 0.014929 | 0.001049 | 2 | 4 | 1 | F6RJ39; Q8 | Apoptotic | F6RJ39 [44 | 0 | 1823.992 | 0 | -0.24 | 0.182182 | 8.36 | 105.2 | 94.8 | 18.03 | 7.54 | Shared | High | High | High | High | High | High | High | High | High | High | High | 0.000555 | 0.004901 | 2.59 |
| FALSE | High | VSDFGGR | [R].VSDFG | 1xPhosphc | Q3U8B3 1x | 0.015603 | 0.001124 | 1 | 3 | 2 | Q3U8B3 | Putative ur | Q3U8B3 [4 | 0 | 1046.487 | 0 | -1.41 | 0.00036 | 0.36 | 144.2 | 55.8 | 2.56 | 12.67 | | High | High | High | High | High | High | High | High | High | High | High | 0.001027 | 0.01109 | 1.94 |
| FALSE | High | VSDPVSSSE | [R].VSDPVS | 1xPhosphc | H3BK88 1x | 0.006713 | 0.00062 | 2 | 8 | 2 | H3BK88; H | Treacle prc | H3BK88 [6 | 0 | 2118.984 | 0 | -0.79 | 0.054411 | 10.15 | 124.1 | 75.9 | 14.12 | 26.37 | Shared | High | High | High | High | High | High | High | High | High | High | High | 0.000968 | 0.01003 | 2.04 |
| FALSE | High | VQSDLEAK | [K].VQSDLI | 1xPhosphc | E9PV87 1xl | 0.002272 | 0.000109 | 1 | 4 | 1 | E9PV87 | Protein TA | E9PV87 [1 | 0 | 1427.755 | 0 | 0.63 | 0.032307 | 26.91 | 77.2 | 122.8 | 9.28 | 19.41 | | High | High | High | High | High | High | High | High | High | High | High | 4.94E-05 | 0.0006 | 3.2 |
| FALSE | High | SPGTATLAI | [R].SPGTAT | 1xPhosphc | Q9CS00 1x | 1.57E-08 | 0 | 1 | 1 | 7 | Q9CS00 | Cactin OS= | Q9CS00 [1 | 0 | 2224.117 | 0 | 0.09 | 0.298148 | 4.56 | 97.9 | 102.1 | 5.58 | 6.58 | | High | High | High | High | High | High | High | High | High | High | High | 0 | 9.49E-10 | 4.71 |
| FALSE | High | TLSVNTDF | [R].TLSVNT | 1xPhosphc | Q91ZS8-6 | 0.003962 | 0.000311 | 1 | 7 | 4 | Q91ZS8-6 | Isoform 6 c | Q91ZS8-6 [ | 0 | 3660.703 | 0 | 6.64 | | 0 | 9.5 | 190.5 | | 10.85 | | High | High | High | High | High | High | High | High | High | High | High | 0.000571 | 0.004964 | 3.57 |
| FALSE | High | TLSVAAAFI | [K].TLSVAA | 1xPhosphc | F8WI88 1xl | 1.91E-07 | 0 | 1 | 5 | 25 | F8WI88 | PEST prote | F8WI88 [1 | 0 | 3160.432 | 0 | -0.15 | 0.041051 | 3.34 | 105.4 | 94.6 | 3.71 | 5.57 | | High | High | High | High | High | High | High | High | High | High | High | 0 | 1.57E-08 | 5.23 |
| FALSE | High | SSGSETEQ | [R].SSGSET | 1xPhosphc | Q5SVQ0 1x | 1.54E-05 | 0 | 1 | 5 | 5 | Q5SVQ0 | Histone ac | Q5SVQ0 [1 | 0 | 1951.848 | 0 | 0.29 | 0.013717 | 8.4 | 89.9 | 110.1 | 3.28 | 5.98 | | High | High | High | High | High | High | High | High | High | High | High | 0 | 4.03E-05 | 4.09 |
| FALSE | High | SSGSETEQ | [R].SSGSET | 2xPhosphc | Q5SVQ0 2x | 1.56E-05 | 0 | 1 | 5 | 5 | Q5SVQ0 | Histone ac | Q5SVQ0 [1 | 0 | 2031.815 | 0 | -0.51 | 0.020122 | 6.06 | 118.4 | 81.6 | 7.99 | 13.1 | | High | High | High | High | High | High | High | High | High | High | High | 0 | 2.22E-06 | 4.64 |
| FALSE | High | SSGSETEQ | [R].SSGSET | 1xPhosphc | Q5SVQ0 1x | 0.038346 | 0.002563 | 1 | 5 | 1 | Q5SVQ0 | Histone ac | Q5SVQ0 [1 | 1 | 2539.197 | 0 | 0.06 | 0.633433 | 10.85 | 108.5 | 91.5 | 13.37 | 11.1 | | High | High | High | High | High | High | High | High | High | High | High | 0.001355 | 0.01423 | 2.54 |
| FALSE | High | SSGSLSPGI | [K].SSGSLS | 1xPhosphc | B9EKJ7 1xP | 9.53E-08 | 0 | 1 | 3 | 8 | B9EKJ7 | Tankyrase | B9EKJ7 [13 | 0 | 2154.016 | 0 | -0.27 | 0.019149 | 8.27 | 108.4 | 91.6 | 5.09 | 6.94 | | High | High | High | High | High | High | High | High | High | High | High | 0 | 7.22E-09 | 4.17 |
| FALSE | High | SSGSLSPGI | [K].SSGSLS | 2xPhosphc | B9EKJ7 2xP | 4.61E-06 | 0 | 1 | 3 | 4 | B9EKJ7 | Tankyrase | B9EKJ7 [13 | 0 | 2233.983 | 0 | 1.09 | 0.001008 | 13.42 | 63 | 137 | 7.85 | 7.84 | | High | High | High | High | High | High | High | High | High | High | High | 0 | 1.13E-06 | 4.78 |
| FALSE | High | SSGSPK | [RK].SSGSF | 1xPhosphc | M1VMF6 1 | 0.096181 | 0.007535 | 1 | 9 | 1 | M1VMF6 | Metastasis | M1VMF6 [ | 0 | 1100.575 | 0 | 0 | 0.970045 | 6.29 | 100.4 | 99.6 | 4.81 | 6.33 | | High | High | High | High | High | High | High | High | High | High | High | 0.003838 | 0.04128 | 1.55 |
| FALSE | High | SSGSPYGG | [R].SSGSPY | 1xPhosphc | Q3UZG3 1x | 8.16E-11 | 0 | 1 | 10 | 57 | Q3UZG3 | Putative ur | Q3UZG3 [3 | 0 | 2219.919 | 0 | -0.39 | 0.002475 | 4.89 | 116.2 | 83.8 | 2.76 | 7.26 | | High | High | High | High | High | High | High | High | High | High | High | 0 | 1.61E-11 | 5.36 |
| FALSE | High | SSGSPYGG | [R].SSGSPY | 2xPhosphc | Q3UZG3 2x | 8.63E-08 | 0 | 1 | 10 | 6 | Q3UZG3 | Putative ur | Q3UZG3 [3 | 0 | 2299.886 | 0 | -0.69 | 0.000749 | 2.97 | 124.4 | 75.6 | 2.68 | 7.14 | | High | High | High | High | High | High | High | High | High | High | High | 0 | 1.16E-05 | 4.74 |
| FALSE | High | SSGSSSSGI | [R].SSGSSS | 1xPhosphc | Q80TJ7 1xl | 3.66E-08 | 0 | 1 | 3 | 2 | Q80TJ7 | Histone lys | Q80TJ7 [75 | 0 | 2134.982 | 0 | 0.02 | 0.606903 | 5.08 | 99.4 | 100.6 | 3.32 | 2.33 | | High | High | High | High | High | High | High | High | High | High | High | 0 | 2.47E-09 | 3.9 |
| FALSE | High | SSGSTPVT | [K].SSGSTP | 1xPhosphc | E9PVX6 1xl | 0.005717 | 0.00062 | 1 | 2 | 2 | E9PVX6 | Proliferatic | E9PVX6 [2 | 0 | 1585.767 | 0 | 0.35 | 0.782726 | 36.71 | 87.8 | 112.2 | 50.26 | 34.64 | | High | High | High | High | High | High | High | High | High | High | High | 0.000275 | 0.001684 | 2.46 |
| FALSE | High | SSGSTPVT | [K].SSGSTP | 1xPhosphc | E9PVX6 1xl | 4.25E-06 | 0 | 1 | 2 | 1 | E9PVX6 | Proliferatic | E9PVX6 [2 | 0 | 1814.93 | 0 | -0.11 | 0.98349 | 20.27 | 99.9 | 100.1 | 5.57 | 17.37 | | High | High | High | High | High | High | High | High | High | High | High | 0 | 5.15E-07 | 2.04 |
| FALSE | High | SSGSTPVT | [K].SSGSTP | 2xPhosphc | E9PVX6 2xl | 5.91E-05 | 0 | 1 | 2 | 2 | E9PVX6 | Proliferatic | E9PVX6 [2 | 0 | 1894.896 | 0 | -0.15 | 0.210499 | 5.26 | 101.9 | 98.1 | 6.15 | 5.2 | | High | High | High | High | High | High | High | High | High | High | High | 0 | 9.97E-06 | 4.49 |
| FALSE | High | SSGTAPAP | [R].SSGTAF | 1xPhosphc | Q9QYY8 1x | 2.41E-06 | 0 | 1 | 2 | 1 | Q9QYY8 | Spastin OS | Q9QYY8 [9 | 0 | 2698.256 | 0 | 1.18 | 0.000567 | 4.3 | 63.6 | 136.4 | 15.17 | 9.83 | | High | High | High | High | High | High | High | High | High | High | High | 0 | 2.72E-07 | 3.85 |
| FALSE | High | SSGSEGSSI | [K].SSGSEC | 1xPhosphc | B9EKJ7 1xP | 0.007673 | 0.000664 | 1 | 3 | 2 | B9EKJ7 | Tankyrase | B9EKJ7 [16 | 0 | 2186.089 | 0 | 0.01 | 0.456469 | 5.69 | 100.7 | 99.3 | 5.9 | 18.2 | | High | High | High | High | High | High | High | High | High | High | High | 0.000801 | 0.007176 | 3.15 |
| FALSE | High | SSGTASSV | [R].SSGTAS | 1xPhosphc | Q8CCF0 1x | 3.39E-06 | 0 | 1 | 2 | 1 | Q8CCF0 | U4/U6 sma | Q8CCF0 [4 | 0 | 3140.628 | 0 | -0.42 | 0.239735 | 2.26 | 108.9 | 91.1 | 9.91 | 15.54 | | High | High | High | High | High | High | High | High | High | High | High | 0 | 4.01E-07 | 4.05 |
| FALSE | High | SSGTFLSGI | [K].SSGTFL | 1xCarbami | Q6P5B0 1x | 1.11E-05 | 0 | 1 | 1 | 4 | Q6P5B0 | RRP12-like | Q6P5B0 [9 | 0 | 2546.225 | 0 | -0.07 | 0.853025 | 16.78 | 97.8 | 102.2 | 5.63 | 16.74 | | High | High | High | High | High | High | High | High | High | High | High | 0 | 2.74E-06 | 3.59 |
| FALSE | High | SSGTNFQG | [K].SSGTNI | 1xPhosphc | Q3UIA2 1x | 0.001968 | 5.66E-05 | 1 | 6 | 4 | Q3UIA2 | Rho GTPas | Q3UIA2 [1 | 0 | 1760.898 | 0 | 1.31 | 0.002728 | 21.63 | 59.2 | 140.8 | 9.28 | 14.18 | | High | High | High | High | High | High | High | High | High | High | High | 2.58E-05 | 0.00051 | 3.05 |
| FALSE | High | SSGTTPQG | [K].SSGTTF | 1xPhosphc | Q542Q3 1x | 0.001278 | 0 | 1 | 1 | 1 | Q542Q3 | G two S ph | Q542Q3 [4 | 0 | 1700.788 | 0 | 0.36 | 0.129259 | 31.89 | 89 | 111 | 9.43 | 20.89 | | High | High | High | High | High | High | High | High | High | High | High | 0 | 0.000313 | 2.16 |
| FALSE | High | SSGVSYK | [R].SSGVSY | 1xPhosphc | B7ZWC4 1x | 0.021989 | 0.001542 | 1 | 8 | 2 | B7ZWC4 | Insulin-like | B7ZWC4 [2 | 0 | 1265.654 | 0 | -0.19 | 0.037231 | 10.47 | 108.3 | 91.7 | 6.11 | 4.55 | | High | High | High | High | High | High | High | High | High | High | High | 0.000814 | 0.007617 | 1.05 |
| FALSE | High | SSGYGVYP | [R].SSGYG\ | 1xPhosphc | Q3U2Z5 1x | 1.38E-05 | 0 | 1 | 7 | 2 | Q3U2Z5 | Putative ur | Q3U2Z5 [3 | 0 | 3022.313 | 0 | 0.94 | 0.002229 | 12.8 | 66.3 | 133.7 | 11.17 | 6.18 | | High | High | High | High | High | High | High | High | High | High | High | 0 | 2E-06 | 4.28 |
| FALSE | High | SSGYGVYP | [R].SSGYG\ | 1xDeamida | Q3U2Z5 1x | 2.34E-05 | 0 | 1 | 7 | 6 | Q3U2Z5 | Putative ur | Q3U2Z5 [3 | 0 | 3023.297 | 0 | 1.03 | 0.002655 | 17.78 | 65.7 | 134.3 | 11.13 | 10.83 | | High | High | High | High | High | High | High | High | High | High | High | 0 | 4.87E-05 | 3.31 |
| FALSE | High | SSHCDSPPI | [K].SSHCDS | 1xCarbami | O55003 1x | 0.065405 | 0.004734 | 1 | 2 | 2 | O55003 | BCL2/aden | O55003 [5 | 0 | 1351.566 | 0 | -0.64 | 0.012625 | 6.02 | 120.2 | 79.8 | 5.54 | 15.06 | | High | High | High | High | High | High | High | High | High | High | High | 0.004193 | 0.04512 | 2.7 |
| FALSE | High | SSHDLLK | [K].SSHDLL | 1xPhosphc | Q3TKY6 1x | 0.050826 | 0.003676 | 1 | 1 | 1 | Q3TKY6 | Peptidyl-pr | Q3TKY6 [2 | 0 | 1337.723 | 0 | -0.44 | 0.325169 | 24.62 | 115.2 | 84.8 | 8.67 | 32.57 | | High | High | High | High | High | High | High | High | High | High | High | 0.001948 | 0.01971 | 1.5 |
| FALSE | High | SSHETLNV | [K].SSHETL | 1xPhosphc | Q80UE5 1x | 0.000349 | 0 | 1 | 8 | 4 | Q80UE5 | Band 4.1-li | Q80UE5 [5 | 0 | 1909.967 | 0 | -0.76 | 0.044662 | 9.05 | 125.7 | 74.3 | 21.49 | 16 | | High | High | High | High | High | High | High | High | High | High | High | 0.000756 | 0.006588 | 2.83 |
| FALSE | High | SSHSGPAE | [R].SSHSGF | 1xPhosphc | Q14AX9 1x | 8.15E-07 | 0 | 1 | 2 | 3 | Q14AX9 | Mannose r | Q14AX9 [1 | 0 | 1738.841 | 0 | 0.07 | 0.220563 | 6.41 | 100.8 | 99.2 | 6.67 | 11.82 | | High | High | High | High | High | High | High | High | High | High | High | 0 | 8.1E-08 | 4.01 |
| FALSE | High | SSHSICNVC | [K].SSHSICI | 1xCarbami | Q9JMB0 1x | 0.11108 | 0.008991 | 1 | 1 | 1 | Q9JMB0 | G kinase-a | Q9JMB0 [8 | 0 | 2758.338 | 0 | 0.66 | 0.001682 | 8.6 | 77.4 | 122.6 | 5.96 | 4.18 | | High | High | High | High | High | High | High | High | High | High | High | 0.004503 | 0.04879 | 2.03 |
| FALSE | High | SSLAETLDS | [K].SSLAET | 1xPhosphc | Q9EPR4 1x | 1.71E-06 | 0 | 1 | 3 | 2 | Q9EPR4 | Solute carr | Q9EPR4 [7 | 0 | 2085.99 | 0 | 1.35 | 0.001992 | 11.59 | 56.5 | 143.5 | 8.91 | 13.22 | | High | High | High | High | High | High | High | High | High | High | High | 0 | 1.86E-07 | 3.8 |
| FALSE | High | SSLAETLDS | [K].SSLAET | 2xPhosphc | Q9EPR4 2x | 9.65E-06 | 0 | 1 | 3 | 3 | Q9EPR4 | Solute carr | Q9EPR4 [7 | 0 | 2165.957 | 0 | 1.38 | 0.000798 | 2.03 | 58.7 | 141.3 | 8.27 | 10.33 | | High | High | High | High | High | High | High | High | High | High | High | 0 | 1.3E-06 | 3.76 |
| FALSE | High | SSGTFCYCI | [R].SSGTFC | 2xCarbami | Q3UID6 2x | 0.004256 | 0.00036 | 1 | 5 | 1 | Q3UID6 | Putative ur | Q3UID6 [5 | 0 | 3468.203 | 0 | -0.12 | 0.842503 | 29.31 | 94.1 | 105.9 | 10.5 | 23.41 | | High | High | High | High | High | High | High | High | High | High | High | 0.000161 | 0.001212 | 3.38 |
| FALSE | High | SSLAETLDS | [K].SSLAET | 3xPhosphc | Q9EPR4 3x | 0.001916 | 5.66E-05 | 1 | 3 | 1 | Q9EPR4 | Solute carr | Q9EPR4 [7 | 0 | 2245.923 | 0 | 1.44 | 0.002159 | 40.74 | 54.1 | 145.9 | 30.7 | 18.14 | | High | High | High | High | High | High | High | High | High | High | High | 2.58E-05 | 0.000496 | 2.75 |
| FALSE | High | SSGSAEQSI | [R].SSGSAE | 1xPhosphc | Q9D2D7 1x | 0.052576 | 0.00384 | 1 | 1 | 1 | Q9D2D7 | Zinc finger | Q9D2D7 [1 | 0 | 1599.794 | 0 | 0.25 | 0.131519 | 6.36 | 92.1 | 107.9 | 5.97 | 25.22 | | High | High | High | High | High | High | High | High | High | High | High | 0.001999 | 0.02044 | 2.58 |
| FALSE | High | SSGRNSPS | [R].SSGRN | 1xPhosphc | Q80TZ9 1x | 0.00129 | 0 | 1 | 2 | 2 | Q80TZ9 | Arginine-gl | Q80TZ9 [6 | 1 | 2278.071 | 0 | 0.31 | 0.002152 | 1.83 | 90.1 | 109.9 | 2.55 | 2.23 | | High | High | High | High | High | High | High | High | High | High | High | 2.58E-05 | 0.00034 | 3.88 |
| FALSE | High | SSFSMEEG | [R].SSFSMI | 1xPhosphc | Q3UKG7 1x | 0.059657 | 0.004365 | 1 | 6 | 1 | Q3UKG7 | Putative ur | Q3UKG7 [5 | 0 | 1509.638 | 0 | -0.96 | 0.001401 | 1.9 | 132.3 | 67.7 | 7.22 | 6.07 | | High | High | High | High | High | High | High | High | High | High | High | 0.002249 | 0.02357 | 1.74 |
| FALSE | High | SSFSMEEG | [R].SSFSMI | 1xPhosphc | Q3UKG7 1x | 0.000219 | 0 | 1 | 6 | 2 | Q3UKG7 | Putative ur | Q3UKG7 [5 | 0 | 1525.633 | 0 | -1.35 | 0.040348 | 6.49 | 143.7 | 56.3 | 11.96 | 44.41 | | High | High | High | High | High | High | High | High | High | High | High | 0 | 4.31E-05 | 2.07 |
| FALSE | High | SSFSNSADI | [K].SSFSNS | 1xPhosphc | Q61686 1x | 0.000316 | 0 | 1 | 1 | 4 | Q61686 | Chromobo | Q61686 [9 | 0 | 1479.657 | 0 | -0.37 | 0.736168 | 28.07 | 109.9 | 90.1 | 70.41 | 64.93 | | High | High | High | High | High | High | High | High | High | High | High | 0 | 6.58E-05 | 2.48 |
| FALSE | High | SSFSNSADI | [K].SSFSNS | 1xPhosphc | Q61686 1x | 3.9E-05 | 0 | 1 | 1 | 6 | Q61686 | Chromobo | Q61686 [9 | 0 | 1708.82 | 0 | -0.58 | 0.000556 | 0.96 | 119.1 | 80.9 | 3.87 | 6.01 | | High | High | High | High | High | High | High | High | High | High | High | 0 | 1.4E-05 | 3.6 |
| FALSE | High | SSFSNSADI | [K].SSFSNS | 1xDeamida | Q61686 1x | 1.48E-06 | 0 | 1 | 1 | 4 | Q61686 | Chromobo | Q61686 [9 | 0 | 1709.804 | 0 | -0.47 | 0.011785 | 7.19 | 116.1 | 83.9 | 7.97 | 6.78 | | High | High | High | High | High | High | High | High | High | High | High | 0 | 1.57E-07 | 3.41 |
| FALSE | High | SSFSNSADI | [K].SSFSNS | 2xPhosphc | Q61686 2x | 0.006957 | 0.00062 | 1 | 1 | 2 | Q61686 | Chromobo | Q61686 [9 | 0 | 1788.786 | 0 | -1.01 | 0.000333 | 2.22 | 133.1 | 66.9 | 4.12 | 6.24 | | High | High | High | High | High | High | High | High | High | High | High | 0.000275 | 0.002094 | 3.16 |
| FALSE | High | SSFSQHAR | [R].SSFSQH | 1xPhosphc | P48678 1xl | 0.013426 | 0.000976 | 1 | 3 | 1 | P48678 | Prelamin-A | P48678 [42 | 0 | 1228.567 | 0 | 0.65 | 0.087035 | 27.99 | 81.1 | 118.9 | 14.13 | 18.61 | | High | High | High | High | High | High | High | High | High | High | High | 0.000505 | 0.004357 | 1.29 |
| FALSE | High | SSGACDISC | [R].SSGACI | 1xCarbami | G3UWZ0 1 | 5.23E-06 | 0 | 1 | 4 | 1 | G3UWZ0 | Bromodom | G3UWZ0 [ | 0 | 1807.755 | 0 | 1.72 | 0.000896 | 16.61 | 45.5 | 154.5 | 10.13 | 9.79 | | High | High | High | High | High | High | High | High | High | High | High | 0 | 6.51E-07 | 3.26 |
| FALSE | High | SSGACDISC | [R].SSGACI | 1xCarbami | G3UWZ0 1 | 0.077825 | 0.005963 | 1 | 4 | 1 | G3UWZ0 | Bromodom | G3UWZ0 [ | 0 | 1823.75 | 0 | 1.77 | 0.001481 | 23.27 | 39.4 | 160.6 | 27.8 | 12.8 | | High | High | High | High | High | High | High | High | High | High | High | 0.003079 | 0.03203 | 2.05 |
| FALSE | High | SSGADGSL | [K].SSGAD( | 1xPhosphc | P97287 1xl | 1.48E-06 | 0 | 1 | 2 | 8 | P97287 | Induced m | P97287 [13 | 0 | 2854.251 | 0 | -0.43 | 0.009977 | 4.37 | 113.3 | 86.7 | 7.17 | 6.41 | | High | High | High | High | High | High | High | High | High | High | High | 0 | 6.81E-06 | 4.24 |
| FALSE | High | SSGAGQGS | [K].SSGAG( | 1xPhosphc | Q5XJV7-2 | 1.77E-05 | 0 | 1 | 3 | 2 | Q5XJV7-2 | Isoform 2 c | Q5XJV7-2 [ | 0 | 3521.629 | 0 | -0.58 | 0.088548 | 17.82 | 130.7 | 69.3 | 25.69 | 26.86 | | High | High | High | High | High | High | High | High | High | High | High | 0 | 2.57E-06 | 4.54 |
| FALSE | High | SSGALSVDI | [R].SSGALS | 1xPhosphc | P27546 1xl | 0.007339 | 0.00062 | 1 | 10 | 1 | P27546 | Microtubu | P27546 [87 | 0 | 1401.739 | 0 | 1.44 | 0.00066 | 5.75 | 53.8 | 146.2 | 7.39 | 8.82 | | High | High | High | High | High | High | High | High | High | High | High | 0.000275 | 0.002229 | 1.85 |
| FALSE | High | SSGDGTEW | [R].SSGDG | 1xPhosphc | B2RQS1 1x | 1.69E-06 | 0 | 1 | 2 | 2 | B2RQS1 | Striatin-3 C | B2RQS1 [3 | 0 | 2518.19 | 0 | 0.58 | 0.00646 | 12.94 | 79.1 | 120.9 | 12.14 | 9.64 | | High | High | High | High | High | High | High | High | High | High | High | 0 | 1.83E-07 | 2.97 |
| FALSE | High | SSGSADSTS | [R].SSGSAD | 1xPhosphc | E9Q286 1x | 0.008313 | 0.000736 | 1 | 1 | 1 | E9Q286 | Little elong | E9Q286 [7 | 0 | 1586.68 | 0 | -0.53 | 0.030551 | 7.28 | 114.1 | 85.9 | 10.62 | 16.35 | | High | High | High | High | High | High | High | High | High | High | High | 0.000327 | 0.002561 | 2.81 |
| FALSE | High | SSGDVHLP | [K].SSGDVI | 1xPhosphc | Q570Z8 1x | 0.000235 | 0 | 1 | 5 | 2 | Q570Z8 | MKIAA411 | Q570Z8 [4 | 0 | 2299.117 | 0 | 0.68 | 0.004784 | 13.29 | 74 | 126 | 5.79 | 11.5 | | High | High | High | High | High | High | High | High | High | High | High | 7.19E-05 | 0.000809 | 2.88 |
| FALSE | High | SSGGEDLSI | [K].SSGGEI | 1xPhosphc | B2RY61 1xl | 6.73E-06 | 0 | 1 | 4 | 2 | B2RY61 | AT rich inte | B2RY61 [7 | 0 | 1728.846 | 0 | -0.22 | 0.029236 | 9.75 | 106.1 | 93.9 | 4.89 | 9.42 | | High | High | High | High | High | High | High | High | High | High | High | 0 | 6.38E-06 | 2.43 |

| | | | | | | | | | | | | | | | | | | | | | | | | | | | | | | | | | | | | | | |
|---|---|---|---|---|---|---|---|---|---|---|---|---|---|---|---|---|---|---|---|---|---|---|---|---|---|---|---|---|---|---|---|---|---|---|---|---|---|---|
| FALSE | High | SSGGGSEA | [R].SSGGG | 1xPhospho | Q9QXY1 1x | 2.18E-06 | 0 | 1 | 3 | 1 | Q9QXY1 | Tight junct | Q9QXY1 [1 | 0 | 2236.09 | 0 | -1.49 | 0.000927 | 2.1 | 147.5 | 52.5 | 5.43 | 12.86 | | High | High | High | High | High | High | High | High | High | High | High | 0 | 2.43E-07 | 3.62 |
| FALSE | High | SSGGGSEA | [R].SSGGG | 1xDeamida | Q9QXY1 1x | 2.31E-05 | 0 | 1 | 3 | 1 | Q9QXY1 | Tight junct | Q9QXY1 [1 | 0 | 2237.074 | 0 | -0.98 | 0.001142 | 6.29 | 132.8 | 67.2 | 8.48 | 18.51 | | High | High | High | High | High | High | High | High | High | High | High | 0 | 3.48E-06 | 3.8 |
| FALSE | High | SSGLQPQG | [R].SSGLQI | 1xPhospho | E9Q4K7 1x | 1.28E-05 | 0 | 1 | 2 | 2 | E9Q4K7 | Kinesin fan | E9Q4K7 [1 | 0 | 1634.81 | 0 | 0.19 | 0.008662 | 6.27 | 94.1 | 105.9 | 1.45 | 5.82 | | High | High | High | High | High | High | High | High | High | High | High | 0 | 1.78E-06 | 3.49 |
| FALSE | High | SSGLQTPE | [R].SSGLQT | 1xCarbami | P42128 1xl | 0.000584 | 0 | 1 | 1 | 2 | P42128 | Forkhead b | P42128 [41 | 0 | 1643.766 | 0 | 1.35 | 0.000199 | 6.52 | 56.3 | 143.7 | 6.63 | 3.39 | | High | High | High | High | High | High | High | High | High | High | High | 2.58E-05 | 0.000341 | 2.75 |
| FALSE | High | SSGLQTPE | [R].SSGLQT | 1xCarbami | P42128 2xl | 0.100129 | 0.008067 | 1 | 1 | 1 | P42128 | Forkhead b | P42128 [41 | 0 | 1723.732 | 0 | 1.61 | 0.000574 | 6.84 | 50.5 | 149.5 | 11.62 | 10.94 | | High | High | High | High | High | High | High | High | High | High | High | 0.004046 | 0.04318 | 2.07 |
| FALSE | High | SSGIQVGE | [R].SSGIQV | 1xPhospho | A2AH22 1x | 6.82E-08 | 0 | 1 | 4 | 2 | A2AH22 | Activating | A2AH22 [2 | 0 | 3720.807 | 0 | 0.68 | 0.196321 | 31.01 | 84.7 | 115.3 | 33.63 | 20.76 | | High | High | High | High | High | High | High | High | High | High | High | 0 | 4.96E-09 | 5.48 |
| FALSE | High | SSGLVQR | [R].SSGLVC | 1xPhospho | Q9CRA5 1x | 0.112856 | 0.009146 | 1 | 4 | 1 | Q9CRA5 | Golgi phos | Q9CRA5 [8 | 0 | 1055.545 | 0 | 0.72 | 0.019018 | 24.21 | 73.2 | 126.8 | 17.67 | 8.66 | | High | High | High | High | High | High | High | High | High | High | High | 0.004583 | 0.04975 | 1.99 |
| FALSE | High | SSGIWEHG | [R].SSGIWI | 1xPhospho | E9Q3P4 1x | 0.013666 | 0.001011 | 1 | 2 | 3 | E9Q3P4 | Centromer | E9Q3P4 [2 | 0 | 1538.777 | 0 | -0.97 | 0.000464 | 2.79 | 133.5 | 66.5 | 2.94 | 9.49 | | High | High | High | High | High | High | High | High | High | High | High | 0.001665 | 0.01662 | 2.5 |
| FALSE | High | SSGPALTES | [R].SSGPAL | 1xPhospho | Q14AX6 1x | 1.7E-08 | 0 | 2 | 2 | 2 | Q14AX6; Q | Cyclin-dep | Q14AX6 [1 | 0 | 2152.167 | 0 | -0.73 | 0.000536 | 2.7 | 124.4 | 75.6 | 6.38 | 6.96 | Shared | High | High | High | High | High | High | High | High | High | High | High | 0 | 1.1E-07 | 4.4 |
| FALSE | High | SSGPPPPS | [K].SSGPPF | 1xPhospho | Q60865 1x | 5.62E-06 | 0 | 1 | 3 | 7 | Q60865 | Caprin-1 O | Q60865 [1 | 0 | 4129.954 | 0 | -0.62 | 0.055523 | 1.14 | 118.2 | 81.8 | 7.99 | 18.38 | | High | High | High | High | High | High | High | High | High | High | High | 0 | 7.07E-07 | 4.86 |
| FALSE | High | SSGPQRPH | [R].SSGPQI | 1xPhospho | Q80UW5 1 | 0.102567 | 0.008249 | 1 | 2 | 1 | Q80UW5 | Serine/thre | Q80UW5 [ | 0 | 1998.938 | 0 | 0.52 | 0.032671 | 26 | 82.3 | 117.7 | 8.37 | 15.39 | | High | High | High | High | High | High | High | High | High | High | High | 0.004138 | 0.04442 | 1.62 |
| FALSE | High | SSGPTSLFA | [K].SSGPTS | 1xPhospho | Q3TXW2 1 | 0.013545 | 0.000976 | 1 | 8 | 4 | Q3TXW2 | Putative ur | Q3TXW2 [ | 0 | 2024.042 | 0 | -0.15 | 0.104379 | 12.72 | 107.3 | 92.7 | 8.3 | 16.07 | | High | High | High | High | High | High | High | High | High | High | High | 0.00074 | 0.006437 | 3.11 |
| FALSE | High | SSGFSGK | [K].SSGFSG | 1xPhospho | Q569Z5 1x | 0.038677 | 0.002563 | 1 | 3 | 1 | Q569Z5 | Probable A | Q569Z5 [7 | 0 | 1207.612 | 0 | -0.49 | 0.0119 | 3.06 | 116.5 | 83.5 | 9.66 | 4.62 | | High | High | High | High | High | High | High | High | High | High | High | 0.001372 | 0.01432 | 1.64 |
| FALSE | High | SSLAGSDG | [K].SSLAGS | 1xPhospho | Q64511 1x | 0.017344 | 0.001204 | 1 | 3 | 2 | Q64511 | DNA topois | Q64511 [4 | 0 | 2330.179 | 0 | -0.17 | 0.017537 | 1.6 | 105.7 | 94.3 | 7.23 | 12.39 | | High | High | High | High | High | High | High | High | High | High | High | 0.000723 | 0.006267 | 2.93 |
| FALSE | High | SSLAIEDGK | [R].SSLAIEI | 1xPhospho | Q4VAC9 1x | 0.000548 | 0 | 1 | 3 | 1 | Q4VAC9 | Pleckstrin I | Q4VAC9 [7 | 0 | 1457.765 | 0 | 1.11 | 0.005969 | 11.6 | 63.4 | 136.6 | 20.12 | 8.59 | | High | High | High | High | High | High | High | High | High | High | High | 0 | 0.000122 | 1.68 |
| FALSE | High | SSLDDLNSI | [K].SSLDDL | 1xCarbami | G3UYN0 1x | 3.43E-07 | 0 | 1 | 2 | 2 | G3UYN0 | L1TD1 OS= | G3UYN0 [6 | 0 | 2073.973 | 0 | -1.45 | 0.001448 | 3.24 | 146.8 | 53.2 | 12.98 | 17.47 | | High | High | High | High | High | High | High | High | High | High | High | 0 | 3.06E-08 | 4.59 |
| FALSE | High | SSIQEDGVS | [K].SSIQED | 1xPhospho | Q3UFY8 1x | 5.75E-05 | 0 | 1 | 1 | 1 | Q3UFY8 | Mitochond | Q3UFY8 [6 | 1 | 2763.241 | 0 | 0.79 | 0.547553 | 44.18 | 73.4 | 126.6 | 59.36 | 55.01 | | High | High | High | High | High | High | High | High | High | High | High | 0 | 9.65E-06 | 3.49 |
| FALSE | High | SSIQEDGVS | [K].SSIQED | 1xPhospho | Q3UFY8 1x | 9.74E-06 | 0 | 1 | 1 | 2 | Q3UFY8 | Mitochond | Q3UFY8 [6 | 1 | 2992.404 | 0 | 0.63 | 0.003002 | 8.9 | 80.3 | 119.7 | 6.47 | 9.18 | | High | High | High | High | High | High | High | High | High | High | High | 0 | 1.31E-06 | 4.7 |
| FALSE | High | SSLQESSSP | [K].SSLQES | 1xPhospho | B9EKI3 1xP | 5.31E-05 | 0 | 1 | 2 | 1 | B9EKI3 | TATA elem | B9EKI3 [13 | 0 | 1658.758 | 0 | 1.16 | 0.003393 | 14.42 | 62.3 | 137.7 | 5.64 | 14.01 | | High | High | High | High | High | High | High | High | High | High | High | 0 | 8.82E-06 | 2.75 |
| FALSE | High | SSLQSMFS | [R].SSLQSM | 1xPhospho | F8VPX2 1xl | 0.000184 | 0 | 1 | 1 | 1 | F8VPX2 | F-box and ' | F8VPX2 [5C | 0 | 2004.966 | 0 | 0.44 | 0.028391 | 20.66 | 82.1 | 117.9 | 9.36 | 15.47 | | High | High | High | High | High | High | High | High | High | High | High | 0 | 3.56E-05 | 2.4 |
| FALSE | High | SSLRGEAEI | [R].SSLRGE | 1xPhospho | P70268-2 1 | 5.63E-07 | 0 | 1 | 3 | 2 | P70268-2 | Isoform 2 c | P70268-2 [ | 1 | 2429.269 | 0 | 1.17 | 0.000484 | 6.51 | 61.8 | 138.2 | 10.27 | 3.42 | | High | High | High | High | High | High | High | High | High | High | High | 0 | 5.32E-08 | 3.46 |
| FALSE | High | SSLSALAPA | [R].SSLSAL | 1xDeamida | Q6J1H4 1xl | 8.63E-05 | 0 | 1 | 2 | 5 | Q6J1H4 | Undifferen | Q6J1H4 [1 | 0 | 3634.809 | 0 | -3.32 | | 0.56 | 182.1 | 17.9 | 26.17 | 23.46 | | High | High | High | High | High | High | High | High | High | High | High | 0 | 1.53E-05 | 4.07 |
| FALSE | High | SSISAQPPC | [R].SSISAQ | 1xCarbami | A2A9C3 1x | 0.004143 | 0.00036 | 1 | 2 | 2 | A2A9C3 | KICSTOR co | A2A9C3 [1 | 0 | 2096.957 | 0 | 1.84 | 0.002665 | 17.35 | 44 | 156 | 6.98 | 15 | | High | High | High | High | High | High | High | High | High | High | High | 0.001388 | 0.01443 | 2 |
| FALSE | High | SSLSASHPN | [R].SSLSAS | 1xPhospho | A5D6P2 1x | 0.002818 | 0.000109 | 1 | 18 | 1 | A5D6P2 | Par-3 (Part | A5D6P2 [2 | 0 | 1611.74 | 0 | -0.76 | 0.026966 | 2.98 | 123.7 | 76.3 | 8.01 | 23.23 | | High | High | High | High | High | High | High | High | High | High | High | 7.19E-05 | 0.000765 | 2.54 |
| FALSE | High | SSLSDSPLN | [R].SSLSDS | 1xPhospho | Q5PPR1 1x | 0.001558 | 5.66E-05 | 1 | 5 | 1 | Q5PPR1 | Ccnc prote | Q5PPR1 [3 | 0 | 1897.893 | 0 | 0.25 | 0.028779 | 10.42 | 89.5 | 110.5 | 6.24 | 7.66 | | High | High | High | High | High | High | High | High | High | High | High | 2.58E-05 | 0.000391 | 2.92 |
| FALSE | High | SSLSETPPG | [K].SSLSETI | 1xCarbami | Q14B71 1x | 1.41E-05 | 0 | 1 | 4 | 4 | Q14B71 | Cell divisio | Q14B71 [4 | 0 | 2418.162 | 0 | -0.69 | 0.000932 | 2 | 123.1 | 76.9 | 3.43 | 6.06 | | High | High | High | High | High | High | High | High | High | High | High | 0 | 2.12E-05 | 3.88 |
| FALSE | High | SSISGTEEE | [R].SSISGTI | 1xPhospho | G5E8W8 1 | 0.001087 | 0 | 2 | 7 | 6 | G5E8W8; A | Zinc finger | G5E8W8 [1 | 0 | 2647.243 | 0 | -1.39 | 0.000241 | 1.13 | 142.9 | 57.1 | 3.42 | 10.48 | Shared | High | High | High | High | High | High | High | High | High | High | High | 0 | 0.000263 | 3.65 |
| FALSE | High | SSLSPGR | [R].SSLSPG | 2xPhospho | D3Z0M9 2x | 0.097743 | 0.007804 | 1 | 1 | 1 | D3Z0M9 | DEAD (Asp | D3Z0M9 [1 | 0 | 1092.469 | 0 | 0.41 | 0.034129 | 18.05 | 87.3 | 112.7 | 9.87 | 12.09 | | High | High | High | High | High | High | High | High | High | High | High | 0.00395 | 0.04196 | 1.45 |
| FALSE | High | SSLSSHSHC | [R].SSLSSH | 1xPhospho | A2AIX1 1xP | 0.00172 | 5.66E-05 | 1 | 2 | 3 | A2AIX1 | SEC16 hor | A2AIX1 [13 | 0 | 1925.907 | 0 | 0.28 | 0.401429 | 44.9 | 85.3 | 114.7 | 29.74 | 21.82 | | High | High | High | High | High | High | High | High | High | High | High | 0.001812 | 0.01784 | 2.9 |
| FALSE | High | SSIQEDGVS | [K].SSIQED | 1xPhospho | Q3UFY8 1x | 7.54E-05 | 0 | 1 | 1 | 1 | Q3UFY8 | Mitochond | Q3UFY8 [6 | 0 | 2017.973 | 0 | -0.09 | 0.688871 | 14.3 | 99.1 | 100.9 | 20.85 | 13.19 | | High | High | High | High | High | High | High | High | High | High | High | 0 | 1.31E-05 | 2.92 |
| FALSE | High | SSISSISGR | [K].SSISSIS | 1xPhospho | Q8K1N2-2 | 0.04913 | 0.003576 | 1 | 3 | 2 | Q8K1N2-2 | Isoform 2 c | Q8K1N2-2 | 0 | 1202.598 | 0 | 2.26 | 0.000356 | 4.14 | 31.7 | 168.3 | 8.76 | 8.05 | | High | High | High | High | High | High | High | High | High | High | High | 0.001889 | 0.01884 | 0.98 |
| FALSE | High | SSITEPEGP | [R].SSITEPI | 1xPhospho | Q62415 1x | 3.02E-07 | 0 | 1 | 2 | 4 | Q62415 | Apoptosis- | Q62415 [7 | 0 | 2149.094 | 0 | -0.97 | 0.001522 | 3.5 | 132.4 | 67.6 | 4.16 | 10.58 | | High | High | High | High | High | High | High | High | High | High | High | 0 | 1.17E-06 | 3.91 |
| FALSE | High | SSITEPEGP | [R].SSITEPI | 1xPhospho | E9QJU8 1x | 0.000114 | 0 | 1 | 2 | 2 | E9QJU8 | Transform | E9QJU8 [7 | 0 | 2206.116 | 0 | -0.28 | 0.036713 | 10.81 | 109.6 | 90.4 | 3.66 | 9.22 | | High | High | High | High | High | High | High | High | High | High | High | 0 | 2.09E-05 | 3.27 |
| FALSE | High | SSITPGTVL | [R].SSITPG | 1xPhospho | P47911 1xl | 0.000651 | 0 | 1 | 3 | 3 | P47911 | 60S riboso | P47911 [15 | 0 | 1837.04 | 0 | -0.06 | 0.777362 | 12.29 | 101.6 | 98.4 | 6.82 | 16.24 | | High | High | High | High | High | High | High | High | High | High | High | 7.19E-05 | 0.000926 | 2.94 |
| FALSE | High | SSITSAASSI | [R].SSITSA | 1xPhospho | F2Z3U3 1xl | 0.001704 | 5.66E-05 | 1 | 3 | 3 | F2Z3U3 | Ras associa | F2Z3U3 [2 | 0 | 2282.088 | 0 | 0.14 | 0.245767 | 32.42 | 98.1 | 101.9 | 6.96 | 28.72 | | High | High | High | High | High | High | High | High | High | High | High | 2.58E-05 | 0.000436 | 2.59 |
| FALSE | High | SSLVANGE | [R].SSLVAN | 1xDeamida | Q810L3 1xl | 0.000346 | 0 | 1 | 5 | 2 | Q810L3 | E3 ubiquiti | Q810L3 [2 | 0 | 2517.289 | 0 | -0.17 | 0.289167 | 8.5 | 110.8 | 89.2 | 17.34 | 9.74 | | High | High | High | High | High | High | High | High | High | High | High | 0 | 7.26E-05 | 3.28 |
| FALSE | High | SSLVDEAPI | [R].SSLVDE | 1xPhospho | Q9R0L6-2 | 2E-05 | 0 | 1 | 3 | 1 | Q9R0L6-2 | Isoform 2 c | Q9R0L6-2 | 0 | 2390.105 | 0 | 0.42 | 0.047695 | 29.64 | 82.7 | 117.3 | 12.36 | 21.16 | | High | High | High | High | High | High | High | High | High | High | High | 0 | 2.96E-06 | 4.01 |
| FALSE | High | SSLVNESEN | [K].SSLVNE | 1xPhospho | A1A549 1x | 0.000314 | 0 | 1 | 3 | 4 | A1A549 | Transcripti | A1A549 [6 | 0 | 2451.036 | 0 | -0.93 | 0.002761 | 7.35 | 130 | 70 | 6.52 | 17.28 | | High | High | High | High | High | High | High | High | High | High | High | 0 | 6.49E-05 | 3.35 |
| FALSE | High | SSLVNESET | [K].SSLVNE | 1xPhospho | E9QQ89 1x | 9.25E-07 | 0 | 1 | 16 | 5 | E9QQ89 | Transcripti | E9QQ89 [4 | 0 | 2579.094 | 0 | 0.18 | 0.044794 | 4.21 | 93.6 | 106.4 | 12.42 | 11.37 | | High | High | High | High | High | High | High | High | High | High | High | 0 | 4.13E-07 | 4.56 |
| FALSE | High | SSLVPADD | [R].SSLVPA | 1xPhospho | Q6PIU9 1xl | 0.037369 | 0.00249 | 1 | 1 | 1 | Q6PIU9 | Uncharacte | Q6PIU9 [13 | 0 | 2031.045 | 0 | 0.15 | 0.535505 | 18.43 | 94.6 | 105.4 | 8.12 | 16.51 | | High | High | High | High | High | High | High | High | High | High | High | 0.001308 | 0.01378 | 2.78 |
| FALSE | High | SSLVTSK | [K].SSLVTS | 1xPhospho | P60840 1xl | 0.017192 | 0.001204 | 1 | 2 | 2 | P60840 | Alpha-endc | P60840 [1C | 0 | 1259.701 | 0 | -0.08 | 0.724765 | 30.6 | 107.2 | 92.8 | 18.07 | 11.72 | | High | High | High | High | High | High | High | High | High | High | High | 0.004462 | 0.04832 | 2.06 |
| FALSE | High | SSMSEQILI | [R].SSMSE( | 1xCarbami | Q8K3Z9 1x | 1E-06 | 0 | 1 | 2 | 3 | Q8K3Z9 | Nuclear en | Q8K3Z9 [2 | 0 | 3177.473 | 0 | -0.6 | 0.00491 | 4.71 | 119.1 | 80.9 | 7.42 | 11.65 | | High | High | High | High | High | High | High | High | High | High | High | 0 | 2.97E-07 | 3.6 |
| FALSE | High | SSMSSCGS | [R].SSMSS( | 2xCarbami | B2ZRS3 1xl | 0.003962 | 0.000311 | 1 | 6 | 3 | B2ZRS3 | Depdc6-00 | B2ZRS3 [27 | 0 | 3320.436 | 0 | 3.37 | 0.000698 | 46.96 | 17.8 | 182.2 | 45.27 | 12.36 | | High | High | High | High | High | High | High | High | High | High | High | 0.00014 | 0.00112 | 2.99 |
| FALSE | High | SSMSSCGS | [R].SSMSS( | 2xCarbami | B2ZRS3 1xl | 3.85E-06 | 0 | 1 | 6 | 6 | B2ZRS3 | Depdc6-00 | B2ZRS3 [27 | 0 | 3533.604 | 0 | 1.17 | 0.002981 | 11.43 | 61 | 139 | 13.11 | 11.24 | | High | High | High | High | High | High | High | High | High | High | High | 0 | 4.63E-07 | 4.45 |
| FALSE | High | SSLTDEEAT | [K].SSLTDE | 1xPhospho | B2RUG8 1x | 0.000193 | 0 | 1 | 5 | 2 | B2RUG8 | Asxl2 prote | B2RUG8 [5 | 0 | 2107.984 | 0 | 0.15 | 0.160238 | 13.83 | 94.7 | 105.3 | 8.55 | 12.92 | | High | High | High | High | High | High | High | High | High | High | High | 0 | 3.75E-05 | 3.12 |
| FALSE | High | SSLPSPMS | [R].SSLPSPI | 1xPhospho | A0A0N4SV | 0.040373 | 0.002982 | 1 | 9 | 1 | A0A0N4SV | Eukaryotic | A0A0N4SV | 0 | 2818.399 | 0 | 0.53 | 0.045798 | 27.25 | 82.8 | 117.2 | 16.98 | 15.95 | | High | High | High | High | High | High | High | High | High | High | High | 0.001566 | 0.01507 | 2.95 |
| FALSE | High | SSLPSPMS | [R].SSLPSPI | 1xPhospho | A0A0N4SV | 0.001476 | 5.66E-05 | 1 | 9 | 2 | A0A0N4SV | Eukaryotic | A0A0N4SV | 0 | 2802.404 | 0 | 1.1 | 0.023494 | 7.06 | 64.1 | 135.9 | 10.64 | 21.85 | | High | High | High | High | High | High | High | High | High | High | High | 2.58E-05 | 0.000371 | 4.07 |
| FALSE | High | SSLPPLVISI | [R].SSLPPL' | 1xPhospho | Q3TJN8 1x | 0.117405 | 0.009523 | 1 | 2 | 1 | Q3TJN8 | Putative ur | Q3TJN8 [8 | 0 | 2050.042 | 0 | 0.18 | 0.562407 | 25.15 | 93 | 107 | 9.18 | 13.71 | | High | High | High | High | High | High | High | High | High | High | High | 0.004782 | 0.05235 | 2.33 |
| FALSE | High | SSIDSQASL | [R].SSIDSQ | 1xPhospho | E9Q6B2 1x | 1.25E-07 | 0 | 1 | 1 | 3 | E9Q6B2 | Coiled-coil | E9Q6B2 [1 | 0 | 2356.134 | 0 | -0.08 | 0.666169 | 9.74 | 102.7 | 97.3 | 14.33 | 6.93 | | High | High | High | High | High | High | High | High | High | High | High | 0 | 3.38E-08 | 4.96 |
| FALSE | High | SSIDSQASL | [R].SSIDSQ | 2xPhospho | E9Q6B2 2x | 1.07E-05 | 0 | 1 | 1 | 1 | E9Q6B2 | Coiled-coil | E9Q6B2 [1 | 0 | 2436.101 | 0 | -0.58 | 0.002895 | 7.18 | 119.5 | 80.5 | 4.25 | 14.74 | | High | High | High | High | High | High | High | High | High | High | High | 0 | 1.45E-06 | 4.35 |
| FALSE | High | SSLELHSR | [R].SSLELH | 1xPhospho | G3X963 1x | 0.028327 | 0.001791 | 1 | 2 | 1 | G3X963 | ATPase fan | G3X963 [6 | 0 | 1237.614 | 0 | -0.25 | 0.188585 | 9.25 | 108.8 | 91.2 | 15.57 | 9.57 | | High | High | High | High | High | High | High | High | High | High | High | 0.000968 | 0.0101 | 1.49 |
| FALSE | High | SSLENNTPI | [R].SSLENN | 1xDeamida | Q8C4D5 1x | 0.102567 | 0.008249 | 1 | 1 | 1 | Q8C4D5 | Putative ur | Q8C4D5 [1 | 0 | 1528.766 | 0 | -0.21 | 0.162503 | 14.38 | 107.1 | 92.9 | 11.49 | 8.67 | | High | High | High | High | High | High | High | High | High | High | High | 0.004138 | 0.04427 | 1.76 |
| FALSE | High | SSLEQCSD | [R].SSLEQC | 1xCarbami | Q9JI57 1xP | 9.14E-06 | 0 | 1 | 16 | 1 | Q9JI57 | General tra | Q9JI57 [11 | 0 | 2208.102 | 0 | 0.08 | 0.188318 | 12.39 | 90.7 | 109.3 | 12.02 | 14.8 | | High | High | High | High | High | High | High | High | High | High | High | 0 | 1.23E-06 | 3.8 |
| FALSE | High | SSLEVLPSQ | [K].SSLEVL | 1xPhospho | O88904 1x | 2.13E-08 | 0 | 1 | 10 | 1 | O88904 | Homeodor | O88904 [8 | 0 | 2427.273 | 0 | 0.1 | 0.215077 | 9.02 | 96.5 | 103.5 | 7.41 | 8.89 | | High | High | High | High | High | High | High | High | High | High | High | 0 | 1.35E-09 | 3.74 |
| FALSE | High | SSIEYSQR | [R].SSIEYS( | 1xPhospho | P20263 1xl | 0.009843 | 0.000736 | 1 | 1 | 2 | P20263 | POU doma | P20263 [28 | 0 | 1278.593 | 0 | 0.3 | 0.060802 | 12.03 | 88.8 | 111.2 | 6.83 | 13.89 | | High | High | High | High | High | High | High | High | High | High | High | 0.000344 | 0.003072 | 2.49 |
| FALSE | High | SSLFPFEDA | [R].SSLFPFI | 1xPhospho | A0A0R4J1C | 7.65E-07 | 0 | 1 | 9 | 2 | A0A0R4J1C | Cytoplasmi | A0A0R4J1C | 0 | 3392.552 | 0 | -0.83 | 0.000953 | 5.06 | 123.6 | 76.4 | 7.77 | 12.55 | | High | High | High | High | High | High | High | High | High | High | High | 0 | 7.51E-08 | 5.78 |
| FALSE | High | SSLGESVTL | [K].SSLGES | 1xPhospho | Q5SYL3 1xl | 0.005565 | 0.00062 | 1 | 1 | 1 | Q5SYL3 | Protein KIA | Q5SYL3 [5 | 0 | 1858.013 | 0 | 0.72 | 0.004956 | 10.39 | 72.7 | 127.3 | 18.87 | 14.22 | | High | High | High | High | High | High | High | High | High | High | High | 0.000275 | 0.001628 | 2.7 |
| FALSE | High | SSLGLSYYP | [R].SSLGLS | 1xPhospho | A2AM95 1 | 2.06E-05 | 0 | 1 | 2 | 3 | A2AM95 | Emerin OS | A2AM95 [1 | 0 | 3294.525 | 0 | 0.39 | 0.034575 | 8.09 | 86.1 | 113.9 | 6.33 | 9.52 | | High | High | High | High | High | High | High | High | High | High | High | 0 | 3.05E-06 | 4.97 |
| FALSE | High | SSLGNDETI | [K].SSLGNI | 1xPhospho | B2RWU4 1 | 1.77E-05 | 0 | 2 | 11 | 5 | B2RWU4; ( | Cordon-ble | B2RWU4 [ | 0 | 1603.762 | 0 | -0.17 | 0.017981 | 4.9 | 106.1 | 93.9 | 4.09 | 1.81 | Shared | High | High | High | High | High | High | High | High | High | High | High | 0 | 5.39E-06 | 2.57 |
| FALSE | High | SSLGQSASI | [R].SSLGQS | 1xPhospho | Q3UAI4 1x | 1.8E-05 | 0 | 1 | 5 | 9 | Q3UAI4 | Putative ur | Q3UAI4 [2 | 0 | 2493.201 | 0 | -0.28 | 0.01642 | 4.12 | 110.6 | 89.4 | 4.62 | 6.37 | | High | High | High | High | High | High | High | High | High | High | High | 0 | 2.6E-06 | 3.21 |
| FALSE | High | SSLGQSASI | [R].SSLGQS | 2xPhospho | Q3UAI4 2x | 1.74E-06 | 0 | 1 | 5 | 13 | Q3UAI4 | Putative ur | Q3UAI4 [2 | 0 | 2573.167 | 0 | -0.73 | 0.000209 | 1.13 | 124.6 | 75.4 | 1.84 | 2.9 | | High | High | High | High | High | High | High | High | High | High | High | 0 | 1.89E-07 | 3.29 |
| FALSE | High | SSLGSHQLI | [R].SSLGSH | 1xPhospho | A6H663 1x | 0.00103 | 0 | 1 | 3 | 2 | A6H663 | BCL2-assoc | A6H663 [2 | 0 | 1390.704 | 0 | 1.47 | 0.000576 | 14.39 | 49.7 | 150.3 | 13.06 | 9.68 | | High | High | High | High | High | High | High | High | High | High | High | 0.001665 | 0.01675 | 1.29 |
| FALSE | High | SSLGWESL | [K].SSLGW | 1xPhospho | G5E8N7 1x | 0.000126 | 0 | 1 | 8 | 2 | G5E8N7 | Bifunction | G5E8N7 [1 | 0 | 1544.813 | 0 | 0.58 | 0.002525 | 13.56 | 82.2 | 117.8 | 6.42 | 4.27 | | High | High | High | High | High | High | High | High | High | High | High | 0 | 3.11E-05 | 2.48 |
| FALSE | High | SSIHNFMTI | [R].SSIHNF | 1xPhospho | G5E829 1x | 0.009332 | 0.000736 | 1 | 5 | 1 | G5E829 | Plasma me | G5E829 [1 | 0 | 1911.877 | 0 | 0.19 | 0.097023 | 18.07 | 88.9 | 111.1 | 6.86 | 12.4 | | High | High | High | High | High | High | High | High | High | High | High | 0.000327 | 0.002891 | 3.13 |
| FALSE | High | SSIHNFMTI | [R].SSIHNF | 1xPhospho | G5E829 1x | 0.004491 | 0.000409 | 1 | 5 | 3 | G5E829 | Plasma me | G5E829 [1 | 0 | 1927.872 | 0 | 0.71 | 0.002296 | 4.07 | 70.1 | 129.9 | 8.24 | 9.92 | | High | High | High | High | High | High | High | High | High | High | High | 0.005205 | 0.05656 | 2.4 |
| FALSE | High | SSIKTPESV | [K].SSIKTPI | 1xPhospho | E9QK89 1x | 7.1E-06 | 0 | 1 | 2 | 2 | E9QK89 | Mediator c | E9QK89 [1 | 1 | 3133.736 | 0 | -0.64 | 0.048343 | 12.35 | 120.1 | 79.9 | 13.58 | 21.57 | | High | High | High | High | High | High | High | High | High | High | High | 0 | 1.3E-06 | 3.79 |
| FALSE | High | SSLIETGQG | [R].SSLIET( | 1xPhospho | Q9DC04 1x | 0.028575 | 0.001829 | 1 | 3 | 1 | Q9DC04 | Regulator c | Q9DC04 [6 | 0 | 2017.979 | 0 | -0.15 | 0.437691 | 4.59 | 104.7 | 95.3 | 7.3 | 20.17 | | High | High | High | High | High | High | High | High | High | High | High | 0.000984 | 0.01017 | 1.18 |
| FALSE | High | SSLLFVK | [K].SSLLFVI | 1xPhospho | Q80ZV0 1x | 0.017809 | 0.001282 | 1 | 2 | 1 | Q80ZV0 | Ribonuclea | Q80ZV0 [3 | 0 | 1331.774 | 0 | -0.09 | 0.06718 | 4.57 | 104.3 | 95.7 | 8.13 | 4.13 | | High | High | High | High | High | High | High | High | High | High | High | 0.000689 | 0.005962 | 1.89 |
| FALSE | High | SSIIIVNR | [K].SSIIIVN | 1xPhospho | Q8BU33 1x | 0.008167 | 0.000736 | 1 | 6 | 1 | Q8BU33 | Acetolactat | Q8BU33 [3 | 0 | 1210.676 | 0 | 2.01 | 0.00179 | 26.27 | 38 | 162 | 16.4 | 16.48 | | High | High | High | High | High | High | High | High | High | High | High | 0.000327 | 0.002508 | 2.19 |
| FALSE | High | SSLISSLEEE | [K].SSLISSL | 1xPhospho | E9Q7G0 1x | 0.000699 | 0 | 1 | 1 | 3 | E9Q7G0 | Nuclear mi | E9Q7G0 [1 | 0 | 2085.068 | 0 | 3.57 | 0.001381 | 21.52 | 14.3 | 185.7 | 15.2 | 18.32 | | High | High | High | High | High | High | High | High | High | High | High | 2.58E-05 | 0.000371 | 2.93 |
| FALSE | High | SSLLTLK | [K].SSLLTLI | 1xPhospho | Q8C9G3 1x | 0.006895 | 0.00062 | 1 | 6 | 1 | Q8C9G3 | Putative ur | Q8C9G3 [3 | 0 | 1299.769 | 0 | -0.59 | 0.032358 | 14.82 | 120.3 | 79.7 | 16.18 | 7.28 | | High | High | High | High | High | High | High | High | High | High | High | 0.000275 | 0.002077 | 1.73 |
| FALSE | High | SSLNSSPW | [K].SSLNSS | 1xPhospho | A0A0R4J0D | 2.76E-07 | 0 | 1 | 2 | 4 | A0A0R4J0D | Dolichyl-di | A0A0R4J0D | 0 | 2173.031 | 0 | 0.03 | 0.409053 | 2.85 | 96.8 | 103.2 | 5.57 | 8.32 | | High | High | High | High | High | High | High | High | High | High | High | 0 | 2.4E-08 | 4.37 |
| FALSE | High | SSLNSSPW | [K].SSLNSS | 1xPhospho | A0A0R4J0D | 2.16E-06 | 0 | 1 | 2 | 2 | A0A0R4J0D | Dolichyl-di | A0A0R4J0D | 0 | 2189.026 | 0 | -0.05 | 0.853025 | 19.98 | 98.1 | 101.9 | 12.4 | 13.93 | | High | High | High | High | High | High | High | High | High | High | High | 0 | 2.42E-07 | 4.63 |
| FALSE | High | SSLPHSQEI | [K].SSLPHS | 1xPhospho | Q3U9K6 1x | 0.006594 | 0.00062 | 1 | 4 | 1 | Q3U9K6 | Interferon | Q3U9K6 [1 | 0 | 1874.978 | 0 | 1.15 | 0.001942 | 3.33 | 62.6 | 137.4 | 7.03 | 11.21 | | High | High | High | High | High | High | High | High | High | High | High | 0.000275 | 0.001964 | 1.61 |
| FALSE | High | SSLPHSQEI | [K].SSLPHS | 2xPhospho | Q3U9K6 2x | 4.51E-05 | 0 | 1 | 4 | 1 | Q3U9K6 | Interferon | Q3U9K6 [1 | 0 | 1954.944 | 0 | 1.84 | 0.001756 | 13.3 | 43.7 | 156.3 | 13.67 | 12.8 | | High | High | High | High | High | High | High | High | High | High | High | 0 | 7.35E-06 | 3.76 |
| FALSE | High | SSLPLELPIC | [R].SSLPLEI | 1xPhospho | A0A0J9YUS | 8.2E-06 | 0 | 1 | 1 | 1 | A0A0J9YUS | Protein fur | A0A0J9YUS | 0 | 2042.145 | 0 | -0.56 | 0.014206 | 1.02 | 119 | 81 | 6.08 | 11.4 | | High | High | High | High | High | High | High | High | High | High | High | 0 | 1.08E-06 | 3.33 |
| FALSE | High | SSLPITEDEI | [R].SSLPITE | 1xPhospho | B2RY61 1xl | 0.019791 | 0.001438 | 1 | 6 | 1 | B2RY61 | AT rich inte | B2RY61 [1 | 0 | 1656.85 | 0 | 0.13 | 0.043063 | 4.13 | 94.6 | 105.4 | 2.91 | 8.59 | | High | High | High | High | High | High | High | High | High | High | High | 0.000773 | 0.006766 | 2.17 |
| FALSE | High | SSFSMEEE | [R].SSFSMI | 1xPhospho | A2AEW1 1 | 0.029586 | 0.001857 | 1 | 7 | 3 | A2AEW1 | Transcripti | A2AEW1 [5 | 0 | 1357.507 | 0 | -0.37 | 0.132842 | 19.46 | 114.7 | 85.3 | 11.13 | 18.56 | | High | High | High | High | High | High | High | High | High | High | High | 0.001199 | 0.01291 | 1.09 |
| FALSE | High | SSFSHYSGL | [R].SSFSHY | 1xPhospho | Q9CY58 1x | 1.17E-05 | 0 | 2 | 4 | 1 | Q9CY58; Q | Plasminoge | Q9CY58 [2 | 0 | 1650.829 | 0 | -0.55 | 0.046233 | 12.16 | 118.7 | 81.3 | 16.37 | 11.21 | Shared | High | High | High | High | High | High | High | High | High | High | High | 0 | 1.6E-06 | 2.16 |
| FALSE | High | SSFSEGQT | [R].SSFSEG | 1xPhospho | Q80U62 1x | 8.22E-07 | 0 | 1 | 5 | 2 | Q80U62 | Run domai | Q80U62 [3 | 0 | 2092.036 | 0 | 0.7 | 0.000211 | 2.98 | 75.3 | 124.7 | 7.89 | 4.4 | | High | High | High | High | High | High | High | High | High | High | High | 0 | 9.81E-08 | 2.88 |
| FALSE | High | SSFQITSVT | [K].SSFQIT | 1xPhospho | Q3TCX6 1x | 0.015741 | 0.001124 | 1 | 7 | 1 | Q3TCX6 | Putative ur | Q3TCX6 [8 | 0 | 3748.794 | 0 | 1.26 | 0.00291 | 3.28 | 58.1 | 141.9 | 16.7 | 10.19 | | High | High | High | High | High | High | High | High | High | High | High | 0.000585 | 0.005233 | 2.45 |
| FALSE | High | SRTLSQSSE | [R].SRTLSQ | 1xCarbami | Q9QYG0 2 | 0.023584 | 0.001578 | 1 | 3 | 2 | Q9QYG0 | Protein ND | Q9QYG0 [3 | 1 | 3078.315 | 0 | -1.97 | 0.001203 | 1.12 | 155.2 | 44.8 | 13.87 | 19.27 | | High | High | High | High | High | High | High | High | High | High | High | 0.002717 | 0.02913 | 2.05 |
| FALSE | High | SRTPPSAPS | [R].SRTPPS | 1xPhospho | Q8BTI8 1xP | 0.08727 | 0.006537 | 1 | 3 | 3 | Q8BTI8 | Serine/argi | Q8BTI8 [23 | 1 | 1579.779 | 0 | -0.36 | 0.000684 | 3.72 | 112.3 | 87.7 | 2.99 | 6.14 | | High | High | High | High | High | High | High | High | High | High | High | 0.009784 | 0.1058 | 2.44 |
| FALSE | High | SRTSPAPW | [R].SRTSPA | 2xPhospho | Q8BTI8 2xP | 0.018125 | 0.001282 | 1 | 3 | 2 | Q8BTI8 | Serine/argi | Q8BTI8 [18 | 1 | 1647.806 | 0 | -0.67 | 0.00743 | 7.35 | 122.8 | 77.2 | 7.89 | 11.08 | | High | High | High | High | High | High | High | High | High | High | High | 0.000689 | 0.006085 | 2.42 |
| FALSE | High | SRTSVQTEI | [R].SRTSVC | 1xPhospho | P26231 1xl | 5.8E-08 | 0 | 1 | 7 | 4 | P26231 | Catenin alp | P26231 [65 | 1 | 2371.145 | 0 | -0.75 | 0.006213 | 6.94 | 125.5 | 74.5 | 15.02 | 3.37 | | High | High | High | High | High | High | High | High | High | High | High | 0 | 4.19E-07 | 3.99 |
| FALSE | High | SRTSVQTEI | [R].SRTSVC | 2xPhospho | P26231 2xl | 0.001398 | 5.66E-05 | 1 | 7 | 2 | P26231 | Catenin alp | P26231 [65 | 1 | 2451.112 | 0 | -0.8 | 0.004118 | 2.21 | 125.1 | 74.9 | 7.86 | 10.89 | | High | High | High | High | High | High | High | High | High | High | High | 2.58E-05 | 0.000348 | 3.19 |
| FALSE | High | SRVSPVPPS | [K].SRVSPV | 1xPhospho | A2SW42 1x | 8E-07 | 0 | 1 | 7 | 3 | A2SW42 | Zinc finger | A2SW42 [2 | 1 | 2307.211 | 0 | -1.52 | 0.000182 | 0.56 | 148.2 | 51.8 | 3.76 | 2.26 | | High | High | High | High | High | High | High | High | High | High | High | 0 | 4.6E-06 | 3.16 |
| FALSE | High | SSAAAAAA | [K].SSAAAA | 1xPhospho | Q9ERD6 1x | 4.84E-08 | 0 | 1 | 7 | 1 | Q9ERD6 | Ras-specifi | Q9ERD6 [3 | 0 | 2485.265 | 0 | -0.63 | 0.001032 | 1 | 121.4 | 78.6 | 3.57 | 7.16 | | High | High | High | High | High | High | High | High | High | High | High | 0 | 3.39E-09 | 3.5 |
| FALSE | High | SSAAAAAA | [K].SSAAAA | 2xPhospho | Q9ERD6 2x | 1.29E-05 | 0 | 1 | 7 | 3 | Q9ERD6 | Ras-specifi | Q9ERD6 [3 | 0 | 2565.231 | 0 | 0.16 | 0.191077 | 5.66 | 96 | 104 | 3.84 | 5.92 | | High | High | High | High | High | High | High | High | High | High | High | 0 | 1.79E-06 | 3.1 |
| FALSE | High | SSAAIQNE | [K].SSAAIQ | 1xPhospho | Q80XP8 1x | 0.001916 | 5.66E-05 | 1 | 2 | 1 | Q80XP8 | Protein FAI | Q80XP8 [2 | 0 | 1454.709 | 0 | -0.61 | 0.393337 | 29.64 | 120.6 | 79.4 | 54.22 | 51.68 | | High | High | High | High | High | High | High | High | High | High | High | 2.58E-05 | 0.000494 | 2.85 |
| FALSE | High | SSAAIQNE | [K].SSAAIQ | 1xDeamida | Q80XP8 1x | 0.013426 | 0.000976 | 1 | 2 | 1 | Q80XP8 | Protein FAI | Q80XP8 [2 | 0 | 1455.693 | 0 | 0.1 | 0.547638 | 38.2 | 98.9 | 101.1 | 18.94 | 12.7 | | High | High | High | High | High | High | High | High | High | High | High | 0.000505 | 0.00437 | 1.42 |
| FALSE | High | SSAAIQNE | [K].SSAAIQ | 1xPhospho | Q80XP8 1x | 0.000496 | 0 | 1 | 2 | 2 | Q80XP8 | Protein FAI | Q80XP8 [2 | 0 | 1683.872 | 0 | -0.57 | 0.001675 | 5.06 | 119.6 | 80.4 | 3.51 | 7.83 | | High | High | High | High | High | High | High | High | High | High | High | 0 | 0.000108 | 3.29 |
| FALSE | High | SSAATANA | [R].SSAATA | 1xCarbami | Q8BMK4 1 | 5.32E-06 | 0 | 1 | 1 | 2 | Q8BMK4 | Cytoskelet | Q8BMK4 [6 | 0 | 1736.747 | 0 | 0.41 | 0.956024 | 60.31 | 89.9 | 110.1 | 34.18 | 37.9 | | High | High | High | High | High | High | High | High | High | High | High | 0 | 6.67E-07 | 2.63 |
| FALSE | High | SSAAVETSI | [K].SSAAVE | 1xPhospho | Q4VAC9 1x | 0.006536 | 0.00062 | 1 | 3 | 2 | Q4VAC9 | Pleckstrin I | Q4VAC9 [1 | 0 | 1313.63 | 0 | 1.83 | 0.00029 | 7.31 | 44.3 | 155.7 | 3.76 | 6.17 | | High | High | High | High | High | High | High | High | High | High | High | 0.000275 | 0.00195 | 1.06 |
| FALSE | High | SRTASLTSA | [R].SRTASL | 2xPhospho | Q9QYG0 2 | 0.009585 | 0.000736 | 1 | 3 | 1 | Q9QYG0 | Protein ND | Q9QYG0 [3 | 1 | 1968.899 | 0 | -1.36 | 0.000308 | 2.72 | 139 | 61 | 6.65 | 16.07 | | High | High | High | High | High | High | High | High | High | High | High | 0.000344 | 0.002992 | 2.21 |
| FALSE | High | SSADSLPGI | [R].SSADSL | 2xPhospho | B7ZC24 2xl | 0.003997 | 0.000311 | 1 | 1 | 3 | B7ZC24 | Nuclear re | B7ZC24 [37 | 0 | 1520.66 | 0 | 0.04 | 0.478281 | 7.09 | 100.9 | 99.1 | 4.09 | 13.25 | | High | High | High | High | High | High | High | High | High | High | High | 0.000399 | 0.003362 | 2.65 |
| FALSE | High | SSADSLPGI | [R].SSADSL | 2xPhospho | Q91W39 2 | 0.002508 | 0.000109 | 1 | 4 | 2 | Q91W39 | Nuclear re | Q91W39 [3 | 0 | 1575.702 | 0 | -0.42 | 0.003744 | 4.78 | 113.5 | 86.5 | 2.12 | 6.32 | | High | High | High | High | High | High | High | High | High | High | High | 4.94E-05 | 0.000667 | 2.6 |
| FALSE | High | SSAELRKIL | [K].SSAELR | 1xDeamida | Q9WVS7 1 | 0.051694 | 0.003745 | 1 | 4 | 2 | Q9WVS7 | Dual specif | Q9WVS7 [ | 2 | 2599.275 | 0 | -0.01 | 0.6853 | 6.74 | 97.1 | 102.9 | 7.41 | 6.27 | | High | High | High | High | High | High | High | High | High | High | High | 0.007722 | 0.0836 | 2.94 |
| FALSE | High | SSAIGIENIC | [R].SSAIGIE | 1xPhospho | A2ALS7 1xl | 0.002948 | 0.00016 | 2 | 6 | 1 | A2ALS7; A | Rap1 GTPa | A2ALS7 [2 | 0 | 2183.136 | 0 | 1.35 | 0.000983 | 9.98 | 53.3 | 146.7 | 9.8 | 7.97 | Shared | High | High | High | High | High | High | High | High | High | High | High | 7.19E-05 | 0.000801 | 2.65 |
| FALSE | High | SSANSTPS | [K].SSANST | 1xPhospho | Q66JX5 1xl | 0.001386 | 5.66E-05 | 1 | 3 | 3 | Q66JX5 | FGFR1 onc | Q66JX5 [1 | 0 | 1416.714 | 0 | -0.88 | 0.001377 | 2 | 129.3 | 70.7 | 6.1 | 4.22 | | High | High | High | High | High | High | High | High | High | High | High | 2.58E-05 | 0.000345 | 1.94 |
| FALSE | High | SSAQLQTN | [R].SSAQL( | 1xPhospho | B2RS19 1xl | 0.001147 | 0 | 1 | 4 | 4 | B2RS19 | Max protei | B2RS19 [1C | 0 | 2827.355 | 0 | 0.02 | 0.914068 | 9.24 | 101 | 99 | 4.76 | 13.85 | | High | High | High | High | High | High | High | High | High | High | High | 2.58E-05 | 0.000561 | 3.22 |
| FALSE | High | SSAQPSST | [K].SSAQPS | 1xPhospho | E9Q1K1 1x | 0.000262 | 0 | 1 | 6 | 1 | E9Q1K1 | Cilia- and fl | E9Q1K1 [2 | 0 | 1430.729 | 0 | 0.82 | 0.020552 | 6.53 | 76.9 | 123.1 | 14.86 | 8.87 | | High | High | High | High | High | High | High | High | High | High | High | 0 | 5.28E-05 | 2.15 |
| FALSE | High | SSASAPDVI | [K].SSASAP | 1xPhospho | Q9CY58 1x | 0.000284 | 0 | 2 | 8 | 8 | Q9CY58; Q | Plasminoge | Q9CY58 [3 | 0 | 2068.931 | 0 | -0.49 | 0.000618 | 1.5 | 118.7 | 81.3 | 6.98 | 11.45 | Shared | High | High | High | High | High | High | High | High | High | High | High | 0.000638 | 0.005666 | 2.75 |
| FALSE | High | SSASFSPGI | [R].SSASFS | 1xPhospho | Q4QRL3-2 | 0.002192 | 0.000109 | 1 | 3 | 3 | Q4QRL3-2 | Isoform 2 c | Q4QRL3-2 | 0 | 1517.683 | 0 | 0.47 | 0.017282 | 25.34 | 81 | 119 | 3.9 | 12.49 | | High | High | High | High | High | High | High | High | High | High | High | 0.000449 | 0.003756 | 1.41 |
| FALSE | High | SSASGIAGI | [K].SSASGI | 1xPhospho | A3KFM7 1x | 3.66E-08 | 0 | 1 | 3 | 2 | A3KFM7 | Chromodo | A3KFM7 [2 | 0 | 2084.97 | 0 | 0.69 | 0.107908 | 10.42 | 80.8 | 119.2 | 13.16 | 17.28 | | High | High | High | High | High | High | High | High | High | High | High | 0 | 2.46E-09 | 4.02 |
| FALSE | High | SSASGTHSI | [R].SSASGT | 1xPhospho | Q8K1J5 1xl | 4.63E-05 | 0 | 1 | 1 | 4 | Q8K1J5 | Protein SDI | Q8K1J5 [2 | 0 | 1841.868 | 0 | -1.34 | 0.002129 | 0.47 | 143.8 | 56.2 | 10.84 | 8.69 | | High | High | High | High | High | High | High | High | High | High | High | 0.001308 | 0.0138 | 2.69 |
| FALSE | High | SSASGYGTI | [R].SSASGY | 1xPhospho | Q8CH25-2 | 0.077188 | 0.005893 | 1 | 2 | 1 | Q8CH25-2 | Isoform 2 c | Q8CH25-2 | 0 | 1194.535 | 0 | 0.12 | 0.223462 | 3.47 | 95.8 | 104.2 | 11.73 | 17.32 | | High | High | High | High | High | High | High | High | High | High | High | 0.003064 | 0.03198 | 1.29 |
| FALSE | High | SSASVTGEI | [K].SSASVT | 1xPhospho | Q80U93 1x | 0.001147 | 0 | 1 | 1 | 1 | Q80U93 | Nuclear po | Q80U93 [4 | 0 | 2304.059 | 0 | 2.05 | 0.004341 | 11.02 | 38.5 | 161.5 | 10.25 | 23.28 | | High | High | High | High | High | High | High | High | High | High | High | 0 | 0.000279 | 3.16 |
| FALSE | High | SSATLWNS | [K].SSATLW | 1xPhospho | A2AJI0 1xP | 0.00311 | 0.00016 | 1 | 2 | 1 | A2AJI0 | MAP7 dom | A2AJI0 [27 | 0 | 1514.72 | 0 | 0.94 | 0.006842 | 10.24 | 67 | 133 | 13.51 | 11.78 | | High | High | High | High | High | High | High | High | High | High | High | 7.19E-05 | 0.000852 | 2.27 |
| FALSE | High | SSADSLPGI | [R].SSADSL | 1xPhospho | Q91W39 1 | 0.002869 | 0.00016 | 1 | 4 | 2 | Q91W39 | Nuclear re | Q91W39 [3 | 0 | 1495.736 | 0 | -0.13 | 0.081246 | 13.52 | 105.4 | 94.6 | 3.32 | 7.26 | | High | High | High | High | High | High | High | High | High | High | High | 7.19E-05 | 0.00078 | 2.09 |

| | | | | | | | | | | | | | | | | | | | | | | | | | | | | | | | | | | | | | | |
|---|---|---|---|---|---|---|---|---|---|---|---|---|---|---|---|---|---|---|---|---|---|---|---|---|---|---|---|---|---|---|---|---|---|---|---|---|---|---|
| FALSE | High | SRSTTPAPK | [R].SRSTTP | 2xPhospho | Q8BTI8 2xP | 0.005135 | 0.000551 | 1 | 3 | 3 | Q8BTI8 | Serine/argi | Q8BTI8 [24 | 1 | 1562.775 | 0 | -1.07 | 0.000912 | 1.34 | 132.9 | 67.1 | 4.04 | 9.63 | | High | High | High | High | High | High | High | High | High | High | High | 0.000538 | 0.004499 | 1.89 |
| FALSE | High | SRSTTELNI | [K].SRSTTE | 1xPhospho | Q6ZQ32 1x | 0.000382 | 0 | 2 | 7 | 1 | Q6ZQ32; F | MKIAA085 | Q6ZQ32 [1 | 1 | 2141.126 | 0 | 0.64 | 0.001603 | 12.52 | 78.4 | 121.6 | 13.51 | 6.02 | Shared | High | High | High | High | High | High | High | High | High | High | High | 0 | 0.000081 | 2.83 |
| FALSE | High | SRSSSVGSS | [R].SRSSSV | 3xPhospho | Q9QXS1-3 | 0.000479 | 0 | 3 | 23 | 1 | Q9QXS1-3; | Isoform PL | Q9QXS1-3 | 1 | 2425.004 | 0 | 0.56 | 0.014715 | 11.99 | 81 | 119 | 8.25 | 12.74 | Shared | High | High | High | High | High | High | High | High | High | High | High | 0 | 0.000104 | 4.78 |
| FALSE | High | SRSDIDVN | [R].SRSDID | 1xDeamida | B9EJA4 1xP | 0.002553 | 0.000109 | 1 | 7 | 1 | B9EJA4 | Clasp2 pro | B9EJA4 [37 | 1 | 1913.973 | 0 | -0.43 | 0.063363 | 13.73 | 114.9 | 85.1 | 14.99 | 10.54 | | High | High | High | High | High | High | High | High | High | High | High | 4.94E-05 | 0.000684 | 3.21 |
| FALSE | High | SRSDIDVN | [R].SRSDID | 1xPhospho | B2RY22 1x | 1.59E-07 | 0 | 2 | 16 | 5 | B2RY22; A | Clasp1 pro | B2RY22 [5 | 1 | 1943 | 0 | -1 | 0.000226 | 1.56 | 133.4 | 66.6 | 5.39 | 14.93 | Shared | High | High | High | High | High | High | High | High | High | High | High | 0 | 2.82E-08 | 4.71 |
| FALSE | High | SRSDIDVN | [R].SRSDID | 1xDeamida | B2RY22 1x | 7.07E-08 | 0 | 2 | 16 | 1 | B2RY22; A | Clasp1 pro | B2RY22 [5 | 1 | 1943.984 | 0 | -0.73 | 0.015035 | 6.88 | 124.7 | 75.3 | 10.71 | 9.94 | Shared | High | High | High | High | High | High | High | High | High | High | High | 0 | 5.17E-09 | 3.98 |
| FALSE | High | SRSDVDMI | [R].SRSDVI | 1xPhospho | Q68FM7 1 | 2.7E-06 | 0 | 1 | 3 | 1 | Q68FM7 | Rho guanir | Q68FM7 [6 | 1 | 1903.842 | 0 | -0.24 | 0.021488 | 1.96 | 105.4 | 94.6 | 5.38 | 6.85 | | High | High | High | High | High | High | High | High | High | High | High | 0 | 3.09E-07 | 3.27 |
| FALSE | High | SRSDVDMI | [R].SRSDVI | 1xPhospho | Q68FM7 1 | 0.000822 | 0 | 1 | 3 | 1 | Q68FM7 | Rho guanir | Q68FM7 [6 | 1 | 1919.837 | 0 | 0.09 | 0.268371 | 10.07 | 94 | 106 | 5.15 | 14.91 | | High | High | High | High | High | High | High | High | High | High | High | 0 | 0.000191 | 2.93 |
| FALSE | High | SRSDVDMI | [R].SRSDVI | 1xPhospho | Q61210-5 | 2.56E-05 | 0 | 1 | 3 | 2 | Q61210-5 | Isoform 5 c | Q61210-5 | 1 | 2357.101 | 0 | 0.62 | 0.012372 | 16.95 | 79.5 | 120.5 | 10.68 | 9.47 | | High | High | High | High | High | High | High | High | High | High | High | 0.000801 | 0.007207 | 3.53 |
| FALSE | High | SRSELCLDL | [R].SRSELC | 1xCarbami | Q8BX02 1x | 1.86E-07 | 0 | 1 | 2 | 1 | Q8BX02 | KN motif a | Q8BX02 [2 | 1 | 2764.306 | 0 | 0.06 | 0.676297 | 36.03 | 96.5 | 103.5 | 9.56 | 17.8 | | High | High | High | High | High | High | High | High | High | High | High | 0 | 1.53E-08 | 4.16 |
| FALSE | High | SRSESDASI | [R].SRSESD | 1xPhospho | Q9CZX2 1x | 1.6E-05 | 0 | 1 | 3 | 2 | Q9CZX2 | Centrosom | Q9CZX2 [7 | 1 | 1861.894 | 0 | 0.13 | 0.399176 | 5.65 | 96.6 | 103.4 | 5.29 | 5.01 | | High | High | High | High | High | High | High | High | High | High | High | 0 | 2.3E-06 | 3.63 |
| FALSE | High | SRSESDLSC | [K].SRSESD | 1xPhospho | E9PZP8 1xP | 2.52E-07 | 0 | 1 | 1 | 3 | E9PZP8 | HECT and | E9PZP8 [15 | 1 | 2708.189 | 0 | -0.8 | 0.00144 | 1.43 | 126.8 | 73.2 | 4.7 | 5.58 | | High | High | High | High | High | High | High | High | High | High | High | 0 | 2.17E-08 | 5.54 |
| FALSE | High | SRSESDSEK | [R].SRSESD | 1xPhospho | Q69ZF8 1x | 0.016743 | 0.001166 | 1 | 1 | 1 | Q69ZF8 | E3 ubiquiti | Q69ZF8 [3 | 1 | 1562.746 | 0 | -0.27 | 0.02772 | 8.6 | 109.8 | 90.2 | 7.64 | 6.85 | | High | High | High | High | High | High | High | High | High | High | High | 0.000638 | 0.005609 | 2.4 |
| FALSE | High | SRSESETST | [R].SRSESE | 1xPhospho | Q0VAW6 1 | 1.04E-05 | 0 | 1 | 5 | 1 | Q0VAW6 | RIKEN cDN | Q0VAW6 [ | 1 | 1922.929 | 0 | -1.14 | 0.00122 | 0.92 | 139.5 | 60.5 | 8.96 | 14.61 | | High | High | High | High | High | High | High | High | High | High | High | 0 | 1.4E-06 | 2.9 |
| FALSE | High | SRSESETST | [R].SRSESE | 1xPhospho | Q0VAW6 1 | 2.1E-06 | 0 | 1 | 5 | 14 | Q0VAW6 | RIKEN cDN | Q0VAW6 [ | 1 | 1938.924 | 0 | -0.79 | 0.001428 | 2.51 | 124.4 | 75.6 | 3.44 | 9.35 | | High | High | High | High | High | High | High | High | High | High | High | 0 | 2.35E-07 | 3.53 |
| FALSE | High | SRSEVQSEY | [R].SRSEVC | 1xPhospho | E9PYK3 1xP | 8.18E-05 | 0 | 1 | 2 | 1 | E9PYK3 | Poly [ADP- | E9PYK3 [10 | 1 | 2604.259 | 0 | 2.16 | 0.000191 | 14.28 | 36.6 | 163.4 | 9.08 | 3.18 | | High | High | High | High | High | High | High | High | High | High | High | 0 | 1.43E-05 | 3.9 |
| FALSE | High | SRSHSDND | [R].SRSHSD | 1xCarbami | Q9R0U0-3 | 8.03E-05 | 0 | 1 | 2 | 3 | Q9R0U0-3 | Isoform 3 c | Q9R0U0-3 | 1 | 3528.426 | 0 | -0.38 | 0.017495 | 9.65 | 115.2 | 84.8 | 4.06 | 13.64 | | High | High | High | High | High | High | High | High | High | High | High | 0 | 0.000014 | 3.97 |
| FALSE | High | SRSHSDND | [R].SRSHSD | 1xCarbami | Q9R0U0-3 | 0.02933 | 0.001857 | 1 | 2 | 1 | Q9R0U0-3 | Isoform 3 c | Q9R0U0-3 | 1 | 3608.393 | 0 | -0.48 | 0.008797 | 7.19 | 112.7 | 87.3 | 10.08 | 12.02 | | High | High | High | High | High | High | High | High | High | High | High | 0.000997 | 0.01054 | 2.85 |
| FALSE | High | SRSPGTATI | [R].SRSPGT | 2xPhospho | Q9CS00 2x | 0.027839 | 0.001755 | 1 | 1 | 2 | Q9CS00 | Cactin OS= | Q9CS00 [1 | 1 | 2547.217 | 0 | -0.12 | 0.011144 | 3.27 | 105.3 | 94.7 | 3.16 | 3.58 | | High | High | High | High | High | High | High | High | High | High | High | 0.000953 | 0.009906 | 3.05 |
| FALSE | High | SRSPIIHSPK | [K].SRSPIIH | 2xPhospho | F6UFZ5 2xP | 0.000415 | 0 | 2 | 7 | 1 | F6UFZ5; Q | PCF11 clea | F6UFZ5 [28 | 1 | 1739.901 | 0 | -0.58 | 0.000854 | 3.16 | 119.8 | 80.2 | 1.55 | 6.77 | Shared | High | High | High | High | High | High | High | High | High | High | High | 0 | 8.88E-05 | 1.56 |
| FALSE | High | SRSPLLNDF | [R].SRSPLL | 1xPhospho | C7G3P2 1x | 0.046687 | 0.00354 | 1 | 3 | 1 | C7G3P2 | MKIAA053 | C7G3P2 [3 | 1 | 1366.704 | 0 | -0.12 | 0.953464 | 12.92 | 103.6 | 96.4 | 17.3 | 20.23 | | High | High | High | High | High | High | High | High | High | High | High | 0.001812 | 0.01789 | 1.32 |
| FALSE | High | SRSPLQPTS | [R].SRSPLQ | 1xPhospho | Q9WTV7 1 | 0.00051 | 0 | 1 | 1 | 2 | Q9WTV7 | E3 ubiquiti | Q9WTV7 [ | 1 | 1776.921 | 0 | 0.73 | 0.000546 | 7.42 | 74.9 | 125.1 | 6.25 | 3.25 | | High | High | High | High | High | High | High | High | High | High | High | 2.58E-05 | 0.00053 | 3.09 |
| FALSE | High | SRSPPDRD | [R].SRSPPD | 1xPhospho | Q8BIA4 1xP | 8.92E-07 | 0 | 1 | 2 | 2 | Q8BIA4 | F-box/WD | Q8BIA4 [84 | 2 | 2700.356 | 0 | -0.45 | 0.005385 | 4.31 | 119.1 | 80.9 | 3.44 | 14.32 | | High | High | High | High | High | High | High | High | High | High | High | 0 | 5.9E-06 | 2.42 |
| FALSE | High | SRSPPPVSK | [R-].SRSPPI | 1xPhospho | Q8C671 1x | 0.098533 | 0.007902 | 1 | 4 | 1 | Q8C671 | Putative ur | Q8C671 [1 | 1 | 1492.829 | 0 | -0.55 | 0.004111 | 6.23 | 120.9 | 79.1 | 7.03 | 7.81 | | High | High | High | High | High | High | High | High | High | High | High | 0.003977 | 0.0424 | 1.96 |
| FALSE | High | SRSPSPAPE | [K].SRSPSP | 2xPhospho | Q52KI8 2xP | 0.071071 | 0.005247 | 1 | 7 | 1 | Q52KI8 | Serine/argi | Q52KI8 [20 | 1 | 1673.807 | 0 | -0.65 | 0.037335 | 18.25 | 124.3 | 75.7 | 16.27 | 22.2 | | High | High | High | High | High | High | High | High | High | High | High | 0.002703 | 0.02903 | 1.98 |
| FALSE | High | SRSPTVNG | [R].SRSPTV | 1xPhospho | Q3UG20 1 | 0.005565 | 0.00062 | 1 | 2 | 1 | Q3UG20 | Histone-lys | Q3UG20 [8 | 1 | 1726.889 | 0 | 1.36 | 0.000253 | 4.97 | 56.8 | 143.2 | 7.25 | 7.15 | | High | High | High | High | High | High | High | High | High | High | High | 0.000275 | 0.001636 | 2.2 |
| FALSE | High | SRSPVDSP | [R].SRSPVD | 1xPhospho | O08582 1x | 6.15E-06 | 0 | 1 | 1 | 1 | O08582 | GTP-bindir | O08582 [6 | 1 | 2806.343 | 0 | -0.81 | 0.001018 | 4.99 | 127.2 | 72.8 | 3.87 | 12.51 | | High | High | High | High | High | High | High | High | High | High | High | 0 | 7.83E-07 | 4.83 |
| FALSE | High | SRSPVDSP | [R].SRSPVD | 1xPhospho | O08582 1x | 2.86E-07 | 0 | 1 | 1 | 5 | O08582 | GTP-bindir | O08582 [6 | 1 | 2822.338 | 0 | -0.54 | 0.006523 | 2.64 | 117.5 | 82.5 | 3.97 | 14.68 | | High | High | High | High | High | High | High | High | High | High | High | 0 | 2.48E-08 | 4.46 |
| FALSE | High | SRSPVDSP | [R].SRSPVD | 2xPhospho | O08582 2x | 0.019107 | 0.001402 | 1 | 1 | 3 | O08582 | GTP-bindir | O08582 [6 | 1 | 2886.31 | 0 | -0.28 | 0.00298 | 5.2 | 111.2 | 88.8 | 5.01 | 6.7 | | High | High | High | High | High | High | High | High | High | High | High | 0.005374 | 0.05901 | 3.53 |
| FALSE | High | SRSPVDSP | [R].SRSPVD | 2xPhospho | O08582 2x | 0.048303 | 0.00354 | 1 | 1 | 5 | O08582 | GTP-bindir | O08582 [6 | 1 | 2902.305 | 0 | -0.37 | 0.539332 | 6.63 | 111.1 | 88.9 | 5.9 | 29.79 | | High | High | High | High | High | High | High | High | High | High | High | 0.001812 | 0.01845 | 3.53 |
| FALSE | High | SRSPVDSP | [R].SRSPVD | 3xPhospho | O08582 3x | 0.039011 | 0.002718 | 1 | 1 | 1 | O08582 | GTP-bindir | O08582 [6 | 1 | 2966.276 | 0 | 0.04 | 0.690861 | 1.6 | 99.1 | 100.9 | 2.49 | 8.17 | | High | High | High | High | High | High | High | High | High | High | High | 0.001453 | 0.01458 | 3.46 |
| FALSE | High | SRSSPSTDH | [R].SRSSPS | 2xPhospho | A7TU71 2x | 4.55E-05 | 0 | 1 | 2 | 2 | A7TU71 | Protein Shi | A7TU71 [9 | 1 | 2661.154 | 0 | 2.8 | 0.003062 | 30.11 | 22.1 | 177.9 | 25.13 | 20.9 | | High | High | High | High | High | High | High | High | High | High | High | 0 | 7.45E-06 | 3.5 |
| FALSE | High | SSCDSLDY | [K].SSCDSL | 1xCarbami | Q8C9G3 2x | 0.109329 | 0.008865 | 1 | 5 | 1 | Q8C9G3 | Putative ur | Q8C9G3 [2 | 0 | 1820.758 | 0 | -1.52 | 0.009282 | 3.26 | 148.3 | 51.7 | 18.73 | 13.13 | | High | High | High | High | High | High | High | High | High | High | High | 0.004462 | 0.04805 | 2.6 |
| FALSE | High | SSMSSCGS | [R].SSMSS | 2xCarbami | B2ZRS3 1xP | 7.89E-05 | 0 | 1 | 6 | 4 | B2ZRS3 | Depdc6-00 | B2ZRS3 [27 | 0 | 3549.599 | 0 | 2.74 | 0.000552 | 2.21 | 25.7 | 174.3 | 18.67 | 12.24 | | High | High | High | High | High | High | High | High | High | High | High | 0 | 1.38E-05 | 3.97 |
| FALSE | High | SSCGSPPDI | [K].SSCGSF | 1xCarbami | A0A140T8 | 0.010568 | 0.000772 | 1 | 1 | 4 | A0A140T8 | Chromoso | A0A140T8 | 0 | 1771.849 | 0 | -0.63 | 0.005179 | 3.73 | 118.6 | 81.4 | 6 | 12.63 | | High | High | High | High | High | High | High | High | High | High | High | 0.002455 | 0.02651 | 2.52 |
| FALSE | High | SSCSLSVAD | [K].SSCSLS | 1xCarbami | E9PWI3 1x | 0.00011 | 0 | 1 | 3 | 1 | E9PWI3 | Mutated in | E9PWI3 [3 | 0 | 1604.719 | 0 | 1.01 | 0.00095 | 5.02 | 67.2 | 132.8 | 6.43 | 7.71 | | High | High | High | High | High | High | High | High | High | High | High | 0 | 2.01E-05 | 2.66 |
| FALSE | High | SSDVPLPLE | [K].SSDVPL | 1xPhospho | B2RXX0 1x | 8.48E-05 | 0 | 1 | 5 | 2 | B2RXX0 | Nek1 prote | B2RXX0 [6 | 0 | 2464.299 | 0 | -0.96 | 0.003222 | 4.96 | 132.1 | 67.9 | 6.83 | 15.13 | | High | High | High | High | High | High | High | High | High | High | High | 0 | 1.49E-05 | 3.91 |
| FALSE | High | SSECLSQAS | [R].SSECLS | 1xCarbami | Q1W617 1 | 1.72E-05 | 0 | 1 | 2 | 1 | Q1W617 | Protein Shi | Q1W617 [6 | 0 | 1937.893 | 0 | 2.89 | 0.002321 | 34.27 | 27 | 173 | 12.42 | 19.9 | | High | High | High | High | High | High | High | High | High | High | High | 0 | 2.49E-06 | 2.84 |
| FALSE | High | SSEDVSAH | [R].SSEDVS | 1xPhospho | Q9JID9 1xP | 0.000161 | 0 | 1 | 2 | 1 | Q9JID9 | SH2B adap | Q9JID9 [10 | 0 | 1740.857 | 0 | 1.05 | 0.002741 | 4.03 | 64.4 | 135.6 | 2.27 | 13.4 | | High | High | High | High | High | High | High | High | High | High | High | 0 | 3.06E-05 | 2.43 |
| FALSE | High | SSEEDVQL | [K].SSEEDV | 1xDeamida | H7BX21 1x | 0.039347 | 0.002793 | 1 | 2 | 1 | H7BX21 | Thioredoxi | H7BX21 [4 | 0 | 1876.883 | 0 | -1.07 | 0.015674 | 7.75 | 135.7 | 64.3 | 17.78 | 13.89 | | High | High | High | High | High | High | High | High | High | High | High | 0.001468 | 0.01467 | 1.9 |
| FALSE | High | SSEEGPGD | [R].SSEEGF | 1xPhospho | Q8R1B0 1x | 4.08E-05 | 0 | 1 | 2 | 1 | Q8R1B0 | SH3 and cy | Q8R1B0 [2 | 0 | 3018.351 | 0 | 0.93 | 0.004569 | 25.65 | 68.7 | 131.3 | 18.07 | 9.1 | | High | High | High | High | High | High | High | High | High | High | High | 0 | 6.56E-06 | 3.9 |
| FALSE | High | SSEEQNLES | [R].SSEEQN | 1xPhospho | Q6P5H2-2 | 0.005467 | 0.00062 | 1 | 2 | 2 | Q6P5H2-2 | Isoform 2 c | Q6P5H2-2 | 0 | 2163.099 | 0 | 2.88 | 0.006139 | 8.34 | 20.9 | 179.1 | 22.33 | 31.15 | | High | High | High | High | High | High | High | High | High | High | High | 0.000275 | 0.001601 | 2.65 |
| FALSE | High | SSEESESDS | [K].SSEESE | 1xPhospho | H3BL37 1xP | 3.7E-05 | 0 | 1 | 7 | 1 | H3BL37 | Treacle prc | H3BL37 [4 | 0 | 2891.345 | 0 | -1.23 | 0.00054 | 0.44 | 140.2 | 59.8 | 10.84 | 9.37 | | High | High | High | High | High | High | High | High | High | High | High | 0 | 5.86E-06 | 3.38 |
| FALSE | High | SSEESESDS | [K].SSEESE | 2xPhospho | H3BL37 2xP | 0.008167 | 0.000736 | 1 | 7 | 3 | H3BL37 | Treacle prc | H3BL37 [4 | 0 | 2971.311 | 0 | 0.01 | 0.992594 | 25.44 | 99.7 | 100.3 | 10.12 | 11.18 | | High | High | High | High | High | High | High | High | High | High | High | 0.000327 | 0.002633 | 2.11 |
| FALSE | High | SSEGDNSV | [K].SSEGDI | 1xPhospho | A0A0R4J07 | 0.00153 | 5.66E-05 | 1 | 5 | 3 | A0A0R4J07 | BRCA1-A c | A0A0R4J07 | 0 | 2531.264 | 0 | 0.06 | 0.823355 | 14.75 | 102.9 | 97.1 | 8.01 | 10.1 | | High | High | High | High | High | High | High | High | High | High | High | 2.58E-05 | 0.000384 | 3.03 |
| FALSE | High | SSEGEQDL | [K].SSEGEC | 1xPhospho | P56183 1xP | 1.57E-05 | 0 | 1 | 3 | 1 | P56183 | Ribosomal | P56183 [30 | 0 | 2068.984 | 0 | 0.15 | 0.066664 | 10.61 | 94.7 | 105.3 | 5.9 | 10.1 | | High | High | High | High | High | High | High | High | High | High | High | 0 | 2.23E-06 | 3.58 |
| FALSE | High | SSELFSVID | [R].SSELFS | 1xPhospho | Q6P1D7 1x | 4.65E-06 | 0 | 1 | 2 | 1 | Q6P1D7 | Structure- | Q6P1D7 [9 | 0 | 3110.512 | 0 | -0.1 | 0.050344 | 3.68 | 105.4 | 94.6 | 7.24 | 3.3 | | High | High | High | High | High | High | High | High | High | High | High | 0 | 5.68E-07 | 3.83 |
| FALSE | High | SSELLVR | [R].SSELLV | 1xPhospho | Q9DBS5 1x | 0.052576 | 0.00384 | 1 | 1 | 1 | Q9DBS5 | Kinesin ligh | Q9DBS5 [5 | 0 | 1112.591 | 0 | 0.54 | 0.01761 | 25.82 | 81.4 | 118.6 | 8.6 | 14.95 | | High | High | High | High | High | High | High | High | High | High | High | 0.001999 | 0.02041 | 2.02 |
| FALSE | High | SSELPLTVP | [K].SSELPL | 1xPhospho | A2APB8 1x | 0.000158 | 0 | 1 | 3 | 1 | A2APB8 | Targeting p | A2APB8 [7 | 0 | 1662.892 | 0 | 0.02 | 0.917061 | 18.72 | 101.5 | 98.5 | 65.13 | 50.97 | | High | High | High | High | High | High | High | High | High | High | High | 0 | 3.01E-05 | 2.97 |
| FALSE | High | SSDTCGSP | [R].SSDTCC | 1xCarbami | B1AYC9 1x | 3.36E-06 | 0 | 1 | 1 | 4 | B1AYC9 | Amyloid be | B1AYC9 [5 | 0 | 1844.887 | 0 | 1.65 | 0.000992 | 11.32 | 46 | 154 | 15.23 | 10 | | High | High | High | High | High | High | High | High | High | High | High | 0 | 3.98E-07 | 3.47 |
| FALSE | High | SSELPLTVP | [K].SSELPL | 1xPhospho | A2APB8 1x | 0.000284 | 0 | 1 | 3 | 3 | A2APB8 | Targeting p | A2APB8 [7 | 0 | 1892.055 | 0 | -0.22 | 0.001253 | 2.84 | 107.4 | 92.6 | 2.59 | 3.56 | | High | High | High | High | High | High | High | High | High | High | High | 0 | 5.81E-05 | 2.36 |
| FALSE | High | SSEQATVA | [R].SSEQAT | 1xCarbami | G3UZM1 1 | 0.000109 | 0 | 1 | 4 | 4 | G3UZM1 | Probable J | G3UZM1 [ | 0 | 3240.406 | 0 | -0.38 | 0.003546 | 6.25 | 113.1 | 86.9 | 5.54 | 4.29 | | High | High | High | High | High | High | High | High | High | High | High | 0 | 0.000294 | 2.51 |
| FALSE | High | SSESGNSVI | [K].SSESGN | 1xPhospho | Q3U9X3 1x | 0.047894 | 0.00354 | 1 | 5 | 1 | Q3U9X3 | Putative ur | Q3U9X3 [1 | 0 | 1432.709 | 0 | -0.24 | 0.143191 | 9.63 | 105.7 | 94.3 | 14.01 | 8.18 | | High | High | High | High | High | High | High | High | High | High | High | 0.001812 | 0.01839 | 1.11 |
| FALSE | High | SSESSDGQ | [K].SSESSD | 1xPhospho | Q6P3Y5 1x | 9.08E-07 | 0 | 1 | 3 | 2 | Q6P3Y5 | Zinc finger | Q6P3Y5 [1 | 0 | 1749.831 | 0 | 1.26 | 0.000412 | 12.47 | 59.5 | 140.5 | 6.21 | 5.35 | | High | High | High | High | High | High | High | High | High | High | High | 0 | 1.28E-07 | 2.17 |
| FALSE | High | SSETENIES | [K].SSETEN | 1xCarbami | Q6P5H2-2 | 1.27E-08 | 0 | 1 | 2 | 3 | Q6P5H2-2 | Isoform 2 c | Q6P5H2-2 | 0 | 2419.09 | 0 | 3.79 | 0.002295 | 3.96 | 13.2 | 186.8 | 20.08 | 23 | | High | High | High | High | High | High | High | High | High | High | High | 0 | 7.58E-10 | 4.77 |
| FALSE | High | SSETLQSIIC | [R].SSETLQ | 1xPhospho | Q8BV49 1x | 0.003167 | 0.00016 | 1 | 2 | 2 | Q8BV49 | Pyrin and H | Q8BV49 [1 | 0 | 1956.045 | 0 | 4.77 | 0.000209 | 27.54 | 7.1 | 192.9 | 24.44 | 7.04 | | High | High | High | High | High | High | High | High | High | High | High | 7.19E-05 | 0.000868 | 2.71 |
| FALSE | High | SSETLTFK | [K].SSETLT | 1xPhospho | Q9JI44 1xP | 0.026652 | 0.001676 | 1 | 2 | 1 | Q9JI44 | DNA methy | Q9JI44 [41 | 0 | 1450.759 | 0 | -0.7 | 0.000965 | 3.76 | 120.8 | 79.2 | 7.6 | 8.25 | | High | High | High | High | High | High | High | High | High | High | High | 0.000903 | 0.009454 | 1.71 |
| FALSE | High | SSEVVLSGI | [R].SSEVVL | 1xPhospho | E9Q616 1x | 2.43E-07 | 0 | 1 | 2 | 11 | E9Q616 | AHNAK nu | E9Q616 [1 | 0 | 2007.875 | 0 | 2.48 | 0.000158 | 9.36 | 29 | 171 | 14.37 | 5.32 | | High | High | High | High | High | High | High | High | High | High | High | 0 | 2.08E-08 | 3.84 |
| FALSE | High | SSEWDPVS | [R].SSEWD | 1xPhospho | Q8VED8 1x | 9.31E-06 | 0 | 1 | 3 | 3 | Q8VED8 | Mitochond | Q8VED8 [3 | 0 | 2184.135 | 0 | 0 | 0.966737 | 11.55 | 98.3 | 101.7 | 5.52 | 6.97 | | High | High | High | High | High | High | High | High | High | High | High | 0 | 1.24E-06 | 2.82 |
| FALSE | High | SSFDGSSLL | [K].SSFDGS | 1xPhospho | Q9D318 1x | 0.004824 | 0.000456 | 1 | 4 | 2 | Q9D318 | Putative ur | Q9D318 [1 | 0 | 1780.877 | 0 | 0.38 | 0.130564 | 8.23 | 93.7 | 106.3 | 19.33 | 10.68 | | High | High | High | High | High | High | High | High | High | High | High | 0.000204 | 0.001386 | 2.37 |
| FALSE | High | SSFESTSSE | [K].SSFEST | 1xPhospho | P26187 1xP | 0.003962 | 0.000311 | 1 | 2 | 2 | P26187 | Methylate | P26187 [19 | 0 | 1666.716 | 0 | 0.8 | 0.02754 | 24.52 | 72.9 | 127.1 | 8.57 | 27.69 | | High | High | High | High | High | High | High | High | High | High | High | 0.003265 | 0.03491 | 1.96 |
| FALSE | High | SSFETEESD | [K].SSFETE | 1xCarbami | Q8BMA5 1 | 0.046291 | 0.003523 | 1 | 1 | 1 | Q8BMA5 | Protein NP | Q8BMA5 [ | 0 | 4159.873 | 0 | -0.43 | 0.030572 | 15.41 | 122.9 | 77.1 | 15.37 | 16.1 | | High | High | High | High | High | High | High | High | High | High | High | 0.001789 | 0.01766 | 2.67 |
| FALSE | High | SSFFGDR | [K].SSFFGD | 1xPhospho | Q3TQX5 1x | 0.01588 | 0.001124 | 1 | 3 | 1 | Q3TQX5 | Putative ur | Q3TQX5 [8 | 0 | 1124.498 | 0 | 0.54 | 0.005814 | 3.12 | 78.5 | 121.5 | 6.72 | 10.22 | | High | High | High | High | High | High | High | High | High | High | High | 0.000585 | 0.00528 | 1.35 |
| FALSE | High | SSFPLTFPE | [K].SSFPLT | 1xPhospho | A0A1W2P6 | 0.000154 | 0 | 1 | 10 | 1 | A0A1W2P6 | Bromodom | A0A1W2P6 | 0 | 1748.809 | 0 | -0.53 | 0.013622 | 8.9 | 120 | 80 | 15.08 | 8.59 | | High | High | High | High | High | High | High | High | High | High | High | 0 | 2.9E-05 | 2.75 |
| FALSE | High | SSENNGSS | [K].SSENN | 1xDeamida | Q6PCM1 1 | 0.004781 | 0.000456 | 1 | 3 | 1 | Q6PCM1 | Lysine-spe | Q6PCM1 [ | 0 | 1634.767 | 0 | -0.69 | 0.001067 | 0.8 | 128.4 | 71.6 | 11.08 | 9.37 | | High | High | High | High | High | High | High | High | High | High | High | 0.000204 | 0.001376 | 1.55 |
| FALSE | High | SSDSWDV | [R].SSDSW | 2xPhospho | V9GXM1 2 | 9.11E-05 | 0 | 1 | 7 | 1 | V9GXM1 | ADP-ribosy | V9GXM1 [ | 0 | 2402.01 | 0 | 0.77 | 0.000743 | 2.98 | 75.5 | 124.5 | 10.22 | 6.74 | | High | High | High | High | High | High | High | High | High | High | High | 0 | 1.63E-05 | 4.05 |
| FALSE | High | SSDSWDV | [R].SSDSW | 1xDeamida | V9GXM1 1 | 2.57E-07 | 0 | 1 | 7 | 8 | V9GXM1 | ADP-ribosy | V9GXM1 [ | 0 | 2323.028 | 0 | 0.62 | 0.005567 | 6.22 | 78.7 | 121.3 | 7.41 | 11.32 | | High | High | High | High | High | High | High | High | High | High | High | 0 | 2.21E-08 | 3.91 |
| FALSE | High | SSDSWDV | [R].SSDSW | 1xPhospho | V9GXM1 1 | 4.08E-08 | 0 | 1 | 7 | 10 | V9GXM1 | ADP-ribosy | V9GXM1 [ | 0 | 2322.044 | 0 | -0.29 | 0.034602 | 4.11 | 108.9 | 91.1 | 4.48 | 12.06 | | High | High | High | High | High | High | High | High | High | High | High | 0 | 1.26E-05 | 4.63 |
| FALSE | High | SSCSLSVAD | [K].SSCSLS | 1xCarbami | E9PWI3 2x | 0.044358 | 0.003264 | 1 | 3 | 1 | E9PWI3 | Mutated in | E9PWI3 [3 | 0 | 1684.685 | 0 | 1.19 | 0.000545 | 2.9 | 64.9 | 135.1 | 15.85 | 9.6 | | High | High | High | High | High | High | High | High | High | High | High | 0.00168 | 0.01678 | 2.11 |
| FALSE | High | SSDAPFGH | [R].SSDAPF | 1xPhospho | A2A6A1 1x | 2.18E-06 | 0 | 1 | 1 | 1 | A2A6A1 | G patch do | A2A6A1 [1 | 0 | 3361.567 | 0 | 0.28 | 0.085609 | 28.9 | 89.6 | 110.4 | 7.39 | 16.91 | | High | High | High | High | High | High | High | High | High | High | High | 0 | 2.43E-07 | 3.47 |
| FALSE | High | SSDAPFGH | [R].SSDAPF | 1xDeamida | A2A6A1 1x | 0.089425 | 0.006807 | 1 | 1 | 1 | A2A6A1 | G patch do | A2A6A1 [1 | 0 | 3362.551 | 0 | 0.19 | 0.798406 | 16.34 | 105.7 | 94.3 | 15.92 | 21.57 | | High | High | High | High | High | High | High | High | High | High | High | 0.003564 | 0.03791 | 2.36 |
| FALSE | High | SSDDEDW | [R].SSDDEI | 1xPhospho | P70270 1xP | 0.014283 | 0.001011 | 1 | 1 | 2 | P70270 | DNA repair | P70270 [19 | 0 | 2200.021 | 0 | -1.18 | 0.00031 | 0.87 | 138.8 | 61.2 | 3.75 | 7.37 | | High | High | High | High | High | High | High | High | High | High | High | 0.000538 | 0.004664 | 2.26 |
| FALSE | High | SSDDEDW | [R].SSDDEI | 2xPhospho | P70270 2xP | 0.0009 | 0 | 1 | 1 | 2 | P70270 | DNA repair | P70270 [19 | 0 | 2279.987 | 0 | -1.43 | 0.000181 | 1.1 | 148.1 | 51.9 | 5.86 | 11.64 | | High | High | High | High | High | High | High | High | High | High | High | 0 | 0.000213 | 2.89 |
| FALSE | High | SSDEDATG | [R].SSDED | 1xPhospho | Q9D483 1x | 0.000393 | 0 | 1 | 2 | 1 | Q9D483 | DNA-direct | Q9D483 [2 | 0 | 1673.767 | 0 | -0.93 | 0.005075 | 2.33 | 131.2 | 68.8 | 17.48 | 11.47 | | High | High | High | High | High | High | High | High | High | High | High | 0 | 8.39E-05 | 2.08 |
| FALSE | High | SSDELDAW | [R].SSDELD | 1xPhospho | Q9WVH4 1 | 3.38E-05 | 0 | 1 | 2 | 2 | Q9WVH4 | Forkhead b | Q9WVH4 [ | 0 | 1750.752 | 0 | -0.93 | 0.001817 | 3.01 | 128.5 | 71.5 | 9.69 | 6.17 | | High | High | High | High | High | High | High | High | High | High | High | 0 | 5.33E-06 | 2.86 |
| FALSE | High | SSDGEDEQ | [K].SSDGEI | 1xPhospho | O08750 1x | 0.053926 | 0.00384 | 1 | 1 | 1 | O08750 | Nuclear fac | O08750 [2 | 0 | 1856.868 | 0 | -0.28 | 0.108696 | 14.11 | 109.4 | 90.6 | 4.99 | 12.92 | | High | High | High | High | High | High | High | High | High | High | High | 0.002045 | 0.02104 | 2.4 |
| FALSE | High | SSDGGQST | [K].SSDGG | 1xPhospho | E9QLG3 1x | 6.47E-05 | 0 | 1 | 4 | 2 | E9QLG3 | MAX gene- | E9QLG3 [2 | 0 | 3249.575 | 0 | -0.74 | 0.011983 | 6.27 | 121.3 | 78.7 | 13.91 | 12.67 | | High | High | High | High | High | High | High | High | High | High | High | 0 | 1.1E-05 | 3.89 |
| FALSE | High | SSDGRPDS | [R].SSDGRI | 1xPhospho | E9Q9C3 1x | 0.006653 | 0.00062 | 1 | 5 | 1 | E9Q9C3 | Afadin OS= | E9Q9C3 [2 | 0 | 1613.748 | 0 | 0.88 | 0.000681 | 7.9 | 70.3 | 129.7 | 8.92 | 7.98 | | High | High | High | High | High | High | High | High | High | High | High | 0.000275 | 0.001983 | 1.69 |
| FALSE | High | SSDGSLSHI | [R].SSDGSL | 1xPhospho | B8QI33 1xP | 0.000244 | 0 | 1 | 3 | 1 | B8QI33 | Liprin-alph | B8QI33 [23 | 0 | 1883.915 | 0 | 0.01 | 0.598825 | 5.49 | 98.5 | 101.5 | 10.84 | 6.47 | | High | High | High | High | High | High | High | High | High | High | High | 0 | 4.9E-05 | 2.85 |
| FALSE | High | SSDGSLSHI | [R].SSDGSL | 2xPhospho | B8QI33 2xP | 0.002974 | 0.00016 | 1 | 3 | 3 | B8QI33 | Liprin-alph | B8QI33 [23 | 0 | 1963.882 | 0 | -0.16 | 0.053157 | 7.24 | 107.9 | 92.1 | 4.45 | 4.39 | | High | High | High | High | High | High | High | High | High | High | High | 7.19E-05 | 0.00081 | 2.69 |
| FALSE | High | SSDLEDNR | [K].SSDLED | 1xCarbami | Q8C0T5 1x | 0.019276 | 0.001402 | 1 | 4 | 3 | Q8C0T5 | Signal-indu | Q8C0T5 [3 | 1 | 2916.36 | 0 | 2.24 | 0.000928 | 14.41 | 31.7 | 168.3 | 12.39 | 12.51 | | High | High | High | High | High | High | High | High | High | High | High | 0.007735 | 0.08363 | 1.95 |
| FALSE | High | SSDIISLLVS | [R].SSDIISL | 1xPhospho | Q6ZPR8 1x | 7.82E-05 | 0 | 1 | 3 | 2 | Q6ZPR8 | MKIAA140 | Q6ZPR8 [5 | 0 | 2120.166 | 0 | 1.27 | 0.000522 | 19.02 | 55.1 | 144.9 | 17.6 | 5.33 | | High | High | High | High | High | High | High | High | High | High | High | 0.001453 | 0.01459 | 3.05 |
| FALSE | High | SSDIVSSVR | [R].SSDIVS | 1xPhospho | Q6PAR5 1x | 0.002647 | 0.000109 | 1 | 9 | 3 | Q6PAR5 | GTPase-act | Q6PAR5 [9 | 0 | 1258.624 | 0 | 0.23 | 0.059875 | 7.77 | 92.2 | 107.8 | 4.63 | 17.24 | | High | High | High | High | High | High | High | High | High | High | High | 0.000275 | 0.001623 | 2.26 |
| FALSE | High | SSDPDLSVI | [K].SSDPDL | 1xCarbami | Q9Z2C9 1x | 5.35E-08 | 0 | 1 | 1 | 7 | Q9Z2C9 | Myotubula | Q9Z2C9 [6 | 0 | 2705.145 | 0 | -0.79 | 0.001212 | 2.78 | 126.9 | 73.1 | 3.27 | 11.37 | | High | High | High | High | High | High | High | High | High | High | High | 0 | 3.77E-09 | 5.4 |
| FALSE | High | SSDPDLSVI | [K].SSDPDL | 1xCarbami | Q9Z2C9 2x | 2.46E-07 | 0 | 1 | 1 | 6 | Q9Z2C9 | Myotubula | Q9Z2C9 [6 | 0 | 2785.111 | 0 | -0.57 | 0.005002 | 2.11 | 121.1 | 78.9 | 4.51 | 14.7 | | High | High | High | High | High | High | High | High | High | High | High | 0 | 2.1E-08 | 5.25 |
| FALSE | High | SSDPDLSVI | [K].SSDPDL | 1xCarbami | Q9Z2C9 2x | 0.007881 | 0.000736 | 1 | 1 | 2 | Q9Z2C9 | Myotubula | Q9Z2C9 [6 | 0 | 2786.095 | 0 | -0.52 | 0.037993 | 8.38 | 117.7 | 82.3 | 6.83 | 20.61 | | High | High | High | High | High | High | High | High | High | High | High | 0.000449 | 0.004104 | 4.05 |
| FALSE | High | SSDPGPDIH | [K].SSDPGF | 1xPhospho | Q8BQ33 1x | 0.000264 | 0 | 1 | 1 | 1 | Q8BQ33 | Treslin OS= | Q8BQ33 [9 | 0 | 2360.142 | 0 | -0.53 | 0.001763 | 3.72 | 117.9 | 82.1 | 7.2 | 6.02 | | High | High | High | High | High | High | High | High | High | High | High | 0 | 5.35E-05 | 3.16 |
| FALSE | High | SSDPLGDT | [K].SSDPLG | 1xPhospho | A2AFQ0 1x | 3.57E-05 | 0 | 1 | 10 | 1 | A2AFQ0 | E3 ubiquiti | A2AFQ0 [6 | 0 | 2444.121 | 0 | 0.47 | 0.011237 | 23.83 | 83.7 | 116.3 | 13.63 | 7.72 | | High | High | High | High | High | High | High | High | High | High | High | 0 | 5.63E-06 | 2.38 |
| FALSE | High | SSDPSLNEK | [R].SSDPSL | 1xPhospho | Q8K296 1x | 0.002022 | 5.66E-05 | 1 | 7 | 2 | Q8K296 | Myotubula | Q8K296 [6 | 0 | 1514.75 | 0 | -0.04 | 0.623693 | 6.95 | 101.7 | 98.3 | 2.49 | 5.75 | | High | High | High | High | High | High | High | High | High | High | High | 7.19E-05 | 0.000801 | 2.39 |
| FALSE | High | SSDPTQDL | [K].SSDPTC | 1xPhospho | Q0VGY8 1x | 0.000128 | 0 | 1 | 4 | 3 | Q0VGY8 | Protein TA | Q0VGY8 [4 | 0 | 3010.482 | 0 | 0.48 | 0.002619 | 3.28 | 82.7 | 117.3 | 7.64 | 8.41 | | High | High | High | High | High | High | High | High | High | High | High | 2.58E-05 | 0.000435 | 3.86 |
| FALSE | High | SSDPWSNS | [R].SSDPW | 1xPhospho | O70472 1x | 0.000173 | 0 | 1 | 3 | 1 | O70472 | Transmem | O70472 [1 | 0 | 1949.802 | 0 | -0.02 | 0.335753 | 7.26 | 101.1 | 98.9 | 10.27 | 15.89 | | High | High | High | High | High | High | High | High | High | High | High | 0 | 3.32E-05 | 3.36 |
| FALSE | High | SSDQAVPA | [R].SSDQA | 1xPhospho | Q571J8 1xP | 8.11E-05 | 0 | 1 | 14 | 2 | Q571J8 | Non-specif | Q571J8 [41 | 0 | 2290.137 | 0 | 1.59 | 0.001713 | 9.83 | 49.8 | 150.2 | 7.85 | 16.52 | | High | High | High | High | High | High | High | High | High | High | High | 0 | 1.42E-05 | 3.79 |
| FALSE | High | SSDQVFVH | [K].SSDQVI | 1xPhospho | B2RX10 1x | 6.65E-05 | 0 | 1 | 3 | 1 | B2RX10 | Fanconi an | B2RX10 [1 | 0 | 1695.805 | 0 | -0.58 | 0.000605 | 2.1 | 121.6 | 78.4 | 3.2 | 5.88 | | High | High | High | High | High | High | High | High | High | High | High | 0 | 1.14E-05 | 2.79 |
| FALSE | High | SSDSDLQE | [K].SSDSDL | 1xPhospho | A0A0R4J0H | 3.49E-07 | 0 | 1 | 2 | 2 | A0A0R4J0H | Fibronectir | A0A0R4J0H | 0 | 2180.041 | 0 | 0.45 | 0.025565 | 21.64 | 84.6 | 115.4 | 8.69 | 10.7 | | High | High | High | High | High | High | High | High | High | High | High | 0 | 3.12E-08 | 3.53 |
| FALSE | High | SSDSEEAFE | [R].SSDSEE | 1xPhospho | E9Q8T1 1x | 1.17E-06 | 0 | 2 | 7 | 1 | E9Q8T1; Q | Transformi | E9Q8T1 [1 | 0 | 2479.153 | 0 | 0 | 0.924754 | 14.79 | 93.5 | 106.5 | 41.82 | 26.99 | Shared | High | High | High | High | High | High | High | High | High | High | High | 0 | 1.21E-07 | 3.52 |
| FALSE | High | SSDSEEAFE | [R].SSDSEE | 2xPhospho | E9Q8T1 2x | 0.000127 | 0 | 2 | 7 | 1 | E9Q8T1; Q | Transformi | E9Q8T1 [1 | 0 | 2559.119 | 0 | -1.61 | 0.001923 | 2.44 | 149.3 | 50.7 | 13.9 | 15.45 | Shared | High | High | High | High | High | High | High | High | High | High | High | 0 | 2.36E-05 | 3.78 |
| FALSE | High | SSDSVDIEE | [R].SSDSVD | 1xPhospho | Q6PR54-3 | 0.000151 | 0 | 2 | 4 | 4 | Q6PR54-3; | Isoform 3 c | Q6PR54-3 | 0 | 2032.936 | 0 | -1.27 | 0.000447 | 1.4 | 141.5 | 58.5 | 5.93 | 8.27 | Shared | High | High | High | High | High | High | High | High | High | High | High | 0 | 2.86E-05 | 3.96 |
| FALSE | High | SSCLSSR | [R].SSCLSS | 1xCarbami | Q3UHK6-3 | 0.096181 | 0.007535 | 1 | 9 | 1 | Q3UHK6-3 | Isoform 3 c | Q3UHK6-3 | 0 | 1105.491 | 0 | -0.93 | 0.003482 | 4.75 | 128.4 | 71.6 | 14.04 | 9.92 | | High | High | High | High | High | High | High | High | High | High | High | 0.003809 | 0.04111 | 1.56 |
| FALSE | High | SRSDIDVN | [R].SRSDID | 1xPhospho | B9EJA4 1xP | 1.56E-07 | 0 | 1 | 7 | 3 | B9EJA4 | Clasp2 pro | B9EJA4 [37 | 1 | 1912.989 | 0 | -0.63 | 0.003826 | 4.31 | 124.3 | 75.7 | 4.21 | 15.44 | | High | High | High | High | High | High | High | High | High | High | High | 0 | 6.44E-08 | 4.93 |
| FALSE | High | SSPADLVTV | [K].SSPADL | 1xPhospho | O55023 1x | 3.46E-07 | 0 | 1 | 8 | 2 | O55023 | Inositol mc | O55023 [3 | 0 | 1898.988 | 0 | -0.6 | 0.000995 | 3.56 | 118.4 | 81.6 | 4.38 | 11.52 | | High | High | High | High | High | High | High | High | High | High | High | 0 | 3.09E-08 | 3.29 |
| FALSE | High | SSPAGFSFC | [K].SSPAGF | 1xPhospho | E9Q3G8 1x | 3.61E-06 | 0 | 1 | 4 | 3 | E9Q3G8 | Nucleopor | E9Q3G8 [1 | 0 | 3360.615 | 0 | -0.35 | 0.010984 | 5.25 | 122.3 | 77.7 | 11.61 | 17.45 | | High | High | High | High | High | High | High | High | High | High | High | 0 | 4.3E-07 | 4.54 |
| FALSE | High | SSSLGNK | [R].SSSLGN | 1xPhospho | P83741-3 1 | 0.009168 | 0.000736 | 1 | 8 | 2 | P83741-3 | Isoform 3 c | P83741-3 [ | 0 | 1230.65 | 0 | 0.26 | 0.009204 | 5.86 | 91.1 | 108.9 | 2.17 | 5.68 | | High | High | High | High | High | High | High | High | High | High | High | 0.000327 | 0.002839 | 1.67 |
| FALSE | High | SSSLGNLK | [K].SSSLGN | 1xPhospho | F6YZ95 1xP | 0.010382 | 0.000772 | 2 | 7 | 2 | F6YZ95; Q8 | Liprin-beta | F6YZ95 [21 | 0 | 1343.734 | 0 | 0.7 | 0.002894 | 6.71 | 71.4 | 128.6 | 7.85 | 9.91 | Shared | High | High | High | High | High | High | High | High | High | High | High | 0.001011 | 0.01084 | 1.95 |
| FALSE | High | SSSLGSTPH | [R].SSSLGS | 1xPhospho | Q9EST3 1x | 0.001659 | 5.66E-05 | 1 | 6 | 3 | Q9EST3 | Eukaryotic | Q9EST3 [3 | 0 | 1837.853 | 0 | 0.11 | 0.497132 | 23.09 | 96.3 | 103.7 | 10.96 | 8.31 | | High | High | High | High | High | High | High | High | High | High | High | 0.000872 | 0.008841 | 1.95 |
| FALSE | High | SSSLGSTPH | [R].SSSLGS | 2xPhospho | Q9EST3 2x | 0.079114 | 0.00613 | 1 | 6 | 1 | Q9EST3 | Eukaryotic | Q9EST3 [3 | 0 | 1917.819 | 0 | -0.01 | 0.570486 | 24.36 | 99.3 | 100.7 | 18.34 | 12.29 | | High | High | High | High | High | High | High | High | High | High | High | 0.003148 | 0.03293 | 2.62 |
| FALSE | High | SSSLIQLTS | [R].SSSLIQI | 1xPhospho | E0CXT7 1xP | 4.94E-09 | 0 | 1 | 1 | 6 | E0CXT7 | Cleavage a | E0CXT7 [2 | 0 | 2284.113 | 0 | 0.23 | 0.006049 | 2.9 | 94.7 | 105.3 | 6.92 | 5.03 | | High | High | High | High | High | High | High | High | High | High | High | 0 | 2.61E-10 | 5.61 |
| FALSE | High | SSSLSDLTH | [R].SSSLSD | 1xPhospho | Q78T81 1x | 0.00407 | 0.00036 | 1 | 2 | 2 | Q78T81 | Protein FA | Q78T81 [2 | 0 | 1411.678 | 0 | -0.89 | 0.000181 | 1.06 | 131 | 69 | 7.68 | 10.41 | | High | High | High | High | High | High | High | High | High | High | High | 0.000161 | 0.001154 | 1.59 |
| FALSE | High | SSSLSDPLE | [R].SSSLSD | 1xPhospho | E9Q557 1x | 7.36E-06 | 0 | 1 | 3 | 12 | E9Q557 | Desmoplak | E9Q557 [2 | 0 | 3217.581 | 0 | 4.44 | 0.000158 | 12.9 | 9.4 | 190.6 | 20.97 | 4.73 | | High | High | High | High | High | High | High | High | High | High | High | 0 | 5.18E-06 | 4.46 |
| FALSE | High | SSSISGEICC | [R].SSSISGI | 1xCarbami | Q14BY7 1x | 7.76E-09 | 0 | 1 | 2 | 3 | Q14BY7 | Regulating | Q14BY7 [2 | 0 | 2750.307 | 0 | -1.9 | 0.002136 | 1.44 | 159.3 | 40.7 | 10.06 | 38.92 | | High | High | High | High | High | High | High | High | High | High | High | 0 | 8.11E-10 | 4.95 |
| FALSE | High | SSSLSVTTD | [K].SSSLSV | 1xPhospho | Q6ZQ32 1x | 3.47E-05 | 0 | 2 | 7 | 2 | Q6ZQ32; F | MKIAA085 | Q6ZQ32 [1 | 0 | 2918.396 | 0 | 0.59 | 0.012596 | 24.99 | 80.1 | 119.9 | 8.54 | 10.44 | Shared | High | High | High | High | High | High | High | High | High | High | High | 0 | 5.45E-06 | 3.49 |

| | | | | | | | | | | | | | | | | | | | | | | | | | | | | | | | | | | | | | | |
|---|---|---|---|---|---|---|---|---|---|---|---|---|---|---|---|---|---|---|---|---|---|---|---|---|---|---|---|---|---|---|---|---|---|---|---|---|---|---|
| FALSE | High | SSSISYSQE! | [R].SSSISYS | 1xCarbami | P50636 1xl | 0.002192 | 0.000109 | 1 | 1 | 2 | P50636 | E3 ubiquiti | P50636 [28 | 0 | 2864.354 | 0 | 0.66 | 0.093672 | 32.92 | 80 | 120 | 15.22 | 28.88 | | High | High | High | High | High | High | High | High | High | High | High | 4.94E-05 | 0.000576 | 3.31 |
| FALSE | High | SSSLTSSPE/ | [K].SSSLTS! | 1xPhospho | Q9D3X5 1x | 0.000121 | 0 | 1 | 3 | 4 | Q9D3X5 | PACRG-like | Q9D3X5 [4 | 0 | 1501.71 | 0 | -0.38 | 0.00323 | 3.68 | 112.6 | 87.4 | 4.3 | 6.07 | | High | High | High | High | High | High | High | High | High | High | High | 0 | 2.21E-05 | 1.93 |
| FALSE | High | SSSLVSPYN | [R].SSSLVS | 1xPhospho | Q3UMG6 1 | 0.000311 | 0 | 1 | 4 | 2 | Q3UMG6 | Non-metas | Q3UMG6 [ | 0 | 1820.927 | 0 | -0.28 | 0.104691 | 18.79 | 113.9 | 86.1 | 9.97 | 18.47 | | High | High | High | High | High | High | High | High | High | High | High | 0 | 6.45E-05 | 2.66 |
| FALSE | High | SSSMAAGL | [R].SSSMA/ | 1xPhospho | Q8BKX1 1x | 0.00925 | 0.000736 | 1 | 5 | 2 | Q8BKX1 | Brain-spec | Q8BKX1 [3 | 0 | 1317.607 | 0 | 1.35 | 0.003874 | 14.92 | 55.7 | 144.3 | 12.37 | 15.89 | | High | High | High | High | High | High | High | High | High | High | High | 0.000327 | 0.002888 | 2.77 |
| FALSE | High | SSSLGDLLR | [R].SSSLGD | 1xPhospho | Q8K124 1x | 0.00582 | 0.00062 | 1 | 1 | 4 | Q8K124 | Pleckstrin I | Q8K124 [3! | 0 | 1256.645 | 0 | -0.17 | 0.778463 | 20.1 | 105.7 | 94.3 | 6.82 | 35.48 | | High | High | High | High | High | High | High | High | High | High | High | 0.000275 | 0.00171 | 1.91 |
| FALSE | High | SSSMAAGL | [R].SSSMA/ | 1xPhospho | Q8BKX1 1x | 0.000184 | 0 | 1 | 5 | 5 | Q8BKX1 | Brain-spec | Q8BKX1 [3 | 0 | 1333.602 | 0 | 1.6 | 0.002128 | 18.43 | 53.3 | 146.7 | 12.46 | 15.09 | | High | High | High | High | High | High | High | High | High | High | High | 0 | 3.55E-05 | 1.9 |
| FALSE | High | SSSMSSIDL | [R].SSSMS! | 1xPhospho | Q9EQG9 1: | 3.3E-06 | 0 | 1 | 1 | 4 | Q9EQG9 | Collagen ty | Q9EQG9 [3 | 0 | 2217.99 | 0 | 0.04 | 0.263667 | 14.16 | 97 | 103 | 4.18 | 15.8 | | High | High | High | High | High | High | High | High | High | High | High | 0 | 6.09E-07 | 3.64 |
| FALSE | High | SSSMSSIDL | [R].SSSMS! | 2xPhospho | Q9EQG9 2: | 0.002576 | 0.000109 | 1 | 1 | 3 | Q9EQG9 | Collagen ty | Q9EQG9 [3 | 0 | 2281.961 | 0 | 0.04 | 0.368515 | 24.87 | 97.4 | 102.6 | 21.02 | 10.52 | | High | High | High | High | High | High | High | High | High | High | High | 0.000161 | 0.001167 | 3.91 |
| FALSE | High | SSSMSYVD | [R].SSSMSY | 1xPhospho | H7BX08 1x | 0.008691 | 0.000736 | 1 | 5 | 2 | H7BX08 | Calmodulir | H7BX08 [4( | 0 | 2300.107 | 0 | 0.81 | 0.002605 | 14.99 | 72.6 | 127.4 | 5.33 | 11.15 | | High | High | High | High | High | High | High | High | High | High | High | 0.000327 | 0.002681 | 2.54 |
| FALSE | High | SSSMSYVD | [R].SSSMSY | 1xPhospho | H7BX08 1x | 0.049971 | 0.003641 | 1 | 5 | 2 | H7BX08 | Calmodulir | H7BX08 [4( | 0 | 2316.102 | 0 | 1.04 | 0.00367 | 21.74 | 61.7 | 138.3 | 14.23 | 11.5 | | High | High | High | High | High | High | High | High | High | High | High | 0.001902 | 0.01917 | 2.21 |
| FALSE | High | SSSNVNGG | [R].SSSNVN | 1xDeamida | Q6PGN3 1: | 0.00625 | 0.00062 | 1 | 9 | 6 | Q6PGN3 | Serine/thre | Q6PGN3 [3 | 0 | 1641.732 | 0 | -1.83 | 0.000962 | 1.5 | 156.2 | 43.8 | 8.48 | 13.64 | | High | High | High | High | High | High | High | High | High | High | High | 0.000968 | 0.01004 | 2.59 |
| FALSE | High | SSSPAELEL | [R].SSSPAE | 1xPhospho | A0A0A0M( | 0.000731 | 0 | 1 | 4 | 1 | A0A0A0M( | Ral GTPase | A0A0A0M( | 0 | 1598.844 | 0 | 1.1 | 0.000795 | 10.21 | 64.4 | 135.6 | 4.73 | 7.03 | | High | High | High | High | High | High | High | High | High | High | High | 0 | 0.000167 | 2.28 |
| FALSE | High | SSSPDLSSS | [R].SSSPDL | 1xPhospho | E9Q8I9-2 1 | 0.001058 | 0 | 1 | 2 | 3 | E9Q8I9-2 | Isoform 2 ( | E9Q8I9-2 [ | 0 | 1619.793 | 0 | -0.8 | 0.000888 | 4.4 | 126.7 | 73.3 | 6.05 | 8.89 | | High | High | High | High | High | High | High | High | High | High | High | 0 | 0.000254 | 2.37 |
| FALSE | High | SSSPDSK | [R].SSSPDS | 1xPhospho | Q8BTI8 1xP | 0.101749 | 0.008194 | 1 | 3 | 1 | Q8BTI8 | Serine/argi | Q8BTI8 [94 | 0 | 1245.613 | 0 | -1.17 | 0.00203 | 3.45 | 138.7 | 61.3 | 12.79 | 5.83 | | High | High | High | High | High | High | High | High | High | High | High | 0.004126 | 0.04415 | 1.13 |
| FALSE | High | SSSPEPVTH | [K].SSSPEP | 1xPhospho | A2AH25 1x | 0.007274 | 0.00062 | 1 | 3 | 7 | A2AH25 | Rho GTPas | A2AH25 [8 | 0 | 1719.908 | 0 | 0.15 | 0.016296 | 5.42 | 93.8 | 106.2 | 8 | 3.34 | | High | High | High | High | High | High | High | High | High | High | High | 0.000275 | 0.002188 | 3.03 |
| FALSE | High | SSSPEPVTH | [K].SSSPEP | 2xPhospho | A2AH25 2x | 0.004824 | 0.000456 | 1 | 3 | 2 | A2AH25 | Rho GTPas | A2AH25 [8 | 0 | 1799.875 | 0 | 0.67 | 0.002999 | 2.55 | 83.2 | 116.8 | 7.12 | 7.48 | | High | High | High | High | High | High | High | High | High | High | High | 0.000204 | 0.001392 | 2.36 |
| FALSE | High | SSSPFLSK | [R].SSSPFL! | 2xPhospho | Q14AX6-2 | 0.074071 | 0.0057 | 3 | 5 | 2 | Q14AX6-2; | Isoform 2 ( | Q14AX6-2 | 0 | 1241.542 | 0 | -0.33 | 0.342423 | 17.18 | 111.4 | 88.6 | 14.45 | 20.8 | Shared | High | High | High | High | High | High | High | High | High | High | High | 0.002924 | 0.03034 | 2.35 |
| FALSE | High | SSSPFLSK | [R].SSSPFL! | 1xPhospho | Q14AX6-2 | 0.04913 | 0.003576 | 3 | 5 | 1 | Q14AX6-2; | Isoform 2 ( | Q14AX6-2 | 0 | 1390.738 | 0 | -0.53 | 0.010569 | 7.57 | 119.3 | 80.7 | 6.86 | 11.71 | Shared | High | High | High | High | High | High | High | High | High | High | High | 0.001889 | 0.01888 | 2.09 |
| FALSE | High | SSSPFLSK | [R].SSSPFL! | 2xPhospho | Q14AX6-2 | 0.035181 | 0.002269 | 3 | 5 | 2 | Q14AX6-2; | Isoform 2 ( | Q14AX6-2 | 0 | 1470.705 | 0 | -0.01 | 0.929149 | 8.75 | 100.5 | 99.5 | 2.09 | 7.83 | Shared | High | High | High | High | High | High | High | High | High | High | High | 0.001199 | 0.01288 | 2.15 |
| FALSE | High | SSSMLGSP( | [R].SSSML( | 1xPhospho | Q80WQ8 1 | 0.058173 | 0.004227 | 1 | 4 | 2 | Q80WQ8 | Mis18-binc | Q80WQ8 [ | 0 | 1999.956 | 0 | -0.43 | 0.042075 | 12.25 | 115.5 | 84.5 | 12.22 | 12.05 | | High | High | High | High | High | High | High | High | High | High | High | 0.004707 | 0.05053 | 2.33 |
| FALSE | High | SSSPGKPQ/ | [R].SSSPGK | 1xPhospho | Q9ESC8 1x | 0.000117 | 0 | 1 | 2 | 2 | Q9ESC8 | AF4/FMR2 | Q9ESC8 [1 | 0 | 2424.228 | 0 | -0.4 | 0.100553 | 10.05 | 122.2 | 77.8 | 29.14 | 23.27 | | High | High | High | High | High | High | High | High | High | High | High | 0 | 2.15E-05 | 3.8 |
| FALSE | High | SSSLESQGI | [R].SSSLES( | 1xPhospho | Q8BIE6-2 1 | 0.001463 | 5.66E-05 | 1 | 4 | 2 | Q8BIE6-2 | Isoform 2 ( | Q8BIE6-2 [ | 0 | 1460.74 | 0 | 0.22 | 0.010811 | 2.84 | 88.6 | 111.4 | 4.48 | 8.54 | | High | High | High | High | High | High | High | High | High | High | High | 0.000275 | 0.001712 | 2.04 |
| FALSE | High | SSSLDMNR | [R].SSSLDM | 1xPhospho | O54916 1x | 0.002818 | 0.000109 | 1 | 5 | 7 | O54916 | RalBP1-ass | O54916 [5! | 0 | 1234.534 | 0 | -0.36 | 0.044386 | 6.27 | 110.3 | 89.7 | 7.67 | 11.9 | | High | High | High | High | High | High | High | High | High | High | High | 0.004317 | 0.04615 | 1.49 |
| FALSE | High | SSSELSPEV | [R].SSSELSI | 1xPhospho | Q8BTI8 1xP | 7.82E-05 | 0 | 1 | 3 | 8 | Q8BTI8 | Serine/argi | Q8BTI8 [13 | 0 | 1828.935 | 0 | -1.12 | 0.000288 | 1.54 | 134.9 | 65.1 | 3.51 | 6.26 | | High | High | High | High | High | High | High | High | High | High | High | 0 | 1.55E-05 | 3.72 |
| FALSE | High | SSSELSPEV | [R].SSSELSI | 2xPhospho | Q8BTI8 2xP | 0.002719 | 0.000109 | 1 | 3 | 3 | Q8BTI8 | Serine/argi | Q8BTI8 [13 | 0 | 1908.901 | 0 | -1.36 | 0.00041 | 0.79 | 144 | 56 | 6.3 | 2.91 | | High | High | High | High | High | High | High | High | High | High | High | 0.001453 | 0.01459 | 2.57 |
| FALSE | High | SSSEITLSE( | [R].SSSEITL | 1xCarbami | G3X9J0 1xI | 4.35E-05 | 0 | 1 | 1 | 7 | G3X9J0 | Signal-indu | G3X9J0 [16 | 0 | 2416.042 | 0 | 0.96 | 0.001529 | 18.15 | 65.9 | 134.1 | 15.08 | 8.29 | | High | High | High | High | High | High | High | High | High | High | High | 0 | 7.02E-06 | 4.65 |
| FALSE | High | SSSESPGST | [K].SSSESP | 1xPhospho | A2A654 1x | 0.003282 | 0.00016 | 1 | 4 | 1 | A2A654 | Bromodom | A2A654 [1: | 0 | 1504.73 | 0 | -0.49 | 0.001217 | 3.75 | 117.1 | 82.9 | 7.89 | 6.33 | | High | High | High | High | High | High | High | High | High | High | High | 7.19E-05 | 0.000903 | 1.4 |
| FALSE | High | SSSESTSST | [R].SSSEST! | 1xDeamida | Q99MY8 1: | 0.051258 | 0.003745 | 1 | 1 | 2 | Q99MY8 | Histone-lys | Q99MY8 [1 | 1 | 2232.034 | 0 | 0.16 | 0.519451 | 20.45 | 95 | 105 | 12.81 | 8.96 | | High | High | High | High | High | High | High | High | High | High | High | 0.00196 | 0.01976 | 2.74 |
| FALSE | High | SSSETALSS | [R].SSSETA | 1xPhospho | P97817 1xl | 1.07E-05 | 0 | 1 | 2 | 6 | P97817 | Cerebellar | P97817 [3( | 0 | 2218.126 | 0 | 0.1 | 0.211213 | 4.34 | 98 | 102 | 7.55 | 7.61 | | High | High | High | High | High | High | High | High | High | High | High | 2.58E-05 | 0.000465 | 2.83 |
| FALSE | High | SSSFGSVST | [R].SSSFGS | 1xPhospho | Q8C080 1x | 7.86E-07 | 0 | 1 | 3 | 3 | Q8C080 | Sorting ne> | Q8C080 [2 | 0 | 1973.947 | 0 | 0.49 | 0.007308 | 14.28 | 81.8 | 118.2 | 12.03 | 5.75 | | High | High | High | High | High | High | High | High | High | High | High | 0 | 7.8E-08 | 2.83 |
| FALSE | High | SSSFSEEK | [K].SSSFSEI | 1xPhospho | F6RJ39 1xP | 0.115566 | 0.009504 | 2 | 5 | 1 | F6RJ39; Q8 | Apoptotic ( | F6RJ39 [15 | 0 | 1438.687 | 0 | -0.41 | 0.00034 | 1.78 | 113.9 | 86.1 | 3.21 | 2.64 | Shared | High | High | High | High | High | High | High | High | High | High | High | 0.004733 | 0.05102 | 2.1 |
| FALSE | High | SSSFSK | [RK].SSSFS | 1xPhospho | Q91W40 1: | 0.109329 | 0.008865 | 1 | 5 | 1 | Q91W40 | Kinesin ligh | Q91W40 [4 | 0 | 1180.602 | 0 | -0.82 | 0.000729 | 1.85 | 129.3 | 70.7 | 3.37 | 14.14 | | High | High | High | High | High | High | High | High | High | High | High | 0.004422 | 0.04786 | 1.38 |
| FALSE | High | SSSFSLPAR | [R].SSSFSLI | 1xPhospho | Q8BGR5 1x | 0.108463 | 0.008839 | 1 | 3 | 1 | Q8BGR5 | Protein FAI | Q8BGR5 [1 | 0 | 1260.619 | 0 | 1.71 | 0.000188 | 2.29 | 47.1 | 152.9 | 2.41 | 4.58 | | High | High | High | High | High | High | High | High | High | High | High | 0.004422 | 0.04764 | 1.6 |
| FALSE | High | SSSFSSWD | [R].SSSFSS\ | 1xPhospho | A0A0R4J0T | 8.79E-05 | 0 | 1 | 2 | 5 | A0A0R4J0T | ADP-ribosy | A0A0R4J0T | 0 | 2363.027 | 0 | 1.11 | 0.002668 | 9.56 | 61.7 | 138.3 | 11.88 | 11.93 | | High | High | High | High | High | High | High | High | High | High | High | 0.004422 | 0.04781 | 2.6 |
| FALSE | High | SSSFTLLQS | [R].SSSFTLI | 1xPhospho | E9Q2E4 1x | 0.005565 | 0.00062 | 1 | 2 | 2 | E9Q2E4 | Uncharacte | E9Q2E4 [1! | 0 | 2063.026 | 0 | 0.9 | 0.003317 | 14.51 | 68.4 | 131.6 | 8.85 | 9.47 | | High | High | High | High | High | High | High | High | High | High | High | 0.000275 | 0.001629 | 3.08 |
| FALSE | High | SSSGESLAC | [R].SSSGES | 1xPhospho | Q3UHI0-2 | 0.003402 | 0.00021 | 1 | 2 | 2 | Q3UHI0-2 | Isoform 2 ( | Q3UHI0-2 | 0 | 2015.969 | 0 | -0.22 | 0.045493 | 5.36 | 107.5 | 92.5 | 4.2 | 11.42 | | High | High | High | High | High | High | High | High | High | High | High | 9.42E-05 | 0.000944 | 2.24 |
| FALSE | High | SSSLEGFHS | [R].SSSLEG | 1xPhospho | A0A1B0GS | 6.18E-05 | 0 | 1 | 2 | 4 | A0A1B0GS | GRB2-asso | A0A1B0GS | 0 | 1907.93 | 0 | -1.89 | 0.000308 | 0.6 | 156.7 | 43.3 | 3.06 | 22.36 | | High | High | High | High | High | High | High | High | High | High | High | 0 | 1.05E-05 | 2.2 |
| FALSE | High | SSSGSEHST | [R].SSSGSE | 1xPhospho | G3X9Q6 1x | 0.002153 | 5.66E-05 | 1 | 4 | 2 | G3X9Q6 | La-related | G3X9Q6 [3 | 0 | 2442.098 | 0 | -0.35 | 0.469387 | 8.29 | 115.9 | 84.1 | 30.34 | 36.57 | | High | High | High | High | High | High | High | High | High | High | High | 2.58E-05 | 0.000565 | 3.62 |
| FALSE | High | SSSGSSVQ! | [K].SSSGSS | 1xPhospho | O54826 1x | 1.63E-06 | 0 | 1 | 4 | 2 | O54826 | Protein AF- | O54826 [3( | 0 | 2656.198 | 0 | -0.48 | 0.003192 | 3.41 | 118.7 | 81.3 | 6.38 | 9.79 | | High | High | High | High | High | High | High | High | High | High | High | 0 | 1.77E-07 | 4.06 |
| FALSE | High | SSSGSTHS( | [R].SSSGST | 1xPhospho | Q9R020 1x | 0.008167 | 0.000736 | 1 | 4 | 1 | Q9R020 | Zinc finger | Q9R020 [3 | 0 | 1358.59 | 0 | -0.21 | 0.589582 | 6 | 108.2 | 91.8 | 17.79 | 10.05 | | High | High | High | High | High | High | High | High | High | High | High | 0.000327 | 0.002513 | 1.57 |
| FALSE | High | SSSGSWGT | [R].SSSGSV | 1xPhospho | E9Q8G4 1x | 0.016163 | 0.001124 | 1 | 18 | 1 | E9Q8G4 | Transcripti | E9Q8G4 [7 | 0 | 1852.818 | 0 | -0.8 | 0.065701 | 4.09 | 128.2 | 71.8 | 25.35 | 18.63 | | High | High | High | High | High | High | High | High | High | High | High | 0.000585 | 0.005364 | 2.28 |
| FALSE | High | SSSGYFSFD | [R].SSSGYF | 1xPhospho | Q542N5 1x | 0.010475 | 0.000772 | 1 | 1 | 2 | Q542N5 | Bcl-2-like p | Q542N5 [8 | 0 | 1677.7 | 0 | 0.24 | 0.15062 | 27.98 | 85.3 | 114.7 | 15.79 | 9.52 | | High | High | High | High | High | High | High | High | High | High | High | 0.000382 | 0.003289 | 2.41 |
| FALSE | High | SSSHDFDPT | [R].SSSHDF | 1xPhospho | Q8R4H2 1x | 6.02E-05 | 0 | 2 | 3 | 2 | Q8R4H2; F: | Rho guanir | Q8R4H2 [3 | 0 | 2034.906 | 0 | 2.63 | 0.00532 | 30.34 | 29 | 171 | 13.8 | 25.63 | Shared | High | High | High | High | High | High | High | High | High | High | High | 0 | 1.02E-05 | 3.49 |
| FALSE | High | SSSHDFDPT | [R].SSSHDF | 1xPhospho | A0A1L1SUI | 0.005044 | 0.000504 | 1 | 1 | 2 | A0A1L1SUI | Rho guanir | A0A1L1SUI | 0 | 1854.858 | 0 | 0.32 | 0.170826 | 12.65 | 89.1 | 110.9 | 13.97 | 24.06 | | High | High | High | High | High | High | High | High | High | High | High | 0.000246 | 0.001464 | 2.72 |
| FALSE | High | SSSHLSSEE | [R].SSSHLS | 1xPhospho | Q80WQ8 1 | 0.004491 | 0.000409 | 1 | 3 | 1 | Q80WQ8 | Mis18-binc | Q80WQ8 [ | 0 | 2213.074 | 0 | -0.04 | 0.749576 | 25.19 | 105.1 | 94.9 | 19.13 | 16.93 | | High | High | High | High | High | High | High | High | High | High | High | 0.000183 | 0.001277 | 2.45 |
| FALSE | High | SSSHLSSEE | [R].SSSHLS | 2xPhospho | Q80WQ8 2 | 9.42E-07 | 0 | 1 | 3 | 1 | Q80WQ8 | Mis18-binc | Q80WQ8 [ | 0 | 2293.04 | 0 | -0.31 | 0.292224 | 25.18 | 110.7 | 89.3 | 12.69 | 13.32 | | High | High | High | High | High | High | High | High | High | High | High | 0 | 9.52E-08 | 2.7 |
| FALSE | High | SSSHSFCSV | [R].SSSHSF | 1xCarbami | Q8BP27 1x | 0.000319 | 0 | 1 | 2 | 8 | Q8BP27 | Swi5-depe | Q8BP27 [1: | 0 | 1762.86 | 0 | -0.51 | 0.022497 | 11.34 | 117.4 | 82.6 | 10.49 | 13.75 | | High | High | High | High | High | High | High | High | High | High | High | 0.001011 | 0.01102 | 2.47 |
| FALSE | High | SSSLDALGF | [R].SSSLDA | 1xPhospho | Q8C3W1 1 | 0.012072 | 0.000889 | 1 | 1 | 2 | Q8C3W1 | Uncharacte | Q8C3W1 [: | 0 | 1382.688 | 0 | 0.41 | 0.043323 | 3.25 | 82.8 | 117.2 | 10.19 | 19.05 | | High | High | High | High | High | High | High | High | High | High | High | 0.000449 | 0.003882 | 1.86 |
| FALSE | High | SSSLDGDH | [R].SSSLDG | 1xPhospho | Q8K4P0 1x | 0.001068 | 0 | 1 | 2 | 3 | Q8K4P0 | pre-mRNA | Q8K4P0 [1: | 0 | 1891.792 | 0 | 0.18 | 0.011774 | 3.83 | 95 | 105 | 10.39 | 7.45 | | High | High | High | High | High | High | High | High | High | High | High | 0.004115 | 0.0439 | 1.71 |
| FALSE | High | SSSIDGSN\ | [R].SSSIDG | 1xPhospho | Q8BXJ2-4 1 | 0.004955 | 0.000504 | 2 | 3 | 4 | Q8BXJ2-4; | Isoform 4 ( | Q8BXJ2-4 [ | 0 | 2837.34 | 0 | 0.3 | 0.072777 | 32.99 | 88.7 | 111.3 | 12.53 | 10.66 | Shared | High | High | High | High | High | High | High | High | High | High | High | 0.000327 | 0.002958 | 3.94 |
| FALSE | High | SSSLDMNR | [R].SSSLDM | 1xPhospho | O54916 1x | 0.008388 | 0.000736 | 1 | 5 | 2 | O54916 | RalBP1-ass | O54916 [5! | 0 | 1218.539 | 0 | -0.2 | 0.103533 | 13.22 | 109.8 | 90.2 | 8.37 | 12.13 | | High | High | High | High | High | High | High | High | High | High | High | 0.000327 | 0.002566 | 1.53 |
| FALSE | High | SSSGSSDSI | [K].SSSGSS | 1xPhospho | P97440 1xl | 0.002355 | 0.000109 | 1 | 3 | 1 | P97440 | Histone RN | P97440 [11 | 0 | 1379.646 | 0 | 0.02 | 0.435481 | 11.75 | 98.3 | 101.7 | 9.99 | 6.24 | | High | High | High | High | High | High | High | High | High | High | High | 4.94E-05 | 0.000623 | 2.02 |
| FALSE | High | SSSPGPIER | [K].SSSPGP | 1xPhospho | J3QNH8 1x | 0.045508 | 0.003373 | 1 | 1 | 2 | J3QNH8 | Uncharacte | J3QNH8 [2 | 0 | 1238.598 | 0 | 4.24 | 0.000201 | 26.21 | 9.9 | 190.1 | 25.46 | 6.96 | | High | High | High | High | High | High | High | High | High | High | High | 0.001741 | 0.01732 | 1.61 |
| FALSE | High | SSSPLPTVC | [K].SSSPLP | 1xPhospho | A0A0A6YY! | 0.016596 | 0.001124 | 1 | 13 | 2 | A0A0A6YY! | Protocadh | A0A0A6YY! | 0 | 2470.311 | 0 | -0.31 | 0.313653 | 31.17 | 117.4 | 82.6 | 16.7 | 24.38 | | High | High | High | High | High | High | High | High | High | High | High | 0.004399 | 0.04715 | 2.38 |
| FALSE | High | SSSPPIEIQ( | [R].SSSPPIE | 1xPhospho | B9EKB3 1xl | 3.57E-05 | 0 | 1 | 3 | 2 | B9EKB3 | E3 ubiquiti | B9EKB3 [1! | 0 | 2842.321 | 0 | 0.06 | 0.253264 | 15.36 | 96.9 | 103.1 | 3.54 | 7.42 | | High | High | High | High | High | High | High | High | High | High | High | 0 | 5.63E-06 | 3.64 |
| FALSE | High | SSSSPEVVK | [K].SSSSPE | 1xPhospho | G3UZM1 1 | 0.005135 | 0.000551 | 1 | 4 | 6 | G3UZM1 | Probable J( | G3UZM1 [( | 0 | 1912.076 | 0 | -0.81 | 0.003196 | 4.73 | 126.4 | 73.6 | 11.19 | 5 | | High | High | High | High | High | High | High | High | High | High | High | 0.000246 | 0.001493 | 2.46 |
| FALSE | High | SSSSQTLTC | [R].SSSSQT | 1xPhospho | O54916 1x | 2.43E-06 | 0 | 1 | 3 | 6 | O54916 | RalBP1-ass | O54916 [4: | 0 | 3473.715 | 0 | -0.04 | 0.83735 | 17.04 | 102.1 | 97.9 | 13.96 | 18.84 | | High | High | High | High | High | High | High | High | High | High | High | 0 | 2.74E-07 | 4.51 |
| FALSE | High | SSSSSASPS | [R].SSSSSA | 1xPhospho | Q8K019 1x | 0.020142 | 0.001472 | 1 | 6 | 4 | Q8K019 | Bcl-2-assoc | Q8K019 [7: | 0 | 1619.711 | 0 | 0.32 | 0.308451 | 30.94 | 87.9 | 112.1 | 19.55 | 18.56 | | High | High | High | High | High | High | High | High | High | High | High | 0.003188 | 0.03329 | 1.55 |
| FALSE | High | SSSSSGDPS | [K].SSSSSG | 1xPhospho | Q3UMA0 1 | 0.002134 | 5.66E-05 | 1 | 3 | 2 | Q3UMA0 | Putative ur | Q3UMA0 [ | 0 | 2284.079 | 0 | 6.64 | | 0 | | 200 | | 20.4 | | Not Found | Not Found | Not Found | Not Found | Not Found | High | High | High | High | High | High | 2.58E-05 | 0.000559 | 2.29 |
| FALSE | High | SSSSSGVPY | [K].SSSSSG | 1xPhospho | O35250 1x | 2.34E-06 | 0 | 1 | 7 | 7 | O35250 | Exocyst co | O35250 [2 | 0 | 2116.073 | 0 | 0.33 | 0.014555 | 12.31 | 88.6 | 111.4 | 2.27 | 9.95 | | High | High | High | High | High | High | High | High | High | High | High | 0 | 3.68E-05 | 2.55 |
| FALSE | High | SSSSSPTSP | [K].SSSSSP | 1xPhospho | Q6PGN3 1: | 0.000162 | 0 | 1 | 9 | 2 | Q6PGN3 | Serine/thre | Q6PGN3 [3 | 0 | 1592.716 | 0 | -1.99 | 0.000727 | 0.83 | 160.3 | 39.7 | 10.53 | 6.74 | | High | High | High | High | High | High | High | High | High | High | High | 0 | 3.09E-05 | 1.35 |
| FALSE | High | SSSSSSQSS | [R].SSSSSS( | 1xPhospho | Q569Z6 1x | 0.009416 | 0.000736 | 1 | 3 | 3 | Q569Z6 | Thyroid ho | Q569Z6 [7 | 0 | 1896.874 | 0 | 0.35 | 0.632185 | 28.25 | 90.1 | 109.9 | 23.41 | 19.63 | | High | High | High | High | High | High | High | High | High | High | High | 0.000449 | 0.004037 | 2.26 |
| FALSE | High | SSSSSSSSS | [R].SSSSSS! | 2xPhospho | Q8BTI8 2xP | 0.117405 | 0.009523 | 1 | 3 | 4 | Q8BTI8 | Serine/argi | Q8BTI8 [24 | 0 | 4827.026 | 0 | -0.51 | 0.06648 | 22.42 | 123 | 77 | 8.08 | 28.87 | | High | High | High | High | High | High | High | High | High | High | High | 0.006033 | 0.06526 | 3.73 |
| FALSE | High | SSSSSTTPT | [R].SSSSST | 1xPhospho | B2RY19 1xl | 0.012845 | 0.000889 | 1 | 7 | 6 | B2RY19 | Cgnl1 prot( | B2RY19 [2! | 0 | 2143.057 | 0 | 1.58 | 0.001004 | 6.64 | 49.4 | 150.6 | 15.07 | 15.54 | | High | High | High | High | High | High | High | High | High | High | High | 0.001151 | 0.01248 | 2.7 |
| FALSE | High | SSSSVLSSL | [K].SSSSVL! | 1xPhospho | M1VMF6 1 | 1.64E-05 | 0 | 1 | 8 | 2 | M1VMF6 | Metastasis | M1VMF6 [ | 0 | 1801.971 | 0 | -0.17 | 0.128754 | 6.84 | 103 | 97 | 7.34 | 7.11 | | High | High | High | High | High | High | High | High | High | High | High | 0 | 2.36E-06 | 3.52 |
| FALSE | High | SSSSVSPSS | [K].SSSSVS | 1xCarbami | A2ALK8 1xl | 1.31E-07 | 0 | 1 | 1 | 7 | A2ALK8 | Tyrosine-p | A2ALK8 [4! | 0 | 2574.098 | 0 | 0.62 | 0.01329 | 7.43 | 76.9 | 123.1 | 9.56 | 12.86 | | High | High | High | High | High | High | High | High | High | High | High | 0 | 1.03E-08 | 3.83 |
| FALSE | High | SSSSVSPSS | [K].SSSSVS | 1xCarbami | A2ALK8 2xl | 0.002058 | 5.66E-05 | 1 | 1 | 1 | A2ALK8 | Tyrosine-p | A2ALK8 [4! | 0 | 2654.064 | 0 | 0.29 | 0.072217 | 8 | 91.9 | 108.1 | 6.25 | 12.47 | | High | High | High | High | High | High | High | High | High | High | High | 2.58E-05 | 0.000538 | 3.14 |
| FALSE | High | SSSSVTTSE | [K].SSSSVT | 1xCarbami | P70460 1xl | 6.62E-07 | 0 | 1 | 1 | 6 | P70460 | Vasodilato | P70460 [31 | 0 | 2948.23 | 0 | 1.19 | 0.002676 | 22.73 | 65.9 | 134.1 | 15.92 | 11.42 | | High | High | High | High | High | High | High | High | High | High | High | 0 | 4.44E-07 | 4.47 |
| FALSE | High | SSSSPDGSI | [R].SSSSPD | 1xPhospho | E9QLN8 1x | 0.004333 | 0.00036 | 1 | 2 | 3 | E9QLN8 | Ribonuclea | E9QLN8 [7 | 0 | 1389.666 | 0 | 0.08 | 0.030716 | 4.8 | 95.8 | 104.2 | 2.17 | 3.85 | | High | High | High | High | High | High | High | High | High | High | High | 0.000327 | 0.002782 | 2.03 |
| FALSE | High | SSSSWIESL | [R].SSSSWI | 1xPhospho | D3YXQ5 1x | 0.01186 | 0.000889 | 1 | 4 | 2 | D3YXQ5 | Phospholip | D3YXQ5 [1 | 0 | 1460.698 | 0 | 1.67 | 0.000167 | 8.71 | 45.5 | 154.5 | 12.49 | 6.66 | | High | High | High | High | High | High | High | High | High | High | High | 0.000847 | 0.008043 | 2.53 |
| FALSE | High | SSSTGNLLL | [K].SSSTGN | 1xPhospho | Q8BKX1 1x | 0.002355 | 0.000109 | 1 | 5 | 3 | Q8BKX1 | Brain-spec | Q8BKX1 [4 | 0 | 1559.808 | 0 | 0.59 | 0.099516 | 24.86 | 80.8 | 119.2 | 5.29 | 51.12 | | High | High | High | High | High | High | High | High | High | High | High | 4.94E-05 | 0.00062 | 2.4 |
| FALSE | High | SSSTLFSTA | [R].SSSTLF! | 1xPhospho | Q91ZX6 1x | 1.06E-05 | 0 | 1 | 1 | 2 | Q91ZX6 | Sentrin-spe | Q91ZX6 [3: | 0 | 2407.205 | 0 | -0.15 | 0.948156 | 3.54 | 101.5 | 98.5 | 6.82 | 11.48 | | High | High | High | High | High | High | High | High | High | High | High | 0 | 1.44E-06 | 3.7 |
| FALSE | High | SSSTRSIDT( | [R].SSSTRS | 2xPhospho | Q3U3E2 2x | 0.012072 | 0.000889 | 1 | 1 | 1 | Q3U3E2 | Protein FAI | Q3U3E2 [3 | 1 | 2326.073 | 0 | 1.08 | 0.000181 | 7.02 | 65.2 | 134.8 | 10.54 | 6.02 | | High | High | High | High | High | High | High | High | High | High | High | 0.000449 | 0.003882 | 2.36 |
| FALSE | High | SSSTSDILEI | [R].SSSTSD | 1xPhospho | A0A1W2P7 | 0.002096 | 5.66E-05 | 1 | 7 | 4 | A0A1W2P7 | Ral GTPase | A0A1W2P7 | 0 | 1976.942 | 0 | 1.28 | 0.002116 | 12.02 | 58.3 | 141.7 | 8.89 | 10.99 | | High | High | High | High | High | High | High | High | High | High | High | 0.000161 | 0.001206 | 2.6 |
| FALSE | High | SSSTSSSTV | [R].SSSTSS! | 1xPhospho | Q8CCI5 1xl | 2.51E-08 | 0 | 1 | 1 | 26 | Q8CCI5 | RING1 and | Q8CCI5 [17 | 0 | 3463.523 | 0 | -1.43 | 0.000547 | 1.65 | 147.3 | 52.7 | 5.64 | 9.81 | | High | High | High | High | High | High | High | High | High | High | High | 0 | 1.49E-08 | 6.23 |
| FALSE | High | SSSTSSSTV | [R].SSSTSS! | 1xDeamida | Q8CCI5 1xl | 6.21E-10 | 0 | 1 | 1 | 27 | Q8CCI5 | RING1 and | Q8CCI5 [17 | 0 | 3464.507 | 0 | -1.59 | 0.002393 | 1.1 | 150 | 50 | 13.96 | 8.29 | | High | High | High | High | High | High | High | High | High | High | High | 0 | 7.26E-11 | 7.22 |
| FALSE | High | SSSTSSSTV | [R].SSSTSS! | 2xPhospho | Q8CCI5 2xl | 2.47E-06 | 0 | 1 | 1 | 5 | Q8CCI5 | RING1 and | Q8CCI5 [17 | 0 | 3543.49 | 0 | -1.18 | 0.019804 | 1.54 | 132.5 | 67.5 | 18.22 | 15.17 | | High | High | High | High | High | High | High | High | High | High | High | 0 | 2.8E-07 | 4.9 |
| FALSE | High | SSSVGSSSS | [R].SSSVGS | 1xPhospho | Q9QXS1-3 | 5.21E-08 | 0 | 3 | 23 | 13 | Q9QXS1-3; | Isoform PL | Q9QXS1-3 | 0 | 2021.938 | 0 | 2.11 | 0.001146 | 10.42 | 33.5 | 166.5 | 8.86 | 14.37 | Shared | High | High | High | High | High | High | High | High | High | High | High | 0 | 8.91E-08 | 3.39 |
| FALSE | High | SSSVGSSSS | [R].SSSVGS | 2xPhospho | Q9QXS1-3 | 0.000668 | 0 | 3 | 23 | 6 | Q9QXS1-3; | Isoform PL | Q9QXS1-3 | 0 | 2101.904 | 0 | 1.47 | 0.002468 | 6.51 | 52.9 | 147.1 | 9.11 | 18.04 | Shared | High | High | High | High | High | High | High | High | High | High | High | 0 | 0.000297 | 3.97 |
| FALSE | High | SSSVLSLEG | [R].SSSVLS | 1xPhospho | Q4VAC9 1x | 0.000738 | 0 | 1 | 3 | 4 | Q4VAC9 | Pleckstrin I | Q4VAC9 [7 | 0 | 1746.893 | 0 | 0.94 | 0.000348 | 8.87 | 68.1 | 131.9 | 5.3 | 3.32 | | High | High | High | High | High | High | High | High | High | High | High | 0.000204 | 0.001318 | 2.31 |
| FALSE | High | SSSVLSLEG | [R].SSSVLS | 2xPhospho | Q4VAC9 2x | 3.33E-06 | 0 | 1 | 3 | 1 | Q4VAC9 | Pleckstrin I | Q4VAC9 [7 | 0 | 1826.859 | 0 | 1.3 | 0.000822 | 15.07 | 58.9 | 141.1 | 3.46 | 8.76 | | High | High | High | High | High | High | High | High | High | High | High | 0 | 3.91E-07 | 3.14 |
| FALSE | High | SSSVSPSAS | [K].SSSVSP | 1xPhospho | Q3UY41 1x | 0.002441 | 0.000109 | 1 | 3 | 3 | Q3UY41 | Putative ur | Q3UY41 [2 | 0 | 1420.667 | 0 | -0.33 | 0.001151 | 3.64 | 109.9 | 90.1 | 4.53 | 4.28 | | High | High | High | High | High | High | High | High | High | High | High | 0.000327 | 0.002471 | 1.77 |
| FALSE | High | SSSVSPSAS | [K].SSSVSP | 2xPhospho | Q3UY41 2x | 0.007209 | 0.00062 | 1 | 3 | 2 | Q3UY41 | Putative ur | Q3UY41 [2 | 0 | 1500.633 | 0 | 0.6 | 0.002969 | 1.67 | 73.1 | 126.9 | 10.45 | 8.26 | | High | High | High | High | High | High | High | High | High | High | High | 0.000275 | 0.002178 | 2.1 |
| FALSE | High | SSSTEPPPP | [R].SSSTEP | 1xPhospho | Q8BTV2 1x | 0.000933 | 0 | 1 | 2 | 3 | Q8BTV2 | Cleavage a | Q8BTV2 [4 | 0 | 1462.714 | 0 | 0.37 | 0.003208 | 5.2 | 88.4 | 111.6 | 4.43 | 3.89 | | High | High | High | High | High | High | High | High | High | High | High | 0.000449 | 0.004078 | 2.28 |
| FALSE | High | SSSSLLASP! | [R].SSSSLL/ | 2xPhospho | Q3TZZ7 2xl | 2.98E-05 | 0 | 1 | 2 | 4 | Q3TZZ7 | Extended s | Q3TZZ7 [6( | 0 | 2074.039 | 0 | 0.89 | 0.002708 | 3.61 | 68.2 | 131.8 | 4.09 | 11.61 | | High | High | High | High | High | High | High | High | High | High | High | 0 | 4.59E-06 | 2.99 |
| FALSE | High | SSSSLLASP! | [R].SSSSLL/ | 1xPhospho | Q3TZZ7 1xl | 3.15E-06 | 0 | 1 | 2 | 3 | Q3TZZ7 | Extended s | Q3TZZ7 [6( | 0 | 1994.072 | 0 | 0.64 | 0.043229 | 11.88 | 78.2 | 121.8 | 12.52 | 15.79 | | High | High | High | High | High | High | High | High | High | High | High | 0 | 3.71E-07 | 3.2 |
| FALSE | High | SSSSLGSTS | [K].SSSSLG | 1xCarbami | Q8BW72 1 | 3.56E-11 | 0 | 1 | 3 | 4 | Q8BW72 | Lysine-spe | Q8BW72 [! | 0 | 3676.606 | 0 | -0.68 | 0.000597 | 1.48 | 123.2 | 76.8 | 1.74 | 7.14 | | High | High | High | High | High | High | High | High | High | High | High | 0 | 1.02E-12 | 5.52 |
| FALSE | High | SSSPQVVP! | [R].SSSPQV | 1xPhospho | Q3TZW5 1: | 1.97E-06 | 0 | 1 | 5 | 3 | Q3TZW5 | Putative ur | Q3TZW5 [1 | 0 | 1814.93 | 0 | -1.47 | 0.000389 | 0.31 | 145.9 | 54.1 | 8.42 | 7.62 | | High | High | High | High | High | High | High | High | High | High | High | 2.58E-05 | 0.000521 | 2.34 |
| FALSE | High | SSSPSASLT | [K].SSSPSA | 1xPhospho | D3Z3E0 1xl | 2.63E-05 | 0 | 1 | 3 | 5 | D3Z3E0 | Histone-lys | D3Z3E0 [6: | 0 | 2823.422 | 0 | 2.12 | 0.001004 | 7.32 | 36.5 | 163.5 | 6.55 | 12.44 | | High | High | High | High | High | High | High | High | High | High | High | 0 | 5.03E-06 | 3.77 |
| FALSE | High | SSSPSASLT | [K].SSSPSA | 1xPhospho | Q8BVE8-2 | 0.0003 | 0 | 1 | 3 | 2 | Q8BVE8-2 | Isoform 2 ( | Q8BVE8-2 | 0 | 4766.089 | 0 | -0.83 | 0.051498 | 7.19 | 118.7 | 81.3 | 13.25 | 22.56 | | High | High | High | High | High | High | High | High | High | High | High | 0 | 6.57E-05 | 4.1 |
| FALSE | High | SSSPSDFM | [K].SSSPSD | 1xPhospho | Q8BL06 1x | 0.002192 | 0.000109 | 1 | 1 | 5 | Q8BL06 | Inactive ub | Q8BL06 [1: | 0 | 2071.936 | 0 | -0.26 | 0.036468 | 9.17 | 109.1 | 90.9 | 10.69 | 8.49 | | High | High | High | High | High | High | High | High | High | High | High | 4.94E-05 | 0.000572 | 3.19 |
| FALSE | High | SSSPSFNSS | [R].SSSPSFI | 1xPhospho | E9Q1M1 1: | 6.94E-11 | 0 | 1 | 1 | 3 | E9Q1M1 | PDZ domai | E9Q1M1 [( | 0 | 2790.217 | 0 | 1.01 | 0.006548 | 22.46 | 67.2 | 132.8 | 20.88 | 7.84 | | High | High | High | High | High | High | High | High | High | High | High | 0 | 5.38E-12 | 4.68 |
| FALSE | High | SSSPSSPAA | [K].SSSPSSI | 1xPhospho | Q9R0A0 1x | 3.77E-05 | 0 | 1 | 2 | 1 | Q9R0A0 | Peroxisom; | Q9R0A0 [2 | 0 | 3649.702 | 0 | 0.57 | 0.459208 | 36.52 | 85.3 | 114.7 | 22.22 | 29.77 | | High | High | High | High | High | High | High | High | High | High | High | 0 | 5.99E-06 | 4.06 |
| FALSE | High | SSSPSSPAA | [K].SSSPSSI | 2xPhospho | Q9R0A0 2x | 0.0008 | 0 | 1 | 2 | 1 | Q9R0A0 | Peroxisom; | Q9R0A0 [2 | 0 | 3729.668 | 0 | 1.2 | 0.02961 | 56.27 | 63.9 | 136.1 | 18.89 | 23.68 | | High | High | High | High | High | High | High | High | High | High | High | 0 | 0.000186 | 4.61 |
| FALSE | High | SSSPTAGP! | [R].SSSPTA | 1xPhospho | Q6P1G2 1x | 5.73E-07 | 0 | 1 | 6 | 6 | Q6P1G2 | Lysine-spe | Q6P1G2 [8 | 0 | 2356.096 | 0 | -1.35 | 0.000663 | 2.99 | 145.8 | 54.2 | 6.93 | 9.09 | | High | High | High | High | High | High | High | High | High | High | High | 0 | 1.18E-07 | 4.51 |
| FALSE | High | SSSPTNSLT | [R].SSSPTN | 1xPhospho | B9EKS2 1xl | 0.000538 | 0 | 1 | 3 | 3 | B9EKS2 | Jumonji do | B9EKS2 [7: | 0 | 4659.327 | 0 | -0.05 | 0.824405 | 19.31 | 102.1 | 97.9 | 11.47 | 9.24 | | High | High | High | High | High | High | High | High | High | High | High | 0 | 0.000119 | 3.71 |
| FALSE | High | SSSPTQYGI | [R].SSSPTQ | 1xPhospho | Q9ESL4 1xl | 0.000852 | 0 | 1 | 1 | 4 | Q9ESL4 | Mitogen-a( | Q9ESL4 [6: | 0 | 1491.704 | 0 | 1.09 | 0.000661 | 4.67 | 63.6 | 136.4 | 2.23 | 7.19 | | High | High | High | High | High | High | High | High | High | High | High | 0 | 0.0002 | 2.39 |
| FALSE | High | SSSPVGLAI | [R].SSSPVG | 1xPhospho | Q9EST3 1x | 0.000779 | 0 | 1 | 5 | 4 | Q9EST3 | Eukaryotic | Q9EST3 [9 | 0 | 1383.765 | 0 | 0.46 | 0.00752 | 7.6 | 87.9 | 112.1 | 3.99 | 8.72 | | High | High | High | High | High | High | High | High | High | High | High | 0 | 0.000181 | 1.83 |
| FALSE | High | SSSPVLVEE | [R].SSSPVL | 1xPhospho | Q501J7-3 1 | 0.003195 | 0.00016 | 1 | 5 | 2 | Q501J7-3 | Isoform 3 ( | Q501J7-3 [ | 0 | 1766.841 | 0 | 0.3 | 0.014963 | 13 | 90.2 | 109.8 | 5.95 | 6.02 | | High | High | High | High | High | High | High | High | High | High | High | 0.000204 | 0.001355 | 2.6 |
| FALSE | High | SSSPVTELT | [R].SSSPVT | 1xPhospho | Q8BTI8 1xP | 0.000194 | 0 | 1 | 3 | 5 | Q8BTI8 | Serine/argi | Q8BTI8 [1( | 0 | 1456.725 | 0 | -0.53 | 0.002566 | 4.51 | 118.6 | 81.4 | 6.24 | 3.85 | | High | High | High | High | High | High | High | High | High | High | High | 0.000486 | 0.004338 | 2.82 |
| FALSE | High | SSSPVTELT | [R].SSSPVT | 2xPhospho | Q8BTI8 2xP | 0.009843 | 0.000736 | 1 | 3 | 4 | Q8BTI8 | Serine/argi | Q8BTI8 [1( | 0 | 1536.691 | 0 | -0.58 | 0.000448 | 1.68 | 120.1 | 79.9 | 3.64 | 6.46 | | High | High | High | High | High | High | High | High | High | High | High | 0.000555 | 0.00486 | 3.36 |
| FALSE | High | SSSPVTELT | [R].SSSPVT | 2xPhospho | Q8BTI8 2xP | 5.75E-05 | 0 | 1 | 3 | 1 | Q8BTI8 | Serine/argi | Q8BTI8 [1( | 1 | 2177.102 | 0 | -0.89 | 0.007633 | 6.26 | 132 | 68 | 14.23 | 7.07 | | High | High | High | High | High | High | High | High | High | High | High | 0 | 9.65E-06 | 1.91 |
| FALSE | High | SSSQENLLI | [K].SSSQEN | 1xPhospho | Q5SNZ0 1x | 0.014797 | 0.001049 | 1 | 4 | 4 | Q5SNZ0 | Girdin OS= | Q5SNZ0 [1 | 0 | 2017.992 | 0 | 0.34 | 0.068583 | 5.17 | 88.3 | 111.7 | 12.01 | 20.09 | | High | High | High | High | High | High | High | High | High | High | High | 0.000585 | 0.005441 | 2.72 |
| FALSE | High | SSSQENLLI | [K].SSSQEN | 1xPhospho | Q5SNZ0 1x | 9.36E-05 | 0 | 1 | 4 | 4 | Q5SNZ0 | Girdin OS= | Q5SNZ0 [1 | 0 | 2033.987 | 0 | 0.71 | 0.052538 | 46.56 | 77.3 | 122.7 | 12.52 | 31.24 | | High | High | High | High | High | High | High | High | High | High | High | 0 | 0.000104 | 3.92 |
| FALSE | High | SSSQIPEGF | [R].SSSQIPI | 1xPhospho | Q920I9 1xP | 1.77E-06 | 0 | 1 | 2 | 4 | Q920I9 | WD repeat | Q920I9 [11 | 0 | 2524.192 | 0 | 0.38 | 0.008473 | 8.86 | 87 | 113 | 5.53 | 3.43 | | High | High | High | High | High | High | High | High | High | High | High | 0 | 1.93E-07 | 4.92 |
| FALSE | High | SSSQPAAK | [R].SSSQP/ | 1xPhospho | Q3TZP3 1x | 0.019618 | 0.001438 | 1 | 2 | 1 | Q3TZP3 | Putative ur | Q3TZP3 [5! | 0 | 1313.687 | 0 | 0.97 | 0.006408 | 34.86 | 63.9 | 136.1 | 12.45 | 13.13 | | High | High | High | High | High | High | High | High | High | High | High | 0.000756 | 0.006659 | 1.24 |
| FALSE | High | SSSQSSPN! | [R].SSSQSS | 1xPhospho | Q9CYA6 1x | 3.35E-05 | 0 | 1 | 2 | 5 | Q9CYA6 | Zinc finger | Q9CYA6 [4 | 0 | 1643.804 | 0 | -0.12 | 0.148755 | 9.58 | 101.5 | 98.5 | 4.57 | 4.42 | | High | High | High | High | High | High | High | High | High | High | High | 0 | 3.55E-05 | 2.78 |
| FALSE | High | SSSSAGSE\ | [R].SSSSAG | 1xPhospho | Q8BKI2 1xl | 1.91E-07 | 0 | 1 | 2 | 4 | Q8BKI2 | Trinucleoti | Q8BKI2 [5! | 0 | 2302.071 | 0 | 0.19 | 0.271866 | 21.8 | 93.8 | 106.2 | 10.34 | 11.15 | | High | High | High | High | High | High | High | High | High | High | High | 0 | 1.58E-08 | 3.43 |
| FALSE | High | SSSSAPSTS | [R].SSSSAP | 1xPhospho | Q8K2E5 1x | 0.043981 | 0.003228 | 1 | 4 | 2 | Q8K2E5 | Budding ur | Q8K2E5 [6 | 0 | 1676.851 | 0 | 1.02 | 0.002529 | 20.32 | 62 | 138 | 6.66 | 11.89 | | High | High | High | High | High | High | High | High | High | High | High | 0.002159 | 0.0225 | 2.12 |
| FALSE | High | SSSSAVGG | [K].SSSSAV | 1xPhospho | Q03141 1x | 0.019791 | 0.001438 | 1 | 5 | 1 | Q03141 | MAP/micr( | Q03141 [4! | 0 | 1317.682 | 0 | -0.91 | 0.00016 | 2.3 | 131 | 69 | 3.85 | 10.58 | | High | High | High | High | High | High | High | High | High | High | High | 0.000773 | 0.006727 | 2.21 |

| | | | | | | | | | | | | | | | | | | | | | | | | | | | | | | | | | | | | | | |
|---|---|---|---|---|---|---|---|---|---|---|---|---|---|---|---|---|---|---|---|---|---|---|---|---|---|---|---|---|---|---|---|---|---|---|---|---|---|---|
| FALSE | High | SSSSDSIHS' | [R].SSSSDS | 1xPhospho | Q76I79 1xF | 0.010853 | 0.000772 | 1 | 3 | 4 | Q76I79 | Protein ph | Q76I79 [92 | 0 | 1470.679 | 0 | 0.78 | 0.059149 | 26.63 | 75.4 | 124.6 | 18.42 | 22.16 | | High | High | High | High | High | High | High | High | High | High | High | 0.002403 | 0.02504 | 1.55 |
| FALSE | High | SSSSERPEP | [K].SSSSER | 1xPhospho | A0A0R4J12 | 0.01029 | 0.000772 | 1 | 7 | 4 | A0A0R4J12 | SRSF prote | A0A0R4J12 | 0 | 1897.942 | 0 | 2.36 | 0.00025 | 10.16 | 32.9 | 167.1 | 8.04 | 6.45 | | High | High | High | High | High | High | High | High | High | High | High | 0.000382 | 0.003251 | 1.72 |
| FALSE | High | SSSSESLSEI | [K].SSSSESI | 2xPhospho | Q9ERD6 2x | 0.001476 | 5.66E-05 | 1 | 9 | 1 | Q9ERD6 | Ras-specifi | Q9ERD6 [2 | 0 | 1658.733 | 0 | -0.37 | 0.002599 | 3.68 | 111.8 | 88.2 | 4.65 | 7.64 | | High | High | High | High | High | High | High | High | High | High | High | 2.58E-05 | 0.00037 | 3.42 |
| FALSE | High | SSSSFTPFIL | [K].SSSSFTI | 1xPhospho | A0A0A6YW | 0.00153 | 5.66E-05 | 2 | 10 | 4 | A0A0A6YW | Mitogen-ac | A0A0A6YW | 0 | 1649.777 | 0 | -0.33 | 0.006677 | 4.29 | 114.3 | 85.7 | 5.32 | 11.84 | Shared | High | High | High | High | High | High | High | High | High | High | High | 4.94E-05 | 0.000687 | 3.37 |
| FALSE | High | SSSSFTPFIL | [K].SSSSFTI | 2xPhospho | A0A0A6YW | 0.111965 | 0.009053 | 2 | 10 | 1 | A0A0A6YW | Mitogen-ac | A0A0A6YW | 0 | 1729.744 | 0 | -0.42 | 0.020741 | 4.3 | 114.3 | 85.7 | 7.46 | 15.39 | Shared | High | High | High | High | High | High | High | High | High | High | High | 0.004543 | 0.04943 | 2.85 |
| FALSE | High | SSSSGSDD' | [R].SSSSGS | 1xPhospho | Q571K4 1x | 1.05E-06 | 0 | 1 | 1 | 9 | Q571K4 | TGF-beta-a | Q571K4 [5 | 0 | 2508.161 | 0 | 0.66 | 0.001527 | 8.09 | 76.1 | 123.9 | 8.57 | 4.52 | | High | High | High | High | High | High | High | High | High | High | High | 0 | 1.07E-07 | 4.35 |
| FALSE | High | SSSELSK | [R].SSSELSI | 1xPhospho | F7D6J5 1xF | 0.032548 | 0.002167 | 1 | 11 | 1 | F7D6J5 | Sodium ch | F7D6J5 [47 | 0 | 1275.66 | 0 | 2.23 | 0.000207 | 5.62 | 34 | 166 | 14.27 | 7.61 | | High | High | High | High | High | High | High | High | High | High | High | 0.001151 | 0.01187 | 1.3 |
| FALSE | High | SSSEDDIEC | [R].SSSEDD | 1xPhospho | Q3UFM5 1 | 0.000276 | 0 | 1 | 1 | 2 | Q3UFM5 | Nucleolar I | Q3UFM5 [3 | 0 | 2745.214 | 0 | -0.35 | 0.061835 | 13.94 | 122.3 | 77.7 | 15.25 | 20.79 | | High | High | High | High | High | High | High | High | High | High | High | 0 | 5.63E-05 | 3.14 |
| FALSE | High | SSSEDDDLI | [R].SSSEDD | 1xDeamida | Q6PDQ2 1 | 1.35E-06 | 0 | 1 | 6 | 7 | Q6PDQ2 | Chromodo | Q6PDQ2 [3 | 0 | 3611.439 | 0 | 0.09 | 0.72558 | 1 | 99 | 101 | 18.66 | 5.37 | | High | High | High | High | High | High | High | High | High | High | High | 0 | 6.9E-05 | 4.37 |
| FALSE | High | SSSEDDDLI | [R].SSSEDD | 1xPhospho | Q6PDQ2 1 | 3.1E-07 | 0 | 1 | 6 | 2 | Q6PDQ2 | Chromodo | Q6PDQ2 [3 | 0 | 3610.455 | 0 | -0.29 | 0.039652 | 5.21 | 110.1 | 89.9 | 12.45 | 4.24 | | High | High | High | High | High | High | High | High | High | High | High | 0 | 2.74E-08 | 4.99 |
| FALSE | High | SSPPATDP( | [R].SSPPAT | 1xPhospho | Q922Y1 1x | 0.017041 | 0.001204 | 1 | 1 | 2 | Q922Y1 | UBX domai | Q922Y1 [1 | 1 | 2854.439 | 0 | 0.25 | 0.545109 | 22.95 | 91.8 | 108.2 | 40.53 | 70.73 | | High | High | High | High | High | High | High | High | High | High | High | 0.000638 | 0.005689 | 2.39 |
| FALSE | High | SSPPPGYIP | [R].SSPPPG | 1xPhospho | G5E8L8 1xl | 0.000102 | 0 | 1 | 1 | 1 | G5E8L8 | Mitogen ac | G5E8L8 [16 | 0 | 2186.084 | 0 | -0.07 | 0.826237 | 4.53 | 102.5 | 97.5 | 15.44 | 8.4 | | High | High | High | High | High | High | High | High | High | High | High | 0 | 1.83E-05 | 3.57 |
| FALSE | High | SSPPPGYVI | [R].SSPPPG | 1xPhospho | Q3UVM5 1 | 0.009416 | 0.000736 | 1 | 3 | 2 | Q3UVM5 | Putative ur | Q3UVM5 [ | 0 | 1494.719 | 0 | 0.39 | 0.024667 | 18.74 | 90.8 | 109.2 | 7.95 | 10.77 | | High | High | High | High | High | High | High | High | High | High | High | 0.000327 | 0.002937 | 2.26 |
| FALSE | High | SSPPPLSGA | [R].SSPPPL | 1xPhospho | A6X919 1x | 3.77E-05 | 0 | 1 | 1 | 2 | A6X919 | Probable C | A6X919 [22 | 0 | 2351.097 | 0 | 0.05 | 0.294997 | 4.75 | 98.3 | 101.7 | 3.32 | 6.79 | | High | High | High | High | High | High | High | High | High | High | High | 0 | 7.08E-06 | 3.81 |
| FALSE | High | SSPPRPQPI | [R].SSPPRP | 1xDeamida | Q3U3E2 2x | 0.054844 | 0.003982 | 1 | 1 | 1 | Q3U3E2 | Protein FAI | Q3U3E2 [1 | 0 | 4372.162 | 0 | 0.28 | 0.209982 | 7.88 | 95.9 | 104.1 | 13.38 | 15.61 | | High | High | High | High | High | High | High | High | High | High | High | 0.002059 | 0.02136 | 2.99 |
| FALSE | High | SSPPVEHPA | [K].SSPPVE | 1xPhospho | Q9EP71 1x | 2.43E-07 | 0 | 1 | 1 | 2 | Q9EP71 | Ankycorbin | Q9EP71 [4 | 0 | 2502.207 | 0 | 0.34 | 0.098045 | 28.24 | 93.9 | 106.1 | 11.01 | 7.8 | | High | High | High | High | High | High | High | High | High | High | High | 0 | 2.07E-08 | 2.94 |
| FALSE | High | SSPPVEHPA | [K].SSPPVE | 1xDeamida | Q9EP71 1x | 0.000488 | 0 | 1 | 1 | 1 | Q9EP71 | Ankycorbin | Q9EP71 [4 | 0 | 2503.191 | 0 | 0.12 | 0.647976 | 13.84 | 95.8 | 104.2 | 2.73 | 11.52 | | High | High | High | High | High | High | High | High | High | High | High | 0 | 0.000106 | 2.44 |
| FALSE | High | SSPPVEHPA | [K].SSPPVE | 2xPhospho | Q9EP71 2x | 0.00083 | 0 | 1 | 1 | 1 | Q9EP71 | Ankycorbin | Q9EP71 [4 | 0 | 2582.174 | 0 | 0.45 | 0.043797 | 10.45 | 87 | 113 | 5.08 | 10.63 | | High | High | High | High | High | High | High | High | High | High | High | 0 | 0.000193 | 3.73 |
| FALSE | High | SSPQHSLSN | [R].SSPQHS | 1xPhospho | B1ASU9 1x | 1.33E-05 | 0 | 1 | 3 | 3 | B1ASU9 | Serine/thre | B1ASU9 [1 | 0 | 1728.863 | 0 | 0.05 | 0.585492 | 8.53 | 96.3 | 103.7 | 8.26 | 4.73 | | High | High | High | High | High | High | High | High | High | High | High | 0 | 1.86E-06 | 2.69 |
| FALSE | High | SSPQHSLSN | [R].SSPQHS | 3xPhospho | B1ASU9 3x | 0.071662 | 0.005278 | 1 | 3 | 1 | B1ASU9 | Serine/thre | B1ASU9 [1 | 0 | 1888.796 | 0 | 1.27 | 0.000252 | 4.8 | 59.3 | 140.7 | 5.4 | 6.49 | | High | High | High | High | High | High | High | High | High | High | High | 0.002717 | 0.02919 | 3.03 |
| FALSE | High | SSPQLDPLF | [R].SSPQLD | 1xPhospho | Q8K2F8 1x | 0.063258 | 0.004664 | 1 | 2 | 1 | Q8K2F8 | Protein LSN | Q8K2F8 [1 | 1 | 1678.929 | 0 | -0.41 | 0.008856 | 1.05 | 113 | 87 | 8.08 | 11.79 | | High | High | High | High | High | High | High | High | High | High | High | 0.002403 | 0.02523 | 1.81 |
| FALSE | High | SSPQIPHK | [K].SSPQIP | 1xPhospho | Q61532 1x | 0.028824 | 0.001829 | 1 | 2 | 2 | Q61532 | Mitogen-ac | Q61532 [7 | 0 | 1431.776 | 0 | 1.28 | 0.014586 | 20.5 | 58.7 | 141.3 | 14.63 | 21.86 | | High | High | High | High | High | High | High | High | High | High | High | 0.000997 | 0.01028 | 2.12 |
| FALSE | High | SSPQNYGP | [R].SSPQNY | 1xPhospho | Q32P12 1x | 1.95E-06 | 0 | 1 | 2 | 1 | Q32P12 | Uncharacte | Q32P12 [2 | 0 | 2021.964 | 0 | 0.38 | 0.120978 | 6.46 | 86.9 | 113.1 | 14.19 | 11.87 | | High | High | High | High | High | High | High | High | High | High | High | 0 | 2.16E-07 | 3.63 |
| FALSE | High | SSPPATDP( | [R].SSPPAT | 2xPhospho | Q922Y1 2x | 3.05E-07 | 0 | 1 | 1 | 4 | Q922Y1 | UBX domai | Q922Y1 [1 | 0 | 2778.304 | 0 | -0.7 | 0.002696 | 2.48 | 121.6 | 78.4 | 5.21 | 7.93 | | High | High | High | High | High | High | High | High | High | High | High | 0 | 2.68E-08 | 4.08 |
| FALSE | High | SSPQPLVA' | [K].SSPQPL | 1xPhospho | Q3UIZ4 1xl | 0.00407 | 0.00036 | 1 | 2 | 1 | Q3UIZ4 | Putative ur | Q3UIZ4 [14 | 0 | 3554.802 | 0 | -0.5 | 0.176618 | 9.6 | 117.2 | 82.8 | 14.16 | 12.74 | | High | High | High | High | High | High | High | High | High | High | High | 0.000161 | 0.00115 | 3.63 |
| FALSE | High | SSPSATSGF | [K].SSPSAT | 1xPhospho | A0A140T8 | 6.77E-05 | 0 | 1 | 1 | 4 | A0A140T8 | Chromosor | A0A140T8 | 0 | 1642.824 | 0 | -0.36 | 0.011758 | 4.73 | 114 | 86 | 5.27 | 10.21 | | High | High | High | High | High | High | High | High | High | High | High | 0 | 1.16E-05 | 2.76 |
| FALSE | High | SSPSESPLN | [K].SSPSES | 1xPhospho | Q61609 1x | 0.01441 | 0.001011 | 1 | 1 | 2 | Q61609 | Sodium-de | Q61609 [2 | 0 | 1745.843 | 0 | 0.72 | 0.002578 | 16.33 | 77.2 | 122.8 | 6.84 | 6.1 | | High | High | High | High | High | High | High | High | High | High | High | 0.000538 | 0.004735 | 2.77 |
| FALSE | High | SSPSESPLN | [K].SSPSES | 2xPhospho | Q61609 2x | 0.023379 | 0.001578 | 1 | 1 | 1 | Q61609 | Sodium-de | Q61609 [2 | 0 | 1809.815 | 0 | -0.43 | 0.004668 | 3.21 | 112.1 | 87.9 | 9.83 | 9.68 | | High | High | High | High | High | High | High | High | High | High | High | 0.000847 | 0.008101 | 3.45 |
| FALSE | High | SSPSGNTQ | [R].SSPSGN | 1xPhospho | Q9D219 1x | 7.68E-05 | 0 | 1 | 1 | 3 | Q9D219 | B-cell CLL/l | Q9D219 [1 | 0 | 1714.841 | 0 | -0.48 | 0.000779 | 1.92 | 117 | 83 | 2.79 | 2.8 | | High | High | High | High | High | High | High | High | High | High | High | 0 | 1.34E-05 | 2.92 |
| FALSE | High | SSPSILAVQ | [K].SSPSILA | 1xPhospho | P40201 1xl | 0.001572 | 5.66E-05 | 1 | 6 | 3 | P40201 | Chromodo | P40201 [8 | 0 | 1366.729 | 0 | 0.33 | 0.010065 | 3.3 | 88.4 | 111.6 | 6.42 | 4.78 | | High | High | High | High | High | High | High | High | High | High | High | 2.58E-05 | 0.000397 | 2.52 |
| FALSE | High | SSPSIICMP | [R].SSPSIIC | 1xCarbami | A0A1S6GW | 0.003083 | 0.00016 | 1 | 8 | 1 | A0A1S6GW | Uncharacte | A0A1S6GW | 0 | 1657.846 | 0 | 0.14 | 0.124965 | 11.35 | 91.6 | 108.4 | 8.54 | 12.34 | | High | High | High | High | High | High | High | High | High | High | High | 7.19E-05 | 0.000839 | 2.63 |
| FALSE | High | SSPSIICMP | [R].SSPSIIC | 1xCarbami | A0A1S6GW | 0.012288 | 0.000889 | 1 | 8 | 1 | A0A1S6GW | Uncharacte | A0A1S6GW | 0 | 1673.841 | 0 | 0.27 | 0.383876 | 37.95 | 86.6 | 113.4 | 10.36 | 56.73 | | High | High | High | High | High | High | High | High | High | High | High | 0.000449 | 0.003962 | 1.71 |
| FALSE | High | SSPSPQK | [K].SSPSPQ | 1xPhospho | O54774 1x | 0.026652 | 0.001676 | 1 | 5 | 3 | O54774 | AP-3 comp | O54774 [6 | 0 | 1268.665 | 0 | 0.14 | 0.094211 | 15.11 | 95.2 | 104.8 | 4.19 | 8.4 | | High | High | High | High | High | High | High | High | High | High | High | 0.006059 | 0.06563 | 1.51 |
| FALSE | High | SSPSSIMR | [K].SSPSSIN | 1xPhospho | Q6ZWQ0 1 | 0.099328 | 0.008003 | 1 | 2 | 1 | Q6ZWQ0 | Nesprin-2 | Q6ZWQ0 [ | 0 | 1173.554 | 0 | 1.95 | 0.00024 | 3.66 | 39.2 | 160.8 | 15.98 | 8.95 | | High | High | High | High | High | High | High | High | High | High | High | 0.004032 | 0.04291 | 1.76 |
| FALSE | High | SSPSSPAVF | [R].SSPSSP | 1xPhospho | E9Q2V5 1x | 0.002355 | 0.000109 | 1 | 3 | 1 | E9Q2V5 | CDNA sequ | E9Q2V5 [4 | 0 | 1196.587 | 0 | 1.62 | 0.00024 | 9.99 | 49.2 | 150.8 | 16.73 | 4.76 | | High | High | High | High | High | High | High | High | High | High | High | 4.94E-05 | 0.000622 | 2.16 |
| FALSE | High | SSPSSPSPT | [R].SSPSSP | 1xPhospho | D3YWD3 1 | 0.058664 | 0.004294 | 1 | 2 | 1 | D3YWD3 | Transmem | D3YWD3 [ | 0 | 1481.72 | 0 | 0.35 | 0.018819 | 17.58 | 87.9 | 112.1 | 8.34 | 5.15 | | High | High | High | High | High | High | High | High | High | High | High | 0.002187 | 0.02323 | 1.55 |
| FALSE | High | SSPSSPSPT | [R].SSPSSP | 2xPhospho | D3YWD3 2 | 0.014797 | 0.001049 | 1 | 2 | 2 | D3YWD3 | Transmem | D3YWD3 [ | 0 | 1561.686 | 0 | 2.02 | 0.00024 | 16.37 | 37.8 | 162.2 | 9.77 | 5.4 | | High | High | High | High | High | High | High | High | High | High | High | 0.000555 | 0.004856 | 2.57 |
| FALSE | High | SSPSTDHYS | [R].SSPSTD | 1xPhospho | A7TU71 1x | 0.000693 | 0 | 1 | 2 | 3 | A7TU71 | Protein Shi | A7TU71 [9 | 0 | 2338.055 | 0 | 1.08 | 0.016974 | 12.79 | 64.4 | 135.6 | 10.08 | 18.3 | | High | High | High | High | High | High | High | High | High | High | High | 0.000161 | 0.001189 | 2.84 |
| FALSE | High | SSPSALHQ( | [R].SSPSAL | 1xPhospho | Q6PG95 1x | 0.112856 | 0.009146 | 1 | 1 | 1 | Q6PG95 | Protein cra | Q6PG95 [7 | 0 | 1419.694 | 0 | -0.84 | 0.024203 | 8.67 | 124.2 | 75.8 | 16.44 | 19.3 | | High | High | High | High | High | High | High | High | High | High | High | 0.004569 | 0.0497 | 2.84 |
| FALSE | High | SSPPATDP( | [R].SSPPAT | 1xPhospho | Q922Y1 1x | 4.23E-11 | 0 | 1 | 1 | 12 | Q922Y1 | UBX domai | Q922Y1 [1 | 0 | 2698.338 | 0 | -0.36 | 0.000635 | 0.29 | 111.9 | 88.1 | 3.52 | 4.96 | | High | High | High | High | High | High | High | High | High | High | High | 0 | 2.19E-08 | 4.76 |
| FALSE | High | SSPPASSPC | [R].SSPPAS | 1xPhospho | O70293-2 | 2.95E-05 | 0 | 1 | 3 | 3 | O70293-2 | Isoform GF | O70293-2 | 0 | 2091.97 | 0 | -0.95 | 0.001522 | 1.53 | 130.1 | 69.9 | 8.46 | 6.44 | | High | High | High | High | High | High | High | High | High | High | High | 0 | 4.54E-06 | 3.77 |
| FALSE | High | SSPPAPSLP | [R].SSPPAP | 2xPhospho | A2A5V4 2x | 0.062209 | 0.004646 | 1 | 2 | 1 | A2A5V4 | SH3 domai | A2A5V4 [5 | 0 | 2586.257 | 0 | -0.44 | 0.016084 | 2.27 | 115.3 | 84.7 | 8.94 | 5.74 | | High | High | High | High | High | High | High | High | High | High | High | 0.002381 | 0.02475 | 2.9 |
| FALSE | High | SSPAHCTG | [K].SSPAHC | 1xCarbami | Q8BP97 1x | 0.021989 | 0.001542 | 1 | 2 | 1 | Q8BP97 | Rhomboid | Q8BP97 [3 | 0 | 1441.598 | 0 | 0.74 | 0.023663 | 12.7 | 77.4 | 122.6 | 11.23 | 21.69 | | High | High | High | High | High | High | High | High | High | High | High | 0.000814 | 0.007574 | 2.23 |
| FALSE | High | SSPALAAPI | [K].SSPALA | 1xPhospho | E9Q4N7 1x | 1.12E-05 | 0 | 1 | 4 | 2 | E9Q4N7 | AT-rich inte | E9Q4N7 [1 | 0 | 1964.014 | 0 | -0.32 | 0.00126 | 2.83 | 110.5 | 89.5 | 2.6 | 3.85 | | High | High | High | High | High | High | High | High | High | High | High | 0 | 1.54E-06 | 2.5 |
| FALSE | High | SSPAVSSSK | [K].SSPAVS | 1xPhospho | A6H8H2-2 | 0.022378 | 0.001542 | 1 | 4 | 1 | A6H8H2-2 | Isoform 2 c | A6H8H2-2 | 0 | 1387.723 | 0 | 0.83 | 0.000661 | 4.01 | 73.2 | 126.8 | 5.93 | 4.62 | | High | High | High | High | High | High | High | High | High | High | High | 0.000814 | 0.007737 | 1.51 |
| FALSE | High | SSPCLSTTS | [R].SSPCLS | 1xCarbami | A0A0A0MC | 0.003464 | 0.00021 | 1 | 4 | 3 | A0A0A0MC | Calmodulir | A0A0A0MC | 0 | 2787.302 | 0 | 0.03 | 0.635843 | 15.07 | 97.4 | 102.6 | 14.76 | 8.37 | | High | High | High | High | High | High | High | High | High | High | High | 0.000327 | 0.002466 | 3.32 |
| FALSE | High | SSPCSTGEI | [K].SSPCST | 1xCarbami | U5LK19 1x | 0.027121 | 0.001676 | 2 | 9 | 1 | U5LK19; E | Marshalin- | U5LK19 [5 | 0 | 2919.514 | 0 | -1.85 | 0.000647 | 1.89 | 156.9 | 43.1 | 4.05 | 27.68 | Shared | High | High | High | High | High | High | High | High | High | High | High | 0.00092 | 0.009592 | 2.11 |
| FALSE | High | SSPDDGND | [R].SSPDD( | 1xPhospho | Q3TQ38 1x | 6.65E-05 | 0 | 1 | 4 | 1 | Q3TQ38 | Putative ur | Q3TQ38 [1 | 0 | 2603.228 | 0 | 0.43 | 0.044662 | 26.4 | 82.8 | 117.2 | 11.31 | 14.62 | | High | High | High | High | High | High | High | High | High | High | High | 0 | 1.14E-05 | 3.47 |
| FALSE | High | SSPDGGLK | [R].SSPDG( | 1xPhospho | Q7TSC1 1x | 0.028327 | 0.001791 | 1 | 3 | 1 | Q7TSC1 | Protein PRI | Q7TSC1 [1 | 0 | 1298.676 | 0 | -0.3 | 0.051924 | 4.01 | 107.5 | 92.5 | 4.39 | 8.61 | | High | High | High | High | High | High | High | High | High | High | High | 0.000968 | 0.01009 | 1.56 |
| FALSE | High | SSPDQFLFS | [R].SSPDQI | 1xPhospho | E9QMX4 1 | 0.000442 | 0 | 1 | 4 | 1 | E9QMX4 | Ribosomal | E9QMX4 [ | 0 | 1692.82 | 0 | 0.21 | 0.056336 | 3.59 | 93 | 107 | 4.25 | 5.15 | | High | High | High | High | High | High | High | High | High | High | High | 0 | 9.57E-05 | 2.63 |
| FALSE | High | SSPDQFLFS | [R].SSPDQI | 3xPhospho | E9QMX4 3 | 0.021231 | 0.001507 | 1 | 4 | 1 | E9QMX4 | Ribosomal | E9QMX4 [ | 1 | 2426.987 | 0 | 0.04 | 0.290832 | 11.32 | 97.8 | 102.2 | 7.08 | 8.08 | | High | High | High | High | High | High | High | High | High | High | High | 0.000801 | 0.007326 | 2.93 |
| FALSE | High | SSPDSPEN' | [K].SSPDSP | 1xPhospho | B0QZW4 1 | 0.000349 | 0 | 1 | 4 | 2 | B0QZW4 | Suppressor | B0QZW4 [2 | 0 | 2884.295 | 0 | -0.18 | 0.108781 | 13.85 | 111.4 | 88.6 | 15.86 | 17.29 | | High | High | High | High | High | High | High | High | High | High | High | 0 | 7.29E-05 | 3.94 |
| FALSE | High | SSPDSPEN' | [K].SSPDSP | 2xPhospho | B0QZW4 2 | 0.001011 | 0 | 1 | 4 | 1 | B0QZW4 | Suppressor | B0QZW4 [2 | 0 | 2964.261 | 0 | 0.54 | 0.091002 | 36.38 | 73 | 127 | 17.07 | 21.84 | | High | High | High | High | High | High | High | High | High | High | High | 0 | 0.000242 | 2.15 |
| FALSE | High | SSPEEVK | [K].SSPEEV | 1xPhospho | Q75NR7-2 | 0.106748 | 0.008634 | 1 | 4 | 1 | Q75NR7-2 | Isoform 2 c | Q75NR7-2 | 0 | 1313.675 | 0 | -0.98 | 0.000661 | 2.59 | 132.2 | 67.8 | 6.12 | 5.38 | | High | High | High | High | High | High | High | High | High | High | High | 0.004327 | 0.0465 | 1.2 |
| FALSE | High | SSPEIAPSD | [R].SSPEIAI | 1xPhospho | H3BK65 1x | 8.66E-06 | 0 | 1 | 6 | 3 | H3BK65 | Epidermal | H3BK65 [5 | 0 | 2970.423 | 0 | -0.27 | 0.025406 | 6.04 | 107.4 | 92.6 | 7.82 | 4.38 | | High | High | High | High | High | High | High | High | High | High | High | 0 | 1.15E-06 | 3.91 |
| FALSE | High | SSPELTR | [R].SSPELT | 1xPhospho | Q8BTI8 1xF | 0.063258 | 0.004664 | 1 | 3 | 2 | Q8BTI8 | Serine/argi | Q8BTI8 [16 | 0 | 1098.539 | 0 | -1.09 | 0.000406 | 1.97 | 136.1 | 63.9 | 5.6 | 7.35 | | High | High | High | High | High | High | High | High | High | High | High | 0.002403 | 0.02516 | 2.12 |
| FALSE | High | SSPERNDW | [K].SSPERN | 1xPhospho | A0A1L7NR | 0.027597 | 0.001716 | 2 | 5 | 1 | A0A1L7NR | C-Cbl assoc | A0A1L7NR | 1 | 2094.99 | 0 | -1.58 | 0.001284 | 1.05 | 148.2 | 51.8 | 6.74 | 14.53 | Shared | High | High | High | High | High | High | High | High | High | High | High | 0.000953 | 0.009825 | 2.38 |
| FALSE | High | SSPEVGSF( | [K].SSPEVG | 1xCarbami | Q6P9L6 1x | 1.75E-06 | 0 | 1 | 2 | 2 | Q6P9L6 | Kinesin-like | Q6P9L6 [6 | 0 | 2242.989 | 0 | -0.14 | 0.318462 | 15.2 | 104.9 | 95.1 | 9.2 | 8.11 | | High | High | High | High | High | High | High | High | High | High | High | 0 | 1.9E-07 | 4.72 |
| FALSE | High | SSPGAGPSI | [R].SSPGAC | 1xPhospho | F6YRW4 1x | 9.25E-07 | 0 | 1 | 3 | 1 | F6YRW4 | F-box and I | F6YRW4 [7 | 0 | 1758.776 | 0 | 1.12 | 0.003242 | 14.64 | 62.1 | 137.9 | 22.15 | 3.72 | | High | High | High | High | High | High | High | High | High | High | High | 0 | 9.32E-08 | 3.86 |
| FALSE | High | SSPGDTLVI | [K].SSPGDT | 1xPhospho | Q69ZT1 1x | 0.075927 | 0.005826 | 1 | 4 | 1 | Q69ZT1 | Fanconi-as | Q69ZT1 [2 | 0 | 1441.77 | 0 | -0.59 | 0.001551 | 3.55 | 120.2 | 79.8 | 2.98 | 7.58 | | High | High | High | High | High | High | High | High | High | High | High | 0.002976 | 0.03126 | 1.52 |
| FALSE | High | SSPHFQHA | [R].SSPHFC | 1xPhospho | B8JK87 1xF | 0.021797 | 0.001507 | 1 | 4 | 1 | B8JK87 | Angiomoti | B8JK87 [26 | 0 | 1603.758 | 0 | 0.01 | 0.418092 | 7.22 | 98.7 | 101.3 | 4.36 | 10.3 | | High | High | High | High | High | High | High | High | High | High | High | 0.000814 | 0.007507 | 2.24 |
| FALSE | High | SSPIPAGDC | [R].SSPIPA( | 1xCarbami | E9Q4S3 1x | 0.006653 | 0.00062 | 1 | 7 | 1 | E9Q4S3 | Ninein OS= | E9Q4S3 [1 | 0 | 2078.977 | 0 | 2.52 | 0.000606 | 11.41 | 28.7 | 171.3 | 9.75 | 10.86 | | High | High | High | High | High | High | High | High | High | High | High | 0.000275 | 0.001981 | 2.63 |
| FALSE | High | SSPITVQTS | [R].SSPITV( | 1xPhospho | Q9D168 1x | 0.028327 | 0.001791 | 1 | 1 | 2 | Q9D168 | Integrator | Q9D168 [1 | 0 | 1585.86 | 0 | -0.43 | 0.021794 | 2.51 | 110.8 | 89.2 | 10.81 | 13.54 | | High | High | High | High | High | High | High | High | High | High | High | 0.000968 | 0.01009 | 2.34 |
| FALSE | High | SSPMLQHK | [R].SSPML( | 1xPhospho | Q5SNZ0 1x | 0.024424 | 0.001612 | 1 | 4 | 4 | Q5SNZ0 | Girdin OS= | Q5SNZ0 [1 | 0 | 1481.759 | 0 | 1.09 | 0.000614 | 4.29 | 64 | 136 | 9.74 | 6.86 | | High | High | High | High | High | High | High | High | High | High | High | 0.002403 | 0.0252 | 2.89 |
| FALSE | High | SSPNDTRPI | [R].SSPNDT | 1xPhospho | Q9D4J7 1x | 0.000462 | 0 | 1 | 3 | 2 | Q9D4J7 | PHD finger | Q9D4J7 [2 | 0 | 1539.793 | 0 | 1.45 | 0.000193 | 7.4 | 55.1 | 144.9 | 4.66 | 4.33 | | High | High | High | High | High | High | High | High | High | High | High | 0 | 9.98E-05 | 2.08 |
| FALSE | High | SSPNPFVG: | [K].SSPNPF | 1xPhospho | A0A0R4J1C | 0.000194 | 0 | 1 | 5 | 2 | A0A0R4J1C | Disabled h | A0A0R4J1C | 0 | 1751.913 | 0 | 1.63 | 0.003361 | 43.44 | 48.9 | 151.1 | 24.23 | 12.49 | | High | High | High | High | High | High | High | High | High | High | High | 0 | 3.8E-05 | 2.77 |
| FALSE | High | SSPNPFVG: | [K].SSPNPF | 2xPhospho | A0A0R4J1C | 0.002695 | 0.000109 | 1 | 5 | 1 | A0A0R4J1C | Disabled h | A0A0R4J1C | 0 | 1831.88 | 0 | 3.06 | 0.001327 | 34.11 | 23.5 | 176.5 | 29.89 | 14.44 | | High | High | High | High | High | High | High | High | High | High | High | 4.94E-05 | 0.000724 | 3 |
| FALSE | High | SSPNSSLGS | [R].SSPNSS | 1xPhospho | A6H8H2-2 | 1.92E-06 | 0 | 1 | 4 | 2 | A6H8H2-2 | Isoform 2 c | A6H8H2-2 | 0 | 2271.2 | 0 | 1.38 | 0.000928 | 13.26 | 52.3 | 147.7 | 9.44 | 8.73 | | High | High | High | High | High | High | High | High | High | High | High | 0 | 2.11E-07 | 3.27 |
| FALSE | High | SSPNVANC | [R].SSPNVA | 1xPhospho | E9Q9C3 1x | 5.58E-07 | 0 | 1 | 5 | 6 | E9Q9C3 | Afadin OS= | E9Q9C3 [1 | 0 | 1975.005 | 0 | 0.06 | 0.158103 | 4.44 | 97.3 | 102.7 | 4.72 | 2.02 | | High | High | High | High | High | High | High | High | High | High | High | 0 | 7.22E-05 | 3.12 |
| FALSE | High | SSPNVANC | [R].SSPNVA | 2xPhospho | E9Q9C3 2x | 3.09E-05 | 0 | 1 | 5 | 4 | E9Q9C3 | Afadin OS= | E9Q9C3 [1 | 0 | 2054.971 | 0 | -0.25 | 0.023103 | 7.45 | 110.3 | 89.7 | 3.33 | 7.47 | | High | High | High | High | High | High | High | High | High | High | High | 0 | 4.8E-06 | 2.66 |
| FALSE | High | SSPPAPSLP | [R].SSPPAP | 1xPhospho | A2A5V4 1x | 1.05E-05 | 0 | 1 | 2 | 7 | A2A5V4 | SH3 domai | A2A5V4 [5 | 0 | 2506.29 | 0 | -0.41 | 0.002332 | 1.49 | 113.7 | 86.3 | 5.04 | 2.34 | | High | High | High | High | High | High | High | High | High | High | High | 0 | 2.95E-06 | 4.29 |
| FALSE | High | SSPSTGSLE | [R].SSPSTG | 1xPhospho | D4AFX6 1x | 1.6E-06 | 0 | 1 | 3 | 4 | D4AFX6 | Centaurin | D4AFX6 [3 | 0 | 1990.937 | 0 | 0.88 | 0.000177 | 1.85 | 70.7 | 129.3 | 3.57 | 3.7 | | High | High | High | High | High | High | High | High | High | High | High | 0 | 1.73E-07 | 3.34 |
| FALSE | High | SSPAEPELN | [R].SSPAEP | 1xPhospho | B7ZWN9 1 | 0.000217 | 0 | 2 | 6 | 2 | B7ZWN9; F | SUMO/sen | B7ZWN9 [4 | 0 | 1932.004 | 0 | -0.26 | 0.027113 | 6.98 | 107.7 | 92.3 | 5.47 | 5.97 | Shared | High | High | High | High | High | High | High | High | High | High | High | 0 | 4.26E-05 | 2.99 |
| FALSE | High | SSPSTPACF | [R].SSPSTP | 1xCarbami | Q9QWF0 1 | 0.057686 | 0.004227 | 1 | 2 | 1 | Q9QWF0 | Chromatin | Q9QWF0 [ | 0 | 1271.565 | 0 | -0.03 | 0.435845 | 3.6 | 102.3 | 97.7 | 5.45 | 5.83 | | High | High | High | High | High | High | High | High | High | High | High | 0.002159 | 0.02272 | 2.25 |
| FALSE | High | SSPTPHSGF | [R].SSPTPH | 1xCarbami | Q812A5 1x | 0.024001 | 0.001612 | 1 | 2 | 2 | Q812A5 | Proline-ricl | Q812A5 [3 | 0 | 1675.746 | 0 | -0.9 | 0.003279 | 5.13 | 128.5 | 71.5 | 6.96 | 10.8 | | High | High | High | High | High | High | High | High | High | High | High | 0.005336 | 0.05814 | 2.69 |
| FALSE | High | SSRSFSLDE | [R].SSRSFS | 2xPhospho | B2RQG2 2x | 0.003495 | 0.00021 | 2 | 3 | 1 | B2RQG2; A | PHD finger | B2RQG2 [6 | 1 | 3051.525 | 0 | 0.4 | 0.067105 | 12.63 | 90.2 | 109.8 | 7.67 | 14.81 | Shared | High | High | High | High | High | High | High | High | High | High | High | 9.42E-05 | 0.000969 | 2.91 |
| FALSE | High | SSSADFGTI | [K].SSSADF | 1xPhospho | A0A087WF | 0.000181 | 0 | 1 | 7 | 4 | A0A087WF | Arf-GAP dc | A0A087WF | 0 | 2886.356 | 0 | 0.26 | 0.997457 | 30.73 | 94.8 | 105.2 | 26.93 | 24.51 | | High | High | High | High | High | High | High | High | High | High | High | 0 | 3.5E-05 | 3.4 |
| FALSE | High | SSSAEFSAA | [R].SSSAEF | 1xPhospho | Q5DU37 1x | 4.28E-06 | 0 | 1 | 4 | 4 | Q5DU37 | Zinc finger | Q5DU37 [1 | 0 | 1987.969 | 0 | 0.43 | 0.057024 | 20.42 | 84.1 | 115.9 | 8.18 | 15.55 | | High | High | High | High | High | High | High | High | High | High | High | 0.000204 | 0.001341 | 3.73 |
| FALSE | High | SSSAENQE( | [R].SSSAEN | 1xPhospho | Q8BVE8-2 | 1.19E-05 | 0 | 2 | 6 | 2 | Q8BVE8-2; | Isoform 2 c | Q8BVE8-2 | 0 | 2059.922 | 0 | 0.99 | 0.003647 | 18.17 | 64.3 | 135.7 | 17.72 | 13.63 | Shared | High | High | High | High | High | High | High | High | High | High | High | 0 | 1.64E-06 | 3.43 |
| FALSE | High | SSSAPNVH | [R].SSSAPN | 1xPhospho | F6SZ47 1xF | 1.65E-06 | 0 | 1 | 2 | 4 | F6SZ47 | Serine/thre | F6SZ47 [34 | 0 | 2713.376 | 0 | -0.49 | 0.01331 | 1.51 | 116.9 | 83.1 | 4.5 | 9.49 | | High | High | High | High | High | High | High | High | High | High | High | 0 | 7.64E-07 | 4.46 |
| FALSE | High | SSSAPNVH | [R].SSSAPN | 1xDeamida | F6SZ47 1xF | 0.040721 | 0.003058 | 1 | 2 | 3 | F6SZ47 | Serine/thre | F6SZ47 [34 | 0 | 2714.36 | 0 | 0.14 | 0.866261 | 1.14 | 97.2 | 102.8 | 12.92 | 10.88 | | High | High | High | High | High | High | High | High | High | High | High | 0.001581 | 0.01517 | 3.13 |
| FALSE | High | SSSAPNVH | [R].SSSAPN | 1xPhospho | P28028 1xl | 4.13E-09 | 0 | 1 | 1 | 2 | P28028 | Serine/thre | P28028 [34 | 0 | 2702.38 | 0 | -0.26 | 0.201992 | 8.99 | 109.1 | 90.9 | 9.24 | 14.64 | | High | High | High | High | High | High | High | High | High | High | High | 0 | 2.13E-10 | 4.54 |
| FALSE | High | SSSASGAS/ | [K].SSSASG | 2xCarbami | Q8K298 1x | 3.72E-11 | 0 | 1 | 2 | 6 | Q8K298 | Anillin OS= | Q8K298 [2 | 0 | 2925.255 | 0 | 0.32 | 0.032216 | 4.26 | 88.5 | 111.5 | 7.56 | 11.31 | | High | High | High | High | High | High | High | High | High | High | High | 0 | 4.33E-12 | 6.19 |
| FALSE | High | SSSASPELK | [R].SSSASP | 1xPhospho | Q8BTI8 1xF | 0.000255 | 0 | 1 | 3 | 4 | Q8BTI8 | Serine/argi | Q8BTI8 [13 | 0 | 1443.75 | 0 | -0.57 | 0.005374 | 2.38 | 119.5 | 80.5 | 7.8 | 4.35 | | High | High | High | High | High | High | High | High | High | High | High | 0 | 5.13E-05 | 2.61 |
| FALSE | High | SSSASPELK | [R].SSSASP | 2xPhospho | Q8BTI8 2xF | 0.000984 | 0 | 1 | 3 | 5 | Q8BTI8 | Serine/argi | Q8BTI8 [13 | 0 | 1523.716 | 0 | -0.71 | 0.000941 | 2.5 | 123 | 77 | 4.34 | 3.01 | | High | High | High | High | High | High | High | High | High | High | High | 0 | 0.000234 | 3.51 |
| FALSE | High | SSSASSPPK | [R].SSSASS | 1xPhospho | Q9Z2D6-2 | 0.00083 | 0 | 1 | 2 | 2 | Q9Z2D6-2 | Isoform B c | Q9Z2D6-2 | 0 | 1385.708 | 0 | 1.4 | 0.000405 | 3.5 | 56.3 | 143.7 | 9.64 | 6.49 | | High | High | High | High | High | High | High | High | High | High | High | 0 | 0.000193 | 1.57 |
| FALSE | High | SSSCESLGA | [K].SSSCES | 1xCarbami | Q9Z2D1 1x | 1.11E-07 | 0 | 1 | 2 | 4 | Q9Z2D1 | Myotubula | Q9Z2D1 [4 | 0 | 1842.862 | 0 | 0.11 | 0.273798 | 10.48 | 97.1 | 102.9 | 7.64 | 12.98 | | High | High | High | High | High | High | High | High | High | High | High | 0 | 1.88E-08 | 3.76 |
| FALSE | High | SSSCESLGA | [K].SSSCES | 1xCarbami | Q9Z2D1 2x | 0.00311 | 0.00016 | 1 | 2 | 4 | Q9Z2D1 | Myotubula | Q9Z2D1 [4 | 0 | 1922.828 | 0 | 0.38 | 0.003364 | 6.77 | 88.4 | 111.6 | 5.89 | 3.36 | | High | High | High | High | High | High | High | High | High | High | High | 7.19E-05 | 0.000853 | 3.46 |
| FALSE | High | SSQYYSQG | [R].SSQYYS | 1xCarbami | Q8CAF4-3 | 1.49E-06 | 0 | 1 | 4 | 1 | Q8CAF4-3 | Isoform 3 c | Q8CAF4-3 | 1 | 4014.69 | 0 | 1.26 | 0.003748 | 2.65 | 59.7 | 140.3 | 8.64 | 15.9 | | High | High | High | High | High | High | High | High | High | High | High | 0 | 1.59E-07 | 5.62 |
| FALSE | High | SSSDALSAS | [R].SSSDAL | 1xPhospho | Q811P8 1x | 1.94E-07 | 0 | 1 | 3 | 7 | Q811P8 | Rho GTPas | Q811P8 [7 | 0 | 2105.97 | 0 | -1.26 | 0.000181 | 1.09 | 140.8 | 59.2 | 1.3 | 9.8 | | High | High | High | High | High | High | High | High | High | High | High | 0 | 6.34E-07 | 4.55 |
| FALSE | High | SSSDALSAS | [R].SSSDAL | 2xPhospho | Q811P8 2x | 0.000207 | 0 | 1 | 3 | 2 | Q811P8 | Rho GTPas | Q811P8 [7 | 0 | 2185.937 | 0 | -0.56 | 0.004435 | 2.97 | 119.2 | 80.8 | 6.31 | 7.72 | | High | High | High | High | High | High | High | High | High | High | High | 0 | 4.06E-05 | 4.19 |
| FALSE | High | SSSDAQLSI | [K].SSSDAC | 1xPhospho | Q80VE5 1x | 0.009756 | 0.000736 | 1 | 1 | 1 | Q80VE5 | TBC1 doma | Q80VE5 [1 | 0 | 1259.583 | 0 | 0.12 | 0.110792 | 10.99 | 95.4 | 104.6 | 9.7 | 7.33 | | High | High | High | High | High | High | High | High | High | High | High | 0.000344 | 0.003053 | 1.93 |
| FALSE | High | SSSDLEK | [K].SSSDLE | 1xPhospho | E9QLH9 1x | 0.030898 | 0.001896 | 1 | 2 | 2 | E9QLH9 | PEAK1-rela | E9QLH9 [7 | 0 | 1303.655 | 0 | 0.97 | 0.000651 | 3.8 | 66.4 | 133.6 | 9.58 | 5.61 | | High | High | High | High | High | High | High | High | High | High | High | 0.002937 | 0.03049 | 2.2 |
| FALSE | High | SSSDLPAVI | [R].SSSDLP | 1xPhospho | E9PX52 1xl | 0.003892 | 0.000261 | 1 | 2 | 2 | E9PX52 | Arf-GAP wi | E9PX52 [8 | 0 | 1992.052 | 0 | 0.25 | 0.047049 | 11.43 | 91.4 | 108.6 | 5.51 | 15.52 | | High | High | High | High | High | High | High | High | High | High | High | 0.000505 | 0.00434 | 2.2 |
| FALSE | High | SSSDMDLC | [R].SSSDM | 1xPhospho | Q91X43 1x | 0.000142 | 0 | 1 | 1 | 1 | Q91X43 | SH3 domai | Q91X43 [3 | 0 | 1548.738 | 0 | 0.95 | 0.003994 | 16.2 | 70 | 130 | 10.36 | 13.2 | | High | High | High | High | High | High | High | High | High | High | High | 0 | 2.65E-05 | 2.37 |
| FALSE | High | SSSDMDLC | [R].SSSDM | 1xPhospho | Q91X43 1x | 0.000568 | 0 | 1 | 1 | 3 | Q91X43 | SH3 domai | Q91X43 [3 | 0 | 1564.733 | 0 | 0.8 | 0.025158 | 20.3 | 70.8 | 129.2 | 8.36 | 24.01 | | High | High | High | High | High | High | High | High | High | High | High | 0.000467 | 0.004234 | 2.74 |
| FALSE | High | SSSDNNTN | [R].SSSDNI | 1xPhospho | Q69ZR2 1x | 7.15E-05 | 0 | 1 | 3 | 3 | Q69ZR2 | E3 ubiquiti | Q69ZR2 [1 | 0 | 1574.701 | 0 | 1.94 | 0.000395 | 9.91 | 42.6 | 157.4 | 11.93 | 5.6 | | High | High | High | High | High | High | High | High | High | High | High | 0 | 1.23E-05 | 2.22 |
| FALSE | High | SSSDSVEEE | [K].SSSDSV | 1xPhospho | E9PZM7 1x | 3.13E-07 | 0 | 1 | 3 | 9 | E9PZM7 | SR-related | E9PZM7 [3 | 0 | 2788.307 | 0 | -0.23 | 0.007293 | 3.13 | 108.6 | 91.4 | 2.8 | 4.65 | | High | High | High | High | High | High | High | High | High | High | High | 0 | 2.54E-07 | 5.72 |
| FALSE | High | SSSDSVEEE | [K].SSSDSV | 2xPhospho | E9PZM7 2x | 4.41E-07 | 0 | 1 | 3 | 5 | E9PZM7 | SR-related | E9PZM7 [3 | 0 | 2868.273 | 0 | -0.09 | 0.438409 | 7.04 | 101.4 | 98.6 | 4.55 | 7.06 | | High | High | High | High | High | High | High | High | High | High | High | 0 | 1.2E-07 | 5.08 |
| FALSE | High | SSSDSWEL' | [K].SSSDSV | 1xPhospho | B2RXQ3 1x | 0.007951 | 0.000736 | 1 | 5 | 2 | B2RXQ3 | Family with | B2RXQ3 [3 | 0 | 1807.852 | 0 | -0.3 | 0.006653 | 4.23 | 110.4 | 89.6 | 4.32 | 9.78 | | High | High | High | High | High | High | High | High | High | High | High | 0.000327 | 0.002439 | 3.25 |
| FALSE | High | SSSDTAAYI | [R].SSSDTA | 1xCarbami | E9PYH6 1x | 9.39E-06 | 0 | 1 | 1 | 3 | E9PYH6 | SET domai | E9PYH6 [2 | 0 | 3644.565 | 0 | -0.99 | 0.056513 | 4.58 | 131 | 69 | 14.27 | 25.81 | | High | High | High | High | High | High | High | High | High | High | High | 0 | 1.26E-06 | 3.56 |
| FALSE | High | SSSDTAAYI | [R].SSSDTA | 1xCarbami | E9PYH6 1x | 1.15E-07 | 0 | 1 | 1 | 2 | E9PYH6 | SET domai | E9PYH6 [2 | 0 | 3645.549 | 0 | -1.45 | 0.001632 | 2.28 | 146.5 | 53.5 | 11.56 | 4.48 | | High | High | High | High | High | High | High | High | High | High | High | 0 | 8.95E-09 | 5.02 |
| FALSE | High | SSSDTSTPE | [R].SSSDTS | 1xPhospho | A0A1D5RL | 0.111965 | 0.009053 | 1 | 5 | 1 | A0A1D5RL | Uncharacte | A0A1D5RL | 0 | 1818.877 | 0 | 0.56 | 0.006898 | 9.36 | 82.7 | 117.3 | 10.63 | 12.28 | | High | High | High | High | High | High | High | High | High | High | High | 0.004569 | 0.04956 | 2.1 |
| FALSE | High | SSSDALSAS | [R].SSSDAL | 1xDeamida | Q811P8 1x | 2.85E-05 | 0 | 1 | 3 | 4 | Q811P8 | Rho GTPas | Q811P8 [7 | 0 | 2106.954 | 0 | -0.38 | 0.054432 | 10.06 | 108.1 | 91.9 | 10.47 | 10.26 | | High | High | High | High | High | High | High | High | High | High | High | 0 | 4.37E-06 | 3.8 |
| FALSE | High | SSQVRPVS' | [K].SSQVRI | 2xPhospho | H3BL37 2xl | 0.006363 | 0.00062 | 1 | 7 | 1 | H3BL37 | Treacle pro | H3BL37 [4 | 0 | 2292.14 | 0 | -0.38 | 0.008386 | 9.09 | 114.8 | 85.2 | 7 | 4.77 | | High | High | High | High | High | High | High | High | High | High | High | 0.000275 | 0.00189 | 1.57 |
| FALSE | High | SSQVRPVS' | [K].SSQVRI | 1xPhospho | H3BL37 1xl | 1.46E-06 | 0 | 1 | 7 | 4 | H3BL37 | Treacle pro | H3BL37 [4 | 0 | 2212.174 | 0 | -0.56 | 0.000757 | 3.28 | 119 | 81 | 2.72 | 6.6 | | High | High | High | High | High | High | High | High | High | High | High | 0.005703 | 0.06287 | 2.56 |
| FALSE | High | SSQTQNK | [R].SSQTQI | 1xPhospho | Q9EQU5-2 | 0.005228 | 0.000551 | 2 | 3 | 1 | Q9EQU5-2 | Isoform 2 c | Q9EQU5-2 | 0 | 1330.677 | 0 | -0.46 | 0.003067 | 6.86 | 119 | 81 | 7.45 | 4.48 | Shared | High | High | High | High | High | High | High | High | High | High | High | 0.000246 | 0.001516 | 1.93 |
| FALSE | High | SSPTPYQDI | [K].SSPTPY | 1xPhospho | H9LAN6 1x | 0.000639 | 0 | 1 | 8 | 2 | H9LAN6 | Histone-lys | H9LAN6 [1 | 0 | 2146.017 | 0 | -0.33 | 0.026204 | 10.26 | 112.7 | 87.3 | 6.7 | 13.4 | | High | High | High | High | High | High | High | High | High | High | High | 0 | 0.000144 | 2.71 |
| FALSE | High | SSPTQSLSN | [K].SSPTQS | 1xPhospho | E9PVX6 1xl | 1.16E-06 | 0 | 1 | 2 | 4 | E9PVX6 | Proliferatic | E9PVX6 [2 | 0 | 1916.937 | 0 | -0.22 | 0.796121 | 14.25 | 105.4 | 94.6 | 5.7 | 21 | | High | High | High | High | High | High | High | High | High | High | High | 0 | 4.1E-07 | 3.19 |
| FALSE | High | SSPTQSLSN | [K].SSPTQS | 1xDeamida | E9PVX6 1xl | 0.002974 | 0.00016 | 1 | 2 | 3 | E9PVX6 | Proliferatic | E9PVX6 [2 | 0 | 1917.921 | 0 | -0.13 | 0.18154 | 13.81 | 108.3 | 91.7 | 7.08 | 13.18 | | High | High | High | High | High | High | High | High | High | High | High | 0.001151 | 0.0121 | 2.72 |

| | | | | | | | | | | | | | | | | | | | | | | | | | | | | | | | | | | | | | | |
|---|---|---|---|---|---|---|---|---|---|---|---|---|---|---|---|---|---|---|---|---|---|---|---|---|---|---|---|---|---|---|---|---|---|---|---|---|---|---|
| FALSE | High | SSPTSASAC | [K].SSPTSA | 1xCarbami | E9QMF5 1: | 0.000152 | 0 | 1 | 2 | 1 | E9QMF5 | Neuron na | E9QMF5 [1 | 0 | 1788.815 | 0 | 3.66 | 0.00026 | 17.08 | 13.2 | 186.8 | 31.85 | 6.44 | | High | High | High | High | High | High | High | High | High | High | High | 0 | 2.88E-05 | 2.48 |
| FALSE | High | SSPTSPAPT | [K].SSPTSP | 1xPhospho | Q7TQC7 1x | 0.019276 | 0.001402 | 1 | 4 | 1 | Q7TQC7 | G patch do | Q7TQC7 [5 | 0 | 1579.72 | 0 | 0.31 | 0.001664 | 1.78 | 91.1 | 108.9 | 5.16 | 3.72 | | High | High | High | High | High | High | High | High | High | High | High | 0.000756 | 0.006532 | 2.94 |
| FALSE | High | SSPVCSTAF | [K].SSPVCS | 1xCarbami | Q80XK6 1x | 0.000845 | 0 | 1 | 2 | 2 | Q80XK6 | Autophagy | Q80XK6 [2 | 0 | 2127.045 | 0 | 0.26 | 0.0162 | 5.41 | 88.4 | 111.6 | 7.5 | 6.97 | | High | High | High | High | High | High | High | High | High | High | High | 0 | 0.000198 | 1.74 |
| FALSE | High | SSPVESLK | [R].SSPVES | 1xPhospho | Q9EST3 1x | 0.002153 | 5.66E-05 | 1 | 5 | 2 | Q9EST3 | Eukaryotic | Q9EST3 [7 | 0 | 1384.749 | 0 | 0.11 | 0.137377 | 9.36 | 97.9 | 102.1 | 2.51 | 9.47 | | High | High | High | High | High | High | High | High | High | High | High | 2.58E-05 | 0.000566 | 1.79 |
| FALSE | High | SSPVNNHS | [K].SSPVNN | 1xCarbami | Q6A065 1x | 1.49E-06 | 0 | 1 | 5 | 3 | Q6A065 | Centrosom | Q6A065 [1 | 0 | 2672.234 | 0 | 0.44 | 0.017726 | 21.55 | 83.4 | 116.6 | 5.85 | 15.77 | | High | High | High | High | High | High | High | High | High | High | High | 0 | 1.59E-07 | 4.18 |
| FALSE | High | SSPVNNHS | [K].SSPVNN | 1xCarbami | Q6A065 2x | 9.39E-06 | 0 | 1 | 5 | 1 | Q6A065 | Centrosom | Q6A065 [1 | 0 | 2752.2 | 0 | 1.12 | 0.000457 | 1.93 | 63.2 | 136.8 | 7.17 | 4.24 | | High | High | High | High | High | High | High | High | High | High | High | 0 | 1.25E-06 | 4.44 |
| FALSE | High | SSPVNRPSS | [R].SSPVNF | 1xPhospho | Q8CHI8 1xI | 0.018447 | 0.001324 | 1 | 5 | 1 | Q8CHI8 | E1A-bindin | Q8CHI8 [72 | 0 | 1882.979 | 0 | -0.55 | 0.003629 | 3.07 | 116.5 | 83.5 | 7.71 | 4.58 | | High | High | High | High | High | High | High | High | High | High | High | 0.000723 | 0.006247 | 1.2 |
| FALSE | High | SSPVNSHLS | [R].SSPVNS | 1xPhospho | Q6NXJ0 1x | 4.21E-06 | 0 | 1 | 1 | 2 | Q6NXJ0 | Protein WV | Q6NXJ0 [2 | 0 | 1592.8 | 0 | 0.25 | 0.070972 | 4.41 | 88.7 | 111.3 | 8.97 | 13.45 | | High | High | High | High | High | High | High | High | High | High | High | 0 | 5.1E-07 | 2.89 |
| FALSE | High | SSPVTVNQ | [R].SSPVTV | 1xPhospho | E9QMF5 1: | 0.100129 | 0.008067 | 1 | 2 | 1 | E9QMF5 | Neuron na | E9QMF5 [1 | 0 | 1713.882 | 0 | 2.07 | 0.001139 | 31.88 | 37.7 | 162.3 | 23.71 | 9.97 | | High | High | High | High | High | High | High | High | High | High | High | 0.004032 | 0.04307 | 1.95 |
| FALSE | High | SSPYGSSLD | [R].SSPYGS | 1xPhospho | A0A1N9PT | 0.098533 | 0.007902 | 1 | 1 | 1 | A0A1N9PT | Probable fi | A0A1N9PT | 0 | 2645.322 | 0 | 1.63 | 0.000396 | 10.43 | 48.7 | 151.3 | 1.01 | 6.99 | | High | High | High | High | High | High | High | High | High | High | High | 0.003963 | 0.04237 | 1.7 |
| FALSE | High | SSQCHSGS | [K].SSQCHS | 1xCarbami | Q8BY02 1x | 0.003028 | 0.00016 | 1 | 1 | 3 | Q8BY02 | NF-kappa-I | Q8BY02 [4 | 0 | 1699.788 | 0 | 0.17 | 0.376443 | 8.35 | 97.6 | 102.4 | 5.78 | 9.67 | | High | High | High | High | High | High | High | High | High | High | High | 0.000449 | 0.003874 | 1.78 |
| FALSE | High | SSQDSDTP. | [K].SSQDSI | 1xPhospho | A2AB59 1x | 0.000529 | 0 | 1 | 3 | 2 | A2AB59 | Rho GTPas | A2AB59 [4 | 0 | 2312.07 | 0 | -0.02 | 0.708765 | 15.93 | 100.8 | 99.2 | 10.82 | 5.6 | | High | High | High | High | High | High | High | High | High | High | High | 0 | 0.000116 | 2.56 |
| FALSE | High | SSQEAPSAI | [R].SSQEAF | 1xPhospho | Q9CQ13 1x | 1.82E-06 | 0 | 1 | 1 | 2 | Q9CQ13 | Coordinatc | Q9CQ13 [1 | 0 | 2261.028 | 0 | -0.43 | 0.001665 | 4 | 115.1 | 84.9 | 9.07 | 6.52 | | High | High | High | High | High | High | High | High | High | High | High | 0 | 1.99E-07 | 4 |
| FALSE | High | SSQEIETSS | [R].SSQEIE | 1xCarbami | Q8C0T5 1x | 0.065405 | 0.004734 | 1 | 4 | 1 | Q8C0T5 | Signal-indu | Q8C0T5 [9 | 0 | 2410.146 | 0 | 0.1 | 0.164884 | 12.44 | 93 | 107 | 12.18 | 6.44 | | High | High | High | High | High | High | High | High | High | High | High | 0.002446 | 0.02635 | 2.76 |
| FALSE | High | SSQESLDIK | [R].SSQESL | 1xPhospho | E9QMX4 1: | 0.000183 | 0 | 1 | 4 | 2 | E9QMX4 | Ribosomal | E9QMX4 [4 | 0 | 1544.797 | 0 | 0.59 | 0.001183 | 3.26 | 79.2 | 120.8 | 3.26 | 6.82 | | High | High | High | High | High | High | High | High | High | High | High | 0 | 3.53E-05 | 2.11 |
| FALSE | High | SSQESLDIK | [R].SSQESL | 2xPhospho | E9QMX4 2: | 0.00095 | 0 | 1 | 4 | 2 | E9QMX4 | Ribosomal | E9QMX4 [4 | 0 | 1624.764 | 0 | 0.5 | 0.00502 | 11.99 | 83 | 117 | 4.14 | 7.8 | | High | High | High | High | High | High | High | High | High | High | High | 0 | 0.000224 | 3.03 |
| FALSE | High | SSQFEGTFI | [K].SSQFEG | 2xPhospho | E9Q5F9 2x | 0.0026 | 0.000109 | 1 | 3 | 3 | E9Q5F9 | Histone-lys | E9Q5F9 [3 | 0 | 2465.995 | 0 | 0 | 0.506477 | 5.52 | 100.5 | 99.5 | 3.02 | 2.84 | | High | High | High | High | High | High | High | High | High | High | High | 4.94E-05 | 0.000696 | 3.69 |
| FALSE | High | SSQGASNF | [R].SSQGA: | 1xPhospho | B7ZP47 1xI | 2.42E-05 | 0 | 1 | 2 | 1 | B7ZP47 | Wapal prot | B7ZP47 [31 | 0 | 1578.757 | 0 | -1.32 | 0.001678 | 2.76 | 141.9 | 58.1 | 10.05 | 6.85 | | High | High | High | High | High | High | High | High | High | High | High | 0 | 3.64E-06 | 2.92 |
| FALSE | High | SSQPPSHSS | [R].SSQPPS | 1xPhospho | Q3U2K8 1x | 0.119269 | 0.00979 | 1 | 4 | 1 | Q3U2K8 | Arf-GAP dc | Q3U2K8 [2 | 0 | 1680.836 | 0 | 1.27 | 0.004388 | 20.56 | 57.8 | 142.2 | 22.16 | 17.37 | | High | High | High | High | High | High | High | High | High | High | High | 0.004866 | 0.053 | 1.61 |
| FALSE | High | SSQPSPTTV | [R].SSQPSF | 1xPhospho | A2AJI0 1xP | 0.00013 | 0 | 1 | 4 | 2 | A2AJI0 | MAP7 dom | A2AJI0 [11 | 0 | 2292.152 | 0 | 0.55 | 0.002433 | 9.78 | 80.5 | 119.5 | 10.01 | 6.26 | | High | High | High | High | High | High | High | High | High | High | High | 0 | 2.4E-05 | 3.24 |
| FALSE | High | SSQQVVEC | [K].SSQQV' | 1xPhospho | Q6NS46 1x | 1.23E-09 | 0 | 1 | 2 | 4 | Q6NS46 | Protein RRI | Q6NS46 [2 | 0 | 2987.465 | 0 | -0.28 | 0.015518 | 5.41 | 108.9 | 91.1 | 9.08 | 8.06 | | High | High | High | High | High | High | High | High | High | High | High | 0 | 7.8E-10 | 5.39 |
| FALSE | High | SSQSESEIV | [R].SSQSES | 1xPhospho | Q69ZS7 1xI | 0.109329 | 0.008865 | 1 | 4 | 2 | Q69ZS7 | HBS1-like p | Q69ZS7 [1 | 0 | 1728.882 | 0 | -1.18 | 0.000489 | 2.5 | 135.1 | 64.9 | 5.03 | 11.3 | | High | High | High | High | High | High | High | High | High | High | High | 0.004757 | 0.05145 | 2.75 |
| FALSE | High | SSQSLEEK | [K].SSQSLE | 1xPhospho | F6RJ39 1xP | 0.009006 | 0.000736 | 1 | 4 | 1 | F6RJ39 | Apoptotic | F6RJ39 [32 | 0 | 1445.729 | 0 | -1.8 | 0.000163 | 0.67 | 156.5 | 43.5 | 4.51 | 10.77 | | High | High | High | High | High | High | High | High | High | High | High | 0.000327 | 0.002797 | 1.62 |
| FALSE | High | SSQSLPSDS | [R].SSQSLP | 1xPhospho | D3Z656 1xI | 2.62E-08 | 0 | 1 | 1 | 6 | D3Z656 | Synaptojar | D3Z656 [13 | 0 | 2226.142 | 0 | 1.23 | 0.000975 | 9.91 | 60.5 | 139.5 | 6.11 | 7.41 | | High | High | High | High | High | High | High | High | High | High | High | 0 | 0.000122 | 3.1 |
| FALSE | High | SSQSPPLA/ | [K].SSQSPF | 1xPhospho | G5E8T2 1x | 0.001189 | 0 | 1 | 8 | 1 | G5E8T2 | E3 ubiquiti | G5E8T2 [1 | 0 | 1705.847 | 0 | -0.86 | 0.000417 | 1.65 | 129 | 71 | 5.96 | 6.47 | | High | High | High | High | High | High | High | High | High | High | High | 0 | 0.00029 | 1.92 |
| FALSE | High | SSQTLSPLV | [R].SSQTLS | 1xPhospho | E9Q286 1x | 0.009168 | 0.000736 | 1 | 1 | 3 | E9Q286 | Little elong | E9Q286 [1 | 0 | 2023.006 | 0 | -0.96 | 0.000477 | 2.03 | 130.5 | 69.5 | 5.82 | 3.23 | | High | High | High | High | High | High | High | High | High | High | High | 0.000467 | 0.004236 | 1.92 |
| FALSE | High | SSPTGFGIK | [K].SSPTGF | 1xPhospho | D3Z5I1 1xP | 0.00509 | 0.000551 | 1 | 3 | 2 | D3Z5I1 | Zinc finger | D3Z5I1 [48 | 0 | 1431.765 | 0 | 1.66 | 0.000209 | 7.31 | 48.2 | 151.8 | 4.85 | 3.32 | | High | High | High | High | High | High | High | High | High | High | High | 0.000246 | 0.001472 | 1.96 |
| FALSE | High | SSSVTSIDK | [R].SSSVTS | 1xPhospho | Q8K400 1x | 0.013545 | 0.000976 | 1 | 5 | 1 | Q8K400 | Syntaxin-b | Q8K400 [7 | 0 | 1461.76 | 0 | 1.81 | 0.001785 | 13.2 | 44.2 | 155.8 | 8.8 | 15.43 | | High | High | High | High | High | High | High | High | High | High | High | 0.000538 | 0.00442 | 2.03 |
| FALSE | High | SRPSSPAVF | [R].SRPSSP | 2xPhospho | Q9R020 2x | 0.038677 | 0.002563 | 1 | 5 | 1 | Q9R020 | Zinc finger | Q9R020 [2 | 0 | 1345.623 | 0 | -0.67 | 0.002355 | 2.95 | 124.7 | 75.3 | 6.82 | 11.42 | | High | High | High | High | High | High | High | High | High | High | High | 0.001405 | 0.01443 | 2.06 |
| FALSE | High | SRPNSGDE | [R].SRPNS( | 1xPhospho | D3YYT1 1xI | 0.113753 | 0.009241 | 1 | 2 | 1 | D3YYT1 | Putative ox | D3YYT1 [1 | 0 | 1527.757 | 0 | -0.59 | 0.002928 | 6.76 | 120.4 | 79.6 | 4.39 | 8.12 | | High | High | High | High | High | High | High | High | High | High | High | 0.004625 | 0.05011 | 0.93 |
| FALSE | High | SPSHSSSNF | [K].SPSHSS | 2xPhospho | Q925J9 2xI | 0.017192 | 0.001204 | 1 | 2 | 1 | Q925J9 | Mediator c | Q925J9 [1C | 0 | 2734.264 | 0 | 1.12 | 0.00292 | 17.09 | 63.8 | 136.2 | 12.48 | 9.81 | | High | High | High | High | High | High | High | High | High | High | High | 0.000638 | 0.005773 | 2.39 |
| FALSE | High | SPSIDSIQK | [R].SPSIDSI | 1xPhospho | Q68FH0 1x | 0.012731 | 0.000889 | 1 | 12 | 2 | Q68FH0 | Plakophilin | Q68FH0 [5 | 0 | 1512.808 | 0 | -0.89 | 0.007426 | 4.06 | 128.9 | 71.1 | 11.39 | 10.21 | | High | High | High | High | High | High | High | High | High | High | High | 0.003633 | 0.03871 | 2.06 |
| FALSE | High | SPSLGNVPI | [R].SPSLGN | 1xPhospho | Q80U49 1x | 0.000674 | 0 | 1 | 3 | 1 | Q80U49 | Centrosom | Q80U49 [1 | 0 | 2078.048 | 0 | 1.75 | 0.000618 | 4.14 | 44.8 | 155.2 | 10.97 | 10.48 | | High | High | High | High | High | High | High | High | High | High | High | 0 | 0.000153 | 1.88 |
| FALSE | High | SPSLGSDLT | [R].SPSLGS | 1xPhospho | B7ZNU9 1x | 2.67E-05 | 0 | 1 | 3 | 3 | B7ZNU9 | Beta-2-syn | B7ZNU9 [3 | 0 | 1660.814 | 0 | -0.41 | 0.001331 | 3.86 | 112.5 | 87.5 | 4.89 | 4.41 | | High | High | High | High | High | High | High | High | High | High | High | 0 | 5.67E-05 | 2.75 |
| FALSE | High | SPSLGSEDI | [R].SPSLGS | 1xPhospho | B2RWU2 1 | 0.000122 | 0 | 1 | 2 | 2 | B2RWU2 | Immunode | B2RWU2 [ | 0 | 2250.131 | 0 | 3.05 | 0.000207 | 5.53 | 22.6 | 177.4 | 9.66 | 6.78 | | High | High | High | High | High | High | High | High | High | High | High | 0 | 2.23E-05 | 3.1 |
| FALSE | High | SPSLGSEDI | [R].SPSLGS | 2xPhospho | B2RWU2 2 | 0.000845 | 0 | 1 | 2 | 1 | B2RWU2 | Immunode | B2RWU2 [ | 0 | 2330.097 | 0 | 1.29 | 0.001147 | 20.81 | 56.2 | 143.8 | 6.77 | 10.2 | | High | High | High | High | High | High | High | High | High | High | High | 0 | 0.000198 | 2.63 |
| FALSE | High | SPSLPGLLS | [R].SPSLPG | 1xPhospho | Q6PCQ0 1x | 0.001255 | 0 | 1 | 3 | 1 | Q6PCQ0 | IQ domain- | Q6PCQ0 [5 | 0 | 2227.169 | 0 | 2.43 | 0.000321 | 20.31 | 32.1 | 167.9 | 12.5 | 6.86 | | High | High | High | High | High | High | High | High | High | High | High | 0 | 0.000308 | 3.11 |
| FALSE | High | SPSLPTGGT | [K].SPSLPT | 1xPhospho | Q1W617 1: | 0.005228 | 0.000551 | 1 | 2 | 2 | Q1W617 | Protein Shi | Q1W617 [4 | 0 | 1753.832 | 0 | 2.43 | 0.000233 | 8.34 | 32.1 | 167.9 | 8.62 | 6.94 | | High | High | High | High | High | High | High | High | High | High | High | 0.000246 | 0.001517 | 1.73 |
| FALSE | High | SPSISNMA/ | [R].SPSISNI | 1xPhospho | B7ZNX2 1x | 0.0095 | 0.000736 | 1 | 5 | 2 | B7ZNX2 | Anaphase | B7ZNX2 [3 | 0 | 1558.75 | 0 | 0.46 | 0.008251 | 11.5 | 82.9 | 117.1 | 8.25 | 9.64 | | High | High | High | High | High | High | High | High | High | High | High | 0.000327 | 0.002965 | 2.03 |
| FALSE | High | SPSLSPSPP | [K].SPSLSP | 1xPhospho | P14873 1xI | 1.45E-05 | 0 | 2 | 2 | 2 | P14873; B2 | Microtubu | P14873 [12 | 0 | 1961.04 | 0 | -0.94 | 0.002101 | 4.44 | 130.3 | 69.7 | 8.36 | 7.09 | Shared | High | High | High | High | High | High | High | High | High | High | High | 0 | 2.04E-06 | 2.75 |
| FALSE | High | SPSLSPSPP | [K].SPSLSP | 2xPhospho | P14873 2xI | 0.103392 | 0.008249 | 2 | 2 | 2 | P14873; B2 | Microtubu | P14873 [12 | 0 | 2041.006 | 0 | -0.9 | 0.003321 | 6.51 | 128.1 | 71.9 | 8.91 | 9.39 | Shared | High | High | High | High | High | High | High | High | High | High | High | 0.004164 | 0.04503 | 2.31 |
| FALSE | High | SPSMNISEI | [K].SPSMN | 1xPhospho | Q811P8 1x | 0.015329 | 0.001089 | 1 | 3 | 2 | Q811P8 | Rho GTPas | Q811P8 [9 | 0 | 2150.077 | 0 | 0.17 | 0.511886 | 30.11 | 100.6 | 99.4 | 6.37 | 18.79 | | High | High | High | High | High | High | High | High | High | High | High | 0.00196 | 0.01973 | 2.73 |
| FALSE | High | SPSNISVQK | [R].SPSNIS' | 1xPhospho | Q6PCQ0 1x | 0.000311 | 0 | 1 | 3 | 2 | Q6PCQ0 | IQ domain- | Q6PCQ0 [3 | 0 | 1497.808 | 0 | 1.98 | 0.000207 | 5.15 | 40.3 | 159.7 | 5.13 | 5.41 | | High | High | High | High | High | High | High | High | High | High | High | 0 | 6.44E-05 | 2.09 |
| FALSE | High | SPSHSGNA | [R].SPSHS( | 2xPhospho | Q99MQ1 2 | 0.003312 | 0.00016 | 1 | 3 | 1 | Q99MQ1 | Protein bic | Q99MQ1 [ | 0 | 1787.813 | 0 | -0.13 | 0.578145 | 34.76 | 107.8 | 92.2 | 11.73 | 36.57 | | High | High | High | High | High | High | High | High | High | High | High | 7.19E-05 | 0.000915 | 2.57 |
| FALSE | High | SPSPAHIVS | [R].SPSPAH | 1xPhospho | E9QQ89 1x | 0.001168 | 0 | 1 | 22 | 1 | E9QQ89 | Transcripti | E9QQ89 [1 | 0 | 1674.898 | 0 | 1.35 | 0.002834 | 38.61 | 56.4 | 143.6 | 14.93 | 8.48 | | High | High | High | High | High | High | High | High | High | High | High | 0 | 0.000285 | 2.44 |
| FALSE | High | SPSPALSGS | [K].SPSPAL | 2xPhospho | G5E8L6 2xI | 0.041072 | 0.003094 | 1 | 3 | 1 | G5E8L6 | Killer cell le | G5E8L6 [4 | 0 | 1444.644 | 0 | 1.46 | 0.002137 | 6.68 | 57.6 | 142.4 | 11.73 | 13.76 | | High | High | High | High | High | High | High | High | High | High | High | 0.001609 | 0.01547 | 2.96 |
| FALSE | High | SPSPAPAAI | [K].SPSPAP | 1xPhospho | B9EKB5 1xI | 0.002695 | 0.000109 | 1 | 2 | 1 | B9EKB5 | Baz2b prot | B9EKB5 [1 | 0 | 3662.996 | 0 | -0.74 | 0.008024 | 8.96 | 124.7 | 75.3 | 10.56 | 15.12 | | High | High | High | High | High | High | High | High | High | High | High | 4.94E-05 | 0.000722 | 3.86 |
| FALSE | High | SPSPAPPPF | [R].SPSPAP | 1xPhospho | Q52KI8 1xI | 0.000343 | 0 | 1 | 7 | 2 | Q52KI8 | Serine/argi | Q52KI8 [57 | 0 | 1893.983 | 0 | -0.15 | 0.369178 | 19.5 | 105.7 | 94.3 | 2.76 | 18.39 | | High | High | High | High | High | High | High | High | High | High | High | 0 | 7.21E-05 | 2 |
| FALSE | High | SPSPAVPLF | [R].SPSPAV | 1xPhospho | Q6PIJ4 1xP | 0.059159 | 0.004294 | 2 | 3 | 2 | Q6PIJ4; A0 | Nuclear fac | Q6PIJ4 [22 | 0 | 1232.66 | 0 | -0.05 | 0.467909 | 15.33 | 103.8 | 96.2 | 8.48 | 7 | Shared | High | High | High | High | High | High | High | High | High | High | High | 0.002202 | 0.02346 | 2.07 |
| FALSE | High | SPSPDSAAS | [R].SPSPDS | 1xPhospho | Q80TU6 1x | 0.075927 | 0.005826 | 1 | 1 | 2 | Q80TU6 | MKIAA067 | Q80TU6 [5 | 0 | 2281.123 | 0 | 2.23 | 0.000719 | 2.21 | 36 | 164 | 7.14 | 11.51 | | High | High | High | High | High | High | High | High | High | High | High | 0.002976 | 0.03128 | 1.82 |
| FALSE | High | SPSPEPIYN | [R].SPSPEP | 1xPhospho | D3YZC9 1xI | 0.000466 | 0 | 1 | 9 | 2 | D3YZC9 | Splicing fac | D3YZC9 [8 | 0 | 1942.956 | 0 | -1.05 | 0.000954 | 1.19 | 134 | 66 | 4.3 | 8.48 | | High | High | High | High | High | High | High | High | High | High | High | 0 | 0.000146 | 3.03 |
| FALSE | High | SPSPFDCSF | [K].SPSPFD | 2xCarbami | A3KFM7 1x | 1.21E-06 | 0 | 1 | 5 | 1 | A3KFM7 | Chromodo | A3KFM7 [3 | 0 | 3474.55 | 0 | 0.24 | 0.001698 | 1.15 | 91.6 | 108.4 | 13.16 | 9.88 | | High | High | High | High | High | High | High | High | High | High | High | 0 | 1.26E-07 | 4.57 |
| FALSE | High | SPSPGLFR | [R].SPSPGL | 1xPhospho | Q80YR7 1x | 0.015741 | 0.001124 | 1 | 2 | 2 | Q80YR7 | Claspin OS | Q80YR7 [7 | 0 | 1169.592 | 0 | -0.41 | 0.001822 | 3.99 | 114.4 | 85.6 | 2.44 | 6.24 | | High | High | High | High | High | High | High | High | High | High | High | 0.000585 | 0.005216 | 2.16 |
| FALSE | High | SPSPGSSPS | [R].SPSPGS | 1xPhospho | P53349 1xI | 0.004256 | 0.00036 | 2 | 3 | 1 | P53349; F8 | Mitogen-a | P53349 [24 | 0 | 1324.61 | 0 | -1.45 | 0.003768 | 0.79 | 144.6 | 55.4 | 12.33 | 22.22 | Shared | High | High | High | High | High | High | High | High | High | High | High | 0.000161 | 0.001211 | 2.01 |
| FALSE | High | SPSPLGVGS | [R].SPSPLG | 1xPhospho | Q3UHU5 1: | 8.43E-06 | 0 | 1 | 5 | 2 | Q3UHU5 | Microtubu | Q3UHU5 [ | 0 | 1642.804 | 0 | -1.58 | 0.000572 | 0.69 | 148 | 52 | 5.69 | 9.18 | | High | High | High | High | High | High | High | High | High | High | High | 0 | 1.12E-06 | 2.38 |
| FALSE | High | SPSPISISSS | [R].SPSPISI | 1xPhospho | E9Q309 1x | 9.2E-05 | 0 | 1 | 1 | 2 | E9Q309 | Centrosom | E9Q309 [1 | 0 | 2420.203 | 0 | 0.19 | 0.876693 | 21.7 | 97.5 | 102.5 | 14.1 | 13.18 | | High | High | High | High | High | High | High | High | High | High | High | 0 | 2.27E-05 | 3.63 |
| FALSE | High | SPSPLVSLQ | [R].SPSPLV | 1xPhospho | P59438 1xI | 0.00095 | 0 | 1 | 7 | 1 | P59438 | Hermansky | P59438 [53 | 0 | 1764.007 | 0 | 2.06 | 0.00135 | 30.62 | 44.5 | 155.5 | 14.49 | 12.52 | | High | High | High | High | High | High | High | High | High | High | High | 0 | 0.000226 | 3.03 |
| FALSE | High | SPSPPPDGS | [K].SPSPPP | 1xPhospho | O08539 1x | 3.61E-06 | 0 | 1 | 3 | 5 | O08539 | Myc box-d | O08539 [2 | 0 | 1984.958 | 0 | 1.19 | 0.000534 | 1.93 | 60.8 | 139.2 | 4.76 | 7.36 | | High | High | High | High | High | High | High | High | High | High | High | 0 | 3.64E-06 | 4.09 |
| FALSE | High | SPSPALHPF | [K].SPSPAL | 1xPhospho | B2RR83 1x | 0.024424 | 0.001612 | 1 | 1 | 1 | B2RR83 | Probable A | B2RR83 [1 | 0 | 1696.919 | 0 | 0.06 | 0.157148 | 2.49 | 96.9 | 103.1 | 4.18 | 4.6 | | High | High | High | High | High | High | High | High | High | High | High | 0.000859 | 0.008506 | 2.06 |
| FALSE | High | SPSPPPDGS | [K].SPSPPP | 2xPhospho | O08539 2x | 9.2E-05 | 0 | 1 | 3 | 3 | O08539 | Myc box-d | O08539 [2 | 0 | 2064.924 | 0 | 1.54 | 0.000784 | 19.38 | 52.4 | 147.6 | 8.51 | 9.59 | | High | High | High | High | High | High | High | High | High | High | High | 0 | 3.07E-05 | 2.96 |
| FALSE | High | SPSGSAFGS | [R-].SPSGS | 2xPhospho | A2A9M4 2: | 0.057203 | 0.004227 | 1 | 7 | 2 | A2A9M4 | Dedicator | A2A9M4 [1 | 0 | 1825.772 | 0 | 0.34 | 0.055 | 17.48 | 87.3 | 112.7 | 7.96 | 11.51 | | High | High | High | High | High | High | High | High | High | High | High | 0.003303 | 0.03615 | 1.86 |
| FALSE | High | SPSGPPGSI | [R].SPSGPF | 1xPhospho | E9Q2Z1 1x | 9.36E-05 | 0 | 1 | 4 | 1 | E9Q2Z1 | CECR2, hist | E9Q2Z1 [1 | 0 | 1481.71 | 0 | -0.94 | 0.002027 | 2.88 | 132.2 | 67.8 | 9.12 | 2.92 | | High | High | High | High | High | High | High | High | High | High | High | 0 | 1.67E-05 | 2.58 |
| FALSE | High | SPSAQELK | [K].SPSAQE | 1xPhospho | Q3TYJ0 1xI | 0.007404 | 0.00062 | 1 | 2 | 1 | Q3TYJ0 | Putative ur | Q3TYJ0 [24 | 0 | 1397.744 | 0 | -0.68 | 0.012129 | 5.59 | 121.5 | 78.5 | 8.26 | 14.24 | | High | High | High | High | High | High | High | High | High | High | High | 0.000275 | 0.002246 | 1.34 |
| FALSE | High | SPSASITDE | [R].SPSASI | 1xPhospho | Q8C8U0-3 | 0.000117 | 0 | 1 | 6 | 2 | Q8C8U0-3 | Isoform 3 | Q8C8U0-3 | 0 | 1630.705 | 0 | 0.34 | 0.018066 | 7.98 | 88.2 | 111.8 | 5.81 | 8.82 | | High | High | High | High | High | High | High | High | High | High | High | 0 | 2.15E-05 | 3.09 |
| FALSE | High | SPSASSVGS | [R].SPSASS | 1xPhospho | Q64127 1x | 2.82E-05 | 0 | 1 | 5 | 1 | Q64127 | Transcripti | Q64127 [6 | 0 | 1243.588 | 0 | -0.53 | 0.058415 | 1.48 | 116.5 | 83.5 | 7.83 | 14.02 | | High | High | High | High | High | High | High | High | High | High | High | 0 | 4.34E-06 | 2.11 |
| FALSE | High | SPSAVSASH | [R].SPSAVS | 1xPhospho | Q6NSV7-3 | 0.000152 | 0 | 1 | 5 | 2 | Q6NSV7-3 | Isoform 3 | Q6NSV7-3 | 0 | 2065.975 | 0 | 0.17 | 0.645217 | 28.22 | 96.9 | 103.1 | 17.36 | 16.58 | | High | High | High | High | High | High | High | High | High | High | High | 0 | 2.88E-05 | 2.99 |
| FALSE | High | SPSDLHISP | [R].SPSDLH | 1xPhospho | F6ZBR8 1xI | 0.000815 | 0 | 2 | 5 | 2 | F6ZBR8; O | Protein Wi | F6ZBR8 [44 | 0 | 1832.992 | 0 | 0.45 | 0.035417 | 27.11 | 83.6 | 116.4 | 13.18 | 10.16 | Shared | High | High | High | High | High | High | High | High | High | High | High | 0 | 0.00019 | 2.68 |
| FALSE | High | SPSDLHISP | [R].SPSDLH | 2xPhospho | F6ZBR8 2xI | 0.043607 | 0.003228 | 2 | 5 | 2 | F6ZBR8; O | Protein Wi | F6ZBR8 [44 | 0 | 1912.959 | 0 | -0.12 | 0.153083 | 7.88 | 105.8 | 94.2 | 4.46 | 9.05 | Shared | High | High | High | High | High | High | High | High | High | High | High | 0.001651 | 0.01644 | 3.03 |
| FALSE | High | SPSDITISA( | [K].SPSDITI | 1xPhospho | Q69ZR9-2 | 0.003495 | 0.00021 | 1 | 2 | 2 | Q69ZR9-2 | Isoform 2 | Q69ZR9-2 | 0 | 3769.971 | 0 | -1.34 | 0.000459 | 1.31 | 139.9 | 60.1 | 6.68 | 7.63 | | High | High | High | High | High | High | High | High | High | High | High | 9.42E-05 | 0.000968 | 3.59 |
| FALSE | High | SPSDLWK | [K].SPSDLV | 1xPhospho | Q01320 1x | 0.005467 | 0.00062 | 3 | 6 | 1 | Q01320; Q | DNA topois | Q01320 [1 | 0 | 1370.712 | 0 | -1.81 | 0.00076 | 1.01 | 154.5 | 45.5 | 9.15 | 6.63 | Shared | High | High | High | High | High | High | High | High | High | High | High | 0.000275 | 0.001594 | 2.16 |
| FALSE | High | SPSDMATS | [R].SPSDM | 1xCarbami | Q8BYY4 1x | 0.000267 | 0 | 1 | 3 | 1 | Q8BYY4 | Tetratricop | Q8BYY4 [2 | 0 | 2536.165 | 0 | -0.77 | 0.008619 | 5.43 | 126.3 | 73.7 | 13.75 | 8.65 | | High | High | High | High | High | High | High | High | High | High | High | 0 | 5.39E-05 | 2.84 |
| FALSE | High | SPSDMATS | [R].SPSDM | 1xCarbami | Q8BYY4 1x | 0.053473 | 0.00384 | 1 | 3 | 1 | Q8BYY4 | Tetratricop | Q8BYY4 [2 | 0 | 2552.16 | 0 | -0.88 | 0.005556 | 6.15 | 127.5 | 72.5 | 16.36 | 5.69 | | High | High | High | High | High | High | High | High | High | High | High | 0.001999 | 0.02086 | 2.2 |
| FALSE | High | SPSDNSLN | [R].SPSDNS | 1xCarbami | Q3TLI0 1xP | 0.000259 | 0 | 1 | 3 | 2 | Q3TLI0 | Trafficking | Q3TLI0 [68 | 0 | 1943.91 | 0 | 0.71 | 0.002165 | 16.73 | 75.7 | 124.3 | 6.64 | 4.77 | | High | High | High | High | High | High | High | High | High | High | High | 0 | 5.24E-05 | 3.15 |
| FALSE | High | SPSDPGGE | [R].SPSDPG | 1xPhospho | A0A0A0M( | 0.000331 | 0 | 1 | 2 | 2 | A0A0A0M( | Calmodulir | A0A0A0M( | 0 | 1750.866 | 0 | 0.27 | 0.006251 | 2.92 | 90.8 | 109.2 | 3.55 | 2 | | High | High | High | High | High | High | High | High | High | High | High | 0 | 6.92E-05 | 2.18 |
| FALSE | High | SPSDSSTAS | [R].SPSDSS | 1xPhospho | Q9QXS6 1x | 4.91E-06 | 0 | 2 | 3 | 7 | Q9QXS6; A | Drebrin OS | Q9QXS6 [3 | 0 | 2185.022 | 0 | -0.96 | 0.007167 | 5.25 | 130.6 | 69.4 | 10.08 | 13.56 | Shared | High | High | High | High | High | High | High | High | High | High | High | 0 | 6.04E-07 | 3.51 |
| FALSE | High | SPSGSAFGS | [R-].SPSGS | 1xPhospho | A2A9M4 1: | 4.44E-06 | 0 | 1 | 7 | 5 | A2A9M4 | Dedicator | A2A9M4 [1 | 0 | 1745.806 | 0 | 0.78 | 0.00108 | 9.01 | 74.1 | 125.9 | 1.98 | 6.44 | | High | High | High | High | High | High | High | High | High | High | High | 0 | 5.44E-07 | 3.28 |
| FALSE | High | SPSEESTTT | [K].SPSEES | 1xPhospho | P54726 1xI | 2.36E-06 | 0 | 2 | 4 | 11 | P54726; Q | UV excisior | P54726 [12 | 0 | 2907.294 | 0 | 0.21 | 0.030305 | 4.68 | 92.9 | 107.1 | 3.95 | 5.41 | Shared | High | High | High | High | High | High | High | High | High | High | High | 0 | 2.37E-06 | 3.57 |
| FALSE | High | SPSELFAQH | [R].SPSELF | 1xPhospho | Q569Z6 1x | 0.000772 | 0 | 1 | 4 | 1 | Q569Z6 | Thyroid ho | Q569Z6 [6 | 0 | 2580.41 | 0 | -0.32 | 0.007082 | 6.58 | 110.1 | 89.9 | 7.07 | 10.76 | | High | High | High | High | High | High | High | High | High | High | High | 0 | 0.000178 | 2.42 |
| FALSE | High | SPSEMAQF | [R].SPSEM | 1xPhospho | B9EKC1 1xI | 0.095408 | 0.007437 | 1 | 2 | 1 | B9EKC1 | PAS domai | B9EKC1 [1 | 0 | 1230.539 | 0 | -0.4 | 0.010551 | 1.52 | 115 | 85 | 5.94 | 12.72 | | High | High | High | High | High | High | High | High | High | High | High | 0.003768 | 0.04066 | 1.24 |
| FALSE | High | SPSESAEET | [R].SPSESA | 1xPhospho | Q9CT10 1x | 0.090888 | 0.007081 | 1 | 1 | 2 | Q9CT10 | Ran-bindin | Q9CT10 [1 | 0 | 1982.879 | 0 | 0.35 | 0.206223 | 42.7 | 84.7 | 115.3 | 38.44 | 8.8 | | High | High | High | High | High | High | High | High | High | High | High | 0.003633 | 0.03856 | 1.98 |
| FALSE | High | SPSESAEET | [R].SPSESA | 1xPhospho | Q9CT10 1x | 5.91E-05 | 0 | 1 | 1 | 13 | Q9CT10 | Ran-bindin | Q9CT10 [1 | 0 | 2212.042 | 0 | 0.51 | 0.000223 | 4.42 | 81.8 | 118.2 | 5.96 | 3.94 | | High | High | High | High | High | High | High | High | High | High | High | 0.000344 | 0.003056 | 2.72 |
| FALSE | High | SPSESPQHI | [R].SPSESP | 1xPhospho | Q5DU37 1x | 0.000375 | 0 | 1 | 2 | 1 | Q5DU37 | Zinc finger | Q5DU37 [6 | 0 | 1818.858 | 0 | 0.33 | 0.162506 | 6.28 | 88.5 | 111.5 | 16.32 | 3.64 | | High | High | High | High | High | High | High | High | High | High | High | 0 | 7.93E-05 | 3.39 |
| FALSE | High | SPSFASEW | [K].SPSFAS | 1xPhospho | Q3UHP6 1x | 4.27E-05 | 0 | 1 | 4 | 3 | Q3UHP6 | Putative ur | Q3UHP6 [6 | 0 | 2062.977 | 0 | 0.33 | 0.005611 | 5.47 | 88.6 | 111.4 | 4.22 | 6.35 | | High | High | High | High | High | High | High | High | High | High | High | 2.58E-05 | 0.000443 | 3.51 |
| FALSE | High | SPSFGEDC | [R].SPSFGE | 1xCarbami | O35492 1x | 0.100936 | 0.008097 | 1 | 4 | 1 | O35492 | Dual specif | O35492 [2 | 0 | 1757.704 | 0 | -0.55 | 0.07248 | 12.83 | 120.5 | 79.5 | 14.97 | 16.61 | | High | High | High | High | High | High | High | High | High | High | High | 0.004072 | 0.04354 | 1.78 |
| FALSE | High | SPSFPIHEE | [R].SPSFPIH | 1xPhospho | Q9D074-5 | 0.029586 | 0.001857 | 1 | 5 | 1 | Q9D074-5 | Isoform 5 | Q9D074-5 | 0 | 2081.983 | 0 | 1.41 | 0.000949 | 9.64 | 55.4 | 144.6 | 7.58 | 12.23 | | High | High | High | High | High | High | High | High | High | High | High | 0.000997 | 0.01058 | 1.77 |
| FALSE | High | SPSFSSTSE | [K].SPSFSS | 1xPhospho | Q3UGE1 1: | 7.18E-07 | 0 | 1 | 7 | 4 | Q3UGE1 | Putative ur | Q3UGE1 [2 | 0 | 1853.893 | 0 | 1.02 | 0.000507 | 9.46 | 64.8 | 135.2 | 2.97 | 5.88 | | High | High | High | High | High | High | High | High | High | High | High | 0 | 3.44E-06 | 2.51 |
| FALSE | High | SPSGGAAG | [R].SPSGG/ | 1xPhospho | A0A0R4J2E | 1.08E-06 | 0 | 1 | 4 | 4 | A0A0R4J2E | BTB/POZ d | A0A0R4J2E | 0 | 2264.112 | 0 | -0.41 | 0.175945 | 17.33 | 110.4 | 89.6 | 16.82 | 17.98 | | High | High | High | High | High | High | High | High | High | High | High | 0 | 1.1E-07 | 4.22 |
| FALSE | High | SPSGGAAG | [R].SPSGG/ | 2xPhospho | A0A0R4J2E | 0.001831 | 5.66E-05 | 1 | 4 | 1 | A0A0R4J2E | BTB/POZ d | A0A0R4J2E | 0 | 2344.078 | 0 | 0.56 | 0.001006 | 4.47 | 81.7 | 118.3 | 10.72 | 5.67 | | High | High | High | High | High | High | High | High | High | High | High | 2.58E-05 | 0.000472 | 3.16 |
| FALSE | High | SPSGGEHA | [R].SPSGGI | 1xPhospho | Q9Z2C9 1x | 0.000167 | 0 | 1 | 1 | 3 | Q9Z2C9 | Myotubula | Q9Z2C9 [6 | 0 | 1979.911 | 0 | -0.72 | 0.011719 | 5.2 | 124.5 | 75.5 | 6.04 | 13.12 | | High | High | High | High | High | High | High | High | High | High | High | 0 | 3.19E-05 | 2.6 |
| FALSE | High | SPSGPHGL | [R].SPSGPH | 1xCarbami | Q61982 1x | 0.008313 | 0.000736 | 1 | 1 | 1 | Q61982 | Neurogeni | Q61982 [2 | 0 | 2807.508 | 0 | -1.58 | 0.000158 | 1.08 | 148.9 | 51.1 | 6.48 | 18.26 | | High | High | High | High | High | High | High | High | High | High | High | 0.000327 | 0.002541 | 3.48 |
| FALSE | High | SPSEGPGA | [K].SPSEGF | 1xPhospho | B2RUG9 1x | 0.000188 | 0 | 1 | 6 | 1 | B2RUG9 | Adenomat | B2RUG9 [2 | 0 | 1552.721 | 0 | 0.01 | 0.841595 | 26.82 | 99.5 | 100.5 | 10.77 | 9.58 | | High | High | High | High | High | High | High | High | High | High | High | 0 | 3.65E-05 | 2.53 |
| FALSE | High | SPSPPQQQ | [R].SPSPPC | 1xPhospho | Q3UIA2 1x | 8.34E-09 | 0 | 1 | 6 | 13 | Q3UIA2 | Rho GTPas | Q3UIA2 [6 | 0 | 3528.624 | 0 | -0.79 | 0.087869 | 3.49 | 138.1 | 61.9 | 55.77 | 22.45 | | High | High | High | High | High | High | High | High | High | High | High | 0 | 9.83E-07 | 5.46 |
| FALSE | High | SPSPPQQQ | [R].SPSPPC | 1xDeamida | Q3UIA2 1x | 5.36E-09 | 0 | 1 | 6 | 56 | Q3UIA2 | Rho GTPas | Q3UIA2 [6 | 0 | 3529.608 | 0 | -2.62 | | 2.96 | 175 | 25 | 94.84 | 40.1 | | High | High | High | High | High | High | High | High | High | High | High | 0 | 1.11E-09 | 5.9 |
| FALSE | High | SPSPQFAP( | [R].SPSPQF | 1xPhospho | A7TU71 1x | 0.031439 | 0.002011 | 1 | 3 | 1 | A7TU71 | Protein Shi | A7TU71 [1 | 0 | 1624.85 | 0 | 1.13 | 0.002521 | 6.63 | 63.4 | 136.6 | 8.03 | 13.92 | | High | High | High | High | High | High | High | High | High | High | High | 0.001076 | 0.01139 | 2.6 |
| FALSE | High | SPSSPIVSIL | [K].SPSSPI\ | 1xPhospho | P97929 1xI | 0.000248 | 0 | 1 | 4 | 1 | P97929 | Breast canc | P97929 [23 | 0 | 2243.163 | 0 | 0.46 | 0.001927 | 2.65 | 81.2 | 118.8 | 7.61 | 4.46 | | High | High | High | High | High | High | High | High | High | High | High | 0 | 4.99E-05 | 3.75 |
| FALSE | High | SPSSPYSVE | [R].SPSSPY | 1xPhospho | A0A068BIR | 0.000233 | 0 | 1 | 3 | 1 | A0A068BIR | KN motif a | A0A068BIR | 0 | 3037.5 | 0 | -0.99 | 0.000405 | 4.27 | 128.8 | 71.2 | 9.76 | 18.34 | | High | High | High | High | High | High | High | High | High | High | High | 0 | 4.65E-05 | 3.11 |
| FALSE | High | SPSSTESLQ | [K].SPSSTE | 1xPhospho | B0V2M3 1: | 0.001147 | 0 | 1 | 5 | 6 | B0V2M3 | Zinc finger | B0V2M3 [1 | 0 | 1898.988 | 0 | -0.56 | 0.006529 | 2.91 | 119.2 | 80.8 | 2.13 | 10.87 | | High | High | High | High | High | High | High | High | High | High | High | 2.58E-05 | 0.000341 | 3.38 |
| FALSE | High | SPSSVTGN/ | [R].SPSSVT | 1xPhospho | Q91VL8 1x | 6.41E-05 | 0 | 1 | 2 | 1 | Q91VL8 | Telomeric | Q91VL8 [1 | 0 | 1555.772 | 0 | -1 | 0.004523 | 2.77 | 130.5 | 69.5 | 15.39 | 13.4 | | High | High | High | High | High | High | High | High | High | High | High | 0 | 1.09E-05 | 2.38 |
| FALSE | High | SPSSVTGN/ | [R].SPSSVT | 1xPhospho | Q91VL8 1x | 6.74E-07 | 0 | 1 | 2 | 2 | Q91VL8 | Telomeric | Q91VL8 [1 | 0 | 1784.935 | 0 | -0.94 | 0.001143 | 3.36 | 130.3 | 69.7 | 9.24 | 7.65 | | High | High | High | High | High | High | High | High | High | High | High | 0 | 6.53E-08 | 2.52 |
| FALSE | High | SPSTAYFLS | [K].SPSTAY | 1xPhospho | P47810 1xI | 0.000407 | 0 | 1 | 1 | 4 | P47810 | Wee1-like | P47810 [12 | 0 | 2052.004 | 0 | -1.04 | 0.00034 | 2.74 | 134.6 | 65.4 | 4.62 | 7.03 | | High | High | High | High | High | High | High | High | High | High | High | 0 | 0.000142 | 3.82 |
| FALSE | High | SPSTAYFLS | [K].SPSTAY | 2xPhospho | P47810 2xI | 0.000422 | 0 | 1 | 1 | 2 | P47810 | Wee1-like | P47810 [12 | 0 | 2131.97 | 0 | -0.78 | 0.001386 | 3.39 | 127.7 | 72.3 | 6.07 | 13.24 | | High | High | High | High | High | High | High | High | High | High | High | 0 | 9.07E-05 | 4.03 |
| FALSE | High | SPSTLSNTP | [K].SPSTLSI | 1xCarbami | B2RX81 1xI | 1.73E-10 | 0 | 1 | 1 | 2 | B2RX81 | BC032203 | B2RX81 [5 | 0 | 2893.396 | 0 | -0.29 | 0.02533 | 6.32 | 107.9 | 92.1 | 7.47 | 8.57 | | High | High | High | High | High | High | High | High | High | High | High | 0 | 6.03E-12 | 4.42 |
| FALSE | High | SPSVASHG | [K].SPSVAS | 1xPhospho | Q5SXJ3 1xI | 0.00153 | 5.66E-05 | 1 | 1 | 1 | Q5SXJ3 | Fanconi an | Q5SXJ3 [9 | 0 | 1392.683 | 0 | 0.07 | 0.910645 | 20.38 | 102.4 | 97.6 | 10.16 | 12.27 | | High | High | High | High | High | High | High | High | High | High | High | 2.58E-05 | 0.000386 | 2.39 |
| FALSE | High | SPSVISLQV | [K].SPSVISI | 1xCarbami | Q68FE8 1x | 0.047488 | 0.00354 | 1 | 7 | 1 | Q68FE8 | Zinc finger | Q68FE8 [2 | 0 | 2885.448 | 0 | 0.91 | 0.003534 | 5.94 | 73.1 | 126.9 | 14.91 | 19.23 | | High | High | High | High | High | High | High | High | High | High | High | 0.001812 | 0.01813 | 3.36 |
| FALSE | High | SPSVPQDA | [K].SPSVPC | 1xPhospho | Q9DB00 1x | 5.35E-05 | 0 | 2 | 7 | 2 | Q9DB00; C | GON-4-like | Q9DB00 [1 | 0 | 1593.747 | 0 | -1.16 | 0.000245 | 1.6 | 138.7 | 61.3 | 3.67 | 3.04 | Shared | High | High | High | High | High | High | High | High | High | High | High | 0 | 8.91E-06 | 2.05 |
| FALSE | High | SPSVSQSQ | [K].SPSVSC | 1xPhospho | Q3UYC8 1x | 4.94E-05 | 0 | 1 | 5 | 2 | Q3UYC8 | Putative ur | Q3UYC8 [4 | 0 | 1442.684 | 0 | -0.07 | 0.897099 | 14.62 | 102.3 | 97.7 | 6.04 | 6.65 | | High | High | High | High | High | High | High | High | High | High | High | 0 | 8.09E-06 | 1.66 |
| FALSE | High | SPSWEPFR | [R].SPSWEI | 1xPhospho | Q545F4 1x | 0.012731 | 0.000889 | 1 | 4 | 14 | Q545F4 | Hspb1 prot | Q545F4 [1 | 0 | 1314.608 | 0 | 0.22 | 0.031024 | 12.37 | 94 | 106 | 7.01 | 6.57 | | High | High | High | High | High | High | High | High | High | High | High | 0.000449 | 0.004118 | 1.69 |
| FALSE | High | SPSSLSANV | [K].SPSSLS/ | 2xPhospho | F8VPZ3 2xI | 0.000158 | 0 | 1 | 3 | 2 | F8VPZ3 | Ubiquitin-s | F8VPZ3 [1 | 0 | 1979.949 | 0 | -0.2 | 0.156553 | 5.13 | 105 | 95 | 6.8 | 8.58 | | High | High | High | High | High | High | High | High | High | High | High | 0 | 2.99E-05 | 3.64 |
| FALSE | High | SPSWISASM | [R].SPSWIS | 1xPhospho | B2RRE0 1x | 0.001503 | 5.66E-05 | 1 | 2 | 3 | B2RRE0 | A kinase (P | B2RRE0 [8 | 0 | 3674.807 | 0 | -1.17 | 0.006474 | 4.69 | 137.4 | 62.6 | 14.19 | 26.5 | | High | High | High | High | High | High | High | High | High | High | High | 2.58E-05 | 0.000377 | 4.82 |
| FALSE | High | SPSWTK | [R].SPSWTI | 1xPhospho | H3BKU0 1x | 0.056724 | 0.004124 | 1 | 4 | 1 | H3BKU0 | Dysbindin | H3BKU0 [1 | 0 | 1243.649 | 0 | 0.81 | 0.000537 | 9.36 | 74.5 | 125.5 | 14.06 | 9.87 | | High | High | High | High | High | High | High | High | High | High | High | 0.002159 | 0.02238 | 1.8 |

| | | | | | | | | | | | | | | | | | | | | | | | | | | | | | | | | | | | | | | |
|---|---|---|---|---|---|---|---|---|---|---|---|---|---|---|---|---|---|---|---|---|---|---|---|---|---|---|---|---|---|---|---|---|---|---|---|---|---|---|
| FALSE | High | SPSYGVPS/ | [K].SPSYGV | 1xPhosphc | Q9CQS4 1x | 0.064323 | 0.004664 | 1 | 1 | 1 | Q9CQS4 | Solute carr | Q9CQS4 [6 | 0 | 4200.971 | 0 | 0.07 | 0.284605 | 13.59 | 96.8 | 103.2 | 12.12 | 14.26 | | High | High | High | High | High | High | High | High | High | High | High | 0.002403 | 0.02577 | 2.35 |
| FALSE | High | SPSYQIK | [K].SPSYQI | 1xPhosphc | B7ZNX2 1x | 0.023379 | 0.001578 | 1 | 2 | 1 | B7ZNX2 | Anaphase | B7ZNX2 [1 | 0 | 1360.728 | 0 | -0.53 | 0.00098 | 3.55 | 119.1 | 80.9 | 4.76 | 5.92 | | High | High | High | High | High | High | High | High | High | High | High | 0.000847 | 0.008152 | 1.24 |
| FALSE | High | SPSYTGSK | [R].SPSYTG | 1xPhosphc | Q2TGE7 1x | 0.009416 | 0.000736 | 1 | 3 | 1 | Q2TGE7 | Palmitoyltr | Q2TGE7 [6 | 0 | 1364.686 | 0 | 1.5 | 0.000154 | 9.06 | 50.2 | 149.8 | 9.6 | 3.78 | | High | High | High | High | High | High | High | High | High | High | High | 0.000327 | 0.002934 | 1.55 |
| FALSE | High | SPTAPSPQ/ | [K].SPTAPS | 1xPhosphc | B2RY19 1x | 2.85E-05 | 0 | 1 | 7 | 1 | B2RY19 | Cgnl1 prot | B2RY19 [2 | 0 | 2001.993 | 0 | 1.82 | 0.00215 | 6.63 | 43.3 | 156.7 | 13.4 | 18.25 | | High | High | High | High | High | High | High | High | High | High | High | 0 | 4.36E-06 | 1.94 |
| FALSE | High | SPTAQSPTS | [K].SPTAQS | 1xPhosphc | Q9D318 1x | 8.11E-05 | 0 | 1 | 4 | 2 | Q9D318 | Putative ur | Q9D318 [1 | 0 | 2484.191 | 0 | 0.31 | 0.001471 | 3.53 | 87.7 | 112.3 | 8.18 | 8.45 | | High | High | High | High | High | High | High | High | High | High | High | 0 | 7.06E-05 | 3.69 |
| FALSE | High | SPTDFLELE | [R].SPTDFL | 1xPhosphc | B7ZW89 1x | 0.045122 | 0.003373 | 1 | 3 | 1 | B7ZW89 | Uncharacte | B7ZW89 [9 | 0 | 1740.841 | 0 | -0.4 | 0.004683 | 0.8 | 114.1 | 85.9 | 6.84 | 8.01 | | High | High | High | High | High | High | High | High | High | High | High | 0.001727 | 0.0172 | 2 |
| FALSE | High | SPTDQPTV | [K].SPTDQI | 1xPhosphc | B7ZDF5 1x | 0.043607 | 0.003228 | 1 | 8 | 1 | B7ZDF5 | Histone de | B7ZDF5 [6 | 0 | 1609.86 | 0 | 0.77 | 0.033311 | 11.15 | 76.7 | 123.3 | 29.57 | 4.18 | | High | High | High | High | High | High | High | High | High | High | High | 0.001651 | 0.01656 | 1.72 |
| FALSE | High | SPTDSDVSI | [R].SPTDSD | 1xPhosphc | E9Q3E2 1x | 2.15E-05 | 0 | 2 | 4 | 2 | E9Q3E2; Q | Synaptopo | E9Q3E2 [1 | 0 | 2290.074 | 0 | 0.23 | 0.616432 | 9.3 | 101.4 | 98.6 | 12.84 | 12 | Shared | High | High | High | High | High | High | High | High | High | High | High | 0 | 3.21E-06 | 3.51 |
| FALSE | High | SPTDWALF | [K].SPTDW | 1xPhosphc | Q62418 1x | 0.000479 | 0 | 1 | 3 | 1 | Q62418 | Drebrin-lik | Q62418 [2 | 0 | 2395.08 | 0 | -0.6 | 0.751924 | 18.71 | 108.2 | 91.8 | 32.4 | 108.23 | | High | High | High | High | High | High | High | High | High | High | High | 0 | 0.000105 | 2.44 |
| FALSE | High | SPTEEKPAS | [K].SPTEEK | 1xPhosphc | G3X8X0 1x | 0.045898 | 0.003408 | 1 | 4 | 2 | G3X8X0 | DEAH (Asp | G3X8X0 [1 | 0 | 2328.08 | 0 | 0.04 | 0.831269 | 2.96 | 99.6 | 100.4 | 4.45 | 10.02 | | High | High | High | High | High | High | High | High | High | High | High | 0.004707 | 0.05073 | 2.55 |
| FALSE | High | SPTEEKPAS | [K].SPTEEK | 2xPhosphc | G3X8X0 2x | 0.029844 | 0.001857 | 1 | 4 | 1 | G3X8X0 | DEAH (Asp | G3X8X0 [1 | 0 | 2408.046 | 0 | -0.11 | 0.101785 | 4.27 | 104.1 | 95.9 | 6.43 | 7.19 | | High | High | High | High | High | High | High | High | High | High | High | 0.001011 | 0.01072 | 1.16 |
| FALSE | High | SPTFEEGS( | [R].SPTFEE | 1xPhosphc | Q62388 1x | 0.006713 | 0.00062 | 1 | 2 | 1 | Q62388 | Serine-pro | Q62388 [1 | 0 | 2523.227 | 0 | 0.64 | 0.012613 | 6.39 | 79.6 | 120.4 | 7.69 | 17.13 | | High | High | High | High | High | High | High | High | High | High | High | 0.000275 | 0.002009 | 2 |
| FALSE | High | SPSWISASI | [R].SPSWIS | 1xPhosphc | B2RRE0 1x | 6.3E-05 | 0 | 1 | 2 | 2 | B2RRE0 | A kinase (P | B2RRE0 [8 | 0 | 3690.801 | 0 | -1.12 | 0.007449 | 4.92 | 135.4 | 64.6 | 21 | 19.39 | | High | High | High | High | High | High | High | High | High | High | High | 0 | 1.06E-05 | 4.1 |
| FALSE | High | SPSSLSANV | [K].SPSSLS | 1xPhosphc | F8VPZ3 1x | 4.17E-06 | 0 | 1 | 3 | 5 | F8VPZ3 | Ubiquitin-s | F8VPZ3 [1 | 0 | 1899.983 | 0 | 0.29 | 0.023056 | 7.07 | 93.8 | 106.2 | 6.63 | 8.47 | | High | High | High | High | High | High | High | High | High | High | High | 0 | 7.28E-07 | 4.5 |
| FALSE | High | SPSSLLEDA | [K].SPSSLLI | 1xPhosphc | F8VQH0 1x | 5.75E-05 | 0 | 1 | 5 | 3 | F8VQH0 | Tyrosine-p | F8VQH0 [6 | 0 | 1584.829 | 0 | -0.68 | 0.000927 | 4.54 | 124.9 | 75.1 | 4.13 | 11.82 | | High | High | High | High | High | High | High | High | High | High | High | 0 | 9.69E-06 | 2.6 |
| FALSE | High | SPSSESSPQ | [R].SPSSES | 2xPhosphc | Q8CE90-7 | 0.01861 | 0.001362 | 1 | 6 | 3 | Q8CE90-7 | Isoform 7 c | Q8CE90-7 | 0 | 2334.048 | 0 | 0.54 | 0.118293 | 35.22 | 81.5 | 118.5 | 22.27 | 15.17 | | High | High | High | High | High | High | High | High | High | High | High | 0.000773 | 0.006714 | 2.53 |
| FALSE | High | SPSPQGTK | [R].SPSPQC | 1xPhosphc | E9PYL2 1x | 0.079114 | 0.00613 | 1 | 2 | 1 | E9PYL2 | Proline-ricl | E9PYL2 [9 | 0 | 1339.702 | 0 | -0.63 | 0.00091 | 2.91 | 121.6 | 78.4 | 2.21 | 8.2 | | High | High | High | High | High | High | High | High | High | High | High | 0.003148 | 0.03263 | 1.67 |
| FALSE | High | SPSPQLPLP | [R].SPSPQL | 1xPhosphc | P59178 1x | 4.43E-05 | 0 | 2 | 2 | 1 | P59178; Q | Lethal(3)m | P59178 [6 | 0 | 2044.15 | 0 | -0.87 | 0.000442 | 2.35 | 129.7 | 70.3 | 3.37 | 5.91 | Shared | High | High | High | High | High | High | High | High | High | High | High | 0 | 7.16E-06 | 2.33 |
| FALSE | High | SPSPSPTPE | [R].SPSPSP | 1xPhosphc | Q6PDG5 1x | 0.021418 | 0.001507 | 1 | 3 | 1 | Q6PDG5 | SWI/SNF c | Q6PDG5 [3 | 0 | 1635.84 | 0 | 1.31 | 0.00869 | 23.63 | 58.1 | 141.9 | 15.34 | 18.86 | | High | High | High | High | High | High | High | High | High | High | High | 0.000801 | 0.007364 | 1.67 |
| FALSE | High | SPSPSPTSP | [K].SPSPSP | 1xPhosphc | Q3UGP0 1x | 0.001968 | 5.66E-05 | 2 | 19 | 5 | Q3UGP0; C | Putative ur | Q3UGP0 [3 | 0 | 1578.773 | 0 | 0.55 | 0.003703 | 9.31 | 78.3 | 121.7 | 13.22 | 11.23 | Shared | High | High | High | High | High | High | High | High | High | High | High | 0.000275 | 0.001937 | 2.39 |
| FALSE | High | SPSPSPTSP | [K].SPSPSP | 2xPhosphc | Q3UGP0 2x | 0.026191 | 0.001676 | 2 | 19 | 1 | Q3UGP0; C | Putative ur | Q3UGP0 [3 | 0 | 1658.739 | 0 | -0.5 | 0.111818 | 10.94 | 113.6 | 86.4 | 14.23 | 16.13 | Shared | High | High | High | High | High | High | High | High | High | High | High | 0.000903 | 0.00927 | 2.6 |
| FALSE | High | SPSPTFSWI | [R].SPSPTF | 1xPhosphc | Q811Q9 1x | 0.000852 | 0 | 1 | 2 | 3 | Q811Q9 | Choline-ph | Q811Q9 [3 | 0 | 1898.982 | 0 | -0.57 | 0.001364 | 4.76 | 119.5 | 80.5 | 1.46 | 9.45 | | High | High | High | High | High | High | High | High | High | High | High | 0 | 0.0002 | 1.96 |
| FALSE | High | SPSPTLGES | [R].SPSPTL | 1xPhosphc | D3Z0X5 1x | 0.0095 | 0.000736 | 1 | 7 | 1 | D3Z0X5 | Pleckstrin I | D3Z0X5 [5 | 0 | 1620.82 | 0 | -0.81 | 0.00079 | 2.59 | 128.6 | 71.4 | 4.2 | 6.69 | | High | High | High | High | High | High | High | High | High | High | High | 0.000344 | 0.002971 | 1.94 |
| FALSE | High | SPSPVAPSF | [R].SPSPVA | 1xPhosphc | Q8BRM2 1 | 6.45E-07 | 0 | 1 | 2 | 3 | Q8BRM2 | RAB6-inter | Q8BRM2 [ | 0 | 2407.216 | 0 | 1.27 | 0.090787 | 7.36 | 59.8 | 140.2 | 16.27 | 44.41 | | High | High | High | High | High | High | High | High | High | High | High | 0 | 5.26E-05 | 4.42 |
| FALSE | High | SPSPVAPSF | [R].SPSPVA | 2xPhosphc | Q8BRM2 2 | 0.002232 | 0.000109 | 1 | 2 | 1 | Q8BRM2 | RAB6-inter | Q8BRM2 [ | 0 | 2487.182 | 0 | 2.12 | 0.024739 | 13.92 | 35.7 | 164.3 | 9.31 | 40.32 | | High | High | High | High | High | High | High | High | High | High | High | 4.94E-05 | 0.000584 | 3.4 |
| FALSE | High | SPSPVHLPE | [R].SPSPVH | 1xPhosphc | Q6A0A2 1x | 0.000589 | 0 | 1 | 3 | 4 | Q6A0A2 | La-related | Q6A0A2 [6 | 0 | 1840.961 | 0 | 0.54 | 0.0007 | 3.94 | 81.6 | 118.4 | 2.47 | 5.17 | | High | High | High | High | High | High | High | High | High | High | High | 0.000275 | 0.001678 | 3 |
| FALSE | High | SPSPVQR | [K].SPSPVC | 1xPhosphc | A0A0R4J0Y | 0.088701 | 0.006669 | 1 | 6 | 1 | A0A0R4J0Y | Probable h | A0A0R4J0Y | 0 | 1079.545 | 0 | 0.36 | 0.01265 | 12.04 | 84.7 | 115.3 | 5.15 | 8.15 | | High | High | High | High | High | High | High | High | High | High | High | 0.003432 | 0.03737 | 2.28 |
| FALSE | High | SPSPYYSR | [R].SPSPYY | 1xPhosphc | E9QP00 1x | 0.039011 | 0.002718 | 2 | 6 | 1 | E9QP00; P | Transforme | E9QP00 [2 | 0 | 1265.576 | 0 | -0.19 | 0.088063 | 10.33 | 107.8 | 92.2 | 5.46 | 19.96 | Shared | High | High | High | High | High | High | High | High | High | High | High | 0.001453 | 0.01458 | 1.95 |
| FALSE | High | SPSQCSPIC | [K].SPSQCS | 1xCarbami | Q3UB74 1x | 0.00195 | 5.66E-05 | 1 | 2 | 3 | Q3UB74 | Transformi | Q3UB74 [3 | 0 | 1601.672 | 0 | -1.04 | 0.000846 | 2.71 | 134.6 | 65.4 | 4.38 | 8.18 | | High | High | High | High | High | High | High | High | High | High | High | 0.000418 | 0.003595 | 2.64 |
| FALSE | High | SPSQEPSAF | [R].SPSQEF | 1xPhosphc | Q8CIJ3 1xP | 0.000916 | 0 | 1 | 2 | 2 | Q8CIJ3 | Eukaryotic | Q8CIJ3 [88 | 0 | 1393.656 | 0 | -0.23 | 0.062295 | 5.15 | 107.8 | 92.2 | 6.43 | 5.62 | | High | High | High | High | High | High | High | High | High | High | High | 0 | 0.000217 | 2.19 |
| FALSE | High | SPSQEPSAF | [R].SPSQEF | 1xPhosphc | Q8CIJ3 1xP | 3.38E-05 | 0 | 1 | 2 | 4 | Q8CIJ3 | Eukaryotic | Q8CIJ3 [88 | 0 | 1622.819 | 0 | -0.1 | 0.21827 | 6.44 | 105 | 95 | 2.81 | 9.31 | | High | High | High | High | High | High | High | High | High | High | High | 0 | 5.33E-06 | 2.96 |
| FALSE | High | SPSQESIGA | [R].SPSQES | 1xPhosphc | Q8VHK1 1x | 0.000496 | 0 | 1 | 1 | 1 | Q8VHK1 | Caskin-2 O | Q8VHK1 [6 | 0 | 1340.641 | 0 | 0.34 | 0.00187 | 5.1 | 88.1 | 111.9 | 2.57 | 4.9 | | High | High | High | High | High | High | High | High | High | High | High | 0 | 0.000109 | 1.6 |
| FALSE | High | SPSQGDED | [R].SPSQGI | 1xPhosphc | Q9Z2C9 1x | 1.37E-08 | 0 | 1 | 1 | 6 | Q9Z2C9 | Myotubula | Q9Z2C9 [5 | 0 | 2569.28 | 0 | -1.1 | 0.000346 | 1.12 | 136.8 | 63.2 | 3.26 | 12.95 | | High | High | High | High | High | High | High | High | High | High | High | 0 | 7.52E-07 | 4.22 |
| FALSE | High | SPSQGDED | [R].SPSQGI | 2xPhosphc | Q9Z2C9 2x | 0.000529 | 0 | 1 | 1 | 1 | Q9Z2C9 | Myotubula | Q9Z2C9 [5 | 0 | 2649.246 | 0 | -0.47 | 0.004899 | 7.79 | 115.6 | 84.4 | 11.13 | 12.81 | | High | High | High | High | High | High | High | High | High | High | High | 0 | 0.000116 | 2.94 |
| FALSE | High | SPSQGSYSI | [R].SPSQGS | 1xPhosphc | Q7TQG1 1x | 0.013909 | 0.001011 | 1 | 6 | 2 | Q7TQG1 | Pleckstrin I | Q7TQG1 [4 | 0 | 1277.572 | 0 | 1.85 | 0.000154 | 5.51 | 42.3 | 157.7 | 5.88 | 2.58 | | High | High | High | High | High | High | High | High | High | High | High | 0.001876 | 0.01879 | 1.19 |
| FALSE | High | SPSQNSQC | [R].SPSQNS | 1xCarbami | Q6PAJ1 1x | 0.003755 | 0.00021 | 1 | 2 | 4 | Q6PAJ1 | Breakpoint | Q6PAJ1 [3 | 0 | 2960.35 | 0 | 0.75 | 0.001236 | 1.24 | 75.6 | 124.4 | 13.06 | 6.4 | | High | High | High | High | High | High | High | High | High | High | High | 0.000801 | 0.007323 | 2.23 |
| FALSE | High | SPSQQEILL | [K].SPSQQI | 1xPhosphc | Q6WQJ1 1 | 0.014033 | 0.001011 | 1 | 2 | 1 | Q6WQJ1 | Sn1-specifi | Q6WQJ1 [ | 0 | 1479.777 | 0 | 1.52 | 0.000621 | 17.15 | 50.5 | 149.5 | 12.05 | 6.7 | | High | High | High | High | High | High | High | High | High | High | High | 0.000538 | 0.004602 | 2.15 |
| FALSE | High | SPSRLDTFL | [K].SPSRLD | 1xPhosphc | Q6P5D4 1x | 0.039347 | 0.002793 | 1 | 2 | 1 | Q6P5D4 | Centrosom | Q6P5D4 [4 | 1 | 1701.934 | 0 | -0.63 | 0.012441 | 5.69 | 122.8 | 77.2 | 15.18 | 3.83 | | High | High | High | High | High | High | High | High | High | High | High | 0.001468 | 0.01473 | 1.65 |
| FALSE | High | SPSRPLPEV | [R].SPSRPL | 1xPhosphc | P32067 1x | 0.000164 | 0 | 1 | 6 | 2 | P32067 | Lupus La p | P32067 [9 | 0 | 2156.104 | 0 | -0.53 | 0.001059 | 4.2 | 116.5 | 83.5 | 3.84 | 8.32 | | High | High | High | High | High | High | High | High | High | High | High | 0 | 3.12E-05 | 2.13 |
| FALSE | High | SPSRPVSD( | [K].SPSRPV | 2xCarbami | Q8C4C7 1x | 3.15E-09 | 0 | 1 | 2 | 2 | Q8C4C7 | Putative ur | Q8C4C7 [6 | 0 | 2685.185 | 0 | 2.25 | 0.000154 | 0.84 | 34.3 | 165.7 | 12.46 | 2.52 | | High | High | High | High | High | High | High | High | High | High | High | 0 | 4.84E-07 | 3.46 |
| FALSE | High | SPSSDSWT | [K].SPSSDS | 1xCarbami | V9GXM1 1 | 2.67E-08 | 0 | 1 | 7 | 5 | V9GXM1 | ADP-ribosy | V9GXM1 [ | 0 | 1993.816 | 0 | 1.33 | 0.00131 | 19.29 | 54.3 | 145.7 | 5.78 | 10.88 | | High | High | High | High | High | High | High | High | High | High | High | 0 | 1.74E-09 | 3.28 |
| FALSE | High | SPSSESEAE | [K].SPSSES | 1xPhosphc | A2AR02 1x | 5.97E-05 | 0 | 1 | 5 | 2 | A2AR02 | Peptidyl-pr | A2AR02 [2 | 0 | 3698.738 | 0 | 0.14 | 0.763371 | 19.84 | 93.5 | 106.5 | 19.64 | 8.48 | | High | High | High | High | High | High | High | High | High | High | High | 0 | 1E-05 | 3.49 |
| FALSE | High | SPSSESEAE | [K].SPSSES | 2xPhosphc | A2AR02 2x | 0.008927 | 0.000736 | 1 | 5 | 2 | A2AR02 | Peptidyl-pr | A2AR02 [2 | 0 | 3778.705 | 0 | -0.13 | 0.056375 | 3.06 | 104.3 | 95.7 | 6.81 | 9.41 | | High | High | High | High | High | High | High | High | High | High | High | 0.001889 | 0.01903 | 3.86 |
| FALSE | High | SPSSESEAE | [K].SPSSES | 3xPhosphc | A2AR02 3x | 7.47E-05 | 0 | 1 | 5 | 5 | A2AR02 | Peptidyl-pr | A2AR02 [2 | 0 | 3858.671 | 0 | -0.32 | 0.00637 | 4.95 | 112.5 | 87.5 | 4.88 | 10.2 | | High | High | High | High | High | High | High | High | High | High | High | 0 | 1.29E-05 | 5.36 |
| FALSE | High | SPSSESSPQ | [R].SPSSES | 1xPhosphc | Q8CE90-7 | 0.013909 | 0.001011 | 1 | 6 | 5 | Q8CE90-7 | Isoform 7 c | Q8CE90-7 | 0 | 2254.081 | 0 | 0.25 | 0.392729 | 24.18 | 91.3 | 108.7 | 20.73 | 19.73 | | High | High | High | High | High | High | High | High | High | High | High | 0.003937 | 0.04192 | 1.51 |
| FALSE | High | SPSAPVCK | [R].SPSAPV | 1xCarbami | Q3U3E2 1x | 0.096181 | 0.007535 | 1 | 1 | 1 | Q3U3E2 | Protein FAI | Q3U3E2 [1 | 0 | 1383.711 | 0 | -0.53 | 0.006058 | 8.2 | 126.5 | 73.5 | 7.76 | 16.93 | | High | High | High | High | High | High | High | High | High | High | High | 0.003823 | 0.04127 | 1.16 |
| FALSE | High | SPSANENSI | [R].SPSANE | 1xPhosphc | Q3USJ8-2 | 0.004411 | 0.000409 | 1 | 3 | 1 | Q3USJ8-2 | Isoform 2 c | Q3USJ8-2 [ | 0 | 2482.12 | 0 | 0.29 | 0.183329 | 17.52 | 89.9 | 110.1 | 12.27 | 15.94 | | High | High | High | High | High | High | High | High | High | High | High | 0.000183 | 0.001258 | 2.49 |
| FALSE | High | SPSAGIQR | [R].SPSAGI | 1xPhosphc | Q99M54 1 | 0.028082 | 0.001791 | 1 | 5 | 1 | Q99M54 | Cell divisio | Q99M54 [ | 0 | 1124.566 | 0 | -0.6 | 0.001363 | 4.31 | 122.4 | 77.6 | 4.1 | 6.63 | | High | High | High | High | High | High | High | High | High | High | High | 0.000968 | 0.01003 | 1.75 |
| FALSE | High | SPSADPAP( | [R].SPSADF | 1xCarbami | Q9CWP9 2 | 0.022378 | 0.001542 | 1 | 1 | 1 | Q9CWP9 | Proline-ricl | Q9CWP9 [ | 0 | 2767.26 | 0 | -1.06 | 0.00183 | 6.03 | 141.1 | 58.9 | 7.06 | 17.49 | | High | High | High | High | High | High | High | High | High | High | High | 0.000814 | 0.007709 | 3.31 |
| FALSE | High | SPLSGIPVR | [R].SPLSGI | 1xPhosphc | Q5SXY1 1x | 0.028327 | 0.001791 | 1 | 5 | 2 | Q5SXY1 | Cytospin-B | Q5SXY1 [8 | 0 | 1234.676 | 0 | 2.76 | 0.000181 | 5.14 | 25.8 | 174.2 | 15.52 | 6.18 | | High | High | High | High | High | High | High | High | High | High | High | 0.000968 | 0.01007 | 2.06 |
| FALSE | High | SPISNGTFG | [K].SPISNG | 1xPhosphc | Q80UP8 1x | 3.16E-07 | 0 | 1 | 1 | 3 | Q80UP8 | Sodium-de | Q80UP8 [3 | 0 | 1961.878 | 0 | 1.22 | 0.001836 | 12.63 | 59.9 | 140.1 | 5.59 | 13.69 | | High | High | High | High | High | High | High | High | High | High | High | 0 | 2.8E-08 | 4.48 |
| FALSE | High | SPISNGTFG | [K].SPISNG | 1xDeamida | Q80UP8 1x | 2.27E-05 | 0 | 1 | 1 | 3 | Q80UP8 | Sodium-de | Q80UP8 [3 | 0 | 1962.862 | 0 | 0.54 | 0.01711 | 2.02 | 80.7 | 119.3 | 8.64 | 12.74 | | High | High | High | High | High | High | High | High | High | High | High | 0 | 3.39E-06 | 2.5 |
| FALSE | High | SPISPELHS/ | [K].SPISPEI | 1xPhosphc | A2A5Y4 1x | 0.000112 | 0 | 1 | 3 | 3 | A2A5Y4 | KAT8 regul | A2A5Y4 [9 | 0 | 2081.099 | 0 | -0.79 | 0.002254 | 4.05 | 124.5 | 75.5 | 5.05 | 11.18 | | High | High | High | High | High | High | High | High | High | High | High | 0 | 2.04E-05 | 2.44 |
| FALSE | High | SPLSPFDTE | [R].SPLSPF | 1xPhosphc | Q4VAC9 1x | 2.91E-06 | 0 | 1 | 3 | 3 | Q4VAC9 | Pleckstrin I | Q4VAC9 [1 | 0 | 2317.074 | 0 | 0.81 | 0.005184 | 9.41 | 69.8 | 130.2 | 9.3 | 12.83 | | High | High | High | High | High | High | High | High | High | High | High | 0 | 3.37E-07 | 3.69 |
| FALSE | High | SPISSDPTA | [K].SPISSDI | 1xPhosphc | Q925J9 1x | 0.01186 | 0.000889 | 1 | 2 | 1 | Q925J9 | Mediator c | Q925J9 [1 | 0 | 2791.382 | 0 | 1.37 | 0.001513 | 12.07 | 56.9 | 143.1 | 12.99 | 7.96 | | High | High | High | High | High | High | High | High | High | High | High | 0.000449 | 0.003789 | 2.3 |
| FALSE | High | SPISSGSSG | [K].SPISSGS | 1xPhosphc | Q925J9 1x | 0.000165 | 0 | 1 | 2 | 3 | Q925J9 | Mediator c | Q925J9 [1 | 0 | 2565.19 | 0 | 1.36 | 0.162257 | 22.13 | 58.2 | 141.8 | 42.82 | 49.82 | | High | High | High | High | High | High | High | High | High | High | High | 0 | 3.16E-05 | 2.39 |
| FALSE | High | SPISVPGGS | [K].SPISVP( | 1xPhosphc | Q61191 1x | 5.42E-06 | 0 | 1 | 2 | 2 | Q61191 | Host cell fa | Q61191 [6 | 0 | 2135.188 | 0 | -0.66 | 0.005613 | 3.42 | 117.5 | 82.5 | 8.35 | 8.1 | | High | High | High | High | High | High | High | High | High | High | High | 0 | 6.78E-07 | 2.54 |
| FALSE | High | SPLTILQDD | [R].SPLTILC | 1xPhosphc | Q99M54 1 | 0.008847 | 0.000736 | 1 | 1 | 2 | Q99M54 | Cell divisio | Q99M54 [1 | 0 | 2250.158 | 0 | 0.46 | 0.003648 | 3.21 | 83.2 | 116.8 | 4.05 | 8.54 | | High | High | High | High | High | High | High | High | High | High | High | 0.000327 | 0.002742 | 3.17 |
| FALSE | High | SPLTILQDD | [R].SPLTILC | 2xPhosphc | Q99M54 2 | 0.002376 | 0.000109 | 1 | 1 | 2 | Q99M54 | Cell divisio | Q99M54 [1 | 0 | 2330.124 | 0 | -0.17 | 0.045047 | 6.5 | 106.6 | 93.4 | 7.94 | 2.91 | | High | High | High | High | High | High | High | High | High | High | High | 4.94E-05 | 0.000628 | 2.73 |
| FALSE | High | SPLTQEQLI | [R].SPLTQE | 1xPhosphc | Q3TYJ0 1x | 0.01029 | 0.000772 | 1 | 2 | 1 | Q3TYJ0 | Putative ur | Q3TYJ0 [2 | 0 | 2221.207 | 0 | -0.45 | 0.013466 | 7.96 | 115.1 | 84.9 | 4.19 | 12.38 | | High | High | High | High | High | High | High | High | High | High | High | 0.000382 | 0.003246 | 2.37 |
| FALSE | High | SPLVSYAAS | [K].SPLVSY | 1xPhosphc | D3YYY8 1x | 1.93E-05 | 0 | 1 | 1 | 1 | D3YYY8 | Rho guanir | D3YYY8 [1 | 0 | 1858.972 | 0 | 1.97 | 0.00047 | 6.44 | 38.8 | 161.2 | 14.48 | 8.36 | | High | High | High | High | High | High | High | High | High | High | High | 0 | 2.84E-06 | 3.18 |
| FALSE | High | SPLVSYAAS | [K].SPLVSY | 2xPhosphc | D3YYY8 2x | 0.000779 | 0 | 1 | 1 | 2 | D3YYY8 | Rho guanir | D3YYY8 [1 | 0 | 1938.938 | 0 | 2.68 | 0.00021 | 5.03 | 27.1 | 172.9 | 10 | 7.13 | | High | High | High | High | High | High | High | High | High | High | High | 0.000872 | 0.008752 | 4.46 |
| FALSE | High | SPLSAEELT | [R].SPLSAE | 1xPhosphc | D3Z3Q3 1x | 1.95E-06 | 0 | 1 | 7 | 1 | D3Z3Q3 | Smoothelir | D3Z3Q3 [5 | 0 | 2554.294 | 0 | -1.07 | 0.002556 | 4.33 | 138.5 | 61.5 | 11.83 | 8.08 | | High | High | High | High | High | High | High | High | High | High | High | 0 | 2.16E-07 | 3.37 |
| FALSE | High | SPIYPLSDSI | [R].SPIYPLS | 1xCarbami | Q924W7 1 | 6.22E-07 | 0 | 1 | 5 | 1 | Q924W7 | Suppressio | Q924W7 [4 | 0 | 1991.898 | 0 | -0.55 | 0.000647 | 5.13 | 117.8 | 82.2 | 3.5 | 11.2 | | High | High | High | High | High | High | High | High | High | High | High | 0 | 5.96E-08 | 4.19 |
| FALSE | High | SPMAESLD | [K].SPMAE | 1xCarbami | Q9EPQ8 1x | 8.2E-06 | 0 | 1 | 3 | 5 | Q9EPQ8 | Transcripti | Q9EPQ8 [5 | 0 | 2510.985 | 0 | -0.1 | 0.877824 | 27.71 | 101.1 | 98.9 | 11.45 | 22.12 | | High | High | High | High | High | High | High | High | High | High | High | 0 | 1.08E-06 | 4.08 |
| FALSE | High | SPNGISDYF | [K].SPNGIS | 1xDeamida | Q91ZX6 1x | 0.097743 | 0.007804 | 1 | 3 | 1 | Q91ZX6 | Sentrin-spe | Q91ZX6 [1 | 0 | 1616.797 | 0 | 1.09 | 0.000534 | 3.71 | 62.7 | 137.3 | 7.46 | 8.73 | | High | High | High | High | High | High | High | High | High | High | High | 0.003963 | 0.04219 | 2.04 |
| FALSE | High | SPNHGTVE | [K].SPNHG | 1xPhosphc | Q68FH0 1x | 0.001386 | 5.66E-05 | 1 | 8 | 1 | Q68FH0 | Plakophilin | Q68FH0 [4 | 0 | 2267.102 | 0 | -0.03 | 0.378474 | 16.74 | 100.7 | 99.3 | 9.92 | 2.28 | | High | High | High | High | High | High | High | High | High | High | High | 2.58E-05 | 0.000346 | 3.45 |
| FALSE | High | SPNKPTFAY | [R].SPNKPT | 1xPhosphc | Q8CEE0 1x | 0.001158 | 0 | 1 | 6 | 1 | Q8CEE0 | Centrosom | Q8CEE0 [5 | 0 | 2233.105 | 0 | -0.92 | 0.000626 | 0.81 | 133.4 | 66.6 | 7.11 | 7.67 | | High | High | High | High | High | High | High | High | High | High | High | 0 | 0.00028 | 1.67 |
| FALSE | High | SPNPSDGG | [R].SPNPSD | 1xCarbami | Q330P7 1x | 0.007471 | 0.00062 | 1 | 1 | 2 | Q330P7 | SCF apopto | Q330P7 [1 | 0 | 3396.317 | 0 | 0.13 | 0.245948 | 27.76 | 87.5 | 112.5 | 13.12 | 9.53 | | High | High | High | High | High | High | High | High | High | High | High | 0.000275 | 0.002263 | 3.32 |
| FALSE | High | SPPAAAAY( | [K].SPPAAA | 1xPhosphc | Q3U1I6 1x | 0.002463 | 0.000109 | 1 | 4 | 2 | Q3U1I6 | Putative ur | Q3U1I6 [8 | 0 | 3894.004 | 0 | 0.38 | 0.230615 | 14.99 | 90.6 | 109.4 | 11.45 | 13.15 | | High | High | High | High | High | High | High | High | High | High | High | 4.94E-05 | 0.000657 | 3.93 |
| FALSE | High | SPPACSSSS | [R].SPPACS | 1xCarbami | A0A0R4J0I | 1.93E-05 | 0 | 1 | 2 | 2 | A0A0R4J0I | ETS domain | A0A0R4J0I | 0 | 2206.041 | 0 | -0.59 | 0.006252 | 5.07 | 120.3 | 79.7 | 15.2 | 4.79 | | High | High | High | High | High | High | High | High | High | High | High | 0 | 2.19E-05 | 2.43 |
| FALSE | High | SPPATGSAY | [K].SPPATC | 1xPhosphc | Q569Z6 1x | 1.56E-07 | 0 | 1 | 4 | 1 | Q569Z6 | Thyroid ho | Q569Z6 [3 | 0 | 1646.762 | 0 | -1 | 0.353176 | 9.12 | 109 | 91 | 42.91 | 43.41 | | High | High | High | High | High | High | High | High | High | High | High | 0 | 1.27E-08 | 4.3 |
| FALSE | High | SPPATGSAY | [K].SPPATC | 1xPhosphc | Q569Z6 1x | 6.97E-06 | 0 | 1 | 4 | 6 | Q569Z6 | Thyroid ho | Q569Z6 [3 | 0 | 1875.925 | 0 | -0.1 | 0.064668 | 6.32 | 103.5 | 96.5 | 4.49 | 8.61 | | High | High | High | High | High | High | High | High | High | High | High | 0 | 9.05E-07 | 4.29 |
| FALSE | High | SPPATGSAY | [K].SPPATC | 2xPhosphc | Q569Z6 2x | 0.013545 | 0.000976 | 1 | 4 | 3 | Q569Z6 | Thyroid ho | Q569Z6 [3 | 0 | 1955.892 | 0 | -0.61 | 0.000391 | 0.86 | 119.3 | 80.7 | 5.58 | 4.53 | | High | High | High | High | High | High | High | High | High | High | High | 0.000505 | 0.004412 | 2.18 |
| FALSE | High | SPPAVPSPF | [R].SPPAVF | 1xPhosphc | A0A0R4J0Y | 0.002508 | 0.000109 | 1 | 4 | 1 | A0A0R4J0Y | Probable h | A0A0R4J0Y | 0 | 3025.353 | 0 | -0.68 | 0.155113 | 12.33 | 121.3 | 78.7 | 13.34 | 27.99 | | High | High | High | High | High | High | High | High | High | High | High | 4.94E-05 | 0.00067 | 2.23 |
| FALSE | High | SPPCEDFS\ | [K].SPPCED | 1xCarbami | A2ARP8 1x | 0.006363 | 0.00062 | 1 | 3 | 1 | A2ARP8 | Microtubu | A2ARP8 [1 | 0 | 2206.998 | 0 | -1.47 | 0.002267 | 3.29 | 150.8 | 49.2 | 11.88 | 15.63 | | High | High | High | High | High | High | High | High | High | High | High | 0.000275 | 0.001888 | 2.47 |
| FALSE | High | SPPDQSAV | [K].SPPDQS | 1xPhosphc | Q8K232 1x | 1.64E-05 | 0 | 2 | 8 | 4 | Q8K232; E | Adducin 1 | Q8K232 [6 | 0 | 2357.215 | 0 | 1 | 0.000554 | 6.63 | 66.2 | 133.8 | 4.01 | 6.09 | Shared | High | High | High | High | High | High | High | High | High | High | High | 0 | 2.35E-06 | 3.11 |
| FALSE | High | SPLYVQSM | [K].SPLYVC | 1xPhosphc | Q69Z89 1x | 1.86E-05 | 0 | 1 | 5 | 1 | Q69Z89 | Ras-associa | Q69Z89 [5 | 0 | 2566.24 | 0 | 0.37 | 0.130972 | 43.29 | 82.6 | 117.4 | 19.17 | 18.63 | | High | High | High | High | High | High | High | High | High | High | High | 0 | 2.71E-06 | 2.63 |
| FALSE | High | SPLQPCENI | [R].SPLQPC | 2xCarbami | Q8CJ27 1x | 7.36E-06 | 0 | 1 | 2 | 3 | Q8CJ27 | Abnormal s | Q8CJ27 [1 | 0 | 2933.313 | 0 | 1.18 | 0.001525 | 2.17 | 60.4 | 139.6 | 8.75 | 9.86 | | High | High | High | High | High | High | High | High | High | High | High | 0 | 1.64E-06 | 3.69 |
| FALSE | High | SPLQLTTDI | [R].SPLQLT | 1xPhosphc | Q99LE1 1x | 7.61E-05 | 0 | 1 | 2 | 1 | Q99LE1 | RILP-like pr | Q99LE1 [2 | 0 | 2450.205 | 0 | 1.15 | 0.005604 | 30.36 | 60.6 | 139.4 | 33.14 | 9.07 | | High | High | High | High | High | High | High | High | High | High | High | 0 | 1.32E-05 | 3.5 |
| FALSE | High | SPLQGADT | [R].SPLQG/ | 1xCarbami | Q3URQ7 1 | 0.000786 | 0 | 1 | 2 | 1 | Q3URQ7 | Methenylt | Q3URQ7 [2 | 0 | 2826.406 | 0 | -0.45 | 0.006223 | 8.33 | 117.4 | 82.6 | 15.77 | 21.94 | | High | High | High | High | High | High | High | High | High | High | High | 0 | 0.000182 | 2.51 |
| FALSE | High | SPGTPAGE | [R].SPGTP/ | 1xPhosphc | Q8K232 1x | 7.43E-06 | 0 | 2 | 7 | 4 | Q8K232; E | Adducin 1 | Q8K232 [3 | 0 | 1863.925 | 0 | 1.14 | 0.000625 | 5.97 | 61.4 | 138.6 | 1.55 | 7.37 | Shared | High | High | High | High | High | High | High | High | High | High | High | 0 | 1.46E-06 | 4.11 |
| FALSE | High | SPGTPAGE | [R].SPGTP/ | 2xPhosphc | Q8K232 2x | 3.49E-07 | 0 | 2 | 7 | 1 | Q8K232; E | Adducin 1 | Q8K232 [3 | 0 | 1943.892 | 0 | 1.07 | 0.000841 | 10.66 | 65.9 | 134.1 | 6.2 | 10.04 | Shared | High | High | High | High | High | High | High | High | High | High | High | 0 | 3.11E-08 | 4.83 |
| FALSE | High | SPGVAVSA | [R].SPGVA\ | 1xPhosphc | Q8CHI8 1x | 4.13E-06 | 0 | 1 | 5 | 2 | Q8CHI8 | E1A-bindin | Q8CHI8 [7 | 0 | 3573.815 | 0 | -0.63 | 0.026118 | 4.32 | 121.3 | 78.7 | 8.37 | 11.32 | | High | High | High | High | High | High | High | High | High | High | High | 0 | 4.99E-07 | 2.91 |
| FALSE | High | SPGVQPQL | [R].SPGVQ | 1xPhosphc | E9QAP7 1x | 0.012397 | 0.000889 | 1 | 2 | 1 | E9QAP7 | TATA-box- | E9QAP7 [5 | 0 | 3383.752 | 0 | 0.01 | 0.901411 | 15.58 | 100.5 | 99.5 | 10.28 | 8.97 | | High | High | High | High | High | High | High | High | High | High | High | 0.000449 | 0.003993 | 3.21 |
| FALSE | High | SPGVVSSPI | [R].SPGVVS | 1xPhosphc | A3KGF9 1x | 0.000189 | 0 | 1 | 1 | 4 | A3KGF9 | Uncharacte | A3KGF9 [2 | 0 | 3566.761 | 0 | 6.64 | | 0 | 9.8 | 190.2 | | 18.97 | | Not Found | Not Found | High | High | High | High | High | High | High | High | High | 0 | 0.000324 | 4 |
| FALSE | High | SPHEAGFC | [K].SPHEA( | 1xCarbami | Q8R4X3 1x | 0.081085 | 0.006219 | 1 | 4 | 2 | Q8R4X3 | RNA-bindir | Q8R4X3 [4 | 0 | 1716.802 | 0 | -0.17 | 0.294018 | 3.04 | 105.5 | 94.5 | 4.5 | 7.01 | | High | High | High | High | High | High | High | High | High | High | High | 0.0032 | 0.03376 | 2.56 |
| FALSE | High | SPHNSGLV | [K].SPHNS( | 1xPhosphc | A2SW42 1 | 0.004738 | 0.000456 | 1 | 2 | 2 | A2SW42 | Zinc finger | A2SW42 [3 | 0 | 1732.858 | 0 | -0.6 | 0.00078 | 2.74 | 120.5 | 79.5 | 6.38 | 9.65 | | High | High | High | High | High | High | High | High | High | High | High | 0.000204 | 0.00136 | 2.12 |
| FALSE | High | SPHVQTM( | [R].SPHVQ | 1xCarbami | Q9QXS1 1x | 0.000225 | 0 | 1 | 4 | 1 | Q9QXS1 | Plectin OS= | Q9QXS1 [1 | 0 | 2042.975 | 0 | 2.16 | 0.000716 | 13.33 | 36.5 | 163.5 | 10.03 | 12.39 | | High | High | High | High | High | High | High | High | High | High | High | 0 | 4.46E-05 | 3.29 |
| FALSE | High | SPHVQTM( | [R].SPHVQ | 1xCarbami | Q9QXS1 1x | 4.95E-06 | 0 | 1 | 4 | 11 | Q9QXS1 | Plectin OS= | Q9QXS1 [1 | 0 | 2058.97 | 0 | 2.27 | 0.00034 | 16.55 | 34.4 | 165.6 | 10.73 | 7.37 | | High | High | High | High | High | High | High | High | High | High | High | 0 | 6.1E-07 | 3.99 |
| FALSE | High | SPHVQTM( | [R].SPHVQ | 1xCarbami | Q9QXS1 1x | 0.000123 | 0 | 1 | 4 | 1 | Q9QXS1 | Plectin OS= | Q9QXS1 [1 | 0 | 2272.138 | 0 | 2.17 | 0.000181 | 3.38 | 36.3 | 163.7 | 13.39 | 7.12 | | High | High | High | High | High | High | High | High | High | High | High | 0 | 2.26E-05 | 2.78 |
| FALSE | High | SPHVQTM( | [R].SPHVQ | 1xCarbami | Q9QXS1 1x | 0.000233 | 0 | 1 | 4 | 10 | Q9QXS1 | Plectin OS= | Q9QXS1 [1 | 0 | 2288.133 | 0 | 1.65 | 0.000727 | 8.65 | 48.3 | 151.7 | 9.68 | 10.08 | | High | High | High | High | High | High | High | High | High | High | High | 0.000399 | 0.003368 | 3.67 |
| FALSE | High | SPHVTAPG | [K].SPHVT/ | 1xPhosphc | Q9ERH4 1x | 0.000993 | 0 | 1 | 2 | 5 | Q9ERH4 | Nucleolar a | Q9ERH4 [2 | 0 | 1676.877 | 0 | -0.46 | 0.103628 | 15.24 | 116.4 | 83.6 | 16.94 | 17.51 | | High | High | High | High | High | High | High | High | High | High | High | 0.000449 | 0.00374 | 2.81 |
| FALSE | High | SPIALPVK | [R].SPIALP\ | 1xPhosphc | A2AJT5 1x | 0.043236 | 0.003228 | 1 | 5 | 2 | A2AJT5 | Arginine/se | A2AJT5 [2 | 0 | 1133.653 | 0 | -0.82 | 0.00035 | 1.55 | 129.1 | 70.9 | 3.7 | 2.7 | | High | High | High | High | High | High | High | High | High | High | High | 0.001651 | 0.01624 | 2.4 |
| FALSE | High | SPIAVPETV | [R].SPIAVP | 1xPhosphc | Q6NZN1-2 | 1.2E-06 | 0 | 1 | 4 | 1 | Q6NZN1-2 | Isoform 2 c | Q6NZN1-2 | 0 | 3072.591 | 0 | -0.16 | 0.373712 | 17.95 | 99.8 | 100.2 | 15.43 | 9.97 | | High | High | High | High | High | High | High | High | High | High | High | 0 | 1.26E-07 | 4.03 |
| FALSE | High | SPLCLLSSSI | [R].SPLCLL | 1xCarbami | Q8CJF7 1x | 4.98E-05 | 0 | 1 | 1 | 1 | Q8CJF7 | Protein ELY | Q8CJF7 [1 | 0 | 2402.281 | 0 | 0.18 | 0.238363 | 13.26 | 96.3 | 103.7 | 9.93 | 7.43 | | High | High | High | High | High | High | High | High | High | High | High | 0 | 8.23E-06 | 2.96 |
| FALSE | High | SPLDFTTVT | [K].SPLDFT | 1xPhosphc | Q14B71 1x | 0.017966 | 0.001282 | 1 | 4 | 1 | Q14B71 | Cell divisio | Q14B71 [5 | 0 | 2946.552 | 0 | -1.36 | 0.002315 | 4.46 | 137.3 | 62.7 | 20.15 | 31.88 | | High | High | High | High | High | High | High | High | High | High | High | 0.000689 | 0.006028 | 2.17 |
| FALSE | High | SPLDPDSGI | [K].SPLDPD | 1xCarbami | Q9EP97 1x | 3.14E-08 | 0 | 1 | 1 | 3 | Q9EP97 | Sentrin-spe | Q9EP97 [2 | 0 | 3138.465 | 0 | -1.24 | 0.001229 | 2.09 | 140.6 | 59.4 | 4.48 | 13.18 | | High | High | High | High | High | High | High | High | High | High | High | 0 | 2.07E-09 | 4.84 |
| FALSE | High | SPLDPDSGI | [K].SPLDPD | 1xCarbami | Q9EP97 1x | 1.85E-08 | 0 | 1 | 1 | 8 | Q9EP97 | Sentrin-spe | Q9EP97 [2 | 0 | 3139.449 | 0 | -0.99 | 0.000518 | 1.02 | 136.1 | 63.9 | 5.09 | 6.37 | | High | High | High | High | High | High | High | High | High | High | High | 0 | 1.21E-09 | 3.71 |
| FALSE | High | SPLEPENVI | [R].SPLEPE | 1xPhosphc | A2A8R7 1x | 0.003282 | 0.00016 | 1 | 5 | 1 | A2A8R7 | Terminal u | A2A8R7 [1 | 0 | 1844.997 | 0 | -0.31 | 0.035543 | 10.67 | 110.7 | 89.3 | 9.31 | 6.18 | | High | High | High | High | High | High | High | High | High | High | High | 7.19E-05 | 0.000904 | 2.27 |
| FALSE | High | SPLETSVQ/ | [R].SPLETS | 1xPhosphc | Q542F3 1x | 0.000331 | 0 | 1 | 3 | 2 | Q542F3 | Putative ur | Q542F3 [4 | 0 | 2238.122 | 0 | 1.28 | 0.001453 | 18.21 | 58.9 | 141.1 | 2.95 | 11.03 | | High | High | High | High | High | High | High | High | High | High | High | 0 | 6.87E-05 | 3.41 |
| FALSE | High | SPLGGGGC | [K].SPLGG( | 1xCarbami | Q6PDM1 1 | 9.71E-05 | 0 | 1 | 3 | 1 | Q6PDM1 | Male-speci | Q6PDM1 [ | 0 | 2200.083 | 0 | 0.35 | 0.087732 | 22 | 84 | 116 | 6.81 | 17.61 | | High | High | High | High | High | High | High | High | High | High | High | 0 | 1.74E-05 | 2.99 |
| FALSE | High | SPLGLYCEC | [R].SPLGLY | 2xCarbami | Q05BA5 1x | 6.65E-05 | 0 | 1 | 7 | 2 | Q05BA5 | Nuclear rec | Q05BA5 [5 | 0 | 2820.253 | 0 | -0.13 | 0.143251 | 12.53 | 111.2 | 88.8 | 9.54 | 14.46 | | High | High | High | High | High | High | High | High | High | High | High | 0 | 1.14E-05 | 3.66 |
| FALSE | High | SPIIADLSLS | [K].SPIIADL | 1xPhosphc | E9Q8L9 1x | 9.2E-05 | 0 | 1 | 2 | 1 | E9Q8L9 | Rab11 fam | E9Q8L9 [8 | 1 | 3818.935 | 0 | -0.89 | 0.002982 | 5.37 | 130.1 | 69.9 | 10 | 13.22 | | High | High | High | High | High | High | High | High | High | High | High | 0 | 1.64E-05 | 4.19 |
| FALSE | High | SPLLAGGSI | [R].SPLLAG | 1xPhosphc | Q8R4R6 1x | 2.84E-07 | 0 | 1 | 2 | 1 | Q8R4R6 | Nucleopor | Q8R4R6 [6 | 0 | 2290.272 | 0 | 0.3 | 0.027245 | 15.28 | 89.8 | 110.2 | 4.14 | 8.27 | | High | High | High | High | High | High | High | High | High | High | High | 0 | 2.48E-08 | 3.68 |
| FALSE | High | SPLLLSDSC | [K].SPLLLSI | 1xCarbami | Q3UG20 1 | 1.95E-05 | 0 | 1 | 3 | 1 | Q3UG20 | Histone-lys | Q3UG20 [8 | 0 | 2595.376 | 0 | 1.56 | 0.001416 | 9.32 | 50.6 | 149.4 | 13.21 | 14.38 | | High | High | High | High | High | High | High | High | High | High | High | 0 | 2.87E-06 | 2.67 |
| FALSE | High | SPLLNQPVI | [K].SPLLNC | 1xPhosphc | Q91YD3 1x | 0.000479 | 0 | 1 | 2 | 3 | Q91YD3 | mRNA-dec | Q91YD3 [3 | 0 | 2899.492 | 0 | -0.06 | 0.7542 | 9.44 | 101.9 | 98.1 | 7.92 | 9.11 | | High | High | High | High | High | High | High | High | High | High | High | 0 | 0.000105 | 3.26 |
| FALSE | High | SPLLSASHS | [R].SPLLSA | 1xPhosphc | Q9CZX7 1x | 7.01E-08 | 0 | 1 | 1 | 6 | Q9CZX7 | Type 2 pho | Q9CZX7 [1 | 0 | 3002.482 | 0 | 1.87 | 0.002351 | 26.1 | 42.9 | 157.1 | 12.5 | 13.97 | | High | High | High | High | High | High | High | High | High | High | High | 0 | 5.15E-09 | 4.7 |
| FALSE | High | SPLLSASHS | [R].SPLLSA | 2xPhosphc | Q9CZX7 2x | 0.096959 | 0.007777 | 1 | 1 | 1 | Q9CZX7 | Type 2 pho | Q9CZX7 [1 | 0 | 3082.449 | 0 | 2.5 | 0.001133 | 28.57 | 27.8 | 172.2 | 22.51 | 14.15 | | High | High | High | High | High | High | High | High | High | High | High | 0.003922 | 0.04152 | 2.94 |
| FALSE | High | SPLPPILGSI | [R].SPLPPIL | 1xPhosphc | Q3UFT4 1x | 0.000573 | 0 | 1 | 7 | 1 | Q3UFT4 | Putative ur | Q3UFT4 [6 | 0 | 1745.001 | 0 | -1.6 | 0.000496 | 1.63 | 150.4 | 49.6 | 1.38 | 18.5 | | High | High | High | High | High | High | High | High | High | High | High | 0 | 0.000127 | 2.52 |

| | | | | | | | | | | | | | | | | | | | | | | | | | | | | | | | | | | | | | | |
|---|---|---|---|---|---|---|---|---|---|---|---|---|---|---|---|---|---|---|---|---|---|---|---|---|---|---|---|---|---|---|---|---|---|---|---|---|---|---|
| FALSE | High | SPPDQSAV | [K].SPPDQ! | 2xPhospho | Q8K232 2x | 2.1E-06 | 0 | 2 | 8 | 5 | Q8K232; E! | Adducin 1 | Q8K232 [6 | 0 | 2437.182 | 0 | 1.63 | 0.000181 | 8.77 | 49.1 | 150.9 | 10.54 | 2.6 | Shared | High | High | High | High | High | High | High | High | High | High | High | 0 | 2.17E-06 | 2.99 |
| FALSE | High | SPTGASDH | [K].SPTGA! | 1xPhospho | Q3UHR0 1x | 0.083783 | 0.006345 | 1 | 2 | 1 | Q3UHR0 | BAH and cc | Q3UHR0 [2 | 0 | 1553.731 | 0 | 0.89 | 0.006779 | 16.41 | 74.8 | 125.2 | 16.67 | 6.92 | | High | High | High | High | High | High | High | High | High | High | High | 0.003265 | 0.03497 | 1.71 |
| FALSE | High | SPPDRDAT | [R].SPPDRD | 1xPhospho | Q8BIA4 1xl | 0.003167 | 0.00016 | 1 | 2 | 1 | Q8BIA4 | F-box/WD | Q8BIA4 [8 | 1 | 2457.222 | 0 | -0.62 | 0.009302 | 10.08 | 118.8 | 81.2 | 9.87 | 20.4 | | High | High | High | High | High | High | High | High | High | High | High | 7.19E-05 | 0.000866 | 1.84 |
| FALSE | High | SPPEHYVQ | [K].SPPEHY | 1xPhospho | B2RUG9 1x | 0.00043 | 0 | 1 | 6 | 2 | B2RUG9 | Adenomat | B2RUG9 [1 | 0 | 2195.074 | 0 | -0.41 | 0.0581 | 8.64 | 114.2 | 85.8 | 5.45 | 16.37 | | High | High | High | High | High | High | High | High | High | High | High | 0 | 9.2E-05 | 3.25 |
| FALSE | High | SPPYTAFLG | [K].SPPYTA | 1xDeamida | Q3TSY9 1x | 0.013545 | 0.000976 | 1 | 7 | 2 | Q3TSY9 | Putative ur | Q3TSY9 [9 | 0 | 2866.42 | 0 | -0.02 | 0.924574 | 61.88 | 97.2 | 102.8 | 44.37 | 12.26 | | High | High | High | High | High | High | High | High | High | High | High | 0.000505 | 0.004398 | 2.27 |
| FALSE | High | SPQASGLLI | [K].SPQAS( | 1xPhospho | Q8BZN4 1x | 0.003892 | 0.000261 | 1 | 2 | 1 | Q8BZN4 | NUAK fami | Q8BZN4 [5 | 0 | 1487.793 | 0 | -0.54 | 0.008493 | 6.66 | 115.1 | 84.9 | 10.25 | 8.29 | | High | High | High | High | High | High | High | High | High | High | High | 0.00014 | 0.001091 | 2.19 |
| FALSE | High | SPQELDQG | [R].SPQELD | 1xCarbami | Q921Q7 1x | 7.16E-06 | 0 | 1 | 1 | 1 | Q921Q7 | Ras and Ra | Q921Q7 [1 | 0 | 3286.569 | 0 | 2.24 | 0.000408 | 18.5 | 36.2 | 163.8 | 23.21 | 4.16 | | High | High | High | High | High | High | High | High | High | High | High | 0 | 9.29E-07 | 5.21 |
| FALSE | High | SPQELDQG | [R].SPQELD | 1xCarbami | Q921Q7 1x | 0.007811 | 0.000664 | 1 | 1 | 1 | Q921Q7 | Ras and Ra | Q921Q7 [1 | 0 | 3515.732 | 0 | 1.97 | 0.001044 | 29.49 | 40.7 | 159.3 | 25.22 | 8.5 | | High | High | High | High | High | High | High | High | High | High | High | 0.000327 | 0.002382 | 2.98 |
| FALSE | High | SPQESTGD | [K].SPQEST | 1xPhospho | V9GX74 1x | 3.26E-08 | 0 | 1 | 10 | 1 | V9GX74 | Zinc finger | V9GX74 [1 | 0 | 2374.081 | 0 | -0.36 | 0.025251 | 5.38 | 112 | 88 | 4.91 | 8.93 | | High | High | High | High | High | High | High | High | High | High | High | 0 | 2.17E-09 | 3.12 |
| FALSE | High | SPQLGVQA | [K].SPQLG\ | 1xPhospho | Q6P1H6 1x | 0.021046 | 0.001507 | 1 | 3 | 1 | Q6P1H6 | Ankyrin re | Q6P1H6 [8 | 0 | 2836.511 | 0 | 0.9 | 0.000964 | 5.08 | 68.5 | 131.5 | 12.79 | 10.66 | | High | High | High | High | High | High | High | High | High | High | High | 0.000801 | 0.007201 | 2.34 |
| FALSE | High | NHLSPQEG | [K].NHLSP( | 2xCarbami | Q9EP97 3x | 2.36E-09 | 0 | 1 | 1 | 2 | Q9EP97 | Sentrin-spe | Q9EP97 [1 | 0 | 2661.059 | 0 | 0.38 | 0.009607 | 14.23 | 85 | 115 | 12.27 | 8.09 | | High | High | High | High | High | High | High | High | High | High | High | 0 | 1.48E-08 | 4.46 |
| FALSE | High | SPQLTTPG( | [K].SPQLTT | 1xCarbami | Q5XJV7-2 | 0.005 | 0.000504 | 1 | 3 | 4 | Q5XJV7-2 | Isoform 2 c | Q5XJV7-2 [ | 0 | 2333.043 | 0 | -0.49 | 0.005584 | 4.02 | 115.5 | 84.5 | 3.46 | 11.33 | | High | High | High | High | High | High | High | High | High | High | High | 0.001076 | 0.01142 | 2.76 |
| FALSE | High | SPQMVSAI | [R].SPQMV | 1xPhospho | Q02248 1x | 0.121159 | 0.009984 | 1 | 4 | 1 | Q02248 | Catenin be | Q02248 [1 | 0 | 1412.717 | 0 | 0.35 | 0.007955 | 6.08 | 91.4 | 108.6 | 6.57 | 7.72 | | High | High | High | High | High | High | High | High | High | High | High | 0.004961 | 0.05405 | 1.44 |
| FALSE | High | SPQPPAEEI | [R].SPQPP/ | 1xCarbami | Q8VDM6 1 | 0.001373 | 5.66E-05 | 1 | 3 | 2 | Q8VDM6 | Heterogen | Q8VDM6 [ | 0 | 3805.63 | 0 | 0.17 | 0.579589 | 9.15 | 94.4 | 105.6 | 21.45 | 12.46 | | High | High | High | High | High | High | High | High | High | High | High | 2.58E-05 | 0.000341 | 3.16 |
| FALSE | High | SPQPPAEEI | [R].SPQPP/ | 1xCarbami | Q8VDM6 1 | 1.33E-05 | 0 | 1 | 3 | 1 | Q8VDM6 | Heterogen | Q8VDM6 [ | 0 | 4034.793 | 0 | 0.02 | 0.351833 | 19.37 | 85.4 | 114.6 | 31.14 | 12.7 | | High | High | High | High | High | High | High | High | High | High | High | 0 | 1.87E-06 | 3.58 |
| FALSE | High | SPQPPVEEI | [K].SPQPP\ | 2xCarbami | Q3TXW2 1 | 0.000331 | 0 | 1 | 10 | 3 | Q3TXW2 | Putative ur | Q3TXW2 [2 | 0 | 3944.687 | 0 | -0.63 | 0.478526 | 21.73 | 105.7 | 94.3 | 36.63 | 38.32 | | High | High | High | High | High | High | High | High | High | High | High | 0 | 6.91E-05 | 3.96 |
| FALSE | High | SPQPPVEEI | [K].SPQPP\ | 2xCarbami | Q3TXW2 1 | 1.12E-05 | 0 | 1 | 10 | 77 | Q3TXW2 | Putative ur | Q3TXW2 [2 | 0 | 4173.85 | 0 | -0.14 | 0.054847 | 6.11 | 104.7 | 95.3 | 4.58 | 8.29 | | High | High | High | High | High | High | High | High | High | High | High | 0 | 0.000188 | 4.37 |
| FALSE | High | SPPYTAFLG | [K].SPPYTA | 1xPhospho | Q3TSY9 1x | 5.72E-06 | 0 | 1 | 7 | 3 | Q3TSY9 | Putative ur | Q3TSY9 [9 | 0 | 2865.436 | 0 | -0.03 | 0.368956 | 8.57 | 100.3 | 99.7 | 5.26 | 11.59 | | High | High | High | High | High | High | High | High | High | High | High | 0 | 4.47E-06 | 4.3 |
| FALSE | High | SPQPPVEEI | [K].SPQPP\ | 2xCarbami | Q3TXW2 1 | 0.004294 | 0.00036 | 1 | 10 | 9 | Q3TXW2 | Putative ur | Q3TXW2 [2 | 0 | 4174.834 | 0 | -0.12 | 0.091999 | 3.35 | 103.6 | 96.4 | 5.19 | 6.22 | | High | High | High | High | High | High | High | High | High | High | High | 0.000786 | 0.007031 | 4.79 |
| FALSE | High | SPQQPVCV | [R].SPQQP' | 1xCarbami | Q3UFY0 1x | 0.000751 | 0 | 1 | 2 | 1 | Q3UFY0 | Ribosomal | Q3UFY0 [6 | 0 | 1537.728 | 0 | 0.04 | 0.334688 | 1.76 | 99.8 | 100.2 | 4.37 | 5.8 | | High | High | High | High | High | High | High | High | High | High | High | 0 | 0.000173 | 2.8 |
| FALSE | High | SPQRPFPM | [R].SPQRPI | 1xPhospho | V9GXX7 1x | 0.008313 | 0.000736 | 1 | 3 | 1 | V9GXX7 | DNA repair | V9GXX7 [3 | 0 | 3143.444 | 0 | -0.18 | 0.547578 | 21.62 | 104 | 96 | 5.65 | 19.3 | | High | High | High | High | High | High | High | High | High | High | High | 0.000327 | 0.002551 | 2.63 |
| FALSE | High | SPQRPFPM | [R].SPQRPI | 1xPhospho | V9GXX7 1x | 0.030898 | 0.001896 | 1 | 3 | 1 | V9GXX7 | DNA repair | V9GXX7 [3 | 0 | 3356.612 | 0 | -0.69 | 0.000302 | 3.17 | 126.3 | 73.7 | 9.61 | 14.97 | | High | High | High | High | High | High | High | High | High | High | High | 0.001027 | 0.01115 | 2.13 |
| FALSE | High | SPQRPFPM | [R].SPQRPI | 1xPhospho | V9GXX7 1x | 0.001087 | 0 | 1 | 3 | 2 | V9GXX7 | DNA repair | V9GXX7 [3 | 0 | 3372.607 | 0 | -0.49 | 0.003857 | 0.3 | 117.2 | 82.8 | 8.04 | 10.97 | | High | High | High | High | High | High | High | High | High | High | High | 0 | 0.000261 | 2.54 |
| FALSE | High | SPQSPASK | [R].SPQSP/ | 1xPhospho | A2A7S8-2 | 0.026652 | 0.001676 | 1 | 4 | 1 | A2A7S8-2 | Isoform 2 c | A2A7S8-2 [ | 0 | 1339.702 | 0 | -0.34 | 0.024854 | 8.56 | 109.6 | 90.4 | 7.31 | 9.47 | | High | High | High | High | High | High | High | High | High | High | High | 0.000903 | 0.009396 | 1.91 |
| FALSE | High | SPQTPELVI | [K].SPQTPE | 1xPhospho | Q6A0A9 1x | 1.68E-06 | 0 | 1 | 1 | 2 | Q6A0A9 | Constitutiv | Q6A0A9 [6 | 0 | 1866.956 | 0 | 1.41 | 0.000408 | 19.68 | 56.9 | 143.1 | 11.52 | 3.64 | | High | High | High | High | High | High | High | High | High | High | High | 0 | 1.82E-07 | 3.61 |
| FALSE | High | SPQTPSQL/ | [K].SPQTPS | 1xPhospho | Q9D219 1x | 0.006594 | 0.00062 | 1 | 1 | 2 | Q9D219 | B-cell CLL/I | Q9D219 [8 | 0 | 2621.377 | 0 | 0.86 | 0.004585 | 2.64 | 70.1 | 129.9 | 11.79 | 10.52 | | High | High | High | High | High | High | High | High | High | High | High | 0.000275 | 0.001977 | 2.57 |
| FALSE | High | SPQVLSSSL | [K].SPQVLS | 1xPhospho | Q67FY2 1x | 0.091627 | 0.007146 | 1 | 2 | 1 | Q67FY2 | B-cell CLL/I | Q67FY2 [9 | 0 | 1538.814 | 0 | 0.98 | 0.008548 | 9.81 | 67.7 | 132.3 | 8.76 | 11.72 | | High | High | High | High | High | High | High | High | High | High | High | 0.003633 | 0.03878 | 1.25 |
| FALSE | High | SPRPASPAS | [K].SPRPAS | 2xPhospho | A0A0U1RN | 0.003755 | 0.00021 | 1 | 2 | 1 | A0A0U1RN | BRCA2-inte | A0A0U1RN | 0 | 2903.509 | 0 | -0.76 | 0.018179 | 11.26 | 125.7 | 74.3 | 10.53 | 18.06 | | High | High | High | High | High | High | High | High | High | High | High | 9.42E-05 | 0.001055 | 3.75 |
| FALSE | High | SPRPTSAP/ | [R].SPRPTS | 1xPhospho | A0A068W/ | 9.5E-07 | 0 | 1 | 12 | 1 | A0A068W/ | Protein 4.1 | A0A068W/ | 0 | 2898.549 | 0 | -0.11 | 0.463381 | 7 | 102.3 | 97.7 | 12.17 | 5.77 | | High | High | High | High | High | High | High | High | High | High | High | 0 | 9.63E-08 | 4.88 |
| FALSE | High | SPSAALND! | [K].SPSAAL | 1xCarbami | Q8BT07 1x | 4.15E-08 | 0 | 1 | 2 | 5 | Q8BT07 | Centrosom | Q8BT07 [4 | 0 | 2126.06 | 0 | 1.5 | 0.000158 | 4.19 | 51.3 | 148.7 | 5.53 | 1.4 | | High | High | High | High | High | High | High | High | High | High | High | 0 | 3.18E-09 | 3.69 |
| FALSE | High | SPSAALND! | [K].SPSAAL | 1xCarbami | Q8BT07 1x | 1.36E-06 | 0 | 1 | 2 | 1 | Q8BT07 | Centrosom | Q8BT07 [4 | 0 | 2127.045 | 0 | 1.47 | 0.001043 | 13.34 | 54.3 | 145.7 | 14.83 | 5.58 | | High | High | High | High | High | High | High | High | High | High | High | 0 | 1.44E-07 | 3.8 |
| FALSE | High | SPSAALND! | [K].SPSAAL | 1xCarbami | Q8BT07 2x | 1.1E-05 | 0 | 1 | 2 | 1 | Q8BT07 | Centrosom | Q8BT07 [4 | 0 | 2206.027 | 0 | 1.53 | 0.000275 | 10.48 | 51 | 149 | 7.14 | 4.97 | | High | High | High | High | High | High | High | High | High | High | High | 0 | 1.51E-06 | 4.29 |
| FALSE | High | SPQPVCSP/ | [R].SPQPV( | 1xCarbami | Q6KCD5 1x | 7.09E-09 | 0 | 1 | 5 | 3 | Q6KCD5 | Nipped-B-l | Q6KCD5 [2 | 0 | 2123.024 | 0 | 0.52 | 0.006664 | 11.05 | 82.2 | 117.8 | 1.99 | 11.7 | | High | High | High | High | High | High | High | High | High | High | High | 0 | 3.9E-10 | 4.11 |
| FALSE | High | SPPVLGSA/ | [K].SPPVLG | 2xPhospho | Q9D219 2x | 0.002869 | 0.00016 | 1 | 1 | 2 | Q9D219 | B-cell CLL/I | Q9D219 [9 | 0 | 2149.122 | 0 | 0.35 | 0.082497 | 3.92 | 86.9 | 113.1 | 8.39 | 8.01 | | High | High | High | High | High | High | High | High | High | High | High | 7.19E-05 | 0.000778 | 1.91 |
| FALSE | High | SPPVLGSA/ | [K].SPPVLG | 1xPhospho | Q9D219 1x | 2.42E-08 | 0 | 1 | 1 | 1 | Q9D219 | B-cell CLL/I | Q9D219 [9 | 0 | 2069.156 | 0 | -0.46 | 0.00036 | 0.42 | 115.7 | 84.3 | 4.57 | 8.62 | | High | High | High | High | High | High | High | High | High | High | High | 0 | 1.55E-09 | 2.48 |
| FALSE | High | SPPTTMLLI | [K].SPPTTM | 2xPhospho | Q80Y32 2x | 0.000793 | 0 | 1 | 3 | 1 | Q80Y32 | HMG box c | Q80Y32 [4 | 0 | 2029.025 | 0 | -0.59 | 0.066443 | 20.45 | 120.1 | 79.9 | 12.68 | 17.86 | | High | High | High | High | High | High | High | High | High | High | High | 0 | 0.000183 | 3.33 |
| FALSE | High | SPPEITVTPI | [K].SPPEIT\ | 1xPhospho | G3X8Q1 1x | 8E-07 | 0 | 1 | 6 | 3 | G3X8Q1 | Calcineurir | G3X8Q1 [2 | 0 | 2413.34 | 0 | -0.01 | 0.706056 | 9.48 | 100.5 | 99.5 | 8.07 | 5.34 | | High | High | High | High | High | High | High | High | High | High | High | 0.003685 | 0.03958 | 2.69 |
| FALSE | High | SPPEITVTPI | [K].SPPEIT\ | 2xPhospho | G3X8Q1 2x | 0.016743 | 0.001166 | 1 | 6 | 1 | G3X8Q1 | Calcineurir | G3X8Q1 [2 | 0 | 2493.306 | 0 | -0.47 | 0.002924 | 4.35 | 115.4 | 84.6 | 2.55 | 7.3 | | High | High | High | High | High | High | High | High | High | High | High | 0.000622 | 0.005572 | 2.06 |
| FALSE | High | SPPESLNDI | [K].SPPESL | 1xPhospho | Q8CI71 1xI | 0.000198 | 0 | 1 | 3 | 1 | Q8CI71 | Syndetin O | Q8CI71 [15 | 0 | 2040.984 | 0 | 1.88 | 0.041664 | 54.6 | 42.9 | 157.1 | 30.06 | 43.82 | | High | High | High | High | High | High | High | High | High | High | High | 0 | 3.89E-05 | 3.71 |
| FALSE | High | SPPGENPSI | [R].SPPGEN | 1xPhospho | Q5SXI5 1xI | 0.003402 | 0.00021 | 1 | 2 | 1 | Q5SXI5 | Zinc finger | Q5SXI5 [23 | 0 | 2342.05 | 0 | -0.42 | 0.030926 | 1.15 | 113.3 | 86.7 | 10.73 | 6.24 | | High | High | High | High | High | High | High | High | High | High | High | 9.42E-05 | 0.000936 | 2.24 |
| FALSE | High | SPPGSARP( | [R].SPPGS/ | 1xPhospho | A2AIX1 1xI | 0.027358 | 0.001676 | 1 | 2 | 1 | A2AIX1 | SEC16 hom | A2AIX1 [55 | 0 | 2863.512 | 0 | 0.11 | 0.267979 | 6.77 | 96.8 | 103.2 | 6.19 | 6.76 | | High | High | High | High | High | High | High | High | High | High | High | 0.000936 | 0.009699 | 2.04 |
| FALSE | High | SPPHHSGF( | [K].SPPHHS | 1xPhospho | Q5RJH6-2 | 0.004491 | 0.000409 | 1 | 4 | 4 | Q5RJH6-2 | Isoform 2 c | Q5RJH6-2 [ | 0 | 2551.213 | 0 | -0.89 | 0.000163 | 1.27 | 130.1 | 69.9 | 4.07 | 8.71 | | High | High | High | High | High | High | High | High | High | High | High | 0.000327 | 0.002868 | 2.56 |
| FALSE | High | SPPHHSGF( | [K].SPPHHS | 1xDeamida | Q5RJH6-2 | 0.010382 | 0.000772 | 1 | 4 | 2 | Q5RJH6-2 | Isoform 2 c | Q5RJH6-2 [ | 0 | 2552.197 | 0 | -0.7 | 0.000742 | 6.69 | 126.2 | 73.8 | 9.05 | 20.48 | | High | High | High | High | High | High | High | High | High | High | High | 0.000382 | 0.00327 | 2.55 |
| FALSE | High | SPPIDLTQS | [K].SPPIDL | 1xPhospho | Q6P1D7 1x | 0.003195 | 0.00016 | 1 | 2 | 1 | Q6P1D7 | Structure-s | Q6P1D7 [9 | 0 | 2142.105 | 0 | -0.32 | 0.025818 | 5.84 | 110.6 | 89.4 | 7 | 7.95 | | High | High | High | High | High | High | High | High | High | High | High | 7.19E-05 | 0.000876 | 1.91 |
| FALSE | High | SPPIDLTQS | [K].SPPIDL | 2xPhospho | Q6P1D7 2x | 0.063258 | 0.004664 | 1 | 2 | 1 | Q6P1D7 | Structure-s | Q6P1D7 [9 | 0 | 2222.071 | 0 | -0.12 | 0.027531 | 5.81 | 107.7 | 92.3 | 5.72 | 8.01 | | High | High | High | High | High | High | High | High | High | High | High | 0.002403 | 0.02531 | 2.27 |
| FALSE | High | SPPLLESPD | [K].SPPLLE! | 1xPhospho | Q8BVL3 1x | 8.56E-05 | 0 | 1 | 1 | 1 | Q8BVL3 | Sorting nex | Q8BVL3 [4 | 0 | 1577.777 | 0 | -0.03 | 0.824013 | 5.41 | 100 | 100 | 6.22 | 2.9 | | High | High | High | High | High | High | High | High | High | High | High | 0 | 1.51E-05 | 3.42 |
| FALSE | High | SPPLLGSES | [R].SPPLLG | 1xPhospho | P14873 1xI | 5.16E-05 | 0 | 2 | 2 | 5 | P14873; B2 | Microtubu | P14873 [13 | 0 | 2677.305 | 0 | -1.04 | 0.001548 | 0.88 | 129.8 | 70.2 | 7.21 | 10.48 | Shared | High | High | High | High | High | High | High | High | High | High | High | 0 | 4.22E-05 | 4.08 |
| FALSE | High | SPPLPAVIK | [R].SPPLPA | 1xPhospho | Q69ZX6 1x | 0.017344 | 0.001204 | 2 | 2 | 2 | Q69ZX6; Q | MORC fam | Q69ZX6 [6 | 0 | 1230.706 | 0 | -0.54 | 0.021448 | 9.61 | 120 | 80 | 10.69 | 16.85 | Shared | High | High | High | High | High | High | High | High | High | High | High | 0.000656 | 0.005811 | 1.65 |
| FALSE | High | SPPLSPVGT | [K].SPPLSP | 1xPhospho | Q922M7 1: | 2.04E-05 | 0 | 1 | 1 | 3 | Q922M7 | Ashwin OS | Q922M7 [1 | 0 | 1818.018 | 0 | -1.36 | 0.000456 | 0.76 | 143.6 | 56.4 | 4.17 | 11.23 | | High | High | High | High | High | High | High | High | High | High | High | 0 | 8.85E-05 | 3.89 |
| FALSE | High | SPPLSPVGT | [K].SPPLSP | 2xPhospho | Q922M7 2: | 0.025738 | 0.001645 | 1 | 1 | 2 | Q922M7 | Ashwin OS | Q922M7 [1 | 0 | 1897.984 | 0 | -1.37 | 0.000377 | 1.73 | 142.2 | 57.8 | 8.1 | 14.62 | | High | High | High | High | High | High | High | High | High | High | High | 0.000872 | 0.009067 | 2.39 |
| FALSE | High | SPPMEMQ | [R].SPPMEI | 1xCarbami | P97473 1xI | 0.021046 | 0.001507 | 1 | 4 | 5 | P97473 | RISC-loadir | P97473 [14 | 0 | 3858.888 | 0 | -0.47 | 0.000774 | 2.37 | 114.6 | 85.4 | 3.35 | 4.43 | | High | High | High | High | High | High | High | High | High | High | High | 0.001151 | 0.01235 | 3.19 |
| FALSE | High | SPPPETSSS | [R].SPPPET | 1xPhospho | Q2MHN3 1 | 4.26E-07 | 0 | 1 | 2 | 3 | Q2MHN3 | ZFH-5 OS=I | Q2MHN3 [ | 0 | 2242.089 | 0 | -0.19 | 0.097569 | 6.28 | 106.6 | 93.4 | 5.02 | 6.09 | | High | High | High | High | High | High | High | High | High | High | High | 0 | 3.88E-08 | 3.71 |
| FALSE | High | SPPPPEEEA | [K].SPPPPE | 1xPhospho | Q00PI9 1xI | 0.013666 | 0.001011 | 1 | 1 | 1 | Q00PI9 | Heterogen | Q00PI9 [22 | 1 | 4669.269 | 0 | 0.09 | 0.360874 | 30.6 | 91.8 | 108.2 | 9.5 | 16.48 | | High | High | High | High | High | High | High | High | High | High | High | 0.000538 | 0.004452 | 3.4 |
| FALSE | High | SPPPPTHN | [K].SPPPPT | 1xPhospho | Q63943-2 | 0.000165 | 0 | 2 | 6 | 1 | Q63943-2; | Isoform Mi | Q63943-2 [ | 0 | 1965.975 | 0 | 1.41 | 0.02023 | 25.07 | 58.4 | 141.6 | 21.98 | 18.6 | Shared | High | High | High | High | High | High | High | High | High | High | High | 0 | 3.15E-05 | 2.55 |
| FALSE | High | SPPPTVDES | [R].SPPPTV | 1xPhospho | Q3UV57 1x | 1.91E-05 | 0 | 1 | 1 | 2 | Q3UV57 | Putative ur | Q3UV57 [4 | 0 | 2429.17 | 0 | 0.29 | 0.016511 | 8.44 | 87.6 | 112.4 | 7.82 | 6.49 | | High | High | High | High | High | High | High | High | High | High | High | 0.000204 | 0.001379 | 2.59 |
| FALSE | High | SPPQYACD | [K].SPPQY/ | 1xCarbami | Q8BHK9 1x | 1.43E-08 | 0 | 1 | 3 | 7 | Q8BHK9 | DNA excisi | Q8BHK9 [9 | 0 | 2655.164 | 0 | -0.61 | 0.01132 | 9.92 | 121 | 79 | 7.76 | 12.8 | | High | High | High | High | High | High | High | High | High | High | High | 0 | 8.25E-09 | 4.42 |
| FALSE | High | SPPSGSQSI | [R].SPPSGS | 1xPhospho | B9EKJ7 1xI | 0.003372 | 0.00016 | 1 | 3 | 2 | B9EKJ7 | Tankyrase | B9EKJ7 [14 | 0 | 2502.256 | 0 | -0.04 | 0.959108 | 29.33 | 101.5 | 98.5 | 9.11 | 18.53 | | High | High | High | High | High | High | High | High | High | High | High | 9.42E-05 | 0.000934 | 2.43 |
| FALSE | High | SPPSGSQSI | [R].SPPSGS | 1xPhospho | B9EKJ7 1xI | 2.77E-05 | 0 | 1 | 3 | 5 | B9EKJ7 | Tankyrase | B9EKJ7 [14 | 0 | 2518.251 | 0 | 0.33 | 0.118675 | 25.82 | 90.5 | 109.5 | 8.87 | 20.28 | | High | High | High | High | High | High | High | High | High | High | High | 0 | 7.91E-06 | 3.07 |
| FALSE | High | SPPSSAEQS | [K].SPPSSA | 1xPhospho | P57016 1xI | 0.000305 | 0 | 1 | 2 | 4 | P57016 | Ladinin-1 C | P57016 [32 | 0 | 2120.068 | 0 | -1.05 | 0.001295 | 2.41 | 135 | 65 | 7.58 | 5.1 | | High | High | High | High | High | High | High | High | High | High | High | 0 | 6.27E-05 | 2.76 |
| FALSE | High | SPPSSPSSP | [K].SPPSSP | 1xPhospho | Q8BKR5 1x | 0.001068 | 0 | 1 | 2 | 3 | Q8BKR5 | Protein ph | Q8BKR5 [5 | 0 | 1665.768 | 0 | 0.53 | 0.004952 | 6.9 | 80.3 | 119.7 | 5.12 | 7.94 | | High | High | High | High | High | High | High | High | High | High | High | 0 | 0.000256 | 2.34 |
| FALSE | High | SPPSVQSL/ | [K].SPPSVC | 1xPhospho | A2CG76 1x | 0.000379 | 0 | 1 | 5 | 1 | A2CG76 | Histone-lys | A2CG76 [1 | 0 | 1497.733 | 0 | 0.26 | 0.294976 | 13.98 | 87.5 | 112.5 | 24.41 | 21.83 | | High | High | High | High | High | High | High | High | High | High | High | 0 | 8.03E-05 | 3.21 |
| FALSE | High | SPPTASLQI | [K].SPPTAS | 1xPhospho | O35711-4 | 0.109329 | 0.008865 | 1 | 4 | 1 | O35711-4 | Isoform 4 c | O35711-4 [ | 0 | 2309.143 | 0 | -0.47 | 0.023334 | 11.87 | 116.3 | 83.7 | 7 | 14.46 | | High | High | High | High | High | High | High | High | High | High | High | 0.004462 | 0.04798 | 2.44 |
| FALSE | High | SPPTPSVPI | [R].SPPTPS | 2xCarbami | Q3TZX8 1x | 0.007209 | 0.00062 | 1 | 4 | 2 | Q3TZX8 | Polynucleo | Q3TZX8 [8 | 0 | 2787.341 | 0 | -0.45 | 0.100917 | 17.96 | 115.4 | 84.6 | 11.89 | 15.87 | | High | High | High | High | High | High | High | High | High | High | High | 0.000327 | 0.002565 | 2.91 |
| FALSE | High | SPPTTMLLI | [K].SPPTTM | 1xPhospho | Q80Y32 1x | 0.002553 | 0.000109 | 1 | 3 | 1 | Q80Y32 | HMG box c | Q80Y32 [4 | 0 | 1949.058 | 0 | 0 | 0.956149 | 19.77 | 99.8 | 100.2 | 8.58 | 10.6 | | High | High | High | High | High | High | High | High | High | High | High | 4.94E-05 | 0.000679 | 2.26 |
| FALSE | High | SPPTTMLLI | [K].SPPTTM | 1xPhospho | Q80Y32 1x | 5.6E-05 | 0 | 1 | 3 | 3 | Q80Y32 | HMG box c | Q80Y32 [4 | 0 | 1965.053 | 0 | 0.02 | 0.984827 | 10.17 | 99.3 | 100.7 | 2.95 | 7.56 | | High | High | High | High | High | High | High | High | High | High | High | 0 | 9.35E-06 | 2.74 |
| FALSE | High | SPPEDAIPK | [R].SPPED/ | 1xPhospho | Q6PFB2 1x | 0.010663 | 0.000772 | 1 | 4 | 3 | Q6PFB2 | Rcc1 prote | Q6PFB2 [1 | 0 | 1262.623 | 0 | -0.68 | 0.592321 | 8.32 | 126 | 74 | 13.15 | 59.3 | | High | High | High | High | High | High | High | High | High | High | High | 0.000723 | 0.006329 | 1.88 |
| FALSE | High | SRPPEQDG | [R].SRPPEC | 1xPhospho | Q3URQ4 1: | 0.000125 | 0 | 1 | 2 | 2 | Q3URQ4 | Family with | Q3URQ4 [1 | 0 | 3735.745 | 0 | 1.21 | 0.009326 | 28.02 | 60.5 | 139.5 | 15.17 | 16.25 | | High | High | High | High | High | High | High | High | High | High | High | 0 | 2.3E-05 | 4.01 |
| FALSE | High | SPTGNTPP' | [R].SPTGN | 1xPhospho | B2RUG9 1x | 2.29E-05 | 0 | 1 | 6 | 2 | B2RUG9 | Adenomat | B2RUG9 [2 | 0 | 2166.11 | 0 | 0.25 | 0.004368 | 4.51 | 92.7 | 107.3 | 5.91 | 2.01 | | High | High | High | High | High | High | High | High | High | High | High | 0 | 3.43E-06 | 2.68 |
| FALSE | High | SPTGPSNSI | [R].SPTGPS | 1xPhospho | Q8JZX4 1xI | 3.73E-08 | 0 | 1 | 1 | 1 | Q8JZX4 | Splicing fac | Q8JZX4 [22 | 0 | 2511.247 | 0 | -0.83 | 0.008105 | 5.73 | 128.1 | 71.9 | 10.94 | 8.44 | | High | High | High | High | High | High | High | High | High | High | High | 0 | 2.52E-09 | 5.01 |
| FALSE | High | SQSESSDE\ | [R].SQSESS | 1xPhospho | Q7TPS5 1x | 2.18E-06 | 0 | 2 | 6 | 6 | Q7TPS5; A( | C2 domain | Q7TPS5 [6 | 0 | 2486.17 | 0 | -0.03 | 0.744565 | 34.75 | 106.8 | 93.2 | 7.27 | 52.83 | Shared | High | High | High | High | High | High | High | High | High | High | High | 0 | 2.44E-07 | 3.79 |
| FALSE | High | SQSESSDE\ | [R].SQSESS | 2xPhospho | Q7TPS5 2x | 4.4E-06 | 0 | 2 | 6 | 2 | Q7TPS5; A( | C2 domain | Q7TPS5 [6 | 0 | 2566.136 | 0 | -0.53 | 0.02249 | 5.88 | 116.9 | 83.1 | 6.37 | 10.47 | Shared | High | High | High | High | High | High | High | High | High | High | High | 0 | 5.37E-07 | 3.97 |
| FALSE | High | SQSETSLLG | [R].SQSETS | 1xPhospho | Q0PHV7 1x | 0.093879 | 0.007306 | 1 | 1 | 2 | Q0PHV7 | Dapper ho | Q0PHV7 [4 | 0 | 1386.683 | 0 | -1.4 | 0.000231 | 1.49 | 142.7 | 57.3 | 4.44 | 11.93 | | High | High | High | High | High | High | High | High | High | High | High | 0.003698 | 0.03992 | 2.25 |
| FALSE | High | SQSEYCNV | [R].SQSEY( | 1xCarbami | G3UYW1 1 | 0.063788 | 0.004664 | 1 | 3 | 1 | G3UYW1 | WD repeat | G3UYW1 [ | 0 | 1810.845 | 0 | 0.97 | 0.000458 | 2.95 | 67.5 | 132.5 | 4 | 6.44 | | High | High | High | High | High | High | High | High | High | High | High | 0.002403 | 0.02543 | 2.29 |
| FALSE | High | SQSFAGVL( | [R].SQSFA( | 1xPhospho | Q68FE6 1x | 0.000501 | 0 | 1 | 1 | 1 | Q68FE6 | Rho family | Q68FE6 [2 | 0 | 1683.805 | 0 | 1.07 | 0.001931 | 3.95 | 66.4 | 133.6 | 14.64 | 7.69 | | High | High | High | High | High | High | High | High | High | High | High | 0 | 0.00011 | 2.04 |
| FALSE | High | SQSFGVAS. | [R].SQSFG\ | 1xPhospho | Q8CI12 1xI | 0.00509 | 0.000551 | 1 | 2 | 2 | Q8CI12 | Smoothelir | Q8CI12 [33 | 0 | 1577.777 | 0 | -1.38 | 0.015675 | 3.87 | 143 | 57 | 18.77 | 20.35 | | High | High | High | High | High | High | High | High | High | High | High | 0.000246 | 0.001473 | 2.61 |
| FALSE | High | SQSFNNYD | [R].SQSFNI | 1xPhospho | E9Q842 1x | 0.004451 | 0.000409 | 1 | 7 | 1 | E9Q842 | Neuron na | E9Q842 [2 | 0 | 1640.772 | 0 | 1.78 | 0.003512 | 35.48 | 45 | 155 | 17.66 | 13.79 | | High | High | High | High | High | High | High | High | High | High | High | 0.000183 | 0.001274 | 2.78 |
| FALSE | High | SQSFNNYD | [R].SQSFNI | 1xDeamida | E9Q842 1x | 0.004738 | 0.000456 | 1 | 7 | 1 | E9Q842 | Neuron na | E9Q842 [2 | 0 | 1641.756 | 0 | 1.29 | 0.002108 | 21.62 | 57 | 143 | 17.07 | 8.86 | | High | High | High | High | High | High | High | High | High | High | High | 0.000204 | 0.001363 | 1.85 |
| FALSE | High | SQSFNTDT | [R].SQSFN | 1xPhospho | Q6QI06 1xI | 1.03E-05 | 0 | 1 | 2 | 3 | Q6QI06 | Rapamycin | Q6QI06 [1 | 0 | 2574.123 | 0 | 0.41 | 0.03787 | 7.06 | 84 | 116 | 19.33 | 2.97 | | High | High | High | High | High | High | High | High | High | High | High | 0 | 1.39E-06 | 3.84 |
| FALSE | High | SQSFNTDT | [R].SQSFN | 1xPhospho | Q6QI06 1xI | 2.82E-11 | 0 | 1 | 2 | 8 | Q6QI06 | Rapamycin | Q6QI06 [1 | 0 | 2590.118 | 0 | 0.5 | 0.009796 | 4.97 | 82.3 | 117.7 | 4.4 | 9.46 | | High | High | High | High | High | High | High | High | High | High | High | 0 | 7.81E-13 | 5.08 |
| FALSE | High | SQSFNTDT | [R].SQSFN | 2xPhospho | Q6QI06 2xI | 4.52E-06 | 0 | 1 | 2 | 1 | Q6QI06 | Rapamycin | Q6QI06 [1 | 0 | 2670.084 | 0 | -0.06 | 0.908769 | 57.15 | 97.1 | 102.9 | 29.19 | 19.33 | | High | High | High | High | High | High | High | High | High | High | High | 0 | 5.54E-07 | 3.93 |
| FALSE | High | SQSFPDVEI | [R].SQSFPD | 1xPhospho | E9Q0S6 1x | 0.000751 | 0 | 1 | 6 | 1 | E9Q0S6 | Tensin 1 O | E9Q0S6 [7 | 0 | 2206.074 | 0 | 0.66 | 0.005994 | 0.75 | 79.7 | 120.3 | 12.26 | 4.31 | | High | High | High | High | High | High | High | High | High | High | High | 0 | 0.000173 | 3.1 |
| FALSE | High | SQSFSHSIC | [R].SQSFSH | 1xPhospho | Q3UHI0-2 | 0.005135 | 0.000551 | 1 | 2 | 3 | Q3UHI0-2 | Isoform 2 c | Q3UHI0-2 [ | 0 | 2541.27 | 0 | -0.01 | 0.72558 | 25.93 | 101.9 | 98.1 | 8.25 | 13.26 | | High | High | High | High | High | High | High | High | High | High | High | 0.000275 | 0.001598 | 3.57 |
| FALSE | High | SQSENLSSI | [K].SQSENI | 1xPhospho | Q9Z2U2 1x | 0.004738 | 0.000456 | 1 | 3 | 1 | Q9Z2U2 | Zinc finger | Q9Z2U2 [1 | 0 | 2198.085 | 0 | -0.36 | 0.002264 | 3.01 | 113.1 | 86.9 | 3.35 | 5.68 | | High | High | High | High | High | High | High | High | High | High | High | 0.000204 | 0.001356 | 2.67 |
| FALSE | High | SQSFTHTPI | [R].SQSFTH | 1xPhospho | Q80U49 1x | 0.014667 | 0.001049 | 1 | 2 | 2 | Q80U49 | Centrosom | Q80U49 [4 | 0 | 1950.973 | 0 | 0.63 | 0.000422 | 4.26 | 77.1 | 122.9 | 3.53 | 4.51 | | High | High | High | High | High | High | High | High | High | High | High | 0.001076 | 0.01127 | 2.64 |
| FALSE | High | SQSGILDGI | [R].SQSGIL | 1xPhospho | Q61037 1x | 8.58E-06 | 0 | 1 | 14 | 1 | Q61037 | Tuberin OS | Q61037 [1 | 0 | 2461.108 | 0 | -0.58 | 0.009533 | 5.46 | 115.9 | 84.1 | 9.16 | 19.67 | | High | High | High | High | High | High | High | High | High | High | High | 0 | 1.14E-06 | 4.04 |
| FALSE | High | SQSGSPAA | [R].SQSGSI | 1xPhospho | E9QK89 1x | 5.53E-07 | 0 | 1 | 2 | 4 | E9QK89 | Mediator c | E9QK89 [5 | 0 | 3183.552 | 0 | -0.76 | 0.004034 | 6.18 | 128.9 | 71.1 | 10.76 | 10.31 | | High | High | High | High | High | High | High | High | High | High | High | 0 | 2.91E-07 | 4.89 |
| FALSE | High | SQSGSPAA | [R].SQSGSI | 2xPhospho | E9QK89 2x | 3.94E-05 | 0 | 1 | 2 | 1 | E9QK89 | Mediator c | E9QK89 [5 | 0 | 3263.518 | 0 | -0.87 | 0.00116 | 1.2 | 130.5 | 69.5 | 12.01 | 17.7 | | High | High | High | High | High | High | High | High | High | High | High | 0 | 6.32E-06 | 3.58 |
| FALSE | High | SQSLIIR | [R].SQSLIIR | 1xPhospho | D3Z6J1 1xI | 0.025738 | 0.001645 | 2 | 8 | 2 | D3Z6J1; F8 | DEP domai | D3Z6J1 [51 | 0 | 1125.623 | 0 | 1.25 | 0.00034 | 1.73 | 58.8 | 141.2 | 7.14 | 8.5 | Shared | High | High | High | High | High | High | High | High | High | High | High | 0.000872 | 0.009016 | 1.74 |
| FALSE | High | SQSLLTFST | [K].SQSLLT | 1xPhospho | Q3U821 1x | 0.012959 | 0.000889 | 1 | 7 | 1 | Q3U821 | WD repeat | Q3U821 [6 | 0 | 1649.892 | 0 | -1.59 | 0.000209 | 0.43 | 150.4 | 49.6 | 5.8 | 10.63 | | High | High | High | High | High | High | High | High | High | High | High | 0.000449 | 0.00421 | 2.7 |
| FALSE | High | SQSLPTTLL | [R].SQSLPT | 1xPhospho | Q922B9 1x | 2.11E-05 | 0 | 1 | 5 | 1 | Q922B9 | Sperm-spe | Q922B9 [7 | 0 | 1707.924 | 0 | 0.36 | 0.004331 | 6.79 | 86 | 114 | 4.68 | 6.97 | | High | High | High | High | High | High | High | High | High | High | High | 0 | 3.13E-06 | 2.92 |
| FALSE | High | SQSLPTTLL | [R].SQSLPT | 2xPhospho | Q922B9 2x | 0.000628 | 0 | 1 | 5 | 1 | Q922B9 | Sperm-spe | Q922B9 [7 | 0 | 1787.891 | 0 | 0.34 | 0.081942 | 29.63 | 79.7 | 120.3 | 17.43 | 19.25 | | High | High | High | High | High | High | High | High | High | High | High | 0 | 0.000141 | 2.56 |
| FALSE | High | SQSLQDQF | [R].SQSLQI | 1xPhospho | Q9JM52-3 | 0.005418 | 0.00062 | 1 | 8 | 2 | Q9JM52-3 | Isoform 3 c | Q9JM52-3 | 0 | 1468.699 | 0 | 0.96 | 0.000461 | 3.67 | 68 | 132 | 3.66 | 6.04 | | High | High | High | High | High | High | High | High | High | High | High | 0.000275 | 0.00158 | 1.65 |
| FALSE | High | SQSLSLQM | [R].SQSLSL | 1xPhospho | A0A1L1SR) | 4.01E-05 | 0 | 1 | 5 | 3 | A0A1L1SR) | AMP deam | A0A1L1SR) | 0 | 2331.146 | 0 | 2.05 | 0.001324 | 13.13 | 40.6 | 159.4 | 8.79 | 15.24 | | High | High | High | High | High | High | High | High | High | High | High | 0 | 3.46E-05 | 3.46 |
| FALSE | High | SQSMDIDG | [R].SQSMD | 1xCarbami | Q80TU3 1x | 0.000141 | 0 | 1 | 6 | 2 | Q80TU3 | MKIAA068 | Q80TU3 [1 | 0 | 1993.901 | 0 | -0.22 | 0.018587 | 6.61 | 107.6 | 92.4 | 4.25 | 8.69 | | High | High | High | High | High | High | High | High | High | High | High | 0 | 2.64E-05 | 3.55 |
| FALSE | High | SQSMDIDG | [R].SQSMD | 1xCarbami | Q80TU3 1x | 5.91E-05 | 0 | 1 | 6 | 6 | Q80TU3 | MKIAA068 | Q80TU3 [1 | 0 | 2009.896 | 0 | -0.27 | 0.151428 | 0.97 | 108.5 | 91.5 | 2.32 | 9.92 | | High | High | High | High | High | High | High | High | High | High | High | 0 | 6.38E-05 | 3.15 |
| FALSE | High | SQSNPILGS | [R].SQSNPI | 1xPhospho | Q9ESL4 1xI | 0.000274 | 0 | 1 | 1 | 1 | Q9ESL4 | Mitogen-a | Q9ESL4 [5 | 0 | 3156.503 | 0 | 1.55 | 0.002056 | 15.77 | 50.1 | 149.9 | 10.84 | 12.32 | | High | High | High | High | High | High | High | High | High | High | High | 0 | 5.58E-05 | 4.77 |
| FALSE | High | SQSPAASD | [R].SQSPA/ | 1xCarbami | A6H663 1x | 9.99E-09 | 0 | 1 | 3 | 5 | A6H663 | BCL2-assoc | A6H663 [1 | 0 | 2509.071 | 0 | 2.52 | 0.000313 | 12.57 | 29.7 | 170.3 | 12.35 | 7.59 | | High | High | High | High | High | High | High | High | High | High | High | 0 | 5.76E-10 | 4.12 |
| FALSE | High | SQSGEDES | [R].SQSGEI | 1xPhospho | Q5BL07 1x | 8.55E-08 | 0 | 1 | 2 | 2 | Q5BL07 | Peroxisom | Q5BL07 [1 | 0 | 2224.09 | 0 | -0.14 | 0.068512 | 4.15 | 104.3 | 95.7 | 5.48 | 5.81 | | High | High | High | High | High | High | High | High | High | High | High | 0 | 6.4E-09 | 5.15 |
| FALSE | High | SQSPEDLS( | [R].SQSPED | 1xPhospho | E9PVX6 1xI | 2.5E-11 | 0 | 1 | 2 | 1 | E9PVX6 | Proliferatic | E9PVX6 [2 | 0 | 2812.355 | 0 | 0.22 | 0.18287 | 35.72 | 92.6 | 107.4 | 7.56 | 20.93 | | High | High | High | High | High | High | High | High | High | High | High | 0 | 6.88E-13 | 6.09 |
| FALSE | High | SQSELDDQ | [R].SQSELD | 2xPhospho | Q68FF6 2xI | 0.00925 | 0.000736 | 1 | 2 | 1 | Q68FF6 | ARF GTPas | Q68FF6 [3 | 0 | 3867.559 | 0 | -0.36 | 0.369237 | 15.08 | 111.4 | 88.6 | 24.05 | 12.89 | | High | High | High | High | High | High | High | High | High | High | High | 0.000327 | 0.002888 | 3.11 |
| FALSE | High | SQSELDDQ | [R].SQSELD | 1xPhospho | Q68FF6 1xI | 1.45E-06 | 0 | 1 | 2 | 4 | Q68FF6 | ARF GTPas | Q68FF6 [3 | 0 | 3787.593 | 0 | -0.87 | 0.011915 | 2.61 | 125.2 | 74.8 | 10.16 | 16.4 | | High | High | High | High | High | High | High | High | High | High | High | 0 | 0.000112 | 3.98 |
| FALSE | High | SQNNFVAI | [R].SQNNF | 1xDeamida | Q3U5I9 1xI | 0.041781 | 0.003126 | 1 | 3 | 1 | Q3U5I9 | Putative ur | Q3U5I9 [1 | 0 | 2741.359 | 0 | 1.03 | 0.000704 | 12.57 | 65.8 | 134.2 | 10.43 | 7.51 | | High | High | High | High | High | High | High | High | High | High | High | 0.001625 | 0.01574 | 3.02 |
| FALSE | High | SQNREDSF | [R].SQNREI | 1xPhospho | Q9Z0U1 1x | 0.021797 | 0.001507 | 1 | 1 | 2 | Q9Z0U1 | Tight junct | Q9Z0U1 [9 | 1 | 2013.932 | 0 | 1.21 | 0.001124 | 8.59 | 60.8 | 139.2 | 7.54 | 7.26 | | High | High | High | High | High | High | High | High | High | High | High | 0.000801 | 0.00748 | 1.9 |
| FALSE | High | SQPAPSSP\ | [R].SQPAPS | 1xPhospho | O89091 1x | 0.000779 | 0 | 1 | 4 | 3 | O89091 | Krueppel-li | O89091 [2 | 0 | 1663.882 | 0 | -0.62 | 0.000922 | 2.94 | 121.7 | 78.3 | 3.55 | 6.04 | | High | High | High | High | High | High | High | High | High | High | High | 0 | 0.000181 | 3.05 |
| FALSE | High | SQPASIPGS | [R].SQPASI | 1xPhospho | D3Z0X5 1xI | 0.041781 | 0.003126 | 1 | 6 | 1 | D3Z0X5 | Pleckstrin I | D3Z0X5 [3 | 0 | 1377.698 | 0 | -0.36 | 0.902037 | 28.24 | 91.1 | 108.9 | 35.76 | 42.68 | | High | High | High | High | High | High | High | High | High | High | High | 0.001609 | 0.01563 | 2.33 |
| FALSE | High | SQPASIPGS | [R].SQPASI | 1xPhospho | D3Z0X5 1xI | 8.25E-05 | 0 | 1 | 6 | 2 | D3Z0X5 | Pleckstrin I | D3Z0X5 [3 | 0 | 1606.861 | 0 | 1 | 0.000492 | 5.88 | 67.2 | 132.8 | 3.82 | 6.29 | | High | High | High | High | High | High | High | High | High | High | High | 0 | 0.000163 | 2.45 |
| FALSE | High | SQPEPAETI | [R].SQPEP/ | 1xCarbami | Q4LDD4 1x | 0.001087 | 0 | 1 | 2 | 1 | Q4LDD4 | Arf-GAP wi | Q4LDD4 [1 | 0 | 4654.455 | 0 | 0.84 | 0.002099 | 5.74 | 73.3 | 126.7 | 6.69 | 10.46 | | High | High | High | High | High | High | High | High | High | High | High | 0 | 0.000262 | 3.61 |
| FALSE | High | SQPEPSPVI | [K].SQPEPS | 1xPhospho | Q91VX2 1x | 0.007471 | 0.00062 | 1 | 3 | 1 | Q91VX2 | Ubiquitin-a | Q91VX2 [4 | 0 | 1961.989 | 0 | 0.68 | 0.004938 | 12.63 | 76.7 | 123.3 | 13.59 | 11.19 | | High | High | High | High | High | High | High | High | High | High | High | 0.000275 | 0.002254 | 2.55 |
| FALSE | High | SQPPTSPE( | [R].SQPPTS | 1xPhospho | E9Q6R7 1x | 0.1059 | 0.008582 | 1 | 2 | 1 | E9Q6R7 | Utrophin C | E9Q6R7 [1 | 0 | 1364.641 | 0 | 1.16 | 0.000542 | 10.68 | 60.8 | 139.2 | 2.22 | 6.54 | | High | High | High | High | High | High | High | High | High | High | High | 0.004291 | 0.04598 | 1.44 |
| FALSE | High | SQPQVLVE | [K].SQPQV | 1xPhospho | E9QK89 1x | 3.35E-05 | 0 | 1 | 2 | 4 | E9QK89 | Mediator c | E9QK89 [4 | 1 | 3416.529 | 0 | -0.9 | 0.004369 | 1.21 | 136.4 | 63.6 | 13.22 | 9.49 | | High | High | High | High | High | High | High | High | High | High | High | 0 | 5.25E-06 | 4.35 |

| | | | | | | | | | | | | | | | | | | | | | | | | | | | | | | | | | | | | | | |
|---|---|---|---|---|---|---|---|---|---|---|---|---|---|---|---|---|---|---|---|---|---|---|---|---|---|---|---|---|---|---|---|---|---|---|---|---|---|---|
| FALSE | High | SQPQVLVE | [K].SQPQV | 2xPhosphc | E9QK89 2x | 0.00509 | 0.000551 | 1 | 2 | 3 | E9QK89 | Mediator c | E9QK89 [4 | 1 | 3496.495 | 0 | -0.87 | 0.000333 | 1.26 | 129 | 71 | 4.28 | 2.8 | | High | High | High | High | High | High | High | High | High | High | High | 0.000246 | 0.001481 | 4.29 |
| FALSE | High | SQPQVLVE | [K].SQPQV | 1xDeamida | E9QK89 2x | 0.000622 | 0 | 1 | 2 | 5 | E9QK89 | Mediator c | E9QK89 [4 | 1 | 3497.479 | 0 | -0.87 | 0.004517 | 6.33 | 125.7 | 74.3 | 8.38 | 13.45 | | High | High | High | High | High | High | High | High | High | High | High | 0 | 0.000141 | 4.72 |
| FALSE | High | SQPQVLVE | [K].SQPQV | 3xPhosphc | E9QK89 3x | 0.00047 | 0 | 1 | 2 | 1 | E9QK89 | Mediator c | E9QK89 [4 | 1 | 3576.461 | 0 | -0.47 | 0.014759 | 4.57 | 119.1 | 80.9 | 10.06 | 21.36 | | High | High | High | High | High | High | High | High | High | High | High | 0 | 0.000103 | 4.22 |
| FALSE | High | SQPSPTLPL | [R].SQPSPT | 1xCarbami | Q99PT3 1x | 0.000415 | 0 | 1 | 3 | 2 | Q99PT3 | INO80 com | Q99PT3 [2 | 0 | 3298.723 | 0 | 0.01 | 0.746967 | 16.99 | 102 | 98 | 9.58 | 13.57 | | High | High | High | High | High | High | High | High | High | High | High | 0 | 8.91E-05 | 3.74 |
| FALSE | High | SQSELDDQ | [R].SQSELD | 1xDeamida | Q68FF6 1x | 0.010568 | 0.000772 | 1 | 2 | 3 | Q68FF6 | ARF GTPas | Q68FF6 [3 | 0 | 3788.577 | 0 | -0.11 | 0.108792 | 9.81 | 107.4 | 92.6 | 15.65 | 8.54 | | High | High | High | High | High | High | High | High | High | High | High | 0.001355 | 0.01419 | 3.81 |
| FALSE | High | SQPSPTLPL | [R].SQPSPT | 1xCarbami | Q99PT3 1x | 0.001689 | 5.66E-05 | 1 | 3 | 2 | Q99PT3 | INO80 com | Q99PT3 [2 | 0 | 3314.718 | 0 | -0.27 | 0.048925 | 8.55 | 112 | 88 | 3.47 | 14.56 | | High | High | High | High | High | High | High | High | High | High | High | 2.58E-05 | 0.000429 | 3.18 |
| FALSE | High | SQQQPTPV | [R].SQQQP | 1xPhosphc | Q5SVQ0 1x | 1.17E-06 | 0 | 1 | 5 | 4 | Q5SVQ0 | Histone ac | Q5SVQ0 [8 | 0 | 1748.935 | 0 | -1.45 | 0.000158 | 1.36 | 146.5 | 53.5 | 3.44 | 11 | | High | High | High | High | High | High | High | High | High | High | High | 0 | 1.21E-07 | 3.69 |
| FALSE | High | SQQQPTPV | [R].SQQQP | 2xPhosphc | Q5SVQ0 2x | 0.000595 | 0 | 1 | 5 | 1 | Q5SVQ0 | Histone ac | Q5SVQ0 [8 | 0 | 1828.901 | 0 | -1.12 | 0.000307 | 2.48 | 136.6 | 63.4 | 4.04 | 9.55 | | High | High | High | High | High | High | High | High | High | High | High | 0 | 0.000133 | 2.6 |
| FALSE | High | SQQVAQS/ | [R].SQQVA | 1xCarbami | E9Q3P4 1x | 5.21E-08 | 0 | 1 | 6 | 2 | E9Q3P4 | Centromer | E9Q3P4 [1 | 0 | 2683.316 | 0 | -0.26 | 0.025917 | 8.82 | 110 | 90 | 3.8 | 12.88 | | High | High | High | High | High | High | High | High | High | High | High | 0 | 3.68E-09 | 3.68 |
| FALSE | High | SQSAAVTP | [R].SQSAA | 1xPhosphc | A0A1S6GW | 1.34E-06 | 0 | 1 | 3 | 10 | A0A1S6GW | Uncharacte | A0A1S6GW | 0 | 1832.859 | 0 | -0.1 | 0.094824 | 3.59 | 103.3 | 96.7 | 3.47 | 3.02 | | High | High | High | High | High | High | High | High | High | High | High | 0 | 1.41E-07 | 2.94 |
| FALSE | High | SQSADTPG | [R].SQSAD | 1xPhosphc | Q99JP6-2 1 | 0.008388 | 0.000736 | 1 | 2 | 2 | Q99JP6-2 | Isoform 2 c | Q99JP6-2 [ | 0 | 1554.7 | 0 | 0.39 | 0.014586 | 11.05 | 86.6 | 113.4 | 9.31 | 4.76 | | High | High | High | High | High | High | High | High | High | High | High | 0.000327 | 0.002577 | 1.86 |
| FALSE | High | SQSASSIDV | [R].SQSAS | 1xPhosphc | Q8CDA1 1x | 1.67E-05 | 0 | 1 | 3 | 2 | Q8CDA1 | Phosphatic | Q8CDA1 [9 | 0 | 3079.499 | 0 | 1.54 | 0.001841 | 18.18 | 49.1 | 150.9 | 6.7 | 13.24 | | High | High | High | High | High | High | High | High | High | High | High | 0 | 2.4E-06 | 3.7 |
| FALSE | High | SQSASYYP | [R].SQSAS | 1xCarbami | A2A690-2 | 0.000142 | 0 | 1 | 3 | 2 | A2A690-2 | Isoform 2 c | A2A690-2 | 0 | 2051.966 | 0 | 1.9 | 0.000233 | 2.7 | 41.3 | 158.7 | 9.58 | 7.04 | | High | High | High | High | High | High | High | High | High | High | High | 0 | 2.65E-05 | 2.48 |
| FALSE | High | SQSATQPA | [K].SQSAT | 1xPhosphc | Q8K1E6 1x | 0.000852 | 0 | 1 | 1 | 2 | Q8K1E6 | Alpha-keto | Q8K1E6 [2 | 0 | 1509.726 | 0 | 0.39 | 0.006374 | 0.76 | 86.8 | 113.2 | 5.02 | 11.14 | | High | High | High | High | High | High | High | High | High | High | High | 0.000183 | 0.001276 | 1.31 |
| FALSE | High | SQSDIFSHK | [R].SQSDIF | 1xPhosphc | Q8C0B6 1x | 0.028824 | 0.001829 | 1 | 4 | 1 | Q8C0B6 | Putative ur | Q8C0B6 [3 | 0 | 1937.005 | 0 | 0.66 | 0.012796 | 21.94 | 77.6 | 122.4 | 26.88 | 14.18 | | High | High | High | High | High | High | High | High | High | High | High | 0.000997 | 0.0103 | 1.36 |
| FALSE | High | SQSDLSSR | [R].SQSDLS | 1xPhosphc | Q8VE94 1x | 0.111965 | 0.009053 | 1 | 1 | 1 | Q8VE94 | Protein FAI | Q8VE94 [3 | 0 | 1188.546 | 0 | 0.64 | 0.001022 | 6.93 | 76 | 124 | 7.76 | 7.41 | | High | High | High | High | High | High | High | High | High | High | High | 0.004569 | 0.04954 | 1.63 |
| FALSE | High | SQSDVDFC | [K].SQSDV | 2xCarbami | Q8R5K2 1x | 1.84E-06 | 0 | 1 | 4 | 1 | Q8R5K2 | Ubiquitin c | Q8R5K2 [2 | 0 | 2605.084 | 0 | -0.2 | 0.21828 | 5.1 | 105.5 | 94.5 | 8.39 | 3.42 | | High | High | High | High | High | High | High | High | High | High | High | 0 | 2E-07 | 5.07 |
| FALSE | High | SQSEEQSE | [K].SQSEE | 1xPhosphc | Q5XG69 1x | 4.73E-06 | 0 | 1 | 1 | 5 | Q5XG69 | Soluble lan | Q5XG69 [6 | 0 | 3134.42 | 0 | -1.79 | 0.001075 | 2.78 | 155.5 | 44.5 | 8.33 | 18.22 | | High | High | High | High | High | High | High | High | High | High | High | 0 | 5.84E-07 | 3.96 |
| FALSE | High | SQSEEQSE | [K].SQSEE | 2xPhosphc | Q5XG69 2x | 0.000765 | 0 | 1 | 1 | 1 | Q5XG69 | Soluble lan | Q5XG69 [6 | 0 | 3214.387 | 0 | -1.27 | 0.000562 | 1.13 | 140.5 | 59.5 | 9.75 | 12.15 | | High | High | High | High | High | High | High | High | High | High | High | 0 | 0.000176 | 2.43 |
| FALSE | High | SQQQPTPV | [R].SQQQP | 1xPhosphc | Q5SVQ0 1x | 0.014667 | 0.001049 | 1 | 5 | 2 | Q5SVQ0 | Histone ac | Q5SVQ0 [8 | 0 | 1519.772 | 0 | -1.08 | 0.191514 | 1.35 | 135 | 65 | 58.69 | 46.87 | | High | High | High | High | High | High | High | High | High | High | High | 0.000555 | 0.004805 | 2.28 |
| FALSE | High | SQSPHYFQ | [R].SQSPH | 1xPhosphc | A2A6A1 1x | 0.001463 | 5.66E-05 | 1 | 1 | 2 | A2A6A1 | G patch do | A2A6A1 [1 | 0 | 1602.726 | 0 | 0.08 | 0.964236 | 22.29 | 98.2 | 101.8 | 11.33 | 16.23 | | High | High | High | High | High | High | High | High | High | High | High | 2.58E-05 | 0.000366 | 1.51 |
| FALSE | High | SQSPHYPM | [R].SQSPH | 1xPhosphc | Q8BIZ6 1xF | 0.002768 | 0.000109 | 1 | 1 | 5 | Q8BIZ6 | Smad nucl | Q8BIZ6 [89 | 0 | 1727.859 | 0 | -0.27 | 0.168442 | 12.85 | 110.4 | 89.6 | 8.84 | 15.03 | | High | High | High | High | High | High | High | High | High | High | High | 0.009336 | 0.1012 | 2.58 |
| FALSE | High | SQSPNWPI | [R].SQSPN | 2xPhosphc | Q8CBR1 2x | 0.020862 | 0.001472 | 1 | 3 | 1 | Q8CBR1 | Immunoglo | Q8CBR1 [4 | 0 | 2190.946 | 0 | -1.92 | 0.000627 | 1.11 | 160.1 | 39.9 | 6.7 | 21.51 | | High | High | High | High | High | High | High | High | High | High | High | 0.000801 | 0.007182 | 2.23 |
| FALSE | High | SQSYIPTSG | [R].SQSYIP | 1xCarbami | Q9ERS5 1x | 0.001644 | 5.66E-05 | 1 | 2 | 1 | Q9ERS5 | Pleckstrin h | Q9ERS5 [1 | 0 | 2684.388 | 0 | 0.39 | 0.015874 | 10.06 | 79 | 121 | 10.75 | 12.09 | | High | High | High | High | High | High | High | High | High | High | High | 2.58E-05 | 0.000419 | 1.71 |
| FALSE | High | SQTDVYNE | [K].SQTDV | 1xCarbami | Q9QYY8 1x | 0.001572 | 5.66E-05 | 1 | 2 | 1 | Q9QYY8 | Spastin OS | Q9QYY8 [2 | 0 | 2096.916 | 0 | 1.42 | 0.001283 | 5.68 | 54.6 | 145.4 | 5.8 | 13.48 | | High | High | High | High | High | High | High | High | High | High | High | 2.58E-05 | 0.000395 | 3.12 |
| FALSE | High | SQTEPGSSI | [R].SQTEP | 1xPhosphc | Q3UGN6 1 | 1.89E-06 | 0 | 1 | 13 | 2 | Q3UGN6 | Putative ur | Q3UGN6 [ | 0 | 1951.932 | 0 | 0.87 | 0.002016 | 10.82 | 69.5 | 130.5 | 10.63 | 5.62 | | High | High | High | High | High | High | High | High | High | High | High | 0 | 2.08E-07 | 2.66 |
| FALSE | High | SQTPPGVA | [R].SQTPP | 1xPhosphc | Q925J9 1x | 0.00025 | 0 | 1 | 2 | 2 | Q925J9 | Mediator c | Q925J9 [10 | 0 | 1928.066 | 0 | 1.23 | 0.000635 | 12.18 | 58.4 | 141.6 | 3.79 | 8.29 | | High | High | High | High | High | High | High | High | High | High | High | 0 | 5.02E-05 | 2.57 |
| FALSE | High | SQTPPGVA | [R].SQTPP | 2xPhosphc | Q925J9 2x | 0.003167 | 0.00016 | 1 | 2 | 1 | Q925J9 | Mediator c | Q925J9 [10 | 0 | 2008.032 | 0 | 1.04 | 0.000605 | 3.91 | 64.2 | 135.8 | 6.25 | 8.05 | | High | High | High | High | High | High | High | High | High | High | High | 7.19E-05 | 0.000866 | 2.25 |
| FALSE | High | SQTPQLHF | [K].SQTPQ | 1xPhosphc | P61406 1x | 0.005182 | 0.000551 | 1 | 1 | 1 | P61406 | Telomeras | P61406 [46 | 0 | 3041.373 | 0 | -0.04 | 0.459598 | 1.78 | 102.7 | 97.3 | 10.62 | 14.24 | | High | High | High | High | High | High | High | High | High | High | High | 0.000246 | 0.001508 | 2.58 |
| FALSE | High | SQTPSPSTL | [R].SQTPSP | 1xPhosphc | A0A1W2P7 | 0.007881 | 0.000736 | 1 | 7 | 1 | A0A1W2P7 | Ral GTPase | A0A1W2P7 | 0 | 1469.72 | 0 | 1.42 | 0.000357 | 7.48 | 56.3 | 143.7 | 8.59 | 4.91 | | High | High | High | High | High | High | High | High | High | High | High | 0.000327 | 0.002393 | 1.98 |
| FALSE | High | SQTSFDNL | [K].SQTSFD | 1xPhosphc | A2ATK9 1x | 0.022973 | 0.001578 | 1 | 3 | 3 | A2ATK9 | Family with | A2ATK9 [4 | 0 | 2191.105 | 0 | 1.3 | 0.000246 | 2.87 | 57.7 | 142.3 | 9.46 | 1.51 | | High | High | High | High | High | High | High | High | High | High | High | 0.009697 | 0.1047 | 2.57 |
| FALSE | High | SQTTSLEN | [R].SQTTSL | 1xPhosphc | Q68FD7 1x | 8.25E-05 | 0 | 1 | 2 | 4 | Q68FD7 | Folliculin-ir | Q68FD7 [2 | 0 | 1744.847 | 0 | -0.19 | 0.11987 | 14.98 | 110.8 | 89.2 | 9.62 | 9.53 | | High | High | High | High | High | High | High | High | High | High | High | 0 | 1.45E-05 | 3.29 |
| FALSE | High | SQVEDAED | [R].SQVEDA | 1xDeamida | E9QPB3 1x | 0.053023 | 0.00384 | 1 | 4 | 5 | E9QPB3 | WD repeat | E9QPB3 [8 | 1 | 4965.27 | 0 | -1.22 | 0.0067 | 5.45 | 142.4 | 57.6 | 17.57 | 26.91 | | High | High | High | High | High | High | High | High | High | High | High | 0.001999 | 0.02053 | 4.19 |
| FALSE | High | SQVEDPLPI | [K].SQVEDI | 1xPhosphc | Q9WVS8 1 | 0.000239 | 0 | 1 | 5 | 1 | Q9WVS8 | Mitogen-ac | Q9WVS8 [ | 0 | 2236.167 | 0 | -0.65 | 0.030198 | 10.68 | 122.1 | 77.9 | 15.66 | 8.26 | | High | High | High | High | High | High | High | High | High | High | High | 0 | 4.79E-05 | 2.49 |
| FALSE | High | SQVGGQV | [R].SQVGG | 1xPhosphc | P19001 1x | 0.000418 | 0 | 1 | 1 | 1 | P19001 | Keratin, typ | P19001 [23 | 0 | 2610.343 | 0 | 2.32 | 0.000154 | 8.05 | 33.5 | 166.5 | 13.26 | 1.23 | | High | High | High | High | High | High | High | High | High | High | High | 0 | 8.96E-05 | 2.79 |
| FALSE | High | SQVGPEDP | [K].SQVGP | 1xPhosphc | Q80UU6 1x | 0.004613 | 0.000456 | 1 | 3 | 1 | Q80UU6 | Opioid gro | Q80UU6 [5 | 0 | 2577.26 | 0 | 0.41 | 0.007885 | 12.02 | 79.3 | 120.7 | 12.39 | 6.32 | | High | High | High | High | High | High | High | High | High | High | High | 0.000204 | 0.001318 | 2.87 |
| FALSE | High | SQSWLSNI | [R].SQSWL | 1xPhosphc | E9QPA1 1x | 0.000745 | 0 | 1 | 4 | 1 | E9QPA1 | RAB6A-GEI | E9QPA1 [1 | 0 | 1791.874 | 0 | 0.23 | 0.236757 | 18.28 | 94.8 | 105.2 | 6.14 | 11.34 | | High | High | High | High | High | High | High | High | High | High | High | 0 | 0.000171 | 2.42 |
| FALSE | High | SQVPPFDSI | [R].SQVPPI | 2xPhosphc | E9QKL0 2x | 0.0008 | 0 | 1 | 2 | 1 | E9QKL0 | Helicase AI | E9QKL0 [1 | 0 | 3253.402 | 0 | -0.13 | 0.103725 | 8.94 | 109.4 | 90.6 | 9.42 | 12.85 | | High | High | High | High | High | High | High | High | High | High | High | 0 | 0.000185 | 3.1 |
| FALSE | High | SREDLSAQI | [K].SREDLS | 1xPhosphc | P39447 1x | 0.002022 | 5.66E-05 | 1 | 3 | 1 | P39447 | Tight junct | P39447 [61 | 1 | 1767.884 | 0 | -0.66 | 0.437802 | 5.01 | 122.6 | 77.4 | 50.5 | 57.85 | | High | High | High | High | High | High | High | High | High | High | High | 2.58E-05 | 0.000525 | 2.54 |
| FALSE | High | SREDLSAQI | [K].SREDLS | 1xPhosphc | P39447 1x | 0.005615 | 0.00062 | 1 | 3 | 3 | P39447 | Tight junct | P39447 [61 | 1 | 1997.047 | 0 | -0.5 | 0.002487 | 4.48 | 119.9 | 80.1 | 4.3 | 12.9 | | High | High | High | High | High | High | High | High | High | High | High | 0.000275 | 0.001642 | 2.78 |
| FALSE | High | SREDSPELN | [R].SREDSF | 1xPhosphc | P55200 1x | 9.36E-05 | 0 | 1 | 3 | 2 | P55200 | Histone-lys | P55200 [18 | 1 | 2317.066 | 0 | -0.66 | 0.000729 | 3.71 | 118.9 | 81.1 | 4.33 | 7.86 | | High | High | High | High | High | High | High | High | High | High | High | 0 | 4.42E-05 | 4.31 |
| FALSE | High | SRESANQP | [K].SRESAN | 1xPhosphc | Q80W49-2 | 8.18E-05 | 0 | 1 | 2 | 1 | Q80W49-2 | Isoform 2 c | Q80W49-2 | 1 | 2583.298 | 0 | 2.16 | 0.000325 | 6.85 | 34.4 | 165.6 | 14.33 | 9.73 | | High | High | High | High | High | High | High | High | High | High | High | 0 | 1.43E-05 | 2.77 |
| FALSE | High | SRGSTNLFS | [R].SRGSTN | 1xPhosphc | Q03141 1x | 0.050826 | 0.003676 | 1 | 5 | 1 | Q03141 | MAP/micrc | Q03141 [5 | 1 | 1634.867 | 0 | -0.95 | 0.002004 | 3.85 | 133.4 | 66.6 | 8.83 | 19.74 | | High | High | High | High | High | High | High | High | High | High | High | 0.001916 | 0.01963 | 1.99 |
| FALSE | High | SRLNVEWL | [K].SRLNVE | 1xPhosphc | F2Z446 1xF | 0.000466 | 0 | 1 | 1 | 1 | F2Z446 | Vacuolar A | F2Z446 [6- | 1 | 3049.571 | 0 | 4.37 | 0.004123 | 64.07 | 7.3 | 192.7 | 43 | 26.92 | | High | High | High | High | High | High | High | High | High | High | High | 0 | 0.000102 | 3.43 |
| FALSE | High | SRLSAIEIDI | [R].SRLSAII | 1xPhosphc | Q9ET43 1x | 0.00051 | 0 | 1 | 1 | 1 | Q9ET43 | Claudin-12 | Q9ET43 [2 | 1 | 2133.115 | 0 | -0.99 | 0.001512 | 1.37 | 132.8 | 67.2 | 7.55 | 6.01 | | High | High | High | High | High | High | High | High | High | High | High | 0 | 0.000112 | 4.57 |
| FALSE | High | SRLSLQESC | [R].SRLSLQ | 1xPhosphc | Q8BUL6 1x | 1.86E-05 | 0 | 1 | 2 | 2 | Q8BUL6 | Pleckstrin h | Q8BUL6 [3 | 1 | 1924.067 | 0 | -0.42 | 0.005792 | 7.14 | 114.4 | 85.6 | 3.53 | 9.54 | | High | High | High | High | High | High | High | High | High | High | High | 0 | 1.12E-05 | 2.21 |
| FALSE | High | SRLSSSSSN | [K].SRLSSS | 2xPhosphc | E9Q616 2x | 0.047086 | 0.00354 | 1 | 4 | 3 | E9Q616 | AHNAK nuc | E9Q616 [5 | 1 | 2030.92 | 0 | 1.68 | 0.000125 | 2.87 | 47.6 | 152.4 | 7.97 | 1.81 | | High | High | High | High | High | High | High | High | High | High | High | 0.004543 | 0.04945 | 3.23 |
| FALSE | High | SRLTPTTPE | [R].SRLTPT | 1xPhosphc | Q64337 1x | 0.011652 | 0.000889 | 1 | 3 | 2 | Q64337 | Sequestoso | Q64337 [2 | 1 | 2432.196 | 0 | 1.01 | 0.000565 | 3.51 | 65.2 | 134.8 | 10.22 | 3.64 | | High | High | High | High | High | High | High | High | High | High | High | 0.002133 | 0.02223 | 2.81 |
| FALSE | High | SRPALSPLG | [R].SRPALS | 1xCarbami | E9PYL2 1xF | 1.39E-07 | 0 | 1 | 2 | 1 | E9PYL2 | Proline-ricl | E9PYL2 [11 | 0 | 2841.359 | 0 | 0.49 | 0.024133 | 9.57 | 83.3 | 116.7 | 8.03 | 9.97 | | High | High | High | High | High | High | High | High | High | High | High | 0 | 1.11E-08 | 3.42 |
| FALSE | High | SRPLNAVS | [R].SRPLNA | 1xPhosphc | Q9JKB3 1x | 0.000553 | 0 | 1 | 5 | 2 | Q9JKB3 | Y-box-bind | Q9JKB3 [3 | 0 | 1580.8 | 0 | 0.93 | 0.034264 | 18.93 | 67.1 | 132.9 | 13.13 | 27.67 | | High | High | High | High | High | High | High | High | High | High | High | 0 | 0.000122 | 3.04 |
| FALSE | High | SRPLNAVS | [R].SRPLNA | 1xPhosphc | Q9JKB3 1x | 0.000941 | 0 | 1 | 5 | 1 | Q9JKB3 | Y-box-bind | Q9JKB3 [3 | 0 | 1809.962 | 0 | 1.56 | 0.003824 | 31.06 | 49.5 | 150.5 | 12.73 | 16.15 | | High | High | High | High | High | High | High | High | High | High | High | 0 | 0.000223 | 1.04 |
| FALSE | High | SRDVPESP | [K].SRDVPE | 1xCarbami | Q9D516 1x | 0.000225 | 0 | 1 | 3 | 2 | Q9D516 | Coiled-coil | Q9D516 [3 | 1 | 2684.182 | 0 | 0.61 | 0.002246 | 8.12 | 79.8 | 120.2 | 7.57 | 7.39 | | High | High | High | High | High | High | High | High | High | High | High | 0 | 8.67E-05 | 3.38 |
| FALSE | High | SQSWDDR | [R].SQSWD | 1xPhosphc | Q99MR1 1 | 0.064323 | 0.004664 | 1 | 2 | 2 | Q99MR1 | GRB10-inte | Q99MR1 [ | 0 | 1202.504 | 0 | 0.55 | 0.003294 | 6.94 | 79.7 | 120.3 | 6.04 | 5.78 | | High | High | High | High | High | High | High | High | High | High | High | 0.004032 | 0.04279 | 1.83 |
| FALSE | High | SQSVPLTVI | [R].SQSVPL | 1xPhosphc | F8VPJ6 1xF | 0.007471 | 0.00062 | 1 | 2 | 1 | F8VPJ6 | Regulatory | F8VPJ6 [12 | 0 | 2729.417 | 0 | -0.8 | 0.017258 | 8.9 | 119.8 | 80.2 | 14.08 | 17.98 | | High | High | High | High | High | High | High | High | High | High | High | 0.000275 | 0.002265 | 3.1 |
| FALSE | High | SQSVPGAV | [K].SQSVP | 1xPhosphc | E9Q0S6 1x | 1.28E-07 | 0 | 1 | 6 | 1 | E9Q0S6 | Tensin 1 O | E9Q0S6 [8 | 0 | 2675.339 | 0 | 0.35 | 0.44256 | 14.96 | 90.6 | 109.4 | 16.12 | 22.6 | | High | High | High | High | High | High | High | High | High | High | High | 0 | 1.02E-08 | 4.45 |
| FALSE | High | SQSPSPPPL | [K].SQSPSF | 1xPhosphc | F6RJ39 1xF | 1.85E-05 | 0 | 2 | 5 | 7 | F6RJ39; Q8 | Apoptotic | F6RJ39 [33 | 0 | 2159.104 | 0 | 0.05 | 0.771516 | 5.67 | 98.4 | 101.6 | 4.67 | 12.2 | Shared | High | High | High | High | High | High | High | High | High | High | High | 0 | 2.7E-06 | 2.7 |
| FALSE | High | SQSPSPPPL | [K].SQSPSF | 2xPhosphc | F6RJ39 2xF | 1.43E-05 | 0 | 2 | 5 | 9 | F6RJ39; Q8 | Apoptotic | F6RJ39 [33 | 0 | 2239.07 | 0 | 0.25 | 0.172198 | 2.55 | 91.5 | 108.5 | 6.59 | 14.05 | Shared | High | High | High | High | High | High | High | High | High | High | High | 0 | 0.000171 | 3.57 |
| FALSE | High | SQSPVHLR | [R].SQSPVI | 1xPhosphc | A2ADB0 1x | 0.058173 | 0.004227 | 1 | 5 | 2 | A2ADB0 | Msx2-inter | A2ADB0 [7 | 0 | 1232.635 | 0 | 0.65 | 0.001184 | 2.67 | 78.4 | 121.6 | 3.25 | 7.52 | | High | High | High | High | High | High | High | High | High | High | High | 0.002159 | 0.02288 | 2 |
| FALSE | High | SQSPVK | [R].SQSPVI | 1xPhosphc | Q6NZE7-2 | 0.081085 | 0.006219 | 1 | 4 | 2 | Q6NZE7-2 | Isoform 2 c | Q6NZE7-2 | 0 | 1183.649 | 0 | -1.48 | 0.000231 | 1.33 | 148.2 | 51.8 | 7.62 | 18.48 | | High | High | High | High | High | High | High | High | High | High | High | 0.0032 | 0.0338 | 1.51 |
| FALSE | High | SQSQASLT | [R].SQSQA | 1xPhosphc | D3YUW7 1 | 2.5E-07 | 0 | 1 | 8 | 2 | D3YUW7 | Cingulin OS | D3YUW7 [ | 0 | 2207.037 | 0 | -0.5 | 0.08949 | 7.38 | 113.3 | 86.7 | 4.2 | 24.27 | | High | High | High | High | High | High | High | High | High | High | High | 0 | 2.14E-08 | 4.34 |
| FALSE | High | SQSQASLT | [R].SQSQA | 2xPhosphc | D3YUW7 2 | 0.000144 | 0 | 1 | 8 | 2 | D3YUW7 | Cingulin OS | D3YUW7 [ | 0 | 2287.003 | 0 | 0.8 | 0.001813 | 3.76 | 71.9 | 128.1 | 4.98 | 6.79 | | High | High | High | High | High | High | High | High | High | High | High | 0 | 2.71E-05 | 3.57 |
| FALSE | High | SQSQSSPS | [R].SQSQS | 1xPhosphc | P97868 1x | 0.000337 | 0 | 1 | 3 | 1 | P97868 | E3 ubiquiti | P97868 [16 | 0 | 1642.764 | 0 | -0.76 | 0.011718 | 12.93 | 127.7 | 72.3 | 9.03 | 21.29 | | High | High | High | High | High | High | High | High | High | High | High | 0 | 7.06E-05 | 2.23 |
| FALSE | High | SQSRSHSPI | [R].SQSRSH | 3xPhosphc | Q3TWW8 | 0.002272 | 0.000109 | 1 | 2 | 2 | Q3TWW8 | Serine/argi | Q3TWW8 | 1 | 2044.886 | 0 | -0.41 | 0.013974 | 3.61 | 111 | 89 | 11.23 | 5.18 | | High | High | High | High | High | High | High | High | High | High | High | 4.94E-05 | 0.0006 | 3 |
| FALSE | High | SQSSDLLEF | [K].SQSSDI | 1xPhosphc | Q9D279 1x | 0.020862 | 0.001472 | 1 | 1 | 4 | Q9D279 | Mitotic inte | Q9D279 [5 | 0 | 1343.641 | 0 | 3.6 | 0.000361 | 23.2 | 13.9 | 186.1 | 17.79 | 9.28 | | High | High | High | High | High | High | High | High | High | High | High | 0.002662 | 0.02831 | 2.38 |
| FALSE | High | SQSSDTEQ | [R].SQSSDT | 1xPhosphc | Q3TSY9 1x | 4.15E-11 | 0 | 1 | 6 | 29 | Q3TSY9 | Putative ur | Q3TSY9 [4 | 0 | 2188.017 | 0 | -0.12 | 0.427641 | 1.9 | 104.2 | 95.8 | 1.97 | 9.16 | | High | High | High | High | High | High | High | High | High | High | High | 0 | 1.22E-12 | 4.01 |
| FALSE | High | SQSSDTEQ | [R].SQSSDT | 2xPhosphc | Q3TSY9 2x | 8.92E-07 | 0 | 1 | 6 | 5 | Q3TSY9 | Putative ur | Q3TSY9 [4 | 0 | 2267.983 | 0 | 0.33 | 0.008175 | 10.46 | 87.8 | 112.2 | 3.38 | 8.03 | | High | High | High | High | High | High | High | High | High | High | High | 0 | 8.9E-08 | 4.53 |
| FALSE | High | SQSSEGVS | [R].SQSSE | 1xPhosphc | Q80TU3 1x | 1.52E-10 | 0 | 1 | 6 | 8 | Q80TU3 | MKIAA068 | Q80TU3 [8 | 0 | 3065.411 | 0 | -0.48 | 0.001455 | 2.35 | 115 | 85 | 3.4 | 5.53 | | High | High | High | High | High | High | High | High | High | High | High | 0 | 1.94E-08 | 6.49 |
| FALSE | High | SQSSEGVS | [R].SQSSE | 1xDeamida | Q80TU3 1x | 2.41E-10 | 0 | 1 | 6 | 8 | Q80TU3 | MKIAA068 | Q80TU3 [8 | 0 | 3066.395 | 0 | 0.05 | 0.729754 | 10.39 | 98.4 | 101.6 | 8.69 | 2.02 | | High | High | High | High | High | High | High | High | High | High | High | 0 | 1.16E-09 | 7 |
| FALSE | High | SQSSEGVS | [R].SQSSE | 2xPhosphc | Q80TU3 2x | 4.47E-05 | 0 | 1 | 6 | 5 | Q80TU3 | MKIAA068 | Q80TU3 [8 | 0 | 3145.377 | 0 | -0.48 | 0.005697 | 7.97 | 118 | 82 | 2.97 | 10.36 | | High | High | High | High | High | High | High | High | High | High | High | 0 | 7.3E-06 | 3.92 |
| FALSE | High | SQSSGSSA | [K].SQSSGS | 1xPhosphc | P81122 1x | 0.000211 | 0 | 1 | 2 | 3 | P81122 | Insulin rece | P81122 [3 | 0 | 2034.981 | 0 | -0.66 | 0.015574 | 2.61 | 122.9 | 77.1 | 8.24 | 19.3 | | High | High | High | High | High | High | High | High | High | High | High | 0 | 4.15E-05 | 1.84 |
| FALSE | High | SQSSHSYDI | [R].SQSSHS | 1xPhosphc | E9Q8Z8 1x | 1.04E-05 | 0 | 1 | 19 | 3 | E9Q8Z8 | Catenin de | E9Q8Z8 [8 | 0 | 2230.023 | 0 | -0.37 | 0.17897 | 7.77 | 111.6 | 88.4 | 12.67 | 19.81 | | High | High | High | High | High | High | High | High | High | High | High | 0 | 5.14E-05 | 3.66 |
| FALSE | High | SQSSHSYDI | [R].SQSSHS | 2xPhosphc | E9Q8Z8 2x | 0.000148 | 0 | 1 | 19 | 3 | E9Q8Z8 | Catenin de | E9Q8Z8 [8 | 0 | 2309.989 | 0 | -0.16 | 0.010234 | 4.57 | 107.1 | 92.9 | 5.71 | 4.77 | | High | High | High | High | High | High | High | High | High | High | High | 0 | 2.8E-05 | 3.72 |
| FALSE | High | SQSSSLLDN | [R].SQSSSL | 1xPhosphc | Q3U1G4 1x | 0.000718 | 0 | 1 | 4 | 6 | Q3U1G4 | Putative ur | Q3U1G4 [4 | 0 | 2624.142 | 0 | 0.11 | 0.247211 | 11.44 | 95.9 | 104.1 | 9.17 | 15.81 | | High | High | High | High | High | High | High | High | High | High | High | 2.58E-05 | 0.000366 | 3.57 |
| FALSE | High | SQSSSQFR | [R].SQSSSC | 1xPhosphc | Q61151 1x | 0.035181 | 0.002269 | 1 | 3 | 5 | Q61151 | Serine/thre | Q61151 [3 | 0 | 1235.562 | 0 | 0.95 | 0.002018 | 4.63 | 68.4 | 131.6 | 3.02 | 11.32 | | High | High | High | High | High | High | High | High | High | High | High | 0.004961 | 0.0539 | 1.41 |
| FALSE | High | SQSSTPK | [R].SQSSTF | 1xPhosphc | P98195 1x | 0.044358 | 0.003264 | 1 | 3 | 1 | P98195 | Probable p | P98195 [52 | 0 | 1272.66 | 0 | 0.88 | 0.000834 | 14.78 | 70.1 | 129.9 | 9.41 | 5.17 | | High | High | High | High | High | High | High | High | High | High | High | 0.00168 | 0.01684 | 1.42 |
| FALSE | High | SQSSTSLES | [R].SQSSTS | 1xPhosphc | Q9Z131 1x | 3.81E-06 | 0 | 1 | 3 | 3 | Q9Z131 | SH3 domai | Q9Z131 [4 | 0 | 1932.911 | 0 | 1.11 | 0.001896 | 13.29 | 55 | 145 | 16.17 | 10.28 | | High | High | High | High | High | High | High | High | High | High | High | 0 | 2.67E-06 | 3.57 |
| FALSE | High | SQSSTSLES | [R].SQSSTS | 2xPhosphc | Q9Z131 2x | 0.000822 | 0 | 1 | 3 | 2 | Q9Z131 | SH3 domai | Q9Z131 [4 | 0 | 2012.878 | 0 | 2.03 | 0.000282 | 12.63 | 41.5 | 158.5 | 17.01 | 8.22 | | High | High | High | High | High | High | High | High | High | High | High | 0 | 0.00025 | 3.18 |
| FALSE | High | SQSSVSTA | [K].SQSSVS | 1xPhosphc | Q01320 1x | 0.003622 | 0.00021 | 2 | 4 | 3 | Q01320; Q | DNA topois | Q01320 [1 | 0 | 1590.814 | 0 | 0.13 | 0.144638 | 18.76 | 89.4 | 110.6 | 9.08 | 15.47 | Shared | High | High | High | High | High | High | High | High | High | High | High | 0.006945 | 0.07453 | 1.99 |
| FALSE | High | SQSTLDLLH | [K].SQSTLD | 1xPhosphc | H9LAN6 1x | 9.36E-05 | 0 | 1 | 6 | 3 | H9LAN6 | Histone-lys | H9LAN6 [4 | 0 | 2941.429 | 0 | -0.5 | 0.002105 | 1.37 | 116.7 | 83.3 | 7.45 | 5.96 | | High | High | High | High | High | High | High | High | High | High | High | 0 | 1.66E-05 | 3.61 |
| FALSE | High | SQSTLDLLH | [K].SQSTLD | 2xPhosphc | H9LAN6 2x | 0.000103 | 0 | 1 | 6 | 2 | H9LAN6 | Histone-lys | H9LAN6 [4 | 0 | 3021.396 | 0 | -0.58 | 0.001524 | 2.54 | 120.5 | 79.5 | 6.9 | 8.2 | | High | High | High | High | High | High | High | High | High | High | High | 0 | 1.86E-05 | 3.48 |
| FALSE | High | SQSTSEQEI | [R].SQSTSE | 1xPhosphc | Q6PAJ1 1x | 0.012288 | 0.000889 | 1 | 2 | 2 | Q6PAJ1 | Breakpoint | Q6PAJ1 [3 | 0 | 1561.751 | 0 | 1.82 | 0.00114 | 4.05 | 42.5 | 157.5 | 2.69 | 13.21 | | High | High | High | High | High | High | High | High | High | High | High | 0.000638 | 0.005699 | 2.31 |
| FALSE | High | SQSTTFNPI | [R].SQSTTF | 1xPhosphc | F6YZ95 1xF | 3.39E-06 | 0 | 2 | 5 | 3 | F6YZ95; Q8 | Liprin-beta | F6YZ95 [15 | 0 | 2498.12 | 0 | 0.86 | 0.001839 | 14.54 | 71.4 | 128.6 | 15.08 | 3.11 | Shared | High | High | High | High | High | High | High | High | High | High | High | 0 | 4.02E-07 | 3.69 |
| FALSE | High | SQSTTFNPI | [R].SQSTTF | 1xPhosphc | F6YZ95 1xF | 0.00015 | 0 | 2 | 5 | 4 | F6YZ95; Q8 | Liprin-beta | F6YZ95 [15 | 0 | 2514.115 | 0 | 0.93 | 0.004429 | 12.03 | 68.8 | 131.2 | 7.88 | 16.14 | Shared | High | High | High | High | High | High | High | High | High | High | High | 2.58E-05 | 0.000534 | 2.29 |
| FALSE | High | SQSVDITAI | [K].SQSVDI | 1xPhosphc | Q4KMM3 | 3.12E-05 | 0 | 1 | 2 | 1 | Q4KMM3 | Oxidation r | Q4KMM3 [ | 0 | 2354.216 | 0 | 2.28 | 0.011726 | 36.64 | 39.7 | 160.3 | 12.96 | 32.23 | | High | High | High | High | High | High | High | High | High | High | High | 0 | 4.86E-06 | 3.21 |
| FALSE | High | SQMSTPTR | [R].SQMST | 1xPhosphc | A0A0J9YU | 0.0016 | 5.66E-05 | 1 | 4 | 1 | A0A0J9YU | Mediator c | A0A0J9YU | 1 | 3353.51 | 0 | -0.11 | 0.589185 | 12.69 | 104.6 | 95.4 | 14.26 | 8.97 | | High | High | High | High | High | High | High | High | High | High | High | 2.58E-05 | 0.000404 | 2.91 |
| FALSE | High | SQLSPVK | [R].SQLSPV | 1xPhosphc | Q91YJ5 1x | 0.006478 | 0.00062 | 1 | 4 | 1 | Q91YJ5 | Translation | Q91YJ5 [8 | 0 | 1296.733 | 0 | -1.53 | 0.000809 | 1.87 | 151.1 | 48.9 | 8.94 | 7.49 | | High | High | High | High | High | High | High | High | High | High | High | 0.000275 | 0.001933 | 1.84 |
| FALSE | High | SQLQALHI | [R].SQLQAI | 1xPhosphc | F8VPK5 1x | 0.071662 | 0.005278 | 1 | 4 | 2 | F8VPK5 | Rho-associ | F8VPK5 [1 | 0 | 3582.589 | 0 | 1.26 | 0.010953 | 27.44 | 61.3 | 138.7 | 10.22 | 21.57 | | High | High | High | High | High | High | High | High | High | High | High | 0.008008 | 0.0867 | 2.92 |
| FALSE | High | SQLPTLEQI | [R].SQLPTL | 2xPhosphc | P46938 2x | 0.014929 | 0.001049 | 1 | 3 | 1 | P46938 | Transcripti | P46938 [3 | 0 | 3104.385 | 0 | 0.14 | 0.301242 | 38.83 | 95.3 | 104.7 | 22.78 | 19.48 | | High | High | High | High | High | High | High | High | High | High | High | 0.000555 | 0.004922 | 2.93 |
| FALSE | High | SPTTVLFTS | [R].SPTTVL | 2xPhosphc | F8VPJ6 2xF | 8.58E-06 | 0 | 1 | 1 | 2 | F8VPJ6 | Regulatory | F8VPJ6 [44 | 0 | 1996.021 | 0 | -0.49 | 0.002019 | 2.19 | 117 | 83 | 2.62 | 6.81 | | High | High | High | High | High | High | High | High | High | High | High | 0 | 1.14E-06 | 3.31 |
| FALSE | High | SPTVPTPG | [K].SPTVPT | 1xPhosphc | A0A0A0M | 0.028327 | 0.001791 | 1 | 4 | 1 | A0A0A0M | Calmodulir | A0A0A0M | 0 | 1522.828 | 0 | 1.12 | 0.000486 | 9.69 | 63 | 137 | 6.79 | 3.62 | | High | High | High | High | High | High | High | High | High | High | High | 0.000968 | 0.01006 | 1.61 |
| FALSE | High | SPVDESGD | [R].SPVDES | 1xDeamida | Q9ET54 1x | 0.015603 | 0.001124 | 1 | 8 | 2 | Q9ET54 | Palladin OS | Q9ET54 [1 | 0 | 3588.638 | 0 | -0.52 | 0.020754 | 9.96 | 126.3 | 73.7 | 15.03 | 19.08 | | High | High | High | High | High | High | High | High | High | High | High | 0.001308 | 0.01392 | 3.33 |
| FALSE | High | SPVDSPVP | [R].SPVDSF | 1xPhosphc | O08582 1x | 1.15E-05 | 0 | 1 | 1 | 4 | O08582 | GTP-bindir | O08582 [8 | 0 | 2563.21 | 0 | -0.22 | 0.004397 | 4.75 | 107.4 | 92.6 | 6.56 | 8.87 | | High | High | High | High | High | High | High | High | High | High | High | 0 | 3.15E-06 | 4.17 |
| FALSE | High | SPVDSPVP | [R].SPVDSF | 1xPhosphc | O08582 1x | 2.45E-06 | 0 | 1 | 1 | 8 | O08582 | GTP-bindir | O08582 [8 | 0 | 2579.205 | 0 | -0.04 | 0.74147 | 5.16 | 101.8 | 98.2 | 3.11 | 19.59 | | High | High | High | High | High | High | High | High | High | High | High | 0 | 2.77E-07 | 3.66 |
| FALSE | High | SPVDSPVP | [R].SPVDSF | 2xPhosphc | O08582 2x | 0.001349 | 0 | 1 | 1 | 1 | O08582 | GTP-bindir | O08582 [8 | 0 | 2643.176 | 0 | -0.4 | 0.014073 | 5.22 | 116.5 | 83.5 | 7.53 | 5.82 | | High | High | High | High | High | High | High | High | High | High | High | 0 | 0.000335 | 3.1 |
| FALSE | High | SPVDSPVP | [R].SPVDSF | 2xPhosphc | O08582 2x | 0.007538 | 0.00062 | 1 | 1 | 1 | O08582 | GTP-bindir | O08582 [8 | 0 | 2659.171 | 0 | -0.26 | 0.85344 | 17.73 | 104.5 | 95.5 | 18.52 | 25.62 | | High | High | High | High | High | High | High | High | High | High | High | 0.000275 | 0.002278 | 2.55 |
| FALSE | High | SPVFSDED | [K].SPVFSD | 1xPhosphc | Q810V0 1x | 0.000375 | 0 | 1 | 3 | 4 | Q810V0 | U3 small nu | Q810V0 [1 | 0 | 2424.126 | 0 | -0.37 | 0.044401 | 4.41 | 114 | 86 | 9.47 | 14.38 | | High | High | High | High | High | High | High | High | High | High | High | 0 | 7.94E-05 | 3.64 |
| FALSE | High | SPVFSDED | [K].SPVFSD | 2xPhosphc | Q810V0 2x | 1.07E-06 | 0 | 1 | 3 | 6 | Q810V0 | U3 small nu | Q810V0 [1 | 0 | 2504.092 | 0 | -0.41 | 0.052638 | 2.14 | 113.4 | 86.6 | 2.64 | 16.13 | | High | High | High | High | High | High | High | High | High | High | High | 0 | 6.25E-06 | 4.23 |
| FALSE | High | SPVGDTGL | [R].SPVGD | 1xPhosphc | B9EKJ7 1xF | 0.009756 | 0.000736 | 1 | 3 | 1 | B9EKJ7 | Tankyrase | B9EKJ7 [76 | 0 | 1468.781 | 0 | 0.2 | 0.096574 | 3.91 | 93 | 107 | 7.71 | 6.73 | | High | High | High | High | High | High | High | High | High | High | High | 0.000344 | 0.00306 | 1.02 |
| FALSE | High | SPVGKSPP | [K].SPVGKS | 1xPhosphc | Q569Z6 1x | 0.067059 | 0.004796 | 1 | 4 | 1 | Q569Z6 | Thyroid ho | Q569Z6 [3 | 1 | 2573.358 | 0 | -0.17 | 0.716448 | 19.36 | 105.8 | 94.2 | 15.29 | 19.2 | | High | High | High | High | High | High | High | High | High | High | High | 0.002485 | 0.02715 | 2.27 |
| FALSE | High | SPVLFPSPN | [R].SPVLFP | 1xPhosphc | B2RPS6 1x | 7.54E-05 | 0 | 1 | 3 | 1 | B2RPS6 | Putative ur | B2RPS6 [4 | 0 | 2772.406 | 0 | 0.82 | 0.025422 | 33.55 | 63.2 | 136.8 | 23.73 | 17.75 | | High | High | High | High | High | High | High | High | High | High | High | 0 | 1.31E-05 | 3.43 |
| FALSE | High | SPVNGEAG | [K].SPVNGI | 1xPhosphc | Q8BG15 1x | 2.56E-05 | 0 | 1 | 4 | 2 | Q8BG15 | CTD small p | Q8BG15 [1 | 0 | 2500.194 | 0 | 0.12 | 0.892429 | 14.4 | 100.3 | 99.7 | 8.48 | 15.09 | | High | High | High | High | High | High | High | High | High | High | High | 0 | 3.88E-06 | 2.79 |
| FALSE | High | SPTTVLFTS | [R].SPTTVL | 1xPhosphc | F8VPJ6 1xF | 0.000693 | 0 | 1 | 1 | 3 | F8VPJ6 | Regulatory | F8VPJ6 [44 | 0 | 1916.055 | 0 | -0.47 | 0.000611 | 2.1 | 115.3 | 84.7 | 3.33 | 5.08 | | High | High | High | High | High | High | High | High | High | High | High | 0 | 0.000158 | 3.5 |
| FALSE | High | SPVNGEAG | [K].SPVNGI | 1xPhosphc | Q8BG15 1x | 9.54E-05 | 0 | 1 | 4 | 3 | Q8BG15 | CTD small p | Q8BG15 [1 | 0 | 2516.189 | 0 | -0.07 | 0.632822 | 12 | 100.3 | 99.7 | 6.07 | 10.56 | | High | High | High | High | High | High | High | High | High | High | High | 4.94E-05 | 0.000711 | 2.43 |
| FALSE | High | SPVPSAFSD | [R].SPVPSA | 1xPhosphc | Q8BTI8 1x | 0.000786 | 0 | 1 | 3 | 3 | Q8BTI8 | Serine/argi | Q8BTI8 [24 | 0 | 1586.741 | 0 | -0.5 | 0.001972 | 4.52 | 116.9 | 83.1 | 2.96 | 4.74 | | High | High | High | High | High | High | High | High | High | High | High | 2.58E-05 | 0.000497 | 1.52 |
| FALSE | High | SPVQEASS | [R].SPVQE | 1xPhosphc | Q3UVG3 1 | 6.73E-06 | 0 | 1 | 1 | 6 | Q3UVG3 | Protein FAI | Q3UVG3 [ | 0 | 4285.976 | 0 | 0.61 | 0.007633 | 2.6 | 73.7 | 126.3 | 10.13 | 12.34 | | High | High | High | High | High | High | High | High | High | High | High | 0 | 8.69E-07 | 4.94 |
| FALSE | High | SPVQTPDH | [K].SPVQTI | 1xPhosphc | Q6P5D4 1x | 0.106748 | 0.008634 | 1 | 1 | 1 | Q6P5D4 | Centrosom | Q6P5D4 [1 | 0 | 1487.709 | 0 | -0.15 | 0.161791 | 14.09 | 105.4 | 94.6 | 8.7 | 12.34 | | High | High | High | High | High | High | High | High | High | High | High | 0.004327 | 0.04667 | 1.22 |
| FALSE | High | SPVSFTEEL | [K].SPVSFT | 1xPhosphc | Q9JIM3 1x | 0.000331 | 0 | 1 | 3 | 1 | Q9JIM3 | DNA excisi | Q9JIM3 [1 | 0 | 1826.852 | 0 | -0.3 | 0.898104 | 8.9 | 105.7 | 94.3 | 14.73 | 24.41 | | High | High | High | High | High | High | High | High | High | High | High | 0 | 6.91E-05 | 2.65 |
| FALSE | High | SPVSNYSE | [K].SPVSNY | 1xCarbami | E9PZM7 1x | 1.24E-08 | 0 | 1 | 3 | 4 | E9PZM7 | SR-related | E9PZM7 [3 | 0 | 3534.598 | 0 | -0.4 | 0.005378 | 8.84 | 114.1 | 85.9 | 5.2 | 6.56 | | High | High | High | High | High | High | High | High | High | High | High | 0 | 7.36E-10 | 5.84 |
| FALSE | High | SPVSNYSE | [K].SPVSNY | 1xCarbami | E9PZM7 1x | 0.004218 | 0.00036 | 1 | 3 | 3 | E9PZM7 | SR-related | E9PZM7 [3 | 0 | 3535.582 | 0 | 0.16 | 0.072802 | 6.3 | 94.1 | 105.9 | 11.16 | 3.97 | | High | High | High | High | High | High | High | High | High | High | High | 0.000801 | 0.007198 | 3.23 |
| FALSE | High | SPVSNYSE | [K].SPVSNY | 1xCarbami | E9PZM7 2x | 0.005978 | 0.00062 | 1 | 3 | 3 | E9PZM7 | SR-related | E9PZM7 [3 | 0 | 3614.565 | 0 | 0.36 | 0.002376 | 5.16 | 90 | 110 | 5.33 | 6.61 | | High | High | High | High | High | High | High | High | High | High | High | 0.006329 | 0.06782 | 3.95 |

| | | | | | | | | | | | | | | | | | | | | | | | | | | | | | | | | | | | | | | |
|---|---|---|---|---|---|---|---|---|---|---|---|---|---|---|---|---|---|---|---|---|---|---|---|---|---|---|---|---|---|---|---|---|---|---|---|---|---|---|
| FALSE | High | SPVSPLAAC | [R].SPVSPL | 1xPhospho | Q8K0E1 1x | 0.00129 | 0 | 1 | 1 | 2 | Q8K0E1 | BTB/POZ d | Q8K0E1 [3 | 0 | 2426.382 | 0 | 1.16 | 0.00032 | 5.39 | 62.9 | 137.1 | 7.57 | 7.45 | | High | High | High | High | High | High | High | High | High | High | High | 2.58E-05 | 0.000363 | 3.53 |
| FALSE | High | SPVSPQLQ | [R].SPVSPQ | 1xPhospho | E9Q969 1x | 0.098533 | 0.007902 | 1 | 4 | 1 | E9Q969 | Cytoplasmi | E9Q969 [5 | 0 | 2713.377 | 0 | 0.37 | 0.131737 | 32.16 | 89.7 | 110.3 | 18.44 | 9.65 | | High | High | High | High | High | High | High | High | High | High | High | 0.003977 | 0.04243 | 2.83 |
| FALSE | High | SPVVELSK | [R].SPVVEL | 1xPhospho | Q3U1V3 1x | 0.010568 | 0.000772 | 1 | 3 | 2 | Q3U1V3 | Protein-ser | Q3U1V3 [6 | 0 | 1167.622 | 0 | 1.22 | 0.000523 | 9.32 | 60.3 | 139.7 | 5.32 | 4.94 | | High | High | High | High | High | High | High | High | High | High | High | 0.000382 | 0.003337 | 2.21 |
| FALSE | High | SPVVITIDSI | [R].SPVVIT | 2xPhospho | Q80Z37 2x | 0.009168 | 0.000736 | 1 | 1 | 1 | Q80Z37 | E3 ubiquiti | Q80Z37 [9 | 0 | 2495.161 | 0 | -0.35 | 0.025 | 6.42 | 112 | 88 | 14.26 | 11.01 | | High | High | High | High | High | High | High | High | High | High | High | 0.000327 | 0.002846 | 3.05 |
| FALSE | High | SPVVSGDT | [K].SPVVSG | 1xPhospho | A2A5Y6 1x | 0.033116 | 0.002195 | 1 | 11 | 1 | A2A5Y6 | Microtubu | A2A5Y6 [7 | 0 | 1410.683 | 0 | -1.04 | 0.001335 | 2.3 | 134.7 | 65.3 | 4.23 | 18.95 | | High | High | High | High | High | High | High | High | High | High | High | 0.001151 | 0.01201 | 1.5 |
| FALSE | High | SPWQPPTT | [R].SPWQP | 1xPhospho | Q9JLF6 1xP | 0.010568 | 0.000772 | 1 | 1 | 3 | Q9JLF6 | Protein-glu | Q9JLF6 [14 | 0 | 2679.218 | 0 | 2.74 | 0.000461 | 5.34 | 24.7 | 175.3 | 10.11 | 11.47 | | High | High | High | High | High | High | High | High | High | High | High | 0.000847 | 0.008106 | 3.46 |
| FALSE | High | SPVPEQFQ | [K].SPVPEQ | 1xPhospho | Q921I6 1xP | 0.01002 | 0.000736 | 1 | 1 | 1 | Q921I6 | SH3 domai | Q921I6 [27 | 0 | 1483.714 | 0 | -1.01 | 0.013381 | 4.73 | 133.8 | 66.2 | 13.92 | 21.27 | | High | High | High | High | High | High | High | High | High | High | High | 0.000344 | 0.003133 | 3.15 |
| FALSE | High | SPTTTQSPK | [K].SPTTTQ | 2xPhospho | G5E870 2x | 0.009931 | 0.000736 | 1 | 2 | 1 | G5E870 | E3 ubiquiti | G5E870 [1 | 0 | 1564.743 | 0 | -1.14 | 0.000676 | 3.47 | 137.7 | 62.3 | 5.37 | 9.79 | | High | High | High | High | High | High | High | High | High | High | High | 0.000344 | 0.003105 | 2.45 |
| FALSE | High | SPTTTQSPK | [K].SPTTTQ | 1xPhospho | G5E870 1x | 0.00824 | 0.000736 | 1 | 2 | 3 | G5E870 | E3 ubiquiti | G5E870 [1 | 0 | 1484.776 | 0 | -1.48 | 0.000864 | 2.04 | 148.2 | 51.8 | 5.41 | 13.3 | | High | High | High | High | High | High | High | High | High | High | High | 0.000327 | 0.002529 | 2.15 |
| FALSE | High | SPTTTAPVS | [R].SPTTTA | 1xPhospho | Q9JHC9 1x | 0.000262 | 0 | 1 | 7 | 2 | Q9JHC9 | ETS-related | Q9JHC9 [3 | 0 | 1706.868 | 0 | -0.09 | 0.361542 | 22.15 | 112.4 | 87.6 | 9.55 | 17.24 | | High | High | High | High | High | High | High | High | High | High | High | 0 | 5.31E-05 | 2.84 |
| FALSE | High | SPTGPSNSI | [R].SPTGPS | 1xPhospho | Q8JZX4 1xP | 9.05E-09 | 0 | 1 | 1 | 13 | Q8JZX4 | Splicing fac | Q8JZX4 [22 | 0 | 2527.242 | 0 | -0.48 | 0.011841 | 10.2 | 116.5 | 83.5 | 5.36 | 10.42 | | High | High | High | High | High | High | High | High | High | High | High | 0 | 5.13E-10 | 4.63 |
| FALSE | High | SPTGSLEQI | [K].SPTGSL | 1xPhospho | A2A7F4 1xP | 9.56E-06 | 0 | 1 | 3 | 2 | A2A7F4 | MCG12287 | A2A7F4 [6 | 0 | 1822.853 | 0 | -0.46 | 0.003523 | 1.73 | 119.2 | 80.8 | 5.1 | 8.15 | | High | High | High | High | High | High | High | High | High | High | High | 0.000327 | 0.002615 | 2.92 |
| FALSE | High | SPTHASPQ | [K].SPTHAS | 1xPhospho | A2RSJ4 1xP | 3.34E-07 | 0 | 1 | 3 | 1 | A2RSJ4 | UHRF1-bin | A2RSJ4 [41 | 0 | 2087.032 | 0 | 0.76 | 0.003047 | 10.03 | 73.9 | 126.1 | 5.19 | 9.47 | | High | High | High | High | High | High | High | High | High | High | High | 0 | 2.97E-08 | 2.67 |
| FALSE | High | SPTHLTVSS | [R].SPTHLT | 1xPhospho | G3UZM1 1 | 3.26E-05 | 0 | 1 | 3 | 2 | G3UZM1 | Probable Ju | G3UZM1 [ | 0 | 1792.916 | 0 | -0.18 | 0.288036 | 27.67 | 107.2 | 92.8 | 17.09 | 13.4 | | High | High | High | High | High | High | High | High | High | High | High | 0 | 5.1E-06 | 2.98 |
| FALSE | High | SPTLGEDSD | [R].SPTLGE | 1xPhospho | E9QK89 1x | 2.38E-06 | 0 | 1 | 2 | 1 | E9QK89 | Mediator c | E9QK89 [2 | 0 | 3973.739 | 0 | -1.44 | 0.000744 | 1.61 | 146.8 | 53.2 | 9.13 | 10.43 | | High | High | High | High | High | High | High | High | High | High | High | 0 | 2.69E-07 | 4.84 |
| FALSE | High | SPTLGEDSD | [R].SPTLGE | 2xPhospho | E9QK89 2x | 0.021231 | 0.001507 | 1 | 2 | 1 | E9QK89 | Mediator c | E9QK89 [2 | 0 | 4053.705 | 0 | -1.25 | 0.00026 | 0.57 | 140.9 | 59.1 | 5.1 | 4.01 | | High | High | High | High | High | High | High | High | High | High | High | 0.000801 | 0.007257 | 3.93 |
| FALSE | High | SPTIGFNNL | [R].SPTIGF | 1xPhospho | B9EKK3 1xP | 3.53E-08 | 0 | 1 | 2 | 1 | B9EKK3 | IQ motif co | B9EKK3 [3 | 0 | 2093.985 | 0 | 0.86 | 0.000983 | 3.68 | 73.1 | 126.9 | 8.38 | 9.58 | | High | High | High | High | High | High | High | High | High | High | High | 0 | 2.38E-09 | 5.38 |
| FALSE | High | SPTLGIAR | [R].SPTLGI | 1xPhospho | Q99M54 1x | 0.103392 | 0.008249 | 1 | 5 | 2 | Q99M54 | Cell divisio | Q99M54 [6 | 0 | 1123.607 | 0 | -1.13 | 0.000661 | 3.67 | 139.1 | 60.9 | 5.28 | 12.1 | | High | High | High | High | High | High | High | High | High | High | High | 0.009784 | 0.1057 | 1.2 |
| FALSE | High | SPTLSQVHS | [K].SPTLSQ | 1xPhospho | Q67FY2 1x | 0.000193 | 0 | 1 | 2 | 2 | Q67FY2 | B-cell CLL/l | Q67FY2 [9 | 0 | 2601.405 | 0 | 0.36 | 0.234139 | 7.47 | 85.6 | 114.4 | 18.47 | 13.34 | | High | High | High | High | High | High | High | High | High | High | High | 0 | 3.75E-05 | 3.07 |
| FALSE | High | SPTMEQAV | [K].SPTME | 1xPhospho | Q8K2F8 1x | 2.65E-05 | 0 | 1 | 2 | 1 | Q8K2F8 | Protein LSM | Q8K2F8 [1 | 0 | 2625.306 | 0 | -0.21 | 0.239364 | 15.48 | 108.5 | 91.5 | 8.97 | 18.55 | | High | High | High | High | High | High | High | High | High | High | High | 0 | 4.03E-06 | 3.16 |
| FALSE | High | SPTMEQAV | [K].SPTME | 1xPhospho | Q8K2F8 1x | 1.66E-05 | 0 | 1 | 2 | 7 | Q8K2F8 | Protein LSM | Q8K2F8 [1 | 0 | 2641.301 | 0 | -0.14 | 0.916737 | 5.59 | 105 | 95 | 8.53 | 24.16 | | High | High | High | High | High | High | High | High | High | High | High | 0 | 9.99E-06 | 4.3 |
| FALSE | High | SPTNGGAG | [R].SPTNG | 1xPhospho | B2RRP1 1xP | 0.000529 | 0 | 1 | 1 | 1 | B2RRP1 | Gene trap | B2RRP1 [9 | 0 | 2357.181 | 0 | 0.22 | 0.088543 | 15.26 | 93.8 | 106.2 | 7.89 | 3.86 | | High | High | High | High | High | High | High | High | High | High | High | 0 | 0.000117 | 2.7 |
| FALSE | High | SPTNSSEIF | [R].SPTNSS | 1xPhospho | Q3UGS4 1x | 2.81E-07 | 0 | 1 | 1 | 2 | Q3UGS4 | Mapk-regu | Q3UGS4 [2 | 0 | 2324.076 | 0 | 0.24 | 0.001744 | 2.23 | 89.1 | 110.9 | 7.02 | 4.28 | | High | High | High | High | High | High | High | High | High | High | High | 0 | 5.91E-07 | 3.96 |
| FALSE | High | SPTNSSEIF | [R].SPTNSS | 1xDeamida | Q3UGS4 1x | 0.025073 | 0.001645 | 1 | 1 | 3 | Q3UGS4 | Mapk-regu | Q3UGS4 [2 | 0 | 2325.06 | 0 | -0.23 | 0.084172 | 10.33 | 108.2 | 91.8 | 5.97 | 12.48 | | High | High | High | High | High | High | High | High | High | High | High | 0.000872 | 0.008792 | 2.74 |
| FALSE | High | SPTNVSHN | [K].SPTNVS | 1xCarbami | Q8C0D9 1x | 0.024638 | 0.001612 | 1 | 5 | 1 | Q8C0D9 | Centrosom | Q8C0D9 [3 | 0 | 2724.265 | 0 | 0.48 | 0.027037 | 22.6 | 88 | 112 | 12.07 | 12.89 | | High | High | High | High | High | High | High | High | High | High | High | 0.000859 | 0.008595 | 3.2 |
| FALSE | High | SPTPESSTI | [K].SPTPES | 1xPhospho | E9Q6H8 1x | 4.95E-06 | 0 | 1 | 12 | 1 | E9Q6H8 | Pleckstrin | E9Q6H8 [1 | 0 | 2018.004 | 0 | -0.49 | 0.007538 | 3.18 | 116.8 | 83.2 | 9.17 | 6.03 | | High | High | High | High | High | High | High | High | High | High | High | 0 | 6.12E-07 | 2.45 |
| FALSE | High | SPTPPQPPI | [K].SPTPPQ | 1xPhospho | Q640N2 1x | 4.44E-06 | 0 | 1 | 1 | 1 | Q640N2 | ADP-ribosy | Q640N2 [3 | 0 | 2278.204 | 0 | 1.58 | 0.002604 | 37.78 | 50.2 | 149.8 | 10.12 | 14.22 | | High | High | High | High | High | High | High | High | High | High | High | 0 | 5.43E-07 | 2.39 |
| FALSE | High | SPTPPQPPI | [K].SPTPPQ | 2xPhospho | Q640N2 2x | 0.1059 | 0.008582 | 1 | 1 | 1 | Q640N2 | ADP-ribosy | Q640N2 [3 | 0 | 2358.17 | 0 | 1.86 | 0.000746 | 19.31 | 44.2 | 155.8 | 14.19 | 7.28 | | High | High | High | High | High | High | High | High | High | High | High | 0.004317 | 0.04634 | 2 |
| FALSE | High | SPTPSAASS | [R].SPTPSA | 1xPhospho | Q76I79 1xP | 0.001222 | 0 | 1 | 3 | 2 | Q76I79 | Protein ph | Q76I79 [9- | 1 | 3032.41 | 0 | 0.86 | 0.001712 | 4.95 | 68.6 | 131.4 | 6.08 | 9.41 | | High | High | High | High | High | High | High | High | High | High | High | 0 | 0.000298 | 2.86 |
| FALSE | High | SPTPVKPTE | [K].SPTPVK | 1xCarbami | A2AQH4 1x | 0.002948 | 0.00016 | 1 | 1 | 1 | A2AQH4 | BCL-6 core | A2AQH4 [1 | 0 | 2393.275 | 0 | 0.99 | 0.003889 | 11.77 | 67.8 | 132.2 | 10.19 | 11.15 | | High | High | High | High | High | High | High | High | High | High | High | 7.19E-05 | 0.000798 | 1.59 |
| FALSE | High | SPTQAPFSI | [R].SPTQA | 1xPhospho | Q32MF0 1x | 0.002531 | 0.000109 | 1 | 3 | 1 | Q32MF0 | Predicted g | Q32MF0 [6 | 0 | 2370.251 | 0 | 1.93 | 0.0014 | 5.06 | 41.2 | 158.8 | 8.19 | 14.3 | | High | High | High | High | High | High | High | High | High | High | High | 4.94E-05 | 0.000676 | 3.36 |
| FALSE | High | SPTQLQLG | [R].SPTQLQ | 1xPhospho | Q811P8 1x | 0.007811 | 0.000664 | 1 | 3 | 2 | Q811P8 | Rho GTPas | Q811P8 [9 | 0 | 1509.844 | 0 | -1.05 | 0.000466 | 0.89 | 135.7 | 64.3 | 3.28 | 9.5 | | High | High | High | High | High | High | High | High | High | High | High | 0.000327 | 0.002387 | 1.84 |
| FALSE | High | SPTSPLNM | [K].SPTSPL | 1xPhospho | Q6PGB8-2 | 0.008022 | 0.000736 | 1 | 5 | 1 | Q6PGB8-2 | Isoform 2 c | Q6PGB8-2 | 0 | 1528.785 | 0 | -1.92 | 0.001252 | 0.49 | 161.8 | 38.2 | 10.69 | 21.57 | | High | High | High | High | High | High | High | High | High | High | High | 0.000327 | 0.002446 | 1.84 |
| FALSE | High | SPTSPVTPE | [R].SPTSPV | 1xPhospho | A0A0A0M | 3.47E-05 | 0 | 1 | 2 | 16 | A0A0A0M | F-box only | A0A0A0M | 0 | 2626.296 | 0 | 4.04 | 0.000396 | 33.74 | 10.2 | 189.8 | 29.13 | 9.76 | | High | High | High | High | High | High | High | High | High | High | High | 0 | 0.000164 | 3.44 |
| FALSE | High | SPTSPVTPE | [R].SPTSPV | 2xPhospho | A0A0A0M | 2.01E-06 | 0 | 1 | 2 | 15 | A0A0A0M | F-box only | A0A0A0M | 0 | 2706.263 | 0 | 2.18 | 0.000281 | 2.85 | 36.3 | 163.7 | 5.97 | 6.11 | | High | High | High | High | High | High | High | High | High | High | High | 0 | 1.15E-05 | 3.27 |
| FALSE | High | SPTSPVTPE | [R].SPTSPV | 3xPhospho | A0A0A0M | 0.000118 | 0 | 1 | 2 | 15 | A0A0A0M | F-box only | A0A0A0M | 0 | 2786.229 | 0 | 4.13 | 0.000264 | 51.09 | 12.2 | 187.8 | 52.14 | 7.83 | | High | High | High | High | High | High | High | High | High | High | High | 0.002896 | 0.03003 | 3.67 |
| FALSE | High | SPTSTNSSV | [R].SPTSTN | 1xPhospho | Q80Y82 1x | 5.32E-06 | 0 | 1 | 14 | 5 | Q80Y82 | Protein kin | Q80Y82 [4 | 0 | 2740.271 | 0 | -1.16 | 0.024903 | 6.48 | 141.1 | 58.9 | 19.64 | 23.56 | | High | High | High | High | High | High | High | High | High | High | High | 0 | 6.62E-07 | 2.93 |
| FALSE | High | SPTSTNSSV | [R].SPTSTN | 2xPhospho | Q80Y82 2x | 5.31E-05 | 0 | 1 | 14 | 4 | Q80Y82 | Protein kin | Q80Y82 [4 | 0 | 2820.238 | 0 | -0.39 | 0.041676 | 13.92 | 114.3 | 85.7 | 8.74 | 18.82 | | High | High | High | High | High | High | High | High | High | High | High | 0 | 8.85E-06 | 2.38 |
| FALSE | High | SPTTPSVFS | [R].SPTTPS | 1xPhospho | Q8BG30 1x | 3.8E-05 | 0 | 1 | 2 | 3 | Q8BG30 | Negative e | Q8BG30 [2 | 0 | 1742.831 | 0 | 0.24 | 0.077409 | 20.42 | 91.4 | 108.6 | 8.03 | 15.09 | | High | High | High | High | High | High | High | High | High | High | High | 0 | 1.48E-05 | 3.16 |
| FALSE | High | SPWQPPTT | [R].SPWQP | 1xPhospho | Q9JLF6 1xP | 0.008769 | 0.000736 | 1 | 1 | 6 | Q9JLF6 | Protein-glu | Q9JLF6 [14 | 1 | 2835.319 | 0 | 2.9 | 0.000637 | 7.04 | 19.4 | 180.6 | 17.65 | 13.58 | | High | High | High | High | High | High | High | High | High | High | High | 0.000638 | 0.005613 | 4.47 |
| FALSE | High | SPTGPSNSI | [R].SPTGPS | 1xPhospho | Q8JZX4 1xP | 0.011755 | 0.000889 | 1 | 1 | 2 | Q8JZX4 | Splicing fac | Q8JZX4 [22 | 0 | 2282.084 | 0 | -0.65 | 0.003262 | 3.46 | 124.4 | 75.6 | 10.98 | 8.27 | | High | High | High | High | High | High | High | High | High | High | High | 0.000449 | 0.003764 | 2.03 |
| FALSE | High | SPWQPPTT | [R].SPWQP | 1xPhospho | A0A0R4J29 | 0.000138 | 0 | 1 | 1 | 2 | A0A0R4J29 | Protein-glu | A0A0R4J29 | 0 | 2649.244 | 0 | -1 | 0.00041 | 0.77 | 134.1 | 65.9 | 5.89 | 2.79 | | High | High | High | High | High | High | High | High | High | High | High | 0 | 5.39E-05 | 2.9 |
| FALSE | High | SPYIDGCSP | [R].SPYIDG | 1xCarbami | Q8BS90 1x | 9.62E-05 | 0 | 1 | 1 | 2 | Q8BS90 | Protein au | Q8BS90 [2 | 0 | 1774.885 | 0 | -1.94 | 0.000207 | 0.35 | 159 | 41 | 5.02 | 14.45 | | High | High | High | High | High | High | High | High | High | High | High | 0 | 1.72E-05 | 3.07 |
| FALSE | High | SQETPAAK | [K].SQETPA | 1xPhospho | Q62189 1x | 0.085859 | 0.006406 | 1 | 4 | 2 | Q62189 | U1 small nu | Q62189 [1 | 0 | 1369.713 | 0 | -0.5 | 0.019623 | 10.64 | 114.2 | 85.8 | 11.31 | 16.95 | | High | High | High | High | High | High | High | High | High | High | High | 0.006781 | 0.07356 | 1.57 |
| FALSE | High | SQETSCPSI | [R].SQETSC | 1xCarbami | V9GX40 1x | 3.17E-08 | 0 | 1 | 5 | 3 | V9GX40 | Synergin ga | V9GX40 [1 | 0 | 2625.227 | 0 | 0.75 | 0.02098 | 23.82 | 78.7 | 121.3 | 15.72 | 16.65 | | High | High | High | High | High | High | High | High | High | High | High | 0 | 2.09E-09 | 3.53 |
| FALSE | High | SQEVTAGG | [R].SQEVTA | 1xPhospho | Q09XV5 1x | 0.000548 | 0 | 1 | 2 | 6 | Q09XV5 | Chromodo | Q09XV5 [2 | 0 | 3876.885 | 0 | -0.32 | 0.057117 | 12.2 | 109.9 | 90.1 | 8.9 | 6.81 | | High | High | High | High | High | High | High | High | High | High | High | 0 | 0.000146 | 4.02 |
| FALSE | High | SQFLDEGM | [K].SQFLDE | 1xDeamida | Q8BWB6 1 | 0.071662 | 0.005278 | 1 | 1 | 1 | Q8BWB6 | Metallored | Q8BWB6 [ | 0 | 2353.073 | 0 | 0.08 | 0.137398 | 0.35 | 91.8 | 108.2 | 16.82 | 16.35 | | High | High | High | High | High | High | High | High | High | High | High | 0.002717 | 0.02912 | 4.02 |
| FALSE | High | SQGAIQGR | [K].SQGAIQ | 1xPhospho | Q6A0A9 1x | 0.005 | 0.000504 | 1 | 3 | 1 | Q6A0A9 | Constitutiv | Q6A0A9 [1 | 0 | 2568.322 | 0 | 1.71 | 0.000324 | 8.48 | 43.8 | 156.2 | 5.61 | 6.05 | | High | High | High | High | High | High | High | High | High | High | High | 0.000225 | 0.001451 | 1.59 |
| FALSE | High | SQGCHPES | [K].SQGCH | 2xCarbami | Q9D842 1x | 7.91E-06 | 0 | 1 | 3 | 2 | Q9D842 | Aprataxin a | Q9D842 [3 | 0 | 4078.805 | 0 | 0.08 | 0.885239 | 27.23 | 107.4 | 92.6 | 21.41 | 27.3 | | High | High | High | High | High | High | High | High | High | High | High | 0 | 1.04E-06 | 5.28 |
| FALSE | High | SQGCHPES | [K].SQGCH | 2xCarbami | Q9D842 2x | 4.82E-06 | 0 | 1 | 3 | 3 | Q9D842 | Aprataxin a | Q9D842 [3 | 0 | 4158.771 | 0 | 0.69 | 0.015727 | 3.95 | 79.3 | 120.7 | 8.88 | 11.52 | | High | High | High | High | High | High | High | High | High | High | High | 0 | 2.02E-06 | 5.33 |
| FALSE | High | SQGGGDSV | [R].SQGGG | 1xPhospho | Q5SSL4-2 1 | 6.73E-06 | 0 | 1 | 6 | 8 | Q5SSL4-2 | Isoform 2 c | Q5SSL4-2 [ | 0 | 2732.354 | 0 | 1.13 | 0.000877 | 13.6 | 60.6 | 139.4 | 7.51 | 6.93 | | High | High | High | High | High | High | High | High | High | High | High | 0 | 4.51E-06 | 3.27 |
| FALSE | High | SQGGSSPL | [K].SQGGS | 1xPhospho | Q8C0T5 1x | 0.000229 | 0 | 1 | 3 | 3 | Q8C0T5 | Signal-indu | Q8C0T5 [1 | 0 | 1284.615 | 0 | 1.76 | 0.000417 | 5.4 | 45.5 | 154.5 | 4.04 | 7.75 | | High | High | High | High | High | High | High | High | High | High | High | 0 | 4.57E-05 | 2 |
| FALSE | High | SQGPVNPS | [K].SQGPV | 1xPhospho | P48754 1xP | 0.007471 | 0.00062 | 1 | 4 | 1 | P48754 | Breast canc | P48754 [71 | 0 | 1475.721 | 0 | -0.7 | 0.000656 | 5 | 124.7 | 75.3 | 3.17 | 7.21 | | High | High | High | High | High | High | High | High | High | High | High | 0.000275 | 0.002267 | 0.71 |
| FALSE | High | SQGSGLSS | [K].SQGSG | 1xPhospho | Q8CGC7 1x | 9E-07 | 0 | 1 | 2 | 5 | Q8CGC7 | Bifunctiona | Q8CGC7 [9 | 0 | 2099.017 | 0 | 0 | 0.461487 | 9.45 | 100.1 | 99.9 | 7.25 | 9.53 | | High | High | High | High | High | High | High | High | High | High | High | 0 | 9.02E-08 | 3.84 |
| FALSE | High | SQGSPEDP | [R].SQGSP | 1xPhospho | Q6PGG2 1x | 2.53E-05 | 0 | 1 | 2 | 2 | Q6PGG2 | GEM-intera | Q6PGG2 [2 | 0 | 2308.086 | 0 | 2.13 | 0.000255 | 9.39 | 36.8 | 163.2 | 13.76 | 4.11 | | High | High | High | High | High | High | High | High | High | High | High | 2.58E-05 | 0.000404 | 3.08 |
| FALSE | High | SQGVGSQI | [K].SQGVG | 1xPhospho | A2AUM9 1 | 0.001222 | 0 | 1 | 1 | 1 | A2AUM9 | Centrosom | A2AUM9 [ | 0 | 1726.878 | 0 | 0.2 | 0.099492 | 14.27 | 92.2 | 107.8 | 7.82 | 8.48 | | High | High | High | High | High | High | High | High | High | High | High | 0 | 0.000298 | 1.67 |
| FALSE | High | SQESDGVE | [R].SQESD | 1xPhospho | G5E8S8 1xP | 0.00095 | 0 | 1 | 2 | 1 | G5E8S8 | MAGUK p5 | G5E8S8 [4 | 0 | 2140.062 | 0 | -0.09 | 0.269248 | 10.13 | 105.3 | 94.7 | 9.04 | 9.71 | | High | High | High | High | High | High | High | High | High | High | High | 0 | 0.000225 | 2.27 |
| FALSE | High | SQHQSPPP | [R].SQHQS | 1xPhospho | F6Y616 1xP | 1.68E-06 | 0 | 1 | 2 | 1 | F6Y616 | Polyadenyl | F6Y616 [43 | 0 | 1865.968 | 0 | 1.76 | 0.002038 | 24.69 | 48.1 | 151.9 | 16.54 | 11.69 | | High | High | High | High | High | High | High | High | High | High | High | 0 | 1.82E-07 | 2.4 |
| FALSE | High | SQHSSGNG | [R].SQHSS | 1xDeamida | Q8BU14 1x | 0.001848 | 5.66E-05 | 1 | 3 | 2 | Q8BU14 | Translocati | Q8BU14 [3 | 0 | 2291.042 | 0 | -0.11 | 0.55171 | 24.96 | 102.2 | 97.8 | 12.46 | 10.59 | | High | High | High | High | High | High | High | High | High | High | High | 2.58E-05 | 0.000476 | 2.81 |
| FALSE | High | SQHTVDTT | [R].SQHTV | 1xPhospho | Q91VX2 1x | 0.00172 | 5.66E-05 | 1 | 2 | 1 | Q91VX2 | Ubiquitin-a | Q91VX2 [6 | 0 | 2093.068 | 0 | 0.27 | 0.145768 | 25.19 | 90.4 | 109.6 | 10.91 | 9.86 | | High | High | High | High | High | High | High | High | High | High | High | 2.58E-05 | 0.000438 | 1.41 |
| FALSE | High | SQLDDHPE | [R].SQLDD | 1xPhospho | A0A0R4J04 | 3.16E-07 | 0 | 1 | 2 | 3 | A0A0R4J04 | CCAAT/enh | A0A0R4J04 | 0 | 3374.411 | 0 | -0.74 | 0.001311 | 6.18 | 124.7 | 75.3 | 3.36 | 9.6 | | High | High | High | High | High | High | High | High | High | High | High | 0 | 2.8E-08 | 5.31 |
| FALSE | High | SQLDDHPE | [R].SQLDD | 2xPhospho | A0A0R4J04 | 1.04E-06 | 0 | 1 | 2 | 7 | A0A0R4J04 | CCAAT/enh | A0A0R4J04 | 0 | 3454.377 | 0 | -0.21 | 0.037628 | 7.86 | 107.3 | 92.7 | 3.41 | 9.1 | | High | High | High | High | High | High | High | High | High | High | High | 0 | 8.49E-07 | 5.45 |
| FALSE | High | SQLDLFDD | [R].SQLDLF | 1xPhospho | Q99K28-2 | 1.47E-05 | 0 | 1 | 2 | 1 | Q99K28-2 | Isoform 2 c | Q99K28-2 | 0 | 2203.052 | 0 | 0.31 | 0.090878 | 29.21 | 90.8 | 109.2 | 17.93 | 10.28 | | High | High | High | High | High | High | High | High | High | High | High | 0 | 2.09E-06 | 3.63 |
| FALSE | High | SQLDLFDD | [R].SQLDLF | 1xPhospho | Q99K28-2 | 0.001516 | 5.66E-05 | 1 | 2 | 1 | Q99K28-2 | Isoform 2 c | Q99K28-2 | 0 | 2432.215 | 0 | 0.74 | 0.036276 | 10.97 | 67.3 | 132.7 | 14.88 | 29.6 | | High | High | High | High | High | High | High | High | High | High | High | 2.58E-05 | 0.000379 | 2.6 |
| FALSE | High | SQIDVALSC | [K].SQIDVA | 1xPhospho | Q9EPU0 1x | 3.64E-06 | 0 | 1 | 2 | 3 | Q9EPU0 | Regulator c | Q9EPU0 [1 | 0 | 2206.023 | 0 | 0.34 | 0.14525 | 14.5 | 94 | 106 | 5.6 | 14.74 | | High | High | High | High | High | High | High | High | High | High | High | 0 | 9.67E-06 | 3.4 |
| FALSE | High | SQIDVALSC | [K].SQIDVA | 2xPhospho | Q9EPU0 2x | 0.004106 | 0.00036 | 1 | 2 | 1 | Q9EPU0 | Regulator c | Q9EPU0 [1 | 0 | 2285.989 | 0 | 0.45 | 0.087414 | 28.29 | 69.2 | 130.8 | 24.12 | 28.23 | | High | High | High | High | High | High | High | High | High | High | High | 0.000161 | 0.001156 | 3.39 |
| FALSE | High | SQIHDIVLV | [K].SQIHDI | 1xPhospho | P63017 1xP | 0.007082 | 0.00062 | 1 | 16 | 1 | P63017 | Heat shock | P63017 [32 | 0 | 1790.936 | 0 | -0.32 | 0.094387 | 7.57 | 111.1 | 88.9 | 2.01 | 20.04 | | High | High | High | High | High | High | High | High | High | High | High | 0.000275 | 0.002126 | 2.27 |
| FALSE | High | SQLPSPGEI | [R].SQLPSP | 1xPhospho | Q8BIJ7 1xP | 0.054844 | 0.003982 | 1 | 1 | 1 | Q8BIJ7 | RUN and F | Q8BIJ7 [48 | 0 | 1392.709 | 0 | 1.77 | 0.000191 | 3.99 | 46.1 | 153.9 | 12.85 | 7.16 | | High | High | High | High | High | High | High | High | High | High | High | 0.002059 | 0.02148 | 1.4 |
| FALSE | High | SQLPTLEQI | [R].SQLPTL | 1xPhospho | P46938 1xP | 2.48E-07 | 0 | 1 | 3 | 2 | P46938 | Transcripti | P46938 [33 | 0 | 3024.418 | 0 | 0.57 | 0.019891 | 19.67 | 79.5 | 120.5 | 11.74 | 12.27 | | High | High | High | High | High | High | High | High | High | High | High | 0 | 2.11E-08 | 5.59 |
| FALSE | High | SQLPTLEQI | [R].SQLPTL | 1xDeamida | P46938 1xP | 0.00083 | 0 | 1 | 3 | 3 | P46938 | Transcripti | P46938 [33 | 0 | 3025.402 | 0 | 0.01 | 0.453309 | 11.37 | 98.7 | 101.3 | 16.98 | 9.99 | | High | High | High | High | High | High | High | High | High | High | High | 2.58E-05 | 0.000387 | 4.1 |
| FALSE | High | SQLPTLEQI | [R].SQLPTL | 2xPhospho | P46938 2xP | 0.009416 | 0.000736 | 1 | 3 | 1 | P46938 | Transcripti | P46938 [33 | 0 | 3088.39 | 0 | 0.02 | 0.612816 | 43.66 | 89.4 | 110.6 | 15.25 | 25.32 | | High | High | High | High | High | High | High | High | High | High | High | 0.000327 | 0.002934 | 2.69 |
| FALSE | High | SQHSSGNG | [R].SQHSS | 1xPhospho | Q8BU14 1x | 0.000362 | 0 | 1 | 3 | 2 | Q8BU14 | Translocati | Q8BU14 [3 | 0 | 2290.058 | 0 | 0.07 | 0.648309 | 1.31 | 99.5 | 100.5 | 25.83 | 8.12 | | High | High | High | High | High | High | High | High | High | High | High | 0 | 7.66E-05 | 2.86 |
| FALSE | High | SQERPPETI | [R].SQERPI | 2xPhospho | B2RWW1 2 | 0.003083 | 0.00016 | 1 | 3 | 2 | B2RWW1 | Microtubu | B2RWW1 [ | 0 | 4685.05 | 0 | 0.14 | 0.127309 | 0.85 | 92.8 | 107.2 | 9.12 | 8.46 | | High | High | High | High | High | High | High | High | High | High | High | 7.19E-05 | 0.000846 | 4.24 |
| FALSE | High | SQERPPETI | [R].SQERPI | 1xPhospho | B2RWW1 1 | 3.71E-06 | 0 | 1 | 3 | 4 | B2RWW1 | Microtubu | B2RWW1 [ | 0 | 4605.084 | 0 | 0.08 | 0.600993 | 5.46 | 97.2 | 102.8 | 6.33 | 7.34 | | High | High | High | High | High | High | High | High | High | High | High | 0 | 4.43E-07 | 3.87 |
| FALSE | High | SQEREETES | [K].SQEREE | 2xCarbami | B2RY61 2xP | 0.014797 | 0.001049 | 1 | 4 | 2 | B2RY61 | AT rich inte | B2RY61 [5 | 2 | 4136.651 | 0 | 0.12 | 0.305871 | 20.41 | 96.1 | 103.9 | 16.25 | 5.33 | | High | High | High | High | High | High | High | High | High | High | High | 0.000814 | 0.007595 | 3.79 |
| FALSE | High | SPYIDGCSP | [R].SPYIDG | 1xCarbami | Q8BS90 2x | 0.045898 | 0.003408 | 1 | 1 | 1 | Q8BS90 | Protein au | Q8BS90 [2 | 0 | 1854.851 | 0 | -0.64 | 0.000926 | 3.74 | 122 | 78 | 6.7 | 10.8 | | High | High | High | High | High | High | High | High | High | High | High | 0.001774 | 0.01754 | 2.59 |
| FALSE | High | SPYLPAEED | [R].SPYLPA | 1xPhospho | Q3UIL6-2 1 | 9.97E-08 | 0 | 1 | 8 | 2 | Q3UIL6-2 | Isoform 2 c | Q3UIL6-2 [ | 0 | 2452.2 | 0 | -1.06 | 0.001142 | 3.46 | 138.4 | 61.6 | 7.32 | 13.87 | | High | High | High | High | High | High | High | High | High | High | High | 0 | 7.59E-09 | 3.97 |
| FALSE | High | SPYNGVR | [K].SPYNGV | 1xPhospho | Q52KE7 1x | 0.015329 | 0.001089 | 1 | 1 | 2 | Q52KE7 | Cyclin-L1 C | Q52KE7 [3 | 0 | 1101.529 | 0 | 0.02 | 0.555299 | 4.87 | 103.8 | 96.2 | 6.76 | 6.01 | | High | High | High | High | High | High | High | High | High | High | High | 0.000571 | 0.005044 | 2.08 |
| FALSE | High | SPYQEFTDI | [K].SPYQEF | 1xPhospho | P25444 1xP | 9.06E-06 | 0 | 1 | 9 | 2 | P25444 | 40S riboso | P25444 [26 | 0 | 2002.009 | 0 | -0.53 | 0.000711 | 1.75 | 118 | 82 | 4.61 | 8.92 | | High | High | High | High | High | High | High | High | High | High | High | 0 | 1.21E-06 | 3.08 |
| FALSE | High | SPYSGPK | [R].SPYSGP | 1xPhospho | Q8VH51 1x | 0.020679 | 0.001472 | 1 | 5 | 2 | Q8VH51 | RNA-bindir | Q8VH51 [9 | 0 | 1273.659 | 0 | -0.25 | 0.009217 | 3.04 | 110.7 | 89.3 | 6.05 | 8.8 | | High | High | High | High | High | High | High | High | High | High | High | 0.000786 | 0.00705 | 1.26 |
| FALSE | High | SQALGNQI | [K].SQALGI | 1xPhospho | E9Q3Y4 1x | 1.1E-08 | 0 | 1 | 7 | 6 | E9Q3Y4 | Lipopolysa | E9Q3Y4 [2 | 0 | 3516.627 | 0 | 0.32 | 0.006344 | 7.86 | 86.4 | 113.6 | 6.37 | 9.57 | | High | High | High | High | High | High | High | High | High | High | High | 0 | 2.83E-09 | 5.86 |
| FALSE | High | SQALGNQI | [K].SQALGI | 2xPhospho | E9Q3Y4 2x | 3.57E-05 | 0 | 1 | 7 | 4 | E9Q3Y4 | Lipopolysa | E9Q3Y4 [2 | 0 | 3596.593 | 0 | 1.35 | 0.000881 | 6.05 | 58.1 | 141.9 | 15.26 | 10.73 | | High | High | High | High | High | High | High | High | High | High | High | 0 | 4.17E-05 | 4.66 |
| FALSE | High | SQAPETTM | [K].SQAPET | 1xPhospho | Q6P5B0 1x | 2.42E-08 | 0 | 1 | 1 | 1 | Q6P5B0 | RRP12-like | Q6P5B0 [7 | 0 | 2662.236 | 0 | 0.11 | 0.210479 | 15.82 | 75.6 | 124.4 | 29.01 | 26.84 | | High | High | High | High | High | High | High | High | High | High | High | 0 | 1.55E-09 | 4.59 |
| FALSE | High | SQAPETTM | [K].SQAPET | 1xPhospho | Q6P5B0 1x | 4.11E-07 | 0 | 1 | 1 | 1 | Q6P5B0 | RRP12-like | Q6P5B0 [7 | 0 | 2678.231 | 0 | 0.24 | 0.132413 | 21.3 | 69 | 131 | 24.76 | 30.36 | | High | High | High | High | High | High | High | High | High | High | High | 0 | 3.75E-08 | 3.54 |
| FALSE | High | SQASEASET | [K].SQASEA | 1xPhospho | Q9NWG9 1 | 0.002508 | 0.000109 | 1 | 1 | 4 | Q9NWG9 | Melanoma | Q9NWG9 [ | 0 | 4280.957 | 0 | -0.58 | 0.022574 | 9.09 | 122.5 | 77.5 | 5.21 | 17.63 | | High | High | High | High | High | High | High | High | High | High | High | 7.19E-05 | 0.000883 | 3.92 |
| FALSE | High | SQASKPTP | [R].SQASKI | 1xPhospho | Q8BI84 1xP | 0.106748 | 0.008634 | 1 | 4 | 1 | Q8BI84 | Melanoma | Q8BI84 [19 | 0 | 1353.718 | 0 | -0.31 | 0.000899 | 2.77 | 110.1 | 89.9 | 5.21 | 3.88 | | High | High | High | High | High | High | High | High | High | High | High | 0.004327 | 0.04654 | 1.14 |
| FALSE | High | SQASPSEDI | [R].SQASPS | 1xPhospho | O70479 1x | 0.006895 | 0.00062 | 1 | 1 | 1 | O70479 | BTB/POZ d | O70479 [2 | 0 | 2019.875 | 0 | 0.88 | 0.003227 | 11.63 | 73.3 | 126.7 | 13.66 | 9.88 | | High | High | High | High | High | High | High | High | High | High | High | 0.000275 | 0.002062 | 1.75 |
| FALSE | High | SQAWAEG | [K].SQAWA | 1xPhospho | A0A0R4J0V | 4.08E-05 | 0 | 1 | 4 | 2 | A0A0R4J0V | Junction-m | A0A0R4J0V | 0 | 1454.663 | 0 | 0.67 | 0.00654 | 15.91 | 77.1 | 122.9 | 5.89 | 9.26 | | High | High | High | High | High | High | High | High | High | High | High | 0 | 6.54E-06 | 1.86 |
| FALSE | High | SQDAPATF | [R].SQDAP | 1xPhospho | Q5FWI3 1x | 0.007404 | 0.00062 | 1 | 1 | 1 | Q5FWI3 | Cell surface | Q5FWI3 [5 | 0 | 2089.943 | 0 | -0.63 | 0.027796 | 6.71 | 119.1 | 80.9 | 17.87 | 9.63 | | High | High | High | High | High | High | High | High | High | High | High | 0.000275 | 0.002248 | 3.03 |
| FALSE | High | SQDATVSP | [R].SQDAT | 1xPhospho | Q80YV2 1x | 5.96E-08 | 0 | 1 | 4 | 5 | Q80YV2 | Nuclear-int | Q80YV2 [3 | 0 | 2087.99 | 0 | -0.57 | 0.002194 | 4.61 | 118.5 | 81.5 | 6.66 | 7.6 | | High | High | High | High | High | High | High | High | High | High | High | 0 | 4.28E-09 | 3.34 |
| FALSE | High | SQDFGNLF | [K].SQDFG | 1xPhospho | Q64511 1x | 5.65E-05 | 0 | 1 | 2 | 4 | Q64511 | DNA topois | Q64511 [1 | 0 | 2390.147 | 0 | 1.59 | 0.002698 | 6.47 | 49.1 | 150.9 | 9.03 | 17.4 | | High | High | High | High | High | High | High | High | High | High | High | 7.19E-05 | 0.000777 | 3.63 |
| FALSE | High | SQDNSISFS | [K].SQDNS | 1xPhospho | B2RXS2 1xP | 0.016743 | 0.001166 | 1 | 5 | 1 | B2RXS2 | Fam179b p | B2RXS2 [84 | 0 | 2248.105 | 0 | -0.54 | 0.000472 | 3.58 | 119.3 | 80.7 | 5.07 | 10.63 | | High | High | High | High | High | High | High | High | High | High | High | 0.000638 | 0.005609 | 2.5 |
| FALSE | High | SQDSYPGS | [K].SQDSY | 1xPhospho | A0A0R4J0G | 2.09E-07 | 0 | 1 | 2 | 2 | A0A0R4J0G | Ran-bindin | A0A0R4J0G | 0 | 1786.821 | 0 | -0.31 | 0.001617 | 2.23 | 109.9 | 90.1 | 1.71 | 3.23 | | High | High | High | High | High | High | High | High | High | High | High | 0.000437 | 0.003647 | 2.84 |
| FALSE | High | SQDSYPGS | [K].SQDSY | 2xPhospho | A0A0R4J0G | 2.16E-06 | 0 | 1 | 2 | 3 | A0A0R4J0G | Ran-bindin | A0A0R4J0G | 0 | 1866.787 | 0 | 0.35 | 0.000484 | 1.73 | 88.5 | 111.5 | 4.42 | 2.54 | | High | High | High | High | High | High | High | High | High | High | High | 0.000161 | 0.001201 | 3.06 |
| FALSE | High | SQDSYPGS | [K].SQDSY | 3xPhospho | A0A0R4J0G | 0.065405 | 0.004734 | 1 | 2 | 2 | A0A0R4J0G | Ran-bindin | A0A0R4J0G | 0 | 1946.754 | 0 | 0.67 | 0.000999 | 6.65 | 78.9 | 121.1 | 8.06 | 4.95 | | High | High | High | High | High | High | High | High | High | High | High | 0.003606 | 0.03841 | 3.52 |
| FALSE | High | SQDVAISP | [K].SQDVA | 1xCarbami | Q8K2D3 1x | 7.63E-06 | 0 | 1 | 1 | 2 | Q8K2D3 | Enhancer c | Q8K2D3 [1 | 0 | 2242.094 | 0 | -0.73 | 0.004771 | 5.57 | 124.8 | 75.2 | 11.49 | 8.67 | | High | High | High | High | High | High | High | High | High | High | High | 0 | 9.96E-07 | 3.82 |
| FALSE | High | SQDVESVC | [K].SQDVE | 1xPhospho | G3X9B1 1x | 0.000382 | 0 | 1 | 1 | 1 | G3X9B1 | HEAT repe | G3X9B1 [1 | 0 | 2345.065 | 0 | -0.1 | 0.353382 | 23.1 | 104.2 | 95.8 | 16.36 | 5.96 | | High | High | High | High | High | High | High | High | High | High | High | 0 | 8.13E-05 | 3.59 |
| FALSE | High | SQEDEEEIS | [K].SQEDE | 1xCarbami | Q9D5V6 1x | 2.39E-07 | 0 | 1 | 1 | 6 | Q9D5V6 | Synapse-as | Q9D5V6 [2 | 0 | 4029.727 | 0 | 0.57 | 0.015977 | 21.86 | 84.2 | 115.8 | 6.17 | 13.08 | | High | High | High | High | High | High | High | High | High | High | High | 0 | 2.04E-08 | 4.15 |
| FALSE | High | SQEDEISPV | [R].SQEDE | 1xPhospho | Q6PR54-3 | 0.000765 | 0 | 2 | 4 | 2 | Q6PR54-3; | Isoform 3 c | Q6PR54-3 | 0 | 1783.888 | 0 | -0.57 | 0.001448 | 5.41 | 119.5 | 80.5 | 4.8 | 4.15 | Shared | High | High | High | High | High | High | High | High | High | High | High | 2.58E-05 | 0.000432 | 2.39 |
| FALSE | High | SQEFSFPVD | [K].SQEFSF | 1xPhospho | E9PUR0 1x | 1.35E-05 | 0 | 1 | 4 | 2 | E9PUR0 | Ankyrin re | E9PUR0 [1 | 0 | 2305.115 | 0 | -0.88 | 0.002001 | 4.16 | 130.8 | 69.2 | 4.05 | 13.44 | | High | High | High | High | High | High | High | High | High | High | High | 0 | 1.79E-05 | 3.51 |
| FALSE | High | SQEGTAEIF | [R].SQEGTA | 1xCarbami | B9EKJ7 1xP | 0.000109 | 0 | 1 | 2 | 2 | B9EKJ7 | Tankyrase | B9EKJ7 [22 | 1 | 3180.456 | 0 | 0.02 | 0.736731 | 10.41 | 98.9 | 101.1 | 9.75 | 6.33 | | High | High | High | High | High | High | High | High | High | High | High | 0 | 1.98E-05 | 2.27 |
| FALSE | High | SQELEPIPD | [R].SQELEP | 1xPhospho | Q62187 1x | 0.0016 | 5.66E-05 | 1 | 2 | 3 | Q62187 | Transcripti | Q62187 [3 | 0 | 2569.176 | 0 | -0.31 | 0.006362 | 0.75 | 110.8 | 89.2 | 7.22 | 7.07 | | High | High | High | High | High | High | High | High | High | High | High | 0.000204 | 0.001335 | 3.82 |
| FALSE | High | SQEPISNDC | [K].SQEPIS | 1xPhospho | Q80WJ7 1x | 0.071071 | 0.005247 | 1 | 4 | 1 | Q80WJ7 | Protein LYF | Q80WJ7 [4 | 1 | 2343.112 | 0 | 0.98 | 0.427509 | 32.5 | 71.7 | 128.3 | 60.48 | 58.7 | | High | High | High | High | High | High | High | High | High | High | High | 0.002703 | 0.02897 | 1.74 |
| FALSE | High | SQEPISNDC | [K].SQEPIS | 1xPhospho | Q80WJ7 1x | 0.002441 | 0.000109 | 1 | 4 | 3 | Q80WJ7 | Protein LYF | Q80WJ7 [4 | 1 | 2572.275 | 0 | 0.55 | 0.004771 | 9.63 | 79.9 | 120.1 | 5.06 | 9.24 | | High | High | High | High | High | High | High | High | High | High | High | 4.94E-05 | 0.000646 | 3.51 |
| FALSE | High | SPYGSRSPF | [R].SPYGSR | 1xPhospho | P40201 1xP | 0.003857 | 0.000261 | 1 | 1 | 1 | P40201 | Chromodo | P40201 [16 | 1 | 2152.976 | 0 | 0.07 | 0.41116 | 12.98 | 96.6 | 103.4 | 11.45 | 12.96 | | High | High | High | High | High | High | High | High | High | High | High | 0.000117 | 0.001079 | 3.15 |
| FALSE | High | SSSWGSTE | [R].SSSWG | 1xPhospho | Q3U3E2 1x | 5.91E-05 | 0 | 1 | 1 | 6 | Q3U3E2 | Protein FA | Q3U3E2 [2 | 0 | 1747.867 | 0 | -0.48 | 0.003925 | 5.37 | 115.8 | 84.2 | 7.66 | 8.91 | | High | High | High | High | High | High | High | High | High | High | High | 0 | 9.97E-06 | 3.67 |
| FALSE | High | SSSWGSTE | [R].SSSWG | 2xPhospho | Q3U3E2 2x | 0.000173 | 0 | 1 | 1 | 1 | Q3U3E2 | Protein FA | Q3U3E2 [2 | 0 | 1827.833 | 0 | -0.17 | 0.257424 | 5.49 | 105.3 | 94.7 | 3.04 | 14.09 | | High | High | High | High | High | High | High | High | High | High | High | 0 | 3.31E-05 | 3.43 |
| FALSE | High | SSSYSEANE | [R].SSSYSE | 1xPhospho | Q3U2G7 1x | 0.000244 | 0 | 1 | 11 | 4 | Q3U2G7 | Putative ur | Q3U2G7 [7 | 0 | 2553.158 | 0 | -0.6 | 0.000554 | 2.52 | 120.2 | 79.8 | 6.7 | 10.69 | | High | High | High | High | High | High | High | High | High | High | High | 0 | 5.72E-05 | 3.22 |
| FALSE | High | TEENVSDG | [R].TEENVS | 1xPhospho | P46414 1xP | 8.4E-08 | 0 | 1 | 2 | 1 | P46414 | Cyclin-dep | P46414 [17 | 0 | 2598.234 | 0 | 2.87 | 0.00031 | 16.29 | 23.8 | 176.2 | 8.92 | 7.68 | | High | High | High | High | High | High | High | High | High | High | High | 0 | 6.28E-09 | 3.77 |
| FALSE | High | TEEPAPIPIS | [K].TEEPAP | 1xPhospho | Q6P5E3 1x | 1.19E-07 | 0 | 1 | 3 | 2 | Q6P5E3 | Sal-like 1 (I | Q6P5E3 [5 | 0 | 2641.364 | 0 | -0.64 | 0.003746 | 4.28 | 122.4 | 77.6 | 5.61 | 9.32 | | High | High | High | High | High | High | High | High | High | High | High | 0 | 9.34E-09 | 3.87 |

| | | | | | | | | | | | | | | | | | | | | | | | | | | | | | | | | | | | | | | |
|---|---|---|---|---|---|---|---|---|---|---|---|---|---|---|---|---|---|---|---|---|---|---|---|---|---|---|---|---|---|---|---|---|---|---|---|---|---|---|
| FALSE | High | TEEPAPIPIS | [K].TEEPAF | 2xPhosphc | Q6P5E3 2x | 1.77E-05 | 0 | 1 | 3 | 1 | Q6P5E3 | Sal-like 1 (I | Q6P5E3 [5 | 0 | 2721.33 | 0 | -1.33 | 0.000158 | 0.45 | 143.3 | 56.7 | 5.58 | 10.53 | | High | High | High | High | High | High | High | High | High | High | High | 0 | 2.56E-06 | 3.43 |
| FALSE | High | TEEPESPES | [R].TEEPES | 1xPhosphc | Q9WVL3 1: | 1.74E-06 | 0 | 1 | 2 | 1 | Q9WVL3 | Solute carr | Q9WVL3 [2 | 0 | 2735.188 | 0 | 0.53 | 0.009796 | 21.9 | 78.6 | 121.4 | 18.77 | 13.26 | | High | High | High | High | High | High | High | High | High | High | High | 0 | 1.89E-07 | 4.35 |
| FALSE | High | TEEPVSPLT | [R].TEEPVS | 1xPhosphc | G5E8K3 1x | 3.36E-06 | 0 | 2 | 26 | 2 | G5E8K3; A | Ankyrin 3, | G5E8K3 [1 | 0 | 2001.035 | 0 | 1.42 | 0.000325 | 6.99 | 54.4 | 145.6 | 5.32 | 6.32 | Shared | High | High | High | High | High | High | High | High | High | High | High | 0 | 3.96E-07 | 2.84 |
| FALSE | High | TEESLESTE | [K].TEESLE | 1xPhosphc | G3X8Q1 1x | 0.000916 | 0 | 1 | 3 | 2 | G3X8Q1 | Calcineurir | G3X8Q1 [1 | 0 | 1707.768 | 0 | 0.63 | 0.005887 | 18.48 | 78.6 | 121.4 | 4.98 | 10.33 | | High | High | High | High | High | High | High | High | High | High | High | 0 | 0.000217 | 2.83 |
| FALSE | High | TEEVEVESE | [K].TEEVEV | 1xPhosphc | O55091 1x | 1.46E-06 | 0 | 1 | 1 | 3 | O55091 | Protein IM | O55091 [1 | 0 | 3183.539 | 0 | 0.89 | 0.002413 | 14.45 | 70.6 | 129.4 | 6.83 | 8.72 | | High | High | High | High | High | High | High | High | High | High | High | 0 | 1.8E-06 | 4.57 |
| FALSE | High | TEEVLSPDC | [R].TEEVLS | 1xPhosphc | Q9QYB5 1x | 5.93E-06 | 0 | 1 | 7 | 4 | Q9QYB5 | Gamma-ad | Q9QYB5 [6 | 0 | 1883.94 | 0 | 0.98 | 0.000384 | 5.73 | 67.2 | 132.8 | 3.16 | 4.96 | | High | High | High | High | High | High | High | High | High | High | High | 0 | 1.04E-06 | 3.59 |
| FALSE | High | TEEVLSPDC | [R].TEEVLS | 2xPhosphc | Q9QYB5 2x | 2.38E-05 | 0 | 1 | 7 | 2 | Q9QYB5 | Gamma-ad | Q9QYB5 [6 | 0 | 1963.907 | 0 | 0.63 | 0.00109 | 9.4 | 77.4 | 122.6 | 6.95 | 6.37 | | High | High | High | High | High | High | High | High | High | High | High | 0.001199 | 0.0129 | 2.87 |
| FALSE | High | TEFSPAAFE | [K].TEFSPA | 1xPhosphc | Q6T264 1x | 0.000117 | 0 | 1 | 2 | 1 | Q6T264 | Mastermin | Q6T264 [3 | 0 | 2430.154 | 0 | 0.41 | 0.004351 | 5.52 | 83.5 | 116.5 | 5.22 | 8.63 | | High | High | High | High | High | High | High | High | High | High | High | 0 | 2.15E-05 | 2.66 |
| FALSE | High | TEFSPAAFE | [K].TEFSPA | 2xPhosphc | Q6T264 2x | 0.001881 | 5.66E-05 | 1 | 2 | 1 | Q6T264 | Mastermin | Q6T264 [3 | 0 | 2510.12 | 0 | 0.73 | 0.002306 | 14.54 | 71.1 | 128.9 | 13.68 | 8.18 | | High | High | High | High | High | High | High | High | High | High | High | 2.58E-05 | 0.000484 | 3.69 |
| FALSE | High | TEGYAAFQ | [R].TEGYA | 2xPhosphc | A6H6B3 2x | 0.005467 | 0.00062 | 1 | 8 | 1 | A6H6B3 | RalA bindir | A6H6B3 [8 | 0 | 2594.006 | 0 | 0.13 | 0.071087 | 11.71 | 93.9 | 106.1 | 12.2 | 12.58 | | High | High | High | High | High | High | High | High | High | High | High | 0.000275 | 0.001605 | 3.04 |
| FALSE | High | TEGYAAFQ | [R].TEGYA | 2xPhosphc | A6H6B3 2x | 0.00145 | 5.66E-05 | 1 | 8 | 1 | A6H6B3 | RalA bindir | A6H6B3 [8 | 0 | 2823.169 | 0 | 0.19 | 0.667996 | 37.48 | 93.3 | 106.7 | 20.84 | 4.88 | | High | High | High | High | High | High | High | High | High | High | High | 2.58E-05 | 0.00036 | 2.32 |
| FALSE | High | TEEIKTPEP | [R].TEEIKT | 1xPhosphc | Q6EDY6 1x | 0.001503 | 5.66E-05 | 1 | 4 | 1 | Q6EDY6 | F-actin-unc | Q6EDY6 [1 | 1 | 2720.343 | 0 | 0.09 | 0.126428 | 8.05 | 94.4 | 105.6 | 7.16 | 6.05 | | High | High | High | High | High | High | High | High | High | High | High | 2.58E-05 | 0.000378 | 2.87 |
| FALSE | High | TEHAPSPSS | [K].TEHAPS | 1xPhosphc | Q69ZA1 1x | 0.001158 | 0 | 1 | 3 | 2 | Q69ZA1 | Cyclin-dep | Q69ZA1 [5 | 0 | 1892.952 | 0 | 0.86 | 0.003512 | 18.67 | 69.6 | 130.4 | 5.35 | 10.28 | | High | High | High | High | High | High | High | High | High | High | High | 0 | 0.000313 | 2.61 |
| FALSE | High | TEHSHSGT | [R].TEHSHS | 1xPhosphc | A2A863-3 | 0.052576 | 0.00384 | 1 | 6 | 3 | A2A863-3 | Isoform 3 c | A2A863-3 | 0 | 1530.726 | 0 | 3.19 | 0.000647 | 13.53 | 20.1 | 179.9 | 13.22 | 13.27 | | High | High | High | High | High | High | High | High | High | High | High | 0.001999 | 0.02044 | 2.1 |
| FALSE | High | TEHTLQHS | [R].TEHTLC | 1xPhosphc | P97929 1xl | 3.9E-05 | 0 | 1 | 2 | 2 | P97929 | Breast can | P97929 [2C | 0 | 2356.195 | 0 | -1.1 | 0.000385 | 1.48 | 138.3 | 61.7 | 6.04 | 6.08 | | High | High | High | High | High | High | High | High | High | High | High | 0 | 6.27E-06 | 3.72 |
| FALSE | High | TELEDTLDS | [K].TELEDT | 1xPhosphc | Q8VDD5 1x | 1.7E-05 | 0 | 1 | 2 | 1 | Q8VDD5 | Myosin-9 ( | Q8VDD5 [1 | 0 | 2229.049 | 0 | 0.97 | 0.008068 | 14.81 | 57.7 | 142.3 | 18.39 | 23.46 | | High | High | High | High | High | High | High | High | High | High | High | 0 | 2.45E-06 | 4.46 |
| FALSE | High | TELGSQTPI | [K].TELGSC | 1xPhosphc | Q8BQR4 1x | 0.001349 | 0 | 1 | 4 | 1 | Q8BQR4 | KAT8 regul | Q8BQR4 [1 | 0 | 1600.742 | 0 | 0.05 | 0.395734 | 3.61 | 98.1 | 101.9 | 7.02 | 3.92 | | High | High | High | High | High | High | High | High | High | High | High | 0 | 0.000333 | 1.86 |
| FALSE | High | TELGSWSS | [R].TELGSV | 1xCarbami | E9PWU7 1: | 0.002115 | 5.66E-05 | 2 | 4 | 4 | E9PWU7; C | Kinesin-like | E9PWU7 [7 | 0 | 3334.504 | 0 | 0.42 | 0.006628 | 8.28 | 82.5 | 117.5 | 12.3 | 5.97 | Shared | High | High | High | High | High | High | High | High | High | High | High | 2.58E-05 | 0.000555 | 3.51 |
| FALSE | High | TELLAADSL | [R].TELLAA | 1xPhosphc | Q6WQJ1 1: | 0.000786 | 0 | 1 | 2 | 1 | Q6WQJ1 | Sn1-specifi | Q6WQJ1 [8 | 0 | 1685.913 | 0 | 1.78 | 0.00062 | 29.1 | 47.2 | 152.8 | 17.63 | 6.13 | | High | High | High | High | High | High | High | High | High | High | High | 0 | 0.000182 | 1.79 |
| FALSE | High | TENAGDQI | [K].TENAGI | 1xPhosphc | G5E924 1x | 6.81E-07 | 0 | 1 | 4 | 2 | G5E924 | Heterogen | G5E924 [5 | 0 | 3378.441 | 0 | -0.57 | 0.009159 | 7.02 | 117.9 | 82.1 | 7.07 | 8.63 | | High | High | High | High | High | High | High | High | High | High | High | 0 | 1.41E-06 | 3.28 |
| FALSE | High | TENDGQPC | [R].TENDG | 1xCarbami | Q9CQS5 2x | 0.001313 | 0 | 1 | 2 | 1 | Q9CQS5 | Serine/thre | Q9CQS5 [4 | 1 | 3937.69 | 0 | -1.31 | 0.001238 | 4.85 | 143.2 | 56.8 | 7.36 | 17.9 | | High | High | High | High | High | High | High | High | High | High | High | 0 | 0.000325 | 2.87 |
| FALSE | High | TEPAPAEE( | [K].TEPAPA | 1xPhosphc | B2RRE0 1x | 0.001068 | 0 | 1 | 3 | 1 | B2RRE0 | A kinase (P | B2RRE0 [3 | 1 | 3141.41 | 0 | -2.56 | 0.205405 | 1.57 | 170 | 30 | 56.38 | 127.33 | | High | High | High | High | High | High | High | High | High | High | High | 0 | 0.000256 | 2.97 |
| FALSE | High | TEPAPAEE( | [K].TEPAPA | 1xDeamida | B2RRE0 1x | 0.014158 | 0.001011 | 1 | 3 | 2 | B2RRE0 | A kinase (P | B2RRE0 [3 | 1 | 3142.394 | 0 | -4.14 | | 0.39 | 171.3 | 28.7 | 64.65 | 125.41 | | High | High | High | High | High | High | High | Not Found | Not Found | High | High | 0.000538 | 0.00463 | 2.97 |
| FALSE | High | TEPAPAEE( | [K].TEPAPA | 1xPhosphc | B2RRE0 1x | 1.18E-08 | 0 | 1 | 3 | 16 | B2RRE0 | A kinase (P | B2RRE0 [3 | 1 | 3370.573 | 0 | -1.86 | 0.000687 | 1.69 | 154.6 | 45.4 | 7.03 | 16.09 | | High | High | High | High | High | High | High | High | High | High | High | 0 | 6.27E-09 | 4.23 |
| FALSE | High | TEPAPAEE( | [K].TEPAPA | 1xDeamida | B2RRE0 1x | 6.43E-06 | 0 | 1 | 3 | 4 | B2RRE0 | A kinase (P | B2RRE0 [3 | 1 | 3371.557 | 0 | -1.91 | 0.000456 | 0.62 | 155.3 | 44.7 | 8.48 | 15.15 | | High | High | High | High | High | High | High | High | High | High | High | 0 | 8.22E-07 | 4.45 |
| FALSE | High | TEPAQDSS | [K].TEPAQI | 1xPhosphc | Q6PGL7 1x | 1.51E-06 | 0 | 1 | 2 | 1 | Q6PGL7 | WASH com | Q6PGL7 [9 | 0 | 2016.009 | 0 | -0.2 | 0.068202 | 4.42 | 107.8 | 92.2 | 5.01 | 10.01 | | High | High | High | High | High | High | High | High | High | High | High | 0 | 1.61E-07 | 3.76 |
| FALSE | High | TEHGLTPK | [K].TEHGLT | 1xPhosphc | F6VAN0 1x | 0.055308 | 0.003982 | 1 | 2 | 1 | F6VAN0 | Cyclic AMP | F6VAN0 [1 | 0 | 1420.76 | 0 | -0.24 | 0.003574 | 1.41 | 108.3 | 91.7 | 3.18 | 6.94 | | High | High | High | High | High | High | High | High | High | High | High | 0.002074 | 0.02159 | 2.7 |
| FALSE | High | TEPEQGTL' | [R].TEPEQ( | 1xDeamida | Q68FH0 1x | 0.003342 | 0.00016 | 1 | 6 | 3 | Q68FH0 | Plakophilin | Q68FH0 [1 | 0 | 3243.453 | 0 | -0.5 | 0.018163 | 7.45 | 120.3 | 79.7 | 13.75 | 10.19 | | High | High | High | High | High | High | High | High | High | High | High | 7.19E-05 | 0.000927 | 4.18 |
| FALSE | High | TEEDRENT | [R].TEEDRE | 2xPhosphc | P70399 2x | 1.43E-06 | 0 | 1 | 4 | 1 | P70399 | TP53-bindi | P70399 [5 | 1 | 3222.449 | 0 | 0.31 | 0.061182 | 12.35 | 87.8 | 112.2 | 10.91 | 10.34 | | High | High | High | High | High | High | High | High | High | High | High | 0 | 1.51E-07 | 4.13 |
| FALSE | High | TEEDRENT | [R].TEEDRE | 1xPhosphc | P70399 1x | 3.14E-08 | 0 | 1 | 4 | 2 | P70399 | TP53-bindi | P70399 [5 | 1 | 3142.483 | 0 | 0.46 | 0.024467 | 9.98 | 87.5 | 112.5 | 10.38 | 12.62 | | High | High | High | High | High | High | High | High | High | High | High | 0 | 2.08E-09 | 3.72 |
| FALSE | High | TEAEVLAEI | [K].TEAEVL | 2xPhosphc | Q9QXX8 2x | 0.000628 | 0 | 1 | 1 | 2 | Q9QXX8 | Nuclear fra | Q9QXX8 [3 | 1 | 2888.436 | 0 | -1.22 | 0.148553 | 5.32 | 142.3 | 57.7 | 51.13 | 42.81 | | High | High | High | High | High | High | High | High | High | High | High | 0 | 0.000142 | 2.67 |
| FALSE | High | TEALEDSEI | [R].TEALED | 1xPhosphc | Q9D479 1x | 0.000458 | 0 | 1 | 2 | 1 | Q9D479 | UV-stimula | Q9D479 [3 | 0 | 2862.286 | 0 | 2.01 | 0.000225 | 11.12 | 39.8 | 160.2 | 14.1 | 7.37 | | High | High | High | High | High | High | High | High | High | High | High | 0 | 9.97E-05 | 2.38 |
| FALSE | High | TEALSPYSK | [K].TEALSP | 1xPhosphc | F8WI37 1x | 0.004033 | 0.000311 | 1 | 2 | 2 | F8WI37 | Histone-lys | F8WI37 [2 | 0 | 1533.797 | 0 | -0.37 | 0.012104 | 1.2 | 113.7 | 86.3 | 7.9 | 6.84 | | High | High | High | High | High | High | High | High | High | High | High | 0.000183 | 0.00125 | 2.31 |
| FALSE | High | TEANSEEAI | [R].TEANSE | 1xPhosphc | Q9CZ62 1x | 0.001986 | 5.66E-05 | 1 | 3 | 1 | Q9CZ62 | Centrosom | Q9CZ62 [5 | 0 | 2236.009 | 0 | 0.21 | 0.034149 | 9.21 | 88.6 | 111.4 | 9.15 | 10.38 | | High | High | High | High | High | High | High | High | High | High | High | 2.58E-05 | 0.000516 | 3.72 |
| FALSE | High | TEATQGLD | [R].TEATQ( | 1xPhosphc | A2ARP8 1x | 2.81E-10 | 0 | 1 | 3 | 1 | A2ARP8 | Microtubu | A2ARP8 [1 | 0 | 3234.571 | 0 | -1.89 | 0.00227 | 1.54 | 159.9 | 40.1 | 16.52 | 11.52 | | High | High | High | High | High | High | High | High | High | High | High | 0 | 1.04E-11 | 4.76 |
| FALSE | High | TEATQPSTI | [K].TEATQI | 1xPhosphc | A2AUZ7 1x | 5.67E-06 | 0 | 2 | 4 | 4 | A2AUZ7; Q | Sal-like pro | A2AUZ7 [1 | 0 | 2414.226 | 0 | -1.74 | 0.00047 | 1.18 | 152.8 | 47.2 | 8.76 | 5.27 | Shared | High | High | High | High | High | High | High | High | High | High | High | 0 | 0.000019 | 2.4 |
| FALSE | High | TEAVSPTVS | [R].TEAVSF | 1xPhosphc | H3BJD6 1x | 2.23E-05 | 0 | 1 | 10 | 2 | H3BJD6 | Protein ph | H3BJD6 [1 | 0 | 3341.692 | 0 | -0.75 | 0.003068 | 6.68 | 126.1 | 73.9 | 9.21 | 8.63 | | High | High | High | High | High | High | High | High | High | High | High | 0 | 4.93E-06 | 3.38 |
| FALSE | High | TECQAYPG | [R].TECQA' | 2xCarbami | Q921Q9 1x | 4.05E-05 | 0 | 2 | 4 | 3 | Q921Q9; C | Mapkbp1 p | Q921Q9 [5 | 0 | 4551.209 | 0 | -0.58 | 0.008081 | 6.58 | 120.6 | 79.4 | 1.52 | 15.13 | Shared | High | High | High | High | High | High | High | High | High | High | High | 0 | 6.52E-06 | 3.26 |
| FALSE | High | TEDCSPEH | [K].TEDCSF | 1xCarbami | Q3U8K7 1x | 0.001337 | 0 | 1 | 3 | 1 | Q3U8K7 | Histone-lys | Q3U8K7 [6 | 0 | 2028.914 | 0 | -0.71 | 0.000384 | 0.43 | 124 | 76 | 2.79 | 1.95 | | High | High | High | High | High | High | High | High | High | High | High | 0 | 0.000331 | 2.46 |
| FALSE | High | TEDEEFLIQ | [R].TEDEEF | 1xPhosphc | E9PYL2 1x | 7.86E-07 | 0 | 1 | 2 | 3 | E9PYL2 | Proline-ricl | E9PYL2 [63 | 0 | 2529.259 | 0 | -0.4 | 0.016183 | 4.39 | 113.4 | 86.6 | 7.11 | 13.43 | | High | High | High | High | High | High | High | High | High | High | High | 0 | 7.78E-08 | 3.27 |
| FALSE | High | TEDGDWE' | [K].TEDGD' | 1xPhosphc | A2AQL0 1x | 0.000194 | 0 | 1 | 3 | 2 | A2AQL0 | Serine/thre | A2AQL0 [3 | 0 | 2525.01 | 0 | 1.48 | 0.00205 | 20.57 | 52.5 | 147.5 | 15.05 | 8.63 | | High | High | High | High | High | High | High | High | High | High | High | 0 | 3.81E-05 | 3.07 |
| FALSE | High | TEDGDWE' | [K].TEDGD' | 1xPhosphc | A2AQL0 1x | 1.57E-06 | 0 | 1 | 3 | 10 | A2AQL0 | Serine/thre | A2AQL0 [3 | 0 | 2541.005 | 0 | 1.98 | 0.000377 | 9.71 | 40.5 | 159.5 | 6.56 | 7.35 | | High | High | High | High | High | High | High | High | High | High | High | 4.94E-05 | 0.000619 | 3.37 |
| FALSE | High | TEDGGHA\ | [K].TEDGG | 1xPhosphc | Q6PB37 1x | 1.4E-06 | 0 | 1 | 2 | 3 | Q6PB37 | 5830417I1( | Q6PB37 [1 | 0 | 3044.498 | 0 | -0.9 | 0.000375 | 0.48 | 130.1 | 69.9 | 7.1 | 4.68 | | High | High | High | High | High | High | High | High | High | High | High | 0 | 4.39E-07 | 3.7 |
| FALSE | High | TEEDRENT | [R].TEEDRE | 1xDeamida | P70399 1x | 2.38E-08 | 0 | 1 | 4 | 6 | P70399 | TP53-bindi | P70399 [5 | 1 | 3143.467 | 0 | 0.43 | 0.106257 | 15.22 | 85.3 | 114.7 | 10.35 | 16.68 | | High | High | High | High | High | High | High | High | High | High | High | 0 | 1.53E-09 | 4.44 |
| FALSE | High | TEDGGWE' | [K].TEDGG | 1xPhosphc | Q6P9R2 1x | 6.35E-08 | 0 | 1 | 3 | 5 | Q6P9R2 | Serine/thre | Q6P9R2 [3 | 0 | 2813.057 | 0 | -0.26 | 0.000396 | 0.28 | 109 | 91 | 2.68 | 1.77 | | High | High | High | High | High | High | High | High | High | High | High | 0 | 4.6E-09 | 4.4 |
| FALSE | High | TEDQHLGY | [K].TEDQH | 1xPhosphc | A2AUM9 1 | 0.003962 | 0.000311 | 1 | 1 | 2 | A2AUM9 | Centrosom | A2AUM9 [ | 0 | 3166.416 | 0 | -0.47 | 0.004114 | 6.31 | 116.7 | 83.3 | 4.27 | 9.06 | | High | High | High | High | High | High | High | High | High | High | High | 0.00014 | 0.001116 | 2.61 |
| FALSE | High | TEDSDDIHI | [K].TEDSDI | 1xPhosphc | Q9ERU9 1x | 7.65E-07 | 0 | 1 | 1 | 2 | Q9ERU9 | E3 SUMO-p | Q9ERU9 [1 | 0 | 2654.246 | 0 | 0.61 | 0.025753 | 27.33 | 75.9 | 124.1 | 7.38 | 15.63 | | High | High | High | High | High | High | High | High | High | High | High | 0 | 7.49E-08 | 3.59 |
| FALSE | High | TEDSDDIHI | [K].TEDSDI | 1xPhosphc | Q9ERU9 1x | 2.87E-05 | 0 | 1 | 1 | 2 | Q9ERU9 | E3 SUMO-p | Q9ERU9 [1 | 0 | 2670.241 | 0 | 0.47 | 0.035441 | 6.18 | 77.4 | 122.6 | 14.29 | 21.14 | | High | High | High | High | High | High | High | High | High | High | High | 0 | 4.43E-06 | 3.55 |
| FALSE | High | TEDSEQDG | [R].TEDSEC | 1xPhosphc | Q5BLK4 1x | 0.000442 | 0 | 1 | 2 | 1 | Q5BLK4 | Terminal u | Q5BLK4 [1 | 0 | 2987.402 | 0 | 0.79 | 0.001007 | 13.46 | 74.6 | 125.4 | 10 | 4.49 | | High | High | High | High | High | High | High | High | High | High | High | 0 | 9.52E-05 | 3.17 |
| FALSE | High | TEDTLQMI | [R].TEDTLC | 1xPhosphc | Q6ZWQ0 1 | 7.35E-09 | 0 | 1 | 2 | 2 | Q6ZWQ0 | Nesprin-2 ( | Q6ZWQ0 [ | 0 | 2970.412 | 0 | 2.69 | 0.002083 | 17.13 | 29.8 | 170.2 | 11.06 | 20.04 | | High | High | High | High | High | High | High | High | High | High | High | 0 | 4.07E-10 | 5.34 |
| FALSE | High | TEDTLQMI | [R].TEDTLC | 1xPhosphc | Q6ZWQ0 1 | 0.000108 | 0 | 1 | 2 | 8 | Q6ZWQ0 | Nesprin-2 ( | Q6ZWQ0 [ | 0 | 2986.407 | 0 | 1.91 | 0.000797 | 6.02 | 42.1 | 157.9 | 10.82 | 10.93 | | High | High | High | High | High | High | High | High | High | High | High | 0 | 5.33E-05 | 4.22 |
| FALSE | High | TEEAADGG | [R].TEEAAI | 1xPhosphc | Q8BP71 1x | 1.09E-05 | 0 | 1 | 3 | 1 | Q8BP71 | RNA bindir | Q8BP71 [5 | 0 | 2854.27 | 0 | -0.42 | 0.165885 | 25.51 | 106.5 | 93.5 | 14.65 | 21.76 | | High | High | High | High | High | High | High | High | High | High | High | 0 | 1.49E-06 | 4.71 |
| FALSE | High | TEEAADGG | [R].TEEAAI | 1xPhosphc | Q8BP71 1x | 1.24E-05 | 0 | 1 | 3 | 1 | Q8BP71 | RNA bindir | Q8BP71 [5 | 0 | 2870.265 | 0 | -0.14 | 0.799466 | 11.46 | 98.5 | 101.5 | 17.07 | 25.44 | | High | High | High | High | High | High | High | High | High | High | High | 0 | 1.72E-06 | 4.51 |
| FALSE | High | TEEAGALA( | [R].TEEAG/ | 2xPhosphc | P97452 2x | 3.23E-08 | 0 | 1 | 4 | 12 | P97452 | Ribosome I | P97452 [97 | 0 | 2990.155 | 0 | -0.5 | 0.010801 | 4.09 | 118 | 82 | 8.64 | 9.25 | | High | High | High | High | High | High | High | High | High | High | High | 0 | 2.14E-09 | 5.38 |
| FALSE | High | TEEAILISLA | [R].TEEAILI | 1xPhosphc | Q3UHX9 1x | 0.015329 | 0.001089 | 1 | 2 | 4 | Q3UHX9 | Putative m | Q3UHX9 [3 | 0 | 3438.856 | 0 | -1.6 | 0.001501 | 1.87 | 150.4 | 49.6 | 10.4 | 14.52 | | High | High | High | High | High | High | High | High | High | High | High | 0.002293 | 0.02385 | 1.91 |
| FALSE | High | TEEAILISLA | [R].TEEAILI | 2xPhosphc | Q3UHX9 2x | 0.000167 | 0 | 1 | 2 | 2 | Q3UHX9 | Putative m | Q3UHX9 [3 | 0 | 3518.822 | 0 | -0.57 | 0.010909 | 6.19 | 120.5 | 79.5 | 5.45 | 14.59 | | High | High | High | High | High | High | High | High | High | High | High | 0 | 3.18E-05 | 2.48 |
| FALSE | High | TEEDETSEI | [K].TEEDET | 1xCarbami | Q3V141 1x | 3.45E-06 | 0 | 1 | 7 | 1 | Q3V141 | Putative ur | Q3V141 [6 | 0 | 2759.212 | 0 | -0.44 | 0.005481 | 1.43 | 114.4 | 85.6 | 4.34 | 6.05 | | High | High | High | High | High | High | High | High | High | High | High | 0 | 4.09E-07 | 4.22 |
| FALSE | High | TEEDRENT | [R].TEEDRE | 1xPhosphc | P70399 1x | 1.06E-06 | 0 | 1 | 4 | 1 | P70399 | TP53-bindi | P70399 [5 | 1 | 2913.32 | 0 | 0.11 | 0.570598 | 107.96 | 87.8 | 112.2 | 49.95 | 74.42 | | High | High | High | High | High | High | High | High | High | High | High | 0 | 1.08E-07 | 4.78 |
| FALSE | High | TEDLEATSE | [K].TEDLEA | 1xPhosphc | A0A0R4J0F | 0.081752 | 0.006219 | 1 | 3 | 1 | A0A0R4J0F | Vesicle-ass | A0A0R4J0F | 0 | 1944.936 | 0 | 1.3 | 0.000706 | 3.75 | 58.9 | 141.1 | 4.58 | 8.54 | | High | High | High | High | High | High | High | High | High | High | High | 0.0032 | 0.03395 | 2.58 |
| FALSE | High | TEPGTATN | [R].TEPGT/ | 1xPhosphc | G5E8R8 1x | 0.000101 | 0 | 1 | 3 | 1 | G5E8R8 | UBX domai | G5E8R8 [3 | 0 | 3323.558 | 0 | -0.37 | 0.050207 | 8.98 | 110.9 | 89.1 | 7.11 | 23.7 | | High | High | High | High | High | High | High | High | High | High | High | 0 | 1.8E-05 | 2.27 |
| FALSE | High | TEPGTATN | [R].TEPGT/ | 1xPhosphc | G5E8R8 1x | 7.36E-06 | 0 | 1 | 3 | 3 | G5E8R8 | UBX domai | G5E8R8 [3 | 0 | 3339.553 | 0 | -0.3 | 0.078517 | 14.12 | 108.2 | 91.8 | 5.36 | 15.6 | | High | High | High | High | High | High | High | High | High | High | High | 0 | 9.62E-07 | 4.14 |
| FALSE | High | TEPGTLFFT | [K].TEPGTL | 1xPhosphc | F8WI37 1x | 3.33E-06 | 0 | 1 | 7 | 2 | F8WI37 | Histone-lys | F8WI37 [4 | 0 | 2405.138 | 0 | -0.48 | 0.031654 | 2.24 | 108.7 | 91.3 | 9.26 | 9.88 | | High | High | High | High | High | High | High | High | High | High | High | 0 | 3.91E-07 | 3.72 |
| FALSE | High | TETDITTVE | [K].TETDIT | 1xPhosphc | E9Q6J5-3 1 | 9.74E-06 | 0 | 1 | 3 | 2 | E9Q6J5-3 | Isoform Ne | E9Q6J5-3 [ | 0 | 2218.089 | 0 | -0.22 | 0.011388 | 6.21 | 107.3 | 92.7 | 3.99 | 6.67 | | High | High | High | High | High | High | High | High | High | High | High | 0 | 1.32E-06 | 4.05 |
| FALSE | High | TETPPPLAS | [R].TETPPP | 1xPhosphc | Q9D0R8 1x | 0.013074 | 0.000889 | 1 | 1 | 3 | Q9D0R8 | Protein LSI | Q9D0R8 [7 | 0 | 1762.919 | 0 | 0.33 | 0.641342 | 16.92 | 94.8 | 105.2 | 36.42 | 56.79 | | High | High | High | High | High | High | High | High | High | High | High | 0.003922 | 0.04164 | 2.09 |
| FALSE | High | TETPPPLAS | [R].TETPPP | 1xPhosphc | Q9D0R8 1x | 8.92E-07 | 0 | 1 | 1 | 3 | Q9D0R8 | Protein LSI | Q9D0R8 [7 | 0 | 1992.082 | 0 | 0.38 | 0.006105 | 2.55 | 86.8 | 113.2 | 7.47 | 3.16 | | High | High | High | High | High | High | High | High | High | High | High | 0 | 1.67E-06 | 3.41 |
| FALSE | High | TETPTEELK | [K].TETPTE | 1xPhosphc | Q6KCD5 1x | 0.020142 | 0.001472 | 1 | 3 | 1 | Q6KCD5 | Nipped-B-I | Q6KCD5 [6 | 0 | 1585.813 | 0 | -0.45 | 0.031401 | 1.69 | 117 | 83 | 16.83 | 5.2 | | High | High | High | High | High | High | High | High | High | High | High | 0.000786 | 0.006852 | 1.57 |
| FALSE | High | TETSLDTPL | [K].TETSLD | 1xPhosphc | Q3TI53-7 1 | 7.98E-06 | 0 | 1 | 8 | 1 | Q3TI53-7 | Isoform 7 c | Q3TI53-7 [ | 0 | 2061.023 | 0 | -1.88 | 0.00075 | 1.05 | 156.8 | 43.2 | 7.31 | 11.81 | | High | High | High | High | High | High | High | High | High | High | High | 0 | 1.05E-06 | 2.49 |
| FALSE | High | TETYVEAN | [K].TETYVE | 1xCarbami | P28700 1x | 0.000125 | 0 | 1 | 3 | 1 | P28700 | Retinoic ac | P28700 [2 | 0 | 3690.707 | 0 | 0.27 | 0.137374 | 10.93 | 90.8 | 109.2 | 8.09 | 29.92 | | High | High | High | High | High | High | High | High | High | High | High | 0 | 2.31E-05 | 4.03 |
| FALSE | High | TEVAQVDE | [K].TEVAQ' | 1xCarbami | A0A0G2JG | 0.001058 | 0 | 1 | 2 | 2 | A0A0G2JG | Crystallin b | A0A0G2JG | 0 | 3478.48 | 0 | 2.34 | 0.002238 | 26.62 | 30.7 | 169.3 | 38.28 | 14.51 | | High | High | High | High | High | High | High | High | High | High | High | 0 | 0.000254 | 2.68 |
| FALSE | High | TEVNGFVH | [K].TEVNGI | 1xPhosphc | Q6P5E3 1x | 6.37E-06 | 0 | 1 | 3 | 1 | Q6P5E3 | Sal-like 1 (I | Q6P5E3 [1 | 0 | 2182.095 | 0 | -1.28 | 0.001332 | 2.1 | 141.5 | 58.5 | 9.03 | 7.85 | | High | High | High | High | High | High | High | High | High | High | High | 0 | 8.13E-07 | 3.29 |
| FALSE | High | TEVNGFVH | [K].TEVNGI | 1xDeamida | Q6P5E3 1x | 0.000573 | 0 | 1 | 3 | 2 | Q6P5E3 | Sal-like 1 (I | Q6P5E3 [1 | 0 | 2183.079 | 0 | -1.11 | 0.000797 | 2.71 | 136.6 | 63.4 | 6.58 | 9.31 | | High | High | High | High | High | High | High | High | High | High | High | 0 | 0.000128 | 2.2 |
| FALSE | High | TEVSGCPEI | [K].TEVSG( | 1xCarbami | Q3UH85 1x | 0.002252 | 0.000109 | 1 | 5 | 1 | Q3UH85 | Nucleolar p | Q3UH85 [3 | 0 | 1828.835 | 0 | -1.47 | 0.000436 | 2.35 | 148.6 | 51.4 | 4.88 | 11.77 | | High | High | High | High | High | High | High | High | High | High | High | 4.94E-05 | 0.00059 | 2.37 |
| FALSE | High | TEVTDSSSI | [K].TEVTDS | 1xCarbami | A2A654 1x | 0.004491 | 0.000409 | 1 | 4 | 1 | A2A654 | Bromodom | A2A654 [1 | 0 | 3407.674 | 0 | -0.3 | 0.01265 | 8.08 | 109.9 | 90.1 | 6.03 | 5.71 | | High | High | High | High | High | High | High | High | High | High | High | 0.000183 | 0.001284 | 2.91 |
| FALSE | High | TFAPSGGE( | [R].TFAPSG | 1xPhosphc | O35185 1x | 4.94E-05 | 0 | 1 | 2 | 3 | O35185 | Class E bas | O35185 [2 | 0 | 3130.378 | 0 | 0.12 | 0.382937 | 42.89 | 93.3 | 106.7 | 9.69 | 17.74 | | High | High | High | High | High | High | High | High | High | High | High | 0 | 8.09E-06 | 4.28 |
| FALSE | High | TFDEESNA | [K].TFDEES | 2xPhosphc | A2AJK6 2x | 0.000407 | 0 | 1 | 2 | 1 | A2AJK6 | Chromodo | A2AJK6 [2 | 0 | 1916.788 | 0 | -0.88 | 0.005355 | 6.98 | 127.5 | 72.5 | 12.03 | 17.18 | | High | High | High | High | High | High | High | High | High | High | High | 0 | 8.71E-05 | 2.49 |
| FALSE | High | TETDITTVE | [K].TETDIT | 1xPhosphc | E9Q6J5-3 1 | 0.000104 | 0 | 1 | 3 | 2 | E9Q6J5-3 | Isoform Ne | E9Q6J5-3 [ | 0 | 1988.926 | 0 | 0.51 | 0.578502 | 13.61 | 80.7 | 119.3 | 63.52 | 69.36 | | High | High | High | High | High | High | High | High | High | High | High | 0 | 1.89E-05 | 3.6 |
| FALSE | High | TFEISEPVIF | [K].TFEISEI | 1xPhosphc | G3XA30 1x | 0.000346 | 0 | 1 | 2 | 3 | G3XA30 | MCG1618, | G3XA30 [3 | 0 | 2268.147 | 0 | -0.65 | 0.000757 | 3.49 | 120.4 | 79.6 | 8.36 | 8.48 | | High | High | High | High | High | High | High | High | High | High | High | 0 | 7.24E-05 | 3.05 |
| FALSE | High | TFPAGSSLE | [K].TFPAGS | 1xPhosphc | Q6P1G0 1x | 0.024001 | 0.001612 | 1 | 1 | 1 | Q6P1G0 | HEAT repe | Q6P1G0 [6 | 0 | 2016.973 | 0 | 2.36 | 0.003594 | 55.08 | 32.8 | 167.2 | 41.19 | 18.7 | | High | High | High | High | High | High | High | High | High | High | High | 0.000859 | 0.008393 | 2.46 |
| FALSE | High | TFPAGSSLE | [K].TFPAGS | 1xPhosphc | Q6P1G0 1x | 9.31E-06 | 0 | 1 | 1 | 2 | Q6P1G0 | HEAT repe | Q6P1G0 [6 | 0 | 2246.136 | 0 | 0.7 | 0.001154 | 6.34 | 72.2 | 127.8 | 7.4 | 7.62 | | High | High | High | High | High | High | High | High | High | High | High | 0 | 1.25E-06 | 4.18 |
| FALSE | High | TFQLQLLSF | [K].TFQLQI | 1xPhosphc | Q3UKX8 1x | 0.001799 | 5.66E-05 | 1 | 5 | 4 | Q3UKX8 | Putative ur | Q3UKX8 [7 | 0 | 3414.869 | 0 | -1.22 | 0.012737 | 6.31 | 134.2 | 65.8 | 11.66 | 34.19 | | High | High | High | High | High | High | High | High | High | High | High | 2.58E-05 | 0.000463 | 3.43 |
| FALSE | High | TFQIVPEVE | [K].TFQIVP | 1xPhosphc | Q501J7-3 1 | 0.000418 | 0 | 1 | 3 | 3 | Q501J7-3 | Isoform 3 c | Q501J7-3 [ | 0 | 3669.761 | 0 | 1.05 | 0.037043 | 43.22 | 65 | 135 | 28.49 | 23.94 | | High | High | High | High | High | High | High | High | High | High | High | 0 | 8.99E-05 | 3.28 |
| FALSE | High | TFQIVPEVE | [K].TFQIVP | 1xDeamida | Q501J7-3 1 | 1.47E-05 | 0 | 1 | 3 | 4 | Q501J7-3 | Isoform 3 c | Q501J7-3 [ | 0 | 3670.745 | 0 | 0.82 | 0.056336 | 15.78 | 70.6 | 129.4 | 11.84 | 32 | | High | High | High | High | High | High | High | High | High | High | High | 0 | 4.07E-06 | 4.78 |
| FALSE | High | TFQQIQEEI | [R].TFQQIC | 1xPhosphc | Q9CSU0 1x | 2.8E-06 | 0 | 1 | 5 | 16 | Q9CSU0 | Regulation | Q9CSU0 [1 | 0 | 4378.058 | 0 | 0.59 | 0.093555 | 24.06 | 86.1 | 113.9 | 13.48 | 15.96 | | High | High | High | High | High | High | High | High | High | High | High | 0 | 4.35E-05 | 4.37 |
| FALSE | High | TFSATVR | [R].TFSATV | 1xPhosphc | Q3U434 1x | 0.015329 | 0.001089 | 1 | 3 | 4 | Q3U434 | Putative ur | Q3U434 [5 | 0 | 1090.55 | 0 | 1.08 | 0.000447 | 3.46 | 64.1 | 135.9 | 5.18 | 6.67 | | High | High | High | High | High | High | High | High | High | High | High | 0.000571 | 0.005048 | 1.34 |
| FALSE | High | TFSDGLAA' | [R].TFSDGI | 1xPhosphc | D3Z0X5 1x | 0.000289 | 0 | 1 | 6 | 4 | D3Z0X5 | Pleckstrin I | D3Z0X5 [4 | 0 | 1347.651 | 0 | 1.08 | 0.001124 | 4.47 | 62 | 138 | 6.15 | 8.85 | | High | High | High | High | High | High | High | High | High | High | High | 0 | 5.94E-05 | 2.38 |
| FALSE | High | TFSDGLESF | [R].TFSDGI | 1xPhosphc | A0A0R4J0( | 0.081085 | 0.006219 | 1 | 6 | 1 | A0A0R4J0( | Disks large | A0A0R4J0( | 0 | 1320.603 | 0 | -0.23 | 0.056032 | 13.8 | 112.8 | 87.2 | 9.77 | 12.26 | | High | High | High | High | High | High | High | High | High | High | High | 0.0032 | 0.03373 | 1.9 |
| FALSE | High | TFSEASIWS | [R].TFSEAS | 2xPhosphc | B9EKC2 2x | 0.006713 | 0.00062 | 1 | 6 | 1 | B9EKC2 | ATP-bindin | B9EKC2 [6 | 0 | 2428.115 | 0 | 0.6 | 0.047532 | 21.71 | 84.9 | 115.1 | 5.58 | 19.83 | | High | High | High | High | High | High | High | High | High | High | High | 0.000275 | 0.002003 | 2.45 |
| FALSE | High | TFSEASIWS | [R].TFSEAS | 1xPhosphc | B9EKC2 1x | 0.001916 | 5.66E-05 | 1 | 6 | 1 | B9EKC2 | ATP-bindin | B9EKC2 [6 | 0 | 2577.311 | 0 | 0.65 | 0.123202 | 8.82 | 80.5 | 119.5 | 11.21 | 17.18 | | High | High | High | High | High | High | High | High | High | High | High | 2.58E-05 | 0.000497 | 2.93 |
| FALSE | High | TFSLDAAP/ | [R].TFSLDA | 1xPhosphc | Q8C6B2 1x | 1.31E-07 | 0 | 1 | 2 | 2 | Q8C6B2 | Rhotekin C | Q8C6B2 [5 | 0 | 2050.98 | 0 | 0.97 | 0.008468 | 23.2 | 70.5 | 129.5 | 10.97 | 18.08 | | High | High | High | High | High | High | High | High | High | High | High | 0 | 1.03E-08 | 4.07 |
| FALSE | High | TFSLDTILSS | [R].TFSLDT | 1xPhosphc | Q7TNF9 1x | 8.56E-05 | 0 | 1 | 2 | 2 | Q7TNF9 | Protein FAI | Q7TNF9 [8 | 0 | 2406.231 | 0 | -0.86 | 0.00149 | 5.89 | 128.4 | 71.6 | 8.25 | 9.69 | | High | High | High | High | High | High | High | High | High | High | High | 0 | 1.51E-05 | 3.23 |
| FALSE | High | TFGGAPGF | [R].TFGGAI | 1xPhosphc | P31001 1x | 0.093879 | 0.007306 | 1 | 3 | 1 | P31001 | Desmin OS | P31001 [17 | 0 | 2387.2 | 0 | -1.6 | 0.000965 | 1.83 | 146.7 | 53.3 | 10.78 | 23.38 | | High | High | High | High | High | High | High | High | High | High | High | 0.003727 | 0.04009 | 3.37 |
| FALSE | High | TESVPSDIN | [R].TESVPS | 1xPhosphc | Q61097 1x | 7.2E-08 | 0 | 1 | 5 | 1 | Q61097 | Kinase sup | Q61097 [3 | 0 | 1851.869 | 0 | -0.88 | 0.003583 | 7.51 | 135.5 | 64.5 | 7.05 | 18.4 | | High | High | High | High | High | High | High | High | High | High | High | 0 | 5.3E-09 | 4.39 |
| FALSE | High | TESVDEAE' | [K].TESVDE | 1xPhosphc | Q3TQQ9-2 | 2.86E-06 | 0 | 1 | 3 | 2 | Q3TQQ9-2 | Isoform 2 c | Q3TQQ9-2 | 0 | 2244.105 | 0 | -0.26 | 0.215815 | 19.05 | 108.9 | 91.1 | 10.06 | 11.63 | | High | High | High | High | High | High | High | High | High | High | High | 0 | 0.000234 | 3.24 |
| FALSE | High | TESTYLAEII | [K].TESTYL | 1xPhosphc | Q80TI0-4 1 | 6.07E-05 | 0 | 1 | 8 | 2 | Q80TI0-4 | Isoform 4 c | Q80TI0-4 [ | 1 | 2298.19 | 0 | -1.49 | 0.000492 | 0.71 | 147.7 | 52.3 | 6.47 | 10.22 | | High | High | High | High | High | High | High | High | High | High | High | 0.00196 | 0.01996 | 3.19 |
| FALSE | High | TEPLRPSVF | [K].TEPLRP | 1xPhosphc | P30415 1x | 0.000203 | 0 | 1 | 2 | 2 | P30415 | NK-tumor r | P30415 [5 | 0 | 4194.216 | 0 | -0.26 | 0.21382 | 22.54 | 109 | 91 | 8.14 | 23.27 | | High | High | High | High | High | High | High | High | High | High | High | 0 | 4E-05 | 3.1 |
| FALSE | High | TEPLRPSVF | [K].TEPLRP | 1xDeamida | P30415 1x | 4.63E-05 | 0 | 1 | 2 | 8 | P30415 | NK-tumor r | P30415 [5 | 0 | 4195.2 | 0 | -0.18 | 0.343166 | 16.65 | 111.2 | 88.8 | 15.24 | 15.13 | | High | High | High | High | High | High | High | High | High | High | High | 0 | 5.24E-05 | 3.7 |
| FALSE | High | TEPPAEGLI | [R].TEPPAE | 1xPhosphc | Q8R5K4 1x | 0.002623 | 0.000109 | 1 | 3 | 2 | Q8R5K4 | Nucleolar p | Q8R5K4 [5 | 1 | 2921.543 | 0 | 0.01 | 0.345289 | 5.1 | 99.3 | 100.7 | 6.78 | 12.36 | | High | High | High | High | High | High | High | High | High | High | High | 4.94E-05 | 0.000705 | 2.53 |
| FALSE | High | TEPSSPLSD | [K].TEPSSP | 1xPhosphc | E9Q444 1x | 0.046291 | 0.003523 | 1 | 9 | 2 | E9Q444 | Zinc finger | E9Q444 [4 | 0 | 1922.931 | 0 | 0.15 | 0.066391 | 4.9 | 99.4 | 100.6 | 8.76 | 6.97 | | High | High | High | High | High | High | High | High | High | High | High | 0.001789 | 0.01764 | 2.74 |
| FALSE | High | TEPSSPLSD | [K].TEPSSP | 2xPhosphc | E9Q444 2x | 0.061176 | 0.004646 | 1 | 9 | 1 | E9Q444 | Zinc finger | E9Q444 [4 | 0 | 2002.897 | 0 | 0.17 | 0.074847 | 13.03 | 91.8 | 108.2 | 6.72 | 9.05 | | High | High | High | High | High | High | High | High | High | High | High | 0.002354 | 0.02431 | 2.09 |
| FALSE | High | TEPTEDSDI | [K].TEPTED | 1xPhosphc | G3UYD0 1x | 0.003892 | 0.000261 | 1 | 1 | 1 | G3UYD0 | General tra | G3UYD0 [2 | 0 | 2335.027 | 0 | -0.4 | 0.013273 | 9.82 | 113 | 87 | 7.78 | 7.01 | | High | High | High | High | High | High | High | High | High | High | High | 0.00014 | 0.001092 | 2.69 |
| FALSE | High | TEPVSPLAE | [K].TEPVSP | 1xPhosphc | Q6PAR7 1x | 1.75E-08 | 0 | 1 | 3 | 3 | Q6PAR7 | Phf20 prot | Q6PAR7 [4 | 0 | 3004.59 | 0 | -1.67 | 0.000209 | 1.09 | 151.6 | 48.4 | 3.26 | 6.69 | | High | High | High | High | High | High | High | High | High | High | High | 0 | 1.08E-09 | 4.24 |
| FALSE | High | TEQDPNCS | [K].TEQDPI | 1xCarbami | O70477 2x | 0.013308 | 0.000889 | 1 | 1 | 1 | O70477 | Homeobox | O70477 [2 | 0 | 3845.621 | 0 | 0.1 | 0.389781 | 26.32 | 96.3 | 103.7 | 11.81 | 5.5 | | High | High | High | High | High | High | High | High | High | High | High | 0.000467 | 0.004297 | 3 |
| FALSE | High | TEQELPRP( | [K].TEQELF | 1xPhosphc | O35126 1x | 0.039011 | 0.002718 | 1 | 2 | 2 | O35126 | Atrophin-1 | O35126 [9 | 0 | 2549.208 | 0 | 1.15 | 0.001754 | 5.16 | 62.4 | 137.6 | 7.66 | 9.24 | | High | High | High | High | High | High | High | High | High | High | High | 0.001453 | 0.01455 | 2.51 |
| FALSE | High | TEQELPRP( | [K].TEQELF | 2xPhosphc | O35126 2x | 0.047488 | 0.00354 | 1 | 2 | 1 | O35126 | Atrophin-1 | O35126 [9 | 0 | 2629.175 | 0 | 0.54 | 0.005145 | 11.94 | 80.6 | 119.4 | 8.91 | 4.92 | | High | High | High | High | High | High | High | High | High | High | High | 0.001812 | 0.01815 | 1.72 |
| FALSE | High | TEQPDDEY | [K].TEQPDI | 1xPhosphc | Q4VBD9 1x | 0.001815 | 5.66E-05 | 1 | 2 | 1 | Q4VBD9 | GDNF-indu | Q4VBD9 [6 | 0 | 1917.888 | 0 | -0.11 | 0.879201 | 7.47 | 100 | 100 | 15.67 | 7.5 | | High | High | High | High | High | High | High | High | High | High | High | 2.58E-05 | 0.000465 | 3.05 |
| FALSE | High | TEQPSPVV | [K].TEQPSF | 1xPhosphc | Q8CJF7 1x | 0.0095 | 0.000736 | 1 | 2 | 1 | Q8CJF7 | Protein ELY | Q8CJF7 [1 | 0 | 3694.81 | 0 | -0.83 | 0.006175 | 0.56 | 124.5 | 75.5 | 11.44 | 8.93 | | High | High | High | High | High | High | High | High | High | High | High | 0.000344 | 0.002974 | 3.32 |
| FALSE | High | TERDTPGH | [K].TERDTF | 1xPhosphc | G5E866 1x | 0.005 | 0.000504 | 1 | 5 | 2 | G5E866 | Splicing fac | G5E866 [2 | 1 | 2162.982 | 0 | 0.09 | 0.733361 | 17.59 | 97 | 103 | 4.41 | 18.71 | | High | High | High | High | High | High | High | High | High | High | High | 0.000327 | 0.00279 | 2.7 |
| FALSE | High | TERDTPGH | [K].TERDTF | 2xPhosphc | G5E866 2x | 0.000183 | 0 | 1 | 5 | 1 | G5E866 | Splicing fac | G5E866 [2 | 1 | 2242.948 | 0 | 0.5 | 0.116655 | 6.61 | 83.3 | 116.7 | 5.48 | 19.73 | | High | High | High | High | High | High | High | High | High | High | High | 0 | 3.54E-05 | 1.79 |
| FALSE | High | TESDSGLK | [R].TESDSG | 1xPhosphc | A0A140LJJ | 0.011549 | 0.00086 | 1 | 3 | 1 | A0A140LJJ | A-kinase ar | A0A140LJJ | 0 | 1374.692 | 0 | -0.23 | 0.044459 | 6.14 | 108.3 | 91.7 | 6.83 | 9.56 | | High | High | High | High | High | High | High | High | High | High | High | 0.000437 | 0.003679 | 2.02 |
| FALSE | High | TESESSSLD | [K].TESESS | 1xPhosphc | Q3UM18 1 | 8.76E-07 | 0 | 1 | 3 | 4 | Q3UM18 | Large subu | Q3UM18 [ | 0 | 2080.939 | 0 | 0.5 | 0.002884 | 11.1 | 83.3 | 116.7 | 3.41 | 7.08 | | High | High | High | High | High | High | High | High | High | High | High | 0 | 4.62E-05 | 3.61 |

| | | | | | | | | | | | | | | | | | | | | | | | | | | | | | | | | | | | | | | |
|---|---|---|---|---|---|---|---|---|---|---|---|---|---|---|---|---|---|---|---|---|---|---|---|---|---|---|---|---|---|---|---|---|---|---|---|---|---|---|
| FALSE | High | TESGETLAC | [K].TESGET | 1xPhospho | Q3ULZ6 1x | 2.9E-05 | 0 | 2 | 3 | 2 | Q3ULZ6; Q | Putative ur | Q3ULZ6 [2 | 0 | 2778.405 | 0 | 2.36 | 0.003083 | 24.66 | 29.3 | 170.7 | 11.18 | 21.95 | Shared | High | High | High | High | High | High | High | High | High | High | High | 0 | 4.48E-06 | 3.29 |
| FALSE | High | TESGYGSE | [K].TESGYG | 1xPhospho | Q9EQG9 1 | 0.049971 | 0.003641 | 1 | 4 | 1 | Q9EQG9 | Collagen ty | Q9EQG9 [1 | 0 | 1581.7 | 0 | 0.27 | 0.070975 | 8.84 | 89.1 | 110.9 | 9.76 | 8.72 | | High | High | High | High | High | High | High | High | High | High | High | 0.001902 | 0.0192 | 1 |
| FALSE | High | TESGYGSE | [K].TESGYG | 2xPhospho | Q9EQG9 2 | 0.115566 | 0.009504 | 1 | 4 | 1 | Q9EQG9 | Collagen ty | Q9EQG9 [1 | 1 | 1817.767 | 0 | 0.53 | 0.014374 | 6.87 | 81.9 | 118.1 | 12.83 | 8.85 | | High | High | High | High | High | High | High | High | High | High | High | 0.004757 | 0.05144 | 1.53 |
| FALSE | High | TESLEGDD | [R].TESLEG | 1xPhospho | E9Q784 1x | 0.003224 | 0.00016 | 1 | 1 | 1 | E9Q784 | Zinc finger | E9Q784 [1 | 0 | 1747.804 | 0 | -1 | 0.00499 | 7.05 | 134.7 | 65.3 | 8.14 | 23 | | High | High | High | High | High | High | High | High | High | High | High | 7.19E-05 | 0.000888 | 3.15 |
| FALSE | High | TESLPSAQ | [K].TESLPS | 1xPhospho | G3UWG2 1 | 2.27E-05 | 0 | 1 | 11 | 1 | G3UWG2 | T-cell lymp | G3UWG2 [ | 0 | 2247.146 | 0 | -1.58 | 0.003969 | 4.58 | 148.5 | 51.5 | 12.93 | 19.36 | | High | High | High | High | High | High | High | High | High | High | High | 0 | 3.39E-06 | 3.88 |
| FALSE | High | TESNQEVA | [K].TESNQ | 1xPhospho | Q3TK95 1x | 2.45E-06 | 0 | 1 | 5 | 2 | Q3TK95 | Putative ur | Q3TK95 [2 | 0 | 2297.122 | 0 | 0.46 | 0.033799 | 22.11 | 86.3 | 113.7 | 13.55 | 7.01 | | High | High | High | High | High | High | High | High | High | High | High | 0 | 2.79E-07 | 4.1 |
| FALSE | High | TESPAEGK | [K].TESPAE | 1xPhospho | Q6ZQ61 1x | 0.019618 | 0.001438 | 1 | 8 | 1 | Q6ZQ61 | MCG12197 | Q6ZQ61 [6 | 0 | 1356.681 | 0 | -0.53 | 0.00545 | 0.82 | 118.6 | 81.4 | 6.81 | 6.24 | | High | High | High | High | High | High | High | High | High | High | High | 0.000756 | 0.00669 | 1.8 |
| FALSE | High | TESPPLPAS | [R].TESPPL | 1xPhospho | Q8VID5 1x | 0.022773 | 0.001542 | 1 | 2 | 1 | Q8VID5 | ATP-depen | Q8VID5 [7 | 0 | 1559.803 | 0 | 0.29 | 0.022408 | 6.86 | 91.1 | 108.9 | 6.09 | 5.79 | | High | High | High | High | High | High | High | High | High | High | High | 0.000832 | 0.007898 | 2.26 |
| FALSE | High | TESPSESCP | [R].TESPSE | 1xCarbami | B7ZP47 1x | 0.090888 | 0.007081 | 1 | 2 | 1 | B7ZP47 | Wapal prot | B7ZP47 [22 | 0 | 1529.676 | 0 | 0.2 | 0.454881 | 25.16 | 94.9 | 105.1 | 13.12 | 6.28 | | High | High | High | High | High | High | High | High | High | High | High | 0.003606 | 0.03844 | 1.83 |
| FALSE | High | TESPSESCP | [R].TESPSE | 1xCarbami | B7ZP47 1x | 6.89E-05 | 0 | 1 | 2 | 3 | B7ZP47 | Wapal prot | B7ZP47 [22 | 0 | 1758.839 | 0 | -0.86 | 0.001008 | 1.83 | 127.8 | 72.2 | 6.15 | 2.68 | | High | High | High | High | High | High | High | High | High | High | High | 0 | 0.000216 | 2.82 |
| FALSE | High | TESPTQPS | [K].TESPTC | 1xPhospho | Q3TN42 1x | 0.000281 | 0 | 1 | 2 | 3 | Q3TN42 | Putative ur | Q3TN42 [5 | 0 | 1843.873 | 0 | -0.22 | 0.038494 | 8.53 | 108.8 | 91.2 | 3.7 | 8.77 | | High | High | High | High | High | High | High | High | High | High | High | 0 | 5.75E-05 | 2.75 |
| FALSE | High | TESRDVDS | [R].TESRD\ | 1xPhospho | Q8CJ61 1x | 0.079114 | 0.00613 | 1 | 1 | 1 | Q8CJ61 | CKLF-like M | Q8CJ61 [1 | 1 | 1996.965 | 0 | -1.01 | 0.000391 | 1.06 | 131.4 | 68.6 | 6.21 | 10.07 | | High | High | High | High | High | High | High | High | High | High | High | 0.003148 | 0.0329 | 1.98 |
| FALSE | High | TESSEALK | [R].TESSEA | 1xPhospho | A2ARP8 1x | 0.022773 | 0.001542 | 1 | 3 | 1 | A2ARP8 | Microtubu | A2ARP8 [6 | 0 | 1402.723 | 0 | -0.86 | 0.002996 | 3.56 | 128.2 | 71.8 | 9.64 | 4.66 | | High | High | High | High | High | High | High | High | High | High | High | 0.000814 | 0.00789 | 2.02 |
| FALSE | High | TEAEVLAE | [K].TEAEVL | 1xDeamida | Q9QXX8 1 | 0.000662 | 0 | 1 | 1 | 2 | Q9QXX8 | Nuclear fra | Q9QXX8 [3 | 0 | 2497.274 | 0 | -0.69 | 0.015593 | 13.39 | 123.5 | 76.5 | 12.95 | 17.88 | | High | High | High | High | High | High | High | High | High | High | High | 9.42E-05 | 0.000982 | 3.17 |
| FALSE | High | TEAEVLAE | [K].TEAEVL | 1xPhospho | Q9QXX8 1 | 9.13E-09 | 0 | 1 | 1 | 4 | Q9QXX8 | Nuclear fra | Q9QXX8 [3 | 0 | 2496.29 | 0 | -1.23 | 0.000222 | 0.24 | 138.9 | 61.1 | 5.38 | 6.01 | | High | High | High | High | High | High | High | High | High | High | High | 0 | 2.23E-07 | 3.81 |
| FALSE | High | TEAEGLRP | [K].TEAEGI | 1xPhospho | Q0VGT2 1 | 0.002334 | 0.000109 | 1 | 7 | 1 | Q0VGT2 | Zinc finger | Q0VGT2 [3 | 0 | 2746.423 | 0 | -0.83 | 0.153089 | 12.45 | 128 | 72 | 47.5 | 20.31 | | High | High | High | High | High | High | High | High | High | High | High | 4.94E-05 | 0.000615 | 4.46 |
| FALSE | High | TEADLPKP | [R].TEADLF | 1xPhospho | A2A5V4 1x | 0.00359 | 0.00021 | 1 | 5 | 3 | A2A5V4 | SH3 domai | A2A5V4 [5 | 0 | 2051.139 | 0 | -0.25 | 0.064607 | 8.46 | 107.7 | 92.3 | 6.27 | 8.99 | | High | High | High | High | High | High | High | High | High | High | High | 9.42E-05 | 0.001 | 2.73 |
| FALSE | High | TATYNGPP | [R].TATYN | 1xPhospho | Q03141 1x | 7.41E-05 | 0 | 1 | 5 | 3 | Q03141 | MAP/micro | Q03141 [5 | 0 | 2792.345 | 0 | -0.57 | 0.073297 | 2.51 | 119.2 | 80.8 | 13.11 | 16.49 | | High | High | High | High | High | High | High | High | High | High | High | 0 | 1.28E-05 | 3.37 |
| FALSE | High | TATYNGPP | [R].TATYN | 1xDeamida | Q03141 1x | 0.030898 | 0.001896 | 1 | 5 | 1 | Q03141 | MAP/micro | Q03141 [5 | 0 | 2793.329 | 0 | -0.04 | 0.237885 | 10.36 | 100 | 100 | 7.66 | 9.63 | | High | High | High | High | High | High | High | High | High | High | High | 0.001027 | 0.01109 | 1.9 |
| FALSE | High | TAVAPSAV | [R].TAVAPS | 1xPhospho | Q8BTI8 1x | 0.01861 | 0.001362 | 1 | 3 | 1 | Q8BTI8 | Serine/argi | Q8BTI8 [22 | 1 | 2870.509 | 0 | -0.8 | 0.004362 | 1.92 | 122 | 78 | 12.98 | 13.67 | | High | High | High | High | High | High | High | High | High | High | High | 0.000723 | 0.006263 | 2.22 |
| FALSE | High | TAVPQLSE | [K-].TAVPC | 1xPhospho | E9Q5F9 1x | 0.001398 | 5.66E-05 | 1 | 4 | 4 | E9Q5F9 | Histone-lys | E9Q5F9 [1 | 0 | 2307.034 | 0 | -0.3 | 0.007769 | 1.06 | 110.2 | 89.8 | 5.39 | 6.21 | | High | High | High | High | High | High | High | High | High | High | High | 2.58E-05 | 0.000347 | 3.06 |
| FALSE | High | TAVPQLSE | [K-].TAVPC | 2xPhospho | E9Q5F9 2x | 9.11E-05 | 0 | 1 | 4 | 2 | E9Q5F9 | Histone-lys | E9Q5F9 [1 | 0 | 2387 | 0 | -0.2 | 0.177983 | 15.88 | 107.3 | 92.7 | 12.47 | 18.47 | | High | High | High | High | High | High | High | High | High | High | High | 0 | 1.62E-05 | 4.2 |
| FALSE | High | TAVSVAQG | [K].TAVSV | 1xPhospho | Q3TWM8 1 | 1.21E-05 | 0 | 1 | 6 | 2 | Q3TWM8 | Putative ur | Q3TWM8 [ | 0 | 1478.731 | 0 | 0.05 | 0.968584 | 12.88 | 98.3 | 101.7 | 6.25 | 5.54 | | High | High | High | High | High | High | High | High | High | High | High | 2.58E-05 | 0.000496 | 2.99 |
| FALSE | High | TAYCDIHT | [K].TAYCDI | 1xCarbami | A0A0N4SU | 0.031713 | 0.002011 | 1 | 10 | 1 | A0A0N4SU | Peregrin O | A0A0N4SU | 0 | 1854.841 | 0 | -0.44 | 0.00815 | 5.69 | 119.3 | 80.7 | 8.44 | 10.1 | | High | High | High | High | High | High | High | High | High | High | High | 0.001093 | 0.0115 | 2.45 |
| FALSE | High | TCDSPQNP | [R].TCDSP | 1xCarbami | P52479-2 1 | 5.84E-07 | 0 | 1 | 2 | 5 | P52479-2 | Isoform 2 c | P52479-2 [ | 0 | 2838.301 | 0 | 0.05 | 0.589532 | 4.02 | 97.8 | 102.2 | 12.34 | 6.53 | | High | High | High | High | High | High | High | High | High | High | High | 0 | 2.68E-06 | 4.43 |
| FALSE | High | TCDSPQNP | [R].TCDSP | 1xCarbami | P52479-2 2 | 0.115566 | 0.009504 | 1 | 2 | 1 | P52479-2 | Isoform 2 c | P52479-2 [ | 0 | 2918.267 | 0 | 0.56 | 0.044958 | 15.89 | 84.6 | 115.4 | 6.11 | 20.79 | | High | High | High | High | High | High | High | High | High | High | High | 0.004733 | 0.05114 | 2.29 |
| FALSE | High | TCGSPVQS | [R].TCGSP\ | 1xCarbami | Q9QZ41 1x | 9.11E-08 | 0 | 1 | 2 | 1 | Q9QZ41 | Protein DB | Q9QZ41 [5 | 0 | 2677.32 | 0 | -0.47 | 0.04524 | 2.96 | 113.3 | 86.7 | 8.37 | 12.85 | | High | High | High | High | High | High | High | High | High | High | High | 0 | 6.86E-09 | 3.77 |
| FALSE | High | TCLSEGNP | [K].TCLSEG | 1xCarbami | Q8K1N2-2 | 5.55E-05 | 0 | 1 | 3 | 1 | Q8K1N2-2 | Isoform 2 c | Q8K1N2-2 | 0 | 2501.194 | 0 | 2.47 | 0.000683 | 25.66 | 33.3 | 166.7 | 10.86 | 11.28 | | High | High | High | High | High | High | High | High | High | High | High | 0 | 9.28E-06 | 3.67 |
| FALSE | High | TCNVLVAL | [K].TCNVL\ | 1xCarbami | Q3TED1 1x | 0.007404 | 0.00062 | 1 | 9 | 1 | Q3TED1 | S-adenosyl | Q3TED1 [1 | 0 | 2872.423 | 0 | -1.32 | 0.002081 | 4.79 | 146.8 | 53.2 | 8.79 | 15.26 | | High | High | High | High | High | High | High | High | High | High | High | 0.000275 | 0.002247 | 3.44 |
| FALSE | High | TCPSLSPTS | [R].TCPSLS | 1xCarbami | Q2M4G9 1 | 0.000132 | 0 | 1 | 2 | 4 | Q2M4G9 | Protein inh | Q2M4G9 [ | 0 | 2027.028 | 0 | 0.2 | 0.184084 | 9.6 | 93.3 | 106.7 | 4.95 | 13.17 | | High | High | High | High | High | High | High | High | High | High | High | 0 | 0.000149 | 2.77 |
| FALSE | High | TATSSPAST | [K].TATSSP | 1xPhospho | Q91YK2 1x | 3.21E-06 | 0 | 1 | 2 | 11 | Q91YK2 | Ribosomal | Q91YK2 [6 | 0 | 1828.871 | 0 | 0.03 | 0.510453 | 2.28 | 98.8 | 101.2 | 5.48 | 2.44 | | High | High | High | High | High | High | High | High | High | High | High | 2.58E-05 | 0.000435 | 2.51 |
| FALSE | High | TCPSLSPTS | [R].TCPSLS | 1xCarbami | Q2M4G9 2 | 0.000725 | 0 | 1 | 2 | 2 | Q2M4G9 | Protein inh | Q2M4G9 [ | 0 | 2106.995 | 0 | 0.39 | 0.041563 | 10.93 | 85.3 | 114.7 | 7.63 | 14.36 | | High | High | High | High | High | High | High | High | High | High | High | 0 | 0.000167 | 2.35 |
| FALSE | High | TCSFGGFD | [R].TCSFG | 1xCarbami | F8VQK5 1x | 2.38E-05 | 0 | 1 | 4 | 2 | F8VQK5 | SAM and S | F8VQK5 [3 | 0 | 1683.74 | 0 | 0.44 | 0.007444 | 3.95 | 84.6 | 115.4 | 10.41 | 5.16 | | High | High | High | High | High | High | High | High | High | High | High | 0 | 3.56E-06 | 2.56 |
| FALSE | High | TCSIGSTVD | [K].TCSIGS | 1xCarbami | Q9WVM1 | 5.77E-06 | 0 | 1 | 2 | 3 | Q9WVM1 | Rac GTPase | Q9WVM1 | 0 | 2418.199 | 0 | -1.07 | 0.000795 | 3.59 | 139 | 61 | 5.36 | 9.29 | | High | High | High | High | High | High | High | High | High | High | High | 0 | 3.22E-06 | 3.17 |
| FALSE | High | TCSLPDLSK | [R].TCSLPD | 1xCarbami | B2RXX0 1x | 0.000305 | 0 | 1 | 6 | 3 | B2RXX0 | Nek1 prote | B2RXX0 [1 | 0 | 1558.795 | 0 | -0.58 | 0.001518 | 2.18 | 121.5 | 78.5 | 4.57 | 7.78 | | High | High | High | High | High | High | High | High | High | High | High | 0 | 6.3E-05 | 2.72 |
| FALSE | High | TCSMVGN | [R].TCSMV | 2xCarbami | B2WTI2 1x | 2.66E-06 | 0 | 1 | 3 | 5 | B2WTI2 | Phosphatic | B2WTI2 [3 | 0 | 2616.103 | 0 | -0.58 | 0.015266 | 3.85 | 118.8 | 81.2 | 6.83 | 17.94 | | High | High | High | High | High | High | High | High | High | High | High | 0 | 3.04E-07 | 3.5 |
| FALSE | High | TCSPTTLSC | [K].TCSPTT | 1xCarbami | Q3U3X7 1x | 0.000993 | 0 | 1 | 8 | 3 | Q3U3X7 | Putative ur | Q3U3X7 [1 | 0 | 2410.161 | 0 | 0.54 | 0.014367 | 22.97 | 79.5 | 120.5 | 6.08 | 11.18 | | High | High | High | High | High | High | High | High | High | High | High | 7.19E-05 | 0.0009 | 2.4 |
| FALSE | High | TCSVPGTP | [K].TCSVP | 1xCarbami | Q9CPY3 1x | 0.002818 | 0.000109 | 1 | 1 | 4 | Q9CPY3 | Sororin OS | Q9CPY3 [1 | 0 | 3603.625 | 0 | 0.52 | 0.066291 | 25.73 | 87.8 | 112.2 | 28.66 | 28.04 | | High | High | High | High | High | High | High | High | High | High | High | 7.19E-05 | 0.000763 | 4.58 |
| FALSE | High | TCYDSEDS | [K].TCYDSE | 2xCarbami | Q8BL06 2x | 0.009585 | 0.000736 | 1 | 1 | 2 | Q8BL06 | Inactive ub | Q8BL06 [3 | 0 | 2864.119 | 0 | -1.18 | 0.003674 | 6.54 | 134.8 | 65.2 | 11.45 | 14.61 | | High | High | High | High | High | High | High | High | High | High | High | 0.000344 | 0.002996 | 2.9 |
| FALSE | High | TDACQSGE | [R].TDACQ | 2xCarbami | O88665 1x | 9.64E-09 | 0 | 1 | 1 | 2 | O88665 | Bromodom | O88665 [2 | 0 | 2065.794 | 0 | -0.16 | 0.483523 | 19.37 | 104 | 96 | 5.98 | 16.85 | | High | High | High | High | High | High | High | High | High | High | High | 0 | 2.16E-09 | 5.16 |
| FALSE | High | TDALSPTM | [R].TDALSF | 1xPhospho | Q3U0X6 1x | 0.008538 | 0.000736 | 1 | 1 | 1 | Q3U0X6 | Putative ur | Q3U0X6 [4 | 0 | 1656.787 | 0 | -0.19 | 0.078697 | 9.43 | 106.6 | 93.4 | 4.25 | 13.61 | | High | High | High | High | High | High | High | High | High | High | High | 0.000327 | 0.002636 | 3.08 |
| FALSE | High | TDALSPTM | [R].TDALSF | 1xPhospho | Q3U0X6 1x | 8.4E-05 | 0 | 1 | 1 | 2 | Q3U0X6 | Putative ur | Q3U0X6 [4 | 0 | 1672.781 | 0 | -0.1 | 0.554195 | 3.62 | 103.3 | 96.7 | 8.6 | 7.51 | | High | High | High | High | High | High | High | High | High | High | High | 0 | 1.48E-05 | 3.19 |
| FALSE | High | TDALTAHS | [R].TDALT | 1xPhospho | Q9D666 1x | 0.043607 | 0.003228 | 1 | 15 | 1 | Q9D666 | SUN doma | Q9D666 [1 | 0 | 1947.949 | 0 | -0.08 | 0.658248 | 27.68 | 100.8 | 99.2 | 8.47 | 13.37 | | High | High | High | High | High | High | High | High | High | High | High | 0.001651 | 0.01641 | 2.4 |
| FALSE | High | TDAPDSPQ | [K].TDAPD | 1xPhospho | Q8CFA1 1x | 0.019791 | 0.001438 | 1 | 4 | 1 | Q8CFA1 | Interleukin | Q8CFA1 [1 | 0 | 1583.772 | 0 | 1.63 | 0.000437 | 14.56 | 47.1 | 152.9 | 21.15 | 10.05 | | High | High | High | High | High | High | High | High | High | High | High | 0.000773 | 0.006768 | 2.32 |
| FALSE | High | TDASSASS | [R].TDASSA | 1xPhospho | Q8VH51 1x | 6.97E-06 | 0 | 1 | 6 | 2 | Q8VH51 | RNA-bindir | Q8VH51 [3 | 0 | 2138.933 | 0 | -0.54 | 0.018661 | 3.26 | 114.9 | 85.1 | 7.74 | 14.1 | | High | High | High | High | High | High | High | High | High | High | High | 0 | 9.38E-05 | 3.93 |
| FALSE | High | TCSASPGC | [R].TCSASP | 3xCarbami | Q8BYN3 1x | 0.045508 | 0.003373 | 1 | 1 | 1 | Q8BYN3 | Inositol-tet | Q8BYN3 [3 | 0 | 1956.809 | 0 | 0.66 | 0.001467 | 3.95 | 77.8 | 122.2 | 18.44 | 14.42 | | High | High | High | High | High | High | High | High | High | High | High | 0.001741 | 0.01735 | 1.46 |
| FALSE | High | TATSSPAST | [K].TATSSP | 1xPhospho | Q91YK2 1x | 4.2E-05 | 0 | 1 | 2 | 1 | Q91YK2 | Ribosomal | Q91YK2 [6 | 0 | 1812.876 | 0 | -0.43 | 0.021759 | 11.09 | 112.8 | 87.2 | 2.79 | 18.72 | | High | High | High | High | High | High | High | High | High | High | High | 0 | 6.79E-06 | 2.23 |
| FALSE | High | TATPQSSH | [R].TATPQ | 2xPhospho | Q3UPW2 2 | 0.001068 | 0 | 1 | 2 | 1 | Q3UPW2 | ETS-relatec | Q3UPW2 [ | 0 | 3123.381 | 0 | -1.55 | 0.006998 | 2.88 | 144.3 | 55.7 | 15.01 | 30.73 | | High | High | High | High | High | High | High | High | High | High | High | 0 | 0.000257 | 4.01 |
| FALSE | High | TATADDK | [R].TATADI | 1xPhospho | Q64152 1x | 0.080423 | 0.006219 | 1 | 4 | 1 | Q64152 | Transcripti | Q64152 [7 | 0 | 1259.628 | 0 | -0.21 | 0.009217 | 5.69 | 108.9 | 91.1 | 7.92 | 6.76 | | High | High | High | High | High | High | High | High | High | High | High | 0.003188 | 0.0334 | 2.01 |
| FALSE | High | TASIDSNQ | [R].TASIDS | 1xDeamida | Q99KE1 1x | 0.003997 | 0.000311 | 1 | 1 | 2 | Q99KE1 | NAD-deper | Q99KE1 [3 | 0 | 3612.715 | 0 | -0.78 | 0.01387 | 7.48 | 128.4 | 71.6 | 19.56 | 10.06 | | High | High | High | High | High | High | High | High | High | High | High | 0.000183 | 0.001268 | 4.23 |
| FALSE | High | TASLNLPP | [R].TASLNL | 1xPhospho | P59729 1x | 0.045122 | 0.003373 | 1 | 5 | 1 | P59729 | Ras and Ra | P59729 [4 | 0 | 2502.291 | 0 | 3.21 | 0.000223 | 17.15 | 20.4 | 179.6 | 20.13 | 8.35 | | High | High | High | High | High | High | High | High | High | High | High | 0.001727 | 0.01715 | 1.54 |
| FALSE | High | TASISSSPS | [R].TASISS | 1xCarbami | Q6ZQ58 2x | 0.001916 | 5.66E-05 | 1 | 2 | 2 | Q6ZQ58 | La-related | Q6ZQ58 [8 | 0 | 3055.357 | 0 | 0.02 | 0.945245 | 111.17 | 99.5 | 100.5 | 80.1 | 71.17 | | High | High | High | High | High | High | High | High | High | High | High | 2.58E-05 | 0.000493 | 2.76 |
| FALSE | High | TASISSSPS | [R].TASISS | 1xCarbami | Q6ZQ58 1x | 2.26E-07 | 0 | 1 | 2 | 9 | Q6ZQ58 | La-related | Q6ZQ58 [8 | 0 | 3204.554 | 0 | 0.83 | 0.000262 | 2.82 | 71.2 | 128.8 | 8.74 | 6.98 | | High | High | High | High | High | High | High | High | High | High | High | 0 | 5.48E-08 | 5.04 |
| FALSE | High | TASISSSPS | [R].TASISS | 1xCarbami | Q6ZQ58 2x | 8.4E-05 | 0 | 1 | 2 | 19 | Q6ZQ58 | La-related | Q6ZQ58 [8 | 0 | 3284.52 | 0 | -0.09 | 0.749692 | 7.48 | 103.1 | 96.9 | 4.85 | 5.05 | | High | High | High | High | High | High | High | High | High | High | High | 0 | 4.57E-05 | 5.17 |
| FALSE | High | TASLTSAAS | [R].TASLTS | 2xPhospho | Q9QYG0 2 | 0.002058 | 5.66E-05 | 1 | 3 | 1 | Q9QYG0 | Protein ND | Q9QYG0 [3 | 0 | 1725.766 | 0 | -1.91 | 0.003074 | 2.89 | 160.2 | 39.8 | 11.8 | 31.01 | | High | High | High | High | High | High | High | High | High | High | High | 2.58E-05 | 0.000534 | 2.16 |
| FALSE | High | TASLTSAAS | [R].TASLTS | 3xPhospho | Q9QYG0 3 | 0.044358 | 0.003264 | 1 | 3 | 1 | Q9QYG0 | Protein ND | Q9QYG0 [3 | 0 | 1805.732 | 0 | -1.15 | 0.000834 | 0.54 | 142.1 | 57.9 | 8.04 | 5.53 | | High | High | High | High | High | High | High | High | High | High | High | 0.00168 | 0.0168 | 2.82 |
| FALSE | High | TASNAEQY | [R].TASNAI | 1xPhospho | Q80YS6 1x | 0.052133 | 0.00384 | 1 | 2 | 1 | Q80YS6 | Actin filam | Q80YS6 [5 | 0 | 1320.603 | 0 | 0.38 | 0.241127 | 30.31 | 89.4 | 110.6 | 14.27 | 18.92 | | High | High | High | High | High | High | High | High | High | High | High | 0.001999 | 0.02026 | 2.57 |
| FALSE | High | TASNAEQY | [R].TASNAI | 1xPhospho | Q80YS6 1x | 0.000322 | 0 | 1 | 2 | 3 | Q80YS6 | Actin filam | Q80YS6 [5 | 0 | 1549.766 | 0 | 0.42 | 0.025494 | 1.81 | 88 | 112 | 10.74 | 5.78 | | High | High | High | High | High | High | High | High | High | High | High | 0 | 6.71E-05 | 2.48 |
| FALSE | High | TASPLEHIL | [K].TASPLE | 1xPhospho | Q8C9B9 1x | 0.015741 | 0.001124 | 1 | 1 | 1 | Q8C9B9 | Death-indu | Q8C9B9 [1 | 0 | 2193.208 | 0 | -0.25 | 0.208802 | 10.19 | 108.3 | 91.7 | 4.58 | 12.08 | | High | High | High | High | High | High | High | High | High | High | High | 0.000585 | 0.005223 | 2.3 |
| FALSE | High | TASPPGPP | [K].TASPPG | 1xPhospho | O35126 1x | 0.001189 | 0 | 1 | 1 | 2 | O35126 | Atrophin-1 | O35126 [6 | 0 | 1767.908 | 0 | 0.41 | 0.00351 | 6.06 | 84.6 | 115.4 | 4.5 | 4.71 | | High | High | High | High | High | High | High | High | High | High | High | 0.000505 | 0.004393 | 1.31 |
| FALSE | High | TASPPPPPK | [R].TASPPP | 2xPhospho | Q52KI8 2x | 0.036416 | 0.002419 | 1 | 7 | 2 | Q52KI8 | Serine/argi | Q52KI8 [63 | 0 | 1280.589 | 0 | -0.39 | 0.049459 | 3.6 | 107.7 | 92.3 | 9.23 | 7.98 | | High | High | High | High | High | High | High | High | High | High | High | 0.001279 | 0.0134 | 0.76 |
| FALSE | High | TASPPPPPK | [R].TASPPP | 1xPhospho | Q52KI8 1x | 0.005666 | 0.00062 | 1 | 7 | 5 | Q52KI8 | Serine/argi | Q52KI8 [63 | 0 | 1429.786 | 0 | -0.49 | 0.001431 | 3.97 | 117 | 83 | 3.68 | 3.7 | | High | High | High | High | High | High | High | High | High | High | High | 0.000344 | 0.002976 | 2.51 |
| FALSE | High | TASPPVLPK | [R].TASPPV | 1xPhospho | A2A7B5 1x | 0.003997 | 0.000311 | 1 | 2 | 2 | A2A7B5 | PR domain | A2A7B5 [6 | 0 | 1447.833 | 0 | -0.87 | 0.000442 | 2.42 | 127.6 | 72.4 | 5.21 | 3.91 | | High | High | High | High | High | High | High | High | High | High | High | 0.00014 | 0.001123 | 2.87 |
| FALSE | High | TASPSPLRP | [R].TASPSP | 1xPhospho | Q09XV5 1x | 0.000656 | 0 | 1 | 2 | 3 | Q09XV5 | Chromodo | Q09XV5 [1 | 0 | 2103.125 | 0 | -0.98 | 0.000427 | 1.56 | 132.5 | 67.5 | 5.99 | 7.11 | | High | High | High | High | High | High | High | High | High | High | High | 0.000204 | 0.001324 | 1.48 |
| FALSE | High | TASPSPLRP | [R].TASPSP | 2xPhospho | Q09XV5 2x | 0.005044 | 0.000504 | 1 | 2 | 2 | Q09XV5 | Chromodo | Q09XV5 [1 | 0 | 2183.091 | 0 | -0.94 | 0.000391 | 2.37 | 132.1 | 67.9 | 2.61 | 7.64 | | High | High | High | High | High | High | High | High | High | High | High | 0.000246 | 0.001461 | 2.19 |
| FALSE | High | TASPSTSSI | [R].TASPST | 1xPhospho | Q3UH98 1 | 0.012619 | 0.000889 | 1 | 4 | 5 | Q3UH98 | Putative ur | Q3UH98 [3 | 0 | 1921.859 | 0 | 0.59 | 0.001214 | 5.86 | 78.2 | 121.8 | 5.84 | 7.61 | | High | High | High | High | High | High | High | High | High | High | High | 0.000571 | 0.00509 | 2.87 |
| FALSE | High | TASPVAPVI | [R].TASPVA | 1xCarbami | A0A158SIT | 2.78E-06 | 0 | 1 | 2 | 1 | A0A158SIT | Proline, glu | A0A158SIT | 0 | 3167.661 | 0 | -0.6 | 0.016002 | 10.98 | 120.4 | 79.6 | 12.13 | 6.42 | | High | High | High | High | High | High | High | High | High | High | High | 0 | 3.2E-07 | 3.05 |
| FALSE | High | TASPVAPVI | [R].TASPVA | 1xCarbami | A0A158SIT | 0.055308 | 0.003982 | 1 | 2 | 1 | A0A158SIT | Proline, glu | A0A158SIT | 0 | 3183.656 | 0 | -0.21 | 0.068597 | 12.5 | 112.1 | 87.9 | 10.13 | 7.53 | | High | High | High | High | High | High | High | High | High | High | High | 0.002074 | 0.02156 | 2.2 |
| FALSE | High | TASPVIALC | [R].TASPVI | 1xPhospho | Q91X58 1x | 6.97E-09 | 0 | 1 | 3 | 1 | Q91X58 | AN1-type z | Q91X58 [1 | 0 | 2421.222 | 0 | -0.21 | 0.096033 | 6.21 | 107.1 | 92.9 | 5.31 | 7.29 | | High | High | High | High | High | High | High | High | High | High | High | 0 | 3.82E-10 | 5.39 |
| FALSE | High | TASPVIALC | [R].TASPVI | 1xDeamida | Q91X58 1x | 2.03E-06 | 0 | 1 | 3 | 1 | Q91X58 | AN1-type z | Q91X58 [1 | 0 | 2422.206 | 0 | -0.36 | 0.013326 | 5.62 | 114.1 | 85.9 | 5.64 | 9.92 | | High | High | High | High | High | High | High | High | High | High | High | 0 | 2.24E-07 | 4.03 |
| FALSE | High | TASRPEDTI | [R].TASRPE | 1xPhospho | Q6EDY6 1x | 3.57E-05 | 0 | 1 | 4 | 2 | Q6EDY6 | F-actin-unc | Q6EDY6 [1 | 0 | 2451.18 | 0 | -0.11 | 0.407829 | 22.99 | 107.3 | 92.7 | 19.76 | 42.68 | | High | High | High | High | High | High | High | High | High | High | High | 0 | 2.95E-05 | 3.52 |
| FALSE | High | TASRPEDTI | [R].TASRPE | 2xPhospho | Q6EDY6 2x | 0.058664 | 0.004294 | 1 | 4 | 1 | Q6EDY6 | F-actin-unc | Q6EDY6 [1 | 0 | 2531.147 | 0 | 0.78 | 0.097101 | 14.69 | 79.8 | 120.2 | 8.41 | 27.55 | | High | High | High | High | High | High | High | High | High | High | High | 0.002187 | 0.02322 | 2.18 |
| FALSE | High | TASSNEIGH | [R].TASSNE | 1xPhospho | A0A1B0GR | 0.00925 | 0.000736 | 1 | 1 | 4 | A0A1B0GR | RIKEN cDN | A0A1B0GR | 0 | 1710.846 | 0 | 2.45 | 0.00157 | 29.24 | 27.6 | 172.4 | 20.15 | 14.94 | | High | High | High | High | High | High | High | High | High | High | High | 0.002159 | 0.02254 | 2.63 |
| FALSE | High | TASSSTEPS | [R].TASSST | 1xPhospho | Q5DTM2 1 | 0.000515 | 0 | 1 | 4 | 2 | Q5DTM2 | Oxysterol-l | Q5DTM2 [ | 0 | 1517.705 | 0 | 2.82 | 0.000285 | 5.74 | 23.9 | 176.1 | 10.45 | 6.52 | | High | High | High | High | High | High | High | High | High | High | High | 0 | 0.000113 | 2.02 |
| FALSE | High | TASTNSIAC | [K].TASTNS | 1xPhospho | A0A0N4SW | 0.015061 | 0.001049 | 1 | 3 | 4 | A0A0N4SW | Guanine nu | A0A0N4SW | 0 | 1428.705 | 0 | 1.38 | 0.003957 | 24.48 | 50 | 150 | 10.11 | 19.34 | | High | High | High | High | High | High | High | High | High | High | High | 0.000689 | 0.005948 | 2.28 |
| FALSE | High | TASTPTPPC | [R].TASTPT | 1xPhospho | Q6NZJ6 1x | 0.002869 | 0.00016 | 1 | 14 | 6 | Q6NZJ6 | Eukaryotic | Q6NZJ6 [2 | 0 | 3821.891 | 0 | 1.94 | 0.001238 | 13.85 | 42.3 | 157.7 | 16 | 12.03 | | High | High | High | High | High | High | High | High | High | High | High | 7.19E-05 | 0.000778 | 3.53 |
| FALSE | High | TASTPTPPC | [R].TASTPT | 1xDeamida | Q6NZJ6 1x | 4.27E-05 | 0 | 1 | 14 | 10 | Q6NZJ6 | Eukaryotic | Q6NZJ6 [2 | 0 | 3822.875 | 0 | 1.2 | 0.00085 | 10.39 | 59.9 | 140.1 | 13.86 | 7.28 | | High | High | High | High | High | High | High | High | High | High | High | 0 | 4.1E-05 | 4.48 |
| FALSE | High | TATAASEQ | [R].TATAAS | 1xPhospho | I3RSH5 1x | 8.28E-06 | 0 | 1 | 3 | 1 | I3RSH5 | E3 ubiquiti | I3RSH5 [20 | 0 | 1555.731 | 0 | -0.35 | 0.001416 | 2.78 | 112.8 | 87.2 | 6.08 | 5.17 | | High | High | High | High | High | High | High | High | High | High | High | 0 | 1.09E-06 | 1.62 |
| FALSE | High | TDCTSESV | [R].TDCTSE | 2xCarbami | E9Q9M8 2 | 0.000868 | 0 | 1 | 3 | 1 | E9Q9M8 | PWWP dor | E9Q9M8 [4 | 0 | 2665.106 | 0 | 0.81 | 0.001826 | 3.14 | 73.5 | 126.5 | 6.97 | 12.33 | | High | High | High | High | High | High | High | High | High | High | High | 0 | 0.000203 | 3.51 |
| FALSE | High | TFSTCSDN | [R].TFSTCS | 1xCarbami | Q9WUF3 1 | 0.004955 | 0.000504 | 1 | 1 | 1 | Q9WUF3 | CASP8-asso | Q9WUF3 [ | 0 | 1698.781 | 0 | -0.23 | 0.155738 | 8.25 | 107.6 | 92.4 | 12.7 | 7.19 | | High | High | High | High | High | High | High | High | High | High | High | 0.000225 | 0.001429 | 2.19 |
| FALSE | High | TDDSQEK | [K].TDDSQ | 1xPhospho | P25206 1x | 0.019276 | 0.001402 | 2 | 3 | 1 | P25206; Q | DNA replic | P25206 [72 | 0 | 1360.64 | 0 | -1.48 | 9.77E-05 | 0.37 | 148 | 52 | 1.91 | 5.76 | Shared | High | High | High | High | High | High | High | High | High | High | High | 0.000756 | 0.006515 | 1.98 |
| FALSE | High | TDDSQETC | [K].TDDSQ | 1xPhospho | P25206 1x | 0.023792 | 0.001612 | 2 | 3 | 2 | P25206; Q | DNA replic | P25206 [72 | 1 | 2730.287 | 0 | -1.01 | 0.032983 | 10.08 | 128.4 | 71.6 | 16.61 | 20.1 | Shared | High | High | High | High | High | High | High | High | High | High | High | 0.000859 | 0.008273 | 2.49 |
| FALSE | High | TDSHDGET | [R].TDSHD | 1xPhospho | P97465 1x | 0.002768 | 0.000109 | 1 | 2 | 3 | P97465 | Docking pr | P97465 [2 | 0 | 1713.773 | 0 | 3.14 | 0.000963 | 31.76 | 20.4 | 179.6 | 17.2 | 13.41 | | High | High | High | High | High | High | High | High | High | High | High | 0.000814 | 0.0075 | 2.37 |
| FALSE | High | TDSIGEK | [R].TDSIGE | 1xPhospho | Q80TL7 1x | 0.007339 | 0.00062 | 1 | 3 | 3 | Q80TL7 | Protein MC | Q80TL7 [1 | 0 | 1287.66 | 0 | 0.51 | 0.014708 | 17.9 | 83.8 | 116.2 | 4.14 | 11.98 | | High | High | High | High | High | High | High | High | High | High | High | 0.000275 | 0.002219 | 1.99 |
| FALSE | High | TDSPLPLEE | [K].TDSPLP | 1xPhospho | Q3U4J6 1x | 2.06E-05 | 0 | 1 | 2 | 3 | Q3U4J6 | Putative ur | Q3U4J6 [3 | 0 | 1947.951 | 0 | 1.03 | 0.141496 | 55.44 | 65.9 | 134.1 | 20.62 | 41.62 | | High | High | High | High | High | High | High | High | High | High | High | 0 | 6.9E-06 | 3.13 |
| FALSE | High | TDSPLPLEE | [K].TDSPLP | 1xPhospho | Q3U4J6 1x | 4.62E-07 | 0 | 1 | 2 | 4 | Q3U4J6 | Putative ur | Q3U4J6 [3 | 0 | 2177.114 | 0 | 1.4 | 0.001627 | 8.66 | 54.3 | 145.7 | 7.09 | 10.07 | | High | High | High | High | High | High | High | High | High | High | High | 0 | 6.15E-06 | 2.72 |
| FALSE | High | TDSPLPLEE | [K].TDSPLP | 2xPhospho | Q3U4J6 2x | 7.86E-07 | 0 | 1 | 2 | 2 | Q3U4J6 | Putative ur | Q3U4J6 [3 | 0 | 2257.081 | 0 | 1.56 | 0.004922 | 7.93 | 51.3 | 148.7 | 17.97 | 14.78 | | High | High | High | High | High | High | High | High | High | High | High | 0 | 7.74E-08 | 3.84 |
| FALSE | High | TDSQTIGD | [R].TDSQTI | 1xPhospho | A0A1B0GS | 2.87E-05 | 0 | 1 | 1 | 2 | A0A1B0GS | GRB2-asso | A0A1B0GS | 0 | 1620.747 | 0 | -1.54 | 0.001723 | 2.58 | 150 | 50 | 7.26 | 20.3 | | High | High | High | High | High | High | High | High | High | High | High | 0 | 2.44E-05 | 3.2 |
| FALSE | High | TDSREDEIS | [R].TDSRED | 1xPhospho | Q3TYK4 1x | 2.93E-05 | 0 | 1 | 5 | 1 | Q3TYK4 | Putative ur | Q3TYK4 [2 | 1 | 2286.122 | 0 | -0.23 | 0.722952 | 75.41 | 107.8 | 92.2 | 52.3 | 91.73 | | High | High | High | High | High | High | High | High | High | High | High | 0 | 4.51E-06 | 3.65 |
| FALSE | High | TDSREDEIS | [R].TDSRED | 1xPhospho | Q3TYK4 1x | 5.09E-06 | 0 | 1 | 5 | 4 | Q3TYK4 | Putative ur | Q3TYK4 [2 | 1 | 2515.285 | 0 | 0.98 | 0.000362 | 4.98 | 67.6 | 132.4 | 2.67 | 4.83 | | High | High | High | High | High | High | High | High | High | High | High | 0 | 2.61E-06 | 3.29 |
| FALSE | High | TDSREDEIS | [R].TDSRED | 2xPhospho | Q3TYK4 2x | 4.55E-08 | 0 | 1 | 5 | 2 | Q3TYK4 | Putative ur | Q3TYK4 [2 | 1 | 2595.251 | 0 | 0.68 | 0.019999 | 7.05 | 78.9 | 121.1 | 9.27 | 15.07 | | High | High | High | High | High | High | High | High | High | High | High | 0 | 1.14E-05 | 4.27 |
| FALSE | High | TDSSGPVS | [K].TDSSGI | 1xPhospho | D3YYY8 1x | 0.000725 | 0 | 1 | 1 | 2 | D3YYY8 | Rho guanir | D3YYY8 [1 | 1 | 3651.878 | 0 | 2.03 | 0.000414 | 6.7 | 37.9 | 162.1 | 5.39 | 9.33 | | High | High | High | High | High | High | High | High | High | High | High | 0 | 0.000167 | 3.48 |
| FALSE | High | TDSSPNQA | [K].TDSSPN | 1xPhospho | Q5RJV5 1x | 0.00509 | 0.000551 | 1 | 15 | 2 | Q5RJV5 | Polypyrimi | Q5RJV5 [1 | 0 | 1284.578 | 0 | -0.44 | 0.007977 | 1.87 | 118.6 | 81.4 | 11.75 | 8.03 | | High | High | High | High | High | High | High | High | High | High | High | 0.000814 | 0.007511 | 1.66 |
| FALSE | High | TDSTESSG | [K].TDSTES | 1xPhospho | B2RUG9 1x | 0.001516 | 5.66E-05 | 1 | 2 | 1 | B2RUG9 | Adenomat | B2RUG9 [2 | 0 | 1832.868 | 0 | -0.09 | 0.620389 | 12.1 | 106.6 | 93.4 | 14.74 | 14.82 | | High | High | High | High | High | High | High | High | High | High | High | 2.58E-05 | 0.000382 | 1.82 |
| FALSE | High | TDSTSDGR | [R].TDSTSD | 1xPhospho | Q9JHU4 1x | 0.006031 | 0.00062 | 1 | 3 | 1 | Q9JHU4 | Cytoplasmi | Q9JHU4 [4 | 0 | 1788.794 | 0 | -0.22 | 0.052281 | 10.39 | 105.3 | 94.7 | 5.31 | 9.59 | | High | High | High | High | High | High | High | High | High | High | High | 0.000275 | 0.001785 | 1.5 |
| FALSE | High | TDSGYLFS | [R].TDSGYI | 1xPhospho | A2A8U2 1x | 0.121159 | 0.009984 | 1 | 2 | 1 | A2A8U2 | Transmem | A2A8U2 [4 | 0 | 1935.916 | 0 | 0.69 | 0.032992 | 5.97 | 76.6 | 123.4 | 18.76 | 10.85 | | High | High | High | High | High | High | High | High | High | High | High | 0.004974 | 0.05423 | 2.1 |
| FALSE | High | TDSVIIADQ | [R].TDSVII | 1xPhospho | Q3TZR9 1x | 4.8E-05 | 0 | 1 | 5 | 2 | Q3TZR9 | Activating | Q3TZR9 [4 | 0 | 1923.963 | 0 | 0.74 | 0.001108 | 6.96 | 75.2 | 124.8 | 10 | 10.32 | | High | High | High | High | High | High | High | High | High | High | High | 0 | 7.87E-06 | 3.14 |
| FALSE | High | TDSVRTPES | [R].TDSVRT | 1xPhospho | P97875 1x | 0.000157 | 0 | 1 | 3 | 1 | P97875 | Jun dimeri | P97875 [14 | 1 | 2766.396 | 0 | -1.39 | 0.000382 | 2.26 | 146.8 | 53.2 | 6.46 | 16.7 | | High | High | High | High | High | High | High | High | High | High | High | 0 | 2.98E-05 | 4.97 |
| FALSE | High | TDSYSAGQ | [R].TDSYSA | 1xPhospho | Q9JLN9 1x | 5.92E-09 | 0 | 1 | 1 | 7 | Q9JLN9 | Serine/thre | Q9JLN9 [2 | 0 | 3040.492 | 0 | -0.37 | 0.016552 | 5.28 | 111.3 | 88.7 | 5.9 | 5.73 | | High | High | High | High | High | High | High | High | High | High | High | 0 | 3.19E-10 | 6.45 |
| FALSE | High | TDTAADGE | [R].TDTAAI | 1xPhospho | Q9WV92-7 | 5.53E-07 | 0 | 1 | 5 | 2 | Q9WV92-7 | Isoform 7 c | Q9WV92-7 | 0 | 2278.038 | 0 | -0.59 | 0.063968 | 19.26 | 118 | 82 | 13.9 | 17.18 | | High | High | High | High | High | High | High | High | High | High | High | 0 | 5.24E-08 | 3.78 |
| FALSE | High | TDTDNLGT | [R].TDTDN | 1xPhospho | Q9WVM1 | 0.002022 | 5.66E-05 | 1 | 1 | 1 | Q9WVM1 | Rac GTPase | Q9WVM1 | 0 | 2002.874 | 0 | -0.59 | 0.024604 | 8.81 | 121.7 | 78.3 | 13.25 | 13.19 | | High | High | High | High | High | High | High | High | High | High | High | 2.58E-05 | 0.000525 | 2.36 |
| FALSE | High | TDTLCLNN | [R].TDTLCL | 1xCarbami | Q9EP71 1x | 5.58E-07 | 0 | 1 | 1 | 1 | Q9EP71 | Ankycorbir | Q9EP71 [4 | 0 | 2877.359 | 0 | 0.85 | 0.000632 | 12.33 | 70.2 | 129.8 | 9.72 | 7.07 | | High | High | High | High | High | High | High | High | High | High | High | 0 | 5.27E-08 | 4.6 |
| FALSE | High | TDTLCLNN | [R].TDTLCL | 1xCarbami | Q9EP71 1x | 6.74E-07 | 0 | 1 | 1 | 3 | Q9EP71 | Ankycorbir | Q9EP71 [4 | 0 | 2878.343 | 0 | 0.84 | 0.001429 | 7.32 | 71.2 | 128.8 | 5.77 | 8.86 | | High | High | High | High | High | High | High | High | High | High | High | 0 | 9.99E-08 | 4.18 |
| FALSE | High | TDTLCLNN | [R].TDTLCL | 1xCarbami | Q9EP71 2x | 0.000294 | 0 | 1 | 1 | 1 | Q9EP71 | Ankycorbir | Q9EP71 [4 | 0 | 2957.325 | 0 | -0.1 | 0.543497 | 2.49 | 102.3 | 97.7 | 8.31 | 6.29 | | High | High | High | High | High | High | High | High | High | High | High | 0 | 6.04E-05 | 3.95 |
| FALSE | High | TDVFAPDY | [K].TDVFAI | 1xPhospho | Q148S3 1x | 1.65E-06 | 0 | 1 | 1 | 1 | Q148S3 | NIN1/RPN1 | Q148S3 [3 | 0 | 2780.302 | 0 | -0.47 | 0.003676 | 4.25 | 116.4 | 83.6 | 7.54 | 6.6 | | High | High | High | High | High | High | High | High | High | High | High | 0 | 1.77E-07 | 4.49 |
| FALSE | High | TDVLGVHE | [K].TDVLG\ | 1xPhospho | Q61097 1x | 0.000454 | 0 | 1 | 4 | 2 | Q61097 | Kinase sup | Q61097 [5 | 1 | 4436.967 | 0 | -0.61 | 0.006027 | 3.92 | 122.6 | 77.4 | 13.01 | 5.79 | | High | High | High | High | High | High | High | High | High | High | High | 0.000204 | 0.001367 | 4.11 |
| FALSE | High | TDVSNFDE | [R].TDVSNI | 1xPhospho | P70268-2 1 | 2.62E-07 | 0 | 1 | 2 | 4 | P70268-2 | Isoform 2 c | P70268-2 [ | 0 | 2618.186 | 0 | 0.7 | 0.001095 | 3.27 | 78.7 | 121.3 | 2.82 | 6.06 | | High | High | High | High | High | High | High | High | High | High | High | 0 | 1.26E-07 | 3.45 |
| FALSE | High | TDWSSGDA | [R].TDWSS | 1xPhospho | Q8R4H2 1x | 0.027358 | 0.001676 | 2 | 2 | 1 | Q8R4H2; F | Rho guanir | Q8R4H2 [2 | 0 | 2362.003 | 0 | 0.82 | 0.136373 | 7.32 | 72.2 | 127.8 | 30.74 | 27.79 | Shared | High | High | High | High | High | High | High | High | High | High | High | 0.000936 | 0.009721 | 2.64 |
| FALSE | High | TDWSSGDA | [R].TDWSS | 1xPhospho | Q8R4H2 1x | 0.001865 | 5.66E-05 | 2 | 2 | 1 | Q8R4H2; F | Rho guanir | Q8R4H2 [2 | 0 | 2591.166 | 0 | 1.25 | 0.0054 | 24.72 | 59.1 | 140.9 | 10.59 | 13.97 | Shared | High | High | High | High | High | High | High | High | High | High | High | 2.58E-05 | 0.000481 | 1.37 |
| FALSE | High | TEADLPKPT | [R].TEADLF | 1xPhospho | A2A5V4 1x | 0.001361 | 0 | 1 | 5 | 4 | A2A5V4 | SH3 domai | A2A5V4 [5 | 0 | 1821.976 | 0 | -0.23 | 0.370304 | 2.25 | 106.8 | 93.2 | 15.63 | 19.05 | | High | High | High | High | High | High | High | High | High | High | High | 2.58E-05 | 0.000338 | 3.43 |

| | | | | | | | | | | | | | | | | | | | | | | | | | | | | | | | | | | | | | | |
|---|---|---|---|---|---|---|---|---|---|---|---|---|---|---|---|---|---|---|---|---|---|---|---|---|---|---|---|---|---|---|---|---|---|---|---|---|---|---|
| FALSE | High | TDSVIIADQ | [R].TDSVII/ | 2xPhospho | Q3TZR9 2x | 0.002272 | 0.000109 | 1 | 5 | 2 | Q3TZR9 | Activating | Q3TZR9 [4 | 0 | 2003.929 | 0 | 0.97 | 0.000305 | 4.71 | 67.9 | 132.1 | 15.45 | 5.95 | | High | High | High | High | High | High | High | High | High | High | High | 4.94E-05 | 0.000598 | 3.26 |
| FALSE | High | TDSDSDLQ | [K].TDSDSI | 1xPhospho | A2AE27 1x | 2.83E-06 | 0 | 1 | 2 | 4 | A2AE27 | AMP deam | A2AE27 [1 | 0 | 1822.888 | 0 | 1.17 | 0.000957 | 11.3 | 61.6 | 138.4 | 5.73 | 10.46 | | High | High | High | High | High | High | High | High | High | High | High | 2.58E-05 | 0.000396 | 2.98 |
| FALSE | High | TDSDSDLQ | [K].TDSDSI | 1xPhospho | A2AE27 1x | 0.062209 | 0.004646 | 1 | 2 | 2 | A2AE27 | AMP deam | A2AE27 [1 | 0 | 1593.725 | 0 | 0.14 | 0.993752 | 27.26 | 98 | 102 | 13.17 | 19.41 | | High | High | High | High | High | High | High | High | High | High | High | 0.002381 | 0.02477 | 2.49 |
| FALSE | High | TDSCSSGQ | [R].TDSCSS | 1xCarbami | Q91ZV3 1x | 0.001002 | 0 | 1 | 2 | 1 | Q91ZV3 | Discoidin, | Q91ZV3 [7 | 0 | 2182.973 | 0 | 0.99 | 0.001717 | 18.69 | 72.2 | 127.8 | 15.48 | 10.74 | | High | High | High | High | High | High | High | High | High | High | High | 0 | 0.00024 | 2.97 |
| FALSE | High | TDDSQPPT | [R].TDDSQ | 1xPhospho | Q921K9 1x | 0.000181 | 0 | 1 | 2 | 1 | Q921K9 | B-cell CLL/l | Q921K9 [1 | 0 | 4043.004 | 0 | 1.17 | 0.003497 | 15.95 | 58.3 | 141.7 | 13.64 | 12.07 | | High | High | High | High | High | High | High | High | High | High | High | 0 | 3.5E-05 | 3.24 |
| FALSE | High | TDEDAVLM | [R].TDEDA | 1xPhospho | Q61584-3 1 | 0.000142 | 0 | 1 | 5 | 3 | Q61584-3 | Isoform B | Q61584-3 [ | 0 | 3219.502 | 0 | -0.54 | 0.053345 | 3.5 | 118.5 | 81.5 | 22.67 | 19.47 | | High | High | High | High | High | High | High | High | High | High | High | 0 | 3.84E-05 | 3.79 |
| FALSE | High | TDEGPDTP | [K].TDEGPI | 1xPhospho | Q6A065 1x | 0.003755 | 0.00021 | 1 | 3 | 1 | Q6A065 | Centrosom | Q6A065 [9 | 0 | 1660.705 | 0 | 0.59 | 0.003976 | 16.15 | 80.4 | 119.6 | 12.43 | 4.5 | | High | High | High | High | High | High | High | High | High | High | High | 9.42E-05 | 0.001055 | 2.73 |
| FALSE | High | TDEPVSGA | [K].TDEPVS | 1xPhospho | E9Q0S6 1x | 1.11E-08 | 0 | 1 | 6 | 1 | E9Q0S6 | Tensin 1 O | E9Q0S6 [5 | 0 | 2508.263 | 0 | -0.15 | 0.583735 | 16.61 | 104.5 | 95.5 | 11.7 | 5.26 | | High | High | High | High | High | High | High | High | High | High | High | 0 | 6.52E-10 | 3.92 |
| FALSE | High | TDESLDWD | [K].TDESLD | 1xPhospho | Q9WVM1 1 | 0.005322 | 0.000599 | 1 | 2 | 1 | Q9WVM1 | Rac GTPase | Q9WVM1 [ | 0 | 2032.988 | 0 | -0.78 | 0.027591 | 8.87 | 124.6 | 75.4 | 11.9 | 15.11 | | High | High | High | High | High | High | High | High | High | High | High | 0.000267 | 0.001546 | 1.84 |
| FALSE | High | TDFYDVSV | [K].TDFYDV | 1xPhospho | F6QK86 1x | 0.000369 | 0 | 1 | 1 | 1 | F6QK86 | COP9 signa | F6QK86 [1 | 0 | 2776.27 | 0 | 0.63 | 0.000829 | 8.94 | 80.3 | 119.7 | 19.21 | 14.7 | | High | High | High | High | High | High | High | High | High | High | High | 0 | 7.82E-05 | 3.07 |
| FALSE | High | TDGFAEAIH | [R].TDGFAI | 1xPhospho | F6ZDS4 1xl | 0.000316 | 0 | 1 | 5 | 2 | F6ZDS4 | Nucleoprot | F6ZDS4 [22 | 0 | 2161.064 | 0 | 0.73 | 0.083847 | 4.61 | 82 | 118 | 26.71 | 9.59 | | High | High | High | High | High | High | High | High | High | High | High | 0 | 6.53E-05 | 3.44 |
| FALSE | High | TDGKEEEE | [R].TDGKEI | 1xPhospho | B2RWS6 1x | 8.3E-07 | 0 | 1 | 1 | 6 | B2RWS6 | Histone ac | B2RWS6 [1 | 1 | 3343.615 | 0 | -0.26 | 0.043134 | 6.82 | 112.9 | 87.1 | 8.74 | 10.9 | | High | High | High | High | High | High | High | High | High | High | High | 0 | 8.21E-08 | 4.19 |
| FALSE | High | TDGSISGDI | [R].TDGSIS | 2xPhospho | Q6P5B5 2x | 0.071071 | 0.005247 | 1 | 2 | 1 | Q6P5B5 | Fragile X m | Q6P5B5 [6 | 1 | 2526.148 | 0 | -0.11 | 0.274161 | 4.73 | 104.4 | 95.6 | 11.9 | 6.24 | | High | High | High | High | High | High | High | High | High | High | High | 0.002703 | 0.02896 | 3.31 |
| FALSE | High | TDLANGSL | [R].TDLAN( | 1xPhospho | Q8N7N5 1x | 3.97E-08 | 0 | 1 | 2 | 1 | Q8N7N5 | DDB1- and | Q8N7N5 [1 | 0 | 2533.096 | 0 | 0.04 | 0.941827 | 8.43 | 98.8 | 101.2 | 20.14 | 5.05 | | High | High | High | High | High | High | High | High | High | High | High | 0 | 2.7E-09 | 4.16 |
| FALSE | High | TDLANGSL | [R].TDLAN( | 1xDeamida | Q8N7N5 1x | 3.01E-05 | 0 | 1 | 2 | 1 | Q8N7N5 | DDB1- and | Q8N7N5 [1 | 0 | 2534.08 | 0 | -0.18 | 0.084089 | 9.72 | 104.9 | 95.1 | 9.91 | 5.95 | | High | High | High | High | High | High | High | High | High | High | High | 0 | 4.64E-06 | 3.11 |
| FALSE | High | TDLANGSL | [R].TDLAN( | 1xPhospho | Q8N7N5 1x | 3.9E-05 | 0 | 1 | 2 | 2 | Q8N7N5 | DDB1- and | Q8N7N5 [1 | 0 | 2549.091 | 0 | 1.56 | 0.022491 | 7.72 | 52.3 | 147.7 | 10.08 | 30.65 | | High | High | High | High | High | High | High | High | High | High | High | 0 | 6.27E-06 | 3.28 |
| FALSE | High | TDLANGSL | [R].TDLAN( | 1xDeamida | Q8N7N5 1x | 0.009843 | 0.000736 | 1 | 2 | 1 | Q8N7N5 | DDB1- and | Q8N7N5 [1 | 0 | 2550.075 | 0 | 0.09 | 0.929891 | 44.86 | 98.7 | 101.3 | 22.05 | 10.76 | | High | High | High | High | High | High | High | High | High | High | High | 0.000344 | 0.003084 | 4.28 |
| FALSE | High | TDLSEQVSI | [K].TDLSEC | 1xPhospho | P58404 1xl | 0.035181 | 0.002269 | 1 | 4 | 1 | P58404 | Striatin-4 C | P58404 [14 | 0 | 3266.66 | 0 | -0.33 | 0.166613 | 5.87 | 108.3 | 91.7 | 13.7 | 16.85 | | High | High | High | High | High | High | High | High | High | High | High | 0.001214 | 0.01293 | 3.29 |
| FALSE | High | TDLSSPPPT | [K].TDLSSP | 1xPhospho | Q80WQ8 1 | 0.014538 | 0.001011 | 1 | 3 | 1 | Q80WQ8 | Mis18-binc | Q80WQ8 [ | 0 | 1379.677 | 0 | -1.28 | 0.000271 | 0.28 | 141.8 | 58.2 | 4.76 | 10.23 | | High | High | High | High | High | High | High | High | High | High | High | 0.000555 | 0.004786 | 1.9 |
| FALSE | High | TDLTPPSPE | [K].TDLTPP | 1xCarbami | E9PZ97 1xl | 0.001704 | 5.66E-05 | 1 | 2 | 1 | E9PZ97 | Bloom synd | E9PZ97 [2C | 0 | 2452.245 | 0 | -1.97 | 0.000275 | 0.34 | 160 | 40 | 6.23 | 7.8 | | High | High | High | High | High | High | High | High | High | High | High | 2.58E-05 | 0.000436 | 2.3 |
| FALSE | High | TDLTQGDE | [R].TDLTQ( | 1xPhospho | Q9ERU9 1x | 1.54E-07 | 0 | 1 | 1 | 2 | Q9ERU9 | E3 SUMO- | Q9ERU9 [2 | 0 | 3209.431 | 0 | -0.12 | 0.896758 | 27.39 | 114 | 86 | 31.58 | 33.7 | | High | High | High | High | High | High | High | High | High | High | High | 0 | 1.24E-08 | 5.44 |
| FALSE | High | TDLTQGDE | [R].TDLTQ( | 1xPhospho | Q9ERU9 1x | 4.15E-08 | 0 | 1 | 1 | 9 | Q9ERU9 | E3 SUMO- | Q9ERU9 [2 | 0 | 3438.594 | 0 | -0.2 | 0.010346 | 3.48 | 108.1 | 91.9 | 7.05 | 5.35 | | High | High | High | High | High | High | High | High | High | High | High | 0 | 2.84E-09 | 4.27 |
| FALSE | High | TDLTQGDE | [R].TDLTQ( | 2xPhospho | Q9ERU9 2x | 1.24E-05 | 0 | 1 | 1 | 7 | Q9ERU9 | E3 SUMO- | Q9ERU9 [2 | 0 | 3518.56 | 0 | -0.34 | 0.032735 | 4.56 | 112.7 | 87.3 | 7.69 | 11.07 | | High | High | High | High | High | High | High | High | High | High | High | 0 | 0.000208 | 4.39 |
| FALSE | High | TDNSVASS | [K].TDNSV/ | 1xPhospho | O70546-2 1 | 7.4E-08 | 0 | 1 | 4 | 2 | O70546-2 | Isoform 2 | O70546-2 [ | 0 | 2543.264 | 0 | -0.27 | 0.111567 | 9.91 | 110.9 | 89.1 | 4.24 | 14.01 | | High | High | High | High | High | High | High | High | High | High | High | 0 | 5.48E-09 | 3.51 |
| FALSE | High | TDNSVASS | [K].TDNSV/ | 2xPhospho | O70546-2 2 | 0.013787 | 0.001011 | 1 | 4 | 2 | O70546-2 | Isoform 2 | O70546-2 [ | 0 | 2623.231 | 0 | 0.28 | 0.062235 | 10.14 | 90.2 | 109.8 | 4.54 | 10.65 | | High | High | High | High | High | High | High | High | High | High | High | 0.000538 | 0.004478 | 2.9 |
| FALSE | High | TDPACLETC | [K].TDPACI | 1xCarbami | Q6ZPR8 1x | 0.01441 | 0.001011 | 1 | 3 | 2 | Q6ZPR8 | MKIAA140 | Q6ZPR8 [4 | 0 | 3894.626 | 0 | -1.52 | 0.930502 | 5.07 | 144.7 | 55.3 | 63.57 | 166.71 | | High | High | High | High | High | High | High | High | High | High | High | 0.000786 | 0.007034 | 3.42 |
| FALSE | High | TDPACLETC | [K].TDPACI | 1xCarbami | Q6ZPR8 1x | 0.000442 | 0 | 1 | 3 | 7 | Q6ZPR8 | MKIAA140 | Q6ZPR8 [4 | 0 | 4123.789 | 0 | -0.27 | 0.201214 | 16.63 | 108.1 | 91.9 | 12.76 | 6.65 | | High | High | High | High | High | High | High | High | High | High | High | 0 | 9.48E-05 | 4.1 |
| FALSE | High | TDPACLETC | [K].TDPACI | 1xCarbami | Q6ZPR8 1x | 4.47E-05 | 0 | 1 | 3 | 4 | Q6ZPR8 | MKIAA140 | Q6ZPR8 [4 | 0 | 4124.773 | 0 | -0.23 | 0.00062 | 0.89 | 109 | 91 | 6.98 | 6.33 | | High | High | High | High | High | High | High | High | High | High | High | 0 | 7.28E-06 | 5.21 |
| FALSE | High | TDPFYHSP( | [K].TDPFYH | 1xPhospho | E9QQ10 1x | 0.00013 | 0 | 1 | 1 | 2 | E9QQ10 | A-kinase ar | E9QQ10 [3 | 0 | 2343.101 | 0 | 0.28 | 0.03348 | 13.66 | 90.5 | 109.5 | 9.7 | 9.11 | | High | High | High | High | High | High | High | High | High | High | High | 0.00196 | 0.01997 | 2.93 |
| FALSE | High | TDPGSIENI | [K].TDPGSI | 1xCarbami | B9EHS3 1x | 0.062731 | 0.004664 | 1 | 7 | 1 | B9EHS3 | Hisppd2a p | B9EHS3 [9 | 0 | 1925.944 | 0 | 0.63 | 0.000609 | 5.59 | 78.8 | 121.2 | 6.27 | 5.38 | | High | High | High | High | High | High | High | High | High | High | High | 0.002403 | 0.02492 | 2.21 |
| FALSE | High | TDPISLSNC | [K].TDPISL | 1xCarbami | Q8K3K8 1x | 7.91E-06 | 0 | 1 | 1 | 1 | Q8K3K8 | Optineurin | Q8K3K8 [2 | 0 | 1887.836 | 0 | 2.34 | 0.001438 | 11.69 | 31 | 169 | 8.75 | 16.24 | | High | High | High | High | High | High | High | High | High | High | High | 0 | 1.04E-06 | 3.93 |
| FALSE | High | TDQEGLNA | [K].TDQEG | 1xPhospho | Q69Z38 1x | 1.29E-05 | 0 | 1 | 2 | 1 | Q69Z38 | Pseudopod | Q69Z38 [1 | 0 | 2428.253 | 0 | -0.57 | 0.040926 | 3.21 | 116.9 | 83.1 | 11.85 | 10.18 | | High | High | High | High | High | High | High | High | High | High | High | 0 | 1.79E-06 | 3.07 |
| FALSE | High | TDSASADP | [R].TDSAS/ | 1xPhospho | Q8BKX6 1x | 5.89E-07 | 0 | 1 | 4 | 2 | Q8BKX6 | Serine/thre | Q8BKX6 [3 | 0 | 1758.856 | 0 | 1.4 | 0.002664 | 30.66 | 55.1 | 144.9 | 13.77 | 11.04 | | High | High | High | High | High | High | High | High | High | High | High | 0 | 5.59E-08 | 3.64 |
| FALSE | High | TDDSQETC | [K].TDDSQ | 1xPhospho | P25206 1xl | 0.000633 | 0 | 2 | 3 | 2 | P25206; Q | DNA replic | P25206 [72 | 0 | 1919.864 | 0 | -0.73 | 0.016927 | 4.58 | 122.7 | 77.3 | 9.34 | 17.97 | Shared | High | High | High | High | High | High | High | High | High | High | High | 0 | 0.000143 | 2.4 |
| FALSE | High | TASGYGGT | [R].TASGY( | 1xPhospho | A0A0R4J0E | 0.115566 | 0.009504 | 1 | 2 | 1 | A0A0R4J0E | Store-oper | A0A0R4J0E | 0 | 1178.54 | 0 | -0.62 | 0.049442 | 6.36 | 121.3 | 78.7 | 7.46 | 20.75 | | High | High | High | High | High | High | High | High | High | High | High | 0.004757 | 0.05131 | 0.93 |
| FALSE | High | TFSYDEIHG | [R].TFSYDE | 1xPhospho | P33174 1xl | 0.034281 | 0.002195 | 1 | 2 | 3 | P33174 | Chromosor | P33174 [8C | 0 | 2479.086 | 0 | 1.07 | 0.01772 | 35.18 | 65.1 | 134.9 | 19.91 | 17.12 | | High | High | High | High | High | High | High | High | High | High | High | 0.001168 | 0.01255 | 2.21 |
| FALSE | High | TFTEPDSLK | [K].TFTEPD | 1xPhospho | B2RY84 1xl | 0.029844 | 0.001857 | 1 | 4 | 1 | B2RY84 | Kif14 prote | B2RY84 [3 | 0 | 1575.807 | 0 | -0.21 | 0.338616 | 8.63 | 99.5 | 100.5 | 18.03 | 9.83 | | High | High | High | High | High | High | High | High | High | High | High | 0.001011 | 0.01072 | 1.8 |
| FALSE | High | TLGTPTQS( | [R].TLGTPT | 1xPhospho | Q8R4R6 1x | 2.14E-06 | 0 | 1 | 1 | 3 | Q8R4R6 | Nucleopor | Q8R4R6 [2 | 0 | 1611.794 | 0 | 0.01 | 0.89226 | 3.81 | 101.5 | 98.5 | 4.31 | 8.19 | | High | High | High | High | High | High | High | High | High | High | High | 0 | 0.000266 | 2.77 |
| FALSE | High | TIGYEQINS | [K].TIGYEQ | 1xPhospho | Q8BMD7 1 | 4.06E-06 | 0 | 1 | 4 | 1 | Q8BMD7 | MORC fam | Q8BMD7 [ | 0 | 2183.131 | 0 | 0.24 | 0.018753 | 2.23 | 89.9 | 110.1 | 8.96 | 12.81 | | High | High | High | High | High | High | High | High | High | High | High | 0 | 4.93E-07 | 3.3 |
| FALSE | High | TIHLDSEED | [R].TIHLDS | 1xPhospho | Q8K4J0 1xl | 1.05E-08 | 0 | 1 | 4 | 1 | Q8K4J0 | Protein art | Q8K4J0 [3 | 0 | 2862.328 | 0 | -0.94 | 0.006573 | 7.47 | 128.6 | 71.4 | 13.85 | 15.62 | | High | High | High | High | High | High | High | High | High | High | High | 0 | 6.05E-10 | 5.06 |
| FALSE | High | TIHTFPCLK | [R].TIHTFP | 1xCarbami | B7ZMV4 1x | 0.040373 | 0.002982 | 1 | 3 | 2 | B7ZMV4 | Cdr2l prote | B7ZMV4 [1 | 0 | 1425.716 | 0 | -0.11 | 0.154323 | 3.79 | 106.3 | 93.7 | 9.69 | 9.95 | | High | High | High | High | High | High | High | High | High | High | High | 0.007451 | 0.0822 | 1.59 |
| FALSE | High | TIHTFPCLK | [R].TIHTFP | 1xCarbami | B7ZMV4 1x | 0.009756 | 0.000736 | 1 | 3 | 1 | B7ZMV4 | Cdr2l prote | B7ZMV4 [1 | 0 | 1654.879 | 0 | -0.17 | 0.148604 | 23.55 | 108.1 | 91.9 | 18.91 | 5.2 | | High | High | High | High | High | High | High | High | High | High | High | 0.000344 | 0.003046 | 1.97 |
| FALSE | High | TLKESDSAE | [R].TLKESD | 1xPhospho | Q8K382 1x | 0.028082 | 0.001791 | 1 | 3 | 2 | Q8K382 | DENN dom | Q8K382 [5 | 1 | 2858.371 | 0 | 0.48 | 0.002107 | 3.76 | 85 | 115 | 4.42 | 7.63 | | High | High | High | High | High | High | High | High | High | High | High | 0.000968 | 0.009949 | 3.13 |
| FALSE | High | TIKTPEDG( | [K].TIKTPEI | 1xCarbami | P14873 1xl | 0.094641 | 0.007408 | 1 | 1 | 2 | P14873 | Microtubu | P14873 [1 | 1 | 2709.366 | 0 | -0.98 | 0.00179 | 0.63 | 130.4 | 69.6 | 9.83 | 6.21 | | High | High | High | High | High | High | High | High | High | High | High | 0.004221 | 0.04536 | 3.13 |
| FALSE | High | TLLAQGNC | [K].TLLAQ( | 1xPhospho | A0A1D5RM | 2.56E-05 | 0 | 1 | 3 | 1 | A0A1D5RM | Kinesin-like | A0A1D5RM | 0 | 2981.567 | 0 | 0.85 | 0.002468 | 9.39 | 71.3 | 128.7 | 10.36 | 9.01 | | High | High | High | High | High | High | High | High | High | High | High | 0 | 3.88E-06 | 4.5 |
| FALSE | High | TLLAQGNC | [K].TLLAQ( | 1xDeamida | A0A1D5RM | 3.87E-05 | 0 | 1 | 3 | 2 | A0A1D5RM | Kinesin-like | A0A1D5RM | 0 | 2982.551 | 0 | 1.11 | 0.007034 | 33.53 | 64.5 | 135.5 | 10.54 | 15.84 | | High | High | High | High | High | High | High | High | High | High | High | 0 | 6.18E-06 | 4.12 |
| FALSE | High | TLLFGALSE | [K].TLLFGA | 1xPhospho | Q8BQ33 1x | 0.001002 | 0 | 1 | 1 | 2 | Q8BQ33 | Treslin OS= | Q8BQ33 [1 | 0 | 2088.139 | 0 | -0.96 | 0.008505 | 9.06 | 134.6 | 65.4 | 14.21 | 13.07 | | High | High | High | High | High | High | High | High | High | High | High | 0 | 0.000238 | 3.01 |
| FALSE | High | TLLGSSDN | [K].TLLGSS | 1xPhospho | Q5XG69 1x | 0.000246 | 0 | 1 | 1 | 3 | Q5XG69 | Soluble lan | Q5XG69 [5 | 0 | 2286.199 | 0 | -1.27 | 0.001017 | 0.54 | 142.3 | 57.7 | 6.79 | 12.74 | | High | High | High | High | High | High | High | High | High | High | High | 0 | 4.92E-05 | 3.45 |
| FALSE | High | TLLLSSDDE | [K].TLLLSSI | 1xPhospho | Q8BG09 1x | 0.001361 | 0 | 1 | 1 | 5 | Q8BG09 | Transmem | Q8BG09 [3 | 0 | 1448.676 | 0 | 1.24 | 0.003531 | 25.71 | 58.7 | 141.3 | 7.03 | 13.36 | | High | High | High | High | High | High | High | High | High | High | High | 2.58E-05 | 0.000339 | 2.84 |
| FALSE | High | TLLLSSDDE | [K].TLLLSSI | 2xPhospho | Q8BG09 2x | 0.001516 | 5.66E-05 | 1 | 1 | 15 | Q8BG09 | Transmem | Q8BG09 [3 | 0 | 1528.642 | 0 | 2.16 | 0.00014 | 6.96 | 36.5 | 163.5 | 6.06 | 3.22 | | High | High | High | High | High | High | High | High | High | High | High | 2.58E-05 | 0.000379 | 2.66 |
| FALSE | High | TLGSDSTGI | [R].TLGSDS | 1xPhospho | P98203-3 1 | 9.5E-07 | 0 | 1 | 6 | 2 | P98203-3 | Isoform 3 | P98203-3 [ | 0 | 1835.868 | 0 | -1.17 | 0.000193 | 1.09 | 139.3 | 60.7 | 3.05 | 7 | | High | High | High | High | High | High | High | High | High | High | High | 0 | 9.58E-08 | 3.11 |
| FALSE | High | TLLPWDSD | [K].TLLPWI | 1xPhospho | F8WJ40 1x | 0.001349 | 0 | 1 | 5 | 2 | F8WJ40 | Histone-lys | F8WJ40 [1 | 0 | 2514.175 | 0 | 0.04 | 0.682276 | 17.63 | 100.7 | 99.3 | 5.62 | 22.88 | | High | High | High | High | High | High | High | High | High | High | High | 0 | 0.000334 | 2.89 |
| FALSE | High | TLIVSEASPI | [K].TLIVSE/ | 1xDeamida | Q5U4C1 1x | 6.71E-05 | 0 | 1 | 3 | 1 | Q5U4C1 | G-protein | Q5U4C1 [9 | 0 | 1883.977 | 0 | -1.33 | 0.000716 | 2.18 | 144.5 | 55.5 | 5.26 | 11.07 | | High | High | High | High | High | High | High | High | High | High | High | 0 | 1.15E-05 | 2.02 |
| FALSE | High | TLMLPLTE( | [K].TLMLPI | 1xPhospho | Q8BK03 1x | 6.83E-05 | 0 | 1 | 2 | 1 | Q8BK03 | Mitoguard | Q8BK03 [2 | 0 | 1655.864 | 0 | 1.17 | 0.002174 | 7.15 | 70.2 | 129.8 | 11.23 | 15.83 | | High | High | High | High | High | High | High | High | High | High | High | 0 | 1.18E-05 | 2.91 |
| FALSE | High | TLNEWSSC | [K].TLNEW | 1xPhospho | O09172 1x | 8.79E-05 | 0 | 1 | 4 | 1 | O09172 | Glutamate- | O09172 [5 | 0 | 2054.016 | 0 | 1.39 | 0.000684 | 3.94 | 55.9 | 144.1 | 5.49 | 8.77 | | High | High | High | High | High | High | High | High | High | High | High | 0 | 1.55E-05 | 3.45 |
| FALSE | High | TINHQMES | [K].TINHQI | 1xPhospho | D3YU22 1x | 0.024001 | 0.001612 | 1 | 7 | 1 | D3YU22 | LIM and ca | D3YU22 [9 | 0 | 1723.767 | 0 | 0.29 | 0.205521 | 14.27 | 86.1 | 113.9 | 14.67 | 45.11 | | High | High | High | High | High | High | High | High | High | High | High | 0.000859 | 0.008398 | 1.94 |
| FALSE | High | TLNLGLSPS | [K].TLNLGI | 1xPhospho | Q7TME2 1 | 0.047086 | 0.00354 | 1 | 2 | 1 | Q7TME2 | Sperm-ass | Q7TME2 [6 | 0 | 1792.997 | 0 | -0.6 | 0.003447 | 1.7 | 120.3 | 79.7 | 9.42 | 5.22 | | High | High | High | High | High | High | High | High | High | High | High | 0.001812 | 0.01795 | 1.89 |
| FALSE | High | TLNMTTSP | [K].TLNMT | 1xPhospho | Q3THK7 1x | 0.000375 | 0 | 1 | 1 | 4 | Q3THK7 | GMP synth | Q3THK7 [3 | 0 | 1788.885 | 0 | -0.48 | 0.000565 | 2.08 | 118.6 | 81.4 | 3.97 | 5.19 | | High | High | High | High | High | High | High | High | High | High | High | 0 | 7.94E-05 | 3.15 |
| FALSE | High | TLNMTTSP | [K].TLNMT | 1xPhospho | Q3THK7 1x | 0.012619 | 0.000889 | 1 | 1 | 15 | Q3THK7 | GMP synth | Q3THK7 [3 | 0 | 1804.88 | 0 | -0.45 | 0.024821 | 2.91 | 115.6 | 84.4 | 2.9 | 13.6 | | High | High | High | High | High | High | High | High | High | High | High | 0.000449 | 0.004172 | 2.5 |
| FALSE | High | TLNQLGTP | [R].TLNQL( | 1xPhospho | O70439 1x | 0.000213 | 0 | 1 | 4 | 1 | O70439 | Syntaxin-7 | O70439 [3 | 0 | 1977.984 | 0 | 0.06 | 0.726273 | 21.58 | 99.3 | 100.7 | 19.31 | 9.47 | | High | High | High | High | High | High | High | High | High | High | High | 0 | 4.2E-05 | 3.11 |
| FALSE | High | TLPDSQQV | [R].TLPDSC | 1xPhospho | E9Q652 1x | 5.38E-07 | 0 | 1 | 9 | 4 | E9Q652 | Regulator | E9Q652 [8 | 0 | 2080.073 | 0 | 1.68 | 0.000193 | 12.09 | 46.3 | 153.7 | 5.6 | 3.79 | | High | High | High | High | High | High | High | High | High | High | High | 0 | 5.07E-08 | 3.64 |
| FALSE | High | TLPFSPSQF | [K].TLPFSP | 1xPhospho | P48972 1xl | 8.71E-05 | 0 | 1 | 2 | 3 | P48972 | Myb-relate | P48972 [4 | 0 | 2364.219 | 0 | -0.65 | 0.001182 | 3.12 | 123 | 77 | 8.86 | 7.79 | | High | High | High | High | High | High | High | High | High | High | High | 0 | 1.57E-05 | 3.57 |
| FALSE | High | TLPGISPPV | [R].TLPGISI | 1xPhospho | Q9CPY3 1x | 0.038346 | 0.002563 | 1 | 2 | 1 | Q9CPY3 | Sororin OS | Q9CPY3 [2 | 0 | 1645.969 | 0 | -0.21 | 0.017274 | 4.23 | 108.6 | 91.4 | 5.43 | 6.15 | | High | High | High | High | High | High | High | High | High | High | High | 0.001355 | 0.01428 | 1.49 |
| FALSE | High | TLPSEDEEC | [R].TLPSED | 1xPhospho | A0A1L1STC | 2.78E-06 | 0 | 1 | 7 | 1 | A0A1L1STC | Nesprin-1 | A0A1L1STC | 0 | 2015.91 | 0 | 3.72 | | 59.49 | 15.8 | 184.2 | 20.41 | 14.8 | | High | Not Found | High | High | High | High | High | High | High | High | High | 0 | 3.2E-07 | 3.91 |
| FALSE | High | TIPSLTLTST | [R].TIPSLTL | 1xPhospho | A0A0R4J2D | 0.059159 | 0.004294 | 1 | 3 | 1 | A0A0R4J2D | Ubiquitin c | A0A0R4J2D | 0 | 1708.945 | 0 | -1.04 | 0.002215 | 2.45 | 131.2 | 68.8 | 12.84 | 18.12 | | High | High | High | High | High | High | High | High | High | High | High | 0.002187 | 0.02329 | 2.6 |
| FALSE | High | TLLSPSSIK | [R].TLLSPS | 1xPhospho | E9QMD3 1 | 0.007209 | 0.00062 | 1 | 5 | 1 | E9QMD3 | Zinc finger | E9QMD3 [ | 0 | 1483.854 | 0 | 2.92 | 0.000374 | 10.8 | 25 | 175 | 13.4 | 8.58 | | High | High | High | High | High | High | High | High | High | High | High | 0.000275 | 0.002171 | 2.05 |
| FALSE | High | TLPSPIDAK | [K].TLPSPII | 1xPhospho | F8VPJ6 1xF | 0.001783 | 5.66E-05 | 1 | 2 | 2 | F8VPJ6 | Regulatory | F8VPJ6 [29 | 0 | 1479.822 | 0 | -0.19 | 0.005318 | 1.07 | 106.8 | 93.2 | 2.87 | 6.51 | | High | High | High | High | High | High | High | High | High | High | High | 2.58E-05 | 0.000455 | 2.49 |
| FALSE | High | TIGGGDDS | [K].TIGGGI | 1xCarbami | A0A0A0MC | 3.47E-05 | 0 | 1 | 7 | 3 | A0A0A0MC | Tubulin alp | A0A0A0MC | 0 | 2606.189 | 0 | -0.46 | 0.01543 | 6.33 | 115.9 | 84.1 | 12.22 | 3.56 | | High | High | High | High | High | High | High | High | High | High | High | 0.004974 | 0.05428 | 3.54 |
| FALSE | High | TIGGGDDS | [K].TIGGGI | 1xPhospho | P68369 1xl | 0.01218 | 0.000889 | 2 | 6 | 1 | P68369; Q | Tubulin alp | P68369 [4 | 0 | 2317.022 | 0 | 0.27 | 0.092995 | 14.74 | 88.3 | 111.7 | 13.35 | 18.3 | Shared | High | High | High | High | High | High | High | High | High | High | High | 0.000449 | 0.003892 | 3.16 |
| FALSE | High | TLASPK | [R].TLASPK | 1xPhospho | Q9CZ46 1x | 0.049971 | 0.003641 | 1 | 2 | 2 | Q9CZ46 | Putative ur | Q9CZ46 [7 | 0 | 1154.659 | 0 | -0.67 | 0.01036 | 2.15 | 122.2 | 77.8 | 5.99 | 14.46 | | High | High | High | High | High | High | High | High | High | High | High | 0.001902 | 0.01917 | 1.45 |
| FALSE | High | TLCGTPNY | [K].TLCGTF | 1xCarbami | Q3TPZ2 1x | 0.009931 | 0.000736 | 1 | 2 | 3 | Q3TPZ2 | Serine/thre | Q3TPZ2 [2 | 0 | 2301.197 | 0 | -0.53 | 0.001778 | 3.36 | 116.4 | 83.6 | 7.14 | 9.16 | | High | High | High | High | High | High | High | High | High | High | High | 0.000344 | 0.003099 | 3.06 |
| FALSE | High | TLDFDPLLS | [R].TLDFDF | 1xPhospho | B1AXD8 1x | 9.82E-06 | 0 | 1 | 1 | 2 | B1AXD8 | Akirin-2 OS | B1AXD8 [1 | 0 | 1809.924 | 0 | -0.56 | 0.526275 | 11.91 | 122.6 | 77.4 | 61.75 | 62.58 | | High | High | High | High | High | High | High | High | High | High | High | 0 | 1.33E-06 | 3.9 |
| FALSE | High | TLDFDPLLS | [R].TLDFDF | 1xPhospho | B1AXD8 1x | 0.000297 | 0 | 1 | 1 | 3 | B1AXD8 | Akirin-2 OS | B1AXD8 [1 | 0 | 2039.087 | 0 | -0.45 | 0.016451 | 5.1 | 114.2 | 85.8 | 2.73 | 12.65 | | High | High | High | High | High | High | High | High | High | High | High | 0 | 6.09E-05 | 2.89 |
| FALSE | High | TLDFDPLLS | [R].TLDFDF | 1xPhospho | B1AXD8 1x | 0.000633 | 0 | 1 | 1 | 1 | B1AXD8 | Akirin-2 OS | B1AXD8 [1 | 1 | 2195.188 | 0 | -0.38 | 0.015674 | 5.57 | 113 | 87 | 9.87 | 4.15 | | High | High | High | High | High | High | High | High | High | High | High | 0 | 0.000143 | 2.32 |
| FALSE | High | TIDIEPDIET | [R].TIDIEPI | 1xPhospho | P11352 1xl | 6.13E-05 | 0 | 1 | 3 | 1 | P11352 | Glutathion | P11352 [1 | 0 | 2369.132 | 0 | 0 | 0.579589 | 36.95 | 104.5 | 95.5 | 14.36 | 18.9 | | High | High | High | High | High | High | High | High | High | High | High | 0 | 1.04E-05 | 3.15 |
| FALSE | High | TLDRSGDL( | [R].TLDRSC | 1xPhospho | E9Q8Z8 1x | 1.06E-07 | 0 | 1 | 19 | 1 | E9Q8Z8 | Catenin de | E9Q8Z8 [9 | 1 | 2185.098 | 0 | -0.15 | 0.228968 | 12.79 | 107 | 93 | 2.38 | 15.63 | | High | High | High | High | High | High | High | High | High | High | High | 0 | 8.2E-09 | 2.85 |
| FALSE | High | TLDRSGDL( | [R].TLDRSC | 1xPhospho | E9Q8Z8 1x | 6.79E-06 | 0 | 1 | 19 | 1 | E9Q8Z8 | Catenin de | E9Q8Z8 [9 | 1 | 2201.093 | 0 | -0.2 | 0.129833 | 1.97 | 107.8 | 92.2 | 8.7 | 13.23 | | High | High | High | High | High | High | High | High | High | High | High | 0 | 8.78E-07 | 3.45 |
| FALSE | High | TLDSEEEQI | [R].TLDSEE | 1xPhospho | Q8BLH7 1x | 0.002553 | 0.000109 | 1 | 1 | 1 | Q8BLH7 | HIRA-intera | Q8BLH7 [5 | 0 | 1512.678 | 0 | -1.65 | 0.000452 | 3.01 | 154.4 | 45.6 | 6.87 | 19.73 | | High | High | High | High | High | High | High | High | High | High | High | 4.94E-05 | 0.000684 | 2.61 |
| FALSE | High | TIEAAEALL | [K].TIEAAE | 1xCarbami | Q9JHC9 1x | 3.44E-05 | 0 | 1 | 9 | 2 | Q9JHC9 | ETS-related | Q9JHC9 [9 | 0 | 2367.129 | 0 | 0.34 | 0.034496 | 18.48 | 87 | 113 | 10.13 | 12.2 | | High | High | High | High | High | High | High | High | High | High | High | 0 | 5.43E-06 | 3.31 |
| FALSE | High | TLEDEEEM | [R].TLEDEE | 1xPhospho | E9Q481 1x | 5.53E-07 | 0 | 1 | 3 | 9 | E9Q481 | Serine/thre | E9Q481 [6 | 0 | 3916.623 | 3 | -0.6 | 0.009986 | 8.83 | 118.1 | 81.9 | 8.83 | 12.99 | | High | High | High | High | High | High | High | High | High | High | High | 0 | 5.25E-08 | 6.26 |
| FALSE | High | TLEDEEEM | [R].TLEDEE | 1xPhospho | E9Q481 1x | 3.98E-05 | 0 | 1 | 3 | 11 | E9Q481 | Serine/thre | E9Q481 [6 | 1 | 3932.598 | 0 | -0.34 | 0.035947 | 11.93 | 111.1 | 88.9 | 7.25 | 10.32 | | High | High | High | High | High | High | High | High | High | High | High | 0 | 6.37E-06 | 5.59 |
| FALSE | High | TLENSPISS( | [R].TLENSP | 1xCarbami | Q80YS6 1x | 2.31E-07 | 0 | 1 | 1 | 3 | Q80YS6 | Actin filam | Q80YS6 [6 | 0 | 3410.68 | 0 | 0.61 | 0.012334 | 20.52 | 78.9 | 121.1 | 21.08 | 9.35 | | High | High | High | High | High | High | High | High | High | High | High | 0 | 1.96E-08 | 5.92 |
| FALSE | High | TIGGGDDS | [K].TIGGGI | 1xPhospho | P68369 1xl | 7.77E-06 | 0 | 2 | 6 | 4 | P68369; Q | Tubulin alp | P68369 [4 | 0 | 2546.185 | 0 | -0.7 | 0.001281 | 3.56 | 120.5 | 79.5 | 6.71 | 11.04 | Shared | High | High | High | High | High | High | High | High | High | High | High | 0 | 1.02E-06 | 3.71 |
| FALSE | High | TLEPVDTEE | [R].TLEPVD | 2xPhospho | Q8BU03 2x | 3.29E-05 | 0 | 1 | 4 | 6 | Q8BU03 | Periodic tr | Q8BU03 [8 | 0 | 2975.265 | 0 | -0.66 | 0.006208 | 4.65 | 123 | 77 | 9.75 | 10.17 | | High | High | High | High | High | High | High | High | High | High | High | 0 | 1.25E-05 | 4.78 |
| FALSE | High | TLEPVKPPT | [K].TLEPVK | 1xPhospho | J3QNK8 1x | 0.006306 | 0.00062 | 1 | 15 | 3 | J3QNK8 | Abl interac | J3QNK8 [2 | 0 | 2767.414 | 0 | -0.27 | 0.080328 | 21.2 | 110.8 | 89.2 | 5.8 | 19 | | High | High | High | High | High | High | High | High | High | High | High | 0.000275 | 0.00187 | 3.49 |
| FALSE | High | TLEQESEQ | [R].TLEQES | 1xPhospho | Q3U2K0 1x | 3.83E-05 | 0 | 1 | 3 | 1 | Q3U2K0 | Protein FAI | Q3U2K0 [7 | 0 | 2906.466 | 0 | 0.82 | 0.00109 | 4.86 | 73.9 | 126.1 | 9.84 | 12.83 | | High | High | High | High | High | High | High | High | High | High | High | 0 | 6.09E-06 | 3.54 |
| FALSE | High | TLEQLSSFS | [R].TLEQLS | 1xCarbami | A2BIE1 2xF | 0.017192 | 0.001204 | 1 | 2 | 1 | A2BIE1 | Glutamine | A2BIE1 [11 | 0 | 3319.416 | 0 | -0.31 | 0.012596 | 2.42 | 112.5 | 87.5 | 13.6 | 15.43 | | High | High | High | High | High | High | High | High | High | High | High | 0.000638 | 0.00574 | 3.55 |
| FALSE | High | TLESLPATN | [K].TLESLP/ | 1xPhospho | Q3U9W6 1 | 0.00311 | 0.00016 | 1 | 3 | 1 | Q3U9W6 | Putative ur | Q3U9W6 [ | 0 | 2971.425 | 0 | -0.79 | 0.005206 | 11.6 | 125.2 | 74.8 | 4.73 | 20.06 | | High | High | High | High | High | High | High | High | High | High | High | 7.19E-05 | 0.000852 | 2.99 |
| FALSE | High | TLEVVSPSC | [K].TLEVVS | 1xPhospho | P14873 1xl | 0.000302 | 0 | 2 | 2 | 4 | P14873; B2 | Microtubu | P14873 [1 | 0 | 3503.698 | 0 | -1.19 | 0.006858 | 0.97 | 135.3 | 64.7 | 16.65 | 18.5 | Shared | High | High | High | High | High | High | High | High | High | High | High | 0 | 6.25E-05 | 3.41 |
| FALSE | High | TLEVVSPSC | [K].TLEVVS | 2xPhospho | P14873 2xl | 0.011652 | 0.000889 | 2 | 2 | 1 | P14873; B2 | Microtubu | P14873 [1 | 0 | 3583.665 | 0 | -1.05 | 0.000779 | 3.24 | 132.8 | 67.2 | 5.17 | 7.55 | Shared | High | High | High | High | High | High | High | High | High | High | High | 0.000449 | 0.003718 | 3.57 |
| FALSE | High | TLFLGIPDE | [R].TLFLGII | 1xPhospho | O35099 1x | 1.72E-05 | 0 | 1 | 3 | 1 | O35099 | Mitogen-ac | O35099 [1 | 0 | 3107.501 | 0 | 1.61 | 0.00185 | 18.21 | 53.4 | 146.6 | 8.71 | 12.19 | | High | High | High | High | High | High | High | High | High | High | High | 0 | 2.49E-06 | 3.35 |
| FALSE | High | TLFLGIPDE | [R].TLFLGII | 2xPhospho | O35099 2x | 0.007339 | 0.00062 | 1 | 3 | 2 | O35099 | Mitogen-ac | O35099 [1 | 0 | 3187.468 | 0 | 1.09 | 0.00036 | 11.58 | 64 | 136 | 13.45 | 5.66 | | High | High | High | High | High | High | High | High | High | High | High | 0.007212 | 0.07825 | 2.68 |
| FALSE | High | TLFTEHYLS | [R].TLFTEH | 1xPhospho | Q9D7I8 1xl | 0.009168 | 0.000736 | 1 | 3 | 2 | Q9D7I8 | Protein FAI | Q9D7I8 [5 | 0 | 2315.095 | 0 | 0.64 | 0.007339 | 5.43 | 78.3 | 121.7 | 7.18 | 13.31 | | High | High | High | High | High | High | High | High | High | High | High | 0.002217 | 0.0235 | 1.85 |
| FALSE | High | TIGDSPPVL | [K].TIGDSP | 1xPhospho | Q91VS8 1x | 1.99E-05 | 0 | 1 | 1 | 1 | Q91VS8 | FERM, Rho | Q91VS8 [8 | 0 | 2051.078 | 0 | 0.99 | 0.001726 | 4.75 | 71.1 | 128.9 | 14.49 | 13.31 | | High | High | High | High | High | High | High | High | High | High | High | 0 | 2.9E-06 | 3.5 |
| FALSE | High | TIGENDEVI | [R].TIGEND | 1xPhospho | Q921N6 1x | 0.01861 | 0.001362 | 1 | 3 | 2 | Q921N6 | Probable A | Q921N6 [1 | 0 | 3406.526 | 0 | -0.52 | 0.01912 | 8.39 | 123.5 | 76.5 | 12.41 | 18.46 | | High | High | High | High | High | High | High | High | High | High | High | 0.000723 | 0.006296 | 3.18 |
| FALSE | High | TIGENDEVI | [R].TIGEND | 2xPhospho | Q921N6 2x | 6.95E-05 | 0 | 1 | 3 | 18 | Q921N6 | Probable A | Q921N6 [1 | 0 | 3486.492 | 0 | 0.01 | 0.963302 | 15.03 | 100 | 100 | 2.02 | 11.43 | | High | High | High | High | High | High | High | High | High | High | High | 0 | 1.95E-05 | 4.05 |
| FALSE | High | TIGENDEVI | [R].TIGEND | 1xDeamida | Q921N6 2x | 0.000255 | 0 | 1 | 3 | 4 | Q921N6 | Probable A | Q921N6 [1 | 0 | 3487.476 | 0 | -0.22 | 0.174114 | 7.68 | 108.4 | 91.6 | 14.76 | 11.88 | | High | High | High | High | High | High | High | High | High | High | High | 0 | 5.14E-05 | 4.4 |
| FALSE | High | TLEPVDTEE | [R].TLEPVD | 3xPhospho | Q8BU03 3x | 4.63E-05 | 0 | 1 | 4 | 4 | Q8BU03 | Periodic tr | Q8BU03 [8 | 0 | 3055.231 | 0 | -0.55 | 0.001675 | 3.95 | 119.7 | 80.3 | 8.05 | 7.74 | | High | High | High | High | High | High | High | High | High | High | High | 0 | 7.54E-06 | 4.66 |
| FALSE | High | TLPSTSSSG | [R].TLPSTS | 1xPhospho | P70255 1xl | 0.011755 | 0.000889 | 1 | 5 | 2 | P70255 | Nuclear fac | P70255 [27 | 0 | 1589.819 | 0 | 2.64 | 0.001084 | 18.39 | 25.7 | 174.3 | 11.59 | 14.3 | | High | High | High | High | High | High | High | High | High | High | High | 0.001566 | 0.01508 | 1.65 |
| FALSE | High | TLPVVFDSF | [R].TLPVVF | 1xPhospho | Q5DTJ8 1x | 0.094641 | 0.007408 | 1 | 5 | 1 | Q5DTJ8 | MKIAA417 | Q5DTJ8 [3 | 0 | 1439.75 | 0 | 1.15 | 0.001021 | 9.59 | 63.7 | 136.3 | 7.99 | 10.49 | | High | High | High | High | High | High | High | High | High | High | High | 0.003754 | 0.04048 | 1.74 |
| FALSE | High | TLQAISPK | [R].TLQAIS | 1xDeamida | B0V2N1-3 | 0.035181 | 0.002269 | 1 | 2 | 1 | B0V2N1-3 | Isoform 3 | B0V2N1-3 | 0 | 1167.622 | 0 | -1.39 | 0.001711 | 1.11 | 145.4 | 54.6 | 9.94 | 8.98 | | High | High | High | High | High | High | High | High | High | High | High | 0.001214 | 0.01294 | 1.87 |
| FALSE | High | TLSNESEES | [R].TLSNES | 1xPhospho | Q6P1G2 1x | 0.042503 | 0.003197 | 1 | 7 | 1 | Q6P1G2 | Lysine-spe | Q6P1G2 [4 | 0 | 1760.872 | 0 | -1.13 | 0.002994 | 4.3 | 137.7 | 62.3 | 10.69 | 14.59 | | High | High | High | High | High | High | High | High | High | High | High | 0.001651 | 0.01608 | 2.37 |
| FALSE | High | TISNPEVVN | [R].TISNPE | 1xPhospho | E9Q9C3 1x | 0.000765 | 0 | 1 | 6 | 5 | E9Q9C3 | Afadin OS= | E9Q9C3 [2 | 0 | 1671.879 | 0 | 0.23 | 0.078556 | 14.39 | 94.7 | 105.3 | 11.21 | 15.71 | | High | High | High | High | High | High | High | High | High | High | High | 0 | 0.000177 | 3.66 |
| FALSE | High | TISPPSAVV | [R].TISPPS/ | 1xPhospho | B2RWU4 1 | 9.27E-08 | 0 | 2 | 6 | 4 | B2RWU4; ( | Cordon-ble | B2RWU4 [ | 0 | 2347.256 | 0 | -0.24 | 0.003877 | 1.98 | 109.1 | 90.9 | 2.89 | 3.3 | Shared | High | High | High | High | High | High | High | High | High | High | High | 0 | 2.71E-07 | 4.53 |
| FALSE | High | TLSPSEPTV | [R].TLSPSE | 1xPhospho | B9EKK8 1x | 0.031713 | 0.002011 | 1 | 4 | 2 | B9EKK8 | Uncharacte | B9EKK8 [9 | 0 | 1496.756 | 0 | 1.05 | 0.000662 | 2.02 | 65.5 | 134.5 | 7.61 | 10.5 | | High | High | High | High | High | High | High | High | High | High | High | 0.001639 | 0.01593 | 2.83 |
| FALSE | High | TLSPSNLK | [R].TLSPSN | 1xPhospho | E9QMU3 1 | 0.004824 | 0.000456 | 1 | 4 | 2 | E9QMU3 | Ensconsin | E9QMU3 [ | 0 | 1397.781 | 0 | -0.42 | 0.001542 | 4.31 | 116.2 | 83.8 | 2.53 | 8.17 | | High | High | High | High | High | High | High | High | High | High | High | 0.000225 | 0.001396 | 2.43 |
| FALSE | High | TISPSTSNN | [KR].TISPST | 1xPhospho | Q5FWX6 1 | 0.002647 | 0.000109 | 1 | 4 | 1 | Q5FWX6 | Serine/thre | Q5FWX6 [3 | 0 | 1839.924 | 0 | 0.66 | 0.000605 | 2.43 | 75.8 | 124.2 | 5.38 | 5.98 | | High | High | High | High | High | High | High | High | High | High | High | 4.94E-05 | 0.000712 | 1.85 |
| FALSE | High | TISPSTSNN | [KR].TISPST | 1xPhospho | Q5FWX6 1 | 0.013666 | 0.001011 | 1 | 4 | 1 | Q5FWX6 | Serine/thre | Q5FWX6 [3 | 0 | 1855.919 | 0 | 0.95 | 0.000793 | 9.08 | 68.3 | 131.7 | 4.46 | 6.96 | | High | High | High | High | High | High | High | High | High | High | High | 0.000538 | 0.00444 | 2.02 |
| FALSE | High | TISPSTSNN | [KR].TISPST | 2xPhospho | Q5FWX6 2 | 0.003526 | 0.00021 | 1 | 4 | 1 | Q5FWX6 | Serine/thre | Q5FWX6 [3 | 0 | 1935.885 | 0 | 0.92 | 0.000236 | 3.25 | 68 | 132 | 9.4 | 7.98 | | High | High | High | High | High | High | High | High | High | High | High | 9.42E-05 | 0.000977 | 2.91 |
| FALSE | High | TLSPTPSAE | [R].TLSPTP | 1xPhospho | E9QK41 1x | 7.31E-07 | 0 | 1 | 17 | 1 | E9QK41 | Actin-bindi | E9QK41 [4 | 0 | 1929.916 | 0 | 1.86 | 0.000313 | 7.16 | 42.5 | 157.5 | 6.66 | 6.07 | | High | High | High | High | High | High | High | High | High | High | High | 0 | 7.12E-08 | 2.7 |

| | | | | | | | | | | | | | | | | | | | | | | | | | | | | | | | | | | | | | | |
|---|---|---|---|---|---|---|---|---|---|---|---|---|---|---|---|---|---|---|---|---|---|---|---|---|---|---|---|---|---|---|---|---|---|---|---|---|---|---|
| FALSE | High | TLSPTPSAE | [R].TLSPTP | 2xPhospho | E9QK41 2x | 0.003892 | 0.000261 | 1 | 17 | 2 | E9QK41 | Actin-bindi | E9QK41 [4 | 0 | 2009.882 | 0 | 2.21 | 0.000552 | 15.11 | 34.2 | 165.8 | 19.21 | 8.63 | | High | High | High | High | High | High | High | High | High | High | High | 0.000275 | 0.001627 | 3.04 |
| FALSE | High | TLSQGESQ | [K].TLSQGE | 1xPhospho | A2A791 1x | 5.65E-05 | 0 | 1 | 3 | 4 | A2A791 | Zinc finger | A2A791 [1( | 0 | 2679.317 | 0 | -0.76 | 0.000213 | 2.06 | 125.4 | 74.6 | 4.5 | 10.27 | | High | High | High | High | High | High | High | High | High | High | High | 0 | 9.42E-06 | 3.33 |
| FALSE | High | TLSQGESQ | [K].TLSQGE | 2xPhospho | A2A791 2x | 9.62E-05 | 0 | 1 | 3 | 1 | A2A791 | Zinc finger | A2A791 [1( | 0 | 2759.283 | 0 | -0.57 | 0.003644 | 4.5 | 121.7 | 78.3 | 4.82 | 8.06 | | High | High | High | High | High | High | High | High | High | High | High | 0 | 1.72E-05 | 3.2 |
| FALSE | High | TLSQSSESG | [R].TLSQSS | 1xCarbami | Q9QYG0 1x | 0.000284 | 0 | 1 | 3 | 3 | Q9QYG0 | Protein ND | Q9QYG0 [3 | 0 | 2755.215 | 0 | -1.24 | 0.007704 | 4.68 | 140.5 | 59.5 | 16.77 | 23.79 | | High | High | High | High | High | High | High | High | High | High | High | 0 | 7.27E-05 | 3.25 |
| FALSE | High | TLSNAEDYI | [R].TLSNAE | 2xPhospho | Q62422 2x | 8.17E-08 | 0 | 1 | 1 | 34 | Q62422 | Osteoclast | Q62422 [2( | 0 | 2090.757 | 0 | 1.27 | 0.002666 | 18.44 | 60.1 | 139.9 | 7.69 | 12.72 | | High | High | High | High | High | High | High | High | High | High | High | 0 | 6.11E-09 | 5.08 |
| FALSE | High | TLSQSSESG | [R].TLSQSS | 1xCarbami | Q9QYG0 2x | 0.001267 | 0 | 1 | 3 | 2 | Q9QYG0 | Protein ND | Q9QYG0 [3 | 0 | 2835.182 | 0 | -0.81 | 0.007004 | 3.09 | 130.3 | 69.7 | 15.89 | 10.35 | | High | High | High | High | High | High | High | High | High | High | High | 0 | 0.000312 | 3.1 |
| FALSE | High | TISSDDDV( | [K].TISSDD | 1xPhospho | Q05BA5 1x | 3.04E-06 | 0 | 1 | 7 | 2 | Q05BA5 | Nuclear re | Q05BA5 [8 | 0 | 1645.809 | 0 | -0.02 | 0.816617 | 4.16 | 99.2 | 100.8 | 7.33 | 5.59 | | High | High | High | High | High | High | High | High | High | High | High | 0 | 3.74E-07 | 3.6 |
| FALSE | High | TLSSFFTPR | [K].TLSSFF | 1xPhospho | Q3U4X8 1x | 0.114657 | 0.009411 | 1 | 6 | 1 | Q3U4X8 | DNA ligase | Q3U4X8 [2 | 0 | 1364.681 | 0 | -0.1 | 0.396455 | 12.41 | 108.3 | 91.7 | 16.67 | 9.3 | | High | High | High | High | High | High | High | High | High | High | High | 0.004707 | 0.05069 | 1.92 |
| FALSE | High | TLSSPSLQT | [R].TLSSPS | 1xPhospho | B2RY04 1x | 0.000633 | 0 | 1 | 2 | 4 | B2RY04 | Dedicator ( | B2RY04 [17 | 0 | 2825.51 | 0 | 0.35 | 0.007781 | 1.37 | 88 | 112 | 4.05 | 7.44 | | High | High | High | High | High | High | High | High | High | High | High | 4.94E-05 | 0.000683 | 3.87 |
| FALSE | High | TLSSPSNRF | [K].TLSSPS | 1xPhospho | A0A0U1RP | 0.082424 | 0.006288 | 1 | 3 | 1 | A0A0U1RP | Ataxin-2-lik | A0A0U1RP | 0 | 2563.271 | 0 | 0.36 | 0.010221 | 7.98 | 90.6 | 109.4 | 5.56 | 8.44 | | High | High | High | High | High | High | High | High | High | High | High | 0.003258 | 0.03435 | 1.85 |
| FALSE | High | TLSSPTGTE | [R].TLSSPT | 1xPhospho | Q3UMF0 1 | 0.00051 | 0 | 1 | 6 | 3 | Q3UMF0 | Cordon-ble | Q3UMF0 [8 | 0 | 1968.009 | 0 | 0.28 | 0.119278 | 13.65 | 84.6 | 115.4 | 13.02 | 18.57 | | High | High | High | High | High | High | High | High | High | High | High | 0 | 0.000112 | 3.52 |
| FALSE | High | TISSPTSCEI | [R].TISSPTS | 1xCarbami | Q5SRF8 1x | 0.071662 | 0.005278 | 1 | 2 | 2 | Q5SRF8 | Sprouty-re | Q5SRF8 [19 | 0 | 1583.709 | 0 | 0.57 | 0.035454 | 13.75 | 81.5 | 118.5 | 9.16 | 13.5 | | High | High | High | High | High | High | High | High | High | High | High | 0.002717 | 0.02923 | 1.68 |
| FALSE | High | TISSSPSIES | [K].TISSSPS | 1xPhospho | Q3U325 1x | 8.1E-08 | 0 | 1 | 4 | 3 | Q3U325 | Putative ur | Q3U325 [5 | 0 | 1796.899 | 0 | 2.02 | 0.00024 | 7.26 | 38.6 | 161.4 | 8.69 | 4.81 | | High | High | High | High | High | High | High | High | High | High | High | 0 | 6.03E-09 | 3.1 |
| FALSE | High | TISSSPSIES | [K].TISSSPS | 2xPhospho | Q3U325 2x | 0.000284 | 0 | 1 | 4 | 1 | Q3U325 | Putative ur | Q3U325 [5 | 0 | 1876.866 | 0 | 1.44 | 0.001121 | 6.02 | 54.9 | 145.1 | 18.94 | 11.32 | | High | High | High | High | High | High | High | High | High | High | High | 0 | 5.8E-05 | 2.87 |
| FALSE | High | TLSSSSMDI | [R].TLSSSSI | 1xPhospho | Q91YS4 1x | 0.012072 | 0.000889 | 1 | 2 | 1 | Q91YS4 | Kinesin ligh | Q91YS4 [6( | 0 | 1492.692 | 0 | 0.52 | 0.03411 | 15.81 | 83.5 | 116.5 | 6.36 | 12.82 | | High | High | High | High | High | High | High | High | High | High | High | 0.000449 | 0.003864 | 2.16 |
| FALSE | High | TLSSSSMDI | [R].TLSSSSI | 1xPhospho | Q91YS4 1x | 0.0016 | 5.66E-05 | 1 | 2 | 3 | Q91YS4 | Kinesin ligh | Q91YS4 [6( | 0 | 1508.687 | 0 | 0.34 | 0.0462 | 20.94 | 83.6 | 116.4 | 7.48 | 14.65 | | High | High | High | High | High | High | High | High | High | High | High | 2.58E-05 | 0.000404 | 2.29 |
| FALSE | High | TLSVAAAFI | [K].TLSVAA | 1xPhospho | F8WI88 1x | 0.004333 | 0.00036 | 1 | 5 | 1 | F8WI88 | PEST prote | F8WI88 [1 | 0 | 2931.269 | 0 | -0.14 | 0.723772 | 5.25 | 106.9 | 93.1 | 14.82 | 53.53 | | High | High | High | High | High | High | High | High | High | High | High | 0.000161 | 0.001227 | 3.61 |
| FALSE | High | TLSVAAAFI | [K].TLSVAA | 1xPhospho | F8WI88 1x | 1.35E-06 | 0 | 1 | 5 | 7 | F8WI88 | PEST prote | F8WI88 [1 | 0 | 3144.437 | 0 | -0.1 | 0.139125 | 6.68 | 101.1 | 98.9 | 5.33 | 10.63 | | High | High | High | High | High | High | High | High | High | High | High | 0 | 9.97E-07 | 4.4 |
| FALSE | High | TLSVAAAFI | [K].TLSVAA | 1xDeamida | F8WI88 1x | 0.004256 | 0.00036 | 1 | 5 | 1 | F8WI88 | PEST prote | F8WI88 [1 | 0 | 3145.421 | 0 | -0.13 | 0.679473 | 11.67 | 100.9 | 99.1 | 12.28 | 16.15 | | High | High | High | High | High | High | High | High | High | High | High | 0.000161 | 0.001206 | 3.66 |
| FALSE | High | TLSQSSESG | [R].TLSQSS | 1xCarbami | Q9QYG0 2x | 7.08E-05 | 0 | 1 | 3 | 2 | Q9QYG0 | Protein ND | Q9QYG0 [3 | 0 | 2851.177 | 0 | -0.7 | 0.004962 | 7.59 | 123.9 | 76.1 | 4.2 | 15.76 | | High | High | High | High | High | High | High | High | High | High | High | 0 | 1.22E-05 | 3.76 |
| FALSE | High | TLSNAEDYI | [R].TLSNAE | 1xDeamida | Q62422 1x | 8.32E-08 | 0 | 1 | 1 | 7 | Q62422 | Osteoclast | Q62422 [2( | 0 | 2011.774 | 0 | 0.7 | 0.024209 | 17.8 | 80.2 | 119.8 | 6.34 | 18.56 | | High | High | High | High | High | High | High | High | High | High | High | 0 | 1.83E-07 | 4.57 |
| FALSE | High | TLSNAEDYI | [R].TLSNAE | 1xPhospho | Q62422 1x | 3.64E-09 | 0 | 1 | 1 | 26 | Q62422 | Osteoclast | Q62422 [2( | 0 | 2010.79 | 0 | 0.79 | 0.001238 | 9.07 | 71.2 | 128.8 | 7.24 | 8.94 | | High | High | High | High | High | High | High | High | High | High | High | 0 | 3.26E-10 | 4.55 |
| FALSE | High | TLSMIEEEII | [R].TLSMIE | 1xPhospho | F7AA26 1x | 0.003253 | 0.00016 | 1 | 10 | 5 | F7AA26 | Paralemmi | F7AA26 [98 | 0 | 1545.743 | 0 | 0.52 | 0.011227 | 7.54 | 81.5 | 118.5 | 4.34 | 13.97 | | High | High | High | High | High | High | High | High | High | High | High | 7.19E-05 | 0.000895 | 3.02 |
| FALSE | High | TLQNPLPTT | [K].TLQNPI | 1xPhospho | O88466 1x | 0.000248 | 0 | 2 | 7 | 2 | O88466; R | Zinc finger | O88466 [5( | 0 | 1747.976 | 0 | -1.41 | 0.0003 | 0.7 | 145.1 | 54.9 | 4.2 | 9.6 | Shared | High | High | High | High | High | High | High | High | High | High | High | 0 | 4.96E-05 | 2.61 |
| FALSE | High | TLQNSSDQ | [R].TLQNSS | 1xPhospho | Q7TSY8 1x | 5.12E-05 | 0 | 1 | 1 | 2 | Q7TSY8 | Shugoshin | Q7TSY8 [4( | 0 | 2258.107 | 0 | -0.42 | 0.010925 | 4.54 | 120.3 | 79.7 | 12.76 | 7.36 | | High | High | High | High | High | High | High | High | High | High | High | 0 | 8.44E-06 | 2.79 |
| FALSE | High | TLQPSNPTI | [K].TLQPSN | 1xPhospho | Q8C050 1x | 1.1E-05 | 0 | 1 | 3 | 4 | Q8C050 | Ribosomal | Q8C050 [8 | 0 | 2604.17 | 0 | 1.08 | 0.00268 | 26.65 | 65.3 | 134.7 | 15.83 | 13.8 | | High | High | High | High | High | High | High | High | High | High | High | 0 | 6.87E-06 | 3.99 |
| FALSE | High | TLQPSNPTI | [K].TLQPSN | 1xDeamida | Q8C050 1x | 0.028824 | 0.001829 | 1 | 3 | 1 | Q8C050 | Ribosomal | Q8C050 [8 | 0 | 2605.154 | 0 | 0.91 | 0.020756 | 46.61 | 63.1 | 136.9 | 31.92 | 17.02 | | High | High | High | High | High | High | High | High | High | High | High | 0.000997 | 0.01031 | 2.84 |
| FALSE | High | TLQTEDSN | [R].TLQTED | 2xPhospho | Q8CJ76 2x | 7.02E-05 | 0 | 1 | 3 | 1 | Q8CJ76 | Leukemia-a | Q8CJ76 [34 | 0 | 2217.879 | 0 | 1.25 | 0.018661 | 30.99 | 57.3 | 142.7 | 16.15 | 21.21 | | High | High | High | High | High | High | High | High | High | High | High | 0 | 1.21E-05 | 3.45 |
| FALSE | High | TLSALPQTF | [R].TLSALP | 1xPhospho | Q5DW34 1 | 0.060666 | 0.00451 | 1 | 13 | 2 | Q5DW34 | Histone-lys | Q5DW34 [ | 0 | 2921.398 | 0 | 1.02 | 0.008771 | 37.51 | 67.2 | 132.8 | 28.16 | 12.61 | | High | High | High | High | High | High | High | High | High | High | High | 0.002516 | 0.0273 | 3.1 |
| FALSE | High | TISAQDTLA | [R].TISAQD | 1xPhospho | E9PV45 1x | 0.033694 | 0.002195 | 1 | 2 | 1 | E9PV45 | Ubiquitin c | E9PV45 [25 | 0 | 2232.136 | 0 | -0.08 | 0.587431 | 16.97 | 101.7 | 98.3 | 17.75 | 57.88 | | High | High | High | High | High | High | High | High | High | High | High | 0.001151 | 0.01227 | 2.31 |
| FALSE | High | TISAQDTLA | [R].TISAQD | 1xPhospho | E9PV45 1x | 0.000262 | 0 | 1 | 2 | 2 | E9PV45 | Ubiquitin c | E9PV45 [25 | 0 | 2461.299 | 0 | 0.68 | 0.001524 | 8.74 | 79.1 | 120.9 | 10.47 | 3.43 | | High | High | High | High | High | High | High | High | High | High | High | 2.58E-05 | 0.000401 | 2.74 |
| FALSE | High | TLSDDLDE | [K].TLSDDL | 1xPhospho | Q8K224 1x | 8.3E-07 | 0 | 1 | 1 | 2 | Q8K224 | RNA cytidir | Q8K224 [9 | 0 | 1715.85 | 0 | -0.55 | 0.000513 | 2.79 | 120.5 | 79.5 | 2.45 | 4.55 | | High | High | High | High | High | High | High | High | High | High | High | 0 | 8.25E-08 | 3.17 |
| FALSE | High | TLSDESLCS | [R].TLSDES | 1xCarbami | G3X9J0 1x | 8.56E-05 | 0 | 2 | 12 | 2 | G3X9J0; Q | Signal-indu | G3X9J0 [15 | 0 | 1533.682 | 0 | 1.03 | 0.000158 | 1.41 | 65.7 | 134.3 | 2.12 | 1.67 | Shared | High | High | High | High | High | High | High | High | High | High | High | 2.58E-05 | 0.000561 | 2.1 |
| FALSE | High | TLSDESLCS | [R].TLSDES | 1xCarbami | G3X9J0 2x | 0.103392 | 0.008249 | 2 | 12 | 1 | G3X9J0; Q | Signal-indu | G3X9J0 [15 | 0 | 1613.648 | 0 | 0.7 | 0.027437 | 22.12 | 69.8 | 130.2 | 20.56 | 21.98 | Shared | High | High | High | High | High | High | High | High | High | High | High | 0.004152 | 0.0448 | 2.28 |
| FALSE | High | TLSDESIYS | [R].TLSDES | 1xPhospho | Q8C0T5 1x | 1.16E-05 | 0 | 1 | 3 | 3 | Q8C0T5 | Signal-indu | Q8C0T5 [1 | 0 | 1694.784 | 0 | 1.66 | 0.000438 | 7.24 | 50.4 | 149.6 | 7.22 | 6.7 | | High | High | High | High | High | High | High | High | High | High | High | 2.58E-05 | 0.000449 | 3.74 |
| FALSE | High | TLSDESVCS | [R].TLSDES | 1xCarbami | B9EK76 1x | 0.000183 | 0 | 1 | 5 | 2 | B9EK76 | Signal-indu | B9EK76 [14 | 0 | 1576.688 | 0 | -0.75 | 0.000289 | 2.34 | 124.6 | 75.4 | 1.98 | 4.92 | | High | High | High | High | High | High | High | High | High | High | High | 0 | 3.54E-05 | 1.78 |
| FALSE | High | TISDGTISA | [R].TISDGT | 1xPhospho | E9PUI5 1xP | 0.021231 | 0.001507 | 1 | 5 | 2 | E9PUI5 | Formin-bin | E9PUI5 [29 | 0 | 1601.855 | 0 | -1.05 | 0.000364 | 1.89 | 134.8 | 65.2 | 6.32 | 12.9 | | High | High | High | High | High | High | High | High | High | High | High | 0.000801 | 0.0073 | 2.45 |
| FALSE | High | TLSDISNAS | [R].TLSDISI | 1xDeamida | P22682 1x | 0.000203 | 0 | 1 | 5 | 2 | P22682 | E3 ubiquiti | P22682 [79 | 0 | 4886.371 | 0 | -0.3 | 0.961829 | 15.43 | 100.5 | 99.5 | 15.65 | 25.05 | | High | High | High | High | High | High | High | High | High | High | High | 0 | 3.98E-05 | 4.99 |
| FALSE | High | TLSDPPSPL | [R].TLSDPP | 2xPhospho | E9QN63 2x | 0.006085 | 0.00062 | 1 | 7 | 1 | E9QN63 | Arf-GAP wi | E9QN63 [8 | 0 | 2042.955 | 0 | 0.79 | 0.01254 | 16.09 | 73.8 | 126.2 | 13.57 | 10.07 | | High | High | High | High | High | High | High | High | High | High | High | 0.000275 | 0.001805 | 1.53 |
| FALSE | High | TLSDYNIQK | [R].TLSDYN | 1xPhospho | E9QNP0 1x | 0.001049 | 0 | 1 | 19 | 1 | E9QNP0 | KxDL motif | E9QNP0 [1 | 0 | 1619.845 | 0 | -0.21 | 0.009637 | 3.74 | 107.2 | 92.8 | 3.72 | 2.99 | | High | High | High | High | High | High | High | High | High | High | High | 0 | 0.000251 | 3.1 |
| FALSE | High | TISEDSIEVL | [R].TISEDSI | 1xCarbami | Q3UMB5 1 | 0.000924 | 0 | 1 | 2 | 4 | Q3UMB5 | Guanine nu | Q3UMB5 [ | 0 | 3204.568 | 0 | 2.77 | 0.000746 | 26.24 | 25.5 | 174.5 | 18.99 | 10.11 | | High | High | High | High | High | High | High | High | High | High | High | 0 | 0.000219 | 3.63 |
| FALSE | High | TLSEENVYE | [R].TLSEEN | 1xPhospho | Q8C4S8 1x | 0.000183 | 0 | 1 | 3 | 2 | Q8C4S8 | DENN dom | Q8C4S8 [3 | 0 | 2531.239 | 0 | -1.14 | 0.002729 | 5.11 | 139.3 | 60.7 | 11.26 | 19.25 | | High | High | High | High | High | High | High | High | High | High | High | 0 | 3.55E-05 | 2.33 |
| FALSE | High | TLSEENVYE | [R].TLSEEN | 1xPhospho | Q8C4S8 1x | 5.93E-06 | 0 | 1 | 3 | 2 | Q8C4S8 | DENN dom | Q8C4S8 [3 | 0 | 2547.234 | 0 | -1.6 | 0.000644 | 3.1 | 152.5 | 47.5 | 5.54 | 19.37 | | High | High | High | High | High | High | High | High | High | High | High | 0 | 7.48E-07 | 3.48 |
| FALSE | High | TLSEPTGPS | [R].TLSEPT | 2xPhospho | Q8VHK1 2x | 0.11108 | 0.008991 | 1 | 1 | 1 | Q8VHK1 | Caskin-2 O | Q8VHK1 [9 | 0 | 4019.752 | 0 | 0.05 | 0.879736 | 38.78 | 109.9 | 90.1 | 19.73 | 17.81 | | High | High | High | High | High | High | High | High | High | High | High | 0.004517 | 0.04895 | 3.21 |
| FALSE | High | TLSFDTSK | [R].TLSFDT | 1xPhospho | G3XA57-2 | 0.077188 | 0.005893 | 1 | 3 | 1 | G3XA57-2 | Isoform 2 ( | G3XA57-2 | 0 | 1436.744 | 0 | 1.44 | 0.001148 | 10.93 | 56.9 | 143.1 | 7.23 | 11.51 | | High | High | High | High | High | High | High | High | High | High | High | 0.003035 | 0.0318 | 2.07 |
| FALSE | High | TLSFGSDLN | [R].TLSFGS | 1xPhospho | P23116 1x | 0.000815 | 0 | 1 | 6 | 2 | P23116 | Eukaryotic | P23116 [49 | 0 | 1753.836 | 0 | 0.14 | 0.072248 | 7.59 | 92.1 | 107.9 | 7.79 | 5.92 | | High | High | High | High | High | High | High | High | High | High | High | 0.000204 | 0.001367 | 2.98 |
| FALSE | High | TLSGDSSSD | [K].TLSGDS | 1xPhospho | F6YRW4 1x | 0.015603 | 0.001124 | 1 | 5 | 2 | F6YRW4 | F-box and l | F6YRW4 [4 | 0 | 1521.663 | 0 | 1.3 | 0.004328 | 5.23 | 59.2 | 140.8 | 10.06 | 13.24 | | High | High | High | High | High | High | High | High | High | High | High | 0.000585 | 0.005165 | 1.88 |
| FALSE | High | TISGIPDIIQ | [K].TISGIPD | 1xPhospho | Q8CAF4-3 | 0.000908 | 0 | 1 | 2 | 1 | Q8CAF4-3 | Isoform 3 ( | Q8CAF4-3 | 0 | 1722.981 | 0 | 1.03 | 0.001691 | 11.33 | 65.6 | 134.4 | 10.11 | 7.13 | | High | High | High | High | High | High | High | High | High | High | High | 0 | 0.000215 | 3.38 |
| FALSE | High | TISINSPK | [R].TISINSP | 1xPhospho | A6H619 1x | 0.002818 | 0.000109 | 1 | 2 | 1 | A6H619 | PHD and RI | A6H619 [8( | 0 | 1397.781 | 0 | 0.41 | 0.029552 | 4.2 | 89.5 | 110.5 | 6.82 | 6.42 | | High | High | High | High | High | High | High | High | High | High | High | 7.19E-05 | 0.000765 | 1.42 |
| FALSE | High | TLSISLSED( | [R].TLSISLS | 2xPhospho | Q922B9 2x | 0.114657 | 0.009411 | 1 | 5 | 1 | Q922B9 | Sperm-spe | Q922B9 [3 | 0 | 1938.923 | 0 | 0.76 | 0.000877 | 4.93 | 75.4 | 124.6 | 11.38 | 6.36 | | High | High | High | High | High | High | High | High | High | High | High | 0.004707 | 0.05067 | 2.34 |
| FALSE | High | TLSISTQDL | [R].TLSIST( | 1xPhospho | Q91W40 1 | 0.009843 | 0.000736 | 1 | 2 | 5 | Q91W40 | Kinesin ligh | Q91W40 [4 | 0 | 1626.83 | 0 | -0.71 | 0.000434 | 1.62 | 125.5 | 74.5 | 5.93 | 10.29 | | High | High | High | High | High | High | High | High | High | High | High | 0.000344 | 0.003091 | 2.09 |
| FALSE | High | TLSMIEEEII | [R].TLSMIE | 1xPhospho | F7AA26 1x | 0.002818 | 0.000109 | 1 | 10 | 2 | F7AA26 | Paralemmi | F7AA26 [98 | 0 | 1529.748 | 0 | 0.52 | 0.004529 | 15.34 | 82.9 | 117.1 | 7.36 | 8.79 | | High | High | High | High | High | High | High | High | High | High | High | 7.19E-05 | 0.000763 | 3.62 |
| FALSE | High | TLAMNATT | [R].TLAMN | 2xPhospho | Q2TAX1 2x | 0.019446 | 0.001438 | 1 | 3 | 1 | Q2TAX1 | Glutamate | Q2TAX1 [3 | 0 | 2751.381 | 0 | -0.23 | 0.023809 | 4.35 | 110.7 | 89.3 | 4.78 | 10.21 | | High | High | High | High | High | High | High | High | High | High | High | 0.000756 | 0.006582 | 2.94 |
| FALSE | High | TIADSEEED | [K].TIADSE | 2xPhospho | Q6NV99 2x | 7.49E-06 | 0 | 1 | 6 | 1 | Q6NV99 | HAUS augr | Q6NV99 [4 | 0 | 2426.067 | 0 | 0.36 | 0.009249 | 7.86 | 87.8 | 112.2 | 7.14 | 8.81 | | High | High | High | High | High | High | High | High | High | High | High | 0 | 9.82E-07 | 3.3 |
| FALSE | High | TIADSEEED | [K].TIADSE | 1xPhospho | Q6NV99 1x | 3.46E-07 | 0 | 1 | 6 | 2 | Q6NV99 | HAUS augr | Q6NV99 [4 | 0 | 2346.1 | 0 | 0.45 | 0.008504 | 15.52 | 80.6 | 119.4 | 5.05 | 10.12 | | High | High | High | High | High | High | High | High | High | High | High | 0 | 4.31E-06 | 3.67 |
| FALSE | High | TIAAASVQT | [R].TIAAAS | 1xPhospho | A0A0R4J1Z | 3.83E-08 | 0 | 1 | 18 | 1 | A0A0R4J1Z | POU doma | A0A0R4J1Z | 0 | 2366.273 | 0 | -0.34 | 0.053394 | 1.63 | 111.6 | 88.4 | 8.06 | 4.98 | | High | High | High | High | High | High | High | High | High | High | High | 0 | 2.61E-09 | 3.86 |
| FALSE | High | TGQGDAPL | [K].TGQGD | 1xPhospho | Q5SUR0 1x | 0.00925 | 0.000736 | 1 | 1 | 2 | Q5SUR0 | Phosphoril | Q5SUR0 [6 | 0 | 3043.579 | 0 | 1.05 | 0.002442 | 5.47 | 66.6 | 133.4 | 6.24 | 8.39 | | High | High | High | High | High | High | High | High | High | High | High | 0.000756 | 0.006609 | 2.63 |
| FALSE | High | TGQPPQPA | [R].TGQPP | 1xPhospho | A2ACV6 1x | 8.84E-07 | 0 | 1 | 1 | 1 | A2ACV6 | Protein SO | A2ACV6 [1 | 0 | 3761.808 | 0 | 1.7 | 0.000426 | 6.2 | 47.3 | 152.7 | 18.44 | 7.26 | | High | High | High | High | High | High | High | High | High | High | High | 0 | 8.87E-08 | 3.82 |
| FALSE | High | TGRDSPEN | [K].TGRDSI | 1xPhospho | A0A0R4IZZ | 3.02E-07 | 0 | 1 | 4 | 2 | A0A0R4IZZ | Formin bin | A0A0R4IZZ | 1 | 2601.208 | 0 | -0.67 | 0.001154 | 1.9 | 123.3 | 76.7 | 6.32 | 5.91 | | High | High | High | High | High | High | High | High | High | High | High | 0 | 2.64E-08 | 3.47 |
| FALSE | High | TGRDSPEN | [K].TGRDSI | 1xDeamida | A0A0R4IZZ | 0.002232 | 0.000109 | 1 | 4 | 1 | A0A0R4IZZ | Formin bin | A0A0R4IZZ | 1 | 2602.192 | 0 | 0 | 0.557493 | 26.11 | 103.8 | 96.2 | 13.48 | 29.02 | | High | High | High | High | High | High | High | High | High | High | High | 4.94E-05 | 0.000586 | 3.59 |
| FALSE | High | TGSEPTLSP | [R].TGSEPT | 1xPhospho | Q9QZK2 1x | 0.001799 | 5.66E-05 | 1 | 4 | 3 | Q9QZK2 | Breast can | Q9QZK2 [3 | 0 | 1636.851 | 0 | 0.33 | 0.018656 | 7.17 | 90 | 110 | 3.43 | 10.43 | | High | High | High | High | High | High | High | High | High | High | High | 2.58E-05 | 0.000462 | 2.9 |
| FALSE | High | TGSESSQT( | [R].TGSESS | 1xPhospho | Q3TSY9 1x | 1.4E-07 | 0 | 1 | 6 | 4 | Q3TSY9 | Putative ur | Q3TSY9 [4 | 0 | 1792.791 | 0 | -0.41 | 0.072038 | 8.23 | 114.1 | 85.9 | 12.89 | 10.63 | | High | High | High | High | High | High | High | High | High | High | High | 0 | 0.000116 | 2.56 |
| FALSE | High | TGSESSQT( | [R].TGSESS | 2xPhospho | Q3TSY9 2x | 0.000102 | 0 | 1 | 6 | 2 | Q3TSY9 | Putative ur | Q3TSY9 [4 | 0 | 1872.758 | 0 | -0.79 | 0.080033 | 11.48 | 124.3 | 75.7 | 18.85 | 17.92 | | High | High | High | High | High | High | High | High | High | High | High | 0 | 1.84E-05 | 3.16 |
| FALSE | High | TGSFHGPG | [R].TGSFH( | 1xPhospho | E9PXD2 1x | 0.028082 | 0.001791 | 1 | 7 | 1 | E9PXD2 | Rho GTPas | E9PXD2 [8 | 0 | 1674.831 | 0 | -1.33 | 0.001069 | 0.95 | 145.7 | 54.3 | 5.56 | 21.53 | | High | High | High | High | High | High | High | High | High | High | High | 0.000968 | 0.01001 | 1.43 |
| FALSE | High | TGSGGVAS | [K].TGSGG | 1xPhospho | G3UVU2 1 | 0.057686 | 0.004227 | 1 | 5 | 1 | G3UVU2 | Splicing fac | G3UVU2 [1 | 0 | 1604.711 | 0 | -0.3 | 0.203779 | 10.81 | 111.7 | 88.3 | 5.17 | 26.41 | | High | High | High | High | High | High | High | High | High | High | High | 0.002159 | 0.02269 | 1.26 |
| FALSE | High | TGSLDESLS | [R].TGSLDE | 1xPhospho | A0A087WF | 0.000611 | 0 | 1 | 2 | 2 | A0A087WF | Proline-rich | A0A087WF | 0 | 1373.651 | 0 | 4.55 | 0.000365 | 30.29 | 8.4 | 191.6 | 34.33 | 10.56 | | High | High | High | High | High | High | High | High | High | High | High | 0 | 0.000137 | 1.35 |
| FALSE | High | TGSLQLSST | [R].TGSLQI | 1xPhospho | Q3UMF0 1 | 1E-05 | 0 | 1 | 6 | 3 | Q3UMF0 | Cordon-ble | Q3UMF0 [ | 0 | 2205.178 | 0 | -0.08 | 0.904676 | 31.44 | 110.4 | 89.6 | 11.3 | 23.73 | | High | High | High | High | High | High | High | High | High | High | High | 0 | 2.07E-06 | 3.62 |
| FALSE | High | TGSNLGLL( | [R].TGSNL( | 1xCarbami | D3YUC5 1x | 0.002743 | 0.000109 | 1 | 1 | 1 | D3YUC5 | Folliculin-ir | D3YUC5 [2 | 0 | 1972.034 | 0 | 0.06 | 0.793145 | 28.52 | 99.2 | 100.8 | 13.98 | 10.55 | | High | High | High | High | High | High | High | High | High | High | High | 4.94E-05 | 0.000736 | 2.82 |
| FALSE | High | TGSNISGAS | [R].TGSNIS | 1xPhospho | Q3B7Z2 1x | 4.51E-08 | 0 | 1 | 2 | 2 | Q3B7Z2 | Oxysterol-l | Q3B7Z2 [3 | 0 | 2267.028 | 0 | 0.61 | 0.209868 | 44.69 | 78.6 | 121.4 | 19.82 | 36.4 | | High | High | High | High | High | High | High | High | High | High | High | 0 | 3.12E-09 | 3.98 |
| FALSE | High | TGPPPISPS | [R].TGPPPI | 1xPhospho | Q80V24 1x | 0.000322 | 0 | 1 | 4 | 2 | Q80V24 | Transcripti | Q80V24 [5 | 0 | 1518.833 | 0 | 0.59 | 0.011632 | 5.41 | 80.1 | 119.9 | 5.29 | 10.27 | | High | High | High | High | High | High | High | High | High | High | High | 0.000275 | 0.001947 | 2.15 |
| FALSE | High | TGSNISGAS | [R].TGSNIS | 3xPhospho | Q3B7Z2 3x | 0.000124 | 0 | 1 | 2 | 2 | Q3B7Z2 | Oxysterol-l | Q3B7Z2 [3 | 0 | 2426.96 | 0 | 0.52 | 0.092851 | 15.51 | 82.2 | 117.8 | 17.16 | 16.95 | | High | High | High | High | High | High | High | High | High | High | High | 0 | 2.27E-05 | 4.96 |
| FALSE | High | TGSNISGAS | [R].TGSNIS | 2xPhospho | Q3B7Z2 2x | 2.6E-08 | 0 | 1 | 2 | 2 | Q3B7Z2 | Oxysterol-l | Q3B7Z2 [3 | 0 | 2576.157 | 0 | 0.43 | 0.007777 | 9.71 | 82.4 | 117.6 | 5.67 | 8.05 | | High | High | High | High | High | High | High | High | High | High | High | 0 | 1.69E-09 | 4.25 |
| FALSE | High | TGSPEIK | [K].TGSPEI | 1xPhospho | D3Z357 1x | 0.057686 | 0.004227 | 1 | 6 | 1 | D3Z357 | Histone-lys | D3Z357 [1 | 0 | 1269.686 | 0 | 1.21 | 0.001792 | 14.4 | 58.2 | 141.8 | 6.38 | 11.14 | | High | High | High | High | High | High | High | High | High | High | High | 0.002159 | 0.0228 | 1.71 |
| FALSE | High | TGSPGAK | [R].TGSPG | 1xPhospho | Q9D8C4 1x | 0.030898 | 0.001896 | 1 | 2 | 1 | Q9D8C4 | Interferon- | Q9D8C4 [3 | 0 | 1155.618 | 0 | 2.33 | 0.000378 | 12.33 | 33.7 | 166.3 | 7.77 | 7.69 | | High | High | High | High | High | High | High | High | High | High | High | 0.001027 | 0.01115 | 1.79 |
| FALSE | High | TGSPGSPS | [K].TGSPG | 1xPhospho | Q6PDL0 1x | 1.72E-06 | 0 | 1 | 3 | 2 | Q6PDL0 | Cytoplasmi | Q6PDL0 [4 | 0 | 2027.021 | 0 | 0.87 | 0.000625 | 1.21 | 70.2 | 129.8 | 7.76 | 6.64 | | High | High | High | High | High | High | High | High | High | High | High | 0 | 1.88E-07 | 3.56 |
| FALSE | High | TGSPGSPS | [K].TGSPG | 2xPhospho | Q6PDL0 2x | 0.00204 | 5.66E-05 | 1 | 3 | 1 | Q6PDL0 | Cytoplasmi | Q6PDL0 [4 | 0 | 2106.987 | 0 | 0.88 | 0.089844 | 56.96 | 67.3 | 132.7 | 17.4 | 30.16 | | High | High | High | High | High | High | High | High | High | High | High | 2.58E-05 | 0.00053 | 3.25 |
| FALSE | High | TGSPLAGG | [K].TGSPLA | 1xPhospho | Q6IRU7 1x | 0.003997 | 0.000311 | 1 | 2 | 1 | Q6IRU7 | Centrosom | Q6IRU7 [6 | 0 | 1448.746 | 0 | -0.86 | 0.005665 | 3.38 | 124.5 | 75.5 | 15.21 | 4.7 | | High | High | High | High | High | High | High | High | High | High | High | 0.00014 | 0.00113 | 1.47 |
| FALSE | High | TGSPTPSN( | [K].TGSPTF | 2xPhospho | E9Q5F9 2x | 0.000651 | 0 | 1 | 3 | 2 | E9Q5F9 | Histone-lys | E9Q5F9 [6 | 0 | 2179.951 | 0 | -0.05 | 0.667891 | 29.46 | 101.8 | 98.2 | 31.88 | 32.91 | | High | High | High | High | High | High | High | High | High | High | High | 0 | 0.000147 | 3.62 |
| FALSE | High | TGSPTPSN( | [K].TGSPTF | 1xPhospho | E9Q5F9 1x | 7.29E-06 | 0 | 1 | 3 | 2 | E9Q5F9 | Histone-lys | E9Q5F9 [6 | 0 | 2329.148 | 0 | 0.25 | 0.199576 | 4.53 | 94.5 | 105.5 | 9.46 | 16.34 | | High | High | High | High | High | High | High | High | High | High | High | 0 | 9.45E-07 | 3.79 |
| FALSE | High | TGSPTPSN( | [K].TGSPTF | 2xPhospho | E9Q5F9 2x | 1.26E-06 | 0 | 1 | 3 | 3 | E9Q5F9 | Histone-lys | E9Q5F9 [6 | 0 | 2409.114 | 0 | -0.09 | 0.170069 | 5.1 | 104.5 | 95.5 | 6.5 | 6.79 | | High | High | High | High | High | High | High | High | High | High | High | 0 | 1.32E-07 | 4.26 |
| FALSE | High | TGSQGQCT | [R].TGSQG | 1xCarbami | P62858 1x | 8.25E-05 | 0 | 1 | 3 | 2 | P62858 | 40S riboso | P62858 [21 | 0 | 1530.693 | 0 | 0.18 | 0.207156 | 10.74 | 85.4 | 114.6 | 18.13 | 29.25 | | High | High | High | High | High | High | High | High | High | High | High | 0 | 1.44E-05 | 1.8 |
| FALSE | High | TGSRVSVQ | [R].TGSRVS | 2xPhospho | Q9Z2A7 2x | 0.014797 | 0.001049 | 1 | 1 | 1 | Q9Z2A7 | Diacylglyce | Q9Z2A7 [1 | 1 | 1934.95 | 0 | -0.24 | 0.010409 | 3.14 | 108.2 | 91.8 | 4.38 | 8.83 | | High | High | High | High | High | High | High | High | High | High | High | 0.000555 | 0.004841 | 3.14 |
| FALSE | High | TGSSPTQG | [R].TGSSPT | 1xPhospho | K3W4S4 1x | 4.69E-06 | 0 | 1 | 14 | 3 | K3W4S4 | C-Jun-amir | K3W4S4 [3 | 0 | 1726.914 | 0 | -0.34 | 0.058349 | 4.46 | 107.9 | 92.1 | 11.86 | 10.25 | | High | High | High | High | High | High | High | High | High | High | High | 9.42E-05 | 0.001038 | 3.38 |
| FALSE | High | TGSSSSILS | [R].TGSSSS | 1xPhospho | B2RUG9 1x | 0.001301 | 0 | 1 | 6 | 4 | B2RUG9 | Adenomat | B2RUG9 [2 | 0 | 1952.89 | 0 | 0.64 | 0.041177 | 31.15 | 76 | 124 | 31.05 | 10.48 | | High | High | High | High | High | High | High | High | High | High | High | 9.42E-05 | 0.000948 | 2.78 |
| FALSE | High | TGSNISGAS | [R].TGSNIS | 1xPhospho | Q3B7Z2 1x | 1.26E-05 | 0 | 1 | 2 | 6 | Q3B7Z2 | Oxysterol-l | Q3B7Z2 [3 | 0 | 2496.191 | 0 | 0.02 | 0.659045 | 2.88 | 99.4 | 100.6 | 4.63 | 6.07 | | High | High | High | High | High | High | High | High | High | High | High | 0 | 0.000244 | 3.42 |
| FALSE | High | TGPGEDED | [R].TGPGEI | 1xPhospho | Q0GNC1 1 | 7.29E-06 | 0 | 1 | 3 | 4 | Q0GNC1 | Inverted fc | Q0GNC1 [1 | 0 | 2786.173 | 0 | 0.05 | 0.985795 | 13.45 | 100.2 | 99.8 | 3.52 | 11.6 | | High | High | High | High | High | High | High | High | High | High | High | 0 | 9.52E-07 | 4.62 |
| FALSE | High | TGPEDVDD | [R].TGPED | 1xCarbami | A0A1L1SQ | 0.002647 | 0.000109 | 1 | 5 | 5 | A0A1L1SQ | Dedicator ( | A0A1L1SQ | 0 | 2605.071 | 0 | -1.37 | 0.001303 | 0.07 | 139.9 | 60.1 | 10.6 | 9.17 | | High | High | High | High | High | High | High | High | High | High | High | 0.000538 | 0.004425 | 3.18 |
| FALSE | High | TGNVSPSG | [R].TGNVSI | 1xCarbami | Q3T9G9 1x | 1.03E-05 | 0 | 1 | 6 | 1 | Q3T9G9 | Putative ur | Q3T9G9 [5 | 0 | 1813.847 | 0 | 1.51 | 0.000363 | 13.75 | 47.8 | 152.2 | 12.68 | 6.16 | | High | High | High | High | High | High | High | High | High | High | High | 0 | 1.39E-06 | 1.5 |
| FALSE | High | TFTPECESC | [K].TFTPEC | 1xCarbami | A2SW42 1x | 0.003312 | 0.00016 | 1 | 2 | 1 | A2SW42 | Zinc finger | A2SW42 [1 | 0 | 1764.828 | 0 | -1.12 | 0.004421 | 2.54 | 140.4 | 59.6 | 10.87 | 19.92 | | High | High | High | High | High | High | High | High | High | High | High | 7.19E-05 | 0.000916 | 3.09 |
| FALSE | High | TFTPLPEFE | [R].TFTPLP | 1xPhospho | Q99PP7 1x | 1.09E-06 | 0 | 1 | 4 | 6 | Q99PP7 | E3 ubiquiti | Q99PP7 [1 | 0 | 3709.579 | 0 | -0.41 | 0.102085 | 16.02 | 112.4 | 87.6 | 9.62 | 26.94 | | High | High | High | High | High | High | High | High | High | High | High | 0 | 1.11E-07 | 4.89 |
| FALSE | High | TFVINTSDS | [R].TFVINT | 1xPhospho | Q9CWJ3 1x | 4.37E-07 | 0 | 1 | 1 | 1 | Q9CWJ3 | KATNB1-lik | Q9CWJ3 [1 | 0 | 2597.238 | 0 | -0.07 | 0.078127 | 1.33 | 109 | 91 | 11.28 | 9.66 | | High | High | High | High | High | High | High | High | High | High | High | 0 | 4E-08 | 4.93 |
| FALSE | High | TFWSPELK | [K].TFWSPI | 1xPhospho | F8VPM7 1x | 0.002695 | 0.000109 | 1 | 18 | 1 | F8VPM7 | ELKS/Rab6 | F8VPM7 [1 | 0 | 1545.812 | 0 | 0.18 | 0.066982 | 4.37 | 94 | 106 | 1.34 | 7.56 | | High | High | High | High | High | High | High | High | High | High | High | 4.94E-05 | 0.000725 | 2.5 |
| FALSE | High | TGANAQFL | [K].TGANA | 1xPhospho | Q8R4R6 1x | 2.98E-08 | 0 | 1 | 2 | 1 | Q8R4R6 | Nucleopor | Q8R4R6 [2 | 0 | 2904.468 | 0 | 0.03 | 0.983605 | 12 | 97.7 | 102.3 | 6.64 | 19.14 | | High | High | High | High | High | High | High | High | High | High | High | 0 | 1.96E-09 | 5.29 |
| FALSE | High | TGANAQFL | [K].TGANA | 1xPhospho | Q8R4R6 1x | 2.91E-07 | 0 | 1 | 2 | 1 | Q8R4R6 | Nucleopor | Q8R4R6 [2 | 0 | 2920.463 | 0 | 0.09 | 0.982035 | 36.3 | 102.6 | 97.4 | 14.9 | 15.71 | | High | High | High | High | High | High | High | High | High | High | High | 0 | 2.56E-08 | 5.21 |
| FALSE | High | TGASQGRD | [R].TGASQ | 2xPhospho | Q8CD14 2x | 0.008094 | 0.000736 | 1 | 3 | 1 | Q8CD14 | Putative ur | Q8CD14 [3 | 1 | 3060.194 | 0 | -0.54 | 0.041766 | 17.01 | 118.5 | 81.5 | 11.7 | 13.95 | | High | High | High | High | High | High | High | High | High | High | High | 0.000327 | 0.002475 | 2.74 |
| FALSE | High | TGATASPS | [R].TGATA | 1xPhospho | O08715-4 | 5.03E-05 | 0 | 1 | 6 | 2 | O08715-4 | Isoform 4 ( | O08715-4 | 0 | 1936.019 | 0 | 0.55 | 0.010095 | 6.63 | 83.8 | 116.2 | 5.96 | 7.84 | | High | High | High | High | High | High | High | High | High | High | High | 7.19E-05 | 0.00091 | 4 |
| FALSE | High | TGDCQAGS | [R].TGDCQ | 2xCarbami | Q9QWF0 1 | 2.54E-10 | 0 | 1 | 2 | 1 | Q9QWF0 | Chromatin | Q9QWF0 [ | 0 | 2430.063 | 0 | -0.11 | 0.111798 | 12.34 | 103 | 97 | 9.33 | 9.56 | | High | High | High | High | High | High | High | High | High | High | High | 0 | 9.3E-12 | 4.53 |
| FALSE | High | TGDLGIPPI | [R].TGDLGI | 2xPhospho | D3YZC9 2x | 3.64E-06 | 0 | 1 | 9 | 6 | D3YZC9 | Splicing fac | D3YZC9 [6 | 1 | 3155.417 | 0 | -0.16 | 0.994524 | 58.63 | 105.5 | 94.5 | 48.32 | 57.37 | | High | High | High | High | High | High | High | High | High | High | High | 0 | 4.36E-07 | 3.61 |
| FALSE | High | TGDLGIPPI | [R].TGDLGI | 2xPhospho | D3YZC9 2x | 0.000543 | 0 | 1 | 9 | 8 | D3YZC9 | Splicing fac | D3YZC9 [6 | 1 | 3384.58 | 0 | -0.14 | 0.067062 | 1.36 | 105.3 | 94.7 | 3.31 | 7.86 | | High | High | High | High | High | High | High | High | High | High | High | 0 | 0.000121 | 4.5 |
| FALSE | High | TGDQDSTD | [R].TGDQD | 1xPhospho | B2RXQ3 1x | 0.003892 | 0.000261 | 1 | 6 | 1 | B2RXQ3 | Family with | B2RXQ3 [6 | 0 | 2268.946 | 0 | 0.13 | 0.912542 | 32.19 | 104.1 | 95.9 | 16.22 | 19.14 | | High | High | High | High | High | High | High | High | High | High | High | 0.00014 | 0.001091 | 2.21 |
| FALSE | High | TGEPSPPHI | [R].TGEPSF | 1xPhospho | A6H6B3 1x | 0.006478 | 0.00062 | 1 | 9 | 1 | A6H6B3 | RalA bindir | A6H6B3 [4 | 0 | 3022.408 | 0 | 0.11 | 0.88729 | 5.39 | 97 | 103 | 13.78 | 5.46 | | High | High | High | High | High | High | High | High | High | High | High | 0.000275 | 0.001938 | 1.93 |
| FALSE | High | TGEPSPPHI | [R].TGEPSF | 2xPhospho | A6H6B3 2x | 0.001424 | 5.66E-05 | 1 | 9 | 3 | A6H6B3 | RalA bindir | A6H6B3 [4 | 0 | 3102.375 | 0 | -0.29 | 0.068567 | 5.36 | 107.6 | 92.4 | 8.56 | 6.66 | | High | High | High | High | High | High | High | High | High | High | High | 0.000161 | 0.001162 | 3.17 |
| FALSE | High | TGESLTVES | [R].TGESLT | 1xPhospho | E9PZM7 1x | 4.12E-05 | 0 | 1 | 3 | 2 | E9PZM7 | SR-related | E9PZM7 [5 | 0 | 3390.661 | 0 | 0.36 | 0.069679 | 18.28 | 87.2 | 112.8 | 7.74 | 15.08 | | High | High | High | High | High | High | High | High | High | High | High | 0 | 6.61E-06 | 3.4 |
| FALSE | High | TGFSTSPES | [R].TGFSTS | 1xPhospho | Q9JIF0 1xP | 0.013308 | 0.000889 | 1 | 13 | 2 | Q9JIF0 | Protein arg | Q9JIF0 [29 | 0 | 2263.084 | 0 | -0.89 | 0.001207 | 3.56 | 126.4 | 73.6 | 7.26 | 3.93 | | High | High | High | High | High | High | High | High | High | High | High | 0.008697 | 0.09279 | 1.71 |
| FALSE | High | TGGFGAAF | [K].TGGFG | 1xPhospho | Q80U93 1x | 0.011246 | 0.000772 | 1 | 3 | 2 | Q80U93 | Nuclear po | Q80U93 [1 | 0 | 4494.221 | 0 | -1 | 0.006085 | 8.03 | 128.9 | 71.1 | 11.23 | 20.07 | | High | High | High | High | High | High | High | High | High | High | High | 0.000399 | 0.003565 | 3.62 |
| FALSE | High | TGGPAYGF | [R].TGGPA | 1xPhospho | Q6P5B5 1x | 2.23E-05 | 0 | 2 | 3 | 2 | Q6P5B5; Q | Fragile X m | Q6P5B5 [4 | 0 | 2464.06 | 0 | -0.03 | 0.713033 | 12.02 | 101 | 99 | 32.59 | 64.26 | Shared | High | High | High | High | High | High | High | High | High | High | High | 0 | 3.31E-06 | 3.88 |
| FALSE | High | TGGPAYGF | [R].TGGPA | 1xPhospho | Q6P5B5 1x | 8.17E-08 | 0 | 2 | 3 | 6 | Q6P5B5; Q | Fragile X m | Q6P5B5 [4 | 0 | 2693.223 | 0 | 0.11 | 0.009982 | 3.34 | 96.3 | 103.7 | 3.52 | 2.04 | Shared | High | High | High | High | High | High | High | High | High | High | High | 0 | 2.77E-08 | 4.4 |
| FALSE | High | TGGPAYGF | [R].TGGPA | 2xPhospho | Q6P5B5 2x | 0.000102 | 0 | 2 | 3 | 4 | Q6P5B5; Q | Fragile X m | Q6P5B5 [4 | 0 | 2773.189 | 0 | 0.28 | 0.043534 | 11.2 | 88 | 112 | 12.81 | 6.38 | Shared | High | High | High | High | High | High | High | High | High | High | High | 0 | 1.86E-05 | 4.28 |
| FALSE | High | TGGPAYGF | [R].TGGPA | 1xPhospho | Q6P5B5 1x | 0.076555 | 0.005826 | 2 | 3 | 1 | Q6P5B5; Q | Fragile X m | Q6P5B5 [4 | 1 | 2849.324 | 0 | -0.08 | 0.476678 | 26.73 | 102.7 | 97.3 | 14.42 | 9.99 | Shared | High | High | High | High | High | High | High | High | High | High | High | 0.00302 | 0.03169 | 1.65 |
| FALSE | High | TGGSSPEVI | [K].TGGSSI | 1xCarbami | F8VQH0 1x | 1.18E-06 | 0 | 1 | 5 | 4 | F8VQH0 | Tyrosine-p | F8VQH0 [6 | 0 | 3207.509 | 0 | -0.53 | 0.003531 | 7.49 | 118.3 | 81.7 | 4.92 | 9 | | High | High | High | High | High | High | High | High | High | High | High | 0 | 1.22E-07 | 4.02 |

| | | | | | | | | | | | | | | | | | | | | | | | | | | | | | | | | | | | | | | |
|---|---|---|---|---|---|---|---|---|---|---|---|---|---|---|---|---|---|---|---|---|---|---|---|---|---|---|---|---|---|---|---|---|---|---|---|---|---|---|
| FALSE | High | TGLEELVLS | [R].TGLEEL | 1xPhospho | A2AEW8 1 | 0.028082 | 0.001791 | 1 | 3 | 1 | A2AEW8 | GRIP1-assc | A2AEW8 [6 | 0 | 2086.993 | 0 | 0.3 | 0.084948 | 16.22 | 89.6 | 110.4 | 8.15 | 7.64 | | High | High | High | High | High | High | High | High | High | High | High | 0.000968 | 0.01002 | 2.41 |
| FALSE | High | TGLGLSSPT | [R].TGLGLS | 1xPhospho | Q8CH02 1x | 0.060159 | 0.004476 | 1 | 2 | 1 | Q8CH02 | SURP and | Q8CH02 [1 | 0 | 1522.808 | 0 | -0.49 | 0.703845 | 14.9 | 108 | 92 | 24.42 | 49.01 | | High | High | High | High | High | High | High | High | High | High | High | 0.002293 | 0.02383 | 1.29 |
| FALSE | High | TGLGLSSPT | [R].TGLGLS | 1xPhospho | Q8CH02 1x | 0.001386 | 5.66E-05 | 1 | 2 | 2 | Q8CH02 | SURP and | Q8CH02 [1 | 0 | 1751.971 | 0 | -0.63 | 0.000676 | 0.71 | 121.6 | 78.4 | 2.04 | 8.85 | | High | High | High | High | High | High | High | High | High | High | High | 2.58E-05 | 0.000343 | 2.27 |
| FALSE | High | TGLLFSPDI | [K].TGLLFS | 1xPhospho | Q80Y19 1x | 0.047086 | 0.00354 | 1 | 3 | 1 | Q80Y19 | Rho GTPas | Q80Y19 [4 | 0 | 1671.819 | 0 | -0.2 | 0.183252 | 5.66 | 104.8 | 95.2 | 7.64 | 7.74 | | High | High | High | High | High | High | High | High | High | High | High | 0.001812 | 0.01792 | 2.62 |
| FALSE | High | TGLSDMLA | [R].TGLSDM | 1xPhospho | B2RS91 1x | 0.000102 | 0 | 1 | 1 | 1 | B2RS91 | RNA polym | B2RS91 [27 | 1 | 2964.54 | 0 | -1.05 | 0.00508 | 1.39 | 134.8 | 65.2 | 10.27 | 13.8 | | High | High | High | High | High | High | High | High | High | High | High | 0 | 1.84E-05 | 2.69 |
| FALSE | High | TGMGSGS | [K].TGMGS | 1xPhospho | Q7TPS5 1x | 0.033404 | 0.002195 | 1 | 4 | 2 | Q7TPS5 | C2 domain | Q7TPS5 [3 | 0 | 1390.68 | 0 | -0.12 | 0.040323 | 5.65 | 103.1 | 96.9 | 4.98 | 8.34 | | High | High | High | High | High | High | High | High | High | High | High | 0.001151 | 0.01221 | 1.52 |
| FALSE | High | TGMGSGS | [K].TGMGS | 1xPhospho | Q7TPS5 1x | 0.000362 | 0 | 1 | 4 | 2 | Q7TPS5 | C2 domain | Q7TPS5 [3 | 0 | 1406.675 | 0 | -0.39 | 0.004552 | 1.59 | 112.1 | 87.9 | 4.08 | 7.03 | | High | High | High | High | High | High | High | High | High | High | High | 0 | 7.66E-05 | 1.34 |
| FALSE | High | TGSTGALG | [K].TGSTG | 1xPhospho | Q3UMT1 1 | 0.002355 | 0.000109 | 1 | 2 | 4 | Q3UMT1 | Protein ph | Q3UMT1 [ | 0 | 1441.689 | 0 | -0.1 | 0.401189 | 14.78 | 103.5 | 96.5 | 5.13 | 12.77 | | High | High | High | High | High | High | High | High | High | High | High | 0.000275 | 0.001954 | 2.63 |
| FALSE | High | TFSYDEIHG | [R].TFSYDE | 1xPhospho | P33174 1x | 0.000369 | 0 | 1 | 2 | 5 | P33174 | Chromoso | P33174 [80 | 0 | 2708.249 | 0 | 0.32 | 0.006504 | 9.21 | 91.9 | 108.1 | 7.19 | 4.48 | | High | High | High | High | High | High | High | High | High | High | High | 0.000327 | 0.002517 | 3.61 |
| FALSE | High | TGSTSSK | [R].TGSTSS | 1xPhospho | A2AN08-5 | 0.01186 | 0.000889 | 1 | 4 | 3 | A2AN08-5 | Isoform 5 | A2AN08-5 | 0 | 1205.618 | 0 | 0.11 | 0.254137 | 7.64 | 96.2 | 103.8 | 4.83 | 11.39 | | High | High | High | High | High | High | High | High | High | High | High | 0.005588 | 0.06106 | 1.36 |
| FALSE | High | TGSTSSSS | [R].TGSTSS | 1xPhospho | Q05BC3-3 | 0.000264 | 0 | 1 | 6 | 2 | Q05BC3-3 | Isoform 3 | Q05BC3-3 | 0 | 1523.735 | 0 | -0.81 | 0.007175 | 8.76 | 131.6 | 68.4 | 13.92 | 20.4 | | High | High | High | High | High | High | High | High | High | High | High | 0 | 0.000194 | 1.42 |
| FALSE | High | THSIESSGK | [R].THSIES | 1xPhospho | Q543X6 1x | 0.003755 | 0.00021 | 1 | 2 | 2 | Q543X6 | Putative ur | Q543X6 [7 | 0 | 1483.756 | 0 | 0.06 | 0.287997 | 14.47 | 96.9 | 103.1 | 10.14 | 12.87 | | High | High | High | High | High | High | High | High | High | High | High | 0.000814 | 0.007706 | 2.46 |
| FALSE | High | THSLLYQP | [R].THSLLY | 1xPhospho | Q8JZZ7-2 1 | 0.010853 | 0.000772 | 1 | 11 | 1 | Q8JZZ7-2 | Isoform 2 | Q8JZZ7-2 [ | 0 | 1752.945 | 0 | 0.07 | 0.757695 | 22.74 | 97.7 | 102.3 | 8.6 | 9.68 | | High | High | High | High | High | High | High | High | High | High | High | 0.000399 | 0.003434 | 3.18 |
| FALSE | High | THSLSNAD | [R].THSLSN | 1xPhospho | E9Q842 1x | 5.15E-07 | 0 | 1 | 3 | 1 | E9Q842 | Neuron na | E9Q842 [1 | 0 | 2336.003 | 0 | 0.85 | 0.000442 | 6.06 | 72.6 | 127.4 | 9.63 | 5.76 | | High | High | High | High | High | High | High | High | High | High | High | 0 | 4.84E-08 | 3.94 |
| FALSE | High | THSNEPGK | [R].THSNEP | 1xPhospho | Q9ERV1 1x | 0.069903 | 0.005181 | 1 | 2 | 1 | Q9ERV1 | Probable E | Q9ERV1 [9 | 0 | 1407.703 | 0 | 1.32 | 0.000555 | 15.67 | 55.8 | 144.2 | 12.39 | 5.58 | | High | High | High | High | High | High | High | High | High | High | High | 0.002662 | 0.02826 | 1.6 |
| FALSE | High | THSPPQWI | [R].THSPPQ | 1xPhospho | E9Q9Q2 1x | 0.032267 | 0.002129 | 1 | 3 | 2 | E9Q9Q2 | R3H domai | E9Q9Q2 [8 | 0 | 1518.787 | 0 | 0.44 | 0.016509 | 2.32 | 82.8 | 117.2 | 4.27 | 11.8 | | High | High | High | High | High | High | High | High | High | High | High | 0.001142 | 0.01175 | 1.47 |
| FALSE | High | THSPVPDM | [K].THSPVP | 1xPhospho | Q9CYA6 1x | 0.004333 | 0.00036 | 1 | 2 | 1 | Q9CYA6 | Zinc finger | Q9CYA6 [6 | 0 | 1636.817 | 0 | 0.43 | 0.173126 | 7.62 | 85.2 | 114.8 | 7.8 | 22.85 | | High | High | High | High | High | High | High | High | High | High | High | 0.000161 | 0.001237 | 1.82 |
| FALSE | High | THSPVPDM | [K].THSPVP | 1xPhospho | Q9CYA6 1x | 0.001783 | 5.66E-05 | 1 | 2 | 2 | Q9CYA6 | Zinc finger | Q9CYA6 [6 | 0 | 1652.812 | 0 | 0.36 | 0.100268 | 3.22 | 87.6 | 112.4 | 10.66 | 14.14 | | High | High | High | High | High | High | High | High | High | High | High | 0.000706 | 0.00624 | 2.21 |
| FALSE | High | THSQGGY | [R].THSQG | 1xPhospho | Q8C0C7 1x | 6.45E-07 | 0 | 1 | 2 | 2 | Q8C0C7 | Phenylalan | Q8C0C7 [2 | 0 | 1907.905 | 0 | -0.85 | 0.000273 | 2.74 | 127.9 | 72.1 | 4.97 | 7.39 | | High | High | High | High | High | High | High | High | High | High | High | 0 | 6.2E-08 | 3.74 |
| FALSE | High | THSSSPQLI | [R].THSSSP | 1xPhospho | A6H619 1x | 0.000372 | 0 | 1 | 2 | 1 | A6H619 | PHD and RI | A6H619 [6 | 0 | 1792.854 | 0 | 0.17 | 0.472252 | 18.77 | 94.3 | 105.7 | 6.55 | 12.78 | | High | High | High | High | High | High | High | High | High | High | High | 0 | 7.84E-05 | 2.78 |
| FALSE | High | THSTSSSIG | [R].THSTSS | 1xPhospho | Q8VCV2 1x | 6.51E-07 | 0 | 1 | 2 | 5 | Q8VCV2 | Ndrg3 prot | Q8VCV2 [3 | 0 | 2032.917 | 0 | 0.91 | 0.020332 | 32.61 | 69.4 | 130.6 | 20.6 | 16.05 | | High | High | High | High | High | High | High | High | High | High | High | 0 | 6.28E-08 | 3.1 |
| FALSE | High | THSTSSSIG | [R].THSTSS | 2xPhospho | Q8VCV2 2x | 0.00418 | 0.00036 | 1 | 2 | 3 | Q8VCV2 | Ndrg3 prot | Q8VCV2 [3 | 0 | 2112.884 | 0 | 0.56 | 0.009386 | 14.26 | 80.1 | 119.9 | 7.38 | 9.84 | | High | High | High | High | High | High | High | High | High | High | High | 0.000267 | 0.001566 | 2.59 |
| FALSE | High | THSTSSSIG | [R].THSTSS | 3xPhospho | Q8VCV2 3x | 0.058664 | 0.004294 | 1 | 2 | 1 | Q8VCV2 | Ndrg3 prot | Q8VCV2 [3 | 0 | 2192.85 | 0 | 0.67 | 0.032983 | 6.42 | 79 | 121 | 12.73 | 12.84 | | High | High | High | High | High | High | High | High | High | High | High | 0.002187 | 0.02328 | 2.67 |
| FALSE | High | THSVSADP | [R].THSVSA | 1xCarbami | S0DHL8 1x | 6.53E-05 | 0 | 1 | 4 | 1 | S0DHL8 | Capping pr | S0DHL8 [1 | 0 | 3104.466 | 0 | -0.57 | 0.08107 | 17.04 | 124.2 | 75.8 | 17.07 | 19.38 | | High | High | High | High | High | High | High | High | High | High | High | 0 | 1.11E-05 | 3.01 |
| FALSE | High | THSLCNGD | [R].THSLCN | 1xCarbami | Q69ZS8 1x | 3.67E-05 | 0 | 1 | 8 | 3 | Q69ZS8 | Kazrin OS= | Q69ZS8 [3 | 0 | 2135.036 | 0 | -1.32 | 0.000348 | 0.98 | 144.8 | 55.2 | 7.2 | 4.73 | | High | High | High | High | High | High | High | High | High | High | High | 0 | 5.83E-06 | 2.49 |
| FALSE | High | THSYSPLET | [K].THSYSP | 1xPhospho | S4R1P5 1x | 1.61E-07 | 0 | 1 | 6 | 6 | S4R1P5 | Dystonin O | S4R1P5 [2 | 0 | 2765.288 | 0 | 0.64 | 0.003309 | 13.23 | 78.6 | 121.4 | 6.9 | 9.08 | | High | High | High | High | High | High | High | High | High | High | High | 0 | 2.25E-08 | 3.97 |
| FALSE | High | THTGERPY | [R].THTGE | 2xCarbami | O70230 1x | 0.005924 | 0.00062 | 1 | 3 | 1 | O70230 | Zinc finger | O70230 [3 | 0 | 2349.994 | 0 | -0.83 | 0.066125 | 5.94 | 126.4 | 73.6 | 22.32 | 9.41 | | High | High | High | High | High | High | High | High | High | High | High | 0.000275 | 0.001745 | 1.37 |
| FALSE | High | THTHSDSA | [K].THTHS | 1xPhospho | A2AAE1-3 | 0.011048 | 0.000772 | 1 | 6 | 1 | A2AAE1-3 | Isoform 3 | A2AAE1-3 | 0 | 1634.83 | 0 | 0.17 | 0.091086 | 7.21 | 94.2 | 105.8 | 18.33 | 9.4 | | High | High | High | High | High | High | High | High | High | High | High | 0.000399 | 0.003517 | 2.09 |
| FALSE | High | THTLPPESA | [R].THTLPP | 1xPhospho | A0A0R4J0C | 3.49E-07 | 0 | 1 | 2 | 4 | A0A0R4J0C | BRF2, subu | A0A0R4J0C | 0 | 3211.488 | 0 | -0.51 | 0.018922 | 12.64 | 118.9 | 81.1 | 13.72 | 14.58 | | High | High | High | High | High | High | High | High | High | High | High | 0 | 3.11E-08 | 4.96 |
| FALSE | High | THVLTEQS | [R].THVLTE | 1xPhospho | Q3UHU5 1 | 1.76E-07 | 0 | 1 | 5 | 1 | Q3UHU5 | Microtubu | Q3UHU5 [ | 0 | 2487.307 | 0 | -1.47 | 0.000193 | 1.34 | 145.2 | 54.8 | 6.67 | 14.04 | | High | High | High | High | High | High | High | High | High | High | High | 0 | 1.44E-08 | 3.55 |
| FALSE | High | TKDYDWTI | [K].TKDYD | 1xPhospho | Q8BVG4 1x | 0.024424 | 0.001612 | 1 | 2 | 2 | Q8BVG4 | Dipeptidyl | Q8BVG4 [4 | 1 | 2510.121 | 1 | 1.05 | 0.005099 | 21.56 | 68.9 | 131.1 | 32.67 | 10.28 | | High | High | High | High | High | High | High | High | High | High | High | 0.001712 | 0.0171 | 3.26 |
| FALSE | High | TKGDSPTQ | [K].TKGDS | 1xCarbami | A3KFM7 1x | 1.99E-05 | 0 | 1 | 3 | 1 | A3KFM7 | Chromodo | A3KFM7 [2 | 1 | 2501.256 | 0 | -0.67 | 0.0409 | 10.53 | 115.1 | 84.9 | 16.13 | 10.24 | | High | High | High | High | High | High | High | High | High | High | High | 0 | 2.91E-06 | 3.98 |
| FALSE | High | TKPLHLVE | [K].TKPLHL | 1xCarbami | Q7TSZ8 1x | 5.4E-05 | 0 | 1 | 1 | 1 | Q7TSZ8 | Nucleus ac | Q7TSZ8 [4 | 0 | 2720.284 | 0 | -1.76 | 0.006003 | 2.4 | 153.1 | 46.9 | 25.91 | 24.87 | | High | High | High | High | High | High | High | High | High | High | High | 0 | 8.97E-06 | 3.95 |
| FALSE | High | TKPPPTYES | [K].TKPPPT | 1xPhospho | Q3UH70 1x | 3.79E-07 | 0 | 1 | 4 | 1 | Q3UH70 | Bromodon | Q3UH70 [5 | 0 | 2188.046 | 0 | -0.77 | 0.87387 | 18.51 | 125.2 | 74.8 | 35.11 | 83.93 | | High | High | High | High | High | High | High | High | High | High | High | 0 | 3.42E-08 | 3.2 |
| FALSE | High | TKPPPTYES | [K].TKPPPT | 1xPhospho | Q3UH70 1x | 2.08E-05 | 0 | 1 | 4 | 5 | Q3UH70 | Bromodon | Q3UH70 [5 | 0 | 2417.209 | 0 | -0.69 | 0.002261 | 1.31 | 124.3 | 75.7 | 4.64 | 6.95 | | High | High | High | High | High | High | High | High | High | High | High | 0 | 3.09E-06 | 2.33 |
| FALSE | High | TKPPRPDSI | [K].TKPPRP | 1xPhospho | Q3V1H4 1x | 0.000462 | 0 | 1 | 3 | 3 | Q3V1H4 | E74-like fa | Q3V1H4 [1 | 0 | 2703.505 | 0 | 0.62 | 0.007702 | 12.17 | 75.8 | 124.2 | 4.54 | 13.13 | | High | High | High | High | High | High | High | High | High | High | High | 0 | 0.000101 | 2.09 |
| FALSE | High | TKSENGLEI | [K].TKSEN | 1xPhospho | Q60932 1x | 0.003253 | 0.00016 | 1 | 6 | 1 | Q60932 | Voltage-de | Q60932 [4 | 1 | 2956.476 | 0 | -0.03 | 0.619019 | 28.93 | 101.3 | 98.7 | 8.44 | 12.79 | | High | High | High | High | High | High | High | High | High | High | High | 7.19E-05 | 0.000892 | 4.2 |
| FALSE | High | TKSNDDLL | [K].TKSNDI | 1xPhospho | A0A0R4J0J | 0.002743 | 0.000109 | 1 | 4 | 2 | A0A0R4J0J | Cytospin-A | A0A0R4J0J | 1 | 2484.237 | 0 | 1.86 | 0.004842 | 35.23 | 43.6 | 156.4 | 29.37 | 17.12 | | High | High | High | High | High | High | High | High | High | High | High | 4.94E-05 | 0.000743 | 3.92 |
| FALSE | High | TKSNDDLL | [K].TKSNDI | 1xPhospho | A0A0R4J0J | 0.001127 | 0 | 1 | 4 | 1 | A0A0R4J0J | Cytospin-A | A0A0R4J0J | 1 | 2500.232 | 0 | 1.31 | 0.007293 | 10.08 | 54.6 | 145.4 | 14.59 | 21.11 | | High | High | High | High | High | High | High | High | High | High | High | 0 | 0.000274 | 3.49 |
| FALSE | High | THTGDKPY | [R].THTGD | 2xCarbami | A2ART0 1x | 0.011048 | 0.000772 | 2 | 5 | 1 | A2ART0; A | Uncharacte | A2ART0 [2 | 0 | 2562.209 | 0 | -0.55 | 0.005355 | 1.38 | 118.2 | 81.8 | 16.82 | 14.72 | Shared | High | High | High | High | High | High | High | High | High | High | High | 0.000399 | 0.003496 | 2.71 |
| FALSE | High | THSGLTDG | [R].THSGLT | 1xPhospho | Q6ZQ08-4 | 2.34E-05 | 0 | 1 | 7 | 1 | Q6ZQ08-4 | Isoform 4 | Q6ZQ08-4 | 0 | 3119.581 | 0 | 1.43 | 0.012364 | 44.59 | 54.2 | 145.8 | 45.39 | 28.51 | | High | High | High | High | High | High | High | High | High | High | High | 0 | 3.51E-06 | 4.23 |
| FALSE | High | THSGERPYI | [R].THSGE | 1xCarbami | Q3UH06-3 | 0.060666 | 0.00451 | 1 | 6 | 1 | Q3UH06-3 | Isoform 3 | Q3UH06-3 | 0 | 1885.94 | 0 | 0.19 | 0.042826 | 5.47 | 91.5 | 108.5 | 10.97 | 4.83 | | High | High | High | High | High | High | High | High | High | High | High | 0.002354 | 0.02416 | 1.85 |
| FALSE | High | THSGENPY | [R].THSGE | 2xCarbami | A2BE20 1x | 5.16E-05 | 0 | 1 | 1 | 1 | A2BE20 | Novel KRA | A2BE20 [1 | 0 | 2327.994 | 0 | -1.16 | 0.006355 | 4.46 | 141 | 59 | 16.7 | 6.3 | | High | High | High | High | High | High | High | High | High | High | High | 0 | 8.58E-06 | 2.76 |
| FALSE | High | TGSVEK | [R].TGSVEK | 1xPhospho | Q80SY5 1x | 0.117405 | 0.009523 | 1 | 1 | 2 | Q80SY5 | Pre-mRNA- | Q80SY5 [4 | 0 | 1158.617 | 0 | -0.16 | 0.191859 | 5.67 | 103.5 | 96.5 | 4.94 | 5.51 | | High | High | High | High | High | High | High | High | High | High | High | 0.004769 | 0.0522 | 1.5 |
| FALSE | High | TGSVGIGN | [R].TGSVGI | 1xPhospho | Q68FH0 1x | 0.002974 | 0.00016 | 1 | 8 | 1 | Q68FH0 | Plakophilin | Q68FH0 [4 | 0 | 1410.73 | 0 | -0.25 | 0.792969 | 12.77 | 106.4 | 93.6 | 22 | 10.21 | | High | High | High | High | High | High | High | High | High | High | High | 7.19E-05 | 0.000809 | 2 |
| FALSE | High | TGSVGSDL | [R].TGSVGS | 1xDeamida | B2RXS2 1x | 5.27E-09 | 0 | 1 | 5 | 5 | B2RXS2 | Fam179b p | B2RXS2 [7 | 0 | 2571.249 | 0 | 0.47 | 0.006541 | 12.34 | 83.8 | 116.2 | 10.23 | 1.49 | | High | High | High | High | High | High | High | High | High | High | High | 0 | 2.79E-10 | 6.36 |
| FALSE | High | TGSVQTSP | [R].TGSVQ | 1xPhospho | D9HP81 1x | 6.15E-06 | 0 | 1 | 5 | 3 | D9HP81 | Regulating | D9HP81 [1 | 0 | 1828.864 | 0 | 0.72 | 0.002031 | 11.39 | 73.2 | 126.8 | 7.22 | 7.9 | | High | High | High | High | High | High | High | High | High | High | High | 0 | 7.85E-07 | 2.51 |
| FALSE | High | TGSYGALA | [K].TGSYG | 1xPhospho | Q9DBR7 1x | 1.13E-05 | 0 | 1 | 2 | 6 | Q9DBR7 | Protein ph | Q9DBR7 [4 | 0 | 1663.814 | 0 | -1.04 | 0.183894 | 4.43 | 135 | 65 | 36.59 | 49.3 | | High | High | High | High | High | High | High | High | High | High | High | 0 | 1.56E-06 | 3.7 |
| FALSE | High | TGSYGALA | [K].TGSYG | 1xPhospho | Q9DBR7 1x | 5.03E-05 | 0 | 1 | 2 | 7 | Q9DBR7 | Protein ph | Q9DBR7 [4 | 0 | 1892.977 | 0 | -1.14 | 0.002624 | 5.84 | 137.7 | 62.3 | 8.14 | 23.07 | | High | High | High | High | High | High | High | High | High | High | High | 0 | 7.22E-05 | 3.24 |
| FALSE | High | TGTGDIAT | [K].TGTGD | 1xCarbami | Q8R4H2 1x | 9.91E-06 | 0 | 2 | 2 | 2 | Q8R4H2; F | Rho guanir | Q8R4H2 [1 | 0 | 1659.725 | 0 | 1.67 | 0.000363 | 6.11 | 47.9 | 152.1 | 6.41 | 7.78 | Shared | High | High | High | High | High | High | High | High | High | High | High | 0 | 1.34E-06 | 2.49 |
| FALSE | High | TGTGSPFA | [K].TGTGS | 1xPhospho | Q6ZPZ3 1x | 2.55E-07 | 0 | 1 | 5 | 7 | Q6ZPZ3 | Zinc finger | Q6ZPZ3 [1 | 0 | 1628.763 | 0 | -0.3 | 0.004611 | 1.28 | 110 | 90 | 4.25 | 5.51 | | High | High | High | High | High | High | High | High | High | High | High | 0 | 2.19E-08 | 2.94 |
| FALSE | High | TGTGSPFA | [K].TGTGS | 2xPhospho | Q6ZPZ3 2x | 6.26E-06 | 0 | 1 | 5 | 2 | Q6ZPZ3 | Zinc finger | Q6ZPZ3 [1 | 0 | 1708.729 | 0 | 0.53 | 0.008233 | 6.65 | 83.2 | 116.8 | 4.94 | 9.11 | | High | High | High | High | High | High | High | High | High | High | High | 0 | 7.94E-07 | 2.67 |
| FALSE | High | TGVLAAVL | [K].TGVLA | 1xPhospho | O54916 1x | 2.02E-05 | 0 | 1 | 6 | 1 | O54916 | RalBP1-ass | O54916 [7 | 0 | 1919.056 | 0 | 0.09 | 0.474918 | 12.34 | 100.8 | 99.2 | 16.32 | 8.34 | | High | High | High | High | High | High | High | High | High | High | High | 0 | 2.97E-06 | 3.59 |
| FALSE | High | TGWDTSES | [K].TGWDT | 2xPhospho | A0A0R4J1S | 0.062731 | 0.004664 | 1 | 6 | 2 | A0A0R4J1S | PRP40 pre- | A0A0R4J1S | 0 | 2313.936 | 0 | -1.03 | 0.001637 | 1 | 125.6 | 74.4 | 15.02 | 23.51 | | High | High | High | High | High | High | High | High | High | High | High | 0.002867 | 0.0297 | 3.42 |
| FALSE | High | TGYESGDY | [R].TGYES | 1xPhospho | Q3TPZ5 1x | 0.016021 | 0.001124 | 1 | 1 | 2 | Q3TPZ5 | Dynactin 2 | Q3TPZ5 [7 | 0 | 2459.145 | 0 | 0.01 | 0.453105 | 32.71 | 95.6 | 104.4 | 15.83 | 26.37 | | High | High | High | High | High | High | High | High | High | High | High | 0.001214 | 0.01303 | 2.78 |
| FALSE | High | THAVSVSET | [R].THAVS | 1xPhospho | P34152-4 1 | 0.002077 | 5.66E-05 | 1 | 3 | 1 | P34152-4 | Isoform 4 | P34152-4 [ | 0 | 3484.519 | 0 | 0.37 | 0.086568 | 11.12 | 82.8 | 117.2 | 18.19 | 17.37 | | High | High | High | High | High | High | High | High | High | High | High | 2.58E-05 | 0.00054 | 2.52 |
| FALSE | High | THAVSVSET | [R].THAVS | 1xPhospho | P34152-4 1 | 9.8E-05 | 0 | 1 | 3 | 3 | P34152-4 | Isoform 4 | P34152-4 [ | 0 | 3500.514 | 0 | 0.34 | 0.588905 | 17.46 | 104.5 | 95.5 | 26.57 | 20.2 | | High | High | High | High | High | High | High | High | High | High | High | 0 | 5.34E-05 | 3.15 |
| FALSE | High | THDILEDVI | [K].THDILE | 1xPhospho | Q05BA5 1x | 7.74E-08 | 0 | 1 | 7 | 4 | Q05BA5 | Nuclear re | Q05BA5 [2 | 0 | 2005.943 | 0 | -0.2 | 0.075259 | 7.54 | 106.6 | 93.4 | 7.4 | 5.35 | | High | High | High | High | High | High | High | High | High | High | High | 0 | 5.76E-09 | 4.46 |
| FALSE | High | THEILGSLE | [K].THEILG | 1xPhospho | Q6P5H2-2 | 0.001373 | 5.66E-05 | 1 | 2 | 2 | Q6P5H2-2 | Isoform 2 | Q6P5H2-2 | 0 | 1578.773 | 0 | 2.34 | 0.001387 | 6.72 | 30.4 | 169.6 | 9.8 | 16.8 | | High | High | High | High | High | High | High | High | High | High | High | 2.58E-05 | 0.000342 | 3.65 |
| FALSE | High | THGHLIPT | [K].THGHLI | 1xPhospho | Q8VE42 1x | 1.11E-05 | 0 | 1 | 2 | 3 | Q8VE42 | Ankyrin re | Q8VE42 [3 | 0 | 3574.674 | 0 | -0.25 | 0.054738 | 5.38 | 109.1 | 90.9 | 9.73 | 9.7 | | High | High | High | High | High | High | High | High | High | High | High | 0 | 1.52E-06 | 4.64 |
| FALSE | High | THGHLIPT | [K].THGHLI | 1xDeamida | Q8VE42 1x | 4.85E-05 | 0 | 1 | 2 | 1 | Q8VE42 | Ankyrin re | Q8VE42 [3 | 0 | 3575.658 | 0 | 0.25 | 0.238038 | 19.3 | 96.4 | 103.6 | 23.52 | 14.71 | | High | High | High | High | High | High | High | High | High | High | High | 0 | 0.000008 | 4.73 |
| FALSE | High | THHDLTTP | [R].THHDL | 1xPhospho | B7ZMS8 1x | 0.000365 | 0 | 1 | 9 | 2 | B7ZMS8 | Mdm1 pro | B7ZMS8 [5 | 0 | 2525.353 | 0 | -1.1 | 0.000475 | 2.32 | 135.8 | 64.2 | 4.42 | 7.08 | | High | High | High | High | High | High | High | High | High | High | High | 0 | 7.72E-05 | 2.16 |
| FALSE | High | THLLQFALI | [R].THLLQF | 1xPhospho | E9Q392 1x | 1.06E-08 | 0 | 1 | 7 | 2 | E9Q392 | E3 ubiquiti | E9Q392 [9 | 0 | 2777.489 | 0 | 0.72 | 0.017207 | 36.6 | 73.6 | 126.4 | 10.74 | 15.43 | | High | High | High | High | High | High | High | High | High | High | High | 0 | 6.18E-10 | 4.26 |
| FALSE | High | THLLQFALI | [R].THLLQF | 2xPhospho | E9Q392 2x | 9.16E-07 | 0 | 1 | 7 | 2 | E9Q392 | E3 ubiquiti | E9Q392 [9 | 0 | 2857.455 | 0 | 0.69 | 0.01071 | 16.06 | 74.4 | 125.6 | 5.97 | 13.01 | | High | High | High | High | High | High | High | High | High | High | High | 0 | 9.19E-08 | 4.27 |
| FALSE | High | THIVFASDN | [K].THIVFA | 1xCarbami | E9QKD1 1x | 5.21E-08 | 0 | 1 | 2 | 3 | E9QKD1 | Nucleolar | E9QKD1 [7 | 0 | 3522.551 | 0 | -1.16 | 0.001914 | 2.29 | 141.6 | 58.4 | 13.63 | 7.92 | | High | High | High | High | High | High | High | High | High | High | High | 0 | 1.05E-07 | 5.97 |
| FALSE | High | THIVFASDN | [K].THIVFA | 1xCarbami | E9QKD1 2x | 7.68E-05 | 0 | 1 | 2 | 3 | E9QKD1 | Nucleolar | E9QKD1 [7 | 0 | 3601.533 | 0 | -0.6 | 0.007348 | 2.71 | 119 | 81 | 5.47 | 12.2 | | High | High | High | High | High | High | High | High | High | High | High | 0 | 1.34E-05 | 4.76 |
| FALSE | High | THSAGTSP | [R].THSAG | 1xPhospho | P35569 1x | 0.043236 | 0.003228 | 1 | 2 | 1 | P35569 | Insulin rec | P35569 [5 | 0 | 1990.016 | 0 | -0.68 | 0.003454 | 6.25 | 127.1 | 72.9 | 9.89 | 5.26 | | High | High | High | High | High | High | High | High | High | High | High | 0.001651 | 0.01638 | 2.12 |
| FALSE | High | THSDASDD | [R].THSDA | 1xPhospho | B9EKC3 1x | 0.011146 | 0.000772 | 1 | 2 | 3 | B9EKC3 | Rho GTPas | B9EKC3 [1 | 0 | 1920.806 | 0 | -0.26 | 0.987611 | 9.37 | 107.6 | 92.4 | 13.65 | 27.31 | | High | High | High | High | High | High | High | High | High | High | High | 0.000399 | 0.003538 | 1.72 |
| FALSE | High | THSDASDD | [R].THSDA | 1xPhospho | B9EKC3 1x | 0.004654 | 0.000456 | 1 | 2 | 2 | B9EKC3 | Rho GTPas | B9EKC3 [1 | 0 | 2149.969 | 0 | -0.34 | 0.00917 | 8.02 | 115.3 | 84.7 | 6.59 | 10.44 | | High | High | High | High | High | High | High | High | High | High | High | 0.000225 | 0.001433 | 2.83 |
| FALSE | High | THSDASDD | [R].THSDA | 2xPhospho | B9EKC3 2x | 9.14E-06 | 0 | 1 | 2 | 3 | B9EKC3 | Rho GTPas | B9EKC3 [1 | 0 | 2229.935 | 0 | -0.26 | 0.043439 | 3.4 | 107.3 | 92.7 | 2.48 | 8.77 | | High | High | High | High | High | High | High | High | High | High | High | 0 | 1.22E-06 | 3.65 |
| FALSE | High | THSDQFLV | [R].THSDQ | 1xPhospho | Q3UD06 1x | 0.002004 | 5.66E-05 | 1 | 7 | 1 | Q3UD06 | ATP syntha | Q3UD06 [1 | 0 | 1846.95 | 0 | -1.86 | 0.000202 | 0.33 | 156.5 | 43.5 | 6.49 | 10.22 | | High | High | High | High | High | High | High | High | High | High | High | 2.58E-05 | 0.000518 | 2.03 |
| FALSE | High | THSFENVN | [R].THSFEN | 1xCarbami | A6H8H2-2 | 2.34E-06 | 0 | 1 | 3 | 2 | A6H8H2-2 | Isoform 2 | A6H8H2-2 | 0 | 2095.922 | 0 | 0.87 | 0.003101 | 15.62 | 70.6 | 129.4 | 3.25 | 11.76 | | High | High | High | High | High | High | High | High | High | High | High | 0 | 1.62E-05 | 3.33 |
| FALSE | High | TGSTSSKEI | [R].TGSTSS | 1xPhospho | A2AN08-5 | 0.000311 | 0 | 1 | 4 | 8 | A2AN08-5 | Isoform 5 | A2AN08-5 | 1 | 2770.307 | 0 | 0.17 | 0.794741 | 27.6 | 90.5 | 109.5 | 29.39 | 25.09 | | High | High | High | High | High | High | High | High | High | High | High | 0 | 0.000132 | 3.29 |
| FALSE | High | TASGSSVT | [R].TASGSS | 2xPhospho | B7ZWC0 2x | 0.000466 | 0 | 1 | 4 | 3 | B7ZWC0 | N-myc dov | B7ZWC0 [3 | 0 | 1741.761 | 0 | 0.54 | 0.036339 | 18.78 | 81.7 | 118.3 | 4.7 | 20.51 | | High | High | High | High | High | High | High | High | High | High | High | 0 | 0.000109 | 3.02 |
| FALSE | High | TASGSSVT | [R].TASGSS | 1xPhospho | B7ZWC0 1x | 5.67E-06 | 0 | 1 | 4 | 7 | B7ZWC0 | N-myc dov | B7ZWC0 [3 | 0 | 1661.794 | 0 | 0.63 | 0.025677 | 14.07 | 79.3 | 120.7 | 4.83 | 23.7 | | High | High | High | High | High | High | High | High | High | High | High | 0 | 7.11E-07 | 2.75 |
| FALSE | High | TASFSESR | [R].TASFSE | 1xPhospho | Q3V117 1x | 0.002192 | 0.000109 | 1 | 5 | 3 | Q3V117 | ATP-citrate | Q3V117 [4 | 0 | 1193.54 | 0 | -0.7 | 0.000825 | 1.87 | 123.1 | 76.9 | 2 | 12.23 | | High | High | High | High | High | High | High | High | High | High | High | 4.94E-05 | 0.000575 | 1.56 |
| FALSE | High | STRPECGQ | [R].STRPEC | 2xCarbami | Q9EQM6 1 | 0.017966 | 0.001282 | 1 | 1 | 2 | Q9EQM6 | Microproc | Q9EQM6 [ | 0 | 4395.079 | 0 | -0.87 | 0.011181 | 6.82 | 129.3 | 70.7 | 13.41 | 9.79 | | High | High | High | High | High | High | High | High | High | High | High | 0.000801 | 0.007437 | 3.14 |
| FALSE | High | STSAGNK | [R].STSAGN | 1xPhospho | A0A0R4J0J | 0.073462 | 0.005631 | 1 | 5 | 1 | A0A0R4J0J | Cytospin-A | A0A0R4J0J | 0 | 1202.618 | 0 | 0.68 | 0.001558 | 7.32 | 74.7 | 125.3 | 4.81 | 8.2 | | High | High | High | High | High | High | High | High | High | High | High | 0.002896 | 0.03013 | 0.68 |
| FALSE | High | STSAPQMS | [K].STSAPC | 1xPhospho | Q80X50-5 | 4.92E-07 | 0 | 1 | 10 | 2 | Q80X50-5 | Isoform 5 | Q80X50-5 | 0 | 3070.419 | 0 | -0.04 | 0.554509 | 20.28 | 103.6 | 96.4 | 9.81 | 17.57 | | High | High | High | High | High | High | High | High | High | High | High | 0 | 4.58E-08 | 3.93 |
| FALSE | High | STSAPQMS | [K].STSAPC | 1xPhospho | Q80X50-5 | 4.42E-11 | 0 | 1 | 10 | 42 | Q80X50-5 | Isoform 5 | Q80X50-5 | 0 | 3086.414 | 0 | -0.2 | 0.121156 | 4.08 | 106.6 | 93.4 | 2.56 | 8.8 | | High | High | High | High | High | High | High | High | High | High | High | 0 | 3.17E-08 | 5.17 |
| FALSE | High | STSAPQMS | [K].STSAPC | 2xPhospho | Q80X50-5 | 3.06E-05 | 0 | 1 | 10 | 3 | Q80X50-5 | Isoform 5 | Q80X50-5 | 0 | 3150.385 | 0 | -0.19 | 0.134146 | 4.63 | 105.5 | 94.5 | 7.55 | 18.4 | | High | High | High | High | High | High | High | High | High | High | High | 0 | 4.73E-06 | 3.92 |
| FALSE | High | STSASSHA | [R].STSASS | 1xCarbami | S4R1P5 1x | 4.89E-05 | 0 | 1 | 11 | 2 | S4R1P5 | Dystonin O | S4R1P5 [7 | 0 | 2819.38 | 0 | 0.32 | 0.103207 | 39.33 | 88.1 | 111.9 | 3.09 | 20.86 | | High | High | High | High | High | High | High | High | High | High | High | 0 | 8.05E-06 | 2.6 |
| FALSE | High | STSATDTHI | [R].STSATD | 1xPhospho | O55028 1x | 8.56E-05 | 0 | 1 | 7 | 10 | O55028 | [3-methyl- | O55028 [3 | 0 | 1833.869 | 0 | -0.14 | 0.272978 | 11.4 | 104.9 | 95.1 | 6.05 | 12.55 | | High | High | High | High | High | High | High | High | High | High | High | 0 | 3.65E-05 | 2.47 |
| FALSE | High | STSCTTSLA | [R].STSCTT | 1xCarbami | Q6A065 1x | 0.035181 | 0.002269 | 1 | 2 | 1 | Q6A065 | Centrosom | Q6A065 [6 | 0 | 1793.794 | 0 | 0.81 | 0.001007 | 12.22 | 72.7 | 127.3 | 9.87 | 7.25 | | High | High | High | High | High | High | High | High | High | High | High | 0.001199 | 0.0129 | 1.57 |
| FALSE | High | STSENKEEI | [K].STSENK | 1xCarbami | E9PV22 1x | 0.00051 | 0 | 1 | 4 | 1 | E9PV22 | Leucine-ric | E9PV22 [5 | 1 | 4167.955 | 0 | -0.36 | 0.096033 | 17 | 112.7 | 87.3 | 11.16 | 13.9 | | High | High | High | High | High | High | High | High | High | High | High | 0 | 0.000112 | 4.41 |
| FALSE | High | STSEPEEAE | [R].STSEPE | 1xPhospho | G3X915 1x | 0.003001 | 0.00016 | 1 | 3 | 2 | G3X915 | LysM and | G3X915 [3 | 0 | 2027.974 | 0 | -1.09 | 0.039737 | 15.37 | 140.2 | 59.8 | 16.06 | 41.63 | | High | High | High | High | High | High | High | High | High | High | High | 7.19E-05 | 0.000816 | 3.81 |
| FALSE | High | STSESSASF | [R].STSESS | 1xPhospho | A2A9C3 1x | 0.019107 | 0.001402 | 1 | 2 | 2 | A2A9C3 | KICSTOR c | A2A9C3 [1 | 0 | 1464.657 | 0 | -0.2 | 0.026397 | 4.71 | 109.5 | 90.5 | 10.66 | 8.42 | | High | High | High | High | High | High | High | High | High | High | High | 0.00074 | 0.006498 | 1.54 |
| FALSE | High | STSFGVPN | [R].STSFGV | 1xPhospho | D3Z3Q3 1x | 0.009843 | 0.000736 | 1 | 7 | 2 | D3Z3Q3 | Smoothelir | D3Z3Q3 [7 | 0 | 1859.967 | 0 | -0.59 | 0.000561 | 4.04 | 116.5 | 83.5 | 7.16 | 9.53 | | High | High | High | High | High | High | High | High | High | High | High | 0.000344 | 0.003082 | 2.15 |
| FALSE | High | STSFISCDK | [K].STSFIS | 1xCarbami | Q99L00 1x | 0.00925 | 0.000736 | 1 | 2 | 1 | Q99L00 | HAUS augr | Q99L00 [1 | 0 | 1582.759 | 0 | 0.27 | 0.019025 | 8.21 | 91.3 | 108.7 | 5.1 | 10.76 | | High | High | High | High | High | High | High | High | High | High | High | 0.000327 | 0.002886 | 1.69 |
| FALSE | High | STRPECGQ | [R].STRPEC | 2xCarbami | Q9EQM6 1 | 6.3E-05 | 0 | 1 | 1 | 2 | Q9EQM6 | Microproc | Q9EQM6 [ | 0 | 4394.095 | 0 | -0.41 | 0.045798 | 6.65 | 114.3 | 85.7 | 6.97 | 15.83 | | High | High | High | High | High | High | High | High | High | High | High | 0 | 1.7E-05 | 3.41 |
| FALSE | High | STSFMSVSI | [K].STSFMS | 1xPhospho | B2ZRS3 1x | 0.018447 | 0.001324 | 1 | 6 | 1 | B2ZRS3 | Depdc6-00 | B2ZRS3 [2 | 0 | 1695.843 | 0 | 1.84 | 0.000478 | 6.5 | 42.3 | 157.7 | 6.99 | 10.27 | | High | High | High | High | High | High | High | High | High | High | High | 0.000723 | 0.006254 | 2.5 |
| FALSE | High | STSFSLSGP | [R].STSFSL | 1xPhospho | Q80U62 1x | 0.000693 | 0 | 1 | 5 | 1 | Q80U62 | Run domai | Q80U62 [2 | 0 | 2218.002 | 0 | 0.03 | 0.220074 | 5.72 | 98.3 | 101.7 | 5.33 | 10.25 | | High | High | High | High | High | High | High | High | High | High | High | 0 | 0.000158 | 3.21 |
| FALSE | High | STSFTENDI | [R].STSFTE | 1xPhospho | E9Q3E2 1x | 0.00172 | 5.66E-05 | 1 | 3 | 2 | E9Q3E2 | Synaptopo | E9Q3E2 [1 | 0 | 1679.829 | 0 | -0.58 | 0.026118 | 8.61 | 116.3 | 83.7 | 12.75 | 18.37 | | High | High | High | High | High | High | High | High | High | High | High | 2.58E-05 | 0.000439 | 2.54 |
| FALSE | High | STSGNDDS | [K].STSGNI | 1xPhospho | E9PYF7 1x | 8.84E-07 | 0 | 1 | 2 | 3 | E9PYF7 | AP1AR OS= | E9PYF7 [1 | 0 | 2912.161 | 1 | -0.23 | 0.015285 | 5.02 | 108.4 | 91.6 | 11.93 | 11.18 | | High | High | High | High | High | High | High | High | High | High | High | 0 | 8.86E-08 | 4.44 |
| FALSE | High | STSIDTGPT | [K].STSIDT | 1xPhospho | E9Q0N0 1x | 1.72E-06 | 0 | 1 | 12 | 6 | E9Q0N0 | Intersectin | E9Q0N0 [9 | 0 | 2129.078 | 0 | 0.05 | 0.474859 | 17.55 | 98.3 | 101.7 | 4.58 | 8.91 | | High | High | High | High | High | High | High | High | High | High | High | 0 | 1.87E-07 | 3.96 |
| FALSE | High | STSIDTGPT | [K].STSIDT | 2xPhospho | E9Q0N0 2x | 0.014283 | 0.001011 | 1 | 12 | 2 | E9Q0N0 | Intersectin | E9Q0N0 [9 | 0 | 2209.044 | 0 | 0.31 | 0.039817 | 14.34 | 87.8 | 112.2 | 11.78 | 6.3 | | High | High | High | High | High | High | High | High | High | High | High | 0.000538 | 0.00469 | 3.55 |
| FALSE | High | STSLPAPR | [R].STSLPA | 1xPhospho | Q80Z37 1x | 0.06169 | 0.004646 | 1 | 2 | 1 | Q80Z37 | E3 ubiquiti | Q80Z37 [5 | 0 | 1137.587 | 0 | 0.14 | 0.028615 | 5.34 | 96.1 | 103.9 | 4 | 6.01 | | High | High | High | High | High | High | High | High | High | High | High | 0.002381 | 0.02455 | 1.02 |
| FALSE | High | STSLQPEDI | [R].STSLQP | 1xPhospho | P48754 1x | 0.00204 | 5.66E-05 | 1 | 5 | 3 | P48754 | Breast can | P48754 [5 | 0 | 1573.771 | 0 | -0.75 | 0.243045 | 11.29 | 125.4 | 74.6 | 21.77 | 31.97 | | High | High | High | High | High | High | High | High | High | High | High | 2.58E-05 | 0.000531 | 2.51 |
| FALSE | High | STSLQPEDI | [R].STSLQP | 1xPhospho | P48754 1x | 0.000496 | 0 | 1 | 5 | 4 | P48754 | Breast can | P48754 [5 | 0 | 1802.934 | 0 | -0.97 | 0.000344 | 1.18 | 131.9 | 68.1 | 4.57 | 4.9 | | High | High | High | High | High | High | High | High | High | High | High | 0 | 0.000109 | 2.73 |
| FALSE | High | STSLQPEDI | [R].STSLQP | 2xPhospho | P48754 2x | 0.004824 | 0.000456 | 1 | 5 | 1 | P48754 | Breast can | P48754 [5 | 0 | 1882.9 | 0 | 0.07 | 0.41864 | 4.62 | 103.3 | 96.7 | 13.64 | 9.81 | | High | High | High | High | High | High | High | High | High | High | High | 0.000225 | 0.001396 | 2.41 |
| FALSE | High | STSLVEGR | [R].STSLVE | 1xPhospho | P22893 1x | 0.031713 | 0.002011 | 1 | 2 | 2 | P22893 | mRNA dec | P22893 [5 | 0 | 1157.576 | 0 | 0.27 | 0.015217 | 13.19 | 91.9 | 108.1 | 11.08 | 10.33 | | High | High | High | High | High | High | High | High | High | High | High | 0.001093 | 0.01152 | 1.37 |
| FALSE | High | STSMLISSA | [R].STSMLI | 1xPhospho | F8WGX5 1 | 0.0009 | 0 | 1 | 2 | 1 | F8WGX5 | DmX-like p | F8WGX5 [5 | 0 | 1802.912 | 0 | 0.39 | 0.307414 | 19.97 | 86.4 | 113.6 | 16.97 | 23.64 | | High | High | High | High | High | High | High | High | High | High | High | 0 | 0.000213 | 2.19 |
| FALSE | High | STSNGTVA | [K].STSNGT | 1xPhospho | D3YYY8 1x | 0.008094 | 0.000736 | 1 | 1 | 1 | D3YYY8 | Rho guanir | D3YYY8 [1 | 0 | 1840.937 | 0 | 0.9 | 0.004947 | 21.15 | 76.9 | 123.1 | 20.99 | 17.88 | | High | High | High | High | High | High | High | High | High | High | High | 0.000327 | 0.002471 | 3.73 |
| FALSE | High | STSPAPAD | [K].STSPAP | 1xPhospho | Q571F9 1x | 1.22E-07 | 0 | 1 | 3 | 22 | Q571F9 | MKIAA411 | Q571F9 [2 | 0 | 2033.974 | 0 | -0.19 | 0.005936 | 4.59 | 106.8 | 93.2 | 0.89 | 4.22 | | High | High | High | High | High | High | High | High | High | High | High | 0 | 6.83E-06 | 3.82 |
| FALSE | High | STSFMSVSI | [K].STSFMS | 1xPhospho | B2ZRS3 1x | 0.002553 | 0.000109 | 1 | 6 | 8 | B2ZRS3 | Depdc6-00 | B2ZRS3 [2 | 0 | 1711.838 | 0 | 1.99 | 0.000781 | 9.35 | 38.9 | 161.1 | 7.64 | 10.49 | | High | High | High | High | High | High | High | High | High | High | High | 0.001011 | 0.01082 | 2.38 |
| FALSE | High | STSPAPAD | [K].STSPAP | 2xPhospho | Q571F9 2x | 0.000731 | 0 | 1 | 3 | 1 | Q571F9 | MKIAA411 | Q571F9 [2 | 0 | 2113.941 | 0 | -0.11 | 0.110186 | 10.86 | 106 | 94 | 6.46 | 9.35 | | High | High | High | High | High | High | High | High | High | High | High | 0 | 0.000168 | 3.46 |
| FALSE | High | STQTAPSS | [R-].STQTA | 1xPhospho | Q80U93 1x | 0.00407 | 0.00036 | 1 | 3 | 2 | Q80U93 | Nuclear po | Q80U93 [6 | 0 | 1985.995 | 0 | -0.06 | 0.977321 | 3.55 | 101.7 | 98.3 | 13.08 | 3.01 | | High | High | High | High | High | High | High | High | High | High | High | 0.000585 | 0.005517 | 3.01 |
| FALSE | High | STQSENQH | [R].STQSEN | 1xPhospho | A0A0R4J0V | 1.12E-06 | 0 | 1 | 5 | 2 | A0A0R4J0V | Integrator | A0A0R4J0V | 0 | 2914.329 | 0 | -0.1 | 0.302415 | 3.13 | 103.9 | 96.1 | 10.29 | 6.04 | | High | High | High | High | High | High | High | High | High | High | High | 0 | 1.16E-07 | 3.25 |
| FALSE | High | STPDTELVT | [K].STPDTE | 1xPhospho | Q14AX6-2 | 6.45E-07 | 0 | 3 | 4 | 2 | Q14AX6-2; | Isoform 2 | Q14AX6-2 | 0 | 2462.258 | 0 | -0.37 | 0.003466 | 3.67 | 113.1 | 86.9 | 4.82 | 2.16 | Shared | High | High | High | High | High | High | High | High | High | High | High | 0 | 6.2E-08 | 3.95 |
| FALSE | High | STPDTQVA | [R].STPDTC | 1xPhospho | Q8BV49 1x | 8.22E-07 | 0 | 1 | 7 | 3 | Q8BV49 | Pyrin and H | Q8BV49 [1 | 0 | 2081.048 | 0 | 2.33 | 0.000406 | 11.41 | 33.9 | 166.1 | 12.33 | 10.29 | | High | High | High | High | High | High | High | High | High | High | High | 0 | 2.43E-07 | 4.26 |

| | | | | | | | | | | | | | | | | | | | | | | | | | | | | | | | | | | | | | | |
|---|---|---|---|---|---|---|---|---|---|---|---|---|---|---|---|---|---|---|---|---|---|---|---|---|---|---|---|---|---|---|---|---|---|---|---|---|---|---|
| FALSE | High | STPEHTWS | [R].STPEHT | 1xPhospho | P40201 1xP | 0.017809 | 0.001282 | 1 | 1 | 1 | P40201 | Chromodo | P40201 [17 | 0 | 1496.673 | 0 | 0.74 | 0.077372 | 17.64 | 75.7 | 124.3 | 20.47 | 13.53 | | High | High | High | High | High | High | High | High | High | High | High | 0.000689 | 0.005971 | 1.66 |
| FALSE | High | STPESGESD | [R].STPESG | 1xPhospho | B2RUR8 1x | 0.050397 | 0.003641 | 1 | 1 | 2 | B2RUR8 | OTU doma | B2RUR8 [4 | 0 | 1574.735 | 0 | 0.7 | 0.000442 | 0.18 | 76.8 | 123.2 | 5.89 | 2.84 | | High | High | High | High | High | High | High | High | High | High | High | 0.001916 | 0.01941 | 2.23 |
| FALSE | High | STPFIVPSSI | [R].STPFIVI | 1xPhospho | P70399 1xP | 5.33E-07 | 0 | 1 | 5 | 5 | P70399 | TP53-bindi | P70399 [37 | 0 | 1968.963 | 0 | 0.43 | 0.048205 | 16.35 | 88.1 | 111.9 | 6.52 | 11.5 | | High | High | High | High | High | High | High | High | High | High | High | 0 | 4.99E-08 | 3.5 |
| FALSE | High | STPGTPLAI | [K].STPGTP | 1xPhospho | Q8CI43 1xP | 0.0102 | 0.000736 | 1 | 1 | 1 | Q8CI43 | Myosin ligh | Q8CI43 [24 | 0 | 1409.781 | 0 | 0.08 | 0.292312 | 5.79 | 97 | 103 | 8.39 | 2.97 | | High | High | High | High | High | High | High | High | High | High | High | 0.000344 | 0.003204 | 2.58 |
| FALSE | High | STPHLDGA | [R].STPHLD | 1xPhospho | Q2WF71 1: | 3.79E-11 | 0 | 1 | 1 | 1 | Q2WF71 | Leucine-ric | Q2WF71 [7 | 0 | 2547.167 | 0 | -0.5 | 0.003619 | 8.04 | 122.5 | 77.5 | 18.7 | 23.1 | | High | High | High | High | High | High | High | High | High | High | High | 0 | 1.09E-12 | 5.94 |
| FALSE | High | STPLVPEVA | [K].STPLVP | 1xPhospho | Q61146 1x | 0.000155 | 0 | 1 | 3 | 1 | Q61146 | Occludin O | Q61146 [3 | 0 | 2735.411 | 0 | 0.61 | 0.056614 | 30.05 | 79.9 | 120.1 | 13.62 | 13.54 | | High | High | High | High | High | High | High | High | High | High | High | 0 | 2.94E-05 | 3.57 |
| FALSE | High | STPPQHPPI | [R].STPPQH | 1xPhospho | H9LAN6 1x | 0.000815 | 0 | 1 | 7 | 2 | H9LAN6 | Histone-lys | H9LAN6 [9 | 0 | 1853.947 | 0 | 0.01 | 0.949411 | 6.74 | 101.4 | 98.6 | 6.59 | 2.94 | | High | High | High | High | High | High | High | High | High | High | High | 0 | 0.000189 | 2.95 |
| FALSE | High | STPSGPVR | [R].STPSGP | 1xPhospho | Q91VM5 1 | 0.002768 | 0.000109 | 1 | 1 | 2 | Q91VM5 | RNA bindir | Q91VM5 [1 | 0 | 1109.555 | 0 | -0.42 | 0.006323 | 5.51 | 114.1 | 85.9 | 2.07 | 8.02 | | High | High | High | High | High | High | High | High | High | High | High | 0.007212 | 0.07821 | 1.92 |
| FALSE | High | STPSHGSVS | [R].STPSHG | 1xPhospho | Q60902 1x | 4.49E-10 | 0 | 1 | 5 | 6 | Q60902 | Epidermal | Q60902 [2 | 0 | 2468.243 | 0 | 0.58 | 0.1343 | 33.91 | 77.9 | 122.1 | 20.87 | 26.47 | | High | High | High | High | High | High | High | High | High | High | High | 0 | 1.77E-11 | 4.6 |
| FALSE | High | STPSHGSVS | [R].STPSHG | 2xPhospho | Q60902 2x | 7.68E-05 | 0 | 1 | 5 | 3 | Q60902 | Epidermal | Q60902 [2 | 0 | 2548.21 | 0 | 0.67 | 0.057609 | 16.91 | 78.3 | 121.7 | 14.08 | 21.2 | | High | High | High | High | High | High | High | High | High | High | High | 0.000275 | 0.00164 | 2.68 |
| FALSE | High | STPSPVPQI | [R].STPSPV | 1xPhospho | H9LAN6 1x | 0.01861 | 0.001362 | 1 | 6 | 1 | H9LAN6 | Histone-lys | H9LAN6 [8 | 0 | 1374.698 | 0 | -0.79 | 0.000805 | 2.06 | 126.9 | 73.1 | 6.34 | 2.71 | | High | High | High | High | High | High | High | High | High | High | High | 0.000723 | 0.006271 | 2.1 |
| FALSE | High | STQSENQH | [R].STQSEN | 1xPhospho | A0A0R4J0V | 7.86E-07 | 0 | 1 | 5 | 2 | A0A0R4J0V | Integrator | A0A0R4J0V | 0 | 2930.324 | 0 | -0.03 | 0.792592 | 12.45 | 101 | 99 | 7.99 | 17.93 | | High | High | High | High | High | High | High | High | High | High | High | 0 | 7.78E-08 | 2.65 |
| FALSE | High | STPSPVPQI | [R].STPSPV | 2xPhospho | H9LAN6 2x | 0.017498 | 0.001245 | 1 | 6 | 1 | H9LAN6 | Histone-lys | H9LAN6 [8 | 0 | 1454.664 | 0 | -0.01 | 0.450942 | 6 | 100.7 | 99.3 | 2.83 | 10.46 | | High | High | High | High | High | High | High | High | High | High | High | 0.000689 | 0.005885 | 2.21 |
| FALSE | High | STPSSGDV | [K].STPSSG | 1xDeamida | Q3THK3 1x | 0.00011 | 0 | 1 | 1 | 2 | Q3THK3 | General tra | Q3THK3 [4 | 0 | 1957.895 | 0 | 0.44 | 0.001008 | 3.24 | 83.8 | 116.2 | 6.78 | 5.09 | | High | High | High | High | High | High | High | High | High | High | High | 0 | 2E-05 | 4.4 |
| FALSE | High | STPSSQPSA | [K].STPSSQ | 1xPhospho | E9QLZ9 1xP | 5.13E-09 | 0 | 1 | 12 | 6 | E9QLZ9 | Protein ena | E9QLZ9 [75 | 0 | 2518.13 | 0 | -1.51 | 0.000311 | 1.25 | 147.9 | 52.1 | 4.9 | 11.49 | | High | High | High | High | High | High | High | High | High | High | High | 0 | 9.2E-07 | 5.38 |
| FALSE | High | STPSSQPSA | [K].STPSSQ | 1xDeamida | E9QLZ9 1xP | 5.52E-10 | 0 | 1 | 12 | 4 | E9QLZ9 | Protein ena | E9QLZ9 [75 | 0 | 2519.114 | 0 | -0.47 | 0.01596 | 5.55 | 113 | 87 | 9.51 | 10.94 | | High | High | High | High | High | High | High | High | High | High | High | 0 | 2.02E-10 | 4.59 |
| FALSE | High | STPSTSTTP | [K].STPSTS | 1xPhospho | B9EIG6 1xP | 0.000884 | 0 | 1 | 5 | 3 | B9EIG6 | Pogo trans | B9EIG6 [25 | 0 | 4005.981 | 0 | -1.16 | 0.00286 | 4.06 | 138.1 | 61.9 | 11.16 | 8.86 | | High | High | High | High | High | High | High | High | High | High | High | 0 | 0.000207 | 2.53 |
| FALSE | High | STPTAEDV | [K].STPTAE | 1xPhospho | A2A5Y6 1x | 0.110202 | 0.008961 | 1 | 5 | 1 | A2A5Y6 | Microtubu | A2A5Y6 [5 | 0 | 2009.963 | 0 | 1.55 | 0.000928 | 0.62 | 49.9 | 150.1 | 14.9 | 15.57 | | High | High | High | High | High | High | High | High | High | High | High | 0.004462 | 0.04839 | 1.77 |
| FALSE | High | STPTNCSPI | [R].STPTNC | 1xCarbami | E9Q286 1x | 6.77E-05 | 0 | 1 | 1 | 4 | E9Q286 | Little elong | E9Q286 [1 | 0 | 1945.934 | 0 | -1.12 | 0.000816 | 0.9 | 136.7 | 63.3 | 6.12 | 8.38 | | High | High | High | High | High | High | High | High | High | High | High | 0 | 1.16E-05 | 2.69 |
| FALSE | High | STPTNCSPI | [R].STPTNC | 1xCarbami | E9Q286 2x | 0.011447 | 0.000816 | 1 | 1 | 1 | E9Q286 | Little elong | E9Q286 [1 | 0 | 2025.901 | 0 | -1.29 | 0.000679 | 2.06 | 140.3 | 59.7 | 6.41 | 8.67 | | High | High | High | High | High | High | High | High | High | High | High | 0.000437 | 0.003643 | 2.2 |
| FALSE | High | STPVFTPGS | [R].STPVFT | 1xPhospho | P70399 1xP | 1.73E-05 | 0 | 1 | 5 | 1 | P70399 | TP53-bindi | P70399 [43 | 0 | 4532.257 | 0 | 0.1 | 0.976976 | 10.23 | 97.3 | 102.7 | 19.01 | 13.3 | | High | High | High | High | High | High | High | High | High | High | High | 0 | 2.52E-06 | 4.18 |
| FALSE | High | STQASSGSI | [R].STQASS | 1xPhospho | F8VQH0 1x | 0.002948 | 0.00016 | 1 | 5 | 2 | F8VQH0 | Tyrosine-p | F8VQH0 [6 | 0 | 1567.768 | 0 | -0.49 | 0.011023 | 1.67 | 117.6 | 82.4 | 2.74 | 14.57 | | High | High | High | High | High | High | High | High | High | High | High | 7.19E-05 | 0.000804 | 1.74 |
| FALSE | High | STQGVTLTI | [R].STQGV | 1xPhospho | A6H644 1x | 0.053473 | 0.00384 | 2 | 6 | 2 | A6H644; Q | MCG13049 | A6H644 [6 | 0 | 1928.942 | 0 | -0.56 | 0.127572 | 10.99 | 119.3 | 80.7 | 17.43 | 19.97 | Shared | High | High | High | High | High | High | High | High | High | High | High | 0.001999 | 0.02072 | 2.66 |
| FALSE | High | STQGVTLTI | [R].STQGV | 1xPhospho | A6H644 1x | 9.48E-06 | 0 | 2 | 6 | 1 | A6H644; Q | MCG13049 | A6H644 [6 | 0 | 2158.104 | 0 | -0.47 | 0.216313 | 16.68 | 111.2 | 88.8 | 27.48 | 20.59 | Shared | High | High | High | High | High | High | High | High | High | High | High | 0 | 1.27E-06 | 3.51 |
| FALSE | High | STQLLQAA | [K].STQLLC | 1xPhospho | Q3UHJ0 1x | 0.000292 | 0 | 1 | 2 | 1 | Q3UHJ0 | AP2-associ | Q3UHJ0 [6 | 0 | 2154.157 | 0 | 1.5 | 0.009741 | 25.42 | 52.7 | 147.3 | 2.62 | 24.56 | | High | High | High | High | High | High | High | High | High | High | High | 0 | 5.98E-05 | 3.85 |
| FALSE | High | STQMDVSV | [K].STQMD | 1xPhospho | Q6PR54-3 1 | 0.000105 | 0 | 2 | 4 | 1 | Q6PR54-3; | Isoform 3 c | Q6PR54-3 | 0 | 2422.11 | 0 | -0.4 | 0.127698 | 15.59 | 113.7 | 86.3 | 23.1 | 14.67 | Shared | High | High | High | High | High | High | High | High | High | High | High | 0 | 1.91E-05 | 2.94 |
| FALSE | High | STPSSGDV | [K].STPSSG | 1xPhospho | Q3THK3 1x | 4.32E-06 | 0 | 1 | 1 | 2 | Q3THK3 | General tra | Q3THK3 [4 | 0 | 1956.911 | 0 | 0.12 | 0.213721 | 10.97 | 92.1 | 107.9 | 13.91 | 13.16 | | High | High | High | High | High | High | High | High | High | High | High | 0 | 3.69E-06 | 3.09 |
| FALSE | High | STSPCQTPI | [R].STSPCC | 1xCarbami | Q8C6M1 1: | 6.35E-05 | 0 | 1 | 1 | 4 | Q8C6M1 | Ubiquitin c | Q8C6M1 [3 | 0 | 2247.99 | 0 | 0.06 | 0.318338 | 9.06 | 101 | 99 | 10.24 | 12.34 | | High | High | High | High | High | High | High | High | High | High | High | 0 | 1.08E-05 | 3.9 |
| FALSE | High | STSPHDVD | [R].STSPHD | 2xCarbami | B2RWW1 1 | 5.21E-08 | 0 | 1 | 3 | 7 | B2RWW1 | Microtubu | B2RWW1 [ | 0 | 2652.179 | 0 | -0.01 | 0.951345 | 20.24 | 99.9 | 100.1 | 9.85 | 9.09 | | High | High | High | High | High | High | High | High | High | High | High | 0 | 8.34E-09 | 4.47 |
| FALSE | High | STSPHDVD | [R].STSPHD | 2xCarbami | B2RWW1 2 | 1.09E-06 | 0 | 1 | 3 | 2 | B2RWW1 | Microtubu | B2RWW1 [ | 0 | 2732.145 | 0 | 0.87 | 0.0175 | 3.51 | 76.1 | 123.9 | 10.6 | 14.01 | | High | High | High | High | High | High | High | High | High | High | High | 0 | 1.12E-07 | 3.89 |
| FALSE | High | STTELNDPI | [R].STTELN | 1xPhospho | Q6ZQ32 1x | 0.000808 | 0 | 2 | 7 | 5 | Q6ZQ32; F | MKIAA085 | Q6ZQ32 [1 | 0 | 1897.992 | 0 | -0.52 | 0.001826 | 3.33 | 117.8 | 82.2 | 2.87 | 6.45 | Shared | High | High | High | High | High | High | High | High | High | High | High | 0 | 0.000187 | 3.17 |
| FALSE | High | STTPQCQH | [R].STTPQC | 1xCarbami | Q80WQ8 1 | 0.000146 | 0 | 1 | 2 | 2 | Q80WQ8 | Mis18-binc | Q80WQ8 [ | 0 | 2486.141 | 0 | -0.23 | 0.012069 | 5.3 | 108.5 | 91.5 | 3.54 | 6.45 | | High | High | High | High | High | High | High | High | High | High | High | 0.003346 | 0.03657 | 2.77 |
| FALSE | High | STTPTPSVI | [R].STTPTP | 1xPhospho | Q7TSH3 1x | 0.004491 | 0.000409 | 1 | 1 | 2 | Q7TSH3 | Zinc finger | Q7TSH3 [9 | 0 | 1468.761 | 0 | -0.32 | 0.003528 | 3.89 | 115.1 | 84.9 | 3.86 | 8.78 | | High | High | High | High | High | High | High | High | High | High | High | 0.000275 | 0.001886 | 2.22 |
| FALSE | High | STTPTSSPF | [R].STTPTS | 1xPhospho | Q3TLH4 1x | 0.030367 | 0.001857 | 1 | 7 | 3 | Q3TLH4 | Protein PRI | Q3TLH4 [2 | 0 | 1389.661 | 0 | 0.72 | 0.002838 | 8.54 | 75.7 | 124.3 | 6.17 | 7.64 | | High | High | High | High | High | High | High | High | High | High | High | 0.008618 | 0.09203 | 2.16 |
| FALSE | High | STTPTSSPF | [R].STTPTS | 2xPhospho | Q3TLH4 2x | 0.096181 | 0.007535 | 1 | 7 | 1 | Q3TLH4 | Protein PRI | Q3TLH4 [2 | 0 | 1469.628 | 0 | -0.12 | 0.030348 | 7.49 | 105.8 | 94.2 | 7.55 | 6.95 | | High | High | High | High | High | High | High | High | High | High | High | 0.003809 | 0.04106 | 1.53 |
| FALSE | High | STTPVK | [K].STTPVK | 1xPhospho | Q8BZX4-2 1 | 0.08447 | 0.006345 | 1 | 2 | 2 | Q8BZX4-2 | Isoform 2 c | Q8BZX4-2 | 0 | 1170.654 | 0 | -0.2 | 0.012856 | 3.24 | 107.7 | 92.3 | 2.49 | 4.64 | | High | High | High | High | High | High | High | High | High | High | High | 0.003265 | 0.03536 | 2.24 |
| FALSE | High | STTPVSK | [K].STTPVS | 1xPhospho | Q3UY41 1x | 0.081085 | 0.006219 | 1 | 4 | 2 | Q3UY41 | Putative ur | Q3UY41 [3 | 0 | 1257.686 | 0 | 1.03 | 0.00049 | 6.41 | 63.4 | 136.6 | 10.77 | 5.59 | | High | High | High | High | High | High | High | High | High | High | High | 0.0032 | 0.03384 | 1.84 |
| FALSE | High | STTSDCPSL | [R].STTSDC | 2xCarbami | Q9D8U0 1x | 0.000426 | 0 | 1 | 2 | 4 | Q9D8U0 | Interferon- | Q9D8U0 [4 | 0 | 4366.947 | 0 | 0.2 | 0.446185 | 14.09 | 93.1 | 106.9 | 20.95 | 25.5 | | High | High | High | High | High | High | High | High | High | High | High | 0 | 0.000121 | 5.03 |
| FALSE | High | STTSNNLSV | [R].STTSNN | 1xPhospho | Q3URU2 1: | 0.058664 | 0.004294 | 1 | 3 | 2 | Q3URU2 | Paternally- | Q3URU2 [9 | 0 | 1716.893 | 0 | 1.85 | 0.008931 | 36.24 | 37.7 | 162.3 | 42.2 | 23.91 | | High | High | High | High | High | High | High | High | High | High | High | 0.005664 | 0.06251 | 1.81 |
| FALSE | High | STTSSATDS | [R].STTSSA | 1xPhospho | G3X963 1x | 1.33E-09 | 0 | 1 | 2 | 8 | G3X963 | ATPase fan | G3X963 [1 | 0 | 2870.314 | 0 | 0.27 | 0.01357 | 6.06 | 90.9 | 109.1 | 8.76 | 3.91 | | High | High | High | High | High | High | High | High | High | High | High | 0 | 5.63E-10 | 5.23 |
| FALSE | High | STTTGHLIY | [K].STTTGH | 1xPhospho | Q3TII3 1xP | 0.085859 | 0.006406 | 1 | 7 | 1 | Q3TII3 | Elongation | Q3TII3 [21 | 0 | 1658.892 | 0 | -0.33 | 0.022911 | 8.35 | 111.5 | 88.5 | 7.16 | 10.96 | | High | High | High | High | High | High | High | High | High | High | High | 0.003291 | 0.03598 | 1.46 |
| FALSE | High | STTTQTEAI | [K].STTTQT | 1xPhospho | Q9D7I8 1xP | 7.74E-08 | 0 | 1 | 3 | 1 | Q9D7I8 | Protein FAI | Q9D7I8 [43 | 0 | 2664.281 | 0 | 1.08 | 0.000191 | 1.44 | 63.6 | 136.4 | 3.28 | 3.37 | | High | High | High | High | High | High | High | High | High | High | High | 0 | 5.72E-09 | 3.12 |
| FALSE | High | STTVYVGQ | [K].STTVYV | 1xPhospho | Q8R4E9 1x | 0.009843 | 0.000736 | 1 | 1 | 1 | Q8R4E9 | DNA replic | Q8R4E9 [1 | 0 | 1802.945 | 0 | 0.75 | 0.028665 | 29.54 | 74.7 | 125.3 | 18.16 | 17.87 | | High | High | High | High | High | High | High | High | High | High | High | 0.000344 | 0.003085 | 1.74 |
| FALSE | High | STTELNDPI | [R].STTELN | 1xPhospho | Q6ZQ32 1x | 0.001021 | 0 | 2 | 7 | 10 | Q6ZQ32; F | MKIAA085 | Q6ZQ32 [1 | 0 | 1668.829 | 0 | 1.26 | 0.000671 | 1.47 | 59.3 | 140.7 | 3.92 | 7.95 | Shared | High | High | High | High | High | High | High | High | High | High | High | 0 | 0.000244 | 3.06 |
| FALSE | High | STVCINGNI | [R].STVCIN | 1xCarbami | E9Q8L9 1xP | 0.00407 | 0.00036 | 1 | 3 | 1 | E9Q8L9 | Rab11 fam | E9Q8L9 [3 | 0 | 2430.078 | 0 | -0.32 | 0.051651 | 11.77 | 110.9 | 89.1 | 9.03 | 15.2 | | High | High | High | High | High | High | High | High | High | High | High | 0.000161 | 0.001149 | 2.56 |
| FALSE | High | STVEDEDSI | [K].STVEDE | 1xPhospho | F8WI37 1xP | 0.045508 | 0.003373 | 1 | 5 | 2 | F8WI37 | Histone-lys | F8WI37 [6 | 0 | 3468.575 | 0 | -0.22 | 0.17885 | 11.6 | 107.4 | 92.6 | 19.05 | 8.88 | | High | High | High | High | High | High | High | High | High | High | High | 0.001741 | 0.01735 | 3.01 |
| FALSE | High | STVLGLPQI | [R].STVLGL | 1xPhospho | A2A7S8-2 1 | 0.001158 | 0 | 1 | 4 | 1 | A2A7S8-2 | Isoform 2 c | A2A7S8-2 [ | 0 | 1845.04 | 0 | -0.04 | 0.297556 | 3.82 | 102 | 98 | 9.64 | 3.02 | | High | High | High | High | High | High | High | High | High | High | High | 0 | 0.000281 | 1.64 |
| FALSE | High | STVTLAGD | [K].STVTLA | 1xCarbami | A7TU71 1x | 2.3E-09 | 0 | 1 | 2 | 2 | A7TU71 | Protein Shr | A7TU71 [6 | 0 | 2348.99 | 0 | 1.76 | 0.00032 | 16.92 | 42.7 | 157.3 | 18.82 | 5.98 | | High | High | High | High | High | High | High | High | High | High | High | 0 | 1.11E-10 | 4.69 |
| FALSE | High | STVTLAGD | [K].STVTLA | 1xCarbami | A7TU71 2x | 1.14E-06 | 0 | 1 | 2 | 1 | A7TU71 | Protein Shr | A7TU71 [6 | 0 | 2428.957 | 0 | 0.48 | 0.032198 | 27.29 | 78.1 | 121.9 | 10.67 | 16.76 | | High | High | High | High | High | High | High | High | High | High | High | 0 | 1.18E-07 | 3.77 |
| FALSE | High | STYQDKPST | [R].STYQDI | 1xPhospho | P51125-3 1 | 0.044739 | 0.003302 | 1 | 12 | 1 | P51125-3 | Isoform 3 c | P51125-3 [ | 0 | 2219.156 | 0 | 1.94 | 0.000806 | 17.71 | 46 | 154 | 12.8 | 10.27 | | High | High | High | High | High | High | High | High | High | High | High | 0.001712 | 0.01702 | 1.97 |
| FALSE | High | SVALSPTPT | [K].SVALSP | 1xPhospho | Q9Z2U2 1x | 0.110202 | 0.008961 | 1 | 3 | 1 | Q9Z2U2 | Zinc finger | Q9Z2U2 [1 | 0 | 1538.86 | 0 | -0.66 | 0.002614 | 0.95 | 122.6 | 77.4 | 7.36 | 4.87 | | High | High | High | High | High | High | High | High | High | High | High | 0.00449 | 0.04864 | 1.95 |
| FALSE | High | SVASNQSE | [R].SVASNC | 1xPhospho | Q8VDP4 1x | 0.001267 | 0 | 1 | 1 | 4 | Q8VDP4 | Cell cycle a | Q8VDP4 [6 | 0 | 2669.224 | 0 | 0.19 | 0.127601 | 16.86 | 94.1 | 105.9 | 10.32 | 26.1 | | High | High | High | High | High | High | High | High | High | High | High | 0 | 0.00031 | 3.81 |
| FALSE | High | SVASNQSE | [R].SVASNC | 1xPhospho | Q8VDP4 1x | 2.39E-07 | 0 | 1 | 1 | 6 | Q8VDP4 | Cell cycle a | Q8VDP4 [6 | 0 | 2685.219 | 0 | 0.37 | 0.052661 | 23.4 | 83.4 | 116.6 | 7.1 | 15.38 | | High | High | High | High | High | High | High | High | High | High | High | 0 | 2.03E-08 | 4.03 |
| FALSE | High | SVASNQSE | [R].SVASNC | 2xPhospho | Q8VDP4 2x | 0.000289 | 0 | 1 | 1 | 1 | Q8VDP4 | Cell cycle a | Q8VDP4 [6 | 0 | 2749.19 | 0 | 0.47 | 0.118736 | 10.45 | 83.1 | 116.9 | 6.02 | 17.74 | | High | High | High | High | High | High | High | High | High | High | High | 0 | 5.93E-05 | 4.35 |
| FALSE | High | SVCPGTSG | [K].SVCPGT | 1xCarbami | A2A432 1x | 3.39E-10 | 0 | 1 | 2 | 6 | A2A432 | Cullin-4B C | A2A432 [7 | 0 | 2744.255 | 0 | -0.84 | 0.001594 | 1.92 | 126.2 | 73.8 | 6.82 | 4.26 | | High | High | High | High | High | High | High | High | High | High | High | 0 | 1.29E-11 | 4.81 |
| FALSE | High | SVDEVNYW | [R].SVDEVI | 1xPhospho | Q3U3J1 1x | 0.004867 | 0.000504 | 1 | 3 | 1 | Q3U3J1 | 2-oxoisova | Q3U3J1 [3 | 0 | 1563.693 | 0 | -1.48 | 0.00032 | 1.24 | 146.5 | 53.5 | 5.81 | 2.47 | | High | High | High | High | High | High | High | High | High | High | High | 0.000225 | 0.001398 | 3.3 |
| FALSE | High | SVDISLGGS | [R].SVDISL | 1xPhospho | Q3UIL6-2 1 | 6.77E-05 | 0 | 1 | 8 | 2 | Q3UIL6-2 | Isoform 2 c | Q3UIL6-2 [ | 0 | 1396.703 | 0 | -1.83 | 0.000231 | 0.36 | 155.9 | 44.1 | 1.38 | 14.29 | | High | High | High | High | High | High | High | High | High | High | High | 0 | 1.16E-05 | 1.82 |
| FALSE | High | SVDISLGGS | [R].SVDISL | 2xPhospho | Q3UIL6-2 2 | 0.019446 | 0.001438 | 1 | 8 | 1 | Q3UIL6-2 | Isoform 2 c | Q3UIL6-2 [ | 0 | 1476.67 | 0 | -1.84 | 0.00037 | 1.34 | 156.4 | 43.6 | 6.41 | 20.57 | | High | High | High | High | High | High | High | High | High | High | High | 0.000756 | 0.006612 | 2.23 |
| FALSE | High | STVDNTGL | [R].STVDNT | 1xPhospho | Q5SSZ5 1xP | 0.000205 | 0 | 1 | 1 | 1 | Q5SSZ5 | Tensin-3 O | Q5SSZ5 [6 | 0 | 2520.281 | 0 | -1.47 | 0.000707 | 2.12 | 146.5 | 53.5 | 4.61 | 14.38 | | High | High | High | High | High | High | High | High | High | High | High | 0 | 4.01E-05 | 2.77 |
| FALSE | High | STTDSPPW | [K].STTDSP | 1xPhospho | Q6VGS5-2 | 2.48E-08 | 0 | 1 | 4 | 2 | Q6VGS5-2 | Isoform 2 c | Q6VGS5-2 | 0 | 2960.388 | 0 | 0.05 | 0.692969 | 28.01 | 100.2 | 99.8 | 12.07 | 17.8 | | High | High | High | High | High | High | High | High | High | High | High | 0 | 1.6E-09 | 4.78 |
| FALSE | High | STTATGQT | [K].STTATG | 1xCarbami | Q9CQN9 1: | 1.08E-05 | 0 | 1 | 2 | 2 | Q9CQN9 | Putative ur | Q9CQN9 [2 | 0 | 2787.291 | 0 | -0.03 | 0.413882 | 27.19 | 101.1 | 98.9 | 17.05 | 11.62 | | High | High | High | High | High | High | High | High | High | High | High | 0 | 4.38E-05 | 2.73 |
| FALSE | High | STTAHSWC | [R].STTAHS | 2xPhospho | A2A6A1 2x | 0.104222 | 0.008348 | 1 | 1 | 1 | A2A6A1 | G patch do | A2A6A1 [9 | 0 | 1462.608 | 0 | 0.13 | 0.520777 | 20.68 | 96.9 | 103.1 | 7.77 | 8.96 | | High | High | High | High | High | High | High | High | High | High | High | 0.004193 | 0.04511 | 1.74 |
| FALSE | High | STSPIIGSPF | [R].STSPIIG | 1xPhospho | Q3TC46 1x | 1.31E-05 | 0 | 1 | 1 | 3 | Q3TC46 | Protein PA | Q3TC46 [1 | 0 | 1519.808 | 0 | 0.01 | 0.400752 | 2.53 | 99.2 | 100.8 | 4.22 | 6.55 | | High | High | High | High | High | High | High | High | High | High | High | 0 | 1.84E-06 | 2.16 |
| FALSE | High | STSPIIGSPF | [R].STSPIIG | 2xPhospho | Q3TC46 2x | 0.066503 | 0.004796 | 1 | 1 | 2 | Q3TC46 | Protein PA | Q3TC46 [1 | 0 | 1599.775 | 0 | -0.54 | 0.131189 | 17.26 | 114.9 | 85.1 | 18.99 | 19.3 | | High | High | High | High | High | High | High | High | High | High | High | 0.002455 | 0.02682 | 1.93 |
| FALSE | High | STSPTSDSI | [R].STSPTS | 1xPhospho | Q499D0 1x | 1.02E-05 | 0 | 1 | 1 | 7 | Q499D0 | Forkhead b | Q499D0 [3 | 0 | 3132.393 | 0 | -1.1 | 0.003164 | 3.85 | 136.7 | 63.3 | 9.69 | 11.73 | | High | High | High | High | High | High | High | High | High | High | High | 0 | 1.38E-06 | 3.6 |
| FALSE | High | STSPVTDPS | [R].STSPVT | 1xPhospho | Q32MF0 1: | 0.022773 | 0.001542 | 1 | 4 | 2 | Q32MF0 | Predicted g | Q32MF0 [1 | 0 | 1696.818 | 0 | 1.16 | 0.006653 | 15.79 | 61.9 | 138.1 | 10.11 | 12.84 | | High | High | High | High | High | High | High | High | High | High | High | 0.000814 | 0.007873 | 2.14 |
| FALSE | High | STSPVTDPS | [R].STSPVT | 1xPhospho | Q32MF0 1: | 0.001586 | 5.66E-05 | 1 | 4 | 1 | Q32MF0 | Predicted g | Q32MF0 [1 | 0 | 1712.813 | 0 | 1.84 | 0.00041 | 9.64 | 41.7 | 158.3 | 7.99 | 8.38 | | High | High | High | High | High | High | High | High | High | High | High | 2.58E-05 | 0.000401 | 2.32 |
| FALSE | High | STSQGSINS | [R].STSQGS | 1xPhospho | E9QK41 1x | 2.06E-06 | 0 | 1 | 18 | 2 | E9QK41 | Actin-bindi | E9QK41 [4 | 0 | 1791.848 | 0 | 2.15 | 0.000167 | 5.13 | 35.9 | 164.1 | 7.16 | 3.27 | | High | High | High | High | High | High | High | High | High | High | High | 0 | 4.57E-07 | 2.89 |
| FALSE | High | STSSAMGG | [K].STSSAM | 1xPhospho | A0A1W2P7 | 0.000221 | 0 | 1 | 2 | 3 | A0A1W2P7 | RAB3A inte | A0A1W2P7 | 0 | 2696.373 | 0 | -0.54 | 0.185198 | 3.17 | 118.4 | 81.6 | 9.21 | 27.75 | | High | High | High | High | High | High | High | High | High | High | High | 0 | 4.37E-05 | 2.99 |
| FALSE | High | STSSANPGI | [R].STSSAN | 1xPhospho | Q8BU22 1x | 0.000984 | 0 | 1 | 4 | 2 | Q8BU22 | Kinesin-like | Q8BU22 [1 | 0 | 1485.69 | 0 | -0.07 | 0.425076 | 13.66 | 103.1 | 96.9 | 12.17 | 10.67 | | High | High | High | High | High | High | High | High | High | High | High | 0.000656 | 0.005834 | 1.94 |
| FALSE | High | STSSCSSK | [R].STSSCS | 1xCarbami | D3YTT6 1xP | 0.007209 | 0.00062 | 1 | 8 | 2 | D3YTT6 | Oxysterol-l | D3YTT6 [1 | 0 | 1381.643 | 0 | 2.01 | 0.00024 | 0.49 | 37.5 | 162.5 | 5.41 | 5.37 | | High | High | High | High | High | High | High | High | High | High | High | 0.000275 | 0.002167 | 1.35 |
| FALSE | High | STSSEEAVA | [R].STSSEE | 1xPhospho | A0A0U1RN | 0.000104 | 0 | 1 | 5 | 6 | A0A0U1RN | Unconvent | A0A0U1RN | 0 | 3211.504 | 0 | 0.24 | 0.02906 | 12.61 | 91.8 | 108.2 | 7.55 | 4.14 | | High | High | High | High | High | High | High | High | High | High | High | 2.58E-05 | 0.000563 | 3.62 |
| FALSE | High | STSSEEAVA | [R].STSSEE | 2xPhospho | A0A0U1RN | 0.011346 | 0.000816 | 1 | 5 | 1 | A0A0U1RN | Unconvent | A0A0U1RN | 0 | 3291.471 | 0 | -0.32 | 0.182026 | 40.92 | 117.7 | 82.3 | 15.41 | 34 | | High | High | High | High | High | High | High | High | High | High | High | 0.000418 | 0.00363 | 3.25 |
| FALSE | High | STSSGETSS | [R].STSSGE | 1xPhospho | Q3TRR0 1x | 0.008769 | 0.000736 | 1 | 5 | 2 | Q3TRR0 | Microtubu | Q3TRR0 [1 | 0 | 2983.42 | 0 | -1.08 | 0.0007 | 1.58 | 137.9 | 62.1 | 12.16 | 12.41 | | High | High | High | High | High | High | High | High | High | High | High | 0.000327 | 0.002719 | 2.48 |
| FALSE | High | STSSIDQLG | [K].STSSID | 1xPhospho | B7ZNA0 1x | 0.014929 | 0.001049 | 2 | 9 | 3 | B7ZNA0; Q | Shroom3 p | B7ZNA0 [2 | 0 | 2075.069 | 0 | -0.04 | 0.245938 | 9.52 | 104.9 | 95.1 | 11.89 | 9.78 | Shared | High | High | High | High | High | High | High | High | High | High | High | 0.001214 | 0.01295 | 2.98 |
| FALSE | High | STSSIPPSPL | [K].STSSIPF | 1xCarbami | E9Q8S5 1x | 0.005322 | 0.000599 | 1 | 2 | 3 | E9Q8S5 | Microtubu | E9Q8S5 [1 | 0 | 2464.251 | 0 | -0.96 | 0.011316 | 3.85 | 131 | 69 | 8.34 | 20.03 | | High | High | High | High | High | High | High | High | High | High | High | 0.000327 | 0.002384 | 2.65 |
| FALSE | High | STSSSPAPT | [K].STSSSP | 1xPhospho | Q3TAB2 1x | 9.85E-07 | 0 | 2 | 6 | 20 | Q3TAB2; H | Putative ur | Q3TAB2 [4 | 0 | 2282.123 | 0 | -0.14 | 0.022408 | 0.52 | 104.2 | 95.8 | 4.4 | 4.37 | Shared | High | High | High | High | High | High | High | High | High | High | High | 0 | 1.41E-05 | 4.87 |
| FALSE | High | STSSSPAPT | [K].STSSSP | 1xDeamida | Q3TAB2 1x | 0.001476 | 5.66E-05 | 2 | 6 | 16 | Q3TAB2; H | Putative ur | Q3TAB2 [4 | 0 | 2283.107 | 0 | -0.25 | 0.013446 | 3.73 | 107.9 | 92.1 | 7.19 | 7.58 | Shared | High | High | High | High | High | High | High | High | High | High | High | 0.000585 | 0.005146 | 3.6 |
| FALSE | High | STSSSPAPT | [K].STSSSP | 2xPhospho | Q3TAB2 2x | 3.83E-05 | 0 | 2 | 6 | 1 | Q3TAB2; H | Putative ur | Q3TAB2 [4 | 0 | 2362.089 | 0 | 0.28 | 0.01039 | 9.85 | 90.2 | 109.8 | 12.22 | 6.5 | Shared | High | High | High | High | High | High | High | High | High | High | High | 0 | 6.1E-06 | 3.18 |
| FALSE | High | STSSTPGM | [R].STSSTP | 1xPhospho | Q8C8U0-3 | 0.003997 | 0.000311 | 1 | 5 | 3 | Q8C8U0-3 | Isoform 3 c | Q8C8U0-3 | 0 | 1576.688 | 0 | 0.99 | 0.009682 | 15.28 | 67.3 | 132.7 | 7.89 | 13.61 | | High | High | High | High | High | High | High | High | High | High | High | 0.006342 | 0.06814 | 2.18 |
| FALSE | High | STSSTPVK | [R].STSSTP | 1xPhospho | Q8CID0 1xP | 0.001751 | 5.66E-05 | 1 | 2 | 2 | Q8CID0 | Cysteine-ri | Q8CID0 [2 | 0 | 1344.718 | 0 | -0.51 | 0.004056 | 4.46 | 116.1 | 83.9 | 9.86 | 6.07 | | High | High | High | High | High | High | High | High | High | High | High | 2.58E-05 | 0.000449 | 1.69 |
| FALSE | High | STSSTSSGS | [R].STSSTS | 1xPhospho | F8VQ05 1x | 1.42E-05 | 0 | 1 | 5 | 6 | F8VQ05 | FRY-like tra | F8VQ05 [2 | 0 | 2521.259 | 0 | 1.33 | 0.000924 | 19.08 | 56.9 | 143.1 | 9.63 | 8.41 | | High | High | High | High | High | High | High | High | High | High | High | 0 | 2.54E-06 | 3.96 |
| FALSE | High | STSTLYSQF | [R].STSTLY | 1xPhospho | Q6ZPE2-2 1 | 0.000558 | 0 | 1 | 3 | 5 | Q6ZPE2-2 | Isoform 2 c | Q6ZPE2-2 [ | 0 | 2258.018 | 0 | 0.67 | 0.023638 | 4.13 | 79.7 | 120.3 | 7.03 | 12.22 | | High | High | High | High | High | High | High | High | High | High | High | 0.000275 | 0.002186 | 2.76 |
| FALSE | High | STSTPNVHI | [R].STSTPN | 1xPhospho | Q99N57-2 | 0.002419 | 0.000109 | 1 | 3 | 4 | Q99N57-2 | Isoform 2 c | Q99N57-2 | 0 | 2378.137 | 0 | -0.5 | 0.167091 | 3.95 | 111.4 | 88.6 | 15.14 | 14.94 | | High | High | High | High | High | High | High | High | High | High | High | 0.00014 | 0.001111 | 2.67 |
| FALSE | High | STSTPNVHI | [R].STSTPN | 1xPhospho | Q99N57-2 | 0.0004 | 0 | 1 | 3 | 3 | Q99N57-2 | Isoform 2 c | Q99N57-2 | 0 | 2394.132 | 0 | -0.32 | 0.423819 | 5.18 | 111.9 | 88.1 | 13.03 | 19.02 | | High | High | High | High | High | High | High | High | High | High | High | 0.002354 | 0.02424 | 2.88 |
| FALSE | High | STSTPTSPG | [K].STSTPT | 1xPhospho | A0A0U1RP | 0.049549 | 0.00361 | 1 | 7 | 5 | A0A0U1RP | Ataxin-2-lik | A0A0U1RP | 0 | 1396.667 | 0 | 0.78 | 0.000704 | 7.91 | 73.3 | 126.7 | 6.07 | 6.15 | | High | High | High | High | High | High | High | High | High | High | High | 0.005455 | 0.05983 | 1.57 |
| FALSE | High | STSTPTSPG | [K].STSTPT | 2xPhospho | A0A0U1RP | 0.000745 | 0 | 1 | 7 | 2 | A0A0U1RP | Ataxin-2-lik | A0A0U1RP | 0 | 1476.633 | 0 | 1.5 | 0.000669 | 5.04 | 52.3 | 147.7 | 4.61 | 8.14 | | High | High | High | High | High | High | High | High | High | High | High | 0 | 0.000171 | 2.49 |
| FALSE | High | STSVPAPYI | [R].STSVPA | 1xPhospho | E9PUJ6 1xP | 3.18E-06 | 0 | 1 | 5 | 4 | E9PUJ6 | E3 ubiquiti | E9PUJ6 [34 | 0 | 3547.696 | 0 | 1.67 | 0.002152 | 3.1 | 48.7 | 151.3 | 16.58 | 11.8 | | High | High | High | High | High | High | High | High | High | High | High | 0 | 6.74E-07 | 5.59 |
| FALSE | High | STSYGYSR | [R].STSYGY | 1xPhospho | Q3TXM9 1: | 0.001147 | 0 | 1 | 3 | 4 | Q3TXM9 | Putative ur | Q3TXM9 [1 | 0 | 1229.54 | 0 | 0.06 | 0.357413 | 7.2 | 96.3 | 103.7 | 5.66 | 7.12 | | High | High | High | High | High | High | High | High | High | High | High | 0.000903 | 0.009235 | 1.75 |
| FALSE | High | STSYGYSR | [R].STSYGY | 2xPhospho | Q3TXM9 2: | 0.08447 | 0.006345 | 1 | 3 | 1 | Q3TXM9 | Putative ur | Q3TXM9 [1 | 0 | 1309.506 | 0 | 0.06 | 0.948003 | 19.61 | 97 | 103 | 2.19 | 16.36 | | High | High | High | High | High | High | High | High | High | High | High | 0.003265 | 0.03534 | 1.98 |
| FALSE | High | STTAHSWC | [R].STTAHS | 1xPhospho | A2A6A1 1x | 0.014283 | 0.001011 | 1 | 1 | 2 | A2A6A1 | G patch do | A2A6A1 [9 | 0 | 1382.642 | 0 | -0.18 | 0.217699 | 23.93 | 106.3 | 93.7 | 4.98 | 21.45 | | High | High | High | High | High | High | High | High | High | High | High | 0.000538 | 0.004685 | 1.95 |
| FALSE | High | STPASPEPG | [K].STPASP | 1xCarbami | Q6VGS5-2 | 2.7E-06 | 0 | 1 | 4 | 2 | Q6VGS5-2 | Isoform 2 c | Q6VGS5-2 | 0 | 2706.163 | 0 | -0.91 | 0.005136 | 6.72 | 126.7 | 73.3 | 6.51 | 17.34 | | High | High | High | High | High | High | High | High | High | High | High | 0 | 9.86E-07 | 4.79 |
| FALSE | High | STNSVFGG | [K].STNSVF | 1xPhospho | B2RT41 1xP | 0.000255 | 0 | 1 | 2 | 1 | B2RT41 | Zinc finger, | B2RT41 [72 | 0 | 2024.018 | 0 | -0.14 | 0.209518 | 17.27 | 105.2 | 94.8 | 14.51 | 16.26 | | High | High | High | High | High | High | High | High | High | High | High | 0 | 5.11E-05 | 2.15 |
| FALSE | High | STNSVFGG | [K].STNSVF | 1xPhospho | B2RT41 1xP | 0.011549 | 0.00086 | 1 | 2 | 1 | B2RT41 | Zinc finger, | B2RT41 [72 | 0 | 2008.023 | 0 | -1.24 | 0.006694 | 6.26 | 145.5 | 54.5 | 16.26 | 29.49 | | High | High | High | High | High | High | High | High | High | High | High | 0.000437 | 0.003693 | 2.79 |
| FALSE | High | STNAAASR | [R].STNAA | 1xPhospho | Q3UL36 1x | 0.053926 | 0.00384 | 1 | 2 | 2 | Q3UL36 | Arginine ar | Q3UL36 [6 | 0 | 1086.514 | 0 | -0.26 | 0.172888 | 19.61 | 109.1 | 90.9 | 6.92 | 13.78 | | High | High | High | High | High | High | High | High | High | High | High | 0.002045 | 0.0211 | 1.47 |
| FALSE | High | SSTSSEPTP | [R].SSTSSE | 1xPhospho | A0A0R4J0J | 0.011246 | 0.000772 | 1 | 3 | 4 | A0A0R4J0J | Cytospin-A | A0A0R4J0J | 0 | 1758.893 | 0 | 1.62 | 0.00018 | 5.97 | 47.8 | 152.2 | 8.42 | 4.17 | | High | High | High | High | High | High | High | High | High | High | High | 0.000399 | 0.003559 | 2.92 |
| FALSE | High | SSTSVVSEV | [R].SSTSVV | 1xPhospho | E9QP99 1x | 1.71E-06 | 0 | 1 | 5 | 5 | E9QP99 | Golgin sub | E9QP99 [2 | 0 | 2176.981 | 0 | 1.27 | 0.001603 | 13.23 | 59.4 | 140.6 | 14.18 | 13.63 | | High | High | High | High | High | High | High | High | High | High | High | 0 | 1.85E-07 | 4.31 |
| FALSE | High | SSTTTTPVC | [K].SSTTTT | 1xPhospho | Q3U0T7 1x | 0.007404 | 0.00062 | 1 | 7 | 4 | Q3U0T7 | Putative ur | Q3U0T7 [1 | 0 | 2115.11 | 0 | 0.5 | 0.114034 | 27.74 | 78.1 | 121.9 | 21.51 | 28.47 | | High | High | High | High | High | High | High | High | High | High | High | 0.000275 | 0.002246 | 2.15 |
| FALSE | High | SSTVASLDT | [R].SSTVAS | 1xPhospho | O35671 1x | 0.020142 | 0.001472 | 1 | 2 | 3 | O35671 | Integrin be | O35671 [3 | 0 | 1849.92 | 0 | 0.35 | 0.015871 | 5.43 | 86.8 | 113.2 | 7.74 | 7.6 | | High | High | High | High | High | High | High | High | High | High | High | 0.000786 | 0.006906 | 2.63 |
| FALSE | High | SSTVTEAPI | [R].SSTVTE | 1xPhospho | Q69Z43 1x | 0.00086 | 0 | 1 | 2 | 3 | Q69Z43 | MKIAA199 | Q69Z43 [3 | 0 | 1826.946 | 0 | 0.99 | 0.002055 | 15.96 | 62.7 | 137.3 | 14.05 | 12.37 | | High | High | High | High | High | High | High | High | High | High | High | 4.94E-05 | 0.000658 | 2.51 |
| FALSE | High | SSVASSAVA | [K].SSVASS | 1xCarbami | Q6NZQ4 1: | 5.27E-06 | 0 | 1 | 1 | 1 | Q6NZQ4 | PAX-intera | Q6NZQ4 [2 | 0 | 2651.326 | 0 | -1.25 | 0.019272 | 1.01 | 136.4 | 63.6 | 25.24 | 13.01 | | High | High | High | High | High | High | High | High | High | High | High | 0 | 6.6E-07 | 4.33 |
| FALSE | High | SSVDLSQVS | [K].SSVDLS | 1xPhospho | A0A0G2JG | 4.68E-05 | 0 | 1 | 4 | 1 | A0A0G2JG | Suppresso | A0A0G2JG | 0 | 2714.327 | 0 | -0.87 | 0.02272 | 10.09 | 131 | 69 | 19.4 | 15.17 | | High | High | High | High | High | High | High | High | High | High | High | 0 | 7.64E-06 | 3.95 |
| FALSE | High | SSVDSQGP | [K].SSVDSC | 1xCarbami | Q91VE6 1x | 8.06E-06 | 0 | 1 | 2 | 7 | Q91VE6 | MKI67 FHA | Q91VE6 [2 | 0 | 2387.115 | 0 | -0.55 | 0.072065 | 2.65 | 116.6 | 83.4 | 6.15 | 18.3 | | High | High | High | High | High | High | High | High | High | High | High | 0 | 7.95E-06 | 4.63 |
| FALSE | High | SSVDSQGP | [K].SSVDSC | 1xCarbami | Q91VE6 2x | 0.000264 | 0 | 1 | 2 | 1 | Q91VE6 | MKI67 FHA | Q91VE6 [2 | 0 | 2467.081 | 0 | -0.57 | 0.00135 | 1.34 | 119.4 | 80.6 | 5.92 | 4.97 | | High | High | High | High | High | High | High | High | High | High | High | 0 | 5.37E-05 | 2.77 |
| FALSE | High | SSVDSQGP | [K].SSVDSC | 1xCarbami | Q91VE6 3x | 0.000132 | 0 | 1 | 2 | 1 | Q91VE6 | MKI67 FHA | Q91VE6 [2 | 0 | 2547.048 | 0 | -0.43 | 0.034292 | 12.1 | 115 | 85 | 11.29 | 6.68 | | High | High | High | High | High | High | High | High | High | High | High | 0 | 2.45E-05 | 3.99 |
| FALSE | High | SSVEGLEPA | [K].SSVEGL | 1xPhospho | P16254 1xP | 3.47E-05 | 0 | 1 | 2 | 3 | P16254 | Signal reco | P16254 [44 | 0 | 1797.904 | 0 | 1.46 | 0.000331 | 3.22 | 53.2 | 146.8 | 6.82 | 5.79 | | High | High | High | High | High | High | High | High | High | High | High | 0 | 8.5E-06 | 3.84 |
| FALSE | High | SSVNGEAG | [R].SSVNGI | 1xPhospho | E9QP49 1x | 0.009585 | 0.000736 | 1 | 7 | 1 | E9QP49 | EH domain | E9QP49 [1 | 0 | 1672.826 | 0 | 0.29 | 0.01436 | 1.75 | 90.1 | 109.9 | 3 | 7.09 | | High | High | High | High | High | High | High | High | High | High | High | 0.000344 | 0.002979 | 2.61 |
| FALSE | High | SSVNGEAG | [R].SSVNGI | 1xDeamida | E9QP49 1x | 0.050397 | 0.003641 | 1 | 7 | 3 | E9QP49 | EH domain | E9QP49 [1 | 0 | 1673.81 | 0 | 1.29 | 0.001475 | 20.09 | 62.2 | 137.8 | 13.67 | 9.87 | | High | High | High | High | High | High | High | High | High | High | High | 0.004451 | 0.04798 | 2.35 |
| FALSE | High | SSTSQTVST | [R].SSTSQT | 2xPhospho | F6ZDS4 2xP | 0.053926 | 0.00384 | 1 | 7 | 2 | F6ZDS4 | Nucleopro | F6ZDS4 [71 | 0 | 3163.51 | 0 | -0.11 | 0.2962 | 7.75 | 106 | 94 | 17.39 | 14.51 | | High | High | High | High | High | High | High | High | High | High | High | 0.002031 | 0.02095 | 2.97 |
| FALSE | High | SSVPETGLI | [K].SSVPET | 1xCarbami | Q61089 1x | 0.036731 | 0.00249 | 1 | 3 | 2 | Q61089 | Frizzled-6 ( | Q61089 [6 | 0 | 3188.532 | 0 | 0.6 | 0.015925 | 17.68 | 81.2 | 118.8 | 35.24 | 19.53 | | High | High | High | High | High | High | High | High | High | High | High | 0.001308 | 0.01356 | 2.96 |

| | | | | | | | | | | | | | | | | | | | | | | | | | | | | | | | | | | | | | | |
|---|---|---|---|---|---|---|---|---|---|---|---|---|---|---|---|---|---|---|---|---|---|---|---|---|---|---|---|---|---|---|---|---|---|---|---|---|---|---|
| FALSE | High | SSVPSGGSI | [R].SSVPSG | 1xPhospho | Q6PGL7 1x | 4.33E-07 | 0 | 1 | 4 | 3 | Q6PGL7 | WASH com | Q6PGL7 [8 | 0 | 2870.302 | 0 | -0.28 | 0.273239 | 15.43 | 100.9 | 99.1 | 15.04 | 11.43 | | High | High | High | High | High | High | High | High | High | High | High | 0 | 8.37E-08 | 3.77 |
| FALSE | High | SSVQDFIC | [R].SSVQDI | 1xCarbami | P59729 1xI | 0.012619 | 0.000889 | 1 | 6 | 1 | P59729 | Ras and Ra | P59729 [87 | 0 | 2809.4 | 0 | 3.11 | 0.000961 | 5.94 | 20.9 | 179.1 | 8.63 | 13.88 | | High | High | High | High | High | High | High | High | High | High | High | 0.000449 | 0.004062 | 2.96 |
| FALSE | High | SSVSGISLEI | [R].SSVSGI | 1xPhospho | P70271 1xI | 0.003962 | 0.000311 | 1 | 3 | 1 | P70271 | PDZ and LII | P70271 [11 | 0 | 1572.747 | 0 | 0.43 | 0.01048 | 14.96 | 81.7 | 118.3 | 10.14 | 8.14 | | High | High | High | High | High | High | High | High | High | High | High | 0.00014 | 0.00111 | 2.7 |
| FALSE | High | SSVSPSPAA | [K].SSVSPS | 1xPhospho | Q80U93 1x | 0.00145 | 5.66E-05 | 1 | 2 | 2 | Q80U93 | Nuclear po | Q80U93 [6 | 0 | 1324.646 | 0 | -0.31 | 0.004603 | 2.45 | 110.1 | 89.9 | 3.86 | 6.25 | | High | High | High | High | High | High | High | High | High | High | High | 2.58E-05 | 0.00036 | 1.53 |
| FALSE | High | SSVSSEIK | [K].SSVSSE | 1xPhospho | F8WI37 1xI | 0.001644 | 5.66E-05 | 1 | 2 | 1 | F8WI37 | Histone-lys | F8WI37 [28 | 0 | 1374.728 | 0 | -0.31 | 0.003482 | 5.24 | 109.4 | 90.6 | 3.57 | 8.12 | | High | High | High | High | High | High | High | High | High | High | High | 2.58E-05 | 0.000417 | 2.34 |
| FALSE | High | SSVSSGSW | [K].SSVSSG | 1xPhospho | A0A140T8S | 0.026652 | 0.001676 | 1 | 1 | 2 | A0A140T8S | Chromosoı | A0A140T8S | 1 | 2802.432 | 0 | -0.08 | 0.274781 | 4.71 | 102.5 | 97.5 | 1.81 | 16.67 | | High | High | High | High | High | High | High | High | High | High | High | 0.000903 | 0.009453 | 3.26 |
| FALSE | High | SSVSSVGAI | [R].SSVSSV | 1xPhospho | Q3UFK8 1x | 0.003927 | 0.000311 | 1 | 2 | 1 | Q3UFK8 | FERM dom | Q3UFK8 [2 | 0 | 1158.572 | 0 | 0.2 | 0.071798 | 2.57 | 94.1 | 105.9 | 4.52 | 5.32 | | High | High | High | High | High | High | High | High | High | High | High | 0.00014 | 0.001109 | 2.25 |
| FALSE | High | SSVSSVGAI | [R].SSVSSV | 2xPhospho | Q3UFK8 2x | 0.0095 | 0.000736 | 1 | 2 | 1 | Q3UFK8 | FERM dom | Q3UFK8 [2 | 0 | 1238.538 | 0 | 0.71 | 0.026497 | 31.86 | 76 | 124 | 11.14 | 17.19 | | High | High | High | High | High | High | High | High | High | High | High | 0.000327 | 0.002965 | 1.89 |
| FALSE | High | SSVTFEDE | [R].SSVTFE | 1xPhospho | B2RPY5-2 1 | 0.002463 | 0.000109 | 1 | 4 | 2 | B2RPY5-2 | Isoform 2 c | B2RPY5-2 [ | 0 | 2049.019 | 0 | 2.49 | 0.000313 | 10.36 | 32.5 | 167.5 | 15.12 | 10.02 | | High | High | High | High | High | High | High | High | High | High | High | 4.94E-05 | 0.000658 | 2.7 |
| FALSE | High | SSVTPADSI | [R].SSVTPA | 1xPhospho | P97929 1xI | 0.000958 | 0 | 1 | 3 | 1 | P97929 | Breast canc | P97929 [21 | 0 | 1811.956 | 0 | -1.21 | 0.001359 | 3.88 | 140.3 | 59.7 | 9.54 | 8.82 | | High | High | High | High | High | High | High | High | High | High | High | 0 | 0.000228 | 1.95 |
| FALSE | High | SSVTSCLDS | [K].SSVTSC | 1xCarbami | A0A0R4J2D | 7.72E-07 | 0 | 1 | 3 | 2 | A0A0R4J2D | Ubiquitin c | A0A0R4J2D | 0 | 2310.046 | 0 | -1.15 | 0.000264 | 1.64 | 137.8 | 62.2 | 3.37 | 5.28 | | High | High | High | High | High | High | High | High | High | High | High | 0 | 7.57E-08 | 3.6 |
| FALSE | High | SSVTSCLDS | [K].SSVTSC | 1xCarbami | A0A0R4J2D | 3.81E-06 | 0 | 1 | 3 | 3 | A0A0R4J2D | Ubiquitin c | A0A0R4J2D | 0 | 2390.012 | 0 | -1.22 | 0.000932 | 0.21 | 138.8 | 61.2 | 6.75 | 5.13 | | High | High | High | High | High | High | High | High | High | High | High | 0 | 4.59E-07 | 4.59 |
| FALSE | High | SSVVIEELSI | [K].SSVVIEI | 1xPhospho | Q8BWW9 1 | 1.76E-07 | 0 | 1 | 6 | 2 | Q8BWW9 | Serine/thre | Q8BWW9 | 0 | 2293.225 | 0 | 1.37 | 0.000181 | 4.54 | 56.1 | 143.9 | 9.38 | 4.62 | | High | High | High | High | High | High | High | High | High | High | High | 0 | 8.48E-08 | 4.23 |
| FALSE | High | SSVPGVR | [R].SSVPGV | 1xPhospho | P20152 1xI | 0.013666 | 0.001011 | 1 | 10 | 4 | P20152 | Vimentin C | P20152 [72 | 0 | 1010.523 | 0 | 0.41 | 0.120754 | 27.92 | 79.4 | 120.6 | 14.25 | 18.41 | | High | High | High | High | High | High | High | High | High | High | High | 0.003122 | 0.03242 | 1.74 |
| FALSE | High | SSTSQTVST | [R].SSTSQT | 1xPhospho | F6ZDS4 1xI | 5.82E-06 | 0 | 1 | 7 | 7 | F6ZDS4 | Nucleopro | F6ZDS4 [71 | 0 | 3083.544 | 0 | 0.16 | 0.025422 | 7.05 | 93 | 107 | 8.08 | 6.72 | | High | High | High | High | High | High | High | High | High | High | High | 0 | 1.03E-06 | 4.26 |
| FALSE | High | SSTSPCSTS | [R].SSTSPC | 1xCarbami | Q5RIM6 1x | 0.000396 | 0 | 1 | 7 | 3 | Q5RIM6 | Nuclear rec | Q5RIM6 [1 | 0 | 1579.744 | 0 | 0.11 | 0.517709 | 17.68 | 97.3 | 102.7 | 3.46 | 8.17 | | High | High | High | High | High | High | High | High | High | High | High | 0.000505 | 0.004362 | 2.12 |
| FALSE | High | SSTSIDK | [K].SSTSIDI | 1xPhospho | E9Q5F9 1x | 0.049971 | 0.003641 | 1 | 3 | 1 | E9Q5F9 | Histone-lys | E9Q5F9 [34 | 0 | 1275.66 | 0 | 0.47 | 0.00125 | 3.42 | 83.9 | 116.1 | 3.78 | 6.07 | | High | High | High | High | High | High | High | High | High | High | High | 0.001902 | 0.01935 | 1.89 |
| FALSE | High | SSSYSEANE | [R].SSSYSE. | 1xDeamida | Q3U2G7 1x | 0.018125 | 0.001282 | 1 | 11 | 2 | Q3U2G7 | Putative ur | Q3U2G7 [7 | 0 | 2554.142 | 0 | -0.26 | 0.086759 | 6.62 | 109.1 | 90.9 | 10.1 | 5.49 | | High | High | High | High | High | High | High | High | High | High | High | 0.000689 | 0.006117 | 3 |
| FALSE | High | SSSYSEANE | [R].SSSYSE. | 1xPhospho | Q3U2G7 1x | 0.002793 | 0.000109 | 1 | 11 | 2 | Q3U2G7 | Putative ur | Q3U2G7 [7 | 0 | 2569.153 | 0 | -0.27 | 0.075456 | 7.14 | 106 | 94 | 9.35 | 12.13 | | High | High | High | High | High | High | High | High | High | High | High | 0.000161 | 0.001227 | 2.84 |
| FALSE | High | SSTADAEL | [K].SSTADA | 1xPhospho | Q80U04 1x | 0.003822 | 0.000261 | 1 | 2 | 1 | Q80U04 | E3 ubiquiti | Q80U04 [2 | 0 | 2326.066 | 0 | -0.07 | 0.461445 | 11.23 | 102.3 | 97.7 | 31.1 | 11.95 | | High | High | High | High | High | High | High | High | High | High | High | 0.000117 | 0.00107 | 1.51 |
| FALSE | High | SSTFPQTD | [R].SSTFPQ | 1xPhospho | Q6NXH3-2 | 3.29E-08 | 0 | 1 | 3 | 4 | Q6NXH3-2 | Isoform 2 c | Q6NXH3-2 | 0 | 2400.128 | 0 | -0.97 | 0.000839 | 2.54 | 131.1 | 68.9 | 7.9 | 6.32 | | High | High | High | High | High | High | High | High | High | High | High | 0 | 2.19E-09 | 5.46 |
| FALSE | High | SSTFQGYCI | [K].SSTFQG | 1xCarbami | Q8BQ33 1x | 0.012508 | 0.000889 | 1 | 1 | 1 | Q8BQ33 | Treslin OS= | Q8BQ33 [9 | 0 | 1615.759 | 0 | -1.39 | 0.000887 | 1.02 | 143.4 | 56.6 | 4.66 | 17.54 | | High | High | High | High | High | High | High | High | High | High | High | 0.000449 | 0.004035 | 1.58 |
| FALSE | High | SSTFSSFDN | [R].SSTFSSI | 1xPhospho | Q60949 1x | 0.016892 | 0.001204 | 1 | 4 | 2 | Q60949 | TBC1 doma | Q60949 [2 | 0 | 3814.685 | 0 | -0.65 | 0.032803 | 5.98 | 122.1 | 77.9 | 10.12 | 22.18 | | High | High | High | High | High | High | High | High | High | High | High | 0.000638 | 0.00562 | 4.12 |
| FALSE | High | SSTGPELPA | [R].SSTGPE | 1xPhospho | Q8BTI8 1xI | 0.013074 | 0.000889 | 1 | 3 | 3 | Q8BTI8 | Serine/argi | Q8BTI8 [35 | 0 | 4952.488 | 0 | -1.23 | 0.000375 | 2.51 | 140.8 | 59.2 | 10.02 | 17.81 | | High | High | High | High | High | High | High | High | High | High | High | 0.000467 | 0.004241 | 5.14 |
| FALSE | High | SSTHASVSI | [K].SSTHAS | 1xPhospho | D3Z5I1 1xP | 0.000178 | 0 | 1 | 3 | 1 | D3Z5I1 | Zinc finger | D3Z5I1 [55 | 0 | 1808.838 | 0 | 1.06 | 0.021004 | 28.05 | 65.1 | 134.9 | 13.93 | 19.54 | | High | High | High | High | High | High | High | High | High | High | High | 0 | 3.43E-05 | 2.34 |
| FALSE | High | SSTHGEGA | [K].SSTHGE | 1xPhospho | Q8VDF2 1x | 1.71E-07 | 0 | 1 | 3 | 10 | Q8VDF2 | E3 ubiquiti | Q8VDF2 [1 | 0 | 1912.869 | 0 | 0.35 | 0.005999 | 12.35 | 89.3 | 110.7 | 11.08 | 7.08 | | High | High | High | High | High | High | High | High | High | High | High | 0 | 1.4E-08 | 4.73 |
| FALSE | High | SSTLAPAQI | [R].SSTLAP | 1xPhospho | E9Q5G3 1x | 2.57E-07 | 0 | 1 | 1 | 8 | E9Q5G3 | Kinesin-like | E9Q5G3 [9 | 0 | 2686.208 | 0 | -1.15 | 0.000192 | 0.97 | 138.4 | 61.6 | 3.06 | 9.51 | | High | High | High | High | High | High | High | High | High | High | High | 0 | 3.35E-08 | 3.82 |
| FALSE | High | SSTLAWDR | [R].SSTLAW | 1xPhospho | G5E880 1x | 0.000738 | 0 | 1 | 6 | 1 | G5E880 | Lymphoid- | G5E880 [4 | 0 | 1244.587 | 0 | 2.66 | 0.000538 | 13.61 | 25.3 | 174.7 | 13.31 | 11.45 | | High | High | High | High | High | High | High | High | High | High | High | 0 | 0.000169 | 2.26 |
| FALSE | High | SSTPAAGC. | [K].SSTPAA | 1xCarbami | E9PYV1 1xI | 1.15E-06 | 0 | 1 | 12 | 2 | E9PYV1 | Glucocortic | E9PYV1 [15 | 0 | 1915.924 | 0 | 1.17 | 0.000359 | 16.96 | 61.5 | 138.5 | 13.05 | 6.4 | | High | High | High | High | High | High | High | High | High | High | High | 0 | 1.18E-07 | 3.9 |
| FALSE | High | SSTPASVTA | [R].SSTPAS | 1xPhospho | O70374 1x | 0.008769 | 0.000736 | 1 | 4 | 4 | O70374 | Protein CBI | O70374 [5 | 0 | 1712.831 | 0 | -0.52 | 0.044698 | 10.76 | 112.7 | 87.3 | 24.78 | 8.03 | | High | High | High | High | High | High | High | High | High | High | High | 0.000327 | 0.002701 | 3.57 |
| FALSE | High | SSTPASVTA | [R].SSTPAS | 1xDeamida | O70374 1x | 0.001373 | 5.66E-05 | 1 | 4 | 4 | O70374 | Protein CBI | O70374 [5 | 0 | 1713.815 | 0 | -0.54 | 0.010701 | 9.56 | 124.1 | 75.9 | 9.83 | 15.66 | | High | High | High | High | High | High | High | High | High | High | High | 2.58E-05 | 0.00034 | 2.81 |
| FALSE | High | SSTPEEVK | [K].SSTPEE | 1xPhospho | F8WGL3 1x | 0.024211 | 0.001612 | 1 | 3 | 3 | F8WGL3 | Cofilin-1 O | F8WGL3 [2 | 0 | 1414.723 | 0 | -0.77 | 0.006518 | 1.82 | 126.3 | 73.7 | 4.23 | 16.18 | | High | High | High | High | High | High | High | High | High | High | High | 0.002432 | 0.02613 | 1.99 |
| FALSE | High | SSTPEGND | [R].SSTPEG | 1xPhospho | M0QWG1 | 0.030367 | 0.001857 | 1 | 1 | 2 | M0QWG1 | Genetic su | M0QWG1 | 0 | 2649.235 | 0 | -0.4 | 0.00445 | 4.41 | 115 | 85 | 10.03 | 12.29 | | High | High | High | High | High | High | High | High | High | High | High | 0.001011 | 0.01091 | 3.36 |
| FALSE | High | SSTPGLK | [K].SSTPGL | 1xPhospho | Q80TC1 1x | 0.068184 | 0.004935 | 1 | 14 | 2 | Q80TC1 | MKIAA154 | Q80TC1 [4 | 0 | 1227.675 | 0 | -1.29 | 0.00036 | 2.47 | 141.9 | 58.1 | 4.27 | 10.89 | | High | High | High | High | High | High | High | High | High | High | High | 0.003243 | 0.03426 | 1.98 |
| FALSE | High | SSTPLHVHT | [R].SSTPLH | 1xPhospho | P97360 1xI | 0.00095 | 0 | 1 | 3 | 3 | P97360 | Transcripti | P97360 [29 | 0 | 1639.852 | 0 | 0.7 | 0.002243 | 12.03 | 77.7 | 122.3 | 9.16 | 10.16 | | High | High | High | High | High | High | High | High | High | High | High | 0 | 0.000226 | 2.55 |
| FALSE | High | SSTPLPTVS | [R].SSTPLP | 1xPhospho | Q3U955 1x | 2.2E-06 | 0 | 2 | 10 | 14 | Q3U955; Q | Putative ur | Q3U955 [1 | 0 | 1942.932 | 0 | 1.65 | 0.000569 | 16.65 | 46.8 | 153.2 | 7.22 | 8.16 | Shared | High | High | High | High | High | High | High | High | High | High | High | 0 | 1.75E-06 | 3.37 |
| FALSE | High | SSTPLPTVS | [R].SSTPLP | 2xPhospho | Q3U955 2x | 3.06E-05 | 0 | 2 | 10 | 5 | Q3U955; Q | Putative ur | Q3U955 [1 | 0 | 2022.898 | 0 | 2.6 | 0.001722 | 31.99 | 28.4 | 171.6 | 14.16 | 15.46 | Shared | High | High | High | High | High | High | High | High | High | High | High | 0 | 9.43E-06 | 3.57 |
| FALSE | High | SSTPPLHVH | [R].SSTPPL | 1xPhospho | A3KGV1-2 | 0.000584 | 0 | 1 | 11 | 2 | A3KGV1-2 | Isoform 2 c | A3KGV1-2 | 0 | 2881.513 | 0 | -0.45 | 0.010114 | 8.64 | 118.6 | 81.4 | 10.39 | 7.46 | | High | High | High | High | High | High | High | High | High | High | High | 0 | 0.00013 | 2.17 |
| FALSE | High | SSTPQELHI | [R].SSTPQE | 1xPhospho | Q6PFY9 1x | 0.090153 | 0.00698 | 1 | 1 | 4 | Q6PFY9 | Dual specif | Q6PFY9 [3 | 0 | 1363.657 | 0 | 0.06 | 0.694866 | 7.02 | 100.5 | 99.5 | 8.18 | 12.96 | | High | High | High | High | High | High | High | High | High | High | High | 0.004907 | 0.05338 | 2.25 |
| FALSE | High | SSTPSPQA | [R].SSTPSP | 1xPhospho | Q0VBD2 1x | 1.47E-05 | 0 | 1 | 1 | 1 | Q0VBD2 | Protein MC | Q0VBD2 [2 | 0 | 1807.936 | 0 | -0.93 | 0.00998 | 5.57 | 131 | 69 | 12.67 | 17.85 | | High | High | High | High | High | High | High | High | High | High | High | 0 | 2.09E-06 | 2.91 |
| FALSE | High | SSTPSPQA | [R].SSTPSP | 2xPhospho | Q0VBD2 2x | 0.010758 | 0.000772 | 1 | 1 | 2 | Q0VBD2 | Protein MC | Q0VBD2 [2 | 0 | 1887.902 | 0 | -0.48 | 0.004444 | 3.73 | 114.1 | 85.9 | 4.3 | 8.74 | | High | High | High | High | High | High | High | High | High | High | High | 0.000399 | 0.003415 | 2.67 |
| FALSE | High | SSTPSSPTG | [R].SSTPSS | 1xPhospho | Q8BWJ3 1x | 0.000674 | 0 | 1 | 2 | 5 | Q8BWJ3 | Phosphory | Q8BWJ3 [1 | 0 | 3525.587 | 0 | -0.88 | 0.056113 | 8.08 | 128.8 | 71.2 | 14.37 | 30.62 | | High | High | High | High | High | High | High | High | High | High | High | 4.94E-05 | 0.000739 | 2.54 |
| FALSE | High | SSTSDWPT | [K].SSTSDV | 1xPhospho | G3X940 1x | 1.38E-05 | 0 | 1 | 4 | 3 | G3X940 | Histone ac | G3X940 [4 | 0 | 2478.086 | 0 | 1.49 | 0.000807 | 8.81 | 52.4 | 147.6 | 9.66 | 11.92 | | High | High | High | High | High | High | High | High | High | High | High | 0 | 1.95E-06 | 3.15 |
| FALSE | High | SSTSFANIQ | [K].SSTSFA | 1xPhospho | Q8BGT0 1x | 1.02E-05 | 0 | 1 | 1 | 6 | Q8BGT0 | Osteopetro | Q8BGT0 [3 | 0 | 1678.752 | 0 | 0.31 | 0.092744 | 17.15 | 89.3 | 110.7 | 2.22 | 14.58 | | High | High | High | High | High | High | High | High | High | High | High | 0 | 1.71E-06 | 2.91 |
| FALSE | High | SSTSFANIQ | [K].SSTSFA | 2xPhospho | Q8BGT0 2x | 0.000645 | 0 | 1 | 1 | 1 | Q8BGT0 | Osteopetro | Q8BGT0 [3 | 0 | 1758.719 | 0 | 1.6 | 0.003152 | 6.58 | 49.7 | 150.3 | 18.02 | 23.26 | | High | High | High | High | High | High | High | High | High | High | High | 0 | 0.000146 | 4.02 |
| FALSE | High | SSTSGSASS | [K].SSTSGS | 1xPhospho | Q542Q3 1x | 2.57E-07 | 0 | 1 | 1 | 3 | Q542Q3 | G two S ph | Q542Q3 [3 | 0 | 1883.859 | 0 | -0.33 | 0.042071 | 7.23 | 112.2 | 87.8 | 15.42 | 10.33 | | High | High | High | High | High | High | High | High | High | High | High | 0 | 2.2E-08 | 3.06 |
| FALSE | High | SSVVPSGG | [R].SSVVPS | 1xPhospho | Q8C9G3 1x | 0.001933 | 5.66E-05 | 1 | 6 | 1 | Q8C9G3 | Putative ur | Q8C9G3 [3 | 0 | 1840.977 | 0 | -0.6 | 0.00462 | 2.64 | 120.5 | 79.5 | 7.7 | 8.12 | | High | High | High | High | High | High | High | High | High | High | High | 2.58E-05 | 0.000498 | 3.16 |
| FALSE | High | SVDNLSCPS | [R].SVDNLS | 1xCarbami | Q922B9 1x | 0.001039 | 0 | 1 | 3 | 1 | Q922B9 | Sperm-spe | Q922B9 [1 | 0 | 2779.361 | 0 | 0.74 | 0.006335 | 26.68 | 73.3 | 126.7 | 12.71 | 12.44 | | High | High | High | High | High | High | High | High | High | High | High | 0 | 0.000248 | 3.46 |
| FALSE | High | SSWSLSPSI | [R].SSWSLS | 1xCarbami | Q3TAA7 1x | 0.000107 | 0 | 1 | 1 | 3 | Q3TAA7 | Serine/thre | Q3TAA7 [7 | 0 | 1920.851 | 0 | -0.74 | 0.000693 | 1.83 | 124.4 | 75.6 | 5.87 | 8.22 | | High | High | High | High | High | High | High | High | High | High | High | 0 | 1.94E-05 | 2.02 |
| FALSE | High | SSYASQHS( | [R].SSYASC | 1xPhospho | Q68FH0 1x | 0.008313 | 0.000736 | 1 | 8 | 1 | Q68FH0 | Plakophilin | Q68FH0 [3 | 0 | 1999.944 | 0 | -1.26 | 0.003083 | 3.69 | 138.4 | 61.6 | 13.38 | 9.85 | | High | High | High | High | High | High | High | High | High | High | High | 0.000327 | 0.002551 | 2.9 |
| FALSE | High | STESLPLTSI | [R].STESLP | 1xPhospho | E9Q7D5 1x | 0.028327 | 0.001791 | 1 | 2 | 1 | E9Q7D5 | Rho guanir | E9Q7D5 [8 | 0 | 1399.703 | 0 | 1.2 | 0.004362 | 12.03 | 59.1 | 140.9 | 9.88 | 14.65 | | High | High | High | High | High | High | High | High | High | High | High | 0.000968 | 0.01008 | 1.26 |
| FALSE | High | STESLQAN | [K].STESLQ | 1xPhospho | P47963 1xI | 0.000134 | 0 | 1 | 3 | 1 | P47963 | 60S riboso | P47963 [10 | 0 | 1541.752 | 0 | -0.39 | 0.008624 | 3.52 | 113.3 | 86.7 | 1.06 | 9.58 | | High | High | High | High | High | High | High | High | High | High | High | 0 | 2.51E-05 | 2.95 |
| FALSE | High | STESVPPSK | [K].STESVP | 1xPhospho | E9Q8T1 1x | 0.007209 | 0.00062 | 2 | 10 | 1 | E9Q8T1; Q | Transformi | E9Q8T1 [1 | 0 | 1469.765 | 0 | -0.99 | 0.000461 | 3.17 | 132.9 | 67.1 | 4.08 | 9.96 | Shared | High | High | High | High | High | High | High | High | High | High | High | 0.000275 | 0.002185 | 1.35 |
| FALSE | High | STETDNGY | [K].STETDN | 1xPhospho | A0A0R4J1C | 0.062731 | 0.004664 | 1 | 3 | 1 | A0A0R4J1C | Putative hc | A0A0R4J1C | 0 | 2023.963 | 0 | 0.36 | 0.001633 | 4.48 | 88.9 | 111.1 | 10.15 | 5.62 | | High | High | High | High | High | High | High | High | High | High | High | 0.002403 | 0.02498 | 2.95 |
| FALSE | High | STETSDFEN | [K].STETSD | 1xPhospho | A0A1L1SV7 | 2.99E-06 | 0 | 1 | 3 | 2 | A0A1L1SV7 | Ubiquitin c | A0A1L1SV7 | 0 | 2277.012 | 0 | -0.01 | 0.612655 | 9.04 | 97.5 | 102.5 | 12.2 | 6.83 | | High | High | High | High | High | High | High | High | High | High | High | 0 | 3.49E-07 | 3.71 |
| FALSE | High | STETSDFEN | [K].STETSD | 2xPhospho | A0A1L1SV7 | 0.000411 | 0 | 1 | 3 | 1 | A0A1L1SV7 | Ubiquitin c | A0A1L1SV7 | 0 | 2356.978 | 0 | -0.36 | 0.243923 | 16.58 | 112.3 | 87.7 | 6.74 | 19.44 | | High | High | High | High | High | High | High | High | High | High | High | 0 | 8.76E-05 | 4.08 |
| FALSE | High | STFSLPEEE | [R].STFSLPI | 1xPhospho | F8WJF7 1x | 2.73E-06 | 0 | 1 | 4 | 6 | F8WJF7 | E3 ubiquiti | F8WJF7 [1 | 0 | 3256.595 | 0 | 0.15 | 0.332136 | 5.73 | 95.4 | 104.6 | 12.22 | 8 | | High | High | High | High | High | High | High | High | High | High | High | 0 | 3.15E-07 | 5.12 |
| FALSE | High | STFSPISDL( | [K].STFSPIS | 1xPhospho | H9LAN6 1x | 0.021231 | 0.001507 | 1 | 5 | 2 | H9LAN6 | Histone-lys | H9LAN6 [1 | 0 | 1874.008 | 0 | -0.23 | 0.418661 | 16.2 | 108.1 | 91.9 | 9.04 | 13.54 | | High | High | High | High | High | High | High | High | High | High | High | 0.000801 | 0.007277 | 2.29 |
| FALSE | High | STGASPPS( | [R].STGASI | 1xCarbami | Q7TNC6 1x | 1.66E-06 | 0 | 1 | 2 | 1 | Q7TNC6 | Kinesin-like | Q7TNC6 [1 | 0 | 2429.178 | 0 | 0.36 | 0.077567 | 10.12 | 84.2 | 115.8 | 17.99 | 9.11 | | High | High | High | High | High | High | High | High | High | High | High | 0 | 1.8E-07 | 4.15 |
| FALSE | High | STGDITAA( | [K].STGDIT | 1xPhospho | Q9Z0U1 1x | 5.3E-08 | 0 | 1 | 1 | 2 | Q9Z0U1 | Tight junct | Q9Z0U1 [4 | 0 | 1744.832 | 0 | -0.84 | 0.010809 | 3.79 | 130.5 | 69.5 | 6.27 | 28.54 | | High | High | High | High | High | High | High | High | High | High | High | 0 | 3.76E-09 | 4.64 |
| FALSE | High | STGNFLTLS | [K].STGNFI | 1xPhospho | Q6ZPT2 1x | 0.043236 | 0.003228 | 1 | 2 | 2 | Q6ZPT2 | MKIAA135 | Q6ZPT2 [7 | 0 | 2019.056 | 0 | -0.27 | 0.009572 | 0.69 | 109.1 | 90.9 | 7.58 | 4.68 | | High | High | High | High | High | High | High | High | High | High | High | 0.002675 | 0.02858 | 2.36 |
| FALSE | High | STGPTLSSP | [K].STGPTL | 1xPhospho | Q8VDF2 1x | 0.006895 | 0.00062 | 1 | 2 | 3 | Q8VDF2 | E3 ubiquiti | Q8VDF2 [6 | 0 | 1311.651 | 0 | -0.04 | 0.223652 | 1.27 | 100.8 | 99.2 | 5.58 | 5.25 | | High | High | High | High | High | High | High | High | High | High | High | 0.000275 | 0.002077 | 1.69 |
| FALSE | High | STGQLNLS | [K].STGQLI | 1xCarbami | F8VQ05 1x | 3.85E-06 | 0 | 1 | 5 | 2 | F8VQ05 | FRY-like tra | F8VQ05 [1 | 0 | 2882.33 | 0 | 1 | 0.01621 | 25.33 | 71.7 | 128.3 | 11.95 | 19.77 | | High | High | High | High | High | High | High | High | High | High | High | 0 | 4.61E-07 | 5.08 |
| FALSE | High | STESEVYDI | [R].STESEV | 1xPhospho | Q9Z1X2 1x | 0.006713 | 0.00062 | 1 | 2 | 1 | Q9Z1X2 | Phosphatic | Q9Z1X2 [2 | 0 | 2363.007 | 0 | -0.44 | 0.055671 | 11.86 | 112.1 | 87.9 | 16.19 | 11.06 | | High | High | High | High | High | High | High | High | High | High | High | 0.000275 | 0.002016 | 3.15 |
| FALSE | High | STGQLNLS | [K].STGQLI | 1xCarbami | F8VQ05 1x | 1.57E-05 | 0 | 1 | 5 | 2 | F8VQ05 | FRY-like tra | F8VQ05 [1 | 0 | 2883.314 | 0 | 0.43 | 0.023491 | 13.92 | 82.9 | 117.1 | 6.89 | 11.96 | | High | High | High | High | High | High | High | High | High | High | High | 0 | 2.79E-05 | 4.09 |
| FALSE | High | STGQLNLS | [K].STGQLI | 1xCarbami | F8VQ05 2x | 0.011146 | 0.000772 | 1 | 5 | 1 | F8VQ05 | FRY-like tra | F8VQ05 [1 | 0 | 2963.28 | 0 | 0.47 | 0.036155 | 12.36 | 81 | 119 | 6.35 | 21.32 | | High | High | High | High | High | High | High | High | High | High | High | 0.000399 | 0.003533 | 3.94 |
| FALSE | High | STGQQAAA | [K].STGQQ | 2xPhospho | F7BJB9 2xP | 0.000205 | 0 | 1 | 2 | 2 | F7BJB9 | MORC fam | F7BJB9 [86 | 0 | 3487.556 | 0 | -0.97 | 0.004507 | 2.45 | 132.7 | 67.3 | 10.11 | 17.43 | | High | High | High | High | High | High | High | High | High | High | High | 0 | 4.03E-05 | 4.1 |
| FALSE | High | STGSLLTPS | [K].STGSLL | 1xPhospho | B7ZCJ1 1xP | 0.034281 | 0.002195 | 1 | 5 | 2 | B7ZCJ1 | Rho GTPas | B7ZCJ1 [17 | 0 | 1327.682 | 0 | -0.29 | 0.07529 | 7.26 | 114.4 | 85.6 | 7.3 | 15.68 | | High | High | High | High | High | High | High | High | High | High | High | 0.001151 | 0.01248 | 1.58 |
| FALSE | High | STGSPSSQS | [K].STGSPS | 1xPhospho | Q3TBZ6 1x | 0.007274 | 0.00062 | 1 | 2 | 4 | Q3TBZ6 | Putative ur | Q3TBZ6 [2 | 0 | 2146.964 | 0 | 0.24 | 0.129392 | 14.17 | 91.6 | 108.4 | 11.04 | 11.48 | | High | High | High | High | High | High | High | High | High | High | High | 0.009162 | 0.09914 | 3.43 |
| FALSE | High | STGVFQDE | [K].STGVFC | 1xPhospho | Q6PGL7 1x | 5.34E-11 | 0 | 1 | 5 | 2 | Q6PGL7 | WASH com | Q6PGL7 [8 | 0 | 2275.141 | 0 | 0.57 | 0.061561 | 18.95 | 88.2 | 111.8 | 9.86 | 18.19 | | High | High | High | High | High | High | High | High | High | High | High | 0 | 1.61E-12 | 5.72 |
| FALSE | High | STGVSFWT | [R].STGVSI | 1xPhospho | Q9DBR7 1x | 6.02E-05 | 0 | 1 | 4 | 1 | Q9DBR7 | Protein ph | Q9DBR7 [8 | 0 | 2424.019 | 0 | -0.36 | 0.007868 | 3.83 | 117.7 | 82.3 | 6.49 | 12.7 | | High | High | High | High | High | High | High | High | High | High | High | 0 | 1.02E-05 | 3.64 |
| FALSE | High | STGVSFWT | [R].STGVSI | 1xPhospho | Q9DBR7 1x | 0.001689 | 5.66E-05 | 1 | 4 | 3 | Q9DBR7 | Protein ph | Q9DBR7 [8 | 1 | 3687.597 | 0 | -0.45 | 0.001964 | 3.25 | 115.5 | 84.5 | 4.21 | 10.68 | | High | High | High | High | High | High | High | High | High | High | High | 2.58E-05 | 0.00043 | 4.01 |
| FALSE | High | STGVSFWT | [R].STGVSI | 1xDeamida | Q9DBR7 1x | 0.000712 | 0 | 1 | 4 | 2 | Q9DBR7 | Protein ph | Q9DBR7 [8 | 1 | 3688.581 | 0 | -0.25 | 0.037048 | 10.06 | 106.1 | 93.9 | 10.82 | 4.61 | | High | High | High | High | High | High | High | High | High | High | High | 0 | 0.000163 | 5.58 |
| FALSE | High | STGVSFWT | [R].STGVSI | 2xPhospho | Q9DBR7 2x | 0.002921 | 0.00016 | 1 | 4 | 5 | Q9DBR7 | Protein ph | Q9DBR7 [8 | 1 | 3767.564 | 0 | -0.24 | 0.057979 | 10.39 | 108.1 | 91.9 | 7.6 | 11.34 | | High | High | High | High | High | High | High | High | High | High | High | 7.19E-05 | 0.000796 | 4.09 |
| FALSE | High | STIGVMVT | [R].STIGVM | 1xPhospho | Q8BME1 1 | 0.001168 | 0 | 1 | 7 | 1 | Q8BME1 | Phosphoac | Q8BME1 [5 | 0 | 2623.284 | 0 | -1.04 | 0.025138 | 8.79 | 132.6 | 67.4 | 11.85 | 22.83 | | High | High | High | High | High | High | High | High | High | High | High | 0 | 0.000284 | 2.79 |
| FALSE | High | STIGVMVT | [R].STIGVM | 1xPhospho | Q8BME1 1 | 0.000294 | 0 | 1 | 7 | 2 | Q8BME1 | Phosphoac | Q8BME1 [5 | 0 | 2639.279 | 0 | -1.97 | 0.00222 | 1.11 | 155.2 | 44.8 | 13.98 | 14.55 | | High | High | High | High | High | High | High | High | High | High | High | 0 | 6.02E-05 | 2.49 |
| FALSE | High | STLNPQWI | [R].STLNPC | 1xPhospho | Q4VA93 1x | 0.000687 | 0 | 1 | 1 | 2 | Q4VA93 | Protein kin | Q4VA93 [2 | 0 | 2237.104 | 0 | 2.06 | 0.00425 | 7.5 | 34.3 | 165.7 | 19.17 | 25.15 | | High | High | High | High | High | High | High | High | High | High | High | 0 | 0.000157 | 3.27 |
| FALSE | High | STLTTLPLP | [R].STLTTLI | 1xPhospho | Q8VEB3 1x | 0.000118 | 0 | 1 | 2 | 1 | Q8VEB3 | UNC119-bi | Q8VEB3 [1 | 0 | 2293.246 | 0 | 0.89 | 0.00034 | 3.85 | 69.2 | 130.8 | 3.16 | 4.88 | | High | High | High | High | High | High | High | High | High | High | High | 0 | 2.18E-05 | 4.01 |
| FALSE | High | STGQLNLS | [K].STGQLI | 1xCarbami | F8VQ05 2x | 0.000178 | 0 | 1 | 5 | 1 | F8VQ05 | FRY-like tra | F8VQ05 [1 | 0 | 2962.296 | 0 | 0.56 | 0.012104 | 16.48 | 80.7 | 119.3 | 12.59 | 11.35 | | High | High | High | High | High | High | High | High | High | High | High | 0 | 3.42E-05 | 4.5 |
| FALSE | High | STEPPPPPP | [K].STEPPP | 1xPhospho | Q9ESZ8-5 1 | 2.65E-08 | 0 | 2 | 9 | 2 | Q9ESZ8-5; | Isoform 5 c | Q9ESZ8-5 [ | 0 | 2320.199 | 0 | 0.79 | 0.003629 | 14.89 | 73.1 | 126.9 | 11.19 | 8.82 | Shared | High | High | High | High | High | High | High | High | High | High | High | 0 | 1.71E-09 | 3.59 |
| FALSE | High | STENVNK | [R].STENVI | 1xPhospho | S4R221 1xI | 0.044358 | 0.003264 | 1 | 8 | 2 | S4R221 | Rho GTPas | S4R221 [10 | 0 | 1329.682 | 0 | -0.97 | 0.002826 | 0.92 | 132.9 | 67.1 | 5.6 | 22.71 | | High | High | High | High | High | High | High | High | High | High | High | 0.001665 | 0.01676 | 1.54 |
| FALSE | High | STELVLSPD | [R].STELVL | 1xPhospho | Q9ESK9 1x | 2.02E-05 | 0 | 1 | 6 | 1 | Q9ESK9 | RB1-induci | Q9ESK9 [2 | 0 | 1653.812 | 0 | -0.48 | 0.00481 | 2.68 | 116 | 84 | 7.62 | 6.95 | | High | High | High | High | High | High | High | High | High | High | High | 0 | 2.97E-06 | 2.53 |
| FALSE | High | SSYTYSK | [R].SSYTYS | 1xPhospho | P97868 1xI | 0.017809 | 0.001282 | 1 | 2 | 1 | P97868 | E3 ubiquiti | P97868 [69 | 0 | 1373.675 | 0 | -0.12 | 0.33687 | 9.06 | 100.4 | 99.6 | 6.38 | 7.33 | | High | High | High | High | High | High | High | High | High | High | High | 0.000689 | 0.006002 | 1.24 |
| FALSE | High | STAACGVE | [K].STAAC( | 2xCarbami | B0QZW4 1 | 9.94E-07 | 0 | 1 | 4 | 2 | B0QZW4 | Suppresso | B0QZW4 [3 | 0 | 2489.126 | 0 | -0.15 | 0.038264 | 7.93 | 105.7 | 94.3 | 5.01 | 3.7 | | High | High | High | High | High | High | High | High | High | High | High | 0 | 1.01E-07 | 3.48 |
| FALSE | High | STAACGVE | [K].STAAC( | 2xCarbami | B0QZW4 1 | 4.02E-06 | 0 | 1 | 4 | 1 | B0QZW4 | Suppresso | B0QZW4 [3 | 0 | 2490.11 | 0 | -0.11 | 0.924907 | 9.48 | 103.4 | 96.6 | 7.78 | 14.16 | | High | High | High | High | High | High | High | High | High | High | High | 0 | 4.86E-07 | 5.18 |
| FALSE | High | STAALSGD | [K].STAALS | 1xCarbami | E9QMU3 1 | 1.66E-05 | 0 | 1 | 2 | 1 | E9QMU3 | Ensconsin | E9QMU3 [ | 0 | 2592.285 | 0 | -0.49 | 0.022755 | 5.49 | 116 | 84 | 10.25 | 11.93 | | High | High | High | High | High | High | High | High | High | High | High | 0 | 2.4E-06 | 2.65 |
| FALSE | High | STAALSGD | [K].STAALS | 1xCarbami | E9QMU3 1 | 1.09E-07 | 0 | 1 | 2 | 1 | E9QMU3 | Ensconsin | E9QMU3 [ | 0 | 2608.28 | 0 | -0.39 | 0.138594 | 19.09 | 113.4 | 86.6 | 11.67 | 16.59 | | High | High | High | High | High | High | High | High | High | High | High | 0 | 8.44E-09 | 4.63 |
| FALSE | High | STAGSPPA | [R].STAGSI | 1xCarbami | E9QKK4 1x | 7.49E-09 | 0 | 1 | 2 | 3 | E9QKK4 | Glucocortic | E9QKK4 [2 | 0 | 2621.198 | 0 | 0.67 | 0.005912 | 8.68 | 77.9 | 122.1 | 11.34 | 12.83 | | High | High | High | High | High | High | High | High | High | High | High | 0 | 6.69E-08 | 6.42 |
| FALSE | High | STANNVEII | [R].STANN | 1xPhospho | P35585 1xI | 0.000183 | 0 | 1 | 1 | 1 | P35585 | AP-1 comp | P35585 [30 | 0 | 2656.338 | 0 | 0.37 | 0.015263 | 12.42 | 87 | 113 | 6.14 | 7.02 | | High | High | High | High | High | High | High | High | High | High | High | 0 | 3.54E-05 | 3.47 |
| FALSE | High | STAQWFG | [RK].STAQ | 1xPhospho | Q80WQ6 1 | 6.07E-05 | 0 | 1 | 2 | 1 | Q80WQ6 | Inactive rh | Q80WQ6 [ | 0 | 2193.042 | 0 | 0.27 | 0.166853 | 27.03 | 90.5 | 109.5 | 18.35 | 5.4 | | High | High | High | High | High | High | High | High | High | High | High | 0 | 1.02E-05 | 3.57 |
| FALSE | High | STASPASTK | [K].STASPA | 1xPhospho | Q80Y82 1x | 0.081752 | 0.006219 | 1 | 11 | 1 | Q80Y82 | Protein kin | Q80Y82 [4 | 0 | 1387.723 | 0 | 0.6 | 0.011926 | 21.33 | 80.2 | 119.8 | 4.34 | 11.66 | | High | High | High | High | High | High | High | High | High | High | High | 0.0032 | 0.03399 | 2.2 |
| FALSE | High | STASPVPCE | [K].STASPV | 2xCarbami | A0A0R4J0V | 1.77E-05 | 0 | 1 | 3 | 4 | A0A0R4J0V | Junction-m | A0A0R4J0V | 0 | 3220.542 | 0 | 0.13 | 0.967434 | 8.79 | 102 | 98 | 8.64 | 10.2 | | High | High | High | High | High | High | High | High | High | High | High | 0 | 2.56E-06 | 2.9 |
| FALSE | High | STAVTTGS. | [R].STAVTT | 1xPhospho | Q64012 1x | 0.002768 | 0.000109 | 1 | 5 | 2 | Q64012 | RNA-bindir | Q64012 [1 | 0 | 1460.776 | 0 | -0.32 | 0.00058 | 2.3 | 111.1 | 88.9 | 3.66 | 4.53 | | High | High | High | High | High | High | High | High | High | High | High | 0.000984 | 0.01019 | 1.34 |
| FALSE | High | STCAPSSPF | [R].STCAPS | 1xCarbami | Q3UFT3 1x | 0.000446 | 0 | 1 | 2 | 2 | Q3UFT3 | GRB2-asso | Q3UFT3 [4 | 0 | 2395.234 | 0 | -0.08 | 0.929149 | 14.64 | 104.7 | 95.3 | 16.68 | 3.05 | | High | High | High | High | High | High | High | High | High | High | High | 0 | 0.000162 | 3.22 |
| FALSE | High | STDFVFIPP | [K].STDFVI | 1xPhospho | Q6PR54-3 | 5.65E-05 | 0 | 2 | 4 | 4 | Q6PR54-3; | Isoform 3 c | Q6PR54-3 | 0 | 1874.971 | 0 | 0.42 | 0.303249 | 29.79 | 85.6 | 114.4 | 23.81 | 10.66 | Shared | High | High | High | High | High | High | High | High | High | High | High | 0 | 9.48E-06 | 2.31 |
| FALSE | High | STDIFSK | [K].STDIFSI | 1xPhospho | P33215 1xI | 0.093123 | 0.007276 | 1 | 2 | 3 | P33215 | Protein NE | P33215 [37 | 0 | 1335.696 | 0 | 1.7 | 0.000442 | 12.1 | 51.1 | 148.9 | 10.43 | 7.77 | | High | High | High | High | High | High | High | High | High | High | High | 0.003698 | 0.03984 | 2.23 |
| FALSE | High | STDPGTEG | [R].STDPGT | 1xPhospho | Q6NV66 1x | 0.004911 | 0.000504 | 1 | 4 | 2 | Q6NV66 | Zinc finger | Q6NV66 [5 | 0 | 1896.936 | 0 | -0.66 | 0.001855 | 5.42 | 122.1 | 77.9 | 4.58 | 7.36 | | High | High | High | High | High | High | High | High | High | High | High | 0.000246 | 0.001521 | 2.39 |
| FALSE | High | STDQDYRP | [R].STDQD | 1xPhospho | E9PYH6 1xI | 3.01E-06 | 0 | 1 | 1 | 1 | E9PYH6 | SET domain | E9PYH6 [4 | 0 | 4393.918 | 0 | -0.22 | 0.06577 | 13.23 | 110.6 | 89.4 | 4.32 | 18.37 | | High | High | High | High | High | High | High | High | High | High | High | 0 | 3.52E-07 | 3.21 |
| FALSE | High | STDQDYRP | [R].STDQD | 1xDeamida | E9PYH6 1xI | 6.41E-08 | 0 | 1 | 1 | 1 | E9PYH6 | SET domain | E9PYH6 [4 | 0 | 4394.902 | 0 | -0.69 | 0.012057 | 9.1 | 124.3 | 75.7 | 5.93 | 18.15 | | High | High | High | High | High | High | High | High | High | High | High | 0 | 4.61E-09 | 3.64 |
| FALSE | High | STDQDYRP | [R].STDQD | 1xDeamida | E9PYH6 1xI | 0.005565 | 0.00062 | 1 | 1 | 1 | E9PYH6 | SET domain | E9PYH6 [4 | 1 | 4880.126 | 0 | -0.65 | 0.004721 | 1.19 | 118.3 | 81.7 | 6.32 | 6.77 | | High | High | High | High | High | High | High | High | High | High | High | 0.000275 | 0.00163 | 2.52 |
| FALSE | High | STDSEVSQ | [R].STDSEV | 1xPhospho | Q8K0V4 1x | 9.11E-05 | 0 | 1 | 3 | 3 | Q8K0V4 | CCR4-NOT | Q8K0V4 [2 | 0 | 1773.867 | 0 | -0.79 | 0.000793 | 3.18 | 126.7 | 73.3 | 3.1 | 5.59 | | High | High | High | High | High | High | High | High | High | High | High | 0 | 2.42E-05 | 3.52 |
| FALSE | High | STDSPGAD | [K].STDSPG | 1xPhospho | A0A0G2JG | 5.27E-09 | 0 | 1 | 3 | 2 | A0A0G2JG | Crystallin b | A0A0G2JG | 0 | 2112.026 | 0 | 5.58 | 0.000253 | 74.97 | 4.3 | 195.7 | 59.84 | 8.76 | | High | High | High | High | High | High | High | High | High | High | High | 0 | 1.2E-09 | 3.01 |
| FALSE | High | STDSPIAIEI | [R].STDSPI. | 1xPhospho | Q3TZW5 1 | 8.18E-05 | 0 | 1 | 5 | 2 | Q3TZW5 | Putative ur | Q3TZW5 [1 | 0 | 2189.021 | 0 | -0.9 | 0.002082 | 5.47 | 136.4 | 63.6 | 10.08 | 7.17 | | High | High | High | High | High | High | High | High | High | High | High | 0 | 2.43E-05 | 4.11 |
| FALSE | High | STDSSSSVS | [R].STDSSS | 1xPhospho | Q8VDY9 1x | 3.36E-06 | 0 | 1 | 1 | 6 | Q8VDY9 | Caspase ac | Q8VDY9 [6 | 0 | 2266.085 | 0 | -0.05 | 0.256668 | 10.53 | 103.5 | 96.5 | 7.57 | 6.16 | | High | High | High | High | High | High | High | High | High | High | High | 0 | 4.16E-07 | 4.17 |
| FALSE | High | STDSSSYPS | [K].STDSSS | 1xCarbami | B2RR83 1xI | 0.006713 | 0.00062 | 1 | 1 | 1 | B2RR83 | Probable A | B2RR83 [1 | 0 | 2621.184 | 0 | 0.19 | 0.080373 | 17.45 | 93.4 | 106.6 | 8.27 | 4.18 | | High | High | High | High | High | High | High | High | High | High | High | 0.000275 | 0.002007 | 1.64 |
| FALSE | High | STDSSSYPS | [K].STDSSS | 1xCarbami | B2RR83 2xI | 0.000305 | 0 | 1 | 1 | 1 | B2RR83 | Probable A | B2RR83 [1 | 0 | 2701.151 | 0 | 0.94 | 0.000489 | 12.14 | 70 | 130 | 7.58 | 5.61 | | High | High | High | High | High | High | High | High | High | High | High | 0 | 6.27E-05 | 2.76 |
| FALSE | High | STECLLLSLI | [K].STECLI | 1xCarbami | D3Z6J1 1xP | 0.027121 | 0.001676 | 1 | 2 | 1 | D3Z6J1 | DEP domai | D3Z6J1 [34 | 0 | 1978.074 | 0 | -0.91 | 0.000574 | 0.45 | 130.4 | 69.6 | 2.77 | 11.42 | | High | High | High | High | High | High | High | High | High | High | High | 0.000903 | 0.009564 | 2.43 |
| FALSE | High | STEDGSHD | [K].STEDGS | 1xPhospho | B1AX39 1x | 0.040028 | 0.002867 | 1 | 2 | 2 | B1AX39 | Zinc finger | B1AX39 [5 | 0 | 2115.041 | 0 | 0.24 | 0.039989 | 9.12 | 88.6 | 111.4 | 8.81 | 9.5 | | High | High | High | High | High | High | High | High | High | High | High | 0.0015 | 0.01494 | 2.38 |
| FALSE | High | STEDLSPQI | [K].STEDLS | 1xPhospho | Q80TH2-2 | 0.022182 | 0.001542 | 1 | 6 | 2 | Q80TH2-2 | Isoform 2 c | Q80TH2-2 | 0 | 1341.625 | 0 | 0.38 | 0.016865 | 10.72 | 85.7 | 114.3 | 6.33 | 15.89 | | High | High | High | High | High | High | High | High | High | High | High | 0.001999 | 0.02075 | 1.98 |
| FALSE | High | STEEDDQG | [K].STEEDI | 1xPhospho | F6ZMJ4 1xI | 4.01E-08 | 0 | 1 | 5 | 2 | F6ZMJ4 | Trinucleoti | F6ZMJ4 [6 | 0 | 3050.363 | 0 | 0.65 | 0.022823 | 9.71 | 78.7 | 121.3 | 10.02 | 19.13 | | High | High | High | High | High | High | High | High | High | High | High | 0 | 2.73E-09 | 4.91 |

| | | | | | | | | | | | | | | | | | | | | | | | | | | | | | | | | | | | | | | |
|---|---|---|---|---|---|---|---|---|---|---|---|---|---|---|---|---|---|---|---|---|---|---|---|---|---|---|---|---|---|---|---|---|---|---|---|---|---|---|
| FALSE | High | STELLLGVK | [K].STELLL | 1xPhosphc | Q6QI06 1x | 0.003655 | 0.00021 | 1 | 2 | 2 | Q6QI06 | Rapamycin | Q6QI06 [1 | 0 | 1497.869 | 0 | 0.01 | 0.505963 | 22.03 | 101.2 | 98.8 | 3.13 | 17.03 | | High | High | High | High | High | High | High | High | High | High | High | 0.000275 | 0.001636 | 2.57 |
| FALSE | High | SSWSSDEG | [R].SSWSSI | 1xPhosphc | Q7TSH8 1x | 0.000705 | 0 | 1 | 2 | 5 | Q7TSH8 | Transmem | Q7TSH8 [8 | 0 | 2161.047 | 0 | 0.53 | 0.014308 | 13.22 | 83.6 | 116.4 | 8.85 | 14.96 | | High | High | High | High | High | High | High | High | High | High | High | 2.58E-05 | 0.000338 | 3.18 |
| FALSE | High | SVDSGIYLD | [K].SVDSGI | 1xPhosphc | P70677 1x | 1.46E-05 | 0 | 1 | 2 | 3 | P70677 | Caspase-3 | P70677 [26 | 0 | 1971.972 | 0 | 1.54 | 0.00252 | 28.91 | 51.9 | 148.1 | 12.27 | 13.75 | | High | High | High | High | High | High | High | High | High | High | High | 4.94E-05 | 0.000696 | 2.58 |
| FALSE | High | SVEAGGST | [K].SVEAG | 1xPhosphc | Q9EPR4 1x | 0.000173 | 0 | 1 | 4 | 2 | Q9EPR4 | Solute carr | Q9EPR4 [1 | 0 | 1559.772 | 0 | 2.07 | 0.000243 | 11.91 | 39.4 | 160.6 | 16.01 | 4.67 | | High | High | High | High | High | High | High | High | High | High | High | 0.000183 | 0.001245 | 2.84 |
| FALSE | High | SVEDEMDS | [K].SVEDEN | 1xPhosphc | Q02780-7 | 2.25E-05 | 0 | 1 | 14 | 3 | Q02780-7 | Isoform 7 c | Q02780-7 [ | 0 | 2555.048 | 0 | 0.73 | 0.00041 | 6.51 | 76 | 124 | 10.95 | 5.38 | | High | High | High | High | High | High | High | High | High | High | High | 0 | 3.36E-06 | 2.73 |
| FALSE | High | SYSALSEPE | [K].SYSALS | 2xPhosphc | E9QK89 2x | 3.12E-06 | 0 | 1 | 2 | 1 | E9QK89 | Mediator c | E9QK89 [1 | 0 | 3101.282 | 0 | -1.69 | 0.002847 | 1.25 | 149.6 | 50.4 | 14.12 | 26.47 | | High | High | High | High | High | High | High | High | High | High | High | 0 | 3.65E-07 | 4.18 |
| FALSE | High | SYSDGSEA | [K].SYSDGS | 2xPhosphc | D3YTT6 2x | 0.015465 | 0.001089 | 1 | 7 | 1 | D3YTT6 | Oxysterol-l | D3YTT6 [3 | 0 | 2098.866 | 0 | 0.37 | 0.087964 | 11.23 | 89.8 | 110.2 | 16.03 | 5.37 | | High | High | High | High | High | High | High | High | High | High | High | 0.000571 | 0.005107 | 2.86 |
| FALSE | High | SYSFEASEE | [R].SYSFEA | 1xPhosphc | H3BKF8 1x | 3.29E-05 | 0 | 1 | 11 | 5 | H3BKF8 | Homeobox | H3BKF8 [1 | 0 | 2387.058 | 0 | -0.26 | 0.042872 | 4.91 | 106.8 | 93.2 | 8.89 | 5.58 | | High | High | High | High | High | High | High | High | High | High | High | 0 | 9E-06 | 4.59 |
| FALSE | High | SYSGIEYHL | [K].SYSGIE | 1xPhosphc | A0A0N4SU | 0.001361 | 0 | 1 | 10 | 1 | A0A0N4SU | Peregrin O | A0A0N4SU | 0 | 3406.573 | 0 | -0.55 | 0.001118 | 3.14 | 117.5 | 82.5 | 2.87 | 7.73 | | High | High | High | High | High | High | High | High | High | High | High | 0 | 0.000337 | 3.1 |
| FALSE | High | SYSLSELSV | [R].SYSLSE | 1xPhosphc | Q921I6 1x | 0.009416 | 0.000736 | 1 | 1 | 3 | Q921I6 | SH3 domai | Q921I6 [24 | 0 | 1963.055 | 0 | -0.63 | 0.001672 | 3.79 | 122.1 | 77.9 | 3.77 | 9.6 | | High | High | High | High | High | High | High | High | High | High | High | 0.000327 | 0.002934 | 2.53 |
| FALSE | High | SYSPAHNR | [R].SYSPAH | 1xPhosphc | Q569Z6 1x | 0.003755 | 0.00021 | 1 | 4 | 6 | Q569Z6 | Thyroid ho | Q569Z6 [5 | 0 | 1240.567 | 0 | -0.24 | 0.215257 | 24.74 | 108.3 | 91.7 | 5.29 | 19.7 | | High | High | High | High | High | High | High | High | High | High | High | 0.000275 | 0.002153 | 2.49 |
| FALSE | High | SYSPDGKE | [R].SYSPDG | 2xPhosphc | Q6ZQ61 2x | 0.033404 | 0.002195 | 1 | 8 | 1 | Q6ZQ61 | MCG12197 | Q6ZQ61 [6 | 1 | 2014.881 | 0 | -0.05 | 0.678765 | 34.41 | 110.7 | 89.3 | 85.76 | 66.81 | | High | High | High | High | High | High | High | High | High | High | High | 0.001151 | 0.01211 | 2.69 |
| FALSE | High | SYSPPLPSG | [R].SYSPPL | 2xPhosphc | Q8CBR1 2x | 0.104222 | 0.008348 | 1 | 3 | 1 | Q8CBR1 | Immunoglc | Q8CBR1 [4 | 0 | 2669.183 | 0 | -1.51 | 0.000546 | 1.33 | 146.6 | 53.4 | 8.4 | 9.03 | | High | High | High | High | High | High | High | High | High | High | High | 0.004206 | 0.04524 | 2.06 |
| FALSE | High | SYSSATEED | [R].SYSSAT | 1xPhosphc | Q4V9W2 1 | 0.000196 | 0 | 1 | 1 | 2 | Q4V9W2 | Protein SRI | Q4V9W2 [ | 0 | 1812.83 | 0 | -0.17 | 0.156886 | 5.23 | 103.7 | 96.3 | 5.99 | 10.01 | | High | High | High | High | High | High | High | High | High | High | High | 7.19E-05 | 0.000888 | 3.1 |
| FALSE | High | SYSSATEED | [R].SYSSAT | 2xPhosphc | Q4V9W2 2 | 8.63E-05 | 0 | 1 | 1 | 1 | Q4V9W2 | Protein SRI | Q4V9W2 [ | 0 | 1892.797 | 0 | 0.48 | 0.002408 | 6.41 | 81.6 | 118.4 | 3.05 | 6.58 | | High | High | High | High | High | High | High | High | High | High | High | 0 | 1.52E-05 | 3.08 |
| FALSE | High | SYSSPDITQ | [R].SYSSPD | 1xPhosphc | A2AI52 1x | 1.57E-05 | 0 | 1 | 5 | 7 | A2AI52 | Ubiquitin c | A2AI52 [68 | 0 | 2133.979 | 0 | -0.95 | 0.006295 | 5.22 | 129.8 | 70.2 | 10.62 | 17.6 | | High | High | High | High | High | High | High | High | High | High | High | 0 | 2.25E-06 | 3.58 |
| FALSE | High | SYSSPDITQ | [R].SYSSPD | 1xPhosphc | A2AI52 1x | 2.03E-07 | 0 | 1 | 5 | 8 | A2AI52 | Ubiquitin c | A2AI52 [68 | 0 | 2363.142 | 0 | -0.66 | 0.003368 | 3.8 | 122.2 | 77.8 | 5.38 | 8.33 | | High | High | High | High | High | High | High | High | High | High | High | 0 | 3.82E-06 | 3.62 |
| FALSE | High | SYSSPDITQ | [R].SYSSPD | 1xPhosphc | A2AI52 1x | 0.000162 | 0 | 1 | 5 | 3 | A2AI52 | Ubiquitin c | A2AI52 [68 | 1 | 2519.243 | 0 | -1.06 | 0.002633 | 5.16 | 135.2 | 64.8 | 10.83 | 11.14 | | High | High | High | High | High | High | High | High | High | High | High | 0 | 3.08E-05 | 1.92 |
| FALSE | High | SYSALSEPE | [K].SYSALS | 1xPhosphc | E9QK89 1x | 3.22E-10 | 0 | 1 | 2 | 7 | E9QK89 | Mediator c | E9QK89 [1 | 0 | 3021.316 | 0 | -0.6 | 0.000443 | 3.65 | 121.1 | 78.9 | 3.29 | 9.76 | | High | High | High | High | High | High | High | High | High | High | High | 0 | 1.21E-11 | 5.65 |
| FALSE | High | SYSSSSGSL | [R].SYSSSS | 1xPhosphc | P08551 1x | 0.006306 | 0.00062 | 1 | 2 | 8 | P08551 | Neurofilan | P08551 [56 | 0 | 3551.759 | 0 | 3.53 | 0.003634 | 12.44 | 15.9 | 184.1 | 15.29 | 26.48 | | High | High | High | High | High | High | High | High | High | High | High | 0.000638 | 0.005752 | 3.06 |
| FALSE | High | SYTMDDGV | [R].SYTMD | 1xPhosphc | Q6ZQ32 1x | 0.000808 | 0 | 2 | 7 | 1 | Q6ZQ32; F | MKIAA085 | Q6ZQ32 [8 | 0 | 1656.738 | 0 | 2.48 | 0.001763 | 14.34 | 31.1 | 168.9 | 22.39 | 19.07 | Shared | High | High | High | High | High | High | High | High | High | High | High | 0 | 0.000187 | 1.87 |
| FALSE | High | SYTSLMPPI | [R].SYTSLN | 1xPhosphc | F2Z3U3 1x | 0.061176 | 0.004646 | 1 | 1 | 1 | F2Z3U3 | Ras associa | F2Z3U3 [6 | 0 | 2088.085 | 0 | 1.25 | 0.000457 | 8.23 | 56.5 | 143.5 | 11.9 | 6.97 | | High | High | High | High | High | High | High | High | High | High | High | 0.00237 | 0.02437 | 2.15 |
| FALSE | High | SYTSLMPPI | [R].SYTSLN | 1xPhosphc | F2Z3U3 1x | 0.01002 | 0.000736 | 1 | 1 | 1 | F2Z3U3 | Ras associa | F2Z3U3 [6 | 0 | 2104.08 | 0 | 1.26 | 0.003924 | 22.15 | 58 | 142 | 13.01 | 13.73 | | High | High | High | High | High | High | High | High | High | High | High | 0.000344 | 0.00314 | 2.86 |
| FALSE | High | SYTYMESD | [K].SYTYMI | 1xPhosphc | Q99N96 1x | 2.69E-07 | 0 | 1 | 3 | 3 | Q99N96 | 39S riboso | Q99N96 [7 | 0 | 2493.118 | 0 | -0.15 | 0.240387 | 4.5 | 103.7 | 96.3 | 6.5 | 7.83 | | High | High | High | High | High | High | High | High | High | High | High | 0 | 2.33E-08 | 3.83 |
| FALSE | High | SYTYMESD | [K].SYTYMI | 1xPhosphc | Q99N96 1x | 0.000369 | 0 | 1 | 3 | 2 | Q99N96 | 39S riboso | Q99N96 [7 | 0 | 2509.113 | 0 | -0.13 | 0.158799 | 9.16 | 99.8 | 100.2 | 8.09 | 8.5 | | High | High | High | High | High | High | High | High | High | High | High | 0 | 7.77E-05 | 2.83 |
| FALSE | High | TAADVVSP | [K].TAADV | 1xPhosphc | B2RRE7 1x | 0.005872 | 0.00062 | 1 | 2 | 2 | B2RRE7 | OTU doma | B2RRE7 [9 | 0 | 1767.848 | 0 | -0.15 | 0.182196 | 17.45 | 105.2 | 94.8 | 9.17 | 9.55 | | High | High | High | High | High | High | High | High | High | High | High | 0.000756 | 0.00665 | 3.3 |
| FALSE | High | TAADVVSP | [K].TAADV | 2xPhosphc | B2RRE7 2x | 7.53E-08 | 0 | 1 | 2 | 1 | B2RRE7 | OTU doma | B2RRE7 [9 | 0 | 1847.814 | 0 | 0.57 | 0.003301 | 8.81 | 81.2 | 118.8 | 2.5 | 9.7 | | High | High | High | High | High | High | High | High | High | High | High | 0 | 5.54E-09 | 4 |
| FALSE | High | TAAIPINGS | [K].TAAIPIN | 1xPhosphc | P13405 1x | 0.00967 | 0.000736 | 1 | 4 | 4 | P13405 | Retinoblast | P13405 [23 | 0 | 1405.74 | 0 | 1.44 | 0.000243 | 9.95 | 54 | 146 | 5.7 | 4.37 | | High | High | High | High | High | High | High | High | High | High | High | 0.000344 | 0.003017 | 1.48 |
| FALSE | High | TAAIPINGS | [K].TAAIPIN | 1xDeamida | P13405 1x | 7.68E-05 | 0 | 1 | 4 | 6 | P13405 | Retinoblast | P13405 [23 | 0 | 1406.724 | 0 | 1.37 | 0.000237 | 6.17 | 55.7 | 144.3 | 3.06 | 4.58 | | High | High | High | High | High | High | High | High | High | High | High | 0 | 1.33E-05 | 2.29 |
| FALSE | High | TAAPESEVS | [K].TAAPES | 1xPhosphc | Q5XJV7-2 | 0.003372 | 0.00016 | 1 | 2 | 3 | Q5XJV7-2 | Isoform 2 c | Q5XJV7-2 [ | 0 | 3192.562 | 0 | -0.05 | 0.428573 | 13.25 | 101.8 | 98.2 | 14.95 | 10.78 | | High | High | High | High | High | High | High | High | High | High | High | 0.009004 | 0.0962 | 3.52 |
| FALSE | High | TAASPGAE | [K].TAASPC | 1xPhosphc | A2A6A1 1x | 0.01186 | 0.000889 | 1 | 2 | 1 | A2A6A1 | G patch do | A2A6A1 [4 | 0 | 1168.556 | 0 | -0.06 | 0.201417 | 8.38 | 103.1 | 96.9 | 9.78 | 4.01 | | High | High | High | High | High | High | High | High | High | High | High | 0.000449 | 0.003783 | 2.18 |
| FALSE | High | TAAYGHFG | [R].TAAYGI | 1xPhosphc | Q3TED1 1x | 0.064862 | 0.004664 | 1 | 2 | 1 | Q3TED1 | S-adenosyl | Q3TED1 [3 | 1 | 2603.285 | 0 | -0.56 | 0.000931 | 3.41 | 117.3 | 82.7 | 2.72 | 6.78 | | High | High | High | High | High | High | High | High | High | High | High | 0.002403 | 0.02593 | 1.9 |
| FALSE | High | TADGRVSP | [R].TADGR | 1xPhosphc | Q8VBW5 1 | 0.036731 | 0.00249 | 1 | 4 | 1 | Q8VBW5 | HMG box t | Q8VBW5 [ | 1 | 2552.369 | 0 | 0.11 | 0.190915 | 19.21 | 96.2 | 103.8 | 7.43 | 7.6 | | High | High | High | High | High | High | High | High | High | High | High | 0.001308 | 0.01355 | 1.4 |
| FALSE | High | SYSSSSGSL | [R].SYSSSS | 1xPhosphc | P08551 1x | 0.000458 | 0 | 1 | 2 | 3 | P08551 | Neurofilan | P08551 [56 | 0 | 3567.754 | 0 | 3.86 | 0.000485 | 17.32 | 12.1 | 187.9 | 22.97 | 12.66 | | High | High | High | High | High | High | High | High | High | High | High | 0 | 9.97E-05 | 2.9 |
| FALSE | High | SYSAGNAS | [R].SYSAGN | 1xPhosphc | E9Q238 1x | 0.047894 | 0.00354 | 1 | 3 | 1 | E9Q238 | KN motif a | E9Q238 [3 | 0 | 1888.937 | 0 | 1.88 | 0.000348 | 15.17 | 44.7 | 155.3 | 12.23 | 4.29 | | High | High | High | High | High | High | High | High | High | High | High | 0.001812 | 0.01832 | 1.5 |
| FALSE | High | SYQVAPLSI | [K].SYQVAI | 1xPhosphc | A0A0R4J0C | 0.00582 | 0.00062 | 1 | 7 | 4 | A0A0R4J0C | Disks large | A0A0R4J0C | 0 | 1426.729 | 0 | -1.22 | 0.000244 | 3.21 | 139.9 | 60.1 | 2.1 | 13.31 | | High | High | High | High | High | High | High | High | High | High | High | 0.000327 | 0.002496 | 1.25 |
| FALSE | High | SYQNSPSSI | [R].SYQNSI | 2xPhosphc | Q925J9 2x | 0.022574 | 0.001542 | 1 | 2 | 2 | Q925J9 | Mediator c | Q925J9 [14 | 0 | 3102.411 | 0 | 1.11 | 0.001015 | 12.31 | 64 | 136 | 11.37 | 7.26 | | High | High | High | High | High | High | High | High | High | High | High | 0.000814 | 0.007814 | 2.22 |
| FALSE | High | SYEDCTSGI | [K].SYEDCT | 1xCarbami | G5E8P8 1x | 0.00025 | 0 | 1 | 6 | 2 | G5E8P8 | Estrogen re | G5E8P8 [9 | 0 | 2360.044 | 0 | -1.14 | 0.000333 | 1.95 | 136 | 64 | 5.54 | 8.32 | | High | High | High | High | High | High | High | High | High | High | High | 0 | 5.02E-05 | 3.07 |
| FALSE | High | SYEDLTELE | [R].SYEDLT | 1xPhosphc | A0A0R4J0D | 0.000579 | 0 | 1 | 4 | 2 | A0A0R4J0D | Vacuolar fu | A0A0R4J0D | 0 | 1678.741 | 0 | 0.46 | 0.008431 | 8.22 | 83.6 | 116.4 | 4.6 | 9.54 | | High | High | High | High | High | High | High | High | High | High | High | 0 | 0.000195 | 2.9 |
| FALSE | High | SYELPDGQ | [RK-].SYELI | 1xPhosphc | Q8BFZ3 1x | 7.43E-06 | 0 | 3 | 35 | 1 | Q8BFZ3; P | Beta-actin- | Q8BFZ3 [2 | 0 | 2100.021 | 0 | 0.68 | 0.000489 | 8.14 | 75.3 | 124.7 | 8.62 | 5.41 | Shared | High | High | High | High | High | High | High | High | High | High | High | 0 | 9.63E-07 | 3.89 |
| FALSE | High | SYESSEDCP | [K].SYESSE | 1xCarbami | Q3TSG4 1x | 5.05E-07 | 0 | 1 | 1 | 6 | Q3TSG4 | RNA deme | Q3TSG4 [3 | 0 | 2181.884 | 0 | -0.55 | 0.015758 | 5.35 | 119.5 | 80.5 | 11.49 | 5.98 | | High | High | High | High | High | High | High | High | High | High | High | 0 | 4.73E-08 | 3.8 |
| FALSE | High | SYESSEDCP | [K].SYESSE | 1xCarbami | Q3TSG4 1x | 5.5E-05 | 0 | 1 | 1 | 5 | Q3TSG4 | RNA deme | Q3TSG4 [3 | 1 | 2539.142 | 0 | -0.22 | 0.137948 | 3.68 | 107 | 93 | 8.67 | 6.36 | | High | High | High | High | High | High | High | High | High | High | High | 0.000225 | 0.001438 | 2.38 |
| FALSE | High | SYESSEDCP | [K].SYESSE | 1xCarbami | Q3TSG4 2x | 0.021231 | 0.001507 | 1 | 1 | 2 | Q3TSG4 | RNA deme | Q3TSG4 [3 | 1 | 2619.109 | 0 | -0.24 | 0.032315 | 2.34 | 105.6 | 94.4 | 6.48 | 6.89 | | High | High | High | High | High | High | High | High | High | High | High | 0.000801 | 0.007322 | 2.97 |
| FALSE | High | SYGSQPNS | [K].SYGSQI | 1xPhosphc | B2RY19 1x | 0.00043 | 0 | 1 | 7 | 1 | B2RY19 | Cgnl1 prot | B2RY19 [1 | 0 | 2219.063 | 0 | 1.98 | 0.003497 | 27.12 | 38.5 | 161.5 | 9.85 | 22.17 | | High | High | High | High | High | High | High | High | High | High | High | 0 | 9.26E-05 | 2.56 |
| FALSE | High | SYGSQPNS | [K].SYGSQI | 2xPhosphc | B2RY19 2x | 0.000104 | 0 | 1 | 7 | 2 | B2RY19 | Cgnl1 prot | B2RY19 [1 | 0 | 2299.03 | 0 | 1.76 | 0.000427 | 7.21 | 47.1 | 152.9 | 10.82 | 5.99 | | High | High | High | High | High | High | High | High | High | High | High | 0 | 1.88E-05 | 2.86 |
| FALSE | High | SYHDLSQA | [K].SYHDLS | 1xCarbami | Q64512 1x | 1.6E-06 | 0 | 1 | 4 | 1 | Q64512 | Tyrosine-p | Q64512 [9 | 0 | 1979.9 | 0 | 0.53 | 0.009677 | 10.74 | 85.3 | 114.7 | 10.88 | 8.39 | | High | High | High | High | High | High | High | High | High | High | High | 0 | 1.72E-07 | 4.49 |
| FALSE | High | SYHEAYEPI | [R].SYHEAY | 1xPhosphc | Q9Z0U1 1x | 3.61E-09 | 0 | 1 | 1 | 24 | Q9Z0U1 | Tight junct | Q9Z0U1 [2 | 0 | 2622.066 | 0 | 0.08 | 0.255505 | 7.1 | 96.5 | 103.5 | 6.53 | 6.98 | | High | High | High | High | High | High | High | High | High | High | High | 0 | 1.13E-08 | 4.78 |
| FALSE | High | SYHEAYEPI | [R].SYHEAY | 2xPhosphc | Q9Z0U1 2x | 8.4E-08 | 0 | 1 | 1 | 1 | Q9Z0U1 | Tight junct | Q9Z0U1 [2 | 0 | 2702.032 | 0 | -0.18 | 0.368752 | 17.89 | 104.6 | 95.4 | 12.3 | 18.02 | | High | High | High | High | High | High | High | High | High | High | High | 0 | 6.26E-09 | 4.79 |
| FALSE | High | SYHEAYEPI | [R].SYHEAY | 1xPhosphc | Q9Z0U1 1x | 6.58E-08 | 0 | 1 | 1 | 4 | Q9Z0U1 | Tight junct | Q9Z0U1 [2 | 1 | 2778.167 | 0 | 0.26 | 0.378549 | 12.34 | 90 | 110 | 11.45 | 14.4 | | High | High | High | High | High | High | High | High | High | High | High | 0.005374 | 0.05854 | 2.87 |
| FALSE | High | SYHEAYEPI | [R].SYHEAY | 2xPhosphc | Q9Z0U1 2x | 0.001021 | 0 | 1 | 1 | 1 | Q9Z0U1 | Tight junct | Q9Z0U1 [2 | 1 | 2858.133 | 0 | -0.26 | 0.007624 | 3.66 | 108.1 | 91.9 | 7.55 | 7.33 | | High | High | High | High | High | High | High | High | High | High | High | 0 | 0.000243 | 3.96 |
| FALSE | High | SYLEGSSDT | [K].SYLEGS | 1xPhosphc | Q8R4Z6 1x | 0.000302 | 0 | 1 | 3 | 3 | Q8R4Z6 | DEAD-box | Q8R4Z6 [6 | 0 | 1865.93 | 0 | -1.07 | 0.000362 | 0.38 | 136.3 | 63.7 | 3.84 | 8.03 | | High | High | High | High | High | High | High | High | High | High | High | 0 | 9.78E-05 | 3.03 |
| FALSE | High | SYLEGSSDT | [K].SYLEGS | 2xPhosphc | Q8R4Z6 2x | 0.030898 | 0.001896 | 1 | 3 | 2 | Q8R4Z6 | DEAD-box | Q8R4Z6 [6 | 0 | 1945.896 | 0 | -0.96 | 0.000503 | 3.11 | 131 | 69 | 4.12 | 8.33 | | High | High | High | High | High | High | High | High | High | High | High | 0.00488 | 0.05318 | 2.26 |
| FALSE | High | SYLGGGAG | [K].SYLGG | 1xPhosphc | Q00899 1x | 7.07E-11 | 0 | 1 | 2 | 1 | Q00899 | Transcripti | Q00899 [1 | 0 | 2172.049 | 0 | -1.14 | 0.000673 | 2.24 | 139.4 | 60.6 | 7.29 | 14.49 | | High | High | High | High | High | High | High | High | High | High | High | 0 | 2.21E-12 | 4.34 |
| FALSE | High | SYLLGNSSF | [R].SYLLGN | 1xPhosphc | P48678 1x | 0.016306 | 0.001124 | 1 | 2 | 2 | P48678 | Prelamin-A | P48678 [64 | 0 | 1402.693 | 0 | 1.55 | 0.000241 | 12.39 | 49.4 | 150.6 | 7.77 | 5.2 | | High | High | High | High | High | High | High | High | High | High | High | 0.000585 | 0.005402 | 1.29 |
| FALSE | High | SYLNFVSPL | [K].SYLNFV | 1xPhosphc | Q99LI5 1x | 0.000731 | 0 | 1 | 1 | 1 | Q99LI5 | Zinc finger | Q99LI5 [47 | 0 | 2301.23 | 0 | -1.13 | 0.002335 | 4.07 | 140.6 | 59.4 | 13.81 | 7.48 | | High | High | High | High | High | High | High | High | High | High | High | 0 | 0.000168 | 2.82 |
| FALSE | High | SYIPHSSPS | [K].SYIPHS | 1xPhosphc | Q80XK6 1x | 0.012072 | 0.000889 | 1 | 1 | 1 | Q80XK6 | Autophagy | Q80XK6 [1 | 0 | 1866.895 | 0 | -0.05 | 0.915429 | 31.08 | 96.6 | 103.4 | 16.46 | 20.63 | | High | High | High | High | High | High | High | High | High | High | High | 0.000449 | 0.003884 | 2.27 |
| FALSE | High | SYIQSLPQS | [K].SYIQSLI | 1xPhosphc | Q3UIB2 1x | 0.004911 | 0.000504 | 1 | 1 | 1 | Q3UIB2 | Mitogen-a | Q3UIB2 [2 | 0 | 1785.955 | 0 | 1.03 | 0.000755 | 11.39 | 65.7 | 134.3 | 9.82 | 6.7 | | High | High | High | High | High | High | High | High | High | High | High | 0.000225 | 0.001414 | 2.07 |
| FALSE | High | SYIVMSPES | [K].SYIVMS | 1xPhosphc | A0A1L1SSL | 0.00011 | 0 | 1 | 4 | 2 | A0A1L1SSL | Zinc finger | A0A1L1SSL | 0 | 1890.969 | 0 | -1.1 | 0.000765 | 1.96 | 136.3 | 63.7 | 5.11 | 7.54 | | High | High | High | High | High | High | High | High | High | High | High | 0 | 1.99E-05 | 3.31 |
| FALSE | High | SYMNPGD | [K].SYMNP | 1xPhosphc | A2APA8 1x | 0.000123 | 0 | 1 | 4 | 1 | A2APA8 | Transcripti | A2APA8 [4 | 0 | 2121.956 | 0 | -0.53 | 0.00331 | 1.27 | 119.1 | 80.9 | 8.42 | 9.34 | | High | High | High | High | High | High | High | High | High | High | High | 0 | 2.25E-05 | 2.95 |
| FALSE | High | SYMNPGD | [K].SYMNP | 1xPhosphc | A2APA8 1x | 0.000191 | 0 | 1 | 4 | 4 | A2APA8 | Transcripti | A2APA8 [4 | 0 | 2137.951 | 0 | -0.42 | 0.003079 | 3.43 | 115.3 | 84.7 | 4.78 | 5.5 | | High | High | High | High | High | High | High | High | High | High | High | 0.000344 | 0.003177 | 3.24 |
| FALSE | High | SYNLQER | [R].SYNLQE | 1xPhosphc | A0A0R4J1C | 0.01319 | 0.000889 | 1 | 11 | 2 | A0A0R4J1C | Anion exch | A0A0R4J1C | 0 | 1218.572 | 0 | 2.48 | 0.005733 | 36.79 | 31.9 | 168.1 | 16.22 | 24.92 | | High | High | High | High | High | High | High | High | High | High | High | 0.000689 | 0.006051 | 1.42 |
| FALSE | High | SYNLVVAK | [R].SYNLVV | 1xPhosphc | B6ZHD0 1x | 0.012845 | 0.000889 | 2 | 16 | 6 | B6ZHD0; Q | Erythrocyt | B6ZHD0 [8 | 0 | 1202.638 | 0 | 1.38 | 0.003575 | 30.73 | 55.9 | 144.1 | 12.23 | 12.32 | Shared | High | High | High | High | High | High | High | High | High | High | High | 0.000449 | 0.004148 | 2.51 |
| FALSE | High | SYNLVVAK | [R].SYNLVV | 1xPhosphc | B6ZHD0 1x | 0.005872 | 0.00062 | 2 | 16 | 1 | B6ZHD0; Q | Erythrocyt | B6ZHD0 [8 | 0 | 1431.801 | 0 | 1.37 | 0.002682 | 5.32 | 55.5 | 144.5 | 19.35 | 10.72 | Shared | High | High | High | High | High | High | High | High | High | High | High | 0.000275 | 0.001738 | 2.13 |
| FALSE | High | SYNMPSLD | [R].SYNMP | 1xPhosphc | Q91ZV0 1x | 0.006653 | 0.00062 | 1 | 12 | 5 | Q91ZV0 | Melanoma | Q91ZV0 [1 | 0 | 1608.774 | 0 | 1.24 | 0.039904 | 53.14 | 58.2 | 141.8 | 25.52 | 36.2 | | High | High | High | High | High | High | High | High | High | High | High | 0.001151 | 0.01207 | 1.8 |
| FALSE | High | SYNTPGPA | [R].SYNTP | 1xPhosphc | D3Z482 1x | 3.17E-05 | 0 | 1 | 3 | 2 | D3Z482 | Ankyrin re | D3Z482 [24 | 0 | 3287.409 | 0 | 0.39 | 0.014788 | 16.91 | 80.3 | 119.7 | 12.87 | 11.58 | | High | High | High | High | High | High | High | High | High | High | High | 0 | 4.94E-06 | 3.57 |
| FALSE | High | SYQNSPSSI | [R].SYQNSI | 1xPhosphc | Q925J9 1x | 0.008167 | 0.000736 | 1 | 2 | 3 | Q925J9 | Mediator c | Q925J9 [14 | 0 | 3022.445 | 0 | 1.19 | 0.000689 | 8.8 | 60.2 | 139.8 | 2.66 | 7.84 | | High | High | High | High | High | High | High | High | High | High | High | 0.00074 | 0.006476 | 2.26 |
| FALSE | High | TADSSSSED | [R].TADSSS | 1xPhosphc | Q61686 1x | 4.27E-05 | 0 | 1 | 2 | 9 | Q61686 | Chromobo | Q61686 [8 | 0 | 2442.085 | 0 | 0.63 | 0.009421 | 18.81 | 80.7 | 119.3 | 10.39 | 8.9 | | High | High | High | High | High | High | High | High | High | High | High | 0 | 7.04E-05 | 3.32 |
| FALSE | High | SYDYDAWA | [K].SYDYDA | 1xPhosphc | Q9D706 1x | 0.005 | 0.000504 | 1 | 1 | 1 | Q9D706 | RNA polym | Q9D706 [8 | 0 | 1656.771 | 0 | 0.66 | 0.025206 | 26.18 | 78.8 | 121.2 | 12.02 | 15.31 | | High | High | High | High | High | High | High | High | High | High | High | 0.000225 | 0.001444 | 2.22 |
| FALSE | High | TADSSSSED | [R].TADSSS | 2xPhosphc | Q61686 2x | 0.000183 | 0 | 1 | 2 | 2 | Q61686 | Chromobo | Q61686 [8 | 0 | 2522.051 | 0 | 0.49 | 0.058658 | 20.9 | 83.7 | 116.3 | 14 | 8.94 | | High | High | High | High | High | High | High | High | High | High | High | 0 | 0.000109 | 3.03 |
| FALSE | High | TAEDAGVA | [R].TAEDA | 1xCarbami | Q91YI1-2 1 | 0.020499 | 0.001472 | 1 | 1 | 1 | Q91YI1-2 | Isoform 2 c | Q91YI1-2 [ | 0 | 4327.008 | 0 | -1.16 | 0.000668 | 3.31 | 134 | 66 | 4.49 | 15.13 | | High | High | High | High | High | High | High | High | High | High | High | 0.000786 | 0.006976 | 3.12 |
| FALSE | High | TAPVTSTPS | [K].TAPVTS | 2xPhosphc | B9EIG6 2x | 0.004824 | 0.000456 | 1 | 5 | 1 | B9EIG6 | Pogo trans | B9EIG6 [42 | 0 | 2668.365 | 0 | -0.18 | 0.025779 | 3.96 | 102 | 98 | 13.31 | 16.38 | | High | High | High | High | High | High | High | High | High | High | High | 0.000204 | 0.001394 | 2.46 |
| FALSE | High | TAQASASP | [R].TAQAS | 2xPhosphc | Q80Y19 2x | 0.011246 | 0.000772 | 1 | 3 | 1 | Q80Y19 | Rho GTPas | Q80Y19 [3 | 0 | 2202.159 | 0 | 0.25 | 0.032726 | 12.44 | 92.9 | 107.1 | 8.27 | 3.07 | | High | High | High | High | High | High | High | High | High | High | High | 0.000418 | 0.003589 | 2.79 |
| FALSE | High | TAQASPVP | [R].TAQASI | 1xPhosphc | A2AEG2 1x | 0.024638 | 0.001612 | 1 | 4 | 1 | A2AEG2 | Oral-facial- | A2AEG2 [7 | 0 | 1322.667 | 0 | 1.05 | 0.000383 | 5.54 | 65.3 | 134.7 | 4.15 | 3.98 | | High | High | High | High | High | High | High | High | High | High | High | 0.000859 | 0.008587 | 1.28 |
| FALSE | High | TAQDLETSI | [R].TAQDLI | 1xPhosphc | A2AIX1 1x | 1.07E-05 | 0 | 1 | 7 | 8 | A2AIX1 | SEC16 hom | A2AIX1 [20 | 0 | 2472.186 | 0 | 0.36 | 0.042344 | 7.04 | 87.7 | 112.3 | 11.51 | 12.17 | | High | High | High | High | High | High | High | High | High | High | High | 0 | 7.33E-05 | 3.15 |
| FALSE | High | TAQQPAGI | [R].TAQQP | 1xPhosphc | Q5DTX6 1x | 0.052133 | 0.00384 | 1 | 1 | 1 | Q5DTX6 | Junctional | Q5DTX6 [8 | 1 | 2303.171 | 0 | 1.56 | 0.000353 | 6.26 | 50.7 | 149.3 | 5.7 | 8.26 | | High | High | High | High | High | High | High | High | High | High | High | 0.001999 | 0.02015 | 1.89 |
| FALSE | High | TAQTLLYTF | [R].TAQTLI | 1xPhosphc | Q8BQ33 1x | 0.030632 | 0.001857 | 1 | 1 | 1 | Q8BQ33 | Treslin OS= | Q8BQ33 [1 | 0 | 1601.814 | 0 | -1.02 | 0.000555 | 2.36 | 133.9 | 66.1 | 6.26 | 5.78 | | High | High | High | High | High | High | High | High | High | High | High | 0.001011 | 0.01107 | 1.44 |
| FALSE | High | TARPNSEAI | [K].TARPNS | 1xPhosphc | Q05D44 1x | 0.000292 | 0 | 1 | 1 | 4 | Q05D44 | Eukaryotic | Q05D44 [1 | 0 | 2470.093 | 0 | -0.76 | 0.482659 | 12.23 | 136.1 | 63.9 | 70.75 | 83.36 | | High | High | High | High | High | High | High | High | High | High | High | 0 | 5.99E-05 | 3.62 |
| FALSE | High | TARPNSEAI | [K].TARPNS | 1xPhosphc | Q05D44 1x | 1.52E-07 | 0 | 1 | 1 | 11 | Q05D44 | Eukaryotic | Q05D44 [1 | 0 | 2699.256 | 0 | -0.86 | 0.004406 | 0.64 | 129.4 | 70.6 | 8.05 | 8.99 | | High | High | High | High | High | High | High | High | High | High | High | 0.000275 | 0.00158 | 2.65 |
| FALSE | High | TARPNSEAI | [K].TARPNS | 1xDeamida | Q05D44 1x | 9.39E-06 | 0 | 1 | 1 | 7 | Q05D44 | Eukaryotic | Q05D44 [1 | 0 | 2700.24 | 0 | -0.69 | 0.018897 | 4.98 | 123 | 77 | 10.78 | 9.91 | | High | High | High | High | High | High | High | High | High | High | High | 0 | 2.71E-05 | 3.49 |
| FALSE | High | TARPNSEAI | [K].TARPNS | 2xPhosphc | Q05D44 2x | 1.43E-06 | 0 | 1 | 1 | 7 | Q05D44 | Eukaryotic | Q05D44 [1 | 0 | 2779.223 | 0 | -0.45 | 0.025158 | 6.39 | 118.9 | 81.1 | 8.2 | 8.16 | | High | High | High | High | High | High | High | High | High | High | High | 0 | 5.46E-05 | 3.79 |
| FALSE | High | TASAGTVS | [R].TASAG | 1xPhosphc | F2Z3U3 1x | 0.000118 | 0 | 1 | 4 | 2 | F2Z3U3 | Ras associa | F2Z3U3 [1 | 0 | 1544.716 | 0 | 0.3 | 0.04162 | 15.54 | 88 | 112 | 4.85 | 13.71 | | High | High | High | High | High | High | High | High | High | High | High | 0 | 2.16E-05 | 2.15 |
| FALSE | High | TASAPNLAI | [R].TASAPN | 1xPhosphc | Q8C8U0-3 | 0.000297 | 0 | 1 | 6 | 3 | Q8C8U0-3 | Isoform 3 c | Q8C8U0-3 | 0 | 1540.746 | 0 | 0.78 | 0.083129 | 17.29 | 75.8 | 124.2 | 11.23 | 35.23 | | High | High | High | High | High | High | High | High | High | High | High | 0.000204 | 0.001334 | 2.41 |
| FALSE | High | TASAPNLAI | [R].TASAPN | 1xPhosphc | Q8C8U0-3 | 1.23E-05 | 0 | 1 | 6 | 1 | Q8C8U0-3 | Isoform 3 c | Q8C8U0-3 | 0 | 1769.909 | 0 | 1.06 | 0.008295 | 28.43 | 66 | 134 | 10.2 | 20.97 | | High | High | High | High | High | High | High | High | High | High | High | 0 | 1.71E-06 | 2.93 |
| FALSE | High | TAPVTSTPS | [K].TAPVTS | 1xPhosphc | B9EIG6 1x | 7.7E-06 | 0 | 1 | 5 | 2 | B9EIG6 | Pogo trans | B9EIG6 [42 | 0 | 2588.399 | 0 | -0.39 | 0.01808 | 3.91 | 114.9 | 85.1 | 7.27 | 9.1 | | High | High | High | High | High | High | High | High | High | High | High | 2.58E-05 | 0.000338 | 3.16 |
| FALSE | High | TASAPTK | [R].TASAPT | 1xPhosphc | A2AP18 1x | 0.080423 | 0.006219 | 1 | 10 | 1 | A2AP18 | 1-phospha | A2AP18 [1 | 0 | 1213.659 | 0 | 0.37 | 0.041941 | 29.1 | 84 | 116 | 19.79 | 7.07 | | High | High | High | High | High | High | High | High | High | High | High | 0.003188 | 0.03348 | 1.35 |
| FALSE | High | TASCSELDV | [R].TASCSE | 1xCarbami | Q5DTN0 1x | 5.45E-05 | 0 | 1 | 6 | 2 | Q5DTN0 | MKIAA410 | Q5DTN0 [7 | 0 | 1888.913 | 0 | 0.69 | 0.002106 | 0.63 | 75.3 | 124.7 | 3.25 | 8.02 | | High | High | High | High | High | High | High | High | High | High | High | 0 | 1.75E-05 | 3.68 |
| FALSE | High | TASDSDEQ | [R].TASDSD | 1xPhosphc | F8WHU5 1 | 1.56E-06 | 0 | 1 | 3 | 13 | F8WHU5 | Sister chro | F8WHU5 [ | 0 | 2187.985 | 0 | 0.16 | 0.194543 | 1.86 | 94.8 | 105.2 | 3.54 | 9.23 | | High | High | High | High | High | High | High | High | High | High | High | 0 | 9.09E-05 | 3.58 |
| FALSE | High | TASDSDEQ | [R].TASDSD | 2xPhosphc | F8WHU5 2 | 7.7E-06 | 0 | 1 | 3 | 6 | F8WHU5 | Sister chro | F8WHU5 [ | 0 | 2267.951 | 0 | 0.85 | 0.00939 | 1.64 | 73.2 | 126.8 | 6.71 | 13.56 | | High | High | High | High | High | High | High | High | High | High | High | 0 | 1.01E-06 | 4.11 |
| FALSE | High | TASEGSEAI | [R].TASEGS | 1xPhosphc | Q05CL8 1x | 3.14E-05 | 0 | 1 | 2 | 7 | Q05CL8 | La-related | Q05CL8 [2 | 0 | 2041.973 | 0 | -0.48 | 0.000452 | 2.62 | 116.9 | 83.1 | 2.42 | 4.9 | | High | High | High | High | High | High | High | High | High | High | High | 0 | 1.41E-05 | 3.25 |
| FALSE | High | TASEGSEAI | [R].TASEGS | 2xPhosphc | Q05CL8 2x | 0.000772 | 0 | 1 | 2 | 4 | Q05CL8 | La-related | Q05CL8 [2 | 0 | 2121.939 | 0 | -0.48 | 0.002691 | 2.26 | 117.3 | 82.7 | 3.16 | 5.93 | | High | High | High | High | High | High | High | High | High | High | High | 4.94E-05 | 0.00075 | 2.68 |
| FALSE | High | TASELLLDR | [R].TASELL | 1xPhosphc | E2JF22 1x | 0.003138 | 0.00016 | 1 | 8 | 3 | E2JF22 | Piezo-type | E2JF22 [16 | 0 | 1326.687 | 0 | 0.76 | 0.000552 | 6.05 | 76.3 | 123.7 | 5.9 | 5.13 | | High | High | High | High | High | High | High | High | High | High | High | 7.19E-05 | 0.000856 | 2.62 |
| FALSE | High | TASEINFDK | [R].TASEIN | 1xPhosphc | E9PX30 1x | 0.000131 | 0 | 1 | 5 | 2 | E9PX30 | Mitogen-a | E9PX30 [3 | 0 | 1562.787 | 0 | 2.06 | 0.001173 | 18.76 | 39.1 | 160.9 | 3.71 | 14.19 | | High | High | High | High | High | High | High | High | High | High | High | 4.94E-05 | 0.000684 | 2.53 |
| FALSE | High | TASESISNL | [K].TASESIS | 1xPhosphc | F8WIA1 1x | 1.87E-07 | 0 | 1 | 6 | 4 | F8WIA1 | CAP-Gly dc | F8WIA1 [1 | 0 | 2118.073 | 0 | 1.36 | 0.00142 | 7.25 | 52.6 | 147.4 | 11.73 | 9.35 | | High | High | High | High | High | High | High | High | High | High | High | 0 | 1.55E-08 | 3.13 |
| FALSE | High | TASESISNL | [K].TASESIS | 2xPhosphc | F8WIA1 2x | 0.000382 | 0 | 1 | 6 | 2 | F8WIA1 | CAP-Gly dc | F8WIA1 [1 | 0 | 2198.04 | 0 | 1.55 | 0.001467 | 17.54 | 48.8 | 151.2 | 11.45 | 12.41 | | High | High | High | High | High | High | High | High | High | High | High | 0 | 8.13E-05 | 2.46 |
| FALSE | High | TASESNVYI | [R].TASESN | 1xPhosphc | Q8K1N2-2 | 3.07E-06 | 0 | 1 | 5 | 5 | Q8K1N2-2 | Isoform 2 c | Q8K1N2-2 | 0 | 2309.044 | 0 | 1.93 | 0.000385 | 6.49 | 44.3 | 155.7 | 11.1 | 9.33 | | High | High | High | High | High | High | High | High | High | High | High | 0 | 3.58E-07 | 4.77 |
| FALSE | High | TASETRSEC | [R].TASETR | 1xPhosphc | Q7TSC1 1x | 0.000645 | 0 | 1 | 3 | 2 | Q7TSC1 | Protein PRI | Q7TSC1 [1 | 1 | 2451.169 | 0 | 0.39 | 0.009386 | 5.36 | 85.2 | 114.8 | 7.65 | 6.92 | | High | High | High | High | High | High | High | High | High | High | High | 0.001183 | 0.01282 | 2.07 |
| FALSE | High | TASETRSEC | [R].TASETR | 2xPhosphc | Q7TSC1 2x | 0.012845 | 0.000889 | 1 | 3 | 3 | Q7TSC1 | Protein PRI | Q7TSC1 [1 | 1 | 2531.136 | 0 | 0.2 | 0.022153 | 3.94 | 92.2 | 107.8 | 5.85 | 4.22 | | High | High | High | High | High | High | High | High | High | High | High | 0.001484 | 0.0149 | 2.91 |
| FALSE | High | TASFGNILD | [K].TASFGN | 1xPhosphc | Q923A8 1x | 3.96E-07 | 0 | 1 | 2 | 1 | Q923A8 | Map3k7 pr | Q923A8 [4 | 0 | 2593.322 | 0 | -0.82 | 0.003925 | 6.85 | 130 | 70 | 16.55 | 25.97 | | High | High | High | High | High | High | High | High | High | High | High | 0 | 3.59E-08 | 4.42 |
| FALSE | High | TASAVAGK | [K].TASAVA | 1xPhosphc | Q3TXS7 1x | 0.093879 | 0.007306 | 1 | 5 | 1 | Q3TXS7 | 26S protea | Q3TXS7 [3 | 1 | 2394.288 | 0 | 0.36 | 0.027624 | 7.67 | 87.7 | 112.3 | 7.53 | 7.25 | | High | High | High | High | High | High | High | High | High | High | High | 0.003741 | 0.04021 | 1.88 |
| FALSE | High | TAPTSTIAP | [R].TAPTST | 1xPhosphc | Q547S9 1x | 1.45E-05 | 0 | 1 | 6 | 4 | Q547S9 | CAMP resp | Q547S9 [2 | 0 | 3146.564 | 0 | -0.05 | 0.688602 | 12.43 | 101.8 | 98.2 | 10.63 | 31.61 | | High | High | High | High | High | High | High | High | High | High | High | 0 | 2.05E-06 | 4.24 |
| FALSE | High | TAPTSTIAP | [R].TAPTST | 1xPhosphc | Q547S9 1x | 9.2E-05 | 0 | 1 | 6 | 2 | Q547S9 | CAMP resp | Q547S9 [2 | 0 | 3130.569 | 0 | 0 | 0.66309 | 18.01 | 100.2 | 99.8 | 6.47 | 21.78 | | High | High | High | High | High | High | High | High | High | High | High | 0 | 9.51E-05 | 4.16 |
| FALSE | High | TAPSSPLTS | [K].TAPSSP | 2xPhosphc | Q6ZPJ0 2x | 2.65E-05 | 0 | 1 | 2 | 2 | Q6ZPJ0 | Testis-expr | Q6ZPJ0 [26 | 0 | 1805.792 | 0 | 0.38 | 0.021108 | 0.7 | 85.7 | 114.3 | 8.98 | 14.07 | | High | High | High | High | High | High | High | High | High | High | High | 0 | 4.02E-06 | 3.35 |
| FALSE | High | TAFDDAIAE | [K].TAFDDA | 1xPhosphc | A8IP69 1x | 3.1E-07 | 0 | 1 | 1 | 2 | A8IP69 | 14-3-3 pro | A8IP69 [19 | 0 | 2669.264 | 0 | 0.1 | 0.253535 | 9.67 | 93.4 | 106.6 | 8.6 | 10.39 | | High | High | High | High | High | High | High | High | High | High | High | 0 | 2.73E-08 | 3.74 |
| FALSE | High | TAFDEAIAE | [K].TAFDEA | 1xPhosphc | F6VW30 1x | 6.81E-07 | 0 | 1 | 4 | 3 | F6VW30 | 14-3-3 pro | F6VW30 [2 | 0 | 2683.279 | 0 | -0.51 | 0.002805 | 4.65 | 118.2 | 81.8 | 5.3 | 10.29 | | High | High | High | High | High | High | High | High | High | High | High | 0 | 6.57E-08 | 3.99 |
| FALSE | High | TAFDEAIAE | [K].TAFDEA | 1xPhosphc | P63101 1x | 4.41E-07 | 0 | 1 | 2 | 8 | P63101 | 14-3-3 pro | P63101 [19 | 0 | 2670.284 | 0 | 0.61 | 0.000623 | 3.23 | 78.6 | 121.4 | 2.89 | 5.8 | | High | High | High | High | High | High | High | High | High | High | High | 0 | 4.05E-08 | 4.02 |
| FALSE | High | TAFYNEDD | [R].TAFYNE | 1xPhosphc | F8VQ93 1x | 4.08E-08 | 0 | 1 | 2 | 9 | F8VQ93 | PH-interact | F8VQ93 [1 | 0 | 2041.823 | 0 | 0.26 | 0.051617 | 2.92 | 91.6 | 108.4 | 3.74 | 8.67 | | High | High | High | High | High | High | High | High | High | High | High | 0 | 2.8E-09 | 4.86 |
| FALSE | High | TAGYRPQA | [K].TAGYRI | 1xPhosphc | P42703 1x | 5.82E-06 | 0 | 1 | 2 | 1 | P42703 | Leukemia i | P42703 [10 | 0 | 2752.341 | 0 | 0.51 | 0.02492 | 8.26 | 77.4 | 122.6 | 14.46 | 20.84 | | High | High | High | High | High | High | High | High | High | High | High | 0 | 7.36E-07 | 3.71 |
| FALSE | High | TAHNSEAD | [K].TAHNSI | 1xPhosphc | Q9D4J7 1x | 0.000186 | 0 | 1 | 3 | 2 | Q9D4J7 | PHD finger | Q9D4J7 [1 | 0 | 3006.32 | 0 | 0.61 | 0.165093 | 24.81 | 84.8 | 115.2 | 15.71 | 38.27 | | High | High | High | High | High | High | High | High | High | High | High | 0 | 8.29E-05 | 3.37 |
| FALSE | High | TAHNSEAD | [K].TAHNSI | 1xPhosphc | Q9D4J7 1x | 4.33E-07 | 0 | 1 | 3 | 6 | Q9D4J7 | PHD finger | Q9D4J7 [1 | 0 | 3235.483 | 3 | 1.35 | 0.000347 | 5.6 | 55.3 | 144.7 | 4.93 | 7.05 | | High | High | High | High | High | High | High | High | High | High | High | 0 | 1.85E-06 | 3.61 |
| FALSE | High | TAHNSEAD | [K].TAHNSI | 1xDeamida | Q9D4J7 1x | 8.2E-06 | 0 | 1 | 3 | 2 | Q9D4J7 | PHD finger | Q9D4J7 [1 | 0 | 3236.467 | 1 | 0.49 | 0.008457 | 10.44 | 81.7 | 118.3 | 12.03 | 7.34 | | High | High | High | High | High | High | High | High | High | High | High | 0 | 1.09E-06 | 2.93 |

| | | | | | | | | | | | | | | | | | | | | | | | | | | | | | | | | | | | | | | |
|---|---|---|---|---|---|---|---|---|---|---|---|---|---|---|---|---|---|---|---|---|---|---|---|---|---|---|---|---|---|---|---|---|---|---|---|---|---|---|
| FALSE | High | TAHNSEAD | [K].TAHNS | 2xPhospho | Q9D4J7 2x | 0.00145 | 5.66E-05 | 1 | 3 | 3 | Q9D4J7 | PHD finger | Q9D4J7 [1 | 0 | 3315.45 | 0 | 0.84 | 0.000912 | 5.53 | 72.5 | 127.5 | 5.67 | 9.21 | | High | High | High | High | High | High | High | High | High | High | High | 2.58E-05 | 0.000363 | 2.62 |
| FALSE | High | TAHNSEAD | [K].TAHNS | 3xPhospho | Q9D4J7 3x | 0.042868 | 0.003228 | 1 | 3 | 1 | Q9D4J7 | PHD finger | Q9D4J7 [1 | 1 | 3395.396 | 0 | 0.7 | 0.025755 | 28.47 | 77.9 | 122.1 | 10.85 | 13.29 | | High | High | High | High | High | High | High | High | High | High | High | 0.001651 | 0.01623 | 3.68 |
| FALSE | High | TAHNSEAD | [K].TAHNS | 1xPhospho | Q9D4J7 1x | 0.000483 | 0 | 1 | 3 | 2 | Q9D4J7 | PHD finger | Q9D4J7 [1 | 1 | 3464.626 | 0 | 1.87 | 0.001703 | 23.68 | 43.9 | 156.1 | 14.62 | 13.92 | | High | High | High | High | High | High | High | High | High | High | High | 2.58E-05 | 0.000431 | 4.06 |
| FALSE | High | TAHYTPLP | [R].TAHYT | 1xCarbami | Q3TI79 1x | 0.000264 | 0 | 1 | 6 | 2 | Q3TI79 | Putative ur | Q3TI79 [66 | 0 | 2194.113 | 0 | -0.43 | 0.044983 | 6.6 | 113.8 | 86.2 | 12.37 | 8.5 | | High | High | High | High | High | High | High | High | High | High | High | 0 | 0.000129 | 3.39 |
| FALSE | High | TALINSTGE | [R].TALINS | 1xPhospho | P48678 1x | 0.055777 | 0.004093 | 1 | 4 | 1 | P48678 | Prelamin-A | P48678 [52 | 0 | 1816.871 | 0 | 0.71 | 0.000646 | 6.75 | 75.2 | 124.8 | 6.08 | 4.18 | | High | High | High | High | High | High | High | High | High | High | High | 0.002117 | 0.02189 | 2.24 |
| FALSE | High | TAIPLPSSP | [R].TAIPLP | 1xPhospho | Q641K5 1x | 0.006536 | 0.00062 | 1 | 1 | 2 | Q641K5 | NUAK fami | Q641K5 [4 | 0 | 2155.076 | 0 | -1.26 | 0.000217 | 1.77 | 143.1 | 56.9 | 4.83 | 9.51 | | High | High | High | High | High | High | High | High | High | High | High | 0.000275 | 0.001944 | 2.66 |
| FALSE | High | TALPLSPEH | [K].TALPLS | 1xPhospho | A0A140T8 | 0.00045 | 0 | 1 | 1 | 1 | A0A140T8 | Chromoso | A0A140T8 | 0 | 1816.976 | 0 | -0.86 | 0.002254 | 3.59 | 129.1 | 70.9 | 5.71 | 11.31 | | High | High | High | High | High | High | High | High | High | High | High | 0 | 9.73E-05 | 1.83 |
| FALSE | High | TALPTSGSS | [R].TALPTS | 1xPhospho | Q80VP1-2 | 0.000171 | 0 | 1 | 2 | 10 | Q80VP1-2 | Isoform 2 c | Q80VP1-2 | 0 | 2565.301 | 0 | -0.64 | 0.092475 | 15.65 | 119.2 | 80.8 | 25.83 | 11.83 | | High | High | High | High | High | High | High | High | High | High | High | 0.000294 | 0.002341 | 3.33 |
| FALSE | High | TALSDLYLE | [R].TALSDL | 1xPhospho | Q542P4 1x | 0.000984 | 0 | 1 | 9 | 2 | Q542P4 | Membrane | Q542P4 [1 | 0 | 2182.192 | 0 | 1.82 | 0.000975 | 14.93 | 40.2 | 159.8 | 10.45 | 13.3 | | High | High | High | High | High | High | High | High | High | High | High | 0 | 0.000236 | 3.23 |
| FALSE | High | TAMGSNQ | [K].TAMGS | 1xPhospho | Q3UW53 1 | 0.000191 | 0 | 1 | 1 | 8 | Q3UW53 | Protein Nik | Q3UW53 [ | 0 | 1515.682 | 0 | -0.02 | 0.480983 | 15.86 | 105.9 | 94.1 | 6.68 | 14.82 | | High | High | High | High | High | High | High | High | High | High | High | 0.004277 | 0.0458 | 1.75 |
| FALSE | High | TAMPSPGV | [K].TAMPS | 1xPhospho | Q67FY2 1x | 0.000815 | 0 | 1 | 2 | 3 | Q67FY2 | B-cell CLL/l | Q67FY2 [1 | 0 | 1770.886 | 0 | 1.1 | 0.001617 | 13.5 | 61.2 | 138.8 | 9.67 | 12.63 | | High | High | High | High | High | High | High | High | High | High | High | 0 | 0.000189 | 2.01 |
| FALSE | High | TAMSSPGN | [K].TAMSS | 1xPhospho | Q6PIJ4 1xP | 0.008388 | 0.000736 | 1 | 1 | 4 | Q6PIJ4 | Nuclear fac | Q6PIJ4 [88 | 0 | 3383.728 | 0 | -0.37 | 0.042948 | 12.34 | 115.3 | 84.7 | 13.49 | 9.29 | | High | High | High | High | High | High | High | High | High | High | High | 0.000872 | 0.009055 | 2.89 |
| FALSE | High | TAMSSPGN | [K].TAMSS | 1xPhospho | Q6PIJ4 1xP | 0.000868 | 0 | 1 | 1 | 2 | Q6PIJ4 | Nuclear fac | Q6PIJ4 [88 | 0 | 3399.723 | 0 | -0.54 | 0.004033 | 5.58 | 116.9 | 83.1 | 5.98 | 4.45 | | High | High | High | High | High | High | High | High | High | High | High | 0 | 0.000203 | 3.46 |
| FALSE | High | TAMSTPHV | [K].TAMST | 1xPhospho | P26041 1x | 0.017344 | 0.001204 | 1 | 3 | 2 | P26041 | Moesin OS | P26041 [46 | 0 | 3660.559 | 0 | 0.46 | 0.173477 | 34.97 | 80.1 | 119.9 | 19.82 | 27.69 | | High | High | High | High | High | High | High | High | High | High | High | 0.000903 | 0.009187 | 3.27 |
| FALSE | High | TAMYLEQE | [K].TAMYL | 1xPhospho | O88466 1x | 0.000492 | 0 | 2 | 5 | 1 | O88466; R | Zinc finger | O88466 [9 | 0 | 2574.236 | 0 | -0.35 | 0.094089 | 5.06 | 115.9 | 84.1 | 7.66 | 26.86 | Shared | High | High | High | High | High | High | High | High | High | High | High | 0 | 0.000108 | 3.18 |
| FALSE | High | TAPNGGQI | [K].TAPNG | 2xPhospho | Q8C796 2x | 0.002022 | 5.66E-05 | 1 | 2 | 2 | Q8C796 | REST corep | Q8C796 [2 | 0 | 3238.262 | 0 | -1.26 | 0.001018 | 0.19 | 142.1 | 57.9 | 7.76 | 10.58 | | High | High | High | High | High | High | High | High | High | High | High | 0.000814 | 0.007543 | 3.57 |
| FALSE | High | TAPNGGQI | [K].TAPNG | 3xPhospho | Q8C796 3x | 0.037692 | 0.00249 | 1 | 2 | 1 | Q8C796 | REST corep | Q8C796 [2 | 0 | 3318.228 | 0 | -1.85 | 0.001275 | 0.36 | 156.7 | 43.3 | 13.15 | 13.17 | | High | High | High | High | High | High | High | High | High | High | High | 0.001308 | 0.0139 | 1.95 |
| FALSE | High | TAPPVLPTC | [K].TAPPVL | 1xPhospho | Q7JJ13 1xP | 0.001325 | 0 | 1 | 3 | 4 | Q7JJ13 | Bromodom | Q7JJ13 [62 | 0 | 3494.596 | 0 | -0.84 | 0.000757 | 1.47 | 128.2 | 71.8 | 3.57 | 10.73 | | High | High | High | High | High | High | High | High | High | High | High | 0 | 0.000326 | 3.38 |
| FALSE | High | TAPPVLPTC | [K].TAPPVL | 1xPhospho | Q7JJ13 1xP | 4.43E-05 | 0 | 1 | 3 | 10 | Q7JJ13 | Bromodom | Q7JJ13 [62 | 0 | 3510.591 | 0 | -0.71 | 0.000222 | 0.82 | 123.6 | 76.4 | 3.34 | 6.91 | | High | High | High | High | High | High | High | High | High | High | High | 0 | 0.000247 | 3.54 |
| FALSE | High | TAPSPSLR | [K].TAPSPS | 1xPhospho | Q7TSY8 1x | 0.018125 | 0.001282 | 1 | 1 | 1 | Q7TSY8 | Shugoshin | Q7TSY8 [2 | 0 | 1137.587 | 0 | -0.51 | 0.010543 | 8.66 | 118.8 | 81.2 | 7.36 | 9.91 | | High | High | High | High | High | High | High | High | High | High | High | 0.000689 | 0.006077 | 1.66 |
| FALSE | High | TAPSSPLTS | [K].TAPSSP | 1xPhospho | Q6ZPJ0 1x | 0.00129 | 0 | 1 | 2 | 2 | Q6ZPJ0 | Testis-expr | Q6ZPJ0 [26 | 0 | 1725.826 | 0 | 1.09 | 0.000718 | 7.94 | 62.9 | 137.1 | 5.88 | 8.83 | | High | High | High | High | High | High | High | High | High | High | High | 0 | 0.000316 | 2.69 |
| FALSE | High | TADSVSPLE | [K].TADSV | 1xPhospho | V9GX40 1x | 0.000411 | 0 | 1 | 4 | 3 | V9GX40 | Synergin ga | V9GX40 [6 | 0 | 1879.982 | 0 | 0.92 | 0.007276 | 28.58 | 66.1 | 133.9 | 19.04 | 8.55 | | High | High | High | High | High | High | High | High | High | High | High | 0.002264 | 0.02368 | 2.59 |
| FALSE | High | TLSVNSSEC | [R].TLSVNS | 1xPhospho | A0A0R4IZX | 0.001222 | 0 | 1 | 3 | 1 | A0A0R4IZX | Differentia | A0A0R4IZX | 0 | 1630.845 | 0 | 2.19 | 0.001269 | 28.91 | 37.4 | 162.6 | 11.4 | 13.46 | | High | High | High | High | High | High | High | High | High | High | High | 0 | 0.000297 | 3.64 |
| FALSE | High | SYDVPPPPI | [R].SYDVPF | 1xPhospho | Q3U7Z6 1x | 0.000146 | 0 | 1 | 2 | 14 | Q3U7Z6 | Phosphogly | Q3U7Z6 [1 | 0 | 2971.399 | 0 | 0.31 | 0.003313 | 2.49 | 89.2 | 110.8 | 3.28 | 5.6 | | High | High | High | High | High | High | High | High | High | High | High | 0.001011 | 0.01105 | 3.52 |
| FALSE | High | SYDEVPQN | [R].SYDEVF | 1xPhospho | Q9CZ70 1x | 5.5E-05 | 0 | 1 | 1 | 1 | Q9CZ70 | MCG12980 | Q9CZ70 [2 | 0 | 2235.002 | 0 | -1.3 | 0.001091 | 2.52 | 143.7 | 56.3 | 7.27 | 15 | | High | High | High | High | High | High | High | High | High | High | High | 0 | 9.2E-06 | 2.44 |
| FALSE | High | SVPTVDSG | [K].SVPTVD | 1xDeamida | Q05D44 1x | 5.48E-07 | 0 | 1 | 1 | 3 | Q05D44 | Eukaryotic | Q05D44 [2 | 0 | 2338.038 | 0 | -0.42 | 0.011611 | 1.27 | 114.6 | 85.4 | 9.26 | 2.37 | | High | High | High | High | High | High | High | High | High | High | High | 0 | 5.15E-08 | 4.14 |
| FALSE | High | SVPTVDSG | [K].SVPTVD | 2xPhospho | Q05D44 2x | 0.08447 | 0.006345 | 1 | 1 | 1 | Q05D44 | Eukaryotic | Q05D44 [2 | 0 | 2417.02 | 0 | -0.51 | 0.415442 | 12.26 | 111.7 | 88.3 | 20.68 | 24.3 | | High | High | High | High | High | High | High | High | High | High | High | 0.003265 | 0.03537 | 2.89 |
| FALSE | High | SVQPEVELS | [K].SVQPEV | 2xPhospho | Q8CH77 2x | 0.005135 | 0.000551 | 1 | 3 | 2 | Q8CH77 | Neuron na | Q8CH77 [8 | 0 | 2695.097 | 0 | 0.17 | 0.265532 | 12.57 | 90.9 | 109.1 | 13.04 | 15.59 | | High | High | High | High | High | High | High | High | High | High | High | 0.000246 | 0.001497 | 3.22 |
| FALSE | High | SVQSAAVP | [R].SVQSA | 1xPhospho | A2ADB0 1x | 0.00195 | 5.66E-05 | 1 | 6 | 1 | A2ADB0 | Msx2-inter | A2ADB0 [3 | 0 | 2551.378 | 0 | 0.05 | 0.80014 | 28.42 | 99 | 101 | 11.69 | 10.74 | | High | High | High | High | High | High | High | High | High | High | High | 2.58E-05 | 0.000503 | 2.75 |
| FALSE | High | SVQTNSVC | [K].SVQTN | 1xPhospho | Q0GNC1 1 | 5E-10 | 0 | 1 | 3 | 2 | Q0GNC1 | Inverted fo | Q0GNC1 [3 | 0 | 2933.425 | 0 | 0.87 | 0.031654 | 5.65 | 76.3 | 123.7 | 26.48 | 9.2 | | High | High | High | High | High | High | High | High | High | High | High | 0 | 1.98E-11 | 6.23 |
| FALSE | High | SVSAITESE | [R].SVSAIT | 1xPhospho | Q8BQ33 1 | 0.004696 | 0.000456 | 1 | 2 | 1 | Q8BQ33 | Treslin OS= | Q8BQ33 [8 | 0 | 1999.979 | 0 | -0.68 | 0.001778 | 1.77 | 127 | 73 | 7.85 | 5.78 | | High | High | High | High | High | High | High | High | High | High | High | 0.000204 | 0.001343 | 2.31 |
| FALSE | High | SVSANPK | [R].SVSANI | 1xPhospho | Q571J8 1x | 0.016743 | 0.001166 | 1 | 14 | 3 | Q571J8 | Non-specif | Q571J8 [40 | 0 | 1240.67 | 0 | 0.54 | 0.003222 | 6.29 | 82.3 | 117.7 | 4.43 | 8.57 | | High | High | High | High | High | High | High | High | High | High | High | 0.000622 | 0.005578 | 1.63 |
| FALSE | High | SVSASHEG | [R].SVSASH | 1xPhospho | Q3U9G9 1 | 0.003224 | 0.00016 | 1 | 3 | 4 | Q3U9G9 | Lamin-B re | Q3U9G9 [1 | 0 | 1653.825 | 0 | -0.16 | 0.516316 | 3.41 | 105.6 | 94.4 | 6.01 | 11.95 | | High | High | High | High | High | High | High | High | High | High | High | 0.000638 | 0.005756 | 1.69 |
| FALSE | High | SVSCDNVS | [R].SVSCDI | 1xCarbami | Q9D168 1x | 0.00017 | 0 | 1 | 1 | 3 | Q9D168 | Integrator | Q9D168 [3 | 0 | 1533.738 | 0 | -0.22 | 0.002894 | 1.8 | 109.2 | 90.8 | 3.25 | 7.09 | | High | High | High | High | High | High | High | High | High | High | High | 0 | 3.25E-05 | 2.78 |
| FALSE | High | SVSCDSALF | [K].SVSCDS | 2xCarbami | Q80Y19 1x | 0.00129 | 0 | 1 | 2 | 2 | Q80Y19 | Rho GTPas | Q80Y19 [8 | 0 | 2378.113 | 0 | 0.11 | 0.966995 | 5.03 | 100.5 | 99.5 | 14.19 | 9.7 | | High | High | High | High | High | High | High | High | High | High | High | 0 | 0.000319 | 2.73 |
| FALSE | High | SVSCSDLTE | [K].SVSCSD | 1xCarbami | Q5SYL3 1x | 2.56E-06 | 0 | 1 | 2 | 2 | Q5SYL3 | Protein KIA | Q5SYL3 [2 | 0 | 3019.469 | 0 | 0.49 | 0.008331 | 15.47 | 91.4 | 108.6 | 16.3 | 15.01 | | High | High | High | High | High | High | High | High | High | High | High | 0 | 2.92E-07 | 4.9 |
| FALSE | High | SVSDNTLV | [R].SVSDN | 1xPhospho | Q03160 1x | 0.00013 | 0 | 1 | 2 | 1 | Q03160 | Growth fac | Q03160 [3 | 0 | 2188.99 | 0 | -0.62 | 0.006548 | 9.08 | 120.9 | 79.1 | 7.9 | 6.59 | | High | High | High | High | High | High | High | High | High | High | High | 0 | 2.4E-05 | 3.95 |
| FALSE | High | SVSEAALA | [R].SVSEA | 1xPhospho | E9Q4M2 1 | 2.57E-07 | 0 | 1 | 3 | 3 | E9Q4M2 | Hormone-s | E9Q4M2 [8 | 0 | 2538.347 | 0 | -0.78 | 0.002525 | 3.72 | 125.8 | 74.2 | 6.58 | 7.68 | | High | High | High | High | High | High | High | High | High | High | High | 0 | 1.65E-05 | 3.47 |
| FALSE | High | SVPTVDSG | [K].SVPTVD | 1xPhospho | Q05D44 1x | 1.43E-07 | 0 | 1 | 1 | 12 | Q05D44 | Eukaryotic | Q05D44 [2 | 0 | 2337.054 | 0 | -0.59 | 0.004499 | 4.88 | 122.9 | 77.1 | 5.32 | 8.28 | | High | High | High | High | High | High | High | High | High | High | High | 0 | 1.92E-08 | 4.6 |
| FALSE | High | SVSEDSPLG | [R].SVSEDS | 1xPhospho | Q3UQI9 1x | 3.33E-06 | 0 | 1 | 3 | 1 | Q3UQI9 | Probable u | Q3UQI9 [3 | 0 | 2230.982 | 0 | -0.02 | 0.500983 | 15.43 | 104.5 | 95.5 | 9.4 | 7.15 | | High | High | High | High | High | High | High | High | High | High | High | 0 | 3.94E-07 | 3.53 |
| FALSE | High | SVSEINSDD | [R].SVSEIN | 1xPhospho | B9EKI3 1xP | 4.68E-05 | 0 | 1 | 2 | 1 | B9EKI3 | TATA elem | B9EKI3 [33 | 0 | 2027.994 | 0 | 0.84 | 0.000855 | 5.62 | 73 | 127 | 7.11 | 5.69 | | High | High | High | High | High | High | High | High | High | High | High | 0 | 7.61E-06 | 2.44 |
| FALSE | High | SVSEINSDD | [R].SVSEIN | 2xPhospho | B9EKI3 2xP | 0.008167 | 0.000736 | 1 | 2 | 2 | B9EKI3 | TATA elem | B9EKI3 [33 | 0 | 2107.96 | 0 | 0.92 | 0.004623 | 3.35 | 69 | 131 | 4.88 | 13.52 | | High | High | High | High | High | High | High | High | High | High | High | 0.000872 | 0.009153 | 3.22 |
| FALSE | High | SVSENSLV | [R].SVSENS | 1xPhospho | Q60760 1x | 6.85E-06 | 0 | 1 | 5 | 2 | Q60760 | Growth fac | Q60760 [4 | 0 | 2206.041 | 0 | 0.97 | 0.012631 | 23.64 | 66.4 | 133.6 | 9.11 | 20.11 | | High | High | High | High | High | High | High | High | High | High | High | 0 | 8.82E-07 | 4.17 |
| FALSE | High | SVSENSLV | [R].SVSENS | 1xPhospho | Q60760 1x | 5.23E-06 | 0 | 1 | 5 | 4 | Q60760 | Growth fac | Q60760 [4 | 0 | 2222.036 | 0 | -0.71 | 0.067218 | 9.95 | 124.1 | 75.9 | 13.45 | 19.36 | | High | High | High | High | High | High | High | High | High | High | High | 0 | 1.57E-06 | 3.55 |
| FALSE | High | SVSENSLV | [R].SVSENS | 2xPhospho | Q60760 2x | 0.053023 | 0.00384 | 1 | 5 | 1 | Q60760 | Growth fac | Q60760 [4 | 0 | 2302.003 | 0 | -0.04 | 0.787709 | 12.92 | 102.3 | 97.7 | 8.09 | 11.67 | | High | High | High | High | High | High | High | High | High | High | High | 0.001999 | 0.02067 | 3.44 |
| FALSE | High | SVSESHTPC | [K].SVSESH | 1xCarbami | Q8R5A6-2 | 0.006653 | 0.00062 | 1 | 2 | 2 | Q8R5A6-2 | Isoform 2 c | Q8R5A6-2 | 0 | 2680.212 | 0 | -0.04 | 0.892723 | 5.94 | 102.3 | 97.7 | 16.36 | 2.85 | | High | High | High | High | High | High | High | High | High | High | High | 0.000275 | 0.001992 | 4.23 |
| FALSE | High | SVSETSEDK | [K].SVSETS | 1xPhospho | E9Q6J5-3 1 | 0.058664 | 0.004294 | 1 | 3 | 1 | E9Q6J5-3 | Isoform Ne | E9Q6J5-3 [ | 0 | 1519.729 | 0 | -0.26 | 0.154868 | 5.89 | 109.1 | 90.9 | 3.23 | 12.77 | | High | High | High | High | High | High | High | High | High | High | High | 0.002187 | 0.02319 | 1.33 |
| FALSE | High | SVSGFLHFD | [R].SVSGFL | 1xPhospho | Q6ZQG1 1 | 0.007404 | 0.00062 | 1 | 5 | 2 | Q6ZQG1 | MCG21756 | Q6ZQG1 [1 | 0 | 1947.998 | 0 | -0.01 | 0.703446 | 20.23 | 95.5 | 104.5 | 10.94 | 20.75 | | High | High | High | High | High | High | High | High | High | High | High | 0.003564 | 0.03804 | 1.7 |
| FALSE | High | SVSGSPEPA | [RK].SVSG | 1xPhospho | Q52KI8 1x | 1.13E-05 | 0 | 1 | 7 | 3 | Q52KI8 | Serine/argi | Q52KI8 [79 | 0 | 1567.813 | 0 | -0.26 | 0.088735 | 10.12 | 107.8 | 92.2 | 7.18 | 6.49 | | High | High | High | High | High | High | High | High | High | High | High | 0 | 1.56E-06 | 2.19 |
| FALSE | High | SVSGSPEPA | [RK].SVSG | 2xPhospho | Q52KI8 2x | 0.001361 | 0 | 1 | 7 | 3 | Q52KI8 | Serine/argi | Q52KI8 [79 | 0 | 1647.78 | 0 | 0 | 0.28459 | 5.61 | 104.9 | 95.1 | 4.89 | 12.38 | | High | High | High | High | High | High | High | High | High | High | High | 0.000267 | 0.001563 | 2.73 |
| FALSE | High | SVSHGSNH | [R].SVSHG | 1xPhospho | Q3TLH4 1x | 0.022182 | 0.001542 | 1 | 5 | 7 | Q3TLH4 | Protein PRI | Q3TLH4 [8 | 0 | 2059.914 | 0 | -0.76 | 0.005498 | 4.13 | 126.7 | 73.3 | 7.1 | 14.52 | | High | High | High | High | High | High | High | High | High | High | High | 0.001876 | 0.01876 | 2.21 |
| FALSE | High | SVSHGSNH | [R].SVSHG | 2xPhospho | Q3TLH4 2x | 0.056248 | 0.004124 | 1 | 5 | 1 | Q3TLH4 | Protein PRI | Q3TLH4 [8 | 0 | 2139.881 | 0 | -1.82 | 0.006395 | 2.8 | 155.8 | 44.2 | 19.16 | 16.41 | | High | High | High | High | High | High | High | High | High | High | High | 0.002117 | 0.02208 | 1.96 |
| FALSE | High | SVSLDSQM | [R].SVSLDS | 1xPhospho | Q67FY2 1x | 0.028327 | 0.001791 | 1 | 3 | 2 | Q67FY2 | B-cell CLL/l | Q67FY2 [1 | 0 | 2653.213 | 0 | 0.81 | 0.152903 | 29.83 | 73.8 | 126.2 | 14.61 | 60.79 | | High | High | High | High | High | High | High | High | High | High | High | 0.000968 | 0.01008 | 3.4 |
| FALSE | High | SVSEESANS | [R].SVSEES | 1xPhospho | F8WI37 1x | 0.002463 | 0.000109 | 1 | 5 | 2 | F8WI37 | Histone-lys | F8WI37 [1 | 0 | 2330.189 | 0 | 0.13 | 0.651542 | 19.74 | 97.4 | 102.6 | 15.07 | 12.61 | | High | High | High | High | High | High | High | High | High | High | High | 4.94E-05 | 0.000658 | 2.67 |
| FALSE | High | SVPTVDSG | [K].SVPTVD | 1xDeamida | Q05D44 1x | 0.045122 | 0.003373 | 1 | 1 | 1 | Q05D44 | Eukaryotic | Q05D44 [2 | 0 | 2108.875 | 0 | -1.29 | 0.580893 | 14.06 | 142 | 58 | 60.38 | 89.74 | | High | High | High | High | High | High | High | High | High | High | High | 0.001727 | 0.01716 | 2.62 |
| FALSE | High | SVPTVDSG | [K].SVPTVD | 1xPhospho | Q05D44 1x | 4.2E-05 | 0 | 1 | 1 | 2 | Q05D44 | Eukaryotic | Q05D44 [2 | 0 | 2107.891 | 0 | -0.72 | 0.582997 | 11.36 | 124.4 | 75.6 | 60.5 | 88.79 | | High | High | High | High | High | High | High | High | High | High | High | 0.000449 | 0.003826 | 3.21 |
| FALSE | High | SVPTETSEG | [R].SVPTET | 2xPhospho | A2A7B5 2x | 0.000131 | 0 | 1 | 2 | 4 | A2A7B5 | PR domain | A2A7B5 [3 | 0 | 2461.1 | 0 | -0.53 | 0.001594 | 3.63 | 118.1 | 81.9 | 4.87 | 10.08 | | High | High | High | High | High | High | High | High | High | High | High | 0.000327 | 0.002498 | 2.81 |
| FALSE | High | SVEEEMEL | [K].SVEEEN | 1xPhospho | P70423 1x | 0.096181 | 0.007535 | 1 | 2 | 1 | P70423 | Cationic an | P70423 [43 | 0 | 2577.193 | 0 | 1.28 | 0.003156 | 24.8 | 55.9 | 144.1 | 38.87 | 13.97 | | High | High | High | High | High | High | High | High | High | High | High | 0.003911 | 0.0414 | 2.65 |
| FALSE | High | SVEVTHGV | [K].SVEVTH | 1xPhospho | Q8R2X8-2 | 0.000563 | 0 | 1 | 4 | 1 | Q8R2X8-2 | Isoform 2 c | Q8R2X8-2 | 0 | 2503.302 | 0 | 0.11 | 0.273821 | 27.42 | 98.2 | 101.8 | 8.11 | 20.18 | | High | High | High | High | High | High | High | High | High | High | High | 0 | 0.000126 | 3.2 |
| FALSE | High | SVFPATVSA | [K].SVFPAT | 1xCarbami | Q5FWX6 1 | 0.000108 | 0 | 1 | 3 | 3 | Q5FWX6 | Serine/thre | Q5FWX6 [1 | 0 | 2637.376 | 0 | 0.86 | 0.003503 | 14.94 | 67.7 | 132.3 | 9.06 | 9.87 | | High | High | High | High | High | High | High | High | High | High | High | 0 | 1.96E-05 | 2.83 |
| FALSE | High | SVFPATVSA | [K].SVFPAT | 1xCarbami | Q5FWX6 2 | 0.000257 | 0 | 1 | 3 | 3 | Q5FWX6 | Serine/thre | Q5FWX6 [1 | 0 | 2717.343 | 0 | 0.97 | 0.009145 | 32.82 | 65.9 | 134.1 | 18.43 | 13.49 | | High | High | High | High | High | High | High | High | High | High | High | 0 | 5.18E-05 | 3.05 |
| FALSE | High | SVFSQSGN | [K].SVFSQS | 1xPhospho | A2A6Q5-3 | 0.034579 | 0.002233 | 1 | 6 | 1 | A2A6Q5-3 | Isoform 3 c | A2A6Q5-3 | 0 | 1377.636 | 0 | 0.34 | 0.015353 | 9.88 | 84.6 | 115.4 | 8.45 | 8.43 | | High | High | High | High | High | High | High | High | High | High | High | 0.001183 | 0.01268 | 0.9 |
| FALSE | High | SVGDGETV | [R].SVGDG | 1xPhospho | Q9JKB3 1x | 3.83E-05 | 0 | 2 | 14 | 3 | Q9JKB3; Q | Y-box-bind | Q9JKB3 [1 | 0 | 2334.115 | 0 | 0.25 | 0.00935 | 4.16 | 92.7 | 107.3 | 7.02 | 5.28 | Shared | High | High | High | High | High | High | High | High | High | High | High | 0 | 6.09E-06 | 3.02 |
| FALSE | High | SVGGSGGC | [R].SVGGS | 1xPhospho | P48678 1x | 1.56E-07 | 0 | 1 | 2 | 6 | P48678 | Prelamin-A | P48678 [62 | 0 | 1875.88 | 0 | 1.38 | 0.000775 | 14.2 | 54.4 | 145.6 | 3.03 | 8.67 | | High | High | High | High | High | High | High | High | High | High | High | 0 | 1.7E-06 | 2.5 |
| FALSE | High | SVICDISPLF | [R].SVICDIS | 1xCarbami | Q9EPQ8 1 | 0.007339 | 0.00062 | 1 | 3 | 1 | Q9EPQ8 | Transcripti | Q9EPQ8 [8 | 0 | 1468.743 | 0 | -0.13 | 0.019522 | 6.34 | 106.7 | 93.3 | 3.8 | 7.63 | | High | High | High | High | High | High | High | High | High | High | High | 0.000275 | 0.00222 | 1.32 |
| FALSE | High | SVLCSTPCV | [R].SVLCST | 2xCarbami | Q9JJ78 1xP | 0.010382 | 0.000772 | 1 | 2 | 1 | Q9JJ78 | Lymphokin | Q9JJ78 [18 | 0 | 2690.316 | 0 | -0.65 | 0.002998 | 4.6 | 120.4 | 79.6 | 9.44 | 8.31 | | High | High | High | High | High | High | High | High | High | High | High | 0.000382 | 0.003258 | 2.16 |
| FALSE | High | SVLEPQAES | [R].SVLEPC | 1xPhospho | Q4VA36 1x | 5.66E-09 | 0 | 1 | 1 | 1 | Q4VA36 | Coiled-coil | Q4VA36 [1 | 0 | 2897.349 | 0 | 0.56 | 0.002799 | 12.38 | 80.9 | 119.1 | 3.52 | 6.13 | | High | High | High | High | High | High | High | High | High | High | High | 0 | 3.05E-10 | 4.43 |
| FALSE | High | SVLTAGGV | [K].SVLTAG | 1xPhospho | A0A1B0GS | 2.87E-08 | 0 | 1 | 2 | 6 | A0A1B0GS | GRB2-asso | A0A1B0GS | 0 | 3154.46 | 0 | -1.92 | 0.00022 | 1.39 | 156.3 | 43.7 | 3.69 | 20.73 | | High | High | High | High | High | High | High | High | High | High | High | 0 | 1.88E-09 | 5.25 |
| FALSE | High | SVLTAGGV | [K].SVLTAG | 2xPhospho | A0A1B0GS | 0.029844 | 0.001857 | 1 | 2 | 1 | A0A1B0GS | GRB2-asso | A0A1B0GS | 0 | 3218.432 | 0 | -0.75 | 0.01863 | 14.57 | 124.7 | 75.3 | 9.95 | 27.41 | | High | High | High | High | High | High | High | High | High | High | High | 0.001011 | 0.0107 | 2.52 |
| FALSE | High | SVLTSPVAN | [R].SVLTSP | 1xPhospho | F6YRW4 1 | 1.9E-08 | 0 | 1 | 5 | 1 | F6YRW4 | F-box and l | F6YRW4 [3 | 0 | 2557.332 | 0 | 1.09 | 0.003257 | 22.54 | 64 | 136 | 7.08 | 12.19 | | High | High | High | High | High | High | High | High | High | High | High | 0 | 1.18E-09 | 5.5 |
| FALSE | High | SVNEENGE | [K].SVNEEI | 1xPhospho | C7G3P2 1x | 5.52E-06 | 0 | 1 | 4 | 3 | C7G3P2 | MKIAA053 | C7G3P2 [2 | 0 | 2431.103 | 0 | 0.23 | 0.029985 | 7.02 | 92.5 | 107.5 | 4.85 | 4.42 | | High | High | High | High | High | High | High | High | High | High | High | 0 | 6.91E-07 | 4.48 |
| FALSE | High | SVNEILGLA | [K].SVNEIL | 1xPhospho | Q8VDY9 1 | 0.000656 | 0 | 1 | 2 | 2 | Q8VDY9 | Caspase ac | Q8VDY9 [2 | 0 | 1752.898 | 0 | 0.09 | 0.644753 | 81.49 | 94.4 | 105.6 | 29.14 | 34 | | High | High | High | High | High | High | High | High | High | High | High | 0 | 0.000149 | 3.53 |
| FALSE | High | SVNEILGLA | [K].SVNEIL | 1xPhospho | Q8VDY9 1 | 0.000174 | 0 | 1 | 2 | 2 | Q8VDY9 | Caspase ac | Q8VDY9 [2 | 0 | 1982.061 | 0 | 0.24 | 0.07782 | 17.14 | 93 | 107 | 6.43 | 7.69 | | High | High | High | High | High | High | High | High | High | High | High | 0 | 3.36E-05 | 2.8 |
| FALSE | High | SVNFSLTPN | [R].SVNFSL | 1xPhospho | P14873 1x | 0.002895 | 0.00016 | 2 | 2 | 1 | P14873; B2 | Microtubu | P14873 [1 | 0 | 1887.003 | 0 | -0.44 | 0.005955 | 7.5 | 115.6 | 84.4 | 8.29 | 7.8 | Shared | High | High | High | High | High | High | High | High | High | High | High | 7.19E-05 | 0.000782 | 2.74 |
| FALSE | High | SVNGGPGS | [R].SVNGG | 1xPhospho | A0A0A6YY | 0.002272 | 0.000109 | 1 | 13 | 1 | A0A0A6YY | Protocadh | A0A0A6YY | 0 | 1535.742 | 0 | -0.59 | 0.005406 | 4.3 | 117.6 | 82.4 | 8.51 | 16.31 | | High | High | High | High | High | High | High | High | High | High | High | 4.94E-05 | 0.000598 | 2.39 |
| FALSE | High | SVNGQQST | [K].SVNGQ | 1xPhospho | P70371 1x | 0.017041 | 0.001204 | 1 | 3 | 2 | P70371 | Telomeric | P70371 [28 | 0 | 2556.239 | 0 | -1.05 | 0.000505 | 4.79 | 136.6 | 63.4 | 5.02 | 15.52 | | High | High | High | High | High | High | High | High | High | High | High | 0.000638 | 0.005678 | 3.4 |
| FALSE | High | SVNPVLSPT | [K].SVNPVI | 1xPhospho | Q8CJ27 1x | 7.89E-05 | 0 | 1 | 2 | 1 | Q8CJ27 | Abnormal | Q8CJ27 [3 | 0 | 1954.081 | 0 | 0.41 | 0.019018 | 17.34 | 87.8 | 112.2 | 10.44 | 7.54 | | High | High | High | High | High | High | High | High | High | High | High | 0 | 1.37E-05 | 2.93 |
| FALSE | High | SVNTTECV | [K].SVNTTE | 1xCarbami | Q05A36 1x | 4.21E-06 | 0 | 1 | 4 | 1 | Q05A36 | RNA-bindir | Q05A36 [2 | 0 | 2676.29 | 0 | -0.08 | 0.183903 | 9.4 | 107.9 | 92.1 | 10.41 | 13.04 | | High | High | High | High | High | High | High | High | High | High | High | 0 | 5.12E-07 | 3.54 |
| FALSE | High | SVNVSLTQ | [R].SVNVSI | 1xPhospho | F7AA26 1x | 1.28E-10 | 0 | 1 | 10 | 4 | F7AA26 | Paralemmi | F7AA26 [7 | 0 | 2599.27 | 0 | 1.13 | 0.001015 | 1.42 | 68.4 | 131.6 | 8.79 | 7.46 | | High | High | High | High | High | High | High | High | High | High | High | 0 | 4.29E-12 | 6.72 |
| FALSE | High | SVPGVTSTI | [K].SVPGV | 1xPhospho | F7AA26 1x | 0.010109 | 0.000736 | 1 | 10 | 2 | F7AA26 | Paralemmi | F7AA26 [2 | 0 | 1734.919 | 0 | 0.24 | 0.025855 | 0.59 | 93.6 | 106.4 | 3.67 | 4.6 | | High | High | High | High | High | High | High | High | High | High | High | 0.000872 | 0.008752 | 2.02 |
| FALSE | High | SVPNTPLRI | [K].SVPNTI | 1xPhospho | B2RY51 1x | 0.021989 | 0.001542 | 1 | 4 | 1 | B2RY51 | Thyroid ho | B2RY51 [1 | 0 | 3360.739 | 0 | 1.26 | 0.001068 | 15.4 | 56 | 144 | 14.75 | 5.87 | | High | High | High | High | High | High | High | High | High | High | High | 0.000814 | 0.007608 | 2.92 |
| FALSE | High | SVPQFTEPT | [K].SVPQF | 1xPhospho | Q5SV77-3 | 2.27E-05 | 0 | 1 | 6 | 1 | Q5SV77-3 | Isoform 3 c | Q5SV77-3 | 0 | 2595.209 | 0 | 0.25 | 0.118894 | 8.97 | 92.1 | 107.9 | 7.23 | 8.09 | | High | High | High | High | High | High | High | High | High | High | High | 0 | 3.38E-06 | 4.74 |
| FALSE | High | SVPSIAAAT | [R].SVPSIA | 1xPhospho | Q9D4H4 1 | 2.58E-05 | 0 | 1 | 2 | 3 | Q9D4H4 | Angiomoti | Q9D4H4 [8 | 0 | 1663.837 | 0 | 1.53 | 0.000831 | 8.99 | 51 | 149 | 5.78 | 8.47 | | High | High | High | High | High | High | High | High | High | High | High | 0 | 3.93E-06 | 3.82 |
| FALSE | High | SVPSIAAAT | [R].SVPSIA | 2xPhospho | Q9D4H4 2 | 0.000579 | 0 | 1 | 2 | 1 | Q9D4H4 | Angiomoti | Q9D4H4 [8 | 0 | 1743.803 | 0 | 0.91 | 0.000378 | 6.09 | 71.2 | 128.8 | 4.36 | 3.58 | | High | High | High | High | High | High | High | High | High | High | High | 0 | 0.000129 | 2.93 |
| FALSE | High | SVPSIFAAA | [K].SVPSIF | 1xPhospho | B8JK87 1xP | 7.38E-07 | 0 | 1 | 4 | 1 | B8JK87 | Angiomoti | B8JK87 [69 | 0 | 2656.333 | 0 | 0.59 | 0.186514 | 50.37 | 92.4 | 107.6 | 20.92 | 29.52 | | High | High | High | High | High | High | High | High | High | High | High | 0 | 7.25E-08 | 3.44 |
| FALSE | High | SVPTETSEG | [R].SVPTET | 1xPhospho | A2A7B5 1x | 1.43E-05 | 0 | 1 | 2 | 2 | A2A7B5 | PR domain | A2A7B5 [3 | 0 | 2381.134 | 0 | -0.43 | 0.001759 | 1.74 | 113.2 | 86.8 | 4.87 | 4.49 | | High | High | High | High | High | High | High | High | High | High | High | 0 | 2.02E-06 | 3.28 |
| FALSE | High | SVSLLSLSTI | [K].SVSLLS | 1xPhospho | Q07139 1x | 1.28E-05 | 0 | 1 | 3 | 2 | Q07139 | Protein EC | Q07139 [3 | 0 | 1783.915 | 0 | -0.34 | 0.075797 | 18 | 111.1 | 88.9 | 11.21 | 18.75 | | High | High | High | High | High | High | High | High | High | High | High | 0 | 1.78E-06 | 4.07 |
| FALSE | High | SYDVPPPPI | [R].SYDVPF | 1xPhospho | Q3U7Z6 1x | 1.16E-06 | 0 | 1 | 2 | 3 | Q3U7Z6 | Phosphogly | Q3U7Z6 [1 | 0 | 2955.404 | 0 | 0.27 | 0.011675 | 8.96 | 87.9 | 112.1 | 5.78 | 8.76 | | High | High | High | High | High | High | High | High | High | High | High | 0.004866 | 0.05302 | 2.88 |
| FALSE | High | SVSLQEPR | [K].SVSLQE | 1xPhospho | Q80WQ6 1 | 0.005924 | 0.00062 | 1 | 1 | 1 | Q80WQ6 | Inactive rh | Q80WQ6 [ | 0 | 1224.619 | 0 | -0.09 | 0.971111 | 13.03 | 103.2 | 96.8 | 8.98 | 12.14 | | High | High | High | High | High | High | High | High | High | High | High | 0.000275 | 0.001751 | 1.51 |
| FALSE | High | SVSIQNITG | [K].SVSIQN | 1xPhospho | A0A140LJJ | 5.16E-05 | 0 | 1 | 5 | 1 | A0A140LJJ | A-kinase ar | A0A140LJJ | 0 | 2848.359 | 0 | -0.26 | 0.120674 | 14.12 | 110.2 | 89.8 | 10.29 | 11.36 | | High | High | High | High | High | High | High | High | High | High | High | 0 | 8.52E-06 | 3.43 |
| FALSE | High | SVTPASTLT | [K].SVTPAS | 1xPhospho | E9Q9C3 1x | 0.008614 | 0.000736 | 1 | 5 | 1 | E9Q9C3 | Afadin OS= | E9Q9C3 [1 | 0 | 1542.854 | 0 | -0.28 | 0.740139 | 1.51 | 109.2 | 90.8 | 7.73 | 36.05 | | High | High | High | High | High | High | High | High | High | High | High | 0.000327 | 0.002644 | 2.12 |
| FALSE | High | SVTPLEEQ | [K].SVTPLE | 1xPhospho | P30306 1x | 4.05E-05 | 0 | 1 | 3 | 2 | P30306 | M-phase ir | P30306 [3 | 0 | 2165.114 | 0 | 1.02 | 0.001 | 8.7 | 62.9 | 137.1 | 11.2 | 8.8 | | High | High | High | High | High | High | High | High | High | High | High | 0 | 6.51E-06 | 3.28 |
| FALSE | High | SVTPPPPPF | [R].SVTPPP | 1xPhospho | Q8BYR2 1x | 0.08447 | 0.006345 | 1 | 1 | 1 | Q8BYR2 | Serine/thre | Q8BYR2 [2 | 0 | 1256.66 | 0 | 1.7 | 0.000676 | 12.56 | 43.6 | 156.4 | 12.26 | 10.91 | | High | High | High | High | High | High | High | High | High | High | High | 0.003279 | 0.03552 | 1.64 |
| FALSE | High | SVTSESEILE | [R].SVTSES | 1xCarbami | Q9QWF0 1 | 1.08E-05 | 0 | 1 | 1 | 2 | Q9QWF0 | Chromatin | Q9QWF0 [ | 0 | 4851.115 | 0 | 0.47 | 0.659875 | 21.99 | 91.9 | 108.1 | 25.01 | 46.09 | | High | High | High | High | High | High | High | High | High | High | High | 0 | 1.48E-06 | 5.7 |
| FALSE | High | SVTSNQSD | [R].SVTSN | 1xCarbami | Q8VCV2 1 | 3.3E-10 | 0 | 1 | 2 | 5 | Q8VCV2 | Ndrg3 prot | Q8VCV2 [3 | 0 | 2720.156 | 0 | 0.73 | 0.000827 | 3.03 | 75.9 | 124.1 | 6.74 | 4.02 | | High | High | High | High | High | High | High | High | High | High | High | 0 | 1.25E-11 | 4.68 |
| FALSE | High | SVVDMDLE | [R].SVVDM | 1xDeamida | E9Q2H1 1x | 0.027121 | 0.001676 | 1 | 2 | 1 | E9Q2H1 | E3 ubiquiti | E9Q2H1 [2 | 0 | 3082.296 | 0 | -0.2 | 0.211905 | 10.65 | 104.2 | 95.8 | 8.55 | 7.11 | | High | High | High | High | High | High | High | High | High | High | High | 0.00092 | 0.009583 | 3.11 |
| FALSE | High | SVVDMDLE | [R].SVVDM | 1xPhospho | E9Q2H1 1x | 0.005924 | 0.00062 | 1 | 2 | 2 | E9Q2H1 | E3 ubiquiti | E9Q2H1 [2 | 0 | 3294.48 | 0 | -0.59 | 0.011295 | 4.67 | 119.1 | 80.9 | 10.82 | 2.92 | | High | High | High | High | High | High | High | High | High | High | High | 0.000275 | 0.001742 | 3.5 |
| FALSE | High | SVVDMDLE | [R].SVVDM | 1xPhospho | E9Q2H1 1x | 0.001158 | 0 | 1 | 2 | 3 | E9Q2H1 | E3 ubiquiti | E9Q2H1 [2 | 0 | 3310.475 | 0 | -0.39 | 0.089579 | 16.77 | 122.9 | 77.1 | 9.8 | 26.2 | | High | High | High | High | High | High | High | High | High | High | High | 0 | 0.00028 | 3.22 |
| FALSE | High | SVVGIECSF | [R].SVVGIE | 1xCarbami | B1ARU1 2x | 0.000674 | 0 | 1 | 6 | 1 | B1ARU1 | Microtubu | B1ARU1 [5 | 0 | 2005.854 | 0 | -0.86 | 0.011227 | 7.26 | 123.9 | 76.1 | 10.83 | 15.35 | | High | High | High | High | High | High | High | High | High | High | High | 0 | 0.000153 | 2.76 |
| FALSE | High | SVVGTPAY | [R].SVVGTI | 1xPhospho | Q8BZ03 1x | 4.16E-05 | 0 | 1 | 1 | 1 | Q8BZ03 | Serine/thre | Q8BZ03 [7 | 0 | 2570.322 | 0 | 1.2 | 0.002168 | 14.69 | 60.3 | 139.7 | 23.91 | 11.83 | | High | High | High | High | High | High | High | High | High | High | High | 0 | 6.7E-06 | 3.84 |
| FALSE | High | SVVGTPAY | [R].SVVGTI | 1xPhospho | Q5FWX6 1 | 8.79E-05 | 0 | 1 | 4 | 2 | Q5FWX6 | Serine/thre | Q5FWX6 [ | 0 | 1881.008 | 0 | 1.18 | 0.001291 | 6.46 | 61.7 | 138.3 | 6.86 | 11.62 | | High | High | High | High | High | High | High | High | High | High | High | 0.001355 | 0.01431 | 2.08 |
| FALSE | High | SVVQTTPV | [R].SVVQT | 1xPhospho | Q8BTI8 1x | 5.33E-10 | 0 | 1 | 3 | 1 | Q8BTI8 | Serine/argi | Q8BTI8 [24 | 0 | 2542.353 | 0 | -0.82 | 0.000858 | 1.14 | 127 | 73 | 7.54 | 11.43 | | High | High | High | High | High | High | High | High | High | High | High | 0 | 2.14E-11 | 3.67 |
| FALSE | High | SVVTSLCSC | [K].SVVTSL | 1xCarbami | G3UWQ7 1 | 9.11E-05 | 0 | 1 | 10 | 1 | G3UWQ7 | Protein reg | G3UWQ7 [ | 0 | 1575.822 | 0 | -0.55 | 0.024882 | 4.28 | 114.5 | 85.5 | 10.4 | 10.29 | | High | High | High | High | High | High | High | High | High | High | High | 0 | 1.61E-05 | 2.55 |
| FALSE | High | SVTNSSSDI | [R].SVTNSS | 1xPhospho | Q9EPK5-2 | 1.31E-06 | 0 | 1 | 2 | 2 | Q9EPK5-2 | Isoform 2 c | Q9EPK5-2 | 0 | 2331.06 | 0 | 0.62 | 0.003684 | 2.02 | 81.8 | 118.2 | 10.07 | 13.65 | | High | High | High | High | High | High | High | High | High | High | High | 0 | 1.39E-07 | 2.64 |
| FALSE | High | SVWEESGF | [R].SVWEE | 1xPhospho | P49452 1x | 0.002441 | 0.000109 | 1 | 1 | 2 | P49452 | Centromer | P49452 [64 | 0 | 1442.651 | 0 | -0.99 | 0.000476 | 2.36 | 130.4 | 69.6 | 6.96 | 6.09 | | High | High | High | High | High | High | High | High | High | High | High | 4.94E-05 | 0.000649 | 1.15 |
| FALSE | High | SVYLPGQTS | [K].SVYLPG | 1xPhospho | Q8CFE5 1x | 0.00537 | 0.000599 | 1 | 1 | 1 | Q8CFE5 | BTB/POZ d | Q8CFE5 [9 | 0 | 1714.918 | 0 | 0.37 | 0.006265 | 2.96 | 85.9 | 114.1 | 6.94 | 4.03 | | High | High | High | High | High | High | High | High | High | High | High | 0.000267 | 0.001566 | 2.63 |
| FALSE | High | SWASPVYT | [R].SWASP | 1xPhospho | F6TFN2 1x | 4.91E-06 | 0 | 1 | 2 | 2 | F6TFN2 | LIM domai | F6TFN2 [1 | 0 | 2112.948 | 0 | 0.85 | 0.004996 | 20.7 | 68.9 | 131.1 | 10.82 | 8.93 | | High | High | High | High | High | High | High | High | High | High | High | 0 | 6.07E-07 | 3.72 |
| FALSE | High | SWDSEEILE | [R].SWDSE | 1xPhospho | Q8BP47 1x | 0.010758 | 0.000772 | 1 | 1 | 2 | Q8BP47 | Asparagine | Q8BP47 [4 | 0 | 1993.956 | 0 | -0.23 | 0.085519 | 10.7 | 109.6 | 90.4 | 6.15 | 12.33 | | High | High | High | High | High | High | High | High | High | High | High | 0.000399 | 0.003408 | 2.47 |
| FALSE | High | SWESSSPV | [R].SWESS | 1xPhospho | Q80YV2 1x | 0.00086 | 0 | 1 | 3 | 4 | Q80YV2 | Nuclear-int | Q80YV2 [3 | 0 | 2611.224 | 0 | -0.44 | 0.000986 | 4.16 | 113.4 | 86.6 | 3.07 | 6.68 | | High | High | High | High | High | High | High | High | High | High | High | 0 | 0.000317 | 2.13 |
| FALSE | High | SWESSSPV | [R].SWESS | 3xPhospho | Q80YV2 3x | 0.028082 | 0.001791 | 1 | 3 | 1 | Q80YV2 | Nuclear-int | Q80YV2 [3 | 0 | 2771.157 | 0 | -0.49 | 0.011789 | 4.83 | 118.3 | 81.7 | 9.6 | 10.93 | | High | High | High | High | High | High | High | High | High | High | High | 0.000968 | 0.01002 | 2.9 |

| | | | | | | | | | | | | | | | | | | | | | | | | | | | | | | | | | | | | | | |
|---|---|---|---|---|---|---|---|---|---|---|---|---|---|---|---|---|---|---|---|---|---|---|---|---|---|---|---|---|---|---|---|---|---|---|---|---|---|---|
| FALSE | High | SWLSPSVSI | [R].SWLSPS | 1xPhospho | Q9CZD0 1x | 0.003558 | 0.00021 | 1 | 1 | 1 | Q9CZD0 | Methylmal | Q9CZD0 [2 | 0 | 1802.893 | 0 | -0.59 | 0.109562 | 5.76 | 114.4 | 85.6 | 12.96 | 17.46 | | High | High | High | High | High | High | High | High | High | High | High | 9.42E-05 | 0.000993 | 2.61 |
| FALSE | High | SWPSPK | [R].SWPSP | 1xPhospho | Q7TSC1 1x | 0.064323 | 0.004664 | 1 | 3 | 1 | Q7TSC1 | Protein PR | Q7TSC1 [1 | 0 | 1239.654 | 0 | 0.53 | 0.045601 | 8.68 | 81.2 | 118.8 | 14.17 | 16.34 | | High | High | High | High | High | High | High | High | High | High | High | 0.002403 | 0.0258 | 1.66 |
| FALSE | High | SWSPSAPP | [R].SWSPS | 1xPhospho | D9HP81 1x | 0.090153 | 0.00698 | 1 | 5 | 1 | D9HP81 | Regulating | D9HP81 [9 | 0 | 1518.73 | 0 | 0.13 | 0.11186 | 12.83 | 91.9 | 108.1 | 10.26 | 5.57 | | High | High | High | High | High | High | High | High | High | High | High | 0.003606 | 0.03823 | 2.15 |
| FALSE | High | SWTALLSA | [R].SWTAL | 1xPhospho | Q8BGR5 1x | 0.002252 | 0.000109 | 1 | 2 | 1 | Q8BGR5 | Protein FA | Q8BGR5 [3 | 0 | 1698.841 | 0 | 1.59 | 0.001974 | 19.08 | 45.2 | 154.8 | 19.66 | 13.31 | | High | High | High | High | High | High | High | High | High | High | High | 4.94E-05 | 0.000592 | 3.12 |
| FALSE | High | SWVIGSPE | [R].SWVIG | 1xPhospho | E9Q1Q1 1x | 0.008614 | 0.000736 | 1 | 6 | 1 | E9Q1Q1 | Microtubu | E9Q1Q1 [9 | 0 | 1565.829 | 0 | -1.26 | 0.000231 | 2.89 | 140.7 | 59.3 | 4.62 | 13.17 | | High | High | High | High | High | High | High | High | High | High | High | 0.000327 | 0.002654 | 2.64 |
| FALSE | High | SWYIADIFT | [K].SWYIAD | 2xPhospho | E9Q4F8 2x | 3.32E-05 | 0 | 1 | 4 | 8 | E9Q4F8 | Ankyrin re | E9Q4F8 [1 | 0 | 2566.108 | 0 | -0.36 | 0.015353 | 3.97 | 108 | 92 | 9.34 | 12.64 | | High | High | High | High | High | High | High | High | High | High | High | 0 | 5.2E-06 | 3.78 |
| FALSE | High | SYAMDPDI | [K].SYAMD | 1xPhospho | Q9Z1T1 1x | 0.015329 | 0.001089 | 1 | 3 | 1 | Q9Z1T1 | AP-3 comp | Q9Z1T1 [2 | 0 | 1416.582 | 0 | 0.01 | 0.679993 | 8.98 | 94.2 | 105.8 | 16.81 | 8.44 | | High | High | High | High | High | High | High | High | High | High | High | 0.000571 | 0.005077 | 3.57 |
| FALSE | High | SYDEGLDD | [K].SYDEGI | 1xPhospho | B7ZCJ1 1xF | 0.007673 | 0.000664 | 1 | 6 | 1 | B7ZCJ1 | Rho GTPas | B7ZCJ1 [88 | 1 | 2214 | 0 | 0.4 | 0.003932 | 8.98 | 86.2 | 113.8 | 8.77 | 3.77 | | High | High | High | High | High | High | High | High | High | High | High | 0.000294 | 0.002328 | 2.27 |
| FALSE | High | SVWGSLAV | [K].SVWGS | 1xPhospho | P19426 1xI | 5.21E-05 | 0 | 1 | 2 | 1 | P19426 | Negative e | P19426 [34 | 0 | 1911.014 | 0 | 0.06 | 0.574996 | 15.06 | 99.4 | 100.6 | 7.03 | 12.21 | | High | High | High | High | High | High | High | High | High | High | High | 0 | 8.59E-06 | 3.35 |
| FALSE | High | SVTIQAPGI | [K].SVTIQA | 1xPhospho | Q6ZPV8 1x | 0.000314 | 0 | 1 | 5 | 2 | Q6ZPV8 | MKIAA121 | Q6ZPV8 [9 | 0 | 2236.106 | 0 | -1.48 | 0.001083 | 1.05 | 154.6 | 45.4 | 9.55 | 19.16 | | High | High | High | High | High | High | High | High | High | High | High | 0 | 6.48E-05 | 2.62 |
| FALSE | High | SVTIQAPA( | [K].SVTIQA | 1xPhospho | Q80Z96 1x | 6.88E-08 | 0 | 1 | 4 | 1 | Q80Z96 | Vang-like p | Q80Z96 [4 | 0 | 2219.127 | 0 | -0.8 | 0.006474 | 3.59 | 124.4 | 75.6 | 7.99 | 13.49 | | High | High | High | High | High | High | High | High | High | High | High | 0 | 5.03E-09 | 4.22 |
| FALSE | High | SVTLLEVC( | [R].SVTLLE | 1xCarbami | Q8CCN5 1x | 3.78E-06 | 0 | 1 | 6 | 1 | Q8CCN5 | Breast carc | Q8CCN5 [8 | 0 | 2316.13 | 0 | -1.3 | 0.002066 | 6.41 | 140.7 | 59.3 | 4.83 | 24.8 | | High | High | High | High | High | High | High | High | High | High | High | 0 | 4.52E-07 | 4.17 |
| FALSE | High | SVSLTGAPI | [R].SVSLTG | 1xPhospho | Q3U0V1 1x | 0.006478 | 0.00062 | 1 | 1 | 1 | Q3U0V1 | Far upstre | Q3U0V1 [1 | 0 | 1840.982 | 0 | 1.29 | 0.003993 | 23.98 | 60.1 | 139.9 | 20.44 | 18.42 | | High | High | High | High | High | High | High | High | High | High | High | 0.000275 | 0.001937 | 2.56 |
| FALSE | High | SVSLYYTGE | [R].SVSLYY | 1xPhospho | P09405 1xI | 0.090153 | 0.00698 | 1 | 6 | 1 | P09405 | Nucleolin ( | P09405 [46 | 0 | 1684.86 | 0 | -0.05 | 0.679761 | 44.61 | 89.7 | 110.3 | 20.98 | 27.99 | | High | High | High | High | High | High | High | High | High | High | High | 0.003606 | 0.03817 | 2.09 |
| FALSE | High | SVSNEGLTI | [R].SVSNE( | 1xPhospho | H7BX08 1x | 0.000458 | 0 | 1 | 5 | 2 | H7BX08 | Calmodulir | H7BX08 [4 | 0 | 1699.821 | 0 | 0.83 | 0.004896 | 12.46 | 74.8 | 125.2 | 4.99 | 13.3 | | High | High | High | High | High | High | High | High | High | High | High | 0 | 9.97E-05 | 2.62 |
| FALSE | High | SVSNGALR | [K].SVSNG | 1xPhospho | Q62130 1x | 0.047488 | 0.00354 | 1 | 4 | 2 | Q62130 | Tyrosine-p | Q62130 [7 | 0 | 1112.566 | 0 | 0.03 | 0.617977 | 2.22 | 98.8 | 101.2 | 7.54 | 8.53 | | High | High | High | High | High | High | High | High | High | High | High | 0.001812 | 0.01813 | 1.7 |
| FALSE | High | SVSPAVHR | [R].SVSPA\ | 1xPhospho | F8VPJ6 1xF | 0.008463 | 0.000736 | 1 | 3 | 1 | F8VPJ6 | Regulatory | F8VPJ6 [11 | 0 | 1161.598 | 0 | -0.59 | 0.001964 | 4.82 | 117.3 | 82.7 | 12.13 | 15.25 | | High | High | High | High | High | High | High | High | High | High | High | 0.000327 | 0.002602 | 1.7 |
| FALSE | High | SVSPGVTQ | [R].SVSPG\ | 1xCarbami | P14873 1xI | 4.65E-06 | 0 | 2 | 2 | 4 | P14873; B2 | Microtubu | P14873 [12 | 0 | 2778.342 | 0 | -0.78 | 0.004363 | 3.47 | 124.7 | 75.3 | 6.7 | 10.36 | Shared | High | High | High | High | High | High | High | High | High | High | High | 0 | 5.69E-07 | 4.68 |
| FALSE | High | SVSPGVTQ | [R].SVSPG\ | 1xCarbami | P14873 2xI | 4.31E-05 | 0 | 2 | 2 | 3 | P14873; B2 | Microtubu | P14873 [12 | 0 | 2858.308 | 0 | -0.63 | 0.00524 | 4.13 | 120.8 | 79.2 | 5.64 | 7.38 | Shared | High | High | High | High | High | High | High | High | High | High | High | 0.000161 | 0.0012 | 3.44 |
| FALSE | High | SVSPLVWC | [K].SVSPLV | 1xCarbami | Q8CI08-2 1 | 0.047894 | 0.00354 | 1 | 3 | 1 | Q8CI08-2 | Isoform 2 ( | Q8CI08-2 [ | 0 | 1412.696 | 0 | 0.8 | 0.008175 | 11.68 | 72.3 | 127.7 | 9.03 | 14.19 | | High | High | High | High | High | High | High | High | High | High | High | 0.001812 | 0.0184 | 1.7 |
| FALSE | High | SVSPSDSLP | [R].SVSPSD | 1xCarbami | P58334 1xI | 3.71E-06 | 0 | 1 | 1 | 4 | P58334 | Krueppel-li | P58334 [22 | 0 | 3041.474 | 0 | -0.12 | 0.286072 | 16.31 | 103.4 | 96.6 | 6.5 | 13.13 | | High | High | High | High | High | High | High | High | High | High | High | 0 | 4.41E-07 | 4.36 |
| FALSE | High | SVSPSPLPL | [R].SVSPSP | 1xPhospho | E9Q5G3 1x | 5.48E-07 | 0 | 1 | 1 | 2 | E9Q5G3 | Kinesin-like | E9Q5G3 [6 | 0 | 3637.836 | 0 | -1.2 | 0.013157 | 2.7 | 150.6 | 49.4 | 25.42 | 16.28 | | High | High | High | High | High | High | High | High | High | High | High | 0 | 5.15E-08 | 3.69 |
| FALSE | High | SVSPSPLPL | [R].SVSPSP | 1xDeamida | E9Q5G3 1x | 0.000211 | 0 | 1 | 1 | 4 | E9Q5G3 | Kinesin-like | E9Q5G3 [6 | 0 | 3638.82 | 0 | -1.36 | 0.002433 | 2.37 | 143.6 | 56.4 | 13.14 | 11 | | High | High | High | High | High | High | High | High | High | High | High | 0 | 4.16E-05 | 4.15 |
| FALSE | High | SVSPSPLPL | [R].SVSPSP | 1xPhospho | E9Q5G3 1x | 0.003927 | 0.000311 | 1 | 1 | 1 | E9Q5G3 | Kinesin-like | E9Q5G3 [6 | 0 | 3653.831 | 0 | -1.29 | 0.006586 | 3.66 | 142.6 | 57.4 | 13.17 | 26.1 | | High | High | High | High | High | High | High | High | High | High | High | 0.00014 | 0.001104 | 2.99 |
| FALSE | High | SVSPSPLPL | [R].SVSPSP | 1xDeamida | E9Q5G3 1x | 0.023175 | 0.001578 | 1 | 1 | 5 | E9Q5G3 | Kinesin-like | E9Q5G3 [6 | 0 | 3654.815 | 0 | -0.72 | 0.001927 | 3.56 | 126.2 | 73.8 | 10.68 | 14.04 | | High | High | High | High | High | High | High | High | High | High | High | 0.001712 | 0.0171 | 3.54 |
| FALSE | High | SVSPVQDLI | [R].SVSPV( | 1xPhospho | Q499D0 1x | 0.002172 | 5.66E-05 | 1 | 4 | 1 | Q499D0 | Forkhead b | Q499D0 [8 | 0 | 3121.366 | 0 | -0.86 | 0.075456 | 15.04 | 128.8 | 71.2 | 22.88 | 23.07 | | High | High | High | High | High | High | High | High | High | High | High | 4.94E-05 | 0.000572 | 2.93 |
| FALSE | High | SVSPVQDLI | [R].SVSPV( | 2xPhospho | Q499D0 2x | 0.000415 | 0 | 1 | 4 | 1 | Q499D0 | Forkhead b | Q499D0 [8 | 0 | 3201.332 | 0 | -1.06 | 0.020975 | 8.76 | 133.5 | 66.5 | 13.29 | 19 | | High | High | High | High | High | High | High | High | High | High | High | 0 | 8.85E-05 | 3.63 |
| FALSE | High | SVSPVQDLI | [R].SVSPV( | 2xPhospho | Q499D0 2x | 0.001361 | 0 | 1 | 4 | 1 | Q499D0 | Forkhead b | Q499D0 [8 | 0 | 3217.327 | 0 | -0.77 | 0.003631 | 0.71 | 124.4 | 75.6 | 5.12 | 14.41 | | High | High | High | High | High | High | High | High | High | High | High | 0 | 0.000337 | 3.17 |
| FALSE | High | SVSSIPVPQ | [R].SVSSIP\ | 1xPhospho | Q8C0J2-3 1 | 1.73E-05 | 0 | 1 | 6 | 2 | Q8C0J2-3 | Isoform 3 ( | Q8C0J2-3 [ | 0 | 2604.314 | 0 | -0.29 | 0.018265 | 4.1 | 109.7 | 90.3 | 6.38 | 8.46 | | High | High | High | High | High | High | High | High | High | High | High | 0 | 2.52E-06 | 3.11 |
| FALSE | High | SVSSNVAS\ | [R].SVSSN\ | 1xPhospho | Q3TWG5 1 | 1.27E-08 | 0 | 1 | 2 | 4 | Q3TWG5 | Dynein cyt | Q3TWG5 [ | 0 | 2125.131 | 0 | 0.7 | 0.00146 | 11.06 | 75.3 | 124.7 | 7.3 | 5.18 | | High | High | High | High | High | High | High | High | High | High | High | 0 | 7.56E-10 | 4.47 |
| FALSE | High | SVSSSSSFD | [K].SVSSSS | 1xPhospho | Q8CGF1 1x | 0.028824 | 0.001829 | 1 | 3 | 2 | Q8CGF1 | Rho GTPas | Q8CGF1 [3 | 0 | 2071.004 | 0 | 0.37 | 0.0162 | 4.24 | 91.7 | 108.3 | 12.96 | 13.84 | | High | High | High | High | High | High | High | High | High | High | High | 0.000997 | 0.01031 | 2.91 |
| FALSE | High | SVSSYGNIF | [R].SVSSYG | 1xPhospho | A2ACM0 1 | 0.01588 | 0.001124 | 1 | 4 | 3 | A2ACM0 | Putative ur | A2ACM0 [7 | 0 | 1291.625 | 0 | 0.43 | 0.04347 | 24.75 | 84.6 | 115.4 | 9.28 | 12.61 | | High | High | High | High | High | High | High | High | High | High | High | 0.000585 | 0.005252 | 2.28 |
| FALSE | High | SVSTMNLS | [R].SVSTM | 1xPhospho | E9QMU3 1 | 0.108463 | 0.008839 | 1 | 5 | 1 | E9QMU3 | Ensconsin | E9QMU3 [ | 0 | 1520.78 | 0 | -0.25 | 0.44771 | 22.46 | 107 | 93 | 26.87 | 14.17 | | High | High | High | High | High | High | High | High | High | High | High | 0.004422 | 0.04764 | 1.02 |
| FALSE | High | SVSTPSEA( | [K].SVSTPS | 1xPhospho | Q3TPJ8 1xI | 5.94E-10 | 0 | 1 | 9 | 9 | Q3TPJ8 | Cytoplasmi | Q3TPJ8 [86 | 0 | 2259.993 | 0 | 0.33 | 0.013926 | 3.96 | 89.4 | 110.6 | 6.01 | 4.91 | | High | High | High | High | High | High | High | High | High | High | High | 0 | 3.89E-09 | 4.82 |
| FALSE | High | SVSTPSVDI | [K].SVSTPS | 1xPhospho | Q3KR45 1x | 6.73E-06 | 0 | 1 | 5 | 7 | Q3KR45 | DNA (cytos | Q3KR45 [1 | 0 | 3263.41 | 0 | -1.28 | 0.000261 | 1.03 | 141.1 | 58.9 | 3.1 | 5.55 | | High | High | High | High | High | High | High | High | High | High | High | 0 | 8.64E-07 | 4.86 |
| FALSE | High | SVSTPSVDI | [K].SVSTPS | 1xPhospho | Q3KR45 1x | 5.96E-08 | 0 | 1 | 5 | 6 | Q3KR45 | DNA (cytos | Q3KR45 [1 | 0 | 3279.405 | 0 | -1.52 | 0.006184 | 1.14 | 145 | 55 | 14.37 | 16.37 | | High | High | High | High | High | High | High | High | High | High | High | 0 | 4.26E-09 | 5.21 |
| FALSE | High | SVSTSSPA( | [K].SVSTSS | 1xPhospho | B1ASU9 1x | 2.51E-09 | 0 | 1 | 5 | 6 | B1ASU9 | Serine/thre | B1ASU9 [7 | 0 | 2521.125 | 0 | -0.27 | 0.088603 | 10.59 | 108 | 92 | 6.17 | 6.97 | | High | High | High | High | High | High | High | High | High | High | High | 0 | 1.22E-10 | 5 |
| FALSE | High | SVSTSSPA( | [K].SVSTSS | 1xDeamida | B1ASU9 1x | 0.005322 | 0.000599 | 1 | 5 | 4 | B1ASU9 | Serine/thre | B1ASU9 [7 | 0 | 2522.109 | 0 | -0.22 | 0.01165 | 6.93 | 108.3 | 91.7 | 8.7 | 9.78 | | High | High | High | High | High | High | High | High | High | High | High | 0.000814 | 0.007577 | 2.8 |
| FALSE | High | SVSTTNIA( | [K].SVSTTN | 1xPhospho | Q60875 1x | 0.00011 | 0 | 1 | 14 | 2 | Q60875 | Rho guanir | Q60875 [1 | 0 | 2424.176 | 0 | 0.14 | 0.2329 | 7.77 | 97.3 | 102.7 | 9.79 | 11.54 | | High | High | High | High | High | High | High | High | High | High | High | 0 | 1.99E-05 | 2.81 |
| FALSE | High | SVSVDLNV | [R].SVSVDI | 1xPhospho | Q3TJN6 1x | 6.46E-08 | 0 | 1 | 3 | 1 | Q3TJN6 | Sorting ne | Q3TJN6 [2 | 0 | 2791.396 | 0 | -1.12 | 0.010214 | 1.3 | 132.9 | 67.1 | 9.84 | 20.08 | | High | High | High | High | High | High | High | High | High | High | High | 0 | 4.69E-09 | 4.78 |
| FALSE | High | SVTLEEAIA | [K].SVTLEE | 1xPhospho | Q8CGN4 1 | 2.35E-07 | 0 | 1 | 4 | 1 | Q8CGN4 | BCL-6 core | Q8CGN4 [1 | 0 | 2085.954 | 0 | 0.51 | 0.005976 | 17.92 | 78.7 | 121.3 | 19.02 | 8.06 | | High | High | High | High | High | High | High | High | High | High | High | 0 | 2E-08 | 4.63 |
| FALSE | High | SVSIQNITG | [K].SVSIQN | 1xPhospho | A0A140LJJ | 0.000136 | 0 | 1 | 5 | 2 | A0A140LJJ | A-kinase ar | A0A140LJJ | 0 | 2832.364 | 0 | -0.15 | 0.456862 | 15.43 | 104.6 | 95.4 | 15.19 | 13.87 | | High | High | High | High | High | High | High | High | High | High | High | 0 | 2.53E-05 | 3.92 |
| FALSE | High | NHLSPQEG | [K].NHLSP( | 2xCarbami | Q9EP97 2x | 1.61E-07 | 0 | 1 | 1 | 1 | Q9EP97 | Sentrin-spe | Q9EP97 [1 | 0 | 2582.077 | 0 | -0.81 | 0.002434 | 1.61 | 127.8 | 72.2 | 8.56 | 9.61 | | High | High | High | High | High | High | High | High | High | High | High | 0 | 1.31E-08 | 5.19 |
| FALSE | High | NEESASHSI | [R].NEESAS | 1xPhospho | P59438 1xI | 0.001168 | 0 | 1 | 6 | 2 | P59438 | Hermansky | P59438 [5 | 0 | 2300.052 | 0 | 0 | 0.911229 | 22.98 | 102.3 | 97.7 | 18.59 | 16.14 | | High | High | High | High | High | High | High | High | High | High | High | 2.58E-05 | 0.000393 | 3.75 |
| FALSE | High | NHLSPQEG | [K].NHLSP( | 2xCarbami | Q9EP97 1x | 3.01E-06 | 0 | 1 | 1 | 3 | Q9EP97 | Sentrin-spe | Q9EP97 [1 | 0 | 2502.11 | 0 | -0.72 | 0.003502 | 4.45 | 125.1 | 74.9 | 4.47 | 14.6 | | High | High | High | High | High | High | High | High | High | High | High | 9.42E-05 | 0.001037 | 3.35 |
| FALSE | High | EADMSSIQ | [K].EADMS | 2xPhospho | B2RUA5 2x | 0.00614 | 0.00062 | 1 | 2 | 2 | B2RUA5 | SAPS doma | B2RUA5 [7 | 0 | 2710.215 | 0 | -0.29 | 0.792363 | 10.14 | 110.5 | 89.5 | 10.44 | 25.85 | | High | High | High | High | High | High | High | High | High | High | High | 0.002403 | 0.02575 | 3.49 |
| FALSE | High | EADPADES | [K].EADPAI | 2xCarbami | Q9CQS5 1x | 2.69E-08 | 0 | 1 | 1 | 3 | Q9CQS5 | Serine/thre | Q9CQS5 [3 | 0 | 2684.089 | 0 | -1.04 | 0.001289 | 3.03 | 131.1 | 68.9 | 6.24 | 17.05 | | High | High | High | High | High | High | High | High | High | High | High | 0 | 1.75E-09 | 5.83 |
| FALSE | High | EAEALLQSI | [R].EAEALL | 1xPhospho | Q3TPJ8 1xI | 3.39E-09 | 0 | 1 | 3 | 3 | Q3TPJ8 | Cytoplasmi | Q3TPJ8 [5 | 0 | 3321.676 | 0 | 0.59 | 0.426324 | 27.97 | 79.8 | 120.2 | 13.3 | 32.51 | | High | High | High | High | High | High | High | High | High | High | High | 0 | 1.7E-10 | 4.8 |
| FALSE | High | EAEALLQSI | [R].EAEALL | 1xPhospho | Q3TPJ8 1xI | 1.16E-06 | 0 | 1 | 3 | 9 | Q3TPJ8 | Cytoplasmi | Q3TPJ8 [5 | 0 | 3337.671 | 0 | 0.5 | 0.005584 | 9.99 | 82.9 | 117.1 | 6.14 | 7.12 | | High | High | High | High | High | High | High | High | High | High | High | 0.000859 | 0.008612 | 3.72 |
| FALSE | High | EAEDVIWE | [K].EAEDVI | 1xPhospho | Q9CWY8 1 | 4.41E-07 | 0 | 1 | 1 | 4 | Q9CWY8 | Ribonuclea | Q9CWY8 [ | 0 | 2968.35 | 0 | -0.79 | 0.001339 | 4.16 | 125.1 | 74.9 | 6.64 | 6.38 | | High | High | High | High | High | High | High | High | High | High | High | 0 | 4.07E-08 | 4.11 |
| FALSE | High | EAEEELAAS | [R].EAEEEL | 1xPhospho | B1ARU1 1x | 1.97E-06 | 0 | 2 | 12 | 3 | B1ARU1; F | Microtubu | B1ARU1 [4 | 0 | 2996.384 | 0 | 0.74 | 0.000378 | 1.93 | 73.7 | 126.3 | 3.72 | 4.81 | Shared | High | High | High | High | High | High | High | High | High | High | High | 0 | 7.38E-06 | 4.27 |
| FALSE | High | EAEEESSG( | [K].EAEEES | 1xPhospho | P59325 1xI | 9.2E-05 | 0 | 1 | 4 | 4 | P59325 | Eukaryotic | P59325 [3 | 0 | 3240.388 | 0 | -0.48 | 0.022914 | 10.38 | 116.4 | 83.6 | 13.29 | 17.71 | | High | High | High | High | High | High | High | High | High | High | High | 0 | 1.64E-05 | 3.25 |
| FALSE | High | EAEEESSG( | [K].EAEEES | 1xDeamida | P59325 1xI | 0.000529 | 0 | 1 | 4 | 1 | P59325 | Eukaryotic | P59325 [3 | 0 | 3241.372 | 0 | -0.28 | 0.01224 | 5.7 | 107.7 | 92.3 | 9.29 | 7.11 | | High | High | High | High | High | High | High | High | High | High | High | 0 | 0.000116 | 3.89 |
| FALSE | High | EAEEESSG( | [K].EAEEES | 2xPhospho | P59325 2xI | 3.41E-05 | 0 | 1 | 4 | 13 | P59325 | Eukaryotic | P59325 [3 | 0 | 3320.354 | 0 | -0.26 | 0.004295 | 4.23 | 110.3 | 89.7 | 5.38 | 6.52 | | High | High | High | High | High | High | High | High | High | High | High | 0 | 1.23E-05 | 5.36 |
| FALSE | High | EAEENPVG | [K].EAEENI | 1xCarbami | O89032 1x | 1.16E-08 | 0 | 1 | 3 | 3 | O89032 | SH3 and P | O89032 [5 | 0 | 2440.147 | 0 | 0.73 | 0.001283 | 7.24 | 75.2 | 124.8 | 8.23 | 5.87 | | High | High | High | High | High | High | High | High | High | High | High | 0 | 3.26E-06 | 4.24 |
| FALSE | High | EAEEQHPE | [R].EAEEQI | 1xPhospho | Q8C080 1x | 0.020499 | 0.001472 | 1 | 3 | 3 | Q8C080 | Sorting ne | Q8C080 [9 | 0 | 3850.747 | 0 | 0.43 | 0.013928 | 11.08 | 82.6 | 117.4 | 4.92 | 9.78 | | High | High | High | High | High | High | High | High | High | High | High | 0.000786 | 0.006991 | 3.26 |
| FALSE | High | EAEESAVC | [K].EAEESA | 2xCarbami | Q80X41 1x | 1.08E-05 | 0 | 1 | 6 | 2 | Q80X41 | Serine/thre | Q80X41 [3 | 0 | 3352.385 | 0 | -0.42 | 0.042071 | 11.63 | 117.5 | 82.5 | 21.93 | 12.49 | | High | High | High | High | High | High | High | High | High | High | High | 0 | 2.54E-06 | 4.89 |
| FALSE | High | EAEESAVC | [K].EAEESA | 2xCarbami | Q80X41 1x | 1.41E-06 | 0 | 1 | 6 | 4 | Q80X41 | Serine/thre | Q80X41 [3 | 0 | 3368.38 | 0 | -0.22 | 0.299789 | 23.34 | 109.4 | 90.6 | 11.96 | 27.54 | | High | High | High | High | High | High | High | High | High | High | High | 0 | 1.5E-07 | 5.51 |
| FALSE | High | EADMSSIQ | [K].EADMS | 1xPhospho | B2RUA5 1x | 6.82E-08 | 0 | 1 | 2 | 4 | B2RUA5 | SAPS doma | B2RUA5 [7 | 0 | 2630.248 | 0 | -0.53 | 0.06354 | 1.12 | 117.2 | 82.8 | 6.72 | 12.49 | | High | High | High | High | High | High | High | High | High | High | High | 0 | 4.31E-07 | 4.41 |
| FALSE | High | EAEGSFTE\ | [K].EAEGSI | 1xPhospho | B1AU76 1x | 2.68E-06 | 0 | 2 | 4 | 3 | B1AU76; B | Nuclear au | B1AU76 [2 | 0 | 1827.845 | 0 | -1.97 | 0.000125 | 0.72 | 159.9 | 40.1 | 3.06 | 14.12 | Shared | High | High | High | High | High | High | High | High | High | High | High | 0 | 3.88E-05 | 3.29 |
| FALSE | High | EAELLTSAK | [R].EAELLT | 1xPhospho | Q8K2E5 1x | 0.012731 | 0.000889 | 1 | 3 | 1 | Q8K2E5 | Budding ur | Q8K2E5 [4 | 0 | 1499.812 | 0 | -0.81 | 0.000576 | 1.45 | 126.6 | 73.4 | 5.92 | 4.31 | | High | High | High | High | High | High | High | High | High | High | High | 0.000449 | 0.004105 | 1.82 |
| FALSE | High | EAEISLK | [R].EAEISLI | 1xPhospho | Q99JF8 1xI | 0.112856 | 0.009146 | 1 | 1 | 1 | Q99JF8 | PC4 and SF | Q99JF8 [5 | 0 | 1327.727 | 0 | -1.26 | 0.000559 | 1.36 | 141 | 59 | 8.74 | 5.43 | | High | High | High | High | High | High | High | High | High | High | High | 0.004569 | 0.04961 | 1.22 |
| FALSE | High | EAEQGSGE | [K].EAEQG | 1xPhospho | Q6P542 1x | 0.00407 | 0.00036 | 2 | 3 | 2 | Q6P542; Q | ATP-bindin | Q6P542 [2 | 0 | 1601.746 | 0 | -0.17 | 0.025375 | 2.74 | 103.1 | 96.9 | 3.71 | 5.69 | Shared | High | High | High | High | High | High | High | High | High | High | High | 0.000161 | 0.001153 | 2.11 |
| FALSE | High | EAEVAPTSI | [R].EAEVAI | 1xPhospho | A2A5Y4 1x | 3.52E-07 | 0 | 1 | 3 | 2 | A2A5Y4 | KAT8 regul | A2A5Y4 [1 | 0 | 2072.144 | 0 | -0.57 | 0.000325 | 1.76 | 118.8 | 81.2 | 1.31 | 3.07 | | High | High | High | High | High | High | High | High | High | High | High | 0 | 1.5E-05 | 3.68 |
| FALSE | High | EAEVQVSA | [K].EAEVQ | 1xPhospho | O88466 1x | 3.88E-06 | 0 | 1 | 4 | 2 | O88466 | Zinc finger | O88466 [6 | 0 | 2703.437 | 2 | -0.88 | 0.000872 | 3.02 | 131.1 | 68.9 | 4.8 | 7.15 | | High | High | High | High | High | High | High | High | High | High | High | 0 | 4.65E-07 | 3.66 |
| FALSE | High | EAEVQVSA | [K].EAEVQ | 1xPhospho | R4GML0 1 | 9.2E-05 | 0 | 1 | 3 | 3 | R4GML0 | Zinc finger | R4GML0 [6 | 0 | 2702.453 | 0 | -1.43 | 0.000847 | 0.78 | 146 | 54 | 9.68 | 5.18 | | High | High | High | High | High | High | High | High | High | High | High | 0 | 1.63E-05 | 3.42 |
| FALSE | High | EAFEEMEG | [K].EAFEEN | 1xPhospho | E9Q0S6 1x | 1.85E-08 | 0 | 1 | 7 | 1 | E9Q0S6 | Tensin 1 O | E9Q0S6 [1 | 0 | 2454.085 | 0 | -0.09 | 0.380242 | 14.34 | 102.9 | 97.1 | 10.39 | 12.12 | | High | High | High | High | High | High | High | High | High | High | High | 0 | 1.14E-09 | 4.15 |
| FALSE | High | EAFEEMEG | [K].EAFEEN | 1xPhospho | E9Q0S6 1x | 1.86E-07 | 0 | 1 | 7 | 3 | E9Q0S6 | Tensin 1 O | E9Q0S6 [1 | 0 | 2470.08 | 0 | -0.6 | 0.102491 | 3.03 | 121.5 | 78.5 | 5.57 | 32.02 | | High | High | High | High | High | High | High | High | High | High | High | 0 | 4.16E-07 | 4.53 |
| FALSE | High | EAFEQPQT | [K].EAFEQI | 1xPhospho | P70236 1xI | 0.000246 | 0 | 1 | 5 | 2 | P70236 | Dual specif | P70236 [1 | 0 | 1883.874 | 0 | 0.54 | 0.004087 | 5.62 | 86.1 | 113.9 | 10.2 | 10.31 | | High | High | High | High | High | High | High | High | High | High | High | 0 | 4.95E-05 | 2.69 |
| FALSE | High | EAFEQPQT | [K].EAFEQI | 1xPhospho | P70236 1xI | 0.022182 | 0.001542 | 1 | 5 | 1 | P70236 | Dual specif | P70236 [1 | 1 | 2671.302 | 0 | 0.22 | 0.021822 | 11.33 | 91.1 | 108.9 | 6.74 | 4.55 | | High | High | High | High | High | High | High | High | High | High | High | 0.000814 | 0.007652 | 2.99 |
| FALSE | High | EAGAGESE | [K].EAGAG | 1xCarbami | E9Q286 1x | 0.000786 | 0 | 1 | 1 | 1 | E9Q286 | Little elong | E9Q286 [5 | 0 | 2180.035 | 0 | -0.07 | 0.501149 | 28.39 | 102.8 | 97.2 | 24.8 | 10.7 | | High | High | High | High | High | High | High | High | High | High | High | 0 | 0.000181 | 2.04 |
| FALSE | High | EAGDSPNS | [R].EAGDSI | 1xPhospho | E9QLK3 1x | 2.23E-08 | 0 | 1 | 3 | 1 | E9QLK3 | BRCA1-ass | E9QLK3 [7 | 0 | 2010.933 | 0 | -0.08 | 0.892939 | 9.3 | 101.9 | 98.1 | 14.68 | 3.11 | | High | High | High | High | High | High | High | High | High | High | High | 0 | 1.42E-09 | 4.83 |
| FALSE | High | EAGELPTSI | [R].EAGELI | 1xPhospho | P59178 1xI | 0.001313 | 0 | 2 | 6 | 4 | P59178; Q | Lethal(3)m | P59178 [6 | 0 | 2349.144 | 0 | -0.93 | 0.00545 | 2.11 | 131.1 | 68.9 | 11.87 | 8.03 | Shared | High | High | High | High | High | High | High | High | High | High | High | 2.58E-05 | 0.000551 | 2.57 |
| FALSE | High | EAELENSGI | [K].EAELEN | 1xPhospho | Q9CYA6 1x | 6.04E-06 | 0 | 1 | 2 | 1 | Q9CYA6 | Zinc finger | Q9CYA6 [3 | 0 | 3735.68 | 0 | -0.2 | 0.137056 | 20.35 | 105 | 95 | 14.37 | 17.96 | | High | High | High | High | High | High | High | High | High | High | High | 0 | 7.69E-07 | 4.92 |
| FALSE | High | EAGELPTSI | [R].EAGELI | 1xDeamida | P59178 1xI | 7.91E-06 | 0 | 2 | 6 | 3 | P59178; Q | Lethal(3)m | P59178 [6 | 0 | 2350.128 | 0 | -0.9 | 0.005337 | 4.87 | 130.3 | 69.7 | 7.49 | 10.88 | Shared | High | High | High | High | High | High | High | High | High | High | High | 0 | 1.04E-06 | 2.95 |
| FALSE | High | EADMSSIQ | [K].EADMS | 1xPhospho | B2RUA5 1x | 0.000148 | 0 | 1 | 2 | 1 | B2RUA5 | SAPS doma | B2RUA5 [7 | 0 | 2614.253 | 0 | -0.27 | 0.162736 | 11.22 | 111.9 | 88.1 | 19.35 | 9.67 | | High | High | High | High | High | High | High | High | High | High | High | 0 | 2.81E-05 | 2.78 |
| FALSE | High | EADIDSSDE | [K].EADIDS | 3xPhospho | O35218 3x | 0.000356 | 0 | 1 | 2 | 2 | O35218 | Cleavage a | O35218 [4 | 1 | 3215.243 | 0 | -0.28 | 0.001957 | 0.91 | 110.9 | 89.1 | 6.38 | 6.62 | | High | High | High | High | High | High | High | High | High | High | High | 0 | 7.48E-05 | 4.15 |
| FALSE | High | DYSGSQG( | [R].DYSGS( | 1xPhospho | Q545K5 1x | 0.005978 | 0.00062 | 1 | 1 | 3 | Q545K5 | MCG11638 | Q545K5 [1 | 0 | 1513.616 | 0 | 1.15 | 0.004298 | 15.43 | 64 | 136 | 8.86 | 14.26 | | High | High | High | High | High | High | High | High | High | High | High | 0.000275 | 0.002191 | 2.1 |
| FALSE | High | DYSGSQG( | [R].DYSGS( | 1xPhospho | Q545K5 1x | 0.0026 | 0.000109 | 1 | 1 | 2 | Q545K5 | MCG11638 | Q545K5 [1 | 1 | 2310.961 | 0 | 0.73 | 0.001134 | 13.17 | 75.3 | 124.7 | 9.53 | 2.09 | | High | High | High | High | High | High | High | High | High | High | High | 4.94E-05 | 0.000696 | 1.17 |
| FALSE | High | DYSHSSSR | [R].DYSHSS | 1xPhospho | Q91VM5 1 | 0.094641 | 0.007408 | 1 | 1 | 2 | Q91VM5 | RNA bindir | Q91VM5 [ | 0 | 1247.526 | 0 | -0.66 | 0.024509 | 1.51 | 121.9 | 78.1 | 17.55 | 3.64 | | High | High | High | High | High | High | High | High | High | High | High | 0.003754 | 0.04041 | 1.87 |
| FALSE | High | DYSPPYAPS | [R].DYSPPY | 1xPhospho | Q6ZPZ3 1x | 0.000718 | 0 | 1 | 6 | 2 | Q6ZPZ3 | Zinc finger | Q6ZPZ3 [1 | 0 | 3109.482 | 0 | -0.01 | 0.415288 | 16.86 | 102.8 | 97.2 | 14.09 | 16.74 | | High | High | High | High | High | High | High | High | High | High | High | 0 | 0.000234 | 3.27 |
| FALSE | High | DYSYER | [R].DYSYEF | 1xPhospho | Q7TSC1 1x | 0.116482 | 0.009523 | 1 | 3 | 2 | Q7TSC1 | Protein PR | Q7TSC1 [1 | 0 | 1141.476 | 0 | -0.16 | 0.147979 | 9.49 | 105.5 | 94.5 | 2.57 | 11.47 | | High | High | High | High | High | High | High | High | High | High | High | 0.007094 | 0.07709 | 1.12 |
| FALSE | High | DYTSHLSP( | [R].DYTSHI | 1xPhospho | F8VQL9 1x | 0.047086 | 0.00354 | 1 | 4 | 1 | F8VQL9 | Nuclear re | F8VQL9 [4 | 0 | 2108.038 | 0 | 1.12 | 0.000444 | 12.31 | 62.1 | 137.9 | 12.81 | 6.64 | | High | High | High | High | High | High | High | High | High | High | High | 0.001812 | 0.01801 | 1.87 |
| FALSE | High | DYYFSNSD | [K].DYYFSN | 1xPhospho | Q80WQ8 1 | 0.007881 | 0.000736 | 1 | 3 | 4 | Q80WQ8 | Mis18-binc | Q80WQ8 [ | 0 | 1319.503 | 0 | -0.75 | 0.002082 | 2.7 | 126.5 | 73.5 | 9.57 | 8.44 | | High | High | High | High | High | High | High | High | High | High | High | 0.000674 | 0.005868 | 2.08 |
| FALSE | High | DYYFSNSD | [K].DYYFSN | 2xPhospho | Q80WQ8 2 | 0.057203 | 0.004227 | 1 | 3 | 1 | Q80WQ8 | Mis18-binc | Q80WQ8 [ | 0 | 1399.469 | 0 | -0.82 | 0.003951 | 5.3 | 127.2 | 72.8 | 6.34 | 11.15 | | High | High | High | High | High | High | High | High | High | High | High | 0.002159 | 0.02241 | 1.17 |
| FALSE | High | EAAANTNF | [R].EAAAN | 1xPhospho | Q9WTK0 1 | 0.075304 | 0.005727 | 1 | 1 | 1 | Q9WTK0 | Nuclear pr | Q9WTK0 [ | 0 | 1972.919 | 0 | 3.19 | | 53.76 | 22.2 | 177.8 | 33.93 | 22.27 | | Not Found | High | High | High | High | High | High | High | High | High | High | 0.002951 | 0.03083 | 1.51 |
| FALSE | High | EAADTDVE | [K].EAADTI | 1xPhospho | Q9CZX5 1x | 9.54E-05 | 0 | 1 | 1 | 2 | Q9CZX5 | PIN2/TERF | Q9CZX5 [2 | 0 | 2078.98 | 0 | -0.71 | 0.00092 | 1.58 | 127 | 73 | 7.22 | 7.04 | | High | High | High | High | High | High | High | High | High | High | High | 0 | 1.7E-05 | 2.62 |
| FALSE | High | EAADTDVE | [K].EAADTI | 2xPhospho | Q9CZX5 2x | 0.003001 | 0.00016 | 1 | 1 | 1 | Q9CZX5 | PIN2/TERF | Q9CZX5 [2 | 0 | 2158.946 | 0 | -0.3 | 0.098153 | 2.69 | 106.8 | 93.2 | 12.45 | 10.17 | | High | High | High | High | High | High | High | High | High | High | High | 7.19E-05 | 0.000819 | 2.82 |
| FALSE | High | EAAEAEPA | [K].EAAEAI | 1xPhospho | P26645 1xI | 7.56E-06 | 0 | 1 | 3 | 17 | P26645 | Myristoyla | P26645 [1 | 0 | 3254.499 | 0 | 1.6 | 0.000897 | 12.57 | 49.3 | 150.7 | 6.19 | 10.68 | | High | High | High | High | High | High | High | High | High | High | High | 0 | 4.55E-06 | 4.23 |
| FALSE | High | EAAEAEPA | [K].EAAEAI | 2xPhospho | P26645 2xI | 0.001933 | 5.66E-05 | 1 | 3 | 1 | P26645 | Myristoyla | P26645 [1 | 0 | 3334.465 | 0 | 1.57 | 0.002301 | 6.75 | 50.8 | 149.2 | 14.63 | 13.24 | | High | High | High | High | High | High | High | High | High | High | High | 2.58E-05 | 0.000498 | 3.83 |
| FALSE | High | EADLVFISV | [R].EADLVI | 1xPhospho | Q3TJE8 1xI | 5.82E-06 | 0 | 1 | 6 | 2 | Q3TJE8 | UDP-gluco | Q3TJE8 [8 | 0 | 2072.108 | 0 | -0.79 | 0.001137 | 3.64 | 126.7 | 73.3 | 4.79 | 8.78 | | High | High | High | High | High | High | High | High | High | High | High | 0 | 1.19E-05 | 3.87 |
| FALSE | High | EAAFGGGL | [R].EAAFG( | 1xPhospho | Q04863 1x | 1.62E-06 | 0 | 1 | 3 | 2 | Q04863 | Transcripti | Q04863 [5 | 0 | 1782.851 | 0 | 0.31 | 0.033733 | 15.81 | 89.1 | 110.9 | 7.46 | 10.98 | | High | High | High | High | High | High | High | High | High | High | High | 0.000859 | 0.008641 | 2.9 |
| FALSE | High | EAALSPEQI | [K].EAALSP | 1xPhospho | Q3URL4 1x | 0.046291 | 0.003523 | 1 | 3 | 1 | Q3URL4 | BTB and C | Q3URL4 [6 | 0 | 2049.14 | 0 | -0.18 | 0.656908 | 18.08 | 104.7 | 95.3 | 12.73 | 11.65 | | High | High | High | High | High | High | High | High | High | High | High | 0.001789 | 0.0177 | 2.32 |
| FALSE | High | EAAQSPVP | [K].EAAQSI | 1xPhospho | Q8VC19 1x | 0.026191 | 0.001676 | 1 | 4 | 1 | Q8VC19 | 5-aminolev | Q8VC19 [1 | 0 | 2074.135 | 0 | -0.67 | 0.014914 | 4.77 | 122.9 | 77.1 | 8.91 | 18.69 | | High | High | High | High | High | High | High | High | High | High | High | 0.000903 | 0.009229 | 2.94 |
| FALSE | High | EAAQTAPS | [R].EAAQT | 1xPhospho | Q3TSW4 1 | 7.34E-05 | 0 | 1 | 4 | 1 | Q3TSW4 | MCG1557( | Q3TSW4 [2 | 0 | 1733.924 | 0 | 1.64 | 0.000342 | 9.44 | 48.5 | 151.5 | 8.72 | 6.42 | | High | High | High | High | High | High | High | High | High | High | High | 0 | 1.27E-05 | 1.99 |
| FALSE | High | EAASGGAD | [R].EAASG( | 1xPhospho | Q9DBY0 1x | 0.046687 | 0.00354 | 1 | 6 | 1 | Q9DBY0 | Forkhead b | Q9DBY0 [5 | 0 | 3549.663 | 0 | -0.12 | 0.352521 | 15.73 | 110.2 | 89.8 | 22.03 | 16.52 | | High | High | High | High | High | High | High | High | High | High | High | 0.001812 | 0.01782 | 2.44 |
| FALSE | High | EAAVLSEEE | [R].EAAVLS | 1xPhospho | B1ARU1 1x | 9.61E-08 | 0 | 1 | 4 | 1 | B1ARU1 | Microtubu | B1ARU1 [1 | 0 | 3287.586 | 0 | -1.49 | 0.000546 | 1.27 | 147.2 | 52.8 | 6.25 | 12.28 | | High | High | High | High | High | High | High | High | High | High | High | 0 | 7.29E-09 | 4.96 |
| FALSE | High | EAAVPTPEI | [K].EAAVPT | 1xPhospho | Q2MHN3 1 | 0.001386 | 5.66E-05 | 1 | 1 | 2 | Q2MHN3 | ZFH-5 OS=I | Q2MHN3 [ | 0 | 3512.749 | 0 | -0.42 | 0.026173 | 11.53 | 118.2 | 81.8 | 14.94 | 6.4 | | High | High | High | High | High | High | High | High | High | High | High | 2.58E-05 | 0.000345 | 2.92 |
| FALSE | High | EADALPAST | [K].EADALI | 1xPhospho | B2RRE0 1xI | 0.000386 | 0 | 1 | 3 | 4 | B2RRE0 | A kinase (P | B2RRE0 [7 | 0 | 4626.083 | 0 | -1.71 | 0.000242 | 1.04 | 153.3 | 46.7 | 4.31 | 14.04 | | High | High | High | High | High | High | High | High | High | High | High | 0 | 0.000123 | 4.71 |
| FALSE | High | EADALPAST | [K].EADALI | 1xDeamida | B2RRE0 1xI | 8.98E-06 | 0 | 1 | 3 | 17 | B2RRE0 | A kinase (P | B2RRE0 [7 | 0 | 4627.067 | 0 | -1.57 | 0.000975 | 2.26 | 150.2 | 49.8 | 10.73 | 4.3 | | High | High | High | High | High | High | High | High | High | High | High | 0 | 1.32E-06 | 4.87 |
| FALSE | High | EADALPAST | [K].EADALI | 2xPhospho | B2RRE0 2xI | 0.002252 | 0.000109 | 1 | 3 | 1 | B2RRE0 | A kinase (P | B2RRE0 [7 | 0 | 4706.05 | 0 | -1.52 | 0.001023 | 1.25 | 148.9 | 51.1 | 10.48 | 5.11 | | High | High | High | High | High | High | High | High | High | High | High | 4.94E-05 | 0.00059 | 3.52 |
| FALSE | High | EADDDEEA | [K].EADDD | 1xPhospho | Q3UHZ3 1x | 2.33E-07 | 0 | 3 | 6 | 9 | Q3UHZ3; C | DNA (cytos | Q3UHZ3 [7 | 0 | 2648.085 | 0 | -0.14 | 0.147804 | 15.6 | 107.1 | 92.9 | 9.62 | 10.98 | Shared | High | High | High | High | High | High | High | High | High | High | High | 0 | 4.95E-08 | 4.72 |
| FALSE | High | EADDDEEA | [K].EADDD | 1xPhospho | Q3UHZ3 1x | 6.91E-06 | 0 | 3 | 6 | 3 | Q3UHZ3; C | DNA (cytos | Q3UHZ3 [7 | 1 | 2989.348 | 0 | -0.24 | 0.039089 | 12.63 | 108.2 | 91.8 | 9.53 | 10.86 | Shared | High | High | High | High | High | High | High | High | High | High | High | 0 | 2.03E-05 | 4.11 |
| FALSE | High | EADDESLDI | [R].EADDE | 1xPhospho | Q3TH99 1x | 0.055777 | 0.004093 | 1 | 4 | 1 | Q3TH99 | Putative ur | Q3TH99 [5 | 0 | 2132.964 | 0 | -0.52 | 0.029032 | 11.47 | 117.8 | 82.2 | 12.95 | 18.74 | | High | High | High | High | High | High | High | High | High | High | High | 0.002117 | 0.02183 | 1.67 |
| FALSE | High | EADGTEEV | [K].EADGTI | 1xPhospho | Q76KJ5 1xI | 1.51E-05 | 0 | 1 | 2 | 4 | Q76KJ5 | DNA-direct | Q76KJ5 [2 | 0 | 4149.078 | 0 | 0.22 | 0.026169 | 10.23 | 86.3 | 113.7 | 18.52 | 8.83 | | High | High | High | High | High | High | High | High | High | High | High | 0 | 2.15E-06 | 4.3 |
| FALSE | High | EAALILGVS | [R].EAALIL | 1xPhospho | G3X8S1 1x | 0.009168 | 0.000736 | 1 | 5 | 1 | G3X8S1 | DnaJ (Hsp4 | G3X8S1 [6 | 0 | 1922.076 | 0 | -0.97 | 0.002086 | 3.9 | 131.5 | 68.5 | 8.5 | 5.82 | | High | High | High | High | High | High | High | High | High | High | High | 0.000327 | 0.002856 | 2.63 |
| FALSE | High | EAGENAPV | [K].EAGENA | 1xPhospho | Q3U568 1x | 4.62E-07 | 0 | 1 | 2 | 2 | Q3U568 | Putative ur | Q3U568 [1 | 0 | 2427.131 | 0 | 0.58 | 0.002141 | 9.94 | 80.3 | 119.7 | 12.26 | 3.41 | | High | High | High | High | High | High | High | High | High | High | High | 0 | 4.27E-08 | 3.65 |
| FALSE | High | EAGENAPV | [K].EAGENA | 1xPhospho | Q3U568 1x | 3.93E-07 | 0 | 1 | 2 | 2 | Q3U568 | Putative ur | Q3U568 [1 | 0 | 2443.126 | 0 | 1.6 | 0.000741 | 17.87 | 50.5 | 149.5 | 9.77 | 9.72 | | High | High | High | High | High | High | High | High | High | High | High | 0 | 3.54E-08 | 4.91 |
| FALSE | High | EAGLQHGE | [K].EAGLQ | 1xCarbami | Q6P1D7 1x | 0.000815 | 0 | 1 | 1 | 1 | Q6P1D7 | Structure-s | Q6P1D7 [1 | 0 | 2520.15 | 0 | -0.35 | 0.037815 | 1.51 | 110 | 90 | 18.95 | 7.65 | | High | High | High | High | High | High | High | High | High | High | High | 0 | 0.00019 | 3.62 |
| FALSE | High | EAPGSPPLS | [R].EAPGSI | 2xPhospho | Q67FY2 2x | 0.093879 | 0.007306 | 1 | 2 | 1 | Q67FY2 | B-cell CLL/ | Q67FY2 [1 | 0 | 1496.675 | 0 | 1.56 | 0.000348 | 12.32 | 51.2 | 148.8 | 6.56 | 5.79 | | High | High | High | High | High | High | High | High | High | High | High | 0.003727 | 0.04006 | 2.12 |

| | | | | | | | | | | | | | | | | | | | | | | | | | | | | | | | | | | | | | | |
|---|---|---|---|---|---|---|---|---|---|---|---|---|---|---|---|---|---|---|---|---|---|---|---|---|---|---|---|---|---|---|---|---|---|---|---|---|---|---|
| FALSE | High | EAPSPLMIH | [R].EAPSPL | 1xPhospho | Q3UH70 1x | 0.003138 | 0.00016 | 1 | 2 | 1 | Q3UH70 | Bromodom | Q3UH70 [1 | 0 | 4162.066 | 0 | -0.74 | 0.045496 | 8.65 | 121.4 | 78.6 | 24.3 | 17.16 | | High | High | High | High | High | High | High | High | High | High | High | 7.19E-05 | 0.000861 | 2.5 |
| FALSE | High | EAPSPSIAT | [R].EAPSPS | 1xPhospho | P33215 1x | 0.000115 | 0 | 1 | 2 | 2 | P33215 | Protein NE | P33215 [34 | 0 | 2615.446 | 0 | 1.03 | 0.000975 | 11.37 | 66.8 | 133.2 | 7.27 | 5.29 | | High | High | High | High | High | High | High | High | High | High | High | 0 | 5.85E-05 | 3.61 |
| FALSE | High | EAQELGSP | [R].EAQEL | 1xPhospho | Q60875 1x | 0.014667 | 0.001049 | 1 | 8 | 1 | Q60875 | Rho guanir | Q60875 [9 | 0 | 1539.689 | 0 | -0.01 | 0.859345 | 10.06 | 100.4 | 99.6 | 14.24 | 8.51 | | High | High | High | High | High | High | High | High | High | High | High | 0.000555 | 0.004799 | 1.86 |
| FALSE | High | EAQLPSCH | [R].EAQLPS | 2xCarbami | A0A0G2JG | 8.71E-05 | 0 | 1 | 2 | 4 | A0A0G2JG | Crystallin b | A0A0G2JG | 0 | 3955.756 | 0 | 1.64 | 0.026389 | 30.99 | 48.2 | 151.8 | 19.48 | 35.56 | | High | High | High | High | High | High | High | High | High | High | High | 0 | 1.87E-05 | 5.57 |
| FALSE | High | EAQLPSCH | [R].EAQLPS | 2xCarbami | A0A0G2JG | 4.53E-07 | 0 | 1 | 2 | 2 | A0A0G2JG | Crystallin b | A0A0G2JG | 0 | 4183.935 | 0 | 6.45 | | 21.12 | 4 | 196 | 69.29 | 16.26 | | Not Found | Not Found | High | High | High | High | High | High | High | High | High | 0 | 4.16E-08 | 4.54 |
| FALSE | High | EASAEDLEI | [K].EASAEI | 1xCarbami | Q8R2M2 1 | 2.31E-07 | 0 | 1 | 1 | 4 | Q8R2M2 | Deoxynucl | Q8R2M2 [4 | 0 | 2807.31 | 0 | -0.96 | 0.000944 | 2.14 | 132.9 | 67.1 | 5.16 | 10.88 | | High | High | High | High | High | High | High | High | High | High | High | 0 | 1.96E-08 | 3.9 |
| FALSE | High | EASANGPV | [K].EASAN | 1xDeamida | Q80YR4 1x | 0.048303 | 0.00354 | 1 | 3 | 1 | Q80YR4 | E3 ubiquiti | Q80YR4 [4 | 0 | 4066.083 | 0 | -0.44 | 0.00132 | 3.67 | 116.4 | 83.6 | 4.78 | 2.7 | | High | High | High | High | High | High | High | High | High | High | High | 0.001844 | 0.01858 | 2.73 |
| FALSE | High | EASFEK | [K].EASFEK | 1xPhospho | Q91ZV0 1x | 0.101749 | 0.008194 | 1 | 10 | 1 | Q91ZV0 | Melanoma | Q91ZV0 [7 | 0 | 1248.628 | 0 | -0.12 | 0.330082 | 6.02 | 105.4 | 94.6 | 3.45 | 7.12 | | High | High | High | High | High | High | High | High | High | High | High | 0.004115 | 0.04388 | 1.67 |
| FALSE | High | EASFSLLES | [K].EASFSL | 1xPhospho | Q810L3 1x | 0.000738 | 0 | 1 | 5 | 2 | Q810L3 | E3 ubiquiti | Q810L3 [2 | 0 | 1648.86 | 0 | -0.59 | 0.001617 | 5.45 | 117 | 83 | 7.68 | 10.32 | | High | High | High | High | High | High | High | High | High | High | High | 4.94E-05 | 0.000655 | 2.33 |
| FALSE | High | EASIDILHSI | [R].EASIDIL | 1xPhospho | Q497E8 1x | 0.116482 | 0.009523 | 1 | 11 | 1 | Q497E8 | Proteasom | Q497E8 [2 | 0 | 1863.039 | 0 | 0.67 | 0.001928 | 10.27 | 78 | 122 | 4.45 | 5.37 | | High | High | High | High | High | High | High | High | High | High | High | 0.004757 | 0.05182 | 2.19 |
| FALSE | High | EASPPGGS | [K].EASPPG | 2xPhospho | Q8BUH8 2x | 0.000475 | 0 | 1 | 2 | 1 | Q8BUH8 | Sentrin-spe | Q8BUH8 [4 | 0 | 4108.929 | 0 | 0.48 | 0.004913 | 8.95 | 80.8 | 119.2 | 3.62 | 9.66 | | High | High | High | High | High | High | High | High | High | High | High | 0 | 0.000103 | 3.95 |
| FALSE | High | EASQMGH | [K].EASQM | 1xPhospho | Q61687 1x | 0.015329 | 0.001089 | 1 | 4 | 3 | Q61687 | Transcripti | Q61687 [7 | 0 | 3084.344 | 0 | -0.29 | 0.025767 | 10.24 | 118.1 | 81.9 | 7.27 | 13.83 | | High | High | High | High | High | High | High | High | High | High | High | 0.000571 | 0.00505 | 2.94 |
| FALSE | High | EAPAGGSG | [R].EAPAG | 2xPhospho | Q8CJ40 2x | 0.000139 | 0 | 1 | 3 | 1 | Q8CJ40 | Rootletin C | Q8CJ40 [14 | 0 | 2520.089 | 0 | -1.35 | 0.000918 | 2.09 | 143.8 | 56.2 | 6.6 | 8.94 | | High | High | High | High | High | High | High | High | High | High | High | 0 | 2.59E-05 | 3.81 |
| FALSE | High | EASRPPEEF | [R].EASRPF | 1xPhospho | Q8BG30 1x | 0.003857 | 0.000261 | 1 | 2 | 1 | Q8BG30 | Negative e | Q8BG30 [3 | 0 | 2804.427 | 0 | 0.15 | 0.086233 | 9.31 | 95.1 | 104.9 | 7.05 | 6.66 | | High | High | High | High | High | High | High | High | High | High | High | 0.000117 | 0.001079 | 2.17 |
| FALSE | High | EASSPPTHI | [K].EASSPP | 1xPhospho | F6RJ39 1xP | 0.032831 | 0.002195 | 2 | 4 | 1 | F6RJ39; Q8 | Apoptotic | F6RJ39 [37 | 0 | 4412.168 | 0 | -0.32 | 0.216698 | 23.97 | 123.8 | 76.2 | 18.89 | 31.34 | Shared | High | High | High | High | High | High | High | High | High | High | High | 0.001151 | 0.01198 | 3.68 |
| FALSE | High | EASSPPTHI | [K].EASSPP | 1xPhospho | F6RJ39 1xP | 0.00311 | 0.00016 | 2 | 4 | 1 | F6RJ39; Q8 | Apoptotic | F6RJ39 [37 | 0 | 4428.163 | 0 | -0.11 | 0.269505 | 34.98 | 109 | 91 | 25.56 | 25.89 | Shared | High | High | High | High | High | High | High | High | High | High | High | 7.19E-05 | 0.000851 | 2.95 |
| FALSE | High | EASSPPTHI | [K].EASSPP | 2xPhospho | F6RJ39 2xP | 7.28E-05 | 0 | 2 | 4 | 3 | F6RJ39; Q8 | Apoptotic | F6RJ39 [37 | 0 | 4492.135 | 0 | 0.22 | 0.806648 | 21.59 | 97.6 | 102.4 | 16.61 | 35.89 | Shared | High | High | High | High | High | High | High | High | High | High | High | 0 | 0.000171 | 3.78 |
| FALSE | High | EASSPPTHI | [K].EASSPP | 2xPhospho | F6RJ39 2xP | 1.3E-05 | 0 | 2 | 4 | 12 | F6RJ39; Q8 | Apoptotic | F6RJ39 [37 | 0 | 4508.13 | 0 | -0.21 | 0.17874 | 6.03 | 104 | 96 | 8.56 | 14.19 | Shared | High | High | High | High | High | High | High | High | High | High | High | 0 | 1.81E-06 | 4.82 |
| FALSE | High | EATAQKPT | [K].EATAQ | 1xPhospho | E9QMN5 1 | 0.011346 | 0.000816 | 1 | 3 | 3 | E9QMN5 | Transcripti | E9QMN5 [ | 0 | 2834.507 | 0 | -0.73 | 0.034624 | 7.68 | 124.8 | 75.2 | 14.08 | 16.17 | | High | High | High | High | High | High | High | High | High | High | High | 0.000418 | 0.003617 | 1.53 |
| FALSE | High | EATGPAPSI | [K].EATGP | 1xPhospho | Q9JJ66 1xP | 0.033116 | 0.002195 | 1 | 1 | 1 | Q9JJ66 | Cell divisio | Q9JJ66 [34 | 0 | 1438.66 | 0 | -0.28 | 0.058759 | 9.87 | 109.3 | 90.7 | 8.97 | 16.69 | | High | High | High | High | High | High | High | High | High | High | High | 0.001151 | 0.01208 | 1.35 |
| FALSE | High | EATGSQK | [K].EATGS | 1xPhospho | Q9R1X5 1x | 0.008538 | 0.000736 | 1 | 1 | 1 | Q9R1X5 | Multidrug | Q9R1X5 [7 | 0 | 1258.644 | 0 | 2.06 | 0.000622 | 7.65 | 38.7 | 161.3 | 8.96 | 11.81 | | High | High | High | High | High | High | High | High | High | High | High | 0.000327 | 0.002641 | 1.31 |
| FALSE | High | EATLPPVSF | [R].EATLPP | 1xPhospho | Q6NZJ6 1x | 0.016021 | 0.001124 | 1 | 9 | 6 | Q6NZJ6 | Eukaryotic | Q6NZJ6 [1 | 0 | 1444.765 | 0 | 0.11 | 0.397168 | 1.93 | 96.5 | 103.5 | 7.99 | 9.17 | | High | High | High | High | High | High | High | High | High | High | High | 0.000585 | 0.005296 | 1.87 |
| FALSE | High | EATLPPVSF | [R].EATLPP | 1xPhospho | Q6NZJ6 1x | 5.12E-05 | 0 | 1 | 9 | 6 | Q6NZJ6 | Eukaryotic | Q6NZJ6 [1 | 0 | 1673.928 | 0 | 0.13 | 0.23528 | 6.39 | 97.6 | 102.4 | 7.49 | 8.71 | | High | High | High | High | High | High | High | High | High | High | High | 0 | 6.35E-05 | 3.39 |
| FALSE | High | EATPLPPPE | [K].EATPLP | 1xPhospho | Q99MR1 1 | 6.15E-06 | 0 | 1 | 1 | 4 | Q99MR1 | GRB10-inte | Q99MR1 [3 | 0 | 4227.079 | 0 | 0.21 | 0.413376 | 30.76 | 91.5 | 108.5 | 7.68 | 25.95 | | High | High | High | High | High | High | High | High | High | High | High | 0 | 7.82E-07 | 4.62 |
| FALSE | High | EATSSLVLP | [R].EATSSL | 1xPhospho | Q91Z58 1x | 0.013074 | 0.000889 | 1 | 1 | 2 | Q91Z58 | Uncharacte | Q91Z58 [5 | 0 | 2835.438 | 0 | 3.88 | 0.002848 | 45.29 | 13.2 | 186.8 | 24.75 | 21.96 | | High | High | High | High | High | High | High | High | High | High | High | 0.000467 | 0.00423 | 3.27 |
| FALSE | High | EATTSTPLL | [R].EATTST | 1xPhospho | Q9DBY8 1x | 0.038017 | 0.002528 | 1 | 2 | 4 | Q9DBY8 | Nuclear va | Q9DBY8 [1 | 0 | 1786.96 | 0 | -0.36 | 0.007528 | 6.31 | 113.3 | 86.7 | 5.76 | 5.7 | | High | High | High | High | High | High | High | High | High | High | High | 0.001355 | 0.01416 | 2.25 |
| FALSE | High | EAVAVAPP | [K].EAVAV | 1xPhospho | A2ARV4 1x | 0.001313 | 0 | 1 | 1 | 1 | A2ARV4 | Low-densit | A2ARV4 [4 | 0 | 2069.145 | 0 | 0.11 | 0.780005 | 4.75 | 99.8 | 100.2 | 15.44 | 9.72 | | High | High | High | High | High | High | High | High | High | High | High | 0 | 0.000324 | 2.14 |
| FALSE | High | EASSGSTPQ | [K].EASSGS | 1xPhospho | Q3TAB2 1x | 0.076555 | 0.005826 | 2 | 6 | 1 | Q3TAB2; H | Putative ur | Q3TAB2 [4 | 0 | 1984.072 | 0 | -0.78 | 0.027288 | 10.11 | 127.1 | 72.9 | 23.57 | 5.97 | Shared | High | High | High | High | High | High | High | High | High | High | High | 0.002976 | 0.03145 | 1.84 |
| FALSE | High | EAPAGGSG | [R].EAPAG | 1xPhospho | Q8CJ40 1x | 4.76E-05 | 0 | 1 | 3 | 1 | Q8CJ40 | Rootletin C | Q8CJ40 [14 | 0 | 2440.123 | 0 | -0.39 | 0.030903 | 9.46 | 108.1 | 91.9 | 8.82 | 9.34 | | High | High | High | High | High | High | High | High | High | High | High | 0 | 7.83E-06 | 4.24 |
| FALSE | High | EANNCSSP | [R].EANNC | 1xCarbami | P42230 1x | 2.06E-06 | 0 | 1 | 4 | 1 | P42230 | Signal tran | P42230 [12 | 0 | 2532.188 | 0 | 1.39 | 0.000779 | 11.2 | 53.3 | 146.7 | 10.29 | 7.47 | | High | High | High | High | High | High | High | High | High | High | High | 0 | 2.29E-07 | 3.22 |
| FALSE | High | EANNCSSP | [R].EANNC | 1xCarbami | P42230 1x | 5.45E-05 | 0 | 1 | 4 | 1 | P42230 | Signal tran | P42230 [12 | 0 | 2516.193 | 0 | 0.15 | 0.290332 | 18.29 | 97.9 | 102.1 | 18.75 | 15.38 | | High | High | High | High | High | High | High | High | High | High | High | 0 | 9.04E-06 | 3 |
| FALSE | High | EAGLSQSH | [K].EAGLSQ | 1xPhospho | Q80X80 1x | 8.63E-05 | 0 | 1 | 5 | 2 | Q80X80 | Phospholip | Q80X80 [6 | 0 | 2495.161 | 0 | 0.68 | 0.01545 | 3.85 | 76.9 | 123.1 | 7.98 | 15.49 | | High | High | High | High | High | High | High | High | High | High | High | 0 | 0.000177 | 1.89 |
| FALSE | High | EAGIVHSDA | [K].EAGIVH | 1xPhospho | Q3TUQ5 1 | 0.000279 | 0 | 1 | 2 | 11 | Q3TUQ5 | Pinin OS=M | Q3TUQ5 [3 | 0 | 1693.856 | 0 | -0.41 | 0.010609 | 7.28 | 113.6 | 86.4 | 2.8 | 7.98 | | High | High | High | High | High | High | High | High | High | High | High | 9.42E-05 | 0.000989 | 2.65 |
| FALSE | High | EAGPAAAS | [K].EAGPA | 1xPhospho | A2ADB0 1x | 7.62E-09 | 0 | 1 | 5 | 2 | A2ADB0 | Msx2-inter | A2ADB0 [2 | 0 | 2222.074 | 0 | 0.02 | 0.500716 | 1.78 | 98.7 | 101.3 | 6.95 | 12.17 | | High | High | High | High | High | High | High | High | High | High | High | 0 | 1.06E-07 | 4.32 |
| FALSE | High | EAGPAAAS | [K].EAGPA | 2xPhospho | A2ADB0 2x | 0.000772 | 0 | 1 | 5 | 1 | A2ADB0 | Msx2-inter | A2ADB0 [2 | 0 | 2302.041 | 0 | 0.04 | 0.832202 | 11.69 | 101.3 | 98.7 | 7.81 | 10.06 | | High | High | High | High | High | High | High | High | High | High | High | 0 | 0.000178 | 3 |
| FALSE | High | EAGTPSLDS | [R].EAGTPS | 1xPhospho | Q67FY2 1x | 0.00582 | 0.00062 | 1 | 2 | 1 | Q67FY2 | B-cell CLL/l | Q67FY2 [1 | 0 | 1742.861 | 0 | 1.66 | 0.00044 | 4.37 | 47.7 | 152.3 | 6.65 | 8.59 | | High | High | High | High | High | High | High | High | High | High | High | 0.000275 | 0.001719 | 2.12 |
| FALSE | High | EALEQCTEI | [R].EALEQ | 2xCarbami | Q6P9P6 1x | 0.001674 | 5.66E-05 | 1 | 1 | 2 | Q6P9P6 | Kinesin-like | Q6P9P6 [9 | 0 | 3358.429 | 0 | -0.91 | 0.028021 | 6.73 | 124.2 | 75.8 | 29.15 | 8.19 | | High | High | High | High | High | High | High | High | High | High | High | 2.58E-05 | 0.000427 | 3.3 |
| FALSE | High | EALEVEECS | [R].EALEVE | 1xCarbami | Q80U04 1x | 6.02E-05 | 0 | 1 | 3 | 2 | Q80U04 | E3 ubiquiti | Q80U04 [3 | 0 | 2372.134 | 0 | 0.62 | 0.002011 | 16.59 | 81.7 | 118.3 | 8.85 | 4.28 | | High | High | High | High | High | High | High | High | High | High | High | 0 | 1.02E-05 | 2.51 |
| FALSE | High | EALGAPEA | [K].EALGAF | 1xPhospho | E9QK83 1x | 0.097743 | 0.007804 | 1 | 2 | 1 | E9QK83 | Small subu | E9QK83 [1 | 0 | 2110.083 | 0 | -0.31 | 0.04537 | 6.2 | 112.2 | 87.8 | 4.86 | 10.18 | | High | High | High | High | High | High | High | High | High | High | High | 0.003922 | 0.04187 | 2.58 |
| FALSE | High | EAILAPQCL | [R].EAILAP | 1xCarbami | Q921C3 1x | 0.08447 | 0.006345 | 1 | 5 | 1 | Q921C3 | Bromodom | Q921C3 [1 | 0 | 1703.85 | 0 | -1.51 | 0.001999 | 2.35 | 148 | 52 | 13.15 | 16.55 | | High | High | High | High | High | High | High | High | High | High | High | 0.003279 | 0.0355 | 2.44 |
| FALSE | High | EAILAPQCL | [R].EAILAP | 1xCarbami | Q921C3 2x | 0.00153 | 5.66E-05 | 1 | 5 | 1 | Q921C3 | Bromodom | Q921C3 [1 | 1 | 3129.391 | 0 | -0.9 | 0.003643 | 5.94 | 133.9 | 66.1 | 11.7 | 5.67 | | High | High | High | High | High | High | High | High | High | High | High | 2.58E-05 | 0.000384 | 2.92 |
| FALSE | High | EAILAPQCL | [R].EAILAP | 1xCarbami | Q921C3 3x | 2.22E-06 | 0 | 1 | 5 | 1 | Q921C3 | Bromodom | Q921C3 [1 | 1 | 3209.357 | 0 | -1.29 | 0.000734 | 2.12 | 142 | 58 | 9.44 | 5.46 | | High | High | High | High | High | High | High | High | High | High | High | 0 | 2.5E-07 | 3.9 |
| FALSE | High | EALLESLGS | [R].EALLES | 1xPhospho | Q3TYD6 1x | 0.000132 | 0 | 1 | 1 | 3 | Q3TYD6 | Serine/thre | Q3TYD6 [9 | 0 | 1718.868 | 0 | -0.66 | 0.000516 | 1.08 | 122.5 | 77.5 | 6.51 | 6.48 | | High | High | High | High | High | High | High | High | High | High | High | 4.94E-05 | 0.000666 | 2.56 |
| FALSE | High | EALNIIGDIS | [K].EALNII | 2xDeamida | F8WIV5 1x | 0.014929 | 0.001049 | 1 | 9 | 3 | F8WIV5 | Dynamin-2 | F8WIV5 [7 | 0 | 4439.149 | 0 | 0.68 | 0.008943 | 22.3 | 75.3 | 124.7 | 12.67 | 12.1 | | High | High | High | High | High | High | High | High | High | High | High | 0.000555 | 0.004924 | 3.42 |
| FALSE | High | EALPDLPSF | [K].EALPDL | 1xPhospho | O54692 1x | 0.000579 | 0 | 1 | 2 | 2 | O54692 | Centromer | O54692 [4 | 0 | 2043.02 | 0 | -0.03 | 0.98349 | 2.14 | 100.7 | 99.3 | 4.16 | 5.9 | | High | High | High | High | High | High | High | High | High | High | High | 0 | 0.000129 | 1.74 |
| FALSE | High | EALPSIWDS | [K].EALPSI | 1xPhospho | Q9CTE8 1x | 0.006478 | 0.00062 | 1 | 4 | 2 | Q9CTE8 | Putative ur | Q9CTE8 [5 | 0 | 1881.976 | 0 | 0.43 | 0.033011 | 24.52 | 81.8 | 118.2 | 12.91 | 12.35 | | High | High | High | High | High | High | High | High | High | High | High | 0.000275 | 0.001929 | 3.31 |
| FALSE | High | EAIQAPSEE | [K].EAIQAP | 1xPhospho | Q8BHZ4 1x | 2.09E-07 | 0 | 1 | 2 | 2 | Q8BHZ4 | Zinc finger | Q8BHZ4 [2 | 0 | 2180.089 | 0 | -0.25 | 0.001694 | 3.72 | 108.8 | 91.2 | 1.18 | 3.93 | | High | High | High | High | High | High | High | High | High | High | High | 0 | 1.6E-07 | 4.18 |
| FALSE | High | EAISDDEDE | [K].EAISDD | 1xPhospho | Q8C3S2 1x | 3.42E-06 | 0 | 1 | 3 | 3 | Q8C3S2 | Transport | Q8C3S2 [5 | 0 | 2164.01 | 0 | -0.56 | 0.001757 | 6.29 | 119.6 | 80.4 | 4.24 | 9.22 | | High | High | High | High | High | High | High | High | High | High | High | 0 | 1.03E-06 | 3.53 |
| FALSE | High | EALSQQLD | [R].EALSQ | 1xPhospho | Q640N2 1x | 1.67E-07 | 0 | 1 | 1 | 3 | Q640N2 | ADP-ribosy | Q640N2 [3 | 0 | 2199.96 | 0 | 0.4 | 0.03699 | 11.29 | 87.1 | 112.9 | 11.33 | 7.99 | | High | High | High | High | High | High | High | High | High | High | High | 0 | 1.36E-08 | 4.27 |
| FALSE | High | EALVEPASE | [R].EALVEP | 1xPhospho | P70441 1x | 1.27E-08 | 0 | 1 | 1 | 3 | P70441 | Na(+)/H(+) | P70441 [26 | 0 | 2102.084 | 0 | 0.92 | 0.000656 | 3.61 | 69.2 | 130.8 | 2.84 | 5.7 | | High | High | High | High | High | High | High | High | High | High | High | 0 | 3.42E-08 | 2.13 |
| FALSE | High | EALVEPASE | [R].EALVEP | 2xPhospho | P70441 2x | 0.001179 | 0 | 1 | 1 | 1 | P70441 | Na(+)/H(+) | P70441 [26 | 0 | 2182.051 | 0 | 0.27 | 0.004112 | 5.61 | 88.1 | 111.9 | 5.94 | 4.45 | | High | High | High | High | High | High | High | High | High | High | High | 0 | 0.000287 | 2.04 |
| FALSE | High | EAIVESDSE | [K].EAIVES | 1xPhospho | Q5H8C4 1x | 2.72E-05 | 0 | 1 | 2 | 1 | Q5H8C4 | Vacuolar p | Q5H8C4 [1 | 0 | 2121.999 | 0 | 1.24 | 0.000769 | 13.75 | 61.6 | 138.4 | 10.29 | 7.89 | | High | High | High | High | High | High | High | High | High | High | High | 0 | 4.18E-06 | 3.57 |
| FALSE | High | EALVSQPSS | [R].EALVSQ | 1xPhospho | P41230 1x | 1.56E-06 | 0 | 1 | 3 | 1 | P41230 | Lysine-spe | P41230 [88 | 0 | 2333.232 | 0 | -0.35 | 0.018989 | 4.54 | 110.8 | 89.2 | 8.43 | 5.57 | | High | High | High | High | High | High | High | High | High | High | High | 0 | 1.67E-07 | 4.84 |
| FALSE | High | EAMEATEA | [R].EAMEA | 1xPhospho | Q52KE1 1x | 0.000132 | 0 | 1 | 2 | 1 | Q52KE1 | THAP dom | Q52KE1 [1 | 0 | 2410.234 | 0 | 0.62 | 0.005735 | 15.79 | 79.5 | 120.5 | 14.63 | 5.29 | | High | High | High | High | High | High | High | High | High | High | High | 0 | 2.44E-05 | 2.23 |
| FALSE | High | EANELSDSA | [K].EANELS | 1xPhospho | Q69ZX6 1x | 0.000117 | 0 | 1 | 1 | 2 | Q69ZX6 | MORC fam | Q69ZX6 [7 | 0 | 2248.992 | 0 | -0.22 | 0.023429 | 7.54 | 112.8 | 87.2 | 8.34 | 8.28 | | High | High | High | High | High | High | High | High | High | High | High | 0 | 0.000219 | 2.48 |
| FALSE | High | EANELSDSA | [K].EANELS | 1xPhospho | Q69ZX6 1x | 2.67E-05 | 0 | 1 | 1 | 2 | Q69ZX6 | MORC fam | Q69ZX6 [7 | 1 | 2606.25 | 0 | -0.52 | 0.015001 | 8.05 | 119.8 | 80.2 | 11.3 | 6.22 | | High | High | High | High | High | High | High | High | High | High | High | 0 | 3.9E-05 | 2.95 |
| FALSE | High | EANELSDSA | [K].EANELS | 2xPhospho | Q69ZX6 2x | 0.019276 | 0.001402 | 1 | 1 | 2 | Q69ZX6 | MORC fam | Q69ZX6 [7 | 1 | 2686.216 | 0 | -0.4 | 0.015235 | 6.53 | 116.5 | 83.5 | 9 | 5.64 | | High | High | High | High | High | High | High | High | High | High | High | 0.000756 | 0.006536 | 1.67 |
| FALSE | High | EANEVLSDS | [K].EANEVI | 1xPhospho | Q6PCN6 1x | 1.16E-08 | 0 | 1 | 1 | 4 | Q6PCN6 | Microrchid | Q6PCN6 [7 | 0 | 2348.06 | 0 | -0.75 | 0.000551 | 1.94 | 125.2 | 74.8 | 5.64 | 3.84 | | High | High | High | High | High | High | High | High | High | High | High | 0 | 6.82E-10 | 4.08 |
| FALSE | High | EANEVLSDS | [K].EANEVI | 2xPhospho | Q6PCN6 2x | 3.63E-05 | 0 | 1 | 1 | 1 | Q6PCN6 | Microrchid | Q6PCN6 [7 | 0 | 2428.027 | 0 | -0.64 | 0.025103 | 16.41 | 122.8 | 77.2 | 11.96 | 17.46 | | High | High | High | High | High | High | High | High | High | High | High | 0 | 5.75E-06 | 2.06 |
| FALSE | High | EANEVLSDS | [K].EANEVI | 2xPhospho | Q6PCN6 2x | 0.01186 | 0.000889 | 1 | 1 | 1 | Q6PCN6 | Microrchid | Q6PCN6 [7 | 1 | 2785.285 | 0 | -0.6 | 0.004954 | 4.4 | 118.6 | 81.4 | 8.71 | 6.77 | | High | High | High | High | High | High | High | High | High | High | High | 0.000449 | 0.003788 | 1.75 |
| FALSE | High | DYSFEEPNT | [R].DYSFEE | 1xPhospho | Q8C0P0 1x | 0.000174 | 0 | 1 | 1 | 4 | Q8C0P0 | Serine/thre | Q8C0P0 [5 | 0 | 2581.252 | 0 | -0.47 | 0.007099 | 4.57 | 116.5 | 83.5 | 8.49 | 6.79 | | High | High | High | High | High | High | High | High | High | High | High | 0 | 3.36E-05 | 2.74 |
| FALSE | High | EAVVDLHA | [K].EAVVDI | 1xPhospho | F6RJ39 1xP | 7.54E-05 | 0 | 2 | 4 | 1 | F6RJ39; Q8 | Apoptotic | F6RJ39 [81 | 1 | 2595.177 | 0 | -0.43 | 0.009739 | 5.74 | 114.5 | 85.5 | 5.63 | 7.9 | Shared | High | High | High | High | High | High | High | High | High | High | High | 0 | 1.31E-05 | 1.33 |
| FALSE | High | DYSDITPGK | [K].DYSDIT | 1xPhospho | Q9CWC4 1 | 0.019966 | 0.001438 | 1 | 2 | 1 | Q9CWC4 | Transcripti | Q9CWC4 [ | 0 | 1533.76 | 0 | -0.55 | 0.022893 | 5.37 | 114.9 | 85.1 | 19.87 | 9.4 | | High | High | High | High | High | High | High | High | High | High | High | 0.000786 | 0.006828 | 1.87 |
| FALSE | High | DYRPIPELD | [R].DYRPIP | 1xPhospho | Q3UJN1 1x | 2.45E-06 | 0 | 1 | 3 | 10 | Q3UJN1 | DNA helica | Q3UJN1 [8 | 0 | 3859.8 | 0 | -0.73 | 0.003074 | 5.95 | 120.5 | 79.5 | 13.83 | 13.76 | | High | High | High | High | High | High | High | High | High | High | High | 0 | 2.78E-07 | 3.78 |
| FALSE | High | DTPSSEGEI | [K].DTPSSE | 1xPhospho | E9QAE4 1x | 0.000765 | 0 | 1 | 2 | 2 | E9QAE4 | Histone-lys | E9QAE4 [1 | 0 | 1762.857 | 0 | -1.35 | 0.000191 | 1.61 | 143.7 | 56.3 | 5.14 | 8.72 | | High | High | High | High | High | High | High | High | High | High | High | 0 | 0.000177 | 3.03 |
| FALSE | High | DTPTSAGP | [R].DTPTSA | 1xPhospho | F8WI88 1x | 0.000579 | 0 | 1 | 4 | 2 | F8WI88 | PEST prote | F8WI88 [1 | 0 | 1873.91 | 0 | 1.15 | 0.000865 | 14.48 | 62.3 | 137.7 | 14.98 | 9.85 | | High | High | High | High | High | High | High | High | High | High | High | 0 | 0.000128 | 2.91 |
| FALSE | High | DTSASPDSA | [K].DTSASF | 1xPhospho | Q6PCS8 1x | 0.000579 | 0 | 1 | 2 | 1 | Q6PCS8 | Zinc finger | Q6PCS8 [7 | 0 | 1516.73 | 0 | -0.18 | 0.012729 | 5.52 | 103.8 | 96.2 | 8.61 | 7.48 | | High | High | High | High | High | High | High | High | High | High | High | 0 | 0.000128 | 1.89 |
| FALSE | High | DTSLSSEGN | [K].DTSLSS | 1xPhospho | Q9EPQ8 1x | 0.001489 | 5.66E-05 | 1 | 3 | 5 | Q9EPQ8 | Transcripti | Q9EPQ8 [6 | 0 | 1676.814 | 0 | -0.15 | 0.01967 | 1.87 | 105 | 95 | 13.35 | 10.78 | | High | High | High | High | High | High | High | High | High | High | High | 2.58E-05 | 0.000374 | 2.93 |
| FALSE | High | DTSSNDIN | [R].DTSSNI | 1xPhospho | P17426 1x | 2.26E-06 | 0 | 1 | 2 | 9 | P17426 | AP-2 comp | P17426 [63 | 0 | 3393.626 | 0 | 0.16 | 0.58577 | 29.3 | 95.6 | 104.4 | 20.08 | 7.25 | | High | High | High | High | High | High | High | High | High | High | High | 0 | 2.54E-07 | 5.58 |
| FALSE | High | DTSSNDIN | [R].DTSSNI | 1xDeamida | P17426 2x | 0.00582 | 0.00062 | 1 | 2 | 2 | P17426 | AP-2 comp | P17426 [63 | 0 | 3474.576 | 0 | 0.14 | 0.391043 | 15.54 | 95.2 | 104.8 | 8.25 | 14.58 | | High | High | High | High | High | High | High | High | High | High | High | 0.000275 | 0.001708 | 3.3 |
| FALSE | High | DTSTEALLK | [K].DTSTEA | 1xPhospho | E9QMN5 1 | 0.000845 | 0 | 1 | 4 | 1 | E9QMN5 | Transcripti | E9QMN5 [ | 0 | 1515.807 | 0 | -1.12 | 0.000647 | 2.15 | 136.1 | 63.9 | 10.4 | 9.41 | | High | High | High | High | High | High | High | High | High | High | High | 0 | 0.000198 | 2.64 |
| FALSE | High | DTSVGTLS | [R].DTSVG | 1xPhospho | E9PVV1 1x | 0.000241 | 0 | 2 | 11 | 1 | E9PVV1; E | Transcripti | E9PVV1 [1 | 0 | 3537.849 | 0 | 0.47 | 0.002813 | 7.7 | 88.1 | 111.9 | 8.71 | 6.89 | Shared | High | High | High | High | High | High | High | High | High | High | High | 0 | 4.82E-05 | 4.72 |
| FALSE | High | DTTLEPYA | [R].DTTLEP | 1xPhospho | G3X8T2 1x | 5.37E-06 | 0 | 1 | 3 | 3 | G3X8T2 | RIKEN cDN | G3X8T2 [3 | 0 | 2562.116 | 0 | 0.53 | 0.101353 | 21.04 | 81.7 | 118.3 | 20.12 | 8.44 | | High | High | High | High | High | High | High | High | High | High | High | 0 | 6.7E-07 | 3.58 |
| FALSE | High | DTTSFEDIS | [K].DTTSFE | 1xPhospho | Q5SNZ0 1x | 0.000108 | 0 | 1 | 6 | 5 | Q5SNZ0 | Girdin OS= | Q5SNZ0 [1 | 0 | 2611.125 | 0 | -0.24 | 0.613473 | 17.08 | 102.7 | 97.3 | 18.19 | 11.54 | | High | High | High | High | High | High | High | High | High | High | High | 0 | 5.69E-05 | 4.49 |
| FALSE | High | DTTSPMEL | [K].DTTSPN | 1xPhospho | Q8K212 1x | 4.68E-05 | 0 | 1 | 3 | 4 | Q8K212 | Phosphofu | Q8K212 [4 | 0 | 1943.98 | 0 | 2.55 | 0.000872 | 2.88 | 30 | 170 | 3.71 | 12.64 | | High | High | High | High | High | High | High | High | High | High | High | 0 | 7.62E-06 | 2.84 |
| FALSE | High | DTTSPMEL | [K].DTTSPN | 1xPhospho | Q8K212 1x | 1.5E-05 | 0 | 1 | 3 | 6 | Q8K212 | Phosphofu | Q8K212 [4 | 0 | 1959.975 | 0 | 3.01 | 0.00086 | 13.95 | 24.6 | 175.4 | 14.34 | 13.9 | | High | High | High | High | High | High | High | High | High | High | High | 0 | 2.14E-06 | 2.62 |
| FALSE | High | DTTTVFCPI | [K].DTTTVF | 1xCarbami | Q9JIM3 1x | 0.052133 | 0.00384 | 1 | 3 | 1 | Q9JIM3 | DNA excisi | Q9JIM3 [1 | 0 | 2975.329 | 0 | -0.07 | 0.426847 | 23.89 | 99.9 | 100.1 | 13.3 | 11.82 | | High | High | High | High | High | High | High | High | High | High | High | 0.001999 | 0.02026 | 3.07 |
| FALSE | High | DTPQTPSR | [K].DTPQTI | 1xPhospho | Q8BTI8 1x | 0.059159 | 0.004294 | 1 | 3 | 1 | Q8BTI8 | Serine/argi | Q8BTI8 [14 | 0 | 1210.567 | 0 | -0.63 | 0.01132 | 7.43 | 120.3 | 79.7 | 12.38 | 6.16 | | High | High | High | High | High | High | High | High | High | High | High | 0.002202 | 0.0234 | 1.36 |
| FALSE | High | DTVDSAGT | [R].DTVDSA | 1xPhospho | Q9DBS2 1x | 0.006713 | 0.00062 | 1 | 2 | 1 | Q9DBS2 | Tumor pro | Q9DBS2 [6 | 0 | 3247.579 | 0 | -0.01 | 0.404224 | 7.54 | 99.4 | 100.6 | 6.65 | 12.31 | | High | High | High | High | High | High | High | High | High | High | High | 0.000275 | 0.002018 | 2.32 |
| FALSE | High | DTVIIVSEPS | [K].DTVIIVS | 2xPhospho | Q04692 2x | 1.51E-06 | 0 | 1 | 1 | 7 | Q04692 | SWI/SNF-r | Q04692 [1 | 0 | 2943.311 | 0 | -1.33 | 0.000313 | 1.72 | 144.1 | 55.9 | 4.68 | 5.12 | | High | High | High | High | High | High | High | High | High | High | High | 0 | 5.29E-06 | 3.51 |
| FALSE | High | DTVIIVSEPS | [K].DTVIIVS | 3xPhospho | Q04692 3x | 0.01441 | 0.001011 | 1 | 1 | 4 | Q04692 | SWI/SNF-r | Q04692 [1 | 0 | 3023.277 | 0 | -0.4 | 0.007077 | 4.42 | 115.5 | 84.5 | 8.49 | 2.74 | | High | High | High | High | High | High | High | High | High | High | High | 0.000538 | 0.00471 | 3.08 |
| FALSE | High | DTVNNVDT | [K].DTVNN | 1xPhospho | S4R1M2 1x | 2.01E-07 | 0 | 1 | 2 | 7 | S4R1M2 | Scaffold at | S4R1M2 [1 | 0 | 3927.744 | 0 | 0.43 | 0.87882 | 9.33 | 89.2 | 110.8 | 11.18 | 38.86 | | High | High | High | High | High | High | High | High | High | High | High | 0 | 1.68E-08 | 5.08 |
| FALSE | High | DTVNNVDT | [K].DTVNN | 1xDeamida | S4R1M2 1x | 0.001373 | 5.66E-05 | 1 | 2 | 13 | S4R1M2 | Scaffold at | S4R1M2 [1 | 0 | 3928.728 | 0 | -0.01 | 0.723629 | 23.74 | 100.4 | 99.6 | 15.2 | 38.94 | | High | High | High | High | High | High | High | High | High | High | High | 0.001372 | 0.01435 | 3.34 |
| FALSE | High | DTVNNVDT | [K].DTVNN | 1xPhospho | S4R1M2 1x | 3.32E-08 | 0 | 1 | 2 | 39 | S4R1M2 | Scaffold at | S4R1M2 [1 | 0 | 3943.739 | 0 | -0.08 | 0.397276 | 9.28 | 99.4 | 100.6 | 6.64 | 17.5 | | High | High | High | High | High | High | High | High | High | High | High | 0 | 4.33E-08 | 4.76 |
| FALSE | High | DTYSDRSG | [R].DTYSDF | 2xPhospho | P70698 2x | 0.00047 | 0 | 1 | 1 | 1 | P70698 | CTP syntha | P70698 [5 | 1 | 2792.232 | 0 | -0.58 | 0.036919 | 5.07 | 116.2 | 83.8 | 7.34 | 13.09 | | High | High | High | High | High | High | High | High | High | High | High | 0 | 0.000103 | 4.36 |
| FALSE | High | DVAEGNVS | [K].DVAEG | 1xPhospho | Q91V83 1x | 1.16E-05 | 0 | 1 | 1 | 1 | Q91V83 | TELO2-inte | Q91V83 [8 | 0 | 2851.365 | 0 | -0.11 | 0.139827 | 6.73 | 103.9 | 96.1 | 8.47 | 5.07 | | High | High | High | High | High | High | High | High | High | High | High | 0 | 1.6E-06 | 3.09 |
| FALSE | High | DVAPPMEE | [K].DVAPPI | 1xPhospho | P27546 1x | 2.32E-06 | 0 | 1 | 3 | 3 | P27546 | Microtubu | P27546 [32 | 0 | 2664.288 | 0 | -1.69 | 0.011828 | 1.71 | 152.7 | 47.3 | 23.68 | 35.15 | | High | High | High | High | High | High | High | High | High | High | High | 0 | 2.62E-07 | 4.12 |
| FALSE | High | DVAPPMEE | [K].DVAPPI | 1xPhospho | P27546 1x | 1.19E-05 | 0 | 1 | 3 | 5 | P27546 | Microtubu | P27546 [32 | 0 | 2680.283 | 0 | -1.38 | 0.002602 | 3.44 | 144.4 | 55.6 | 11.65 | 22.35 | | High | High | High | High | High | High | High | High | High | High | High | 0 | 1.64E-06 | 3.76 |
| FALSE | High | DVASATPE | [R].DVASA | 1xPhospho | Q91VA6 1x | 0.012508 | 0.000889 | 1 | 2 | 2 | Q91VA6 | Polymeras | Q91VA6 [1 | 0 | 1455.75 | 0 | -0.02 | 0.472252 | 2.61 | 101.5 | 98.5 | 6.28 | 5.56 | | High | High | High | High | High | High | High | High | High | High | High | 0.009016 | 0.09636 | 2.05 |
| FALSE | High | DVATSPISP | [R].DVATSI | 1xPhospho | G5E884 1x | 1.75E-05 | 0 | 1 | 2 | 6 | G5E884 | Serine/thre | G5E884 [2 | 0 | 2606.255 | 0 | -0.71 | 0.001224 | 4.41 | 124.2 | 75.8 | 6.5 | 8.03 | | High | High | High | High | High | High | High | High | High | High | High | 0 | 7.02E-05 | 4.67 |
| FALSE | High | DVATSPISP | [R].DVATSI | 1xDeamida | G5E884 1x | 0.000553 | 0 | 1 | 2 | 2 | G5E884 | Serine/thre | G5E884 [2 | 0 | 2607.239 | 0 | -0.61 | 0.000945 | 3.58 | 118.8 | 81.2 | 5.14 | 6.56 | | High | High | High | High | High | High | High | High | High | High | High | 0 | 0.000123 | 3.52 |
| FALSE | High | DVATSPISP | [R].DVATSI | 2xPhospho | G5E884 2x | 0.000379 | 0 | 1 | 2 | 4 | G5E884 | Serine/thre | G5E884 [2 | 0 | 2686.221 | 0 | -0.61 | 0.000877 | 1.01 | 120.9 | 79.1 | 5.18 | 4.87 | | High | High | High | High | High | High | High | High | High | High | High | 0 | 0.000117 | 3.84 |
| FALSE | High | DTVIIVSEPS | [K].DTVIIVS | 1xPhospho | Q04692 1x | 5.7E-05 | 0 | 1 | 1 | 6 | Q04692 | SWI/SNF-r | Q04692 [1 | 0 | 2863.345 | 0 | -1.33 | 0.000401 | 1.07 | 143.1 | 56.9 | 7.32 | 7.3 | | High | High | High | High | High | High | High | High | High | High | High | 0 | 0.000146 | 3.52 |
| FALSE | High | DVDELPSL | [R].DVDELI | 1xPhospho | O55106 1x | 0.000422 | 0 | 1 | 4 | 3 | O55106 | Striatin OS | O55106 [3 | 0 | 2606.266 | 0 | 0.53 | 0.000729 | 6.71 | 81.3 | 118.7 | 7.15 | 3.92 | | High | High | High | High | High | High | High | High | High | High | High | 0 | 9.03E-05 | 2.01 |
| FALSE | High | DTPPPLVPF | [K].DTPPPL | 1xPhospho | Q99J09 1x | 0.04913 | 0.003576 | 1 | 2 | 1 | Q99J09 | Methyloso | Q99J09 [4- | 0 | 1539.813 | 0 | -1.1 | 0.002341 | 5.34 | 137 | 63 | 10.78 | 16.56 | | High | High | High | High | High | High | High | High | High | High | High | 0.001889 | 0.01881 | 1.59 |
| FALSE | High | DTPLPGPA | [R].DTPLPG | 1xPhospho | Q3UZ64 1x | 0.01319 | 0.000889 | 1 | 4 | 1 | Q3UZ64 | Nuclear fac | Q3UZ64 [1 | 0 | 2231.025 | 0 | 0.13 | 0.984827 | 11.61 | 99.7 | 100.3 | 12.52 | 26.94 | | High | High | High | High | High | High | High | High | High | High | High | 0.000467 | 0.004272 | 1.87 |
| FALSE | High | DTDSGTPF | [R].DTDSG | 1xPhospho | Q6PCL9 1x | 0.012397 | 0.000889 | 1 | 4 | 2 | Q6PCL9 | Poly(A) pol | Q6PCL9 [5 | 0 | 3554.669 | 0 | -0.84 | 0.002536 | 5.06 | 130.3 | 69.7 | 10.49 | 9.06 | | High | High | High | High | High | High | High | High | High | High | High | 0.002045 | 0.02114 | 3.66 |
| FALSE | High | DTECLLQD | [R].DTECLL | 1xCarbami | B2RUC3 1x | 1.96E-07 | 0 | 1 | 8 | 2 | B2RUC3 | Excision re | B2RUC3 [6 | 0 | 2515.068 | 0 | -0.26 | 0.098604 | 11.1 | 108.7 | 91.3 | 7.51 | 14.04 | | High | High | High | High | High | High | High | High | High | High | High | 0 | 1.63E-08 | 5.03 |
| FALSE | High | DTECLLQD | [R].DTECLL | 1xCarbami | B2RUC3 1x | 5.72E-06 | 0 | 1 | 8 | 2 | B2RUC3 | Excision re | B2RUC3 [6 | 0 | 2531.063 | 0 | -0.17 | 0.180676 | 17.79 | 109.7 | 90.3 | 7.16 | 14.48 | | High | High | High | High | High | High | High | High | High | High | High | 0 | 7.19E-07 | 5.06 |
| FALSE | High | DTECTPVG | [K].DTECTI | 1xCarbami | Q8R4Z6 1x | 0.000138 | 0 | 1 | 3 | 3 | Q8R4Z6 | DEAD-box | Q8R4Z6 [6 | 0 | 4597.169 | 0 | 0.62 | 0.017137 | 18.36 | 79.4 | 120.6 | 9.07 | 8.99 | | High | High | High | High | High | High | High | High | High | High | High | 0 | 2.59E-05 | 4.75 |
| FALSE | High | DTECTPVG | [K].DTECTI | 1xCarbami | Q8R4Z6 1x | 3.48E-06 | 0 | 1 | 3 | 12 | Q8R4Z6 | DEAD-box | Q8R4Z6 [6 | 0 | 4598.153 | 0 | 0.51 | 0.004373 | 15.15 | 83.6 | 116.4 | 8.37 | 7.46 | | High | High | High | High | High | High | High | High | High | High | High | 0 | 4.12E-07 | 5.31 |
| FALSE | High | DTEDTSSL | [R].DTEDTS | 1xPhospho | Q9D6X5 1x | 5.38E-07 | 0 | 1 | 1 | 8 | Q9D6X5 | Solute carr | Q9D6X5 [2 | 0 | 2186.063 | 0 | 0.93 | 0.001422 | 12.42 | 69.5 | 130.5 | 2.56 | 10.57 | | High | High | High | High | High | High | High | High | High | High | High | 0 | 1.89E-07 | 3.45 |
| FALSE | High | DTESSPGS | [K].DTESSP | 1xPhospho | Q3SYK5 1x | 0.000529 | 0 | 1 | 4 | 2 | Q3SYK5 | Tyrosine-p | Q3SYK5 [8 | 0 | 2125.047 | 0 | -0.11 | 0.247888 | 2.49 | 100.8 | 99.2 | 5.81 | 6.94 | | High | High | High | High | High | High | High | High | High | High | High | 0 | 0.000117 | 2.41 |
| FALSE | High | DTESSPGS | [K].DTESSP | 2xPhospho | Q3SYK5 2x | 0.001398 | 5.66E-05 | 1 | 4 | 1 | Q3SYK5 | Tyrosine-p | Q3SYK5 [8 | 0 | 2205.013 | 0 | 0.23 | 0.139874 | 14.97 | 91.9 | 108.1 | 7.36 | 13.13 | | High | High | High | High | High | High | High | High | High | High | High | 2.58E-05 | 0.000349 | 2.36 |
| FALSE | High | DTETGTEA | [R].DTETGT | 1xPhospho | Q8R2Y8 1x | 0.000411 | 0 | 1 | 1 | 1 | Q8R2Y8 | Peptidyl-tR | Q8R2Y8 [5 | 0 | 2425.142 | 0 | 1.14 | 0.021563 | 1.29 | 64.8 | 135.2 | 11.67 | 25.05 | | High | High | High | High | High | High | High | High | High | High | High | 0 | 8.81E-05 | 3.39 |
| FALSE | High | DTEVFICTT | [K].DTEVFI | 1xCarbami | Q9JM14 1x | 0.002818 | 0.000109 | 1 | 2 | 2 | Q9JM14 | 5'(3')-deox | Q9JM14 [9 | 0 | 2075.09 | 0 | 0.82 | 0.009495 | 20.18 | 72.2 | 127.8 | 10.83 | 9.77 | | High | High | High | High | High | High | High | High | High | High | High | 0.000275 | 0.00202 | 2.76 |
| FALSE | High | DTFEHDPSI | [K].DTFEHI | 1xPhospho | Q8K019 1x | 3.39E-06 | 0 | 1 | 10 | 5 | Q8K019 | Bcl-2-asso | Q8K019 [1 | 0 | 2448.101 | 0 | -1.02 | 0.000218 | 0.6 | 133.8 | 66.2 | 3.35 | 2.06 | | High | High | High | High | High | High | High | High | High | High | High | 0 | 8.13E-07 | 3.31 |
| FALSE | High | DTFEHDPSI | [K].DTFEHI | 2xPhospho | Q8K019 2x | 0.000605 | 0 | 1 | 10 | 1 | Q8K019 | Bcl-2-asso | Q8K019 [1 | 0 | 2528.067 | 0 | -0.92 | 0.003272 | 7.61 | 134.1 | 65.9 | 10.28 | 10.74 | | High | High | High | High | High | High | High | High | High | High | High | 0 | 0.000136 | 2.49 |
| FALSE | High | DTGKTPVE | [K].DTGKTI | 1xPhospho | Q5BLK2 1x | 0.019618 | 0.001438 | 1 | 1 | 2 | Q5BLK2 | MCG3574 | Q5BLK2 [5 | 1 | 2187.158 | 0 | 0.71 | 0.006321 | 12.29 | 77.7 | 122.3 | 7.12 | 10.48 | | High | High | High | High | High | High | High | High | High | High | High | 0.001372 | 0.01431 | 2.75 |
| FALSE | High | DTPLPGPA | [R].DTPLPG | 1xPhospho | Q3UZ64 1x | 0.003195 | 0.00016 | 1 | 4 | 1 | Q3UZ64 | Nuclear fac | Q3UZ64 [1 | 0 | 2247.02 | 0 | -0.34 | 0.470764 | 15.25 | 111.8 | 88.2 | 9.15 | 29.38 | | High | High | High | High | High | High | High | High | High | High | High | 7.19E-05 | 0.000873 | 2.13 |

| | | | | | | | | | | | | | | | | | | | | | | | | | | | | | | | | | | | | | | |
|---|---|---|---|---|---|---|---|---|---|---|---|---|---|---|---|---|---|---|---|---|---|---|---|---|---|---|---|---|---|---|---|---|---|---|---|---|---|---|
| FALSE | High | DTLCSVENI | [R].DTLCSV | 1xCarbami | Q8R2M2 1 | 0.000264 | 0 | 1 | 1 | 1 | Q8R2M2 | Deoxynucl | Q8R2M2 [4 | 0 | 1902.903 | 0 | -0.39 | 0.014107 | 0.64 | 112.2 | 87.8 | 4.69 | 7.98 | | High | High | High | High | High | High | High | High | High | High | High | 0 | 5.37E-05 | 3.1 |
| FALSE | High | DTLEGTGD | [K].DTLEGT | 1xPhospho | Q91X20 1x | 1.86E-11 | 0 | 1 | 6 | 3 | Q91X20 | Set1/Ash2 | Q91X20 [7 | 0 | 3022.267 | 0 | -0.24 | 0.103236 | 4.43 | 109.9 | 90.1 | 16.13 | 12.12 | | High | High | High | High | High | High | High | High | High | High | High | 0 | 4.9E-13 | 5.98 |
| FALSE | High | DTLEGTGD | [K].DTLEGT | 1xDeamida | Q91X20 1x | 1.13E-05 | 0 | 1 | 6 | 2 | Q91X20 | Set1/Ash2 | Q91X20 [7 | 0 | 3023.251 | 0 | 0.05 | 0.472254 | 25.16 | 99.4 | 100.6 | 15.71 | 17.47 | | High | High | High | High | High | High | High | High | High | High | High | 0 | 6.72E-06 | 4.99 |
| FALSE | High | DTLEGTGD | [K].DTLEGT | 1xPhospho | Q91X20 1x | 3.65E-07 | 0 | 1 | 6 | 13 | Q91X20 | Set1/Ash2 | Q91X20 [7 | 0 | 3038.262 | 0 | -0.56 | 0.000858 | 3.81 | 119.5 | 80.5 | 3.53 | 4.14 | | High | High | High | High | High | High | High | High | High | High | High | 0 | 1.02E-05 | 5.07 |
| FALSE | High | DTLEGTGD | [K].DTLEGT | 1xDeamida | Q91X20 1x | 2.42E-08 | 0 | 1 | 6 | 3 | Q91X20 | Set1/Ash2 | Q91X20 [7 | 0 | 3039.246 | 0 | 0 | 0.758934 | 38.91 | 97.7 | 102.3 | 25.84 | 45.58 | | High | High | High | High | High | High | High | High | High | High | High | 0 | 1.56E-09 | 5.54 |
| FALSE | High | DTIFSTSPK | [R].DTIFST | 1xPhospho | B1AQJ2 1x | 0.001783 | 5.66E-05 | 1 | 2 | 3 | B1AQJ2 | Ubiquitin c | B1AQJ2 [5 | 0 | 1533.797 | 0 | -0.44 | 0.007045 | 3.9 | 115.9 | 84.1 | 5.81 | 8.92 | | High | High | High | High | High | High | High | High | High | High | High | 2.58E-05 | 0.000458 | 2.52 |
| FALSE | High | DTLLALHQ | [R].DTLLAL | 1xPhospho | E9Q1P8 1x | 0.013074 | 0.000889 | 1 | 1 | 1 | E9Q1P8 | Interferon | E9Q1P8 [2 | 0 | 2512.275 | 0 | 1.68 | 0.000808 | 2.97 | 46.8 | 153.2 | 14.57 | 12.91 | | High | High | High | High | High | High | High | High | High | High | High | 0.000467 | 0.004254 | 1.32 |
| FALSE | High | DTIIDVVGA | [K].DTIIDV | 1xPhospho | Q9JI44 1xP | 4.13E-06 | 0 | 1 | 1 | 5 | Q9JI44 | DNA methy | Q9JI44 [43 | 0 | 1977.026 | 0 | -0.39 | 0.001364 | 0.94 | 116.3 | 83.7 | 5.88 | 4.62 | | High | High | High | High | High | High | High | High | High | High | High | 0 | 5E-07 | 3.6 |
| FALSE | High | DTIIDVVGA | [K].DTIIDV | 1xDeamida | Q9JI44 1xP | 0.00311 | 0.00016 | 1 | 1 | 3 | Q9JI44 | DNA methy | Q9JI44 [43 | 0 | 1978.01 | 0 | -0.37 | 0.225793 | 11.8 | 106 | 94 | 17.92 | 14.33 | | High | High | High | High | High | High | High | High | High | High | High | 9.42E-05 | 0.000951 | 2.94 |
| FALSE | High | DTLSCGLD | [R].DTLSCG | 1xCarbami | Q7TNL7 1x | 0.000699 | 0 | 1 | 2 | 2 | Q7TNL7 | Dual specif | Q7TNL7 [2 | 0 | 4220.119 | 0 | -1.73 | 0.00082 | 1 | 153.6 | 46.4 | 11.02 | 10.37 | | High | High | High | High | High | High | High | High | High | High | High | 0 | 0.000159 | 3.23 |
| FALSE | High | DTNSTTAA | [K].DTNST | 1xPhospho | Q3UH06-3 | 1.62E-06 | 0 | 1 | 4 | 2 | Q3UH06-3 | Isoform 3 c | Q3UH06-3 | 0 | 2009.036 | 0 | -0.47 | 0.007524 | 3.14 | 113.8 | 86.2 | 3.87 | 7.42 | | High | High | High | High | High | High | High | High | High | High | High | 0 | 0.000117 | 2.91 |
| FALSE | High | DTPDSLSPE | [K].DTPDSL | 1xPhospho | Q7TMK6 1 | 3.17E-05 | 0 | 1 | 5 | 1 | Q7TMK6 | Protein Ho | Q7TMK6 [1 | 0 | 2452.05 | 0 | -0.05 | 0.201156 | 9.54 | 103.4 | 96.6 | 6 | 5.64 | | High | High | High | High | High | High | High | High | High | High | High | 0 | 4.94E-06 | 3.81 |
| FALSE | High | DTPGHGSG | [R].DTPGH | 2xPhospho | G5E866 2x | 0.000111 | 0 | 1 | 5 | 2 | G5E866 | Splicing fac | G5E866 [3 | 0 | 1856.757 | 0 | 0.13 | 0.133338 | 19.03 | 90.5 | 109.5 | 7.06 | 13.51 | | High | High | High | High | High | High | High | High | High | High | High | 0 | 2.02E-05 | 2.88 |
| FALSE | High | DTPGQATT | [K].DTPGQ | 1xPhospho | Q91Z58 1x | 0.029076 | 0.001857 | 1 | 1 | 2 | Q91Z58 | Uncharacte | Q91Z58 [6 | 0 | 1553.798 | 0 | 1.65 | 0.000515 | 21.77 | 48.4 | 151.6 | 11.16 | 5.48 | | High | High | High | High | High | High | High | High | High | High | High | 0.000997 | 0.0104 | 1.83 |
| FALSE | High | DTLCSVENI | [R].DTLCSV | 1xCarbami | Q8R2M2 1 | 0.000553 | 0 | 1 | 1 | 2 | Q8R2M2 | Deoxynucl | Q8R2M2 [4 | 0 | 1903.887 | 0 | -0.37 | 0.127572 | 5.27 | 106.3 | 93.7 | 17.52 | 10.55 | | High | High | High | High | High | High | High | High | High | High | High | 7.19E-05 | 0.000834 | 2.78 |
| FALSE | High | DVDQDVSI | [K].DVDQD | 1xPhospho | Q8BGC0 1x | 0.000223 | 0 | 1 | 1 | 3 | Q8BGC0 | HIV Tat-spe | Q8BGC0 [5 | 0 | 1558.74 | 0 | -0.35 | 0.001311 | 4.18 | 111.7 | 88.3 | 2.25 | 3.71 | | High | High | High | High | High | High | High | High | High | High | High | 0 | 4.41E-05 | 2.02 |
| FALSE | High | DVEDMELS | [R].DVEDM | 1xPhospho | Q6NXI6 1x | 1.6E-05 | 0 | 1 | 5 | 4 | Q6NXI6 | Regulation | Q6NXI6 [3 | 0 | 2321.014 | 0 | -0.35 | 0.018628 | 6.84 | 113.5 | 86.5 | 2.43 | 15.98 | | High | High | High | High | High | High | High | High | High | High | High | 0 | 6.48E-06 | 4.01 |
| FALSE | High | DVEDMELS | [R].DVEDM | 1xPhospho | Q6NXI6 1x | 1.91E-07 | 0 | 1 | 5 | 14 | Q6NXI6 | Regulation | Q6NXI6 [3 | 0 | 2337.009 | 0 | -0.23 | 0.032602 | 2.32 | 108.2 | 91.8 | 2.5 | 12.18 | | High | High | High | High | High | High | High | High | High | High | High | 0 | 1.87E-08 | 4.7 |
| FALSE | High | DVSSPDESI | [R].DVSSPD | 1xPhospho | Q544H8 1x | 0.049971 | 0.003641 | 1 | 1 | 2 | Q544H8 | Putative ur | Q544H8 [1 | 0 | 1887.874 | 0 | -0.06 | 0.236212 | 5.26 | 106.6 | 93.4 | 8.65 | 12.12 | | High | High | High | High | High | High | High | High | High | High | High | 0.001902 | 0.01934 | 2.28 |
| FALSE | High | DVTNLTVG | [R].DVTNL | 1xPhospho | O55201 1x | 1.17E-06 | 0 | 1 | 2 | 1 | O55201 | Transcripti | O55201 [6 | 0 | 2000.971 | 0 | -0.44 | 0.326319 | 14.41 | 113.7 | 86.3 | 8.1 | 29.82 | | High | High | High | High | High | High | High | High | High | High | High | 0 | 1.22E-07 | 4.07 |
| FALSE | High | DVTNLTVG | [R].DVTNL | 1xPhospho | O55201 1x | 2.16E-06 | 0 | 1 | 2 | 6 | O55201 | Transcripti | O55201 [6 | 0 | 2016.966 | 0 | -0.04 | 0.968584 | 5.89 | 102.8 | 97.2 | 4.85 | 11.1 | | High | High | High | High | High | High | High | High | High | High | High | 0 | 2.41E-07 | 3.51 |
| FALSE | High | DVVVEFPSI | [K].DVVVEI | 1xDeamida | F2Z3U3 1x | 1.78E-05 | 0 | 1 | 1 | 2 | F2Z3U3 | Ras associa | F2Z3U3 [1 | 0 | 4990.535 | 0 | 1.16 | 0.009223 | 24.06 | 64.1 | 135.9 | 17.9 | 12.64 | | High | High | High | High | High | High | High | High | High | High | High | 0 | 2.6E-06 | 4.51 |
| FALSE | High | DVYLSPR | [R].DVYLSP | 1xPhospho | Q91VM5 1 | 0.008463 | 0.000736 | 1 | 5 | 4 | Q91VM5 | RNA bindir | Q91VM5 [2 | 0 | 1158.576 | 0 | 0.13 | 0.098134 | 13.17 | 96.7 | 103.3 | 4.73 | 6.49 | | High | High | High | High | High | High | High | High | High | High | High | 0.000327 | 0.002595 | 1.78 |
| FALSE | High | DVYPSCSSS | [K].DVYPS | 1xCarbami | Q8BMI0 1x | 2.37E-07 | 0 | 1 | 2 | 8 | Q8BMI0 | F-box only | Q8BMI0 [7 | 0 | 3734.596 | 0 | 0.88 | 0.001166 | 4.08 | 71.6 | 128.4 | 7.13 | 10.73 | | High | High | High | High | High | High | High | High | High | High | High | 0 | 2.02E-08 | 5.45 |
| FALSE | High | DVYTIFDAE | [K].DVYTIF | 1xPhospho | A6H5Z3 1x | 5.07E-05 | 0 | 1 | 3 | 5 | A6H5Z3 | Exocyst co | A6H5Z3 [2 | 0 | 2339.146 | 0 | -0.01 | 0.597529 | 7.27 | 99.6 | 100.4 | 8.23 | 7.06 | | High | High | High | High | High | High | High | High | High | High | High | 0 | 8.34E-06 | 2.98 |
| FALSE | High | DVYVTTHT | [K].DVYVT | 1xPhospho | Q3ULB8 1x | 0.078467 | 0.006104 | 1 | 3 | 1 | Q3ULB8 | Putative ur | Q3ULB8 [1 | 0 | 1497.73 | 0 | 0.42 | 0.043199 | 24.6 | 85.7 | 114.3 | 17.56 | 5.33 | | High | High | High | High | High | High | High | High | High | High | High | 0.003148 | 0.03258 | 2.54 |
| FALSE | High | DWDKESE( | [K].DWDKE | 1xPhospho | Q9CYX7 1x | 0.000365 | 0 | 1 | 1 | 2 | Q9CYX7 | RRP15-like | Q9CYX7 [2 | 1 | 2199.996 | 0 | -0.61 | 0.004673 | 6.86 | 117.7 | 82.3 | 7.51 | 5.3 | | High | High | High | High | High | High | High | High | High | High | High | 0 | 7.67E-05 | 2.11 |
| FALSE | High | DWEDDSD | [K].DWEDD | 1xPhospho | Q9R0Q7 1x | 5.58E-07 | 0 | 1 | 2 | 24 | Q9R0Q7 | Prostaglan | Q9R0Q7 [1 | 0 | 2184.79 | 0 | -0.3 | 0.016752 | 6.43 | 112.5 | 87.5 | 2.75 | 9.98 | | High | High | High | High | High | High | High | High | High | High | High | 0 | 5.26E-08 | 4.71 |
| FALSE | High | DWEDDSD | [K].DWEDD | 1xPhospho | Q9R0Q7 1x | 2.41E-07 | 0 | 1 | 2 | 76 | Q9R0Q7 | Prostaglan | Q9R0Q7 [1 | 0 | 2200.785 | 0 | -0.32 | 0.016128 | 5.75 | 111.2 | 88.8 | 1.36 | 9.55 | | High | High | High | High | High | High | High | High | High | High | High | 0 | 8.92E-08 | 4.6 |
| FALSE | High | DWEDDSD | [R].DWEDD | 1xDeamida | Q9CRH1 1x | 0.006594 | 0.00062 | 1 | 1 | 1 | Q9CRH1 | Putative ur | Q9CRH1 [1 | 0 | 2215.785 | 0 | -0.33 | 0.043486 | 9.02 | 115.6 | 84.4 | 15.25 | 9.94 | | High | High | High | High | High | High | High | High | High | High | High | 0.000275 | 0.001971 | 2.07 |
| FALSE | High | DYDDMSPI | [R].DYDDM | 1xPhospho | Q5FWJ5 1x | 0.006594 | 0.00062 | 2 | 13 | 13 | Q5FWJ5; C | Hnrpk prot | Q5FWJ5 [2 | 0 | 1323.513 | 0 | -0.09 | 0.031262 | 1.64 | 102.7 | 97.3 | 2.36 | 6.83 | Shared | High | High | High | High | High | High | High | High | High | High | High | 0.000327 | 0.002465 | 1.85 |
| FALSE | High | DVSQGTLP | [K].DVSQG | 1xPhospho | Q8CJF7 1x | 1.5E-08 | 0 | 1 | 1 | 2 | Q8CJF7 | Protein ELY | Q8CJF7 [1 | 0 | 2801.319 | 0 | -0.58 | 0.001977 | 6.01 | 120.1 | 79.9 | 2.15 | 12.3 | | High | High | High | High | High | High | High | High | High | High | High | 0 | 9.1E-10 | 5.42 |
| FALSE | High | DYDEEEQG | [R].DYDEE | 1xPhospho | F6RDI8 1xP | 1.23E-06 | 0 | 1 | 7 | 5 | F6RDI8 | Serine/argi | F6RDI8 [44 | 0 | 2144.895 | 0 | -0.42 | 0.009388 | 7.55 | 114.5 | 85.5 | 4.79 | 6.61 | | High | High | High | High | High | High | High | High | High | High | High | 0 | 7.8E-06 | 4.09 |
| FALSE | High | DYDPYSDG | [K].DYDPY | 1xCarbami | Q9JJ48 2xP | 0.014158 | 0.001011 | 1 | 3 | 3 | Q9JJ48 | Zinc finger | Q9JJ48 [80 | 0 | 3047.158 | 0 | -0.23 | 0.282765 | 20.7 | 101.9 | 98.1 | 9.85 | 14.02 | | High | High | High | High | High | High | High | High | High | High | High | 0.000538 | 0.004647 | 3.46 |
| FALSE | High | DYEGPAISI | [R].DYEGP | 1xPhospho | Q80U27 1x | 0.01186 | 0.000889 | 1 | 4 | 1 | Q80U27 | Kinesin-like | Q80U27 [6 | 0 | 1614.818 | 0 | 0.02 | 0.459917 | 2.81 | 98.4 | 101.6 | 8.58 | 10.07 | | High | High | High | High | High | High | High | High | High | High | High | 0.000449 | 0.003795 | 2.95 |
| FALSE | High | DYEIENQN | [K].DYEIEN | 1xPhospho | Q5F2E7-2 | 2.82E-08 | 0 | 1 | 2 | 3 | Q5F2E7-2 | Isoform 2 c | Q5F2E7-2 | 0 | 2913.465 | 0 | 0.63 | 0.002408 | 8.32 | 78.6 | 121.4 | 6.65 | 5.44 | | High | High | High | High | High | High | High | High | High | High | High | 0 | 1.85E-09 | 3.99 |
| FALSE | High | DYEIENQN | [K].DYEIEN | 1xDeamida | Q5F2E7-2 | 0.002058 | 5.66E-05 | 1 | 2 | 4 | Q5F2E7-2 | Isoform 2 c | Q5F2E7-2 | 0 | 2914.449 | 0 | 0.27 | 0.080492 | 11.1 | 88.8 | 111.2 | 18.41 | 11.29 | | High | High | High | High | High | High | High | High | High | High | High | 2.58E-05 | 0.000536 | 3.5 |
| FALSE | High | DYEIENQN | [K].DYEIEN | 2xPhospho | Q5F2E7-2 | 0.000475 | 0 | 1 | 2 | 2 | Q5F2E7-2 | Isoform 2 c | Q5F2E7-2 | 0 | 2993.431 | 0 | 0.22 | 0.317066 | 19.4 | 92.3 | 107.7 | 13.74 | 13.83 | | High | High | High | High | High | High | High | High | High | High | High | 0 | 0.000103 | 3.24 |
| FALSE | High | DYETTAPTY | [K].DYETTA | 1xPhospho | Q9D8L0-2 | 0.100129 | 0.008067 | 1 | 2 | 1 | Q9D8L0-2 | Isoform 2 c | Q9D8L0-2 | 0 | 2839.384 | 0 | 2.19 | 0.00252 | 9.68 | 37.9 | 162.1 | 22.74 | 22.82 | | High | High | High | High | High | High | High | High | High | High | High | 0.004046 | 0.04319 | 2.13 |
| FALSE | High | DYGEPGPS | [R].DYGEP | 1xPhospho | Q3UMA0 1 | 1.34E-06 | 0 | 1 | 3 | 2 | Q3UMA0 | Putative ur | Q3UMA0 [ | 0 | 3892.866 | 0 | 2.14 | 0.001911 | 30.92 | 36.3 | 163.7 | 41.69 | 11.16 | | High | High | High | High | High | High | High | High | High | High | High | 0 | 1.41E-07 | 4.79 |
| FALSE | High | DYGSSPLHI | [K].DYGSS | 1xPhospho | Q5RJV5 1x | 0.002768 | 0.000109 | 1 | 13 | 1 | Q5RJV5 | Polypyrimi | Q5RJV5 [4 | 0 | 1340.62 | 0 | 0.17 | 0.255507 | 9.28 | 93.4 | 106.6 | 8.46 | 8.16 | | High | High | High | High | High | High | High | High | High | High | High | 4.94E-05 | 0.000745 | 1.54 |
| FALSE | High | DYHSLFSPF | [K].DYHSLF | 1xPhospho | F8VQ28 1x | 0.096959 | 0.007777 | 1 | 6 | 1 | F8VQ28 | Paxillin OS | F8VQ28 [4 | 0 | 1430.667 | 0 | 0.19 | 0.064476 | 3.31 | 91.5 | 108.5 | 15.95 | 18.02 | | High | High | High | High | High | High | High | High | High | High | High | 0.003922 | 0.04164 | 2.12 |
| FALSE | High | DYLQTQQF | [R].DYLQT | 1xPhospho | Q6VGS5-2 | 6.59E-05 | 0 | 1 | 3 | 2 | Q6VGS5-2 | Isoform 2 c | Q6VGS5-2 | 0 | 1997.015 | 0 | -0.88 | 0.000188 | 2.41 | 130.4 | 69.6 | 1.84 | 6.82 | | High | High | High | High | High | High | High | High | High | High | High | 0 | 1.13E-05 | 2.58 |
| FALSE | High | DYNASAST | [K].DYNAS | 1xPhospho | P14873 1x | 0.000534 | 0 | 2 | 2 | 2 | P14873; B2 | Microtubu | P14873 [11 | 0 | 2567.162 | 0 | -0.68 | 0.019505 | 2.46 | 125 | 75 | 15.33 | 17.05 | Shared | High | High | High | High | High | High | High | High | High | High | High | 2.58E-05 | 0.000527 | 3.28 |
| FALSE | High | DYNPYNYS | [R].DYNPY | 1xPhospho | B9EJA4 1x | 1.72E-08 | 0 | 1 | 7 | 1 | B9EJA4 | Clasp2 pro | B9EJA4 [10 | 0 | 2462.132 | 0 | -0.14 | 0.549056 | 11.68 | 102.1 | 97.9 | 11.41 | 6.76 | | High | High | High | High | High | High | High | High | High | High | High | 0 | 1.05E-09 | 4.13 |
| FALSE | High | DYQSEADL | [K].DYQSE | 1xPhospho | E9QML5 1x | 0.035486 | 0.002304 | 1 | 16 | 2 | E9QML5 | Zinc finger | E9QML5 [3 | 0 | 1615.757 | 0 | -0.34 | 0.187366 | 3.87 | 107.1 | 92.9 | 10.21 | 10.24 | | High | High | High | High | High | High | High | High | High | High | High | 0.001214 | 0.01304 | 1.81 |
| FALSE | High | DYDPYSDG | [K].DYDPY | 1xCarbami | Q9JJ48 1xP | 1.02E-07 | 0 | 1 | 3 | 7 | Q9JJ48 | Zinc finger | Q9JJ48 [80 | 0 | 2967.192 | 0 | -0.86 | 0.000642 | 2.21 | 130.4 | 69.6 | 4.49 | 4.35 | | High | High | High | High | High | High | High | High | High | High | High | 0 | 7.85E-09 | 5.31 |
| FALSE | High | DVSPIVEIPI | [R].DVSPIV | 1xPhospho | Q8BKT3 1x | 9.08E-07 | 0 | 1 | 3 | 3 | Q8BKT3 | GC-rich sec | Q8BKT3 [1 | 0 | 2221.147 | 0 | 0.22 | 0.041421 | 12.04 | 92.8 | 107.2 | 1.32 | 8.75 | | High | High | High | High | High | High | High | High | High | High | High | 0 | 3.52E-07 | 3.88 |
| FALSE | High | DVSDERLSI | [K].DVSDE | 1xPhospho | P14873 1x | 0.089425 | 0.006807 | 2 | 2 | 1 | P14873; B2 | Microtubu | P14873 [12 | 1 | 1754.909 | 0 | -0.95 | 0.000786 | 5.23 | 133.4 | 66.6 | 4.03 | 16.63 | Shared | High | High | High | High | High | High | High | High | High | High | High | 0.003564 | 0.03791 | 1.32 |
| FALSE | High | DVQDAIAC | [K].DVQDA | 1xPhospho | Q8CH25-2 | 2.71E-07 | 0 | 1 | 4 | 4 | Q8CH25-2 | Isoform 2 c | Q8CH25-2 | 0 | 1838.93 | 0 | -0.05 | 0.495081 | 11.25 | 102.2 | 97.8 | 4.13 | 6.16 | | High | High | High | High | High | High | High | High | High | High | High | 0 | 2.35E-08 | 3.45 |
| FALSE | High | DVEETGIT\ | [R].DVEET | 1xPhospho | Q3TIV5 1x | 0.029586 | 0.001857 | 1 | 2 | 1 | Q3TIV5 | Zinc finger | Q3TIV5 [34 | 0 | 1813.878 | 0 | -0.33 | 0.118143 | 11.13 | 106.1 | 93.9 | 13.39 | 14.43 | | High | High | High | High | High | High | High | High | High | High | High | 0.000997 | 0.01061 | 2.48 |
| FALSE | High | DVEFEVVG | [K].DVEFE | 1xPhospho | Q9Z1F9 1x | 0.000426 | 0 | 1 | 2 | 1 | Q9Z1F9 | SUMO-acti | Q9Z1F9 [5 | 0 | 1758.804 | 0 | -0.41 | 0.620081 | 6.89 | 114.3 | 85.7 | 73.43 | 72.82 | | High | High | High | High | High | High | High | High | High | High | High | 0 | 9.19E-05 | 2.34 |
| FALSE | High | DVEFEVVG | [K].DVEFE | 1xPhospho | Q9Z1F9 1x | 4.32E-06 | 0 | 1 | 2 | 5 | Q9Z1F9 | SUMO-acti | Q9Z1F9 [5 | 0 | 1987.967 | 0 | -0.56 | 0.002101 | 6.66 | 117.2 | 82.8 | 4.22 | 7.15 | | High | High | High | High | High | High | High | High | High | High | High | 0 | 4.72E-06 | 4.18 |
| FALSE | High | DVEGSTSP | [K].DVEGS | 1xPhospho | Q3U1S6 1x | 0.101749 | 0.008194 | 1 | 10 | 1 | Q3U1S6 | Putative ur | Q3U1S6 [4 | 0 | 1870.92 | 0 | -0.35 | 0.008566 | 8.34 | 113.3 | 86.7 | 5.21 | 9.28 | | High | High | High | High | High | High | High | High | High | High | High | 0.004115 | 0.04389 | 1.55 |
| FALSE | High | DVETEHAA | [K].DVETEH | 1xCarbami | Q6PR54-3 | 0.000125 | 0 | 2 | 4 | 1 | Q6PR54-3; | Isoform 3 c | Q6PR54-3 | 0 | 3665.514 | 0 | -0.94 | 0.003903 | 3.44 | 135.3 | 64.7 | 8.21 | 24.54 | Shared | High | High | High | High | High | High | High | High | High | High | High | 0 | 2.31E-05 | 3.24 |
| FALSE | High | DVETEHAA | [K].DVETEH | 1xCarbami | Q6PR54-3 | 1.05E-05 | 0 | 2 | 4 | 3 | Q6PR54-3; | Isoform 3 c | Q6PR54-3 | 0 | 3681.509 | 0 | -1.33 | 0.00207 | 4.06 | 142.6 | 57.4 | 7.72 | 16.83 | Shared | High | High | High | High | High | High | High | High | High | High | High | 0 | 1.43E-06 | 5.1 |
| FALSE | High | DVFASYLN | [K].DVFAS | 1xPhospho | Q9CYA6 1x | 4.65E-06 | 0 | 1 | 2 | 1 | Q9CYA6 | Zinc finger | Q9CYA6 [3 | 0 | 2238.046 | 0 | 0.2 | 0.061494 | 4.57 | 92.9 | 107.1 | 8.2 | 15.71 | | High | High | High | High | High | High | High | High | High | High | High | 0 | 5.74E-07 | 4.6 |
| FALSE | High | DVFASYLN | [K].DVFAS | 1xPhospho | Q9CYA6 1x | 4.65E-06 | 0 | 1 | 2 | 1 | Q9CYA6 | Zinc finger | Q9CYA6 [3 | 0 | 2254.041 | 0 | -0.34 | 0.024109 | 5.39 | 109.9 | 90.1 | 6.52 | 8.42 | | High | High | High | High | High | High | High | High | High | High | High | 0 | 5.71E-07 | 4.13 |
| FALSE | High | DVFYYSPS( | [R].DVFYY | 1xPhospho | Q9Z2D8 1x | 0.026191 | 0.001676 | 1 | 1 | 1 | Q9Z2D8 | Methyl-Cp | Q9Z2D8 [3 | 0 | 1700.834 | 0 | -0.22 | 0.084959 | 17.85 | 108.1 | 91.9 | 12.37 | 8.93 | | High | High | High | High | High | High | High | High | High | High | High | 0.000903 | 0.009275 | 1.55 |
| FALSE | High | DVGHLEEG | [R].DVGHL | 1xPhospho | B9EKJ7 1x | 8.91E-10 | 0 | 1 | 3 | 4 | B9EKJ7 | Tankyrase | B9EKJ7 [10 | 0 | 2512.203 | 0 | -0.04 | 0.349971 | 4.11 | 102.1 | 97.9 | 8.04 | 13.05 | | High | High | High | High | High | High | High | High | High | High | High | 0 | 3.78E-11 | 4.47 |
| FALSE | High | DVGHLEEG | [R].DVGHL | 2xPhospho | B9EKJ7 2x | 5.48E-07 | 0 | 1 | 3 | 1 | B9EKJ7 | Tankyrase | B9EKJ7 [10 | 0 | 2592.169 | 0 | 0 | 0.938397 | 16.02 | 99.9 | 100.1 | 13.3 | 8.34 | | High | High | High | High | High | High | High | High | High | High | High | 0 | 5.17E-08 | 4.48 |
| FALSE | High | DVGTEPRS | [K].DVGTE | 1xPhospho | P97412 1x | 0.050397 | 0.003641 | 1 | 2 | 1 | P97412 | Lysosomal- | P97412 [21 | 1 | 2503.046 | 0 | 0.68 | 0.008431 | 7.6 | 72.8 | 127.2 | 13.62 | 14.36 | | High | High | High | High | High | High | High | High | High | High | High | 0.001916 | 0.0194 | 2.14 |
| FALSE | High | DVHSPNAV | [R].DVHSP | 1xPhospho | Q8R0T2 1x | 0.000305 | 0 | 1 | 3 | 2 | Q8R0T2 | Zinc finger | Q8R0T2 [1 | 0 | 2132.079 | 0 | 1.01 | 0.000532 | 6.65 | 64.6 | 135.4 | 5.56 | 7.35 | | High | High | High | High | High | High | High | High | High | High | High | 0 | 6.3E-05 | 3.87 |
| FALSE | High | DVICIDNDS | [R].DVICID | 1xCarbami | E9PZ97 1x | 2.84E-08 | 0 | 1 | 2 | 3 | E9PZ97 | Bloom sync | E9PZ97 [24 | 0 | 2476.157 | 0 | -1.98 | 0.000223 | 0.91 | 159.4 | 40.6 | 3.62 | 17.86 | | High | High | High | High | High | High | High | High | High | High | High | 0 | 1.86E-09 | 3.72 |
| FALSE | High | DVILPEQSF | [R].DVILPE | 1xPhospho | Q9EST3 1x | 6.04E-06 | 0 | 1 | 5 | 1 | Q9EST3 | Eukaryotic | Q9EST3 [2 | 0 | 3138.511 | 0 | 0.43 | 0.002082 | 1.41 | 85.4 | 114.6 | 12.51 | 6.63 | | High | High | High | High | High | High | High | High | High | High | High | 0 | 7.65E-07 | 3.67 |
| FALSE | High | DVINRSPT( | [R].DVINRS | 1xPhospho | Q811P8 1x | 0.058664 | 0.004294 | 1 | 3 | 1 | Q811P8 | Rho GTPas | Q811P8 [9 | 1 | 2107.168 | 0 | -1.22 | 0.000443 | 0.84 | 143.6 | 56.4 | 6.39 | 8.04 | | High | High | High | High | High | High | High | High | High | High | High | 0.002187 | 0.02307 | 1.89 |
| FALSE | High | DVLQAAAA | [K].DVLQA | 1xDeamida | E9QP99 1x | 0.026885 | 0.001676 | 1 | 5 | 2 | E9QP99 | Golgin sub | E9QP99 [3 | 1 | 2945.37 | 0 | 0.83 | 0.012003 | 11.8 | 78.7 | 121.3 | 11.47 | 14.01 | | High | High | High | High | High | High | High | High | High | High | High | 0.000903 | 0.009497 | 2.28 |
| FALSE | High | DVMLQNG | [K].DVMLC | 1xPhospho | Q6PDQ2 1x | 0.005516 | 0.00062 | 1 | 5 | 1 | Q6PDQ2 | Chromodo | Q6PDQ2 [1 | 0 | 1769.891 | 0 | -0.72 | 0.00126 | 0.64 | 124.8 | 75.2 | 5.01 | 9.29 | | High | High | High | High | High | High | High | High | High | High | High | 0.000275 | 0.001613 | 3.48 |
| FALSE | High | DVMLQNG | [K].DVMLC | 1xPhospho | Q6PDQ2 1x | 1.33E-05 | 0 | 1 | 5 | 1 | Q6PDQ2 | Chromodo | Q6PDQ2 [1 | 0 | 1785.886 | 0 | -0.35 | 0.122336 | 7.81 | 110.2 | 89.8 | 5.11 | 16.42 | | High | High | High | High | High | High | High | High | High | High | High | 0 | 1.88E-06 | 3.4 |
| FALSE | High | DVMSDETI | [R].DVMSD | 1xPhospho | P14873 1x | 0.000693 | 0 | 2 | 2 | 1 | P14873; B2 | Microtubu | P14873 [11 | 0 | 3281.481 | 0 | -0.18 | 0.146804 | 13.77 | 106.2 | 93.8 | 8.89 | 23.94 | Shared | High | High | High | High | High | High | High | High | High | High | High | 0 | 0.000158 | 3.12 |
| FALSE | High | DVMSDETI | [R].DVMSD | 1xPhospho | P14873 1x | 0.006085 | 0.00062 | 2 | 2 | 2 | P14873; B2 | Microtubu | P14873 [11 | 0 | 3297.476 | 0 | -0.01 | 0.930474 | 29.78 | 99.9 | 100.1 | 14.82 | 14.57 | Shared | High | High | High | High | High | High | High | High | High | High | High | 0.000275 | 0.001798 | 3.29 |
| FALSE | High | DVNDTGFL | [K].DVNDT | 2xPhospho | Q6NS45 2x | 0.000297 | 0 | 1 | 3 | 1 | Q6NS45 | Coiled-coil | Q6NS45 [1 | 0 | 3279.431 | 0 | -0.18 | 0.340349 | 6.9 | 105.9 | 94.1 | 10.73 | 6.74 | | High | High | High | High | High | High | High | High | High | High | High | 0 | 6.12E-05 | 3.49 |
| FALSE | High | DVNGSPFS | [K].DVNGS | 1xPhospho | Q8BJL1 1x | 0.079766 | 0.00613 | 1 | 1 | 1 | Q8BJL1 | F-box only | Q8BJL1 [37 | 0 | 2027.999 | 0 | 0.36 | 0.009109 | 7.14 | 85.2 | 114.8 | 6.5 | 8.99 | | High | High | High | High | High | High | High | High | High | High | High | 0.003163 | 0.03311 | 2.2 |
| FALSE | High | DVPCNNHS | [R].DVPCN | 1xCarbami | A0A0A0M( | 0.000286 | 0 | 1 | 4 | 6 | A0A0A0M( | Anoctamin | A0A0A0M( | 0 | 3629.548 | 0 | 6.04 | 0.004886 | 13.79 | 2.4 | 197.6 | 75.32 | 30.9 | | High | High | High | High | High | High | High | High | High | High | High | 7.19E-05 | 0.000766 | 3.66 |
| FALSE | High | DVPCNNHS | [R].DVPCN | 1xCarbami | A0A0A0M( | 0.007742 | 0.000664 | 1 | 4 | 2 | A0A0A0M( | Anoctamin | A0A0A0M( | 0 | 3709.514 | 0 | 2.97 | 0.00502 | 20.67 | 20.2 | 179.8 | 11.67 | 27.4 | | High | High | High | High | High | High | High | High | High | High | High | 0.002896 | 0.03013 | 3.17 |
| FALSE | High | DVPCNNHS | [R].DVPCN | 1xCarbami | A0A0A0M( | 0.000793 | 0 | 1 | 4 | 1 | A0A0A0M( | Anoctamin | A0A0A0M( | 0 | 3789.48 | 0 | 3.08 | 0.005538 | 34.69 | 20.6 | 179.4 | 10.23 | 29 | | High | High | High | High | High | High | High | High | High | High | High | 0 | 0.000183 | 3.88 |
| FALSE | High | DVPPDILLD | [R].DVPPD | 1xPhospho | Q6KCD5 1x | 3.17E-05 | 0 | 1 | 5 | 3 | Q6KCD5 | Nipped-B-l | Q6KCD5 [3 | 0 | 1774.883 | 0 | 0.02 | 0.838781 | 7.5 | 100.1 | 99.9 | 9.05 | 8.93 | | High | High | High | High | High | High | High | High | High | High | High | 0 | 4.94E-06 | 3.62 |
| FALSE | High | DVPPDILLD | [R].DVPPD | 1xPhospho | Q6KCD5 1x | 0.000359 | 0 | 1 | 5 | 6 | Q6KCD5 | Nipped-B-l | Q6KCD5 [3 | 1 | 2132.14 | 0 | 0.16 | 0.179968 | 12.16 | 94.5 | 105.5 | 3.06 | 10.6 | | High | High | High | High | High | High | High | High | High | High | High | 0 | 7.53E-05 | 3.38 |
| FALSE | High | DVQDAIAC | [K].DVQDA | 1xPhospho | Q8CH25-2 | 0.002623 | 0.000109 | 1 | 4 | 1 | Q8CH25-2 | Isoform 2 c | Q8CH25-2 | 0 | 1609.767 | 0 | -0.07 | 0.770294 | 21.25 | 106 | 94 | 72.21 | 68.08 | | High | High | High | High | High | High | High | High | High | High | High | 4.94E-05 | 0.000705 | 2.25 |
| FALSE | High | DYSDHPSG | [R].DYSDH | 1xPhospho | Q91VM5 1 | 0.120211 | 0.009851 | 1 | 5 | 1 | Q91VM5 | RNA bindir | Q91VM5 [2 | 0 | 1649.679 | 0 | -0.03 | 0.418416 | 11.95 | 94.8 | 105.2 | 15.81 | 9.02 | | High | High | High | High | High | High | High | High | High | High | High | 0.004907 | 0.05357 | 1.68 |
| FALSE | High | DTDSDREA | [K].DTDSD | 1xPhospho | Q571L5 1x | 0.018285 | 0.001282 | 1 | 7 | 1 | Q571L5 | MKIAA406 | Q571L5 [4 | 2 | 3422.532 | 0 | -0.94 | 0.000988 | 2.26 | 131 | 69 | 1.52 | 12.78 | | High | High | High | High | High | High | High | High | High | High | High | 0.000689 | 0.006181 | 2.23 |
| FALSE | High | EAWTDGR( | [R].EAWTD | 1xPhospho | A0A1B0GS | 0.001674 | 5.66E-05 | 1 | 2 | 1 | A0A1B0GS | GRB2-asso | A0A1B0GS | 1 | 2461.165 | 0 | -1.56 | 0.000396 | 2.18 | 149.5 | 50.5 | 1.68 | 20.51 | | High | High | High | High | High | High | High | High | High | High | High | 2.58E-05 | 0.000424 | 2.75 |
| FALSE | High | EAYSGCSG | [K].EAYSG | 2xCarbami | Q80YS6 1x | 3.82E-07 | 0 | 1 | 4 | 1 | Q80YS6 | Actin filam | Q80YS6 [2 | 0 | 3228.463 | 0 | 1.03 | 0.035701 | 16.16 | 65.7 | 134.3 | 29.14 | 14.84 | | High | High | High | High | High | High | High | High | High | High | High | 0 | 3.44E-08 | 3.27 |
| FALSE | High | EEGTVGSG | [K].EEGTV | 2xPhospho | Q8BLH7 2x | 0.001386 | 5.66E-05 | 1 | 1 | 1 | Q8BLH7 | HIRA-intera | Q8BLH7 [3 | 0 | 2667.111 | 0 | -1.3 | 0.006362 | 4.39 | 140.5 | 59.5 | 15.25 | 9.76 | | High | High | High | High | High | High | High | High | High | High | High | 2.58E-05 | 0.000345 | 3.41 |
| FALSE | High | EEHGGLIRS | [K].EEHGG | 1xPhospho | P26369 1x | 0.000426 | 0 | 1 | 4 | 6 | P26369 | Splicing fac | P26369 [71 | 1 | 1559.789 | 0 | -0.21 | 0.004527 | 1.9 | 108.1 | 91.9 | 2.18 | 4.3 | | High | High | High | High | High | High | High | High | High | High | High | 0 | 9.11E-05 | 2.42 |
| FALSE | High | EEHQDISSL | [K].EEHQD | 2xPhospho | P59178 2x | 0.00204 | 5.66E-05 | 1 | 1 | 1 | P59178 | Lethal(3)m | P59178 [67 | 1 | 3433.706 | 0 | -0.03 | 0.632875 | 15.08 | 104.6 | 95.4 | 8.84 | 13.03 | | High | High | High | High | High | High | High | High | High | High | High | 2.58E-05 | 0.000529 | 3.35 |
| FALSE | High | EEKPEVQSI | [K].EEKPEV | 1xPhospho | Q6ZPJ3 1x | 0.000699 | 0 | 1 | 1 | 3 | Q6ZPJ3 | (E3-indepe | Q6ZPJ3 [88 | 0 | 2037.124 | 0 | -0.15 | 0.090026 | 6.1 | 104.4 | 95.6 | 4.96 | 7.64 | | High | High | High | High | High | High | High | High | High | High | High | 0 | 0.00016 | 2.32 |
| FALSE | High | EELADISTD | [K].EELADI | 1xPhospho | E9Q784 1x | 0.000219 | 0 | 1 | 1 | 2 | E9Q784 | Zinc finger | E9Q784 [1 | 0 | 1658.829 | 0 | -1.21 | 0.000444 | 1.94 | 141.8 | 58.2 | 5.08 | 6.3 | | High | High | High | High | High | High | High | High | High | High | High | 0 | 4.33E-05 | 2.82 |
| FALSE | High | EELASDLEE | [K].EELASD | 1xPhospho | Q9D6Z1 1x | 1.11E-05 | 0 | 1 | 2 | 14 | Q9D6Z1 | Nucleolar p | Q9D6Z1 [5 | 0 | 2249.066 | 0 | -0.23 | 0.056703 | 2.57 | 107.3 | 92.7 | 0.97 | 16.8 | | High | High | High | High | High | High | High | High | High | High | High | 0 | 1.3E-05 | 3.23 |
| FALSE | High | EELASDLEE | [K].EELASD | 1xPhospho | Q9D6Z1 1x | 2.35E-07 | 0 | 1 | 2 | 76 | Q9D6Z1 | Nucleolar p | Q9D6Z1 [5 | 0 | 2265.061 | 0 | -0.14 | 0.008972 | 3.47 | 105.2 | 94.8 | 3.36 | 5.54 | | High | High | High | High | High | High | High | High | High | High | High | 0 | 6.31E-07 | 3.93 |
| FALSE | High | EELASDLEE | [K].EELASD | 2xPhospho | Q9D6Z1 2x | 0.00614 | 0.00062 | 1 | 2 | 3 | Q9D6Z1 | Nucleolar p | Q9D6Z1 [5 | 0 | 2329.032 | 0 | -0.3 | 0.08644 | 11.4 | 109.6 | 90.4 | 4.64 | 19.61 | | High | High | High | High | High | High | High | High | High | High | High | 0.004317 | 0.0462 | 2.55 |
| FALSE | High | EELDVSVRI | [K].EELDVS | 1xCarbami | Q3THU3 1x | 0.001011 | 0 | 1 | 12 | 1 | Q3THU3 | Putative ur | Q3THU3 [1 | 1 | 2915.411 | 0 | -0.44 | 0.002737 | 2.63 | 115.8 | 84.2 | 7.19 | 8.03 | | High | High | High | High | High | High | High | High | High | High | High | 0 | 0.000241 | 2.67 |
| FALSE | High | EELEQQTD | [K].EELEQ | 1xCarbami | Q8BU14 1x | 2.49E-05 | 0 | 1 | 3 | 1 | Q8BU14 | Translocati | Q8BU14 [3 | 1 | 3562.551 | 0 | -0.35 | 0.237131 | 14.38 | 112.5 | 87.5 | 16.03 | 16.52 | | High | High | High | High | High | High | High | High | High | High | High | 0 | 3.78E-06 | 4.44 |
| FALSE | High | EELIAQGLL | [K].EELIAQ | 1xPhospho | Q8R0L9 1x | 0.000279 | 0 | 1 | 3 | 1 | Q8R0L9 | Transcripti | Q8R0L9 [3 | 0 | 3464.652 | 0 | 0.19 | 0.027388 | 8.39 | 92.9 | 107.1 | 5.81 | 6.04 | | High | High | High | High | High | High | High | High | High | High | High | 0 | 5.69E-05 | 3.82 |
| FALSE | High | EELLDGED' | [K].EELLDG | 1xPhospho | Q6P5G3 1x | 0.028824 | 0.001829 | 1 | 3 | 3 | Q6P5G3 | MBT doma | Q6P5G3 [6 | 0 | 3712.731 | 0 | -0.88 | 0.046634 | 10.03 | 129.7 | 70.3 | 19.11 | 13.05 | | High | High | High | High | High | High | High | High | High | High | High | 0.003346 | 0.03654 | 2.69 |
| FALSE | High | EELLDGED' | [K].EELLDG | 1xDeamida | Q6P5G3 1x | 1.73E-07 | 0 | 1 | 3 | 3 | Q6P5G3 | MBT doma | Q6P5G3 [6 | 0 | 3713.715 | 0 | -0.71 | 0.009731 | 8.19 | 124.4 | 75.6 | 7.09 | 12.89 | | High | High | High | High | High | High | High | High | High | High | High | 0 | 1.41E-08 | 4.42 |
| FALSE | High | EEGTVGSG | [K].EEGTV | 1xPhospho | Q8BLH7 1x | 0.068184 | 0.004935 | 1 | 1 | 2 | Q8BLH7 | HIRA-intera | Q8BLH7 [3 | 0 | 2587.145 | 0 | -0.25 | 0.135524 | 17.6 | 108.7 | 91.3 | 12.24 | 12.93 | | High | High | High | High | High | High | High | High | High | High | High | 0.002559 | 0.02756 | 2.25 |
| FALSE | High | EELLENEDE | [K].EELLEN | 1xPhospho | F8WHU5 1 | 6.2E-09 | 0 | 1 | 3 | 3 | F8WHU5 | Sister chro | F8WHU5 [1 | 0 | 2309.095 | 0 | 0.5 | 0.001274 | 9.21 | 82.8 | 117.2 | 6.75 | 4.53 | | High | High | High | High | High | High | High | High | High | High | High | 0 | 3.37E-10 | 3.94 |
| FALSE | High | EELLENEDE | [K].EELLEN | 1xPhospho | F8WHU5 1 | 2.36E-06 | 0 | 1 | 3 | 3 | F8WHU5 | Sister chro | F8WHU5 [1 | 1 | 2666.353 | 0 | 0.58 | 0.009585 | 6.49 | 80 | 120 | 8 | 9.42 | | High | High | High | High | High | High | High | High | High | High | High | 0 | 4.47E-06 | 4.03 |
| FALSE | High | EELLLPSVS | [K].EELLLP | 1xPhospho | Q4LDD4 1x | 0.016892 | 0.001204 | 1 | 2 | 1 | Q4LDD4 | Arf-GAP wi | Q4LDD4 [1 | 0 | 1548.824 | 0 | 0.31 | 0.027241 | 5.54 | 89.9 | 110.1 | 7.29 | 8.98 | | High | High | High | High | High | High | High | High | High | High | High | 0.000638 | 0.005645 | 1.46 |
| FALSE | High | EEILQPPEP | [K].EEILQP | 1xPhospho | P35831 1x | 0.000745 | 0 | 1 | 1 | 2 | P35831 | Tyrosine-p | P35831 [3 | 0 | 4284.147 | 0 | 1.05 | 0.002897 | 18.06 | 65.1 | 134.9 | 15.54 | 13.36 | | High | High | High | High | High | High | High | High | High | High | High | 0 | 0.000171 | 3.04 |
| FALSE | High | EELNGAMI | [R].EELNG | 1xPhospho | P58462-5 1 | 0.004372 | 0.000409 | 1 | 14 | 4 | P58462-5 | Isoform 5 c | P58462-5 [ | 0 | 2586.061 | 0 | -0.16 | 0.694866 | 5.32 | 100.3 | 99.7 | 8.36 | 12.03 | | High | High | High | High | High | High | High | High | High | High | High | 0.000183 | 0.001243 | 3.27 |
| FALSE | High | EELNGSFLN | [K].EELNGS | 1xPhospho | Q8BTI8 1x | 1.73E-08 | 0 | 1 | 3 | 2 | Q8BTI8 | Serine/argi | Q8BTI8 [13 | 0 | 2762.3 | 0 | -1.02 | 0.003775 | 3.51 | 134.7 | 65.3 | 6.9 | 24.07 | | High | High | High | High | High | High | High | High | High | High | High | 0 | 1.07E-09 | 5.39 |
| FALSE | High | EELNGSFLN | [K].EELNGS | 1xDeamida | Q8BTI8 1x | 9.7E-08 | 0 | 1 | 3 | 6 | Q8BTI8 | Serine/argi | Q8BTI8 [13 | 0 | 2763.284 | 0 | -0.79 | 0.003522 | 4.99 | 127.6 | 72.4 | 6.25 | 19.14 | | High | High | High | High | High | High | High | High | High | High | High | 0 | 7.41E-09 | 5.27 |
| FALSE | High | EELNGSFLN | [K].EELNGS | 1xPhospho | Q8BTI8 1x | 1.13E-07 | 0 | 1 | 3 | 11 | Q8BTI8 | Serine/argi | Q8BTI8 [13 | 0 | 2778.295 | 0 | -1.18 | 0.004775 | 0.98 | 134.6 | 65.4 | 12.95 | 10.73 | | High | High | High | High | High | High | High | High | High | High | High | 0 | 8.78E-09 | 5.31 |
| FALSE | High | EELNSPVAI | [K].EELNSP | 1xPhospho | Q497V5 1x | 4.47E-08 | 0 | 1 | 3 | 5 | Q497V5 | S1 RNA-bir | Q497V5 [8 | 0 | 2409.22 | 0 | 0.5 | 0.005808 | 6.41 | 84.3 | 115.7 | 3.05 | 8.3 | | High | High | High | High | High | High | High | High | High | High | High | 0 | 1.32E-07 | 4.8 |
| FALSE | High | EEIPTHSYS | [K].EEIPTH | 1xPhospho | Q9CYV5 1x | 0.033116 | 0.002195 | 1 | 2 | 1 | Q9CYV5 | Transmem | Q9CYV5 [1 | 0 | 2361.142 | 0 | -0.52 | 0.004408 | 4.44 | 118.3 | 81.7 | 4.08 | 9.52 | | High | High | High | High | High | High | High | High | High | High | High | 0.001151 | 0.01208 | 3.13 |
| FALSE | High | EELQANGS | [K].EELQAN | 1xPhospho | P26645 1x | 0.000237 | 0 | 1 | 3 | 1 | P26645 | Myristoyla | P26645 [5 | 0 | 1938.957 | 0 | 1.38 | 0.000712 | 7.02 | 52.4 | 147.6 | 9.37 | 10.97 | | High | High | High | High | High | High | High | High | High | High | High | 0 | 4.74E-05 | 2.79 |
| FALSE | High | EELQSAPA | [R].EELQSA | 1xPhospho | Q8CIL4 1x | 0.000118 | 0 | 1 | 3 | 2 | Q8CIL4 | Uncharacte | Q8CIL4 [85 | 0 | 2814.438 | 0 | -0.47 | 0.018373 | 8.4 | 118.6 | 81.4 | 10.48 | 11.3 | | High | High | High | High | High | High | High | High | High | High | High | 0 | 2.18E-05 | 3.63 |
| FALSE | High | EELQSAPA | [R].EELQSA | 1xPhospho | Q8CIL4 1x | 0.008769 | 0.000736 | 1 | 3 | 2 | Q8CIL4 | Uncharacte | Q8CIL4 [85 | 0 | 3043.6 | 0 | -0.74 | 0.003655 | 3.88 | 125.1 | 74.9 | 6.6 | 8.56 | | High | High | High | High | High | High | High | High | High | High | High | 0.000327 | 0.002721 | 2.39 |

| | | | | | | | | | | | | | | | | | | | | | | | | | | | | | | | | | | | | | | |
|---|---|---|---|---|---|---|---|---|---|---|---|---|---|---|---|---|---|---|---|---|---|---|---|---|---|---|---|---|---|---|---|---|---|---|---|---|---|---|
| FALSE | High | EEISDHENN | [R].EEISDH | 1xPhospho | Q61687 1x | 7.56E-06 | 0 | 1 | 4 | 7 | Q61687 | Transcripti | Q61687 [6 | 0 | 2650.245 | 0 | 0.11 | 0.284827 | 4.42 | 96.4 | 103.6 | 3.57 | 8.38 | | High | High | High | High | High | High | High | High | High | High | High | 0 | 9.9E-07 | 3.84 |
| FALSE | High | EELLENEDE | [K].EELLEN | 1xPhospho | F8WHU5 1 | 0.000699 | 0 | 1 | 3 | 2 | F8WHU5 | Sister chro | F8WHU5 [1 | 1 | 2437.19 | 0 | 0.45 | 0.523891 | 22.63 | 84 | 116 | 28.53 | 37.32 | | High | High | High | High | High | High | High | High | High | High | High | 0 | 0.000216 | 2.83 |
| FALSE | High | EEISDHENN | [R].EEISDH | 1xDeamida | Q61687 1x | 0.003028 | 0.00016 | 1 | 4 | 4 | Q61687 | Transcripti | Q61687 [6 | 0 | 2651.229 | 0 | 0.08 | 0.058816 | 3.21 | 97.1 | 102.9 | 6.04 | 3.14 | | High | High | High | High | High | High | High | High | High | High | High | 0.000538 | 0.00444 | 3.11 |
| FALSE | High | EEGSSTPDC | [K].EEGSST | 1xPhospho | Q9D6K7 1x | 0.002974 | 0.00016 | 1 | 1 | 3 | Q9D6K7 | Tetratricop | Q9D6K7 [2 | 0 | 1990.977 | 0 | 0.46 | 0.000669 | 2.49 | 84.1 | 115.9 | 4.59 | 4.57 | | High | High | High | High | High | High | High | High | High | High | High | 9.42E-05 | 0.000955 | 2.74 |
| FALSE | High | EEGSELEQI | [K].EEGSEL | 1xPhospho | S4R1M2 1x | 4.23E-08 | 0 | 2 | 5 | 5 | S4R1M2; C | Scaffold att | S4R1M2 [2 | 0 | 2543.15 | 0 | -0.4 | 0.006738 | 4.34 | 113.7 | 86.3 | 4.3 | 8.72 | Shared | High | High | High | High | High | High | High | High | High | High | High | 0 | 4.63E-08 | 3.93 |
| FALSE | High | EEDAPPVPS | [K].EEDAPP | 2xCarbami | A0A0N4SV | 1.53E-06 | 0 | 1 | 9 | 5 | A0A0N4SV | Eukaryotic | A0A0N4SV | 0 | 3541.612 | 0 | 0.86 | 0.005503 | 6.71 | 72.6 | 127.4 | 11.72 | 8.32 | | High | High | High | High | High | High | High | High | High | High | High | 0 | 1.64E-07 | 3.61 |
| FALSE | High | EEDCHSPTS | [K].EEDCHS | 1xCarbami | Q3TSY9 1x | 0.041425 | 0.003126 | 1 | 6 | 2 | Q3TSY9 | Putative ur | Q3TSY9 [4 | 0 | 2957.505 | 0 | -0.13 | 0.929136 | 72.6 | 90.2 | 109.8 | 42.92 | 43.69 | | High | High | High | High | High | High | High | High | High | High | High | 0.001916 | 0.01943 | 2.26 |
| FALSE | High | EEDEEAESI | [K].EEDEEA | 1xPhospho | Q80XU3 1x | 0.00407 | 0.00036 | 1 | 2 | 1 | Q80XU3 | Nuclear ub | Q80XU3 [2 | 1 | 2284.12 | 0 | 0.25 | 0.098734 | 16.24 | 96.7 | 103.3 | 15.03 | 7.82 | | High | High | High | High | High | High | High | High | High | High | High | 0.000161 | 0.001145 | 3.09 |
| FALSE | High | EEDEELLGI | [R].EEDEEL | 1xPhospho | Q91VU1 1x | 0.000184 | 0 | 1 | 2 | 1 | Q91VU1 | Protein YIP | Q91VU1 [5 | 0 | 2986.382 | 0 | 0.76 | 0.009479 | 19.74 | 74.8 | 125.2 | 21.22 | 7.73 | | High | High | High | High | High | High | High | High | High | High | High | 0 | 3.58E-05 | 3.1 |
| FALSE | High | EEDGSLSLD | [K].EEDGSI | 1xPhospho | Q62318 1x | 2.17E-08 | 0 | 1 | 2 | 10 | Q62318 | Transcripti | Q62318 [6 | 0 | 2388.158 | 0 | -0.56 | 0.00397 | 7.44 | 121.2 | 78.8 | 6.77 | 5.27 | | High | High | High | High | High | High | High | High | High | High | High | 0 | 1.09E-06 | 3.96 |
| FALSE | High | EEDGSVELI | [K].EEDGSV | 1xPhospho | Q0VAW6 1 | 3.64E-06 | 0 | 1 | 3 | 2 | Q0VAW6 | RIKEN cDN | Q0VAW6 [ | 0 | 2115.026 | 0 | -0.78 | 0.000181 | 0.15 | 126.3 | 73.7 | 4.66 | 10.47 | | High | High | High | High | High | High | High | High | High | High | High | 0 | 1.73E-06 | 3.09 |
| FALSE | High | EEDLSDSTD | [K].EEDLSD | 1xPhospho | Z4YKQ9 1x | 0.051694 | 0.003745 | 1 | 1 | 1 | Z4YKQ9 | ATPase fan | Z4YKQ9 [1 | 0 | 1823.835 | 0 | -0.25 | 0.030027 | 7.66 | 111.3 | 88.7 | 4.06 | 10.97 | | High | High | High | High | High | High | High | High | High | High | High | 0.001999 | 0.0201 | 2.2 |
| FALSE | High | EEDMLSGT | [K].EEDML | 1xCarbami | E9PV22 1x | 1.6E-08 | 0 | 1 | 4 | 3 | E9PV22 | Leucine-ric | E9PV22 [5 | 0 | 3292.5 | 0 | -0.33 | 0.033847 | 11.08 | 114.2 | 85.8 | 11.45 | 5.81 | | High | High | High | High | High | High | High | High | High | High | High | 0 | 1.12E-09 | 5.57 |
| FALSE | High | EEDMLSGT | [K].EEDML | 1xCarbami | E9PV22 1x | 1.24E-08 | 0 | 1 | 4 | 9 | E9PV22 | Leucine-ric | E9PV22 [5 | 0 | 3308.495 | 0 | -0.19 | 0.296625 | 10.35 | 107.1 | 92.9 | 8.22 | 11.91 | | High | High | High | High | High | High | High | High | High | High | High | 0 | 1.71E-09 | 6.2 |
| FALSE | High | EEDVPHRE | [K].EEDVPH | 1xPhospho | E9PZY8 1x | 0.003788 | 0.00021 | 1 | 4 | 1 | E9PZY8 | RIKEN cDN | E9PZY8 [2 | 1 | 2539.134 | 0 | -0.26 | 0.002175 | 2.42 | 112.1 | 87.9 | 4.35 | 4.63 | | High | High | High | High | High | High | High | High | High | High | High | 0.000117 | 0.001063 | 1.51 |
| FALSE | High | EEECLGTA | [K].EEECLG | 1xCarbami | Q7M6U3 1 | 0.050397 | 0.003641 | 1 | 2 | 2 | Q7M6U3 | Inactive se | Q7M6U3 [ | 0 | 3953.789 | 0 | -0.33 | 0.051085 | 10.22 | 120.4 | 79.6 | 14.37 | 15.28 | | High | High | High | High | High | High | High | High | High | High | High | 0.001916 | 0.01947 | 3.35 |
| FALSE | High | EEECLSDFS | [R].EEECLS | 1xCarbami | Q9CQK3 2x | 0.000745 | 0 | 1 | 1 | 1 | Q9CQK3 | RAD52 mot | Q9CQK3 [2 | 0 | 2590.096 | 0 | 0.5 | 0.448961 | 67.04 | 82.8 | 117.2 | 25.94 | 21.96 | | High | High | High | High | High | High | High | High | High | High | High | 0 | 0.000171 | 3.91 |
| FALSE | High | EEEDSCSEA | [K].EEEDSC | 2xCarbami | Q3UTB7 1x | 1.75E-05 | 0 | 1 | 3 | 2 | Q3UTB7 | Forkhead b | Q3UTB7 [8 | 0 | 2656.111 | 0 | -0.91 | 0.03621 | 8.02 | 124.2 | 75.8 | 12.24 | 29.94 | | High | High | High | High | High | High | High | High | High | High | High | 0 | 2.54E-06 | 4.5 |
| FALSE | High | EEGSELEQI | [K].EEGSEL | 1xPhospho | S4R1M2 1x | 0.000492 | 0 | 2 | 5 | 3 | S4R1M2; C | Scaffold att | S4R1M2 [2 | 1 | 2900.408 | 0 | -0.34 | 0.015716 | 7.49 | 111.8 | 88.2 | 5.07 | 13.49 | Shared | High | High | High | High | High | High | High | High | High | High | High | 0 | 0.000108 | 3.38 |
| FALSE | High | EEEEEEEEC | [R].EEEEEE | 1xCarbami | Q148S3 1x | 0.000352 | 0 | 1 | 1 | 1 | Q148S3 | NIN1/RPN1 | Q148S3 [1 | 0 | 3307.283 | 0 | -0.85 | 0.012741 | 1.74 | 130.5 | 69.5 | 19.2 | 12.82 | | High | High | High | High | High | High | High | High | High | High | High | 0 | 7.42E-05 | 2.68 |
| FALSE | High | EEEEQPSTS | [K].EEEEQF | 1xPhospho | B2RWS6 1x | 3.36E-06 | 0 | 1 | 1 | 1 | B2RWS6 | Histone ac | B2RWS6 [1 | 0 | 2713.261 | 0 | -0.79 | 0.000822 | 1.72 | 126.8 | 73.2 | 8.72 | 6.42 | | High | High | High | High | High | High | High | High | High | High | High | 0 | 3.98E-07 | 2.93 |
| FALSE | High | EEEKESDSD | [R].EEEKES | 1xPhospho | Q501J7-3 1 | 0.030632 | 0.001857 | 1 | 3 | 2 | Q501J7-3 | Isoform 3 c | Q501J7-3 [ | 1 | 2446.184 | 0 | 1.02 | 0.014483 | 13.91 | 65.1 | 134.9 | 6.04 | 20.65 | | High | High | High | High | High | High | High | High | High | High | High | 0.001027 | 0.01108 | 1.74 |
| FALSE | High | EEELNDSSE | [R].EEELND | 1xPhospho | A2A7B5 1x | 8.63E-05 | 0 | 1 | 3 | 3 | A2A7B5 | PR domain | A2A7B5 [1 | 0 | 2438.163 | 0 | -0.57 | 0.000823 | 4.2 | 120.5 | 79.5 | 2.82 | 5.27 | | High | High | High | High | High | High | High | High | High | High | High | 0 | 1.53E-05 | 3.56 |
| FALSE | High | EEEPCQQS | [R].EEEPCC | 1xCarbami | Q8K3Z9 1x | 0.035486 | 0.002304 | 1 | 2 | 1 | Q8K3Z9 | Nuclear en | Q8K3Z9 [4 | 1 | 3183.48 | 0 | 0.66 | 0.010718 | 21.51 | 73.9 | 126.1 | 10.03 | 18.21 | | High | High | High | High | High | High | High | High | High | High | High | 0.001214 | 0.01302 | 3.76 |
| FALSE | High | EEEPDPEE | [K].EEEPDF | 1xPhospho | O70494 1x | 0.047488 | 0.00354 | 1 | 4 | 1 | O70494 | Transcripti | O70494 [5 | 0 | 3334.495 | 0 | -0.8 | 0.020161 | 8.28 | 125.1 | 74.9 | 10.94 | 14.67 | | High | High | High | High | High | High | High | High | High | High | High | 0.001812 | 0.0181 | 1.78 |
| FALSE | High | EEEPTGWE | [K].EEEPTG | 1xPhospho | F6ZMJ4 1x | 1.08E-05 | 0 | 1 | 6 | 5 | F6ZMJ4 | Trinucleoti | F6ZMJ4 [9 | 0 | 2180.959 | 0 | 1.06 | 0.014254 | 9.6 | 65.9 | 134.1 | 9.01 | 17.35 | | High | High | High | High | High | High | High | High | High | High | High | 2.58E-05 | 0.0004 | 2.64 |
| FALSE | High | EEERPSSED | [K].EEERPS | 1xPhospho | P41183 1x | 0.001087 | 0 | 1 | 2 | 2 | P41183 | B-cell lymp | P41183 [3 | 0 | 3085.447 | 0 | 2.47 | 0.000517 | 16.52 | 29.9 | 170.1 | 13.55 | 11.56 | | High | High | High | High | High | High | High | High | High | High | High | 0 | 0.000262 | 3.36 |
| FALSE | High | EEESQPLYH | [R].EEESQF | 1xDeamida | Q3UGE1 1x | 0.000142 | 0 | 1 | 5 | 2 | Q3UGE1 | Putative ur | Q3UGE1 [3 | 0 | 4792.062 | 0 | -0.1 | 0.999774 | 9.75 | 98.2 | 101.8 | 24.62 | 17.46 | | High | High | High | High | High | High | High | High | High | High | High | 0 | 3.56E-05 | 4.3 |
| FALSE | High | EEETDENE | [K].EEETDE | 1xPhospho | B7ZWC4 1x | 0.058173 | 0.004227 | 1 | 8 | 1 | B7ZWC4 | Insulin-like | B7ZWC4 [2 | 0 | 2999.307 | 0 | 0.09 | 0.734272 | 26.1 | 106.7 | 93.3 | 16.6 | 22.83 | | High | High | High | High | High | High | High | High | High | High | High | 0.002187 | 0.02296 | 3.14 |
| FALSE | High | EEEWDPEY | [RK].EEEW | 1xPhospho | Q569Z6 1x | 0.000205 | 0 | 2 | 6 | 2 | Q569Z6; Q | Thyroid ho | Q569Z6 [8 | 0 | 1731.735 | 0 | -0.01 | 0.425076 | 12.49 | 100.3 | 99.7 | 17.02 | 13.8 | Shared | High | High | High | High | High | High | High | High | High | High | High | 0 | 4.04E-05 | 3.14 |
| FALSE | High | EEEWDPEY | [RK].EEEW | 1xPhospho | Q569Z6 1x | 8.53E-07 | 0 | 2 | 6 | 7 | Q569Z6; Q | Thyroid ho | Q569Z6 [8 | 0 | 1960.898 | 0 | -0.08 | 0.108696 | 6.41 | 103.2 | 96.8 | 8.15 | 8.46 | Shared | High | High | High | High | High | High | High | High | High | High | High | 0 | 8.69E-07 | 3.42 |
| FALSE | High | EEGPLSPFL | [R].EEGPLS | 1xPhospho | B0V2M3 1x | 0.002974 | 0.00016 | 1 | 5 | 1 | B0V2M3 | Zinc finger | B0V2M3 [2 | 0 | 2274.053 | 0 | -0.09 | 0.531112 | 16.24 | 101.6 | 98.4 | 8.62 | 15.32 | | High | High | High | High | High | High | High | High | High | High | High | 7.19E-05 | 0.000806 | 2.37 |
| FALSE | High | EEGSDEVL | [K].EEGSDI | 1xPhospho | Q8CAF4-3 | 0.041072 | 0.003094 | 1 | 3 | 1 | Q8CAF4-3 | Isoform 3 c | Q8CAF4-3 | 0 | 2096.068 | 0 | 2.17 | 0.001223 | 33.53 | 33.2 | 166.8 | 17.72 | 13.33 | | High | High | High | High | High | High | High | High | High | High | High | 0.001597 | 0.01533 | 2.33 |
| FALSE | High | EEEEEEEEC | [R].EEEEEE | 1xCarbami | Q148S3 1x | 7.63E-06 | 0 | 1 | 1 | 7 | Q148S3 | NIN1/RPN1 | Q148S3 [1 | 0 | 3536.446 | 0 | -0.59 | 0.002783 | 6.54 | 120.1 | 79.9 | 5.16 | 6.84 | | High | High | High | High | High | High | High | High | High | High | High | 0 | 1.22E-05 | 4.22 |
| FALSE | High | EELSHSPEP | [K].EELSHS | 1xCarbami | P41230 1x | 0.000132 | 0 | 1 | 3 | 3 | P41230 | Lysine-spe | P41230 [2 | 0 | 1951.924 | 0 | -0.36 | 0.006795 | 6.75 | 113.4 | 86.6 | 3.87 | 5.83 | | High | High | High | High | High | High | High | High | High | High | High | 0 | 2.45E-05 | 2.25 |
| FALSE | High | EELSSGDSL | [K].EELSSG | 1xPhospho | E9Q9C3 1x | 7.72E-07 | 0 | 1 | 6 | 1 | E9Q9C3 | Afadin OS= | E9Q9C3 [1 | 0 | 2185.047 | 0 | -0.6 | 0.082383 | 11.47 | 119.2 | 80.8 | 6.75 | 28.7 | | High | High | High | High | High | High | High | High | High | High | High | 0 | 7.58E-08 | 3.16 |
| FALSE | High | EELVSSEDE | [K].EELVSS | 1xPhospho | Q3U319 1x | 1.67E-07 | 0 | 1 | 3 | 3 | Q3U319 | E3 ubiquiti | Q3U319 [5 | 0 | 2751.329 | 0 | 1.35 | 0.001883 | 17.82 | 58 | 142 | 7.74 | 13.04 | | High | High | High | High | High | High | High | High | High | High | High | 0 | 1.35E-08 | 4.09 |
| FALSE | High | EEQTDTSD | [R].EEQTD | 2xPhospho | P36916 2x | 0.065405 | 0.004734 | 1 | 1 | 1 | P36916 | Guanine nu | P36916 [4 | 0 | 2329.954 | 0 | -1.29 | 0.00318 | 3.05 | 138 | 62 | 7.02 | 24.79 | | High | High | High | High | High | High | High | High | High | High | High | 0.002446 | 0.02632 | 2.68 |
| FALSE | High | EESAASGG | [K].EESAAS | 1xPhospho | Q569Z6 1x | 0.002818 | 0.000109 | 1 | 4 | 2 | Q569Z6 | Thyroid ho | Q569Z6 [3 | 0 | 1765.841 | 0 | -0.47 | 0.003145 | 3.46 | 115.7 | 84.3 | 6.6 | 6.63 | | High | High | High | High | High | High | High | High | High | High | High | 0.000204 | 0.001289 | 3.38 |
| FALSE | High | EESDGEYD | [R].EESDGI | 1xPhospho | Q9R020 1x | 0.064323 | 0.004664 | 1 | 5 | 3 | Q9R020 | Zinc finger | Q9R020 [1 | 1 | 2098.937 | 0 | -0.34 | 0.010855 | 5.85 | 111.9 | 88.1 | 6.13 | 5.64 | | High | High | High | High | High | High | High | High | High | High | High | 0.002403 | 0.02573 | 2.01 |
| FALSE | High | EESDTDLEV | [K].EESDTD | 1xPhospho | H7BX94 1x | 4.94E-05 | 0 | 1 | 4 | 8 | H7BX94 | Lysine-rich | H7BX94 [3 | 0 | 2044.014 | 0 | -0.23 | 0.011023 | 5.7 | 106.9 | 93.1 | 3.21 | 5.98 | | High | High | High | High | High | High | High | High | High | High | High | 0 | 4.47E-05 | 3.65 |
| FALSE | High | EESDTDLEV | [K].EESDTD | 2xPhospho | H7BX94 2x | 0.000693 | 0 | 1 | 4 | 11 | H7BX94 | Lysine-rich | H7BX94 [3 | 0 | 2123.98 | 0 | 0.4 | 0.025943 | 19.82 | 81.9 | 118.1 | 12.93 | 14.8 | | High | High | High | High | High | High | High | High | High | High | High | 0 | 0.000159 | 3.38 |
| FALSE | High | EESDWPAS | [K].EESDW | 1xPhospho | Q8BSN5 1x | 1.74E-06 | 0 | 1 | 2 | 4 | Q8BSN5 | PHD finger | Q8BSN5 [5 | 0 | 1913.848 | 0 | -1.19 | 0.000158 | 1.14 | 139.5 | 60.5 | 1.73 | 5.74 | | High | High | High | High | High | High | High | High | High | High | High | 0.002187 | 0.02334 | 3.48 |
| FALSE | High | EESDWPAS | [K].EESDW | 1xPhospho | Q8BSN5 1x | 0.008769 | 0.000736 | 1 | 2 | 4 | Q8BSN5 | PHD finger | Q8BSN5 [5 | 1 | 3730.598 | 0 | -1.06 | 0.004626 | 4.46 | 143.6 | 56.4 | 12.13 | 19.57 | | High | High | High | High | High | High | High | High | High | High | High | 0.000449 | 0.004089 | 3.02 |
| FALSE | High | EESDWPAS | [K].EESDW | 2xPhospho | Q8BSN5 2x | 0.000633 | 0 | 1 | 2 | 2 | Q8BSN5 | PHD finger | Q8BSN5 [5 | 1 | 3810.564 | 0 | -0.09 | 0.142289 | 1.33 | 103.9 | 96.1 | 7.69 | 10.69 | | High | High | High | High | High | High | High | High | High | High | High | 0 | 0.000143 | 3.13 |
| FALSE | High | EESDWPAS | [K].EESDW | 3xPhospho | Q8BSN5 3x | 0.000349 | 0 | 1 | 2 | 1 | Q8BSN5 | PHD finger | Q8BSN5 [5 | 1 | 3890.531 | 0 | -0.58 | 0.021162 | 10.49 | 121.8 | 78.2 | 17.71 | 11.11 | | High | High | High | High | High | High | High | High | High | High | High | 0 | 7.32E-05 | 4.53 |
| FALSE | High | EESEDELEE | [R].EESEDE | 1xPhospho | Q8CFE3 1x | 2.5E-10 | 0 | 1 | 2 | 2 | Q8CFE3 | REST corep | Q8CFE3 [2 | 0 | 3011.309 | 0 | 0.69 | 0.017586 | 8.13 | 78.6 | 121.4 | 10.22 | 9.41 | | High | High | High | High | High | High | High | High | High | High | High | 0 | 9.11E-12 | 5.86 |
| FALSE | High | EESEDELEE | [R].EESEDE | 1xPhospho | Q8CFE3 1x | 1.37E-05 | 0 | 1 | 2 | 2 | Q8CFE3 | REST corep | Q8CFE3 [2 | 0 | 3240.472 | 0 | 0.39 | 0.050261 | 6.5 | 84.3 | 115.7 | 6.66 | 16.48 | | High | High | High | High | High | High | High | High | High | High | High | 0 | 1.93E-06 | 5.83 |
| FALSE | High | EESEDELEE | [R].EESEDE | 1xDeamida | Q8CFE3 1x | 7.49E-09 | 0 | 1 | 2 | 8 | Q8CFE3 | REST corep | Q8CFE3 [2 | 0 | 3241.456 | 0 | 0.31 | 0.013116 | 13.82 | 85.2 | 114.8 | 8.94 | 5.3 | | High | High | High | High | High | High | High | High | High | High | High | 0 | 8.54E-10 | 6.07 |
| FALSE | High | EESEDPDY | [K].EESEDF | 1xPhospho | Q9ESZ8-5 | 3.75E-07 | 0 | 1 | 1 | 3 | Q9ESZ8-5 | Isoform 5 c | Q9ESZ8-5 [ | 0 | 3349.435 | 0 | -0.24 | 0.157791 | 5.42 | 106.5 | 93.5 | 11.31 | 12.3 | | High | High | High | High | High | High | High | High | High | High | High | 0 | 3.39E-08 | 4.37 |
| FALSE | High | EEQTDTSD | [R].EEQTD | 1xPhospho | P36916 1x | 7.41E-05 | 0 | 1 | 1 | 6 | P36916 | Guanine nu | P36916 [4 | 0 | 2249.987 | 0 | -1.28 | 0.009628 | 2.2 | 139.2 | 60.8 | 14.98 | 19.87 | | High | High | High | High | High | High | High | High | High | High | High | 0.001279 | 0.01336 | 3.02 |
| FALSE | High | EESEDSFTF | [K].EESEDS | 1xPhospho | Q497V5 1x | 0.000239 | 0 | 1 | 2 | 2 | Q497V5 | S1 RNA-bir | Q497V5 [1 | 0 | 2225.042 | 0 | 0.13 | 0.124049 | 9.72 | 97.3 | 102.7 | 6.95 | 6.21 | | High | High | High | High | High | High | High | High | High | High | High | 0 | 0.000177 | 4.18 |
| FALSE | High | EESEESDEE | [K].EESEES | 1xPhospho | Q5M8R8 1 | 3.98E-09 | 0 | 1 | 2 | 25 | Q5M8R8 | 60S acidic r | Q5M8R8 [3 | 0 | 2160.804 | 3 | 0.09 | 0.651178 | 5.76 | 95.6 | 104.4 | 10.05 | 18.88 | | High | High | High | High | High | High | High | High | High | High | High | 0 | 2.04E-10 | 3.95 |
| FALSE | High | EESEESDEE | [K].EESEES | 2xPhospho | Q5M8R8 2 | 6.78E-09 | 0 | 1 | 2 | 115 | Q5M8R8 | 60S acidic r | Q5M8R8 [3 | 0 | 2240.771 | 5 | -0.27 | 0.011534 | 6.65 | 110.6 | 89.4 | 5.39 | 5.44 | | High | High | High | High | High | High | High | High | High | High | High | 0 | 4.66E-10 | 4.38 |
| FALSE | High | EESEESEDE | [K].EESEES | 1xPhospho | E9Q3T0 1x | 1.44E-07 | 0 | 1 | 2 | 41 | E9Q3T0 | Uncharacte | E9Q3T0 [9 | 0 | 2160.804 | 0 | 0.27 | 0.253769 | 12.52 | 87.2 | 112.8 | 10.49 | 21.73 | | High | High | High | High | High | High | High | High | High | High | High | 0 | 1.61E-06 | 3.85 |
| FALSE | High | EESETRSES | [R].EESETR | 1xPhospho | Q3TLH4 1x | 0.007605 | 0.000664 | 1 | 5 | 1 | Q3TLH4 | Protein PRI | Q3TLH4 [1 | 1 | 2493.18 | 0 | -0.32 | 0.018722 | 3.69 | 113.6 | 86.4 | 10.67 | 7.22 | | High | High | High | High | High | High | High | High | High | High | High | 0.000294 | 0.00232 | 2.36 |
| FALSE | High | EESETRSES | [R].EESETR | 2xPhospho | Q3TLH4 2x | 2.27E-05 | 0 | 1 | 5 | 2 | Q3TLH4 | Protein PRI | Q3TLH4 [1 | 1 | 2573.146 | 0 | 0.03 | 0.911229 | 5.54 | 98.9 | 101.1 | 9.1 | 5.15 | | High | High | High | High | High | High | High | High | High | High | High | 0 | 3.39E-06 | 3.71 |
| FALSE | High | EESFRDPSE | [K].EESFRD | 1xPhospho | Q9JJ11 1xP | 0.000462 | 0 | 1 | 5 | 1 | Q9JJ11 | Transformi | Q9JJ11 [31 | 1 | 3762.783 | 0 | 0.16 | 0.545179 | 29.81 | 97.3 | 102.7 | 37.67 | 28.98 | | High | High | High | High | High | High | High | High | High | High | High | 0 | 0.000101 | 4.41 |
| FALSE | High | EESFRDPSE | [K].EESFRD | 1xDeamida | Q9JJ11 1xP | 0.032548 | 0.002167 | 1 | 5 | 1 | Q9JJ11 | Transformi | Q9JJ11 [31 | 1 | 3763.767 | 0 | 0.3 | 0.024228 | 12.31 | 89.5 | 110.5 | 10.69 | 10.88 | | High | High | High | High | High | High | High | High | High | High | High | 0.001142 | 0.01182 | 4.56 |
| FALSE | High | EESPEPYFF | [R].EESPEP | 1xPhospho | Q148W8 1 | 0.008538 | 0.000736 | 1 | 1 | 1 | Q148W8 | Inactive du | Q148W8 [1 | 0 | 1609.714 | 0 | -1.81 | 0.001041 | 1.48 | 153.5 | 46.5 | 10.18 | 15 | | High | High | High | High | High | High | High | High | High | High | High | 0.000327 | 0.002631 | 2.34 |
| FALSE | High | EESPMDVD | [R].EESPMI | 1xPhospho | A2AFQ0 1x | 2.32E-06 | 0 | 1 | 4 | 2 | A2AFQ0 | E3 ubiquiti | A2AFQ0 [3 | 0 | 3810.767 | 0 | -0.01 | 0.67052 | 10.17 | 98.8 | 101.2 | 4.22 | 14.07 | | High | High | High | High | High | High | High | High | High | High | High | 0 | 2.61E-07 | 5.26 |
| FALSE | High | EESPMDVD | [R].EESPMI | 1xPhospho | A2AFQ0 1x | 0.008538 | 0.000736 | 1 | 4 | 1 | A2AFQ0 | E3 ubiquiti | A2AFQ0 [3 | 0 | 3826.762 | 0 | 0.2 | 0.288901 | 16.2 | 95.6 | 104.4 | 8.78 | 8.85 | | High | High | High | High | High | High | High | High | High | High | High | 0.000327 | 0.00263 | 3.95 |
| FALSE | High | EESRTINLY | [K].EESRTII | 1xPhospho | B8JK39 1x | 0.001039 | 0 | 1 | 5 | 1 | B8JK39 | Integrin alp | B8JK39 [9 | 1 | 2734.45 | 0 | -0.93 | 0.000824 | 4.3 | 130.4 | 69.6 | 4.67 | 11.87 | | High | High | High | High | High | High | High | High | High | High | High | 0 | 0.000249 | 2.49 |
| FALSE | High | EESSEDENI | [K].EESSED | 1xPhospho | B2RRE7 1x | 1.1E-06 | 0 | 1 | 2 | 7 | B2RRE7 | OTU doma | B2RRE7 [1 | 0 | 2058.907 | 0 | -0.34 | 0.002004 | 0.9 | 109.4 | 90.6 | 4.72 | 4.41 | | High | High | High | High | High | High | High | High | High | High | High | 0 | 4.38E-07 | 4.61 |
| FALSE | High | EESSEDENI | [K].EESSED | 2xPhospho | B2RRE7 2x | 4.73E-06 | 0 | 1 | 2 | 20 | B2RRE7 | OTU doma | B2RRE7 [1 | 0 | 2138.873 | 0 | -0.25 | 0.009865 | 7.48 | 109 | 91 | 3.16 | 5.76 | | High | High | High | High | High | High | High | High | High | High | High | 0 | 1.45E-05 | 4.38 |
| FALSE | High | EESEDSFTF | [K].EESEDS | 2xPhospho | Q497V5 2x | 0.000786 | 0 | 1 | 2 | 1 | Q497V5 | S1 RNA-bir | Q497V5 [1 | 0 | 2305.008 | 0 | 0.1 | 0.290011 | 2.63 | 96.9 | 103.1 | 13.42 | 4.57 | | High | High | High | High | High | High | High | High | High | High | High | 0 | 0.000182 | 2.68 |
| FALSE | High | EEQSSPSLL | [K].EEQSSF | 1xPhospho | A2CG63 1x | 7.56E-06 | 0 | 1 | 4 | 4 | A2CG63 | AT-rich inte | A2CG63 [7 | 0 | 2027.994 | 0 | -0.42 | 0.005911 | 2.31 | 113.7 | 86.3 | 3.73 | 5.04 | | High | High | High | High | High | High | High | High | High | High | High | 0 | 9.88E-07 | 3.7 |
| FALSE | High | EEQSQITSC | [K].EEQSQI | 1xPhospho | Q6A0C9 1x | 1.77E-07 | 0 | 1 | 5 | 1 | Q6A0C9 | MKIAA010 | Q6A0C9 [1 | 0 | 2256.086 | 0 | 0.35 | 0.146558 | 45.72 | 68.5 | 131.5 | 28.26 | 32.77 | | High | High | High | High | High | High | High | High | High | High | High | 0 | 1.46E-08 | 3.31 |
| FALSE | High | EEQSPVK | [K].EEQSPV | 1xPhospho | P14873 1x | 0.008022 | 0.000736 | 2 | 6 | 1 | P14873; B2 | Microtubu | P14873 [6 | 0 | 1354.702 | 0 | -1.13 | 0.000361 | 1.82 | 137.8 | 62.2 | 1.74 | 12.98 | Shared | High | High | High | High | High | High | High | High | High | High | High | 0.000327 | 0.002462 | 1.48 |
| FALSE | High | EENFCSPEN | [K].EENFCS | 1xCarbami | Q924W5 1 | 0.025073 | 0.001645 | 1 | 5 | 1 | Q924W5 | Structural | Q924W5 [6 | 0 | 1862.84 | 0 | -0.28 | 0.010443 | 7.57 | 107 | 93 | 15.32 | 16.64 | | High | High | High | High | High | High | High | High | High | High | High | 0.000872 | 0.0088 | 2.71 |
| FALSE | High | EENPVFKTI | [K].EENPVF | 1xCarbami | Q80YR6 1x | 0.026885 | 0.001676 | 1 | 3 | 1 | Q80YR6 | DNA endor | Q80YR6 [5 | 1 | 3761.859 | 0 | -1.57 | 0.000662 | 0.89 | 151.6 | 48.4 | 7.7 | 12.24 | | High | High | High | High | High | High | High | High | High | High | High | 0.000903 | 0.009496 | 2.35 |
| FALSE | High | EENQTGPT | [K].EENQT | 1xPhospho | Q8CB16 1x | 2.83E-06 | 0 | 1 | 6 | 2 | Q8CB16 | Putative ur | Q8CB16 [3 | 0 | 2762.248 | 0 | 0.69 | 0.006327 | 9.91 | 74.5 | 125.5 | 11.46 | 9.07 | | High | High | High | High | High | High | High | High | High | High | High | 0 | 3.28E-07 | 3.68 |
| FALSE | High | EENSGDVE | [K].EENSGI | 1xPhospho | E9Q6J5-3 1 | 2.75E-09 | 0 | 1 | 3 | 2 | E9Q6J5-3 | Isoform Ne | E9Q6J5-3 [ | 0 | 2282.048 | 0 | 0.32 | 0.006456 | 3.8 | 87.1 | 112.9 | 6.33 | 9.42 | | High | High | High | High | High | High | High | High | High | High | High | 0 | 1.36E-10 | 4.27 |
| FALSE | High | EENVNSPE | [K].EENVN | 1xPhospho | Q61687 1x | 6.91E-06 | 0 | 1 | 4 | 1 | Q61687 | Transcripti | Q61687 [9 | 0 | 1698.799 | 0 | -0.05 | 0.427919 | 18.93 | 105 | 95 | 11.76 | 8.2 | | High | High | High | High | High | High | High | High | High | High | High | 0 | 8.95E-07 | 2.46 |
| FALSE | High | EEPASGSA | [K].EEPASG | 1xPhospho | P26645 1x | 0.013074 | 0.000889 | 1 | 3 | 1 | P26645 | Myristoyla | P26645 [7 | 0 | 2024.994 | 0 | 1.1 | 0.006647 | 29.57 | 66.3 | 133.7 | 12.9 | 17.52 | | High | High | High | High | High | High | High | High | High | High | High | 0.000467 | 0.004233 | 1.97 |
| FALSE | High | EEPEDVGM | [R].EEPEDV | 1xPhospho | Q6ZPJ3 1x | 0.004218 | 0.00036 | 1 | 1 | 3 | Q6ZPJ3 | (E3-indepe | Q6ZPJ3 [1 | 0 | 3269.374 | 0 | -0.82 | 0.008022 | 8.31 | 129.2 | 70.8 | 12.84 | 17.51 | | High | High | High | High | High | High | High | High | High | High | High | 0.000275 | 0.001877 | 4.1 |
| FALSE | High | EEPEDVGM | [R].EEPEDV | 1xPhospho | Q6ZPJ3 1x | 1.5E-05 | 0 | 1 | 1 | 5 | Q6ZPJ3 | (E3-indepe | Q6ZPJ3 [1 | 0 | 3285.369 | 0 | -1.01 | 0.008466 | 4.77 | 132.4 | 67.6 | 11.02 | 37.69 | | High | High | High | High | High | High | High | High | High | High | High | 0 | 2.13E-06 | 4.64 |
| FALSE | High | EEPEPLSPE | [K].EEPEPL | 1xPhospho | Q69ZP3-3 | 0.020499 | 0.001472 | 1 | 4 | 1 | Q69ZP3-3 | Isoform 3 c | Q69ZP3-3 | 0 | 2107.02 | 0 | -0.23 | 0.3003 | 16.2 | 106.6 | 93.4 | 14.77 | 21.66 | | High | High | High | High | High | High | High | High | High | High | High | 0.000786 | 0.006979 | 2.13 |
| FALSE | High | EEPGESSPC | [K].EEPGES | 1xPhospho | Q5DTL2 1x | 0.007673 | 0.000664 | 1 | 2 | 2 | Q5DTL2 | MKIAA414 | Q5DTL2 [6 | 0 | 2167.021 | 0 | 1 | 0.013621 | 26.57 | 63.5 | 136.5 | 14.61 | 21.02 | | High | High | High | High | High | High | High | High | High | High | High | 0.000801 | 0.007207 | 2.13 |
| FALSE | High | EEPLDPEEA | [K].EEPLDP | 1xPhospho | P58462-5 1 | 0.035793 | 0.002304 | 1 | 14 | 1 | P58462-5 | Isoform 5 c | P58462-5 [ | 0 | 3526.585 | 0 | 0.43 | 0.055487 | 14.86 | 80.7 | 119.3 | 15.31 | 14.72 | | High | High | High | High | High | High | High | High | High | High | High | 0.00123 | 0.01319 | 3.79 |
| FALSE | High | EEPLDPEEA | [K].EEPLDP | 1xPhospho | P58462-5 1 | 0.104222 | 0.008348 | 1 | 14 | 1 | P58462-5 | Isoform 5 c | P58462-5 [ | 1 | 4992.104 | 0 | 0.67 | 0.064031 | 8.78 | 75.7 | 124.3 | 9.5 | 24.97 | | High | High | High | High | High | High | High | High | High | High | High | 0.004206 | 0.04526 | 3.51 |
| FALSE | High | EEPLSEEEA | [K].EEPLSE | 1xCarbami | Q3TWE8 1 | 3.27E-06 | 0 | 1 | 4 | 2 | Q3TWE8 | Putative ur | Q3TWE8 [4 | 0 | 4161.818 | 0 | -0.54 | 0.073029 | 7.68 | 112.7 | 87.3 | 10.39 | 16.41 | | High | High | High | High | High | High | High | High | High | High | High | 0 | 3.85E-07 | 5.51 |
| FALSE | High | EEPLSEEEA | [K].EEPLSE | 1xCarbami | Q3TWE8 1 | 1.34E-07 | 0 | 1 | 4 | 3 | Q3TWE8 | Putative ur | Q3TWE8 [4 | 0 | 4177.813 | 0 | -0.74 | 0.066227 | 6.55 | 122.9 | 77.1 | 9.74 | 22.87 | | High | High | High | High | High | High | High | High | High | High | High | 0 | 5.7E-08 | 5.74 |
| FALSE | High | EEPLSEEEP | [K].EEPLSE | 1xCarbami | Q6DFW4 1 | 1.52E-06 | 0 | 1 | 1 | 11 | Q6DFW4 | Nucleolar p | Q6DFW4 [ | 0 | 2754.283 | 0 | -0.33 | 0.00854 | 0.84 | 110.8 | 89.2 | 5.5 | 4.25 | | High | High | High | High | High | High | High | High | High | High | High | 0 | 1.63E-07 | 4.69 |
| FALSE | High | EEPLSEEEP | [K].EEPLSE | 1xCarbami | Q6DFW4 2 | 2.17E-05 | 0 | 1 | 1 | 14 | Q6DFW4 | Nucleolar p | Q6DFW4 [ | 0 | 2834.25 | 0 | -0.58 | 0.00114 | 3.98 | 119.4 | 80.6 | 3.76 | 5.84 | | High | High | High | High | High | High | High | High | High | High | High | 0 | 7.27E-06 | 3.83 |
| FALSE | High | EEPMEVDI | [K].EEPME | 1xPhospho | Q505G8 1x | 0.001097 | 0 | 1 | 2 | 1 | Q505G8 | Zinc finger | Q505G8 [6 | 0 | 2396.089 | 0 | 0.81 | 0.001799 | 6.92 | 67.7 | 132.3 | 14.36 | 10.95 | | High | High | High | High | High | High | High | High | High | High | High | 0 | 0.000265 | 3.06 |
| FALSE | High | EEPPEPVPS | [K].EEPPEP | 1xPhospho | F8WI37 1x | 0.002313 | 0.000109 | 1 | 6 | 1 | F8WI37 | Histone-lys | F8WI37 [4 | 0 | 2734.508 | 0 | -0.41 | 0.009155 | 7.84 | 114.4 | 85.6 | 3.02 | 9.1 | | High | High | High | High | High | High | High | High | High | High | High | 4.94E-05 | 0.000609 | 2.95 |
| FALSE | High | EEPPGSQD | [K].EEPPGS | 1xPhospho | B0FTY3 1x | 0.000157 | 0 | 1 | 4 | 1 | B0FTY3 | NudC dom | B0FTY3 [1 | 0 | 2625.26 | 0 | -0.03 | 0.600106 | 6.29 | 101.7 | 98.3 | 4.87 | 3.72 | | High | High | High | High | High | High | High | High | High | High | High | 0 | 2.98E-05 | 2.94 |
| FALSE | High | EEPQETAIP | [K].EEPQET | 1xPhospho | E9QK89 1x | 2.58E-05 | 0 | 1 | 2 | 3 | E9QK89 | Mediator c | E9QK89 [1 | 0 | 2007.009 | 0 | -1.17 | 0.000176 | 0.32 | 137.4 | 62.6 | 3.51 | 2.34 | | High | High | High | High | High | High | High | High | High | High | High | 0 | 3.93E-06 | 2.34 |
| FALSE | High | EEPQGCAG | [K].EEPQG | 1xCarbami | Q8R3F9 1x | 2.31E-05 | 0 | 1 | 1 | 2 | Q8R3F9 | Speckle tar | Q8R3F9 [6 | 0 | 2952.259 | 0 | -0.23 | 0.081248 | 9.37 | 110.2 | 89.8 | 9.44 | 13.56 | | High | High | High | High | High | High | High | High | High | High | High | 0 | 6.78E-05 | 3.98 |
| FALSE | High | EEPQGCAG | [K].EEPQG | 1xCarbami | Q8R3F9 1x | 1.68E-07 | 0 | 1 | 1 | 2 | Q8R3F9 | Speckle tar | Q8R3F9 [6 | 0 | 2968.254 | 0 | -0.34 | 0.512102 | 7.27 | 106.1 | 93.9 | 8.11 | 25.42 | | High | High | High | High | High | High | High | High | High | High | High | 0 | 1.38E-08 | 4.77 |
| FALSE | High | EEPSPISK | [K].EEPSPIS | 1xPhospho | G3X8R8 1x | 0.015603 | 0.001124 | 1 | 4 | 1 | G3X8R8 | MCG12388 | G3X8R8 [9 | 0 | 1424.744 | 0 | 0.33 | 0.035862 | 12.96 | 90.2 | 109.8 | 8.23 | 8.28 | | High | High | High | High | High | High | High | High | High | High | High | 0.000585 | 0.005137 | 1.55 |
| FALSE | High | EEPVLTPID | [K].EEPVLT | 1xPhospho | P23298 1x | 0.005275 | 0.000599 | 1 | 3 | 2 | P23298 | Protein kin | P23298 [6 | 0 | 2888.374 | 0 | 4.11 | 0.000926 | 40.03 | 10.7 | 189.3 | 41.18 | 16.3 | | High | High | High | High | High | High | High | High | High | High | High | 0.000267 | 0.001533 | 3.52 |
| FALSE | High | EEPVLTPID | [K].EEPVLT | 1xPhospho | P23298 1x | 0.000442 | 0 | 1 | 3 | 1 | P23298 | Protein kin | P23298 [6 | 0 | 2904.369 | 0 | 2.57 | 0.001153 | 14.76 | 26.9 | 173.1 | 5.29 | 14.86 | | High | High | High | High | High | High | High | High | High | High | High | 0 | 9.52E-05 | 2.8 |
| FALSE | High | EEQAASPV | [K].EEQAA | 1xPhospho | E9PWN0 1 | 1.4E-10 | 0 | 1 | 5 | 4 | E9PWN0 | Phosphatic | E9PWN0 [2 | 0 | 2871.318 | 0 | 1.76 | 0.000181 | 6.63 | 44 | 156 | 21.71 | 5.62 | | High | High | High | High | High | High | High | High | High | High | High | 0 | 1.39E-08 | 5.01 |
| FALSE | High | EEQAASPV | [K].EEQAA | 2xPhospho | E9PWN0 2 | 9.64E-09 | 0 | 1 | 5 | 2 | E9PWN0 | Phosphatic | E9PWN0 [2 | 0 | 2951.285 | 0 | 1.12 | 0.000487 | 4.19 | 62.8 | 137.2 | 7.23 | 8.51 | | High | High | High | High | High | High | High | High | High | High | High | 0 | 5.5E-10 | 6.75 |
| FALSE | High | EEQIDAPPA | [K].EEQIDA | 1xCarbami | F6UFZ5 1x | 2.39E-07 | 0 | 2 | 7 | 4 | F6UFZ5; Q | PCF11 clea | F6UFZ5 [1 | 0 | 2939.4 | 0 | -0.55 | 0.012854 | 5.52 | 117.9 | 82.1 | 10.91 | 7.41 | Shared | High | High | High | High | High | High | High | High | High | High | High | 0 | 2.03E-08 | 4.22 |
| FALSE | High | EEQIDAPPA | [K].EEQIDA | 1xCarbami | F6UFZ5 1x | 0.033694 | 0.002195 | 2 | 7 | 1 | F6UFZ5; Q | PCF11 clea | F6UFZ5 [1 | 0 | 2940.384 | 0 | -0.48 | 0.050373 | 3.17 | 115.4 | 84.6 | 6.63 | 18.7 | Shared | High | High | High | High | High | High | High | High | High | High | High | 0.001151 | 0.01227 | 2.81 |
| FALSE | High | EECPEAME | [K].EECPEA | 1xCarbami | P70399 2x | 0.000198 | 0 | 1 | 4 | 2 | P70399 | TP53-bindi | P70399 [7 | 0 | 2996.332 | 0 | -0.15 | 0.294976 | 2.28 | 104.5 | 95.5 | 7.11 | 26 | | High | High | High | High | High | High | High | High | High | High | High | 4.94E-05 | 0.000642 | 3.73 |
| FALSE | High | EAYNDSSD | [R].EAYND | 1xCarbami | Q9QYJ0 2x | 0.079114 | 0.00613 | 1 | 2 | 1 | Q9QYJ0 | DnaJ homo | Q9QYJ0 [3 | 0 | 3017.124 | 0 | -0.33 | 0.138545 | 25.7 | 111.4 | 88.6 | 25.34 | 12.87 | | High | High | High | High | High | High | High | High | High | High | High | 0.003148 | 0.03267 | 3.45 |
| FALSE | High | EECPEAME | [K].EECPEA | 1xCarbami | P70399 1x | 7.33E-08 | 0 | 1 | 4 | 3 | P70399 | TP53-bindi | P70399 [7 | 0 | 2932.361 | 0 | 0.63 | 0.055019 | 19.1 | 79.2 | 120.8 | 17.11 | 24.91 | | High | High | High | High | High | High | High | High | High | High | High | 0 | 2.36E-07 | 5.33 |
| FALSE | High | EECGDTAL | [K].EECGD | 1xCarbami | A0A1L1SSC | 2.24E-06 | 0 | 1 | 1 | 4 | A0A1L1SSC | Nuclear fac | A0A1L1SSC | 0 | 2958.262 | 0 | 0.35 | 0.02431 | 3.1 | 91.7 | 108.3 | 8.01 | 10.92 | | High | High | High | High | High | High | High | High | High | High | High | 0 | 2.51E-07 | 4.38 |
| FALSE | High | EDELSIGSA | [K].EDELSI | 1xPhospho | Q80TY5 1x | 0.01002 | 0.000736 | 1 | 2 | 1 | Q80TY5 | Vacuolar p | Q80TY5 [9 | 0 | 1853.966 | 0 | 0.16 | 0.026274 | 6.86 | 97.2 | 102.8 | 5.6 | 5.52 | | High | High | High | High | High | High | High | High | High | High | High | 0.000344 | 0.00313 | 2.57 |
| FALSE | High | EDELSIGSA | [K].EDELSI | 2xPhospho | Q80TY5 2x | 0.000328 | 0 | 1 | 2 | 1 | Q80TY5 | Vacuolar p | Q80TY5 [9 | 0 | 1933.933 | 0 | 0.59 | 0.008626 | 14.69 | 72.8 | 127.2 | 10.09 | 16.8 | | High | High | High | High | High | High | High | High | High | High | High | 0 | 6.81E-05 | 2.89 |
| FALSE | High | EDEISPPPP | [R].EDEISP | 1xPhospho | Q3TYK4 1x | 1.23E-05 | 0 | 1 | 5 | 3 | Q3TYK4 | Putative ur | Q3TYK4 [2 | 0 | 1826.914 | 0 | 0.27 | 0.005452 | 4.36 | 90.9 | 109.1 | 3.24 | 6.61 | | High | High | High | High | High | High | High | High | High | High | High | 0 | 1.71E-06 | 3.48 |
| FALSE | High | EDEISPPPP | [R].EDEISP | 1xPhospho | Q3TYK4 1x | 2.26E-06 | 0 | 1 | 5 | 4 | Q3TYK4 | Putative ur | Q3TYK4 [2 | 0 | 2056.077 | 0 | 0.18 | 0.058087 | 9.05 | 92.4 | 107.6 | 6.33 | 9.36 | | High | High | High | High | High | High | High | High | High | High | High | 0 | 2.55E-07 | 2.91 |
| FALSE | High | EDELWASF | [K].EDELW | 1xPhospho | O88271 1x | 3.87E-05 | 0 | 1 | 1 | 1 | O88271 | Craniofacia | O88271 [1 | 0 | 2258.115 | 0 | -0.2 | 0.128286 | 8.38 | 111.7 | 88.3 | 27.26 | 19.03 | | High | High | High | High | High | High | High | High | High | High | High | 0 | 6.17E-06 | 3.24 |
| FALSE | High | EDENTEYS | [R].EDENTI | 1xPhospho | B2RSC8 1x | 0.000369 | 0 | 1 | 2 | 4 | B2RSC8 | E3 ubiquiti | B2RSC8 [3 | 0 | 3994.793 | 0 | -0.89 | 0.000406 | 1.99 | 126.5 | 73.5 | 7.12 | 9.2 | | High | High | High | High | High | High | High | High | High | High | High | 0 | 7.8E-05 | 3.75 |
| FALSE | High | EDEQSDLV | [R].EDEQSI | 1xPhospho | B2RQG2 1x | 0.000213 | 0 | 2 | 2 | 1 | B2RQG2; A | PHD finger | B2RQG2 [2 | 0 | 2616.233 | 0 | -0.68 | 0.058087 | 10.07 | 124.5 | 75.5 | 19.94 | 18.01 | Shared | High | High | High | High | High | High | High | High | High | High | High | 0 | 4.21E-05 | 3.52 |
| FALSE | High | EDFDSTSSS | [R].EDFDST | 1xPhospho | A2A432 1x | 8.76E-07 | 0 | 1 | 4 | 6 | A2A432 | Cullin-4B O | A2A432 [1 | 0 | 2046.885 | 0 | -1.27 | 0.00194 | 4.81 | 141.4 | 58.6 | 7.71 | 12.13 | | High | High | High | High | High | High | High | High | High | High | High | 7.19E-05 | 0.000817 | 2.93 |

| | | | | | | | | | | | | | | | | | | | | | | | | | | | | | | | | | | | | | | |
|---|---|---|---|---|---|---|---|---|---|---|---|---|---|---|---|---|---|---|---|---|---|---|---|---|---|---|---|---|---|---|---|---|---|---|---|---|---|---|
| FALSE | High | EDGGTVGV | [R].EDGGT | 1xPhospho | O35071 1x | 4.16E-05 | 0 | 1 | 4 | 1 | O35071 | Kinesin-like | O35071 [4 | 0 | 1730.877 | 0 | 1.03 | 0.000389 | 8.54 | 65.2 | 134.8 | 6.09 | 3.51 | | High | High | High | High | High | High | High | High | High | High | High | 0 | 6.72E-06 | 3.35 |
| FALSE | High | EDGHSESST | [K].EDGHS | 1xCarbami | Q6PCN7 1x | 1.36E-09 | 0 | 1 | 5 | 1 | Q6PCN7 | Helicase-lik | Q6PCN7 [3 | 0 | 2858.244 | 0 | -0.08 | 0.590539 | 18.52 | 99.9 | 100.1 | 7.02 | 11.45 | | High | High | High | High | High | High | High | High | High | High | High | 0 | 6.13E-11 | 4.37 |
| FALSE | High | EDGIDAVE | [R].EDGIDA | 1xPhospho | Q8K1J5 1x | 1.59E-06 | 0 | 1 | 1 | 9 | Q8K1J5 | Protein SD | Q8K1J5 [25 | 0 | 2163.028 | 0 | -1.5 | 0.000282 | 0.82 | 148 | 52 | 2.93 | 8.5 | | High | High | High | High | High | High | High | High | High | High | High | 0 | 2.23E-06 | 3.19 |
| FALSE | High | EDGQSMD | [R].EDGQS | 1xPhospho | E2JF22 1xP | 1.38E-06 | 0 | 1 | 6 | 1 | E2JF22 | Piezo-type | E2JF22 [75 | 0 | 2824.286 | 0 | 0.76 | 0.065986 | 44.73 | 78.1 | 121.9 | 19.04 | 25.9 | | High | High | High | High | High | High | High | High | High | High | High | 0 | 1.46E-07 | 4.54 |
| FALSE | High | EDGRESPS | [K].EDGRES | 1xPhospho | I1VGT4 1x | 0.001503 | 5.66E-05 | 2 | 19 | 1 | I1VGT4; I1 | Anion exch | I1VGT4 [84 | 1 | 2032.881 | 0 | 1.13 | 0.001255 | 0.4 | 63.3 | 136.7 | 8.58 | 8.16 | Shared | High | High | High | High | High | High | High | High | High | High | High | 2.58E-05 | 0.000376 | 2.58 |
| FALSE | High | EDEEEEDD | [K].EDEEEE | 1xPhospho | Q3TPJ8 1x | 0.000868 | 0 | 1 | 9 | 1 | Q3TPJ8 | Cytoplasmi | Q3TPJ8 [16 | 0 | 3307.56 | 0 | 0.3 | 0.061522 | 8.26 | 90.6 | 109.4 | 9.4 | 5.39 | | High | High | High | High | High | High | High | High | High | High | High | 0 | 0.000203 | 2.42 |
| FALSE | High | EDGVMSP | [K].EDGVM | 1xPhospho | Q99P72 1x | 0.105058 | 0.008518 | 1 | 2 | 2 | Q99P72 | Reticulon-4 | Q99P72 [3 | 0 | 1545.727 | 0 | -0.32 | 0.135083 | 17.43 | 112.4 | 87.6 | 20.86 | 25.57 | | High | High | High | High | High | High | High | High | High | High | High | 0.006622 | 0.07133 | 1.86 |
| FALSE | High | EDLAGPDI | [R].EDLAGI | 1xPhospho | Q9ERD6 1x | 8.76E-07 | 0 | 1 | 7 | 1 | Q9ERD6 | Ras-specifi | Q9ERD6 [2 | 0 | 1848.869 | 0 | -0.42 | 0.013347 | 5.69 | 114.4 | 85.6 | 6.35 | 7.69 | | High | High | High | High | High | High | High | High | High | High | High | 0 | 8.75E-08 | 4.4 |
| FALSE | High | EDLEAEEE | [K].EDLEAE | 1xCarbami | Q5QGZ6 1x | 1.13E-07 | 0 | 1 | 4 | 1 | Q5QGZ6 | Retinoic ac | Q5QGZ6 [8 | 0 | 4184.003 | 0 | 1.62 | 0.001134 | 18.5 | 48.4 | 151.6 | 18.02 | 14.3 | | High | High | High | High | High | High | High | High | High | High | High | 0 | 8.77E-09 | 6.2 |
| FALSE | High | EDLPPLTPP | [K].EDLPPL | 1xPhospho | B9EKJ7 1xP | 0.000529 | 0 | 1 | 2 | 4 | B9EKJ7 | Tankyrase | B9EKJ7 [12 | 0 | 1514.782 | 0 | 0.2 | 0.23021 | 11.25 | 95.8 | 104.2 | 12.16 | 8.07 | | High | High | High | High | High | High | High | High | High | High | High | 0 | 0.000117 | 3.82 |
| FALSE | High | EDLSDVTD | [R].EDLSDV | 2xPhospho | Q5U3K5 2x | 0.001558 | 5.66E-05 | 1 | 1 | 2 | Q5U3K5 | Rab-like pr | Q5U3K5 [5 | 0 | 2813.257 | 0 | 0 | 0.546191 | 5.07 | 100.4 | 99.6 | 1.74 | 5.6 | | High | High | High | High | High | High | High | High | High | High | High | 2.58E-05 | 0.000394 | 3.17 |
| FALSE | High | EDLSPICDH | [K].EDLSPI | 1xCarbami | Q06180 1x | 0.017192 | 0.001204 | 1 | 2 | 3 | Q06180 | Tyrosine-p | Q06180 [2 | 0 | 1879.821 | 0 | -0.31 | 0.004164 | 3.35 | 111.5 | 88.5 | 4.25 | 3.92 | | High | High | High | High | High | High | High | High | High | High | High | 0.000847 | 0.007977 | 2.52 |
| FALSE | High | EDLTDLEQ | [K].EDLTDL | 1xPhospho | Q6DFZ1 1x | 5.07E-05 | 0 | 1 | 6 | 2 | Q6DFZ1 | Golgi-speci | Q6DFZ1 [3 | 0 | 1739.805 | 0 | 0 | 0.634289 | 2.78 | 102.1 | 97.9 | 5.8 | 4.75 | | High | High | High | High | High | High | High | High | High | High | High | 0 | 8.34E-06 | 3.16 |
| FALSE | High | EDLTYCEHI | [R].EDLTYC | 1xCarbami | A0A1W2P6 | 4.6E-09 | 0 | 1 | 9 | 2 | A0A1W2P6 | Bromodom | A0A1W2P6 | 0 | 2681.179 | 0 | -1.66 | 0.000191 | 1.35 | 151.7 | 48.3 | 4.56 | 12.23 | | High | High | High | High | High | High | High | High | High | High | High | 0 | 2.42E-10 | 4.93 |
| FALSE | High | EDLVCSAA | [K].EDLVCS | 1xCarbami | Q99P72 1x | 0.000589 | 0 | 1 | 2 | 4 | Q99P72 | Reticulon-4 | Q99P72 [3 | 0 | 2819.405 | 0 | -0.1 | 0.067556 | 5.04 | 102.4 | 97.6 | 7.72 | 8.95 | | High | High | High | High | High | High | High | High | High | High | High | 0 | 0.000132 | 4.82 |
| FALSE | High | EDNLSLPG | [K].EDNLSL | 1xPhospho | Q571N6 1x | 1.56E-05 | 0 | 1 | 6 | 2 | Q571N6 | MFLJ00274 | Q571N6 [6 | 0 | 2231.054 | 0 | -1.43 | 0.002733 | 5.43 | 146.8 | 53.2 | 11.77 | 29.77 | | High | High | High | High | High | High | High | High | High | High | High | 0 | 2.22E-06 | 4.26 |
| FALSE | High | EDNSEGEE | [K].EDNSE | 1xCarbami | Q6PDQ2 1x | 1.94E-06 | 0 | 1 | 6 | 6 | Q6PDQ2 | Chromodo | Q6PDQ2 [4 | 0 | 3741.468 | 0 | -0.16 | 0.253264 | 36.05 | 103.2 | 96.8 | 16.25 | 28.11 | | High | High | High | High | High | High | High | High | High | High | High | 0 | 2.14E-07 | 4.73 |
| FALSE | High | EDNSEGEE | [K].EDNSE | 1xCarbami | Q6PDQ2 1x | 2.01E-07 | 0 | 1 | 6 | 14 | Q6PDQ2 | Chromodo | Q6PDQ2 [4 | 0 | 3757.463 | 0 | -0.4 | 0.005221 | 4.75 | 113.1 | 86.9 | 7.18 | 7.65 | | High | High | High | High | High | High | High | High | High | High | High | 0 | 1.57E-06 | 5.76 |
| FALSE | High | EDNSSSVE | [K].EDNSS | 1xPhospho | Q6ZQH0 1x | 0.000237 | 0 | 1 | 5 | 6 | Q6ZQH0 | MKIAA018 | Q6ZQH0 [6 | 0 | 1961.947 | 0 | 0.48 | 0.024998 | 10.75 | 83.7 | 116.3 | 11.37 | 12.29 | | High | High | High | High | High | High | High | High | High | High | High | 0 | 4.74E-05 | 3.18 |
| FALSE | High | EDNSSSVE | [K].EDNSS | 2xPhospho | Q6ZQH0 2x | 4.21E-06 | 0 | 1 | 5 | 3 | Q6ZQH0 | MKIAA018 | Q6ZQH0 [6 | 0 | 2041.913 | 0 | 0.09 | 0.181258 | 6.08 | 95.7 | 104.3 | 7.22 | 6.82 | | High | High | High | High | High | High | High | High | High | High | High | 0 | 5.1E-07 | 3.64 |
| FALSE | High | EDKSPSEES | [R].EDKSPS | 1xPhospho | P54726 1x | 0.057203 | 0.004227 | 2 | 4 | 1 | P54726; Q | UV excision | P54726 [12 | 1 | 3508.622 | 0 | -0.06 | 0.722999 | 12.75 | 99.2 | 100.8 | 9.23 | 12.99 | Shared | High | High | High | High | High | High | High | High | High | High | High | 0.002159 | 0.02251 | 3.62 |
| FALSE | High | EDPGQVSS | [R].EDPGQ | 1xPhospho | Q69ZC8 1x | 1.57E-06 | 0 | 1 | 1 | 1 | Q69ZC8 | GPALPP mo | Q69ZC8 [1 | 0 | 3506.503 | 0 | -0.5 | 0.131088 | 26.54 | 114.8 | 85.2 | 16.86 | 18.75 | | High | High | High | High | High | High | High | High | High | High | High | 0 | 1.69E-07 | 3.78 |
| FALSE | High | EDEDDAED | [K].EDEDD | 1xPhospho | Q571C7 1x | 1.08E-05 | 0 | 1 | 3 | 2 | Q571C7 | Transcripti | Q571C7 [1 | 0 | 3758.568 | 0 | -0.4 | 0.022596 | 4.92 | 113.4 | 86.6 | 4.6 | 10.57 | | High | High | High | High | High | High | High | High | High | High | High | 0 | 1.48E-06 | 4.73 |
| FALSE | High | EDDPLKSP | [K].EDDPLK | 1xCarbami | Q8BHK9 1x | 6.93E-07 | 0 | 1 | 3 | 1 | Q8BHK9 | DNA excisi | Q8BHK9 [9 | 1 | 3581.655 | 0 | -0.86 | 0.001006 | 3.1 | 127.9 | 72.1 | 8.8 | 7.84 | | High | High | High | High | High | High | High | High | High | High | High | 0 | 6.73E-08 | 3.63 |
| FALSE | High | ECADAASP | [K].ECADA | 1xCarbami | E9PWI3 1x | 0.001107 | 0 | 1 | 3 | 1 | E9PWI3 | Mutated in | E9PWI3 [8 | 0 | 1982.946 | 0 | 0.82 | 0.008712 | 28.16 | 72.4 | 127.6 | 16.65 | 10.28 | | High | High | High | High | High | High | High | High | High | High | High | 0 | 0.000268 | 2.51 |
| FALSE | High | ECDSFDLE | [K].ECDSFD | 1xCarbami | A8C756 1x | 3.08E-07 | 0 | 1 | 1 | 4 | A8C756 | Thyroid ad | A8C756 [1 | 0 | 3185.449 | 0 | 0.88 | 0.003071 | 12.8 | 69.2 | 130.8 | 9.24 | 8.76 | | High | High | High | High | High | High | High | High | High | High | High | 0 | 2.71E-08 | 5.02 |
| FALSE | High | ECDSVSAV | [K].ECDSVS | 1xCarbami | B2RRE7 1x | 0.017809 | 0.001282 | 1 | 2 | 1 | B2RRE7 | OTU doma | B2RRE7 [8 | 0 | 2005.841 | 0 | 0.66 | 0.122158 | 12.58 | 77.2 | 122.8 | 11.33 | 22.08 | | High | High | High | High | High | High | High | High | High | High | High | 0.000689 | 0.006 | 3.85 |
| FALSE | High | ECELSPGV | [R].ECELSP | 1xCarbami | Q99MR6 1 | 0.000589 | 0 | 1 | 5 | 1 | Q99MR6 | Serrate RN | Q99MR6 [4 | 0 | 1469.666 | 0 | -0.47 | 0.049634 | 6.6 | 112.6 | 87.4 | 8.66 | 10.66 | | High | High | High | High | High | High | High | High | High | High | High | 0 | 0.000132 | 1.73 |
| FALSE | High | ECESPEVD | [R].ECESPE | 1xCarbami | Q63943-2 | 0.002623 | 0.000109 | 1 | 2 | 1 | Q63943-2 | Isoform M | Q63943-2 | 0 | 2847.305 | 0 | -0.31 | 0.943683 | 10.28 | 99.7 | 100.3 | 17.82 | 28.45 | | High | High | High | High | High | High | High | High | High | High | High | 4.94E-05 | 0.000706 | 3.82 |
| FALSE | High | ECLGRPDS | [R].ECLGRI | 1xCarbami | Q9ESK9 1x | 0.005565 | 0.00062 | 1 | 6 | 1 | Q9ESK9 | RB1-induci | Q9ESK9 [2 | 0 | 2495.164 | 0 | 0.15 | 0.94233 | 13.42 | 97.5 | 102.5 | 9.66 | 11.49 | | High | High | High | High | High | High | High | High | High | High | High | 0.000275 | 0.001637 | 2.88 |
| FALSE | High | ECLSGLDS | [R].ECLSGL | 1xCarbami | Q3USC0 1x | 1.21E-05 | 0 | 1 | 3 | 2 | Q3USC0 | Putative ur | Q3USC0 [1 | 0 | 2362.071 | 0 | -0.77 | 0.000565 | 0.92 | 126 | 74 | 3.94 | 4 | | High | High | High | High | High | High | High | High | High | High | High | 0 | 1.67E-06 | 4.46 |
| FALSE | High | ECLSGLDS | [R].ECLSGL | 1xCarbami | Q3USC0 1x | 1.49E-05 | 0 | 1 | 3 | 3 | Q3USC0 | Putative ur | Q3USC0 [1 | 0 | 2378.066 | 0 | -0.24 | 0.008872 | 2.09 | 107.3 | 92.7 | 4.03 | 2.15 | | High | High | High | High | High | High | High | High | High | High | High | 0 | 2.12E-06 | 4.16 |
| FALSE | High | ECISSPSGD | [R].ECISSPS | 1xCarbami | G3X8S2 1x | 0.004867 | 0.000504 | 1 | 2 | 3 | G3X8S2 | BRF1 homo | G3X8S2 [4 | 0 | 1714.812 | 0 | 0.28 | 0.076546 | 4.76 | 91.9 | 108.1 | 7.43 | 6.61 | | High | High | High | High | High | High | High | High | High | High | High | 0.000246 | 0.001458 | 2.22 |
| FALSE | High | ECPDQLGP | [K].ECPDQ | 1xCarbami | Q61026 1x | 0.001158 | 0 | 1 | 8 | 1 | Q61026 | Nuclear rec | Q61026 [2 | 0 | 1765.86 | 0 | -0.38 | 0.001811 | 3.71 | 111.4 | 88.6 | 2.6 | 5.2 | | High | High | High | High | High | High | High | High | High | High | High | 0 | 0.000283 | 3.48 |
| FALSE | High | ECPDVSPE | [R].ECPDVS | 1xCarbami | D0VLQ4 1x | 0.004824 | 0.000456 | 1 | 4 | 1 | D0VLQ4 | Tet oncoge | D0VLQ4 [1 | 0 | 2345.079 | 0 | -1.01 | 0.000285 | 3.88 | 133.6 | 66.4 | 2.93 | 11.83 | | High | High | High | High | High | High | High | High | High | High | High | 0.000204 | 0.001387 | 1.69 |
| FALSE | High | ECPGIQDTI | [K].ECPGIC | 1xCarbami | Q8BHB4 1x | 1.41E-06 | 0 | 1 | 1 | 3 | Q8BHB4 | WD repeat | Q8BHB4 [2 | 0 | 3003.307 | 0 | -0.4 | 0.122113 | 18.32 | 112 | 88 | 3.36 | 26.96 | | High | High | High | High | High | High | High | High | High | High | High | 0 | 1.5E-07 | 4.13 |
| FALSE | High | ECPGIQDTI | [K].ECPGIC | 1xCarbami | Q8BHB4 2x | 0.001021 | 0 | 1 | 1 | 1 | Q8BHB4 | WD repeat | Q8BHB4 [2 | 0 | 3083.273 | 0 | -0.67 | 0.003309 | 5.37 | 122.7 | 77.3 | 6.34 | 7.75 | | High | High | High | High | High | High | High | High | High | High | High | 0 | 0.000245 | 3.4 |
| FALSE | High | EDEASTDV | [K].EDEAST | 1xPhospho | P53986 1x | 3.06E-05 | 0 | 1 | 5 | 7 | P53986 | Monocarbo | P53986 [4 | 0 | 2230.11 | 0 | -0.01 | 0.567198 | 13.12 | 99.6 | 100.4 | 11.82 | 13.23 | | High | High | High | High | High | High | High | High | High | High | High | 0.000382 | 0.003292 | 3.4 |
| FALSE | High | ECSSDESVI | [K].ECSSDE | 1xCarbami | Q569L8 2x | 3.73E-05 | 0 | 1 | 1 | 3 | Q569L8 | Centromer | Q569L8 [7 | 0 | 2428.076 | 0 | -0.56 | 0.005351 | 7.47 | 118.9 | 81.1 | 9.57 | 4.44 | | High | High | High | High | High | High | High | High | High | High | High | 0 | 5.94E-06 | 2.99 |
| FALSE | High | ECSTDLSI | [K].ECSTDL | 1xCarbami | Q6RHR9 1x | 0.012959 | 0.000889 | 1 | 4 | 1 | Q6RHR9 | Membrane | Q6RHR9 [1 | 0 | 1233.527 | 0 | 0.07 | 0.848529 | 22.51 | 96.5 | 103.5 | 10.3 | 13.17 | | High | High | High | High | High | High | High | High | High | High | High | 0.000449 | 0.004191 | 1.87 |
| FALSE | High | ECSVEK | [K].ECSVEK | 1xCarbami | F6RDI8 1x | 0.091627 | 0.007146 | 1 | 7 | 1 | F6RDI8 | Serine/argi | F6RDI8 [47 | 0 | 1289.621 | 0 | -0.49 | 0.006536 | 7.21 | 116.7 | 83.3 | 6.75 | 9 | | High | High | High | High | High | High | High | High | High | High | High | 0.003659 | 0.03909 | 1.51 |
| FALSE | High | EDALDDSV | [R].EDALDI | 1xPhospho | Q9ERU9 1x | 3.93E-07 | 0 | 1 | 1 | 1 | Q9ERU9 | E3 SUMO-p | Q9ERU9 [2 | 0 | 2722.277 | 0 | 0.33 | 0.15054 | 28.31 | 88.2 | 111.8 | 10.23 | 18.89 | | High | High | High | High | High | High | High | High | High | High | High | 0 | 3.54E-08 | 5.51 |
| FALSE | High | EDALDDSV | [R].EDALDI | 3xPhospho | Q9ERU9 3x | 0.000712 | 0 | 1 | 1 | 1 | Q9ERU9 | E3 SUMO-p | Q9ERU9 [2 | 0 | 2882.21 | 0 | 0.47 | 0.055104 | 24.43 | 84.6 | 115.4 | 9.52 | 12.32 | | High | High | High | High | High | High | High | High | High | High | High | 0 | 0.000163 | 5.92 |
| FALSE | High | EDALDDSV | [R].EDALDI | 1xPhospho | Q9ERU9 1x | 0.008691 | 0.000736 | 1 | 1 | 2 | Q9ERU9 | E3 SUMO-p | Q9ERU9 [2 | 1 | 3079.535 | 0 | -0.22 | 0.145945 | 21.3 | 107.1 | 92.9 | 10.13 | 16.24 | | High | High | High | High | High | High | High | High | High | High | High | 0.000327 | 0.00268 | 3.2 |
| FALSE | High | EDALDDSV | [R].EDALDI | 2xPhospho | Q9ERU9 2x | 0.087983 | 0.006639 | 1 | 1 | 1 | Q9ERU9 | E3 SUMO-p | Q9ERU9 [2 | 1 | 3159.501 | 0 | 0.01 | 0.702264 | 22.41 | 98.1 | 101.9 | 5.07 | 17.1 | | High | High | High | High | High | High | High | High | High | High | High | 0.003402 | 0.0372 | 3.26 |
| FALSE | High | EDALLYQSI | [K].EDALLY | 1xPhospho | A2AM95 1 | 0.016743 | 0.001166 | 1 | 4 | 1 | A2AM95 | Emerin OS | A2AM95 [8 | 0 | 1604.834 | 0 | 0.11 | 0.127232 | 2.92 | 95.8 | 104.2 | 14.45 | 5.7 | | High | High | High | High | High | High | High | High | High | High | High | 0.000638 | 0.005611 | 2.15 |
| FALSE | High | EDAPPEDK | [R].EDAPPE | 1xPhospho | Q99L43 1x | 0.072258 | 0.005346 | 1 | 2 | 1 | Q99L43 | Phosphatic | Q99L43 [1 | 1 | 2428.174 | 0 | 1.07 | 0.000533 | 1.76 | 64 | 136 | 9.49 | 9.73 | | High | High | High | High | High | High | High | High | High | High | High | 0.002748 | 0.02938 | 1.92 |
| FALSE | High | EDASQSIQ | [R].EDASQ | 1xCarbami | Q0VGT4 1x | 4.87E-10 | 0 | 1 | 3 | 2 | Q0VGT4 | Protein ZG | Q0VGT4 [6 | 0 | 2657.264 | 0 | 0.12 | 0.430334 | 15.85 | 96 | 104 | 7.94 | 6.06 | | High | High | High | High | High | High | High | High | High | High | High | 0 | 1.94E-11 | 5.15 |
| FALSE | High | EDDDVIVN | [K].EDDDV | 1xPhospho | H9KV00 1x | 1.64E-05 | 0 | 1 | 9 | 2 | H9KV00 | Protein SO | H9KV00 [2 | 0 | 4144.978 | 0 | 0.04 | 0.947039 | 16.47 | 97.3 | 102.7 | 4.37 | 12.77 | | High | High | High | High | High | High | High | High | High | High | High | 0 | 2.37E-06 | 4.79 |
| FALSE | High | EDDGDSCV | [K].EDDGD | 2xCarbami | Q5SV77-3 | 3.89E-07 | 0 | 1 | 6 | 4 | Q5SV77-3 | Isoform 3 o | Q5SV77-3 | 0 | 2909.201 | 0 | -0.04 | 0.372338 | 7.4 | 103.1 | 96.9 | 4.42 | 5.3 | | High | High | High | High | High | High | High | High | High | High | High | 0 | 3.53E-08 | 4.39 |
| FALSE | High | EDDLDVVL | [K].EDDLD | 1xPhospho | Q9EST3 1x | 0.019618 | 0.001438 | 1 | 4 | 1 | Q9EST3 | Eukaryotic | Q9EST3 [1 | 0 | 1694.82 | 0 | -0.02 | 0.468272 | 16.28 | 101 | 99 | 5.16 | 15.71 | | High | High | High | High | High | High | High | High | High | High | High | 0.000756 | 0.006658 | 2.73 |
| FALSE | High | EDDPAQDS | [R].EDDPA | 1xPhospho | P13405 1x | 1.32E-05 | 0 | 1 | 4 | 2 | P13405 | Retinoblast | P13405 [24 | 0 | 2147.97 | 0 | 1.7 | 0.000177 | 4.73 | 47.2 | 152.8 | 8.71 | 6.04 | | High | High | High | High | High | High | High | High | High | High | High | 0 | 1.84E-06 | 2.89 |
| FALSE | High | ECSSSSPVA | [R].ECSSSS | 1xCarbami | Q3UHE0 1x | 0.002844 | 0.00016 | 1 | 4 | 3 | Q3UHE0 | Putative ur | Q3UHE0 [1 | 0 | 2568.068 | 0 | -0.38 | 0.01835 | 6.61 | 112.7 | 87.3 | 5.65 | 13.66 | | High | High | High | High | High | High | High | High | High | High | High | 7.19E-05 | 0.000773 | 3.16 |
| FALSE | High | EDPGQVSS | [R].EDPGQ | 1xDeamida | Q69ZC8 1x | 0.002096 | 5.66E-05 | 1 | 1 | 5 | Q69ZC8 | GPALPP mo | Q69ZC8 [1 | 0 | 3507.487 | 0 | -0.17 | 0.703596 | 31.84 | 102.2 | 97.8 | 17 | 20.44 | | High | High | High | High | High | High | High | High | High | High | High | 2.58E-05 | 0.000545 | 4.74 |
| FALSE | High | EDPGQVSS | [R].EDPGQ | 1xPhospho | Q69ZC8 1x | 1.3E-05 | 0 | 1 | 1 | 1 | Q69ZC8 | GPALPP mo | Q69ZC8 [1 | 0 | 3522.498 | 0 | -0.37 | 0.589352 | 18.32 | 107.3 | 92.7 | 10.89 | 37.14 | | High | High | High | High | High | High | High | High | High | High | High | 0 | 1.81E-06 | 3.22 |
| FALSE | High | EDPGQVSS | [R].EDPGQ | 1xPhospho | Q69ZC8 1x | 0.002719 | 0.000109 | 1 | 1 | 1 | Q69ZC8 | GPALPP mo | Q69ZC8 [1 | 0 | 3735.666 | 0 | -0.02 | 0.332338 | 18.47 | 113.9 | 86.1 | 19.61 | 27.72 | | High | High | High | High | High | High | High | High | High | High | High | 4.94E-05 | 0.00073 | 4.36 |
| FALSE | High | EDTAFSDS | [K].EDTAFS | 1xCarbami | E9Q444 1x | 3.49E-07 | 0 | 1 | 10 | 6 | E9Q444 | Zinc finger | E9Q444 [8 | 0 | 3876.693 | 0 | 1.09 | 0.007281 | 8.46 | 64.6 | 135.4 | 9.7 | 14.68 | | High | High | High | High | High | High | High | High | High | High | High | 0 | 1.38E-06 | 4.12 |
| FALSE | High | EDTDQEEK | [K].EDTDQ | 1xPhospho | Q99JF8 1x | 0.042141 | 0.003162 | 1 | 4 | 1 | Q99JF8 | PC4 and SF | Q99JF8 [12 | 0 | 1531.693 | 0 | -0.64 | 0.039094 | 5.84 | 116.4 | 83.6 | 16.63 | 13.74 | | High | High | High | High | High | High | High | High | High | High | High | 0.001639 | 0.01589 | 2.07 |
| FALSE | High | EDTVVDNT | [K].EDTVVI | 1xPhospho | Q7TT18 1x | 2.36E-05 | 0 | 1 | 4 | 3 | Q7TT18 | Activating | Q7TT18 [4 | 0 | 3060.462 | 0 | -0.41 | 0.198731 | 11.68 | 114 | 86 | 19.96 | 27.25 | | High | High | High | High | High | High | High | High | High | High | High | 0 | 3.56E-06 | 3.88 |
| FALSE | High | EDVDPHSG | [K].EDVDPI | 1xPhospho | Q9DAT2 1x | 5.33E-07 | 0 | 1 | 2 | 1 | Q9DAT2 | MRG/MOR | Q9DAT2 [1 | 0 | 2644.218 | 0 | 0.2 | 0.20676 | 14.02 | 94.3 | 105.7 | 3.96 | 11.6 | | High | High | High | High | High | High | High | High | High | High | High | 0 | 5.02E-08 | 4.18 |
| FALSE | High | EDVEGVGT | [K].EDVEG | 1xPhospho | Q99JP4 1x | 0.000389 | 0 | 1 | 1 | 2 | Q99JP4 | Anaphase- | Q99JP4 [34 | 0 | 2615.213 | 0 | -0.14 | 0.147947 | 9.14 | 105.5 | 94.5 | 3.09 | 14.68 | | High | High | High | High | High | High | High | High | High | High | High | 0 | 8.29E-05 | 3.62 |
| FALSE | High | EDVGSLILD | [K].EDVGSI | 1xCarbami | A0A1W2P7 | 0.001898 | 5.66E-05 | 1 | 1 | 2 | A0A1W2P7 | Uncharacte | A0A1W2P7 | 0 | 3849.749 | 0 | -1.34 | 0.015997 | 9.4 | 141.1 | 58.9 | 12.45 | 38.33 | | High | High | High | High | High | High | High | High | High | High | High | 2.58E-05 | 0.000488 | 3.2 |
| FALSE | High | EDVPAFGP | [R].EDVPAI | 1xPhospho | Q5SVL6 1x | 0.000616 | 0 | 1 | 3 | 1 | Q5SVL6 | Rap1 GTPa | Q5SVL6 [3 | 0 | 2460.298 | 0 | 3.1 | 0.00034 | 23.31 | 19.9 | 180.1 | 15.86 | 8.68 | | High | High | High | High | High | High | High | High | High | High | High | 0 | 0.000138 | 3.79 |
| FALSE | High | EDVSNFDD | [-R].EDVSN | 1xPhospho | Q8BWW9 | 0.054844 | 0.003982 | 1 | 6 | 1 | Q8BWW9 | Serine/thre | Q8BWW9 | 0 | 2688.228 | 0 | -0.74 | 0.002367 | 2.31 | 125.1 | 74.9 | 7.62 | 6 | | High | High | High | High | High | High | High | High | High | High | High | 0.002059 | 0.02139 | 2.25 |
| FALSE | High | EDVSNFDD | [-R].EDVSN | 1xPhospho | Q8BWW9 | 0.000284 | 0 | 1 | 6 | 1 | Q8BWW9 | Serine/thre | Q8BWW9 | 1 | 3070.424 | 0 | -0.04 | 0.729662 | 19.56 | 101.9 | 98.1 | 15.72 | 8.88 | | High | High | High | High | High | High | High | High | High | High | High | 0 | 5.81E-05 | 3.3 |
| FALSE | High | EDVVQVLS | [R].EDVVQ | 1xPhospho | Q8CJ26 1x | 0.001689 | 5.66E-05 | 1 | 2 | 2 | Q8CJ26 | Death dom | Q8CJ26 [2 | 0 | 2040.994 | 0 | 0.62 | 0.057609 | 42.92 | 73.7 | 126.3 | 13.98 | 22.24 | | High | High | High | High | High | High | High | High | High | High | High | 2.58E-05 | 0.000431 | 3.47 |
| FALSE | High | EDYFEPISP | [R].EDYFEF | 1xPhospho | E9PZY8 1x | 0.007881 | 0.000736 | 1 | 4 | 2 | E9PZY8 | RIKEN cDN | E9PZY8 [2 | 0 | 1676.741 | 0 | -0.4 | 0.00096 | 4.11 | 115.2 | 84.8 | 7 | 7.76 | | High | High | High | High | High | High | High | High | High | High | High | 0.000706 | 0.006229 | 2.22 |
| FALSE | High | EDYPHTHG | [K].EDYPHT | 1xDeamida | G3X8R8 1x | 3.01E-05 | 0 | 1 | 4 | 4 | G3X8R8 | MCG12388 | G3X8R8 [1 | 0 | 4903.106 | 0 | 1.63 | 0.008365 | 56.15 | 48.3 | 151.7 | 37.8 | 18.04 | | High | High | High | High | High | High | High | High | High | High | High | 0 | 7.06E-06 | 6.24 |
| FALSE | High | EEAAEAAA | [K].EEAAEA | 1xDeamida | Q9WV35 1 | 0.017041 | 0.001204 | 1 | 1 | 1 | Q9WV35 | C->U-editir | Q9WV35 [ | 0 | 3152.42 | 0 | -2.87 | | 0.52 | 173.4 | 26.6 | 14.81 | 8.49 | | High | High | High | High | High | High | High | Not Found | High | High | High | 0.000638 | 0.005705 | 3.02 |
| FALSE | High | EDSVVAET | [R].EDSVVA | 1xDeamida | Q6ZQ32 1x | 0.000259 | 0 | 2 | 7 | 1 | Q6ZQ32; F | MKIAA085 | Q6ZQ32 [7 | 0 | 2422.097 | 0 | 0.34 | 0.024213 | 8.39 | 87 | 113 | 4.08 | 10.44 | Shared | High | High | High | High | High | High | High | High | High | High | High | 0 | 5.26E-05 | 3.67 |
| FALSE | High | EEAAEVAA | [R].EEAAEV | 1xPhospho | Q8BLH7 1x | 4.55E-05 | 0 | 1 | 1 | 2 | Q8BLH7 | HIRA-intera | Q8BLH7 [5 | 0 | 2578.308 | 0 | -1.25 | 0.000445 | 0.75 | 140.6 | 59.4 | 7.54 | 7.31 | | High | High | High | High | High | High | High | High | High | High | High | 0.001183 | 0.01277 | 3.13 |
| FALSE | High | EEAAPPTPA | [R].EEAAPF | 1xPhospho | Q80WJ7 1x | 3.03E-05 | 0 | 1 | 3 | 1 | Q80WJ7 | Protein LYF | Q80WJ7 [9 | 0 | 2301.178 | 0 | 1.77 | 0.00319 | 29.97 | 49.3 | 150.7 | 9.26 | 17.25 | | High | High | High | High | High | High | High | High | High | High | High | 0 | 4.7E-06 | 2.47 |
| FALSE | High | EEAASPLK | [K].EEAASF | 1xPhospho | Q80TZ9 1x | 0.008388 | 0.000736 | 1 | 2 | 2 | Q80TZ9 | Arginine-gl | Q80TZ9 [6 | 0 | 1382.733 | 0 | -0.15 | 0.037194 | 2.43 | 106.8 | 93.2 | 5.28 | 5.57 | | High | High | High | High | High | High | High | High | High | High | High | 0.000399 | 0.003458 | 2.1 |
| FALSE | High | EEALASWE | [K].EEALAS | 1xCarbami | Q6P1D7 1x | 1.81E-05 | 0 | 1 | 2 | 2 | Q6P1D7 | Structure-s | Q6P1D7 [7 | 0 | 3507.593 | 0 | -0.4 | 0.156736 | 14.19 | 113.4 | 86.6 | 13.86 | 13.32 | | High | High | High | High | High | High | High | High | High | High | High | 0 | 2.63E-06 | 3.69 |
| FALSE | High | EEALDEAE | [K].EEALDE | 3xPhospho | Q80TZ9 3x | 0.002441 | 0.000109 | 1 | 3 | 2 | Q80TZ9 | Arginine-gl | Q80TZ9 [1 | 1 | 4488.936 | 0 | 0.17 | 0.432379 | 9.38 | 97.8 | 102.2 | 11.58 | 14.84 | | High | High | High | High | High | High | High | High | High | High | High | 4.94E-05 | 0.000652 | 4.73 |
| FALSE | High | EEAIHPSLG | [R].EEAIHP | 1xPhospho | Q6P5H2-2 | 0.026191 | 0.001676 | 1 | 2 | 1 | Q6P5H2-2 | Isoform 2 o | Q6P5H2-2 | 0 | 2063.046 | 0 | 2.71 | 0.007032 | 26.05 | 23.1 | 176.9 | 20.71 | 32.3 | | High | High | High | High | High | High | High | High | High | High | High | 0.000903 | 0.009254 | 2.39 |
| FALSE | High | EEALQNISA | [R].EEALQI | 1xPhospho | Q6PAR5 1x | 0.000171 | 0 | 1 | 7 | 1 | Q6PAR5 | GTPase-act | Q6PAR5 [1 | 0 | 3542.591 | 0 | 0.13 | 0.209669 | 19.18 | 92.5 | 107.5 | 11.92 | 18.99 | | High | High | High | High | High | High | High | High | High | High | High | 0 | 3.28E-05 | 3.66 |
| FALSE | High | EEALSDGD | [K].EEALSD | 1xCarbami | P51612 1x | 0.000529 | 0 | 1 | 3 | 1 | P51612 | DNA repair | P51612 [8 | 1 | 2565.122 | 0 | 0.12 | 0.787847 | 22.36 | 97.7 | 102.3 | 7.71 | 11.34 | | High | High | High | High | High | High | High | High | High | High | High | 0 | 0.000117 | 1.22 |
| FALSE | High | EEAPASPLF | [K].EEAPAS | 1xPhospho | P42128 1x | 5.12E-05 | 0 | 1 | 1 | 2 | P42128 | Forkhead b | P42128 [1 | 0 | 2644.452 | 0 | 1.26 | 0.000546 | 4.13 | 56.6 | 143.4 | 2.94 | 7.62 | | High | High | High | High | High | High | High | High | High | High | High | 0 | 8.44E-06 | 3.02 |
| FALSE | High | EEAPASPLF | [K].EEAPAS | 2xPhospho | P42128 2x | 1.47E-05 | 0 | 1 | 1 | 2 | P42128 | Forkhead b | P42128 [1 | 0 | 2724.418 | 0 | 0.92 | 0.000439 | 7.04 | 67.2 | 132.8 | 4.46 | 5.86 | | High | High | High | High | High | High | High | High | High | High | High | 0 | 2.09E-06 | 2.15 |
| FALSE | High | EEATSNSP | [K].EEATSN | 1xPhospho | D3YZK4 1x | 0.000718 | 0 | 1 | 5 | 2 | D3YZK4 | Transcripti | D3YZK4 [5 | 0 | 1587.767 | 0 | -0.77 | 0.001341 | 0.84 | 124.1 | 75.9 | 2.72 | 9.24 | | High | High | High | High | High | High | High | High | High | High | High | 0 | 0.000165 | 2.37 |
| FALSE | High | EEATSTSEI | [R].EEATST | 1xPhospho | A0A068F1 | 5.68E-07 | 0 | 1 | 11 | 2 | A0A068F1 | Glyco-gag | A0A068F1 | 0 | 2180.069 | 0 | 3.18 | 0.00014 | 33.33 | 19.1 | 180.9 | 26.22 | 3.09 | | High | High | High | High | High | High | High | High | High | High | High | 0 | 5.39E-08 | 3.01 |
| FALSE | High | EEATYGVW | [K].EEATYG | 1xPhospho | Q3UGE9 1x | 0.001222 | 0 | 1 | 4 | 2 | Q3UGE9 | Tuftelin-in | Q3UGE9 [4 | 1 | 3153.457 | 0 | -0.03 | 0.604514 | 5.85 | 100.2 | 99.8 | 6.15 | 11.71 | | High | High | High | High | High | High | High | High | High | High | High | 0 | 0.000299 | 3.01 |
| FALSE | High | EEAVQIPSF | [K].EEAVQI | 1xPhospho | B2RUG8 1x | 0.002869 | 0.00016 | 1 | 5 | 1 | B2RUG8 | Asxl2 prote | B2RUG8 [4 | 0 | 1850.967 | 0 | 0.19 | 0.035417 | 8.23 | 94.9 | 105.1 | 3.82 | 9.46 | | High | High | High | High | High | High | High | High | High | High | High | 7.19E-05 | 0.000778 | 2.06 |
| FALSE | High | EEAAGQQ | [R].EEAAG | 1xDeamida | D6RI76 1x | 0.028824 | 0.001829 | 1 | 1 | 1 | D6RI76 | Pre-rRNA-p | D6RI76 [37 | 0 | 2977.28 | 0 | -0.62 | 0.23258 | 30.4 | 149.8 | 50.2 | 104.2 | 131.16 | | High | High | High | High | High | High | High | High | High | High | High | 0.000997 | 0.01029 | 2.04 |
| FALSE | High | EDSVVAET | [R].EDSVVA | 1xPhospho | Q6ZQ32 1x | 2.77E-08 | 0 | 2 | 7 | 5 | Q6ZQ32; F | MKIAA085 | Q6ZQ32 [7 | 0 | 2421.113 | 0 | 0.6 | 0.035062 | 2.4 | 82 | 118 | 7.88 | 10.79 | Shared | High | High | High | High | High | High | High | High | High | High | High | 0 | 1.82E-09 | 4.6 |
| FALSE | High | EDSVSSASF | [K].EDSVSS | 1xPhospho | F8VQG7 1x | 2.72E-05 | 0 | 1 | 4 | 2 | F8VQG7 | NEDD4-bin | F8VQG7 [5 | 0 | 1544.761 | 0 | -0.46 | 0.001741 | 2.91 | 113.9 | 86.1 | 2.97 | 5.39 | | High | High | High | High | High | High | High | High | High | High | High | 0 | 4.16E-06 | 3.48 |
| FALSE | High | EDSSSTEFV | [K].EDSSST | 1xPhospho | Q3U5L3 1x | 0.001211 | 0 | 1 | 3 | 2 | Q3U5L3 | Putative ur | Q3U5L3 [2 | 0 | 1795.84 | 0 | 1.44 | 0.000609 | 2.99 | 54.8 | 145.2 | 8.85 | 9.46 | | High | High | High | High | High | High | High | High | High | High | High | 4.94E-05 | 0.00058 | 2.63 |
| FALSE | High | EDPGQVSS | [R].EDPGQ | 1xPhospho | Q69ZC8 1x | 6.56E-07 | 0 | 1 | 1 | 1 | Q69ZC8 | GPALPP mo | Q69ZC8 [1 | 0 | 3751.661 | 0 | -0.23 | 0.953807 | 29.46 | 107.3 | 92.7 | 25.47 | 54.7 | | High | High | High | High | High | High | High | High | High | High | High | 0 | 6.36E-08 | 3.85 |
| FALSE | High | EDPLLTPVF | [K].EDPLLT | 1xPhospho | P63213 1x | 0.054844 | 0.003982 | 1 | 1 | 1 | P63213 | Guanine nu | P63213 [47 | 0 | 2091.036 | 0 | -1.61 | 0.000209 | 0.87 | 150.2 | 49.8 | 5.96 | 8.32 | | High | High | High | High | High | High | High | High | High | High | High | 0.002059 | 0.02142 | 2.04 |
| FALSE | High | EDPWQEEI | [K].EDPWC | 1xPhospho | H7BX94 1x | 0.007471 | 0.00062 | 1 | 2 | 1 | H7BX94 | Lysine-rich | H7BX94 [3 | 1 | 3314.617 | 0 | -0.47 | 0.026625 | 5.59 | 117.6 | 82.4 | 10.68 | 14.7 | | High | High | High | High | High | High | High | High | High | High | High | 0.000275 | 0.002255 | 2.88 |
| FALSE | High | EDQCLTPE | [K].EDQCL | 1xCarbami | O55106 1x | 2.43E-07 | 0 | 1 | 5 | 1 | O55106 | Striatin OS | O55106 [3 | 0 | 2727.31 | 0 | 2.02 | 0.001225 | 18.52 | 36.7 | 163.3 | 11.85 | 12.32 | | High | High | High | High | High | High | High | High | High | High | High | 0 | 2.07E-08 | 4.11 |
| FALSE | High | EDQFNGSP | [K].EDQFN | 1xPhospho | E9PZY8 1x | 3.73E-05 | 0 | 1 | 4 | 1 | E9PZY8 | RIKEN cDN | E9PZY8 [16 | 0 | 1933.912 | 0 | -0.6 | 0.000297 | 2.7 | 120.7 | 79.3 | 1.54 | 5.08 | | High | High | High | High | High | High | High | High | High | High | High | 0 | 5.95E-06 | 0.99 |
| FALSE | High | EDQFNGSP | [K].EDQFN | 1xDeamida | E9PZY8 1x | 0.043607 | 0.003228 | 1 | 4 | 2 | E9PZY8 | RIKEN cDN | E9PZY8 [16 | 0 | 1934.896 | 0 | -0.48 | 0.000618 | 2.53 | 114.8 | 85.2 | 3.49 | 4.8 | | High | High | High | High | High | High | High | High | High | High | High | 0.001651 | 0.01643 | 1.48 |
| FALSE | High | EDSARPGA | [R].EDSARI | 1xPhospho | Q9CX86 1x | 0.001659 | 5.66E-05 | 1 | 1 | 1 | Q9CX86 | Heterogen | Q9CX86 [8 | 0 | 1676.852 | 0 | 0.5 | 0.040129 | 17.48 | 83.8 | 116.2 | 4.08 | 16.91 | | High | High | High | High | High | High | High | High | High | High | High | 2.58E-05 | 0.000423 | 1.24 |
| FALSE | High | EDSCGEGN | [K].EDSCGI | 1xCarbami | G3X9T5 1x | 0.006363 | 0.00062 | 1 | 7 | 1 | G3X9T5 | MCG12342 | G3X9T5 [7 | 1 | 2804.227 | 0 | 0.24 | 0.417962 | 3.18 | 93.1 | 106.9 | 2.87 | 16.78 | | High | High | High | High | High | High | High | High | High | High | High | 0.000275 | 0.001888 | 2.12 |
| FALSE | High | EDSCSGSD | [K].EDSCSG | 1xCarbami | Q6P1G0 2x | 1.95E-05 | 0 | 1 | 1 | 1 | Q6P1G0 | HEAT repe | Q6P1G0 [6 | 0 | 2766.051 | 0 | 0.19 | 0.213721 | 34.6 | 97.8 | 102.2 | 39.72 | 11.47 | | High | High | High | High | High | High | High | High | High | High | High | 0 | 2.87E-06 | 4.52 |
| FALSE | High | EDSEADEL | [R].EDSEAI | 1xPhospho | Q6P5H2-2 | 0.001039 | 0 | 1 | 2 | 2 | Q6P5H2-2 | Isoform 2 o | Q6P5H2-2 | 0 | 3030.485 | 0 | 1.63 | 0.054277 | 68.51 | 49.6 | 150.4 | 36.01 | 35.3 | | High | High | High | High | High | High | High | High | High | High | High | 0 | 0.000249 | 2.95 |
| FALSE | High | EDSEDLLCF | [R].EDSEDI | 1xCarbami | F6RGI3 1x | 0.000967 | 0 | 1 | 1 | 2 | F6RGI3 | Uncharacte | F6RGI3 [37 | 0 | 2299.104 | 0 | 0.1 | 0.800956 | 7.51 | 97 | 103 | 5.64 | 11.01 | | High | High | High | High | High | High | High | High | High | High | High | 0.000183 | 0.001278 | 1.5 |
| FALSE | High | EDSEPEDV | [K].EDSEPE | 1xPhospho | A2ANY6 1x | 2.2E-06 | 0 | 1 | 2 | 1 | A2ANY6 | Midasin OS | A2ANY6 [4 | 0 | 2156.964 | 0 | -1.22 | 0.027215 | 13.34 | 126.4 | 73.6 | 18.87 | 40.16 | | High | High | High | High | High | High | High | High | High | High | High | 0 | 2.46E-07 | 4.06 |
| FALSE | High | EDSFDYSK | [R].EDSFDY | 1xPhospho | Q9Z0U1 1x | 0.01441 | 0.001011 | 1 | 1 | 3 | Q9Z0U1 | Tight junct | Q9Z0U1 [9 | 0 | 1299.534 | 0 | 1.28 | 0.003083 | 10.54 | 58.2 | 141.8 | 10.09 | 12.62 | | High | High | High | High | High | High | High | High | High | High | High | 0.000638 | 0.005603 | 2.6 |
| FALSE | High | EDSFDYSK | [R].EDSFDY | 1xPhospho | Q9Z0U1 1x | 0.009416 | 0.000736 | 1 | 1 | 3 | Q9Z0U1 | Tight junct | Q9Z0U1 [9 | 0 | 1528.697 | 0 | 1.17 | 0.002809 | 16.48 | 59.9 | 140.1 | 13.59 | 11.52 | | High | High | High | High | High | High | High | High | High | High | High | 0.000449 | 0.003934 | 2.91 |
| FALSE | High | EDSFESLDS | [R].EDSFES | 1xPhospho | Q6ZQ32 1x | 0.001137 | 0 | 2 | 5 | 1 | Q6ZQ32; F | MKIAA085 | Q6ZQ32 [1 | 0 | 1750.773 | 0 | 0.7 | 0.00079 | 7.24 | 75.7 | 124.3 | 3.5 | 4.56 | Shared | High | High | High | High | High | High | High | High | High | High | High | 0 | 0.000275 | 2.2 |
| FALSE | High | EDSGDAET | [R].EDSGD | 1xPhospho | O88665 1x | 0.00311 | 0.00016 | 1 | 1 | 1 | O88665 | Bromodom | O88665 [2 | 1 | 2247.069 | 0 | -0.45 | 0.00318 | 4.9 | 115.4 | 84.6 | 5.73 | 6.53 | | High | High | High | High | High | High | High | High | High | High | High | 7.19E-05 | 0.000848 | 1.3 |
| FALSE | High | EDSGTFSL | [R].EDSGTI | 1xPhospho | Q9WTX2 1 | 0.00153 | 5.66E-05 | 1 | 1 | 2 | Q9WTX2 | Interferon- | Q9WTX2 [ | 0 | 1578.782 | 0 | 1.04 | 0.000509 | 3.14 | 67.6 | 132.4 | 4.55 | 4.63 | | High | High | High | High | High | High | High | High | High | High | High | 2.58E-05 | 0.000386 | 1.71 |
| FALSE | High | EDSLESTEI | [R].EDSLES | 1xCarbami | F8VPV0 1x | 1.9E-06 | 0 | 1 | 6 | 4 | F8VPV0 | Pericentrin | F8VPV0 [4 | 0 | 2721.201 | 0 | 0.46 | 0.01951 | 16.97 | 84.9 | 115.1 | 5.14 | 13.05 | | High | High | High | High | High | High | High | High | High | High | High | 0 | 2.08E-07 | 4.1 |
| FALSE | High | EDSLGDEV | [R].EDSLGI | 1xPhospho | A2AJN7 1x | 0.006773 | 0.00062 | 1 | 2 | 1 | A2AJN7 | Sodium bic | A2AJN7 [3 | 0 | 2764.276 | 0 | 0.67 | 0.001262 | 14.91 | 77.1 | 122.9 | 13.9 | 8.99 | | High | High | High | High | High | High | High | High | High | High | High | 0.000275 | 0.00203 | 3.22 |
| FALSE | High | EDSLGDEV | [R].EDSLGI | 2xPhospho | A2AJN7 2x | 7.16E-06 | 0 | 1 | 2 | 1 | A2AJN7 | Sodium bic | A2AJN7 [3 | 0 | 2844.243 | 0 | 1.52 | 0.000925 | 3.69 | 49.3 | 150.7 | 32.1 | 4.79 | | High | High | High | High | High | High | High | High | High | High | High | 0 | 9.27E-07 | 3.91 |
| FALSE | High | EDSLLTHK | [R].EDSLLT | 1xPhospho | Q8BGS1 1x | 0.00407 | 0.00036 | 1 | 2 | 1 | Q8BGS1 | Band 4.1-li | Q8BGS1 [5 | 0 | 1480.781 | 0 | -0.33 | 0.101642 | 1.27 | 112.2 | 87.8 | 18.38 | 24.89 | | High | High | High | High | High | High | High | High | High | High | High | 0.000161 | 0.001152 | 1.57 |
| FALSE | High | EDSLPSAPF | [R].EDSLPS | 1xPhospho | F8WJ40 1x | 9.5E-07 | 0 | 1 | 5 | 5 | F8WJ40 | Histone-lys | F8WJ40 [2 | 0 | 3226.547 | 0 | -1.31 | 0.001469 | 1.75 | 143.1 | 56.9 | 8.57 | 12.08 | | High | High | High | High | High | High | High | High | High | High | High | 0 | 9.57E-08 | 3.33 |

| | | | | | | | | | | | | | | | | | | | | | | | | | | | | | | | | | | | | | | |
|---|---|---|---|---|---|---|---|---|---|---|---|---|---|---|---|---|---|---|---|---|---|---|---|---|---|---|---|---|---|---|---|---|---|---|---|---|---|---|
| FALSE | High | EDSLPSAPF | [R].EDSLPS | 1xDeamida | F8WJ40 1x | 2.51E-05 | 0 | 1 | 5 | 10 | F8WJ40 | Histone-lys | F8WJ40 [2 | 0 | 3227.531 | 0 | -1.23 | 0.001361 | 2.82 | 142.2 | 57.8 | 8.68 | 8.54 | | High | High | High | High | High | High | High | High | High | High | High | 0 | 4.37E-05 | 3.82 |
| FALSE | High | EDSLSRPQI | [R].EDSLSR | 1xPhospho | A2AKX3 1x | 0.017809 | 0.001282 | 1 | 2 | 1 | A2AKX3 | Probable h | A2AKX3 [1 | 0 | 2341.242 | 0 | 0.95 | 0.000757 | 7.22 | 68.3 | 131.7 | 5.14 | 7.04 | | High | High | High | High | High | High | High | High | High | High | High | 0.000689 | 0.005963 | 2.58 |
| FALSE | High | EDSQRPGA | [R].EDSQRI | 1xPhospho | Q3UK83 1x | 0.075304 | 0.005727 | 4 | 7 | 1 | Q3UK83; P | Putative ur | Q3UK83 [9 | 0 | 1976.037 | 0 | 0.32 | 0.280522 | 63.06 | 84.3 | 115.7 | 12.52 | 31.02 | Shared | High | High | High | High | High | High | High | High | High | High | High | 0.002976 | 0.03109 | 2.15 |
| FALSE | High | EDSRPPTP\ | [K].EDSRPF | 1xPhospho | D3Z5N6 1x | 0.00537 | 0.000599 | 1 | 5 | 4 | D3Z5N6 | Zinc finger | D3Z5N6 [2 | 0 | 1677.816 | 0 | -0.83 | 0.000291 | 1.11 | 128 | 72 | 3.68 | 3.09 | | High | High | High | High | High | High | High | High | High | High | High | 0.006563 | 0.07053 | 2.3 |
| FALSE | High | EDSSEEPGT | [K].EDSSEE | 1xPhospho | Q3U4Z0 1x | 0.000246 | 0 | 1 | 3 | 2 | Q3U4Z0 | Putative ur | Q3U4Z0 [5 | 0 | 2970.295 | 0 | 0.75 | 0.005561 | 13.19 | 70.4 | 129.6 | 12.74 | 11.27 | | High | High | High | High | High | High | High | High | High | High | High | 0 | 4.92E-05 | 3.79 |
| FALSE | High | EDSSGMHI | [R].EDSSGI | 1xPhospho | P39447 1xI | 1.5E-05 | 0 | 1 | 3 | 5 | P39447 | Tight junct | P39447 [88 | 0 | 3414.516 | 0 | 0.47 | 0.224087 | 23.78 | 80.6 | 119.4 | 12.18 | 21.17 | | High | High | High | High | High | High | High | High | High | High | High | 0 | 4.08E-06 | 3.47 |
| FALSE | High | EDSSSLVAE | [R].EDSSSL | 1xPhospho | Q9ESK9 1x | 0.000125 | 0 | 1 | 5 | 5 | Q9ESK9 | RB1-induci | Q9ESK9 [1 | 0 | 1972.988 | 0 | 0.16 | 0.126435 | 11.5 | 94.7 | 105.3 | 17.79 | 9.25 | | High | High | High | High | High | High | High | High | High | High | High | 0 | 0.000267 | 3.53 |
| FALSE | High | EECPEAME | [K].EECPEA | 1xCarbami | P70399 1xI | 8.9E-06 | 0 | 1 | 4 | 1 | P70399 | TP53-bindi | P70399 [70 | 0 | 2916.366 | 0 | 0.79 | 0.152997 | 17.18 | 74.4 | 125.6 | 10.3 | 28.42 | | High | High | High | High | High | High | High | High | High | High | High | 0 | 1.18E-06 | 4.03 |
| FALSE | High | DTDAGEG/ | [R].DTDAG | 1xPhospho | Q3ULT8 1x | 0.000147 | 0 | 1 | 6 | 2 | Q3ULT8 | Putative ur | Q3ULT8 [4 | 1 | 3484.607 | 0 | 1.1 | 0.000634 | 8.38 | 60.9 | 139.1 | 11.94 | 6.55 | | High | High | High | High | High | High | High | High | High | High | High | 0 | 2.78E-05 | 3.09 |
| FALSE | High | DTASLSTTF | [R].DTASLS | 1xPhospho | A0JNY3 1xI | 0.000389 | 0 | 1 | 2 | 1 | A0JNY3 | Gephyrin C | A0JNY3 [29 | 0 | 1757.816 | 0 | 1.63 | 0.000503 | 5.36 | 50.8 | 149.2 | 16.17 | 10.14 | | High | High | High | High | High | High | High | High | High | High | High | 0 | 8.3E-05 | 2.25 |
| FALSE | High | DTAEVPSLF | [R].DTAEVI | 1xPhospho | Q9ERH4 1x | 0.00095 | 0 | 1 | 1 | 5 | Q9ERH4 | Nucleolar a | Q9ERH4 [1 | 0 | 3649.727 | 0 | -0.52 | 0.00099 | 1.61 | 117.7 | 82.3 | 4 | 5.52 | | High | High | High | High | High | High | High | High | High | High | High | 0 | 0.000271 | 3.17 |
| FALSE | High | DLILPNGGT | [K].DLILPN | 1xPhospho | Q8CE50 1x | 1.08E-05 | 0 | 1 | 1 | 4 | Q8CE50 | Sorting nex | Q8CE50 [5 | 0 | 2705.371 | 0 | 0.66 | 0.009012 | 13.47 | 79.5 | 120.5 | 7.5 | 8.69 | | High | High | High | High | High | High | High | High | High | High | High | 0 | 1.48E-06 | 3.78 |
| FALSE | High | DLLPDTELS | [K].DLLPDT | 1xPhospho | Q5SXJ3 1xI | 2.9E-05 | 0 | 1 | 2 | 2 | Q5SXJ3 | Fanconi an | Q5SXJ3 [10 | 0 | 2253.13 | 0 | 0.32 | 0.01224 | 12.08 | 89.5 | 110.5 | 5.67 | 4.92 | | High | High | High | High | High | High | High | High | High | High | High | 0 | 4.48E-06 | 3.64 |
| FALSE | High | DLLPSPAGI | [K].DLLPSP | 1xPhospho | Q9ESC8 1x | 8.25E-05 | 0 | 1 | 2 | 1 | Q9ESC8 | AF4/FMR2 | Q9ESC8 [8 | 0 | 1846.049 | 0 | -0.04 | 0.199519 | 6.19 | 101.4 | 98.6 | 3.8 | 2.4 | | High | High | High | High | High | High | High | High | High | High | High | 0 | 1.45E-05 | 2.6 |
| FALSE | High | DLLPTVSSV | [R].DLLPTV | 1xPhospho | Q9D0P7 1x | 2.21E-05 | 0 | 1 | 3 | 1 | Q9D0P7 | Proline/ser | Q9D0P7 [1 | 0 | 2490.284 | 0 | -0.82 | 0.002072 | 3.02 | 127.9 | 72.1 | 10.39 | 11.35 | | High | High | High | High | High | High | High | High | High | High | High | 0 | 3.3E-06 | 2.32 |
| FALSE | High | DLLSDCSVI | [K].DLLSDC | 1xCarbami | P97329 1xI | 0.000121 | 0 | 1 | 1 | 1 | P97329 | Kinesin-like | P97329 [38 | 0 | 2290.129 | 0 | -0.18 | 0.750626 | 33.72 | 107.6 | 92.4 | 5.76 | 32.26 | | High | High | High | High | High | High | High | High | High | High | High | 0 | 2.23E-05 | 3.87 |
| FALSE | High | DLLSDLQDI | [R].DLLSDL | 1xPhospho | P24788 1xI | 2.41E-07 | 0 | 1 | 4 | 6 | P24788 | Cyclin-dep | P24788 [26 | 0 | 1914.89 | 0 | -0.51 | 0.011252 | 7.15 | 118.1 | 81.9 | 9.15 | 4.98 | | High | High | High | High | High | High | High | High | High | High | High | 0 | 4.17E-07 | 3.78 |
| FALSE | High | DLLSDLQDI | [R].DLLSDL | 2xPhospho | P24788 2xI | 6.02E-05 | 0 | 1 | 4 | 3 | P24788 | Cyclin-dep | P24788 [26 | 0 | 1994.856 | 0 | -0.84 | 0.005685 | 8.64 | 128.6 | 71.4 | 10.48 | 9.2 | | High | High | High | High | High | High | High | High | High | High | High | 0 | 3.27E-05 | 3.97 |
| FALSE | High | DLISLDSSP/ | [K].DLISLD! | 1xPhospho | Q04891 1x | 0.000183 | 0 | 1 | 2 | 2 | Q04891 | Transcripti | Q04891 [3 | 0 | 1683.897 | 0 | 0.55 | 0.000632 | 2.42 | 80.1 | 119.9 | 5.82 | 6.01 | | High | High | High | High | High | High | High | High | High | High | High | 0 | 3.54E-05 | 2.8 |
| FALSE | High | DLISNSPR | [R].DLISNS | 1xPhospho | A0A1D5RM | 0.054383 | 0.003951 | 1 | 6 | 3 | A0A1D5RM | DNA-direct | A0A1D5RM | 0 | 1210.603 | 0 | -0.5 | 0.006792 | 6.2 | 116.7 | 83.3 | 5.4 | 5.55 | | High | High | High | High | High | High | High | High | High | High | High | 0.0032 | 0.03381 | 1.56 |
| FALSE | High | DIISSDGDS | [R].DIISSD( | 1xPhospho | Q8BLH7 1x | 0.003892 | 0.000261 | 1 | 1 | 5 | Q8BLH7 | HIRA-inter | Q8BLH7 [5 | 0 | 1304.546 | 0 | -1.39 | 0.005605 | 2.18 | 140.6 | 59.4 | 16.07 | 30.32 | | High | High | High | High | High | High | High | High | High | High | High | 0.002937 | 0.03065 | 1.59 |
| FALSE | High | DLMQTLPS | [K].DLMQT | 1xPhospho | A0A0R4J0I | 0.038677 | 0.002563 | 1 | 4 | 2 | A0A0R4J0I | CTF18, chr | A0A0R4J0I | 0 | 1466.728 | 0 | -0.39 | 0.004633 | 6.24 | 111.9 | 88.1 | 5.34 | 9.44 | | High | High | High | High | High | High | High | High | High | High | High | 0.009869 | 0.1071 | 1.62 |
| FALSE | High | DLMQTLPS | [K].DLMQT | 1xPhospho | A0A0R4J0I | 0.093123 | 0.007276 | 1 | 4 | 1 | A0A0R4J0I | CTF18, chr | A0A0R4J0I | 0 | 1482.723 | 0 | -0.5 | 0.000381 | 2.61 | 116.2 | 83.8 | 2.76 | 4.7 | | High | High | High | High | High | High | High | High | High | High | High | 0.003685 | 0.03964 | 1.35 |
| FALSE | High | DLNFSVEQ | [K].DLNFS\ | 1xPhospho | O88466 1x | 0.002252 | 0.000109 | 2 | 4 | 1 | O88466; R | Zinc finger | O88466 [1 | 0 | 1674.769 | 0 | -0.1 | 0.483825 | 12.59 | 103.3 | 96.7 | 12.83 | 4.71 | Shared | High | High | High | High | High | High | High | High | High | High | High | 4.94E-05 | 0.000592 | 2.97 |
| FALSE | High | DILLISETEG | [K].DILLISE | 1xPhospho | Q80WQ8 1 | 0.001831 | 5.66E-05 | 1 | 3 | 1 | Q80WQ8 | Mis18-binc | Q80WQ8 [ | 0 | 1683.84 | 0 | -1.17 | 0.000154 | 0.89 | 138.3 | 61.7 | 2.32 | 6.11 | | High | High | High | High | High | High | High | High | High | High | High | 2.58E-05 | 0.000473 | 2.89 |
| FALSE | High | DLNLVQAV | [K].DLNLV( | 1xCarbami | B2RXX0 1x | 1.8E-05 | 0 | 1 | 6 | 1 | B2RXX0 | Nek1 prote | B2RXX0 [1 | 0 | 2496.252 | 0 | -0.64 | 0.001726 | 0.93 | 117.5 | 82.5 | 7.96 | 9.92 | | High | High | High | High | High | High | High | High | High | High | High | 0 | 2.62E-06 | 3.36 |
| FALSE | High | DLNSSFETS | [R].DLNSSF | 1xPhospho | Q8K3G5 1x | 0.000193 | 0 | 1 | 5 | 3 | Q8K3G5 | Inactive se | Q8K3G5 [5 | 0 | 1762.866 | 0 | 0.02 | 0.740299 | 9.95 | 100.9 | 99.1 | 5.41 | 3.19 | | High | High | High | High | High | High | High | High | High | High | High | 0 | 8.83E-05 | 2.7 |
| FALSE | High | DLNSSFETS | [R].DLNSSF | 2xPhospho | Q8K3G5 2x | 0.01218 | 0.000889 | 1 | 5 | 1 | Q8K3G5 | Inactive se | Q8K3G5 [5 | 0 | 1842.833 | 0 | -0.49 | 0.0162 | 10.42 | 115.7 | 84.3 | 7.68 | 7.47 | | High | High | High | High | High | High | High | High | High | High | High | 0.000449 | 0.00391 | 2.87 |
| FALSE | High | DLNYCFSGI | [R].DLNYCI | 1xCarbami | Q8C2Q7 1x | 6.89E-05 | 0 | 2 | 4 | 1 | Q8C2Q7; P | Heterogen | Q8C2Q7 [2 | 0 | 1926.771 | 0 | 0.6 | 0.000614 | 7.97 | 78.2 | 121.8 | 7.36 | 5.86 | Shared | High | High | High | High | High | High | High | High | High | High | High | 0 | 1.18E-05 | 2.99 |
| FALSE | High | DLPAIPGVT | [K].DLPAIP | 1xPhospho | Q3TII9 1xP | 3.63E-05 | 0 | 1 | 2 | 7 | Q3TII9 | Putative ur | Q3TII9 [73 | 0 | 3844.86 | 0 | -0.24 | 0.246537 | 19.06 | 117.9 | 82.1 | 24.76 | 30.31 | | High | High | High | High | High | High | High | High | High | High | High | 0.0015 | 0.01497 | 3.78 |
| FALSE | High | DLPAIPGVT | [K].DLPAIP | 1xDeamida | Q3TII9 1xP | 4.83E-07 | 0 | 1 | 2 | 4 | Q3TII9 | Putative ur | Q3TII9 [73 | 0 | 3845.844 | 0 | -0.14 | 0.258695 | 24.29 | 119.8 | 80.2 | 25.51 | 28.71 | | High | High | High | High | High | High | High | High | High | High | High | 0 | 4.48E-08 | 4.19 |
| FALSE | High | DLPAIPGVT | [K].DLPAIP | 1xPhospho | Q3TII9 1xP | 9.74E-06 | 0 | 1 | 2 | 7 | Q3TII9 | Putative ur | Q3TII9 [73 | 0 | 3860.855 | 0 | -0.66 | 0.329053 | 6.77 | 117.3 | 82.7 | 12.09 | 38.69 | | High | High | High | High | High | High | High | High | High | High | High | 0 | 1.83E-05 | 3.67 |
| FALSE | High | DLPAIPGVT | [K].DLPAIP | 2xPhospho | Q3TII9 2xP | 0.012959 | 0.000889 | 1 | 2 | 2 | Q3TII9 | Putative ur | Q3TII9 [73 | 0 | 3924.827 | 0 | -0.3 | 0.183293 | 18.91 | 119.8 | 80.2 | 20.55 | 32.95 | | High | High | High | High | High | High | High | High | High | High | High | 0.000449 | 0.004188 | 3.03 |
| FALSE | High | DLPIPGTIS | [R].DLPIPG | 1xPhospho | A0A087WC | 0.000316 | 0 | 1 | 6 | 2 | A0A087WC | Snf2-relate | A0A087WC | 0 | 2176.085 | 0 | 0.49 | 0.006141 | 12.26 | 82 | 118 | 10.52 | 7.53 | | High | High | High | High | High | High | High | High | High | High | High | 0 | 6.53E-05 | 2.61 |
| FALSE | High | DLPPFGINS | [K].DLPPFG | 1xPhospho | A0A0A6YW | 0.016163 | 0.001124 | 1 | 4 | 1 | A0A0A6YW | Rap guanin | A0A0A6YW | 0 | 2035.103 | 0 | 0.45 | 0.003285 | 2.48 | 85.2 | 114.8 | 6.27 | 5.48 | | High | High | High | High | High | High | High | High | High | High | High | 0.000585 | 0.005365 | 2.81 |
| FALSE | High | DLPQPSLSF | [R].DLPQPS | 1xPhospho | B9EJ86 1xP | 1.49E-06 | 0 | 1 | 2 | 1 | B9EJ86 | Oxysterol-l | B9EJ86 [58 | 0 | 2375.159 | 0 | 0.22 | 0.231296 | 17.31 | 92.2 | 107.8 | 5.17 | 11.29 | | High | High | High | High | High | High | High | High | High | High | High | 0 | 1.59E-07 | 1.98 |
| FALSE | High | DIPSPLHVG | [K].DIPSPLI | 1xPhospho | Q8BJ34-6 1 | 0.036103 | 0.002342 | 1 | 5 | 1 | Q8BJ34-6 | Isoform 6 c | Q8BJ34-6 [ | 0 | 1687.918 | 0 | 2.36 | 0.00183 | 8.94 | 34.7 | 165.3 | 12.45 | 15.87 | | High | High | High | High | High | High | High | High | High | High | High | 0.001279 | 0.01335 | 1.65 |
| FALSE | High | DIQQDLGL | [K].DIQQDI | 1xPhospho | E9Q3P4 1x | 7.83E-09 | 0 | 1 | 4 | 2 | E9Q3P4 | Centromer | E9Q3P4 [1 | 0 | 2431.144 | 0 | -0.14 | 0.086098 | 4.52 | 103.2 | 96.8 | 11.01 | 9.75 | | High | High | High | High | High | High | High | High | High | High | High | 0 | 4.38E-10 | 4.64 |
| FALSE | High | DLQQSIEA/ | [R].DLQQSI | 1xPhospho | Q543N7 1x | 9.59E-07 | 0 | 1 | 1 | 1 | Q543N7 | Protein kin | Q543N7 [2 | 0 | 2127.964 | 0 | -0.04 | 0.811815 | 6.83 | 98 | 102 | 8.65 | 7.07 | | High | High | High | High | High | High | High | High | High | High | High | 0 | 9.65E-08 | 4.24 |
| FALSE | High | DLNSSFETS | [R].DLNSSF | 1xPhospho | Q8K3G5 1x | 0.004256 | 0.00036 | 1 | 5 | 1 | Q8K3G5 | Inactive se | Q8K3G5 [5 | 0 | 1533.704 | 0 | -0.09 | 0.79069 | 15.47 | 103.3 | 96.7 | 37.77 | 44.1 | | High | High | High | High | High | High | High | High | High | High | High | 0.000161 | 0.001203 | 1.84 |
| FALSE | High | DLQSPDFT/ | [R].DLQSPI | 1xPhospho | Q6ZPR6 1x | 2.58E-05 | 0 | 1 | 2 | 3 | Q6ZPR6 | Inhibitor of | Q6ZPR6 [1 | 0 | 2203.047 | 0 | -0.65 | 0.006587 | 5.79 | 121.8 | 78.2 | 5.58 | 8.65 | | High | High | High | High | High | High | High | High | High | High | High | 0 | 5.52E-06 | 4.03 |
| FALSE | High | DIILAPESCE | [R].DIILAPE | 1xCarbami | A0A0R4J0S | 1.96E-07 | 0 | 1 | 4 | 1 | A0A0R4J0S | Lethal(2) gi | A0A0R4J0S | 0 | 2603.187 | 0 | -0.4 | 0.059274 | 19.86 | 113.8 | 86.2 | 14.46 | 9.82 | | High | High | High | High | High | High | High | High | High | High | High | 0 | 1.63E-08 | 3.9 |
| FALSE | High | DLLHPSPEE | [K].DLLHPS | 1xPhospho | Q6ZWU9 1 | 0.000144 | 0 | 1 | 3 | 4 | Q6ZWU9 | 40S riboso | Q6ZWU9 [ | 0 | 1831.924 | 0 | 0.01 | 0.419573 | 6.31 | 99.5 | 100.5 | 2.03 | 8.59 | | High | High | High | High | High | High | High | High | High | High | High | 0 | 2.71E-05 | 2.99 |
| FALSE | High | DLEGLASAI | [K].DLEGL/ | 1xPhospho | E9QK89 1x | 6.29E-08 | 0 | 1 | 2 | 1 | E9QK89 | Mediator c | E9QK89 [9 | 0 | 2438.258 | 0 | -0.2 | 0.971553 | 17.02 | 93 | 107 | 19.18 | 29.74 | | High | High | High | High | High | High | High | High | High | High | High | 0 | 4.53E-09 | 4.11 |
| FALSE | High | DLELPLSPS | [R].DLELPL | 1xPhospho | P41969 1xI | 0.000314 | 0 | 1 | 2 | 1 | P41969 | ETS domain | P41969 [31 | 0 | 2187.126 | 0 | -0.07 | 0.740299 | 10.64 | 101.6 | 98.4 | 18.88 | 29.81 | | High | High | High | High | High | High | High | High | High | High | High | 0 | 6.48E-05 | 2.79 |
| FALSE | High | DLETQPNN | [K].DLETQI | 1xPhospho | Q3UZ01 1x | 1.07E-08 | 0 | 1 | 3 | 5 | Q3UZ01 | RNA-bindir | Q3UZ01 [3 | 0 | 3548.559 | 0 | -0.4 | 0.000647 | 2.94 | 112.9 | 87.1 | 2.8 | 4.92 | | High | High | High | High | High | High | High | High | High | High | High | 0 | 6.51E-10 | 5.81 |
| FALSE | High | DLETQPNN | [K].DLETQI | 2xPhospho | Q3UZ01 2x | 1.62E-06 | 0 | 1 | 3 | 1 | Q3UZ01 | RNA-bindir | Q3UZ01 [3 | 0 | 3628.525 | 0 | 0.37 | 0.051977 | 11.69 | 92.9 | 107.1 | 9.9 | 11.44 | | High | High | High | High | High | High | High | High | High | High | High | 0 | 1.75E-07 | 4.31 |
| FALSE | High | DIFDSPK | [K].DIFDSP | 1xPhospho | Q3UHI0-2 1 | 0.002869 | 0.00016 | 1 | 2 | 1 | Q3UHI0-2 | Isoform 2 c | Q3UHI0-2 | 0 | 1359.696 | 0 | -0.14 | 0.483129 | 8.8 | 105.5 | 94.5 | 6.86 | 9.45 | | High | High | High | High | High | High | High | High | High | High | High | 7.19E-05 | 0.000774 | 2.17 |
| FALSE | High | DLFELDSSE | [K].DLFELD | 1xPhospho | J3QK52 1xI | 1.77E-06 | 0 | 1 | 4 | 3 | J3QK52 | Nucleolar c | J3QK52 [67 | 0 | 2547.065 | 0 | -0.45 | 0.007981 | 8.8 | 113 | 87 | 3.87 | 10.99 | | High | High | High | High | High | High | High | High | High | High | High | 0 | 2.02E-06 | 3.54 |
| FALSE | High | DLFELDSSE | [K].DLFELD | 2xPhospho | J3QK52 2xI | 8.7E-08 | 0 | 1 | 4 | 24 | J3QK52 | Nucleolar c | J3QK52 [67 | 0 | 2627.031 | 0 | -0.3 | 0.009175 | 7.99 | 110 | 90 | 1.81 | 11.33 | | High | High | High | High | High | High | High | High | High | High | High | 0 | 7.35E-07 | 4.3 |
| FALSE | High | DIFHCDDD | [K].DIFHCD | 1xCarbami | Q62187 1x | 3.44E-05 | 0 | 1 | 1 | 5 | Q62187 | Transcripti | Q62187 [8 | 0 | 2860.098 | 0 | -0.27 | 0.049801 | 8.79 | 109 | 91 | 5.12 | 11.82 | | High | High | High | High | High | High | High | High | High | High | High | 0 | 5.42E-06 | 3.47 |
| FALSE | High | DLFNGLDS | [R].DLFNGI | 1xPhospho | A6H624 1x | 4.25E-06 | 0 | 1 | 1 | 1 | A6H624 | LysM, puta | A6H624 [9 | 0 | 3046.393 | 0 | -0.07 | 0.576865 | 4.48 | 99.1 | 100.9 | 7.78 | 15.77 | | High | High | High | High | High | High | High | High | High | High | High | 0 | 5.17E-07 | 3.13 |
| FALSE | High | DLFSLDSEG | [R].DLFSLD | 1xPhospho | Q99L02 1xI | 1.7E-07 | 0 | 1 | 3 | 3 | Q99L02 | PAXIP1-ass | Q99L02 [2 | 0 | 2224.074 | 0 | 0.05 | 0.338616 | 7.19 | 98.4 | 101.6 | 5.71 | 5.23 | | High | High | High | High | High | High | High | High | High | High | High | 0 | 1.38E-08 | 4.71 |
| FALSE | High | DLGELELSG | [K].DLGELE | 1xPhospho | Q9CR02 1x | 0.000758 | 0 | 1 | 2 | 2 | Q9CR02 | Translation | Q9CR02 [1 | 0 | 2299.147 | 0 | -0.34 | 0.012639 | 6.41 | 115.4 | 84.6 | 7.8 | 6.74 | | High | High | High | High | High | High | High | High | High | High | High | 0 | 0.000175 | 2.47 |
| FALSE | High | DIGEQLYTT | [K].DIGEQL | 1xPhospho | J3QPN0 1x | 4.59E-05 | 0 | 1 | 3 | 2 | J3QPN0 | MCG14131 | J3QPN0 [8 | 0 | 2961.468 | 0 | -1.29 | 0.004203 | 1.1 | 132.5 | 67.5 | 14.62 | 18.27 | | High | High | High | High | High | High | High | High | High | High | High | 0 | 7.48E-06 | 3.34 |
| FALSE | High | DIGEQLYTT | [K].DIGEQL | 1xDeamida | J3QPN0 1x | 1.51E-05 | 0 | 1 | 3 | 2 | J3QPN0 | MCG14131 | J3QPN0 [8 | 0 | 2962.452 | 0 | -0.89 | 0.001694 | 2.98 | 131.3 | 68.7 | 10.79 | 5.1 | | High | High | High | High | High | High | High | High | High | High | High | 0 | 7.46E-06 | 3.54 |
| FALSE | High | DIILAPESCE | [R].DIILAPE | 1xCarbami | A0A0R4J0S | 1.13E-05 | 0 | 1 | 4 | 3 | A0A0R4J0S | Lethal(2) gi | A0A0R4J0S | 0 | 2523.22 | 0 | -0.72 | 0.022408 | 4.98 | 127.8 | 72.2 | 14.02 | 13.96 | | High | High | High | High | High | High | High | High | High | High | High | 0 | 1.56E-06 | 2.83 |
| FALSE | High | DIGEQLYTT | [K].DIGEQL | 1xPhospho | J3QPN0 1x | 3.07E-10 | 0 | 1 | 3 | 1 | J3QPN0 | MCG14131 | J3QPN0 [8 | 0 | 2977.463 | 0 | -1.65 | 0.000154 | 1.09 | 151.8 | 48.2 | 1.79 | 10.27 | | High | High | High | High | High | High | High | High | High | High | High | 0 | 1.15E-11 | 6.24 |
| FALSE | High | DLGHPVEE | [K].DLGHP\ | 1xDeamida | Q9Z1Z0 1xI | 7.09E-09 | 0 | 1 | 3 | 4 | Q9Z1Z0 | General ve | Q9Z1Z0 [9 | 0 | 3556.417 | 0 | 1.05 | 0.004627 | 37.64 | 68.3 | 131.7 | 30.73 | 12.36 | | High | High | High | High | High | High | High | High | High | High | High | 0 | 3.9E-10 | 6.26 |
| FALSE | High | DLGNCSDV | [R].DLGNC | 1xCarbami | Q3TFK5 1x | 0.007209 | 0.00062 | 1 | 3 | 2 | Q3TFK5 | G patch do | Q3TFK5 [1 | 0 | 2433.116 | 0 | -0.02 | 0.999012 | 7.26 | 100.8 | 99.2 | 4.31 | 9.02 | | High | High | High | High | High | High | High | High | High | High | High | 0.000275 | 0.002166 | 2.24 |
| FALSE | High | DLGNCSDV | [R].DLGNC | 1xCarbami | Q3TFK5 2x | 0.000908 | 0 | 1 | 3 | 1 | Q3TFK5 | G patch do | Q3TFK5 [1 | 0 | 2513.082 | 0 | -0.34 | 0.045066 | 10.21 | 111.8 | 88.2 | 11.12 | 8.53 | | High | High | High | High | High | High | High | High | High | High | High | 0 | 0.000215 | 3.21 |
| FALSE | High | DLGQSNSD | [R].DLGQSI | 1xPhospho | Q8C8U0-3 | 0.000162 | 0 | 1 | 5 | 1 | Q8C8U0-3 | Isoform 3 c | Q8C8U0-3 | 0 | 2192.035 | 0 | 1.09 | 0.027619 | 35.38 | 58.7 | 141.3 | 17.32 | 26.82 | | High | High | High | High | High | High | High | High | High | High | High | 0 | 3.11E-05 | 3.99 |
| FALSE | High | DLGSVQGF | [R].DLGSV( | 1xPhospho | E9Q784 1x | 0.001373 | 5.66E-05 | 1 | 2 | 1 | E9Q784 | Zinc finger | E9Q784 [1 | 0 | 1918.956 | 0 | -0.17 | 0.138789 | 13.37 | 104.7 | 95.3 | 9.68 | 4.79 | | High | High | High | High | High | High | High | High | High | High | High | 2.58E-05 | 0.000342 | 2.81 |
| FALSE | High | DIHHIDYYK | [R].DIHHID | 1xPhospho | J3QN85 1x | 5.6E-05 | 0 | 1 | 10 | 1 | J3QN85 | Fibroblast | J3QN85 [6 | 0 | 1741.871 | 0 | 2.38 | 0.001271 | 33.67 | 37.2 | 162.8 | 27.04 | 17.84 | | High | High | High | High | High | High | High | High | High | High | High | 0 | 9.39E-06 | 2.8 |
| FALSE | High | DIHPYSDGI | [K].DIHPYS | 1xCarbami | Q571G1 1x | 0.00509 | 0.000551 | 1 | 5 | 1 | Q571G1 | MKIAA402 | Q571G1 [2 | 0 | 3257.49 | 0 | -0.57 | 0.001434 | 6.55 | 115.4 | 84.6 | 5.46 | 9.94 | | High | High | High | High | High | High | High | High | High | High | High | 0.000246 | 0.001473 | 2.96 |
| FALSE | High | DIHPYSDGI | [K].DIHPYS | 1xCarbami | Q571G1 1x | 0.016596 | 0.001124 | 1 | 5 | 4 | Q571G1 | MKIAA402 | Q571G1 [2 | 0 | 3258.474 | 0 | -0.58 | 0.018079 | 8.96 | 120.9 | 79.1 | 9.84 | 15.32 | | High | High | High | High | High | High | High | High | High | High | High | 0.000585 | 0.005506 | 2.94 |
| FALSE | High | DIHPYSDGI | [K].DIHPYS | 1xCarbami | Q571G1 1x | 5.89E-07 | 0 | 1 | 5 | 2 | Q571G1 | MKIAA402 | Q571G1 [2 | 0 | 3273.485 | 0 | -1.29 | 0.000242 | 1.05 | 141.8 | 58.2 | 4.74 | 6.28 | | High | High | High | High | High | High | High | High | High | High | High | 0 | 5.59E-08 | 3.56 |
| FALSE | High | DIKEESDEE | [K].DIKEESI | 1xPhospho | Q9CXK9 1x | 2.38E-05 | 0 | 1 | 4 | 2 | Q9CXK9 | RNA-bindir | Q9CXK9 [2 | 1 | 2650.093 | 0 | -0.19 | 0.150827 | 1.3 | 106.4 | 93.6 | 6.21 | 10.61 | | High | High | High | High | High | High | High | High | High | High | High | 0 | 3.59E-06 | 4.05 |
| FALSE | High | DLKPSTPR | [K].DLKPST | 1xPhospho | Q5SQB7 1x | 0.118334 | 0.009621 | 1 | 7 | 1 | Q5SQB7 | MCG68069 | Q5SQB7 [2 | 0 | 1222.639 | 0 | -0.99 | 0.482824 | 21.87 | 139.5 | 60.5 | 112.8 | 114.21 | | High | High | High | High | High | High | High | High | High | High | High | 0.004866 | 0.05283 | 1.86 |
| FALSE | High | DLLGGEESI | [R].DLLGGI | 1xPhospho | B7ZMV4 1x | 0.051694 | 0.003745 | 1 | 3 | 1 | B7ZMV4 | Cdr2l prote | B7ZMV4 [4 | 0 | 1825.899 | 0 | -0.07 | 0.722482 | 5.87 | 99.2 | 100.8 | 9.6 | 6.73 | | High | High | High | High | High | High | High | High | High | High | High | 0.00196 | 0.02005 | 2.23 |
| FALSE | High | DLLHPSPEE | [K].DLLHPS | 1xPhospho | Q6ZWU9 1 | 0.036731 | 0.00249 | 1 | 3 | 1 | Q6ZWU9 | 40S riboso | Q6ZWU9 [ | 0 | 1602.761 | 0 | -0.35 | 0.8096 | 27.19 | 111.9 | 88.1 | 50.39 | 54.25 | | High | High | High | High | High | High | High | High | High | High | High | 0.001308 | 0.01355 | 2.73 |
| FALSE | High | DLGHPVEE | [K].DLGHP\ | 1xPhospho | Q9Z1Z0 1xI | 1.18E-11 | 0 | 1 | 3 | 9 | Q9Z1Z0 | General ve | Q9Z1Z0 [9 | 0 | 3555.433 | 0 | 0.75 | 0.000592 | 2.62 | 78.7 | 121.3 | 9.45 | 9.39 | | High | High | High | High | High | High | High | High | High | High | High | 0 | 2.96E-13 | 6.5 |
| FALSE | High | DLQYITISK | [K].DLQYIT | 1xPhospho | E9Q9C3 1x | 0.004033 | 0.000311 | 1 | 6 | 1 | E9Q9C3 | Afadin OS= | E9Q9C3 [1 | 0 | 1618.886 | 0 | -0.86 | 0.073783 | 11.95 | 129.2 | 70.8 | 4.84 | 47.12 | | High | High | High | High | High | High | High | High | High | High | High | 0.000161 | 0.001135 | 2.43 |
| FALSE | High | DLRSPLAAT | [K].DLRSPL | 1xPhospho | Q505F1 1x | 9.99E-09 | 0 | 1 | 2 | 2 | Q505F1 | Nuclear re | Q505F1 [2 | 1 | 2457.222 | 0 | 1.34 | 0.000372 | 11.95 | 54.1 | 145.9 | 7.43 | 6.96 | | High | High | High | High | High | High | High | High | High | High | High | 0 | 2.36E-07 | 4.8 |
| FALSE | High | DLSDFPSLE | [R].DLSDFF | 1xPhospho | Q8VE91 1x | 3.77E-05 | 0 | 1 | 4 | 1 | Q8VE91 | Reticuloph | Q8VE91 [3 | 0 | 3644.751 | 0 | 6.64 | | 0 | 4.9 | 195.1 | 32.65 | 15.34 | | Not Found | Not Found | High | High | Not Found | High | High | High | High | High | High | 0 | 5.99E-06 | 4.72 |
| FALSE | High | DMPGGFLI | [R].DMPG( | 1xPhospho | Q52L66 1xI | 1.33E-05 | 0 | 1 | 2 | 14 | Q52L66 | Putative ur | Q52L66 [2 | 0 | 2895.202 | 0 | -0.56 | 0.009319 | 3.36 | 120.2 | 79.8 | 5.74 | 8.84 | | High | High | High | High | High | High | High | High | High | High | High | 2.58E-05 | 0.000392 | 4.05 |
| FALSE | High | DMSPLPES | [K].DMSPL | 1xPhospho | P27546 1xI | 2.22E-06 | 0 | 1 | 3 | 2 | P27546 | Microtubu | P27546 [47 | 0 | 2041.033 | 0 | 1.06 | 0.083857 | 6.59 | 69.2 | 130.8 | 18.27 | 28.77 | | High | High | High | High | High | High | High | High | High | High | High | 0 | 2.48E-07 | 3.11 |
| FALSE | High | DMSPLPES | [K].DMSPL | 1xPhospho | P27546 1xI | 1.11E-05 | 0 | 1 | 3 | 6 | P27546 | Microtubu | P27546 [47 | 0 | 2057.028 | 0 | 0.98 | 0.001419 | 19.93 | 71.2 | 128.8 | 11.72 | 9.28 | | High | High | High | High | High | High | High | High | High | High | High | 0 | 1.52E-06 | 2.74 |
| FALSE | High | DMSPSAET | [K].DMSPS | 1xPhospho | P27546 1xI | 0.000151 | 0 | 1 | 3 | 4 | P27546 | Microtubu | P27546 [51 | 0 | 1984.97 | 0 | -1.33 | 0.014184 | 3.4 | 147.1 | 52.9 | 25.73 | 23.13 | | High | High | High | High | High | High | High | High | High | High | High | 0 | 2.87E-05 | 3.23 |
| FALSE | High | DMSPSAET | [K].DMSPS | 1xPhospho | P27546 1xI | 3.55E-06 | 0 | 1 | 3 | 2 | P27546 | Microtubu | P27546 [51 | 0 | 2000.965 | 0 | -1.27 | 0.002259 | 3.71 | 141.7 | 58.3 | 12.53 | 13.26 | | High | High | High | High | High | High | High | High | High | High | High | 0 | 4.2E-07 | 4.29 |
| FALSE | High | DMSPTSTD | [R].DMSPT | 1xPhospho | E9PYB0 1xI | 1.55E-08 | 0 | 1 | 2 | 11 | E9PYB0 | AHNAK nu | E9PYB0 [28 | 0 | 1800.767 | 0 | 1.13 | 0.000154 | 2.06 | 61.6 | 138.4 | 5.34 | 4.14 | | High | High | High | High | High | High | High | High | High | High | High | 0 | 3.23E-06 | 3.86 |
| FALSE | High | DMTECSTP | [R].DMTEC | 2xCarbami | A0A140LJJ | 0.002004 | 5.66E-05 | 1 | 3 | 2 | A0A140LJJ | A-kinase ar | A0A140LJJ | 0 | 2722.043 | 0 | 0.12 | 0.183384 | 6.2 | 93.6 | 106.4 | 5.16 | 12.18 | | High | High | High | High | High | High | High | High | High | High | High | 0.000246 | 0.001466 | 3.55 |
| FALSE | High | DNAVLFPD | [K].DNAVLI | 1xPhospho | A0A0R4J0Y | 0.007742 | 0.000664 | 1 | 5 | 1 | A0A0R4J0Y | Probable h | A0A0R4J0Y | 0 | 2680.304 | 0 | -0.35 | 0.32481 | 4.03 | 112 | 88 | 18.03 | 19.88 | | High | High | High | High | High | High | High | High | High | High | High | 0.000294 | 0.002365 | 3.36 |
| FALSE | High | DNDDDQSI | [K].DNDDD | 2xPhospho | Q6A065 2x | 0.042141 | 0.003162 | 1 | 2 | 1 | Q6A065 | Centrosom | Q6A065 [4 | 1 | 3701.578 | 0 | 2.33 | 0.030855 | 29.04 | 35.8 | 164.2 | 43.82 | 42.57 | | High | High | High | High | High | High | High | High | High | High | High | 0.001639 | 0.01582 | 2.84 |
| FALSE | High | DNFFQVFS | [K].DNFFQ | 1xPhospho | P54103 1xI | 0.000269 | 0 | 1 | 7 | 3 | P54103 | DnaJ homo | P54103 [17 | 0 | 1940.915 | 0 | -0.4 | 0.003183 | 5.38 | 113.3 | 86.7 | 3.94 | 4.64 | | High | High | High | High | High | High | High | High | High | High | High | 0.000786 | 0.00707 | 2.84 |
| FALSE | High | DNLACASA | [R].DNLAC/ | 1xCarbami | Q24JP2 1xI | 0.063788 | 0.004664 | 1 | 1 | 1 | Q24JP2 | Additional | Q24JP2 [49 | 0 | 1585.688 | 0 | -0.19 | 0.042673 | 3.59 | 112.5 | 87.5 | 10.53 | 10.86 | | High | High | High | High | High | High | High | High | High | High | High | 0.002403 | 0.0254 | 2.03 |
| FALSE | High | DNLACASA | [R].DNLAC/ | 1xCarbami | Q24JP2 1xI | 0.004613 | 0.000456 | 1 | 1 | 2 | Q24JP2 | Additional | Q24JP2 [49 | 1 | 3380.645 | 0 | -0.24 | 0.013699 | 2.87 | 107.5 | 92.5 | 8.06 | 6.4 | | High | High | High | High | High | High | High | High | High | High | High | 0.000275 | 0.001734 | 3.18 |
| FALSE | High | DNLGEAPN | [R].DNLGE. | 1xPhospho | B2RXS1 1xI | 5.88E-06 | 0 | 1 | 8 | 7 | B2RXS1 | Nbeal2 pro | B2RXS1 [18 | 0 | 2852.44 | 0 | 2.33 | 0.00058 | 11.18 | 33.3 | 166.7 | 7.32 | 9.41 | | High | High | High | High | High | High | High | High | High | High | High | 0 | 1.25E-06 | 3.76 |
| FALSE | High | DMPGGFLI | [R].DMPG( | 1xPhospho | Q52L66 1xI | 1.52E-06 | 0 | 1 | 2 | 3 | Q52L66 | Putative ur | Q52L66 [2 | 0 | 2879.207 | 0 | -0.75 | 0.001066 | 1.1 | 122.5 | 77.5 | 9.5 | 7.95 | | High | High | High | High | High | High | High | High | High | High | High | 0 | 1.63E-07 | 3.62 |
| FALSE | High | DNLGEAPN | [R].DNLGE. | 1xPhospho | B2RXS1 1xI | 1.64E-05 | 0 | 1 | 8 | 1 | B2RXS1 | Nbeal2 pro | B2RXS1 [18 | 0 | 2868.435 | 0 | 2.33 | 0.001539 | 18.52 | 32.9 | 167.1 | 14.43 | 14.57 | | High | High | High | High | High | High | High | High | High | High | High | 0 | 2.36E-06 | 3.82 |
| FALSE | High | DNLGNSN\ | [K].DNLGN | 1xCarbami | Q91ZV0 1x | 0.001068 | 0 | 1 | 12 | 4 | Q91ZV0 | Melanoma | Q91ZV0 [1 | 0 | 2593.144 | 0 | 0.91 | 0.003213 | 9.71 | 69.4 | 130.6 | 2.53 | 11.32 | | High | High | High | High | High | High | High | High | High | High | High | 0.000275 | 0.001823 | 3.02 |
| FALSE | High | DNLLLTDG | [R].DNLLLT | 1xCarbami | Q53Z05 1x | 1.03E-05 | 0 | 1 | 3 | 1 | Q53Z05 | Sir2alpha C | Q53Z05 [1 | 0 | 3031.319 | 0 | -1.07 | 0.00061 | 1.38 | 136.2 | 63.8 | 7.91 | 10.99 | | High | High | High | High | High | High | High | High | High | High | High | 0 | 1.39E-06 | 4.54 |
| FALSE | High | DNLLLTDG | [R].DNLLLT | 1xCarbami | Q53Z05 1x | 0.004955 | 0.000504 | 1 | 3 | 1 | Q53Z05 | Sir2alpha C | Q53Z05 [1 | 0 | 3032.303 | 0 | -0.44 | 0.068108 | 13.18 | 113.2 | 86.8 | 21.17 | 16.95 | | High | High | High | High | High | High | High | High | High | High | High | 0.000225 | 0.001429 | 2.9 |
| FALSE | High | DNLTLWT/ | [R].DNLTL\ | 1xPhospho | O70456 1x | 1.85E-06 | 0 | 1 | 1 | 13 | O70456 | 14-3-3 pro | O70456 [2 | 0 | 2812.204 | 0 | 0.61 | 0.000633 | 2.42 | 80.4 | 119.6 | 6.79 | 7.9 | | High | High | High | High | High | High | High | High | High | High | High | 0 | 3.35E-05 | 4.6 |
| FALSE | High | DNLTLWTS | [R].DNLTL\ | 1xPhospho | Q5SS40 1xI | 3.81E-06 | 0 | 1 | 3 | 2 | Q5SS40 | Putative ur | Q5SS40 [2 | 0 | 2719.232 | 0 | -1.27 | 0.000226 | 0.63 | 140.5 | 59.5 | 4.5 | 6.63 | | High | High | High | High | High | High | High | High | High | High | High | 0 | 4.57E-07 | 4.31 |
| FALSE | High | DNLTLWTS | [R].DNLTL\ | 1xPhospho | Q5SS40 1xI | 9.93E-10 | 0 | 1 | 3 | 1 | Q5SS40 | Putative ur | Q5SS40 [2 | 0 | 2735.227 | 0 | -1.24 | 0.000333 | 2.84 | 140.6 | 59.4 | 4.96 | 15.67 | | High | High | High | High | High | High | High | High | High | High | High | 0 | 4.27E-11 | 4.16 |
| FALSE | High | DNLTLWTS | [R].DNLTL\ | 1xPhospho | A8IP69 1xP | 0.001301 | 0 | 1 | 1 | 2 | A8IP69 | 14-3-3 pro | A8IP69 [22 | 0 | 2503.01 | 0 | -0.28 | 0.150641 | 4.1 | 107.8 | 92.2 | 3.95 | 17.91 | | High | High | High | High | High | High | High | High | High | High | High | 0 | 0.000319 | 3.07 |
| FALSE | High | DNLTLWTS | [R].DNLTL\ | 1xCarbami | F6VW30 1x | 2.44E-08 | 0 | 1 | 2 | 24 | F6VW30 | 14-3-3 pro | F6VW30 [2 | 0 | 2764.113 | 0 | -0.29 | 0.011438 | 5.69 | 110.4 | 89.6 | 3.36 | 6.12 | | High | High | High | High | High | High | High | High | High | High | High | 0 | 2.99E-06 | 5.43 |
| FALSE | High | DNLTLWTS | [R].DNLTL\ | 1xPhospho | P63101 1xI | 1.69E-06 | 0 | 1 | 2 | 16 | P63101 | 14-3-3 pro | P63101 [22 | 0 | 2718.125 | 0 | 0.48 | 0.084865 | 41.93 | 83.4 | 116.6 | 26.6 | 18.23 | | High | High | High | High | High | High | High | High | High | High | High | 0 | 6.36E-07 | 5.19 |
| FALSE | High | DNMVLIDS | [R].DNMVI | 2xPhospho | Q6ZPR8 2x | 0.003558 | 0.00021 | 1 | 4 | 1 | Q6ZPR8 | MKIAA140 | Q6ZPR8 [7 | 0 | 3089.277 | 0 | 0.09 | 0.109991 | 6.39 | 99.9 | 100.1 | 6.3 | 8.88 | | High | High | High | High | High | High | High | High | High | High | High | 9.42E-05 | 0.000989 | 3.4 |
| FALSE | High | DNPAQDFS | [K].DNPAQ | 1xPhospho | Q925J9 1xI | 6.95E-05 | 0 | 1 | 2 | 1 | Q925J9 | Mediator c | Q925J9 [65 | 0 | 2306.054 | 0 | 1.68 | 0.000729 | 14.71 | 48.5 | 151.5 | 9.15 | 9.16 | | High | High | High | High | High | High | High | High | High | High | High | 0 | 1.19E-05 | 3.28 |
| FALSE | High | DNPLGQNS | [R].DNPLG( | 1xCarbami | Q8R3I2 1xI | 0.000248 | 0 | 1 | 3 | 1 | Q8R3I2 | Lysophosp | Q8R3I2 [48 | 0 | 2635.123 | 0 | 0.93 | 0.007147 | 29.28 | 72.1 | 127.9 | 32.58 | 21.13 | | High | High | High | High | High | High | High | High | High | High | High | 0 | 4.98E-05 | 3.05 |
| FALSE | High | DNPSPEPQ | [K].DNPSPI | 1xPhospho | Q9D4C5 1x | 2.93E-06 | 0 | 1 | 1 | 4 | Q9D4C5 | ELL-associa | Q9D4C5 [1 | 0 | 2005.988 | 0 | -0.64 | 0.000245 | 1.21 | 122.2 | 77.8 | 1.98 | 2.15 | | High | High | High | High | High | High | High | High | High | High | High | 0 | 1.7E-06 | 2.16 |
| FALSE | High | DNLGNSN\ | [K].DNLGN | 1xCarbami | Q91ZV0 1x | 5.48E-07 | 0 | 1 | 12 | 2 | Q91ZV0 | Melanoma | Q91ZV0 [1 | 0 | 2592.16 | 0 | 1.54 | 0.002238 | 22.97 | 50.4 | 149.6 | 4.32 | 14.93 | | High | High | High | High | High | High | High | High | High | High | High | 0 | 9.56E-07 | 3.91 |
| FALSE | High | DMDEPSP\ | [K].DMDEF | 1xPhospho | Q6PDG5 1x | 0.002376 | 0.000109 | 1 | 3 | 4 | Q6PDG5 | SWI/SNF c | Q6PDG5 [3 | 0 | 2550.245 | 0 | 1.73 | 0.000693 | 9.05 | 46.3 | 153.7 | 10 | 9.63 | | High | High | High | High | High | High | High | High | High | High | High | 0.000161 | 0.001199 | 2.86 |
| FALSE | High | DMDEPSP\ | [K].DMDEF | 1xPhospho | Q6PDG5 1x | 0.000134 | 0 | 1 | 3 | 2 | Q6PDG5 | SWI/SNF c | Q6PDG5 [3 | 0 | 2534.25 | 0 | 2.16 | 0.003696 | 32.11 | 33.8 | 166.2 | 4.57 | 21.58 | | High | High | High | High | High | High | High | High | High | High | High | 7.19E-05 | 0.000845 | 2.04 |
| FALSE | High | DMDDGSD | [R].DMDD( | 2xPhospho | Q4G0F8-3 | 0.019446 | 0.001438 | 1 | 3 | 2 | Q4G0F8-3 | Isoform 3 c | Q4G0F8-3 | 0 | 2360.919 | 0 | 0.66 | 0.015383 | 25.25 | 80.6 | 119.4 | 8.07 | 16.96 | | High | High | High | High | High | High | High | High | High | High | High | 0.001453 | 0.01459 | 3.29 |
| FALSE | High | DLSDFPSLE | [R].DLSDFF | 1xDeamida | Q8VE91 1x | 3.85E-06 | 0 | 1 | 4 | 2 | Q8VE91 | Reticuloph | Q8VE91 [3 | 0 | 3645.735 | 0 | 6.64 | | 0 | 5.4 | 194.6 | | 14.34 | | Not Found | Not Found | Not Found | Not Found | High | High | High | High | High | High | High | 0 | 1.15E-06 | 4.87 |
| FALSE | High | DLSELSFSE | [R].DLSELS | 1xPhospho | B9EKJ9 1xP | 0.001361 | 0 | 1 | 3 | 1 | B9EKJ9 | RNA bindir | B9EKJ9 [74 | 0 | 2049.896 | 0 | -1.71 | 0.00062 | 2.05 | 152.4 | 47.6 | 7.6 | 17.53 | | High | High | High | High | High | High | High | High | High | High | High | 0 | 0.000336 | 2.77 |
| FALSE | High | DLSGPYDSI | [K].DLSGP\ | 1xPhospho | Q9WVB2 1 | 0.001629 | 5.66E-05 | 1 | 4 | 1 | Q9WVB2 | Transducin | Q9WVB2 [ | 0 | 1892.857 | 0 | 0.72 | 0.001489 | 12.24 | 72.4 | 127.6 | 10.04 | 8.09 | | High | High | High | High | High | High | High | High | High | High | High | 2.58E-05 | 0.000412 | 2.63 |

| | | | | | | | | | | | | | | | | | | | | | | | | | | | | | | | | | | | | | | |
|---|---|---|---|---|---|---|---|---|---|---|---|---|---|---|---|---|---|---|---|---|---|---|---|---|---|---|---|---|---|---|---|---|---|---|---|---|---|---|
| FALSE | High | DLSIVAETG | [R].DLSIVA | 1xCarbami | E9Q2Z1 1x | 1.26E-07 | 0 | 1 | 4 | 2 | E9Q2Z1 | CECR2, hist | E9Q2Z1 [1 | 0 | 2794.317 | 0 | -1.46 | 0.003791 | 2.23 | 145.7 | 54.3 | 11.25 | 24.42 | | High | High | High | High | High | High | High | High | High | High | High | 0 | 1.49E-08 | 3.94 |
| FALSE | High | DLSPEGPAI | [R].DLSPEG | 1xPhospho | Q6PGG2 1x | 3.21E-06 | 0 | 1 | 2 | 2 | Q6PGG2 | GEM-intera | Q6PGG2 [7 | 0 | 2290.105 | 0 | 1.89 | 0.00022 | 11.77 | 41.2 | 158.8 | 8.14 | 4.94 | | High | High | High | High | High | High | High | High | High | High | High | 0 | 3.75E-07 | 3.95 |
| FALSE | High | DLSPELAPD | [R].DLSPEL | 1xPhospho | Q497V6 1x | 4.59E-05 | 0 | 1 | 4 | 2 | Q497V6 | Bromo adja | Q497V6 [1 | 0 | 1678.789 | 0 | 0.63 | 0.016809 | 6.05 | 79.4 | 120.6 | 10.35 | 13.53 | | High | High | High | High | High | High | High | High | High | High | High | 0 | 7.5E-06 | 3.65 |
| FALSE | High | DLSPPDNH | [R].DLSPPD | 1xPhospho | Q6PCX9 1x | 0.00086 | 0 | 1 | 1 | 1 | Q6PCX9 | E3 ubiquiti | Q6PCX9 [4 | 0 | 2543.125 | 0 | 0.54 | 0.011135 | 4.86 | 86.8 | 113.2 | 10.07 | 7.7 | | High | High | High | High | High | High | High | High | High | High | High | 0 | 0.000202 | 2.52 |
| FALSE | High | DLSPQQMV | [R].DLSPQ( | 1xPhospho | Q99KK9 1x | 0.006536 | 0.00062 | 1 | 2 | 1 | Q99KK9 | Probable h | Q99KK9 [6 | 0 | 1481.738 | 0 | 0.77 | 0.002359 | 11.97 | 75.4 | 124.6 | 1.9 | 10.04 | | High | High | High | High | High | High | High | High | High | High | High | 0.000275 | 0.001942 | 2.48 |
| FALSE | High | DISPSVTK | [K].DISPSV | 1xPhospho | Q8CCI5 1xP | 0.032267 | 0.002129 | 1 | 1 | 3 | Q8CCI5 | RING1 and | Q8CCI5 [97 | 0 | 1384.749 | 0 | -1.16 | 0.000209 | 0.63 | 138.6 | 61.4 | 5.45 | 7 | | High | High | High | High | High | High | High | High | High | High | High | 0.001142 | 0.01172 | 1.9 |
| FALSE | High | DLSPTAPCF | [R].DLSPTA | 1xCarbami | Q6EBV9 1x | 0.001325 | 0 | 1 | 1 | 2 | Q6EBV9 | Autophagy | Q6EBV9 [7 | 0 | 2479.247 | 0 | 2.59 | 0.000405 | 8.3 | 32.7 | 167.3 | 36.91 | 11.35 | | High | High | High | High | High | High | High | High | High | High | High | 0 | 0.000329 | 2.83 |
| FALSE | High | DLSPTLLDN | [R].DLSPTL | 1xPhospho | Q99LJ0 1xP | 9.91E-06 | 0 | 1 | 1 | 3 | Q99LJ0 | CTTNBP2 N | Q99LJ0 [48 | 0 | 1882.993 | 0 | 0.57 | 0.003365 | 4.23 | 82.5 | 117.5 | 6.48 | 4.4 | | High | High | High | High | High | High | High | High | High | High | High | 0 | 1.34E-06 | 3.62 |
| FALSE | High | DISSSLNSL | [R].DISSSLN | 1xPhospho | Q3TZP3 1x | 1.27E-06 | 0 | 1 | 3 | 3 | Q3TZP3 | Putative ur | Q3TZP3 [50 | 0 | 1858.875 | 0 | -0.24 | 0.094925 | 4.6 | 106.7 | 93.3 | 5.45 | 6.27 | | High | High | High | High | High | High | High | High | High | High | High | 0 | 2.66E-06 | 3.69 |
| FALSE | High | DLSSSPPGF | [R].DLSSSP | 1xPhospho | Q6PAM0 1 | 1.95E-06 | 0 | 1 | 1 | 2 | Q6PAM0 | 5'-AMP-act | Q6PAM0 [1 | 0 | 2339.062 | 0 | -0.27 | 0.001693 | 2.68 | 108.7 | 91.3 | 5.53 | 3.76 | | High | High | High | High | High | High | High | High | High | High | High | 0 | 2.17E-07 | 3.98 |
| FALSE | High | DLSSSPPGF | [R].DLSSSP | 1xPhospho | Q6PAM0 1 | 4.96E-07 | 0 | 1 | 1 | 4 | Q6PAM0 | 5'-AMP-act | Q6PAM0 [1 | 0 | 2355.057 | 0 | -0.3 | 0.024959 | 8.19 | 109.6 | 90.4 | 7.7 | 6.65 | | High | High | High | High | High | High | High | High | High | High | High | 0 | 4.62E-08 | 3.85 |
| FALSE | High | DITAELSTP | [K].DITAELS | 1xPhospho | Q80WT5 1 | 0.001659 | 5.66E-05 | 1 | 5 | 1 | Q80WT5 | Aftiphilin C | Q80WT5 [9 | 0 | 1711.928 | 0 | 0.63 | 0.012979 | 5.65 | 78.4 | 121.6 | 18.84 | 9.15 | | High | High | High | High | High | High | High | High | High | High | High | 2.58E-05 | 0.000419 | 2.71 |
| FALSE | High | DITDPLSLN | [K].DITDPL | 1xCarbami | Q8K3A9 1x | 7.08E-05 | 0 | 1 | 1 | 2 | Q8K3A9 | 7SK snRNA | Q8K3A9 [2 | 0 | 3147.605 | 0 | -0.36 | 0.003021 | 0.78 | 112.8 | 87.2 | 8.29 | 5.06 | | High | High | High | High | High | High | High | High | High | High | High | 0 | 1.21E-05 | 2.77 |
| FALSE | High | DLTEHKPSA | [R].DLTEHK | 1xPhospho | Q925P3 1x | 4.65E-06 | 0 | 1 | 4 | 1 | Q925P3 | CEA-relate | Q925P3 [4 | 0 | 3240.625 | 0 | 4.62 | | 72.85 | 8.1 | 191.9 | 8.52 | 35.57 | | Not Found | High | Not Found | High | High | High | High | High | High | High | High | 0 | 5.73E-07 | 3.93 |
| FALSE | High | DITEINNLT | [R].DITEINI | 1xPhospho | Q8BHX3-2 | 1.94E-06 | 0 | 1 | 3 | 1 | Q8BHX3-2 | Isoform 2 c | Q8BHX3-2 | 0 | 2522.352 | 0 | -1.04 | 0.003529 | 2.59 | 135.7 | 64.3 | 11.35 | 9.26 | | High | High | High | High | High | High | High | High | High | High | High | 0 | 2.14E-07 | 2.9 |
| FALSE | High | DLTGQVPT | [K].DLTGQ | 1xPhospho | P14873 1xP | 0.031713 | 0.002011 | 2 | 6 | 1 | P14873; B2 | Microtubu | P14873 [52 | 0 | 1789.987 | 0 | -0.24 | 0.039334 | 11.59 | 107 | 93 | 9.49 | 6.1 | Shared | High | High | High | High | High | High | High | High | High | High | High | 0.001076 | 0.01144 | 1.88 |
| FALSE | High | DLTNNNIPI | [K].DLTNNI | 1xPhospho | A0A0G2JG | 6.53E-05 | 0 | 1 | 4 | 1 | A0A0G2JG | Cyclin-dep | A0A0G2JG | 0 | 2114.141 | 0 | 0.39 | 0.02797 | 14.2 | 87.3 | 112.7 | 7.93 | 10.08 | | High | High | High | High | High | High | High | High | High | High | High | 0 | 1.11E-05 | 2.89 |
| FALSE | High | DLTPETSGF | [R].DLTPET | 1xPhospho | Q80TS5 1x | 0.083783 | 0.006345 | 1 | 4 | 1 | Q80TS5 | Zinc finger | Q80TS5 [1 | 0 | 1381.656 | 0 | -1.61 | 0.018838 | 6.69 | 158.5 | 41.5 | 23.84 | 37.23 | | High | High | High | High | High | High | High | High | High | High | High | 0.003265 | 0.03522 | 1.76 |
| FALSE | High | DLTTSQEA | [K].DLTTSC | 1xPhospho | Q6NXY1 1x | 3.35E-05 | 0 | 1 | 2 | 2 | Q6NXY1 | TBC1 doma | Q6NXY1 [8 | 0 | 1700.887 | 0 | -0.62 | 0.006247 | 0.09 | 120.6 | 79.4 | 1.31 | 12.31 | | High | High | High | High | High | High | High | High | High | High | High | 0 | 5.26E-06 | 3.71 |
| FALSE | High | DLVLANLA | [K].DLVLAN | 1xPhospho | O35144 1x | 9.85E-07 | 0 | 1 | 5 | 1 | O35144 | Telomeric | O35144 [3 | 0 | 2158.167 | 0 | -1.06 | 0.001989 | 4.05 | 134.2 | 65.8 | 9.69 | 5.38 | | High | High | High | High | High | High | High | High | High | High | High | 0 | 1E-07 | 3.35 |
| FALSE | High | DLVLGSSP( | [R].DLVLGS | 1xPhospho | Q8C0J6 1xP | 0.005322 | 0.000599 | 1 | 1 | 1 | Q8C0J6 | Ankyrin re | Q8C0J6 [19 | 0 | 1694.949 | 0 | 3.01 | 0.000181 | 6.85 | 22.9 | 177.1 | 17.73 | 2.86 | | High | High | High | High | High | High | High | High | High | High | High | 0.000267 | 0.001552 | 1.75 |
| FALSE | High | DLVQPDKP | [R].DLVQP | 1xPhospho | Q8BJ05 1xP | 0.04913 | 0.003576 | 1 | 1 | 2 | Q8BJ05 | Zinc finger | Q8BJ05 [50 | 0 | 2062.155 | 0 | 0.23 | 0.135412 | 21.51 | 91.3 | 108.7 | 11.01 | 9.61 | | High | High | High | High | High | High | High | High | High | High | High | 0.004596 | 0.04989 | 1.29 |
| FALSE | High | DIVSIQELIE | [R].DIVSIQ | 1xPhospho | P33611 1xP | 0.000203 | 0 | 1 | 6 | 5 | P33611 | DNA polym | P33611 [92 | 0 | 3591.786 | 0 | -0.12 | 0.114926 | 2.89 | 107.7 | 92.3 | 17.91 | 16.87 | | High | High | High | High | High | High | High | High | High | High | High | 0 | 3.99E-05 | 3.85 |
| FALSE | High | DLWPMVS | [K].DLWPM | 1xPhospho | A2ARP8 1x | 0.004654 | 0.000456 | 1 | 3 | 3 | A2ARP8 | Microtubu | A2ARP8 [1 | 0 | 3150.463 | 0 | -1.82 | 0.000702 | 0.79 | 155.9 | 44.1 | 12.62 | 13.52 | | High | High | High | High | High | High | High | High | High | High | High | 0.000872 | 0.008753 | 4.21 |
| FALSE | High | DIYMSPPA | [R].DIYMSF | 1xPhospho | Q793C9 1x | 4.52E-06 | 0 | 1 | 2 | 1 | Q793C9 | U2 small nu | Q793C9 [3 | 0 | 2151.023 | 0 | 0.15 | 0.103581 | 12.27 | 94.7 | 105.3 | 5.16 | 7 | | High | High | High | High | High | High | High | High | High | High | High | 0 | 5.52E-07 | 3.35 |
| FALSE | High | DLYRPLSSD | [R].DLYRPL | 1xPhospho | Q3UGP0 1x | 0.107603 | 0.008634 | 1 | 5 | 1 | Q3UGP0 | Putative ur | Q3UGP0 [3 | 0 | 2262.038 | 0 | -0.98 | 0.01048 | 1.03 | 130.9 | 69.1 | 16.65 | 21.93 | | High | High | High | High | High | High | High | High | High | High | High | 0.004385 | 0.04707 | 2.4 |
| FALSE | High | DLEFCSTEE | [R].DLEFCS | 1xCarbami | Q05CL8 2x | 9.98E-05 | 0 | 1 | 3 | 7 | Q05CL8 | La-related | Q05CL8 [3 | 0 | 2004.831 | 0 | 0.09 | 0.786517 | 7.62 | 95.1 | 104.9 | 13.32 | 9.69 | | High | High | High | High | High | High | High | High | High | High | High | 4.94E-05 | 0.000583 | 2.63 |
| FALSE | High | DNPSPEPQ | [K].DNPSPE | 1xPhospho | Q9D4C5 1x | 0.007145 | 0.00062 | 1 | 1 | 1 | Q9D4C5 | ELL-associa | Q9D4C5 [1 | 1 | 2162.089 | 0 | -0.69 | 0.01055 | 8.3 | 129 | 71 | 11.37 | 16.21 | | High | High | High | High | High | High | High | High | High | High | High | 0.000275 | 0.002158 | 2.14 |
| FALSE | High | DLEFCSTEE | [R].DLEFCS | 1xCarbami | Q05CL8 1x | 3.7E-05 | 0 | 1 | 3 | 8 | Q05CL8 | La-related | Q05CL8 [3 | 0 | 1924.865 | 0 | 0 | 0.686036 | 6.66 | 98.3 | 101.7 | 6.58 | 6.83 | | High | High | High | High | High | High | High | High | High | High | High | 0 | 1.26E-05 | 3.84 |
| FALSE | High | DLEEPNLD | [R].DLEEPN | 1xPhospho | Q3TVI4 1xP | 1.68E-07 | 0 | 1 | 1 | 2 | Q3TVI4 | Protein HE | Q3TVI4 [15 | 0 | 3541.53 | 0 | -0.49 | 0.003103 | 4.73 | 116.9 | 83.1 | 7.28 | 4.2 | | High | High | High | High | High | High | High | High | High | High | High | 0 | 1.37E-08 | 5.34 |
| FALSE | High | DGPTSPK | [K].DGPTSP | 1xPhospho | F8VQ93 1x | 0.030898 | 0.001896 | 1 | 3 | 2 | F8VQ93 | PH-interact | F8VQ93 [9 | 0 | 1239.639 | 0 | 0.06 | 0.813937 | 7.6 | 101.1 | 98.9 | 6.9 | 11.74 | | High | High | High | High | High | High | High | High | High | High | High | 0.001027 | 0.01112 | 1.35 |
| FALSE | High | DGQAEPA( | [R].DGQAE | 1xPhospho | Q8K0L9 1x | 4.67E-08 | 0 | 1 | 2 | 8 | Q8K0L9 | Zinc finger | Q8K0L9 [4 | 0 | 3853.735 | 0 | 4.03 | 0.000781 | 13.19 | 9.9 | 190.1 | 49.75 | 15.54 | | High | High | High | High | High | High | High | High | High | High | High | 0 | 3.25E-09 | 5.06 |
| FALSE | High | DGQTVSPC | [R].DGQTV | 1xPhospho | Q8JZR6 1xP | 3.62E-07 | 0 | 1 | 2 | 2 | Q8JZR6 | Electroneu | Q8JZR6 [25 | 0 | 2212.126 | 0 | 1.55 | 0.000631 | 1.71 | 50.7 | 149.3 | 9.59 | 11.3 | | High | High | High | High | High | High | High | High | High | High | High | 0 | 3.24E-08 | 3.86 |
| FALSE | High | DGQVINET | [R].DGQVII | 1xPhospho | P20152 1xP | 3.1E-06 | 0 | 1 | 6 | 3 | P20152 | Vimentin C | P20152 [45 | 0 | 2145.929 | 0 | 2.7 | 0.000847 | 8.35 | 25 | 175 | 8 | 14.3 | | High | High | High | High | High | High | High | High | High | High | High | 0 | 3.63E-07 | 4.74 |
| FALSE | High | DGQWTPD | [K].DGQW | 1xPhospho | Q9CXF7 1x | 0.006653 | 0.00062 | 1 | 2 | 2 | Q9CXF7 | Chromodo | Q9CXF7 [5 | 0 | 3377.537 | 0 | -0.54 | 0.213075 | 14.24 | 104.5 | 95.5 | 14.97 | 26.88 | | High | High | High | High | High | High | High | High | High | High | High | 0.000275 | 0.001992 | 3.04 |
| FALSE | High | DGSAAHFL | [R].DGSAA | 1xPhospho | E9Q3P4 1x | 7.08E-05 | 0 | 1 | 6 | 1 | E9Q3P4 | Centromer | E9Q3P4 [2 | 0 | 2196.074 | 0 | -0.12 | 0.373625 | 7.89 | 103.2 | 96.8 | 8.78 | 9.13 | | High | High | High | High | High | High | High | High | High | High | High | 0 | 1.22E-05 | 2.56 |
| FALSE | High | DGSEDPST | [R].DGSED | 1xPhospho | Q9DB90 1x | 0.0095 | 0.000736 | 1 | 3 | 1 | Q9DB90 | Protein SM | Q9DB90 [5 | 0 | 1732.73 | 0 | -0.01 | 0.952052 | 51.9 | 100.3 | 99.7 | 16.1 | 32.64 | | High | High | High | High | High | High | High | High | High | High | High | 0.000327 | 0.002962 | 2.42 |
| FALSE | High | DGSEDPST | [R].DGSED | 1xPhospho | Q9DB90 1x | 6.59E-05 | 0 | 1 | 3 | 1 | Q9DB90 | Protein SM | Q9DB90 [5 | 0 | 1945.898 | 0 | -0.09 | 0.849172 | 23.15 | 106.4 | 93.6 | 12.98 | 20.14 | | High | High | High | High | High | High | High | High | High | High | High | 0 | 1.13E-05 | 3.47 |
| FALSE | High | DGSEDPST | [R].DGSED | 1xPhospho | Q9DB90 1x | 0.031439 | 0.002011 | 1 | 3 | 1 | Q9DB90 | Protein SM | Q9DB90 [5 | 0 | 1961.893 | 0 | -0.03 | 0.902065 | 36.57 | 105.1 | 94.9 | 10.81 | 25.86 | | High | High | High | High | High | High | High | High | High | High | High | 0.001076 | 0.01138 | 3.49 |
| FALSE | High | DGSEQTDE | [K].DGSEQ | 1xPhospho | A0A1W2P8 | 2.04E-05 | 0 | 1 | 2 | 2 | A0A1W2P8 | Receptor e | A0A1W2P8 | 0 | 3091.313 | 0 | 0.33 | 0.137078 | 46.61 | 90.2 | 109.8 | 17.25 | 16.15 | | High | High | High | High | High | High | High | High | High | High | High | 0 | 3.02E-06 | 3.84 |
| FALSE | High | DGSEQTDE | [K].DGSEQ | 1xPhospho | A0A1W2P8 | 3.9E-05 | 0 | 1 | 2 | 3 | A0A1W2P8 | Receptor e | A0A1W2P8 | 0 | 3107.308 | 0 | 1.15 | 0.000995 | 4.87 | 60.9 | 139.1 | 3.44 | 10.69 | | High | High | High | High | High | High | High | High | High | High | High | 0 | 2.15E-05 | 3.6 |
| FALSE | High | DGSEVLSP | [R].DGSEVI | 1xPhospho | A2A5K6 1x | 0.002022 | 5.66E-05 | 1 | 1 | 2 | A2A5K6 | Zinc finger | A2A5K6 [9 | 0 | 1570.813 | 0 | 0.05 | 0.588258 | 1.71 | 99.4 | 100.6 | 3.37 | 3.72 | | High | High | High | High | High | High | High | High | High | High | High | 0.000449 | 0.003924 | 1.31 |
| FALSE | High | DGSFEGLT | [K].DGSFE( | 1xPhospho | Q8BGB2 1x | 0.003688 | 0.00021 | 1 | 4 | 1 | Q8BGB2 | Tetratricop | Q8BGB2 [6 | 0 | 1361.655 | 0 | 1.75 | 0.037955 | 28.07 | 45.9 | 154.1 | 30.35 | 36.64 | | High | High | High | High | High | High | High | High | High | High | High | 9.42E-05 | 0.00103 | 2.66 |
| FALSE | High | DGPSSAPA | [R].DGPSS/ | 2xPhospho | Q80T83 2x | 2.06E-05 | 0 | 1 | 4 | 2 | Q80T83 | MKIAA185 | Q80T83 [7 | 0 | 1746.812 | 0 | 0.14 | 0.340059 | 1.46 | 101.8 | 98.2 | 6.06 | 7.14 | | High | High | High | High | High | High | High | High | High | High | High | 0 | 3.04E-06 | 2.69 |
| FALSE | High | DGSFEGLT | [K].DGSFE( | 1xPhospho | Q8BGB2 1x | 0.001189 | 0 | 1 | 4 | 1 | Q8BGB2 | Tetratricop | Q8BGB2 [6 | 0 | 1590.818 | 0 | 2.33 | 0.000483 | 15.39 | 34.6 | 165.4 | 10.4 | 7.99 | | High | High | High | High | High | High | High | High | High | High | High | 0 | 0.00029 | 2.35 |
| FALSE | High | DGSGLGLE | [R].DGSGL( | 1xPhospho | B7ZNK9 1x | 0.000132 | 0 | 1 | 4 | 1 | B7ZNK9 | Spata5 pro | B7ZNK9 [2 | 0 | 1900.967 | 0 | -0.88 | 0.007505 | 2.53 | 129.5 | 70.5 | 7.95 | 12.52 | | High | High | High | High | High | High | High | High | High | High | High | 0 | 2.46E-05 | 2.73 |
| FALSE | High | DGSLASNP | [R].DGSLAS | 1xPhospho | Q80X50-5 | 8.4E-08 | 0 | 1 | 14 | 2 | Q80X50-5 | Isoform 5 c | Q80X50-5 | 0 | 2063.01 | 0 | 1.87 | 0.000711 | 8.94 | 41.6 | 158.4 | 13.82 | 11.42 | | High | High | High | High | High | High | High | High | High | High | High | 0 | 6.3E-09 | 3.75 |
| FALSE | High | DGSLETVA | [R].DGSLET | 1xPhospho | Q5SS00 1xP | 0.000292 | 0 | 1 | 1 | 1 | Q5SS00 | DBF4-type | Q5SS00 [6 | 0 | 1763.853 | 0 | -0.94 | 0.000439 | 1.31 | 132.6 | 67.4 | 7.19 | 6.39 | | High | High | High | High | High | High | High | High | High | High | High | 0 | 5.99E-05 | 3.11 |
| FALSE | High | DGSLNQSS | [R].DGSLN( | 1xPhospho | P39688 1xP | 0.001933 | 5.66E-05 | 1 | 5 | 3 | P39688 | Tyrosine-p | P39688 [19 | 0 | 1492.663 | 0 | -0.35 | 0.000927 | 1.97 | 111.9 | 88.1 | 2.45 | 2.47 | | High | High | High | High | High | High | High | High | High | High | High | 4.94E-05 | 0.000739 | 2.09 |
| FALSE | High | DGSISPVSS | [R].DGSISP | 1xCarbami | A0A1L1SSL | 7.65E-07 | 0 | 1 | 4 | 2 | A0A1L1SSL | Zinc finger | A0A1L1SSL | 0 | 1988.883 | 0 | -0.33 | 0.013082 | 7.25 | 111.5 | 88.5 | 3.13 | 9.74 | | High | High | High | High | High | High | High | High | High | High | High | 0 | 7.52E-08 | 2.09 |
| FALSE | High | DGSISPVSS | [R].DGSISP | 1xCarbami | A0A1L1SSL | 2.72E-05 | 0 | 1 | 4 | 1 | A0A1L1SSL | Zinc finger | A0A1L1SSL | 0 | 2068.85 | 0 | -1.08 | 0.00571 | 0.63 | 135 | 65 | 9.37 | 16.41 | | High | High | High | High | High | High | High | High | High | High | High | 0 | 4.15E-06 | 2.66 |
| FALSE | High | DGSPELAEI | [R].DGSPEI | 1xPhospho | E9Q8X5 1x | 0.072858 | 0.005599 | 1 | 1 | 1 | E9Q8X5 | Zinc finger | E9Q8X5 [5 | 0 | 1998.942 | 0 | 1.66 | 0.000193 | 6.08 | 45.2 | 154.8 | 8.01 | 5.44 | | High | High | High | High | High | High | High | High | High | High | High | 0.002867 | 0.0297 | 2.47 |
| FALSE | High | DGSPSQAK | [K].DGSPS( | 1xPhospho | Q99PL5 1xP | 0.090153 | 0.00698 | 2 | 10 | 1 | Q99PL5; Q | Ribosome- | Q99PL5 [78 | 0 | 1327.666 | 0 | -0.56 | 0.010801 | 8.48 | 119.2 | 80.8 | 7.03 | 9.71 | Shared | High | High | High | High | High | High | High | High | High | High | High | 0.003564 | 0.03805 | 2.3 |
| FALSE | High | DGSTLTLQ | [K].DGSTLT | 1xCarbami | Q9WV30-2 | 0.000284 | 0 | 1 | 8 | 3 | Q9WV30-2 | Isoform 2 c | Q9WV30-2 | 0 | 3361.773 | 0 | 1.25 | 0.000313 | 5.54 | 60.5 | 139.5 | 7.67 | 6.95 | | High | High | High | High | High | High | High | High | High | High | High | 0 | 5.82E-05 | 3.76 |
| FALSE | High | DGTAPPPQ | [R].DGTAPI | 1xPhospho | Q543N7 1x | 0.000438 | 0 | 1 | 2 | 4 | Q543N7 | Protein kin | Q543N7 [3 | 0 | 3221.338 | 0 | 0.17 | 0.365123 | 21.2 | 92.3 | 107.7 | 17.92 | 7.4 | | High | High | High | High | High | High | High | High | High | High | High | 0 | 9.42E-05 | 3.62 |
| FALSE | High | DGTAPPPQ | [R].DGTAPI | 1xPhospho | Q543N7 1x | 0.000411 | 0 | 1 | 2 | 2 | Q543N7 | Protein kin | Q543N7 [3 | 1 | 3578.596 | 0 | -0.15 | 0.56001 | 1.26 | 104.1 | 95.9 | 11.64 | 9.18 | | High | High | High | High | High | High | High | High | High | High | High | 2.58E-05 | 0.000363 | 3.76 |
| FALSE | High | DGTAPPPQ | [R].DGTAPI | 2xPhospho | Q543N7 2x | 0.001349 | 0 | 1 | 2 | 1 | Q543N7 | Protein kin | Q543N7 [3 | 1 | 3658.562 | 0 | -0.32 | 0.01881 | 4.47 | 109.2 | 90.8 | 11.2 | 13.66 | | High | High | High | High | High | High | High | High | High | High | High | 0 | 0.000332 | 3.93 |
| FALSE | High | DGTILNVA | [K].DGTILN | 1xPhospho | P39654 1xP | 0.002768 | 0.000109 | 1 | 1 | 2 | P39654 | Arachidona | P39654 [14 | 0 | 2394.212 | 0 | -0.73 | 0.76172 | 15.81 | 114 | 86 | 37.77 | 116.53 | | High | High | High | High | High | High | High | High | High | High | High | 4.94E-05 | 0.00075 | 1.47 |
| FALSE | High | DGSFFTNP | [K].DGSFFT | 1xCarbami | Q69Z45 1x | 1.95E-05 | 0 | 1 | 2 | 1 | Q69Z45 | MKIAA199 | Q69Z45 [1 | 0 | 2619.215 | 0 | -1.31 | 0.000228 | 1.17 | 141.5 | 58.5 | 4.31 | 6.67 | | High | High | High | High | High | High | High | High | High | High | High | 0 | 2.87E-06 | 2.42 |
| FALSE | High | DGWNQNI | [K].DGWN( | 1xPhospho | Q68FH0 1x | 0.000725 | 0 | 1 | 4 | 2 | Q68FH0 | Plakophilin | Q68FH0 [1 | 0 | 2323.107 | 0 | -1.12 | 0.001986 | 3.93 | 137 | 63 | 9.73 | 10.66 | | High | High | High | High | High | High | High | High | High | High | High | 0 | 0.000167 | 2 |
| FALSE | High | DGPSSAPA | [R].DGPSS/ | 1xPhospho | Q80T83 1x | 0.0004 | 0 | 1 | 4 | 4 | Q80T83 | MKIAA185 | Q80T83 [7 | 0 | 1666.845 | 0 | -0.62 | 0.001728 | 5.74 | 120.9 | 79.1 | 3.95 | 5.9 | | High | High | High | High | High | High | High | High | High | High | High | 0.001665 | 0.01672 | 2.42 |
| FALSE | High | DGLTNAGE | [R].DGLTN/ | 2xPhospho | Q2LC58 2x | 0.019107 | 0.001402 | 1 | 2 | 2 | Q2LC58 | MCG9550, | Q2LC58 [2 | 0 | 2312.994 | 0 | 1.23 | 0.000375 | 1.63 | 57 | 143 | 12.58 | 11.01 | | High | High | High | High | High | High | High | High | High | High | High | 0.00074 | 0.006479 | 2.33 |
| FALSE | High | DGEESPTV | [K].DGEESI | 1xPhospho | Q3USA0 1x | 0.001916 | 5.66E-05 | 1 | 2 | 1 | Q3USA0 | Putative ur | Q3USA0 [5 | 0 | 1612.824 | 0 | 0.09 | 0.396981 | 13.66 | 97.9 | 102.1 | 4.7 | 9.23 | | High | High | High | High | High | High | High | High | High | High | High | 2.58E-05 | 0.000496 | 2.35 |
| FALSE | High | DGEGPDNI | [K].DGEGP | 1xCarbami | P13405 2xP | 0.014797 | 0.001049 | 1 | 4 | 1 | P13405 | Retinoblast | P13405 [57 | 1 | 4479.117 | 0 | 0.73 | 0.003738 | 18.47 | 78.9 | 121.1 | 13.2 | 9.15 | | High | High | High | High | High | High | High | High | High | High | High | 0.000555 | 0.004856 | 5.11 |
| FALSE | High | DGELPVED | [K].DGELP\ | 1xPhospho | Q3THH1 1x | 2.59E-06 | 0 | 1 | 4 | 13 | Q3THH1 | Putative ur | Q3THH1 [4 | 0 | 3011.427 | 0 | -1.18 | 0.000443 | 2.28 | 140.5 | 59.5 | 4.38 | 8.27 | | High | High | High | High | High | High | High | High | High | High | High | 0 | 1.93E-06 | 4.04 |
| FALSE | High | DGEQSPN\ | [R].DGEQS | 1xPhospho | Q9DC22 1x | 0.00407 | 0.00036 | 1 | 1 | 1 | Q9DC22 | DDB1- and | Q9DC22 [3 | 0 | 1785.804 | 0 | 0.3 | 0.059884 | 6.25 | 90 | 110 | 3.6 | 11.53 | | High | High | High | High | High | High | High | High | High | High | High | 0.000161 | 0.001147 | 2.06 |
| FALSE | High | DGEQSPSE | [K].DGEQS | 1xPhospho | Q3UGE1 1x | 0.012288 | 0.000889 | 1 | 6 | 2 | Q3UGE1 | Putative ur | Q3UGE1 [6 | 0 | 2977.308 | 0 | 0.42 | 0.244307 | 15.16 | 85.8 | 114.2 | 10.58 | 22.41 | | High | High | High | High | High | High | High | High | High | High | High | 0.000638 | 0.005593 | 3.16 |
| FALSE | High | DGEQSPSE | [K].DGEQS | 1xPhospho | Q3UGE1 1x | 0.000725 | 0 | 1 | 6 | 7 | Q3UGE1 | Putative ur | Q3UGE1 [6 | 0 | 2993.303 | 0 | 0.39 | 0.248537 | 21.17 | 88.4 | 111.6 | 16.02 | 28.12 | | High | High | High | High | High | High | High | High | High | High | High | 4.94E-05 | 0.000638 | 3.53 |
| FALSE | High | DGEQSPSE | [K].DGEQS | 2xPhospho | Q3UGE1 2x | 0.002974 | 0.00016 | 1 | 6 | 1 | Q3UGE1 | Putative ur | Q3UGE1 [6 | 0 | 3073.27 | 0 | -0.39 | 0.787429 | 6.78 | 110.4 | 89.6 | 23.11 | 20.09 | | High | High | High | High | High | High | High | High | High | High | High | 7.19E-05 | 0.000811 | 3.9 |
| FALSE | High | DGESYDPY | [K].DGESYI | 1xPhospho | P25206 1xP | 6.89E-05 | 0 | 2 | 3 | 1 | P25206; Q | DNA replic | P25206 [69 | 0 | 3080.307 | 0 | -0.48 | 0.478963 | 15.16 | 114.1 | 85.9 | 85.36 | 62.24 | Shared | High | High | High | High | High | High | High | High | High | High | High | 0 | 1.19E-05 | 4.38 |
| FALSE | High | DGESYDPY | [K].DGESYI | 1xPhospho | P25206 1xP | 0.000209 | 0 | 2 | 3 | 1 | P25206; Q | DNA replic | P25206 [69 | 0 | 3096.302 | 0 | -1.27 | 0.743085 | 19 | 147.8 | 52.2 | 80.34 | 107.88 | Shared | High | High | High | High | High | High | High | High | High | High | High | 0 | 4.1E-05 | 3.22 |
| FALSE | High | DGESYDPY | [K].DGESYI | 1xPhospho | P25206 1xP | 1.4E-07 | 0 | 2 | 3 | 12 | P25206; Q | DNA replic | P25206 [69 | 0 | 3309.47 | 0 | -0.44 | 0.00378 | 2.03 | 115.6 | 84.4 | 2.01 | 10.23 | Shared | High | High | High | High | High | High | High | High | High | High | High | 0 | 1.12E-08 | 4.97 |
| FALSE | High | DGESYDPY | [K].DGESYI | 1xPhospho | P25206 1xP | 6.01E-08 | 0 | 2 | 3 | 28 | P25206; Q | DNA replic | P25206 [69 | 0 | 3325.465 | 0 | -0.34 | 0.001541 | 2.96 | 112 | 88 | 3.99 | 2.26 | Shared | High | High | High | High | High | High | High | High | High | High | High | 0 | 4.85E-06 | 4.47 |
| FALSE | High | DGESYDPY | [K].DGESYI | 2xPhospho | P25206 2xP | 0.002077 | 5.66E-05 | 2 | 3 | 1 | P25206; Q | DNA replic | P25206 [69 | 0 | 3389.436 | 0 | 0.39 | 0.139039 | 40.77 | 74.8 | 125.2 | 18.37 | 24.81 | Shared | High | High | High | High | High | High | High | High | High | High | High | 2.58E-05 | 0.000542 | 3.04 |
| FALSE | High | DGETPGQI | [K].DGETP( | 1xPhospho | O88995 1x | 9.8E-05 | 0 | 1 | 2 | 2 | O88995 | NF-kappa-I | O88995 [1 | 0 | 3624.631 | 0 | -0.09 | 0.971941 | 36.33 | 101.4 | 98.6 | 14.87 | 22.74 | | High | High | High | High | High | High | High | High | High | High | High | 0 | 1.76E-05 | 3.34 |
| FALSE | High | DGPLGEDS | [R].DGPLGI | 1xPhospho | Q9Z1E4 1x | 0.000121 | 0 | 1 | 2 | 2 | Q9Z1E4 | Glycogen [ | Q9Z1E4 [6 | 1 | 2677.192 | 0 | 0.28 | 0.00687 | 2.35 | 90.1 | 109.9 | 6.89 | 9.82 | | High | High | High | High | High | High | High | High | High | High | High | 0 | 2.21E-05 | 2.61 |
| FALSE | High | DGFGDTC( | [R].DGFGD | 1xCarbami | A0A1B0GR | 0.000152 | 0 | 1 | 1 | 1 | A0A1B0GR | RIKEN cDN | A0A1B0GR | 0 | 1749.792 | 0 | 1.49 | 0.000413 | 12.03 | 54.3 | 145.7 | 10.44 | 7.78 | | High | High | High | High | High | High | High | High | High | High | High | 0 | 2.89E-05 | 3 |
| FALSE | High | DGFPSGTP | [K].DGFPS( | 1xPhospho | Q8CIN4 1xP | 0.032267 | 0.002129 | 1 | 1 | 2 | Q8CIN4 | Serine/thre | Q8CIN4 [1 | 0 | 1842.94 | 0 | 1.52 | 0.000484 | 6.92 | 48.1 | 151.9 | 12.27 | 9.15 | | High | High | High | High | High | High | High | High | High | High | High | 0.001812 | 0.01828 | 2.23 |
| FALSE | High | DGGAASPA | [R].DGGAA | 1xPhospho | B2RY22 1xP | 1.35E-05 | 0 | 2 | 18 | 1 | B2RY22; A( | Clasp1 pro | B2RY22 [1 | 0 | 1397.626 | 0 | 0.24 | 0.020354 | 4.71 | 92.6 | 107.4 | 9.66 | 10.37 | Shared | High | High | High | High | High | High | High | High | High | High | High | 0 | 1.9E-06 | 2.84 |
| FALSE | High | DGGEALVS | [R].DGGEA | 1xPhospho | Q68FL4 1xP | 5.82E-06 | 0 | 1 | 6 | 2 | Q68FL4 | Putative ac | Q68FL4 [1 | 0 | 2079.98 | 0 | 0.76 | 0.000534 | 7.09 | 74.1 | 125.9 | 11.48 | 6.17 | | High | High | High | High | High | High | High | High | High | High | High | 0 | 7.37E-07 | 3.03 |
| FALSE | High | DGGGENTI | [K].DGGGE | 1xPhospho | Q7TNL5 1x | 1.35E-08 | 0 | 1 | 3 | 7 | Q7TNL5 | Protein ph | Q7TNL5 [2 | 0 | 2896.3 | 0 | 0.7 | 0.001963 | 11.96 | 77.3 | 122.7 | 5.93 | 7.59 | | High | High | High | High | High | High | High | High | High | High | High | 0 | 1.26E-08 | 4.61 |
| FALSE | High | DGGSGNST | [R].DGGSG | 1xPhospho | A2AFQ0 1x | 0.000346 | 0 | 1 | 6 | 5 | A2AFQ0 | E3 ubiquiti | A2AFQ0 [2 | 0 | 1571.763 | 0 | 0.07 | 0.550554 | 13.6 | 99.4 | 100.6 | 6.37 | 7.33 | | High | High | High | High | High | High | High | High | High | High | High | 0.000117 | 0.001082 | 2.68 |
| FALSE | High | DGGSGNST | [R].DGGSG | 2xPhospho | A2AFQ0 2x | 0.00051 | 0 | 1 | 6 | 3 | A2AFQ0 | E3 ubiquiti | A2AFQ0 [2 | 0 | 1651.729 | 0 | 0.33 | 0.060071 | 9.49 | 88.6 | 111.4 | 12.99 | 9.06 | | High | High | High | High | High | High | High | High | High | High | High | 0 | 0.000112 | 3.24 |
| FALSE | High | DGGSHIDS | [K].DGGSH | 1xPhospho | Q9D3A8 1x | 0.0016 | 5.66E-05 | 1 | 6 | 4 | Q9D3A8 | Carboxyl-te | Q9D3A8 [2 | 0 | 2841.308 | 0 | 0.62 | 0.006323 | 15.9 | 72.1 | 127.9 | 9.93 | 11.66 | | High | High | High | High | High | High | High | High | High | High | High | 2.58E-05 | 0.000406 | 2.56 |
| FALSE | High | DGGSPPLLI | [K].DGGSP | 1xPhospho | Q8CGN4 1x | 0.027597 | 0.001716 | 1 | 5 | 2 | Q8CGN4 | BCL-6 core | Q8CGN4 [4 | 0 | 1550.823 | 0 | 0.81 | 0.003069 | 4.94 | 74.7 | 125.3 | 3.31 | 9.4 | | High | High | High | High | High | High | High | High | High | High | High | 0.000953 | 0.009844 | 2.31 |
| FALSE | High | DGLAPPSP | [K].DGLAPI | 1xPhospho | Q921C5-2 | 0.000822 | 0 | 1 | 4 | 2 | Q921C5-2 | Isoform 2 c | Q921C5-2 | 0 | 3183.695 | 0 | 2.44 | 0.001377 | 19.89 | 28.3 | 171.7 | 28 | 17.06 | | High | High | High | High | High | High | High | High | High | High | High | 0 | 0.000196 | 3.42 |
| FALSE | High | DGLLAAAA | [R].DGLLA/ | 1xPhospho | Q8CJF7 1xP | 0.000119 | 0 | 1 | 1 | 3 | Q8CJF7 | Protein EL | Q8CJF7 [2 | 0 | 1921.995 | 0 | -1.12 | 0.001693 | 1.88 | 134.1 | 65.9 | 9.59 | 8.64 | | High | High | High | High | High | High | High | High | High | High | High | 0 | 0.000118 | 3.54 |
| FALSE | High | DGLLTPSM | [R].DGLLTF | 1xPhospho | G5E8J4 1xP | 0.00311 | 0.00016 | 1 | 5 | 2 | G5E8J4 | Tyrosine-p | G5E8J4 [79 | 0 | 2176.004 | 0 | 0.85 | 0.003094 | 24.37 | 69.4 | 130.6 | 13.91 | 8.38 | | High | High | High | High | High | High | High | High | High | High | High | 7.19E-05 | 0.00085 | 2.63 |
| FALSE | High | DGLLTPSM | [R].DGLLTF | 1xPhospho | G5E8J4 1xP | 0.000112 | 0 | 1 | 5 | 5 | G5E8J4 | Tyrosine-p | G5E8J4 [79 | 0 | 2191.999 | 0 | 0.81 | 0.005818 | 8.98 | 65.1 | 134.9 | 12.35 | 15.44 | | High | High | High | High | High | High | High | High | High | High | High | 0 | 4.52E-05 | 2.57 |
| FALSE | High | DGLTNAGE | [R].DGLTN/ | 1xPhospho | Q2LC58 1x | 0.0004 | 0 | 1 | 2 | 1 | Q2LC58 | MCG9550, | Q2LC58 [2 | 0 | 2233.027 | 0 | 1.11 | 0.007155 | 2.13 | 67.1 | 132.9 | 16.19 | 11.02 | | High | High | High | High | High | High | High | High | High | High | High | 0 | 8.55E-05 | 2.91 |
| FALSE | High | DGFGLSPG | [R].DGFGL! | 1xPhospho | E9QQ10 1x | 4.35E-05 | 0 | 1 | 4 | 1 | E9QQ10 | A-kinase ar | E9QQ10 [3 | 0 | 1657.86 | 0 | 1.1 | 0.000725 | 13.28 | 63.7 | 136.3 | 1.91 | 7.29 | | High | High | High | High | High | High | High | High | High | High | High | 0 | 7.05E-06 | 2.74 |
| FALSE | High | DGWSLDR | [R].DGWSL | 1xPhospho | Q0VBL3 1x | 0.007538 | 0.00062 | 1 | 6 | 1 | Q0VBL3 | RNA bindir | Q0VBL3 [6 | 0 | 1157.519 | 0 | -0.64 | 0.000656 | 2.61 | 122.2 | 77.8 | 2.67 | 4.46 | | High | High | High | High | High | High | High | High | High | High | High | 0.000275 | 0.002283 | 1.82 |
| FALSE | High | DHDSDAEA | [R].DHDSD. | 1xCarbami | Q91YN0 1x | 2.86E-06 | 0 | 1 | 1 | 1 | Q91YN0 | Protein C1 | Q91YN0 [7 | 0 | 2097.956 | 0 | 1 | 0.001836 | 18.18 | 68.2 | 131.8 | 5.58 | 9.31 | | High | High | High | High | High | High | High | High | High | High | High | 0 | 3.32E-07 | 2.97 |
| FALSE | High | DHFGLEGD | [K].DHFGLI | 1xPhospho | Q80TM2 1 | 6.41E-08 | 0 | 1 | 3 | 1 | Q80TM2 | MKIAA102 | Q80TM2 [4 | 0 | 2860.3 | 0 | 0.31 | 0.11127 | 24.71 | 89.3 | 110.7 | 9.45 | 15.22 | | High | High | High | High | High | High | High | High | High | High | High | 0 | 4.62E-09 | 4.33 |
| FALSE | High | DLAATQISF | [R].DLAAT( | 1xPhospho | Q8BRG8 1x | 0.000229 | 0 | 1 | 5 | 3 | Q8BRG8 | Transmem | Q8BRG8 [1 | 0 | 3179.582 | 0 | 0.05 | 0.50415 | 2.68 | 100.2 | 99.8 | 10.08 | 15.18 | | High | High | High | High | High | High | High | High | High | High | High | 0 | 4.55E-05 | 3.8 |
| FALSE | High | DIAETESNF | [K].DIAETE | 1xPhospho | Q69Z38 1x | 0.000105 | 0 | 1 | 1 | 2 | Q69Z38 | Pseudopod | Q69Z38 [3 | 0 | 2645.202 | 0 | -0.51 | 0.007714 | 1.58 | 116 | 84 | 8.06 | 2.18 | | High | High | High | High | High | High | High | High | High | High | High | 0 | 1.9E-05 | 3.12 |
| FALSE | High | DLAFGITTS | [K].DLAFGI | 1xPhospho | B2RX88 1x | 0.031439 | 0.002011 | 1 | 3 | 1 | B2RX88 | Centrosom | B2RX88 [3 | 1 | 2671.338 | 0 | -0.86 | 0.00466 | 4.31 | 128.5 | 71.5 | 10.13 | 6.49 | | High | High | High | High | High | High | High | High | High | High | High | 0.001076 | 0.01139 | 1.89 |
| FALSE | High | DIASETDEH | [K].DIASET | 1xPhospho | A2A3V1 1x | 0.00086 | 0 | 1 | 1 | 1 | A2A3V1 | A-kinase ar | A2A3V1 [6 | 0 | 2957.371 | 0 | -1.2 | 0.000378 | 1.91 | 136.4 | 63.6 | 5.34 | 6.21 | | High | High | High | High | High | High | High | High | High | High | High | 0 | 0.000202 | 3.3 |
| FALSE | High | DIASETDEH | [K].DIASET | 2xPhospho | A2A3V1 2x | 4.62E-07 | 0 | 1 | 1 | 2 | A2A3V1 | A-kinase ar | A2A3V1 [6 | 0 | 3037.337 | 0 | -1.12 | 0.00053 | 1.83 | 137.9 | 62.1 | 4.49 | 7.83 | | High | High | High | High | High | High | High | High | High | High | High | 0 | 1.5E-05 | 3.79 |
| FALSE | High | DLASPLLGS | [K].DLASPL | 1xPhospho | P30276 1xP | 0.000322 | 0 | 1 | 2 | 3 | P30276 | G2/mitotic | P30276 [3 | 0 | 1278.654 | 0 | -0.89 | 0.000668 | 3.43 | 127.1 | 72.9 | 4.03 | 10.66 | | High | High | High | High | High | High | High | High | High | High | High | 0 | 6.71E-05 | 2.8 |
| FALSE | High | DIAVFPGTI | [R].DIAVFP | 1xPhospho | A2AD39 1x | 0.090153 | 0.00698 | 1 | 4 | 1 | A2AD39 | SCL-interru | A2AD39 [1 | 0 | 2223.208 | 0 | 0.37 | 0.00867 | 4.36 | 85.4 | 114.6 | 6.91 | 6.42 | | High | High | High | High | High | High | High | High | High | High | High | 0.003564 | 0.03806 | 1.97 |
| FALSE | High | DLCPSEAES | [K].DLCPSE | 1xCarbami | P70324 1xP | 0.000227 | 0 | 1 | 3 | 2 | P70324 | T-box trans | P70324 [3 | 0 | 2276.004 | 0 | -1.78 | 0.000261 | 1.13 | 154.8 | 45.2 | 6.2 | 13.41 | | High | High | High | High | High | High | High | High | High | High | High | 0 | 4.48E-05 | 2.88 |
| FALSE | High | DLCPSEAES | [K].DLCPSE | 1xCarbami | P70324 2xP | 0.001068 | 0 | 1 | 3 | 3 | P70324 | T-box trans | P70324 [3 | 0 | 2355.97 | 0 | -0.84 | 0.003985 | 3.77 | 128.2 | 71.8 | 8.9 | 8.32 | | High | High | High | High | High | High | High | High | High | High | High | 0 | 0.000258 | 2.89 |
| FALSE | High | DIDASPSPL | [R].DIDASP | 1xPhospho | Q8CH02 1x | 1.74E-07 | 0 | 1 | 1 | 5 | Q8CH02 | SURP and ( | Q8CH02 [4 | 0 | 2123.104 | 0 | 0.09 | 0.060345 | 5.02 | 96.6 | 103.4 | 7.1 | 1.95 | | High | High | High | High | High | High | High | High | High | High | High | 0 | 1.3E-07 | 3.84 |
| FALSE | High | DLDAYLVQ | [R].DLDAYI | 1xCarbami | P48754 1xP | 1.18E-05 | 0 | 1 | 3 | 7 | P48754 | Breast canc | P48754 [17 | 0 | 2808.176 | 0 | -0.5 | 0.004837 | 3.67 | 117.6 | 82.4 | 9.95 | 14.85 | | High | High | High | High | High | High | High | High | High | High | High | 0 | 1.62E-06 | 4.23 |
| FALSE | High | DLDDSDK | [K].DLDDSI | 1xPhospho | E9PZ97 1xP | 0.007538 | 0.00062 | 1 | 2 | 2 | E9PZ97 | Bloom syn | E9PZ97 [33 | 0 | 1345.629 | 0 | -1.73 | 0.00057 | 1.98 | 153.6 | 46.4 | 6.65 | 10.18 | | High | High | High | High | High | High | High | High | High | High | High | 0.000275 | 0.002286 | 1.89 |
| FALSE | High | DLDEDELL( | [K].DLDEDI | 1xPhospho | Q3TFA9 1x | 0.027839 | 0.001755 | 1 | 4 | 1 | Q3TFA9 | Putative ur | Q3TFA9 [1 | 1 | 2700.42 | 0 | 1.55 | 0.000223 | 3.95 | 49.3 | 150.7 | 6.7 | 6.67 | | High | High | High | High | High | High | High | High | High | High | High | 0.000953 | 0.009929 | 3.24 |
| FALSE | High | DIAADGLCI | [R].DIAAD( | 1xCarbami | P0CB42 1xP | 1.54E-07 | 0 | 1 | 1 | 1 | P0CB42 | Nucleic aci | P0CB42 [3 | 0 | 2497.231 | 0 | -1.23 | 0.001697 | 3.88 | 135.8 | 64.2 | 11.13 | 10.04 | | High | High | High | High | High | High | High | High | High | High | High | 0 | 1.24E-08 | 2.42 |
| FALSE | High | DLDGSPEA | [R].DLDGSI | 1xDeamida | Q3UFB2 1x | 0.011246 | 0.000772 | 1 | 1 | 1 | Q3UFB2 | Box C/D sn | Q3UFB2 [3 | 1 | 2627.224 | 0 | -0.7 | 0.000251 | 1 | 123.3 | 76.7 | 6.92 | 7.42 | | High | High | High | High | High | High | High | High | High | High | High | 0.000399 | 0.003566 | 1.89 |
| FALSE | High | DIDHETVVI | [K].DIDHET | 1xDeamida | Q00899 1x | 3.94E-08 | 0 | 1 | 3 | 2 | Q00899 | Transcripti | Q00899 [2 | 0 | 3734.707 | 0 | -0.67 | 0.070942 | 11.36 | 122.7 | 77.3 | 17.62 | 33.72 | | High | High | High | High | High | High | High | High | High | High | High | 0 | 2.68E-09 | 6.15 |

| | | | | | | | | | | | | | | | | | | | | | | | | | | | | | | | | | | | | | | |
|---|---|---|---|---|---|---|---|---|---|---|---|---|---|---|---|---|---|---|---|---|---|---|---|---|---|---|---|---|---|---|---|---|---|---|---|---|---|---|
| FALSE | High | DIDHETVVI | [K].DIDHET | 1xPhospho | Q00899 1x | 4.3E-07 | 0 | 1 | 3 | 3 | Q00899 | Transcripti | Q00899 [2 | 0 | 3749.718 | 0 | -0.86 | 0.010413 | 6.88 | 123.7 | 76.3 | 13.75 | 14.52 | | High | High | High | High | High | High | High | High | High | High | High | 0 | 3.92E-08 | 4.73 |
| FALSE | High | DLDLEPLSD | [K].DLDLEP | 1xPhospho | Q3UXZ9 1x | 1.18E-06 | 0 | 1 | 2 | 4 | Q3UXZ9 | Lysine-spe | Q3UXZ9 [1 | 0 | 2368.101 | 0 | 0.04 | 0.864762 | 13.41 | 97.4 | 102.6 | 5.75 | 9.7 | | High | High | High | High | High | High | High | High | High | High | High | 0 | 1.23E-07 | 3.7 |
| FALSE | High | DIDLFGSDE | [K].DIDLFG | 1xPhospho | F6ZFU0 1xP | 3.08E-07 | 0 | 2 | 12 | 12 | F6ZFU0; A0 | Elongation | F6ZFU0 [42 | 0 | 1949.81 | 0 | 0.04 | 0.971891 | 29.32 | 107.3 | 92.7 | 73.49 | 81.68 | Shared | High | High | High | High | High | High | High | High | High | High | High | 0 | 2.69E-08 | 4.47 |
| FALSE | High | DIDLFGSDE | [K].DIDLFG | 1xPhospho | F6ZFU0 1xP | 1.72E-08 | 0 | 2 | 12 | 38 | F6ZFU0; A0 | Elongation | F6ZFU0 [42 | 0 | 2178.973 | 0 | 0 | 0.546191 | 4.18 | 99 | 101 | 2.57 | 6.65 | Shared | High | High | High | High | High | High | High | High | High | High | High | 0 | 2.37E-06 | 4.12 |
| FALSE | High | DIDLFGSDE | [K].DIDLFG | 1xPhospho | F6ZFU0 1xP | 0.06169 | 0.004646 | 2 | 12 | 1 | F6ZFU0; A0 | Elongation | F6ZFU0 [42 | 1 | 2606.191 | 0 | -0.53 | 0.766373 | 27.25 | 106.4 | 93.6 | 40.25 | 115.28 | Shared | High | High | High | High | High | High | High | High | High | High | High | 0.002381 | 0.02458 | 2.87 |
| FALSE | High | DLDLLASVF | [K].DLDLLA | 1xPhospho | Q7TN29 1x | 0.003857 | 0.000261 | 1 | 2 | 2 | Q7TN29 | Stromal me | Q7TN29 [2 | 0 | 1951.994 | 0 | 0.62 | 0.001438 | 3.84 | 77.7 | 122.3 | 8.76 | 6.24 | | High | High | High | High | High | High | High | High | High | High | High | 0.000117 | 0.001078 | 2.38 |
| FALSE | High | DLDLMLLD | [K].DLDLM | 1xPhospho | Q3UKG7 1x | 2.93E-06 | 0 | 1 | 6 | 1 | Q3UKG7 | Putative ur | Q3UKG7 [4 | 0 | 3688.839 | 0 | -0.75 | 0.008783 | 9 | 123.5 | 76.5 | 16.44 | 22.35 | | High | High | High | High | High | High | High | High | High | High | High | 0 | 3.42E-07 | 4.23 |
| FALSE | High | DIDITSPEFI | [K].DIDITSP | 1xPhospho | E9Q616 1x | 0.000118 | 0 | 1 | 2 | 5 | E9Q616 | AHNAK nuc | E9Q616 [1 | 0 | 1962.99 | 0 | 2.71 | 0.000191 | 7.13 | 27.4 | 172.6 | 9.22 | 5.79 | | High | High | High | High | High | High | High | High | High | High | High | 0 | 2.17E-05 | 3.33 |
| FALSE | High | DLDPDVVT | [K].DLDPDV | 1xPhospho | G3UWZ0 1 | 1.49E-09 | 0 | 1 | 3 | 4 | G3UWZ0 | Bromodom | G3UWZ0 [ | 0 | 2521.138 | 0 | 1.62 | 0.000692 | 11.03 | 50.9 | 149.1 | 3.82 | 9.73 | | High | High | High | High | High | High | High | High | High | High | High | 0 | 1.66E-08 | 3.66 |
| FALSE | High | DLEEFGEN | [K].DLEEFG | 1xPhospho | Q8CH25-2 | 1.72E-05 | 0 | 1 | 3 | 2 | Q8CH25-2 | Isoform 2 o | Q8CH25-2 | 0 | 3485.625 | 0 | -0.47 | 0.001406 | 4.15 | 118.4 | 81.6 | 5.46 | 5.49 | | High | High | High | High | High | High | High | High | High | High | High | 0 | 2.47E-06 | 5.11 |
| FALSE | High | DLEEFGEN | [K].DLEEFG | 1xDeamida | Q8CH25-2 | 0.001476 | 5.66E-05 | 1 | 3 | 1 | Q8CH25-2 | Isoform 2 o | Q8CH25-2 | 0 | 3486.609 | 0 | -0.21 | 0.093499 | 9.85 | 109.3 | 90.7 | 11.1 | 7.28 | | High | High | High | High | High | High | High | High | High | High | High | 2.58E-05 | 0.00037 | 3.2 |
| FALSE | High | DLEELGAN | [K].DLEELG | 1xPhospho | A0A1S6GW | 0.003464 | 0.00021 | 1 | 6 | 2 | A0A1S6GW | Uncharacte | A0A1S6GW | 0 | 2126.078 | 0 | 2.3 | 0.002107 | 19.36 | 32.6 | 167.4 | 10.52 | 19.55 | | High | High | High | High | High | High | High | High | High | High | High | 9.42E-05 | 0.000956 | 2.68 |
| FALSE | High | DIDHETVVI | [K].DIDHET | 1xPhospho | Q00899 1x | 7.35E-09 | 0 | 1 | 3 | 4 | Q00899 | Transcripti | Q00899 [2 | 0 | 3733.723 | 0 | -0.79 | 0.0054 | 2.6 | 126.6 | 73.4 | 12.29 | 7.39 | | High | High | High | High | High | High | High | High | High | High | High | 0 | 4.08E-10 | 5.59 |
| FALSE | High | DIAADGLCI | [R].DIAADC | 1xCarbami | P0CB42 1x | 0.050397 | 0.003641 | 1 | 1 | 1 | P0CB42 | Nucleic aci | P0CB42 [36 | 0 | 2268.068 | 0 | -0.38 | 0.018987 | 3.25 | 112.5 | 87.5 | 9.77 | 13.86 | | High | High | High | High | High | High | High | High | High | High | High | 0.001916 | 0.01952 | 2.53 |
| FALSE | High | DKTPVQSC | [R].DKTPVC | 1xPhospho | Q3TI61 1xP | 5.25E-08 | 0 | 1 | 8 | 2 | Q3TI61 | Putative ur | Q3TI61 [7- | 1 | 2939.497 | 0 | 0.02 | 0.872418 | 18.36 | 99.1 | 100.9 | 6.47 | 8.91 | | High | High | High | High | High | High | High | High | High | High | High | 0 | 3.89E-09 | 5.05 |
| FALSE | High | DKTPVQSC | [R].DKTPVC | 1xPhospho | Q3TI61 1xP | 3.13E-07 | 0 | 1 | 8 | 2 | Q3TI61 | Putative ur | Q3TI61 [7- | 1 | 2710.334 | 0 | 0.26 | 0.928357 | 45.56 | 98.1 | 101.9 | 42.57 | 39.56 | | High | High | High | High | High | High | High | High | High | High | High | 0 | 2.75E-08 | 4.7 |
| FALSE | High | DHFGLEGD | [K].DHFGLE | 1xPhospho | Q80TM2 1x | 0.000215 | 0 | 1 | 3 | 2 | Q80TM2 | MKIAA102 | Q80TM2 [4 | 0 | 2876.295 | 0 | 0.41 | 0.044432 | 14.52 | 84.4 | 115.6 | 14.01 | 11.55 | | High | High | High | High | High | High | High | High | High | High | High | 0 | 4.23E-05 | 3.73 |
| FALSE | High | DHFGLEGD | [K].DHFGLE | 1xPhospho | Q80TM2 1x | 1.21E-05 | 0 | 1 | 3 | 1 | Q80TM2 | MKIAA102 | Q80TM2 [4 | 1 | 3217.558 | 0 | 0.28 | 0.241644 | 16.85 | 90.2 | 109.8 | 9.88 | 16.34 | | High | High | High | High | High | High | High | High | High | High | High | 0 | 1.68E-06 | 4.59 |
| FALSE | High | DHLYGVHN | [K].DHLYGV | 1xPhospho | G5E8G5 1x | 0.005467 | 0.00062 | 1 | 4 | 4 | G5E8G5 | Prominin 1 | G5E8G5 [8 | 1 | 2298.058 | 0 | 2.94 | 0.001494 | 7.11 | 27.2 | 172.8 | 46.68 | 14.88 | | High | High | High | High | High | High | High | High | High | High | High | 0.000275 | 0.001592 | 3.35 |
| FALSE | High | DHMPSQD | [R].DHMPS | 1xPhospho | B9EKC1 1x | 0.043236 | 0.003228 | 1 | 2 | 1 | B9EKC1 | PAS domai | B9EKC1 [44 | 0 | 1854.871 | 0 | -0.62 | 0.002342 | 2.7 | 121.3 | 78.7 | 12.75 | 13.37 | | High | High | High | High | High | High | High | High | High | High | High | 0.001651 | 0.01634 | 2.08 |
| FALSE | High | DHPAPVPG | [K].DHPAPV | 1xPhospho | Q3TW36 1x | 0.00824 | 0.000736 | 1 | 4 | 4 | Q3TW36 | Putative ur | Q3TW36 [2 | 0 | 1767.936 | 0 | 0.12 | 0.159882 | 8.55 | 94.1 | 105.9 | 6.05 | 7.31 | | High | High | High | High | High | High | High | High | High | High | High | 0.000327 | 0.002518 | 2.59 |
| FALSE | High | DHQSSPYA | [R].DHQSSI | 1xPhospho | O35690 1x | 0.002252 | 0.000109 | 1 | 2 | 2 | O35690 | Paired mes | O35690 [7 | 0 | 2000.988 | 0 | 4.27 | 0.00046 | 46.58 | 9.2 | 190.8 | 33.75 | 10.15 | | High | High | High | High | High | High | High | High | High | High | High | 4.94E-05 | 0.00059 | 2.84 |
| FALSE | High | DHSDSEST | [K].DHSDSI | 1xPhospho | O70546-2 1 | 1.6E-06 | 0 | 1 | 5 | 3 | O70546-2 | Isoform 2 o | O70546-2 | | 0 | 2090.892 | 0 | 0.43 | 0.045626 | 16.12 | 92 | 108 | 7.35 | 13.25 | | High | High | High | High | High | High | High | High | High | High | High | 0 | 1.73E-07 | 3.44 |
| FALSE | High | DHSPGPWS | [K].DHSPGI | 1xPhospho | B1B0E8 1x | 5.23E-06 | 0 | 1 | 1 | 2 | B1B0E8 | U2 small nu | B1B0E8 [43 | 0 | 1562.695 | 0 | -0.46 | 0.010182 | 11.32 | 117.1 | 82.9 | 7.55 | 8.98 | | High | High | High | High | High | High | High | High | High | High | High | 0 | 6.51E-07 | 2.36 |
| FALSE | High | DHSPTPSVI | [R].DHSPTF | 1xPhospho | Q9D824 1x | 1.18E-06 | 0 | 1 | 3 | 13 | Q9D824 | Pre-mRNA | Q9D824 [4 | 0 | 2025.875 | 0 | -0.06 | 0.321015 | 4.78 | 101.8 | 98.2 | 3.01 | 7.39 | | High | High | High | High | High | High | High | High | High | High | High | 0.000437 | 0.003646 | 3.52 |
| FALSE | High | DHSPTPSVI | [R].DHSPTF | 1xDeamida | Q9D824 1x | 1.3E-05 | 0 | 1 | 3 | 4 | Q9D824 | Pre-mRNA | Q9D824 [4 | 0 | 2026.859 | 0 | 0.3 | 0.160225 | 19.58 | 91.5 | 108.5 | 10.68 | 8.71 | | High | High | High | High | High | High | High | High | High | High | High | 0.000814 | 0.007749 | 2.96 |
| FALSE | High | DHSPTPSVI | [R].DHSPTF | 2xPhospho | Q9D824 2x | 1.09E-05 | 0 | 1 | 3 | 10 | Q9D824 | Pre-mRNA | Q9D824 [4 | 0 | 2105.842 | 0 | 0.09 | 0.591416 | 16.77 | 99.1 | 100.9 | 3.48 | 8.12 | | High | High | High | High | High | High | High | High | High | High | High | 0 | 0.00026 | 2.94 |
| FALSE | High | DHSPTPSVI | [R].DHSPTF | 1xDeamida | Q9D824 2x | 0.003083 | 0.00016 | 1 | 3 | 2 | Q9D824 | Pre-mRNA | Q9D824 [4 | 0 | 2106.826 | 0 | -0.11 | 0.850474 | 19.22 | 103.7 | 96.3 | 10.85 | 16.43 | | High | High | High | High | High | High | High | High | High | High | High | 0.000275 | 0.001623 | 2.75 |
| FALSE | High | DHSPTPSVI | [R].DHSPTF | 2xPhospho | Q9D824 2x | 0.006478 | 0.00062 | 1 | 3 | 1 | Q9D824 | Pre-mRNA | Q9D824 [4 | 1 | 2425.006 | 0 | -0.09 | 0.216698 | 12.06 | 107.9 | 92.1 | 11.6 | 13.04 | | High | High | High | High | High | High | High | High | High | High | High | 0.000275 | 0.001936 | 1.55 |
| FALSE | High | DHSSQSEE | [R].DHSSQ | 1xPhospho | Q3TEW0 1x | 0.000538 | 0 | 1 | 3 | 6 | Q3TEW0 | Putative ur | Q3TEW0 [1 | 0 | 2227.017 | 0 | 0.06 | 0.956024 | 10.66 | 98.9 | 101.1 | 5.51 | 10.27 | | High | High | High | High | High | High | High | High | High | High | High | 0 | 0.00012 | 3.54 |
| FALSE | High | DHTLSPPR | [R].DHTLSP | 1xPhospho | B7ZNG4 1x | 0.077825 | 0.005963 | 1 | 1 | 1 | B7ZNG4 | Tastin OS= | B7ZNG4 [1 | 0 | 1231.603 | 0 | 0.39 | 0.056383 | 22.67 | 87 | 113 | 15.78 | 8.93 | | High | High | High | High | High | High | High | High | High | High | High | 0.003109 | 0.03219 | 2.2 |
| FALSE | High | DHVLGPQE | [K].DHVLGI | 1xPhospho | A0A140LJ6 | 0.006536 | 0.00062 | 1 | 6 | 2 | A0A140LJ6 | Protein un | A0A140LJ6 | 0 | 2805.329 | 0 | -0.23 | 0.108264 | 10.7 | 107.8 | 92.2 | 12.12 | 7.19 | | High | High | High | High | High | High | High | High | High | High | High | 0.000275 | 0.001948 | 3.28 |
| FALSE | High | DHVPSTPTS | [R].DHVPST | 1xPhospho | Q3UHU5 1x | 1.12E-05 | 0 | 1 | 5 | 1 | Q3UHU5 | Microtubu | Q3UHU5 [6 | 0 | 2702.185 | 0 | -1.23 | 0.003583 | 5.23 | 135.8 | 64.2 | 8.09 | 18.9 | | High | High | High | High | High | High | High | High | High | High | High | 0 | 1.54E-06 | 2.66 |
| FALSE | High | DHVPSTPTS | [R].DHVPST | 2xPhospho | Q3UHU5 2x | 0.012619 | 0.000889 | 1 | 5 | 2 | Q3UHU5 | Microtubu | Q3UHU5 [6 | 0 | 2766.157 | 0 | -1.23 | 0.001846 | 4.97 | 141.7 | 58.3 | 10.56 | 7.92 | | High | High | High | High | High | High | High | High | High | High | High | 0.000449 | 0.004053 | 3.12 |
| FALSE | High | DHVPSTPTS | [R].DHVPST | 2xPhospho | Q3UHU5 2x | 0.005228 | 0.000551 | 1 | 5 | 2 | Q3UHU5 | Microtubu | Q3UHU5 [6 | 0 | 2782.152 | 0 | -1.63 | 0.000336 | 1.72 | 152.7 | 47.3 | 6.86 | 11.9 | | High | High | High | High | High | High | High | High | High | High | High | 0.000399 | 0.003456 | 3 |
| FALSE | High | DHYAYSDR | [K].DHYAYS | 1xPhospho | A2AIX1 1xP | 0.076555 | 0.005826 | 1 | 2 | 1 | A2AIX1 | SEC16 hom | A2AIX1 [13 | 0 | 1918.91 | 0 | -0.02 | 0.89704 | 14.39 | 99.5 | 100.5 | 4.34 | 8.9 | | High | High | High | High | High | High | High | High | High | High | High | 0.00299 | 0.03156 | 2.21 |
| FALSE | High | DHYQDPVF | [K].DHYQD | 1xPhospho | Q62383 1x | 8.35E-06 | 0 | 1 | 2 | 1 | Q62383 | Transcripti | Q62383 [1 | 0 | 2179.002 | 0 | 0.05 | 0.26863 | 3.41 | 93.4 | 106.6 | 9 | 9.37 | | High | High | High | High | High | High | High | High | High | High | High | 0 | 1.1E-06 | 3.55 |
| FALSE | High | DHYQDPVF | [K].DHYQD | 2xPhospho | Q62383 2x | 0.007339 | 0.00062 | 1 | 2 | 3 | Q62383 | Transcripti | Q62383 [1 | 0 | 2258.968 | 0 | 0.09 | 0.146127 | 7.03 | 94.5 | 105.5 | 7.26 | 10.39 | | High | High | High | High | High | High | High | High | High | High | High | 0.000275 | 0.002222 | 3.58 |
| FALSE | High | DKDSEDEQ | [K].DKDSEI | 1xPhospho | E9QNZ2 1x | 2.58E-08 | 0 | 1 | 3 | 1 | E9QNZ2 | E3 ubiquiti | E9QNZ2 [6 | 1 | 2569.235 | 0 | -0.41 | 0.01841 | 9.08 | 114.2 | 85.8 | 6.07 | 10.41 | | High | High | High | High | High | High | High | High | High | High | High | 0 | 1.66E-09 | 5.35 |
| FALSE | High | DKEVSDDE | [R].DKEVSI | 1xPhospho | Q3UIF3 1x | 0.005615 | 0.00062 | 1 | 9 | 1 | Q3UIF3 | Putative ur | Q3UIF3 [22 | 1 | 2161.052 | 0 | -1.84 | 0.019763 | 2.99 | 146.7 | 53.3 | 21.23 | 49.38 | | High | High | High | High | High | High | High | High | High | High | High | 0.000275 | 0.001651 | 2.39 |
| FALSE | High | DKFSPTQD | [R].DKFSPT | 1xPhospho | Q8BTI8 1xP | 0.002004 | 5.66E-05 | 1 | 3 | 2 | Q8BTI8 | Serine/argi | Q8BTI8 [11 | 1 | 2702.437 | 0 | -1.23 | 0.027964 | 6.71 | 139 | 61 | 26.91 | 15.31 | | High | High | High | High | High | High | High | High | High | High | High | 0.000275 | 0.001789 | 1.04 |
| FALSE | High | DKPTYDEIF | [K].DKPTYI | 1xPhospho | Q8K1X5 1x | 5.93E-06 | 0 | 1 | 6 | 1 | Q8K1X5 | EH-domain | Q8K1X5 [4 | 0 | 2854.488 | 0 | 1.07 | 0.000385 | 8.21 | 65.7 | 134.3 | 9.97 | 7.24 | | High | High | High | High | High | High | High | High | High | High | High | 0 | 7.54E-07 | 3.52 |
| FALSE | High | DKPVYDELI | [K].DKPVYI | 1xPhospho | Q1MWP8 1 | 3.67E-05 | 0 | 1 | 5 | 1 | Q1MWP8 | EH-domain | Q1MWP8 [ | 0 | 2866.525 | 0 | 1.69 | 0.000241 | 10.1 | 50.9 | 149.1 | 10.58 | 5.04 | | High | High | High | High | High | High | High | High | High | High | High | 0 | 5.84E-06 | 3.97 |
| FALSE | High | DKSFDDEE | [K].DKSFDI | 1xPhospho | A0A1L1ST2 | 0.004911 | 0.000504 | 1 | 3 | 2 | A0A1L1ST2 | CLIP-associ | A0A1L1ST2 | 1 | 3226.551 | 0 | -0.69 | 0.017383 | 10.4 | 127.3 | 72.7 | 18.5 | 7.29 | | High | High | High | High | High | High | High | High | High | High | High | 0.002159 | 0.02292 | 3.23 |
| FALSE | High | DKSPATGG | [K].DKSPAT | 1xPhospho | Q8K2E5 1x | 0.00311 | 0.00016 | 1 | 3 | 2 | Q8K2E5 | Budding ur | Q8K2E5 [5 | 1 | 2177.148 | 0 | -0.66 | 0.00817 | 9 | 122.5 | 77.5 | 6.79 | 11.65 | | High | High | High | High | High | High | High | High | High | High | High | 7.19E-05 | 0.000852 | 2.9 |
| FALSE | High | DLEEPNLD | [R].DLEEPN | 2xPhospho | Q3TVI4 2xP | 0.000237 | 0 | 1 | 1 | 1 | Q3TVI4 | Protein HE | Q3TVI4 [15 | 0 | 3621.497 | 0 | -0.66 | 0.003024 | 3.86 | 120.9 | 79.1 | 5.71 | 11.39 | | High | High | High | High | High | High | High | High | High | High | High | 0 | 4.76E-05 | 4.07 |
| FALSE | High | DNPVSPQK | [R].DNPVSI | 1xPhospho | A0A0R4J20 | 0.002077 | 5.66E-05 | 1 | 1 | 2 | A0A0R4J20 | tRNA meth | A0A0R4J20 | 0 | 1422.739 | 0 | 0.2 | 0.008583 | 4.37 | 91.8 | 108.2 | 2.82 | 5.87 | | High | High | High | High | High | High | High | High | High | High | High | 0.000418 | 0.003583 | 1.85 |
| FALSE | High | DNQTLSHS | [K].DNQTL | 1xPhospho | E9Q496 1x | 0.001463 | 5.66E-05 | 1 | 6 | 1 | E9Q496 | Vesicle tra | E9Q496 [2 | 0 | 1680.872 | 0 | -0.01 | 0.892722 | 29.38 | 102.8 | 97.2 | 15.76 | 8.08 | | High | High | High | High | High | High | High | High | High | High | High | 2.58E-05 | 0.000366 | 2.07 |
| FALSE | High | DNSDPAKP | [R].DNSDP | 1xPhospho | Q3UMU9-3 | 0.096959 | 0.007777 | 1 | 4 | 1 | Q3UMU9-3 | Isoform 3 o | Q3UMU9-3 | 0 | 1837.885 | 0 | -0.76 | 0.009505 | 4.96 | 125.7 | 74.3 | 15.88 | 13.63 | | High | High | High | High | High | High | High | High | High | High | High | 0.003922 | 0.04173 | 1.85 |
| FALSE | High | DSLGDFIEH | [R].DSLGDI | 1xPhospho | O35295 1x | 5.98E-06 | 0 | 1 | 1 | 5 | O35295 | Transcripti | O35295 [9 | 0 | 3564.648 | 0 | 0.78 | 0.00192 | 18.15 | 75.9 | 124.1 | 10.34 | 3.71 | | High | High | High | High | High | High | High | High | High | High | High | 0 | 7.61E-07 | 4.47 |
| FALSE | High | DSLGSDSS | [K].DSLGSD | 1xPhospho | Q3U0Z8 1x | 5.07E-08 | 0 | 1 | 5 | 1 | Q3U0Z8 | Suppressor | Q3U0Z8 [3 | 0 | 2220.111 | 0 | -0.36 | 0.084037 | 14.07 | 115.1 | 84.9 | 9.91 | 14.74 | | High | High | High | High | High | High | High | High | High | High | High | 0 | 3.57E-09 | 4.58 |
| FALSE | High | DSLLSTSLN | [R].DSLLST | 1xPhospho | Q8R4Y8 1x | 0.007881 | 0.000736 | 1 | 2 | 1 | Q8R4Y8 | Rotatin OS | Q8R4Y8 [1 | 0 | 1414.714 | 0 | 0.29 | 0.00567 | 2.55 | 88.6 | 111.4 | 12.28 | 11.91 | | High | High | High | High | High | High | High | High | High | High | High | 0.000327 | 0.002396 | 1.68 |
| FALSE | High | DSLITPHVS | [K].DSLITPI | 1xCarbami | Q9ERU9 2x | 0.008167 | 0.000736 | 1 | 1 | 1 | Q9ERU9 | E3 SUMO-p | Q9ERU9 [2 | 1 | 2936.414 | 0 | -0.49 | 0.023817 | 14.49 | 118.5 | 81.5 | 10.14 | 12.79 | | High | High | High | High | High | High | High | High | High | High | High | 0.000327 | 0.002507 | 1.8 |
| FALSE | High | DSLPELQLH | [R].DSLPEL | 2xPhospho | I7H459 2xP | 7.96E-05 | 0 | 1 | 2 | 1 | I7H459 | Protein inc | I7H459 [43 | 0 | 2993.391 | 0 | 0.44 | 0.023123 | 12.72 | 88.1 | 111.9 | 9.33 | 5.98 | | High | High | High | High | High | High | High | High | High | High | High | 0 | 1.39E-05 | 2.34 |
| FALSE | High | DSLPLQEN | [K].DSLPLC | 1xPhospho | Q8BU22 1x | 0.023584 | 0.001578 | 1 | 5 | 2 | Q8BU22 | Kinesin-like | Q8BU22 [7 | 0 | 1978.066 | 0 | -1.07 | 0.003314 | 4.13 | 136.3 | 63.7 | 12.77 | 12.98 | | High | High | High | High | High | High | High | High | High | High | High | 0.001789 | 0.01765 | 1.95 |
| FALSE | High | DSLPSDSPF | [K].DSLPSD | 1xPhospho | Q80U56 1x | 7.84E-06 | 0 | 1 | 1 | 2 | Q80U56 | Late secret | Q80U56 [3 | 0 | 2345.134 | 0 | 0.12 | 0.249318 | 12.05 | 96.3 | 103.7 | 9.17 | 5.78 | | High | High | High | High | High | High | High | High | High | High | High | 0 | 1.17E-06 | 3.71 |
| FALSE | High | DSLQENGS | [K].DSLQEI | 1xPhospho | A2AUZ7 1x | 0.055308 | 0.003982 | 2 | 4 | 1 | A2AUZ7; Q | Sal-like pro | A2AUZ7 [9 | 0 | 1974.942 | 0 | -1.36 | 0.003874 | 2.34 | 142.2 | 57.8 | 12.88 | 15.29 | Shared | High | High | High | High | High | High | High | High | High | High | High | 0.002074 | 0.02156 | 2.04 |
| FALSE | High | DSIQSPK | [R].DSIQSP | 1xPhospho | Q6ZQ33 1x | 0.011652 | 0.000889 | 1 | 4 | 2 | Q6ZQ33 | MKIAA085 | Q6ZQ33 [3 | 0 | 1312.691 | 0 | 1.05 | 0.00045 | 3.5 | 67 | 133 | 8.2 | 6.37 | | High | High | High | High | High | High | High | High | High | High | High | 0.000449 | 0.003707 | 1.5 |
| FALSE | High | DSISEDEM | [K].DSISED | 1xPhospho | Q80ZK9 1x | 0.003892 | 0.000261 | 1 | 3 | 2 | Q80ZK9 | WD and te | Q80ZK9 [5 | 0 | 1618.723 | 0 | -0.44 | 0.170769 | 22.17 | 115.2 | 84.8 | 24.29 | 16.93 | | High | High | High | High | High | High | High | High | High | High | High | 0.00014 | 0.001091 | 3.05 |
| FALSE | High | DSLSMGVS | [K].DSLSM | 1xPhospho | Q80XK6 1x | 0.005516 | 0.00062 | 1 | 2 | 3 | Q80XK6 | Autophagy | Q80XK6 [3 | 0 | 1693.812 | 0 | 1.01 | 0.001485 | 7.43 | 66.6 | 133.4 | 2.36 | 9.13 | | High | High | High | High | High | High | High | High | High | High | High | 0.000275 | 0.00162 | 2.91 |
| FALSE | High | DSLSSQPS | [R].DSLSSC | 1xCarbami | Q9JLM4 1x | 2.14E-06 | 0 | 1 | 4 | 3 | Q9JLM4 | Zinc finger | Q9JLM4 [1 | 0 | 2972.39 | 0 | -0.35 | 0.054915 | 9.57 | 111.9 | 88.1 | 14.96 | 12.76 | | High | High | High | High | High | High | High | High | High | High | High | 0 | 2.39E-07 | 4.71 |
| FALSE | High | DSLSTHTSF | [K].DSLSTH | 2xPhospho | Q80Y56 2x | 0.026421 | 0.001676 | 1 | 2 | 1 | Q80Y56 | Rabenosyn | Q80Y56 [2 | 0 | 2470.06 | 0 | 0.85 | 0.021812 | 34.63 | 69.4 | 130.6 | 21.48 | 14.12 | | High | High | High | High | High | High | High | High | High | High | High | 0.000903 | 0.009286 | 2.48 |
| FALSE | High | DSLESAPSL | [R].DSLESA | 1xPhospho | E9Q309 1x | 4.68E-05 | 0 | 1 | 1 | 1 | E9Q309 | Centrosom | E9Q309 [2 | 0 | 1867.982 | 0 | -0.36 | 0.004737 | 4.5 | 111.7 | 88.3 | 5.29 | 8.46 | | High | High | High | High | High | High | High | High | High | High | High | 0 | 7.62E-06 | 2.8 |
| FALSE | High | DSLTGSSDI | [R].DSLTGS | 1xPhospho | Q3UR00 1x | 0.000699 | 0 | 1 | 8 | 1 | Q3UR00 | Putative ur | Q3UR00 [7 | 0 | 1723.856 | 0 | -0.53 | 0.00276 | 4.39 | 120.1 | 79.9 | 7.26 | 4.31 | | High | High | High | High | High | High | High | High | High | High | High | 0 | 0.00016 | 2.92 |
| FALSE | High | DSLTQAQE | [K].DSLTQA | 1xPhospho | I7H459 1xP | 2.38E-06 | 0 | 1 | 2 | 2 | I7H459 | Protein inc | I7H459 [53 | 0 | 1785.847 | 0 | 0.83 | 0.011434 | 17.88 | 72 | 128 | 10.4 | 14.15 | | High | High | High | High | High | High | High | High | High | High | High | 0 | 2.7E-07 | 2.77 |
| FALSE | High | DSMNATST | [K].DSMNA | 1xPhospho | Q61666 1x | 1.05E-08 | 0 | 1 | 1 | 5 | Q61666 | Protein HIF | Q61666 [5 | 0 | 2704.315 | 0 | -0.25 | 0.2116 | 9.83 | 104.2 | 95.8 | 7.84 | 9.86 | | High | High | High | High | High | High | High | High | High | High | High | 0 | 6.11E-10 | 4.26 |
| FALSE | High | DSNTSYTE | [R].DSNTSY | 1xPhospho | F6RJ39 1xP | 2.21E-05 | 0 | 2 | 4 | 4 | F6RJ39; Q8 | Apoptotic | F6RJ39 [60 | 0 | 1683.788 | 0 | -0.73 | 0.000284 | 0.71 | 124.1 | 75.9 | 5.71 | 6.76 | Shared | High | High | High | High | High | High | High | High | High | High | High | 0 | 9.06E-05 | 2.38 |
| FALSE | High | DSPDSPEPI | [K].DSPDSF | 1xPhospho | Q8QZV7 1x | 0.001222 | 0 | 1 | 3 | 3 | Q8QZV7 | Integrator | Q8QZV7 [6 | 0 | 1720.82 | 0 | -0.06 | 0.542264 | 9.81 | 101 | 99 | 5.26 | 4.23 | | High | High | High | High | High | High | High | High | High | High | High | 7.19E-05 | 0.00077 | 3.25 |
| FALSE | High | DSPEILSR | [K].DSPEILS | 1xPhospho | Q920R0 1x | 0.018774 | 0.001362 | 1 | 2 | 4 | Q920R0 | Alsin OS=M | Q920R0 [1 | 0 | 1225.603 | 0 | -1.03 | 0.001066 | 2.83 | 136.5 | 63.5 | 7.88 | 9.22 | | High | High | High | High | High | High | High | High | High | High | High | 0.002159 | 0.02276 | 2.27 |
| FALSE | High | DSPGIPPSA | [R].DSPGIP | 1xPhospho | Q810V8 1x | 5.88E-06 | 0 | 1 | 8 | 6 | Q810V8 | Rps6ka1 pr | Q810V8 [4 | 0 | 1958.969 | 0 | -0.71 | 0.001651 | 2.96 | 123.4 | 76.6 | 6.53 | 4.29 | | High | High | High | High | High | High | High | High | High | High | High | 0 | 7.47E-07 | 2.82 |
| FALSE | High | DSPGIPPSA | [K].DSPGIP | 1xPhospho | A7E215 1x | 2.06E-05 | 0 | 1 | 3 | 9 | A7E215 | Rps6ka3 pr | A7E215 [36 | 0 | 2015.99 | 0 | 1.81 | 0.001112 | 16.19 | 44.1 | 155.9 | 9.34 | 10.78 | | High | High | High | High | High | High | High | High | High | High | High | 0 | 0.000123 | 2.71 |
| FALSE | High | DSPGNFLS | [R].DSPGN | 1xPhospho | B2RWW9 1 | 0.000779 | 0 | 1 | 2 | 1 | B2RWW9 | Absent in r | B2RWW9 [ | 0 | 1884.951 | 0 | -1.3 | 0.002593 | 2.88 | 142.2 | 57.8 | 10.69 | 10.71 | | High | High | High | High | High | High | High | High | High | High | High | 0 | 0.000179 | 2.61 |
| FALSE | High | DSPLATR | [K].DSPLAT | 1xPhospho | Q80TU3 1x | 0.042868 | 0.003228 | 1 | 4 | 1 | Q80TU3 | MKIAA068 | Q80TU3 [6 | 0 | 1068.529 | 0 | -1.25 | 0.00151 | 4.47 | 142.5 | 57.5 | 8.03 | 21.58 | | High | High | High | High | High | High | High | High | High | High | High | 0.001651 | 0.0162 | 2.49 |
| FALSE | High | DSPLGGVN | [K].DSPLGC | 1xPhospho | Q8R3F9 1x | 0.020142 | 0.001472 | 1 | 1 | 2 | Q8R3F9 | Speckle tar | Q8R3F9 [6 | 0 | 1424.755 | 0 | -0.34 | 0.000445 | 0.59 | 113.6 | 86.4 | 3.19 | 5.01 | | High | High | High | High | High | High | High | High | High | High | High | 0.000786 | 0.00684 | 2.13 |
| FALSE | High | DSPISPHFT | [K].DSPISPI | 1xPhospho | E9Q3Y4 1x | 2.93E-05 | 0 | 1 | 7 | 2 | E9Q3Y4 | Lipopolysa | E9Q3Y4 [9 | 0 | 1465.704 | 0 | 0.18 | 0.174775 | 17.63 | 90.6 | 109.4 | 11.37 | 19.81 | | High | High | High | High | High | High | High | High | High | High | High | 0 | 4.52E-06 | 2.89 |
| FALSE | High | DSPPLVTPI | [K].DSPPLV | 2xPhospho | Q8CI12 2xP | 9.28E-05 | 0 | 1 | 2 | 2 | Q8CI12 | Smoothelin | Q8CI12 [26 | 0 | 2972.383 | 0 | -1.83 | 0.000774 | 0.89 | 155.7 | 44.3 | 5.15 | 36.94 | | High | High | High | High | High | High | High | High | High | High | High | 7.19E-05 | 0.000843 | 3.46 |
| FALSE | High | DSPPLVTPI | [K].DSPPLV | 2xPhospho | Q8CI12 2xP | 0.005666 | 0.00062 | 1 | 2 | 3 | Q8CI12 | Smoothelin | Q8CI12 [26 | 0 | 2988.378 | 0 | -0.78 | 0.002671 | 7.18 | 128 | 72 | 4.62 | 13.13 | | High | High | High | High | High | High | High | High | High | High | High | 0.000275 | 0.00166 | 2.8 |
| FALSE | High | DSLTPVHG | [K].DSLTPV | 1xPhospho | Q8BGS1 1x | 9.28E-05 | 0 | 1 | 2 | 2 | Q8BGS1 | Band 4.1-li | Q8BGS1 [6 | 0 | 2307.2 | 0 | -0.67 | 0.009826 | 7.81 | 126.7 | 73.3 | 9.73 | 14.35 | | High | High | High | High | High | High | High | High | High | High | High | 0 | 6.35E-05 | 2.34 |
| FALSE | High | DSPPPEGL | [R].DSPPPE | 1xPhospho | Q148V8 1x | 0.002671 | 0.000109 | 1 | 1 | 1 | Q148V8 | Protein FAI | Q148V8 [8 | 0 | 2587.285 | 0 | 1.6 | 0.000278 | 20.72 | 49 | 151 | 13.27 | 5.1 | | High | High | High | High | High | High | High | High | High | High | High | 4.94E-05 | 0.000719 | 2.55 |
| FALSE | High | DSLDDLFPI | [K].DSLDDI | 1xPhospho | Q9DBS5 1x | 0.085162 | 0.006376 | 1 | 2 | 1 | Q9DBS5 | Kinesin ligh | Q9DBS5 [1 | 0 | 2705.13 | 0 | -0.91 | 0.01134 | 3.77 | 129 | 71 | 3.24 | 24.23 | | High | High | High | High | High | High | High | High | High | High | High | 0.003291 | 0.03565 | 2.29 |
| FALSE | High | DSGVGASL | [R].DSGVG | 1xPhospho | E9PXD2 1x | 0.000738 | 0 | 1 | 7 | 1 | E9PXD2 | Rho GTPas | E9PXD2 [1 | 0 | 1271.619 | 0 | -1.73 | 0.000799 | 2.64 | 155.7 | 44.3 | 10.02 | 17.18 | | High | High | High | High | High | High | High | High | High | High | High | 0 | 0.000169 | 1.95 |
| FALSE | High | DSDSVILTL | [K].DSDSVI | 1xPhospho | Q8VCH5 1x | 3.87E-05 | 0 | 1 | 3 | 5 | Q8VCH5 | Rab9 effec | Q8VCH5 [3 | 0 | 2001.019 | 0 | 0.39 | 0.001703 | 2.75 | 86.8 | 113.2 | 1.84 | 6.41 | | High | High | High | High | High | High | High | High | High | High | High | 0 | 2.92E-05 | 3.16 |
| FALSE | High | DSDVFTDL | [K].DSDVFT | 1xPhospho | E9PZJ8 1xP | 0.000475 | 0 | 1 | 2 | 2 | E9PZJ8 | Activating | E9PZJ8 [21 | 0 | 1406.629 | 0 | 0.02 | 0.217462 | 4.34 | 96.5 | 103.5 | 8.8 | 4.8 | | High | High | High | High | High | High | High | High | High | High | High | 0 | 0.000103 | 2.74 |
| FALSE | High | DSDYVYPSI | [K].DSDYV | 1xPhospho | Q9WTU0 1 | 1.31E-06 | 0 | 1 | 2 | 2 | Q9WTU0 | Lysine-spe | Q9WTU0 [ | 0 | 2757.258 | 0 | 1.62 | 0.001758 | 8.29 | 50.1 | 149.9 | 6.11 | 11.77 | | High | High | High | High | High | High | High | High | High | High | High | 0 | 1.38E-07 | 3.55 |
| FALSE | High | DSEEGEESI | [K].DSEEGI | 1xPhospho | G3UWX1 1 | 3.38E-08 | 0 | 2 | 9 | 4 | G3UWX1; I | Replication | G3UWX1 [ | 0 | 2526.117 | 0 | 0.1 | 0.048736 | 6.35 | 94.8 | 105.2 | 6.36 | 3.24 | Shared | High | High | High | High | High | High | High | High | High | High | High | 0 | 2.26E-09 | 4.09 |
| FALSE | High | DSEEGEESI | [K].DSEEGI | 2xPhospho | G3UWX1 2 | 4.35E-05 | 0 | 2 | 9 | 3 | G3UWX1; I | Replication | G3UWX1 [ | 0 | 2606.084 | 0 | -0.13 | 0.115535 | 9.2 | 103 | 97 | 15.62 | 7.95 | Shared | High | High | High | High | High | High | High | High | High | High | High | 0 | 8.85E-06 | 3.99 |
| FALSE | High | DSEGDSQG | [R].DSEGD | 1xPhospho | Q6PD29 1x | 0.002192 | 0.000109 | 1 | 2 | 2 | Q6PD29 | Zinc finger | Q6PD29 [6 | 0 | 2888.253 | 0 | 0.04 | 0.416855 | 12.47 | 98.7 | 101.3 | 10.7 | 4.63 | | High | High | High | High | High | High | High | High | High | High | High | 4.94E-05 | 0.000576 | 2.52 |
| FALSE | High | DSEGYSESI | [K].DSEGYS | 1xPhospho | Q3UGC1 1x | 5.8E-08 | 0 | 1 | 4 | 7 | Q3UGC1 | Putative ur | Q3UGC1 [5 | 0 | 2835.217 | 0 | 0.64 | 0.005039 | 7.46 | 78.2 | 121.8 | 11.24 | 6.19 | | High | High | High | High | High | High | High | High | High | High | High | 0 | 4.14E-09 | 4.69 |
| FALSE | High | DSELSDSDS | [R].DSELSD | 1xPhospho | Q8VDF2 1x | 1E-06 | 0 | 1 | 3 | 11 | Q8VDF2 | E3 ubiquiti | Q8VDF2 [8 | 0 | 2379.966 | 0 | -0.04 | 0.47415 | 9.85 | 101.5 | 98.5 | 6.15 | 8.35 | | High | High | High | High | High | High | High | High | High | High | High | 0.000903 | 0.009464 | 3.2 |
| FALSE | High | DSELSDSDS | [R].DSELSD | 2xPhospho | Q8VDF2 2x | 0.008538 | 0.000736 | 1 | 3 | 5 | Q8VDF2 | E3 ubiquiti | Q8VDF2 [8 | 0 | 2459.933 | 0 | 0.22 | 0.464397 | 16.74 | 95.6 | 104.4 | 11.42 | 20.69 | | High | High | High | High | High | High | High | High | High | High | High | 0.003809 | 0.04103 | 2.61 |
| FALSE | High | DSELSDSDS | [R].DSELSD | 1xPhospho | Q8VDF2 1x | 3.14E-08 | 0 | 1 | 3 | 12 | Q8VDF2 | E3 ubiquiti | Q8VDF2 [8 | 0 | 2609.129 | 0 | 0.29 | 0.026679 | 7.55 | 88.2 | 111.8 | 4.67 | 14.41 | | High | High | High | High | High | High | High | High | High | High | High | 0 | 2.09E-09 | 3.11 |
| FALSE | High | DSELSDSDS | [R].DSELSD | 2xPhospho | Q8VDF2 2x | 1.41E-06 | 0 | 1 | 3 | 4 | Q8VDF2 | E3 ubiquiti | Q8VDF2 [8 | 0 | 2689.096 | 0 | 0.62 | 0.035032 | 12.25 | 82.6 | 117.4 | 10.37 | 18.4 | | High | High | High | High | High | High | High | High | High | High | High | 0 | 1.49E-07 | 3.01 |
| FALSE | High | DSELSDSDS | [R].DSELSD | 1xPhospho | Q8VDF2 1x | 0.007274 | 0.00062 | 1 | 3 | 2 | Q8VDF2 | E3 ubiquiti | Q8VDF2 [8 | 1 | 3964.688 | 0 | -0.24 | 0.43661 | 35.75 | 108.4 | 91.6 | 23.03 | 35.79 | | High | High | High | High | High | High | High | High | High | High | High | 0.000275 | 0.002189 | 3.11 |
| FALSE | High | DSELSDSDS | [R].DSELSD | 2xPhospho | Q8VDF2 2x | 0.052576 | 0.00384 | 1 | 3 | 1 | Q8VDF2 | E3 ubiquiti | Q8VDF2 [8 | 1 | 4044.655 | 0 | 0.53 | 0.097653 | 38.56 | 84.5 | 115.5 | 21.17 | 16.16 | | High | High | High | High | High | High | High | High | High | High | High | 0.001999 | 0.02036 | 3.65 |
| FALSE | High | DSHSPEGS | [K].DSHSPE | 1xPhospho | Q9R0A0 1x | 0.000127 | 0 | 1 | 2 | 7 | Q9R0A0 | Peroxisoma | Q9R0A0 [2 | 0 | 3490.678 | 0 | 0.4 | 0.110409 | 46.16 | 86.3 | 113.7 | 11.24 | 21.12 | | High | High | High | High | High | High | High | High | High | High | High | 0 | 2.76E-05 | 3.45 |
| FALSE | High | DSELSPPTS | [K].DSELSP | 1xPhospho | E9QM75 1x | 7.77E-06 | 0 | 1 | 9 | 6 | E9QM75 | TNFAIP3-in | E9QM75 [6 | 0 | 3458.687 | 0 | -0.4 | 0.018074 | 10.84 | 120.8 | 79.2 | 13.42 | 10.58 | | High | High | High | High | High | High | High | High | High | High | High | 0 | 1.01E-06 | 4.26 |
| FALSE | High | DSESESDET | [R].DSESES | 1xPhospho | Q3TI79 1xP | 0.01029 | 0.000772 | 1 | 2 | 1 | Q3TI79 | Putative ur | Q3TI79 [45 | 0 | 2300.095 | 0 | -0.72 | 0.007148 | 6.84 | 122.7 | 77.3 | 3.59 | 15.66 | | High | High | High | High | High | High | High | High | High | High | High | 0.000382 | 0.003239 | 2.97 |
| FALSE | High | DSESESDET | [R].DSESES | 2xPhospho | Q3TI79 2xP | 0.000633 | 0 | 1 | 2 | 2 | Q3TI79 | Putative ur | Q3TI79 [45 | 0 | 2380.061 | 0 | -0.69 | 0.002116 | 0.3 | 123.3 | 76.7 | 4.94 | 8.66 | | High | High | High | High | High | High | High | High | High | High | High | 2.58E-05 | 0.000384 | 3.55 |
| FALSE | High | DSETIEDVA | [R].DSETIEI | 1xPhospho | Q8CIL4 1xP | 1.01E-08 | 0 | 1 | 3 | 4 | Q8CIL4 | Uncharacte | Q8CIL4 [50 | 0 | 2529.269 | 0 | -0.8 | 0.000637 | 3.31 | 126.9 | 73.1 | 5 | 14.13 | | High | High | High | High | High | High | High | High | High | High | High | 0 | 1.42E-06 | 4.23 |
| FALSE | High | DSFEMEEV | [K].DSFEM | 1xCarbami | Q922B9 1x | 0.000386 | 0 | 1 | 5 | 2 | Q922B9 | Sperm-spe | Q922B9 [4 | 0 | 3628.695 | 0 | 0.52 | 0.072288 | 21.99 | 82.2 | 117.8 | 18.17 | 24.05 | | High | High | High | High | High | High | High | High | High | High | High | 0.004411 | 0.04748 | 2.72 |
| FALSE | High | DSFIDSSSS | [K].DSFIDS | 1xCarbami | E9QM77 1x | 7.49E-09 | 0 | 1 | 8 | 2 | E9QM77 | Ataxin-2 O | E9QM77 [8 | 0 | 2478.074 | 0 | 1.18 | 0.00066 | 2.78 | 61.3 | 138.7 | 4.24 | 7.04 | | High | High | High | High | High | High | High | High | High | High | High | 0 | 4.17E-10 | 5.49 |
| FALSE | High | DSFSGSK | [R].DSFSGS | 1xPhospho | Q8BM75 1x | 0.112856 | 0.009146 | 1 | 3 | 1 | Q8BM75 | AT-rich inte | Q8BM75 [2 | 0 | 1265.618 | 0 | 0.71 | 0.000946 | 12.83 | 74.6 | 125.4 | 9.08 | 3.55 | | High | High | High | High | High | High | High | High | High | High | High | 0.004569 | 0.04968 | 1.84 |
| FALSE | High | DSGCYESSI | [R].DSGCYI | 1xCarbami | F8VQK5 1x | 0.037048 | 0.00249 | 1 | 3 | 2 | F8VQK5 | SAM and S | F8VQK5 [7 | 0 | 2240.978 | 0 | 0.35 | 0.067812 | 27.46 | 84.2 | 115.8 | 7.57 | 18.53 | | High | High | High | High | High | High | High | High | High | High | High | 0.001308 | 0.01369 | 2.75 |
| FALSE | High | DSGNWDT | [K].DSGNW | 2xPhospho | Q9R1C7 2x | 0.000176 | 0 | 1 | 3 | 3 | Q9R1C7 | Pre-mRNA- | Q9R1C7 [9 | 0 | 2658.126 | 0 | -0.46 | 0.004581 | 7.42 | 115.7 | 84.3 | 6.27 | 20.17 | | High | High | High | High | High | High | High | High | High | High | High | 0 | 3.39E-05 | 3.35 |
| FALSE | High | DSGSDEDF | [K].DSGSDI | 1xPhospho | Q80XU3 1x | 8.76E-07 | 0 | 1 | 1 | 2 | Q80XU3 | Nuclear ub | Q80XU3 [1 | 0 | 2983.16 | 0 | -0.26 | 0.448961 | 32.97 | 112.4 | 87.6 | 21.63 | 33.65 | | High | High | High | High | High | High | High | High | High | High | High | 0 | 8.77E-08 | 3.74 |
| FALSE | High | DSGSDLSH | [K].DSGSDI | 1xPhospho | Q5SVQ0 1x | 0.003788 | 0.00021 | 1 | 7 | 3 | Q5SVQ0 | Histone ac | Q5SVQ0 [1 | 0 | 1736.873 | 0 | -1.65 | 0.000254 | 1.01 | 149.4 | 50.6 | 5.2 | 7.24 | | High | High | High | High | High | High | High | High | High | High | High | 0.000117 | 0.001065 | 1.72 |
| FALSE | High | DSGSPLQFI | [K].DSGSPL | 1xPhospho | Q91YK2 1x | 0.000112 | 0 | 1 | 3 | 3 | Q91YK2 | Ribosomal | Q91YK2 [2 | 0 | 1794.872 | 0 | -0.93 | 0.004087 | 6.66 | 129.4 | 70.6 | 3.96 | 16.74 | | High | High | High | High | High | High | High | High | High | High | High | 0 | 2.05E-05 | 2.69 |
| FALSE | High | DSGSSSPLF | [K].DSGSSS | 1xPhospho | E9QKK4 1x | 0.001077 | 0 | 1 | 6 | 2 | E9QKK4 | Glucocortic | E9QKK4 [3 | 0 | 1512.771 | 0 | 0.98 | 0.001634 | 7.39 | 67.6 | 132.4 | 11.19 | 7.52 | | High | High | High | High | High | High | High | High | High | High | High | 0 | 0.000259 | 1.72 |
| FALSE | High | DSGSVSLQ | [K].DSGSVS | 1xPhospho | O54988 1x | 0.033694 | 0.002195 | 1 | 2 | 1 | O54988 | STE20-like | O54988 [7 | 0 | 1487.694 | 0 | 0.58 | 0.005551 | 3.51 | 79 | 121 | 5.46 | 14.13 | | High | High | High | High | High | High | High | High | High | High | High | 0.001151 | 0.01232 | 2.19 |
| FALSE | High | DSENTPVK | [K].DSENTI | 1xPhospho | P97496 1xP | 0.097743 | 0.007804 | 1 | 6 | 2 | P97496 | SWI/SNF c | P97496 [3 | 0 | 1427.718 | 0 | -1.2 | 0.000181 | 0.4 | 141 | 59 | 3.45 | 8.33 | | High | High | High | High | High | High | High | High | High | High | High | 0.006195 | 0.06697 | 2.31 |

| | | | | | | | | | | | | | | | | | | | | | | | | | | | | | | | | | | | | | | |
|---|---|---|---|---|---|---|---|---|---|---|---|---|---|---|---|---|---|---|---|---|---|---|---|---|---|---|---|---|---|---|---|---|---|---|---|---|---|---|
| FALSE | High | DSPPVSLSE | [R].DSPPVS | 1xPhosphc | Q3U3C9-2 | 0.003167 | 0.00016 | 1 | 3 | 2 | Q3U3C9-2 | Isoform 2 c | Q3U3C9-2 | 0 | 3964.024 | 0 | -1.42 | 0.000716 | 1.7 | 142.3 | 57.7 | 7.88 | 11.86 | | High | High | High | High | High | High | High | High | High | High | High | 0.000275 | 0.002033 | 3.65 |
| FALSE | High | DSPSAGSPI | [R].DSPSAC | 1xPhosphc | Q80U35 1x | 0.003495 | 0.00021 | 1 | 1 | 1 | Q80U35 | Rho guanir | Q80U35 [4 | 0 | 3642.761 | 0 | 0.03 | 0.030911 | 14.03 | 112.5 | 87.5 | 3.76 | 21.83 | | High | High | High | High | High | High | High | High | High | High | High | 9.42E-05 | 0.000973 | 3.42 |
| FALSE | High | DSPSAIPAC | [K].DSPSAI | 1xPhosphc | Q9D0N7 1x | 0.001147 | 0 | 1 | 1 | 1 | Q9D0N7 | Chromatin | Q9D0N7 [4 | 0 | 1480.781 | 0 | 0.7 | 0.002441 | 7.26 | 75.5 | 124.5 | 14.88 | 3.23 | | High | High | High | High | High | High | High | High | High | High | High | 0 | 0.000278 | 2.27 |
| FALSE | High | DSSPVPSFC | [R].DSSPVF | 1xPhosphc | E9PYD1 1x | 0.037692 | 0.00249 | 1 | 4 | 2 | E9PYD1 | Family wit | E9PYD1 [1 | 0 | 1845.904 | 0 | -1.13 | 0.004327 | 4.53 | 134.4 | 65.6 | 7.22 | 17.57 | | High | High | High | High | High | High | High | High | High | High | High | 0.001308 | 0.0139 | 2.16 |
| FALSE | High | DSSQESQS | [R].DSSQES | 1xCarbami | Q8R2M2 1 | 4.2E-05 | 0 | 1 | 1 | 10 | Q8R2M2 | Deoxynucl | Q8R2M2 [ | 0 | 3348.535 | 0 | 0.07 | 0.875315 | 9.2 | 102.1 | 97.9 | 8.52 | 11.34 | | High | High | High | High | High | High | High | High | High | High | High | 0 | 6.79E-06 | 3.75 |
| FALSE | High | DSSQEYTD | [R].DSSQEY | 1xPhosphc | E9Q9Q2 1x | 9.59E-07 | 0 | 2 | 12 | 2 | E9Q9Q2; C | R3H domai | E9Q9Q2 [1 | 0 | 3150.528 | 0 | 0.55 | 0.005024 | 6.56 | 82.2 | 117.8 | 1.14 | 10.31 | Shared | High | High | High | High | High | High | High | High | High | High | High | 0 | 1.02E-07 | 5 |
| FALSE | High | DSSSLEPM | [K].DSSSLE | 1xPhosphc | Q6ZPJ3 1x | 0.022378 | 0.001542 | 1 | 1 | 1 | Q6ZPJ3 | (E3-indepe | Q6ZPJ3 [11 | 0 | 2131.942 | 0 | -0.46 | 0.045008 | 11.64 | 117.9 | 82.1 | 18.09 | 7.18 | | High | High | High | High | High | High | High | High | High | High | High | 0.000814 | 0.007756 | 3.12 |
| FALSE | High | DSSSLEPM | [K].DSSSLE | 1xPhosphc | Q6ZPJ3 1x | 0.001398 | 5.66E-05 | 1 | 1 | 1 | Q6ZPJ3 | (E3-indepe | Q6ZPJ3 [11 | 0 | 2147.937 | 0 | 0.01 | 0.485791 | 7.19 | 101.8 | 98.2 | 13.84 | 6.17 | | High | High | High | High | High | High | High | High | High | High | High | 2.58E-05 | 0.000347 | 2.58 |
| FALSE | High | DSSSLSSPL | [R].DSSSLS | 1xPhosphc | Q9Z277 1x | 0.028327 | 0.001791 | 1 | 2 | 1 | Q9Z277 | Tyrosine-p | Q9Z277 [3 | 0 | 1769.909 | 0 | 1.39 | 0.003596 | 11.31 | 55.3 | 144.7 | 8.97 | 17.02 | | High | High | High | High | High | High | High | High | High | High | High | 0.000968 | 0.01011 | 2.5 |
| FALSE | High | DSSSLSSYT | [R].DSSSLS | 1xPhosphc | Q3TH99 1x | 2.79E-07 | 0 | 1 | 2 | 2 | Q3TH99 | Putative ur | Q3TH99 [2 | 0 | 2025.922 | 0 | -0.88 | 0.000808 | 7.15 | 132.3 | 67.7 | 9.38 | 23.49 | | High | High | High | High | High | High | High | High | High | High | High | 0 | 2.42E-08 | 3.81 |
| FALSE | High | DSSSGSGS | [K].DSSSGS | 1xPhosphc | Q61687 1x | 0.000209 | 0 | 1 | 3 | 3 | Q61687 | Transcripti | Q61687 [1 | 0 | 2221.027 | 0 | 0.36 | 0.007915 | 4.07 | 87.1 | 112.9 | 7.14 | 7.95 | | High | High | High | High | High | High | High | High | High | High | High | 0 | 0.000103 | 3.56 |
| FALSE | High | DSSSGSGS | [K].DSSSGS | 2xPhosphc | Q61687 2x | 0.007881 | 0.000736 | 1 | 3 | 4 | Q61687 | Transcripti | Q61687 [1 | 0 | 2300.994 | 0 | -0.07 | 0.183774 | 6.78 | 104.9 | 95.1 | 9.39 | 10.18 | | High | High | High | High | High | High | High | High | High | High | High | 0.001011 | 0.01107 | 3.66 |
| FALSE | High | DSSSSLVDF | [K].DSSSSL | 1xPhosphc | Q91VS8 1x | 0.000369 | 0 | 1 | 2 | 4 | Q91VS8 | FERM, Rho | Q91VS8 [4 | 0 | 2081.093 | 0 | 0.67 | 0.008081 | 28.35 | 70.2 | 129.8 | 11.14 | 11.47 | | High | High | High | High | High | High | High | High | High | High | High | 2.58E-05 | 0.000355 | 3.01 |
| FALSE | High | DSSTCPGD | [R].DSSTCF | 1xCarbami | P47941 1x | 0.000284 | 0 | 1 | 2 | 2 | P47941 | Crk-like pr | P47941 [4 | 0 | 2281.985 | 0 | -0.12 | 0.114871 | 2.12 | 108.9 | 91.1 | 15.96 | 5.96 | | High | High | High | High | High | High | High | High | High | High | High | 0 | 5.79E-05 | 3.17 |
| FALSE | High | DSSTETEHC | [K].DSSTET | 1xPhosphc | A0A140T8 | 0.080423 | 0.006219 | 1 | 2 | 3 | A0A140T8 | Protein Jac | A0A140T8 | 0 | 2780.278 | 0 | -1 | 0.094159 | 9.71 | 133.4 | 66.6 | 32.42 | 30.7 | | High | High | High | High | High | High | High | High | High | High | High | 0.005149 | 0.05601 | 1.69 |
| FALSE | High | DSSTETEHC | [K].DSSTET | 2xPhosphc | A0A140T8 | 1.54E-05 | 0 | 1 | 2 | 2 | A0A140T8 | Protein Jac | A0A140T8 | 0 | 2860.244 | 0 | -0.78 | 0.017445 | 11.68 | 126.4 | 73.6 | 13.93 | 18.46 | | High | High | High | High | High | High | High | High | High | High | High | 0 | 2.21E-06 | 3.83 |
| FALSE | High | DSSPTLLEA | [K].DSSPTL | 1xCarbami | A0A0R4J0 | 1.33E-06 | 0 | 1 | 2 | 3 | A0A0R4J0C | E3 ubiquiti | A0A0R4J0C | 0 | 2492.24 | 0 | 2.17 | 0.000485 | 29.86 | 36.4 | 163.6 | 17.19 | 7.46 | | High | High | High | High | High | High | High | High | High | High | High | 0 | 1.78E-07 | 3.78 |
| FALSE | High | DSSTSPGD | [K].DSSTSP | 1xPhosphc | Q64010 1x | 4.95E-06 | 0 | 1 | 7 | 8 | Q64010 | Adapter m | Q64010 [3 | 0 | 2208.986 | 0 | 0.46 | 0.001387 | 5.15 | 84.3 | 115.7 | 3.05 | 5.86 | | High | High | High | High | High | High | High | High | High | High | High | 0 | 6.13E-07 | 3.77 |
| FALSE | High | DSTEAPKPE | [R].DSTEAF | 1xPhosphc | K3W4P2 1x | 0.002895 | 0.00016 | 1 | 4 | 4 | K3W4P2 | Integrator | K3W4P2 [1 | 0 | 2601.26 | 0 | -1.16 | 0.000461 | 0.4 | 138.1 | 61.9 | 5.3 | 8.2 | | High | High | High | High | High | High | High | High | High | High | High | 7.19E-05 | 0.000786 | 2.46 |
| FALSE | High | DSTMNLNI | [K].DSTMN | 1xPhosphc | Q91ZT5 1x | 0.000137 | 0 | 1 | 6 | 1 | Q91ZT5 | FYVE, Rho | Q91ZT5 [4 | 1 | 3106.601 | 0 | -0.04 | 0.534273 | 12.51 | 101.8 | 98.2 | 5.48 | 11.7 | | High | High | High | High | High | High | High | High | High | High | High | 0 | 2.56E-05 | 2.7 |
| FALSE | High | DSTPGSGP | [K].DSTPGS | 1xPhosphc | Q5U4C3 1x | 0.000356 | 0 | 1 | 1 | 2 | Q5U4C3 | Splicing fac | Q5U4C3 [7 | 0 | 1593.829 | 0 | -0.07 | 0.356572 | 2.01 | 99.8 | 100.2 | 6.09 | 5.87 | | High | High | High | High | High | High | High | High | High | High | High | 2.58E-05 | 0.000383 | 3.18 |
| FALSE | High | DSTSQHDD | [K].DSTSQI | 1xPhosphc | E9Q3G8 1x | 4.47E-05 | 0 | 1 | 1 | 6 | E9Q3G8 | Nucleopor | E9Q3G8 [1 | 0 | 2466.026 | 0 | 0.17 | 0.979441 | 18.47 | 96.9 | 103.1 | 15.65 | 8.46 | | High | High | High | High | High | High | High | High | High | High | High | 0 | 7.27E-06 | 3.62 |
| FALSE | High | DSTSQHDD | [K].DSTSQI | 2xPhosphc | E9Q3G8 2x | 0.006031 | 0.00062 | 1 | 1 | 1 | E9Q3G8 | Nucleopor | E9Q3G8 [1 | 0 | 2545.992 | 0 | -0.46 | 0.043057 | 13.81 | 115.3 | 84.7 | 5.66 | 18.07 | | High | High | High | High | High | High | High | High | High | High | High | 0.000275 | 0.001788 | 2.19 |
| FALSE | High | DSVCVSGE | [R].DSVCVS | 1xCarbami | B9EKI3 1x | 0.000538 | 0 | 1 | 2 | 1 | B9EKI3 | TATA elem | B9EKI3 [15 | 0 | 2756.31 | 0 | 1.35 | 0.00172 | 14.26 | 58.3 | 141.7 | 13.51 | 10.12 | | High | High | High | High | High | High | High | High | High | High | High | 0 | 0.00012 | 2.48 |
| FALSE | High | DSVTEDPC | [R].DSVTEI | 1xCarbami | B7ZNH9 1x | 0.000958 | 0 | 1 | 3 | 1 | B7ZNH9 | Phosphoin | B7ZNH9 [1 | 0 | 2419.141 | 0 | 1.23 | 0.005133 | 31.28 | 63.1 | 136.9 | 25.6 | 9.91 | | High | High | High | High | High | High | High | High | High | High | High | 0 | 0.000228 | 3.11 |
| FALSE | High | DSYGGPPR | [R].DSYGG | 1xPhosphc | Q91VM5 1 | 0.000543 | 0 | 1 | 1 | 2 | Q91VM5 | RNA bindir | Q91VM5 [ | 0 | 1157.519 | 0 | -0.58 | 0.015289 | 9.8 | 119.8 | 80.2 | 11.04 | 5.95 | | High | High | High | High | High | High | High | High | High | High | High | 0 | 0.00012 | 3.17 |
| FALSE | High | DSYSSSRSD | [R].DSYSSS | 1xPhosphc | Q91VM5 1 | 0.003138 | 0.00016 | 1 | 3 | 1 | Q91VM5 | RNA bindir | Q91VM5 [ | 1 | 1975.86 | 0 | -0.36 | 0.008011 | 6.87 | 112.4 | 87.6 | 8.56 | 3.92 | | High | High | High | High | High | High | High | High | High | High | High | 7.19E-05 | 0.000861 | 1.39 |
| FALSE | High | DSYVGDEA | [KR].DSYV | 1xPhosphc | P60710 1x | 0.01861 | 0.001362 | 2 | 40 | 2 | P60710; P6 | Actin, cyto | P60710 [51 | 0 | 1736.814 | 0 | 0.55 | 0.012174 | 20.29 | 81.1 | 118.9 | 10.96 | 12.45 | Shared | High | High | High | High | High | High | High | High | High | High | High | 0.000723 | 0.006315 | 2.09 |
| FALSE | High | DSYYVYTQ | [R].DSYYV | 1xPhosphc | B2RY04 1x | 0.000246 | 0 | 1 | 1 | 3 | B2RY04 | Dedicator | B2RY04 [1 | 0 | 2075.014 | 0 | -0.06 | 0.999903 | 36.54 | 98 | 102 | 18.07 | 19.4 | | High | High | High | High | High | High | High | High | High | High | High | 0 | 4.94E-05 | 3.2 |
| FALSE | High | DTAATFQS | [K].DTAATI | 1xPhosphc | Q80YV2 1x | 7.9E-09 | 0 | 1 | 5 | 4 | Q80YV2 | Nuclear-in | Q80YV2 [5 | 0 | 3119.482 | 0 | -0.5 | 0.000231 | 0.65 | 116.7 | 83.3 | 3.22 | 2.33 | | High | High | High | High | High | High | High | High | High | High | High | 0 | 4.43E-10 | 5.24 |
| FALSE | High | DTAATFQS | [K].DTAATI | 2xPhosphc | Q80YV2 2x | 6.37E-06 | 0 | 1 | 5 | 4 | Q80YV2 | Nuclear-in | Q80YV2 [5 | 0 | 3199.449 | 0 | -0.41 | 0.005667 | 3.37 | 116.3 | 83.7 | 6.4 | 8.6 | | High | High | High | High | High | High | High | High | High | High | High | 0 | 8.14E-07 | 4.33 |
| FALSE | High | DSSVQEAT | [K].DSSVQI | 1xPhosphc | Q52KI8 1x | 6.73E-06 | 0 | 1 | 7 | 3 | Q52KI8 | Serine/arg | Q52KI8 [23 | 0 | 2119.057 | 0 | 0.04 | 0.774926 | 6.87 | 99.5 | 100.5 | 8.62 | 5.06 | | High | High | High | High | High | High | High | High | High | High | High | 0 | 8.69E-07 | 3.51 |
| FALSE | High | DSSPGSLPF | [K].DSSPGS | 1xPhosphc | Q8R4Z6 1x | 0.00145 | 5.66E-05 | 1 | 2 | 3 | Q8R4Z6 | DEAD-box | Q8R4Z6 [5 | 0 | 1425.739 | 0 | -0.5 | 0.001951 | 6.68 | 119.6 | 80.4 | 4.05 | 7.15 | | High | High | High | High | High | High | High | High | High | High | High | 2.58E-05 | 0.000363 | 1.85 |
| FALSE | High | DSSLPPQP | [K].DSSLPF | 1xPhosphc | Q6P9Q6-2 | 4.68E-05 | 0 | 1 | 2 | 5 | Q6P9Q6-2 | Isoform B c | Q6P9Q6-2 | 0 | 1851.932 | 0 | 1.81 | 0.002272 | 13.84 | 47.7 | 152.3 | 11.44 | 18.65 | | High | High | High | High | High | High | High | High | High | High | High | 0.000344 | 0.00312 | 2.52 |
| FALSE | High | DSSLLESDK | [K].DSSLLE | 2xPhosphc | Q9DBY8 2x | 0.039686 | 0.00283 | 1 | 2 | 1 | Q9DBY8 | Nuclear va | Q9DBY8 [2 | 0 | 1611.732 | 0 | -0.5 | 0.010986 | 5.3 | 115.6 | 84.4 | 11.06 | 4.8 | | High | High | High | High | High | High | High | High | High | High | High | 0.001468 | 0.01477 | 1.81 |
| FALSE | High | DSSPVPEPE | [K].DSSPVF | 1xPhosphc | Q8K2Z4-2 | 0.000164 | 0 | 1 | 3 | 3 | Q8K2Z4-2 | Isoform 2 c | Q8K2Z4-2 | 0 | 2552.217 | 0 | -0.71 | 0.003083 | 1.85 | 124.2 | 75.8 | 4.49 | 10.25 | | High | High | High | High | High | High | High | High | High | High | High | 0 | 4.73E-05 | 2.94 |
| FALSE | High | DSPSVWA | [K].DSPSVV | 1xPhosphc | P62962 1x | 1.97E-05 | 0 | 1 | 5 | 1 | P62962 | Profilin-1 C | P62962 [27 | 0 | 1751.913 | 0 | 0.05 | 0.872326 | 24.74 | 98.3 | 101.7 | 13.75 | 11.2 | | High | High | High | High | High | High | High | High | High | High | High | 0 | 2.88E-06 | 3.2 |
| FALSE | High | DSPVLSAR | [K].DSPVLS | 1xPhosphc | A6H619 1x | 0.025738 | 0.001645 | 1 | 8 | 2 | A6H619 | PHD and RI | A6H619 [1 | 0 | 1153.582 | 0 | 0.39 | 0.011244 | 9.79 | 88.8 | 111.2 | 4.99 | 9.29 | | High | High | High | High | High | High | High | High | High | High | High | 0.000872 | 0.009084 | 2.7 |
| FALSE | High | DSPVLSNS | [K].DSPVLS | 1xPhosphc | Q9EPE9 1x | 0.000568 | 0 | 1 | 2 | 2 | Q9EPE9 | Manganes | Q9EPE9 [8 | 0 | 1437.694 | 0 | 1.76 | 0.000236 | 3.34 | 47.5 | 152.5 | 6.45 | 6.16 | | High | High | High | High | High | High | High | High | High | High | High | 0 | 0.000126 | 1.83 |
| FALSE | High | DSPVIVR | [R].DSPVIV | 1xPhosphc | Q924A2 1x | 0.050826 | 0.003676 | 1 | 1 | 1 | Q924A2 | Protein cap | Q924A2 [8 | 0 | 1094.581 | 0 | 0.76 | 0.003196 | 4.33 | 74.9 | 125.1 | 8.12 | 7.99 | | High | High | High | High | High | High | High | High | High | High | High | 0.001948 | 0.01967 | 2.36 |
| FALSE | High | DSQDTSAE | [R].DSQDT | 1xPhosphc | Q3UJB9 1x | 1.01E-08 | 0 | 1 | 5 | 6 | Q3UJB9 | Enhancer c | Q3UJB9 [8 | 0 | 3436.507 | 0 | -0.63 | 0.001007 | 3.5 | 121.4 | 78.6 | 5.57 | 4.51 | | High | High | High | High | High | High | High | High | High | High | High | 0 | 1.33E-08 | 5.56 |
| FALSE | High | DSQDTSAE | [R].DSQDT | 2xPhosphc | Q3UJB9 2x | 1.15E-07 | 0 | 1 | 5 | 4 | Q3UJB9 | Enhancer c | Q3UJB9 [8 | 0 | 3516.473 | 0 | 0.61 | 0.002198 | 13.56 | 79.7 | 120.3 | 4.63 | 6.45 | | High | High | High | High | High | High | High | High | High | High | High | 0 | 8.96E-09 | 4.62 |
| FALSE | High | DSQENSDA | [K].DSQEN | 1xPhosphc | A0A140T8 | 1.72E-05 | 0 | 1 | 1 | 7 | A0A140T8 | Chromosor | A0A140T8 | 0 | 2238.96 | 0 | -0.78 | 0.002467 | 1.31 | 126.3 | 73.7 | 8.86 | 7.49 | | High | High | High | High | High | High | High | High | High | High | High | 0 | 2.47E-06 | 4.2 |
| FALSE | High | DSQENSDA | [K].DSQEN | 2xPhosphc | A0A140T8 | 1.81E-05 | 0 | 1 | 1 | 2 | A0A140T8 | Chromosor | A0A140T8 | 0 | 2318.926 | 0 | -0.49 | 0.011822 | 8.39 | 114 | 86 | 12.12 | 5.62 | | High | High | High | High | High | High | High | High | High | High | High | 0 | 2.63E-06 | 4.46 |
| FALSE | High | DSQFTIVLE | [R].DSQFTI | 1xPhosphc | Q3U1J1-2 | 0.01218 | 0.000889 | 1 | 4 | 1 | Q3U1J1-2 | Isoform 2 c | Q3U1J1-2 | 0 | 3824.779 | 0 | -0.12 | 0.033901 | 2.85 | 106.3 | 93.7 | 8.88 | 8.36 | | High | High | High | High | High | High | High | High | High | High | High | 0.000449 | 0.003902 | 3.98 |
| FALSE | High | DSQFTIVLE | [R].DSQFTI | 1xDeamida | Q3U1J1-2 | 5.75E-05 | 0 | 1 | 4 | 1 | Q3U1J1-2 | Isoform 2 c | Q3U1J1-2 | 0 | 3825.763 | 0 | -0.12 | 0.164538 | 10.65 | 107.6 | 92.4 | 5.78 | 10.79 | | High | High | High | High | High | High | High | High | High | High | High | 0 | 9.69E-06 | 5.3 |
| FALSE | High | DSQGLLDS | [K].DSQGL | 1xPhosphc | Q3TYU7 1x | 0.021797 | 0.001507 | 1 | 2 | 1 | Q3TYU7 | Putative ur | Q3TYU7 [4 | 1 | 3523.651 | 0 | -0.5 | 0.024083 | 1.41 | 117 | 83 | 7.1 | 12.69 | | High | High | High | High | High | High | High | High | High | High | High | 0.000801 | 0.007486 | 2.45 |
| FALSE | High | DSQSTAAIL | [K].DSQST | 1xCarbami | P55200 1x | 0.006834 | 0.00062 | 1 | 3 | 3 | P55200 | Histone-lys | P55200 [21 | 0 | 4012.915 | 0 | 0.04 | 0.560961 | 6.06 | 98.9 | 101.1 | 15.27 | 6.83 | | High | High | High | High | High | High | High | High | High | High | High | 0.000275 | 0.002043 | 2.86 |
| FALSE | High | DSRPLSPVI | [R].DSRPLS | 1xPhosphc | Q80ZX0 1x | 0.000386 | 0 | 1 | 2 | 2 | Q80ZX0 | Sec24 relat | Q80ZX0 [1 | 0 | 1999.151 | 0 | 0.11 | 0.076157 | 2.14 | 96.5 | 103.5 | 1.55 | 3.04 | | High | High | High | High | High | High | High | High | High | High | High | 0 | 8.21E-05 | 2.1 |
| FALSE | High | DSSAGSQS | [K].DSSAGS | 1xCarbami | Q8VCI7 1x | 0.002671 | 0.000109 | 1 | 1 | 2 | Q8VCI7 | ATPase SW | Q8VCI7 [2 | 0 | 4042.92 | 0 | -0.35 | 0.305751 | 11.44 | 109.5 | 90.5 | 8.92 | 14.84 | | High | High | High | High | High | High | High | High | High | High | High | 0.000585 | 0.005503 | 3.99 |
| FALSE | High | DSSDSADG | [R].DSSDSA | 1xPhosphc | Q8BTV2 1x | 0.075304 | 0.005727 | 1 | 2 | 2 | Q8BTV2 | Cleavage a | Q8BTV2 [1 | 1 | 2528.146 | 0 | 1.57 | 0.001031 | 6.4 | 54.6 | 145.4 | 14.23 | 10.58 | | High | High | High | High | High | High | High | High | High | High | High | 0.003188 | 0.03329 | 1.9 |
| FALSE | High | DSSEATGW | [K].DSSEAT | 1xPhosphc | B1ATC3 1x | 5.16E-05 | 0 | 1 | 3 | 1 | B1ATC3 | Trinucleoti | B1ATC3 [9 | 0 | 2153.959 | 0 | -1.3 | 0.00289 | 2.53 | 144.3 | 55.7 | 11.76 | 16.16 | | High | High | High | High | High | High | High | High | High | High | High | 0 | 8.57E-06 | 3.09 |
| FALSE | High | DSSESQLA | [R].DSSESC | 1xPhosphc | Q0VAW6 1 | 0.018774 | 0.001362 | 1 | 3 | 1 | Q0VAW6 | RIKEN cDN | Q0VAW6 [ | 0 | 2534.202 | 0 | -0.37 | 0.142677 | 15.68 | 111.4 | 88.6 | 6.94 | 17.1 | | High | High | High | High | High | High | High | High | High | High | High | 0.000723 | 0.006369 | 2.31 |
| FALSE | High | DSSGDGEL | [K].DSSGD | 1xPhosphc | G3X8S2 1x | 0.000191 | 0 | 1 | 2 | 2 | G3X8S2 | BRF1 homo | G3X8S2 [4 | 0 | 2430.076 | 0 | -0.22 | 0.734373 | 18.95 | 99.5 | 100.5 | 23.21 | 15.57 | | High | High | High | High | High | High | High | High | High | High | High | 0 | 3.7E-05 | 2.88 |
| FALSE | High | DSSGDSSE | [K].DSSGD | 1xPhosphc | Q9CX66 1x | 0.000605 | 0 | 1 | 1 | 4 | Q9CX66 | Uncharacte | Q9CX66 [1 | 0 | 3250.46 | 0 | -0.39 | 0.017382 | 2.91 | 113.6 | 86.4 | 8.34 | 11.86 | | High | High | High | High | High | High | High | High | High | High | High | 0 | 0.000192 | 4.37 |
| FALSE | High | DSSGPVSP | [K].DSSGP | 1xCarbami | Q3U0S1 1x | 2.24E-06 | 0 | 1 | 2 | 1 | Q3U0S1 | Putative ur | Q3U0S1 [2 | 0 | 3082.403 | 0 | 1.45 | 0.001791 | 20.55 | 53.3 | 146.7 | 11.39 | 11.24 | | High | High | High | High | High | High | High | High | High | High | High | 0 | 2.51E-07 | 4.14 |
| FALSE | High | DSSGQEDE | [K].DSSGQ | 1xPhosphc | Q5UAK0-1 | 0.000205 | 0 | 1 | 4 | 2 | Q5UAK0-1 | Isoform 1 c | Q5UAK0-1 | 0 | 3815.668 | 0 | -0.21 | 0.368752 | 8.23 | 107.1 | 92.9 | 18.25 | 11.77 | | High | High | High | High | High | High | High | High | High | High | High | 0 | 4.01E-05 | 3.26 |
| FALSE | High | DSSGQEDE | [K].DSSGQ | 1xDeamida | Q5UAK0-1 | 0.013545 | 0.000976 | 1 | 4 | 1 | Q5UAK0-1 | Isoform 1 c | Q5UAK0-1 | 0 | 3896.618 | 0 | -0.08 | 0.320999 | 4.88 | 107.8 | 92.2 | 10.83 | 10.09 | | High | High | High | High | High | High | High | High | High | High | High | 0.000505 | 0.004398 | 4.39 |
| FALSE | High | DSSGSISEC | [K].DSSGSI | 1xPhosphc | F8WI37 1x | 0.000892 | 0 | 1 | 4 | 2 | F8WI37 | Histone-lys | F8WI37 [1 | 0 | 1671.779 | 0 | -0.11 | 0.037124 | 2.77 | 103.8 | 96.2 | 2.81 | 5.96 | | High | High | High | High | High | High | High | High | High | High | High | 0 | 0.00021 | 2.94 |
| FALSE | High | DSSGSISEC | [K].DSSGSI | 2xPhosphc | F8WI37 2x | 0.01861 | 0.001362 | 1 | 4 | 1 | F8WI37 | Histone-lys | F8WI37 [1 | 0 | 1751.745 | 0 | -0.46 | 0.014737 | 6.3 | 114.9 | 85.1 | 6.69 | 10.51 | | High | High | High | High | High | High | High | High | High | High | High | 0.000723 | 0.006271 | 2.47 |
| FALSE | High | DSSGWSSS | [R].DSSGW | 1xPhosphc | Q3TQX5 1x | 0.038017 | 0.002528 | 1 | 3 | 3 | Q3TQX5 | Putative ur | Q3TQX5 [5 | 0 | 1478.693 | 0 | 0.01 | 0.815031 | 5.45 | 97.8 | 102.2 | 3.5 | 6.28 | | High | High | High | High | High | High | High | High | High | High | High | 0.004988 | 0.05454 | 1.76 |
| FALSE | High | DSSIIDPGT | [R].DSSIIDF | 1xPhosphc | F6WJB7 1x | 5.84E-07 | 0 | 1 | 4 | 4 | F6WJB7 | Ubiquitin c | F6WJB7 [3 | 0 | 2853.381 | 0 | 0.48 | 0.006629 | 1.18 | 81.5 | 118.5 | 2.52 | 10.98 | | High | High | High | High | High | High | High | High | High | High | High | 0 | 5.55E-08 | 3.71 |
| FALSE | High | DSSIIDPGT | [R].DSSIIDF | 2xPhosphc | F6WJB7 2x | 0.000308 | 0 | 1 | 4 | 3 | F6WJB7 | Ubiquitin c | F6WJB7 [3 | 0 | 2933.347 | 0 | 0 | 0.782527 | 9.14 | 98.2 | 101.8 | 3.55 | 8.14 | | High | High | High | High | High | High | High | High | High | High | High | 0 | 6.37E-05 | 3.56 |
| FALSE | High | DSSLLESDK | [K].DSSLLE | 1xPhosphc | Q9DBY8 1x | 0.013308 | 0.000889 | 1 | 2 | 2 | Q9DBY8 | Nuclear va | Q9DBY8 [2 | 0 | 1531.766 | 0 | -0.76 | 0.003766 | 5.15 | 125.4 | 74.6 | 11.25 | 5.4 | | High | High | High | High | High | High | High | High | High | High | High | 0.000585 | 0.005275 | 2.49 |
| FALSE | High | DSDQVAQS | [K].DSDQV | 1xPhosphc | Q3UPL0 1x | 1.61E-08 | 0 | 1 | 1 | 9 | Q3UPL0 | Protein tra | Q3UPL0 [5 | 0 | 2869.257 | 0 | 0.55 | 0.019194 | 20.8 | 82.2 | 117.8 | 6.11 | 10.46 | | High | High | High | High | High | High | High | High | High | High | High | 0 | 9.86E-10 | 5.97 |
| FALSE | High | DSDKEDGR | [K].DSDKEI | 1xPhosphc | I1VGT4 1x | 0.041781 | 0.003126 | 2 | 19 | 1 | I1VGT4; I1 | Anion exch | I1VGT4 [8 | 2 | 2707.225 | 0 | 0.53 | 0.088949 | 39.12 | 72.9 | 127.1 | 29.35 | 23.81 | Shared | High | High | High | High | High | High | High | High | High | High | High | 0.001609 | 0.0157 | 2.51 |
| FALSE | High | DSDGVDGI | [R].DSDGV | 1xPhosphc | O88532 1x | 5.6E-08 | 0 | 1 | 2 | 4 | O88532 | Zinc finger | O88532 [1 | 0 | 1863.841 | 0 | 0.03 | 0.251538 | 6.41 | 97 | 103 | 4.63 | 7.42 | | High | High | High | High | High | High | High | High | High | High | High | 0 | 6.98E-09 | 3.69 |
| FALSE | High | DSDGVDGI | [R].DSDGV | 1xPhosphc | O88532 1x | 7.08E-05 | 0 | 1 | 2 | 1 | O88532 | Zinc finger | O88532 [1 | 0 | 1634.678 | 0 | 0.08 | 0.475399 | 14.93 | 100.3 | 99.7 | 9.09 | 12.57 | | High | High | High | High | High | High | High | High | High | High | High | 0 | 1.22E-05 | 2.89 |
| FALSE | High | DSPEPDA | [R].DSPEF | 2xPhosphc | Q8K3W3 2 | 2.68E-09 | 0 | 1 | 1 | 2 | Q8K3W3 | Protein CA | Q8K3W3 [ | 0 | 2267.065 | 0 | -0.85 | 0.003514 | 3.43 | 126.8 | 73.2 | 4.21 | 14.31 | | High | High | High | High | High | High | High | High | High | High | High | 0 | 1.31E-10 | 4.5 |
| FALSE | High | DSPGPLID | [K].DSPGF | 1xPhosphc | Q69ZD1 1x | 1.03E-05 | 0 | 1 | 7 | 4 | Q69ZD1 | MKIAA169 | Q69ZD1 [2 | 0 | 2084.063 | 0 | 0.24 | 0.006126 | 2.76 | 92.1 | 107.9 | 5.07 | 3.29 | | High | High | High | High | High | High | High | High | High | High | High | 0 | 2.64E-05 | 3.36 |
| FALSE | High | DSPGPLID | [K].DSPGF | 2xPhosphc | Q69ZD1 2x | 0.024001 | 0.001612 | 1 | 7 | 1 | Q69ZD1 | MKIAA169 | Q69ZD1 [2 | 0 | 2164.029 | 0 | -0.26 | 0.071879 | 12.78 | 109.8 | 90.2 | 12.21 | 8.19 | | High | High | High | High | High | High | High | High | High | High | High | 0.000859 | 0.008352 | 2.37 |
| FALSE | High | DSPTLASE | [K].DSPTL | 1xPhosphc | Q8C0T5 1x | 0.013545 | 0.000976 | 1 | 2 | 1 | Q8C0T5 | Signal-indu | Q8C0T5 [1 | 0 | 1453.77 | 0 | 1.77 | 0.000676 | 13.84 | 45.3 | 154.7 | 7.93 | 8.07 | | High | High | High | High | High | High | High | High | High | High | High | 0.000538 | 0.004419 | 1.87 |
| FALSE | High | DPSQTSPL | [K].DPSQT | 1xPhosphc | Q7TQG1 1x | 0.003655 | 0.00021 | 1 | 1 | 6 | Q7TQG1 | Pleckstrin | Q7TQG1 [8 | 0 | 1564.757 | 0 | 2.04 | 0.000353 | 4.89 | 39.6 | 160.4 | 2.41 | 7.32 | | High | High | High | High | High | High | High | High | High | High | High | 0.002403 | 0.02507 | 1.93 |
| FALSE | High | DPSSPGPE | [R].DPSSPC | 1xPhosphc | Q8K120 1x | 4.76E-05 | 0 | 1 | 2 | 3 | Q8K120 | Nuclear fac | Q8K120 [3 | 0 | 2824.385 | 0 | -1.21 | 0.000409 | 0.77 | 143.2 | 56.8 | 6.83 | 10.54 | | High | High | High | High | High | High | High | High | High | High | High | 0 | 0.000166 | 4.3 |
| FALSE | High | DPSSSSPL | [K].DPSSSS | 1xPhosphc | Q6A0A2 1x | 0.00034 | 0 | 1 | 3 | 4 | Q6A0A2 | La-related | Q6A0A2 [6 | 0 | 1777.914 | 0 | 0.82 | 0.000484 | 1.58 | 74.1 | 125.9 | 6.01 | 6.51 | | High | High | High | High | High | High | High | High | High | High | High | 0 | 7.08E-05 | 2.78 |
| FALSE | High | DPSVDITQ | [K].DPSVDI | 1xPhosphc | E9Q481 1x | 5.36E-09 | 0 | 1 | 2 | 3 | E9Q481 | Serine/thr | E9Q481 [1 | 0 | 2385.054 | 0 | -0.55 | 0.001626 | 1.24 | 118.7 | 81.3 | 3.32 | 7.72 | | High | High | High | High | High | High | High | High | High | High | High | 0 | 2.85E-10 | 5.66 |
| FALSE | High | DPSVDITQ | [K].DPSVDI | 1xCarbami | E9Q481 1x | 2.29E-05 | 0 | 1 | 2 | 1 | E9Q481 | Serine/thr | E9Q481 [1 | 1 | 4405.941 | 0 | -0.12 | 0.175562 | 10.43 | 111 | 89 | 13.3 | 17.32 | | High | High | High | High | High | High | High | High | High | High | High | 0 | 3.42E-06 | 3.5 |
| FALSE | High | DPSVDITQ | [K].DPSVDI | 1xCarbami | E9Q481 1x | 0.000279 | 0 | 1 | 2 | 9 | E9Q481 | Serine/thr | E9Q481 [1 | 1 | 4421.936 | 0 | -0.23 | 0.057993 | 9.1 | 108.7 | 91.3 | 5.13 | 8.82 | | High | High | High | High | High | High | High | High | High | High | High | 0.000275 | 0.001721 | 4.44 |
| FALSE | High | DPSVDITQ | [K].DPSVDI | 1xCarbami | E9Q481 2x | 0.013545 | 0.000976 | 1 | 2 | 2 | E9Q481 | Serine/thr | E9Q481 [1 | 1 | 4501.903 | 0 | -0.39 | 0.049662 | 15.11 | 108.8 | 91.2 | 13.31 | 10.56 | | High | High | High | High | High | High | High | High | High | High | High | 0.000538 | 0.004423 | 3.9 |
| FALSE | High | DPSVEVTQ | [K].DPSVE | 1xPhosphc | Q922R5 1x | 3.06E-05 | 0 | 1 | 4 | 1 | Q922R5 | Serine/thr | Q922R5 [1 | 0 | 2399.07 | 0 | 0.29 | 0.092821 | 26.16 | 94.2 | 105.8 | 15.86 | 13.63 | | High | High | High | High | High | High | High | High | High | High | High | 0 | 4.73E-06 | 3.41 |
| FALSE | High | DPSVLDDT | [K].DPSVLD | 1xPhosphc | Q3TC93 1x | 0.065405 | 0.004734 | 1 | 2 | 1 | Q3TC93 | HCLS1-binc | Q3TC93 [1 | 0 | 4650.134 | 0 | 0.05 | 0.684445 | 8.95 | 100.6 | 99.4 | 16.12 | 18.46 | | High | High | High | High | High | High | High | High | High | High | High | 0.002432 | 0.02624 | 3.96 |
| FALSE | High | DPSLSDSST | [K].DPSLSD | 2xPhosphc | G3UWQ7 2 | 3.06E-05 | 0 | 1 | 1 | 1 | G3UWQ7 | Protein rep | G3UWQ7 [ | 0 | 1850.814 | 0 | 0.18 | 0.138424 | 12.45 | 93.7 | 106.3 | 10.21 | 4.61 | | High | High | High | High | High | High | High | High | High | High | High | 0 | 4.74E-06 | 2.77 |
| FALSE | High | DPVGEIAV | [R].DPVGE | 1xPhosphc | A2ADL9 1x | 0.000334 | 0 | 1 | 7 | 2 | A2ADL9 | InaD-like ( | A2ADL9 [3 | 0 | 2876.574 | 0 | -0.01 | 0.63281 | 19.94 | 99.1 | 100.9 | 7.66 | 22.61 | | High | High | High | High | High | High | High | High | High | High | High | 0 | 6.93E-05 | 2.95 |
| FALSE | High | DPVTYVSE | [K].DPVTY | 1xCarbami | G3UWX1 1x | 5.7E-08 | 0 | 2 | 9 | 4 | G3UWX1; G | Replication | G3UWX1 [ | 0 | 2672.173 | 0 | -0.13 | 0.76829 | 18.22 | 103.3 | 96.7 | 7.99 | 11.62 | Shared | High | High | High | High | High | High | High | High | High | High | High | 0 | 1.92E-07 | 3.61 |
| FALSE | High | DPVTYVSE | [K].DPVTY | 1xCarbami | G3UWX1 2 | 1.25E-06 | 0 | 2 | 9 | 10 | G3UWX1; G | Replication | G3UWX1 [ | 0 | 2752.139 | 0 | -0.21 | 0.058793 | 1.49 | 106.6 | 93.4 | 6.01 | 6.4 | Shared | High | High | High | High | High | High | High | High | High | High | High | 0 | 1.3E-07 | 4.46 |
| FALSE | High | DQANEGRI | [R].DQANE | 1xCarbami | A0A1B0GS | 0.000475 | 0 | 1 | 5 | 1 | A0A1B0GS | Supervillin | A0A1B0GS | 0 | 3155.478 | 0 | 1.47 | 0.002505 | 24.88 | 52.4 | 147.6 | 12.51 | 13 | | High | High | High | High | High | High | High | High | High | High | High | 0 | 0.000103 | 3.7 |
| FALSE | High | DQATSMSI | [R].DQATS | 1xPhosphc | A2ATK9 1x | 0.000159 | 0 | 1 | 3 | 1 | A2ATK9 | Family wit | A2ATK9 [3 | 0 | 2044.972 | 0 | 1.42 | 0.001079 | 5.54 | 54.4 | 145.6 | 5.29 | 10.55 | | High | High | High | High | High | High | High | High | High | High | High | 0 | 3.03E-05 | 2.4 |
| FALSE | High | DQDELTPG | [K].DQDEL | 1xPhosphc | Q80YR5 1x | 7.84E-06 | 0 | 1 | 2 | 1 | Q80YR5 | Scaffold at | Q80YR5 [5 | 0 | 1762.796 | 0 | -0.9 | 0.00132 | 0.68 | 128.5 | 71.5 | 5.22 | 5.61 | | High | High | High | High | High | High | High | High | High | High | High | 0 | 1.03E-06 | 3.03 |
| FALSE | High | DQEDGALF | [K].DQEDG | 1xPhosphc | P81068 1x | 1.73E-07 | 0 | 1 | 1 | 1 | P81068 | Iroquois-cl | P81068 [1 | 0 | 2448.086 | 0 | -0.13 | 0.668673 | 21.61 | 107.8 | 92.2 | 24.11 | 25.43 | | High | High | High | High | High | High | High | High | High | High | High | 0 | 1.41E-08 | 3 |
| FALSE | High | DQELPPEG | [R].DQELPI | 1xCarbami | B1AS46 1x | 0.000681 | 0 | 1 | 4 | 1 | B1AS46 | Zinc finger | B1AS46 [1 | 0 | 3854.681 | 0 | -0.83 | 0.005483 | 10.68 | 128.1 | 71.9 | 3.02 | 19.01 | | High | High | High | High | High | High | High | High | High | High | High | 0 | 0.000155 | 3.39 |
| FALSE | High | DQEMVSD | [K].DQEMV | 1xPhosphc | Q8VIG1 1x | 1.14E-07 | 0 | 1 | 1 | 2 | Q8VIG1 | RE1-silenci | Q8VIG1 [8 | 0 | 2403.192 | 0 | -0.88 | 0.010214 | 7.71 | 129.7 | 70.3 | 6.51 | 29.22 | | High | High | High | High | High | High | High | High | High | High | High | 0 | 7.06E-08 | 3.54 |
| FALSE | High | DQEMVSD | [K].DQEMV | 2xPhosphc | Q8VIG1 1x | 7.93E-07 | 0 | 1 | 1 | 5 | Q8VIG1 | RE1-silenci | Q8VIG1 [8 | 0 | 2419.187 | 0 | -0.94 | 0.002335 | 4.28 | 134.6 | 65.4 | 8.97 | 8.85 | | High | High | High | High | High | High | High | High | High | High | High | 0 | 7.86E-08 | 4.64 |
| FALSE | High | DQEPDVVC | [R].DQEPD | 1xPhosphc | B1AQJ2 1x | 0.000271 | 0 | 1 | 2 | 2 | B1AQJ2 | Ubiquitin c | B1AQJ2 [9 | 0 | 2425.19 | 0 | -0.66 | 0.002762 | 5.37 | 126.2 | 73.8 | 6.15 | 8.87 | | High | High | High | High | High | High | High | High | High | High | High | 0 | 5.5E-05 | 2.74 |
| FALSE | High | DQGEPVSP | [R].DQGEP | 1xPhosphc | B7ZCJ1 1x | 0.00034 | 0 | 1 | 6 | 5 | B7ZCJ1 | Rho GTPas | B7ZCJ1 [3 | 0 | 2723.264 | 0 | -0.06 | 0.601389 | 11.85 | 102.2 | 97.8 | 4.46 | 4.83 | | High | High | High | High | High | High | High | High | High | High | High | 0 | 7.73E-05 | 3.69 |
| FALSE | High | DQGSTYSG | [K].DQGST | 1xPhosphc | A2A791 1x | 0.0102 | 0.000736 | 1 | 3 | 2 | A2A791 | Zinc finger | A2A791 [1 | 0 | 4017.843 | 0 | -0.45 | 0.061196 | 8.31 | 116.6 | 83.4 | 20.21 | 12.32 | | High | High | High | High | High | High | High | High | High | High | High | 0.000382 | 0.003268 | 4.23 |
| FALSE | High | DQHLNNSS | [R].DQHLN | 1xPhosphc | Q9R0L6-2 | 7.41E-05 | 0 | 1 | 3 | 6 | Q9R0L6-2 | Isoform 2 c | Q9R0L6-2 | 0 | 2239.071 | 0 | -0.42 | 0.025239 | 8.48 | 114.9 | 85.1 | 9.58 | 7.1 | | High | High | High | High | High | High | High | High | High | High | High | 0 | 3.06E-05 | 3.25 |
| FALSE | High | DPVSGSGA | [K].DPVSGS | 1xPhosphc | Q8C180 1x | 8.35E-06 | 0 | 1 | 2 | 2 | Q8C180 | Fibroblast | Q8C180 [2 | 0 | 2780.163 | 0 | 0.81 | 0.008497 | 10.26 | 74.4 | 125.6 | 6.31 | 14.25 | | High | High | High | High | High | High | High | High | High | High | High | 0 | 1.1E-06 | 3.32 |
| FALSE | High | DPRSPEPLE | [K].DPRSPE | 1xPhosphc | A0A1B0GS | 5.26E-05 | 0 | 1 | 3 | 1 | A0A1B0GS | 39S riboso | A0A1B0GS | 0 | 2229.075 | 0 | -0.5 | 0.001972 | 5.1 | 112 | 88 | 5.67 | 7.09 | | High | High | High | High | High | High | High | High | High | High | High | 0 | 8.72E-06 | 3.36 |
| FALSE | High | DPQTPILQ | [R].DPQTP | 1xPhosphc | A2APB8 1x | 0.002376 | 0.000109 | 1 | 3 | 1 | A2APB8 | Targeting p | A2APB8 [3 | 0 | 1678.918 | 0 | 0.79 | 0.04239 | 28.73 | 65.2 | 134.8 | 32.96 | 24.15 | | High | High | High | High | High | High | High | High | High | High | High | 4.94E-05 | 0.000628 | 2.61 |
| FALSE | High | DPQDPSQE | [K].DPQDP | 1xPhosphc | E2JF22 1xP | 0.00034 | 0 | 1 | 6 | 6 | E2JF22 | Piezo-type | E2JF22 [13 | 1 | 2569.152 | 0 | 0.19 | 0.100179 | 12.74 | 98.8 | 101.2 | 5.86 | 10.61 | | High | High | High | High | High | High | High | High | High | High | High | 4.94E-05 | 0.00063 | 3.52 |
| FALSE | High | DNSISPESD | [R].DNSISP | 1xDeamida | Q6A065 1x | 0.046291 | 0.003523 | 1 | 3 | 1 | Q6A065 | Centrosom | Q6A065 [9 | 0 | 3013.301 | 0 | -0.43 | 0.180372 | 5.84 | 116.1 | 83.9 | 6.96 | 19.2 | | High | High | High | High | High | High | High | High | High | High | High | 0.001789 | 0.01765 | 2.21 |
| FALSE | High | DNSPAVFP | [R].DNSPA | 1xPhosphc | Q3U0S4 1x | 0.001881 | 5.66E-05 | 1 | 2 | 3 | Q3U0S4 | Putative ur | Q3U0S4 [7 | 0 | 1426.657 | 0 | -0.26 | 0.015001 | 6.25 | 109.2 | 90.8 | 5.11 | 4.89 | | High | High | High | High | High | High | High | High | High | High | High | 0.000161 | 0.001204 | 2.76 |
| FALSE | High | DNSPDSTP | [R].DNSPD | 1xPhosphc | G5E893 1x | 0.030898 | 0.001896 | 1 | 7 | 1 | G5E893 | Ankyrin re | G5E893 [1 | 0 | 2503.187 | 0 | 0.4 | 0.039504 | 17.05 | 87.8 | 112.2 | 10.18 | 12.26 | | High | High | High | High | High | High | High | High | High | High | High | 0.001044 | 0.01118 | 2.1 |
| FALSE | High | DNSPSLK | [R].DNSPSI | 1xPhosphc | Q9D486 1x | 0.036416 | 0.002419 | 1 | 2 | 2 | Q9D486 | C-Maf-indu | Q9D486 [3 | 0 | 1298.676 | 0 | -0.58 | 0.004156 | 3 | 119.4 | 80.6 | 4.31 | 7.69 | | High | High | High | High | High | High | High | High | High | High | High | 0.001279 | 0.0134 | 2.39 |
| FALSE | High | DNSVLEEK | [K].DNSVLE | 1xPhosphc | B1AS46 1x | 0.014538 | 0.001011 | 1 | 4 | 1 | B1AS46 | Zinc finger | B1AS46 [1 | 0 | 1471.745 | 0 | -1.72 | 0.000721 | 1.1 | 149.7 | 50.3 | 7.07 | 15.73 | | High | High | High | High | High | High | High | High | High | High | High | 0.000555 | 0.004787 | 2.28 |
| FALSE | High | DNTRPGAN | [K].DNTRP | 1xPhosphc | Q3V1Z7 1x | 4.8E-08 | 0 | 1 | 3 | 1 | Q3V1Z7 | Putative ur | Q3V1Z7 [4 | 0 | 2541.221 | 0 | -0.03 | 0.40683 | 18.69 | 101.2 | 98.8 | 2.77 | 16.64 | | High | High | High | High | High | High | High | High | High | High | High | 0 | 3.34E-09 | 3.49 |
| FALSE | High | DPAPPSPT | [K].DPAPPS | 1xPhosphc | Q8BGT6 1x | 0.000279 | 0 | 1 | 1 | 5 | Q8BGT6 | MICAL-like | Q8BGT6 [1 | 0 | 4029.885 | 0 | 1.44 | 0.001272 | 7.35 | 53.8 | 146.2 | 12.04 | 8.76 | | High | High | High | High | High | High | High | High | High | High | High | 0 | 5.71E-05 | 3.54 |
| FALSE | High | DPASPSSIT | [K].DPASPS | 1xPhosphc | Q80WQ8 1 | 0.013308 | 0.000889 | 1 | 2 | 1 | Q80WQ8 | Mis18-binc | Q80WQ8 [ | 0 | 1749.899 | 0 | 0.26 | 0.062763 | 16.76 | 89 | 111 | 7.44 | 12.65 | | High | High | High | High | High | High | High | High | High | High | High | 0.000486 | 0.004315 | 1.97 |
| FALSE | High | DPAVDDSP | [R].DPAVD | 1xCarbami | B2RXX0 1x | 0.003997 | 0.000311 | 1 | 6 | 1 | B2RXX0 | Nek1 prote | B2RXX0 [9 | 0 | 2257.01 | 0 | -0.44 | 0.005072 | 4.31 | 115.1 | 84.9 | 6.17 | 11.67 | | High | High | High | High | High | High | High | High | High | High | High | 0.00014 | 0.001126 | 2.58 |
| FALSE | High | DPDANWD | [R].DPDAN | 1xPhosphc | Q9ESC8 1x | 0.000573 | 0 | 1 | 2 | 2 | Q9ESC8 | AF4/FMR2 | Q9ESC8 [2 | 0 | 1568.658 | 0 | 0.42 | 0.013862 | 12.22 | 86.9 | 113.1 | 4.93 | 6.65 | | High | High | High | High | High | High | High | High | High | High | High | 0 | 0.000128 | 2.31 |

| | | | | | | | | | | | | | | | | | | | | | | | | | | | | | | | | | | | | | | |
|---|---|---|---|---|---|---|---|---|---|---|---|---|---|---|---|---|---|---|---|---|---|---|---|---|---|---|---|---|---|---|---|---|---|---|---|---|---|---|
| FALSE | High | DPDSALGIS | [K].DPDSAI | 2xPhospho | Q3UMF0 2 | 2.02E-05 | 0 | 1 | 6 | 1 | Q3UMF0 | Cordon-ble | Q3UMF0 [5 | 0 | 2381.056 | 0 | 0.24 | 0.087541 | 18.67 | 84.3 | 115.7 | 16.99 | 12.95 | | High | High | High | High | High | High | High | High | High | High | High | 0 | 2.99E-06 | 4.19 |
| FALSE | High | DPETPQSS( | [R].DPETP( | 1xPhospho | Q99M54 1: | 0.041425 | 0.003126 | 1 | 3 | 1 | Q99M54 | Cell divisio | Q99M54 [1 | 0 | 1670.804 | 0 | -0.23 | 0.231855 | 12.43 | 106 | 94 | 18.03 | 11.46 | | High | High | High | High | High | High | High | High | High | High | High | 0.001609 | 0.01552 | 1.71 |
| FALSE | High | DPFSPDK | [K].DPFSPD | 1xPhospho | Q3U1T3 1x | 0.016743 | 0.001166 | 1 | 1 | 2 | Q3U1T3 | Breast canc | Q3U1T3 [1 | 0 | 1343.665 | 0 | -0.59 | 0.000677 | 2.9 | 118.6 | 81.4 | 6.1 | 4.64 | | High | High | High | High | High | High | High | High | High | High | High | 0.002381 | 0.02459 | 1.74 |
| FALSE | High | DPGLSQSH | [K].DPGLS( | 1xPhospho | E9Q3C1 1x | 0.002671 | 0.000109 | 1 | 2 | 2 | E9Q3C1 | C2 domain | E9Q3C1 [6 | 0 | 2546.224 | 0 | 0.11 | 0.278424 | 4.16 | 95.9 | 104.1 | 8.89 | 8.79 | | High | High | High | High | High | High | High | High | High | High | High | 4.94E-05 | 0.00072 | 3.23 |
| FALSE | High | DPGTPNDD | [R].DPGTPI | 1xPhospho | Q9D219 1x | 8.43E-06 | 0 | 1 | 1 | 1 | Q9D219 | B-cell CLL/l | Q9D219 [1 | 0 | 1811.846 | 0 | 0.59 | 0.009873 | 16.57 | 80.2 | 119.8 | 10.44 | 8.67 | | High | High | High | High | High | High | High | High | High | High | High | 0 | 1.12E-06 | 3.36 |
| FALSE | High | DPLEEIQPL | [K].DPLEEI( | 1xPhospho | E9Q8S5 1x | 4.65E-06 | 0 | 1 | 2 | 3 | E9Q8S5 | Microtubu | E9Q8S5 [3 | 0 | 3139.467 | 0 | -0.71 | 0.00346 | 2.39 | 121.2 | 78.8 | 5.87 | 7.12 | | High | High | High | High | High | High | High | High | High | High | High | 0 | 5.72E-07 | 4.81 |
| FALSE | High | DPLHVSPA1 | [K].DPLHVS | 1xPhospho | E9Q921 1x | 4.8E-05 | 0 | 1 | 7 | 1 | E9Q921 | AF4/FMR2 | E9Q921 [3 | 0 | 1946.015 | 0 | -0.68 | 0.003862 | 1.51 | 123.8 | 76.2 | 6.14 | 11.45 | | High | High | High | High | High | High | High | High | High | High | High | 0 | 7.86E-06 | 3.18 |
| FALSE | High | DPLSPFAV( | [R].DPLSPF | 1xCarbami | A0A1S6GW | 2.96E-09 | 0 | 1 | 3 | 5 | A0A1S6GW | Uncharacte | A0A1S6GW | 0 | 2472.11 | 0 | 1.8 | 0.000784 | 7.4 | 44.1 | 155.9 | 10.85 | 11.52 | | High | High | High | High | High | High | High | High | High | High | High | 0 | 1.46E-10 | 5.07 |
| FALSE | High | DPLSSPGGI | [K].DPLSSP | 1xPhospho | Q8VC03 1x | 0.000553 | 0 | 1 | 1 | 2 | Q8VC03 | Echinoderr | Q8VC03 [1 | 0 | 1435.678 | 0 | 2.7 | 0.000373 | 14.27 | 25.8 | 174.2 | 13.61 | 7.05 | | High | High | High | High | High | High | High | High | High | High | High | 0 | 0.000123 | 2.62 |
| FALSE | High | DPLTHASN | [K].DPLTHA | 1xPhospho | Q9QYY8 1x | 0.086562 | 0.006435 | 1 | 2 | 1 | Q9QYY8 | Spastin OS | Q9QYY8 [2 | 0 | 1616.8 | 0 | 1.48 | 0.012083 | 9.13 | 47.7 | 152.3 | 21.32 | 26.41 | | High | High | High | High | High | High | High | High | High | High | High | 0.003303 | 0.03635 | 1.9 |
| FALSE | High | DPLTTTPPF | [K].DPLTTT | 1xPhospho | Q6A074 1x | 0.000271 | 0 | 1 | 5 | 4 | Q6A074 | MKIAA040 | Q6A074 [8 | 0 | 1872.028 | 0 | 2.72 | 0.00034 | 9.83 | 26.6 | 173.4 | 14.05 | 6.39 | | High | High | High | High | High | High | High | High | High | High | High | 0 | 5.81E-05 | 3.32 |
| FALSE | High | DPNASPGD | [R].DPNASI | 1xPhospho | Q64727 1x | 1.72E-06 | 0 | 1 | 1 | 2 | Q64727 | Vinculin OS | Q64727 [2 | 0 | 1806.822 | 0 | 0.27 | 0.008644 | 7 | 90.6 | 109.4 | 13.38 | 11.22 | | High | High | High | High | High | High | High | High | High | High | High | 0 | 1.88E-07 | 2.53 |
| FALSE | High | DPNIEALN( | [R].DPNIEA | 1xPhospho | Q8K0X8 1x | 6.2E-06 | 0 | 1 | 3 | 3 | Q8K0X8 | Fasciculati | Q8K0X8 [1 | 0 | 2633.286 | 0 | 1.81 | 0.00024 | 12.27 | 44.7 | 155.3 | 9.87 | 3.63 | | High | High | High | High | High | High | High | High | High | High | High | 0 | 7.9E-07 | 4.25 |
| FALSE | High | DPNIEALN( | [R].DPNIEA | 1xDeamida | Q8K0X8 1x | 7.25E-07 | 0 | 1 | 3 | 2 | Q8K0X8 | Fasciculati | Q8K0X8 [1 | 0 | 2634.27 | 0 | 2.75 | 0.000466 | 48.8 | 24.5 | 175.5 | 34.18 | 7.45 | | High | High | High | High | High | High | High | High | High | High | High | 0 | 8.73E-07 | 4.31 |
| FALSE | High | DPPAGASA | [R].DPPAG | 1xCarbami | O35245 1x | 0.0003 | 0 | 1 | 5 | 1 | O35245 | Polycystin- | O35245 [6 | 0 | 2149.979 | 0 | -0.09 | 0.228867 | 7.44 | 104.8 | 95.2 | 7.82 | 6.48 | | High | High | High | High | High | High | High | High | High | High | High | 0 | 6.17E-05 | 3.31 |
| FALSE | High | DPPESPVV | [K].DPPESF | 1xPhospho | Q8VE88 1x | 0.038017 | 0.002528 | 1 | 2 | 2 | Q8VE88 | Protein FAI | Q8VE88 [5 | 0 | 2135.14 | 0 | -1.37 | 0.00279 | 4.18 | 144.7 | 55.3 | 8.34 | 19.44 | | High | High | High | High | High | High | High | High | High | High | High | 0.00269 | 0.02876 | 2.43 |
| FALSE | High | DPPSEANSI | [K].DPPSEA | 1xPhospho | Q8CCI5 1xl | 3.14E-05 | 0 | 1 | 1 | 3 | Q8CCI5 | RING1 and | Q8CCI5 [12 | 0 | 2269.111 | 0 | -0.96 | 0.001634 | 2.71 | 129.9 | 70.1 | 8.84 | 7.12 | | High | High | High | High | High | High | High | High | High | High | High | 0 | 1.78E-05 | 3.06 |
| FALSE | High | DPPSEANSI | [K].DPPSEA | 2xPhospho | Q8CCI5 2xl | 0.000442 | 0 | 1 | 1 | 2 | Q8CCI5 | RING1 and | Q8CCI5 [12 | 0 | 2349.078 | 0 | -0.36 | 0.004376 | 4.6 | 114.4 | 85.6 | 3.83 | 6.44 | | High | High | High | High | High | High | High | High | High | High | High | 0 | 9.55E-05 | 3.38 |
| FALSE | High | DPQDPSQE | [K].DPQDP | 1xPhospho | E2JF22 1xP | 1.08E-07 | 0 | 1 | 6 | 2 | E2JF22 | Piezo-type | E2JF22 [13 | 0 | 2413.051 | 0 | 1 | 0.001286 | 13.79 | 67.8 | 132.2 | 11.46 | 4.68 | | High | High | High | High | High | High | High | High | High | High | High | 0 | 8.36E-09 | 3.18 |
| FALSE | High | DQHNGSLT | [K].DQHNG | 1xPhospho | Q99KU0 1x | 2.72E-05 | 0 | 1 | 2 | 2 | Q99KU0 | Vacuole m | Q99KU0 [1 | 0 | 2289.091 | 0 | 1.29 | 0.016284 | 31.64 | 59.6 | 140.4 | 23.22 | 16.02 | | High | High | High | High | High | High | High | High | High | High | High | 0 | 4.15E-06 | 2.71 |
| FALSE | High | EESTFASDL | [R].EESTFA | 1xPhospho | Q80Z37 1x | 4.31E-08 | 0 | 1 | 1 | 3 | Q80Z37 | E3 ubiquiti | Q80Z37 [9 | 0 | 3540.669 | 0 | 0.22 | 0.280464 | 7.09 | 95.3 | 104.7 | 8.2 | 9.49 | | High | High | High | High | High | High | High | High | High | High | High | 0 | 2.97E-09 | 4.19 |
| FALSE | High | DQMEGSPI | [R].DQME( | 1xPhospho | Q3TLH4 1x | 2.64E-07 | 0 | 1 | 5 | 2 | Q3TLH4 | Protein PRI | Q3TLH4 [7 | 0 | 2345.991 | 0 | -0.21 | 0.855727 | 23.58 | 108.3 | 91.7 | 16.57 | 52.89 | | High | High | High | High | High | High | High | High | High | High | High | 0 | 2.29E-08 | 4.64 |
| FALSE | High | DQNGTWE | [K].DQNGT | 2xDeamida | Q059R7 1x | 0.031439 | 0.002011 | 1 | 1 | 1 | Q059R7 | Retinol bin | Q059R7 [4 | 0 | 2620.009 | 0 | -0.17 | 0.153431 | 2.79 | 109.7 | 90.3 | 8.23 | 25.91 | | High | High | High | High | High | High | High | High | High | High | High | 0.001076 | 0.01132 | 2.41 |
| FALSE | High | DRPVSQPS | [K].DRPVS( | 1xPhospho | P83741-3 1 | 8.43E-06 | 0 | 1 | 7 | 3 | P83741-3 | Isoform 3 c | P83741-3 [ | 0 | 1908.036 | 0 | 2.61 | 0.000748 | 35.58 | 28.1 | 171.9 | 15.58 | 9.98 | | High | High | High | High | High | High | High | High | High | High | High | 7.19E-05 | 0.000855 | 2.3 |
| FALSE | High | DRPVSQPS | [K].DRPVS( | 2xPhospho | P83741-3 2 | 0.066503 | 0.004796 | 1 | 7 | 2 | P83741-3 | Isoform 3 c | P83741-3 [ | 0 | 1988.002 | 0 | 3.02 | 0.000474 | 30.28 | 22.7 | 177.3 | 15.29 | 9.53 | | High | High | High | High | High | High | High | High | High | High | High | 0.002455 | 0.02666 | 1.92 |
| FALSE | High | DRQSPSTC | [K].DRQSP | 1xCarbami | B9EKJ7 1xF | 4.61E-06 | 0 | 1 | 3 | 1 | B9EKJ7 | Tankyrase | B9EKJ7 [71 | 1 | 2558.247 | 0 | 0.01 | 0.867244 | 25.2 | 97.4 | 102.6 | 14.23 | 19.04 | | High | High | High | High | High | High | High | High | High | High | High | 0 | 5.66E-07 | 3.5 |
| FALSE | High | DRSDSDDC | [K].DRSDSI | 1xPhospho | Q7TQC7 1x | 0.018447 | 0.001324 | 1 | 7 | 1 | Q7TQC7 | G patch do | Q7TQC7 [1 | 1 | 2033.962 | 0 | 0.41 | 0.095837 | 3.51 | 84.9 | 115.1 | 10.08 | 25.26 | | High | High | High | High | High | High | High | High | High | High | High | 0.000723 | 0.006245 | 2.71 |
| FALSE | High | DRSDSDDC | [K].DRSDSI | 2xPhospho | Q7TQC7 2x | 2.29E-05 | 0 | 1 | 7 | 2 | Q7TQC7 | G patch do | Q7TQC7 [1 | 1 | 2097.933 | 0 | 0.45 | 0.033601 | 12.74 | 84.4 | 115.6 | 6.35 | 13.02 | | High | High | High | High | High | High | High | High | High | High | High | 0 | 3.45E-06 | 3.99 |
| FALSE | High | DRSSPPPGY | [R].DRSSPF | 1xPhospho | G5E8L8 1xl | 1.03E-06 | 0 | 1 | 1 | 4 | G5E8L8 | Mitogen ac | G5E8L8 [16 | 1 | 2457.213 | 0 | 0.91 | 0.000396 | 3.91 | 70.2 | 129.8 | 6.86 | 8.41 | | High | High | High | High | High | High | High | High | High | High | High | 0 | 3.39E-07 | 3.73 |
| FALSE | High | DRTPPHLLY | [R].DRTPPH | 1xPhospho | Q6PHZ5 1x | 0.054844 | 0.003982 | 1 | 2 | 1 | Q6PHZ5 | Putative RI | Q6PHZ5 [5 | 1 | 1778.879 | 0 | 0.55 | 0.002729 | 8.05 | 83.8 | 116.2 | 8.27 | 6.04 | | High | High | High | High | High | High | High | High | High | High | High | 0.002059 | 0.02147 | 1.6 |
| FALSE | High | DRWSDSQ( | [K].DRWSD | 1xPhospho | Q5HZJ0 1xl | 0.000137 | 0 | 1 | 2 | 2 | Q5HZJ0 | Ribonuclea | Q5HZJ0 [36 | 1 | 1760.837 | 0 | -0.4 | 0.025535 | 12.85 | 113.8 | 86.2 | 6.22 | 13.66 | | High | High | High | High | High | High | High | High | High | High | High | 0 | 0.000263 | 2.7 |
| FALSE | High | DSAAEPVSI | [K].DSAAEI | 1xPhospho | Q8VIG1 1x | 4.14E-07 | 0 | 1 | 1 | 4 | Q8VIG1 | RE1-silenci | Q8VIG1 [9 | 0 | 1947.926 | 0 | -0.85 | 0.000793 | 1.18 | 132.9 | 67.1 | 9.97 | 9.34 | | High | High | High | High | High | High | High | High | High | High | High | 0 | 3.8E-08 | 3.91 |
| FALSE | High | DSAAEPVSI | [K].DSAAEI | 2xPhospho | Q8VIG1 2x | 0.010109 | 0.000736 | 1 | 1 | 1 | Q8VIG1 | RE1-silenci | Q8VIG1 [9 | 0 | 2027.893 | 0 | -1.2 | 0.001024 | 5.02 | 139.8 | 60.2 | 5.61 | 11.07 | | High | High | High | High | High | High | High | High | High | High | High | 0.000344 | 0.003181 | 3.37 |
| FALSE | High | DSAAPSPIP | [R].DSAAPS | 1xPhospho | Q6P9Q6-2 | 0.001704 | 5.66E-05 | 1 | 6 | 2 | Q6P9Q6-2 | Isoform B c | Q6P9Q6-2 | 0 | 4000.145 | 0 | 0.97 | 0.005383 | 28.54 | 68.6 | 131.4 | 13.29 | 7.86 | | High | High | High | High | High | High | High | High | High | High | High | 2.58E-05 | 0.000435 | 3.51 |
| FALSE | High | DSALEPVN( | [K].DSALEF | 1xPhospho | Q5XG69 1x | 7.28E-05 | 0 | 1 | 1 | 9 | Q5XG69 | Soluble lan | Q5XG69 [4 | 0 | 4702.267 | 0 | -1.75 | 0.003573 | 2.5 | 151.9 | 48.1 | 17.38 | 13.16 | | High | High | High | High | High | High | High | High | High | High | High | 0 | 9E-05 | 3.95 |
| FALSE | High | DSALLSK | [K].DSALLS | 1xPhospho | F6R6E1 1xl | 0.021797 | 0.001507 | 1 | 1 | 2 | F6R6E1 | Anoctamin | F6R6E1 [14 | 0 | 1271.701 | 0 | 4.87 | 0.000181 | 17.66 | 6.7 | 193.3 | 30.69 | 6.37 | | High | High | High | High | High | High | High | High | High | High | High | 0.000814 | 0.007522 | 2.19 |
| FALSE | High | DRPAAVDL | [K].DRPAA\ | 1xPhospho | Q6NV99 1x | 0.000203 | 0 | 1 | 6 | 2 | Q6NV99 | HAUS augn | Q6NV99 [3 | 0 | 3511.784 | 0 | 0.44 | 0.001678 | 2.73 | 86.4 | 113.6 | 5.18 | 2.89 | | High | High | High | High | High | High | High | High | High | High | High | 0 | 4.42E-05 | 4.54 |
| FALSE | High | DSANLESL/ | [R].DSANLI | 1xPhospho | Q7TNL7 1x | 0.000738 | 0 | 1 | 2 | 1 | Q7TNL7 | Dual specif | Q7TNL7 [2 | 0 | 1585.824 | 0 | -1.84 | 0.000509 | 0.07 | 153.6 | 46.4 | 12.29 | 13.26 | | High | High | High | High | High | High | High | High | High | High | High | 0 | 0.000169 | 3.82 |
| FALSE | High | DSAPELNV | [K].DSAPEL | 1xPhospho | Q3V0C5-2 | 1.44E-06 | 0 | 1 | 5 | 3 | Q3V0C5-2 | Isoform 2 c | Q3V0C5-2 | 0 | 2375.09 | 0 | -1.04 | 0.001021 | 3.88 | 136.8 | 63.2 | 7.12 | 5.97 | | High | High | High | High | High | High | High | High | High | High | High | 0 | 1.53E-07 | 4.06 |
| FALSE | High | DSAPELNV | [K].DSAPEL | 2xPhospho | Q3V0C5-2 | 0.000186 | 0 | 1 | 5 | 6 | Q3V0C5-2 | Isoform 2 c | Q3V0C5-2 | 0 | 2455.057 | 0 | -0.81 | 0.008189 | 5.37 | 127.4 | 72.6 | 12.93 | 12.04 | | High | High | High | High | High | High | High | High | High | High | High | 0 | 3.59E-05 | 2.67 |
| FALSE | High | DSAPTTPTS | [R].DSAPTT | 1xPhospho | E9QNY8 1x | 0.005666 | 0.00062 | 1 | 4 | 3 | E9QNY8 | Sacsin OS= | E9QNY8 [4 | 0 | 2297.126 | 0 | 0.19 | 0.057083 | 6.64 | 93.8 | 106.2 | 5.52 | 11.07 | | High | High | High | High | High | High | High | High | High | High | High | 0.000275 | 0.001662 | 2.75 |
| FALSE | High | DSAQDVQ/ | [K].DSAQD | 1xPhospho | Q9JJ28 1xP | 0.000279 | 0 | 1 | 1 | 3 | Q9JJ28 | Protein flig | Q9JJ28 [43 | 0 | 1499.751 | 0 | 1.18 | 0.001035 | 4.05 | 60.1 | 139.9 | 9.64 | 8.67 | | High | High | High | High | High | High | High | High | High | High | High | 0 | 9.46E-05 | 2.71 |
| FALSE | High | DSASDTAS/ | [K].DSASDT | 1xPhospho | E9Q8L9 1xl | 8.22E-07 | 0 | 1 | 3 | 12 | E9Q8L9 | Rab11 fam | E9Q8L9 [18 | 0 | 3252.508 | 0 | -0.05 | 0.459814 | 3.64 | 101.5 | 98.5 | 3.53 | 6.12 | | High | High | High | High | High | High | High | High | High | High | High | 0 | 2.6E-06 | 4.32 |
| FALSE | High | DSASPIQK | [K].DSASPI | 1xPhospho | H3BL37 1xl | 0.002844 | 0.00016 | 1 | 2 | 6 | H3BL37 | Treacle prc | H3BL37 [13 | 0 | 1383.729 | 0 | -0.59 | 0.000943 | 1.75 | 119.7 | 80.3 | 4.45 | 3.83 | | High | High | High | High | High | High | High | High | High | High | High | 0.005216 | 0.05695 | 2.63 |
| FALSE | High | DSASPSTSS | [R].DSASPS | 1xCarbami | A2A432 1x | 1.74E-06 | 0 | 1 | 4 | 3 | A2A432 | Cullin-4B C | A2A432 [14 | 0 | 3197.595 | 0 | -1.03 | 0.010303 | 8.64 | 134.4 | 65.6 | 15.02 | 16.87 | | High | High | High | High | High | High | High | High | High | High | High | 0 | 5.86E-07 | 3.13 |
| FALSE | High | DSAYFSDN | [R].DSAYFS | 1xPhospho | Q3TYD6 1x | 0.000237 | 0 | 1 | 1 | 4 | Q3TYD6 | Serine/thre | Q3TYD6 [1 | 0 | 2127.916 | 0 | -0.69 | 0.001927 | 4.74 | 122.1 | 77.9 | 5.14 | 6.72 | | High | High | High | High | High | High | High | High | High | High | High | 0 | 4.76E-05 | 3.43 |
| FALSE | High | DSAYFSDN | [R].DSAYFS | 2xPhospho | Q3TYD6 2x | 0.000225 | 0 | 1 | 1 | 1 | Q3TYD6 | Serine/thre | Q3TYD6 [1 | 0 | 2207.882 | 0 | -0.16 | 0.371419 | 6.67 | 105.2 | 94.8 | 5.93 | 8.58 | | High | High | High | High | High | High | High | High | High | High | High | 0 | 4.47E-05 | 2.67 |
| FALSE | High | DSAYGSFS1 | [R].DSAYG | 1xCarbami | A7TU71 1x | 0.098533 | 0.007902 | 1 | 2 | 2 | A7TU71 | Protein Shi | A7TU71 [2 | 0 | 2696.232 | 0 | 1.2 | 0.000336 | 1.71 | 60.2 | 139.8 | 7.31 | 5.66 | | High | High | High | High | High | High | High | High | High | High | High | 0.006366 | 0.06848 | 2.29 |
| FALSE | High | DSAYISSSPI | [R].DSAYIS | 1xPhospho | A2A690-2 | 0.001516 | 5.66E-05 | 1 | 3 | 2 | A2A690-2 | Isoform 2 c | A2A690-2 | 0 | 2422.128 | 0 | 1.59 | 0.000591 | 10.45 | 50.9 | 149.1 | 13.72 | 10.54 | | High | High | High | High | High | High | High | High | High | High | High | 0.000204 | 0.001337 | 3.06 |
| FALSE | High | DSCSSPPSL | [R].DSCSSP | 1xCarbami | A7TU71 1x | 3.39E-06 | 0 | 1 | 3 | 6 | A7TU71 | Protein Shi | A7TU71 [1 | 0 | 1949.908 | 0 | 1.54 | 0.000245 | 7.99 | 52.3 | 147.7 | 4.9 | 5.61 | | High | High | High | High | High | High | High | High | High | High | High | 0 | 5.23E-07 | 2.87 |
| FALSE | High | DSDAGCDS | [K].DSDAG | 1xCarbami | A2BDY3 1x | 0.000113 | 0 | 1 | 2 | 1 | A2BDY3 | Forkhead b | A2BDY3 [3 | 0 | 1938.81 | 0 | -1.95 | 0.357253 | 1.67 | 153.5 | 46.5 | 19.49 | 123.41 | | High | High | High | High | High | High | High | High | High | High | High | 0 | 2.06E-05 | 2.46 |
| FALSE | High | DSAPAPVN | [K].DSAPAI | 1xCarbami | Q80YR6 1x | 1.8E-05 | 0 | 1 | 3 | 1 | Q80YR6 | DNA endor | Q80YR6 [2 | 0 | 3698.741 | 0 | -1.89 | 0.000456 | 0.57 | 157.5 | 42.5 | 8.34 | 6.77 | | High | High | High | High | High | High | High | High | High | High | High | 0 | 2.61E-06 | 5.2 |
| FALSE | High | DRPAAVDL | [K].DRPAA\ | 1xPhospho | Q6NV99 1x | 6.55E-06 | 0 | 1 | 6 | 1 | Q6NV99 | HAUS augn | Q6NV99 [3 | 0 | 3495.789 | 0 | 0.25 | 0.014636 | 7.5 | 91 | 109 | 10.12 | 10.35 | | High | High | High | High | High | High | High | High | High | High | High | 0 | 8.37E-07 | 4.23 |
| FALSE | High | DRNTLPK | [R].DRNTLI | 1xPhospho | Q8CH77 1x | 0.061176 | 0.004646 | 1 | 4 | 1 | Q8CH77 | Neuron na | Q8CH77 [9 | 1 | 1381.76 | 0 | -1.05 | 0.003718 | 4.96 | 136.8 | 63.2 | 6.51 | 25.88 | | High | High | High | High | High | High | High | High | High | High | High | 0.002381 | 0.02442 | 1.2 |
| FALSE | High | DRNSWGG | [R].DRNSW | 1xPhospho | Q8BH66 1x | 0.007605 | 0.000664 | 1 | 1 | 1 | Q8BH66 | Atlastin-1 ( | Q8BH66 [7 | 1 | 1820.873 | 0 | 0.15 | 0.118809 | 16.56 | 94.8 | 105.2 | 8.22 | 12.12 | | High | High | High | High | High | High | High | High | High | High | High | 0.000294 | 0.002317 | 2.38 |
| FALSE | High | DQPAFTPS( | [R].DQPAF1 | 1xPhospho | Q9DBR1 1x | 2.47E-06 | 0 | 1 | 2 | 1 | Q9DBR1 | 5'-3' exorib | Q9DBR1 [4 | 0 | 2274.148 | 0 | 0.65 | 0.028431 | 7.21 | 79.2 | 120.8 | 9.37 | 10.18 | | High | High | High | High | High | High | High | High | High | High | High | 0 | 2.81E-07 | 3.82 |
| FALSE | High | DQPAPSSPI | [R].DQPAP | 1xPhospho | A0A0R4J0I | 0.000349 | 0 | 1 | 4 | 2 | A0A0R4J0I | CTF18, chr | A0A0R4J0I | 0 | 1694.859 | 0 | -0.19 | 0.200005 | 17.73 | 106.2 | 93.8 | 6.83 | 12.71 | | High | High | High | High | High | High | High | High | High | High | High | 0 | 7.33E-05 | 2.65 |
| FALSE | High | DQPAPSSPI | [R].DQPAP | 1xPhospho | A0A0R4J0I | 0.009168 | 0.000736 | 1 | 4 | 3 | A0A0R4J0I | CTF18, chr | A0A0R4J0I | 0 | 1710.854 | 0 | -0.14 | 0.384923 | 2.41 | 105.2 | 94.8 | 4.09 | 11.5 | | High | High | High | High | High | High | High | High | High | High | High | 0.006167 | 0.06681 | 1.87 |
| FALSE | High | DQPDGTSL | [K].DQPDG | 1xCarbami | P97434-2 1 | 1.48E-06 | 0 | 1 | 9 | 2 | P97434-2 | Isoform 2 c | P97434-2 [ | 0 | 2989.356 | 0 | 0.61 | 0.007013 | 19.82 | 79.1 | 120.9 | 12.88 | 7.05 | | High | High | High | High | High | High | High | High | High | High | High | 0 | 1.57E-07 | 5.17 |
| FALSE | High | DQPDGTSL | [K].DQPDG | 1xCarbami | P97434-2 1 | 1.4E-06 | 0 | 1 | 9 | 7 | P97434-2 | Isoform 2 c | P97434-2 [ | 0 | 2990.34 | 0 | 0.22 | 0.041451 | 6.12 | 88.5 | 111.5 | 8.1 | 10.23 | | High | High | High | High | High | High | High | High | High | High | High | 0 | 1.49E-07 | 6.25 |
| FALSE | High | DQPDGTSL | [K].DQPDG | 1xCarbami | P97434-2 2 | 6.83E-05 | 0 | 1 | 9 | 2 | P97434-2 | Isoform 2 c | P97434-2 [ | 0 | 3069.322 | 0 | 0.56 | 0.003596 | 14.92 | 79.8 | 120.2 | 8.82 | 6.64 | | High | High | High | High | High | High | High | High | High | High | High | 0 | 1.16E-05 | 3.73 |
| FALSE | High | DQPPSTGK | [K].DQPPS1 | 1xPhospho | Q3U2W2 1 | 0.095408 | 0.007437 | 1 | 2 | 2 | Q3U2W2 | MYB bindir | Q3U2W2 [ | 0 | 1367.697 | 0 | 0.02 | 0.92593 | 10.84 | 101.6 | 98.4 | 2.92 | 9.3 | | High | High | High | High | High | High | High | High | High | High | High | 0.003796 | 0.04094 | 2.95 |
| FALSE | High | DQQNLPY( | [R].DQQNL | 1xPhospho | Q571J8 1xl | 1.29E-09 | 0 | 1 | 18 | 1 | Q571J8 | Non-specif | Q571J8 [60 | 0 | 2505.172 | 0 | -0.03 | 0.830708 | 18.64 | 100 | 100 | 6.04 | 8.89 | | High | High | High | High | High | High | High | High | High | High | High | 0 | 5.77E-11 | 5.43 |
| FALSE | High | DQQPSGSE | [K].DQQPS | 1xPhospho | Q99J36 1xl | 0.000113 | 0 | 1 | 1 | 3 | Q99J36 | THUMP do | Q99J36 [82 | 0 | 2500.113 | 0 | -0.14 | 0.156553 | 6.11 | 107.1 | 92.9 | 8.62 | 14.45 | | High | High | High | High | High | High | High | High | High | High | High | 0 | 2.07E-05 | 3.1 |
| FALSE | High | DQQPSGSE | [K].DQQPS | 1xDeamida | Q99J36 1xl | 0.00418 | 0.00036 | 1 | 1 | 4 | Q99J36 | THUMP do | Q99J36 [82 | 0 | 2501.097 | 0 | -0.35 | 0.006486 | 2.9 | 114.3 | 85.7 | 6.44 | 8.94 | | High | High | High | High | High | High | High | High | High | High | High | 0.000161 | 0.001179 | 3.78 |
| FALSE | High | DQQPSGSE | [K].DQQPS | 2xPhospho | Q99J36 2xl | 0.000279 | 0 | 1 | 1 | 5 | Q99J36 | THUMP do | Q99J36 [82 | 0 | 2580.079 | 0 | -0.52 | 0.003473 | 4.02 | 118.4 | 81.6 | 6.16 | 17.19 | | High | High | High | High | High | High | High | High | High | High | High | 0 | 5.7E-05 | 3.7 |
| FALSE | High | DQRPSSPIF | [R].DQRPS | 1xPhospho | E9Q784 1x | 0.06169 | 0.004646 | 1 | 2 | 2 | E9Q784 | Zinc finger | E9Q784 [6 | 0 | 1364.688 | 0 | -0.91 | 0.001982 | 2.42 | 128.5 | 71.5 | 6.68 | 6.96 | | High | High | High | High | High | High | High | High | High | High | High | 0.002381 | 0.02458 | 1.5 |
| FALSE | High | DQSEQETS | [K].DQSEQ | 1xPhospho | Q811D0-2 | 9.03E-05 | 0 | 1 | 2 | 3 | Q811D0-2 | Isoform 2 c | Q811D0-2 | 0 | 3182.287 | 0 | 0.03 | 0.624565 | 0.53 | 102.2 | 97.8 | 18 | 7.08 | | High | High | High | High | High | High | High | High | High | High | High | 0 | 3.84E-05 | 2.98 |
| FALSE | High | DQSFPEGC | [R].DQSFPI | 1xPhospho | Q8VID5 1xl | 3.88E-06 | 0 | 1 | 1 | 2 | Q8VID5 | ATP-depen | Q8VID5 [8 | 0 | 1967.988 | 0 | -0.05 | 0.167303 | 7.43 | 106.3 | 93.7 | 7.73 | 4.43 | | High | High | High | High | High | High | High | High | High | High | High | 0.000418 | 0.003587 | 2.81 |
| FALSE | High | DQSPSQM/ | [K].DQSPS( | 1xPhospho | P14901 1xl | 0.070485 | 0.005214 | 1 | 2 | 1 | P14901 | Heme oxyg | P14901 [22 | 0 | 1528.703 | 0 | -0.37 | 0.010061 | 8.57 | 114.2 | 85.8 | 2.63 | 13.13 | | High | High | High | High | High | High | High | High | High | High | High | 0.002675 | 0.02864 | 2.24 |
| FALSE | High | DQSPSQM/ | [K].DQSPS( | 1xPhospho | P14901 1xl | 0.000765 | 0 | 1 | 2 | 4 | P14901 | Heme oxyg | P14901 [22 | 0 | 1544.698 | 0 | -0.38 | 0.002238 | 3.98 | 115 | 85 | 5.97 | 8.47 | | High | High | High | High | High | High | High | High | High | High | High | 0.000997 | 0.01033 | 2.01 |
| FALSE | High | DQTEAFVD | [K].DQTEA | 1xPhospho | A3KGI3 1xl | 5.43E-07 | 0 | 1 | 2 | 1 | A3KGI3 | Kinetochor | A3KGI3 [12 | 0 | 3423.683 | 0 | -0.63 | 0.001231 | 5.83 | 121.9 | 78.1 | 1.87 | 9.83 | | High | High | High | High | High | High | High | High | High | High | High | 0 | 5.13E-08 | 4.29 |
| FALSE | High | DQTPDENI | [R].DQTPD | 1xPhospho | O88477 1x | 0.000475 | 0 | 1 | 2 | 2 | O88477 | Insulin-like | O88477 [5 | 0 | 2039.01 | 0 | -1.25 | 0.000795 | 3.03 | 136.5 | 63.5 | 6.26 | 20.05 | | High | High | High | High | High | High | High | High | High | High | High | 0 | 0.000103 | 3.51 |
| FALSE | High | DQTPDENI | [R].DQTPD | 1xPhospho | Q9CPN8 1x | 0.001986 | 5.66E-05 | 1 | 1 | 1 | Q9CPN8 | Insulin-like | Q9CPN8 [5 | 0 | 2024.994 | 0 | -1.52 | 0.000158 | 0.43 | 150 | 50 | 2.34 | 5.55 | | High | High | High | High | High | High | High | High | High | High | High | 2.58E-05 | 0.000513 | 2.18 |
| FALSE | High | DQVLDDVS | [K].DQVLD | 1xCarbami | A2A654 1x | 2.06E-08 | 0 | 1 | 4 | 1 | A2A654 | Bromodom | A2A654 [1 | 0 | 2337.074 | 0 | 0.12 | 0.797459 | 19.2 | 98.6 | 101.4 | 10.4 | 7.8 | | High | High | High | High | High | High | High | High | High | High | High | 0 | 1.3E-09 | 4.04 |
| FALSE | High | DQVVGGN | [R].DQVVG | 1xPhospho | B0V2M3 1x | 0.000845 | 0 | 1 | 3 | 2 | B0V2M3 | Zinc finger | B0V2M3 [2 | 0 | 1436.71 | 0 | -0.46 | 0.002373 | 5.39 | 116 | 84 | 3.27 | 7.13 | | High | High | High | High | High | High | High | High | High | High | High | 4.94E-05 | 0.0007 | 1.48 |
| FALSE | High | DQWGSEEI | [K].DQWG | 1xPhospho | Q3TH99 1x | 1.1E-05 | 0 | 1 | 2 | 2 | Q3TH99 | Putative ur | Q3TH99 [2 | 0 | 2050.823 | 0 | -0.84 | 0.004473 | 4.36 | 124.1 | 75.9 | 11.07 | 9.56 | | High | High | High | High | High | High | High | High | High | High | High | 0 | 1.52E-06 | 3.23 |
| FALSE | High | DRCPLPCIP | [R].DRCPLF | 2xCarbami | Q9WVM1 | 0.088701 | 0.006669 | 1 | 1 | 1 | Q9WVM1 | Rac GTPase | Q9WVM1 | 1 | 2457.316 | 0 | -0.01 | 0.477332 | 17.62 | 99.5 | 100.5 | 12.94 | 9.57 | | High | High | High | High | High | High | High | High | High | High | High | 0.003461 | 0.03759 | 2.32 |
| FALSE | High | DREEAAAA | [K].DREEA/ | 1xPhospho | Q9D2D7 1x | 0.021797 | 0.001507 | 1 | 6 | 2 | Q9D2D7 | Zinc finger | Q9D2D7 [5 | 1 | 4368.092 | 0 | -0.3 | 0.029037 | 7.9 | 109.3 | 90.7 | 7.17 | 17.87 | | High | High | High | High | High | High | High | High | High | High | High | 0.000814 | 0.007505 | 3.56 |
| FALSE | High | DREPFAFSS | [K].DREPF/ | 1xPhospho | B9EJX5 1xF | 0.001011 | 0 | 1 | 2 | 2 | B9EJX5 | TAF2 RNA | B9EJX5 [11 | 1 | 1832.853 | 0 | -0.38 | 0.023498 | 9.97 | 113.2 | 86.8 | 8.67 | 8.18 | | High | High | High | High | High | High | High | High | High | High | High | 0 | 0.000241 | 2.12 |
| FALSE | High | DRGTPATA | [K].DRGTP/ | 1xPhospho | Q3UI84 1xl | 0.001503 | 5.66E-05 | 1 | 4 | 1 | Q3UI84 | Putative ur | Q3UI84 [19 | 1 | 1972.974 | 0 | 0.51 | 0.006366 | 18.64 | 77 | 123 | 15.04 | 10.24 | | High | High | High | High | High | High | High | High | High | High | High | 2.58E-05 | 0.000377 | 3.48 |
| FALSE | High | DRLEEDSQ | [K].DRLEED | 1xPhospho | Q8C0V9 1x | 1.31E-08 | 0 | 1 | 3 | 1 | Q8C0V9 | FERM dom | Q8C0V9 [4 | 1 | 2742.201 | 0 | 0.86 | 0.007417 | 25.75 | 70.2 | 129.8 | 7.29 | 15.67 | | High | High | High | High | High | High | High | High | High | High | High | 0 | 7.81E-10 | 4.9 |
| FALSE | High | DRLEEDSQ | [K].DRLEED | 1xPhospho | Q8C0V9 1x | 8.13E-06 | 0 | 1 | 3 | 1 | Q8C0V9 | FERM dom | Q8C0V9 [4 | 1 | 2758.196 | 0 | 1.08 | 0.006998 | 26.24 | 72.2 | 127.8 | 21.9 | 19.5 | | High | High | High | High | High | High | High | High | High | High | High | 0 | 1.07E-06 | 3.81 |
| FALSE | High | DRMDGSD | [K].DRMD( | 1xPhospho | Q9D8Y8 1x | 0.008313 | 0.000736 | 1 | 3 | 1 | Q9D8Y8 | Inhibitor of | Q9D8Y8 [1 | 1 | 1868.768 | 0 | -1.51 | 0.000209 | 0.86 | 147.4 | 52.6 | 4.97 | 6.95 | | High | High | High | High | High | High | High | High | High | High | High | 0.000327 | 0.002547 | 1.64 |
| FALSE | High | DQNESLDE | [K].DQNES | 1xPhospho | Q8CIB5 1xl | 5.19E-07 | 0 | 1 | 2 | 2 | Q8CIB5 | Fermitin fa | Q8CIB5 [66 | 0 | 2201.971 | 0 | -0.83 | 0.003114 | 1.85 | 126.8 | 73.2 | 10.49 | 8.41 | | High | High | High | High | High | High | High | High | High | High | High | 0 | 4.86E-08 | 4.38 |
| FALSE | High | EESWEYGF | [R].EESWE | 1xPhospho | Q9CYI4 1xF | 0.105058 | 0.008518 | 1 | 3 | 1 | Q9CYI4 | Putative RI | Q9CYI4 [33 | 0 | 1364.572 | 0 | -1.34 | 0.000278 | 1.33 | 143.3 | 56.7 | 5.2 | 6.88 | | High | High | High | High | High | High | High | High | High | High | High | 0.004291 | 0.04591 | 1.86 |
| FALSE | High | EESWGPPF | [R].EESWG | 1xPhospho | P23116 1xl | 0.101749 | 0.008194 | 1 | 1 | 1 | P23116 | Eukaryotic | P23116 [11 | 0 | 1266.572 | 0 | 0.84 | 0.011295 | 17.33 | 71.8 | 128.2 | 13.41 | 11.45 | | High | High | High | High | High | High | High | High | High | High | High | 0.004126 | 0.04415 | 1.06 |
| FALSE | High | EETDSVVLI | [K].EETDSV | 1xPhospho | O09030 1x | 0.001117 | 0 | 1 | 4 | 1 | O09030 | Interleukin | O09030 [4 | 0 | 2027.071 | 0 | 2.53 | 0.000781 | 17.69 | 27.5 | 172.5 | 21.07 | 15.46 | | High | High | High | High | High | High | High | High | High | High | High | 0 | 0.00027 | 3.06 |
| FALSE | High | ESLSPPGLK | [R].ESLSPP | 1xPhospho | Q569L8 1xl | 0.011755 | 0.000889 | 1 | 1 | 2 | Q569L8 | Centromer | Q569L8 [4 | 0 | 1465.807 | 0 | -1.12 | 0.000311 | 1.69 | 139 | 61 | 5.18 | 7.46 | | High | High | High | High | High | High | High | High | High | High | High | 0.000449 | 0.00374 | 2.75 |
| FALSE | High | ESLSPSVR | [K].ESLSPS\ | 1xPhospho | Q8BRN9 1x | 0.014667 | 0.001049 | 1 | 3 | 2 | Q8BRN9 | Coiled-coil | Q8BRN9 [5 | 0 | 1183.592 | 0 | 0.02 | 0.890708 | 6.38 | 99.8 | 100.2 | 5.54 | 2.8 | | High | High | High | High | High | High | High | High | High | High | High | 0.007083 | 0.07673 | 1.36 |
| FALSE | High | ESLSTSSDL | [R].ESLSTS | 1xPhospho | Q80U58 1x | 5.16E-05 | 0 | 1 | 3 | 1 | Q80U58 | Pumilio ho | Q80U58 [5 | 0 | 1767.882 | 0 | 0.89 | 0.001723 | 3.04 | 66.4 | 133.6 | 14.73 | 11.84 | | High | High | High | High | High | High | High | High | High | High | High | 0 | 8.51E-06 | 2.96 |
| FALSE | High | ESLTEVSET | [K].ESLTEV | 1xPhospho | Q99P72 1x | 0.000396 | 0 | 1 | 2 | 1 | Q99P72 | Reticulon-4 | Q99P72 [7 | 0 | 2126.078 | 0 | -0.15 | 0.01382 | 2.03 | 104.8 | 95.2 | 8.97 | 10.89 | | High | High | High | High | High | High | High | High | High | High | High | 0 | 8.45E-05 | 3.52 |
| FALSE | High | ESITSAVVS | [R].ESITSA\ | 2xPhospho | Q9JL04 2xF | 0.08447 | 0.006345 | 1 | 1 | 1 | Q9JL04 | Formin-2 C | Q9JL04 [36 | 0 | 2241.077 | 0 | 1.87 | 0.000758 | 6.26 | 42.9 | 157.1 | 33.97 | 14.13 | | High | High | High | High | High | High | High | High | High | High | High | 0.003265 | 0.0354 | 2.28 |
| FALSE | High | ESMSPANV | [R].ESMSP/ | 1xPhospho | E9Q8L9 1xl | 0.001689 | 5.66E-05 | 1 | 3 | 1 | E9Q8L9 | Rab11 fam | E9Q8L9 [4 | 0 | 1586.708 | 0 | 0.02 | 0.653478 | 26.72 | 103.6 | 96.4 | 12.25 | 14.75 | | High | High | High | High | High | High | High | High | High | High | High | 2.58E-05 | 0.000429 | 1.87 |
| FALSE | High | ESNQPPED | [R].ESNQPI | 1xPhospho | A2AFQ0 1x | 7.89E-05 | 0 | 1 | 6 | 3 | A2AFQ0 | E3 ubiquiti | A2AFQ0 [2 | 0 | 2485.057 | 0 | 0.07 | 0.968724 | 24.46 | 104.6 | 95.4 | 20.2 | 9.53 | | High | High | High | High | High | High | High | High | High | High | High | 0 | 1.37E-05 | 2.75 |
| FALSE | High | ESPAPPPPF | [R].ESPAPP | 1xPhospho | Q7TSG1 1x | 2.55E-07 | 0 | 1 | 2 | 1 | Q7TSG1 | Centrosom | Q7TSG1 [3 | 0 | 2193.136 | 0 | -1.7 | 0.000539 | 0.46 | 153.4 | 46.6 | 6.41 | 15.11 | | High | High | High | High | High | High | High | High | High | High | High | 0 | 2.19E-08 | 3.47 |
| FALSE | High | ESPDDQQE | [R].ESPDD( | 1xPhospho | Q80U72-4 | 4.43E-05 | 0 | 2 | 5 | 5 | Q80U72-4; | Isoform 4 c | Q80U72-4 | 0 | 4044.856 | 0 | -0.08 | 0.287125 | 14.93 | 104 | 96 | 10.67 | 14.51 | Shared | High | High | High | High | High | High | High | High | High | High | High | 0 | 1.35E-05 | 4.45 |
| FALSE | High | ESPFSTSAS | [R].ESPFST | 1xPhospho | Q6P5F9 1x | 0.000411 | 0 | 1 | 1 | 9 | Q6P5F9 | Exportin-1 | Q6P5F9 [3 | 0 | 2766.334 | 0 | -0.71 | 0.041819 | 3.05 | 124.1 | 75.9 | 10.04 | 14.92 | | High | High | High | High | High | High | High | High | High | High | High | 0.001011 | 0.01082 | 3.26 |
| FALSE | High | ESPFSTSAS | [R].ESPFST | 1xPhospho | Q6P5F9 1x | 0.067619 | 0.004864 | 1 | 1 | 1 | Q6P5F9 | Exportin-1 | Q6P5F9 [3 | 1 | 2922.435 | 0 | 0.35 | 0.05264 | 4.85 | 94.8 | 105.2 | 8.34 | 9.33 | | High | High | High | High | High | High | High | High | High | High | High | 0.002516 | 0.02729 | 2.88 |
| FALSE | High | ESPGGCTSI | [K].ESPGG( | 1xCarbami | Q3TRR0 1x | 3.17E-05 | 0 | 1 | 5 | 3 | Q3TRR0 | Microtubu | Q3TRR0 [2 | 0 | 1958.893 | 0 | 0.52 | 0.002661 | 3.24 | 81.7 | 118.3 | 3.81 | 4.87 | | High | High | High | High | High | High | High | High | High | High | High | 0 | 3.4E-05 | 3.28 |
| FALSE | High | ESPGSSEFF | [K].ESPGSS | 1xPhospho | Q5UAK0-1 | 2.8E-05 | 0 | 1 | 6 | 2 | Q5UAK0-1 | Isoform 1 c | Q5UAK0-1 | 0 | 2361.116 | 0 | -0.08 | 0.161141 | 3.04 | 106.3 | 93.7 | 9.36 | 4.49 | | High | High | High | High | High | High | High | High | High | High | High | 0 | 4.28E-06 | 4.38 |
| FALSE | High | ESLSLPSHP | [R].ESLSLP | 1xPhospho | Q6NXI6 1xl | 0.000579 | 0 | 1 | 3 | 1 | Q6NXI6 | Regulation | Q6NXI6 [1 | 0 | 3852.923 | 0 | 0.11 | 0.467107 | 55.89 | 92 | 108 | 31.07 | 15.45 | | High | High | High | High | High | High | High | High | High | High | High | 0 | 0.000129 | 4 |
| FALSE | High | ESPPEGGA | [K].ESPPEG | 1xPhospho | G3UYN0 1x | 0.000605 | 0 | 1 | 2 | 1 | G3UYN0 | L1TD1 OS= | G3UYN0 [1 | 0 | 1550.741 | 0 | -1.68 | 0.00207 | 0.81 | 152.4 | 47.6 | 12.8 | 8.66 | | High | High | High | High | High | High | High | High | High | High | High | 0 | 0.000136 | 1.46 |
| FALSE | High | ESPRPPAA/ | [R].ESPRPP | 1xPhospho | Q6ZQ58 1x | 0.00204 | 5.66E-05 | 1 | 2 | 4 | Q6ZQ58 | La-related | Q6ZQ58 [6 | 0 | 2403.078 | 0 | 0.65 | 0.001324 | 2.86 | 78.9 | 121.1 | 5.57 | 4.6 | | High | High | High | High | High | High | High | High | High | High | High | 0.000786 | 0.007087 | 2.16 |
| FALSE | High | ESPRPPAA/ | [R].ESPRPP | 2xPhospho | Q6ZQ58 2x | 0.015329 | 0.001089 | 1 | 2 | 1 | Q6ZQ58 | La-related | Q6ZQ58 [6 | 1 | 2639.145 | 0 | 0.8 | 0.006715 | 8.38 | 75.2 | 124.8 | 17.23 | 13.69 | | High | High | High | High | High | High | High | High | High | High | High | 0.000571 | 0.005081 | 2.63 |
| FALSE | High | ESPSLVSR | [R].ESPSLV | 1xPhospho | A0A0U1RP | 0.026191 | 0.001676 | 1 | 7 | 2 | A0A0U1RP | Ataxin-2-lik | A0A0U1RP | 0 | 1183.592 | 0 | 0.41 | 0.016916 | 6.78 | 85.4 | 114.6 | 4.32 | 8.46 | | High | High | High | High | High | High | High | High | High | High | High | 0.000903 | 0.009233 | 1.8 |
| FALSE | High | ESPSSAADI | [K].ESPSSA | 1xPhospho | A2AAE1-3 | 0.001437 | 5.66E-05 | 1 | 6 | 1 | A2AAE1-3 | Isoform 3 c | A2AAE1-3 | 0 | 1995.942 | 0 | -0.56 | 0.015263 | 7.57 | 117 | 83 | 4.5 | 12.09 | | High | High | High | High | High | High | High | High | High | High | High | 2.58E-05 | 0.000359 | 2.95 |
| FALSE | High | ESPSSTEAD | [K].ESPSST | 1xPhospho | A0A1B0GS | 5.68E-07 | 0 | 1 | 2 | 7 | A0A1B0GS | GRB2-asso | A0A1B0GS | 0 | 3371.482 | 0 | -1.09 | 0.005103 | 2.84 | 136.5 | 63.5 | 6.22 | 18.33 | | High | High | High | High | High | High | High | High | High | High | High | 0 | 1.89E-07 | 4.83 |
| FALSE | High | ESPSSTEAD | [K].ESPSST | 1xPhospho | A0A1B0GS | 1.35E-06 | 0 | 1 | 2 | 4 | A0A1B0GS | GRB2-asso | A0A1B0GS | 0 | 3387.477 | 0 | -1.02 | 0.000995 | 1.44 | 136.1 | 63.9 | 6.13 | 11.85 | | High | High | High | High | High | High | High | High | High | High | High | 0 | 2.33E-07 | 5.13 |
| FALSE | High | ESPVTVDA | [K].ESPVTV | 1xPhospho | P70399 1xl | 0.000107 | 0 | 1 | 5 | 3 | P70399 | TP53-bindi | P70399 [8 | 0 | 2012.983 | 0 | 0.53 | 0.00128 | 2.54 | 81.2 | 118.8 | 7.13 | 5.94 | | High | High | High | High | High | High | High | High | High | High | High | 0 | 1.94E-05 | 3.12 |

| | | | | | | | | | | | | | | | | | | | | | | | | | | | | | | | | | | | | | | |
|---|---|---|---|---|---|---|---|---|---|---|---|---|---|---|---|---|---|---|---|---|---|---|---|---|---|---|---|---|---|---|---|---|---|---|---|---|---|---|
| FALSE | High | ESQEFLRSF | [K].ESQEFL | 1xPhosphc | Q6P5H2-2 | 0.008094 | 0.000736 | 1 | 2 | 3 | Q6P5H2-2 | Isoform 2 c | Q6P5H2-2 | 1 | 2860.291 | 0 | 2 | 0.01798 | 29.25 | 39.6 | 160.4 | 11.1 | 32.87 | | High | High | High | High | High | High | High | High | High | High | High | 0.000327 | 0.002464 | 3.23 |
| FALSE | High | ESQEFLRSF | [K].ESQEFL | 1xPhosphc | Q6P5H2-2 | 0.000501 | 0 | 1 | 2 | 2 | Q6P5H2-2 | Isoform 2 c | Q6P5H2-2 | 1 | 2876.286 | 0 | 2.18 | 0.000632 | 1.44 | 36.5 | 163.5 | 12.58 | 12.5 | | High | High | High | High | High | High | High | High | High | High | High | 0 | 0.000109 | 3.7 |
| FALSE | High | ESQEQLAP | [K].ESQEQI | 1xPhosphc | P51125-3 1 | 1.78E-05 | 0 | 1 | 12 | 2 | P51125-3 | Isoform 3 c | P51125-3 [ | 0 | 4595.201 | 0 | 2.81 | 0.00085 | 38.57 | 25.3 | 174.7 | 36.21 | 9.75 | | High | High | High | High | High | High | High | High | High | High | High | 0 | 2.57E-06 | 4.87 |
| FALSE | High | ESQEQLAP | [K].ESQEQI | 1xDeamida | P51125-3 1 | 0.000389 | 0 | 1 | 12 | 1 | P51125-3 | Isoform 3 c | P51125-3 [ | 0 | 4596.185 | 0 | 2.29 | 0.001712 | 18.29 | 35.3 | 164.7 | 14.72 | 14.13 | | High | High | High | High | High | High | High | High | High | High | High | 0 | 8.28E-05 | 4.23 |
| FALSE | High | ESQESADQ | [K].ESQES/ | 1xPhosphc | F8WIP8 1xl | 0.000765 | 0 | 1 | 5 | 1 | F8WIP8 | Osteopont | F8WIP8 [23 | 0 | 2549.166 | 0 | 0.38 | 0.059059 | 21.6 | 81.2 | 118.8 | 17.13 | 29.43 | | High | High | High | High | High | High | High | High | High | High | High | 0 | 0.000177 | 2.95 |
| FALSE | High | ESQESLKSF | [K].ESQESL | 1xPhosphc | Q6P5H2-2 | 0.017041 | 0.001204 | 1 | 2 | 1 | Q6P5H2-2 | Isoform 2 c | Q6P5H2-2 | 1 | 2329.096 | 0 | 1.98 | 0.004914 | 47.61 | 38 | 162 | 19.01 | 20.34 | | High | High | High | High | High | High | High | High | High | High | High | 0.000638 | 0.00568 | 1.6 |
| FALSE | High | ESPPPSAPS | [R].ESPPPS | 1xPhosphc | Q56A08 1x | 0.000379 | 0 | 1 | 1 | 1 | Q56A08 | G patch do | Q56A08 [5 | 0 | 2301.082 | 0 | 0.78 | 0.009341 | 27.81 | 73.5 | 126.5 | 11.22 | 10.52 | | High | High | High | High | High | High | High | High | High | High | High | 0 | 8.04E-05 | 3.82 |
| FALSE | High | ESQMTPSH | [R].ESQMT | 1xPhosphc | A0A0N4SU | 0.003464 | 0.00021 | 1 | 10 | 2 | A0A0N4SU | Peregrin O | A0A0N4SU | 0 | 2901.384 | 0 | 0.05 | 0.604024 | 40.69 | 108.4 | 91.6 | 15.65 | 26.19 | | High | High | High | High | High | High | High | High | High | High | High | 9.42E-05 | 0.00096 | 2.98 |
| FALSE | High | ESLQEAGK | [K].ESLQEA | 1xPhosphc | P53986 1xl | 0.002313 | 0.000109 | 1 | 5 | 3 | P53986 | Monocarb | P53986 [21 | 0 | 1399.723 | 0 | 0.63 | 0.0116 | 10.7 | 78.3 | 121.7 | 12.88 | 12.42 | | High | High | High | High | High | High | High | High | High | High | High | 0.000204 | 0.001335 | 2.79 |
| FALSE | High | ESLPSTVSA | [R].ESLPST | 1xPhosphc | D4AFX6 1x | 0.004911 | 0.000504 | 1 | 3 | 2 | D4AFX6 | Centaurin I | D4AFX6 [5 | 0 | 2374.101 | 0 | 2.68 | 0.001953 | 23.36 | 29.1 | 170.9 | 18.19 | 19.35 | | High | High | High | High | High | High | High | High | High | High | High | 0.000225 | 0.001418 | 2.72 |
| FALSE | High | ESECLAPQ( | [K].ESECLA | 1xCarbami | Q14B71 1x | 2.08E-06 | 0 | 1 | 1 | 2 | Q14B71 | Cell divisio | Q14B71 [7 | 0 | 2715.295 | 0 | -0.94 | 0.000269 | 2.17 | 131.6 | 68.4 | 2.26 | 6.52 | | High | High | High | High | High | High | High | High | High | High | High | 0 | 1.7E-06 | 4.3 |
| FALSE | High | ESEDNDLG | [K].ESEDNI | 1xPhosphc | Q8BGC0 1x | 0.033694 | 0.002195 | 1 | 1 | 3 | Q8BGC0 | HIV Tat-spe | Q8BGC0 [5 | 1 | 2748.307 | 0 | -0.04 | 0.505435 | 9.54 | 101.3 | 98.7 | 4.27 | 6.83 | | High | High | High | High | High | High | High | High | High | High | High | 0.007292 | 0.07918 | 2.71 |
| FALSE | High | ESEEPTSPC | [R].ESEEPT | 1xCarbami | Q6ZWQ0 1 | 0.00129 | 0 | 1 | 9 | 4 | Q6ZWQ0 | Nesprin-2 | Q6ZWQ0 [ | 0 | 2879.36 | 0 | 5 | 0.001095 | 31.69 | 5.7 | 194.3 | 44.87 | 16.69 | | High | High | High | High | High | High | High | High | High | High | High | 0.000161 | 0.00114 | 1.69 |
| FALSE | High | ESEPESPMI | [K].ESEPES | 1xPhosphc | O88622 1x | 0.000126 | 0 | 1 | 4 | 1 | O88622 | Poly(ADP-r | O88622 [2 | 0 | 2101.94 | 0 | -0.58 | 0.013446 | 8.29 | 126 | 74 | 10.42 | 15.04 | | High | High | High | High | High | High | High | High | High | High | High | 0 | 2.34E-05 | 3.27 |
| FALSE | High | ESEPESPMI | [K].ESEPES | 1xPhosphc | O88622 1x | 0.000422 | 0 | 1 | 4 | 2 | O88622 | Poly(ADP-r | O88622 [2 | 0 | 2117.935 | 0 | -0.87 | 0.000323 | 1.43 | 130 | 70 | 3.65 | 3.4 | | High | High | High | High | High | High | High | High | High | High | High | 0.000275 | 0.00205 | 2.85 |
| FALSE | High | ESEPTPAAS | [R].ESEPTP | 1xPhosphc | A0A075F6( | 2.61E-06 | 0 | 1 | 5 | 2 | A0A075F6( | Heat shock | A0A075F6( | 0 | 3703.781 | 0 | 0.31 | 0.58341 | 14.3 | 93.6 | 106.4 | 7.85 | 24.49 | | High | High | High | High | High | High | High | High | High | High | High | 0 | 5.12E-06 | 4.14 |
| FALSE | High | ESEPTPAAS | [R].ESEPTP | 1xPhosphc | A0A075F6( | 7.51E-07 | 0 | 1 | 5 | 3 | A0A075F6( | Heat shock | A0A075F6( | 0 | 3719.776 | 0 | -0.01 | 0.418161 | 8.67 | 97.4 | 102.6 | 12.09 | 35.12 | | High | High | High | High | High | High | High | High | High | High | High | 0 | 9.85E-08 | 4.17 |
| FALSE | High | ESEQEQSSI | [K].ESEQE( | 1xPhosphc | Q8R2U7 1x | 0.000136 | 0 | 1 | 1 | 1 | Q8R2U7 | Leucine-ric | Q8R2U7 [3 | 0 | 1972.927 | 0 | -0.34 | 0.004055 | 3.31 | 111.6 | 88.4 | 4.09 | 3.64 | | High | High | High | High | High | High | High | High | High | High | High | 0 | 2.53E-05 | 2.91 |
| FALSE | High | ESETFSDSS | [K].ESETFS | 1xPhosphc | Q99P72 1x | 1.08E-08 | 0 | 1 | 2 | 19 | Q99P72 | Reticulon-4 | Q99P72 [8 | 0 | 3213.553 | 0 | 0.27 | 0.292214 | 25.49 | 88.6 | 111.4 | 10.42 | 15.16 | | High | High | High | High | High | High | High | High | High | High | High | 0 | 1.86E-08 | 4.13 |
| FALSE | High | ESEVCPVPT | [K].ESEVCF | 1xCarbami | E9Q444 2x | 0.036731 | 0.00249 | 1 | 8 | 2 | E9Q444 | Zinc finger | E9Q444 [9 | 0 | 3248.612 | 0 | 0.23 | 0.033326 | 14.31 | 89.1 | 110.9 | 4.88 | 11.67 | | High | High | High | High | High | High | High | High | High | High | High | 0.001308 | 0.01354 | 2.84 |
| FALSE | High | ESGPGSPA | [K].ESGPGS | 1xCarbami | P55200 1xl | 2.03E-06 | 0 | 1 | 2 | 1 | P55200 | Histone-lys | P55200 [25 | 0 | 2630.212 | 0 | 0.47 | 0.032992 | 17.68 | 83.3 | 116.7 | 15.78 | 16.52 | | High | High | High | High | High | High | High | High | High | High | High | 0 | 2.25E-07 | 4.59 |
| FALSE | High | ESGQEGVD | [K].ESGQE( | 1xPhosphc | O54941 1x | 0.045898 | 0.003408 | 1 | 1 | 4 | O54941 | SWI/SNF-r | O54941 [3 | 0 | 4891.104 | 0 | 0.15 | 0.979024 | 16.15 | 97.3 | 102.7 | 6.53 | 33.49 | | High | High | High | High | High | High | High | High | High | High | High | 0.00449 | 0.04845 | 3.14 |
| FALSE | High | ESGQEGVD | [K].ESGQE( | 1xPhosphc | O54941 1x | 0.007673 | 0.000664 | 1 | 1 | 8 | O54941 | SWI/SNF-r | O54941 [3 | 0 | 4907.099 | 0 | 0.06 | 0.549965 | 15.86 | 103.7 | 96.3 | 15.99 | 40.23 | | High | High | High | High | High | High | High | High | High | High | High | 0.000814 | 0.007787 | 3.52 |
| FALSE | High | ESLPSTVSA | [R].ESLPST | 2xPhosphc | D4AFX6 2x | 0.00172 | 5.66E-05 | 1 | 3 | 1 | D4AFX6 | Centaurin I | D4AFX6 [5 | 0 | 2454.068 | 0 | 2.94 | 0.002674 | 22.41 | 19.5 | 180.5 | 43.75 | 24.84 | | High | High | High | High | High | High | High | High | High | High | High | 2.58E-05 | 0.00044 | 3.27 |
| FALSE | High | ESGSPAQE | [R].ESGSPA | 1xPhosphc | Q7TPM1 1: | 1.49E-06 | 0 | 1 | 3 | 2 | Q7TPM1 | Protein PRI | Q7TPM1 [6 | 0 | 1704.825 | 0 | 0.5 | 0.00359 | 6.86 | 82.7 | 117.3 | 2.22 | 7.87 | | High | High | High | High | High | High | High | High | High | High | High | 0 | 1.6E-07 | 3.56 |
| FALSE | High | ESHSPFGLI | [K].ESHSPF | 1xPhosphc | B9EKS2 1xl | 5.3E-08 | 0 | 1 | 3 | 4 | B9EKS2 | Jumonji do | B9EKS2 [12 | 0 | 2262.084 | 0 | 0.24 | 0.066658 | 18.16 | 93.3 | 106.7 | 6.92 | 13.06 | | High | High | High | High | High | High | High | High | High | High | High | 0 | 1.23E-06 | 3.56 |
| FALSE | High | ESLAQGPD | [R].ESLAQ( | 1xDeamida | Q5SPX8 1x | 2.31E-05 | 0 | 1 | 4 | 4 | Q5SPX8 | TBC1 doma | Q5SPX8 [5 | 0 | 3305.453 | 0 | -0.36 | 0.101015 | 20.22 | 112.3 | 87.7 | 17.51 | 20.3 | | High | High | High | High | High | High | High | High | High | High | High | 0.000183 | 0.001282 | 4.26 |
| FALSE | High | ESLAQGPD | [R].ESLAQ( | 2xPhosphc | Q5SPX8 2x | 4.01E-05 | 0 | 1 | 4 | 2 | Q5SPX8 | TBC1 doma | Q5SPX8 [5 | 0 | 3384.436 | 0 | 0.25 | 0.248012 | 41.83 | 86.3 | 113.7 | 21.16 | 15.27 | | High | High | High | High | High | High | High | High | High | High | High | 0 | 9.53E-05 | 5 |
| FALSE | High | ESLAQGPD | [R].ESLAQ( | 1xDeamida | Q5SPX8 2x | 7.62E-09 | 0 | 1 | 4 | 4 | Q5SPX8 | TBC1 doma | Q5SPX8 [5 | 0 | 3385.42 | 0 | 0.45 | 0.011872 | 11.26 | 83.1 | 116.9 | 7.97 | 11.71 | | High | High | High | High | High | High | High | High | High | High | High | 0 | 4.26E-10 | 6.23 |
| FALSE | High | ESLFLGPES | [K].ESLFLG | 1xPhosphc | Q5DTT3 1x | 0.002355 | 0.000109 | 1 | 1 | 2 | Q5DTT3 | Protein FAI | Q5DTT3 [1 | 0 | 1512.755 | 0 | -0.39 | 0.391247 | 13.49 | 119.3 | 80.7 | 32.01 | 35.49 | | High | High | High | High | High | High | High | High | High | High | High | 9.42E-05 | 0.001039 | 2.7 |
| FALSE | High | ESLFLGPES | [K].ESLFLG | 1xPhosphc | Q5DTT3 1x | 3.83E-05 | 0 | 1 | 1 | 2 | Q5DTT3 | Protein FAI | Q5DTT3 [1 | 0 | 1741.918 | 0 | -0.47 | 0.011496 | 1.92 | 114.9 | 85.1 | 6.18 | 10.26 | | High | High | High | High | High | High | High | High | High | High | High | 0 | 1.2E-05 | 3.12 |
| FALSE | High | ESLGSEEES | [K].ESLGSE | 1xPhosphc | G3X956 1x | 0.001463 | 5.66E-05 | 1 | 2 | 1 | G3X956 | Suppresso | G3X956 [9 | 0 | 1689.798 | 0 | -1.46 | 0.002393 | 1.49 | 147.7 | 52.3 | 15.01 | 6.43 | | High | High | High | High | High | High | High | High | High | High | High | 2.58E-05 | 0.000367 | 2.11 |
| FALSE | High | ESLKEEDES | [K].ESLKEE | 1xPhosphc | Q58EV4 1x | 0.100936 | 0.008097 | 1 | 3 | 2 | Q58EV4 | Proteasom | Q58EV4 [2 | 1 | 2209.91 | 0 | 0.14 | 0.180283 | 4.43 | 97.1 | 102.9 | 18.93 | 6.86 | | High | High | High | High | High | High | High | High | High | High | High | 0.004543 | 0.0494 | 2.2 |
| FALSE | High | ESLLLEDSD | [K].ESLLLEI | 1xCarbami | Q9WV66 1 | 3.06E-08 | 0 | 1 | 1 | 3 | Q9WV66 | E3 ubiquiti | Q9WV66 [ | 0 | 2433.021 | 0 | -0.21 | 0.120762 | 4.72 | 107.7 | 92.3 | 11.26 | 4.87 | | High | High | High | High | High | High | High | High | High | High | High | 0 | 1.97E-08 | 3.83 |
| FALSE | High | ESLPLDLSD | [K].ESLPLD | 1xPhosphc | Q9DBY8 1x | 2E-07 | 0 | 1 | 2 | 5 | Q9DBY8 | Nuclear va | Q9DBY8 [1 | 0 | 2186.063 | 0 | -0.47 | 0.008041 | 4.08 | 114.8 | 85.2 | 3.95 | 7.58 | | High | High | High | High | High | High | High | High | High | High | High | 0 | 2.36E-05 | 4.19 |
| FALSE | High | ESLPLDLSD | [K].ESLPLD | 1xDeamida | Q9DBY8 1x | 3.59E-07 | 0 | 1 | 2 | 5 | Q9DBY8 | Nuclear va | Q9DBY8 [1 | 0 | 2187.047 | 0 | -0.52 | 0.000751 | 2.75 | 118.4 | 81.6 | 1.42 | 6.3 | | High | High | High | High | High | High | High | High | High | High | High | 0 | 3.2E-08 | 3.07 |
| FALSE | High | ESLPLDLSD | [K].ESLPLD | 2xPhosphc | Q9DBY8 2x | 0.000605 | 0 | 1 | 2 | 4 | Q9DBY8 | Nuclear va | Q9DBY8 [1 | 0 | 2266.029 | 0 | -0.3 | 0.00407 | 5.44 | 110.5 | 89.5 | 2.99 | 6.17 | | High | High | High | High | High | High | High | High | High | High | High | 0 | 0.000136 | 2.8 |
| FALSE | High | ESLPSLPQS | [K].ESLPSLI | 1xPhosphc | F8WI37 1xl | 0.073462 | 0.005631 | 1 | 7 | 1 | F8WI37 | Histone-lys | F8WI37 [4 | 0 | 1867.964 | 0 | -0.48 | 0.007798 | 4.19 | 116.6 | 83.4 | 5.06 | 9.29 | | High | High | High | High | High | High | High | High | High | High | High | 0.002882 | 0.02998 | 1.92 |
| FALSE | High | ESGSPYSVL | [R].ESGSPY | 1xPhosphc | Q5DU09 1x | 0.079766 | 0.00613 | 1 | 4 | 1 | Q5DU09 | Zinc finger | Q5DU09 [5 | 0 | 1891.945 | 0 | 0.13 | 0.927127 | 26.52 | 98.7 | 101.3 | 17.94 | 17.6 | | High | High | High | High | High | High | High | High | High | High | High | 0.003163 | 0.03315 | 2 |
| FALSE | High | ESQSVLIAD | [K].ESQSVL | 1xPhosphc | A0A0R4J0F | 2.52E-07 | 0 | 1 | 6 | 2 | A0A0R4J0F | Cyclin-G-as | A0A0R4J0F | 0 | 3307.421 | 0 | 0.51 | 0.069452 | 5.97 | 89.1 | 110.9 | 15.84 | 13.89 | | High | High | High | High | High | High | High | High | High | High | High | 0 | 2.17E-08 | 4.27 |
| FALSE | High | ESQSVLIAD | [K].ESQSVL | 2xPhosphc | A0A0R4J0F | 2.21E-05 | 0 | 1 | 6 | 4 | A0A0R4J0F | Cyclin-G-as | A0A0R4J0F | 0 | 3387.388 | 0 | 0.86 | 0.000525 | 8.95 | 71 | 129 | 9.54 | 6.47 | | High | High | High | High | High | High | High | High | High | High | High | 0 | 3.31E-06 | 4.85 |
| FALSE | High | ESSEDTEVF | [K].ESSEDT | 1xPhosphc | H9KV00 1x | 0.007404 | 0.00062 | 1 | 9 | 4 | H9KV00 | Protein SO | H9KV00 [1 | 0 | 1982.972 | 0 | -0.02 | 0.313178 | 4.7 | 104.5 | 95.5 | 8.86 | 10.1 | | High | High | High | High | High | High | High | High | High | High | High | 0.000275 | 0.002242 | 3.01 |
| FALSE | High | ESTHQSED | [R].ESTHQS | 1xPhosphc | B9EKC3 1xl | 0.004824 | 0.000456 | 1 | 3 | 1 | B9EKC3 | Rho GTPas | B9EKC3 [9 | 0 | 2037.948 | 0 | 0.6 | 0.022785 | 28.31 | 80.9 | 119.1 | 13.24 | 11.14 | | High | High | High | High | High | High | High | High | High | High | High | 0.000204 | 0.001391 | 2.57 |
| FALSE | High | ESTIASPVS | [R].ESTIASI | 1xPhosphc | Q8CGC6 1x | 0.000125 | 0 | 1 | 1 | 1 | Q8CGC6 | RNA-bindir | Q8CGC6 [2 | 0 | 1792.961 | 0 | -0.22 | 0.037499 | 9.39 | 108.1 | 91.9 | 7.02 | 13.74 | | High | High | High | High | High | High | High | High | High | High | High | 0 | 2.3E-05 | 1.7 |
| FALSE | High | ESTNIFSPSI | [K].ESTNIF: | 1xPhosphc | P28658 1xl | 7.21E-05 | 0 | 1 | 4 | 3 | P28658 | Ataxin-10 ( | P28658 [34 | 0 | 1962.983 | 0 | -0.94 | 0.00043 | 2.73 | 133 | 67 | 4.88 | 6.72 | | High | High | High | High | High | High | High | High | High | High | High | 0.000382 | 0.003228 | 3.1 |
| FALSE | High | ESTNSTSPN | [K].ESTNST | 1xCarbami | Q6P5D4 1x | 0.001968 | 5.66E-05 | 1 | 2 | 2 | Q6P5D4 | Centrosom | Q6P5D4 [5 | 0 | 2427.163 | 0 | -0.91 | 0.007035 | 4.66 | 129 | 71 | 13.18 | 4.58 | | High | High | High | High | High | High | High | High | High | High | High | 2.58E-05 | 0.000509 | 2.33 |
| FALSE | High | ESTQISSPG | [K].ESTQIS: | 1xCarbami | Q8VC34 1x | 0.000808 | 0 | 1 | 5 | 1 | Q8VC34 | Putative RI | Q8VC34 [2 | 0 | 1855.846 | 0 | -0.39 | 0.003094 | 3.4 | 113.2 | 86.8 | 5.62 | 3.93 | | High | High | High | High | High | High | High | High | High | High | High | 0 | 0.000187 | 3.45 |
| FALSE | High | ESTSPKPAI | [R].ESTSPK | 1xCarbami | F6Z9A1 1xl | 0.003083 | 0.00016 | 1 | 5 | 3 | F6Z9A1 | Sentrin-spe | F6Z9A1 [37 | 0 | 2857.426 | 0 | -0.26 | 0.688932 | 11.33 | 101.3 | 98.7 | 21.45 | 16.55 | | High | High | High | High | High | High | High | High | High | High | High | 7.19E-05 | 0.000845 | 2.23 |
| FALSE | High | ESTSPQPAI | [R].ESTSPC | 1xCarbami | B7ZWN9 2: | 0.005768 | 0.00062 | 1 | 1 | 1 | B7ZWN9 | SUMO/sen | B7ZWN9 [3 | 0 | 2708.193 | 0 | -1.49 | 0.001158 | 2.64 | 148.4 | 51.6 | 7.55 | 19.43 | | High | High | High | High | High | High | High | High | High | High | High | 0.000275 | 0.0017 | 2.02 |
| FALSE | High | ESVGSTLSF | [K].ESVGST | 1xPhosphc | Q8C9B9 1x | 9.45E-05 | 0 | 1 | 1 | 1 | Q8C9B9 | Death-indu | Q8C9B9 [1 | 0 | 2009.995 | 0 | 0.61 | 0.042368 | 20.04 | 82.1 | 117.9 | 9.04 | 16.57 | | High | High | High | High | High | High | High | High | High | High | High | 0 | 1.69E-05 | 1.83 |
| FALSE | High | ESVPDFPLS | [K].ESVPDF | 1xPhosphc | Q545B6 1x | 0.001021 | 0 | 1 | 5 | 1 | Q545B6 | Stathmin C | Q545B6 [3 | 0 | 1621.808 | 0 | -1.4 | 0.061601 | 3.28 | 143.4 | 56.6 | 42.08 | 38.91 | | High | High | High | High | High | High | High | High | High | High | High | 0 | 0.000244 | 3.31 |
| FALSE | High | ESVPDFPLS | [K].ESVPDF | 1xPhosphc | Q545B6 1x | 1.43E-06 | 0 | 1 | 5 | 9 | Q545B6 | Stathmin C | Q545B6 [3 | 0 | 1850.971 | 0 | -1.47 | 0.002983 | 0.76 | 147.7 | 52.3 | 13.32 | 16.1 | | High | High | High | High | High | High | High | High | High | High | High | 0 | 8.59E-07 | 3.1 |
| FALSE | High | ESVPDFPLS | [K].ESVPDF | 1xPhosphc | Q545B6 1x | 0.028327 | 0.001791 | 1 | 5 | 1 | Q545B6 | Stathmin C | Q545B6 [3 | 1 | 2208.228 | 0 | -1.09 | 0.006898 | 2.75 | 131.3 | 68.7 | 11.76 | 20.06 | | High | High | High | High | High | High | High | High | High | High | High | 0.000968 | 0.01011 | 2.78 |
| FALSE | High | ESVSPEDSE | [R].ESVSPE | 1xPhosphc | G3UWX1 1 | 0.021797 | 0.001507 | 2 | 8 | 1 | G3UWX1; I | Replicatior | G3UWX1 [ | 0 | 1644.777 | 0 | -0.25 | 0.111517 | 20.13 | 111.8 | 88.2 | 12.18 | 20.48 | Shared | High | High | High | High | High | High | High | High | High | High | High | 0.000814 | 0.007537 | 2.12 |
| FALSE | High | ESVTDYTTF | [K].ESVTDY | 1xPhosphc | Q3TLH4 1x | 0.026421 | 0.001676 | 1 | 6 | 2 | Q3TLH4 | Protein PRI | Q3TLH4 [2 | 0 | 2935.434 | 0 | 0.58 | 0.034818 | 33.54 | 75.4 | 124.6 | 14.01 | 18.18 | | High | High | High | High | High | High | High | High | High | High | High | 0.000903 | 0.009288 | 2.84 |
| FALSE | High | ESTGSYSDI | [R].ESTGSY | 2xPhosphc | F6WJB7 2x | 0.014158 | 0.001011 | 1 | 5 | 1 | F6WJB7 | Ubiquitin c | F6WJB7 [1 | 0 | 3372.361 | 0 | -0.05 | 0.384613 | 0.96 | 98.2 | 101.8 | 18.97 | 14.68 | | High | High | High | High | High | High | High | High | High | High | High | 0.000538 | 0.00462 | 3.13 |
| FALSE | High | ESYAFDGS( | [R].ESYAFD | 1xPhosphc | O35387 1x | 2.53E-05 | 0 | 1 | 6 | 3 | O35387 | HCLS1-assc | O35387 [5 | 0 | 2690.165 | 0 | -0.83 | 0.005679 | 8.07 | 127.5 | 72.5 | 9.57 | 11.97 | | High | High | High | High | High | High | High | High | High | High | High | 0 | 3.84E-06 | 3.58 |
| FALSE | High | ESYDDVSSI | [K].ESYDD\ | 1xPhosphc | Q9D903 1x | 0.031713 | 0.002011 | 1 | 1 | 1 | Q9D903 | Probable rI | Q9D903 [2 | 0 | 1513.641 | 0 | -0.15 | 0.079294 | 2.71 | 106.5 | 93.5 | 8.14 | 6.45 | | High | High | High | High | High | High | High | High | High | High | High | 0.001076 | 0.01143 | 1.97 |
| FALSE | High | ESYSVYVYK | [K].ESYSVY | 1xPhosphc | Q8CGP2-2 | 0.021418 | 0.001507 | 1 | 14 | 1 | Q8CGP2-2 | Isoform 2 c | Q8CGP2-2 | 0 | 1675.838 | 0 | -0.1 | 0.508914 | 10.54 | 103.5 | 96.5 | 20.33 | 4.53 | | High | High | High | High | High | High | High | High | High | High | High | 0.000801 | 0.007363 | 1.9 |
| FALSE | High | ESYYFGLSP | [R].ESYYFG | 1xPhosphc | B2RRE7 1xl | 0.000611 | 0 | 1 | 2 | 3 | B2RRE7 | OTU doma | B2RRE7 [4 | 0 | 1785.793 | 0 | 0.16 | 0.019492 | 8.24 | 94.2 | 105.8 | 5.02 | 1.99 | | High | High | High | High | High | High | High | High | High | High | High | 0 | 0.000137 | 2.59 |
| FALSE | High | ETAAAAAG | [K].ETAAA/ | 1xDeamida | A0A0A0M( | 1.08E-05 | 0 | 1 | 2 | 4 | A0A0A0M( | Histone-lys | A0A0A0M( | 0 | 3206.499 | 0 | 1.67 | 0.008671 | 21.52 | 47.9 | 152.1 | 15.08 | 20.83 | | High | High | High | High | High | High | High | High | High | High | High | 0 | 1.6E-06 | 4.93 |
| FALSE | High | ETAAAHQA | [K].ETAAAI | 1xPhosphc | P14873 1xl | 2.15E-07 | 0 | 2 | 2 | 2 | P14873; B2 | Microtubu | P14873 [18 | 0 | 2854.325 | 0 | -0.35 | 0.004529 | 0.9 | 115.8 | 84.2 | 4.28 | 10.14 | Shared | High | High | High | High | High | High | High | High | High | High | High | 0 | 1.81E-08 | 3.97 |
| FALSE | High | ETAATTETI | [K].ETAATT | 1xPhosphc | Q8R2P8 1x | 2.31E-07 | 0 | 1 | 5 | 5 | Q8R2P8 | Lysine--tRI | Q8R2P8 [6 | 0 | 2116.937 | 0 | -0.81 | 0.008699 | 6.74 | 124.9 | 75.1 | 14.55 | 15.92 | | High | High | High | High | High | High | High | High | High | High | High | 0 | 1.96E-08 | 3.7 |
| FALSE | High | ETAEQDRP | [K].ETAEQI | 1xPhosphc | A0A1S6GW | 1.23E-05 | 0 | 1 | 3 | 1 | A0A1S6GW | Uncharacte | A0A1S6GW | 0 | 3190.521 | 0 | -0.16 | 0.07541 | 6.87 | 104.2 | 95.8 | 3.4 | 5.67 | | High | High | High | High | High | High | High | High | High | High | High | 0 | 1.7E-06 | 4.39 |
| FALSE | High | ETAFEFLSS | [K].ETAFEF | 1xPhosphc | P14152 1xl | 0.102567 | 0.008249 | 1 | 1 | 2 | P14152 | Malate del | P14152 [32 | 0 | 1410.639 | 0 | -0.28 | 0.075671 | 2.56 | 111.2 | 88.8 | 12.86 | 16.1 | | High | High | High | High | High | High | High | High | High | High | High | 0.004138 | 0.04442 | 1.74 |
| FALSE | High | ETASTLVQI | [K].ETASTL | 1xPhosphc | Q9CXL3 1x | 8.1E-08 | 0 | 1 | 2 | 7 | Q9CXL3 | Uncharacte | Q9CXL3 [3 | 0 | 2382.128 | 0 | -0.26 | 0.002995 | 3.11 | 110.2 | 89.8 | 4.02 | 3.98 | | High | High | High | High | High | High | High | High | High | High | High | 0 | 6.03E-09 | 4.5 |
| FALSE | High | ETASTLVQI | [K].ETASTL | 2xPhosphc | Q9CXL3 2x | 2.39E-07 | 0 | 1 | 2 | 2 | Q9CXL3 | Uncharacte | Q9CXL3 [3 | 0 | 2462.094 | 0 | -0.34 | 0.011483 | 7.22 | 111.8 | 88.2 | 5.05 | 6.55 | | High | High | High | High | High | High | High | High | High | High | High | 0 | 2.03E-08 | 4.51 |
| FALSE | High | ETAYIVERP | [K].ETAYIV | 1xPhosphc | Q8K019 1x | 0.095408 | 0.007437 | 1 | 10 | 1 | Q8K019 | Bcl-2-assoc | Q8K019 [4 | 0 | 2003.061 | 0 | -0.26 | 0.143477 | 3.14 | 108.8 | 91.2 | 4.39 | 12.48 | | High | High | High | High | High | High | High | High | High | High | High | 0.003782 | 0.04087 | 1.12 |
| FALSE | High | ETCSTLTES | [R].ETCSTL | 1xCarbami | Q3UJB9 1x | 0.0008 | 0 | 1 | 2 | 1 | Q3UJB9 | Enhancer c | Q3UJB9 [8 | 0 | 1589.708 | 0 | 0.43 | 0.134497 | 27.92 | 85.9 | 114.1 | 14.32 | 14.83 | | High | High | High | High | High | High | High | High | High | High | High | 0 | 0.000186 | 2.68 |
| FALSE | High | ETCSTLTES | [R].ETCSTL | 1xCarbami | Q3UJB9 1x | 0.019618 | 0.001438 | 1 | 2 | 1 | Q3UJB9 | Enhancer c | Q3UJB9 [8 | 1 | 2489.2 | 0 | 0.33 | 0.011799 | 12.46 | 86.1 | 113.9 | 5.66 | 8.77 | | High | High | High | High | High | High | High | High | High | High | High | 0.000756 | 0.006684 | 1.35 |
| FALSE | High | ESYDAPPTI | [R].ESYDAF | 1xPhosphc | Q91XU0 1x | 7.63E-06 | 0 | 1 | 2 | 2 | Q91XU0 | ATPase WF | Q91XU0 [1 | 0 | 1656.747 | 0 | 0.08 | 0.54377 | 31.96 | 87.4 | 112.6 | 13.09 | 20.88 | | High | High | High | High | High | High | High | High | High | High | High | 0 | 9.99E-07 | 2.38 |
| FALSE | High | ESTGSYSDI | [R].ESTGSY | 1xPhosphc | F6WJB7 1x | 3.63E-05 | 0 | 1 | 5 | 3 | F6WJB7 | Ubiquitin c | F6WJB7 [1 | 0 | 3292.394 | 0 | 0.08 | 0.481806 | 21.77 | 96.2 | 103.8 | 17.36 | 7.75 | | High | High | High | High | High | High | High | High | High | High | High | 0 | 5.97E-06 | 3.79 |
| FALSE | High | ESTESSPGK | [R].ESTESS | 1xPhosphc | Q9JKX6 1xl | 0.000276 | 0 | 1 | 3 | 6 | Q9JKX6 | ADP-sugar | Q9JKX6 [5- | 0 | 1459.708 | 0 | -0.88 | 0.000574 | 1.79 | 131 | 69 | 3.77 | 13.71 | | High | High | High | High | High | High | High | High | High | High | High | 0.002354 | 0.02421 | 2.09 |
| FALSE | High | ESTDFSSTS | [R].ESTDFS | 1xPhosphc | Q3TW94 1: | 0.00195 | 5.66E-05 | 1 | 2 | 2 | Q3TW94 | Putative ur | Q3TW94 [2 | 0 | 2601.13 | 0 | 0.29 | 0.016358 | 7.61 | 91.3 | 108.7 | 6.83 | 8.81 | | High | High | High | High | High | High | High | High | High | High | High | 2.58E-05 | 0.000504 | 2.65 |
| FALSE | High | ESSEEPALP | [K].ESSEEP | 1xCarbami | Q8CGF1 1x | 0.003558 | 0.00021 | 1 | 3 | 2 | Q8CGF1 | Rho GTPas | Q8CGF1 [1 | 0 | 2450.122 | 0 | 0.31 | 0.086724 | 13.76 | 90.8 | 109.2 | 9.78 | 6.5 | | High | High | High | High | High | High | High | High | High | High | High | 9.42E-05 | 0.000987 | 3.24 |
| FALSE | High | ESSEPPPAA | [K].ESSEPP | 1xPhosphc | Q3UH06-3 | 0.000534 | 0 | 1 | 4 | 2 | Q3UH06-3 | Isoform 3 c | Q3UH06-3 | 0 | 3176.447 | 0 | 0.03 | 0.976209 | 23.07 | 100.6 | 99.4 | 7.02 | 22.89 | | High | High | High | High | High | High | High | High | High | High | High | 0 | 0.000118 | 4.13 |
| FALSE | High | ESSEPTGSF | [K].ESSEPT | 1xPhosphc | Q3TZR9 1x | 9.42E-07 | 0 | 1 | 1 | 3 | Q3TZR9 | Activating t | Q3TZR9 [4 | 0 | 3058.468 | 0 | 1.2 | 0.006746 | 2.62 | 60.2 | 139.8 | 12.43 | 15.58 | | High | High | High | High | High | High | High | High | High | High | High | 0 | 2.58E-07 | 4.52 |
| FALSE | High | ESSEPTGSF | [K].ESSEPT | 1xDeamida | Q3TZR9 1x | 0.004294 | 0.00036 | 1 | 1 | 5 | Q3TZR9 | Activating t | Q3TZR9 [4 | 0 | 3059.452 | 0 | 0.87 | 0.002928 | 17.1 | 70.9 | 129.1 | 5.06 | 11.98 | | High | High | High | High | High | High | High | High | High | High | High | 0.000161 | 0.001222 | 3.62 |
| FALSE | High | ESSIIAPVPT | [K].ESSIIAP | 1xPhosphc | Q571L5 1xl | 0.000328 | 0 | 1 | 6 | 4 | Q571L5 | MKIAA406 | Q571L5 [5 | 0 | 2232.1 | 0 | -1.26 | 0.002106 | 4.39 | 141 | 59 | 9.8 | 8.25 | | High | High | High | High | High | High | High | High | High | High | High | 0 | 6.82E-05 | 2.84 |
| FALSE | High | ESSIIAPVPT | [K].ESSIIAP | 1xPhosphc | Q571L5 1xl | 9.79E-08 | 0 | 1 | 6 | 5 | Q571L5 | MKIAA406 | Q571L5 [5 | 1 | 2589.358 | 0 | -0.9 | 0.0003 | 1.59 | 129.2 | 70.8 | 3.43 | 4.68 | | High | High | High | High | High | High | High | High | High | High | High | 0 | 2.74E-08 | 4.34 |
| FALSE | High | ESSMCLTP: | [K].ESSMCI | 1xCarbami | Q5SSZ5 1xl | 9.45E-05 | 0 | 1 | 1 | 1 | Q5SSZ5 | Tensin-3 O | Q5SSZ5 [8 | 0 | 2794.348 | 0 | -0.53 | 0.052059 | 12.77 | 112.5 | 87.5 | 12.65 | 22.39 | | High | High | High | High | High | High | High | High | High | High | High | 0 | 1.68E-05 | 3.03 |
| FALSE | High | ESSMCLTP: | [K].ESSMCI | 1xCarbami | Q5SSZ5 1xl | 0.000103 | 0 | 1 | 1 | 1 | Q5SSZ5 | Tensin-3 O | Q5SSZ5 [8 | 0 | 2810.343 | 0 | -0.42 | 0.058616 | 1.89 | 110.6 | 89.4 | 8.32 | 18.33 | | High | High | High | High | High | High | High | High | High | High | High | 0 | 1.86E-05 | 3.44 |
| FALSE | High | ESSPDGSYT | [R].ESSPDG | 1xPhosphc | Q80TY0-3 | 1.24E-05 | 0 | 1 | 1 | 2 | Q80TY0-3 | Isoform 3 c | Q80TY0-3 | 0 | 2692.166 | 0 | -0.77 | 0.061494 | 3.39 | 126.2 | 73.8 | 15.43 | 33.13 | | High | High | High | High | High | High | High | High | High | High | High | 0 | 2.27E-06 | 4.02 |
| FALSE | High | ESSPGK | [R].ESSPGK | 1xPhosphc | E9QLG3 1x | 0.073462 | 0.005631 | 1 | 7 | 1 | E9QLG3 | MAX gene- | E9QLG3 [5 | 0 | 1142.586 | 0 | -0.76 | 0.000304 | 2.71 | 124.4 | 75.6 | 3.43 | 5.56 | | High | High | High | High | High | High | High | High | High | High | High | 0.002867 | 0.02995 | 1.86 |
| FALSE | High | ESSPLFDD( | [K].ESSPLFI | 1xPhosphc | Q9D0F1 1x | 2.49E-05 | 0 | 1 | 1 | 2 | Q9D0F1 | Kinetochor | Q9D0F1 [2 | 0 | 2890.344 | 0 | -0.47 | 0.047906 | 9.24 | 114.4 | 85.6 | 5.54 | 15.37 | | High | High | High | High | High | High | High | High | High | High | High | 0 | 6.04E-06 | 3.92 |
| FALSE | High | ESSPLGSR | [K].ESSPLG | 1xPhosphc | A0A0R4J0V | 0.063788 | 0.004664 | 1 | 8 | 1 | A0A0R4J0V | Phosphatic | A0A0R4J0V | 0 | 1141.545 | 0 | -0.34 | 0.026928 | 8.2 | 111.5 | 88.5 | 6.88 | 9.74 | | High | High | High | High | High | High | High | High | High | High | High | 0.002403 | 0.02558 | 1.87 |
| FALSE | High | ESSPLYSPG | [R].ESSPLY: | 1xPhosphc | P14873 1xl | 9.98E-05 | 0 | 2 | 2 | 4 | P14873; B2 | Microtubu | P14873 [17 | 0 | 2369.131 | 0 | -0.79 | 0.004952 | 3.34 | 124.2 | 75.8 | 7.6 | 9.63 | Shared | High | High | High | High | High | High | High | High | High | High | High | 0 | 1.99E-05 | 3.56 |
| FALSE | High | ESSPLYSPG | [R].ESSPLY: | 2xPhosphc | P14873 2xl | 0.00034 | 0 | 2 | 2 | 2 | P14873; B2 | Microtubu | P14873 [17 | 0 | 2449.098 | 0 | -0.25 | 0.254626 | 13.44 | 105.8 | 94.2 | 10.35 | 13.45 | Shared | High | High | High | High | High | High | High | High | High | High | High | 0 | 7.11E-05 | 2.72 |
| FALSE | High | ESSPSHGLI | [R].ESSPSH | 1xPhosphc | Q80T83 1x | 0.085859 | 0.006406 | 1 | 3 | 2 | Q80T83 | MKIAA185 | Q80T83 [1 | 0 | 1592.845 | 0 | -0.06 | 0.780483 | 17.81 | 103.7 | 96.3 | 12.02 | 10.09 | | High | High | High | High | High | High | High | High | High | High | High | 0.003303 | 0.03622 | 2.19 |
| FALSE | High | ESSPSNSPS | [R].ESSPSN | 1xPhosphc | E9Q8L9 1xl | 2.07E-08 | 0 | 1 | 3 | 3 | E9Q8L9 | Rab11 fam | E9Q8L9 [3 | 0 | 1785.801 | 0 | -0.72 | 0.005201 | 3.28 | 121.2 | 78.8 | 5.79 | 11.11 | | High | High | High | High | High | High | High | High | High | High | High | 0 | 1.3E-09 | 3.43 |
| FALSE | High | ESSPSWSTI | [K].ESSPSW | 1xPhosphc | Q9QXD8 1: | 1.21E-10 | 0 | 1 | 1 | 7 | Q9QXD8 | LIM domai | Q9QXD8 [3 | 0 | 2809.313 | 0 | 0.63 | 0.002349 | 6.61 | 80.8 | 119.2 | 5.52 | 9.24 | | High | High | High | High | High | High | High | High | High | High | High | 0 | 4.05E-12 | 4.71 |
| FALSE | High | ESSPVPSPT | [K].ESSPVP | 1xPhosphc | Q62261 1x | 0.000365 | 0 | 1 | 2 | 2 | Q62261 | Spectrin be | Q62261 [2 | 0 | 1593.772 | 0 | 0.4 | 0.006215 | 5.12 | 85.6 | 114.4 | 4.28 | 9.48 | | High | High | High | High | High | High | High | High | High | High | High | 0 | 7.69E-05 | 2.64 |
| FALSE | High | ESSPVPSPT | [K].ESSPVP | 2xPhosphc | Q62261 2x | 0.000231 | 0 | 1 | 2 | 8 | Q62261 | Spectrin be | Q62261 [2 | 0 | 1673.739 | 0 | 1.32 | 0.000311 | 3.39 | 57.5 | 142.5 | 2.35 | 5.13 | | High | High | High | High | High | High | High | High | High | High | High | 0 | 0.000302 | 3.72 |
| FALSE | High | ESSPVPSPT | [K].ESSPVP | 2xPhosphc | Q62261 2x | 0.009332 | 0.000736 | 1 | 2 | 2 | Q62261 | Spectrin be | Q62261 [2 | 1 | 2030.997 | 0 | 1.13 | 0.000726 | 17.52 | 57.7 | 142.3 | 8.27 | 6.43 | | High | High | High | High | High | High | High | High | High | High | High | 0.008836 | 0.09427 | 2.32 |
| FALSE | High | ESSQLWG( | [R].ESSQLV | 1xCarbami | Q3UGJ1 1x | 1.53E-05 | 0 | 1 | 5 | 1 | Q3UGJ1 | Putative ur | Q3UGJ1 [1 | 0 | 2410.151 | 0 | 0.62 | 0.033884 | 22.04 | 78.7 | 121.3 | 14.05 | 12.17 | | High | High | High | High | High | High | High | High | High | High | High | 0 | 2.18E-06 | 3 |
| FALSE | High | ESSRSPSPV | [R].ESSRSP | 2xPhosphc | P48754 2xl | 0.045898 | 0.003408 | 1 | 6 | 1 | P48754 | Breast canc | P48754 [11 | 1 | 2087.993 | 0 | -1.2 | 0.000883 | 1.74 | 137.3 | 62.7 | 13.26 | 16.93 | | High | High | High | High | High | High | High | High | High | High | High | 0.001789 | 0.01754 | 1.42 |
| FALSE | High | ESSSLGSVG | [K].ESSSLG | 1xPhosphc | J3QNH8 1x | 1.06E-07 | 0 | 1 | 1 | 9 | J3QNH8 | Uncharacte | J3QNH8 [7 | 0 | 2057.093 | 0 | 2.61 | 0.000218 | 9.12 | 27.7 | 172.3 | 11.58 | 7.05 | | High | High | High | High | High | High | High | High | High | High | High | 0 | 1.4E-08 | 4.32 |
| FALSE | High | ESSSPESFG | [R].ESSSPE: | 1xPhosphc | Q9D074-5 | 1.97E-08 | 0 | 1 | 5 | 6 | Q9D074-5 | Isoform 5 c | Q9D074-5 | 0 | 2550.154 | 0 | 0.61 | 0.002614 | 4 | 79.2 | 120.8 | 6.65 | 6.42 | | High | High | High | High | High | High | High | High | High | High | High | 0 | 1.24E-09 | 3.9 |
| FALSE | High | ESSSPVALK | [K].ESSSPV | 1xPhosphc | Q4VAC9 1x | 0.002695 | 0.000109 | 1 | 3 | 3 | Q4VAC9 | Pleckstrin I | Q4VAC9 [1 | 0 | 1455.786 | 0 | 1.71 | 0.000609 | 12.54 | 48.1 | 151.9 | 7.13 | 8.82 | | High | High | High | High | High | High | High | High | High | High | High | 0.000437 | 0.003682 | 2.27 |
| FALSE | High | ESSTNNSV: | [R].ESSTNN | 1xDeamida | Q80TC1 1x | 0.003464 | 0.00021 | 1 | 14 | 6 | Q80TC1 | MKIAA154 | Q80TC1 [3 | 0 | 1903.848 | 0 | -1.1 | 0.002215 | 5.14 | 134.9 | 65.1 | 10.04 | 21.12 | | High | High | High | High | High | High | High | High | High | High | High | 0.001812 | 0.01806 | 2.01 |
| FALSE | High | ESSTVGGG | [R].ESSTVG | 1xPhosphc | Q3TW94 1: | 2.2E-07 | 0 | 1 | 3 | 7 | Q3TW94 | Putative ur | Q3TW94 [5 | 0 | 2409.052 | 0 | 0.06 | 0.439515 | 24.32 | 97.8 | 102.2 | 2.36 | 19.46 | | High | High | High | High | High | High | High | High | High | High | High | 0 | 1.87E-08 | 4.68 |
| FALSE | High | ESSYSSLSPI | [K].ESSYSS | 1xPhosphc | G3UZM1 1 | 1.7E-07 | 0 | 1 | 3 | 2 | G3UZM1 | Probable Ji | G3UZM1 [ | 0 | 2857.446 | 0 | 0.28 | 0.638265 | 13.08 | 91.6 | 108.4 | 9.7 | 21.37 | | High | High | High | High | High | High | High | High | High | High | High | 0 | 1.39E-08 | 4.82 |
| FALSE | High | ESSYSSLSPI | [K].ESSYSS | 1xPhosphc | G3UZM1 1 | 3.5E-05 | 0 | 1 | 3 | 2 | G3UZM1 | Probable Ji | G3UZM1 [ | 0 | 2873.441 | 0 | -0.28 | 0.054984 | 14.35 | 106.8 | 93.2 | 5.69 | 15.57 | | High | High | High | High | High | High | High | High | High | High | High | 0 | 5.54E-06 | 4.16 |
| FALSE | High | ESEASTPTA | [K].ESEAST | 1xPhosphc | A2AR56 1x | 3.95E-06 | 0 | 1 | 10 | 3 | A2AR56 | FERM dom | A2AR56 [4 | 0 | 2328.081 | 0 | 0.98 | 0.00286 | 19.68 | 64 | 136 | 6.26 | 12.13 | | High | High | High | High | High | High | High | High | High | High | High | 0 | 4.09E-06 | 3.84 |
| FALSE | High | ETDEAAFEI | [R].ETDEA/ | 1xPhosphc | P97868 1xl | 0.000331 | 0 | 1 | 3 | 1 | P97868 | E3 ubiquiti | P97868 [16 | 0 | 3100.356 | 0 | 0.46 | 0.031183 | 3.92 | 83.4 | 116.6 | 6.15 | 11.37 | | High | High | High | High | High | High | High | High | High | High | High | 0 | 6.92E-05 | 3.43 |
| FALSE | High | ESDSDSEGI | [K].ESDSDS | 1xPhosphc | Q501J7-3 1 | 0.002115 | 5.66E-05 | 1 | 3 | 2 | Q501J7-3 | Isoform 3 c | Q501J7-3 [ | 0 | 1701.798 | 0 | 0.58 | 0.000245 | 0.62 | 80.2 | 119.8 | 7.65 | 7.62 | | High | High | High | High | High | High | High | High | High | High | High | 2.58E-05 | 0.000554 | 2.46 |
| FALSE | High | ESDPFHSSS | [K].ESDPFH | 1xPhosphc | Q60902 1x | 0.000379 | 0 | 1 | 3 | 2 | Q60902 | Epidermal | Q60902 [6 | 0 | 2270.005 | 0 | 0.22 | 0.13164 | 21.7 | 90.3 | 109.7 | 4.44 | 15.11 | | High | High | High | High | High | High | High | High | High | High | High | 0 | 7.98E-05 | 2.61 |
| FALSE | High | EQAEMDD. | [R].EQAEM | 1xPhosphc | C7G3P2 1x | 3.39E-06 | 0 | 1 | 4 | 3 | C7G3P2 | MKIAA053 | C7G3P2 [1 | 0 | 2035.857 | 0 | -0.07 | 0.21174 | 5.07 | 101.8 | 98.2 | 8.21 | 12.87 | | High | High | High | High | High | High | High | High | High | High | High | 0 | 4.01E-07 | 4.12 |
| FALSE | High | EQAGQMP | [K].EQAGQ | 1xPhosphc | A0A0N4SV | 3.1E-06 | 0 | 1 | 9 | 1 | A0A0N4SV | Eukaryotic | A0A0N4SV | 0 | 2418.085 | 0 | 0.76 | 0.006784 | 8.6 | 76.9 | 123.1 | 13.08 | 7.8 | | High | High | High | High | High | High | High | High | High | High | High | 0 | 3.6E-07 | 4.36 |
| FALSE | High | EQAGQMP | [K].EQAGQ | 2xPhosphc | A0A0N4SV | 0.04913 | 0.003576 | 1 | 9 | 1 | A0A0N4SV | Eukaryotic | A0A0N4SV | 1 | 3705.707 | 0 | 1.02 | 0.008577 | 30.98 | 63.7 | 136.3 | 6.16 | 17.52 | | High | High | High | High | High | High | High | High | High | High | High | 0.001889 | 0.01885 | 3.02 |

| | | | | | | | | | | | | | | | | | | | | | | | | | | | | | | | | | | | | | | |
|---|---|---|---|---|---|---|---|---|---|---|---|---|---|---|---|---|---|---|---|---|---|---|---|---|---|---|---|---|---|---|---|---|---|---|---|---|---|---|
| FALSE | High | EQAQWTP | [K].EQAQV | 1xPhospho | Q6A065 1x | 0.002096 | 5.66E-05 | 1 | 2 | 1 | Q6A065 | Centrosom | Q6A065 [7 | 0 | 1626.829 | 0 | 0.78 | 0.011739 | 15.37 | 75.6 | 124.4 | 6.12 | 12.43 | | High | High | High | High | High | High | High | High | High | High | High | 2.58E-05 | 0.000547 | 2.12 |
| FALSE | High | EQASERLSI | [K].EQASEI | 1xPhospho | E9QK83 1x | 0.06169 | 0.004646 | 1 | 2 | 1 | E9QK83 | Small subu | E9QK83 [1 | 1 | 1787.894 | 0 | -0.78 | 0.001125 | 2.54 | 126.4 | 73.6 | 3.35 | 7.8 | | High | High | High | High | High | High | High | High | High | High | High | 0.002381 | 0.02463 | 1.19 |
| FALSE | High | EQASPGET | [K].EQASP( | 1xCarbami | Q3UH06-3 | 2E-07 | 0 | 1 | 4 | 3 | Q3UH06-3 | Isoform 3 c | Q3UH06-3 | 0 | 3291.622 | 0 | 0.11 | 0.430088 | 24.34 | 94.9 | 105.1 | 7.64 | 21.26 | | High | High | High | High | High | High | High | High | High | High | High | 0 | 1.66E-08 | 3.8 |
| FALSE | High | EQASPGET | [K].EQASP( | 1xCarbami | Q3UH06-3 | 6.54E-09 | 0 | 1 | 4 | 3 | Q3UH06-3 | Isoform 3 c | Q3UH06-3 | 0 | 3371.588 | 0 | 0.06 | 0.63307 | 10.2 | 100.8 | 99.2 | 11.89 | 9.35 | | High | High | High | High | High | High | High | High | High | High | High | 0 | 3.59E-10 | 5.35 |
| FALSE | High | EQATHHGI | [K].EQATH | 1xPhospho | Q8K2J4 1xl | 0.042141 | 0.003162 | 1 | 1 | 1 | Q8K2J4 | Coiled-coil | Q8K2J4 [8! | 0 | 2471.26 | 0 | -0.17 | 0.380242 | 23.94 | 109.3 | 90.7 | 15.27 | 22.01 | | High | High | High | High | High | High | High | High | High | High | High | 0.001625 | 0.01578 | 2.69 |
| FALSE | High | EQDSDVSE | [K].EQDSD' | 1xPhospho | P97496 1xl | 5.47E-06 | 0 | 1 | 5 | 4 | P97496 | SWI/SNF c | P97496 [81 | 0 | 2501.226 | 0 | -0.92 | 0.000311 | 0.69 | 131.5 | 68.5 | 4.58 | 3.31 | | High | High | High | High | High | High | High | High | High | High | High | 0 | 6.84E-07 | 3.3 |
| FALSE | High | EQESSSGG | [K].EQESSS | 1xPhospho | Q8BXZ1 1x | 0.007951 | 0.000736 | 1 | 1 | 3 | Q8BXZ1 | Protein dis | Q8BXZ1 [4 | 0 | 2369.164 | 0 | -0.03 | 0.533817 | 5.44 | 105.7 | 94.3 | 9.95 | 8.77 | | High | High | High | High | High | High | High | High | High | High | High | 0.000327 | 0.00242 | 2.94 |
| FALSE | High | EQEWDSSS | [R].EQEWD | 2xPhospho | Q61687 2x | 0.061176 | 0.004646 | 1 | 6 | 2 | Q61687 | Transcripti | Q61687 [9 | 1 | 3270.36 | 0 | -0.09 | 0.258909 | 4.49 | 104.4 | 95.6 | 9.65 | 10.48 | | High | High | High | High | High | High | High | High | High | High | High | 0.002381 | 0.0244 | 2.78 |
| FALSE | High | EQEWDSSS | [R].EQEWD | 2xPhospho | Q61687 2x | 0.052133 | 0.00384 | 1 | 6 | 2 | Q61687 | Transcripti | Q61687 [9 | 1 | 3499.523 | 0 | -0.37 | 0.021715 | 13.09 | 111.3 | 88.7 | 5.14 | 13.28 | | High | High | High | High | High | High | High | High | High | High | High | 0.001999 | 0.02027 | 2.76 |
| FALSE | High | EQFDTLTPI | [R].EQFDTI | 1xPhospho | Q6ZQ58 1x | 0.000101 | 0 | 1 | 3 | 5 | Q6ZQ58 | La-related | Q6ZQ58 [6 | 0 | 2965.418 | 0 | 1.3 | 0.001034 | 3.05 | 56.8 | 143.2 | 13.17 | 6.66 | | High | High | High | High | High | High | High | High | High | High | High | 0 | 1.81E-05 | 4.25 |
| FALSE | High | EPVVGGTL | [K].EPVVG( | 1xPhospho | Q6A060 1x | 0.00043 | 0 | 1 | 7 | 1 | Q6A060 | MKIAA048 | Q6A060 [7 | 0 | 2104.182 | 0 | 0.7 | 0.003483 | 15.68 | 76.9 | 123.1 | 4.65 | 7.9 | | High | High | High | High | High | High | High | High | High | High | High | 0 | 9.25E-05 | 3.18 |
| FALSE | High | EQGSQIPG | [K].EQGSQ | 1xCarbami | R4GML0 1: | 5.57E-06 | 0 | 1 | 3 | 4 | R4GML0 | Zinc finger | R4GML0 [6 | 0 | 3180.565 | 0 | -2 | 0.000406 | 1.1 | 159.7 | 40.3 | 6.53 | 7.4 | | High | High | High | High | High | High | High | High | High | High | High | 0 | 6.98E-07 | 4.12 |
| FALSE | High | EQGSQIPG | [K].EQGSQ | 1xPhospho | O88466 1x | 5.09E-06 | 0 | 1 | 4 | 4 | O88466 | Zinc finger | O88466 [6 | 0 | 2207.054 | 0 | -0.88 | 0.000328 | 2.39 | 129.8 | 70.2 | 5.95 | 7.73 | | High | High | High | High | High | High | High | High | High | High | High | 0 | 6.35E-07 | 2.89 |
| FALSE | High | EQGSQIPG | [K].EQGSQ | 2xPhospho | O88466 2x | 0.000124 | 0 | 1 | 4 | 3 | O88466 | Zinc finger | O88466 [6 | 0 | 2287.021 | 0 | -0.79 | 0.000255 | 1.15 | 124.7 | 75.3 | 4.8 | 5.69 | | High | High | High | High | High | High | High | High | High | High | High | 0 | 4.28E-05 | 3.54 |
| FALSE | High | EQGSSGLG | [R].EQGSS( | 1xPhospho | Q3UHJ0 1x | 2.02E-09 | 0 | 1 | 3 | 2 | Q3UHJ0 | AP2-associ | Q3UHJ0 [1 | 0 | 2711.211 | 0 | 0.22 | 0.066181 | 9.91 | 92.9 | 107.1 | 5.06 | 5.76 | | High | High | High | High | High | High | High | High | High | High | High | 0 | 9.58E-11 | 4.78 |
| FALSE | High | EQGSTSFS( | [K].EQGST! | 1xPhospho | Q9EQD6 1: | 6.35E-05 | 0 | 1 | 4 | 2 | Q9EQD6 | Keratin inte | Q9EQD6 [4 | 1 | 2154.914 | 0 | 1.6 | 0.000949 | 7.13 | 51.1 | 148.9 | 16.63 | 12.92 | | High | High | High | High | High | High | High | High | High | High | High | 0 | 5.3E-05 | 3.6 |
| FALSE | High | EQGTESRS | [R].EQGTE! | 2xPhospho | Q3U955 2x | 0.000404 | 0 | 2 | 10 | 2 | Q3U955; C | Putative ur | Q3U955 [1 | 1 | 2810.244 | 0 | 2.16 | 0.002109 | 13.32 | 32.1 | 167.9 | 13.46 | 18.03 | Shared | High | High | High | High | High | High | High | High | High | High | High | 0 | 8.6E-05 | 2.85 |
| FALSE | High | EQGVESPG | [K].EQGVE | 1xPhospho | Q3UD58 1x | 2.24E-07 | 0 | 1 | 1 | 7 | Q3UD58 | Putative ur | Q3UD58 [5 | 0 | 1905.891 | 0 | -0.4 | 0.003085 | 5.21 | 114.8 | 85.2 | 3.41 | 5.37 | | High | High | High | High | High | High | High | High | High | High | High | 0 | 1.9E-08 | 3.58 |
| FALSE | High | EQLAVNQ\ | [R].EQLAVI | 2xPhospho | Q61687 2x | 0.06169 | 0.004646 | 1 | 3 | 1 | Q61687 | Transcripti | Q61687 [1 | 0 | 2813.217 | 0 | 0.13 | 0.646446 | 13.14 | 95.5 | 104.5 | 10.99 | 13.35 | | High | High | High | High | High | High | High | High | High | High | High | 0.002381 | 0.02447 | 2.64 |
| FALSE | High | EQIEVIAPS | [K].EQIEVI/ | 1xPhospho | E9QQ30 1x | 0.080423 | 0.006219 | 1 | 3 | 1 | E9QQ30 | Eukaryotic | E9QQ30 [7 | 0 | 1676.846 | 0 | -0.9 | 0.00059 | 3.42 | 129.1 | 70.9 | 3.5 | 9.85 | | High | High | High | High | High | High | High | High | High | High | High | 0.003188 | 0.03326 | 2.01 |
| FALSE | High | EQIEVIAPS | [K].EQIEVI/ | 2xPhospho | E9QQ30 2x | 0.000141 | 0 | 1 | 3 | 1 | E9QQ30 | Eukaryotic | E9QQ30 [7 | 1 | 1999.945 | 0 | -0.74 | 0.006391 | 3.57 | 120.2 | 79.8 | 10.69 | 6.73 | | High | High | High | High | High | High | High | High | High | High | High | 0 | 2.62E-05 | 3.34 |
| FALSE | High | EQIMGPPI' | [K].EQIMG | 1xPhospho | F8WJE0 1x | 1.33E-06 | 0 | 1 | 9 | 3 | F8WJE0 | Deoxynucl | F8WJE0 [3 | 0 | 1848.011 | 0 | 0.37 | 0.008486 | 3.52 | 86.5 | 113.5 | 4.73 | 5.26 | | High | High | High | High | High | High | High | High | High | High | High | 0 | 4.87E-06 | 3.4 |
| FALSE | High | EQIMGPPI' | [K].EQIMG | 1xPhospho | F8WJE0 1x | 1.02E-07 | 0 | 1 | 9 | 32 | F8WJE0 | Deoxynucl | F8WJE0 [3 | 0 | 1864.006 | 0 | 0.42 | 0.000797 | 2.11 | 86.2 | 113.8 | 3 | 2.9 | | High | High | High | High | High | High | High | High | High | High | High | 0 | 2.22E-07 | 4.29 |
| FALSE | High | EQLSEVEG | [K].EQLSEV | 1xPhospho | P35123 1xl | 1.68E-07 | 0 | 1 | 9 | 4 | P35123 | Ubiquitin c | P35123 [67 | 0 | 3114.342 | 0 | 0.22 | 0.044633 | 10.69 | 93.1 | 106.9 | 10.13 | 1.79 | | High | High | High | High | High | High | High | High | High | High | High | 0 | 7.94E-07 | 4.37 |
| FALSE | High | EQLTEHNG | [R].EQLTEH | 1xPhospho | Q9JMB0 1x | 9.03E-05 | 0 | 1 | 1 | 1 | Q9JMB0 | G kinase-a | Q9JMB0 [2 | 0 | 3182.516 | 0 | 0.17 | 0.340349 | 6.19 | 93.8 | 106.2 | 7.95 | 11.26 | | High | High | High | High | High | High | High | High | High | High | High | 0 | 1.61E-05 | 2.7 |
| FALSE | High | EQGSQIPG | [K].EQGSQ | 1xCarbami | R4GML0 2: | 0.001437 | 5.66E-05 | 1 | 3 | 1 | R4GML0 | Zinc finger | R4GML0 [6 | 0 | 3260.531 | 0 | -1.7 | 0.001282 | 2.24 | 152.8 | 47.2 | 8.75 | 16.9 | | High | High | High | High | High | High | High | High | High | High | High | 2.58E-05 | 0.000359 | 3.47 |
| FALSE | High | EQMMNSS | [R].EQMM | 1xPhospho | Q811D0-2 | 0.050826 | 0.003676 | 2 | 11 | 3 | Q811D0-2; | Isoform 2 c | Q811D0-2 | 0 | 1997.851 | 0 | 0.85 | 0.001347 | 10.32 | 69.8 | 130.2 | 5.85 | 8.91 | Shared | High | High | High | High | High | High | High | High | High | High | High | 0.001916 | 0.01956 | 2.08 |
| FALSE | High | EPVRADSP' | [K].EPVRAI | 1xPhospho | Q7TN02 1x | 0.002272 | 0.000109 | 1 | 1 | 1 | Q7TN02 | Mediator c | Q7TN02 [4 | 1 | 2100.069 | 0 | 0.09 | 0.502075 | 3.31 | 98 | 102 | 6.56 | 7.42 | | High | High | High | High | High | High | High | High | High | High | High | 4.94E-05 | 0.000597 | 2.46 |
| FALSE | High | EPVLSPEDE | [K].EPVLSP | 1xPhospho | Q02526 1x | 7.23E-06 | 0 | 1 | 2 | 1 | Q02526 | Zinc finger | Q02526 [4 | 0 | 2902.395 | 0 | -0.74 | 0.013079 | 2.33 | 127.7 | 72.3 | 17.21 | 10.45 | | High | High | High | High | High | High | High | High | High | High | High | 0 | 9.4E-07 | 4.51 |
| FALSE | High | EPSESPDQI | [R].EPSESP | 1xPhospho | Q2MHN3 1 | 2.54E-06 | 0 | 1 | 1 | 1 | Q2MHN3 | ZFH-5 OS=I | Q2MHN3 [ | 0 | 2252.043 | 0 | -0.29 | 0.052661 | 12.32 | 110.2 | 89.8 | 8.92 | 9.92 | | High | High | High | High | High | High | High | High | High | High | High | 0 | 2.91E-07 | 3.56 |
| FALSE | High | EPSEVPTPK | [K].EPSEVP | 1xPhospho | Q3TE85 1x | 0.000837 | 0 | 1 | 3 | 5 | Q3TE85 | Putative ur | Q3TE85 [4 | 0 | 1521.797 | 0 | -0.18 | 0.120848 | 10.06 | 104.6 | 95.4 | 8.47 | 5.18 | | High | High | High | High | High | High | High | High | High | High | High | 0.001388 | 0.01441 | 2.4 |
| FALSE | High | EPSGGGG( | [K].EPSGG( | 1xPhospho | A0A1S6GW | 4.26E-07 | 0 | 1 | 2 | 1 | A0A1S6GW | Uncharacte | A0A1S6GW | 0 | 2289.095 | 0 | -1.2 | 0.001114 | 3.54 | 139.3 | 60.7 | 7.74 | 10.3 | | High | High | High | High | High | High | High | High | High | High | High | 0 | 3.89E-08 | 4.01 |
| FALSE | High | EPSGQPED | [K].EPSGQI | 1xPhospho | E9Q7E9 1x | 0.011549 | 0.00086 | 1 | 2 | 1 | E9Q7E9 | Phosphofu | E9Q7E9 [3 | 0 | 3031.407 | 0 | -0.22 | 0.136273 | 13.5 | 110.8 | 89.2 | 10.27 | 20.13 | | High | High | High | High | High | High | High | High | High | High | High | 0.000437 | 0.003694 | 2.8 |
| FALSE | High | EPSLFHSGI | [K].EPSLFH | 1xCarbami | E9PWN0 1: | 0.033404 | 0.002195 | 1 | 4 | 3 | E9PWN0 | Phosphatic | E9PWN0 [2 | 0 | 4146.908 | 0 | 0.98 | 0.002445 | 15.84 | 67.4 | 132.6 | 7.69 | 10.19 | | High | High | High | High | High | High | High | High | High | High | High | 0.002403 | 0.02561 | 3.94 |
| FALSE | High | EPSLHLDTF | [K].EPSLHL | 1xPhospho | Q6ZQF0 1x | 9.71E-05 | 0 | 1 | 1 | 2 | Q6ZQF0 | DNA topois | Q6ZQF0 [8 | 0 | 1761.919 | 0 | 0.29 | 0.050837 | 8.46 | 91.2 | 108.8 | 4.55 | 19.05 | | High | High | High | High | High | High | High | High | High | High | High | 0.000399 | 0.003363 | 2.21 |
| FALSE | High | EPSPEGSK | [R].EPSPEG | 1xPhospho | A2APA8 1x | 0.007538 | 0.00062 | 1 | 2 | 1 | A2APA8 | Transcripti | A2APA8 [6 | 0 | 1368.681 | 0 | 0.11 | 0.146635 | 8.74 | 96.2 | 103.8 | 4.93 | 9.26 | | High | High | High | High | High | High | High | High | High | High | High | 0.000275 | 0.002277 | 1.73 |
| FALSE | High | EPSQPLQLI | [K].EPSQPL | 1xPhospho | E9PYU8 1x | 0.000134 | 0 | 1 | 4 | 1 | E9PYU8 | E3 ubiquiti | E9PYU8 [3 | 0 | 2844.35 | 0 | 3.15 | 0.000747 | 19.58 | 20 | 180 | 13.44 | 13.71 | | High | High | High | High | High | High | High | High | High | High | High | 0 | 2.49E-05 | 2.87 |
| FALSE | High | EPSSPGTDI | [K].EPSSPG | 1xPhospho | Q3U3A6 1x | 1.01E-05 | 0 | 1 | 5 | 8 | Q3U3A6 | Sodium/hy | Q3U3A6 [7 | 0 | 2646.141 | 0 | -0.04 | 0.578272 | 7.85 | 101.7 | 98.3 | 8.39 | 10.76 | | High | High | High | High | High | High | High | High | High | High | High | 0 | 1.18E-05 | 4.45 |
| FALSE | High | EPSSPGTDI | [K].EPSSPG | 2xPhospho | Q3U3A6 2x | 0.0003 | 0 | 1 | 5 | 4 | Q3U3A6 | Sodium/hy | Q3U3A6 [7 | 0 | 2726.107 | 0 | 0.4 | 0.004516 | 12.41 | 88.6 | 111.4 | 9.64 | 5.53 | | High | High | High | High | High | High | High | High | High | High | High | 0 | 6.18E-05 | 4.42 |
| FALSE | High | EPSSPYTQI | [R].EPSSPY | 1xPhospho | A0A0G2JD | 0.041072 | 0.003094 | 1 | 3 | 3 | A0A0G2JD | Non-canor | A0A0G2JD | 0 | 1373.63 | 0 | 1.42 | 9.77E-05 | 4.1 | 55.2 | 144.8 | 6.29 | 2.11 | | High | High | High | High | High | High | High | High | High | High | High | 0.001597 | 0.01533 | 1.73 |
| FALSE | High | EPSSSQPV\ | [R].EPSSSQ | 1xPhospho | Q3TVI8 1xl | 1.49E-05 | 0 | 1 | 2 | 2 | Q3TVI8 | Pre-B-cell l | Q3TVI8 [16 | 0 | 2449.226 | 0 | 0.14 | 0.386033 | 8.31 | 95.9 | 104.1 | 12.92 | 12.55 | | High | High | High | High | High | High | High | High | High | High | High | 0 | 2.12E-06 | 4.49 |
| FALSE | High | EPSSSQPV\ | [R].EPSSSQ | 1xDeamida | Q3TVI8 1xl | 0.00925 | 0.000736 | 1 | 2 | 3 | Q3TVI8 | Pre-B-cell l | Q3TVI8 [16 | 0 | 2450.21 | 0 | -0.05 | 0.400201 | 3.01 | 101 | 99 | 10.95 | 11.36 | | High | High | High | High | High | High | High | High | High | High | High | 0.000689 | 0.00614 | 4.12 |
| FALSE | High | EPVQTAQS | [K].EPVQT/ | 1xPhospho | Q6DID3 1x | 6.62E-07 | 0 | 1 | 1 | 3 | Q6DID3 | Protein SC/ | Q6DID3 [6 | 0 | 2019.056 | 0 | -0.21 | 0.00534 | 1.85 | 107 | 93 | 3.41 | 1.36 | | High | High | High | High | High | High | High | High | High | High | High | 0 | 6.37E-08 | 3.81 |
| FALSE | High | EPSTTSGG( | [R].EPSTTS | 1xPhospho | E9QM90 1: | 3.78E-06 | 0 | 1 | 7 | 7 | E9QM90 | RIKEN cDN | E9QM90 [1 | 0 | 1612.753 | 0 | 2.04 | 0.000462 | 18.79 | 38.1 | 161.9 | 5.84 | 8.08 | | High | High | High | High | High | High | High | High | High | High | High | 0 | 0.000312 | 0.99 |
| FALSE | High | EPTPSEQV( | [R].EPTPSE | 1xPhospho | Q922Y1 1x | 0.000396 | 0 | 1 | 1 | 2 | Q922Y1 | UBX domai | Q922Y1 [6 | 0 | 3305.537 | 0 | 0.22 | 0.140024 | 21.52 | 91.4 | 108.6 | 12.37 | 8.02 | | High | High | High | High | High | High | High | High | High | High | High | 0.000449 | 0.00394 | 2.32 |
| FALSE | High | EPTPSEQV( | [R].EPTPSE | 1xDeamida | Q922Y1 1x | 0.001361 | 0 | 1 | 1 | 1 | Q922Y1 | UBX domai | Q922Y1 [6 | 0 | 3306.521 | 0 | 0.11 | 0.140552 | 17.39 | 96.2 | 103.8 | 7.76 | 6.52 | | High | High | High | High | High | High | High | High | High | High | High | 0 | 0.000336 | 2.48 |
| FALSE | High | EPTPSIASD | [K].EPTPSI/ | 1xPhospho | Q3TZZ7 1xl | 8.28E-06 | 0 | 1 | 2 | 10 | Q3TZZ7 | Extended s | Q3TZZ7 [6 | 0 | 2447.263 | 0 | -0.03 | 0.132198 | 1.74 | 103.1 | 96.9 | 6.72 | 8.15 | | High | High | High | High | High | High | High | High | High | High | High | 0 | 2.21E-06 | 4.12 |
| FALSE | High | EPTPSIASD | [K].EPTPSI/ | 2xPhospho | Q3TZZ7 2xl | 2.42E-05 | 0 | 1 | 2 | 9 | Q3TZZ7 | Extended s | Q3TZZ7 [6 | 0 | 2527.23 | 0 | 0.08 | 0.493184 | 32.3 | 98 | 102 | 12.43 | 10.51 | | High | High | High | High | High | High | High | High | High | High | High | 0 | 1.32E-05 | 3.38 |
| FALSE | High | EPTPSIASD | [K].EPTPSI/ | 3xPhospho | Q3TZZ7 3xl | 0.001799 | 5.66E-05 | 1 | 2 | 2 | Q3TZZ7 | Extended s | Q3TZZ7 [6 | 0 | 2607.196 | 0 | -0.01 | 0.48049 | 14.47 | 104.5 | 95.5 | 12.56 | 21 | | High | High | High | High | High | High | High | High | High | High | High | 2.58E-05 | 0.000462 | 3.3 |
| FALSE | High | EPTQADVE | [K].EPTQAI | 1xPhospho | B6ZHD0 1x | 1.49E-05 | 0 | 2 | 14 | 1 | B6ZHD0; Q | Erythrocyt | B6ZHD0 [1 | 1 | 3266.572 | 0 | 1.14 | 0.007497 | 25.51 | 63.9 | 136.1 | 14.81 | 13.48 | Shared | High | High | High | High | High | High | High | High | High | High | High | 0 | 2.11E-06 | 4.12 |
| FALSE | High | EPTSVSDSE | [K].EPTSVS | 1xPhospho | F8WIJ5 1xF | 3.29E-05 | 0 | 1 | 9 | 5 | F8WIJ5 | Phospholip | F8WIJ5 [72 | 0 | 2862.434 | 0 | 0.68 | 0.00198 | 1.67 | 80.2 | 119.8 | 13.6 | 7.03 | | High | High | High | High | High | High | High | High | High | High | High | 0 | 1.61E-05 | 3.37 |
| FALSE | High | EPTSVSDSE | [K].EPTSVS | 2xPhospho | F8WIJ5 2xF | 5.16E-05 | 0 | 1 | 9 | 2 | F8WIJ5 | Phospholip | F8WIJ5 [72 | 0 | 2942.4 | 0 | 1.12 | 0.001119 | 17 | 63 | 137 | 6.58 | 6.46 | | High | High | High | High | High | High | High | High | High | High | High | 0 | 8.54E-06 | 4.16 |
| FALSE | High | EPVAATWE | [R].EPVAAT | 2xPhospho | Q6ZQ32 2x | 0.000616 | 0 | 2 | 7 | 2 | Q6ZQ32; F | MKIAA085 | Q6ZQ32 [1 | 0 | 2367.071 | 0 | 0.94 | 0.010173 | 16.3 | 69.7 | 130.3 | 10.31 | 20.48 | Shared | High | High | High | High | High | High | High | High | High | High | High | 0 | 0.000138 | 2.1 |
| FALSE | High | EPVAESPK | [R].EPVAES | 1xPhospho | D3YWS8 1: | 0.039347 | 0.002793 | 1 | 2 | 2 | D3YWS8 | Leukocyte | D3YWS8 [3 | 0 | 1394.733 | 0 | -0.81 | 0.003089 | 1.1 | 127.9 | 72.1 | 7.17 | 8.4 | | High | High | High | High | High | High | High | High | High | High | High | 0.001999 | 0.02038 | 1.45 |
| FALSE | High | EPVCEELSF | [K].EPVCEE | 1xCarbami | E9Q7D5 1x | 1.03E-05 | 0 | 1 | 1 | 5 | E9Q7D5 | Rho guanir | E9Q7D5 [4 | 0 | 4651.154 | 0 | 3.3 | 0.002712 | 24.42 | 19.4 | 180.6 | 24.21 | 20.14 | | High | High | High | High | High | High | High | High | High | High | High | 0 | 1.39E-06 | 4.2 |
| FALSE | High | EPVEELSDS | [R].EPVEEL | 1xPhospho | Q9DB28 1x | 8.92E-07 | 0 | 1 | 1 | 2 | Q9DB28 | Ribonuclea | Q9DB28 [1 | 0 | 2345.978 | 0 | -0.37 | 0.07199 | 12.66 | 112.2 | 87.8 | 10.63 | 13.2 | | High | High | High | High | High | High | High | High | High | High | High | 0 | 3.42E-06 | 4.23 |
| FALSE | High | EPVGLEEP\ | [R].EPVGLE | 1xPhospho | S4R1M2 1x | 0.000365 | 0 | 1 | 2 | 2 | S4R1M2 | Scaffold at | S4R1M2 [3 | 0 | 3303.619 | 0 | -0.25 | 0.06626 | 6.76 | 107.4 | 92.6 | 6.9 | 6.91 | | High | High | High | High | High | High | High | High | High | High | High | 0 | 7.73E-05 | 2.93 |
| FALSE | High | EPSVPAPC/ | [R].EPSVPA | 1xCarbami | Q6A0A2 2x | 0.005182 | 0.000551 | 1 | 3 | 1 | Q6A0A2 | La-related | Q6A0A2 [5 | 1 | 3589.72 | 0 | -0.12 | 0.323487 | 10.94 | 101.5 | 98.5 | 7.05 | 8.47 | | High | High | High | High | High | High | High | High | High | High | High | 0.000246 | 0.001511 | 2.66 |
| FALSE | High | EQPGDEYS | [R].EQPGD | 1xPhospho | B2RXX0 1x | 1.66E-05 | 0 | 1 | 7 | 2 | B2RXX0 | Nek1 prote | B2RXX0 [1 | 0 | 2406.1 | 0 | -0.39 | 0.023468 | 11.87 | 110.8 | 89.2 | 5.4 | 12.72 | | High | High | High | High | High | High | High | High | High | High | High | 0 | 2.38E-06 | 3.29 |
| FALSE | High | EQPPAEGA | [K].EQPPAI | 1xPhospho | Q3UH06-3 | 7.4E-08 | 0 | 1 | 3 | 1 | Q3UH06-3 | Isoform 3 c | Q3UH06-3 | 0 | 2676.353 | 0 | -0.04 | 0.976184 | 17.32 | 101.3 | 98.7 | 9.21 | 12.48 | | High | High | High | High | High | High | High | High | High | High | High | 0 | 5.45E-09 | 3.85 |
| FALSE | High | EQPPPLQSI | [K].EQPPPI | 1xPhospho | E9Q2I4 1xF | 1.51E-05 | 0 | 1 | 3 | 1 | E9Q2I4 | ELM2 and | E9Q2I4 [27 | 0 | 1882.963 | 0 | 0.9 | 0.023689 | 21.35 | 73.7 | 126.3 | 24.11 | 12.2 | | High | High | High | High | High | High | High | High | High | High | High | 0 | 2.16E-06 | 4.53 |
| FALSE | High | ERQESESE( | [R].ERQESE | 1xPhospho | Q6NS46 1x | 1.69E-05 | 0 | 1 | 1 | 1 | Q6NS46 | Protein RR | Q6NS46 [1 | 1 | 2243.096 | 0 | -0.67 | 0.108653 | 4.54 | 122.8 | 77.2 | 9.89 | 29.91 | | High | High | High | High | High | High | High | High | High | High | High | 0 | 2.43E-06 | 2.63 |
| FALSE | High | ERQSPFHG | [K].ERQSPI | 1xPhospho | Q3U3E2 1x | 0.008769 | 0.000736 | 1 | 1 | 1 | Q3U3E2 | Protein FAI | Q3U3E2 [2 | 1 | 2923.473 | 0 | -0.5 | 0.038131 | 6.34 | 119.5 | 80.5 | 14.85 | 23.58 | | High | High | High | High | High | High | High | High | High | High | High | 0.000327 | 0.00271 | 2.36 |
| FALSE | High | ERSCDYQF | [R].ERSCDY | 1xCarbami | Q69ZR9-2 | 0.003167 | 0.00016 | 1 | 1 | 2 | Q69ZR9-2 | Isoform 2 c | Q69ZR9-2 | 1 | 2736.203 | 0 | 0.18 | 0.069096 | 14.3 | 92.2 | 107.8 | 9.83 | 9.19 | | High | High | High | High | High | High | High | High | High | High | High | 7.19E-05 | 0.000869 | 4.22 |
| FALSE | High | ERSPEVLSC | [R].ERSPEV | 1xPhospho | Q5U4D9 1x | 0.010475 | 0.000772 | 1 | 4 | 1 | Q5U4D9 | THO comp | Q5U4D9 [1 | 1 | 1966.943 | 0 | -0.4 | 0.020774 | 9.08 | 116.6 | 83.4 | 7.64 | 10.43 | | High | High | High | High | High | High | High | High | High | High | High | 0.000382 | 0.003295 | 3.29 |
| FALSE | High | ERSPFGNQ | [R].ERSPFG | 1xPhospho | A0A1L1SQI | 0.00145 | 5.66E-05 | 1 | 4 | 2 | A0A1L1SQI | Dedicator c | A0A1L1SQI | 1 | 1741.822 | 0 | -1.43 | 0.001501 | 1.13 | 145 | 55 | 11.81 | 9.5 | | High | High | High | High | High | High | High | High | High | High | High | 2.58E-05 | 0.000363 | 3.26 |
| FALSE | High | ERSPPLTPK | [K].ERSPPL | 1xPhospho | E9QP54 1x | 0.015465 | 0.001089 | 1 | 6 | 2 | E9QP54 | Joubertin O | E9QP54 [9 | 1 | 1562.871 | 0 | 0.6 | 0.000236 | 2.76 | 79.8 | 120.2 | 2.6 | 3.42 | | High | High | High | High | High | High | High | High | High | High | High | 0.000571 | 0.005092 | 2.2 |
| FALSE | High | ERSPSPPA\ | [R].ERSPSP | 1xCarbami | Q3U3C9-2 | 0.000207 | 0 | 1 | 4 | 2 | Q3U3C9-2 | Isoform 2 c | Q3U3C9-2 | 1 | 1973.932 | 0 | -1.72 | 0.000299 | 1.5 | 153.6 | 46.4 | 4.32 | 8.08 | | High | High | High | High | High | High | High | High | High | High | High | 0 | 4.05E-05 | 2.55 |
| FALSE | High | ESAALLAS/ | [K].ESAALL | 1xPhospho | Q8K136 1x | 5.5E-05 | 0 | 1 | 2 | 8 | Q8K136 | Sodium ch | Q8K136 [1 | 0 | 2028.049 | 0 | -1.62 | 0.001817 | 1.81 | 150.3 | 49.7 | 6.63 | 21.19 | | High | High | High | High | High | High | High | High | High | High | High | 0 | 0.000332 | 2.79 |
| FALSE | High | ESAEFSVLE | [R].ESAEFS | 1xPhospho | Q99P72 1x | 6.68E-07 | 0 | 1 | 2 | 2 | Q99P72 | Reticulon-4 | Q99P72 [2 | 0 | 2936.331 | 0 | -0.31 | 0.562148 | 16.83 | 110.8 | 89.2 | 12.46 | 31.12 | | High | High | High | High | High | High | High | High | High | High | High | 0 | 7.8E-05 | 4.59 |
| FALSE | High | ESAEQEGF | [R].ESAEQI | 1xPhospho | A2AJ15 1xF | 3.54E-05 | 0 | 1 | 4 | 2 | A2AJ15 | Endoplasm | A2AJ15 [1C | 0 | 2190.073 | 0 | 1.59 | 0.001008 | 23.24 | 49.8 | 150.2 | 11.3 | 8.9 | | High | High | High | High | High | High | High | High | High | High | High | 0 | 5.57E-06 | 3.09 |
| FALSE | High | ESAEQSSG | [R].ESAEQ! | 1xPhospho | Q7TSC1 1x | 0.06169 | 0.004646 | 1 | 6 | 1 | Q7TSC1 | Protein PRI | Q7TSC1 [1 | 0 | 2681.252 | 0 | 1.6 | 0.001371 | 18.9 | 49.5 | 150.5 | 9.4 | 10.94 | | High | High | High | High | High | High | High | High | High | High | High | 0.002381 | 0.02454 | 1.53 |
| FALSE | High | ESAPQVLLI | [K].ESAPQ\ | 1xPhospho | Q9ES28 1x | 0.003055 | 0.00016 | 1 | 8 | 1 | Q9ES28 | Rho guanir | Q9ES28 [7 | 0 | 2007.045 | 0 | 0.38 | 0.054842 | 18.97 | 86.7 | 113.3 | 12.78 | 13.23 | | High | High | High | High | High | High | High | High | High | High | High | 7.19E-05 | 0.000832 | 2.64 |
| FALSE | High | ESAQAVAV | [K].ESAQA\ | 1xPhospho | H9KV00 1x | 0.037692 | 0.00249 | 1 | 8 | 1 | H9KV00 | Protein SO | H9KV00 [1 | 0 | 1579.829 | 0 | -0.14 | 0.798275 | 24.65 | 105.7 | 94.3 | 37.92 | 47.32 | | High | High | High | High | High | High | High | High | High | High | High | 0.001324 | 0.01396 | 2.52 |
| FALSE | High | ERPTSAIYP | [R].ERPTSA | 1xPhospho | Q60875 1x | 0.029586 | 0.001857 | 1 | 16 | 1 | Q60875 | Rho guanir | Q60875 [1 | 0 | 1934.921 | 0 | -1.33 | 0.001052 | 1.85 | 144.9 | 55.1 | 16.17 | 19.73 | | High | High | High | High | High | High | High | High | High | High | High | 0.000997 | 0.01058 | 2.55 |
| FALSE | High | ESAQAVAV | [K].ESAQA\ | 1xPhospho | H9KV00 1x | 1.26E-05 | 0 | 1 | 8 | 6 | H9KV00 | Protein SO | H9KV00 [1 | 0 | 1808.992 | 0 | 0.2 | 0.080641 | 21.63 | 93.1 | 106.9 | 2.49 | 11.34 | | High | High | High | High | High | High | High | High | High | High | High | 2.58E-05 | 0.000415 | 3.54 |
| FALSE | High | ESASPTIPN | [K].ESASPT | 1xPhospho | Q9WTU0 1 | 4.35E-05 | 0 | 1 | 1 | 2 | Q9WTU0 | Lysine-spe | Q9WTU0 [ | 0 | 2474.294 | 0 | 1.35 | 0.00311 | 11.88 | 53.4 | 146.6 | 16.15 | 10.04 | | High | High | High | High | High | High | High | High | High | High | High | 0 | 1.77E-05 | 2.81 |
| FALSE | High | ESASVLQQ | [K].ESASVL | 1xPhospho | Q9EQH4 1x | 0.000434 | 0 | 1 | 2 | 3 | Q9EQH4 | Transcripti | Q9EQH4 [2 | 0 | 1931.927 | 0 | -0.39 | 0.041446 | 4.55 | 112.4 | 87.6 | 12.65 | 6.66 | | High | High | High | High | High | High | High | High | High | High | High | 0 | 9.36E-05 | 2.96 |
| FALSE | High | ESAVDSSSS | [K].ESAVD! | 1xCarbami | Q05CL8 1x | 0.005768 | 0.00062 | 1 | 2 | 2 | Q05CL8 | La-related | Q05CL8 [2 | 0 | 1850.861 | 0 | -0.1 | 0.439433 | 6.25 | 101.6 | 98.4 | 8.08 | 9.1 | | High | High | High | High | High | High | High | High | High | High | High | 0.000275 | 0.001698 | 2.47 |
| FALSE | High | ESCEDLQV | [K].ESCEDL | 1xCarbami | Q6GTE9 1x | 0.000319 | 0 | 1 | 4 | 1 | Q6GTE9 | NIMA (Nev | Q6GTE9 [6 | 0 | 2188.978 | 0 | 0.11 | 0.607767 | 18.33 | 91.1 | 108.9 | 14.63 | 18.91 | | High | High | High | High | High | High | High | High | High | High | High | 0 | 6.62E-05 | 2.84 |
| FALSE | High | ESCSSPSAV | [R].ESCSSP | 1xCarbami | Q8BMB0 1 | 2.21E-05 | 0 | 1 | 3 | 3 | Q8BMB0 | BRCA2-inte | Q8BMB0 [ | 0 | 1954.914 | 0 | 0.01 | 0.544611 | 9.25 | 103.9 | 96.1 | 11.11 | 7.65 | | High | High | High | High | High | High | High | High | High | High | High | 0 | 3.31E-06 | 3.77 |
| FALSE | High | ESDDALTVI | [K].ESDDAI | 1xPhospho | Q0VGB7 1x | 0.000422 | 0 | 2 | 4 | 1 | Q0VGB7; A | Serine/thre | Q0VGB7 [3 | 0 | 4048.692 | 0 | -0.29 | 0.042275 | 16.9 | 110.1 | 89.9 | 12.31 | 19.09 | Shared | High | High | High | High | High | High | High | High | High | High | High | 0 | 9.02E-05 | 4.75 |
| FALSE | High | ESDDKPEIE | [K].ESDDKI | 1xPhospho | Q3UIF3 1xl | 1.06E-07 | 0 | 1 | 9 | 9 | Q3UIF3 | Putative ur | Q3UIF3 [2 | 0 | 2846.239 | 0 | -1.18 | 0.269767 | 2.87 | 138.7 | 61.3 | 60.69 | 65.14 | | High | High | High | High | High | High | High | High | High | High | High | 0 | 8.21E-09 | 5.29 |
| FALSE | High | ESDDKPEIE | [K].ESDDKI | 1xPhospho | Q3UIF3 1xl | 7.25E-07 | 0 | 1 | 9 | 18 | Q3UIF3 | Putative ur | Q3UIF3 [2 | 0 | 3075.402 | 0 | -1.16 | 0.000951 | 4.55 | 140.5 | 59.5 | 5.75 | 11.03 | | High | High | High | High | High | High | High | High | High | High | High | 0 | 0.000216 | 3.63 |
| FALSE | High | ESDDKPEIE | [K].ESDDKI | 1xPhospho | Q3UIF3 1xl | 0.007082 | 0.00062 | 1 | 9 | 1 | Q3UIF3 | Putative ur | Q3UIF3 [2 | 1 | 3203.497 | 0 | -1.36 | 0.221356 | 2.25 | 147.2 | 52.8 | 84.75 | 54.3 | | High | High | High | High | High | High | High | High | High | High | High | 0.000275 | 0.002136 | 2.82 |
| FALSE | High | ESDDSDVD | [R].ESDDSI | 1xPhospho | Q62187 1x | 0.020862 | 0.001472 | 1 | 2 | 2 | Q62187 | Transcripti | Q62187 [4 | 0 | 1773.774 | 0 | -0.54 | 0.546997 | 16.54 | 101.9 | 98.1 | 28.18 | 29.58 | | High | High | High | High | High | High | High | High | High | High | High | 0.002446 | 0.02635 | 2.37 |
| FALSE | High | ESDHEFDN | [K].ESDHEI | 1xPhospho | Q8BGF7 1x | 0.005275 | 0.000599 | 1 | 4 | 1 | Q8BGF7 | PAB-deper | Q8BGF7 [4 | 0 | 2491.061 | 0 | -0.01 | 0.714829 | 13.41 | 100.4 | 99.6 | 8.51 | 7.54 | | High | High | High | High | High | High | High | High | High | High | High | 0.000267 | 0.001533 | 3.62 |
| FALSE | High | ESDLSDTLS | [K].ESDLSD | 1xPhospho | Q3URD3 1x | 1.26E-05 | 0 | 1 | 10 | 2 | Q3URD3 | Sarcolemm | Q3URD3 [4 | 0 | 1816.898 | 0 | 0.79 | 0.010441 | 6.86 | 76.6 | 123.4 | 6.13 | 13.43 | | High | High | High | High | High | High | High | High | High | High | High | 0 | 1.76E-06 | 2.62 |
| FALSE | High | ESDLSDTLS | [K].ESDLSD | 2xPhospho | Q3URD3 2x | 0.090888 | 0.007081 | 1 | 10 | 1 | Q3URD3 | Sarcolemm | Q3URD3 [4 | 0 | 1896.864 | 0 | 0.25 | 0.332394 | 13.81 | 94.2 | 105.8 | 12.13 | 13.63 | | High | High | High | High | High | High | High | High | High | High | High | 0.003633 | 0.0386 | 2.85 |
| FALSE | High | ESASEGPN' | [R].ESASEG | 2xPhospho | S4R1N2 2x | 2.14E-06 | 0 | 1 | 2 | 1 | S4R1N2 | DNA cross- | S4R1N2 [5 | 0 | 2444.151 | 0 | -0.02 | 0.573685 | 5.14 | 102.3 | 97.7 | 7.25 | 8.67 | | High | High | High | High | High | High | High | High | High | High | High | 0 | 2.4E-07 | 4.69 |
| FALSE | High | ERPSSWSS | [R].ERPSSV | 1xPhospho | B7ZNA0 1x | 0.003055 | 0.00016 | 2 | 8 | 1 | B7ZNA0; Q | Shroom3 p | B7ZNA0 [3 | 0 | 1957.978 | 0 | -0.41 | 0.015365 | 5.38 | 116.8 | 83.2 | 5.39 | 15.71 | Shared | High | High | High | High | High | High | High | High | High | High | High | 7.19E-05 | 0.00083 | 1.75 |
| FALSE | High | ERPQSPGK | [K].ERPQSI | 1xPhospho | E9PVX6 1xl | 0.008094 | 0.000736 | 1 | 3 | 2 | E9PVX6 | Proliferatic | E9PVX6 [5 | 0 | 1436.766 | 0 | -0.14 | 0.565547 | 23.39 | 106.1 | 93.9 | 4.94 | 17.81 | | High | High | High | High | High | High | High | High | High | High | High | 0.000327 | 0.002489 | 1.23 |
| FALSE | High | ERPQLTECI | [K].ERPQLT | 1xCarbami | O35954 1x | 0.001097 | 0 | 1 | 1 | 2 | O35954 | Membrane | O35954 [8 | 0 | 2715.269 | 0 | 2.31 | 0.000347 | 24.45 | 34.7 | 165.3 | 19.83 | 5.66 | | High | High | High | High | High | High | High | High | High | High | High | 0 | 0.000264 | 2.72 |
| FALSE | High | EQPQSSSG | [R].EQPQS! | 1xPhospho | O55176 1x | 2.25E-05 | 0 | 1 | 6 | 1 | O55176 | E3 ubiquiti | O55176 [3 | 0 | 2397.174 | 0 | 0.56 | 0.002231 | 7.71 | 82.6 | 117.4 | 14.69 | 11.23 | | High | High | High | High | High | High | High | High | High | High | High | 0 | 3.35E-06 | 3.12 |
| FALSE | High | EQQEDSSA | [K].EQQED | 1xPhospho | E9QAD8 1x | 1.23E-07 | 0 | 1 | 5 | 1 | E9QAD8 | IQ motif ar | E9QAD8 [5 | 0 | 3525.615 | 0 | 1.36 | 0.011287 | 49.48 | 56 | 144 | 45.43 | 11.61 | | High | High | High | High | High | High | High | High | High | High | High | 0 | 9.61E-09 | 3.62 |
| FALSE | High | EQSALESEE | [R].EQSALE | 1xPhospho | Q8VDX6 1x | 2.08E-05 | 0 | 1 | 1 | 1 | Q8VDX6 | Transmem | Q8VDX6 [5 | 0 | 2701.207 | 0 | 1.94 | 0.000922 | 6.76 | 41.3 | 158.7 | 13.68 | 10.78 | | High | High | High | High | High | High | High | High | High | High | High | 0 | 3.06E-06 | 2.9 |
| FALSE | High | EQSDISISPI | [K].EQSDIS | 1xPhospho | S4R2L4 1xF | 0.000687 | 0 | 1 | 4 | 3 | S4R2L4 | Arginine/s | S4R2L4 [25 | 0 | 1440.693 | 0 | -0.75 | 0.000256 | 1.27 | 124.3 | 75.7 | 2.09 | 3.02 | | High | High | High | High | High | High | High | High | High | High | High | 0.000161 | 0.001163 | 2.16 |
| FALSE | High | EQSESVNT | [R].EQSES\ | 1xPhospho | Q99MP8 1: | 4.44E-06 | 0 | 1 | 3 | 3 | Q99MP8 | BRCA1-ass | Q99MP8 [1 | 0 | 2128.021 | 0 | 0.72 | 0.001085 | 10.28 | 77.1 | 122.9 | 2.09 | 7.03 | | High | High | High | High | High | High | High | High | High | High | High | 0 | 5.41E-07 | 3.04 |
| FALSE | High | EQSETSELH | [R].EQSETS | 1xPhospho | E9QK89 1x | 0.020142 | 0.001472 | 1 | 2 | 1 | E9QK89 | Mediator c | E9QK89 [7 | 0 | 2528.162 | 0 | -0.42 | 0.008329 | 2.89 | 115.4 | 84.6 | 8.99 | 10.23 | | High | High | High | High | High | High | High | High | High | High | High | 0.000786 | 0.00685 | 2.24 |
| FALSE | High | EQSMLTPK | [R].EQSML | 1xPhospho | G3UVV3 1x | 0.074685 | 0.005727 | 1 | 1 | 1 | G3UVV3 | MCG1556, | G3UVV3 [8 | 0 | 1487.758 | 0 | -0.81 | 0.997457 | 14.29 | 118.1 | 81.9 | 32.83 | 88.88 | | High | High | High | High | High | High | High | High | High | High | High | 0.002937 | 0.03077 | 0.86 |
| FALSE | High | EQSSEASE1 | [R].EQSSEA | 1xPhospho | E9QPI5 1xF | 1.4E-06 | 0 | 1 | 3 | 2 | E9QPI5 | Sister chro | E9QPI5 [11 | 0 | 2487.072 | 0 | -0.1 | 0.979952 | 10.76 | 99.1 | 100.9 | 11.08 | 14.86 | | High | High | High | High | High | High | High | High | High | High | High | 0 | 1.49E-07 | 2.86 |
| FALSE | High | EQTASAPA | [R].EQTAS/ | 1xPhospho | Q3UMF0 1 | 0.000259 | 0 | 1 | 7 | 2 | Q3UMF0 | Cordon-ble | Q3UMF0 [ | 0 | 1938.035 | 0 | 0.03 | 0.385059 | 13.29 | 94.2 | 105.8 | 16.06 | 10.33 | | High | High | High | High | High | High | High | High | High | High | High | 0 | 0.000068 | 3.05 |
| FALSE | High | EQTFSGGT | [K].EQTFS( | 1xPhospho | Q569Z6 1x | 0.000815 | 0 | 1 | 4 | 2 | Q569Z6 | Thyroid ho | Q569Z6 [2 | 0 | 1935.947 | 0 | -0.26 | 0.033416 | 4.74 | 108.1 | 91.9 | 4.28 | 9.81 | | High | High | High | High | High | High | High | High | High | High | High | 4.94E-05 | 0.000662 | 3.05 |
| FALSE | High | EQTSPQK | [K].EQTSP( | 1xPhospho | Q8CJF7 1xl | 0.040373 | 0.002982 | 1 | 1 | 2 | Q8CJF7 | Protein EL\ | Q8CJF7 [1 | 0 | 1355.697 | 0 | -0.32 | 0.107341 | 11.66 | 111 | 89 | 9.08 | 14.33 | | High | High | High | High | High | High | High | High | High | High | High | 0.001566 | 0.01512 | 1.91 |
| FALSE | High | EQVYNISIN | [K].EQVYN | 1xPhospho | Q8BHX3-2 | 5.3E-08 | 0 | 1 | 2 | 3 | Q8BHX3-2 | Isoform 2 c | Q8BHX3-2 | 0 | 2373.21 | 0 | -0.76 | 0.000781 | 3.03 | 123.6 | 76.4 | 5.22 | 7.44 | | High | High | High | High | High | High | High | High | High | High | High | 0 | 2.58E-08 | 3.95 |
| FALSE | High | EQVYNISIN | [K].EQVYN | 1xDeamida | Q8BHX3-2 | 6.23E-08 | 0 | 1 | 2 | 5 | Q8BHX3-2 | Isoform 2 c | Q8BHX3-2 | 0 | 2374.194 | 0 | -0.67 | 0.000758 | 3.27 | 122.3 | 77.7 | 4.83 | 7.35 | | High | High | High | High | High | High | High | High | High | High | High | 0 | 4.49E-09 | 4.05 |
| FALSE | High | EQYGLGPY | [R].EQYGL( | 1xPhospho | P70399 1xl | 3.17E-05 | 0 | 1 | 7 | 1 | P70399 | TP53-bindi | P70399 [15 | 0 | 2304.193 | 0 | -0.14 | 0.206847 | 12.3 | 105.9 | 94.1 | 7.87 | 8.94 | | High | High | High | High | High | High | High | High | High | High | High | 0 | 4.96E-06 | 3.06 |
| FALSE | High | EQYSDAPE | [K].EQYSD/ | 1xPhospho | Q9DB96 1x | 0.000462 | 0 | 1 | 1 | 4 | Q9DB96 | Neuroguid | Q9DB96 [2 | 0 | 1645.731 | 0 | 0.43 | 0.018308 | 16.23 | 84.3 | 115.7 | 6.63 | 12.27 | | High | High | High | High | High | High | High | High | High | High | High | 0.001609 | 0.01572 | 2.79 |
| FALSE | High | EQYVPPRSI | [K].EQYVPI | 1xPhospho | G3X9Q6 1x | 0.000876 | 0 | 1 | 4 | 4 | G3X9Q6 | La-related | G3X9Q6 [7 | 1 | 1738.929 | 0 | -0.32 | 0.002185 | 3.12 | 112 | 88 | 3.04 | 5.94 | | High | High | High | High | High | High | High | High | High | High | High | 0.006342 | 0.06802 | 1.56 |
| FALSE | High | ERDHSPTP! | [R].ERDHSI | 1xPhospho | Q9D824 1x | 0.001021 | 0 | 1 | 3 | 5 | Q9D824 | Pre-mRNA | Q9D824 [4 | 1 | 2311.019 | 0 | -0.14 | 0.104985 | 5.42 | 102.8 | 97.2 | 4.47 | 5.98 | | High | High | High | High | High | High | High | High | High | High | High | 0.005745 | 0.0633 | 3.56 |

| | | | | | | | | | | | | | | | | | | | | | | | | | | | | | | | | | | | | | | |
|---|---|---|---|---|---|---|---|---|---|---|---|---|---|---|---|---|---|---|---|---|---|---|---|---|---|---|---|---|---|---|---|---|---|---|---|---|---|---|
| FALSE | High | ERDHSPTP! | [R].ERDHSI | 2xPhosphc | Q9D824 2x | 0.000993 | 0 | 1 | 3 | 2 | Q9D824 | Pre-mRNA | Q9D824 [4 | 1 | 2390.985 | 0 | -0.2 | 0.120495 | 13.55 | 109.5 | 90.5 | 7.28 | 11.04 | | High | High | High | High | High | High | High | High | High | High | High | 0 | 0.000238 | 3.49 |
| FALSE | High | ERDSELSD! | [K].ERDSEL | 1xPhosphc | Q8VDF2 1x | 0.002272 | 0.000109 | 1 | 3 | 6 | Q8VDF2 | E3 ubiquiti | Q8VDF2 [8 | 1 | 2894.273 | 0 | 0.26 | 0.002762 | 1.73 | 93.3 | 106.7 | 8.79 | 6.86 | | High | High | High | High | High | High | High | High | High | High | High | 0.00014 | 0.001097 | 3.19 |
| FALSE | High | ERDSYHGS | [R].ERDSYH | 1xPhosphc | Q8R0F5 1x | 0.090888 | 0.007081 | 1 | 1 | 1 | Q8R0F5 | RNA-bindir | Q8R0F5 [3 | 1 | 1530.654 | 0 | 0.5 | 0.006842 | 12.9 | 83 | 117 | 5.59 | 8.66 | | High | High | High | High | High | High | High | High | High | High | High | 0.003606 | 0.03841 | 1.95 |
| FALSE | High | EREDSGDA | [R].EREDSC | 1xPhosphc | O88665 1x | 0.020142 | 0.001472 | 1 | 1 | 1 | O88665 | Bromodom | O88665 [2 | 1 | 1919.833 | 0 | 0.04 | 0.454322 | 7.43 | 98.6 | 101.4 | 11.5 | 5.4 | | High | High | High | High | High | High | High | High | High | High | High | 0.000786 | 0.00688 | 2.11 |
| FALSE | High | EREDSGDA | [R].EREDSC | 1xPhosphc | O88665 1x | 0.112856 | 0.009146 | 1 | 1 | 1 | O88665 | Bromodom | O88665 [2 | 2 | 2532.213 | 0 | -0.53 | 0.001148 | 3.04 | 121.7 | 78.3 | 4.24 | 7 | | High | High | High | High | High | High | High | High | High | High | High | 0.004569 | 0.0497 | 1.42 |
| FALSE | High | EREESEDEL | [R].EREESE | 1xPhosphc | Q8CFE3 1x | 2.88E-06 | 0 | 1 | 2 | 3 | Q8CFE3 | REST corep | Q8CFE3 [2 | 1 | 3525.616 | 1 | 0.53 | 0.013009 | 18.08 | 81.9 | 118.1 | 5.93 | 11.73 | | High | High | High | High | High | High | High | High | High | High | High | 0 | 7.85E-06 | 3.54 |
| FALSE | High | EREESEDEL | [R].EREESE | 1xDeamida | Q8CFE3 1x | 1.7E-05 | 0 | 1 | 2 | 4 | Q8CFE3 | REST corep | Q8CFE3 [2 | 1 | 3526.6 | 0 | 0.5 | 0.003185 | 3.85 | 83.1 | 116.9 | 4.54 | 6.27 | | High | High | High | High | High | High | High | High | High | High | High | 0 | 6.18E-06 | 3.92 |
| FALSE | High | EREVSVSS\ | [R].EREVSV | 2xPhosphc | Q8CJF7 2xI | 0.068752 | 0.005042 | 1 | 1 | 1 | Q8CJF7 | Protein EL\ | Q8CJF7 [1 | 1 | 2194.045 | 0 | -0.13 | 0.084994 | 1.58 | 110.2 | 89.8 | 17.1 | 9.41 | | High | High | High | High | High | High | High | High | High | High | High | 0.002618 | 0.02796 | 2.11 |
| FALSE | High | ERNQVVPE | [K].ERNQV | 1xDeamida | Q3U4X8 1x | 0.004033 | 0.000311 | 1 | 7 | 2 | Q3U4X8 | DNA ligase | Q3U4X8 [5 | 1 | 2123.079 | 0 | -0.79 | 0.004823 | 7.32 | 128 | 72 | 12.15 | 5.32 | | High | High | High | High | High | High | High | High | High | High | High | 0.00014 | 0.001134 | 3.5 |
| FALSE | High | ERPESCDD/ | [R].ERPESC | 1xCarbami | Q8CH77 1x | 0.000105 | 0 | 1 | 4 | 1 | Q8CH77 | Neuron na | Q8CH77 [5 | 0 | 1831.83 | 0 | 1.13 | 0.003314 | 29.21 | 61.2 | 138.8 | 12.15 | 11.15 | | High | High | High | High | High | High | High | High | High | High | High | 0 | 1.9E-05 | 2 |
| FALSE | High | ERPISLGIFF | [K].ERPISL( | 1xPhosphc | B8X349 1xI | 3.2E-05 | 0 | 2 | 12 | 2 | B8X349; F6 | JNK-interac | B8X349 [1 | 0 | 3073.674 | 0 | 3.11 | 0.00041 | 19.1 | 20.7 | 179.3 | 18.41 | 8.21 | Shared | High | High | High | High | High | High | High | High | High | High | High | 0 | 5E-06 | 4.02 |
| FALSE | High | ERPPETLTP | [K].ERPPET | 1xPhosphc | Q62219 1x | 0.00011 | 0 | 1 | 7 | 4 | Q62219 | Transformi | Q62219 [2 | 0 | 3743.795 | 0 | -0.08 | 0.229226 | 14.64 | 107.4 | 92.6 | 5.97 | 19.59 | | High | High | High | High | High | High | High | High | High | High | High | 0.000344 | 0.00315 | 3.68 |
| FALSE | High | ESDPGVVT | [K].ESDPG\ | 1xPhosphc | E9PWG6 1: | 1.53E-06 | 0 | 1 | 2 | 2 | E9PWG6 | Non-SMC c | E9PWG6 [8 | 0 | 3532.635 | 0 | -0.17 | 0.060888 | 8.36 | 105 | 95 | 8.75 | 10.35 | | High | High | High | High | High | High | High | High | High | High | High | 0 | 1.64E-07 | 4.48 |
| FALSE | High | ETDEAAFEI | [R].ETDEA/ | 2xPhosphc | P97868 2xI | 3.44E-05 | 0 | 1 | 3 | 5 | P97868 | E3 ubiquiti | P97868 [16 | 0 | 3180.322 | 0 | -0.08 | 0.633629 | 7.91 | 102 | 98 | 7.3 | 7.65 | | High | High | High | High | High | High | High | High | High | High | High | 0 | 8.28E-06 | 4.34 |
| FALSE | High | ETDGSETPI | [R].ETDGSI | 1xPhosphc | Q3TRW3 1 | 0.000165 | 0 | 1 | 5 | 2 | Q3TRW3 | Putative ur | Q3TRW3 [2 | 0 | 2217.036 | 0 | 0.57 | 0.001789 | 0.86 | 75.2 | 124.8 | 6.93 | 6.39 | | High | High | High | High | High | High | High | High | High | High | High | 0 | 0.000144 | 3.31 |
| FALSE | High | ETDILSDDD | [K].ETDILSI | 1xCarbami | G3UWG2 1 | 1.63E-06 | 0 | 1 | 9 | 3 | G3UWG2 | T-cell lymp | G3UWG2 [ | 0 | 2454.104 | 0 | -1.63 | 0.001113 | 3.66 | 151.2 | 48.8 | 8.06 | 14.66 | | High | High | High | High | High | High | High | High | High | High | High | 0 | 1.75E-07 | 4.08 |
| FALSE | High | FDHSPTPS/ | [R].FDHSPT | 1xPhosphc | Q6ZQD6 1x | 0.008463 | 0.000736 | 1 | 5 | 1 | Q6ZQD6 | MKIAA033 | Q6ZQD6 [4 | 0 | 1597.721 | 0 | 1.59 | 0.000688 | 7.36 | 47.4 | 152.6 | 13.37 | 8.68 | | High | High | High | High | High | High | High | High | High | High | High | 0.000327 | 0.002598 | 1.75 |
| FALSE | High | FDIYDPFHF | [R].FDIYDP | 1xPhosphc | Q5U4C3 1x | 0.008388 | 0.000736 | 1 | 1 | 3 | Q5U4C3 | Splicing fac | Q5U4C3 [2 | 0 | 3199.507 | 0 | 0 | 0.688127 | 10.74 | 98.4 | 101.6 | 9.44 | 8.35 | | High | High | High | High | High | High | High | High | High | High | High | 0.003265 | 0.0345 | 2.16 |
| FALSE | High | FDPAMFNI | [R].FDPAM | 1xPhosphc | B0FTY3 1xI | 9.03E-05 | 0 | 1 | 4 | 1 | B0FTY3 | NudC dom | B0FTY3 [34 | 0 | 1949.907 | 0 | 0.59 | 0.109554 | 40.83 | 82.9 | 117.1 | 21.15 | 19.83 | | High | High | High | High | High | High | High | High | High | High | High | 0 | 1.6E-05 | 4.07 |
| FALSE | High | FDPFGPIGT | [R].FDPFGI | 1xPhosphc | A0A1S6GW | 0.000231 | 0 | 1 | 3 | 3 | A0A1S6GW | Uncharacte | A0A1S6GW | 0 | 3425.527 | 0 | 0.67 | 0.029354 | 26.19 | 75.8 | 124.2 | 16.86 | 15.95 | | High | High | High | High | High | High | High | High | High | High | High | 0 | 4.62E-05 | 4.37 |
| FALSE | High | FDQLFDDE | [R].FDQLFI | 1xPhosphc | Q9CY58 1x | 6.61E-06 | 0 | 2 | 11 | 3 | Q9CY58; Q | Plasminoge | Q9CY58 [1 | 0 | 2482.183 | 0 | 0.29 | 0.003479 | 6.24 | 92 | 108 | 5.38 | 3.96 | Shared | High | High | High | High | High | High | High | High | High | High | High | 0 | 8.47E-07 | 3.66 |
| FALSE | High | FDSDEEDEI | [K].FDSDEE | 1xPhosphc | A2AJT5 1xI | 5.5E-05 | 0 | 1 | 5 | 6 | A2AJT5 | Arginine/se | A2AJT5 [28 | 0 | 2724.181 | 0 | -0.93 | 0.001718 | 6.32 | 131.1 | 68.9 | 7.03 | 9.6 | | High | High | High | High | High | High | High | High | High | High | High | 0 | 1.07E-05 | 3.5 |
| FALSE | High | FDSDEEDEI | [K].FDSDEE | 2xPhosphc | A2AJT5 2xI | 0.007538 | 0.00062 | 1 | 5 | 1 | A2AJT5 | Arginine/se | A2AJT5 [28 | 1 | 3160.365 | 0 | -0.13 | 0.282765 | 9.66 | 102.4 | 97.6 | 5.36 | 14.46 | | High | High | High | High | High | High | High | High | High | High | High | 0.000275 | 0.002286 | 3.4 |
| FALSE | High | FDSNEEDT. | [K].FDSNEE | 1xPhosphc | Q64511 1x | 7.25E-07 | 0 | 1 | 2 | 3 | Q64511 | DNA topois | Q64511 [1 | 0 | 2370.074 | 0 | 1.03 | 0.386237 | 48.34 | 72.4 | 127.6 | 47.33 | 58.97 | | High | High | High | High | High | High | High | High | High | High | High | 0 | 5.66E-06 | 3.66 |
| FALSE | High | FDSNEEDT. | [K].FDSNEE | 1xPhosphc | Q64511 1x | 1.01E-06 | 0 | 1 | 2 | 4 | Q64511 | DNA topois | Q64511 [1 | 0 | 2599.237 | 0 | 1.13 | 0.001453 | 6.38 | 66.2 | 133.8 | 6.45 | 8.85 | | High | High | High | High | High | High | High | High | High | High | High | 0 | 7.36E-07 | 4.22 |
| FALSE | High | FDSQLFHTI | [K].FDSQLF | 1xPhosphc | A2AFP5 1xI | 0.000808 | 0 | 1 | 3 | 1 | A2AFP5 | RAB9, men | A2AFP5 [4 | 0 | 2433.262 | 0 | -0.77 | 0.00151 | 4.37 | 128.3 | 71.7 | 8.57 | 11.06 | | High | High | High | High | High | High | High | High | High | High | High | 0 | 0.000187 | 2.99 |
| FALSE | High | FEDEDSDD | [K].FEDEDS | 1xPhosphc | B2RY56 1xI | 0.069903 | 0.005181 | 1 | 2 | 2 | B2RY56 | RNA-bindir | B2RY56 [6 | 1 | 1989.921 | 0 | 0.09 | 0.378474 | 13.38 | 97.6 | 102.4 | 4.18 | 6.01 | | High | High | High | High | High | High | High | High | High | High | High | 0.005336 | 0.0583 | 2.6 |
| FALSE | High | FEDEGAGF | [K].FEDEG/ | 1xPhosphc | P14873 1xI | 1.41E-08 | 0 | 2 | 2 | 9 | P14873; B2 | Microtubu | P14873 [9 | 0 | 2793.17 | 0 | -0.52 | 0.000707 | 3.58 | 118.2 | 81.8 | 1.11 | 5.79 | Shared | High | High | High | High | High | High | High | High | High | High | High | 0 | 8.47E-10 | 5.44 |
| FALSE | High | FEDILSPAE | [R].FEDILSI | 1xPhosphc | Q8K202 1x | 2.67E-05 | 0 | 1 | 4 | 4 | Q8K202 | DNA-direct | Q8K202 [2 | 0 | 2695.238 | 0 | -0.14 | 0.139619 | 2.72 | 103.4 | 96.6 | 9.73 | 11 | | High | High | High | High | High | High | High | High | High | High | High | 0 | 0.000261 | 3.03 |
| FALSE | High | FDGSETRPI | [K].FDGSET | 1xPhosphc | J7I1C3 1xPI | 0.002004 | 5.66E-05 | 1 | 1 | 4 | J7I1C3 | Rif1 OS=M | J7I1C3 [12 | 0 | 2951.381 | 0 | -1.03 | 0.001413 | 1.56 | 136.1 | 63.9 | 5.2 | 13.82 | | High | High | High | High | High | High | High | High | High | High | High | 2.58E-05 | 0.000518 | 2.46 |
| FALSE | High | FEDVYEPYI | [K].FEDVYE | 1xPhosphc | Q9R0X5-5 | 1.69E-05 | 0 | 1 | 11 | 1 | Q9R0X5-5 | Isoform 5 c | Q9R0X5-5 | 0 | 2845.317 | 0 | -0.8 | 0.001549 | 4.26 | 124.3 | 75.7 | 8.06 | 5.1 | | High | High | High | High | High | High | High | High | High | High | High | 0 | 2.43E-06 | 4.08 |
| FALSE | High | FEDYLREP/ | [R].FEDYLR | 1xCarbami | E9Q0A7 1x | 0.023792 | 0.001612 | 1 | 1 | 1 | E9Q0A7 | Oxidation r | E9Q0A7 [3 | 1 | 4389.042 | 0 | 0.38 | 0.020078 | 9.16 | 82.8 | 117.2 | 14.08 | 15.75 | | High | High | High | High | High | High | High | High | High | High | High | 0.000859 | 0.008282 | 3.02 |
| FALSE | High | FEEVEEEPE | [R].FEEVEE | 1xPhosphc | Q3UT19 1x | 0.000141 | 0 | 1 | 2 | 3 | Q3UT19 | Putative ur | Q3UT19 [1 | 0 | 3165.535 | 0 | 0.05 | 0.812375 | 16.26 | 98.9 | 101.1 | 7.63 | 8.39 | | High | High | High | High | High | High | High | High | High | High | High | 0.002432 | 0.02607 | 2.69 |
| FALSE | High | FEEVEEEPE | [R].FEEVEE | 1xPhosphc | Q3UT19 1x | 0.0009 | 0 | 1 | 2 | 3 | Q3UT19 | Putative ur | Q3UT19 [1 | 0 | 3181.53 | 0 | 0.13 | 0.109538 | 1.6 | 97.6 | 102.4 | 6.55 | 6.13 | | High | High | High | High | High | High | High | High | High | High | High | 0 | 0.000212 | 2.89 |
| FALSE | High | FELDHSSPT | [R].FELDHS | 1xPhosphc | Q8VDF2 1x | 0.005275 | 0.000599 | 1 | 2 | 2 | Q8VDF2 | E3 ubiquiti | Q8VDF2 [7 | 0 | 1497.694 | 0 | -0.75 | 0.005316 | 6.4 | 125 | 75 | 8.11 | 9.54 | | High | High | High | High | High | High | High | High | High | High | High | 0.007424 | 0.08199 | 2.74 |
| FALSE | High | FELLDDLPA | [K].FELLDD | 2xCarbami | Q9D7I8 1xI | 0.000731 | 0 | 1 | 1 | 1 | Q9D7I8 | Protein FAI | Q9D7I8 [5- | 0 | 3425.6 | 0 | 0.79 | 0.001211 | 9.4 | 71.9 | 128.1 | 12.34 | 10.89 | | High | High | High | High | High | High | High | High | High | High | High | 0 | 0.000169 | 4.14 |
| FALSE | High | FELLPTPPL! | [K].FELLPTI | 1xPhosphc | B2RSN1 1x | 0.016892 | 0.001204 | 1 | 10 | 2 | B2RSN1 | Myc proto- | B2RSN1 [6 | 0 | 1762.934 | 0 | 0.3 | 0.08086 | 9.69 | 89.5 | 110.5 | 8.53 | 9.34 | | High | High | High | High | High | High | High | High | High | High | High | 0.001011 | 0.01069 | 2.14 |
| FALSE | High | FEINAYDP/ | [K].FEINAY | 1xPhosphc | A0A1S6GW | 2.67E-05 | 0 | 1 | 6 | 1 | A0A1S6GW | Uncharacte | A0A1S6GW | 0 | 3200.571 | 0 | 2.82 | 0.020379 | 83.04 | 20.5 | 179.5 | 70.58 | 37.83 | | High | High | High | High | High | High | High | High | High | High | High | 0 | 4.08E-06 | 4.16 |
| FALSE | High | FELPHGSP( | [K].FELPHG | 1xPhosphc | Q61097 1x | 4.68E-05 | 0 | 1 | 5 | 2 | Q61097 | Kinase sup | Q61097 [3 | 0 | 1688.872 | 0 | -0.78 | 0.002539 | 5.99 | 126.5 | 73.5 | 6.58 | 17.81 | | High | High | High | High | High | High | High | High | High | High | High | 0 | 7.68E-06 | 2.81 |
| FALSE | High | FEPESPGFE | [K].FEPESP | 1xPhosphc | Q8R0T2 1x | 0.001313 | 0 | 1 | 3 | 1 | Q8R0T2 | Zinc finger | Q8R0T2 [8 | 0 | 1590.704 | 0 | 2.68 | 0.000374 | 22.19 | 27.4 | 172.6 | 14.18 | 8.21 | | High | High | High | High | High | High | High | High | High | High | High | 0 | 0.000323 | 2.43 |
| FALSE | High | FEPPQSDSI | [R].FEPPQS | 1xPhosphc | Q6NV83 1x | 0.00614 | 0.00062 | 1 | 4 | 1 | Q6NV83 | U2 snRNP-; | Q6NV83 [2 | 0 | 1671.721 | 0 | -0.24 | 0.089432 | 3.66 | 106.2 | 93.8 | 7.64 | 4.64 | | High | High | High | High | High | High | High | High | High | High | High | 0.000275 | 0.001824 | 2.44 |
| FALSE | High | FESPATGIL | [R].FESPAT | 1xPhosphc | Q9ERU9 1x | 1.75E-05 | 0 | 1 | 1 | 2 | Q9ERU9 | E3 SUMO- | Q9ERU9 [9 | 0 | 1583.803 | 0 | 0.97 | 0.002365 | 8.09 | 69.2 | 130.8 | 2.94 | 10.26 | | High | High | High | High | High | High | High | High | High | High | High | 0.006552 | 0.0703 | 2.18 |
| FALSE | High | FESPDSR | [R].FESPDS | 1xPhosphc | Q9D1R2 1x | 0.022378 | 0.001542 | 1 | 1 | 1 | Q9D1R2 | Protein KTI | Q9D1R2 [2 | 0 | 1146.503 | 0 | -0.01 | 0.993116 | 5.59 | 100.3 | 99.7 | 7.9 | 12.2 | | High | High | High | High | High | High | High | High | High | High | High | 0.000814 | 0.007759 | 1.75 |
| FALSE | High | FESPEVAEF | [K].FESPEV | 1xPhosphc | A0A1S6GW | 0.028824 | 0.001829 | 1 | 7 | 2 | A0A1S6GW | Uncharacte | A0A1S6GW | 0 | 1372.635 | 0 | -0.4 | 0.005918 | 2.67 | 112.6 | 87.4 | 3.97 | 6.05 | | High | High | High | High | High | High | High | High | High | High | High | 0.000997 | 0.01028 | 2.44 |
| FALSE | High | FEDYLREP/ | [R].FEDYLR | 1xCarbami | E9Q0A7 1x | 0.027358 | 0.001676 | 1 | 1 | 2 | E9Q0A7 | Oxidation r | E9Q0A7 [3 | 1 | 4388.058 | 0 | 0.36 | 0.014383 | 9.78 | 84.3 | 115.7 | 7.92 | 9.55 | | High | High | High | High | High | High | High | High | High | High | High | 0.000936 | 0.009719 | 2.91 |
| FALSE | High | FEVNVAELI | [R].FEVNV/ | 1xPhosphc | Q3T9L0 1xI | 2.78E-09 | 0 | 1 | 8 | 2 | Q3T9L0 | Putative ur | Q3T9L0 [4 | 0 | 2890.396 | 0 | -0.31 | 0.065342 | 10.42 | 110.8 | 89.2 | 5.74 | 11.21 | | High | High | High | High | High | High | High | High | High | High | High | 0 | 1.36E-10 | 4.41 |
| FALSE | High | FDGSETRPI | [K].FDGSET | 1xPhosphc | J7I1C3 1xPI | 0.00925 | 0.000736 | 1 | 1 | 1 | J7I1C3 | Rif1 OS=M | J7I1C3 [12 | 0 | 2935.386 | 0 | -1.29 | 0.000716 | 1.23 | 142.6 | 57.4 | 9.7 | 4.06 | | High | High | High | High | High | High | High | High | High | High | High | 0.000327 | 0.00288 | 1.77 |
| FALSE | High | FDFGEDSEI | [R].FDFGEI | 2xPhosphc | Q6DID3 2x | 0.013909 | 0.001011 | 1 | 2 | 2 | Q6DID3 | Protein SC/ | Q6DID3 [2 | 0 | 2260.909 | 0 | 0.24 | 0.001147 | 4.66 | 91.4 | 108.6 | 5.4 | 3.7 | | High | High | High | High | High | High | High | High | High | High | High | 0.002937 | 0.0307 | 2.36 |
| FALSE | High | EYVPPTAT! | [R].EYVPPT | 1xPhosphc | E9QAE4 1x | 1.54E-05 | 0 | 1 | 2 | 3 | E9QAE4 | Histone-lys | E9QAE4 [2 | 0 | 2392.22 | 0 | -1.43 | 0.000192 | 0.87 | 145.6 | 54.4 | 2.83 | 7.21 | | High | High | High | High | High | High | High | High | High | High | High | 0 | 7.39E-06 | 3.61 |
| FALSE | High | EYVPPTAT! | [R].EYVPPT | 2xPhosphc | E9QAE4 2x | 0.034281 | 0.002195 | 1 | 2 | 1 | E9QAE4 | Histone-lys | E9QAE4 [2 | 0 | 2472.187 | 0 | -1.55 | 0.000179 | 0.82 | 149 | 51 | 3.51 | 6.5 | | High | High | High | High | High | High | High | High | High | High | High | 0.001151 | 0.01251 | 2.19 |
| FALSE | High | EYVSNDAT | [K].EYVSNI | 1xPhosphc | Q6NVE8 1x | 4.98E-08 | 0 | 1 | 1 | 5 | Q6NVE8 | WD repeat | Q6NVE8 [3 | 0 | 2267.996 | 0 | 0.31 | 0.027796 | 14.31 | 89.2 | 110.8 | 7.09 | 6.47 | | High | High | High | High | High | High | High | High | High | High | High | 0 | 3.48E-09 | 3.42 |
| FALSE | High | FACLSPGY\ | [K].FACLSP | 1xCarbami | E9Q4F8 1x | 5.47E-06 | 0 | 1 | 3 | 2 | E9Q4F8 | Ankyrin re | E9Q4F8 [1 | 0 | 2657.276 | 0 | -0.48 | 0.038698 | 9.08 | 117.2 | 82.8 | 13.95 | 13.26 | | High | High | High | High | High | High | High | High | High | High | High | 0 | 7.6E-07 | 3.18 |
| FALSE | High | FACLSPGY\ | [K].FACLSP | 1xCarbami | E9Q4F8 2x | 0.071662 | 0.005278 | 1 | 3 | 1 | E9Q4F8 | Ankyrin re | E9Q4F8 [1 | 0 | 2737.243 | 0 | -0.07 | 0.154702 | 5.21 | 112.1 | 87.9 | 12.15 | 18.05 | | High | High | High | High | High | High | High | High | High | High | High | 0.002717 | 0.0292 | 2.1 |
| FALSE | High | FADQDDIG | [K].FADQD | 1xPhosphc | G5E8R4 1x | 1.39E-06 | 0 | 1 | 3 | 1 | G5E8R4 | SAPS doma | G5E8R4 [5 | 0 | 1907.837 | 0 | -0.61 | 0.045114 | 8.72 | 115.2 | 84.8 | 8.63 | 17.2 | | High | High | High | High | High | High | High | High | High | High | High | 0 | 1.46E-07 | 2.41 |
| FALSE | High | FADQHVP( | [K].FADQH | 1xPhosphc | Q8BTM8 1: | 0.001039 | 0 | 1 | 5 | 1 | Q8BTM8 | Filamin-A C | Q8BTM8 [2 | 0 | 2054.051 | 0 | 0.4 | 0.017754 | 6.5 | 86.3 | 113.7 | 12.68 | 4.76 | | High | High | High | High | High | High | High | High | High | High | High | 0 | 0.00025 | 2.58 |
| FALSE | High | FAELPEFRP | [R].FAELPE | 1xPhosphc | Q924A2 1x | 0.011549 | 0.00086 | 1 | 3 | 2 | Q924A2 | Protein cap | Q924A2 [2 | 0 | 3220.649 | 0 | -0.54 | 0.023241 | 11.63 | 118.4 | 81.6 | 11.38 | 10.19 | | High | High | High | High | High | High | High | High | High | High | High | 0.000571 | 0.005013 | 2.67 |
| FALSE | High | FAESPK | [R].FAESPK | 1xPhosphc | E9Q4F8 1x | 0.079114 | 0.00613 | 1 | 3 | 3 | E9Q4F8 | Ankyrin re | E9Q4F8 [1 | 0 | 1216.638 | 0 | -0.08 | 0.803747 | 19.34 | 101.8 | 98.2 | 7.4 | 8.48 | | High | High | High | High | High | High | High | High | High | High | High | 0.005149 | 0.05614 | 1.97 |
| FALSE | High | FAFSPLSEE | [R].FAFSPL | 1xPhosphc | A2AQJ6 1x | 2.31E-08 | 0 | 1 | 5 | 4 | A2AQJ6 | Transmem | A2AQJ6 [4 | 0 | 2438.105 | 0 | 0.49 | 0.002023 | 5.96 | 85 | 115 | 7.69 | 4.8 | | High | High | High | High | High | High | High | High | High | High | High | 0 | 1.47E-09 | 3.86 |
| FALSE | High | FAIQDISVE | [R].FAIQDI | 1xPhosphc | A0A1L1SV2 | 5.42E-06 | 0 | 2 | 13 | 2 | A0A1L1SV2 | Alpha-actir | A0A1L1SV2 | 0 | 2076.067 | 0 | 0.01 | 0.90967 | 11.11 | 99 | 101 | 10.44 | 14.71 | Shared | High | High | High | High | High | High | High | High | High | High | High | 0 | 6.82E-07 | 2.71 |
| FALSE | High | FAPPAESG! | [K].FAPPAE | 1xPhosphc | F2Z3U3 1xI | 0.001629 | 5.66E-05 | 1 | 1 | 2 | F2Z3U3 | Ras associa | F2Z3U3 [1 | 0 | 1712.866 | 0 | 0.98 | 0.000774 | 11.88 | 69.9 | 130.1 | 7.5 | 5.14 | | High | High | High | High | High | High | High | High | High | High | High | 2.58E-05 | 0.000414 | 2.55 |
| FALSE | High | FAQSAPGS | [R].FAQSAI | 1xPhosphc | Q3UCQ1 1: | 1.35E-05 | 0 | 1 | 2 | 1 | Q3UCQ1 | Forkhead k | Q3UCQ1 [4 | 0 | 2492.311 | 0 | 1.56 | 0.000756 | 9.96 | 51.9 | 148.1 | 8.84 | 7.61 | | High | High | High | High | High | High | High | High | High | High | High | 0 | 1.88E-06 | 4.09 |
| FALSE | High | FDGSETRPI | [K].FDGSET | 1xPhosphc | Q6PR54-3 | 0.000137 | 0 | 1 | 3 | 2 | Q6PR54-3 | Isoform 3 c | Q6PR54-3 | 0 | 2917.429 | 0 | -0.44 | 0.002933 | 3.48 | 114.6 | 85.4 | 8.03 | 3.05 | | High | High | High | High | High | High | High | High | High | High | High | 0 | 2.57E-05 | 2.32 |
| FALSE | High | FASENDLPI | [R].FASENI | 1xPhosphc | P34022 1xI | 8.63E-05 | 0 | 1 | 4 | 1 | P34022 | Ran-specifi | P34022 [5 | 0 | 1873.914 | 0 | 0 | 0.997457 | 10.64 | 99.4 | 100.6 | 4.87 | 8.03 | | High | High | High | High | High | High | High | High | High | High | High | 0 | 1.52E-05 | 3.36 |
| FALSE | High | FATLSLHDI | [K].FATLSLI | 1xPhosphc | Q3UWE6 1 | 0.071071 | 0.005247 | 1 | 3 | 1 | Q3UWE6 | MCG14935 | Q3UWE6 [ | 0 | 1368.687 | 0 | 1.45 | 0.00747 | 6.35 | 54.2 | 145.8 | 24.77 | 24.63 | | High | High | High | High | High | High | High | High | High | High | High | 0.002703 | 0.02884 | 1.59 |
| FALSE | High | FCDSPTSDI | [R].FCDSPT | 1xCarbami | Q8BZ47 1x | 0.000168 | 0 | 1 | 2 | 2 | Q8BZ47 | Zinc finger | Q8BZ47 [3 | 0 | 1766.733 | 0 | -0.4 | 0.004715 | 0.74 | 112.5 | 87.5 | 2.58 | 9.8 | | High | High | High | High | High | High | High | High | High | High | High | 0 | 4.24E-05 | 2.93 |
| FALSE | High | FCDSPTSDI | [R].FCDSPT | 1xCarbami | Q8BZ47 1x | 0.000475 | 0 | 1 | 2 | 1 | Q8BZ47 | Zinc finger | Q8BZ47 [3 | 0 | 1782.728 | 0 | -0.41 | 0.021844 | 2.15 | 114 | 86 | 4.52 | 9.22 | | High | High | High | High | High | High | High | High | High | High | High | 0 | 0.000104 | 2.88 |
| FALSE | High | FCDSPTSDI | [R].FCDSPT | 1xCarbami | Q8BZ47 2x | 5.31E-05 | 0 | 1 | 2 | 2 | Q8BZ47 | Zinc finger | Q8BZ47 [3 | 0 | 1862.694 | 0 | 0 | 0.404404 | 4.63 | 99.9 | 100.1 | 5.04 | 9.55 | | High | High | High | High | High | High | High | High | High | High | High | 0 | 0.000154 | 4.22 |
| FALSE | High | FCPSPSTSS | [R].FCPSPS | 1xCarbami | Q545E8 1x | 0.000209 | 0 | 1 | 2 | 3 | Q545E8 | Deoxycytid | Q545E8 [8 | 0 | 1721.74 | 0 | -0.09 | 0.019183 | 2.44 | 104.9 | 95.1 | 4.85 | 5.28 | | High | High | High | High | High | High | High | High | High | High | High | 0 | 4.12E-05 | 2.38 |
| FALSE | High | FCPSPSTSS | [R].FCPSPS | 1xCarbami | Q545E8 2x | 0.055777 | 0.004093 | 1 | 2 | 1 | Q545E8 | Deoxycytid | Q545E8 [8 | 0 | 1801.707 | 0 | -0.41 | 0.001361 | 3.04 | 112.5 | 87.5 | 4.56 | 3.06 | | High | High | High | High | High | High | High | High | High | High | High | 0.002117 | 0.02187 | 2.62 |
| FALSE | High | FCSPISSEL/ | [K].FCSPISS | 1xCarbami | Q9DBY0 1x | 4.17E-06 | 0 | 1 | 6 | 1 | Q9DBY0 | Forkhead k | Q9DBY0 [4 | 0 | 2595.236 | 0 | -0.46 | 0.000334 | 2.22 | 115.8 | 84.2 | 6.14 | 8.1 | | High | High | High | High | High | High | High | High | High | High | High | 0 | 5.05E-07 | 4.27 |
| FALSE | High | FCTGDSPPI | [R].FCTGDS | 1xCarbami | Q8C0J6 1xI | 5.16E-05 | 0 | 1 | 1 | 3 | Q8C0J6 | Ankyrin re | Q8C0J6 [7 | 0 | 1859.901 | 0 | 2.45 | 0.000247 | 1.18 | 30.8 | 169.2 | 5.8 | 6.25 | | High | High | High | High | High | High | High | High | High | High | High | 0 | 0.000103 | 3.28 |
| FALSE | High | FDDMSSPG | [R].FDDMS | 1xCarbami | E9Q481 1x | 7.28E-05 | 0 | 1 | 2 | 4 | E9Q481 | Serine/thre | E9Q481 [1 | 0 | 2365.029 | 0 | -0.21 | 0.147878 | 3.04 | 107.3 | 92.7 | 7.91 | 6.41 | | High | High | High | High | High | High | High | High | High | High | High | 0 | 1.25E-05 | 3.39 |
| FALSE | High | FDDSDTEQ | [K].FDDSD1 | 1xPhosphc | D3Z5N5 1x | 3.75E-07 | 0 | 1 | 1 | 4 | D3Z5N5 | 1-phospha | D3Z5N5 [4 | 0 | 2650.181 | 0 | -0.47 | 0.001421 | 3.51 | 116.3 | 83.7 | 2.05 | 6.04 | | High | High | High | High | High | High | High | High | High | High | High | 0 | 3.37E-08 | 4.22 |
| FALSE | High | FDDSDTEQ | [K].FDDSD1 | 1xPhosphc | Q9Z1T6 1x | 3.46E-07 | 0 | 1 | 1 | 7 | Q9Z1T6 | 1-phospha | Q9Z1T6 [4 | 0 | 3193.41 | 0 | 0 | 0.466265 | 5.43 | 96.6 | 103.4 | 6.94 | 5.07 | | High | High | High | High | High | High | High | High | High | High | High | 0 | 5.12E-07 | 4.06 |
| FALSE | High | FDDSDTEQ | [K].FDDSD1 | 2xPhosphc | Q9Z1T6 2x | 3.21E-06 | 0 | 1 | 1 | 3 | Q9Z1T6 | 1-phospha | Q9Z1T6 [4 | 0 | 3273.376 | 0 | -0.51 | 0.004941 | 4.19 | 117.2 | 82.8 | 5.87 | 10.86 | | High | High | High | High | High | High | High | High | High | High | High | 0 | 3.87E-06 | 3.04 |
| FALSE | High | FDFGEDSEI | [R].FDFGEI | 1xPhosphc | Q6DID3 1x | 0.004033 | 0.000311 | 1 | 2 | 1 | Q6DID3 | Protein SC/ | Q6DID3 [2 | 0 | 2180.943 | 0 | 0.71 | 0.000932 | 9.95 | 77.2 | 122.8 | 9.82 | 8.6 | | High | High | High | High | High | High | High | High | High | High | High | 0.00014 | 0.001134 | 1.94 |
| FALSE | High | FASGTFLSF | [K].FASGTF | 1xCarbami | A0A1B0GS | 4.06E-06 | 0 | 1 | 6 | 2 | A0A1B0GS | Nuclear po | A0A1B0GS | 0 | 2152.994 | 0 | -0.06 | 0.289616 | 14.46 | 103.5 | 96.5 | 5.79 | 8.92 | | High | High | High | High | High | High | High | High | High | High | High | 0 | 6.52E-06 | 3.03 |
| FALSE | High | FFDSEDEDI | [R].FFDSED | 1xPhosphc | A0A0R4J06 | 0.000233 | 0 | 1 | 3 | 3 | A0A0R4J06 | Cysteine pi | A0A0R4J06 | 0 | 2014.877 | 0 | -0.25 | 0.047093 | 2.85 | 109.4 | 90.6 | 5.12 | 11.09 | | High | High | High | High | High | High | High | High | High | High | High | 2.58E-05 | 0.000521 | 2.36 |
| FALSE | High | FFHHAEEEI | [R].FFHHAI | 1xPhosphc | E9Q9J4 1xI | 9.11E-08 | 0 | 1 | 4 | 4 | E9Q9J4 | Inositol he: | E9Q9J4 [24 | 0 | 2681.088 | 0 | 0.65 | 0.001441 | 6.26 | 78.3 | 121.7 | 5.08 | 4.85 | | High | High | High | High | High | High | High | High | High | High | High | 0 | 6.86E-09 | 4.11 |
| FALSE | High | FFLSSEDPP | [R].FFLSSEI | 1xPhosphc | Q5DU30 1x | 0.004824 | 0.000456 | 1 | 4 | 3 | Q5DU30 | Apoptotic | Q5DU30 [7 | 0 | 2488.266 | 0 | 1.37 | 0.001147 | 31.73 | 52.3 | 147.7 | 27.21 | 9.59 | | High | High | High | High | High | High | High | High | High | High | High | 0.000344 | 0.003126 | 2.66 |
| FALSE | High | FGTFGGLG | [K].FGTFG( | 1xPhosphc | E9Q616 1x | 0.019446 | 0.001438 | 1 | 4 | 1 | E9Q616 | AHNAK nu( | E9Q616 [5 | 0 | 1508.791 | 0 | 2.25 | 0.024714 | 30.09 | 34.1 | 165.9 | 15.07 | 44.11 | | High | High | High | High | High | High | High | High | High | High | High | 0.000756 | 0.006607 | 1.19 |
| FALSE | High | FGTPLSASK | [R].FGTPLS | 1xPhosphc | A0A0R4J0( | 0.00967 | 0.000736 | 1 | 5 | 1 | A0A0R4J0( | Disks large- | A0A0R4J0( | 0 | 1445.781 | 0 | -1.24 | 0.000304 | 0.9 | 139.2 | 60.8 | 5.63 | 3.08 | | High | High | High | High | High | High | High | High | High | High | High | 0.000344 | 0.003023 | 1.34 |
| FALSE | High | FGVPAEGR | [R].FGVPAI | 1xPhosphc | Q9JK92 1xI | 4.01E-05 | 0 | 1 | 1 | 1 | Q9JK92 | Heat shock | Q9JK92 [7 | 1 | 2590.326 | 0 | 1.95 | 0.002179 | 14.74 | 38.7 | 161.3 | 12.79 | 18.84 | | High | High | High | High | High | High | High | High | High | High | High | 0 | 6.45E-06 | 2.54 |
| FALSE | High | FHALSSPQ! | [K].FHALSS | 1xPhosphc | Q8C0T5 1x | 4.68E-05 | 0 | 1 | 3 | 5 | Q8C0T5 | Signal-indu | Q8C0T5 [1 | 0 | 2267.106 | 0 | 1.57 | 0.001591 | 14.75 | 50.3 | 149.7 | 3.44 | 13.43 | | High | High | High | High | High | High | High | High | High | High | High | 4.94E-05 | 0.000719 | 3.57 |
| FALSE | High | FHAQNLSC | [R].FHAQN | 1xPhosphc | P18762 1xI | 4.51E-05 | 0 | 1 | 3 | 1 | P18762 | Beta-2 adre | P18762 [24 | 0 | 1937.907 | 0 | 1.79 | 0.005133 | 7.69 | 45 | 155 | 14.9 | 25.95 | | High | High | High | High | High | High | High | High | High | High | High | 0 | 7.36E-06 | 2.96 |
| FALSE | High | FHDSEGDD | [K].FHDSE( | 1xPhosphc | Q8K019 1x | 3.79E-07 | 0 | 1 | 11 | 10 | Q8K019 | Bcl-2-assoc | Q8K019 [3 | 0 | 2253.866 | 0 | -0.3 | 0.006074 | 4.38 | 107.9 | 92.1 | 5.18 | 10.08 | | High | High | High | High | High | High | High | High | High | High | High | 0 | 3.4E-08 | 4.38 |
| FALSE | High | FHDTSSPLI | [K].FHDTSS | 1xPhosphc | Q60902 1x | 7.83E-09 | 0 | 1 | 5 | 3 | Q60902 | Epidermal | Q60902 [1 | 0 | 2768.27 | 0 | -0.4 | 0.151243 | 24.35 | 113.8 | 86.2 | 31.95 | 15.87 | | High | High | High | High | High | High | High | High | High | High | High | 0 | 4.39E-10 | 4.99 |
| FALSE | High | FHDTSSPLI | [K].FHDTSS | 1xPhosphc | Q60902 1x | 6.87E-07 | 0 | 1 | 5 | 3 | Q60902 | Epidermal | Q60902 [1 | 0 | 2784.265 | 0 | -0.32 | 0.513901 | 4.99 | 111 | 89 | 26.53 | 33.16 | | High | High | High | High | High | High | High | High | High | High | High | 0 | 1.18E-07 | 4.21 |
| FALSE | High | FHFPPLDSI | [R].FHFPPL | 1xPhosphc | Q3U1H6 1x | 0.003001 | 0.00016 | 1 | 3 | 1 | Q3U1H6 | Putative ur | Q3U1H6 [1 | 0 | 2400.134 | 0 | 0.62 | 0.001361 | 2.42 | 81.6 | 118.4 | 9.09 | 6.69 | | High | High | High | High | High | High | High | High | High | High | High | 7.19E-05 | 0.00082 | 3.33 |
| FALSE | High | FHHNSLEVI | [R].FHHNSI | 1xCarbami | D3Z5I1 2xP | 9.14E-06 | 0 | 1 | 5 | 2 | D3Z5I1 | Zinc finger | D3Z5I1 [25 | 0 | 3537.689 | 0 | 0.54 | 0.005085 | 5.24 | 85.5 | 114.5 | 5.44 | 7.76 | | High | High | High | High | High | High | High | High | High | High | High | 0 | 1.23E-06 | 4.68 |
| FALSE | High | FHLDQNM( | [R].FHLDQI | 1xPhosphc | Q6NXJ0 1x | 2.43E-07 | 0 | 1 | 1 | 2 | Q6NXJ0 | Protein W\ | Q6NXJ0 [2 | 0 | 2070.915 | 0 | 0.06 | 0.804643 | 19.83 | 101.8 | 98.2 | 11.9 | 10.9 | | High | High | High | High | High | High | High | High | High | High | High | 0 | 2.08E-08 | 3.36 |
| FALSE | High | FHSLPFSLT | [R].FHSLPF | 1xPhosphc | Q80TQ2-2 | 0.081752 | 0.006219 | 1 | 4 | 1 | Q80TQ2-2 | Isoform 1 c | Q80TQ2-2 | 0 | 1714.933 | 0 | 2.21 | 0.000391 | 3.9 | 34.7 | 165.3 | 6.44 | 8.94 | | High | High | High | High | High | High | High | High | High | High | High | 0.003229 | 0.03414 | 1.39 |
| FALSE | High | FHSPSTTW | [R].FHSPST | 1xPhosphc | E9PZM7 1x | 0.042503 | 0.003197 | 1 | 3 | 1 | E9PZM7 | SR-related | E9PZM7 [8 | 0 | 1697.789 | 0 | 0.46 | 0.974473 | 39.49 | 86.7 | 113.3 | 56.69 | 31.82 | | High | High | High | High | High | High | High | High | High | High | High | 0.001639 | 0.01593 | 2.31 |
| FALSE | High | FGSTGSTPI | [K].FGSTGS | 2xPhosphc | Q3U1I6 2xI | 0.005182 | 0.000551 | 1 | 6 | 1 | Q3U1I6 | Putative ur | Q3U1I6 [4 | 0 | 2092.919 | 0 | 0.28 | 0.030934 | 10.93 | 89.1 | 110.9 | 8.44 | 7.67 | | High | High | High | High | High | High | High | High | High | High | High | 0.000246 | 0.001502 | 3.19 |
| FALSE | High | FHSPSTTW | [R].FHSPST | 2xPhosphc | E9PZM7 2x | 0.045898 | 0.003408 | 1 | 3 | 1 | E9PZM7 | SR-related | E9PZM7 [8 | 0 | 1777.755 | 0 | -0.04 | 0.738776 | 24.92 | 101.5 | 98.5 | 42.85 | 31.32 | | High | High | High | High | High | High | High | High | High | High | High | 0.001774 | 0.01752 | 2.22 |
| FALSE | High | FHSPSTTW | [R].FHSPST | 2xPhosphc | E9PZM7 2x | 3.81E-06 | 0 | 1 | 3 | 3 | E9PZM7 | SR-related | E9PZM7 [8 | 0 | 2006.918 | 0 | 0.17 | 0.956884 | 13.88 | 97.5 | 102.5 | 9.25 | 22.04 | | High | High | High | High | High | High | High | High | High | High | High | 0.002187 | 0.02302 | 3 |
| FALSE | High | FLAAVAHSI | [R].FLAAV/ | 1xPhosphc | D3YVL0 1xI | 0.053473 | 0.00384 | 1 | 7 | 1 | D3YVL0 | Putative he | D3YVL0 [3 | 0 | 2975.709 | 0 | -0.78 | 0.003207 | 0.17 | 126.3 | 73.7 | 9.89 | 8.64 | | High | High | High | High | High | High | High | High | High | High | High | 0.001999 | 0.02087 | 1.83 |
| FALSE | High | FIAFSGEG( | [R].FIAFSGI | 1xPhosphc | P70362 1xI | 0.093879 | 0.007306 | 1 | 4 | 1 | P70362 | Ubiquitin r | P70362 [2 | 1 | 1978.056 | 0 | 1.53 | 0.006625 | 13.93 | 46.8 | 153.2 | 11.83 | 24.62 | | High | High | High | High | High | High | High | High | High | High | High | 0.003727 | 0.04014 | 1.52 |
| FALSE | High | FLAGDNGF | [R].FLAGDI | 1xPhosphc | Q3U3N0 1x | 0.000279 | 0 | 1 | 3 | 3 | Q3U3N0 | Mesoderm | Q3U3N0 [1 | 0 | 2988.388 | 0 | -0.91 | 0.001391 | 3.21 | 131.2 | 68.8 | 4.17 | 10.25 | | High | High | High | High | High | High | High | High | High | High | High | 0 | 0.000057 | 3.47 |
| FALSE | High | FLAGDNGF | [R].FLAGDI | 2xPhosphc | Q3U3N0 2x | 0.001986 | 5.66E-05 | 1 | 3 | 3 | Q3U3N0 | Mesoderm | Q3U3N0 [1 | 0 | 3068.354 | 0 | -0.6 | 0.014034 | 3.76 | 118 | 82 | 6.76 | 9.61 | | High | High | High | High | High | High | High | High | High | High | High | 0.000246 | 0.001457 | 2.89 |
| FALSE | High | FLAVSQNP | [K].FLAVSC | 1xPhosphc | E9PUQ5 1x | 5.45E-05 | 0 | 1 | 5 | 2 | E9PUQ5 | Golgin sub | E9PUQ5 [9 | 0 | 3681.764 | 0 | 1.4 | 0.00756 | 25.62 | 52 | 148 | 13.38 | 26.01 | | High | High | High | High | High | High | High | High | High | High | High | 0 | 9.08E-06 | 4.04 |
| FALSE | High | FICVTPTTC | [K].FICVTP | 2xCarbami | P42227-3 1 | 2.03E-06 | 0 | 2 | 8 | 1 | P42227-3; | Isoform De | P42227-3 [ | 0 | 2619.222 | 1 | 0.93 | 0.000247 | 4.39 | 69 | 131 | 8.31 | 6.29 | Shared | High | High | High | High | High | High | High | High | High | High | High | 0 | 2.24E-07 | 3.87 |
| FALSE | High | FICVTPTTC | [K].FICVTP | 2xCarbami | P42227-3 1 | 0.000257 | 0 | 2 | 8 | 4 | P42227-3; | Isoform De | P42227-3 [ | 0 | 2635.217 | 1 | 0.51 | 0.000555 | 5.4 | 81.9 | 118.1 | 3.71 | 3.34 | Shared | High | High | High | High | High | High | High | High | High | High | High | 0 | 5.2E-05 | 3.37 |
| FALSE | High | FICVTPTTC | [K].FICVTP | 2xCarbami | P42227-3 2 | 8.18E-05 | 0 | 2 | 8 | 2 | P42227-3; | Isoform De | P42227-3 [ | 0 | 2715.183 | 0 | 2.22 | 0.002501 | 50.19 | 35.4 | 164.6 | 25.83 | 13.86 | Shared | High | High | High | High | High | High | High | High | High | High | High | 0 | 1.55E-05 | 3.43 |
| FALSE | High | FLDFSDNSI | [K].FLDFSD | 1xPhosphc | E9Q777 1x | 0.051694 | 0.003745 | 1 | 3 | 1 | E9Q777 | A kinase (P | E9Q777 [9 | 0 | 1643.742 | 0 | 0.32 | 0.017374 | 13.25 | 91.2 | 108.8 | 10.47 | 12.87 | | High | High | High | High | High | High | High | High | High | High | High | 0.00196 | 0.02004 | 2.23 |
| FALSE | High | FLEENSSDF | [R].FLEENS | 1xPhosphc | Q8C8K1 1x | 2.34E-05 | 0 | 1 | 7 | 3 | Q8C8K1 | Ephrin typ( | Q8C8K1 [7 | 0 | 2470.179 | 0 | 0.97 | 0.00049 | 3.47 | 66.9 | 133.1 | 13.45 | 9.04 | | High | High | High | High | High | High | High | High | High | High | High | 0 | 1.15E-05 | 3.25 |
| FALSE | High | FLEESVSM! | [K].FLEESV | 1xPhosphc | Q9JKB1 1xI | 0.004411 | 0.000409 | 1 | 3 | 2 | Q9JKB1 | Ubiquitin c | Q9JKB1 [1 | 0 | 1864.824 | 0 | -1.03 | 0.004077 | 1.46 | 134.1 | 65.9 | 7.13 | 14.9 | | High | High | High | High | High | High | High | High | High | High | High | 0.000183 | 0.001254 | 1.69 |
| FALSE | High | FLEQSRSPS | [K].FLEQSR | 1xPhosphc | Q8CI71 1xI | 0.005666 | 0.00062 | 1 | 3 | 1 | Q8CI71 | Syndetin O | Q8CI71 [48 | 1 | 2087.094 | 0 | 0.51 | 0.001602 | 3.55 | 82.8 | 117.2 | 3.56 | 7.85 | | High | High | High | High | High | High | High | High | High | High | High | 0.000275 | 0.00167 | 1.01 |
| FALSE | High | FHSPSTTW | [R].FHSPST | 1xPhosphc | E9PZM7 1x | 3.87E-05 | 0 | 1 | 3 | 1 | E9PZM7 | SR-related | E9PZM7 [8 | 0 | 1926.952 | 0 | 0.11 | 0.79523 | 22.16 | 103.1 | 96.9 | 15.91 | 19.4 | | High | High | High | High | High | High | High | High | High | High | High | 0 | 6.16E-06 | 2.09 |
| FALSE | High | FGSNINLE/ | [K].FGSNIN | 1xPhosphc | Q3UWL8 1 | 0.0002 | 0 | 1 | 3 | 2 | Q3UWL8 | Prefoldin s | Q3UWL8 [ | 0 | 1604.704 | 0 | 0.03 | 0.944517 | 17.76 | 99.1 | 100.9 | 8.91 | 11.02 | | High | High | High | High | High | High | High | High | High | High | High | 0 | 3.91E-05 | 2.7 |

| | | | | | | | | | | | | | | | | | | | | | | | | | | | | | | | | | | | | | | |
|---|---|---|---|---|---|---|---|---|---|---|---|---|---|---|---|---|---|---|---|---|---|---|---|---|---|---|---|---|---|---|---|---|---|---|---|---|---|---|
| FALSE | High | FGSADNIPI | [K].FGSADI | 1xPhospho | F8WJ98 1x | 0.000167 | 0 | 1 | 6 | 1 | F8WJ98 | Transmem | F8WJ98 [4 | 0 | 1713.898 | 0 | 0.37 | 0.067639 | 3.35 | 88.5 | 111.5 | 8.45 | 8.22 | | High | High | High | High | High | High | High | High | High | High | High | 0 | 3.18E-05 | 2.56 |
| FALSE | High | FGSADNIPI | [K].FGSADI | 1xPhospho | F8WJ98 1x | 0.011966 | 0.000889 | 1 | 6 | 1 | F8WJ98 | Transmem | F8WJ98 [4 | 0 | 1484.735 | 0 | 0.36 | 0.765648 | 35.91 | 89.5 | 110.5 | 23.24 | 35.25 | | High | High | High | High | High | High | High | High | High | High | High | 0.000449 | 0.003824 | 3.01 |
| FALSE | High | FFTSSTSLTI | [R].FFTSST | 1xPhospho | Q8CEC0 1x | 0.086562 | 0.006435 | 1 | 7 | 1 | Q8CEC0 | Nuclear po | Q8CEC0 [1 | 0 | 1769.949 | 0 | 0.02 | 0.608912 | 33.21 | 99.8 | 100.2 | 8.45 | 22.27 | | High | High | High | High | High | High | High | High | High | High | High | 0.003317 | 0.03644 | 2.3 |
| FALSE | High | FGATAHLG | [R].FGATAI | 1xCarbami | F8WJA0 1x | 0.000359 | 0 | 1 | 5 | 1 | F8WJA0 | ATP-depen | F8WJA0 [3 | 0 | 1783.897 | 0 | -0.05 | 0.33997 | 3.25 | 101.5 | 98.5 | 5.11 | 6.41 | | High | High | High | High | High | High | High | High | High | High | High | 0 | 7.56E-05 | 2.15 |
| FALSE | High | FGDFLQHS | [K].FGDFLC | 1xPhospho | Q80WE4 1 | 0.001799 | 5.66E-05 | 1 | 4 | 1 | Q80WE4 | Kinesin-like | Q80WE4 [1 | 0 | 2256.183 | 0 | -0.03 | 0.129471 | 1.07 | 101.1 | 98.9 | 5.7 | 7.19 | | High | High | High | High | High | High | High | High | High | High | High | 2.58E-05 | 0.000462 | 2.33 |
| FALSE | High | FGDISEGEV | [R].FGDISE | 1xPhospho | E9Q1K1 1x | 1.1E-06 | 0 | 1 | 8 | 1 | E9Q1K1 | Cilia- and fl | E9Q1K1 [4 | 0 | 3031.329 | 0 | 0.38 | 0.023446 | 19.32 | 88.3 | 111.7 | 10.38 | 5.67 | | High | High | High | High | High | High | High | High | High | High | High | 0 | 1.13E-07 | 3.56 |
| FALSE | High | FGEAATME | [R].FGEAAT | 1xPhospho | Q3TMM5 1 | 0.002212 | 0.000109 | 1 | 1 | 1 | Q3TMM5 | Putative ur | Q3TMM5 [ | 0 | 3797.876 | 0 | 0.11 | 0.161462 | 10.4 | 96.6 | 103.4 | 13.41 | 15.28 | | High | High | High | High | High | High | High | High | High | High | High | 4.94E-05 | 0.000582 | 2.67 |
| FALSE | High | FGESEEVEI | [K].FGESEE | 1xPhospho | Q3TVM1 1 | 1.53E-09 | 0 | 1 | 2 | 14 | Q3TVM1 | Putative ur | Q3TVM1 [3 | 0 | 2912.232 | 0 | -0.26 | 0.035035 | 4.13 | 108 | 92 | 3.61 | 14.45 | | High | High | High | High | High | High | High | High | High | High | High | 0 | 6.99E-11 | 4.82 |
| FALSE | High | FGESEEVEI | [K].FGESEE | 1xDeamida | Q3TVM1 1 | 4.26E-07 | 0 | 1 | 2 | 3 | Q3TVM1 | Putative ur | Q3TVM1 [3 | 0 | 2913.216 | 0 | -0.32 | 0.014441 | 5.18 | 111 | 89 | 2.34 | 14.42 | | High | High | High | High | High | High | High | High | High | High | High | 0 | 3.9E-08 | 4.49 |
| FALSE | High | FGESEEVEI | [K].FGESEE | 1xPhospho | Q3TVM1 1 | 1.23E-08 | 0 | 1 | 2 | 18 | Q3TVM1 | Putative ur | Q3TVM1 [3 | 0 | 2928.227 | 0 | -0.27 | 0.010481 | 5.7 | 108.8 | 91.2 | 1.38 | 7.72 | | High | High | High | High | High | High | High | High | High | High | High | 0 | 7.25E-10 | 5.38 |
| FALSE | High | FGESTTGFI | [R].FGESTT | 1xPhospho | Q9ERU9 1x | 2.1E-05 | 0 | 1 | 1 | 1 | Q9ERU9 | E3 SUMO-p | Q9ERU9 [2 | 0 | 2006.967 | 0 | 0.3 | 0.061077 | 21.46 | 88.3 | 111.7 | 13.32 | 9.2 | | High | High | High | High | High | High | High | High | High | High | High | 0 | 3.1E-06 | 3.88 |
| FALSE | High | FGEYNSNIS | [R].FGEYNS | 1xPhospho | Q8R3N1 1x | 8.25E-05 | 0 | 1 | 4 | 1 | Q8R3N1 | Nucleolar p | Q8R3N1 [8 | 0 | 1822.81 | 0 | -0.19 | 0.115051 | 8.65 | 108.9 | 91.1 | 9.38 | 10.86 | | High | High | High | High | High | High | High | High | High | High | High | 0 | 1.45E-05 | 3.23 |
| FALSE | High | FGEYNSNIS | [R].FGEYNS | 1xPhospho | Q8R3N1 1x | 9.82E-06 | 0 | 1 | 4 | 2 | Q8R3N1 | Nucleolar p | Q8R3N1 [8 | 0 | 2051.973 | 0 | -0.18 | 0.055783 | 0.85 | 106.2 | 93.8 | 5.64 | 11.01 | | High | High | High | High | High | High | High | High | High | High | High | 0.001142 | 0.01183 | 3.2 |
| FALSE | High | FGEYNSNIS | [R].FGEYNS | 1xDeamida | Q8R3N1 1x | 0.00359 | 0.00021 | 1 | 4 | 2 | Q8R3N1 | Nucleolar p | Q8R3N1 [8 | 0 | 2052.957 | 0 | -0.15 | 0.647754 | 7.22 | 105.7 | 94.3 | 16.96 | 12.84 | | High | High | High | High | High | High | High | High | High | High | High | 9.42E-05 | 0.000996 | 3.14 |
| FALSE | High | FGGEHVPN | [R].FGGEH | 1xPhospho | Q8BTM8 1 | 0.000483 | 0 | 1 | 6 | 2 | Q8BTM8 | Filamin-A C | Q8BTM8 [1 | 0 | 3272.63 | 0 | 1.43 | 0.004299 | 2.26 | 46.8 | 153.2 | 11.54 | 18.51 | | High | High | High | High | High | High | High | High | High | High | High | 0 | 0.000244 | 4.52 |
| FALSE | High | FGGFVAWI | [K].FGGFV/ | 1xPhospho | B9EHW0 1 | 3.8E-05 | 0 | 1 | 2 | 1 | B9EHW0 | Protein ph | B9EHW0 [2 | 0 | 2252.096 | 0 | 1.37 | 0.008254 | 28.92 | 49.6 | 150.4 | 10.48 | 24.17 | | High | High | High | High | High | High | High | High | High | High | High | 0 | 6.08E-06 | 3.64 |
| FALSE | High | FGGFVAWI | [K].FGGFV/ | 1xDeamida | B9EHW0 1 | 0.010568 | 0.000772 | 1 | 2 | 2 | B9EHW0 | Protein ph | B9EHW0 [2 | 0 | 2253.08 | 0 | 0.99 | 0.010401 | 21.78 | 56.4 | 143.6 | 15.15 | 23.23 | | High | High | High | High | High | High | High | High | High | High | High | 0.000399 | 0.003524 | 3.34 |
| FALSE | High | FGGNPGGI | [R].FGGNP | 1xPhospho | Q921F2 1x | 0.001301 | 0 | 1 | 3 | 1 | Q921F2 | TAR DNA-b | Q921F2 [2 | 0 | 2035.897 | 0 | 0.21 | 0.319984 | 45.2 | 84.8 | 115.2 | 11.72 | 23.02 | | High | High | High | High | High | High | High | High | High | High | High | 0 | 0.000322 | 1.62 |
| FALSE | High | FGGSGSQV | [R].FGGSG | 1xPhospho | Q3TI61 1xP | 1.86E-05 | 0 | 1 | 8 | 4 | Q3TI61 | Putative ur | Q3TI61 [35 | 0 | 1476.668 | 0 | -0.39 | 0.00037 | 1.11 | 113.8 | 86.2 | 5.34 | 5.6 | | High | High | High | High | High | High | High | High | High | High | High | 0 | 3.58E-06 | 2.12 |
| FALSE | High | FGGTSLPPV | [K].FGGTSI | 1xPhospho | Q6P9R4 1x | 3.01E-05 | 0 | 1 | 1 | 2 | Q6P9R4 | Rho guanir | Q6P9R4 [9 | 0 | 3295.711 | 0 | 2.61 | 0.000994 | 25.09 | 31.7 | 168.3 | 23.61 | 11.66 | | High | High | High | High | High | High | High | High | High | High | High | 0 | 8.78E-06 | 3.37 |
| FALSE | High | FGLGQSGR | [R].FGLGQ | 1xPhospho | B2RY22 1xI | 0.010109 | 0.000736 | 2 | 18 | 1 | B2RY22; A( | Clasp1 pro | B2RY22 [72 | 0 | 1130.556 | 0 | -0.05 | 0.384303 | 22.52 | 100.5 | 99.5 | 10.34 | 18.62 | Shared | High | High | High | High | High | High | High | High | High | High | High | 0.000344 | 0.003189 | 0.66 |
| FALSE | High | FGLHGSSG | [K].FGLHG | 1xPhospho | Q3U1G4 1x | 0.070485 | 0.005214 | 1 | 2 | 1 | Q3U1G4 | Putative ur | Q3U1G4 [4 | 0 | 1427.745 | 0 | -0.26 | 0.048094 | 17.5 | 115.1 | 84.9 | 7.36 | 18.93 | | High | High | High | High | High | High | High | High | High | High | High | 0.00269 | 0.02871 | 1.93 |
| FALSE | High | FGIHVYQFI | [K].FGIHVY | 1xCarbami | B7ZNM7 1 | 7.14E-08 | 0 | 1 | 4 | 5 | B7ZNM7 | Sept5 prot | B7ZNM7 [2 | 0 | 3015.316 | 0 | 1.32 | 0.000209 | 2.79 | 57.8 | 142.2 | 8.02 | 2.53 | | High | High | High | High | High | High | High | High | High | High | High | 0 | 6.64E-08 | 4.74 |
| FALSE | High | FGIHVYQFI | [K].FGIHVY | 1xCarbami | B7ZNM7 1 | 0.004613 | 0.000456 | 1 | 4 | 1 | B7ZNM7 | Sept5 prot | B7ZNM7 [2 | 0 | 3016.3 | 0 | 1.46 | 0.001221 | 16.12 | 49.7 | 150.3 | 17.14 | 13.24 | | High | High | High | High | High | High | High | High | High | High | High | 0.000204 | 0.001328 | 3.4 |
| FALSE | High | FGINIPSPEI | [R].FGINIPS | 1xCarbami | Q69ZF8 1xI | 0.0012 | 0 | 1 | 1 | 1 | Q69ZF8 | E3 ubiquiti | Q69ZF8 [2( | 0 | 3118.568 | 0 | -0.01 | 0.779352 | 12.04 | 100.6 | 99.4 | 1.27 | 8.17 | | High | High | High | High | High | High | High | High | High | High | High | 0 | 0.000294 | 3.83 |
| FALSE | High | FGIVTSSAG | [R].FGIVTS | 1xPhospho | B2RXM7 1x | 0.018125 | 0.001282 | 2 | 3 | 2 | B2RXM7; E | MCG11369 | B2RXM7 [1 | 0 | 2410.179 | 0 | 0.1 | 0.160895 | 16.56 | 96.2 | 103.8 | 13.63 | 14.98 | Shared | High | High | High | High | High | High | High | High | High | High | High | 0.000847 | 0.008064 | 2.12 |
| FALSE | High | FGNVDADS | [K].FGNVD | 1xPhospho | B0V2P5 1xI | 5.33E-07 | 0 | 1 | 5 | 1 | B0V2P5 | DmX-like p | B0V2P5 [12 | 0 | 2810.365 | 0 | 1.74 | 0.000402 | 11.89 | 46 | 154 | 11.68 | 5.29 | | High | High | High | High | High | High | High | High | High | High | High | 0 | 5.01E-08 | 4.59 |
| FALSE | High | FGQAATMI | [R].FGQAA | 1xPhospho | Q3TF40 1x | 6.37E-06 | 0 | 1 | 4 | 1 | Q3TF40 | Putative ur | Q3TF40 [4 | 0 | 3780.897 | 0 | 0 | 0.700633 | 15.44 | 96.7 | 103.3 | 6.97 | 15.86 | | High | High | High | High | High | High | High | High | High | High | High | 0 | 8.14E-07 | 4.63 |
| FALSE | High | FGQAATMI | [R].FGQAA | 1xPhospho | Q3TF40 1x | 6.73E-06 | 0 | 1 | 4 | 3 | Q3TF40 | Putative ur | Q3TF40 [4 | 0 | 3796.892 | 0 | 0.2 | 0.030529 | 8.76 | 92.6 | 107.4 | 10.47 | 8.62 | | High | High | High | High | High | High | High | High | High | High | High | 0 | 1.17E-05 | 3.49 |
| FALSE | High | FGSADNIAI | [K].FGSADI | 1xPhospho | Q8C440 1x | 0.002004 | 5.66E-05 | 1 | 8 | 1 | Q8C440 | Putative ur | Q8C440 [2 | 0 | 1481.735 | 0 | 1.24 | 0.04423 | 7.99 | 58.6 | 141.4 | 11.71 | 32.75 | | High | High | High | High | High | High | High | High | High | High | High | 2.58E-05 | 0.000521 | 1.75 |
| FALSE | High | FGSADNIAI | [K].FGSADI | 1xPhospho | Q8C440 1x | 0.002895 | 0.00016 | 1 | 8 | 2 | Q8C440 | Putative ur | Q8C440 [2 | 0 | 1710.898 | 0 | 1.92 | 0.000534 | 10.04 | 42.9 | 157.1 | 6.96 | 8.82 | | High | High | High | High | High | High | High | High | High | High | High | 7.19E-05 | 0.000784 | 3.15 |
| FALSE | High | EYVNSTSEI | [R].EYVNST | 1xPhospho | Q9D5T0 1x | 0.000123 | 0 | 1 | 2 | 2 | Q9D5T0 | ATPase fan | Q9D5T0 [3 | 0 | 3449.544 | 0 | 1.27 | 0.001627 | 16.29 | 57.2 | 142.8 | 5.03 | 11.11 | | High | High | High | High | High | High | High | High | High | High | High | 0.001651 | 0.01615 | 2.27 |
| FALSE | High | EYSSELNAF | [R].EYSSEL | 1xPhospho | B2RY56 1xI | 0.000155 | 0 | 1 | 2 | 1 | B2RY56 | RNA-bindir | B2RY56 [2 | 0 | 2342.014 | 0 | -0.3 | 0.018734 | 7.37 | 115.4 | 84.6 | 8.48 | 12.73 | | High | High | High | High | High | High | High | High | High | High | High | 0 | 2.94E-05 | 3.19 |
| FALSE | High | EYQEFSSPE | [R].EYQEFS | 1xPhospho | P48972 1xI | 0.003857 | 0.000261 | 1 | 1 | 2 | P48972 | Myb-relate | P48972 [27 | 0 | 2343.12 | 0 | -1.33 | 0.000356 | 0.33 | 141.9 | 58.1 | 5.58 | 5.43 | | High | High | High | High | High | High | High | High | High | High | High | 0.000449 | 0.003943 | 2.75 |
| FALSE | High | EYPDQSPV | [K].EYPDQ | 1xPhospho | Q60855 1x | 0.000227 | 0 | 1 | 2 | 3 | Q60855 | Receptor-i | Q60855 [3 | 0 | 1640.788 | 0 | 0.97 | 0.000724 | 10.35 | 66.9 | 133.1 | 3.47 | 7.2 | | High | High | High | High | High | High | High | High | High | High | High | 0 | 4.53E-05 | 2.86 |
| FALSE | High | ETSQPSSSV | [K].ETSQPS | 1xPhospho | Q64512 1x | 2.24E-07 | 0 | 1 | 2 | 2 | Q64512 | Tyrosine-p | Q64512 [1 | 0 | 2791.267 | 1 | 2.36 | 0.001735 | 36.46 | 30.3 | 169.7 | 20.12 | 15.94 | | High | High | High | High | High | High | High | High | High | High | High | 0 | 1.89E-08 | 3.92 |
| FALSE | High | ETSSLQELE | [R].ETSSLQ | 1xPhospho | Q60902 1x | 0.012072 | 0.000889 | 1 | 5 | 2 | Q60902 | Epidermal | Q60902 [4 | 0 | 1900.967 | 0 | 0.55 | 0.042717 | 12.53 | 85.8 | 114.2 | 10.78 | 11.14 | | High | High | High | High | High | High | High | High | High | High | High | 0.000449 | 0.00388 | 2.59 |
| FALSE | High | ETSVEGPG | [K].ETSVEC | 1xPhospho | P51612 1xI | 8.1E-08 | 0 | 1 | 3 | 4 | P51612 | DNA repair | P51612 [33 | 0 | 3396.596 | 0 | -0.16 | 0.60881 | 12.71 | 107.5 | 92.5 | 10.63 | 15.73 | | High | High | High | High | High | High | High | High | High | High | High | 0 | 6.04E-09 | 5.88 |
| FALSE | High | ETTGLSPQ | [R].ETTGLS | 1xPhospho | Q8BGS3 1x | 6.87E-07 | 0 | 1 | 2 | 2 | Q8BGS3 | Zinc finger | Q8BGS3 [8 | 0 | 1897.967 | 0 | 2.56 | 0.000192 | 6.92 | 29.5 | 170.5 | 9.33 | 6.65 | | High | High | High | High | High | High | High | High | High | High | High | 0 | 6.64E-08 | 3.57 |
| FALSE | High | ETTGSESD( | [R].ETTGSE | 1xPhospho | Q8R550-3 | 0.027358 | 0.001676 | 1 | 6 | 2 | Q8R550-3 | Isoform 3 c | Q8R550-3 | 0 | 2082.912 | 0 | 2.84 | 0.001296 | 44.55 | 22.7 | 177.3 | 38.05 | 12.2 | | High | High | High | High | High | High | High | High | High | High | High | 0.00092 | 0.009692 | 2.99 |
| FALSE | High | ETTHAQSG | [R].ETTHA( | 1xPhospho | Q9CT10 1x | 3.68E-10 | 0 | 1 | 1 | 11 | Q9CT10 | Ran-bindin | Q9CT10 [2 | 0 | 2470.15 | 0 | 0.4 | 0.005126 | 5.71 | 85.7 | 114.3 | 4.29 | 4.9 | | High | High | High | High | High | High | High | High | High | High | High | 0 | 1.41E-11 | 4.31 |
| FALSE | High | ETTSDPSRT | [K].ETTSDF | 1xPhospho | E9Q0S6 1x | 0.0002 | 0 | 1 | 6 | 8 | E9Q0S6 | Tensin 1 O | E9Q0S6 [1 | 1 | 2986.436 | 0 | -0.55 | 0.011318 | 4.91 | 122.5 | 77.5 | 11.71 | 10.14 | | High | High | High | High | High | High | High | High | High | High | High | 0 | 0.000115 | 3.94 |
| FALSE | High | ETVELSPTG | [K].ETVELS | 1xPhospho | Q60848 1x | 0.029844 | 0.001857 | 1 | 2 | 1 | Q60848 | Lymphocyt | Q60848 [4 | 0 | 1851.998 | 0 | -0.54 | 0.001151 | 4.1 | 118.7 | 81.3 | 4.95 | 10.06 | | High | High | High | High | High | High | High | High | High | High | High | 0.001011 | 0.01072 | 1.11 |
| FALSE | High | ETVNSPAIY | [K].ETVNSI | 1xPhospho | Q9CZJ1 1xI | 0.020679 | 0.001472 | 1 | 2 | 1 | Q9CZJ1 | Probable U | Q9CZJ1 [23 | 0 | 1458.719 | 0 | -0.33 | 0.009215 | 0.79 | 110.7 | 89.3 | 8.18 | 6.27 | | High | High | High | High | High | High | High | High | High | High | High | 0.000786 | 0.007099 | 2.87 |
| FALSE | High | ETVSLTPSG | [R].ETVSLT | 1xPhospho | Q6PCQ0 1x | 1.57E-05 | 0 | 1 | 3 | 3 | Q6PCQ0 | IQ domain- | Q6PCQ0 [6 | 0 | 1995 | 0 | 1.26 | 0.000209 | 2.77 | 58.6 | 141.4 | 5.79 | 2.99 | | High | High | High | High | High | High | High | High | High | High | High | 0 | 2.25E-06 | 4.04 |
| FALSE | High | ETVVESQS | [R].ETVVES | 1xPhospho | E9PZM7 1x | 1.87E-07 | 0 | 1 | 3 | 5 | E9PZM7 | SR-related | E9PZM7 [8 | 0 | 2128.058 | 0 | 0.34 | 0.026629 | 21.14 | 83.3 | 116.7 | 6.91 | 9.08 | | High | High | High | High | High | High | High | High | High | High | High | 0 | 1.55E-08 | 4.38 |
| FALSE | High | ETVVESQS | [R].ETVVES | 2xPhospho | E9PZM7 2x | 1.79E-07 | 0 | 1 | 3 | 2 | E9PZM7 | SR-related | E9PZM7 [8 | 0 | 2208.024 | 0 | 0.09 | 0.506736 | 11.32 | 95.9 | 104.1 | 6.78 | 8.1 | | High | High | High | High | High | High | High | High | High | High | High | 0 | 1.48E-08 | 4.14 |
| FALSE | High | ETVVSGPL( | [R].ETVVS( | 1xPhospho | P70399 1xI | 9.6E-11 | 0 | 1 | 6 | 3 | P70399 | TP53-bindi | P70399 [14 | 0 | 2797.362 | 0 | 0.65 | 0.049257 | 22.95 | 78.4 | 121.6 | 8.33 | 17.01 | | High | High | High | High | High | High | High | High | High | High | High | 0 | 1.71E-07 | 5.92 |
| FALSE | High | ETSQGLAV | [R].ETSQGI | 1xCarbami | Q14B71 1x | 0.000389 | 0 | 1 | 4 | 1 | Q14B71 | Cell divisio | Q14B71 [2 | 0 | 3115.47 | 0 | -1.08 | 0.000816 | 2.6 | 131.4 | 68.6 | 12.9 | 14.47 | | High | High | High | High | High | High | High | High | High | High | High | 0 | 8.27E-05 | 5.25 |
| FALSE | High | ETVVSGPL( | [R].ETVVS( | 1xPhospho | P70399 1xI | 1.29E-06 | 0 | 1 | 6 | 1 | P70399 | TP53-bindi | P70399 [14 | 0 | 2813.357 | 0 | 0.45 | 0.081107 | 24.78 | 83.9 | 116.1 | 11.12 | 13.88 | | High | High | High | High | High | High | High | High | High | High | High | 0 | 1.36E-07 | 4.6 |
| FALSE | High | EVAGGAAA | [R].EVAGG | 1xPhospho | Q61321 1x | 6.47E-05 | 0 | 1 | 3 | 1 | Q61321 | Homeobox | Q61321 [3 | 0 | 3024.503 | 0 | -1.06 | 0.001414 | 1.38 | 135.6 | 64.4 | 6.01 | 11.86 | | High | High | High | High | High | High | High | High | High | High | High | 0 | 1.1E-05 | 4.58 |
| FALSE | High | EVCVSGGD | [K].EVCVS( | 1xCarbami | B2RRE0 1xI | 5.8E-08 | 0 | 1 | 3 | 3 | B2RRE0 | A kinase (P | B2RRE0 [4 | 0 | 2739.295 | 2 | -1.76 | 0.003589 | 1.37 | 156.1 | 43.9 | 20.35 | 9.3 | | High | High | High | High | High | High | High | High | High | High | High | 0 | 4.14E-09 | 4.63 |
| FALSE | High | EVDDILGE( | [K].EVDDIL | 1xPhospho | Q3U0S4 1x | 5.11E-08 | 0 | 1 | 4 | 5 | Q3U0S4 | Putative ur | Q3U0S4 [8 | 0 | 2374.095 | 0 | -0.56 | 0.003629 | 4.6 | 119 | 81 | 4.05 | 9.86 | | High | High | High | High | High | High | High | High | High | High | High | 0 | 3.58E-09 | 4.3 |
| FALSE | High | EVDDILGE( | [K].EVDDIL | 2xPhospho | Q3U0S4 2x | 3.94E-05 | 0 | 1 | 4 | 5 | Q3U0S4 | Putative ur | Q3U0S4 [8 | 0 | 2454.061 | 0 | -0.61 | 0.000273 | 1.94 | 121.7 | 78.3 | 3.73 | 2.8 | | High | High | High | High | High | High | High | High | High | High | High | 0 | 5.78E-05 | 3.59 |
| FALSE | High | EVDGLLTSI | [K].EVDGLI | 1xPhospho | A0A0U1RP | 7.29E-09 | 0 | 1 | 7 | 5 | A0A0U1RP | Ataxin-2-lik | A0A0U1RP | 0 | 2357.171 | 0 | 0.47 | 0.346713 | 20.54 | 83.6 | 116.4 | 12.49 | 28.72 | | High | High | High | High | High | High | High | High | High | High | High | 0 | 4.03E-10 | 4.21 |
| FALSE | High | EVDGLLTSI | [K].EVDGLI | 1xPhospho | A0A0U1RP | 2.37E-10 | 0 | 1 | 7 | 38 | A0A0U1RP | Ataxin-2-lik | A0A0U1RP | 0 | 2373.166 | 0 | 0.48 | 0.040272 | 3.42 | 84.3 | 115.7 | 7.19 | 10.66 | | High | High | High | High | High | High | High | High | High | High | High | 0 | 1.96E-09 | 4.15 |
| FALSE | High | EVDGLLTSI | [K].EVDGLI | 2xPhospho | A0A0U1RP | 0.001097 | 0 | 1 | 7 | 1 | A0A0U1RP | Ataxin-2-lik | A0A0U1RP | 0 | 2437.138 | 0 | 0.19 | 0.581853 | 33.98 | 95.6 | 104.4 | 10.27 | 33.29 | | High | High | High | High | High | High | High | High | High | High | High | 0 | 0.000265 | 4.55 |
| FALSE | High | EVDPSTGEI | [K].EVDPST | 1xPhospho | Q64337 1x | 2.32E-06 | 0 | 1 | 1 | 4 | Q64337 | Sequestoso | Q64337 [3 | 0 | 4064.93 | 0 | 2.04 | 0.004795 | 8.35 | 40.2 | 159.8 | 2.86 | 21.99 | | High | High | High | High | High | High | High | High | High | High | High | 0 | 2.99E-06 | 4.59 |
| FALSE | High | EVDPSTGEI | [K].EVDPST | 1xPhospho | Q64337 1x | 9.76E-07 | 0 | 1 | 1 | 21 | Q64337 | Sequestoso | Q64337 [3 | 0 | 4080.925 | 0 | 1.84 | 0.001558 | 20.08 | 44.9 | 155.1 | 15.47 | 13.74 | | High | High | High | High | High | High | High | High | High | High | High | 0 | 1.28E-06 | 5.98 |
| FALSE | High | EVDPSTGEI | [K].EVDPST | 2xPhospho | Q64337 2x | 6.95E-05 | 0 | 1 | 1 | 2 | Q64337 | Sequestoso | Q64337 [3 | 0 | 4144.896 | 0 | 1.91 | 0.010182 | 23.56 | 43.9 | 156.1 | 17.61 | 25.5 | | High | High | High | High | High | High | High | High | High | High | High | 0 | 0.000165 | 4.36 |
| FALSE | High | EVDSLGSIH | [R].EVDSL( | 1xPhospho | B2RY27 1xI | 0.015061 | 0.001049 | 1 | 4 | 2 | B2RY27 | MCG12578 | B2RY27 [3 | 0 | 2425.181 | 0 | 0.68 | 0.056091 | 8.6 | 76.9 | 123.1 | 13.37 | 19.8 | | High | High | High | High | High | High | High | High | High | High | High | 0.000571 | 0.004982 | 2.55 |
| FALSE | High | EVDTPGSV | [R].EVDTP( | 2xCarbami | Q6NV99 1x | 3.77E-05 | 0 | 1 | 5 | 2 | Q6NV99 | HAUS augn | Q6NV99 [4 | 0 | 4289.003 | 0 | 0.59 | 0.01502 | 18.1 | 84.1 | 115.9 | 16.67 | 11.54 | | High | High | High | High | High | High | High | High | High | High | High | 0 | 6E-06 | 4.25 |
| FALSE | High | EVDYSDSLT | [K].EVDYSI | 1xPhospho | A0A0R4J17 | 3.94E-05 | 0 | 1 | 8 | 7 | A0A0R4J17 | Transcripti | A0A0R4J17 | 0 | 1823.872 | 0 | -0.95 | 0.007825 | 5.48 | 129.5 | 70.5 | 7.11 | 16.2 | | High | High | High | High | High | High | High | High | High | High | High | 0 | 0.000145 | 3.02 |
| FALSE | High | ETVVSGPL( | [R].ETVVS( | 2xPhospho | P70399 2xI | 5.7E-05 | 0 | 1 | 6 | 2 | P70399 | TP53-bindi | P70399 [14 | 0 | 2877.328 | 0 | 0.79 | 0.027177 | 19.68 | 71.7 | 128.3 | 5.54 | 19.94 | | High | High | High | High | High | High | High | High | High | High | High | 0 | 1.15E-05 | 4.1 |
| FALSE | High | ETSQGLAV | [R].ETSQGI | 1xCarbami | Q14B71 1x | 3.33E-06 | 0 | 1 | 4 | 4 | Q14B71 | Cell divisio | Q14B71 [2 | 0 | 3114.486 | 0 | -0.86 | 0.000469 | 3.3 | 130.3 | 69.7 | 7.58 | 9.76 | | High | High | High | High | High | High | High | High | High | High | High | 0 | 1.87E-06 | 4.89 |
| FALSE | High | ETSLLAVPG | [K].ETSLLA | 1xPhospho | Q91Z31-2 1 | 9.02E-08 | 0 | 1 | 3 | 1 | Q91Z31-2 | Isoform 2 c | Q91Z31-2 [ | 0 | 2995.618 | 0 | -1.36 | 0.000241 | 1.38 | 143.9 | 56.1 | 5.72 | 14.46 | | High | High | High | High | High | High | High | High | High | High | High | 0 | 6.84E-09 | 3.88 |
| FALSE | High | ETSEELLPS | [K].ETSEEL | 1xPhospho | Q3TLI6 1xP | 0.0026 | 0.000109 | 1 | 2 | 1 | Q3TLI6 | Putative ur | Q3TLI6 [11 | 0 | 2268.178 | 0 | 0.67 | 0.003503 | 6.79 | 77.2 | 122.8 | 8.15 | 15.22 | | High | High | High | High | High | High | High | High | High | High | High | 4.94E-05 | 0.000699 | 2.19 |
| FALSE | High | ETDSSSAPA | [R].ETDSSS | 1xPhospho | Q9JMA1 1x | 0.021607 | 0.001507 | 1 | 3 | 1 | Q9JMA1 | Ubiquitin c | Q9JMA1 [2 | 0 | 1914.946 | 0 | 0.38 | 0.020522 | 11.61 | 91.3 | 108.7 | 9.62 | 6.7 | | High | High | High | High | High | High | High | High | High | High | High | 0.000801 | 0.007471 | 1.64 |
| FALSE | High | ETDVAEDD | [K].ETDVAI | 1xPhospho | H7BWY8 1 | 0.000524 | 0 | 1 | 1 | 1 | H7BWY8 | Pre-rRNA-p | H7BWY8 [1 | 0 | 2577.157 | 0 | 0.04 | 0.918194 | 10.29 | 97.6 | 102.4 | 6.67 | 17.08 | | High | High | High | High | High | High | High | High | High | High | High | 0 | 0.000115 | 3.21 |
| FALSE | High | ETDVAEDD | [K].ETDVAI | 1xPhospho | H7BWY8 1 | 1.75E-06 | 0 | 1 | 1 | 3 | H7BWY8 | Pre-rRNA-p | H7BWY8 [1 | 0 | 2593.152 | 0 | 0.26 | 0.040051 | 17.58 | 92.5 | 107.5 | 11.83 | 13 | | High | High | High | High | High | High | High | High | High | High | High | 0 | 3.03E-07 | 3.62 |
| FALSE | High | ETEHISSPW | [R].ETEHIS | 1xPhospho | Q8C0M0-3 | 0.002077 | 5.66E-05 | 1 | 4 | 2 | Q8C0M0-3 | Isoform 3 c | Q8C0M0-3 | 0 | 2379.07 | 0 | -0.35 | 0.030897 | 8.13 | 112.7 | 87.3 | 5.89 | 8.15 | | High | High | High | High | High | High | High | High | High | High | High | 2.58E-05 | 0.000541 | 2.4 |
| FALSE | High | ETENQGLD | [K].ETENQ | 1xPhospho | Q9R1C0 1x | 9.24E-10 | 0 | 1 | 2 | 2 | Q9R1C0 | Transcripti | Q9R1C0 [1 | 0 | 2223.99 | 0 | -1.04 | 0.001295 | 0.44 | 139.6 | 60.4 | 10.12 | 11.13 | | High | High | High | High | High | High | High | High | High | High | High | 0 | 3.95E-11 | 5.49 |
| FALSE | High | ETENQGLD | [K].ETENQ | 1xPhospho | Q9R1C0 1x | 4.08E-08 | 0 | 1 | 2 | 2 | Q9R1C0 | Transcripti | Q9R1C0 [1 | 0 | 2239.985 | 0 | -0.88 | 0.0103 | 1.55 | 126.5 | 73.5 | 9.79 | 11.98 | | High | High | High | High | High | High | High | High | High | High | High | 0 | 5.64E-06 | 4.22 |
| FALSE | High | ETENQGLD | [K].ETENQ | 2xPhospho | Q9R1C0 2x | 1.4E-05 | 0 | 1 | 2 | 1 | Q9R1C0 | Transcripti | Q9R1C0 [1 | 0 | 2303.957 | 0 | -1.12 | 0.000945 | 1.37 | 135.9 | 64.1 | 10.03 | 8.12 | | High | High | High | High | High | High | High | High | High | High | High | 0 | 1.96E-06 | 4.36 |
| FALSE | High | ETESAPGSI | [K].ETESAP | 1xPhospho | Q6ZQ58 1x | 0.007019 | 0.00062 | 1 | 3 | 1 | Q6ZQ58 | La-related | Q6ZQ58 [4 | 0 | 1339.609 | 0 | 0.94 | 0.00598 | 12.4 | 70.2 | 129.8 | 21.09 | 9.5 | | High | High | High | High | High | High | High | High | High | High | High | 0.000275 | 0.002119 | 1.43 |
| FALSE | High | ETFASDTNI | [R].ETFASD | 1xPhospho | Q9JJT9 1xP | 3.88E-06 | 0 | 1 | 2 | 2 | Q9JJT9 | Phosphory | Q9JJT9 [34 | 0 | 3148.461 | 0 | -0.87 | 0.000716 | 2.94 | 129.1 | 70.9 | 3.91 | 7.34 | | High | High | High | High | High | High | High | High | High | High | High | 0 | 4.66E-07 | 4.07 |
| FALSE | High | ETFLHRPEF | [R].ETFLHR | 1xCarbami | B0V2M3 1x | 0.031989 | 0.00205 | 1 | 5 | 1 | B0V2M3 | Zinc finger | B0V2M3 [2 | 0 | 2894.349 | 0 | -0.44 | 0.016428 | 9.26 | 112.2 | 87.8 | 8.31 | 8.66 | | High | High | High | High | High | High | High | High | High | High | High | 0.001126 | 0.01163 | 2.26 |
| FALSE | High | ETFPRTPPG | [R].ETFPRT | 1xPhospho | Q9D279 1x | 0.003312 | 0.00016 | 1 | 2 | 1 | Q9D279 | Mitotic inte | Q9D279 [1 | 1 | 2191.168 | 0 | 3.1 | 0.00054 | 19.34 | 22.2 | 177.8 | 16.33 | 11.12 | | High | High | High | High | High | High | High | High | High | High | High | 7.19E-05 | 0.000911 | 1.99 |
| FALSE | High | ETGSPPCFI | [R].ETGSPF | 1xCarbami | A2A7B5 1x | 6.02E-05 | 0 | 1 | 2 | 2 | A2A7B5 | PR domain | A2A7B5 [7 | 0 | 1967.886 | 0 | -0.77 | 0.000384 | 2.59 | 125.2 | 74.8 | 3.86 | 3.81 | | High | High | High | High | High | High | High | High | High | High | High | 0 | 1.01E-05 | 1.95 |
| FALSE | High | ETKPQSPK | [K].ETKPQS | 1xPhospho | D3YWH1 1 | 0.053023 | 0.00384 | 1 | 1 | 1 | D3YWH1 | Sorting ne | D3YWH1 [5 | 0 | 1681.949 | 0 | -0.18 | 0.004274 | 4.34 | 105.7 | 94.3 | 3.09 | 2.43 | | High | High | High | High | High | High | High | High | High | High | High | 0.001999 | 0.02054 | 1.31 |
| FALSE | High | ETIQGTGS\ | [K].ETIQGT | 1xPhospho | Q9EP97 1x | 2.46E-08 | 0 | 1 | 1 | 2 | Q9EP97 | Sentrin-spe | Q9EP97 [3 | 0 | 2795.288 | 0 | 0.08 | 0.100462 | 9.43 | 93.1 | 106.9 | 5.17 | 6.75 | | High | High | High | High | High | High | High | High | High | High | High | 0 | 9.88E-06 | 4.19 |
| FALSE | High | ETISAIDTSI | [R].ETISAIC | 1xPhospho | A0A172Q4 | 0.000244 | 0 | 1 | 3 | 1 | A0A172Q4 | Cysteine-k | A0A172Q4 | 0 | 1699.892 | 0 | 5.22 | 0.000606 | 13 | 5.8 | 194.2 | 24.29 | 13.6 | | High | High | High | High | High | High | High | High | High | High | High | 0 | 4.91E-05 | 2.96 |
| FALSE | High | ETNLESLPL | [R].ETNLES | 1xPhospho | P20152 1xI | 3.38E-08 | 0 | 1 | 7 | 9 | P20152 | Vimentin C | P20152 [42 | 0 | 2221.152 | 0 | 2.83 | 0.009458 | 17.06 | 24.8 | 175.2 | 8.12 | 33.28 | | High | High | High | High | High | High | High | High | High | High | High | 0 | 2.25E-09 | 5.03 |
| FALSE | High | ETNNNTDL | [K].ETNNN | 1xPhospho | Q3U821 1x | 2.45E-09 | 0 | 1 | 2 | 1 | Q3U821 | WD repeat | Q3U821 [7 | 0 | 2537.229 | 0 | -1.31 | 0.000363 | 1.12 | 142.4 | 57.6 | 6.27 | 3.34 | | High | High | High | High | High | High | High | High | High | High | High | 0 | 1.18E-10 | 3.61 |
| FALSE | High | ETPPGGNL | [K].ETPPG( | 1xPhospho | Q7TSC1 1x | 3.99E-06 | 0 | 1 | 3 | 1 | Q7TSC1 | Protein PRI | Q7TSC1 [9 | 0 | 1601.789 | 0 | 0.93 | 0.000664 | 3.23 | 69.4 | 130.6 | 6.4 | 6.05 | | High | High | High | High | High | High | High | High | High | High | High | 0 | 4.79E-07 | 4.34 |
| FALSE | High | ETPPSQDQ | [R].ETPPSC | 1xPhospho | Q5DTP8 1x | 0.000462 | 0 | 1 | 8 | 4 | Q5DTP8 | MKIAA407 | Q5DTP8 [9 | 0 | 3036.29 | 0 | 0.41 | 0.065156 | 25.06 | 85.9 | 114.1 | 15.22 | 20.6 | | High | High | High | High | High | High | High | High | High | High | High | 2.58E-05 | 0.000545 | 3.61 |
| FALSE | High | ETPSPEME | [K].ETPSPE | 1xPhospho | Q9QY06-2 | 0.000779 | 0 | 2 | 9 | 1 | Q9QY06-2; | Isoform 2 c | Q9QY06-2 | 0 | 1956.939 | 0 | -0.17 | 0.16138 | 6.57 | 107.6 | 92.4 | 14.45 | 9.06 | Shared | High | High | High | High | High | High | High | High | High | High | High | 0 | 0.000181 | 3.13 |
| FALSE | High | ETPSPEME | [K].ETPSPE | 1xPhospho | Q9QY06-2 | 1.63E-05 | 0 | 2 | 9 | 3 | Q9QY06-2; | Isoform 2 c | Q9QY06-2 | 0 | 1972.934 | 0 | -0.32 | 0.002016 | 3.75 | 110.9 | 89.1 | 7.54 | 11.28 | Shared | High | High | High | High | High | High | High | High | High | High | High | 0 | 2.33E-06 | 3.51 |
| FALSE | High | ETPSPEVSV | [K].ETPSPE | 1xPhospho | E9Q286 1x | 0.00359 | 0.00021 | 1 | 1 | 1 | E9Q286 | Little elong | E9Q286 [8 | 0 | 1682.799 | 0 | -1.15 | 0.003128 | 3.41 | 137.9 | 62.1 | 8.47 | 19.51 | | High | High | High | High | High | High | High | High | High | High | High | 9.42E-05 | 0.000999 | 1.9 |
| FALSE | High | ETPSQENG | [R].ETPSQE | 1xPhospho | Q61074 1x | 0.001255 | 0 | 1 | 3 | 2 | Q61074 | Protein ph | Q61074 [1 | 0 | 1796.883 | 0 | -0.21 | 0.178078 | 9.73 | 106.1 | 93.9 | 14.81 | 3.08 | | High | High | High | High | High | High | High | High | High | High | High | 0.008101 | 0.08746 | 2.22 |
| FALSE | High | ETQDSGSG | [R].ETQDS( | 1xPhospho | Q3UYK2 1x | 6.29E-08 | 0 | 1 | 2 | 8 | Q3UYK2 | Presenilin ( | Q3UYK2 [3 | 0 | 2710.074 | 0 | 0.66 | 0.00998 | 20.02 | 81.7 | 118.3 | 24.5 | 9.82 | | High | High | High | High | High | High | High | High | High | High | High | 0 | 4.54E-09 | 6.56 |
| FALSE | High | ETQPWSSP | [K].ETQPW | 1xPhospho | Q04690 1x | 0.000852 | 0 | 1 | 6 | 3 | Q04690 | Neurofibro | Q04690 [2 | 0 | 1597.803 | 0 | 1.42 | 0.000704 | 5.92 | 54.5 | 145.5 | 8.55 | 9.18 | | High | High | High | High | High | High | High | High | High | High | High | 0 | 0.000198 | 2.92 |
| FALSE | High | ETQQEPGE | [K].ETQQE | 1xPhospho | E9Q8T1 1x | 0.009168 | 0.000736 | 2 | 10 | 2 | E9Q8T1; Q | Transformi | E9Q8T1 [2 | 0 | 3086.461 | 0 | -1.14 | 0.000534 | 2.64 | 138 | 62 | 3.91 | 10.3 | Shared | High | High | High | High | High | High | High | High | High | High | High | 0.000327 | 0.002852 | 2.81 |
| FALSE | High | ETQSPADQ | [K].ETQSPA | 1xPhospho | O88466 1x | 3.32E-09 | 0 | 2 | 4 | 3 | O88466; R | Zinc finger | O88466 [1 | 0 | 2783.257 | 0 | -1.19 | 0.003497 | 2.72 | 135.4 | 64.6 | 14.34 | 6.06 | Shared | High | High | High | High | High | High | High | High | High | High | High | 0 | 1.67E-10 | 5.45 |
| FALSE | High | ETQVILLDT | [K].ETQVIL | 1xPhospho | Q9CZU4 1x | 8.86E-08 | 0 | 1 | 3 | 1 | Q9CZU4 | GTPase Era | Q9CZU4 [1 | 0 | 2361.345 | 0 | -1.93 | 0.001751 | 1.07 | 154.8 | 45.2 | 14.81 | 6.69 | | High | High | High | High | High | High | High | High | High | High | High | 0 | 6.7E-09 | 3.17 |
| FALSE | High | ETSATTLSP | [R].ETSATT | 1xPhospho | E9Q137 1x | 9.16E-07 | 0 | 1 | 4 | 3 | E9Q137 | Testis-expr | E9Q137 [2 | 0 | 1670.795 | 0 | 1.46 | 0.00085 | 2.3 | 53.9 | 146.1 | 7.5 | 10.92 | | High | High | High | High | High | High | High | High | High | High | High | 0 | 9.25E-08 | 2.76 |
| FALSE | High | EVEKPDSEI | [K].EVEKPD | 1xPhospho | B2RUG9 1x | 0.023379 | 0.001578 | 1 | 6 | 1 | B2RUG9 | Adenomat | B2RUG9 [1 | 0 | 1827.971 | 0 | 0.26 | 0.069384 | 8.19 | 90.9 | 109.1 | 7.34 | 9.31 | | High | High | High | High | High | High | High | High | High | High | High | 0.000847 | 0.008152 | 1.64 |
| FALSE | High | EPQTLDSQ | [R].EPQTLD | 1xPhospho | P27546 1xI | 5.88E-06 | 0 | 1 | 4 | 5 | P27546 | Microtubu | P27546 [11 | 0 | 1897.899 | 0 | 0.74 | 0.000715 | 4.62 | 73.6 | 126.4 | 8.35 | 3.67 | | High | High | High | High | High | High | High | High | High | High | High | 0 | 7.46E-07 | 3.38 |
| FALSE | High | EVEMSVDD | [R].EVEMS | 1xPhospho | Q8K298 1x | 0.096959 | 0.007777 | 1 | 1 | 2 | Q8K298 | Anillin OS= | Q8K298 [5 | 0 | 2033.857 | 0 | -0.07 | 0.532508 | 3.74 | 98 | 102 | 8.98 | 19.18 | | High | High | High | High | High | High | High | High | High | High | High | 0.003922 | 0.04174 | 2.71 |
| FALSE | High | EVETPEK | [K].EVETPE | 1xPhospho | B1ASU9 1x | 0.091627 | 0.007146 | 1 | 7 | 2 | B1ASU9 | Serine/thre | B1ASU9 [4 | 0 | 1369.702 | 0 | -0.45 | 0.005368 | 1.49 | 114.4 | 85.6 | 9.48 | 3.21 | | High | High | High | High | High | High | High | High | High | High | High | 0.004667 | 0.0503 | 2.05 |
| FALSE | High | EVTLDGSFI | [K].EVTLD( | 1xPhospho | Q80WT5 1 | 0.005666 | 0.00062 | 1 | 6 | 1 | Q80WT5 | Aftiphilin C | Q80WT5 [2 | 0 | 2078.912 | 0 | 0.6 | 0.019051 | 10.89 | 83.2 | 116.8 | 12.55 | 16.44 | | High | High | High | High | High | High | High | High | High | High | High | 0.000275 | 0.001657 | 2.25 |
| FALSE | High | EVTPEGLQ | [K].EVTPE( | 1xPhospho | Q9DCL9 1x | 0.003083 | 0.00016 | 1 | 1 | 1 | Q9DCL9 | Multifunct | Q9DCL9 [2 | 0 | 1784.927 | 0 | -0.46 | 0.009804 | 2.35 | 113.9 | 86.1 | 16.84 | 16.19 | | High | High | High | High | High | High | High | High | High | High | High | 7.19E-05 | 0.000842 | 2.76 |
| FALSE | High | EVTPVLVA( | [R].EVTPVL | 1xPhospho | A2A6Q5-3 | 4.98E-05 | 0 | 1 | 6 | 5 | A2A6Q5-3 | Isoform 3 c | A2A6Q5-3 | 0 | 4109.137 | 0 | 0.76 | 0.001682 | 14.5 | 73.4 | 126.6 | 8.73 | 5.85 | | High | High | High | High | High | High | High | High | High | High | High | 0 | 1.38E-05 | 4.47 |
| FALSE | High | EVVDTLSN | [K].EVVDTI | 1xPhospho | Q3UGE1 1x | 2.78E-06 | 0 | 1 | 7 | 2 | Q3UGE1 | Putative ur | Q3UGE1 [1 | 0 | 2316.008 | 0 | -0.31 | 0.310707 | 29.44 | 104.3 | 95.7 | 7.7 | 30.2 | | High | High | High | High | High | High | High | High | High | High | High | 0 | 3.2E-07 | 3.53 |
| FALSE | High | EVVKPVPIT | [K].EVVKP\ | 1xPhospho | Q8CI51 1xP | 5.79E-07 | 0 | 2 | 3 | 2 | Q8CI51; Q2 | PDZ and LII | Q8CI51 [10 | 0 | 2318.37 | 0 | 0.61 | 0.005469 | 8.47 | 79.8 | 120.2 | 7.89 | 7.27 | Shared | High | High | High | High | High | High | High | High | High | High | High | 0 | 5.52E-08 | 3.7 |
| FALSE | High | EVVVHEAS | [K].EVVVHI | 1xPhospho | Q5SS00 1xI | 4.53E-07 | 0 | 1 | 1 | 1 | Q5SS00 | DBF4-type | Q5SS00 [18 | 0 | 2567.363 | 0 | -1.84 | 0.000445 | 0.55 | 157.1 | 42.9 | 5.96 | 14.72 | | High | High | High | High | High | High | High | High | High | High | High | 0 | 4.18E-08 | 4.14 |
| FALSE | High | EVYFAER | [K].EVYFAE | 1xPhospho | A0A0G2JEI | 0.08447 | 0.006345 | 1 | 4 | 1 | A0A0G2JEI | Cysteine-ri | A0A0G2JEI | 0 | 1222.571 | 0 | 4.18 | 0.000361 | 30.45 | 10.6 | 189.4 | 18.99 | 8.62 | | High | High | High | High | High | High | High | High | High | High | High | 0.003279 | 0.0355 | 2.25 |
| FALSE | High | EVYQQQQ | [K].EVYQQ | 1xPhospho | Q99020 1x | 9.62E-05 | 0 | 1 | 6 | 2 | Q99020 | Heterogen | Q99020 [2 | 0 | 1808.817 | 0 | 0.05 | 0.99483 | 8.73 | 98.9 | 101.1 | 5.81 | 8 | | High | High | High | High | High | High | High | High | High | High | High | 0 | 1.72E-05 | 3.27 |
| FALSE | High | EVYSSDSES | [R].EVYSSD | 1xPhospho | E9QQ10 1x | 0.002313 | 0.000109 | 1 | 4 | 5 | E9QQ10 | A-kinase ar | E9QQ10 [2 | 0 | 2959.247 | 0 | -0.08 | 0.251967 | 13.41 | 106.4 | 93.6 | 13.27 | 9.24 | | High | High | High | High | High | High | High | High | High | High | High | 0.000275 | 0.00164 | 3.5 |
| FALSE | High | EWPSAESP | [K].EWPSA | 1xPhospho | H3BJQ9 1x | 1.73E-05 | 0 | 1 | 10 | 6 | H3BJQ9 | Homeobox | H3BJQ9 [8 | 0 | 2678.218 | 0 | 1.25 | 0.000921 | 19.52 | 62.8 | 137.2 | 26.31 | 10.55 | | High | High | High | High | High | High | High | High | High | High | High | 0 | 2.52E-06 | 4.24 |

| | | | | | | | | | | | | | | | | | | | | | | | | | | | | | | | | | | | | | | |
|---|---|---|---|---|---|---|---|---|---|---|---|---|---|---|---|---|---|---|---|---|---|---|---|---|---|---|---|---|---|---|---|---|---|---|---|---|---|---|
| FALSE | High | EWPVSSFN | [K].EWPVS | 1xPhosphc | Q9CR78 1x | 0.016451 | 0.001124 | 1 | 1 | 1 | Q9CR78 | Myb/SANT | Q9CR78 [2 | 0 | 2069.968 | 0 | -0.87 | 0.00127 | 2.49 | 129.4 | 70.6 | 5.98 | 7.02 | | High | High | High | High | High | High | High | High | High | High | High | 0.000585 | 0.005449 | 2.09 |
| FALSE | High | EWSLESSP/ | [K].EWSLE! | 1xPhosphc | V9GXM1 1 | 2.38E-06 | 0 | 1 | 8 | 1 | V9GXM1 | ADP-ribosy | V9GXM1 [: | 0 | 2620.285 | 0 | 0.5 | 0.005396 | 4.24 | 88.3 | 111.7 | 6.54 | 7.61 | | High | High | High | High | High | High | High | High | High | High | High | 0 | 2.69E-07 | 4.08 |
| FALSE | High | EWTPLGK | [K].EWTPL | 1xPhosphc | Q9Z1T1 1x | 0.041781 | 0.003126 | 1 | 3 | 1 | Q9Z1T1 | AP-3 comp | Q9Z1T1 [6 | 0 | 1368.733 | 0 | 0.09 | 0.521683 | 5.38 | 97 | 103 | 4.96 | 11.92 | | High | High | High | High | High | High | High | High | High | High | High | 0.001609 | 0.01569 | 1.95 |
| FALSE | High | EVTDITADS | [K].EVTDIT | 1xPhosphc | E9PUQ3 1x | 0.007673 | 0.000664 | 1 | 4 | 1 | E9PUQ3 | Expressed | E9PUQ3 [1 | 0 | 2312.002 | 0 | 0.3 | 0.013074 | 8 | 88.6 | 111.4 | 15.02 | 10.47 | | High | High | High | High | High | High | High | High | High | High | High | 0.000294 | 0.00233 | 3.14 |
| FALSE | High | EWYQSTIP | [R].EWYQS | 1xPhosphc | E9PVU7 1x | 0.000118 | 0 | 1 | 11 | 1 | E9PVU7 | Phosphodi | E9PVU7 [7 | 0 | 2826.246 | 0 | 0.25 | 0.058619 | 19.08 | 87.7 | 112.3 | 11.33 | 11.2 | | High | High | High | High | High | High | High | High | High | High | High | 0 | 2.17E-05 | 2.29 |
| FALSE | High | EYAENIGD | [K].EYAENI | 1xPhosphc | Q3UJX2 1x | 0.029076 | 0.001857 | 1 | 3 | 2 | Q3UJX2 | Putative ur | Q3UJX2 [5 | 1 | 2048.928 | 0 | -0.18 | 0.006243 | 4.92 | 106.2 | 93.8 | 1.73 | 5.69 | | High | High | High | High | High | High | High | High | High | High | High | 0.000997 | 0.01044 | 1.7 |
| FALSE | High | EYDFGTQL | [K].EYDFGT | 1xPhosphc | Q6PDX6 1x | 2.6E-05 | 0 | 1 | 2 | 2 | Q6PDX6 | E3 ubiquiti | Q6PDX6 [1 | 0 | 2351.157 | 0 | -1.38 | 0.001684 | 3.2 | 144.6 | 55.4 | 11.02 | 5.91 | | High | High | High | High | High | High | High | High | High | High | High | 2.58E-05 | 0.000448 | 2.56 |
| FALSE | High | EYDNEGTV | [K].EYDNE | 1xCarbami | E9QMD3 1 | 6.95E-08 | 0 | 1 | 5 | 1 | E9QMD3 | Zinc finger | E9QMD3 [ | 0 | 3773.641 | 0 | 3.65 | 0.003621 | 61.59 | 21.7 | 178.3 | 43.93 | 24.8 | | High | High | High | High | High | High | High | High | High | High | High | 0 | 5.1E-09 | 4.12 |
| FALSE | High | EYDQLYEE | [R].EYDQL | 1xPhosphc | O08908 1x | 0.000181 | 0 | 1 | 2 | 2 | O08908 | Phosphatic | O08908 [4 | 0 | 1817.783 | 0 | 0.53 | 0.079495 | 15.81 | 89.7 | 110.3 | 11.32 | 15.41 | | High | High | High | High | High | High | High | High | High | High | High | 0 | 3.51E-05 | 2.5 |
| FALSE | High | EYEVMYSP | [K].EYEVM | 1xCarbami | G3X9Q6 1x | 3.41E-05 | 0 | 1 | 5 | 3 | G3X9Q6 | La-related | G3X9Q6 [6 | 0 | 2060.854 | 0 | 0.43 | 0.051987 | 3.65 | 88.5 | 111.5 | 3.84 | 14.27 | | High | High | High | High | High | High | High | High | High | High | High | 0 | 5.34E-06 | 3.04 |
| FALSE | High | EYEVMYSP | [K].EYEVM | 1xCarbami | G3X9Q6 1x | 3.44E-05 | 0 | 1 | 5 | 6 | G3X9Q6 | La-related | G3X9Q6 [6 | 0 | 2076.849 | 0 | 0.43 | 0.000858 | 2.94 | 86.9 | 113.1 | 6.76 | 5.07 | | High | High | High | High | High | High | High | High | High | High | High | 0 | 5.4E-06 | 3.37 |
| FALSE | High | EYFDDSTEI | [K].EYFDDS | 1xPhosphc | Q9CZ91 1x | 0.000346 | 0 | 1 | 1 | 2 | Q9CZ91 | Serum resp | Q9CZ91 [2 | 0 | 1599.641 | 0 | -0.47 | 0.004685 | 2.01 | 117.2 | 82.8 | 3.98 | 12.01 | | High | High | High | High | High | High | High | High | High | High | High | 0 | 7.25E-05 | 2.71 |
| FALSE | High | EYGSPLK | [R].EYGSPL | 1xPhosphc | P24788 1x | 0.056724 | 0.004124 | 1 | 4 | 4 | P24788 | Cyclin-dep | P24788 [57 | 0 | 1331.701 | 0 | -0.5 | 0.000537 | 0.67 | 118.8 | 81.2 | 4.65 | 3.64 | | High | High | High | High | High | High | High | High | High | High | High | 0.002117 | 0.0222 | 1.52 |
| FALSE | High | EYGSQEGK | [R].EYGSQ | 1xPhosphc | Q99KG3 1x | 0.068752 | 0.005042 | 2 | 5 | 1 | Q99KG3; C | RNA-bindir | Q99KG3 [4 | 0 | 1435.687 | 0 | -0.37 | 0.00611 | 1.35 | 112.5 | 87.5 | 1.59 | 7.39 | Shared | High | High | High | High | High | High | High | High | High | High | High | 0.002604 | 0.02784 | 1.16 |
| FALSE | High | EYGSTSSID | [R].EYGSTS | 1xPhosphc | Q8C0T5 1x | 0.000793 | 0 | 1 | 4 | 5 | Q8C0T5 | Signal-indu | Q8C0T5 [2 | 0 | 1624.787 | 0 | 1.79 | 0.001574 | 17.7 | 43.5 | 156.5 | 4.97 | 15 | | High | High | High | High | High | High | High | High | High | High | High | 4.94E-05 | 0.000582 | 2.68 |
| FALSE | High | EYGSTSSID | [R].EYGSTS | 2xPhosphc | Q8C0T5 2x | 0.043981 | 0.003228 | 1 | 4 | 1 | Q8C0T5 | Signal-indu | Q8C0T5 [2 | 0 | 1704.753 | 0 | 1.83 | 0.000158 | 1.38 | 42 | 158 | 9.95 | 4.44 | | High | High | High | High | High | High | High | High | High | High | High | 0.001665 | 0.0166 | 2.79 |
| FALSE | High | EYGSTSSID | [R].EYGSTS | 1xPhosphc | B9EK76 1x | 0.004572 | 0.000456 | 1 | 4 | 2 | B9EK76 | Signal-indu | B9EK76 [1 | 0 | 1423.63 | 0 | -0.82 | 0.001493 | 3.51 | 122.5 | 77.5 | 7.9 | 8.79 | | High | High | High | High | High | High | High | High | High | High | High | 0.000204 | 0.001314 | 1.5 |
| FALSE | High | EYINECDSD | [R].EYINEC | 1xCarbami | P70398 1x | 0.022182 | 0.001542 | 1 | 2 | 1 | P70398 | Probable u | P70398 [85 | 0 | 2167.848 | 0 | 0.36 | 0.146637 | 36.16 | 87.4 | 112.6 | 31.41 | 9.79 | | High | High | High | High | High | High | High | High | High | High | High | 0.000814 | 0.007635 | 2.41 |
| FALSE | High | EWYQSTIP | [R].EWYQS | 2xPhosphc | E9PVU7 2x | 0.009931 | 0.000736 | 1 | 11 | 1 | E9PVU7 | Phosphodi | E9PVU7 [7 | 0 | 2906.212 | 0 | 0.13 | 0.067806 | 8.38 | 92.7 | 107.3 | 7.38 | 6.41 | | High | High | High | High | High | High | High | High | High | High | High | 0.000344 | 0.003116 | 2.75 |
| FALSE | High | EVSVSSVTE | [R].EVSVSS | 2xPhosphc | Q8CJF7 2x | 0.000286 | 0 | 1 | 1 | 5 | Q8CJF7 | Protein EL | Q8CJF7 [1 | 0 | 1908.901 | 0 | 0.07 | 0.705484 | 14.2 | 103.3 | 96.7 | 14.64 | 7.58 | | High | High | High | High | High | High | High | High | High | High | High | 0 | 5.84E-05 | 3.1 |
| FALSE | High | EVSVSSVTE | [R].EVSVSS | 1xPhosphc | Q8CJF7 1x | 2.38E-05 | 0 | 1 | 1 | 6 | Q8CJF7 | Protein EL | Q8CJF7 [1 | 0 | 1828.935 | 0 | -0.36 | 0.070069 | 15.96 | 112.7 | 87.3 | 12.43 | 9.71 | | High | High | High | High | High | High | High | High | High | High | High | 0 | 8.18E-06 | 3.44 |
| FALSE | High | EVSSRPSTF | [R].EVSSRP | 2xPhosphc | Q6PAR5 2x | 0.001058 | 0 | 1 | 8 | 5 | Q6PAR5 | GTPase-act | Q6PAR5 [7 | 0 | 3332.643 | 0 | 0.97 | 0.000485 | 11.8 | 68.2 | 131.8 | 8.67 | 4.45 | | High | High | High | High | High | High | High | High | High | High | High | 0 | 0.000319 | 3.82 |
| FALSE | High | EVGSSGLLS | [R].EVGSS( | 1xPhosphc | Q80U38 1x | 0.000142 | 0 | 1 | 2 | 1 | Q80U38 | Protein KH | Q80U38 [2 | 0 | 1969.995 | 0 | 0.7 | 0.033552 | 27.26 | 79.3 | 120.7 | 7.86 | 16.29 | | High | High | High | High | High | High | High | High | High | High | High | 0 | 2.65E-05 | 3.34 |
| FALSE | High | EVHDELED | [K].EVHDEI | 2xPhosphc | A0JNY3 2x | 0.079766 | 0.00613 | 1 | 2 | 1 | A0JNY3 | Gephyrin ( | A0JNY3 [17 | 0 | 3431.658 | 0 | 2.07 | 0.000181 | 9.71 | 39.4 | 160.6 | 8.26 | 4.83 | | High | High | High | High | High | High | High | High | High | High | High | 0.003178 | 0.03319 | 1.38 |
| FALSE | High | EVIATEDV/ | [K].EVIATE | 1xPhosphc | Q6ZQ32 1x | 2.45E-06 | 0 | 2 | 7 | 3 | Q6ZQ32; F | MKIAA085 | Q6ZQ32 [9 | 0 | 2907.419 | 0 | 1.91 | 0.003074 | 15.37 | 42.5 | 157.5 | 8.53 | 18.55 | Shared | High | High | High | High | High | High | High | High | High | High | High | 0 | 1.23E-05 | 4.37 |
| FALSE | High | EVIEASDK | [K].EVIEAS | 1xPhosphc | C7G3P2 1x | 0.001558 | 5.66E-05 | 1 | 3 | 4 | C7G3P2 | MKIAA053 | C7G3P2 [8 | 0 | 1428.739 | 0 | 0.08 | 0.101693 | 6.88 | 97.2 | 102.8 | 3.47 | 6.53 | | High | High | High | High | High | High | High | High | High | High | High | 0.000801 | 0.007457 | 2.27 |
| FALSE | High | EVIEIEDASI | [R].EVIEIED | 1xPhosphc | Q61687 1x | 2.26E-07 | 0 | 1 | 3 | 5 | Q61687 | Transcripti | Q61687 [1 | 0 | 1868.966 | 0 | -0.16 | 0.029835 | 5.52 | 105.2 | 94.8 | 2.99 | 8 | | High | High | High | High | High | High | High | High | High | High | High | 0 | 1.92E-08 | 3.87 |
| FALSE | High | EVLESILSP( | [K].EVLESIL | 1xPhosphc | Q8BQ30-2 | 3.58E-06 | 0 | 1 | 4 | 3 | Q8BQ30-2 | Isoform 2 ( | Q8BQ30-2 | 0 | 2236.151 | 0 | 0.97 | 0.000434 | 7.89 | 68.9 | 131.1 | 4.78 | 4.31 | | High | High | High | High | High | High | High | High | High | High | High | 0 | 4.27E-07 | 3.2 |
| FALSE | High | EVLFDGGG | [R].EVLFD( | 1xPhosphc | Q6PG95 1x | 1.11E-07 | 0 | 1 | 1 | 1 | Q6PG95 | Protein cra | Q6PG95 [1 | 0 | 2026.932 | 0 | -0.33 | 0.155355 | 4.61 | 111.9 | 88.1 | 15.56 | 4.57 | | High | High | High | High | High | High | High | High | High | High | High | 0 | 8.57E-09 | 4.17 |
| FALSE | High | EVLPASPR | [K].EVLPAS | 1xPhosphc | E9Q4F8 1x | 0.001933 | 5.66E-05 | 1 | 3 | 2 | E9Q4F8 | Ankyrin re | E9Q4F8 [1 | 0 | 1177.618 | 0 | -0.06 | 0.914003 | 2.52 | 99.8 | 100.2 | 2.55 | 6.17 | | High | High | High | High | High | High | High | High | High | High | High | 2.58E-05 | 0.000498 | 1.61 |
| FALSE | High | EVLPSTAPT | [K].EVLPST | 1xPhosphc | Q9CXH7 1x | 0.000109 | 0 | 1 | 1 | 7 | Q9CXH7 | Shugoshin | Q9CXH7 [4 | 0 | 2415.201 | 0 | 0.19 | 0.052933 | 14.84 | 92.4 | 107.6 | 7.57 | 6.07 | | High | High | High | High | High | High | High | High | High | High | High | 0.000246 | 0.001474 | 3.65 |
| FALSE | High | EVLTPTSTS | [R].EVLTPT | 1xPhosphc | F6WJB7 1x | 1.19E-05 | 0 | 1 | 4 | 5 | F6WJB7 | Ubiquitin c | F6WJB7 [3 | 0 | 1858.863 | 0 | 0.22 | 0.070924 | 16.38 | 92.4 | 107.6 | 5.13 | 5.78 | | High | High | High | High | High | High | High | High | High | High | High | 0 | 1.64E-06 | 3.42 |
| FALSE | High | EVLYDSEGI | [R].EVLYDS | 2xPhosphc | Q5U4C3 2x | 0.000488 | 0 | 1 | 1 | 3 | Q5U4C3 | Splicing fac | Q5U4C3 [6 | 0 | 1971.819 | 0 | -0.05 | 0.357166 | 10.41 | 103.3 | 96.7 | 7.86 | 10.8 | | High | High | High | High | High | High | High | High | High | High | High | 0 | 0.000106 | 3.19 |
| FALSE | High | EVMESAV( | [R].EVMES. | 1xPhosphc | Q8BIG4 1x | 0.001424 | 5.66E-05 | 1 | 2 | 2 | Q8BIG4 | F-box only | Q8BIG4 [3 | 1 | 2777.306 | 0 | 0.3 | 0.015907 | 5.4 | 88.8 | 111.2 | 1.99 | 9.07 | | High | High | High | High | High | High | High | High | High | High | High | 2.58E-05 | 0.000354 | 3.22 |
| FALSE | High | EVMESAV( | [R].EVMES. | 1xPhosphc | Q8BIG4 1x | 0.011755 | 0.000889 | 1 | 2 | 2 | Q8BIG4 | F-box only | Q8BIG4 [3 | 1 | 2793.301 | 0 | 0.25 | 0.005831 | 6.61 | 90.6 | 109.4 | 3.71 | 5.85 | | High | High | High | High | High | High | High | High | High | High | High | 0.000832 | 0.007953 | 2.42 |
| FALSE | High | EVNDSPK | [K].EVNDSI | 1xPhosphc | S4R2I9 1xP | 0.011755 | 0.000889 | 1 | 4 | 1 | S4R2I9 | Zinc finger | S4R2I9 [30 | 0 | 1326.671 | 0 | -1.22 | 0.00016 | 1.39 | 139.5 | 60.5 | 4.32 | 10.61 | | High | High | High | High | High | High | High | High | High | High | High | 0.000449 | 0.00377 | 1.87 |
| FALSE | High | EVPRSPGIE | [K].EVPRSF | 1xPhosphc | Q7TPM1 1: | 0.013426 | 0.000976 | 1 | 3 | 2 | Q7TPM1 | Protein PR | Q7TPM1 [5 | 1 | 1649.903 | 0 | -0.03 | 0.687129 | 19.27 | 102.2 | 97.8 | 4.83 | 11.44 | | High | High | High | High | High | High | High | High | High | High | High | 0.007639 | 0.08329 | 2.32 |
| FALSE | High | EVQNDLM | [R].EVQND | 1xPhosphc | Q3UP01 1x | 0.000738 | 0 | 1 | 5 | 2 | Q3UP01 | Putative ur | Q3UP01 [5 | 0 | 2731.26 | 0 | -0.11 | 0.177383 | 1.58 | 104 | 96 | 6.9 | 5.04 | | High | High | High | High | High | High | High | High | High | High | High | 9.42E-05 | 0.000975 | 2.86 |
| FALSE | High | EVQSPEQV | [K].EVQSP | 1xPhosphc | Q8K019 1x | 0.000257 | 0 | 1 | 9 | 8 | Q8K019 | Bcl-2-asso | Q8K019 [4 | 0 | 1581.829 | 0 | -0.39 | 0.006676 | 6.95 | 113.6 | 86.4 | 5.49 | 5.88 | | High | High | High | High | High | High | High | High | High | High | High | 2.58E-05 | 0.000404 | 3.42 |
| FALSE | High | EVRFSEEP/ | [R].EVRFSE | 1xPhosphc | Q921T2 1x | 0.037048 | 0.00249 | 1 | 5 | 1 | Q921T2 | Torsin-1A-i | Q921T2 [5 | 1 | 2465.122 | 0 | 0.48 | 0.013303 | 24.15 | 78 | 122 | 17.05 | 5.92 | | High | High | High | High | High | High | High | High | High | High | High | 0.001308 | 0.01363 | 2.45 |
| FALSE | High | EVSASQK | [R].EVSAS( | 1xPhosphc | B2RRE0 1x | 0.013666 | 0.001011 | 1 | 2 | 2 | B2RRE0 | A kinase (P | B2RRE0 [1 | 0 | 1286.676 | 0 | -1.52 | 0.000805 | 0.86 | 148.8 | 51.2 | 6.48 | 9.79 | | High | High | High | High | High | High | High | High | High | High | High | 0.006993 | 0.07555 | 1.74 |
| FALSE | High | EVSDDEAE | [K].EVSDDI | 1xPhosphc | Q3UIF3 1x | 2.77E-05 | 0 | 1 | 9 | 4 | Q3UIF3 | Putative ur | Q3UIF3 [22 | 0 | 1688.767 | 0 | -1.1 | 0.000154 | 2 | 137.6 | 62.4 | 4.67 | 10.36 | | High | High | High | High | High | High | High | High | High | High | High | 0 | 3.42E-05 | 3.68 |
| FALSE | High | EVSDSQAG | [R].EVSDS( | 1xPhosphc | Q8BLH7 1x | 2.64E-07 | 0 | 1 | 1 | 3 | Q8BLH7 | HIRA-intera | Q8BLH7 [3 | 0 | 1801.873 | 0 | -1.11 | 0.001145 | 1.56 | 137 | 63 | 9.47 | 6.32 | | High | High | High | High | High | High | High | High | High | High | High | 0 | 1.69E-05 | 4.05 |
| FALSE | High | EVSGHTVR | [R].EVSGH | 1xPhosphc | Q8CH25-2 | 0.008463 | 0.000736 | 1 | 2 | 1 | Q8CH25-2 | Isoform 2 ( | Q8CH25-2 | 0 | 1193.588 | 0 | -0.16 | 0.289093 | 6.41 | 102.5 | 97.5 | 9.6 | 8.14 | | High | High | High | High | High | High | High | High | High | High | High | 0.000327 | 0.002602 | 0.43 |
| FALSE | High | EVSPAPAT( | [R].EVSPAF | 1xPhosphc | A6H619 1x | 0.000302 | 0 | 1 | 5 | 1 | A6H619 | PHD and RI | A6H619 [1 | 0 | 1637.773 | 0 | 0.13 | 0.018639 | 2.39 | 95 | 105 | 2 | 6.48 | | High | High | High | High | High | High | High | High | High | High | High | 0 | 6.24E-05 | 2.39 |
| FALSE | High | EVSPAPVV/ | [R].EVSPAF | 1xPhosphc | G3X9J0 1x | 1.56E-06 | 0 | 2 | 10 | 2 | G3X9J0; Q | Signal-indu | G3X9J0 [13 | 0 | 1833.988 | 0 | 1.06 | 0.000261 | 5.07 | 67.3 | 132.7 | 4.34 | 4.22 | Shared | High | High | High | High | High | High | High | High | High | High | High | 0 | 1.02E-06 | 4.82 |
| FALSE | High | EVSPDPSTT | [K].EVSPDF | 1xPhosphc | O09100 1x | 3.04E-09 | 0 | 1 | 4 | 2 | O09100 | Endothelia | O09100 [1 | 0 | 2555.182 | 0 | 0.99 | 0.002351 | 24.78 | 67.1 | 132.9 | 13.8 | 11.4 | | High | High | High | High | High | High | High | High | High | High | High | 0 | 1.5E-10 | 5.58 |
| FALSE | High | EVSPEVVR | [R].EVSPEV | 1xPhosphc | E9Q784 1x | 0.080423 | 0.006219 | 1 | 3 | 3 | E9Q784 | Zinc finger | E9Q784 [1 | 0 | 1223.623 | 0 | -0.79 | 0.000489 | 1.43 | 125.2 | 74.8 | 3.03 | 5.85 | | High | High | High | High | High | High | High | High | High | High | High | 0.004707 | 0.05068 | 2.81 |
| FALSE | High | EVSPPGAR | [R].EVSPP( | 1xPhosphc | E9Q6J5-3 1 | 0.013666 | 0.001011 | 1 | 3 | 2 | E9Q6J5-3 | Isoform Ne | E9Q6J5-3 [ | 0 | 1121.555 | 0 | -0.12 | 0.169116 | 12.19 | 107 | 93 | 7.39 | 6.95 | | High | High | High | High | High | High | High | High | High | High | High | 0.000538 | 0.004678 | 1.84 |
| FALSE | High | EVSQGEPW | [K].EVSQG | 1xPhosphc | Q58FA4 1x | 0.101749 | 0.008194 | 1 | 1 | 1 | Q58FA4 | Transcripti | Q58FA4 [5 | 0 | 2195.115 | 0 | -0.26 | 0.375087 | 10.76 | 104.5 | 95.5 | 14.33 | 10.96 | | High | High | High | High | High | High | High | High | High | High | High | 0.004126 | 0.04403 | 2.69 |
| FALSE | High | EVSSRPSTF | [R].EVSSRP | 1xPhosphc | Q6PAR5 1x | 0.001349 | 0 | 1 | 8 | 2 | Q6PAR5 | GTPase-act | Q6PAR5 [7 | 0 | 3252.676 | 0 | 0.13 | 0.684462 | 12.08 | 98.9 | 101.1 | 14.55 | 11.13 | | High | High | High | High | High | High | High | High | High | High | High | 0 | 0.000333 | 3.14 |
| FALSE | High | EVENEQTP | [K].EVENE( | 1xPhosphc | B6ZHD0 1x | 2.59E-07 | 0 | 2 | 16 | 2 | B6ZHD0; Q | Erythrocyt | B6ZHD0 [2 | 0 | 2411.127 | 0 | 0.2 | 0.03685 | 8.5 | 92.9 | 107.1 | 7.85 | 7.6 | Shared | High | High | High | High | High | High | High | High | High | High | High | 0 | 2.22E-08 | 2.57 |
| FALSE | High | DGEEPQLT | [R].DGEEP( | 2xPhosphc | Q8CHC8 2x | 5.37E-06 | 0 | 1 | 1 | 1 | Q8CHC8 | Activity-de | Q8CHC8 [1 | 0 | 2567.12 | 0 | -0.52 | 0.005547 | 6.24 | 121.7 | 78.3 | 14.01 | 15.19 | | High | High | High | High | High | High | High | High | High | High | High | 0 | 6.73E-07 | 4.76 |
| FALSE | High | EPQSPSSQ | [K].EPQSPS | 1xCarbami | B7ZNK9 1x | 0.090153 | 0.00698 | 1 | 4 | 1 | B7ZNK9 | Spata5 pro | B7ZNK9 [2 | 0 | 2555.185 | 0 | -0.21 | 0.073771 | 9.4 | 106.3 | 93.7 | 10.32 | 5.66 | | High | High | High | High | High | High | High | High | High | High | High | 0.003606 | 0.03829 | 1.72 |
| FALSE | High | EPPVGTTST | [K].EPPVGT | 1xPhosphc | A2A3Z3 1x | 0.024001 | 0.001612 | 1 | 1 | 1 | A2A3Z3 | Zinc finger | A2A3Z3 [6 | 0 | 1991.025 | 0 | 0.95 | 0.000396 | 4.8 | 66.4 | 133.6 | 8.53 | 4.05 | | High | High | High | High | High | High | High | High | High | High | High | 0.000859 | 0.008412 | 2.07 |
| FALSE | High | EHQTSPSV | [K].EHQTSI | 1xCarbami | Q8BQ33 3x | 0.005872 | 0.00062 | 1 | 1 | 1 | Q8BQ33 | Treslin OS= | Q8BQ33 [1 | 1 | 3288.436 | 0 | -0.29 | 0.053378 | 5.97 | 111.6 | 88.4 | 7.87 | 13.2 | | High | High | High | High | High | High | High | High | High | High | High | 0.000275 | 0.001723 | 2.61 |
| FALSE | High | EHQTSPYH | [K].EHQTSI | 1xPhosphc | Q9DAZ9 1x | 0.007339 | 0.00062 | 1 | 1 | 1 | Q9DAZ9 | Abscission, | Q9DAZ9 [3 | 0 | 1560.716 | 0 | -0.41 | 0.098762 | 7.94 | 114.4 | 85.6 | 13.45 | 8.23 | | High | High | High | High | High | High | High | High | High | High | High | 0.000275 | 0.002217 | 2.12 |
| FALSE | High | EHRPALLPT | [K].EHRPAI | 1xPhosphc | Q9CXK9 1x | 5.35E-05 | 0 | 1 | 2 | 1 | Q9CXK9 | RNA-bindir | Q9CXK9 [3 | 0 | 2401.27 | 0 | 0.11 | 0.910712 | 16.88 | 98.8 | 101.2 | 17.54 | 7.43 | | High | High | High | High | High | High | High | High | High | High | High | 0 | 8.88E-06 | 3.1 |
| FALSE | High | EHRPSLEG( | [R].EHRPSL | 1xPhosphc | P40201 1x | 0.001158 | 0 | 1 | 1 | 2 | P40201 | Chromodo | P40201 [16 | 0 | 1760.946 | 0 | 1.32 | 0.001636 | 15.39 | 52.6 | 147.4 | 10.05 | 10.67 | | High | High | High | High | High | High | High | High | High | High | High | 9.42E-05 | 0.000961 | 2.1 |
| FALSE | High | EHSGLSPQ | [R].EHSGLS | 1xPhosphc | P70365 1x | 0.007673 | 0.000664 | 1 | 5 | 1 | P70365 | Nuclear re | P70365 [36 | 0 | 2339.017 | 0 | 0.56 | 0.003397 | 4.57 | 78.2 | 121.8 | 7.52 | 6.19 | | High | High | High | High | High | High | High | High | High | High | High | 0.000294 | 0.002334 | 3.37 |
| FALSE | High | EHSLASPR | [K].EHSLAS | 1xPhosphc | F8QZ85 1x | 0.040028 | 0.002867 | 1 | 2 | 1 | F8QZ85 | Ovum mut | F8QZ85 [1 | 0 | 1205.588 | 0 | 0.2 | 0.433819 | 5.95 | 93.1 | 106.9 | 9.94 | 8.44 | | High | High | High | High | High | High | High | High | High | High | High | 0.001484 | 0.0149 | 1.93 |
| FALSE | High | EHSPDSDT | [R].EHSPDS | 1xPhosphc | E9Q784 1x | 0.000422 | 0 | 1 | 2 | 1 | E9Q784 | Zinc finger | E9Q784 [9 | 0 | 2227.955 | 0 | -0.78 | 0.003332 | 2.56 | 127 | 73 | 7.52 | 10.21 | | High | High | High | High | High | High | High | High | High | High | High | 0 | 9.06E-05 | 2.96 |
| FALSE | High | EHSPSESEK | [R].EHSPSE | 1xPhosphc | B1AY10 1x | 0.007605 | 0.000664 | 1 | 3 | 1 | B1AY10 | Transcripti | B1AY10 [1 | 0 | 1567.741 | 0 | -0.2 | 0.510502 | 7.28 | 107.4 | 92.6 | 6.8 | 14.38 | | High | High | High | High | High | High | High | High | High | High | High | 0.000294 | 0.002308 | 1.48 |
| FALSE | High | EHSPYGPSI | [R].EHSPY( | 2xPhosphc | Q91ZV0 2x | 0.000365 | 0 | 1 | 10 | 1 | Q91ZV0 | Melanoma | Q91ZV0 [1 | 0 | 2353.058 | 0 | 0.11 | 0.923765 | 10.91 | 96.5 | 103.5 | 8.78 | 15.79 | | High | High | High | High | High | High | High | High | High | High | High | 0 | 7.7E-05 | 1.93 |
| FALSE | High | EHVPFHVS | [R].EHVPFH | 1xPhosphc | E9QMU3 1 | 1.08E-05 | 0 | 1 | 4 | 2 | E9QMU3 | Ensconsin | E9QMU3 [ | 0 | 1641.81 | 0 | -0.53 | 0.000747 | 1.39 | 116.1 | 83.9 | 2.43 | 5.67 | | High | High | High | High | High | High | High | High | High | High | High | 0 | 1.49E-06 | 2.47 |
| FALSE | High | EHYPVSSP | [K].EHYPV | 1xPhosphc | Q99NH0 1x | 3.76E-11 | 0 | 1 | 4 | 7 | Q99NH0 | Ankyrin re | Q99NH0 [2 | 0 | 2529.197 | 0 | -0.08 | 0.155811 | 10.2 | 102.4 | 97.6 | 4.69 | 9.48 | | High | High | High | High | High | High | High | High | High | High | High | 0 | 6.03E-05 | 3.86 |
| FALSE | High | EHYPVSSP | [K].EHYPV | 2xPhosphc | Q99NH0 2x | 1.08E-06 | 0 | 1 | 4 | 4 | Q99NH0 | Ankyrin re | Q99NH0 [2 | 0 | 2609.164 | 0 | 0.14 | 0.340459 | 15.74 | 97.1 | 102.9 | 5.38 | 9.49 | | High | High | High | High | High | High | High | High | High | High | High | 0 | 1.11E-07 | 4.8 |
| FALSE | High | EHYPVSSP | [K].EHYPV | 3xPhosphc | Q99NH0 3x | 0.037048 | 0.00249 | 1 | 4 | 3 | Q99NH0 | Ankyrin re | Q99NH0 [2 | 0 | 2689.13 | 0 | 0.37 | 0.012242 | 11.89 | 86.7 | 113.3 | 7.97 | 4.78 | | High | High | High | High | High | High | High | High | High | High | High | 0.001308 | 0.01372 | 3.02 |
| FALSE | High | EHQSQNAT | [K].EHQSQ | 1xPhosphc | Q9R207 1x | 0.082424 | 0.006288 | 1 | 2 | 1 | Q9R207 | Nibrin OS= | Q9R207 [5 | 0 | 1607.738 | 0 | 0.06 | 0.367911 | 27.74 | 95.3 | 104.7 | 13.16 | 17.06 | | High | High | High | High | High | High | High | High | High | High | High | 0.003243 | 0.03426 | 3.03 |
| FALSE | High | EKDECTPTI | [K].EKDECT | 1xCarbami | Q6NV83 1x | 0.014283 | 0.001011 | 1 | 3 | 1 | Q6NV83 | U2 snRNP- | Q6NV83 [9 | 2 | 2031.055 | 0 | -0.99 | 0.005665 | 5.32 | 132.7 | 67.3 | 12.95 | 5.24 | | High | High | High | High | High | High | High | High | High | High | High | 0.000538 | 0.004679 | 1.46 |
| FALSE | High | EKEEEETSF | [K].EKEEEE | 1xPhosphc | E9Q7D5 1x | 0.007881 | 0.000736 | 1 | 1 | 4 | E9Q7D5 | Rho guanir | E9Q7D5 [1 | 1 | 2285.095 | 0 | 3.13 | 0.000163 | 3.29 | 21.3 | 178.7 | 8.53 | 4.87 | | High | High | High | High | High | High | High | High | High | High | High | 0.005664 | 0.06238 | 2.65 |
| FALSE | High | EKEISDDEA | [R].EKEISD | 1xPhosphc | P11499 1x | 0.001097 | 0 | 1 | 1 | 1 | P11499 | Heat shock | P11499 [22 | 1 | 2318.126 | 0 | -0.78 | 0.197296 | 23.84 | 131.1 | 68.9 | 29.5 | 49.28 | | High | High | High | High | High | High | High | High | High | High | High | 0 | 0.000265 | 1.87 |
| FALSE | High | EKPDSDDD | [R].EKPDSI | 1xPhosphc | Q8R4U7 1x | 1.6E-08 | 0 | 2 | 4 | 5 | Q8R4U7; B | Leucine zip | Q8R4U7 [6 | 0 | 2791.389 | 0 | -0.34 | 0.011867 | 6.54 | 111.8 | 88.2 | 3.77 | 6.72 | Shared | High | High | High | High | High | High | High | High | High | High | High | 0 | 1.93E-07 | 4.29 |
| FALSE | High | EKPDSDDS | [K].EKPDSI | 1xPhosphc | Q8C1D8 1x | 7.06E-07 | 0 | 1 | 3 | 2 | Q8C1D8 | Protein IW | Q8C1D8 [2 | 0 | 2203.037 | 0 | 0.77 | 0.137056 | 28.93 | 75.1 | 124.9 | 8.75 | 41.04 | | High | High | High | High | High | High | High | High | High | High | High | 0 | 6.88E-08 | 2.64 |
| FALSE | High | EKPEGHGC | [R].EKPEGH | 1xPhosphc | Q5NCR9 1x | 0.020319 | 0.001472 | 1 | 1 | 2 | Q5NCR9 | Nuclear sp | Q5NCR9 [4 | 0 | 1878.911 | 0 | 0.05 | 0.639617 | 14.97 | 99.9 | 100.1 | 8.52 | 11.84 | | High | High | High | High | High | High | High | High | High | High | High | 0.000936 | 0.009771 | 1.23 |
| FALSE | High | EKPELAEPS | [K].EKPELA | 1xPhosphc | P83741-3 1 | 4.51E-05 | 0 | 1 | 8 | 2 | P83741-3 | Isoform 3 ( | P83741-3 [ | 0 | 3232.629 | 0 | 1.66 | 0.000181 | 18.28 | 45.2 | 154.8 | 20.03 | 6.59 | | High | High | High | High | High | High | High | High | High | High | High | 0 | 7.31E-06 | 3.57 |
| FALSE | High | EKPELAEPS | [K].EKPELA | 1xDeamida | P83741-3 1 | 3.04E-06 | 0 | 1 | 8 | 2 | P83741-3 | Isoform 3 ( | P83741-3 [ | 0 | 3233.613 | 0 | 1.23 | 0.001983 | 16.18 | 59.5 | 140.5 | 4.6 | 11.26 | | High | High | High | High | High | High | High | High | High | High | High | 0 | 3.54E-07 | 3.13 |
| FALSE | High | EKPQALVT | [R].EKPQAI | 1xPhosphc | O88271 1x | 0.001898 | 5.66E-05 | 1 | 1 | 2 | O88271 | Craniofacia | O88271 [1 | 0 | 2829.609 | 0 | -0.4 | 0.105025 | 26.78 | 122.8 | 77.2 | 20.58 | 35.6 | | High | High | High | High | High | High | High | High | High | High | High | 2.58E-05 | 0.000487 | 1.76 |
| FALSE | High | EKSPEIDDS | [K].EKSPEII | 1xPhosphc | Z4YKQ9 1x | 0.004333 | 0.00036 | 1 | 3 | 1 | Z4YKQ9 | ATPase fan | Z4YKQ9 [7 | 2 | 3537.854 | 0 | -0.66 | 0.016431 | 7.71 | 122.6 | 77.4 | 7.17 | 12.81 | | High | High | High | High | High | High | High | High | High | High | High | 0.000161 | 0.00123 | 2.71 |
| FALSE | High | EKSPELPEP | [K].EKSPEL | 1xPhosphc | Q52KI8 1x | 0.042503 | 0.003197 | 1 | 7 | 3 | Q52KI8 | Serine/argi | Q52KI8 [21 | 1 | 1906.009 | 0 | -0.4 | 0.008602 | 1.83 | 113.9 | 86.1 | 4.76 | 6.33 | | High | High | High | High | High | High | High | High | High | High | High | 0.002432 | 0.02623 | 1.87 |
| FALSE | High | EKVEEDTSI | [K].EKVEED | 1xPhosphc | Q9QWF0 1 | 0.107603 | 0.008634 | 1 | 2 | 1 | Q9QWF0 | Chromatin | Q9QWF0 [ | 1 | 1929.019 | 0 | -0.71 | 0.243908 | 13.21 | 124.1 | 75.9 | 34.45 | 14.28 | | High | High | High | High | High | High | High | High | High | High | High | 0.004341 | 0.04694 | 1.42 |
| FALSE | High | ELAEATDT( | [R].ELAEAT | 1xPhosphc | E9Q137 1x | 0.000104 | 0 | 1 | 4 | 1 | E9Q137 | Testis-expr | E9Q137 [2 | 0 | 3518.597 | 0 | 0.82 | 0.002821 | 9.05 | 72.2 | 127.8 | 8.66 | 12.47 | | High | High | High | High | High | High | High | High | High | High | High | 0 | 1.88E-05 | 4.89 |
| FALSE | High | ELAEDEPST | [R].ELAEDE | 1xPhosphc | Q3UMU9-3 | 0.003138 | 0.00016 | 1 | 4 | 3 | Q3UMU9-3 | Isoform 3 ( | Q3UMU9-3 | 1 | 2969.414 | 0 | 0.09 | 0.676132 | 13.61 | 95 | 105 | 11.47 | 11.34 | | High | High | High | High | High | High | High | High | High | High | High | 7.19E-05 | 0.000863 | 3.24 |
| FALSE | High | EKDSPHM( | [K].EKDSPH | 1xPhosphc | Q9DBV7 1x | 1.58E-05 | 0 | 1 | 8 | 1 | Q9DBV7 | Nfya prote | Q9DBV7 [3 | 1 | 3221.501 | 0 | -0.71 | 0.026866 | 16.28 | 118.9 | 81.1 | 11.66 | 21.6 | | High | High | High | High | High | High | High | High | High | High | High | 0 | 2.27E-06 | 4.12 |
| FALSE | High | ELAEDEPST | [R].ELAEDE | 1xDeamida | Q3UMU9-3 | 0.011755 | 0.000889 | 1 | 4 | 1 | Q3UMU9-3 | Isoform 3 ( | Q3UMU9-3 | 1 | 2970.398 | 0 | -0.68 | 0.002994 | 4.19 | 124.6 | 75.4 | 5.92 | 11.04 | | High | High | High | High | High | High | High | High | High | High | High | 0.000449 | 0.003759 | 2.17 |
| FALSE | High | EHQGYSPE | [R].EHQGY | 1xCarbami | A2A7F4 1x | 1.25E-06 | 0 | 1 | 2 | 4 | A2A7F4 | MCG12287 | A2A7F4 [1 | 0 | 2228.046 | 0 | -0.42 | 0.009748 | 6.95 | 114.9 | 85.1 | 4.44 | 8.93 | | High | High | High | High | High | High | High | High | High | High | High | 0 | 1.33E-06 | 3.18 |
| FALSE | High | EHLYHLESE | [K].EHLYHL | 1xPhosphc | Q8R038 1x | 9.08E-07 | 0 | 1 | 4 | 1 | Q8R038 | Melanocyt | Q8R038 [2 | 0 | 2120.083 | 0 | -0.31 | 0.006931 | 2.04 | 111.7 | 88.3 | 5.77 | 9.15 | | High | High | High | High | High | High | High | High | High | High | High | 0 | 9.08E-08 | 3.55 |
| FALSE | High | EGTQASEG | [K].EGTQA | 1xCarbami | Q9QXS6 1x | 1.25E-06 | 0 | 1 | 2 | 2 | Q9QXS6 | Drebrin OS | Q9QXS6 [6 | 0 | 3591.551 | 0 | -0.73 | 0.011939 | 10.88 | 124.8 | 75.2 | 10.57 | 12.05 | | High | High | High | High | High | High | High | High | High | High | High | 0 | 1.3E-07 | 4.13 |
| FALSE | High | EGTQASEG | [K].EGTQA | 1xCarbami | Q9QXS6 1x | 1.35E-08 | 0 | 1 | 2 | 3 | Q9QXS6 | Drebrin OS | Q9QXS6 [6 | 0 | 3592.535 | 0 | -0.84 | 0.043173 | 6.39 | 122.9 | 77.1 | 17.09 | 15.98 | | High | High | High | High | High | High | High | High | High | High | High | 0 | 1.49E-09 | 5.4 |
| FALSE | High | EGTQQASE | [K].EGTQQ | 1xCarbami | A0A0R4J1E | 1.1E-06 | 0 | 1 | 1 | 2 | A0A0R4J1E | Drebrin OS | A0A0R4J1E | 0 | 3719.61 | 0 | -1.51 | 0.020561 | 5.8 | 154.7 | 45.3 | 17.54 | 56.03 | | High | High | High | High | High | High | High | High | High | High | High | 0 | 4.64E-07 | 3.82 |
| FALSE | High | EGTSSSIVF | [R].EGTSSS | 1xCarbami | E9QQ30 1x | 0.002096 | 5.66E-05 | 1 | 3 | 1 | E9QQ30 | Eukaryotic | E9QQ30 [7 | 0 | 2557.153 | 0 | -0.22 | 0.112697 | 6.68 | 101.9 | 98.1 | 8.36 | 7.87 | | High | High | High | High | High | High | High | High | High | High | High | 2.58E-05 | 0.000546 | 3.26 |
| FALSE | High | EGTTQQLC | [R].EGTTQ | 1xPhosphc | Q6NZN0 1x | 0.000179 | 0 | 1 | 7 | 4 | Q6NZN0 | RNA-bindir | Q6NZN0 [6 | 0 | 1957.004 | 0 | -0.63 | 0.000289 | 0.63 | 122.9 | 77.1 | 2.71 | 4.55 | | High | High | High | High | High | High | High | High | High | High | High | 0 | 3.46E-05 | 3.46 |
| FALSE | High | EGVAPLLG | [K].EGVAPI | 1xPhosphc | B2RXC2 1x | 0.0095 | 0.000736 | 1 | 2 | 1 | B2RXC2 | Kinase OS= | B2RXC2 [2 | 0 | 1673.883 | 0 | 1.32 | 0.001095 | 14.34 | 53 | 147 | 11.27 | 8.25 | | High | High | High | High | High | High | High | High | High | High | High | 0.000327 | 0.002966 | 2.52 |
| FALSE | High | EGVGEETL | [K].EGVGEI | 1xPhosphc | B2RY19 1x | 0.013666 | 0.001011 | 1 | 7 | 2 | B2RY19 | Cgnl1 prot | B2RY19 [24 | 0 | 1482.704 | 0 | 1.4 | 0.000434 | 9.73 | 56 | 144 | 6.03 | 5.55 | | High | High | High | High | High | High | High | High | High | High | High | 0.000538 | 0.004461 | 1.47 |
| FALSE | High | EGVLGQVF | [K].EGVLG( | 1xPhosphc | Q7TSC1 1x | 1.26E-05 | 0 | 1 | 3 | 1 | Q7TSC1 | Protein PR | Q7TSC1 [1 | 0 | 2612.38 | 0 | 1.07 | 0.000924 | 10.27 | 60 | 140 | 16.34 | 10.94 | | High | High | High | High | High | High | High | High | High | High | High | 0 | 1.75E-06 | 3.23 |
| FALSE | High | EGVILTNEI | [K].EGVILT | 1xPhosphc | A0A1S6GW | 2.15E-05 | 0 | 1 | 4 | 2 | A0A1S6GW | Uncharacte | A0A1S6GW | 0 | 2593.187 | 0 | -0.44 | 0.682466 | 20.16 | 120 | 80 | 64.33 | 65.97 | | High | High | High | High | High | High | High | High | High | High | High | 0 | 3.18E-06 | 3.98 |
| FALSE | High | EGVILTNEI | [K].EGVILT | 1xPhosphc | A0A1S6GW | 2.42E-09 | 0 | 1 | 4 | 10 | A0A1S6GW | Uncharacte | A0A1S6GW | 0 | 2822.35 | 0 | -0.17 | 0.003285 | 1.72 | 105.5 | 94.5 | 1.22 | 2.92 | | High | High | High | High | High | High | High | High | High | High | High | 0 | 1.17E-10 | 4.53 |
| FALSE | High | EGVILTNEI | [K].EGVILT | 1xDeamida | A0A1S6GW | 3.25E-07 | 0 | 1 | 4 | 9 | A0A1S6GW | Uncharacte | A0A1S6GW | 0 | 2823.334 | 0 | -0.09 | 0.212745 | 3.14 | 101.4 | 98.6 | 8.02 | 4.26 | | High | High | High | High | High | High | High | High | High | High | High | 0 | 2.88E-08 | 4.42 |
| FALSE | High | EGVSSTTP/ | [R].EGVSST | 1xPhosphc | Q5U4C3 1x | 0.008388 | 0.000736 | 1 | 1 | 4 | Q5U4C3 | Splicing fac | Q5U4C3 [7 | 0 | 1514.787 | 0 | 0.2 | 0.185523 | 18.89 | 93.5 | 106.5 | 6.96 | 14.62 | | High | High | High | High | High | High | High | High | High | High | High | 0.000327 | 0.002566 | 2.58 |
| FALSE | High | EGVTEAGA | [K].EGVTE/ | 1xCarbami | G5E8V9 1x | 1.72E-08 | 0 | 1 | 2 | 4 | G5E8V9 | ADP-ribosy | G5E8V9 [5 | 0 | 2773.343 | 0 | 1.29 | 0.002264 | 13.3 | 58.2 | 141.8 | 4.87 | 11.07 | | High | High | High | High | High | High | High | High | High | High | High | 0 | 1.18E-09 | 4.36 |
| FALSE | High | EHQEQEPE | [R].EHQEQ | 1xPhosphc | Q8VCF0 1x | 0.004738 | 0.000456 | 1 | 2 | 1 | Q8VCF0 | Mitochond | Q8VCF0 [1 | 0 | 4390.171 | 0 | 4.56 | | 29.37 | 10.4 | 189.6 | 68.73 | 19.14 | | Not Found | High | High | High | High | High | High | High | High | High | High | 0.000204 | 0.001358 | 3.71 |
| FALSE | High | EGWTDSD | [R].EGWTD | 1xCarbami | P70371 1x | 0.056248 | 0.004124 | 1 | 3 | 3 | P70371 | Telomeric | P70371 [16 | 0 | 3812.668 | 0 | 0.03 | 0.738776 | 23.69 | 101.6 | 98.4 | 23.26 | 51.34 | | High | High | High | High | High | High | High | High | High | High | High | 0.005703 | 0.06284 | 4.15 |
| FALSE | High | EHANIDAQ | [R].EHANII | 1xPhosphc | Q8VDQ8 1: | 3.8E-05 | 0 | 1 | 5 | 1 | Q8VDQ8 | NAD-depe | Q8VDQ8 [3 | 0 | 2945.441 | 0 | 0.06 | 0.126406 | 0.82 | 97.3 | 102.7 | 6.89 | 4.84 | | High | High | High | High | High | High | High | High | High | High | High | 0 | 6.05E-06 | 3.36 |
| FALSE | High | EHANIDAQ | [R].EHANII | 1xPhosphc | Q8VDQ8 1: | 2.06E-05 | 0 | 1 | 5 | 1 | Q8VDQ8 | NAD-depe | Q8VDQ8 [3 | 1 | 3751.926 | 0 | 0.13 | 0.262885 | 16.09 | 93.7 | 106.3 | 12.44 | 11.15 | | High | High | High | High | High | High | High | High | High | High | High | 0 | 3.04E-06 | 2.93 |
| FALSE | High | EHATVEDE | [K].EHATVI | 1xPhosphc | Q69ZC8 1x | 0.000308 | 0 | 1 | 1 | 3 | Q69ZC8 | GPALPP mo | Q69ZC8 [1 | 1 | 3055.436 | 0 | -0.27 | 0.018401 | 2.26 | 109.6 | 90.4 | 5.88 | 6.2 | | High | High | High | High | High | High | High | High | High | High | High | 0 | 0.000211 | 3.45 |
| FALSE | High | EHCDSEGE | [K].EHCDSI | 1xCarbami | A0A0R4J17 | 6.46E-08 | 0 | 1 | 3 | 5 | A0A0R4J17 | Bromodom | A0A0R4J17 | 0 | 2301.974 | 0 | -0.88 | 0.000202 | 2.2 | 129.7 | 70.3 | 0.35 | 5.15 | | High | High | High | High | High | High | High | High | High | High | High | 0 | 4.68E-09 | 3.68 |
| FALSE | High | EHDSAEGS | [K].EHDSAI | 1xPhosphc | Q8CJG0 1x | 0.000103 | 0 | 1 | 4 | 1 | Q8CJG0 | Protein arg | Q8CJG0 [8 | 0 | 2064.857 | 0 | -1.66 | 0.000934 | 0.56 | 155.6 | 44.4 | 8.73 | 15.76 | | High | High | High | High | High | High | High | High | High | High | High | 0 | 1.86E-05 | 1.56 |
| FALSE | High | EHEADTAN | [K].EHEAD | 1xPhosphc | G3X8Y3 1x | 0.000205 | 0 | 1 | 4 | 1 | G3X8Y3 | N-alpha-ac | G3X8Y3 [5 | 0 | 1885.84 | 0 | 0.19 | 0.116092 | 12.42 | 93.1 | 106.9 | 10.59 | 9.33 | | High | High | High | High | High | High | High | High | High | High | High | 0 | 4.03E-05 | 3.51 |
| FALSE | High | EHEADTAN | [K].EHEAD | 1xPhosphc | G3X8Y3 1x | 1.55E-07 | 0 | 1 | 4 | 1 | G3X8Y3 | N-alpha-ac | G3X8Y3 [5 | 0 | 1901.835 | 0 | 0.7 | 0.035739 | 5.29 | 77.5 | 122.5 | 14 | 14.35 | | High | High | High | High | High | High | High | High | High | High | High | 0 | 1.26E-08 | 4.78 |
| FALSE | High | EHELVTGL | [R].EHELVT | 1xPhosphc | Q3ULZ2 1x | 1.81E-08 | 0 | 1 | 1 | 1 | Q3ULZ2 | FH2 domai | Q3ULZ2 [6 | 0 | 2480.284 | 0 | 0.7 | 0.000248 | 2.35 | 75.5 | 124.5 | 6.11 | 4.14 | | High | High | High | High | High | High | High | High | High | High | High | 0 | 1.12E-09 | 2.47 |

| | | | | | | | | | | | | | | | | | | | | | | | | | | | | | | | | | | | | | | |
|---|---|---|---|---|---|---|---|---|---|---|---|---|---|---|---|---|---|---|---|---|---|---|---|---|---|---|---|---|---|---|---|---|---|---|---|---|---|---|
| FALSE | High | EHFETPNPI | [K].EHFETF | 1xPhosphc | E9PVX6 1xI | 2.29E-05 | 0 | 1 | 2 | 1 | E9PVX6 | Proliferatic | E9PVX6 [97 | 0 | 1636.814 | 0 | 0.29 | 0.023759 | 9.92 | 90.2 | 109.8 | 4.87 | 7.4 | | High | High | High | High | High | High | High | High | High | High | High | 0 | 3.45E-06 | 2.34 |
| FALSE | High | EHHPEEGF | [K].EHHPEI | 1xCarbami | P70399 1xI | 1.61E-07 | 0 | 1 | 4 | 6 | P70399 | TP53-bindi | P70399 [64 | 0 | 3128.325 | 0 | 0.32 | 0.050373 | 17.74 | 86.8 | 113.2 | 8.28 | 14.9 | | High | High | High | High | High | High | High | High | High | High | High | 0 | 5.05E-08 | 4.47 |
| FALSE | High | EHHPEEGF | [K].EHHPEI | 1xCarbami | P70399 2xI | 0.000178 | 0 | 1 | 4 | 4 | P70399 | TP53-bindi | P70399 [64 | 0 | 3208.292 | 0 | 0.68 | 0.026798 | 15.56 | 75 | 125 | 10.15 | 19.01 | | High | High | High | High | High | High | High | High | High | High | High | 0 | 3.41E-05 | 3.96 |
| FALSE | High | EHLESANSI | [K].EHLESA | 1xPhosphc | Q7TPS5 1x | 0.000162 | 0 | 2 | 3 | 2 | Q7TPS5; A( | C2 domain | Q7TPS5 [84 | 0 | 2089.987 | 0 | 0.36 | 0.005547 | 10.01 | 85.7 | 114.3 | 5.31 | 6.66 | Shared | High | High | High | High | High | High | High | High | High | High | High | 0 | 0.000156 | 3.57 |
| FALSE | High | EHLVLGSDI | [R].EHLVLG | 1xPhosphc | P58659 1xI | 0.005 | 0.000504 | 1 | 5 | 1 | P58659 | Protein eva | P58659 [35 | 0 | 1535.823 | 0 | 3.27 | 0.000656 | 13.34 | 18.7 | 181.3 | 11.34 | 13.02 | | High | High | High | High | High | High | High | High | High | High | High | 0.000225 | 0.001443 | 3.21 |
| FALSE | High | EHAFDYVN | [K].EHAFDY | 1xPhosphc | Q99M03 1: | 1.63E-06 | 0 | 1 | 1 | 1 | Q99M03 | RWD doma | Q99M03 [1 | 0 | 2238.079 | 0 | -0.05 | 0.995746 | 5.93 | 97.1 | 102.9 | 8.4 | 5.68 | | High | High | High | High | High | High | High | High | High | High | High | 0 | 1.76E-07 | 3.09 |
| FALSE | High | ELAEQLGLS | [K].ELAEQL | 1xPhosphc | Q9Z1D1 1x | 0.012508 | 0.000889 | 1 | 2 | 1 | Q9Z1D1 | Eukaryotic | Q9Z1D1 [1 | 0 | 1913.003 | 0 | -0.22 | 0.033452 | 5.79 | 110.2 | 89.8 | 6.31 | 8.58 | | High | High | High | High | High | High | High | High | High | High | High | 0.000449 | 0.004037 | 2.38 |
| FALSE | High | ELAGDTGT | [K].ELAGDT | 1xPhosphc | Q8BRB7 1x | 0.001211 | 0 | 1 | 3 | 2 | Q8BRB7 | Histone ac | Q8BRB7 [1 | 0 | 2938.361 | 0 | -1.84 | 0.000314 | 0.74 | 155.9 | 44.1 | 5.79 | 12.16 | | High | High | High | High | High | High | High | High | High | High | High | 0 | 0.000297 | 3.75 |
| FALSE | High | EIAGTSASS | [R].EIAGTS | 1xPhosphc | Q14B19 1x | 2.12E-06 | 0 | 1 | 1 | 2 | Q14B19 | Ventral ant | Q14B19 [4 | 0 | 1599.758 | 0 | 3.52 | 0.000191 | 13.12 | 16.1 | 183.9 | 16.8 | 6.22 | | High | High | High | High | High | High | High | High | High | High | High | 0 | 2.37E-07 | 2.56 |
| FALSE | High | ELFASALSN | [K].ELFASA | 1xCarbami | Q9R0L6-2 | 4.89E-05 | 0 | 1 | 3 | 1 | Q9R0L6-2 | Isoform 2 c | Q9R0L6-2 | | 0 | 2904.328 | 0 | -0.18 | 0.246105 | 20.69 | 106.2 | 93.8 | 14.41 | 9.8 | | High | High | High | High | High | High | High | High | High | High | High | 0 | 8.04E-06 | 3.39 |
| FALSE | High | ELFDDPSY\ | [R].ELFDDF | 1xPhosphc | P98083 1xI | 6.41E-05 | 0 | 1 | 3 | 4 | P98083 | SHC-transf | P98083 [41 | 0 | 2448.21 | 0 | 1.2 | 0.002558 | 13.76 | 59.9 | 140.1 | 9.61 | 11.23 | | High | High | High | High | High | High | High | High | High | High | High | 0 | 8.44E-05 | 3.55 |
| FALSE | High | ELFDYSPPL | [K].ELFDYS | 1xPhosphc | Q8K019 1x | 0.08727 | 0.006537 | 1 | 9 | 2 | Q8K019 | Bcl-2-assoc | Q8K019 [5( | 0 | 1883.971 | 0 | -0.53 | 0.004914 | 4.79 | 115.9 | 84.1 | 4.45 | 7.11 | | High | High | High | High | High | High | High | High | High | High | High | 0.003633 | 0.0386 | 1.57 |
| FALSE | High | ELFGDDSEI | [K].ELFGDI | 1xPhosphc | Q5XJE5 1xI | 4.94E-05 | 0 | 1 | 2 | 2 | Q5XJE5 | RNA polym | Q5XJE5 [60 | 0 | 2937.176 | 0 | -0.5 | 0.140338 | 19.84 | 117.3 | 82.7 | 24.48 | 6.11 | | High | High | High | High | High | High | High | High | High | High | High | 0 | 8.16E-06 | 3.53 |
| FALSE | High | ELFIEEIQA( | [R].ELFIEEI | 1xPhosphc | H3BJQ9 1x | 0.001799 | 5.66E-05 | 1 | 12 | 3 | H3BJQ9 | Homeobox | H3BJQ9 [1 | 0 | 2757.293 | 0 | 1.71 | 0.001302 | 24.48 | 44.9 | 155.1 | 30.92 | 7.73 | | High | High | High | High | High | High | High | High | High | High | High | 2.58E-05 | 0.000461 | 4 |
| FALSE | High | ELFQHVHN | [R].ELFQH\ | 1xPhosphc | Q8BRH3 1x | 0.003927 | 0.000311 | 1 | 4 | 1 | Q8BRH3 | Rho GTPas | Q8BRH3 [3 | 0 | 2246.116 | 0 | -0.11 | 0.393337 | 6.17 | 98.8 | 101.2 | 7.09 | 8.54 | | High | High | High | High | High | High | High | High | High | High | High | 0.00014 | 0.001105 | 2.1 |
| FALSE | High | ELFSSHQW | [K].ELFSSH | 1xCarbami | Q3TZW9 1: | 3.52E-07 | 0 | 1 | 2 | 3 | Q3TZW9 | Tyrosine-p | Q3TZW9 [3 | 0 | 2891.239 | 0 | 1.24 | 0.00163 | 20.29 | 59.6 | 140.4 | 7.82 | 10.07 | | High | High | High | High | High | High | High | High | High | High | High | 9.42E-05 | 0.000937 | 4.47 |
| FALSE | High | ELGDSSGE( | [K].ELGDSS | 1xPhosphc | Q6PDQ2 1: | 0.028575 | 0.001829 | 1 | 4 | 1 | Q6PDQ2 | Chromodo | Q6PDQ2 [7 | 1 | 4168.865 | 0 | 0.16 | 0.398532 | 36.21 | 101.9 | 98.1 | 21.81 | 10.84 | | High | High | High | High | High | High | High | High | High | High | High | 0.000984 | 0.01018 | 3.91 |
| FALSE | High | ELGHCGGE | [K].ELGHC( | 1xCarbami | Q3TYD6 1x | 3.81E-06 | 0 | 1 | 1 | 2 | Q3TYD6 | Serine/thre | Q3TYD6 [1 | 0 | 3031.357 | 0 | -0.63 | 0.011397 | 6.05 | 123 | 77 | 10.26 | 13.14 | | High | High | High | High | High | High | High | High | High | High | High | 0 | 4.55E-07 | 3.12 |
| FALSE | High | ELGHCGGE | [K].ELGHC( | 1xCarbami | Q3TYD6 2x | 0.000205 | 0 | 1 | 1 | 1 | Q3TYD6 | Serine/thre | Q3TYD6 [1 | 0 | 3111.323 | 0 | -0.58 | 0.006596 | 4.68 | 121.7 | 78.3 | 9.04 | 8.3 | | High | High | High | High | High | High | High | High | High | High | High | 0 | 4.01E-05 | 4.53 |
| FALSE | High | ELGHEPVP/ | [K].ELGHEI | 1xPhosphc | P48972 1xI | 1.75E-05 | 0 | 1 | 1 | 2 | P48972 | Myb-relate | P48972 [25 | 1 | 3134.618 | 0 | -0.94 | 0.000209 | 1.11 | 132.8 | 67.2 | 4.03 | 2.28 | | High | High | High | High | High | High | High | High | High | High | High | 0 | 2.27E-05 | 3.3 |
| FALSE | High | EIGINEDQF | [K].EIGINEI | 1xCarbami | Q8C6E0 1x | 1.82E-09 | 0 | 1 | 3 | 1 | Q8C6E0 | Cilia- and fl | Q8C6E0 [7 | 0 | 2688.299 | 0 | -0.39 | 0.016327 | 10.18 | 114.6 | 85.4 | 11.28 | 8.38 | | High | High | High | High | High | High | High | High | High | High | High | 0 | 8.5E-11 | 4.22 |
| FALSE | High | ELGLVDDSI | [R].ELGLVD | 1xPhosphc | Q8VDL9 1x | 0.000501 | 0 | 1 | 2 | 1 | Q8VDL9 | Zinc finger | Q8VDL9 [3 | 0 | 3096.606 | 0 | 0.17 | 0.461527 | 13.16 | 99.5 | 100.5 | 14.32 | 17.13 | | High | High | High | High | High | High | High | High | High | High | High | 0 | 0.00011 | 2.63 |
| FALSE | High | EIESSPQYR | [R].EIESSP( | 1xPhosphc | A0A0U1RP | 0.095408 | 0.007437 | 1 | 7 | 2 | A0A0U1RP | Ataxin-2-lik | A0A0U1RP | 0 | 1417.656 | 0 | 0.13 | 0.034005 | 5.05 | 95.9 | 104.1 | 6 | 6.16 | | High | High | High | High | High | High | High | High | High | High | High | 0.005981 | 0.0647 | 2.16 |
| FALSE | High | ELGNIHETS | [R].ELGNIH | 1xPhosphc | A0A0R4J0( | 6.13E-05 | 0 | 1 | 6 | 2 | A0A0R4J0( | Disks large | A0A0R4J0( | 0 | 2335.158 | 0 | -1.94 | 0.002449 | 0.96 | 157.3 | 42.7 | 13.7 | 24.96 | | High | High | High | High | High | High | High | High | High | High | High | 0 | 1.04E-05 | 3.24 |
| FALSE | High | ELGPLPDDI | [K].ELGPLP | 1xPhosphc | F8VQD1 1x | 0.000173 | 0 | 1 | 13 | 1 | F8VQD1 | Protein pol | F8VQD1 [6 | 0 | 2138.013 | 0 | 0.22 | 0.456949 | 18.59 | 93.6 | 106.4 | 5.68 | 13.02 | | High | High | High | High | High | High | High | High | High | High | High | 0 | 3.32E-05 | 2.57 |
| FALSE | High | ELGPLPDDI | [K].ELGPLP | 1xPhosphc | F8VQD1 1x | 8.22E-07 | 0 | 1 | 13 | 2 | F8VQD1 | Protein pol | F8VQD1 [6 | 0 | 2154.008 | 0 | 0.21 | 0.128234 | 10.44 | 95 | 105 | 5.79 | 8.98 | | High | High | High | High | High | High | High | High | High | High | High | 0 | 8.12E-08 | 3.76 |
| FALSE | High | ELGSPYTGI | [R].ELGSPY | 1xPhosphc | Q5DTI8 1xI | 0.039347 | 0.002793 | 1 | 2 | 1 | Q5DTI8 | Extended s | Q5DTI8 [59 | 0 | 1885.971 | 0 | 1.72 | 0.008085 | 43.24 | 46.6 | 153.4 | 12.31 | 24.06 | | High | High | High | High | High | High | High | High | High | High | High | 0.001453 | 0.01459 | 2.06 |
| FALSE | High | EIGSSTSSSS | [K].EIGSST | 1xPhosphc | Q8C0V9 1x | 0.01029 | 0.000772 | 1 | 2 | 3 | Q8C0V9 | FERM dom | Q8C0V9 [5 | 0 | 1834.863 | 0 | 0.82 | 0.000338 | 1.66 | 72.9 | 127.1 | 2.94 | 3.27 | | High | High | High | High | High | High | High | High | High | High | High | 0.000382 | 0.003241 | 1.91 |
| FALSE | High | EIHDVAGD | [R].EIHDVA | 1xPhosphc | Q80U49 1x | 8.29E-10 | 0 | 1 | 3 | 1 | Q80U49 | Centrosom | Q80U49 [1 | 0 | 2189.007 | 0 | 1.4 | 0.000246 | 16.96 | 56.3 | 143.7 | 13.78 | 6.09 | | High | High | High | High | High | High | High | High | High | High | High | 0 | 3.49E-11 | 4.63 |
| FALSE | High | EIHDVAGD | [R].EIHDVA | 2xPhosphc | Q80U49 2x | 0.052133 | 0.00384 | 1 | 3 | 1 | Q80U49 | Centrosom | Q80U49 [1 | 0 | 2268.974 | 0 | -0.08 | 0.851285 | 15.17 | 103.8 | 96.2 | 14.34 | 11.16 | | High | High | High | High | High | High | High | High | High | High | High | 0.001999 | 0.02029 | 2.7 |
| FALSE | High | ELLAFQTPE | [K].ELLAFQ | 1xPhosphc | P70398 1xI | 0.010109 | 0.000736 | 1 | 2 | 1 | P70398 | Probable u | P70398 [14 | 0 | 1713.923 | 0 | -0.55 | 0.009994 | 6.53 | 120.6 | 79.4 | 8.99 | 11.75 | | High | High | High | High | High | High | High | High | High | High | High | 0.000344 | 0.00317 | 2.19 |
| FALSE | High | ELLALATPE | [R].ELLALA | 1xCarbami | Q9DC40 1x | 0.000852 | 0 | 2 | 3 | 1 | Q9DC40; C | Telomere l | Q9DC40 [4 | 0 | 2282.094 | 0 | -0.44 | 0.025984 | 7.37 | 114.2 | 85.8 | 12.83 | 12.69 | Shared | High | High | High | High | High | High | High | High | High | High | High | 0 | 0.000199 | 3.03 |
| FALSE | High | EILASPDR | [K].EILASPI | 1xPhosphc | B9EKJ7 1xF | 0.050826 | 0.003676 | 1 | 2 | 1 | B9EKJ7 | Tankyrase | B9EKJ7 [17 | 0 | 1209.608 | 0 | -0.07 | 0.322787 | 11.02 | 101.8 | 98.2 | 6.16 | 7.25 | | High | High | High | High | High | High | High | High | High | High | High | 0.001932 | 0.01965 | 1.3 |
| FALSE | High | ELLDSDNDI | [K].ELLDSD | 1xPhosphc | A2A654 1x | 0.003342 | 0.00016 | 1 | 4 | 1 | A2A654 | Bromodom | A2A654 [1 | 0 | 1586.772 | 0 | -0.27 | 0.002408 | 1.22 | 109.1 | 90.9 | 2.57 | 6.06 | | High | High | High | High | High | High | High | High | High | High | High | 7.19E-05 | 0.000921 | 2.61 |
| FALSE | High | ELLDSDNDI | [K].ELLDSD | 1xDeamida | A2A654 1x | 0.001558 | 5.66E-05 | 1 | 4 | 1 | A2A654 | Bromodom | A2A654 [1 | 0 | 1587.756 | 0 | -0.11 | 0.062866 | 8.77 | 100.4 | 99.6 | 10.56 | 8.6 | | High | High | High | High | High | High | High | High | High | High | High | 2.58E-05 | 0.000393 | 2.84 |
| FALSE | High | ELLEEGPQ\ | [R].ELLEEG | 1xPhosphc | P70399 1xI | 2.46E-07 | 0 | 1 | 6 | 12 | P70399 | TP53-bindi | P70399 [28 | 0 | 3857.826 | 0 | 0.75 | 0.05647 | 32.15 | 71.9 | 128.1 | 11.46 | 21.21 | | High | High | High | High | High | High | High | High | High | High | High | 0 | 3.19E-08 | 4.9 |
| FALSE | High | ELLEEGPQ\ | [R].ELLEEG | 2xPhosphc | P70399 2xI | 0.001848 | 5.66E-05 | 1 | 6 | 2 | P70399 | TP53-bindi | P70399 [28 | 0 | 3937.792 | 0 | 0.69 | 0.008673 | 6.77 | 74.3 | 125.7 | 7.18 | 12.79 | | High | High | High | High | High | High | High | High | High | High | High | 0.008718 | 0.09353 | 4.33 |
| FALSE | High | ELGNIHETS | [R].ELGNIH | 2xPhosphc | A0A0R4J0( | 0.001767 | 5.66E-05 | 1 | 6 | 1 | A0A0R4J0( | Disks large | A0A0R4J0( | 0 | 2415.125 | 0 | -0.66 | 0.003855 | 5.32 | 122.6 | 77.4 | 3.36 | 11.55 | | High | High | High | High | High | High | High | High | High | High | High | 2.58E-05 | 0.00045 | 1.6 |
| FALSE | High | ELESPLTPG | [K].ELESPL | 2xPhosphc | Q8BZ47 2x | 0.012288 | 0.000889 | 1 | 2 | 1 | Q8BZ47 | Zinc finger | Q8BZ47 [7 | 0 | 1688.831 | 0 | -0.5 | 0.006795 | 2.82 | 116.9 | 83.1 | 9.55 | 2.76 | | High | High | High | High | High | High | High | High | High | High | High | 0.000449 | 0.003945 | 2.94 |
| FALSE | High | ELESPLTPG | [K].ELESPL | 1xPhosphc | Q8BZ47 1x | 0.001349 | 0 | 1 | 2 | 2 | Q8BZ47 | Zinc finger | Q8BZ47 [7 | 0 | 1608.865 | 0 | -0.01 | 0.698105 | 4.35 | 100.4 | 99.6 | 3.38 | 0.58 | | High | High | High | High | High | High | High | High | High | High | High | 0.000984 | 0.01024 | 2.99 |
| FALSE | High | EIESEIDSEE | [K].EIESEID | 1xPhosphc | Q3UC10 1x | 0.025514 | 0.001645 | 1 | 4 | 1 | Q3UC10 | DNA helica | Q3UC10 [7 | 1 | 2672.389 | 0 | -0.22 | 0.086501 | 1.2 | 110.3 | 89.7 | 12.05 | 9.85 | | High | High | High | High | High | High | High | High | High | High | High | 0.000872 | 0.008949 | 2.48 |
| FALSE | High | ELAPDQVS | [K].ELAPD( | 1xPhosphc | A0A140LJJ | 0.003495 | 0.00021 | 1 | 3 | 3 | A0A140LJJ | A-kinase ar | A0A140LJJ | 0 | 3285.618 | 0 | -0.78 | 0.006842 | 4.86 | 124.7 | 75.3 | 13.87 | 3.44 | | High | High | High | High | High | High | High | High | High | High | High | 9.42E-05 | 0.000971 | 3.64 |
| FALSE | High | ELAQIAGRI | [K].ELAQIA | 1xPhosphc | Q61074 1x | 0.009087 | 0.000736 | 1 | 3 | 1 | Q61074 | Protein ph | Q61074 [1 | 0 | 2325.138 | 0 | -0.06 | 0.512612 | 13.56 | 102.4 | 97.6 | 6.87 | 9.91 | | High | High | High | High | High | High | High | High | High | High | High | 0.000327 | 0.002821 | 1.86 |
| FALSE | High | ELASPLALT | [R].ELASPL | 1xPhosphc | Q9QY81 1x | 0.000639 | 0 | 1 | 4 | 3 | Q9QY81 | Nuclear po | Q9QY81 [1 | 0 | 1476.802 | 0 | -0.04 | 0.573998 | 18.44 | 100.4 | 99.6 | 10.11 | 8.29 | | High | High | High | High | High | High | High | High | High | High | High | 0.000382 | 0.003245 | 2.45 |
| FALSE | High | ELASPLALT | [R].ELASPL | 2xPhosphc | Q9QY81 2x | 0.000446 | 0 | 1 | 4 | 2 | Q9QY81 | Nuclear po | Q9QY81 [1 | 0 | 1556.769 | 0 | 0.33 | 0.162494 | 13.43 | 86.2 | 113.8 | 10.3 | 12.63 | | High | High | High | High | High | High | High | High | High | High | High | 0 | 9.58E-05 | 2.78 |
| FALSE | High | ELASPVSPE | [K].ELASPV | 1xPhosphc | A2AN08-5 | 0.001629 | 5.66E-05 | 1 | 4 | 3 | A2AN08-5 | Isoform 5 c | A2AN08-5 | 0 | 1506.777 | 0 | -0.03 | 0.850221 | 4.5 | 102.2 | 97.8 | 3.03 | 6.86 | | High | High | High | High | High | High | High | High | High | High | High | 0.000449 | 0.004034 | 2.91 |
| FALSE | High | EIATPTHW | [K].EIATPTI | 1xPhosphc | Q8CH18 1x | 0.0095 | 0.000736 | 1 | 2 | 3 | Q8CH18 | Cell divisio | Q8CH18 [6 | 0 | 1707.887 | 0 | 0.37 | 0.3502 | 12.84 | 86.9 | 113.1 | 11.24 | 16.15 | | High | High | High | High | High | High | High | High | High | High | High | 0.000327 | 0.002954 | 1.89 |
| FALSE | High | EICQLSNEA | [K].EICQLSI | 1xCarbami | Q8K205 1x | 1.79E-06 | 0 | 1 | 2 | 1 | Q8K205 | Processing | Q8K205 [7 | 0 | 2238.042 | 0 | -0.06 | 0.26794 | 0.43 | 102.2 | 97.8 | 5.95 | 8.06 | | High | High | High | High | High | High | High | High | High | High | High | 0 | 1.94E-07 | 4.16 |
| FALSE | High | ELCQVQSN | [K].ELCQV( | 1xCarbami | Q8C156 1x | 7.62E-09 | 0 | 1 | 2 | 1 | Q8C156 | Condensin | Q8C156 [3 | 0 | 2485.148 | 0 | -0.28 | 0.049529 | 11.1 | 110.3 | 89.7 | 5.57 | 12.46 | | High | High | High | High | High | High | High | High | High | High | High | 0 | 4.24E-10 | 4.35 |
| FALSE | High | EICSLFGEA | [R].EICSLF( | 1xCarbami | P70398 1xI | 7.58E-07 | 0 | 1 | 4 | 2 | P70398 | Probable u | P70398 [57 | 0 | 2373.111 | 0 | 0.21 | 0.209866 | 36.04 | 89.3 | 110.7 | 9.37 | 20.59 | | High | High | High | High | High | High | High | High | High | High | High | 0 | 7.43E-08 | 3.7 |
| FALSE | High | EICSLFGEA | [R].EICSLF( | 1xCarbami | G3UY52 1x | 2.13E-07 | 0 | 1 | 1 | 6 | G3UY52 | Probable u | G3UY52 [6 | 0 | 2204.026 | 0 | 0.24 | 0.118736 | 6.72 | 91.8 | 108.2 | 7.74 | 7.74 | | High | High | High | High | High | High | High | High | High | High | High | 0 | 1.78E-08 | 4.17 |
| FALSE | High | EIDCLSPEA | [K].EIDCLSI | 1xCarbami | Q3UZ39 1x | 0.002818 | 0.000109 | 1 | 6 | 2 | Q3UZ39 | Leucine-ric | Q3UZ39 [1 | 0 | 1626.74 | 0 | 1.56 | 0.000192 | 2.12 | 48.8 | 151.2 | 7.07 | 5.3 | | High | High | High | High | High | High | High | High | High | High | High | 4.94E-05 | 0.000761 | 2.63 |
| FALSE | High | ELDDALDEI | [K].ELDDAI | 1xPhosphc | P51125-3 1 | 0.119269 | 0.00979 | 1 | 12 | 1 | P51125-3 | Isoform 3 c | P51125-3 [ | 0 | 4003.963 | 0 | 1.71 | 0.003556 | 11.36 | 51.8 | 148.2 | 19.73 | 15.48 | | High | High | High | High | High | High | High | High | High | High | High | 0.004866 | 0.05306 | 2.3 |
| FALSE | High | ELDENVEYI | [K].ELDEN\ | 1xPhosphc | Q8BX09 1x | 0.027358 | 0.001676 | 1 | 4 | 3 | Q8BX09 | Retinoblast | Q8BX09 [3 | 1 | 3299.44 | 0 | -0.27 | 0.027415 | 4.18 | 109.2 | 90.8 | 6.73 | 7.28 | | High | High | High | High | High | High | High | High | High | High | High | 0.001214 | 0.01308 | 3.61 |
| FALSE | High | ELDPEEILPS | [K].ELDPEE | 1xPhosphc | Q9D1R2 1x | 0.004218 | 0.00036 | 1 | 1 | 1 | Q9D1R2 | Protein KTI | Q9D1R2 [1 | 0 | 2878.42 | 0 | 0.31 | 0.123612 | 8.73 | 86 | 114 | 20.5 | 13.6 | | High | High | High | High | High | High | High | High | High | High | High | 0.000161 | 0.001191 | 2.57 |
| FALSE | High | EIDVLEFGE | [K].EIDVLEI | 1xPhosphc | Q8CB16 1x | 0.045898 | 0.003408 | 1 | 7 | 1 | Q8CB16 | Putative ur | Q8CB16 [9 | 0 | 2201.114 | 0 | 2.29 | 0.001382 | 32.75 | 31.4 | 168.6 | 48.48 | 16.79 | | High | High | High | High | High | High | High | High | High | High | High | 0.001774 | 0.01749 | 2.71 |
| FALSE | High | EIEAIESDSI | [K].EIEAIES | 1xPhosphc | Q80XR9 1x | 0.001424 | 5.66E-05 | 1 | 6 | 1 | Q80XR9 | Rsrc1 prote | Q80XR9 [2 | 0 | 2208.042 | 0 | -0.1 | 0.03716 | 4.14 | 101.7 | 98.3 | 16.5 | 16.38 | | High | High | High | High | High | High | High | High | High | High | High | 2.58E-05 | 0.000355 | 3.05 |
| FALSE | High | ELEDLLSPL | [K].ELEDLL | 1xPhosphc | Q3TFE2 1x | 0.001222 | 0 | 1 | 6 | 2 | Q3TFE2 | Putative ur | Q3TFE2 [4( | 0 | 2139.124 | 0 | -0.39 | 0.124926 | 21.99 | 117 | 83 | 19.42 | 35.64 | | High | High | High | High | High | High | High | High | High | High | High | 0 | 0.000299 | 2.16 |
| FALSE | High | ELEEEEENS | [R].ELEEEE | 1xPhosphc | Q9JI10 1xP | 0.000396 | 0 | 1 | 3 | 2 | Q9JI10 | Serine/thre | Q9JI10 [30 | 0 | 3045.317 | 0 | 1.01 | 0.000426 | 4.71 | 63.7 | 136.3 | 7.68 | 6.26 | | High | High | High | High | High | High | High | High | High | High | High | 0 | 8.46E-05 | 4.17 |
| FALSE | High | ELEEEEENS | [R].ELEEEE | 1xPhosphc | Q9JI10 1xP | 3.11E-08 | 0 | 1 | 3 | 14 | Q9JI10 | Serine/thre | Q9JI10 [30 | 0 | 3061.312 | 0 | 0.99 | 0.000928 | 10.26 | 66.1 | 133.9 | 8.95 | 4.93 | | High | High | High | High | High | High | High | High | High | High | High | 0 | 2.05E-09 | 5.13 |
| FALSE | High | ELEELSLEA | [R].ELEELSI | 1xPhosphc | Q05D44 1x | 3.54E-05 | 0 | 1 | 1 | 2 | Q05D44 | Eukaryotic | Q05D44 [6 | 0 | 1795.904 | 0 | 0.58 | 0.008776 | 13.34 | 78.9 | 121.1 | 8.52 | 8.32 | | High | High | High | High | High | High | High | High | High | High | High | 0 | 5.58E-06 | 2.69 |
| FALSE | High | ELELAEEAS | [K].ELELAE | 1xPhosphc | Q8R3F9 1x | 0.023175 | 0.001578 | 1 | 2 | 1 | Q8R3F9 | Speckle tar | Q8R3F9 [3 | 0 | 1656.85 | 0 | -0.55 | 0.014476 | 9.71 | 123.5 | 76.5 | 10.97 | 11.54 | | High | High | High | High | High | High | High | High | High | High | High | 0.000847 | 0.00802 | 2.4 |
| FALSE | High | ELELLFGCH | [K].ELELLF( | 1xCarbami | P31938 1xI | 0.002293 | 0.000109 | 1 | 5 | 2 | P31938 | Dual specif | P31938 [27 | 0 | 2802.348 | 0 | 0.02 | 0.945722 | 4.65 | 100.5 | 99.5 | 9.09 | 9.08 | | High | High | High | High | High | High | High | High | High | High | High | 0.004032 | 0.04309 | 2.65 |
| FALSE | High | ELEILNSTSI | [R].ELEILNS | 1xCarbami | Q80WQ9 1 | 2.17E-05 | 0 | 1 | 1 | 4 | Q80WQ9 | Zinc finger | Q80WQ9 [ | 0 | 3163.475 | 0 | -0.59 | 0.004634 | 3.14 | 126.3 | 73.7 | 11.79 | 13.47 | | High | High | High | High | High | High | High | High | High | High | High | 0 | 3.2E-05 | 3.99 |
| FALSE | High | ELELPSQE\ | [R].ELELPS( | 1xCarbami | Q80WQ6 1 | 0.001137 | 0 | 1 | 2 | 1 | Q80WQ6 | Inactive rh | Q80WQ6 [ | 0 | 3254.662 | 0 | -0.44 | 0.017753 | 9.16 | 114.8 | 85.2 | 5.57 | 10.9 | | High | High | High | High | High | High | High | High | High | High | High | 0 | 0.000275 | 2.71 |
| FALSE | High | ELENWAGS | [K].ELENW | 1xPhosphc | Q9D8B3 1x | 0.099328 | 0.008003 | 1 | 1 | 1 | Q9D8B3 | Charged m | Q9D8B3 [2 | 0 | 1345.57 | 0 | 0.17 | 0.968584 | 40.92 | 103.2 | 96.8 | 25.47 | 29.95 | | High | High | High | High | High | High | High | High | High | High | High | 0.004032 | 0.04275 | 1.87 |
| FALSE | High | ELENWAGS | [K].ELENW | 1xPhosphc | Q9D8B3 1x | 0.002252 | 0.000109 | 1 | 1 | 2 | Q9D8B3 | Charged m | Q9D8B3 [2 | 0 | 1361.565 | 0 | -0.21 | 0.137535 | 11.61 | 111.2 | 88.8 | 9.49 | 13.19 | | High | High | High | High | High | High | High | High | High | High | High | 9.42E-05 | 0.000974 | 2.41 |
| FALSE | High | EIEQEETLS | [K].EIEQEE | 2xPhosphc | P11862 2xI | 1.27E-06 | 0 | 1 | 2 | 1 | P11862 | Growth arr | P11862 [17 | 0 | 2718.256 | 0 | -0.98 | 0.000712 | 1.02 | 131.6 | 68.4 | 7.11 | 8.95 | | High | High | High | High | High | High | High | High | High | High | High | 0 | 1.32E-07 | 3.27 |
| FALSE | High | ELEQHIQTS | [R].ELEQHI | 1xPhosphc | A0A1L1SV | 3.24E-06 | 0 | 1 | 3 | 4 | A0A1L1SV | Ubiquitin c | A0A1L1SV | 0 | 2625.167 | 0 | 0.24 | 0.004696 | 6.03 | 92.9 | 107.1 | 9.86 | 10.47 | | High | High | High | High | High | High | High | High | High | High | High | 0 | 3.8E-07 | 3.7 |
| FALSE | High | EIESEIDSEE | [K].EIESEID | 1xPhosphc | Q3UC10 1x | 2.6E-05 | 0 | 1 | 4 | 4 | Q3UC10 | DNA helica | Q3UC10 [7 | 0 | 2315.131 | 0 | -0.13 | 0.371419 | 18.83 | 104.4 | 95.6 | 9.02 | 7.94 | | High | High | High | High | High | High | High | High | High | High | High | 0 | 3.94E-06 | 3.1 |
| FALSE | High | EGTLSSPAL | [R].EGTLSS | 1xPhosphc | Q62431 1x | 0.005322 | 0.000599 | 1 | 2 | 2 | Q62431 | AT-rich inte | Q62431 [9 | 0 | 2656.257 | 0 | -0.8 | 0.001756 | 1.5 | 125.5 | 74.5 | 10.73 | 13.68 | | High | High | High | High | High | High | High | High | High | High | High | 0.000638 | 0.005664 | 3 |
| FALSE | High | ELLELGIDN | [R].ELLELG | 1xPhosphc | E9Q2H1 1x | 6.49E-06 | 0 | 1 | 2 | 4 | E9Q2H1 | E3 ubiquiti | E9Q2H1 [1 | 0 | 3784.696 | 0 | -0.26 | 0.00481 | 0.98 | 108.8 | 91.2 | 1.13 | 7.49 | | High | High | High | High | High | High | High | High | High | High | High | 0 | 1.61E-05 | 4.75 |
| FALSE | High | EGSVLDILK | [R].EGSVLI | 1xPhosphc | E9Q3G8 1x | 0.001386 | 5.66E-05 | 1 | 3 | 2 | E9Q3G8 | Nucleopor | E9Q3G8 [6 | 0 | 1511.849 | 0 | 1.18 | 0.011019 | 17.04 | 59.5 | 140.5 | 26.79 | 24.86 | | High | High | High | High | High | High | High | High | High | High | High | 2.58E-05 | 0.00053 | 2.79 |
| FALSE | High | EGSTPEDD | [R].EGSTPE | 1xDeamida | E0CYH0 1x | 1.01E-07 | 0 | 1 | 2 | 11 | E0CYH0 | MCG16685 | E0CYH0 [2 | 0 | 2364.028 | 0 | -0.38 | 0.002673 | 0.54 | 114.6 | 85.4 | 5.94 | 5.78 | | High | High | High | High | High | High | High | High | High | High | High | 0 | 7.79E-09 | 3.79 |
| FALSE | High | EFESPLLFTI | [K].EFESPLI | 1xPhosphc | Q7TSY8 1x | 0.011048 | 0.000772 | 1 | 1 | 1 | Q7TSY8 | Shugoshin | Q7TSY8 [6 | 0 | 1547.771 | 0 | 0.25 | 0.023243 | 6.5 | 91.5 | 108.5 | 11.06 | 9.02 | | High | High | High | High | High | High | High | High | High | High | High | 0.000399 | 0.003512 | 2.03 |
| FALSE | High | EFETPEK | [K].EFETPE | 1xPhosphc | Q6DFW4 1 | 0.025293 | 0.001645 | 1 | 3 | 2 | Q6DFW4 | Nucleolar p | Q6DFW4 [ | 0 | 1417.702 | 0 | -0.9 | 0.000209 | 0.67 | 128.3 | 71.7 | 3.64 | 7.08 | | High | High | High | High | High | High | High | High | High | High | High | 0.000872 | 0.008859 | 2.18 |
| FALSE | High | EFFVTSAP\ | [K].EFFVTS | 1xPhosphc | P56546-2 1 | 1.67E-05 | 0 | 1 | 4 | 2 | P56546-2 | Isoform 2 c | P56546-2 [ | 0 | 3385.646 | 0 | -1.89 | 0.002027 | 0.79 | 160.5 | 39.5 | 18.1 | 14.16 | | High | High | High | High | High | High | High | High | High | High | High | 0 | 2.42E-06 | 4.42 |
| FALSE | High | EFFVTSAP\ | [K].EFFVTS | 1xDeamida | P56546-2 1 | 0.040373 | 0.002982 | 1 | 4 | 1 | P56546-2 | Isoform 2 c | P56546-2 [ | 0 | 3386.63 | 0 | -1.81 | 0.004697 | 3.41 | 159.9 | 40.1 | 16.73 | 25.5 | | High | High | High | High | High | High | High | High | High | High | High | 0.00155 | 0.01503 | 1.97 |
| FALSE | High | EFGEDLVE( | [K].EFGEDI | 1xPhosphc | Q9D0L8 1x | 4.07E-10 | 0 | 1 | 6 | 14 | Q9D0L8 | mRNA cap | Q9D0L8 [4 | 0 | 2796.302 | 0 | -0.65 | 9.77E-05 | 0.03 | 121.6 | 78.4 | 1.88 | 2.56 | | High | High | High | High | High | High | High | High | High | High | High | 0 | 1.58E-11 | 5.16 |
| FALSE | High | EFGEDLVE( | [K].EFGEDI | 1xDeamida | Q9D0L8 1x | 0.002252 | 0.000109 | 1 | 6 | 2 | Q9D0L8 | mRNA cap | Q9D0L8 [4 | 0 | 2797.286 | 0 | -0.53 | 0.015617 | 10.95 | 118.5 | 81.5 | 12.91 | 4.73 | | High | High | High | High | High | High | High | High | High | High | High | 4.94E-05 | 0.000592 | 3.72 |
| FALSE | High | EFGEDLVE( | [K].EFGEDI | 2xPhosphc | Q9D0L8 2x | 0.007274 | 0.00062 | 1 | 6 | 2 | Q9D0L8 | mRNA cap | Q9D0L8 [4 | 0 | 2876.268 | 0 | -0.33 | 0.015758 | 8.1 | 117.1 | 82.9 | 9.09 | 11.28 | | High | High | High | High | High | High | High | High | High | High | High | 0.001916 | 0.01957 | 3.95 |
| FALSE | High | EFGSLPTTF | [K].EFGSLP | 1xPhosphc | O70439 1x | 0.001783 | 5.66E-05 | 1 | 4 | 2 | O70439 | Syntaxin-7 | O70439 [7 | 0 | 1757.831 | 0 | 0.48 | 0.007249 | 9.75 | 82.4 | 117.6 | 7.23 | 6.7 | | High | High | High | High | High | High | High | High | High | High | High | 2.58E-05 | 0.000455 | 2.78 |
| FALSE | High | EFHLNESGI | [K].EFHLNE | 1xPhosphc | Q9EQU5-2 | 0.053926 | 0.00384 | 2 | 4 | 1 | Q9EQU5-2 | Isoform 2 c | Q9EQU5-2 | 0 | 1984.942 | 0 | 0.6 | 0.083267 | 42.18 | 78.8 | 121.2 | 5.61 | 40.38 | Shared | High | High | High | High | High | High | High | High | High | High | High | 0.002031 | 0.02095 | 2.4 |
| FALSE | High | EFLEPDSW | [K].EFLEPD | 1xPhosphc | P81117 1xI | 3.55E-07 | 0 | 1 | 2 | 4 | P81117 | Nucleobin( | P81117 [32 | 0 | 3378.607 | 0 | 3.8 | 0.010904 | 75.03 | 10.7 | 189.3 | 40.84 | 37.74 | | High | High | High | High | High | High | High | High | High | High | High | 0 | 3.17E-08 | 4.74 |
| FALSE | High | EFITGDVEF | [R].EFITGD | 1xPhosphc | Q8C1W9 1 | 1.23E-08 | 0 | 1 | 4 | 12 | Q8C1W9 | Putative ur | Q8C1W9 [ | 0 | 3402.494 | 0 | 0.09 | 0.820711 | 3.18 | 96.6 | 103.4 | 15.85 | 14.79 | | High | High | High | High | High | High | High | High | High | High | High | 0 | 6.83E-07 | 5.01 |
| FALSE | High | EFITGDVEF | [R].EFITGD | 1xDeamida | Q8C1W9 1 | 3.8E-05 | 0 | 1 | 4 | 1 | Q8C1W9 | Putative ur | Q8C1W9 [ | 0 | 3403.478 | 0 | 0.11 | 0.728878 | 25.37 | 99 | 101 | 8.06 | 11.67 | | High | High | High | High | High | High | High | High | High | High | High | 0 | 6.07E-06 | 4.81 |
| FALSE | High | EFNIPEAGS | [K].EFNIPE. | 1xCarbami | B9EK83 1xI | 4.8E-05 | 0 | 2 | 3 | 2 | B9EK83; A2 | Transient r | B9EK83 [13 | 0 | 2941.4 | 0 | 2.2 | 0.000232 | 1.9 | 37.1 | 162.9 | 9.45 | 4.68 | Shared | High | High | High | High | High | High | High | High | High | High | High | 0 | 7.85E-06 | 4.07 |
| FALSE | High | EFESGDFQ | [K].EFESGI | 1xPhosphc | V9GX40 1x | 0.001021 | 0 | 1 | 5 | 2 | V9GX40 | Synergin ga | V9GX40 [9 | 0 | 1786.752 | 0 | 2.69 | 0.000176 | 10.59 | 25.7 | 174.3 | 9.65 | 4.3 | | High | High | High | High | High | High | High | High | High | High | High | 0.000275 | 0.002145 | 2.09 |
| FALSE | High | EFQDYIEPE | [K].EFQDYI | 1xCarbami | Q6PDL0 1x | 4.09E-09 | 0 | 1 | 3 | 5 | Q6PDL0 | Cytoplasmi | Q6PDL0 [1 | 0 | 2478.048 | 0 | 0.05 | 0.121001 | 1.88 | 98.4 | 101.6 | 0.72 | 5.01 | | High | High | High | High | High | High | High | High | High | High | High | 0 | 2.01E-09 | 3.95 |
| FALSE | High | EFSAPSTPT | [R].EFSAPS | 1xPhosphc | E9Q616 1x | 2.1E-06 | 0 | 1 | 4 | 1 | E9Q616 | AHNAK nu | E9Q616 [5 | 0 | 2292.063 | 0 | 1.27 | 0.034611 | 44.65 | 58.6 | 141.4 | 13.5 | 28.7 | | High | High | High | High | High | High | High | High | High | High | High | 0 | 2.35E-07 | 2.85 |
| FALSE | High | EFSAPSTPT | [R].EFSAPS | 2xPhosphc | E9Q616 2x | 0.065952 | 0.004766 | 1 | 4 | 1 | E9Q616 | AHNAK nu | E9Q616 [5 | 0 | 2372.03 | 0 | 3.01 | 0.003866 | 52.53 | 20.1 | 179.9 | 35.71 | 23.18 | | High | High | High | High | High | High | High | High | High | High | High | 0.002455 | 0.02651 | 2.81 |
| FALSE | High | EFSAPSTPT | [R].EFSAPS | 1xPhosphc | E9Q616 1x | 2.87E-08 | 0 | 1 | 4 | 14 | E9Q616 | AHNAK nu | E9Q616 [5 | 0 | 2521.226 | 0 | 3.16 | 0.000605 | 18.16 | 19.6 | 180.4 | 14.42 | 11.42 | | High | High | High | High | High | High | High | High | High | High | High | 0 | 1.21E-06 | 4.37 |
| FALSE | High | EFSAPSTPT | [R].EFSAPS | 2xPhosphc | E9Q616 2x | 4.02E-06 | 0 | 1 | 4 | 4 | E9Q616 | AHNAK nu | E9Q616 [5 | 0 | 2601.193 | 0 | 2.68 | 0.001312 | 17.86 | 29 | 171 | 19.92 | 17.17 | | High | High | High | High | High | High | High | High | High | High | High | 0 | 4.86E-07 | 4.21 |
| FALSE | High | EFSGGSYN | [R].EFSGGS | 1xPhosphc | A0A1D5RL | 0.020142 | 0.001472 | 1 | 9 | 1 | A0A1D5RL | Serine/thre | A0A1D5RL | 0 | 1627.777 | 0 | 1.56 | 0.000513 | 4.91 | 49.5 | 150.5 | 7.98 | 6.47 | | High | High | High | High | High | High | High | High | High | High | High | 0.000786 | 0.006849 | 2.26 |
| FALSE | High | EFSLAIFSPI | [R].EFSLAIF | 1xPhosphc | Q6NV99 1x | 0.036103 | 0.002342 | 1 | 5 | 1 | Q6NV99 | HAUS augn | Q6NV99 [6 | 0 | 1852.872 | 0 | -0.37 | 0.057154 | 20.6 | 112.9 | 87.1 | 14.65 | 9 | | High | High | High | High | High | High | High | High | High | High | High | 0.001263 | 0.01332 | 1.87 |
| FALSE | High | EFTGSPPPS | [R].EFTGSF | 1xPhosphc | Q3U325 1x | 0.000148 | 0 | 1 | 4 | 3 | Q3U325 | Putative ur | Q3U325 [5 | 0 | 1756.892 | 0 | 1.84 | 0.000269 | 6.35 | 43 | 157 | 4.69 | 5.17 | | High | High | High | High | High | High | High | High | High | High | High | 0 | 2.79E-05 | 2.61 |
| FALSE | High | EFVSSDESS | [K].EFVSSD | 1xPhosphc | Q08943-2 | 0.096181 | 0.007535 | 1 | 2 | 1 | Q08943-2 | Isoform 2 c | Q08943-2 | 0 | 2039.921 | 0 | -0.96 | 0.000631 | 3.06 | 132.1 | 67.9 | 7.47 | 14.29 | | High | High | High | High | High | High | High | High | High | High | High | 0.003823 | 0.04127 | 2.06 |
| FALSE | High | EGAAHAFA | [R].EGAAH. | 1xPhosphc | P47753 1xI | 4.06E-06 | 0 | 1 | 5 | 1 | P47753 | F-actin-cap | P47753 [48 | 0 | 2663.273 | 0 | 0.18 | 0.02274 | 5.84 | 93.2 | 106.8 | 4.13 | 5.13 | | High | High | High | High | High | High | High | High | High | High | High | 0 | 4.91E-07 | 3.64 |
| FALSE | High | EGADSLGV | [R].EGADSI | 1xPhosphc | Q8VBT6 1x | 4.22E-07 | 0 | 1 | 1 | 1 | Q8VBT6 | Apolipopro | Q8VBT6 [2 | 0 | 2993.3 | 0 | 2.11 | 0.00231 | 16.84 | 44.6 | 155.4 | 19.09 | 18.26 | | High | High | High | High | High | High | High | High | High | High | High | 0 | 3.86E-08 | 4.75 |
| FALSE | High | EGAEGASP | [K].EGAEG. | 1xPhosphc | B7ZNK9 1x | 7.86E-07 | 0 | 1 | 5 | 3 | B7ZNK9 | Spata5 pro | B7ZNK9 [1 | 0 | 1795.842 | 0 | -0.4 | 0.002202 | 5.48 | 112.6 | 87.4 | 5.01 | 6.36 | | High | High | High | High | High | High | High | High | High | High | High | 0 | 4.88E-05 | 3.27 |
| FALSE | High | EGAHTSGA | [K].EGAHT! | 1xPhosphc | A2AGQ4 1: | 0.001349 | 0 | 1 | 14 | 2 | A2AGQ4 | MAP kinase | A2AGQ4 [1 | 0 | 4078.795 | 0 | -0.48 | 0.157601 | 9 | 113.9 | 86.1 | 18.83 | 36.74 | | High | High | High | High | High | High | High | High | High | High | High | 0.002187 | 0.02339 | 3.32 |
| FALSE | High | EGCAISSET | [R].EGCAIS | 1xCarbami | E9Q6J5-3 1 | 5.7E-05 | 0 | 1 | 3 | 3 | E9Q6J5-3 | Isoform Ne | E9Q6J5-3 [ | 0 | 2957.268 | 0 | -0.16 | 0.371817 | 22.93 | 102.1 | 97.9 | 14.98 | 18.13 | | High | High | High | High | High | High | High | High | High | High | High | 0 | 0.000311 | 2.59 |
| FALSE | High | EFQSFMPV | [K].EFQSFN | 1xCarbami | Q9JHI4 2xF | 0.096959 | 0.007777 | 1 | 2 | 1 | Q9JHI4 | Solute carr | Q9JHI4 [42 | 0 | 4262.957 | 0 | -0.51 | 0.194485 | 10.37 | 116.3 | 83.7 | 16.09 | 22.29 | | High | High | High | High | High | High | High | High | High | High | High | 0.003922 | 0.04177 | 2.54 |
| FALSE | High | EGCAISSET | [R].EGCAIS | 1xCarbami | E9Q6J5-3 1 | 9.11E-08 | 0 | 1 | 3 | 4 | E9Q6J5-3 | Isoform Ne | E9Q6J5-3 [ | 0 | 2973.263 | 0 | 0.03 | 0.278236 | 7.02 | 96.5 | 103.5 | 10.23 | 6.43 | | High | High | High | High | High | High | High | High | High | High | High | 0 | 6.85E-09 | 4.64 |
| FALSE | High | EFDYDSPH( | [K].EFDYDS | 2xPhosphc | Q7TPE5 2x | 2.58E-05 | 0 | 1 | 1 | 2 | Q7TPE5 | Probable R | Q7TPE5 [2 | 0 | 1885.677 | 0 | -0.28 | 0.022392 | 2.86 | 108.1 | 91.9 | 5.56 | 6.78 | | High | High | High | High | High | High | High | High | High | High | High | 0 | 3.9E-06 | 2.73 |
| FALSE | High | EFDSDSLR | [R].EFDSDS | 1xPhosphc | G5E8K3 1x | 0.059657 | 0.004365 | 2 | 24 | 1 | G5E8K3; A( | Ankyrin 3, | G5E8K3 [9 | 0 | 1277.561 | 0 | 1.99 | 0.000313 | 9.28 | 37 | 163 | 13.35 | 8.5 | Shared | High | High | High | High | High | High | High | High | High | High | High | 0.002233 | 0.02352 | 1.7 |
| FALSE | High | EETPGTPK | [K].EETPGT | 1xPhosphc | Q8BU14 1x | 0.043981 | 0.003228 | 1 | 2 | 2 | Q8BU14 | Translocati | Q8BU14 [1 | 0 | 1396.713 | 0 | 0.2 | 0.026937 | 5.26 | 93 | 107 | 3.72 | 4.35 | | High | High | High | High | High | High | High | High | High | High | High | 0.001665 | 0.01672 | 1.89 |
| FALSE | High | EETPPSVEF | [K].EETPPS | 1xPhosphc | Q05D44 1x | 1.08E-05 | 0 | 1 | 1 | 4 | Q05D44 | Eukaryotic | Q05D44 [4 | 1 | 4431.875 | 0 | -0.4 | 0.055784 | 17.19 | 111.2 | 88.8 | 10.04 | 13.12 | | High | High | High | High | High | High | High | High | High | High | High | 0 | 1.47E-06 | 4.82 |
| FALSE | High | EETPPSVEF | [K].EETPPS | 1xPhosphc | Q05D44 1x | 4.8E-05 | 0 | 1 | 1 | 7 | Q05D44 | Eukaryotic | Q05D44 [4 | 1 | 4447.87 | 0 | -1.29 | 0.004016 | 2.75 | 142.2 | 57.8 | 8.49 | 32.79 | | High | High | High | High | High | High | High | High | High | High | High | 0 | 4.46E-05 | 4.66 |
| FALSE | High | EETQPLSHS | [K].EETQPL | 1xPhosphc | B9EJ80 1xF | 2.13E-05 | 0 | 1 | 2 | 2 | B9EJ80 | PDZ domai | B9EJ80 [50 | 0 | 1790.909 | 0 | 0.82 | 0.001284 | 5.83 | 73 | 127 | 3.81 | 8.08 | | High | High | High | High | High | High | High | High | High | High | High | 0 | 3.17E-06 | 3.47 |

| | | | | | | | | | | | | | | | | | | | | | | | | | | | | | | | | | | | | | | |
|---|---|---|---|---|---|---|---|---|---|---|---|---|---|---|---|---|---|---|---|---|---|---|---|---|---|---|---|---|---|---|---|---|---|---|---|---|---|---|
| FALSE | High | EETSNESAI | [R].EETSNE | 1xPhospho | Q8BUY8 1x | 0.000519 | 0 | 1 | 1 | 1 | Q8BUY8 | G-protein c | Q8BUY8 [2 | 0 | 2633.055 | 0 | -0.71 | 0.001009 | 4.29 | 125.5 | 74.5 | 4.86 | 12.05 | | High | High | High | High | High | High | High | High | High | High | High | 0 | 0.000115 | 4.05 |
| FALSE | High | EETSPPLTK | [K].EETSPP | 1xPhospho | Q8BQ33 1x | 0.004955 | 0.000504 | 1 | 1 | 2 | Q8BQ33 | Treslin OS= | Q8BQ33 [1 | 0 | 1539.807 | 0 | -0.78 | 0.000313 | 1.4 | 126.5 | 73.5 | 3.78 | 3.69 | | High | High | High | High | High | High | High | High | High | High | High | 0.000225 | 0.001437 | 3.11 |
| FALSE | High | EEVASEPEE | [K].EEVASE | 1xPhospho | Q9D6Z1 1x | 2.44E-08 | 0 | 1 | 3 | 12 | Q9D6Z1 | Nucleolar p | Q9D6Z1 [5 | 0 | 2310.115 | 0 | -0.36 | 0.001945 | 2.27 | 112.5 | 87.5 | 2.95 | 4.6 | | High | High | High | High | High | High | High | High | High | High | High | 0 | 1.56E-05 | 5.27 |
| FALSE | High | EEVASEPEE | [K].EEVASE | 2xPhospho | Q9D6Z1 2x | 0.000164 | 0 | 1 | 3 | 3 | Q9D6Z1 | Nucleolar p | Q9D6Z1 [5 | 0 | 2390.082 | 0 | -0.5 | 0.007259 | 2.88 | 117.3 | 82.7 | 5.07 | 8.55 | | High | High | High | High | High | High | High | High | High | High | High | 0 | 3.14E-05 | 3.9 |
| FALSE | High | EEVASEPEE | [K].EEVASE | 1xPhospho | Q9D6Z1 1x | 4.11E-07 | 0 | 1 | 3 | 4 | Q9D6Z1 | Nucleolar p | Q9D6Z1 [5 | 1 | 2667.373 | 0 | -1.45 | 0.000678 | 0.39 | 147.5 | 52.5 | 9.39 | 7.9 | | High | High | High | High | High | High | High | High | High | High | High | 0 | 4.57E-08 | 4.26 |
| FALSE | High | EEVASETPA | [K].EEVASE | 1xPhospho | Q8K3W3 1 | 0.014797 | 0.001049 | 1 | 1 | 1 | Q8K3W3 | Protein CA | Q8K3W3 [3 | 0 | 3167.628 | 0 | -0.43 | 0.003168 | 1.84 | 116.3 | 83.7 | 7.63 | 10.72 | | High | High | High | High | High | High | High | High | High | High | High | 0.000555 | 0.004852 | 2.27 |
| FALSE | High | EEVDSENIF | [K].EEVDSE | 1xPhospho | Q569Z9 1x | 5.75E-05 | 0 | 1 | 4 | 1 | Q569Z9 | Putative ur | Q569Z9 [4 | 0 | 2931.42 | 0 | -0.23 | 0.136642 | 7.59 | 110.3 | 89.7 | 12.35 | 15.42 | | High | High | High | High | High | High | High | High | High | High | High | 0 | 9.6E-06 | 2.49 |
| FALSE | High | EEVDSVAG | [K].EEVDSV | 1xPhospho | Q3UM18 1 | 0.000213 | 0 | 1 | 3 | 1 | Q3UM18 | Large subu | Q3UM18 [ | 0 | 1801.862 | 0 | 0.32 | 0.30957 | 39.4 | 85.2 | 114.8 | 16.15 | 29.95 | | High | High | High | High | High | High | High | High | High | High | High | 0 | 4.2E-05 | 2.95 |
| FALSE | High | EEVGAGYN | [R].EEVGA | 1xPhospho | A2AES4 1x | 4.02E-09 | 0 | 1 | 6 | 3 | A2AES4 | OTU doma | A2AES4 [1 | 0 | 2339.987 | 0 | -0.02 | 0.767123 | 16.34 | 98.5 | 101.5 | 8.83 | 12.34 | | High | High | High | High | High | High | High | High | High | High | High | 0 | 2.06E-10 | 4 |
| FALSE | High | EFDYDSPH | [K].EFDYDS | 1xPhospho | Q7TPE5 1x | 5.75E-05 | 0 | 1 | 1 | 4 | Q7TPE5 | Probable R | Q7TPE5 [2 | 0 | 1805.71 | 0 | -1.11 | 0.000583 | 1.12 | 137.2 | 62.8 | 8.53 | 6.29 | | High | High | High | High | High | High | High | High | High | High | High | 0.003535 | 0.03784 | 2.8 |
| FALSE | High | EEVGNISIL | [R].EEVGN | 1xPhospho | A1L366 1xl | 1.26E-09 | 0 | 1 | 5 | 2 | A1L366 | RAD21 hor | A1L366 [14 | 0 | 2809.223 | 0 | 0.15 | 0.204315 | 17.43 | 84.5 | 115.5 | 18.59 | 22.89 | | High | High | High | High | High | High | High | High | High | High | High | 0 | 4.43E-07 | 5.04 |
| FALSE | High | EEVIPGSPE | [K].EEVIPG | 1xCarbami | Q9JJ89 1xP | 0.004411 | 0.000409 | 1 | 1 | 2 | Q9JJ89 | Coiled-coil | Q9JJ89 [21 | 0 | 4287.011 | 0 | 0.16 | 0.618281 | 34.03 | 93.5 | 106.5 | 8.74 | 23.89 | | High | High | High | High | High | High | High | High | High | High | High | 0.000246 | 0.00149 | 2.92 |
| FALSE | High | EEVLSESET | [K].EEVLSE | 1xPhospho | Q3TI31 1xP | 0.000505 | 0 | 1 | 3 | 3 | Q3TI31 | Putative ur | Q3TI31 [77 | 0 | 3233.587 | 0 | -0.35 | 0.042352 | 7.09 | 111.9 | 88.1 | 9.99 | 5.21 | | High | High | High | High | High | High | High | High | High | High | High | 0 | 0.000111 | 2.93 |
| FALSE | High | EEVLSESET | [K].EEVLSE | 2xPhospho | Q3TI31 2xP | 5.37E-06 | 0 | 1 | 3 | 3 | Q3TI31 | Putative ur | Q3TI31 [77 | 0 | 3313.553 | 0 | 0.13 | 0.294792 | 13.55 | 95.4 | 104.6 | 11.03 | 4.24 | | High | High | High | High | High | High | High | High | High | High | High | 0 | 6.71E-07 | 4.56 |
| FALSE | High | EEYEVAEM | [K].EEYEVA | 1xPhospho | Q9CQW9 1 | 0.005666 | 0.00062 | 1 | 1 | 1 | Q9CQW9 | Interferon- | Q9CQW9 [ | 0 | 2227.989 | 0 | 0.83 | 0.015233 | 18.23 | 71.9 | 128.1 | 23.47 | 15.43 | | High | High | High | High | High | High | High | High | High | High | High | 0.000275 | 0.001669 | 3.48 |
| FALSE | High | EEYEVAEM | [K].EEYEVA | 1xPhospho | Q9CQW9 1 | 0.062209 | 0.004646 | 1 | 1 | 2 | Q9CQW9 | Interferon- | Q9CQW9 [ | 0 | 2243.984 | 0 | 0.95 | 0.010801 | 22.52 | 71.8 | 128.2 | 19.54 | 13.65 | | High | High | High | High | High | High | High | High | High | High | High | 0.002381 | 0.02481 | 1.29 |
| FALSE | High | EEYGVTEL | [K].EEYGVT | 1xPhospho | A0A1B0GT | 1.89E-06 | 0 | 1 | 2 | 3 | A0A1B0GT | Uncharacte | A0A1B0GT | 0 | 2245.059 | 0 | 0.91 | 0.006112 | 14.5 | 72.1 | 127.9 | 10.7 | 18.5 | | High | High | High | High | High | High | High | High | High | High | High | 0 | 6.23E-07 | 3.62 |
| FALSE | High | EFAGEDTSI | [K].EFAGEI | 1xPhospho | Q6P5F9 1x | 0.041425 | 0.003126 | 1 | 3 | 2 | Q6P5F9 | Exportin-1 | Q6P5F9 [1 | 0 | 2066.916 | 0 | -0.91 | 0.004139 | 6.4 | 130.7 | 69.3 | 9.53 | 8.73 | | High | High | High | High | High | High | High | High | High | High | High | 0.001609 | 0.01551 | 1.92 |
| FALSE | High | EFAPSDEEI | [K].EFAPSD | 1xPhospho | Q7TNE3 1x | 3.93E-07 | 0 | 1 | 2 | 3 | Q7TNE3 | Sperm-ass | Q7TNE3 [1 | 0 | 1866.8 | 0 | -0.04 | 0.740174 | 16.24 | 100.1 | 99.9 | 6.2 | 9.83 | | High | High | High | High | High | High | High | High | High | High | High | 0 | 3.55E-08 | 3.33 |
| FALSE | High | EFCADHDIS | [K].EFCADH | 1xCarbami | E9PZM7 1x | 0.002974 | 0.00016 | 1 | 3 | 2 | E9PZM7 | SR-related | E9PZM7 [4 | 0 | 2071.899 | 0 | 0.16 | 0.368922 | 10.25 | 96.2 | 103.8 | 3.32 | 8.26 | | High | High | High | High | High | High | High | High | High | High | High | 0.000275 | 0.002244 | 2.86 |
| FALSE | High | EFCADHDIS | [K].EFCADH | 1xCarbami | E9PZM7 2x | 0.010475 | 0.000772 | 1 | 3 | 4 | E9PZM7 | SR-related | E9PZM7 [4 | 0 | 2151.866 | 0 | -0.15 | 0.016892 | 5.5 | 106.5 | 93.5 | 2.71 | 6.59 | | High | High | High | High | High | High | High | High | High | High | High | 0.007335 | 0.08016 | 2.67 |
| FALSE | High | EFCADHDIS | [K].EFCADH | 1xCarbami | E9PZM7 2x | 0.001968 | 5.66E-05 | 1 | 3 | 2 | E9PZM7 | SR-related | E9PZM7 [4 | 1 | 2364.988 | 0 | -0.3 | 0.052102 | 4.55 | 108.7 | 91.3 | 5.62 | 11.57 | | High | High | High | High | High | High | High | High | High | High | High | 0.000437 | 0.00366 | 3.29 |
| FALSE | High | EFCNSPECS | [R].EFCNSF | 2xCarbami | Q6ZQE4 1x | 0.000334 | 0 | 1 | 3 | 1 | Q6ZQE4 | Nuclear en | Q6ZQE4 [3 | 0 | 2052.896 | 0 | 0.2 | 0.253907 | 35.96 | 91.8 | 108.2 | 14.75 | 11.95 | | High | High | High | High | High | High | High | High | High | High | High | 0 | 6.96E-05 | 2.96 |
| FALSE | High | EFDELSPSA | [R].EFDELS | 1xPhospho | Q5DTI2 1xl | 9.71E-05 | 0 | 1 | 3 | 3 | Q5DTI2 | ATPase, Ca | Q5DTI2 [67 | 0 | 1587.725 | 0 | 0.38 | 0.009539 | 11.03 | 86.8 | 113.2 | 3.86 | 6.27 | | High | High | High | High | High | High | High | High | High | High | High | 0 | 1.73E-05 | 1.66 |
| FALSE | High | EEVGNISIL | [R].EEVGN | 1xPhospho | A1L366 1xl | 1.13E-07 | 0 | 1 | 5 | 2 | A1L366 | RAD21 hor | A1L366 [14 | 0 | 2825.217 | 0 | 0.07 | 0.916792 | 12.45 | 97.6 | 102.4 | 26.98 | 13.48 | | High | High | High | High | High | High | High | High | High | High | High | 0 | 9.55E-09 | 5.72 |
| FALSE | High | EGCPLSEAE | [R].EGCPLS | 1xCarbami | Q8BPI1 1xl | 1.78E-08 | 0 | 1 | 2 | 1 | Q8BPI1 | Protein kin | Q8BPI1 [66 | 0 | 3542.661 | 0 | -0.06 | 0.650159 | 13.77 | 100.7 | 99.3 | 16.08 | 7.12 | | High | High | High | High | High | High | High | High | High | High | High | 0 | 1.1E-09 | 5.6 |
| FALSE | High | EGCPSENP | [K].EGCPSE | 1xCarbami | Q5DTN0 1x | 5.45E-05 | 0 | 1 | 5 | 1 | Q5DTN0 | MKIAA410 | Q5DTN0 [4 | 0 | 2408.172 | 0 | -0.36 | 0.001181 | 4.08 | 114.9 | 85.1 | 4.37 | 8.09 | | High | High | High | High | High | High | High | High | High | High | High | 0 | 9.05E-06 | 1.74 |
| FALSE | High | EGCPSPPG | [R].EGCPSF | 1xCarbami | Q62219 1x | 1.66E-05 | 0 | 1 | 9 | 2 | Q62219 | Transformi | Q62219 [1 | 0 | 1782.85 | 0 | -0.53 | 0.234921 | 12.63 | 113.6 | 86.4 | 16.52 | 24.35 | | High | High | High | High | High | High | High | High | High | High | High | 0 | 2.39E-06 | 2.7 |
| FALSE | High | EGRPQEPT | [R].EGRPQ | 1xPhospho | Q8BTI8 1xP | 0.004738 | 0.000456 | 1 | 2 | 2 | Q8BTI8 | Serine/argi | Q8BTI8 [25 | 0 | 1747.914 | 0 | -0.76 | 0.000905 | 3.23 | 124.3 | 75.7 | 6.58 | 1.6 | | High | High | High | High | High | High | High | High | High | High | High | 0.000204 | 0.001357 | 1.58 |
| FALSE | High | EGRPSGEA | [R].EGRPSC | 1xPhospho | Q8C2Q7 1x | 5.13E-06 | 0 | 2 | 4 | 2 | Q8C2Q7; P | Heterogen | Q8C2Q7 [5 | 0 | 2645.275 | 0 | 0.4 | 0.010452 | 10.22 | 86.4 | 113.6 | 6.82 | 9.46 | Shared | High | High | High | High | High | High | High | High | High | High | High | 0 | 6.41E-07 | 3.18 |
| FALSE | High | EGSAFEDD | [R].EGSAFE | 1xPhospho | A1L366 1xl | 0.000207 | 0 | 1 | 5 | 6 | A1L366 | RAD21 hor | A1L366 [17 | 0 | 4124.915 | 0 | 0.56 | 0.026386 | 17.33 | 86.4 | 113.6 | 16.14 | 11.05 | | High | High | High | High | High | High | High | High | High | High | High | 0 | 4.07E-05 | 5.16 |
| FALSE | High | EGSGLYFN | [R].EGSGLY | 1xPhospho | E9Q2I4 1xP | 8.11E-05 | 0 | 1 | 3 | 1 | E9Q2I4 | ELM2 and I | E9Q2I4 [65 | 0 | 3122.658 | 0 | 0.43 | 0.008916 | 19.19 | 86.7 | 113.3 | 8.1 | 8.11 | | High | High | High | High | High | High | High | High | High | High | High | 0 | 1.43E-05 | 2.48 |
| FALSE | High | EGSGSSGT | [R].EGSGSS | 1xPhospho | P38647 1xl | 0.011755 | 0.000889 | 1 | 1 | 1 | P38647 | Stress-70 p | P38647 [66 | 0 | 1661.778 | 0 | -0.2 | 0.450942 | 13.45 | 106.9 | 93.1 | 8.23 | 22.95 | | High | High | High | High | High | High | High | High | High | High | High | 0.000449 | 0.003765 | 2.21 |
| FALSE | High | EGSHSPLDS | [R].EGSHSF | 1xPhospho | Q6IR34-5 1 | 3.67E-05 | 0 | 1 | 5 | 1 | Q6IR34-5 | Isoform 5 c | Q6IR34-5 [ | 0 | 1678.764 | 0 | -0.38 | 0.013055 | 4.33 | 109.6 | 90.4 | 7.15 | 10.26 | | High | High | High | High | High | High | High | High | High | High | High | 0 | 5.8E-06 | 1.97 |
| FALSE | High | EGSLAEAD | [K].EGSLAE | 1xPhospho | Q8CH25-2 | 0.010382 | 0.000772 | 1 | 1 | 2 | Q8CH25-2 | Isoform 2 c | Q8CH25-2 | 0 | 2836.286 | 0 | -0.07 | 0.629462 | 6.96 | 102.3 | 97.7 | 13.5 | 15.11 | | High | High | High | High | High | High | High | High | High | High | High | 0.000382 | 0.003269 | 2.92 |
| FALSE | High | EGSLAEAD | [K].EGSLAE | 1xPhospho | Q8CH25-2 | 0.000203 | 0 | 1 | 1 | 3 | Q8CH25-2 | Isoform 2 c | Q8CH25-2 | 0 | 2852.281 | 0 | -0.22 | 0.292077 | 1.75 | 102.4 | 97.6 | 10.09 | 14.45 | | High | High | High | High | High | High | High | High | High | High | High | 0 | 3.99E-05 | 3.41 |
| FALSE | High | EGSIETLSN | [R].EGSIET | 1xPhospho | Q8CCN5 1x | 8.28E-06 | 0 | 1 | 5 | 3 | Q8CCN5 | Breast carc | Q8CCN5 [8 | 0 | 2275.065 | 0 | -0.11 | 0.148786 | 5.09 | 101.5 | 98.5 | 3.51 | 4.74 | | High | High | High | High | High | High | High | High | High | High | High | 0 | 1.09E-06 | 4.54 |
| FALSE | High | EGSLGDELI | [K].EGSLGI | 1xPhospho | A0A0R4J2E | 4.18E-07 | 0 | 1 | 4 | 4 | A0A0R4J2E | BTB/POZ d | A0A0R4J2E | 0 | 3917.89 | 0 | -0.72 | 0.217762 | 16 | 109.9 | 90.1 | 22.99 | 32.87 | | High | High | High | High | High | High | High | High | High | High | High | 0 | 3.83E-08 | 5.48 |
| FALSE | High | EGSLSPAR | [R].EGSLSP | 1xPhospho | Q3TDH0 1x | 0.045122 | 0.003373 | 1 | 7 | 1 | Q3TDH0 | Putative ur | Q3TDH0 [4 | 0 | 1125.55 | 0 | 0.2 | 0.111266 | 14.92 | 93.8 | 106.2 | 0.79 | 11.09 | | High | High | High | High | High | High | High | High | High | High | High | 0.001727 | 0.0172 | 1.28 |
| FALSE | High | EGSLSPASV | [K].EGSLSP | 1xPhospho | Q5SWU9-2 | 0.01441 | 0.001011 | 1 | 6 | 4 | Q5SWU9-2 | Isoform 2 c | Q5SWU9-2 | 0 | 3456.656 | 0 | 1.08 | 0.014011 | 10.27 | 63.1 | 136.9 | 22.11 | 10.67 | | High | High | High | High | High | High | High | High | High | High | High | 0.001597 | 0.01538 | 3.67 |
| FALSE | High | EGSLSPASV | [K].EGSLSP | 1xPhospho | Q5SWU9-2 | 0.019276 | 0.001402 | 1 | 6 | 1 | Q5SWU9-2 | Isoform 2 c | Q5SWU9-2 | 0 | 3472.65 | 0 | 0.73 | 0.024604 | 30.01 | 74.4 | 125.6 | 16.52 | 16.45 | | High | High | High | High | High | High | High | High | High | High | High | 0.000756 | 0.00653 | 3.91 |
| FALSE | High | EGRLSPDIV | [K].EGRLSF | 1xPhospho | Q64669 1x | 0.004451 | 0.000409 | 1 | 2 | 1 | Q64669 | NAD(P)H de | Q64669 [7 | 1 | 1989.057 | 0 | 1.99 | 0.000356 | 4.81 | 38.7 | 161.3 | 5.43 | 7.89 | | High | High | High | High | High | High | High | High | High | High | High | 0.000183 | 0.001269 | 1.29 |
| FALSE | High | EGSLSTEDL | [R].EGSLST | 1xPhospho | Q80TX5 1x | 0.096959 | 0.007777 | 1 | 3 | 2 | Q80TX5 | MKIAA057 | Q80TX5 [1 | 0 | 1528.746 | 0 | -0.19 | 0.003881 | 3.63 | 107 | 93 | 2.81 | 4.78 | | High | High | High | High | High | High | High | High | High | High | High | 0.003922 | 0.04166 | 2.28 |
| FALSE | High | EGSPANWI | [R].EGSPAI | 1xPhospho | Q3TF62 1x | 0.012845 | 0.000889 | 1 | 4 | 3 | Q3TF62 | Putative ur | Q3TF62 [2 | 0 | 1426.713 | 0 | -1.74 | 0.000207 | 0.66 | 156 | 44 | 5.06 | 10.51 | | High | High | High | High | High | High | High | High | High | High | High | 0.000585 | 0.005502 | 1.93 |
| FALSE | High | EGSPAPLEF | [R].EGSPAF | 1xPhospho | Q3UCQ1 1 | 0.003822 | 0.000261 | 1 | 3 | 2 | Q3UCQ1 | Forkhead b | Q3UCQ1 [3 | 0 | 1999.958 | 0 | 0.99 | 0.064278 | 46.59 | 59 | 141 | 24.8 | 31.2 | | High | High | High | High | High | High | High | High | High | High | High | 0.000117 | 0.001076 | 3.51 |
| FALSE | High | EGSPAPLEF | [R].EGSPAF | 1xPhospho | Q3UCQ1 1 | 3.89E-07 | 0 | 1 | 3 | 3 | Q3UCQ1 | Forkhead b | Q3UCQ1 [3 | 0 | 2229.12 | 0 | 1.35 | 0.000466 | 6.57 | 56.8 | 143.2 | 6.28 | 8.72 | | High | High | High | High | High | High | High | High | High | High | High | 0 | 3.53E-08 | 3.98 |
| FALSE | High | EGSPIPHDF | [R].EGSPIP | 1xPhospho | P42128 1xl | 0.002623 | 0.000109 | 1 | 1 | 1 | P42128 | Forkhead b | P42128 [42 | 0 | 1757.831 | 0 | 1.59 | 0.09118 | 52.56 | 49.9 | 150.1 | 33.98 | 40.63 | | High | High | High | High | High | High | High | High | High | High | High | 4.94E-05 | 0.000705 | 2.74 |
| FALSE | High | EGSPIPHDF | [R].EGSPIP | 1xPhospho | P42128 1xl | 5.77E-06 | 0 | 1 | 1 | 3 | P42128 | Forkhead b | P42128 [42 | 0 | 1986.994 | 0 | 1.4 | 0.000222 | 4.16 | 55.2 | 144.8 | 5.78 | 4.71 | | High | High | High | High | High | High | High | High | High | High | High | 0 | 3.43E-06 | 3.22 |
| FALSE | High | EGSPQTCA | [R].EGSPQ | 1xCarbami | Q5D1E7 1x | 0.000941 | 0 | 1 | 1 | 1 | Q5D1E7 | Endoribon | Q5D1E7 [3 | 0 | 1539.682 | 0 | 0.14 | 0.197078 | 11.49 | 96.6 | 103.4 | 7.4 | 9.37 | | High | High | High | High | High | High | High | High | High | High | High | 0 | 0.000222 | 2.22 |
| FALSE | High | EGSQGADA | [R].EGSQG | 1xPhospho | I3PQW4 1x | 0.002793 | 0.000109 | 1 | 1 | 1 | I3PQW4 | Uncharacte | I3PQW4 [4 | 0 | 3487.751 | 0 | -0.99 | 0.000931 | 2.24 | 133.1 | 66.9 | 10.77 | 5.48 | | High | High | High | High | High | High | High | High | High | High | High | 4.94E-05 | 0.000756 | 2.93 |
| FALSE | High | EGSQGADA | [R].EGSQG | 1xPhospho | Q0VGT4 1x | 2.99E-07 | 0 | 1 | 3 | 1 | Q0VGT4 | Protein ZG | Q0VGT4 [4 | 0 | 3471.72 | 0 | 0.21 | 0.452373 | 29.7 | 97.2 | 102.8 | 15.52 | 14.82 | | High | High | High | High | High | High | High | High | High | High | High | 0 | 2.62E-08 | 3.97 |
| FALSE | High | EGSQGELT | [R].EGSQG | 1xPhospho | G3UXW9 1 | 8.22E-07 | 0 | 1 | 4 | 4 | G3UXW9 | COP9 signa | G3UXW9 [ | 0 | 1869.854 | 0 | -0.1 | 0.169103 | 9.24 | 103.6 | 96.4 | 4.99 | 5.17 | | High | High | High | High | High | High | High | High | High | High | High | 0 | 0.000183 | 2.46 |
| FALSE | High | EGSRPEEAI | [R].EGSRPE | 1xPhospho | Q99L02 1xl | 0.001211 | 0 | 1 | 3 | 1 | Q99L02 | PAXIP1-ass | Q99L02 [6 | 0 | 2310.045 | 0 | -0.09 | 0.210627 | 6.19 | 103.9 | 96.1 | 8.47 | 5.79 | | High | High | High | High | High | High | High | High | High | High | High | 0 | 0.000296 | 2.44 |
| FALSE | High | EGSSIPELA | [K].EGSSIP | 1xPhospho | E9QQ10 1x | 2.06E-05 | 0 | 1 | 4 | 2 | E9QQ10 | A-kinase ar | E9QQ10 [2 | 0 | 2117.995 | 0 | 0.8 | 0.001333 | 11.65 | 75 | 125 | 6.22 | 8.34 | | High | High | High | High | High | High | High | High | High | High | High | 0 | 3.03E-06 | 3.88 |
| FALSE | High | EGSTEFDW | [R].EGSTEF | 1xPhospho | Q924A2 1x | 0.033116 | 0.002195 | 1 | 1 | 1 | Q924A2 | Protein cap | Q924A2 [5 | 0 | 1924.78 | 0 | -0.16 | 0.042286 | 4.18 | 107.7 | 92.3 | 10.6 | 8.47 | | High | High | High | High | High | High | High | High | High | High | High | 0.001151 | 0.01209 | 2.65 |
| FALSE | High | EGSTPEDD | [R].EGSTPE | 1xPhospho | E0CYH0 1x | 4.74E-10 | 0 | 1 | 2 | 7 | E0CYH0 | MCG16685 | E0CYH0 [2 | 0 | 2363.044 | 0 | -0.36 | 0.009502 | 11.56 | 115.9 | 84.1 | 6.82 | 9.87 | | High | High | High | High | High | High | High | High | High | High | High | 0 | 7.73E-11 | 4.48 |
| FALSE | High | EGSLTGTK | [R].EGSLTG | 1xPhospho | G3X9K3 1x | 0.021231 | 0.001507 | 1 | 1 | 2 | G3X9K3 | Brefeldin A | G3X9K3 [1 | 0 | 1330.702 | 0 | 0.68 | 0.00212 | 8.64 | 76.2 | 123.8 | 9.57 | 4.56 | | High | High | High | High | High | High | High | High | High | High | High | 0.000801 | 0.00729 | 1.43 |
| FALSE | High | EGQDPAEL | [K].EGQDP | 1xPhospho | Q3U343 1x | 5.7E-05 | 0 | 1 | 2 | 3 | Q3U343 | Putative ur | Q3U343 [3 | 0 | 2141.042 | 0 | 0.24 | 0.029268 | 9.25 | 88.2 | 111.8 | 6.63 | 8.96 | | High | High | High | High | High | High | High | High | High | High | High | 0 | 9.55E-06 | 2.43 |
| FALSE | High | EGPVGGES | [R].EGPVG | 1xPhospho | Q3UHU5 1 | 5.81E-05 | 0 | 1 | 5 | 2 | Q3UHU5 | Microtubu | Q3UHU5 [ | 0 | 2250.988 | 0 | -1.86 | 0.000402 | 0.49 | 155.2 | 44.8 | 5.51 | 12.65 | | High | High | High | High | High | High | High | High | High | High | High | 9.42E-05 | 0.001041 | 3.04 |
| FALSE | High | EGPSSLNL | [K].EGPSSL | 1xPhospho | A0A0R4J14 | 0.066503 | 0.004796 | 1 | 6 | 1 | A0A0R4J14 | Clustered r | A0A0R4J14 | 0 | 1310.619 | 0 | 2.14 | 0.000688 | 16.42 | 36.3 | 163.7 | 10.84 | 10.3 | | High | High | High | High | High | High | High | High | High | High | High | 0.002455 | 0.0268 | 2.03 |
| FALSE | High | EGDTEEEA | [K].EGDTEI | 1xPhospho | P28667 1xl | 0.004572 | 0.000456 | 1 | 1 | 6 | P28667 | MARCKS-re | P28667 [1 | 0 | 3662.534 | 0 | -1.29 | 0.002146 | 4.74 | 141.9 | 58.1 | 4.52 | 31.59 | | High | High | High | High | High | High | High | High | High | High | High | 0.000327 | 0.002424 | 3.32 |
| FALSE | High | EGEATKDS | [K].EGEATI | 1xPhospho | Q9DBS5 1x | 0.003788 | 0.00021 | 1 | 2 | 1 | Q9DBS5 | Kinesin ligh | Q9DBS5 [1 | 1 | 3549.579 | 0 | 0.77 | 0.021899 | 18.46 | 73.9 | 126.1 | 15.19 | 25.52 | | High | High | High | High | High | High | High | High | High | High | High | 0.000117 | 0.001065 | 3.77 |
| FALSE | High | EGEDSSVIH | [K].EGEDSS | 1xPhospho | Q6PDQ2 1 | 0.000589 | 0 | 1 | 8 | 3 | Q6PDQ2 | Chromodo | Q6PDQ2 [1 | 0 | 2031.931 | 0 | -0.3 | 0.033452 | 10.39 | 107.1 | 92.9 | 12.97 | 19.94 | | High | High | High | High | High | High | High | High | High | High | High | 0 | 0.000274 | 2.3 |
| FALSE | High | EGEEPTVYS | [K].EGEEPT | 1xPhospho | O55022 1x | 6E-07 | 0 | 1 | 3 | 4 | O55022 | Membrane | O55022 [1 | 0 | 2262.01 | 0 | 0.28 | 0.02895 | 6.78 | 90.5 | 109.5 | 3.16 | 8.37 | | High | High | High | High | High | High | High | High | High | High | High | 0 | 5.74E-08 | 3.39 |
| FALSE | High | EGEEPTVYS | [K].EGEEPT | 1xPhospho | O55022 1x | 0.001659 | 5.66E-05 | 1 | 3 | 3 | O55022 | Membrane | O55022 [1 | 1 | 2834.266 | 0 | -0.28 | 0.72894 | 21.36 | 109.5 | 90.5 | 24.02 | 18.54 | | High | High | High | High | High | High | High | High | High | High | High | 2.58E-05 | 0.000423 | 2 |
| FALSE | High | EGEGSLRPS | [K].EGEGSI | 1xPhospho | Q9DBN9 1 | 0.080423 | 0.006219 | 1 | 2 | 2 | Q9DBN9 | Probable A | Q9DBN9 [1 | 0 | 1708.822 | 0 | -0.36 | 0.002004 | 5.16 | 111.9 | 88.1 | 3.69 | 3.42 | | High | High | High | High | High | High | High | High | High | High | High | 0.003188 | 0.0335 | 1.29 |
| FALSE | High | EGEPAAEG | [K].EGEPA | 1xCarbami | E9Q3P4 1x | 0.001255 | 0 | 1 | 3 | 4 | E9Q3P4 | Centromer | E9Q3P4 [1 | 0 | 4067.877 | 0 | -0.44 | 0.050261 | 11.63 | 116.3 | 83.7 | 13.74 | 13.7 | | High | High | High | High | High | High | High | High | High | High | High | 0.000161 | 0.001216 | 3.16 |
| FALSE | High | EGETQSGT | [R].EGETQ | 1xPhospho | S4R1W5 1x | 5.91E-05 | 0 | 1 | 4 | 2 | S4R1W5 | RNA-bindir | S4R1W5 [3 | 0 | 4275.992 | 0 | -0.8 | 0.074601 | 17.35 | 126.9 | 73.1 | 19.18 | 22.09 | | High | High | High | High | High | High | High | High | High | High | High | 0 | 0.00011 | 3.71 |
| FALSE | High | EGGASSLK | [R].EGGAS | 1xPhospho | Q3U9K6 1x | 0.119269 | 0.00979 | 1 | 4 | 1 | Q3U9K6 | Interferon | Q3U9K6 [3 | 0 | 1286.676 | 0 | 1.37 | 0.000261 | 13.03 | 57.6 | 142.4 | 10.3 | 5.01 | | High | High | High | High | High | High | High | High | High | High | High | 0.004866 | 0.05292 | 1.57 |
| FALSE | High | EGGIDVSD | [K].EGGID | 1xCarbami | B2RS91 2xl | 8.5E-06 | 0 | 1 | 1 | 4 | B2RS91 | RNA polym | B2RS91 [1 | 0 | 3181.254 | 0 | -0.33 | 0.020774 | 10.83 | 116 | 84 | 6.52 | 10.38 | | High | High | High | High | High | High | High | High | High | High | High | 0 | 1.13E-06 | 3.49 |
| FALSE | High | EGGSEDGS | [R].EGGSEI | 2xPhospho | Q3U0X6 2x | 0.004106 | 0.00036 | 1 | 1 | 1 | Q3U0X6 | Putative ur | Q3U0X6 [4 | 0 | 2415.979 | 0 | -0.56 | 0.02226 | 8.27 | 119 | 81 | 15.29 | 9.04 | | High | High | High | High | High | High | High | High | High | High | High | 0.000161 | 0.001158 | 2.67 |
| FALSE | High | EGGSEDGS | [R].EGGSEI | 1xPhospho | Q3U0X6 1x | 8.79E-05 | 0 | 1 | 1 | 4 | Q3U0X6 | Putative ur | Q3U0X6 [4 | 0 | 2565.176 | 0 | -0.26 | 0.002954 | 4.54 | 107.9 | 92.1 | 5.11 | 6.1 | | High | High | High | High | High | High | High | High | High | High | High | 0 | 0.000283 | 3.75 |
| FALSE | High | EGGSEDGS | [R].EGGSEI | 2xPhospho | Q3U0X6 2x | 3.8E-05 | 0 | 1 | 1 | 3 | Q3U0X6 | Putative ur | Q3U0X6 [4 | 0 | 2645.142 | 0 | -0.67 | 0.000487 | 3.59 | 123.1 | 76.9 | 2.39 | 4.35 | | High | High | High | High | High | High | High | High | High | High | High | 0 | 6.08E-06 | 4.05 |
| FALSE | High | EGGTVGSG | [K].EGGTV | 1xPhospho | Q8BLH7 1x | 2.1E-06 | 0 | 1 | 1 | 4 | Q8BLH7 | HIRA-intera | Q8BLH7 [3 | 0 | 1747.815 | 0 | -1.23 | 0.000306 | 1.71 | 138.6 | 61.4 | 2.03 | 9.33 | | High | High | High | High | High | High | High | High | High | High | High | 0 | 2.35E-07 | 3.85 |
| FALSE | High | EGHGSSSFI | [R].EGHGS | 1xPhospho | Q80SY5 1x | 0.042868 | 0.003228 | 1 | 1 | 3 | Q80SY5 | Pre-mRNA- | Q80SY5 [2 | 0 | 1387.584 | 0 | -0.57 | 0.005665 | 7.27 | 118.1 | 81.9 | 5.28 | 11.75 | | High | High | High | High | High | High | High | High | High | High | High | 0.002159 | 0.02259 | 2.41 |
| FALSE | High | EGHPAPPP | [R].EGHPAI | 1xPhospho | Q6PEV3 1x | 0.008388 | 0.000736 | 1 | 1 | 1 | Q6PEV3 | WAS/WAS | Q6PEV3 [2 | 0 | 2524.384 | 0 | 0.61 | 0.00116 | 13.02 | 79.1 | 120.9 | 6.91 | 5.17 | | High | High | High | High | High | High | High | High | High | High | High | 0.000327 | 0.002574 | 1.76 |
| FALSE | High | EGLCGQPS | [K].EGLCG | 1xCarbami | E9QM75 1 | 0.0016 | 5.66E-05 | 1 | 8 | 2 | E9QM75 | TNFAIP3-in | E9QM75 [2 | 0 | 2466.267 | 0 | -0.06 | 0.916669 | 10.47 | 102 | 98 | 11 | 3.36 | | High | High | High | High | High | High | High | High | High | High | High | 2.58E-05 | 0.000405 | 2.23 |
| FALSE | High | EGLLPASD | [K].EGLLPA | 1xPhospho | Q6PGL7 1x | 6.26E-06 | 0 | 1 | 5 | 4 | Q6PGL7 | WASH com | Q6PGL7 [7 | 0 | 3142.523 | 0 | 0 | 0.780612 | 12.14 | 101.5 | 98.5 | 9.29 | 9.43 | | High | High | High | High | High | High | High | High | High | High | High | 0 | 1.93E-06 | 4.51 |
| FALSE | High | EGLLPISPA | [K].EGLLPIS | 1xPhospho | B9EHK8 1x | 0.001386 | 5.66E-05 | 1 | 4 | 2 | B9EHK8 | Homeo bo | B9EHK8 [2 | 0 | 2246.172 | 0 | 2.56 | 0.000464 | 24.78 | 31.5 | 168.5 | 15.97 | 8.18 | | High | High | High | High | High | High | High | High | High | High | High | 2.58E-05 | 0.000344 | 3.02 |
| FALSE | High | EGLISQDGS | [K].EGLISQ | 1xPhospho | D3Z5N6 1x | 0.007881 | 0.000736 | 1 | 4 | 2 | D3Z5N6 | Zinc finger | D3Z5N6 [1 | 0 | 1997.015 | 0 | -0.26 | 0.007837 | 2.49 | 110.4 | 89.6 | 4.96 | 3.52 | | High | High | High | High | High | High | High | High | High | High | High | 0.000382 | 0.003292 | 2.03 |
| FALSE | High | EGINPGYD | [K].EGINPG | 1xPhospho | Q08943-2 | 1.3E-09 | 0 | 1 | 4 | 21 | Q08943-2 | Isoform 2 c | Q08943-2 | 0 | 3096.258 | 0 | -0.95 | 0.000163 | 0.22 | 132.1 | 67.9 | 1.59 | 3.08 | | High | High | High | High | High | High | High | High | High | High | High | 0 | 2.55E-09 | 5.68 |
| FALSE | High | EGINPGYD | [K].EGINPG | 1xDeamida | Q08943-2 | 0.012397 | 0.000889 | 1 | 4 | 3 | Q08943-2 | Isoform 2 c | Q08943-2 | 0 | 3097.242 | 0 | -0.8 | 0.000176 | 1.21 | 127.5 | 72.5 | 4.25 | 7.82 | | High | High | High | High | High | High | High | High | High | High | High | 0.000449 | 0.004002 | 3.69 |
| FALSE | High | EGLPVNNII | [K].EGLPVN | 1xPhospho | G3UW74 1 | 3.71E-06 | 0 | 1 | 4 | 1 | G3UW74 | Mediator c | G3UW74 [ | 0 | 2592.384 | 0 | 0.55 | 0.020327 | 20.09 | 78.6 | 121.4 | 7.55 | 14.21 | | High | High | High | High | High | High | High | High | High | High | High | 0 | 4.42E-07 | 3.35 |
| FALSE | High | EGLRFSDEF | [K].EGLRFS | 1xPhospho | Q99MJ9 1x | 0.068752 | 0.005042 | 1 | 2 | 1 | Q99MJ9 | ATP-depen | Q99MJ9 [7 | 1 | 2174.068 | 0 | -0.19 | 0.116307 | 11.1 | 106 | 94 | 5.55 | 16.55 | | High | High | High | High | High | High | High | High | High | High | High | 0.002589 | 0.02777 | 1.93 |
| FALSE | High | EGLSPAK | [K].EGLSPA | 1xPhospho | C7G3P2 1x | 0.00824 | 0.000736 | 1 | 3 | 2 | C7G3P2 | MKIAA053 | C7G3P2 [9 | 0 | 1239.675 | 0 | -0.02 | 0.761625 | 4.12 | 98.9 | 101.1 | 4.94 | 4.51 | | High | High | High | High | High | High | High | High | High | High | High | 0.004115 | 0.04392 | 1.63 |
| FALSE | High | EGLSPSQR | [K].EGLSPS | 1xPhospho | Q3U1J1-2 | 0.104222 | 0.008348 | 1 | 4 | 1 | Q3U1J1-2 | Isoform 2 c | Q3U1J1-2 | 0 | 1182.572 | 0 | -0.2 | 0.033217 | 4.92 | 107.7 | 92.3 | 3.05 | 13.84 | | High | High | High | High | High | High | High | High | High | High | High | 0.004206 | 0.04529 | 1.55 |
| FALSE | High | EGPEPPEE | [K].EGPEPF | 1xPhospho | Q7TSC1 1x | 0.000611 | 0 | 1 | 3 | 4 | Q7TSC1 | Protein PRI | Q7TSC1 [5 | 0 | 2504.273 | 0 | 0.53 | 0.001842 | 2.79 | 82.1 | 117.9 | 6.19 | 5.35 | | High | High | High | High | High | High | High | High | High | High | High | 0 | 0.000137 | 2.31 |
| FALSE | High | EGPGPGPG | [K].EGPGP | 1xPhospho | F8WJ40 1x | 0.000302 | 0 | 1 | 2 | 1 | F8WJ40 | Histone-lys | F8WJ40 [2 | 0 | 1531.792 | 0 | 0.08 | 0.737014 | 1.24 | 98.4 | 101.6 | 14.45 | 2.68 | | High | High | High | High | High | High | High | High | High | High | High | 0 | 6.21E-05 | 2.58 |
| FALSE | High | EGPRDSLTI | [K].EGPRDS | 1xPhospho | Q8BJS8 1xl | 0.000712 | 0 | 1 | 3 | 2 | Q8BJS8 | Mdm2-bin | Q8BJS8 [5 | 1 | 1953.046 | 0 | -0.67 | 0.00341 | 3.22 | 119.2 | 80.8 | 6.78 | 6.8 | | High | High | High | High | High | High | High | High | High | High | High | 0 | 0.000163 | 3.06 |
| FALSE | High | EGSVLDILK | [R].EGSVLD | 1xPhospho | E9Q3G8 1x | 0.058664 | 0.004294 | 1 | 3 | 1 | E9Q3G8 | Nucleopor | E9Q3G8 [6 | 0 | 1282.686 | 0 | 0.89 | 0.153371 | 30.95 | 72.7 | 127.3 | 19.18 | 29.68 | | High | High | High | High | High | High | High | High | High | High | High | 0.002187 | 0.02325 | 1.94 |
| FALSE | High | ELLELGIDN | [R].ELLELG | 2xPhospho | E9Q2H1 2x | 3.71E-06 | 0 | 1 | 2 | 1 | E9Q2H1 | E3 ubiquiti | E9Q2H1 [1 | 0 | 3864.663 | 0 | 0.85 | 0.02662 | 21.19 | 70.6 | 129.4 | 36.82 | 20.04 | | High | High | High | High | High | High | High | High | High | High | High | 0 | 4.45E-07 | 4.6 |
| FALSE | High | ELILGSETL | [R].ELILGSI | 1xPhospho | B2RS36 1xl | 0.004572 | 0.000456 | 1 | 4 | 2 | B2RS36 | Adrenocor | B2RS36 [1 | 0 | 1710.888 | 0 | -0.79 | 0.002496 | 0.61 | 126.1 | 73.9 | 2.99 | 10.03 | | High | High | High | High | High | High | High | High | High | High | High | 0.000538 | 0.004447 | 2.38 |
| FALSE | High | ELILSSEPSF | [R].ELILSSE | 1xPhospho | Q3U5L3 1x | 3.78E-06 | 0 | 1 | 3 | 3 | Q3U5L3 | Putative ur | Q3U5L3 [8 | 0 | 2798.515 | 0 | 0.33 | 0.038891 | 23.25 | 90.3 | 109.7 | 14.47 | 6.91 | | High | High | High | High | High | High | High | High | High | High | High | 0 | 4.53E-07 | 4.17 |
| FALSE | High | ENQESLVSI | [K].ENQESI | 1xPhospho | Q6P5H2-2 | 0.007811 | 0.000664 | 1 | 2 | 1 | Q6P5H2-2 | Isoform 2 c | Q6P5H2-2 | 0 | 2518.215 | 0 | 0.81 | 0.001737 | 10.2 | 73.9 | 126.1 | 5.86 | 10.67 | | High | High | High | High | High | High | High | High | High | High | High | 0.000327 | 0.002374 | 3.38 |
| FALSE | High | ENQGSTLP | [K].ENQGS | 1xPhospho | Q3TBP4 1x | 2.58E-08 | 0 | 1 | 2 | 2 | Q3TBP4 | Putative ur | Q3TBP4 [1 | 0 | 3387.523 | 0 | -0.06 | 0.150758 | 10.7 | 102.8 | 97.2 | 6.81 | 4.96 | | High | High | High | High | High | High | High | High | High | High | High | 0 | 9.93E-08 | 5.28 |
| FALSE | High | ENQGSTLP | [K].ENQGS | 2xPhospho | Q3TBP4 2x | 6.66E-09 | 0 | 1 | 2 | 2 | Q3TBP4 | Putative ur | Q3TBP4 [1 | 0 | 3467.489 | 0 | 0.1 | 0.423823 | 5.17 | 98.4 | 101.6 | 7.96 | 5.28 | | High | High | High | High | High | High | High | High | High | High | High | 0 | 4.67E-05 | 4.74 |
| FALSE | High | ENQVVELH | [R].ENQVV | 1xCarbami | Q56A08 1x | 2.49E-05 | 0 | 1 | 1 | 1 | Q56A08 | G patch do | Q56A08 [4 | 0 | 3108.28 | 0 | 1.11 | 0.007244 | 30.74 | 62.1 | 137.9 | 10.38 | 16.26 | | High | High | High | High | High | High | High | High | High | High | High | 0 | 3.76E-06 | 3.11 |
| FALSE | High | ENRQSIINP | [K].ENRQSI | 1xPhospho | P46460 1xl | 6.71E-05 | 0 | 1 | 2 | 2 | P46460 | Vesicle-fus | P46460 [2 | 1 | 2428.206 | 0 | 0.47 | 0.047232 | 24.19 | 77.4 | 122.6 | 9 | 22.57 | | High | High | High | High | High | High | High | High | High | High | High | 0 | 1.15E-05 | 3.76 |
| FALSE | High | ENSAAPSP | [K].ENSAAI | 1xPhospho | Q8CB16 1x | 0.014797 | 0.001049 | 1 | 7 | 1 | Q8CB16 | Putative ur | Q8CB16 [1 | 0 | 2290.159 | 0 | 1.09 | 0.010932 | 17.38 | 66 | 134 | 10.06 | 14.96 | | High | High | High | High | High | High | High | High | High | High | High | 0.000555 | 0.004873 | 1.34 |
| FALSE | High | ENSAAPSP | [K].ENSAAI | 2xPhospho | Q8CB16 2x | 0.030367 | 0.001857 | 1 | 7 | 2 | Q8CB16 | Putative ur | Q8CB16 [1 | 0 | 2370.126 | 0 | 1.68 | 0.005644 | 9.4 | 48.9 | 151.1 | 6.77 | 20.41 | | High | High | High | High | High | High | High | High | High | High | High | 0.002159 | 0.02286 | 1.74 |
| FALSE | High | ENSEGEK | [K].ENSEGI | 1xPhospho | A2AR02 1x | 0.117405 | 0.009523 | 1 | 2 | 1 | A2AR02 | Peptidyl-pr | A2AR02 [6 | 0 | 1330.629 | 0 | -0.98 | 0.001301 | 1.23 | 130.5 | 69.5 | 5.95 | 11.25 | | High | High | High | High | High | High | High | High | High | High | High | 0.004769 | 0.05197 | 1.73 |
| FALSE | High | ENSFESSLE | [R].ENSFES | 1xPhospho | Q8BTI8 1xP | 0.053023 | 0.00384 | 1 | 3 | 1 | Q8BTI8 | Serine/argi | Q8BTI8 [12 | 0 | 1625.73 | 0 | -0.09 | 0.276869 | 10.11 | 102.4 | 97.6 | 9.55 | 3.33 | | High | High | High | High | High | High | High | High | High | High | High | 0.001999 | 0.02056 | 1.98 |
| FALSE | High | ENSFESSLE | [R].ENSFES | 1xPhospho | Q8BTI8 1xP | 0.000483 | 0 | 1 | 3 | 4 | Q8BTI8 | Serine/argi | Q8BTI8 [12 | 0 | 1854.893 | 0 | -0.77 | 0.000682 | 1.45 | 128.5 | 71.5 | 5.95 | 8.77 | | High | High | High | High | High | High | High | High | High | High | High | 9.42E-05 | 0.001026 | 3.14 |
| FALSE | High | ENSFESSLE | [R].ENSFES | 2xPhospho | Q8BTI8 2xP | 0.033986 | 0.002195 | 1 | 3 | 1 | Q8BTI8 | Serine/argi | Q8BTI8 [12 | 0 | 1934.859 | 0 | -0.98 | 0.000456 | 1.91 | 134.7 | 65.3 | 6.95 | 8.59 | | High | High | High | High | High | High | High | High | High | High | High | 0.001151 | 0.0124 | 2.79 |
| FALSE | High | ENSPFINNV | [R].ENSPFI | 1xPhospho | Q9WVL3 1 | 0.001361 | 0 | 1 | 3 | 1 | Q9WVL3 | Solute carr | Q9WVL3 [4 | 0 | 1855.879 | 0 | -0.07 | 0.836574 | 6.81 | 101.1 | 98.9 | 6.67 | 4.95 | | High | High | High | High | High | High | High | High | High | High | High | 0 | 0.000336 | 2.47 |
| FALSE | High | ENSPFINNV | [R].ENSPFI | 1xPhospho | Q9WVL3 1 | 0.106748 | 0.008634 | 1 | 3 | 2 | Q9WVL3 | Solute carr | Q9WVL3 [4 | 1 | 2925.407 | 0 | 0.41 | 0.119278 | 28.49 | 81.7 | 118.3 | 7.16 | 26.91 | | High | High | High | High | High | High | High | High | High | High | High | 0.004462 | 0.04817 | 2.01 |
| FALSE | High | ENPVLDIVS | [K].ENPVLD | 1xPhospho | P70399 1xP | 1.02E-05 | 0 | 1 | 5 | 8 | P70399 | TP53-bindi | P70399 [6 | 0 | 3821.827 | 0 | -0.14 | 0.044819 | 5.08 | 106.4 | 93.6 | 4.43 | 12.26 | | High | High | High | High | High | High | High | High | High | High | High | 0 | 3.09E-05 | 4.17 |
| FALSE | High | ENSSLEGG | [K].ENSSLE | 1xCarbami | Q8BX22 1x | 2.22E-07 | 0 | 1 | 1 | 11 | Q8BX22 | Sal-like pro | Q8BX22 [7 | 0 | 3231.395 | 0 | -1.58 | 0.001245 | 2.42 | 149.3 | 50.7 | 9.36 | 10.59 | | High | High | High | High | High | High | High | High | High | High | High | 0 | 1.88E-08 | 4.18 |
| FALSE | High | ENSSTSYTF | [K].ENSSTS | 1xPhospho | Q9ERU9 1x | 0.118334 | 0.009621 | 1 | 1 | 1 | Q9ERU9 | E3 SUMO-p | Q9ERU9 [2 | 1 | 2386.215 | 0 | -0.55 | 0.142758 | 14.79 | 116.9 | 83.1 | 19.12 | 14.41 | | High | High | High | High | High | High | High | High | High | High | High | 0.004866 | 0.05284 | 1.91 |
| FALSE | High | ENSTVDFSI | [R].ENSTVI | 1xPhospho | I1VGT4 1xP | 0.006895 | 0.00062 | 2 | 34 | 2 | I1VGT4; I1 | Anion exch | I1VGT4 [28 | 0 | 1335.603 | 0 | 0.54 | 0.042372 | 9.62 | 83.8 | 116.2 | 7.66 | 14.17 | Shared | High | High | High | High | High | High | High | High | High | High | High | 0.005336 | 0.05805 | 2.45 |

| | | | | | | | | | | | | | | | | | | | | | | | | | | | | | | | | | | | | | | |
|---|---|---|---|---|---|---|---|---|---|---|---|---|---|---|---|---|---|---|---|---|---|---|---|---|---|---|---|---|---|---|---|---|---|---|---|---|---|---|
| FALSE | High | ENSTVDFSI | [R].ENSTVI | 1xPhospho | I1VGT4 1xP | 0.002719 | 0.000109 | 2 | 34 | 2 | I1VGT4; I1V | Anion exch | I1VGT4 [28 | 0 | 1564.766 | 0 | 0.98 | 0.002927 | 7.43 | 66.3 | 133.7 | 6.38 | 15.07 | Shared | High | High | High | High | High | High | High | High | High | High | High | 0.000275 | 0.002069 | 2.62 |
| FALSE | High | ENTAFSDW | [K].ENTAFS | 1xPhospho | E9Q784 1x | 2.63E-05 | 0 | 1 | 1 | 1 | E9Q784 | Zinc finger | E9Q784 [1 | 0 | 2191.902 | 0 | -0.75 | 0.02113 | 0.61 | 125.5 | 74.5 | 14.06 | 17.06 | | High | High | High | High | High | High | High | High | High | High | High | 0 | 3.99E-06 | 3.3 |
| FALSE | High | ENTAFSDW | [K].ENTAFS | 2xPhospho | E9Q784 2x | 1.73E-07 | 0 | 1 | 1 | 2 | E9Q784 | Zinc finger | E9Q784 [1 | 0 | 2271.868 | 0 | -0.48 | 0.041628 | 10.38 | 126.4 | 73.6 | 15.18 | 26.99 | | High | High | High | High | High | High | High | High | High | High | High | 0 | 1.41E-08 | 5.04 |
| FALSE | High | ENTETPAA | [K].ENTETI | 1xPhospho | A0A0N4SU | 1.6E-06 | 0 | 1 | 10 | 1 | A0A0N4SU | Peregrin O | A0A0N4SU | 0 | 1696.856 | 0 | 0.68 | 0.001309 | 5.85 | 75.7 | 124.3 | 7.98 | 6.78 | | High | High | High | High | High | High | High | High | High | High | High | 0 | 1.73E-07 | 3.6 |
| FALSE | High | ENTGVLHA | [K].ENTGV | 1xCarbami | Q0VBL3 1x | 0.017041 | 0.001204 | 1 | 8 | 1 | Q0VBL3 | RNA bindir | Q0VBL3 [9 | 0 | 3257.574 | 0 | -0.21 | 0.149084 | 9.74 | 101.2 | 98.8 | 9.33 | 9.71 | | High | High | High | High | High | High | High | High | High | High | High | 0.000638 | 0.005713 | 2.12 |
| FALSE | High | ENTPEELQ | [K].ENTPEE | 1xPhospho | B2RY19 1xI | 2.74E-07 | 0 | 1 | 7 | 2 | B2RY19 | Cgnl1 prot | B2RY19 [96 | 0 | 2777.36 | 0 | 0.52 | 0.023738 | 25.97 | 85.8 | 114.2 | 14.06 | 19.96 | | High | High | High | High | High | High | High | High | High | High | High | 0 | 2.36E-08 | 4.28 |
| FALSE | High | ENTQIDDT | [R].ENTQID | 1xPhospho | P70399 1xI | 9.25E-07 | 0 | 1 | 4 | 2 | P70399 | TP53-bindi | P70399 [54 | 0 | 2512.222 | 0 | 0.31 | 0.014325 | 9.97 | 89.7 | 110.3 | 3 | 9.39 | | High | High | High | High | High | High | High | High | High | High | High | 0 | 9.32E-08 | 3.96 |
| FALSE | High | ENVALQLG | [R].ENVAL( | 1xPhospho | O54928 1x | 0.003622 | 0.00021 | 1 | 2 | 1 | O54928 | Suppresso | O54928 [6 | 0 | 1894.045 | 0 | 1.69 | 0.000457 | 16.86 | 46.5 | 153.5 | 9.63 | 8.19 | | High | High | High | High | High | High | High | High | High | High | High | 9.42E-05 | 0.001012 | 2.55 |
| FALSE | High | ENVDSPLLI | [K].ENVDSI | 1xPhospho | Q542Q3 1x | 0.00047 | 0 | 1 | 1 | 1 | Q542Q3 | G two S ph | Q542Q3 [7 | 0 | 1552.839 | 0 | 0.38 | 0.020793 | 10.23 | 85.4 | 114.6 | 3.92 | 10.55 | | High | High | High | High | High | High | High | High | High | High | High | 0 | 0.000102 | 2.3 |
| FALSE | High | ENVELSPTA | [K].ENVELS | 1xCarbami | Q04690 1x | 1.24E-05 | 0 | 1 | 5 | 1 | Q04690 | Neurofibro | Q04690 [2 | 0 | 2036.906 | 0 | 1.42 | 0.000605 | 19.62 | 54.5 | 145.5 | 11.66 | 7.38 | | High | High | High | High | High | High | High | High | High | High | High | 0 | 1.73E-06 | 1.77 |
| FALSE | High | ENVEYIERE | [R].ENVEYI | 1xPhospho | Q9R020 1x | 0.005717 | 0.00062 | 1 | 5 | 6 | Q9R020 | Zinc finger | Q9R020 [1 | 1 | 2774.167 | 0 | -0.4 | 0.000364 | 1.59 | 115.2 | 84.8 | 2.29 | 4.21 | | High | High | High | High | High | High | High | High | High | High | High | 0.00196 | 0.01993 | 2.75 |
| FALSE | High | ENSSQMA( | [K].ENSSQI | 1xPhospho | Q9Z277 1x | 0.005182 | 0.000551 | 1 | 1 | 2 | Q9Z277 | Tyrosine-p | Q9Z277 [1 | 0 | 1932.914 | 0 | 0.32 | 0.239595 | 25.64 | 91.2 | 108.8 | 13.22 | 13.15 | | High | High | High | High | High | High | High | High | High | High | High | 0.000246 | 0.001507 | 3.41 |
| FALSE | High | ENVEYIERE | [R].ENVEYI | 1xPhospho | Q9R020 1x | 0.046687 | 0.00354 | 1 | 5 | 2 | Q9R020 | Zinc finger | Q9R020 [1 | 2 | 3131.425 | 0 | -0.38 | 0.004815 | 3.94 | 111.6 | 88.4 | 5.73 | 5.63 | | High | High | High | High | High | High | High | High | High | High | High | 0.001805 | 0.01778 | 2.57 |
| FALSE | High | ENPSSQSTY | [K].ENPSS( | 1xPhospho | Q8BJ34-6 1 | 1.35E-08 | 0 | 1 | 7 | 1 | Q8BJ34-6 | Isoform 6 c | Q8BJ34-6 [ | 0 | 2141.053 | 0 | 1.6 | 0.00038 | 9.55 | 48.9 | 151.1 | 9.49 | 7.13 | | High | High | High | High | High | High | High | High | High | High | High | 0 | 8.06E-10 | 4.02 |
| FALSE | High | ENPPSPPTS | [SR].ENPPS | 3xPhospho | Q8BP27 3x | 1.07E-05 | 0 | 1 | 3 | 4 | Q8BP27 | Swi5-depe | Q8BP27 [6 | 0 | 2111.88 | 0 | -0.49 | 0.004808 | 4.88 | 116.7 | 83.3 | 12.45 | 6.46 | | High | High | High | High | High | High | High | High | High | High | High | 0.000418 | 0.003582 | 2.87 |
| FALSE | High | ENFTPLK | [K].ENFTPL | 1xPhospho | Q8VDR2 1x | 0.026885 | 0.001676 | 1 | 3 | 1 | Q8VDR2 | Hmmr prot | Q8VDR2 [8 | 0 | 1386.743 | 0 | -0.35 | 0.018795 | 5.71 | 112 | 88 | 4.98 | 11.17 | | High | High | High | High | High | High | High | High | High | High | High | 0.000903 | 0.009507 | 1.82 |
| FALSE | High | ENGADSILI | [R].ENGAD | 1xPhospho | E9Q2H1 1x | 8.63E-08 | 0 | 1 | 2 | 4 | E9Q2H1 | E3 ubiquiti | E9Q2H1 [2 | 0 | 2414.21 | 0 | -0.67 | 0.000475 | 1.37 | 123.3 | 76.7 | 5.09 | 6.47 | | High | High | High | High | High | High | High | High | High | High | High | 0 | 6.5E-09 | 3.91 |
| FALSE | High | ENGDLSPF | [K].ENGDL | 1xPhospho | Q62077 1x | 0.015465 | 0.001089 | 1 | 5 | 1 | Q62077 | 1-phospha | Q62077 [1 | 0 | 1800.873 | 0 | -0.27 | 0.042998 | 14.96 | 109.2 | 90.8 | 9.78 | 14.66 | | High | High | High | High | High | High | High | High | High | High | High | 0.000571 | 0.005133 | 1.83 |
| FALSE | High | ENGFSSPPI | [K].ENGFS | 1xPhospho | Q04750 1x | 0.031168 | 0.001935 | 1 | 3 | 2 | Q04750 | DNA topoi | Q04750 [1 | 0 | 1299.593 | 0 | -0.27 | 0.080237 | 11.2 | 109.3 | 90.7 | 8.77 | 14.5 | | High | High | High | High | High | High | High | High | High | High | High | 0.001308 | 0.01366 | 0.75 |
| FALSE | High | ENGFSSPPI | [K].ENGFS | 1xDeamida | Q04750 1x | 0.010109 | 0.000736 | 1 | 3 | 1 | Q04750 | DNA topoi | Q04750 [1 | 0 | 1300.577 | 0 | -0.3 | 0.019183 | 8.25 | 110.3 | 89.7 | 7.64 | 3.91 | | High | High | High | High | High | High | High | High | High | High | High | 0.000344 | 0.003173 | 1.39 |
| FALSE | High | ENGNNSPL | [K].ENGNN | 1xPhospho | Q3U3E2 1x | 0.058664 | 0.004294 | 1 | 2 | 2 | Q3U3E2 | Protein FA | Q3U3E2 [4 | 0 | 1607.819 | 0 | 0.16 | 0.136449 | 19.21 | 93.8 | 106.2 | 8.73 | 10.26 | | High | High | High | High | High | High | High | High | High | High | High | 0.002855 | 0.0296 | 2.15 |
| FALSE | High | ENGNNSPL | [K].ENGNN | 1xDeamida | Q3U3E2 1x | 0.037048 | 0.00249 | 1 | 2 | 1 | Q3U3E2 | Protein FA | Q3U3E2 [4 | 0 | 1608.803 | 0 | 0.39 | 0.015862 | 12.29 | 87.6 | 112.4 | 6.03 | 8.25 | | High | High | High | High | High | High | High | High | High | High | High | 0.001308 | 0.01364 | 2.82 |
| FALSE | High | ENGSDNLP | [R].ENGSD | 1xPhospho | A2AUD5 1x | 6.41E-05 | 0 | 1 | 5 | 6 | A2AUD5 | Tumor pro | A2AUD5 [2 | 0 | 2758.22 | 0 | 0.88 | 0.00022 | 4.41 | 71.6 | 128.4 | 5.29 | 2.79 | | High | High | High | High | High | High | High | High | High | High | High | 0 | 1.09E-05 | 3.53 |
| FALSE | High | ENGSDNLP | [R].ENGSD | 1xDeamida | A2AUD5 1x | 0.000346 | 0 | 1 | 5 | 19 | A2AUD5 | Tumor pro | A2AUD5 [2 | 0 | 2759.204 | 0 | 1 | 0.000313 | 3.46 | 67 | 133 | 5.51 | 2.85 | | High | High | High | High | High | High | High | High | High | High | High | 0 | 0.00017 | 3.38 |
| FALSE | High | ENGSDNLP | [R].ENGSD | 2xPhospho | A2AUD5 2x | 0.0002 | 0 | 1 | 5 | 6 | A2AUD5 | Tumor pro | A2AUD5 [2 | 0 | 2838.186 | 0 | 2.02 | 0.000317 | 5.11 | 40.8 | 159.2 | 8.04 | 6.54 | | High | High | High | High | High | High | High | High | High | High | High | 0 | 0.00017 | 3.61 |
| FALSE | High | ENGSDNLP | [R].ENGSD | 1xDeamida | A2AUD5 2x | 0.001361 | 0 | 1 | 5 | 9 | A2AUD5 | Tumor pro | A2AUD5 [2 | 0 | 2839.17 | 0 | 2 | 0.000217 | 0.16 | 39.1 | 160.9 | 8.52 | 4.94 | | High | High | High | High | High | High | High | High | High | High | High | 2.58E-05 | 0.000338 | 3.53 |
| FALSE | High | ENLESLRSL | [K].ENLESL | 2xPhospho | Q6P5H2-2 | 0.004613 | 0.000456 | 1 | 2 | 1 | Q6P5H2-2 | Isoform 2 c | Q6P5H2-2 | 1 | 2692.263 | 0 | 3.2 | 0.020295 | 23.69 | 19.9 | 180.1 | 14.56 | 43.9 | | High | High | High | High | High | High | High | High | High | High | High | 0.000204 | 0.001319 | 3.17 |
| FALSE | High | ENLILQETS | [K].ENLILQ | 1xPhospho | Q5RJH6-2 | 2.59E-07 | 0 | 1 | 3 | 3 | Q5RJH6-2 | Isoform 2 c | Q5RJH6-2 | 0 | 2938.543 | 0 | -0.14 | 0.561446 | 11.61 | 103.3 | 96.7 | 10.99 | 7.65 | | High | High | High | High | High | High | High | High | High | High | High | 0 | 2.22E-08 | 4.18 |
| FALSE | High | ENPQLDISF | [R].ENPQLI | 1xCarbami | Q9D8S9 1x | 0.000808 | 0 | 1 | 1 | 4 | Q9D8S9 | BolA-like p | Q9D8S9 [1 | 0 | 2250.124 | 0 | -0.36 | 0.016487 | 3.89 | 112.5 | 87.5 | 3.21 | 9.75 | | High | High | High | High | High | High | High | High | High | High | High | 0.001789 | 0.0176 | 2.16 |
| FALSE | High | ENINSQEVI | [K].ENINSC | 1xPhospho | Q3TUQ5 1 | 0.002272 | 0.000109 | 1 | 2 | 2 | Q3TUQ5 | Pinin OS=N | Q3TUQ5 [4 | 0 | 1727.862 | 0 | 0.27 | 0.434039 | 10.98 | 89 | 111 | 9.99 | 13.4 | | High | High | High | High | High | High | High | High | High | High | High | 4.94E-05 | 0.000596 | 3.31 |
| FALSE | High | ENLPVSPD | [K].ENLPVS | 1xPhospho | E9QJU8 1x | 2.29E-05 | 0 | 1 | 2 | 1 | E9QJU8 | Transform | E9QJU8 [3 | 0 | 2485.229 | 0 | 1.38 | 0.001923 | 12.03 | 54.5 | 145.5 | 13.91 | 12.1 | | High | High | High | High | High | High | High | High | High | High | High | 0 | 3.43E-06 | 4.3 |
| FALSE | High | ENMAQEN | [K].ENMAC | 1xPhospho | A2BEF2 1xI | 2.07E-08 | 0 | 1 | 1 | 4 | A2BEF2 | Disks large | A2BEF2 [2 | 0 | 3325.478 | 0 | 0.85 | 0.022853 | 24.85 | 72 | 128 | 5.37 | 19.56 | | High | High | High | High | High | High | High | High | High | High | High | 0 | 1.3E-09 | 5.47 |
| FALSE | High | ENMAQEN | [K].ENMAC | 1xPhospho | A2BEF2 1xI | 8.02E-11 | 0 | 1 | 1 | 5 | A2BEF2 | Disks large | A2BEF2 [2 | 0 | 3341.473 | 0 | 0.62 | 0.012803 | 26.91 | 77.5 | 122.5 | 10.74 | 10.46 | | High | High | High | High | High | High | High | High | High | High | High | 0 | 3.84E-11 | 6.78 |
| FALSE | High | ENNSPVVY | [K].ENNSP\ | 1xPhospho | P49452 1xI | 3.64E-06 | 0 | 1 | 1 | 2 | P49452 | Centromer | P49452 [51 | 0 | 1798.914 | 0 | -0.91 | 0.002708 | 5.17 | 128.8 | 71.2 | 6.64 | 13.57 | | High | High | High | High | High | High | High | High | High | High | High | 0 | 3.68E-06 | 3.42 |
| FALSE | High | ENNVSQPA | [K].ENNVS | 1xPhospho | K4DI62 1xF | 6.57E-11 | 0 | 1 | 3 | 2 | K4DI62 | Transcripti | K4DI62 [11 | 0 | 3052.374 | 1 | -0.44 | 0.06099 | 18.04 | 115 | 85 | 9.98 | 16.39 | | High | High | High | High | High | High | High | High | High | High | High | 0 | 2.04E-12 | 5.87 |
| FALSE | High | ENNVSQPA | [K].ENNVS | 1xDeamida | K4DI62 1xF | 2.74E-11 | 0 | 1 | 3 | 7 | K4DI62 | Transcripti | K4DI62 [11 | 0 | 3053.358 | 2 | -0.29 | 0.081674 | 1.75 | 110.1 | 89.9 | 9.07 | 16.83 | | High | High | High | High | High | High | High | High | High | High | High | 0 | 7.59E-13 | 5.87 |
| FALSE | High | ENPEPEISN | [K].ENPEPE | 1xPhospho | Q8BRB7 1x | 0.034281 | 0.002195 | 1 | 3 | 2 | Q8BRB7 | Histone ac | Q8BRB7 [1 | 0 | 1856.875 | 0 | -1.84 | 0.000361 | 1.18 | 157.8 | 42.2 | 6.77 | 12.01 | | High | High | High | High | High | High | High | High | High | High | High | 0.007957 | 0.0859 | 1.77 |
| FALSE | High | ENPPGPPIA | [R].ENPPGI | 1xDeamida | Q80TU3 1x | 0.032267 | 0.002129 | 1 | 6 | 3 | Q80TU3 | MKIAA068 | Q80TU3 [4 | 0 | 4265.101 | 0 | -0.93 | 0.086427 | 6.58 | 132 | 68 | 19.72 | 31.31 | | High | High | High | High | High | High | High | High | High | High | High | 0.001142 | 0.01171 | 2.89 |
| FALSE | High | ENPPSPHSI | [R].ENPPSF | 1xPhospho | Q8BP27 1x | 0.000125 | 0 | 1 | 3 | 6 | Q8BP27 | Swi5-depe | Q8BP27 [1 | 0 | 1875.9 | 0 | -0.81 | 0.012872 | 9.75 | 126 | 74 | 13.81 | 9.76 | | High | High | High | High | High | High | High | High | High | High | High | 0 | 0.00021 | 2.17 |
| FALSE | High | ENPPSPHSI | [R].ENPPSF | 1xDeamida | Q8BP27 1x | 0.00145 | 5.66E-05 | 1 | 3 | 4 | Q8BP27 | Swi5-depe | Q8BP27 [1 | 0 | 1876.884 | 0 | -0.67 | 0.008837 | 3.16 | 121.7 | 78.3 | 8.59 | 9.38 | | High | High | High | High | High | High | High | High | High | High | High | 2.58E-05 | 0.000529 | 1.88 |
| FALSE | High | ENPPSPHSI | [R].ENPPSF | 2xPhospho | Q8BP27 2x | 0.019966 | 0.001438 | 1 | 3 | 1 | Q8BP27 | Swi5-depe | Q8BP27 [1 | 0 | 1955.867 | 0 | -0.55 | 0.000985 | 1.05 | 119.9 | 80.1 | 2.61 | 5.68 | | High | High | High | High | High | High | High | High | High | High | High | 0.000786 | 0.006834 | 1.83 |
| FALSE | High | ENPPSPPTS | [SR].ENPPS | 1xPhospho | Q8BP27 1x | 2.38E-06 | 0 | 1 | 3 | 10 | Q8BP27 | Swi5-depe | Q8BP27 [6 | 0 | 1951.948 | 0 | -0.45 | 0.005104 | 4.38 | 116.8 | 83.2 | 4.99 | 4.94 | | High | High | High | High | High | High | High | High | High | High | High | 0 | 5.01E-05 | 3.36 |
| FALSE | High | ENPPSPPTS | [SR].ENPPS | 2xPhospho | Q8BP27 2x | 3.59E-07 | 0 | 1 | 3 | 9 | Q8BP27 | Swi5-depe | Q8BP27 [6 | 0 | 2031.914 | 0 | -0.32 | 0.000445 | 2.11 | 110.4 | 89.6 | 0.71 | 2.18 | | High | High | High | High | High | High | High | High | High | High | High | 0 | 9.27E-05 | 3.06 |
| FALSE | High | ENINSQEVI | [K].ENINSC | 1xPhospho | Q3TUQ5 1 | 0.001799 | 5.66E-05 | 1 | 2 | 1 | Q3TUQ5 | Pinin OS=N | Q3TUQ5 [4 | 1 | 2559.279 | 0 | 0.23 | 0.461797 | 10.45 | 92.7 | 107.3 | 6.25 | 16.8 | | High | High | High | High | High | High | High | High | High | High | High | 2.58E-05 | 0.00046 | 2.6 |
| FALSE | High | ENVGLPEE | [K].ENVGLI | 1xPhospho | Q8CJF7 1xI | 0.000302 | 0 | 1 | 2 | 4 | Q8CJF7 | Protein EL | Q8CJF7 [14 | 0 | 1535.73 | 0 | -0.68 | 0.002755 | 2.33 | 124.8 | 75.2 | 6.8 | 7.29 | | High | High | High | High | High | High | High | High | High | High | High | 0 | 6.21E-05 | 2.66 |
| FALSE | High | ENVIPLLVT | [K].ENVIPL | 1xPhospho | Q4VBD2 1x | 2.8E-05 | 0 | 1 | 1 | 1 | Q4VBD2 | Transmem | Q4VBD2 [5 | 0 | 3170.591 | 0 | 2.08 | 0.003212 | 16.46 | 35.3 | 164.7 | 50.44 | 24.59 | | High | High | High | High | High | High | High | High | High | High | High | 0 | 4.31E-06 | 4.23 |
| FALSE | High | ENVQYSSV | [R].ENVQY | 1xCarbami | Q60977 1x | 6.09E-06 | 0 | 1 | 4 | 10 | Q60977 | Tumor-ass | Q60977 [3 | 0 | 1836.779 | 0 | 1.03 | 0.000154 | 3.57 | 66.8 | 133.2 | 4.31 | 3.35 | | High | High | High | High | High | High | High | High | High | High | High | 0 | 5.28E-05 | 3.63 |
| FALSE | High | EPGSDQNC | [K].EPGSD( | 1xCarbami | A3KFM7 1x | 5.93E-06 | 0 | 1 | 3 | 4 | A3KFM7 | Chromodo | A3KFM7 [2 | 1 | 3151.395 | 0 | -0.36 | 0.022633 | 4.4 | 113.6 | 86.4 | 9.37 | 14.09 | | High | High | High | High | High | High | High | High | High | High | High | 0 | 7.53E-07 | 3.96 |
| FALSE | High | EPGSSGPQ | [K].EPGSS( | 1xPhospho | Q9DB90 1x | 1.28E-05 | 0 | 1 | 3 | 2 | Q9DB90 | Protein SM | Q9DB90 [2 | 0 | 1805.838 | 0 | 0.22 | 0.017677 | 11.54 | 93.5 | 106.5 | 1.67 | 7.35 | | High | High | High | High | High | High | High | High | High | High | High | 0 | 1.78E-06 | 3.33 |
| FALSE | High | EPHLNYSPT | [K].EPHLNY | 1xCarbami | A0A0A6YX | 0.01002 | 0.000736 | 1 | 2 | 2 | A0A0A6YX | Immunogl | A0A0A6YX | 0 | 2965.437 | 0 | -1.5 | 0.003131 | 3.71 | 147.6 | 52.4 | 14.9 | 6.8 | | High | High | High | High | High | High | High | High | High | High | High | 0.000638 | 0.005779 | 2.71 |
| FALSE | High | EPHSPDGN | [R].EPHSPD | 1xCarbami | Q8BG80 1x | 0.007881 | 0.000736 | 1 | 1 | 1 | Q8BG80 | F-box only | Q8BG80 [2 | 0 | 3224.453 | 0 | -0.1 | 0.642546 | 10.19 | 103.4 | 96.6 | 8.93 | 10.53 | | High | High | High | High | High | High | High | High | High | High | High | 0.000327 | 0.002395 | 2.17 |
| FALSE | High | EPHSPDGN | [R].EPHSPD | 1xCarbami | Q8BG80 1x | 0.058664 | 0.004294 | 1 | 1 | 1 | Q8BG80 | F-box only | Q8BG80 [2 | 0 | 3225.437 | 0 | 0.02 | 0.303257 | 7.28 | 93.8 | 106.2 | 8.03 | 12.65 | | High | High | High | High | High | High | High | High | High | High | High | 0.002187 | 0.02313 | 2.95 |
| FALSE | High | EPHTESVVS | [R].EPHTES | 1xPhospho | Q80Y44 1x | 0.000111 | 0 | 1 | 6 | 1 | Q80Y44 | Probable A | Q80Y44 [6 | 0 | 2221.115 | 0 | -0.36 | 0.011982 | 4.42 | 112.4 | 87.6 | 10.52 | 1.97 | | High | High | High | High | High | High | High | High | High | High | High | 0 | 2.03E-05 | 2.22 |
| FALSE | High | EPLASPHPA | [R].EPLASP | 1xPhospho | Q80TU6 1x | 0.002895 | 0.00016 | 1 | 1 | 3 | Q80TU6 | MKIAA067 | Q80TU6 [4 | 0 | 1723.909 | 0 | 1.62 | 0.000824 | 13.09 | 50.6 | 149.4 | 8.6 | 11.35 | | High | High | High | High | High | High | High | High | High | High | High | 0.001168 | 0.01256 | 2.22 |
| FALSE | High | EPLKPDLSP | [K].EPLKPD | 1xCarbami | A0A1D5RM | 0.098533 | 0.007902 | 1 | 1 | 1 | A0A1D5RM | Alstrom sy | A0A1D5RM | 0 | 2278.249 | 0 | -1.13 | 0.001525 | 3.89 | 140 | 60 | 9.12 | 6.35 | | High | High | High | High | High | High | High | High | High | High | High | 0.003977 | 0.04247 | 2 |
| FALSE | High | EPLLSSSEN | [R].EPLLSS | 1xPhospho | Q61210-5 | 1.65E-06 | 0 | 1 | 7 | 3 | Q61210-5 | Isoform 5 c | Q61210-5 | 0 | 2485.112 | 0 | 0.16 | 0.124926 | 6.18 | 94.2 | 105.8 | 10.63 | 13.35 | | High | High | High | High | High | High | High | High | High | High | High | 0 | 1.78E-07 | 5.63 |
| FALSE | High | EPLLSSSEN | [R].EPLLSS | 1xDeamida | Q61210-5 | 0.000699 | 0 | 1 | 7 | 3 | Q61210-5 | Isoform 5 c | Q61210-5 | 0 | 2486.096 | 0 | 0.15 | 0.373852 | 28.01 | 92.6 | 107.4 | 15.82 | 9.84 | | High | High | High | High | High | High | High | High | High | High | High | 0 | 0.000315 | 4.06 |
| FALSE | High | EPIPEEQEN | [K].EPIPEE( | 2xPhospho | Q6Y7W8 2 | 0.117405 | 0.009523 | 1 | 10 | 1 | Q6Y7W8 | GRB10-inte | Q6Y7W8 [3 | 1 | 4349.847 | 0 | -0.67 | 0.002161 | 2.55 | 120.9 | 79.1 | 8.09 | 7.47 | | High | High | High | High | High | High | High | High | High | High | High | 0.004769 | 0.05201 | 2.5 |
| FALSE | High | EPLPVEQD | [K].EPLPVE | 1xPhospho | Q8C0J2-3 1 | 9.82E-06 | 0 | 1 | 6 | 2 | Q8C0J2-3 | Isoform 3 c | Q8C0J2-3 [ | 0 | 3602.647 | 0 | -0.74 | 0.008406 | 7.28 | 125.2 | 74.8 | 4.84 | 19.16 | | High | High | High | High | High | High | High | High | High | High | High | 0 | 1.32E-06 | 5.69 |
| FALSE | High | EPLSDEDEE | [K].EPLSDE | 1xPhospho | Q923F1 1x | 2.73E-06 | 0 | 1 | 1 | 7 | Q923F1 | Chloride ch | Q923F1 [9 | 0 | 2788.156 | 0 | -0.72 | 0.003771 | 8.87 | 124.2 | 75.8 | 9.04 | 14.47 | | High | High | High | High | High | High | High | High | High | High | High | 0 | 3.13E-07 | 4.46 |
| FALSE | High | EPGQASPK | [R].EPGQA | 1xPhospho | Q9D2Q2 1 | 0.035793 | 0.002304 | 1 | 2 | 1 | Q9D2Q2 | Probable t | Q9D2Q2 [7 | 0 | 1351.702 | 0 | 0.55 | 0.003427 | 5.13 | 84.7 | 115.3 | 5.59 | 10.29 | | High | High | High | High | High | High | High | High | High | High | High | 0.00123 | 0.01321 | 1.47 |
| FALSE | High | EPLSPSSPP | [K].EPLSPS | 2xPhospho | Q9CWR8 2 | 0.055777 | 0.004093 | 1 | 1 | 1 | Q9CWR8 | DNA (cytos | Q9CWR8 [ | 0 | 1613.717 | 0 | -1.72 | 0.000954 | 0.85 | 152.1 | 47.9 | 10.13 | 12.32 | | High | High | High | High | High | High | High | High | High | High | High | 0.002117 | 0.02178 | 1.81 |
| FALSE | High | EPMAQQD | [K].EPMAC | 1xPhospho | Q3UIV3 1x | 4.59E-05 | 0 | 1 | 12 | 3 | Q3UIV3 | Putative ur | Q3UIV3 [6 | 0 | 3779.746 | 0 | -0.66 | 0.041116 | 18.55 | 122.4 | 77.6 | 11.82 | 25.96 | | High | High | High | High | High | High | High | High | High | High | High | 0 | 9.4E-06 | 4.61 |
| FALSE | High | EPMAQQD | [K].EPMAC | 1xPhospho | Q3UIV3 1x | 8.33E-05 | 0 | 1 | 12 | 5 | Q3UIV3 | Putative ur | Q3UIV3 [6 | 0 | 3795.74 | 0 | -1.03 | 0.001413 | 1.42 | 143.1 | 56.9 | 11.19 | 16.87 | | High | High | High | High | High | High | High | High | High | High | High | 0 | 2.39E-05 | 4.24 |
| FALSE | High | EPMAQQD | [K].EPMAC | 1xPhospho | Q3UIV3 1x | 3.33E-06 | 0 | 1 | 12 | 5 | Q3UIV3 | Putative ur | Q3UIV3 [6 | 0 | 3811.735 | 0 | -0.4 | 0.031911 | 10.52 | 111.9 | 88.1 | 10.16 | 12.06 | | High | High | High | High | High | High | High | High | High | High | High | 0 | 5.25E-06 | 4.71 |
| FALSE | High | EPMGLGPF | [R].EPMGL | 1xPhospho | Q9CW46 1 | 1.75E-06 | 0 | 1 | 2 | 1 | Q9CW46 | Ribonuclec | Q9CW46 [ | 0 | 2505.314 | 0 | -0.33 | 0.013451 | 5.53 | 113 | 87 | 6.08 | 8.29 | | High | High | High | High | High | High | High | High | High | High | High | 0 | 1.92E-07 | 3.28 |
| FALSE | High | EPMGLGPF | [R].EPMGL | 1xPhospho | Q9CW46 1 | 6.39E-07 | 0 | 1 | 2 | 1 | Q9CW46 | Ribonuclec | Q9CW46 [ | 0 | 2521.309 | 0 | -0.06 | 0.326701 | 8.63 | 103 | 97 | 13.23 | 13.92 | | High | High | High | High | High | High | High | High | High | High | High | 0 | 6.12E-08 | 3.66 |
| FALSE | High | EPPGHQHL | [R].EPPGH( | 1xPhospho | E9QK89 1x | 1.56E-05 | 0 | 1 | 2 | 4 | E9QK89 | Mediator c | E9QK89 [7 | 0 | 2593.324 | 0 | -0.81 | 0.002427 | 1.41 | 127.7 | 72.3 | 7.87 | 8.48 | | High | High | High | High | High | High | High | High | High | High | High | 0 | 2.23E-06 | 5 |
| FALSE | High | EPPPPQTL( | [R].EPPPPC | 1xPhospho | Q5U4C3 1x | 6.52E-08 | 0 | 1 | 1 | 1 | Q5U4C3 | Splicing fac | Q5U4C3 [3 | 0 | 2423.181 | 0 | 0.11 | 0.086638 | 12.94 | 96.4 | 103.6 | 7.43 | 5.42 | | High | High | High | High | High | High | High | High | High | High | High | 0 | 4.73E-09 | 3.59 |
| FALSE | High | EPPQGCST | [R].EPPQG | 1xCarbami | Q80WR5 1 | 3.23E-05 | 0 | 1 | 3 | 3 | Q80WR5 | UPF0688 p | Q80WR5 [ | 0 | 2338.104 | 0 | 0.1 | 0.16905 | 11.2 | 92.8 | 107.2 | 5.04 | 11.75 | | High | High | High | High | High | High | High | High | High | High | High | 0 | 5.03E-06 | 3.44 |
| FALSE | High | EPPRASPP( | [K].EPPRAS | 1xPhospho | Q6ZQ88 1x | 0.001313 | 0 | 1 | 2 | 1 | Q6ZQ88 | Lysine-spe | Q6ZQ88 [6 | 1 | 4514.15 | 0 | -1.31 | 0.002034 | 5.2 | 136.1 | 63.9 | 9.57 | 14.91 | | High | High | High | High | High | High | High | High | High | High | High | 0 | 0.000322 | 4.83 |
| FALSE | High | EPPRASPP( | [K].EPPRAS | 1xPhospho | Q6ZQ88 1x | 8.48E-05 | 0 | 1 | 2 | 5 | Q6ZQ88 | Lysine-spe | Q6ZQ88 [6 | 1 | 4530.145 | 0 | -1.13 | 0.002608 | 0.79 | 136.3 | 63.7 | 8.27 | 15.34 | | High | High | High | High | High | High | High | High | High | High | High | 0 | 3.33E-05 | 5.13 |
| FALSE | High | EPPRPEPPF | [R].EPPRPE | 1xPhospho | Q80WC1-4 | 0.000102 | 0 | 1 | 4 | 1 | Q80WC1-4 | Isoform 4 c | Q80WC1-4 | 0 | 2518.342 | 0 | -0.34 | 0.026292 | 12.3 | 112.1 | 87.9 | 11.93 | 3.51 | | High | High | High | High | High | High | High | High | High | High | High | 0 | 1.84E-05 | 2.48 |
| FALSE | High | EPPSPSEA( | [R].EPPSPS | 1xPhospho | Q8CGF1 1x | 0.00172 | 5.66E-05 | 1 | 3 | 1 | Q8CGF1 | Rho GTPas | Q8CGF1 [5 | 0 | 2223.11 | 0 | 0.06 | 0.384841 | 9 | 98.2 | 101.8 | 4.15 | 4.91 | | High | High | High | High | High | High | High | High | High | High | High | 2.58E-05 | 0.00044 | 2.92 |
| FALSE | High | EPPTSWSP | [R].EPPTSV | 1xPhospho | Q8C7U1 1x | 0.055308 | 0.003982 | 1 | 2 | 1 | Q8C7U1 | NEDD4-bin | Q8C7U1 [5 | 0 | 1365.64 | 0 | -0.14 | 0.023857 | 6.83 | 105.4 | 94.6 | 5.4 | 8.28 | | High | High | High | High | High | High | High | High | High | High | High | 0.002105 | 0.02164 | 1.8 |
| FALSE | High | EPIYTLPAII | [K].EPIYTLF | 1xPhospho | B2RQE8 1x | 0.01441 | 0.001011 | 1 | 1 | 2 | B2RQE8 | Rho GTPas | B2RQE8 [3 | 0 | 1883.07 | 0 | 0.36 | 0.025825 | 20.54 | 88.9 | 111.1 | 15.64 | 4.31 | | High | High | High | High | High | High | High | High | High | High | High | 0.000997 | 0.01032 | 2.34 |
| FALSE | High | EPGPPLASS | [K].EPGPPL | 1xCarbami | Q6PDM1 1 | 6.27E-07 | 0 | 1 | 3 | 3 | Q6PDM1 | Male-speci | Q6PDM1 [ | 0 | 2774.322 | 0 | 0.66 | 0.002203 | 9.79 | 81.9 | 118.1 | 12.47 | 9 | | High | High | High | High | High | High | High | High | High | High | High | 0 | 2.37E-07 | 4.02 |
| FALSE | High | EPGFEAGP | [K].EPGFEA | 1xPhospho | Q80WC3 1 | 8.57E-09 | 0 | 1 | 2 | 1 | Q80WC3 | Trinucleoti | Q80WC3 [ | 0 | 2300.991 | 0 | 1.21 | 0.008231 | 1.48 | 64.8 | 135.2 | 22.64 | 14.56 | | High | High | High | High | High | High | High | High | High | High | High | 0 | 4.83E-10 | 3.37 |
| FALSE | High | EPGEQAST | [K].EPGEQ, | 1xPhospho | G3X9H5 1x | 0.005516 | 0.00062 | 1 | 3 | 1 | G3X9H5 | Huntingtin | G3X9H5 [1 | 0 | 1912.913 | 0 | 0.59 | 0.017101 | 3.44 | 79 | 121 | 14.65 | 9.65 | | High | High | High | High | High | High | High | High | High | High | High | 0.000275 | 0.001615 | 2.04 |
| FALSE | High | ENVQYSSV | [R].ENVQY | 1xCarbami | Q60977 1x | 0.08447 | 0.006345 | 1 | 4 | 6 | Q60977 | Tumor-ass | Q60977 [3 | 0 | 1837.763 | 0 | 0.89 | 0.000307 | 11.17 | 70.1 | 129.9 | 12.27 | 6.26 | | High | High | High | High | High | High | High | High | High | High | High | 0.003265 | 0.03541 | 2.66 |
| FALSE | High | ENVSAVCL | [R].ENVSA\ | 1xCarbami | G3X939 2x | 0.042141 | 0.003162 | 1 | 2 | 1 | G3X939 | Sodium/hy | G3X939 [5 | 0 | 2283.959 | 0 | -0.6 | 0.004501 | 3.09 | 116.7 | 83.3 | 8.41 | 17.33 | | High | High | High | High | High | High | High | High | High | High | High | 0.001639 | 0.01592 | 3.19 |
| FALSE | High | ENVSDLTL( | [R].ENVSDI | 1xPhospho | Q3V0N3 1x | 2.83E-09 | 0 | 1 | 2 | 2 | Q3V0N3 | Putative ur | Q3V0N3 [1 | 0 | 2079.032 | 0 | 0.58 | 0.005058 | 15.44 | 80.6 | 119.4 | 9.38 | 6.82 | | High | High | High | High | High | High | High | High | High | High | High | 0 | 1.39E-10 | 4.28 |
| FALSE | High | ENVSLLAEI | [R].ENVSLL | 1xPhospho | Q8CDZ5 1x | 3.83E-08 | 0 | 1 | 2 | 10 | Q8CDZ5 | Putative ur | Q8CDZ5 [4 | 0 | 2514.217 | 0 | -0.38 | 0.00472 | 8.01 | 118.8 | 81.2 | 6.13 | 11.96 | | High | High | High | High | High | High | High | High | High | High | High | 0 | 3.59E-07 | 3.56 |
| FALSE | High | ENVTNASC | [K].ENVTN | 1xPhospho | G5E8B9 1x | 5.58E-07 | 0 | 1 | 3 | 5 | G5E8B9 | MCG13089 | G5E8B9 [4 | 0 | 3132.389 | 0 | -0.67 | 0.002716 | 1.33 | 123.1 | 76.9 | 4.77 | 8.81 | | High | High | High | High | High | High | High | High | High | High | High | 0 | 1.29E-07 | 5.3 |
| FALSE | High | ENVTNASC | [K].ENVTN | 2xPhospho | G5E8B9 2x | 0.001398 | 5.66E-05 | 1 | 3 | 3 | G5E8B9 | MCG13089 | G5E8B9 [4 | 0 | 3212.356 | 0 | -0.42 | 0.030211 | 7.13 | 112.4 | 87.6 | 8.35 | 12.57 | | High | High | High | High | High | High | High | High | High | High | High | 2.58E-05 | 0.000349 | 2.98 |
| FALSE | High | ENYEDQEP | [K].ENYED( | 1xPhospho | B9EKJ7 1xF | 2.77E-08 | 0 | 1 | 2 | 4 | B9EKJ7 | Tankyrase | B9EKJ7 [58 | 0 | 2677.271 | 0 | 0.37 | 0.061453 | 12.6 | 87.1 | 112.9 | 5.36 | 14.31 | | High | High | High | High | High | High | High | High | High | High | High | 0 | 1.81E-09 | 5.4 |
| FALSE | High | EPAAGGSL | [R].EPAAG( | 1xPhospho | Q5SPX8 1x | 0.017809 | 0.001282 | 1 | 3 | 2 | Q5SPX8 | TBC1 doma | Q5SPX8 [4 | 0 | 1411.678 | 0 | 0.32 | 0.025762 | 2.44 | 91.6 | 108.4 | 7.69 | 6.72 | | High | High | High | High | High | High | High | High | High | High | High | 0.001044 | 0.01118 | 2.04 |
| FALSE | High | EPAAPASP | [K].EPAAP | 1xPhospho | A2AJI0 1xP | 1.17E-05 | 0 | 1 | 5 | 2 | A2AJI0 | MAP7 dom | A2AJI0 [53 | 0 | 2760.437 | 0 | 1.74 | 0.000162 | 6.75 | 44.9 | 155.1 | 8.8 | 3.94 | | High | High | High | High | High | High | High | High | High | High | High | 0 | 1.61E-06 | 4.54 |
| FALSE | High | EPAAPASP | [K].EPAAP | 2xPhospho | A2AJI0 2xP | 0.012288 | 0.000889 | 1 | 5 | 1 | A2AJI0 | MAP7 dom | A2AJI0 [53 | 0 | 2840.404 | 0 | 1.12 | 0.000216 | 0.53 | 62 | 138 | 6.99 | 4.35 | | High | High | High | High | High | High | High | High | High | High | High | 0.000449 | 0.003953 | 2.54 |
| FALSE | High | EPACTYGN | [K].EPACTY | 1xCarbami | Q3UMF0 1 | 7.7E-06 | 0 | 1 | 6 | 2 | Q3UMF0 | Cordon-ble | Q3UMF0 [ | 0 | 2657.268 | 0 | 0.35 | 0.038394 | 13.48 | 86.5 | 113.5 | 13.83 | 9.78 | | High | High | High | High | High | High | High | High | High | High | High | 0 | 1.01E-06 | 3.49 |
| FALSE | High | EPAEPPLQI | [K].EPAEPF | 1xPhospho | Q80UE6 1x | 0.000758 | 0 | 1 | 2 | 2 | Q80UE6 | Serine/thre | Q80UE6 [1 | 0 | 1999.01 | 0 | 1.47 | 0.000985 | 15.93 | 51.3 | 148.7 | 10.64 | 8.2 | | High | High | High | High | High | High | High | High | High | High | High | 0 | 0.000176 | 1.7 |
| FALSE | High | EPALAVAG | [R].EPALAV | 1xPhospho | Q8CEC6 1x | 0.000438 | 0 | 1 | 2 | 1 | Q8CEC6 | Peptidylpr | Q8CEC6 [3 | 0 | 2383.014 | 0 | -0.34 | 0.035273 | 4.02 | 114.4 | 85.6 | 12.26 | 13.19 | | High | High | High | High | High | High | High | High | High | High | High | 0 | 9.4E-05 | 3.94 |
| FALSE | High | EPAISSQNS | [K].EPAISS( | 1xPhospho | E9PYD5 1xI | 1.19E-05 | 0 | 1 | 4 | 14 | E9PYD5 | Transcripti | E9PYD5 [1 | 0 | 1694.795 | 0 | -1.07 | 0.000163 | 0.67 | 136.1 | 63.9 | 2.44 | 8.2 | | High | High | High | High | High | High | High | High | High | High | High | 0 | 3.74E-06 | 3.76 |
| FALSE | High | EPAISSQNS | [K].EPAISS( | 1xDeamida | E9PYD5 1xI | 0.022973 | 0.001578 | 1 | 4 | 3 | E9PYD5 | Transcripti | E9PYD5 [1 | 0 | 1695.779 | 0 | -0.42 | 0.005856 | 6.7 | 118.9 | 81.1 | 6.15 | 14.22 | | High | High | High | High | High | High | High | High | High | High | High | 0.000832 | 0.007948 | 2.53 |
| FALSE | High | EPAPGDPG | [R].EPAPGI | 1xCarbami | E9Q0A7 1x | 0.004613 | 0.000456 | 1 | 1 | 1 | E9Q0A7 | Oxidation r | E9Q0A7 [4 | 0 | 3564.672 | 0 | 0.75 | 0.007381 | 25.4 | 74.6 | 125.4 | 8.81 | 9.04 | | High | High | High | High | High | High | High | High | High | High | High | 0.000204 | 0.001318 | 3.49 |
| FALSE | High | EPAPLLSAC | [R].EPAPLL | 1xPhospho | Q5RIM6 1x | 1.9E-05 | 0 | 1 | 12 | 4 | Q5RIM6 | Nuclear re | Q5RIM6 [2 | 0 | 2260.011 | 0 | -0.92 | 0.064086 | 4.71 | 122.1 | 77.9 | 24.88 | 14.63 | | High | High | High | High | High | High | High | High | High | High | High | 0 | 2.76E-06 | 3.78 |
| FALSE | High | EPAPLNGS | [R].EPAPLN | 1xPhospho | Q8CB16 1x | 0.00013 | 0 | 1 | 8 | 2 | Q8CB16 | Putative ur | Q8CB16 [3 | 0 | 2104.109 | 0 | 1.39 | 0.002186 | 14.04 | 54.7 | 145.3 | 3.32 | 15.22 | | High | High | High | High | High | High | High | High | High | High | High | 0 | 9.39E-05 | 3.08 |
| FALSE | High | EPAPLNGS | [R].EPAPLN | 1xDeamida | Q8CB16 1x | 1.33E-05 | 0 | 1 | 8 | 2 | Q8CB16 | Putative ur | Q8CB16 [3 | 0 | 2105.093 | 0 | 1.71 | 0.001711 | 11.7 | 46.9 | 153.1 | 3.75 | 14.67 | | High | High | High | High | High | High | High | High | High | High | High | 0 | 1.86E-06 | 4.44 |
| FALSE | High | EPDSPSMP | [K].EPDSPS | 1xPhospho | Q5QGZ6 1 | 0.032831 | 0.002195 | 1 | 4 | 1 | Q5QGZ6 | Retinoic ac | Q5QGZ6 [1 | 0 | 1598.754 | 0 | 0.96 | 0.00724 | 11.82 | 71.5 | 128.5 | 14.09 | 20.45 | | High | High | High | High | High | High | High | High | High | High | High | 0.001151 | 0.01199 | 2.49 |
| FALSE | High | EPEAAPPSF | [K].EPEAAF | 1xPhospho | B2RUC3 1x | 0.002743 | 0.000109 | 1 | 8 | 1 | B2RUC3 | Excision re | B2RUC3 [3 | 0 | 1359.651 | 0 | -0.36 | 0.016883 | 6.37 | 113.7 | 86.3 | 8.75 | 7.03 | | High | High | High | High | High | High | High | High | High | High | High | 4.94E-05 | 0.000741 | 1.41 |
| FALSE | High | EPEERPTLE | [K].EPEERP | 1xPhospho | Q3UY28 1x | 0.001386 | 5.66E-05 | 1 | 1 | 2 | Q3UY28 | Putative ur | Q3UY28 [2 | 0 | 4435.159 | 0 | 0.05 | 0.430782 | 3.78 | 97.2 | 102.8 | 5.7 | 9.4 | | High | High | High | High | High | High | High | High | High | High | High | 2.58E-05 | 0.000343 | 3.82 |
| FALSE | High | EPEQEGPS | [K].EPEQE( | 1xPhospho | Q62415 1x | 8.2E-06 | 0 | 1 | 2 | 1 | Q62415 | Apoptosis- | Q62415 [6 | 0 | 3342.687 | 0 | -0.39 | 0.00252 | 4.97 | 114.2 | 85.8 | 4.94 | 7.68 | | High | High | High | High | High | High | High | High | High | High | High | 0 | 1.09E-06 | 2.7 |
| FALSE | High | EPESPPAPI | [R].EPESPP | 1xPhospho | A0A0A0M( | 0.036731 | 0.00249 | 1 | 2 | 1 | A0A0A0M( | Tetratricop | A0A0A0M( | 0 | 1886.961 | 0 | -0.07 | 0.638265 | 6.72 | 103 | 97 | 11.84 | 10.54 | | High | High | High | High | High | High | High | High | High | High | High | 0.001308 | 0.01357 | 2.5 |
| FALSE | High | EPETGLDN | [K].EPETGL | 1xPhospho | A0A0R4J0F | 4.66E-07 | 0 | 1 | 7 | 3 | A0A0R4J0F | Cyclin-G-as | A0A0R4J0F | 0 | 1825.899 | 0 | 0.93 | 0.002355 | 1.23 | 68.7 | 131.3 | 9.78 | 6.84 | | High | High | High | High | High | High | High | High | High | High | High | 0 | 4.29E-08 | 2.89 |
| FALSE | High | EPETVGAQ | [K].EPETV( | 1xPhospho | Q9R0L6-2 | 0.002232 | 0.000109 | 1 | 3 | 2 | Q9R0L6-2 | Isoform 2 c | Q9R0L6-2 | 0 | 1339.634 | 0 | 1.25 | 0.000328 | 6.03 | 59.1 | 140.9 | 6.55 | 6.53 | | High | High | High | High | High | High | High | High | High | High | High | 4.94E-05 | 0.000587 | 2.21 |
| FALSE | High | EPFEDGFAI | [K].EPFED( | 1xPhospho | E9QM38 1 | 0.000164 | 0 | 1 | 2 | 5 | E9QM38 | Solute carr | E9QM38 [2 | 0 | 2191.938 | 0 | -0.39 | 0.014006 | 2.32 | 112 | 88 | 8.43 | 4.12 | | High | High | High | High | High | High | High | High | High | High | High | 0 | 5.29E-05 | 3.15 |
| FALSE | High | EPFEDGFAI | [K].EPFED( | 1xDeamida | E9QM38 1 | 4.36E-06 | 0 | 1 | 2 | 6 | E9QM38 | Solute carr | E9QM38 [2 | 0 | 2192.922 | 0 | -0.09 | 0.580186 | 13.6 | 104.7 | 95.3 | 14.93 | 8.12 | | High | High | High | High | High | High | High | High | High | High | High | 0.000294 | 0.002303 | 3.62 |
| FALSE | High | EPGEQAST | [K].EPGEQ, | 1xPhospho | G3X9H5 1x | 0.000543 | 0 | 1 | 3 | 1 | G3X9H5 | Huntingtin | G3X9H5 [1 | 0 | 1896.918 | 0 | 0.45 | 0.008837 | 14.4 | 83.8 | 116.2 | 11.59 | 5.43 | | High | High | High | High | High | High | High | High | High | High | High | 0 | 0.000121 | 2.12 |
| FALSE | High | ENFPVQSI( | [R].ENFPV( | 1xPhospho | Q8BQ33 1x | 0.000141 | 0 | 1 | 1 | 2 | Q8BQ33 | Treslin OS= | Q8BQ33 [1 | 0 | 1911.998 | 0 | -0.75 | 0.001718 | 1.81 | 125.6 | 74.4 | 3.97 | 9.5 | | High | High | High | High | High | High | High | High | High | High | High | 7.19E-05 | 0.00085 | 2.22 |
| FALSE | High | ENFPSISTY | [K].ENFPSI | 1xPhospho | A0A0R4J0V | 2.75E-05 | 0 | 1 | 4 | 2 | A0A0R4J0V | Phosphatic | A0A0R4J0V | 0 | 2542.134 | 0 | -0.38 | 0.055806 | 10.49 | 112.5 | 87.5 | 16.42 | 13.2 | | High | High | High | High | High | High | High | High | High | High | High | 0 | 4.2E-06 | 2.89 |

| | | | | | | | | | | | | | | | | | | | | | | | | | | | | | | | | | | | | | | |
|---|---|---|---|---|---|---|---|---|---|---|---|---|---|---|---|---|---|---|---|---|---|---|---|---|---|---|---|---|---|---|---|---|---|---|---|---|---|---|
| FALSE | High | ENETDEEN | [K].ENETDI | 1xPhospho | A2AT37 1x | 1.74E-06 | 0 | 1 | 2 | 4 | A2AT37 | UPF2 regul | A2AT37 [1 | 0 | 2205.003 | 0 | -0.59 | 0.000289 | 2.16 | 119.1 | 80.9 | 2.3 | 6.61 | | High | High | High | High | High | High | High | High | High | High | High | 0 | 1.8E-06 | 3.6 |
| FALSE | High | ENETDEEN | [K].ENETDI | 1xPhospho | A2AT37 1x | 1.45E-05 | 0 | 1 | 2 | 1 | A2AT37 | UPF2 regul | A2AT37 [1 | 0 | 2189.009 | 0 | -0.69 | 0.001066 | 4.31 | 123.4 | 76.6 | 5.05 | 5.86 | | High | High | High | High | High | High | High | High | High | High | High | 0 | 2.06E-06 | 3.28 |
| FALSE | High | EIQNGTLRI | [K].EIQNGT | 1xDeamida | F8VQC7 1x | 0.007339 | 0.00062 | 1 | 35 | 1 | F8VQC7 | Kinectin O | F8VQC7 [6 | 1 | 2277.071 | 0 | 1.13 | 0.005391 | 10.05 | 66.6 | 133.4 | 10.77 | 14.54 | | High | High | High | High | High | High | High | High | High | High | High | 0.000275 | 0.002223 | 2.33 |
| FALSE | High | ELQPIVYTP | [K].ELQPIV | 1xCarbami | Q91WD7 1 | 0.000434 | 0 | 1 | 1 | 1 | Q91WD7 | Kinesin-like | Q91WD7 [ | 0 | 2130.059 | 0 | -0.27 | 0.177102 | 11.23 | 105.1 | 94.9 | 12.65 | 11.81 | | High | High | High | High | High | High | High | High | High | High | High | 0 | 9.33E-05 | 3.48 |
| FALSE | High | ELSAPAR | [R].ELSAPA | 1xPhospho | D3Z7W0 1x | 0.028575 | 0.001829 | 1 | 3 | 1 | D3Z7W0 | Iron-sulfur | D3Z7W0 [2 | 0 | 1052.534 | 0 | -0.54 | 0.003932 | 5.95 | 122 | 78 | 5.11 | 14.61 | | High | High | High | High | High | High | High | High | High | High | High | 0.000984 | 0.01015 | 1.72 |
| FALSE | High | EISDDEAEE | [K].EISDDE | 1xPhospho | P11499 1xP | 2.4E-05 | 0 | 1 | 1 | 5 | P11499 | Heat shock | P11499 [22 | 0 | 1831.825 | 0 | -0.61 | 0.001364 | 2.63 | 120.9 | 79.1 | 3.05 | 12.08 | | High | High | High | High | High | High | High | High | High | High | High | 0 | 3.6E-06 | 3.49 |
| FALSE | High | ELSDDSFEN | [K].ELSDDS | 1xPhospho | B9EKT8 1xP | 0.000128 | 0 | 1 | 2 | 1 | B9EKT8 | Uncharacte | B9EKT8 [19 | 0 | 1922.878 | 0 | -0.52 | 0.004531 | 2.43 | 119.5 | 80.5 | 7.3 | 12.01 | | High | High | High | High | High | High | High | High | High | High | High | 0 | 2.38E-05 | 2.42 |
| FALSE | High | ELSDPCVPS | [K].ELSDPC | 1xCarbami | Q8C263 1x | 0.08447 | 0.006345 | 1 | 1 | 1 | Q8C263 | Spindle and | Q8C263 [1 | 1 | 3495.666 | 0 | -1.61 | 0.000477 | 1.66 | 150.9 | 49.1 | 9.68 | 10.76 | | High | High | High | High | High | High | High | High | High | High | High | 0.003265 | 0.03529 | 2.99 |
| FALSE | High | ELSEESEDE | [K].ELSEES | 1xPhospho | P54103 1xP | 2.34E-06 | 0 | 1 | 6 | 4 | P54103 | DnaJ homo | P54103 [58 | 0 | 2962.393 | 0 | -1.22 | 0.00025 | 1.31 | 138.9 | 61.1 | 6.14 | 10.21 | | High | High | High | High | High | High | High | High | High | High | High | 0 | 2.64E-07 | 4.16 |
| FALSE | High | ELSEESEDE | [K].ELSEES | 1xPhospho | P54103 1xP | 1.25E-06 | 0 | 1 | 6 | 12 | P54103 | DnaJ homo | P54103 [58 | 0 | 2978.388 | 0 | -1.19 | 0.001718 | 3.05 | 138.1 | 61.9 | 3.74 | 16.91 | | High | High | High | High | High | High | High | High | High | High | High | 0 | 1.3E-07 | 4.77 |
| FALSE | High | ELSEESEDE | [K].ELSEES | 2xPhospho | P54103 2xP | 2.78E-06 | 0 | 1 | 6 | 12 | P54103 | DnaJ homo | P54103 [58 | 0 | 3042.36 | 0 | -0.35 | 0.016499 | 7.97 | 113.9 | 86.1 | 10.76 | 9.28 | | High | High | High | High | High | High | High | High | High | High | High | 0 | 1.15E-06 | 4.67 |
| FALSE | High | ELSESVQQ | [K].ELSESV | 1xPhospho | Q80Y82 1x | 1.37E-08 | 0 | 1 | 11 | 2 | Q80Y82 | Protein kin | Q80Y82 [5 | 0 | 2575.378 | 0 | -0.04 | 0.308005 | 6 | 102.2 | 97.8 | 6.16 | 5.43 | | High | High | High | High | High | High | High | High | High | High | High | 0 | 8.22E-10 | 5.69 |
| FALSE | High | ELSGESDEL | [K].ELSGES | 1xPhospho | Q8R550-3 | 1.91E-07 | 0 | 1 | 6 | 2 | Q8R550-3 | Isoform 3 o | Q8R550-3 | 0 | 2602.254 | 0 | 1.13 | 0.00024 | 18.52 | 62.8 | 137.2 | 22.11 | 7.83 | | High | High | High | High | High | High | High | High | High | High | High | 0 | 2.04E-08 | 4.52 |
| FALSE | High | ELSGLPSGF | [K].ELSGLP | 1xPhospho | Q3TC53 1x | 0.000259 | 0 | 1 | 3 | 5 | Q3TC53 | Adenylyl cy | Q3TC53 [2 | 0 | 3937.906 | 0 | 0.15 | 0.415902 | 13.43 | 92.3 | 107.7 | 22.06 | 22.12 | | High | High | High | High | High | High | High | High | High | High | High | 0 | 5.22E-05 | 4.34 |
| FALSE | High | ELSGSCDH | [R].ELSGSC | 1xCarbami | Q8BKT3 2x | 0.037369 | 0.00249 | 1 | 1 | 1 | Q8BKT3 | GC-rich se | Q8BKT3 [4 | 0 | 3756.531 | 0 | 0.6 | 0.01611 | 3.5 | 79.8 | 120.2 | 6.25 | 13.37 | | High | High | High | High | High | High | High | High | High | High | High | 0.001308 | 0.0138 | 3.41 |
| FALSE | High | ELQLEELES | [K].ELQLEE | 1xPhospho | Q80TL6 1xP | 0.007471 | 0.00062 | 1 | 8 | 1 | Q80TL6 | MKIAA104 | Q80TL6 [3 | 0 | 2447.247 | 0 | 0.26 | 0.676202 | 11.43 | 96.1 | 103.9 | 20.51 | 11.01 | | High | High | High | High | High | High | High | High | High | High | High | 0.000275 | 0.00227 | 2.3 |
| FALSE | High | ELSHSPPR | [K].ELSHSP | 1xPhospho | Q8BTI8 1xP | 0.005924 | 0.00062 | 1 | 3 | 4 | Q8BTI8 | Serine/argi | Q8BTI8 [12 | 0 | 1231.603 | 0 | -0.45 | 0.008538 | 2.76 | 117.2 | 82.8 | 7.84 | 6.9 | | High | High | High | High | High | High | High | High | High | High | High | 0.000656 | 0.005822 | 2.34 |
| FALSE | High | ELSHSPPRE | [K].ELSHSP | 2xPhospho | Q8BTI8 2xP | 0.001424 | 5.66E-05 | 1 | 3 | 2 | Q8BTI8 | Serine/argi | Q8BTI8 [12 | 1 | 2838.315 | 0 | -1.01 | 0.000885 | 1.95 | 133.8 | 66.2 | 6.05 | 8.29 | | High | High | High | High | High | High | High | High | High | High | High | 0.001214 | 0.01296 | 2.34 |
| FALSE | High | ELSISK | [R].ELSISK. | 1xPhospho | B9EK83 1xP | 0.100129 | 0.008067 | 2 | 3 | 2 | B9EK83; A2 | Transient r | B9EK83 [12 | 0 | 1214.68 | 0 | 2.34 | 0.006213 | 19.17 | 32 | 168 | 13.93 | 27.9 | Shared | High | High | High | High | High | High | High | High | High | High | High | 0.004059 | 0.0433 | 1.66 |
| FALSE | High | ELSISQSPE | [R].ELSISQ | 1xPhospho | Q99MU3 1 | 0.000793 | 0 | 1 | 3 | 1 | Q99MU3 | Double-str | Q99MU3 [ | 0 | 1783.924 | 0 | -0.7 | 0.000404 | 2.41 | 125.3 | 74.7 | 3.76 | 4.24 | | High | High | High | High | High | High | High | High | High | High | High | 0 | 0.000184 | 2.27 |
| FALSE | High | ELSLSTEEG | [K].ELSLSTI | 2xPhospho | B2RWU2 2 | 0.000259 | 0 | 1 | 2 | 1 | B2RWU2 | Immunode | B2RWU2 [ | 0 | 2364.066 | 0 | 1.79 | 0.00093 | 21.91 | 44.9 | 155.1 | 18.04 | 8.35 | | High | High | High | High | High | High | High | High | High | High | High | 0 | 5.26E-05 | 2.97 |
| FALSE | High | EISPAENIL | [R].EISPAEI | 1xPhospho | Q8K2E5 1x | 2.93E-06 | 0 | 1 | 2 | 1 | Q8K2E5 | Budding ur | Q8K2E5 [5 | 0 | 2336.179 | 1 | -1.21 | 0.003265 | 2.26 | 139.7 | 60.3 | 11.59 | 19.54 | | High | High | High | High | High | High | High | High | High | High | High | 0 | 3.42E-07 | 4.62 |
| FALSE | High | ELSPEGPGI | [R].ELSPEG | 1xPhospho | Q80U72-4 | 0.003892 | 0.000261 | 2 | 5 | 1 | Q80U72-4; | Isoform 4 c | Q80U72-4 | 0 | 1451.755 | 0 | 0.22 | 0.155644 | 4.88 | 92.9 | 107.1 | 3.16 | 8.67 | Shared | High | High | High | High | High | High | High | High | High | High | High | 0.00014 | 0.001098 | 1.85 |
| FALSE | High | ELSPEQDD | [R].ELSPEQ | 1xPhospho | Q6P1G0 1x | 0.026885 | 0.001676 | 1 | 1 | 2 | Q6P1G0 | HEAT repe | Q6P1G0 [2 | 0 | 1601.705 | 0 | 0.42 | 0.031728 | 22.18 | 86.7 | 113.3 | 9.43 | 7.92 | | High | High | High | High | High | High | High | High | High | High | High | 0.003317 | 0.03647 | 2.24 |
| FALSE | High | EISPLAAITF | [K].EISPLA | 1xPhospho | L7N209 1xP | 1.43E-05 | 0 | 1 | 2 | 1 | L7N209 | HBS1-like p | L7N209 [58 | 0 | 2087.123 | 0 | -0.34 | 0.069157 | 13.2 | 111.8 | 88.2 | 6.88 | 12.22 | | High | High | High | High | High | High | High | High | High | High | High | 0 | 2.03E-06 | 2.93 |
| FALSE | High | ELSPQHK | [R].ELSPQH | 1xPhospho | Q8R4U7 1x | 0.007605 | 0.000664 | 2 | 4 | 1 | Q8R4U7; B | Leucine zip | Q8R4U7 [3 | 0 | 1376.734 | 0 | -0.59 | 0.000733 | 5.19 | 121.9 | 78.1 | 3.69 | 11.01 | Shared | High | High | High | High | High | High | High | High | High | High | High | 0.000294 | 0.002303 | 1.68 |
| FALSE | High | ELSPSNSSS | [K].ELSPSN | 1xPhospho | B0V2M3 1x | 0.001881 | 5.66E-05 | 1 | 5 | 2 | B0V2M3 | Zinc finger | B0V2M3 [1 | 0 | 1774.862 | 0 | -0.74 | 0.001716 | 1.5 | 123.7 | 76.3 | 5.93 | 5.86 | | High | High | High | High | High | High | High | High | High | High | High | 2.58E-05 | 0.000483 | 2.68 |
| FALSE | High | EISPVGEK | [R].EISPVG | 1xPhospho | Q3UZ39 1x | 0.024638 | 0.001612 | 1 | 2 | 1 | Q3UZ39 | Leucine-ric | Q3UZ39 [6 | 0 | 1396.749 | 0 | 1.36 | 0.001385 | 14.06 | 57.4 | 142.6 | 5.79 | 12.6 | | High | High | High | High | High | High | High | High | High | High | High | 0.000859 | 0.008593 | 2.34 |
| FALSE | High | ELSREDLDL | [K].ELSRED | 1xPhospho | Q3TZW9 1 | 0.044358 | 0.003264 | 1 | 4 | 1 | Q3TZW9 | Tyrosine-p | Q3TZW9 [2 | 1 | 3053.623 | 0 | 0.41 | 0.004591 | 9.67 | 83.7 | 116.3 | 6.26 | 8.18 | | High | High | High | High | High | High | High | High | High | High | High | 0.00168 | 0.01682 | 1.29 |
| FALSE | High | ELSSGQDV | [K].ELSSGC | 2xCarbami | Q5SW15 1 | 4.32E-09 | 0 | 1 | 3 | 2 | Q5SW15 | Methyltran | Q5SW15 [4 | 0 | 4373.022 | 0 | -0.01 | 0.720958 | 5.88 | 98.5 | 101.5 | 9.4 | 9.18 | | High | High | High | High | High | High | High | High | High | High | High | 0 | 1.08E-06 | 4.59 |
| FALSE | High | ELSHSPPRE | [K].ELSHSP | 1xPhospho | Q8BTI8 1xP | 0.010475 | 0.000772 | 1 | 3 | 1 | Q8BTI8 | Serine/argi | Q8BTI8 [12 | 1 | 2758.349 | 0 | -0.66 | 0.023663 | 8.28 | 122.4 | 77.6 | 15.44 | 12.43 | | High | High | High | High | High | High | High | High | High | High | High | 0.000382 | 0.003316 | 2.23 |
| FALSE | High | ELQELLIDG | [R].ELQELL | 1xCarbami | Q148S3 1x | 6.71E-05 | 0 | 1 | 1 | 4 | Q148S3 | NIN1/RPN1 | Q148S3 [1 | 1 | 4703.076 | 0 | -0.84 | 0.000344 | 1.88 | 128.9 | 71.1 | 6.7 | 6.88 | | High | High | High | High | High | High | High | High | High | High | High | 0 | 1.15E-05 | 4.85 |
| FALSE | High | ELPTLSPAP | [R].ELPTLS | 1xPhospho | E9Q8A3 1x | 1.35E-05 | 0 | 1 | 3 | 1 | E9Q8A3 | Phosphatic | E9Q8A3 [2 | 0 | 2248.188 | 0 | -0.05 | 0.943661 | 3.96 | 102.7 | 97.3 | 8.54 | 13.3 | | High | High | High | High | High | High | High | High | High | High | High | 0 | 1.9E-06 | 3.39 |
| FALSE | High | ELPSPVEPA | [K].ELPSPV | 1xPhospho | Q8VIG1 1x | 4.62E-07 | 0 | 1 | 1 | 2 | Q8VIG1 | RE1-silenci | Q8VIG1 [7 | 0 | 3855.926 | 0 | -1.2 | 0.004034 | 2.45 | 142.7 | 57.3 | 18.46 | 17.39 | | High | High | High | High | High | High | High | High | High | High | High | 0 | 4.27E-08 | 3.87 |
| FALSE | High | ELLNSEPFD | [R].ELLNSE | 1xDeamida | O08696 1x | 1.11E-05 | 0 | 1 | 6 | 4 | O08696 | Forkhead b | O08696 [6 | 0 | 4959.486 | 0 | 2.01 | 0.000466 | 23.59 | 35.3 | 164.7 | 37.46 | 16.89 | | High | High | High | High | High | High | High | High | High | High | High | 0 | 1.41E-05 | 4.75 |
| FALSE | High | ELLSNDTPE | [R].ELLSND | 1xPhospho | B9EK83 1xP | 0.016021 | 0.001124 | 2 | 3 | 2 | B9EK83; A2 | Transient r | B9EK83 [14 | 0 | 2012.035 | 0 | 3.01 | 0.000264 | 11.62 | 21.7 | 178.3 | 6.31 | 7.07 | Shared | High | High | High | High | High | High | High | High | High | High | High | 0.000585 | 0.005336 | 2.64 |
| FALSE | High | ELLSPGIK | [R].ELLSPG | 1xPhospho | P52479-2 1 | 0.016596 | 0.001124 | 1 | 2 | 1 | P52479-2 | Isoform 2 o | P52479-2 [ | 0 | 1394.806 | 0 | -1.1 | 0.000209 | 1.77 | 136.5 | 63.5 | 4.67 | 8.52 | | High | High | High | High | High | High | High | High | High | High | High | 0.000585 | 0.005517 | 1.73 |
| FALSE | High | ELLSPLSEPI | [K].ELLSPLS | 1xPhospho | Q9ESC8 1x | 0.001373 | 5.66E-05 | 1 | 2 | 1 | Q9ESC8 | AF4/FMR2 | Q9ESC8 [6 | 0 | 1679.809 | 0 | -0.41 | 0.040784 | 8.83 | 112.2 | 87.8 | 11.46 | 8.51 | | High | High | High | High | High | High | High | High | High | High | High | 2.58E-05 | 0.000341 | 3.34 |
| FALSE | High | ELLSPLSEPI | [K].ELLSPLS | 2xPhospho | Q9ESC8 2x | 0.001881 | 5.66E-05 | 1 | 2 | 1 | Q9ESC8 | AF4/FMR2 | Q9ESC8 [6 | 0 | 1759.775 | 0 | -0.44 | 0.006454 | 8.56 | 115 | 85 | 6.31 | 12.31 | | High | High | High | High | High | High | High | High | High | High | High | 2.58E-05 | 0.000486 | 2.84 |
| FALSE | High | ELLSPLSEPI | [K].ELLSPLS | 1xPhospho | Q9ESC8 1x | 0.038017 | 0.002528 | 1 | 2 | 2 | Q9ESC8 | AF4/FMR2 | Q9ESC8 [6 | 1 | 2622.42 | 0 | -0.1 | 0.416137 | 2.93 | 101 | 99 | 8.68 | 9.54 | | High | High | High | High | High | High | High | High | High | High | High | 0.004596 | 0.05001 | 2.55 |
| FALSE | High | ELLSPLSEPI | [K].ELLSPLS | 2xPhospho | Q9ESC8 2x | 0.00537 | 0.000599 | 1 | 2 | 3 | Q9ESC8 | AF4/FMR2 | Q9ESC8 [6 | 1 | 2702.386 | 0 | -0.46 | 0.028732 | 9.25 | 113.9 | 86.1 | 7.85 | 11.73 | | High | High | High | High | High | High | High | High | High | High | High | 0.000267 | 0.001572 | 2.76 |
| FALSE | High | EILSPVDIID | [R].EILSPVI | 1xPhospho | Q99JY8 1xP | 0.004824 | 0.000456 | 1 | 1 | 1 | Q99JY8 | Phospholip | Q99JY8 [2 | 0 | 1578.834 | 0 | -1.68 | 0.000525 | 1.2 | 153 | 47 | 8.99 | 10.3 | | High | High | High | High | High | High | High | High | High | High | High | 0.000204 | 0.001384 | 2.79 |
| FALSE | High | EIITEEPSEE | [K].EIITEEP | 1xPhospho | Q3ULC7 1x | 0.000237 | 0 | 1 | 4 | 2 | Q3ULC7 | Putative ur | Q3ULC7 [1 | 0 | 2839.429 | 0 | -0.64 | 0.218193 | 23.12 | 120.8 | 79.2 | 32.11 | 18.41 | | High | High | High | High | High | High | High | High | High | High | High | 0 | 4.74E-05 | 2.77 |
| FALSE | High | ELLTSDSPP | [K].ELLTSD | 1xPhospho | Q9R1E0 1x | 4.98E-05 | 0 | 1 | 3 | 5 | Q9R1E0 | Forkhead b | Q9R1E0 [4 | 0 | 3198.498 | 0 | 0.04 | 0.614299 | 15.88 | 101 | 99 | 10.46 | 27.53 | | High | High | High | High | High | High | High | High | High | High | High | 0 | 1E-05 | 4.45 |
| FALSE | High | ELLVSQQEI | [R].ELLVSQ | 1xPhospho | E9PV87 1xP | 0.009931 | 0.000736 | 1 | 6 | 1 | E9PV87 | Protein TAI | E9PV87 [13 | 0 | 2883.346 | 0 | -0.13 | 0.219398 | 9.14 | 107.6 | 92.4 | 7.57 | 10.63 | | High | High | High | High | High | High | High | High | High | High | High | 0.000344 | 0.003099 | 3.16 |
| FALSE | High | ELNFVTDS | [K].ELNFVT | 1xPhospho | Q0VEJ0 1xP | 8.48E-05 | 0 | 1 | 1 | 2 | Q0VEJ0 | Centrosom | Q0VEJ0 [68 | 0 | 2314.105 | 0 | 0.22 | 0.793472 | 7.29 | 92 | 108 | 30.1 | 20.91 | | High | High | High | High | High | High | High | High | High | High | High | 0 | 1.49E-05 | 2.99 |
| FALSE | High | ELNFVTDS | [K].ELNFVT | 1xPhospho | Q0VEJ0 1xP | 6.48E-09 | 0 | 1 | 1 | 6 | Q0VEJ0 | Centrosom | Q0VEJ0 [68 | 0 | 2543.268 | 0 | -0.09 | 0.580974 | 6.07 | 102.4 | 97.6 | 5.18 | 7.66 | | High | High | High | High | High | High | High | High | High | High | High | 0 | 4.09E-10 | 4.46 |
| FALSE | High | EINSDQST | [K].EINSDC | 1xPhospho | E9QPI5 1xP | 0.000109 | 0 | 1 | 2 | 2 | E9QPI5 | Sister chro | E9QPI5 [12 | 0 | 2059.913 | 0 | -0.14 | 0.521043 | 22.07 | 104.7 | 95.3 | 15.43 | 18.4 | | High | High | High | High | High | High | High | High | High | High | High | 0 | 1.98E-05 | 3.34 |
| FALSE | High | ELNSNHDG | [K].ELNSNH | 1xPhospho | Q9EQU5-2 | 0.000158 | 0 | 1 | 1 | 8 | Q9EQU5-2 | Isoform 2 o | Q9EQU5-2 | 0 | 2183.986 | 0 | -1.08 | 0.000311 | 0.99 | 133.4 | 66.6 | 5.72 | 6.57 | | High | High | High | High | High | High | High | High | High | High | High | 0 | 3E-05 | 3.28 |
| FALSE | High | ELNSNHDG | [K].ELNSNH | 1xDeamida | Q9EQU5-2 | 0.010758 | 0.000772 | 1 | 1 | 4 | Q9EQU5-2 | Isoform 2 o | Q9EQU5-2 | 0 | 2184.97 | 0 | -0.96 | 0.003629 | 2.31 | 129.7 | 70.3 | 10.84 | 9.44 | | High | High | High | High | High | High | High | High | High | High | High | 0.000399 | 0.003389 | 2.86 |
| FALSE | High | ELNSPYLLS | [K].ELNSPY | 1xCarbami | E9PZ97 1xP | 2.38E-05 | 0 | 1 | 2 | 2 | E9PZ97 | Bloom sync | E9PZ97 [46 | 0 | 3296.616 | 0 | -1.77 | 0.000494 | 0.27 | 151.7 | 48.3 | 8.42 | 7.19 | | High | High | High | High | High | High | High | High | High | High | High | 0 | 3.59E-06 | 2.61 |
| FALSE | High | ELNVSFINE | [K].ELNVSF | 1xPhospho | P49586 1xP | 4.89E-05 | 0 | 2 | 3 | 1 | P49586; Q8 | Choline-ph | P49586 [22 | 0 | 1730.913 | 0 | 0.66 | 0.010455 | 19.93 | 73.1 | 126.9 | 17.53 | 16.08 | Shared | High | High | High | High | High | High | High | High | High | High | High | 0 | 8.05E-06 | 4.05 |
| FALSE | High | EIPAAPALP | [K].EIPAAP | 1xPhospho | Q7TSC1 1x | 0.000294 | 0 | 1 | 3 | 1 | Q7TSC1 | Protein PRI | Q7TSC1 [5 | 0 | 2365.282 | 0 | 1.2 | 0.000696 | 15.59 | 62.3 | 137.7 | 13.27 | 10.86 | | High | High | High | High | High | High | High | High | High | High | High | 0 | 6.07E-05 | 2.42 |
| FALSE | High | EIPASPTRF | [R].EIPASPT | 1xPhospho | Q3TU15 1x | 0.007082 | 0.00062 | 1 | 3 | 1 | Q3TU15 | Non-specif | Q3TU15 [4 | 1 | 2050.114 | 0 | -0.79 | 0.000941 | 4.65 | 129.6 | 70.4 | 4.27 | 9.34 | | High | High | High | High | High | High | High | High | High | High | High | 0.000275 | 0.002126 | 1.49 |
| FALSE | High | ELPDSSSPV | [K].ELPDSS | 1xPhospho | Q6PCL9 1x | 0.001881 | 5.66E-05 | 1 | 2 | 1 | Q6PCL9 | Poly(A) pol | Q6PCL9 [7 | 0 | 1904.932 | 0 | -0.06 | 0.809605 | 20.24 | 96.3 | 103.7 | 13.55 | 11.32 | | High | High | High | High | High | High | High | High | High | High | High | 2.58E-05 | 0.000483 | 2.49 |
| FALSE | High | EIPFNEDPN | [K].EIPFNE | 1xPhospho | Q7TPS5 1x | 0.004738 | 0.000456 | 1 | 4 | 3 | Q7TPS5 | C2 domain | Q7TPS5 [2 | 0 | 2916.418 | 0 | 0.14 | 0.163178 | 12.56 | 95.7 | 104.3 | 4.37 | 9 | | High | High | High | High | High | High | High | High | High | High | High | 0.000275 | 0.001616 | 2.4 |
| FALSE | High | EIPFNEDPN | [K].EIPFNE | 2xPhospho | Q7TPS5 2x | 2.77E-05 | 0 | 1 | 4 | 2 | Q7TPS5 | C2 domain | Q7TPS5 [2 | 0 | 2996.384 | 0 | 0.25 | 0.073671 | 13.57 | 91 | 109 | 8.46 | 9.52 | | High | High | High | High | High | High | High | High | High | High | High | 0 | 4.26E-06 | 3.64 |
| FALSE | High | EIPGTLQD | [K].EIPGTL | 1xPhospho | G5E8W8 1 | 0.000356 | 0 | 2 | 6 | 2 | G5E8W8; A | Zinc finger | G5E8W8 [2 | 0 | 4671.344 | 0 | -1.66 | 0.001546 | 4.4 | 148.5 | 51.5 | 10.27 | 41.3 | Shared | High | High | High | High | High | High | High | High | High | High | High | 0 | 0.000132 | 4.04 |
| FALSE | High | ELPPAEPVL | [K].ELPPAE | 1xPhospho | Q571G8 1x | 0.002334 | 0.000109 | 1 | 5 | 1 | Q571G8 | MKIAA164 | Q571G8 [2 | 0 | 2215.203 | 0 | -0.06 | 0.300442 | 2.24 | 101.4 | 98.6 | 11.68 | 9.09 | | High | High | High | High | High | High | High | High | High | High | High | 4.94E-05 | 0.000619 | 3.36 |
| FALSE | High | ELPPELSHL | [K].ELPPEL | 1xPhospho | E9Q2Z1 1x | 3.54E-05 | 0 | 1 | 3 | 1 | E9Q2Z1 | CECR2, hist | E9Q2Z1 [3 | 0 | 2157.061 | 0 | -0.95 | 0.002503 | 1.14 | 131.5 | 68.5 | 7.05 | 7.24 | | High | High | High | High | High | High | High | High | High | High | High | 0 | 5.61E-06 | 1.55 |
| FALSE | High | ELPPLSPSL | [R].ELPPLS | 1xPhospho | D3Z0X5 1xP | 0.008927 | 0.000736 | 1 | 6 | 4 | D3Z0X5 | Pleckstrin h | D3Z0X5 [4 | 0 | 1504.797 | 0 | 0.64 | 0.001495 | 3.23 | 79.3 | 120.7 | 6.2 | 10.36 | | High | High | High | High | High | High | High | High | High | High | High | 0.003963 | 0.04235 | 1.99 |
| FALSE | High | ELPPPPPAP | [R].ELPPPP | 1xPhospho | F8WJ40 1x | 4.96E-07 | 0 | 1 | 2 | 3 | F8WJ40 | Histone-lys | F8WJ40 [6 | 0 | 2644.374 | 0 | -0.63 | 0.000485 | 3.17 | 123.5 | 76.5 | 4.67 | 6.9 | | High | High | High | High | High | High | High | High | High | High | High | 0 | 1.51E-07 | 3.72 |
| FALSE | High | ELPRPGASI | [R].ELPRPG | 1xPhospho | Q05915 1x | 0.000219 | 0 | 1 | 2 | 1 | Q05915 | GTP cycloh | Q05915 [1 | 0 | 1887.014 | 0 | 1.09 | 0.000672 | 6.46 | 65.2 | 134.8 | 11.78 | 8.1 | | High | High | High | High | High | High | High | High | High | High | High | 0 | 4.32E-05 | 1.91 |
| FALSE | High | EISSPARPC | [K].EISSPAI | 1xCarbami | Q9WV30-2 | 0.083783 | 0.006345 | 1 | 6 | 2 | Q9WV30-2 | Isoform 2 o | Q9WV30-2 | 0 | 2393.128 | 0 | 0.92 | 0.008321 | 24.85 | 77.2 | 122.8 | 13.42 | 13.08 | | High | High | High | High | High | High | High | High | High | High | High | 0.004072 | 0.04349 | 2.57 |
| FALSE | High | EPQSPSSQ | [K].EPQSPS | 1xCarbami | B7ZNK9 1x | 0.001799 | 5.66E-05 | 1 | 4 | 3 | B7ZNK9 | Spata5 pro | B7ZNK9 [2 | 0 | 2554.201 | 0 | -0.5 | 0.026093 | 4.59 | 120.4 | 79.6 | 14.79 | 7.4 | | High | High | High | High | High | High | High | High | High | High | High | 9.42E-05 | 0.000941 | 1.86 |
| FALSE | High | EISSSPTSK | [R].EISSSPT | 1xPhospho | Q8BTI8 1xP | 5.7E-05 | 0 | 1 | 3 | 5 | Q8BTI8 | Serine/argi | Q8BTI8 [45 | 0 | 1473.76 | 0 | -0.38 | 0.037937 | 5.54 | 116.4 | 83.6 | 10.55 | 7.96 | | High | High | High | High | High | High | High | High | High | High | High | 0 | 9.59E-06 | 3.47 |
| FALSE | High | ELSVEEQIK | [R].ELSVEE | 1xPhospho | Q8C7S2 1x | 0.001659 | 5.66E-05 | 1 | 5 | 2 | Q8C7S2 | Putative ur | Q8C7S2 [7 | 0 | 1612.86 | 0 | 2.01 | 0.001499 | 30.15 | 39.8 | 160.2 | 11.17 | 12.03 | | High | High | High | High | High | High | High | High | High | High | High | 2.58E-05 | 0.000423 | 2.8 |
| FALSE | High | EMFPYEAS | [K].EMFPYI | 1xCarbami | Q3U955 1x | 5.46E-09 | 0 | 1 | 7 | 3 | Q3U955 | Putative ur | Q3U955 [3 | 0 | 2329.008 | 0 | 0.17 | 0.122467 | 8.44 | 94.8 | 105.2 | 8.1 | 5.1 | | High | High | High | High | High | High | High | High | High | High | High | 0 | 3.06E-07 | 3.63 |
| FALSE | High | EMFPYEAS | [K].EMFPYI | 1xCarbami | Q3U955 2x | 0.001503 | 5.66E-05 | 1 | 7 | 1 | Q3U955 | Putative ur | Q3U955 [3 | 0 | 2392.979 | 0 | -0.38 | 0.004615 | 1.86 | 112.2 | 87.8 | 7.38 | 7.63 | | High | High | High | High | High | High | High | High | High | High | High | 2.58E-05 | 0.000375 | 4.06 |
| FALSE | High | EMGWQED | [K].EMGW | 1xCarbami | Q6NXH3-2 | 1.35E-06 | 0 | 1 | 3 | 1 | Q6NXH3-2 | Isoform 2 o | Q6NXH3-2 | 0 | 3097.195 | 0 | -0.52 | 0.017912 | 10.23 | 117.7 | 82.3 | 6.67 | 15.51 | | High | High | High | High | High | High | High | High | High | High | High | 0 | 1.42E-07 | 4.74 |
| FALSE | High | EMGWQED | [K].EMGW | 1xCarbami | Q6NXH3-2 | 7.67E-08 | 0 | 1 | 3 | 2 | Q6NXH3-2 | Isoform 2 o | Q6NXH3-2 | 0 | 3113.19 | 0 | -0.27 | 0.056779 | 11.99 | 105.8 | 94.2 | 7.95 | 11.69 | | High | High | High | High | High | High | High | High | High | High | High | 0 | 5.68E-09 | 5.65 |
| FALSE | High | EMLLEDVG | [R].EMLLEI | 1xPhospho | Q80XU3 1x | 4.25E-06 | 0 | 1 | 1 | 7 | Q80XU3 | Nuclear ub | Q80XU3 [1 | 0 | 3332.469 | 0 | -0.26 | 0.22474 | 18.26 | 108.9 | 91.1 | 16.21 | 13.05 | | High | High | High | High | High | High | High | High | High | High | High | 0 | 5.18E-07 | 4.88 |
| FALSE | High | EMLLEDVG | [R].EMLLEI | 1xPhospho | Q80XU3 1x | 2.34E-09 | 0 | 1 | 1 | 37 | Q80XU3 | Nuclear ub | Q80XU3 [1 | 0 | 3348.464 | 0 | -0.17 | 0.123306 | 6.55 | 104.3 | 95.7 | 7.94 | 1.33 | | High | High | High | High | High | High | High | High | High | High | High | 0 | 1.51E-08 | 5.65 |
| FALSE | High | EMNDMKD | [R].EMNDN | 1xCarbami | Q9JM14 1x | 0.08447 | 0.006345 | 1 | 2 | 1 | Q9JM14 | 5'(3')-deox | Q9JM14 [8 | 1 | 2611.194 | 0 | -1.24 | 0.009158 | 2.77 | 143.2 | 56.8 | 14.91 | 35.53 | | High | High | High | High | High | High | High | High | High | High | High | 0.003265 | 0.03523 | 3.01 |
| FALSE | High | EMNPPSPS | [K].EMNPP | 1xPhospho | Q6P2K3 1x | 0.012731 | 0.000889 | 1 | 1 | 1 | Q6P2K3 | Uncharacte | Q6P2K3 [5 | 0 | 1550.687 | 0 | -0.71 | 0.000486 | 2.96 | 124.2 | 75.8 | 5.73 | 7.67 | | High | High | High | High | High | High | High | High | High | High | High | 0.000449 | 0.004094 | 2.24 |
| FALSE | High | EMPGSNIE | [K].EMPGS | 1xPhospho | Q8BTI8 1xP | 0.004738 | 0.000456 | 1 | 3 | 2 | Q8BTI8 | Serine/argi | Q8BTI8 [12 | 0 | 3964.925 | 0 | -0.8 | 0.017061 | 3.18 | 128.4 | 71.6 | 14.66 | 18.67 | | High | High | High | High | High | High | High | High | High | High | High | 0.003606 | 0.03816 | 3.59 |
| FALSE | High | EMPGSNIE | [K].EMPGS | 1xPhospho | Q8BTI8 1xP | 0.008167 | 0.000736 | 1 | 3 | 3 | Q8BTI8 | Serine/argi | Q8BTI8 [12 | 0 | 3980.92 | 0 | -0.84 | 0.083111 | 14.29 | 134 | 66 | 16.84 | 42.08 | | High | High | High | High | High | High | High | High | High | High | High | 0.000756 | 0.006675 | 2.86 |
| FALSE | High | EMPGSNIE | [K].EMPGS | 2xPhospho | Q8BTI8 2xP | 0.020319 | 0.001472 | 1 | 3 | 2 | Q8BTI8 | Serine/argi | Q8BTI8 [12 | 0 | 4044.891 | 0 | -0.52 | 0.069972 | 21.24 | 117.8 | 82.2 | 14.76 | 29.41 | | High | High | High | High | High | High | High | High | High | High | High | 0.005428 | 0.0596 | 3.63 |
| FALSE | High | EMPSVFQS | [K].EMPSV | 1xPhospho | Q3U0X6 1x | 5.11E-08 | 0 | 1 | 1 | 2 | Q3U0X6 | Putative ur | Q3U0X6 [5 | 0 | 3139.496 | 0 | -0.48 | 0.015285 | 11.13 | 116.4 | 83.6 | 9.88 | 10.61 | | High | High | High | High | High | High | High | High | High | High | High | 0 | 3.62E-09 | 4.33 |
| FALSE | High | EMQSTLSC | [R].EMQST | 1xPhospho | A0A0R4J07 | 7.44E-10 | 0 | 1 | 5 | 3 | A0A0R4J07 | BRCA1-A co | A0A0R4J07 | 0 | 2486.261 | 0 | 0.32 | 0.070098 | 5.39 | 88.3 | 111.7 | 9.1 | 20.56 | | High | High | High | High | High | High | High | High | High | High | High | 0 | 3.1E-11 | 4.4 |
| FALSE | High | EMFPYEAS | [K].EMFPYI | 1xCarbami | Q3U955 1x | 4.12E-05 | 0 | 1 | 7 | 1 | Q3U955 | Putative ur | Q3U955 [3 | 0 | 2313.013 | 0 | 0.15 | 0.167621 | 10.76 | 92.3 | 107.7 | 9.85 | 14.73 | | High | High | High | High | High | High | High | High | High | High | High | 0 | 6.6E-06 | 3.7 |
| FALSE | High | EMSPGHGI | [K].EMSPG | 1xPhospho | B6ZHD0 1x | 0.000738 | 0 | 2 | 12 | 9 | B6ZHD0; Q | Erythrocyt | B6ZHD0 [6 | 0 | 1579.677 | 0 | -0.07 | 0.252799 | 13.39 | 102.7 | 97.3 | 3.4 | 15.05 | Shared | High | High | High | High | High | High | High | High | High | High | High | 0.009931 | 0.108 | 2.89 |
| FALSE | High | EMTHSWP | [K].EMTHS | 1xCarbami | Q9ESC8 1x | 0.000651 | 0 | 1 | 1 | 1 | Q9ESC8 | AF4/FMR2 | Q9ESC8 [3 | 0 | 2657.302 | 0 | -0.15 | 0.096737 | 10.57 | 107.7 | 92.3 | 5.16 | 9.69 | | High | High | High | High | High | High | High | High | High | High | High | 0 | 0.000147 | 3.18 |
| FALSE | High | EMTTSADC | [K].EMTTS | 1xPhospho | Q5SW75 1 | 0.113753 | 0.009241 | 1 | 4 | 1 | Q5SW75 | Protein ph | Q5SW75 [4 | 0 | 1960.97 | 0 | 1.26 | 0.00475 | 28.54 | 60.9 | 139.1 | 15.3 | 15.26 | | High | High | High | High | High | High | High | High | High | High | High | 0.00464 | 0.05013 | 2.85 |
| FALSE | High | EMTTSADC | [K].EMTTS | 1xPhospho | Q5SW75 1 | 4.59E-05 | 0 | 1 | 4 | 4 | Q5SW75 | Protein ph | Q5SW75 [4 | 0 | 1976.965 | 0 | 1.12 | 0.000555 | 4.82 | 61.8 | 138.2 | 2.96 | 7.86 | | High | High | High | High | High | High | High | High | High | High | High | 0 | 3.11E-05 | 4.04 |
| FALSE | High | ENASPAPG | [R].ENASP | 1xPhospho | Q9Z1R2 1x | 8.63E-05 | 0 | 1 | 7 | 3 | Q9Z1R2 | Large proli | Q9Z1R2 [9 | 0 | 2027.894 | 0 | -0.15 | 0.141153 | 3.4 | 105.4 | 94.6 | 5.38 | 10.58 | | High | High | High | High | High | High | High | High | High | High | High | 0.002867 | 0.02981 | 3.16 |
| FALSE | High | ENASPAPG | [R].ENASP | 1xPhospho | Q9Z1R2 1x | 1.49E-09 | 0 | 1 | 7 | 20 | Q9Z1R2 | Large proli | Q9Z1R2 [9 | 0 | 2043.889 | 0 | -0.05 | 0.503027 | 5.43 | 103.2 | 96.8 | 2.96 | 9.01 | | High | High | High | High | High | High | High | High | High | High | High | 0 | 2.23E-07 | 4.42 |
| FALSE | High | ENDPENSL | [K].ENDPEI | 1xPhospho | E9Q286 1x | 2.52E-07 | 0 | 1 | 1 | 2 | E9Q286 | Little elong | E9Q286 [6 | 0 | 2069.02 | 0 | -1.29 | 0.00035 | 1.39 | 140.7 | 59.3 | 3.7 | 9.67 | | High | High | High | High | High | High | High | High | High | High | High | 0 | 3.95E-08 | 3.7 |
| FALSE | High | ENDPLRTPI | [K].ENDPLF | 1xPhospho | B2RT41 1xP | 0.000426 | 0 | 1 | 2 | 1 | B2RT41 | Zinc finger, | B2RT41 [7 | 1 | 2276.158 | 0 | -0.47 | 0.022691 | 8.54 | 116.2 | 83.8 | 9.69 | 9.51 | | High | High | High | High | High | High | High | High | High | High | High | 0 | 9.15E-05 | 2.43 |
| FALSE | High | ENEAQLPH | [K].ENEAQ | 1xPhospho | F8VQ87 1x | 0.026191 | 0.001676 | 1 | 5 | 1 | F8VQ87 | 5'-3' exorib | F8VQ87 [1 | 0 | 3412.592 | 0 | 0.7 | 0.055154 | 23.04 | 76.2 | 123.8 | 10.49 | 19.29 | | High | High | High | High | High | High | High | High | High | High | High | 0.000903 | 0.00928 | 2.55 |
| FALSE | High | ENEAQQSF | [K].ENEAQ | 1xPhospho | O88466 1x | 1.86E-12 | 0 | 2 | 6 | 3 | O88466; R | Zinc finger | O88466 [9 | 0 | 2724.324 | 0 | -1.14 | 0.000445 | 2.97 | 136 | 64 | 5.94 | 11.78 | Shared | High | High | High | High | High | High | High | High | High | High | High | 0 | 3.72E-14 | 4.51 |
| FALSE | High | ENEAQQSF | [K].ENEAQ | 2xPhospho | O88466 2x | 6.58E-08 | 0 | 2 | 6 | 1 | O88466; R | Zinc finger | O88466 [9 | 0 | 2804.291 | 0 | -1.3 | 0.002461 | 4.84 | 140.6 | 59.4 | 8.81 | 15.77 | Shared | High | High | High | High | High | High | High | High | High | High | High | 0 | 4.79E-09 | 3.52 |
| FALSE | High | ENEDSSDA | [K].ENEDS | 1xPhospho | Q6PIJ4 1xP | 0.011652 | 0.000889 | 1 | 1 | 2 | Q6PIJ4 | Nuclear fac | Q6PIJ4 [48 | 0 | 1872.788 | 0 | 0.26 | 0.223553 | 20.45 | 91.4 | 108.6 | 9.12 | 18.43 | | High | High | High | High | High | High | High | High | High | High | High | 0.007994 | 0.08644 | 2.72 |
| FALSE | High | ENEEAEPST | [K].ENEEAI | 1xPhospho | B9EKC2 1xP | 4.94E-05 | 0 | 1 | 6 | 4 | B9EKC2 | ATP-bindin | B9EKC2 [6 | 0 | 2107.975 | 0 | 0.93 | 0.000833 | 7.3 | 65.5 | 134.5 | 8.2 | 10.33 | | High | High | High | High | High | High | High | High | High | High | High | 0 | 0.00017 | 2.57 |
| FALSE | High | ENEEGDTG | [R].ENEEGI | 2xPhospho | Q6ZPZ3 2xP | 4.89E-05 | 0 | 1 | 2 | 6 | Q6ZPZ3 | Zinc finger | Q6ZPZ3 [8 | 0 | 3523.472 | 0 | -0.28 | 0.0325 | 7.54 | 113.4 | 86.6 | 8.58 | 14.83 | | High | High | High | High | High | High | High | High | High | High | High | 0 | 8.01E-06 | 4.41 |
| FALSE | High | EMSVDESK | [R].EMSVD | 1xPhospho | Q3V0B5 1x | 0.054383 | 0.003951 | 1 | 4 | 1 | Q3V0B5 | Putative ur | Q3V0B5 [8 | 0 | 1478.685 | 0 | -0.51 | 0.058826 | 17.33 | 121.5 | 78.5 | 12.33 | 27.53 | | High | High | High | High | High | High | High | High | High | High | High | 0.002059 | 0.02122 | 1.37 |
| FALSE | High | EMEPLLSPI | [R].EMEPLI | 1xPhospho | E9Q5F9 1x | 0.011652 | 0.000889 | 1 | 3 | 3 | E9Q5F9 | Histone-lys | E9Q5F9 [7 | 0 | 1986.976 | 0 | -0.07 | 0.577811 | 9.19 | 101.9 | 98.1 | 5.91 | 7.78 | | High | High | High | High | High | High | High | High | High | High | High | 0.000449 | 0.003736 | 1.78 |
| FALSE | High | EMEPLLSPI | [R].EMEPLI | 1xPhospho | E9Q5F9 1x | 8.48E-05 | 0 | 1 | 3 | 1 | E9Q5F9 | Histone-lys | E9Q5F9 [7 | 0 | 1970.981 | 0 | -0.12 | 0.248575 | 6.98 | 104 | 96 | 4.69 | 21.16 | | High | High | High | High | High | High | High | High | High | High | High | 0 | 1.49E-05 | 1.76 |
| FALSE | High | EMDTARTF | [R].EMDTA | 1xPhospho | H3BJW3 1x | 0.004531 | 0.000456 | 1 | 4 | 1 | H3BJW3 | Cleavage a | H3BJW3 [4 | 1 | 2694.199 | 0 | -0.05 | 0.502298 | 7.83 | 101.2 | 98.8 | 4.54 | 5.68 | | High | High | High | High | High | High | High | High | High | High | High | 0.000204 | 0.001294 | 2.47 |
| FALSE | High | ELSVQDQP | [K].ELSVQD | 1xPhospho | Q8BT14-3 | 1.46E-05 | 0 | 1 | 6 | 1 | Q8BT14-3 | Isoform 3 o | Q8BT14-3 | 0 | 3265.635 | 0 | -0.39 | 0.53828 | 39.47 | 111.7 | 88.3 | 34.32 | 33.84 | | High | High | High | High | High | High | High | High | High | High | High | 0 | 2.07E-06 | 3.72 |
| FALSE | High | EISWEAR | [R].EISWEA | 1xPhospho | Q8C0P0 1x | 0.071662 | 0.005278 | 1 | 1 | 1 | Q8C0P0 | Serine/thre | Q8C0P0 [3 | 0 | 1199.566 | 0 | 0.4 | 0.013787 | 20.51 | 86.2 | 113.8 | 10.08 | 4.25 | | High | High | High | High | High | High | High | High | High | High | High | 0.002717 | 0.02912 | 1.19 |
| FALSE | High | ELTDSPATT | [K].ELTDSP | 1xPhospho | E9QMD3 1 | 5.35E-05 | 0 | 1 | 2 | 2 | E9QMD3 | Zinc finger | E9QMD3 [ | 0 | 1600.824 | 0 | 1.8 | 0.000401 | 11.74 | 41 | 159 | 12.88 | 8.33 | | High | High | High | High | High | High | High | High | High | High | High | 0.004769 | 0.05213 | 2.06 |
| FALSE | High | ELTGSNAST | [R].ELTGSN | 1xPhospho | A0A1W2P6 | 1.13E-06 | 0 | 1 | 6 | 3 | A0A1W2P6 | Bromodom | A0A1W2P6 | 0 | 1599.758 | 0 | -0.91 | 0.000542 | 4.11 | 129.6 | 70.4 | 3.74 | 7.74 | | High | High | High | High | High | High | High | High | High | High | High | 0 | 1.16E-07 | 2.13 |
| FALSE | High | ELTNRHSLF | [R].ELTNRH | 1xPhospho | P22682 1xP | 0.011652 | 0.000889 | 1 | 5 | 1 | P22682 | E3 ubiquiti | P22682 [61 | 1 | 2564.253 | 0 | 0.66 | 0.04512 | 14.29 | 77.4 | 122.6 | 9.53 | 27.09 | | High | High | High | High | High | High | High | High | High | High | High | 0.000449 | 0.00373 | 1.73 |
| FALSE | High | ELTVGSQ | [R].ELTVGS | 1xPhospho | A0A1B0GS | 0.047086 | 0.00354 | 1 | 2 | 1 | A0A1B0GS | Nuclear po | A0A1B0GS | 0 | 1042.502 | 0 | -0.35 | 0.02597 | 13.3 | 116.7 | 83.3 | 8.28 | 14.39 | | High | High | High | High | High | High | High | High | High | High | High | 0.001812 | 0.01798 | 1.76 |
| FALSE | High | ELTVNPAA | [K].ELTVNF | 1xPhospho | E0CZ72 1xP | 0.063258 | 0.004664 | 1 | 5 | 1 | E0CZ72 | Kinesin-like | E0CZ72 [58 | 0 | 4412.097 | 0 | 0.28 | 0.348383 | 20.51 | 95.8 | 104.2 | 16.85 | 8.79 | | High | High | High | High | High | High | High | High | High | High | High | 0.002403 | 0.02526 | 2.98 |
| FALSE | High | ELVEPLTPS | [R].ELVEPL | 1xPhospho | Q01279 1x | 2.98E-05 | 0 | 1 | 2 | 1 | Q01279 | Epidermal | Q01279 [6 | 0 | 2367.191 | 0 | 2.64 | 0.000848 | 12.89 | 27.2 | 172.8 | 10.22 | 13.12 | | High | High | High | High | High | High | High | High | High | High | High | 0 | 4.6E-06 | 3.47 |
| FALSE | High | ELVGDTGS | [R].ELVGD | 1xPhospho | E9Q942 1x | 1.82E-06 | 0 | 1 | 1 | 5 | E9Q942 | Small integ | E9Q942 [4 | 0 | 3742.649 | 0 | -0.71 | 0.103338 | 12.25 | 120.8 | 79.2 | 18.1 | 17.33 | | High | High | High | High | High | High | High | High | High | High | High | 0 | 1.99E-07 | 5.72 |
| FALSE | High | ELVGDTGS | [R].ELVGD | 2xPhospho | E9Q942 2x | 8.33E-05 | 0 | 1 | 1 | 3 | E9Q942 | Small integ | E9Q942 [4 | 0 | 3822.616 | 0 | 0.04 | 0.621058 | 1.14 | 101.1 | 98.9 | 7.49 | 7.81 | | High | High | High | High | High | High | High | High | High | High | High | 0 | 1.46E-05 | 3.93 |
| FALSE | High | ELVGYPLSF | [R].ELVGYF | 1xPhospho | Q924H5 1x | 0.005 | 0.000504 | 1 | 3 | 1 | Q924H5 | DNA repair | Q924H5 [4 | 0 | 1609.855 | 0 | -0.69 | 0.004921 | 3.69 | 123.4 | 76.6 | 7.05 | 7.59 | | High | High | High | High | High | High | High | High | High | High | High | 0.000225 | 0.001439 | 1.96 |

| | | | | | | | | | | | | | | | | | | | | | | | | | | | | | | | | | | | | | | |
|---|---|---|---|---|---|---|---|---|---|---|---|---|---|---|---|---|---|---|---|---|---|---|---|---|---|---|---|---|---|---|---|---|---|---|---|---|---|---|
| FALSE | High | ELVHLSELP | [K].ELVHLS | 2xPhospho | Q53Z05 2x | 0.001689 | 5.66E-05 | 1 | 4 | 2 | Q53Z05 | Sir2alpha C | Q53Z05 [5 | 0 | 3058.437 | 0 | -0.15 | 0.164523 | 10.34 | 104.6 | 95.4 | 9.49 | 6.35 | | High | High | High | High | High | High | High | High | High | High | High | 2.58E-05 | 0.000432 | 2.73 |
| FALSE | High | ELVSDISSQ | [K].ELVSDI | 1xPhospho | Q8CJF7 1x | 8.48E-05 | 0 | 1 | 3 | 2 | Q8CJF7 | Protein ELY | Q8CJF7 [21 | 0 | 2253.173 | 0 | -0.64 | 0.005364 | 5.93 | 116.5 | 83.5 | 9.97 | 6.09 | | High | High | High | High | High | High | High | High | High | High | High | 0 | 1.49E-05 | 3.39 |
| FALSE | High | EIVSPNQGI | [R].EIVSPN | 1xPhospho | E9PV87 1x | 0.042503 | 0.003197 | 1 | 4 | 1 | E9PV87 | Protein TA | E9PV87 [2 | 0 | 1623.851 | 0 | -0.23 | 0.17872 | 11.77 | 104.7 | 95.3 | 12.18 | 16.79 | | High | High | High | High | High | High | High | High | High | High | High | 0.001639 | 0.01603 | 2.65 |
| FALSE | High | ELVSSSSSG | [K].ELVSSS | 1xPhospho | P11031 1x | 8.56E-05 | 0 | 1 | 1 | 1 | P11031 | Activated F | P11031 [6- | 0 | 2367.085 | 0 | -1.6 | 0.001184 | 0.76 | 151 | 49 | 6.55 | 30.81 | | High | High | High | High | High | High | High | High | High | High | High | 0 | 1.5E-05 | 2.57 |
| FALSE | High | ELVSSSSSG | [K].ELVSSS | 2xPhospho | P11031 2x | 2.42E-05 | 0 | 1 | 1 | 1 | P11031 | Activated F | P11031 [6- | 0 | 2447.052 | 0 | -0.49 | 0.050305 | 11.77 | 116.9 | 83.1 | 9.42 | 15.78 | | High | High | High | High | High | High | High | High | High | High | High | 0 | 3.67E-06 | 3.24 |
| FALSE | High | ELWDSPEP | [R].ELWDS | 1xPhospho | Q6P9Q4 1x | 0.006713 | 0.00062 | 1 | 1 | 2 | Q6P9Q4 | FH1/FH2 d | Q6P9Q4 [4 | 0 | 1763.82 | 0 | 0.43 | 0.006529 | 10.53 | 83.1 | 116.9 | 11.31 | 3.5 | | High | High | High | High | High | High | High | High | High | High | High | 0.001308 | 0.01377 | 2.44 |
| FALSE | High | ELYEITPPE | [K].ELYEITF | 1xPhospho | Q7TPS5 1x | 1.39E-06 | 0 | 2 | 6 | 1 | Q7TPS5; A | C2 domain | Q7TPS5 [6 | 0 | 2921.409 | 0 | 0.43 | 0.000668 | 4.38 | 86.6 | 113.4 | 11.18 | 4.59 | Shared | High | High | High | High | High | High | High | High | High | High | High | 0 | 1.47E-07 | 4.53 |
| FALSE | High | ELYEITPPE | [K].ELYEITF | 1xPhospho | Q7TPS5 1x | 2.18E-07 | 0 | 2 | 6 | 2 | Q7TPS5; A | C2 domain | Q7TPS5 [6 | 0 | 2937.404 | 0 | 0.59 | 0.019727 | 24.81 | 84.1 | 115.9 | 9.07 | 13.21 | Shared | High | High | High | High | High | High | High | High | High | High | High | 0 | 1.83E-08 | 3.58 |
| FALSE | High | ELYPEEFET | [K].ELYPEE | 2xPhospho | E9PUQ3 2x | 0.002293 | 0.000109 | 1 | 4 | 1 | E9PUQ3 | Expressed | E9PUQ3 [1 | 0 | 3063.316 | 0 | -0.26 | 0.003379 | 2.71 | 110.8 | 89.2 | 3.94 | 6.3 | | High | High | High | High | High | High | High | High | High | High | High | 4.94E-05 | 0.000603 | 3.64 |
| FALSE | High | EMASPSSE | [K].EMASP | 1xPhospho | G3X928 1x | 0.012959 | 0.000889 | 1 | 4 | 1 | G3X928 | SEC23-inte | G3X928 [7 | 0 | 1920.866 | 0 | -0.14 | 0.08508 | 11.37 | 106.9 | 93.1 | 11.42 | 3.9 | | High | High | High | High | High | High | High | High | High | High | High | 0.000449 | 0.004199 | 2.71 |
| FALSE | High | EMASPSSE | [K].EMASP | 1xPhospho | G3X928 1x | 6.13E-05 | 0 | 1 | 4 | 2 | G3X928 | SEC23-inte | G3X928 [7 | 0 | 1936.861 | 0 | -0.26 | 0.322951 | 8.81 | 108.2 | 91.8 | 12.45 | 7.01 | | High | High | High | High | High | High | High | High | High | High | High | 0 | 1.03E-05 | 2.27 |
| FALSE | High | EMCEGLLE | [R].EMCEG | 1xCarbami | Q8BK35 1x | 4E-07 | 0 | 1 | 3 | 2 | Q8BK35 | Ribosome | Q8BK35 [2 | 0 | 2601.032 | 0 | 0.01 | 0.707436 | 23.23 | 97.3 | 102.7 | 4.84 | 9.8 | | High | High | High | High | High | High | High | High | High | High | High | 0 | 3.63E-08 | 4.38 |
| FALSE | High | EMCEGLLE | [R].EMCEG | 1xCarbami | Q8BK35 1x | 5.87E-09 | 0 | 1 | 3 | 19 | Q8BK35 | Ribosome | Q8BK35 [2 | 0 | 2617.027 | 0 | 0.1 | 0.671564 | 7.04 | 96.1 | 103.9 | 4.02 | 13.94 | | High | High | High | High | High | High | High | High | High | High | High | 0 | 2.86E-09 | 4.67 |
| FALSE | High | EMDESLAN | [R].EMDES | 1xPhospho | Q6ZQ88 1x | 1.04E-07 | 0 | 1 | 3 | 1 | Q6ZQ88 | Lysine-spe | Q6ZQ88 [1 | 0 | 2747.112 | 0 | -0.71 | 0.003838 | 2.73 | 126.4 | 73.6 | 11.58 | 14.82 | | High | High | High | High | High | High | High | High | High | High | High | 0 | 8.05E-09 | 4.53 |
| FALSE | High | EMDESLAN | [R].EMDES | 1xPhospho | Q6ZQ88 1x | 3.22E-07 | 0 | 1 | 3 | 7 | Q6ZQ88 | Lysine-spe | Q6ZQ88 [1 | 0 | 2763.107 | 0 | -0.61 | 0.000434 | 2.34 | 119.1 | 80.9 | 5.3 | 7.27 | | High | High | High | High | High | High | High | High | High | High | High | 0 | 2.84E-08 | 3.57 |
| FALSE | High | EMDESLAN | [R].EMDES | 2xPhospho | Q6ZQ88 2x | 1.21E-09 | 0 | 1 | 3 | 2 | Q6ZQ88 | Lysine-spe | Q6ZQ88 [1 | 0 | 2827.078 | 0 | -0.84 | 0.000724 | 2.43 | 128.3 | 71.7 | 3.17 | 11.22 | | High | High | High | High | High | High | High | High | High | High | High | 0 | 5.36E-11 | 4.82 |
| FALSE | High | EMDESLAN | [R].EMDES | 2xPhospho | Q6ZQ88 2x | 0.000233 | 0 | 1 | 3 | 17 | Q6ZQ88 | Lysine-spe | Q6ZQ88 [1 | 0 | 2843.073 | 0 | -0.67 | 0.002588 | 2.07 | 122.2 | 77.8 | 6.85 | 10.52 | | High | High | High | High | High | High | High | High | High | High | High | 0 | 4.65E-05 | 3.74 |
| FALSE | High | EMDTARTF | [R].EMDTA | 1xPhospho | H3BJW3 1x | 0.002553 | 0.000109 | 1 | 4 | 2 | H3BJW3 | Cleavage a | H3BJW3 [4 | 1 | 2678.204 | 0 | -0.09 | 0.450438 | 14.92 | 101 | 99 | 3.3 | 18.28 | | High | High | High | High | High | High | High | High | High | High | High | 4.94E-05 | 0.00068 | 2.46 |
| FALSE | High | EISSSPTSK | [R].EISSSPT | 2xPhospho | Q8BTI8 2x | 0.003892 | 0.000261 | 1 | 3 | 7 | Q8BTI8 | Serine/argi | Q8BTI8 [45 | 0 | 1553.727 | 0 | -0.58 | 0.001063 | 2.66 | 120.5 | 79.5 | 2.62 | 5.65 | | High | High | High | High | High | High | High | High | High | High | High | 0.000689 | 0.006022 | 2.53 |
| FALSE | High | FLESAAADV | [K].FLESAA | 1xPhospho | O55106 1x | 2.77E-11 | 0 | 1 | 5 | 8 | O55106 | Striatin OS | O55106 [2 | 0 | 2595.046 | 0 | 0.85 | 0.00311 | 20.25 | 71.7 | 128.3 | 7.76 | 7.4 | | High | High | High | High | High | High | High | High | High | High | High | 0 | 7.66E-13 | 6.4 |
| FALSE | High | DGEEPQLT | [R].DGEEP | 1xPhospho | Q8CHC8 1x | 8.89E-09 | 0 | 1 | 1 | 3 | Q8CHC8 | Activity-de | Q8CHC8 [1 | 0 | 2487.154 | 0 | -0.13 | 0.351316 | 8.72 | 105.1 | 94.9 | 7.1 | 8.13 | | High | High | High | High | High | High | High | High | High | High | High | 0 | 3.4E-08 | 4.52 |
| FALSE | High | DGASPFQR | [K].DGASP | 1xPhospho | F8WI37 1x | 0.01894 | 0.001362 | 1 | 5 | 1 | F8WI37 | Histone-lys | F8WI37 [4 | 0 | 1186.546 | 0 | -0.4 | 0.001746 | 1.79 | 114.2 | 85.8 | 6.46 | 4.52 | | High | High | High | High | High | High | High | High | High | High | High | 0.000723 | 0.006406 | 1.54 |
| FALSE | High | ALPSFPTSE | [R].ALPSFP | 1xCarbami | Q8CD14 1x | 2.37E-07 | 0 | 1 | 3 | 4 | Q8CD14 | Putative ur | Q8CD14 [6 | 0 | 2416.094 | 0 | 0.15 | 0.095163 | 14.85 | 94.7 | 105.3 | 3.69 | 8.14 | | High | High | High | High | High | High | High | High | High | High | High | 0 | 2.01E-08 | 3.65 |
| FALSE | High | ALPSFWIPS | [K].ALPSFW | 1xPhospho | Q9D6T0 1x | 0.001349 | 0 | 1 | 5 | 1 | Q9D6T0 | Nitric oxide | Q9D6T0 [1 | 0 | 2195.192 | 0 | -0.22 | 0.009672 | 3.2 | 107.6 | 92.4 | 2.51 | 7.18 | | High | High | High | High | High | High | High | High | High | High | High | 0 | 0.000334 | 2.9 |
| FALSE | High | ALPTCTSAS | [R].ALPTCT | 1xCarbami | Q91XH8 1x | 0.000584 | 0 | 1 | 10 | 3 | Q91XH8 | Cellular tur | Q91XH8 [3 | 0 | 1895.97 | 0 | -0.28 | 0.053388 | 8.95 | 111 | 89 | 8.03 | 6.79 | | High | High | High | High | High | High | High | High | High | High | High | 0 | 0.00013 | 2.68 |
| FALSE | High | AIPTVNHS( | [R].AIPTVN | 1xPhospho | Q8BWW9 | 0.111965 | 0.009053 | 1 | 5 | 1 | Q8BWW9 | Serine/thre | Q8BWW9 | 0 | 3067.582 | 0 | 0.3 | 0.230045 | 30.26 | 83 | 117 | 20.02 | 17.63 | | High | High | High | High | High | High | High | High | High | High | High | 0.004543 | 0.04934 | 2.37 |
| FALSE | High | ALPVPEQG | [K].ALPVPE | 1xPhospho | D0VLQ5 1x | 4.84E-08 | 0 | 1 | 3 | 2 | D0VLQ5 | Tet oncoge | D0VLQ5 [8 | 0 | 2401.253 | 0 | -1.82 | 0.000469 | 1.91 | 154.2 | 45.8 | 6.34 | 15.72 | | High | High | High | High | High | High | High | High | High | High | High | 0 | 3.37E-09 | 3.83 |
| FALSE | High | ALQDSHVI | [K].ALQDSI | 1xPhospho | Q5SVR0 1x | 1.64E-05 | 0 | 1 | 2 | 1 | Q5SVR0 | TBC1 dom | Q5SVR0 [1 | 0 | 3563.752 | 0 | 0.81 | 0.006504 | 17.63 | 72.1 | 127.9 | 16.32 | 9.3 | | High | High | High | High | High | High | High | High | High | High | High | 0 | 2.37E-06 | 3.5 |
| FALSE | High | ALQEGEGD | [K].ALQEGI | 1xPhospho | Q3UXN3 1 | 9.1E-11 | 0 | 1 | 3 | 2 | Q3UXN3 | Putative ur | Q3UXN3 [3 | 0 | 1869.879 | 0 | -1.08 | 0.000886 | 2.58 | 135.2 | 64.8 | 1.44 | 17.08 | | High | High | High | High | High | High | High | High | High | High | High | 0 | 2.94E-12 | 4.95 |
| FALSE | High | AIQEYIDLS | [R].AIQEYII | 1xCarbami | Q04692 1x | 2.35E-07 | 0 | 1 | 3 | 8 | Q04692 | SWI/SNF-r | Q04692 [8 | 0 | 3453.602 | 0 | -1.43 | 0.000417 | 1.33 | 145.7 | 54.3 | 4.73 | 6.3 | | High | High | High | High | High | High | High | High | High | High | High | 0 | 6.06E-07 | 4.29 |
| FALSE | High | AIQEYIDLS | [R].AIQEYII | 1xCarbami | Q04692 2x | 1.04E-05 | 0 | 1 | 3 | 9 | Q04692 | SWI/SNF-r | Q04692 [8 | 0 | 3533.568 | 0 | -1.13 | 0.00104 | 4.03 | 137.6 | 62.4 | 6.48 | 7.47 | | High | High | High | High | High | High | High | High | High | High | High | 0 | 1.41E-06 | 4.3 |
| FALSE | High | ALQQHQH | [K].ALQQH | 1xPhospho | Q8CH02 1x | 1.35E-10 | 0 | 1 | 1 | 11 | Q8CH02 | SURP and ( | Q8CH02 [4 | 0 | 3516.565 | 0 | -0.46 | 0.003246 | 4.5 | 116.3 | 83.7 | 5.1 | 7.53 | | High | High | High | High | High | High | High | High | High | High | High | 0 | 7.44E-09 | 6.03 |
| FALSE | High | ALQQHQH | [K].ALQQH | 1xDeamida | Q8CH02 1x | 2.9E-05 | 0 | 1 | 1 | 7 | Q8CH02 | SURP and ( | Q8CH02 [4 | 0 | 3517.55 | 0 | -0.48 | 0.002253 | 3.47 | 119 | 81 | 7.42 | 10.45 | | High | High | High | High | High | High | High | High | High | High | High | 0 | 4.46E-06 | 4.99 |
| FALSE | High | ALQSNPSE | [R].ALQSNI | 1xPhospho | Q60977 1x | 0.053473 | 0.00384 | 1 | 4 | 2 | Q60977 | Tumor-ass | Q60977 [3 | 0 | 1310.63 | 0 | 1.4 | 0.000191 | 2.78 | 56.8 | 143.2 | 4.36 | 4.63 | | High | High | High | High | High | High | High | High | High | High | High | 0.00749 | 0.08254 | 0.72 |
| FALSE | High | ALQSNPSE | [R].ALQSNI | 1xCarbami | Q60977 1x | 0.000264 | 0 | 1 | 4 | 1 | Q60977 | Tumor-ass | Q60977 [3 | 1 | 2819.262 | 0 | 1.14 | 0.008465 | 30.02 | 62.3 | 137.7 | 10.66 | 15.12 | | High | High | High | High | High | High | High | High | High | High | High | 0 | 5.37E-05 | 1.27 |
| FALSE | High | ALPQTPR | [K].ALPQTF | 1xPhospho | Q8BTI8 1x | 0.090153 | 0.00698 | 1 | 3 | 1 | Q8BTI8 | Serine/argi | Q8BTI8 [14 | 0 | 1091.581 | 0 | -0.65 | 0.000954 | 4.59 | 122.4 | 77.6 | 4.98 | 3.85 | | High | High | High | High | High | High | High | High | High | High | High | 0.003606 | 0.03837 | 1.7 |
| FALSE | High | ALQSNPSE | [R].ALQSNI | 1xCarbami | Q60977 2x | 0.042141 | 0.003162 | 1 | 4 | 1 | Q60977 | Tumor-ass | Q60977 [3 | 1 | 2899.228 | 0 | 1.62 | 0.002055 | 15.62 | 52.1 | 147.9 | 20.7 | 18.47 | | High | High | High | High | High | High | High | High | High | High | High | 0.001639 | 0.01586 | 2.5 |
| FALSE | High | ALRPGDLP | [R].ALRPGI | 1xPhospho | D3YUY7 1x | 0.008022 | 0.000736 | 1 | 2 | 4 | D3YUY7 | Tripartite r | D3YUY7 [4 | 0 | 2115.125 | 0 | -0.11 | 0.027104 | 1.8 | 103.7 | 96.3 | 5.64 | 3.11 | | High | High | High | High | High | High | High | High | High | High | High | 0.000786 | 0.00697 | 2.74 |
| FALSE | High | ALRPPSSPG | [K].ALRPPS | 1xPhospho | Q9WTU0 1 | 1.59E-06 | 0 | 1 | 1 | 2 | Q9WTU0 | Lysine-spe | Q9WTU0 | 0 | 2397.31 | 0 | 2.49 | 0.000361 | 16.38 | 30.3 | 169.7 | 12.12 | 9.04 | | High | High | High | High | High | High | High | High | High | High | High | 0 | 1.71E-07 | 2.72 |
| FALSE | High | ALRSEDSDS | [R].ALRSED | 1xPhospho | Q9JJF3 1xP | 0.012845 | 0.000889 | 1 | 1 | 1 | Q9JJF3 | Ribosomal | Q9JJF3 [62 | 1 | 1645.82 | 0 | 1.92 | 0.001102 | 20.82 | 45.1 | 154.9 | 8.13 | 12.36 | | High | High | High | High | High | High | High | High | High | High | High | 0.000449 | 0.004136 | 2.25 |
| FALSE | High | ALSAGNIDI | [K].ALSAGI | 1xCarbami | Q3THQ5 1 | 0.000207 | 0 | 1 | 2 | 1 | Q3THQ5 | Putative ur | Q3THQ5 [1 | 0 | 2577.267 | 0 | -0.2 | 0.136725 | 19.83 | 107.7 | 92.3 | 11.82 | 19.76 | | High | High | High | High | High | High | High | High | High | High | High | 0 | 4.08E-05 | 3.57 |
| FALSE | High | AISAPTSPT | [R].AISAPT | 1xPhospho | B2RPW2 1 | 0.020679 | 0.001472 | 1 | 1 | 2 | B2RPW2 | Smith-Mag | B2RPW2 [5 | 0 | 1309.671 | 0 | -0.05 | 0.627143 | 15.57 | 101.7 | 98.3 | 7.91 | 4.34 | | High | High | High | High | High | High | High | High | High | High | High | 0.000786 | 0.007057 | 1.04 |
| FALSE | High | ALSDVTDEI | [R].ALSDVT | 1xPhospho | Q61532 1x | 1.31E-05 | 0 | 1 | 2 | 4 | Q61532 | Mitogen-a | Q61532 [3 | 0 | 2110.974 | 0 | 0.81 | 0.040926 | 36.97 | 72.8 | 127.2 | 23.4 | 11.34 | | High | High | High | High | High | High | High | High | High | High | High | 0 | 1.83E-06 | 3.72 |
| FALSE | High | ALSDVTDEI | [R].ALSDVT | 2xPhospho | Q61532 2x | 3.7E-05 | 0 | 1 | 2 | 1 | Q61532 | Mitogen-a | Q61532 [3 | 0 | 2190.941 | 0 | 1.08 | 0.023662 | 26.59 | 66.2 | 133.8 | 14.74 | 18.5 | | High | High | High | High | High | High | High | High | High | High | High | 0 | 5.9E-06 | 3.31 |
| FALSE | High | ALSEDEPCS | [R].ALSEDE | 1xCarbami | Q8VDF2 1x | 0.000139 | 0 | 1 | 2 | 1 | Q8VDF2 | E3 ubiquiti | Q8VDF2 [1 | 0 | 2017.955 | 0 | -0.26 | 0.011721 | 3.17 | 111.7 | 88.3 | 4.67 | 10.49 | | High | High | High | High | High | High | High | High | High | High | High | 0 | 2.62E-05 | 2.15 |
| FALSE | High | ALSFSEK | [R].ALSFSE | 1xPhospho | Q61884 1x | 0.087983 | 0.006639 | 1 | 3 | 1 | Q61884 | Meiosis-sp | Q61884 [7 | 0 | 1319.701 | 0 | -0.55 | 0.003667 | 7.3 | 118.9 | 81.1 | 4.01 | 8.57 | | High | High | High | High | High | High | High | High | High | High | High | 0.003402 | 0.03716 | 1.39 |
| FALSE | High | AISGPLPAP | [K].AISGPL | 1xPhospho | P0C7L0 1x | 0.0095 | 0.000736 | 1 | 4 | 1 | P0C7L0 | WAS/WAS | P0C7L0 [15 | 0 | 1542.824 | 0 | -1.01 | 0.000181 | 1.98 | 133.5 | 66.5 | 3.07 | 9.76 | | High | High | High | High | High | High | High | High | High | High | High | 0.000327 | 0.002965 | 2.46 |
| FALSE | High | ALSHAELF( | [R].ALSHAE | 1xPhospho | Q7TT28 1x | 3.65E-07 | 0 | 1 | 1 | 4 | Q7TT28 | RNA exonu | Q7TT28 [5 | 0 | 2883.288 | 0 | -0.39 | 0.006141 | 3.06 | 117.2 | 82.8 | 8.68 | 12.28 | | High | High | High | High | High | High | High | High | High | High | High | 0 | 3.29E-08 | 5.49 |
| FALSE | High | ALSHAELF( | [R].ALSHAE | 2xPhospho | Q7TT28 2x | 6.72E-09 | 0 | 1 | 1 | 2 | Q7TT28 | RNA exonu | Q7TT28 [5 | 0 | 2963.255 | 0 | -0.1 | 0.704255 | 19.31 | 103.5 | 96.5 | 8.81 | 17.54 | | High | High | High | High | High | High | High | High | High | High | High | 0 | 3.68E-10 | 7.44 |
| FALSE | High | AISHTDVF | [R].AISHTD | 1xPhospho | A0A172Q4 | 0.002671 | 0.000109 | 1 | 4 | 3 | A0A172Q4 | Cysteine-ki | A0A172Q4 | 0 | 2066.052 | 0 | 3.03 | 0.000533 | 20.18 | 21.5 | 178.5 | 13.36 | 11.37 | | High | High | High | High | High | High | High | High | High | High | High | 0.003633 | 0.03877 | 2.83 |
| FALSE | High | ALQTQNSE | [R].ALQTQ | 1xPhospho | Q02395 1x | 0.000568 | 0 | 1 | 3 | 2 | Q02395 | Metal-resp | Q02395 [5 | 0 | 1754.945 | 0 | -0.77 | 0.005702 | 5.02 | 122.4 | 77.6 | 8.32 | 10.82 | | High | High | High | High | High | High | High | High | High | High | High | 0 | 0.000127 | 2.48 |
| FALSE | High | AISHTDVF | [R].AISHTD | 2xPhospho | A0A172Q4 | 0.02933 | 0.001857 | 1 | 4 | 1 | A0A172Q4 | Cysteine-ki | A0A172Q4 | 0 | 2146.018 | 0 | 1.94 | 0.001953 | 57.72 | 37.7 | 162.3 | 45.77 | 9.04 | | High | High | High | High | High | High | High | High | High | High | High | 0.000997 | 0.0105 | 2.56 |
| FALSE | High | ALPLSSAEK | [K].ALPLSS | 1xPhospho | F8VQ54 1x | 0.030367 | 0.001857 | 1 | 4 | 2 | F8VQ54 | Endoribon | F8VQ54 [1 | 0 | 1453.807 | 0 | -1.09 | 0.001593 | 1.51 | 136.1 | 63.9 | 6.22 | 9.53 | | High | High | High | High | High | High | High | High | High | High | High | 0.001011 | 0.01093 | 2.01 |
| FALSE | High | ALPLPLSSP | [R].ALPLPL | 1xPhospho | Q91XC0 1x | 0.031989 | 0.00205 | 1 | 1 | 1 | Q91XC0 | LIM domai | Q91XC0 [1 | 0 | 1718.908 | 0 | -1.73 | 0.002231 | 1.51 | 149.5 | 50.5 | 11.26 | 16.28 | | High | High | High | High | High | High | High | High | High | High | High | 0.001093 | 0.01154 | 2.49 |
| FALSE | High | ALGSLEGS | [K].ALGSLE | 2xPhospho | B2RRE0 2x | 1.7E-05 | 0 | 1 | 2 | 1 | B2RRE0 | A kinase (P | B2RRE0 [1 | 0 | 2232.06 | 0 | -1.47 | 0.000739 | 2.75 | 147.1 | 52.9 | 5.71 | 13.07 | | High | High | High | High | High | High | High | High | High | High | High | 0 | 2.47E-06 | 3.81 |
| FALSE | High | ALGSPLK | [R].ALGSPL | 1xPhospho | A2AG50 1x | 0.00582 | 0.00062 | 1 | 5 | 1 | A2AG50 | MAP7 dom | A2AG50 [2 | 0 | 1223.717 | 0 | -0.57 | 0.001068 | 2.3 | 121.3 | 78.7 | 6.03 | 11.35 | | High | High | High | High | High | High | High | High | High | High | High | 0.000275 | 0.001711 | 1.17 |
| FALSE | High | ALGSSVNC | [K].ALGSS | 1xCarbami | Q8VID5 1x | 1.86E-07 | 0 | 1 | 2 | 1 | Q8VID5 | ATP-depen | Q8VID5 [7 | 0 | 2055.982 | 0 | -0.14 | 0.129886 | 7.16 | 104.2 | 95.8 | 3.41 | 6.89 | | High | High | High | High | High | High | High | High | High | High | High | 0 | 1.54E-08 | 2.81 |
| FALSE | High | ALGVISNF( | [K].ALGVIS | 1xPhospho | F8WJ98 1x | 3.89E-07 | 0 | 1 | 6 | 2 | F8WJ98 | Transmem | F8WJ98 [4 | 0 | 1886.019 | 0 | 0.85 | 0.001518 | 9.36 | 72.2 | 127.8 | 6.19 | 8.4 | | High | High | High | High | High | High | High | High | High | High | High | 0 | 3.53E-08 | 3.22 |
| FALSE | High | ALGYWGD | [K].ALGYW | 1xPhospho | Q3U2C5 1x | 7.47E-05 | 0 | 1 | 2 | 2 | Q3U2C5 | E3 ubiquiti | Q3U2C5 [3 | 0 | 2779.281 | 0 | -0.09 | 0.653054 | 16.19 | 107.4 | 92.6 | 14.58 | 16.01 | | High | High | High | High | High | High | High | High | High | High | High | 0 | 1.39E-05 | 3.73 |
| FALSE | High | ALHGAQTS | [K].ALHGA | 1xPhospho | Q14AR0 1x | 5.65E-05 | 0 | 1 | 2 | 21 | Q14AR0 | RIKEN cDN | Q14AR0 [9 | 0 | 1622.737 | 0 | 0.57 | 0.005001 | 6.4 | 80.4 | 119.6 | 4.3 | 7.7 | | High | High | High | High | High | High | High | High | High | High | High | 0 | 9.47E-06 | 3.51 |
| FALSE | High | ALHGAQTS | [K].ALHGA | 2xPhospho | Q14AR0 2x | 4.21E-06 | 0 | 1 | 2 | 4 | Q14AR0 | RIKEN cDN | Q14AR0 [9 | 0 | 1702.704 | 0 | 0.92 | 0.001472 | 10.53 | 65.5 | 134.5 | 8.06 | 10.99 | | High | High | High | High | High | High | High | High | High | High | High | 0 | 5.94E-05 | 3.54 |
| FALSE | High | ALHSDSP | [R].ALHSD | 1xPhospho | Q6PD29 1x | 0.112856 | 0.009146 | 1 | 2 | 1 | Q6PD29 | Zinc finger | Q6PD29 [5 | 0 | 1035.471 | 0 | 0.03 | 0.944517 | 4.96 | 99.4 | 100.6 | 3.58 | 5.27 | | High | High | High | High | High | High | High | High | High | High | High | 0.004583 | 0.04982 | 1.26 |
| FALSE | High | ALHSFITPP | [R].ALHSFI | 1xPhospho | Q61097 1x | 0.003857 | 0.000261 | 1 | 6 | 1 | Q61097 | Kinase sup | Q61097 [2 | 0 | 1988.057 | 0 | -0.25 | 0.046192 | 3.64 | 108.7 | 91.3 | 16.3 | 7.38 | | High | High | High | High | High | High | High | High | High | High | High | 0.000117 | 0.001078 | 1.83 |
| FALSE | High | ALHSPGPSI | [K].ALHSPC | 1xPhospho | A0A1B0GR | 0.104222 | 0.008348 | 1 | 1 | 2 | A0A1B0GR | RIKEN cDN | A0A1B0GR | 0 | 1431.776 | 0 | 1.22 | 0.000449 | 11.22 | 61.1 | 138.9 | 11.45 | 6.29 | | High | High | High | High | High | High | High | High | High | High | High | 0.008572 | 0.09108 | 1.2 |
| FALSE | High | ALLAPEELI | [K].ALLAPE | 2xCarbami | A2SW42 1 | 0.00017 | 0 | 1 | 4 | 1 | A2SW42 | Zinc finger | A2SW42 [1 | 1 | 4864.383 | 0 | -0.77 | 0.003383 | 0.9 | 130 | 70 | 10.5 | 8.54 | | High | High | High | High | High | High | High | High | High | High | High | 0 | 3.27E-05 | 4.53 |
| FALSE | High | ALLATETPE | [K].ALLATE | 1xPhospho | Q3UL54 1x | 8.11E-05 | 0 | 1 | 8 | 7 | Q3UL54 | Putative ur | Q3UL54 [4 | 0 | 2239.162 | 0 | -0.58 | 0.010377 | 4.61 | 116.5 | 83.5 | 8.44 | 7.66 | | High | High | High | High | High | High | High | High | High | High | High | 0 | 4.35E-05 | 3.49 |
| FALSE | High | ALLATETPE | [K].ALLATE | 2xPhospho | Q3UL54 2x | 0.00625 | 0.00062 | 1 | 8 | 3 | Q3UL54 | Putative ur | Q3UL54 [4 | 0 | 2319.129 | 0 | 0.25 | 0.041308 | 0.21 | 91.5 | 108.5 | 3.96 | 8.81 | | High | High | High | High | High | High | High | High | High | High | High | 0.001999 | 0.02084 | 2.75 |
| FALSE | High | ALPLPQSST | [R].ALPLPC | 1xCarbami | Q60751 1x | 0.047086 | 0.00354 | 1 | 4 | 1 | Q60751 | Insulin-like | Q60751 [1 | 0 | 1382.659 | 0 | 0.67 | 0.004188 | 10.31 | 69.8 | 130.2 | 10.43 | 15.85 | | High | High | High | High | High | High | High | High | High | High | High | 0.001812 | 0.01801 | 2.53 |
| FALSE | High | ALLFSNNS( | [K].ALLFSN | 1xPhospho | Q60902 1x | 1.3E-05 | 0 | 1 | 3 | 1 | Q60902 | Epidermal | Q60902 [6 | 0 | 3182.56 | 0 | -0.13 | 0.067952 | 9.11 | 107.5 | 92.5 | 4.89 | 9.89 | | High | High | High | High | High | High | High | High | High | High | High | 0 | 1.81E-06 | 3.29 |
| FALSE | High | ALIQTCLNS | [K].ALIQTC | 1xCarbami | Q9JM73 1x | 0.000259 | 0 | 1 | 1 | 1 | Q9JM73 | Serum resp | Q9JM73 [2 | 1 | 2878.36 | 0 | -0.48 | 0.01807 | 5.53 | 117.4 | 82.6 | 10.78 | 7.9 | | High | High | High | High | High | High | High | High | High | High | High | 0 | 5.22E-05 | 2.77 |
| FALSE | High | ALLSSPEGE | [K].ALLSSP | 2xPhospho | E9PUJ6 2x | 0.002486 | 0.000109 | 1 | 5 | 2 | E9PUJ6 | E3 ubiquiti | E9PUJ6 [39 | 0 | 1777.843 | 0 | 0.45 | 0.003085 | 13.02 | 82.4 | 117.6 | 7.25 | 6.06 | | High | High | High | High | High | High | High | High | High | High | High | 4.94E-05 | 0.000665 | 2.21 |
| FALSE | High | ALLSTPNG | [K].ALLSTP | 1xPhospho | A0A1W2P6 | 8.3E-07 | 0 | 1 | 9 | 2 | A0A1W2P6 | Bromodom | A0A1W2P6 | 0 | 2799.365 | 0 | -0.81 | 0.001349 | 3.63 | 128 | 72 | 5.92 | 5.59 | | High | High | High | High | High | High | High | High | High | High | High | 0 | 8.22E-08 | 4.84 |
| FALSE | High | ALMGSPQI | [R].ALMGS | 1xPhospho | Q02257 1x | 0.000235 | 0 | 1 | 1 | 1 | Q02257 | Junction pl | Q02257 [1 | 0 | 1736.933 | 0 | 0.93 | 0.003818 | 22.18 | 69 | 131 | 12.06 | 8.22 | | High | High | High | High | High | High | High | High | High | High | High | 0 | 4.67E-05 | 2.99 |
| FALSE | High | ALNAETPK | [R].ALNAET | 1xPhospho | Q8K3A9 1x | 0.006031 | 0.00062 | 1 | 1 | 1 | Q8K3A9 | 7SK snRNA | Q8K3A9 [1 | 0 | 1381.749 | 0 | -0.12 | 0.10969 | 5.08 | 102.9 | 97.1 | 5.73 | 3.48 | | High | High | High | High | High | High | High | High | High | High | High | 0.000275 | 0.001791 | 2.04 |
| FALSE | High | AINGPTSAS | [K].AINGPT | 1xPhospho | Q61584-3 | 6.81E-07 | 0 | 2 | 6 | 4 | Q61584-3; | Isoform B c | Q61584-3 | 0 | 1995.02 | 0 | 0.19 | 0.004809 | 3.96 | 92.4 | 107.6 | 4.4 | 5.66 | Shared | High | High | High | High | High | High | High | High | High | High | High | 0 | 6.62E-08 | 3.22 |
| FALSE | High | ALNGVSSP | [R].ALNGV | 1xPhospho | Q3UMT1 1 | 0.000311 | 0 | 1 | 2 | 3 | Q3UMT1 | Protein ph | Q3UMT1 [ | 0 | 1895.004 | 0 | 0.12 | 0.028718 | 7.21 | 96 | 104 | 7.92 | 4.37 | | High | High | High | High | High | High | High | High | High | High | High | 7.19E-05 | 0.00079 | 3.21 |
| FALSE | High | ALNGVSSP | [R].ALNGV | 1xDeamida | Q3UMT1 1 | 2.47E-06 | 0 | 1 | 2 | 4 | Q3UMT1 | Protein ph | Q3UMT1 [ | 0 | 1895.988 | 0 | 0.08 | 0.270912 | 20.05 | 94.7 | 105.3 | 7.53 | 10.3 | | High | High | High | High | High | High | High | High | High | High | High | 0 | 2.8E-07 | 2.74 |
| FALSE | High | ALNHSPMS | [R].ALNHSI | 1xPhospho | A3KFM7 1 | 0.000568 | 0 | 1 | 5 | 1 | A3KFM7 | Chromodo | A3KFM7 [1 | 0 | 2434.151 | 0 | -0.23 | 0.173895 | 13.25 | 114.5 | 85.5 | 12.49 | 21 | | High | High | High | High | High | High | High | High | High | High | High | 0 | 0.000126 | 2.36 |
| FALSE | High | AIPEQHSPI | [R].AIPEQH | 1xPhospho | F8VQL9 1x | 0.036731 | 0.00249 | 1 | 8 | 1 | F8VQL9 | Nuclear re | F8VQL9 [1 | 0 | 1794.967 | 0 | 1.19 | 0.000485 | 14.3 | 59.3 | 140.7 | 9.96 | 4.23 | | High | High | High | High | High | High | High | High | High | High | High | 0.001308 | 0.01359 | 2.99 |
| FALSE | High | ALPGAAFC | [R].ALPGA | 1xCarbami | Q3TWC8 1 | 0.064862 | 0.004664 | 1 | 2 | 2 | Q3TWC8 | Putative ur | Q3TWC8 [ | 0 | 2396.167 | 0 | 0 | 0.64213 | 7.67 | 100.5 | 99.5 | 5.39 | 4.75 | | High | High | High | High | High | High | High | High | High | High | High | 0.009584 | 0.1033 | 1.86 |
| FALSE | High | AIPGEAGD | [R].AIPGEA | 1xPhospho | Q3UJU9 1x | 6.73E-06 | 0 | 1 | 1 | 1 | Q3UJU9 | Regulator | Q3UJU9 [6 | 0 | 2719.396 | 0 | -0.24 | 0.242421 | 7.49 | 103.5 | 96.5 | 13.76 | 14.55 | | High | High | High | High | High | High | High | High | High | High | High | 0 | 8.65E-07 | 3.58 |
| FALSE | High | ALPGPSTQ | [R].ALPGPS | 1xPhospho | E9QQ25 1x | 0.00204 | 5.66E-05 | 1 | 5 | 2 | E9QQ25 | Striated mu | E9QQ25 [3 | 0 | 2121.069 | 0 | -0.82 | 0.003114 | 6.23 | 127.6 | 72.4 | 6.81 | 15.79 | | High | High | High | High | High | High | High | High | High | High | High | 2.58E-05 | 0.000531 | 2.92 |
| FALSE | High | ALIQTCLNS | [K].ALIQTC | 1xCarbami | Q9JM73 1x | 1.6E-06 | 0 | 1 | 1 | 3 | Q9JM73 | Serum resp | Q9JM73 [2 | 0 | 1977.967 | 0 | 0.89 | 0.006095 | 19.66 | 68.2 | 131.8 | 2.17 | 15.44 | | High | High | High | High | High | High | High | High | High | High | High | 0 | 1.72E-07 | 4.49 |
| FALSE | High | ALSIESLSPT | [K].ALSIESL | 1xPhospho | F8WIY7 1x | 0.000884 | 0 | 1 | 3 | 2 | F8WIY7 | StAR-relate | F8WIY7 [4 | 0 | 2369.134 | 0 | 0.08 | 0.907335 | 20.39 | 104.6 | 95.4 | 16.86 | 15.75 | | High | High | High | High | High | High | High | High | High | High | High | 0 | 0.000208 | 2.79 |
| FALSE | High | ALSIESLSPT | [K].ALSIESL | 1xDeamida | F8WIY7 1x | 0.000422 | 0 | 1 | 3 | 3 | F8WIY7 | StAR-relate | F8WIY7 [4 | 0 | 2370.118 | 0 | 0.23 | 0.103034 | 15.26 | 92.8 | 107.2 | 10.72 | 5.14 | | High | High | High | High | High | High | High | High | High | High | High | 7.19E-05 | 0.00086 | 3.8 |
| FALSE | High | ALSINSGNT | [K].ALSINS | 1xPhospho | Q8JZR6 1x | 0.002096 | 5.66E-05 | 1 | 2 | 4 | Q8JZR6 | Electroneu | Q8JZR6 [1 | 0 | 1542.829 | 0 | 2.03 | 0.000209 | 7.46 | 39.3 | 160.7 | 15.44 | 7.66 | | High | High | High | High | High | High | High | High | High | High | High | 0.001263 | 0.01332 | 2.34 |
| FALSE | High | ALSYASLDK | [K].ALSYAS | 1xPhospho | Q6QI06 1x | 0.000269 | 0 | 1 | 2 | 3 | Q6QI06 | Rapamycin | Q6QI06 [1 | 0 | 1505.802 | 0 | 1.2 | 0.003962 | 13.36 | 60.6 | 139.4 | 5.58 | 12.9 | | High | High | High | High | High | High | High | High | High | High | High | 0 | 5.48E-05 | 1.88 |
| FALSE | High | ALSYASLDK | [K].ALSYAS | 2xPhospho | Q6QI06 2x | 0.005135 | 0.000551 | 1 | 2 | 1 | Q6QI06 | Rapamycin | Q6QI06 [1 | 0 | 1585.768 | 0 | -0.06 | 0.727314 | 32.39 | 102.1 | 97.9 | 2.94 | 23.57 | | High | High | High | High | High | High | High | High | High | High | High | 0.000246 | 0.001486 | 2.58 |
| FALSE | High | ALTLAPGLS | [R].ALTLAP | 1xPhospho | Q8R3Y8 1x | 0.004333 | 0.00036 | 1 | 2 | 1 | Q8R3Y8 | Interferon | Q8R3Y8 [1 | 0 | 2725.473 | 0 | 0.34 | 0.045254 | 13.58 | 87.7 | 112.3 | 11.27 | 12.25 | | High | High | High | High | High | High | High | High | High | High | High | 0.000161 | 0.001237 | 1.5 |
| FALSE | High | AITLPCGPG | [R].AITLPC | 1xCarbami | Q7TNB8 1x | 6.13E-05 | 0 | 1 | 2 | 1 | Q7TNB8 | Protein str | Q7TNB8 [1 | 0 | 3275.626 | 0 | 0.68 | 0.009335 | 17.09 | 75.8 | 124.2 | 14.12 | 8.37 | | High | High | High | High | High | High | High | High | High | High | High | 0 | 1.04E-05 | 4.15 |
| FALSE | High | ALTPASLQI | [K].ALTPAS | 1xPhospho | Q3THU3 1 | 0.002553 | 0.000109 | 1 | 12 | 2 | Q3THU3 | Putative ur | Q3THU3 [2 | 0 | 1466.838 | 0 | -0.27 | 0.005261 | 5.74 | 111.1 | 88.9 | 5.19 | 2.3 | | High | High | High | High | High | High | High | High | High | High | High | 0.002896 | 0.03008 | 1.85 |
| FALSE | High | ALTPDSASS | [R].ALTPDS | 1xPhospho | Q8K124 1x | 1.56E-07 | 0 | 1 | 1 | 5 | Q8K124 | Pleckstrin I | Q8K124 [2 | 0 | 2911.361 | 0 | 0.21 | 0.012031 | 6.14 | 92.6 | 107.4 | 7.77 | 6.55 | | High | High | High | High | High | High | High | High | High | High | High | 0 | 1.27E-08 | 4.68 |
| FALSE | High | ALTPESPDS | [K].ALTPES | 1xCarbami | Q8CCE9 1x | 0.001137 | 0 | 1 | 5 | 3 | Q8CCE9 | Transcripti | Q8CCE9 [3 | 0 | 2514.09 | 0 | -0.36 | 0.169809 | 5.68 | 111.9 | 88.1 | 9.89 | 11.32 | | High | High | High | High | High | High | High | High | High | High | High | 0 | 0.000277 | 4.01 |
| FALSE | High | ALTPESPDS | [K].ALTPES | 1xCarbami | Q8CCE9 1x | 0.014158 | 0.001011 | 1 | 5 | 1 | Q8CCE9 | Transcripti | Q8CCE9 [3 | 0 | 2515.074 | 0 | -0.59 | 0.002035 | 2.79 | 118.5 | 81.5 | 4.7 | 9.81 | | High | High | High | High | High | High | High | High | High | High | High | 0.000538 | 0.004635 | 3.58 |
| FALSE | High | AITPPQQP | [K].AITPPQ | 1xPhospho | Q14AX6-2 | 0.001373 | 5.66E-05 | 3 | 4 | 4 | Q14AX6-2; | Isoform 2 c | Q14AX6-2 | 0 | 1680.913 | 0 | -0.25 | 0.046466 | 5.89 | 109.5 | 90.5 | 4.38 | 8.6 | Shared | High | High | High | High | High | High | High | High | High | High | High | 2.58E-05 | 0.000342 | 2.74 |
| FALSE | High | ALTPSIEAK | [R].ALTPSII | 1xPhospho | E9QM77 1 | 0.00418 | 0.00036 | 1 | 9 | 3 | E9QM77 | Ataxin-2 O | E9QM77 [7 | 0 | 1467.822 | 0 | 0.31 | 0.03405 | 16.19 | 92.1 | 107.9 | 4.33 | 11.62 | | High | High | High | High | High | High | High | High | High | High | High | 0.000161 | 0.001182 | 2.37 |
| FALSE | High | AITSLLGGC | [K].AITSLL( | 1xPhospho | Q6KCD5 1x | 0.0004 | 0 | 1 | 2 | 1 | Q6KCD5 | Nipped-B-l | Q6KCD5 [2 | 0 | 1409.76 | 0 | -0.04 | 0.654046 | 20.94 | 104.4 | 95.6 | 42.35 | 36.48 | | High | High | High | High | High | High | High | High | High | High | High | 0 | 8.52E-05 | 1.92 |
| FALSE | High | AITSLLGGC | [K].AITSLL( | 1xPhospho | Q6KCD5 1x | 1.69E-05 | 0 | 1 | 2 | 1 | Q6KCD5 | Nipped-B-l | Q6KCD5 [2 | 0 | 1638.923 | 0 | -0.06 | 0.840115 | 8.61 | 99.6 | 100.4 | 6.11 | 6.48 | | High | High | High | High | High | High | High | High | High | High | High | 0 | 2.43E-06 | 4.11 |
| FALSE | High | AITSPLAAG | [K].AITSPL | 1xPhospho | Q80Y32 1x | 0.000515 | 0 | 1 | 5 | 2 | Q80Y32 | HMG box c | Q80Y32 [1 | 0 | 1537.876 | 0 | -0.77 | 0.000291 | 1.02 | 126.5 | 73.5 | 4.75 | 5.44 | | High | High | High | High | High | High | High | High | High | High | High | 7.19E-05 | 0.000898 | 2.56 |
| FALSE | High | ALSVTAAT( | [R].ALSVTA | 1xPhospho | Q811L6 1x | 0.000209 | 0 | 1 | 4 | 1 | Q811L6 | Microtubu | Q811L6 [2 | 0 | 1682.913 | 0 | 0.63 | 0.007789 | 11.87 | 77.1 | 122.9 | 8.08 | 12.25 | | High | High | High | High | High | High | High | High | High | High | High | 0 | 4.12E-05 | 1.8 |
| FALSE | High | ALTTPTHY | [K].ALTTPT | 1xPhospho | A0A1B0GS | 0.007145 | 0.00062 | 1 | 9 | 2 | A0A1B0GS | Nuclear po | A0A1B0GS | 0 | 1569.844 | 0 | -0.93 | 0.000363 | 0.5 | 131.5 | 68.5 | 2.15 | 11.38 | | High | High | High | High | High | High | High | High | High | High | High | 0.001296 | 0.01341 | 1.81 |
| FALSE | High | ALVHAASP | [R].ALVHA | 1xDeamida | Q9JLY0 1x | 0.000171 | 0 | 1 | 2 | 2 | Q9JLY0 | Suppresso | Q9JLY0 [15 | 0 | 1754.915 | 0 | 0.88 | 0.001489 | 19.86 | 70.7 | 129.3 | 9.25 | 5.77 | | High | High | High | High | High | High | High | High | High | High | High | 0 | 3.28E-05 | 2.87 |
| FALSE | High | ALVHQLSN | [R].ALVHQ | 1xPhospho | D3YTT6 1x | 0.008927 | 0.000736 | 1 | 7 | 2 | D3YTT6 | Oxysterol-I | D3YTT6 [4 | 0 | 1562.789 | 0 | 1.57 | 0.001363 | 12.38 | 52.8 | 147.2 | 5.08 | 13.05 | | High | High | High | High | High | High | High | High | High | High | High | 0.000968 | 0.01006 | 1.72 |
| FALSE | High | ALVIQESES | [K].ALVIQE | 1xPhospho | A2CG77 1x | 7.09E-09 | 0 | 1 | 1 | 1 | A2CG77 | Histone-lys | A2CG77 [1 | 0 | 2312.137 | 0 | -0.45 | 0.026403 | 13.42 | 119.7 | 80.3 | 12.34 | 14.42 | | High | High | High | High | High | High | High | High | High | High | High | 0 | 3.91E-10 | 4.78 |
| FALSE | High | ALVIQESES | [K].ALVIQE | 2xPhospho | A2CG77 2x | 4.07E-07 | 0 | 1 | 1 | 2 | A2CG77 | Histone-lys | A2CG77 [1 | 0 | 2392.104 | 0 | -0.09 | 0.078888 | 3.91 | 104.8 | 95.2 | 5.6 | 5.38 | | High | High | High | High | High | High | High | High | High | High | High | 0 | 3.69E-08 | 4.31 |
| FALSE | High | ALVPTPGK | [K].ALVPTF | 1xPhospho | P09405 1x | 0.007811 | 0.000664 | 1 | 7 | 2 | P09405 | Nucleolin ( | P09405 [11 | 0 | 1320.769 | 0 | -0.91 | 0.00238 | 5.08 | 130.6 | 69.4 | 7.8 | 11.17 | | High | High | High | High | High | High | High | High | High | High | High | 0.000327 | 0.00239 | 2.3 |
| FALSE | High | ALVSPETPF | [R].ALVSPE | 1xPhospho | Q80U35 1x | 0.029844 | 0.001857 | 1 | 1 | 2 | Q80U35 | Rho guanir | Q80U35 [6 | 0 | 1814.961 | 0 | -0.22 | 0.106756 | 12.49 | 108.7 | 91.3 | 12.27 | 10.86 | | High | High | High | High | High | High | High | High | High | High | High | 0.007121 | 0.07735 | 2.05 |
| FALSE | High | ALVSPLCVF | [R].ALVSPL | 1xCarbami | Q542Q3 1x | 0.011346 | 0.000816 | 1 | 1 | 2 | Q542Q3 | G two S ph | Q542Q3 [5 | 0 | 1491.796 | 0 | -0.13 | 0.051833 | 3.19 | 105.1 | 94.9 | 5.15 | 2.66 | | High | High | High | High | High | High | High | High | High | High | High | 0.000638 | 0.005612 | 3.1 |
| FALSE | High | ALVVPEPEI | [K].ALVVPE | 1xPhospho | Q5DTM8 1 | 1.5E-05 | 0 | 1 | 3 | 10 | Q5DTM8 | E3 ubiquiti | Q5DTM8 [ | 0 | 2191.012 | 0 | 0.34 | 0.003916 | 8.27 | 88.2 | 111.8 | 3.69 | 3.92 | | High | High | High | High | High | High | High | High | High | High | High | 0 | 0.000151 | 3.86 |
| FALSE | High | ALVVPEPEI | [K].ALVVPE | 2xPhospho | Q5DTM8 2 | 0.000167 | 0 | 1 | 3 | 2 | Q5DTM8 | E3 ubiquiti | Q5DTM8 [ | 0 | 2270.978 | 0 | 0.82 | 0.004216 | 7.71 | 74.8 | 125.2 | 13.06 | 7.08 | | High | High | High | High | High | High | High | High | High | High | High | 2.58E-05 | 0.000526 | 2.99 |

| | | | | | | | | | | | | | | | | | | | | | | | | | | | | | | | | | | | | | | |
|---|---|---|---|---|---|---|---|---|---|---|---|---|---|---|---|---|---|---|---|---|---|---|---|---|---|---|---|---|---|---|---|---|---|---|---|---|---|---|
| FALSE | High | ALVVPEPEI | [K].ALVVPE | 1xPhospho | Q5DTM8 1 | 0.003857 | 0.000261 | 1 | 3 | 2 | Q5DTM8 | E3 ubiquiti | Q5DTM8 [ | 1 | 2548.27 | 0 | 0.49 | 0.007276 | 9.92 | 84 | 116 | 5.96 | 5.16 | | High | High | High | High | High | High | High | High | High | High | High | 0.000117 | 0.001079 | 3.07 |
| FALSE | High | ALVVPEPEI | [K].ALVVPE | 1xDeamida | Q5DTM8 1 | 0.035486 | 0.002304 | 1 | 3 | 2 | Q5DTM8 | E3 ubiquiti | Q5DTM8 [ | 1 | 2549.254 | 0 | 0.39 | 0.00621 | 2.97 | 85.6 | 114.4 | 7.1 | 3.49 | | High | High | High | High | High | High | High | High | High | High | High | 0.001324 | 0.01403 | 3.06 |
| FALSE | High | ALVVPEPEI | [K].ALVVPE | 2xPhospho | Q5DTM8 2 | 0.009756 | 0.000736 | 1 | 3 | 1 | Q5DTM8 | E3 ubiquiti | Q5DTM8 [ | 1 | 2628.236 | 0 | 0.56 | 0.004615 | 11.55 | 80.8 | 119.2 | 9.12 | 5.59 | | High | High | High | High | High | High | High | High | High | High | High | 0.000344 | 0.003045 | 3.35 |
| FALSE | High | ALYDYAGQ | [R].ALYDYA | 1xPhospho | Q543N7 1x | 0.014538 | 0.001011 | 1 | 2 | 1 | Q543N7 | Protein kin | Q543N7 [3 | 0 | 2156.974 | 0 | 0.26 | 0.151866 | 9.4 | 92.5 | 107.5 | 10.85 | 6.83 | | High | High | High | High | High | High | High | High | High | High | High | 0.000555 | 0.004781 | 2.7 |
| FALSE | High | ALVHAASP | [K].ALVHA | 1xPhospho | Q9JLY0 1xP | 0.000255 | 0 | 1 | 2 | 1 | Q9JLY0 | Suppressor | Q9JLY0 [15 | 0 | 1753.931 | 0 | 1.49 | 0.001723 | 8.44 | 47.9 | 152.1 | 14.13 | 12.77 | | High | High | High | High | High | High | High | High | High | High | High | 0 | 5.15E-05 | 2.9 |
| FALSE | High | ALSVQFEG | [R].ALSVQF | 1xPhospho | Q8BSD5 1x | 0.003224 | 0.00016 | 1 | 2 | 1 | Q8BSD5 | SH2 domai | Q8BSD5 [3 | 0 | 1924.948 | 0 | 1.52 | 0.006784 | 34.6 | 50.6 | 149.4 | 19.18 | 20.48 | | High | High | High | High | High | High | High | High | High | High | High | 7.19E-05 | 0.000884 | 2.29 |
| FALSE | High | ALSVDWPG | [R].ALSVDV | 1xPhospho | O35231 1x | 0.047086 | 0.00354 | 1 | 4 | 1 | O35231 | Kinesin-like | O35231 [8 | 0 | 1406.703 | 0 | 1.2 | 0.000669 | 17 | 61.1 | 138.9 | 12.09 | 3.4 | | High | High | High | High | High | High | High | High | High | High | High | 0.001812 | 0.01798 | 2.3 |
| FALSE | High | ALSTVGPAS | [R].ALSTVG | 1xPhospho | E9Q842 1x | 0.000492 | 0 | 1 | 7 | 3 | E9Q842 | Neuron na | E9Q842 [4 | 0 | 1739.934 | 0 | 0.11 | 0.070153 | 6.17 | 96.4 | 103.6 | 1.81 | 6.8 | | High | High | High | High | High | High | High | High | High | High | High | 0 | 0.000108 | 2.69 |
| FALSE | High | ALSLSSADS | [R].ALSLSS | 1xPhospho | Q7TPM1 1 | 2.1E-06 | 0 | 1 | 3 | 4 | Q7TPM1 | Protein PRI | Q7TPM1 [4 | 0 | 1803.914 | 0 | -0.34 | 0.002439 | 2.79 | 111.7 | 88.3 | 1.64 | 7.38 | | High | High | High | High | High | High | High | High | High | High | High | 0 | 2.33E-07 | 3.05 |
| FALSE | High | ALSLSSADS | [R].ALSLSS | 2xPhospho | Q7TPM1 2 | 0.005182 | 0.000551 | 1 | 3 | 2 | Q7TPM1 | Protein PRI | Q7TPM1 [4 | 0 | 1883.88 | 0 | 0.28 | 0.16228 | 21.78 | 86.6 | 113.4 | 12.34 | 13.14 | | High | High | High | High | High | High | High | High | High | High | High | 0.000382 | 0.003297 | 2.45 |
| FALSE | High | ALSMNTEN | [K].ALSMN | 1xPhospho | I1VGT4 1xP | 0.096959 | 0.007777 | 2 | 13 | 1 | I1VGT4; I1V | Anion exch | I1VGT4 [11 | 0 | 1632.807 | 0 | 2.22 | 0.004023 | 24.02 | 32.8 | 167.2 | 11.24 | 24.1 | Shared | High | High | High | High | High | High | High | High | High | High | High | 0.003911 | 0.04148 | 1.71 |
| FALSE | High | ALSMTHGS | [R].ALSMTI | 1xPhospho | Q80UP8 1x | 0.112856 | 0.009146 | 1 | 1 | 1 | Q80UP8 | Sodium-de | Q80UP8 [3 | 0 | 1556.791 | 0 | 1.96 | 0.001212 | 26.35 | 41 | 159 | 33.85 | 8.73 | | High | High | High | High | High | High | High | High | High | High | High | 0.004583 | 0.04985 | 1.66 |
| FALSE | High | AISNDALDS | [R].AISNDA | 1xPhospho | Q3TRR0 1x | 0.000108 | 0 | 1 | 5 | 1 | Q3TRR0 | Microtubu | Q3TRR0 [1 | 0 | 3685.701 | 0 | 1.64 | 0.000709 | 18.5 | 51 | 149 | 15.06 | 5.72 | | High | High | High | High | High | High | High | High | High | High | High | 0 | 1.97E-05 | 3.9 |
| FALSE | High | AISNDALDS | [R].AISNDA | 1xDeamida | Q3TRR0 1x | 0.001097 | 0 | 1 | 5 | 3 | Q3TRR0 | Microtubu | Q3TRR0 [1 | 0 | 3686.685 | 0 | -0.06 | 0.734069 | 10.02 | 104.6 | 95.4 | 15.76 | 21.83 | | High | High | High | High | High | High | High | High | High | High | High | 0 | 0.000264 | 3.57 |
| FALSE | High | ALSPAER | [R].ALSPAE | 1xPhospho | Q80U72-4 | 0.118334 | 0.009621 | 2 | 5 | 2 | Q80U72-4; | Isoform 4 c | Q80U72-4 | 0 | 1052.534 | 0 | 0.34 | 0.009319 | 8.24 | 90 | 110 | 5.78 | 4.77 | Shared | High | High | High | High | High | High | High | High | High | High | High | 0.004812 | 0.05246 | 2.37 |
| FALSE | High | ALSPEGALF | [R].ALSPEG | 1xPhospho | Q5NCX5 1x | 0.037369 | 0.00249 | 1 | 8 | 2 | Q5NCX5 | Neuralized | Q5NCX5 [5 | 0 | 1222.639 | 0 | 0.11 | 0.102329 | 1.6 | 96.6 | 103.4 | 1.75 | 6.6 | | High | High | High | High | High | High | High | High | High | High | High | 0.001308 | 0.01386 | 2.46 |
| FALSE | High | ALSPLPAR | [R].ALSPLP | 1xPhospho | D3Z0X5 1x | 0.108463 | 0.008839 | 1 | 6 | 1 | D3Z0X5 | Pleckstrin I | D3Z0X5 [4 | 0 | 1133.628 | 0 | 0.89 | 0.001479 | 5.05 | 69.8 | 130.2 | 7.37 | 11.09 | | High | High | High | High | High | High | High | High | High | High | High | 0.004411 | 0.04733 | 1.46 |
| FALSE | High | AISPIPTETA | [R].AISPIPT | 1xPhospho | B2RX10 1x | 0.001179 | 0 | 1 | 3 | 2 | B2RX10 | Fanconi an | B2RX10 [1 | 0 | 2663.363 | 0 | -0.56 | 0.027531 | 13.19 | 116.3 | 83.7 | 10.02 | 14.83 | | High | High | High | High | High | High | High | High | High | High | High | 0 | 0.000286 | 3.04 |
| FALSE | High | ALSPLQSTL | [R].ALSPLQ | 1xCarbami | E4NKG5 1x | 0.004867 | 0.000504 | 1 | 4 | 1 | E4NKG5 | MCG11736 | E4NKG5 [1 | 0 | 3836.943 | 0 | -1.32 | 0.001745 | 1.86 | 150.7 | 49.3 | 13.41 | 23.09 | | High | High | High | High | High | High | High | High | High | High | High | 0.000225 | 0.0014 | 3.39 |
| FALSE | High | AISPPVK | [K].AISPPV | 1xPhospho | E9Q309 1x | 0.048715 | 0.00354 | 1 | 1 | 1 | E9Q309 | Centrosom | E9Q309 [8 | 0 | 1249.732 | 0 | -0.08 | 0.398943 | 9.14 | 102.9 | 97.1 | 3.46 | 9.11 | | High | High | High | High | High | High | High | High | High | High | High | 0.00186 | 0.01871 | 1.43 |
| FALSE | High | ALSPQVEPI | [K].ALSPQV | 1xPhospho | Q80W88 1 | 3.9E-05 | 0 | 1 | 3 | 2 | Q80W88 | Homeobox | Q80W88 [3 | 0 | 1829.951 | 0 | -0.05 | 0.979201 | 12.58 | 101 | 99 | 6.5 | 11.7 | | High | High | High | High | High | High | High | High | High | High | High | 0 | 6.27E-06 | 4.05 |
| FALSE | High | AISPSIK | [R].AISPSIK | 1xPhospho | K3W4P2 1x | 0.083101 | 0.006345 | 1 | 5 | 1 | K3W4P2 | Integrator | K3W4P2 [1 | 0 | 1253.727 | 0 | 0.09 | 0.857827 | 16.58 | 97.2 | 102.8 | 5.6 | 8.66 | | High | High | High | High | High | High | High | High | High | High | High | 0.003265 | 0.03479 | 1.43 |
| FALSE | High | ALSPVIPIIP | [R].ALSPVII | 1xPhospho | A0A0A0M | 0.010475 | 0.000772 | 1 | 2 | 1 | A0A0A0M | Histone-lys | A0A0A0M | 0 | 1484.88 | 0 | 0.31 | 0.035229 | 13.75 | 90.4 | 109.6 | 8.3 | 4.62 | | High | High | High | High | High | High | High | High | High | High | High | 0.000382 | 0.003295 | 3.09 |
| FALSE | High | ALSQAAVE | [R].ALSQA | 1xPhospho | Q6P5B0 1x | 2.74E-07 | 0 | 1 | 1 | 3 | Q6P5B0 | RRP12-like | Q6P5B0 [1 | 0 | 3156.44 | 0 | -0.95 | 0.001121 | 4.72 | 131.7 | 68.3 | 6.79 | 8.68 | | High | High | High | High | High | High | High | High | High | High | High | 0 | 2.36E-08 | 4.67 |
| FALSE | High | AISSDMFF | [K].AISSDM | 1xPhospho | Q99K28-2 | 0.005275 | 0.000599 | 1 | 2 | 1 | Q99K28-2 | Isoform 2 c | Q99K28-2 | 0 | 1439.659 | 0 | 1.45 | 0.016406 | 28.15 | 53.6 | 146.4 | 19.22 | 23.47 | | High | High | High | High | High | High | High | High | High | High | High | 0.000267 | 0.001532 | 2.31 |
| FALSE | High | AISSDMFF | [K].AISSDM | 1xPhospho | Q99K28-2 | 0.001881 | 5.66E-05 | 1 | 2 | 5 | Q99K28-2 | Isoform 2 c | Q99K28-2 | 0 | 1455.654 | 0 | 1.08 | 0.027544 | 32.54 | 61.5 | 138.5 | 11.91 | 31.87 | | High | High | High | High | High | High | High | High | High | High | High | 0.000538 | 0.004467 | 2.12 |
| FALSE | High | ALSSEWDL | [R].ALSSEW | 1xPhospho | B2RXC5 1x | 0.00086 | 0 | 1 | 1 | 2 | B2RXC5 | Zinc finger | B2RXC5 [2 | 0 | 2051.036 | 0 | -1.16 | 0.006997 | 5.64 | 135.9 | 64.1 | 7.69 | 25.92 | | High | High | High | High | High | High | High | High | High | High | High | 0 | 0.000202 | 3.27 |
| FALSE | High | ALSSGGSIT | [R].ALSSGG | 1xPhospho | Q6Y7W8 1 | 1.41E-06 | 0 | 1 | 7 | 6 | Q6Y7W8 | GRB10-inte | Q6Y7W8 [1 | 0 | 2318.277 | 0 | 0.08 | 0.04121 | 1.28 | 96.3 | 103.7 | 5.23 | 4.42 | | High | High | High | High | High | High | High | High | High | High | High | 0 | 1.5E-07 | 3.41 |
| FALSE | High | ALSSGGSIT | [R].ALSSGG | 2xPhospho | Q6Y7W8 2 | 0.000319 | 0 | 1 | 7 | 4 | Q6Y7W8 | GRB10-inte | Q6Y7W8 [1 | 0 | 2398.244 | 0 | 0.13 | 0.116483 | 8.59 | 93.4 | 106.6 | 4.41 | 10.57 | | High | High | High | High | High | High | High | High | High | High | High | 0 | 0.000287 | 3.15 |
| FALSE | High | ALSSGGSV | [R].ALSSGG | 1xPhospho | Q99MR1 1 | 0.021607 | 0.001507 | 1 | 2 | 1 | Q99MR1 | GRB10-inte | Q99MR1 [1 | 0 | 2276.176 | 0 | 0.77 | 0.004638 | 5.94 | 72.3 | 127.7 | 13.04 | 8.41 | | High | High | High | High | High | High | High | High | High | High | High | 0.000801 | 0.007422 | 1.69 |
| FALSE | High | ALSSGGSV | [R].ALSSGG | 1xPhospho | Q99MR1 1 | 0.000128 | 0 | 1 | 2 | 1 | Q99MR1 | GRB10-inte | Q99MR1 [1 | 0 | 2292.171 | 0 | 0.99 | 0.019572 | 35.37 | 70.9 | 129.1 | 11.82 | 21.79 | | High | High | High | High | High | High | High | High | High | High | High | 0 | 2.38E-05 | 2.38 |
| FALSE | High | ALSSGPASI | [R].ALSSGP | 1xPhospho | Q9R194 1x | 0.000611 | 0 | 1 | 2 | 2 | Q9R194 | Cryptochro | Q9R194 [5 | 0 | 1452.786 | 0 | -0.84 | 0.000483 | 1.39 | 125.2 | 74.8 | 4.8 | 5.31 | | High | High | High | High | High | High | High | High | High | High | High | 0.000859 | 0.008623 | 2.11 |
| FALSE | High | ALSTDPASI | [K].ALSTDP | 1xPhospho | Q80TM2 1 | 0.000305 | 0 | 1 | 3 | 2 | Q80TM2 | MKIAA102 | Q80TM2 [1 | 0 | 1751.934 | 0 | -0.71 | 0.002257 | 3.73 | 121.9 | 78.1 | 8.08 | 11.07 | | High | High | High | High | High | High | High | High | High | High | High | 0 | 6.27E-05 | 2.82 |
| FALSE | High | AISTGSLAS | [R].AISTGS | 1xPhospho | Q64512 1x | 1.5E-05 | 0 | 1 | 4 | 2 | Q64512 | Tyrosine-p | Q64512 [8 | 0 | 1888.019 | 0 | 1.57 | 0.001117 | 9.36 | 50 | 150 | 6.86 | 11.96 | | High | High | High | High | High | High | High | High | High | High | High | 0 | 1.75E-05 | 2.14 |
| FALSE | High | ALSTTSTST | [R].ALSTTS | 1xPhospho | Q8K2Y9 1x | 0.003167 | 0.00016 | 1 | 4 | 3 | Q8K2Y9 | Cerebral ca | Q8K2Y9 [4 | 0 | 1731.848 | 0 | -0.17 | 0.062444 | 13.6 | 105.1 | 94.9 | 3.2 | 11.67 | | High | High | High | High | High | High | High | High | High | High | High | 7.19E-05 | 0.000872 | 1.25 |
| FALSE | High | ALSTTSTST | [R].ALSTTS | 1xDeamida | Q8K2Y9 1x | 0.082424 | 0.006288 | 1 | 4 | 1 | Q8K2Y9 | Cerebral ca | Q8K2Y9 [4 | 0 | 1732.832 | 0 | 0.68 | 0.008669 | 14.78 | 77.6 | 122.4 | 4.81 | 12.17 | | High | High | High | High | High | High | High | High | High | High | High | 0.003258 | 0.03436 | 1.96 |
| FALSE | High | ALSTVGPAS | [R].ALSTVG | 1xPhospho | E9Q842 1x | 0.086562 | 0.006435 | 1 | 7 | 1 | E9Q842 | Neuron na | E9Q842 [4 | 0 | 1510.772 | 0 | 0.71 | 0.243703 | 72.84 | 75.6 | 124.4 | 36.92 | 54.55 | | High | High | High | High | High | High | High | High | High | High | High | 0.003303 | 0.03626 | 1.89 |
| FALSE | High | ALGSLEGSI | [K].ALGSLE | 1xPhospho | B2RRE0 1x | 1.54E-07 | 0 | 1 | 2 | 7 | B2RRE0 | A kinase (P | B2RRE0 [1 | 0 | 2152.094 | 0 | -1.34 | 0.000975 | 3.4 | 143.1 | 56.9 | 7.49 | 13.07 | | High | High | High | High | High | High | High | High | High | High | High | 0 | 2.78E-05 | 3.13 |
| FALSE | High | ALYSPLFPT | [K].ALYSPL | 2xCarbami | E9QPJ5 1x | 1.08E-05 | 0 | 1 | 2 | 2 | E9QPJ5 | DNA segme | E9QPJ5 [12 | 0 | 2918.339 | 0 | -0.83 | 0.004563 | 2.53 | 127.9 | 72.1 | 11.44 | 6.37 | | High | High | High | High | High | High | High | High | High | High | High | 0 | 1.48E-06 | 3.62 |
| FALSE | High | AIGSGESET | [K].AIGSGE | 2xPhospho | E9QPA1 2x | 2.47E-06 | 0 | 1 | 4 | 2 | E9QPA1 | RAB6A-GEI | E9QPA1 [9 | 0 | 3362.434 | 0 | 0.72 | 0.000761 | 2.66 | 75.9 | 124.1 | 5.5 | 6.62 | | High | High | High | High | High | High | High | High | High | High | High | 0 | 2.82E-07 | 4.21 |
| FALSE | High | ALGSEGEA | [R].ALGSEG | 1xPhospho | O54988 1x | 3.11E-08 | 0 | 1 | 2 | 2 | O54988 | STE20-like | O54988 [6 | 0 | 2026.953 | 0 | 0.97 | 0.000537 | 6.22 | 68.7 | 131.3 | 9.79 | 2.83 | | High | High | High | High | High | High | High | High | High | High | High | 0 | 2.06E-09 | 5.4 |
| FALSE | High | AHDASSPS | [K].AHDAS | 1xPhospho | A2ANY6 1x | 0.001615 | 5.66E-05 | 1 | 3 | 3 | A2ANY6 | Midasin OS | A2ANY6 [2 | 0 | 2427.187 | 0 | -0.17 | 0.30535 | 6.9 | 103.3 | 96.7 | 9.56 | 7.24 | | High | High | High | High | High | High | High | High | High | High | High | 2.58E-05 | 0.000484 | 3.22 |
| FALSE | High | AHEEQDEE | [K].AHEEQ | 1xPhospho | B2RY84 1x | 0.017809 | 0.001282 | 1 | 4 | 1 | B2RY84 | Kif14 prote | B2RY84 [1 | 0 | 2357.972 | 0 | -0.25 | 0.100753 | 2.48 | 108.7 | 91.3 | 11.97 | 17.4 | | High | High | High | High | High | High | High | High | High | High | High | 0.000689 | 0.005999 | 2.16 |
| FALSE | High | AHGCFQD | [R].AHGCF | 1xCarbami | Q8K3X4 1x | 3.2E-05 | 0 | 1 | 1 | 1 | Q8K3X4 | Interferon | Q8K3X4 [6 | 1 | 2598.305 | 0 | -0.65 | 0.002442 | 2.78 | 122.2 | 77.8 | 10.13 | 10.09 | | High | High | High | High | High | High | High | High | High | High | High | 0 | 4.99E-06 | 2.91 |
| FALSE | High | AHHESPDL | [R].AHHESI | 1xPhospho | Q8R149 1x | 0.006031 | 0.00062 | 1 | 2 | 3 | Q8R149 | BUD13 hor | Q8R149 [2 | 0 | 1950.984 | 0 | -0.19 | 0.084157 | 11.41 | 109.3 | 90.7 | 3.54 | 9.46 | | High | High | High | High | High | High | High | High | High | High | High | 0.005257 | 0.05727 | 2.22 |
| FALSE | High | AHLDHIPD | [R].AHLDHI | 1xPhospho | Q8CFC7 1x | 1.95E-06 | 0 | 1 | 5 | 2 | Q8CFC7 | CLK4-asso | Q8CFC7 [8 | 0 | 3283.572 | 0 | -1.13 | 0.000154 | 0.49 | 137.4 | 62.6 | 5.94 | 12.43 | | High | High | High | High | High | High | High | High | High | High | High | 0 | 2.17E-07 | 3.72 |
| FALSE | High | AHLNAQAA | [R].AHLNA | 1xCarbami | Q6P5H2-2 | 8.6E-07 | 0 | 1 | 2 | 1 | Q6P5H2-2 | Isoform 2 c | Q6P5H2-2 | 0 | 1618.772 | 0 | 1.05 | 0.004534 | 19.57 | 59.8 | 140.2 | 16.98 | 12.89 | | High | High | High | High | High | High | High | High | High | High | High | 0 | 8.63E-08 | 3.64 |
| FALSE | High | AHLPLGQS | [K].AHLPLG | 1xCarbami | Q8K1N2-2 | 0.053473 | 0.00384 | 1 | 5 | 2 | Q8K1N2-2 | Isoform 2 c | Q8K1N2-2 | 0 | 3060.541 | 0 | 0.95 | 0.011372 | 12.9 | 73.7 | 126.3 | 7.43 | 18.76 | | High | High | High | High | High | High | High | High | High | High | High | 0.001999 | 0.02088 | 3.9 |
| FALSE | High | AHLSENELI | [R].AHLSEN | 1xPhospho | Q99KG3 1x | 2.11E-05 | 0 | 2 | 3 | 1 | Q99KG3; C | RNA-bindi | Q99KG3 [7 | 0 | 2021.036 | 0 | 0.16 | 0.044747 | 11.82 | 94.4 | 105.6 | 1.58 | 8.52 | Shared | High | High | High | High | High | High | High | High | High | High | High | 0 | 3.15E-06 | 3.16 |
| FALSE | High | AHNLQDG | [K].AHNLQ | 1xPhospho | E9Q0J5 1xP | 3.33E-06 | 0 | 1 | 7 | 2 | E9Q0J5 | Kinesin-like | E9Q0J5 [16 | 0 | 2579.146 | 0 | 1.24 | 0.000382 | 16.25 | 60.2 | 139.8 | 20.04 | 10.11 | | High | High | High | High | High | High | High | High | High | High | High | 0 | 1.38E-05 | 3.22 |
| FALSE | High | AHNLQDG | [K].AHNLQ | 2xPhospho | E9Q0J5 2xP | 0.008463 | 0.000736 | 1 | 7 | 2 | E9Q0J5 | Kinesin-like | E9Q0J5 [16 | 0 | 2659.112 | 0 | 0.94 | 0.028892 | 26.55 | 70.8 | 129.2 | 27.03 | 18.47 | | High | High | High | High | High | High | High | High | High | High | High | 0.003883 | 0.04135 | 3.75 |
| FALSE | High | AHNSGEDS | [K].AHNSG | 1xPhospho | P52963 1xP | 4.79E-07 | 0 | 1 | 2 | 1 | P52963 | Band 4.1-li | P52963 [46 | 0 | 1710.81 | 0 | -1.89 | 0.000571 | 0.88 | 155.7 | 44.3 | 4.84 | 17.37 | | High | High | High | High | High | High | High | High | High | High | High | 0 | 4.46E-08 | 3.2 |
| FALSE | High | AHPIEEAQI | [K].AHPIEE | 2xCarbami | P49452 1xP | 0.000119 | 0 | 1 | 3 | 1 | P49452 | Centromer | P49452 [33 | 1 | 4321.963 | 0 | -0.85 | 0.000702 | 1.53 | 129.2 | 70.8 | 4.59 | 5.74 | | High | High | High | High | High | High | High | High | High | High | High | 0 | 2.19E-05 | 4.82 |
| FALSE | High | AHPIEEAQI | [K].AHPIEE | 2xCarbami | P49452 2xP | 0.00043 | 0 | 1 | 3 | 2 | P49452 | Centromer | P49452 [33 | 1 | 4401.929 | 0 | -0.79 | 0.003782 | 3.31 | 126.7 | 73.3 | 6.73 | 8.84 | | High | High | High | High | High | High | High | High | High | High | High | 0 | 9.22E-05 | 4.58 |
| FALSE | High | AGYTTDES | [K].AGYTTI | 1xPhospho | Q6P5B5 1x | 2.91E-06 | 0 | 2 | 3 | 6 | Q6P5B5; Q | Fragile X m | Q6P5B5 [4 | 0 | 2108.933 | 0 | -0.3 | 0.65751 | 30.64 | 110.2 | 89.8 | 32.25 | 31.79 | Shared | High | High | High | High | High | High | High | High | High | High | High | 0 | 3.39E-07 | 3.09 |
| FALSE | High | AHPLSPETF | [R].AHPLSF | 1xPhospho | E9Q1Q1 1x | 0.074071 | 0.0057 | 1 | 6 | 1 | E9Q1Q1 | Microtubu | E9Q1Q1 [1 | 0 | 2270.246 | 0 | -0.9 | 0.002628 | 1.22 | 126.2 | 73.8 | 7.38 | 10.35 | | High | High | High | High | High | High | High | High | High | High | High | 0.002911 | 0.03024 | 1.75 |
| FALSE | High | AHQVTDES | [R].AHQVT | 1xPhospho | Q9D3L3 1x | 7.96E-05 | 0 | 1 | 5 | 2 | Q9D3L3 | Synaptoso | Q9D3L3 [1 | 0 | 1781.827 | 0 | -0.07 | 0.251971 | 6.48 | 103.8 | 96.2 | 2.42 | 9.34 | | High | High | High | High | High | High | High | High | High | High | High | 0 | 1.4E-05 | 3.72 |
| FALSE | High | AHSAENCS | [K].AHSAE | 1xCarbami | B2RSV7 1x | 5.93E-06 | 0 | 1 | 2 | 2 | B2RSV7 | Fgd6 prote | B2RSV7 [6 | 0 | 1998.936 | 0 | 1.42 | 0.001524 | 11.25 | 52.4 | 147.6 | 7.61 | 13.86 | | High | High | High | High | High | High | High | High | High | High | High | 0 | 7.54E-07 | 3.88 |
| FALSE | High | AHSDSNLS | [R].AHSDSI | 1xPhospho | Q8K2D0 1x | 1.95E-06 | 0 | 1 | 1 | 1 | Q8K2D0 | Protein FAI | Q8K2D0 [3 | 0 | 1724.78 | 0 | -0.42 | 0.097454 | 2.38 | 110.6 | 89.4 | 8.99 | 9.98 | | High | High | High | High | High | High | High | High | High | High | High | 0 | 2.16E-07 | 3.22 |
| FALSE | High | AHSGQFPG | [K].AHSGQ | 1xPhospho | Q9CQC5 1x | 0.009756 | 0.000736 | 1 | 1 | 1 | Q9CQC5 | Cdc42 effe | Q9CQC5 [7 | 0 | 1925.865 | 0 | 1.71 | 0.001566 | 15.25 | 46.9 | 153.1 | 19.7 | 10.76 | | High | High | High | High | High | High | High | High | High | High | High | 0.000344 | 0.003052 | 2.21 |
| FALSE | High | AHSLGTLDI | [R].AHSLGT | 1xCarbami | G3X9J0 1xP | 5.03E-05 | 0 | 1 | 1 | 2 | G3X9J0 | Signal-indu | G3X9J0 [38 | 0 | 2715.31 | 0 | 0.83 | 0.00198 | 14.24 | 73.9 | 126.1 | 12.48 | 6.56 | | High | High | High | High | High | High | High | High | High | High | High | 0 | 8.33E-06 | 3.8 |
| FALSE | High | AHSLGTLDI | [R].AHSLGT | 1xCarbami | G3X9J0 2xP | 3.83E-05 | 0 | 1 | 1 | 1 | G3X9J0 | Signal-indu | G3X9J0 [38 | 0 | 2795.276 | 0 | 1.04 | 0.022035 | 6.89 | 70.3 | 129.7 | 11.44 | 18.6 | | High | High | High | High | High | High | High | High | High | High | High | 0 | 6.09E-06 | 3.9 |
| FALSE | High | AHSLLFENS | [R].AHSLLF | 1xPhospho | Q501J7-3 1 | 6.8E-10 | 0 | 1 | 3 | 5 | Q501J7-3 | Isoform 3 c | Q501J7-3 [ | 0 | 2592.182 | 0 | 1.24 | 0.00014 | 5.73 | 59.1 | 140.9 | 8.74 | 5.08 | | High | High | High | High | High | High | High | High | High | High | High | 0 | 2.81E-11 | 6.1 |
| FALSE | High | AHSIQIMK | [R].AHSIQI | 1xPhospho | A0A1D5RM | 0.021607 | 0.001507 | 1 | 6 | 3 | A0A1D5RM | 60S riboso | A0A1D5RM | 0 | 1481.795 | 0 | -0.64 | 0.005619 | 2.67 | 123.4 | 76.6 | 6.92 | 11.86 | | High | High | High | High | High | High | High | High | High | High | High | 0.000801 | 0.007453 | 2.77 |
| FALSE | High | AHSPAEGA | [R].AHSPAI | 1xPhospho | Q8QZS3 1x | 1.07E-08 | 0 | 1 | 2 | 2 | Q8QZS3 | Folliculin O | Q8QZS3 [6 | 0 | 2221.054 | 0 | -0.02 | 0.955015 | 1.21 | 100.9 | 99.1 | 3.67 | 5.92 | | High | High | High | High | High | High | High | High | High | High | High | 0 | 6.25E-10 | 4.02 |
| FALSE | High | AHSPAEGA | [R].AHSPAI | 2xPhospho | Q8QZS3 2x | 0.080423 | 0.006219 | 1 | 2 | 1 | Q8QZS3 | Folliculin O | Q8QZS3 [6 | 0 | 2301.02 | 0 | -0.6 | 0.034391 | 10.71 | 118.9 | 81.1 | 13.52 | 8.67 | | High | High | High | High | High | High | High | High | High | High | High | 0.003188 | 0.03344 | 2.45 |
| FALSE | High | AHSPALGV | [R].AHSPAI | 1xPhospho | B7ZNX2 1x | 7.77E-06 | 0 | 1 | 5 | 1 | B7ZNX2 | Anaphase | B7ZNX2 [3 | 0 | 1930.949 | 0 | -0.23 | 0.209028 | 26.57 | 111.7 | 88.3 | 14.78 | 14.18 | | High | High | High | High | High | High | High | High | High | High | High | 0 | 1.02E-06 | 3 |
| FALSE | High | AHSSGGAA | [R].AHSSG | 2xCarbami | Q3TIU4 1x | 1.76E-11 | 0 | 1 | 1 | 2 | Q3TIU4 | 2',5'-phosp | Q3TIU4 [9 | 0 | 3020.426 | 0 | -0.27 | 0.024837 | 6.2 | 116 | 84 | 6.59 | 16.91 | | High | High | High | High | High | High | High | High | High | High | High | 0 | 4.62E-13 | 6.63 |
| FALSE | High | AHSSPASL | [R].AHSSPA | 1xPhospho | P46938 1xP | 5.81E-05 | 0 | 1 | 7 | 5 | P46938 | Transcripti | P46938 [11 | 0 | 3743.943 | 0 | 0.19 | 0.357051 | 32.51 | 95.3 | 104.7 | 20.67 | 9.86 | | High | High | High | High | High | High | High | High | High | High | High | 0 | 1.32E-05 | 3.82 |
| FALSE | High | AHQSPQTN | [K].AHQSP | 1xPhospho | B2RUA5 1x | 0.001147 | 0 | 1 | 2 | 1 | B2RUA5 | SAPS doma | B2RUA5 [8 | 0 | 1767.85 | 0 | 0.01 | 0.425076 | 10.2 | 103.5 | 96.5 | 11.02 | 5.69 | | High | High | High | High | High | High | High | High | High | High | High | 0 | 0.000278 | 2.62 |
| FALSE | High | AHSSPASL | [R].AHSSPA | 1xDeamida | P46938 1xP | 0.000375 | 0 | 1 | 7 | 2 | P46938 | Transcripti | P46938 [11 | 0 | 3744.927 | 0 | -0.37 | 0.049067 | 12.22 | 112.8 | 87.2 | 17.93 | 14.86 | | High | High | High | High | High | High | High | High | High | High | High | 0 | 0.000109 | 3.2 |
| FALSE | High | AGWVSPCI | [K].AGWVS | 1xCarbami | Q5QGZ6 1x | 5.67E-06 | 0 | 1 | 4 | 1 | Q5QGZ6 | Retinoic ac | Q5QGZ6 [8 | 0 | 3142.616 | 0 | 1.2 | 0.012005 | 11.91 | 61.6 | 138.4 | 7.58 | 23.53 | | High | High | High | High | High | High | High | High | High | High | High | 0 | 7.1E-07 | 4.38 |
| FALSE | High | AGVLTPPD | [K].AGVLTI | 1xPhospho | Q924A2 1x | 0.000157 | 0 | 1 | 1 | 1 | Q924A2 | Protein cap | Q924A2 [6 | 0 | 2195.158 | 0 | 0.68 | 0.001084 | 11.91 | 77.3 | 122.7 | 9.62 | 4.43 | | High | High | High | High | High | High | High | High | High | High | High | 0 | 2.99E-05 | 3.25 |
| FALSE | High | AGSGEDPV | [R].AGSGEI | 2xPhospho | A0A158SIT | 0.015061 | 0.001049 | 1 | 1 | 1 | A0A158SIT | Proline, glu | A0A158SIT | 0 | 3094.401 | 0 | 0.05 | 0.349526 | 25.64 | 98.2 | 101.8 | 21.56 | 18.4 | | High | High | High | High | High | High | High | High | High | High | High | 0.000571 | 0.00495 | 2.94 |
| FALSE | High | AGSLKDPEI | [R].AGSLKI | 1xPhospho | B2RX66 1x | 0.014283 | 0.001011 | 1 | 2 | 1 | B2RX66 | MCG12481 | B2RX66 [7- | 1 | 2432.345 | 0 | 1.05 | 0.000696 | 6.53 | 63.6 | 136.4 | 6.09 | 5.42 | | High | High | High | High | High | High | High | High | High | High | High | 0.000538 | 0.004664 | 2.84 |
| FALSE | High | AGSLPNYA | [R].AGSLPN | 1xPhospho | E9Q0S6 1x | 0.0012 | 0 | 1 | 10 | 1 | E9Q0S6 | Tensin 1 O | E9Q0S6 [1 | 0 | 1843.972 | 0 | 0.3 | 0.040272 | 12.87 | 89.9 | 110.1 | 12.56 | 13.99 | | High | High | High | High | High | High | High | High | High | High | High | 0 | 0.000293 | 2.74 |
| FALSE | High | AGSISTLDS | [R].AGSIST | 1xPhospho | E9QM90 1 | 4.56E-06 | 0 | 1 | 7 | 3 | E9QM90 | RIKEN cDN | E9QM90 [1 | 0 | 1761.862 | 0 | -1.4 | 0.000487 | 2.13 | 146.7 | 53.3 | 4.38 | 10.35 | | High | High | High | High | High | High | High | High | High | High | High | 0 | 5.57E-07 | 2.71 |
| FALSE | High | AGSISTLDS | [R].AGSIST | 2xPhospho | E9QM90 2 | 0.000793 | 0 | 1 | 7 | 1 | E9QM90 | RIKEN cDN | E9QM90 [1 | 0 | 1841.828 | 0 | 0.41 | 0.005514 | 5.86 | 81.5 | 118.5 | 8.5 | 9.53 | | High | High | High | High | High | High | High | High | High | High | High | 0 | 0.000185 | 3.22 |
| FALSE | High | AGSISTLDS | [R].AGSIST | 3xPhospho | E9QM90 3 | 0.08727 | 0.006537 | 1 | 7 | 1 | E9QM90 | RIKEN cDN | E9QM90 [1 | 0 | 1921.795 | 0 | 0.52 | 0.077039 | 18.73 | 91.3 | 108.7 | 15.24 | 12.31 | | High | High | High | High | High | High | High | High | High | High | High | 0.003375 | 0.03684 | 1.34 |
| FALSE | High | AGSPASPPS | [R].AGSPAS | 1xPhospho | Q8BUV6 1x | 0.079114 | 0.00613 | 1 | 1 | 1 | Q8BUV6 | U7 snRNA- | Q8BUV6 [1 | 0 | 1332.651 | 0 | 0.03 | 0.514645 | 7.33 | 100.5 | 99.5 | 3.02 | 6.28 | | High | High | High | High | High | High | High | High | High | High | High | 0.003148 | 0.03264 | 1.28 |
| FALSE | High | AGSPESPH | [R].AGSPES | 1xPhospho | E9Q8L9 1xP | 9.98E-05 | 0 | 1 | 2 | 1 | E9Q8L9 | Rab11 fam | E9Q8L9 [7 | 0 | 3530.669 | 0 | -0.61 | 0.032475 | 11.72 | 123.2 | 76.8 | 13.24 | 13.4 | | High | High | High | High | High | High | High | High | High | High | High | 0 | 1.8E-05 | 4.11 |
| FALSE | High | AGSPESPH | [R].AGSPES | 2xPhospho | E9Q8L9 2xP | 6.22E-07 | 0 | 1 | 2 | 1 | E9Q8L9 | Rab11 fam | E9Q8L9 [7 | 0 | 3610.635 | 0 | 0.4 | 0.098507 | 13.08 | 86.9 | 113.1 | 9.18 | 12.5 | | High | High | High | High | High | High | High | High | High | High | High | 0 | 5.98E-08 | 3.68 |
| FALSE | High | AGSPGSEV | [K].AGSPGS | 1xPhospho | Q5SNZ0 1x | 4.43E-08 | 0 | 1 | 4 | 3 | Q5SNZ0 | Girdin OS= | Q5SNZ0 [1 | 0 | 2658.343 | 0 | 0.54 | 0.009236 | 18.71 | 78.3 | 121.7 | 11.02 | 8.09 | | High | High | High | High | High | High | High | High | High | High | High | 0 | 3.07E-09 | 5.31 |
| FALSE | High | AGSPHSAV | [R].AGSPHS | 1xPhospho | Q5NCI3 1xP | 0.00043 | 0 | 1 | 4 | 3 | Q5NCI3 | DNA-direct | Q5NCI3 [1 | 0 | 2016.011 | 0 | 0.5 | 0.007289 | 15.19 | 83 | 117 | 5.04 | 6.02 | | High | High | High | High | High | High | High | High | High | High | High | 0 | 9.23E-05 | 2.78 |
| FALSE | High | AGSPKPEAI | [R].AGSPKI | 1xPhospho | A2ADL9 1x | 0.000235 | 0 | 1 | 7 | 1 | A2ADL9 | InaD-like (I | A2ADL9 [4 | 0 | 2621.322 | 0 | 0.38 | 0.020612 | 19.01 | 83.7 | 116.3 | 5.41 | 9.98 | | High | High | High | High | High | High | High | High | High | High | High | 0 | 4.7E-05 | 3.07 |
| FALSE | High | AGSPKPEAI | [R].AGSPKI | 2xPhospho | A2ADL9 2x | 0.086562 | 0.006435 | 1 | 7 | 1 | A2ADL9 | InaD-like (I | A2ADL9 [4 | 0 | 2701.289 | 0 | 0.24 | 0.124873 | 14.61 | 93.7 | 106.3 | 13.4 | 10.62 | | High | High | High | High | High | High | High | High | High | High | High | 0.003317 | 0.03646 | 2.92 |
| FALSE | High | AGWSDSS | [R].AGWSD | 1xPhospho | E4NKG5 1x | 7.1E-06 | 0 | 1 | 2 | 2 | E4NKG5 | MCG11736 | E4NKG5 [8 | 0 | 1801.796 | 0 | -1.5 | 0.000448 | 0.75 | 146.3 | 53.7 | 6.73 | 12.31 | | High | High | High | High | High | High | High | High | High | High | High | 0 | 2.73E-05 | 2.01 |
| FALSE | High | AGSPQLDD | [R].AGSPQ | 1xPhospho | Q8CHW4 1 | 0.004372 | 0.000409 | 1 | 1 | 7 | Q8CHW4 | Translation | Q8CHW4 [ | 0 | 1380.672 | 0 | -0.32 | 0.011332 | 5.63 | 110.2 | 89.8 | 1.38 | 8.82 | | High | High | High | High | High | High | High | High | High | High | High | 0.00168 | 0.01684 | 2.22 |
| FALSE | High | AGSREPVE | [R].AGSREI | 1xPhospho | Q3UZ39 1x | 0.000325 | 0 | 1 | 2 | 2 | Q3UZ39 | Leucine-ric | Q3UZ39 [6 | 1 | 2226.08 | 0 | -0.18 | 0.051403 | 9.8 | 106.8 | 93.2 | 5.38 | 7.39 | | High | High | High | High | High | High | High | High | High | High | High | 0 | 9.56E-05 | 2.9 |
| FALSE | High | AGSSDAGS | [R].AGSSD | 1xPhospho | Q5NCT2 1x | 0.000225 | 0 | 1 | 3 | 1 | Q5NCT2 | DNA topois | Q5NCT2 [9 | 0 | 1517.761 | 0 | -0.1 | 0.647484 | 8.87 | 104.8 | 95.2 | 12.05 | 7.1 | | High | High | High | High | High | High | High | High | High | High | High | 0 | 4.43E-05 | 2.61 |
| FALSE | High | AGSSDSHN | [K].AGSSD | 1xPhospho | Q8BKI2 1xP | 0.027597 | 0.001716 | 1 | 2 | 1 | Q8BKI2 | Trinucleoti | Q8BKI2 [61 | 0 | 1383.585 | 0 | -0.39 | 0.118714 | 12.9 | 113.1 | 86.9 | 22.98 | 2.54 | | High | High | High | High | High | High | High | High | High | High | High | 0.000936 | 0.009772 | 1.37 |
| FALSE | High | AGSSPCLGS | [K].AGSSP | 1xCarbami | A0A087W | 1.1E-08 | 0 | 1 | 8 | 6 | A0A087W | Snf2-relate | A0A087W | 0 | 2365.106 | 0 | 0.37 | 0.03508 | 8.29 | 88.2 | 111.8 | 6.47 | 9.84 | | High | High | High | High | High | High | High | High | High | High | High | 0 | 6.45E-10 | 5.53 |
| FALSE | High | AGSSPCLGS | [K].AGSSP | 1xCarbami | A0A087W | 0.037048 | 0.00249 | 1 | 8 | 1 | A0A087W | Snf2-relate | A0A087W | 0 | 2445.072 | 0 | 0.11 | 0.66842 | 18.57 | 95.1 | 104.9 | 12.41 | 12.74 | | High | High | High | High | High | High | High | High | High | High | High | 0.001308 | 0.01364 | 2.11 |
| FALSE | High | AGSSPTQG | [K].AGSSPT | 1xPhospho | Q9EPQ8 1x | 5.48E-07 | 0 | 1 | 3 | 4 | Q9EPQ8 | Transcripti | Q9EPQ8 [5 | 0 | 1779.822 | 0 | -0.18 | 0.039354 | 8.86 | 105.7 | 94.3 | 2.73 | 9.54 | | High | High | High | High | High | High | High | High | High | High | High | 0 | 5.18E-08 | 3.38 |
| FALSE | High | AGSVSLEN | [K].AGSVSI | 1xCarbami | Q921C3 1x | 0.000284 | 0 | 1 | 3 | 1 | Q921C3 | Bromodom | Q921C3 [1 | 0 | 1515.682 | 0 | -1.46 | 0.00046 | 0.87 | 147.1 | 52.9 | 5.42 | 6.46 | | High | High | High | High | High | High | High | High | High | High | High | 0 | 5.8E-05 | 1.54 |
| FALSE | High | AGSVSLEN | [K].AGSVSI | 1xCarbami | Q921C3 1x | 0.000589 | 0 | 1 | 3 | 1 | Q921C3 | Bromodom | Q921C3 [1 | 0 | 1516.666 | 0 | -0.81 | 0.000495 | 2.27 | 126.6 | 73.4 | 5.17 | 4.5 | | High | High | High | High | High | High | High | High | High | High | High | 0 | 0.000132 | 2.47 |
| FALSE | High | AGSVSPGS | [R].AGSVSI | 1xPhospho | Q8CGB6-3 | 1.75E-05 | 0 | 1 | 4 | 2 | Q8CGB6-3 | Isoform 3 c | Q8CGB6-3 | 0 | 1821.91 | 0 | -0.35 | 0.056375 | 15.26 | 112.1 | 87.9 | 11.66 | 13.96 | | High | High | High | High | High | High | High | High | High | High | High | 0 | 2.54E-06 | 2.55 |
| FALSE | High | AGTGYMSS | [R].AGTGYI | 1xPhospho | E9QK36 1x | 0.001361 | 0 | 1 | 5 | 2 | E9QK36 | WD repeat | E9QK36 [1 | 0 | 2327.099 | 0 | -0.22 | 0.486579 | 9.93 | 104.2 | 95.8 | 5.93 | 17.36 | | High | High | High | High | High | High | High | High | High | High | High | 0 | 0.000336 | 3.26 |
| FALSE | High | AGTQVPVL | [R].AGTQV | 1xPhospho | K3W4P2 1x | 0.001147 | 0 | 1 | 4 | 1 | K3W4P2 | Integrator | K3W4P2 [1 | 0 | 3371.749 | 0 | 0.52 | 0.007508 | 8.29 | 83.4 | 116.6 | 8.52 | 6.67 | | High | High | High | High | High | High | High | High | High | High | High | 0 | 0.000279 | 3.69 |
| FALSE | High | AGTSPAYH | [R].AGTSPA | 1xPhospho | Q4VAC9 1x | 0.004867 | 0.000504 | 1 | 5 | 2 | Q4VAC9 | Pleckstrin I | Q4VAC9 [8 | 0 | 1469.755 | 0 | 1.59 | 0.001448 | 20.04 | 47.2 | 152.8 | 16.4 | 10.31 | | High | High | High | High | High | High | High | High | High | High | High | 0.000225 | 0.001397 | 1.8 |
| FALSE | High | AGVDPAPS | [K].AGVDP | 1xPhospho | Q5SSZ5 1xP | 6E-07 | 0 | 1 | 1 | 1 | Q5SSZ5 | Tensin-3 O | Q5SSZ5 [7 | 0 | 2412.21 | 0 | -0.9 | 0.015688 | 5.71 | 130.3 | 69.7 | 13.47 | 15.11 | | High | High | High | High | High | High | High | High | High | High | High | 0 | 5.71E-08 | 3.91 |
| FALSE | High | AGSQLGPG | [R].AGSQL | 1xPhospho | E9Q5G3 1x | 0.003857 | 0.000261 | 1 | 1 | 2 | E9Q5G3 | Kinesin-like | E9Q5G3 [9 | 0 | 2214.122 | 0 | -0.76 | 0.00868 | 6.92 | 130.3 | 69.7 | 13.37 | 14.45 | | High | High | High | High | High | High | High | High | High | High | High | 0.000117 | 0.001085 | 2.31 |
| FALSE | High | AHSVDVEK | [R].AHSVD | 1xPhospho | Q8QZT2 1x | 0.017041 | 0.001204 | 1 | 1 | 1 | Q8QZT2 | Centriole, | Q8QZT2 [2 | 0 | 1422.739 | 0 | 0.62 | 0.00605 | 15.01 | 81 | 119 | 8.76 | 7.35 | | High | High | High | High | High | High | High | High | High | High | High | 0.000638 | 0.005706 | 1.92 |
| FALSE | High | AHSVNGITI | [K].AHSVN | 1xPhospho | Q3U1S6 1x | 5.37E-06 | 0 | 1 | 10 | 1 | Q3U1S6 | Putative ur | Q3U1S6 [5 | 0 | 2526.264 | 0 | -0.53 | 0.000696 | 1.85 | 119.1 | 80.9 | 6.68 | 9.64 | | High | High | High | High | High | High | High | High | High | High | High | 0 | 6.73E-07 | 3.88 |
| FALSE | High | AHTDEPHS | [R].AHTDEI | 1xPhospho | Q3U0M8 1 | 0.000159 | 0 | 1 | 3 | 6 | Q3U0M8 | Putative ur | Q3U0M8 [ | 0 | 1871.905 | 0 | -0.74 | 0.001883 | 3.28 | 123.9 | 76.1 | 6.28 | 7.35 | | High | High | High | High | High | High | High | High | High | High | High | 0 | 3.04E-05 | 3.58 |
| FALSE | High | ALEEGDGS | [R].ALEEGI | 1xPhospho | A0A0R4IZZ | 2.93E-06 | 0 | 1 | 4 | 9 | A0A0R4IZZ | Formin bin | A0A0R4IZZ | 0 | 1756.795 | 0 | -0.13 | 0.021793 | 5.18 | 105.9 | 94.1 | 6.16 | 2 | | High | High | High | High | High | High | High | High | High | High | High | 0 | 4.28E-05 | 3.26 |
| FALSE | High | ALEEGDGS | [R].ALEEGI | 2xPhospho | A0A0R4IZZ | 0.002334 | 0.000109 | 1 | 4 | 1 | A0A0R4IZZ | Formin bin | A0A0R4IZZ | 1 | 3177.394 | 0 | -0.35 | 0.050745 | 7.88 | 108.8 | 91.2 | 7.38 | 13.85 | | High | High | High | High | High | High | High | High | High | High | High | 4.94E-05 | 0.000616 | 3.42 |
| FALSE | High | ALEEGDGS | [R].ALEEGI | 3xPhospho | A0A0R4IZZ | 0.020862 | 0.001472 | 1 | 4 | 3 | A0A0R4IZZ | Formin bin | A0A0R4IZZ | 1 | 3257.36 | 0 | -0.47 | 0.00467 | 6.52 | 115.4 | 84.6 | 4.29 | 11.39 | | High | High | High | High | High | High | High | High | High | High | High | 0.001812 | 0.01836 | 3.43 |
| FALSE | High | ALEETPPDS | [R].ALEETP | 1xCarbami | Q3UMZ4 1 | 1.32E-07 | 0 | 1 | 3 | 4 | Q3UMZ4 | Putative ur | Q3UMZ4 [ | 0 | 3047.387 | 0 | -0.75 | 0.000278 | 2.7 | 125.5 | 74.5 | 2.19 | 6.83 | | High | High | High | High | High | High | High | High | High | High | High | 0 | 1.04E-08 | 5.78 |
| FALSE | High | ALEETPPDS | [R].ALEETP | 1xCarbami | Q3UMZ4 2 | 0.003788 | 0.00021 | 1 | 3 | 4 | Q3UMZ4 | Putative ur | Q3UMZ4 [ | 0 | 3127.353 | 0 | -0.72 | 0.006794 | 4.7 | 124.9 | 75.1 | 7.65 | 14.94 | | High | High | High | High | High | High | High | High | High | High | High | 0.000117 | 0.001062 | 3.21 |

| | | | | | | | | | | | | | | | | | | | | | | | | | | | | | | | | | | | | | | |
|---|---|---|---|---|---|---|---|---|---|---|---|---|---|---|---|---|---|---|---|---|---|---|---|---|---|---|---|---|---|---|---|---|---|---|---|---|---|---|
| FALSE | High | ALENGEAD | [R].ALENGI | 1xDeamida | Q8CIJ3 1xP | 0.000109 | 0 | 1 | 2 | 16 | Q8CIJ3 | Eukaryotic | Q8CIJ3 [13 | 0 | 4148.889 | 0 | -1.2 | 0.000245 | 1.09 | 141 | 59 | 5.5 | 11.41 | | High | High | High | High | High | High | High | High | High | High | High | 0 | 1.97E-05 | 4.93 |
| FALSE | High | ALEQASSPS | [K].ALEQAS | 1xPhospho | Q91XB0 1x | 2.27E-05 | 0 | 1 | 1 | 2 | Q91XB0 | Three-prim | Q91XB0 [1 | 0 | 1669.774 | 0 | 1.64 | 0.000518 | 4.99 | 47 | 153 | 9.86 | 8.37 | | High | High | High | High | High | High | High | High | High | High | High | 0 | 6.74E-06 | 2.3 |
| FALSE | High | ALEQASSPS | [K].ALEQAS | 1xDeamida | Q91XB0 1x | 3.26E-05 | 0 | 1 | 1 | 5 | Q91XB0 | Three-prim | Q91XB0 [1 | 0 | 1670.758 | 0 | 1.27 | 0.00059 | 4.18 | 58.1 | 141.9 | 5.94 | 7.67 | | High | High | High | High | High | High | High | High | High | High | High | 0 | 5.11E-06 | 2.8 |
| FALSE | High | ALEQSHSLF | [R].ALEQSH | 1xPhospho | E9PXE2 1xI | 7.04E-06 | 0 | 1 | 19 | 4 | E9PXE2 | Guanine nu | E9PXE2 [93 | 0 | 2616.369 | 0 | -1.76 | 0.000719 | 2.39 | 154.5 | 45.5 | 7.79 | 11.71 | | High | High | High | High | High | High | High | High | High | High | High | 0 | 9.12E-07 | 3.81 |
| FALSE | High | ALESPERPF | [K].ALESPE | 1xPhospho | P09411 1xI | 6.05E-07 | 0 | 1 | 2 | 2 | P09411 | Phosphogly | P09411 [20 | 0 | 2307.288 | 0 | 0.62 | 0.001684 | 3.78 | 81.9 | 118.1 | 4.89 | 9.3 | | High | High | High | High | High | High | High | High | High | High | High | 0 | 0.000175 | 2.31 |
| FALSE | High | ALETIPITPII | [K].ALETIPI | 1xPhospho | A2AAE1-3 | 0.000144 | 0 | 1 | 8 | 1 | A2AAE1-3 | Isoform 3 c | A2AAE1-3 | 0 | 1661.908 | 0 | -0.69 | 0.001656 | 4.87 | 124.6 | 75.4 | 5.95 | 6.04 | | High | High | High | High | High | High | High | High | High | High | High | 0 | 0.000027 | 2.31 |
| FALSE | High | ALEYSNGIF | [K].ALEYSN | 1xCarbami | Q8C8U0-3 | 8.56E-05 | 0 | 1 | 5 | 1 | Q8C8U0-3 | Isoform 3 c | Q8C8U0-3 | 0 | 2886.304 | 0 | 0.74 | 0.008009 | 19.8 | 77.4 | 122.6 | 10.12 | 11.25 | | High | High | High | High | High | High | High | High | High | High | High | 0 | 1.51E-05 | 4.7 |
| FALSE | High | ALEYSNGIF | [K].ALEYSN | 1xCarbami | Q8C8U0-3 | 2.6E-05 | 0 | 1 | 5 | 5 | Q8C8U0-3 | Isoform 3 c | Q8C8U0-3 | 0 | 2887.288 | 0 | 0.36 | 0.057163 | 16.28 | 88.6 | 111.4 | 16.22 | 7.07 | | High | High | High | High | High | High | High | High | High | High | High | 0 | 3.97E-06 | 4.22 |
| FALSE | High | ALEAPPEED | [R].ALEAPP | 1xPhospho | Q793C9 1x | 4.84E-08 | 0 | 1 | 3 | 2 | Q793C9 | U2 small nu | Q793C9 [3 | 0 | 2608.15 | 0 | 0.1 | 0.479593 | 16.02 | 98.2 | 101.8 | 11.07 | 2.44 | | High | High | High | High | High | High | High | High | High | High | High | 0 | 3.4E-09 | 3.87 |
| FALSE | High | ALEYSNGIF | [K].ALEYSN | 1xCarbami | Q8C8U0-3 | 1.67E-08 | 0 | 1 | 5 | 5 | Q8C8U0-3 | Isoform 3 c | Q8C8U0-3 | 0 | 2902.299 | 0 | 0.7 | 0.004667 | 7.94 | 76.5 | 123.5 | 6.35 | 10.13 | | High | High | High | High | High | High | High | High | High | High | High | 0 | 1.02E-09 | 4.86 |
| FALSE | High | ALFHQFSLK | [K].ALFHQI | 1xPhospho | Q8C0Q9 1x | 0.090888 | 0.007081 | 1 | 3 | 1 | Q8C0Q9 | Rap guanin | Q8C0Q9 [4 | 0 | 1628.897 | 0 | 0.68 | 0.003502 | 12.14 | 75.9 | 124.1 | 13.05 | 6.45 | | High | High | High | High | High | High | High | High | High | High | High | 0.003633 | 0.03853 | 0.95 |
| FALSE | High | ALFQDVQK | [R].ALFQDV | 1xPhospho | P29391 1xI | 0.00311 | 0.00016 | 1 | 10 | 1 | P29391 | Ferritin ligh | P29391 [77 | 0 | 2643.379 | 0 | 0.15 | 0.806589 | 33.33 | 95.6 | 104.4 | 10.81 | 18.26 | | High | High | High | High | High | High | High | High | High | High | High | 7.19E-05 | 0.000849 | 1.61 |
| FALSE | High | ALFTLEDGI | [K].ALFTLE | 1xPhospho | E9QMX4 1: | 0.000302 | 0 | 1 | 4 | 2 | E9QMX4 | Ribosomal | E9QMX4 [6 | 0 | 2731.312 | 0 | 1.44 | 0.000466 | 19.14 | 53.9 | 146.1 | 18.88 | 5 | | High | High | High | High | High | High | High | High | High | High | High | 0 | 6.26E-05 | 3.05 |
| FALSE | High | ALGDSSPS( | [K].ALGDSS | 1xPhospho | Q14BV7 1x | 0.00642 | 0.00062 | 1 | 3 | 2 | Q14BV7 | Acyl-Coenz | Q14BV7 [1 | 0 | 2504.251 | 0 | -0.28 | 0.30957 | 2.62 | 104.8 | 95.2 | 15.95 | 11.75 | | High | High | High | High | High | High | High | High | High | High | High | 0.000275 | 0.001904 | 2.63 |
| FALSE | High | AIGGGLSSS | [R].AIGGGI | 1xPhospho | P50446 1xI | 0.001489 | 5.66E-05 | 1 | 7 | 4 | P50446 | Keratin, typ | P50446 [52 | 0 | 2117.126 | 0 | 2.25 | 0.002235 | 5.54 | 36 | 164 | 6.13 | 18.34 | | High | High | High | High | High | High | High | High | High | High | High | 0.000327 | 0.002459 | 2.42 |
| FALSE | High | AIGGIILTAS | [K].AIGGIIL | 1xPhospho | Q66JR7 1xI | 2.77E-05 | 0 | 1 | 6 | 3 | Q66JR7 | Pgm2 prote | Q66JR7 [13 | 0 | 2744.454 | 0 | 0.51 | 0.019515 | 14.61 | 81.4 | 118.6 | 15.53 | 3.01 | | High | High | High | High | High | High | High | High | High | High | High | 0 | 4.27E-06 | 3.24 |
| FALSE | High | AIGGIILTAS | [K].AIGGIIL | 1xDeamida | Q66JR7 1xI | 2.32E-06 | 0 | 1 | 6 | 5 | Q66JR7 | Pgm2 prote | Q66JR7 [13 | 0 | 2745.438 | 0 | 0.53 | 0.027704 | 18.52 | 82.8 | 117.2 | 17.42 | 8.77 | | High | High | High | High | High | High | High | High | High | High | High | 0 | 1.12E-05 | 3.27 |
| FALSE | High | ALGLPATPT | [R].ALGLPA | 1xPhospho | Q3TZJ9 1xP | 0.000103 | 0 | 1 | 6 | 2 | Q3TZJ9 | Putative ur | Q3TZJ9 [11 | 0 | 2217.23 | 0 | 1.84 | 0.000586 | 22.96 | 43.7 | 156.3 | 15.11 | 9.89 | | High | High | High | High | High | High | High | High | High | High | High | 0 | 1.87E-05 | 3.34 |
| FALSE | High | AIGLSPMSI | [R].AIGLSPI | 1xPhospho | Q7TPM1 1: | 0.023792 | 0.001612 | 1 | 5 | 1 | Q7TPM1 | Protein PRI | Q7TPM1 [1 | 0 | 2316.233 | 0 | 0.68 | 0.00128 | 7.99 | 79.2 | 120.8 | 7.72 | 7.1 | | High | High | High | High | High | High | High | High | High | High | High | 0.000859 | 0.008267 | 2.73 |
| FALSE | High | ALGLSSSG/ | [K].ALGLSS | 1xPhospho | Q69ZR9-2 1 | 4.52E-06 | 0 | 1 | 2 | 2 | Q69ZR9-2 | Isoform 2 c | Q69ZR9-2 | 0 | 1963.019 | 0 | -1.36 | 0.000612 | 4.12 | 145.7 | 54.3 | 2.95 | 17.68 | | High | High | High | High | High | High | High | High | High | High | High | 0 | 1.19E-05 | 2.98 |
| FALSE | High | ALGPASEQ | [K].ALGPAS | 1xPhospho | Q62315 1x | 6.07E-05 | 0 | 1 | 3 | 5 | Q62315 | Protein Jur | Q62315 [70 | 0 | 1897.931 | 0 | -1.76 | 0.000434 | 2.01 | 154.1 | 45.9 | 3.38 | 24.7 | | High | High | High | High | High | High | High | High | High | High | High | 0 | 1.03E-05 | 2.96 |
| FALSE | High | ALGPASEQ | [K].ALGPAS | 2xPhospho | Q62315 2x | 0.028327 | 0.001791 | 1 | 3 | 1 | Q62315 | Protein Jur | Q62315 [70 | 0 | 1977.897 | 0 | -1.47 | 0.000374 | 0.85 | 147.9 | 52.1 | 6.23 | 6.84 | | High | High | High | High | High | High | High | High | High | High | High | 0.000968 | 0.01005 | 1.82 |
| FALSE | High | ALGQAASD | [R].ALGQA | 1xPhospho | Q6GQT9 1: | 1.56E-05 | 0 | 1 | 1 | 1 | Q6GQT9 | Nodal mod | Q6GQT9 [1 | 0 | 2117.983 | 0 | 1.13 | 0.00562 | 13.22 | 71.4 | 128.6 | 9.98 | 18.81 | | High | High | High | High | High | High | High | High | High | High | High | 0 | 2.21E-06 | 3.87 |
| FALSE | High | ALEYSNGIF | [K].ALEYSN | 1xCarbami | Q8C8U0-3 | 9.91E-06 | 0 | 1 | 5 | 1 | Q8C8U0-3 | Isoform 3 c | Q8C8U0-3 | 0 | 2903.283 | 0 | 0.98 | 0.001481 | 14.74 | 71.8 | 128.2 | 8.83 | 7.59 | | High | High | High | High | High | High | High | High | High | High | High | 0 | 1.34E-06 | 3.93 |
| FALSE | High | ALEAGGST | [K].ALEAG( | 2xPhospho | P15261 2xI | 0.003253 | 0.00016 | 1 | 4 | 1 | P15261 | Interferon | P15261 [35 | 0 | 2561.119 | 0 | 0.36 | 0.105269 | 15.95 | 87.5 | 112.5 | 11.68 | 14.76 | | High | High | High | High | High | High | High | High | High | High | High | 7.19E-05 | 0.000893 | 2.79 |
| FALSE | High | ALEAGGST | [K].ALEAG( | 1xPhospho | P15261 1xI | 7.68E-05 | 0 | 1 | 4 | 2 | P15261 | Interferon | P15261 [35 | 0 | 2497.148 | 0 | 1.85 | 0.002282 | 21.13 | 43.4 | 156.6 | 22.9 | 17.78 | | High | High | High | High | High | High | High | High | High | High | High | 0 | 1.33E-05 | 3.78 |
| FALSE | High | ALEAGGST | [K].ALEAG( | 1xPhospho | P15261 1xI | 0.000105 | 0 | 1 | 4 | 3 | P15261 | Interferon | P15261 [35 | 0 | 2481.153 | 0 | 0.72 | 0.005211 | 12.49 | 71.7 | 128.3 | 7.43 | 13.58 | | High | High | High | High | High | High | High | High | High | High | High | 0 | 2.03E-05 | 3.83 |
| FALSE | High | AKALHGAC | [R].AKALH( | 1xPhospho | Q14AR0 1x | 0.064862 | 0.004664 | 1 | 2 | 2 | Q14AR0 | RIKEN cDN | Q14AR0 [9 | 1 | 1821.869 | 0 | 0.58 | 0.373883 | 10.59 | 93.4 | 106.6 | 16.28 | 27.64 | | High | High | High | High | High | High | High | High | High | High | High | 0.002432 | 0.02613 | 2.86 |
| FALSE | High | AKFSPQPQ | [K].AKFSPC | 1xPhospho | Q7TSH3 1x | 0.000112 | 0 | 1 | 1 | 1 | Q7TSH3 | Zinc finger | Q7TSH3 [9 | 1 | 2290.293 | 0 | -0.04 | 0.704419 | 12.52 | 100.3 | 99.7 | 5.1 | 7.79 | | High | High | High | High | High | High | High | High | High | High | High | 0 | 2.04E-05 | 2.85 |
| FALSE | High | AKPAAQSE | [R].AKPAA( | 2xPhospho | Q8CIJ3 2xP | 0.001068 | 0 | 1 | 2 | 2 | Q8CIJ3 | Eukaryotic | Q8CIJ3 [62 | 0 | 3230.502 | 0 | -0.19 | 0.73068 | 29.1 | 113.5 | 86.5 | 16.9 | 93.3 | | High | High | High | High | High | High | High | High | High | High | High | 0 | 0.000258 | 3.04 |
| FALSE | High | AKPLSPEA( | [K].AKPLSP | 1xPhospho | Q9CXK9 1x | 1.06E-05 | 0 | 1 | 2 | 2 | Q9CXK9 | RNA-bindir | Q9CXK9 [8 | 0 | 1933.113 | 0 | -0.27 | 0.018867 | 5.27 | 110.5 | 89.5 | 7.86 | 2.56 | | High | High | High | High | High | High | High | High | High | High | High | 0 | 1.43E-06 | 1.8 |
| FALSE | High | AKPQAEDE | [K].AKPQAI | 1xDeamida | Q3UIL6-2 1 | 0.005182 | 0.000551 | 1 | 7 | 2 | Q3UIL6-2 | Isoform 2 c | Q3UIL6-2 [ | 0 | 4138.13 | 0 | -1.4 | 0.00469 | 4.56 | 146.5 | 53.5 | 16.5 | 22.05 | | High | High | High | High | High | High | High | High | High | High | High | 0.000246 | 0.00151 | 3.84 |
| FALSE | High | AKPSPEHAI | [K].AKPSPE | 1xPhospho | Q3TDI5 1xI | 8.98E-06 | 0 | 1 | 2 | 1 | Q3TDI5 | Peroxisom | Q3TDI5 [32 | 0 | 2853.511 | 0 | -0.84 | 0.005061 | 4.47 | 127.8 | 72.2 | 12.99 | 14.61 | | High | High | High | High | High | High | High | High | High | High | High | 0 | 1.2E-06 | 3 |
| FALSE | High | AKPTSDKP( | [K].AKPTSD | 1xPhospho | Q9QUJ7 1x | 0.012845 | 0.000889 | 1 | 3 | 1 | Q9QUJ7 | Long-chain | Q9QUJ7 [4 | 0 | 2171.183 | 0 | 2.71 | 0.001018 | 10.66 | 25.9 | 174.1 | 13.73 | 14.47 | | High | High | High | High | High | High | High | High | High | High | High | 0.000449 | 0.004163 | 1.24 |
| FALSE | High | AKSEENTK | [K].AKSEEN | 1xPhospho | Q3U3N7 1: | 0.085162 | 0.006376 | 1 | 7 | 1 | Q3U3N7 | Putative ur | Q3U3N7 [4 | 1 | 1673.908 | 0 | -0.5 | 0.007339 | 6.78 | 119 | 81 | 5.36 | 8.94 | | High | High | High | High | High | High | High | High | High | High | High | 0.003279 | 0.03561 | 1.19 |
| FALSE | High | AKSELVLAF | [R].AKSELV | 1xPhospho | Q3U912 1x | 0.000244 | 0 | 1 | 2 | 1 | Q3U912 | Putative ur | Q3U912 [3 | 1 | 2772.533 | 0 | -0.55 | 0.011762 | 12.35 | 119.9 | 80.1 | 7.14 | 18.53 | | High | High | High | High | High | High | High | High | High | High | High | 0 | 4.87E-05 | 2.89 |
| FALSE | High | AKSPAPGL/ | [K].AKSPAF | 1xCarbami | F6YAU7 1x | 0.120211 | 0.009851 | 2 | 9 | 1 | F6YAU7; F8 | Nuclear re | F6YAU7 [2 | 1 | 3373.636 | 0 | 0.97 | 0.024853 | 15.85 | 67.6 | 132.4 | 17.63 | 16.15 | Shared | High | High | High | High | High | High | High | High | High | High | High | 0.004907 | 0.0535 | 2.54 |
| FALSE | High | AKSPQPPV | [R].AKSPQI | 2xCarbami | Q3TXW2 1 | 0.000725 | 0 | 1 | 10 | 2 | Q3TXW2 | Putative ur | Q3TXW2 [2 | 1 | 4373.966 | 0 | -0.28 | 0.260481 | 30.75 | 102.8 | 97.2 | 17.13 | 22.61 | | High | High | High | High | High | High | High | High | High | High | High | 0 | 0.000167 | 5.3 |
| FALSE | High | AKSPQPPV | [R].AKSPQI | 2xCarbami | Q3TXW2 1 | 0.000616 | 0 | 1 | 10 | 8 | Q3TXW2 | Putative ur | Q3TXW2 [2 | 1 | 4602.145 | 0 | -0.25 | 0.303653 | 14.39 | 108.6 | 91.4 | 10.7 | 10.74 | | High | High | High | High | High | High | High | High | High | High | High | 0 | 0.000273 | 5.71 |
| FALSE | High | AKSPSSDS\ | [K].AKSPSS | 1xCarbami | V9GXM1 1 | 0.0095 | 0.000736 | 1 | 7 | 1 | V9GXM1 | ADP-ribosy | V9GXM1 [3 | 1 | 2422.111 | 0 | 1.13 | 0.003893 | 16.92 | 64.9 | 135.1 | 11.56 | 11.58 | | High | High | High | High | High | High | High | High | High | High | High | 0.000327 | 0.002958 | 1.99 |
| FALSE | High | ALAEEDSEI | [K].ALAEED | 1xPhospho | B2RXU2 1x | 7.23E-06 | 0 | 1 | 2 | 5 | B2RXU2 | ESF1, nucle | B2RXU2 [6 | 0 | 3611.624 | 0 | -0.86 | 0.001077 | 5.58 | 128.2 | 71.8 | 3.78 | 9.09 | | High | High | High | High | High | High | High | High | High | High | High | 0 | 9.41E-07 | 5.46 |
| FALSE | High | ALAEEDSEI | [K].ALAEED | 1xDeamida | B2RXU2 1x | 1.65E-09 | 0 | 1 | 2 | 2 | B2RXU2 | ESF1, nucle | B2RXU2 [6 | 0 | 3612.608 | 0 | -1.11 | 0.002273 | 4.62 | 136.8 | 63.2 | 10.8 | 8.82 | | High | High | High | High | High | High | High | High | High | High | High | 0 | 4.08E-10 | 6.25 |
| FALSE | High | ALAEEDSEI | [K].ALAEED | 2xPhospho | B2RXU2 2x | 3.12E-06 | 0 | 1 | 2 | 17 | B2RXU2 | ESF1, nucle | B2RXU2 [6 | 0 | 3691.591 | 0 | -0.43 | 0.000987 | 2.07 | 113.9 | 86.1 | 5.38 | 3.5 | | High | High | High | High | High | High | High | High | High | High | High | 0 | 9.47E-06 | 4.71 |
| FALSE | High | AIAEPESPG | [R].AIAEPE | 1xPhospho | Q9QXY1 1x | 0.072258 | 0.005346 | 1 | 3 | 1 | Q9QXY1 | Tight junct | Q9QXY1 [3 | 0 | 1551.725 | 0 | -0.84 | 0.003131 | 4.55 | 128.3 | 71.7 | 7.88 | 7.91 | | High | High | High | High | High | High | High | High | High | High | High | 0.002855 | 0.02953 | 2.4 |
| FALSE | High | AIAIDPTEE' | [K].AIAIDP | 1xPhospho | F8WGX5 1: | 0.074685 | 0.005727 | 1 | 2 | 1 | F8WGX5 | DmX-like p | F8WGX5 [2 | 0 | 2534.226 | 0 | 1.54 | 0.096071 | 22.35 | 56.9 | 143.1 | 19.49 | 68.69 | | High | High | High | High | High | High | High | High | High | High | High | 0.002937 | 0.03061 | 2.73 |
| FALSE | High | ALALNCHSI | [R].ALALN( | 1xCarbami | Q8VDF2 1x | 1.3E-06 | 0 | 1 | 2 | 2 | Q8VDF2 | E3 ubiquiti | Q8VDF2 [5 | 0 | 2005.034 | 0 | -0.51 | 0.013491 | 4.56 | 117.4 | 82.6 | 8.96 | 9.57 | | High | High | High | High | High | High | High | High | High | High | High | 0 | 4.51E-05 | 2.62 |
| FALSE | High | ALANDGTS | [R].ALAND( | 1xPhospho | Q91ZM2 1: | 0.011652 | 0.000889 | 1 | 7 | 2 | Q91ZM2 | SH2B adap | Q91ZM2 [2 | 0 | 1496.694 | 0 | -0.02 | 0.725015 | 8.65 | 100.8 | 99.2 | 3.82 | 6.43 | | High | High | High | High | High | High | High | High | High | High | High | 0.000449 | 0.003738 | 2.06 |
| FALSE | High | ALASNTSFF | [R].ALASNT | 1xCarbami | P33174 1xI | 3.93E-07 | 0 | 1 | 2 | 4 | P33174 | Chromosor | P33174 [12 | 0 | 2478.085 | 0 | 0.4 | 0.01045 | 8.55 | 86.3 | 113.7 | 3.13 | 8.1 | | High | High | High | High | High | High | High | High | High | High | High | 0 | 3.54E-08 | 4.97 |
| FALSE | High | ALDESLAEF | [K].ALDESL | 1xPhospho | Q60953 1x | 2.08E-06 | 0 | 2 | 6 | 2 | Q60953; Q | Protein PM | Q60953 [60 | 0 | 1903.9 | 0 | -0.21 | 0.886284 | 50.78 | 95.5 | 104.5 | 24.57 | 37.79 | Shared | High | High | High | High | High | High | High | High | High | High | High | 0 | 2.32E-07 | 3.13 |
| FALSE | High | ALDLAGDL' | [R].ALDLA( | 1xPhospho | B7ZMS8 1x | 0.018125 | 0.001282 | 1 | 9 | 1 | B7ZMS8 | Mdm1 pro | B7ZMS8 [4 | 1 | 2288.216 | 0 | -1.25 | 0.000333 | 0.88 | 140.3 | 59.7 | 6.71 | 7.45 | | High | High | High | High | High | High | High | High | High | High | High | 0.000689 | 0.006128 | 2.55 |
| FALSE | High | ALDIDSDEE | [K].ALDIDS | 1xPhospho | D3YYY8 1xI | 0.000302 | 0 | 1 | 1 | 2 | D3YYY8 | Rho guanir | D3YYY8 [38 | 0 | 1969.941 | 0 | 2.55 | 0.000162 | 6.94 | 29.6 | 170.4 | 5.96 | 3.77 | | High | High | High | High | High | High | High | High | High | High | High | 0 | 6.24E-05 | 2.97 |
| FALSE | High | ALDISLSSEI | [K].ALDISL | 2xPhospho | Q9DBE9 2x | 8.87E-05 | 0 | 1 | 1 | 12 | Q9DBE9 | pre-rRNA p | Q9DBE9 [3 | 0 | 3227.406 | 0 | -0.17 | 0.18411 | 3.4 | 99.8 | 100.2 | 13.24 | 10.94 | | High | High | High | High | High | High | High | High | High | High | High | 0 | 4.27E-05 | 4.5 |
| FALSE | High | ALDLSSWS | [R].ALDLSS | 1xPhospho | F8VPV0 1xI | 4.35E-05 | 0 | 1 | 8 | 3 | F8VPV0 | Pericentrin | F8VPV0 [18 | 0 | 1854.92 | 0 | 0.37 | 0.44178 | 15.5 | 89.4 | 110.6 | 10.53 | 23.87 | | High | High | High | High | High | High | High | High | High | High | High | 0 | 7.02E-06 | 3.7 |
| FALSE | High | ALEAASLSC | [R].ALEAAS | 2xPhospho | E9PVA2 2x | 1.16E-06 | 0 | 1 | 3 | 4 | E9PVA2 | Ribosomal | E9PVA2 [9 | 0 | 3205.402 | 0 | 0 | 0.847927 | 19.21 | 102.1 | 97.9 | 12.74 | 6.66 | | High | High | High | High | High | High | High | High | High | High | High | 0 | 1.2E-07 | 5.45 |
| FALSE | High | ALEAASLSC | [R].ALEAAS | 2xPhospho | E9PVA2 2x | 0.01861 | 0.001362 | 1 | 3 | 1 | E9PVA2 | Ribosomal | E9PVA2 [9 | 1 | 3562.66 | 0 | 0.18 | 0.429458 | 17.28 | 93.9 | 106.1 | 10.34 | 19.01 | | High | High | High | High | High | High | High | High | High | High | High | 0.000723 | 0.006277 | 2.8 |
| FALSE | High | ALEAELDA( | [K].ALEAEL | 1xPhospho | B2RUG9 1x | 0.003495 | 0.00021 | 1 | 8 | 1 | B2RUG9 | Adenomat | B2RUG9 [7 | 0 | 3208.582 | 0 | 0.04 | 0.833454 | 16.13 | 104.4 | 95.6 | 17.44 | 22.59 | | High | High | High | High | High | High | High | High | High | High | High | 9.42E-05 | 0.000968 | 3.68 |
| FALSE | High | AIGSGESET | [K].AIGSGE | 1xPhospho | E9QPA1 1x | 9.8E-05 | 0 | 1 | 4 | 2 | E9QPA1 | RAB6A-GEI | E9QPA1 [9 | 0 | 3282.468 | 0 | -0.99 | 0.012404 | 7.53 | 135.5 | 64.5 | 11.91 | 18.97 | | High | High | High | High | High | High | High | High | High | High | High | 0 | 6.02E-05 | 4.23 |
| FALSE | High | AGSGEDPV | [R].AGSGEI | 1xPhospho | A0A158SIT | 6.09E-06 | 0 | 1 | 1 | 8 | A0A158SIT | Proline, glu | A0A158SIT | 0 | 3014.435 | 0 | -0.45 | 0.010545 | 6.78 | 114.7 | 85.3 | 4.5 | 10.77 | | High | High | High | High | High | High | High | High | High | High | High | 0 | 5.56E-06 | 3.6 |
| FALSE | High | AMALGDSS | [R].AMALG | 1xPhospho | A2APF7 1xI | 0.005768 | 0.00062 | 1 | 2 | 1 | A2APF7 | Z-DNA-bin( | A2APF7 [37 | 0 | 2098.009 | 0 | 0.98 | 0.059204 | 14.7 | 70.9 | 129.1 | 7.32 | 40.66 | | High | High | High | High | High | High | High | High | High | High | High | 0.000275 | 0.001688 | 3.15 |
| FALSE | High | AMDATLSP | [K].AMDAT | 1xPhospho | B1ARU1 1x | 0.00925 | 0.000736 | 2 | 13 | 3 | B1ARU1; F' | Microtubu | B1ARU1 [2 | 0 | 1588.806 | 0 | -0.22 | 0.037937 | 3.31 | 107.9 | 92.1 | 5.5 | 5.2 | Shared | High | High | High | High | High | High | High | High | High | High | High | 0.000327 | 0.002886 | 2.7 |
| FALSE | High | APSIDGKEI | [K].APSIDG | 1xPhospho | D3YZC8 1xI | 0.043981 | 0.003228 | 1 | 4 | 1 | D3YZC8 | Bromodom | D3YZC8 [45 | 1 | 2368.262 | 0 | -1.37 | 0.023392 | 6.57 | 144.3 | 55.7 | 23.5 | 19.99 | | High | High | High | High | High | High | High | High | High | High | High | 0.001665 | 0.01666 | 1.81 |
| FALSE | High | APSIHGGS( | [R].APSIHG | 1xPhospho | P19001 1xI | 0.001039 | 0 | 1 | 1 | 4 | P19001 | Keratin, typ | P19001 [33 | 0 | 1304.631 | 0 | 4.89 | 0.001361 | 30.62 | 7.6 | 192.4 | 32.77 | 18.81 | | High | High | High | High | High | High | High | High | High | High | High | 0.001812 | 0.01852 | 1.65 |
| FALSE | High | APSLPAPPF | [R].APSLPA | 1xPhospho | B2RW11 1: | 0.007742 | 0.000664 | 1 | 1 | 1 | B2RW11 | Ankyrin rep | B2RW11 [4 | 0 | 1864.952 | 0 | -0.86 | 0.007202 | 2.7 | 127.2 | 72.8 | 6.21 | 15.04 | | High | High | High | High | High | High | High | High | High | High | High | 0.000294 | 0.002357 | 3.02 |
| FALSE | High | APSLPNDE/ | [R].APSLPN | 1xPhospho | Q80Y44 1x | 4.02E-06 | 0 | 1 | 5 | 2 | Q80Y44 | Probable A | Q80Y44 [5 | 0 | 2557.211 | 0 | -0.38 | 0.001302 | 4.31 | 112.6 | 87.4 | 1.86 | 3.59 | | High | High | High | High | High | High | High | High | High | High | High | 0 | 4.87E-07 | 4.03 |
| FALSE | High | APSLTDLVK | [K].APSLTD | 1xPhospho | Q8BZ47 1x | 0.002115 | 5.66E-05 | 1 | 2 | 1 | Q8BZ47 | Zinc finger | Q8BZ47 [1( | 0 | 1481.838 | 0 | -0.24 | 0.021335 | 6.66 | 108.2 | 91.8 | 5.57 | 5.95 | | High | High | High | High | High | High | High | High | High | High | High | 2.58E-05 | 0.000555 | 2.24 |
| FALSE | High | APSLTSDSE | [R].APSLTS | 1xPhospho | A2AIX1 1xP | 0.000104 | 0 | 1 | 7 | 2 | A2AIX1 | SEC16 hom | A2AIX1 [20 | 0 | 1629.814 | 0 | 0.4 | 0.000654 | 3.7 | 85.2 | 114.8 | 3.78 | 2.89 | | High | High | High | High | High | High | High | High | High | High | High | 0 | 1.89E-05 | 2.92 |
| FALSE | High | APSLTSDSE | [R].APSLTS | 2xPhospho | A2AIX1 2xP | 8.79E-05 | 0 | 1 | 7 | 2 | A2AIX1 | SEC16 hom | A2AIX1 [20 | 0 | 1709.78 | 0 | 0.84 | 0.00252 | 15.99 | 71.6 | 128.4 | 14.26 | 6.74 | | High | High | High | High | High | High | High | High | High | High | High | 0 | 1.55E-05 | 3.12 |
| FALSE | High | APSPAPAVS | [R].APSPAF | 2xPhospho | Q5U4C3 2x | 0.004411 | 0.000409 | 1 | 1 | 2 | Q5U4C3 | Splicing fac | Q5U4C3 [6 | 0 | 1639.826 | 0 | -0.04 | 0.084197 | 4.26 | 101.9 | 98.1 | 4.84 | 3.21 | | High | High | High | High | High | High | High | High | High | High | High | 0.000246 | 0.001496 | 3.43 |
| FALSE | High | APSPAPSTF | [R].APSPAF | 1xPhospho | G5E8L6 1xI | 0.042868 | 0.003228 | 1 | 3 | 1 | G5E8L6 | Killer cell le | G5E8L6 [12 | 0 | 1192.592 | 0 | 3.65 | 0.000269 | 13.99 | 14.4 | 185.6 | 19.45 | 7.37 | | High | High | High | High | High | High | High | High | High | High | High | 0.001651 | 0.01616 | 1.57 |
| FALSE | High | APSPASFQI | [R].APSPAS | 1xPhospho | E9Q4N7 1x | 0.079766 | 0.00613 | 1 | 3 | 1 | E9Q4N7 | AT-rich inte | E9Q4N7 [1 | 0 | 1269.619 | 0 | 1.37 | 0.000992 | 7.92 | 55.9 | 144.1 | 13.92 | 8.13 | | High | High | High | High | High | High | High | High | High | High | High | 0.003148 | 0.03299 | 2.01 |
| FALSE | High | APSPASR | [R].APSPAS | 1xPhospho | Q8BTI8 1xP | 0.109329 | 0.008865 | 1 | 3 | 1 | Q8BTI8 | Serine/argi | Q8BTI8 [23 | 0 | 994.492 | 0 | -0.59 | 0.000823 | 2.24 | 118.2 | 81.8 | 5.66 | 4.16 | | High | High | High | High | High | High | High | High | High | High | High | 0.004437 | 0.04793 | 1.88 |
| FALSE | High | APSPEPPTE | [K].APSPEP | 1xPhospho | Q9ESJ4 1xP | 0.0012 | 0 | 1 | 1 | 2 | Q9ESJ4 | NCK-intera | Q9ESJ4 [25 | 0 | 4334.001 | 0 | -0.25 | 0.014955 | 0.81 | 110.2 | 89.8 | 4.55 | 8.6 | | High | High | High | High | High | High | High | High | High | High | High | 0 | 0.000324 | 3.67 |
| FALSE | High | APSPGDYK | [R].APSPGI | 1xPhospho | B1AR09 1x | 0.003253 | 0.00016 | 1 | 4 | 1 | B1AR09 | Myeloid/ly | B1AR09 [40 | 0 | 1372.691 | 0 | 0.33 | 0.004784 | 4.49 | 87.1 | 112.9 | 2.68 | 6.15 | | High | High | High | High | High | High | High | High | High | High | High | 7.19E-05 | 0.000896 | 2.02 |
| FALSE | High | APSIDGK | [K].APSIDG | 1xPhospho | D3YZC8 1xI | 0.050397 | 0.003641 | 1 | 4 | 2 | D3YZC8 | Bromodom | D3YZC8 [45 | 0 | 1225.659 | 0 | -0.92 | 0.000181 | 0.86 | 129.7 | 70.3 | 2.25 | 5.15 | | High | High | High | High | High | High | High | High | High | High | High | 0.001916 | 0.0195 | 1.79 |
| FALSE | High | APSPGEVS/ | [R].APSPGE | 1xPhospho | A2AIV6 1xP | 8.37E-07 | 0 | 1 | 4 | 3 | A2AIV6 | snRNA-acti | A2AIV6 [12 | 0 | 2901.408 | 0 | -1.51 | 0.000295 | 1.94 | 149 | 51 | 5.03 | 15.52 | | High | High | High | High | High | High | High | High | High | High | High | 0 | 8.35E-08 | 4.64 |
| FALSE | High | APSPHVVQ | [R].APSPH\ | 1xCarbami | E9PUJ6 1xP | 8.79E-05 | 0 | 1 | 5 | 2 | E9PUJ6 | E3 ubiquiti | E9PUJ6 [29 | 0 | 3285.659 | 0 | 0.28 | 0.002604 | 6.73 | 90.3 | 109.7 | 4.02 | 3.65 | | High | High | High | High | High | High | High | High | High | High | High | 0 | 0.00015 | 2.08 |
| FALSE | High | APSPHVVQ | [R].APSPH\ | 1xCarbami | E9PUJ6 1xP | 0.000595 | 0 | 1 | 5 | 3 | E9PUJ6 | E3 ubiquiti | E9PUJ6 [29 | 0 | 3286.643 | 0 | 0.32 | 0.042483 | 17.74 | 88.6 | 111.4 | 12.88 | 9.67 | | High | High | High | High | High | High | High | High | High | High | High | 4.94E-05 | 0.000702 | 2.99 |
| FALSE | High | APSPPMSC | [R].APSPPN | 1xPhospho | Q9EST3 1x | 0.026652 | 0.001676 | 1 | 5 | 3 | Q9EST3 | Eukaryotic | Q9EST3 [50 | 0 | 1541.738 | 0 | 0.18 | 0.199496 | 14.03 | 96 | 104 | 7.91 | 18.77 | | High | High | High | High | High | High | High | High | High | High | High | 0.000903 | 0.00943 | 2.02 |
| FALSE | High | APSPPPLGL | [K].APSPPP | 1xPhospho | A7TU71 1x | 0.090153 | 0.00698 | 1 | 2 | 1 | A7TU71 | Protein Shr | A7TU71 [3 | 0 | 1523.855 | 0 | 2.02 | 0.000158 | 8.81 | 38.4 | 161.6 | 9.69 | 3.01 | | High | High | High | High | High | High | High | High | High | High | High | 0.003606 | 0.03825 | 2.36 |
| FALSE | High | APSPPPQP( | [R].APSPPP | 1xPhospho | D3Z5N8 1x | 0.017498 | 0.001245 | 1 | 6 | 2 | D3Z5N8 | Pleckstrin I | D3Z5N8 [1 | 0 | 1913.984 | 0 | -0.36 | 0.04807 | 6.33 | 113.1 | 86.9 | 13.33 | 10.45 | | High | High | High | High | High | High | High | High | High | High | High | 0.000674 | 0.005851 | 2.82 |
| FALSE | High | APSPPPTAS | [R].APSPPP | 1xPhospho | Q5UAK0-1 | 1.07E-09 | 0 | 1 | 6 | 5 | Q5UAK0-1 | Isoform 1 c | Q5UAK0-1 | 0 | 2324.117 | 0 | -0.1 | 0.035562 | 5.4 | 104 | 96 | 4.38 | 2.07 | | High | High | High | High | High | High | High | High | High | High | High | 0 | 5.25E-11 | 5.9 |
| FALSE | High | APSPPPTAS | [R].APSPPP | 1xDeamida | Q5UAK0-1 | 2.65E-08 | 0 | 1 | 6 | 2 | Q5UAK0-1 | Isoform 1 c | Q5UAK0-1 | 0 | 2325.101 | 0 | -0.03 | 0.597503 | 24.39 | 99.1 | 100.9 | 17.39 | 7.86 | | High | High | High | High | High | High | High | High | High | High | High | 0 | 1.72E-09 | 4.52 |
| FALSE | High | APSPPPTAS | [R].APSPPP | 2xPhospho | Q5UAK0-1 | 0.000294 | 0 | 1 | 6 | 1 | Q5UAK0-1 | Isoform 1 c | Q5UAK0-1 | 0 | 2404.083 | 0 | -0.24 | 0.043679 | 10.32 | 108 | 92 | 7.66 | 11.51 | | High | High | High | High | High | High | High | High | High | High | High | 0 | 6.06E-05 | 4.2 |
| FALSE | High | APSPPWPP | [R].APSPPV | 1xPhospho | Q3UR85 1x | 4.2E-05 | 0 | 1 | 4 | 1 | Q3UR85 | Myelin reg | Q3UR85 [2 | 0 | 2789.459 | 0 | -0.61 | 0.01565 | 8.54 | 121 | 79 | 8.6 | 22.21 | | High | High | High | High | High | High | High | High | High | High | High | 0 | 6.79E-06 | 3.73 |
| FALSE | High | APSPQGR | [R].APSPQ( | 1xPhospho | Q76LS9 1xI | 0.015329 | 0.001089 | 1 | 5 | 1 | Q76LS9 | Ubiquitin c | Q76LS9 [43 | 0 | 1021.503 | 0 | 1.53 | 0.000191 | 9.69 | 51.5 | 148.5 | 7.31 | 2.23 | | High | High | High | High | High | High | High | High | High | High | High | 0.000571 | 0.005079 | 1.7 |
| FALSE | High | APSPSGLM | [R].APSPSG | 1xPhospho | U5LK19 1xI | 0.077825 | 0.005963 | 2 | 16 | 1 | U5LK19; E9 | Marshalin- | U5LK19 [10 | 0 | 1495.718 | 0 | -1.58 | 0.000392 | 0.72 | 148.1 | 51.9 | 7.04 | 8.33 | Shared | High | High | High | High | High | High | High | High | High | High | High | 0.003094 | 0.03208 | 1.77 |
| FALSE | High | APSPSINR | [K].APSPSII | 1xPhospho | Q9JM13 1x | 0.030104 | 0.001857 | 1 | 3 | 1 | Q9JM13 | Rab5 GDP/ | Q9JM13 [1 | 0 | 1150.582 | 0 | 0.95 | 0.004746 | 10.1 | 74.6 | 125.4 | 7.31 | 15.78 | | High | High | High | High | High | High | High | High | High | High | High | 0.001011 | 0.01079 | 1.42 |
| FALSE | High | APSPSQPPI | [K].APSPSC | 1xPhospho | F8WHU5 1 | 0.021046 | 0.001507 | 1 | 3 | 2 | F8WHU5 | Sister chro | F8WHU5 [1 | 0 | 1446.776 | 0 | 0.28 | 0.00114 | 1.48 | 90.7 | 109.3 | 5.65 | 3.95 | | High | High | High | High | High | High | High | High | High | High | High | 0.000801 | 0.007299 | 1.98 |
| FALSE | High | APSPGSYK | [R].APSPGS | 1xPhospho | O35126 1x | 0.01894 | 0.001362 | 1 | 1 | 2 | O35126 | Atrophin-1 | O35126 [6 | 0 | 1344.696 | 0 | 0.72 | 0.000293 | 1.93 | 74.2 | 125.8 | 4.01 | 4.47 | | High | High | High | High | High | High | High | High | High | High | High | 0.00074 | 0.006421 | 1.57 |
| FALSE | High | APSPSQPPI | [K].APSPSC | 1xPhospho | F8WHU5 1 | 0.032831 | 0.002195 | 1 | 3 | 1 | F8WHU5 | Sister chro | F8WHU5 [1 | 1 | 1804.034 | 0 | -0.05 | 0.830093 | 27.17 | 101.9 | 98.1 | 16.25 | 13.77 | | High | High | High | High | High | High | High | High | High | High | High | 0.001151 | 0.01194 | 1.69 |
| FALSE | High | APSKESSEE | [R].APSKES | 2xPhospho | Q9JHS4 2xI | 0.00153 | 5.66E-05 | 1 | 2 | 1 | Q9JHS4 | ATP-depen | Q9JHS4 [60 | 1 | 3101.307 | 0 | 0.95 | 0.002226 | 8.18 | 70.2 | 129.8 | 7.55 | 12.73 | | High | High | High | High | High | High | High | High | High | High | High | 2.58E-05 | 0.000386 | 3.2 |
| FALSE | High | APSGSGQP | [R].APSGS( | 1xPhospho | Q80U49 1x | 0.008769 | 0.000736 | 1 | 3 | 1 | Q80U49 | Centrosom | Q80U49 [1 | 0 | 1593.84 | 0 | 0.48 | 0.007034 | 6.62 | 77.9 | 122.1 | 14.26 | 6.42 | | High | High | High | High | High | High | High | High | High | High | High | 0.000327 | 0.002719 | 2.06 |
| FALSE | High | APPIQSSSG | [K].APPIQS | 1xPhospho | A0A0G2JG! | 1.21E-05 | 0 | 1 | 2 | 2 | A0A0G2JG! | Crystallin b | A0A0G2JG! | 0 | 2935.422 | 0 | 2.15 | 0.003119 | 12.63 | 39.4 | 160.6 | 31.51 | 24 | | High | High | High | High | High | High | High | High | High | High | High | 0 | 1.68E-06 | 4.55 |
| FALSE | High | APPPPISPT( | [K].APPPPI! | 1xPhospho | Q9EP71 1x | 1.31E-07 | 0 | 1 | 1 | 7 | Q9EP71 | Ankycorbir | Q9EP71 [2 | 0 | 2155.1 | 0 | 0.84 | 0.000251 | 1.77 | 73.9 | 126.1 | 4.82 | 3.49 | | High | High | High | High | High | High | High | High | High | High | High | 0 | 3.49E-06 | 3.72 |
| FALSE | High | APPPPISPT( | [K].APPPPI! | 2xPhospho | Q9EP71 2x | 0.000595 | 0 | 1 | 1 | 3 | Q9EP71 | Ankycorbir | Q9EP71 [2 | 0 | 2235.066 | 0 | 0.08 | 0.064607 | 3.48 | 94.4 | 105.6 | 3.43 | 7.52 | | High | High | High | High | High | High | High | High | High | High | High | 0.001812 | 0.01838 | 3.71 |
| FALSE | High | APPPPISPT( | [K].APPPPI! | 3xPhospho | Q9EP71 3x | 0.000292 | 0 | 1 | 1 | 7 | Q9EP71 | Ankycorbir | Q9EP71 [2 | 0 | 2315.032 | 0 | 1.11 | 0.000562 | 14.89 | 62.4 | 137.6 | 9.93 | 5.52 | | High | High | High | High | High | High | High | High | High | High | High | 0 | 6E-05 | 3.57 |
| FALSE | High | APPPPSPK | [R].APPPPS | 1xPhospho | B2RWU4 1 | 0.016021 | 0.001124 | 2 | 11 | 3 | B2RWU4; ( | Cordon-ble | B2RWU4 [3 | 0 | 1328.738 | 0 | 0.01 | 0.63569 | 13.34 | 98.3 | 101.7 | 5.57 | 4.09 | Shared | High | High | High | High | High | High | High | High | High | High | High | 0.005205 | 0.05654 | 1.7 |
| FALSE | High | APPPSRPPF | [R].APPPSR | 1xPhospho | Q9CZ52 1x | 0.090888 | 0.007081 | 1 | 1 | 1 | Q9CZ52 | Anthrax to: | Q9CZ52 [5 | 0 | 1663.888 | 0 | 1.43 | 0.000812 | 23.14 | 55 | 145 | 13.86 | 6.67 | | High | High | High | High | High | High | High | High | High | High | High | 0.003606 | 0.03843 | 1.18 |
| FALSE | High | APPQDCAS | [K].APPQD( | 1xCarbami | Q04891 1x | 2.56E-06 | 0 | 1 | 2 | 1 | Q04891 | Transcripti | Q04891 [6 | 0 | 2379.094 | 0 | 0.9 | 0.000919 | 4.83 | 67.6 | 132.4 | 5.43 | 7.1 | | High | High | High | High | High | High | High | High | High | High | High | 0 | 2.92E-07 | 3.77 |
| FALSE | High | APPQDCAS | [K].APPQD( | 1xCarbami | Q04891 2x | 0.000483 | 0 | 1 | 2 | 1 | Q04891 | Transcripti | Q04891 [6 | 0 | 2459.06 | 0 | 0.72 | 0.002652 | 14.16 | 77.6 | 122.4 | 5.61 | 10.52 | | High | High | High | High | High | High | High | High | High | High | High | 0 | 0.000106 | 2.98 |
| FALSE | High | APPQPGSS' | [R].APPQP( | 1xPhospho | F8WJA0 1x | 0.006085 | 0.00062 | 1 | 3 | 6 | F8WJA0 | ATP-depen | F8WJA0 [8 | 0 | 1237.566 | 0 | -0.41 | 0.137755 | 13.36 | 112.6 | 87.4 | 8.44 | 15.42 | | High | High | High | High | High | High | High | High | High | High | High | 0.002381 | 0.02442 | 1.71 |
| FALSE | High | APPQQPSP | [R].APPQQ | 1xPhospho | A0A0R4J10 | 9.71E-05 | 0 | 1 | 9 | 3 | A0A0R4J10 | Anion exch | A0A0R4J10 | 0 | 3207.52 | 0 | 1.43 | 0.001845 | 8.9 | 51.8 | 148.2 | 10.48 | 13.11 | | High | High | High | High | High | High | High | High | High | High | High | 0 | 1.74E-05 | 3.75 |
| FALSE | High | APQASICSI | [R].APQASI | 1xCarbami | Q5DTX6 1x | 1.67E-08 | 0 | 1 | 1 | 3 | Q5DTX6 | Junctional | Q5DTX6 [1 | 0 | 3085.459 | 0 | 1.69 | 0.001507 | 15.05 | 46.1 | 153.9 | 8.91 | 13.3 | | High | High | High | High | High | High | High | High | High | High | High | 0 | 2.35E-05 | 3.85 |
| FALSE | High | APQQQPPF | [R].APQQC | 1xPhospho | Q8VDM6 1 | 0.001211 | 0 | 1 | 2 | 4 | Q8VDM6 | Heterogen | Q8VDM6 [ | 0 | 3637.815 | 0 | 0.4 | 0.000552 | 2.42 | 87.3 | 112.7 | 9.6 | 5.94 | | High | High | High | High | High | High | High | High | High | High | High | 2.58E-05 | 0.00047 | 3 |
| FALSE | High | APQSPTLAI | [R].APQSPT | 1xPhospho | P97792 1xI | 0.000107 | 0 | 1 | 3 | 3 | P97792 | Coxsackiev | P97792 [32 | 0 | 1618.897 | 0 | 1.55 | 0.000434 | 9.01 | 50 | 150 | 5.06 | 7.65 | | High | High | High | High | High | High | High | High | High | High | High | 0.000161 | 0.001197 | 2.93 |
| FALSE | High | APSGVSETI | [K].APSGV! | 1xPhospho | Q8VDZ4 1x | 5.4E-08 | 0 | 1 | 1 | 1 | Q8VDZ4 | Palmitoyltr | Q8VDZ4 [6 | 0 | 2604.394 | 0 | 1.91 | 0.000416 | 31.14 | 47.5 | 152.5 | 24.06 | 9.39 | | High | High | High | High | High | High | High | High | High | High | High | 0 | 3.83E-09 | 2.73 |
| FALSE | High | APQTSSPPI | [R].APQTS! | 1xPhospho | Q52KI8 1xP | 0.000188 | 0 | 1 | 7 | 17 | Q52KI8 | Serine/argi | Q52KI8 [70 | 0 | 1445.735 | 0 | -0.25 | 0.001426 | 2.59 | 108.6 | 91.4 | 1.37 | 3.76 | | High | High | High | High | High | High | High | High | High | High | High | 0.000294 | 0.002309 | 2.52 |
| FALSE | High | APSAPPGT( | [R].APSAPF | 1xPhospho | Q8K3X4 1x | 0.027121 | 0.001676 | 1 | 1 | 2 | Q8K3X4 | Interferon | Q8K3X4 [4 | 0 | 1995.026 | 0 | 0.99 | 0.000949 | 13.13 | 66.2 | 133.8 | 8.57 | 7.25 | | High | High | High | High | High | High | High | High | High | High | High | 0.002559 | 0.0276 | 2.48 |
| FALSE | High | APSASPLAI | [R].APSASF | 1xPhospho | Q8BGT6 1x | 1.81E-05 | 0 | 1 | 2 | 4 | Q8BGT6 | MICAL-like | Q8BGT6 [4 | 0 | 1586.825 | 0 | 1.74 | 0.001341 | 2.15 | 43.1 | 156.9 | 8.62 | 13.87 | | High | High | High | High | High | High | High | High | High | High | High | 0 | 2.65E-06 | 2.35 |
| FALSE | High | APSASPLAI | [R].APSASF | 2xPhospho | Q8BGT6 2x | 0.010109 | 0.000736 | 1 | 2 | 1 | Q8BGT6 | MICAL-like | Q8BGT6 [4 | 0 | 1666.792 | 0 | 1.13 | 0.001041 | 20.93 | 63.5 | 136.5 | 12.21 | 4.25 | | High | High | High | High | High | High | High | High | High | High | High | 0.000344 | 0.003177 | 2.46 |

| | | | | | | | | | | | | | | | | | | | | | | | | | | | | | | | | | | | | | | |
|---|---|---|---|---|---|---|---|---|---|---|---|---|---|---|---|---|---|---|---|---|---|---|---|---|---|---|---|---|---|---|---|---|---|---|---|---|---|---|
| FALSE | High | APSDDGSL | [K].APSDD( | 2xPhospho | Q9JMH9-6 | 8.4E-05 | 0 | 2 | 15 | 1 | Q9JMH9-6 | Isoform 6 ( | Q9JMH9-6 | 0 | 1507.685 | 0 | 0.29 | 0.024172 | 11.82 | 90.1 | 109.9 | 6.03 | 5.9 | Shared | High | High | High | High | High | High | High | High | High | High | High | 0 | 1.47E-05 | 3.13 |
| FALSE | High | APSDRSLSI | [R].APSDRS | 2xPhospho | Q91ZV0 2x | 0.011246 | 0.000772 | 1 | 12 | 1 | Q91ZV0 | Melanoma | Q91ZV0 [1 | 1 | 2129.926 | 0 | 0.3 | 0.456511 | 11.14 | 92.6 | 107.4 | 13.03 | 12.84 | | High | High | High | High | High | High | High | High | High | High | High | 0.000418 | 0.003571 | 2.17 |
| FALSE | High | APSENEK | [R].APSENE | 1xPhospho | E9Q6J5-3 1 | 0.088701 | 0.006669 | 1 | 4 | 1 | E9Q6J5-3 | Isoform Ne | E9Q6J5-3 [ | 0 | 1312.655 | 0 | 1.02 | 0.000968 | 6.6 | 64.3 | 135.7 | 4.69 | 9.21 | | High | High | High | High | High | High | High | High | High | High | High | 0.003402 | 0.03731 | 1.28 |
| FALSE | High | APSEVAIEE | [K].APSEVA | 1xPhospho | Q8R2M2 1 | 4.05E-05 | 0 | 1 | 1 | 3 | Q8R2M2 | Deoxynucl | Q8R2M2 [5 | 0 | 1739.887 | 0 | -0.03 | 0.548534 | 3.96 | 99.4 | 100.6 | 8.82 | 5.2 | | High | High | High | High | High | High | High | High | High | High | High | 0 | 0.000233 | 3.56 |
| FALSE | High | APSEVAIEE | [K].APSEVA | 1xPhospho | Q8R2M2 1 | 0.000148 | 0 | 1 | 1 | 1 | Q8R2M2 | Deoxynucl | Q8R2M2 [5 | 1 | 2384.152 | 0 | 0.29 | 0.402939 | 13.72 | 86.4 | 113.6 | 29.65 | 26.23 | | High | High | High | High | High | High | High | High | High | High | High | 0 | 2.8E-05 | 3.78 |
| FALSE | High | APSEVAIEE | [K].APSEVA | 1xPhospho | Q8R2M2 1 | 0.000253 | 0 | 1 | 1 | 1 | Q8R2M2 | Deoxynucl | Q8R2M2 [5 | 1 | 2613.315 | 0 | -0.18 | 0.034904 | 6.06 | 105.4 | 94.6 | 5.39 | 11.09 | | High | High | High | High | High | High | High | High | High | High | High | 0 | 5.08E-05 | 3.37 |
| FALSE | High | APSFSVSAI | [K].APSFSV | 1xPhospho | E9Q616 1x | 2.1E-06 | 0 | 1 | 4 | 4 | E9Q616 | AHNAK nu | E9Q616 [4 | 0 | 2707.447 | 0 | 4.07 | 0.004676 | 14.52 | 8.8 | 191.2 | 30.38 | 29.26 | | High | High | High | High | High | High | High | High | High | High | High | 0 | 2.35E-07 | 2.68 |
| FALSE | High | APSGETPPS | [R].APSGET | 1xPhospho | Q6IQX8 1x | 0.000584 | 0 | 1 | 5 | 2 | Q6IQX8 | Zinc finger | Q6IQX8 [6 | 0 | 2500.181 | 0 | -1.7 | 0.000425 | 2.15 | 153.2 | 46.8 | 4.23 | 12.65 | | High | High | High | High | High | High | High | High | High | High | High | 0 | 0.000131 | 3.06 |
| FALSE | High | APSGETPPS | [R].APSGET | 2xPhospho | Q6IQX8 2x | 3.78E-06 | 0 | 1 | 5 | 4 | Q6IQX8 | Zinc finger | Q6IQX8 [6 | 0 | 2580.147 | 0 | -1.31 | 0.00098 | 2.4 | 140.7 | 59.3 | 7.58 | 8.6 | | High | High | High | High | High | High | High | High | High | High | High | 0 | 4.51E-07 | 4.13 |
| FALSE | High | APSGETPPS | [R].APSGET | 3xPhospho | Q6IQX8 3x | 5.21E-05 | 0 | 1 | 5 | 4 | Q6IQX8 | Zinc finger | Q6IQX8 [6 | 0 | 2660.113 | 0 | -0.95 | 0.000879 | 1.49 | 131.9 | 68.1 | 5.09 | 9.23 | | High | High | High | High | High | High | High | High | High | High | High | 0 | 9.17E-06 | 4.8 |
| FALSE | High | APQTSSPPI | [R].APQTSS | 2xPhospho | Q52KI8 2xI | 0.022973 | 0.001578 | 1 | 7 | 1 | Q52KI8 | Serine/argi | Q52KI8 [70 | 0 | 1525.701 | 0 | -0.36 | 0.445496 | 4.31 | 110.6 | 89.4 | 12.03 | 14.35 | | High | High | High | High | High | High | High | High | High | High | High | 0.000847 | 0.007997 | 2.5 |
| FALSE | High | APSPTAEQ | [R].APSPTA | 1xPhospho | Q61127 1x | 0.054844 | 0.003982 | 1 | 5 | 1 | Q61127 | NGFI-A-bin | Q61127 [4 | 0 | 1516.736 | 0 | -0.07 | 0.510184 | 7.28 | 100.4 | 99.6 | 2.08 | 7.61 | | High | High | High | High | High | High | High | High | High | High | High | 0.002059 | 0.02136 | 2.47 |
| FALSE | High | APSPTDLPE | [K].APSPTD | 1xPhospho | Q8R550-3 | 2.96E-06 | 0 | 1 | 2 | 1 | Q8R550-3 | Isoform 3 ( | Q8R550-3 | 0 | 1921.992 | 0 | 2.55 | 0.000229 | 27.75 | 29.2 | 170.8 | 29.05 | 6.73 | | High | High | High | High | High | High | High | High | High | High | High | 0 | 3.45E-07 | 3.12 |
| FALSE | High | APSPTTAA( | [R].APSPTT | 1xCarbami | A2ACV6 1x | 8.55E-08 | 0 | 1 | 5 | 5 | A2ACV6 | Protein SO | A2ACV6 [1 | 0 | 1943.919 | 0 | 1.79 | 0.000417 | 13.76 | 43.9 | 156.1 | 5.46 | 6.96 | | High | High | High | High | High | High | High | High | High | High | High | 0 | 6.41E-09 | 3.94 |
| FALSE | High | APVSPSSPS | [R].APVSPS | 1xPhospho | Q9QYE6 1x | 8.22E-07 | 0 | 1 | 1 | 4 | Q9QYE6 | Golgin sub | Q9QYE6 [1 | 0 | 2628.353 | 0 | 0.02 | 0.443788 | 8.84 | 102.5 | 97.5 | 14.79 | 6.68 | | High | High | High | High | High | High | High | High | High | High | High | 0 | 2.16E-06 | 4.27 |
| FALSE | High | APVSPSSPS | [R].APVSPS | 2xPhospho | Q9QYE6 2x | 0.036731 | 0.00249 | 1 | 1 | 1 | Q9QYE6 | Golgin sub | Q9QYE6 [1 | 0 | 2708.32 | 0 | 0.01 | 0.407188 | 9.81 | 94.8 | 105.2 | 10.97 | 8.23 | | High | High | High | High | High | High | High | High | High | High | High | 0.001308 | 0.01351 | 2.81 |
| FALSE | High | APVSSVESI | [R].APVSSV | 1xPhospho | Q32P12 1x | 0.000458 | 0 | 1 | 2 | 2 | Q32P12 | Uncharacte | Q32P12 [3 | 0 | 1721.867 | 0 | -0.55 | 0.002279 | 4.98 | 120.1 | 79.9 | 2.04 | 10.24 | | High | High | High | High | High | High | High | High | High | High | High | 0 | 9.9E-05 | 2.97 |
| FALSE | High | APVSVACG | [K].APVSV/ | 2xCarbami | F8VQ95 1x | 1.35E-07 | 0 | 1 | 5 | 1 | F8VQ95 | Transformi | F8VQ95 [5 | 0 | 2870.372 | 0 | 0.04 | 0.740572 | 28.88 | 99.9 | 100.1 | 18.85 | 19.82 | | High | High | High | High | High | High | High | High | High | High | High | 0 | 1.08E-08 | 3.52 |
| FALSE | High | APVTPEGA | [K].APVTPE | 1xPhospho | Q8QZS3 1x | 0.051694 | 0.003745 | 1 | 1 | 1 | Q8QZS3 | Folliculin O | Q8QZS3 [3 | 0 | 1392.672 | 0 | 0.43 | 0.005426 | 6.06 | 84.9 | 115.1 | 7.15 | 7.16 | | High | High | High | High | High | High | High | High | High | High | High | 0.00196 | 0.02001 | 2.5 |
| FALSE | High | APVVLQPE | [K].APVVL( | 1xPhospho | F6RJ39 1xP | 1.04E-06 | 0 | 2 | 4 | 3 | F6RJ39; Q8 | Apoptotic | F6RJ39 [34 | 0 | 3052.662 | 0 | 0.22 | 0.083764 | 13.2 | 93.1 | 106.9 | 5.69 | 10.23 | Shared | High | High | High | High | High | High | High | High | High | High | High | 0 | 1.06E-07 | 3.34 |
| FALSE | High | APVVLQPE | [K].APVVL( | 1xDeamida | F6RJ39 1xP | 0.01588 | 0.001124 | 2 | 4 | 2 | F6RJ39; Q8 | Apoptotic | F6RJ39 [34 | 0 | 3053.646 | 0 | 0.93 | 0.01807 | 20.61 | 68.9 | 131.1 | 18.19 | 9.48 | Shared | High | High | High | High | High | High | High | High | High | High | High | 0.000638 | 0.0056 | 1.89 |
| FALSE | High | APVVLQPE | [K].APVVL( | 2xPhospho | F6RJ39 2xP | 0.000319 | 0 | 2 | 4 | 3 | F6RJ39; Q8 | Apoptotic | F6RJ39 [34 | 0 | 3132.629 | 0 | 0.52 | 0.077726 | 12.64 | 81.5 | 118.5 | 14.06 | 20.35 | Shared | High | High | High | High | High | High | High | High | High | High | High | 0.000585 | 0.005513 | 4.02 |
| FALSE | High | APWIPPNP | [R].APWIPI | 1xPhospho | A0A0R4J1C | 0.052133 | 0.00384 | 1 | 3 | 1 | A0A0R4J1C | RIKEN cDN | A0A0R4J1C | 0 | 2195.167 | 0 | 1.02 | 0.001147 | 1.42 | 66 | 134 | 4.15 | 8.71 | | High | High | High | High | High | High | High | High | High | High | High | 0.001999 | 0.02017 | 2.83 |
| FALSE | High | APWWDAS | [R].APWW | 1xPhospho | Q53ZY9 1x | 0.000543 | 0 | 1 | 1 | 1 | Q53ZY9 | Adrenal hy | Q53ZY9 [2 | 0 | 1678.794 | 0 | -1.32 | 0.002432 | 1.31 | 141.5 | 58.5 | 18.76 | 15.05 | | High | High | High | High | High | High | High | High | High | High | High | 0 | 0.00012 | 3.16 |
| FALSE | High | APYSPTTSK | [K].APYSPT | 1xPhospho | Q80T83 1x | 0.014158 | 0.001011 | 1 | 4 | 2 | Q80T83 | MKIAA185 | Q80T83 [7 | 0 | 1489.77 | 0 | 0.42 | 0.007008 | 7.63 | 87.2 | 112.8 | 8.08 | 3.14 | | High | High | High | High | High | High | High | High | High | High | High | 0.000538 | 0.00463 | 2.25 |
| FALSE | High | APYTLDTLI | [K].APYTLD | 1xCarbami | A2BE28 1x | 0.002172 | 5.66E-05 | 1 | 2 | 1 | A2BE28 | Ribosomal | A2BE28 [5 | 0 | 3738.761 | 0 | -0.54 | 0.006462 | 4.11 | 120 | 80 | 7.32 | 4.92 | | High | High | High | High | High | High | High | High | High | High | High | 2.58E-05 | 0.000568 | 2.46 |
| FALSE | High | AQAGLTAT | [R].AQAGL | 1xPhospho | Q9JI13 1xP | 9.85E-07 | 0 | 1 | 4 | 15 | Q9JI13 | Something | Q9JI13 [18 | 0 | 4446.936 | 0 | -0.28 | 0.046215 | 7.27 | 109.8 | 90.2 | 3.81 | 9.45 | | High | High | High | High | High | High | High | High | High | High | High | 0 | 1E-07 | 4.45 |
| FALSE | High | APVSICGG( | [R].APVSIC | 1xCarbami | O55176 1x | 5.16E-05 | 0 | 1 | 2 | 1 | O55176 | E3 ubiquiti | O55176 [1 | 0 | 1924.96 | 0 | 1.1 | 0.00198 | 12.4 | 64.6 | 135.4 | 7.97 | 12.04 | | High | High | High | High | High | High | High | High | High | High | High | 0 | 8.51E-06 | 4.05 |
| FALSE | High | AQAPASPY | [R].AQAPA | 1xPhospho | G5E8C3 1x | 1.53E-05 | 0 | 1 | 2 | 4 | G5E8C3 | G protein-( | G5E8C3 [3 | 0 | 1847.816 | 0 | 1.92 | 0.000264 | 8.14 | 42.5 | 157.5 | 3.3 | 5.77 | | High | High | High | High | High | High | High | High | High | High | High | 0 | 2.18E-06 | 3.35 |
| FALSE | High | AQASAPAC | [K].AQASA | 1xPhospho | P47915 1xI | 0.000122 | 0 | 1 | 3 | 2 | P47915 | 60S riboso | P47915 [13 | 0 | 1577.845 | 0 | 0.19 | 0.165719 | 0.84 | 94.3 | 105.7 | 11.35 | 5.14 | | High | High | High | High | High | High | High | High | High | High | High | 0 | 5.39E-05 | 2.64 |
| FALSE | High | AQASTAAG | [R].AQAST/ | 1xPhospho | Q8VBZ3 1x | 3.47E-05 | 0 | 1 | 1 | 2 | Q8VBZ3 | Cleft lip an | Q8VBZ3 [6 | 0 | 3094.571 | 0 | -0.54 | 0.007384 | 8.89 | 115.9 | 84.1 | 3.79 | 11.48 | | High | High | High | High | High | High | High | High | High | High | High | 0 | 5.48E-06 | 3.12 |
| FALSE | High | AQAVSEEE | [K].AQAVS | 1xPhospho | F8WJA0 1x | 0.000718 | 0 | 1 | 6 | 2 | F8WJA0 | ATP-depen | F8WJA0 [1 | 0 | 2588.165 | 0 | -0.35 | 0.057514 | 16.81 | 112.1 | 87.9 | 12.84 | 14.06 | | High | High | High | High | High | High | High | High | High | High | High | 0 | 0.000164 | 2.6 |
| FALSE | High | AQAVSEEE | [K].AQAVS | 2xPhospho | F8WJA0 2x | 0.000343 | 0 | 1 | 6 | 1 | F8WJA0 | ATP-depen | F8WJA0 [1 | 0 | 2668.132 | 0 | -0.44 | 0.008294 | 1.99 | 114.8 | 85.2 | 17.98 | 16.86 | | High | High | High | High | High | High | High | High | High | High | High | 0 | 7.18E-05 | 2.03 |
| FALSE | High | AQAVSEEE | [K].AQAVS | 1xPhospho | F8WJA0 1x | 0.007471 | 0.00062 | 1 | 6 | 3 | F8WJA0 | ATP-depen | F8WJA0 [1 | 1 | 2945.423 | 0 | 0.9 | 0.045644 | 18.86 | 73.4 | 126.6 | 11.26 | 22.91 | | High | High | High | High | High | High | High | High | High | High | High | 0.000275 | 0.002273 | 2.87 |
| FALSE | High | AQCETLSPI | [R].AQCETI | 1xCarbami | A2AFQ0 1x | 6.15E-06 | 0 | 1 | 5 | 2 | A2AFQ0 | E3 ubiquiti | A2AFQ0 [3 | 0 | 2767.326 | 0 | 0.11 | 0.15132 | 7.51 | 94 | 106 | 7.05 | 5.97 | | High | High | High | High | High | High | High | High | High | High | High | 0 | 1.26E-06 | 3.96 |
| FALSE | High | AQDSAMA | [K].AQDSA | 1xCarbami | B2RQC2 1x | 0.006773 | 0.00062 | 1 | 2 | 1 | B2RQC2 | Ubiquitin c | B2RQC2 [8 | 1 | 3483.69 | 0 | 0.18 | 0.706954 | 37.06 | 93.8 | 106.2 | 15.62 | 26.05 | | High | High | High | High | High | High | High | High | High | High | High | 0.000275 | 0.002028 | 2.86 |
| FALSE | High | AQDSAMA | [K].AQDSA | 1xCarbami | B2RQC2 1x | 0.007145 | 0.00062 | 1 | 2 | 1 | B2RQC2 | Ubiquitin c | B2RQC2 [8 | 1 | 3499.685 | 0 | 0.08 | 0.261799 | 18.39 | 94.7 | 105.3 | 6.2 | 14.15 | | High | High | High | High | High | High | High | High | High | High | High | 0.000275 | 0.002165 | 2.86 |
| FALSE | High | AQEAAAEE | [R].AQEAA | 1xPhospho | Q9CXS4 1x | 2.52E-06 | 0 | 1 | 1 | 1 | Q9CXS4 | Centromer | Q9CXS4 [6 | 0 | 3084.509 | 0 | 0.34 | 0.085571 | 20.95 | 87.2 | 112.8 | 15.86 | 14.1 | | High | High | High | High | High | High | High | High | High | High | High | 0 | 2.86E-07 | 4.83 |
| FALSE | High | AQEAAAEE | [R].AQEAA | 1xDeamida | Q9CXS4 1x | 9.08E-07 | 0 | 1 | 1 | 1 | Q9CXS4 | Centromer | Q9CXS4 [6 | 0 | 3085.493 | 0 | 0.41 | 0.007525 | 4.61 | 86.9 | 113.1 | 14.97 | 8.99 | | High | High | High | High | High | High | High | High | High | High | High | 0 | 9.08E-08 | 4.75 |
| FALSE | High | AQENYEGQ | [R].AQENY | 1xPhospho | P70398 1xI | 9.08E-07 | 0 | 1 | 5 | 3 | P70398 | Probable u | P70398 [2 | 0 | 2401.132 | 0 | -0.34 | 0.004775 | 3.47 | 111.9 | 88.1 | 1.88 | 5.86 | | High | High | High | High | High | High | High | High | High | High | High | 0 | 4.44E-05 | 2.96 |
| FALSE | High | AQEPPPAQ | [R].AQEPPI | 1xPhospho | A0A0A0MC | 2.03E-07 | 0 | 1 | 2 | 1 | A0A0A0MC | Histone-lys | A0A0A0MC | 0 | 2618.297 | 0 | 0.43 | 0.046034 | 15.19 | 79.4 | 120.6 | 6.63 | 19.63 | | High | High | High | High | High | High | High | High | High | High | High | 0 | 1.71E-08 | 5.08 |
| FALSE | High | AQEPSPAK | [R].AQEPSI | 1xPhospho | B2RQC2 1x | 0.01029 | 0.000772 | 1 | 3 | 1 | B2RQC2 | Ubiquitin c | B2RQC2 [9 | 0 | 1365.718 | 0 | 0.12 | 0.176975 | 12.89 | 96.4 | 103.6 | 2.13 | 7.76 | | High | High | High | High | High | High | High | High | High | High | High | 0.000382 | 0.003232 | 1.82 |
| FALSE | High | AQAPASPY | [R].AQAPA | 1xDeamida | G5E8C3 1x | 0.003433 | 0.00021 | 1 | 2 | 1 | G5E8C3 | G protein-( | G5E8C3 [3 | 0 | 1848.8 | 0 | 1.73 | 0.000547 | 25.62 | 46.4 | 153.6 | 10.1 | 5.97 | | High | High | High | High | High | High | High | High | High | High | High | 9.42E-05 | 0.000954 | 2.89 |
| FALSE | High | APVHFVEP | [K].APVHF | 1xPhospho | A0A0A0MC | 1.2E-05 | 0 | 1 | 2 | 2 | A0A0A0MC | Calmodulir | A0A0A0MC | 0 | 2206.137 | 0 | 0.59 | 0.00356 | 5.91 | 77.6 | 122.4 | 7.4 | 7.31 | | High | High | High | High | High | High | High | High | High | High | High | 7.19E-05 | 0.000773 | 2.74 |
| FALSE | High | APVGSVVS | [K].APVGS | 1xPhospho | A0A087WF | 0.011447 | 0.000816 | 1 | 6 | 1 | A0A087WF | Arf-GAP dc | A0A087WF | 0 | 2336.19 | 0 | 1.02 | 0.008642 | 18.98 | 67.7 | 132.3 | 20.96 | 10.5 | | High | High | High | High | High | High | High | High | High | High | High | 0.000437 | 0.003651 | 1.39 |
| FALSE | High | APTVSISCL | [K].APTVSI | 1xCarbami | Q60738 1x | 9.5E-07 | 0 | 1 | 1 | 1 | Q60738 | Zinc transp | Q60738 [4 | 0 | 2428.245 | 0 | -0.3 | 0.144166 | 3.96 | 110.3 | 89.7 | 15.15 | 4.39 | | High | High | High | High | High | High | High | High | High | High | High | 0 | 9.6E-08 | 3.36 |
| FALSE | High | APSPVLLSP | [R].APSPVL | 1xPhospho | P41971 1xI | 0.000237 | 0 | 1 | 4 | 1 | P41971 | ETS domai | P41971 [3 | 0 | 1849.024 | 0 | 1 | 0.003603 | 18.76 | 69.1 | 130.9 | 13.61 | 9.57 | | High | High | High | High | High | High | High | High | High | High | High | 0 | 4.76E-05 | 2.86 |
| FALSE | High | APSPVLLSP | [R].APSPVL | 2xPhospho | P41971 2xI | 0.004033 | 0.000311 | 1 | 4 | 1 | P41971 | ETS domai | P41971 [3 | 0 | 1928.99 | 0 | 1.03 | 0.004341 | 25.93 | 69.1 | 130.9 | 16.85 | 7.02 | | High | High | High | High | High | High | High | High | High | High | High | 0.000161 | 0.001141 | 3.21 |
| FALSE | High | APSQDSLD | [R].APSQD | 1xPhospho | A0A0R4J29 | 0.117405 | 0.009523 | 1 | 5 | 1 | A0A0R4J29 | Eukaryotic | A0A0R4J29 | 0 | 3007.521 | 0 | -0.88 | 0.000885 | 0.89 | 127.3 | 72.7 | 6.86 | 9.61 | | High | High | High | High | High | High | High | High | High | High | High | 0.004782 | 0.05226 | 2.29 |
| FALSE | High | APSQPPSPT | [R].APSQPI | 1xPhospho | B2RQ83 1x | 0.005516 | 0.00062 | 1 | 4 | 3 | B2RQ83 | Gem (Nucl | B2RQ83 [1 | 0 | 1604.752 | 0 | -0.7 | 0.015144 | 5.44 | 124 | 76 | 10.29 | 11.26 | | High | High | High | High | High | High | High | High | High | High | High | 0.003122 | 0.03229 | 1.8 |
| FALSE | High | APSQPPSPT | [R].APSQPI | 2xPhospho | B2RQ83 2x | 0.002948 | 0.00016 | 1 | 4 | 1 | B2RQ83 | Gem (Nucl | B2RQ83 [1 | 0 | 1684.718 | 0 | -0.21 | 0.08465 | 15.91 | 112.9 | 87.1 | 8.88 | 16.51 | | High | High | High | High | High | High | High | High | High | High | High | 7.19E-05 | 0.000802 | 2.16 |
| FALSE | High | APSQPPSPT | [R].APSQPI | 2xPhospho | B7ZN04 2x | 0.012508 | 0.000889 | 1 | 3 | 2 | B7ZN04 | Gemin5 pr | B7ZN04 [1 | 0 | 1426.633 | 0 | -0.38 | 0.000474 | 2.34 | 113.6 | 86.4 | 2.09 | 5.84 | | High | High | High | High | High | High | High | High | High | High | High | 0.000538 | 0.004419 | 2.12 |
| FALSE | High | APSQPPSPT | [R].APSQPI | 1xPhospho | B7ZN04 1x | 0.000142 | 0 | 1 | 3 | 1 | B7ZN04 | Gemin5 pr | B7ZN04 [1 | 1 | 1760.853 | 0 | -0.54 | 0.012867 | 5.53 | 116.3 | 83.7 | 6.77 | 11.29 | | High | High | High | High | High | High | High | High | High | High | High | 0 | 2.66E-05 | 2.09 |
| FALSE | High | APSQPPSPT | [R].APSQPI | 2xPhospho | B7ZN04 2x | 0.044358 | 0.003264 | 1 | 3 | 1 | B7ZN04 | Gemin5 pr | B7ZN04 [1 | 1 | 1840.819 | 0 | -0.36 | 0.000746 | 2.43 | 113.5 | 86.5 | 3.94 | 7.62 | | High | High | High | High | High | High | High | High | High | High | High | 0.001665 | 0.01673 | 1.74 |
| FALSE | High | APSQVSISA | [R].APSQV | 1xPhospho | E9Q842 1x | 0.005978 | 0.00062 | 1 | 3 | 2 | E9Q842 | Neuron na | E9Q842 [1 | 0 | 1508.767 | 0 | 1.34 | 0.001307 | 16.6 | 60.6 | 139.4 | 8.07 | 11.35 | | High | High | High | High | High | High | High | High | High | High | High | 0.000275 | 0.001758 | 1.8 |
| FALSE | High | APSSDEECI | [R].APSSDE | 1xCarbami | Q6IR34-5 1 | 0.009168 | 0.000736 | 1 | 7 | 4 | Q6IR34-5 | Isoform 5 ( | Q6IR34-5 [ | 0 | 2283.066 | 0 | -0.25 | 0.059607 | 13.73 | 108.9 | 91.1 | 8.29 | 8.91 | | High | High | High | High | High | High | High | High | High | High | High | 0.000449 | 0.003981 | 2.45 |
| FALSE | High | APSSEVK | [R].APSSEV | 1xPhospho | Q6P6I6 1xI | 0.059159 | 0.004294 | 1 | 2 | 2 | Q6P6I6 | DNA-direct | Q6P6I6 [16 | 0 | 1255.67 | 0 | -1.25 | 0.00131 | 2.41 | 137 | 63 | 6 | 15.16 | | High | High | High | High | High | High | High | High | High | High | High | 0.002937 | 0.03077 | 1.68 |
| FALSE | High | APSSHFSAS | [K].APSSHI | 1xPhospho | Q80Y82 1x | 0.000375 | 0 | 1 | 14 | 1 | Q80Y82 | Protein kin | Q80Y82 [4 | 0 | 2523.151 | 0 | 0.32 | 0.11516 | 25.59 | 86.5 | 113.5 | 11.94 | 17.38 | | High | High | High | High | High | High | High | High | High | High | High | 0 | 7.92E-05 | 3.79 |
| FALSE | High | APSSHFSAS | [K].APSSHI | 1xPhospho | Q80Y82 1x | 0.000144 | 0 | 1 | 14 | 1 | Q80Y82 | Protein kin | Q80Y82 [4 | 0 | 2539.146 | 0 | 0.1 | 0.233903 | 4.17 | 99.7 | 100.3 | 12.82 | 5.74 | | High | High | High | High | High | High | High | High | High | High | High | 0 | 2.71E-05 | 3.79 |
| FALSE | High | APSSSSPGS | [R].APSSSS | 1xPhospho | Q6DFZ1 1x | 3.32E-09 | 0 | 1 | 4 | 14 | Q6DFZ1 | Golgi-speci | Q6DFZ1 [1 | 0 | 2116.003 | 0 | 0.29 | 0.009489 | 4.01 | 90.1 | 109.9 | 5.86 | 5.65 | | High | High | High | High | High | High | High | High | High | High | High | 0 | 7.05E-09 | 4.3 |
| FALSE | High | APSSSSPGS | [R].APSSSS | 2xPhospho | Q6DFZ1 2x | 0.120211 | 0.009851 | 1 | 4 | 1 | Q6DFZ1 | Golgi-speci | Q6DFZ1 [1 | 0 | 2179.975 | 0 | 0.48 | 0.011061 | 11.96 | 83.5 | 116.5 | 8.6 | 12.82 | | High | High | High | High | High | High | High | High | High | High | High | 0.004961 | 0.05389 | 2.65 |
| FALSE | High | APSTPVPPS | [K].APSTPV | 1xPhospho | Q3TYU7 1x | 0.000524 | 0 | 1 | 3 | 5 | Q3TYU7 | Putative ur | Q3TYU7 [1 | 0 | 2253.23 | 0 | -0.13 | 0.04068 | 4.84 | 106.4 | 93.6 | 4.16 | 3.53 | | High | High | High | High | High | High | High | High | High | High | High | 0 | 0.000115 | 3.19 |
| FALSE | High | APSTPVPPS | [K].APSTPV | 2xPhospho | Q3TYU7 2x | 0.030367 | 0.001857 | 1 | 3 | 1 | Q3TYU7 | Putative ur | Q3TYU7 [1 | 0 | 2333.196 | 0 | 0.02 | 0.585128 | 7.08 | 98.4 | 101.6 | 18.25 | 5.92 | | High | High | High | High | High | High | High | High | High | High | High | 0.001011 | 0.01095 | 2.33 |
| FALSE | High | APSTSDSVI | [R].APSTSD | 1xPhospho | E9PYD5 1xI | 0.012731 | 0.000889 | 1 | 4 | 1 | E9PYD5 | Transcripti | E9PYD5 [1 | 0 | 1228.577 | 0 | -1.58 | 0.000154 | 0.14 | 149 | 51 | 3.28 | 3.03 | | High | High | High | High | High | High | High | High | High | High | High | 0.000449 | 0.00411 | 1.54 |
| FALSE | High | APSVANIGS | [R].APSVAI | 1xCarbami | Q8BTM8 1 | 1.07E-07 | 0 | 1 | 5 | 3 | Q8BTM8 | Filamin-A ( | Q8BTM8 [2 | 0 | 1977.967 | 0 | 0.4 | 0.006947 | 8.93 | 87.3 | 112.7 | 3.94 | 9.01 | | High | High | High | High | High | High | High | High | High | High | High | 0 | 8.3E-09 | 3.94 |
| FALSE | High | APSVANIGS | [R].APSVAI | 1xCarbami | Q8BTM8 1 | 1.07E-05 | 0 | 1 | 5 | 3 | Q8BTM8 | Filamin-A ( | Q8BTM8 [2 | 0 | 2207.13 | 0 | 0.73 | 0.000361 | 4.68 | 73 | 127 | 6.94 | 4.31 | | High | High | High | High | High | High | High | High | High | High | High | 0.000275 | 0.00185 | 2.4 |
| FALSE | High | APSVSVVSI | [K].APSVSV | 1xPhospho | Q8VDU0 1 | 0.000718 | 0 | 1 | 1 | 1 | Q8VDU0 | G-protein- | Q8VDU0 [5 | 0 | 2828.392 | 0 | 1.1 | 0.008624 | 7.61 | 68.3 | 131.7 | 20.32 | 13.23 | | High | High | High | High | High | High | High | High | High | High | High | 0 | 0.000165 | 3.15 |
| FALSE | High | APSWIDTG | [R].APSWII | 1xPhospho | Q52L78 1xI | 0.003433 | 0.00021 | 1 | 2 | 2 | Q52L78 | Cryab prot | Q52L78 [5 | 0 | 1771.829 | 0 | 2.84 | 0.000707 | 4.83 | 22.8 | 177.2 | 9.39 | 12.94 | | High | High | High | High | High | High | High | High | High | High | High | 0.000786 | 0.006831 | 2.07 |
| FALSE | High | APSWIDTG | [R].APSWII | 1xPhospho | Q52L78 1xI | 1.39E-06 | 0 | 1 | 2 | 3 | Q52L78 | Cryab prot | Q52L78 [5 | 0 | 1787.824 | 0 | 1.54 | 0.001302 | 16.15 | 49.3 | 150.7 | 9.52 | 10.48 | | High | High | High | High | High | High | High | High | High | High | High | 0 | 1.47E-07 | 4.5 |
| FALSE | High | APTAALSPE | [R].APTAAI | 1xPhospho | S4R1M2 1x | 0.003495 | 0.00021 | 2 | 4 | 1 | S4R1M2; C | Scaffold at | S4R1M2 [3 | 0 | 1720.836 | 0 | -0.12 | 0.404498 | 48.66 | 104.3 | 95.7 | 30.09 | 19 | Shared | High | High | High | High | High | High | High | High | High | High | High | 9.42E-05 | 0.000968 | 1.95 |
| FALSE | High | APTAALSPE | [R].APTAAI | 1xPhospho | S4R1M2 1x | 3.38E-05 | 0 | 2 | 4 | 6 | S4R1M2; C | Scaffold at | S4R1M2 [3 | 0 | 1949.999 | 0 | -0.04 | 0.15757 | 5.45 | 103.3 | 96.7 | 3.1 | 7 | Shared | High | High | High | High | High | High | High | High | High | High | High | 0.000225 | 0.001449 | 1.94 |
| FALSE | High | APTAGSGC | [K].APTAG | 1xCarbami | Q05CL8 1x | 0.000112 | 0 | 1 | 3 | 2 | Q05CL8 | La-related | Q05CL8 [4 | 0 | 2088.967 | 0 | -0.33 | 0.011672 | 7.68 | 117.4 | 82.6 | 6.63 | 8.58 | | High | High | High | High | High | High | High | High | High | High | High | 0 | 2.04E-05 | 3.42 |
| FALSE | High | APTGSTPG | [R].APTGST | 1xPhospho | Q8C9B9 1x | 0.031989 | 0.00205 | 1 | 1 | 2 | Q8C9B9 | Death-indu | Q8C9B9 [1 | 0 | 1816.916 | 0 | -0.4 | 0.021595 | 8.41 | 109.2 | 90.8 | 17.27 | 12.41 | | High | High | High | High | High | High | High | High | High | High | High | 0.002485 | 0.02718 | 1.56 |
| FALSE | High | APTNPSVEI | [K].APTNPS | 1xPhospho | P11157 1xI | 0.021046 | 0.001507 | 1 | 3 | 1 | P11157 | Ribonuclec | P11157 [62 | 0 | 1846.915 | 0 | 0.44 | 0.00125 | 8.37 | 88 | 112 | 10.6 | 9.06 | | High | High | High | High | High | High | High | High | High | High | High | 0.000801 | 0.007216 | 2.71 |
| FALSE | High | APTPQEK | [K].APTPQI | 1xPhospho | Q3THK3 1x | 0.091627 | 0.007146 | 1 | 1 | 1 | Q3THK3 | General tr | Q3THK3 [3 | 0 | 1308.696 | 0 | -0.1 | 0.607616 | 20.58 | 104 | 96 | 12.6 | 12.34 | | High | High | High | High | High | High | High | High | High | High | High | 0.003659 | 0.03901 | 1.82 |
| FALSE | High | APPGAPGP | [R].APPGAI | 2xPhospho | Q8CCS6 2x | 0.000379 | 0 | 1 | 3 | 1 | Q8CCS6 | Polyadenyl | Q8CCS6 [7 | 0 | 4938.254 | 0 | -0.59 | 0.1363 | 2.79 | 117.7 | 82.3 | 17.09 | 14.99 | | High | High | High | High | High | High | High | High | High | High | High | 0 | 8.04E-05 | 4.27 |
| FALSE | High | AMALPSSG | [K].AMALP | 1xPhospho | P14901 1xI | 2.36E-05 | 0 | 1 | 2 | 1 | P14901 | Heme oxyg | P14901 [1 | 0 | 3052.514 | 0 | -1.45 | 0.000756 | 2.25 | 142 | 58 | 9.18 | 14.74 | | High | High | High | High | High | High | High | High | High | High | High | 0 | 3.54E-06 | 4.7 |
| FALSE | High | APPGAPGP | [R].APPGAI | 1xDeamida | Q8CCS6 1x | 1.15E-06 | 0 | 1 | 3 | 2 | Q8CCS6 | Polyadenyl | Q8CCS6 [7 | 0 | 4859.271 | 0 | -1.26 | 0.002068 | 5 | 141.1 | 58.9 | 16.87 | 29.99 | | High | High | High | High | High | High | High | High | High | High | High | 0 | 1.18E-07 | 5.4 |
| FALSE | High | APPAAPAA | [R].APPAAI | 1xPhospho | A2A7S8-2 | 9.79E-08 | 0 | 1 | 1 | 3 | A2A7S8-2 | Isoform 2 ( | A2A7S8-2 [ | 0 | 2020.969 | 0 | 1.22 | 0.00041 | 7.37 | 59.2 | 140.8 | 5.12 | 4.8 | | High | High | High | High | High | High | High | High | High | High | High | 0 | 7.45E-09 | 4.14 |
| FALSE | High | ANGQENG | [K].ANGQE | 1xDeamida | P28667 1xI | 0.047488 | 0.00354 | 1 | 1 | 2 | P28667 | MARCKS-re | P28667 [2 | 1 | 2433.192 | 0 | -1.29 | 0.000255 | 0.97 | 144.8 | 55.2 | 5.09 | 7.46 | | High | High | High | High | High | High | High | High | High | High | High | 0.003291 | 0.03599 | 3.52 |
| FALSE | High | ANGSQPTC | [K].ANGSQ | 1xPhospho | Q62311 1x | 2.6E-05 | 0 | 1 | 1 | 2 | Q62311 | Transcripti | Q62311 [6 | 0 | 1787.853 | 0 | -0.2 | 0.058378 | 5.84 | 106.4 | 93.6 | 6.62 | 13.29 | | High | High | High | High | High | High | High | High | High | High | High | 0 | 3.94E-06 | 3.95 |
| FALSE | High | ANGSQPTC | [K].ANGSQ | 1xDeamida | Q62311 1x | 2.95E-05 | 0 | 1 | 1 | 3 | Q62311 | Transcripti | Q62311 [6 | 0 | 1788.837 | 0 | -0.42 | 0.004633 | 5.01 | 113.7 | 86.3 | 6.4 | 2.41 | | High | High | High | High | High | High | High | High | High | High | High | 0 | 4.58E-06 | 3.47 |
| FALSE | High | ANGTPASL | [K].ANGTP | 1xPhospho | E9Q5C9 1x | 2.8E-05 | 0 | 1 | 2 | 2 | E9Q5C9 | Nucleolar a | E9Q5C9 [5 | 0 | 1567.825 | 0 | -0.25 | 0.076909 | 9.96 | 104.8 | 95.2 | 7.35 | 9.45 | | High | High | High | High | High | High | High | High | High | High | High | 0.000571 | 0.005075 | 2.1 |
| FALSE | High | ANGTPASL | [K].ANGTP | 1xDeamida | E9Q5C9 1x | 0.013426 | 0.000976 | 1 | 2 | 2 | E9Q5C9 | Nucleolar a | E9Q5C9 [5 | 0 | 1568.809 | 0 | -0.21 | 0.054842 | 6.75 | 107.8 | 92.2 | 10.09 | 5.81 | | High | High | High | High | High | High | High | High | High | High | High | 0.000505 | 0.004378 | 2.28 |
| FALSE | High | ANGTPASL | [K].ANGTP | 2xPhospho | E9Q5C9 2x | 0.120211 | 0.009851 | 1 | 2 | 1 | E9Q5C9 | Nucleolar a | E9Q5C9 [5 | 0 | 1647.791 | 0 | -0.31 | 0.027267 | 7.88 | 111.6 | 88.4 | 7.05 | 8.94 | | High | High | High | High | High | High | High | High | High | High | High | 0.004907 | 0.05355 | 1.55 |
| FALSE | High | ANHLGDTD | [K].ANHLG | 1xPhospho | Q60949 1x | 1.93E-05 | 0 | 1 | 5 | 1 | Q60949 | TBC1 doma | Q60949 [7 | 0 | 1862.941 | 0 | -1.08 | 0.000158 | 0.57 | 136.6 | 63.4 | 6.42 | 8.93 | | High | High | High | High | High | High | High | High | High | High | High | 0 | 2.84E-06 | 2.37 |
| FALSE | High | ANIAQEQC | [K].ANIAQI | 1xPhospho | P28229 1xI | 4.94E-05 | 0 | 1 | 2 | 1 | P28229 | Gap juncti | P28229 [3 | 0 | 3186.505 | 0 | -1.42 | 0.000158 | 1.45 | 145 | 55 | 5.79 | 20.67 | | High | High | High | High | High | High | High | High | High | High | High | 0 | 8.14E-06 | 2.42 |
| FALSE | High | ANIEPCDSC | [R].ANIEPC | 1xCarbami | A0A1B0GR | 2.45E-09 | 0 | 1 | 1 | 2 | A0A1B0GR | RIKEN cDN | A0A1B0GR | 0 | 2681.289 | 0 | 0.82 | 0.004356 | 11.04 | 71.5 | 128.5 | 4.48 | 11.07 | | High | High | High | High | High | High | High | High | High | High | High | 0 | 1.18E-10 | 6.04 |
| FALSE | High | ANLSPDR | [K].ANLSPI | 1xPhospho | G3X8T2 1x | 0.046687 | 0.00354 | 1 | 4 | 1 | G3X8T2 | RIKEN cDN | G3X8T2 [8 | 0 | 1081.524 | 0 | -0.54 | 0.001033 | 2.58 | 117 | 83 | 4.69 | 5.2 | | High | High | High | High | High | High | High | High | High | High | High | 0.001812 | 0.01785 | 1.01 |
| FALSE | High | ANNLHSGI | [R].ANNLH | 1xPhospho | B9EJ86 1xI | 0.002232 | 0.000109 | 1 | 2 | 1 | B9EJ86 | Oxysterol-l | B9EJ86 [3 | 0 | 2482.12 | 0 | 0.55 | 0.00759 | 23.52 | 83.4 | 116.6 | 12.05 | 9.4 | | High | High | High | High | High | High | High | High | High | High | High | 4.94E-05 | 0.00059 | 3.12 |
| FALSE | High | ANNSGVSL | [K].ANNSG | 1xCarbami | S4R1P5 1xI | 3.54E-05 | 0 | 1 | 7 | 1 | S4R1P5 | Dystonin O | S4R1P5 [4 | 0 | 2567.367 | 0 | -0.31 | 0.056967 | 7.9 | 117.1 | 82.9 | 11.78 | 23.98 | | High | High | High | High | High | High | High | High | High | High | High | 0 | 5.58E-06 | 2.42 |
| FALSE | High | ANPDSLEP/ | [K].ANPDSI | 1xDeamida | A2CG76 1x | 0.002153 | 5.66E-05 | 1 | 4 | 5 | A2CG76 | Histone-lys | A2CG76 [1 | 0 | 4958.279 | 0 | -0.12 | 0.362023 | 15.18 | 110.5 | 89.5 | 14.06 | 12.81 | | High | High | High | High | High | High | High | High | High | High | High | 0.000183 | 0.001257 | 4.85 |
| FALSE | High | ANGPLSPS | [R].ANGPL | 1xPhospho | Q9Z2B9 1x | 0.071071 | 0.005247 | 1 | 1 | 1 | Q9Z2B9 | Ribosomal | Q9Z2B9 [7 | 0 | 1051.502 | 0 | 2.99 | 0.000616 | 20.53 | 21.8 | 178.2 | 7.52 | 11.42 | | High | High | High | High | High | High | High | High | High | High | High | 0.002703 | 0.02886 | 1.51 |
| FALSE | High | ANPTTGSLI | [R].ANPTT( | 1xDeamida | Q922X9 1x | 0.016451 | 0.001124 | 1 | 3 | 1 | Q922X9 | Protein arg | Q922X9 [8 | 0 | 3313.452 | 0 | 0.19 | 0.058592 | 13.86 | 97.1 | 102.9 | 10.89 | 7.58 | | High | High | High | High | High | High | High | High | High | High | High | 0.000585 | 0.005468 | 3.89 |
| FALSE | High | ANSAEGGN | [R].ANSAE( | 1xDeamida | Q80XK6 1x | 4.45E-07 | 0 | 1 | 1 | 1 | Q80XK6 | Autophagy | Q80XK6 [1 | 0 | 3399.474 | 0 | 0.07 | 0.806051 | 7.1 | 101 | 99 | 12.12 | 15.73 | | High | High | High | High | High | High | High | High | High | High | High | 0 | 4.11E-08 | 6.02 |
| FALSE | High | ANSDTDLV | [R].ANSDTI | 1xPhospho | Q6I6G8 1xI | 0.000132 | 0 | 1 | 3 | 1 | Q6I6G8 | E3 ubiquiti | Q6I6G8 [4 | 0 | 1703.769 | 0 | 0.97 | 0.00102 | 8.39 | 70.6 | 129.4 | 7.76 | 7.42 | | High | High | High | High | High | High | High | High | High | High | High | 0 | 2.44E-05 | 2.92 |
| FALSE | High | ANSEASSSE | [R].ANSEA | 1xPhospho | Q6PAL7 1x | 0.000745 | 0 | 1 | 1 | 1 | Q6PAL7 | AT-hook D | Q6PAL7 [1 | 0 | 2436.154 | 0 | -0.62 | 0.010673 | 2.05 | 119.4 | 80.6 | 9.86 | 5.99 | | High | High | High | High | High | High | High | High | High | High | High | 0 | 0.000172 | 2.87 |
| FALSE | High | ANSFVGTA | [R].ANSFV( | 1xPhospho | Q3UGN6 1 | 9.76E-07 | 0 | 1 | 10 | 2 | Q3UGN6 | Putative ur | Q3UGN6 [ | 0 | 2592.336 | 0 | 0.56 | 0.002625 | 4.3 | 81.8 | 118.2 | 5.92 | 4.87 | | High | High | High | High | High | High | High | High | High | High | High | 0 | 9.9E-08 | 5.53 |
| FALSE | High | ANSFVGTA | [R].ANSFV( | 1xDeamida | Q3UGN6 1 | 2.63E-05 | 0 | 1 | 10 | 4 | Q3UGN6 | Putative ur | Q3UGN6 [ | 0 | 2593.32 | 0 | 0.41 | 0.051602 | 30.24 | 75.1 | 124.9 | 30.14 | 10.96 | | High | High | High | High | High | High | High | High | High | High | High | 0 | 4E-06 | 4.14 |
| FALSE | High | ANSGGGGI | [R].ANSGG | 1xPhospho | Q9CXK9 1x | 4.73E-06 | 0 | 1 | 3 | 1 | Q9CXK9 | RNA-bindir | Q9CXK9 [1 | 0 | 1864.895 | 0 | 0.48 | 0.033571 | 18.14 | 83.4 | 116.6 | 6.7 | 10.06 | | High | High | High | High | High | High | High | High | High | High | High | 0 | 5.85E-07 | 1.8 |
| FALSE | High | ANSLASEK | [R].ANSLAS | 2xPhospho | Q8BYJ6 2xI | 0.016021 | 0.001124 | 1 | 9 | 1 | Q8BYJ6 | TBC1 doma | Q8BYJ6 [6 | 0 | 1437.679 | 0 | 1.56 | 0.004121 | 6.16 | 49.6 | 150.4 | 6.01 | 19.36 | | High | High | High | High | High | High | High | High | High | High | High | 0.000585 | 0.005313 | 1.95 |
| FALSE | High | ANSLPSTP/ | [K].ANSLPS | 1xPhospho | B0V2M3 1 | 0.006478 | 0.00062 | 1 | 5 | 1 | B0V2M3 | Zinc finger | B0V2M3 [5 | 0 | 1622.892 | 0 | -0.38 | 0.114618 | 10.26 | 110.5 | 89.5 | 5.51 | 17.68 | | High | High | High | High | High | High | High | High | High | High | High | 0.000275 | 0.001922 | 2.37 |
| FALSE | High | ANSLVTLGS | [R].ANSLVT | 1xPhospho | E9Q1M1 1 | 0.054383 | 0.003951 | 1 | 1 | 1 | E9Q1M1 | PDZ domai | E9Q1M1 [9 | 0 | 1454.757 | 0 | 0.95 | 0.004156 | 17.74 | 63.4 | 136.6 | 12.54 | 11.92 | | High | High | High | High | High | High | High | High | High | High | High | 0.002045 | 0.0212 | 1.62 |
| FALSE | High | ANSPLESIV | [R].ANSPLE | 2xCarbami | Q9WUF3 1 | 7.22E-09 | 0 | 1 | 1 | 5 | Q9WUF3 | CASP8-ass( | Q9WUF3 [ | 0 | 3503.657 | 0 | -0.69 | 0.011547 | 2.9 | 123 | 77 | 10.15 | 13.98 | | High | High | High | High | High | High | High | High | High | High | High | 0 | 3.97E-10 | 5.39 |
| FALSE | High | ANSQDSLA | [R].ANSQD | 1xPhospho | B2RWW1 1 | 0.000289 | 0 | 1 | 3 | 2 | B2RWW1 | Microtubu | B2RWW1 [ | 0 | 1357.631 | 0 | -0.21 | 0.053096 | 11.86 | 107.9 | 92.1 | 2.97 | 13.29 | | High | High | High | High | High | High | High | High | High | High | High | 0 | 5.89E-05 | 2.48 |
| FALSE | High | ANSSGLYK | [R].ANSSGI | 1xPhospho | Q99KZ6 1x | 0.001313 | 0 | 1 | 2 | 2 | Q99KZ6 | Zinc finger | Q99KZ6 [1 | 0 | 1377.718 | 0 | -0.6 | 0.004102 | 5.65 | 117.7 | 82.3 | 7.69 | 4.53 | | High | High | High | High | High | High | High | High | High | High | High | 0 | 0.000325 | 2.15 |
| FALSE | High | ANSTQEPS | [K].ANSTQ | 1xPhospho | Q2Q7P0 1x | 9.91E-06 | 0 | 1 | 1 | 2 | Q2Q7P0 | ENH1 OS=I | Q2Q7P0 [3 | 0 | 4058.925 | 0 | 0.27 | 0.341735 | 1.79 | 99.9 | 100.1 | 9.8 | 13.06 | | High | High | High | High | High | High | High | High | High | High | High | 0 | 1.33E-06 | 3.48 |
| FALSE | High | ANQISPSNS | [K].ANQISI | 1xPhospho | Q67FY2 1x | 0.00642 | 0.00062 | 1 | 2 | 1 | Q67FY2 | B-cell CLL/l | Q67FY2 [8 | 0 | 1783.936 | 0 | 0.6 | 0.04534 | 7.77 | 79.5 | 120.5 | 12.11 | 12.27 | | High | High | High | High | High | High | High | High | High | High | High | 0.000275 | 0.001905 | 2.6 |
| FALSE | High | ANSTQEPS | [K].ANSTQ | 2xPhospho | Q8CI51 2xI | 2.87E-05 | 0 | 1 | 2 | 3 | Q8CI51 | PDZ and LI | Q8CI51 [3 | 0 | 4154.887 | 0 | 1.29 | 0.002798 | 34.11 | 58 | 142 | 14.31 | 8.82 | | High | High | High | High | High | High | High | High | High | High | High | 0.000204 | 0.001357 | 4.1 |
| FALSE | High | ANGPHLYS | [K].ANGPH | 1xPhospho | Q3U2C9 1x | 0.002313 | 0.000109 | 1 | 5 | 1 | Q3U2C9 | PHD finger | Q3U2C9 [5 | 0 | 2123.96 | 0 | 0.06 | 0.649576 | 5.11 | 98.3 | 101.7 | 7.71 | 7.03 | | High | High | High | High | High | High | High | High | High | High | High | 4.94E-05 | 0.000612 | 2.36 |

| | | | | | | | | | | | | | | | | | | | | | | | | | | | | | | | | | | | | | | |
|---|---|---|---|---|---|---|---|---|---|---|---|---|---|---|---|---|---|---|---|---|---|---|---|---|---|---|---|---|---|---|---|---|---|---|---|---|---|---|
| FALSE | High | ANGHSTDC | [R].ANGHS | 1xCarbami | F7AA26 1x | 0.004911 | 0.000504 | 1 | 11 | 2 | F7AA26 | Paralemmi | F7AA26 [3 | 0 | 3792.661 | 0 | -0.17 | 0.072512 | 12.11 | 106.2 | 93.8 | 7.57 | 7.46 | | High | High | High | High | High | High | High | High | High | High | High | 0.000225 | 0.001415 | 3.26 |
| FALSE | High | AMDDPCSI | [R].AMDDF | 1xCarbami | P47810 1x | 1.43E-07 | 0 | 1 | 1 | 2 | P47810 | Wee1-like | P47810 [15 | 0 | 2694.198 | 0 | -1.27 | 0.002403 | 3.53 | 143.6 | 56.4 | 12.42 | 5.9 | | High | High | High | High | High | High | High | High | High | High | High | 0 | 7.19E-08 | 4.65 |
| FALSE | High | AMDDPCSI | [R].AMDDF | 1xCarbami | P47810 2x | 0.001158 | 0 | 1 | 1 | 1 | P47810 | Wee1-like | P47810 [15 | 0 | 2758.17 | 0 | -0.66 | 0.004486 | 7.61 | 122.6 | 77.4 | 9.29 | 14.13 | | High | High | High | High | High | High | High | High | High | High | High | 0 | 0.000281 | 2.76 |
| FALSE | High | AMEVLPLA | [R].AMEVL | 1xPhospho | E9Q166 1x | 0.000924 | 0 | 1 | 1 | 1 | E9Q166 | ATPase fan | E9Q166 [9 | 0 | 1799.969 | 0 | -0.91 | 0.002127 | 1.81 | 128.2 | 71.8 | 6.61 | 10.29 | | High | High | High | High | High | High | High | High | High | High | High | 0 | 0.000219 | 2.48 |
| FALSE | High | AMEVLPLA | [R].AMEVL | 1xPhospho | E9Q166 1x | 5.35E-05 | 0 | 1 | 1 | 1 | E9Q166 | ATPase fan | E9Q166 [9 | 0 | 1815.964 | 0 | -0.66 | 0.010089 | 4.99 | 121.2 | 78.8 | 10.58 | 9.56 | | High | High | High | High | High | High | High | High | High | High | High | 0 | 8.91E-06 | 3.32 |
| FALSE | High | AMGGNAG | [K].AMGGI | 1xPhospho | Q9QYE6 1x | 9.25E-07 | 0 | 1 | 1 | 2 | Q9QYE6 | Golgin sub | Q9QYE6 [1 | 0 | 2866.344 | 0 | 0.31 | 0.011772 | 2.7 | 85.3 | 114.7 | 8.88 | 8.28 | | High | High | High | High | High | High | High | High | High | High | High | 0 | 9.27E-08 | 4.56 |
| FALSE | High | AMLDSGV | [R].AMLDS | 1xPhospho | Q3UY32 1x | 0.040373 | 0.002982 | 1 | 3 | 1 | Q3UY32 | SNARE-ass | Q3UY32 [1 | 0 | 2044.023 | 0 | -0.1 | 0.264816 | 3.76 | 104.3 | 95.7 | 4.44 | 10.03 | | High | High | High | High | High | High | High | High | High | High | High | 0.00155 | 0.01503 | 2.04 |
| FALSE | High | AMLDSGV | [R].AMLDS | 1xPhospho | Q3UY32 1x | 0.000442 | 0 | 1 | 3 | 3 | Q3UY32 | SNARE-ass | Q3UY32 [1 | 0 | 2060.018 | 0 | -0.08 | 0.690622 | 3.06 | 102.6 | 97.4 | 2.86 | 25 | | High | High | High | High | High | High | High | High | High | High | High | 0.000275 | 0.002169 | 2.15 |
| FALSE | High | AMPSNSPV | [K].AMPSN | 1xPhospho | Q8BH80 1x | 0.062209 | 0.004646 | 1 | 4 | 2 | Q8BH80 | Putative ur | Q8BH80 [2 | 0 | 2040.06 | 0 | -0.11 | 0.594602 | 18.99 | 102.1 | 97.9 | 11.76 | 9.08 | | High | High | High | High | High | High | High | High | High | High | High | 0.002403 | 0.02538 | 2.56 |
| FALSE | High | AMQAAGC | [K].AMQA/ | 1xPhospho | P26369 1x | 0.011447 | 0.000816 | 1 | 4 | 2 | P26369 | Splicing fac | P26369 [10 | 0 | 4130.094 | 0 | 0.24 | 0.411041 | 10.21 | 95.6 | 104.4 | 12.22 | 13.27 | | High | High | High | High | High | High | High | High | High | High | High | 0.000656 | 0.005815 | 3.33 |
| FALSE | High | AMSEVTSL | [R].AMSEV | 1xPhospho | Q99KG5 1x | 0.016163 | 0.001124 | 1 | 4 | 1 | Q99KG5 | Lipolysis-st | Q99KG5 [3 | 0 | 1984.867 | 0 | -1.84 | 0.000986 | 1.76 | 157.4 | 42.6 | 9.51 | 12.54 | | High | High | High | High | High | High | High | High | High | High | High | 0.000585 | 0.005342 | 1.74 |
| FALSE | High | AMSGSWE | [R].AMSGS | 1xPhospho | Q9CQV4 1x | 0.041425 | 0.003126 | 1 | 1 | 2 | Q9CQV4 | Reticuloph | Q9CQV4 [2 | 0 | 1232.533 | 0 | 0.36 | 0.02937 | 11.98 | 87.4 | 112.6 | 1.68 | 10.83 | | High | High | High | High | High | High | High | High | High | High | High | 0.001609 | 0.01555 | 1.24 |
| FALSE | High | AMSGSWE | [R].AMSGS | 1xPhospho | Q9CQV4 1x | 0.017192 | 0.001204 | 1 | 1 | 3 | Q9CQV4 | Reticuloph | Q9CQV4 [2 | 0 | 1248.528 | 0 | 0.5 | 0.004236 | 1.85 | 82.8 | 117.2 | 4.15 | 6.74 | | High | High | High | High | High | High | High | High | High | High | High | 0.004411 | 0.04726 | 2.06 |
| FALSE | High | AMSQGEIT | [K].AMSQG | 1xPhospho | D3Z3A8 1x | 0.006306 | 0.00062 | 1 | 3 | 1 | D3Z3A8 | Unconvent | D3Z3A8 [1 | 0 | 1518.764 | 0 | 0.27 | 0.149303 | 20 | 95.9 | 104.1 | 6.3 | 18.5 | | High | High | High | High | High | High | High | High | High | High | High | 0.000275 | 0.001879 | 1.51 |
| FALSE | High | ANGLPSGP | [K].ANGLP | 2xPhospho | Q7TQG1 2x | 0.001544 | 5.66E-05 | 1 | 6 | 4 | Q7TQG1 | Pleckstrin I | Q7TQG1 [2 | 0 | 2642.137 | 0 | 0.88 | 0.000699 | 9.12 | 69 | 131 | 16.28 | 8.23 | | High | High | High | High | High | High | High | High | High | High | High | 2.58E-05 | 0.000387 | 3.43 |
| FALSE | High | AMSTTSVT | [R].AMSTT | 1xPhospho | Q62418 1x | 0.000311 | 0 | 1 | 3 | 3 | Q62418 | Drebrin-lik | Q62418 [2 | 0 | 1919.955 | 0 | -0.1 | 0.024344 | 5.68 | 102.3 | 97.7 | 4.98 | 9.88 | | High | High | High | High | High | High | High | High | High | High | High | 0 | 6.45E-05 | 2.7 |
| FALSE | High | AMTPPLPV | [K].AMTPP | 1xPhospho | O08715-4 | 0.0026 | 0.000109 | 1 | 7 | 1 | O08715-4 | Isoform 4 c | O08715-4 | 0 | 2522.349 | 0 | 0.86 | 0.029016 | 1.66 | 73.6 | 126.4 | 6.16 | 23.78 | | High | High | High | High | High | High | High | High | High | High | High | 4.94E-05 | 0.000694 | 3.72 |
| FALSE | High | AMTPPLPV | [K].AMTPP | 1xDeamida | O08715-4 | 0.001544 | 5.66E-05 | 1 | 7 | 2 | O08715-4 | Isoform 4 c | O08715-4 | 0 | 2523.333 | 0 | 0.75 | 0.037177 | 9.66 | 72.7 | 127.3 | 7.08 | 20.11 | | High | High | High | High | High | High | High | High | High | High | High | 2.58E-05 | 0.000387 | 3.25 |
| FALSE | High | AMTPPLPV | [K].AMTPP | 1xPhospho | O08715-4 | 0.000372 | 0 | 1 | 7 | 1 | O08715-4 | Isoform 4 c | O08715-4 | 0 | 2538.344 | 0 | 0.88 | 0.004153 | 6.77 | 70.7 | 129.3 | 4.75 | 13.28 | | High | High | High | High | High | High | High | High | High | High | High | 0 | 7.85E-05 | 4.27 |
| FALSE | High | ANATNSPE | [K].ANATN | 1xPhospho | Q0VEE6 1x | 0.001021 | 0 | 1 | 1 | 1 | Q0VEE6 | Zinc finger | Q0VEE6 [6 | 0 | 1640.805 | 0 | 2.18 | 0.00032 | 5.9 | 35.9 | 164.1 | 4.94 | 7.65 | | High | High | High | High | High | High | High | High | High | High | High | 0 | 0.000245 | 1.42 |
| FALSE | High | ANAVDQSI | [K].ANAVD | 1xPhospho | B2RXR2 1x | 0.032548 | 0.002167 | 1 | 4 | 1 | B2RXR2 | Phosphatic | B2RXR2 [8 | 0 | 2175.06 | 0 | 0.21 | 0.167585 | 11.53 | 90.4 | 109.6 | 10.57 | 21.04 | | High | High | High | High | High | High | High | High | High | High | High | 0.001151 | 0.01186 | 2.94 |
| FALSE | High | ANDQDSLI | [K].ANDQD | 1xPhospho | Q4TVN0 1x | 0.000876 | 0 | 1 | 4 | 1 | Q4TVN0 | TBC1 doma | Q4TVN0 [2 | 0 | 1710.863 | 0 | 0.8 | 0.006042 | 16.68 | 73 | 127 | 18.16 | 7.22 | | High | High | High | High | High | High | High | High | High | High | High | 0 | 0.000206 | 2.77 |
| FALSE | High | ANEHGPAV | [K].ANEHG | 1xCarbami | E9Q444 1x | 0.005615 | 0.00062 | 1 | 9 | 1 | E9Q444 | Zinc finger | E9Q444 [5 | 1 | 3935.698 | 0 | 0.12 | 0.759311 | 12.41 | 99.4 | 100.6 | 8.87 | 15.32 | | High | High | High | High | High | High | High | High | High | High | High | 0.000275 | 0.001646 | 4.89 |
| FALSE | High | ANEISVIQN | [K].ANEISV | 1xPhospho | Q6P5E3 1x | 0.000573 | 0 | 1 | 1 | 3 | Q6P5E3 | Sal-like 1 (I | Q6P5E3 [1 | 0 | 3972.073 | 0 | -1.14 | 0.001334 | 4.01 | 139.1 | 60.9 | 8.25 | 10.99 | | High | High | High | High | High | High | High | High | High | High | High | 0 | 0.000128 | 3.93 |
| FALSE | High | ANFDEENA | [R].ANFDE | 1xPhospho | Q810A7 1x | 2.78E-06 | 0 | 1 | 2 | 57 | Q810A7 | ATP-depen | Q810A7 [6 | 0 | 4115.739 | 0 | -0.09 | 0.448961 | 10.63 | 103.5 | 96.5 | 7.25 | 4.72 | | High | High | High | High | High | High | High | High | High | High | High | 0 | 1.47E-06 | 4.83 |
| FALSE | High | ANFNAMF | [R].ANFNA | 1xCarbami | D3Z7I0 1xP | 0.049971 | 0.003641 | 1 | 2 | 3 | D3Z7I0 | General tra | D3Z7I0 [92 | 0 | 3457.558 | 0 | 0.18 | 0.749046 | 13.85 | 98.5 | 101.5 | 14.62 | 8.7 | | High | High | High | High | High | High | High | High | High | High | High | 0.002187 | 0.02296 | 2.97 |
| FALSE | High | ANFNAMF | [R].ANFNA | 1xCarbami | D3Z7I0 1xP | 0.001629 | 5.66E-05 | 1 | 2 | 2 | D3Z7I0 | General tra | D3Z7I0 [92 | 0 | 3473.553 | 0 | 0.48 | 0.078092 | 39.36 | 83.5 | 116.5 | 19.67 | 16.16 | | High | High | High | High | High | High | High | High | High | High | High | 2.58E-05 | 0.000511 | 4.75 |
| FALSE | High | ANGHSTDC | [R].ANGHS | 1xCarbami | F7AA26 1x | 5.47E-06 | 0 | 1 | 11 | 2 | F7AA26 | Paralemmi | F7AA26 [3 | 0 | 3790.693 | 0 | 0.54 | 0.387054 | 81.13 | 65.7 | 134.3 | 26.62 | 48.39 | | High | High | High | High | High | High | High | High | High | High | High | 0 | 5.86E-06 | 4.33 |
| FALSE | High | ANGHSTDC | [R].ANGHS | 1xCarbami | F7AA26 1x | 0.000628 | 0 | 1 | 11 | 4 | F7AA26 | Paralemmi | F7AA26 [3 | 0 | 3791.677 | 0 | -0.23 | 0.448961 | 35.79 | 107.9 | 92.1 | 27.32 | 12.97 | | High | High | High | High | High | High | High | High | High | High | High | 0 | 0.000198 | 4.28 |
| FALSE | High | AMSTTSVT | [R].AMSTT | 1xPhospho | Q62418 1x | 0.003526 | 0.00021 | 1 | 3 | 1 | Q62418 | Drebrin-lik | Q62418 [2 | 0 | 1935.95 | 0 | 0.61 | 0.149743 | 24.81 | 79.8 | 120.2 | 14.15 | 62.19 | | High | High | High | High | High | High | High | High | High | High | High | 9.42E-05 | 0.000981 | 1.88 |
| FALSE | High | ANSTSDSM | [R].ANSTS | 1xPhospho | Q9CQC5 1x | 0.061176 | 0.004646 | 1 | 1 | 3 | Q9CQC5 | Cdc42 effe | Q9CQC5 [8 | 0 | 2450.193 | 0 | 1.11 | 0.013397 | 20.54 | 67.7 | 132.3 | 10.72 | 22.36 | | High | High | High | High | High | High | High | High | High | High | High | 0.002354 | 0.02433 | 2.76 |
| FALSE | High | ANSTSDSM | [R].ANSTS | 1xPhospho | Q9CQC5 1x | 6.46E-08 | 0 | 1 | 1 | 2 | Q9CQC5 | Cdc42 effe | Q9CQC5 [8 | 0 | 2466.188 | 0 | 2.52 | 0.00118 | 11.75 | 29 | 171 | 9.74 | 15.69 | | High | High | High | High | High | High | High | High | High | High | High | 0 | 4.7E-09 | 4.51 |
| FALSE | High | ANSWQLVI | [K].ANSWC | 1xPhospho | B1AZP2 1x | 8.35E-06 | 0 | 1 | 9 | 5 | B1AZP2 | Disks large | B1AZP2 [9 | 0 | 1939.993 | 0 | -0.58 | 0.001618 | 1.82 | 120.6 | 79.4 | 2.05 | 9.28 | | High | High | High | High | High | High | High | High | High | High | High | 0 | 3.42E-06 | 4.17 |
| FALSE | High | APGELEEQ | [K].APGELE | 1xPhospho | B0V2M3 1x | 0.000176 | 0 | 1 | 3 | 1 | B0V2M3 | Zinc finger | B0V2M3 [1 | 0 | 2424.158 | 0 | -0.48 | 0.013178 | 6.24 | 112 | 88 | 4.97 | 14.5 | | High | High | High | High | High | High | High | High | High | High | High | 0 | 3.39E-05 | 2.66 |
| FALSE | High | APGEQDGI | [R].APGEQ | 1xPhospho | P25206 1x | 8.9E-06 | 0 | 2 | 3 | 3 | P25206; Q | DNA replic | P25206 [5 | 0 | 4011.782 | 0 | -0.45 | 0.003498 | 0.6 | 117.9 | 82.1 | 6.67 | 13.43 | Shared | High | High | High | High | High | High | High | High | High | High | High | 0 | 7.42E-05 | 4.46 |
| FALSE | High | APGESSAIS | [R].APGES | 1xPhospho | Q69ZR2 1x | 2.75E-05 | 0 | 1 | 2 | 1 | Q69ZR2 | E3 ubiquiti | Q69ZR2 [1 | 0 | 3373.685 | 0 | 2.16 | 0.001455 | 15.15 | 36.2 | 163.8 | 8.6 | 15.31 | | High | High | High | High | High | High | High | High | High | High | High | 0 | 4.21E-06 | 5.26 |
| FALSE | High | APGPPTPG | [R].APGPP | 1xPhospho | Q8C708 1x | 0.002844 | 0.00016 | 1 | 1 | 1 | Q8C708 | Transmem | Q8C708 [1 | 0 | 1615.804 | 0 | 0.76 | 0.001067 | 3.53 | 75.8 | 124.2 | 6.72 | 8.07 | | High | High | High | High | High | High | High | High | High | High | High | 7.19E-05 | 0.000772 | 2.37 |
| FALSE | High | APGPPTPG | [R].APGPP | 2xPhospho | Q8C708 2x | 0.013074 | 0.000889 | 1 | 1 | 1 | Q8C708 | Transmem | Q8C708 [1 | 0 | 1695.771 | 0 | 0.6 | 0.022767 | 14.49 | 79.4 | 120.6 | 14.98 | 7.9 | | High | High | High | High | High | High | High | High | High | High | High | 0.000467 | 0.004237 | 2.99 |
| FALSE | High | APGSPGAG | [K].APGSP | 1xPhospho | Q56A10 1x | 0.001704 | 5.66E-05 | 1 | 2 | 1 | Q56A10 | Zinc finger | Q56A10 [6 | 0 | 1842.952 | 0 | 1.79 | 0.002145 | 13.77 | 44.3 | 155.7 | 4.12 | 16.79 | | High | High | High | High | High | High | High | High | High | High | High | 2.58E-05 | 0.000433 | 2.36 |
| FALSE | High | APGSPVSSI | [K].APGSP | 1xPhospho | Q8VBT9 1x | 0.003167 | 0.00016 | 1 | 8 | 4 | Q8VBT9 | Tether con | Q8VBT9 [1 | 0 | 3171.541 | 0 | -0.26 | 0.014436 | 1.9 | 109.6 | 90.4 | 13.29 | 12.33 | | High | High | High | High | High | High | High | High | High | High | High | 0.007598 | 0.08293 | 3.49 |
| FALSE | High | APGSPVSSI | [K].APGSP | 1xDeamida | Q8VBT9 1x | 0.011447 | 0.000816 | 1 | 8 | 4 | Q8VBT9 | Tether con | Q8VBT9 [1 | 0 | 3172.525 | 0 | -0.22 | 0.415677 | 22.61 | 107.7 | 92.3 | 19.16 | 14.81 | | High | High | High | High | High | High | High | High | High | High | High | 0.000437 | 0.003658 | 3.85 |
| FALSE | High | APGTPAPV | [R].APGTP/ | 2xPhospho | E9PVX6 2x | 0.002844 | 0.00016 | 1 | 2 | 1 | E9PVX6 | Proliferatic | E9PVX6 [1 | 0 | 2722.156 | 0 | 0 | 0.55351 | 36.49 | 102.5 | 97.5 | 21.03 | 80.65 | | High | High | High | High | High | High | High | High | High | High | High | 7.19E-05 | 0.000769 | 3.24 |
| FALSE | High | APGTPAPV | [R].APGTP/ | 1xPhospho | E9PVX6 1x | 2.54E-06 | 0 | 1 | 2 | 1 | E9PVX6 | Proliferatic | E9PVX6 [1 | 0 | 2855.357 | 0 | -0.22 | 0.038097 | 6.38 | 109 | 91 | 8.19 | 16.43 | | High | High | High | High | High | High | High | High | High | High | High | 0 | 2.9E-07 | 4.74 |
| FALSE | High | APGTPAPV | [R].APGTP/ | 1xPhospho | E9PVX6 1x | 1.41E-08 | 0 | 1 | 2 | 1 | E9PVX6 | Proliferatic | E9PVX6 [1 | 0 | 2871.352 | 0 | -0.4 | 0.053706 | 10 | 111.1 | 88.9 | 9.23 | 20.68 | | High | High | High | High | High | High | High | High | High | High | High | 0 | 8.45E-10 | 4.68 |
| FALSE | High | APGTPAPV | [R].APGTP/ | 2xPhospho | E9PVX6 2x | 0.002463 | 0.000109 | 1 | 2 | 1 | E9PVX6 | Proliferatic | E9PVX6 [1 | 0 | 2935.324 | 0 | 0.12 | 0.660137 | 21.89 | 95.6 | 104.4 | 9.04 | 20.53 | | High | High | High | High | High | High | High | High | High | High | High | 4.94E-05 | 0.000655 | 3.05 |
| FALSE | High | APHLSGTSI | [R].APHLS | 1xPhospho | Q3U0T7 1x | 0.052576 | 0.00384 | 1 | 7 | 1 | Q3U0T7 | Putative ur | Q3U0T7 [3 | 0 | 1435.771 | 0 | -1.55 | 0.000424 | 1.58 | 148.8 | 51.2 | 2.47 | 15.42 | | High | High | High | High | High | High | High | High | High | High | High | 0.001999 | 0.02044 | 1.76 |
| FALSE | High | APFKDEIDE | [R].APFKDE | 1xPhospho | A2AJK6 1x | 0.002419 | 0.000109 | 1 | 2 | 1 | A2AJK6 | Chromodo | A2AJK6 [18 | 1 | 2790.384 | 0 | -1.03 | 0.005673 | 7.46 | 137.3 | 62.7 | 10.76 | 16.5 | | High | High | High | High | High | High | High | High | High | High | High | 4.94E-05 | 0.000642 | 2.27 |
| FALSE | High | APHSPFYQ | [R].APHSPF | 2xPhospho | H9LAN6 2x | 0.016306 | 0.001124 | 1 | 6 | 1 | H9LAN6 | Histone-lys | H9LAN6 [4 | 0 | 4062.106 | 0 | -1.27 | 0.000161 | 1.34 | 141.2 | 58.8 | 5.27 | 13.12 | | High | High | High | High | High | High | High | High | High | High | High | 0.000585 | 0.005395 | 3.7 |
| FALSE | High | APLATGED | [K].APLATG | 1xPhospho | Q64152 1x | 5.38E-07 | 0 | 1 | 2 | 2 | Q64152 | Transcripti | Q64152 [1 | 0 | 3229.471 | 0 | -0.44 | 0.008052 | 6.5 | 113 | 87 | 3.93 | 9.97 | | High | High | High | High | High | High | High | High | High | High | High | 0 | 5.07E-08 | 5.6 |
| FALSE | High | APLATGED | [K].APLATG | 1xDeamida | Q64152 1x | 0.007605 | 0.000664 | 1 | 2 | 2 | Q64152 | Transcripti | Q64152 [1 | 1 | 3658.621 | 0 | -1.09 | 0.004079 | 8.44 | 133 | 67 | 7.89 | 17.14 | | High | High | High | High | High | High | High | High | High | High | High | 0.000344 | 0.00318 | 2.95 |
| FALSE | High | APLLSEPAS | [R].APLLSE | 1xPhospho | E9Q4K7 1x | 0.011447 | 0.000816 | 1 | 2 | 3 | E9Q4K7 | Kinesin fan | E9Q4K7 [1 | 0 | 2049.062 | 0 | 0.23 | 0.034731 | 7.28 | 93.7 | 106.3 | 2.37 | 8.48 | | High | High | High | High | High | High | High | High | High | High | High | 0.000449 | 0.003768 | 2.97 |
| FALSE | High | APLPPPLPE | [R].APLPPP | 1xPhospho | Q3U5C0 1x | 0.001489 | 5.66E-05 | 1 | 2 | 8 | Q3U5C0 | Putative ur | Q3U5C0 [6 | 0 | 3373.654 | 0 | 0.21 | 0.244688 | 20.8 | 100.1 | 99.9 | 13.45 | 19.7 | | High | High | High | High | High | High | High | High | High | High | High | 0.000275 | 0.002131 | 3.18 |
| FALSE | High | APISLSQDP | [R].APISLS | 1xCarbami | A2ARP8 1x | 0.005275 | 0.000599 | 1 | 3 | 1 | A2ARP8 | Microtubu | A2ARP8 [2 | 0 | 2763.286 | 0 | -1.2 | 0.001193 | 0.43 | 145 | 55 | 13.81 | 11.42 | | High | High | High | High | High | High | High | High | High | High | High | 0.000267 | 0.001535 | 2.85 |
| FALSE | High | APLSPEGEE | [R].APLSPE | 1xPhospho | G3X8Q1 1x | 2.31E-05 | 0 | 1 | 4 | 1 | G3X8Q1 | Calcineurir | G3X8Q1 [2 | 0 | 2860.381 | 0 | 0.03 | 0.311444 | 18.31 | 92.2 | 107.8 | 14.01 | 18.66 | | High | High | High | High | High | High | High | High | High | High | High | 0 | 3.45E-06 | 3.38 |
| FALSE | High | APLSPGVTT | [R].APLSPG | 1xPhospho | A3KGF9 1x | 0.000127 | 0 | 1 | 1 | 1 | A3KGF9 | Uncharacte | A3KGF9 [1 | 0 | 3255.471 | 0 | 0.56 | 0.027796 | 30.34 | 77.9 | 122.1 | 14.67 | 14.13 | | High | High | High | High | High | High | High | High | High | High | High | 0 | 2.35E-05 | 4.01 |
| FALSE | High | APLTLAGSI | [K].APLTLA | 1xPhospho | F6ZBR8 1x | 0.000302 | 0 | 2 | 5 | 1 | F6ZBR8; O | Protein Wi | F6ZBR8 [18 | 0 | 1690.954 | 0 | 0.07 | 0.345814 | 3.88 | 98.4 | 101.6 | 6.92 | 4.14 | Shared | High | High | High | High | High | High | High | High | High | High | High | 0 | 6.25E-05 | 3.24 |
| FALSE | High | APLVGSPVI | [K].APLVGS | 1xPhospho | Q6P5H2-2 | 7.82E-05 | 0 | 1 | 3 | 1 | Q6P5H2-2 | Isoform 2 c | Q6P5H2-2 | 0 | 2006.104 | 0 | 1.65 | 0.034837 | 40.95 | 43.8 | 156.2 | 23.41 | 41.78 | | High | High | High | High | High | High | High | High | High | High | High | 0 | 1.36E-05 | 3.61 |
| FALSE | High | APLVGSPVI | [K].APLVGS | 1xPhospho | Q6P5H2-2 | 6.09E-06 | 0 | 1 | 3 | 2 | Q6P5H2-2 | Isoform 2 c | Q6P5H2-2 | 0 | 2235.267 | 0 | 2.56 | 0.000411 | 14.06 | 28 | 172 | 10.13 | 9.53 | | High | High | High | High | High | High | High | High | High | High | High | 0 | 7.72E-07 | 3.28 |
| FALSE | High | APLVPHSG | [K].APLVPH | 1xPhospho | F8QZ85 1x | 0.014929 | 0.001049 | 1 | 2 | 3 | F8QZ85 | Ovum mut | F8QZ85 [2 | 0 | 1772.971 | 0 | 1.21 | 0.009897 | 21.32 | 58.8 | 141.2 | 7.31 | 20.53 | | High | High | High | High | High | High | High | High | High | High | High | 0.000555 | 0.004921 | 2.18 |
| FALSE | High | APLVPPGSI | [K].APLVPP | 1xPhospho | Q9JHU9 1x | 0.087983 | 0.006639 | 1 | 1 | 1 | Q9JHU9 | Inositol-3-p | Q9JHU9 [4 | 0 | 1943.072 | 0 | -1.59 | 0.001395 | 2.91 | 147 | 53 | 7.53 | 17.57 | | High | High | High | High | High | High | High | High | High | High | High | 0.003389 | 0.03695 | 1.53 |
| FALSE | High | APNSPDVLI | [K].APNSPI | 1xPhospho | P16460 1x | 0.001361 | 0 | 1 | 2 | 2 | P16460 | Argininosu | P16460 [21 | 0 | 1997.04 | 0 | -1.5 | 0.000432 | 3.76 | 146.4 | 53.6 | 5.19 | 29.16 | | High | High | High | High | High | High | High | High | High | High | High | 0 | 0.000336 | 3.04 |
| FALSE | High | APHWTSAS | [K].APHWT | 1xPhospho | P20357 1x | 0.00153 | 5.66E-05 | 1 | 9 | 1 | P20357 | Microtubu | P20357 [11 | 0 | 2910.401 | 0 | 0.57 | 0.049053 | 32.21 | 80.4 | 119.6 | 16.27 | 20.02 | | High | High | High | High | High | High | High | High | High | High | High | 2.58E-05 | 0.000385 | 2.66 |
| FALSE | High | APEPEPGLI | [R].APEPEF | 1xPhospho | Q9D8C8 1x | 8.46E-11 | 0 | 1 | 1 | 3 | Q9D8C8 | Protein ph | Q9D8C8 [3 | 0 | 2795.452 | 0 | 0.99 | 0.002936 | 20.32 | 66.8 | 133.2 | 7.14 | 9.44 | | High | High | High | High | High | High | High | High | High | High | High | 0 | 2.69E-12 | 5.42 |
| FALSE | High | APEPEPGLI | [R].APEPEF | 1xPhospho | Q9D8C8 1x | 4.17E-06 | 0 | 1 | 1 | 1 | Q9D8C8 | Protein ph | Q9D8C8 [3 | 0 | 2566.289 | 0 | 1.13 | 0.186556 | 18.88 | 64.1 | 135.9 | 41.13 | 47.39 | | High | High | High | High | High | High | High | High | High | High | High | 0 | 5.03E-07 | 4.52 |
| FALSE | High | APEPDVMS | [K].APEPD\ | 1xPhospho | B2RY04 1x | 7.47E-05 | 0 | 1 | 2 | 1 | B2RY04 | Dedicator ( | B2RY04 [1 | 0 | 1681.827 | 0 | 0.03 | 0.664303 | 20.18 | 97.3 | 102.7 | 12.26 | 13.42 | | High | High | High | High | High | High | High | High | High | High | High | 0 | 1.3E-05 | 2.89 |
| FALSE | High | ANTFPCTP | [R].ANTFP | 1xCarbami | A2A9C3 1x | 2.49E-05 | 0 | 1 | 3 | 1 | A2A9C3 | KICSTOR cc | A2A9C3 [2 | 0 | 2524.256 | 0 | 0.43 | 0.00041 | 0.84 | 85 | 115 | 4.5 | 1.52 | | High | High | High | High | High | High | High | High | High | High | High | 0 | 3.77E-06 | 3.7 |
| FALSE | High | ANTLSHFP | [R].ANTLSH | 1xCarbami | Q60949 1x | 7.08E-05 | 0 | 1 | 5 | 4 | Q60949 | TBC1 doma | Q60949 [5 | 0 | 3169.498 | 0 | -0.97 | 0.004113 | 4.43 | 130.9 | 69.1 | 9.73 | 11.94 | | High | High | High | High | High | High | High | High | High | High | High | 0 | 1.22E-05 | 3.2 |
| FALSE | High | ANTLSHFP | [R].ANTLSH | 1xCarbami | Q60949 1x | 1.35E-05 | 0 | 1 | 5 | 2 | Q60949 | TBC1 doma | Q60949 [5 | 0 | 3170.482 | 0 | -1.6 | 0.000434 | 3.29 | 148.1 | 51.9 | 13.99 | 32 | | High | High | High | High | High | High | High | High | High | High | High | 0 | 3.36E-05 | 4.3 |
| FALSE | High | ANTPDSDV | [R].ANTPD | 1xPhospho | Q04692 1x | 3.03E-05 | 0 | 1 | 5 | 4 | Q04692 | SWI/SNF-r | Q04692 [5 | 0 | 1714.83 | 0 | -1.1 | 0.000158 | 0.32 | 136.3 | 63.7 | 2.15 | 2.31 | | High | High | High | High | High | High | High | High | High | High | High | 0 | 1.37E-05 | 3.01 |
| FALSE | High | ANTSSDLEI | [K].ANTSSI | 1xPhospho | Q9ERU9 1x | 0.000337 | 0 | 1 | 1 | 8 | Q9ERU9 | E3 SUMO-p | Q9ERU9 [1 | 0 | 1502.75 | 0 | -0.16 | 0.009393 | 5 | 104.9 | 95.1 | 3.72 | 5.78 | | High | High | High | High | High | High | High | High | High | High | High | 0 | 7.01E-05 | 2.62 |
| FALSE | High | ANTSSDLEI | [K].ANTSSI | 2xPhospho | Q9ERU9 2x | 0.003688 | 0.00021 | 1 | 1 | 3 | Q9ERU9 | E3 SUMO-p | Q9ERU9 [1 | 0 | 1582.717 | 0 | 0.24 | 0.060647 | 18.58 | 87.6 | 112.4 | 6.64 | 11.71 | | High | High | High | High | High | High | High | High | High | High | High | 0.001372 | 0.01432 | 3.56 |
| FALSE | High | ANVVTTPS | [R].ANVVT | 1xPhospho | E9PUR0 1x | 0.032831 | 0.002195 | 1 | 4 | 1 | E9PUR0 | Ankyrin rep | E9PUR0 [1 | 0 | 1468.736 | 0 | -0.48 | 0.000571 | 1.11 | 116.4 | 83.6 | 1.48 | 4.43 | | High | High | High | High | High | High | High | High | High | High | High | 0.001151 | 0.01192 | 2.3 |
| FALSE | High | ANYSQSSSI | [K].ANYSQ | 1xPhospho | P49452 1x | 0.008691 | 0.000736 | 1 | 4 | 4 | P49452 | Centromer | P49452 [66 | 0 | 1509.735 | 0 | -1.04 | 0.000528 | 0.76 | 139.4 | 60.6 | 7.06 | 8.77 | | High | High | High | High | High | High | High | High | High | High | High | 0.007957 | 0.08555 | 1.12 |
| FALSE | High | APAAQPPA | [R].APAAQ | 1xPhospho | B2RPU8 1x | 3.75E-07 | 0 | 1 | 4 | 2 | B2RPU8 | MCG13067 | B2RPU8 [2 | 0 | 2306.186 | 0 | -0.68 | 0.008902 | 8.63 | 123.2 | 76.8 | 8.35 | 10.51 | | High | High | High | High | High | High | High | High | High | High | High | 0 | 3.37E-08 | 3.7 |
| FALSE | High | APACSPLLF | [R].APACSF | 1xCarbami | Q3U325 1x | 0.005 | 0.000504 | 1 | 4 | 3 | Q3U325 | Putative ur | Q3U325 [7 | 0 | 4091.85 | 0 | 0.37 | 0.040051 | 16.7 | 85.5 | 114.5 | 17.05 | 6.65 | | High | High | High | High | High | High | High | High | High | High | High | 0.000225 | 0.001452 | 3.37 |
| FALSE | High | APAPPPPQ | [R].APAPPI | 1xPhospho | B2RWU4 1 | 5.31E-05 | 0 | 2 | 10 | 2 | B2RWU4; ( | Cordon-ble | B2RWU4 [3 | 0 | 2252.179 | 0 | -0.21 | 0.020314 | 2.51 | 105.4 | 94.6 | 6.69 | 6.01 | Shared | High | High | High | High | High | High | High | High | High | High | High | 0 | 1.18E-05 | 2.72 |
| FALSE | High | APAPPPPQ | [R].APAPPI | 1xPhospho | B2RWU4 1 | 0.035486 | 0.002304 | 2 | 10 | 1 | B2RWU4; ( | Cordon-ble | B2RWU4 [3 | 1 | 2609.437 | 0 | -0.06 | 0.384923 | 2.94 | 100.9 | 99.1 | 8.39 | 5.52 | Shared | High | High | High | High | High | High | High | High | High | High | High | 0.001214 | 0.01301 | 2.12 |
| FALSE | High | APAPVVLG | [K].APAPV | 1xPhospho | Q8BY46 1x | 4.69E-06 | 0 | 1 | 1 | 1 | Q8BY46 | Zinc finger | Q8BY46 [1 | 0 | 2290.289 | 0 | -0.28 | 0.09717 | 12.12 | 111.8 | 88.2 | 8.78 | 9.31 | | High | High | High | High | High | High | High | High | High | High | High | 0 | 5.75E-07 | 3.81 |
| FALSE | High | APARPSSA | [K].APARPS | 1xPhospho | B2RWW1 1 | 0.071071 | 0.005247 | 1 | 3 | 3 | B2RWW1 | Microtubu | B2RWW1 [ | 0 | 1577.8 | 0 | -1.66 | 0.000308 | 1.21 | 153.5 | 46.5 | 3.76 | 19.91 | | High | High | High | High | High | High | High | High | High | High | High | 0.002703 | 0.02898 | 1.27 |
| FALSE | High | APASDYGT | [R].APASD | 1xDeamida | Q80W68 1 | 0.000605 | 0 | 1 | 1 | 2 | Q80W68 | Kin of IRRE | Q80W68 [6 | 0 | 4014.806 | 0 | 0.77 | 0.000679 | 3.49 | 71.5 | 128.5 | 14 | 10.8 | | High | High | High | High | High | High | High | High | High | High | High | 0 | 0.000276 | 4.34 |
| FALSE | High | APASPTAAI | [R].APASPT | 1xCarbami | Q8BJ56 1x | 0.003755 | 0.00021 | 1 | 3 | 2 | Q8BJ56 | Patatin-like | Q8BJ56 [4 | 0 | 2437.131 | 0 | 0.7 | 0.002007 | 17.75 | 73.9 | 126.1 | 13.18 | 2.49 | | High | High | High | High | High | High | High | High | High | High | High | 0.000327 | 0.002451 | 3.6 |
| FALSE | High | APASTLVP | [R].APASTL | 1xPhospho | A0A087W | 6.51E-07 | 0 | 1 | 5 | 1 | A0A087W | Snf2-relate | A0A087W | 0 | 2034.047 | 0 | 0.35 | 0.030573 | 16.63 | 87.3 | 112.7 | 8.67 | 7.29 | | High | High | High | High | High | High | High | High | High | High | High | 0 | 6.3E-08 | 2.88 |
| FALSE | High | APCPDRNS | [R].APCPD | 1xCarbami | A2A654 1x | 1.26E-05 | 0 | 1 | 5 | 1 | A2A654 | Bromodon | A2A654 [1 | 1 | 3156.463 | 0 | -0.47 | 0.003032 | 6.71 | 120.6 | 79.4 | 9.74 | 20.24 | | High | High | High | High | High | High | High | High | High | High | High | 0 | 1.75E-06 | 3.62 |
| FALSE | High | APCPDRNS | [R].APCPD | 1xCarbami | A2A654 1x | 3.14E-05 | 0 | 1 | 5 | 2 | A2A654 | Bromodon | A2A654 [1 | 1 | 3157.447 | 0 | -0.41 | 0.032184 | 3.15 | 112.1 | 87.9 | 6.62 | 19.46 | | High | High | High | High | High | High | High | High | High | High | High | 0 | 4.89E-06 | 4.03 |
| FALSE | High | APCPDRNS | [R].APCPD | 1xCarbami | A2A654 1x | 0.015329 | 0.001089 | 1 | 5 | 1 | A2A654 | Bromodon | A2A654 [1 | 1 | 3172.458 | 0 | -0.16 | 0.867757 | 12.4 | 101.5 | 98.5 | 10.38 | 18.7 | | High | High | High | High | High | High | High | High | High | High | High | 0.000571 | 0.00507 | 3.85 |
| FALSE | High | APCYAPSSI | [K].APCYAI | 1xCarbami | Q9QYP6 1x | 0.111965 | 0.009053 | 1 | 2 | 2 | Q9QYP6 | 5-azacytidi | Q9QYP6 [3 | 1 | 2154.082 | 0 | 0.74 | 0.001698 | 12.92 | 72.7 | 127.3 | 12.02 | 13.46 | | High | High | High | High | High | High | High | High | High | High | High | 0.004569 | 0.04954 | 1.98 |
| FALSE | High | APDPAITES | [R].APDPAI | 1xPhospho | F8VQH0 1x | 0.007404 | 0.00062 | 1 | 5 | 2 | F8VQH0 | Tyrosine-p | F8VQH0 [8 | 0 | 2715.401 | 0 | -0.18 | 0.306697 | 14.35 | 106 | 94 | 11.84 | 10.21 | | High | High | High | High | High | High | High | High | High | High | High | 0.000344 | 0.003084 | 3.15 |
| FALSE | High | APDRTPPSI | [R].APDRTI | 1xPhospho | Q8BJU0 1x | 0.010758 | 0.000772 | 1 | 4 | 1 | Q8BJU0 | Small gluta | Q8BJU0 [7 | 1 | 2165.955 | 0 | 0.24 | 0.022477 | 3.01 | 89.3 | 110.7 | 11.48 | 8.52 | | High | High | High | High | High | High | High | High | High | High | High | 0.000399 | 0.003395 | 2.77 |
| FALSE | High | APDSPSK | [K].APDSPS | 1xPhospho | Q7TT79 1x | 0.045508 | 0.003373 | 1 | 4 | 1 | Q7TT79 | Microceph | Q7TT79 [2 | 0 | 1239.639 | 0 | -1.02 | 0.000378 | 2.31 | 136.5 | 63.5 | 6.82 | 7.54 | | High | High | High | High | High | High | High | High | High | High | High | 0.001727 | 0.01725 | 1.61 |
| FALSE | High | APDVLQVD | [K].APDVL( | 1xPhospho | H3BLJ7 1xP | 4.85E-05 | 0 | 1 | 2 | 1 | H3BLJ7 | Protein CA | H3BLJ7 [54 | 0 | 1920.927 | 0 | 0.13 | 0.078585 | 7.66 | 94.6 | 105.4 | 4.7 | 5.43 | | High | High | High | High | High | High | High | High | High | High | High | 0 | 7.96E-06 | 3.48 |
| FALSE | High | APDVLQVD | [K].APDVL( | 2xPhospho | H3BLJ7 2xP | 3.73E-05 | 0 | 1 | 2 | 3 | H3BLJ7 | Protein CA | H3BLJ7 [54 | 0 | 2000.893 | 0 | 2.05 | 0.00083 | 13.79 | 40.6 | 159.4 | 8.74 | 12.59 | | High | High | High | High | High | High | High | High | High | High | High | 0 | 4.84E-05 | 3.29 |
| FALSE | High | APEAAVSD | [K].APEAA\ | 1xPhospho | Q8CH77 1x | 0.061176 | 0.004646 | 1 | 3 | 1 | Q8CH77 | Neuron na | Q8CH77 [1 | 0 | 1597.787 | 0 | -1.17 | 0.000384 | 1.76 | 139.2 | 60.8 | 3.66 | 12.61 | | High | High | High | High | High | High | High | High | High | High | High | 0.002381 | 0.02438 | 2.3 |
| FALSE | High | APEASEAA | [K].APEASE | 1xPhospho | Q3TFU3 1x | 4.18E-07 | 0 | 1 | 6 | 2 | Q3TFU3 | Putative ur | Q3TFU3 [7 | 0 | 3482.73 | 0 | -0.29 | 0.003141 | 1.09 | 111 | 89 | 4.85 | 9.17 | | High | High | High | High | High | High | High | High | High | High | High | 0 | 3.83E-08 | 5.3 |
| FALSE | High | APEATVTLI | [K].APEAT\ | 1xCarbami | Q99KN1 1x | 2.17E-05 | 0 | 1 | 2 | 1 | Q99KN1 | Arrestin dc | Q99KN1 [2 | 0 | 3228.669 | 0 | 0.68 | 0.000505 | 4.05 | 78.1 | 121.9 | 4.47 | 5.54 | | High | High | High | High | High | High | High | High | High | High | High | 0 | 3.24E-06 | 4.21 |
| FALSE | High | APPGAPGP | [R].APPGAI | 1xPhospho | Q8CCS6 1x | 4.17E-06 | 0 | 1 | 3 | 4 | Q8CCS6 | Polyadenyl | Q8CCS6 [7 | 0 | 4858.287 | 0 | -1.15 | 0.00041 | 2.6 | 138.1 | 61.9 | 4.11 | 9.39 | | High | High | High | High | High | High | High | High | High | High | High | 0 | 1.43E-06 | 4.3 |
| FALSE | High | AGSGDQG | [K].AGSGD | 1xCarbami | D3YYU8 1x | 0.045122 | 0.003373 | 1 | 2 | 1 | D3YYU8 | Obscurin-li | D3YYU8 [3 | 0 | 1757.788 | 0 | -0.64 | 0.003299 | 1.22 | 124.6 | 75.4 | 9.99 | 8.43 | | High | High | High | High | High | High | High | High | High | High | High | 0.001727 | 0.01716 | 2.06 |
| FALSE | High | AGSFTGPSI | [R].AGSFT( | 1xPhospho | Q80U49 1x | 9.28E-05 | 0 | 1 | 2 | 1 | Q80U49 | Centrosom | Q80U49 [1 | 0 | 1859.837 | 0 | 1.2 | 0.020498 | 28.8 | 58.8 | 141.2 | 15.08 | 25.4 | | High | High | High | High | High | High | High | High | High | High | High | 0 | 1.66E-05 | 2.1 |
| FALSE | High | AGSEECVF | [R].AGSEE( | 1xCarbami | Q6P9S0 1x | 3.42E-06 | 0 | 1 | 2 | 2 | Q6P9S0 | MTSS1-like | Q6P9S0 [5 | 0 | 3007.441 | 0 | -1.71 | 0.002847 | 2.48 | 152.3 | 47.7 | 10.51 | 19.65 | | High | High | High | High | High | High | High | High | High | High | High | 0 | 4.07E-07 | 5.2 |
| FALSE | High | AASPSKPPI | [K].AASPSK | 1xDeamida | Q8K298 1x | 0.007339 | 0.00062 | 1 | 2 | 3 | Q8K298 | Anillin OS= | Q8K298 [1 | 0 | 2347.242 | 0 | -0.25 | 0.066822 | 7.85 | 108.5 | 91.5 | 7.24 | 6.45 | | High | High | High | High | High | High | High | High | High | High | High | 0.002976 | 0.03145 | 2.45 |
| FALSE | High | AASPSKPPI | [K].AASPSK | 2xPhospho | Q8K298 2x | 0.007019 | 0.00062 | 1 | 2 | 2 | Q8K298 | Anillin OS= | Q8K298 [1 | 0 | 2426.225 | 0 | 0.26 | 0.011004 | 8.89 | 94.5 | 105.5 | 9.04 | 4.02 | | High | High | High | High | High | High | High | High | High | High | High | 0.000275 | 0.002109 | 2.92 |
| FALSE | High | AASPSPQS | [KR].AASPS | 1xPhospho | Q52KI8 1xP | 0.000418 | 0 | 1 | 6 | 6 | Q52KI8 | Serine/argi | Q52KI8 [77 | 0 | 1308.651 | 0 | 0.02 | 0.703602 | 12.98 | 101.1 | 98.9 | 5.65 | 6.04 | | High | High | High | High | High | High | High | High | High | High | High | 0.000872 | 0.008686 | 2.32 |
| FALSE | High | AASPSPQS | [KR].AASPS | 2xPhospho | Q52KI8 2xP | 0.00086 | 0 | 1 | 6 | 6 | Q52KI8 | Serine/argi | Q52KI8 [77 | 0 | 1388.617 | 0 | -0.57 | 0.000445 | 2.17 | 119.9 | 80.1 | 1.74 | 3.55 | | High | High | High | High | High | High | High | High | High | High | High | 0.000399 | 0.003492 | 2.35 |
| FALSE | High | AASPTLPR | [R].AASPTL | 1xPhospho | Q9D952 1x | 0.016892 | 0.001204 | 1 | 3 | 1 | Q9D952 | Envoplakin | Q9D952 [2 | 0 | 1121.592 | 0 | 2.11 | 0.00036 | 2.91 | 38.5 | 161.5 | 5.29 | 8.34 | | High | High | High | High | High | High | High | High | High | High | High | 0.000638 | 0.005634 | 1.55 |
| FALSE | High | AASPYSQR | [R].AASPYS | 1xPhospho | Q68FH0 1x | 4.43E-05 | 0 | 1 | 6 | 3 | Q68FH0 | Plakophilin | Q68FH0 [2 | 0 | 1967.99 | 0 | -1.16 | 0.00036 | 2.48 | 138.2 | 61.8 | 3.9 | 6.47 | | High | High | High | High | High | High | High | High | High | High | High | 0 | 7.23E-06 | 1.91 |
| FALSE | High | AASQGSES | [R].AASQG | 1xPhospho | O89032 1x | 0.001325 | 0 | 1 | 3 | 2 | O89032 | SH3 and PX | O89032 [1 | 0 | 1850.921 | 0 | -0.14 | 0.574175 | 11.83 | 104.1 | 95.9 | 2.46 | 14.54 | | High | High | High | High | High | High | High | High | High | High | High | 0 | 0.000328 | 2.77 |
| FALSE | High | AASQPSK | [R].AASQP | 1xPhospho | Q3TAB2 1x | 0.094641 | 0.007408 | 2 | 8 | 1 | Q3TAB2; H | Putative ur | Q3TAB2 [2 | 0 | 1226.655 | 0 | 1.46 | 0.001294 | 13.8 | 50.3 | 149.7 | 8.27 | 12.14 | Shared | High | High | High | High | High | High | High | High | High | High | High | 0.003754 | 0.0404 | 1.64 |
| FALSE | High | AASREEPIE | [K].AASREE | 1xPhospho | H3BJQ9 1x | 0.01029 | 0.000772 | 1 | 12 | 1 | H3BJQ9 | Homeobox | H3BJQ9 [1 | 1 | 1772.809 | 0 | 2.73 | 0.002624 | 23.42 | 24.8 | 175.2 | 13.02 | 21.62 | | High | High | High | High | High | High | High | High | High | High | High | 0.000382 | 0.003253 | 1.76 |
| FALSE | High | AASSANLLI | [K].AASSAN | 1xPhospho | Q8VC03 1x | 0.000884 | 0 | 1 | 1 | 3 | Q8VC03 | Echinoderr | Q8VC03 [1 | 0 | 1324.719 | 0 | 2.03 | 0.000652 | 20.49 | 39.2 | 160.8 | 14.14 | 7.96 | | High | High | High | High | High | High | High | High | High | High | High | 0.000275 | 0.001675 | 2.61 |
| FALSE | High | AASSPNLSI | [R].AASSPN | 1xPhospho | A0A0A6YW | 1.87E-07 | 0 | 2 | 11 | 6 | A0A0A6YW | Mitogen-ac | A0A0A6YW | 0 | 2129.053 | 0 | -0.75 | 0.000438 | 1.08 | 125.4 | 74.6 | 3.99 | 3.94 | Shared | High | High | High | High | High | High | High | High | High | High | High | 0 | 1.55E-08 | 4.04 |
| FALSE | High | AASSPNLSI | [R].AASSPN | 1xDeamida | A0A0A6YW | 4.82E-06 | 0 | 2 | 11 | 6 | A0A0A6YW | Mitogen-ac | A0A0A6YW | 0 | 2130.037 | 0 | -0.68 | 0.004072 | 2.95 | 117.9 | 82.1 | 8.05 | 9.28 | Shared | High | High | High | High | High | High | High | High | High | High | High | 0 | 7.79E-07 | 3.51 |

| | | | | | | | | | | | | | | | | | | | | | | | | | | | | | | | | | | | | | | |
|---|---|---|---|---|---|---|---|---|---|---|---|---|---|---|---|---|---|---|---|---|---|---|---|---|---|---|---|---|---|---|---|---|---|---|---|---|---|---|
| FALSE | High | AASSPSNP/ | [K].AASSPS | 1xPhospho | Q5QGZ6 1x | 1.66E-05 | 0 | 1 | 4 | 1 | Q5QGZ6 | Retinoic ac | Q5QGZ6 [1 | 0 | 2683.305 | 0 | 1.06 | 0.004821 | 27.15 | 69.4 | 130.6 | 17.89 | 16.65 | | High | High | High | High | High | High | High | High | High | High | High | 0 | 2.39E-06 | 4.31 |
| FALSE | High | AASPSKPPI | [K].AASPSK | 1xPhospho | Q8K298 1x | 0.056248 | 0.004124 | 1 | 2 | 3 | Q8K298 | Anillin OS= | Q8K298 [1 | 0 | 2346.258 | 0 | -0.21 | 0.025604 | 7.17 | 107.2 | 92.8 | 3.15 | 6.76 | | High | High | High | High | High | High | High | High | High | High | High | 0.002117 | 0.02217 | 2.51 |
| FALSE | High | AASSYSAK | [R].AASSYS | 1xPhospho | G3UY93 1x | 0.028082 | 0.001791 | 1 | 5 | 1 | G3UY93 | Valine--tRN | G3UY93 [1 | 0 | 1322.676 | 0 | 1.56 | 0.000574 | 6.07 | 49.4 | 150.6 | 3.73 | 9.91 | | High | High | High | High | High | High | High | High | High | High | High | 0.000968 | 0.009992 | 1.73 |
| FALSE | High | AASTDLGA | [K].AASTDI | 1xPhospho | E9Q9Q2 1x | 0.000121 | 0 | 1 | 3 | 4 | E9Q9Q2 | R3H domai | E9Q9Q2 [1 | 0 | 1883.988 | 0 | 0.93 | 0.01882 | 20.38 | 68.8 | 131.2 | 9.07 | 23.6 | | High | High | High | High | High | High | High | High | High | High | High | 0 | 2.21E-05 | 2.95 |
| FALSE | High | AASVPCSV/ | [R].AASVPC | 1xCarbami | Q7TNF9 1x | 0.001986 | 5.66E-05 | 1 | 2 | 1 | Q7TNF9 | Protein FAI | Q7TNF9 [6 | 0 | 1753.896 | 0 | -0.15 | 0.466741 | 14.46 | 102.7 | 97.3 | 5.23 | 10.51 | | High | High | High | High | High | High | High | High | High | High | High | 2.58E-05 | 0.000514 | 2.81 |
| FALSE | High | AATATRPP( | [R].AATATI | 1xPhospho | E9QKI4 1xF | 0.01441 | 0.001011 | 1 | 6 | 1 | E9QKI4 | Proline-ricl | E9QKI4 [14 | 0 | 3326.772 | 0 | 0.21 | 0.198479 | 30.25 | 92.6 | 107.4 | 7.94 | 13.46 | | High | High | High | High | High | High | High | High | High | High | High | 0.000538 | 0.004742 | 1.56 |
| FALSE | High | AATATRPP( | [R].AATATI | 2xPhospho | E9QKI4 2xF | 0.006195 | 0.00062 | 1 | 6 | 1 | E9QKI4 | Proline-ricl | E9QKI4 [14 | 0 | 3406.739 | 0 | 0.41 | 0.002262 | 2.51 | 84.8 | 115.2 | 4.06 | 5.12 | | High | High | High | High | High | High | High | High | High | High | High | 0.000275 | 0.001829 | 1.63 |
| FALSE | High | AATENSEN | [K].AATENS | 1xPhospho | Q61687 1x | 0.000128 | 0 | 1 | 13 | 1 | Q61687 | Transcripti | Q61687 [1( | 0 | 2387.157 | 0 | -0.26 | 0.067537 | 12.42 | 104.9 | 95.1 | 10.31 | 14.62 | | High | High | High | High | High | High | High | High | High | High | High | 0 | 2.39E-05 | 3.1 |
| FALSE | High | AATESVPCI | [K].AATESV | 1xCarbami | G5E8E3 1x | 2.23E-05 | 0 | 1 | 7 | 1 | G5E8E3 | RIKEN cDN | G5E8E3 [1( | 0 | 2235.068 | 0 | -0.6 | 0.000464 | 2.98 | 120.1 | 79.9 | 3.69 | 3.14 | | High | High | High | High | High | High | High | High | High | High | High | 0 | 3.34E-06 | 3.88 |
| FALSE | High | AATPHLLHI | [R].AATPHI | 1xPhospho | Q6ZPZ3 1x | 0.048303 | 0.00354 | 1 | 2 | 1 | Q6ZPZ3 | Zinc finger | Q6ZPZ3 [3 | 0 | 1461.768 | 0 | -0.12 | 0.643462 | 8.27 | 103 | 97 | 15.69 | 14.25 | | High | High | High | High | High | High | High | High | High | High | High | 0.001812 | 0.01849 | 1.8 |
| FALSE | High | AATPPNPSI | [K].AATPPN | 1xPhospho | A6H663 1x | 0.001783 | 5.66E-05 | 1 | 3 | 3 | A6H663 | BCL2-assoc | A6H663 [5 | 0 | 2170.038 | 0 | 1.24 | 0.001468 | 11.95 | 60.7 | 139.3 | 6.59 | 11.39 | | High | High | High | High | High | High | High | High | High | High | High | 0.001011 | 0.01092 | 2.84 |
| FALSE | High | AATPSPSG/ | [R].AATPSF | 1xPhospho | Q8VEL2 1x | 5.62E-06 | 0 | 1 | 3 | 3 | Q8VEL2 | Myotubula | Q8VEL2 [6 | 0 | 2152.1 | 0 | -0.15 | 0.897998 | 22.97 | 97.2 | 102.8 | 24.12 | 13.82 | | High | High | High | High | High | High | High | High | High | High | High | 0 | 6.18E-06 | 4.46 |
| FALSE | High | AATPVVVT | [K].AATPVV | 1xPhospho | A0A0R4J03 | 9.76E-07 | 0 | 1 | 1 | 4 | A0A0R4J03 | Tripartite r | A0A0R4J03 | 0 | 2742.374 | 0 | -0.28 | 0.207856 | 7.13 | 109.6 | 90.4 | 12.49 | 12.17 | | High | High | High | High | High | High | High | High | High | High | High | 0 | 9.84E-08 | 3.46 |
| FALSE | High | AATSGVPS | [K].AATSGV | 1xPhospho | Q99KG5 1x | 0.000483 | 0 | 1 | 4 | 1 | Q99KG5 | Lipolysis-st | Q99KG5 [2 | 0 | 2769.463 | 0 | -1.67 | 0.001218 | 0.25 | 150.2 | 49.8 | 10.42 | 15.07 | | High | High | High | High | High | High | High | High | High | High | High | 0 | 0.000106 | 2.89 |
| FALSE | High | AAVAASSS | [R].AAVAA | 1xPhospho | Q9D823 1x | 0.002313 | 0.000109 | 1 | 1 | 4 | Q9D823 | 60S riboso | Q9D823 [8 | 0 | 1059.492 | 0 | -0.22 | 0.048061 | 2.12 | 106.7 | 93.3 | 6.21 | 9.04 | | High | High | High | High | High | High | High | High | High | High | High | 4.94E-05 | 0.000687 | 2.23 |
| FALSE | High | AAVASLLQ | [R].AAVASI | 1xPhospho | A0A1S6GW | 6.95E-05 | 0 | 1 | 2 | 1 | A0A1S6GW | Uncharacte | A0A1S6GW | 0 | 2423.335 | 0 | 0.52 | 0.011384 | 16.73 | 80 | 120 | 10.29 | 12.87 | | High | High | High | High | High | High | High | High | High | High | High | 0 | 1.19E-05 | 3.85 |
| FALSE | High | AASTAPSST | [R].AASTAF | 1xPhospho | E9Q2H1 1x | 0.001313 | 0 | 1 | 2 | 2 | E9Q2H1 | E3 ubiquiti | E9Q2H1 [1 | 0 | 3015.487 | 0 | -0.47 | 0.008322 | 3.98 | 116.3 | 83.7 | 8.85 | 4.15 | | High | High | High | High | High | High | High | High | High | High | High | 0 | 0.000323 | 3.15 |
| FALSE | High | AAVASVTP | [R].AAVASV | 1xPhospho | Q60875 1x | 0.017652 | 0.001282 | 1 | 8 | 1 | Q60875 | Rho guanir | Q60875 [7 | 0 | 1510.828 | 0 | 0.46 | 0.001621 | 3.61 | 83.4 | 116.6 | 2.85 | 5.9 | | High | High | High | High | High | High | High | High | High | High | High | 0.000689 | 0.005903 | 1.89 |
| FALSE | High | AASPQEVT | [R].AASPQI | 1xPhospho | E9QAE4 1x | 1.56E-06 | 0 | 1 | 2 | 2 | E9QAE4 | Histone-lys | E9QAE4 [2 | 0 | 2008.999 | 0 | -1.58 | 0.000158 | 0.98 | 148.7 | 51.3 | 2.25 | 8.23 | | High | High | High | High | High | High | High | High | High | High | High | 0 | 1.67E-07 | 3.1 |
| FALSE | High | AASPPASA | [R].AASPPA | 2xPhospho | Q3UGC1 2 | 0.000116 | 0 | 1 | 5 | 1 | Q3UGC1 | Putative ur | Q3UGC1 [3 | 0 | 2359.135 | 0 | -0.06 | 0.115456 | 3.97 | 106.1 | 93.9 | 4.72 | 6.8 | | High | High | High | High | High | High | High | High | High | High | High | 0 | 2.13E-05 | 3.88 |
| FALSE | High | AASASISSN | [K].AASASI | 1xPhospho | Q811Q9 1x | 4.37E-07 | 0 | 1 | 2 | 1 | Q811Q9 | Choline-ph | Q811Q9 [3 | 0 | 2252.004 | 0 | -0.82 | 0.000941 | 4.54 | 123.4 | 76.6 | 5.52 | 9.6 | | High | High | High | High | High | High | High | High | High | High | High | 0 | 4E-08 | 3.4 |
| FALSE | High | AASASISSN | [K].AASASI | 1xPhospho | Q811Q9 1x | 2.59E-06 | 0 | 1 | 2 | 1 | Q811Q9 | Choline-ph | Q811Q9 [3 | 0 | 2267.999 | 0 | -0.75 | 0.009564 | 5.65 | 123.2 | 76.8 | 13.05 | 6.95 | | High | High | High | High | High | High | High | High | High | High | High | 0 | 2.95E-07 | 3.68 |
| FALSE | High | AASDGQYE | [R].AASDG | 2xPhospho | E9Q0S6 2x | 0.003001 | 0.00016 | 1 | 6 | 3 | E9Q0S6 | Tensin 1 O | E9Q0S6 [1( | 1 | 2793.208 | 0 | -0.46 | 0.009682 | 5.86 | 115.7 | 84.3 | 10.63 | 6.83 | | High | High | High | High | High | High | High | High | High | High | High | 7.19E-05 | 0.000821 | 3.7 |
| FALSE | High | AASDPLLSS | [K].AASDPI | 1xPhospho | A2AEW1 1 | 2.4E-05 | 0 | 1 | 7 | 3 | A2AEW1 | Transcripti | A2AEW1 [5 | 0 | 2067.114 | 0 | 0.17 | 0.044797 | 8.63 | 91.9 | 108.1 | 3.67 | 8.55 | | High | High | High | High | High | High | High | High | High | High | High | 0 | 3.6E-06 | 3.34 |
| FALSE | High | AASDPLLSS | [K].AASDPI | 2xPhospho | A2AEW1 2 | 1.06E-05 | 0 | 1 | 7 | 5 | A2AEW1 | Transcripti | A2AEW1 [5 | 0 | 2147.08 | 0 | 0.37 | 0.001042 | 2.68 | 86.8 | 113.2 | 5.5 | 2.7 | | High | High | High | High | High | High | High | High | High | High | High | 0 | 1.43E-06 | 5.39 |
| FALSE | High | AASEELVEK | [K].AASEEL | 1xPhospho | B2RSV7 1x | 0.000131 | 0 | 1 | 2 | 2 | B2RSV7 | Fgd6 prote | B2RSV7 [4 | 0 | 1513.792 | 0 | 1.17 | 0.000384 | 8.99 | 63.2 | 136.8 | 6.58 | 6.36 | | High | High | High | High | High | High | High | High | High | High | High | 0 | 2.43E-05 | 2.7 |
| FALSE | High | AASEQQAD | [K].AASEQ | 1xPhospho | P42128 1x | 3.48E-09 | 0 | 1 | 1 | 2 | P42128 | Forkhead b | P42128 [27 | 0 | 2056.959 | 0 | 1.14 | 0.000879 | 14.03 | 60.2 | 139.8 | 6.5 | 7.59 | | High | High | High | High | High | High | High | High | High | High | High | 0 | 1.76E-10 | 3.7 |
| FALSE | High | AASEQQAD | [K].AASEQ | 2xPhospho | P42128 2x | 3.83E-05 | 0 | 1 | 1 | 1 | P42128 | Forkhead b | P42128 [27 | 0 | 2136.925 | 0 | 0.66 | 0.00337 | 5.54 | 73.6 | 126.4 | 16.9 | 5.12 | | High | High | High | High | High | High | High | High | High | High | High | 0 | 6.12E-06 | 3.3 |
| FALSE | High | AASGGSSG | [R].AASGG | 1xPhospho | Q6ZQ29-2 | 7.38E-07 | 0 | 1 | 2 | 4 | Q6ZQ29-2 | Isoform 2 c | Q6ZQ29-2 | 0 | 2344.15 | 0 | 0.7 | 0.008166 | 13.12 | 76.9 | 123.1 | 13.52 | 5.88 | | High | High | High | High | High | High | High | High | High | High | High | 0 | 7.21E-08 | 3.59 |
| FALSE | High | AASGPEESI | [R].AASGPI | 1xPhospho | Q7TPN9 1x | 0.000221 | 0 | 1 | 2 | 3 | Q7TPN9 | Proline-ricl | Q7TPN9 [1 | 0 | 1597.787 | 0 | -0.42 | 0.00138 | 3.85 | 113 | 87 | 5.23 | 7.84 | | High | High | High | High | High | High | High | High | High | High | High | 0 | 8.4E-05 | 2.45 |
| FALSE | High | AASGSQPE | [R].AASGS( | 1xPhospho | Q9Z2F6 1x | 5.22E-09 | 0 | 1 | 3 | 2 | Q9Z2F6 | B-cell lymp | Q9Z2F6 [3 | 0 | 2683.168 | 0 | -0.27 | 0.156496 | 14.08 | 112 | 88 | 6.73 | 22.6 | | High | High | High | High | High | High | High | High | High | High | High | 0 | 3.61E-09 | 4.4 |
| FALSE | High | AASLDGAE | [R].AASLD( | 1xPhospho | Q99PU5 1x | 0.002974 | 0.00016 | 1 | 3 | 1 | Q99PU5 | Long-chain | Q99PU5 [6 | 0 | 1899.905 | 0 | 0.61 | 0.046407 | 20.67 | 86.2 | 113.8 | 18.9 | 9.99 | | High | High | High | High | High | High | High | High | High | High | High | 7.19E-05 | 0.000812 | 2.65 |
| FALSE | High | AASIDLLED | [R].AASIDL | 1xPhospho | Q8K382 1x | 0.000611 | 0 | 1 | 3 | 1 | Q8K382 | DENN dom | Q8K382 [5 | 0 | 3364.769 | 0 | 0.23 | 0.203938 | 34.18 | 87.1 | 112.9 | 17.62 | 25.88 | | High | High | High | High | High | High | High | High | High | High | High | 0 | 0.000137 | 3.08 |
| FALSE | High | AASPQDLA | [R].AASPQI | 1xCarbami | P51949 1x | 5.85E-08 | 0 | 1 | 1 | 2 | P51949 | CDK-activa | P51949 [27 | 0 | 2271.043 | 0 | -0.56 | 0.010312 | 5.62 | 119.1 | 80.9 | 11.82 | 7.91 | | High | High | High | High | High | High | High | High | High | High | High | 0 | 4.21E-09 | 4.87 |
| FALSE | High | AASLNYLN( | [R].AASLNY | 1xPhospho | Q9DBS5 1x | 3.51E-06 | 0 | 1 | 1 | 1 | Q9DBS5 | Kinesin ligh | Q9DBS5 [5 | 0 | 2336.196 | 0 | 0.21 | 0.14416 | 11.51 | 91.7 | 108.3 | 9.44 | 13.58 | | High | High | High | High | High | High | High | High | High | High | High | 0 | 4.17E-07 | 3.06 |
| FALSE | High | AASLSPAA/ | [K].AASLSP | 1xPhospho | Q8BRB7 1x | 0.002441 | 0.000109 | 1 | 3 | 3 | Q8BRB7 | Histone ac | Q8BRB7 [1 | 0 | 2138.059 | 0 | -0.73 | 0.000629 | 0.54 | 124.7 | 75.3 | 8.45 | 5.99 | | High | High | High | High | High | High | High | High | High | High | High | 0.00014 | 0.001117 | 3.57 |
| FALSE | High | AASLTEDR | [K].AASLTE | 1xPhospho | Q6NZJ6 1x | 0.000553 | 0 | 1 | 9 | 3 | Q6NZJ6 | Eukaryotic | Q6NZJ6 [1 | 0 | 1171.556 | 0 | 1.43 | 0.001312 | 9.14 | 53.3 | 146.7 | 18.27 | 6.31 | | High | High | High | High | High | High | High | High | High | High | High | 0 | 0.000122 | 2.25 |
| FALSE | High | AASITSEVF | [K].AASITS | 1xPhospho | G3X956 1x | 0.028824 | 0.001829 | 1 | 2 | 2 | G3X956 | Suppresso | G3X956 [1 | 0 | 1704.897 | 0 | -1.14 | 0.000738 | 1.38 | 137.8 | 62.2 | 6.14 | 4.99 | | High | High | High | High | High | High | High | High | High | High | High | 0.001597 | 0.01538 | 2.77 |
| FALSE | High | AASMDNN | [R].AASMD | 1xPhospho | Q9R1E0 1x | 0.001933 | 5.66E-05 | 1 | 2 | 3 | Q9R1E0 | Forkhead b | Q9R1E0 [2 | 0 | 1491.691 | 0 | -1.49 | 0.000167 | 1.15 | 150.2 | 49.8 | 4.54 | 9.81 | | High | High | High | High | High | High | High | High | High | High | High | 2.58E-05 | 0.000501 | 1.31 |
| FALSE | High | AASMDSSS | [R].AASMD | 1xPhospho | Q9WVH3 1 | 0.001117 | 0 | 1 | 2 | 1 | Q9WVH3 | Forkhead b | Q9WVH3 [ | 0 | 1437.67 | 0 | -1.37 | 0.000605 | 1.55 | 143.5 | 56.5 | 3.71 | 10.82 | | High | High | High | High | High | High | High | High | High | High | High | 0 | 0.000271 | 1.92 |
| FALSE | High | AASPAGQK | [K].AASPA( | 1xPhospho | P49452 1x | 0.023584 | 0.001578 | 1 | 3 | 1 | P49452 | Centromer | P49452 [14 | 0 | 1267.681 | 0 | -0.93 | 0.001728 | 2.31 | 132.7 | 67.3 | 3.8 | 14 | | High | High | High | High | High | High | High | High | High | High | High | 0.000859 | 0.008241 | 1.8 |
| FALSE | High | AASPFRSP | [K].AASPFF | 1xPhospho | A6H663 1x | 0.037369 | 0.00249 | 1 | 3 | 1 | A6H663 | BCL2-assoc | A6H663 [2 | 1 | 1396.73 | 0 | 2.45 | 0.000338 | 20.15 | 33.2 | 166.8 | 13.41 | 6.36 | | High | High | High | High | High | High | High | High | High | High | High | 0.001308 | 0.01386 | 1.89 |
| FALSE | High | AASPGAAN | [R].AASPG/ | 1xPhospho | E9QMQ3 1 | 0.003655 | 0.00021 | 1 | 1 | 1 | E9QMQ3 | TRIO and F | E9QMQ3 [ | 0 | 2142.107 | 0 | 0.43 | 0.1656 | 33.41 | 85.2 | 114.8 | 15.58 | 10.7 | | High | High | High | High | High | High | High | High | High | High | High | 9.42E-05 | 0.001022 | 3.22 |
| FALSE | High | AASPGAAN | [R].AASPG/ | 1xPhospho | E9QMQ3 1 | 9.65E-06 | 0 | 1 | 1 | 1 | E9QMQ3 | TRIO and F | E9QMQ3 [ | 0 | 2158.102 | 0 | 1.01 | 0.00462 | 16.26 | 66.4 | 133.6 | 8.46 | 15.47 | | High | High | High | High | High | High | High | High | High | High | High | 0 | 1.3E-06 | 3.95 |
| FALSE | High | AASPGPPG | [K].AASPGI | 1xPhospho | E9Q309 1x | 0.010568 | 0.000772 | 1 | 1 | 1 | E9Q309 | Centrosom | E9Q309 [1 | 0 | 1461.73 | 0 | 0.23 | 0.01106 | 7.05 | 93 | 107 | 2.74 | 5.13 | | High | High | High | High | High | High | High | High | High | High | High | 0.000382 | 0.003337 | 1.37 |
| FALSE | High | AASPLELQ( | [R].AASPLE | 1xCarbami | Q9Z206 1x | 2.29E-05 | 0 | 1 | 2 | 2 | Q9Z206 | Neuroepitl | Q9Z206 [5 | 0 | 3369.562 | 0 | -0.51 | 0.001422 | 2.9 | 116.6 | 83.4 | 7.88 | 4.68 | | High | High | High | High | High | High | High | High | High | High | High | 0 | 3.62E-05 | 4.43 |
| FALSE | High | AASPLELQ( | [R].AASPLE | 1xCarbami | Q9Z206 1x | 1.72E-06 | 0 | 1 | 2 | 2 | Q9Z206 | Neuroepitl | Q9Z206 [5 | 0 | 3370.546 | 0 | -0.46 | 0.001176 | 0.89 | 116 | 84 | 8.86 | 6.13 | | High | High | High | High | High | High | High | High | High | High | High | 0 | 1.87E-07 | 4.17 |
| FALSE | High | AASPPASA | [R].AASPPA | 1xPhospho | Q3UGC1 1 | 1.44E-07 | 0 | 1 | 5 | 7 | Q3UGC1 | Putative ur | Q3UGC1 [3 | 0 | 2279.168 | 0 | 0.3 | 0.011663 | 6.05 | 89.6 | 110.4 | 2.16 | 6.91 | | High | High | High | High | High | High | High | High | High | High | High | 0 | 7.27E-07 | 4.26 |
| FALSE | High | AASLNYLN( | [R].AASLNY | 1xDeamida | Q9DBS5 1x | 3.9E-05 | 0 | 1 | 1 | 5 | Q9DBS5 | Kinesin ligh | Q9DBS5 [5 | 0 | 2337.18 | 0 | 0.18 | 0.134183 | 7.57 | 94.9 | 105.1 | 6.94 | 19.02 | | High | High | High | High | High | High | High | High | High | High | High | 0 | 6.23E-06 | 3.59 |
| FALSE | High | AAVEDINP/ | [R].AAVEDI | 1xPhospho | D3YVW2 1 | 3.54E-05 | 0 | 1 | 2 | 3 | D3YVW2 | Golgi integ | D3YVW2 [5 | 1 | 4981.206 | 0 | 0.2 | 0.38161 | 44.61 | 97.7 | 102.3 | 19.88 | 20.67 | | High | High | High | High | High | High | High | High | High | High | High | 0 | 5.59E-06 | 4.79 |
| FALSE | High | AAVEDINP/ | [R].AAVEDI | 1xDeamida | D3YVW2 1 | 6.55E-06 | 0 | 1 | 2 | 5 | D3YVW2 | Golgi integ | D3YVW2 [5 | 1 | 4982.19 | 0 | 0.26 | 0.396169 | 6.89 | 92.2 | 107.8 | 20.14 | 12.84 | | High | High | High | High | High | High | High | High | High | High | High | 0 | 3.89E-06 | 5.49 |
| FALSE | High | AAVGTQEF | [K].AAVGT | 1xPhospho | B1AQY2 1x | 0.082424 | 0.006288 | 1 | 4 | 1 | B1AQY2 | Rho GTPas | B1AQY2 [1 | 1 | 4313.104 | 0 | 0.56 | 0.002602 | 9.01 | 79.6 | 120.4 | 4.37 | 6.6 | | High | High | High | High | High | High | High | High | High | High | High | 0.003243 | 0.03432 | 3.03 |
| FALSE | High | ACYITPK | [R].ACYITP | 1xCarbami | Q80WQ8 1 | 0.022182 | 0.001542 | 1 | 3 | 1 | Q80WQ8 | Mis18-binc | Q80WQ8 [ | 0 | 1390.721 | 0 | -1.41 | 0.000177 | 0.75 | 146.2 | 53.8 | 4.33 | 7.22 | | High | High | High | High | High | High | High | High | High | High | High | 0.000814 | 0.007691 | 1.19 |
| FALSE | High | ADADAGP/ | [R].ADADA | 1xPhospho | A0A1B0GC | 3.23E-05 | 0 | 1 | 1 | 1 | A0A1B0GC | Cytohesin- | A0A1B0GC | 0 | 1834.817 | 0 | -0.5 | 0.003389 | 3.5 | 117 | 83 | 7.59 | 6.15 | | High | High | High | High | High | High | High | High | High | High | High | 0 | 5.04E-06 | 2.99 |
| FALSE | High | ADALQAG/ | [R].ADALQ | 1xPhospho | B0QZN5 1x | 2.32E-06 | 0 | 1 | 4 | 2 | B0QZN5 | Vesicle-ass | B0QZN5 [6 | 0 | 2204.1 | 0 | 0.17 | 0.285453 | 20.08 | 94.3 | 105.7 | 12.39 | 7.89 | | High | High | High | High | High | High | High | High | High | High | High | 0 | 2.61E-07 | 3.01 |
| FALSE | High | ADALTSSP( | [R].ADALTS | 1xPhospho | Q3UJN1 1x | 0.003253 | 0.00016 | 1 | 3 | 6 | Q3UJN1 | DNA helica | Q3UJN1 [3 | 0 | 1283.619 | 0 | -0.33 | 0.003414 | 3.74 | 109.4 | 90.6 | 2.35 | 5.15 | | High | High | High | High | High | High | High | High | High | High | High | 0.00155 | 0.01504 | 1.21 |
| FALSE | High | ADALTSSP( | [R].ADALTS | 2xPhospho | Q3UJN1 2x | 0.009843 | 0.000736 | 1 | 3 | 4 | Q3UJN1 | DNA helica | Q3UJN1 [3 | 0 | 1363.586 | 0 | -0.78 | 0.000214 | 0.34 | 125.7 | 74.3 | 4.64 | 3.46 | | High | High | High | High | High | High | High | High | High | High | High | 0.005745 | 0.06329 | 1.66 |
| FALSE | High | ADANLLTD | [R].ADANL | 1xPhospho | Q3U2K4 1x | 4.56E-06 | 0 | 1 | 4 | 2 | Q3U2K4 | Putative ur | Q3U2K4 [7 | 0 | 1956.911 | 0 | 0.06 | 0.628172 | 0.85 | 97.9 | 102.1 | 4.44 | 9.89 | | High | High | High | High | High | High | High | High | High | High | High | 0 | 5.57E-07 | 4.05 |
| FALSE | High | ADANLLTD | [R].ADANL | 1xCarbami | Q3U2K4 1x | 0.000183 | 0 | 1 | 4 | 3 | Q3U2K4 | Putative ur | Q3U2K4 [7 | 1 | 2756.312 | 0 | -0.09 | 0.545793 | 3.46 | 102.9 | 97.1 | 4.91 | 5.27 | | High | High | High | High | High | High | High | High | High | High | High | 0 | 3.52E-05 | 3.05 |
| FALSE | High | ADANLLTD | [R].ADANL | 1xCarbami | Q3U2K4 2x | 2.56E-05 | 0 | 1 | 4 | 1 | Q3U2K4 | Putative ur | Q3U2K4 [7 | 1 | 2836.279 | 0 | 0.22 | 0.038787 | 16.64 | 92.5 | 107.5 | 5 | 5.92 | | High | High | High | High | High | High | High | High | High | High | High | 0 | 3.89E-06 | 3.22 |
| FALSE | High | ADAPDAG/ | [R].ADAPD | 1xPhospho | Q6DFZ1 1x | 4.9E-09 | 0 | 1 | 4 | 4 | Q6DFZ1 | Golgi-speci | Q6DFZ1 [1 | 0 | 3067.348 | 0 | 0.32 | 0.10088 | 14.73 | 89.1 | 110.9 | 7.93 | 13.14 | | High | High | High | High | High | High | High | High | High | High | High | 0 | 2.58E-10 | 4.59 |
| FALSE | High | ADAPDAG/ | [R].ADAPD | 1xDeamida | Q6DFZ1 1x | 0.000359 | 0 | 1 | 4 | 2 | Q6DFZ1 | Golgi-speci | Q6DFZ1 [1 | 0 | 3068.332 | 0 | 0.16 | 0.302188 | 7.38 | 98.9 | 101.1 | 7.97 | 16.47 | | High | High | High | High | High | High | High | High | High | High | High | 0 | 7.56E-05 | 3.74 |
| FALSE | High | ADASSLTVI | [K].ADASSL | 1xPhospho | Q3TIK1 1x | 8.3E-07 | 0 | 1 | 7 | 7 | Q3TIK1 | Putative ur | Q3TIK1 [4( | 0 | 2087.067 | 0 | 0.12 | 0.119091 | 7.74 | 94.5 | 105.5 | 5.39 | 11.58 | | High | High | High | High | High | High | High | High | High | High | High | 0 | 1.31E-07 | 4.57 |
| FALSE | High | ADASSTEN | [R].ADASST | 1xPhospho | A7TU71 1x | 0.00034 | 0 | 1 | 2 | 6 | A7TU71 | Protein Shi | A7TU71 [2 | 0 | 1849.935 | 0 | 1.84 | 0.00049 | 9.81 | 46.2 | 153.8 | 5.64 | 9.76 | | High | High | High | High | High | High | High | High | High | High | High | 2.58E-05 | 0.000468 | 2.64 |
| FALSE | High | ADDADEFG | [R].ADDAD | 1xPhospho | Q3TRW3 1 | 0.041072 | 0.003094 | 1 | 4 | 2 | Q3TRW3 | Putative ur | Q3TRW3 [9 | 0 | 1554.631 | 0 | 0.4 | 0.016638 | 10.58 | 90.9 | 109.1 | 10.55 | 5.97 | | High | High | High | High | High | High | High | High | High | High | High | 0.0025 | 0.02725 | 1.56 |
| FALSE | High | ACVAVEGT | [R].ACVAV | 1xCarbami | O88495 1x | 0.011146 | 0.000772 | 1 | 1 | 2 | O88495 | Melatonin- | O88495 [3 | 0 | 1368.654 | 0 | 2.58 | 0.001773 | 31.2 | 29.8 | 170.2 | 11.82 | 15.93 | | High | High | High | High | High | High | High | High | High | High | High | 0.000399 | 0.003543 | 1.59 |
| FALSE | High | ADDDVDLE | [K].ADDDV | 1xPhospho | E9Q842 1x | 0.000173 | 0 | 1 | 4 | 2 | E9Q842 | Neuron na | E9Q842 [8 | 0 | 3269.452 | 0 | 0.89 | 0.00537 | 12.53 | 70.2 | 129.8 | 14.82 | 19.46 | | High | High | High | High | High | High | High | High | High | High | High | 0 | 3.31E-05 | 3.5 |
| FALSE | High | ADDTFEALI | [R].ADDTFI | 1xPhospho | P62869 1x | 0.00145 | 5.66E-05 | 1 | 1 | 1 | P62869 | Elongin-B ( | P62869 [8 | 0 | 1346.619 | 0 | 0.28 | 0.01329 | 10.98 | 84.6 | 115.4 | 20.11 | 12.58 | | High | High | High | High | High | High | High | High | High | High | High | 2.58E-05 | 0.000364 | 1.75 |
| FALSE | High | ADEATAGC | [R].ADEAT/ | 1xPhospho | A0A1S6GW | 0.028082 | 0.001791 | 1 | 4 | 2 | A0A1S6GW | DEAD (Asp | A0A1S6GW | 1 | 3409.443 | 0 | -0.73 | 0.01702 | 6.66 | 122.7 | 77.3 | 12.52 | 18.85 | | High | High | High | High | High | High | High | High | High | High | High | 0.000968 | 0.00997 | 3.69 |
| FALSE | High | ADEATAGC | [R].ADEAT/ | 2xPhospho | A0A1S6GW | 0.000225 | 0 | 1 | 4 | 5 | A0A1S6GW | DEAD (Asp | A0A1S6GW | 1 | 3489.409 | 0 | 0.18 | 0.436521 | 10.73 | 96.5 | 103.5 | 8.93 | 6.24 | | High | High | High | High | High | High | High | High | High | High | High | 0 | 4.8E-05 | 3.52 |
| FALSE | High | ADEDLIEHL | [R].ADEDLI | 1xPhospho | Q3UTC9 1x | 0.001158 | 0 | 1 | 4 | 1 | Q3UTC9 | Protein Mc | Q3UTC9 [1 | 0 | 2166.975 | 0 | 0.16 | 0.028717 | 9.83 | 95.9 | 104.1 | 7.67 | 3.28 | | High | High | High | High | High | High | High | High | High | High | High | 0 | 0.00028 | 2.4 |
| FALSE | High | ADEENDSC | [K].ADEENI | 1xPhospho | A0A0G2JG | 0.000237 | 0 | 1 | 4 | 1 | A0A0G2JG | Suppresso | A0A0G2JG | 0 | 3615.574 | 0 | -0.34 | 0.029743 | 11.67 | 108.8 | 91.2 | 6.89 | 11.17 | | High | High | High | High | High | High | High | High | High | High | High | 0 | 4.72E-05 | 2.73 |
| FALSE | High | ADEENDSC | [K].ADEENI | 1xPhospho | A0A0G2JG | 0.01441 | 0.001011 | 1 | 4 | 1 | A0A0G2JG | Suppresso | A0A0G2JG | 0 | 3631.568 | 0 | -0.9 | 0.009526 | 5.22 | 132.5 | 67.5 | 8.77 | 15.73 | | High | High | High | High | High | High | High | High | High | High | High | 0.000538 | 0.004718 | 2.09 |
| FALSE | High | ADFAVVAG | [K].ADFAV | 1xCarbami | K4DI59 1xF | 7.07E-08 | 0 | 1 | 9 | 8 | K4DI59 | DNA ligase | K4DI59 [84 | 0 | 3592.615 | 0 | 0.22 | 0.086968 | 4.23 | 94.9 | 105.1 | 4.41 | 10.2 | | High | High | High | High | High | High | High | High | High | High | High | 0 | 1.47E-07 | 5.82 |
| FALSE | High | ADFAVVAG | [K].ADFAV | 1xCarbami | K4DI59 1xF | 6.07E-05 | 0 | 1 | 9 | 2 | K4DI59 | DNA ligase | K4DI59 [84 | 0 | 3593.599 | 0 | -0.02 | 0.718729 | 22.96 | 99.2 | 100.8 | 17.54 | 13.61 | | High | High | High | High | High | High | High | High | High | High | High | 0 | 1.03E-05 | 4.47 |
| FALSE | High | ADGATSDD | [K].ADGAT | 1xPhospho | P39447 1x | 0.002508 | 0.000109 | 1 | 3 | 2 | P39447 | Tight junct | P39447 [8 | 0 | 1924.812 | 0 | -0.8 | 0.013457 | 5.92 | 127.2 | 72.8 | 2.28 | 22.14 | | High | High | High | High | High | High | High | High | High | High | High | 4.94E-05 | 0.000671 | 1.91 |
| FALSE | High | ADGDNAGT | [K].ADGDN | 1xPhospho | Q3TXH6 1x | 9.41E-10 | 0 | 1 | 3 | 1 | Q3TXH6 | Putative ur | Q3TXH6 [2 | 0 | 2282.07 | 0 | 0.85 | 0.001274 | 6.53 | 71.9 | 128.1 | 4.66 | 7.86 | | High | High | High | High | High | High | High | High | High | High | High | 0 | 4.06E-11 | 4.7 |
| FALSE | High | ADGDSDAE | [R].ADGDS | 1xPhospho | Q91YU8 1x | 1.06E-05 | 0 | 1 | 1 | 5 | Q91YU8 | Suppresso | Q91YU8 [3 | 0 | 3730.503 | 0 | -0.71 | 0.00024 | 0.96 | 124.6 | 75.4 | 3.71 | 6.75 | | High | High | High | High | High | High | High | High | High | High | High | 7.19E-05 | 0.000913 | 3.85 |
| FALSE | High | ADGDWDD | [R].ADGDV | 1xPhospho | Q8K019 1x | 1.45E-06 | 0 | 1 | 11 | 5 | Q8K019 | Bcl-2-assoc | Q8K019 [3 | 0 | 2326.959 | 0 | -0.13 | 0.592459 | 25.71 | 115.8 | 84.2 | 67.99 | 52.5 | | High | High | High | High | High | High | High | High | High | High | High | 0 | 1.54E-07 | 4.23 |
| FALSE | High | ADGDWDD | [R].ADGDV | 1xPhospho | Q8K019 1x | 5.25E-08 | 0 | 1 | 11 | 18 | Q8K019 | Bcl-2-assoc | Q8K019 [3 | 0 | 2556.122 | 0 | -0.34 | 0.027619 | 3.98 | 110.5 | 89.5 | 5.89 | 14.88 | | High | High | High | High | High | High | High | High | High | High | High | 0 | 9.04E-09 | 5.29 |
| FALSE | High | ADDEYETS( | [R].ADDEYI | 1xPhospho | B8JJC1 1xP | 3.99E-06 | 0 | 1 | 3 | 3 | B8JJC1 | Tyrosyl-DN | B8JJC1 [14 | 0 | 2855.237 | 0 | -1.03 | 0.004179 | 5.62 | 134.4 | 65.6 | 7.07 | 17.13 | | High | High | High | High | High | High | High | High | High | High | High | 0 | 4.79E-07 | 4.17 |
| FALSE | High | ACTLELGPI | [K].ACTLEL | 1xCarbami | F8VPU2 1x | 0.001544 | 5.66E-05 | 1 | 2 | 1 | F8VPU2 | FERM, Rho | F8VPU2 [4 | 0 | 2257.182 | 0 | 0.95 | 0.005749 | 20.92 | 70.5 | 129.5 | 5.72 | 12.39 | | High | High | High | High | High | High | High | High | High | High | High | 2.58E-05 | 0.000387 | 1.75 |
| FALSE | High | ACSVEELY/ | [K].ACSVEE | 1xCarbami | Q69Z38 1x | 0.014667 | 0.001049 | 1 | 1 | 1 | Q69Z38 | Pseudopoc | Q69Z38 [7 | 0 | 2504.203 | 0 | -0.24 | 0.060872 | 5.82 | 108.2 | 91.8 | 8.54 | 11.16 | | High | High | High | High | High | High | High | High | High | High | High | 0.000555 | 0.00481 | 2.45 |
| FALSE | High | ACSPSLCD( | [K].ACSPSL | 2xCarbami | E9Q777 1x | 3.54E-05 | 0 | 1 | 3 | 2 | E9Q777 | A kinase (P | E9Q777 [9 | 0 | 2314.061 | 0 | -0.16 | 0.33427 | 4.15 | 104.6 | 95.4 | 2.59 | 12.69 | | High | High | High | High | High | High | High | High | High | High | High | 0 | 6.53E-06 | 4.34 |
| FALSE | High | AAVGTQEF | [K].AAVGT | 1xDeamida | B1AQY2 1x | 8.5E-06 | 0 | 1 | 4 | 1 | B1AQY2 | Rho GTPas | B1AQY2 [1 | 1 | 4314.088 | 0 | 0.44 | 0.004179 | 13.44 | 81.2 | 118.8 | 9.91 | 10.23 | | High | High | High | High | High | High | High | High | High | High | High | 0 | 1.13E-06 | 4.24 |
| FALSE | High | AAVGVTGN | [K].AAVGV | 1xPhospho | P27546 1x | 1.97E-06 | 0 | 1 | 7 | 5 | P27546 | Microtubu | P27546 [64 | 0 | 2093.104 | 0 | 0.05 | 0.164808 | 5.88 | 97.9 | 102.1 | 4.96 | 2 | | High | High | High | High | High | High | High | High | High | High | High | 0 | 6.53E-07 | 3.3 |
| FALSE | High | AAVGVTGN | [K].AAVGV | 1xDeamida | P27546 1x | 2.75E-05 | 0 | 1 | 7 | 2 | P27546 | Microtubu | P27546 [64 | 0 | 2094.088 | 0 | -0.41 | 0.000798 | 1.18 | 114.1 | 85.9 | 5.13 | 2.63 | | High | High | High | High | High | High | High | High | High | High | High | 0 | 4.21E-06 | 4.24 |
| FALSE | High | AAVLSDSEI | [K].AAVLSD | 1xPhospho | Q8C1D8 1x | 0.000496 | 0 | 1 | 3 | 3 | Q8C1D8 | Protein IW | Q8C1D8 [3 | 0 | 2159.027 | 0 | 0.17 | 0.085742 | 13.14 | 90.1 | 109.9 | 8.16 | 13.01 | | High | High | High | High | High | High | High | High | High | High | High | 0 | 0.000108 | 3.39 |
| FALSE | High | AAVLSDSEI | [K].AAVLSD | 2xPhospho | Q8C1D8 2x | 0.000128 | 0 | 1 | 3 | 3 | Q8C1D8 | Protein IW | Q8C1D8 [3 | 0 | 2238.993 | 0 | -0.39 | 0.01334 | 10.45 | 113.6 | 86.4 | 5.14 | 8.22 | | High | High | High | High | High | High | High | High | High | High | High | 0.000275 | 0.001746 | 3.57 |
| FALSE | High | AAVPSGAS | [R].AAVPS( | 1xPhospho | Q5FW97 1 | 1E-05 | 0 | 1 | 18 | 5 | Q5FW97 | Enolase 1, | Q5FW97 [3 | 0 | 2114.073 | 0 | -0.07 | 0.772383 | 16.17 | 102.4 | 97.6 | 7.88 | 7.76 | | High | High | High | High | High | High | High | High | High | High | High | 0 | 1.35E-06 | 3.49 |
| FALSE | High | AAVPTPAK | [K].AAVPTI | 1xPhospho | P09405 1x | 0.002869 | 0.00016 | 1 | 8 | 4 | P09405 | Nucleolin ( | P09405 [72 | 0 | 1292.738 | 0 | -1.04 | 0.000173 | 1.05 | 135.3 | 64.7 | 5.94 | 10.59 | | High | High | High | High | High | High | High | High | High | High | High | 7.19E-05 | 0.000778 | 2.16 |
| FALSE | High | AAVPTPAK | [K].AAVPTI | 1xPhospho | P09405 1x | 0.019446 | 0.001438 | 1 | 8 | 1 | P09405 | Nucleolin ( | P09405 [72 | 1 | 1649.996 | 0 | -1.45 | 0.008985 | 2.31 | 144.1 | 55.9 | 22.14 | 6.74 | | High | High | High | High | High | High | High | High | High | High | High | 0.000756 | 0.006576 | 1.39 |
| FALSE | High | AAVQELSG | [R].AAVQE | 1xPhospho | Q3UYK2 1x | 5.54E-08 | 0 | 1 | 2 | 2 | Q3UYK2 | Presenilin ( | Q3UYK2 [3 | 0 | 2340.117 | 0 | 0.64 | 0.003856 | 3.2 | 79.7 | 120.3 | 3.55 | 7.02 | | High | High | High | High | High | High | High | High | High | High | High | 0 | 3.95E-09 | 3.75 |
| FALSE | High | AAVQELSG | [R].AAVQE | 2xPhospho | Q3UYK2 2x | 4.96E-07 | 0 | 1 | 2 | 8 | Q3UYK2 | Presenilin ( | Q3UYK2 [3 | 0 | 2420.083 | 0 | -0.13 | 0.070863 | 3.95 | 104.6 | 95.4 | 4.26 | 3.7 | | High | High | High | High | High | High | High | High | High | High | High | 0 | 6.59E-06 | 4.53 |
| FALSE | High | AAVQGLQ/ | [K].AAVQG | 1xPhospho | B2RX03 1x | 1.3E-05 | 0 | 1 | 3 | 1 | B2RX03 | MCG8063 ( | B2RX03 [1 | 0 | 1894.02 | 0 | -1.39 | 0.001723 | 3.46 | 144.3 | 55.7 | 6.98 | 16.47 | | High | High | High | High | High | High | High | High | High | High | High | 0 | 1.82E-06 | 3.14 |
| FALSE | High | AAVSHWQ | [K].AAVSH | 1xPhospho | Q02248 1x | 0.000178 | 0 | 1 | 6 | 1 | Q02248 | Catenin be | Q02248 [2 | 0 | 3623.79 | 0 | -1.16 | 0.002648 | 1.98 | 133.4 | 66.6 | 10.94 | 22.05 | | High | High | High | High | High | High | High | High | High | High | High | 0 | 3.42E-05 | 3.91 |
| FALSE | High | AAVSPQK | [R].AAVSP( | 1xPhospho | Q547N0 1x | 0.017966 | 0.001282 | 1 | 2 | 2 | Q547N0 | BRMS1 OS | Q547N0 [2 | 0 | 1238.691 | 0 | -1.04 | 0.000456 | 2.04 | 133.2 | 66.8 | 5.28 | 6.95 | | High | High | High | High | High | High | High | High | High | High | High | 0.008046 | 0.08712 | 2.21 |
| FALSE | High | AAVTPGK | [K].AAVTP( | 1xPhospho | P09405 1x | 0.033986 | 0.002195 | 1 | 7 | 2 | P09405 | Nucleolin ( | P09405 [8 | 0 | 1181.67 | 0 | -1.12 | 0.000554 | 2.14 | 136.9 | 63.1 | 5.29 | 10.33 | | High | High | High | High | High | High | High | High | High | High | High | 0.001151 | 0.01242 | 1.91 |
| FALSE | High | AAVVASPLI | [R].AAVVA | 1xPhospho | E9Q784 1x | 3.48E-06 | 0 | 1 | 3 | 3 | E9Q784 | Zinc finger | E9Q784 [2 | 0 | 1676.893 | 0 | -0.53 | 0.006492 | 8.09 | 116 | 84 | 7.7 | 8.6 | | High | High | High | High | High | High | High | High | High | High | High | 0 | 4.12E-07 | 3.78 |
| FALSE | High | AAVVTSPP | [R].AAVVT | 1xPhospho | Q8K232 1x | 0.003495 | 0.00021 | 2 | 8 | 2 | Q8K232; E | Adducin 1 | Q8K232 [7 | 0 | 2012.098 | 0 | 1.65 | 0.000364 | 6.02 | 47.6 | 152.4 | 6.08 | 7.19 | Shared | High | High | High | High | High | High | High | High | High | High | High | 9.42E-05 | 0.00097 | 1.94 |
| FALSE | High | AAYPPSYSI | [R].AAYPPS | 1xPhospho | Q80XC3-2 | 4.28E-06 | 0 | 1 | 2 | 1 | Q80XC3-2 | Isoform 2 c | Q80XC3-2 | 0 | 2584.264 | 0 | 0.9 | 0.006438 | 32.23 | 69.3 | 130.7 | 11.78 | 10.11 | | High | High | High | High | High | High | High | High | High | High | High | 0 | 5.21E-07 | 3 |
| FALSE | High | ACEPHGDA | [R].ACEPH( | 2xCarbami | B2RY84 1x | 0.009756 | 0.000736 | 1 | 4 | 1 | B2RY84 | Kif14 prote | B2RY84 [1 | 0 | 2508.048 | 0 | 0.1 | 0.865205 | 9.15 | 96.6 | 103.4 | 14.87 | 22.5 | | High | High | High | High | High | High | High | High | High | High | High | 0.000344 | 0.003045 | 2.63 |
| FALSE | High | ACESSPQD | [R].ACESSP | 2xCarbami | B9EIT8 2xP | 0.002096 | 5.66E-05 | 2 | 3 | 1 | B9EIT8; A0 | Rg9mtd2 p | B9EIT8 [28 | 0 | 3295.254 | 0 | -0.11 | 0.652621 | 7.69 | 106.3 | 93.7 | 8.74 | 19.81 | Shared | High | High | High | High | High | High | High | High | High | High | High | 2.58E-05 | 0.000548 | 2.56 |
| FALSE | High | ACESSPQD | [K].ACESSP | 2xCarbami | B9EIT8 3xP | 0.108463 | 0.008839 | 2 | 3 | 1 | B9EIT8; A0 | Rg9mtd2 p | B9EIT8 [28 | 0 | 3375.221 | 0 | -0.94 | 0.001827 | 1.85 | 129.1 | 70.9 | 7.23 | 10.03 | Shared | High | High | High | High | High | High | High | High | High | High | High | 0.004411 | 0.04746 | 2.56 |
| FALSE | High | ACPDLQLS/ | [K].ACPDL( | 1xCarbami | Q9R0L6-2 | 3.1E-06 | 0 | 1 | 3 | 4 | Q9R0L6-2 | Isoform 2 c | Q9R0L6-2 | 0 | 2371.198 | 0 | 0.13 | 0.173652 | 7.7 | 96 | 104 | 6.6 | 10.32 | | High | High | High | High | High | High | High | High | High | High | High | 0 | 3.61E-07 | 4.58 |
| FALSE | High | ACPDLQLS/ | [K].ACPDL( | 1xCarbami | Q9R0L6-2 | 0.000126 | 0 | 1 | 3 | 8 | Q9R0L6-2 | Isoform 2 c | Q9R0L6-2 | 0 | 2451.164 | 0 | -0.2 | 0.060688 | 12.68 | 106.8 | 93.2 | 8.22 | 8.58 | | High | High | High | High | High | High | High | High | High | High | High | 0 | 2.5E-05 | 3.6 |
| FALSE | High | ACPGQSAL | [K].ACPGQ | 2xCarbami | E9Q156 1x | 0.014283 | 0.001011 | 1 | 2 | 1 | E9Q156 | tRNA (guar | E9Q156 [4 | 0 | 1359.564 | 0 | -0.04 | 0.912392 | 18.49 | 100.7 | 99.3 | 4.72 | 8.05 | | High | High | High | High | High | High | High | High | High | High | High | 0.000538 | 0.004654 | 2.35 |
| FALSE | High | ACPVTVPS | [K].ACPVT | 1xCarbami | Q3TLR7 1x | 8.22E-07 | 0 | 1 | 2 | 1 | Q3TLR7 | Denticleles | Q3TLR7 [4 | 0 | 2068.055 | 0 | 0.52 | 0.02799 | 6.54 | 85.2 | 114.8 | 6.16 | 10.18 | | High | High | High | High | High | High | High | High | High | High | High | 0 | 8.15E-08 | 3.96 |
| FALSE | High | ACQMVNS | [R].ACQM | 2xCarbami | A0A0R4J2D | 0.001301 | 0 | 1 | 1 | 1 | A0A0R4J2D | Ubiquitin c | A0A0R4J2D | 0 | 2392.111 | 0 | -0.3 | 0.017472 | 4.17 | 112.4 | 87.6 | 19.61 | 16.03 | | High | High | High | High | High | High | High | High | High | High | High | 0 | 0.000319 | 1.87 |
| FALSE | High | ACSDSLESI | [R].ACSDSL | 1xCarbami | Q80TM9 1 | 4.01E-08 | 0 | 1 | 5 | 7 | Q80TM9 | Nischarin ( | Q80TM9 [5 | 0 | 2400.095 | 0 | -0.18 | 0.152785 | 14.13 | 107.3 | 92.7 | 6.85 | 9.07 | | High | High | High | High | High | High | High | High | High | High | High | 0 | 2.74E-09 | 4.02 |

| | | | | | | | | | | | | | | | | | | | | | | | | | | | | | | | | | | | | | | |
|---|---|---|---|---|---|---|---|---|---|---|---|---|---|---|---|---|---|---|---|---|---|---|---|---|---|---|---|---|---|---|---|---|---|---|---|---|---|---|
| FALSE | High | ACSDSLESI | [R].ACSDSL | 1xCarbami | Q80TM9 2: | 1.15E-05 | 0 | 1 | 5 | 3 | Q80TM9 | Nischarin ( | Q80TM9 [5 | 0 | 2480.061 | 0 | 0.28 | 0.240448 | 37.97 | 88.3 | 111.7 | 15.61 | 13.77 | | High | High | High | High | High | High | High | High | High | High | High | 0 | 1.57E-06 | 4.1 |
| FALSE | High | ACSPDPLR | [R].ACSPDF | 1xCarbami | Q64512 1x | 0.043236 | 0.003228 | 1 | 3 | 1 | Q64512 | Tyrosine-p | Q64512 [2( | 0 | 1224.565 | 0 | 2.07 | 0.000348 | 12.26 | 39.2 | 160.8 | 5.92 | 6.4 | | High | High | High | High | High | High | High | High | High | High | High | 0.001651 | 0.01639 | 2.1 |
| FALSE | High | ACSPGASEI | [K].ACSPG/ | 1xCarbami | Q91XB0 1x | 0.000933 | 0 | 1 | 2 | 1 | Q91XB0 | Three-prim | Q91XB0 [7 | 0 | 1915.96 | 0 | 1.13 | 0.017746 | 21.17 | 62.8 | 137.2 | 14 | 18.65 | | High | High | High | High | High | High | High | High | High | High | High | 0 | 0.00022 | 2.25 |
| FALSE | High | AASAPAKE | [R].AASAP/ | 1xPhospho | Q3TAB2 1x | 0.116482 | 0.009523 | 2 | 9 | 1 | Q3TAB2; H | Putative ur | Q3TAB2 [1 | 1 | 1594.861 | 0 | -0.32 | 0.181916 | 12.48 | 112.5 | 87.5 | 28.2 | 19.54 | Shared | High | High | High | High | High | High | High | High | High | High | High | 0.004757 | 0.05185 | 2.16 |
| FALSE | High | ADGDWDD | [R].ADGDV | 1xDeamida | Q8K019 1x | 0.000589 | 0 | 1 | 11 | 2 | Q8K019 | Bcl-2-assoc | Q8K019 [3( | 0 | 2557.106 | 0 | -0.44 | 0.021284 | 8.47 | 117.6 | 82.4 | 11.51 | 7.47 | | High | High | High | High | High | High | High | High | High | High | High | 0 | 0.000132 | 4.09 |
| FALSE | High | AASALLLR | [R].AASALL | 1xPhospho | D3Z7W0 1: | 0.008691 | 0.000736 | 1 | 3 | 3 | D3Z7W0 | Iron-sulfur | D3Z7W0 [1 | 0 | 1123.644 | 0 | -0.33 | 0.02872 | 5.23 | 111.5 | 88.5 | 7.31 | 10.56 | | High | High | High | High | High | High | High | High | High | High | High | 0.000327 | 0.002694 | 2.29 |
| FALSE | High | AASADSTTI | [K].AASADS | 2xPhospho | Q9QZQ8 2: | 2.72E-05 | 0 | 1 | 3 | 5 | Q9QZQ8 | Core histor | Q9QZQ8 [1 | 0 | 2775.278 | 0 | 0.45 | 0.016522 | 14.5 | 81.8 | 118.2 | 5.34 | 13.33 | | High | High | High | High | High | High | High | High | High | High | High | 0 | 4.16E-06 | 3.73 |
| FALSE | High | AAEAHPQD | [K].AAEAH | 2xCarbami | Q3TDF3 1x | 0.000308 | 0 | 1 | 4 | 1 | Q3TDF3 | Putative ur | Q3TDF3 [4 | 0 | 4546.923 | 0 | 0.26 | 0.005171 | 6.82 | 88 | 112 | 6.26 | 6.63 | | High | High | High | High | High | High | High | High | High | High | High | 0 | 6.35E-05 | 2.95 |
| FALSE | High | AAEAHPQD | [K].AAEAH | 2xCarbami | Q3TDF3 2x | 0.001968 | 5.66E-05 | 1 | 4 | 1 | Q3TDF3 | Putative ur | Q3TDF3 [4 | 0 | 4625.905 | 0 | -0.03 | 0.499891 | 6.37 | 99.4 | 100.6 | 6.98 | 6.06 | | High | High | High | High | High | High | High | High | High | High | High | 2.58E-05 | 0.00051 | 2.67 |
| FALSE | High | AAEDISAGI | [K].AAEDIS | 1xPhospho | E9PV87 1xl | 0.000705 | 0 | 1 | 6 | 2 | E9PV87 | Protein TAI | E9PV87 [13 | 0 | 3386.556 | 0 | 0.08 | 0.480088 | 7.95 | 97.6 | 102.4 | 20.61 | 23 | | High | High | High | High | High | High | High | High | High | High | High | 0 | 0.000162 | 3.26 |
| FALSE | High | AAEDNPYV | [K].AAEDN | 1xPhospho | G3X934 1x | 0.030632 | 0.001857 | 1 | 2 | 1 | G3X934 | MCG11596 | G3X934 [6 | 0 | 2236.073 | 0 | -0.05 | 0.81087 | 3.51 | 100.4 | 99.6 | 7.36 | 7.69 | | High | High | High | High | High | High | High | High | High | High | High | 0.001011 | 0.01102 | 2.85 |
| FALSE | High | AAEEAFVN | [R].AAEEAI | 1xPhospho | Q3THU7 1: | 9.02E-08 | 0 | 1 | 3 | 3 | Q3THU7 | Clathrin lig | Q3THU7 [1 | 0 | 2661.156 | 0 | 0.51 | 0.003645 | 9.52 | 82 | 118 | 3.52 | 7.24 | | High | High | High | High | High | High | High | High | High | High | High | 0 | 6.78E-09 | 5.14 |
| FALSE | High | AAEGAAAC | [R].AAEGA | 1xCarbami | F8WJA0 1x | 2.59E-07 | 0 | 1 | 4 | 1 | F8WJA0 | ATP-depen | F8WJA0 [1 | 1 | 4005.899 | 0 | -0.71 | 0.005145 | 5.48 | 125.2 | 74.8 | 13.98 | 8.05 | | High | High | High | High | High | High | High | High | High | High | High | 0 | 2.23E-08 | 5.02 |
| FALSE | High | AAEGLPQP | [K].AAEGLI | 1xPhospho | Q05AH6 1: | 0.000102 | 0 | 1 | 1 | 2 | Q05AH6 | Uncharacte | Q05AH6 [2 | 0 | 2283.133 | 0 | -1.1 | 0.000291 | 0.39 | 136.2 | 63.8 | 2.37 | 6.83 | | High | High | High | High | High | High | High | High | High | High | High | 0 | 1.83E-05 | 2.83 |
| FALSE | High | AAEGISSDS | [K].AAEGIS | 1xPhospho | B0V2P5 1xl | 0.001127 | 0 | 1 | 5 | 2 | B0V2P5 | DmX-like p | B0V2P5 [92 | 0 | 2197.152 | 0 | 2.13 | 0.001117 | 21.22 | 38.3 | 161.7 | 8.57 | 11.67 | | High | High | High | High | High | High | High | High | High | High | High | 0 | 0.000272 | 2.68 |
| FALSE | High | AAELAETQ | [K].AAELAE | 1xPhospho | E9QPB3 1x | 2.41E-07 | 0 | 1 | 4 | 5 | E9QPB3 | WD repeat | E9QPB3 [8 | 0 | 1972.952 | 0 | -1.25 | 0.00032 | 1.27 | 139.3 | 60.7 | 4.98 | 6.18 | | High | High | High | High | High | High | High | High | High | High | High | 0 | 2.05E-08 | 3.6 |
| FALSE | High | AAELAETQ | [K].AAELAE | 2xPhospho | E9QPB3 2x | 0.047894 | 0.00354 | 1 | 4 | 1 | E9QPB3 | WD repeat | E9QPB3 [8 | 0 | 2052.918 | 0 | -0.59 | 0.007509 | 5.97 | 121.7 | 78.3 | 9.52 | 9.38 | | High | High | High | High | High | High | High | High | High | High | High | 0.001812 | 0.01828 | 2.25 |
| FALSE | High | AAELAETQ | [K].AAELAE | 1xPhospho | E9QPB3 1x | 0.000369 | 0 | 1 | 4 | 2 | E9QPB3 | WD repeat | E9QPB3 [8 | 1 | 2649.233 | 0 | -0.97 | 0.003196 | 1.69 | 137.7 | 62.3 | 14.51 | 12.64 | | High | High | High | High | High | High | High | High | High | High | High | 0 | 7.81E-05 | 3.34 |
| FALSE | High | AAEQMSTI | [K].AAEQM | 1xPhospho | P27546 1xl | 5.5E-05 | 0 | 1 | 4 | 1 | P27546 | Microtubu | P27546 [6( | 0 | 2920.478 | 0 | -1.55 | 0.011559 | 4.52 | 152.7 | 47.3 | 22.31 | 28.46 | | High | High | High | High | High | High | High | High | High | High | High | 0 | 9.21E-06 | 3.2 |
| FALSE | High | AAEQMSTI | [K].AAEQM | 1xPhospho | P27546 1xl | 0.00025 | 0 | 1 | 4 | 2 | P27546 | Microtubu | P27546 [6( | 0 | 2936.473 | 0 | -1.52 | 0.015144 | 6.06 | 148.4 | 51.6 | 11.46 | 43.82 | | High | High | High | High | High | High | High | High | High | High | High | 0 | 5.04E-05 | 2.59 |
| FALSE | High | AAEAHPQD | [K].AAEAH | 2xCarbami | Q3TDF3 1x | 0.022378 | 0.001542 | 1 | 4 | 2 | Q3TDF3 | Putative ur | Q3TDF3 [4 | 0 | 4545.939 | 0 | 0.02 | 0.97806 | 2.28 | 98.8 | 101.2 | 11.14 | 6.67 | | High | High | High | High | High | High | High | High | High | High | High | 0.000814 | 0.007734 | 3.94 |
| FALSE | High | AAESSEESC | [R].AAESSE | 1xCarbami | P35831 1xl | 3.09E-05 | 0 | 1 | 1 | 1 | P35831 | Tyrosine-p | P35831 [5( | 0 | 3290.54 | 0 | 0.97 | 0.001282 | 1.81 | 66.9 | 133.1 | 6.64 | 8.57 | | High | High | High | High | High | High | High | High | High | High | High | 0 | 4.81E-06 | 4.91 |
| FALSE | High | AAETPAVA | [K].AAETP/ | 1xCarbami | Q8BTI8 2xl | 2.51E-05 | 0 | 1 | 3 | 1 | Q8BTI8 | Serine/argi | Q8BTI8 [12 | 0 | 2827.293 | 0 | -0.66 | 0.048984 | 15.9 | 123.7 | 76.3 | 14.64 | 23.25 | | High | High | High | High | High | High | High | High | High | High | High | 0 | 3.78E-06 | 4.15 |
| FALSE | High | AAETVPDLI | [K].AAETVI | 1xPhospho | Q5SRX1 1x | 0.000515 | 0 | 1 | 6 | 1 | Q5SRX1 | TOM1-like | Q5SRX1 [4 | 0 | 2786.345 | 0 | 1.66 | 0.000363 | 6.82 | 49.6 | 150.4 | 22.24 | 3.07 | | High | High | High | High | High | High | High | High | High | High | High | 0 | 0.000113 | 3.23 |
| FALSE | High | AAFDDAIAI | [K].AAFDD/ | 1xPhospho | Q5SS40 1xl | 9.27E-08 | 0 | 1 | 3 | 7 | Q5SS40 | Putative ur | Q5SS40 [1! | 0 | 2626.258 | 0 | -0.43 | 0.008372 | 6.32 | 114.7 | 85.3 | 3.59 | 8.52 | | High | High | High | High | High | High | High | High | High | High | High | 0 | 1.1E-06 | 3.7 |
| FALSE | High | AAFSETTSF | [R].AAFSET | 1xPhospho | Q8C9G3 1: | 0.000352 | 0 | 1 | 5 | 3 | Q8C9G3 | Putative ur | Q8C9G3 [2 | 0 | 1673.855 | 0 | -0.56 | 0.001394 | 1.93 | 120.1 | 79.9 | 2.39 | 8.12 | | High | High | High | High | High | High | High | High | High | High | High | 0.000344 | 0.003116 | 2.33 |
| FALSE | High | AAFTTPDH | [R].AAFTTF | 1xPhospho | G5E8A7 1x | 2.8E-06 | 0 | 1 | 2 | 4 | G5E8A7 | Protein Sm | G5E8A7 [2 | 0 | 3555.607 | 0 | 1.15 | 0.001088 | 12.77 | 62.1 | 137.9 | 5.6 | 8.76 | | High | High | High | High | High | High | High | High | High | High | High | 0 | 3.25E-07 | 4.99 |
| FALSE | High | AAGADPLV | [K].AAGAD | 1xPhospho | P25799 1xl | 1.16E-09 | 0 | 1 | 8 | 1 | P25799 | Nuclear fac | P25799 [73 | 0 | 3132.486 | 0 | -0.94 | 0.002281 | 4.18 | 134.5 | 65.5 | 6.49 | 14.52 | | High | High | High | High | High | High | High | High | High | High | High | 0 | 5.11E-11 | 5.16 |
| FALSE | High | AAGALGSA | [R].AAGAL( | 1xPhospho | Q8BIE6-2 1 | 3.96E-07 | 0 | 1 | 4 | 5 | Q8BIE6-2 | Isoform 2 c | Q8BIE6-2 [ | 0 | 2013.963 | 0 | -0.04 | 0.627714 | 28.1 | 95.9 | 104.1 | 10.6 | 29.38 | | High | High | High | High | High | High | High | High | High | High | High | 0 | 3.58E-08 | 3.58 |
| FALSE | High | AAGDTVVI | [K].AAGDT | 1xPhospho | F8VQC7 1x | 1.2E-08 | 0 | 1 | 17 | 1 | F8VQC7 | Kinectin O! | F8VQC7 [1 | 0 | 2856.363 | 0 | 0.96 | 0.014934 | 3.7 | 69.4 | 130.6 | 5.54 | 15.99 | | High | High | High | High | High | High | High | High | High | High | High | 0 | 7.07E-10 | 4.3 |
| FALSE | High | AAGDTVVI | [K].AAGDT | 1xPhospho | F8VQC7 1x | 2.29E-08 | 0 | 1 | 17 | 3 | F8VQC7 | Kinectin O! | F8VQC7 [1 | 0 | 2872.358 | 0 | 0.98 | 0.010665 | 18.01 | 67.4 | 132.6 | 8.56 | 16.58 | | High | High | High | High | High | High | High | High | High | High | High | 0 | 1.46E-09 | 4.86 |
| FALSE | High | AAGGAGA( | [K].AAGGA | 1xPhospho | P48678 1xl | 3.55E-06 | 0 | 1 | 2 | 9 | P48678 | Prelamin-A | P48678 [6( | 0 | 2531.23 | 0 | 1.42 | 0.000478 | 18.57 | 56.2 | 143.8 | 11.64 | 7.18 | | High | High | High | High | High | High | High | High | High | High | High | 0 | 5.26E-06 | 3.17 |
| FALSE | High | AAGGAVST | [K].AAGGA | 1xPhospho | E9Q5C9 1x | 0.000699 | 0 | 1 | 2 | 2 | E9Q5C9 | Nucleolar a | E9Q5C9 [4 | 0 | 1621.871 | 0 | 0.18 | 0.015601 | 4.07 | 94.8 | 105.2 | 2.56 | 4.44 | | High | High | High | High | High | High | High | High | High | High | High | 0 | 0.000159 | 2.21 |
| FALSE | High | AAGHQELV | [R].AAGHC | 1xPhospho | Q9D279 1x | 0.099328 | 0.008003 | 1 | 1 | 1 | Q9D279 | Mitotic inte | Q9D279 [2 | 0 | 2482.395 | 0 | 2.92 | 0.000489 | 3.52 | 20.7 | 179.3 | 17.71 | 9.48 | | High | High | High | High | High | High | High | High | High | High | High | 0.004032 | 0.04276 | 2.39 |
| FALSE | High | AAGSAQGS | [R].AAGSA( | 1xPhospho | B2RY19 1xl | 7.28E-12 | 0 | 1 | 7 | 2 | B2RY19 | Cgnl1 prote | B2RY19 [5E | 0 | 2966.426 | 0 | 1.67 | 0.000975 | 21.14 | 46.5 | 153.5 | 6.78 | 11.51 | | High | High | High | High | High | High | High | High | High | High | High | 0 | 1.71E-13 | 4.73 |
| FALSE | High | AAETPAVA | [K].AAETP/ | 1xCarbami | Q8BTI8 1x | 1.31E-09 | 0 | 1 | 3 | 5 | Q8BTI8 | Serine/argi | Q8BTI8 [12 | 0 | 2747.326 | 0 | -0.64 | 0.040631 | 10.39 | 125.5 | 74.5 | 18.81 | 24.48 | | High | High | High | High | High | High | High | High | High | High | High | 0 | 5.86E-11 | 4.92 |
| FALSE | High | AAGSQESL | [R].AAGSQ | 1xPhospho | E9QPI5 1xF | 9.74E-06 | 0 | 1 | 2 | 3 | E9QPI5 | Sister chro | E9QPI5 [12 | 0 | 1870.931 | 0 | 0.01 | 0.531112 | 8.47 | 100.6 | 99.4 | 6.3 | 8.5 | | High | High | High | High | High | High | High | High | High | High | High | 0 | 1.32E-06 | 3.28 |
| FALSE | High | AADSQNSC | [R].AADSQ | 2xPhospho | F6ZDS4 2xl | 1.05E-05 | 0 | 1 | 5 | 2 | F6ZDS4 | Nucleoprot | F6ZDS4 [2( | 0 | 2889.177 | 0 | -0.93 | 0.000468 | 2.72 | 130.8 | 69.2 | 3.74 | 5.92 | | High | High | High | High | High | High | High | High | High | High | High | 0 | 1.42E-06 | 5.34 |
| FALSE | High | AADSQNSC | [R].AADSQ | 1xPhospho | F6ZDS4 1xl | 4.51E-12 | 0 | 1 | 5 | 13 | F6ZDS4 | Nucleoprot | F6ZDS4 [2( | 0 | 2809.211 | 0 | -0.43 | 0.002458 | 6.87 | 112.1 | 87.9 | 4.11 | 9.38 | | High | High | High | High | High | High | High | High | High | High | High | 0 | 1.44E-08 | 5.75 |
| FALSE | High | AAAAGLGH | [R].AAAAG | 1xPhospho | Q62431 1x | 9.84E-10 | 0 | 1 | 3 | 4 | Q62431 | AT-rich inte | Q62431 [6 | 0 | 2985.342 | 0 | -1 | 0.000721 | 1.37 | 136.4 | 63.6 | 6.38 | 11.83 | | High | High | High | High | High | High | High | High | High | High | High | 0 | 1.52E-10 | 5.75 |
| FALSE | High | AAAAGLGH | [R].AAAAG | 2xPhospho | Q62431 2x | 1.25E-08 | 0 | 1 | 3 | 1 | Q62431 | AT-rich inte | Q62431 [6 | 0 | 3065.309 | 0 | -1.88 | 0.000985 | 1.23 | 157.2 | 42.8 | 11.13 | 19.97 | | High | High | High | High | High | High | High | High | High | High | High | 0 | 7.39E-10 | 5.78 |
| FALSE | High | AAAAGLGH | [R].AAAAG | 3xPhospho | Q62431 3x | 0.000386 | 0 | 1 | 3 | 1 | Q62431 | AT-rich inte | Q62431 [6 | 0 | 3145.275 | 0 | -0.99 | 0.002056 | 6.31 | 133 | 67 | 6.14 | 23.86 | | High | High | High | High | High | High | High | High | High | High | High | 0 | 8.21E-05 | 3.74 |
| FALSE | High | AAAASAAE | [R].AAAAS/ | 1xPhospho | O35226 1x | 2.83E-09 | 0 | 1 | 3 | 5 | O35226 | 26S protea | O35226 [2 | 0 | 2854.412 | 0 | 0.37 | 0.004531 | 0.73 | 88 | 112 | 7.84 | 2.94 | | High | High | High | High | High | High | High | High | High | High | High | 0 | 3.34E-09 | 5.15 |
| FALSE | High | AAAASAAE | [R].AAAAS/ | 1xPhospho | Q3ULG4 1x | 1.54E-08 | 0 | 1 | 4 | 2 | Q3ULG4 | Putative ur | Q3ULG4 [2 | 0 | 2110.002 | 0 | 0.55 | 0.000525 | 4.89 | 81.5 | 118.5 | 3.12 | 3.09 | | High | High | High | High | High | High | High | High | High | High | High | 0 | 9.39E-10 | 3.99 |
| FALSE | High | AAAASAAE | [R].AAAAS/ | 1xPhospho | Q3ULG4 1x | 8.89E-09 | 0 | 1 | 3 | 2 | Q3ULG4 | Putative ur | Q3ULG4 [2 | 1 | 3196.578 | 0 | 0.25 | 0.005279 | 3.14 | 92.4 | 107.6 | 4.93 | 7.76 | | High | High | High | High | High | High | High | High | High | High | High | 0 | 1.18E-08 | 4.73 |
| FALSE | High | AAAASSPV | [R].AAAAS! | 1xPhospho | Q9ERQ3 1x | 4.39E-08 | 0 | 1 | 1 | 2 | Q9ERQ3 | Zinc finger | Q9ERQ3 [6 | 0 | 2708.287 | 0 | 0.72 | 0.037468 | 13.64 | 77.7 | 122.3 | 22.58 | 17.07 | | High | High | High | High | High | High | High | High | High | High | High | 0 | 9.63E-09 | 4.28 |
| FALSE | High | AAAATAPP | [R].AAAAT/ | 1xPhospho | D3YXK1 1x | 6.97E-06 | 0 | 1 | 1 | 2 | D3YXK1 | Atherin OS | D3YXK1 [1 | 0 | 1922.969 | 0 | -0.52 | 0.001522 | 3.35 | 118.4 | 81.6 | 4.72 | 3.08 | | High | High | High | High | High | High | High | High | High | High | High | 0 | 8.99E-07 | 2.89 |
| FALSE | High | AAADDGEE | [K].AAADD | 1xPhospho | Q544Z7 1x | 0.003055 | 0.00016 | 1 | 4 | 1 | Q544Z7 | DNA-(apur | Q544Z7 [8 | 1 | 2441.205 | 0 | -1.48 | 0.000328 | 1.13 | 147.9 | 52.1 | 3.6 | 7.42 | | High | High | High | High | High | High | High | High | High | High | High | 7.19E-05 | 0.000832 | 1.54 |
| FALSE | High | AAAEEPQE | [K].AAAEEI | 1xPhospho | Q7TND5 1x | 3.44E-08 | 0 | 1 | 2 | 2 | Q7TND5 | Ribosome | Q7TND5 [2 | 0 | 2894.382 | 0 | -0.12 | 0.147797 | 9.08 | 103.2 | 96.8 | 10.36 | 4.94 | | High | High | High | High | High | High | High | High | High | High | High | 0 | 6.79E-09 | 3.81 |
| FALSE | High | AAAGAGGS | [R].AAAGA | 1xPhospho | Q91YU6 1x | 0.018125 | 0.001282 | 1 | 1 | 1 | Q91YU6 | Leucine zip | Q91YU6 [5 | 0 | 1139.577 | 0 | 0.04 | 0.838781 | 13.78 | 100.1 | 99.9 | 5.64 | 11.45 | | High | High | High | High | High | High | High | High | High | High | High | 0.000689 | 0.00611 | 0.9 |
| FALSE | High | AAAPAPQP | [R].AAAPAI | 1xPhospho | Q1PSW8 1: | 0.000158 | 0 | 1 | 1 | 2 | Q1PSW8 | E3 ubiquiti | Q1PSW8 [1 | 0 | 2379.209 | 0 | -1.98 | 0.00129 | 1.1 | 159.9 | 40.1 | 12.08 | 9.36 | | High | High | High | High | High | High | High | High | High | High | High | 0 | 0.000108 | 2.51 |
| FALSE | High | AAAPPQPS | [R].AAAPP( | 1xPhospho | Q62311 1x | 0.025293 | 0.001645 | 1 | 2 | 2 | Q62311 | Transcripti | Q62311 [5 | 0 | 1796.971 | 0 | 0.09 | 0.162366 | 10.13 | 95.9 | 104.1 | 10.85 | 9.62 | | High | High | High | High | High | High | High | High | High | High | High | 0.000872 | 0.00886 | 2.3 |
| FALSE | High | AADSQNSC | [R].AADSQ | 1xDeamida | F6ZDS4 1xl | 5.61E-09 | 0 | 1 | 5 | 3 | F6ZDS4 | Nucleoprot | F6ZDS4 [2( | 0 | 2810.195 | 0 | -0.75 | 0.000533 | 3.2 | 125.1 | 74.9 | 2.67 | 5.41 | | High | High | High | High | High | High | High | High | High | High | High | 0 | 3E-10 | 5.82 |
| FALSE | High | AAAPSAAS | [R].AAAPS/ | 1xPhospho | B2RWW2 1 | 0.0006 | 0 | 1 | 7 | 4 | B2RWW2 | Golgb1 prc | B2RWW2 [ | 0 | 2123.115 | 0 | 0.8 | 0.002897 | 6.22 | 75.7 | 124.3 | 5.06 | 11.36 | | High | High | High | High | High | High | High | High | High | High | High | 0.000327 | 0.002927 | 3.65 |
| FALSE | High | AAASFSPE( | [K].AAASFS | 1xPhospho | Q06831 1x | 0.041781 | 0.003126 | 1 | 2 | 1 | Q06831 | Transcripti | Q06831 [2( | 0 | 2869.515 | 0 | 0.41 | 0.011785 | 2.64 | 86.2 | 113.8 | 8.87 | 9.92 | | High | High | High | High | High | High | High | High | High | High | High | 0.001625 | 0.01575 | 2.53 |
| FALSE | High | AAASGPAS | [K].AAASGI | 1xPhospho | Q3U0F3 1x | 1.13E-06 | 0 | 1 | 3 | 1 | Q3U0F3 | Putative ur | Q3U0F3 [2 | 0 | 2302.201 | 0 | 0.03 | 0.628926 | 27.89 | 98.9 | 101.1 | 15.73 | 8.26 | | High | High | High | High | High | High | High | High | High | High | High | 0 | 1.17E-07 | 3.59 |
| FALSE | High | AAASPLGE! | [R].AAASPI | 1xPhospho | Q3U0S4 1x | 0.007209 | 0.00062 | 1 | 2 | 2 | Q3U0S4 | Putative ur | Q3U0S4 [4 | 0 | 2309.169 | 0 | -0.9 | 0.002704 | 4.27 | 129.5 | 70.5 | 7.2 | 15.39 | | High | High | High | High | High | High | High | High | High | High | High | 0.000275 | 0.00218 | 2.93 |
| FALSE | High | AAATEAPA | [K].AAATE/ | 1xPhospho | H9LAN6 1x | 1.86E-07 | 0 | 1 | 6 | 2 | H9LAN6 | Histone-lys | H9LAN6 [3 | 0 | 2084.979 | 0 | -0.8 | 0.000717 | 2.73 | 125.9 | 74.1 | 5.76 | 5.52 | | High | High | High | High | High | High | High | High | High | High | High | 0 | 1.53E-08 | 4.02 |
| FALSE | High | AAATPESQ | [R].AAATPI | 1xPhospho | P28667 1xl | 6.07E-05 | 0 | 1 | 1 | 3 | P28667 | MARCKS-re | P28667 [14 | 0 | 1865.941 | 0 | -1.93 | 0.001438 | 0.19 | 160.7 | 39.3 | 17.91 | 13.79 | | High | High | High | High | High | High | High | High | High | High | High | 0 | 1.03E-05 | 3.69 |
| FALSE | High | AAATPTDT | [K].AAATPT | 1xPhospho | Q3UH06-3 | 0.000442 | 0 | 1 | 4 | 1 | Q3UH06-3 | Isoform 3 c | Q3UH06-3 | 0 | 1688.851 | 0 | -0.62 | 0.009769 | 8.8 | 124.2 | 75.8 | 13.74 | 9.75 | | High | High | High | High | High | High | High | High | High | High | High | 0 | 9.48E-05 | 2.56 |
| FALSE | High | AADEDWD | [R].AADED | 1xPhospho | Q9CXK9 1x | 1.95E-06 | 0 | 1 | 4 | 4 | Q9CXK9 | RNA-bindir | Q9CXK9 [3 | 0 | 3073.418 | 0 | 0.16 | 0.500705 | 3.02 | 97.1 | 102.9 | 4.19 | 15.13 | | High | High | High | High | High | High | High | High | High | High | High | 0 | 2.16E-07 | 5.34 |
| FALSE | High | AADGDLVS | [K].AADGD | 1xCarbami | Q8BHN1 1: | 0.0102 | 0.000736 | 1 | 2 | 2 | Q8BHN1 | Gamma-ta: | Q8BHN1 [4 | 0 | 2700.336 | 0 | 0.74 | 0.004388 | 6.08 | 69.8 | 130.2 | 10.52 | 13.32 | | High | High | High | High | High | High | High | High | High | High | High | 0.001889 | 0.0189 | 2.89 |
| FALSE | High | AADLGSSPI | [R].AADLG! | 1xPhospho | Q7TT79 1x | 0.002921 | 0.00016 | 1 | 4 | 1 | Q7TT79 | Microceph | Q7TT79 [3! | 0 | 1154.566 | 0 | -0.75 | 0.370874 | 19.92 | 118.6 | 81.4 | 61.52 | 49.75 | | High | High | High | High | High | High | High | High | High | High | High | 7.19E-05 | 0.000792 | 1.33 |
| FALSE | High | AADLGSSPI | [R].AADLG! | 1xPhospho | Q7TT79 1x | 0.016743 | 0.001166 | 1 | 4 | 2 | Q7TT79 | Microceph | Q7TT79 [3! | 0 | 1383.729 | 0 | -0.86 | 0.00087 | 3.11 | 129.5 | 70.5 | 6.31 | 5.04 | | High | High | High | High | High | High | High | High | High | High | High | 0.000638 | 0.005605 | 1.88 |
| FALSE | High | AADISLDNI | [K].AADISL | 1xPhospho | P70399 1xl | 0.000705 | 0 | 1 | 7 | 2 | P70399 | TP53-bindi | P70399 [16 | 0 | 1882.993 | 0 | 0.02 | 0.459588 | 5.79 | 97.5 | 102.5 | 7.65 | 4.43 | | High | High | High | High | High | High | High | High | High | High | High | 0 | 0.000161 | 3.71 |
| FALSE | High | AADPDLSP | [K].AADPD | 1xPhospho | Q9Z2E2 1x | 0.002743 | 0.000109 | 1 | 6 | 1 | Q9Z2E2 | Methyl-Cp | Q9Z2E2 [5! | 0 | 1550.823 | 0 | 0.85 | 0.001006 | 9.71 | 72.4 | 127.6 | 3.45 | 6.07 | | High | High | High | High | High | High | High | High | High | High | High | 4.94E-05 | 0.000743 | 2.06 |
| FALSE | High | AADSGERP | [R].AADSG | 1xPhospho | F6ZBR8 1xl | 3.1E-06 | 0 | 2 | 5 | 3 | F6ZBR8; Ol | Protein Wi | F6ZBR8 [82 | 0 | 2292.2 | 0 | 0.27 | 0.043258 | 4.75 | 92.9 | 107.1 | 4.32 | 7.02 | Shared | High | High | High | High | High | High | High | High | High | High | High | 0 | 3.6E-07 | 2.7 |
| FALSE | High | AAAQYSSQ | [R].AAAQY | 1xPhospho | Q3TIY5 1xF | 0.008769 | 0.000736 | 1 | 4 | 5 | Q3TIY5 | Putative ur | Q3TIY5 [12 | 0 | 1491.761 | 0 | 0.16 | 0.192505 | 23.44 | 90.1 | 109.9 | 6.43 | 13.77 | | High | High | High | High | High | High | High | High | High | High | High | 0.000327 | 0.00272 | 1.95 |
| FALSE | High | AAGTLLNG | [K].AAGTLI | 1xDeamida | Q8K0V4 1x | 7.41E-05 | 0 | 1 | 2 | 1 | Q8K0V4 | CCR4-NOT | Q8K0V4 [5 | 0 | 2481.304 | 0 | 0.13 | 0.017423 | 6.12 | 95.6 | 104.4 | 10.61 | 8.64 | | High | High | High | High | High | High | High | High | High | High | High | 0 | 1.29E-05 | 4.46 |
| FALSE | High | AAGVSVEP | [K].AAGVS | 1xPhospho | E9Q3T0 1x | 0.001097 | 0 | 1 | 2 | 1 | E9Q3T0 | Uncharacte | E9Q3T0 [3 | 0 | 2214.176 | 0 | -0.57 | 0.006328 | 8.88 | 118.4 | 81.6 | 7.79 | 16.59 | | High | High | High | High | High | High | High | High | High | High | High | 0 | 0.000265 | 2.66 |
| FALSE | High | AAGYDLFS/ | [K].AAGYD | 1xPhospho | Q8VCG1 1: | 7.98E-06 | 0 | 1 | 3 | 1 | Q8VCG1 | Deoxyuridi | Q8VCG1 [8 | 0 | 2680.266 | 0 | -1.52 | 0.000652 | 0.67 | 146.2 | 53.8 | 12.28 | 13.85 | | High | High | High | High | High | High | High | High | High | High | High | 0 | 1.05E-06 | 4.32 |
| FALSE | High | AAPPPPPPI | [K].AAPPPF | 1xPhospho | A2AN08-5 | 1.05E-06 | 0 | 1 | 3 | 20 | A2AN08-5 | Isoform 5 c | A2AN08-5 | 0 | 2013.041 | 0 | 0.2 | 0.023859 | 6.13 | 89.7 | 110.3 | 5.44 | 7.51 | | High | High | High | High | High | High | High | High | High | High | High | 0 | 1.07E-07 | 4.6 |
| FALSE | High | AAPQSPSV | [K].AAPQSI | 1xPhospho | Q6ZQ45 1x | 0.001815 | 5.66E-05 | 1 | 4 | 5 | Q6ZQ45 | MKIAA079 | Q6ZQ45 [1 | 0 | 1519.829 | 0 | -0.02 | 0.97851 | 11.71 | 100.5 | 99.5 | 3.35 | 7.68 | | High | High | High | High | High | High | High | High | High | High | High | 7.19E-05 | 0.000832 | 2.98 |
| FALSE | High | AAPSPAPAI | [K].AAPSP/ | 1xPhospho | A6H663 1x | 9.11E-05 | 0 | 1 | 2 | 2 | A6H663 | BCL2-assoc | A6H663 [4( | 0 | 1812.966 | 0 | 2.23 | 0.000292 | 1.16 | 33.7 | 166.3 | 6.86 | 7.01 | | High | High | High | High | High | High | High | High | High | High | High | 0 | 1.62E-05 | 3.4 |
| FALSE | High | AAPSSEGD | [R].AAPSSE | 1xCarbami | H3BJQ9 2x | 5.4E-05 | 0 | 1 | 12 | 1 | H3BJQ9 | Homeobox | H3BJQ9 [1 | 0 | 3452.485 | 0 | 1.48 | 0.003177 | 5.66 | 53.1 | 146.9 | 14.72 | 13.58 | | High | High | High | High | High | High | High | High | High | High | High | 0 | 9E-06 | 4.2 |
| FALSE | High | AAPTSSPG! | [K].AAPTSS | 1xPhospho | Q8VDI4 1x | 0.000687 | 0 | 1 | 4 | 5 | Q8VDI4 | MutL homo | Q8VDI4 [4 | 0 | 1326.625 | 0 | -0.95 | 0.000492 | 3.18 | 131.8 | 68.2 | 4.95 | 6.89 | | High | High | High | High | High | High | High | High | High | High | High | 0 | 0.000157 | 1.73 |
| FALSE | High | AAPTTPPPI | [R].AAPTTF | 1xPhospho | Q9ESJ4 1xl | 0.000524 | 0 | 1 | 1 | 2 | Q9ESJ4 | NCK-intera | Q9ESJ4 [17 | 0 | 1613.907 | 0 | -0.17 | 0.043165 | 7.57 | 109.5 | 90.5 | 4.09 | 9.87 | | High | High | High | High | High | High | High | High | High | High | High | 0 | 0.000115 | 2.37 |
| FALSE | High | AAPTTSSSS | [R].AAPTTS | 1xPhospho | E9QAU4 1x | 4.05E-09 | 0 | 1 | 9 | 21 | E9QAU4 | Enhancer t | E9QAU4 [1 | 0 | 2665.267 | 0 | -0.19 | 0.073446 | 6.72 | 107.7 | 92.3 | 6.12 | 5.31 | | High | High | High | High | High | High | High | High | High | High | High | 0 | 2.08E-10 | 4.28 |
| FALSE | High | AAPTTSSSS | [R].AAPTTS | 2xPhospho | E9QAU4 2x | 0.003083 | 0.00016 | 1 | 9 | 3 | E9QAU4 | Enhancer t | E9QAU4 [1 | 0 | 2745.233 | 0 | 0 | 0.857726 | 13.81 | 100.2 | 99.8 | 6.44 | 5.63 | | High | High | High | High | High | High | High | High | High | High | High | 0.001651 | 0.0164 | 3.22 |
| FALSE | High | AAPWGSN | [K].AAPWC | 1xPhospho | A2A9M4 1: | 0.014158 | 0.001011 | 1 | 4 | 2 | A2A9M4 | Dedicator | A2A9M4 [9 | 0 | 2369.043 | 0 | 0.08 | 0.148342 | 5.29 | 98.4 | 101.6 | 3.38 | 4.94 | | High | High | High | High | High | High | High | High | High | High | High | 0.001279 | 0.01339 | 1.99 |
| FALSE | High | AAQASLNA | [K].AAQAS | 1xPhospho | E9Q6R7 1x | 9.88E-08 | 0 | 1 | 2 | 2 | E9Q6R7 | Utrophin O | E9Q6R7 [9 | 0 | 2832.491 | 0 | 1.58 | 0.003285 | 21.83 | 48.8 | 151.2 | 5.25 | 17.69 | | High | High | High | High | High | High | High | High | High | High | High | 0 | 7.54E-09 | 4.39 |
| FALSE | High | AAQCSVAT | [K].AAQCS | 1xCarbami | E0CZ26 1x | 0.000205 | 0 | 1 | 2 | 4 | E0CZ26 | Potassium | E0CZ26 [33 | 0 | 3836.806 | 0 | 0.71 | 0.050436 | 16.52 | 78.6 | 121.4 | 10.9 | 15.35 | | High | High | High | High | High | High | High | High | High | High | High | 0.000801 | 0.007233 | 3.76 |
| FALSE | High | AAQCSVAT | [K].AAQCS | 1xCarbami | E0CZ26 1xl | 1.71E-07 | 0 | 1 | 2 | 5 | E0CZ26 | Potassium | E0CZ26 [33 | 0 | 3837.79 | 0 | 0.46 | 0.098549 | 21.55 | 84.7 | 115.3 | 4.32 | 19.08 | | High | High | High | High | High | High | High | High | High | High | High | 0 | 4.2E-08 | 3.62 |
| FALSE | High | AAQQAASS | [K].AAQQA | 1xPhospho | E9Q2U6 1x | 9.89E-05 | 0 | 1 | 4 | 1 | E9Q2U6 | Casein kina | E9Q2U6 [3 | 0 | 2330.061 | 0 | -0.48 | 0.022761 | 4.55 | 114.8 | 85.2 | 6.31 | 9.21 | | High | High | High | High | High | High | High | High | High | High | High | 0 | 1.77E-05 | 4.28 |
| FALSE | High | AAPPACSSI | [K].AAPPA( | 1xCarbami | B2RRF6 1xl | 0.058173 | 0.004227 | 1 | 1 | 2 | B2RRF6 | Zinc finger | B2RRF6 [5( | 0 | 1605.802 | 0 | -0.74 | 0.001559 | 2.07 | 123.3 | 76.7 | 7.69 | 5.63 | | High | High | High | High | High | High | High | High | High | High | High | 0.002187 | 0.02298 | 2.46 |
| FALSE | High | AAQQAASS | [K].AAQQA | 1xPhospho | E9Q3W1 1: | 2.88E-09 | 0 | 1 | 6 | 5 | E9Q3W1 | Casein kina | E9Q3W1 [3 | 0 | 2540.251 | 0 | 0.01 | 0.739769 | 4.02 | 96.8 | 103.2 | 6.54 | 10.53 | | High | High | High | High | High | High | High | High | High | High | High | 0 | 4.1E-09 | 4.45 |
| FALSE | High | AAQQQQP | [R].AAQQC | 1xPhospho | F8VQD1 1x | 0.000227 | 0 | 1 | 3 | 4 | F8VQD1 | Protein pol | F8VQD1 [1 | 0 | 1577.763 | 0 | 0.06 | 0.719591 | 8.58 | 99.8 | 100.2 | 2.65 | 8.92 | | High | High | High | High | High | High | High | High | High | High | High | 0.001651 | 0.01656 | 2.34 |
| FALSE | High | AAQSLDEN | [K].AAQSLI | 1xPhospho | Q9ESK9 1x | 0.000143 | 0 | 1 | 6 | 1 | Q9ESK9 | RB1-induci | Q9ESK9 [6 | 0 | 2773.373 | 0 | 0.56 | 0.019497 | 15.24 | 77.6 | 122.4 | 12.14 | 9.64 | | High | High | High | High | High | High | High | High | High | High | High | 0 | 2.69E-05 | 4.47 |
| FALSE | High | AAQSNENL | [K].AAQSN | 1xPhospho | D3YWX2 1: | 3.56E-08 | 0 | 1 | 2 | 1 | D3YWX2 | YLP motif-c | D3YWX2 [8 | 0 | 2385.134 | 0 | 0.17 | 0.032606 | 4.92 | 96.5 | 103.5 | 6.16 | 5.57 | | High | High | High | High | High | High | High | High | High | High | High | 0 | 2.41E-09 | 5.58 |
| FALSE | High | AAQSPQQI | [K].AAQSP( | 1xPhospho | P53986 1xl | 0.000217 | 0 | 1 | 6 | 6 | P53986 | Monocarbo | P53986 [47 | 0 | 2390.035 | 0 | 0.8 | 0.003526 | 10.69 | 76.3 | 123.7 | 7.37 | 9.03 | | High | High | High | High | High | High | High | High | High | High | High | 0 | 4.27E-05 | 2.75 |
| FALSE | High | AAQSSVTA | [R].AAQSS\ | 1xPhospho | Q6IR34-5 1 | 0.016306 | 0.001124 | 1 | 7 | 2 | Q6IR34-5 | Isoform 5 c | Q6IR34-5 [ | 0 | 2966.398 | 0 | -0.04 | 0.922271 | 10.12 | 97.3 | 102.7 | 14.87 | 18.62 | | High | High | High | High | High | High | High | High | High | High | High | 0.000585 | 0.005415 | 2.57 |
| FALSE | High | AAQSSVTA | [R].AAQSS\ | 1xDeamida | Q6IR34-5 1 | 0.057203 | 0.004227 | 1 | 7 | 4 | Q6IR34-5 | Isoform 5 c | Q6IR34-5 [ | 0 | 2967.382 | 0 | 0.01 | 0.72762 | 26.62 | 96.6 | 103.4 | 24.89 | 27.33 | | High | High | High | High | High | High | High | High | High | High | High | 0.004769 | 0.05203 | 3.09 |
| FALSE | High | AAQTCGGT | [K].AAQTC | 1xCarbami | B2RS91 1xl | 0.001881 | 5.66E-05 | 1 | 1 | 1 | B2RS91 | RNA polym | B2RS91 [26 | 0 | 3485.465 | 0 | -0.2 | 0.49955 | 5.99 | 106.5 | 93.5 | 10.54 | 6.16 | | High | High | High | High | High | High | High | High | High | High | High | 2.58E-05 | 0.000483 | 2.21 |
| FALSE | High | AAQTCGGT | [K].AAQTC | 1xCarbami | B2RS91 1xl | 0.000138 | 0 | 1 | 1 | 1 | B2RS91 | RNA polym | B2RS91 [26 | 0 | 3486.449 | 0 | -0.2 | 0.58159 | 16.72 | 101.6 | 98.4 | 23.89 | 14.47 | | High | High | High | High | High | High | High | High | High | High | High | 0 | 2.58E-05 | 3.46 |
| FALSE | High | AAQTCGGT | [K].AAQTC | 1xCarbami | B2RS91 1xl | 3.9E-05 | 0 | 1 | 1 | 1 | B2RS91 | RNA polym | B2RS91 [26 | 0 | 3501.46 | 0 | -0.01 | 0.820023 | 30.72 | 98.8 | 101.2 | 13.58 | 19.19 | | High | High | High | High | High | High | High | High | High | High | High | 0 | 6.24E-06 | 4.15 |
| FALSE | High | AASAAAAT | [R].AASAA/ | 1xPhospho | Q5DU09 1: | 4.28E-06 | 0 | 1 | 3 | 4 | Q5DU09 | Zinc finger | Q5DU09 [1 | 0 | 1551.773 | 0 | -0.09 | 0.062049 | 2.92 | 101.9 | 98.1 | 3.37 | 3.8 | | High | High | High | High | High | High | High | High | High | High | High | 0 | 5.23E-07 | 3.42 |
| FALSE | High | AASAAGAS | [R].AASAA( | 1xPhospho | A2AL50 1xl | 6.18E-05 | 0 | 1 | 3 | 3 | A2AL50 | Alkylglycer | A2AL50 [7( | 0 | 2532.311 | 0 | 0.34 | 0.015391 | 5.35 | 88.4 | 111.6 | 4.78 | 11.72 | | High | High | High | High | High | High | High | High | High | High | High | 0.000327 | 0.002911 | 3.68 |
| FALSE | High | AASACEAQ | [R].AASACI | 1xCarbami | D3Z3A8 1x | 4.66E-07 | 0 | 1 | 3 | 1 | D3Z3A8 | Unconvent | D3Z3A8 [2! | 0 | 2284.104 | 0 | 0.67 | 0.016956 | 28.77 | 77.1 | 122.9 | 8.72 | 15.96 | | High | High | High | High | High | High | High | High | High | High | High | 0 | 4.3E-08 | 3.54 |
| FALSE | High | AASADSTTI | [K].AASADS | 1xPhospho | Q9QZQ8 1: | 3.19E-07 | 0 | 1 | 3 | 3 | Q9QZQ8 | Core histor | Q9QZQ8 [1 | 0 | 2695.312 | 0 | 1.36 | 0.002367 | 13.42 | 59 | 141 | 9.6 | 14.51 | | High | High | High | High | High | High | High | High | High | High | High | 0 | 2.83E-08 | 4.23 |
| FALSE | High | AAQQLQQ | [K].AAQQL | 1xPhospho | Q9EQM3 1 | 4.33E-07 | 0 | 1 | 2 | 1 | Q9EQM3 | Homeobox | Q9EQM3 [ | 0 | 2625.339 | 0 | 0.74 | 0.107596 | 18.46 | 80.5 | 119.5 | 27.23 | 12.22 | | High | High | High | High | High | High | High | High | High | High | High | 0 | 3.97E-08 | 3.98 |
| FALSE | High | AAPGSQTK | [K].AAPGS( | 1xPhospho | O88271 1x | 0.049971 | 0.003641 | 1 | 1 | 1 | O88271 | Craniofacia | O88271 [1 | 0 | 1297.692 | 0 | -1.29 | 0.004098 | 2.9 | 141.9 | 58.1 | 7.7 | 18.57 | | High | High | High | High | High | High | High | High | High | High | High | 0.001902 | 0.01924 | 1.22 |
| FALSE | High | AANEGHPF | [K].AANEG | 1xPhospho | Q3UHR0 1: | 0.000822 | 0 | 1 | 2 | 1 | Q3UHR0 | BAH and cc | Q3UHR0 [2 | 0 | 2425.216 | 0 | -0.21 | 0.188616 | 5.54 | 104.9 | 95.1 | 13.05 | 9.39 | | High | High | High | High | High | High | High | High | High | High | High | 0 | 0.000192 | 2.3 |
| FALSE | High | AAMYDIISS | [K].AAMYD | 1xPhospho | Q6KCD5 1x | 0.000164 | 0 | 1 | 4 | 6 | Q6KCD5 | Nipped-B-I | Q6KCD5 [3 | 0 | 1850.938 | 0 | 0.11 | 0.232124 | 6.04 | 92.9 | 107.1 | 3.69 | 20.27 | | High | High | High | High | High | High | High | High | High | High | High | 0 | 0.000233 | 3.6 |
| FALSE | High | AAGYDLFS/ | [R].AAGYD | 1xPhospho | Q8VCG1 1x | 2.13E-07 | 0 | 1 | 3 | 1 | Q8VCG1 | Deoxyuridi | Q8VCG1 [8 | 0 | 2696.261 | 0 | -1.59 | 0.000695 | 1.25 | 148.4 | 51.6 | 6.81 | 13.84 | | High | High | High | High | High | High | High | High | High | High | High | 0 | 1.78E-08 | 4.41 |
| FALSE | High | AAHDESPG | [R].AAHDE | 1xDeamida | Q78ZJ8 1xl | 0.001337 | 0 | 1 | 3 | 2 | Q78ZJ8 | MCG22989 | Q78ZJ8 [18 | 0 | 3354.637 | 0 | 0.33 | 0.052243 | 18.04 | 85.2 | 114.8 | 6.74 | 13.92 | | High | High | High | High | High | High | High | High | High | High | High | 0 | 0.00033 | 3.41 |
| FALSE | High | AAHQESDN | [R].AAHQE | 1xPhospho | A2A791 1x | 2.5E-10 | 0 | 1 | 2 | 4 | A2A791 | Zinc finger | A2A791 [1: | 0 | 2747.34 | 0 | -0.39 | 0.002252 | 2.99 | 112.3 | 87.7 | 1.66 | 5.97 | | High | High | High | High | High | High | High | High | High | High | High | 0 | 9.08E-12 | 5.37 |
| FALSE | High | AAHQESDN | [R].AAHQE | 1xDeamida | A2A791 1x | 1.28E-05 | 0 | 1 | 2 | 4 | A2A791 | Zinc finger | A2A791 [1: | 0 | 2748.324 | 0 | -0.22 | 0.17681 | 5.37 | 108.5 | 91.5 | 8.13 | 12.51 | | High | High | High | High | High | High | High | High | High | High | High | 0 | 6.24E-06 | 3.57 |
| FALSE | High | AAHTEDIN/ | [R].AAHTEI | 1xCarbami | Q99N57-2 | 2.78E-06 | 0 | 1 | 4 | 5 | Q99N57-2 | Isoform 2 c | Q99N57-2 | 0 | 2167.005 | 0 | -0.2 | 0.028156 | 6.64 | 108.8 | 91.2 | 3.53 | 10.67 | | High | High | High | High | High | High | High | High | High | High | High | 0 | 5.38E-07 | 3.8 |
| FALSE | High | AAKESEEEI | [K].AAKESE | 1xPhospho | E9Q5C9 1x | 1.52E-07 | 0 | 1 | 2 | 12 | E9Q5C9 | Nucleolar a | E9Q5C9 [5 | 1 | 2792.285 | 0 | -0.09 | 0.077511 | 7.94 | 102.8 | 97.2 | 2.43 | 6.73 | | High | High | High | High | High | High | High | High | High | High | High | 0 | 1.5E-06 | 4.92 |
| FALSE | High | AALDDDVP | [K].AALDDI | 1xPhospho | Q7TT79 1x | 0.000113 | 0 | 1 | 5 | 2 | Q7TT79 | Microceph | Q7TT79 [1 | 0 | 2066.041 | 0 | -1.15 | 0.000432 | 2.49 | 137.8 | 62.2 | 7.75 | 13.03 | | High | High | High | High | High | High | High | High | High | High | High | 0 | 2.08E-05 | 4.25 |

| | | | | | | | | | | | | | | | | | | | | | | | | | | | | | | | | | | | | | | |
|---|---|---|---|---|---|---|---|---|---|---|---|---|---|---|---|---|---|---|---|---|---|---|---|---|---|---|---|---|---|---|---|---|---|---|---|---|---|---|
| FALSE | High | AALEDSNG | [K].AALEDS | 1xDeamida | Q3U2Z5 1x | 3.12E-05 | 0 | 1 | 3 | 1 | Q3U2Z5 | Putative ur | Q3U2Z5 [8 | 0 | 2175.968 | 0 | 0 | 0.848529 | 24.87 | 99.9 | 100.1 | 6.64 | 12.96 | | High | High | High | High | High | High | High | High | High | High | High | 0 | 4.85E-06 | 3.42 |
| FALSE | High | AALFLLQHT | [K].AALFLL | 1xPhospho | E9PZJ2 1xP | 0.113753 | 0.009241 | 1 | 3 | 1 | E9PZJ2 | Interferon | E9PZJ2 [44 | 0 | 2229.127 | 0 | 0.21 | 0.037743 | 12.96 | 90.3 | 109.7 | 3.27 | 9.09 | | High | High | High | High | High | High | High | High | High | High | High | 0.004654 | 0.05013 | 1.71 |
| FALSE | High | AALGLQDS | [K].AALGL( | 1xPhospho | Q569Z5 1x | 2.36E-05 | 0 | 1 | 3 | 6 | Q569Z5 | Probable A | Q569Z5 [7 | 0 | 3564.634 | 0 | -0.81 | 0.009277 | 6.31 | 128.8 | 71.2 | 13 | 21.14 | | High | High | High | High | High | High | High | High | High | High | High | 0 | 3.54E-06 | 4.04 |
| FALSE | High | AALGLQDS | [K].AALGL( | 1xPhospho | Q569Z5 1x | 1.09E-07 | 0 | 1 | 3 | 20 | Q569Z5 | Probable A | Q569Z5 [7 | 0 | 3580.629 | 0 | -0.47 | 0.210155 | 9.26 | 116.1 | 83.9 | 12.1 | 23.89 | | High | High | High | High | High | High | High | High | High | High | High | 0 | 8.42E-09 | 6.53 |
| FALSE | High | AALLAAFQ | [R].AALLA/ | 1xPhospho | Q3UHQ6 1 | 9.11E-05 | 0 | 1 | 3 | 1 | Q3UHQ6 | Protein do | Q3UHQ6 [ | 0 | 1679.872 | 0 | 1.24 | 0.0019 | 20.44 | 62 | 138 | 7.62 | 12.48 | | High | High | High | High | High | High | High | High | High | High | High | 0 | 1.62E-05 | 2.76 |
| FALSE | High | AALLAQYA | [K].AALLA( | 1xPhospho | Q9CR29 1x | 4.68E-09 | 0 | 1 | 2 | 1 | Q9CR29 | Coiled-coil | Q9CR29 [1 | 0 | 2376.069 | 0 | -0.02 | 0.774201 | 17.68 | 107.1 | 92.9 | 54.39 | 45.82 | | High | High | High | High | High | High | High | High | High | High | High | 0 | 2.45E-10 | 4.09 |
| FALSE | High | AALLAQYA | [K].AALLA( | 1xPhospho | Q9CR29 1x | 1.18E-09 | 0 | 1 | 2 | 8 | Q9CR29 | Coiled-coil | Q9CR29 [1 | 0 | 2605.232 | 0 | 0.05 | 0.722999 | 17.14 | 100.2 | 99.8 | 3.52 | 10.71 | | High | High | High | High | High | High | High | High | High | High | High | 0 | 3.23E-09 | 5.29 |
| FALSE | High | AALLAQYA | [K].AALLA( | 1xPhospho | Q9CR29 1x | 3.57E-05 | 0 | 1 | 2 | 1 | Q9CR29 | Coiled-coil | Q9CR29 [1 | 1 | 2733.327 | 0 | 0.25 | 0.998391 | 48.54 | 91.5 | 108.5 | 71.33 | 76.04 | | High | High | High | High | High | High | High | High | High | High | High | 0 | 5.64E-06 | 4.25 |
| FALSE | High | AALLAQYA | [K].AALLA( | 1xPhospho | Q9CR29 1x | 1.25E-06 | 0 | 1 | 2 | 4 | Q9CR29 | Coiled-coil | Q9CR29 [1 | 1 | 2962.49 | 0 | -0.15 | 0.037628 | 4.73 | 105.1 | 94.9 | 6.26 | 6.5 | | High | High | High | High | High | High | High | High | High | High | High | 0 | 2.76E-06 | 3.46 |
| FALSE | High | AALLTQQA | [R].AALLTC | 1xPhospho | Q3U0M8 1 | 0.000109 | 0 | 1 | 3 | 1 | Q3U0M8 | Putative ur | Q3U0M8 [ | 0 | 1823.874 | 0 | -0.28 | 0.111673 | 14.89 | 104.3 | 95.7 | 11.39 | 15 | | High | High | High | High | High | High | High | High | High | High | High | 0 | 1.99E-05 | 3.74 |
| FALSE | High | AALLTQQA | [R].AALLTC | 1xPhospho | Q3U0M8 1 | 4.26E-07 | 0 | 1 | 3 | 2 | Q3U0M8 | Putative ur | Q3U0M8 [ | 1 | 2663.344 | 0 | -0.57 | 0.000629 | 3.24 | 119 | 81 | 2.14 | 8.38 | | High | High | High | High | High | High | High | High | High | High | High | 0 | 3.88E-08 | 3.52 |
| FALSE | High | AAIPPPVYE | [K].AAIPPP | 1xPhospho | Q8JZX4 1xI | 0.000616 | 0 | 1 | 1 | 4 | Q8JZX4 | Splicing fac | Q8JZX4 [20 | 1 | 4199.115 | 0 | -0.87 | 0.005619 | 4.8 | 127.6 | 72.4 | 8.27 | 10.91 | | High | High | High | High | High | High | High | High | High | High | High | 0.002187 | 0.02307 | 3.05 |
| FALSE | High | AAIPVFDLD | [R].AAIPVF | 1xCarbami | Z4YKQ9 1x | 6.41E-05 | 0 | 1 | 3 | 8 | Z4YKQ9 | ATPase fan | Z4YKQ9 [7 | 0 | 3597.634 | 0 | 0.13 | 0.89327 | 7.49 | 98.7 | 101.3 | 13.03 | 7.05 | | High | High | High | High | High | High | High | High | High | High | High | 0 | 1.09E-05 | 4.38 |
| FALSE | High | AALQAAPE | [K].AALQA/ | 1xPhospho | Q9WVH4 1 | 5.23E-10 | 0 | 1 | 2 | 2 | Q9WVH4 | Forkhead b | Q9WVH4 [ | 0 | 2424.206 | 0 | -0.75 | 0.000348 | 2.45 | 127.2 | 72.8 | 3.93 | 7.16 | | High | High | High | High | High | High | High | High | High | High | High | 0 | 2.1E-11 | 3.67 |
| FALSE | High | AALSPGSVI | [R].AALSPG | 1xPhospho | B2RXC2 1x | 0.000128 | 0 | 1 | 2 | 2 | B2RXC2 | Kinase OS= | B2RXC2 [3 | 0 | 1554.788 | 0 | 2.32 | 0.000864 | 28.96 | 36.6 | 163.4 | 22.12 | 10.01 | | High | High | High | High | High | High | High | High | High | High | High | 0.000538 | 0.004623 | 2.46 |
| FALSE | High | AALSSQQC | [K].AALSSC | 1xPhospho | Q6Y7W8 1: | 5.93E-06 | 0 | 1 | 8 | 2 | Q6Y7W8 | GRB10-inte | Q6Y7W8 [6 | 0 | 2993.659 | 0 | -0.85 | 0.034448 | 14.52 | 118.9 | 81.1 | 19.79 | 26.73 | | High | High | High | High | High | High | High | High | High | High | High | 0 | 1.35E-06 | 4.35 |
| FALSE | High | AAMHTYSS | [K].AAMHT | 1xPhospho | Q8R4Z6 1x | 0.000174 | 0 | 1 | 2 | 3 | Q8R4Z6 | DEAD-box | Q8R4Z6 [4 | 0 | 2101.983 | 0 | -1.11 | 0.112706 | 11.09 | 122.8 | 77.2 | 37.65 | 28.24 | | High | High | High | High | High | High | High | High | High | High | High | 0 | 3.35E-05 | 3.4 |
| FALSE | High | AAMHTYSS | [K].AAMHT | 1xPhospho | Q8R4Z6 1x | 1.41E-05 | 0 | 1 | 2 | 2 | Q8R4Z6 | DEAD-box | Q8R4Z6 [4 | 0 | 2315.151 | 0 | -1.58 | 0.000209 | 1.02 | 149.9 | 50.1 | 5.52 | 13.18 | | High | High | High | High | High | High | High | High | High | High | High | 0 | 5.56E-06 | 3.96 |
| FALSE | High | AAMHTYSS | [K].AAMHT | 1xPhospho | Q8R4Z6 1x | 1.14E-08 | 0 | 1 | 2 | 12 | Q8R4Z6 | DEAD-box | Q8R4Z6 [4 | 0 | 2331.146 | 0 | -1.08 | 0.002445 | 2.3 | 136.2 | 63.8 | 5.88 | 13.91 | | High | High | High | High | High | High | High | High | High | High | High | 0 | 2.41E-08 | 3.92 |
| FALSE | High | AAMHTYSS | [K].AAMHT | 2xPhospho | Q8R4Z6 2x | 7.7E-06 | 0 | 1 | 2 | 2 | Q8R4Z6 | DEAD-box | Q8R4Z6 [4 | 0 | 2395.117 | 0 | -0.34 | 0.030282 | 12.88 | 111.3 | 88.7 | 4.01 | 12.75 | | High | High | High | High | High | High | High | High | High | High | High | 0 | 1E-06 | 2.74 |
| FALSE | High | AAMHTYSS | [K].AAMHT | 1xPhospho | Q8R4Z6 1x | 0.000916 | 0 | 1 | 2 | 2 | Q8R4Z6 | DEAD-box | Q8R4Z6 [4 | 1 | 2672.409 | 0 | -1.29 | 0.002205 | 3.01 | 138.3 | 61.7 | 12.48 | 6.52 | | High | High | High | High | High | High | High | High | High | High | High | 0 | 0.000216 | 2.84 |
| FALSE | High | AAMYDIISS | [K].AAMYD | 1xPhospho | Q6KCD5 1x | 0.000186 | 0 | 1 | 4 | 1 | Q6KCD5 | Nipped-B-l | Q6KCD5 [3 | 0 | 1834.943 | 0 | 0.15 | 0.972151 | 12.05 | 93.6 | 106.4 | 8.02 | 22.44 | | High | High | High | High | High | High | High | High | High | High | High | 0 | 3.61E-05 | 3.61 |
| FALSE | High | AASAEAGD | [R].AASAE/ | 1xPhospho | Q9WVA7 1 | 0.002313 | 0.000109 | 1 | 1 | 2 | Q9WVA7 | FRAT3 OS= | Q9WVA7 [ | 0 | 1745.842 | 0 | -0.81 | 0.000562 | 2.72 | 125.1 | 74.9 | 4.66 | 7.98 | | High | High | High | High | High | High | High | High | High | High | High | 4.94E-05 | 0.000613 | 2.45 |
| FALSE | High | ADGDWDD | [R].ADGDV | 1xPhospho | Q8K019 1x | 0.000112 | 0 | 1 | 11 | 1 | Q8K019 | Bcl-2-assoc | Q8K019 [3 | 1 | 3200.492 | 0 | 0.03 | 0.872326 | 22.07 | 102.6 | 97.4 | 20.07 | 26.38 | | High | High | High | High | High | High | High | High | High | High | High | 0 | 2.04E-05 | 2.81 |
| FALSE | High | ADGGTGDI | [R].ADGGT | 1xPhospho | Q921C5-2 | 4.31E-05 | 0 | 1 | 4 | 4 | Q921C5-2 | Isoform 2 c | Q921C5-2 | 0 | 2607.177 | 0 | 1.55 | 0.000812 | 19.11 | 50.5 | 149.5 | 14.03 | 7.19 | | High | High | High | High | High | High | High | High | High | High | High | 0 | 7E-06 | 4.62 |
| FALSE | High | ADGPRPSLI | [R].ADGPR | 1xPhospho | E9QMQ3 1 | 2.29E-05 | 0 | 2 | 7 | 6 | E9QMQ3; ( | TRIO and F | E9QMQ3 [ | 0 | 2761.376 | 0 | 1 | 0.000364 | 2.61 | 66.4 | 133.6 | 5.63 | 6.54 | Shared | High | High | High | High | High | High | High | High | High | High | High | 0 | 3.45E-06 | 3.55 |
| FALSE | High | AETFGGYD | [R].AETFG( | 1xPhospho | Q6P9R4 1x | 3.34E-07 | 0 | 1 | 2 | 2 | Q6P9R4 | Rho guanir | Q6P9R4 [5 | 0 | 1810.81 | 0 | 0.78 | 0.001337 | 2.79 | 73.3 | 126.7 | 5.8 | 5.45 | | High | High | High | High | High | High | High | High | High | High | High | 0 | 2.95E-08 | 3.68 |
| FALSE | High | AETFGGYD | [R].AETFG( | 1xPhospho | Q6P9R4 1x | 1.53E-06 | 0 | 1 | 2 | 2 | Q6P9R4 | Rho guanir | Q6P9R4 [5 | 0 | 2039.973 | 0 | 0.81 | 0.000546 | 13.15 | 70.7 | 129.3 | 9.18 | 2.15 | | High | High | High | High | High | High | High | High | High | High | High | 0 | 0.00021 | 3.15 |
| FALSE | High | AETPPSHQ | [R].AETPPS | 1xPhospho | B1ATC3 1x | 4.89E-05 | 0 | 1 | 3 | 2 | B1ATC3 | Trinucleoti | B1ATC3 [9 | 0 | 1915.922 | 0 | -1.06 | 0.000309 | 1.68 | 134.2 | 65.8 | 2.77 | 6.01 | | High | High | High | High | High | High | High | High | High | High | High | 0 | 8.02E-06 | 2.3 |
| FALSE | High | AETSESSGS | [K].AETSES | 1xPhospho | Q9Z2D6-2 | 0.003857 | 0.000261 | 1 | 4 | 1 | Q9Z2D6-2 | Isoform B ( | Q9Z2D6-2 | 0 | 2269.043 | 0 | 0.3 | 0.412897 | 32.73 | 89.8 | 110.2 | 19.33 | 41.89 | | High | High | High | High | High | High | High | High | High | High | High | 0.000117 | 0.001084 | 2.67 |
| FALSE | High | AETSESSGS | [K].AETSES | 1xPhospho | Q9Z2D6-2 | 2.91E-07 | 0 | 1 | 4 | 2 | Q9Z2D6-2 | Isoform B ( | Q9Z2D6-2 | 0 | 2498.206 | 0 | 1.27 | 0.000558 | 8.44 | 59.7 | 140.3 | 2.24 | 7.19 | | High | High | High | High | High | High | High | High | High | High | High | 0 | 2.54E-08 | 3.49 |
| FALSE | High | AETSLGIPS | [K].AETSLG | 1xPhospho | Q8VE88 1x | 3.1E-06 | 0 | 1 | 2 | 3 | Q8VE88 | Protein FAI | Q8VE88 [1 | 0 | 2367.246 | 0 | 1.33 | 0.001718 | 7.88 | 56.6 | 143.4 | 10.79 | 13.83 | | High | High | High | High | High | High | High | High | High | High | High | 0 | 3.6E-07 | 2.54 |
| FALSE | High | AEVDGSLSI | [K].AEVDG | 2xPhospho | E9Q166 2x | 0.087983 | 0.006639 | 1 | 1 | 3 | E9Q166 | ATPase fan | E9Q166 [7 | 0 | 2314.077 | 0 | -0.46 | 0.000226 | 1.33 | 116.4 | 83.6 | 3.04 | 2.87 | | High | High | High | High | High | High | High | High | High | High | High | 0.005324 | 0.05783 | 2.93 |
| FALSE | High | AEVEQETV | [K].AEVEQI | 1xPhospho | Q8R3C6 1x | 0.024001 | 0.001612 | 1 | 2 | 1 | Q8R3C6 | Probable R | Q8R3C6 [6 | 1 | 3527.761 | 0 | -0.92 | 0.019318 | 3.2 | 127.8 | 72.2 | 12.76 | 14.87 | | High | High | High | High | High | High | High | High | High | High | High | 0.000859 | 0.008407 | 2.97 |
| FALSE | High | AEVESPLK | [-].AEVESP | 1xAcetyl [N | G3UWH9 1 | 0.00614 | 0.00062 | 1 | 1 | 2 | G3UWH9 | Conserved | G3UWH9 [ | 0 | 1223.612 | 0 | -1.07 | 0.000241 | 1.31 | 134.2 | 65.8 | 2.51 | 8.82 | | High | High | High | High | High | High | High | High | High | High | High | 0.000538 | 0.004627 | 2.41 |
| FALSE | High | AEVTAPSTS | [R].AEVTAI | 1xPhospho | A0A087W( | 0.004411 | 0.000409 | 1 | 5 | 2 | A0A087W( | Snf2-relate | A0A087W( | 0 | 2859.409 | 0 | -0.64 | 0.001285 | 3.54 | 121.8 | 78.2 | 6.39 | 6.54 | | High | High | High | High | High | High | High | High | High | High | High | 0.000183 | 0.001257 | 2.82 |
| FALSE | High | AFAMSAPT | [R].AFAMS | 1xDeamida | Q8BP60 1x | 2.9E-05 | 0 | 1 | 3 | 9 | Q8BP60 | Putative ur | Q8BP60 [5 | 0 | 4965.344 | 0 | -0.46 | 0.068366 | 9.3 | 110.1 | 89.9 | 16.06 | 11.53 | | High | High | High | High | High | High | High | High | High | High | High | 0 | 9.03E-05 | 5.65 |
| FALSE | High | AFDGAGSP | [R].AFDGA( | 1xPhospho | Q5I1X5 1xI | 0.003688 | 0.00021 | 1 | 1 | 2 | Q5I1X5 | RelA-assoc | Q5I1X5 [14 | 0 | 1380.626 | 0 | 1.7 | 0.000278 | 15.26 | 47 | 153 | 6.57 | 4.3 | | High | High | High | High | High | High | High | High | High | High | High | 9.42E-05 | 0.001029 | 2.53 |
| FALSE | High | AFEEDIETP | [K].AFEEDI | 1xPhospho | A2AJK6 1xI | 4.12E-05 | 0 | 1 | 1 | 1 | A2AJK6 | Chromodo | A2AJK6 [25 | 0 | 1725.83 | 0 | 0.23 | 0.571249 | 14.62 | 92 | 108 | 28.73 | 17.8 | | High | High | High | High | High | High | High | High | High | High | High | 0 | 6.64E-06 | 2.98 |
| FALSE | High | AESVSSDEI | [K].AESVSS | 2xPhospho | Q5DTL2 2x | 0.000173 | 0 | 1 | 3 | 2 | Q5DTL2 | MKIAA414 | Q5DTL2 [7 | 0 | 2979.266 | 0 | 2.77 | 0.000581 | 42.3 | 25.6 | 174.4 | 43.86 | 8.37 | | High | High | High | High | High | High | High | High | High | High | High | 0 | 3.34E-05 | 4.94 |
| FALSE | High | AFEGQAHS | [R].AFEGQ. | 1xPhospho | P32037 1xI | 0.000543 | 0 | 1 | 3 | 2 | P32037 | Solute carr | P32037 [46 | 0 | 1569.783 | 0 | -1.75 | 0.01 | 4.68 | 156.9 | 43.1 | 20.99 | 36.14 | | High | High | High | High | High | High | High | High | High | High | High | 0 | 0.00012 | 2.11 |
| FALSE | High | AFGPGLQG | [K].AFGPGI | 1xPhospho | Q8BTM8 1: | 1.08E-06 | 0 | 1 | 6 | 2 | Q8BTM8 | Filamin-A ( | Q8BTM8 [ | 0 | 1765.858 | 0 | 0.64 | 0.000816 | 5.16 | 79.1 | 120.9 | 6.66 | 4.39 | | High | High | High | High | High | High | High | High | High | High | High | 0 | 1.1E-07 | 4.05 |
| FALSE | High | AFKPEETSS | [R].AFKPEE | 1xPhospho | B2RT41 1xI | 0.090153 | 0.00698 | 1 | 2 | 1 | B2RT41 | Zinc finger, | B2RT41 [64 | 0 | 3433.704 | 0 | -0.44 | 0.028848 | 12.31 | 115 | 85 | 6.07 | 14.29 | | High | High | High | High | High | High | High | High | High | High | High | 0.003606 | 0.03822 | 2.57 |
| FALSE | High | AFKPEETSS | [R].AFKPEE | 1xDeamida | B2RT41 1xI | 2.15E-05 | 0 | 1 | 2 | 3 | B2RT41 | Zinc finger, | B2RT41 [64 | 0 | 3434.688 | 0 | -0.42 | 0.017524 | 1.24 | 116.3 | 83.7 | 9 | 11.35 | | High | High | High | High | High | High | High | High | High | High | High | 0 | 3.19E-06 | 3.4 |
| FALSE | High | AFLAELEQI | [K].AFLAEL | 1xPhospho | B1ARU1 1x | 3.18E-06 | 0 | 2 | 13 | 2 | B1ARU1; F | Microtubu | B1ARU1 [2 | 0 | 1884.987 | 0 | -0.06 | 0.916669 | 5.11 | 101.1 | 98.9 | 3.15 | 7.8 | Shared | High | High | High | High | High | High | High | High | High | High | High | 0 | 3.72E-07 | 2.67 |
| FALSE | High | AFLSPPTLL | [R].AFLSPP | 1xPhospho | Q91ZV0 1x | 0.004143 | 0.00036 | 1 | 9 | 2 | Q91ZV0 | Melanoma | Q91ZV0 [1 | 0 | 1819.992 | 0 | 0.6 | 0.003019 | 4.45 | 79.2 | 120.8 | 5.79 | 11.58 | | High | High | High | High | High | High | High | High | High | High | High | 0.000161 | 0.001168 | 2.3 |
| FALSE | High | AFLSPPTLL | [R].AFLSPP | 1xPhospho | Q91ZV0 1x | 0.004143 | 0.00036 | 1 | 9 | 1 | Q91ZV0 | Melanoma | Q91ZV0 [1 | 1 | 2824.569 | 0 | 0.06 | 0.873799 | 17.74 | 101.7 | 98.3 | 9.48 | 9.03 | | High | High | High | High | High | High | High | High | High | High | High | 0.000161 | 0.001177 | 2.43 |
| FALSE | High | AFQFQPPL | [K].AFQFQ | 1xPhospho | Q9CYA6 1x | 0.076555 | 0.005826 | 1 | 2 | 3 | Q9CYA6 | Zinc finger | Q9CYA6 [4 | 0 | 3810.084 | 0 | 0.83 | 0.028177 | 22.78 | 65 | 135 | 10.41 | 25.26 | | High | High | High | High | High | High | High | High | High | High | High | 0.007121 | 0.07726 | 3.16 |
| FALSE | High | AFQFQPPL | [K].AFQFQ | 2xPhospho | Q9CYA6 2x | 0.002719 | 0.000109 | 1 | 2 | 1 | Q9CYA6 | Zinc finger | Q9CYA6 [4 | 0 | 3890.05 | 0 | -0.69 | 0.027495 | 12.18 | 116 | 84 | 9.76 | 25.33 | | High | High | High | High | High | High | High | High | High | High | High | 4.94E-05 | 0.000733 | 2.96 |
| FALSE | High | AFQTSTSEI | [K].AFQTST | 1xPhospho | Q569L8 1xI | 1.67E-05 | 0 | 1 | 1 | 3 | Q569L8 | Centromer | Q569L8 [6 | 0 | 2163.991 | 0 | -0.38 | 0.030639 | 5.25 | 114.6 | 85.4 | 6.01 | 18.92 | | High | High | High | High | High | High | High | High | High | High | High | 0 | 2.49E-06 | 3.75 |
| FALSE | High | AFSDPFVE/ | [R].AFSDPF | 1xPhospho | Q571M2 1: | 0.000337 | 0 | 1 | 5 | 2 | Q571M2 | MKIAA402 | Q571M2 [1 | 0 | 1777.881 | 0 | 1.47 | 0.000562 | 11.87 | 52.9 | 147.1 | 4.49 | 7.57 | | High | High | High | High | High | High | High | High | High | High | High | 0.000294 | 0.002366 | 3.9 |
| FALSE | High | AFSEDEAL/ | [R].AFSEDE | 1xPhospho | G3X932 1x | 4.31E-05 | 0 | 1 | 5 | 1 | G3X932 | DNA segm | G3X932 [5 | 0 | 2076.989 | 0 | -0.12 | 0.364859 | 15.72 | 105 | 95 | 4.87 | 12.74 | | High | High | High | High | High | High | High | High | High | High | High | 0 | 6.97E-06 | 4.02 |
| FALSE | High | AFSEQGPLI | [R].AFSEQ( | 1xPhospho | Q68FD7 1x | 0.108463 | 0.008839 | 1 | 2 | 1 | Q68FD7 | Folliculin-ir | Q68FD7 [2 | 0 | 1313.645 | 0 | -0.13 | 0.227279 | 1.06 | 103.6 | 96.4 | 6.19 | 3.42 | | High | High | High | High | High | High | High | High | High | High | High | 0.004422 | 0.0476 | 1.86 |
| FALSE | High | AFSESHISLI | [R].AFSESH | 1xPhospho | Q1W617 1: | 3.21E-06 | 0 | 1 | 2 | 2 | Q1W617 | Protein Shr | Q1W617 [ | 0 | 2668.317 | 0 | 2.11 | 0.000405 | 12.26 | 36.8 | 163.2 | 4.07 | 8.23 | | High | High | High | High | High | High | High | High | High | High | High | 0 | 5.57E-07 | 4.25 |
| FALSE | High | AFGESSTES | [R].AFGESS | 1xCarbami | A5A4Y9 1x | 1.39E-06 | 0 | 2 | 4 | 4 | A5A4Y9; Q | Protein ph | A5A4Y9 [7 | 0 | 2768.128 | 0 | -0.18 | 0.275412 | 13.25 | 105.4 | 94.6 | 11.93 | 27.99 | Shared | High | High | High | High | High | High | High | High | High | High | High | 0 | 1.47E-07 | 3.44 |
| FALSE | High | AFSSPQEEI | [K].AFSSPC | 1xPhospho | Q8CB77 1x | 0.000179 | 0 | 1 | 1 | 3 | Q8CB77 | Elongin-A ( | Q8CB77 [5 | 0 | 2050.896 | 0 | -0.7 | 0.001775 | 3.51 | 125.6 | 74.4 | 7.82 | 4.46 | | High | High | High | High | High | High | High | High | High | High | High | 0 | 0.000109 | 2.99 |
| FALSE | High | AESSSSISSS | [R].AESSSS | 2xPhospho | Q3UQN2 2 | 0.012731 | 0.000889 | 1 | 3 | 1 | Q3UQN2 | F-BAR dom | Q3UQN2 [ | 0 | 2742.243 | 0 | 0.07 | 0.421291 | 10.77 | 96.9 | 103.1 | 8.26 | 11.17 | | High | High | High | High | High | High | High | High | High | High | High | 0.000449 | 0.004096 | 2.75 |
| FALSE | High | AESSQNSP | [K].AESSQI | 1xPhospho | Q58FA4 1x | 0.000225 | 0 | 1 | 2 | 1 | Q58FA4 | Transcripti | Q58FA4 [4 | 0 | 1892.952 | 0 | -0.79 | 0.000354 | 1.73 | 127.7 | 72.3 | 3.72 | 3.23 | | High | High | High | High | High | High | High | High | High | High | High | 0 | 4.46E-05 | 2.58 |
| FALSE | High | AENYDISPA | [R].AENYDI | 1xPhospho | A0A1S6GW | 0.006031 | 0.00062 | 1 | 2 | 1 | A0A1S6GW | Uncharacte | A0A1S6GW | 0 | 1559.694 | 0 | -0.52 | 0.021713 | 6.49 | 115.3 | 84.7 | 5.64 | 11.77 | | High | High | High | High | High | High | High | High | High | High | High | 0.000275 | 0.001776 | 1.84 |
| FALSE | High | AEPATPAE/ | [K].AEPATI | 1xPhospho | Q3U2W2 1 | 1.08E-05 | 0 | 1 | 3 | 2 | Q3U2W2 | MYB bindir | Q3U2W2 [ | 1 | 2682.283 | 0 | -0.24 | 0.098754 | 4.22 | 108.6 | 91.4 | 9.93 | 12.79 | | High | High | High | High | High | High | High | High | High | High | High | 0 | 5.53E-06 | 2.26 |
| FALSE | High | AEPLTPTED | [K].AEPLTP | 1xCarbami | Q8VBW5 1 | 8.48E-05 | 0 | 1 | 5 | 4 | Q8VBW5 | HMG box t | Q8VBW5 [ | 0 | 3566.724 | 0 | 0.16 | 0.166849 | 26.44 | 93 | 107 | 7.29 | 11.54 | | High | High | High | High | High | High | High | High | High | High | High | 0.006415 | 0.06909 | 2.85 |
| FALSE | High | AEPPKPEV\ | [R].AEPPKF | 2xPhospho | Q9JLM4 2x | 0.006031 | 0.00062 | 1 | 7 | 2 | Q9JLM4 | Zinc finger | Q9JLM4 [2 | 0 | 4848.1 | 0 | 0.01 | 0.983921 | 18.15 | 100.4 | 99.6 | 9.96 | 10 | | High | High | High | High | High | High | High | High | High | High | High | 0.000275 | 0.001784 | 4.77 |
| FALSE | High | AEPQPLSP/ | [K].AEPQPI | 1xCarbami | F6VAN0 1x | 0.001278 | 0 | 1 | 2 | 1 | F6VAN0 | Cyclic AMP | F6VAN0 [7 | 0 | 2267.058 | 0 | 1.53 | 0.001212 | 7.2 | 50.6 | 149.4 | 8.72 | 9.02 | | High | High | High | High | High | High | High | High | High | High | High | 0 | 0.000313 | 3.27 |
| FALSE | High | AEPTPLASS | [K].AEPTPL | 1xPhospho | Q3UMA3 1 | 4.47E-05 | 0 | 1 | 6 | 4 | Q3UMA3 | Hepatocyte | Q3UMA3 [ | 0 | 3986.947 | 0 | 1.4 | 0.000847 | 11.17 | 51 | 149 | 14.56 | 10.58 | | High | High | High | High | High | High | High | High | High | High | High | 0 | 7.29E-06 | 4.22 |
| FALSE | High | AEPYDTSQ | [K].AEPYDT | 1xPhospho | B1ASU9 1x | 0.001107 | 0 | 1 | 7 | 1 | B1ASU9 | Serine/thre | B1ASU9 [6 | 0 | 1633.787 | 0 | 0.78 | 0.00491 | 14.9 | 72.4 | 127.6 | 13.87 | 8.2 | | High | High | High | High | High | High | High | High | High | High | High | 0 | 0.000268 | 1.57 |
| FALSE | High | AEPYVASE\ | [K].AEPYV/ | 1xPhospho | F8VPN7 1x | 0.110202 | 0.008961 | 1 | 11 | 1 | F8VPN7 | Dedicator ( | F8VPN7 [9 | 0 | 1694.844 | 0 | 0.86 | 0.039417 | 17.81 | 71 | 129 | 17.33 | 22.21 | | High | High | High | High | High | High | High | High | High | High | High | 0.00449 | 0.04851 | 1.58 |
| FALSE | High | AEQAPEAL | [R].AEQAPI | 1xPhospho | Q3U480 1x | 1.74E-06 | 0 | 1 | 3 | 1 | Q3U480 | Putative ur | Q3U480 [4 | 0 | 2505.238 | 0 | -1.17 | 0.004532 | 0.36 | 133.5 | 66.5 | 9.95 | 12.47 | | High | High | High | High | High | High | High | High | High | High | High | 0 | 1.9E-07 | 4.24 |
| FALSE | High | AEQLAVNG | [R].AEQLA\ | 1xPhospho | Q62130 1x | 1.24E-08 | 0 | 1 | 3 | 1 | Q62130 | Tyrosine-p | Q62130 [7 | 0 | 2921.527 | 0 | 0.8 | 0.000892 | 8.49 | 74.9 | 125.1 | 7.1 | 5.04 | | High | High | High | High | High | High | High | High | High | High | High | 0 | 7.35E-10 | 4.75 |
| FALSE | High | AEQLAVNG | [R].AEQLA\ | 1xDeamida | Q62130 1x | 3.79E-07 | 0 | 1 | 3 | 4 | Q62130 | Tyrosine-p | Q62130 [7 | 0 | 2922.511 | 0 | 0.78 | 0.008671 | 17.75 | 73.7 | 126.3 | 8.64 | 14.64 | | High | High | High | High | High | High | High | High | High | High | High | 0 | 3.63E-06 | 4.39 |
| FALSE | High | AEQSLHDLI | [R].AEQSLI | 1xPhospho | A0A1B0GR | 0.004451 | 0.000409 | 1 | 3 | 2 | A0A1B0GR | Stromal int | A0A1B0GR | 0 | 1634.774 | 0 | 1.37 | 0.058587 | 41.63 | 52.2 | 147.8 | 11.17 | 40.05 | | High | High | High | High | High | High | High | High | High | High | High | 0.000183 | 0.001267 | 1.95 |
| FALSE | High | AEQSPVSSI | [K].AEQSP\ | 1xPhospho | Q80TS5 1x | 8.95E-05 | 0 | 1 | 4 | 1 | Q80TS5 | Zinc finger | Q80TS5 [6 | 0 | 2184.084 | 0 | -1.97 | 0.00022 | 1.32 | 158.6 | 41.4 | 3.53 | 16.59 | | High | High | High | High | High | High | High | High | High | High | High | 0 | 1.59E-05 | 3.26 |
| FALSE | High | AESSSSISSS | [R].AESSSS | 1xPhospho | Q3UQN2 1 | 2.68E-09 | 0 | 1 | 3 | 11 | Q3UQN2 | F-BAR dom | Q3UQN2 [ | 0 | 2662.277 | 0 | 0.78 | 0.015277 | 27.57 | 77.4 | 122.6 | 11.09 | 13.37 | | High | High | High | High | High | High | High | High | High | High | High | 0 | 2.34E-10 | 4.82 |
| FALSE | High | AERNSAEA | [R].AERNS/ | 1xPhospho | Q9WTV7 1 | 0.002232 | 0.000109 | 1 | 1 | 1 | Q9WTV7 | E3 ubiquiti | Q9WTV7 [ | 1 | 2039.996 | 0 | 1.85 | 0.001802 | 21.42 | 44.2 | 155.8 | 6.18 | 15.12 | | High | High | High | High | High | High | High | High | High | High | High | 4.94E-05 | 0.000586 | 2.64 |
| FALSE | High | AESCGSFDI | [K].AESCGS | 1xCarbami | Q9CRB0 1x | 0.003028 | 0.00016 | 1 | 1 | 2 | Q9CRB0 | Sorting nex | Q9CRB0 [1 | 0 | 2417.01 | 0 | 1.09 | 0.001091 | 10.98 | 64.5 | 135.5 | 13.02 | 7.28 | | High | High | High | High | High | High | High | High | High | High | High | 0.00607 | 0.06592 | 2.74 |
| FALSE | High | AESEEERDI | [K].AESEEE | 1xPhospho | Q64028 1x | 0.075927 | 0.005826 | 1 | 7 | 1 | Q64028 | Polyhomec | Q64028 [6 | 1 | 2698.359 | 0 | -1.32 | 0.004104 | 6.47 | 142.9 | 57.1 | 11.91 | 23.85 | | High | High | High | High | High | High | High | High | High | High | High | 0.002976 | 0.03113 | 2.23 |
| FALSE | High | AESEEMET | [K].AESEEN | 1xPhospho | Q571M2 1: | 0.000868 | 0 | 1 | 4 | 1 | Q571M2 | MKIAA402 | Q571M2 [6 | 0 | 2037.909 | 0 | -0.37 | 0.011088 | 6.34 | 111.2 | 88.8 | 4.66 | 11.66 | | High | High | High | High | High | High | High | High | High | High | High | 0 | 0.000204 | 2.36 |
| FALSE | High | AESESDGQ | [K].AESESD | 1xPhospho | A0A0N4SV | 6E-07 | 0 | 1 | 9 | 4 | A0A0N4SV | Eukaryotic | A0A0N4SV | 0 | 2610.115 | 0 | 0.82 | 0.006931 | 14.03 | 75.7 | 124.3 | 15.3 | 17.25 | | High | High | High | High | High | High | High | High | High | High | High | 0 | 5.75E-08 | 4.75 |
| FALSE | High | AESFPGHS( | [R].AESFPG | 1xCarbami | Q811P8 1x | 0.040028 | 0.002867 | 1 | 3 | 2 | Q811P8 | Rho GTPas | Q811P8 [1 | 0 | 2294.063 | 0 | -0.11 | 0.151118 | 9.27 | 106.6 | 93.4 | 8.58 | 8.15 | | High | High | High | High | High | High | High | High | High | High | High | 0.008216 | 0.08879 | 2.68 |
| FALSE | High | AESPAGRV | [K].AESPA( | 2xPhospho | F6RJ39 2xP | 0.045898 | 0.003408 | 2 | 4 | 1 | F6RJ39; Q8 | Apoptotic ( | F6RJ39 [43 | 1 | 2572.283 | 0 | 0 | 0.574996 | 9.26 | 100 | 100 | 1.71 | 9.85 | Shared | High | High | High | High | High | High | High | High | High | High | High | 0.001774 | 0.01753 | 2.11 |
| FALSE | High | AESPEMEV | [K].AESPEN | 1xPhospho | E9Q616 1x | 0.000121 | 0 | 1 | 3 | 2 | E9Q616 | AHNAK nu( | E9Q616 [8 | 0 | 1881.943 | 0 | 3.06 | 0.003981 | 2.7 | 22.9 | 177.1 | 14.25 | 22.75 | | High | High | High | High | High | High | High | High | High | High | High | 0 | 3.7E-05 | 4.06 |
| FALSE | High | AESPEMEV | [K].AESPEN | 1xPhospho | E9Q616 1x | 0.000568 | 0 | 1 | 3 | 4 | E9Q616 | AHNAK nu( | E9Q616 [8 | 0 | 1897.938 | 0 | 2.71 | 0.001309 | 11.38 | 27.1 | 172.9 | 9.56 | 16.02 | | High | High | High | High | High | High | High | High | High | High | High | 0.000183 | 0.001242 | 3 |
| FALSE | High | AESPETSA\ | [R].AESPET | 1xPhospho | F8WHU5 1 | 1.09E-08 | 0 | 1 | 3 | 5 | F8WHU5 | Sister chro | F8WHU5 [ | 0 | 2415.169 | 0 | 0.42 | 0.000609 | 4.49 | 86.2 | 113.8 | 3.49 | 3.07 | | High | High | High | High | High | High | High | High | High | High | High | 0 | 6.25E-09 | 4.65 |
| FALSE | High | AESPETSA\ | [R].AESPET | 2xPhospho | F8WHU5 2 | 6.89E-05 | 0 | 1 | 3 | 5 | F8WHU5 | Sister chro | F8WHU5 [ | 0 | 2495.136 | 0 | 0.03 | 0.928237 | 11.05 | 99.6 | 100.4 | 3.99 | 5.91 | | High | High | High | High | High | High | High | High | High | High | High | 0 | 1.18E-05 | 4.05 |
| FALSE | High | AESPTPGLT | [R].AESPTP | 1xPhospho | A0A0R4J0D | 3.59E-07 | 0 | 1 | 4 | 2 | A0A0R4J0D | Vacuolar fu | A0A0R4J0D | 0 | 3035.413 | 0 | 0.5 | 0.007362 | 2.72 | 77.2 | 122.8 | 12.04 | 14.7 | | High | High | High | High | High | High | High | High | High | High | High | 0 | 3.22E-08 | 3.65 |
| FALSE | High | AESPTPGLT | [R].AESPTP | 1xPhospho | A0A0R4J0D | 5.81E-05 | 0 | 1 | 4 | 3 | A0A0R4J0D | Vacuolar fu | A0A0R4J0D | 0 | 3051.408 | 0 | 0.26 | 0.127319 | 2.76 | 86.9 | 113.1 | 5.24 | 28.3 | | High | High | High | High | High | High | High | High | High | High | High | 0 | 9.72E-06 | 4.1 |
| FALSE | High | AESQNESS | [K].AESQNI | 1xPhospho | Q8C0V0 1x | 0.002232 | 0.000109 | 1 | 2 | 1 | Q8C0V0 | Serine/thre | Q8C0V0 [1 | 0 | 1702.805 | 0 | 0.39 | 0.04239 | 6.74 | 90.8 | 109.2 | 5.07 | 11.49 | | High | High | High | High | High | High | High | High | High | High | High | 4.94E-05 | 0.000586 | 2.3 |
| FALSE | High | AESCGNGN | [K].AESCGI | 1xCarbami | Q8VHK9 1) | 0.103392 | 0.008249 | 1 | 3 | 1 | Q8VHK9 | ATP-depen | Q8VHK9 [2 | 0 | 1922.827 | 0 | -0.4 | 0.128286 | 10.34 | 109.9 | 90.1 | 14.49 | 13.14 | | High | High | High | High | High | High | High | High | High | High | High | 0.004164 | 0.04488 | 1.53 |
| FALSE | High | AFTFDDED | [K].AFTFDI | 1xPhospho | Q3UM95 1 | 0.000579 | 0 | 1 | 3 | 1 | Q3UM95 | Putative ur | Q3UM95 [ | 0 | 2311.078 | 0 | 0.28 | 0.199328 | 40.63 | 86.7 | 113.3 | 10.7 | 30.32 | | High | High | High | High | High | High | High | High | High | High | High | 0 | 0.000129 | 2.48 |
| FALSE | High | AFVEDSSDI | [R].AFVEDS | 2xPhospho | F6WIU1 2x | 2.78E-06 | 0 | 1 | 3 | 4 | F6WIU1 | Protein KRI | F6WIU1 [1 | 0 | 2816.232 | 0 | -0.24 | 0.099941 | 14.96 | 109.2 | 90.8 | 10.39 | 7.94 | | High | High | High | High | High | High | High | High | High | High | High | 0 | 3.21E-07 | 3.43 |
| FALSE | High | AFVGPLSPS | [R].AFVGPI | 1xPhospho | Q6P1H6 1x | 0.002623 | 0.000109 | 1 | 3 | 1 | Q6P1H6 | Ankyrin re | Q6P1H6 [5 | 0 | 1540.854 | 0 | 0.31 | 0.048258 | 14.14 | 91.7 | 108.3 | 9.1 | 7.4 | | High | High | High | High | High | High | High | High | High | High | High | 4.94E-05 | 0.000705 | 2.4 |
| FALSE | High | AGGAGVP/ | [R].AGGAG | 1xPhospho | Q3U9P7 1x | 0.120211 | 0.009851 | 1 | 6 | 1 | Q3U9P7 | Succinyl-C( | Q3U9P7 [1 | 0 | 3123.58 | 0 | -0.82 | 0.004383 | 5.82 | 125.7 | 74.3 | 15.92 | 17.8 | | High | High | High | High | High | High | High | High | High | High | High | 0.004949 | 0.05381 | 2.67 |
| FALSE | High | AGGANYD/ | [K].AGGAN | 1xPhospho | Q9CQI6 1x | 1.38E-05 | 0 | 1 | 1 | 2 | Q9CQI6 | Coactosin- | Q9CQI6 [1 | 0 | 1391.568 | 0 | -1.42 | 0.000949 | 1.04 | 147.2 | 52.8 | 8.8 | 17.5 | | High | High | High | High | High | High | High | High | High | High | High | 0 | 1.94E-06 | 3.02 |
| FALSE | High | AGGASPAA | [R].AGGAS | 1xPhospho | Q8R3Y8 1x | 2.3E-06 | 0 | 1 | 2 | 3 | Q8R3Y8 | Interferon | Q8R3Y8 [4 | 0 | 2101.003 | 0 | 0.73 | 0.001678 | 4.8 | 75.6 | 124.4 | 3.39 | 6.02 | | High | High | High | High | High | High | High | High | High | High | High | 0 | 2.58E-07 | 5.24 |
| FALSE | High | AGGGGSN/ | [K].AGGG( | 1xPhospho | Q7JJ13 1xP | 0.000916 | 0 | 1 | 3 | 3 | Q7JJ13 | Bromodom | Q7JJ13 [58 | 0 | 2744.304 | 0 | -0.51 | 0.158121 | 13.43 | 117.5 | 82.5 | 19.4 | 24.42 | | High | High | High | High | High | High | High | High | High | High | High | 0 | 0.000217 | 3.74 |
| FALSE | High | AGGGSQDI | [K].AGGGS | 1xPhospho | Q543E3 1x | 0.000984 | 0 | 1 | 3 | 2 | Q543E3 | Suppresso | Q543E3 [9 | 0 | 1702.857 | 0 | -1.84 | 0.000163 | 1.47 | 156.4 | 43.6 | 1.5 | 13.73 | | High | High | High | High | High | High | High | High | High | High | High | 0 | 0.000234 | 2.7 |
| FALSE | High | AGGQAPSS | [R].AGGQA | 1xPhospho | Q61286 1x | 0.000808 | 0 | 1 | 4 | 2 | Q61286 | Transcripti | Q61286 [3 | 0 | 2468.222 | 0 | -0.6 | 0.026931 | 6.46 | 121.1 | 78.9 | 15.36 | 17.69 | | High | High | High | High | High | High | High | High | High | High | High | 0 | 0.000283 | 2.5 |
| FALSE | High | AGGQAPSS | [R].AGGQA | 1xPhospho | Q61286-2 | 6.08E-09 | 0 | 1 | 5 | 6 | Q61286-2 | Isoform AL | Q61286-2 | 0 | 2482.156 | 0 | -0.43 | 0.1636 | 10.42 | 115.5 | 84.5 | 27.37 | 18.76 | | High | High | High | High | High | High | High | High | High | High | High | 0 | 4.69E-09 | 3.75 |
| FALSE | High | AGGQAPSS | [R].AGGQA | 1xDeamida | Q61286-2 | 0.000579 | 0 | 1 | 5 | 5 | Q61286-2 | Isoform AL | Q61286-2 | 0 | 2483.14 | 0 | -0.54 | 0.085068 | 10.24 | 118.5 | 81.5 | 19.63 | 17.05 | | High | High | High | High | High | High | High | High | High | High | High | 4.94E-05 | 0.000605 | 3.28 |
| FALSE | High | AGGSPASY | [K].AGGSP. | 1xPhospho | Q8CHU3 1: | 0.118334 | 0.009621 | 1 | 3 | 1 | Q8CHU3 | Epsin-2 OS | Q8CHU3 [1 | 0 | 1740.79 | 0 | 1.08 | 0.035756 | 48.03 | 63.6 | 136.4 | 26.37 | 29.07 | | High | High | High | High | High | High | High | High | High | High | High | 0.004866 | 0.05287 | 3.12 |
| FALSE | High | AGGSPASY | [K].AGGSP. | 2xPhospho | Q8CHU3 2: | 0.000434 | 0 | 1 | 3 | 2 | Q8CHU3 | Epsin-2 OS | Q8CHU3 [1 | 0 | 1820.757 | 0 | 1.16 | 0.000983 | 6.25 | 61.2 | 138.8 | 6.63 | 9.62 | | High | High | High | High | High | High | High | High | High | High | High | 0.009953 | 0.1085 | 2.25 |
| FALSE | High | AGGSPASY | [K].AGGSP. | 3xPhospho | Q8CHU3 3: | 0.011755 | 0.000889 | 1 | 3 | 1 | Q8CHU3 | Epsin-2 OS | Q8CHU3 [1 | 0 | 1900.723 | 0 | 3.12 | 0.002167 | 51.87 | 20.6 | 179.4 | 67.58 | 20.05 | | High | High | High | High | High | High | High | High | High | High | High | 0.000449 | 0.00374 | 2.36 |
| FALSE | High | AGGSVESLI | [K].AGGSV | 1xPhospho | F8VQK5 1x | 0.005717 | 0.00062 | 1 | 3 | 2 | F8VQK5 | SAM and S | F8VQK5 [4 | 0 | 1184.587 | 0 | 0.22 | 0.016374 | 3.81 | 90.5 | 109.5 | 2.97 | 8.24 | | High | High | High | High | High | High | High | High | High | High | High | 0.000538 | 0.004523 | 1.62 |
| FALSE | High | AGLLTSCSS | [K].AGLLTS | 1xCarbami | P48754 1xI | 0.022378 | 0.001542 | 1 | 4 | 1 | P48754 | Breast can | P48754 [69 | 0 | 1457.702 | 0 | -0.84 | 0.002245 | 2.27 | 124.3 | 75.7 | 6.24 | 11.03 | | High | High | High | High | High | High | High | High | High | High | High | 0.000814 | 0.007765 | 2 |
| FALSE | High | AGFSPDSSI | [K].AGFSPI | 2xPhospho | Q7TN02 2x | 0.008927 | 0.000736 | 1 | 1 | 1 | Q7TN02 | Mediator c | Q7TN02 [3 | 0 | 1513.674 | 0 | 0.45 | 0.000277 | 4.43 | 87.1 | 112.9 | 8.53 | 6.43 | | High | High | High | High | High | High | High | High | High | High | High | 0.000327 | 0.002772 | 1.71 |
| FALSE | High | AGLLTSCSS | [K].AGLLTS | 1xCarbami | P48754 1xI | 0.102567 | 0.008249 | 1 | 4 | 1 | P48754 | Breast can | P48754 [69 | 1 | 1814.96 | 0 | -0.98 | 0.003421 | 4.38 | 131.6 | 68.4 | 10.83 | 5.94 | | High | High | High | High | High | High | High | High | High | High | High | 0.004138 | 0.04429 | 1.28 |
| FALSE | High | AGIVGSPAI | [R].AGIVGS | 1xPhospho | O88852 1x | 0.013787 | 0.001011 | 1 | 1 | 2 | O88852 | Testis-spec | O88852 [4 | 0 | 1233.655 | 0 | 1.32 | 0.000158 | 3.68 | 57.2 | 142.8 | 9.1 | 4.53 | | High | High | High | High | High | High | High | High | High | High | High | 0.000538 | 0.00448 | 1.64 |
| FALSE | High | AGMTSSPD | [R].AGMTS | 1xPhospho | Q3UJ69 1x | 7.16E-06 | 0 | 1 | 5 | 2 | Q3UJ69 | Putative ur | Q3UJ69 [7 | 0 | 1853.783 | 0 | 1.72 | 0.002652 | 3.64 | 46.6 | 153.4 | 4.37 | 17.59 | | High | High | High | High | High | High | High | High | High | High | High | 0 | 9.28E-07 | 3.91 |
| FALSE | High | AGNSDSEE | [K].AGNSD | 2xPhospho | Q6PGL7 2x | 0.037048 | 0.00249 | 1 | 5 | 1 | Q6PGL7 | WASH com | Q6PGL7 [1 | 1 | 3034.37 | 0 | -0.26 | 0.051322 | 5.99 | 110.6 | 89.4 | 7.49 | 10.67 | | High | High | High | High | High | High | High | High | High | High | High | 0.001308 | 0.01369 | 2.53 |
| FALSE | High | AGNWPGS | [R].AGNWI | 1xPhospho | A2ABV5 1x | 1.07E-05 | 0 | 1 | 2 | 1 | A2ABV5 | Mediator c | A2ABV5 [1 | 0 | 2074.991 | 0 | 1.3 | 0.000681 | 8.78 | 57.7 | 142.3 | 5.35 | 9.32 | | High | High | High | High | High | High | High | High | High | High | High | 0 | 1.46E-06 | 3.23 |
| FALSE | High | AGNWPGS | [R].AGNWI | 2xPhospho | A2ABV5 2x | 0.008167 | 0.000736 | 1 | 2 | 2 | A2ABV5 | Mediator c | A2ABV5 [1 | 0 | 2154.957 | 0 | 1.14 | 0.000242 | 5.28 | 62.2 | 137.8 | 4.07 | 5.39 | | High | High | High | High | High | High | High | High | High | High | High | 0.000327 | 0.002511 | 3.72 |

| | | | | | | | | | | | | | | | | | | | | | | | | | | | | | | | | | | | | | | |
|---|---|---|---|---|---|---|---|---|---|---|---|---|---|---|---|---|---|---|---|---|---|---|---|---|---|---|---|---|---|---|---|---|---|---|---|---|---|---|
| FALSE | High | AGPGSSPLI | [R].AGPGS | 1xPhospho | Q9DB90 1x | 0.005717 | 0.00062 | 1 | 1 | 2 | Q9DB90 | Protein SM | Q9DB90 [4 | 0 | 1927.988 | 0 | 0.63 | 0.113234 | 4.18 | 72 | 128 | 44.25 | 10.24 | | High | High | High | High | High | High | High | High | High | High | High | 0.000294 | 0.002352 | 3.78 |
| FALSE | High | AGPPAPAP | [R].AGPPAI | 1xPhospho | Q5I1X5 1xF | 0.000189 | 0 | 1 | 1 | 2 | Q5I1X5 | RelA-assoc | Q5I1X5 [57 | 0 | 3565.706 | 0 | 0.81 | 0.031921 | 22.24 | 73.3 | 126.7 | 20.74 | 19.24 | | High | High | High | High | High | High | High | High | High | High | High | 0 | 3.68E-05 | 3.36 |
| FALSE | High | AGPPGSPS | [K].AGPPG | 1xPhospho | O88466 1x | 0.001898 | 5.66E-05 | 2 | 7 | 3 | O88466; R | Zinc finger | O88466 [4 | 0 | 1472.766 | 0 | -1.5 | 0.000295 | 1.47 | 147.5 | 52.5 | 4.46 | 5.91 | Shared | High | High | High | High | High | High | High | High | High | High | High | 2.58E-05 | 0.000564 | 2.08 |
| FALSE | High | AGQTQPNI | [K].AGQTQ | 1xPhospho | Q3UHJ0 1x | 4.89E-05 | 0 | 1 | 2 | 1 | Q3UHJ0 | AP2-associ | Q3UHJ0 [3 | 0 | 2378.279 | 0 | 0.27 | 0.023389 | 11.49 | 90 | 110 | 8.96 | 5.27 | | High | High | High | High | High | High | High | High | High | High | High | 0 | 8.03E-06 | 3.14 |
| FALSE | High | AGQTVTVV | [R].AGQTV | 1xPhospho | A0A0R4J0C | 1.62E-09 | 0 | 1 | 3 | 1 | A0A0R4J0C | Lamin-B2 C | A0A0R4J0C | 0 | 3031.581 | 0 | -0.11 | 0.196955 | 16.98 | 103.4 | 96.6 | 5.85 | 13.13 | | High | High | High | High | High | High | High | High | High | High | High | 0 | 7.42E-11 | 4.62 |
| FALSE | High | AGQTVTVV | [R].AGQTV | 1xDeamida | A0A0R4J0C | 0.007605 | 0.000664 | 1 | 3 | 2 | A0A0R4J0C | Lamin-B2 C | A0A0R4J0C | 0 | 3032.565 | 0 | 0.07 | 0.163016 | 5.89 | 97.9 | 102.1 | 15.02 | 11.04 | | High | High | High | High | High | High | High | High | High | High | High | 0.000294 | 0.002321 | 3.04 |
| FALSE | High | AGSAPAFG | [R].AGSAP | 1xPhospho | Q60692 1x | 5.52E-06 | 0 | 1 | 1 | 2 | Q60692 | Proteasom | Q60692 [1 | 0 | 2295.064 | 0 | 1.26 | 0.004778 | 21.94 | 58.5 | 141.5 | 8.04 | 16.85 | | High | High | High | High | High | High | High | High | High | High | High | 0 | 6.94E-07 | 4.56 |
| FALSE | High | AGSDAAAS | [K].AGSDA | 1xPhospho | P47911 1xI | 0.015061 | 0.001049 | 1 | 2 | 1 | P47911 | 60S riboso | P47911 [19 | 0 | 1367.663 | 0 | -0.12 | 0.223224 | 7.7 | 104.2 | 95.8 | 5.1 | 6.9 | | High | High | High | High | High | High | High | High | High | High | High | 0.000571 | 0.004962 | 0.78 |
| FALSE | High | AGIPQHHP | [K].AGIPQH | 1xPhospho | Q9JKF6 1xI | 0.000227 | 0 | 1 | 1 | 3 | Q9JKF6 | Nectin-1 O | Q9JKF6 [40 | 0 | 3257.497 | 0 | -1.83 | 0.000816 | 3.06 | 156 | 44 | 4.94 | 21.56 | | High | High | High | High | High | High | High | High | High | High | High | 0 | 4.5E-05 | 2.62 |
| FALSE | High | AGFSPDSSI | [R].AGFSPD | 1xPhospho | Q7TN02 1x | 0.055777 | 0.004093 | 1 | 1 | 2 | Q7TN02 | Mediator c | Q7TN02 [3 | 0 | 1433.708 | 0 | -0.14 | 0.009104 | 3.95 | 104.3 | 95.7 | 4 | 4.08 | | High | High | High | High | High | High | High | High | High | High | High | 0.002117 | 0.02192 | 2.52 |
| FALSE | High | AGFQTEDF | [R].AGFQT | 1xCarbami | P28028 1xI | 0.000415 | 0 | 2 | 3 | 1 | P28028; F6 | Serine/thre | P28028 [77 | 0 | 2430.145 | 0 | -0.19 | 0.118727 | 14.7 | 106.5 | 93.5 | 4.04 | 12.06 | Shared | High | High | High | High | High | High | High | High | High | High | High | 0 | 8.85E-05 | 4.11 |
| FALSE | High | AGFNYESP | [K].AGFNY | 1xPhospho | V9GX74 1x | 0.058173 | 0.004227 | 1 | 8 | 1 | V9GX74 | Zinc finger | V9GX74 [8 | 0 | 1637.798 | 0 | -0.66 | 0.001698 | 4.41 | 122.8 | 77.2 | 5.34 | 6.16 | | High | High | High | High | High | High | High | High | High | High | High | 0.002187 | 0.02298 | 1.57 |
| FALSE | High | AGAEASEE | [R].AGAEA | 1xPhospho | Q6Y7W8 1 | 0.002463 | 0.000109 | 1 | 8 | 9 | Q6Y7W8 | GRB10-inte | Q6Y7W8 [3 | 0 | 4004.846 | 0 | -0.28 | 0.07664 | 12.3 | 109.7 | 90.3 | 10.21 | 14.73 | | High | High | High | High | High | High | High | High | High | High | High | 4.94E-05 | 0.000654 | 4.09 |
| FALSE | High | AGAGMITC | [R].AGAGM | 1xPhospho | Q8CH25-2 | 3.6E-05 | 0 | 1 | 2 | 14 | Q8CH25-2 | Isoform 2 c | Q8CH25-2 | 0 | 2109.995 | 0 | 0.7 | 0.040364 | 23 | 76.2 | 123.8 | 9.19 | 19.85 | | High | High | High | High | High | High | High | High | High | High | High | 0 | 5.7E-06 | 2.86 |
| FALSE | High | AGAISASGI | [R].AGAISA | 1xPhospho | E9Q616 1x | 0.000884 | 0 | 1 | 3 | 2 | E9Q616 | AHNAK nu | E9Q616 [2 | 0 | 2177.1 | 0 | 2.29 | 0.001573 | 16.17 | 34.7 | 165.3 | 7.22 | 15.24 | | High | High | High | High | High | High | High | High | High | High | High | 0 | 0.000207 | 1.98 |
| FALSE | High | AGAPGPAS | [K].AGAPG | 1xPhospho | E9Q7F2 1x | 0.001189 | 0 | 1 | 1 | 2 | E9Q7F2 | E3 ubiquiti | E9Q7F2 [3 | 0 | 1785.983 | 0 | 0.01 | 0.764555 | 17.37 | 97 | 103 | 15.38 | 18.22 | | High | High | High | High | High | High | High | High | High | High | High | 4.94E-05 | 0.000658 | 3.75 |
| FALSE | High | AGAPGPGL | [R].AGAPG | 1xCarbami | Q8CHI8 1xI | 1.22E-06 | 0 | 1 | 7 | 3 | Q8CHI8 | E1A-bindin | Q8CHI8 [1 | 0 | 2865.417 | 0 | 0.77 | 0.049053 | 37.18 | 76.3 | 123.7 | 14.21 | 19.44 | | High | High | High | High | High | High | High | High | High | High | High | 0 | 3.44E-07 | 4.35 |
| FALSE | High | AGAPSALSI | [R].AGAPS | 1xPhospho | E9PVV1 1x | 4.04E-08 | 0 | 1 | 4 | 3 | E9PVV1 | Transcripti | E9PVV1 [3 | 0 | 2592.322 | 0 | -0.29 | 0.001353 | 1.54 | 110.4 | 89.6 | 3.98 | 6.75 | | High | High | High | High | High | High | High | High | High | High | High | 0 | 2.76E-09 | 2.42 |
| FALSE | High | AGAPSALSI | [R].AGAPS | 1xPhospho | E9PVV2 1x | 6.52E-08 | 0 | 1 | 7 | 2 | E9PVV2 | Transcripti | E9PVV2 [3 | 0 | 2464.264 | 0 | -0.42 | 0.00589 | 3.72 | 114.4 | 85.6 | 7.29 | 2.75 | | High | High | High | High | High | High | High | High | High | High | High | 0 | 4.71E-09 | 2.75 |
| FALSE | High | AGASPPPV | [R].AGASPI | 1xPhospho | Q9D938 1x | 0.013545 | 0.000976 | 1 | 2 | 1 | Q9D938 | Transmem | Q9D938 [4 | 0 | 1604.788 | 0 | -0.65 | 0.000609 | 3.19 | 120.1 | 79.9 | 5.51 | 6.1 | | High | High | High | High | High | High | High | High | High | High | High | 0.000505 | 0.004401 | 2.43 |
| FALSE | High | AGATCVSC | [R].AGATC | 2xCarbami | Q99J10 1xI | 0.000209 | 0 | 1 | 1 | 2 | Q99J10 | Cytoplasmi | Q99J10 [36 | 0 | 2302.963 | 0 | -0.16 | 0.337344 | 20.16 | 107.4 | 92.6 | 6.15 | 13.13 | | High | High | High | High | High | High | High | High | High | High | High | 0 | 4.12E-05 | 2.68 |
| FALSE | High | AGATCVSC | [R].AGATC | 2xCarbami | Q99J10 1xI | 0.000161 | 0 | 1 | 1 | 1 | Q99J10 | Cytoplasmi | Q99J10 [36 | 0 | 2303.947 | 0 | -0.1 | 0.681384 | 7.4 | 104.5 | 95.5 | 9 | 14.64 | | High | High | High | High | High | High | High | High | High | High | High | 0 | 3.07E-05 | 4.27 |
| FALSE | High | AGATCVSC | [R].AGATC | 2xCarbami | Q99J10 1xI | 1.07E-07 | 0 | 1 | 1 | 2 | Q99J10 | Cytoplasmi | Q99J10 [40 | 0 | 1973.855 | 0 | 0.18 | 0.056967 | 14.5 | 88.2 | 111.8 | 9.91 | 9 | | High | High | High | High | High | High | High | High | High | High | High | 0 | 4.07E-05 | 3.85 |
| FALSE | High | AGAVTSSA | [K].AGAVT | 1xPhospho | H3BL37 1xI | 1.9E-05 | 0 | 1 | 7 | 3 | H3BL37 | Treacle pro | H3BL37 [58 | 0 | 2056.109 | 0 | -0.22 | 0.019942 | 5.85 | 109.8 | 90.2 | 5.11 | 8.52 | | High | High | High | High | High | High | High | High | High | High | High | 0 | 2.77E-06 | 3.24 |
| FALSE | High | AGCFSPK | [K].AGCFSI | 1xCarbami | Q80Y19 1x | 0.083101 | 0.006345 | 1 | 4 | 1 | Q80Y19 | Rho GTPas | Q80Y19 [4 | 0 | 1304.647 | 0 | -0.1 | 0.194947 | 4.31 | 103.7 | 96.3 | 6.87 | 7.75 | | High | High | High | High | High | High | High | High | High | High | High | 0.003265 | 0.03456 | 1.2 |
| FALSE | High | AGDELAYN | [R].AGDEL | 1xCarbami | Q8R0X2 1x | 0.044739 | 0.003302 | 1 | 4 | 2 | Q8R0X2 | CDK2-asso | Q8R0X2 [3 | 0 | 2064.89 | 0 | 0.07 | 0.065406 | 0.94 | 95.2 | 104.8 | 9.24 | 5.57 | | High | High | High | High | High | High | High | High | High | High | High | 0.001697 | 0.01694 | 1.46 |
| FALSE | High | AGDIQGQL | [R].AGDIQ | 1xPhospho | Q3U0P1 1x | 6.05E-07 | 0 | 1 | 4 | 3 | Q3U0P1 | Partner an | Q3U0P1 [1 | 0 | 2349.185 | 0 | -1 | 0.004623 | 0.16 | 133.3 | 66.7 | 10.73 | 8.63 | | High | High | High | High | High | High | High | High | High | High | High | 0 | 3.63E-06 | 2.62 |
| FALSE | High | AGDSDEES | [R].AGDSD | 1xPhospho | B7ZWA9 1x | 0.06169 | 0.004646 | 1 | 4 | 1 | B7ZWA9 | Zbtb7a pro | B7ZWA9 [3 | 1 | 1962.869 | 0 | 1.99 | 0.001189 | 9.11 | 40.2 | 159.8 | 20.04 | 10.26 | | High | High | High | High | High | High | High | High | High | High | High | 0.002381 | 0.02448 | 2.1 |
| FALSE | High | AGDVLEDS | [R].AGDVLI | 1xPhospho | P51859 1xI | 0.000229 | 0 | 1 | 2 | 8 | P51859 | Hepatoma- | P51859 [15 | 0 | 1339.634 | 0 | -0.04 | 0.768475 | 17.7 | 105.2 | 94.8 | 34.19 | 37.21 | | High | High | High | High | High | High | High | High | High | High | High | 0 | 0.00016 | 2.75 |
| FALSE | High | AGDVLEDS | [R].AGDVLI | 1xPhospho | P51859 1xI | 5.4E-05 | 0 | 1 | 2 | 10 | P51859 | Hepatoma- | P51859 [15 | 0 | 1568.797 | 0 | -0.18 | 0.123492 | 7.9 | 106.9 | 93.1 | 5.99 | 7.15 | | High | High | High | High | High | High | High | High | High | High | High | 0 | 9.56E-05 | 3.12 |
| FALSE | High | AGEAPTEN | [K].AGEAP | 1xPhospho | Q9JKB3 1xI | 0.000219 | 0 | 1 | 3 | 2 | Q9JKB3 | Y-box-bind | Q9JKB3 [3 | 0 | 2165.944 | 0 | -0.83 | 0.019243 | 9.17 | 128.1 | 71.9 | 15.4 | 9.59 | | High | High | High | High | High | High | High | High | High | High | High | 0 | 4.35E-05 | 2.24 |
| FALSE | High | AGEPQASS | [K].AGEPQ | 1xCarbami | Q3U0X6 1x | 0.01186 | 0.000889 | 1 | 1 | 3 | Q3U0X6 | Putative ur | Q3U0X6 [6 | 0 | 1567.714 | 0 | 0.15 | 0.016364 | 0.71 | 94.2 | 105.8 | 6.05 | 7.75 | | High | High | High | High | High | High | High | High | High | High | High | 0.000449 | 0.0038 | 2.19 |
| FALSE | High | AGEQDPVP | [R].AGEQD | 1xPhospho | A0A1L7NR | 0.002376 | 0.000109 | 2 | 12 | 4 | A0A1L7NR | C-Cbl asso | A0A1L7NR | 0 | 2131.027 | 0 | 1.41 | 0.000213 | 3.39 | 55.4 | 144.6 | 7.94 | 3.03 | Shared | High | High | High | High | High | High | High | High | High | High | High | 4.94E-05 | 0.000628 | 2.68 |
| FALSE | High | AGEQDPVP | [R].AGEQD | 1xDeamida | A0A1L7NR | 0.048715 | 0.00354 | 2 | 12 | 2 | A0A1L7NR | C-Cbl asso | A0A1L7NR | 0 | 2132.011 | 0 | 1.27 | 0.000414 | 6.15 | 58.7 | 141.3 | 11.54 | 7.16 | Shared | High | High | High | High | High | High | High | High | High | High | High | 0.00186 | 0.01873 | 2.73 |
| FALSE | High | AGEQQLSE | [K].AGEQQ | 1xPhospho | F8VPX1 1xI | 4.39E-08 | 0 | 1 | 4 | 1 | F8VPX1 | Ubiquitin c | F8VPX1 [1 | 0 | 2956.206 | 0 | -0.43 | 0.053992 | 12.51 | 110.7 | 89.3 | 18.46 | 19.58 | | High | High | High | High | High | High | High | High | High | High | High | 0 | 3.03E-09 | 5.8 |
| FALSE | High | AGEQQLSE | [K].AGEQQ | 1xPhospho | F8VPX1 1xI | 4.85E-05 | 0 | 1 | 4 | 5 | F8VPX1 | Ubiquitin c | F8VPX1 [1 | 0 | 2972.201 | 0 | -0.31 | 0.050849 | 12.45 | 110.6 | 89.4 | 5.57 | 14.2 | | High | High | High | High | High | High | High | High | High | High | High | 0 | 7.94E-06 | 3.95 |
| FALSE | High | AGEQQLSE | [K].AGEQQ | 1xDeamida | F8VPX1 1xI | 0.016021 | 0.001124 | 1 | 4 | 4 | F8VPX1 | Ubiquitin c | F8VPX1 [1 | 0 | 2973.185 | 0 | -0.3 | 0.193984 | 8.45 | 115.1 | 84.9 | 13.92 | 30.41 | | High | High | High | High | High | High | High | High | High | High | High | 0.001142 | 0.01182 | 3.21 |
| FALSE | High | AGEQQLSE | [K].AGEQQ | 1xPhospho | F8VPX1 1xI | 2.61E-06 | 0 | 1 | 4 | 16 | F8VPX1 | Ubiquitin c | F8VPX1 [1 | 0 | 2988.196 | 0 | -0.27 | 0.024601 | 7.93 | 114 | 86 | 10.36 | 10.64 | | High | High | High | High | High | High | High | High | High | High | High | 0 | 3.48E-05 | 4.14 |
| FALSE | High | AGEVLLDV | [K].AGEVLI | 1xPhospho | B7ZNK9 1x | 0.002096 | 5.66E-05 | 1 | 4 | 4 | B7ZNK9 | Spata5 pro | B7ZNK9 [2 | 0 | 1693.872 | 0 | -0.46 | 0.002009 | 4.38 | 115.9 | 84.1 | 4.41 | 9.8 | | High | High | High | High | High | High | High | High | High | High | High | 2.58E-05 | 0.000547 | 3.44 |
| FALSE | High | AGEVLLDV | [K].AGEVLI | 2xPhospho | B7ZNK9 2x | 0.000235 | 0 | 1 | 4 | 1 | B7ZNK9 | Spata5 pro | B7ZNK9 [2 | 0 | 1773.839 | 0 | -1.15 | 0.001041 | 3.02 | 140.8 | 59.2 | 5.6 | 14.2 | | High | High | High | High | High | High | High | High | High | High | High | 0 | 4.69E-05 | 2.84 |
| FALSE | High | AGFDSDYD | [K].AGFDSI | 1xPhospho | P54276 1xI | 9.71E-05 | 0 | 1 | 3 | 3 | P54276 | DNA mism | P54276 [92 | 0 | 1965.879 | 0 | -0.29 | 0.021681 | 7.23 | 109.9 | 90.1 | 4.98 | 5.63 | | High | High | High | High | High | High | High | High | High | High | High | 0 | 1.73E-05 | 3.49 |
| FALSE | High | AENQRPAE | [K].AENQR | 1xPhospho | Q3UZ39 1x | 4.81E-09 | 0 | 1 | 2 | 4 | Q3UZ39 | Leucine-ric | Q3UZ39 [2 | 0 | 2617.339 | 0 | 1.26 | 0.001078 | 8.95 | 58.8 | 141.2 | 10.52 | 6.13 | | High | High | High | High | High | High | High | High | High | High | High | 0 | 7.41E-08 | 3.47 |
| FALSE | High | AENPLTPTI | [K].AENPLT | 1xPhospho | L0HN04 1x | 1.87E-07 | 0 | 1 | 4 | 3 | L0HN04 | Methylcyto | L0HN04 [7 | 0 | 2452.314 | 0 | 1.89 | 0.000777 | 3.9 | 44.5 | 155.5 | 4.85 | 11.19 | | High | High | High | High | High | High | High | High | High | High | High | 0 | 9.91E-08 | 4.03 |
| FALSE | High | AENGLSTEI | [K].AENGL | 1xPhospho | Q80WR5 1 | 1.31E-06 | 0 | 1 | 3 | 2 | Q80WR5 | UPF0688 p | Q80WR5 [ | 0 | 3348.507 | 0 | 0.34 | 0.01956 | 8.1 | 90.2 | 109.8 | 23.74 | 13.52 | | High | High | High | High | High | High | High | High | High | High | High | 0 | 1.38E-07 | 4.31 |
| FALSE | High | AENGLLMT | [R].AENGLI | 1xCarbami | A7E215 1xI | 0.000411 | 0 | 2 | 15 | 3 | A7E215; Q | Rps6ka3 pr | A7E215 [57 | 0 | 2992.497 | 0 | 2.64 | 0.040745 | 64.52 | 31.4 | 168.6 | 25.51 | 45.36 | Shared | High | High | High | High | High | High | High | High | High | High | High | 0 | 8.8E-05 | 4.47 |
| FALSE | High | ADSPAGLE | [R].ADSPA | 1xPhospho | Q80U49 1x | 0.003083 | 0.00016 | 1 | 2 | 1 | Q80U49 | Centrosom | Q80U49 [7 | 0 | 1366.657 | 0 | 0.9 | 0.001609 | 6.58 | 68.7 | 131.3 | 7.74 | 8.14 | | High | High | High | High | High | High | High | High | High | High | High | 7.19E-05 | 0.000845 | 2.48 |
| FALSE | High | ADSPELDP | [K].ADSPEL | 1xPhospho | Q14B71 1x | 2.9E-05 | 0 | 1 | 2 | 2 | Q14B71 | Cell divisio | Q14B71 [6 | 0 | 2499.192 | 0 | -1.13 | 0.000159 | 0.39 | 136.6 | 63.4 | 2.59 | 4.34 | | High | High | High | High | High | High | High | High | High | High | High | 0 | 4.49E-06 | 3.65 |
| FALSE | High | ADSPVPTE | [R].ADSPVI | 1xPhospho | Q7TN02 1x | 0.00967 | 0.000736 | 1 | 1 | 3 | Q7TN02 | Mediator c | Q7TN02 [4 | 0 | 1618.804 | 0 | 0.07 | 0.225897 | 3.15 | 98.6 | 101.4 | 3.9 | 3.9 | | High | High | High | High | High | High | High | High | High | High | High | 0.001774 | 0.01752 | 2.44 |
| FALSE | High | ADSSSGGG | [R].ADSSS | 1xPhospho | Q3UIA2 1x | 8.98E-06 | 0 | 1 | 2 | 2 | Q3UIA2 | Rho GTPas | Q3UIA2 [5 | 0 | 3810.941 | 0 | -0.5 | 0.023117 | 0.26 | 114.5 | 85.5 | 9.13 | 20.77 | | High | High | High | High | High | High | High | High | High | High | High | 0 | 1.2E-06 | 2.85 |
| FALSE | High | ADSVDIQD | [R].ADSVDI | 1xPhospho | A0A0R4J21 | 0.000147 | 0 | 1 | 4 | 1 | A0A0R4J21 | Probable A | A0A0R4J21 | 0 | 1627.834 | 0 | 0.75 | 0.014387 | 19.33 | 74.5 | 125.5 | 8.26 | 14.89 | | High | High | High | High | High | High | High | High | High | High | High | 0 | 2.77E-05 | 2.47 |
| FALSE | High | ADSVPNLE | [R].ADSVPI | 1xPhospho | Q7TQ95 1x | 0.015603 | 0.001124 | 1 | 1 | 2 | Q7TQ95 | Protein lun | Q7TQ95 [4 | 0 | 2353.194 | 0 | 1.62 | 0.00079 | 11.83 | 49.1 | 150.9 | 5.08 | 10.99 | | High | High | High | High | High | High | High | High | High | High | High | 0.000585 | 0.005139 | 2.08 |
| FALSE | High | ADSVVEAG | [R].ADSVVI | 1xPhospho | Q9DBU5 1x | 0.000205 | 0 | 1 | 1 | 1 | Q9DBU5 | E3 ubiquiti | Q9DBU5 [5 | 0 | 1749.935 | 0 | 1.78 | 0.002105 | 11.19 | 46.4 | 153.6 | 13.5 | 17.67 | | High | High | High | High | High | High | High | High | High | High | High | 0 | 4.04E-05 | 2.75 |
| FALSE | High | ADTFTEEN | [R].ADTFTE | 1xPhospho | E9QAU4 1x | 2.56E-06 | 0 | 1 | 3 | 2 | E9QAU4 | Enhancer t | E9QAU4 [1 | 0 | 1975.941 | 0 | -0.97 | 0.000434 | 2.08 | 131.3 | 68.7 | 5.16 | 4.28 | | High | High | High | High | High | High | High | High | High | High | High | 4.94E-05 | 0.000587 | 3.36 |
| FALSE | High | ADTPSGDE | [R].ADTPS | 1xPhospho | O70551 1x | 1.26E-06 | 0 | 1 | 2 | 7 | O70551 | SRSF prote | O70551 [4 | 0 | 2369.091 | 0 | -1.12 | 0.000344 | 1.12 | 138.2 | 61.8 | 2.77 | 8.57 | | High | High | High | High | High | High | High | High | High | High | High | 0 | 8.7E-06 | 4.34 |
| FALSE | High | ADTPSGDE | [R].ADTPS | 1xDeamida | O70551 1x | 0.000211 | 0 | 1 | 2 | 7 | O70551 | SRSF prote | O70551 [4 | 0 | 2370.075 | 0 | -0.96 | 0.0012 | 3.99 | 133.1 | 66.9 | 7.01 | 8.73 | | High | High | High | High | High | High | High | High | High | High | High | 0 | 4.16E-05 | 3.66 |
| FALSE | High | ADTQAVSS | [K].ADTQA | 1xPhospho | E9QMX4 1 | 6.51E-07 | 0 | 1 | 4 | 3 | E9QMX4 | Ribosomal | E9QMX4 [5 | 0 | 2649.308 | 0 | 0.19 | 0.314745 | 51.62 | 93.2 | 106.8 | 29.1 | 13.97 | | High | High | High | High | High | High | High | High | High | High | High | 0 | 6.25E-08 | 5.04 |
| FALSE | High | ADTQAVSS | [K].ADTQA | 2xPhospho | E9QMX4 2 | 0.023379 | 0.001578 | 1 | 4 | 2 | E9QMX4 | Ribosomal | E9QMX4 [5 | 0 | 2729.275 | 0 | 0.27 | 0.114145 | 8.27 | 95.1 | 104.9 | 12.02 | 12.73 | | High | High | High | High | High | High | High | High | High | High | High | 0.000847 | 0.008116 | 3.98 |
| FALSE | High | ADTSPTYVI | [R].ADTSPT | 1xPhospho | Q6IQX8 1x | 0.005228 | 0.000551 | 1 | 5 | 5 | Q6IQX8 | Zinc finger | Q6IQX8 [6 | 0 | 1318.624 | 0 | -1.27 | 0.000389 | 0.77 | 141.7 | 58.3 | 5.42 | 7.99 | | High | High | High | High | High | High | High | High | High | High | High | 0.002159 | 0.02249 | 2.66 |
| FALSE | High | ADSLTTSPI | [K].ADSLTT | 1xPhospho | Q059U7 1x | 0.000976 | 0 | 1 | 1 | 3 | Q059U7 | Protein int | Q059U7 [6 | 0 | 1569.809 | 0 | -0.4 | 0.046276 | 15.12 | 113.9 | 86.1 | 7.38 | 12.3 | | High | High | High | High | High | High | High | High | High | High | High | 0 | 0.000232 | 2.13 |
| FALSE | High | ADTTTPTPT | [K].ADTTTI | 1xPhospho | Q7JJ13 1xP | 0.000411 | 0 | 1 | 3 | 2 | Q7JJ13 | Bromodom | Q7JJ13 [28 | 0 | 3112.622 | 0 | -0.39 | 0.054177 | 6.17 | 115.9 | 84.1 | 11.49 | 19.84 | | High | High | High | High | High | High | High | High | High | High | High | 0 | 8.83E-05 | 3 |
| FALSE | High | ADTTTPTTI | [K].ADTTTI | 1xPhospho | Q3UH70 1x | 6.97E-06 | 0 | 1 | 8 | 7 | Q3UH70 | Bromodom | Q3UH70 [2 | 0 | 2966.516 | 0 | 0.27 | 0.252181 | 22.32 | 91.4 | 108.6 | 2.82 | 16.01 | | High | High | High | High | High | High | High | High | High | High | High | 0.000327 | 0.002868 | 3.51 |
| FALSE | High | ADTTTPTTS | [K].ADTTTI | 1xPhospho | Q8K2F0-2 | 0.006363 | 0.00062 | 1 | 6 | 3 | Q8K2F0-2 | Isoform 2 c | Q8K2F0-2 [ | 0 | 1802.873 | 0 | -0.02 | 0.624972 | 10.12 | 101.9 | 98.1 | 8.82 | 6.7 | | High | High | High | High | High | High | High | High | High | High | High | 0.000275 | 0.001892 | 2.22 |
| FALSE | High | ADVQLFMA | [K].ADVQLI | 2xPhospho | Q9WVC3 2 | 0.000519 | 0 | 1 | 4 | 3 | Q9WVC3 | Caveolin-2 | Q9WVC3 [ | 0 | 3399.468 | 0 | 0.69 | 0.03537 | 26.81 | 72.2 | 127.8 | 17.29 | 19.69 | | High | High | High | High | High | High | High | High | High | High | High | 0.00014 | 0.001124 | 2.72 |
| FALSE | High | ADVSCEPLI | [R].ADVSCI | 1xCarbami | P70399 1xI | 0.000501 | 0 | 1 | 5 | 2 | P70399 | TP53-bindi | P70399 [7 | 0 | 2042.976 | 0 | -0.12 | 0.085118 | 2.1 | 107.6 | 92.4 | 7.19 | 9.06 | | High | High | High | High | High | High | High | High | High | High | High | 0 | 0.00011 | 3.52 |
| FALSE | High | ADVSSPK | [K].ADVSSI | 1xPhospho | Q7TT28 1x | 0.00824 | 0.000736 | 1 | 4 | 1 | Q7TT28 | RNA exonu | Q7TT28 [2 | 0 | 1241.654 | 0 | -0.26 | 0.009544 | 5.03 | 109 | 91 | 2.46 | 5.87 | | High | High | High | High | High | High | High | High | High | High | High | 0.000327 | 0.002522 | 1.5 |
| FALSE | High | ADVTLSALI | [K].ADVTLS | 1xPhospho | Q9DCT6-3 | 1.72E-06 | 0 | 1 | 4 | 2 | Q9DCT6-3 | Isoform 3 c | Q9DCT6-3 | 0 | 3551.741 | 0 | 0.07 | 0.157978 | 6.21 | 98.3 | 101.7 | 4.46 | 3.64 | | High | High | High | High | High | High | High | High | High | High | High | 0 | 1.87E-07 | 4.57 |
| FALSE | High | AEAAAAPA | [K].AEAAA | 1xPhospho | O08919 1x | 1.43E-06 | 0 | 1 | 2 | 3 | O08919 | Numb-like | O08919 [2 | 0 | 3373.719 | 0 | -0.58 | 0.006829 | 7.19 | 118.7 | 81.3 | 8.82 | 8.05 | | High | High | High | High | High | High | High | High | High | High | High | 0 | 2.49E-07 | 4.67 |
| FALSE | High | AEADASGN | [K].AEADA | 1xPhospho | B2RRE0 1xI | 0.003822 | 0.000261 | 1 | 2 | 1 | B2RRE0 | A kinase (P | B2RRE0 [1 | 1 | 2769.391 | 0 | -1.34 | 0.00574 | 2.35 | 147.1 | 52.9 | 19.34 | 14.08 | | High | High | High | High | High | High | High | High | High | High | High | 0.000117 | 0.001069 | 2.38 |
| FALSE | High | AEAENNSG | [K].AEAENI | 1xPhospho | E9QNN1 1x | 4.39E-12 | 0 | 1 | 4 | 11 | E9QNN1 | ATP-depen | E9QNN1 [1 | 0 | 2676.141 | 0 | -0.01 | 0.269441 | 1.97 | 104.1 | 95.9 | 5.04 | 7.51 | | High | High | High | High | High | High | High | High | High | High | High | 0 | 3.29E-09 | 6.39 |
| FALSE | High | AEAENNSG | [K].AEAENI | 1xDeamida | E9QNN1 1x | 1.33E-09 | 0 | 1 | 4 | 19 | E9QNN1 | ATP-depen | E9QNN1 [1 | 0 | 2677.125 | 0 | -0.11 | 0.018764 | 2.07 | 103.5 | 96.5 | 4.29 | 3.68 | | High | High | High | High | High | High | High | High | High | High | High | 0 | 5.95E-11 | 5.68 |
| FALSE | High | AEAENNSG | [K].AEAENI | 2xPhospho | E9QNN1 2x | 1.81E-07 | 0 | 1 | 4 | 1 | E9QNN1 | ATP-depen | E9QNN1 [1 | 0 | 2756.108 | 0 | -0.17 | 0.261796 | 16.76 | 110.2 | 89.8 | 14.14 | 9.08 | | High | High | High | High | High | High | High | High | High | High | High | 0 | 1.49E-08 | 5.38 |
| FALSE | High | AEAGAGSA | [K].AEAGA | 1xPhospho | Q5M9K7 1 | 4.95E-06 | 0 | 1 | 2 | 1 | Q5M9K7 | MCG21688 | Q5M9K7 [ | 0 | 1750.8 | 0 | -0.13 | 0.445314 | 7.48 | 101.5 | 98.5 | 9.79 | 3.5 | | High | High | High | High | High | High | High | High | High | High | High | 0 | 6.12E-07 | 3.08 |
| FALSE | High | AEAGSPTG | [R].AEAGSI | 1xPhospho | Q8C0Y1 1x | 6.24E-05 | 0 | 1 | 2 | 2 | Q8C0Y1 | Homeobox | Q8C0Y1 [1 | 0 | 2846.401 | 0 | -0.55 | 0.004235 | 4.94 | 117.9 | 82.1 | 7.16 | 11.42 | | High | High | High | High | High | High | High | High | High | High | High | 0 | 1.06E-05 | 3.83 |
| FALSE | High | ADTTTPTPT | [K].ADTTTI | 2xPhospho | Q7JJ13 2xP | 3.95E-06 | 0 | 1 | 3 | 8 | Q7JJ13 | Bromodom | Q7JJ13 [28 | 0 | 3192.588 | 0 | -0.34 | 0.001286 | 1.77 | 110.9 | 89.1 | 3.63 | 4.29 | | High | High | High | High | High | High | High | High | High | High | High | 0 | 0.000103 | 4.3 |
| FALSE | High | ADSLTDDIN | [K].ADSLTD | 1xPhospho | P50446 1xI | 0.000115 | 0 | 1 | 1 | 2 | P50446 | Keratin, ty | P50446 [27 | 0 | 1688.809 | 0 | 4.16 | 0.000415 | 38.25 | 10.7 | 189.3 | 38.28 | 12.62 | | High | High | High | High | High | High | High | High | High | High | High | 0 | 2.11E-05 | 2.73 |
| FALSE | High | ADSISEEAD | [R].ADSISE | 1xCarbami | Q921C3 2x | 3.24E-06 | 0 | 1 | 5 | 4 | Q921C3 | Bromodom | Q921C3 [1 | 0 | 2671.15 | 0 | -0.89 | 0.000813 | 2.72 | 129.6 | 70.4 | 7.32 | 10.18 | | High | High | High | High | High | High | High | High | High | High | High | 0 | 3.79E-07 | 3.66 |
| FALSE | High | ADSEYTDK | [R].ADSEYT | 1xPhospho | P54763-2 1 | 0.015465 | 0.001089 | 1 | 4 | 1 | P54763-2 | Isoform 2 c | P54763-2 [ | 0 | 1466.682 | 0 | -2 | 0.001851 | 0.36 | 160.1 | 39.9 | 12.89 | 15.68 | | High | High | High | High | High | High | High | High | High | High | High | 0.000571 | 0.005097 | 2.07 |
| FALSE | High | ADGPRPSLI | [K].ADGPR | 2xPhospho | E9QMQ3 2 | 1.29E-05 | 0 | 2 | 7 | 3 | E9QMQ3; ( | TRIO and F | E9QMQ3 [ | 0 | 2841.342 | 0 | 1.1 | 0.000689 | 12.37 | 63.4 | 136.6 | 7.18 | 4.85 | Shared | High | High | High | High | High | High | High | High | High | High | High | 0 | 1.79E-06 | 4.34 |
| FALSE | High | ADGYEPPV | [R].ADGYEI | 1xPhospho | P97351 1xI | 1.37E-05 | 0 | 1 | 5 | 4 | P97351 | 40S riboso | P97351 [25 | 0 | 1599.714 | 0 | -0.22 | 0.023047 | 11.77 | 107.9 | 92.1 | 2.19 | 10.05 | | High | High | High | High | High | High | High | High | High | High | High | 0 | 1.93E-06 | 2.95 |
| FALSE | High | ADHGVPLP | [R].ADHGV | 1xPhospho | A0A0R4J1F | 2.36E-06 | 0 | 1 | 6 | 1 | A0A0R4J1F | Ubiquitin c | A0A0R4J1F | 0 | 2454.311 | 0 | -0.52 | 0.053053 | 10.72 | 117.4 | 82.6 | 12.74 | 9.99 | | High | High | High | High | High | High | High | High | High | High | High | 0 | 2.67E-07 | 3.95 |
| FALSE | High | ADLAPPSP | [R].ADLAPI | 1xPhospho | A2ASY1 1xI | 6.61E-06 | 0 | 1 | 5 | 1 | A2ASY1 | TAF3 RNA | A2ASY1 [2 | 0 | 2386.265 | 0 | -0.15 | 0.154898 | 12.66 | 107.3 | 92.7 | 9.49 | 14.85 | | High | High | High | High | High | High | High | High | High | High | High | 0 | 8.45E-07 | 3.47 |
| FALSE | High | ADLAPPSP | [R].ADLAPI | 1xDeamida | A2ASY1 1xI | 0.057686 | 0.004227 | 1 | 5 | 1 | A2ASY1 | TAF3 RNA | A2ASY1 [2 | 0 | 2387.249 | 0 | -0.2 | 0.150477 | 20.3 | 102.5 | 97.5 | 6.49 | 14.98 | | High | High | High | High | High | High | High | High | High | High | High | 0.002159 | 0.02263 | 2.08 |
| FALSE | High | ADLAPPSP | [R].ADLAPI | 1xPhospho | A2ASY1 1xI | 0.000115 | 0 | 1 | 5 | 1 | A2ASY1 | TAF3 RNA | A2ASY1 [2 | 0 | 2402.26 | 0 | -0.29 | 0.081231 | 9.24 | 108.1 | 91.9 | 3.43 | 11.97 | | High | High | High | High | High | High | High | High | High | High | High | 0 | 2.11E-05 | 2.08 |
| FALSE | High | ADLDISGTK | [K].ADLDIS | 1xPhospho | E9Q616 1x | 0.018447 | 0.001324 | 1 | 3 | 1 | E9Q616 | AHNAK nu | E9Q616 [7 | 0 | 1457.765 | 0 | 2.47 | 0.000505 | 4.53 | 26.2 | 173.8 | 15.69 | 11.99 | | High | High | High | High | High | High | High | High | High | High | High | 0.000706 | 0.006225 | 1.66 |
| FALSE | High | ADIDVSGPI | [KN].ADIDV | 1xPhospho | E9Q616 1x | 0.001068 | 0 | 1 | 6 | 6 | E9Q616 | AHNAK nu | E9Q616 [1 | 0 | 1439.755 | 3 | 3.19 | 0.000281 | 7.7 | 18.9 | 181.1 | 12.25 | 8.42 | | High | High | High | High | High | High | High | High | High | High | High | 2.58E-05 | 0.000477 | 1.2 |
| FALSE | High | ADLIPYQDS | [K].ADLIPY | 1xPhospho | D3Z7I0 1xP | 0.000984 | 0 | 1 | 3 | 4 | D3Z7I0 | General tra | D3Z7I0 [79 | 0 | 3519.609 | 0 | -0.22 | 0.119735 | 6.61 | 106.9 | 93.1 | 9.06 | 12.41 | | High | High | High | High | High | High | High | High | High | High | High | 0 | 0.000235 | 3.67 |
| FALSE | High | ADLIPYQDS | [K].ADLIPY | 1xDeamida | D3Z7I0 1xP | 0.028082 | 0.001791 | 1 | 3 | 3 | D3Z7I0 | General tra | D3Z7I0 [79 | 0 | 3520.593 | 0 | -0.12 | 0.634657 | 5.42 | 101 | 99 | 13.41 | 16.26 | | High | High | High | High | High | High | High | High | High | High | High | 0.00196 | 0.01982 | 3.49 |
| FALSE | High | ADLIPYQDS | [K].ADLIPY | 2xPhospho | D3Z7I0 2xP | 0.002671 | 0.000109 | 1 | 3 | 3 | D3Z7I0 | General tra | D3Z7I0 [79 | 0 | 3599.576 | 0 | 0.07 | 0.636612 | 15.33 | 101.5 | 98.5 | 13.7 | 12.81 | | High | High | High | High | High | High | High | High | High | High | High | 0.000449 | 0.003994 | 3.79 |
| FALSE | High | ADLNQGIG | [R].ADLNQ | 1xPhospho | Q8C845 1x | 6.33E-07 | 0 | 1 | 2 | 3 | Q8C845 | EF-hand dc | Q8C845 [6 | 0 | 1877.896 | 0 | 1.37 | 0.000879 | 8.88 | 58.1 | 141.9 | 5.51 | 9.63 | | High | High | High | High | High | High | High | High | High | High | High | 0 | 2.57E-07 | 3.96 |
| FALSE | High | ADLNQGIG | [R].ADLNQ | 1xPhospho | Q8C845 1x | 0.001516 | 5.66E-05 | 1 | 2 | 1 | Q8C845 | EF-hand dc | Q8C845 [6 | 1 | 2033.997 | 0 | 1.35 | 0.000154 | 7.58 | 56.4 | 143.6 | 5.9 | 2.25 | | High | High | High | High | High | High | High | High | High | High | High | 2.58E-05 | 0.000381 | 1.94 |
| FALSE | High | ADLQGSAS | [K].ADLQG | 1xPhospho | Q8K212 1x | 0.00359 | 0.00021 | 1 | 3 | 3 | Q8K212 | Phosphofu | Q8K212 [4 | 0 | 1598.819 | 0 | 1.7 | 0.000181 | 11.33 | 47.9 | 152.1 | 12.83 | 4.4 | | High | High | High | High | High | High | High | High | High | High | High | 9.42E-05 | 0.000996 | 2.79 |
| FALSE | High | ADISASVLP | [R].ADISAS | 1xPhospho | O88495 1x | 0.000124 | 0 | 1 | 1 | 2 | O88495 | Melatonin- | O88495 [5 | 0 | 3651.84 | 0 | 6.64 | | 0 | 4.5 | 195.5 | | 10.22 | | Not Found | Not Found | Not Found | High | Not Found | High | High | High | High | High | High | 0 | 2.29E-05 | 3.63 |
| FALSE | High | ADLSEPSVE | [K].ADLSEP | 1xDeamida | F8VQG7 1x | 0.000253 | 0 | 1 | 4 | 1 | F8VQG7 | NEDD4-bin | F8VQG7 [8 | 0 | 4615.185 | 0 | -0.44 | 0.046245 | 3.49 | 116.1 | 83.9 | 16.51 | 14.81 | | High | High | High | High | High | High | High | High | High | High | High | 0 | 5.1E-05 | 4.4 |
| FALSE | High | ADIVESLCS | [R].ADIVES | 1xCarbami | Q8CFT3 1x | 1.59E-06 | 0 | 1 | 4 | 3 | Q8CFT3 | Nerve grov | Q8CFT3 [4 | 0 | 2074.945 | 0 | 2.9 | 0.00123 | 12.05 | 23 | 177 | 5.79 | 15.78 | | High | High | High | High | High | High | High | High | High | High | High | 4.94E-05 | 0.000662 | 3.38 |
| FALSE | High | ADNCSPVA | [R].ADNCSI | 1xCarbami | Q0VGY8 1x | 0.000213 | 0 | 1 | 4 | 1 | Q0VGY8 | Protein TA | Q0VGY8 [2 | 0 | 2600.164 | 0 | -0.53 | 0.046407 | 6.9 | 118.3 | 81.7 | 7.47 | 20.63 | | High | High | High | High | High | High | High | High | High | High | High | 0 | 4.21E-05 | 2.97 |
| FALSE | High | ADNCSPVA | [R].ADNCSI | 1xCarbami | Q0VGY8 1x | 0.013545 | 0.000976 | 1 | 4 | 1 | Q0VGY8 | Protein TA | Q0VGY8 [2 | 0 | 2829.327 | 0 | -0.06 | 0.48072 | 15.25 | 90.9 | 109.1 | 14.13 | 24 | | High | High | High | High | High | High | High | High | High | High | High | 0.000505 | 0.004392 | 2.47 |
| FALSE | High | ADNEPSCS | [K].ADNEP | 1xCarbami | Q0VGY8 1x | 9.2E-05 | 0 | 1 | 4 | 2 | Q0VGY8 | Protein TA | Q0VGY8 [1 | 0 | 2167.989 | 0 | -0.12 | 0.125134 | 6.58 | 106.1 | 93.9 | 6.99 | 6.06 | | High | High | High | High | High | High | High | High | High | High | High | 0 | 1.63E-05 | 2.75 |
| FALSE | High | ADNPTPQT | [K].ADNPTI | 2xPhospho | Q9D6J3 2xI | 4.16E-05 | 0 | 1 | 1 | 4 | Q9D6J3 | Coiled-coil | Q9D6J3 [2 | 0 | 3186.287 | 0 | 0.14 | 0.025338 | 2.33 | 92.9 | 107.1 | 5.08 | 6.14 | | High | High | High | High | High | High | High | High | High | High | High | 0 | 6.7E-06 | 4.11 |
| FALSE | High | ADPLILQAF | [K].ADPLIL | 1xPhospho | Q80Y19 1x | 5.65E-08 | 0 | 1 | 2 | 2 | Q80Y19 | Rho GTPas | Q80Y19 [6 | 0 | 2700.312 | 0 | -0.15 | 0.422651 | 7.53 | 102.4 | 97.6 | 5.77 | 10.01 | | High | High | High | High | High | High | High | High | High | High | High | 0 | 4.01E-09 | 3.75 |
| FALSE | High | ADPSLLGFS | [R].ADPSLL | 1xPhospho | Q6Y7W8 1 | 5.32E-06 | 0 | 1 | 3 | 2 | Q6Y7W8 | GRB10-inte | Q6Y7W8 [1 | 0 | 1958.942 | 0 | 0.25 | 0.03388 | 11.93 | 90.5 | 109.5 | 3.84 | 6.66 | | High | High | High | High | High | High | High | High | High | High | High | 0 | 6.66E-07 | 4.47 |
| FALSE | High | ADQSISEEF | [R].ADQSIS | 1xPhospho | Q80Y55 1x | 0.002004 | 5.66E-05 | 1 | 1 | 3 | Q80Y55 | BSD domai | Q80Y55 [2 | 0 | 3451.608 | 0 | -0.58 | 0.035585 | 7.47 | 118.8 | 81.2 | 11.41 | 21.01 | | High | High | High | High | High | High | High | High | High | High | High | 2.58E-05 | 0.000518 | 3.21 |
| FALSE | High | ADREQSPV | [K].ADREQ | 1xPhospho | A2AR02 1x | 0.025073 | 0.001645 | 1 | 2 | 1 | A2AR02 | Peptidyl-pr | A2AR02 [6 | 1 | 1654.857 | 0 | -0.48 | 0.001795 | 0.34 | 115.8 | 84.2 | 5.99 | 4.46 | | High | High | High | High | High | High | High | High | High | High | High | 0.000872 | 0.008816 | 1.4 |
| FALSE | High | ADSDGAK | [K].ADSDG | 1xPhospho | Q3UMU9-3 | 0.010758 | 0.000772 | 1 | 4 | 3 | Q3UMU9-3 | Isoform 3 c | Q3UMU9-3 | 0 | 1201.587 | 0 | 0.14 | 0.428926 | 13.16 | 97.2 | 102.8 | 7.56 | 5.35 | | High | High | High | High | High | High | High | High | High | High | High | 0.000449 | 0.003994 | 2.18 |
| FALSE | High | ADSDLEDSI | [K].ADSDLE | 1xPhospho | P97329 1xI | 3.37E-07 | 0 | 1 | 1 | 6 | P97329 | Kinesin-like | P97329 [54 | 0 | 2679.196 | 0 | -0.47 | 0.00047 | 1.64 | 116.5 | 83.5 | 2.06 | 4.95 | | High | High | High | High | High | High | High | High | High | High | High | 0 | 3E-08 | 3.33 |
| FALSE | High | ADSDSEDK | [K].ADSDSI | 1xPhospho | P83917 1xI | 0.008167 | 0.000736 | 1 | 3 | 1 | P83917 | Chromobo | P83917 [87 | 0 | 1404.63 | 0 | -0.95 | 0.000964 | 1.61 | 131.9 | 68.1 | 5.97 | 10.01 | | High | High | High | High | High | High | High | High | High | High | High | 0.000327 | 0.002495 | 1.83 |
| FALSE | High | ADSDSSQK | [R].ADSDS | 1xPhospho | Q60875 1x | 0.002441 | 0.000109 | 1 | 8 | 1 | Q60875 | Rho guanir | Q60875 [7 | 0 | 1375.651 | 0 | 1.28 | 0.000488 | 8.8 | 59.4 | 140.6 | 7.65 | 7.33 | | High | High | High | High | High | High | High | High | High | High | High | 4.94E-05 | 0.000649 | 2.09 |
| FALSE | High | AEAGSPTG | [R].AEAGSI | 2xPhospho | Q8C0Y1 2x | 0.001222 | 0 | 1 | 2 | 1 | Q8C0Y1 | Homeobox | Q8C0Y1 [1 | 0 | 2926.368 | 0 | -0.19 | 0.125495 | 17.87 | 108.7 | 91.3 | 5.97 | 15.6 | | High | High | High | High | High | High | High | High | High | High | High | 0 | 0.0003 | 3.14 |
| FALSE | High | AQGGHEF | [R].AQGGH | 1xPhospho | Q3TFZ4 1xI | 0.000908 | 0 | 1 | 2 | 1 | Q3TFZ4 | Putative ur | Q3TFZ4 [6 | 0 | 2659.292 | 0 | -0.01 | 0.804829 | 5.71 | 101.6 | 98.4 | 11.79 | 6.75 | | High | High | High | High | High | High | High | High | High | High | High | 0 | 0.000213 | 2.61 |
| FALSE | High | AEAKEESEI | [K].AEAKEI | 1xPhospho | Q5M8R8 1 | 8.18E-05 | 0 | 1 | 2 | 3 | Q5M8R8 | 60S acidic | Q5M8R8 [2 | 1 | 2789.179 | 0 | 0.38 | 0.122899 | 7.4 | 85.3 | 114.7 | 8.99 | 19.62 | | High | High | High | High | High | High | High | High | High | High | High | 0 | 0.000103 | 3.68 |
| FALSE | High | AEAKPGTP | [K].AEAKP | 1xPhospho | E9Q5C9 1x | 0.080423 | 0.006219 | 1 | 2 | 2 | E9Q5C9 | Nucleolar a | E9Q5C9 [2 | 0 | 1736.992 | 0 | -0.22 | 0.557496 | 8.5 | 106.5 | 93.5 | 14.3 | 6.95 | | High | High | High | High | High | High | High | High | High | High | High | 0.005231 | 0.05703 | 1.61 |

| | | | | | | | | | | | | | | | | | | | | | | | | | | | | | | | | | | | | | | |
|---|---|---|---|---|---|---|---|---|---|---|---|---|---|---|---|---|---|---|---|---|---|---|---|---|---|---|---|---|---|---|---|---|---|---|---|---|---|---|
| FALSE | High | AEESLTVES | [R].AEESLT | 1xPhosphc | E9PZM7 1x | 8.28E-06 | 0 | 1 | 3 | 3 | E9PZM7 | SR-related | E9PZM7 [5 | 0 | 2116.948 | 0 | -0.19 | 0.138191 | 10.67 | 107.4 | 92.6 | 6.12 | 11.96 | | High | High | High | High | High | High | High | High | High | High | High | 0 | 1.09E-06 | 2.64 |
| FALSE | High | AEETGPGV | [R].AEETGI | 1xPhosphc | A2ANY6 1x | 8.71E-05 | 0 | 1 | 2 | 1 | A2ANY6 | Midasin OS | A2ANY6 [4 | 0 | 2413.142 | 0 | -0.74 | 0.001518 | 4.44 | 123.6 | 76.4 | 9.12 | 10.48 | | High | High | High | High | High | High | High | High | High | High | High | 0 | 1.53E-05 | 2.41 |
| FALSE | High | AEETPNESV | [K].AEETPN | 1xPhosphc | B1AU76 1x | 6.49E-06 | 0 | 2 | 4 | 3 | B1AU76; B | Nuclear au | B1AU76 [1 | 0 | 1883.94 | 0 | -1.93 | 0.000158 | 0.36 | 158.8 | 41.2 | 3.63 | 5.37 | Shared | High | High | High | High | High | High | High | High | High | High | High | 0 | 8.28E-07 | 3.3 |
| FALSE | High | AEETPSQLI | [K].AEETPS | 1xPhosphc | Q3TVM1 1 | 5.45E-08 | 0 | 1 | 2 | 8 | Q3TVM1 | Putative ur | Q3TVM1 [3 | 0 | 3791.6 | 0 | -0.12 | 0.383155 | 9.12 | 103.8 | 96.2 | 8.38 | 24.86 | | High | High | High | High | High | High | High | High | High | High | High | 0 | 3.86E-09 | 6.84 |
| FALSE | High | AEETPSQLI | [K].AEETPS | 1xPhosphc | Q3TVM1 1 | 6.38E-10 | 0 | 1 | 2 | 28 | Q3TVM1 | Putative ur | Q3TVM1 [3 | 0 | 3807.595 | 0 | -0.24 | 0.114561 | 11.53 | 105.4 | 94.6 | 4.75 | 18.45 | | High | High | High | High | High | High | High | High | High | High | High | 0 | 2.62E-11 | 6.86 |
| FALSE | High | AEETSEELA | [R].AEETSE | 1xCarbami | P97452 1xl | 0.000257 | 0 | 1 | 3 | 4 | P97452 | Ribosome | P97452 [8C | 0 | 2170.989 | 0 | 0.06 | 0.724298 | 7.91 | 100.7 | 99.3 | 9.64 | 12.28 | | High | High | High | High | High | High | High | High | High | High | High | 0 | 5.2E-05 | 3.55 |
| FALSE | High | AEFSPAQFS | [R].AEFSPA | 1xPhosphc | Q3U0W6 1 | 0.000349 | 0 | 1 | 6 | 1 | Q3U0W6 | Putative ur | Q3U0W6 [ | 0 | 1803.908 | 0 | -0.04 | 0.713424 | 2.92 | 100.7 | 99.3 | 12.41 | 4.12 | | High | High | High | High | High | High | High | High | High | High | High | 0 | 7.35E-05 | 2.68 |
| FALSE | High | AEFTSPPSL | [K].AEFTSP | 1xPhosphc | Q9D824 1x | 0.000143 | 0 | 1 | 5 | 4 | Q9D824 | Pre-mRNA | Q9D824 [2 | 0 | 1761.923 | 0 | 0.02 | 0.698256 | 3.72 | 99.2 | 100.8 | 2.15 | 8.54 | | High | High | High | High | High | High | High | High | High | High | High | 0 | 2.69E-05 | 3.22 |
| FALSE | High | AEGEPQEE | [K].AEGEP( | 1xPhosphc | Q8K019 1x | 9.11E-05 | 0 | 1 | 10 | 6 | Q8K019 | Bcl-2-assoc | Q8K019 [1 | 0 | 1851.914 | 0 | -0.23 | 0.004625 | 1.42 | 107.3 | 92.7 | 6.65 | 3.55 | | High | High | High | High | High | High | High | High | High | High | High | 0 | 1.62E-05 | 3.99 |
| FALSE | High | AEGHASSP | [K].AEGHA | 1xCarbami | Q5DTX6 1x | 1.13E-05 | 0 | 1 | 1 | 2 | Q5DTX6 | Junctional | Q5DTX6 [5 | 0 | 2309.063 | 0 | 1.97 | 0.000348 | 4.96 | 38.8 | 161.2 | 20.64 | 11.63 | | High | High | High | High | High | High | High | High | High | High | High | 0 | 1.55E-06 | 3.61 |
| FALSE | High | AEGSLHSSI | [K].AEGSLH | 1xPhosphc | Q6ZPZ3 1x | 1.47E-05 | 0 | 1 | 5 | 2 | Q6ZPZ3 | Zinc finger | Q6ZPZ3 [9 | 0 | 2037.005 | 0 | 0.16 | 0.785814 | 25.64 | 96.4 | 103.6 | 16.41 | 11.93 | | High | High | High | High | High | High | High | High | High | High | High | 0 | 2.09E-06 | 2.69 |
| FALSE | High | AEHLGLSG. | [K].AEHLGI | 1xPhosphc | B2RWU4 1 | 3.66E-08 | 0 | 1 | 9 | 3 | B2RWU4 | Cordon-ble | B2RWU4 [ | 0 | 2250.069 | 0 | -0.12 | 0.056444 | 5.54 | 103.6 | 96.4 | 4.88 | 2.06 | | High | High | High | High | High | High | High | High | High | High | High | 0 | 2.47E-09 | 3.42 |
| FALSE | High | AEHLGLSG. | [K].AEHLGI | 2xPhosphc | B2RWU4 2 | 0.00086 | 0 | 1 | 9 | 1 | B2RWU4 | Cordon-ble | B2RWU4 [ | 0 | 2330.035 | 0 | 0.05 | 0.754903 | 14.43 | 98.7 | 101.3 | 7.98 | 10.27 | | High | High | High | High | High | High | High | High | High | High | High | 0 | 0.000201 | 2.19 |
| FALSE | High | AEESEPLPS | [K].AEESEP | 2xCarbami | Q3UMC6 1 | 8.29E-10 | 0 | 1 | 5 | 2 | Q3UMC6 | Putative ur | Q3UMC6 [ | 0 | 3386.594 | 0 | -0.42 | 0.000986 | 4.39 | 114.5 | 85.5 | 1.72 | 6.49 | | High | High | High | High | High | High | High | High | High | High | High | 0 | 3.5E-11 | 5.32 |
| FALSE | High | AELDIATTP | [K].AELDIA | 1xPhosphc | Q6PCM1 1 | 1.29E-05 | 0 | 1 | 5 | 3 | Q6PCM1 | Lysine-spe | Q6PCM1 [4 | 0 | 1967.05 | 0 | -1.04 | 0.003204 | 5.29 | 131.4 | 68.6 | 12.27 | 6.27 | | High | High | High | High | High | High | High | High | High | High | High | 0.000327 | 0.002894 | 2.7 |
| FALSE | High | AELEEMEE | [K].AELEEN | 1xPhosphc | A2ARP8 1x | 2.2E-07 | 0 | 1 | 3 | 2 | A2ARP8 | Microtubu | A2ARP8 [8 | 0 | 2927.279 | 0 | -1.3 | 0.002056 | 1.36 | 141.1 | 58.9 | 11.43 | 13.67 | | High | High | High | High | High | High | High | High | High | High | High | 0 | 1.86E-08 | 5.03 |
| FALSE | High | AELGMNDS | [R].AELGM | 1xPhosphc | A2AQN4 1 | 4.27E-05 | 0 | 1 | 8 | 2 | A2AQN4 | Acetyl-coe | A2AQN4 [2 | 0 | 2195.082 | 0 | 0.7 | 0.001282 | 3.25 | 76.1 | 123.9 | 6.18 | 6.69 | | High | High | High | High | High | High | High | High | High | High | High | 0 | 6.94E-06 | 3.63 |
| FALSE | High | AELGMNDS | [R].AELGM | 1xPhosphc | A2AQN4 1 | 4.55E-05 | 0 | 1 | 8 | 2 | A2AQN4 | Acetyl-coe | A2AQN4 [2 | 0 | 2211.077 | 0 | 0.62 | 0.00713 | 8.47 | 77.8 | 122.2 | 2.15 | 13.89 | | High | High | High | High | High | High | High | High | High | High | High | 0 | 0.000225 | 4.53 |
| FALSE | High | AELGQLIDL | [K].AELGQ | 1xPhosphc | Q542Q3 1x | 5.87E-09 | 0 | 1 | 1 | 3 | Q542Q3 | G two S ph | Q542Q3 [7 | 0 | 2845.536 | 0 | -0.39 | 0.089626 | 11.78 | 114.2 | 85.8 | 16.98 | 6.03 | | High | High | High | High | High | High | High | High | High | High | High | 0 | 3.17E-10 | 4.95 |
| FALSE | High | AELGQLIDL | [K].AELGQ | 2xPhosphc | Q542Q3 2x | 0.003312 | 0.00016 | 1 | 1 | 2 | Q542Q3 | G two S ph | Q542Q3 [7 | 0 | 2925.503 | 0 | 0.35 | 0.041256 | 4.69 | 87.9 | 112.1 | 16.07 | 9.89 | | High | High | High | High | High | High | High | High | High | High | High | 7.19E-05 | 0.000911 | 3.23 |
| FALSE | High | AELPPPYTA | [R].AELPPP | 1xCarbami | Q9CZX7 1x | 0.001968 | 5.66E-05 | 1 | 1 | 1 | Q9CZX7 | Type 2 phc | Q9CZX7 [3 | 0 | 2790.41 | 0 | 0.35 | 0.001136 | 6.23 | 87.8 | 112.2 | 18.62 | 14.82 | | High | High | High | High | High | High | High | High | High | High | High | 2.58E-05 | 0.000508 | 3.18 |
| FALSE | High | AELSDK | [R].AELSDK | 1xPhosphc | Q69ZX3 1x | 0.105058 | 0.008518 | 1 | 4 | 1 | Q69ZX3 | MKIAA086 | Q69ZX3 [1 | 0 | 1200.628 | 0 | -0.09 | 0.353967 | 6.34 | 97.7 | 102.3 | 14.85 | 6.51 | | High | High | High | High | High | High | High | High | High | High | High | 0.004277 | 0.04561 | 1.61 |
| FALSE | High | AELSGNCSI | [R].AELSGN | 1xCarbami | G3X932 1x | 0.008927 | 0.000736 | 1 | 4 | 1 | G3X932 | DNA segm | G3X932 [2 | 0 | 1963.987 | 0 | 0.39 | 0.056806 | 26.3 | 91.6 | 108.4 | 19.52 | 7.07 | | High | High | High | High | High | High | High | High | High | High | High | 0.000327 | 0.002761 | 2 |
| FALSE | High | AEISSIQEN | [K].AEISSIC | 1xPhosphc | Q9EP71 1x | 0.000868 | 0 | 1 | 1 | 1 | Q9EP71 | Ankycorbir | Q9EP71 [3 | 0 | 1656.861 | 0 | 1.22 | 0.000609 | 12.07 | 59.1 | 140.9 | 4.65 | 6.7 | | High | High | High | High | High | High | High | High | High | High | High | 0 | 0.000203 | 2.74 |
| FALSE | High | AELTQESS( | [K].AELTQE | 1xPhosphc | Q542Q3 1x | 3.3E-06 | 0 | 1 | 1 | 5 | Q542Q3 | G two S ph | Q542Q3 [5 | 0 | 4560.13 | 0 | -0.7 | 0.001191 | 1.9 | 124.8 | 75.2 | 7.76 | 9.3 | | High | High | High | High | High | High | High | High | High | High | High | 0 | 1.81E-05 | 5.04 |
| FALSE | High | AELTQESS( | [R].AELTQE | 1xDeamida | Q542Q3 1x | 1.63E-05 | 0 | 1 | 1 | 3 | Q542Q3 | G two S ph | Q542Q3 [5 | 0 | 4561.114 | 0 | -1.38 | 0.001201 | 0.83 | 145.9 | 54.1 | 15.01 | 20.02 | | High | High | High | High | High | High | High | High | High | High | High | 0 | 2.49E-06 | 5.23 |
| FALSE | High | AENAEGQT | [K].AENAE( | 1xPhosphc | A0A0R4J17 | 3.36E-06 | 0 | 1 | 5 | 4 | A0A0R4J17 | Transcripti | A0A0R4J17 | 0 | 3634.724 | 0 | -0.36 | 0.047387 | 5.13 | 113.2 | 86.8 | 5.97 | 24.2 | | High | High | High | High | High | High | High | High | High | High | High | 0 | 3.95E-07 | 4.26 |
| FALSE | High | AENAEGQT | [K].AENAE( | 1xPhosphc | A0A0R4J17 | 3.79E-07 | 0 | 1 | 5 | 9 | A0A0R4J17 | Transcripti | A0A0R4J17 | 0 | 3650.718 | 0 | -0.24 | 0.015809 | 2.49 | 107 | 93 | 9.12 | 6.29 | | High | High | High | High | High | High | High | High | High | High | High | 0 | 3.42E-08 | 4.51 |
| FALSE | High | AEIDSHSFG | [K].AEIDSH | 1xPhosphc | O55003 1x | 0.000217 | 0 | 1 | 2 | 1 | O55003 | BCL2/aden | O55003 [7 | 0 | 1757.851 | 0 | -0.4 | 0.003325 | 2.81 | 113.7 | 86.3 | 4.03 | 9.72 | | High | High | High | High | High | High | High | High | High | High | High | 0 | 4.3E-05 | 2.24 |
| FALSE | High | AEEQLPPLL | [R].AEEQLF | 2xPhosphc | P97434-2 2 | 0.005 | 0.000504 | 1 | 9 | 1 | P97434-2 | Isoform 2 c | P97434-2 [ | 0 | 2529.199 | 0 | 0.69 | 0.0054 | 11.26 | 78 | 122 | 7.3 | 7.3 | | High | High | High | High | High | High | High | High | High | High | High | 0.000225 | 0.001453 | 1.81 |
| FALSE | High | AEEQLPPLL | [R].AEEQLF | 1xPhosphc | P97434-2 1 | 1.98E-07 | 0 | 1 | 9 | 3 | P97434-2 | Isoform 2 c | P97434-2 [ | 0 | 2449.233 | 0 | 0.72 | 0.000408 | 6.42 | 74.2 | 125.8 | 13.53 | 6.63 | | High | High | High | High | High | High | High | High | High | High | High | 0 | 1.65E-08 | 4.65 |
| FALSE | High | AEEPPASP( | [R].AEEPPA | 1xPhosphc | Q3UV57 1x | 0.002671 | 0.000109 | 1 | 1 | 1 | Q3UV57 | Putative ur | Q3UV57 [7 | 0 | 1617.829 | 0 | 1.56 | 0.000222 | 3.83 | 48.9 | 151.1 | 5.3 | 5.39 | | High | High | High | High | High | High | High | High | High | High | High | 4.94E-05 | 0.000716 | 1.74 |
| FALSE | High | AEALSSLHC | [K].AEALSS | 1xPhosphc | B7ZWC4 1 | 9.65E-06 | 0 | 1 | 8 | 2 | B7ZWC4 | Insulin-like | B7ZWC4 [2 | 0 | 3221.55 | 0 | -1.53 | 0.002496 | 2.64 | 150.2 | 49.8 | 11.08 | 16.44 | | High | High | High | High | High | High | High | High | High | High | High | 0 | 1.3E-06 | 3.02 |
| FALSE | High | AEAPAGPA | [K].AEAPA( | 1xPhosphc | Q8BZR9 1x | 2.86E-06 | 0 | 1 | 3 | 4 | Q8BZR9 | Nuclear ca | Q8BZR9 [1 | 0 | 2456.227 | 0 | -0.49 | 0.002929 | 3.64 | 116.9 | 83.1 | 6.78 | 4.15 | | High | High | High | High | High | High | High | High | High | High | High | 0 | 4.02E-06 | 4.45 |
| FALSE | High | AEAPLPSPK | [K].AEAPLF | 1xPhosphc | E9Q616 1x | 0.008769 | 0.000736 | 1 | 5 | 1 | E9Q616 | AHNAK nu | E9Q616 [4 | 0 | 1218.633 | 0 | 3.56 | 0.072631 | 83.82 | 22 | 178 | 53.88 | 66.92 | | High | High | High | High | High | High | High | High | High | High | High | 0.000327 | 0.002706 | 1.31 |
| FALSE | High | AEAPLPSPK | [K].AEAPLF | 1xPhosphc | E9Q616 1x | 0.000302 | 0 | 1 | 5 | 2 | E9Q616 | AHNAK nu | E9Q616 [4 | 0 | 1447.796 | 0 | 3.67 | 0.000236 | 11.47 | 15.2 | 184.8 | 21.61 | 6.75 | | High | High | High | High | High | High | High | High | High | High | High | 0 | 6.23E-05 | 1.79 |
| FALSE | High | AEAQALLQ | [R].AEAQA | 1xPhosphc | A2A4P4 1x | 5.09E-06 | 0 | 1 | 3 | 5 | A2A4P4 | Uncharact | A2A4P4 [5 | 0 | 2262.038 | 0 | 0.33 | 0.018212 | 1.99 | 90.2 | 109.8 | 7.17 | 11.31 | | High | High | High | High | High | High | High | High | High | High | High | 0 | 3.69E-06 | 4.15 |
| FALSE | High | AEASPIIQP. | [R].AEASPI | 1xPhosphc | A2AI52 1xF | 0.020862 | 0.001472 | 1 | 5 | 1 | A2AI52 | Ubiquitin c | A2AI52 [4C | 0 | 1932.061 | 0 | -0.74 | 0.007624 | 2.2 | 118.3 | 81.7 | 9.97 | 9.35 | | High | High | High | High | High | High | High | High | High | High | High | 0.000801 | 0.007157 | 2.3 |
| FALSE | High | AEDCEGSP | [K].AEDCE( | 2xCarbami | Q6ZPR6 1x | 8.35E-06 | 0 | 1 | 2 | 1 | Q6ZPR6 | Inhibitor o | Q6ZPR6 [6 | 0 | 3060.323 | 0 | -0.87 | 0.004486 | 2.4 | 129.3 | 70.7 | 7.56 | 10.94 | | High | High | High | High | High | High | High | High | High | High | High | 0 | 1.1E-06 | 2.01 |
| FALSE | High | AEDGAAPS | [K].AEDGA | 1xPhosphc | P26645 1xl | 3.24E-09 | 0 | 1 | 3 | 12 | P26645 | Myristoyla | P26645 [1 | 0 | 1981.952 | 0 | 1.72 | 0.00054 | 6.76 | 48.2 | 151.8 | 4.67 | 8.62 | | High | High | High | High | High | High | High | High | High | High | High | 0 | 1.63E-10 | 3.82 |
| FALSE | High | AEDGAAPS | [K].AEDGA | 1xPhosphc | P26645 1xl | 7.16E-09 | 0 | 1 | 3 | 1 | P26645 | Myristoyla | P26645 [1 | 1 | 2110.047 | 0 | 1.5 | 0.237536 | 48.26 | 60.6 | 139.4 | 48.49 | 62.2 | | High | High | High | High | High | High | High | High | High | High | High | 0 | 3.97E-10 | 4.15 |
| FALSE | High | AEDGAAPS | [K].AEDGA | 1xPhosphc | P26645 1xl | 2.89E-07 | 0 | 1 | 3 | 4 | P26645 | Myristoyla | P26645 [1 | 1 | 2339.21 | 0 | 1.28 | 0.004921 | 31.74 | 56.2 | 143.8 | 11.46 | 15.26 | | High | High | High | High | High | High | High | High | High | High | High | 0 | 2.52E-08 | 3.1 |
| FALSE | High | AEDGTGTV | [R].AEDGT | 1xPhosphc | Q69ZV4 1x | 2.67E-08 | 0 | 1 | 3 | 2 | Q69ZV4 | MKIAA092 | Q69ZV4 [3 | 0 | 2546.134 | 0 | 0.13 | 0.034287 | 5.59 | 95.4 | 104.6 | 11.64 | 7.33 | | High | High | High | High | High | High | High | High | High | High | High | 0 | 1.74E-09 | 5.01 |
| FALSE | High | AEDLENVS | [K].AEDLEN | 1xPhosphc | S4R1P5 1xl | 4.31E-05 | 0 | 1 | 6 | 1 | S4R1P5 | Dystonin C | S4R1P5 [2 | 0 | 2480.176 | 0 | 0.82 | 0.001789 | 13.7 | 74.5 | 125.5 | 13.89 | 5.01 | | High | High | High | High | High | High | High | High | High | High | High | 0 | 6.98E-06 | 3.87 |
| FALSE | High | AEDLENVS | [K].AEDLEN | 2xPhosphc | S4R1P5 2xl | 5.05E-07 | 0 | 1 | 6 | 2 | S4R1P5 | Dystonin C | S4R1P5 [2 | 0 | 2560.142 | 0 | 0.75 | 0.004584 | 8.94 | 76.2 | 123.8 | 5.87 | 9.96 | | High | High | High | High | High | High | High | High | High | High | High | 0 | 4.71E-08 | 3.98 |
| FALSE | High | AEDPEGTT | [K].AEDPE( | 1xPhosphc | G3X961 1x | 4.68E-05 | 0 | 1 | 5 | 2 | G3X961 | Kelch dom | G3X961 [3 | 0 | 3109.487 | 0 | -0.58 | 0.01576 | 9.04 | 120 | 80 | 14.13 | 10.63 | | High | High | High | High | High | High | High | High | High | High | High | 0 | 7.67E-06 | 3.61 |
| FALSE | High | AEDSGAEC | [K].AEDSG. | 1xPhosphc | B2RRE0 1x | 1.29E-07 | 0 | 1 | 2 | 4 | B2RRE0 | A kinase (P | B2RRE0 [7 | 0 | 2097.954 | 0 | -1.12 | 0.00054 | 3.33 | 138.3 | 61.7 | 4.61 | 14.2 | | High | High | High | High | High | High | High | High | High | High | High | 0 | 1.02E-08 | 4.06 |
| FALSE | High | AEDTSPVIC | [R].AEDTSF | 2xPhosphc | Q6P1H6 2x | 0.00642 | 0.00062 | 1 | 3 | 2 | Q6P1H6 | Ankyrin re | Q6P1H6 [4 | 0 | 2480.14 | 0 | -0.22 | 0.04339 | 3.1 | 107.6 | 92.4 | 15.59 | 17.49 | | High | High | High | High | High | High | High | High | High | High | High | 0.000449 | 0.003943 | 2.37 |
| FALSE | High | AEECEHSPA | [K].AEECEH | 1xCarbami | A2AJK6 1xl | 0.002895 | 0.00016 | 1 | 2 | 1 | A2AJK6 | Chromodo | A2AJK6 [2 | 0 | 1695.781 | 0 | -1.23 | 0.000429 | 2.74 | 138 | 62 | 3.64 | 12.48 | | High | High | High | High | High | High | High | High | High | High | High | 7.19E-05 | 0.000788 | 2.52 |
| FALSE | High | AEEDEILNF | [K].AEEDEI | 1xPhosphc | Q3TXE5 1x | 0.000157 | 0 | 1 | 2 | 16 | Q3TXE5 | Putative ur | Q3TXE5 [5 | 1 | 1737.837 | 0 | 1 | 0.000536 | 4.99 | 67.3 | 132.7 | 2.78 | 5.53 | | High | High | High | High | High | High | High | High | High | High | High | 4.94E-05 | 0.000649 | 2.38 |
| FALSE | High | AEEDLSEEE | [K].AEEDLS | 1xPhosphc | Q9JJG9 1xF | 2.34E-06 | 0 | 1 | 2 | 1 | Q9JJG9 | Nitric oxide | Q9JJG9 [4 | 0 | 2528.217 | 0 | -0.18 | 0.405882 | 13.59 | 103.8 | 96.2 | 7.35 | 12.1 | | High | High | High | High | High | High | High | High | High | High | High | 0 | 2.64E-07 | 3.34 |
| FALSE | High | AEEDSEDES | [R].AEEDSE | 2xPhosphc | Q3UH06-3 | 4.68E-05 | 0 | 1 | 4 | 3 | Q3UH06-3 | Isoform 3 c | Q3UH06-3 | 0 | 4145.693 | 0 | 0 | 0.976029 | 16.24 | 102.6 | 97.4 | 16.2 | 12.89 | | High | High | High | High | High | High | High | High | High | High | High | 0 | 7.61E-06 | 4.18 |
| FALSE | High | AEEDSEDES | [R].AEEDSE | 1xDeamida | Q3UH06-3 | 0.056724 | 0.004124 | 1 | 4 | 3 | Q3UH06-3 | Isoform 3 c | Q3UH06-3 | 0 | 4146.677 | 0 | 0 | 0.90072 | 20.51 | 96.7 | 103.3 | 13.63 | 11.09 | | High | High | High | High | High | High | High | High | High | High | High | 0.002133 | 0.02224 | 3.16 |
| FALSE | High | AEEEDLAAY | [R].AEEEDI | 1xPhosphc | Q4U2R1 1x | 1.28E-06 | 0 | 1 | 3 | 2 | Q4U2R1 | E3 ubiquiti | Q4U2R1 [2 | 0 | 2874.229 | 0 | 0.44 | 0.004846 | 6.3 | 83.8 | 116.2 | 11.03 | 7.75 | | High | High | High | High | High | High | High | High | High | High | High | 0 | 1.35E-07 | 4.62 |
| FALSE | High | AEEEDLAAY | [R].AEEEDI | 1xPhosphc | Q4U2R1 1x | 1.07E-07 | 0 | 1 | 3 | 5 | Q4U2R1 | E3 ubiquiti | Q4U2R1 [2 | 0 | 3103.392 | 0 | 0.55 | 0.001626 | 12.05 | 83.4 | 116.6 | 7.85 | 3.67 | | High | High | High | High | High | High | High | High | High | High | High | 0 | 8.25E-09 | 4.06 |
| FALSE | High | AEEEECENI | [R].AEEEEC | 2xCarbami | Q80XI4 1xl | 1.5E-05 | 0 | 1 | 2 | 13 | Q80XI4 | Phosphatic | Q80XI4 [3C | 0 | 3962.73 | 0 | -1.35 | 0.00191 | 3 | 142.5 | 57.5 | 12.15 | 13.73 | | High | High | High | High | High | High | High | High | High | High | High | 0 | 1.48E-05 | 4.91 |
| FALSE | High | AEEENRSEI | [K].AEEENI | 1xPhosphc | Q6Y7W8 1 | 0.063788 | 0.004664 | 1 | 9 | 2 | Q6Y7W8 | GRB10-inte | Q6Y7W8 [4 | 1 | 2102.017 | 0 | -0.51 | 0.005097 | 1.13 | 118 | 82 | 3.55 | 8.07 | | High | High | High | High | High | High | High | High | High | High | High | 0.005205 | 0.05637 | 1.69 |
| FALSE | High | AEEEQLSSS | [R].AEEEQI | 1xPhosphc | Q8CGF1 1x | 0.001615 | 5.66E-05 | 1 | 3 | 2 | Q8CGF1 | Rho GTPas | Q8CGF1 [3 | 0 | 1986.02 | 0 | -0.16 | 0.173477 | 6.79 | 110 | 90 | 12.19 | 15.75 | | High | High | High | High | High | High | High | High | High | High | High | 2.58E-05 | 0.000407 | 3.2 |
| FALSE | High | AEEGLMLP | [R].AEEGLI | 1xPhosphc | P32507-2 1 | 0.0102 | 0.000736 | 1 | 3 | 1 | P32507-2 | Isoform Al | P32507-2 [ | 0 | 2089.044 | 0 | -0.53 | 0.001897 | 4.29 | 118.1 | 81.9 | 5.91 | 6.91 | | High | High | High | High | High | High | High | High | High | High | High | 0.000382 | 0.003213 | 2.32 |
| FALSE | High | AEENNSGV | [K].AEENN | 1xPhosphc | A0A087WF | 3.36E-10 | 0 | 1 | 4 | 4 | A0A087WF | ATP-depen | A0A087WF | 0 | 2605.104 | 0 | 0.26 | 0.162865 | 10.21 | 91.1 | 108.9 | 6.4 | 9.54 | | High | High | High | High | High | High | High | High | High | High | High | 0 | 1.27E-11 | 5.47 |
| FALSE | High | AEENNSGV | [K].AEENN | 1xDeamida | A0A087WF | 3.98E-09 | 0 | 1 | 4 | 12 | A0A087WF | ATP-depen | A0A087WF | 0 | 2606.088 | 0 | 0.27 | 0.033162 | 7.19 | 90.9 | 109.1 | 6.27 | 5.42 | | High | High | High | High | High | High | High | High | High | High | High | 0 | 2.04E-10 | 5.96 |
| FALSE | High | AEAKEESEI | [K].AEAKEE | 2xPhosphc | Q5M8R8 2 | 5.57E-06 | 0 | 1 | 2 | 18 | Q5M8R8 | 60S acidic | Q5M8R8 [ | 1 | 2853.15 | 0 | -0.48 | 0.003474 | 3.46 | 118.1 | 81.9 | 6.4 | 10.36 | | High | High | High | High | High | High | High | High | High | High | High | 0.001151 | 0.01232 | 2.99 |
| FALSE | High | AQGHSPVN | [K].AQGHS | 1xPhosphc | Q9JJA7 1xF | 0.000466 | 0 | 1 | 1 | 3 | Q9JJA7 | Cyclin-L2 C | Q9JJA7 [3 | 0 | 1758.967 | 0 | -1.26 | 0.000229 | 1.76 | 141.7 | 58.3 | 4.96 | 8.27 | | High | High | High | High | High | High | High | High | High | High | High | 0.000275 | 0.001575 | 2.21 |
| FALSE | High | AQGHSPVN | [K].AQGHS | 1xDeamida | Q9JJA7 1xF | 1.85E-05 | 0 | 1 | 1 | 6 | Q9JJA7 | Cyclin-L2 C | Q9JJA7 [3 | 0 | 1759.951 | 0 | -1.08 | 0.000254 | 1.19 | 136.4 | 63.6 | 2.43 | 5.28 | | High | High | High | High | High | High | High | High | High | High | High | 0.000467 | 0.004301 | 3.17 |
| FALSE | High | AQGLASTS | [R].AQGLA | 1xPhosphc | Q91X58 1x | 0.000274 | 0 | 1 | 3 | 2 | Q91X58 | AN1-type z | Q91X58 [1 | 0 | 1739.853 | 0 | -0.27 | 0.024889 | 8.63 | 109.1 | 90.9 | 7.77 | 5.2 | | High | High | High | High | High | High | High | High | High | High | High | 0.000903 | 0.009346 | 1.96 |
| FALSE | High | CLSETHGEI | [R].CLSETH | 2xCarbami | O55187 1x | 0.051258 | 0.003745 | 1 | 3 | 1 | O55187 | E3 SUMO- | O55187 [4 | 1 | 2140.992 | 0 | 0.87 | 0.001286 | 5.42 | 71 | 129 | 9.83 | 10.63 | | High | High | High | High | High | High | High | High | High | High | High | 0.00196 | 0.01982 | 1 |
| FALSE | High | CISMPVDIS | [R].CISMPV | 1xCarbami | Q3TRM4 1 | 6.85E-06 | 0 | 1 | 6 | 1 | Q3TRM4 | Neuropath | Q3TRM4 [4 | 0 | 1995.959 | 0 | 1.2 | 0.000552 | 10.34 | 57.5 | 142.5 | 18.8 | 13.84 | | High | High | High | High | High | High | High | High | High | High | High | 0 | 8.79E-07 | 4.22 |
| FALSE | High | CISMPVDIS | [R].CISMPV | 1xCarbami | Q3TRM4 1 | 0.000157 | 0 | 1 | 6 | 1 | Q3TRM4 | Neuropath | Q3TRM4 [4 | 0 | 2011.954 | 0 | -0.03 | 0.475877 | 5.44 | 99.2 | 100.8 | 6.34 | 8.97 | | High | High | High | High | High | High | High | High | High | High | High | 0 | 2.97E-05 | 2.02 |
| FALSE | High | CLSPDDST\ | [K].CLSPDD | 1xCarbami | A2ARP8 1x | 0.000837 | 0 | 1 | 3 | 3 | A2ARP8 | Microtubu | A2ARP8 [1 | 0 | 1659.807 | 0 | -1.73 | 0.001856 | 1.3 | 149 | 51 | 13.3 | 17.89 | | High | High | High | High | High | High | High | High | High | High | High | 0 | 0.000195 | 1.99 |
| FALSE | High | CLSPEGVN | [R].CLSPEG | 1xCarbami | Q6PGN3 1 | 0.053473 | 0.00384 | 1 | 9 | 1 | Q6PGN3 | Serine/thre | Q6PGN3 [3 | 0 | 1511.688 | 0 | -0.82 | 0.001633 | 1.16 | 127.5 | 72.5 | 5.97 | 9.48 | | High | High | High | High | High | High | High | High | High | High | High | 0.001999 | 0.02075 | 1.41 |
| FALSE | High | CLSSHDDSI | [K].CLSSHD | 1xCarbami | Q8BS90 1x | 6.16E-07 | 0 | 1 | 2 | 2 | Q8BS90 | Protein aur | Q8BS90 [1 | 0 | 1855.83 | 0 | -1.33 | 0.000445 | 0.89 | 143.2 | 56.8 | 3.72 | 14.56 | | High | High | High | High | High | High | High | High | High | High | High | 0 | 5.89E-08 | 3.38 |
| FALSE | High | CLSYGDTS( | [R].CLSYGD | 1xCarbami | A0A0A0MC | 0.024638 | 0.001612 | 1 | 2 | 2 | A0A0A0MC | Mitogen-a | A0A0A0MC | 0 | 1608.729 | 0 | 1.76 | 0.000381 | 4.82 | 45.4 | 154.6 | 4.82 | 7.65 | | High | High | High | High | High | High | High | High | High | High | High | 0.000859 | 0.008639 | 2.39 |
| FALSE | High | CLVAGLLSI | [R].CLVAGI | 1xCarbami | Q8CJF7 1xl | 0.008167 | 0.000736 | 1 | 1 | 1 | Q8CJF7 | Protein EL' | Q8CJF7 [5 | 0 | 1394.743 | 0 | -0.04 | 0.933157 | 6.71 | 101.2 | 98.8 | 5.2 | 9.83 | | High | High | High | High | High | High | High | High | High | High | High | 0.000327 | 0.002512 | 0.87 |
| FALSE | High | CNESLCHSI | [R].CNESLC | 2xCarbami | Q6P5D8 1x | 0.003312 | 0.00016 | 1 | 1 | 1 | Q6P5D8 | Structural | Q6P5D8 [1 | 0 | 1540.649 | 0 | 1.06 | 0.098008 | 46.61 | 64.9 | 135.1 | 24.38 | 30.37 | | High | High | High | High | High | High | High | High | High | High | High | 7.19E-05 | 0.000909 | 3.06 |
| FALSE | High | CNESLCHSI | [R].CNESLC | 2xCarbami | Q6P5D8 1x | 0.007538 | 0.00062 | 1 | 1 | 3 | Q6P5D8 | Structural | Q6P5D8 [1 | 0 | 1769.812 | 0 | 1.56 | 0.00302 | 21.94 | 51.8 | 148.2 | 13.34 | 14.83 | | High | High | High | High | High | High | High | High | High | High | High | 0.000275 | 0.002279 | 2.22 |
| FALSE | High | CNHSSETTS | [K].CNHSSE | 3xCarbami | A2A7F4 1x | 5.58E-07 | 0 | 1 | 4 | 2 | A2A7F4 | MCG12287 | A2A7F4 [1 | 0 | 3076.358 | 0 | -1.02 | 0.000816 | 3.08 | 133.2 | 66.8 | 5.88 | 9.95 | | High | High | High | High | High | High | High | High | High | High | High | 0 | 5.26E-08 | 4.63 |
| FALSE | High | CNNEVSPG | [R].CNNEV | 1xCarbami | E9Q7D5 1x | 0.002647 | 0.000109 | 1 | 2 | 3 | E9Q7D5 | Rho guanir | E9Q7D5 [9 | 0 | 4625.282 | 0 | 3.8 | 0.000408 | 8.41 | 13.5 | 186.5 | 1.79 | 10.13 | | High | High | High | High | High | High | High | High | High | High | High | 0.00014 | 0.001097 | 3.62 |
| FALSE | High | CNSDESNL | [R].CNSDES | 1xCarbami | Q3THY9 1x | 2.4E-05 | 0 | 1 | 2 | 3 | Q3THY9 | Putative ur | Q3THY9 [1 | 0 | 1820.814 | 0 | -0.04 | 0.970156 | 3.69 | 101 | 99 | 2.98 | 8.23 | | High | High | High | High | High | High | High | High | High | High | High | 0 | 2.31E-05 | 3.88 |
| FALSE | High | CLSDPGPH | [R].CLSDPC | 1xCarbami | Q3U3A6 1x | 4.43E-05 | 0 | 1 | 4 | 2 | Q3U3A6 | Sodium/hy | Q3U3A6 [7 | 0 | 2698.299 | 0 | 0.66 | 0.005736 | 17.97 | 74.6 | 125.4 | 8.25 | 9.64 | | High | High | High | High | High | High | High | High | High | High | High | 0 | 7.19E-06 | 3.24 |
| FALSE | High | CNSSRLSSD | [K].CNSSRL | 1xCarbami | B2RUC3 2x | 3.14E-05 | 0 | 1 | 8 | 1 | B2RUC3 | Excision re | B2RUC3 [5 | 1 | 2867.232 | 0 | -0.3 | 0.098629 | 10.7 | 112.7 | 87.3 | 13.5 | 13.39 | | High | High | High | High | High | High | High | High | High | High | High | 0 | 4.89E-06 | 3.73 |
| FALSE | High | CNTVSPVS( | [K].CNTVSI | 1xCarbami | Q8C804 1x | 0.006957 | 0.00062 | 1 | 1 | 4 | Q8C804 | Spindle an | Q8C804 [8 | 0 | 1673.752 | 0 | -0.51 | 0.010337 | 2.31 | 117.5 | 82.5 | 7.59 | 8.06 | | High | High | High | High | High | High | High | High | High | High | High | 0.000327 | 0.002436 | 1.78 |
| FALSE | High | CPEILSDES | [K].CPEILSI | 1xCarbami | Q7TNV0 2 | 0.002252 | 0.000109 | 1 | 4 | 1 | Q7TNV0 | Protein DE | Q7TNV0 [2 | 0 | 2458.002 | 0 | -0.99 | 0.045341 | 10.43 | 134 | 66 | 23.42 | 21.71 | | High | High | High | High | High | High | High | High | High | High | High | 4.94E-05 | 0.000593 | 3.16 |
| FALSE | High | CPFPAGAA | [R].CPFPA( | 4xCarbami | E9QMB7 1 | 1.2E-08 | 0 | 1 | 4 | 3 | E9QMB7 | N-alpha-ac | E9QMB7 [ | 0 | 4534.846 | 0 | -1.16 | 0.000158 | 1.17 | 138.1 | 61.9 | 2.35 | 6.69 | | High | High | High | High | High | High | High | High | High | High | High | 0 | 7.06E-10 | 7.01 |
| FALSE | High | CPFPAGAA | [R].CPFPA( | 4xCarbami | E9QMB7 2 | 1.41E-08 | 0 | 1 | 4 | 2 | E9QMB7 | N-alpha-ac | E9QMB7 [ | 0 | 4614.812 | 0 | 0.13 | 0.829569 | 13.86 | 98.2 | 101.8 | 11.44 | 5.76 | | High | High | High | High | High | High | High | High | High | High | High | 0 | 8.42E-10 | 5.23 |
| FALSE | High | CPGDGDDI | [R].CPGDG | 1xCarbami | Q80TS5 1x | 4.86E-06 | 0 | 1 | 3 | 3 | Q80TS5 | Zinc finger | Q80TS5 [9 | 0 | 2743.232 | 0 | -1.33 | 0.001179 | 1.73 | 145.5 | 54.5 | 12.99 | 12.22 | | High | High | High | High | High | High | High | High | High | High | High | 0 | 5.98E-07 | 3.74 |
| FALSE | High | CPGDGDDI | [R].CPGDG | 1xCarbami | Q80TS5 2x | 0.00967 | 0.000736 | 1 | 3 | 2 | Q80TS5 | Zinc finger | Q80TS5 [9 | 0 | 2823.199 | 0 | -0.86 | 0.004965 | 3 | 129.1 | 70.9 | 12.16 | 11.53 | | High | High | High | High | High | High | High | High | High | High | High | 0.000344 | 0.003024 | 3.1 |
| FALSE | High | CPHGSPQ( | [R].CPHGSI | 1xCarbami | Q542N5 1x | 0.000259 | 0 | 1 | 1 | 1 | Q542N5 | Bcl-2-like p | Q542N5 [5 | 0 | 2508.206 | 0 | 0.31 | 0.166808 | 17.79 | 92.9 | 107.1 | 9.31 | 15.77 | | High | High | High | High | High | High | High | High | High | High | High | 0 | 5.25E-05 | 3.23 |
| FALSE | High | CPHGSPQ( | [R].CPHGSI | 1xCarbami | Q542N5 2x | 2.02E-05 | 0 | 1 | 1 | 1 | Q542N5 | Bcl-2-like p | Q542N5 [5 | 0 | 2588.172 | 0 | -0.5 | 0.00501 | 3.2 | 118.8 | 81.2 | 14.09 | 15.02 | | High | High | High | High | High | High | High | High | High | High | High | 0 | 2.98E-06 | 3.09 |
| FALSE | High | CPLPCIPPL\ | [R].CPLPCII | 2xCarbami | Q9WVM1 | 0.000718 | 0 | 1 | 1 | 1 | Q9WVM1 | Rac GTPase | Q9WVM1 | 0 | 1957.025 | 0 | 0.25 | 0.224162 | 14.9 | 90.6 | 109.4 | 11.45 | 13.38 | | High | High | High | High | High | High | High | High | High | High | High | 0 | 0.000164 | 3.06 |
| FALSE | High | CPLPCIPPL\ | [R].CPLPCII | 2xCarbami | Q9WVM1 | 1.28E-05 | 0 | 1 | 1 | 1 | Q9WVM1 | Rac GTPase | Q9WVM1 | 0 | 2186.188 | 0 | 0.04 | 0.560516 | 22.19 | 99 | 101 | 14.91 | 0.59 | | High | High | High | High | High | High | High | High | High | High | High | 0 | 1.77E-06 | 3.21 |
| FALSE | High | CPLPPMAP | [K].CPLPPN | 1xCarbami | A0A0R4J0I | 0.00103 | 0 | 1 | 2 | 4 | A0A0R4J0I | ETS domai | A0A0R4J0I | 0 | 3457.631 | 0 | -0.63 | 0.049844 | 5.79 | 119 | 81 | 9.94 | 21.4 | | High | High | High | High | High | High | High | High | High | High | High | 0 | 0.000245 | 3.31 |
| FALSE | High | CPLSGACYS | [K].CPLSGA | 2xCarbami | F8WHL2 1 | 0.009416 | 0.000736 | 1 | 7 | 1 | F8WHL2 | Coatomer | F8WHL2 [1 | 0 | 2053.953 | 0 | -1.01 | 0.000254 | 2.12 | 134.8 | 65.2 | 2.04 | 5.91 | | High | High | High | High | High | High | High | High | High | High | High | 0.000327 | 0.002934 | 2.82 |
| FALSE | High | CPPEEPASA | [K].CPPEEP | 1xCarbami | E9QAU4 1 | 3.07E-06 | 0 | 1 | 11 | 2 | E9QAU4 | Enhancer t | E9QAU4 [1 | 0 | 2894.42 | 0 | 0.33 | 0.209316 | 29.1 | 87.8 | 112.2 | 12.62 | 18.55 | | High | High | High | High | High | High | High | High | High | High | High | 0 | 1.91E-06 | 3.15 |
| FALSE | High | CNTGSEGP | [K].CNTGSI | 1xCarbami | O35954 1x | 0.0003 | 0 | 1 | 1 | 1 | O35954 | Membrane | O35954 [2 | 0 | 2070.957 | 0 | 1.32 | 0.002055 | 14.32 | 54 | 146 | 4.97 | 13.56 | | High | High | High | High | High | High | High | High | High | High | High | 0 | 6.15E-05 | 2.32 |
| FALSE | High | CPPSPAPLS | [R].CPPSPA | 1xCarbami | A2RSJ4 1xF | 0.000605 | 0 | 1 | 2 | 1 | A2RSJ4 | UHRF1-bin | A2RSJ4 [1C | 0 | 1940.996 | 0 | 1.18 | 0.000618 | 4.98 | 64.1 | 135.9 | 9.28 | 5.42 | | High | High | High | High | High | High | High | High | High | High | High | 0 | 0.000136 | 2.16 |
| FALSE | High | CLSAPDLTI | [R].CLSAPD | 1xCarbami | E9Q7F2 1x | 5.16E-05 | 0 | 1 | 1 | 1 | E9Q7F2 | E3 ubiquiti | E9Q7F2 [3 | 0 | 1784.927 | 0 | -0.59 | 0.001923 | 3.82 | 122.3 | 77.7 | 7.75 | 7.44 | | High | High | High | High | High | High | High | High | High | High | High | 0 | 8.56E-06 | 2.97 |
| FALSE | High | CLPNDPEEI | [R].CLPNDI | 1xCarbami | P97929 1xl | 0.0003 | 0 | 1 | 3 | 2 | P97929 | Breast canc | P97929 [6 | 0 | 2803.326 | 0 | -0.94 | 0.003243 | 3.2 | 126.8 | 73.2 | 9.92 | 10.55 | | High | High | High | High | High | High | High | High | High | High | High | 0 | 6.14E-05 | 3.75 |
| FALSE | High | CESPLSK | [K].CESPLS | 1xCarbami | P27641 1xl | 0.115566 | 0.009504 | 1 | 1 | 2 | P27641 | X-ray repai | P27641 [5 | 0 | 1358.679 | 0 | -0.61 | 0.003674 | 2.67 | 119.3 | 80.7 | 5.5 | 6.31 | | High | High | High | High | High | High | High | High | High | High | High | 0.004733 | 0.05113 | 1.83 |
| FALSE | High | CGESSSLVI | [R].CGESSS | 1xCarbami | G3UYN0 1 | 0.003312 | 0.00016 | 1 | 2 | 1 | G3UYN0 | L1TD1 OS= | G3UYN0 [1 | 0 | 1805.855 | 0 | -0.43 | 0.014825 | 8.48 | 115.3 | 84.7 | 10.97 | 8.83 | | High | High | High | High | High | High | High | High | High | High | High | 7.19E-05 | 0.000916 | 2.3 |
| FALSE | High | CGETGHVA | [R].CGETG | 2xCarbami | P53996 1xl | 0.034878 | 0.002269 | 1 | 5 | 1 | P53996 | Cellular nu | P53996 [1 | 0 | 1970.923 | 0 | 0.05 | 0.616311 | 37.64 | 98.2 | 101.8 | 20.74 | 48.16 | | High | High | High | High | High | High | High | High | High | High | High | 0.001183 | 0.01275 | 2.33 |
| FALSE | High | CGETVESG | [K].CGETVI | 1xCarbami | Q923B1 1x | 1.91E-05 | 0 | 1 | 2 | 2 | Q923B1 | Lariat debr | Q923B1 [4 | 0 | 1748.781 | 0 | -0.37 | 0.005547 | 2.11 | 113.1 | 86.9 | 4.07 | 6.21 | | High | High | High | High | High | High | High | High | High | High | High | 0 | 2.79E-06 | 1.76 |
| FALSE | High | CGGGELAS | [K].CGGGE | 3xCarbami | A2A791 1x | 0.000171 | 0 | 1 | 3 | 3 | A2A791 | Zinc finger | A2A791 [1 | 0 | 4005.73 | 0 | -0.59 | 0.008089 | 6.45 | 116.3 | 83.7 | 11.47 | 9.96 | | High | High | High | High | High | High | High | High | High | High | High | 0 | 0.000119 | 3.01 |
| FALSE | High | CGGGELAS | [K].CGGGE | 3xCarbami | A2A791 2x | 1.15E-05 | 0 | 1 | 3 | 1 | A2A791 | Zinc finger | A2A791 [1 | 0 | 4085.697 | 0 | -0.6 | 0.006456 | 8.23 | 120.6 | 79.4 | 7.49 | 9.47 | | High | High | High | High | High | High | High | High | High | High | High | 0 | 1.58E-06 | 4.11 |
| FALSE | High | CGGHSGSF | [R].CGGHS | 1xCarbami | H7BWZ9 1 | 0.000422 | 0 | 1 | 6 | 1 | H7BWZ9 | Protein do | H7BWZ9 [2 | 0 | 2456.183 | 0 | 0.74 | 0.009034 | 9.65 | 75.8 | 124.2 | 12.44 | 10.99 | | High | High | High | High | High | High | High | High | High | High | High | 0 | 9.09E-05 | 2.49 |
| FALSE | High | CGGPGDAS | [R].CGGPG | 2xCarbami | P47810 1xl | 2.14E-09 | 0 | 1 | 1 | 2 | P47810 | Wee1-like | P47810 [1 | 0 | 1852.767 | 0 | -1.16 | 0.003611 | 2.41 | 137.5 | 62.5 | 10.21 | 9.99 | | High | High | High | High | High | High | High | High | High | High | High | 0 | 1.01E-10 | 3.04 |
| FALSE | High | CGMVDDQ | [K].CGMVI | 1xCarbami | E9QAU4 1 | 6.53E-05 | 0 | 1 | 3 | 1 | E9QAU4 | Enhancer t | E9QAU4 [1 | 0 | 1950.874 | 0 | -1.59 | 0.001213 | 1.09 | 150 | 50 | 9.83 | 15.72 | | High | High | High | High | High | High | High | High | High | High | High | 0 | 1.12E-05 | 3.88 |
| FALSE | High | CGMVDDQ | [K].CGMVI | 1xCarbami | E9QAU4 1 | 0.000111 | 0 | 1 | 3 | 1 | E9QAU4 | Enhancer t | E9QAU4 [1 | 0 | 1966.869 | 0 | -1.11 | 0.00212 | 0.59 | 135.8 | 64.2 | 8.46 | 9.51 | | High | High | High | High | High | High | High | High | High | High | High | 0 | 2.02E-05 | 3.71 |
| FALSE | High | CGRESDGG | [R].CGRESI | 1xCarbami | Q8BFX3 1x | 0.00509 | 0.000551 | 1 | 2 | 1 | Q8BFX3 | BTB/POZ d | Q8BFX3 [7 | 1 | 1780.8 | 0 | 0.07 | 0.886053 | 31.6 | 103.8 | 96.2 | 15.06 | 11.44 | | High | High | High | High | High | High | High | High | High | High | High | 0.000246 | 0.001473 | 2.36 |
| FALSE | High | CGSGPVHIS | [K].CGSGP\ | 1xCarbami | Q5SQB7 1 | 2.39E-07 | 0 | 2 | 9 | 18 | Q5SQB7; C | MCG68069 | Q5SQB7 [1 | 0 | 3675.584 | 0 | -0.17 | 0.715043 | 23.88 | 110.3 | 89.7 | 65.13 | 56.6 | Shared | High | High | High | High | High | High | High | High | High | High | High | 0 | 8.6E-06 | 4.73 |
| FALSE | High | CGSGPVHIS | [K].CGSGP\ | 1xCarbami | Q5SQB7 1x | 1.85E-05 | 0 | 2 | 9 | 7 | Q5SQB7; C | MCG68069 | Q5SQB7 [1 | 0 | 3676.568 | 0 | -0.45 | 0.413218 | 25.37 | 108.3 | 91.7 | 24.62 | 34.66 | Shared | High | High | High | High | High | High | High | High | High | High | High | 2.58E-05 | 0.000389 | 4.74 |

| | | | | | | | | | | | | | | | | | | | | | | | | | | | | | | | | | | | | | | |
|---|---|---|---|---|---|---|---|---|---|---|---|---|---|---|---|---|---|---|---|---|---|---|---|---|---|---|---|---|---|---|---|---|---|---|---|---|---|---|
| FALSE | High | CLQVLFPPA | [R].CLQVLF | 1xCarbami | Q3U5F9 1x | 0.099328 | 0.008003 | 1 | 3 | 1 | Q3U5F9 | Putative ur | Q3U5F9 [5 | 0 | 3148.725 | 0 | 0.15 | 0.434294 | 10.22 | 94.9 | 105.1 | 15.61 | 15.17 | | High | High | High | High | High | High | High | High | High | High | High | 0.004006 | 0.04263 | 2.99 |
| FALSE | High | CGSGPVHIS | [K].CGSGPV | 1xCarbami | Q5SQB7 1x | 1.4E-09 | 0 | 2 | 9 | 148 | Q5SQB7; C | MCG68069 | Q5SQB7 [1 | 0 | 3904.747 | 0 | -0.17 | 0.104397 | 10.91 | 104.1 | 95.9 | 3.24 | 11.43 | Shared | High | High | High | High | High | High | High | High | High | High | High | 0 | 5.62E-06 | 5.29 |
| FALSE | High | CHGSDHGF | [R].CHGSD | 1xCarbami | Q921C3 1x | 0.002553 | 0.000109 | 1 | 5 | 5 | Q921C3 | Bromodom | Q921C3 [1 | 0 | 2537.113 | 0 | -1 | 0.001542 | 2.73 | 133.9 | 66.1 | 6.11 | 12.08 | | High | High | High | High | High | High | High | High | High | High | High | 4.94E-05 | 0.000684 | 2.14 |
| FALSE | High | CHGSDHGF | [R].CHGSD | 1xCarbami | Q921C3 2x | 0.000434 | 0 | 1 | 5 | 1 | Q921C3 | Bromodom | Q921C3 [1 | 0 | 2617.079 | 0 | -0.92 | 0.001636 | 3.57 | 130.7 | 69.3 | 3.46 | 12.82 | | High | High | High | High | High | High | High | High | High | High | High | 0 | 9.38E-05 | 3.07 |
| FALSE | High | CHNTPPKP | [K].CHNTP | 1xCarbami | Q9JJT9 1xP | 0.121159 | 0.009984 | 1 | 4 | 1 | Q9JJT9 | Phosphory | Q9JJT9 [77 | 0 | 2661.294 | 0 | -0.29 | 0.15751 | 34.95 | 115 | 85 | 25.96 | 20.68 | | High | High | High | High | High | High | High | High | High | High | High | 0.004961 | 0.05395 | 3.17 |
| FALSE | High | CHNTPPKP | [K].CHNTP | 1xCarbami | Q9JJT9 1xP | 6.65E-05 | 0 | 1 | 4 | 1 | Q9JJT9 | Phosphory | Q9JJT9 [77 | 0 | 2889.473 | 0 | -0.91 | 0.007278 | 3.21 | 130.5 | 69.5 | 7.73 | 14.67 | | High | High | High | High | High | High | High | High | High | High | High | 0 | 1.14E-05 | 2.37 |
| FALSE | High | CHSLGYNF | [K].CHSLGY | 1xCarbami | Q9ERD6 1x | 4.31E-05 | 0 | 1 | 7 | 1 | Q9ERD6 | Ras-specifi | Q9ERD6 [3 | 0 | 1913.95 | 0 | -0.17 | 0.556704 | 16.66 | 101.1 | 98.9 | 12.5 | 8.98 | | High | High | High | High | High | High | High | High | High | High | High | 0 | 7E-06 | 2.24 |
| FALSE | High | CHSPPATTA | [R].CHSPPA | 1xCarbami | Q2KHI9-2 1 | 2.8E-06 | 0 | 1 | 3 | 2 | Q2KHI9-2 | Isoform 2 c | Q2KHI9-2 [ | 0 | 2028.989 | 0 | -1.06 | 0.002988 | 3.89 | 135.3 | 64.7 | 7.85 | 10.89 | | High | High | High | High | High | High | High | High | High | High | High | 0 | 1.12E-05 | 3.58 |
| FALSE | High | CHVEDTPC | [R].CHVEDT | 2xCarbami | A0A0R4J0C | 1.41E-05 | 0 | 1 | 6 | 3 | A0A0R4J0C | Disks large | A0A0R4J0C | 0 | 3206.381 | 0 | -0.79 | 0.003306 | 2.59 | 127 | 73 | 6.91 | 11.48 | | High | High | High | High | High | High | High | High | High | High | High | 0 | 0.000104 | 3.86 |
| FALSE | High | CICPSLSYSF | [R].CICPSLS | 2xCarbami | Q8C078-5 1 | 0.077825 | 0.005963 | 1 | 7 | 1 | Q8C078-5 | Isoform 5 c | Q8C078-5 | 0 | 2470.038 | 0 | -0.06 | 0.689786 | 20.75 | 100.2 | 99.8 | 18.57 | 12.35 | | High | High | High | High | High | High | High | High | High | High | High | 0.003064 | 0.03201 | 2.14 |
| FALSE | High | CLESPVGK | [K].CLESPV | 1xCarbami | E9Q3W1 1: | 0.008463 | 0.000736 | 2 | 2 | 2 | E9Q3W1; E | Casein kina | E9Q3W1 [1 | 0 | 1427.737 | 0 | -0.37 | 0.000406 | 2.27 | 113.3 | 86.7 | 1.54 | 2.87 | Shared | High | High | High | High | High | High | High | High | High | High | High | 0.000327 | 0.002614 | 2.02 |
| FALSE | High | CIFMSETQ | [K].CIFMSE | 1xCarbami | Q8C5D8 1x | 0.000442 | 0 | 1 | 8 | 1 | Q8C5D8 | E3 SUMO-p | Q8C5D8 [4 | 0 | 2053.974 | 0 | -0.8 | 0.000707 | 2.66 | 128.1 | 71.9 | 3.5 | 6.69 | | High | High | High | High | High | High | High | High | High | High | High | 0 | 9.54E-05 | 2.68 |
| FALSE | High | CIFMSETQ | [K].CIFMSE | 1xCarbami | Q8C5D8 1x | 2.62E-08 | 0 | 1 | 8 | 7 | Q8C5D8 | E3 SUMO-p | Q8C5D8 [4 | 0 | 2069.969 | 0 | -1.26 | 0.000217 | 1.92 | 141.9 | 58.1 | 1.8 | 7.53 | | High | High | High | High | High | High | High | High | High | High | High | 0 | 1.69E-09 | 4.01 |
| FALSE | High | CINDPNAM | [K].CINDPN | 1xCarbami | Q8BV79 1x | 0.099328 | 0.008003 | 1 | 1 | 1 | Q8BV79 | TPR and an | Q8BV79 [1 | 0 | 2538.116 | 0 | -0.22 | 0.033459 | 2.34 | 109.5 | 90.5 | 3.86 | 17.68 | | High | High | High | High | High | High | High | High | High | High | High | 0.004032 | 0.043 | 3.11 |
| FALSE | High | CLPLSVDLL | [K].CLPLSV | 1xCarbami | Q8BGS1 1x | 4.69E-06 | 0 | 1 | 2 | 1 | Q8BGS1 | Band 4.1-li | Q8BGS1 [4 | 0 | 2950.505 | 0 | -1.37 | 0.015494 | 9.63 | 152.3 | 47.7 | 23.05 | 36.59 | | High | High | High | High | High | High | High | High | High | High | High | 0 | 5.75E-07 | 4.21 |
| FALSE | High | CGSGPVHIS | [K].CGSGPV | 1xCarbami | Q5SQB7 1x | 4.19E-08 | 0 | 2 | 9 | 81 | Q5SQB7; C | MCG68069 | Q5SQB7 [1 | 0 | 3905.731 | 0 | 0.02 | 0.390184 | 4.34 | 103.9 | 96.1 | 12.2 | 12.32 | Shared | High | High | High | High | High | High | High | High | High | High | High | 0 | 2.88E-09 | 5.87 |
| FALSE | High | CPPSTPSPF | [R].CPPSTP | 1xCarbami | Q8BQ33 1x | 0.000375 | 0 | 1 | 1 | 1 | Q8BQ33 | Treslin OS= | Q8BQ33 [1 | 0 | 2852.373 | 0 | -0.33 | 0.004876 | 4.95 | 111.4 | 88.6 | 11.01 | 18.18 | | High | High | High | High | High | High | High | High | High | High | High | 0 | 7.94E-05 | 2.76 |
| FALSE | High | CPSDTEGLY | [K].CPSDTE | 1xCarbami | Q3TEW6 1: | 0.111965 | 0.009053 | 1 | 2 | 1 | Q3TEW6 | Myelin pro | Q3TEW6 [2 | 0 | 1643.812 | 0 | 0.19 | 0.239516 | 15.13 | 93.3 | 106.7 | 10.17 | 12.6 | | High | High | High | High | High | High | High | High | High | High | High | 0.004543 | 0.04943 | 2.69 |
| FALSE | High | CPSLDNLAY | [R].CPSLDN | 1xCarbami | A0A0A0MC | 8.28E-06 | 0 | 1 | 2 | 2 | A0A0A0MC | Histone-lys | A0A0A0MC | 0 | 2406.214 | 0 | 0.73 | 0.241607 | 8.1 | 73.9 | 126.1 | 14.71 | 44.65 | | High | High | High | High | High | High | High | High | High | High | High | 4.94E-05 | 0.000717 | 3.03 |
| FALSE | High | CSPFQSAR | [R].CSPFQS | 1xCarbami | Q8BPY9 1x | 0.045508 | 0.003373 | 1 | 1 | 2 | Q8BPY9 | Fidgetin-lik | Q8BPY9 [1 | 0 | 1261.56 | 0 | -0.23 | 0.061155 | 5.2 | 107 | 93 | 6.81 | 4.04 | | High | High | High | High | High | High | High | High | High | High | High | 0.002159 | 0.02238 | 2.2 |
| FALSE | High | CSPGSPTDI | [K].CSPGSF | 1xCarbami | Q9ES18 1x | 0.073462 | 0.005631 | 1 | 3 | 2 | Q9ES18 | Forkhead b | Q9ES18 [4 | 0 | 2069.998 | 0 | 0.97 | 0.011021 | 4.04 | 68.7 | 131.3 | 9.52 | 12.52 | | High | High | High | High | High | High | High | High | High | High | High | 0.002911 | 0.0302 | 2.39 |
| FALSE | High | CSPGSSSK | [R].CSPGSS | 1xCarbami | Q6P3Y5 1x | 0.025514 | 0.001645 | 1 | 3 | 2 | Q6P3Y5 | Zinc finger | Q6P3Y5 [7 | 0 | 1347.638 | 0 | 0.56 | 0.006782 | 6.49 | 80.7 | 119.3 | 5.98 | 10.8 | | High | High | High | High | High | High | High | High | High | High | High | 0.000872 | 0.008936 | 1.07 |
| FALSE | High | CSPPETLAS | [R].CSPPET | 1xCarbami | F8VQ05 1x | 6.42E-09 | 0 | 1 | 1 | 4 | F8VQ05 | FRY-like tra | F8VQ05 [8 | 0 | 2650.236 | 0 | 0.39 | 0.025522 | 9.45 | 84.4 | 115.6 | 11.17 | 10.41 | | High | High | High | High | High | High | High | High | High | High | High | 0 | 3.5E-10 | 4.79 |
| FALSE | High | CSPPVPSPL | [K].CSPPVF | 1xCarbami | P52479-2 1 | 1.91E-07 | 0 | 1 | 2 | 3 | P52479-2 | Isoform 2 c | P52479-2 [ | 0 | 1906.975 | 0 | -0.84 | 0.014748 | 5.11 | 126.3 | 73.7 | 10.43 | 12.79 | | High | High | High | High | High | High | High | High | High | High | High | 0 | 4.71E-05 | 3.03 |
| FALSE | High | CSPVPGLSS | [R].CSPVPG | 1xCarbami | Q8CCN5 1x | 2.19E-08 | 0 | 1 | 8 | 4 | Q8CCN5 | Breast carc | Q8CCN5 [4 | 0 | 2389.199 | 0 | -0.54 | 0.016903 | 2.21 | 119.3 | 80.7 | 7 | 10.34 | | High | High | High | High | High | High | High | High | High | High | High | 0 | 1.39E-09 | 4.75 |
| FALSE | High | CSPVPGLSS | [R].CSPVPG | 1xCarbami | Q8CCN5 2x | 0.1059 | 0.008582 | 1 | 8 | 1 | Q8CCN5 | Breast carc | Q8CCN5 [4 | 0 | 2469.165 | 0 | -0.03 | 0.280682 | 2.94 | 105.4 | 94.6 | 9.07 | 7.14 | | High | High | High | High | High | High | High | High | High | High | High | 0.004305 | 0.04602 | 1.99 |
| FALSE | High | CSQPSPAEI | [R].CSQPSF | 1xCarbami | Q8BJL0 1xP | 0.000772 | 0 | 1 | 4 | 1 | Q8BJL0 | SWI/SNF-re | Q8BJL0 [89 | 0 | 2229.066 | 0 | -0.33 | 0.731201 | 9.29 | 107.3 | 92.7 | 17.15 | 28.71 | | High | High | High | High | High | High | High | High | High | High | High | 0 | 0.000178 | 2.5 |
| FALSE | High | CSQQSPDS | [K].CSQQS | 1xCarbami | Q0VGR5 1x | 0.034281 | 0.002195 | 1 | 2 | 1 | Q0VGR5 | CDK5 regul | Q0VGR5 [5 | 0 | 1574.729 | 0 | 0.02 | 0.510502 | 14.52 | 98 | 102 | 10.33 | 10.14 | | High | High | High | High | High | High | High | High | High | High | High | 0.001151 | 0.01251 | 1.89 |
| FALSE | High | CSSAFFEEE | [R].CSSAFF | 1xCarbami | A0A0A6YV | 0.045898 | 0.003408 | 1 | 5 | 2 | A0A0A6YV | Protein lin- | A0A0A6YV | 0 | 1570.645 | 0 | -0.65 | 0.020531 | 6.7 | 120.6 | 79.4 | 10.25 | 10.24 | | High | High | High | High | High | High | High | High | High | High | High | 0.001741 | 0.01741 | 2.59 |
| FALSE | High | CSSCEPFAA | [K].CSSCEP | 2xCarbami | Q3THY9 1x | 1.16E-05 | 0 | 1 | 2 | 1 | Q3THY9 | Putative ur | Q3THY9 [1 | 0 | 2150.92 | 0 | 0.11 | 0.671652 | 11.33 | 96.2 | 103.8 | 8.36 | 11.67 | | High | High | High | High | High | High | High | High | High | High | High | 0 | 1.59E-06 | 2.58 |
| FALSE | High | CSSGLFCSE | [R].CSSGLF | 2xCarbami | Q0VB14 1x | 0.000365 | 0 | 1 | 2 | 2 | Q0VB14 | Interleukin | Q0VB14 [5 | 0 | 2193.971 | 0 | 2.38 | 0.001426 | 27.03 | 29.4 | 170.6 | 20.65 | 13.66 | | High | High | High | High | High | High | High | High | High | High | High | 0 | 7.73E-05 | 2.84 |
| FALSE | High | CSSIADSGI | [R].CSSIAD | 3xCarbami | Q8BMK0 1 | 0.000151 | 0 | 1 | 2 | 2 | Q8BMK0 | Centrosom | Q8BMK0 [4 | 0 | 4279.876 | 0 | -0.57 | 0.157676 | 11.02 | 112.4 | 87.6 | 9.38 | 26.84 | | High | High | High | High | High | High | High | High | High | High | High | 0 | 3.18E-05 | 4.53 |
| FALSE | High | CSMSSSAD | [R].CSMSS | 1xCarbami | S4R1P5 1xl | 1.08E-07 | 0 | 1 | 5 | 2 | S4R1P5 | Dystonin O | S4R1P5 [14 | 0 | 2597.046 | 0 | 0.53 | 0.025199 | 16.11 | 80 | 120 | 3.36 | 17.98 | | High | High | High | High | High | High | High | High | High | High | High | 0 | 8.34E-09 | 3.31 |
| FALSE | High | CSSLEGIVD | [R].CSSLEG | 1xCarbami | D3Z6J1 1xP | 0.023584 | 0.001578 | 1 | 2 | 2 | D3Z6J1 | DEP domai | D3Z6J1 [42 | 0 | 2415.199 | 0 | -0.8 | 0.059306 | 13.16 | 121.6 | 78.4 | 13.3 | 35.18 | | High | High | High | High | High | High | High | High | High | High | High | 0.000859 | 0.008195 | 1.75 |
| FALSE | High | CSSLSGAPC | [R].CSSLSG | 1xCarbami | E9PWU7 1: | 0.064323 | 0.004664 | 1 | 2 | 2 | E9PWU7 | Kinesin-like | E9PWU7 [1 | 0 | 1944.92 | 0 | 2.38 | 0.000262 | 7.79 | 31.7 | 168.3 | 5.94 | 6.84 | | High | High | High | High | High | High | High | High | High | High | High | 0.002403 | 0.02571 | 2.52 |
| FALSE | High | CSSLSSLSQ | [R].CSSLSS | 1xCarbami | A2AIX1 1xP | 0.000426 | 0 | 1 | 3 | 3 | A2AIX1 | SEC16 hom | A2AIX1 [23 | 0 | 1748.809 | 0 | 0.67 | 0.004567 | 18.88 | 79.8 | 120.2 | 5.52 | 10.04 | | High | High | High | High | High | High | High | High | High | High | High | 9.42E-05 | 0.00096 | 2.06 |
| FALSE | High | CSSPESAEF | [R].CSSPES | 2xCarbami | A0A140T8I | 6.62E-07 | 0 | 1 | 2 | 4 | A0A140T8I | Protein Jad | A0A140T8I | 0 | 2837.232 | 0 | -0.96 | 0.002254 | 2.51 | 132.5 | 67.5 | 7.12 | 7.45 | | High | High | High | High | High | High | High | High | High | High | High | 0 | 1.52E-07 | 4.9 |
| FALSE | High | CSSPLQLNI | [R].CSSPLQ | 1xCarbami | Q6A060 1x | 3.34E-07 | 0 | 1 | 5 | 2 | Q6A060 | MKIAA048 | Q6A060 [1 | 0 | 2293.146 | 0 | -0.41 | 0.001684 | 3.58 | 116.1 | 83.9 | 5.76 | 5.5 | | High | High | High | High | High | High | High | High | High | High | High | 0 | 2.97E-08 | 4.58 |
| FALSE | High | CSSPTPGPI | [R].CSSPTP | 1xCarbami | O35711-4 1 | 0.000483 | 0 | 1 | 4 | 2 | O35711-4 | Isoform 4 c | O35711-4 | 0 | 2001.028 | 0 | -0.64 | 0.057277 | 6.46 | 124.9 | 75.1 | 16.24 | 10.58 | | High | High | High | High | High | High | High | High | High | High | High | 0 | 0.000209 | 2.38 |
| FALSE | High | CSSPVDMD | [R].CSSPVD | 1xCarbami | G3X9K4 1x | 3.3E-06 | 0 | 1 | 5 | 2 | G3X9K4 | SAPS doma | G3X9K4 [7 | 0 | 2840.227 | 0 | 0.91 | 0.023158 | 31.37 | 75.2 | 124.8 | 20.77 | 18.87 | | High | High | High | High | High | High | High | High | High | High | High | 0 | 3.9E-07 | 4.21 |
| FALSE | High | CSSSSGGG | [R].CSSSSG | 1xCarbami | Q8BK67 1x | 0.026191 | 0.001676 | 1 | 2 | 2 | Q8BK67 | Protein RC | Q8BK67 [4 | 0 | 2862.214 | 0 | -0.06 | 0.559433 | 15.43 | 102.1 | 97.9 | 7.33 | 14.15 | | High | High | High | High | High | High | High | High | High | High | High | 0.000903 | 0.009243 | 2.3 |
| FALSE | High | CSSSSGGG | [R].CSSSSG | 1xCarbami | Q8BK67 2x | 1.65E-06 | 0 | 1 | 2 | 4 | Q8BK67 | Protein RC | Q8BK67 [4 | 0 | 2942.18 | 0 | -0.57 | 0.00145 | 2.13 | 119.5 | 80.5 | 1.8 | 7.94 | | High | High | High | High | High | High | High | High | High | High | High | 0 | 1.77E-07 | 3.97 |
| FALSE | High | CSSVYLLSP | [R].CSSVYL | 1xCarbami | A0A0R4J0F | 0.001301 | 0 | 1 | 3 | 1 | A0A0R4J0F | Germ cell-s | A0A0R4J0F | 0 | 1804.968 | 0 | 0.2 | 0.079225 | 5.26 | 91 | 109 | 7.95 | 11.35 | | High | High | High | High | High | High | High | High | High | High | High | 0 | 0.00032 | 3.26 |
| FALSE | High | CSTLDLEIQ | [R].CSTLDL | 1xCarbami | A0A1D5RN | 0.000584 | 0 | 1 | 3 | 2 | A0A1D5RN | Kinesin-like | A0A1D5RN | 0 | 1800.84 | 0 | 1.28 | 0.002161 | 11.62 | 61.7 | 138.3 | 19.42 | 7.17 | | High | High | High | High | High | High | High | High | High | High | High | 0 | 0.000329 | 3.56 |
| FALSE | High | CSTPELGLD | [R].CSTPEL | 1xCarbami | A2A5Y4 1x | 0.00025 | 0 | 1 | 3 | 3 | A2A5Y4 | KAT8 regul | A2A5Y4 [1 | 0 | 2440.105 | 0 | -0.16 | 0.017942 | 3.89 | 105.4 | 94.6 | 2.23 | 8.51 | | High | High | High | High | High | High | High | High | High | High | High | 0 | 5.04E-05 | 4.03 |
| FALSE | High | CSTPLLHQ( | [R].CSTPLL | 2xCarbami | Q09XV5 1x | 0.000328 | 0 | 1 | 2 | 2 | Q09XV5 | Chromodo | Q09XV5 [1 | 0 | 1896.866 | 0 | -0.6 | 0.003924 | 4.46 | 120.6 | 79.4 | 6.58 | 5.54 | | High | High | High | High | High | High | High | High | High | High | High | 0 | 0.00014 | 1.97 |
| FALSE | High | CSTQTSSP( | [K].CSTQTS | 1xCarbami | A0A1L1SSL | 5.76E-09 | 0 | 1 | 4 | 3 | A0A1L1SSL | Zinc finger | A0A1L1SSL | 0 | 2596.155 | 0 | -0.47 | 0.001128 | 3.99 | 114.7 | 85.3 | 6.4 | 6.44 | | High | High | High | High | High | High | High | High | High | High | High | 0 | 1.43E-09 | 4.44 |
| FALSE | High | CSSLGNAA | [K].CSSLGN | 1xCarbami | Q9CZ09 1x | 3.36E-10 | 0 | 1 | 1 | 4 | Q9CZ09 | Histidine p | Q9CZ09 [5 | 0 | 2498.107 | 0 | 0.23 | 0.22652 | 12.49 | 94.6 | 105.4 | 11.67 | 8.15 | | High | High | High | High | High | High | High | High | High | High | High | 0 | 1.28E-11 | 4.98 |
| FALSE | High | CSMSSSAD | [R].CSMSS | 1xCarbami | S4R1P5 1xl | 4.93E-08 | 0 | 1 | 5 | 2 | S4R1P5 | Dystonin O | S4R1P5 [14 | 0 | 2581.051 | 0 | 0.36 | 0.041974 | 13.09 | 86.9 | 113.1 | 8.34 | 9.95 | | High | High | High | High | High | High | High | High | High | High | High | 0 | 3.44E-09 | 3.83 |
| FALSE | High | CSLLSASPA | [R].CSLLSA | 1xCarbami | A2A7S8-2 2 | 0.00047 | 0 | 1 | 4 | 1 | A2A7S8-2 | Isoform 2 c | A2A7S8-2 [ | 0 | 1636.737 | 0 | -0.71 | 0.014402 | 12.19 | 131 | 69 | 14.23 | 13.69 | | High | High | High | High | High | High | High | High | High | High | High | 0 | 0.000102 | 2.34 |
| FALSE | High | CSLLSASPA | [R].CSLLSA | 1xCarbami | A2A7S8-2 1 | 0.000225 | 0 | 1 | 4 | 3 | A2A7S8-2 | Isoform 2 c | A2A7S8-2 [ | 0 | 1556.771 | 0 | -0.53 | 0.012693 | 4.82 | 121.7 | 78.3 | 11.46 | 14.27 | | High | High | High | High | High | High | High | High | High | High | High | 0.000327 | 0.002623 | 2.05 |
| FALSE | High | CPSLDNLAY | [R].CPSLDN | 1xCarbami | A0A0A0MC | 0.001077 | 0 | 1 | 2 | 1 | A0A0A0MC | Histone-lys | A0A0A0MC | 0 | 2486.18 | 0 | 1.11 | 0.158064 | 37.7 | 62.2 | 137.8 | 16.06 | 48.87 | | High | High | High | High | High | High | High | High | High | High | High | 0 | 0.000261 | 3.24 |
| FALSE | High | CPSPLFLDN | [R].CPSPLF | 1xCarbami | Q0VB14 1x | 0.012288 | 0.000889 | 1 | 2 | 3 | Q0VB14 | Interleukin | Q0VB14 [4 | 0 | 3796.829 | 0 | 1.5 | 0.000399 | 15.92 | 54.4 | 145.6 | 14.96 | 5.36 | | High | High | High | High | High | High | High | High | High | High | High | 0.000449 | 0.003958 | 4.07 |
| FALSE | High | CPSPLFLDN | [R].CPSPLF | 1xCarbami | Q0VB14 1x | 0.000852 | 0 | 1 | 2 | 6 | Q0VB14 | Interleukin | Q0VB14 [4 | 0 | 3797.813 | 0 | 1.4 | 0.000741 | 13.22 | 55 | 145 | 6.8 | 9.19 | | High | High | High | High | High | High | High | High | High | High | High | 0 | 0.000326 | 4.02 |
| FALSE | High | CPSPISEHN | [K].CPSPISI | 1xCarbami | Q8BX90 1x | 0.000103 | 0 | 1 | 3 | 2 | Q8BX90 | Fibronectir | Q8BX90 [2 | 0 | 1990.023 | 0 | 0.74 | 0.013883 | 12.34 | 75 | 125 | 9.24 | 9.63 | | High | High | High | High | High | High | High | High | High | High | High | 0 | 1.86E-05 | 2.53 |
| FALSE | High | CPSPISEHN | [K].CPSPISI | 1xCarbami | Q8BX90 1x | 0.013308 | 0.000889 | 1 | 3 | 3 | Q8BX90 | Fibronectir | Q8BX90 [2 | 0 | 1991.007 | 0 | 0.57 | 0.002034 | 12.04 | 78.3 | 121.7 | 7.9 | 6.92 | | High | High | High | High | High | High | High | High | High | High | High | 0.000467 | 0.004303 | 2.73 |
| FALSE | High | CPVSINLSK | [R].CPVSIN | 1xCarbami | Q9R0L6-2 1 | 0.018447 | 0.001324 | 1 | 3 | 1 | Q9R0L6-2 | Isoform 2 c | Q9R0L6-2 | 0 | 1555.832 | 0 | 0.16 | 0.214866 | 15.51 | 91.4 | 108.6 | 9.75 | 14.35 | | High | High | High | High | High | High | High | High | High | High | High | 0.000706 | 0.006221 | 1.86 |
| FALSE | High | CQPGGGPI | [R].CQPGG | 1xCarbami | Q64151 1x | 0.075304 | 0.005727 | 1 | 2 | 1 | Q64151 | Semaphori | Q64151 [7 | 0 | 2657.311 | 0 | -0.3 | 0.078245 | 4.66 | 107 | 93 | 6.35 | 10.06 | | High | High | High | High | High | High | High | High | High | High | High | 0.002976 | 0.03108 | 1.81 |
| FALSE | High | CQPGGGPI | [R].CQPGG | 1xCarbami | Q64151 2x | 5.33E-07 | 0 | 1 | 2 | 2 | Q64151 | Semaphori | Q64151 [7 | 0 | 2737.277 | 0 | 0.11 | 0.538337 | 12.43 | 93.4 | 106.6 | 14.61 | 19.36 | | High | High | High | High | High | High | High | High | High | High | High | 0 | 5.01E-08 | 4.35 |
| FALSE | High | CQSLTEDLI | [R].CQSLTE | 1xCarbami | P14733 1xl | 0.026421 | 0.001676 | 1 | 2 | 1 | P14733 | Lamin-B1 ( | P14733 [1 | 0 | 1706.766 | 0 | -0.82 | 0.004104 | 4.89 | 124.5 | 75.5 | 12.61 | 16.85 | | High | High | High | High | High | High | High | High | High | High | High | 0.000903 | 0.009328 | 2.21 |
| FALSE | High | CQSPILHSS | [R].CQSPIL | 1xCarbami | P0DOV2 1x | 4.51E-08 | 0 | 1 | 1 | 1 | P0DOV2 | Interferon- | P0DOV2 [1 | 0 | 2596.284 | 0 | 2.13 | 0.000659 | 12.34 | 32.5 | 167.5 | 18.17 | 13.89 | | High | High | High | High | High | High | High | High | High | High | High | 0 | 3.12E-09 | 2.33 |
| FALSE | High | CQTLFGFD | [K].CQTLFC | 1xCarbami | A0A140LI3 | 3.24E-06 | 0 | 1 | 2 | 3 | A0A140LI3 | Zinc finger | A0A140LI3 | 0 | 1900.751 | 0 | -0.13 | 0.4921 | 27.62 | 99 | 101 | 18.8 | 15.65 | | High | High | High | High | High | High | High | High | High | High | High | 0 | 3.8E-07 | 3.47 |
| FALSE | High | CQWSEAD | [R].CQWSE | 1xCarbami | S4R1M2 1x | 4.63E-05 | 0 | 2 | 5 | 1 | S4R1M2; C | Scaffold att | S4R1M2 [3 | 0 | 2112.06 | 0 | -0.48 | 0.047993 | 6.32 | 120.3 | 79.7 | 11.78 | 19.53 | Shared | High | High | High | High | High | High | High | High | High | High | High | 0 | 7.56E-06 | 2.71 |
| FALSE | High | CRDDSFFG | [R].CRDDSI | 1xCarbami | Q6ZQ61 1x | 0.000876 | 0 | 1 | 6 | 1 | Q6ZQ61 | MCG12197 | Q6ZQ61 [2 | 1 | 2538.127 | 0 | -0.04 | 0.42561 | 5.59 | 99.1 | 100.9 | 13.96 | 18.38 | | High | High | High | High | High | High | High | High | High | High | High | 0 | 0.000206 | 1.86 |
| FALSE | High | CRNSIASCA | [R].CRNSIA | 2xCarbami | Q03141 1x | 0.045508 | 0.003373 | 1 | 6 | 1 | Q03141 | MAP/micro | Q03141 [3 | 1 | 2557.127 | 0 | -1.02 | 0.002357 | 2.81 | 134.7 | 65.3 | 7.7 | 11.56 | | High | High | High | High | High | High | High | High | High | High | High | 0.001741 | 0.01734 | 2.18 |
| FALSE | High | CRPSPSALC | [K].CRPSPS | 1xCarbami | P51432 1xl | 0.007951 | 0.000736 | 1 | 4 | 1 | P51432 | 1-phospha | P51432 [1 | 0 | 1610.849 | 0 | 0.25 | 0.015545 | 3.86 | 89.3 | 110.7 | 6.03 | 6.61 | | High | High | High | High | High | High | High | High | High | High | High | 0.000327 | 0.002418 | 1.62 |
| FALSE | High | CRSPGMLE | [R].CRSPGI | 1xCarbami | Q8BTI8 1xP | 0.002172 | 5.66E-05 | 1 | 3 | 1 | Q8BTI8 | Serine/argi | Q8BTI8 [2 | 1 | 1839.881 | 0 | -0.39 | 0.052703 | 11.37 | 109.7 | 90.3 | 11.49 | 11.33 | | High | High | High | High | High | High | High | High | High | High | High | 2.58E-05 | 0.000569 | 3.61 |
| FALSE | High | CRSPGMLE | [R].CRSPGI | 1xCarbami | Q8BTI8 1xP | 0.000194 | 0 | 1 | 3 | 1 | Q8BTI8 | Serine/argi | Q8BTI8 [2 | 1 | 1855.876 | 0 | -0.57 | 0.030338 | 14.37 | 120.5 | 79.5 | 10.73 | 17.53 | | High | High | High | High | High | High | High | High | High | High | High | 0 | 3.78E-05 | 2.5 |
| FALSE | High | CSAEDEDC | [R].CSAEDI | 2xCarbami | Q05CL8 1x | 0.069903 | 0.005181 | 1 | 3 | 2 | Q05CL8 | La-related | Q05CL8 [2 | 0 | 2212.018 | 0 | -0.27 | 0.011104 | 3.47 | 107.7 | 92.3 | 4.59 | 4.79 | | High | High | High | High | High | High | High | High | High | High | High | 0.002675 | 0.0285 | 1.82 |
| FALSE | High | CSASTTDLF | [K].CSASTT | 1xCarbami | Q3UQN2 1 | 0.016021 | 0.001124 | 1 | 4 | 2 | Q3UQN2 | F-BAR dom | Q3UQN2 [ | 0 | 1319.586 | 0 | 0.39 | 0.130141 | 20.94 | 89.2 | 110.8 | 18.2 | 12.23 | | High | High | High | High | High | High | High | High | High | High | High | 0.005651 | 0.06211 | 1.38 |
| FALSE | High | CSCASPGCI | [K].CSCASF | 3xCarbami | Q8BMK1 1 | 4.01E-05 | 0 | 1 | 1 | 1 | Q8BMK1 | Methyltrar | Q8BMK1 [ | 0 | 3012.392 | 0 | 1.04 | 0.003548 | 12.72 | 65.5 | 134.5 | 4.15 | 11.25 | | High | High | High | High | High | High | High | High | High | High | High | 0 | 6.41E-06 | 2.43 |
| FALSE | High | CSDAAGYP | [R].CSDAA | 1xCarbami | P60335 1xl | 0.003028 | 0.00016 | 1 | 1 | 1 | P60335 | Poly(rC)-bi | P60335 [2 | 0 | 4484.163 | 0 | -0.05 | 0.534561 | 48.89 | 103.6 | 96.4 | 18.9 | 22.05 | | High | High | High | High | High | High | High | High | High | High | High | 7.19E-05 | 0.000825 | 2.66 |
| FALSE | High | CSDNSSYEI | [R].CSDNS | 1xCarbami | B2KFW6 1x | 0.000235 | 0 | 1 | 3 | 1 | B2KFW6 | Polyhomec | B2KFW6 [7 | 0 | 2670.144 | 0 | 0.8 | 0.01581 | 17.79 | 76.7 | 123.3 | 10.48 | 16.53 | | High | High | High | High | High | High | High | High | High | High | High | 0 | 4.67E-05 | 3.64 |
| FALSE | High | CSDSQSWE | [K].CSDSQ | 2xCarbami | P70399 1xl | 1.02E-07 | 0 | 1 | 5 | 4 | P70399 | TP53-bindi | P70399 [7 | 0 | 2746.096 | 0 | 0.68 | 0.135319 | 54.43 | 77.5 | 122.5 | 23.16 | 27.35 | | High | High | High | High | High | High | High | High | High | High | High | 0 | 9.09E-08 | 5.52 |
| FALSE | High | CSDVSELSS | [K].CSDVSE | 1xCarbami | Q3U5I9 1xl | 0.095408 | 0.007437 | 1 | 2 | 2 | Q3U5I9 | Putative ur | Q3U5I9 [1 | 0 | 3647.644 | 0 | 0.54 | 0.156954 | 22.42 | 81.3 | 118.7 | 8.98 | 45.78 | | High | High | High | High | High | High | High | High | High | High | High | 0.003768 | 0.04065 | 2.03 |
| FALSE | High | CSGNSVAC | [R].CSGNS | 1xCarbami | Q6P3Z3 1x | 1.64E-07 | 0 | 1 | 2 | 2 | Q6P3Z3 | THAP dom | Q6P3Z3 [2 | 0 | 2642.29 | 0 | -0.38 | 0.016818 | 10.86 | 117.6 | 82.4 | 9.25 | 12.75 | | High | High | High | High | High | High | High | High | High | High | High | 0 | 1.33E-08 | 3.92 |
| FALSE | High | CSGPGLSP( | [K].CSGPGI | 1xCarbami | Q8BTM8 1: | 0.014797 | 0.001049 | 1 | 5 | 1 | Q8BTM8 | Filamin-A ( | Q8BTM8 [ | 0 | 1526.706 | 0 | 0.44 | 0.283301 | 22.76 | 91.2 | 108.8 | 12.74 | 24.09 | | High | High | High | High | High | High | High | High | High | High | High | 0.000555 | 0.004849 | 2.4 |
| FALSE | High | CSGPGLSP( | [K].CSGPGI | 1xCarbami | Q8BTM8 1: | 8.13E-06 | 0 | 1 | 5 | 9 | Q8BTM8 | Filamin-A ( | Q8BTM8 [ | 0 | 1542.701 | 0 | 0.33 | 0.052754 | 16.65 | 88.8 | 111.2 | 7.4 | 17.69 | | High | High | High | High | High | High | High | High | High | High | High | 0 | 0.00025 | 2.88 |
| FALSE | High | CSGSEPSSS | [R].CSGSEF | 1xCarbami | Q3TYU7 1x | 0.036416 | 0.002419 | 1 | 3 | 1 | Q3TYU7 | Putative ur | Q3TYU7 [8 | 0 | 1692.755 | 0 | -0.68 | 0.0015 | 3.11 | 127 | 73 | 8.13 | 8.79 | | High | High | High | High | High | High | High | High | High | High | High | 0.001296 | 0.01341 | 2.12 |
| FALSE | High | CSLGIFLPK | [R].CSLGIFI | 1xCarbami | A3KGI3 1xl | 0.000543 | 0 | 1 | 2 | 1 | A3KGI3 | Kinetochor | A3KGI3 [14 | 0 | 1572.863 | 0 | -0.7 | 0.057501 | 6.06 | 125 | 75 | 23.42 | 14.84 | | High | High | High | High | High | High | High | High | High | High | High | 0 | 0.00012 | 2.19 |
| FALSE | High | CEESDPESF | [K].CEESDF | 2xCarbami | E9QA22 2x | 0.004696 | 0.000456 | 1 | 4 | 1 | E9QA22 | Zinc finger | E9QA22 [3 | 0 | 3411.413 | 0 | 0.78 | 0.033286 | 37.57 | 74.7 | 125.3 | 15.03 | 20.89 | | High | High | High | High | High | High | High | High | High | High | High | 0.000204 | 0.001343 | 2.3 |
| FALSE | High | CSTSSPSK | [R].CSTSSP | 1xCarbami | Q6A065 1x | 0.019618 | 0.001438 | 1 | 3 | 3 | Q6A065 | Centrosom | Q6A065 [9 | 0 | 1391.664 | 0 | 0.65 | 0.012058 | 14.47 | 79.1 | 120.9 | 8.51 | 10.78 | | High | High | High | High | High | High | High | High | High | High | High | 0.000756 | 0.006683 | 1.44 |
| FALSE | High | CEERSPSFC | [R].CEERSP | 2xCarbami | O35492 2x | 0.014538 | 0.001011 | 1 | 4 | 1 | O35492 | Dual specif | O35492 [2 | 1 | 2411.887 | 0 | -0.14 | 0.066391 | 4.8 | 106.6 | 93.4 | 7.86 | 6.81 | | High | High | High | High | High | High | High | High | High | High | High | 0.000555 | 0.004766 | 2.35 |
| FALSE | High | CEENTSPTI | [R].CEENTS | 1xCarbami | E9Q5F9 1x | 8.43E-06 | 0 | 1 | 3 | 5 | E9Q5F9 | Histone-lys | E9Q5F9 [8 | 0 | 2218.035 | 0 | -0.18 | 0.085001 | 2.25 | 106.3 | 93.7 | 7.24 | 6.46 | | High | High | High | High | High | High | High | High | High | High | High | 0 | 9.16E-06 | 3.69 |
| FALSE | High | AVQQQTPS | [K].AVQQC | 1xPhospho | B2RSV7 1xl | 5.26E-05 | 0 | 1 | 2 | 3 | B2RSV7 | Fgd6 prote | B2RSV7 [3 | 0 | 3502.684 | 0 | 1.54 | 0.002348 | 12.7 | 50.8 | 149.2 | 15.42 | 12.3 | | High | High | High | High | High | High | High | High | High | High | High | 0 | 8.71E-06 | 4.28 |
| FALSE | High | AVSAECLG | [K].AVSAE( | 1xCarbami | Q9QXL1 1x | 1.99E-06 | 0 | 1 | 3 | 2 | Q9QXL1 | Kinesin-like | Q9QXL1 [1 | 0 | 2170.087 | 0 | -0.73 | 0.000851 | 4.48 | 127.3 | 72.7 | 4.13 | 7.86 | | High | High | High | High | High | High | High | High | High | High | High | 0 | 2.21E-07 | 4.14 |
| FALSE | High | AVSETEDL( | [K].AVSETE | 1xPhospho | Q9DCH6 1x | 0.000159 | 0 | 1 | 4 | 1 | Q9DCH6 | AN1-type z | Q9DCH6 [1 | 0 | 1610.762 | 0 | 0.05 | 0.983921 | 12.65 | 96.7 | 103.3 | 11.48 | 14.23 | | High | High | High | High | High | High | High | High | High | High | High | 0 | 3.03E-05 | 3.23 |
| FALSE | High | AVSGNTPV | [R].AVSGN | 1xPhospho | A2APB8 1x | 0.003253 | 0.00016 | 1 | 3 | 4 | A2APB8 | Targeting p | A2APB8 [1 | 0 | 1562.789 | 0 | -0.06 | 0.176572 | 2.54 | 102.7 | 97.3 | 3.62 | 2.9 | | High | High | High | High | High | High | High | High | High | High | High | 9.42E-05 | 0.000951 | 2.78 |
| FALSE | High | AVSGYQSH | [K].AVSGY( | 1xCarbami | Q9CU65 1x | 4.56E-06 | 0 | 1 | 1 | 7 | Q9CU65 | Zinc finger | Q9CU65 [1 | 0 | 3002.292 | 0 | -0.84 | 0.000658 | 2.55 | 128.3 | 71.7 | 5.79 | 7.14 | | High | High | High | High | High | High | High | High | High | High | High | 0 | 1.78E-05 | 3.2 |
| FALSE | High | AVSLDSPVS | [R].AVSLDS | 1xPhospho | Q05BA5 1x | 1.62E-07 | 0 | 1 | 6 | 2 | Q05BA5 | Nuclear re | Q05BA5 [8 | 0 | 2134.156 | 0 | -0.36 | 0.023621 | 1.89 | 111.7 | 88.3 | 6.96 | 5.12 | | High | High | High | High | High | High | High | High | High | High | High | 0 | 1.32E-08 | 4.85 |
| FALSE | High | AVSLNLGN | [R].AVSLNL | 1xPhospho | Q8BFY7 1x | 1.5E-05 | 0 | 2 | 4 | 1 | Q8BFY7; Q | Protein PIN | Q8BFY7 [5 | 0 | 1881.004 | 0 | -1.13 | 0.000435 | 0.69 | 136.7 | 63.3 | 7.24 | 7.13 | Shared | High | High | High | High | High | High | High | High | High | High | High | 0 | 2.13E-06 | 3.69 |
| FALSE | High | AVSLNLGN | [R].AVSLNL | 1xDeamida | Q8BFY7 1x | 0.093123 | 0.007276 | 2 | 4 | 1 | Q8BFY7; Q | Protein PIN | Q8BFY7 [5 | 0 | 1881.988 | 0 | -0.59 | 0.009504 | 8.43 | 121.7 | 78.3 | 16.61 | 14.67 | Shared | High | High | High | High | High | High | High | High | High | High | High | 0.003698 | 0.03986 | 2.25 |
| FALSE | High | AVSMDNSI | [R].AVSMD | 1xPhospho | Q9WVH4 1 | 0.016596 | 0.001124 | 1 | 2 | 1 | Q9WVH4 | Forkhead b | Q9WVH4 [ | 0 | 1503.728 | 0 | -1.83 | 0.00014 | 0.38 | 155.5 | 44.5 | 4.46 | 8.51 | | High | High | High | High | High | High | High | High | High | High | High | 0.000622 | 0.005543 | 1.96 |
| FALSE | High | AVSMDNSI | [R].AVSMD | 1xPhospho | Q9WVH4 1 | 0.007082 | 0.00062 | 1 | 2 | 2 | Q9WVH4 | Forkhead b | Q9WVH4 [ | 0 | 1519.723 | 0 | -1.7 | 0.000965 | 1.95 | 151.9 | 48.1 | 2.93 | 25.7 | | High | High | High | High | High | High | High | High | High | High | High | 0.000275 | 0.002143 | 1.66 |
| FALSE | High | AVSPAATQ | [R].AVSPA/ | 1xPhospho | E9QQ25 1x | 0.040028 | 0.002867 | 1 | 4 | 1 | E9QQ25 | Striated mu | E9QQ25 [5 | 0 | 2071.099 | 0 | -0.53 | 0.09696 | 4.12 | 118.1 | 81.9 | 14.88 | 17.49 | | High | High | High | High | High | High | High | High | High | High | High | 0.001516 | 0.01499 | 1.92 |
| FALSE | High | AVSPLDPAI | [R].AVSPLD | 1xPhospho | Q8BM75 1: | 0.017966 | 0.001282 | 1 | 2 | 1 | Q8BM75 | AT-rich inte | Q8BM75 [ | 0 | 1435.796 | 0 | 1.21 | 0.00047 | 6.6 | 60.7 | 139.3 | 5.18 | 7.82 | | High | High | High | High | High | High | High | High | High | High | High | 0.000689 | 0.006053 | 1.81 |
| FALSE | High | AVSPQPAC | [K].AVSPQI | 1xCarbami | Q52KB6 1x | 1.79E-07 | 0 | 1 | 1 | 2 | Q52KB6 | C2 domain | Q52KB6 [2 | 0 | 2800.338 | 0 | -0.77 | 0.004148 | 4.59 | 128.6 | 71.4 | 13.87 | 7.18 | | High | High | High | High | High | High | High | High | High | High | High | 0 | 1.47E-08 | 5 |
| FALSE | High | AVQQPSSP | [R].AVQQP | 1xPhospho | P53349 1xl | 0.000191 | 0 | 2 | 2 | 2 | P53349; F8 | Mitogen-ac | P53349 [5 | 0 | 2074.028 | 0 | -0.65 | 0.006953 | 2.82 | 122.8 | 77.2 | 5.72 | 12.61 | Shared | High | High | High | High | High | High | High | High | High | High | High | 0 | 3.73E-05 | 4.32 |
| FALSE | High | AVSPQPAC | [K].AVSPQI | 1xCarbami | Q52KB6 2x | 0.072258 | 0.005346 | 1 | 1 | 1 | Q52KB6 | C2 domain | Q52KB6 [2 | 0 | 2880.304 | 0 | -0.18 | 0.502977 | 5.71 | 105.2 | 94.8 | 9.62 | 10.73 | | High | High | High | High | High | High | High | High | High | High | High | 0.002855 | 0.02957 | 2.93 |
| FALSE | High | AVSSLDILG | [R].AVSSLD | 1xPhospho | Q1W617 1: | 0.000155 | 0 | 1 | 2 | 1 | Q1W617 | Protein Sh | Q1W617 [ | 0 | 1802.971 | 0 | 2.92 | 0.000561 | 17.98 | 25.9 | 174.1 | 16.08 | 11.07 | | High | High | High | High | High | High | High | High | High | High | High | 0 | 2.94E-05 | 3.23 |
| FALSE | High | AVSSPPTSF | [K].AVSSPF | 2xPhospho | F8VQC1 2x | 0.103392 | 0.008249 | 1 | 3 | 3 | F8VQC1 | Signal reco | F8VQC1 [6 | 0 | 3080.502 | 0 | 0.26 | 0.000294 | 0.74 | 91.2 | 108.8 | 5.09 | 4.03 | | High | High | High | High | High | High | High | High | High | High | High | 0.004164 | 0.04504 | 2.22 |
| FALSE | High | AVSSPPTSF | [K].AVSSPF | 3xPhospho | F8VQC1 3x | 0.023175 | 0.001578 | 1 | 3 | 1 | F8VQC1 | Signal reco | F8VQC1 [6 | 0 | 3160.468 | 0 | -0.14 | 0.847223 | 12.94 | 101.8 | 98.2 | 17.32 | 14.32 | | High | High | High | High | High | High | High | High | High | High | High | 0.000847 | 0.00805 | 3.06 |
| FALSE | High | AVSTAAAN | [R].AVSTA/ | 1xPhospho | Q3UXJ3 1x | 2.17E-11 | 0 | 1 | 1 | 13 | Q3UXJ3 | Putative ur | Q3UXJ3 [8 | 0 | 2110.014 | 0 | 0.38 | 0.203313 | 29.81 | 86.4 | 113.6 | 16.78 | 44.1 | | High | High | High | High | High | High | High | High | High | High | High | 0 | 5.82E-13 | 3.92 |
| FALSE | High | AVSTVVVT | [K].AVSTV\ | 1xPhospho | Q6PIJ4 1xP | 3.21E-06 | 0 | 1 | 2 | 1 | Q6PIJ4 | Nuclear fac | Q6PIJ4 [12 | 0 | 1895.066 | 0 | -0.09 | 0.234788 | 8.6 | 104.8 | 95.2 | 4.98 | 10.65 | | High | High | High | High | High | High | High | High | High | High | High | 0 | 3.76E-07 | 2.55 |
| FALSE | High | AVSVSCMS | [R].AVSVS( | 1xCarbami | Q3UHU5 1: | 0.002719 | 0.000109 | 1 | 5 | 1 | Q3UHU5 | Microtubu | Q3UHU5 [ | 0 | 1725.754 | 0 | -1.79 | 0.00228 | 0.87 | 155.2 | 44.8 | 12.32 | 18.64 | | High | High | High | High | High | High | High | High | High | High | High | 4.94E-05 | 0.000733 | 2.01 |
| FALSE | High | AVSWTFSE | [R].AVSWT | 1xPhospho | Q9D710 1x | 0.011246 | 0.000772 | 1 | 2 | 3 | Q9D710 | Thioredoxi | Q9D710 [2 | 0 | 1846.894 | 0 | -0.31 | 0.008731 | 5.58 | 110.8 | 89.2 | 3.54 | 5.74 | | High | High | High | High | High | High | High | High | High | High | High | 0.000399 | 0.00356 | 2.7 |
| FALSE | High | AVTENELTI | [K].AVTENI | 1xPhospho | E9Q6E9 1x | 0.001544 | 5.66E-05 | 1 | 9 | 2 | E9Q6E9 | SUMO-inte | E9Q6E9 [1 | 0 | 2422.134 | 0 | -0.19 | 0.030465 | 8.19 | 111.2 | 88.8 | 14.85 | 11.56 | | High | High | High | High | High | High | High | High | High | High | High | 0.000327 | 0.002793 | 3.5 |
| FALSE | High | AVTFPASPV | [R].AVTFP/ | 1xPhospho | P25962-2 1 | 0.002355 | 0.000109 | 1 | 2 | 1 | P25962-2 | Isoform B c | P25962-2 [ | 0 | 1553.793 | 0 | -1.5 | 0.000181 | 1.17 | 145.2 | 54.8 | 4.35 | 14.35 | | High | High | High | High | High | High | High | High | High | High | High | 4.94E-05 | 0.000622 | 2.97 |
| FALSE | High | AVTGQTTC | [K].AVTGQ | 1xPhospho | Q571G4 1x | 3.67E-05 | 0 | 1 | 3 | 1 | Q571G4 | Protein lin- | Q571G4 [2 | 0 | 1979.016 | 0 | -0.56 | 0.00063 | 1.09 | 118.1 | 81.9 | 3.92 | 3.37 | | High | High | High | High | High | High | High | High | High | High | High | 0 | 5.81E-06 | 3.78 |
| FALSE | High | AVTIATPAT | [K].AVTIAT | 1xPhospho | Q52KI8 1xl | 5.8E-08 | 0 | 1 | 7 | 5 | Q52KI8 | Serine/argi | Q52KI8 [87 | 0 | 3499.752 | 0 | -0.21 | 0.191368 | 24.78 | 110.3 | 89.7 | 14.66 | 26.21 | | High | High | High | High | High | High | High | High | High | High | High | 0 | 1.25E-08 | 5.3 |

| | | | | | | | | | | | | | | | | | | | | | | | | | | | | | | | | | | | | | | |
|---|---|---|---|---|---|---|---|---|---|---|---|---|---|---|---|---|---|---|---|---|---|---|---|---|---|---|---|---|---|---|---|---|---|---|---|---|---|---|
| FALSE | High | AVTPGHYV | [K].AVTPGI | 1xPhosphc | Q3UTQ7 1: | 9E-07 | 0 | 1 | 4 | 3 | Q3UTQ7 | PR domain | Q3UTQ7 [1 | 0 | 2651.337 | 0 | -0.43 | 0.007393 | 1.77 | 113.7 | 86.3 | 6.92 | 7.44 | | High | High | High | High | High | High | High | High | High | High | High | 0 | 3.35E-07 | 4.55 |
| FALSE | High | AVTSAEERS | [R].AVTSAE | 1xPhosphc | B2RRE0 1x | 7.98E-06 | 0 | 1 | 2 | 5 | B2RRE0 | A kinase (P | B2RRE0 [8 | 1 | 4518.215 | 0 | -1.27 | 0.026207 | 7.63 | 136.7 | 63.3 | 24.59 | 30.2 | | High | High | High | High | High | High | High | High | High | High | High | 7.19E-05 | 0.000876 | 4.92 |
| FALSE | High | AVSPTEPTI | [R].AVSPTE | 1xPhosphc | Q8C0M0-3 | 0.004572 | 0.000456 | 1 | 4 | 2 | Q8C0M0-3 | Isoform 3 c | Q8C0M0-3 | 0 | 1363.682 | 0 | -0.56 | 0.002931 | 7.31 | 117.9 | 82.1 | 2.51 | 8.94 | | High | High | High | High | High | High | High | High | High | High | High | 0.000204 | 0.001303 | 1.95 |
| FALSE | High | AVTSAEERS | [R].AVTSAE | 1xPhosphc | B2RRE0 1x | 0.000138 | 0 | 1 | 2 | 2 | B2RRE0 | A kinase (P | B2RRE0 [8 | 1 | 4534.21 | 0 | -1.21 | 0.009373 | 7.31 | 147.1 | 52.9 | 19.03 | 35.49 | | High | High | High | High | High | High | High | High | High | High | High | 0 | 4.93E-05 | 4.75 |
| FALSE | High | AVQLMQQ | [R].AVQLM | 1xDeamida | Q8BKX1 1x | 3.48E-06 | 0 | 1 | 6 | 4 | Q8BKX1 | Brain-spec | Q8BKX1 [2 | 0 | 2985.519 | 0 | 0.86 | 0.01298 | 7.34 | 74.9 | 125.1 | 8.13 | 23.16 | | High | High | High | High | High | High | High | High | High | High | High | 0 | 6.67E-07 | 4.22 |
| FALSE | High | AVPVSPSAY | [K].AVPVSI | 1xPhosphc | P39447 1xI | 8.03E-08 | 0 | 1 | 3 | 9 | P39447 | Tight junct | P39447 [16 | 0 | 2889.335 | 0 | -0.44 | 0.008095 | 4.53 | 117.1 | 82.9 | 5.97 | 6.49 | | High | High | High | High | High | High | High | High | High | High | High | 0 | 5.97E-09 | 4.9 |
| FALSE | High | AVEDMGSI | [R].AVEDM | 1xPhosphc | A0A0R4J14 | 6.15E-06 | 0 | 1 | 6 | 1 | A0A0R4J14 | Clustered r | A0A0R4J14 | 0 | 1771.87 | 0 | -0.23 | 0.032669 | 5 | 110.4 | 89.6 | 8.77 | 14.76 | | High | High | High | High | High | High | High | High | High | High | High | 0 | 7.8E-07 | 2.81 |
| FALSE | High | AVEDMGSI | [R].AVEDM | 1xPhosphc | A0A0R4J14 | 3.41E-05 | 0 | 1 | 6 | 4 | A0A0R4J14 | Clustered r | A0A0R4J14 | 0 | 1787.865 | 0 | -0.37 | 0.699355 | 10.9 | 112.9 | 87.1 | 7.56 | 34.74 | | High | High | High | High | High | High | High | High | High | High | High | 0 | 5.36E-06 | 2.92 |
| FALSE | High | AVEEGELL( | [K].AVEEGI | 1xPhosphc | Q6P5H2-2 | 4.22E-07 | 0 | 1 | 2 | 2 | Q6P5H2-2 | Isoform 2 c | Q6P5H2-2 | 0 | 3039.496 | 0 | 2.89 | 0.015085 | 71.15 | 16.5 | 183.5 | 59.33 | 42.13 | | High | High | High | High | High | High | High | High | High | High | High | 0 | 3.87E-08 | 4.66 |
| FALSE | High | AVELTSLSD | [K].AVELTS | 1xPhosphc | Q4FK48 1x | 0.001077 | 0 | 1 | 1 | 1 | Q4FK48 | Snai1 prote | Q4FK48 [8 | 0 | 1988.983 | 0 | -0.65 | 0.000686 | 3.2 | 121.4 | 78.6 | 5.87 | 7.16 | | High | High | High | High | High | High | High | High | High | High | High | 0 | 0.000259 | 2.5 |
| FALSE | High | AVEPALPQ | [R].AVEPAI | 1xPhosphc | P58468 1xI | 0.000475 | 0 | 1 | 3 | 2 | P58468 | Protein FAI | P58468 [3C | 0 | 2059.099 | 0 | 0.04 | 0.442032 | 8.62 | 98.7 | 101.3 | 5.36 | 5.03 | | High | High | High | High | High | High | High | High | High | High | High | 0 | 0.000104 | 2.89 |
| FALSE | High | AVESCPDEI | [R].AVESCF | 1xCarbami | Q6P9R4 1x | 0.000538 | 0 | 1 | 3 | 1 | Q6P9R4 | Rho guanir | Q6P9R4 [4 | 0 | 2737.184 | 0 | 0.55 | 0.072577 | 32.15 | 78.9 | 121.1 | 14.73 | 19.29 | | High | High | High | High | High | High | High | High | High | High | High | 0 | 0.000119 | 3.08 |
| FALSE | High | AVESTDED | [K].AVESTI | 1xPhosphc | Q8BLH7 1x | 0.000207 | 0 | 1 | 2 | 3 | Q8BLH7 | HIRA-intera | Q8BLH7 [1 | 0 | 2312.07 | 0 | -0.97 | 0.004244 | 3.31 | 132.5 | 67.5 | 9.9 | 9.56 | | High | High | High | High | High | High | High | High | High | High | High | 0.000161 | 0.001147 | 2.43 |
| FALSE | High | AVESTDED | [K].AVESTI | 2xPhosphc | Q8BLH7 2x | 0.00582 | 0.00062 | 1 | 2 | 6 | Q8BLH7 | HIRA-intera | Q8BLH7 [1 | 0 | 2392.036 | 0 | -0.95 | 0.000304 | 1.11 | 132.6 | 67.4 | 3.55 | 5.82 | | High | High | High | High | High | High | High | High | High | High | High | 0.000968 | 0.01003 | 3.05 |
| FALSE | High | AVEYLLTGI | [R].AVEYLL | 1xDeamida | P54726 1xI | 2.67E-05 | 0 | 1 | 3 | 1 | P54726 | UV excisior | P54726 [19 | 0 | 4855.319 | 0 | -0.3 | 0.187005 | 6.52 | 109.6 | 90.4 | 10.97 | 14.07 | | High | High | High | High | High | High | High | High | High | High | High | 0 | 4.1E-06 | 5.25 |
| FALSE | High | AVEYLLTGI | [R].AVEYLL | 1xPhosphc | Q8CAP3 1x | 0.000515 | 0 | 1 | 2 | 1 | Q8CAP3 | Putative ur | Q8CAP3 [1 | 0 | 4783.298 | 0 | 0 | 0.800331 | 56.07 | 102.6 | 97.4 | 19.15 | 28.79 | | High | High | High | High | High | High | High | High | High | High | High | 0 | 0.000113 | 4.52 |
| FALSE | High | AVFGESGG | [R].AVFGES | 1xPhosphc | P55200 1xI | 3.65E-07 | 0 | 1 | 2 | 3 | P55200 | Histone-lys | P55200 [12 | 0 | 3085.326 | 0 | 0.28 | 0.048833 | 17.41 | 90.3 | 109.7 | 5.99 | 7.27 | | High | High | High | High | High | High | High | High | High | High | High | 0 | 3.29E-08 | 4.41 |
| FALSE | High | AVGGAPSP | [R].AVGGA | 1xPhosphc | Q8BHL3 1x | 8.11E-05 | 0 | 1 | 1 | 2 | Q8BHL3 | TBC1 doma | Q8BHL3 [6 | 0 | 1510.798 | 0 | -0.05 | 0.3263 | 15.03 | 102.6 | 97.4 | 6.87 | 10.58 | | High | High | High | High | High | High | High | High | High | High | High | 0 | 1.43E-05 | 2.28 |
| FALSE | High | AVGMPSP\ | [K].AVGMF | 1xPhosphc | Q9ESC8 1x | 0.002623 | 0.000109 | 1 | 2 | 2 | Q9ESC8 | AF4/FMR2 | Q9ESC8 [1 | 0 | 1623.858 | 0 | -0.48 | 0.004136 | 4.74 | 116.4 | 83.6 | 11.89 | 7.88 | | High | High | High | High | High | High | High | High | High | High | High | 4.94E-05 | 0.000701 | 2.73 |
| FALSE | High | AVQLMQQ | [R].AVQLM | 1xPhosphc | Q8BKX1 1x | 3.23E-05 | 0 | 1 | 6 | 2 | Q8BKX1 | Brain-spec | Q8BKX1 [2 | 0 | 2984.535 | 0 | 0.41 | 0.065126 | 10.3 | 84.6 | 115.4 | 12.18 | 13.89 | | High | High | High | High | High | High | High | High | High | High | High | 0 | 5.04E-06 | 3.9 |
| FALSE | High | AVGSMPT/ | [K].AVGSM | 1xPhosphc | Q7TN29 1x | 7.47E-05 | 0 | 1 | 1 | 1 | Q7TN29 | Stromal me | Q7TN29 [2 | 0 | 3152.574 | 0 | 0.67 | 0.004117 | 16.05 | 77.8 | 122.2 | 9.52 | 13.4 | | High | High | High | High | High | High | High | High | High | High | High | 0 | 1.3E-05 | 4.17 |
| FALSE | High | AVLFCLSEI | [K].AVLFCL | 1xCarbami | F8WGL3 1x | 0.076555 | 0.005826 | 1 | 3 | 1 | F8WGL3 | Cofilin-1 O | F8WGL3 [3 | 0 | 1719.879 | 0 | 0.2 | 0.109901 | 10.89 | 88.8 | 111.2 | 15.54 | 10.79 | | High | High | High | High | High | High | High | High | High | High | High | 0.002976 | 0.03143 | 1.73 |
| FALSE | High | AVLPASPEI | [K].AVLPAS | 1xPhosphc | A0A140T8S | 0.013787 | 0.001011 | 1 | 1 | 3 | A0A140T8S | Chromosoi | A0A140T8S | 0 | 1345.708 | 0 | -0.53 | 0.000436 | 1.72 | 118.7 | 81.3 | 4.57 | 3.98 | | High | High | High | High | High | High | High | High | High | High | High | 0.006456 | 0.06962 | 2.13 |
| FALSE | High | AVLPGSPIF | [K].AVLPGS | 1xPhosphc | Q5I1X5 1xI | 0.019618 | 0.001438 | 1 | 1 | 1 | Q5I1X5 | RelA-assoc | Q5I1X5 [38 | 0 | 1452.781 | 0 | 1.95 | 0.004352 | 15.31 | 40.4 | 159.6 | 20.1 | 21.16 | | High | High | High | High | High | High | High | High | High | High | High | 0.000756 | 0.006693 | 3.01 |
| FALSE | High | AVITPDFEK | [K].AVITPD | 1xPhosphc | Q8R2M2 1 | 0.007339 | 0.00062 | 1 | 1 | 3 | Q8R2M2 | Deoxynucl | Q8R2M2 [( | 0 | 1557.833 | 0 | -0.93 | 0.003241 | 3.71 | 131.2 | 68.8 | 13.84 | 4.91 | | High | High | High | High | High | High | High | High | High | High | High | 0.000538 | 0.004496 | 2.96 |
| FALSE | High | AVLWQDSI | [K].AVLWC | 1xPhosphc | Q3UR00 1x | 0.051258 | 0.003745 | 1 | 10 | 1 | Q3UR00 | Putative ur | Q3UR00 [1 | 0 | 2066.088 | 0 | -0.71 | 0.001843 | 2.81 | 121.5 | 78.5 | 8.55 | 9.83 | | High | High | High | High | High | High | High | High | High | High | High | 0.00196 | 0.01981 | 2.42 |
| FALSE | High | AVNGESPK | [R].AVNGE | 1xPhosphc | Q6A065 1x | 0.001463 | 5.66E-05 | 1 | 2 | 1 | Q6A065 | Centrosom | Q6A065 [6 | 0 | 1339.702 | 0 | 1.3 | 0.000181 | 4.16 | 57.5 | 142.5 | 8.82 | 2.34 | | High | High | High | High | High | High | High | High | High | High | High | 2.58E-05 | 0.000364 | 1.71 |
| FALSE | High | AVNPTMA/ | [R].AVNPTI | 1xPhosphc | E9Q0S6 1x | 6E-07 | 0 | 1 | 11 | 8 | E9Q0S6 | Tensin 1 O | E9Q0S6 [1: | 0 | 2017.973 | 0 | -0.5 | 0.00408 | 4.52 | 117.2 | 82.8 | 11.91 | 9.8 | | High | High | High | High | High | High | High | High | High | High | High | 0 | 3.9E-06 | 3.49 |
| FALSE | High | AVNSTRET | [R].AVNSTI | 1xPhosphc | Q3TEA8 1x | 0.026652 | 0.001676 | 1 | 10 | 1 | Q3TEA8 | Heterochrc | Q3TEA8 [4 | 1 | 1737.93 | 0 | 1.54 | 0.000154 | 3.5 | 52.3 | 147.7 | 8.31 | 2.36 | | High | High | High | High | High | High | High | High | High | High | High | 0.000903 | 0.009427 | 1.1 |
| FALSE | High | AVNTQALS | [R].AVNTQ | 1xPhosphc | Q3U5L3 1x | 8.63E-05 | 0 | 1 | 3 | 1 | Q3U5L3 | Putative ur | Q3U5L3 [2 | 0 | 1679.904 | 0 | 0.02 | 0.984455 | 3.67 | 99.8 | 100.2 | 9.79 | 11.04 | | High | High | High | High | High | High | High | High | High | High | High | 0 | 1.53E-05 | 2.66 |
| FALSE | High | AVPEIWTP. | [R].AVPEIW | 1xPhosphc | Q7TT37 1x | 0.005467 | 0.00062 | 1 | 1 | 2 | Q7TT37 | Elongator c | Q7TT37 [1: | 0 | 4269.148 | 0 | 0.12 | 0.433106 | 7.12 | 98.2 | 101.8 | 6.16 | 10.25 | | High | High | High | High | High | High | High | High | High | High | High | 0.000275 | 0.001593 | 3.22 |
| FALSE | High | AVPGQVQ/ | [R].AVPGQ | 1xPhosphc | Q8C6B9 1x | 1.08E-06 | 0 | 1 | 1 | 2 | Q8C6B9 | Active regu | Q8C6B9 [2 | 0 | 1848.998 | 0 | 0.75 | 0.014402 | 16.53 | 72.3 | 127.7 | 17.56 | 6.93 | | High | High | High | High | High | High | High | High | High | High | High | 0 | 2.16E-07 | 4.28 |
| FALSE | High | AVPSESPSS | [R].AVPSES | 1xPhosphc | Q9CWC4 1 | 9.45E-05 | 0 | 1 | 5 | 8 | Q9CWC4 | Transcripti | Q9CWC4 [: | 0 | 1787.883 | 0 | -0.46 | 0.008249 | 4.94 | 115.7 | 84.3 | 4.23 | 10.56 | | High | High | High | High | High | High | High | High | High | High | High | 0 | 3.25E-05 | 3.54 |
| FALSE | High | AVPSESPSS | [R].AVPSES | 2xPhosphc | Q9CWC4 2 | 0.068752 | 0.005042 | 1 | 5 | 2 | Q9CWC4 | Transcripti | Q9CWC4 [: | 0 | 1867.849 | 0 | -0.48 | 0.014184 | 7.17 | 117.6 | 82.4 | 12.24 | 12.63 | | High | High | High | High | High | High | High | High | High | High | High | 0.002604 | 0.02787 | 2.2 |
| FALSE | High | AVGTQQS\ | [K].AVGTQ | 1xCarbami | U5KVR9 1x | 0.093123 | 0.007276 | 1 | 6 | 1 | U5KVR9 | Centrosom | U5KVR9 [3 | 0 | 2483.237 | 0 | 0.71 | 0.00409 | 10.06 | 77 | 123 | 5.11 | 10.44 | | High | High | High | High | High | High | High | High | High | High | High | 0.003698 | 0.0398 | 1.12 |
| FALSE | High | AVTTVTQS | [R].AVTTV | 1xPhosphc | Q61191 1x | 2.86E-07 | 0 | 1 | 2 | 2 | Q61191 | Host cell fa | Q61191 [1 | 0 | 3233.666 | 0 | -0.54 | 0.006026 | 6.43 | 121.4 | 78.6 | 9.56 | 4.29 | | High | High | High | High | High | High | High | High | High | High | High | 0 | 2.5E-08 | 3.62 |
| FALSE | High | AVVLPGGN | [K].AVVLP( | 1xPhosphc | Q9D2D7 1x | 1.1E-05 | 0 | 1 | 3 | 1 | Q9D2D7 | Zinc finger | Q9D2D7 [4 | 0 | 1748.971 | 0 | -0.5 | 0.000673 | 4.43 | 117.3 | 82.7 | 1.46 | 5.38 | | High | High | High | High | High | High | High | High | High | High | High | 0 | 1.52E-06 | 3.51 |
| FALSE | High | AVVSGFDF | [R].AVVSGI | 1xPhosphc | D3YYY8 1x | 0.000259 | 0 | 1 | 1 | 1 | D3YYY8 | Rho guanir | D3YYY8 [3 | 0 | 2082.02 | 0 | 1.12 | 0.001161 | 2.53 | 63.6 | 136.4 | 9.29 | 13.28 | | High | High | High | High | High | High | High | High | High | High | High | 0 | 5.23E-05 | 3.3 |
| FALSE | High | CADDNQE/ | [R].CADDN | 2xCarbami | F8VQG7 1x | 5.37E-06 | 0 | 1 | 4 | 2 | F8VQG7 | NEDD4-bin | F8VQG7 [6 | 0 | 3540.48 | 0 | -0.54 | 0.055689 | 15.7 | 117.3 | 82.7 | 8.54 | 24.53 | | High | High | High | High | High | High | High | High | High | High | High | 0 | 6.71E-07 | 4.11 |
| FALSE | High | CAENLFPSI | [R].CAENLI | 1xCarbami | Q8CIB9 1xI | 0.018285 | 0.001282 | 1 | 2 | 2 | Q8CIB9 | N-acetyltra | Q8CIB9 [34 | 0 | 1814.891 | 0 | -1.83 | 0.000477 | 1.31 | 155.9 | 44.1 | 6.67 | 8.66 | | High | High | High | High | High | High | High | High | High | High | High | 0.001061 | 0.01124 | 2.12 |
| FALSE | High | CAPLSAPAS | [R].CAPLSA | 1xCarbami | B1AXD8 1x | 6.68E-07 | 0 | 1 | 1 | 1 | B1AXD8 | Akirin-2 OS | B1AXD8 [2 | 0 | 3289.665 | 0 | -0.44 | 0.029107 | 9.62 | 116.4 | 83.6 | 6.08 | 22.8 | | High | High | High | High | High | High | High | High | High | High | High | 0 | 6.49E-08 | 4 |
| FALSE | High | CAPPASPAI | [R].CAPPAS | 1xCarbami | Q8BQ30-2 | 2.5E-10 | 0 | 1 | 6 | 1 | Q8BQ30-2 | Isoform 2 c | Q8BQ30-2 | 0 | 2786.358 | 0 | 1.37 | 0.003325 | 12.27 | 55.9 | 144.1 | 10.95 | 11.87 | | High | High | High | High | High | High | High | High | High | High | High | 0 | 9.08E-12 | 6.54 |
| FALSE | High | CASESSICS | [R].CASESS | 3xCarbami | E9PZ26 1xI | 2.77E-05 | 0 | 1 | 4 | 3 | E9PZ26 | Bromodom | E9PZ26 [98 | 0 | 3046.325 | 0 | 0 | 0.633273 | 4.38 | 98.7 | 101.3 | 2.3 | 12.71 | | High | High | High | High | High | High | High | High | High | High | High | 0 | 4.23E-06 | 2.76 |
| FALSE | High | CASGPQCE | [R].CASGP( | 2xCarbami | A2AIX1 1xI | 0.057686 | 0.004227 | 1 | 2 | 1 | A2AIX1 | SEC16 hom | A2AIX1 [46 | 0 | 3126.422 | 0 | 0.52 | 0.10088 | 17.69 | 85.8 | 114.2 | 18.46 | 23 | | High | High | High | High | High | High | High | High | High | High | High | 0.002159 | 0.02277 | 2.39 |
| FALSE | High | CASSSSYTA | [K].CASSSS | 1xCarbami | Q6A060 1x | 0.010663 | 0.000772 | 1 | 7 | 4 | Q6A060 | MKIAA048 | Q6A060 [6 | 0 | 2276.963 | 0 | -0.02 | 0.914347 | 10.1 | 102 | 98 | 11.26 | 17.72 | | High | High | High | High | High | High | High | High | High | High | High | 0.000399 | 0.003382 | 2.99 |
| FALSE | High | CAVPHSSPI | [K].CAVPHS | 1xCarbami | Q8CGC3 1x | 1.35E-05 | 0 | 1 | 6 | 3 | Q8CGC3 | Histone de | Q8CGC3 [3 | 0 | 2047.045 | 0 | -0.53 | 0.0161 | 2.99 | 116.9 | 83.1 | 2.3 | 13.31 | | High | High | High | High | High | High | High | High | High | High | High | 0 | 1.88E-06 | 3.1 |
| FALSE | High | CCAADWSI | [K].CCAAD | 2xCarbami | F8WGX5 1: | 1.03E-07 | 0 | 1 | 2 | 2 | F8WGX5 | DmX-like p | F8WGX5 [1 | 0 | 2896.196 | 0 | 0.63 | 0.008671 | 18.34 | 75.5 | 124.5 | 5.15 | 11.09 | | High | High | High | High | High | High | High | High | High | High | High | 0 | 5.87E-07 | 4.22 |
| FALSE | High | CCASATSQ | [K].CCASAT | 2xCarbami | Q6NWW5 | 1.55E-07 | 0 | 1 | 2 | 3 | Q6NWW5 | Kinesin-like | Q6NWW5 | 0 | 2421.094 | 0 | -1.33 | 0.004011 | 5.09 | 143.2 | 56.8 | 12.36 | 13.16 | | High | High | High | High | High | High | High | High | High | High | High | 0 | 1.25E-08 | 4.15 |
| FALSE | High | CCSNADLQ | [R].CCSNAI | 2xCarbami | Q9Z0X0 1x | 0.008769 | 0.000736 | 1 | 1 | 1 | Q9Z0X0 | Cdc42 effe | Q9Z0X0 [1: | 0 | 2001.886 | 0 | 0.84 | 0.036519 | 28.6 | 67.6 | 132.4 | 7.36 | 28.78 | | High | High | High | High | High | High | High | High | High | High | High | 0.000327 | 0.002695 | 1.18 |
| FALSE | High | CCSPLVTLS | [K].CCSPLV | 2xCarbami | Q3TZW5 1: | 1.8E-06 | 0 | 1 | 5 | 1 | Q3TZW5 | Putative ur | Q3TZW5 [1 | 0 | 2403.17 | 0 | -1.34 | 0.001291 | 2.02 | 145.3 | 54.7 | 6.95 | 13.68 | | High | High | High | High | High | High | High | High | High | High | High | 0 | 1.98E-07 | 4.2 |
| FALSE | High | CCSPPNEL( | [R].CCSPPN | 2xCarbami | E9Q5F9 1x | 0.0016 | 5.66E-05 | 1 | 3 | 2 | E9Q5F9 | Histone-lys | E9Q5F9 [5: | 0 | 1645.707 | 0 | -0.1 | 0.184947 | 3.49 | 105.5 | 94.5 | 7.75 | 14.38 | | High | High | High | High | High | High | High | High | High | High | High | 2.58E-05 | 0.000405 | 2.38 |
| FALSE | High | AYTSPLIDN | [K].AYTSPL | 1xPhosphc | Q9DBJ3 1x | 0.000167 | 0 | 1 | 1 | 2 | Q9DBJ3 | Brain-spec | Q9DBJ3 [2 | 0 | 2894.405 | 0 | 0.31 | 0.440727 | 30.96 | 92.9 | 107.1 | 39.87 | 18.96 | | High | High | High | High | High | High | High | High | High | High | High | 0 | 4.64E-05 | 2.78 |
| FALSE | High | CCSSDTNG | [K].CCSSDT | 2xCarbami | E9QAE4 1x | 0.003083 | 0.00016 | 1 | 2 | 1 | E9QAE4 | Histone-lys | E9QAE4 [8 | 0 | 2249.034 | 0 | -0.89 | 0.001006 | 1.65 | 129.9 | 70.1 | 7.82 | 9.82 | | High | High | High | High | High | High | High | High | High | High | High | 7.19E-05 | 0.000843 | 2.21 |
| FALSE | High | CDGSPRTP | [R].CDGSPI | 1xCarbami | F8WJE0 2x | 4.06E-06 | 0 | 1 | 9 | 3 | F8WJE0 | Deoxynucl | F8WJE0 [4 | 1 | 3605.545 | 0 | 0.99 | 0.003048 | 14.96 | 63.4 | 136.6 | 10.42 | 10.92 | | High | High | High | High | High | High | High | High | High | High | High | 0 | 4.88E-07 | 3.43 |
| FALSE | High | CDGSPRTP | [R].CDGSPI | 1xCarbami | F8WJE0 3x | 8.25E-05 | 0 | 1 | 9 | 3 | F8WJE0 | Deoxynucl | F8WJE0 [4 | 1 | 3685.512 | 0 | 0.67 | 0.013797 | 17.6 | 82.6 | 117.4 | 13.92 | 10.28 | | High | High | High | High | High | High | High | High | High | High | High | 0.000555 | 0.004808 | 3.11 |
| FALSE | High | CDHESSPG | [K].CDHESS | 1xCarbami | A2AR02 1x | 0.000179 | 0 | 1 | 2 | 3 | A2AR02 | Peptidyl-pr | A2AR02 [7: | 0 | 2014.847 | 0 | -0.12 | 0.244048 | 7.53 | 101.7 | 98.3 | 10.78 | 9.68 | | High | High | High | High | High | High | High | High | High | High | High | 0 | 3.45E-05 | 2.8 |
| FALSE | High | CDLSLSGGI | [K].CDLSLS | 1xCarbami | A6H630 1x | 1.09E-06 | 0 | 1 | 1 | 1 | A6H630 | Protein-glu | A6H630 [1 | 0 | 1992.935 | 0 | -0.68 | 0.027401 | 10.73 | 123.3 | 76.7 | 16.3 | 6.37 | | High | High | High | High | High | High | High | High | High | High | High | 0 | 1.12E-07 | 2.97 |
| FALSE | High | CDSADLR | [R].CDSADI | 1xCarbami | Q8K019 1x | 0.023379 | 0.001578 | 1 | 9 | 1 | Q8K019 | Bcl-2-assoc | Q8K019 [6 | 0 | 1145.486 | 0 | 0.25 | 0.035106 | 11.07 | 89.7 | 110.3 | 9.6 | 13.58 | | High | High | High | High | High | High | High | High | High | High | High | 0.000847 | 0.008122 | 1.88 |
| FALSE | High | CDSASSTPI | [R].CDSASS | 1xCarbami | Q5SVL6 1x | 0.012731 | 0.000889 | 1 | 3 | 1 | Q5SVL6 | Rap1 GTPa | Q5SVL6 [5 | 0 | 1490.696 | 0 | 3.33 | 0.001472 | 13.56 | 15.9 | 184.1 | 11.23 | 18.31 | | High | High | High | High | High | High | High | High | High | High | High | 0.000449 | 0.004128 | 2.03 |
| FALSE | High | CDSDMWII | [K].CDSDM | 1xCarbami | E9Q9C3 1x | 0.006085 | 0.00062 | 1 | 5 | 2 | E9Q9C3 | Afadin OS= | E9Q9C3 [1 | 0 | 3706.604 | 0 | -0.62 | 0.035629 | 1.17 | 120.4 | 79.6 | 11.62 | 22.57 | | High | High | High | High | High | High | High | High | High | High | High | 0.000275 | 0.001794 | 3.7 |
| FALSE | High | CDSDMWII | [K].CDSDM | 1xCarbami | E9Q9C3 1x | 0.023379 | 0.001578 | 1 | 5 | 1 | E9Q9C3 | Afadin OS= | E9Q9C3 [1 | 0 | 3721.615 | 0 | 0.6 | 0.001378 | 7.74 | 79.7 | 120.3 | 7.3 | 5.28 | | High | High | High | High | High | High | High | High | High | High | High | 0.000847 | 0.008165 | 3.56 |
| FALSE | High | CDSDPASP( | [R].CDSDP/ | 2xCarbami | K4DI61 1xI | 7.72E-07 | 0 | 1 | 4 | 2 | K4DI61 | PHD finger | K4DI61 [15 | 0 | 1954.898 | 0 | 0.11 | 0.386069 | 6.15 | 98.9 | 101.1 | 4.68 | 6.72 | | High | High | High | High | High | High | High | High | High | High | High | 0 | 7.57E-08 | 2.89 |
| FALSE | High | CDSEGEDE | [R].CDSEGI | 2xCarbami | B2RY61 1xI | 1.97E-05 | 0 | 1 | 4 | 2 | B2RY61 | AT rich inte | B2RY61 [5 | 0 | 2825.142 | 0 | 0.25 | 0.087971 | 11.06 | 93.4 | 106.6 | 9.89 | 7.69 | | High | High | High | High | High | High | High | High | High | High | High | 4.94E-05 | 0.000604 | 3.65 |
| FALSE | High | CDSLAQEG | [K].CDSLAC | 1xCarbami | A0A140T8S | 1.63E-06 | 0 | 1 | 1 | 2 | A0A140T8S | Chromosoi | A0A140T8S | 0 | 1811.881 | 0 | -0.36 | 0.117315 | 3.19 | 113.6 | 86.4 | 17.62 | 24.2 | | High | High | High | High | High | High | High | High | High | High | High | 0 | 5.89E-07 | 3.44 |
| FALSE | High | CDSLAQEG | [K].CDSLAC | 1xCarbami | A0A140T8S | 7.67E-08 | 0 | 1 | 1 | 4 | A0A140T8S | Chromosoi | A0A140T8S | 0 | 2041.044 | 0 | -0.97 | 0.004957 | 3.21 | 131.2 | 68.8 | 9.88 | 9.28 | | High | High | High | High | High | High | High | High | High | High | High | 0 | 5.67E-09 | 3.29 |
| FALSE | High | CEEDTEDEI | [R].CEEDTE | 1xCarbami | G3X8S2 1x | 2.84E-08 | 0 | 1 | 2 | 1 | G3X8S2 | BRF1 homc | G3X8S2 [3 | 0 | 2762.157 | 0 | 0.25 | 0.222283 | 20.41 | 92.5 | 107.5 | 12.28 | 10.55 | | High | High | High | High | High | High | High | High | High | High | High | 0 | 1.86E-09 | 5.62 |
| FALSE | High | CCSSSPHRI | [K].CCSSSP | 3xCarbami | Q8BRM2 1 | 0.000563 | 0 | 1 | 2 | 1 | Q8BRM2 | RAB6-inter | Q8BRM2 [: | 1 | 2482.108 | 0 | 1.27 | 0.005398 | 14.16 | 60.6 | 139.4 | 11.8 | 11.75 | | High | High | High | High | High | High | High | High | High | High | High | 0 | 0.000125 | 2.67 |
| FALSE | High | AYTPVVVT | [K].AYTPV\ | 1xPhosphc | P24788 1xI | 6.37E-06 | 0 | 1 | 4 | 3 | P24788 | Cyclin-dep | P24788 [5 | 0 | 1776.929 | 0 | -0.46 | 0.000729 | 1.84 | 114.8 | 85.2 | 6.22 | 6.09 | | High | High | High | High | High | High | High | High | High | High | High | 0 | 8.18E-07 | 3.86 |
| FALSE | High | AYTPPPPL( | [R].AYTPPF | 1xPhosphc | A0A0R4J0Y | 0.000108 | 0 | 1 | 6 | 2 | A0A0R4J0Y | Probable h | A0A0R4J0Y | 0 | 2194.183 | 0 | 0.74 | 0.000642 | 7.19 | 73.5 | 126.5 | 6.25 | 2.73 | | High | High | High | High | High | High | High | High | High | High | High | 0 | 1.96E-05 | 2.4 |
| FALSE | High | AYTHQVVT | [R].AYTHQ' | 1xPhosphc | Q03147 1x | 0.118334 | 0.009621 | 1 | 2 | 2 | Q03147 | Cyclin-dep | Q03147 [1 | 0 | 1383.698 | 0 | -0.44 | 0.093909 | 27.54 | 115.1 | 84.9 | 11.63 | 28.01 | | High | High | High | High | High | High | High | High | High | High | High | 0.004855 | 0.05262 | 1.49 |
| FALSE | High | AVVSPPK | [K].AVVSPI | 1xPhosphc | Q9ERU9 1x | 0.075927 | 0.005826 | 1 | 1 | 3 | Q9ERU9 | E3 SUMO-p | Q9ERU9 [2 | 0 | 1235.717 | 0 | 0.01 | 0.612252 | 3.48 | 100.7 | 99.3 | 7.21 | 10.63 | | High | High | High | High | High | High | High | High | High | High | High | 0.005122 | 0.05567 | 1.32 |
| FALSE | High | AVVVSPK | [R].AVVVSI | 1xPhosphc | Q3U3M5 1 | 0.015329 | 0.001089 | 1 | 2 | 2 | Q3U3M5 | Cyclin-K OS | Q3U3M5 [: | 0 | 1237.732 | 0 | -0.27 | 0.008366 | 6.62 | 106.5 | 93.5 | 4.67 | 4.58 | | High | High | High | High | High | High | High | High | High | High | High | 0.000997 | 0.01051 | 1.76 |
| FALSE | High | AVWPSTAI | [K].AVWPS | 1xCarbami | E9QKG8 1x | 6.41E-08 | 0 | 1 | 5 | 1 | E9QKG8 | Holliday ju | E9QKG8 [5 | 0 | 2481.261 | 0 | -0.78 | 0.000986 | 3.19 | 126.5 | 73.5 | 3.45 | 11.77 | | High | High | High | High | High | High | High | High | High | High | High | 0 | 4.65E-09 | 4.08 |
| FALSE | High | AVYQGPGS | [K].AVYQG | 1xPhosphc | O88587 1x | 0.024638 | 0.001612 | 1 | 2 | 3 | O88587 | Catechol O | O88587 [2 | 0 | 1727.913 | 0 | 0.07 | 0.016932 | 0.77 | 97.9 | 102.1 | 6.47 | 8.56 | | High | High | High | High | High | High | High | High | High | High | High | 0.002432 | 0.02609 | 1.6 |
| FALSE | High | AWDGQSF | [R].AWDG( | 1xPhosphc | A2BE28 1x | 0.031989 | 0.00205 | 1 | 2 | 1 | A2BE28 | Ribosomal | A2BE28 [7- | 0 | 1476.729 | 0 | -0.63 | 0.003407 | 3.62 | 118.6 | 81.4 | 8.9 | 6.7 | | High | High | High | High | High | High | High | High | High | High | High | 0.001126 | 0.0116 | 1.98 |
| FALSE | High | AWESGSSF | [R].AWESG | 1xPhosphc | Q8VIG1 1x | 5.16E-05 | 0 | 1 | 4 | 2 | Q8VIG1 | RE1-silenci | Q8VIG1 [1 | 0 | 2236.021 | 0 | -0.79 | 0.001888 | 4.6 | 128.5 | 71.5 | 4.93 | 14.85 | | High | High | High | High | High | High | High | High | High | High | High | 0 | 8.51E-06 | 2.92 |
| FALSE | High | AWLDEDSN | [R].AWLDE | 1xPhosphc | Q99PT3 1x | 0.000213 | 0 | 1 | 3 | 2 | Q99PT3 | INO80 com | Q99PT3 [1 | 0 | 2008.958 | 0 | -0.32 | 0.017249 | 3.95 | 105.1 | 94.9 | 8.02 | 15.45 | | High | High | High | High | High | High | High | High | High | High | High | 0 | 0.000042 | 2.65 |
| FALSE | High | AWLDEDSN | [R].AWLDE | 2xPhosphc | Q99PT3 2x | 0.00311 | 0.00016 | 1 | 3 | 4 | Q99PT3 | INO80 com | Q99PT3 [1 | 0 | 2088.924 | 0 | -0.16 | 0.226421 | 8.87 | 103.4 | 96.6 | 3.83 | 8.79 | | High | High | High | High | High | High | High | High | High | High | High | 0.000117 | 0.001066 | 3.62 |
| FALSE | High | AWSSQDE\ | [K].AWSSQ | 1xPhosphc | Q4VAC9 1x | 0.000134 | 0 | 1 | 3 | 2 | Q4VAC9 | Pleckstrin I | Q4VAC9 [4 | 0 | 1796.817 | 0 | 1.26 | 0.003731 | 2.41 | 58.3 | 141.7 | 5.92 | 15.49 | | High | High | High | High | High | High | High | High | High | High | High | 0 | 5.16E-05 | 3.41 |
| FALSE | High | AYFNDSSEI | [R].AYFNDS | 1xDeamida | P47810 1xI | 3.09E-05 | 0 | 1 | 1 | 2 | P47810 | Wee1-like | P47810 [2 | 0 | 3485.466 | 0 | -1.28 | 0.000209 | 1.8 | 143.2 | 56.8 | 2.9 | 7 | | High | High | High | High | High | High | High | High | High | High | High | 0 | 8.28E-06 | 3.96 |
| FALSE | High | AYFNDSSEI | [R].AYFNDS | 1xPhosphc | P47810 1xI | 0.014538 | 0.001011 | 1 | 1 | 2 | P47810 | Wee1-like | P47810 [2 | 0 | 3500.477 | 0 | -1.38 | 0.001422 | 0.88 | 141.5 | 58.5 | 8.1 | 14.09 | | High | High | High | High | High | High | High | High | High | High | High | 0.000872 | 0.008911 | 3.39 |
| FALSE | High | AYGLLSPCI | [K].AYGLLS | 1xCarbami | Q8BIX3 1xI | 3.06E-05 | 0 | 1 | 1 | 2 | Q8BIX3 | ARL14 effe | Q8BIX3 [2 | 0 | 2413.239 | 0 | -0.31 | 0.000928 | 2.53 | 110.1 | 89.9 | 5.4 | 9.04 | | High | High | High | High | High | High | High | High | High | High | High | 0 | 4.73E-06 | 2.22 |
| FALSE | High | AYGSQEPL | [R].AYGSQ | 1xPhosphc | A7MCV2 1: | 0.002376 | 0.000109 | 1 | 2 | 2 | A7MCV2 | DEP domai | A7MCV2 [4 | 0 | 2670.368 | 0 | -1.1 | 0.004104 | 9.84 | 133.6 | 66.4 | 7.86 | 51.69 | | High | High | High | High | High | High | High | High | High | High | High | 4.94E-05 | 0.000629 | 2.62 |
| FALSE | High | AYGVDVGS | [K].AYGVD | 1xPhosphc | F8VQ05 1x | 0.001127 | 0 | 1 | 5 | 1 | F8VQ05 | FRY-like tra | F8VQ05 [2 | 0 | 2078.061 | 0 | 1.98 | 0.000213 | 2.27 | 39.9 | 160.1 | 7.11 | 4.74 | | High | High | High | High | High | High | High | High | High | High | High | 0 | 0.000274 | 3.38 |
| FALSE | High | AYLPDGYEI | [R].AYLPD( | 1xCarbami | Q8BHG9 1: | 8.18E-05 | 0 | 1 | 1 | 1 | Q8BHG9 | CGG triplet | Q8BHG9 [1 | 0 | 2525.074 | 0 | 0.05 | 0.490412 | 12.11 | 95 | 105 | 4.32 | 9.05 | | High | High | High | High | High | High | High | High | High | High | High | 0 | 1.43E-05 | 2.33 |
| FALSE | High | AYLQSITPC | [R].AYLQSI | 1xPhosphc | A0A0M3HE | 0.000183 | 0 | 1 | 7 | 2 | A0A0M3HE | Zinc finger | A0A0M3HE | 0 | 2253.027 | 0 | 0.63 | 0.124703 | 31.6 | 78.7 | 121.3 | 20.23 | 17.14 | | High | High | High | High | High | High | High | High | High | High | High | 0 | 3.52E-05 | 3.1 |
| FALSE | High | AYPSYSPPE | [K].AYPSYS | 1xPhosphc | Q7TS75 1x | 0.00204 | 5.66E-05 | 1 | 1 | 2 | Q7TS75 | APC memb | Q7TS75 [7 | 0 | 2677.185 | 0 | 0.42 | 0.009854 | 13.56 | 80.6 | 119.4 | 11.33 | 7.31 | | High | High | High | High | High | High | High | High | High | High | High | 2.58E-05 | 0.000529 | 3.15 |
| FALSE | High | AYQHGGV1 | [R].AYQHG | 1xPhosphc | Q9EPU0 1x | 0.067619 | 0.004864 | 1 | 2 | 1 | Q9EPU0 | Regulator c | Q9EPU0 [1 | 0 | 1689.784 | 0 | 0.07 | 0.907614 | 20.82 | 91.9 | 108.1 | 14.18 | 12.09 | | High | High | High | High | High | High | High | High | High | High | High | 0.00253 | 0.02739 | 2.28 |
| FALSE | High | AYSFCGTVI | [K].AYSFCC | 1xCarbami | E9QK39 1x | 2.36E-06 | 0 | 3 | 20 | 8 | E9QK39; A | Ribosomal | E9QK39 [3 | 0 | 2402.076 | 0 | 0.25 | 0.001727 | 5.02 | 92.5 | 107.5 | 5.98 | 3.16 | Shared | High | High | High | High | High | High | High | High | High | High | High | 0 | 1.75E-06 | 4.55 |
| FALSE | High | AYSFCGTVI | [K].AYSFCC | 1xCarbami | E9QK39 1x | 2.15E-08 | 0 | 3 | 20 | 59 | E9QK39; A | Ribosomal | E9QK39 [3 | 0 | 2418.071 | 0 | -0.12 | 0.053051 | 6.02 | 107 | 93 | 6.36 | 3.6 | Shared | High | High | High | High | High | High | High | High | High | High | High | 0 | 7.43E-09 | 4.83 |
| FALSE | High | AYSFCGTVI | [K].AYSFCC | 1xCarbami | E9QK39 1x | 0.085162 | 0.006376 | 3 | 20 | 2 | E9QK39; A | Ribosomal | E9QK39 [3 | 0 | 2419.055 | 0 | 0.23 | 0.251786 | 24.64 | 89.8 | 110.2 | 18.18 | 11.31 | Shared | High | High | High | High | High | High | High | High | High | High | High | 0.006622 | 0.07148 | 2.46 |
| FALSE | High | AYSGSDLPS | [R].AYSGSI | 1xPhosphc | Q71FD5 1x | 4.09E-09 | 0 | 1 | 1 | 6 | Q71FD5 | E3 ubiquiti | Q71FD5 [1 | 0 | 2205.961 | 0 | -0.98 | 0.000831 | 3.37 | 132.7 | 67.3 | 3.73 | 10.89 | | High | High | High | High | High | High | High | High | High | High | High | 0 | 2.1E-10 | 4.34 |
| FALSE | High | AYSGSQIDI | [R].AYSGS( | 1xPhosphc | H3BKH6 1x | 3.68E-06 | 0 | 1 | 6 | 5 | H3BKH6 | S-formylglu | H3BKH6 [2 | 0 | 2146.12 | 0 | 0.77 | 0.00062 | 7.4 | 73.9 | 126.1 | 4.6 | 4.44 | | High | High | High | High | High | High | High | High | High | High | High | 0 | 3.75E-06 | 3.36 |
| FALSE | High | AYSNSIVR | [R].AYSNSI | 1xPhosphc | Q8K412 1x | 0.113753 | 0.009241 | 1 | 3 | 1 | Q8K412 | Memory-re | Q8K412 [7 | 0 | 1218.608 | 0 | 1.55 | 0.00041 | 15.06 | 49.3 | 150.7 | 8.77 | 7.54 | | High | High | High | High | High | High | High | High | High | High | High | 0.004667 | 0.05032 | 1.71 |
| FALSE | High | AYSPLGGG | [R].AYSPLG | 1xCarbami | A0A0R4J0E | 1.34E-10 | 0 | 1 | 2 | 2 | A0A0R4J0E | Store-oper | A0A0R4J0E | 0 | 2357.007 | 0 | -0.19 | 0.160995 | 11.65 | 101.8 | 98.2 | 9.77 | 12.43 | | High | High | High | High | High | High | High | High | High | High | High | 0 | 4.55E-12 | 4.8 |
| FALSE | High | AYSPTCSPT | [K].AYSPTC | 1xCarbami | Q6PAL7 1x | 0.000198 | 0 | 1 | 1 | 2 | Q6PAL7 | AT-hook D | Q6PAL7 [1: | 0 | 1966.975 | 0 | -1.13 | 0.001653 | 2 | 137.3 | 62.7 | 8.36 | 6.61 | | High | High | High | High | High | High | High | High | High | High | High | 0 | 3.87E-05 | 3.21 |
| FALSE | High | AYSPTCSPT | [K].AYSPTC | 1xCarbami | Q6PAL7 2x | 0.022574 | 0.001542 | 1 | 1 | 3 | Q6PAL7 | AT-hook D | Q6PAL7 [1: | 0 | 2046.941 | 0 | -1.47 | 0.000237 | 0.74 | 146.1 | 53.9 | 3.27 | 8.94 | | High | High | High | High | High | High | High | High | High | High | High | 0.000814 | 0.007834 | 2.76 |
| FALSE | High | AYSQEEITC | [R].AYSQEI | 1xPhosphc | G5E8C3 1x | 0.001437 | 5.66E-05 | 1 | 2 | 1 | G5E8C3 | G protein-c | G5E8C3 [3 | 0 | 4266.026 | 0 | 2.15 | 0.01815 | 52.78 | 36.1 | 163.9 | 29.39 | 28.26 | | High | High | High | High | High | High | High | High | High | High | High | 2.58E-05 | 0.000358 | 3.49 |
| FALSE | High | AYSSPSTTP | [K].AYSSPS | 1xPhosphc | Q8CBY1 1x | 0.0002 | 0 | 1 | 4 | 4 | Q8CBY1 | Protein Sm | Q8CBY1 [4 | 0 | 1603.757 | 0 | 2.26 | 0.001869 | 21.9 | 34.5 | 165.5 | 7 | 17.46 | | High | High | High | High | High | High | High | High | High | High | High | 0.000399 | 0.00338 | 2.7 |
| FALSE | High | CEERSPSFC | [R].CEERSP | 2xCarbami | O35492 1x | 0.001039 | 0 | 1 | 4 | 1 | O35492 | Dual specif | O35492 [2 | 1 | 2331.921 | 0 | -0.34 | 0.017942 | 7.99 | 111.7 | 88.3 | 5.99 | 6.97 | | High | High | High | High | High | High | High | High | High | High | High | 0 | 0.00025 | 2.61 |
| FALSE | High | CSVSAVNLI | [R].CSVSAV | 1xCarbami | A2AJI0 1xP | 0.012397 | 0.000889 | 1 | 2 | 1 | A2AJI0 | MAP7 dom | A2AJI0 [25 | 0 | 1612.853 | 0 | 1.08 | 0.001515 | 6.44 | 66.5 | 133.5 | 12.04 | 11.09 | | High | High | High | High | High | High | High | High | High | High | High | 0.000449 | 0.003972 | 2.21 |
| FALSE | High | CSVSLSNVE | [R].CSVSLS | 1xCarbami | P47713 1xI | 7.41E-05 | 0 | 1 | 2 | 3 | P47713 | Cytosolic p | P47713 [72 | 0 | 1530.718 | 0 | 0.1 | 0.909251 | 50.96 | 106.8 | 93.2 | 16.95 | 29.17 | | High | High | High | High | High | High | High | High | High | High | High | 0.004032 | 0.04275 | 2.35 |
| FALSE | High | CTAPLTPSS | [K].CTAPLT | 1xCarbami | G5E8R4 1x | 4.31E-05 | 0 | 1 | 3 | 7 | G5E8R4 | SAPS doma | G5E8R4 [8 | 0 | 1839.851 | 0 | -0.07 | 0.325228 | 11.51 | 101.7 | 98.3 | 4.81 | 6.77 | | High | High | High | High | High | High | High | High | High | High | High | 0 | 2.59E-05 | 3.19 |
| FALSE | High | DEEADSDN | [K].DEEADS | 1xPhosphc | Q8C167 1x | 4.76E-05 | 0 | 1 | 5 | 2 | Q8C167 | Prolyl endc | Q8C167 [1 | 0 | 2810.306 | 0 | 0.53 | 0.079392 | 19.22 | 88.7 | 111.3 | 17.43 | 15.46 | | High | High | High | High | High | High | High | High | High | High | High | 0 | 7.78E-06 | 4.1 |
| FALSE | High | DEEEAFLD\ | [R].DEEEAF | 1xPhosphc | Q80Y44 1x | 1.99E-06 | 0 | 1 | 3 | 8 | Q80Y44 | Probable A | Q80Y44 [7 | 0 | 3350.362 | 0 | -0.77 | 0.000377 | 2 | 125.1 | 74.9 | 5.31 | 9.82 | | High | High | High | High | High | High | High | High | High | High | High | 0 | 2.2E-07 | 3.88 |
| FALSE | High | DEEEAFLD\ | [R].DEEEAF | 1xPhosphc | Q80Y44 1x | 5.7E-05 | 0 | 1 | 3 | 2 | Q80Y44 | Probable A | Q80Y44 [7 | 0 | 3579.525 | 0 | -0.54 | 0.015784 | 10.34 | 121.1 | 78.9 | 10.04 | 16.76 | | High | High | High | High | High | High | High | High | High | High | High | 0 | 9.59E-06 | 3.58 |
| FALSE | High | DEENDALL( | [K].DEEND/ | 1xPhosphc | P33609 1xI | 0.037048 | 0.00249 | 1 | 1 | 1 | P33609 | DNA polym | P33609 [12 | 0 | 2582.227 | 0 | -0.46 | 0.039038 | 9.31 | 113.9 | 86.1 | 9.49 | 13.42 | | High | High | High | High | High | High | High | High | High | High | High | 0.001308 | 0.01366 | 1.92 |
| FALSE | High | DEEPTCQS | [K].DEEPTC | 1xCarbami | Q3UW53 1 | 6.73E-06 | 0 | 1 | 1 | 4 | Q3UW53 | Protein Nit | Q3UW53 [ | 0 | 2566.215 | 0 | 1.49 | 0.000555 | 4.54 | 52.7 | 147.3 | 10.29 | 10.08 | | High | High | High | High | High | High | High | High | High | High | High | 0 | 8.7E-07 | 3.69 |
| FALSE | High | DEESTGDE | [R].DEESTC | 1xPhosphc | E9QK83 1x | 3.81E-09 | 0 | 1 | 3 | 1 | E9QK83 | Small subu | E9QK83 [7 | 0 | 3836.743 | 0 | -1.09 | 0.003141 | 5.95 | 137.1 | 62.9 | 10.37 | 9.22 | | High | High | High | High | High | High | High | High | High | High | High | 0 | 1.94E-10 | 6.27 |
| FALSE | High | DEETEDVL/ | [K].DEETEI | 1xPhosphc | Q8CI71 1xI | 0.11108 | 0.008991 | 1 | 3 | 1 | Q8CI71 | Syndetin O | Q8CI71 [53 | 0 | 2596.066 | 0 | -0.97 | 0.005976 | 6.2 | 134.9 | 65.1 | 13.71 | 18.17 | | High | High | High | High | High | High | High | High | High | High | High | 0.004503 | 0.04873 | 1.97 |

| | | | | | | | | | | | | | | | | | | | | | | | | | | | | | | | | | | | | | | |
|---|---|---|---|---|---|---|---|---|---|---|---|---|---|---|---|---|---|---|---|---|---|---|---|---|---|---|---|---|---|---|---|---|---|---|---|---|---|---|
| FALSE | High | DEETEDVL | [K].DEETED | 1xPhospho | Q8CI71 1xP | 2.36E-09 | 0 | 1 | 3 | 2 | Q8CI71 | Syndetin O | Q8CI71 [53 | 0 | 2825.229 | 0 | 0.55 | 0.028781 | 6.71 | 81.5 | 118.5 | 13.67 | 8.76 | | High | High | High | High | High | High | High | High | High | High | High | 0 | 6.86E-09 | 5.27 |
| FALSE | High | DEETEDVL | [K].DEETED | 1xDeamida | Q8CI71 1xP | 9.42E-07 | 0 | 1 | 3 | 3 | Q8CI71 | Syndetin O | Q8CI71 [53 | 0 | 2826.213 | 0 | 0.36 | 0.104253 | 20.9 | 87.8 | 112.2 | 20.38 | 6.9 | | High | High | High | High | High | High | High | High | High | High | High | 0 | 1.15E-05 | 4.03 |
| FALSE | High | DEFTTADV | [K].DEFTTA | 1xPhospho | Q9QZM9 1 | 0.000772 | 0 | 1 | 3 | 1 | Q9QZM9 | F-box only | Q9QZM9 [ | 0 | 2598.238 | 0 | -0.41 | 0.039052 | 11.88 | 114 | 86 | 7.87 | 7.77 | | High | High | High | High | High | High | High | High | High | High | High | 0 | 0.000178 | 3.2 |
| FALSE | High | DEGGNDQ | [R].DEGGN | 1xCarbami | Q8BKR5 1x | 2.19E-05 | 0 | 1 | 2 | 10 | Q8BKR5 | Protein ph | Q8BKR5 [5 | 1 | 4665.155 | 0 | -0.22 | 0.936352 | 6.01 | 99.2 | 100.8 | 21.33 | 14.16 | | High | High | High | High | High | High | High | High | High | High | High | 2.58E-05 | 0.00054 | 4.12 |
| FALSE | High | DEGGNDQ | [R].DEGGN | 1xCarbami | Q8BKR5 2x | 2.38E-05 | 0 | 1 | 2 | 2 | Q8BKR5 | Protein ph | Q8BKR5 [5 | 1 | 4744.137 | 0 | 0.87 | 0.012602 | 30.64 | 75.3 | 124.7 | 25.34 | 18.94 | | High | High | High | High | High | High | High | High | High | High | High | 0 | 3.58E-06 | 5.29 |
| FALSE | High | DEGIGSPD | [K].DEGIGS | 1xPhospho | Q9JIZ5 1xP | 2.28E-07 | 0 | 1 | 2 | 2 | Q9JIZ5 | Activator p | Q9JIZ5 [13 | 0 | 2127.989 | 0 | 0.61 | 0.00163 | 6.57 | 78.9 | 121.1 | 8.94 | 11.5 | | High | High | High | High | High | High | High | High | High | High | High | 0 | 1.94E-08 | 2.83 |
| FALSE | High | DEDSSECS | [K].DEDSSE | 1xCarbami | Q80U04 3x | 0.011549 | 0.00086 | 1 | 1 | 1 | Q80U04 | E3 ubiquiti | Q80U04 [4 | 0 | 2632.914 | 0 | 0.03 | 0.452535 | 13.71 | 99.6 | 100.4 | 4.77 | 8.33 | | High | High | High | High | High | High | High | High | High | High | High | 0.000437 | 0.003673 | 3.5 |
| FALSE | High | DEGSDIDD | [K].DEGSDI | 1xPhospho | Q810A4 1x | 2.08E-06 | 0 | 1 | 3 | 14 | Q810A4 | RNA helica | Q810A4 [5 | 0 | 2536.012 | 0 | -0.1 | 0.467428 | 21.09 | 105 | 95 | 17.69 | 9.27 | | High | High | High | High | High | High | High | High | High | High | High | 0 | 2.31E-07 | 4.74 |
| FALSE | High | DEGSYSLE | [K].DEGSYS | 1xPhospho | Q3THY2 1x | 0.000225 | 0 | 1 | 3 | 6 | Q3THY2 | Syndecan ( | Q3THY2 [2 | 0 | 1791.845 | 0 | 1.24 | 0.002195 | 7.33 | 59.3 | 140.7 | 10.04 | 16.57 | | High | High | High | High | High | High | High | High | High | High | High | 0.000327 | 0.002848 | 3.8 |
| FALSE | High | DEHGLEPA | [K].DEHGLI | 1xPhospho | E9PZM4 1x | 0.001865 | 5.66E-05 | 1 | 2 | 1 | E9PZM4 | Chromodo | E9PZM4 [1 | 0 | 1516.699 | 0 | -0.31 | 0.002231 | 2.23 | 110.5 | 89.5 | 4.83 | 2.79 | | High | High | High | High | High | High | High | High | High | High | High | 2.58E-05 | 0.000482 | 3.34 |
| FALSE | High | DELADEIAN | [R].DELADE | 1xPhospho | Q8VDD5 1x | 1.08E-05 | 0 | 1 | 5 | 3 | Q8VDD5 | Myosin-9 ( | Q8VDD5 [1 | 0 | 1886.915 | 0 | 0.5 | 0.002429 | 10.75 | 78.7 | 121.3 | 11.76 | 5.28 | | High | High | High | High | High | High | High | High | High | High | High | 0 | 1.47E-06 | 3.51 |
| FALSE | High | DEIDEFANS | [K].DEIDEF | 1xPhospho | A2AJK6 1xP | 3.61E-06 | 0 | 1 | 2 | 2 | A2AJK6 | Chromodo | A2AJK6 [18 | 0 | 2117.968 | 0 | -0.69 | 0.037488 | 8.74 | 123.6 | 76.4 | 22.04 | 8.69 | | High | High | High | High | High | High | High | High | High | High | High | 0 | 6.07E-07 | 4.19 |
| FALSE | High | DELEMQQ | [K].DELEM | 1xDeamida | Q3UIK4 1x | 0.020319 | 0.001472 | 1 | 4 | 1 | Q3UIK4 | N6-adenos | Q3UIK4 [8 | 0 | 2868.226 | 0 | -0.05 | 0.561785 | 75.15 | 122.2 | 77.8 | 32.73 | 51.56 | | High | High | High | High | High | High | High | High | High | High | High | 0.000786 | 0.006946 | 2.79 |
| FALSE | High | DELHIVEAE | [K].DELHIV | 1xPhospho | Q5SQB7 1x | 2.38E-05 | 0 | 2 | 9 | 3 | Q5SQB7; C | MCG68069 | Q5SQB7 [5 | 0 | 2470.141 | 0 | -0.36 | 0.407594 | 7.44 | 111.5 | 88.5 | 29.82 | 26.98 | Shared | High | High | High | High | High | High | High | High | High | High | High | 0 | 3.57E-06 | 3.44 |
| FALSE | High | DELHIVEAE | [K].DELHIV | 1xPhospho | Q5SQB7 1x | 9.48E-13 | 0 | 2 | 9 | 110 | Q5SQB7; C | MCG68069 | Q5SQB7 [5 | 0 | 2683.309 | 0 | -0.86 | 0.001189 | 2.9 | 127.3 | 72.7 | 4.2 | 12.49 | Shared | High | High | High | High | High | High | High | High | High | High | High | 0 | 4.19E-12 | 5.16 |
| FALSE | High | DELHIVEAE | [K].DELHIV | 1xDeamida | Q5SQB7 1x | 5.56E-09 | 0 | 2 | 9 | 3 | Q5SQB7; C | MCG68069 | Q5SQB7 [5 | 0 | 2684.293 | 0 | -0.61 | 0.005266 | 2.02 | 121.4 | 78.6 | 6.89 | 11.5 | Shared | High | High | High | High | High | High | High | High | High | High | High | 0 | 2.99E-10 | 4.33 |
| FALSE | High | DELHIVEAE | [K].DELHIV | 1xPhospho | Q5SQB7 1x | 3.41E-12 | 0 | 2 | 9 | 52 | Q5SQB7; C | MCG68069 | Q5SQB7 [5 | 0 | 2699.304 | 0 | -0.79 | 0.000273 | 2.24 | 126.9 | 73.1 | 2.6 | 3.87 | Shared | High | High | High | High | High | High | High | High | High | High | High | 0 | 3.57E-11 | 4.91 |
| FALSE | High | DELKPDDL | [K].DELKPD | 1xCarbami | A2AQE2 1x | 0.000221 | 0 | 1 | 2 | 2 | A2AQE2 | SECIS bindi | A2AQE2 [9 | 0 | 3462.577 | 0 | 0.46 | 0.001242 | 4.16 | 83.5 | 116.5 | 9.24 | 4.93 | | High | High | High | High | High | High | High | High | High | High | High | 0 | 4.37E-05 | 3.92 |
| FALSE | High | DELKPDDL | [K].DELKPD | 1xCarbami | A2AQE2 1x | 0.120211 | 0.009851 | 1 | 2 | 2 | A2AQE2 | SECIS bindi | A2AQE2 [9 | 0 | 3463.561 | 0 | 0.62 | 0.003857 | 8.27 | 85.5 | 114.5 | 16.11 | 13.13 | | High | High | High | High | High | High | High | High | High | High | High | 0.004935 | 0.0537 | 3.44 |
| FALSE | High | DEILPTTPIS | [K].DEILPT | 1xPhospho | Q3UCL7 1x | 0.023792 | 0.001612 | 1 | 5 | 4 | Q3UCL7 | Putative ur | Q3UCL7 [2 | 0 | 1779.898 | 0 | 0.01 | 0.870826 | 37.06 | 109.7 | 90.3 | 88.66 | 78.82 | | High | High | High | High | High | High | High | High | High | High | High | 0.001999 | 0.02036 | 3.04 |
| FALSE | High | DEILPTTPIS | [K].DEILPT | 1xPhospho | Q3UCL7 1x | 2.52E-07 | 0 | 1 | 5 | 13 | Q3UCL7 | Putative ur | Q3UCL7 [2 | 0 | 2009.061 | 0 | 0.14 | 0.006832 | 3.07 | 93.7 | 106.3 | 2.6 | 3.2 | | High | High | High | High | High | High | High | High | High | High | High | 0 | 2.17E-08 | 4.03 |
| FALSE | High | DEGSSDLA | [K].DEGSSI | 1xPhospho | Q3U9K6 1x | 0.000492 | 0 | 1 | 4 | 1 | Q3U9K6 | Interferon | Q3U9K6 [1 | 0 | 4370.159 | 0 | 0.65 | 0.010061 | 24.17 | 73.9 | 126.1 | 9.21 | 11.77 | | High | High | High | High | High | High | High | High | High | High | High | 0 | 0.000107 | 3.32 |
| FALSE | High | DEILPTTPIS | [K].DEILPT | 2xPhospho | Q3UCL7 2x | 0.00011 | 0 | 1 | 5 | 1 | Q3UCL7 | Putative ur | Q3UCL7 [2 | 0 | 2089.027 | 0 | 0.12 | 0.657871 | 4.94 | 99.8 | 100.2 | 9.01 | 8.36 | | High | High | High | High | High | High | High | High | High | High | High | 0 | 0.00002 | 2.22 |
| FALSE | High | DEDSDDES | [K].DEDSDI | 1xPhospho | Q8CCJ3 1xP | 3.55E-06 | 0 | 1 | 2 | 4 | Q8CCJ3 | E3 UFM1-p | Q8CCJ3 [45 | 0 | 2130.886 | 0 | -0.08 | 0.259749 | 10.59 | 101.1 | 98.9 | 6.05 | 4.8 | | High | High | High | High | High | High | High | High | High | High | High | 0 | 4.26E-07 | 4.66 |
| FALSE | High | DEDMLYSP | [R].DEDML | 1xPhospho | Q3U6Q1 1x | 0.033986 | 0.002195 | 1 | 4 | 1 | Q3U6Q1 | Putative ur | Q3U6Q1 [2 | 0 | 1875.84 | 0 | -0.17 | 0.158899 | 22.95 | 109.1 | 90.9 | 19.69 | 16.47 | | High | High | High | High | High | High | High | High | High | High | High | 0.001151 | 0.01243 | 2.48 |
| FALSE | High | DDLQAAIA | [K].DDLQA | 1xPhospho | Q5I043 1xP | 2.6E-08 | 0 | 1 | 4 | 2 | Q5I043 | Ubiquitin c | Q5I043 [10 | 0 | 2676.344 | 0 | -1.27 | 0.000409 | 2.16 | 142 | 58 | 7.88 | 14.65 | | High | High | High | High | High | High | High | High | High | High | High | 0 | 1.69E-09 | 4.04 |
| FALSE | High | DDISEIQSL | [R].DDISEI | 1xPhospho | P39447 1xP | 5.79E-07 | 0 | 1 | 3 | 3 | P39447 | Tight junct | P39447 [28 | 0 | 2038.928 | 0 | 0.34 | 0.113817 | 26.33 | 90.9 | 109.1 | 7.41 | 12.46 | | High | High | High | High | High | High | High | High | High | High | High | 0 | 5.47E-08 | 4.4 |
| FALSE | High | DDISEIQSL | [R].DDISEI | 2xPhospho | P39447 2xP | 3.15E-06 | 0 | 1 | 3 | 4 | P39447 | Tight junct | P39447 [28 | 0 | 2118.894 | 0 | 0.53 | 0.012129 | 18.37 | 81.8 | 118.2 | 5.49 | 10.59 | | High | High | High | High | High | High | High | High | High | High | High | 0 | 3.69E-07 | 4.27 |
| FALSE | High | DDISEIQSL | [R].DDISEI | 3xPhospho | P39447 3xP | 8.33E-05 | 0 | 1 | 3 | 2 | P39447 | Tight junct | P39447 [28 | 0 | 2198.861 | 0 | -0.06 | 0.020213 | 2.14 | 102.1 | 97.9 | 2.85 | 2.55 | | High | High | High | High | High | High | High | High | High | High | High | 0 | 1.46E-05 | 2.77 |
| FALSE | High | DDISEIQSL | [R].DDISEI | 3xPhospho | P39447 3xP | 0.002576 | 0.000109 | 1 | 3 | 1 | P39447 | Tight junct | P39447 [28 | 1 | 3044.286 | 0 | 0.17 | 0.096113 | 20.67 | 88.4 | 111.6 | 8.81 | 7.72 | | High | High | High | High | High | High | High | High | High | High | High | 4.94E-05 | 0.000692 | 2.85 |
| FALSE | High | DDLSPTAEV | [K].DDLSPT | 1xPhospho | Q9ERG2 1x | 5.55E-05 | 0 | 1 | 3 | 4 | Q9ERG2 | Striatin-3 C | Q9ERG2 [3 | 0 | 2626.305 | 0 | 1.35 | 0.002667 | 2.14 | 57.4 | 142.6 | 12.92 | 11.33 | | High | High | High | High | High | High | High | High | High | High | High | 0 | 9.27E-06 | 4.1 |
| FALSE | High | DDNEDNSN | [K].DDNED | 1xCarbami | K4DI61 1xP | 0.001011 | 0 | 1 | 4 | 1 | K4DI61 | PHD finger | K4DI61 [29 | 0 | 2255.928 | 0 | 0.23 | 0.044401 | 13.08 | 87.7 | 112.3 | 7.05 | 10.32 | | High | High | High | High | High | High | High | High | High | High | High | 0 | 0.000243 | 1.53 |
| FALSE | High | DDNNNDE | [K].DDNNN | 1xDeamida | P97500 2xP | 0.021231 | 0.001507 | 1 | 3 | 1 | P97500 | Myelin trar | P97500 [18 | 0 | 2435.9 | 0 | -0.12 | 0.497617 | 25.16 | 100.5 | 99.5 | 15.87 | 13.49 | | High | High | High | High | High | High | High | High | High | High | High | 0.000801 | 0.007313 | 2.22 |
| FALSE | High | DDPFAQPD | [R].DDPFA | 1xPhospho | A0A0R4J1C | 0.004333 | 0.00036 | 1 | 5 | 3 | A0A0R4J1C | Disabled h | A0A0R4J1C | 0 | 3118.429 | 0 | 2.23 | 0.002019 | 54.55 | 39.7 | 160.3 | 43.22 | 11.47 | | High | High | High | High | High | High | High | High | High | High | High | 0.000161 | 0.001236 | 3.76 |
| FALSE | High | DDPHLSSVI | [K].DDPHLS | 1xPhospho | Q3TKY6 1x | 0.000233 | 0 | 1 | 1 | 1 | Q3TKY6 | Peptidyl-pr | Q3TKY6 [2 | 0 | 2148.063 | 0 | -0.23 | 0.107945 | 4.72 | 107.3 | 92.7 | 10.49 | 8.41 | | High | High | High | High | High | High | High | High | High | High | High | 0 | 4.66E-05 | 3.39 |
| FALSE | High | DDSELDFS | [K].DDSELD | 1xCarbami | Q8VE91 1x | 0.000958 | 0 | 1 | 4 | 2 | Q8VE91 | Reticuloph | Q8VE91 [2 | 0 | 2034.95 | 0 | 2.56 | 0.024719 | 60.32 | 27.5 | 172.5 | 10.67 | 45.47 | | High | High | High | High | High | High | High | High | High | High | High | 0 | 0.000232 | 2.82 |
| FALSE | High | DDSHSAED | [K].DDSHS | 2xPhospho | Q80XU3 2x | 0.003253 | 0.00016 | 1 | 2 | 1 | Q80XU3 | Nuclear ub | Q80XU3 [7 | 0 | 2081.799 | 0 | -0.13 | 0.445109 | 13.03 | 98.3 | 101.7 | 16.15 | 28.1 | | High | High | High | High | High | High | High | High | High | High | High | 7.19E-05 | 0.000893 | 2.79 |
| FALSE | High | DDSLDAGN | [K].DDSLD | 1xPhospho | A2ANY6 1x | 0.003083 | 0.00016 | 1 | 2 | 2 | A2ANY6 | Midasin OS | A2ANY6 [4 | 0 | 1470.667 | 0 | -0.71 | 0.101692 | 13.38 | 125.1 | 74.9 | 25.59 | 26.64 | | High | High | High | High | High | High | High | High | High | High | High | 0.001151 | 0.01249 | 2.13 |
| FALSE | High | DEDMLYSP | [R].DEDML | 1xPhospho | Q3U6Q1 1x | 5.07E-05 | 0 | 1 | 4 | 2 | Q3U6Q1 | Putative ur | Q3U6Q1 [2 | 0 | 1891.835 | 0 | 0.14 | 0.073936 | 11.37 | 98 | 102 | 9.29 | 9.97 | | High | High | High | High | High | High | High | High | High | High | High | 0 | 9.16E-06 | 3.07 |
| FALSE | High | DDSLDAGN | [K].DDSLD | 1xPhospho | A2ANY6 1x | 3.14E-05 | 0 | 1 | 2 | 2 | A2ANY6 | Midasin OS | A2ANY6 [4 | 0 | 1699.83 | 0 | -0.86 | 0.000162 | 0.89 | 130.1 | 69.9 | 2.86 | 5.94 | | High | High | High | High | High | High | High | High | High | High | High | 0 | 4.91E-06 | 3.66 |
| FALSE | High | DDSLDLSP | [R].DDSLDI | 2xPhospho | G5E870 2x | 0.038677 | 0.002563 | 1 | 3 | 2 | G5E870 | E3 ubiquiti | G5E870 [1 | 1 | 2476.214 | 0 | -1.39 | 0.00043 | 1.92 | 143.8 | 56.2 | 6.33 | 8.21 | | High | High | High | High | High | High | High | High | High | High | High | 0.001388 | 0.01442 | 2.23 |
| FALSE | High | DDSNSTQE | [R].DDSNS | 1xPhospho | Q8BLH7 1x | 1.89E-07 | 0 | 1 | 1 | 6 | Q8BLH7 | HIRA-intera | Q8BLH7 [3 | 0 | 2290.06 | 0 | -0.8 | 0.001125 | 4.57 | 128.3 | 71.7 | 4.07 | 8.79 | | High | High | High | High | High | High | High | High | High | High | High | 0 | 1.57E-07 | 3.11 |
| FALSE | High | DDSPTQTL | [K].DDSPT | 1xCarbami | Q4TVN0 1x | 1.05E-05 | 0 | 1 | 4 | 6 | Q4TVN0 | TBC1 doma | Q4TVN0 [6 | 0 | 1941.857 | 0 | 0.07 | 0.095925 | 3.75 | 97.7 | 102.3 | 4.32 | 4.98 | | High | High | High | High | High | High | High | High | High | High | High | 0 | 3.34E-05 | 3.8 |
| FALSE | High | DDSSEDLS | [K].DDSSED | 1xDeamida | Q640L6 1xP | 0.000319 | 0 | 1 | 8 | 3 | Q640L6 | Atf2 protei | Q640L6 [3 | 0 | 3998.839 | 0 | -0.44 | 0.050652 | 19.02 | 120 | 80 | 14.56 | 26.86 | | High | High | High | High | High | High | High | High | High | High | High | 0 | 8.84E-05 | 3.73 |
| FALSE | High | DEASSPAQ | [R].DEASSF | 1xPhospho | O88888 1x | 0.00025 | 0 | 1 | 2 | 2 | O88888 | Amyloid be | O88888 [1 | 0 | 2361.092 | 0 | 0.45 | 0.006853 | 11.81 | 82.6 | 117.4 | 10.46 | 8.04 | | High | High | High | High | High | High | High | High | High | High | High | 0 | 5.05E-05 | 3.16 |
| FALSE | High | DEAVDFSE | [K].DEAVD | 1xPhospho | G3XA08 1x | 0.025738 | 0.001645 | 1 | 3 | 1 | G3XA08 | MCG67939 | G3XA08 [4 | 0 | 2224.01 | 0 | -0.19 | 0.046669 | 11.5 | 108.6 | 91.4 | 11.53 | 5.06 | | High | High | High | High | High | High | High | High | High | High | High | 0.000872 | 0.00905 | 2.53 |
| FALSE | High | DECTPTRK | [K].DECTPT | 1xCarbami | Q6NV83 1x | 0.040721 | 0.003058 | 1 | 3 | 1 | Q6NV83 | U2 snRNP- | Q6NV83 [9 | 1 | 1544.754 | 0 | -0.52 | 0.000466 | 1.67 | 116.8 | 83.2 | 6.07 | 6.62 | | High | High | High | High | High | High | High | High | High | High | High | 0.001597 | 0.01526 | 1.43 |
| FALSE | High | DEDADTPS | [K].DEDAD | 1xPhospho | P57016 1xP | 1.25E-05 | 0 | 1 | 2 | 6 | P57016 | Ladinin-1 C | P57016 [36 | 0 | 2578.258 | 0 | -0.75 | 0.023056 | 12.76 | 127.2 | 72.8 | 19.1 | 7.63 | | High | High | High | High | High | High | High | High | High | High | High | 0 | 1.74E-06 | 3.81 |
| FALSE | High | DEDDVEQE | [K].DEDDV | 1xPhospho | A0A0R4J12 | 0.000154 | 0 | 1 | 2 | 1 | A0A0R4J12 | SRSF prote | A0A0R4J12 | 0 | 2738.228 | 0 | -0.2 | 0.303709 | 13.38 | 107.7 | 92.3 | 5.13 | 17.28 | | High | High | High | High | High | High | High | High | High | High | High | 0 | 2.92E-05 | 3.55 |
| FALSE | High | DEDDVEQE | [K].DEDDV | 1xPhospho | A0A0R4J12 | 7.53E-11 | 0 | 1 | 2 | 4 | A0A0R4J12 | SRSF prote | A0A0R4J12 | 0 | 2967.391 | 0 | 0.78 | 0.011006 | 8.73 | 74.8 | 125.2 | 9.19 | 13.7 | | High | High | High | High | High | High | High | High | High | High | High | 0 | 2.38E-12 | 5.6 |
| FALSE | High | DEDEDFLPI | [K].DEDEDI | 1xPhospho | Q01320 1x | 7.58E-07 | 0 | 2 | 4 | 5 | Q01320; Q | DNA topois | Q01320 [1 | 0 | 2010.915 | 0 | -0.61 | 0.407188 | 27.37 | 120.3 | 79.7 | 86.51 | 76.14 | Shared | High | High | High | High | High | High | High | High | High | High | High | 0 | 7.44E-08 | 2.88 |
| FALSE | High | DEDEDFLPI | [K].DEDEDI | 1xPhospho | Q01320 1x | 2.32E-06 | 0 | 2 | 4 | 17 | Q01320; Q | DNA topois | Q01320 [1 | 0 | 2240.078 | 0 | -1.02 | 0.000337 | 0.53 | 130.7 | 69.3 | 6.04 | 4.14 | Shared | High | High | High | High | High | High | High | High | High | High | High | 0 | 1.94E-06 | 3.53 |
| FALSE | High | DEDGESGP | [K].DEDGE | 1xPhospho | B7ZDF5 1xP | 5.29E-07 | 0 | 1 | 8 | 2 | B7ZDF5 | Histone de | B7ZDF5 [61 | 0 | 2821.198 | 0 | 0.51 | 0.005174 | 11.1 | 84 | 116 | 5.12 | 7.37 | | High | High | High | High | High | High | High | High | High | High | High | 0 | 8.63E-05 | 2.98 |
| FALSE | High | DDSLDLSP | [R].DDSLDI | 1xPhospho | G5E870 1x | 0.000496 | 0 | 1 | 3 | 2 | G5E870 | E3 ubiquiti | G5E870 [1 | 0 | 1511.694 | 0 | -1.17 | 0.000923 | 3.79 | 138.7 | 61.3 | 4.12 | 12.02 | | High | High | High | High | High | High | High | High | High | High | High | 0 | 0.000108 | 1.87 |
| FALSE | High | DELSDISQ | [K].DELSDI | 1xPhospho | B8X349 1xP | 0.001689 | 5.66E-05 | 1 | 10 | 2 | B8X349 | JNK-intera | B8X349 [24 | 0 | 1773.867 | 0 | 1.67 | 0.000415 | 13.52 | 47.9 | 152.1 | 6.35 | 5.66 | | High | High | High | High | High | High | High | High | High | High | High | 2.58E-05 | 0.000432 | 3.69 |
| FALSE | High | DELSDWSL | [K].DELSDV | 1xPhospho | Q61584-3 1 | 1.94E-06 | 0 | 2 | 9 | 6 | Q61584-3; | Isoform B c | Q61584-3 [ | 0 | 1916.811 | 0 | -0.47 | 0.068746 | 6.88 | 115.7 | 84.3 | 13.75 | 14.37 | Shared | High | High | High | High | High | High | High | High | High | High | High | 0 | 7.42E-06 | 3.04 |
| FALSE | High | DELSDWSL | [K].DELSDV | 2xPhospho | Q61584-3 2 | 0.000183 | 0 | 2 | 9 | 1 | Q61584-3; | Isoform B c | Q61584-3 [ | 0 | 1996.778 | 0 | -0.37 | 0.153392 | 19.63 | 112.8 | 87.2 | 22.06 | 14.12 | Shared | High | High | High | High | High | High | High | High | High | High | High | 0 | 3.54E-05 | 3.71 |
| FALSE | High | DFDISEQNI | [K].DFDISE | 1xPhospho | Q61712 1x | 1.77E-07 | 0 | 1 | 2 | 2 | Q61712 | DnaJ homo | Q61712 [4 | 0 | 2295.991 | 0 | 0.6 | 0.009861 | 6.04 | 80.6 | 119.4 | 8.52 | 7.72 | | High | High | High | High | High | High | High | High | High | High | High | 0 | 1.46E-08 | 3.88 |
| FALSE | High | DFDISEQNI | [K].DFDISE | 2xPhospho | Q61712 2x | 3.1E-06 | 0 | 1 | 2 | 4 | Q61712 | DnaJ homo | Q61712 [4 | 0 | 2375.957 | 0 | 0.73 | 0.006272 | 5.5 | 74.7 | 125.3 | 5.01 | 9.85 | | High | High | High | High | High | High | High | High | High | High | High | 0 | 3.61E-07 | 3.53 |
| FALSE | High | DFEDSMDE | [K].DFEDSN | 1xPhospho | B2RY61 1xP | 1.85E-05 | 0 | 1 | 6 | 2 | B2RY61 | AT rich inte | B2RY61 [42 | 0 | 1853.828 | 0 | 0.04 | 0.456469 | 5.56 | 98.7 | 101.3 | 9.01 | 8.28 | | High | High | High | High | High | High | High | High | High | High | High | 0 | 2.68E-06 | 3.6 |
| FALSE | High | DFFDPVESI | [K].DFFDPV | 1xPhospho | Q810V0 1x | 6.12E-08 | 0 | 1 | 2 | 11 | Q810V0 | U3 small nu | Q810V0 [2 | 0 | 3364.492 | 0 | -0.13 | 0.042098 | 9.28 | 104.7 | 95.3 | 7.96 | 7.06 | | High | High | High | High | High | High | High | High | High | High | High | 0 | 4.41E-09 | 5.71 |
| FALSE | High | DFKLSDASI | [R].DFKLSD | 1xPhospho | F8VQC7 1x | 0.006536 | 0.00062 | 1 | 35 | 1 | F8VQC7 | Kinectin OS | F8VQC7 [8 | 1 | 3744.841 | 0 | 1.37 | 0.00133 | 6.86 | 53.9 | 146.1 | 9.53 | 14.17 | | High | High | High | High | High | High | High | High | High | High | High | 0.000275 | 0.001956 | 3.84 |
| FALSE | High | DFLGSPED | [K].DFLGSF | 1xPhospho | A0A0R4J0S | 0.000356 | 0 | 1 | 1 | 1 | A0A0R4J0S | Lethal(2) gi | A0A0R4J0S | 0 | 1761.835 | 0 | -0.01 | 0.873148 | 4.97 | 97.8 | 102.2 | 6.93 | 7.02 | | High | High | High | High | High | High | High | High | High | High | High | 0 | 7.47E-05 | 2.82 |
| FALSE | High | DFLSGPDYI | [R].DFLSGF | 2xPhospho | G3X960 2x | 0.000159 | 0 | 1 | 2 | 1 | G3X960 | MCG5982 | G3X960 [6 | 0 | 3373.278 | 0 | 1.44 | 0.000374 | 2.49 | 54.2 | 145.8 | 12.13 | 8.02 | | High | High | High | High | High | High | High | High | High | High | High | 0 | 3.04E-05 | 3.9 |
| FALSE | High | DFPDSFQP | [R].DFPDSF | 1xPhospho | Q8VCZ7 1x | 0.000217 | 0 | 1 | 1 | 1 | Q8VCZ7 | Zinc finger | Q8VCZ7 [2 | 0 | 2235.08 | 0 | 1.85 | 0.00344 | 21.14 | 48.6 | 151.4 | 11.73 | 18.54 | | High | High | High | High | High | High | High | High | High | High | High | 0 | 0.000043 | 3.19 |
| FALSE | High | DFPGISNTS | [K].DFPGIS | 1xPhospho | P59438 1xP | 0.000108 | 0 | 1 | 7 | 3 | P59438 | Hermansky | P59438 [65 | 0 | 1988.014 | 0 | 1.73 | 0.000192 | 12.58 | 46.7 | 153.3 | 8.12 | 3.83 | | High | High | High | High | High | High | High | High | High | High | High | 2.58E-05 | 0.000351 | 3.04 |
| FALSE | High | DFQEYVEP | [R].DFQEY | 1xPhospho | Q3TWG5 1 | 1.83E-09 | 0 | 1 | 2 | 6 | Q3TWG5 | Dynein cyt | Q3TWG5 [ | 0 | 2420.065 | 0 | -0.14 | 0.079573 | 11.07 | 105 | 95 | 4.6 | 7.86 | | High | High | High | High | High | High | High | High | High | High | High | 0 | 1.28E-07 | 4.15 |
| FALSE | High | DFQEYVEP | [R].DFQEY | 1xPhospho | Q3TWG5 1 | 2.53E-05 | 0 | 1 | 2 | 3 | Q3TWG5 | Dynein cyt | Q3TWG5 [ | 1 | 2576.166 | 0 | -0.31 | 0.004055 | 3.13 | 107.6 | 92.4 | 3.68 | 6.7 | | High | High | High | High | High | High | High | High | High | High | High | 0.000267 | 0.001543 | 2.7 |
| FALSE | High | DFSDEDQS | [R].DFSDED | 1xPhospho | E9QAK8 1x | 2.78E-06 | 0 | 1 | 5 | 3 | E9QAK8 | Expressed | E9QAK8 [1 | 0 | 2696.193 | 0 | 0.55 | 0.005613 | 5.91 | 80.2 | 119.8 | 8.88 | 15.04 | | High | High | High | High | High | High | High | High | High | High | High | 0 | 3.22E-07 | 4.09 |
| FALSE | High | DFSPEALK | [K].DFSPEA | 1xPhospho | Q3TIK5 1xP | 0.020679 | 0.001472 | 1 | 5 | 2 | Q3TIK5 | Eukaryotic | Q3TIK5 [18 | 0 | 1444.749 | 0 | -0.8 | 0.002085 | 4.13 | 127.3 | 72.7 | 4.11 | 13.48 | | High | High | High | High | High | High | High | High | High | High | High | 0.007994 | 0.0866 | 2.14 |
| FALSE | High | DFATPTLHT | [R].DFATPT | 1xPhospho | E9Q555 1x | 1E-05 | 0 | 1 | 2 | 1 | E9Q555 | E3 ubiquiti | E9Q555 [2 | 0 | 2038.943 | 0 | 1.23 | 0.000707 | 2.91 | 59.3 | 140.7 | 2.86 | 8.97 | | High | High | High | High | High | High | High | High | High | High | High | 0 | 1.36E-06 | 1.98 |
| FALSE | High | DFSPGLFEI | [K].DFSPGL | 1xPhospho | Q5QGZ6 1x | 4.43E-05 | 0 | 1 | 4 | 1 | Q5QGZ6 | Retinoic ac | Q5QGZ6 [6 | 0 | 2679.3 | 0 | 1.27 | 0.003474 | 7.65 | 57.9 | 142.1 | 9.86 | 16.76 | | High | High | High | High | High | High | High | High | High | High | High | 0 | 7.2E-06 | 3.13 |
| FALSE | High | DFTAEPTEI | [R].DFTAEF | 1xPhospho | Q8CAF4-3 | 3.11E-08 | 0 | 1 | 3 | 4 | Q8CAF4-3 | Isoform 3 c | Q8CAF4-3 | 0 | 2615.186 | 0 | 0.89 | 0.000903 | 10.61 | 68.2 | 131.8 | 5.68 | 7.4 | | High | High | High | High | High | High | High | High | High | High | High | 0 | 9.21E-08 | 4.27 |
| FALSE | High | DFTAEPTEI | [R].DFTAEF | 1xDeamida | Q8CAF4-3 | 2.66E-06 | 0 | 1 | 3 | 2 | Q8CAF4-3 | Isoform 3 c | Q8CAF4-3 | 0 | 2616.17 | 0 | 1.46 | 0.000669 | 3.76 | 54.4 | 145.6 | 13.21 | 7.99 | | High | High | High | High | High | High | High | High | High | High | High | 0 | 3.06E-07 | 2.71 |
| FALSE | High | DFTPGR | [K].DFTPGF | 1xPhospho | P97868 1xP | 0.107603 | 0.008634 | 1 | 3 | 1 | P97868 | E3 ubiquiti | P97868 [14 | 0 | 1001.465 | 0 | 0.3 | 0.030321 | 10.32 | 89.5 | 110.5 | 6.75 | 5.8 | | High | High | High | High | High | High | High | High | High | High | High | 0.004327 | 0.04687 | 1.85 |
| FALSE | High | DFTPVCTTI | [RK].DFTPV | 1xCarbami | Q6A0D0 1x | 0.004738 | 0.000456 | 1 | 5 | 1 | Q6A0D0 | MKIAA010 | Q6A0D0 [4 | 0 | 1704.787 | 0 | 0.23 | 0.048085 | 10.13 | 89.5 | 110.5 | 7.54 | 9.86 | | High | High | High | High | High | High | High | High | High | High | High | 0.000204 | 0.001366 | 2.17 |
| FALSE | High | DFVHLGAS | [R].DFVHL | 1xPhospho | Q3U9K6 1x | 0.000114 | 0 | 1 | 4 | 2 | Q3U9K6 | Interferon | Q3U9K6 [1 | 0 | 1995.994 | 0 | 0.71 | 0.000646 | 8.8 | 75.9 | 124.1 | 5.6 | 4.23 | | High | High | High | High | High | High | High | High | High | High | High | 0.000204 | 0.001286 | 2.26 |
| FALSE | High | DFVHLGAS | [R].DFVHL | 1xDeamida | Q3U9K6 1x | 1.76E-07 | 0 | 1 | 4 | 3 | Q3U9K6 | Interferon | Q3U9K6 [1 | 0 | 1996.978 | 0 | 1.06 | 0.001004 | 10.9 | 64.6 | 135.4 | 5.9 | 8.66 | | High | High | High | High | High | High | High | High | High | High | High | 0 | 1.44E-08 | 3.24 |
| FALSE | High | DFVIYQEPN | [K].DFVIYQ | 1xPhospho | B2RRE7 1xP | 0.001179 | 0 | 1 | 2 | 1 | B2RRE7 | OTU doma | B2RRE7 [1 | 0 | 3228.565 | 0 | -0.04 | 0.607456 | 13.73 | 101.5 | 98.5 | 11.81 | 6.85 | | High | High | High | High | High | High | High | High | High | High | High | 0 | 0.000286 | 2.88 |
| FALSE | High | DFVTISPVC | [R].DFVTIS | 1xPhospho | Q80SX8 1x | 0.000924 | 0 | 1 | 4 | 2 | Q80SX8 | ATP-depen | Q80SX8 [1 | 0 | 2067.036 | 0 | -0.02 | 0.830697 | 8.77 | 102.5 | 97.5 | 6.98 | 9.65 | | High | High | High | High | High | High | High | High | High | High | High | 4.94E-05 | 0.000593 | 2.91 |
| FALSE | High | DGAERESP | [R].DGAER | 1xPhospho | Q6ZQ58 1x | 0.000165 | 0 | 1 | 2 | 1 | Q6ZQ58 | La-related | Q6ZQ58 [6 | 1 | 2931.307 | 0 | 0.84 | 0.004291 | 8.88 | 69.9 | 130.1 | 8.01 | 10.59 | | High | High | High | High | High | High | High | High | High | High | High | 0 | 3.17E-05 | 1.77 |
| FALSE | High | DGAERESP | [R].DGAER | 1xPhospho | Q6ZQ58 1x | 0.002647 | 0.000109 | 1 | 2 | 2 | Q6ZQ58 | La-related | Q6ZQ58 [6 | 2 | 3087.408 | 0 | 0.74 | 0.002075 | 2.14 | 75.1 | 124.9 | 7.55 | 7.02 | | High | High | High | High | High | High | High | High | High | High | High | 4.94E-05 | 0.000708 | 3.93 |
| FALSE | High | DGAESSRD | [R].DGAES | 1xPhospho | Q3UGY8 1x | 0.005565 | 0.00062 | 1 | 2 | 1 | Q3UGY8 | Brefeldin A | Q3UGY8 [4 | 1 | 2353.981 | 0 | 2.11 | 0.001015 | 15.85 | 37.4 | 162.6 | 10.67 | 12.97 | | High | High | High | High | High | High | High | High | High | High | High | 0.000275 | 0.001624 | 1.79 |
| FALSE | High | DGAHSGDI | [R].DGAHS | 1xPhospho | Q5SVR0 1x | 0.00625 | 0.00062 | 1 | 2 | 2 | Q5SVR0 | TBC1 doma | Q5SVR0 [1 | 0 | 2689.203 | 0 | 0.73 | 0.004069 | 21.85 | 75.1 | 124.9 | 14.34 | 8.25 | | High | High | High | High | High | High | High | High | High | High | High | 0.000275 | 0.001853 | 3.51 |
| FALSE | High | DGAPHLQA | [R].DGAPH | 1xPhospho | Q8VCQ3 1x | 0.036103 | 0.002342 | 1 | 1 | 1 | Q8VCQ3 | Nuclear rec | Q8VCQ3 [9 | 0 | 3849.865 | 0 | 0.71 | 0.001184 | 0.29 | 79 | 121 | 6.15 | 4.95 | | High | High | High | High | High | High | High | High | High | High | High | 0.001246 | 0.01327 | 3.13 |
| FALSE | High | DFSSQDSFI | [R].DFSSQI | 1xPhospho | P47713 1xP | 3.03E-08 | 0 | 1 | 2 | 9 | P47713 | Cytosolic p | P47713 [50 | 0 | 3029.241 | 0 | -0.69 | 0.01994 | 10.21 | 122 | 78 | 8.82 | 17.66 | | High | High | High | High | High | High | High | High | High | High | High | 0 | 2E-09 | 5.47 |
| FALSE | High | DFATAPPTS | [K].DFATAF | 1xPhospho | Q80XK6 1x | 0.000543 | 0 | 1 | 1 | 3 | Q80XK6 | Autophagy | Q80XK6 [1 | 0 | 1740.897 | 0 | -0.5 | 0.00188 | 1.8 | 118.6 | 81.4 | 3.72 | 9.26 | | High | High | High | High | High | High | High | High | High | High | High | 0 | 0.00012 | 2.65 |
| FALSE | High | DFASEAPTA | [R].DFASEA | 1xCarbami | P97434-2 1 | 0.004372 | 0.000409 | 1 | 6 | 2 | P97434-2 | Isoform 2 c | P97434-2 [ | 1 | 2704.275 | 0 | 0.76 | 0.004392 | 13.82 | 74.3 | 125.7 | 13.15 | 6.61 | | High | High | High | High | High | High | High | High | High | High | High | 0.000204 | 0.001293 | 1.97 |
| FALSE | High | DFASEAPTA | [R].DFASEA | 1xCarbami | P97434-2 1 | 1.49E-10 | 0 | 1 | 6 | 2 | P97434-2 | Isoform 2 c | P97434-2 [ | 0 | 2548.174 | 0 | 0.85 | 0.001161 | 17.89 | 70.7 | 129.3 | 12.64 | 1.44 | | High | High | High | High | High | High | High | High | High | High | High | 0 | 5.12E-12 | 6.2 |
| FALSE | High | DELSDWSL | [K].DELSDV | 1xPhospho | Q61584-3 1 | 0.005615 | 0.00062 | 2 | 9 | 1 | Q61584-3; | Isoform B c | Q61584-3 [ | 1 | 2303.003 | 0 | -0.07 | 0.92272 | 21.34 | 97.2 | 102.8 | 21.24 | 22.13 | Shared | High | High | High | High | High | High | High | High | High | High | High | 0.000275 | 0.001653 | 1.44 |
| FALSE | High | DELTDLDQ | [K].DELTDL | 1xPhospho | Q9CY58-2 1 | 7.2E-08 | 0 | 1 | 3 | 12 | Q9CY58-2 | Isoform 2 c | Q9CY58-2 | 0 | 3779.706 | 0 | -0.3 | 0.145945 | 8.29 | 109.9 | 90.1 | 9.45 | 7.71 | | High | High | High | High | High | High | High | High | High | High | High | 0 | 2.48E-08 | 5.48 |
| FALSE | High | DELTDLDQ | [K].DELTDL | 2xPhospho | Q9CY58-2 2 | 1.3E-05 | 0 | 1 | 3 | 9 | Q9CY58-2 | Isoform 2 c | Q9CY58-2 | 0 | 3859.672 | 0 | -0.77 | 0.00435 | 4.93 | 128 | 72 | 6.34 | 14.7 | | High | High | High | High | High | High | High | High | High | High | High | 0 | 1.8E-06 | 5.36 |
| FALSE | High | DELTESPK | [K].DELTES | 1xPhospho | Q9CY58 1x | 0.004451 | 0.000409 | 1 | 2 | 4 | Q9CY58 | Plasminoge | Q9CY58 [2 | 0 | 1456.734 | 0 | 0.17 | 0.051637 | 2.63 | 93 | 107 | 4.53 | 7.57 | | High | High | High | High | High | High | High | High | High | High | High | 0.000183 | 0.001269 | 2.14 |
| FALSE | High | DEITGECLE | [R].DEITGE | 1xCarbami | Q8K4J0 1xP | 5.35E-05 | 0 | 1 | 1 | 1 | Q8K4J0 | Protein art | Q8K4J0 [49 | 0 | 2880.317 | 0 | -0.18 | 0.053147 | 9.55 | 103.5 | 96.5 | 8.82 | 8.29 | | High | High | High | High | High | High | High | High | High | High | High | 0 | 8.92E-06 | 5.05 |
| FALSE | High | DEITGECLE | [R].DEITGE | 1xCarbami | Q8K4J0 1xP | 0.001117 | 0 | 1 | 1 | 2 | Q8K4J0 | Protein art | Q8K4J0 [49 | 0 | 3109.48 | 0 | -0.45 | 0.046113 | 13.01 | 113.9 | 86.1 | 13.59 | 9.23 | | High | High | High | High | High | High | High | High | High | High | High | 0 | 0.000271 | 3.04 |
| FALSE | High | DEPDWEDS | [K].DEPDW | 1xPhospho | Q8VBT6 1x | 5.21E-08 | 0 | 1 | 1 | 3 | Q8VBT6 | Apolipoprc | Q8VBT6 [4 | 0 | 2774.192 | 0 | 2.03 | 0.003174 | 31.23 | 34.8 | 165.2 | 24.06 | 21.74 | | High | High | High | High | High | High | High | High | High | High | High | 0 | 3.67E-09 | 4.3 |
| FALSE | High | DEPPPLSPA | [K].DEPPPL | 1xPhospho | A2AFQ0 1x | 1.98E-07 | 0 | 1 | 4 | 3 | A2AFQ0 | E3 ubiquiti | A2AFQ0 [3 | 0 | 2945.454 | 0 | -0.35 | 0.129807 | 3.72 | 110 | 90 | 10.26 | 11.89 | | High | High | High | High | High | High | High | High | High | High | High | 0 | 1.65E-08 | 4.57 |
| FALSE | High | DEPPPLSPA | [K].DEPPPL | 2xPhospho | A2AFQ0 2x | 8.79E-05 | 0 | 1 | 4 | 11 | A2AFQ0 | E3 ubiquiti | A2AFQ0 [3 | 0 | 3025.42 | 0 | -0.29 | 0.001187 | 0.9 | 110.2 | 89.8 | 7.47 | 8.1 | | High | High | High | High | High | High | High | High | High | High | High | 0 | 5.79E-05 | 4.33 |
| FALSE | High | DEPQTVPD | [K].DEPQT | 1xPhospho | P15066 1xP | 2.53E-05 | 0 | 1 | 2 | 2 | P15066 | Transcripti | P15066 [23 | 0 | 3378.529 | 0 | -0.32 | 0.12841 | 15.21 | 111.2 | 88.8 | 6.82 | 21.13 | | High | High | High | High | High | High | High | High | High | High | High | 0 | 3.82E-06 | 3.72 |
| FALSE | High | DEPQTVPD | [K].DEPQT | 1xPhospho | P15066 1xP | 0.011146 | 0.000772 | 1 | 2 | 3 | P15066 | Transcripti | P15066 [23 | 0 | 3394.524 | 0 | -0.32 | 0.436564 | 5.86 | 111.3 | 88.7 | 9.21 | 22.25 | | High | High | High | High | High | High | High | High | High | High | High | 0.000571 | 0.005067 | 3.15 |
| FALSE | High | DEPQTVPD | [K].DEPQT | 2xPhospho | P15066 2xP | 6E-07 | 0 | 1 | 2 | 4 | P15066 | Transcripti | P15066 [23 | 0 | 3458.495 | 0 | 1.09 | 0.018181 | 21.22 | 65 | 135 | 4.73 | 22.4 | | High | High | High | High | High | High | High | High | High | High | High | 0 | 5.71E-08 | 5.25 |
| FALSE | High | DEPQTVPD | [K].DEPQT | 2xPhospho | P15066 2xP | 1.87E-06 | 0 | 1 | 2 | 4 | P15066 | Transcripti | P15066 [23 | 0 | 3474.49 | 0 | 1.03 | 0.002718 | 21.73 | 64 | 136 | 3.21 | 12.37 | | High | High | High | High | High | High | High | High | High | High | High | 0 | 2.05E-07 | 4.13 |
| FALSE | High | DEQGQQS | [K].DEQGQ | 1xPhospho | B9EIT8 1xP | 8.15E-07 | 0 | 2 | 3 | 1 | B9EIT8; A0 | Rg9mtd2 p | B9EIT8 [31 | 0 | 1624.742 | 0 | -0.39 | 0.012027 | 4.31 | 111.2 | 88.8 | 8.3 | 12.24 | Shared | High | High | High | High | High | High | High | High | High | High | High | 0 | 8.12E-08 | 3.75 |
| FALSE | High | DERSPQNL | [K].DERSP | 1xPhospho | F8WJD4 1x | 0.083783 | 0.006345 | 1 | 2 | 3 | F8WJD4 | Symplekin | F8WJD4 [1 | 1 | 2461.249 | 0 | 0.63 | 0.002143 | 5.34 | 77.2 | 122.8 | 4.04 | 9.76 | | High | High | High | High | High | High | High | High | High | High | High | 0.007763 | 0.0838 | 3.1 |
| FALSE | High | DESCYDNA | [K].DESCYD | 1xCarbami | Q7TPD0 1x | 1.39E-11 | 0 | 1 | 3 | 12 | Q7TPD0 | Integrator | Q7TPD0 [5 | 0 | 3062.25 | 0 | -0.27 | 0.00402 | 2.74 | 109.4 | 90.6 | 4.27 | 2.53 | | High | High | High | High | High | High | High | High | High | High | High | 0 | 9.1E-11 | 6.02 |
| FALSE | High | DESGGGG | [K].DESGG | 1xPhospho | Q8K3A9 1x | 2.14E-06 | 0 | 1 | 1 | 1 | Q8K3A9 | 7SK snRNA | Q8K3A9 [2 | 0 | 2563.126 | 0 | 1.42 | 0.00128 | 6.69 | 49.6 | 150.4 | 12.7 | 15.76 | | High | High | High | High | High | High | High | High | High | High | High | 0 | 2.39E-07 | 4 |
| FALSE | High | DESPPPSPF | [K].DESPPF | 2xPhospho | A2A7S8-2 2 | 0.00967 | 0.000736 | 1 | 4 | 2 | A2A7S8-2 | Isoform 2 c | A2A7S8-2 [ | 0 | 2739.287 | 0 | -0.12 | 0.105978 | 13.65 | 103.8 | 96.2 | 8.6 | 9.82 | | High | High | High | High | High | High | High | High | High | High | High | 0.000344 | 0.003027 | 2.34 |
| FALSE | High | DESSDEEDI | [K].DESSDE | 2xPhospho | Q8R2M2 2x | 4.47E-08 | 0 | 1 | 1 | 8 | Q8R2M2 | Deoxynucle | Q8R2M2 [5 | 0 | 2199.935 | 0 | -0.18 | 0.015192 | 4.94 | 107.3 | 92.7 | 5.27 | 8.31 | | High | High | High | High | High | High | High | High | High | High | High | 0 | 3.1E-09 | 4.28 |
| FALSE | High | DESSPPQL | [K].DESSPF | 1xCarbami | A2A7B5 1x | 9.67E-07 | 0 | 1 | 2 | 2 | A2A7B5 | PR domain | A2A7B5 [4 | 0 | 2668.294 | 0 | -1.21 | 0.000424 | 0.8 | 142.2 | 57.8 | 6.29 | 8.77 | | High | High | High | High | High | High | High | High | High | High | High | 0 | 9.82E-08 | 4.95 |
| FALSE | High | DESSPSQS | [K].DESSPS | 1xPhospho | Q91VW5 1 | 1.96E-07 | 0 | 1 | 7 | 3 | Q91VW5 | Golgin sub | Q91VW5 [1 | 0 | 2351.08 | 0 | 1.01 | 0.000727 | 5.51 | 64 | 136 | 9.42 | 6.3 | | High | High | High | High | High | High | High | High | High | High | High | 0 | 1.63E-08 | 4.03 |

| | | | | | | | | | | | | | | | | | | | | | | | | | | | | | | | | | | | | | | |
|---|---|---|---|---|---|---|---|---|---|---|---|---|---|---|---|---|---|---|---|---|---|---|---|---|---|---|---|---|---|---|---|---|---|---|---|---|---|---|
| FALSE | High | DESTDSGLS | [R].DESTDS | 2xPhospho | P46938 2xP | 0.065952 | 0.004766 | 1 | 3 | 1 | P46938 | Transcripti | P46938 [38 | 0 | 2236.892 | 0 | 0.11 | 0.743411 | 13.95 | 96.7 | 103.3 | 10.88 | 18.63 | | High | High | High | High | High | High | High | High | High | High | High | 0.002455 | 0.02656 | 1.39 |
| FALSE | High | DESTDWPL | [K].DESTDW | 1xPhospho | E9QQ30 1x | 0.003464 | 0.00021 | 1 | 3 | 1 | E9QQ30 | Eukaryotic | E9QQ30 [6 | 0 | 2684.257 | 0 | -0.49 | 0.011553 | 3.13 | 119.5 | 80.5 | 13.81 | 6.06 | | High | High | High | High | High | High | High | High | High | High | High | 9.42E-05 | 0.000959 | 2.93 |
| FALSE | High | DESVDLEEI | [K].DESVDI | 1xPhospho | Q9EQM6 1 | 0.000292 | 0 | 1 | 1 | 1 | Q9EQM6 | Microproc | Q9EQM6 [ | 0 | 1547.683 | 0 | -1.01 | 0.000489 | 1.52 | 133.7 | 66.3 | 8.59 | 13.7 | | High | High | High | High | High | High | High | High | High | High | High | 0 | 5.98E-05 | 3.14 |
| FALSE | High | DESYPQELI | [R].DESYPC | 1xPhospho | Q6NXI6 1x | 0.000492 | 0 | 1 | 3 | 1 | Q6NXI6 | Regulation | Q6NXI6 [79 | 0 | 3571.703 | 0 | -0.26 | 0.058703 | 4.06 | 108.2 | 91.8 | 6.98 | 8.8 | | High | High | High | High | High | High | High | High | High | High | High | 0 | 0.000108 | 3.34 |
| FALSE | High | DETEYEYSC | [K].DETEYE | 2xPhospho | A0A0A6YW | 0.000225 | 0 | 2 | 11 | 2 | A0A0A6YW | Mitogen-a | A0A0A6YW | 0 | 4547.875 | 0 | -0.98 | 0.011023 | 1.18 | 124.3 | 75.7 | 22.53 | 20.26 | Shared | High | High | High | High | High | High | High | High | High | High | High | 0 | 4.45E-05 | 5.63 |
| FALSE | High | DEVQNEGT | [K].DEVQN | 1xPhospho | Q8VDP4 1x | 4.34E-11 | 0 | 1 | 1 | 6 | Q8VDP4 | Cell cycle a | Q8VDP4 [5 | 0 | 2328.101 | 0 | 0 | 0.979024 | 8.28 | 100 | 100 | 5.84 | 8.08 | | High | High | High | High | High | High | High | High | High | High | High | 0 | 5E-12 | 5.99 |
| FALSE | High | DEVVSPLPS | [R].DEVVSF | 1xPhospho | Q8K3A9 1x | 5.91E-08 | 0 | 1 | 1 | 3 | Q8K3A9 | 7SK snRNA | Q8K3A9 [3 | 0 | 3780.838 | 0 | 0.14 | 0.554195 | 35.65 | 91.2 | 108.8 | 8.74 | 21.95 | | High | High | High | High | High | High | High | High | High | High | High | 0 | 4.22E-09 | 5.02 |
| FALSE | High | DEYEYVSDI | [R].DEYEYV | 1xPhospho | Q9WTU0 1 | 3.79E-07 | 0 | 1 | 1 | 3 | Q9WTU0 | Lysine-spe | Q9WTU0 [ | 0 | 2099.946 | 0 | 1.33 | 0.002286 | 17.52 | 56.8 | 143.2 | 8.94 | 10.23 | | High | High | High | High | High | High | High | High | High | High | High | 0 | 3.43E-08 | 2.79 |
| FALSE | High | DDLLGDILC | [K].DDLLGI | 1xPhospho | P33609 1xP | 8.03E-05 | 0 | 1 | 3 | 1 | P33609 | DNA polym | P33609 [16 | 0 | 3467.869 | 0 | 0.06 | 0.567196 | 17.84 | 97.9 | 102.1 | 17.21 | 15.68 | | High | High | High | High | High | High | High | High | High | High | High | 0 | 1.41E-05 | 4.05 |
| FALSE | High | DDIFSSLEE | [R].DDIFSS | 1xPhospho | A2AM95 1x | 0.000281 | 0 | 1 | 3 | 1 | A2AM95 | Emerin OS | A2AM95 [1 | 0 | 1906.909 | 0 | 1.57 | 0.001851 | 28.8 | 49.9 | 150.1 | 11.57 | 11.02 | | High | High | High | High | High | High | High | High | High | High | High | 0 | 5.75E-05 | 3.17 |
| FALSE | High | DDLETSSDA | [K].DDLETS | 1xPhospho | E9Q3Y4 1x | 0.000154 | 0 | 1 | 7 | 2 | E9Q3Y4 | Lipopolysa | E9Q3Y4 [1 | 0 | 3027.445 | 0 | 0.34 | 0.060252 | 6.63 | 86.9 | 113.1 | 12.26 | 8.54 | | High | High | High | High | High | High | High | High | High | High | High | 0 | 5.48E-05 | 2.84 |
| FALSE | High | DDIEDDVSS | [K].DDIEDD | 1xPhospho | Q8BYH8 1x | 7.81E-08 | 0 | 1 | 4 | 2 | Q8BYH8 | Chromodo | Q8BYH8 [1 | 0 | 3373.528 | 0 | 0.45 | 0.004614 | 8.51 | 84.9 | 115.1 | 6.85 | 5.17 | | High | High | High | High | High | High | High | High | High | High | High | 0 | 5.78E-09 | 5.38 |
| FALSE | High | DALADDDS | [K].DALADI | 1xPhospho | O70472 1x | 0.00047 | 0 | 1 | 2 | 4 | O70472 | Transmem | O70472 [1 | 0 | 2960.31 | 0 | -0.88 | 0.014206 | 6.35 | 128.5 | 71.5 | 14.85 | 13.51 | | High | High | High | High | High | High | High | High | High | High | High | 2.58E-05 | 0.000441 | 3.5 |
| FALSE | High | DALDSLDTS | [K].DALDSL | 1xCarbami | P43120 1xP | 0.000822 | 0 | 1 | 2 | 1 | P43120 | Hematopo | P43120 [20 | 0 | 2988.355 | 0 | 1.22 | 0.00258 | 9.42 | 61.5 | 138.5 | 14.73 | 11.54 | | High | High | High | High | High | High | High | High | High | High | High | 0 | 0.000191 | 3.35 |
| FALSE | High | DALPSPGEY | [K].DALPSF | 1xPhospho | B9EIW5 1x | 0.031168 | 0.001935 | 1 | 2 | 1 | B9EIW5 | Ncoa7 prot | B9EIW5 [2 | 0 | 2282.099 | 0 | 3.42 | | 11.93 | 17.8 | 182.2 | 36.52 | 8.71 | | High | High | Not Found | High | High | High | High | High | High | High | High | 0.001044 | 0.0112 | 2.71 |
| FALSE | High | DALSDSAK | [R].DALSDS | 1xPhospho | B2RWW2 1 | 0.019107 | 0.001402 | 1 | 4 | 2 | B2RWW2 | Golgb1 pro | B2RWW2 [ | 0 | 1344.681 | 0 | 2.04 | 0.001649 | 10.75 | 39.7 | 160.3 | 11.27 | 13.64 | | High | High | High | High | High | High | High | High | High | High | High | 0.00074 | 0.006504 | 1.98 |
| FALSE | High | DALSDSQA | [K].DALSDS | 1xPhospho | Q8K2J4 1xP | 0.002293 | 0.000109 | 1 | 2 | 1 | Q8K2J4 | Coiled-coil | Q8K2J4 [35 | 0 | 3376.635 | 0 | -0.02 | 0.984747 | 11.14 | 97.8 | 102.2 | 7.02 | 10.02 | | High | High | High | High | High | High | High | High | High | High | High | 4.94E-05 | 0.000608 | 3.16 |
| FALSE | High | DAMNSQF | [K].DAMNS | 1xPhospho | Q8R1S9 1x | 9.76E-07 | 0 | 1 | 1 | 2 | Q8R1S9 | Sodium-co | Q8R1S9 [3 | 0 | 2109.863 | 0 | -1.67 | 0.086302 | 0.7 | 148 | 52 | 49.26 | 35.55 | | High | High | High | High | High | High | High | High | High | High | High | 0 | 9.86E-08 | 3.45 |
| FALSE | High | DAMNSQF | [K].DAMNS | 1xPhospho | Q8R1S9 1x | 2.24E-09 | 0 | 1 | 1 | 4 | Q8R1S9 | Sodium-co | Q8R1S9 [3 | 0 | 2339.026 | 0 | -1.87 | 0.002017 | 2.28 | 157.1 | 42.9 | 10.52 | 19.78 | | High | High | High | High | High | High | High | High | High | High | High | 0 | 2.28E-08 | 4.52 |
| FALSE | High | DAPISPASV | [K].DAPISP | 1xPhospho | Q3UY41 1x | 0.000286 | 0 | 1 | 4 | 1 | Q3UY41 | Putative ur | Q3UY41 [2 | 0 | 2314.158 | 0 | -0.43 | 0.008476 | 6.67 | 118.9 | 81.1 | 11.68 | 5.77 | | High | High | High | High | High | High | High | High | High | High | High | 0 | 5.87E-05 | 3.48 |
| FALSE | High | DAPISPASV | [K].DAPISP | 2xPhospho | Q3UY41 2x | 0.009756 | 0.000736 | 1 | 4 | 1 | Q3UY41 | Putative ur | Q3UY41 [2 | 0 | 2394.124 | 0 | -0.32 | 0.057117 | 6.08 | 108.9 | 91.1 | 9.23 | 6.48 | | High | High | High | High | High | High | High | High | High | High | High | 0.000344 | 0.003045 | 3.1 |
| FALSE | High | DAPQTSPV | [R].DAPQT | 1xPhospho | A0A1B0GR | 0.0026 | 0.000109 | 1 | 1 | 5 | A0A1B0GR | RIKEN cDN | A0A1B0GR | 0 | 1392.709 | 0 | 1.24 | 0.000264 | 8.55 | 59.9 | 140.1 | 5.42 | 5.42 | | High | High | High | High | High | High | High | High | High | High | High | 0.003265 | 0.02437 | 2.11 |
| FALSE | High | DAPTSHVP | [K].DAPTSH | 1xPhospho | Q6P5E3 1x | 1.56E-07 | 0 | 1 | 3 | 3 | Q6P5E3 | Sal-like 1 ( | Q6P5E3 [1 | 0 | 3004.571 | 0 | -1.23 | 0.001818 | 2.26 | 144.3 | 55.7 | 10.61 | 9.29 | | High | High | High | High | High | High | High | High | High | High | High | 0 | 2.48E-08 | 4.68 |
| FALSE | High | DAPTTLAES | [K].DAPTTL | 1xPhospho | P70670 1xP | 0.000319 | 0 | 1 | 1 | 2 | P70670 | Nascent po | P70670 [1 | 0 | 1938.983 | 0 | -1.77 | 0.001125 | 0.51 | 154.1 | 45.9 | 11.76 | 16.74 | | High | High | High | High | High | High | High | High | High | High | High | 0 | 0.000199 | 2.92 |
| FALSE | High | DAPWTSNS | [R].DAPWT | 1xPhospho | Q3V1M8 1 | 0.015329 | 0.001089 | 1 | 3 | 2 | Q3V1M8 | Putative ur | Q3V1M8 [ | 0 | 1759.83 | 0 | 0.58 | 0.005132 | 14.21 | 80.2 | 119.8 | 5.39 | 10.27 | | High | High | High | High | High | High | High | High | High | High | High | 0.000571 | 0.005054 | 2.9 |
| FALSE | High | DAGSQSLS | [R].DAGSQ | 1xPhospho | E9Q9R9 1x | 1.24E-05 | 0 | 1 | 5 | 1 | E9Q9R9 | Disks large | E9Q9R9 [1 | 0 | 1962.923 | 0 | 0.69 | 0.038186 | 17.41 | 76.9 | 123.1 | 12.98 | 12.01 | | High | High | High | High | High | High | High | High | High | High | High | 0 | 1.73E-06 | 2.21 |
| FALSE | High | DAQPVATC | [R].DAQPV | 1xCarbami | Q5SXI5 1xP | 0.003312 | 0.00016 | 1 | 3 | 1 | Q5SXI5 | Zinc finger | Q5SXI5 [20 | 1 | 2663.378 | 0 | -0.59 | 0.002631 | 5.33 | 120 | 80 | 7.32 | 8.71 | | High | High | High | High | High | High | High | High | High | High | High | 7.19E-05 | 0.00091 | 2.35 |
| FALSE | High | DARPPSSPI | [K].DARPPS | 1xPhospho | B9EKK8 1x | 0.025964 | 0.001645 | 1 | 4 | 2 | B9EKK8 | Uncharacte | B9EKK8 [4 | 0 | 2835.41 | 0 | 0.56 | 0.033286 | 2.51 | 79.5 | 120.5 | 5.47 | 18.52 | | High | High | High | High | High | High | High | High | High | High | High | 0.001999 | 0.02076 | 2.89 |
| FALSE | High | DARPPSSPI | [K].DARPPS | 1xPhospho | B9EKK8 1x | 0.024855 | 0.001645 | 1 | 4 | 2 | B9EKK8 | Uncharacte | B9EKK8 [4 | 0 | 2851.405 | 0 | 0.15 | 0.101171 | 0.68 | 94.7 | 105.3 | 8.66 | 11.5 | | High | High | High | High | High | High | High | High | High | High | High | 0.000872 | 0.008694 | 3.64 |
| FALSE | High | DASDLDDE | [K].DASDD | 1xPhospho | Q3ULL5 1x | 3.49E-07 | 0 | 1 | 2 | 6 | Q3ULL5 | Eif2s2 prot | Q3ULL5 [6 | 0 | 2295.058 | 0 | -0.67 | 0.001794 | 2.15 | 122.3 | 77.7 | 7.43 | 5.75 | | High | High | High | High | High | High | High | High | High | High | High | 0 | 8.32E-06 | 4.73 |
| FALSE | High | DASDVEAE | [K].DASDV | 1xPhospho | E9Q4F8 1x | 5.21E-05 | 0 | 1 | 3 | 1 | E9Q4F8 | Ankyrin rep | E9Q4F8 [2 | 0 | 1643.793 | 0 | 0.33 | 0.020531 | 9.12 | 91.7 | 108.3 | 9.35 | 7.31 | | High | High | High | High | High | High | High | High | High | High | High | 0 | 8.66E-06 | 3.4 |
| FALSE | High | DASDVEDE | [R].DASDV | 1xPhospho | Q6P549 1x | 0.005872 | 0.00062 | 1 | 2 | 5 | Q6P549 | Phosphatic | Q6P549 [1 | 0 | 2203.105 | 0 | 0.71 | 0.000729 | 1.45 | 75.6 | 124.4 | 4.77 | 4.71 | | High | High | High | High | High | High | High | High | High | High | High | 0.000275 | 0.001734 | 2.81 |
| FALSE | High | DASGGIDL | [R].DASGG | 1xPhospho | B9EKB5 1x | 0.002398 | 0.000109 | 1 | 3 | 1 | B9EKB5 | Baz2b prot | B9EKB5 [1 | 0 | 2513.187 | 0 | -0.33 | 0.023225 | 4.67 | 110.7 | 89.3 | 13.15 | 6.42 | | High | High | High | High | High | High | High | High | High | High | High | 4.94E-05 | 0.000633 | 2.46 |
| FALSE | High | DASGSPAS | [K].DASGS | 1xPhospho | Q5SQA3 1x | 1.57E-08 | 0 | 1 | 12 | 2 | Q5SQA3 | Transducin | Q5SQA3 [2 | 0 | 2222.059 | 0 | 0.08 | 0.450942 | 5.77 | 97 | 103 | 5.6 | 5.65 | | High | High | High | High | High | High | High | High | High | High | High | 0 | 9.57E-10 | 4.83 |
| FALSE | High | DASLESSVE | [R].DASLES | 2xPhospho | Q8K387 2x | 0.00204 | 5.66E-05 | 1 | 2 | 1 | Q8K387 | Ubiquitin c | Q8K387 [4 | 0 | 2855.252 | 0 | -0.1 | 0.539332 | 9.19 | 107 | 93 | 10.51 | 21.05 | | High | High | High | High | High | High | High | High | High | High | High | 2.58E-05 | 0.000532 | 3.68 |
| FALSE | High | DASLIEAN | [K].DASLIE | 1xPhospho | Q3V3X3 1x | 0.000622 | 0 | 1 | 4 | 1 | Q3V3X3 | MCG3705, | Q3V3X3 [4 | 0 | 1797.94 | 0 | -0.2 | 0.161766 | 8.62 | 107.9 | 92.1 | 9.1 | 11.62 | | High | High | High | High | High | High | High | High | High | High | High | 0 | 0.000139 | 3.38 |
| FALSE | High | DASLQDWI | [R].DASLQI | 1xPhospho | B9EKJ7 1xP | 0.002576 | 0.000109 | 1 | 3 | 1 | B9EKJ7 | Tankyrase | B9EKJ7 [87 | 0 | 1833.882 | 0 | 1.48 | 0.010717 | 54.52 | 50 | 150 | 12.09 | 24.54 | | High | High | High | High | High | High | High | High | High | High | High | 4.94E-05 | 0.000692 | 2.78 |
| FALSE | High | DASPPPAFI | [R].DASPPF | 1xPhospho | Q9Z0U1 1x | 0.000227 | 0 | 1 | 1 | 3 | Q9Z0U1 | Tight junct | Q9Z0U1 [9 | 0 | 2245.224 | 0 | 0.2 | 0.00751 | 6.31 | 92.3 | 107.7 | 3.59 | 3.08 | | High | High | High | High | High | High | High | High | High | High | High | 2.58E-05 | 0.000513 | 2.5 |
| FALSE | High | DASPSSSSA | [R].DASPSS | 1xPhospho | G3X959 1x | 0.005666 | 0.00062 | 1 | 7 | 1 | G3X959 | Periphilin 1 | G3X959 [2 | 0 | 1589.726 | 0 | 0.51 | 0.351072 | 9.27 | 84 | 116 | 31.19 | 14.42 | | High | High | High | High | High | High | High | High | High | High | High | 0.000275 | 0.001667 | 3.45 |
| FALSE | High | DASPSSSSA | [R].DASPSS | 1xPhospho | G3X959 1x | 3.24E-06 | 0 | 1 | 7 | 8 | G3X959 | Periphilin 1 | G3X959 [2 | 0 | 1818.889 | 0 | 0.22 | 0.031088 | 12.87 | 92.9 | 107.1 | 3.95 | 6.37 | | High | High | High | High | High | High | High | High | High | High | High | 0 | 9.9E-07 | 3.05 |
| FALSE | High | DAQSPLLEY | [K].DAQSP | 1xPhospho | Q69ZR9-2 | 6.47E-05 | 0 | 1 | 2 | 2 | Q69ZR9-2 | Isoform 2 c | Q69ZR9-2 | 0 | 2081.093 | 0 | -1.42 | 0.000209 | 0.48 | 144.1 | 55.9 | 3.83 | 5.29 | | High | High | High | High | High | High | High | High | High | High | High | 0 | 1.11E-05 | 3.27 |
| FALSE | High | DAGLSPEQ | [R].DAGLSI | 2xPhospho | Q60591 2x | 6.56E-10 | 0 | 1 | 10 | 1 | Q60591 | Nuclear fac | Q60591 [1 | 0 | 2380.151 | 0 | 0.33 | 0.012404 | 4.75 | 83.3 | 116.7 | 11.03 | 7.46 | | High | High | High | High | High | High | High | High | High | High | High | 0 | 2.69E-11 | 4.88 |
| FALSE | High | DAGLSPEQ | [R].DAGLSI | 1xPhospho | Q60591 1x | 3.22E-07 | 0 | 1 | 10 | 5 | Q60591 | Nuclear fac | Q60591 [1 | 0 | 2300.185 | 0 | 1.48 | 0.000284 | 8.6 | 53.8 | 146.2 | 4.86 | 5.36 | | High | High | High | High | High | High | High | High | High | High | High | 0 | 0.000192 | 3.7 |
| FALSE | High | DAGGSLTE | [R].DAGGS | 1xPhospho | Q91YU3 1x | 0.000183 | 0 | 1 | 1 | 1 | Q91YU3 | Myb/SANT | Q91YU3 [1 | 0 | 1514.787 | 0 | -0.82 | 0.001718 | 3.54 | 123.7 | 76.3 | 5.44 | 8.19 | | High | High | High | High | High | High | High | High | High | High | High | 0 | 3.55E-05 | 2.37 |
| FALSE | High | CTAPLTPSS | [K].CTAPLT | 1xCarbami | G5E8R4 2x | 0.006957 | 0.00062 | 1 | 3 | 2 | G5E8R4 | SAPS doma | G5E8R4 [8 | 0 | 1919.817 | 0 | -0.35 | 0.036052 | 1.3 | 110.6 | 89.4 | 3.92 | 12.52 | | High | High | High | High | High | High | High | High | High | High | High | 0.002354 | 0.02414 | 2.26 |
| FALSE | High | CTSTTSSQS | [R].CTSTTS | 1xCarbami | E9QKL0 1x | 0.000758 | 0 | 1 | 3 | 6 | E9QKL0 | Helicase AF | E9QKL0 [7 | 0 | 2106.921 | 0 | -0.22 | 0.286542 | 12.65 | 107.5 | 92.5 | 4.56 | 22.74 | | High | High | High | High | High | High | High | High | High | High | High | 0.004291 | 0.04598 | 3.63 |
| FALSE | High | CVACNNPV | [K].CVACN | 1xCarbami | Q9ERU9 1x | 5.91E-05 | 0 | 1 | 3 | 1 | Q9ERU9 | E3 SUMO-p | Q9ERU9 [1 | 0 | 2685.318 | 0 | 0.7 | 0.050243 | 45.22 | 68.3 | 131.7 | 12.46 | 28.4 | | High | High | High | High | High | High | High | High | High | High | High | 0 | 9.98E-06 | 3.72 |
| FALSE | High | CVGLPSATI | [R].CVGLPS | 1xCarbami | Q8VBT0 1x | 8.56E-05 | 0 | 1 | 1 | 4 | Q8VBT0 | Thioredoxi | Q8VBT0 [2 | 0 | 1416.628 | 0 | 2.02 | 0.000176 | 13.04 | 40 | 160 | 13.08 | 4.61 | | High | High | High | High | High | High | High | High | High | High | High | 0 | 6.01E-05 | 2.66 |
| FALSE | High | CVIDPEDSS | [K].CVIDPE | 1xCarbami | Q9R0L6-2 | 1.43E-05 | 0 | 1 | 3 | 4 | Q9R0L6-2 | Isoform 2 c | Q9R0L6-2 | 0 | 2675.157 | 0 | 0.67 | 0.018867 | 9.32 | 82.2 | 117.8 | 12.97 | 21.18 | | High | High | High | High | High | High | High | High | High | High | High | 0 | 2.02E-06 | 3.64 |
| FALSE | High | CVIDPEDSS | [K].CVIDPE | 1xCarbami | Q9R0L6-2 | 0.032831 | 0.002195 | 1 | 3 | 1 | Q9R0L6-2 | Isoform 2 c | Q9R0L6-2 | 0 | 2676.141 | 0 | 0.2 | 0.18391 | 4.31 | 91.8 | 108.2 | 4.56 | 14.34 | | High | High | High | High | High | High | High | High | High | High | High | 0.001151 | 0.01188 | 3.98 |
| FALSE | High | CVIDPEDSS | [K].CVIDPE | 1xCarbami | Q9R0L6-2 | 3.03E-05 | 0 | 1 | 3 | 2 | Q9R0L6-2 | Isoform 2 c | Q9R0L6-2 | 0 | 2691.152 | 0 | 0.28 | 0.04978 | 21.38 | 91.6 | 108.4 | 17.56 | 11.48 | | High | High | High | High | High | High | High | High | High | High | High | 0 | 4.69E-06 | 3.43 |
| FALSE | High | CVLTFPASF | [K].CVLTFP | 1xCarbami | Q8C4S8 1x | 0.000731 | 0 | 1 | 3 | 2 | Q8C4S8 | DENN dom | Q8C4S8 [3 | 0 | 2547.282 | 0 | -1.48 | 0.001377 | 1.79 | 147.2 | 52.8 | 9.22 | 9.44 | | High | High | High | High | High | High | High | High | High | High | High | 0 | 0.000167 | 2.85 |
| FALSE | High | CVSELEEEK | [R].CVSELE | 1xCarbami | B2RWW2 1 | 0.011048 | 0.000772 | 1 | 5 | 1 | B2RWW2 | Golgb1 pro | B2RWW2 [ | 0 | 1660.791 | 0 | 1.14 | 0.002393 | 20.42 | 61.8 | 138.2 | 19.17 | 17.8 | | High | High | High | High | High | High | High | High | High | High | High | 0.000399 | 0.003495 | 2.71 |
| FALSE | High | CVSSVTPSS | [K].CVSSVT | 1xCarbami | E9Q1K1 1x | 0.000334 | 0 | 1 | 2 | 2 | E9Q1K1 | Cilia- and fl | E9Q1K1 [2 | 0 | 1686.854 | 0 | 0.38 | 0.031407 | 15.39 | 87 | 113 | 9.62 | 5.65 | | High | High | High | High | High | High | High | High | High | High | High | 0 | 6.93E-05 | 3.02 |
| FALSE | High | CVSVQTDP | [R].CVSVQ | 1xCarbami | H9KV00 1x | 2.2E-06 | 0 | 1 | 7 | 1 | H9KV00 | Protein SO | H9KV00 [9 | 0 | 2214.077 | 0 | 0.6 | 0.003152 | 2.28 | 79.2 | 120.8 | 5.23 | 11.64 | | High | High | High | High | High | High | High | High | High | High | High | 0 | 2.46E-07 | 2.8 |
| FALSE | High | CVWSPLAS | [R].CVWSP | 1xCarbami | Q6PR54-3 | 2.7E-05 | 0 | 2 | 4 | 2 | Q6PR54-3; | Isoform 3 c | Q6PR54-3 | 0 | 2184.154 | 0 | -0.72 | 0.000471 | 1.19 | 128.1 | 71.9 | 4.19 | 6.46 | Shared | High | High | High | High | High | High | High | High | High | High | High | 4.94E-05 | 0.000671 | 3.9 |
| FALSE | High | CVWSPLAS | [R].CVWSP | 1xCarbami | Q6PR54-3 | 0.002355 | 0.000109 | 2 | 4 | 2 | Q6PR54-3; | Isoform 3 c | Q6PR54-3 | 0 | 2264.12 | 0 | -1.24 | 0.001191 | 3.35 | 140.4 | 59.6 | 6.32 | 13.56 | Shared | High | High | High | High | High | High | High | High | High | High | High | 0.000161 | 0.001203 | 2.79 |
| FALSE | High | CYSGGSVQ | [R].CYSGG | 1xCarbami | Q8BGR5 1x | 0.056724 | 0.004124 | 1 | 3 | 2 | Q8BGR5 | Protein FAI | Q8BGR5 [1 | 0 | 1322.576 | 0 | 1.34 | 0.000917 | 17.17 | 56.7 | 143.3 | 7.01 | 7.66 | | High | High | High | High | High | High | High | High | High | High | High | 0.004126 | 0.04419 | 1.25 |
| FALSE | High | DAADPSPP | [R].DAADP | 1xPhospho | Q924H7 1x | 0.002376 | 0.000109 | 1 | 10 | 2 | Q924H7 | WW doma | Q924H7 [4 | 0 | 1549.766 | 0 | -0.07 | 0.229431 | 6.24 | 100.7 | 99.3 | 2.54 | 6.15 | | High | High | High | High | High | High | High | High | High | High | High | 0.000467 | 0.004282 | 2.45 |
| FALSE | High | DAAGTAQV | [K].DAAGT | 1xPhospho | A0A0R4J0N | 9.76E-07 | 0 | 1 | 1 | 2 | A0A0R4J0N | Tubulin-sp | A0A0R4J0N | 0 | 2730.412 | 0 | 0.76 | 0.000444 | 8.51 | 74.2 | 125.8 | 7.43 | 3.39 | | High | High | High | High | High | High | High | High | High | High | High | 0 | 9.93E-08 | 3.82 |
| FALSE | High | DAAQSSPA | [K].DAAQS | 1xPhospho | P10518 1xP | 9.74E-06 | 0 | 1 | 2 | 4 | P10518 | Delta-amin | P10518 [21 | 0 | 1530.679 | 0 | -0.12 | 0.191532 | 13.77 | 106.9 | 93.1 | 2.94 | 13.31 | | High | High | High | High | High | High | High | High | High | High | High | 0 | 1.31E-06 | 3.18 |
| FALSE | High | DAASSPQK | [K].DAASSF | 1xPhospho | B9EKX8 1x | 0.039011 | 0.002718 | 1 | 5 | 1 | B9EKX8 | Ell2 protei | B9EKX8 [3 | 0 | 1341.682 | 0 | -1.07 | 0.000458 | 2.04 | 135 | 65 | 6.09 | 8.8 | | High | High | High | High | High | High | High | High | High | High | High | 0.001421 | 0.01449 | 2.34 |
| FALSE | High | DADESPGS | [K].DADESI | 1xPhospho | P52194 1xP | 9.2E-05 | 0 | 1 | 1 | 1 | P52194 | Calmegin C | P52194 [58 | 0 | 1896.899 | 0 | -0.72 | 0.002065 | 4.46 | 130.2 | 69.8 | 10.05 | 10.93 | | High | High | High | High | High | High | High | High | High | High | High | 0 | 1.64E-05 | 3.56 |
| FALSE | High | DADLDLSD | [R].DADLD | 1xPhospho | Q569L8 1x | 0.010853 | 0.000772 | 1 | 1 | 2 | Q569L8 | Centromer | Q569L8 [7 | 0 | 1529.75 | 0 | -0.69 | 0.001264 | 4.68 | 124.1 | 75.9 | 4.82 | 7.15 | | High | High | High | High | High | High | High | High | High | High | High | 0.000997 | 0.01057 | 2.13 |
| FALSE | High | DADSEYLP | [K].DADSE | 1xPhospho | Q7TPQ3 2x | 0.000446 | 0 | 1 | 5 | 4 | Q7TPQ3 | E3 ubiquiti | Q7TPQ3 [5 | 0 | 3428.352 | 0 | 0.06 | 0.515519 | 1.75 | 97.9 | 102.1 | 8.07 | 15.57 | | High | High | High | High | High | High | High | High | High | High | High | 0 | 0.000301 | 3.4 |
| FALSE | High | DADSGDSC | [R].DADSG | 1xPhospho | Q3ULZ6 1x | 1.12E-06 | 0 | 2 | 3 | 1 | Q3ULZ6; Q | Putative ur | Q3ULZ6 [2 | 0 | 2195.002 | 0 | 2.94 | 0.003268 | 22.82 | 24.3 | 175.7 | 6.88 | 22.31 | Shared | High | High | High | High | High | High | High | High | High | High | High | 0 | 1.15E-07 | 2.96 |
| FALSE | High | DADVSEES | [K].DADVS | 1xPhospho | P35831 1x | 2.18E-07 | 0 | 1 | 1 | 5 | P35831 | Tyrosine-p | P35831 [66 | 0 | 1946.895 | 0 | 0.88 | 0.000173 | 1.73 | 70.5 | 129.5 | 6.62 | 5.41 | | High | High | High | High | High | High | High | High | High | High | High | 0 | 0.000146 | 3.01 |
| FALSE | High | DAEAGGSC | [K].DAEAG | 1xPhospho | Q811P8 1x | 1.57E-11 | 0 | 1 | 3 | 2 | Q811P8 | Rho GTPas | Q811P8 [8 | 0 | 3169.494 | 0 | -1.43 | 0.000426 | 1.34 | 147.6 | 52.4 | 7.21 | 3.97 | | High | High | High | High | High | High | High | High | High | High | High | 0 | 4.47E-13 | 6.18 |
| FALSE | High | DAEAGGSC | [K].DAEAG | 2xPhospho | Q811P8 2x | 0.000108 | 0 | 1 | 3 | 2 | Q811P8 | Rho GTPas | Q811P8 [8 | 0 | 3249.46 | 0 | -0.98 | 0.001161 | 3.2 | 132.5 | 67.5 | 5.58 | 6.79 | | High | High | High | High | High | High | High | High | High | High | High | 0 | 1.97E-05 | 4.41 |
| FALSE | High | DAESPSWF | [K].DAESPS | 1xPhospho | Q9WTI7 1x | 0.024001 | 0.001612 | 1 | 4 | 1 | Q9WTI7 | Unconvent | Q9WTI7 [4 | 0 | 1256.551 | 0 | -0.55 | 0.001981 | 1.39 | 120.2 | 79.8 | 10.58 | 9.25 | | High | High | High | High | High | High | High | High | High | High | High | 0.000859 | 0.008395 | 1.2 |
| FALSE | High | DAEYIYPSL | [K].DAEYIY | 1xPhospho | Q80TJ7 1x | 1.51E-08 | 0 | 1 | 2 | 4 | Q80TJ7 | Histone lys | Q80TJ7 [8 | 0 | 2694.248 | 0 | -0.21 | 0.005206 | 3.13 | 107.2 | 92.8 | 3.38 | 3.67 | | High | High | High | High | High | High | High | High | High | High | High | 0 | 9.12E-06 | 4.23 |
| FALSE | High | DAEYIYPSL | [K].DAEYIY | 2xPhospho | Q80TJ7 2x | 0.000352 | 0 | 1 | 2 | 3 | Q80TJ7 | Histone lys | Q80TJ7 [8 | 0 | 2774.214 | 0 | -0.07 | 0.127698 | 6.28 | 101.9 | 98.1 | 4.34 | 8.26 | | High | High | High | High | High | High | High | High | High | High | High | 0.000449 | 0.003802 | 2.65 |
| FALSE | High | DAGAEVPS | [R].DAGAE | 1xPhospho | E9PYH6 1x | 0.002463 | 0.000109 | 1 | 1 | 2 | E9PYH6 | SET domai | E9PYH6 [5 | 1 | 4728.148 | 0 | -0.3 | 0.026608 | 6.52 | 112.1 | 87.9 | 5.9 | 10.36 | | High | High | High | High | High | High | High | High | High | High | High | 4.94E-05 | 0.000655 | 4.23 |
| FALSE | High | DASPTPSTI | [R].DASPTF | 1xPhospho | O55033 1x | 6.97E-06 | 0 | 1 | 3 | 8 | O55033 | Cytoplasm | O55033 [8 | 0 | 2388.019 | 0 | 0.96 | 0.000396 | 8.64 | 68.8 | 131.2 | 10.92 | 2.15 | | High | High | High | High | High | High | High | High | High | High | High | 0 | 8.98E-07 | 4.65 |
| FALSE | High | AVDSSSEEI | [K].AVDSSS | 3xPhospho | Q5XG69 3x | 0.028824 | 0.001829 | 1 | 1 | 1 | Q5XG69 | Soluble lan | Q5XG69 [6 | 1 | 2584.118 | 0 | -0.93 | 0.006192 | 3.95 | 128.1 | 71.9 | 16.37 | 11.54 | | High | High | High | High | High | High | High | High | High | High | High | 0.000997 | 0.01028 | 2.69 |
| FALSE | High | DASPTPSTI | [R].DASPTF | 1xDeamida | O55033 1x | 0.000231 | 0 | 1 | 3 | 2 | O55033 | Cytoplasm | O55033 [8 | 0 | 2389.003 | 0 | -0.2 | 0.049945 | 14.06 | 107.6 | 92.4 | 3.79 | 13.21 | | High | High | High | High | High | High | High | High | High | High | High | 0 | 4.58E-05 | 3.17 |
| FALSE | High | DASVYQDE | [R].DASVY | 1xPhospho | D3YZK4 1x | 6.65E-05 | 0 | 1 | 6 | 3 | D3YZK4 | Transcripti | D3YZK4 [1 | 0 | 3006.46 | 0 | 0.86 | 0.024316 | 52.97 | 71.2 | 128.8 | 37.97 | 18.56 | | High | High | High | High | High | High | High | High | High | High | High | 0 | 4.59E-05 | 2.98 |
| FALSE | High | DDCYDSSSI | [R].DDCYD | 1xCarbami | Q6PAL0 1x | 0.033986 | 0.002195 | 1 | 1 | 4 | Q6PAL0 | BEN domai | Q6PAL0 [4 | 0 | 2453.12 | 0 | -0.57 | 0.002601 | 4.73 | 120.7 | 79.3 | 6.2 | 7.85 | | High | High | High | High | High | High | High | High | High | High | High | 0.008966 | 0.09583 | 2.86 |
| FALSE | High | DDCYDSSSI | [R].DDCYD | 1xCarbami | Q6PAL0 2x | 0.000529 | 0 | 1 | 1 | 3 | Q6PAL0 | BEN domai | Q6PAL0 [4 | 0 | 2533.086 | 0 | -1.46 | 0.000361 | 0.99 | 147.7 | 52.3 | 7.37 | 10.27 | | High | High | High | High | High | High | High | High | High | High | High | 0.002293 | 0.02392 | 2.58 |
| FALSE | High | DDDDIDLF | [K].DDDDII | 1xPhospho | O70251 1x | 6.48E-09 | 0 | 1 | 2 | 120 | O70251 | Elongation | O70251 [9 | 0 | 2811.166 | 0 | -0.23 | 0.042717 | 9.69 | 108.2 | 91.8 | 3.12 | 7.96 | | High | High | High | High | High | High | High | High | High | High | High | 0 | 4.81E-10 | 5.02 |
| FALSE | High | DDDDIDLF | [K].DDDDII | 1xPhospho | O70251 1x | 0.031989 | 0.00205 | 1 | 2 | 4 | O70251 | Elongation | O70251 [9 | 1 | 2939.261 | 0 | 0.35 | 0.992627 | 26.76 | 87.9 | 112.1 | 77.03 | 30.26 | | High | High | High | High | High | High | High | High | High | High | High | 0.001126 | 0.01163 | 3.51 |
| FALSE | High | DDDDIDLF | [K].DDDDII | 1xPhospho | O70251 1x | 7.4E-08 | 0 | 1 | 2 | 6 | O70251 | Elongation | O70251 [9 | 1 | 3168.424 | 0 | -0.86 | 0.025565 | 3.19 | 127.3 | 72.7 | 16.77 | 6.93 | | High | High | High | High | High | High | High | High | High | High | High | 0 | 7.98E-08 | 3.88 |
| FALSE | High | DDDLVEFS | [K].DDDLV | 1xPhospho | Q09XV5 1x | 8.65E-09 | 0 | 1 | 2 | 4 | Q09XV5 | Chromodo | Q09XV5 [1 | 0 | 2590.103 | 0 | -0.59 | 0.002067 | 5.75 | 124 | 76 | 6.55 | 7.55 | | High | High | High | High | High | High | High | High | High | High | High | 0 | 5.23E-08 | 3.44 |
| FALSE | High | DDDLVEFS | [K].DDDLV | 2xPhospho | Q09XV5 2x | 5.84E-07 | 0 | 1 | 2 | 3 | Q09XV5 | Chromodo | Q09XV5 [1 | 0 | 2670.069 | 0 | -0.96 | 0.005584 | 7.83 | 137.3 | 62.7 | 8.99 | 20.05 | | High | High | High | High | High | High | High | High | High | High | High | 0 | 5.53E-08 | 4.86 |
| FALSE | High | DDDSDLILN | [K].DDDSD | 1xPhospho | A1L3P4 1x | 3.09E-05 | 0 | 1 | 5 | 2 | A1L3P4 | Sodium/hy | A1L3P4 [6 | 0 | 3737.675 | 0 | -0.26 | 0.13408 | 10.92 | 106.9 | 93.1 | 12.5 | 3.68 | | High | High | High | High | High | High | High | High | High | High | High | 0 | 4.79E-06 | 4.35 |
| FALSE | High | DDDSLPAE | [K].DDDSLI | 1xPhospho | E9Q2H1 1x | 3.68E-06 | 0 | 1 | 2 | 1 | E9Q2H1 | E3 ubiquiti | E9Q2H1 [2 | 0 | 2254.997 | 0 | 0.71 | 0.004895 | 10.47 | 77.6 | 122.4 | 9.59 | 11.91 | | High | High | High | High | High | High | High | High | High | High | High | 0 | 4.39E-07 | 3.75 |
| FALSE | High | DDDSSVAD | [K].DDDSS | 1xPhospho | Q9CQK1 1 | 2.86E-06 | 0 | 1 | 2 | 4 | Q9CQK1 | Zinc finger | Q9CQK1 [6 | 0 | 2366.935 | 0 | -1.2 | 0.003822 | 3.23 | 139.3 | 60.7 | 12.78 | 6.5 | | High | High | High | High | High | High | High | High | High | High | High | 0 | 3.31E-07 | 5.13 |
| FALSE | High | DDDSSVAD | [K].DDDSS | 1xPhospho | Q9CQK1 1 | 2.38E-05 | 0 | 1 | 2 | 5 | Q9CQK1 | Zinc finger | Q9CQK1 [6 | 1 | 3378.563 | 0 | -1.19 | 0.001468 | 3.5 | 140.5 | 59.5 | 10.07 | 4.67 | | High | High | High | High | High | High | High | High | High | High | High | 0 | 3.58E-06 | 3.37 |
| FALSE | High | DDDTASVT | [R].DDDTA | 1xPhospho | D3YZK4 1x | 0.000157 | 0 | 1 | 6 | 6 | D3YZK4 | Transcripti | D3YZK4 [1 | 0 | 2005.891 | 0 | -0.45 | 0.007864 | 3.92 | 115.6 | 84.4 | 6.95 | 4.8 | | High | High | High | High | High | High | High | High | High | High | High | 0.001355 | 0.01425 | 1.89 |
| FALSE | High | DDEDTPPN | [R].DDEDT | 1xPhospho | P52734 1x | 0.003927 | 0.000311 | 1 | 4 | 1 | P52734 | FYVE, Rhol | P52734 [7 | 0 | 2251.053 | 0 | -0.75 | 0.002244 | 2.33 | 124.9 | 75.1 | 7.5 | 6.61 | | High | High | High | High | High | High | High | High | High | High | High | 0.00014 | 0.001107 | 2.62 |
| FALSE | High | DDATPVRD | [R].DDATP | 1xPhospho | P97868 1x | 0.001674 | 5.66E-05 | 1 | 2 | 2 | P97868 | E3 ubiquiti | P97868 [9 | 1 | 2691.263 | 0 | 0.07 | 0.28526 | 10.45 | 94.8 | 105.2 | 14.56 | 27.36 | | High | High | High | High | High | High | High | High | High | High | High | 2.58E-05 | 0.000426 | 3 |
| FALSE | High | DDEDTPPN | [R].DDEDT | 2xPhospho | P52734 2x | 0.098533 | 0.007902 | 1 | 4 | 1 | P52734 | FYVE, Rhol | P52734 [7 | 0 | 2331.019 | 0 | -0.49 | 0.029032 | 9.2 | 117.2 | 82.8 | 11.07 | 9.82 | | High | High | High | High | High | High | High | High | High | High | High | 0.003963 | 0.04223 | 2.41 |
| FALSE | High | DDEHIVLEF | [K].DDEHIV | 1xPhospho | Q7TSH8 1x | 9.76E-07 | 0 | 1 | 2 | 2 | Q7TSH8 | Transmem | Q7TSH8 [2 | 0 | 2947.352 | 0 | 1.31 | 0.003469 | 16 | 55.6 | 144.4 | 14.09 | 21.24 | | High | High | High | High | High | High | High | High | High | High | High | 0 | 9.92E-08 | 4.61 |
| FALSE | High | DDEPSGWI | [R].DDEPS | 1xPhospho | Q8BKI2 1x | 1.88E-05 | 0 | 1 | 3 | 2 | Q8BKI2 | Trinucleoti | Q8BKI2 [9 | 0 | 2224.96 | 0 | 0.88 | 0.015593 | 6.99 | 71 | 129 | 7.88 | 18.53 | | High | High | High | High | High | High | High | High | High | High | High | 0 | 0.000329 | 3.53 |
| FALSE | High | DDFESEEEI | [K].DDFESE | 1xPhospho | P97868 1x | 0.030104 | 0.001857 | 1 | 3 | 2 | P97868 | E3 ubiquiti | P97868 [1 | 0 | 1879.825 | 0 | 0.02 | 0.758753 | 10.84 | 102.1 | 97.9 | 8.72 | 13.11 | | High | High | High | High | High | High | High | High | High | High | High | 0.001011 | 0.01081 | 2.05 |
| FALSE | High | DDGDEEGL | [R].DDGDE | 1xPhospho | Q64012 1x | 0.000158 | 0 | 1 | 2 | 5 | Q64012 | RNA-bindir | Q64012 [2 | 0 | 3593.476 | 0 | -0.17 | 0.037719 | 3.72 | 112.4 | 87.6 | 12.51 | 10.31 | | High | High | High | High | High | High | High | High | High | High | High | 0 | 3.01E-05 | 4.35 |
| FALSE | High | DDGEMAA | [R].DDGEN | 1xCarbami | A0A1B0GR | 0.095408 | 0.007437 | 1 | 1 | 1 | A0A1B0GR | Single stra | A0A1B0GR | 0 | 3697.65 | 0 | 0.82 | 0.002226 | 7.66 | 72.7 | 127.3 | 11.43 | 13.75 | | High | High | High | High | High | High | High | High | High | High | High | 0.003782 | 0.0408 | 2.93 |
| FALSE | High | DDGESQPF | [R].DDGES | 1xPhospho | B9EKJ7 1x | 0.000725 | 0 | 1 | 2 | 5 | B9EKJ7 | Tankyrase | B9EKJ7 [5 | 0 | 3465.803 | 0 | 0.17 | 0.131366 | 15.37 | 91.8 | 108.2 | 9.22 | 14.09 | | High | High | High | High | High | High | High | High | High | High | High | 0 | 0.000166 | 3.6 |
| FALSE | High | DDGLADLL | [R].DDGLA | 1xPhospho | P27546 1x | 2.72E-05 | 0 | 1 | 3 | 3 | P27546 | Microtubu | P27546 [1 | 0 | 2692.27 | 0 | 0.69 | 0.002786 | 4.86 | 75.4 | 124.6 | 8.34 | 10.64 | | High | High | High | High | High | High | High | High | High | High | High | 0 | 0.000281 | 3.85 |
| FALSE | High | DDGSWEV | [R].DDGSW | 1xPhospho | P26443 1x | 0.000616 | 0 | 1 | 1 | 2 | P26443 | Glutamate | P26443 [1 | 0 | 1734.757 | 0 | -0.09 | 0.21338 | 10.36 | 102.2 | 97.8 | 7.86 | 9.98 | | High | High | High | High | High | High | High | High | High | High | High | 0 | 0.000138 | 3.36 |
| FALSE | High | DDGVFVQI | [R].DDGVF | 1xPhospho | E9Q616 1x | 1.76E-08 | 0 | 1 | 4 | 1 | E9Q616 | AHNAK nu | E9Q616 [3 | 0 | 2171.999 | 0 | 2.19 | 0.002305 | 5.64 | 35.9 | 164.1 | 8.76 | 19.18 | | High | High | High | High | High | High | High | High | High | High | High | 0 | 1.09E-09 | 5.15 |
| FALSE | High | DDGVFVQI | [R].DDGVF | 1xDeamida | E9Q616 1x | 0.007538 | 0.00062 | 1 | 4 | 1 | E9Q616 | AHNAK nu | E9Q616 [3 | 0 | 2172.983 | 0 | 0.07 | 0.277402 | 9.31 | 99.5 | 100.5 | 3.78 | 8 | | High | High | High | High | High | High | High | High | High | High | High | 0.000294 | 0.002295 | 2.73 |
| FALSE | High | DDKDEEDE | [K].DDKDE | 1xPhospho | J3QNU6 1x | 0.007881 | 0.000736 | 1 | 4 | 3 | J3QNU6 | Beta-arres | J3QNU6 [4 | 1 | 2552.13 | 0 | 1.7 | 0.00261 | 8.48 | 47.6 | 152.4 | 7.61 | 5.63 | | High | High | High | High | High | High | High | High | High | High | High | 0.000327 | 0.002402 | 2.72 |
| FALSE | High | DDLDVTEL | [R].DDLDV | 1xDeamida | Q3U955 1x | 0.004613 | 0.000456 | 1 | 1 | 1 | Q3U955 | Putative ur | Q3U955 [1 | 0 | 2525.222 | 0 | 0.64 | 0.013072 | 12.12 | 71.8 | 128.2 | 8.9 | 18.06 | | High | High | High | High | High | High | High | High | High | High | High | 0.000204 | 0.001326 | 2.82 |
| FALSE | High | DDIEDDVSS | [K].DDIEDD | 1xPhospho | Q8BYH8 1x | 5.33E-07 | 0 | 1 | 4 | 1 | Q8BYH8 | Chromodo | Q8BYH8 [1 | 0 | 3357.533 | 0 | 0.71 | 0.034296 | 21.85 | 81.4 | 118.6 | 13.16 | 11.8 | | High | High | High | High | High | High | High | High | High | High | High | 0 | 5.03E-08 | 6.04 |
| FALSE | High | DDEHIVLEF | [K].DDEHIV | 1xPhospho | Q7TSH8 1x | 0.000143 | 0 | 1 | 2 | 4 | Q7TSH8 | Transmem | Q7TSH8 [2 | 0 | 2867.385 | 0 | 0.41 | 0.007436 | 14.32 | 85 | 115 | 11.25 | 7.62 | | High | High | High | High | High | High | High | High | High | High | High | 0 | 2.74E-05 | 3.39 |
| FALSE | High | DDATPVRD | [R].DDATP | 1xPhospho | P97868 1x | 0.004411 | 0.000409 | 1 | 2 | 2 | P97868 | E3 ubiquiti | P97868 [9 | 1 | 2675.268 | 0 | 0.29 | 0.403566 | 3.06 | 91.2 | 108.8 | 7 | 18.68 | | High | High | High | High | High | High | High | High | High | High | High | 0.000555 | 0.004907 | 2.9 |
| FALSE | High | DDATESEV | [R].DDATE | 1xPhospho | Q7TSG1 1x | 0.003055 | 0.00016 | 1 | 2 | 3 | Q7TSG1 | Centrosom | Q7TSG1 [3 | 0 | 2267.037 | 0 | -0.54 | 0.000975 | 0.22 | 119.1 | 80.9 | 5.94 | 2.3 | | High | High | High | High | High | High | High | High | High | High | High | 0.000344 | 0.003177 | 4.3 |
| FALSE | High | DDASQVSS | [K].DDASQ | 1xPhospho | Q62315 1x | 7.67E-08 | 0 | 1 | 3 | 6 | Q62315 | Protein Jur | Q62315 [8 | 0 | 2812.163 | 0 | -1.06 | 0.001007 | 0.81 | 134.2 | 65.8 | 6.17 | 5.69 | | High | High | High | High | High | High | High | High | High | High | High | 0 | 5.7E-09 | 5.3 |
| FALSE | High | DASVYQDE | [R].DASVY | 1xDeamida | D3YZK4 1x | 0.004911 | 0.000504 | 1 | 6 | 1 | D3YZK4 | Transcripti | D3YZK4 [1 | 0 | 3007.444 | 0 | 0.49 | 0.148285 | 16.98 | 83.1 | 116.9 | 30.28 | 11.57 | | High | High | High | High | High | High | High | High | High | High | High | 0.000225 | 0.001421 | 3.62 |

| | | | | | | | | | | | | | | | | | | | | | | | | | | | | | | | | | | | | | | |
|---|---|---|---|---|---|---|---|---|---|---|---|---|---|---|---|---|---|---|---|---|---|---|---|---|---|---|---|---|---|---|---|---|---|---|---|---|---|---|
| FALSE | High | DATIPVESD | [R].DATIPV | 1xPhospho | Q99L48 1x | 1.01E-07 | 0 | 1 | 1 | 3 | Q99L48 | 60S riboso | Q99L48 [4 | 0 | 2096.922 | 0 | -0.19 | 0.097721 | 5.86 | 108.6 | 91.4 | 7.94 | 15.29 | | High | High | High | High | High | High | High | High | High | High | High | 0 | 7.79E-09 | 4.3 |
| FALSE | High | DATPPVSPI | [R].DATPPV | 2xPhospho | P09450 2x | 7.47E-05 | 0 | 1 | 3 | 1 | P09450 | Transcripti | P09450 [25 | 0 | 2187.923 | 0 | 2.19 | 0.007369 | 8.48 | 34.8 | 165.2 | 13.9 | 25.07 | | High | High | High | High | High | High | High | High | High | High | High | 0 | 1.29E-05 | 3.35 |
| FALSE | High | DAVFDGSS | [K].DAVFD | 1xCarbami | Q99N57-2 | 4.27E-05 | 0 | 1 | 3 | 1 | Q99N57-2 | Isoform 2 c | Q99N57-2 | 0 | 2784.29 | 0 | 1.17 | 0.010598 | 18.24 | 58.1 | 141.9 | 9.77 | 24.11 | | High | High | High | High | High | High | High | High | High | High | High | 0 | 6.92E-06 | 3.14 |
| FALSE | High | DAVQAPLN | [K].DAVQA | 1xPhospho | F6ZDS4 1x | 4.87E-07 | 0 | 1 | 3 | 2 | F6ZDS4 | Nucleopro | F6ZDS4 [12 | 0 | 2222.147 | 0 | -1.17 | 0.00046 | 2.4 | 137.9 | 62.1 | 5.55 | 11.01 | | High | High | High | High | High | High | High | High | High | High | High | 0 | 4.53E-08 | 3.88 |
| FALSE | High | DAVSPFSFS | [K].DAVSPI | 1xPhospho | Q8BTI7 1x | 0.002172 | 5.66E-05 | 1 | 2 | 2 | Q8BTI7 | Serine/thre | Q8BTI7 [10 | 0 | 1848.991 | 0 | 2.58 | 0.000278 | 19.04 | 31.6 | 168.4 | 14.78 | 6.81 | | High | High | High | High | High | High | High | High | High | High | High | 4.94E-05 | 0.000572 | 2.53 |
| FALSE | High | DAVTEDSP | [K].DAVTEI | 1xPhospho | P70399 1x | 9.67E-07 | 0 | 1 | 5 | 2 | P70399 | TP53-bindi | P70399 [81 | 0 | 2016.984 | 0 | 0.92 | 0.007979 | 7.35 | 70.5 | 129.5 | 4.67 | 13.52 | | High | High | High | High | High | High | High | High | High | High | High | 0 | 9.82E-08 | 2.28 |
| FALSE | High | DAVYGTPC | [K].DAVYG | 1xPhospho | Q8BY89-2 | 0.119269 | 0.00979 | 1 | 2 | 1 | Q8BY89-2 | Isoform 2 c | Q8BY89-2 | 0 | 1516.781 | 0 | 0.28 | 0.023033 | 5.57 | 91.4 | 108.6 | 5.34 | 8.8 | | High | High | High | High | High | High | High | High | High | High | High | 0.004894 | 0.05335 | 1.61 |
| FALSE | High | DAWIAAEC | [K].DAWIA | 1xPhospho | O88466 1x | 6.07E-05 | 0 | 2 | 3 | 1 | O88466; R | Zinc finger | O88466 [1 | 0 | 2009.953 | 0 | -0.59 | 0.00874 | 7.39 | 120.9 | 79.1 | 5.32 | 14.39 | Shared | High | High | High | High | High | High | High | High | High | High | High | 0 | 1.03E-05 | 3.17 |
| FALSE | High | DAYSSFGSI | [K].DAYSSF | 1xPhospho | Q3TQX5 1x | 0.045122 | 0.003373 | 1 | 6 | 1 | Q3TQX5 | Putative ur | Q3TQX5 [6 | 0 | 1298.562 | 0 | 0.43 | 0.000735 | 6.67 | 86.4 | 113.6 | 8.92 | 6.68 | | High | High | High | High | High | High | High | High | High | High | High | 0.001712 | 0.01706 | 0.8 |
| FALSE | High | DCAESPNP | [K].DCAESI | 2xCarbami | Q7TPQ3 1x | 4.55E-08 | 0 | 1 | 5 | 2 | Q7TPQ3 | E3 ubiquiti | Q7TPQ3 [6 | 0 | 3395.388 | 0 | 0.22 | 0.0997 | 5.81 | 95.3 | 104.7 | 6.17 | 16.39 | | High | High | High | High | High | High | High | High | High | High | High | 0 | 3.16E-09 | 7.49 |
| FALSE | High | DCAPGEGP | [K].DCAPG | 1xCarbami | G3X9I0 1x | 2.56E-05 | 0 | 1 | 5 | 1 | G3X9I0 | Zinc finger | G3X9I0 [44 | 0 | 2664.179 | 0 | -0.66 | 0.01048 | 9.52 | 122.4 | 77.6 | 5.31 | 14.84 | | High | High | High | High | High | High | High | High | High | High | High | 0 | 3.86E-06 | 3.07 |
| FALSE | High | DCDSPVSLI | [K].DCDSPV | 1xCarbami | H3BJL0 1x | 0.013074 | 0.000889 | 1 | 19 | 1 | H3BJL0 | DNA dC->d | H3BJL0 [63 | 0 | 2136.035 | 0 | -1.48 | 0.000181 | 0.61 | 147.6 | 52.4 | 5.58 | 7.05 | | High | High | High | High | High | High | High | High | High | High | High | 0.000467 | 0.004221 | 2.38 |
| FALSE | High | DCDVTALA | [K].DCDVT | 2xCarbami | Q80Z37 1x | 5.38E-07 | 0 | 1 | 1 | 5 | Q80Z37 | E3 ubiquiti | Q80Z37 [9 | 0 | 4029.814 | 0 | 0.86 | 0.012719 | 17.13 | 72.6 | 127.4 | 12.46 | 22.43 | | High | High | High | High | High | High | High | High | High | High | High | 0 | 5.09E-08 | 5.12 |
| FALSE | High | DCEESNMI | [R].DCEESN | 1xCarbami | Q9WV30-2 | 2.17E-05 | 0 | 1 | 7 | 1 | Q9WV30-2 | Isoform 2 c | Q9WV30-2 | 0 | 2385.003 | 0 | 1.45 | 0.007106 | 35.44 | 53.1 | 146.9 | 13.2 | 21.15 | | High | High | High | High | High | High | High | High | High | High | High | 0 | 3.24E-06 | 4.15 |
| FALSE | High | DCEYQQISI | [K].DCEYQ | 1xCarbami | Q5Y5T2 1x | 9E-07 | 0 | 1 | 1 | 1 | Q5Y5T2 | Palmitoyltr | Q5Y5T2 [3 | 0 | 2478.132 | 0 | -0.62 | 0.000488 | 1.78 | 121 | 79 | 6.21 | 4.05 | | High | High | High | High | High | High | High | High | High | High | High | 0 | 9.05E-08 | 2.84 |
| FALSE | High | DCGASLPH | [R].DCGASI | 1xCarbami | E9QAU4 1x | 0.041781 | 0.003126 | 1 | 3 | 1 | E9QAU4 | Enhancer t | E9QAU4 [1 | 0 | 2063.924 | 0 | -0.47 | 0.02918 | 2.89 | 117.5 | 82.5 | 6.24 | 20 | | High | High | High | High | High | High | High | High | High | High | High | 0.001609 | 0.01564 | 1.5 |
| FALSE | High | DCGASLPH | [R].DCGASI | 1xCarbami | E9QAU4 1x | 0.046291 | 0.003523 | 1 | 3 | 1 | E9QAU4 | Enhancer t | E9QAU4 [1 | 0 | 2079.919 | 0 | -0.21 | 0.056462 | 8.72 | 105.2 | 94.8 | 8.06 | 6.13 | | High | High | High | High | High | High | High | High | High | High | High | 0.001789 | 0.01772 | 1.34 |
| FALSE | High | DCGSPK | [K].DCGSPI | 1xCarbami | Q7TMQ1 1 | 0.082424 | 0.006288 | 1 | 2 | 2 | Q7TMQ1 | Gap junctio | Q7TMQ1 [ | 0 | 1201.569 | 0 | 2.95 | 0.001215 | 18.58 | 23 | 177 | 6.27 | 16.42 | | High | High | High | High | High | High | High | High | High | High | High | 0.003243 | 0.03432 | 1.6 |
| FALSE | High | DCLLVYSQI | [K].DCLLVY | 1xCarbami | O55111 1x | 3.81E-09 | 0 | 1 | 2 | 3 | O55111 | Desmoglei | O55111 [7 | 0 | 2078.967 | 0 | -0.23 | 0.137806 | 8.6 | 111 | 89 | 12.18 | 7.24 | | High | High | High | High | High | High | High | High | High | High | High | 0 | 1.94E-10 | 4.72 |
| FALSE | High | DCLQPEDY | [R].DCLQPI | 1xCarbami | S4R1P5 1x | 0.000496 | 0 | 1 | 6 | 1 | S4R1P5 | Dystonin C | S4R1P5 [25 | 1 | 4299.903 | 0 | 1.04 | 0.000436 | 14.24 | 69.3 | 130.7 | 13.49 | 6.67 | | High | High | High | High | High | High | High | High | High | High | High | 0 | 0.000108 | 4.24 |
| FALSE | High | DCLSSNSV | [K].DCLSSN | 2xCarbami | Q8C0P0 1x | 1.19E-05 | 0 | 1 | 1 | 2 | Q8C0P0 | Serine/thre | Q8C0P0 [5 | 0 | 1952.756 | 0 | -0.5 | 0.018207 | 4.03 | 119 | 81 | 7.19 | 10.94 | | High | High | High | High | High | High | High | High | High | High | High | 0 | 1.64E-06 | 3.49 |
| FALSE | High | DCPDHSQS | [K].DCPDH | 1xCarbami | G5E8B9 1x | 0.000147 | 0 | 1 | 3 | 2 | G5E8B9 | MCG13089 | G5E8B9 [4 | 0 | 2077.941 | 0 | -0.13 | 0.032577 | 7.6 | 104 | 96 | 3.3 | 6.01 | | High | High | High | High | High | High | High | High | High | High | High | 0 | 2.76E-05 | 2.21 |
| FALSE | High | DCPNSGQ | [R].DCPNS | 3xCarbami | G5E8D8 1x | 1.47E-05 | 0 | 1 | 4 | 1 | G5E8D8 | MLX intera | G5E8D8 [6 | 0 | 3640.55 | 0 | 0.81 | 0.001659 | 11.22 | 71.1 | 128.9 | 9.45 | 8.92 | | High | High | High | High | High | High | High | High | High | High | High | 0 | 2.1E-06 | 4.13 |
| FALSE | High | DCSYGAVT | [K].DCSYG | 1xCarbami | A2A690-2 | 5.81E-05 | 0 | 1 | 2 | 1 | A2A690-2 | Isoform 2 c | A2A690-2 | 0 | 2139.947 | 0 | 0.26 | 0.129748 | 19.69 | 88.2 | 111.8 | 17.66 | 11.57 | | High | High | High | High | High | High | High | High | High | High | High | 0 | 9.74E-06 | 2.85 |
| FALSE | High | DCTGCQQ | [K].DCTGC | 2xCarbami | O88622 1x | 6.74E-07 | 0 | 1 | 4 | 2 | O88622 | Poly(ADP-r | O88622 [2 | 0 | 3840.76 | 0 | -1.06 | 0.001613 | 1.95 | 133.8 | 66.2 | 7.23 | 11.01 | | High | High | High | High | High | High | High | High | High | High | High | 0 | 0.000197 | 3.98 |
| FALSE | High | DCTGCQQ | [K].DCTGC | 2xCarbami | O88622 1x | 1.84E-06 | 0 | 1 | 4 | 2 | O88622 | Poly(ADP-r | O88622 [2 | 0 | 3841.744 | 0 | -1.23 | 0.002281 | 2.46 | 140.3 | 59.7 | 7.56 | 14.83 | | High | High | High | High | High | High | High | High | High | High | High | 0 | 2.82E-07 | 5.11 |
| FALSE | High | DCTGCQQ | [K].DCTGC | 2xCarbami | O88622 2x | 2.05E-06 | 0 | 1 | 4 | 7 | O88622 | Poly(ADP-r | O88622 [2 | 0 | 3920.726 | 0 | -1.06 | 0.000959 | 2.82 | 133.3 | 66.7 | 4.98 | 7.78 | | High | High | High | High | High | High | High | High | High | High | High | 4.94E-05 | 0.00071 | 5.4 |
| FALSE | High | DDASESTSI | [K].DDASE | 1xDeamida | Q9CRA9 2x | 0.007538 | 0.00062 | 1 | 2 | 1 | Q9CRA9 | FGFR1 onc | Q9CRA9 [2 | 0 | 2571.131 | 0 | -0.15 | 0.7602 | 22.4 | 99.3 | 100.7 | 12.58 | 29.53 | | High | High | High | High | High | High | High | High | High | High | High | 0.000275 | 0.002276 | 2.53 |
| FALSE | High | DASSQDCY | [K].DASSQI | 1xCarbami | A0A1B0GS | 0.012731 | 0.000889 | 1 | 2 | 3 | A0A1B0GS | GRB2-asso | A0A1B0GS | 0 | 1735.72 | 0 | -0.71 | 0.019804 | 15.82 | 124.9 | 75.1 | 10.29 | 22.16 | | High | High | High | High | High | High | High | High | High | High | High | 0.000449 | 0.004118 | 2.65 |
| FALSE | High | DGDESSPIL | [K].DGDES | 1xPhospho | Q6PF93 1x | 0.000297 | 0 | 1 | 5 | 3 | Q6PF93 | Phosphatic | Q6PF93 [2 | 0 | 2275.151 | 0 | 0.2 | 0.175789 | 18.36 | 94.9 | 105.1 | 11.08 | 4.96 | | High | High | High | High | High | High | High | High | High | High | High | 0 | 0.000186 | 3.56 |
| FALSE | High | AVDSSSEEI | [K].AVDSSS | 2xPhospho | Q5XG69 2x | 0.072258 | 0.005346 | 1 | 1 | 1 | Q5XG69 | Soluble lan | Q5XG69 [6 | 1 | 2504.152 | 0 | -1.13 | 0.000291 | 1.95 | 136.8 | 63.2 | 3.32 | 9.23 | | High | High | High | High | High | High | High | High | High | High | High | 0.002855 | 0.02962 | 3.26 |
| FALSE | High | AVDSSSEEI | [K].AVDSSS | 2xPhospho | Q5XG69 2x | 2.39E-07 | 0 | 1 | 1 | 16 | Q5XG69 | Soluble lan | Q5XG69 [6 | 0 | 2348.051 | 0 | -1.32 | 0.001012 | 1.61 | 140.2 | 59.8 | 4.59 | 13.71 | | High | High | High | High | High | High | High | High | High | High | High | 0 | 9.82E-08 | 5.3 |
| FALSE | High | ASIANSDGI | [K].ASIANS | 1xPhospho | Q91VK6 1x | 0.000151 | 0 | 1 | 7 | 2 | Q91VK6 | Phosphata | Q91VK6 [1 | 0 | 2380.195 | 0 | -0.98 | 0.000634 | 4.86 | 134.9 | 65.1 | 8.32 | 15.88 | | High | High | High | High | High | High | High | High | High | High | High | 0 | 2.85E-05 | 4.33 |
| FALSE | High | ASLAPLASP | [R].ASLAPL | 1xPhospho | Q6ZQ33 1x | 0.008538 | 0.000736 | 1 | 2 | 1 | Q6ZQ33 | MKIAA085 | Q6ZQ33 [6 | 0 | 1549.876 | 0 | -0.15 | 0.023554 | 3.42 | 107 | 93 | 5.27 | 10.28 | | High | High | High | High | High | High | High | High | High | High | High | 0.000327 | 0.002624 | 2.3 |
| FALSE | High | ASLDSAGA | [R].ASLDSA | 1xPhospho | Q8C6B2 1x | 0.003962 | 0.000311 | 1 | 3 | 2 | Q8C6B2 | Rhotekin C | Q8C6B2 [2 | 0 | 2600.328 | 0 | 1.3 | 0.000444 | 7.34 | 60.1 | 139.9 | 14.01 | 7.49 | | High | High | High | High | High | High | High | High | High | High | High | 0.000275 | 0.001887 | 3.11 |
| FALSE | High | ASLEAAIAD | [R].ASLEAA | 1xPhospho | P11679 1x | 0.000215 | 0 | 1 | 1 | 1 | P11679 | Keratin, ty | P11679 [33 | 0 | 1653.805 | 0 | 1.86 | 0.00045 | 19.4 | 45.2 | 154.8 | 23.57 | 6.94 | | High | High | High | High | High | High | High | High | High | High | High | 0 | 4.23E-05 | 2.81 |
| FALSE | High | ASLEIGESF | [R].ASLEIG | 1xPhospho | Q9QY06-2 | 0.027358 | 0.001676 | 2 | 8 | 1 | Q9QY06-2; | Isoform 2 c | Q9QY06-2 | 0 | 2003.014 | 0 | 0.27 | 0.055288 | 13.15 | 92.9 | 107.1 | 7.34 | 10.82 | Shared | High | High | High | High | High | High | High | High | High | High | High | 0.00092 | 0.009665 | 2.44 |
| FALSE | High | ASLENVHP | [R].ASLENV | 1xPhospho | Q9JIY2 1xP | 0.012397 | 0.000889 | 1 | 6 | 1 | Q9JIY2 | E3 ubiquiti | Q9JIY2 [20 | 0 | 2445.238 | 0 | 0.1 | 0.929887 | 32.34 | 103.4 | 96.6 | 12.96 | 17.65 | | High | High | High | High | High | High | High | High | High | High | High | 0.000449 | 0.003998 | 1.96 |
| FALSE | High | ASLGPNLD | [K].ASLGPN | 1xPhospho | Q8BHK9 1x | 8.6E-07 | 0 | 1 | 2 | 1 | Q8BHK9 | DNA excisi | Q8BHK9 [8 | 0 | 2306.174 | 0 | 1.28 | 0.0034 | 12.9 | 58.2 | 141.8 | 10.5 | 19.32 | | High | High | High | High | High | High | High | High | High | High | High | 0 | 8.58E-08 | 5.52 |
| FALSE | High | ASLGSLEGI | [K].ASLGSL | 1xPhospho | E9Q616 1x | 0.000231 | 0 | 1 | 3 | 3 | E9Q616 | AHNAK nu | E9Q616 [5 | 0 | 2069.984 | 0 | 2.2 | 0.058931 | 41.48 | 35.7 | 164.3 | 19.57 | 56.02 | | High | High | High | High | High | High | High | High | High | High | High | 0 | 0.000224 | 3.37 |
| FALSE | High | ASLGSLEGI | [K].ASLGSL | 2xPhospho | E9Q616 2x | 0.000186 | 0 | 1 | 3 | 3 | E9Q616 | AHNAK nu | E9Q616 [5 | 0 | 2149.95 | 0 | 1.29 | 0.089857 | 33.69 | 55.9 | 144.1 | 50.17 | 46.3 | | High | High | High | High | High | High | High | High | High | High | High | 0 | 3.61E-05 | 3.57 |
| FALSE | High | ASLGSLEGI | [K].ASLGSL | 1xPhospho | E9Q616 1x | 1.29E-07 | 0 | 1 | 3 | 21 | E9Q616 | AHNAK nu | E9Q616 [5 | 0 | 2299.147 | 0 | 2.37 | 0.00025 | 1.09 | 31.8 | 168.2 | 4.97 | 7.23 | | High | High | High | High | High | High | High | High | High | High | High | 0 | 1.04E-07 | 3.47 |
| FALSE | High | ASLGSLEGI | [K].ASLGSL | 2xPhospho | E9Q616 2x | 2.31E-05 | 0 | 1 | 3 | 17 | E9Q616 | AHNAK nu | E9Q616 [5 | 0 | 2379.113 | 0 | 2 | 0.000348 | 6.1 | 40.5 | 159.5 | 6.7 | 8.6 | | High | High | High | High | High | High | High | High | High | High | High | 0 | 3.23E-05 | 3.97 |
| FALSE | High | ASLGSSLPG | [K].ASLGSS | 1xPhospho | E9Q842 1x | 0.090888 | 0.007081 | 1 | 4 | 2 | E9Q842 | Neuron na | E9Q842 [1 | 0 | 2125.131 | 0 | 1.09 | 0.005925 | 26.57 | 59 | 141 | 17.2 | 11.23 | | High | High | High | High | High | High | High | High | High | High | High | 0.003633 | 0.03872 | 3.13 |
| FALSE | High | ASIHEAWT | [K].ASIHEA | 1xPhospho | A0A1L1SV2 | 0.006478 | 0.00062 | 2 | 7 | 1 | A0A1L1SV2 | Alpha-actir | A0A1L1SV2 | 0 | 1752.872 | 0 | 0.99 | 0.001493 | 17.78 | 66.8 | 133.2 | 5.59 | 7.1 | Shared | High | High | High | High | High | High | High | High | High | High | High | 0.000275 | 0.00194 | 3.27 |
| FALSE | High | ASLADALPI | [R].ASLADA | 1xPhospho | Q91V17 1x | 4.43E-05 | 0 | 1 | 5 | 1 | Q91V17 | E3 ubiquiti | Q91V17 [1 | 0 | 1753.956 | 0 | -0.5 | 0.064786 | 2.94 | 109 | 91 | 10.34 | 16.07 | | High | High | High | High | High | High | High | High | High | High | High | 0 | 7.17E-06 | 3.24 |
| FALSE | High | ASLLDQALI | [R].ASLLDC | 1xPhospho | Q8C0T5 1x | 0.017344 | 0.001204 | 1 | 3 | 1 | Q8C0T5 | Signal-indu | Q8C0T5 [1 | 0 | 3014.589 | 0 | 1.27 | 0.000917 | 6.88 | 55.5 | 144.5 | 9.41 | 9.44 | | High | High | High | High | High | High | High | High | High | High | High | 0.000656 | 0.005837 | 2.68 |
| FALSE | High | ASLLPLQLL | [K].ASLLPL | 1xPhospho | Q3U0L2 1x | 0.051258 | 0.003745 | 1 | 2 | 1 | Q3U0L2 | Ankyrin re | Q3U0L2 [3 | 0 | 1432.849 | 0 | 2.76 | 0.000881 | 18.55 | 27.9 | 172.1 | 17.13 | 13.53 | | High | High | High | High | High | High | High | High | High | High | High | 0.00196 | 0.01982 | 2.46 |
| FALSE | High | ASLLPVETS | [K].ASLLPV | 1xPhospho | Q7TT18 1x | 0.021418 | 0.001507 | 1 | 4 | 2 | Q7TT18 | Activating | Q7TT18 [4 | 0 | 2183.125 | 0 | -0.18 | 0.010598 | 6.24 | 106.2 | 93.8 | 9.06 | 13.21 | | High | High | High | High | High | High | High | High | High | High | High | 0.000801 | 0.00734 | 2.06 |
| FALSE | High | ASINSLLSD | [R].ASINSL | 1xPhospho | A2AG68 1x | 0.000246 | 0 | 1 | 3 | 1 | A2AG68 | Copper-tra | A2AG68 [1 | 0 | 1585.86 | 0 | 0.26 | 0.062406 | 21.12 | 87.5 | 112.5 | 11.75 | 8.47 | | High | High | High | High | High | High | High | High | High | High | High | 0 | 4.93E-05 | 2.32 |
| FALSE | High | ASLPEHLIQ | [R].ASLPEH | 1xPhospho | Q8BW88-4 | 0.010663 | 0.000772 | 1 | 4 | 1 | Q8BW88-4 | Isoform 4 c | Q8BW88-4 | 0 | 1673.939 | 0 | 5.76 | 0.000389 | 90.87 | 3.3 | 196.7 | 67.21 | 9.75 | | High | High | High | High | High | High | High | High | High | High | High | 0.000399 | 0.003377 | 2.66 |
| FALSE | High | ASLPVVR | [R].ASLPVV | 1xPhospho | E9QAU4 1x | 0.022378 | 0.001542 | 1 | 9 | 1 | E9QAU4 | Enhancer t | E9QAU4 [2 | 0 | 1050.591 | 0 | 0.25 | 0.005393 | 6.1 | 92.2 | 107.8 | 8.5 | 5.02 | | High | High | High | High | High | High | High | High | High | High | High | 0.000814 | 0.007716 | 2.31 |
| FALSE | High | ASIQENVLI | [K].ASIQEN | 1xPhospho | Q6ZPR5-4 | 4.14E-07 | 0 | 1 | 7 | 3 | Q6ZPR5-4 | Isoform 4 c | Q6ZPR5-4 | 0 | 2463.269 | 0 | -0.15 | 0.457954 | 11.94 | 106.3 | 93.7 | 9.12 | 9.93 | | High | High | High | High | High | High | High | High | High | High | High | 0 | 3.77E-08 | 4.01 |
| FALSE | High | ASLQSGQE | [K].ASLQSC | 1xPhospho | Q9R1E0 1x | 0.002058 | 5.66E-05 | 1 | 2 | 1 | Q9R1E0 | Forkhead b | Q9R1E0 [2 | 0 | 2502.191 | 0 | 0.2 | 0.257614 | 14.71 | 94.3 | 105.7 | 6.63 | 13.69 | | High | High | High | High | High | High | High | High | High | High | High | 2.58E-05 | 0.000537 | 3.31 |
| FALSE | High | ASLQSGSRI | [R].ASLQSC | 1xPhospho | Q8R409 1x | 0.002313 | 0.000109 | 1 | 1 | 1 | Q8R409 | Protein HE | Q8R409 [5 | 0 | 2296.17 | 0 | -0.28 | 0.023887 | 1.23 | 106.6 | 93.4 | 8.59 | 6.26 | | High | High | High | High | High | High | High | High | High | High | High | 4.94E-05 | 0.00061 | 1.91 |
| FALSE | High | ASLSAPTVC | [R].ASLSAP | 1xPhospho | Q6KAQ0 1x | 0.016306 | 0.001124 | 1 | 2 | 1 | Q6KAQ0 | MFLJ00211 | Q6KAQ0 [7 | 0 | 1364.714 | 0 | -0.4 | 0.043309 | 9.81 | 113.6 | 86.4 | 12.66 | 3.11 | | High | High | High | High | High | High | High | High | High | High | High | 0.000585 | 0.005402 | 1.42 |
| FALSE | High | ASLSATPV | [R].ASLSAT | 1xPhospho | Q8CDB0 1x | 0.014667 | 0.001049 | 1 | 4 | 1 | Q8CDB0 | MAP kinas | Q8CDB0 [4 | 0 | 1693.909 | 0 | 1.34 | 0.004961 | 30.52 | 58.4 | 141.6 | 24.18 | 7.46 | | High | High | High | High | High | High | High | High | High | High | High | 0.000555 | 0.004821 | 2.94 |
| FALSE | High | ASLSCFQN | [K].ASLSCF | 1xCarbami | Q04692 1x | 0.00025 | 0 | 1 | 3 | 2 | Q04692 | SWI/SNF-r | Q04692 [7 | 0 | 1519.693 | 0 | -0.97 | 0.000812 | 1.49 | 132.2 | 67.8 | 9.24 | 5.16 | | High | High | High | High | High | High | High | High | High | High | High | 0 | 5.05E-05 | 2.3 |
| FALSE | High | ASLSDLSSL | [R].ASLSDL | 1xPhospho | Q99KR6 1x | 7.56E-06 | 0 | 1 | 1 | 7 | Q99KR6 | E3 ubiquiti | Q99KR6 [2 | 0 | 2218.051 | 0 | -0.82 | 0.007523 | 7.27 | 124.3 | 75.7 | 16.15 | 16.64 | | High | High | High | High | High | High | High | High | High | High | High | 0 | 9.83E-07 | 4.19 |
| FALSE | High | ASLSDLSSL | [R].ASLSDL | 1xPhospho | Q99KR6 1x | 8.86E-08 | 0 | 1 | 1 | 15 | Q99KR6 | E3 ubiquiti | Q99KR6 [2 | 0 | 2234.046 | 0 | -0.39 | 0.021409 | 8.41 | 111.5 | 88.5 | 9.53 | 6.56 | | High | High | High | High | High | High | High | High | High | High | High | 0 | 7.59E-09 | 5.02 |
| FALSE | High | ASLLEQALF | [K].ASLLEQ | 1xPhospho | B1ARU1 1x | 0.000705 | 0 | 2 | 3 | 1 | B1ARU1; F | Microtubu | B1ARU1 [3 | 0 | 1641.856 | 0 | 2.74 | 0.00028 | 15.04 | 26 | 174 | 19.41 | 9.01 | Shared | High | High | High | High | High | High | High | High | High | High | High | 0 | 0.000162 | 2.38 |
| FALSE | High | ASLSDLTHL | [R].ASLSDL | 1xPhospho | Q3UCG9 1x | 0.000725 | 0 | 1 | 6 | 2 | Q3UCG9 | E3 ubiquiti | Q3UCG9 [2 | 0 | 2278.153 | 0 | 0.71 | 0.001286 | 5.15 | 76.1 | 123.9 | 6.01 | 7.9 | | High | High | High | High | High | High | High | High | High | High | High | 0 | 0.000267 | 2.89 |
| FALSE | High | ASLACSPLG | [R].ASLACS | 1xCarbami | E9PYL2 1x | 3.07E-09 | 0 | 1 | 1 | 1 | E9PYL2 | Proline-ric | E9PYL2 [32 | 0 | 2564.247 | 0 | 0.39 | 0.003819 | 4.92 | 86.1 | 113.9 | 7.42 | 7.3 | | High | High | High | High | High | High | High | High | High | High | High | 0 | 1.52E-10 | 3.86 |
| FALSE | High | ASHYSDQL | [R].ASHYSI | 1xPhospho | A2AIX1 1x | 0.000244 | 0 | 1 | 2 | 3 | A2AIX1 | SEC16 hom | A2AIX1 [12 | 0 | 1553.731 | 0 | 0.07 | 0.270006 | 5.6 | 96.9 | 103.1 | 4.76 | 5.1 | | High | High | High | High | High | High | High | High | High | High | High | 7.19E-05 | 0.000892 | 2.66 |
| FALSE | High | ASFLDDFTI | [K].ASFLDI | 1xPhospho | P49452 1x | 0.000105 | 0 | 1 | 4 | 3 | P49452 | Centromer | P49452 [47 | 0 | 2415.173 | 0 | -0.43 | 0.018686 | 11.66 | 114.4 | 85.6 | 5.46 | 10.35 | | High | High | High | High | High | High | High | High | High | High | High | 0 | 1.9E-05 | 3.49 |
| FALSE | High | ASFSNLPGI | [R].ASFSNL | 1xPhospho | Q8VDU0 1x | 0.032267 | 0.002129 | 1 | 1 | 1 | Q8VDU0 | G-protein- | Q8VDU0 [5 | 0 | 1370.703 | 0 | 1.37 | 0.000191 | 5.79 | 53.4 | 146.6 | 9.6 | 5.4 | | High | High | High | High | High | High | High | High | High | High | High | 0.001142 | 0.01172 | 2.11 |
| FALSE | High | ASGDPGAG | [K].ASGDP | 1xPhospho | Q6PHZ5 1x | 0.000112 | 0 | 1 | 1 | 2 | Q6PHZ5 | Putative RI | Q6PHZ5 [9 | 0 | 1408.642 | 0 | 0.2 | 0.122796 | 19.63 | 89.3 | 110.7 | 6.68 | 11.16 | | High | High | High | High | High | High | High | High | High | High | High | 0 | 2.05E-05 | 1.99 |
| FALSE | High | ASGDPGAG | [K].ASGDP | 2xPhospho | Q6PHZ5 2x | 0.113753 | 0.009241 | 1 | 1 | 1 | Q6PHZ5 | Putative RI | Q6PHZ5 [9 | 1 | 4249.041 | 0 | 0.65 | 0.025602 | 17.21 | 82.3 | 117.7 | 12.61 | 16.35 | | High | High | High | High | High | High | High | High | High | High | High | 0.004654 | 0.05014 | 3.03 |
| FALSE | High | ASGDYDNI | [R].ASGDYI | 2xCarbami | Q8BH57 1x | 1.14E-08 | 0 | 1 | 4 | 4 | Q8BH57 | WD repeat | Q8BH57 [3 | 0 | 3360.553 | 0 | 1.14 | 0.000304 | 7.6 | 62.5 | 137.5 | 6.93 | 5.11 | | High | High | High | High | High | High | High | High | High | High | High | 0 | 6.72E-10 | 4.15 |
| FALSE | High | ASGEMASA | [R].ASGEM | 1xPhospho | Q9JMA1 1x | 1.4E-05 | 0 | 1 | 3 | 3 | Q9JMA1 | Ubiquitin c | Q9JMA1 [1 | 0 | 1948.94 | 0 | 0.02 | 0.240784 | 3.53 | 98.6 | 101.4 | 7.39 | 5.72 | | High | High | High | High | High | High | High | High | High | High | High | 0 | 1.97E-06 | 4.42 |
| FALSE | High | ASGEMASA | [R].ASGEM | 1xPhospho | Q9JMA1 1x | 1.41E-06 | 0 | 1 | 3 | 3 | Q9JMA1 | Ubiquitin c | Q9JMA1 [1 | 0 | 1964.935 | 0 | 0.41 | 0.014138 | 8.11 | 87.8 | 112.2 | 10.4 | 10.2 | | High | High | High | High | High | High | High | High | High | High | High | 0 | 1.31E-05 | 4.3 |
| FALSE | High | ASGGGSAG | [R].ASGGG | 1xPhospho | D3YYI7 1x | 0.001799 | 5.66E-05 | 1 | 1 | 1 | D3YYI7 | Probable E | D3YYI7 [16 | 0 | 1819.865 | 0 | 1.64 | 0.000361 | 11.23 | 50.1 | 149.9 | 3.87 | 6.17 | | High | High | High | High | High | High | High | High | High | High | High | 2.58E-05 | 0.000461 | 3.32 |
| FALSE | High | ASGLPPTES | [K].ASGLPF | 1xCarbami | Q571J8 1x | 0.110202 | 0.008961 | 1 | 10 | 2 | Q571J8 | Non-specif | Q571J8 [54 | 0 | 2660.27 | 0 | 0.6 | 0.003257 | 11.79 | 75.6 | 124.4 | 13.7 | 13.63 | | High | High | High | High | High | High | High | High | High | High | High | 0.004477 | 0.04843 | 2.29 |
| FALSE | High | ASGITGSYS | [R].ASGITG | 1xPhospho | Q9D279 1x | 0.00025 | 0 | 1 | 1 | 1 | Q9D279 | Mitotic inte | Q9D279 [5 | 0 | 3471.797 | 0 | 5.42 | 0.001014 | 27.49 | 4.2 | 195.8 | 60.12 | 16.88 | | High | High | High | High | High | High | High | High | High | High | High | 0 | 5.05E-05 | 3.78 |
| FALSE | High | ASGPASPEI | [K].ASGPAS | 1xPhospho | Q3TAB2 1x | 0.001689 | 5.66E-05 | 2 | 6 | 3 | Q3TAB2; H | Putative ur | Q3TAB2 [1 | 0 | 1381.713 | 0 | -0.47 | 0.009837 | 1.88 | 116.2 | 83.8 | 5.47 | 8.56 | Shared | High | High | High | High | High | High | High | High | High | High | High | 2.58E-05 | 0.000431 | 1.98 |
| FALSE | High | ASGPESPL | [R].ASGPES | 1xPhospho | F8WJ40 1x | 1.63E-06 | 0 | 1 | 4 | 2 | F8WJ40 | Histone-lys | F8WJ40 [8 | 0 | 1504.736 | 0 | -0.43 | 0.000986 | 2.94 | 114.7 | 85.3 | 1.44 | 5 | | High | High | High | High | High | High | High | High | High | High | High | 0 | 1.76E-07 | 3.44 |
| FALSE | High | ASGPESPL | [R].ASGPES | 2xPhospho | F8WJ40 2x | 0.017344 | 0.001204 | 1 | 4 | 1 | F8WJ40 | Histone-lys | F8WJ40 [8 | 0 | 1584.702 | 0 | -0.09 | 0.213681 | 10.64 | 105.3 | 94.7 | 14.16 | 10.3 | | High | High | High | High | High | High | High | High | High | High | High | 0.000638 | 0.005782 | 2.18 |
| FALSE | High | ASHYSDQL | [R].ASHYSI | 2xPhospho | A2AIX1 2x | 0.007538 | 0.00062 | 1 | 2 | 2 | A2AIX1 | SEC16 hom | A2AIX1 [12 | 0 | 1633.697 | 0 | -0.54 | 0.002619 | 2.89 | 117.9 | 82.1 | 8.13 | 6.16 | | High | High | High | High | High | High | High | High | High | High | High | 0.006167 | 0.0666 | 2.17 |
| FALSE | High | ASGPPVSEI | [K].ASGPPV | 1xPhospho | Q149Z9 1x | 5.26E-05 | 0 | 1 | 3 | 2 | Q149Z9 | Histone clu | Q149Z9 [3 | 0 | 1736.96 | 0 | 0.59 | 0.001237 | 3.48 | 78.8 | 121.2 | 4.98 | 4.46 | | High | High | High | High | High | High | High | High | High | High | High | 0 | 8.71E-06 | 2.74 |
| FALSE | High | ASGQAFEL | [R].ASGQA | 1xPhospho | Q9CZ46 1x | 0.006773 | 0.00062 | 1 | 2 | 1 | Q9CZ46 | Putative ur | Q9CZ46 [4 | 0 | 2746.494 | 0 | -0.92 | 0.000209 | 1.24 | 131 | 69 | 3.17 | 8.64 | | High | High | High | High | High | High | High | High | High | High | High | 0.000275 | 0.002024 | 3.08 |
| FALSE | High | ASGQAFEL | [R].ASGQA | 1xPhospho | Q545B6 1x | 6.59E-05 | 0 | 1 | 8 | 3 | Q545B6 | Stathmin C | Q545B6 [1 | 0 | 1697.883 | 0 | -1.62 | 0.003728 | 0.73 | 149.3 | 50.7 | 14.09 | 13.08 | | High | High | High | High | High | High | High | High | High | High | High | 0 | 3.28E-05 | 3.52 |
| FALSE | High | ASGSENEG | [K].ASGSEN | 1xPhospho | Q64511 1x | 8.2E-06 | 0 | 1 | 1 | 4 | Q64511 | DNA topois | Q64511 [1 | 0 | 1761.728 | 0 | 0.62 | 0.002184 | 7.14 | 79.9 | 120.1 | 6.45 | 6.1 | | High | High | High | High | High | High | High | High | High | High | High | 0 | 0.000261 | 3.37 |
| FALSE | High | ASGSPPLLF | [R].ASGSPF | 1xPhospho | Q8VBT9 1x | 0.076555 | 0.005826 | 1 | 3 | 2 | Q8VBT9 | Tether con | Q8VBT9 [4 | 0 | 4567.362 | 0 | -0.69 | 0.001593 | 5.83 | 123.3 | 76.7 | 4.09 | 9.74 | | High | High | High | High | High | High | High | High | High | High | High | 0.00299 | 0.03159 | 3.93 |
| FALSE | High | ASGTYAGP | [K].ASGTY | 1xPhospho | Q6PAL7 1x | 0.005615 | 0.00062 | 1 | 1 | 3 | Q6PAL7 | AT-hook D | Q6PAL7 [8 | 0 | 1939.948 | 0 | 0.43 | 0.01438 | 18.38 | 85.4 | 114.6 | 9.02 | 4.7 | | High | High | High | High | High | High | High | High | High | High | High | 0.000275 | 0.001645 | 2.77 |
| FALSE | High | ASGVSFTY | [K].ASGVSI | 1xPhospho | E9PY43 1x | 9.84E-10 | 0 | 1 | 2 | 10 | E9PY43 | Glutamate- | E9PY43 [17 | 0 | 3711.518 | 0 | 0.14 | 0.636497 | 22.38 | 95.2 | 104.8 | 12.14 | 8.7 | | High | High | High | High | High | High | High | High | High | High | High | 0 | 4.25E-11 | 6.44 |
| FALSE | High | ASGVSFTY | [K].ASGVSI | 1xDeamida | E9PY43 1x | 4.89E-08 | 0 | 1 | 2 | 4 | E9PY43 | Glutamate- | E9PY43 [17 | 0 | 3712.502 | 0 | 0.14 | 0.647586 | 15.59 | 99.5 | 100.5 | 5.03 | 13.74 | | High | High | High | High | High | High | High | High | High | High | High | 0 | 3.42E-09 | 7.05 |
| FALSE | High | ASGVSFTY | [K].ASGVSI | 2xPhospho | E9PY43 2x | 2.52E-06 | 0 | 1 | 2 | 4 | E9PY43 | Glutamate- | E9PY43 [17 | 0 | 3791.484 | 0 | 0.11 | 0.013139 | 3.08 | 94.8 | 105.2 | 5.03 | 2.77 | | High | High | High | High | High | High | High | High | High | High | High | 0 | 2.51E-05 | 6.16 |
| FALSE | High | ASGYQSSQ | [K].ASGYQ | 1xPhospho | P51859 1x | 0.029586 | 0.001857 | 1 | 2 | 1 | P51859 | Hepatoma- | P51859 [9 | 0 | 1493.74 | 0 | 0.36 | 0.007345 | 5.52 | 83.2 | 116.8 | 6.11 | 10.67 | | High | High | High | High | High | High | High | High | High | High | High | 0.000997 | 0.01065 | 1.53 |
| FALSE | High | ASHHESNG | [R].ASHHE | 1xDeamida | A0A0A6YW | 0.041072 | 0.003094 | 1 | 3 | 1 | A0A0A6YW | Mitogen-a | A0A0A6YW | 0 | 1579.685 | 0 | 0.32 | 0.038547 | 18.66 | 86.8 | 113.2 | 11.43 | 17.19 | | High | High | High | High | High | High | High | High | High | High | High | 0.001597 | 0.01537 | 2.22 |
| FALSE | High | ASHHGLEQ | [R].ASHHG | 1xPhospho | P49070 1x | 0.00311 | 0.00016 | 1 | 1 | 1 | P49070 | Calcium sig | P49070 [14 | 0 | 1706.821 | 0 | 0.42 | 0.058452 | 30.91 | 82.6 | 117.4 | 9.24 | 20.54 | | High | High | High | High | High | High | High | High | High | High | High | 7.19E-05 | 0.00085 | 2.26 |
| FALSE | High | ASHPAVES | [R].ASHPAV | 1xCarbami | A2AQE2 1x | 5.88E-10 | 0 | 1 | 5 | 1 | A2AQE2 | SECIS bindi | A2AQE2 [2 | 0 | 3506.531 | 0 | 1.13 | 0.000513 | 19.2 | 62.4 | 137.6 | 16.04 | 7.14 | | High | High | High | High | High | High | High | High | High | High | High | 0 | 2.39E-11 | 5.35 |
| FALSE | High | ASHPGDFP | [R].ASHPGI | 1xPhospho | Q3TM84 1x | 0.000155 | 0 | 1 | 2 | 1 | Q3TM84 | Putative ur | Q3TM84 [7 | 1 | 2267.092 | 0 | 2.69 | 0.000269 | 30.62 | 29.3 | 170.7 | 45.16 | 9.16 | | High | High | High | High | High | High | High | High | High | High | High | 0 | 2.95E-05 | 1.95 |
| FALSE | High | ASGPSSPDI | [R].ASGPSS | 1xPhospho | Q80WC3 1 | 0.003195 | 0.00016 | 1 | 2 | 1 | Q80WC3 | Trinucleoti | Q80WC3 [ | 0 | 1383.692 | 0 | 1.61 | 0.000779 | 15.47 | 46.6 | 153.4 | 4.53 | 9.82 | | High | High | High | High | High | High | High | High | High | High | High | 7.19E-05 | 0.000878 | 1.65 |
| FALSE | High | ASLSPMDE | [R].ASLSPN | 1xPhospho | P14873 1x | 2.47E-06 | 0 | 2 | 2 | 3 | P14873; B2 | Microtubu | P14873 [13 | 0 | 2552.224 | 0 | -0.39 | 0.080046 | 1.21 | 116.8 | 83.2 | 12.81 | 25.99 | Shared | High | High | High | High | High | High | High | High | High | High | High | 0 | 2.81E-07 | 3.78 |
| FALSE | High | ASLSPMDE | [R].ASLSPN | 1xPhospho | P14873 1x | 2.59E-06 | 0 | 2 | 2 | 4 | P14873; B2 | Microtubu | P14873 [13 | 0 | 2568.219 | 0 | -0.33 | 0.074531 | 11.17 | 111.9 | 88.1 | 8.27 | 12.77 | Shared | High | High | High | High | High | High | High | High | High | High | High | 0 | 1.36E-05 | 3.69 |
| FALSE | High | ASLSPMDE | [R].ASLSPN | 2xPhospho | P14873 2x | 0.001337 | 0 | 2 | 2 | 1 | P14873; B2 | Microtubu | P14873 [13 | 0 | 2632.191 | 0 | -0.25 | 0.224236 | 4.78 | 107.6 | 92.4 | 12.93 | 22.19 | Shared | High | High | High | High | High | High | High | High | High | High | High | 0 | 0.00033 | 3.22 |
| FALSE | High | ASPEEPGD | [K].ASPEEP | 1xPhospho | Q6GQT6 1x | 0.113753 | 0.009241 | 1 | 1 | 1 | Q6GQT6 | Sterol regu | Q6GQT6 [8 | 0 | 1660.778 | 0 | 0.42 | 0.011696 | 9.07 | 89.3 | 110.7 | 6 | 7.92 | | High | High | High | High | High | High | High | High | High | High | High | 0.004667 | 0.05019 | 2.35 |
| FALSE | High | ASPEPEGET | [K].ASPEPE | 1xPhospho | Q8R3Y8 1x | 0.000976 | 0 | 1 | 2 | 2 | Q8R3Y8 | Interferon | Q8R3Y8 [3 | 0 | 1710.835 | 0 | 0.34 | 0.028338 | 14.15 | 89 | 111 | 5.64 | 7.07 | | High | High | High | High | High | High | High | High | High | High | High | 0 | 0.000231 | 2.39 |
| FALSE | High | ASPEPPDS | [K].ASPEPP | 1xPhospho | Q8K3X4 1x | 5.91E-05 | 0 | 1 | 1 | 2 | Q8K3X4 | Interferon | Q8K3X4 [5 | 0 | 1936.967 | 0 | -0.45 | 0.001583 | 4 | 116.3 | 83.7 | 5.42 | 4.58 | | High | High | High | High | High | High | High | High | High | High | High | 0 | 1E-05 | 2.51 |
| FALSE | High | ASPEPSALF | [R].ASPEPS | 1xPhospho | O35954 1x | 0.000466 | 0 | 1 | 1 | 1 | O35954 | Membrane | O35954 [6 | 0 | 1532.767 | 0 | 2.79 | 0.000231 | 10.01 | 25.6 | 174.4 | 4.52 | 5.95 | | High | High | High | High | High | High | High | High | High | High | High | 0 | 0.000101 | 2.28 |
| FALSE | High | ASPFEEDQ | [K].ASPFEE | 1xPhospho | Q80U58 1x | 0.007019 | 0.00062 | 1 | 7 | 2 | Q80U58 | Pumilio ho | Q80U58 [1 | 0 | 1501.652 | 0 | 1 | 9.77E-05 | 1.74 | 66.1 | 133.9 | 1.95 | 1.44 | | High | High | High | High | High | High | High | High | High | High | High | 0.000344 | 0.003001 | 2.78 |
| FALSE | High | ASPGGVST | [R].ASPGG | 1xPhospho | O35126 1x | 2.04E-05 | 0 | 1 | 2 | 1 | O35126 | Atrophin-1 | O35126 [3 | 0 | 1545.7 | 0 | 1.05 | 0.000834 | 16.3 | 66.1 | 133.9 | 7.79 | 4.43 | | High | High | High | High | High | High | High | High | High | High | High | 0 | 3E-06 | 2.4 |
| FALSE | High | ASPGGVST | [R].ASPGG | 1xPhospho | O35126 1x | 1.35E-05 | 0 | 1 | 2 | 2 | O35126 | Atrophin-1 | O35126 [3 | 0 | 1774.862 | 0 | 0.98 | 0.000408 | 4.96 | 66.5 | 133.5 | 7.88 | 5.07 | | High | High | High | High | High | High | High | High | High | High | High | 0 | 1.88E-06 | 2.48 |
| FALSE | High | ASPGLLPVA | [R].ASPGLL | 1xPhospho | Q63943-2 | 2.27E-05 | 0 | 2 | 6 | 1 | Q63943-2; | Isoform M | Q63943-2 | 0 | 2090.141 | 0 | 2.19 | 0.000616 | 23.01 | 36 | 164 | 10.06 | 9.54 | Shared | High | High | High | High | High | High | High | High | High | High | High | 0 | 3.39E-06 | 3.29 |
| FALSE | High | ASPGLLPVA | [R].ASPGLL | 1xDeamida | Q63943-2 | 0.001398 | 5.66E-05 | 2 | 6 | 2 | Q63943-2; | Isoform M | Q63943-2 | 0 | 2091.125 | 0 | 2.07 | 0.000308 | 9.28 | 39 | 161 | 9.94 | 5.94 | Shared | High | High | High | High | High | High | High | High | High | High | High | 2.58E-05 | 0.000348 | 3.06 |
| FALSE | High | ASPGLSMP | [R].ASPGLS | 1xPhospho | Q545J5 1x | 4.35E-05 | 0 | 1 | 3 | 3 | Q545J5 | Basic Krup | Q545J5 [9 | 0 | 2010.038 | 0 | -0.14 | 0.368173 | 2.51 | 102.5 | 97.5 | 9.28 | 13.54 | | High | High | High | High | High | High | High | High | High | High | High | 0 | 7.06E-06 | 2.96 |
| FALSE | High | ASPGLSMP | [R].ASPGLS | 2xPhospho | Q545J5 2x | 0.104222 | 0.008348 | 1 | 3 | 1 | Q545J5 | Basic Krup | Q545J5 [9 | 0 | 2074.01 | 0 | 1.08 | 0.008244 | 9.78 | 63.5 | 136.5 | 8.89 | 21.22 | | High | High | High | High | High | High | High | High | High | High | High | 0.004193 | 0.04517 | 2.36 |
| FALSE | High | ASPGPGGL | [R].ASPGP | 1xPhospho | P58404 1x | 0.000418 | 0 | 1 | 4 | 2 | P58404 | Striatin-4 C | P58404 [22 | 0 | 2064.114 | 0 | 0.44 | 0.002655 | 11.85 | 83.2 | 116.8 | 7.09 | 2.61 | | High | High | High | High | High | High | High | High | High | High | High | 0 | 8.96E-05 | 3.09 |
| FALSE | High | ASPHDILET | [R].ASPHDI | 1xPhospho | Q91YI1-2 1 | 0.001865 | 5.66E-05 | 1 | 2 | 1 | Q91YI1-2 | Isoform 2 c | Q91YI1-2 [ | 0 | 1806.935 | 0 | -0.69 | 0.001255 | 1.54 | 123.5 | 76.5 | 4.56 | 7.31 | | High | High | High | High | High | High | High | High | High | High | High | 2.58E-05 | 0.00048 | 2.8 |
| FALSE | High | ASPEDGQE | [K].ASPEDC | 1xPhospho | G3X8Q1 1x | 9.89E-05 | 0 | 1 | 3 | 1 | G3X8Q1 | Calcineurir | G3X8Q1 [1 | 0 | 2044.974 | 0 | -0.22 | 0.0196 | 5.29 | 108 | 92 | 3.6 | 8.25 | | High | High | High | High | High | High | High | High | High | High | High | 0 | 1.77E-05 | 2.9 |

| | | | | | | | | | | | | | | | | | | | | | | | | | | | | | | | | | | | | | | |
|---|---|---|---|---|---|---|---|---|---|---|---|---|---|---|---|---|---|---|---|---|---|---|---|---|---|---|---|---|---|---|---|---|---|---|---|---|---|---|
| FALSE | High | ASPHLHSCI | [R].ASPHLH | 1xCarbami | Q9D6K9 1x | 0.014538 | 0.001011 | 1 | 2 | 2 | Q9D6K9 | Ceramide s | Q9D6K9 [4 | 0 | 1332.561 | 0 | 1.24 | 0.00122 | 2.57 | 58.1 | 141.9 | 4.03 | 10.28 | | High | High | High | High | High | High | High | High | High | High | High | 0.001609 | 0.01556 | 1.81 |
| FALSE | High | ASPINEDIR | [R].ASPINE | 1xPhospho | Q80TZ9 1x | 0.001865 | 5.66E-05 | 1 | 2 | 3 | Q80TZ9 | Arginine-gl | Q80TZ9 [5 | 0 | 1323.651 | 0 | 0.13 | 0.015021 | 3.22 | 97.7 | 102.3 | 4.01 | 5.39 | | High | High | High | High | High | High | High | High | High | High | High | 0.000275 | 0.001865 | 2.16 |
| FALSE | High | ASPLPPPPP | [R].ASPLPP | 1xPhospho | Q6PHZ5 1x | 1.01E-05 | 0 | 1 | 1 | 2 | Q6PHZ5 | Putative RI | Q6PHZ5 [1 | 0 | 3088.58 | 0 | 0.55 | 0.001214 | 3.03 | 84.3 | 115.7 | 6.23 | 5.01 | | High | High | High | High | High | High | High | High | High | High | High | 0 | 7.49E-05 | 4.27 |
| FALSE | High | ASPLPVLNV | [R].ASPLPV | 1xPhospho | B2RS11 1xI | 0.017192 | 0.001204 | 1 | 4 | 1 | B2RS11 | Cyclin E1 C | B2RS11 [1C | 0 | 1632.846 | 0 | 0.17 | 0.044081 | 5 | 90.7 | 109.3 | 5.58 | 8.74 | | High | High | High | High | High | High | High | High | High | High | High | 0.000638 | 0.005769 | 2.14 |
| FALSE | High | ASPITNDGI | [K].ASPITN | 1xPhospho | Q64511 1x | 0.000158 | 0 | 1 | 2 | 4 | Q64511 | DNA topois | Q64511 [1 | 0 | 2644.243 | 0 | 0.74 | 0.000348 | 2.32 | 74.3 | 125.7 | 5.11 | 4.11 | | High | High | High | High | High | High | High | High | High | High | High | 0 | 3.01E-05 | 3.61 |
| FALSE | High | ASPITNDGI | [K].ASPITN | 2xPhospho | Q64511 2x | 3.06E-05 | 0 | 1 | 2 | 4 | Q64511 | DNA topois | Q64511 [1 | 0 | 2724.209 | 0 | 0.73 | 0.002027 | 13.79 | 73.1 | 126.9 | 5.57 | 8.23 | | High | High | High | High | High | High | High | High | High | High | High | 4.94E-05 | 0.000703 | 3.1 |
| FALSE | High | ASPITNDGI | [K].ASPITN | 1xPhospho | Q64511 1x | 0.002134 | 5.66E-05 | 1 | 2 | 1 | Q64511 | DNA topois | Q64511 [1 | 1 | 3857.825 | 0 | 0.49 | 0.080771 | 26.08 | 81.6 | 118.4 | 15.63 | 21.7 | | High | High | High | High | High | High | High | High | High | High | High | 2.58E-05 | 0.000561 | 4.14 |
| FALSE | High | ASPNGSMS | [R].ASPNG | 1xPhospho | E9Q6Y9 1x | 0.006478 | 0.00062 | 1 | 4 | 2 | E9Q6Y9 | BTB/POZ d | E9Q6Y9 [1 | 0 | 1734.732 | 0 | 0.93 | 0.021436 | 21.91 | 74 | 126 | 18.41 | 20.79 | | High | High | High | High | High | High | High | High | High | High | High | 0.001044 | 0.0112 | 2.18 |
| FALSE | High | ASPNLIGNT | [R].ASPNLI | 1xPhospho | Q60929-3 1 | 0.000573 | 0 | 2 | 3 | 1 | Q60929-3; | Isoform 3 c | Q60929-3 [ | 0 | 2052.089 | 0 | 1.5 | 0.000373 | 3.8 | 52.1 | 147.9 | 6.72 | 7.22 | Shared | High | High | High | High | High | High | High | High | High | High | High | 0 | 0.000128 | 2.6 |
| FALSE | High | ASPNPVAM | [R].ASPNPV | 1xPhospho | J3QNK5 1x | 0.001325 | 0 | 1 | 4 | 3 | J3QNK5 | Histone de | J3QNK5 [4 | 0 | 1497.697 | 0 | 0 | 0.779427 | 18.1 | 101.7 | 98.3 | 9.5 | 21.2 | | High | High | High | High | High | High | High | High | High | High | High | 0.001812 | 0.01797 | 1.94 |
| FALSE | High | ASPPGGLA | [R].ASPPGC | 1xPhospho | Q6ZQ88 1x | 0.000501 | 0 | 1 | 2 | 3 | Q6ZQ88 | Lysine-spe | Q6ZQ88 [6 | 0 | 4034.9 | 0 | -0.82 | 0.001879 | 0.89 | 127.2 | 72.8 | 10.07 | 2.32 | | High | High | High | High | High | High | High | High | High | High | High | 0 | 0.000109 | 4.54 |
| FALSE | High | ASPPGGLA | [R].ASPPGC | 1xDeamida | Q6ZQ88 1x | 0.001107 | 0 | 1 | 2 | 2 | Q6ZQ88 | Lysine-spe | Q6ZQ88 [6 | 0 | 4035.884 | 0 | -1.01 | 0.003121 | 2.62 | 129.2 | 70.8 | 12.57 | 8.21 | | High | High | High | High | High | High | High | High | High | High | High | 0 | 0.000266 | 3.73 |
| FALSE | High | ASPPGGLA | [R].ASPPGC | 1xPhospho | Q6ZQ88 1x | 0.010382 | 0.000772 | 1 | 2 | 2 | Q6ZQ88 | Lysine-spe | Q6ZQ88 [6 | 0 | 4050.895 | 0 | -0.92 | 0.003573 | 4.58 | 129.9 | 70.1 | 7.1 | 13.34 | | High | High | High | High | High | High | High | High | High | High | High | 0.000814 | 0.007503 | 4.13 |
| FALSE | High | ASPPLEVSE | [R].ASPPLE | 1xPhospho | Q8R1K1 1x | 0.000779 | 0 | 1 | 2 | 1 | Q8R1K1 | Ubiquitin-a | Q8R1K1 [3 | 0 | 1820.899 | 0 | -0.01 | 0.731081 | 9.01 | 97.8 | 102.2 | 7.13 | 5.67 | | High | High | High | High | High | High | High | High | High | High | High | 0 | 0.000181 | 2.3 |
| FALSE | High | ASPLLLSSA | [R].ASPLLL | 1xPhospho | E9QAU9 1x | 0.007881 | 0.000736 | 1 | 8 | 1 | E9QAU9 | Leucine-ric | E9QAU9 [6 | 0 | 2306.184 | 0 | 0.69 | 0.002897 | 5.12 | 73.7 | 126.3 | 10.13 | 10.98 | | High | High | High | High | High | High | High | High | High | High | High | 0.000327 | 0.002412 | 2.87 |
| FALSE | High | ASPDLWPE | [R].ASPDLV | 1xPhospho | Q8CD94 1x | 0.000418 | 0 | 1 | 1 | 1 | Q8CD94 | Protein lin- | Q8CD94 [2 | 0 | 2798.421 | 0 | 0.99 | 0.053804 | 20.03 | 58 | 142 | 40.48 | 48.3 | | High | High | High | High | High | High | High | High | High | High | High | 0 | 8.98E-05 | 3.44 |
| FALSE | High | ASPATQPP | [R].ASPATC | 1xPhospho | Q60749 1x | 0.008463 | 0.000736 | 1 | 1 | 5 | Q60749 | KH domain | Q60749 [5 | 0 | 4307.334 | 0 | 0.84 | 0.016614 | 36.24 | 77.7 | 122.3 | 14.88 | 16.99 | | High | High | High | High | High | High | High | High | High | High | High | 0.000467 | 0.004277 | 3.75 |
| FALSE | High | ASPATIEM/ | [R].ASPATI | 1xPhospho | P48381 1xI | 0.01319 | 0.000889 | 1 | 2 | 2 | P48381 | Transcripti | P48381 [15 | 0 | 2141.133 | 0 | 0.19 | 0.083748 | 15.26 | 90.4 | 109.6 | 11.26 | 10.37 | | High | High | High | High | High | High | High | High | High | High | High | 0.000467 | 0.004281 | 2.11 |
| FALSE | High | ASLSSSSTP | [K].ASLSSS | 1xPhospho | A0A0R4J1C | 0.000147 | 0 | 1 | 5 | 2 | A0A0R4J1C | Disabled h | A0A0R4J1C | 0 | 1717.877 | 0 | 2.9 | 0.000885 | 30.55 | 23.6 | 176.4 | 19.41 | 13.77 | | High | High | High | High | High | High | High | High | High | High | High | 0.000382 | 0.00324 | 2.74 |
| FALSE | High | ASITEESWL | [R].ASITEE | 1xPhospho | B5B2N2 1x | 0.096181 | 0.007535 | 1 | 11 | 1 | B5B2N2 | Nuclear fac | B5B2N2 [2 | 0 | 1628.788 | 0 | -0.04 | 0.60052 | 7.92 | 101.3 | 98.7 | 7.5 | 6.95 | | High | High | High | High | High | High | High | High | High | High | High | 0.003898 | 0.04138 | 2.24 |
| FALSE | High | ASITPVLGS | [R].ASITPV | 1xPhospho | Q8K2H3 1x | 0.003167 | 0.00016 | 1 | 2 | 1 | Q8K2H3 | Protein FAI | Q8K2H3 [6 | 0 | 1795.997 | 0 | 1.2 | 0.004235 | 17.19 | 62.1 | 137.9 | 7.52 | 13.99 | | High | High | High | High | High | High | High | High | High | High | High | 7.19E-05 | 0.00087 | 2.44 |
| FALSE | High | ASLTTPGSE | [K].ASLTTP | 1xPhospho | Q60793 1x | 1.15E-05 | 0 | 1 | 1 | 2 | Q60793 | Krueppel-li | Q60793 [2 | 0 | 2535.3 | 0 | 1.02 | 0.001045 | 21.9 | 66 | 134 | 16.27 | 4.66 | | High | High | High | High | High | High | High | High | High | High | High | 0 | 2.26E-06 | 3.02 |
| FALSE | High | ASITYDLND | [K].ASITYD | 1xPhospho | Q80WQ8 1 | 2.35E-07 | 0 | 1 | 2 | 1 | Q80WQ8 | Mis18-binc | Q80WQ8 [ | 0 | 1919.895 | 0 | -1 | 0.000644 | 2.95 | 134.9 | 65.1 | 4.56 | 10.39 | | High | High | High | High | High | High | High | High | High | High | High | 0 | 1.99E-08 | 3.52 |
| FALSE | High | ASITYDLND | [K].ASITYD | 2xPhospho | Q80WQ8 2 | 0.002553 | 0.000109 | 1 | 2 | 2 | Q80WQ8 | Mis18-binc | Q80WQ8 [ | 0 | 1999.861 | 0 | -0.67 | 0.003534 | 3.07 | 119.6 | 80.4 | 11.54 | 7.01 | | High | High | High | High | High | High | High | High | High | High | High | 4.94E-05 | 0.000684 | 3.04 |
| FALSE | High | ASLYNAVT | [R].ASLYNA | 1xPhospho | Q3ULZ3 1x | 5.6E-05 | 0 | 1 | 5 | 2 | Q3ULZ3 | Phosphose | Q3ULZ3 [3 | 0 | 2078.046 | 0 | 0.2 | 0.447169 | 27.2 | 88.5 | 111.5 | 25.35 | 14.69 | | High | High | High | High | High | High | High | High | High | High | High | 0 | 1.91E-05 | 2.92 |
| FALSE | High | ASIYQNQN | [K].ASIYQN | 1xPhospho | D3Z5N6 1x | 0.000349 | 0 | 1 | 4 | 2 | D3Z5N6 | Zinc finger | D3Z5N6 [3 | 0 | 1420.631 | 0 | -0.38 | 0.005955 | 8.37 | 113.2 | 86.8 | 4.09 | 8.76 | | High | High | High | High | High | High | High | High | High | High | High | 0 | 7.33E-05 | 2.85 |
| FALSE | High | ASMSEFLES | [K].ASMSE | 1xPhospho | Q80U70 1x | 3.13E-07 | 0 | 1 | 5 | 8 | Q80U70 | Polycomb | Q80U70 [5 | 0 | 2380.021 | 5 | -0.7 | 0.000544 | 3.72 | 125.6 | 74.4 | 2.72 | 5.23 | | High | High | High | High | High | High | High | High | High | High | High | 0 | 3.58E-06 | 4.83 |
| FALSE | High | ASMSEFLES | [K].ASMSE | 1xPhospho | Q80U70 1x | 3.93E-07 | 0 | 1 | 5 | 17 | Q80U70 | Polycomb | Q80U70 [5 | 0 | 2396.016 | 6 | -0.46 | 0.003644 | 7.35 | 115.3 | 84.7 | 3.97 | 5.76 | | High | High | High | High | High | High | High | High | High | High | High | 0 | 2.35E-07 | 5.22 |
| FALSE | High | ASMSEFLES | [K].ASMSE | 1xDeamida | Q80U70 1x | 0.011447 | 0.000816 | 1 | 5 | 1 | Q80U70 | Polycomb | Q80U70 [5 | 0 | 2397 | 0 | -0.48 | 0.084648 | 6.49 | 110.3 | 89.7 | 7.6 | 17.05 | | High | High | High | High | High | High | High | High | High | High | High | 0.000437 | 0.003656 | 3.88 |
| FALSE | High | ASNASTPTI | [K].ASNAS | 1xDeamida | Q69ZA1 1x | 0.030104 | 0.001857 | 1 | 3 | 1 | Q69ZA1 | Cyclin-dep | Q69ZA1 [4 | 0 | 1415.718 | 0 | -0.02 | 0.932262 | 15.94 | 101.6 | 98.4 | 5.62 | 14.67 | | High | High | High | High | High | High | High | High | High | High | High | 0.001011 | 0.01086 | 1.6 |
| FALSE | High | ASNEDVTK | [K].ASNED | 1xPhospho | Q99JF8 1x | 0.000941 | 0 | 1 | 4 | 3 | Q99JF8 | PC4 and SF | Q99JF8 [12 | 0 | 1401.703 | 0 | -0.82 | 0.001701 | 1.71 | 127.2 | 72.8 | 6.14 | 11.79 | | High | High | High | High | High | High | High | High | High | High | High | 0 | 0.000285 | 2.06 |
| FALSE | High | ASNEDVTK | [K].ASNED | 1xDeamida | Q99JF8 1x | 0.090888 | 0.007081 | 1 | 4 | 1 | Q99JF8 | PC4 and SF | Q99JF8 [12 | 0 | 1402.687 | 1 | 0 | 0.455351 | 6.09 | 100 | 100 | 3.72 | 9.33 | | High | High | High | High | High | High | High | High | High | High | High | 0.003633 | 0.03854 | 1.13 |
| FALSE | High | ASNSPEQS | [R].ASNSPE | 1xCarbami | Q3UZ39 2x | 0.01029 | 0.000772 | 1 | 2 | 1 | Q3UZ39 | Leucine-ric | Q3UZ39 [3 | 0 | 3135.332 | 0 | 0.61 | 0.021512 | 4.96 | 79.6 | 120.4 | 17.94 | 4.16 | | High | High | High | High | High | High | High | High | High | High | High | 0.000382 | 0.003225 | 3.26 |
| FALSE | High | ASNSPEQS | [R].ASNSPE | 1xCarbami | Q3UZ39 1x | 8.08E-07 | 0 | 1 | 2 | 3 | Q3UZ39 | Leucine-ric | Q3UZ39 [3 | 0 | 3284.528 | 0 | 1 | 0.001041 | 13.48 | 68 | 132 | 4.71 | 8.77 | | High | High | High | High | High | High | High | High | High | High | High | 0 | 8.01E-08 | 5.06 |
| FALSE | High | ASNSPEQS | [R].ASNSPE | 1xCarbami | Q3UZ39 2x | 0.000107 | 0 | 1 | 2 | 3 | Q3UZ39 | Leucine-ric | Q3UZ39 [3 | 0 | 3364.495 | 0 | 1.36 | 0.000344 | 13.55 | 59.8 | 140.2 | 13.53 | 5.66 | | High | High | High | High | High | High | High | High | High | High | High | 0 | 1.95E-05 | 3.73 |
| FALSE | High | ASPAAESPI | [K].ASPAAE | 1xPhospho | Q64028 1x | 0.001881 | 5.66E-05 | 1 | 7 | 4 | Q64028 | Polyhomec | Q64028 [6 | 0 | 1395.729 | 0 | -1.36 | 0.000209 | 1.21 | 142.5 | 57.5 | 4.43 | 5.1 | | High | High | High | High | High | High | High | High | High | High | High | 0.006342 | 0.068 | 2.79 |
| FALSE | High | ASPAAPLD | [R].ASPAAF | 1xPhospho | U5LK19 1xI | 0.002743 | 0.000109 | 2 | 5 | 1 | U5LK19; E9 | Marshalin- | U5LK19 [1 | 0 | 2015.016 | 0 | -0.9 | 0.006699 | 1.57 | 126.9 | 73.1 | 13.16 | 12.49 | Shared | High | High | High | High | High | High | High | High | High | High | High | 4.94E-05 | 0.000736 | 4.18 |
| FALSE | High | ASPAGTAG | [K].ASPAG | 1xPhospho | Q8K0H5 1x | 0.000155 | 0 | 1 | 1 | 2 | Q8K0H5 | Transcripti | Q8K0H5 [4 | 0 | 2373.213 | 0 | 0.21 | 0.11304 | 12.16 | 90 | 110 | 11.7 | 8.2 | | High | High | High | High | High | High | High | High | High | High | High | 0 | 2.94E-05 | 3.59 |
| FALSE | High | ASPALGSG | [R].ASPALC | 1xPhospho | Q80TT4 1x | 0.007673 | 0.000664 | 1 | 3 | 1 | Q80TT4 | MKIAA071 | Q80TT4 [1 | 0 | 2692.187 | 0 | 1.32 | 0.033561 | 29.61 | 57.2 | 142.8 | 28.7 | 21.12 | | High | High | High | High | High | High | High | High | High | High | High | 0.000294 | 0.002331 | 2.54 |
| FALSE | High | ASPALGSG | [R].ASPALC | 1xPhospho | Q80TT4 1x | 4.14E-07 | 0 | 1 | 3 | 8 | Q80TT4 | MKIAA071 | Q80TT4 [1 | 0 | 2708.182 | 0 | 0.86 | 0.047122 | 20.16 | 71.2 | 128.8 | 14.97 | 25.82 | | High | High | High | High | High | High | High | High | High | High | High | 0 | 0.000148 | 3.51 |
| FALSE | High | ASPALVAK | [K].ASPALV | 1xPhospho | Q3UMC8 1 | 0.002531 | 0.000109 | 1 | 4 | 2 | Q3UMC8 | Putative ur | Q3UMC8 [ | 0 | 1294.754 | 0 | 1.31 | 0.000551 | 11.8 | 56 | 144 | 6.08 | 6.32 | | High | High | High | High | High | High | High | High | High | High | High | 4.94E-05 | 0.000674 | 1.64 |
| FALSE | High | ASPAPAPQ | [R].ASPAPA | 1xPhospho | F8VPZ9 1xI | 2.26E-06 | 0 | 1 | 1 | 1 | F8VPZ9 | BRD4-inter | F8VPZ9 [7 | 0 | 2038.15 | 0 | 1.6 | 0.000429 | 9.36 | 52.3 | 147.7 | 8.91 | 8.71 | | High | High | High | High | High | High | High | High | High | High | High | 0 | 2.53E-07 | 3 |
| FALSE | High | ASPAPAQL | [K].ASPAPA | 1xCarbami | A0A0R4J1C | 5.18E-06 | 0 | 1 | 3 | 1 | A0A0R4J1C | Histone-lys | A0A0R4J1C | 0 | 3436.633 | 0 | 0.81 | 0.005511 | 9.32 | 68.5 | 131.5 | 9.07 | 16.19 | | High | High | High | High | High | High | High | High | High | High | High | 0 | 6.45E-07 | 5.2 |
| FALSE | High | ASPASGHQ | [K].ASPASC | 1xPhospho | Q9QXY1 1x | 1.56E-06 | 0 | 1 | 4 | 3 | Q9QXY1 | Tight junct | Q9QXY1 [1 | 0 | 2301.009 | 0 | -1.65 | 0.004036 | 1.83 | 151.6 | 48.4 | 11.76 | 29.45 | | High | High | High | High | High | High | High | High | High | High | High | 0 | 1.68E-07 | 3.45 |
| FALSE | High | ASPASPVSI | [K].ASPASF | 1xCarbami | F8VQK5 1x | 0.000543 | 0 | 1 | 3 | 2 | F8VQK5 | SAM and S | F8VQK5 [1 | 0 | 1920.872 | 0 | 0.7 | 0.001613 | 4.89 | 73.9 | 126.1 | 13.39 | 5.34 | | High | High | High | High | High | High | High | High | High | High | High | 0 | 0.000172 | 2.73 |
| FALSE | High | ASPASPVSI | [K].ASPASF | 1xCarbami | F8VQK5 2x | 0.000389 | 0 | 1 | 3 | 1 | F8VQK5 | SAM and S | F8VQK5 [1 | 0 | 2000.839 | 0 | -0.08 | 0.567449 | 13.58 | 103 | 97 | 10.48 | 6.02 | | High | High | High | High | High | High | High | High | High | High | High | 0 | 8.25E-05 | 2.57 |
| FALSE | High | ASPATHR | [R].ASPATH | 1xPhospho | Q8BTI8 1xI | 0.076555 | 0.005826 | 1 | 3 | 1 | Q8BTI8 | Serine/argi | Q8BTI8 [18 | 0 | 1048.514 | 0 | 0.08 | 0.846025 | 4.68 | 97.5 | 102.5 | 22.22 | 14.97 | | High | High | High | High | High | High | High | High | High | High | High | 0.00302 | 0.03165 | 1.76 |
| FALSE | High | ASFETLPNI | [R].ASFETL | 1xCarbami | Q9CWE0 1 | 0.000584 | 0 | 1 | 5 | 2 | Q9CWE0 | Mitochond | Q9CWE0 [ | 0 | 2246.154 | 0 | -0.53 | 0.008414 | 8.44 | 117.8 | 82.2 | 8.77 | 5.26 | | High | High | High | High | High | High | High | High | High | High | High | 0 | 0.000131 | 3.51 |
| FALSE | High | ASPPLQDS | [K].ASPPLC | 2xCarbami | Q3URL4 1x | 7.24E-10 | 0 | 1 | 3 | 4 | Q3URL4 | BTB and CI | Q3URL4 [1 | 0 | 2730.288 | 0 | -0.1 | 0.287318 | 8.72 | 104.2 | 95.8 | 8.59 | 6.48 | | High | High | High | High | High | High | High | High | High | High | High | 0 | 3.02E-11 | 5.37 |
| FALSE | High | ASFDQLR | [K].ASFDQI | 1xPhospho | E9Q4F8 1x | 0.007019 | 0.00062 | 1 | 4 | 2 | E9Q4F8 | Ankyrin re | E9Q4F8 [1 | 0 | 1145.555 | 0 | 0.06 | 0.92774 | 17.2 | 99.4 | 100.6 | 9.1 | 8.29 | | High | High | High | High | High | High | High | High | High | High | High | 0.001812 | 0.01855 | 2.45 |
| FALSE | High | ASETSPSFS | [K].ASETSP | 1xPhospho | E9QM77 1 | 0.000563 | 0 | 1 | 9 | 4 | E9QM77 | Ataxin-2 O | E9QM77 [7 | 0 | 1578.782 | 0 | 0.22 | 0.003651 | 0.68 | 93 | 107 | 1.75 | 2.34 | | High | High | High | High | High | High | High | High | High | High | High | 0 | 0.000125 | 2.56 |
| FALSE | High | AQSGSPES | [R].AQSGS | 1xPhospho | Q8BXJ2-2 1 | 2.27E-05 | 0 | 1 | 2 | 2 | Q8BXJ2-2 | Isoform 2 c | Q8BXJ2-2 [ | 0 | 1888.905 | 0 | 0.95 | 0.00212 | 6.96 | 71 | 129 | 6.49 | 10.65 | | High | High | High | High | High | High | High | High | High | High | High | 0 | 3.39E-06 | 2.55 |
| FALSE | High | AQSGTDSS | [R].AQSGT | 1xPhospho | Q8BTI8 1xI | 0.000924 | 0 | 1 | 3 | 4 | Q8BTI8 | Serine/argi | Q8BTI8 [15 | 0 | 1781.847 | 0 | -0.97 | 0.001066 | 2.56 | 132.3 | 67.7 | 7.43 | 4.6 | | High | High | High | High | High | High | High | High | High | High | High | 0 | 0.000219 | 2.23 |
| FALSE | High | AQSLEPYG | [R].AQSLEF | 1xPhospho | Q8BX02 1x | 0.000365 | 0 | 1 | 2 | 2 | Q8BX02 | KN motif a | Q8BX02 [3 | 0 | 1801.95 | 0 | 1.28 | 0.000297 | 8.25 | 61.2 | 138.8 | 8.55 | 6.51 | | High | High | High | High | High | High | High | High | High | High | High | 0.001011 | 0.01074 | 2.45 |
| FALSE | High | AQSLLSTDI | [R].AQSLLS | 1xPhospho | Q497E8 1x | 0.038346 | 0.002563 | 1 | 10 | 2 | Q497E8 | Proteasom | Q497E8 [1 | 0 | 1299.651 | 0 | 1.88 | 0.001527 | 17.68 | 46.2 | 153.8 | 11.15 | 14.67 | | High | High | High | High | High | High | High | High | High | High | High | 0.001355 | 0.01418 | 1.97 |
| FALSE | High | AQSLPSVPI | [R].AQSLPS | 1xCarbami | Q8BJ56 1xI | 0.001783 | 5.66E-05 | 1 | 3 | 1 | Q8BJ56 | Patatin-like | Q8BJ56 [4C | 0 | 2855.4 | 0 | 0.82 | 0.002085 | 0.53 | 77.2 | 122.8 | 8.98 | 9.36 | | High | High | High | High | High | High | High | High | High | High | High | 2.58E-05 | 0.000457 | 3.62 |
| FALSE | High | AQSNGSGI | [K].AQSNG | 1xPhospho | B2RTN5 1x | 3.23E-05 | 0 | 1 | 4 | 1 | B2RTN5 | Glucocortic | B2RTN5 [1 | 0 | 2453.993 | 0 | 0.06 | 0.994373 | 18.55 | 99 | 101 | 10.91 | 5.99 | | High | High | High | High | High | High | High | High | High | High | High | 0 | 5.06E-06 | 3.38 |
| FALSE | High | AQSPSPK | [R].AQSPSF | 1xPhospho | E9PZM7 1x | 0.034579 | 0.002233 | 1 | 3 | 1 | E9PZM7 | SR-related | E9PZM7 [8 | 0 | 1252.67 | 0 | 0.1 | 0.949411 | 4.6 | 96.8 | 103.2 | 8.14 | 5.17 | | High | High | High | High | High | High | High | High | High | High | High | 0.001183 | 0.01269 | 1.76 |
| FALSE | High | AQSPSYVT | [R].AQSPSY | 1xPhospho | Q68FH0 1x | 0.000941 | 0 | 1 | 6 | 2 | Q68FH0 | Plakophilin | Q68FH0 [2 | 0 | 1932.927 | 0 | -0.56 | 0.000407 | 1.22 | 119.3 | 80.7 | 4.51 | 2.15 | | High | High | High | High | High | High | High | High | High | High | High | 0 | 0.000223 | 2.49 |
| FALSE | High | AQSPSYVT | [R].AQSPSY | 1xPhospho | Q68FH0 1x | 0.001301 | 0 | 1 | 6 | 2 | Q68FH0 | Plakophilin | Q68FH0 [2 | 1 | 2346.165 | 0 | -0.31 | 0.154683 | 20.79 | 112.1 | 87.9 | 23.13 | 12.95 | | High | High | High | High | High | High | High | High | High | High | High | 9.42E-05 | 0.001002 | 1.34 |
| FALSE | High | AQSSDLFSI | [R].AQSSDI | 1xPhospho | Q8BWR4 1 | 3.8E-05 | 0 | 1 | 3 | 2 | Q8BWR4 | Ubiquitin c | Q8BWR4 [ | 0 | 1828.879 | 0 | 1.66 | 0.000429 | 13.92 | 48 | 152 | 20.77 | 4.55 | | High | High | High | High | High | High | High | High | High | High | High | 0 | 1.46E-05 | 2.77 |
| FALSE | High | AQSSEVPL | [K].AQSSE | 2xPhospho | V9GWU7 2 | 0.015603 | 0.001124 | 1 | 3 | 1 | V9GWU7 | Serine/thre | V9GWU7 [ | 0 | 1814.829 | 0 | -1.85 | 0.005981 | 0.5 | 156.2 | 43.8 | 5.7 | 41.06 | | High | High | High | High | High | High | High | High | High | High | High | 0.000585 | 0.005168 | 2.29 |
| FALSE | High | AQSSGAVE | [R].AQSSG | 1xPhospho | A0A1B0GR | 0.000108 | 0 | 1 | 7 | 1 | A0A1B0GR | BET1-like p | A0A1B0GR | 0 | 1669.8 | 0 | 1.07 | 0.017613 | 32.84 | 64.4 | 135.6 | 21.58 | 13.83 | | High | High | High | High | High | High | High | High | High | High | High | 0 | 1.97E-05 | 3.72 |
| FALSE | High | AQSSLSDLI | [K].AQSSLS | 1xPhospho | S4R1N2 1x | 0.048715 | 0.00354 | 1 | 2 | 2 | S4R1N2 | DNA cross- | S4R1N2 [6 | 0 | 1948.967 | 0 | -0.13 | 0.237947 | 16 | 104.3 | 95.7 | 10.02 | 8.24 | | High | High | High | High | High | High | High | High | High | High | High | 0.001844 | 0.01865 | 2.93 |
| FALSE | High | AQSGSPES | [R].AQSGS | 1xPhospho | Q8BXJ2-4 1 | 1.18E-05 | 0 | 1 | 2 | 1 | Q8BXJ2-4 | Isoform 4 c | Q8BXJ2-4 [ | 0 | 1801.873 | 0 | 0.81 | 0.013332 | 5.49 | 75.5 | 124.5 | 9.44 | 13.2 | | High | High | High | High | High | High | High | High | High | High | High | 0 | 1.62E-06 | 2.87 |
| FALSE | High | AQSTDSLG | [R].AQSTD | 1xPhospho | O35551-6 1 | 8.98E-06 | 0 | 1 | 6 | 8 | O35551-6 | Isoform 6 c | O35551-6 [ | 0 | 2135.063 | 0 | 0.12 | 0.086519 | 10.15 | 95.8 | 104.2 | 4.43 | 9.99 | | High | High | High | High | High | High | High | High | High | High | High | 0 | 1.2E-06 | 3.34 |
| FALSE | High | AQSTVIQM | [K].AQSTVI | 1xPhospho | E9PWP1 1x | 0.022574 | 0.001542 | 1 | 2 | 1 | E9PWP1 | Zinc finger | E9PWP1 [1 | 0 | 1643.803 | 0 | -0.76 | 0.000364 | 2.33 | 127.8 | 72.2 | 3.52 | 7.8 | | High | High | High | High | High | High | High | High | High | High | High | 0.000814 | 0.007813 | 1.25 |
| FALSE | High | AQSVSPVQ | [K].AQSVSI | 1xPhospho | Q924A2 1x | 1.1E-05 | 0 | 1 | 4 | 1 | Q924A2 | Protein cap | Q924A2 [1 | 0 | 2618.396 | 0 | 0.17 | 0.071917 | 11.75 | 93.1 | 106.9 | 4.76 | 6.78 | | High | High | High | High | High | High | High | High | High | High | High | 0 | 1.51E-06 | 3.38 |
| FALSE | High | AQSYDIQA | [R].AQSYDI | 1xPhospho | F8VQ93 1x | 0.000286 | 0 | 1 | 2 | 3 | F8VQ93 | PH-interact | F8VQ93 [1 | 0 | 1747.882 | 0 | 1.04 | 0.001286 | 15.37 | 64.9 | 135.1 | 3.88 | 9.98 | | High | High | High | High | High | High | High | High | High | High | High | 2.58E-05 | 0.000482 | 3.39 |
| FALSE | High | AQSYPDNH | [R].AQSYPI | 1xPhospho | G5E8L8 1xI | 1.77E-09 | 0 | 1 | 2 | 1 | G5E8L8 | Mitogen ac | G5E8L8 [2 | 0 | 3096.403 | 0 | 1.38 | 0.000144 | 4.54 | 55.7 | 144.3 | 6.61 | 3.13 | | High | High | High | High | High | High | High | High | High | High | High | 0 | 8.18E-11 | 5.04 |
| FALSE | High | AQTEMQF | [K].AQTEM | 1xPhospho | A0A0R4J0V | 7.91E-06 | 0 | 1 | 7 | 1 | A0A0R4J0V | Phosphatic | A0A0R4J0V | 0 | 3754.817 | 0 | 0.1 | 0.883194 | 23.98 | 96.9 | 103.1 | 12.8 | 13.9 | | High | High | High | High | High | High | High | High | High | High | High | 0 | 1.04E-06 | 4.05 |
| FALSE | High | AQTIDLDLT | [R].AQTIDL | 1xCarbami | P70423 1xI | 0.002115 | 5.66E-05 | 1 | 2 | 3 | P70423 | Cationic an | P70423 [6C | 0 | 2082.998 | 0 | 1.14 | 0.002491 | 19.24 | 66.4 | 133.6 | 4.48 | 11.88 | | High | High | High | High | High | High | High | High | High | High | High | 2.58E-05 | 0.000553 | 3.21 |
| FALSE | High | AQTLPTSV | [R].AQTLPT | 1xPhospho | Q62261 1x | 0.006895 | 0.00062 | 1 | 3 | 1 | Q62261 | Spectrin be | Q62261 [2 | 0 | 2212.131 | 0 | 0.31 | 0.241972 | 12.61 | 92 | 108 | 19.89 | 26.54 | | High | High | High | High | High | High | High | High | High | High | High | 0.000275 | 0.002067 | 2.33 |
| FALSE | High | AQTLPTSV | [R].AQTLPT | 1xPhospho | Q62261 1x | 1.71E-06 | 0 | 1 | 3 | 3 | Q62261 | Spectrin be | Q62261 [2 | 0 | 2441.294 | 0 | 1.17 | 0.000897 | 14.32 | 60.6 | 139.4 | 8.32 | 5.82 | | High | High | High | High | High | High | High | High | High | High | High | 0 | 1.85E-07 | 3.15 |
| FALSE | High | AQTLPTSV | [R].AQTLPT | 2xPhospho | Q62261 2x | 0.000101 | 0 | 1 | 3 | 1 | Q62261 | Spectrin be | Q62261 [2 | 0 | 2521.26 | 0 | 0.37 | 0.079048 | 25.83 | 85.7 | 114.3 | 18.93 | 14.26 | | High | High | High | High | High | High | High | High | High | High | High | 0 | 1.82E-05 | 3.78 |
| FALSE | High | AQTPPISSL | [R].AQTPPI | 3xPhospho | E9PZP8 3xI | 0.002022 | 5.66E-05 | 1 | 3 | 1 | E9PZP8 | HECT and I | E9PZP8 [26 | 0 | 2378.028 | 0 | 1.65 | 0.001541 | 41.26 | 54.8 | 145.2 | 39.19 | 13.12 | | High | High | High | High | High | High | High | High | High | High | High | 2.58E-05 | 0.000525 | 3.46 |
| FALSE | High | AQTSPGLG | [R].AQTSPC | 1xPhospho | Q99MD6 1 | 0.120211 | 0.009851 | 1 | 2 | 1 | Q99MD6 | Thioredoxi | Q99MD6 [ | 0 | 1396.76 | 0 | -0.34 | 0.188798 | 3.84 | 111.5 | 88.5 | 5.71 | 14.77 | | High | High | High | High | High | High | High | High | High | High | High | 0.004907 | 0.05346 | 2.15 |
| FALSE | High | AQTSYGFA | [K].AQTSYC | 1xPhospho | Q5HZI2 1xI | 1.86E-05 | 0 | 1 | 1 | 1 | Q5HZI2 | C2 calcium | Q5HZI2 [14 | 0 | 2146.017 | 0 | -0.89 | 0.000381 | 1.18 | 131 | 69 | 4.96 | 7.31 | | High | High | High | High | High | High | High | High | High | High | High | 0 | 2.71E-06 | 3.77 |
| FALSE | High | AQTTNSNS | [R].AQTTN | 1xPhospho | B2RY22 1xI | 5.7E-08 | 0 | 2 | 18 | 6 | B2RY22; AC | Clasp1 pro | B2RY22 [14 | 0 | 2105.882 | 0 | 0.05 | 0.279223 | 8.76 | 98 | 102 | 4.35 | 15.16 | Shared | High | High | High | High | High | High | High | High | High | High | High | 0 | 4.05E-09 | 3.88 |
| FALSE | High | AQSTDSLG | [R].AQSTD | 2xPhospho | O35551-6 1 | 0.000967 | 0 | 1 | 6 | 2 | O35551-6 | Isoform 6 c | O35551-6 [ | 0 | 2215.03 | 0 | 0.67 | 0.000548 | 7.31 | 76.4 | 123.6 | 6.68 | 4.87 | | High | High | High | High | High | High | High | High | High | High | High | 0.000936 | 0.009747 | 2.23 |
| FALSE | High | AQVSPQS | [K].AQVSPC | 1xPhospho | Q52KI8 1xI | 0.024638 | 0.001612 | 1 | 4 | 2 | Q52KI8 | Serine/argi | Q52KI8 [94 | 0 | 1025.487 | 0 | -0.42 | 0.003503 | 5.03 | 116.1 | 83.9 | 4.29 | 5.39 | | High | High | High | High | High | High | High | High | High | High | High | 0.000872 | 0.008655 | 2.11 |
| FALSE | High | AQSFDVAS | [K].AQSFD | 1xPhospho | E9Q0N0 1x | 3.03E-08 | 0 | 1 | 10 | 2 | E9Q0N0 | Intersectin | E9Q0N0 [2 | 0 | 2581.229 | 0 | 0.33 | 0.06283 | 16.82 | 87 | 113 | 9.15 | 10.86 | | High | High | High | High | High | High | High | High | High | High | High | 0 | 9.84E-09 | 3.72 |
| FALSE | High | AQSDDEK | [K].AQSDD | 1xPhospho | Q5XJE5 1xI | 0.083783 | 0.006345 | 1 | 2 | 1 | Q5XJE5 | RNA polym | Q5XJE5 [13 | 0 | 1330.629 | 0 | -0.44 | 0.012741 | 9.21 | 114.3 | 85.7 | 5.36 | 11.7 | | High | High | High | High | High | High | High | High | High | High | High | 0.003265 | 0.03507 | 1.94 |
| FALSE | High | AQGLSEQT | [K].AQGLSI | 3xPhospho | B1AY10 3xI | 0.005044 | 0.000504 | 1 | 3 | 1 | B1AY10 | Transcripti | B1AY10 [1 | 0 | 2434.998 | 0 | 0.6 | 0.013135 | 18.87 | 79.4 | 120.6 | 7.7 | 18.36 | | High | High | High | High | High | High | High | High | High | High | High | 0.000246 | 0.001461 | 2.73 |
| FALSE | High | AQGSHDPT | [R].AQGSH | 1xPhospho | Q69ZP6 1x | 3.99E-06 | 0 | 1 | 2 | 3 | Q69ZP6 | MKIAA117 | Q69ZP6 [2 | 0 | 3859.757 | 0 | -0.11 | 0.724349 | 18.3 | 103.8 | 96.2 | 19.58 | 21.68 | | High | High | High | High | High | High | High | High | High | High | High | 0 | 4.79E-07 | 3.78 |
| FALSE | High | AQGSPEQF | [K].AQGSPI | 1xPhospho | E9PYL1 1xI | 0.058664 | 0.004294 | 1 | 5 | 1 | E9PYL1 | Chromodo | E9PYL1 [16 | 0 | 1592.845 | 0 | -0.01 | 0.83441 | 10.18 | 99.3 | 100.7 | 6.9 | 5.9 | | High | High | High | High | High | High | High | High | High | High | High | 0.002187 | 0.02319 | 1.68 |
| FALSE | High | AQHPDYSF | [K].AQHPD | 1xPhospho | F8VQD1 1x | 0.030104 | 0.001857 | 1 | 10 | 1 | F8VQD1 | Protein pol | F8VQD1 [1 | 0 | 1815.826 | 0 | 0.56 | 0.104308 | 15.63 | 85.1 | 114.9 | 6.2 | 22.43 | | High | High | High | High | High | High | High | High | High | High | High | 0.001011 | 0.01084 | 2.35 |
| FALSE | High | AQHTLPQD | [K].AQHTLI | 1xPhospho | Q8CGC4 1x | 0.002576 | 0.000109 | 1 | 1 | 4 | Q8CGC4 | Protein LSI | Q8CGC4 [8 | 0 | 3926.943 | 0 | 0.81 | 0.0175 | 26.17 | 75.4 | 124.6 | 10.49 | 18.52 | | High | High | High | High | High | High | High | High | High | High | High | 4.94E-05 | 0.000688 | 3.15 |
| FALSE | High | AQKPPSPP/ | [K].AQKPP | 1xPhospho | A0A0R4IZV | 0.070485 | 0.005214 | 1 | 12 | 1 | A0A0R4IZV | tRNA (guar | A0A0R4IZV | 0 | 2119.077 | 0 | -0.16 | 0.067543 | 2.89 | 108 | 92 | 5.25 | 15.22 | | High | High | High | High | High | High | High | High | High | High | High | 0.00269 | 0.02873 | 2.17 |
| FALSE | High | AQKPPSPP/ | [K].AQKPP | 1xPhospho | A0A0R4IZV | 0.00172 | 5.66E-05 | 1 | 12 | 4 | A0A0R4IZV | tRNA (guar | A0A0R4IZV | 0 | 2135.072 | 0 | -0.2 | 0.043381 | 1.42 | 107.2 | 92.8 | 2.78 | 11.56 | | High | High | High | High | High | High | High | High | High | High | High | 0.003265 | 0.035 | 1.94 |
| FALSE | High | AQLEPVAS | [K].AQLEP | 1xPhospho | Q9QXS1-3 | 0.000121 | 0 | 3 | 24 | 4 | Q9QXS1-3; | Isoform PL | Q9QXS1-3 | 0 | 1648.908 | 0 | 0.96 | 0.003329 | 9.06 | 67.9 | 132.1 | 3.81 | 12.47 | Shared | High | High | High | High | High | High | High | High | High | High | High | 0 | 2.22E-05 | 2.38 |
| FALSE | High | AQLSPGLYI | [K].AQLSPC | 1xPhospho | Q5RIM6 1x | 1.31E-05 | 0 | 1 | 10 | 2 | Q5RIM6 | Nuclear re | Q5RIM6 [1 | 0 | 1788.837 | 0 | -0.01 | 0.77479 | 8.61 | 100.5 | 99.5 | 4.04 | 4.87 | | High | High | High | High | High | High | High | High | High | High | High | 0 | 1.82E-06 | 2.43 |
| FALSE | High | AQLSPSVK | [K].AQLSPS | 1xPhospho | E9Q6J5-3 1 | 0.025964 | 0.001645 | 1 | 3 | 2 | E9Q6J5-3 | Isoform Ne | E9Q6J5-3 [ | 0 | 1367.77 | 0 | -0.28 | 0.005793 | 2.18 | 108 | 92 | 5.59 | 4.21 | | High | High | High | High | High | High | High | High | High | High | High | 0.000872 | 0.009167 | 1.55 |
| FALSE | High | AQISSPNLF | [R].AQISSP | 1xPhospho | O89032 1x | 0.067619 | 0.004864 | 1 | 3 | 3 | O89032 | SH3 and PX | O89032 [4 | 0 | 1294.672 | 0 | 0.79 | 0.002043 | 14.33 | 75.5 | 124.5 | 3.71 | 10.42 | | High | High | High | High | High | High | High | High | High | High | High | 0.007451 | 0.08209 | 1.4 |
| FALSE | High | AQLYLAPT | [R].AQLYLA | 1xPhospho | Q8CB62 1x | 0.00011 | 0 | 1 | 2 | 1 | Q8CB62 | Centrobin | Q8CB62 [5 | 0 | 2966.471 | 0 | -0.35 | 0.015526 | 3.98 | 110.1 | 89.9 | 11.03 | 4.29 | | High | High | High | High | High | High | High | High | High | High | High | 0 | 1.99E-05 | 2.73 |
| FALSE | High | AQMCADV | [K].AQMCA | 1xCarbami | Q01320 1x | 0.000168 | 0 | 2 | 3 | 2 | Q01320; Q | DNA topois | Q01320 [1 | 0 | 1653.769 | 0 | -0.46 | 0.001091 | 0.78 | 116.1 | 83.9 | 3.58 | 8.6 | Shared | High | High | High | High | High | High | High | High | High | High | High | 0 | 3.23E-05 | 3.44 |
| FALSE | High | AQSEPANT | [K].AQSEPA | 1xPhospho | Q3V1H1 1x | 0.018447 | 0.001324 | 1 | 1 | 1 | Q3V1H1 | Cytoskelet | Q3V1H1 [2 | 0 | 1768.925 | 0 | -0.04 | 0.653859 | 4.84 | 100.3 | 99.7 | 5.91 | 7.55 | | High | High | High | High | High | High | High | High | High | High | High | 0.000706 | 0.006206 | 1.72 |
| FALSE | High | AQMCADV | [K].AQMCA | 1xCarbami | Q01320 1x | 3.95E-06 | 0 | 2 | 3 | 19 | Q01320; Q | DNA topois | Q01320 [1 | 0 | 1669.764 | 0 | -0.44 | 0.001843 | 1.19 | 115.2 | 84.8 | 4.6 | 2.81 | Shared | High | High | High | High | High | High | High | High | High | High | High | 2.58E-05 | 0.00049 | 3.55 |
| FALSE | High | AQNGLENC | [K].AQNGL | 1xDeamida | Q3UQV4 1 | 0.115566 | 0.009504 | 1 | 3 | 3 | Q3UQV4 | Putative ur | Q3UQV4 [ | 0 | 1928.9 | 0 | -1.57 | 0.000209 | 0.92 | 151.9 | 48.1 | 7.1 | 13.1 | | High | High | High | High | High | High | High | High | High | High | High | 0.00728 | 0.07892 | 2.39 |
| FALSE | High | AQPAGSPS | [R].AQPAG | 1xPhospho | Q64700 1x | 3.09E-05 | 0 | 1 | 5 | 1 | Q64700 | Retinoblast | Q64700 [3 | 0 | 1813.891 | 0 | 1.53 | 0.000236 | 11.15 | 49.4 | 150.6 | 7.81 | 5.23 | | High | High | High | High | High | High | High | High | High | High | High | 0 | 4.79E-06 | 1.79 |
| FALSE | High | AQPAHSPG | [K].AQPAH | 1xPhospho | Q60795 1x | 0.000343 | 0 | 1 | 2 | 3 | Q60795 | Nuclear fac | Q60795 [3 | 0 | 2862.356 | 0 | 0.11 | 0.207807 | 11.95 | 96.1 | 103.9 | 4.6 | 6.86 | | High | High | High | High | High | High | High | High | High | High | High | 0 | 0.000292 | 2.69 |
| FALSE | High | AQPGSPQ/ | [K].AQPGS | 1xPhospho | Q6P9K8 1x | 0.008769 | 0.000736 | 1 | 4 | 1 | Q6P9K8 | Caskin-1 O | Q6P9K8 [7 | 0 | 2280.19 | 0 | -0.88 | 0.077089 | 3.37 | 128 | 72 | 21.51 | 20.14 | | High | High | High | High | High | High | High | High | High | High | High | 0.000327 | 0.002702 | 3.04 |
| FALSE | High | AQPLEDLD | [R].AQPLEI | 1xPhospho | E9PVX6 1xI | 0.003167 | 0.00016 | 1 | 2 | 1 | E9PVX6 | Proliferatic | E9PVX6 [1 | 0 | 3813.851 | 0 | 0.08 | 0.448961 | 25.4 | 97.2 | 102.8 | 17.31 | 26.42 | | High | High | High | High | High | High | High | High | High | High | High | 7.19E-05 | 0.000871 | 4.12 |
| FALSE | High | AQPLEDLT | [K].AQPLEI | 1xCarbami | E9PVX6 1xI | 0.000105 | 0 | 1 | 2 | 1 | E9PVX6 | Proliferatic | E9PVX6 [1 | 0 | 3311.74 | 0 | 0.13 | 0.069479 | 5.28 | 93.7 | 106.3 | 6.24 | 8.76 | | High | High | High | High | High | High | High | High | High | High | High | 0 | 1.91E-05 | 4.27 |
| FALSE | High | AQPSASLG | [R].AQPSA | 1xPhospho | Q8C2Q3 1x | 0.00095 | 0 | 1 | 5 | 2 | Q8C2Q3 | RNA-bindir | Q8C2Q3 [2 | 0 | 1514.757 | 0 | 2.55 | 0.022939 | 30.28 | 29.2 | 170.8 | 16.97 | 44.95 | | High | High | High | High | High | High | High | High | High | High | High | 0 | 0.000226 | 1.61 |
| FALSE | High | AQPSSSEDI | [K].AQPSS | 1xPhospho | B7ZCJ1 1xF | 0.003433 | 0.00021 | 1 | 5 | 2 | B7ZCJ1 | Rho GTPas | B7ZCJ1 [14 | 0 | 2324.11 | 0 | -0.61 | 0.001949 | 7.78 | 129.2 | 70.8 | 10.08 | 19.49 | | High | High | High | High | High | High | High | High | High | High | High | 0.000275 | 0.001989 | 2.71 |
| FALSE | High | AQPSSSEDI | [K].AQPSS | 2xPhospho | B7ZCJ1 2xF | 0.012288 | 0.000889 | 1 | 5 | 2 | B7ZCJ1 | Rho GTPas | B7ZCJ1 [14 | 0 | 2404.076 | 0 | 0.38 | 0.053688 | 18.4 | 81 | 119 | 11.76 | 16.23 | | High | High | High | High | High | High | High | High | High | High | High | 0.000449 | 0.003932 | 2.25 |
| FALSE | High | AQPSTSPK | [K].AQPSTS | 1xPhospho | Q3U343 1x | 0.024424 | 0.001612 | 1 | 3 | 3 | Q3U343 | Putative ur | Q3U343 [1 | 0 | 1353.718 | 0 | -1.44 | 0.000737 | 2.34 | 148.7 | 51.3 | 4.84 | 15.84 | | High | High | High | High | High | High | High | High | High | High | High | 0.00554 | 0.06 | 1.6 |
| FALSE | High | AQQNLCSG | [R].AQQNL | 1xCarbami | Q8BQ30-2 | 0.000558 | 0 | 1 | 4 | 1 | Q8BQ30-2 | Isoform 2 c | Q8BQ30-2 | 0 | 2715.24 | 0 | 0.7 | 0.005755 | 24.37 | 78.8 | 121.2 | 14.36 | 12.32 | | High | High | High | High | High | High | High | High | High | High | High | 0 | 0.000124 | 2.46 |
| FALSE | High | AQQPAPAS | [R].AQQPA | 1xPhospho | Q3U1I6 1xI | 3.57E-05 | 0 | 1 | 5 | 2 | Q3U1I6 | Putative ur | Q3U1I6 [5 | 0 | 1718.924 | 0 | -0.51 | 0.020299 | 2.46 | 116.7 | 83.3 | 7.72 | 8.54 | | High | High | High | High | High | High | High | High | High | High | High | 0 | 3.92E-05 | 2.99 |

| | | | | | | | | | | | | | | | | | | | | | | | | | | | | | | | | | | | | | | |
|---|---|---|---|---|---|---|---|---|---|---|---|---|---|---|---|---|---|---|---|---|---|---|---|---|---|---|---|---|---|---|---|---|---|---|---|---|---|---|
| FALSE | High | AQSASVES | [K].AQSAS | 1xCarbami | Q6DFZ1 2x | 0.033404 | 0.002195 | 1 | 5 | 1 | Q6DFZ1 | Golgi-speci | Q6DFZ1 [3 | 0 | 4434.862 | 0 | 0.03 | 0.387746 | 17.03 | 94.2 | 105.8 | 17.69 | 21.95 | | High | High | High | High | High | High | High | High | High | High | High | 0.001151 | 0.01219 | 2.99 |
| FALSE | High | AQNGLEN | [K].AQNGL | 1xPhospho | Q3UQV4 1 | 7.65E-07 | 0 | 1 | 3 | 2 | Q3UQV4 | Putative ur | Q3UQV4 [ | 0 | 1927.916 | 0 | -1.35 | 0.000125 | 0.24 | 143.6 | 56.4 | 3.58 | 7.18 | | High | High | High | High | High | High | High | High | High | High | High | 0 | 7.56E-08 | 3.1 |
| FALSE | High | AQWPQSE | [K].AQWP | 1xPhospho | F6ZBR8 1x | 5.29E-07 | 0 | 1 | 1 | 4 | F6ZBR8 | Protein Wi | F6ZBR8 [21 | 0 | 2694.225 | 0 | -0.37 | 0.012005 | 3.3 | 113.2 | 86.8 | 6.4 | 5.93 | | High | High | High | High | High | High | High | High | High | High | High | 0 | 7.92E-06 | 4.76 |
| FALSE | High | AQWPQSE | [K].AQWP | 2xPhospho | F6ZBR8 2x | 0.000393 | 0 | 1 | 1 | 2 | F6ZBR8 | Protein Wi | F6ZBR8 [21 | 0 | 2774.191 | 0 | -0.11 | 0.188269 | 15.41 | 103.9 | 96.1 | 15.79 | 13.67 | | High | High | High | High | High | High | High | High | High | High | High | 0 | 8.32E-05 | 4.23 |
| FALSE | High | AQWPQSE | [K].AQWP | 1xPhospho | O88286-3 | 0.001349 | 0 | 1 | 4 | 1 | O88286-3 | Isoform 2 | O88286-3 | 0 | 2718.261 | 0 | -0.13 | 0.250797 | 22.82 | 111.3 | 88.7 | 14.27 | 13.1 | | High | High | High | High | High | High | High | High | High | High | High | 0 | 0.000332 | 3.77 |
| FALSE | High | ASAVSPEK | [R].ASAVSF | 1xPhospho | Q3TAB2 1x | 0.000845 | 0 | 2 | 6 | 5 | Q3TAB2; H | Putative ur | Q3TAB2 [5 | 0 | 1326.707 | 0 | -0.71 | 0.000992 | 1.67 | 124.5 | 75.5 | 5.54 | 3.55 | Shared | High | High | High | High | High | High | High | High | High | High | High | 4.94E-05 | 0.000595 | 2.29 |
| FALSE | High | ASAYVFASI | [K].ASAYVF | 1xCarbami | A0A1B0GR | 0.002172 | 5.66E-05 | 1 | 5 | 1 | A0A1B0GR | 3'(2'),5'-bis | A0A1B0GR | 0 | 1795.885 | 0 | -0.14 | 0.013698 | 1.36 | 105.2 | 94.8 | 7.08 | 4.74 | | High | High | High | High | High | High | High | High | High | High | High | 2.58E-05 | 0.00057 | 2.26 |
| FALSE | High | ASCSDTLLT | [K].ASCSDT | 2xCarbami | Q3UQJ0 1x | 0.00153 | 5.66E-05 | 1 | 5 | 1 | Q3UQJ0 | Putative ur | Q3UQJ0 [8 | 0 | 2712.104 | 0 | -0.94 | 0.002113 | 5.7 | 135 | 65 | 8.81 | 10.9 | | High | High | High | High | High | High | High | High | High | High | High | 2.58E-05 | 0.000383 | 4.21 |
| FALSE | High | ASCSDTLLT | [K].ASCSDT | 2xCarbami | Q3UQJ0 1x | 2.47E-06 | 0 | 1 | 5 | 3 | Q3UQJ0 | Putative ur | Q3UQJ0 [8 | 0 | 2728.099 | 0 | -0.48 | 0.006853 | 3.67 | 113.6 | 86.4 | 5.93 | 5.5 | | High | High | High | High | High | High | High | High | High | High | High | 0 | 2.81E-07 | 4.06 |
| FALSE | High | ASCSSSTG | [R].ASCSSS | 1xCarbami | Q9Z1E4 1x | 0.0016 | 5.66E-05 | 1 | 2 | 5 | Q9Z1E4 | Glycogen [ | Q9Z1E4 [6 | 0 | 1566.724 | 0 | -0.54 | 0.005559 | 4.23 | 117.2 | 82.8 | 11.02 | 7.61 | | High | High | High | High | High | High | High | High | High | High | High | 2.58E-05 | 0.000404 | 2.16 |
| FALSE | High | ASDAFPEEI | [R].ASDAFF | 1xPhospho | P48754 1x | 0.002232 | 0.000109 | 1 | 4 | 1 | P48754 | Breast can | P48754 [68 | 0 | 1531.745 | 0 | -0.6 | 0.001261 | 3.49 | 120.8 | 79.2 | 3.43 | 5.98 | | High | High | High | High | High | High | High | High | High | High | High | 4.94E-05 | 0.000585 | 2.33 |
| FALSE | High | ASDDLGEP | [R].ASDDL | 1xPhospho | Q6PIU9 1x | 2.6E-05 | 0 | 1 | 1 | 2 | Q6PIU9 | Uncharacte | Q6PIU9 [1 | 0 | 2116.979 | 0 | 0.23 | 0.031206 | 10.28 | 93.2 | 106.8 | 11.3 | 13.79 | | High | High | High | High | High | High | High | High | High | High | High | 0 | 3.98E-06 | 3.9 |
| FALSE | High | ASDDSSSEL | [K].ASDDS | 1xPhospho | Q3TDF3 1x | 9.74E-06 | 0 | 1 | 4 | 13 | Q3TDF3 | Putative ur | Q3TDF3 [8 | 0 | 2710.264 | 0 | 0.15 | 0.050186 | 7.27 | 94.3 | 105.7 | 4.94 | 6.01 | | High | High | High | High | High | High | High | High | High | High | High | 0 | 6.99E-06 | 4.09 |
| FALSE | High | ASDEEEQG | [R].ASDEEE | 1xPhospho | Q8VC34 1x | 1.45E-06 | 0 | 1 | 5 | 2 | Q8VC34 | Putative RI | Q8VC34 [2 | 0 | 2698.349 | 0 | -0.45 | 0.000434 | 2.51 | 116 | 84 | 3.22 | 2.22 | | High | High | High | High | High | High | High | High | High | High | High | 0 | 1.54E-07 | 3.43 |
| FALSE | High | ASDGGANI | [R].ASDGG | 1xPhospho | E9PU87 1x | 1.6E-05 | 0 | 1 | 6 | 1 | E9PU87 | Serine/thre | E9PU87 [5 | 0 | 2302.232 | 0 | 0.53 | 0.008808 | 9.91 | 80.2 | 119.8 | 7.5 | 10.88 | | High | High | High | High | High | High | High | High | High | High | High | 0 | 2.29E-06 | 3.51 |
| FALSE | High | ASDLETK | [K].ASDLET | 1xPhospho | E9Q284 1x | 0.007145 | 0.00062 | 1 | 3 | 2 | E9Q284 | Coilin OS= | E9Q284 [1 | 0 | 1301.675 | 2 | -1.31 | 0.000181 | 0.49 | 141.8 | 58.2 | 3.96 | 6.89 | | High | High | High | High | High | High | High | High | High | High | High | 0.000275 | 0.002154 | 1.91 |
| FALSE | High | ASDLSPVSI | [K].ASDLSP | 1xDeamida | E9PUR0 1x | 0.0102 | 0.000736 | 1 | 7 | 1 | E9PUR0 | Ankyrin re | E9PUR0 [2 | 0 | 2956.421 | 0 | -0.08 | 0.927553 | 21.22 | 101.3 | 98.7 | 7.7 | 19.13 | | High | High | High | High | High | High | High | High | High | High | High | 0.000344 | 0.003199 | 3.71 |
| FALSE | High | ASDPGLPA | [R].ASDPG | 1xPhospho | B2RQC6 1x | 0.002869 | 0.00016 | 1 | 6 | 2 | B2RQC6 | CAD protei | B2RQC6 [1 | 0 | 1519.724 | 0 | -0.61 | 0.014652 | 7.29 | 121.1 | 78.9 | 5.78 | 20.95 | | High | High | High | High | High | High | High | High | High | High | High | 0.000449 | 0.004093 | 1.59 |
| FALSE | High | ASAVISPSC | [R].ASAVIS | 1xPhospho | Q8VCD7 1x | 2.6E-05 | 0 | 1 | 2 | 2 | Q8VCD7 | Lysine-spe | Q8VCD7 [4 | 0 | 1638.923 | 0 | -0.7 | 0.001269 | 5.59 | 123.8 | 76.2 | 7 | 8.35 | | High | High | High | High | High | High | High | High | High | High | High | 0 | 3.97E-06 | 3.5 |
| FALSE | High | ASDPGLPA | [R].ASDPG | 1xPhospho | B2RQC6 1x | 3.27E-06 | 0 | 1 | 6 | 5 | B2RQC6 | CAD protei | B2RQC6 [1 | 0 | 1748.887 | 0 | -0.4 | 0.074553 | 19.73 | 116.1 | 83.9 | 10.33 | 22.61 | | High | High | High | High | High | High | High | High | High | High | High | 0.000275 | 0.002135 | 2.8 |
| FALSE | High | ASDSNTHE | [R].ASDSN | 1xPhospho | Q9QZN4 1 | 4.08E-05 | 0 | 1 | 1 | 1 | Q9QZN4 | F-box only | Q9QZN4 [2 | 0 | 2475.108 | 0 | 0.38 | 0.013046 | 8.94 | 88.1 | 111.9 | 7.16 | 6.75 | | High | High | High | High | High | High | High | High | High | High | High | 0 | 6.56E-06 | 3.13 |
| FALSE | High | ASDTSLTQ | [R].ASDTSL | 1xPhospho | Q60670 1x | 0.060666 | 0.00451 | 1 | 3 | 1 | Q60670 | Serine/thre | Q60670 [5 | 0 | 1658.877 | 0 | 0.11 | 0.748423 | 5.7 | 98 | 102 | 9.17 | 11.74 | | High | High | High | High | High | High | High | High | High | High | High | 0.002293 | 0.024 | 2.04 |
| FALSE | High | ASDTSLTQ | [R].ASDTSL | 1xPhospho | F8VPT7 1x | 0.01588 | 0.001124 | 1 | 4 | 1 | F8VPT7 | Serine/thre | F8VPT7 [58 | 0 | 1774.894 | 0 | 0.42 | 0.106958 | 21.2 | 93.2 | 106.8 | 14.09 | 16.01 | | High | High | High | High | High | High | High | High | High | High | High | 0.000585 | 0.005284 | 2.85 |
| FALSE | High | ASEAEQTQ | [R].ASEAE | 1xPhospho | A2ADB0 1x | 4.47E-08 | 0 | 1 | 2 | 1 | A2ADB0 | Msx2-inter | A2ADB0 [2 | 0 | 3525.616 | 0 | 0.05 | 0.878937 | 11.44 | 100.6 | 99.4 | 5.87 | 15.92 | | High | High | High | High | High | High | High | High | High | High | High | 0 | 3.1E-09 | 3.95 |
| FALSE | High | ASEAGPLN | [R].ASEAG | 1xPhospho | B2RWW2 1 | 8.7E-11 | 0 | 1 | 3 | 4 | B2RWW2 | Golgb1 pro | B2RWW2 [ | 0 | 2328.119 | 0 | 0.62 | 0.000667 | 1.65 | 79.4 | 120.6 | 11.17 | 10.12 | | High | High | High | High | High | High | High | High | High | High | High | 0 | 2.79E-12 | 4.43 |
| FALSE | High | ASEATSSP | [R].ASEATS | 1xPhospho | Q8BQ33 1x | 0.096959 | 0.007777 | 1 | 1 | 2 | Q8BQ33 | Treslin OS= | Q8BQ33 [1 | 0 | 3091.495 | 0 | -0.4 | 0.026839 | 7.55 | 116.5 | 83.5 | 11.72 | 18.22 | | High | High | High | High | High | High | High | High | High | High | High | 0.004046 | 0.04315 | 2.54 |
| FALSE | High | ASEEGGGC | [R].ASEEG | 1xPhospho | Q3TIZ8 1x | 3.44E-05 | 0 | 1 | 4 | 2 | Q3TIZ8 | Putative ur | Q3TIZ8 [31 | 0 | 1400.682 | 0 | 0.62 | 0.004106 | 11.52 | 83.6 | 116.4 | 4.49 | 10.01 | | High | High | High | High | High | High | High | High | High | High | High | 0 | 3.2E-05 | 2.31 |
| FALSE | High | ASEGPSSA | [K].ASEGPS | 1xPhospho | E9PU96 1x | 6.13E-05 | 0 | 1 | 3 | 1 | E9PU96 | Nucleolar p | E9PU96 [8 | 0 | 1954.989 | 0 | -1.03 | 0.001237 | 2.56 | 130.1 | 69.9 | 8.18 | 9 | | High | High | High | High | High | High | High | High | High | High | High | 0 | 1.03E-05 | 2.69 |
| FALSE | High | ASEISIDDEI | [K].ASEISID | 1xPhospho | Q9DBE9 1x | 0.000235 | 0 | 1 | 1 | 1 | Q9DBE9 | pre-rRNA p | Q9DBE9 [2 | 0 | 2749.277 | 0 | -0.3 | 0.499925 | 10.64 | 101.6 | 98.4 | 13.32 | 13.96 | | High | High | High | High | High | High | High | High | High | High | High | 0 | 4.7E-05 | 3.13 |
| FALSE | High | ASENQSPS | [R].ASENQ | 1xPhospho | Q3TW94 1 | 6.91E-06 | 0 | 1 | 2 | 4 | Q3TW94 | Putative ur | Q3TW94 [ | 0 | 2066.902 | 0 | 0.66 | 0.002019 | 2.53 | 78.1 | 121.9 | 8.71 | 12.06 | | High | High | High | High | High | High | High | High | High | High | High | 0 | 3.66E-05 | 2.49 |
| FALSE | High | ASEPEAQG | [K].ASEPEA | 1xCarbami | F8WJA0 1x | 3.32E-09 | 0 | 1 | 6 | 3 | F8WJA0 | ATP-depen | F8WJA0 [1 | 0 | 2345.073 | 0 | -0.49 | 0.049004 | 11.33 | 115.5 | 84.5 | 11.63 | 13.13 | | High | High | High | High | High | High | High | High | High | High | High | 0 | 1.66E-10 | 5.58 |
| FALSE | High | ASEPLLSPP | [K].ASEPLL | 1xPhospho | A0A0A0M | 0.010382 | 0.000772 | 1 | 2 | 1 | A0A0A0M | Histone-lys | A0A0A0M | 0 | 1892.936 | 0 | 0.42 | 0.025832 | 19.02 | 83.7 | 116.3 | 10.04 | 15.6 | | High | High | High | High | High | High | High | High | High | High | High | 0.000382 | 0.003271 | 3.12 |
| FALSE | High | ASESSSEEK | [R].ASESSS | 1xPhospho | H9KV00 1x | 0.091627 | 0.007146 | 1 | 9 | 1 | H9KV00 | Protein SO | H9KV00 [1 | 0 | 1491.698 | 0 | 0 | 0.793278 | 30.34 | 94.9 | 105.1 | 10.3 | 14.27 | | High | High | High | High | High | High | High | High | High | High | High | 0.003646 | 0.03892 | 2.17 |
| FALSE | High | ASDPQSPP | [R].ASDPQ | 1xPhospho | B2RX66 1x | 5.65E-05 | 0 | 1 | 1 | 3 | B2RX66 | MCG12481 | B2RX66 [4 | 0 | 1577.752 | 0 | 0.93 | 0.000927 | 11.34 | 68.4 | 131.6 | 2.58 | 6.67 | | High | High | High | High | High | High | High | High | High | High | High | 0 | 6.63E-05 | 2.12 |
| FALSE | High | ASAVLCQIV | [K].ASAVLC | 1xCarbami | Q69Z38 1x | 0.065952 | 0.004766 | 1 | 1 | 2 | Q69Z38 | Pseudopod | Q69Z38 [7 | 0 | 3339.742 | 0 | -0.22 | 0.138801 | 6.95 | 105.4 | 94.6 | 16.58 | 11.26 | | High | High | High | High | High | High | High | High | High | High | High | 0.002455 | 0.02657 | 2.66 |
| FALSE | High | ASAVAQSP | [K].ASAVA | 1xPhospho | Q3TUF7 1x | 3.64E-06 | 0 | 1 | 4 | 2 | Q3TUF7 | YEATS dom | Q3TUF7 [3 | 0 | 3049.435 | 0 | -0.39 | 0.132153 | 11.39 | 111.5 | 88.5 | 16.74 | 16.64 | | High | High | High | High | High | High | High | High | High | High | High | 0 | 0.000155 | 4.84 |
| FALSE | High | ASATSLSGS | [KR].ASATS | 2xPhospho | Q9DBC3 2x | 4.37E-07 | 0 | 2 | 4 | 1 | Q9DBC3; C | Cap-specifi | Q9DBC3 [4 | 0 | 2144.94 | 0 | 0.05 | 0.664073 | 3.39 | 101.1 | 98.9 | 7.51 | 7.76 | Shared | High | High | High | High | High | High | High | High | High | High | High | 0 | 4.02E-08 | 3.6 |
| FALSE | High | ARDEEEAFI | [K].ARDEEE | 1xPhospho | Q80Y44 1x | 0.000568 | 0 | 1 | 3 | 3 | Q80Y44 | Probable A | Q80Y44 [7 | 1 | 3806.663 | 0 | -0.64 | 0.004344 | 7.96 | 122 | 78 | 9.16 | 10.92 | | High | High | High | High | High | High | High | High | High | High | High | 4.94E-05 | 0.000658 | 4.07 |
| FALSE | High | ARDSPPPSS | [K].ARDSPI | 1xPhospho | Q920I9 1x | 4.79E-07 | 0 | 1 | 2 | 2 | Q920I9 | WD repeat | Q920I9 [93 | 1 | 2332.243 | 0 | 1.48 | 0.000304 | 14.03 | 52.8 | 147.2 | 6.12 | 3.89 | | High | High | High | High | High | High | High | High | High | High | High | 0 | 4.46E-08 | 3.97 |
| FALSE | High | ARGSPEDG | [K].ARGSPI | 1xPhospho | Q3UQS8 1 | 0.001039 | 0 | 1 | 3 | 1 | Q3UQS8 | RNA-bindir | Q3UQS8 [1 | 1 | 2362.144 | 0 | 3.42 | 0.000439 | 31.23 | 17.1 | 182.9 | 19.68 | 10.32 | | High | High | High | High | High | High | High | High | High | High | High | 0 | 0.000248 | 2.41 |
| FALSE | High | ARPAAQSS | [K].ARPAA | 1xPhospho | Q80U49 1x | 0.003721 | 0.00021 | 1 | 3 | 3 | Q80U49 | Centrosom | Q80U49 [1 | 0 | 2728.46 | 0 | 1.09 | 0.000757 | 8.17 | 63.8 | 136.2 | 5.34 | 7.41 | | High | High | High | High | High | High | High | High | High | High | High | 9.42E-05 | 0.001038 | 3.24 |
| FALSE | High | ARPGSGST | [K].ARPGS | 1xPhospho | Q3U3Q3 1 | 0.000286 | 0 | 1 | 2 | 1 | Q3U3Q3 | Putative ur | Q3U3Q3 [2 | 0 | 1566.841 | 0 | -0.56 | 0.006137 | 7.81 | 122.9 | 77.1 | 7.67 | 9.28 | | High | High | High | High | High | High | High | High | High | High | High | 0 | 5.83E-05 | 1.63 |
| FALSE | High | ARPSDTGS | [K].ARPSD | 1xPhospho | B7ZNX2 1x | 1.06E-05 | 0 | 1 | 3 | 3 | B7ZNX2 | Anaphase | B7ZNX2 [6 | 0 | 2949.289 | 0 | 0.2 | 0.162233 | 19.95 | 93.2 | 106.8 | 11.25 | 9.51 | | High | High | High | High | High | High | High | High | High | High | High | 0 | 2.19E-06 | 3.85 |
| FALSE | High | ARPSPGHP | [R].ARPSP | 1xPhospho | Q80TE0 1x | 4.1E-06 | 0 | 1 | 5 | 1 | Q80TE0 | RNA polym | Q80TE0 [7 | 0 | 2345.087 | 0 | -1.96 | 0.000547 | 2.04 | 159 | 41 | 5.36 | 18.04 | | High | High | High | High | High | High | High | High | High | High | High | 0 | 4.95E-07 | 2.86 |
| FALSE | High | ARPSPSVIK | [K].ARPSPS | 1xPhospho | Q3TEA8 1x | 0.004143 | 0.00036 | 1 | 5 | 1 | Q3TEA8 | Heterochr | Q3TEA8 [5 | 0 | 1492.865 | 0 | 1.65 | 0.000181 | 2.8 | 49.4 | 150.6 | 5.34 | 3.73 | | High | High | High | High | High | High | High | High | High | High | High | 0.000161 | 0.001174 | 1.59 |
| FALSE | High | ARPSQLPE | [R].ARPSQ | 1xPhospho | E0CZ72 1x | 0.003055 | 0.00016 | 1 | 5 | 1 | E0CZ72 | Kinesin-like | E0CZ72 [11 | 0 | 3505.769 | 0 | -0.86 | 0.012792 | 4.44 | 124.6 | 75.4 | 15.72 | 6.98 | | High | High | High | High | High | High | High | High | High | High | High | 7.19E-05 | 0.000837 | 3.48 |
| FALSE | High | ARPSQLPE | [R].ARPSQ | 1xDeamida | E0CZ72 1x | 9.94E-07 | 0 | 1 | 5 | 7 | E0CZ72 | Kinesin-like | E0CZ72 [11 | 0 | 3506.753 | 0 | -0.39 | 0.013686 | 6.94 | 113 | 87 | 7.45 | 8.33 | | High | High | High | High | High | High | High | High | High | High | High | 0 | 2.63E-07 | 3.55 |
| FALSE | High | ARPTSAGG | [R].ARPTSA | 1xPhospho | Q3TBZ6 1x | 8.03E-05 | 0 | 1 | 2 | 2 | Q3TBZ6 | Putative ur | Q3TBZ6 [1 | 0 | 2311.294 | 0 | -0.15 | 0.237371 | 4.58 | 102.7 | 97.3 | 3.79 | 9.66 | | High | High | High | High | High | High | High | High | High | High | High | 0 | 1.41E-05 | 2.47 |
| FALSE | High | ARSNENLD | [R].ARSNE | 1xPhospho | Q91Z49 1x | 0.026652 | 0.001676 | 1 | 2 | 2 | Q91Z49 | UAP56-inte | Q91Z49 [2 | 1 | 1584.815 | 0 | 0.18 | 0.044873 | 6.32 | 91.8 | 108.2 | 6.16 | 6.11 | | High | High | High | High | High | High | High | High | High | High | High | 0.000903 | 0.009463 | 2.5 |
| FALSE | High | ARSPAEEK | [K].ARSPAE | 1xPhospho | C7G3P2 1x | 0.019791 | 0.001438 | 1 | 3 | 1 | C7G3P2 | MKIAA053 | C7G3P2 [2 | 1 | 1425.75 | 0 | -0.11 | 0.142091 | 7.9 | 104.6 | 95.4 | 8.19 | 2.63 | | High | High | High | High | High | High | High | High | High | High | High | 0.000773 | 0.006719 | 2.12 |
| FALSE | High | ARSPQELD | [K].ARSPQ | 1xCarbami | Q921Q7 1x | 0.085162 | 0.006376 | 1 | 1 | 1 | Q921Q7 | Ras and Ra | Q921Q7 [1 | 1 | 3742.87 | 0 | 1.8 | 0.000718 | 7.02 | 47.3 | 152.7 | 14.83 | 7.73 | | High | High | High | High | High | High | High | High | High | High | High | 0.003291 | 0.03585 | 2.63 |
| FALSE | High | ARSPQELD | [K].ARSPQ | 1xCarbami | Q921Q7 1x | 0.000331 | 0 | 1 | 1 | 2 | Q921Q7 | Ras and Ra | Q921Q7 [1 | 1 | 3743.854 | 0 | 2.72 | 0.000784 | 19.65 | 27.2 | 172.8 | 47.52 | 10.04 | | High | High | High | High | High | High | High | High | High | High | High | 0 | 6.91E-05 | 5.22 |
| FALSE | High | ARSPYLPAE | [K].ARSPYL | 1xPhospho | Q3UIL6-2 1 | 1.08E-06 | 0 | 1 | 8 | 1 | Q3UIL6-2 | Isoform 2 | Q3UIL6-2 [ | 1 | 2679.338 | 0 | -0.85 | 0.008623 | 3.47 | 127.1 | 72.9 | 15.17 | 5 | | High | High | High | High | High | High | High | High | High | High | High | 0 | 1.1E-07 | 3.6 |
| FALSE | High | ARTPLTAA | [R].ARTPLT | 1xPhospho | Q924A2 1x | 0.004256 | 0.00036 | 1 | 1 | 2 | Q924A2 | Protein cap | Q924A2 [4 | 1 | 1722.967 | 0 | 0.33 | 0.026858 | 4.65 | 88 | 112 | 6.99 | 7.09 | | High | High | High | High | High | High | High | High | High | High | High | 0.000997 | 0.01056 | 2.06 |
| FALSE | High | ARTPVSELT | [K].ARTPVS | 1xPhospho | Q6A037 1x | 0.002313 | 0.000109 | 1 | 2 | 2 | Q6A037 | NEDD4-bin | Q6A037 [2 | 1 | 1639.918 | 0 | -0.93 | 0.001445 | 4.95 | 130.2 | 69.8 | 5.11 | 8.6 | | High | High | High | High | High | High | High | High | High | High | High | 0.000418 | 0.003607 | 2.44 |
| FALSE | High | ASAGPSASI | [R].ASAGP | 1xPhospho | E9QJT6 1x | 1.43E-06 | 0 | 1 | 4 | 3 | E9QJT6 | Non-canon | E9QJT6 [9 | 0 | 3999.97 | 0 | -0.6 | 0.008768 | 5.1 | 124.8 | 75.2 | 12.9 | 8.39 | | High | High | High | High | High | High | High | High | High | High | High | 0 | 1.64E-07 | 3.62 |
| FALSE | High | ASAGPSASI | [R].ASAGP | 2xPhospho | E9QJT6 2x | 0.000168 | 0 | 1 | 4 | 2 | E9QJT6 | Non-canon | E9QJT6 [9 | 0 | 4079.936 | 0 | 0.53 | 0.006738 | 14.57 | 79.6 | 120.4 | 12.71 | 9.39 | | High | High | High | High | High | High | High | High | High | High | High | 0 | 3.93E-05 | 4.32 |
| FALSE | High | ASAGTPSLS | [R].ASAGT | 1xPhospho | H9LAN6 1x | 3.74E-09 | 0 | 1 | 7 | 2 | H9LAN6 | Histone-lys | H9LAN6 [1 | 0 | 1898.004 | 0 | -0.51 | 0.000834 | 4.83 | 117.3 | 82.7 | 4.14 | 7.74 | | High | High | High | High | High | High | High | High | High | High | High | 0 | 1.9E-10 | 4.07 |
| FALSE | High | ASAISIR | [K].ASAISIR | 1xPhospho | F8WI88 1x | 0.016021 | 0.001124 | 1 | 5 | 2 | F8WI88 | PEST prote | F8WI88 [9 | 0 | 1026.555 | 0 | 0.24 | 0.467999 | 18.66 | 91.8 | 108.2 | 9.82 | 20.29 | | High | High | High | High | High | High | High | High | High | High | High | 0.000585 | 0.005292 | 1.7 |
| FALSE | High | ASAPATPLS | [R].ASAPA | 1xPhospho | P14733 1x | 0.000343 | 0 | 1 | 2 | 8 | P14733 | Lamin-B1 | P14733 [16 | 0 | 1477.761 | 0 | -0.84 | 0.000546 | 2.2 | 127.4 | 72.6 | 4.82 | 5.07 | | High | High | High | High | High | High | High | High | High | High | High | 2.58E-05 | 0.00042 | 2.42 |
| FALSE | High | ASAPATPLS | [R].ASAPA | 2xPhospho | P14733 2x | 0.000325 | 0 | 1 | 2 | 4 | P14733 | Lamin-B1 | P14733 [16 | 0 | 1557.728 | 0 | -0.42 | 0.005799 | 1.98 | 114 | 86 | 2.95 | 9.64 | | High | High | High | High | High | High | High | High | High | High | High | 0.000275 | 0.001674 | 2.24 |
| FALSE | High | ASAPATPLS | [R].ASAPA | 3xPhospho | P14733 3x | 0.004451 | 0.000409 | 1 | 2 | 2 | P14733 | Lamin-B1 | P14733 [16 | 0 | 1637.694 | 0 | -0.79 | 0.000731 | 0.91 | 127 | 73 | 9.42 | 7.82 | | High | High | High | High | High | High | High | High | High | High | High | 0.000183 | 0.00127 | 3.1 |
| FALSE | High | ASAPRPPP | [R].ASAPRI | 1xPhospho | Q6PAJ1 1x | 0.000884 | 0 | 1 | 1 | 2 | Q6PAJ1 | Breakpoint | Q6PAJ1 [9 | 0 | 3847.854 | 0 | 0.77 | 0.000726 | 6.76 | 75.1 | 124.9 | 3.76 | 4.54 | | High | High | High | High | High | High | High | High | High | High | High | 0 | 0.000207 | 2.91 |
| FALSE | High | ASAQASLA | [K].ASAQA | 1xPhospho | Q9R0L6-2 | 0.003001 | 0.00016 | 1 | 3 | 2 | Q9R0L6-2 | Isoform 2 | Q9R0L6-2 | 0 | 1471.792 | 0 | -0.29 | 0.034689 | 13.8 | 106.6 | 93.4 | 9.99 | 15.89 | | High | High | High | High | High | High | High | High | High | High | High | 7.19E-05 | 0.000819 | 2.33 |
| FALSE | High | ASASPGAG | [K].ASASP | 1xPhospho | Q8BGZ3 1x | 0.01186 | 0.000889 | 1 | 1 | 1 | Q8BGZ3 | DDB1- and | Q8BGZ3 [1 | 0 | 2946.394 | 0 | -0.27 | 0.056758 | 12.93 | 110.7 | 89.3 | 9.36 | 9.43 | | High | High | High | High | High | High | High | High | High | High | High | 0.000449 | 0.003793 | 3.47 |
| FALSE | High | ASATSLSGS | [KR].ASATS | 1xPhospho | Q9DBC3 1x | 0.002077 | 5.66E-05 | 2 | 4 | 3 | Q9DBC3; C | Cap-specifi | Q9DBC3 [4 | 0 | 2064.974 | 0 | -0.11 | 0.298336 | 13.87 | 103.8 | 96.2 | 8.49 | 7.5 | Shared | High | High | High | High | High | High | High | High | High | High | High | 0.000555 | 0.004862 | 2.73 |
| FALSE | High | ASETVSEAS | [R].ASETVS | 1xPhospho | P48377 1x | 2.36E-05 | 0 | 1 | 4 | 2 | P48377 | MHC class | P48377 [1 | 0 | 3682.776 | 0 | 0.01 | 0.587192 | 21.09 | 99.7 | 100.3 | 11.36 | 9.1 | | High | High | High | High | High | High | High | High | High | High | High | 0 | 3.53E-06 | 4.48 |
| FALSE | High | ASPPPQGP | [R].ASPPP | 1xPhospho | A2AES4 1x | 3.09E-08 | 0 | 1 | 6 | 2 | A2AES4 | OTU doma | A2AES4 [6 | 0 | 2055.074 | 0 | -0.45 | 0.004896 | 4.1 | 115.5 | 84.5 | 2.7 | 9.74 | | High | High | High | High | High | High | High | High | High | High | High | 0 | 2.04E-09 | 4.23 |
| FALSE | High | ASPPSGLW | [K].ASPPSG | 1xPhospho | Q9QY81 1x | 0.001189 | 0 | 1 | 3 | 2 | Q9QY81 | Nuclear po | Q9QY81 [1 | 0 | 1836.852 | 0 | -0.26 | 0.328605 | 20.27 | 108.4 | 91.6 | 16.69 | 19.35 | | High | High | High | High | High | High | High | High | High | High | High | 0 | 0.000291 | 3.36 |
| FALSE | High | ASPPTESDL | [K].ASPPTE | 1xCarbami | Q3UQS8 1 | 0.000227 | 0 | 1 | 3 | 1 | Q3UQS8 | RNA-bindir | Q3UQS8 [1 | 0 | 1955.873 | 0 | 2.26 | 0.000275 | 8.16 | 33 | 167 | 19.34 | 10.56 | | High | High | High | High | High | High | High | High | High | High | High | 0 | 4.48E-05 | 2.71 |
| FALSE | High | ATGAALTP | [R].ATGAA | 1xPhospho | Q9QY06-2 | 0.000124 | 0 | 2 | 10 | 4 | Q9QY06-2; | Isoform 2 | Q9QY06-2 | 0 | 1525.746 | 0 | 0.93 | 0.00053 | 2.24 | 68.3 | 131.7 | 7.97 | 3.04 | Shared | High | High | High | High | High | High | High | High | High | High | High | 0 | 6.04E-05 | 2.23 |
| FALSE | High | ATGAALTP | [R].ATGAA | 1xPhospho | Q9QY06-2 | 0.079766 | 0.00613 | 2 | 10 | 1 | Q9QY06-2; | Isoform 2 | Q9QY06-2 | 1 | 1681.847 | 0 | 0.81 | 0.012754 | 23.85 | 75.6 | 124.4 | 16 | 11.51 | Shared | High | High | High | High | High | High | High | High | High | High | High | 0.003148 | 0.03307 | 1.54 |
| FALSE | High | ATGAATPK | [K].ATGAA | 1xPhospho | Q149Z9 1x | 0.002313 | 0.000109 | 1 | 4 | 2 | Q149Z9 | Histone clu | Q149Z9 [1 | 0 | 1254.686 | 0 | -0.25 | 0.246891 | 7.28 | 106.9 | 93.1 | 5.11 | 13.96 | | High | High | High | High | High | High | High | High | High | High | High | 4.94E-05 | 0.000614 | 1.4 |
| FALSE | High | ATGDGSSP | [R].ATGDG | 1xPhospho | O88466 1x | 3.4E-07 | 0 | 2 | 4 | 4 | O88466; R | Zinc finger | O88466 [1 | 0 | 1824.858 | 0 | -1.35 | 0.000743 | 2.19 | 146.1 | 53.9 | 7.32 | 6.92 | Shared | High | High | High | High | High | High | High | High | High | High | High | 0 | 3.01E-08 | 3.8 |
| FALSE | High | ATGDGSSP | [R].ATGDG | 2xPhospho | O88466 2x | 2.29E-05 | 0 | 2 | 4 | 3 | O88466; R | Zinc finger | O88466 [1 | 0 | 1904.824 | 0 | -1.38 | 0.000181 | 0.76 | 144.2 | 55.8 | 1.63 | 8.05 | Shared | High | High | High | High | High | High | High | High | High | High | High | 0 | 3.44E-06 | 4.32 |
| FALSE | High | ATGGLCLL | [K].ATGGL | 1xCarbami | A0A0R4IZZ | 1.16E-06 | 0 | 1 | 5 | 13 | A0A0R4IZZ | Formin bin | A0A0R4IZZ | 0 | 3026.359 | 0 | -0.48 | 0.002468 | 4.07 | 116.6 | 83.4 | 1.83 | 7.81 | | High | High | High | High | High | High | High | High | High | High | High | 0 | 1.49E-07 | 4.43 |
| FALSE | High | ATGNDIGS | [R].ATGND | 1xCarbami | B1AQJ2 1x | 5.29E-07 | 0 | 1 | 2 | 4 | B1AQJ2 | Ubiquitin c | B1AQJ2 [6 | 0 | 2807.303 | 0 | -0.49 | 0.015183 | 5.75 | 114.9 | 85.1 | 6.46 | 13.63 | | High | High | High | High | High | High | High | High | High | High | High | 0 | 4.97E-08 | 4.2 |
| FALSE | High | ATGSGSDE | [R].ATGSG | 1xCarbami | E9Q309 1x | 3.26E-05 | 0 | 1 | 1 | 4 | E9Q309 | Centrosom | E9Q309 [2 | 0 | 2307.046 | 0 | 0.14 | 0.143097 | 3.71 | 94.5 | 105.5 | 6.46 | 13.23 | | High | High | High | High | High | High | High | High | High | High | High | 0 | 5.12E-06 | 3.73 |
| FALSE | High | ATGSGSDE | [R].ATGSG | 1xCarbami | E9Q309 2x | 0.001233 | 0 | 1 | 1 | 1 | E9Q309 | Centrosom | E9Q309 [2 | 0 | 2387.013 | 0 | 0.25 | 0.13738 | 14.69 | 91.3 | 108.7 | 5.93 | 13.78 | | High | High | High | High | High | High | High | High | High | High | High | 0 | 0.000301 | 2.81 |
| FALSE | High | ATLDDLLD | [R].ATLDD | 1xPhospho | J3QMP9 1x | 0.007471 | 0.00062 | 1 | 3 | 2 | J3QMP9 | Centrosom | J3QMP9 [4 | 0 | 1526.792 | 0 | -0.66 | 0.009187 | 7.21 | 124.7 | 75.3 | 16.36 | 7.84 | | High | High | High | High | High | High | High | High | High | High | High | 0.000275 | 0.002263 | 2.6 |
| FALSE | High | ATLEQPETI | [R].ATLEQ | 1xPhospho | O54988 1x | 1.51E-07 | 0 | 1 | 2 | 5 | O54988 | STE20-like | O54988 [6 | 0 | 3170.446 | 0 | 0.6 | 0.001818 | 9.76 | 79.5 | 120.5 | 6.07 | 6.25 | | High | High | High | High | High | High | High | High | High | High | High | 0 | 1.22E-08 | 4.94 |
| FALSE | High | ATLEQPETI | [R].ATLEQ | 2xPhospho | O54988 2x | 0.000164 | 0 | 1 | 2 | 5 | O54988 | STE20-like | O54988 [6 | 0 | 3250.413 | 0 | 0.4 | 0.022037 | 11.2 | 87.4 | 112.6 | 7.88 | 11.87 | | High | High | High | High | High | High | High | High | High | High | High | 0 | 0.000236 | 4.26 |
| FALSE | High | ATLLNVPDI | [R].ATLLN | 1xPhospho | P39447 1x | 4.51E-05 | 0 | 1 | 3 | 2 | P39447 | Tight junct | P39447 [2 | 0 | 2519.234 | 0 | 0.21 | 0.079048 | 14.68 | 92.5 | 107.5 | 10.55 | 6.1 | | High | High | High | High | High | High | High | High | High | High | High | 0 | 7.37E-06 | 3.55 |
| FALSE | High | ATFSSMSG | [K].ATFSSM | 1xPhospho | Q6ZQ32 1x | 0.001898 | 5.66E-05 | 2 | 7 | 2 | Q6ZQ32; F | MKIAA085 | Q6ZQ32 [9 | 1 | 2725.234 | 0 | 2.09 | 0.014011 | 21.09 | 39.2 | 160.8 | 14.98 | 32.58 | Shared | High | High | High | High | High | High | High | High | High | High | High | 0.005651 | 0.06205 | 2.89 |
| FALSE | High | ATLPSPDK | [K].ATLPSP | 1xPhospho | A0A1S6GW | 0.074685 | 0.005727 | 1 | 2 | 2 | A0A1S6GW | Uncharacte | A0A1S6GW | 0 | 1366.738 | 0 | -0.43 | 0.003174 | 7.48 | 117.6 | 82.4 | 4.04 | 7.64 | | High | High | High | High | High | High | High | High | High | High | High | 0.004411 | 0.04756 | 1.89 |
| FALSE | High | ATISDEEIE | [K].ATISDE | 1xPhospho | Q8BJF9 1x | 0.001783 | 5.66E-05 | 1 | 1 | 2 | Q8BJF9 | Charged m | Q8BJF9 [1 | 0 | 1471.688 | 0 | 1.32 | 0.002355 | 2.02 | 58 | 142 | 7.88 | 15.6 | | High | High | High | High | High | High | High | High | High | High | High | 2.58E-05 | 0.000458 | 1.6 |
| FALSE | High | ATISDSAD | [K].ATISDS | 1xCarbami | A5D6P2 1x | 0.000651 | 0 | 1 | 18 | 2 | A5D6P2 | Par-3 (Part | A5D6P2 [7 | 0 | 2773.295 | 0 | 0.35 | 0.246983 | 32.26 | 91.1 | 108.9 | 17.16 | 18.17 | | High | High | High | High | High | High | High | High | High | High | High | 0 | 0.000148 | 4.39 |
| FALSE | High | ATLSDSEIE | [R].ATLSDS | 1xPhospho | A2AGQ4 1 | 5.97E-05 | 0 | 1 | 25 | 7 | A2AGQ4 | MAP kinas | A2AGQ4 [1 | 0 | 2580.285 | 0 | 0.38 | 0.015366 | 5.89 | 89.2 | 110.8 | 5.61 | 9.49 | | High | High | High | High | High | High | High | High | High | High | High | 0.000622 | 0.005558 | 2.43 |
| FALSE | High | ATNLAGET | [K].ATNLA | 1xCarbami | E9Q3Y4 2x | 0.000181 | 0 | 1 | 7 | 2 | E9Q3Y4 | Lipopolysa | E9Q3Y4 [1 | 0 | 3499.523 | 0 | 0.52 | 0.026379 | 8.06 | 86.7 | 113.3 | 12.81 | 6.22 | | High | High | High | High | High | High | High | High | High | High | High | 0.001126 | 0.01163 | 3.32 |
| FALSE | High | ATPGLTLPL | [K].ATPGL | 1xPhospho | Q91Z58 1x | 0.006478 | 0.00062 | 1 | 1 | 3 | Q91Z58 | Uncharacte | Q91Z58 [6 | 0 | 2844.609 | 0 | 2.67 | 0.000223 | 8.51 | 28.1 | 171.9 | 11.05 | 6.36 | | High | High | High | High | High | High | High | High | High | High | High | 0.000275 | 0.001931 | 2.29 |
| FALSE | High | ATPPPSPLL | [K].ATPPPS | 1xPhospho | D3YZC8 1x | 0.000372 | 0 | 1 | 3 | 2 | D3YZC8 | Bromodom | D3YZC8 [2 | 0 | 2001.144 | 0 | -0.41 | 0.034604 | 9.31 | 115.3 | 84.7 | 12.25 | 9.65 | | High | High | High | High | High | High | High | High | High | High | High | 0 | 7.84E-05 | 2.6 |
| FALSE | High | ATPRPDSN | [K].ATPRPD | 1xPhospho | Q3TAB2 1x | 5.98E-06 | 0 | 2 | 7 | 2 | Q3TAB2; H | Putative ur | Q3TAB2 [4 | 0 | 2309.19 | 0 | 0.05 | 0.962931 | 5.02 | 100.5 | 99.5 | 8.53 | 3.29 | Shared | High | High | High | High | High | High | High | High | High | High | High | 0 | 7.55E-07 | 3.02 |
| FALSE | High | ATPRPTPVI | [K].ATPRPT | 1xPhospho | H3BK88 1x | 7.28E-05 | 0 | 2 | 8 | 1 | H3BK88; H | Treacle pro | H3BK88 [8 | 0 | 2317.268 | 0 | 0.19 | 0.078582 | 12.18 | 93.4 | 106.6 | 9.79 | 4.95 | Shared | High | High | High | High | High | High | High | High | High | High | High | 0 | 1.26E-05 | 1.49 |
| FALSE | High | ATPSPCSSE | [R].ATPSPC | 1xCarbami | A0A0R4J01 | 0.000553 | 0 | 1 | 5 | 3 | A0A0R4J01 | Teashirt ho | A0A0R4J01 | 0 | 2156.035 | 0 | -0.04 | 0.399274 | 10.43 | 108.2 | 91.8 | 14.38 | 13.14 | | High | High | High | High | High | High | High | High | High | High | High | 0 | 0.000122 | 2.03 |
| FALSE | High | ATPSTSPGI | [R].ATPSTS | 1xPhospho | Q9WUF3 1 | 0.000183 | 0 | 1 | 1 | 6 | Q9WUF3 | CASP8-ass | Q9WUF3 [ | 0 | 1496.813 | 0 | -0.16 | 0.006654 | 3.24 | 104.5 | 95.5 | 2.35 | 4.19 | | High | High | High | High | High | High | High | High | High | High | High | 0.001789 | 0.01768 | 2.68 |
| FALSE | High | ATPSVPTPI | [R].ATPSVI | 1xPhospho | Q9JLM4 1x | 0.003495 | 0.00021 | 1 | 4 | 1 | Q9JLM4 | Zinc finger | Q9JLM4 [8 | 0 | 2086.094 | 0 | -0.48 | 0.003746 | 1.66 | 117.7 | 82.3 | 7.54 | 9.15 | | High | High | High | High | High | High | High | High | High | High | High | 9.42E-05 | 0.000966 | 3.16 |
| FALSE | High | ATPTESPEK | [K].ATPTES | 1xPhospho | Q8C3W1 1 | 0.029076 | 0.001857 | 1 | 1 | 1 | Q8C3W1 | Uncharacte | Q8C3W1 [ | 0 | 2340.236 | 0 | -0.02 | 0.414929 | 9.13 | 103.2 | 96.8 | 4.21 | 7.06 | | High | High | High | High | High | High | High | High | High | High | High | 0.000997 | 0.01036 | 2.09 |
| FALSE | High | ATPTTAPG | [R].ATPTTA | 1xPhospho | F8VQE9 1x | 0.000305 | 0 | 1 | 4 | 4 | F8VQE9 | Arf-GAP wi | F8VQE9 [4 | 0 | 1465.725 | 0 | 0.35 | 0.013136 | 6.92 | 87.9 | 112.1 | 8.51 | 5.93 | | High | High | High | High | High | High | High | High | High | High | High | 0.000275 | 0.002212 | 2.15 |
| FALSE | High | ATLSDEDD | [K].ATLSDE | 1xPhospho | P48964 1x | 6.15E-06 | 0 | 1 | 3 | 2 | P48964 | M-phase ir | P48964 [2 | 0 | 2752.268 | 0 | -0.8 | 0.084157 | 12.36 | 130.6 | 69.4 | 22.29 | 27.68 | | High | High | High | High | High | High | High | High | High | High | High | 0 | 7.82E-07 | 4.35 |
| FALSE | High | ATPVQLPSI | [R].ATPVQ | 1xCarbami | Q5XJV5 1x | 0.001572 | 5.66E-05 | 1 | 6 | 1 | Q5XJV5 | Nuclear re | Q5XJV5 [1 | 0 | 3323.747 | 0 | 0.98 | 0.020162 | 30.59 | 68.4 | 131.6 | 15.43 | 14.3 | | High | High | High | High | High | High | High | High | High | High | High | 2.58E-05 | 0.000395 | 4.45 |
| FALSE | High | ATFSSMSG | [K].ATFSSM | 1xPhospho | Q6ZQ32 1x | 4.06E-06 | 0 | 2 | 7 | 1 | Q6ZQ32; F | MKIAA085 | Q6ZQ32 [9 | 1 | 2709.239 | 0 | 1.96 | 0.003176 | 15.68 | 39.9 | 160.1 | 7.72 | 17.31 | Shared | High | High | High | High | High | High | High | High | High | High | High | 0 | 4.89E-07 | 3.28 |
| FALSE | High | ATENDIYNI | [R].ATENDI | 1xPhospho | P70333 1x | 5.82E-06 | 0 | 1 | 1 | 1 | P70333 | Heterogen | P70333 [3 | 0 | 2354.073 | 0 | 1.03 | 0.00187 | 5.47 | 65.6 | 134.4 | 5.01 | 11.84 | | High | High | High | High | High | High | High | High | High | High | High | 0 | 7.34E-07 | 4.43 |
| FALSE | High | ASWDQASI | [K].ASWD | 1xDeamida | Q3TH99 1x | 0.01218 | 0.000889 | 1 | 2 | 2 | Q3TH99 | Putative ur | Q3TH99 [2 | 0 | 3460.572 | 0 | -0.65 | 0.054499 | 11.88 | 122.3 | 77.7 | 15.23 | 15.29 | | High | High | High | High | High | High | High | High | High | High | High | 0.001741 | 0.01745 | 4.46 |
| FALSE | High | ASWEEEK | [K].ASWEE | 1xPhospho | A2A6M5-4 | 0.036416 | 0.002419 | 1 | 6 | 1 | A2A6M5-4 | Isoform 4 | A2A6M5-4 | 0 | 1416.681 | 0 | -1.35 | 0.003506 | 1.39 | 139.3 | 60.7 | 11.3 | 20.2 | | High | High | High | High | High | High | High | High | High | High | High | 0.001279 | 0.01338 | 1.99 |
| FALSE | High | ASWGASR | [GR].ASW | 1xPhospho | A0A0R4J0S | 0.021231 | 0.001507 | 1 | 2 | 2 | A0A0R4J0S | Cdc42 effe | A0A0R4J0S | 0 | 1043.487 | 0 | 1.1 | 0.001089 | 10.01 | 62 | 138 | 4.83 | 10.1 | | High | High | High | High | High | High | High | High | High | High | High | 0.000801 | 0.007273 | 1.27 |
| FALSE | High | ASWGLQD | [R].ASWGL | 1xPhospho | Q9WU62 1 | 4.63E-08 | 0 | 1 | 2 | 7 | Q9WU62 | Inner cent | Q9WU62 [ | 0 | 2413.066 | 0 | -0.54 | 0.001177 | 1.25 | 119 | 81 | 3.76 | 5.75 | | High | High | High | High | High | High | High | High | High | High | High | 0 | 3.23E-09 | 5.33 |
| FALSE | High | ASWGLQD | [R].ASWGL | 2xPhospho | Q9WU62 2 | 0.000151 | 0 | 1 | 2 | 1 | Q9WU62 | Inner cent | Q9WU62 [ | 0 | 2493.032 | 0 | -0.97 | 0.000502 | 1.7 | 132.8 | 67.2 | 3.63 | 6.86 | | High | High | High | High | High | High | High | High | High | High | High | 0 | 2.86E-05 | 3.03 |
| FALSE | High | ASYANVFG | [K].ASYAN | 1xPhospho | A3KGV1-2 | 1.98E-07 | 0 | 1 | 4 | 2 | A3KGV1-2 | Isoform 2 | A3KGV1-2 | 0 | 2776.307 | 0 | -1 | 0.006234 | 7.56 | 133.2 | 66.8 | 10.02 | 18.17 | | High | High | High | High | High | High | High | High | High | High | High | 0 | 1.64E-08 | 4.2 |
| FALSE | High | ASYANVFG | [K].ASYAN | 1xDeamida | A3KGV1-2 | 0.003402 | 0.00021 | 1 | 4 | 2 | A3KGV1-2 | Isoform 2 | A3KGV1-2 | 0 | 2777.291 | 0 | -1.47 | 0.001006 | 3.18 | 149.2 | 50.8 | 5.97 | 16.73 | | High | High | High | High | High | High | High | High | High | High | High | 0.000327 | 0.002426 | 3 |
| FALSE | High | ASYQETDG | [R].ASYQE | 1xCarbami | A0A140T8I | 3.16E-07 | 0 | 1 | 2 | 2 | A0A140T8I | Protein Jad | A0A140T8I | 0 | 3359.419 | 0 | -0.59 | 0.006271 | 7.99 | 117.9 | 82.1 | 5.19 | 10.88 | | High | High | High | High | High | High | High | High | High | High | High | 0 | 6.79E-08 | 3.17 |

| | | | | | | | | | | | | | | | | | | | | | | | | | | | | | | | | | | | | | | |
|---|---|---|---|---|---|---|---|---|---|---|---|---|---|---|---|---|---|---|---|---|---|---|---|---|---|---|---|---|---|---|---|---|---|---|---|---|---|---|
| FALSE | High | ASYQETDG | [R].ASYQET | 1xCarbami | A0A140T8I | 4.69E-06 | 0 | 1 | 2 | 5 | A0A140T8I | Protein Jac | A0A140T8I | 0 | 3439.385 | 0 | -0.6 | 0.009509 | 7.6 | 122.1 | 77.9 | 8 | 12.44 | | High | High | High | High | High | High | High | High | High | High | High | 0 | 0.000202 | 4.13 |
| FALSE | High | ASYSGTSPS | [R].ASYSGT | 1xPhospho | Q8K1N2-2 | 0.001629 | 5.66E-05 | 1 | 3 | 6 | Q8K1N2-2 | Isoform 2 c | Q8K1N2-2 | 0 | 2291.018 | 0 | 1.79 | 0.000912 | 36.9 | 43.4 | 156.6 | 30.78 | 6.28 | | High | High | High | High | High | High | High | High | High | High | High | 2.58E-05 | 0.000413 | 2.09 |
| FALSE | High | ASYTPSPAF | [R].ASYTPS | 2xPhospho | Q3UFM6 2 | 0.057686 | 0.004227 | 1 | 3 | 1 | Q3UFM6 | Expressed | Q3UFM6 [ | 0 | 1338.569 | 0 | -0.01 | 0.495097 | 17.91 | 100.9 | 99.1 | 5.01 | 9.81 | | High | High | High | High | High | High | High | High | High | High | High | 0.002159 | 0.02275 | 0.96 |
| FALSE | High | ATAAAAAA | [R].ATAAA | 1xPhospho | Q9WU62 1 | 1.95E-08 | 0 | 1 | 2 | 1 | Q9WU62 | Inner centr | Q9WU62 [ | 0 | 3158.635 | 0 | -0.9 | 0.000642 | 1.59 | 129.4 | 70.6 | 6.55 | 12.62 | | High | High | High | High | High | High | High | High | High | High | High | 0 | 1.22E-09 | 4.7 |
| FALSE | High | ATAPQTQH | [R].ATAPQ | 1xPhospho | F6ZFU0 1x | 2.18E-07 | 0 | 2 | 12 | 2 | F6ZFU0; A( | Elongation | F6ZFU0 [3 | 0 | 1732.84 | 0 | -0.29 | 0.009616 | 1.44 | 114 | 86 | 7.94 | 15.78 | Shared | High | High | High | High | High | High | High | High | High | High | High | 0 | 3.68E-06 | 2.58 |
| FALSE | High | ATENDIYNI | [RK].ATENI | 1xPhospho | Q8C2Q7 1 | 3.74E-06 | 0 | 2 | 6 | 1 | Q8C2Q7; C | Heterogen | Q8C2Q7 [3 | 0 | 2306.106 | 0 | -0.21 | 0.007664 | 3.95 | 109.1 | 90.9 | 6.46 | 6.99 | Shared | High | High | High | High | High | High | High | High | High | High | High | 0 | 4.48E-07 | 3.24 |
| FALSE | High | ATAPQTQH | [R].ATAPQ | 1xPhospho | F6ZFU0 1x | 0.000884 | 0 | 2 | 12 | 40 | F6ZFU0; A( | Elongation | F6ZFU0 [3 | 0 | 1748.835 | 0 | -0.33 | 0.000979 | 3.75 | 112.3 | 87.7 | 3.73 | 8.41 | Shared | High | High | High | High | High | High | High | High | High | High | High | 0 | 0.000309 | 2.92 |
| FALSE | High | ATAQPIIEIL | [R].ATAQP | 1xPhospho | Q8BVK9 1 | 0.017192 | 0.001204 | 1 | 1 | 1 | Q8BVK9 | Sp110 nucl | Q8BVK9 [1 | 0 | 2371.211 | 0 | -0.08 | 0.673241 | 20.14 | 98.2 | 101.8 | 18.89 | 23.73 | | High | High | High | High | High | High | High | High | High | High | High | 0.000638 | 0.005747 | 2.77 |
| FALSE | High | ATASSSAQ | [K].ATASSS | 1xPhospho | Q3TDL0 1x | 5.63E-07 | 0 | 1 | 2 | 4 | Q3TDL0 | Putative ur | Q3TDL0 [2 | 0 | 2145.968 | 0 | 0.85 | 0.015108 | 3.04 | 70.8 | 129.2 | 7.79 | 17.33 | | High | High | High | High | High | High | High | High | High | High | High | 0 | 5.33E-08 | 4.94 |
| FALSE | High | ATDSHSDS | [R].ATDSH | 1xCarbami | Q9CQV4 3 | 3.5E-05 | 0 | 1 | 2 | 7 | Q9CQV4 | Reticuloph | Q9CQV4 [2 | 0 | 3320.294 | 0 | 0.7 | 0.001169 | 9.57 | 76.3 | 123.7 | 7.51 | 9.96 | | High | High | High | High | High | High | High | High | High | High | High | 0 | 5.54E-06 | 4.18 |
| FALSE | High | ATEAAPQP | [K].ATEAA | 1xPhospho | Q8VDP4 1 | 7.28E-05 | 0 | 1 | 1 | 1 | Q8VDP4 | Cell cycle a | Q8VDP4 [4 | 0 | 3790.733 | 0 | -0.45 | 0.054984 | 4.7 | 114.9 | 85.1 | 8.95 | 21.14 | | High | High | High | High | High | High | High | High | High | High | High | 0 | 1.25E-05 | 3.68 |
| FALSE | High | ATEDDSAS | [R].ATEDD | 2xCarbami | P43136 1x | 0.000418 | 0 | 1 | 2 | 2 | P43136 | Nuclear re | P43136 [2 | 0 | 4269.884 | 0 | 1.18 | 0.002167 | 12.64 | 65.9 | 134.1 | 12.44 | 8.41 | | High | High | High | High | High | High | High | High | High | High | High | 0 | 8.99E-05 | 3.91 |
| FALSE | High | ATEDDSAS | [R].ATEDD | 2xCarbami | P43136 1x | 1.36E-08 | 0 | 1 | 2 | 1 | P43136 | Nuclear re | P43136 [2 | 0 | 4270.868 | 0 | 0.57 | 0.046664 | 18.49 | 80.7 | 119.3 | 9.12 | 17.46 | | High | High | High | High | High | High | High | High | High | High | High | 0 | 8.1E-10 | 6.91 |
| FALSE | High | ATEDDSAS | [R].ATEDD | 2xCarbami | P43136 2x | 0.000311 | 0 | 1 | 2 | 1 | P43136 | Nuclear re | P43136 [2 | 0 | 4349.85 | 0 | 0.73 | 0.003273 | 12.74 | 75.9 | 124.1 | 9.89 | 14.35 | | High | High | High | High | High | High | High | High | High | High | High | 0 | 6.4E-05 | 5.92 |
| FALSE | High | ATEDDSPP | [K].ATEDD | 1xPhospho | O88554 1x | 6.07E-05 | 0 | 1 | 1 | 3 | O88554 | Poly [ADP- | O88554 [2 | 0 | 1554.745 | 0 | -1.78 | 0.000364 | 1.09 | 155 | 45 | 6.23 | 9.25 | | High | High | High | High | High | High | High | High | High | High | High | 0 | 1.03E-05 | 2.24 |
| FALSE | High | ATEEEDSDI | [R].ATEEED | 1xPhospho | Q9CT10 1x | 0.000248 | 0 | 1 | 1 | 16 | Q9CT10 | Ran-bindin | Q9CT10 [4 | 0 | 3601.502 | 0 | 0.45 | 0.000261 | 1.5 | 83.4 | 116.6 | 5.35 | 4.1 | | High | High | High | High | High | High | High | High | High | High | High | 0 | 0.000332 | 4.81 |
| FALSE | High | ATEEPSGT | [R].ATEEPS | 1xPhospho | O54724 1x | 2.99E-06 | 0 | 1 | 3 | 3 | O54724 | Caveolae-a | O54724 [3 | 0 | 2072.02 | 0 | -0.69 | 0.016374 | 8.68 | 124.6 | 75.4 | 20.08 | 12.13 | | High | High | High | High | High | High | High | High | High | High | High | 0 | 3.46E-07 | 3.15 |
| FALSE | High | ATEETGPV | [K].ATEETG | 1xCarbami | V9GX74 1x | 8.42E-09 | 0 | 1 | 9 | 2 | V9GX74 | Zinc finger | V9GX74 [1 | 0 | 3140.534 | 0 | -0.33 | 0.039792 | 2.89 | 111.3 | 88.7 | 10.38 | 9.26 | | High | High | High | High | High | High | High | High | High | High | High | 0 | 4.72E-10 | 5.29 |
| FALSE | High | ATENDIAN | [R].ATEND | 1xPhospho | D3Z3N4 1x | 0.000415 | 0 | 1 | 2 | 1 | D3Z3N4 | Heterogen | D3Z3N4 [2 | 0 | 2228.095 | 0 | -1.01 | 0.000609 | 2.12 | 133.7 | 66.3 | 6.62 | 5.92 | | High | High | High | High | High | High | High | High | High | High | High | 0 | 8.91E-05 | 3.64 |
| FALSE | High | ATENDIYNI | [R].ATEND | 1xPhospho | P70333 1x | 0.0008 | 0 | 1 | 1 | 1 | P70333 | Heterogen | P70333 [3 | 0 | 2338.078 | 0 | 0.89 | 0.000177 | 3.35 | 68.1 | 131.9 | 7.99 | 6.2 | | High | High | High | High | High | High | High | High | High | High | High | 0 | 0.000187 | 2.94 |
| FALSE | High | ATAPQTQH | [R].ATAPQ | 1xPhospho | F6ZFU0 1x | 6.59E-05 | 0 | 2 | 12 | 1 | F6ZFU0; A( | Elongation | F6ZFU0 [3 | 1 | 2757.416 | 0 | -0.8 | 0.793145 | 17.07 | 115.3 | 84.7 | 40.11 | 116.95 | Shared | High | High | High | High | High | High | High | High | High | High | High | 0 | 1.13E-05 | 2.59 |
| FALSE | High | ATQNPSTS | [R].ATQNP | 1xPhospho | A2AEG2 1x | 1.19E-07 | 0 | 1 | 2 | 2 | A2AEG2 | Oral-facial- | A2AEG2 [6 | 0 | 1864.957 | 0 | 0.58 | 0.000373 | 1.65 | 80.2 | 119.8 | 3.78 | 4.99 | | High | High | High | High | High | High | High | High | High | High | High | 0 | 9.35E-09 | 2.83 |
| FALSE | High | ATSACAPIS | [R].ATSAC | 1xCarbami | Q8BXK8 1x | 0.030367 | 0.001857 | 1 | 4 | 3 | Q8BXK8 | Arf-GAP wi | Q8BXK8 [4 | 0 | 1727.88 | 0 | -0.65 | 0.010694 | 1.31 | 118.5 | 81.5 | 13.08 | 6.24 | | High | High | High | High | High | High | High | High | High | High | High | 0.001011 | 0.01092 | 1.85 |
| FALSE | High | ATSAETVES | [K].ATSAET | 1xPhospho | Q76LS9 1x | 0.004451 | 0.000409 | 1 | 2 | 1 | Q76LS9 | Ubiquitin c | Q76LS9 [1 | 0 | 2624.249 | 0 | 0.47 | 0.061923 | 28.14 | 85 | 115 | 16.02 | 15.31 | | High | High | High | High | High | High | High | High | High | High | High | 0.000183 | 0.001273 | 1.76 |
| FALSE | High | ATSWYSPY | [R].ATSWY | 1xPhospho | Q8CCS6 1x | 0.002376 | 0.000109 | 1 | 3 | 1 | Q8CCS6 | Polyadenyl | Q8CCS6 [2 | 0 | 1283.555 | 0 | -0.65 | 0.003241 | 2.9 | 122.6 | 77.4 | 9.36 | 4.65 | | High | High | High | High | High | High | High | High | High | High | High | 4.94E-05 | 0.000632 | 2.12 |
| FALSE | High | ATTLSNAV | [R].ATTLSN | 1xPhospho | Q570Z8 1x | 5.5E-05 | 0 | 1 | 9 | 4 | Q570Z8 | MKIAA411 | Q570Z8 [3 | 0 | 2460.336 | 0 | 0.86 | 0.014659 | 3.56 | 70.4 | 129.6 | 7.78 | 22.35 | | High | High | High | High | High | High | High | High | High | High | High | 0 | 9.19E-06 | 3.4 |
| FALSE | High | ATTNNASS | [K].ATTNN | 1xCarbami | Q62315 1x | 2.39E-07 | 0 | 1 | 4 | 5 | Q62315 | Protein Jur | Q62315 [1 | 0 | 1890.821 | 0 | -1.59 | 0.002347 | 4.11 | 153.1 | 46.9 | 15.39 | 26.62 | | High | High | High | High | High | High | High | High | High | High | High | 0 | 2.03E-08 | 3.78 |
| FALSE | High | ATTPASTAI | [K].ATTPAS | 1xPhospho | B8X349 1x | 0.017041 | 0.001204 | 2 | 12 | 1 | B8X349; F6 | JNK-interac | B8X349 [2 | 0 | 2666.333 | 0 | 1.09 | 0.000847 | 13.3 | 60.8 | 139.2 | 18.78 | 6.46 | Shared | High | High | High | High | High | High | High | High | High | High | High | 0.000638 | 0.005722 | 1.86 |
| FALSE | High | ATTPPNQG | [R].ATTPPN | 1xPhospho | S4R221 1x | 0.000496 | 0 | 1 | 8 | 3 | S4R221 | Rho GTPas | S4R221 [2 | 0 | 2934.513 | 0 | -1.26 | 0.000332 | 2.17 | 145.6 | 54.4 | 5 | 12.49 | | High | High | High | High | High | High | High | High | High | High | High | 0 | 0.000108 | 3.05 |
| FALSE | High | ATTPPNQG | [R].ATTPPN | 2xPhospho | S4R221 2x | 0.000699 | 0 | 1 | 8 | 2 | S4R221 | Rho GTPas | S4R221 [2 | 0 | 3014.479 | 0 | -0.54 | 0.009487 | 7.85 | 117.5 | 82.5 | 9.65 | 4.98 | | High | High | High | High | High | High | High | High | High | High | High | 0 | 0.00016 | 3.51 |
| FALSE | High | ATTPQLTD | [R].ATTPQ | 1xPhospho | Q6NS82 1x | 7.15E-05 | 0 | 1 | 4 | 6 | Q6NS82 | Reticuloph | Q6NS82 [3 | 0 | 3392.594 | 0 | -0.19 | 0.038156 | 7.75 | 107.6 | 92.4 | 13.17 | 10.36 | | High | High | High | High | High | High | High | High | High | High | High | 0 | 0.000147 | 4.51 |
| FALSE | High | ATTPQLTD | [R].ATTPQ | 2xPhospho | Q6NS82 2x | 0.002355 | 0.000109 | 1 | 4 | 2 | Q6NS82 | Reticuloph | Q6NS82 [3 | 0 | 3472.561 | 0 | -0.44 | 0.339009 | 22.99 | 115.3 | 84.7 | 19.22 | 24.52 | | High | High | High | High | High | High | High | High | High | High | High | 0.000275 | 0.001641 | 3.84 |
| FALSE | High | ATTTPAK | [K].ATTTPA | 1xPhospho | P09405 1x | 0.008313 | 0.000736 | 1 | 8 | 5 | P09405 | Nucleolin ( | P09405 [5 | 0 | 1227.675 | 0 | -1.3 | 0.000158 | 0.92 | 143.9 | 56.1 | 7.52 | 13.53 | | High | High | High | High | High | High | High | High | High | High | High | 0.000327 | 0.002559 | 2.31 |
| FALSE | High | ATVASSSQ | [R].ATVASS | 1xPhospho | Q80UG5 1 | 0.002463 | 0.000109 | 1 | 3 | 3 | Q80UG5 | Septin-9 O | Q80UG5 [5 | 0 | 1416.75 | 0 | 1.37 | 0.001195 | 14.8 | 57.8 | 142.2 | 11.91 | 10.34 | | High | High | High | High | High | High | High | High | High | High | High | 0.00134 | 0.01407 | 1.5 |
| FALSE | High | ATVSDSAA | [K].ATVSD | 1xPhospho | Q9D4H9-2 | 0.00824 | 0.000736 | 1 | 4 | 2 | Q9D4H9-2 | Isoform 2 c | Q9D4H9-2 | 0 | 4741.323 | 0 | -0.03 | 0.935722 | 11.77 | 97.1 | 102.9 | 14.4 | 18.11 | | High | High | High | High | High | High | High | High | High | High | High | 0.001876 | 0.01877 | 3.37 |
| FALSE | High | ATVSDSAA | [K].ATVSD | 2xPhospho | Q9D4H9-2 | 4.76E-05 | 0 | 1 | 4 | 1 | Q9D4H9-2 | Isoform 2 c | Q9D4H9-2 | 0 | 4821.29 | 0 | 0.75 | 0.005654 | 10.58 | 71.3 | 128.7 | 14.01 | 9.97 | | High | High | High | High | High | High | High | High | High | High | High | 0 | 7.8E-06 | 4.7 |
| FALSE | High | ATVTAATT | [R].ATVTA | 1xPhospho | Q8CIG3 1x | 0.0016 | 5.66E-05 | 1 | 2 | 1 | Q8CIG3 | Lysine-spe | Q8CIG3 [2 | 0 | 2649.273 | 0 | 0.28 | 0.056771 | 17.9 | 92.2 | 107.8 | 10.84 | 9.06 | | High | High | High | High | High | High | High | High | High | High | High | 2.58E-05 | 0.000404 | 3.17 |
| FALSE | High | ATSVDYSSI | [R].ATSVD | 1xPhospho | A0A0N4SW | 0.0008 | 0 | 1 | 1 | 1 | A0A0N4SW | C2 domain | A0A0N4SW | 0 | 1627.72 | 0 | 1.36 | 0.000913 | 5.57 | 59.2 | 140.8 | 10.35 | 6.39 | | High | High | High | High | High | High | High | High | High | High | High | 0 | 0.000187 | 2.33 |
| FALSE | High | ATVTPSPVI | [K].ATVTPS | 1xPhospho | Q80XU3 1 | 0.000379 | 0 | 1 | 2 | 8 | Q80XU3 | Nuclear ub | Q80XU3 [1 | 0 | 1437.812 | 0 | -0.39 | 0.005325 | 3.93 | 118.3 | 81.7 | 7.92 | 12.58 | | High | High | High | High | High | High | High | High | High | High | High | 0 | 8.03E-05 | 2.53 |
| FALSE | High | ATVVNGTG | [K].ATVVN | 1xDeamida | M1VMF6 1 | 0.000984 | 0 | 1 | 10 | 1 | M1VMF6 | Metastasis | M1VMF6 [ | 0 | 1936.933 | 0 | 0.24 | 0.135636 | 12.12 | 92.3 | 107.7 | 8.94 | 5.09 | | High | High | High | High | High | High | High | High | High | High | High | 0 | 0.000235 | 2.73 |
| FALSE | High | ATVVSSPK | [K].ATVVSS | 1xPhospho | E9QML5 1 | 0.001147 | 0 | 1 | 5 | 6 | E9QML5 | Zinc finger | E9QML5 [1 | 0 | 1326.743 | 0 | -0.31 | 0.00062 | 2.41 | 110.4 | 89.6 | 2.2 | 3.6 | | High | High | High | High | High | High | High | High | High | High | High | 0 | 0.000278 | 2.04 |
| FALSE | High | ATWGDGG | [K].ATWGD | 1xPhospho | Q9D3L3 1x | 1.06E-05 | 0 | 1 | 3 | 7 | Q9D3L3 | Synaptoso | Q9D3L3 [1 | 0 | 2229.059 | 0 | -1.13 | 0.000176 | 1.01 | 136.9 | 63.1 | 3.53 | 2.64 | | High | High | High | High | High | High | High | High | High | High | High | 0 | 1.44E-06 | 2.57 |
| FALSE | High | ATYGGLSSI | [K].ATYGG | 1xPhospho | Q3UJW4 1 | 3.6E-05 | 0 | 1 | 2 | 7 | Q3UJW4 | Putative ur | Q3UJW4 [7 | 0 | 2852.364 | 0 | -0.35 | 0.012684 | 7.42 | 104.8 | 95.2 | 12.34 | 17.52 | | High | High | High | High | High | High | High | High | High | High | High | 0 | 5.72E-06 | 4 |
| FALSE | High | ATYGGLSSI | [K].ATYGG | 2xPhospho | Q3UJW4 2 | 0.001137 | 0 | 1 | 2 | 5 | Q3UJW4 | Putative ur | Q3UJW4 [7 | 0 | 2932.331 | 0 | -0.17 | 0.049506 | 4.02 | 104.9 | 95.1 | 6.34 | 7.52 | | High | High | High | High | High | High | High | High | High | High | High | 0 | 0.000327 | 3.51 |
| FALSE | High | AVAGVMIT | [K].AVAGV | 1xPhospho | Q8CAA7 1 | 0.003892 | 0.000261 | 1 | 2 | 2 | Q8CAA7 | Glucose 1,( | Q8CAA7 [1 | 0 | 1651.819 | 0 | -0.06 | 0.576021 | 13.36 | 101.8 | 98.2 | 10.52 | 3.29 | | High | High | High | High | High | High | High | High | High | High | High | 0.00014 | 0.001097 | 2.38 |
| FALSE | High | AVALDSDN | [K].AVALD | 1xPhospho | Q8CH77 1 | 2.13E-05 | 0 | 1 | 4 | 1 | Q8CH77 | Neuron na | Q8CH77 [7 | 0 | 1783.961 | 0 | -0.16 | 0.055827 | 5.16 | 106.1 | 93.9 | 4.85 | 7.95 | | High | High | High | High | High | High | High | High | High | High | High | 0 | 3.17E-06 | 2.84 |
| FALSE | High | AVASVSEE | [R].AVASV | 1xPhospho | B2RR83 1x | 9.08E-07 | 0 | 1 | 1 | 3 | B2RR83 | Probable A | B2RR83 [4 | 0 | 3296.586 | 0 | 1.57 | 0.024694 | 92.7 | 50.4 | 149.6 | 52.67 | 29.55 | | High | High | High | High | High | High | High | High | High | High | High | 0 | 1.12E-07 | 5.07 |
| FALSE | High | AVATPAK | [K].AVATP | 1xPhospho | P09405 1x | 0.008022 | 0.000736 | 1 | 7 | 3 | P09405 | Nucleolin ( | P09405 [8 | 0 | 1195.685 | 0 | -1.1 | 0.000191 | 2.91 | 138.6 | 61.4 | 6.16 | 12.65 | | High | High | High | High | High | High | High | High | High | High | High | 0.000327 | 0.002446 | 2.6 |
| FALSE | High | AVCSTYLQ | [K].AVCST | 1xCarbami | O88904 1x | 0.001222 | 0 | 2 | 10 | 5 | O88904; Q | Homeodor | O88904 [3 | 0 | 1493.702 | 0 | 0.56 | 0.00116 | 4.01 | 80.8 | 119.2 | 5.5 | 5.59 | Shared | High | High | High | High | High | High | High | High | High | High | High | 0 | 0.000327 | 3.12 |
| FALSE | High | AVDITTPK | [K].AVDITT | 1xPhospho | Q99JF8 1x | 0.033404 | 0.002195 | 1 | 4 | 3 | Q99JF8 | PC4 and SF | Q99JF8 [1 | 0 | 1382.77 | 0 | -1.29 | 9.77E-05 | 1.47 | 142.1 | 57.9 | 4.45 | 9.25 | | High | High | High | High | High | High | High | High | High | High | High | 0.003291 | 0.03569 | 2.11 |
| FALSE | High | AVDNPPRG | [K].AVDNP | 1xPhospho | Q9D706 1x | 0.018774 | 0.001362 | 1 | 1 | 2 | Q9D706 | RNA polym | Q9D706 [4 | 1 | 1675.893 | 0 | -0.45 | 0.000885 | 0.83 | 119.5 | 80.5 | 7.74 | 8.4 | | High | High | High | High | High | High | High | High | High | High | High | 0.001324 | 0.01394 | 1.36 |
| FALSE | High | AVDSSSEEI | [K].AVDSS | 1xPhospho | Q5XG69 1 | 1.03E-07 | 0 | 1 | 1 | 13 | Q5XG69 | Soluble lan | Q5XG69 [6 | 0 | 2268.085 | 0 | -1.45 | 0.00126 | 3.75 | 145 | 55 | 7.39 | 15.78 | | High | High | High | High | High | High | High | High | High | High | High | 0 | 7.94E-09 | 4.49 |
| FALSE | High | ATVVNGTG | [K].ATVVN | 1xPhospho | M1VMF6 1 | 1.64E-05 | 0 | 1 | 10 | 2 | M1VMF6 | Metastasis | M1VMF6 [ | 0 | 1935.949 | 0 | -0.08 | 0.193445 | 3.36 | 102.8 | 97.2 | 5.38 | 6 | | High | High | High | High | High | High | High | High | High | High | High | 0 | 3.83E-06 | 2.89 |
| FALSE | High | ATSVALPG | [K].ATSVAL | 1xPhospho | Q8BWW9 | 0.000191 | 0 | 1 | 5 | 4 | Q8BWW9 | Serine/thr | Q8BWW9 | 0 | 1866.895 | 0 | 0.18 | 0.037575 | 9.62 | 94.8 | 105.2 | 8.43 | 5.46 | | High | High | High | High | High | High | High | High | High | High | High | 0 | 3.7E-05 | 2.7 |
| FALSE | High | ATSTSPNS | [R].ATSTSP | 1xDeamida | Q3TLH4 1x | 6.41E-05 | 0 | 1 | 7 | 6 | Q3TLH4 | Protein PR | Q3TLH4 [2 | 0 | 1733.836 | 0 | 0.5 | 0.002227 | 2.9 | 82.3 | 117.7 | 9.4 | 2.79 | | High | High | High | High | High | High | High | High | High | High | High | 0.000756 | 0.006588 | 3 |
| FALSE | High | ATSTSPNS | [R].ATSTSP | 1xPhospho | Q3TLH4 1x | 9.31E-06 | 0 | 1 | 7 | 4 | Q3TLH4 | Protein PR | Q3TLH4 [2 | 0 | 1732.852 | 0 | 0.7 | 0.000944 | 3.05 | 75.6 | 124.4 | 5.09 | 6.89 | | High | High | High | High | High | High | High | High | High | High | High | 0 | 3.83E-05 | 3.51 |
| FALSE | High | ATSAGSSP | [R].ATSAG | 1xCarbami | S4R1M9 1 | 0.000198 | 0 | 1 | 1 | 2 | S4R1M9 | Oxysterol-l | S4R1M9 [2 | 0 | 1717.778 | 0 | -0.32 | 0.046427 | 7.64 | 113.1 | 86.9 | 7.36 | 12.9 | | High | High | High | High | High | High | High | High | High | High | High | 0 | 3.86E-05 | 2.7 |
| FALSE | High | ATSAHLEEI | [K].ATSAHI | 1xPhospho | P81069 1x | 9.39E-06 | 0 | 1 | 1 | 1 | P81069 | GA-binding | P81069 [2 | 0 | 3313.484 | 0 | 0.47 | 0.026231 | 11.97 | 80.4 | 119.6 | 17.04 | 20.49 | | High | High | High | High | High | High | High | High | High | High | High | 0 | 1.26E-06 | 3.97 |
| FALSE | High | ATSAHLEEI | [K].ATSAHI | 1xPhospho | P81069 1x | 3.6E-08 | 0 | 1 | 1 | 1 | P81069 | GA-binding | P81069 [2 | 0 | 3329.479 | 0 | 0.13 | 0.580974 | 4.08 | 97.6 | 102.4 | 14.77 | 29.58 | | High | High | High | High | High | High | High | High | High | High | High | 0 | 2.42E-09 | 4.86 |
| FALSE | High | ATSDSGGD | [K].ATSDS | 1xPhospho | E9Q6J5-3 1 | 0.000319 | 0 | 1 | 3 | 3 | E9Q6J5-3 | Isoform Ne | E9Q6J5-3 [ | 0 | 1929.957 | 0 | -0.18 | 0.609267 | 11.86 | 105 | 95 | 18.52 | 7.14 | | High | High | High | High | High | High | High | High | High | High | High | 2.58E-05 | 0.000391 | 3.12 |
| FALSE | High | ATSFSSVR | [R].ATSFSS | 1xPhospho | G5E8W8 1 | 0.031168 | 0.001935 | 2 | 3 | 1 | G5E8W8; A | Zinc finger | G5E8W8 [6 | 0 | 1163.566 | 0 | -1.97 | 0.000547 | 0.96 | 158.8 | 41.2 | 7.15 | 10.4 | Shared | High | High | High | High | High | High | High | High | High | High | High | 0.001076 | 0.01126 | 1.37 |
| FALSE | High | ATSGGMSI | [R].ATSGG | 1xPhospho | O35280 1x | 1.02E-08 | 0 | 1 | 3 | 10 | O35280 | Serine/thr | O35280 [2 | 0 | 1973.893 | 0 | -1.06 | 0.006684 | 2.21 | 137.4 | 62.6 | 10.9 | 13.43 | | High | High | High | High | High | High | High | High | High | High | High | 0 | 2.09E-05 | 3.79 |
| FALSE | High | ATSGPAAN | [R].ATSGP | 1xPhospho | P58929 1x | 0.058664 | 0.004294 | 1 | 2 | 1 | P58929 | Glucocorti | P58929 [3 | 0 | 4133.179 | 0 | -0.22 | 0.17579 | 14.41 | 111.6 | 88.4 | 16.14 | 11.09 | | High | High | High | High | High | High | High | High | High | High | High | 0.002187 | 0.02313 | 2.56 |
| FALSE | High | ATSGPAAN | [R].ATSGP | 1xPhospho | P58929 1x | 0.018774 | 0.001362 | 1 | 2 | 2 | P58929 | Glucocorti | P58929 [3 | 0 | 4149.174 | 0 | 0.06 | 0.302917 | 11.85 | 96.7 | 103.3 | 17.9 | 16.23 | | High | High | High | High | High | High | High | High | High | High | High | 0.000723 | 0.00634 | 2.62 |
| FALSE | High | ATSGSSLPV | [R].ATSGS | 1xPhospho | Q3UHC1 1 | 5.03E-05 | 0 | 1 | 5 | 4 | Q3UHC1 | Putative ur | Q3UHC1 [3 | 0 | 1567.804 | 0 | -0.14 | 0.072777 | 5 | 105.5 | 94.5 | 2.87 | 6.33 | | High | High | High | High | High | High | High | High | High | High | High | 0 | 1.07E-05 | 2.19 |
| FALSE | High | ATSIPGSPF | [R].ATSIPG | 1xPhospho | Q3TYD6 1x | 0.000112 | 0 | 1 | 1 | 3 | Q3TYD6 | Serine/thr | Q3TYD6 [6 | 0 | 2347.199 | 0 | -0.33 | 0.004997 | 2.39 | 111.4 | 88.6 | 8.59 | 7.18 | | High | High | High | High | High | High | High | High | High | High | High | 0 | 2.04E-05 | 2.41 |
| FALSE | High | ATSNVFAM | [R].ATSNV | 1xPhospho | Q3THE2 1x | 2.76E-07 | 0 | 2 | 7 | 4 | Q3THE2; Q | Myosin re | Q3THE2 [1 | 0 | 2629.277 | 0 | -0.06 | 0.994706 | 24.61 | 101.7 | 98.3 | 14.07 | 13.19 | Shared | High | High | High | High | High | High | High | High | High | High | High | 0 | 3.35E-08 | 4.47 |
| FALSE | High | ATSNVFAM | [R].ATSNV | 1xPhospho | Q3THE2 1x | 5.3E-08 | 0 | 2 | 7 | 49 | Q3THE2; Q | Myosin re | Q3THE2 [1 | 0 | 2645.272 | 0 | -0.03 | 0.806812 | 16.21 | 102.2 | 97.8 | 8.85 | 8.64 | Shared | High | High | High | High | High | High | High | High | High | High | High | 0 | 5.03E-07 | 4.2 |
| FALSE | High | ATSPDAAP | [K].ATSPD | 1xPhospho | Q3U3X7 1 | 0.000165 | 0 | 1 | 4 | 4 | Q3U3X7 | Putative ur | Q3U3X7 [4 | 0 | 1395.729 | 0 | -1.05 | 0.000455 | 1.42 | 136.4 | 63.6 | 8.33 | 10.84 | | High | High | High | High | High | High | High | High | High | High | High | 0 | 4.55E-05 | 2.78 |
| FALSE | High | ATSPESTSR | [R].ATSPES | 1xPhospho | A0A0R4IZZ | 0.107603 | 0.008634 | 1 | 4 | 2 | A0A0R4IZZ | Formin bin | A0A0R4IZZ | 0 | 1244.572 | 0 | -0.63 | 0.000875 | 4.93 | 122.2 | 77.8 | 2.7 | 10.36 | | High | High | High | High | High | High | High | High | High | High | High | 0.008294 | 0.08936 | 1.8 |
| FALSE | High | ATSPLVSLY | [R].ATSPLV | 1xCarbami | A2AES4 1x | 0.118334 | 0.009621 | 1 | 6 | 2 | A2AES4 | OTU doma | A2AES4 [5 | 0 | 1985.997 | 0 | -0.05 | 0.124826 | 4.57 | 106 | 94 | 5.68 | 9.04 | | High | High | High | High | High | High | High | High | High | High | High | 0.006415 | 0.06906 | 2.54 |
| FALSE | High | ATSPPHLD | [K].ATSPP | 1xPhospho | Q60953 1x | 1.91E-07 | 0 | 2 | 6 | 10 | Q60953; Q | Protein PM | Q60953 [5 | 0 | 2602.28 | 0 | -0.14 | 0.086825 | 3.81 | 107.6 | 92.4 | 5.42 | 10.44 | Shared | High | High | High | High | High | High | High | High | High | High | High | 0 | 1.59E-08 | 3.25 |
| FALSE | High | ATSPPHLD | [K].ATSPP | 2xPhospho | Q60953 2x | 0.000375 | 0 | 2 | 6 | 2 | Q60953; Q | Protein PM | Q60953 [5 | 0 | 2682.247 | 0 | 0.31 | 0.049646 | 14.82 | 87 | 113 | 5.34 | 13.99 | Shared | High | High | High | High | High | High | High | High | High | High | High | 0.001516 | 0.01501 | 2.94 |
| FALSE | High | ATSPSSSVS | [R].ATSPSS | 1xPhospho | F8VQD1 1x | 0.002398 | 0.000109 | 1 | 14 | 5 | F8VQD1 | Protein pol | F8VQD1 [8 | 0 | 2890.284 | 0 | -0.22 | 0.523573 | 9.59 | 104.5 | 95.5 | 9.51 | 8.86 | | High | High | High | High | High | High | High | High | High | High | High | 0.00196 | 0.01994 | 2.51 |
| FALSE | High | ATSPSSSVS | [R].ATSPSS | 2xPhospho | F8VQD1 2x | 0.076555 | 0.005826 | 1 | 14 | 2 | F8VQD1 | Protein pol | F8VQD1 [8 | 0 | 2970.251 | 0 | -0.01 | 0.689963 | 7.91 | 97.3 | 102.7 | 8.99 | 10.51 | | High | High | High | High | High | High | High | High | High | High | High | 0.003672 | 0.03943 | 2.56 |
| FALSE | High | ATSPSTLVS | [K].ATSPST | 1xPhospho | P27546 1x | 6.83E-05 | 0 | 1 | 10 | 4 | P27546 | Microtubu | P27546 [7 | 0 | 1756.868 | 0 | 1.04 | 0.000181 | 3.24 | 65.5 | 134.5 | 2.25 | 2.42 | | High | High | High | High | High | High | High | High | High | High | High | 0 | 9.09E-05 | 2.7 |
| FALSE | High | ATSPVANG | [R].ATSPV | 1xPhospho | Q9D1R2 1x | 6.07E-05 | 0 | 1 | 1 | 2 | Q9D1R2 | Protein KT | Q9D1R2 [1 | 0 | 1980.093 | 0 | -0.16 | 0.325932 | 8.68 | 105.5 | 94.5 | 7.84 | 11.11 | | High | High | High | High | High | High | High | High | High | High | High | 0 | 1.03E-05 | 2.59 |
| FALSE | High | ATSPVANG | [R].ATSPV | 1xDeamida | Q9D1R2 1x | 2.07E-07 | 0 | 1 | 1 | 9 | Q9D1R2 | Protein KT | Q9D1R2 [1 | 0 | 1981.077 | 0 | -0.14 | 0.518372 | 10.85 | 104.9 | 95.1 | 3.36 | 9.27 | | High | High | High | High | High | High | High | High | High | High | High | 0 | 1.73E-08 | 3.21 |
| FALSE | High | ATSRPGCS | [K].ATSRP | 1xCarbami | Q3UJP8 1x | 0.003028 | 0.00016 | 1 | 2 | 3 | Q3UJP8 | Trifunction | Q3UJP8 [4 | 0 | 2706.373 | 0 | 0.36 | 0.036377 | 13.11 | 86.4 | 113.6 | 8.13 | 10.69 | | High | High | High | High | High | High | High | High | High | High | High | 0.000538 | 0.004476 | 2.35 |
| FALSE | High | ATSSATTT | [K].ATSSAT | 1xPhospho | G5E8E3 1x | 1.29E-05 | 0 | 1 | 3 | 1 | G5E8E3 | RIKEN cDN | G5E8E3 [3 | 0 | 1775.919 | 0 | -0.36 | 0.000487 | 1.54 | 110.1 | 89.9 | 5.55 | 7.28 | | High | High | High | High | High | High | High | High | High | High | High | 0 | 1.8E-06 | 1.96 |
| FALSE | High | ATSSSSSSS | [R].ATSSSS | 1xPhospho | A0A0R4J0C | 5.32E-06 | 0 | 1 | 3 | 2 | A0A0R4J0C | Lamin-B2 ( | A0A0R4J0C | 0 | 2037.911 | 0 | 0.83 | 0.019942 | 26.21 | 73.9 | 126.1 | 6.54 | 19.88 | | High | High | High | High | High | High | High | High | High | High | High | 0 | 6.64E-07 | 2.63 |
| FALSE | High | ATSSSSSSS | [R].ATSSSS | 1xPhospho | A0A0R4J0C | 1.91E-08 | 0 | 1 | 3 | 18 | A0A0R4J0C | Lamin-B2 ( | A0A0R4J0C | 0 | 2053.906 | 0 | 0.29 | 0.014491 | 8.92 | 86.9 | 113.1 | 4.89 | 9.22 | | High | High | High | High | High | High | High | High | High | High | High | 0 | 5.09E-09 | 4.61 |
| FALSE | High | ATSSSSSSS | [R].ATSSSS | 2xPhospho | A0A0R4J0C | 0.000595 | 0 | 1 | 3 | 3 | A0A0R4J0C | Lamin-B2 ( | A0A0R4J0C | 0 | 2133.872 | 0 | 0.03 | 0.810119 | 10.67 | 99.5 | 100.5 | 4.64 | 10.9 | | High | High | High | High | High | High | High | High | High | High | High | 0 | 0.000133 | 3.44 |
| FALSE | High | ATSSTQSLA | [R].ATSSTC | 1xPhospho | E9Q7G0 1x | 9.54E-05 | 0 | 1 | 4 | 7 | E9Q7G0 | Nuclear mi | E9Q7G0 [1 | 0 | 1330.657 | 0 | 0.43 | 0.012943 | 10.81 | 85.2 | 114.8 | 7.75 | 7 | | High | High | High | High | High | High | High | High | High | High | High | 0 | 1.7E-05 | 3.48 |
| FALSE | High | ATSTSDML | [K].ATSTSD | 1xPhospho | Q8BHS3 1x | 0.004256 | 0.00036 | 1 | 1 | 1 | Q8BHS3 | Pre-mRNA- | Q8BHS3 [1 | 0 | 1620.832 | 0 | 0.28 | 0.155811 | 38.99 | 89.1 | 110.9 | 12.34 | 20.72 | | High | High | High | High | High | High | High | High | High | High | High | 0.000161 | 0.001203 | 1.84 |
| FALSE | High | ASVTIQAR | [K].ASVTIQ | 1xPhospho | D3YZP9 1x | 0.002334 | 0.000109 | 1 | 1 | 1 | D3YZP9 | Coiled-coil | D3YZP9 [9 | 0 | 1154.613 | 1 | -0.16 | 0.161792 | 17.23 | 107.4 | 92.6 | 8.82 | 9.83 | | High | High | High | High | High | High | High | High | High | High | High | 4.94E-05 | 0.000615 | 1.54 |
| FALSE | High | ASVSDLSPF | [R].ASVSDI | 1xPhospho | Q569Z6 1x | 0.000908 | 0 | 1 | 4 | 5 | Q569Z6 | Thyroid ho | Q569Z6 [2 | 0 | 1240.614 | 0 | -0.1 | 0.198374 | 4.29 | 102.6 | 97.4 | 4.42 | 6.4 | | High | High | High | High | High | High | High | High | High | High | High | 0 | 0.000215 | 1.28 |
| FALSE | High | ASVQTEGS | [R].ASVQTI | 1xPhospho | Q3ULW8 1 | 0.001087 | 0 | 1 | 9 | 1 | Q3ULW8 | Poly [ADP- | Q3ULW8 [ | 0 | 1444.745 | 0 | 0.29 | 0.005233 | 9.62 | 86.4 | 113.6 | 11.81 | 6.76 | | High | High | High | High | High | High | High | High | High | High | High | 0 | 0.000262 | 2.18 |
| FALSE | High | ASVLQMDS | [R].ASVLQI | 1xPhospho | Q80WR5 1 | 0.00129 | 0 | 1 | 3 | 1 | Q80WR5 | UPF0688 p | Q80WR5 [ | 0 | 2993.347 | 0 | 0.81 | 0.035713 | 11.88 | 73.7 | 126.3 | 12.53 | 25.36 | | High | High | High | High | High | High | High | High | High | High | High | 0 | 0.000319 | 3.21 |
| FALSE | High | ASQTPVPQ | [K].ASQTP | 2xPhospho | E9QMD3 2 | 0.000976 | 0 | 1 | 4 | 1 | E9QMD3 | Zinc finger | E9QMD3 [ | 0 | 2098.002 | 0 | 1.79 | 0.00034 | 5.36 | 44.7 | 155.3 | 13.18 | 9.14 | | High | High | High | High | High | High | High | High | High | High | High | 0 | 0.000233 | 2.39 |
| FALSE | High | ASQTPVPQ | [K].ASQTP | 1xPhospho | E9QMD3 1 | 0.000496 | 0 | 1 | 4 | 1 | E9QMD3 | Zinc finger | E9QMD3 [ | 1 | 2658.411 | 0 | 2.59 | 0.003291 | 40.06 | 25 | 175 | 21.67 | 20.99 | | High | High | High | High | High | High | High | High | High | High | High | 0 | 0.000108 | 2.49 |
| FALSE | High | ASQVEPQS | [R].ASQVEI | 1xPhospho | A0A0A0M( | 7.18E-07 | 0 | 1 | 2 | 2 | A0A0A0M( | Histone-lys | A0A0A0M( | 0 | 1747.894 | 0 | 0.75 | 0.008152 | 5.56 | 69.9 | 130.1 | 7.98 | 16.42 | | High | High | High | High | High | High | High | High | High | High | High | 0 | 2.48E-05 | 2.86 |
| FALSE | High | ASRDSFCAI | [R].ASRDSF | 1xCarbami | E9PVX6 1x | 1.31E-05 | 0 | 1 | 2 | 2 | E9PVX6 | Proliferatic | E9PVX6 [1 | 1 | 2494.096 | 0 | -0.39 | 0.001742 | 2.5 | 115.3 | 84.7 | 5.28 | 4.3 | | High | High | High | High | High | High | High | High | High | High | High | 0 | 1.82E-06 | 3.27 |
| FALSE | High | ASSAAFLPS | [K].ASSAAF | 1xPhospho | Q9Z0E1 1x | 4.15E-08 | 0 | 1 | 2 | 1 | Q9Z0E1 | Meiosis 1 a | Q9Z0E1 [5 | 0 | 2642.146 | 0 | -1.4 | 0.001041 | 1.45 | 147.5 | 52.5 | 10.43 | 15.29 | | High | High | High | High | High | High | High | High | High | High | High | 0 | 2.85E-09 | 2.66 |
| FALSE | High | ASSAAFLPS | [K].ASSAAF | 2xPhospho | Q9Z0E1 2x | 8.95E-05 | 0 | 1 | 2 | 2 | Q9Z0E1 | Meiosis 1 a | Q9Z0E1 [5 | 0 | 2722.112 | 0 | -1.82 | 0.001112 | 1.67 | 157.9 | 42.1 | 11.67 | 27.45 | | High | High | High | High | High | High | High | High | High | High | High | 0 | 1.59E-05 | 1.94 |
| FALSE | High | ASSAAPSG | [K].ASSAAF | 1xPhospho | Q7M753 1 | 2.51E-05 | 0 | 1 | 4 | 2 | Q7M753 | Pantothen | Q7M753 [5 | 0 | 1771.806 | 0 | -0.85 | 0.003428 | 4.3 | 129 | 71 | 2.79 | 19.02 | | High | High | High | High | High | High | High | High | High | High | High | 0 | 3.78E-06 | 2.67 |
| FALSE | High | ASSAASLPS | [R].ASSAAS | 1xPhospho | B1AQY2 1x | 6.56E-07 | 0 | 1 | 4 | 2 | B1AQY2 | Rho GTPas | B1AQY2 [1 | 0 | 2305.175 | 0 | -0.04 | 0.909421 | 4.23 | 101.3 | 98.7 | 2.19 | 5.95 | | High | High | High | High | High | High | High | High | High | High | High | 0 | 1.27E-07 | 3.89 |
| FALSE | High | ASSCETYEY | [R].ASSCET | 1xCarbami | Q3ZB56 1x | 0.000924 | 0 | 1 | 4 | 3 | Q3ZB56 | Growth fac | Q3ZB56 [4 | 0 | 1742.729 | 0 | 2.41 | 0.000275 | 9.07 | 32.4 | 167.6 | 7.42 | 5.7 | | High | High | High | High | High | High | High | High | High | High | High | 0 | 0.000218 | 3.23 |
| FALSE | High | ASSCSLAVI | [R].ASSCSL | 1xCarbami | P58801 1x | 7.34E-05 | 0 | 1 | 1 | 5 | P58801 | Receptor-i | P58801 [4 | 0 | 2246.191 | 0 | -0.24 | 0.009671 | 7.34 | 107.5 | 92.5 | 8.41 | 6.31 | | High | High | High | High | High | High | High | High | High | High | High | 0 | 1.51E-05 | 3.22 |
| FALSE | High | ASSDLCDL | [K].ASSDLC | 2xCarbami | Q9JLM4 1x | 3.49E-07 | 0 | 1 | 4 | 1 | Q9JLM4 | Zinc finger | Q9JLM4 [9 | 0 | 3352.491 | 0 | -0.25 | 0.135546 | 8.6 | 114.3 | 85.7 | 18.86 | 19.59 | | High | High | High | High | High | High | High | High | High | High | High | 0 | 3.11E-08 | 5.24 |
| FALSE | High | ASSDLSIAS | [R].ASSDLS | 1xPhospho | O54988 1x | 4.79E-07 | 0 | 1 | 2 | 1 | O54988 | STE20-like | O54988 [3 | 0 | 1976.947 | 0 | -0.06 | 0.52028 | 13.51 | 105.7 | 94.3 | 8.29 | 11.42 | | High | High | High | High | High | High | High | High | High | High | High | 0 | 4.43E-08 | 4.16 |
| FALSE | High | ASSDLSIAS | [R].ASSDLS | 2xPhospho | O54988 2x | 0.000693 | 0 | 1 | 2 | 2 | O54988 | STE20-like | O54988 [3 | 0 | 2056.913 | 0 | -0.08 | 0.370152 | 10.06 | 100.4 | 99.6 | 7.63 | 8.63 | | High | High | High | High | High | High | High | High | High | High | High | 0 | 0.000157 | 3.27 |
| FALSE | High | ASQTPAQE | [R].ASQTP | 1xPhospho | Q62187 1x | 3.67E-05 | 0 | 1 | 2 | 2 | Q62187 | Transcripti | Q62187 [5 | 0 | 2468.211 | 0 | 0.54 | 0.001644 | 7.27 | 81.4 | 118.6 | 7.2 | 6.61 | | High | High | High | High | High | High | High | High | High | High | High | 0 | 5.8E-06 | 3.29 |
| FALSE | High | ASSEDEAP | [K].ASSEDE | 1xPhospho | P18911 1x | 0.005322 | 0.000599 | 1 | 3 | 2 | P18911 | Retinoic ac | P18911 [4 | 0 | 1770.831 | 0 | -0.16 | 0.092074 | 7.43 | 111 | 89 | 12.1 | 10.43 | | High | High | High | High | High | High | High | High | High | High | High | 0.000267 | 0.001557 | 2.19 |
| FALSE | High | ASSESGEGI | [K].ASSESG | 1xCarbami | Q3UIV3 1x | 1.5E-08 | 0 | 1 | 9 | 5 | Q3UIV3 | Putative ur | Q3UIV3 [4 | 0 | 2739.189 | 0 | -0.77 | 0.015202 | 2.11 | 130.3 | 69.7 | 20.62 | 8.19 | | High | High | High | High | High | High | High | High | High | High | High | 0 | 9.1E-10 | 5.16 |
| FALSE | High | ASSFADMN | [K].ASSFAD | 1xPhospho | Q9CWE0 1 | 7.82E-05 | 0 | 1 | 1 | 2 | Q9CWE0 | Mitochond | Q9CWE0 [ | 0 | 1808.909 | 0 | 0.21 | 0.699717 | 10.75 | 93.4 | 106.6 | 4.61 | 17.35 | | High | High | High | High | High | High | High | High | High | High | High | 0 | 1.36E-05 | 2.62 |
| FALSE | High | ASSFADMN | [K].ASSFAD | 1xPhospho | Q9CWE0 1 | 3.73E-05 | 0 | 1 | 1 | 5 | Q9CWE0 | Mitochond | Q9CWE0 [ | 0 | 1824.904 | 0 | 0.05 | 0.601546 | 11.87 | 97.6 | 102.4 | 2.45 | 11.14 | | High | High | High | High | High | High | High | High | High | High | High | 0 | 5.94E-06 | 3.22 |
| FALSE | High | ASSFGSISIL | [K].ASSFG | 1xPhospho | Q80TM6 1 | 6.77E-05 | 0 | 1 | 8 | 1 | Q80TM6 | R3H domai | Q80TM6 [3 | 0 | 1547.803 | 0 | 0.91 | 0.001686 | 8.13 | 69.9 | 130.1 | 5.17 | 8.05 | | High | High | High | High | High | High | High | High | High | High | High | 0 | 1.16E-05 | 2.51 |
| FALSE | High | ASSFGSISV | [K].ASSFG | 1xPhospho | E9Q9Q2 1x | 0.000372 | 0 | 1 | 6 | 1 | E9Q9Q2 | R3H domai | E9Q9Q2 [3 | 0 | 1533.788 | 0 | 0.64 | 0.004564 | 8.76 | 80.5 | 119.5 | 5.37 | 8.14 | | High | High | High | High | High | High | High | High | High | High | High | 0 | 7.85E-05 | 1.6 |
| FALSE | High | ASSFYDSPF | [R].ASSFYD | 1xPhospho | A2ADB0 1x | 0.051694 | 0.003745 | 1 | 5 | 1 | A2ADB0 | Msx2-inter | A2ADB0 [1 | 0 | 1338.593 | 0 | 0.58 | 0.013862 | 21.87 | 79.2 | 120.8 | 8.88 | 12.92 | | High | High | High | High | High | High | High | High | High | High | High | 0.00196 | 0.02003 | 1.71 |

| | | | | | | | | | | | | | | | | | | | | | | | | | | | | | | | | | | | | | | |
|---|---|---|---|---|---|---|---|---|---|---|---|---|---|---|---|---|---|---|---|---|---|---|---|---|---|---|---|---|---|---|---|---|---|---|---|---|---|---|
| FALSE | High | ASSFYGSAS | [R].ASSFYG | 1xPhospho | Q7TT79 1x | 1.91E-07 | 0 | 1 | 5 | 2 | Q7TT79 | Microceph | Q7TT79 [2 | 0 | 1802.842 | 0 | -0.9 | 0.006249 | 4.14 | 130.3 | 69.7 | 15.31 | 4.38 | | High | High | High | High | High | High | High | High | High | High | High | 0 | 1.58E-08 | 3.82 |
| FALSE | High | ASSGGSQE | [R].ASSGG | 1xPhospho | Q80T69 1x | 0.001848 | 5.66E-05 | 1 | 2 | 1 | Q80T69 | Round spe | Q80T69 [9 | 0 | 1388.682 | 0 | -0.45 | 0.002272 | 4.5 | 114 | 86 | 7.27 | 12.67 | | High | High | High | High | High | High | High | High | High | High | High | 2.58E-05 | 0.000474 | 1.85 |
| FALSE | High | ASSHSSQS | [R].ASSHSS | 1xPhospho | P48678 1x | 0.0008 | 0 | 1 | 3 | 5 | P48678 | Prelamin-A | P48678 [40 | 0 | 1813.828 | 0 | 0.79 | 0.292423 | 119.58 | 62.5 | 137.5 | 61.88 | 75.26 | | High | High | High | High | High | High | High | High | High | High | High | 0.002279 | 0.02375 | 2.06 |
| FALSE | High | ASSHSSQS | [R].ASSHSS | 2xPhospho | P48678 2x | 0.08727 | 0.006537 | 1 | 3 | 1 | P48678 | Prelamin-A | P48678 [40 | 0 | 1893.794 | 0 | 1.03 | 0.006315 | 15.45 | 71.2 | 128.8 | 17.2 | 10.16 | | High | High | High | High | High | High | High | High | High | High | High | 0.003346 | 0.03665 | 1.78 |
| FALSE | High | ASSHSSQS | [R].ASSHSS | 1xPhospho | P48678 1x | 3.99E-10 | 0 | 1 | 3 | 23 | P48678 | Prelamin-A | P48678 [40 | 0 | 2042.991 | 0 | 1.94 | 0.00943 | 41.16 | 41.3 | 158.7 | 15.89 | 22.53 | | High | High | High | High | High | High | High | High | High | High | High | 0 | 1.55E-11 | 3.24 |
| FALSE | High | ASSHSSQS | [R].ASSHSS | 2xPhospho | P48678 2x | 2.23E-05 | 0 | 1 | 3 | 7 | P48678 | Prelamin-A | P48678 [40 | 0 | 2122.957 | 0 | 1.02 | 0.013875 | 17.74 | 65 | 135 | 10.08 | 21.07 | | High | High | High | High | High | High | High | High | High | High | High | 0.000872 | 0.009168 | 3.03 |
| FALSE | High | ASSIAASR | [R].ASSIAA | 1xPhospho | Q8BGB2 1x | 0.037692 | 0.00249 | 1 | 5 | 2 | Q8BGB2 | Tetratricop | Q8BGB2 [6 | 0 | 1071.54 | 0 | 1.52 | 0.000465 | 3.88 | 50.6 | 149.4 | 8.16 | 5.95 | | High | High | High | High | High | High | High | High | High | High | High | 0.001308 | 0.01391 | 1.5 |
| FALSE | High | ASSESGEGI | [K].ASSESG | 1xCarbami | Q3UIV3 1x | 2.24E-06 | 0 | 1 | 9 | 2 | Q3UIV3 | Putative ur | Q3UIV3 [4 | 0 | 2723.194 | 0 | -1.68 | 0.001599 | 2.35 | 152.3 | 47.7 | 10.6 | 10.63 | | High | High | High | High | High | High | High | High | High | High | High | 0 | 2.52E-07 | 4.83 |
| FALSE | High | ASQQSSPI | [K].ASQQS | 1xPhospho | Q8BG15 1x | 0.000189 | 0 | 1 | 6 | 5 | Q8BG15 | CTD small | Q8BG15 [8 | 0 | 1639.8 | 0 | -0.1 | 0.364279 | 11 | 102.4 | 97.6 | 4.51 | 7.9 | | High | High | High | High | High | High | High | High | High | High | High | 0 | 3.66E-05 | 3.47 |
| FALSE | High | ASQGSNSL | [R].ASQGS | 1xPhospho | E9Q9R9 1x | 0.00083 | 0 | 1 | 5 | 3 | E9Q9R9 | Disks large | E9Q9R9 [1 | 0 | 1570.742 | 0 | 0.44 | 0.02471 | 11.52 | 83.5 | 116.5 | 2.11 | 15.62 | | High | High | High | High | High | High | High | High | High | High | High | 0.000275 | 0.001581 | 1.3 |
| FALSE | High | ASQGLISAV | [R].ASQGLI | 2xPhospho | Q6PR54-3 | 8.63E-08 | 0 | 2 | 4 | 5 | Q6PR54-3; | Isoform 3 c | Q6PR54-3 | 0 | 2627.189 | 0 | -0.69 | 0.000457 | 0.51 | 123 | 77 | 5.28 | 4.25 | Shared | High | High | High | High | High | High | High | High | High | High | High | 0 | 0.000158 | 3.76 |
| FALSE | High | ASPQHSPH | [R].ASPQH | 1xPhospho | Q9QY81 1x | 0.000133 | 0 | 1 | 3 | 1 | Q9QY81 | Nuclear po | Q9QY81 [1 | 0 | 2802.356 | 0 | -0.2 | 0.280409 | 32.47 | 110.9 | 89.1 | 27.79 | 26.58 | | High | High | High | High | High | High | High | High | High | High | High | 0 | 2.49E-05 | 2.75 |
| FALSE | High | ASPQHSPH | [R].ASPQH | 1xDeamida | Q9QY81 1x | 1.11E-06 | 0 | 1 | 3 | 2 | Q9QY81 | Nuclear po | Q9QY81 [1 | 0 | 2803.34 | 0 | -0.44 | 0.281636 | 19.39 | 113.9 | 86.1 | 22.82 | 20.89 | | High | High | High | High | High | High | High | High | High | High | High | 0 | 1.14E-07 | 4.84 |
| FALSE | High | ASPQHSPH | [R].ASPQH | 2xPhospho | Q9QY81 2x | 0.000227 | 0 | 1 | 3 | 1 | Q9QY81 | Nuclear po | Q9QY81 [1 | 0 | 2882.323 | 0 | -0.46 | 0.118547 | 9.2 | 115.9 | 84.1 | 19.93 | 14.75 | | High | High | High | High | High | High | High | High | High | High | High | 0 | 4.5E-05 | 3.68 |
| FALSE | High | ASPQVQLF | [K].ASPQV | 1xPhospho | F6WIU1 1x | 0.000563 | 0 | 1 | 2 | 2 | F6WIU1 | Protein KR | F6WIU1 [6 | 0 | 2239.152 | 0 | 0.24 | 0.116492 | 15.4 | 95.4 | 104.6 | 6.89 | 6.74 | | High | High | High | High | High | High | High | High | High | High | High | 0 | 0.000125 | 3.6 |
| FALSE | High | ASPQVQLF | [K].ASPQV | 1xDeamida | F6WIU1 1x | 8.11E-05 | 0 | 1 | 2 | 10 | F6WIU1 | Protein KR | F6WIU1 [6 | 0 | 2240.136 | 0 | 0.14 | 0.174096 | 5.73 | 94.9 | 105.1 | 5.45 | 8.75 | | High | High | High | High | High | High | High | High | High | High | High | 4.94E-05 | 0.000576 | 3.28 |
| FALSE | High | ASPRPWTP | [R].ASPRPV | 1xPhospho | Q8K120 1x | 0.030898 | 0.001896 | 1 | 2 | 2 | Q8K120 | Nuclear fa | Q8K120 [2 | 0 | 2709.266 | 0 | -1.74 | 0.001154 | 0.67 | 150.6 | 49.4 | 13.52 | 12.05 | | High | High | High | High | High | High | High | High | High | High | High | 0.001027 | 0.01109 | 2.19 |
| FALSE | High | ASPSFLK | [K].ASPSFL | 1xPhospho | Q0VBM2 1 | 0.056248 | 0.004124 | 1 | 1 | 1 | Q0VBM2 | Protein FA | Q0VBM2 [ | 0 | 1287.711 | 0 | 2.27 | 0.000222 | 3.17 | 32.9 | 167.1 | 7.57 | 6.15 | | High | High | High | High | High | High | High | High | High | High | High | 0.002117 | 0.02209 | 1.42 |
| FALSE | High | ASPSPPPK | [R].ASPSPP | 2xPhospho | Q52KI8 2x | 0.015465 | 0.001089 | 1 | 7 | 4 | Q52KI8 | Serine/argi | Q52KI8 [64 | 0 | 1398.684 | 0 | -0.37 | 0.001518 | 3.97 | 113.1 | 86.9 | 3.72 | 5.25 | | High | High | High | High | High | High | High | High | High | High | High | 0.000571 | 0.005108 | 1.42 |
| FALSE | High | ASPSPPSHT | [R].ASPSPP | 1xPhospho | Q3UFT4 1x | 0.001077 | 0 | 1 | 7 | 1 | Q3UFT4 | Putative ur | Q3UFT4 [4 | 0 | 1919.888 | 0 | -0.32 | 0.64213 | 9.18 | 110.7 | 89.3 | 12.3 | 21.79 | | High | High | High | High | High | High | High | High | High | High | High | 0 | 0.00026 | 1.56 |
| FALSE | High | ASPSPQPS | [R].ASPSPC | 1xPhospho | F8WJ93 1x | 9.16E-07 | 0 | 1 | 3 | 3 | F8WJ93 | Echinoderr | F8WJ93 [1 | 0 | 2016.011 | 0 | 0.49 | 0.005217 | 11.13 | 82.6 | 117.4 | 3.89 | 8.12 | | High | High | High | High | High | High | High | High | High | High | High | 0 | 9.23E-08 | 3.46 |
| FALSE | High | ASPSPQPS | [R].ASPSPC | 2xPhospho | F8WJ93 2x | 1.11E-05 | 0 | 1 | 3 | 2 | F8WJ93 | Echinoderr | F8WJ93 [1 | 0 | 2095.978 | 0 | -0.47 | 0.006918 | 7.72 | 116.2 | 83.8 | 4.38 | 12.84 | | High | High | High | High | High | High | High | High | High | High | High | 0 | 1.53E-06 | 3.74 |
| FALSE | High | ASPSSDEDI | [R].ASPSSD | 1xPhospho | Q3TJN8 1x | 0.054383 | 0.003951 | 1 | 2 | 1 | Q3TJN8 | Putative ur | Q3TJN8 [2 | 0 | 2149.058 | 0 | -0.15 | 0.100522 | 3.2 | 105.4 | 94.6 | 12.71 | 9.43 | | High | High | High | High | High | High | High | High | High | High | High | 0.002059 | 0.02132 | 1.71 |
| FALSE | High | ASPTPAELC | [R].ASPTPA | 1xPhospho | Q2VPA6 1x | 1.51E-06 | 0 | 1 | 6 | 2 | Q2VPA6 | Helicase PC | Q2VPA6 [2 | 0 | 2622.177 | 0 | 0.44 | 0.037637 | 23.04 | 85 | 115 | 17.36 | 4.47 | | High | High | High | High | High | High | High | High | High | High | High | 0 | 1.61E-07 | 4.18 |
| FALSE | High | ASPVAVGS | [K].ASPVAV | 1xPhospho | Q6PAL7 1x | 3.09E-08 | 0 | 1 | 1 | 1 | Q6PAL7 | AT-hook D | Q6PAL7 [8 | 0 | 2563.276 | 0 | 0.08 | 0.391857 | 9.15 | 95.4 | 104.6 | 5.49 | 12.72 | | High | High | High | High | High | High | High | High | High | High | High | 0 | 2.04E-09 | 4.16 |
| FALSE | High | ASPVPGPG | [R].ASPVPC | 1xPhospho | Q8BHL3 1x | 0.007019 | 0.00062 | 1 | 1 | 2 | Q8BHL3 | TBC1 doma | Q8BHL3 [1 | 0 | 1445.735 | 0 | 0.24 | 0.1571 | 8.8 | 91.9 | 108.1 | 7.2 | 8.29 | | High | High | High | High | High | High | High | High | High | High | High | 0.000275 | 0.002113 | 2.85 |
| FALSE | High | ASPVPSDPI | [K].ASPVPS | 1xPhospho | A0A0A0M( | 0.000454 | 0 | 1 | 3 | 1 | A0A0A0M( | Histone-lys | A0A0A0M( | 0 | 3045.518 | 0 | 0.33 | 0.007419 | 9.11 | 88.1 | 111.9 | 5.04 | 9.98 | | High | High | High | High | High | High | High | High | High | High | High | 0 | 9.8E-05 | 2.85 |
| FALSE | High | ASPVTAPSS | [R].ASPVTA | 1xPhospho | A2A7S8-2 | 6.59E-05 | 0 | 1 | 4 | 1 | A2A7S8-2 | Isoform 2 c | A2A7S8-2 [ | 0 | 1829.947 | 0 | -0.06 | 0.5692 | 33.8 | 101.7 | 98.3 | 31.75 | 18.98 | | High | High | High | High | High | High | High | High | High | High | High | 0 | 1.12E-05 | 3.81 |
| FALSE | High | ASPVTTSPT | [K].ASPVTT | 1xPhospho | A4FTY7 1x | 1.95E-09 | 0 | 1 | 1 | 3 | A4FTY7 | Lcor protei | A4FTY7 [36 | 0 | 2509.332 | 0 | 0.64 | 0.001821 | 17.48 | 75.5 | 124.5 | 14.08 | 4.75 | | High | High | High | High | High | High | High | High | High | High | High | 0 | 9.13E-11 | 5.25 |
| FALSE | High | ASPVTTSPT | [K].ASPVTT | 2xPhospho | A4FTY7 2x | 0.000916 | 0 | 1 | 1 | 1 | A4FTY7 | Lcor protei | A4FTY7 [36 | 0 | 2589.298 | 0 | 0.35 | 0.04343 | 17.52 | 87.1 | 112.9 | 6.68 | 11.83 | | High | High | High | High | High | High | High | High | High | High | High | 0 | 0.000216 | 3.33 |
| FALSE | High | ASQAQATF | [R].ASQAQ | 1xPhospho | Q3TCN5 1x | 0.000758 | 0 | 1 | 1 | 3 | Q3TCN5 | Putative ur | Q3TCN5 [4 | 0 | 1690.789 | 0 | 0.65 | 0.000522 | 5.59 | 77.9 | 122.1 | 2.68 | 3.04 | | High | High | High | High | High | High | High | High | High | High | High | 0.002748 | 0.0294 | 2.68 |
| FALSE | High | ASQASQEY | [K].ASQAS | 1xPhospho | P27661 1x | 0.001068 | 0 | 1 | 1 | 2 | P27661 | Histone H2 | P27661 [13 | 0 | 1192.508 | 0 | -1.21 | 0.00199 | 3.41 | 138.7 | 61.3 | 9.99 | 10.08 | | High | High | High | High | High | High | High | High | High | High | High | 0.000275 | 0.002148 | 1.62 |
| FALSE | High | ASQASSPK | [K].ASQAS | 1xPhospho | A2A870 1x | 0.030367 | 0.001857 | 1 | 2 | 2 | A2A870 | Fas-binding | A2A870 [4 | 0 | 1313.687 | 0 | 0.62 | 0.006898 | 15.22 | 79.8 | 120.2 | 9 | 7.69 | | High | High | High | High | High | High | High | High | High | High | High | 0.008684 | 0.09274 | 1.52 |
| FALSE | High | ASQDGSGF | [R].ASQDG | 1xPhospho | B0V2M3 1x | 0.017966 | 0.001282 | 1 | 5 | 1 | B0V2M3 | Zinc finger | B0V2M3 [5 | 0 | 1320.578 | 0 | 0.4 | 0.009734 | 9.5 | 88.9 | 111.1 | 9.47 | 9.07 | | High | High | High | High | High | High | High | High | High | High | High | 0.000689 | 0.006025 | 0.98 |
| FALSE | High | ASQEFLED | [R].ASQEFL | 1xPhospho | D3Z5I1 1xP | 5.08E-09 | 0 | 1 | 5 | 2 | D3Z5I1 | Zinc finger | D3Z5I1 [32 | 0 | 2191.975 | 0 | 0.73 | 0.002546 | 5.82 | 77.8 | 122.2 | 7.33 | 10.35 | | High | High | High | High | High | High | High | High | High | High | High | 0 | 2.68E-10 | 4.15 |
| FALSE | High | ASQEFLED | [R].ASQEFL | 1xDeamida | D3Z5I1 1xP | 0.002252 | 0.000109 | 1 | 5 | 1 | D3Z5I1 | Zinc finger | D3Z5I1 [32 | 0 | 2192.959 | 0 | 0.61 | 0.00604 | 15.78 | 78.3 | 121.7 | 13.4 | 16.15 | | High | High | High | High | High | High | High | High | High | High | High | 4.94E-05 | 0.000595 | 4.39 |
| FALSE | High | ASQEPFSPA | [R].ASQEPI | 1xPhospho | P70399 1x | 1.09E-09 | 0 | 1 | 3 | 2 | P70399 | TP53-bindi | P70399 [11 | 0 | 3783.819 | 0 | 0.61 | 0.302767 | 20.89 | 79.1 | 120.9 | 12.8 | 30.59 | | High | High | High | High | High | High | High | High | High | High | High | 0 | 4.75E-11 | 5.33 |
| FALSE | High | ASQEPFSPA | [R].ASQEPI | 1xPhospho | P70399 1x | 7.13E-11 | 0 | 1 | 3 | 3 | P70399 | TP53-bindi | P70399 [11 | 0 | 3799.814 | 0 | 0.34 | 0.015836 | 4.79 | 90.4 | 109.6 | 6.94 | 10.79 | | High | High | High | High | High | High | High | High | High | High | High | 0 | 2.24E-12 | 3.99 |
| FALSE | High | ASQGLISAV | [R].ASQGLI | 1xPhospho | Q6PR54-3 | 1.55E-08 | 0 | 2 | 4 | 6 | Q6PR54-3; | Isoform 3 c | Q6PR54-3 | 0 | 2547.223 | 0 | -0.77 | 0.000928 | 0.46 | 126 | 74 | 5.38 | 3.98 | Shared | High | High | High | High | High | High | High | High | High | High | High | 0 | 1.37E-09 | 5.22 |
| FALSE | High | ASQGLISAV | [R].ASQGLI | 1xDeamida | Q6PR54-3 | 0.00509 | 0.000551 | 2 | 4 | 2 | Q6PR54-3; | Isoform 3 c | Q6PR54-3 | 0 | 2548.207 | 0 | -0.46 | 0.016464 | 13.57 | 120.6 | 79.4 | 7.87 | 15 | Shared | High | High | High | High | High | High | High | High | High | High | High | 0.000246 | 0.001482 | 3.19 |
| FALSE | High | ASSLDLEEP | [R].ASSLDL | 1xPhospho | G3X9F6 1x | 0.002334 | 0.000109 | 1 | 6 | 2 | G3X9F6 | SH3 domai | G3X9F6 [4 | 0 | 1425.682 | 0 | 1.79 | 0.000236 | 6.96 | 44 | 156 | 13.23 | 5.01 | | High | High | High | High | High | High | High | High | High | High | High | 4.94E-05 | 0.000615 | 2.7 |
| FALSE | High | AVDSSSEEI | [K].AVDSSS | 1xPhospho | Q5XG69 1x | 3.23E-05 | 0 | 1 | 1 | 4 | Q5XG69 | Soluble lan | Q5XG69 [6 | 1 | 2424.186 | 0 | -1.91 | 0.000396 | 1.3 | 155.4 | 44.6 | 5.95 | 14.04 | | High | High | High | High | High | High | High | High | High | High | High | 0 | 9E-06 | 3.24 |
| FALSE | High | ASSIDSASN | [R].ASSIDS | 1xPhospho | E9Q3Y4 1x | 4.21E-06 | 0 | 1 | 7 | 4 | E9Q3Y4 | Lipopolysa | E9Q3Y4 [9 | 0 | 2891.302 | 0 | -0.13 | 0.3339 | 3.09 | 101.8 | 98.2 | 6.19 | 6.44 | | High | High | High | High | High | High | High | High | High | High | High | 0 | 5.13E-07 | 5.54 |
| FALSE | High | ASSIDSASN | [R].ASSIDS | 2xPhospho | E9Q3Y4 2x | 3.44E-05 | 0 | 1 | 7 | 1 | E9Q3Y4 | Lipopolysa | E9Q3Y4 [9 | 0 | 2971.268 | 0 | 0.4 | 0.067521 | 4.88 | 89.1 | 110.9 | 6.34 | 17.34 | | High | High | High | High | High | High | High | High | High | High | High | 0 | 5.42E-06 | 4.86 |
| FALSE | High | ASTDNEELI | [R].ASTDNI | 1xCarbami | Q7TPS5 1x | 0.000941 | 0 | 2 | 6 | 2 | Q7TPS5; A | C2 domain | Q7TPS5 [8 | 0 | 3762.838 | 0 | 0.57 | 0.000815 | 9.2 | 82.5 | 117.5 | 9.32 | 5.09 | Shared | High | High | High | High | High | High | High | High | High | High | High | 0 | 0.000223 | 3.35 |
| FALSE | High | ASTDVETA | [R].ASTDVI | 1xCarbami | D3Z357 1x | 8.35E-06 | 0 | 1 | 3 | 1 | D3Z357 | Histone-lys | D3Z357 [64 | 0 | 1769.761 | 0 | 1.9 | 0.000348 | 26.19 | 42.1 | 157.9 | 20.86 | 5.42 | | High | High | High | High | High | High | High | High | High | High | High | 0 | 1.1E-06 | 2.22 |
| FALSE | High | ASTFCGTPI | [R].ASTFCG | 1xCarbami | P28867-2 1 | 0.001544 | 5.66E-05 | 1 | 9 | 2 | P28867-2 | Isoform 2 c | P28867-2 [ | 0 | 2719.382 | 0 | 1.02 | 0.015782 | 53.16 | 58.7 | 141.3 | 27.92 | 24.38 | | High | High | High | High | High | High | High | High | High | High | High | 2.58E-05 | 0.000389 | 3.63 |
| FALSE | High | ASTGPHLA | [K].ASTGPI | 1xPhospho | Q9D279 1x | 1.73E-11 | 0 | 1 | 2 | 1 | Q9D279 | Mitotic inte | Q9D279 [2 | 0 | 2542.278 | 0 | 4.61 | 0.002364 | 43.52 | 7.1 | 192.9 | 59.17 | 23.09 | | High | High | High | High | High | High | High | High | High | High | High | 0 | 4.55E-13 | 5.91 |
| FALSE | High | ASTGPHLA | [K].ASTGPI | 1xPhospho | Q9D279 1x | 1.86E-08 | 0 | 1 | 2 | 2 | Q9D279 | Mitotic inte | Q9D279 [2 | 0 | 2558.273 | 0 | 4.71 | 0.001084 | 64.29 | 5.7 | 194.3 | 46.52 | 16.13 | | High | High | High | High | High | High | High | High | High | High | High | 0 | 8.83E-09 | 3.42 |
| FALSE | High | ASTPDWPS | [R].ASTPD\ | 1xPhospho | Q9D279 1x | 9.25E-07 | 0 | 1 | 1 | 6 | Q9D279 | Mitotic inte | Q9D279 [3 | 0 | 2189.023 | 0 | 3.09 | 0.000406 | 16.59 | 20.2 | 179.8 | 11.29 | 10.21 | | High | High | High | High | High | High | High | High | High | High | High | 0 | 1.75E-07 | 4.82 |
| FALSE | High | ASTPSAPAI | [R].ASTPSA | 1xPhospho | Q9DB43 1x | 0.008691 | 0.000736 | 1 | 3 | 2 | Q9DB43 | Zinc finger | Q9DB43 [1 | 0 | 4741.33 | 0 | 0.26 | 0.115834 | 15.3 | 94.6 | 105.4 | 10.48 | 6.43 | | High | High | High | High | High | High | High | High | High | High | High | 0.000327 | 0.002683 | 2.97 |
| FALSE | High | ASTPSQVN | [K].ASTPSC | 1xPhospho | Q3U2W2 1 | 3.52E-07 | 0 | 1 | 2 | 8 | Q3U2W2 | MYB bindir | Q3U2W2 [ | 0 | 1868.988 | 0 | -0.18 | 0.094655 | 0.54 | 106.4 | 93.6 | 4.55 | 8.49 | | High | High | High | High | High | High | High | High | High | High | High | 0 | 4.42E-06 | 3.17 |
| FALSE | High | ASTPSQVN | [K].ASTPSC | 1xDeamida | Q3U2W2 1 | 9.08E-07 | 0 | 1 | 2 | 22 | Q3U2W2 | MYB bindir | Q3U2W2 [ | 0 | 1869.972 | 0 | -0.03 | 0.438447 | 5.58 | 101.5 | 98.5 | 3 | 8.69 | | High | High | High | High | High | High | High | High | High | High | High | 0 | 4.37E-06 | 3.59 |
| FALSE | High | ASTSDYQV | [K].ASTSDY | 1xPhospho | Q8R4R6 1x | 0.000501 | 0 | 1 | 1 | 2 | Q8R4R6 | Nucleopor | Q8R4R6 [2 | 1 | 2334.174 | 0 | -0.56 | 0.000596 | 3.07 | 119.7 | 80.3 | 5.5 | 4.3 | | High | High | High | High | High | High | High | High | High | High | High | 0 | 0.000109 | 3.22 |
| FALSE | High | ASTSGLSFK | [R].ASTSGL | 1xPhospho | D3YYC3 1x | 0.047894 | 0.00354 | 1 | 5 | 1 | D3YYC3 | Histone-lys | D3YYC3 [1 | 0 | 1435.76 | 0 | -1.29 | 0.003859 | 2.4 | 139 | 61 | 11.22 | 12.56 | | High | High | High | High | High | High | High | High | High | High | High | 0.001812 | 0.01826 | 0.96 |
| FALSE | High | ASTSGLSFK | [R].ASTSGL | 1xPhospho | D3YYC3 1x | 0.072258 | 0.005346 | 1 | 5 | 1 | D3YYC3 | Histone-lys | D3YYC3 [1 | 1 | 2323.179 | 0 | -1.46 | 0.002496 | 0.83 | 146.5 | 53.5 | 13.36 | 4.05 | | High | High | High | High | High | High | High | High | High | High | High | 0.002855 | 0.02957 | 1.07 |
| FALSE | High | ASTSSSSSS | [R].ASTSSS | 1xPhospho | P52482 1x | 1.82E-06 | 0 | 1 | 1 | 6 | P52482 | Ubiquitin-c | P52482 [8- | 0 | 2241.028 | 0 | 0.14 | 0.054984 | 8.03 | 96.7 | 103.3 | 7.02 | 7.99 | | High | High | High | High | High | High | High | High | High | High | High | 0 | 1.99E-07 | 4.01 |
| FALSE | High | ASTDNEELI | [R].ASTDNI | 1xCarbami | Q7TPS5 1x | 0.000454 | 0 | 2 | 6 | 2 | Q7TPS5; A | C2 domain | Q7TPS5 [8 | 0 | 3761.854 | 0 | 0.73 | 0.042631 | 32.79 | 78.3 | 121.7 | 11.94 | 19.29 | Shared | High | High | High | High | High | High | High | High | High | High | High | 0 | 0.00029 | 3.12 |
| FALSE | High | ASTSTEAPC | [K].ASTSTE | 1xPhospho | E9Q3Y4 1x | 0.071662 | 0.005278 | 1 | 7 | 1 | E9Q3Y4 | Lipopolysa | E9Q3Y4 [1 | 0 | 1581.747 | 0 | 0.7 | 0.024338 | 18.2 | 76.2 | 123.8 | 20.59 | 9.08 | | High | High | High | High | High | High | High | High | High | High | High | 0.002703 | 0.0291 | 1.88 |
| FALSE | High | ASTVSPGG | [R].ASTVSF | 1xPhospho | Q6PAL7 1x | 0.007209 | 0.00062 | 1 | 1 | 4 | Q6PAL7 | AT-hook D | Q6PAL7 [1 | 0 | 1847.938 | 0 | -0.85 | 0.035006 | 10.1 | 128.6 | 71.4 | 16.1 | 17.17 | | High | High | High | High | High | High | High | High | High | High | High | 0.000555 | 0.004908 | 2.89 |
| FALSE | High | ASVASDPES | [K].ASVASI | 1xPhospho | Q9D0L8 1x | 1.32E-08 | 0 | 1 | 6 | 7 | Q9D0L8 | mRNA cap | Q9D0L8 [7 | 0 | 2461.119 | 0 | -0.28 | 0.001284 | 3.26 | 108.2 | 91.8 | 3.05 | 2.59 | | High | High | High | High | High | High | High | High | High | High | High | 0 | 5.61E-08 | 4.86 |
| FALSE | High | ASVASDPES | [K].ASVASI | 2xPhospho | Q9D0L8 2x | 2.06E-06 | 0 | 1 | 6 | 2 | Q9D0L8 | mRNA cap | Q9D0L8 [7 | 0 | 2541.086 | 0 | 0.34 | 0.008789 | 9.86 | 88.3 | 111.7 | 4.63 | 5.82 | | High | High | High | High | High | High | High | High | High | High | High | 0 | 2.31E-07 | 4.08 |
| FALSE | High | ASVAVHSP | [K].ASVAVI | 1xPhospho | O88845 1x | 0.003892 | 0.000261 | 1 | 3 | 2 | O88845 | A-kinase ar | O88845 [4 | 0 | 1561.85 | 0 | 0.06 | 0.188409 | 6.95 | 98.1 | 101.9 | 8.37 | 1.66 | | High | High | High | High | High | High | High | High | High | High | High | 0.00014 | 0.001093 | 2.18 |
| FALSE | High | ASVCNLQIS | [K].ASVCNI | 1xCarbami | F8VPL2 1x | 0.000112 | 0 | 1 | 6 | 1 | F8VPL2 | Phosphatic | F8VPL2 [25 | 0 | 1754.928 | 0 | 0.59 | 0.003526 | 9.7 | 81.6 | 118.4 | 5.95 | 6.38 | | High | High | High | High | High | High | High | High | High | High | High | 0 | 2.04E-05 | 3.29 |
| FALSE | High | ASVEGAEA | [K].ASVEG | 1xPhospho | Q8R3C6 1x | 0.00509 | 0.000551 | 1 | 2 | 1 | Q8R3C6 | Probable R | Q8R3C6 [6 | 0 | 1644.825 | 0 | -0.16 | 0.021822 | 3.62 | 105.8 | 94.2 | 4.56 | 7.27 | | High | High | High | High | High | High | High | High | High | High | High | 0.000246 | 0.001473 | 1.9 |
| FALSE | High | ASVGDSSQ | [R].ASVGD | 1xPhospho | Q3V1Z0 1x | 1.11E-06 | 0 | 1 | 5 | 1 | Q3V1Z0 | Iporin OS= | Q3V1Z0 [5 | 0 | 2488.119 | 0 | 0.55 | 0.005533 | 13.12 | 80.7 | 119.3 | 6.92 | 7.86 | | High | High | High | High | High | High | High | High | High | High | High | 0 | 1.14E-07 | 3.88 |
| FALSE | High | ASVGQSPV | [K].ASVGQ | 1xPhospho | Q6NXI6 1x | 1.26E-05 | 0 | 1 | 3 | 1 | Q6NXI6 | Regulation | Q6NXI6 [6 | 0 | 2057.108 | 0 | -0.18 | 0.025579 | 1.08 | 107.2 | 92.8 | 5.14 | 5.43 | | High | High | High | High | High | High | High | High | High | High | High | 0 | 1.76E-06 | 2.66 |
| FALSE | High | ASVHSSGR | [K].ASVHS | 1xPhospho | G3X956 1x | 0.040373 | 0.002982 | 1 | 2 | 1 | G3X956 | Suppressor | G3X956 [1 | 0 | 1109.53 | 0 | 0.06 | 0.049678 | 1.04 | 99.8 | 100.2 | 3.53 | 4.65 | | High | High | High | High | High | High | High | High | High | High | High | 0.001566 | 0.01511 | 0.75 |
| FALSE | High | ASVHSSGR | [K].ASVHS | 2xPhospho | G3X956 2x | 0.028575 | 0.001829 | 1 | 2 | 1 | G3X956 | Suppressor | G3X956 [1 | 0 | 1189.497 | 0 | -0.25 | 0.506206 | 22.19 | 108 | 92 | 14.41 | 21.42 | | High | High | High | High | High | High | High | High | High | High | High | 0.000984 | 0.01023 | 1.48 |
| FALSE | High | ASVHTPVK | [R].ASVHTI | 1xPhospho | D4AFX6 1x | 0.052133 | 0.00384 | 1 | 3 | 1 | D4AFX6 | Centaurin | D4AFX6 [5 | 0 | 1376.77 | 0 | 1.12 | 0.000446 | 11.94 | 63.4 | 136.6 | 7.27 | 3.19 | | High | High | High | High | High | High | High | High | High | High | High | 0.001999 | 0.02026 | 1.68 |
| FALSE | High | ASVLQMDS | [R].ASVLQI | 1xPhospho | Q80WR5 1 | 0.000822 | 0 | 1 | 3 | 1 | Q80WR5 | UPF0688 p | Q80WR5 [ | 0 | 2961.357 | 0 | 0.41 | 0.125108 | 10.47 | 85.6 | 114.4 | 17.93 | 15.91 | | High | High | High | High | High | High | High | High | High | High | High | 0 | 0.000191 | 3.59 |
| FALSE | High | ASVLQMDS | [R].ASVLQI | 1xPhospho | Q80WR5 1 | 0.003857 | 0.000261 | 1 | 3 | 1 | Q80WR5 | UPF0688 p | Q80WR5 [ | 0 | 2977.352 | 0 | 0.97 | 0.008087 | 48.55 | 66.7 | 133.3 | 42 | 13.23 | | High | High | High | High | High | High | High | High | High | High | High | 0.000117 | 0.001079 | 3.13 |
| FALSE | High | ASTTQPED | [K].ASTTQI | 1xPhospho | Q8BGS1 1x | 6.31E-06 | 0 | 1 | 2 | 8 | Q8BGS1 | Band 4.1-li | Q8BGS1 [3 | 0 | 2368.216 | 0 | -0.49 | 0.003121 | 1.62 | 119.5 | 80.5 | 5.83 | 6.85 | | High | High | High | High | High | High | High | High | High | High | High | 0 | 0.000278 | 4.05 |
| FALSE | High | ASTASLPLG | [R].ASTASL | 1xPhospho | P43267 1x | 0.000718 | 0 | 1 | 1 | 1 | P43267 | Protein SO | P43267 [18 | 0 | 3244.65 | 0 | 0.88 | 0.01513 | 15.43 | 72.8 | 127.2 | 16.15 | 13.89 | | High | High | High | High | High | High | High | High | High | High | High | 0 | 0.000164 | 3.15 |
| FALSE | High | ASSVPAGR | [R].ASSVPA | 1xPhospho | Q8CD14 1x | 0.003195 | 0.00016 | 1 | 6 | 2 | Q8CD14 | Putative ur | Q8CD14 [8 | 0 | 1053.529 | 0 | -0.27 | 0.100447 | 19.52 | 108.4 | 91.6 | 12.29 | 16.51 | | High | High | High | High | High | High | High | High | High | High | High | 0.000968 | 0.01005 | 1.09 |
| FALSE | High | ASSVNSPL( | [K].ASSVNS | 1xPhospho | Q8BJH1 1x | 0.000568 | 0 | 1 | 2 | 2 | Q8BJH1 | Zinc finger | Q8BJH1 [2 | 0 | 2615.371 | 0 | -0.85 | 0.003027 | 7.64 | 128.8 | 71.2 | 6.53 | 11.86 | | High | High | High | High | High | High | High | High | High | High | High | 0 | 0.000127 | 3.06 |
| FALSE | High | ASSLEDLVL | [R].ASSLED | 1xPhospho | Q3TW36 1 | 0.000154 | 0 | 1 | 5 | 7 | Q3TW36 | Putative ur | Q3TW36 [2 | 0 | 1612.896 | 0 | 0.31 | 0.003855 | 5.33 | 90.3 | 109.7 | 2.69 | 7.18 | | High | High | High | High | High | High | High | High | High | High | High | 0 | 0.000259 | 3.7 |
| FALSE | High | ASSLGETDI | [R].ASSLGE | 1xPhospho | Q8BWW9 | 6.9E-09 | 0 | 1 | 5 | 3 | Q8BWW9 | Serine/thr | Q8BWW9 | 0 | 1789.805 | 0 | 0.06 | 0.114181 | 5.74 | 94.8 | 105.2 | 9.27 | 11.61 | | High | High | High | High | High | High | High | High | High | High | High | 0 | 3.8E-10 | 3.67 |
| FALSE | High | ASSLIEEAK | [K].ASSLIEE | 1xPhospho | Q80TM2 1 | 0.001783 | 5.66E-05 | 1 | 3 | 1 | Q80TM2 | MKIAA102 | Q80TM2 [1 | 0 | 1485.797 | 0 | 0.01 | 0.577525 | 12.27 | 101.2 | 98.8 | 9.71 | 11.69 | | High | High | High | High | High | High | High | High | High | High | High | 2.58E-05 | 0.000455 | 3.14 |
| FALSE | High | ASSLLTQKF | [K].ASSLLT | 1xPhospho | F8WI37 1x | 0.006306 | 0.00062 | 1 | 6 | 1 | F8WI37 | Histone-lys | F8WI37 [3 | 0 | 2933.532 | 0 | -0.19 | 0.198761 | 12.95 | 109 | 91 | 13 | 14.34 | | High | High | High | High | High | High | High | High | High | High | High | 0.000275 | 0.00187 | 2.44 |
| FALSE | High | ASSLNFLNI | [R].ASSLNF | 1xPhospho | Q91YS4 1x | 0.008538 | 0.000736 | 1 | 3 | 3 | Q91YS4 | Kinesin ligh | Q91YS4 [5 | 0 | 1302.666 | 0 | 0.23 | 0.137398 | 18.15 | 87.4 | 112.6 | 13.94 | 19.54 | | High | High | High | High | High | High | High | High | High | High | High | 0.000327 | 0.002618 | 2.71 |
| FALSE | High | ASSLNFLNI | [R].ASSLNF | 1xPhospho | Q91YS4 1x | 0.003822 | 0.000261 | 1 | 3 | 1 | Q91YS4 | Kinesin ligh | Q91YS4 [5 | 0 | 1531.829 | 0 | 0.22 | 0.059173 | 12.02 | 93.6 | 106.4 | 2.31 | 16.37 | | High | High | High | High | High | High | High | High | High | High | High | 0.000117 | 0.001074 | 2.27 |
| FALSE | High | ASSNESLVA | [K].ASSNES | 1xPhospho | Q3U955 1x | 0.001615 | 5.66E-05 | 1 | 6 | 1 | Q3U955 | Putative ur | Q3U955 [2 | 0 | 1456.699 | 0 | 0.58 | 0.007585 | 10.63 | 79.3 | 120.7 | 2.04 | 11.96 | | High | High | High | High | High | High | High | High | High | High | High | 2.58E-05 | 0.000408 | 1.22 |
| FALSE | High | ASSPAQCV | [R].ASSPAC | 1xCarbami | Q9Z2D1 1x | 0.005135 | 0.000551 | 1 | 4 | 3 | Q9Z2D1 | Myotubula | Q9Z2D1 [6 | 0 | 1852.908 | 0 | 0.44 | 0.060877 | 25.64 | 87.7 | 112.3 | 5.85 | 18.78 | | High | High | High | High | High | High | High | High | High | High | High | 0.000246 | 0.001485 | 2.31 |
| FALSE | High | ASSPATSS( | [K].ASSPAT | 1xPhospho | Q99PL5 1x | 9.82E-06 | 0 | 2 | 11 | 2 | Q99PL5; Q | Ribosome- | Q99PL5 [1 | 0 | 1558.788 | 0 | -0.37 | 0.104771 | 19.22 | 109.4 | 90.6 | 7.78 | 17.27 | Shared | High | High | High | High | High | High | High | High | High | High | High | 0 | 0.000108 | 1.88 |
| FALSE | High | ASSPCQEFI | [R].ASSPCC | 1xCarbami | Q60575 1x | 2.52E-06 | 0 | 1 | 3 | 1 | Q60575 | Kinesin-like | Q60575 [1 | 0 | 3106.479 | 0 | 1.14 | 0.002809 | 4.01 | 62.5 | 137.5 | 4.72 | 13.14 | | High | High | High | High | High | High | High | High | High | High | High | 0 | 2.85E-07 | 3.96 |
| FALSE | High | ASSPDDDV | [K].ASSPDI | 1xCarbami | H7BWZ9 1 | 0.00083 | 0 | 1 | 5 | 2 | H7BWZ9 | Protein do | H7BWZ9 [1 | 0 | 2574.256 | 0 | 1.46 | 0.00171 | 13.79 | 56.5 | 143.5 | 10.17 | 15.31 | | High | High | High | High | High | High | High | High | High | High | High | 0 | 0.000193 | 2.87 |
| FALSE | High | ASSPFR | [R].ASSPFR | 2xPhospho | E9Q5C9 2x | 0.033404 | 0.002195 | 1 | 2 | 1 | E9Q5C9 | Nucleolar a | E9Q5C9 [6 | 0 | 1053.437 | 0 | 0.04 | 0.612576 | 20.83 | 97.3 | 102.7 | 11.45 | 6.19 | | High | High | High | High | High | High | High | High | High | High | High | 0.001151 | 0.01216 | 1.76 |
| FALSE | High | ASSPGPQM | [R].ASSPGI | 1xPhospho | E9QPA1 1x | 1.56E-06 | 0 | 1 | 4 | 2 | E9QPA1 | RAB6A-GEI | E9QPA1 [1 | 0 | 2513.251 | 0 | 0.16 | 0.102234 | 16.57 | 94.7 | 105.3 | 7.78 | 5.63 | | High | High | High | High | High | High | High | High | High | High | High | 0 | 1.73E-07 | 4.5 |
| FALSE | High | ASSPGQK | [K].ASSPG( | 1xPhospho | A2ASY1 1x | 0.012288 | 0.000889 | 1 | 5 | 2 | A2ASY1 | TAF3 RNA | A2ASY1 [2 | 0 | 1212.639 | 0 | -0.49 | 0.006209 | 7.66 | 116.7 | 83.3 | 2.96 | 8.78 | | High | High | High | High | High | High | High | High | High | High | High | 0.000449 | 0.003948 | 1.77 |
| FALSE | High | ASSPPVEV( | [R].ASSPPV | 1xPhospho | Q9Z273 1x | 0.011346 | 0.000816 | 1 | 3 | 2 | Q9Z273 | Tubby-rela | Q9Z273 [2 | 0 | 1533.751 | 0 | -1.1 | 0.001626 | 4.55 | 136.6 | 63.4 | 7.14 | 11.72 | | High | High | High | High | High | High | High | High | High | High | High | 0.000418 | 0.003602 | 2.19 |
| FALSE | High | ASSPQLALI | [R].ASSPQL | 1xPhospho | Q7TSV9 1x | 0.001049 | 0 | 1 | 2 | 2 | Q7TSV9 | Enkurin do | Q7TSV9 [1 | 0 | 1903.045 | 0 | 0.73 | 0.011564 | 20.33 | 75 | 125 | 9.93 | 10.51 | | High | High | High | High | High | High | High | High | High | High | High | 0 | 0.000251 | 1.84 |
| FALSE | High | ASSPSASSS | [K].ASSPSA | 1xPhospho | Q3UUL4 1x | 0.000462 | 0 | 1 | 1 | 1 | Q3UUL4 | Putative ur | Q3UUL4 [3 | 0 | 1566.7 | 0 | 0.04 | 0.105107 | 3.78 | 95.9 | 104.1 | 6.65 | 9.54 | | High | High | High | High | High | High | High | High | High | High | High | 0 | 9.99E-05 | 1.69 |
| FALSE | High | ASSPSDTFS | [K].ASSPSD | 1xPhospho | Q56A10 1x | 0.004033 | 0.000311 | 1 | 3 | 2 | Q56A10 | Zinc finger | Q56A10 [9 | 0 | 1678.809 | 0 | 0.83 | 0.002291 | 13.37 | 70.2 | 129.8 | 4.83 | 10.93 | | High | High | High | High | High | High | High | High | High | High | High | 0.002059 | 0.02149 | 2.91 |
| FALSE | High | ASSPSLR | [K].ASSPSL | 1xPhospho | Q8C7S2 1x | 0.028824 | 0.001829 | 1 | 5 | 2 | Q8C7S2 | Putative ur | Q8C7S2 [6 | 0 | 1026.518 | 0 | 1.84 | 0.000714 | 3.46 | 44.4 | 155.6 | 6.69 | 12.02 | | High | High | High | High | High | High | High | High | High | High | High | 0.006317 | 0.0676 | 1.63 |
| FALSE | High | ASSPSPLTV | [K].ASSPSP | 1xPhospho | Q91YD3 1x | 3.79E-07 | 0 | 1 | 1 | 12 | Q91YD3 | mRNA-dec | Q91YD3 [5 | 0 | 1896.927 | 0 | -0.16 | 0.072838 | 4.09 | 105.4 | 94.6 | 4.53 | 3.78 | | High | High | High | High | High | High | High | High | High | High | High | 0 | 4.08E-08 | 3.86 |
| FALSE | High | ASSPSPLTV | [K].ASSPSP | 2xPhospho | Q91YD3 2x | 5.42E-06 | 0 | 1 | 1 | 3 | Q91YD3 | mRNA-dec | Q91YD3 [5 | 0 | 1976.893 | 0 | 0.23 | 0.010502 | 3.75 | 90.3 | 109.7 | 9.67 | 3.45 | | High | High | High | High | High | High | High | High | High | High | High | 0 | 6.76E-07 | 3.42 |
| FALSE | High | ASSPVTAAT | [K].ASSPVT | 1xPhospho | Q8K298 1x | 1.79E-06 | 0 | 1 | 2 | 8 | Q8K298 | Anillin OS= | Q8K298 [2 | 0 | 1873.926 | 0 | 0.1 | 0.715415 | 10.33 | 96.6 | 103.4 | 8.73 | 6.77 | | High | High | High | High | High | High | High | High | High | High | High | 0 | 5.41E-07 | 3.91 |
| FALSE | High | ASSRVSGG | [R].ASSRVS | 2xPhospho | O70439 2x | 4.35E-05 | 0 | 1 | 4 | 2 | O70439 | Syntaxin-7 | O70439 [1 | 1 | 2128.972 | 0 | -1.2 | 0.000916 | 1.92 | 136.6 | 63.4 | 6.39 | 8.94 | | High | High | High | High | High | High | High | High | High | High | High | 0 | 7.03E-06 | 3.58 |
| FALSE | High | ASSSAQAV | [R].ASSSAC | 1xCarbami | P54276 1x | 0.000113 | 0 | 1 | 2 | 6 | P54276 | DNA mism | P54276 [76 | 0 | 3002.437 | 0 | 0.12 | 0.146715 | 8.7 | 97.7 | 102.3 | 10.33 | 4.35 | | High | High | High | High | High | High | High | High | High | High | High | 0 | 2.06E-05 | 4.07 |
| FALSE | High | ASSSGGSD | [R].ASSSG( | 2xPhospho | Q9WTN3 2 | 0.000751 | 0 | 1 | 6 | 4 | Q9WTN3 | Sterol regu | Q9WTN3 [ | 0 | 2802.18 | 0 | 0.95 | 0.002503 | 37.04 | 66.6 | 133.4 | 25.16 | 3.86 | | High | High | High | High | High | High | High | High | High | High | High | 9.42E-05 | 0.001038 | 3.66 |
| FALSE | High | ASSSPGSPT | [R].ASSSPG | 1xPhospho | B2RQ06 1x | 0.003655 | 0.00021 | 1 | 3 | 2 | B2RQ06 | Zinc finger | B2RQ06 [4 | 0 | 1516.673 | 0 | 2.41 | 0.000181 | 10.59 | 31.9 | 168.1 | 13.71 | 4.93 | | High | High | High | High | High | High | High | High | High | High | High | 9.42E-05 | 0.001017 | 2.07 |
| FALSE | High | ASSTTPK | [K].ASSTTP | 1xPhospho | B1AZI6 1xP | 0.08727 | 0.006537 | 1 | 1 | 1 | B1AZI6 | THO comp | B1AZI6 [12 | 0 | 1229.654 | 0 | -0.21 | 0.001017 | 2.26 | 108.1 | 91.9 | 3.07 | 3.82 | | High | High | High | High | High | High | High | High | High | High | High | 0.003346 | 0.03669 | 1.32 |
| FALSE | High | ASSVGVLSI | [R].ASSVG\ | 1xPhospho | Q80Y32 1x | 0.0009 | 0 | 1 | 3 | 2 | Q80Y32 | HMG box c | Q80Y32 [4 | 0 | 1610.892 | 0 | -0.52 | 0.002872 | 5 | 116.2 | 83.8 | 4.25 | 5.8 | | High | High | High | High | High | High | High | High | High | High | High | 0 | 0.000213 | 2.96 |
| FALSE | High | ASSVGVLSI | [R].ASSVG\ | 2xPhospho | Q80Y32 2x | 0.011246 | 0.000772 | 1 | 3 | 1 | Q80Y32 | HMG box c | Q80Y32 [4 | 0 | 1690.858 | 0 | -0.77 | 0.003671 | 4.26 | 126.1 | 73.9 | 8.06 | 5.08 | | High | High | High | High | High | High | High | High | High | High | High | 0.000418 | 0.003583 | 2.3 |
| FALSE | High | ASSIDSASN | [R].ASSIDS | 1xPhospho | E9Q3Y4 1x | 1.22E-05 | 0 | 1 | 7 | 2 | E9Q3Y4 | Lipopolysa | E9Q3Y4 [9 | 0 | 2907.297 | 0 | -0.03 | 0.467792 | 6.59 | 99.3 | 100.7 | 14.93 | 57.33 | | High | High | High | High | High | High | High | High | High | High | High | 0 | 2.78E-06 | 4.89 |
| FALSE | High | NHLSPQEG | [K].NHLSP( | 2xCarbami | Q9EP97 2x | 5.08E-09 | 0 | 1 | 1 | 4 | Q9EP97 | Sentrin-spe | Q9EP97 [1 | 0 | 2581.093 | 0 | -0.67 | 0.002141 | 2.1 | 122.5 | 77.5 | 6.96 | 2.42 | | High | High | High | High | High | High | High | High | High | High | High | 0 | 1.52E-06 | 5.46 |
| FALSE | High | FLESDSEDE | [R].FLESDS | 1xPhospho | E9QKD1 1x | 6.83E-05 | 0 | 1 | 3 | 3 | E9QKD1 | Nucleolar | E9QKD1 [8 | 0 | 1736.803 | 0 | -0.6 | 0.002153 | 3.89 | 122 | 78 | 6.15 | 4.63 | | High | High | High | High | High | High | High | High | High | High | High | 2.58E-05 | 0.00051 | 2.89 |
| FALSE | High | FLGSDEEDI | [K].FLGSDE | 1xPhospho | Q8CEC0 1x | 0.003464 | 0.00021 | 1 | 6 | 2 | Q8CEC0 | Nuclear po | Q8CEC0 [4 | 0 | 1577.75 | 0 | -0.49 | 0.002811 | 2.51 | 117.2 | 82.8 | 8.41 | 7.14 | | High | High | High | High | High | High | High | High | High | High | High | 9.42E-05 | 0.000964 | 2.97 |
| FALSE | High | LLSSPGK | [K].LLSSPG | 1xPhospho | Q8JZR6 1x | 0.009843 | 0.000736 | 1 | 2 | 1 | Q8JZR6 | Electroneu | Q8JZR6 [1 | 0 | 1239.711 | 0 | 1.2 | 0.000428 | 7.83 | 62.9 | 137.1 | 5.03 | 6.96 | | High | High | High | High | High | High | High | High | High | High | High | 0.000344 | 0.003077 | 2.08 |

| | | | | | | | | | | | | | | | | | | | | | | | | | | | | | | | | | | | | | | |
|---|---|---|---|---|---|---|---|---|---|---|---|---|---|---|---|---|---|---|---|---|---|---|---|---|---|---|---|---|---|---|---|---|---|---|---|---|---|---|
| FALSE | High | LLSSSDDD/ | [R].LLSSSD | 1xPhosphc | A6PW47 1: | 2.26E-06 | 0 | 1 | 3 | 4 | A6PW47 | Mediator c | A6PW47 [8 | 0 | 2215.033 | 0 | 0.11 | 0.01229 | 3.44 | 96.1 | 103.9 | 4.55 | 5.45 | | High | High | High | High | High | High | High | High | High | High | High | 2.58E-05 | 0.000533 | 3.68 |
| FALSE | High | LLSSSDDD/ | [R].LLSSSD | 2xPhosphc | A6PW47 2: | 6.89E-05 | 0 | 1 | 3 | 3 | A6PW47 | Mediator c | A6PW47 [8 | 0 | 2294.999 | 0 | 1.22 | 0.000298 | 11.23 | 60.1 | 139.9 | 6 | 4.89 | | High | High | High | High | High | High | High | High | High | High | High | 0 | 1.19E-05 | 4.18 |
| FALSE | High | LLSSSLTTG | [R].LLSSSLT | 1xCarbami | E9QMB7 1: | 0.000422 | 0 | 1 | 3 | 2 | E9QMB7 | N-alpha-ac | E9QMB7 [1 | 0 | 1703.86 | 0 | -0.88 | 0.003731 | 3.65 | 129.8 | 70.2 | 8.15 | 9.43 | | High | High | High | High | High | High | High | High | High | High | High | 0 | 9.06E-05 | 2.48 |
| FALSE | High | IISSSPNK | [K].IISSSPN | 1xPhosphc | Q6ZPY5 1x | 0.016892 | 0.001204 | 1 | 1 | 4 | Q6ZPY5 | Zinc finger | Q6ZPY5 [3 | 0 | 1383.765 | 0 | -1.05 | 0.000154 | 0.93 | 134.9 | 65.1 | 2.4 | 7.11 | | High | High | High | High | High | High | High | High | High | High | High | 0.000638 | 0.005636 | 2.26 |
| FALSE | High | LLSSSSPQP | [K].LLSSSSI | 1xPhosphc | Q9DC40 1x | 0.008388 | 0.000736 | 1 | 1 | 2 | Q9DC40 | Telomere I | Q9DC40 [8 | 0 | 1224.607 | 0 | -0.18 | 0.043475 | 1.54 | 105.7 | 94.3 | 4.38 | 5.93 | | High | High | High | High | High | High | High | High | High | High | High | 0.000327 | 0.002567 | 2.09 |
| FALSE | High | LISSVDDFT | [R].LISSVDI | 1xPhosphc | Q61037 1x | 5.27E-06 | 0 | 1 | 13 | 3 | Q61037 | Tuberin OS | Q61037 [1: | 0 | 1680.797 | 0 | -0.65 | 0.002468 | 0.96 | 126.8 | 73.2 | 8.86 | 13.54 | | High | High | High | High | High | High | High | High | High | High | High | 0 | 6.56E-07 | 2.49 |
| FALSE | High | LLSTDAEA | [R].LLSTDA | 1xPhosphc | Q9R1C0 1x | 0.007404 | 0.00062 | 1 | 2 | 1 | Q9R1C0 | Transcripti | Q9R1C0 [1 | 0 | 1571.788 | 0 | -1.6 | 0.000824 | 1.29 | 150.9 | 49.1 | 11.09 | 5.77 | | High | High | High | High | High | High | High | High | High | High | High | 0.000275 | 0.002237 | 2.22 |
| FALSE | High | ILSTMDSPS | [R].ILSTMD | 1xPhosphc | Q5SWU9-2 | 0.010382 | 0.000772 | 1 | 2 | 1 | Q5SWU9-2 | Isoform 2 c | Q5SWU9-2 | 0 | 1376.622 | 0 | -0.06 | 0.966301 | 16.8 | 102.2 | 97.8 | 9.7 | 8.97 | | High | High | High | High | High | High | High | High | High | High | High | 0.000382 | 0.003266 | 2.13 |
| FALSE | High | LLSTVDQES | [K].LLSTVD | 1xPhosphc | Q8VDM1 1 | 3.54E-05 | 0 | 1 | 2 | 1 | Q8VDM1 | Zinc finger | Q8VDM1 [ | 0 | 3020.491 | 0 | -0.4 | 0.241445 | 16.79 | 106.5 | 93.5 | 15.45 | 20.23 | | High | High | High | High | High | High | High | High | High | High | High | 0 | 5.58E-06 | 3.61 |
| FALSE | High | IISVTPVK | [R].IISVTPV | 1xPhosphc | E9QPI5 1xF | 0.00129 | 0 | 1 | 3 | 1 | E9QPI5 | Sister chro | E9QPI5 [12 | 0 | 1394.842 | 0 | -0.1 | 0.016746 | 3.05 | 105.8 | 94.2 | 4.25 | 6.81 | | High | High | High | High | High | High | High | High | High | High | High | 0 | 0.000317 | 1.99 |
| FALSE | High | LLSVTSDEG | [R].LLSVTS | 2xPhosphc | Q8BRB7 2x | 0.00195 | 5.66E-05 | 1 | 4 | 1 | Q8BRB7 | Histone acı | Q8BRB7 [3 | 0 | 2162.928 | 0 | -1.37 | 0.000825 | 0.88 | 143.5 | 56.5 | 9.19 | 13.25 | | High | High | High | High | High | High | High | High | High | High | High | 2.58E-05 | 0.000504 | 2.47 |
| FALSE | High | LLSYLEK | [R].LLSYLEI | 1xPhosphc | Q3TFA9 1x | 0.095408 | 0.007437 | 1 | 4 | 1 | Q3TFA9 | Putative ur | Q3TFA9 [6 | 0 | 1403.795 | 0 | 1.81 | 0.020599 | 36.52 | 34.3 | 165.7 | 16.96 | 39.41 | | High | High | High | High | High | High | High | High | High | High | High | 0.003782 | 0.04074 | 2.43 |
| FALSE | High | LLSSPAK | [K].LLSSPA | 1xPhosphc | Q920Q2 1x | 0.071662 | 0.005278 | 1 | 1 | 1 | Q920Q2 | DNA repair | Q920Q2 [1 | 0 | 1253.727 | 0 | -1.19 | 0.000461 | 0.79 | 136.9 | 63.1 | 5.84 | 7.68 | | High | High | High | High | High | High | High | High | High | High | High | 0.002717 | 0.02925 | 1.78 |
| FALSE | High | IITEDELTA( | [K].IITEDEL | 1xCarbami | B2KF05 1xl | 1.07E-07 | 0 | 1 | 1 | 1 | B2KF05 | Bromodom | B2KF05 [6( | 0 | 2732.31 | 0 | -0.28 | 0.035328 | 5.94 | 109 | 91 | 14.7 | 6.58 | | High | High | High | High | High | High | High | High | High | High | High | 0 | 8.24E-09 | 5.19 |
| FALSE | High | LLTPDSQP/ | [K].LLTPDS | 1xPhosphc | B2KFW1 1x | 0.001127 | 0 | 1 | 2 | 1 | B2KFW1 | Zinc finger | B2KFW1 [6 | 0 | 2240.162 | 0 | -0.5 | 0.039817 | 8.36 | 117.3 | 82.7 | 12.59 | 13.37 | | High | High | High | High | High | High | High | High | High | High | High | 0 | 0.000274 | 2.93 |
| FALSE | High | LLTPNPGY( | [R].LLTPNP | 1xPhosphc | Q99J21 1xl | 6.13E-05 | 0 | 1 | 4 | 6 | Q99J21 | Mucolipin- | Q99J21 [1! | 0 | 3250.572 | 0 | 1.13 | 0.003589 | 12.11 | 60.6 | 139.4 | 11.09 | 15.9 | | High | High | High | High | High | High | High | High | High | High | High | 0 | 1.04E-05 | 4 |
| FALSE | High | LLTPNPGY( | [R].LLTPNP | 1xPhosphc | Q99J21 1xl | 4.01E-05 | 0 | 1 | 4 | 3 | Q99J21 | Mucolipin- | Q99J21 [1! | 1 | 3406.673 | 0 | 1.64 | 0.000197 | 4.54 | 49.4 | 150.6 | 5.45 | 5.44 | | High | High | High | High | High | High | High | High | High | High | High | 0 | 6.45E-06 | 3.75 |
| FALSE | High | LLTPTHSFL | [R].LLTPTH | 1xPhosphc | E9QMU3 1 | 0.050397 | 0.003641 | 1 | 4 | 1 | E9QMU3 | Ensconsin | E9QMU3 [ | 0 | 1564.845 | 0 | -0.83 | 0.000374 | 1.87 | 128.8 | 71.2 | 4.83 | 4.8 | | High | High | High | High | High | High | High | High | High | High | High | 0.001916 | 0.01937 | 2.45 |
| FALSE | High | ILTPTLK | [K].ILTPTLK | 1xPhosphc | Q3TAF0 1x | 0.036416 | 0.002419 | 1 | 5 | 1 | Q3TAF0 | Putative ur | Q3TAF0 [5 | 0 | 1323.805 | 0 | 0.67 | 0.00564 | 12.9 | 76.8 | 123.2 | 6.09 | 10.46 | | High | High | High | High | High | High | High | High | High | High | High | 0.001308 | 0.01348 | 2.23 |
| FALSE | High | LLTSITSK | [R].LLTSITS | 1xPhosphc | Q8K298 1x | 0.004696 | 0.000456 | 1 | 2 | 1 | Q8K298 | Anillin OS= | Q8K298 [9 | 0 | 1400.817 | 0 | 1.83 | 0.006488 | 38.77 | 44.1 | 155.9 | 23.72 | 17.74 | | High | High | High | High | High | High | High | High | High | High | High | 0.000204 | 0.001353 | 2.6 |
| FALSE | High | LLVDVDES | [K].LLVDVI | 1xPhosphc | G5E866 1x | 3.63E-08 | 0 | 1 | 4 | 2 | G5E866 | Splicing fac | G5E866 [4 | 0 | 2340.199 | 0 | -1.29 | 0.000242 | 1.34 | 141.4 | 58.6 | 3.68 | 4.6 | | High | High | High | High | High | High | High | High | High | High | High | 0 | 2.45E-09 | 3.64 |
| FALSE | High | ILVGHLEPS | [R].ILVGHL | 1xPhosphc | Q6IQX8 1x | 2.98E-05 | 0 | 1 | 3 | 1 | Q6IQX8 | Zinc finger | Q6IQX8 [8 | 0 | 2612.332 | 0 | -1.42 | 0.00219 | 3.26 | 144.7 | 55.3 | 10.38 | 15.07 | | High | High | High | High | High | High | High | High | High | High | High | 0 | 4.62E-06 | 2.97 |
| FALSE | High | IIVLDQSDL | [R].IIVLDQ | 1xPhosphc | Q7M6U3 1 | 2.72E-05 | 0 | 1 | 3 | 5 | Q7M6U3 | Inactive se | Q7M6U3 [ | 0 | 1526.755 | 0 | -0.76 | 0.00291 | 1.47 | 124.8 | 75.2 | 6.56 | 13.5 | | High | High | High | High | High | High | High | High | High | High | High | 0 | 8.23E-06 | 3.04 |
| FALSE | High | LIVPNSNG | [R].LIVPNS | 1xPhosphc | P09450 1xl | 1.08E-05 | 0 | 1 | 4 | 3 | P09450 | Transcripti | P09450 [9( | 0 | 2941.506 | 0 | 1.23 | 0.005408 | 4.19 | 57.8 | 142.2 | 14.28 | 15.3 | | High | High | High | High | High | High | High | High | High | High | High | 0 | 1.48E-06 | 4.34 |
| FALSE | High | IIYCSPGLVS | [R].IIYCSPG | 1xCarbami | Q8CFJ9 1x | 0.000104 | 0 | 1 | 1 | 1 | Q8CFJ9 | GATOR cor | Q8CFJ9 [4! | 0 | 2654.378 | 0 | -0.96 | 0.006432 | 6.33 | 125.4 | 74.6 | 10.24 | 18.47 | | High | High | High | High | High | High | High | High | High | High | High | 0 | 1.89E-05 | 3.83 |
| FALSE | High | IIYDSDSESI | [R].IIYDSDS | 1xPhosphc | P35601 1xl | 0.00824 | 0.000736 | 1 | 3 | 4 | P35601 | Replication | P35601 [64 | 0 | 2380.157 | 0 | 0.59 | 0.004373 | 12.26 | 84.7 | 115.3 | 7.79 | 9.51 | | High | High | High | High | High | High | High | High | High | High | High | 0.001308 | 0.01386 | 3.3 |
| FALSE | High | IIYDSDSESI | [R].IIYDSDS | 2xPhosphc | P35601 2xl | 9.56E-06 | 0 | 1 | 3 | 5 | P35601 | Replication | P35601 [64 | 0 | 2460.124 | 0 | 0.7 | 0.00078 | 7.23 | 76.9 | 123.1 | 4.7 | 6.84 | | High | High | High | High | High | High | High | High | High | High | High | 0 | 6.04E-05 | 3.35 |
| FALSE | High | ILTFDQLAL | [K].ILTFDQ | 1xPhosphc | Q58EW0 1: | 1.45E-06 | 0 | 1 | 6 | 1 | Q58EW0 | MCG13247 | Q58EW0 [: | 0 | 2013.107 | 0 | -0.77 | 0.001338 | 1.32 | 127.1 | 72.9 | 4.04 | 7.34 | | High | High | High | High | High | High | High | High | High | High | High | 0 | 1.54E-07 | 3.27 |
| FALSE | High | LLYEDPTA/ | [K].LLYEDP | 1xPhosphc | Q8K015 1x | 0.00642 | 0.00062 | 1 | 3 | 2 | Q8K015 | Centromer | Q8K015 [2 | 0 | 3841.9 | 0 | -0.68 | 0.002521 | 0.77 | 123.3 | 76.7 | 6.54 | 12.07 | | High | High | High | High | High | High | High | High | High | High | High | 0.000275 | 0.002159 | 2.55 |
| FALSE | High | LLSQEAAD: | [K].LLSQEA | 1xPhosphc | Q80YR7 1x | 7.61E-05 | 0 | 1 | 4 | 2 | Q80YR7 | Claspin OS | Q80YR7 [1: | 1 | 4550.175 | 0 | -0.04 | 0.436767 | 19.11 | 103.9 | 96.1 | 14.6 | 13.24 | | High | High | High | High | High | High | High | High | High | High | High | 0 | 1.32E-05 | 5.21 |
| FALSE | High | IISPSDITPG | [K].IISPSDI | 1xPhosphc | A3KGI3 1xl | 0.00195 | 5.66E-05 | 1 | 2 | 4 | A3KGI3 | Kinetochor | A3KGI3 [7! | 0 | 2009.061 | 0 | -1.12 | 0.013608 | 3.3 | 137.5 | 62.5 | 25.34 | 1.36 | | High | High | High | High | High | High | High | High | High | High | High | 2.58E-05 | 0.000506 | 2.31 |
| FALSE | High | ILSDEQLLS | [R].ILSDEQ | 1xPhosphc | A0A172Q4 | 6.95E-05 | 0 | 1 | 4 | 1 | A0A172Q4 | Cysteine-ki | A0A172Q4 | 0 | 1712.867 | 0 | 3.03 | 0.002715 | 44.8 | 20.8 | 179.2 | 11.28 | 20.76 | | High | High | High | High | High | High | High | High | High | High | High | 0 | 1.2E-05 | 3.3 |
| FALSE | High | ILSDSEDCE | [K].ILSDSEI | 1xCarbami | Q921C3 1x | 0.001489 | 5.66E-05 | 1 | 5 | 2 | Q921C3 | Bromodom | Q921C3 [1 | 0 | 1862.85 | 0 | -0.94 | 0.003362 | 4.15 | 131.3 | 68.7 | 8.51 | 6.75 | | High | High | High | High | High | High | High | High | High | High | High | 2.58E-05 | 0.000371 | 3.21 |
| FALSE | High | ILSDSEDCE | [K].ILSDSEI | 1xCarbami | Q921C3 2x | 0.004613 | 0.000456 | 1 | 5 | 4 | Q921C3 | Bromodom | Q921C3 [1 | 0 | 1942.816 | 0 | -0.98 | 0.000961 | 1.83 | 131.7 | 68.3 | 6.51 | 5.28 | | High | High | High | High | High | High | High | High | High | High | High | 0.00253 | 0.02744 | 2.41 |
| FALSE | High | LLSDSLPPS | [R].LLSDSLI | 1xPhosphc | Q80TM2 1: | 0.000157 | 0 | 1 | 3 | 5 | Q80TM2 | MKIAA102 | Q80TM2 [1 | 0 | 2343.143 | 0 | 0.81 | 0.023461 | 1.1 | 77.1 | 122.9 | 19.93 | 7.39 | | High | High | High | High | High | High | High | High | High | High | High | 0.000183 | 0.001252 | 2.6 |
| FALSE | High | ILSDVTHSA | [R].ILSDVTI | 1xPhosphc | Q3UHJ0 1x | 0.091627 | 0.007146 | 1 | 2 | 1 | Q3UHJ0 | AP2-associ | Q3UHJ0 [6 | 0 | 2037.062 | 0 | 0.03 | 0.763547 | 55.27 | 97.8 | 102.2 | 28.57 | 60.57 | | High | High | High | High | High | High | High | High | High | High | High | 0.003646 | 0.039 | 2.88 |
| FALSE | High | ILSDVTHSA | [R].ILSDVTI | 1xPhosphc | Q3UHJ0 1x | 2.82E-05 | 0 | 1 | 2 | 4 | Q3UHJ0 | AP2-associ | Q3UHJ0 [6 | 0 | 2266.225 | 0 | 0.74 | 0.00201 | 10.25 | 77.4 | 122.6 | 11.81 | 4.53 | | High | High | High | High | High | High | High | High | High | High | High | 4.94E-05 | 0.000682 | 3.56 |
| FALSE | High | IISENSMDS | [K].IISENSM | 1xCarbami | E9Q5F9 1x | 7.54E-05 | 0 | 1 | 4 | 3 | E9Q5F9 | Histone-lys | E9Q5F9 [1! | 0 | 2853.319 | 0 | 0.37 | 0.212809 | 19.16 | 88.1 | 111.9 | 19.71 | 16.5 | | High | High | High | High | High | High | High | High | High | High | High | 0 | 1.57E-05 | 5.07 |
| FALSE | High | IISENSMDS | [K].IISENSM | 1xCarbami | E9Q5F9 1x | 1.68E-07 | 0 | 1 | 4 | 7 | E9Q5F9 | Histone-lys | E9Q5F9 [1! | 0 | 2868.33 | 0 | 0.43 | 0.005771 | 1.88 | 86.1 | 113.9 | 5.26 | 11.79 | | High | High | High | High | High | High | High | High | High | High | High | 0 | 2.7E-08 | 5.41 |
| FALSE | High | LLSESQK | [R].LLSESQ | 1xPhosphc | Q6PAL0 1x | 0.019791 | 0.001438 | 1 | 1 | 1 | Q6PAL0 | BEN domai | Q6PAL0 [1( | 0 | 1342.738 | 0 | 0.72 | 0.058173 | 4.21 | 75.5 | 124.5 | 17.47 | 22.95 | | High | High | High | High | High | High | High | High | High | High | High | 0.000773 | 0.006746 | 1.92 |
| FALSE | High | LLSFSPEEFI | [R].LLSFSPI | 1xPhosphc | Q7TPM1 1: | 2.1E-05 | 0 | 1 | 3 | 1 | Q7TPM1 | Protein PRI | Q7TPM1 [1 | 0 | 2046.096 | 0 | -0.42 | 0.013787 | 6.16 | 114.2 | 85.8 | 5.84 | 7.83 | | High | High | High | High | High | High | High | High | High | High | High | 0 | 3.11E-06 | 2.79 |
| FALSE | High | LLSGDSAK | [R].LLSGDS | 1xPhosphc | Q8CB77 1x | 0.094641 | 0.007408 | 1 | 1 | 1 | Q8CB77 | Elongin-A ( | Q8CB77 [2 | 0 | 1328.723 | 0 | 0.92 | 0.002264 | 12.88 | 72 | 128 | 8.12 | 7.7 | | High | High | High | High | High | High | High | High | High | High | High | 0.003754 | 0.04049 | 1.79 |
| FALSE | High | LLSGDSIEK | [R].LLSGDS | 1xPhosphc | E9Q1Q1 1x | 0.003402 | 0.00021 | 1 | 6 | 3 | E9Q1Q1 | Microtubu | E9Q1Q1 [1 | 0 | 1499.812 | 0 | -1.59 | 0.000163 | 0.85 | 149.5 | 50.5 | 3.63 | 8.5 | | High | High | High | High | High | High | High | High | High | High | High | 0.000267 | 0.001541 | 2.51 |
| FALSE | High | LLSGGGR | [R].LLSGGG | 1xPhosphc | Q9D309 1x | 0.094641 | 0.007408 | 1 | 1 | 1 | Q9D309 | Protein FAI | Q9D309 [7 | 0 | 968.5128 | 0 | -1.21 | 0.01089 | 6.96 | 139.3 | 60.7 | 15.52 | 21.11 | | High | High | High | High | High | High | High | High | High | High | High | 0.003754 | 0.04057 | 0.78 |
| FALSE | High | LLSPTHEK | [R].LLSPTH | 1xPhosphc | P52479-2 1 | 0.006306 | 0.00062 | 1 | 3 | 1 | P52479-2 | Isoform 2 c | P52479-2 [ | 0 | 1462.807 | 0 | -1.86 | 0.00022 | 0.44 | 156.9 | 43.1 | 6.49 | 17.55 | | High | High | High | High | High | High | High | High | High | High | High | 0.000275 | 0.001872 | 1.63 |
| FALSE | High | LLSGNESSS | [K].LLSGNE | 1xPhosphc | Q9CVI2 1xl | 0.000916 | 0 | 1 | 4 | 2 | Q9CVI2 | Protein FAI | Q9CVI2 [8( | 0 | 1559.808 | 0 | -0.26 | 0.007968 | 5.2 | 108.9 | 91.1 | 4.18 | 10.37 | | High | High | High | High | High | High | High | High | High | High | High | 0 | 0.000217 | 1.91 |
| FALSE | High | LISGPLSPM | [K].LISGPLS | 1xPhosphc | Q7TSC1 1x | 0.016021 | 0.001124 | 1 | 3 | 1 | Q7TSC1 | Protein PRI | Q7TSC1 [1 | 0 | 1466.764 | 0 | 1.12 | 0.011423 | 14.66 | 63 | 137 | 5.51 | 23.53 | | High | High | High | High | High | High | High | High | High | High | High | 0.000585 | 0.005341 | 1.67 |
| FALSE | High | LISGPLSPM | [K].LISGPLS | 1xPhosphc | Q7TSC1 1x | 0.000908 | 0 | 1 | 3 | 4 | Q7TSC1 | Protein PRI | Q7TSC1 [1 | 0 | 1482.759 | 0 | 0.87 | 0.013136 | 14.71 | 69.2 | 130.8 | 9.53 | 20.14 | | High | High | High | High | High | High | High | High | High | High | High | 0.001214 | 0.01313 | 1.94 |
| FALSE | High | IISFSGTEK | [K].IISFSGT | 1xPhosphc | Q8CIN4 1x | 0.003138 | 0.00016 | 1 | 1 | 1 | Q8CIN4 | Serine/thre | Q8CIN4 [5: | 0 | 1632.901 | 0 | 2.54 | 0.003445 | 14.35 | 28.8 | 171.2 | 12.71 | 22.88 | | High | High | High | High | High | High | High | High | High | High | High | 7.19E-05 | 0.000856 | 2.81 |
| FALSE | High | LLSPAEVSS | [K].LLSPAE | 1xPhosphc | Q3U0P1 1x | 0.006031 | 0.00062 | 1 | 3 | 1 | Q3U0P1 | Partner an | Q3U0P1 [4 | 0 | 2045.108 | 0 | -0.66 | 0.00251 | 3.36 | 122.3 | 77.7 | 5.19 | 8.04 | | High | High | High | High | High | High | High | High | High | High | High | 0.000275 | 0.001779 | 2.57 |
| FALSE | High | LLSPAEVSS | [K].LLSPAE | 2xPhosphc | Q3U0P1 2x | 0.044358 | 0.003264 | 1 | 3 | 1 | Q3U0P1 | Partner an | Q3U0P1 [4 | 0 | 2125.075 | 0 | -0.48 | 0.014396 | 10.26 | 120.3 | 79.7 | 12.4 | 9.89 | | High | High | High | High | High | High | High | High | High | High | High | 0.001665 | 0.01676 | 2.78 |
| FALSE | High | IISPGSSTPS | [K].IISPGSS | 1xPhosphc | Q6NXI6 1x | 0.000589 | 0 | 1 | 3 | 1 | Q6NXI6 | Regulation | Q6NXI6 [7 | 0 | 1598.799 | 0 | -0.8 | 0.00916 | 0.92 | 127 | 73 | 13.56 | 11.18 | | High | High | High | High | High | High | High | High | High | High | High | 0 | 0.000131 | 2.22 |
| FALSE | High | LLSPIASNR | [R].LLSPIAS | 1xPhosphc | Q9CW46 1 | 0.018447 | 0.001324 | 1 | 2 | 2 | Q9CW46 | Ribonuclec | Q9CW46 [! | 0 | 1279.697 | 0 | 0.05 | 0.564022 | 6.53 | 98.7 | 101.3 | 5.85 | 3.87 | | High | High | High | High | High | High | High | High | High | High | High | 0.003809 | 0.04107 | 2.06 |
| FALSE | High | LISPNK | [R].LISPNK | 1xPhosphc | Q5DTT3 1x | 0.075304 | 0.005727 | 1 | 2 | 2 | Q5DTT3 | Protein FAI | Q5DTT3 [1 | 0 | 1209.701 | 0 | -0.5 | 0.000719 | 2.22 | 117 | 83 | 3.04 | 5.86 | | High | High | High | High | High | High | High | High | High | High | High | 0.004164 | 0.04493 | 1.79 |
| FALSE | High | ILSPNMTN | [R].ILSPNM | 1xPhosphc | Q9JJ11 1xF | 0.010663 | 0.000772 | 1 | 7 | 1 | Q9JJ11 | Transformi | Q9JJ11 [69 | 0 | 1555.832 | 0 | -0.07 | 0.413376 | 4.77 | 102.5 | 97.5 | 7.28 | 23.49 | | High | High | High | High | High | High | High | High | High | High | High | 0.000399 | 0.003374 | 2.31 |
| FALSE | High | ILSPNMTN | [R].ILSPNM | 1xPhosphc | Q9JJ11 1xP | 0.000505 | 0 | 1 | 7 | 3 | Q9JJ11 | Transformi | Q9JJ11 [69 | 0 | 1571.827 | 0 | -0.37 | 0.809351 | 15.41 | 110.9 | 89.1 | 13.37 | 33.84 | | High | High | High | High | High | High | High | High | High | High | High | 0 | 0.000111 | 3.14 |
| FALSE | High | LLSPPTPK | [R].LLSPPTI | 2xPhosphc | Q8BK35 2x | 0.039011 | 0.002718 | 1 | 3 | 1 | Q8BK35 | Ribosome I | Q8BK35 [1 | 0 | 1470.777 | 0 | 0.06 | 0.90342 | 0.9 | 102.4 | 97.6 | 9.65 | 8.2 | | High | High | High | High | High | High | High | High | High | High | High | 0.001421 | 0.01447 | 2.18 |
| FALSE | High | LLSPQQPAI | [R].LLSPQC | 1xPhosphc | Q63943-2 : | 0.004411 | 0.000409 | 2 | 6 | 2 | Q63943-2; | Isoform Mu | Q63943-2 | 0 | 1559.851 | 0 | 0.97 | 0.001647 | 22.45 | 67.1 | 132.9 | 11.41 | 5.51 | Shared | High | High | High | High | High | High | High | High | High | High | High | 0.000183 | 0.001258 | 1.98 |
| FALSE | High | LISPQTLSIK | [K].LISPQTI | 1xPhosphc | A2AKX3 1x | 0.078467 | 0.006104 | 1 | 2 | 1 | A2AKX3 | Probable h | A2AKX3 [1 | 0 | 1637.964 | 0 | 0.05 | 0.066435 | 2.95 | 96.7 | 103.3 | 6.95 | 5.33 | | High | High | High | High | High | High | High | High | High | High | High | 0.003122 | 0.03232 | 2 |
| FALSE | High | LLSGNESSS | [K].LLSGNE | 2xPhosphc | Q9CVI2 2xl | 0.100936 | 0.008097 | 1 | 4 | 1 | Q9CVI2 | Protein FAI | Q9CVI2 [8( | 0 | 1639.775 | 0 | -0.08 | 0.284441 | 12.95 | 103.3 | 96.7 | 11.47 | 9.14 | | High | High | High | High | High | High | High | High | High | High | High | 0.004072 | 0.04352 | 2.13 |
| FALSE | High | IIYGGSVTG | [R].IIYGGSV | 1xCarbami | P17751 1xl | 4.72E-05 | 0 | 1 | 2 | 1 | P17751 | Triosephos | P17751 [2! | 0 | 1864.964 | 0 | 1.61 | 0.000181 | 3.8 | 49.2 | 150.8 | 8.85 | 3.84 | | High | High | High | High | High | High | High | High | High | High | High | 0 | 7.7E-06 | 3.34 |
| FALSE | High | LMDGQETI | [K].LMDGC | 1xPhosphc | B2RUE3 1x | 0.037048 | 0.00249 | 1 | 3 | 1 | B2RUE3 | UTP14, U3 | B2RUE3 [4 | 0 | 2590.2 | 0 | -1.24 | 0.00028 | 0.92 | 137.7 | 62.3 | 3.82 | 7.18 | | High | High | High | High | High | High | High | High | High | High | High | 0.001308 | 0.01364 | 3.13 |
| FALSE | High | LMDGQETI | [K].LMDGC | 1xPhosphc | B2RUE3 1x | 1.32E-07 | 0 | 1 | 3 | 2 | B2RUE3 | UTP14, U3 | B2RUE3 [4 | 0 | 2606.194 | 0 | -0.72 | 0.001124 | 0.74 | 123.5 | 76.5 | 4.26 | 4.71 | | High | High | High | High | High | High | High | High | High | High | High | 0 | 1.04E-08 | 4.02 |
| FALSE | High | LNESDEQH | [K].LNESDE | 1xPhosphc | Q9R1C0 1x | 6.01E-08 | 0 | 1 | 1 | 2 | Q9R1C0 | Transcripti | Q9R1C0 [2 | 0 | 3179.508 | 0 | -1.85 | 0.000566 | 0.62 | 155.8 | 44.2 | 7.06 | 20.6 | | High | High | High | High | High | High | High | High | High | High | High | 0 | 4.33E-09 | 4.89 |
| FALSE | High | LNESDEQH | [K].LNESDE | 1xDeamida | Q9R1C0 1x | 2.76E-07 | 0 | 1 | 1 | 2 | Q9R1C0 | Transcripti | Q9R1C0 [2 | 0 | 3180.492 | 0 | -1.7 | 0.000392 | 3.56 | 154.4 | 45.6 | 9.14 | 32.6 | | High | High | High | High | High | High | High | High | High | High | High | 0 | 2.4E-08 | 5.34 |
| FALSE | High | LNESDEQH | [K].LNESDE | 1xPhosphc | Q9R1C0 1x | 1.05E-09 | 0 | 1 | 1 | 6 | Q9R1C0 | Transcripti | Q9R1C0 [2 | 0 | 3195.503 | 0 | -1.84 | 0.000429 | 1.76 | 156.3 | 43.7 | 5.52 | 11.95 | | High | High | High | High | High | High | High | High | High | High | High | 0 | 4.58E-11 | 4.85 |
| FALSE | High | LNETELTDI | [R].LNETEL | 1xPhosphc | Q6P5F6 1x | 6.62E-07 | 0 | 1 | 1 | 2 | Q6P5F6 | Zinc transp | Q6P5F6 [5 | 0 | 2580.285 | 0 | -0.25 | 0.051322 | 4.77 | 108.8 | 91.2 | 7.6 | 5.78 | | High | High | High | High | High | High | High | High | High | High | High | 0 | 0.000124 | 3.81 |
| FALSE | High | LNFDMTAS | [K].LNFDM | 1xPhosphc | Q80Y82 1x | 0.004491 | 0.000409 | 1 | 14 | 1 | Q80Y82 | Protein kin | Q80Y82 [4 | 0 | 1661.837 | 0 | 0.35 | 0.23258 | 17.87 | 87.6 | 112.4 | 5.43 | 21.1 | | High | High | High | High | High | High | High | High | High | High | High | 0.000183 | 0.001278 | 1.15 |
| FALSE | High | LNFDMTAS | [K].LNFDM | 1xPhosphc | Q80Y82 1x | 0.001659 | 5.66E-05 | 1 | 14 | 2 | Q80Y82 | Protein kin | Q80Y82 [4 | 0 | 1677.832 | 0 | 0.34 | 0.033021 | 9.89 | 89.5 | 110.5 | 4.38 | 8.53 | | High | High | High | High | High | High | High | High | High | High | High | 2.58E-05 | 0.000553 | 2.6 |
| FALSE | High | INFSDLDQI | [R].INFSDL | 1xPhosphc | Q9R0L6-2 : | 0.121159 | 0.009984 | 1 | 3 | 1 | Q9R0L6-2 | Isoform 2 c | Q9R0L6-2 | 0 | 1416.672 | 0 | 0.24 | 0.036036 | 6.64 | 90.7 | 109.3 | 9.55 | 4.92 | | High | High | High | High | High | High | High | High | High | High | High | 0.004974 | 0.05432 | 1.74 |
| FALSE | High | LNGHSGLI | [K].LNGHS | 1xPhosphc | Q8C0V9 1x | 0.001767 | 5.66E-05 | 1 | 2 | 1 | Q8C0V9 | FERM dom | Q8C0V9 [4 | 0 | 1575.902 | 0 | 0.56 | 0.011089 | 14.91 | 80.8 | 119.2 | 4.59 | 14.14 | | High | High | High | High | High | High | High | High | High | High | High | 2.58E-05 | 0.000452 | 2.24 |
| FALSE | High | LNHSPPQS | [R].LNHSPF | 1xDeamida | Q6NZN0 1x | 0.001898 | 5.66E-05 | 1 | 8 | 1 | Q6NZN0 | RNA-bindir | Q6NZN0 [1 | 0 | 1519.71 | 0 | -1.04 | 0.000246 | 1.77 | 133.9 | 66.1 | 3.85 | 5.24 | | High | High | High | High | High | High | High | High | High | High | High | 2.58E-05 | 0.00049 | 1.53 |
| FALSE | High | INHTVILDD | [R].INHTVI | 1xPhosphc | Q9CXG3 1x | 0.00359 | 0.00021 | 1 | 4 | 1 | Q9CXG3 | Peptidyl-pr | Q9CXG3 [1 | 1 | 3478.734 | 0 | -0.41 | 0.514341 | 14.42 | 118 | 82 | 21.34 | 42.52 | | High | High | High | High | High | High | High | High | High | High | High | 9.42E-05 | 0.001004 | 3.24 |
| FALSE | High | INHTVILDD | [R].INHTVI | 1xPhosphc | Q9CXG3 1x | 0.000107 | 0 | 1 | 4 | 4 | Q9CXG3 | Peptidyl-pr | Q9CXG3 [1 | 1 | 3707.897 | 0 | -0.46 | 0.004833 | 8.05 | 117.2 | 82.8 | 7.2 | 3.86 | | High | High | High | High | High | High | High | High | High | High | High | 0 | 2.53E-05 | 5.25 |
| FALSE | High | LNHVAAGL | [K].LNHVA | 1xPhosphc | F6RDI8 1xF | 1.4E-09 | 0 | 1 | 11 | 2 | F6RDI8 | Serine/argi | F6RDI8 [20 | 0 | 1944.108 | 0 | -0.63 | 0.004077 | 2.42 | 125.2 | 74.8 | 9.01 | 12.49 | | High | High | High | High | High | High | High | High | High | High | High | 0 | 6.28E-11 | 3.73 |
| FALSE | High | LNHVAAGL | [K].LNHVA | 1xDeamida | F6RDI8 1xF | 9.82E-06 | 0 | 1 | 11 | 1 | F6RDI8 | Serine/argi | F6RDI8 [20 | 0 | 1945.092 | 0 | -0.55 | 0.003358 | 3 | 116.9 | 83.1 | 3.71 | 13.65 | | High | High | High | High | High | High | High | High | High | High | High | 0 | 1.32E-06 | 3.7 |
| FALSE | High | LNESCCSSI | [K].LNESCC | 2xCarbami | B9EKI3 2xP | 0.063258 | 0.004664 | 1 | 2 | 1 | B9EKI3 | TATA elem | B9EKI3 [30 | 0 | 1963.717 | 0 | 0.44 | 0.070266 | 5.06 | 83.2 | 116.8 | 3.61 | 23.27 | | High | High | High | High | High | High | High | High | High | High | High | 0.002403 | 0.0252 | 2.23 |
| FALSE | High | LNLAEFLNE | [K].LNLAEF | 1xPhosphc | E9PWG6 1: | 7.77E-06 | 0 | 1 | 2 | 16 | E9PWG6 | Non-SMC c | E9PWG6 [! | 0 | 1674.783 | 0 | 0.04 | 0.310702 | 1.47 | 95.4 | 104.6 | 7.35 | 7.66 | | High | High | High | High | High | High | High | High | High | High | High | 0 | 8.01E-06 | 3.01 |
| FALSE | High | LNLQSLGD | [K].LNLQSL | 2xPhosphc | Q6P1H6 2x | 0.004696 | 0.000456 | 1 | 3 | 2 | Q6P1H6 | Ankyrin re | Q6P1H6 [7 | 0 | 2385.15 | 0 | 0.17 | 0.404359 | 5.94 | 94.1 | 105.9 | 10.26 | 12.46 | | High | High | High | High | High | High | High | High | High | High | High | 0.000204 | 0.001352 | 2.51 |
| FALSE | High | LNLSCIHSP | [K].LNLSCII | 1xCarbami | Q6DFW4 1 | 0.000231 | 0 | 1 | 3 | 1 | Q6DFW4 | Nucleolar ı | Q6DFW4 [ | 0 | 2191.097 | 0 | -0.08 | 0.783367 | 6.19 | 102.1 | 97.9 | 9.81 | 8.6 | | High | High | High | High | High | High | High | High | High | High | High | 0 | 4.59E-05 | 2.69 |
| FALSE | High | LNLSHSETN | [K].LNLSHS | 2xCarbami | E9QLH9 1x | 1.67E-08 | 0 | 1 | 2 | 2 | E9QLH9 | PEAK1-rela | E9QLH9 [8 | 0 | 3378.483 | 0 | 0.86 | 0.010173 | 23.99 | 67.7 | 132.3 | 7.29 | 16.36 | | High | High | High | High | High | High | High | High | High | High | High | 0 | 1.02E-09 | 4.03 |
| FALSE | High | LNLTLSPAK | [K].LNLTLS | 1xPhosphc | Q9WUF3 1 | 0.002058 | 5.66E-05 | 1 | 3 | 2 | Q9WUF3 | CASP8-assc | Q9WUF3 [! | 0 | 1494.87 | 0 | -0.75 | 0.002468 | 5.75 | 125.4 | 74.6 | 5.36 | 11.1 | | High | High | High | High | High | High | High | High | High | High | High | 2.58E-05 | 0.000537 | 2.1 |
| FALSE | High | INLTPLK | [R].INLTPLI | 1xPhosphc | Q9D0N7 1x | 0.087983 | 0.006639 | 1 | 1 | 1 | Q9D0N7 | Chromatin | Q9D0N7 [5 | 0 | 1107.638 | 0 | -0.06 | 0.90008 | 8.35 | 101.6 | 98.4 | 15.54 | 12.81 | | High | High | High | High | High | High | High | High | High | High | High | 0.003402 | 0.03708 | 1.59 |
| FALSE | High | LNITPTK | [K].LNITPTI | 1xPhosphc | Q62383 1x | 0.036103 | 0.002342 | 1 | 1 | 1 | Q62383 | Transcripti | Q62383 [6 | 0 | 1324.764 | 0 | -1.22 | 0.000263 | 0.46 | 140.3 | 59.7 | 5.03 | 10.25 | | High | High | High | High | High | High | High | High | High | High | High | 0.00123 | 0.01324 | 1.33 |
| FALSE | High | LNLTSPK | [K].LNLTSP | 1xPhosphc | E9PUR0 1x | 0.049971 | 0.003641 | 1 | 4 | 1 | E9PUR0 | Ankyrin re | E9PUR0 [1( | 0 | 1081.586 | 0 | -1.01 | 0.199483 | 3.01 | 132.9 | 67.1 | 60.93 | 45.68 | | High | High | High | High | High | High | High | High | High | High | High | 0.001902 | 0.01934 | 1.18 |
| FALSE | High | LNLTSPK | [K].LNLTSP | 1xPhosphc | E9PUR0 1x | 0.045898 | 0.003408 | 1 | 4 | 2 | E9PUR0 | Ankyrin re | E9PUR0 [1( | 0 | 1310.749 | 0 | -1.43 | 0.000519 | 2.1 | 149 | 51 | 7.27 | 13.41 | | High | High | High | High | High | High | High | High | High | High | High | 0.00196 | 0.01992 | 1.58 |
| FALSE | High | LNMTLPDA | [R].LNMTL | 1xDeamida | Q9DC13 2x | 0.093123 | 0.007276 | 1 | 4 | 1 | Q9DC13 | Lysosomal | Q9DC13 [2 | 1 | 3820.884 | 0 | -0.01 | 0.757344 | 17.19 | 98.4 | 101.6 | 20.46 | 9.08 | | High | High | High | High | High | High | High | High | High | High | High | 0.003685 | 0.03963 | 4.32 |
| FALSE | High | LNPDSPQQ | [R].LNPDSF | 1xPhosphc | E9QKG8 1x | 1.86E-05 | 0 | 1 | 9 | 2 | E9QKG8 | Holliday ju | E9QKG8 [5 | 0 | 1866.936 | 0 | -0.84 | 0.000816 | 1.18 | 128.3 | 71.7 | 10.69 | 7.79 | | High | High | High | High | High | High | High | High | High | High | High | 0 | 2.72E-06 | 3.28 |
| FALSE | High | INPPSSGGT | [R].INPPSS | 1xPhosphc | A0A0R4J12 | 0.000117 | 0 | 1 | 19 | 2 | A0A0R4J12 | POU doma | A0A0R4J12 | 0 | 1737.862 | 0 | -0.63 | 0.025643 | 14.24 | 122.7 | 77.3 | 14.7 | 12.77 | | High | High | High | High | High | High | High | High | High | High | High | 0 | 2.14E-05 | 2.41 |
| FALSE | High | INPPSSGGT | [R].INPPSS | 1xPhosphc | A0A0R4J12 | 2.52E-07 | 0 | 1 | 19 | 3 | A0A0R4J12 | POU doma | A0A0R4J12 | 0 | 1967.025 | 0 | -0.58 | 0.000408 | 1.24 | 119.7 | 80.3 | 1.99 | 3.31 | | High | High | High | High | High | High | High | High | High | High | High | 0 | 2.17E-08 | 4.55 |
| FALSE | High | INPSVNPGI | [R].INPSVN | 1xPhosphc | P70365 1xl | 3.98E-05 | 0 | 1 | 5 | 1 | P70365 | Nuclear rec | P70365 [3{ | 0 | 2025.048 | 0 | 0.59 | 0.001593 | 5.23 | 79.4 | 120.6 | 11.79 | 8.15 | | High | High | High | High | High | High | High | High | High | High | High | 0 | 6.37E-06 | 3.37 |
| FALSE | High | LNLESSNSK | [R].LNLESS | 1xPhosphc | A2A6Q5-3 | 0.053473 | 0.00384 | 1 | 6 | 2 | A2A6Q5-3 | Isoform 3 c | A2A6Q5-3 | 0 | 1529.798 | 0 | 0.51 | 0.006569 | 5.42 | 82.1 | 117.9 | 8.12 | 7.29 | | High | High | High | High | High | High | High | High | High | High | High | 0.001999 | 0.02075 | 1.63 |
| FALSE | High | LNESCCSSI | [K].LNESCC | 2xCarbami | B9EKI3 1xP | 0.001049 | 0 | 1 | 2 | 3 | B9EKI3 | TATA elem | B9EKI3 [30 | 0 | 1883.75 | 0 | 0.71 | 0.045035 | 16.85 | 75.8 | 124.2 | 15.49 | 21.98 | | High | High | High | High | High | High | High | High | High | High | High | 0 | 0.000252 | 3.32 |
| FALSE | High | LNDSLYNEI | [K].LNDSLY | 1xCarbami | B7ZNX2 1x | 0.0006 | 0 | 1 | 4 | 1 | B7ZNX2 | Anaphase ı | B7ZNX2 [5( | 0 | 3082.418 | 0 | 0.33 | 0.001325 | 2.9 | 89.2 | 110.8 | 14.55 | 12.2 | | High | High | High | High | High | High | High | High | High | High | High | 0 | 0.000135 | 2.69 |
| FALSE | High | LNDPFQPF | [K].LNDPF( | 1xDeamida | H3BK65 1x | 0.001337 | 0 | 1 | 5 | 1 | H3BK65 | Epidermal | H3BK65 [8 | 0 | 2212.073 | 0 | 0.97 | 0.019804 | 10.46 | 65.4 | 134.6 | 14.13 | 20.31 | | High | High | High | High | High | High | High | High | High | High | High | 0 | 0.000331 | 2.54 |
| FALSE | High | LMDGQETI | [K].LMDGC | 2xPhosphc | B2RUE3 2x | 0.018774 | 0.001362 | 1 | 3 | 1 | B2RUE3 | UTP14, U3 | B2RUE3 [4 | 0 | 2670.166 | 0 | -0.41 | 0.013634 | 8.14 | 115.4 | 84.6 | 6.67 | 13.02 | | High | High | High | High | High | High | High | High | High | High | High | 0.000723 | 0.006348 | 2.57 |
| FALSE | High | LMEALESPI | [K].LMEALI | 1xPhosphc | A0A140T8! | 1E-05 | 0 | 1 | 1 | 1 | A0A140T8! | Chromosoı | A0A140T8! | 0 | 1938.933 | 0 | -0.04 | 0.377166 | 29.65 | 105.3 | 94.7 | 24.14 | 30.21 | | High | High | High | High | High | High | High | High | High | High | High | 0 | 1.35E-06 | 2.86 |
| FALSE | High | IMEETSGIE | [R].IMEETS | 1xPhosphc | A0A1W2P6 | 0.000193 | 0 | 1 | 6 | 4 | A0A1W2P6 | Bromodom | A0A1W2P6 | 0 | 2928.266 | 0 | -0.7 | 0.003062 | 6.29 | 123.7 | 76.3 | 7.91 | 11.61 | | High | High | High | High | High | High | High | High | High | High | High | 0 | 6.1E-05 | 5.07 |
| FALSE | High | IMEETSGIE | [R].IMEETS | 1xPhosphc | A0A1W2P6 | 0.000442 | 0 | 1 | 6 | 17 | A0A1W2P6 | Bromodom | A0A1W2P6 | 0 | 2944.26 | 0 | -0.47 | 0.002479 | 4.36 | 115.5 | 84.5 | 4.63 | 6.22 | | High | High | High | High | High | High | High | High | High | High | High | 0 | 0.000187 | 4.57 |
| FALSE | High | IMEETSGIE | [R].IMEETS | 1xDeamida | A0A1W2P6 | 0.024638 | 0.001612 | 1 | 6 | 2 | A0A1W2P6 | Bromodom | A0A1W2P6 | 0 | 2945.244 | 0 | -0.72 | 0.006901 | 1.25 | 119.6 | 80.4 | 12.44 | 11.31 | | High | High | High | High | High | High | High | High | High | High | High | 0.000872 | 0.008706 | 3.78 |
| FALSE | High | LMELLNET | [R].LMELLN | 1xCarbami | B2ZRS3 1xl | 0.000793 | 0 | 1 | 6 | 1 | B2ZRS3 | Depdc6-00 | B2ZRS3 [22 | 0 | 3000.368 | 0 | 2.5 | 0.00148 | 17.06 | 31.4 | 168.6 | 18.4 | 13.84 | | High | High | High | High | High | High | High | High | High | High | High | 0 | 0.000184 | 3.03 |
| FALSE | High | LMELLNET | [R].LMELLN | 1xCarbami | B2ZRS3 1xl | 2.59E-06 | 0 | 1 | 6 | 2 | B2ZRS3 | Depdc6-00 | B2ZRS3 [22 | 0 | 3016.363 | 0 | 2.38 | 0.004967 | 17.27 | 33 | 167 | 23.53 | 22.4 | | High | High | High | High | High | High | High | High | High | High | High | 0 | 2.96E-07 | 3.86 |
| FALSE | High | LMETNIGF: | [K].LMETN | 1xPhosphc | E9QLG3 1x | 1.22E-06 | 0 | 1 | 5 | 2 | E9QLG3 | MAX gene- | E9QLG3 [8 | 0 | 3132.609 | 0 | -0.16 | 0.453413 | 11.36 | 103 | 97 | 6.75 | 15.97 | | High | High | High | High | High | High | High | High | High | High | High | 0 | 1.27E-07 | 3.37 |
| FALSE | High | LMETNIGF: | [K].LMETN | 1xPhosphc | E9QLG3 1x | 6.12E-08 | 0 | 1 | 5 | 1 | E9QLG3 | MAX gene- | E9QLG3 [8 | 0 | 3148.604 | 0 | -0.3 | 0.108513 | 9.19 | 110.5 | 89.5 | 14.04 | 10.15 | | High | High | High | High | High | High | High | High | High | High | High | 0 | 4.4E-09 | 4.07 |
| FALSE | High | IMETSPTLL | [R].IMETSF | 1xPhosphc | Q6ZQ33 1x | 0.000524 | 0 | 1 | 4 | 1 | Q6ZQ33 | MKIAA085 | Q6ZQ33 [1 | 0 | 2347.275 | 0 | 0.56 | 0.010136 | 17.43 | 80.9 | 119.1 | 7.35 | 7.1 | | High | High | High | High | High | High | High | High | High | High | High | 0 | 0.000115 | 2.63 |
| FALSE | High | IMETSPTLL | [R].IMETSF | 1xPhosphc | Q6ZQ33 1x | 0.077188 | 0.005893 | 1 | 4 | 1 | Q6ZQ33 | MKIAA085 | Q6ZQ33 [1 | 0 | 2363.27 | 0 | 0.15 | 0.396111 | 18.25 | 93.6 | 106.4 | 10.75 | 6.85 | | High | High | High | High | High | High | High | High | High | High | High | 0.003064 | 0.0319 | 2.85 |
| FALSE | High | IMGPNYTP | [R-].IMGPN | 1xPhosphc | P58252 1xl | 0.001916 | 5.66E-05 | 1 | 15 | 3 | P58252 | Elongation | P58252 [42 | 0 | 1631.827 | 0 | -0.67 | 0.016084 | 4.8 | 121.8 | 78.2 | 5.75 | 13.87 | | High | High | High | High | High | High | High | High | High | High | High | 2.58E-05 | 0.000492 | 2.53 |
| FALSE | High | LMHDASDS | [K].LMHDA | 2xPhosphc | Q3UYC8 2x | 0.109329 | 0.008865 | 1 | 6 | 1 | Q3UYC8 | Putative ur | Q3UYC8 [9 | 0 | 2623.051 | 0 | -0.04 | 0.729754 | 9.4 | 98.4 | 101.6 | 12.03 | 13.98 | | High | High | High | High | High | High | High | High | High | High | High | 0.004451 | 0.04797 | 2.68 |
| FALSE | High | LMHDASDS | [K].LMHDA | 1xPhosphc | Q3UYC8 1x | 1.21E-05 | 0 | 1 | 6 | 1 | Q3UYC8 | Putative ur | Q3UYC8 [9 | 0 | 2756.253 | 0 | 0.22 | 0.925932 | 23.83 | 93.4 | 106.6 | 10.21 | 30.78 | | High | High | High | High | High | High | High | High | High | High | High | 0 | 1.67E-06 | 4.87 |
| FALSE | High | LMHDASDS | [K].LMHDA | 1xPhosphc | Q3UYC8 1x | 3.78E-06 | 0 | 1 | 6 | 5 | Q3UYC8 | Putative ur | Q3UYC8 [9 | 0 | 2772.248 | 0 | 0.03 | 0.480197 | 13.58 | 99 | 101 | 9.14 | 14.75 | | High | High | High | High | High | High | High | High | High | High | High | 0 | 4.52E-07 | 3.95 |

| | | | | | | | | | | | | | | | | | | | | | | | | | | | | | | | | | | | | | | |
|---|---|---|---|---|---|---|---|---|---|---|---|---|---|---|---|---|---|---|---|---|---|---|---|---|---|---|---|---|---|---|---|---|---|---|---|---|---|---|
| FALSE | High | LMHDASDS | [K].LMHDA | 2xPhospho | Q3UYC8 2x | 1.31E-06 | 0 | 1 | 6 | 4 | Q3UYC8 | Putative ur | Q3UYC8 [9 | 0 | 2836.219 | 0 | 0.15 | 0.952052 | 20.36 | 94.8 | 105.2 | 6.89 | 27.03 | | High | High | High | High | High | High | High | High | High | High | High | 0 | 1.39E-07 | 4.4 |
| FALSE | High | LMHGLSED | [K].LMHGL | 1xPhospho | Q6GTE9 1x | 6.31E-06 | 0 | 1 | 4 | 2 | Q6GTE9 | NIMA (Nev | Q6GTE9 [6 | 0 | 2578.2 | 0 | -0.36 | 0.851285 | 5.85 | 111.8 | 88.2 | 10.86 | 25.05 | | High | High | High | High | High | High | High | High | High | High | High | 0 | 8.08E-07 | 3.88 |
| FALSE | High | LMKPPSPE | [R].LMKPP | 1xPhospho | Q69ZX6 1x | 1.11E-05 | 0 | 2 | 2 | 4 | Q69ZX6; Q | MORC fam | Q69ZX6 [8 | 0 | 3661.692 | 0 | -0.49 | 0.006251 | 6.65 | 116.4 | 83.6 | 4.12 | 6.54 | Shared | High | High | High | High | High | High | High | High | High | High | High | 0 | 1.52E-06 | 3.58 |
| FALSE | High | IMPDSNDS | [R].IMPDSN | 1xPhospho | Q3UWM7 | 0.01894 | 0.001362 | 1 | 5 | 2 | Q3UWM7 | Putative ur | Q3UWM7 | 1 | 3192.554 | 0 | -1.33 | 0.000742 | 2.45 | 141.6 | 58.4 | 8.18 | 12.21 | | High | High | High | High | High | High | High | High | High | High | High | 0.00074 | 0.006445 | 2.28 |
| FALSE | High | LMTGDTYT | [R].LMTGD | 1xPhospho | Q3SYK5 1x | 0.000651 | 0 | 2 | 11 | 1 | Q3SYK5; F8 | Tyrosine-p | Q3SYK5 [4 | 0 | 1990.971 | 0 | -1.15 | 0.006493 | 2.92 | 136.5 | 63.5 | 17.56 | 11.25 | Shared | High | High | High | High | High | High | High | High | High | High | High | 0 | 0.000148 | 2.72 |
| FALSE | High | IMVGSTDD | [K].IMVGST | 1xPhospho | Q6PAM0 1 | 0.000191 | 0 | 1 | 3 | 2 | Q6PAM0 | 5'-AMP-act | Q6PAM0 [3 | 0 | 2449.197 | 0 | -0.38 | 0.12344 | 5.93 | 112.6 | 87.4 | 2.65 | 16.48 | | High | High | High | High | High | High | High | High | High | High | High | 0 | 3.73E-05 | 3.26 |
| FALSE | High | INAAVASPI | [R].INAAVA | 1xPhospho | Q60953 1x | 2.38E-06 | 0 | 1 | 4 | 4 | Q60953 | Protein PM | Q60953 [3 | 0 | 4097.012 | 0 | 0.13 | 0.659119 | 23.67 | 97 | 103 | 8.41 | 12.04 | | High | High | High | High | High | High | High | High | High | High | High | 0 | 2.7E-07 | 5.88 |
| FALSE | High | INAAVASPI | [R].INAAVA | 1xDeamida | Q60953 1x | 1.02E-05 | 0 | 1 | 4 | 13 | Q60953 | Protein PM | Q60953 [3 | 0 | 4097.996 | 0 | 0.26 | 0.129997 | 25.51 | 91.2 | 108.8 | 10.7 | 9.33 | | High | High | High | High | High | High | High | High | High | High | High | 0 | 1.37E-06 | 4.36 |
| FALSE | High | INAAVASPI | [R].INAAVA | 2xPhospho | Q60953 2x | 0.099328 | 0.008003 | 1 | 4 | 1 | Q60953 | Protein PM | Q60953 [3 | 0 | 4176.978 | 0 | -0.65 | 0.008072 | 8.9 | 119.5 | 80.5 | 5.61 | 16.18 | | High | High | High | High | High | High | High | High | High | High | High | 0.004006 | 0.04263 | 3.11 |
| FALSE | High | LNATASLE( | [R].LNATAS | 1xPhospho | P56695 1xl | 0.002895 | 0.00016 | 1 | 4 | 1 | P56695 | Wolframin | P56695 [27 | 0 | 1727.898 | 0 | 0.73 | 0.005198 | 6.85 | 73.9 | 126.1 | 6.01 | 10.56 | | High | High | High | High | High | High | High | High | High | High | High | 7.19E-05 | 0.000785 | 2.83 |
| FALSE | High | INDELEGL( | [R].INDELE | 1xPhospho | Q6PAL0 1x | 4.09E-09 | 0 | 1 | 1 | 3 | Q6PAL0 | BEN domai | Q6PAL0 [4 | 0 | 2351.097 | 0 | -0.89 | 0.000208 | 0.47 | 128.6 | 71.4 | 4.62 | 5.1 | | High | High | High | High | High | High | High | High | High | High | High | 0 | 2.79E-09 | 4.16 |
| FALSE | High | INDELEGL( | [R].INDELE | 1xCarbami | Q6PAL0 1x | 0.000314 | 0 | 1 | 1 | 2 | Q6PAL0 | BEN domai | Q6PAL0 [4 | 1 | 4477.053 | 0 | -0.18 | 0.211367 | 14.74 | 109.9 | 90.1 | 15.38 | 20.33 | | High | High | High | High | High | High | High | High | High | High | High | 0 | 6.5E-05 | 4.74 |
| FALSE | High | LNDPFQPF | [K].LNDPF( | 1xPhospho | H3BK65 1x | 5.81E-05 | 0 | 1 | 5 | 1 | H3BK65 | Epidermal | H3BK65 [8 | 0 | 1981.926 | 0 | 1.15 | 0.182828 | 29.57 | 66.8 | 133.2 | 40.25 | 36.41 | | High | High | High | High | High | High | High | High | High | High | High | 0 | 9.72E-06 | 2.34 |
| FALSE | High | LNDPFQPF | [K].LNDPF( | 1xPhospho | H3BK65 1x | 2.21E-05 | 0 | 1 | 5 | 2 | H3BK65 | Epidermal | H3BK65 [8 | 0 | 2211.089 | 0 | 1.11 | 0.01588 | 17.76 | 66 | 134 | 10.68 | 19.28 | | High | High | High | High | High | High | High | High | High | High | High | 0 | 3.45E-06 | 2.61 |
| FALSE | High | ILSDDEDEI | [R].ILSDDE | 1xPhospho | Q8BHK9 1x | 3.03E-08 | 0 | 1 | 3 | 11 | Q8BHK9 | DNA excisi | Q8BHK9 [1 | 0 | 2308.016 | 0 | -0.26 | 0.087761 | 9.53 | 111.8 | 88.2 | 3.83 | 12.74 | | High | High | High | High | High | High | High | High | High | High | High | 0 | 7.11E-08 | 4.43 |
| FALSE | High | INPYMSSP( | [R].INPYMS | 1xCarbami | Q80V08 1x | 1.75E-05 | 0 | 1 | 5 | 2 | Q80V08 | Rpl17 prot | Q80V08 [1 | 0 | 2701.32 | 0 | -0.99 | 0.004618 | 3.09 | 134.5 | 65.5 | 15.86 | 6.96 | | High | High | High | High | High | High | High | High | High | High | High | 0 | 2.52E-06 | 2.87 |
| FALSE | High | LLSASEEDE | [K].LLSASE | 1xPhospho | A2AHJ4 1xl | 2.48E-07 | 0 | 1 | 1 | 2 | A2AHJ4 | Bromodom | A2AHJ4 [1 | 0 | 2265.025 | 0 | -0.18 | 0.146469 | 10.1 | 107.4 | 92.6 | 8.33 | 9.87 | | High | High | High | High | High | High | High | High | High | High | High | 0 | 2.12E-08 | 4.04 |
| FALSE | High | LIRPEYAWI | [R].LIRPEY/ | 1xPhospho | Q9Z0W3 1: | 0.001815 | 5.66E-05 | 1 | 1 | 1 | Q9Z0W3 | Nuclear po | Q9Z0W3 [1 | 0 | 3303.766 | 0 | -0.57 | 0.001804 | 3.09 | 118.2 | 81.8 | 4.33 | 14.32 | | High | High | High | High | High | High | High | High | High | High | High | 2.58E-05 | 0.000467 | 2.32 |
| FALSE | High | LLGSENDT( | [K].LLGSEN | 1xPhospho | Q8BIE6-2 1 | 0.000157 | 0 | 1 | 4 | 1 | Q8BIE6-2 | Isoform 2 c | Q8BIE6-2 [ | 0 | 2177.995 | 0 | 0.2 | 0.016134 | 4.57 | 89.1 | 110.9 | 8.1 | 9.21 | | High | High | High | High | High | High | High | High | High | High | High | 0 | 2.98E-05 | 3.52 |
| FALSE | High | ILGSLGALP | [R].ILGSLG | 1xPhospho | Q9D0M2 1 | 5.79E-07 | 0 | 1 | 1 | 1 | Q9D0M2 | Cell divisio | Q9D0M2 [ | 0 | 3142.449 | 0 | -0.42 | 0.012209 | 4.49 | 112.8 | 87.2 | 15.87 | 14.62 | | High | High | High | High | High | High | High | High | High | High | High | 0 | 5.47E-08 | 5.51 |
| FALSE | High | LLGSMDEV | [K].LLGSMI | 1xPhospho | B7ZNX2 1x | 9.28E-05 | 0 | 1 | 4 | 1 | B7ZNX2 | Anaphase | B7ZNX2 [5 | 0 | 2276.217 | 0 | 1.22 | 0.001384 | 12.3 | 59.8 | 140.2 | 6.63 | 11.92 | | High | High | High | High | High | High | High | High | High | High | High | 0 | 1.65E-05 | 3.86 |
| FALSE | High | LLGSMDEV | [K].LLGSMI | 1xPhospho | B7ZNX2 1x | 0.000369 | 0 | 1 | 4 | 3 | B7ZNX2 | Anaphase | B7ZNX2 [5 | 0 | 2292.212 | 0 | 0.85 | 0.004713 | 7.94 | 68.5 | 131.5 | 11.91 | 10.05 | | High | High | High | High | High | High | High | High | High | High | High | 0 | 0.000124 | 4.03 |
| FALSE | High | LLGSQGSQ | [R].LLGSQ( | 1xPhospho | Q3V300 1x | 0.000382 | 0 | 1 | 1 | 1 | Q3V300 | Kinesin-like | Q3V300 [4 | 0 | 2249.231 | 0 | 0.16 | 0.219053 | 16.13 | 96 | 104 | 14.73 | 11.61 | | High | High | High | High | High | High | High | High | High | High | High | 0 | 8.06E-05 | 2.87 |
| FALSE | High | IIGVHQEDE | [K].IIGVHQ | 1xCarbami | F8VPX1 1xl | 2.34E-06 | 0 | 1 | 5 | 2 | F8VPX1 | Ubiquitin c | F8VPX1 [9 | 0 | 2576.263 | 0 | 0.08 | 0.193984 | 4.66 | 95.7 | 104.3 | 5.62 | 4.9 | | High | High | High | High | High | High | High | High | High | High | High | 0 | 2.64E-07 | 4.89 |
| FALSE | High | LLHEDLDES | [R].LLHEDL | 1xPhospho | Q6A074 1x | 9.67E-07 | 0 | 2 | 8 | 2 | Q6A074; E | MKIAA040 | Q6A074 [7 | 0 | 2195.943 | 0 | 0.89 | 0.010548 | 11.46 | 70.1 | 129.9 | 8.47 | 15.1 | Shared | High | High | High | High | High | High | High | High | High | High | High | 0 | 9.77E-08 | 3.58 |
| FALSE | High | LLHEDLDES | [R].LLHEDL | 1xPhospho | Q6A074 1x | 1.68E-09 | 0 | 2 | 8 | 7 | Q6A074; E | MKIAA040 | Q6A074 [7 | 0 | 2425.106 | 0 | 1.94 | 0.000209 | 6.97 | 40.4 | 159.6 | 3.21 | 4.64 | Shared | High | High | High | High | High | High | High | High | High | High | High | 0 | 1.35E-07 | 5.09 |
| FALSE | High | LIHGEDSDS | [R].LIHGED | 2xPhospho | Q6ZPJ3 2xl | 0.000126 | 0 | 1 | 1 | 5 | Q6ZPJ3 | (E3-indepe | Q6ZPJ3 [7 | 0 | 2105.79 | 0 | 0.07 | 0.764475 | 21.94 | 99.2 | 100.8 | 12.57 | 9.57 | | High | High | High | High | High | High | High | High | High | High | High | 0 | 2.33E-05 | 4.1 |
| FALSE | High | LLHLDSDDI | [R].LLHLDS | 1xPhospho | B2RX88 1x | 0.000171 | 0 | 1 | 5 | 1 | B2RX88 | Centrosom | B2RX88 [9 | 0 | 1878.887 | 0 | 0.97 | 0.00486 | 14.86 | 64.8 | 135.2 | 5.93 | 15.76 | | High | High | High | High | High | High | High | High | High | High | High | 0 | 3.3E-05 | 3.2 |
| FALSE | High | LLHLGPPSF | [K].LLHLGP | 1xPhospho | Q9CQJ7 1x | 2.24E-06 | 0 | 1 | 4 | 2 | Q9CQJ7 | Securin OS | Q9CQJ7 [1 | 0 | 1710.012 | 0 | -1.37 | 0.000289 | 0.16 | 144.3 | 55.7 | 7.9 | 14.59 | | High | High | High | High | High | High | High | High | High | High | High | 0 | 2.5E-07 | 3.26 |
| FALSE | High | LLHNSDDQ | [K].LLHNSD | 1xPhospho | Q921C3 1x | 0.072858 | 0.005599 | 1 | 4 | 1 | Q921C3 | Bromodom | Q921C3 [1 | 1 | 2853.485 | 0 | -0.82 | 0.000567 | 4.62 | 127.1 | 72.9 | 5.92 | 10.96 | | High | High | High | High | High | High | High | High | High | High | High | 0.002867 | 0.02978 | 2.12 |
| FALSE | High | LIHPIDNITF | [K].LIHPIDN | 1xPhospho | Q8R1M0 1 | 0.003558 | 0.00021 | 1 | 1 | 1 | Q8R1M0 | Embryonic | Q8R1M0 [2 | 0 | 2678.402 | 0 | -0.26 | 0.01808 | 0.39 | 110.1 | 89.9 | 8.94 | 8.87 | | High | High | High | High | High | High | High | High | High | High | High | 9.42E-05 | 0.000993 | 3.19 |
| FALSE | High | LLGPSAAAI | [R].LLGPSA | 1xPhospho | A2AFQ0 1x | 5.12E-05 | 0 | 1 | 6 | 1 | A2AFQ0 | E3 ubiquiti | A2AFQ0 [2 | 0 | 2546.379 | 0 | -0.74 | 0.013974 | 8.73 | 125.1 | 74.9 | 12.61 | 24.13 | | High | High | High | High | High | High | High | High | High | High | High | 0 | 8.49E-06 | 3.9 |
| FALSE | High | LLHPSPDLV | [R].LLHPSP | 1xPhospho | Q9D486 1x | 0.008847 | 0.000736 | 1 | 4 | 1 | Q9D486 | C-Maf-indu | Q9D486 [3 | 0 | 2398.222 | 0 | -0.03 | 0.266699 | 7.77 | 99.9 | 100.1 | 10.04 | 3.42 | | High | High | High | High | High | High | High | High | High | High | High | 0.000327 | 0.00273 | 1.93 |
| FALSE | High | LLKPGEEPS | [R].LLKPGE | 1xPhospho | Q80UU9 1: | 1.16E-06 | 0 | 1 | 1 | 3 | Q80UU9 | Membrane | Q80UU9 [1 | 0 | 2847.416 | 0 | 0.34 | 0.058697 | 21.41 | 85.8 | 114.2 | 12.09 | 9.8 | | High | High | High | High | High | High | High | High | High | High | High | 0 | 1.2E-07 | 3.52 |
| FALSE | High | LILASSPQP | [R].LILASSF | 1xPhospho | M0QWZ1 1 | 0.001222 | 0 | 1 | 4 | 1 | M0QWZ1 | Protein FAI | M0QWZ1 [ | 0 | 1591.922 | 0 | -0.23 | 0.089626 | 10.2 | 109.4 | 90.6 | 4 | 9.83 | | High | High | High | High | High | High | High | High | High | High | High | 0 | 0.000298 | 1.76 |
| FALSE | High | IILDLISESPI | [R].IILDLISE | 1xPhospho | Q5FWJ5 1x | 6.99E-07 | 0 | 2 | 14 | 2 | Q5FWJ5; C | Hnrpk prot | Q5FWJ5 [2 | 0 | 1879.096 | 0 | -0.99 | 0.001214 | 2.58 | 132.9 | 67.1 | 3.83 | 10.2 | Shared | High | High | High | High | High | High | High | High | High | High | High | 0 | 6.78E-08 | 3.93 |
| FALSE | High | LLLDIPLQTI | [R].LLLDIPL | 1xPhospho | Q9ERU9 1x | 9.03E-05 | 0 | 1 | 1 | 1 | Q9ERU9 | E3 SUMO-| | Q9ERU9 [1 | 0 | 1926.123 | 0 | 0.29 | 0.325336 | 32.85 | 86.4 | 113.6 | 10.22 | 26.37 | | High | High | High | High | High | High | High | High | High | High | High | 0 | 1.61E-05 | 2.88 |
| FALSE | High | LLIDPNCSG | [R].LLIDPN | 1xCarbami | Q9EQM6 1 | 7.58E-07 | 0 | 1 | 1 | 3 | Q9EQM6 | Microproc | Q9EQM6 [ | 0 | 1774.851 | 0 | -0.52 | 0.020576 | 2.93 | 117.1 | 82.9 | 10.8 | 5.76 | | High | High | High | High | High | High | High | High | High | High | High | 0 | 7.45E-08 | 3.35 |
| FALSE | High | LLLDPSSTP | [K].LLLDPS | 1xPhospho | Q9CQ22 1x | 0.001933 | 5.66E-05 | 1 | 2 | 5 | Q9CQ22 | Ragulator c | Q9CQ22 [2 | 0 | 1709.949 | 0 | 0.47 | 0.006784 | 11.95 | 84 | 116 | 5.5 | 5.18 | | High | High | High | High | High | High | High | High | High | High | High | 2.58E-05 | 0.000499 | 2.4 |
| FALSE | High | LIIDSSGSIS | [R].LIIDSSG | 1xPhospho | E9Q284 1x | 8.63E-05 | 0 | 1 | 3 | 3 | E9Q284 | Coilin OS=I | E9Q284 [5 | 0 | 1488.74 | 0 | -1.02 | 0.00323 | 5 | 133.8 | 66.2 | 14.62 | 19.15 | | High | High | High | High | High | High | High | High | High | High | High | 7.19E-05 | 0.000909 | 2.06 |
| FALSE | High | LLLDTTPPG | [R].LLLDTT | 1xPhospho | A0A068BD | 0.000699 | 0 | 1 | 2 | 2 | A0A068BD | Ral guanine | A0A068BD | 0 | 2081.125 | 0 | 0.34 | 0.226722 | 20.43 | 93.4 | 106.6 | 12.09 | 11.53 | | High | High | High | High | High | High | High | High | High | High | High | 2.58E-05 | 0.000392 | 3.03 |
| FALSE | High | LLLGESEDS | [R].LLLGES | 2xPhospho | Q9JIM3 2x | 0.045898 | 0.003408 | 1 | 3 | 1 | Q9JIM3 | DNA excisi | Q9JIM3 [9 | 0 | 2348.108 | 0 | -0.52 | 0.085969 | 9.19 | 117.9 | 82.1 | 13.83 | 14.86 | | High | High | High | High | High | High | High | High | High | High | High | 0.001741 | 0.01746 | 2.54 |
| FALSE | High | ILLLFGESEL | [R].ILLLFGE | 1xPhospho | O09130 1x | 1.3E-05 | 0 | 1 | 1 | 2 | O09130 | NFATC2-in | O09130 [2 | 0 | 2655.466 | 0 | -0.66 | 0.005775 | 7.98 | 120.9 | 79.1 | 7.96 | 8.43 | | High | High | High | High | High | High | High | High | High | High | High | 0 | 3.36E-06 | 4.11 |
| FALSE | High | IILPDDDSSI | [R].IILPDDI | 2xPhospho | B2RX10 2x | 0.011755 | 0.000889 | 1 | 4 | 2 | B2RX10 | Fanconi an | B2RX10 [1 | 0 | 2432.129 | 0 | -0.56 | 0.02533 | 9.42 | 119.1 | 80.9 | 9.98 | 10.36 | | High | High | High | High | High | High | High | High | High | High | High | 0.00196 | 0.01987 | 2.51 |
| FALSE | High | ILLPNNNH\ | [K].ILLPNN | 1xPhospho | Q60953 1x | 2.63E-05 | 0 | 2 | 6 | 3 | Q60953; Q | Protein PM | Q60953 [5 | 0 | 2448.161 | 0 | -0.34 | 0.01055 | 6.87 | 111.9 | 88.1 | 4.07 | 7.54 | Shared | High | High | High | High | High | High | High | High | High | High | High | 0 | 4E-06 | 3.44 |
| FALSE | High | LLLQDQEN | [R].LLLQD( | 1xPhospho | B2RQG2 1x | 0.030104 | 0.001857 | 1 | 2 | 1 | B2RQG2 | PHD finger | B2RQG2 [1 | 0 | 1994.975 | 0 | 0.99 | 0.000937 | 3.42 | 66.8 | 133.2 | 1.68 | 8.59 | | High | High | High | High | High | High | High | High | High | High | High | 0.001011 | 0.01085 | 2.4 |
| FALSE | High | LLKPGEEPS | [R].LLKPGE | 1xPhospho | Q80UU9 1: | 0.002232 | 0.000109 | 1 | 1 | 2 | Q80UU9 | Membrane | Q80UU9 [1 | 0 | 2618.253 | 0 | 0.01 | 0.503204 | 22.06 | 100.4 | 99.6 | 7.21 | 16.69 | | High | High | High | High | High | High | High | High | High | High | High | 7.19E-05 | 0.000921 | 3.6 |
| FALSE | High | LIIQSNGLV | [R].LIIQSN( | 1xPhospho | P15066 1x | 1.97E-05 | 0 | 1 | 2 | 1 | P15066 | Transcripti | P15066 [1 | 0 | 2960.615 | 0 | 0.31 | 0.198168 | 48.25 | 80.8 | 119.2 | 23.89 | 13.34 | | High | High | High | High | High | High | High | High | High | High | High | 0 | 2.88E-06 | 2.97 |
| FALSE | High | LLGMSGK | [K].LLGMS( | 1xPhospho | Q5SQB7 1x | 0.021418 | 0.001507 | 2 | 9 | 3 | Q5SQB7; C | MCG6806 | Q5SQB7 [1 | 0 | 1259.683 | 0 | -0.74 | 0.001482 | 3.68 | 125 | 75 | 2.57 | 13.19 | Shared | High | High | High | High | High | High | High | High | High | High | High | 0.003768 | 0.04058 | 1.8 |
| FALSE | High | LLGGHGED | [K].LLGGH( | 1xPhospho | Q8R149 1x | 0.00824 | 0.000736 | 1 | 2 | 2 | Q8R149 | BUD13 hor | Q8R149 [1 | 1 | 2875.286 | 0 | -0.15 | 0.080075 | 3.8 | 112.6 | 87.4 | 10.24 | 13.25 | | High | High | High | High | High | High | High | High | High | High | High | 0.00074 | 0.006429 | 2.3 |
| FALSE | High | LLEENEAM | [K].LLEENE | 1xPhospho | Q9DBJ3 1x | 3.42E-06 | 0 | 1 | 1 | 2 | Q9DBJ3 | Brain-spec | Q9DBJ3 [4 | 0 | 2362.156 | 0 | 1.44 | 0.016407 | 17.43 | 52.5 | 147.5 | 10.2 | 31.45 | | High | High | High | High | High | High | High | High | High | High | High | 0 | 4.04E-07 | 2.87 |
| FALSE | High | ILEGDHGSI | [R].ILEGDH | 1xPhospho | Q9R0L6-2 | 1.15E-06 | 0 | 1 | 3 | 4 | Q9R0L6-2 | Isoform 2 c | Q9R0L6-2 [ | 0 | 2435.138 | 0 | 0.98 | 0.000249 | 4.34 | 66.7 | 133.3 | 6.47 | 6.29 | | High | High | High | High | High | High | High | High | High | High | High | 0 | 1.18E-07 | 3.23 |
| FALSE | High | ILEGDNGLI | [K].ILEGDN | 1xCarbami | Q8VDY9 1x | 6.95E-08 | 0 | 1 | 2 | 3 | Q8VDY9 | Caspase ac | Q8VDY9 [1 | 0 | 2850.21 | 0 | 0.07 | 0.691005 | 12.67 | 98.7 | 101.3 | 5.9 | 16.97 | | High | High | High | High | High | High | High | High | High | High | High | 0 | 5.08E-09 | 5.01 |
| FALSE | High | ILEGDNGLI | [K].ILEGDN | 1xCarbami | Q8VDY9 1x | 1.22E-08 | 0 | 1 | 2 | 2 | Q8VDY9 | Caspase ac | Q8VDY9 [1 | 0 | 2851.194 | 0 | 0.05 | 0.773766 | 4.37 | 98.2 | 101.8 | 9.58 | 17.34 | | High | High | High | High | High | High | High | High | High | High | High | 0 | 7.22E-10 | 5.17 |
| FALSE | High | ILEGDNGLI | [K].ILEGDN | 1xCarbami | Q8VDY9 1x | 1.01E-07 | 0 | 1 | 2 | 3 | Q8VDY9 | Caspase ac | Q8VDY9 [1 | 0 | 2866.205 | 0 | 0 | 0.691555 | 2.4 | 99.1 | 100.9 | 5.79 | 10.76 | | High | High | High | High | High | High | High | High | High | High | High | 0 | 1.11E-08 | 5.31 |
| FALSE | High | LLEGEESRI | [K].LLEGEE | 1xPhospho | P20152 1xl | 0.090153 | 0.00698 | 1 | 7 | 1 | P20152 | Vimentin C | P20152 [4 | 1 | 2780.48 | 0 | 2.51 | 0.001921 | 18.84 | 24.3 | 175.7 | 18.32 | 19.18 | | High | High | High | High | High | High | High | High | High | High | High | 0.003606 | 0.03822 | 1.79 |
| FALSE | High | LLEGEESRI | [K].LLEGEE | 1xDeamida | P20152 1xl | 0.086562 | 0.006435 | 1 | 7 | 1 | P20152 | Vimentin C | P20152 [4 | 1 | 2781.464 | 0 | 2.37 | 0.002683 | 30.79 | 29.4 | 170.6 | 9.82 | 19.42 | | High | High | High | High | High | High | High | High | High | High | High | 0.003317 | 0.03649 | 2.29 |
| FALSE | High | LLEHAEPAI | [K].LLEHAE | 1xPhospho | Q9ES34 1x | 2.38E-08 | 0 | 1 | 2 | 3 | Q9ES34 | Ubiquitin-| | Q9ES34 [4 | 0 | 3100.685 | 0 | 2.35 | 0.000489 | 38.19 | 33.1 | 166.9 | 28.64 | 8.5 | | High | High | High | High | High | High | High | High | High | High | High | 0 | 1.51E-09 | 4.88 |
| FALSE | High | ILELTPEPDI | [R].ILELTPE | 1xPhospho | Q69ZA1 1x | 0.003372 | 0.00016 | 1 | 2 | 3 | Q69ZA1 | Cyclin-dep | Q69ZA1 [1 | 0 | 1744.92 | 0 | 0.2 | 0.004737 | 4.12 | 93.2 | 106.8 | 3.52 | 3.11 | | High | High | High | High | High | High | High | High | High | High | High | 0.002589 | 0.02777 | 1.74 |
| FALSE | High | LLEPCSPIPI | [K].LLEPCS | 1xCarbami | Q6P5H2-2 | 0.000639 | 0 | 1 | 2 | 1 | Q6P5H2-2 | Isoform 2 c | Q6P5H2-2 | 0 | 4631.269 | 0 | 4.8 | | 105.42 | 10.8 | 189.2 | 21.16 | 40.42 | | High | Not Found | High | High | Not Found | High | High | High | High | High | High | 0 | 0.000144 | 4.08 |
| FALSE | High | LLEPCSPIPI | [K].LLEPCS | 1xCarbami | Q6P5H2-2 | 1.9E-08 | 0 | 1 | 2 | 3 | Q6P5H2-2 | Isoform 2 c | Q6P5H2-2 | 0 | 4632.253 | 0 | 4.32 | 0.006499 | 53.23 | 10.8 | 189.2 | 23.79 | 31.38 | | High | High | High | High | High | High | High | High | High | High | High | 0 | 8.06E-09 | 5.93 |
| FALSE | High | IIEPDDFLDI | [R].IIEPDDI | 1xPhospho | D3Z5N6 1x | 4.03E-10 | 0 | 1 | 4 | 10 | D3Z5N6 | Zinc finger | D3Z5N6 [1 | 0 | 3179.412 | 0 | -0.83 | 0.001772 | 2.12 | 128.4 | 71.6 | 10.16 | 5.84 | | High | High | High | High | High | High | High | High | High | High | High | 0 | 1.57E-11 | 6.07 |
| FALSE | High | LLEPLDLSG | [R].LLEPLD | 1xPhospho | Q3TW36 1 | 2.49E-09 | 0 | 1 | 5 | 2 | Q3TW36 | Putative ur | Q3TW36 [ | 0 | 2506.155 | 0 | 0.55 | 0.001448 | 5.7 | 81 | 119 | 2.28 | 5.93 | | High | High | High | High | High | High | High | High | High | High | High | 0 | 4.71E-10 | 4.42 |
| FALSE | High | LLGLNELDI | [R].LLGLNE | 1xPhospho | Q68FH0 1x | 1.21E-05 | 0 | 1 | 7 | 1 | Q68FH0 | Plakophilin | Q68FH0 [7 | 1 | 2708.509 | 0 | -1.96 | 0.000214 | 0.56 | 160.6 | 39.4 | 3.73 | 10.61 | | High | High | High | High | High | High | High | High | High | High | High | 0 | 1.67E-06 | 3.92 |
| FALSE | High | LLEPLVTQ\ | [R].LLEPLV | 1xPhospho | P26231 1xl | 3.85E-06 | 0 | 1 | 2 | 1 | P26231 | Catenin alp | P26231 [2 | 0 | 2508.409 | 0 | 0.62 | 0.002989 | 14.15 | 79.4 | 120.6 | 6.11 | 8.08 | | High | High | High | High | High | High | High | High | High | High | High | 0 | 4.63E-07 | 3.81 |
| FALSE | High | IIEQPECSSI | [K].IIEQPE( | 1xCarbami | Q80V94 1x | 2.9E-05 | 0 | 1 | 1 | 3 | Q80V94 | AP-4 comp | Q80V94 [9 | 0 | 2196.046 | 0 | 0.23 | 0.045955 | 5.38 | 93.9 | 106.1 | 5.63 | 8.25 | | High | High | High | High | High | High | High | High | High | High | High | 0 | 4.45E-06 | 4.47 |
| FALSE | High | LIESPDSK | [K].LIESPDS | 1xPhospho | Q3U0P1 1x | 0.011146 | 0.000772 | 1 | 3 | 1 | Q3U0P1 | Partner an | Q3U0P1 [5 | 0 | 1426.759 | 0 | -0.94 | 0.000437 | 0.64 | 131.5 | 68.5 | 3.85 | 6.82 | | High | High | High | High | High | High | High | High | High | High | High | 0.000399 | 0.003525 | 1.78 |
| FALSE | High | IIETMSSPK | [K].IIETMS | 1xPhospho | Q80WE4 1 | 0.003526 | 0.00021 | 1 | 4 | 3 | Q80WE4 | Kinesin-like | Q80WE4 [ | 0 | 1559.816 | 0 | -0.07 | 0.229431 | 9.47 | 102.4 | 97.6 | 4.37 | 4.87 | | High | High | High | High | High | High | High | High | High | High | High | 0.000814 | 0.007593 | 1.92 |
| FALSE | High | ILEYNSDGI | [K].ILEYNSI | 1xPhospho | G3UYU0 1x | 1.11E-05 | 0 | 1 | 1 | 5 | G3UYU0 | Mex3 RNA | G3UYU0 [2 | 0 | 2635.213 | 0 | -0.33 | 0.062018 | 16.5 | 109.6 | 90.4 | 8.12 | 16.91 | | High | High | High | High | High | High | High | High | High | High | High | 0 | 0.00016 | 4.39 |
| FALSE | High | LLFDPESG( | [R].LLFDPE | 1xPhospho | A0A087WF | 0.000286 | 0 | 1 | 2 | 1 | A0A087WF | Proline-ricl | A0A087WF | 0 | 2836.437 | 0 | 6.13 | | 63.4 | 6.6 | 193.4 | 57.12 | 7.56 | | High | Not Found | Not Found | High | High | High | High | High | High | High | High | 0 | 5.88E-05 | 3.3 |
| FALSE | High | LIFLDSRVTI | [K].LIFLDSF | 1xDeamida | Q61079 2x | 0.110202 | 0.008961 | 1 | 1 | 1 | Q61079 | Single-min | Q61079 [2 | 1 | 2969.44 | 0 | -0.86 | 0.001359 | 2.01 | 130.2 | 69.8 | 6.08 | 5.56 | | High | High | High | High | High | High | High | High | High | High | High | 0.004462 | 0.04826 | 2.85 |
| FALSE | High | IIFPQDESSI | [K].IIFPQDI | 1xPhospho | B9EK83 1xl | 5.23E-06 | 0 | 2 | 3 | 2 | B9EK83; A | Transient r | B9EK83 [1 | 0 | 2731.359 | 0 | 2.07 | 0.000181 | 6.49 | 38.7 | 161.3 | 8.41 | 5.26 | Shared | High | High | High | High | High | High | High | High | High | High | High | 0 | 6.5E-07 | 5.1 |
| FALSE | High | ILGADTSVI | [R].ILGADT | 1xPhospho | Q03265 1x | 0.056248 | 0.004124 | 1 | 3 | 2 | Q03265 | ATP syntha | Q03265 [5 | 0 | 1884.915 | 0 | 0.12 | 0.306693 | 26.27 | 96.6 | 103.4 | 9.34 | 14.4 | | High | High | High | High | High | High | High | High | High | High | High | 0.002117 | 0.02199 | 2.4 |
| FALSE | High | ILGDHLLLD | [R].ILGDHL | 1xPhospho | Q8BXJ2-4 1 | 0.005 | 0.000504 | 2 | 3 | 3 | Q8BXJ2-4; | Isoform 4 c | Q8BXJ2-4 [ | 0 | 3313.726 | 0 | 0.37 | 0.208654 | 17.84 | 87.2 | 112.8 | 17.99 | 21.67 | Shared | High | High | High | High | High | High | High | High | High | High | High | 0.000225 | 0.001441 | 2.96 |
| FALSE | High | ILGDHLLLD | [R].ILGDHL | 2xPhospho | Q8BXJ2-4 2 | 5.26E-05 | 0 | 2 | 3 | 1 | Q8BXJ2-4; | Isoform 4 c | Q8BXJ2-4 [ | 0 | 3377.697 | 0 | 0.86 | 0.002719 | 1.75 | 78.8 | 121.2 | 14.47 | 14.81 | Shared | High | High | High | High | High | High | High | High | High | High | High | 0 | 8.69E-06 | 3.79 |
| FALSE | High | ILGDHLLLD | [R].ILGDHL | 2xPhospho | Q8BXJ2-4 2 | 0.017041 | 0.001204 | 2 | 3 | 1 | Q8BXJ2-4; | Isoform 4 c | Q8BXJ2-4 [ | 0 | 3393.692 | 0 | 0.53 | 0.043812 | 15.1 | 85.4 | 114.6 | 11.2 | 13.85 | Shared | High | High | High | High | High | High | High | High | High | High | High | 0.000638 | 0.005704 | 3.22 |
| FALSE | High | LIGDLPQTF | [K].LIGDLP | 1xPhospho | Q8VI59 1xl | 0.002058 | 5.66E-05 | 1 | 1 | 1 | Q8VI59 | Pecanex-lik | Q8VI59 [1 | 0 | 2983.486 | 0 | 0.15 | 0.179955 | 24.17 | 88.2 | 111.8 | 20.31 | 7.33 | | High | High | High | High | High | High | High | High | High | High | High | 2.58E-05 | 0.000538 | 3.56 |
| FALSE | High | LLGGHGED | [K].LLGGH( | 1xPhospho | Q8R149 1x | 0.000101 | 0 | 1 | 2 | 3 | Q8R149 | BUD13 hor | Q8R149 [1 | 0 | 2719.185 | 0 | -0.45 | 0.002936 | 5.73 | 115.1 | 84.9 | 4.79 | 9.35 | | High | High | High | High | High | High | High | High | High | High | High | 0 | 1.8E-05 | 2.95 |
| FALSE | High | IIEQLSPQA | [K].IIEQLSF | 1xPhospho | F8VPZ5 1xl | 0.005467 | 0.00062 | 1 | 1 | 1 | F8VPZ5 | Excision re | F8VPZ5 [1 | 0 | 1735.894 | 0 | -1.33 | 0.000784 | 3.1 | 148.5 | 51.5 | 7.82 | 21.57 | | High | High | High | High | High | High | High | High | High | High | High | 0.000275 | 0.001601 | 1.67 |
| FALSE | High | LIIQSNGLV | [R].LIIQSN( | 1xDeamida | P15066 1xl | 0.002115 | 5.66E-05 | 1 | 2 | 3 | P15066 | Transcripti | P15066 [1 | 0 | 2961.599 | 0 | 0.45 | 0.003126 | 8.03 | 85.7 | 114.3 | 12.77 | 8.86 | | High | High | High | High | High | High | High | High | High | High | High | 2.58E-05 | 0.000552 | 2.33 |
| FALSE | High | ILLSQTTGN | [R].ILLSQT | 1xPhospho | F6ZDS4 1xl | 6.77E-05 | 0 | 1 | 7 | 1 | F6ZDS4 | Nucleopro | F6ZDS4 [6 | 0 | 3451.878 | 0 | 0.38 | 0.02885 | 22.32 | 86.2 | 113.8 | 9.98 | 8.55 | | High | High | High | High | High | High | High | High | High | High | High | 0 | 1.16E-05 | 3.71 |
| FALSE | High | ILLSQTTGN | [R].ILLSQT | 1xPhospho | F6ZDS4 1xl | 0.000328 | 0 | 1 | 7 | 3 | F6ZDS4 | Nucleopro | F6ZDS4 [6 | 0 | 3467.872 | 0 | 0.01 | 0.675366 | 26 | 94.9 | 105.1 | 7.31 | 16.82 | | High | High | High | High | High | High | High | High | High | High | High | 0 | 6.81E-05 | 4.29 |
| FALSE | High | ILPQINPELI | [R].ILPQINI | 1xPhospho | F7BJB9 1xF | 1.56E-06 | 0 | 1 | 1 | 3 | F7BJB9 | MORC fam | F7BJB9 [4 | 0 | 3213.721 | 0 | -0.95 | 0.000242 | 0.28 | 133 | 67 | 5.56 | 5.22 | | High | High | High | High | High | High | High | High | High | High | High | 0 | 1.68E-07 | 4.91 |
| FALSE | High | ILPQINPELI | [R].ILPQINI | 1xDeamida | F7BJB9 1xF | 6.79E-06 | 0 | 1 | 1 | 8 | F7BJB9 | MORC fam | F7BJB9 [4 | 0 | 3214.705 | 0 | -0.77 | 0.060452 | 6.86 | 115.7 | 84.3 | 26.04 | 26.21 | | High | High | High | High | High | High | High | High | High | High | High | 0 | 1.73E-06 | 3.21 |
| FALSE | High | LIPSAVATG | [R].LIPSAV/ | 1xPhospho | P70268-2 1 | 0.005565 | 0.00062 | 1 | 3 | 2 | P70268-2 | Isoform 2 c | P70268-2 [ | 0 | 2465.264 | 0 | 1.62 | 0.00046 | 24.94 | 47.8 | 152.2 | 14.12 | 7.33 | | High | High | High | High | High | High | High | High | High | High | High | 0.001076 | 0.01143 | 3.75 |
| FALSE | High | LLPSQSPD( | [R].LLPSQS | 1xPhospho | Q7TSG3 1x | 0.000356 | 0 | 1 | 1 | 2 | Q7TSG3 | F-box only | Q7TSG3 [1 | 0 | 1999.041 | 0 | -1.02 | 0.001401 | 2.62 | 134.5 | 65.5 | 6.34 | 8.68 | | High | High | High | High | High | High | High | High | High | High | High | 0 | 7.44E-05 | 2.49 |
| FALSE | High | LLPSSGPCV | [K].LLPSSG | 1xCarbami | A0A140LJ0 | 7.29E-06 | 0 | 1 | 4 | 4 | A0A140LJ0 | Zinc finger | A0A140LJ0 | 0 | 2975.427 | 0 | -1.03 | 0.003496 | 8.97 | 132.6 | 67.4 | 8.62 | 21.66 | | High | High | High | High | High | High | High | High | High | High | High | 0 | 2.77E-06 | 3.93 |
| FALSE | High | IIPTLEEGL( | [R].IIPTLEE | 1xCarbami | Q5FWJ5 1x | 0.015741 | 0.001124 | 1 | 7 | 7 | Q5FWJ5 | Hnrpk prot | Q5FWJ5 [1 | 0 | 4593.324 | 0 | -0.2 | 0.168246 | 11.38 | 107.6 | 92.4 | 13.89 | 6.75 | | High | High | High | High | High | High | High | High | High | High | High | 0.000786 | 0.006858 | 3.27 |
| FALSE | High | IIPTLEEGL( | [K].IIPTLEE | 1xCarbami | Q5FWJ5 1x | 6.91E-06 | 0 | 1 | 7 | 11 | Q5FWJ5 | Hnrpk prot | Q5FWJ5 [1 | 0 | 4594.308 | 0 | -0.03 | 0.593022 | 11.46 | 101 | 99 | 10.13 | 7.09 | | High | High | High | High | High | High | High | High | High | High | High | 2.58E-05 | 0.00041 | 3.99 |
| FALSE | High | LLPYPTLAS | [K].LLPYPT | 1xPhospho | Q3U1J1-2 | 2.27E-05 | 0 | 1 | 2 | 4 | Q3U1J1-2 | Isoform 2 c | Q3U1J1-2 [ | 0 | 1739.886 | 0 | -0.31 | 0.02194 | 10.45 | 110.6 | 89.4 | 3.76 | 6.99 | | High | High | High | High | High | High | High | High | High | High | High | 0 | 3.39E-06 | 2.98 |
| FALSE | High | LLQAHAPS | [K].LLQAH/ | 1xPhospho | B8JJC1 1xP | 0.004451 | 0.000409 | 1 | 3 | 1 | B8JJC1 | Tyrosyl-DN | B8JJC1 [37 | 0 | 1700.95 | 0 | -0.36 | 0.189702 | 10.49 | 112.4 | 87.6 | 4.56 | 20.58 | | High | High | High | High | High | High | High | High | High | High | High | 0.000183 | 0.001274 | 1.82 |
| FALSE | High | LLQDSSSP\ | [R].LLQDSS | 1xPhospho | Q61026 1x | 1.83E-05 | 0 | 1 | 5 | 2 | Q61026 | Nuclear re | Q61026 [6 | 0 | 1911.024 | 0 | 0.93 | 0.000881 | 2.6 | 68.1 | 131.9 | 8.51 | 3.74 | | High | High | High | High | High | High | High | High | High | High | High | 0 | 2.68E-06 | 3.75 |
| FALSE | High | LLQDSVDF | [R].LLQDSV | 1xPhospho | P20152 1xl | 1.72E-08 | 0 | 1 | 10 | 10 | P20152 | Vimentin C | P20152 [7 | 0 | 2664.357 | 0 | 1.72 | 0.001187 | 4.66 | 46.1 | 153.9 | 13.69 | 15.76 | | High | High | High | High | High | High | High | High | High | High | High | 0 | 1.06E-09 | 4.68 |
| FALSE | High | ILQEAQNLI | [R].ILQEAC | 1xPhospho | A0A1W2P7 | 1.6E-08 | 0 | 1 | 6 | 1 | A0A1W2P7 | Uncharacte | A0A1W2P7 | 0 | 2650.429 | 0 | -0.42 | 0.030592 | 8.89 | 116.3 | 83.7 | 10.62 | 11.11 | | High | High | High | High | High | High | High | High | High | High | High | 0 | 9.73E-10 | 4.15 |
| FALSE | High | ILQEAQNLI | [R].ILQEAC | 1xPhospho | A0A1W2P7 | 3.51E-09 | 0 | 1 | 6 | 2 | A0A1W2P7 | Uncharacte | A0A1W2P7 | 0 | 2666.424 | 0 | -0.29 | 0.240147 | 10.82 | 110.1 | 89.9 | 10.41 | 19.66 | | High | High | High | High | High | High | High | High | High | High | High | 0 | 1.77E-10 | 5.04 |
| FALSE | High | LLPPVGTGI | [R].LLPPVG | 1xPhospho | Q91YL3 1x | 0.004911 | 0.000504 | 1 | 4 | 1 | Q91YL3 | Uridine-cyt | Q91YL3 [4 | 1 | 1558.867 | 0 | 1.11 | 0.00037 | 7.44 | 64.3 | 135.7 | 6.55 | 5.62 | | High | High | High | High | High | High | High | High | High | High | High | 0.000225 | 0.00142 | 2.1 |
| FALSE | High | LLQEGSPSI | [R].LLQEGS | 1xPhospho | P70365 1xl | 1.3E-05 | 0 | 1 | 5 | 2 | P70365 | Nuclear re | P70365 [6 | 0 | 2581.341 | 0 | 1.61 | 0.000696 | 10.92 | 46 | 154 | 6.3 | 9.29 | | High | High | High | High | High | High | High | High | High | High | High | 0 | 1.82E-06 | 2.53 |
| FALSE | High | ILQEYQVQ | [K].ILQEYQ | 1xPhospho | D3Z7P3 1xl | 3.36E-06 | 0 | 1 | 3 | 1 | D3Z7P3 | Glutaminas | D3Z7P3 [6 | 0 | 2722.301 | 0 | 2.97 | 0.001492 | 20.4 | 24.8 | 175.2 | 11.74 | 17.05 | | High | High | High | High | High | High | High | High | High | High | High | 0 | 3.95E-07 | 3.9 |
| FALSE | High | LLQFLAGQ | [K].LLQFLA | 1xPhospho | Q8BQY8-2 | 1.45E-06 | 0 | 1 | 3 | 1 | Q8BQY8-2 | Isoform 2 c | Q8BQY8-2 | 0 | 2082.176 | 0 | -0.74 | 0.006174 | 8.81 | 122.8 | 77.2 | 7.79 | 9.38 | | High | High | High | High | High | High | High | High | High | High | High | 0 | 1.55E-07 | 3.08 |
| FALSE | High | LLQGSPEQ | [K].LLQGSF | 1xPhospho | Q9CZB3 1x | 6.97E-06 | 0 | 1 | 2 | 2 | Q9CZB3 | THUMP do | Q9CZB3 [1 | 0 | 1793.9 | 0 | 0.1 | 0.004853 | 1.32 | 96.3 | 103.7 | 1.04 | 1.94 | | High | High | High | High | High | High | High | High | High | High | High | 0 | 9.02E-07 | 2.88 |
| FALSE | High | LLQHPSTCS | [R].LLQHPS | 1xCarbami | F8VQH0 1x | 0.000356 | 0 | 1 | 4 | 3 | F8VQH0 | Tyrosine-p | F8VQH0 [9 | 0 | 3963.847 | 0 | 0.05 | 0.856445 | 25.99 | 98.8 | 101.2 | 5.78 | 11.08 | | High | High | High | High | High | High | High | High | High | High | High | 0 | 7.48E-05 | 3.62 |
| FALSE | High | LLQNGNSP | [K].LLQNGI | 1xPhospho | Q05BA5 1x | 0.000553 | 0 | 1 | 6 | 1 | Q05BA5 | Nuclear re | Q05BA5 [6 | 0 | 1879.009 | 0 | 0.31 | 0.057117 | 11.62 | 92.5 | 107.5 | 12.15 | 9.88 | | High | High | High | High | High | High | High | High | High | High | High | 0 | 0.000123 | 2.68 |
| FALSE | High | LLQPLSPVT | [K].LLQPLS | 1xPhospho | Q5XJV7-2 | 8.98E-06 | 0 | 1 | 3 | 2 | Q5XJV7-2 | Isoform 2 c | Q5XJV7-2 [ | 0 | 2340.298 | 0 | -0.43 | 0.00135 | 3.65 | 115.7 | 84.3 | 4.79 | 4.25 | | High | High | High | High | High | High | High | High | High | High | High | 0.000859 | 0.008364 | 2.78 |
| FALSE | High | LLQQQEEE | [K].LLQQQ | 1xCarbami | Q9DBT3 2x | 3.76E-08 | 0 | 1 | 1 | 3 | Q9DBT3 | Coiled-coil | Q9DBT3 [2 | 1 | 3962.609 | 0 | 0.01 | 0.976028 | 13.33 | 100.4 | 99.6 | 9.41 | 7.13 | | High | High | High | High | High | High | High | High | High | High | High | 0 | 2.55E-09 | 5.92 |
| FALSE | High | LLQQQQA1 | [K].LLQQQ | 1xPhospho | Q99MR1 1 | 8.92E-07 | 0 | 1 | 1 | 2 | Q99MR1 | GRB10-inte | Q99MR1 [ | 0 | 3443.886 | 0 | 0.94 | 0.002204 | 12.91 | 65.5 | 134.5 | 9.83 | 8.1 | | High | High | High | High | High | High | High | High | High | High | High | 0 | 8.91E-08 | 3.4 |
| FALSE | High | LLQQTSGA | [R].LLQQTS | 1xPhospho | E9PYH6 1xl | 9.42E-07 | 0 | 1 | 5 | 1 | E9PYH6 | SET domai | E9PYH6 [1 | 0 | 3742.899 | 0 | -0.82 | 0.001018 | 3.78 | 127.6 | 72.4 | 6.53 | 7.26 | | High | High | High | High | High | High | High | High | High | High | High | 0 | 9.54E-08 | 3.26 |
| FALSE | High | LLQSPCVD\ | [R].LLQSPC | 1xCarbami | Q8BWH0 1 | 0.047488 | 0.00354 | 1 | 2 | 1 | Q8BWH0 | Putative so | Q8BWH0 [ | 0 | 1893.032 | 0 | 0.51 | 0.025422 | 16.91 | 82.2 | 117.8 | 9.12 | 10.43 | | High | High | High | High | High | High | High | High | High | High | High | 0.001812 | 0.01822 | 2.15 |
| FALSE | High | LLQTCFSSF | [K].LLQTCF | 1xCarbami | Q99P31 1x | 0.002486 | 0.000109 | 1 | 1 | 2 | Q99P31 | Hsp70-bind | Q99P31 [3 | 0 | 2181.94 | 0 | -0.57 | 0.008036 | 5.73 | 121.5 | 78.5 | 6.93 | 17.13 | | High | High | High | High | High | High | High | High | High | High | High | 4.94E-05 | 0.000663 | 2.6 |
| FALSE | High | LLQTCFSSF | [K].LLQTCF | 1xCarbami | Q99P31 1x | 9.76E-07 | 0 | 1 | 1 | 3 | Q99P31 | Hsp70-bind | Q99P31 [3 | 0 | 2197.934 | 0 | -0.26 | 0.049161 | 5.76 | 109.6 | 90.4 | 2.01 | 12.26 | | High | High | High | High | High | High | High | High | High | High | High | 0 | 9.87E-08 | 3.2 |
| FALSE | High | LIRPEYAWI | [R].LIRPEY/ | 1xPhospho | Q9Z0W3 1: | 2.1E-05 | 0 | 1 | 1 | 1 | Q9Z0W3 | Nuclear po | Q9Z0W3 [1 | 0 | 3074.603 | 0 | 0.56 | 0.773942 | 34.03 | 87 | 113 | 59.89 | 33.49 | | High | High | High | High | High | High | High | High | High | High | High | 0 | 3.09E-06 | 3.75 |

| | | | | | | | | | | | | | | | | | | | | | | | | | | | | | | | | | | | | | | |
|---|---|---|---|---|---|---|---|---|---|---|---|---|---|---|---|---|---|---|---|---|---|---|---|---|---|---|---|---|---|---|---|---|---|---|---|---|---|---|
| FALSE | High | LLQETELVE | [R].LLQETE | 1xPhospho | P70424 1xP | 0.002398 | 0.000109 | 1 | 2 | 1 | P70424 | Receptor t | P70424 [69 | 0 | 2946.485 | 0 | 1.32 | 0.000843 | 18.02 | 55 | 145 | 17.78 | 12.89 | | High | High | High | High | High | High | High | High | High | High | High | 4.94E-05 | 0.000636 | 2.93 |
| FALSE | High | LIPPVGSPG | [K].LIPPVG | 1xPhospho | Q9D219 1x | 0.003282 | 0.00016 | 1 | 1 | 3 | Q9D219 | B-cell CLL/I | Q9D219 [2 | 0 | 3035.504 | 0 | 0.91 | 0.004336 | 3.38 | 75.2 | 124.8 | 10.4 | 10.26 | | High | High | High | High | High | High | High | High | High | High | High | 7.19E-05 | 0.000907 | 3.65 |
| FALSE | High | ILPPPLIPTP | [R].ILPPPLI | 1xPhospho | Q91WE1 1 | 0.001127 | 0 | 1 | 2 | 2 | Q91WE1 | Sorting nex | Q91WE1 [1 | 0 | 2032.108 | 0 | 0.51 | 0.002939 | 5.65 | 81.4 | 118.6 | 9.16 | 4.27 | | High | High | High | High | High | High | High | High | High | High | High | 0.000204 | 0.001312 | 2.5 |
| FALSE | High | LIPITGGNA | [K].LIPITGG | 1xPhospho | Q69ZR9-2 | 1.18E-07 | 0 | 1 | 2 | 2 | Q69ZR9-2 | Isoform 2 c | Q69ZR9-2 | 0 | 2329.195 | 0 | -0.94 | 0.000932 | 4.22 | 131.6 | 68.4 | 6.54 | 6.31 | | High | High | High | High | High | High | High | High | High | High | High | 0 | 8.28E-07 | 3.62 |
| FALSE | High | LIISSPSQR | [R].LIISSPS | 1xPhospho | D3Z357 1xP | 0.040028 | 0.002867 | 1 | 5 | 1 | D3Z357 | Histone-lys | D3Z357 [55 | 0 | 1309.708 | 0 | 0.69 | 0.001707 | 6.83 | 75.8 | 124.2 | 7.61 | 5.06 | | High | High | High | High | High | High | High | High | High | High | High | 0.001516 | 0.015 | 1.15 |
| FALSE | High | LILTDNGSA | [R].LILTDN | 1xCarbami | M0QWZ1 1 | 3.1E-06 | 0 | 1 | 3 | 2 | M0QWZ1 | Protein FAI | M0QWZ1 [ | 0 | 3174.532 | 0 | -0.42 | 0.011572 | 2.53 | 113.9 | 86.1 | 4.15 | 8.52 | | High | High | High | High | High | High | High | High | High | High | High | 0 | 3.64E-07 | 3.58 |
| FALSE | High | LILTDNGSA | [R].LILTDN | 1xCarbami | M0QWZ1 1 | 0.000259 | 0 | 1 | 3 | 4 | M0QWZ1 | Protein FAI | M0QWZ1 [ | 0 | 3175.516 | 0 | -0.33 | 0.058703 | 10.44 | 111.4 | 88.6 | 12.32 | 8.44 | | High | High | High | High | High | High | High | High | High | High | High | 0 | 5.26E-05 | 3.65 |
| FALSE | High | LIITPETPEP | [R].LIITPET | 1xPhospho | Q7TS72 1x | 0.000264 | 0 | 1 | 1 | 1 | Q7TS72 | Inositol-tri | Q7TS72 [3 | 0 | 2468.264 | 0 | 0.72 | 0.004376 | 14.03 | 76.2 | 123.8 | 11.58 | 11.03 | | High | High | High | High | High | High | High | High | High | High | High | 0 | 5.37E-05 | 3 |
| FALSE | High | ILLVDSPGN | [K].ILLVDSI | 1xPhospho | Q3UAY3 1x | 1.07E-09 | 0 | 1 | 6 | 3 | Q3UAY3 | Putative ur | Q3UAY3 [6 | 0 | 2972.385 | 0 | -0.01 | 0.748265 | 21.66 | 98.9 | 101.1 | 8.52 | 10.11 | | High | High | High | High | High | High | High | High | High | High | High | 0 | 4.68E-11 | 5.91 |
| FALSE | High | ILLVDSPGN | [K].ILLVDSI | 1xPhospho | Q3UAY3 1x | 5.06E-11 | 0 | 1 | 6 | 4 | Q3UAY3 | Putative ur | Q3UAY3 [6 | 0 | 2988.38 | 0 | 0.06 | 0.5425 | 14.07 | 101.1 | 98.9 | 6.64 | 8.01 | | High | High | High | High | High | High | High | High | High | High | High | 0 | 1.52E-12 | 6.58 |
| FALSE | High | ILIVTQTPPY | [K].ILIVTQT | 1xPhospho | Q6ZQ58 1x | 0.000483 | 0 | 1 | 3 | 3 | Q6ZQ58 | La-related | Q6ZQ58 [6 | 0 | 1740.932 | 0 | 0.3 | 0.054389 | 24.17 | 87.6 | 112.4 | 7.88 | 9.42 | | High | High | High | High | High | High | High | High | High | High | High | 0 | 0.000106 | 2.69 |
| FALSE | High | ILIVTQTPPY | [K].ILIVTQT | 1xPhospho | Q6ZQ58 1x | 0.000916 | 0 | 1 | 3 | 20 | Q6ZQ58 | La-related | Q6ZQ58 [6 | 0 | 1756.927 | 0 | 0.52 | 0.000556 | 2.37 | 82.5 | 117.5 | 3.14 | 2.73 | | High | High | High | High | High | High | High | High | High | High | High | 0.000786 | 0.006829 | 3.02 |
| FALSE | High | ILMDSPEDA | [K].ILMDSP | 1xPhospho | Q3U5I9 1xP | 1.89E-06 | 0 | 1 | 3 | 2 | Q3U5I9 | Putative ur | Q3U5I9 [36 | 0 | 2627.272 | 0 | 1.18 | 0.001294 | 16.51 | 64 | 136 | 11.1 | 8.36 | | High | High | High | High | High | High | High | High | High | High | High | 0 | 2.07E-07 | 5.22 |
| FALSE | High | ILMDSPEDA | [K].ILMDSP | 1xPhospho | Q3U5I9 1xP | 1.92E-06 | 0 | 1 | 3 | 2 | Q3U5I9 | Putative ur | Q3U5I9 [36 | 0 | 2643.267 | 0 | 1.2 | 0.000537 | 12.69 | 61.7 | 138.3 | 5.91 | 7.17 | | High | High | High | High | High | High | High | High | High | High | High | 0 | 2.11E-07 | 5.42 |
| FALSE | High | ILNDLSSDA | [K].ILNDLS | 1xPhospho | Q547S9 1x | 6.53E-05 | 0 | 1 | 5 | 2 | Q547S9 | CAMP resp | Q547S9 [13 | 0 | 1762.894 | 0 | 0.5 | 0.006375 | 7.11 | 81.9 | 118.1 | 6.72 | 5.42 | | High | High | High | High | High | High | High | High | High | High | High | 0 | 1.12E-05 | 2.71 |
| FALSE | High | ILNELSSDV | [K].ILNELS | 1xPhospho | P27699 1xP | 0.016451 | 0.001124 | 1 | 31 | 2 | P27699 | cAMP-resp | P27699 [13 | 0 | 2020.113 | 0 | 1.06 | 0.001713 | 4.67 | 63.7 | 136.3 | 10.57 | 12.37 | | High | High | High | High | High | High | High | High | High | High | High | 0.000585 | 0.005462 | 2.51 |
| FALSE | High | LLNGSHEN | [K].LLNGSH | 1xCarbami | Q7TT18 1x | 6.9E-09 | 0 | 1 | 4 | 2 | Q7TT18 | Activating t | Q7TT18 [4 | 0 | 3219.446 | 0 | 0.23 | 0.033462 | 10.13 | 92 | 108 | 4.83 | 10.26 | | High | High | High | High | High | High | High | High | High | High | High | 0 | 3.78E-10 | 4.37 |
| FALSE | High | LLNGSHEN | [K].LLNGSH | 1xCarbami | Q7TT18 1x | 4.82E-06 | 0 | 1 | 4 | 3 | Q7TT18 | Activating t | Q7TT18 [4 | 0 | 3220.43 | 0 | 0.28 | 0.040002 | 8.55 | 90.3 | 109.7 | 13.58 | 13.3 | | High | High | High | High | High | High | High | High | High | High | High | 0 | 5.96E-07 | 4.03 |
| FALSE | High | IINLGPVHP | [R].IINLGPV | 1xPhospho | Q3TWL2 1 | 0.000235 | 0 | 1 | 3 | 1 | Q3TWL2 | Type 1 pho | Q3TWL2 [1 | 0 | 2614.378 | 0 | -0.28 | 0.009386 | 0.7 | 117 | 83 | 11.59 | 12.91 | | High | High | High | High | High | High | High | High | High | High | High | 0 | 4.69E-05 | 3.91 |
| FALSE | High | IINLGPVHP | [R].IINLGPV | 1xPhospho | Q3TWL2 1 | 4.08E-05 | 0 | 1 | 3 | 1 | Q3TWL2 | Type 1 pho | Q3TWL2 [1 | 0 | 2630.373 | 0 | -0.49 | 0.0642 | 3.24 | 116.5 | 83.5 | 3.8 | 14.79 | | High | High | High | High | High | High | High | High | High | High | High | 0 | 6.57E-06 | 2.03 |
| FALSE | High | IINLNLASPA | [R].IINLNLA | 1xPhospho | F7BJB9 1xP | 0.005182 | 0.000551 | 1 | 2 | 1 | F7BJB9 | MORC fam | F7BJB9 [52 | 0 | 2553.304 | 0 | -0.5 | 0.011345 | 5.48 | 116.6 | 83.4 | 9.76 | 5.49 | | High | High | High | High | High | High | High | High | High | High | High | 0.000246 | 0.001498 | 3.27 |
| FALSE | High | IINLNLASPA | [R].IINLNLA | 1xPhospho | F7BJB9 1xP | 6.79E-06 | 0 | 1 | 2 | 1 | F7BJB9 | MORC fam | F7BJB9 [52 | 0 | 2569.298 | 0 | -0.43 | 0.180092 | 2.95 | 113.3 | 86.7 | 13.32 | 15.1 | | High | High | High | High | High | High | High | High | High | High | High | 0 | 8.77E-07 | 3.9 |
| FALSE | High | LINSELGSP | [R].LINSELC | 1xPhospho | Q3UDW8 1 | 0.01441 | 0.001011 | 1 | 2 | 1 | Q3UDW8 | Heparan-al | Q3UDW8 [ | 0 | 1481.756 | 0 | 0.88 | 0.000997 | 10.89 | 72.2 | 127.8 | 2.87 | 6.5 | | High | High | High | High | High | High | High | High | High | High | High | 0.000538 | 0.004698 | 1.12 |
| FALSE | High | LLPAASSDA | [R].LLPAAS | 1xPhospho | Q3UHU5 1 | 0.057686 | 0.004227 | 1 | 2 | 4 | Q3UHU5 | Microtubu | Q3UHU5 [2 | 1 | 2525.208 | 0 | -1.79 | 0.00041 | 1.93 | 155.3 | 44.7 | 2.05 | 23.78 | | High | High | High | High | High | High | High | High | High | High | High | 0.007374 | 0.08109 | 2.26 |
| FALSE | High | LLPDAVLEC | [K].LLPDAV | 1xPhospho | Q9D7Z3 1x | 8.18E-05 | 0 | 1 | 2 | 1 | Q9D7Z3 | Nucleolar p | Q9D7Z3 [1 | 0 | 2565.383 | 0 | -0.1 | 0.401835 | 20.81 | 98.4 | 101.6 | 13.76 | 16.56 | | High | High | High | High | High | High | High | High | High | High | High | 0 | 1.43E-05 | 3.63 |
| FALSE | High | LLPEFDDSD | [R].LLPEFD | 1xPhospho | Q4LDD4 1x | 0.000217 | 0 | 1 | 2 | 3 | Q4LDD4 | Arf-GAP wi | Q4LDD4 [2 | 0 | 3432.585 | 0 | 0.71 | 0.025512 | 29.48 | 67 | 133 | 15.34 | 20.46 | | High | High | High | High | High | High | High | High | High | High | High | 0 | 4.29E-05 | 2.82 |
| FALSE | High | LLPEFDDSD | [R].LLPEFD | 1xPhospho | Q4LDD4 1x | 0.022773 | 0.001542 | 1 | 2 | 2 | Q4LDD4 | Arf-GAP wi | Q4LDD4 [2 | 0 | 3448.58 | 0 | 0.87 | 0.036576 | 10.94 | 72.9 | 127.1 | 3.19 | 30.94 | | High | High | High | High | High | High | High | High | High | High | High | 0.004115 | 0.04384 | 3.63 |
| FALSE | High | LLPEGEETV | [R].LLPEGE | 1xPhospho | Q9CWU4 1 | 1.09E-06 | 0 | 1 | 1 | 3 | Q9CWU4 | UPF0690 p | Q9CWU4 [ | 0 | 2214.047 | 0 | -0.54 | 0.008939 | 5.4 | 119.6 | 80.4 | 7.77 | 9.62 | | High | High | High | High | High | High | High | High | High | High | High | 0 | 1.11E-07 | 4.11 |
| FALSE | High | LLPELSPEP | [R].LLPELSI | 1xPhospho | B0QZP6 1x | 0.011549 | 0.00086 | 1 | 4 | 1 | B0QZP6 | Transmem | B0QZP6 [6 | 0 | 1924.973 | 0 | 0.78 | 0.007032 | 21.13 | 74.1 | 125.9 | 11.26 | 13.93 | | High | High | High | High | High | High | High | High | High | High | High | 0.000437 | 0.003695 | 2.79 |
| FALSE | High | LIPGTPDVE | [R].LIPGTPI | 1xPhospho | Q7TT79 1x | 6.11E-07 | 0 | 1 | 4 | 1 | Q7TT79 | Microceph | Q7TT79 [3 | 0 | 3095.49 | 0 | 0.28 | 0.230947 | 20.09 | 85.4 | 114.6 | 27.55 | 7.17 | | High | High | High | High | High | High | High | High | High | High | High | 0 | 5.85E-08 | 4.3 |
| FALSE | High | LIPLDEECA | [K].LIPLDEI | 1xCarbami | Q80W00 1 | 4.1E-06 | 0 | 1 | 3 | 4 | Q80W00 | Serine/thre | Q80W00 [ | 0 | 4073.865 | 0 | 0.67 | 0.055165 | 33.87 | 71.2 | 128.8 | 15.06 | 26.53 | | High | High | High | High | High | High | High | High | High | High | High | 0 | 4.97E-07 | 4.76 |
| FALSE | High | LIPLDEECA | [K].LIPLDEI | 1xCarbami | Q80W00 1 | 0.000314 | 0 | 1 | 3 | 10 | Q80W00 | Serine/thre | Q80W00 [ | 0 | 4089.86 | 0 | 0.65 | 0.057117 | 19.62 | 77.8 | 122.2 | 15.52 | 24.35 | | High | High | High | High | High | High | High | High | High | High | High | 0 | 9.03E-05 | 4.48 |
| FALSE | High | LLPIDGAND | [R].LLPIDG. | 1xPhospho | Q9EQQ9-3 | 0.001097 | 0 | 1 | 2 | 1 | Q9EQQ9-3 | Isoform 3 c | Q9EQQ9-3 | 0 | 2919.567 | 0 | 0.17 | 0.168872 | 16.22 | 89.2 | 110.8 | 11.34 | 13.23 | | High | High | High | High | High | High | High | High | High | High | High | 0 | 0.000264 | 3.66 |
| FALSE | High | IISAFPNAN | [R].IISAFPN | 2xDeamida | A0A140LJ6 | 0.030367 | 0.001857 | 1 | 1 | 2 | A0A140LJ6 | Protein un | A0A140LJ6 | 0 | 2177.984 | 0 | -0.18 | 0.888072 | 1.53 | 105.5 | 94.5 | 18.08 | 17.07 | | High | High | High | High | High | High | High | High | High | High | High | 0.0032 | 0.03377 | 3.01 |
| FALSE | High | LLEDSESGE | [K].LLEDSE | 1xPhospho | G3X8X0 1x | 1.59E-06 | 0 | 1 | 4 | 1 | G3X8X0 | DEAH (Asp | G3X8X0 [9 | 0 | 2674.286 | 0 | -0.07 | 0.290998 | 18.52 | 102.3 | 97.7 | 11.86 | 8.5 | | High | High | High | High | High | High | High | High | High | High | High | 0 | 1.71E-07 | 3.25 |
| FALSE | High | INPYMSSP | [R].INPYM | 1xCarbami | Q80V08 1x | 0.002398 | 0.000109 | 1 | 5 | 4 | Q80V08 | Rpl17 prot | Q80V08 [1 | 0 | 2717.315 | 0 | -0.7 | 0.010624 | 11.18 | 123.9 | 76.1 | 11.86 | 10.04 | | High | High | High | High | High | High | High | High | High | High | High | 4.94E-05 | 0.000636 | 2.66 |
| FALSE | High | LNQSGASV | [K].LNQSG | 1xPhospho | Q9ERU9 1x | 4.79E-11 | 0 | 1 | 1 | 9 | Q9ERU9 | E3 SUMO-p | Q9ERU9 [2 | 0 | 2689.168 | 0 | 0.36 | 0.106461 | 11.19 | 89.2 | 110.8 | 9.28 | 20.1 | | High | High | High | High | High | High | High | High | High | High | High | 0 | 1.42E-12 | 5.28 |
| FALSE | High | LPSVEEAEV | [R].LPSVEE | 1xPhospho | P57016 1xP | 0.053926 | 0.00384 | 1 | 2 | 1 | P57016 | Ladinin-1 C | P57016 [60 | 0 | 2619.425 | 0 | -0.91 | 0.017708 | 7.1 | 130.6 | 69.4 | 19.59 | 7.34 | | High | High | High | High | High | High | High | High | High | High | High | 0.002031 | 0.02096 | 1.54 |
| FALSE | High | LPSVGEEST | [R].LPSVGE | 1xDeamida | Q8C180 1x | 0.101749 | 0.008194 | 1 | 1 | 2 | Q8C180 | Fibroblast | Q8C180 [1 | 0 | 3858.864 | 0 | 0.54 | 0.014795 | 5.75 | 81.4 | 118.6 | 12.32 | 10.53 | | High | High | High | High | High | High | High | High | High | High | High | 0.004126 | 0.04398 | 3.35 |
| FALSE | High | LPSVLLAAV | [R].LPSVLL | 1xPhospho | B9EKU4 1x | 0.019791 | 0.001438 | 1 | 4 | 1 | B9EKU4 | URB2 ribos | B9EKU4 [1 | 0 | 2711.477 | 0 | -0.45 | 0.344107 | 17.23 | 113.1 | 86.9 | 18.52 | 18.6 | | High | High | High | High | High | High | High | High | High | High | High | 0.000773 | 0.006706 | 2.66 |
| FALSE | High | LPSVSDENI | [R].LPSVSD | 1xPhospho | Q8CBC4 1x | 1.42E-07 | 0 | 1 | 2 | 3 | Q8CBC4 | Consortin | Q8CBC4 [3 | 0 | 3198.446 | 0 | 0.32 | 0.130583 | 12.27 | 89.6 | 110.4 | 11.99 | 16.21 | | High | High | High | High | High | High | High | High | High | High | High | 0 | 1.13E-08 | 4.9 |
| FALSE | High | LPSYGLCES | [R].LPSYGL | 1xCarbami | Q8CGU1 1 | 9.08E-07 | 0 | 1 | 3 | 3 | Q8CGU1 | Calcium-bi | Q8CGU1 [5 | 0 | 2372.079 | 0 | -0.97 | 0.001142 | 6.31 | 130.4 | 69.6 | 6.69 | 13.64 | | High | High | High | High | High | High | High | High | High | High | High | 0 | 9.16E-08 | 4.46 |
| FALSE | High | IPTDTSTPP | [K].IPTDTS | 1xPhospho | Q6ZQD6 1 | 0.107603 | 0.008634 | 1 | 5 | 2 | Q6ZQD6 | MKIAA033 | Q6ZQD6 [4 | 0 | 1393.693 | 0 | 2.07 | 0.000154 | 1.31 | 37.3 | 162.7 | 7.27 | 4.47 | | High | High | High | High | High | High | High | High | High | High | High | 0.004341 | 0.04695 | 1.62 |
| FALSE | High | LPTKPETSF | [K].LPTKPE | 1xPhospho | Q8C8U0-3 | 0.003495 | 0.00021 | 1 | 6 | 2 | Q8C8U0-3 | Isoform 3 c | Q8C8U0-3 | 0 | 2201.089 | 0 | 1.41 | 0.003575 | 32.35 | 54.6 | 145.4 | 10.66 | 13.93 | | High | High | High | High | High | High | High | High | High | High | High | 9.42E-05 | 0.000969 | 2.17 |
| FALSE | High | LPTPSPEPP | [R].LPTPSP | 1xPhospho | Q61115 1x | 0.081752 | 0.006219 | 1 | 3 | 1 | Q61115 | Protein pat | Q61115 [1 | 0 | 1781.94 | 0 | -1.01 | 0.015347 | 1.47 | 135.7 | 64.3 | 13.59 | 42.53 | | High | High | High | High | High | High | High | High | High | High | High | 0.003243 | 0.03421 | 1.66 |
| FALSE | High | LPTPTASLP | [K].LPTPTA | 1xPhospho | Q924H7 1x | 6.71E-05 | 0 | 1 | 6 | 1 | Q924H7 | WW doma | Q924H7 [2 | 0 | 1761.992 | 0 | 0.41 | 0.025597 | 15.99 | 82.8 | 117.2 | 9.49 | 11.85 | | High | High | High | High | High | High | High | High | High | High | High | 0 | 1.14E-05 | 2.33 |
| FALSE | High | LPTSEEERS | [K].LPTSEE | 1xPhospho | P70399 1xP | 0.052576 | 0.00384 | 1 | 7 | 1 | P70399 | TP53-bindi | P70399 [16 | 1 | 1881.972 | 0 | 0.06 | 0.164511 | 6.57 | 96.8 | 103.2 | 9.79 | 6.22 | | High | High | High | High | High | High | High | High | High | High | High | 0.001999 | 0.02043 | 2.66 |
| FALSE | High | LPTSEEERS | [K].LPTSEE | 2xPhospho | P70399 2xP | 0.114657 | 0.009411 | 1 | 7 | 1 | P70399 | TP53-bindi | P70399 [16 | 1 | 1961.939 | 0 | -0.54 | 0.000905 | 3.44 | 119.1 | 80.9 | 4.38 | 2.96 | | High | High | High | High | High | High | High | High | High | High | High | 0.004707 | 0.05076 | 2.21 |
| FALSE | High | LPTSEPEAD | [K].LPTSEP | 1xPhospho | E9Q5F9 1x | 0.008167 | 0.000736 | 1 | 5 | 2 | E9Q5F9 | Histone-lys | E9Q5F9 [1 | 0 | 2182.057 | 0 | -1.47 | 0.002434 | 4.27 | 143.5 | 56.5 | 7.48 | 33.03 | | High | High | High | High | High | High | High | High | High | High | High | 0.000555 | 0.004773 | 2.1 |
| FALSE | High | LPTSPSGSE | [R].LPTSPS | 1xPhospho | A2RSY1 1xP | 9.91E-06 | 0 | 1 | 3 | 6 | A2RSY1 | KAT8 regul | A2RSY1 [52 | 0 | 2772.359 | 1 | -0.61 | 0.001348 | 4.39 | 120.5 | 79.5 | 5.38 | 8.58 | | High | High | High | High | High | High | High | High | High | High | High | 0 | 3.49E-06 | 4.29 |
| FALSE | High | LPSTSGSEG | [R].LPSTSG | 2xPhospho | Q03145 2x | 0.000656 | 0 | 1 | 1 | 3 | Q03145 | Ephrin type | Q03145 [8 | 0 | 1722.77 | 0 | 1.24 | 0.001214 | 13.25 | 60.4 | 139.6 | 5.11 | 9.02 | | High | High | High | High | High | High | High | High | High | High | High | 4.94E-05 | 0.0006 | 3.37 |
| FALSE | High | LPTSPSGSE | [R].LPTSPS | 2xPhospho | A2RSY1 2xP | 1.57E-05 | 0 | 1 | 3 | 3 | A2RSY1 | KAT8 regul | A2RSY1 [52 | 0 | 2852.326 | 1 | -0.11 | 0.198519 | 8.93 | 100.5 | 99.5 | 6.06 | 11.91 | | High | High | High | High | High | High | High | High | High | High | High | 0 | 5.57E-06 | 4.49 |
| FALSE | High | LPVASQAS | [K].LPVASC | 1xPhospho | Q3TUF7 1x | 9.97E-08 | 0 | 1 | 4 | 1 | Q3TUF7 | YEATS dom | Q3TUF7 [5 | 0 | 2176.178 | 0 | -0.61 | 0.000402 | 0.77 | 120.8 | 79.2 | 3.75 | 4.94 | | High | High | High | High | High | High | High | High | High | High | High | 0 | 7.59E-09 | 3.31 |
| FALSE | High | IPVEGPLSP | [R].IPVEGP | 1xPhospho | Q9D219 1x | 0.096959 | 0.007777 | 1 | 1 | 1 | Q9D219 | B-cell CLL/I | Q9D219 [6 | 0 | 1460.771 | 0 | -0.39 | 0.003356 | 4.03 | 113.2 | 86.8 | 5.12 | 2.05 | | High | High | High | High | High | High | High | High | High | High | High | 0.003922 | 0.04154 | 1.75 |
| FALSE | High | LPVPQGAL | [R].LPVPQ | 1xDeamida | A0A0A0MC | 0.011549 | 0.00086 | 1 | 2 | 2 | A0A0A0MC | Histone-lys | A0A0A0MC | 0 | 3492.854 | 0 | 0.82 | 0.01123 | 15.68 | 72.1 | 127.9 | 8.64 | 18.99 | | High | High | High | High | High | High | High | High | High | High | High | 0.000437 | 0.003669 | 2.95 |
| FALSE | High | IPVPSSPVP | [R].IPVPSSI | 1xPhospho | D3Z183 1xP | 0.017041 | 0.001204 | 1 | 4 | 2 | D3Z183 | Major facil | D3Z183 [64 | 0 | 3139.631 | 0 | 6.64 | | 0 | 24 | 176 | | 21.93 | | Not Found | Not Found | Not Found | Not Found | High | High | High | High | High | High | High | 0.000814 | 0.007818 | 2.62 |
| FALSE | High | LPVSPVAGS | [R].LPVSPV | 1xPhospho | Q91WM3 1 | 0.119269 | 0.00979 | 1 | 1 | 1 | Q91WM3 | U3 small nu | Q91WM3 [ | 0 | 1135.596 | 0 | -0.49 | 0.509764 | 24.48 | 114.4 | 85.6 | 24.75 | 28.81 | | High | High | High | High | High | High | High | High | High | High | High | 0.004866 | 0.05301 | 1.27 |
| FALSE | High | LPVSSSTSN | [K].LPVSSS | 1xPhospho | E9Q6P5 1x | 0.000372 | 0 | 1 | 2 | 4 | E9Q6P5 | Tetratricop | E9Q6P5 [1 | 0 | 1820.911 | 0 | 0 | 0.809082 | 27.17 | 99.9 | 100.1 | 7.16 | 20.96 | | High | High | High | High | High | High | High | High | High | High | High | 4.94E-05 | 0.000755 | 2.39 |
| FALSE | High | LPVSVDSSF | [R].LPVSVD | 1xPhospho | Q5SNZ0 1x | 3.85E-06 | 0 | 1 | 4 | 2 | Q5SNZ0 | Girdin OS= | Q5SNZ0 [1 | 0 | 2841.428 | 0 | 1.99 | 0.000831 | 36.13 | 41.1 | 158.9 | 22.97 | 8.37 | | High | High | High | High | High | High | High | High | High | High | High | 0 | 1.32E-05 | 3.89 |
| FALSE | High | IPVTPPVAP | [R].IPVTPP | 1xPhospho | E9QK89 1x | 0.039686 | 0.00283 | 1 | 2 | 1 | E9QK89 | Mediator c | E9QK89 [3 | 0 | 1426.827 | 0 | -0.4 | 0.133708 | 14.47 | 113.1 | 86.9 | 14.1 | 24.58 | | High | High | High | High | High | High | High | High | High | High | High | 0.001468 | 0.01482 | 2.41 |
| FALSE | High | IPVTPPVAP | [R].IPVTPP | 1xPhospho | E9QK89 1x | 0.000343 | 0 | 1 | 2 | 2 | E9QK89 | Mediator c | E9QK89 [3 | 0 | 1655.99 | 0 | -0.95 | 0.001079 | 4.01 | 132.4 | 67.6 | 6.7 | 6.98 | | High | High | High | High | High | High | High | High | High | High | High | 0 | 7.15E-05 | 3.07 |
| FALSE | High | LPYSDGYSI | [K].LPYSDG | 2xPhospho | P70303 2xP | 0.001077 | 0 | 1 | 3 | 1 | P70303 | CTP syntha | P70303 [56 | 0 | 2682.13 | 0 | -0.58 | 0.005575 | 5.56 | 118.4 | 81.6 | 11.02 | 7.97 | | High | High | High | High | High | High | High | High | High | High | High | 0 | 0.000259 | 3.38 |
| FALSE | High | IPYTPGEIPI | [R].IPYTPG | 1xPhospho | Q3TJE8 1xP | 0.093123 | 0.007276 | 1 | 5 | 1 | Q3TJE8 | UDP-gluco | Q3TJE8 [47 | 0 | 1423.744 | 0 | -0.81 | 0.001139 | 4.99 | 127.2 | 72.8 | 4.95 | 19.6 | | High | High | High | High | High | High | High | High | High | High | High | 0.003698 | 0.03978 | 2.04 |
| FALSE | High | IPYTPGEIPI | [R].IPYTPG | 1xPhospho | Q3TJE8 1xP | 0.00359 | 0.00021 | 1 | 5 | 3 | Q3TJE8 | UDP-gluco | Q3TJE8 [47 | 0 | 1652.906 | 0 | -0.74 | 0.001321 | 6.11 | 127.4 | 72.6 | 7.19 | 11.06 | | High | High | High | High | High | High | High | High | High | High | High | 0.000872 | 0.008711 | 2.68 |
| FALSE | High | IQAAASPPA | [R].IQAAAS | 1xPhospho | E9QAQ5 1 | 6.95E-08 | 0 | 1 | 2 | 1 | E9QAQ5 | Glycogen s | E9QAQ5 [3 | 0 | 2379.114 | 0 | 0.24 | 0.044229 | 11.66 | 92.8 | 107.2 | 11.27 | 2.42 | | High | High | High | High | High | High | High | High | High | High | High | 0 | 5.1E-09 | 4.56 |
| FALSE | High | IPTYEDQW | [R].IPTYED | 1xPhospho | P39447 1xP | 3.41E-05 | 0 | 1 | 4 | 4 | P39447 | Tight junct | P39447 [10 | 0 | 2361.073 | 0 | 1.33 | 0.001039 | 14.95 | 56.6 | 143.4 | 14.07 | 7.48 | | High | High | High | High | High | High | High | High | High | High | High | 0 | 8.75E-05 | 3.33 |
| FALSE | High | LQAADEEE | [R].LQAAD | 1xPhospho | P32507 1xP | 8.55E-08 | 0 | 1 | 2 | 2 | P32507 | Nectin-2 O | P32507 [38 | 0 | 3192.632 | 0 | -1.02 | 0.000845 | 3.06 | 134.8 | 65.2 | 5.97 | 7.65 | | High | High | High | High | High | High | High | High | High | High | High | 0 | 6.41E-09 | 3.45 |
| FALSE | High | LPSTSGSEG | [R].LPSTSG | 1xPhospho | Q03145 1x | 2.82E-05 | 0 | 1 | 1 | 6 | Q03145 | Ephrin type | Q03145 [8 | 0 | 1642.804 | 0 | 0.86 | 0.001842 | 16.25 | 71.3 | 128.7 | 5.6 | 7.08 | | High | High | High | High | High | High | High | High | High | High | High | 0.004059 | 0.04336 | 2.53 |
| FALSE | High | LPSTSDDCF | [K].LPSTSD | 1xCarbami | Q925J9 1xP | 7.34E-05 | 0 | 1 | 2 | 3 | Q925J9 | Mediator c | Q925J9 [79 | 0 | 2020.961 | 0 | 1.16 | 0.001377 | 10.84 | 61.3 | 138.7 | 3.16 | 10.95 | | High | High | High | High | High | High | High | High | High | High | High | 0 | 4.63E-05 | 2.61 |
| FALSE | High | IPSAVSTVS | [K].IPSAVS | 1xPhospho | Q3U3A6 1 | 0.001361 | 0 | 1 | 4 | 3 | Q3U3A6 | Sodium/hy | Q3U3A6 [6 | 0 | 2263.192 | 0 | 0.31 | 0.109569 | 7.05 | 90.7 | 109.3 | 9.47 | 22.08 | | High | High | High | High | High | High | High | High | High | High | High | 2.58E-05 | 0.000339 | 2.42 |
| FALSE | High | LPSDASEDA | [R].LPSDAS | 1xPhospho | Q9JIG7 1xP | 0.028824 | 0.001829 | 1 | 1 | 1 | Q9JIG7 | Coiled-coil | Q9JIG7 [10 | 0 | 2484.149 | 0 | 0.03 | 0.949534 | 10.55 | 99.1 | 100.9 | 10.95 | 21.12 | | High | High | High | High | High | High | High | High | High | High | High | 0.000997 | 0.01034 | 2.6 |
| FALSE | High | LPSDEDESC | [K].LPSDED | 1xPhospho | E9Q9B7 1x | 5.5E-05 | 0 | 1 | 11 | 8 | E9Q9B7 | Kinase D-ir | E9Q9B7 [1 | 0 | 2714.27 | 0 | 0.23 | 0.001891 | 2.87 | 93.1 | 106.9 | 2.95 | 5.19 | | High | High | High | High | High | High | High | High | High | High | High | 0 | 9.19E-06 | 3.57 |
| FALSE | High | LPSDEDESC | [K].LPSDED | 2xPhospho | E9Q9B7 2x | 0.00418 | 0.00036 | 1 | 11 | 1 | E9Q9B7 | Kinase D-ir | E9Q9B7 [1 | 0 | 2794.236 | 0 | 0.31 | 0.106711 | 19.75 | 90.4 | 109.6 | 7.63 | 17.01 | | High | High | High | High | High | High | High | High | High | High | High | 0.000161 | 0.001189 | 2.2 |
| FALSE | High | LPSDGELPD | [R].LPSDGE | 1xCarbami | Q570Z6 1x | 3.31E-07 | 0 | 1 | 2 | 2 | Q570Z6 | MKIAA412 | Q570Z6 [1 | 0 | 2600.283 | 0 | -0.13 | 0.237978 | 6.86 | 104.7 | 95.3 | 6.05 | 6.74 | | High | High | High | High | High | High | High | High | High | High | High | 0 | 2.93E-08 | 3.55 |
| FALSE | High | IPSDNLEEP | [K].IPSDNL | 1xPhospho | Q3UYC8 1 | 9.11E-05 | 0 | 1 | 5 | 2 | Q3UYC8 | Putative ur | Q3UYC8 [9 | 0 | 1875.987 | 0 | -0.68 | 0.000965 | 1.46 | 123 | 77 | 4.83 | 5.4 | | High | High | High | High | High | High | High | High | High | High | High | 0 | 1.61E-05 | 3.04 |
| FALSE | High | LPSDSSASP | [R].LPSDSS | 1xPhospho | Q6P4T3 1x | 1.9E-05 | 0 | 1 | 5 | 10 | Q6P4T3 | Eyes absen | Q6P4T3 [2 | 0 | 2365.205 | 0 | 0.06 | 0.11081 | 1.8 | 98.1 | 101.9 | 2.61 | 2.93 | | High | High | High | High | High | High | High | High | High | High | High | 0 | 2.78E-06 | 3.42 |
| FALSE | High | LPSDSSASP | [R].LPSDSS | 2xPhospho | Q6P4T3 2x | 0.000161 | 0 | 1 | 5 | 1 | Q6P4T3 | Eyes absen | Q6P4T3 [2 | 0 | 2445.172 | 0 | 0.48 | 0.032685 | 17.46 | 82.7 | 117.3 | 7.98 | 18.69 | | High | High | High | High | High | High | High | High | High | High | High | 0 | 3.05E-05 | 2.72 |
| FALSE | High | LPSFNWSP | [K].LPSFNV | 1xPhospho | F7DBB3 1x | 0.014797 | 0.001049 | 1 | 1 | 1 | F7DBB3 | AHNAK nu | F7DBB3 [7 | 0 | 1613.849 | 0 | 2.08 | 0.002007 | 6.74 | 38.7 | 161.3 | 14.74 | 14.64 | | High | High | High | High | High | High | High | High | High | High | High | 0.000555 | 0.004848 | 2.25 |
| FALSE | High | LPSGSGPAS | [R].LPSGSG | 1xPhospho | E9Q616 1x | 1.87E-10 | 0 | 1 | 3 | 14 | E9Q616 | AHNAK nu | E9Q616 [2 | 0 | 2079.032 | 0 | 2.27 | 0.000181 | 1.42 | 35.4 | 164.6 | 4.75 | 5.53 | | High | High | High | High | High | High | High | High | High | High | High | 0 | 6.6E-12 | 4.27 |
| FALSE | High | LPSGSGPAS | [R].LPSGSG | 2xPhospho | E9Q616 2x | 0.000334 | 0 | 1 | 3 | 6 | E9Q616 | AHNAK nu | E9Q616 [2 | 0 | 2158.998 | 0 | 2.35 | 0.000223 | 5.61 | 32.5 | 167.5 | 8.85 | 7.5 | | High | High | High | High | High | High | High | High | High | High | High | 0 | 7.64E-05 | 3.29 |
| FALSE | High | LPSGSGPAS | [R].LPSGSG | 3xPhospho | E9Q616 3x | 4.94E-05 | 0 | 1 | 3 | 6 | E9Q616 | AHNAK nu | E9Q616 [2 | 0 | 2238.965 | 0 | 0.43 | 0.00243 | 5.14 | 86.1 | 113.9 | 4.5 | 8.59 | | High | High | High | High | High | High | High | High | High | High | High | 0 | 4.31E-05 | 3.44 |
| FALSE | High | LPSLSVQPV | [R].LPSLSV | 1xPhospho | Q3UGY8 1 | 4.96E-07 | 0 | 1 | 1 | 1 | Q3UGY8 | Brefeldin A | Q3UGY8 [1 | 0 | 2676.457 | 0 | 5.01 | 0.000218 | 51.74 | 6.4 | 193.6 | 67.45 | 4.41 | | High | High | High | High | High | High | High | High | High | High | High | 0 | 4.63E-08 | 5.08 |
| FALSE | High | LPSTSDDCF | [K].LPSTSD | 1xCarbami | Q925J9 2xP | 4.89E-05 | 0 | 1 | 2 | 2 | Q925J9 | Mediator c | Q925J9 [79 | 0 | 2100.928 | 0 | 0.45 | 0.030095 | 14.29 | 81.8 | 118.2 | 9.7 | 15.35 | | High | High | High | High | High | High | High | High | High | High | High | 0 | 8.03E-06 | 3.22 |
| FALSE | High | LPSPAQTQ | [K].LPSPAC | 1xPhospho | Q3TN34 1x | 0.053023 | 0.00384 | 1 | 4 | 1 | Q3TN34 | MICAL-like | Q3TN34 [1 | 0 | 1306.672 | 0 | 0.1 | 0.510189 | 20.62 | 91.4 | 108.6 | 6.71 | 20.24 | | High | High | High | High | High | High | High | High | High | High | High | 0.001999 | 0.02061 | 1.64 |
| FALSE | High | IPSPIGFPSC | [R].IPSPIGF | 1xPhospho | Q9EST3 1x | 5.85E-08 | 0 | 1 | 6 | 1 | Q9EST3 | Eukaryotic | Q9EST3 [5 | 0 | 2748.378 | 0 | 0.33 | 0.038375 | 19.97 | 86.9 | 113.1 | 8.16 | 10.57 | | High | High | High | High | High | High | High | High | High | High | High | 0 | 4.2E-09 | 3.63 |
| FALSE | High | IPSPIGFPSC | [R].IPSPIGF | 1xPhospho | Q9EST3 1x | 0.001127 | 0 | 1 | 6 | 2 | Q9EST3 | Eukaryotic | Q9EST3 [5 | 0 | 2764.373 | 0 | 0.29 | 0.019917 | 2.59 | 88.8 | 111.2 | 4.3 | 9.49 | | High | High | High | High | High | High | High | High | High | High | High | 0 | 0.000272 | 3.96 |
| FALSE | High | IPSPLQPET | [R].IPSPLQ | 1xPhospho | Q9D8N6 1 | 5.72E-06 | 0 | 1 | 3 | 3 | Q9D8N6 | Protein lin- | Q9D8N6 [1 | 0 | 3587.688 | 0 | 0.21 | 0.084197 | 14.59 | 91.6 | 108.4 | 8.45 | 4.48 | | High | High | High | High | High | High | High | High | High | High | High | 0 | 7.19E-07 | 4.42 |
| FALSE | High | LPSPTAAPC | [R].LPSPTA | 1xPhospho | O88935 1x | 0.001127 | 0 | 1 | 3 | 2 | O88935 | Synapsin-1 | O88935 [5 | 0 | 2687.335 | 0 | -0.8 | 0.000816 | 0.52 | 129.5 | 70.5 | 6.84 | 11.58 | | High | High | High | High | High | High | High | High | High | High | High | 0 | 0.000274 | 3.14 |
| FALSE | High | LPSPTPENK | [R].LPSPTP | 1xPhospho | Q3U2G7 1 | 0.019618 | 0.001438 | 1 | 9 | 2 | Q3U2G7 | Putative ur | Q3U2G7 [7 | 0 | 1520.813 | 0 | -1.2 | 0.000223 | 0.53 | 139.1 | 60.9 | 3.38 | 10.64 | | High | High | High | High | High | High | High | High | High | High | High | 0.000756 | 0.006669 | 1.73 |
| FALSE | High | LPSPVFAQT | [K].LPSPVF | 1xPhospho | Q3TYA6 1x | 0.017809 | 0.001282 | 1 | 2 | 1 | Q3TYA6 | M-phase p | Q3TYA6 [3 | 0 | 1738.991 | 0 | -1.06 | 0.000501 | 0.58 | 135.2 | 64.8 | 9.06 | 6.56 | | High | High | High | High | High | High | High | High | High | High | High | 0.000689 | 0.00596 | 2.87 |
| FALSE | High | LPSPVTQVI | [K].LPSPVT | 1xPhospho | Q9D1F5 1x | 0.00051 | 0 | 1 | 1 | 1 | Q9D1F5 | MRN comp | Q9D1F5 [2 | 0 | 1963.067 | 0 | -0.2 | 0.008364 | 6.23 | 105.8 | 94.2 | 5.96 | 6.9 | | High | High | High | High | High | High | High | High | High | High | High | 0 | 0.000112 | 3.08 |
| FALSE | High | LPSQGNEL | [R].LPSQGI | 1xPhospho | D3YYY8 1xP | 0.003083 | 0.00016 | 1 | 1 | 1 | D3YYY8 | Rho guanir | D3YYY8 [2 | 0 | 2971.591 | 0 | 2.83 | 0.000435 | 20.02 | 25.1 | 174.9 | 10.84 | 8.26 | | High | High | High | High | High | High | High | High | High | High | High | 7.19E-05 | 0.000844 | 3.41 |
| FALSE | High | LPSQTESVT | [R].LPSQTE | 1xPhospho | Q3UHJ0 1x | 0.009087 | 0.000736 | 1 | 2 | 2 | Q3UHJ0 | AP2-associ | Q3UHJ0 [8 | 0 | 1627.789 | 0 | 0.78 | 0.001599 | 8.89 | 68.2 | 131.8 | 13.44 | 5.92 | | High | High | High | High | High | High | High | High | High | High | High | 0.000327 | 0.002821 | 2.77 |
| FALSE | High | LPSSPASPS | [R].LPSSPA | 1xPhospho | A2AUK5 1x | 0.081752 | 0.006219 | 2 | 10 | 2 | A2AUK5; C | Band 4.1-li | A2AUK5 [5 | 0 | 1605.865 | 0 | -0.26 | 0.003357 | 2.01 | 109.4 | 90.6 | 6.97 | 9.74 | Shared | High | High | High | High | High | High | High | High | High | High | High | 0.0032 | 0.0339 | 1.82 |
| FALSE | High | LPSSSPPLEI | [K].LPSSSPI | 1xPhospho | E9PVX6 1xP | 0.004033 | 0.000311 | 1 | 2 | 4 | E9PVX6 | Proliferatic | E9PVX6 [2 | 0 | 2180.069 | 0 | 0.19 | 0.179875 | 20.44 | 93.9 | 106.1 | 9.06 | 15.39 | | High | High | High | High | High | High | High | High | High | High | High | 0.00014 | 0.001134 | 2.34 |
| FALSE | High | LPSTEVK | [K].LPSTEV | 1xPhospho | Q3ULT8 1x | 0.038677 | 0.002563 | 1 | 3 | 1 | Q3ULT8 | Putative ur | Q3ULT8 [8 | 0 | 1311.733 | 0 | 1.08 | 0.001344 | 2.86 | 63.7 | 136.3 | 4.99 | 12.09 | | High | High | High | High | High | High | High | High | High | High | High | 0.001372 | 0.01432 | 1.64 |
| FALSE | High | LPSTPTTTA | [R].LPSTPT | 1xCarbami | Q6A0A2 1x | 0.000164 | 0 | 1 | 3 | 7 | Q6A0A2 | La-related | Q6A0A2 [6 | 0 | 1714.885 | 0 | 2.07 | 0.000348 | 7.06 | 39.8 | 160.2 | 5.21 | 6.82 | | High | High | High | High | High | High | High | High | High | High | High | 2.58E-05 | 0.000456 | 2.84 |
| FALSE | High | LPSPLAHSS | [R].LPSPLA | 1xPhospho | O35185 1x | 4.79E-07 | 0 | 1 | 2 | 1 | O35185 | Class E bas | O35185 [3 | 0 | 2586.431 | 0 | 1.69 | 0.00323 | 38.11 | 48.4 | 151.6 | 16.95 | 12.82 | | High | High | High | High | High | High | High | High | High | High | High | 0 | 4.46E-08 | 4.02 |
| FALSE | High | LQAALDNE | [R].LQAALI | 1xPhospho | Q3TXW2 1 | 0.000725 | 0 | 1 | 8 | 2 | Q3TXW2 | Putative ur | Q3TXW2 [3 | 0 | 3289.547 | 0 | -0.36 | 0.042861 | 15.73 | 118.7 | 81.3 | 6.58 | 18.52 | | High | High | High | High | High | High | High | High | High | High | High | 0 | 0.000167 | 2.05 |
| FALSE | High | LQAALDNE | [R].LQAALI | 1xDeamida | Q3TXW2 1 | 0.000375 | 0 | 1 | 8 | 1 | Q3TXW2 | Putative ur | Q3TXW2 [3 | 0 | 3290.531 | 0 | -0.21 | 0.096071 | 12.11 | 110.4 | 89.6 | 12.48 | 9.36 | | High | High | High | High | High | High | High | High | High | High | High | 0 | 7.93E-05 | 2.84 |
| FALSE | High | LQAALDNE | [R].LQAALI | 1xPhospho | Q3TXW2 1 | 0.002921 | 0.00016 | 1 | 8 | 1 | Q3TXW2 | Putative ur | Q3TXW2 [3 | 0 | 3305.542 | 0 | -0.39 | 0.042295 | 14.25 | 117.7 | 82.3 | 13.44 | 21.3 | | High | High | High | High | High | High | High | High | High | High | High | 7.19E-05 | 0.00079 | 3.19 |
| FALSE | High | LQELNAHIV | [K].LQELNA | 1xCarbami | Q8BWW9 | 0.01029 | 0.000772 | 1 | 6 | 2 | Q8BWW9 | Serine/thre | Q8BWW9 | 1 | 3774.712 | 0 | 0.11 | 0.077139 | 4.97 | 94.4 | 105.6 | 9.81 | 5.65 | | High | High | High | High | High | High | High | High | High | High | High | 0.000382 | 0.003251 | 4.41 |
| FALSE | High | LQESLPILQ | [R].LQESLP | 1xPhospho | Q9DBG5 1 | 0.0004 | 0 | 1 | 2 | 1 | Q9DBG5 | Perilipin-3 | Q9DBG5 [1 | 0 | 2162.187 | 0 | 2.03 | 0.000574 | 8.44 | 38 | 162 | 16.37 | 12.09 | | High | High | High | High | High | High | High | High | High | High | High | 0 | 8.48E-05 | 3.4 |
| FALSE | High | LQETPDPW | [R].LQETPD | 1xPhospho | Q8BSP2-2 | 9.62E-05 | 0 | 1 | 5 | 2 | Q8BSP2-2 | Isoform 2 c | Q8BSP2-2 | 0 | 2870.39 | 0 | -0.39 | 0.089107 | 16.12 | 114 | 86 | 15.76 | 12.78 | | High | High | High | High | High | High | High | High | High | High | High | 0 | 1.71E-05 | 4.13 |
| FALSE | High | LQGHSESQ | [R].LQGHS | 1xPhospho | B2RWU3 1 | 0.000264 | 0 | 1 | 3 | 2 | B2RWU3 | Fanconi an | B2RWU3 [1 | 0 | 2347.242 | 0 | -0.27 | 0.043135 | 8.64 | 110.4 | 89.6 | 7.53 | 8.11 | | High | High | High | High | High | High | High | High | High | High | High | 0 | 5.34E-05 | 3.41 |
| FALSE | High | LQGPPYAE | [R].LQGPPY | 1xCarbami | Q61213 1x | 0.003464 | 0.00021 | 1 | 3 | 1 | Q61213 | Gag OS=M | Q61213 [1 | 0 | 1978.966 | 0 | -0.45 | 0.108021 | 8.33 | 114 | 86 | 17.77 | 8.98 | | High | High | High | High | High | High | High | High | High | High | High | 9.42E-05 | 0.000961 | 2.36 |
| FALSE | High | LQGPQGDS | [R].LQGPQ | 2xCarbami | Q99KW3-2 | 0.001058 | 0 | 1 | 4 | 2 | Q99KW3-2 | Isoform 2 c | Q99KW3-2 | 0 | 3716.594 | 0 | -0.43 | 0.143146 | 16.62 | 111.2 | 88.8 | 20.26 | 23.7 | | High | High | High | High | High | High | High | High | High | High | High | 0.000467 | 0.004298 | 4.41 |
| FALSE | High | LQGSGVSL | [K].LQGSG | 1xPhospho | E9Q616 1x | 0.000133 | 0 | 1 | 4 | 1 | E9Q616 | AHNAK nu | E9Q616 [5 | 0 | 1584.876 | 0 | 2.3 | 0.005007 | 38.99 | 33.7 | 166.3 | 12.56 | 25.43 | | High | High | High | High | High | High | High | High | High | High | High | 0 | 2.48E-05 | 1.89 |
| FALSE | High | LQGSGVSL | [K].LQGSG | 2xPhospho | E9Q616 2x | 0.000712 | 0 | 1 | 4 | 2 | E9Q616 | AHNAK nu | E9Q616 [5 | 0 | 1664.843 | 0 | 2.34 | 0.00072 | 20.09 | 34.8 | 165.2 | 9.97 | 11.86 | | High | High | High | High | High | High | High | High | High | High | High | 0 | 0.000163 | 3.12 |
| FALSE | High | LQHGSTET | [K].LQHGS | 1xPhospho | Q8CH77 1x | 3.15E-06 | 0 | 1 | 2 | 2 | Q8CH77 | Neuron na | Q8CH77 [1 | 0 | 1994.036 | 0 | 0.92 | 0.001148 | 11.03 | 69.1 | 130.9 | 15.05 | 4.28 | | High | High | High | High | High | High | High | High | High | High | High | 0 | 3.71E-07 | 3.51 |

| | | | | | | | | | | | | | | | | | | | | | | | | | | | | | | | | | | | | | | |
|---|---|---|---|---|---|---|---|---|---|---|---|---|---|---|---|---|---|---|---|---|---|---|---|---|---|---|---|---|---|---|---|---|---|---|---|---|---|---|
| FALSE | High | IQHPITALP | [K].IQHPIT | 1xPhospho | E9PV87 1xl | 0.000543 | 0 | 1 | 3 | 1 | E9PV87 | Protein TA | E9PV87 [71 | 0 | 2714.332 | 0 | 0.44 | 0.010091 | 15.75 | 85.3 | 114.7 | 9.75 | 10.72 | | High | High | High | High | High | High | High | High | High | High | High | 0 | 0.000121 | 2.45 |
| FALSE | High | LQHTGPAD | [R].LQHTG | 1xPhospho | Q68FH0 1x | 0.006713 | 0.00062 | 1 | 8 | 1 | Q68FH0 | Plakophilin | Q68FH0 [4 | 1 | 2832.429 | 0 | -0.88 | 0.018977 | 5.87 | 126.1 | 73.9 | 18.45 | 15.82 | | High | High | High | High | High | High | High | High | High | High | High | 0.000275 | 0.002014 | 3.41 |
| FALSE | High | LQLAVCDE | [R].LQLAV | 1xCarbami | P54276 1xl | 0.012959 | 0.000889 | 1 | 2 | 1 | P54276 | DNA misma | P54276 [19 | 0 | 3813.734 | 0 | -1.19 | 0.002546 | 3.66 | 135.7 | 64.3 | 7.24 | 13.28 | | High | High | High | High | High | High | High | High | High | High | High | 0.000449 | 0.00419 | 2.87 |
| FALSE | High | LQLAVCDE | [R].LQLAV | 1xCarbami | P54276 2xl | 0.000125 | 0 | 1 | 2 | 2 | P54276 | DNA misma | P54276 [19 | 0 | 3893.701 | 0 | -0.64 | 0.003695 | 3.99 | 125 | 75 | 7.68 | 8.86 | | High | High | High | High | High | High | High | High | High | High | High | 0 | 2.31E-05 | 4.6 |
| FALSE | High | LQELNAHIV | [K].LQELNA | 1xCarbami | Q8BWW9 | 7.82E-05 | 0 | 1 | 6 | 2 | Q8BWW9 | Serine/thre | Q8BWW9 [ | 0 | 2704.249 | 0 | 0.82 | 0.002592 | 16.46 | 66.5 | 133.5 | 13.62 | 15.39 | | High | High | High | High | High | High | High | High | High | High | High | 0 | 1.36E-05 | 3.5 |
| FALSE | High | LQLDGSLTI | [K].LQLDG | 1xPhospho | Q3UJB9 1x | 0.000237 | 0 | 1 | 7 | 2 | Q3UJB9 | Enhancer c | Q3UJB9 [6 | 0 | 2470.239 | 0 | -0.09 | 0.9447 | 33.18 | 100 | 100 | 23.33 | 13.69 | | High | High | High | High | High | High | High | High | High | High | High | 0 | 4.73E-05 | 3.17 |
| FALSE | High | LQIGPPSPG | [K].LQIGPP | 1xPhospho | Q6GQX2 1 | 0.001049 | 0 | 1 | 1 | 1 | Q6GQX2 | Nck-associ | Q6GQX2 [4 | 0 | 2377.248 | 0 | 0.6 | 0.203403 | 59.5 | 79.3 | 120.7 | 30.59 | 24.53 | | High | High | High | High | High | High | High | High | High | High | High | 0 | 0.000251 | 3.55 |
| FALSE | High | LQLGTPPPI | [R].LQLGTP | 1xPhospho | Q5SFM8 1x | 0.006773 | 0.00062 | 1 | 2 | 1 | Q5SFM8 | RNA-bindir | Q5SFM8 [4 | 0 | 1655.945 | 0 | 0.74 | 0.001972 | 11.72 | 75.6 | 124.4 | 8.45 | 4.87 | | High | High | High | High | High | High | High | High | High | High | High | 0.000275 | 0.002035 | 2.57 |
| FALSE | High | LQLHESQK | [K].LQLHES | 1xPhospho | Q60598 1x | 0.081085 | 0.006219 | 1 | 3 | 1 | Q60598 | Src substra | Q60598 [2 | 0 | 1520.824 | 0 | 0.69 | 0.11415 | 44.5 | 84 | 116 | 22.46 | 24.2 | | High | High | High | High | High | High | High | High | High | High | High | 0.0032 | 0.03382 | 1.7 |
| FALSE | High | LQLLSFSCG | [R].LQLLSF | 1xCarbami | Q3TU15 1x | 7.96E-05 | 0 | 1 | 2 | 3 | Q3TU15 | Non-specif | Q3TU15 [3 | 0 | 2249.165 | 0 | -0.41 | 0.033728 | 12.44 | 116.9 | 83.1 | 12.08 | 8.7 | | High | High | High | High | High | High | High | High | High | High | High | 0 | 0.000277 | 2.47 |
| FALSE | High | IQIPDSQPC | [R].IQIPDS | 1xPhospho | K4DI77 1xP | 0.039011 | 0.002718 | 1 | 6 | 1 | K4DI77 | WD repeat | K4DI77 [15 | 0 | 3101.547 | 0 | 1.45 | 0.001083 | 26.7 | 52.9 | 147.1 | 15.46 | 9.92 | | High | High | High | High | High | High | High | High | High | High | High | 0.001453 | 0.01457 | 3.89 |
| FALSE | High | LQLPSFSCG | [R].LQLPSF | 1xCarbami | Q3UMB0 1 | 9.74E-06 | 0 | 1 | 1 | 1 | Q3UMB0 | Non-specif | Q3UMB0 [ | 0 | 2233.134 | 0 | -1.14 | 0.001124 | 2.6 | 137.7 | 62.3 | 4.41 | 15.68 | | High | High | High | High | High | High | High | High | High | High | High | 0 | 1.31E-06 | 3.11 |
| FALSE | High | IQIQLTQSF | [K].IQIQLT | 1xPhospho | E9Q9Q2 1x | 1.97E-05 | 0 | 1 | 7 | 1 | E9Q9Q2 | R3H domai | E9Q9Q2 [1 | 0 | 1873.024 | 0 | -0.07 | 0.243371 | 7 | 104 | 96 | 4.54 | 8.39 | | High | High | High | High | High | High | High | High | High | High | High | 0 | 2.88E-06 | 2.43 |
| FALSE | High | LQLSDEESV | [K].LQLSDE | 1xCarbami | Q8K1N2-2 | 3.57E-05 | 0 | 1 | 3 | 4 | Q8K1N2-2 | Isoform 2 c | Q8K1N2-2 | 0 | 2602.195 | 0 | -0.05 | 0.863641 | 6.33 | 97.4 | 102.6 | 16.18 | 16.02 | | High | High | High | High | High | High | High | High | High | High | High | 0 | 0.000224 | 3.79 |
| FALSE | High | LQLSPWES | [R].LQLSPV | 1xPhospho | E9QMU3 1 | 0.017809 | 0.001282 | 1 | 4 | 1 | E9QMU3 | Ensconsin | E9QMU3 [ | 0 | 1796.915 | 0 | -0.51 | 0.005076 | 5.56 | 117.3 | 82.7 | 6.32 | 7.33 | | High | High | High | High | High | High | High | High | High | High | High | 0.000689 | 0.005966 | 2.65 |
| FALSE | High | LQISSPPFD | [K].LQISSPI | 1xPhospho | Q8K0D0 1x | 0.020319 | 0.001472 | 1 | 4 | 2 | Q8K0D0 | Cyclin-dep | Q8K0D0 [1 | 0 | 1911.924 | 0 | 0.14 | 0.434334 | 31.92 | 93.6 | 106.4 | 10.48 | 14.13 | | High | High | High | High | High | High | High | High | High | High | High | 0.000786 | 0.006939 | 2.4 |
| FALSE | High | LQISSPPFD | [K].LQISSPI | 1xPhospho | Q8K0D0 1x | 0.067619 | 0.004864 | 1 | 4 | 4 | Q8K0D0 | Cyclin-dep | Q8K0D0 [1 | 0 | 1927.919 | 0 | 0.03 | 0.252012 | 4.91 | 97.9 | 102.1 | 4.06 | 14.78 | | High | High | High | High | High | High | High | High | High | High | High | 0.007374 | 0.0811 | 2.09 |
| FALSE | High | LQLSVTEVC | [R].LQLSVT | 1xPhospho | E9Q9C3 1x | 0.002948 | 0.00016 | 1 | 5 | 1 | E9Q9C3 | Afadin OS= | E9Q9C3 [4 | 0 | 1842.003 | 0 | 0.86 | 0.003869 | 12.32 | 67.7 | 132.3 | 9.44 | 9.9 | | High | High | High | High | High | High | High | High | High | High | High | 7.19E-05 | 0.000804 | 2.98 |
| FALSE | High | LQITPDSNC | [R].LQITPD | 1xPhospho | Q8CFI0 1xP | 0.000662 | 0 | 1 | 8 | 1 | Q8CFI0 | E3 ubiquiti | Q8CFI0 [34 | 0 | 2342.159 | 0 | 0.24 | 0.020742 | 7.79 | 91.8 | 108.2 | 5.08 | 6.58 | | High | High | High | High | High | High | High | High | High | High | High | 0 | 0.000151 | 3.01 |
| FALSE | High | LQLERPVSF | [K].LQLERP | 1xPhospho | A2A7S8-2 1 | 0.000233 | 0 | 1 | 4 | 3 | A2A7S8-2 | Isoform 2 c | A2A7S8-2 [ | 0 | 2259.17 | 0 | -0.57 | 0.000715 | 1.57 | 118.5 | 81.5 | 4.81 | 3.38 | | High | High | High | High | High | High | High | High | High | High | High | 4.94E-05 | 0.000731 | 2.15 |
| FALSE | High | LQELLDPSF | [R].LQELLD | 1xPhospho | Q7TMI6 1x | 0.006306 | 0.00062 | 1 | 3 | 2 | Q7TMI6 | Zinc finger | Q7TMI6 [1 | 0 | 1604.825 | 0 | -0.05 | 0.282725 | 1.51 | 100.8 | 99.2 | 3.97 | 4.73 | | High | High | High | High | High | High | High | High | High | High | High | 0.000275 | 0.001868 | 2.3 |
| FALSE | High | LQELESCSG | [R].LQELES | 1xCarbami | Q6PAR5 2x | 0.000362 | 0 | 1 | 9 | 3 | Q6PAR5 | GTPase-act | Q6PAR5 [7 | 0 | 2558.057 | 0 | 0.07 | 0.606868 | 10.33 | 98.9 | 101.1 | 9.14 | 9.17 | | High | High | High | High | High | High | High | High | High | High | High | 2.58E-05 | 0.000407 | 3.88 |
| FALSE | High | LQELESCSG | [R].LQELES | 1xCarbami | Q6PAR5 1x | 1.71E-06 | 0 | 1 | 9 | 7 | Q6PAR5 | GTPase-act | Q6PAR5 [7 | 0 | 2478.09 | 0 | 0.34 | 0.012767 | 16.67 | 84 | 116 | 9.59 | 4.92 | | High | High | High | High | High | High | High | High | High | High | High | 0 | 1.85E-07 | 4.4 |
| FALSE | High | LQAALDNE | [R].LQAALI | 1xDeamida | Q3TXW2 1 | 0.003433 | 0.00021 | 1 | 8 | 5 | Q3TXW2 | Putative ur | Q3TXW2 [3 | 0 | 3306.526 | 0 | -0.38 | 0.260579 | 8.53 | 109.9 | 90.1 | 9.43 | 20.05 | | High | High | High | High | High | High | High | High | High | High | High | 0.000117 | 0.001059 | 3.14 |
| FALSE | High | LQAEAPHIV | [K].LQAEAI | 1xPhospho | P10630-2 1 | 0.000311 | 0 | 1 | 4 | 1 | P10630-2 | Isoform 2 c | P10630-2 [ | 0 | 1853.984 | 0 | -0.84 | 0.000248 | 1.42 | 128.2 | 71.8 | 3.58 | 5.17 | | High | High | High | High | High | High | High | High | High | High | High | 0 | 6.43E-05 | 1.85 |
| FALSE | High | LQALQSQA | [K].LQALQ | 1xPhospho | A2BIE1 1xP | 1.49E-06 | 0 | 1 | 2 | 2 | A2BIE1 | Glutamine | A2BIE1 [53 | 0 | 2042.012 | 0 | 1.23 | 0.003336 | 16.44 | 61.5 | 138.5 | 13.5 | 10.27 | | High | High | High | High | High | High | High | High | High | High | High | 0 | 1.59E-07 | 4.18 |
| FALSE | High | LQASPVAL | [K].LQASPV | 1xPhospho | Q8N9S3 1x | 0.068184 | 0.004935 | 1 | 5 | 2 | Q8N9S3 | Activator o | Q8N9S3 [1 | 0 | 1419.792 | 0 | 1.09 | 0.000699 | 17.22 | 62.6 | 137.4 | 10.12 | 5.72 | | High | High | High | High | High | High | High | High | High | High | High | 0.002559 | 0.02749 | 2.64 |
| FALSE | High | LQAYLEDH | [R].LQAYLE | 1xPhospho | B2RXM7 1x | 3.92E-06 | 0 | 1 | 1 | 3 | B2RXM7 | MCG11369 | B2RXM7 [3 | 0 | 4904.375 | 0 | 0.28 | 0.02662 | 3.66 | 90.3 | 109.7 | 3.17 | 11.31 | | High | High | High | High | High | High | High | High | High | High | High | 0 | 4.73E-07 | 5.09 |
| FALSE | High | LQAYLEDH | [R].LQAYLE | 1xPhospho | B7ZWI2 1x | 0.000112 | 0 | 1 | 2 | 2 | B7ZWI2 | 1110005A2 | B7ZWI2 [38 | 0 | 4833.337 | 0 | -0.22 | 0.047695 | 3.28 | 107.3 | 92.7 | 7.29 | 1.93 | | High | High | High | High | High | High | High | High | High | High | High | 0 | 2.03E-05 | 5.35 |
| FALSE | High | LQCPVTEP | [K].LQCPVT | 1xCarbami | Q8K1J5 1xl | 0.002172 | 5.66E-05 | 1 | 1 | 2 | Q8K1J5 | Protein SDI | Q8K1J5 [28 | 0 | 3066.418 | 0 | -0.7 | 0.292028 | 6.53 | 124.3 | 75.7 | 50.07 | 44.23 | | High | High | High | High | High | High | High | High | High | High | High | 4.94E-05 | 0.00057 | 3.97 |
| FALSE | High | LQCPVTEP | [K].LQCPVT | 1xCarbami | Q8K1J5 1xl | 2.24E-07 | 0 | 1 | 1 | 3 | Q8K1J5 | Protein SDI | Q8K1J5 [28 | 0 | 3295.581 | 0 | -0.81 | 0.00095 | 2.67 | 126.2 | 73.8 | 4.87 | 7.96 | | High | High | High | High | High | High | High | High | High | High | High | 0 | 1.9E-08 | 5.68 |
| FALSE | High | IQCQAQQE | [R].IQCQA | 1xCarbami | A0A1L1SSL | 6.67E-06 | 0 | 1 | 4 | 1 | A0A1L1SSL | Serine/thre | A0A1L1SSL | 0 | 2568.16 | 0 | -0.22 | 0.590539 | 24.46 | 109.6 | 90.4 | 24.47 | 26.97 | | High | High | High | High | High | High | High | High | High | High | High | 0 | 8.54E-07 | 4.1 |
| FALSE | High | IQCQAQQE | [R].IQCQA | 1xCarbami | A0A1L1SSL | 3.75E-10 | 0 | 1 | 4 | 2 | A0A1L1SSL | Serine/thre | A0A1L1SSL | 0 | 2797.323 | 0 | -0.28 | 0.007402 | 3.47 | 111.1 | 88.9 | 3.83 | 5.99 | | High | High | High | High | High | High | High | High | High | High | High | 0 | 2.12E-11 | 4.98 |
| FALSE | High | IQDECPVEI | [K].IQDECP | 1xCarbami | Z4YKQ9 1x | 5.29E-07 | 0 | 1 | 4 | 1 | Z4YKQ9 | ATPase fan | Z4YKQ9 [1 | 0 | 2286.073 | 0 | 0.24 | 0.045394 | 11.37 | 91.7 | 108.3 | 8.05 | 10.41 | | High | High | High | High | High | High | High | High | High | High | High | 0 | 4.97E-08 | 4.18 |
| FALSE | High | IQDEIPALS | [K].IQDEIPA | 1xPhospho | P54775 1xl | 0.000438 | 0 | 1 | 6 | 2 | P54775 | 26S protea | P54775 [11 | 0 | 3980.973 | 0 | -0.07 | 0.382392 | 8.02 | 100.4 | 99.6 | 18.7 | 13.11 | | High | High | High | High | High | High | High | High | High | High | High | 0 | 0.000133 | 4.62 |
| FALSE | High | LQDISELLA | [R].LQDISE | 1xPhospho | Q91YU8 1x | 0.006085 | 0.00062 | 1 | 1 | 1 | Q91YU8 | Suppressor | Q91YU8 [2 | 0 | 4157.012 | 0 | -0.72 | 0.015392 | 9.18 | 124.6 | 75.4 | 14.21 | 19.83 | | High | High | High | High | High | High | High | High | High | High | High | 0.000275 | 0.001794 | 3.32 |
| FALSE | High | LQDISELLA | [R].LQDISE | 1xDeamida | Q91YU8 1x | 2.28E-06 | 0 | 1 | 1 | 4 | Q91YU8 | Suppressor | Q91YU8 [2 | 0 | 4157.996 | 0 | -0.16 | 0.210823 | 2.79 | 115.5 | 84.5 | 33.9 | 23.09 | | High | High | High | High | High | High | High | High | High | High | High | 0 | 7E-06 | 4.71 |
| FALSE | High | IQDPQEPD | [R].IQDPQE | 1xPhospho | Q924N4-2 | 1.99E-06 | 0 | 1 | 1 | 2 | Q924N4-2 | Isoform 2 c | Q924N4-2 | 0 | 3523.674 | 0 | 1.9 | 0.002072 | 11.86 | 36.6 | 163.4 | 15.32 | 18.07 | | High | High | High | High | High | High | High | High | High | High | High | 0 | 2.19E-07 | 3.33 |
| FALSE | High | IQDQSFGE | [K].IQDQSF | 1xPhospho | Q501J7-3 1 | 0.01441 | 0.001011 | 1 | 3 | 1 | Q501J7-3 | Isoform 3 c | Q501J7-3 [ | 0 | 1909.901 | 0 | 0.79 | 0.012148 | 16.48 | 72.5 | 127.5 | 16.13 | 10.47 | | High | High | High | High | High | High | High | High | High | High | High | 0.000538 | 0.004733 | 1.68 |
| FALSE | High | LQDSSDPD | [R].LQDSSD | 1xPhospho | Q60875 1x | 8.62E-11 | 0 | 1 | 8 | 1 | Q60875 | Rho guanir | Q60875 [9 | 1 | 3118.416 | 0 | -0.88 | 0.004041 | 4.74 | 128.4 | 71.6 | 9.09 | 12.35 | | High | High | High | High | High | High | High | High | High | High | High | 0 | 2.74E-12 | 5.05 |
| FALSE | High | LQDSSDPD | [R].LQDSSD | 2xPhospho | Q60875 2x | 7.79E-07 | 0 | 1 | 8 | 5 | Q60875 | Rho guanir | Q60875 [9 | 1 | 3198.383 | 0 | -0.37 | 0.00838 | 1.73 | 112.5 | 87.5 | 4.66 | 8.3 | | High | High | High | High | High | High | High | High | High | High | High | 0 | 7.7E-08 | 4.03 |
| FALSE | High | LQDSSDPD | [R].LQDSSD | 1xDeamida | Q60875 2x | 0.041781 | 0.003126 | 1 | 8 | 1 | Q60875 | Rho guanir | Q60875 [9 | 1 | 3199.367 | 0 | -0.59 | 0.099819 | 5.23 | 116.9 | 83.1 | 9.69 | 18.14 | | High | High | High | High | High | High | High | High | High | High | High | 0.001609 | 0.01566 | 4.42 |
| FALSE | High | LQDSSDPD | [R].LQDSSD | 3xPhospho | Q60875 3x | 0.081752 | 0.006219 | 1 | 8 | 1 | Q60875 | Rho guanir | Q60875 [9 | 1 | 3278.349 | 0 | 0.31 | 0.003483 | 8.04 | 87.3 | 112.7 | 11.61 | 8.02 | | High | High | High | High | High | High | High | High | High | High | High | 0.003229 | 0.0342 | 3.16 |
| FALSE | High | IQDTGDTIL | [K].IQDTGD | 1xPhospho | E9PYV1 1xl | 1.69E-06 | 0 | 1 | 12 | 1 | E9PYV1 | Glucocortic | E9PYV1 [27 | 0 | 2694.437 | 0 | 1.33 | 0.002447 | 27.73 | 56.6 | 143.4 | 9.95 | 11.26 | | High | High | High | High | High | High | High | High | High | High | High | 0 | 1.84E-07 | 4.96 |
| FALSE | High | LQDVSGQL | [R].LQDVS | 1xPhospho | Q7JJ13 1xP | 0.000645 | 0 | 1 | 3 | 2 | Q7JJ13 | Bromodom | Q7JJ13 [73 | 0 | 1827.962 | 0 | 0.57 | 0.000778 | 4.13 | 79.1 | 120.9 | 7.23 | 6.04 | | High | High | High | High | High | High | High | High | High | High | High | 4.94E-05 | 0.00069 | 2.63 |
| FALSE | High | LQEDPNYS | [R].LQEDPI | 1xPhospho | Q9Z1R2 1x | 0.00509 | 0.000551 | 1 | 6 | 3 | Q9Z1R2 | Large proli | Q9Z1R2 [1 | 0 | 1655.763 | 0 | -0.41 | 0.011602 | 5.32 | 113.8 | 86.2 | 2.97 | 7.38 | | High | High | High | High | High | High | High | High | High | High | High | 0.000246 | 0.001483 | 2.53 |
| FALSE | High | LQEDPNYS | [R].LQEDPI | 1xPhospho | Q9Z1R2 1x | 0.006195 | 0.00062 | 1 | 6 | 2 | Q9Z1R2 | Large proli | Q9Z1R2 [1 | 1 | 2378.124 | 0 | -0.48 | 0.022122 | 10.92 | 116.1 | 83.9 | 10.58 | 15.68 | | High | High | High | High | High | High | High | High | High | High | High | 0.00196 | 0.01979 | 2.14 |
| FALSE | High | LQEEFEATL | [R].LQEEFE | 2xPhospho | B2RRB9 2x | 0.000305 | 0 | 1 | 4 | 1 | B2RRB9 | Family with | B2RRB9 [2 | 0 | 3069.374 | 0 | 0.26 | 0.017036 | 13.69 | 87.9 | 112.1 | 12.98 | 6.35 | | High | High | High | High | High | High | High | High | High | High | High | 0 | 6.29E-05 | 2.95 |
| FALSE | High | LQEEGGSE | [K].LQEEG | 1xPhospho | Q99LL5 1xl | 6.46E-08 | 0 | 1 | 2 | 4 | Q99LL5 | Periodic try | Q99LL5 [43 | 0 | 3422.453 | 0 | 0.02 | 0.973074 | 10.68 | 99.8 | 100.2 | 10.53 | 6.16 | | High | High | High | High | High | High | High | High | High | High | High | 0 | 1.76E-07 | 5.59 |
| FALSE | High | LQEESDLEL | [K].LQEESD | 1xPhospho | Q66JS6 1xP | 0.000292 | 0 | 1 | 3 | 1 | Q66JS6 | Eukaryotic | Q66JS6 [12 | 0 | 1812.94 | 0 | 0.65 | 0.001383 | 7.19 | 77.7 | 122.3 | 7.15 | 7.28 | | High | High | High | High | High | High | High | High | High | High | High | 0 | 6.01E-05 | 3.56 |
| FALSE | High | LQEEVPSEE | [K].LQEEVP | 1xPhospho | Q3TIZ8 1xP | 1.63E-05 | 0 | 1 | 4 | 1 | Q3TIZ8 | Putative ur | Q3TIZ8 [20 | 0 | 2339.124 | 0 | 1.16 | 0.044662 | 16.06 | 63.4 | 136.6 | 7.3 | 29.06 | | High | High | High | High | High | High | High | High | High | High | High | 0 | 2.34E-06 | 4.55 |
| FALSE | High | LQEEVPSEE | [K].LQEEVP | 1xPhospho | Q3TIZ8 1xP | 4.2E-05 | 0 | 1 | 4 | 5 | Q3TIZ8 | Putative ur | Q3TIZ8 [20 | 0 | 2355.119 | 0 | 0.92 | 0.002159 | 11.32 | 66.2 | 133.8 | 7.9 | 10.52 | | High | High | High | High | High | High | High | High | High | High | High | 0 | 9.76E-06 | 3.79 |
| FALSE | High | IPSAVSTVS | [K].IPSAVST | 1xPhospho | Q3U3A6 1x | 0.007811 | 0.000664 | 1 | 4 | 1 | Q3U3A6 | Sodium/hy | Q3U3A6 [6 | 0 | 2247.197 | 0 | 0.36 | 0.002347 | 10.77 | 88.7 | 111.3 | 10.62 | 3.81 | | High | High | High | High | High | High | High | High | High | High | High | 0.000294 | 0.002371 | 2.53 |
| FALSE | High | LNQNNSAC | [R].LNQNN | 1xDeamida | Q99NF3 1x | 0.004696 | 0.000456 | 1 | 4 | 1 | Q99NF3 | Centrosom | Q99NF3 [3 | 0 | 1484.751 | 0 | -0.67 | 0.002013 | 4.54 | 121 | 79 | 4.74 | 8.31 | | High | High | High | High | High | High | High | High | High | High | High | 0.000204 | 0.001342 | 2.54 |
| FALSE | High | IPSAVSSPLI | [R].IPSAVSS | 2xPhospho | E9Q3G8 2x | 2.72E-05 | 0 | 1 | 2 | 2 | E9Q3G8 | Nucleopor | E9Q3G8 [3 | 0 | 1941.929 | 0 | -0.51 | 0.010958 | 3.04 | 117.9 | 82.1 | 10.2 | 11.13 | | High | High | High | High | High | High | High | High | High | High | High | 0 | 4.15E-06 | 3.37 |
| FALSE | High | IPSASPQTC | [K].IPSASP | 1xPhospho | P53349 1xl | 0.038346 | 0.002563 | 2 | 2 | 1 | P53349; F8 | Mitogen-ac | P53349 [99 | 0 | 1393.704 | 0 | -1.15 | 0.000964 | 1.82 | 140.6 | 59.4 | 12.21 | 12.62 | Shared | High | High | High | High | High | High | High | High | High | High | High | 0.001355 | 0.01426 | 1.31 |
| FALSE | High | LNYGDQSA | [R].LNYGD | 1xPhospho | B7ZWN9 1 | 0.012959 | 0.000889 | 2 | 4 | 1 | B7ZWN9; F | SUMO/sen | B7ZWN9 [9 | 0 | 1762.841 | 0 | -0.59 | 0.017349 | 9.59 | 118.9 | 81.1 | 11.95 | 6.59 | Shared | High | High | High | High | High | High | High | High | High | High | High | 0.000449 | 0.004194 | 1.79 |
| FALSE | High | LPAAEDQS | [K].LPAAED | 1xPhospho | Q6ZWQ0 1 | 4.52E-06 | 0 | 1 | 2 | 1 | Q6ZWQ0 | Nesprin-2 c | Q6ZWQ0 [ | 0 | 2935.464 | 0 | 5.33 | 0.00085 | 42.63 | 5.7 | 194.3 | 49.9 | 15.29 | | High | High | High | High | High | High | High | High | High | High | High | 0 | 5.51E-07 | 4.1 |
| FALSE | High | LPADSENV | [K].LPADSE | 1xPhospho | P70399 1xl | 6.05E-07 | 0 | 1 | 4 | 2 | P70399 | TP53-bindi | P70399 [55 | 0 | 3396.628 | 0 | 0.17 | 0.55171 | 40.72 | 94.3 | 105.7 | 9.62 | 28.01 | | High | High | High | High | High | High | High | High | High | High | High | 0 | 5.81E-08 | 4.49 |
| FALSE | High | LPADSENV | [K].LPADSE | 1xPhospho | P70399 1xl | 0.007742 | 0.000664 | 1 | 4 | 1 | P70399 | TP53-bindi | P70399 [55 | 0 | 3412.623 | 0 | 0.23 | 0.145267 | 20.5 | 84.6 | 115.4 | 17.28 | 18.64 | | High | High | High | High | High | High | High | High | High | High | High | 0.000294 | 0.002365 | 2.48 |
| FALSE | High | LPAEVNHV | [K].LPAEVN | 1xPhospho | A2ADB0 1x | 0.101749 | 0.008194 | 1 | 5 | 3 | A2ADB0 | Msx2-inter | A2ADB0 [2 | 0 | 2153.059 | 0 | 0.32 | 0.004686 | 8.71 | 88.8 | 111.2 | 8.71 | 6.65 | | High | High | High | High | High | High | High | High | High | High | High | 0.004126 | 0.04405 | 2.68 |
| FALSE | High | LPAGSSELA | [R].LPAGSS | 1xPhospho | E9Q652 1x | 2.06E-05 | 0 | 1 | 7 | 1 | E9Q652 | Regulator c | E9Q652 [1 | 0 | 2188.203 | 0 | 1.93 | 0.000357 | 8.69 | 41.8 | 158.2 | 7.15 | 8.16 | | High | High | High | High | High | High | High | High | High | High | High | 0 | 3.03E-06 | 3.24 |
| FALSE | High | LPAHSFLGF | [K].LPAHSF | 1xPhospho | Q9WVA7 1 | 4.23E-05 | 0 | 1 | 1 | 1 | Q9WVA7 | FRAT3 OS= | Q9WVA7 [ | 0 | 2782.464 | 0 | 1.3 | 0.000292 | 10.08 | 58.6 | 141.4 | 11.63 | 1.85 | | High | High | High | High | High | High | High | High | High | High | High | 0 | 6.84E-06 | 3.95 |
| FALSE | High | LPALSHSEG | [R].LPALSH | 2xPhospho | A0A0N4SU | 1.58E-08 | 0 | 1 | 10 | 3 | A0A0N4SU | Peregrin O | A0A0N4SU | 0 | 2976.232 | 0 | -0.13 | 0.018705 | 4.21 | 105.7 | 94.3 | 2.71 | 5.68 | | High | High | High | High | High | High | High | High | High | High | High | 0 | 9.61E-10 | 5.66 |
| FALSE | High | IPALSPSSG | [K].IPALSPS | 1xPhospho | E9QAU4 1x | 0.006031 | 0.00062 | 1 | 9 | 1 | E9QAU4 | Enhancer t | E9QAU4 [1 | 0 | 1494.833 | 0 | -0.14 | 0.025631 | 3.13 | 106.1 | 93.9 | 7.47 | 6.26 | | High | High | High | High | High | High | High | High | High | High | High | 0.000275 | 0.001785 | 1.83 |
| FALSE | High | LPAPQEDT | [R].LPAPQE | 1xPhospho | Q64012 1x | 2.45E-09 | 0 | 1 | 3 | 7 | Q64012 | RNA-bindir | Q64012 [2 | 0 | 2591.219 | 0 | -0.02 | 0.471273 | 6.83 | 102.2 | 97.8 | 6.77 | 5.41 | | High | High | High | High | High | High | High | High | High | High | High | 0 | 1.18E-10 | 5.38 |
| FALSE | High | LPAPQEDT | [R].LPAPQE | 2xPhospho | Q64012 2x | 3.52E-07 | 0 | 1 | 3 | 6 | Q64012 | RNA-bindir | Q64012 [2 | 0 | 2671.185 | 0 | 0.14 | 0.113197 | 5.91 | 96.7 | 103.3 | 4.54 | 8.44 | | High | High | High | High | High | High | High | High | High | High | High | 0 | 1.08E-07 | 5.66 |
| FALSE | High | LPAPQEDT | [R].LPAPQE | 3xPhospho | Q64012 3x | 2.91E-07 | 0 | 1 | 3 | 8 | Q64012 | RNA-bindir | Q64012 [2 | 0 | 2751.151 | 0 | 0.43 | 0.039961 | 5.3 | 85.2 | 114.8 | 4.57 | 12.53 | | High | High | High | High | High | High | High | High | High | High | High | 4.94E-05 | 0.000602 | 5.56 |
| FALSE | High | LPAPSLPLM | [K].LPAPSL | 2xCarbami | O70310 1x | 0.00153 | 5.66E-05 | 1 | 1 | 1 | O70310 | Glycylpept | O70310 [1 | 0 | 4056.657 | 0 | -0.13 | 0.947177 | 33.12 | 103.4 | 96.6 | 15.63 | 18.03 | | High | High | High | High | High | High | High | High | High | High | High | 2.58E-05 | 0.000384 | 3.71 |
| FALSE | High | LNVEWLGY | [R].LNVEW | 1xPhospho | F2Z446 1xP | 5.38E-07 | 0 | 1 | 1 | 2 | F2Z446 | Vacuolar A | F2Z446 [8- | 0 | 2806.438 | 0 | 4.21 | 0.000652 | 27.02 | 12.1 | 187.9 | 17.55 | 12.72 | | High | High | High | High | High | High | High | High | High | High | High | 0 | 5.07E-08 | 4.73 |
| FALSE | High | LPAPSLPLM | [K].LPAPSL | 2xCarbami | O70310 1x | 0.018774 | 0.001362 | 1 | 1 | 1 | O70310 | Glycylpept | O70310 [1 | 0 | 4057.641 | 0 | -0.25 | 0.497453 | 42.84 | 108.9 | 91.1 | 19.95 | 21.18 | | High | High | High | High | High | High | High | High | High | High | High | 0.000723 | 0.00633 | 3.97 |
| FALSE | High | LPAPSLPLM | [K].LPAPSL | 2xCarbami | O70310 1x | 0.014158 | 0.001011 | 1 | 1 | 4 | O70310 | Glycylpept | O70310 [1 | 0 | 4072.652 | 0 | -0.1 | 0.358533 | 14.13 | 103.6 | 96.4 | 4.12 | 12.42 | | High | High | High | High | High | High | High | High | High | High | High | 0.000538 | 0.00465 | 3.17 |
| FALSE | High | LPAPSLPLM | [K].LPAPSL | 2xCarbami | O70310 1x | 5.29E-07 | 0 | 1 | 1 | 31 | O70310 | Glycylpept | O70310 [1 | 0 | 4073.636 | 0 | -0.01 | 0.840277 | 2.68 | 99.3 | 100.7 | 3.83 | 1.95 | | High | High | High | High | High | High | High | High | High | High | High | 0 | 4.97E-08 | 5.71 |
| FALSE | High | LPAPSLPLM | [K].LPAPSL | 2xCarbami | O70310 1x | 0.00172 | 5.66E-05 | 1 | 1 | 4 | O70310 | Glycylpept | O70310 [1 | 0 | 4088.647 | 0 | -0.07 | 0.670439 | 36.78 | 109.1 | 90.9 | 10.32 | 90.8 | | High | High | High | High | High | High | High | High | High | High | High | 4.94E-05 | 0.000756 | 3.88 |
| FALSE | High | LPASPGPGT | [R].LPASPG | 1xPhospho | Q66JR8 1xl | 0.016306 | 0.001124 | 1 | 1 | 2 | Q66JR8 | Ptms prote | Q66JR8 [10 | 0 | 1825.917 | 0 | 0.56 | 0.02636 | 8.25 | 77.4 | 122.6 | 12.22 | 18.45 | | High | High | High | High | High | High | High | High | High | High | High | 0.000585 | 0.00542 | 2.64 |
| FALSE | High | LPASPVISN | [R].LPASPV | 1xPhospho | A2A5V4 1x | 0.000112 | 0 | 1 | 2 | 1 | A2A5V4 | SH3 domai | A2A5V4 [6 | 0 | 2720.316 | 0 | 0.7 | 0.164466 | 34.9 | 76.3 | 123.7 | 28.89 | 43.09 | | High | High | High | High | High | High | High | High | High | High | High | 0 | 2.05E-05 | 4.25 |
| FALSE | High | LPATSAESE | [R].LPATSA | 1xPhospho | Q3UKE7 1x | 1.61E-05 | 0 | 1 | 5 | 1 | Q3UKE7 | Hydroxyme | Q3UKE7 [5 | 0 | 2107.975 | 0 | -0.35 | 0.028782 | 9.47 | 107.3 | 92.7 | 11.87 | 7.27 | | High | High | High | High | High | High | High | High | High | High | High | 0 | 2.3E-06 | 2.94 |
| FALSE | High | LPAYLATSP | [K].LPAYLA | 1xPhospho | E9PWN0 1 | 3.7E-05 | 0 | 1 | 5 | 2 | E9PWN0 | Phosphatic | E9PWN0 [1 | 0 | 2576.345 | 0 | 1.62 | 0.000241 | 6.36 | 47.9 | 152.1 | 10.1 | 4.37 | | High | High | High | High | High | High | High | High | High | High | High | 0 | 2.39E-05 | 2.13 |
| FALSE | High | LPCPEGLDI | [K].LPCPEG | 2xCarbami | A0A1S6GW | 3.53E-08 | 0 | 1 | 6 | 1 | A0A1S6GW | Uncharacte | A0A1S6GW | 0 | 2449.098 | 0 | 1 | 0.001718 | 20.09 | 66.1 | 133.9 | 9.91 | 7.81 | | High | High | High | High | High | High | High | High | High | High | High | 0 | 2.37E-09 | 4.65 |
| FALSE | High | IPDPDSDD | [K].IPDPDS | 1xPhospho | A0A0R4J17 | 1E-05 | 0 | 1 | 5 | 3 | A0A0R4J17 | Transcripti | A0A0R4J17 | 0 | 1938.853 | 0 | -0.51 | 0.005476 | 5.91 | 117.9 | 82.1 | 9.62 | 7.84 | | High | High | High | High | High | High | High | High | High | High | High | 0 | 1.35E-06 | 2.87 |
| FALSE | High | IPDPDSDD | [K].IPDPDS | 2xPhospho | A0A0R4J17 | 0.003083 | 0.00016 | 1 | 5 | 1 | A0A0R4J17 | Transcripti | A0A0R4J17 | 0 | 2018.819 | 0 | -0.75 | 0.014476 | 4.53 | 126.1 | 73.9 | 6.44 | 18.3 | | High | High | High | High | High | High | High | High | High | High | High | 7.19E-05 | 0.00084 | 3.1 |
| FALSE | High | LPDSPALAK | [K].LPDSPA | 1xPhospho | A0A1B0GR | 0.003464 | 0.00021 | 1 | 3 | 2 | A0A1B0GR | Stromal int | A0A1B0GR | 0 | 1449.812 | 0 | 1.62 | 0.001286 | 23.44 | 49.3 | 150.7 | 14.02 | 14.18 | | High | High | High | High | High | High | High | High | High | High | High | 9.42E-05 | 0.000956 | 2.99 |
| FALSE | High | IPDTESPVP | [R].IPDTES | 1xPhospho | Q8BYH8 1x | 0.048303 | 0.00354 | 1 | 3 | 2 | Q8BYH8 | Chromodo | Q8BYH8 [2 | 0 | 2060.144 | 0 | 0.31 | 0.054921 | 13.05 | 89 | 111 | 10.66 | 9.29 | | High | High | High | High | High | High | High | High | High | High | High | 0.003265 | 0.03492 | 2.26 |
| FALSE | High | LPEASQSPL | [R].LPEASC | 1xPhospho | M1VMF6 1 | 0.000668 | 0 | 1 | 8 | 2 | M1VMF6 | Metastasis | M1VMF6 [ | 0 | 1820.033 | 0 | -0.22 | 0.093462 | 4.93 | 107.3 | 92.7 | 3.2 | 10.01 | | High | High | High | High | High | High | High | High | High | High | High | 0 | 0.000152 | 2.46 |
| FALSE | High | LPAPSLPLM | [K].LPAPSL | 2xCarbami | O70310 1x | 0.100129 | 0.008067 | 1 | 1 | 1 | O70310 | Glycylpept | O70310 [1 | 0 | 4058.625 | 0 | 0.35 | 0.190824 | 26.71 | 89.5 | 110.5 | 19.57 | 16.43 | | High | High | High | High | High | High | High | High | High | High | High | 0.004059 | 0.04324 | 3.51 |
| FALSE | High | LPEDDEPPA | [K].LPEDDE | 1xPhospho | Q8K3G7 1x | 0.001097 | 0 | 3 | 4 | 4 | Q8K3G7; C | Reticulon C | Q8K3G7 [1 | 0 | 3820.956 | 0 | 0.65 | 0.012849 | 10.55 | 77.1 | 122.9 | 4.02 | 14.45 | Shared | High | High | High | High | High | High | High | High | High | High | High | 0 | 0.000265 | 2.49 |
| FALSE | High | INVDIPEEA | [R].INVDIPI | 1xPhospho | A0A0R4J0V | 0.000134 | 0 | 1 | 5 | 2 | A0A0R4J0V | Leucine ric | A0A0R4J0V | 0 | 2572.213 | 0 | 0.22 | 0.277103 | 20.58 | 91.4 | 108.6 | 19.11 | 6.91 | | High | High | High | High | High | High | High | High | High | High | High | 0 | 2.51E-05 | 3.64 |
| FALSE | High | LNTSDFQK | [R].LNTSDF | 1xPhospho | E9QKI4 1xP | 0.01095 | 0.000772 | 1 | 3 | 1 | E9QKI4 | Proline-rich | E9QKI4 [31 | 0 | 1261.603 | 0 | -1.18 | 0.113424 | 8.56 | 141.8 | 58.2 | 40.55 | 48.04 | | High | High | High | High | High | High | High | High | High | High | High | 0.000399 | 0.003462 | 2.38 |
| FALSE | High | LNQSGASV | [K].LNQSG | 2xPhospho | Q9ERU9 2x | 3.64E-06 | 0 | 1 | 1 | 10 | Q9ERU9 | E3 SUMO-p | Q9ERU9 [2 | 0 | 2769.134 | 0 | 0.35 | 0.040272 | 7.22 | 89 | 111 | 1.91 | 12.82 | | High | High | High | High | High | High | High | High | High | High | High | 0 | 1.54E-06 | 4.31 |
| FALSE | High | LNQSPGLA | [R].LNQSP | 1xPhospho | Q3UYC8 1x | 0.005228 | 0.000551 | 1 | 5 | 1 | Q3UYC8 | Putative ur | Q3UYC8 [7 | 0 | 1661.939 | 0 | -0.06 | 0.382109 | 8.03 | 102.1 | 97.9 | 5.74 | 6.17 | | High | High | High | High | High | High | High | High | High | High | High | 0.000246 | 0.001513 | 2.99 |
| FALSE | High | LNQSSPDN | [K].LNQSSP | 1xPhospho | H7BX08 1x | 3.01E-06 | 0 | 1 | 5 | 2 | H7BX08 | Calmodulir | H7BX08 [5 | 0 | 1970.984 | 0 | 1.27 | 0.002433 | 12.44 | 58.6 | 141.4 | 7.33 | 11.93 | | High | High | High | High | High | High | High | High | High | High | High | 0 | 3.6E-06 | 3.37 |
| FALSE | High | LNQVCFDD | [K].LNQVC | 1xCarbami | Q8BJ71 1xl | 0.017652 | 0.001282 | 1 | 4 | 2 | Q8BJ71 | Nuclear po | Q8BJ71 [41 | 0 | 2262.954 | 0 | -0.68 | 0.011928 | 8.82 | 129.6 | 70.4 | 11.37 | 18.74 | | High | High | High | High | High | High | High | High | High | High | High | 0.001468 | 0.01476 | 2.58 |
| FALSE | High | LNSALSPK | [R].LNSALS | 1xPhospho | A2ADB0 1x | 0.002293 | 0.000109 | 1 | 5 | 1 | A2ADB0 | Msx2-inter | A2ADB0 [1 | 0 | 1367.77 | 0 | 0.26 | 0.24038 | 8.24 | 90.6 | 109.4 | 4.61 | 17.21 | | High | High | High | High | High | High | High | High | High | High | High | 4.94E-05 | 0.000605 | 1.03 |
| FALSE | High | LNSANEEEI | [R].LNSANI | 1xPhospho | Q8BHB4 1x | 0.000164 | 0 | 1 | 1 | 2 | Q8BHB4 | WD repeat | Q8BHB4 [3 | 0 | 3045.381 | 0 | 0.19 | 0.076241 | 1.21 | 87.3 | 112.7 | 21.62 | 11.97 | | High | High | High | High | High | High | High | High | High | High | High | 0 | 3.12E-05 | 4.06 |
| FALSE | High | LNSCETK | [R].LNSCET | 1xCarbami | A0A0A0MC | 0.050826 | 0.003676 | 1 | 4 | 2 | A0A0A0MC | Calmodulir | A0A0A0MC | 0 | 1389.685 | 0 | 0.16 | 0.093492 | 11.45 | 90.2 | 109.8 | 5.61 | 12.68 | | High | High | High | High | High | High | High | High | High | High | High | 0.002117 | 0.02179 | 2.31 |
| FALSE | High | LNSDEEGE | [K].LNSDEE | 1xPhospho | Q5XJE5 1xl | 3.36E-06 | 0 | 1 | 1 | 5 | Q5XJE5 | RNA polym | Q5XJE5 [62 | 0 | 1789.826 | 0 | -0.49 | 0.001937 | 2.46 | 116.6 | 83.4 | 5 | 4.28 | | High | High | High | High | High | High | High | High | High | High | High | 0 | 3.95E-07 | 2.91 |
| FALSE | High | LNSGAGTK | [R].LNSGA | 1xPhospho | E9Q166 1x | 0.022378 | 0.001542 | 1 | 2 | 1 | E9Q166 | ATPase fan | E9Q166 [1 | 0 | 1285.692 | 0 | -0.66 | 0.002822 | 3.46 | 122.4 | 77.6 | 5.07 | 7.62 | | High | High | High | High | High | High | High | High | High | High | High | 0.000814 | 0.00776 | 1.18 |
| FALSE | High | LNSGGSSEI | [K].LNSGG | 1xPhospho | Q8K3K8 1x | 0.000645 | 0 | 1 | 1 | 2 | Q8K3K8 | Optineurin | Q8K3K8 [1 | 0 | 1905.879 | 0 | 3.47 | 0.0175 | 47.77 | 14.6 | 185.4 | 17.15 | 44.2 | | High | High | High | High | High | High | High | High | High | High | High | 0.001581 | 0.01524 | 2.79 |
| FALSE | High | LNSGSLSED | [K].LNSGSL | 1xPhospho | A2CG76 1x | 2.02E-12 | 0 | 1 | 5 | 9 | A2CG76 | Histone-lys | A2CG76 [2 | 0 | 2757.396 | 0 | -0.41 | 0.017176 | 4.54 | 115.3 | 84.7 | 8.64 | 12.28 | | High | High | High | High | High | High | High | High | High | High | High | 0 | 2.36E-09 | 5.81 |
| FALSE | High | INSLLEYGR | [R].INSLLEY | 1xPhospho | Q80Y19 1x | 0.020679 | 0.001472 | 1 | 2 | 1 | Q80Y19 | Rho GTPas | Q80Y19 [8 | 0 | 1373.703 | 0 | 0.12 | 0.228501 | 16.55 | 94 | 106 | 7.13 | 8.27 | | High | High | High | High | High | High | High | High | High | High | High | 0.000786 | 0.007067 | 2.52 |
| FALSE | High | INSLTFK | [R].INSLTFI | 1xPhospho | A0A1L1SQI | 0.009416 | 0.000736 | 1 | 4 | 1 | A0A1L1SQI | Dedicator c | A0A1L1SQI | 0 | 1360.764 | 0 | -1.6 | 0.001541 | 1.12 | 149 | 51 | 11.59 | 9.04 | | High | High | High | High | High | High | High | High | High | High | High | 0.000327 | 0.002943 | 1.77 |
| FALSE | High | INTYVFVGI | [R].INTYVF | 1xPhospho | Q3TM36 1 | 3.92E-06 | 0 | 1 | 5 | 1 | Q3TM36 | Putative ur | Q3TM36 [2 | 0 | 3186.601 | 0 | -0.39 | 0.008768 | 5.53 | 112.4 | 87.6 | 5.77 | 14.08 | | High | High | High | High | High | High | High | High | High | High | High | 0 | 4.72E-07 | 3.25 |
| FALSE | High | INSIYNAK | [R].INSIYNA | 1xPhospho | Q924W7 1 | 0.023584 | 0.001578 | 1 | 4 | 1 | Q924W7 | Suppressio | Q924W7 [6 | 0 | 1460.791 | 0 | -0.22 | 0.470371 | 31.35 | 107.5 | 92.5 | 18.46 | 16.26 | | High | High | High | High | High | High | High | High | High | High | High | 0.000859 | 0.008179 | 2.33 |
| FALSE | High | LNSNNSAS | [K].LNSNN | 1xPhospho | Q9ERU9 1x | 2.25E-05 | 0 | 1 | 1 | 2 | Q9ERU9 | E3 SUMO-p | Q9ERU9 [8 | 0 | 1505.706 | 0 | -0.77 | 0.002925 | 2.56 | 127.7 | 72.3 | 10.36 | 8.1 | | High | High | High | High | High | High | High | High | High | High | High | 0 | 3.36E-06 | 4.19 |
| FALSE | High | LNSPPSTIYI | [R].LNSPPS | 1xPhospho | A0A0R4J0H | 0.00034 | 0 | 1 | 2 | 1 | A0A0R4J0H | Fibronectir | A0A0R4J0H | 0 | 1657.897 | 0 | 1.42 | 0.002667 | 11.79 | 54.6 | 145.4 | 12.6 | 9.53 | | High | High | High | High | High | High | High | High | High | High | High | 0 | 7.09E-05 | 1.72 |
| FALSE | High | LNSPTNSSF | [K].LNSPTN | 1xPhospho | Q9DBX1 1x | 0.000131 | 0 | 1 | 1 | 3 | Q9DBX1 | Regulator c | Q9DBX1 [9 | 0 | 2231.163 | 0 | -0.97 | 0.000559 | 1.49 | 135.5 | 64.5 | 7.12 | 7.24 | | High | High | High | High | High | High | High | High | High | High | High | 0 | 2.42E-05 | 3.18 |
| FALSE | High | INSQLVAQ | [R].INSQLV | 1xPhospho | Q8CCI5 1xl | 0.000568 | 0 | 1 | 1 | 2 | Q8CCI5 | RING1 and | Q8CCI5 [57 | 0 | 2816.511 | 0 | -0.22 | 0.267438 | 20.35 | 102 | 98 | 14.17 | 17.34 | | High | High | High | High | High | High | High | High | High | High | High | 0 | 0.000126 | 2.38 |
| FALSE | High | LNSSENGE | [R].LNSSEN | 1xPhospho | S4R2L4 1xP | 0.008463 | 0.000736 | 1 | 8 | 1 | S4R2L4 | Arginine/se | S4R2L4 [10 | 0 | 1429.616 | 0 | -0.6 | 0.008923 | 3.92 | 123 | 77 | 10.85 | 16.64 | | High | High | High | High | High | High | High | High | High | High | High | 0.000327 | 0.002596 | 2.7 |
| FALSE | High | LNSSENGE | [R].LNSSEN | 1xDeamida | S4R2L4 1xP | 0.037692 | 0.00249 | 1 | 8 | 1 | S4R2L4 | Arginine/se | S4R2L4 [10 | 0 | 1430.6 | 0 | -0.48 | 0.140727 | 13 | 118.1 | 81.9 | 21.89 | 9.35 | | High | High | High | High | High | High | High | High | High | High | High | 0.001324 | 0.01395 | 3.17 |
| FALSE | High | LNSSFDDW | [K].LNSSFD | 1xCarbami | Q5SVT3 1x | 0.001168 | 0 | 1 | 3 | 2 | Q5SVT3 | Ewing's tur | Q5SVT3 [5 | 0 | 3015.322 | 0 | -1.25 | 0.001768 | 6.4 | 140.7 | 59.3 | 9.21 | 24.75 | | High | High | High | High | High | High | High | High | High | High | High | 0 | 0.000285 | 3.36 |

| | | | | | | | | | | | | | | | | | | | | | | | | | | | | | | | | | | | | | | |
|---|---|---|---|---|---|---|---|---|---|---|---|---|---|---|---|---|---|---|---|---|---|---|---|---|---|---|---|---|---|---|---|---|---|---|---|---|---|---|
| FALSE | High | LNSSFDDW | [K].LNSSFD | 1xCarbami | Q5SVT3 1x | 0.003001 | 0.00016 | 1 | 3 | 4 | Q5SVT3 | Ewing's tur | Q5SVT3 [5 | 0 | 3016.306 | 0 | -0.99 | 0.018772 | 2.52 | 128.1 | 71.9 | 13.39 | 18.59 | | High | High | High | High | High | High | High | High | High | High | High | 0.000449 | 0.003942 | 3.43 |
| FALSE | High | INSSGK | [R].INSSGK | 1xPhospho | Q5UAK0-1 | 0.087983 | 0.006639 | 1 | 6 | 1 | Q5UAK0-1 | Isoform 1 c | Q5UAK0-1 | 0 | 1143.618 | 0 | -0.05 | 0.501434 | 9.86 | 99.1 | 100.9 | 4.84 | 4.69 | | High | High | High | High | High | High | High | High | High | High | High | 0.003402 | 0.0372 | 1.33 |
| FALSE | High | LNSSSGTGI | [K].LNSSSG | 1xPhospho | E9Q2H1 1x | 0.037369 | 0.00249 | 1 | 2 | 1 | E9Q2H1 | E3 ubiquiti | E9Q2H1 [9 | 0 | 1187.562 | 0 | -0.67 | 0.012277 | 9.5 | 124.3 | 75.7 | 11.07 | 14.4 | | High | High | High | High | High | High | High | High | High | High | High | 0.001308 | 0.01378 | 0.56 |
| FALSE | High | LNSSSSSSN | [K].LNSSSS | 1xPhospho | A2A432 1x | 0.000103 | 0 | 1 | 4 | 6 | A2A432 | Cullin-4B C | A2A432 [1 | 0 | 1764.76 | 0 | -0.5 | 0.006038 | 5.74 | 118.1 | 81.9 | 4.39 | 10.53 | | High | High | High | High | High | High | High | High | High | High | High | 0 | 5.2E-05 | 3.04 |
| FALSE | High | LNSSSSSSN | [K].LNSSSS | 1xPhospho | A2A432 1x | 0.000279 | 0 | 1 | 4 | 14 | A2A432 | Cullin-4B C | A2A432 [1 | 1 | 3483.498 | 0 | -1.89 | 0.000402 | 0.14 | 157.5 | 42.5 | 9.88 | 7.58 | | High | High | High | High | High | High | High | High | High | High | High | 0.000275 | 0.00158 | 2.4 |
| FALSE | High | LNTDATAE | [K].LNTDAT | 1xPhospho | Q9ERQ3 1x | 0.000793 | 0 | 1 | 2 | 3 | Q9ERQ3 | Zinc finger | Q9ERQ3 [3 | 0 | 3790.874 | 0 | 0.69 | 0.030735 | 12.05 | 78.7 | 121.3 | 19.89 | 16.17 | | High | High | High | High | High | High | High | High | High | High | High | 4.94E-05 | 0.000578 | 2.65 |
| FALSE | High | LNSNNALIE | [R].LNSNN | 1xPhospho | E9QL65 1x | 9.11E-05 | 0 | 1 | 2 | 1 | E9QL65 | Conserved | E9QL65 [6 | 0 | 2452.269 | 0 | 1.13 | 0.004342 | 25.88 | 68.1 | 131.9 | 21.56 | 11.38 | | High | High | High | High | High | High | High | High | High | High | High | 0 | 1.62E-05 | 3.02 |
| FALSE | High | LPEDDEPP | [K].LPEDDE | 2xPhospho | Q8K3G7 2x | 0.065952 | 0.004766 | 3 | 4 | 1 | Q8K3G7; C | Reticulon ( | Q8K3G7 [1 | 0 | 3900.923 | 0 | -0.47 | 0.039448 | 9.11 | 111 | 89 | 10.92 | 19.82 | Shared | High | High | High | High | High | High | High | High | High | High | High | 0.002455 | 0.02664 | 3.11 |
| FALSE | High | LPEEPSSED | [R].LPEEPS | 1xPhospho | E9Q0N0 1x | 0.00153 | 5.66E-05 | 1 | 10 | 3 | E9Q0N0 | Intersectin | E9Q0N0 [3 | 0 | 2280.068 | 0 | 0.18 | 0.016448 | 5.38 | 94.7 | 105.3 | 5.97 | 10.12 | | High | High | High | High | High | High | High | High | High | High | High | 2.58E-05 | 0.000385 | 3.23 |
| FALSE | High | LPEGPVDSI | [R].LPEGPV | 1xPhospho | B2RUJ6 1x | 0.001137 | 0 | 1 | 9 | 1 | B2RUJ6 | Rap guanin | B2RUJ6 [22 | 1 | 3421.501 | 0 | 1.09 | 0.001562 | 7.03 | 61.9 | 138.1 | 11.88 | 8.7 | | High | High | High | High | High | High | High | High | High | High | High | 0 | 0.000275 | 2.93 |
| FALSE | High | LPLVEQSPE | [K].LPLVEC | 1xPhospho | A2AUM0 1 | 0.021607 | 0.001507 | 1 | 9 | 2 | A2AUM0 | eIF-2-alpha | A2AUM0 [ | 0 | 4560.248 | 0 | 0.99 | 0.002795 | 20.08 | 66.9 | 133.1 | 10.61 | 11.67 | | High | High | High | High | High | High | High | High | High | High | High | 0.000801 | 0.00741 | 3.72 |
| FALSE | High | LPMVAGPL | [R].LPMVA | 1xPhospho | Q5NBY9 1x | 0.000257 | 0 | 1 | 6 | 1 | Q5NBY9 | POZ (BTB) a | Q5NBY9 [2 | 0 | 3555.93 | 0 | -0.35 | 0.040916 | 11.87 | 113.8 | 86.2 | 9.43 | 15.65 | | High | High | High | High | High | High | High | High | High | High | High | 0 | 5.17E-05 | 3.5 |
| FALSE | High | LPNLASPSA | [R].LPNLAS | 1xPhospho | Q3UDC3 1x | 0.000311 | 0 | 1 | 3 | 2 | Q3UDC3 | Putative ur | Q3UDC3 [4 | 0 | 2478.27 | 0 | 0.2 | 0.019882 | 7.9 | 93.1 | 106.9 | 1.58 | 7.2 | | High | High | High | High | High | High | High | High | High | High | High | 0 | 6.42E-05 | 1.94 |
| FALSE | High | LPNLASPSA | [R].LPNLAS | 2xPhospho | Q3UDC3 2x | 0.035793 | 0.002304 | 1 | 3 | 2 | Q3UDC3 | Putative ur | Q3UDC3 [4 | 0 | 2558.237 | 0 | 0.37 | 0.01502 | 9.28 | 89.5 | 110.5 | 5.78 | 5.36 | | High | High | High | High | High | High | High | High | High | High | High | 0.00123 | 0.01322 | 2.04 |
| FALSE | High | LPNNSSRPS | [R].LPNNSS | 1xPhospho | Q571L5 1x | 0.005615 | 0.00062 | 1 | 8 | 1 | Q571L5 | MKIAA406 | Q571L5 [3 | 0 | 2336.206 | 0 | -0.46 | 0.17158 | 9.62 | 116.2 | 83.8 | 20.5 | 15.12 | | High | High | High | High | High | High | High | High | High | High | High | 0.000275 | 0.00164 | 2.66 |
| FALSE | High | LPNNSSRPS | [R].LPNNSS | 1xPhospho | Q571L5 1x | 0.000524 | 0 | 1 | 8 | 5 | Q571L5 | MKIAA406 | Q571L5 [3 | 0 | 2565.369 | 0 | -1 | 0.002821 | 3.86 | 131.2 | 68.8 | 8.92 | 8.99 | | High | High | High | High | High | High | High | High | High | High | High | 0 | 0.000291 | 2.18 |
| FALSE | High | LPNNSSRPS | [R].LPNNSS | 1xDeamida | Q571L5 1x | 0.062209 | 0.004646 | 1 | 8 | 2 | Q571L5 | MKIAA406 | Q571L5 [3 | 0 | 2566.353 | 0 | -1.04 | 0.002608 | 1.37 | 133.3 | 66.7 | 10.83 | 10.49 | | High | High | High | High | High | High | High | High | High | High | High | 0.004327 | 0.04674 | 2.19 |
| FALSE | High | LPNNSSRPS | [R].LPNNSS | 2xPhospho | Q571L5 2x | 0.001349 | 0 | 1 | 8 | 2 | Q571L5 | MKIAA406 | Q571L5 [3 | 0 | 2645.335 | 0 | -0.35 | 0.078245 | 13.52 | 111.6 | 88.4 | 12.04 | 8.35 | | High | High | High | High | High | High | High | High | High | High | High | 0 | 0.000335 | 2.75 |
| FALSE | High | LPNTYPNSI | [R].LPNTYP | 1xPhospho | A2A9M4 1 | 0.000718 | 0 | 1 | 9 | 1 | A2A9M4 | Dedicator c | A2A9M4 [8 | 0 | 3007.433 | 0 | 0.72 | 0.021289 | 26.34 | 77.5 | 122.5 | 9.77 | 13.32 | | High | High | High | High | High | High | High | High | High | High | High | 0 | 0.000165 | 4.34 |
| FALSE | High | LPNTYPNSI | [R].LPNTYP | 1xPhospho | A2A9M4 1 | 0.000237 | 0 | 1 | 9 | 2 | A2A9M4 | Dedicator c | A2A9M4 [8 | 0 | 3023.428 | 0 | 0.73 | 0.005355 | 19.54 | 76 | 124 | 9.38 | 8.37 | | High | High | High | High | High | High | High | High | High | High | High | 0 | 4.74E-05 | 3.29 |
| FALSE | High | LPPASTPTS | [K].LPPAST | 1xPhospho | Q6PD28 1x | 0.062209 | 0.004646 | 1 | 1 | 1 | Q6PD28 | Serine/thre | Q6PD28 [5 | 0 | 2764.457 | 0 | 1.22 | 0.00161 | 4.39 | 61.2 | 138.8 | 7.1 | 9.81 | | High | High | High | High | High | High | High | High | High | High | High | 0.002381 | 0.02482 | 1.77 |
| FALSE | High | LPPEAQQL | [R].LPPEAC | 1xCarbami | Q7TT79 1x | 5.31E-05 | 0 | 1 | 4 | 1 | Q7TT79 | Microceph | Q7TT79 [4 | 0 | 2260.115 | 0 | 0.28 | 0.297315 | 24.62 | 87.2 | 112.8 | 15.98 | 11.69 | | High | High | High | High | High | High | High | High | High | High | High | 0 | 8.83E-06 | 3.46 |
| FALSE | High | LPPPPPQAI | [K].LPPPPP | 1xPhospho | Q6ZQ88 1x | 3.06E-05 | 0 | 1 | 2 | 4 | Q6ZQ88 | Lysine-spe | Q6ZQ88 [1 | 0 | 3707.695 | 0 | -0.5 | 0.015951 | 11.2 | 115.4 | 84.6 | 7.63 | 11.99 | | High | High | High | High | High | High | High | High | High | High | High | 0 | 4.76E-06 | 3.96 |
| FALSE | High | IPITPTSSFV | [K].IPITPTS | 2xPhospho | E9Q1P8 2x | 0.016743 | 0.001166 | 1 | 1 | 3 | E9Q1P8 | Interferon | E9Q1P8 [3 | 0 | 2776.331 | 0 | 0.45 | 0.010825 | 6.33 | 85.3 | 114.7 | 15.4 | 8.18 | | High | High | High | High | High | High | High | High | High | High | High | 0.000622 | 0.005564 | 3.8 |
| FALSE | High | LPPPPPQAI | [K].LPPPPP | 1xDeamida | Q6ZQ88 1x | 9.36E-05 | 0 | 1 | 2 | 2 | Q6ZQ88 | Lysine-spe | Q6ZQ88 [1 | 0 | 3708.679 | 0 | -0.03 | 0.532313 | 21.58 | 103.9 | 96.1 | 16.57 | 27.81 | | High | High | High | High | High | High | High | High | High | High | High | 0 | 1.67E-05 | 4.25 |
| FALSE | High | LPPQGEAE | [K].LPPQGI | 1xPhospho | Q8C0J6 1x | 0.065952 | 0.004766 | 1 | 1 | 1 | Q8C0J6 | Ankyrin re | Q8C0J6 [17 | 0 | 2244.05 | 0 | -0.55 | 0.007711 | 8.39 | 119.4 | 80.6 | 2.9 | 11.67 | | High | High | High | High | High | High | High | High | High | High | High | 0.002455 | 0.02665 | 1.29 |
| FALSE | High | LPPQSCLLL | [R].LPPQSC | 1xCarbami | E9QK83 1x | 0.003755 | 0.00021 | 1 | 4 | 1 | E9QK83 | Small subu | E9QK83 [2 | 0 | 1971.07 | 0 | -0.44 | 0.010501 | 7.07 | 115.1 | 84.9 | 7.93 | 7.31 | | High | High | High | High | High | High | High | High | High | High | High | 9.42E-05 | 0.001052 | 2.63 |
| FALSE | High | LPPSPLSSS | [R].LPPSPL | 1xPhospho | D3Z5I1 1xP | 0.002313 | 0.000109 | 1 | 3 | 2 | D3Z5I1 | Zinc finger | D3Z5I1 [53 | 0 | 1674.841 | 0 | 1.18 | 0.011298 | 24.84 | 61.3 | 138.7 | 9.96 | 17.51 | | High | High | High | High | High | High | High | High | High | High | High | 0.000183 | 0.001248 | 2.15 |
| FALSE | High | LPPTFFCG | [K].LPPTFF | 2xCarbami | Q9ERU9 1x | 3.65E-07 | 0 | 1 | 1 | 7 | Q9ERU9 | E3 SUMO-p | Q9ERU9 [2 | 0 | 3515.461 | 0 | 0.07 | 0.299121 | 29.73 | 94 | 106 | 8.08 | 20.52 | | High | High | High | High | High | High | High | High | High | High | High | 0 | 1.17E-06 | 4.72 |
| FALSE | High | LPPYPYSPP | [R].LPPYPY | 1xPhospho | Q3U182 1x | 0.001002 | 0 | 2 | 4 | 1 | Q3U182; C | CREB-regu | Q3U182 [4 | 0 | 2836.545 | 0 | 0.39 | 0.012611 | 9.82 | 89.8 | 110.2 | 5.85 | 8.94 | Shared | High | High | High | High | High | High | High | High | High | High | High | 0 | 0.00024 | 1.76 |
| FALSE | High | LPQCLLVQ | [K].LPQCLL | 1xCarbami | Q6PR54-3 | 3.5E-08 | 0 | 2 | 4 | 4 | Q6PR54-3; | Isoform 3 c | Q6PR54-3 | 0 | 3109.589 | 0 | -1.19 | 0.005083 | 5.7 | 142.1 | 57.9 | 15.44 | 11.48 | Shared | High | High | High | High | High | High | High | High | High | High | High | 0 | 2.36E-09 | 4.86 |
| FALSE | High | IPQCSQLQI | [K].IPQCSC | 1xCarbami | Q14A61 1x | 4.95E-06 | 0 | 1 | 3 | 1 | Q14A61 | Cenpl prot | Q14A61 [5 | 0 | 2337.156 | 0 | -0.33 | 0.026044 | 12.92 | 111.1 | 88.9 | 11.79 | 16.38 | | High | High | High | High | High | High | High | High | High | High | High | 0 | 6.15E-07 | 4.41 |
| FALSE | High | LPQFGISTP | [K].LPQFGI | 1xPhospho | E9Q616 1x | 8.87E-05 | 0 | 1 | 6 | 1 | E9Q616 | AHNAK nuc | E9Q616 [5 | 0 | 2453.309 | 0 | 2.46 | 0.000779 | 8.27 | 29.3 | 170.7 | 12.87 | 14 | | High | High | High | High | High | High | High | High | High | High | High | 0 | 1.56E-05 | 3.55 |
| FALSE | High | IPQMESSPI | [K].IPQME | 1xPhospho | Q5I043 1xP | 8.4E-05 | 0 | 1 | 5 | 2 | Q5I043 | Ubiquitin c | Q5I043 [70 | 0 | 3560.569 | 0 | -1.82 | 0.001075 | 0.93 | 155.8 | 44.2 | 9.35 | 15.77 | | High | High | High | High | High | High | High | High | High | High | High | 0 | 1.48E-05 | 2.91 |
| FALSE | High | LPQPPEGQ | [K].LPQPPE | 1xCarbami | E9PY39 1x | 0.000845 | 0 | 1 | 4 | 2 | E9PY39 | Uncharacte | E9PY39 [36 | 0 | 1698.74 | 0 | 0.86 | 0.000513 | 7.05 | 71.2 | 128.8 | 3.88 | 6.56 | | High | High | High | High | High | High | High | High | High | High | High | 0 | 0.000197 | 2.06 |
| FALSE | High | LPQSALNTI | [R].LPQSAL | 1xPhospho | Q9ESL4 1x | 0.001489 | 5.66E-05 | 1 | 1 | 1 | Q9ESL4 | Mitogen-ac | Q9ESL4 [71 | 0 | 2221.142 | 0 | 1.1 | 0.000399 | 11.46 | 62.7 | 137.3 | 7.2 | 2.94 | | High | High | High | High | High | High | High | High | High | High | High | 2.58E-05 | 0.000373 | 1.62 |
| FALSE | High | LPQSGSQS | [R].LPQSGS | 1xPhospho | A2AWA9 1 | 0.001558 | 5.66E-05 | 1 | 2 | 6 | A2AWA9 | Rab GTPas | A2AWA9 [4 | 0 | 4386.974 | 0 | 1.03 | 0.001078 | 6.86 | 68.2 | 131.8 | 6.84 | 10.49 | | High | High | High | High | High | High | High | High | High | High | High | 0.00014 | 0.001101 | 4.02 |
| FALSE | High | LPQTPLDT | [K].LPQTPL | 1xPhospho | B9EKS2 1x | 0.005182 | 0.000551 | 1 | 2 | 1 | B9EKS2 | Jumonji do | B9EKS2 [13 | 0 | 2994.599 | 0 | 0.84 | 0.050179 | 42.18 | 74.7 | 125.3 | 11.29 | 26.32 | | High | High | High | High | High | High | High | High | High | High | High | 0.000246 | 0.001507 | 2.78 |
| FALSE | High | LPPPVIPGG | [K].LPPPVII | 1xPhospho | A2ADB0 1x | 0.031168 | 0.001935 | 1 | 5 | 2 | A2ADB0 | Msx2-inter | A2ADB0 [2 | 0 | 2352.325 | 0 | 0.25 | 0.135199 | 7.99 | 90.6 | 109.4 | 5.44 | 12.99 | | High | High | High | High | High | High | High | High | High | High | High | 0.001044 | 0.0112 | 2.65 |
| FALSE | High | IPITPTSSFV | [K].IPITPTS | 1xPhospho | E9Q1P8 1x | 1.51E-06 | 0 | 1 | 1 | 3 | E9Q1P8 | Interferon | E9Q1P8 [3 | 0 | 2696.365 | 0 | 0.11 | 0.128234 | 7.12 | 97.3 | 102.7 | 5.37 | 9.82 | | High | High | High | High | High | High | High | High | High | High | High | 0 | 1.61E-07 | 4.56 |
| FALSE | High | IPISTFSPPS | [K].IPISTFS | 1xPhospho | Q7TT18 1x | 7.33E-08 | 0 | 1 | 1 | 2 | Q7TT18 | Activating | Q7TT18 [9 | 0 | 2483.202 | 0 | -0.29 | 0.009845 | 7.92 | 111.5 | 88.5 | 4.11 | 10.63 | | High | High | High | High | High | High | High | High | High | High | High | 0 | 5.42E-09 | 4.25 |
| FALSE | High | LPLRNDSP | [R].LPLRND | 1xDeamida | Q8R1J3 1x | 0.075927 | 0.005826 | 1 | 3 | 1 | Q8R1J3 | Zinc finger | Q8R1J3 [47 | 1 | 1777.961 | 0 | -0.62 | 0.00521 | 7.93 | 123 | 77 | 10.28 | 9.77 | | High | High | High | High | High | High | High | High | High | High | High | 0.002976 | 0.03122 | 2.13 |
| FALSE | High | LPEGSPTW | [K].LPEGSP | 1xCarbami | Q61122 1x | 1.7E-05 | 0 | 1 | 3 | 2 | Q61122 | NGFI-A-bin | Q61122 [9 | 0 | 2375.094 | 0 | 0.73 | 0.001253 | 4.17 | 75.5 | 124.5 | 2.41 | 6.5 | | High | High | High | High | High | High | High | High | High | High | High | 0 | 2.45E-06 | 3.87 |
| FALSE | High | LPELNWRS | [K].LPELNV | 1xPhospho | Q69ZG4 1x | 0.0026 | 0.000109 | 1 | 2 | 1 | Q69ZG4 | MKIAA156 | Q69ZG4 [7 | 1 | 3004.268 | 0 | 0.78 | 0.002367 | 13.52 | 72.9 | 127.1 | 12.51 | 3.61 | | High | High | High | High | High | High | High | High | High | High | High | 4.94E-05 | 0.0007 | 3.92 |
| FALSE | High | LPELNWRS | [K].LPELNV | 2xPhospho | Q69ZG4 2x | 4.2E-05 | 0 | 1 | 2 | 1 | Q69ZG4 | MKIAA156 | Q69ZG4 [7 | 1 | 3084.235 | 0 | 0.74 | 0.000658 | 2.68 | 72 | 128 | 6.19 | 7.23 | | High | High | High | High | High | High | High | High | High | High | High | 0 | 6.77E-06 | 3.87 |
| FALSE | High | IPEISIQDM | [K].IPEISIQ | 1xPhospho | Q8BTM8 1 | 0.091627 | 0.007146 | 1 | 5 | 1 | Q8BTM8 | Filamin-A C | Q8BTM8 [2 | 0 | 2540.308 | 0 | -0.19 | 0.133584 | 10.71 | 105.7 | 94.3 | 12.44 | 6.43 | | High | High | High | High | High | High | High | High | High | High | High | 0.003646 | 0.03895 | 3.06 |
| FALSE | High | LPENEGISP | [K].LPENEC | 1xPhospho | Q80U44 1x | 0.00015 | 0 | 1 | 2 | 1 | Q80U44 | Zinc finger | Q80U44 [3 | 0 | 2079.041 | 0 | 0.35 | 0.002505 | 7.34 | 84.7 | 115.3 | 6.22 | 5.92 | | High | High | High | High | High | High | High | High | High | High | High | 0 | 2.83E-05 | 3.32 |
| FALSE | High | LPENTPPC | [R].LPENTF | 1xCarbami | Q9D0L8 1x | 2.91E-07 | 0 | 1 | 6 | 2 | Q9D0L8 | mRNA cap | Q9D0L8 [3 | 0 | 2289.135 | 0 | -0.5 | 0.001604 | 2.33 | 116.5 | 83.5 | 2.47 | 9.63 | | High | High | High | High | High | High | High | High | High | High | High | 0 | 2.53E-08 | 3.42 |
| FALSE | High | IPESELGSP | [R].IPESEL | 1xPhospho | A0A140LJD | 0.001189 | 0 | 1 | 1 | 1 | A0A140LJD | Protein TA | A0A140LJD | 0 | 2196.157 | 0 | 1.36 | 0.000524 | 9.71 | 59.9 | 140.1 | 19.69 | 9.44 | | High | High | High | High | High | High | High | High | High | High | High | 0 | 0.00029 | 3.06 |
| FALSE | High | LPESEPVEA | [R].LPESEP | 1xPhospho | Q3U5F4 1x | 6.83E-05 | 0 | 1 | 1 | 1 | Q3U5F4 | YrdC doma | Q3U5F4 [5 | 0 | 1819.868 | 0 | -0.24 | 0.054277 | 8.53 | 110 | 90 | 9.25 | 13.7 | | High | High | High | High | High | High | High | High | High | High | High | 0 | 1.17E-05 | 3.21 |
| FALSE | High | IPESETESTA | [R].IPESETI | 1xPhospho | Q62245 1x | 1.93E-07 | 0 | 1 | 4 | 2 | Q62245 | Son of seve | Q62245 [1 | 0 | 2081.959 | 0 | 0.23 | 0.124701 | 13.42 | 93 | 107 | 11.4 | 6.57 | | High | High | High | High | High | High | High | High | High | High | High | 0 | 9.77E-07 | 4.02 |
| FALSE | High | IPESETESTA | [R].IPESETI | 2xPhospho | Q62245 2x | 0.038346 | 0.002563 | 1 | 4 | 2 | Q62245 | Son of seve | Q62245 [1 | 0 | 2161.925 | 0 | -0.95 | 0.001003 | 1.55 | 129.9 | 70.1 | 12.86 | 12.88 | | High | High | High | High | High | High | High | High | High | High | High | 0.001355 | 0.01429 | 3.03 |
| FALSE | High | LPEYSQTAE | [K].LPEYSQ | 1xPhospho | F6RJ39 1xP | 1.07E-07 | 0 | 1 | 4 | 3 | F6RJ39 | Apoptotic | F6RJ39 [18 | 0 | 2547.097 | 0 | -0.87 | 0.005796 | 3.63 | 126.9 | 73.1 | 11.16 | 11.35 | | High | High | High | High | High | High | High | High | High | High | High | 0 | 8.29E-09 | 3.94 |
| FALSE | High | IPFTLEAHS | [R].IPFTLEA | 2xCarbami | Q8BS90 1x | 1.51E-06 | 0 | 1 | 1 | 3 | Q8BS90 | Protein aur | Q8BS90 [3 | 0 | 3840.693 | 0 | -1.16 | 0.000319 | 1.63 | 135.8 | 64.2 | 5.35 | 8.95 | | High | High | High | High | High | High | High | High | High | High | High | 0 | 1.96E-07 | 3.7 |
| FALSE | High | IPFTLEAHS | [R].IPFTLEA | 2xCarbami | Q8BS90 1x | 2.18E-07 | 0 | 1 | 1 | 2 | Q8BS90 | Protein aur | Q8BS90 [3 | 0 | 3841.677 | 0 | -1.24 | 0.001414 | 2.4 | 135.9 | 64.1 | 12.78 | 33.82 | | High | High | High | High | High | High | High | High | High | High | High | 0 | 1.83E-08 | 5.51 |
| FALSE | High | IPGGLQDS | [K].IPGGLC | 1xPhospho | D3YWX2 1 | 0.078467 | 0.006104 | 1 | 2 | 1 | D3YWX2 | YLP motif-c | D3YWX2 [1 | 0 | 1581.747 | 0 | -0.05 | 0.300382 | 14.91 | 105.4 | 94.6 | 14.02 | 9.94 | | High | High | High | High | High | High | High | High | High | High | High | 0.003122 | 0.0324 | 2.45 |
| FALSE | High | IPGGNIYISI | [R].IPGGNI | 1xPhospho | P13405 1x | 1.93E-05 | 0 | 1 | 4 | 2 | P13405 | Retinoblast | P13405 [79 | 0 | 1810.028 | 0 | 1.27 | 0.000154 | 4.15 | 58.7 | 141.3 | 3.23 | 1.52 | | High | High | High | High | High | High | High | High | High | High | High | 0 | 2.84E-06 | 3.18 |
| FALSE | High | LPGGPAVS | [R].LPGGP | 1xPhospho | A2AKB9 1x | 0.025964 | 0.001645 | 1 | 3 | 2 | A2AKB9 | DDB1- and | A2AKB9 [6 | 0 | 1459.751 | 0 | 0.96 | 0.000181 | 1.62 | 68 | 132 | 5.64 | 5.33 | | High | High | High | High | High | High | High | High | High | High | High | 0.000903 | 0.009185 | 1.6 |
| FALSE | High | LPGMSPAN | [R].LPGMS | 1xPhospho | A2ABV5 1x | 0.000112 | 0 | 1 | 2 | 2 | A2ABV5 | Mediator c | A2ABV5 [1 | 0 | 2688.317 | 0 | 0.34 | 0.036498 | 6.32 | 89.1 | 110.9 | 9.97 | 9.83 | | High | High | High | High | High | High | High | High | High | High | High | 0 | 2.04E-05 | 4.16 |
| FALSE | High | LPGMSPAN | [R].LPGMS | 2xPhospho | A2ABV5 2x | 0.032831 | 0.002195 | 1 | 2 | 1 | A2ABV5 | Mediator c | A2ABV5 [1 | 0 | 2784.278 | 0 | 0.06 | 0.522816 | 16.89 | 97.9 | 102.1 | 5.14 | 14.45 | | High | High | High | High | High | High | High | High | High | High | High | 0.001151 | 0.01188 | 2.44 |
| FALSE | High | IPGSPSAAA | [R].IPGSPS | 1xPhospho | E9PUJ6 1x | 0.000595 | 0 | 1 | 4 | 2 | E9PUJ6 | E3 ubiquiti | E9PUJ6 [23 | 0 | 2658.325 | 0 | 0.3 | 0.017307 | 5.37 | 88.4 | 111.6 | 9.82 | 3.26 | | High | High | High | High | High | High | High | High | High | High | High | 0 | 0.000133 | 2.99 |
| FALSE | High | IPHLSEGPK | [R].IPHLSE | 1xPhospho | Q5CZX6 1x | 0.011048 | 0.000772 | 1 | 5 | 1 | Q5CZX6 | Acbd5 prot | Q5CZX6 [3 | 0 | 1515.834 | 0 | -0.71 | 0.002372 | 2.55 | 124.6 | 75.4 | 10.26 | 3.83 | | High | High | High | High | High | High | High | High | High | High | High | 0.000399 | 0.003509 | 1.83 |
| FALSE | High | LPHISSHPL | [K].LPHISS | 1xPhospho | Q80U49 1x | 0.007145 | 0.00062 | 1 | 2 | 1 | Q80U49 | Centrosom | Q80U49 [8 | 0 | 2148.153 | 0 | 0.78 | 0.001006 | 10.78 | 73.2 | 126.8 | 3.99 | 6.12 | | High | High | High | High | High | High | High | High | High | High | High | 0.000275 | 0.00215 | 2.24 |
| FALSE | High | LPHPILPFS | [K].LPHPIL | 1xPhospho | B2RSV7 1x | 1.36E-06 | 0 | 1 | 2 | 2 | B2RSV7 | Fgd6 prote | B2RSV7 [5 | 0 | 2241.245 | 0 | 1.62 | 0.000382 | 11.04 | 49.6 | 150.4 | 2.59 | 7.07 | | High | High | High | High | High | High | High | High | High | High | High | 0 | 1.44E-07 | 4.69 |
| FALSE | High | IPHSSSSEN | [R].IPHSSS | 1xPhospho | Q5SW75 1 | 0.008691 | 0.000736 | 1 | 4 | 1 | Q5SW75 | Protein ph | Q5SW75 [1 | 0 | 1535.742 | 0 | 0.98 | 0.000456 | 10.26 | 67.6 | 132.4 | 7.79 | 4.91 | | High | High | High | High | High | High | High | High | High | High | High | 0.000327 | 0.002682 | 3.38 |
| FALSE | High | IPLDAYQPY | [R].IPLDAY | 1xPhospho | Q9D279 1x | 0.000993 | 0 | 1 | 1 | 1 | Q9D279 | Mitotic inte | Q9D279 [3 | 0 | 2268.208 | 0 | 1.95 | 0.002092 | 16.79 | 41.1 | 158.9 | 11.66 | 15.23 | | High | High | High | High | High | High | High | High | High | High | High | 0 | 0.000236 | 3.48 |
| FALSE | High | LPLEPETPG | [R].LPLEPE | 1xPhospho | A2A9C3 1x | 3.38E-08 | 0 | 1 | 3 | 2 | A2A9C3 | KICSTOR co | A2A9C3 [3 | 0 | 1958.02 | 0 | -0.4 | 0.244374 | 7.52 | 110 | 90 | 15.76 | 12.41 | | High | High | High | High | High | High | High | High | High | High | High | 0 | 2.25E-09 | 3.83 |
| FALSE | High | LPLGCSPET | [K].LPLGCS | 1xCarbami | Q7TSY8 1x | 2.47E-05 | 0 | 1 | 1 | 1 | Q7TSY8 | Shugoshin | Q7TSY8 [9 | 0 | 2988.515 | 0 | -1.04 | 0.023819 | 8.93 | 134.7 | 65.3 | 14.68 | 22.95 | | High | High | High | High | High | High | High | High | High | High | High | 0 | 3.72E-06 | 3.27 |
| FALSE | High | LPLPDDHD | [R].LPLPDD | 1xPhospho | A0A1L1STC | 0.074685 | 0.005727 | 1 | 6 | 1 | A0A1L1STC | Nesprin-1 | A0A1L1STC | 0 | 1701.805 | 0 | 1.12 | 0.000676 | 15.17 | 62.9 | 137.1 | 17.26 | 6.87 | | High | High | High | High | High | High | High | High | High | High | High | 0.002937 | 0.03054 | 1.86 |
| FALSE | High | IPLPTTLAS | [K].IPLPTTI | 1xPhospho | O70472 1x | 0.066503 | 0.004796 | 1 | 2 | 1 | O70472 | Transmem | O70472 [1 | 0 | 1722.981 | 0 | -0.32 | 0.03009 | 3.52 | 108.4 | 91.6 | 10.25 | 11.13 | | High | High | High | High | High | High | High | High | High | High | High | 0.002455 | 0.02691 | 2.2 |
| FALSE | High | LPLQSSPST | [K].LPLQSS | 1xPhospho | Q9CYC5 1x | 0.047488 | 0.00354 | 1 | 1 | 2 | Q9CYC5 | Kinetochor | Q9CYC5 [3 | 0 | 1366.682 | 0 | 0.29 | 0.00421 | 9.29 | 91.2 | 108.8 | 10.09 | 3.72 | | High | High | High | High | High | High | High | High | High | High | High | 0.003922 | 0.04183 | 2.17 |
| FALSE | High | IPSAVSSPLI | [R].IPSAVS | 1xPhospho | E9Q3G8 1x | 5.72E-06 | 0 | 1 | 2 | 4 | E9Q3G8 | Nucleopor | E9Q3G8 [3 | 0 | 1861.962 | 0 | -0.53 | 0.001993 | 5.31 | 118.3 | 81.7 | 2.55 | 8.73 | | High | High | High | High | High | High | High | High | High | High | High | 0 | 4.11E-05 | 3.05 |
| FALSE | High | LLEDSESED | [R].LLEDSE | 2xPhospho | Q9D6J3 2x | 0.000274 | 0 | 1 | 1 | 3 | Q9D6J3 | Coiled-coil | Q9D6J3 [2 | 0 | 2316 | 0 | 0.16 | 0.054927 | 6.47 | 94.1 | 105.9 | 2.74 | 6.39 | | High | High | High | High | High | High | High | High | High | High | High | 0.000467 | 0.004242 | 2.52 |
| FALSE | High | LLEDSESED | [R].LLEDSE | 1xPhospho | Q9D6J3 1x | 0.040028 | 0.002867 | 1 | 1 | 3 | Q9D6J3 | Coiled-coil | Q9D6J3 [2 | 0 | 2236.033 | 0 | 0.14 | 0.275267 | 18.17 | 92.1 | 107.9 | 9.62 | 11.88 | | High | High | High | High | High | High | High | High | High | High | High | 0.001741 | 0.01737 | 2.63 |
| FALSE | High | LIEDNEYTA | [R].LIEDNE | 1xPhospho | Q04736 1x | 6.41E-05 | 0 | 4 | 15 | 8 | Q04736; E | Tyrosine-p | Q04736 [4 | 0 | 1532.72 | 0 | -0.68 | 0.006701 | 3.42 | 123.1 | 76.9 | 9.94 | 11.69 | Shared | High | High | High | High | High | High | High | High | High | High | High | 0 | 4.25E-05 | 3.41 |
| FALSE | High | IEPDMMSN | [K].IEPDMI | 1xPhospho | Q7TSZ8 1x | 0.007339 | 0.00062 | 1 | 1 | 4 | Q7TSZ8 | Nucleus ac | Q7TSZ8 [4 | 0 | 3473.442 | 0 | -1.21 | 0.01858 | 3.16 | 134.1 | 65.9 | 15.78 | 24.83 | | High | High | High | High | High | High | High | High | High | High | High | 0.003094 | 0.03211 | 3.6 |
| FALSE | High | LEPDSEDDI | [R].LEPDSE | 1xPhospho | F6ZGN3 1x | 0.054844 | 0.003982 | 1 | 1 | 1 | F6ZGN3 | Unconvent | F6ZGN3 [3 | 1 | 1856.868 | 0 | -1.29 | 0.00044 | 0.97 | 139.1 | 60.9 | 9.08 | 9.11 | | High | High | High | High | High | High | High | High | High | High | High | 0.002059 | 0.02148 | 2.18 |
| FALSE | High | IEPDSESQE | [R].IEPDSE | 1xPhospho | P70444 1x | 0.000215 | 0 | 1 | 1 | 1 | P70444 | BH3-intera | P70444 [72 | 0 | 2289.096 | 0 | 0.98 | 0.000517 | 4.38 | 66.3 | 133.7 | 5.3 | 5.66 | | High | High | High | High | High | High | High | High | High | High | High | 0 | 4.23E-05 | 2.86 |
| FALSE | High | LEPETLSDG | [R].LEPETL | 1xCarbami | Q8K119 1x | 0.070485 | 0.005214 | 1 | 3 | 2 | Q8K119 | Postmeioti | Q8K119 [5 | 0 | 1769.798 | 0 | -0.59 | 0.013209 | 5.2 | 121.2 | 78.8 | 8.88 | 9.18 | | High | High | High | High | High | High | High | High | High | High | High | 0.005324 | 0.05764 | 1.82 |
| FALSE | High | IEPGASTPR | [K].IEPGAS | 1xPhospho | Q9ERD6 1x | 0.043236 | 0.003228 | 1 | 7 | 1 | Q9ERD6 | Ras-specifi | Q9ERD6 [2 | 0 | 1236.619 | 0 | 0.17 | 0.137095 | 27.72 | 90.9 | 109.1 | 6.29 | 14.55 | | High | High | High | High | High | High | High | High | High | High | High | 0.001651 | 0.0163 | 1.43 |
| FALSE | High | IEPIPGESP | [K].IEPIPGE | 1xPhospho | Q05D44 1x | 0.000751 | 0 | 1 | 5 | 1 | Q05D44 | Eukaryotic | Q05D44 [1 | 0 | 1604.87 | 0 | -1.33 | 0.00074 | 2.04 | 142.8 | 57.2 | 5.1 | 15.08 | | High | High | High | High | High | High | High | High | High | High | High | 0 | 0.000173 | 2.17 |
| FALSE | High | LEPMETER | [K].LEPMET | 1xPhospho | Q7TMI6 1x | 0.064323 | 0.004664 | 1 | 2 | 1 | Q7TMI6 | Zinc finger | Q7TMI6 [1 | 1 | 1823.856 | 0 | -0.56 | 0.004615 | 9.01 | 119.4 | 80.6 | 6.26 | 9.69 | | High | High | High | High | High | High | High | High | High | High | High | 0.002403 | 0.02571 | 1.87 |
| FALSE | High | LEPSPESEA | [K].LEPSPE | 1xPhospho | Q3V300 1x | 3.09E-05 | 0 | 1 | 2 | 2 | Q3V300 | Kinesin-like | Q3V300 [5 | 0 | 1852.935 | 0 | -1.81 | 0.000158 | 0.43 | 155.4 | 44.6 | 4.69 | 7.91 | | High | High | High | High | High | High | High | High | High | High | High | 0 | 0.000118 | 2.98 |
| FALSE | High | LEPVFDSPF | [K].LEPVFD | 1xPhospho | Q9D666 1x | 0.002743 | 0.000109 | 1 | 11 | 2 | Q9D666 | SUN doma | Q9D666 [4 | 0 | 1368.676 | 0 | 0.28 | 0.026075 | 13.15 | 90.6 | 109.4 | 7.13 | 4.92 | | High | High | High | High | High | High | High | High | High | High | High | 0.000689 | 0.00604 | 1.88 |
| FALSE | High | LEPVSPPSP | [K].LEPVSP | 1xPhospho | F8VQL9 1x | 0.024211 | 0.001612 | 1 | 4 | 2 | F8VQL9 | Nuclear re | F8VQL9 [1 | 0 | 2644.322 | 0 | 0.23 | 0.052172 | 7.83 | 90.2 | 109.8 | 3.2 | 10.15 | | High | High | High | High | High | High | High | High | High | High | High | 0.000872 | 0.009025 | 1.94 |
| FALSE | High | LEPVSPPSP | [K].LEPVSP | 2xPhospho | F8VQL9 2x | 8.56E-05 | 0 | 1 | 4 | 3 | F8VQL9 | Nuclear re | F8VQL9 [1 | 0 | 2724.288 | 0 | 0.64 | 0.004613 | 7.02 | 78.3 | 121.7 | 4.69 | 8.38 | | High | High | High | High | High | High | High | High | High | High | High | 9.42E-05 | 0.000999 | 2.93 |
| FALSE | High | LEPVYSPAC | [R].LEPVYS | 1xPhospho | Q9Z2B9 1x | 2.49E-06 | 0 | 1 | 1 | 4 | Q9Z2B9 | Ribosomal | Q9Z2B9 [3 | 0 | 2044.994 | 0 | 1.42 | 0.000226 | 18.69 | 54.4 | 145.6 | 13.53 | 2.68 | | High | High | High | High | High | High | High | High | High | High | High | 0 | 2.84E-07 | 3.61 |
| FALSE | High | LEPVYSPAC | [R].LEPVYS | 2xPhospho | Q9Z2B9 2x | 4.22E-07 | 0 | 1 | 1 | 3 | Q9Z2B9 | Ribosomal | Q9Z2B9 [3 | 0 | 2124.961 | 0 | 2.09 | 0.000583 | 13.24 | 37.3 | 162.7 | 11.52 | 8.74 | | High | High | High | High | High | High | High | High | High | High | High | 0.001812 | 0.01795 | 3.16 |
| FALSE | High | IEPDMMSN | [K].IEPDMI | 1xDeamida | Q7TSZ8 1x | 0.075927 | 0.005826 | 1 | 1 | 1 | Q7TSZ8 | Nucleus ac | Q7TSZ8 [4 | 0 | 3442.436 | 0 | -1.07 | 0.005374 | 7.36 | 137.9 | 62.1 | 9.91 | 19.66 | | High | High | High | High | High | High | High | High | High | High | High | 0.002976 | 0.03112 | 2.72 |
| FALSE | High | LEQALSPE | [K].LEQALS | 1xPhospho | Q9D8S9 1x | 0.009168 | 0.000736 | 1 | 1 | 1 | Q9D8S9 | BolA-like p | Q9D8S9 [4 | 0 | 1805.961 | 0 | -0.01 | 0.774264 | 31.82 | 100.2 | 99.8 | 10.46 | 23.87 | | High | High | High | High | High | High | High | High | High | High | High | 0.000327 | 0.002849 | 2.42 |
| FALSE | High | LEQKPEDG | [R].LEQKPE | 2xCarbami | Q8BGX7 1x | 0.079114 | 0.00613 | 1 | 3 | 1 | Q8BGX7 | DNA repair | Q8BGX7 [8 | 0 | 3588.712 | 0 | -0.05 | 0.90034 | 30.88 | 100.8 | 99.2 | 13.08 | 14.28 | | High | High | High | High | High | High | High | High | High | High | High | 0.003148 | 0.03268 | 2.59 |
| FALSE | High | LEQKPEDG | [R].LEQKPE | 2xCarbami | Q8BGX7 1x | 0.006031 | 0.00062 | 1 | 3 | 3 | Q8BGX7 | DNA repair | Q8BGX7 [8 | 0 | 3589.696 | 0 | 0.04 | 0.8727 | 19.89 | 100.1 | 99.9 | 9.98 | 13.82 | | High | High | High | High | High | High | High | High | High | High | High | 0.000344 | 0.00306 | 3.15 |
| FALSE | High | LEQLCEEFS | [R].LEQLCE | 1xCarbami | Q5SV77-3 | 0.000446 | 0 | 1 | 5 | 3 | Q5SV77-3 | Isoform 3 c | Q5SV77-3 | 0 | 2006.862 | 0 | 0.15 | 0.060525 | 3.53 | 93.1 | 106.9 | 4.95 | 8.4 | | High | High | High | High | High | High | High | High | High | High | High | 0 | 9.66E-05 | 2.55 |
| FALSE | High | IEQLDGVD | [R].IEQLDG | 2xPhospho | F8WJ40 2x | 0.001586 | 5.66E-05 | 1 | 5 | 2 | F8WJ40 | Histone-lys | F8WJ40 [2 | 0 | 2761.18 | 0 | -0.41 | 0.016192 | 6.84 | 114.6 | 85.4 | 6.71 | 12.68 | | High | High | High | High | High | High | High | High | High | High | High | 2.58E-05 | 0.000398 | 3.28 |
| FALSE | High | IEQIEAGTP | [K].IEQIEA | 1xPhospho | Q9JMH6 1 | 0.066503 | 0.004796 | 1 | 3 | 1 | Q9JMH6 | Thioredoxi | Q9JMH6 [3 | 0 | 1479.741 | 0 | 0.27 | 0.020009 | 4.43 | 94.5 | 105.5 | 6.5 | 8.79 | | High | High | High | High | High | High | High | High | High | High | High | 0.002455 | 0.02686 | 1.56 |
| FALSE | High | IEQNLQSP | [R].IEQNLC | 1xPhospho | Q3UFT4 1x | 8.6E-07 | 0 | 1 | 7 | 1 | Q3UFT4 | Putative ur | Q3UFT4 [4 | 0 | 2051.012 | 0 | -1.26 | 0.001598 | 1.86 | 140.8 | 59.2 | 9.58 | 5.14 | | High | High | High | High | High | High | High | High | High | High | High | 0 | 8.62E-08 | 4.29 |
| FALSE | High | LEQTPVRD | [R].LEQTPV | 1xPhospho | Q8K3W3 1 | 0.035486 | 0.002304 | 1 | 1 | 2 | Q8K3W3 | Protein CA | Q8K3W3 [ | 1 | 2781.391 | 0 | -0.43 | 0.095154 | 7.63 | 109.2 | 90.8 | 14.7 | 11.23 | | High | High | High | High | High | High | High | High | High | High | High | 0.005149 | 0.05615 | 2.21 |
| FALSE | High | LEQTPVRD | [R].LEQTPV | 2xPhospho | Q8K3W3 2 | 1.09E-06 | 0 | 1 | 1 | 1 | Q8K3W3 | Protein CA | Q8K3W3 [ | 1 | 2861.357 | 0 | -0.91 | 0.003508 | 3.19 | 132 | 68 | 10.47 | 10.04 | | High | High | High | High | High | High | High | High | High | High | High | 0 | 1.12E-07 | 4.9 |
| FALSE | High | LEQTPVRD | [R].LEQTPV | 1xPhospho | Q8K3W3 1 | 0.088701 | 0.006669 | 1 | 1 | 1 | Q8K3W3 | Protein CA | Q8K3W3 [ | 1 | 3010.554 | 0 | 0.04 | 0.139556 | 6.88 | 98.8 | 101.2 | 6.67 | 6.18 | | High | High | High | High | High | High | High | High | High | High | High | 0.003432 | 0.03732 | 2.37 |
| FALSE | High | LEQTPVRD | [R].LEQTPV | 2xPhospho | Q8K3W3 2 | 0.000466 | 0 | 1 | 1 | 1 | Q8K3W3 | Protein CA | Q8K3W3 [ | 1 | 3090.52 | 0 | -0.81 | 0.008467 | 5.18 | 129.1 | 70.9 | 11.08 | 8.84 | | High | High | High | High | High | High | High | High | High | High | High | 0 | 0.000102 | 2.71 |
| FALSE | High | IERPCCSPK | [R].IERPCC | 2xCarbami | A2ACV6 1x | 0.072858 | 0.005599 | 1 | 4 | 1 | A2ACV6 | Protein SO | A2ACV6 [1 | 0 | 1684.832 | 0 | 1.65 | 0.000241 | 3.66 | 48.6 | 151.4 | 4.59 | 5.91 | | High | High | High | High | High | High | High | High | High | High | High | 0.002855 | 0.02965 | 1.95 |
| FALSE | High | LERPPETPT | [K].LERPPE | 1xPhospho | Q6PDQ2 1 | 6.43E-06 | 0 | 1 | 6 | 2 | Q6PDQ2 | Chromodo | Q6PDQ2 [6 | 0 | 2217.193 | 0 | -0.45 | 0.018931 | 8.72 | 119.3 | 80.7 | 10.66 | 10.63 | | High | High | High | High | High | High | High | High | High | High | High | 0 | 8.22E-07 | 2.24 |
| FALSE | High | IESDAQEP | [K].IESDAQ | 1xPhospho | Q3ULL5 1x | 3.41E-05 | 0 | 1 | 1 | 4 | Q3ULL5 | Eif2s2 prot | Q3ULL5 [1 | 0 | 2967.395 | 0 | 0.28 | 0.087458 | 14.73 | 90.3 | 109.7 | 4.24 | 14.88 | | High | High | High | High | High | High | High | High | High | High | High | 4.94E-05 | 0.000749 | 3.01 |
| FALSE | High | LEQHSQQF | [R].LEQHS | 1xPhospho | Q921T2 1x | 0.000372 | 0 | 1 | 6 | 5 | Q921T2 | Torsin-1A- | Q921T2 [1 | 0 | 2173.06 | 0 | 0.35 | 0.001448 | 2.93 | 87.8 | 112.2 | 2.77 | 6.05 | | High | High | High | High | High | High | High | High | High | High | High | 0.000327 | 0.002691 | 3.61 |
| FALSE | High | IESDAQEP | [K].IESDAQ | 1xPhospho | Q3ULL5 1x | 2.66E-06 | 0 | 1 | 1 | 12 | Q3ULL5 | Eif2s2 prot | Q3ULL5 [1 | 0 | 2983.39 | 0 | 0.35 | 0.003952 | 9.04 | 88.6 | 111.4 | 4.36 | 4.05 | | High | High | High | High | High | High | High | High | High | High | High | 0 | 1.56E-06 | 4.38 |
| FALSE | High | LEPASPPED | [K].LEPASP | 1xCarbami | Q9JI57 1xP | 0.004491 | 0.000409 | 1 | 18 | 1 | Q9JI57 | General tra | Q9JI57 [44 | 0 | 2096.941 | 0 | -0.48 | 0.00273 | 6.37 | 118.8 | 81.2 | 8.49 | 8.41 | | High | High | High | High | High | High | High | High | High | High | High | 0.000183 | 0.001278 | 1.19 |
| FALSE | High | LENSAQAE | [K].LENSAC | 1xPhospho | P23804 1x | 7.75E-05 | 0 | 1 | 4 | 1 | P23804 | E3 ubiquiti | P23804 [34 | 0 | 2310.127 | 0 | 0.13 | 0.810119 | 26.74 | 95.5 | 104.5 | 12.91 | 15.63 | | High | High | High | High | High | High | High | High | High | High | High | 0 | 1.35E-05 | 3.53 |
| FALSE | High | LEGSPEPPE | [K].LEGSPE | 2xPhospho | Q8CFT2 2x | 0.002921 | 0.00016 | 1 | 2 | 2 | Q8CFT2 | Histone-lys | Q8CFT2 [1 | 0 | 3362.51 | 0 | -0.23 | 0.51091 | 27.9 | 103.9 | 96.1 | 25.12 | 12.59 | | High | High | High | High | High | High | High | High | High | High | High | 7.19E-05 | 0.000797 | 4.01 |
| FALSE | High | LEHEQASQ | [R].LEHEQ | 1xPhospho | O70585 1x | 5.97E-05 | 0 | 1 | 10 | 1 | O70585 | Dystrobrev | O70585 [4 | 0 | 2032.015 | 0 | -0.54 | 0.007515 | 7.23 | 118.6 | 81.4 | 9.59 | 5.32 | | High | High | High | High | High | High | High | High | High | High | High | 0 | 1E-05 | 2.84 |
| FALSE | High | IEHQTITTP | [R].IEHQTI | 1xPhospho | Q8R323 1x | 0.000229 | 0 | 1 | 2 | 1 | Q8R323 | Replication | Q8R323 [6 | 0 | 1792.961 | 0 | -0.95 | 0.010356 | 2.02 | 134 | 66 | 16.04 | 9.64 | | High | High | High | High | High | High | High | High | High | High | High | 0 | 4.56E-05 | 3.03 |

| | | | | | | | | | | | | | | | | | | | | | | | | | | | | | | | | | | | | | | |
|---|---|---|---|---|---|---|---|---|---|---|---|---|---|---|---|---|---|---|---|---|---|---|---|---|---|---|---|---|---|---|---|---|---|---|---|---|---|---|
| FALSE | High | LEHSKESQI | [R].LEHSKE | 1xPhospho | F8WIP8 1x | 3.57E-05 | 0 | 1 | 5 | 3 | F8WIP8 | Osteopont | F8WIP8 [2 | 1 | 3372.641 | 0 | 0.14 | 0.921401 | 47.8 | 92.3 | 107.7 | 34.58 | 16.21 | | High | High | High | High | High | High | High | High | High | High | High | 0 | 5.65E-06 | 2.86 |
| FALSE | High | LELASLGIST | [R].LELASL | 1xPhospho | Q3TCX6 1x | 0.042503 | 0.003197 | 1 | 6 | 3 | Q3TCX6 | Putative ur | Q3TCX6 [1 | 0 | 4113.174 | 0 | 1.48 | 0.000461 | 3.9 | 50.7 | 149.3 | 11.35 | 9.12 | | High | High | High | High | High | High | High | High | High | High | High | 0.001639 | 0.01602 | 2.09 |
| FALSE | High | LEICSDDQ | [K].LEICSD | 1xCarbami | Q3UZ64 1x | 5.94E-07 | 0 | 1 | 10 | 3 | Q3UZ64 | Nuclear fac | Q3UZ64 [3 | 0 | 1999.899 | 0 | -0.5 | 0.009193 | 2.12 | 114.3 | 85.7 | 6.39 | 6.96 | | High | High | High | High | High | High | High | High | High | High | High | 0 | 5.69E-08 | 3.81 |
| FALSE | High | LELDSFGSV | [K].LELDSF | 1xPhospho | J7I1C3 1xP | 4.35E-08 | 0 | 1 | 1 | 3 | J7I1C3 | Rif1 OS=M | J7I1C3 [20 | 0 | 2333.068 | 0 | -1.02 | 0.000529 | 3.6 | 131.4 | 68.6 | 5.72 | 14.47 | | High | High | High | High | High | High | High | High | High | High | High | 0 | 2.99E-09 | 4.62 |
| FALSE | High | LELDSFGSV | [K].LELDSF | 1xPhospho | J7I1C3 1xP | 1.11E-07 | 0 | 1 | 1 | 2 | J7I1C3 | Rif1 OS=M | J7I1C3 [20 | 0 | 2349.063 | 0 | -1.08 | 0.009552 | 4.69 | 135.3 | 64.7 | 9.95 | 19.32 | | High | High | High | High | High | High | High | High | High | High | High | 0 | 8.64E-09 | 4.57 |
| FALSE | High | LELDSFGSV | [K].LELDSF | 1xPhospho | Q6PR54-3 | 5.19E-07 | 0 | 1 | 3 | 1 | Q6PR54-3 | Isoform 3 c | Q6PR54-3 | 0 | 2461.127 | 0 | -0.64 | 0.003644 | 6.04 | 123.8 | 76.2 | 4.89 | 13.63 | | High | High | High | High | High | High | High | High | High | High | High | 0 | 4.86E-08 | 3.81 |
| FALSE | High | LELDSFGSV | [K].LELDSF | 1xPhospho | Q6PR54-3 | 5.88E-10 | 0 | 1 | 3 | 2 | Q6PR54-3 | Isoform 3 c | Q6PR54-3 | 0 | 2477.122 | 0 | -0.83 | 0.040926 | 7.43 | 126.7 | 73.3 | 13.33 | 18.64 | | High | High | High | High | High | High | High | High | High | High | High | 0 | 2.39E-11 | 5.35 |
| FALSE | High | IELEDTESEI | [R].IELEDTI | 1xPhospho | Q3UV57 1x | 0.002623 | 0.000109 | 1 | 1 | 3 | Q3UV57 | Putative ur | Q3UV57 [9 | 0 | 2235.032 | 0 | 1.49 | 0.000458 | 13.6 | 50.8 | 149.2 | 7.44 | 6.33 | | High | High | High | High | High | High | High | High | High | High | High | 0.000847 | 0.008088 | 2.6 |
| FALSE | High | LELGLHEGS | [K].LELGLH | 1xPhospho | U5KVR9 1x | 0.000343 | 0 | 1 | 2 | 3 | U5KVR9 | Centrosom | U5KVR9 [5 | 0 | 2457.295 | 0 | -0.39 | 0.033453 | 4.68 | 116.8 | 83.2 | 12.08 | 12.09 | | High | High | High | High | High | High | High | High | High | High | High | 0 | 7.2E-05 | 2.63 |
| FALSE | High | LELGLHEGS | [K].LELGLH | 2xPhospho | U5KVR9 2x | 0.003892 | 0.000261 | 1 | 2 | 1 | U5KVR9 | Centrosom | U5KVR9 [5 | 0 | 2537.262 | 0 | -0.66 | 0.003101 | 2.29 | 118.4 | 81.6 | 7.91 | 7.33 | | High | High | High | High | High | High | High | High | High | High | High | 0.00014 | 0.001094 | 2.86 |
| FALSE | High | LENSPLGEA | [K].LENSPL | 1xPhospho | Q9CRD0 1x | 0.004781 | 0.000456 | 1 | 5 | 1 | Q9CRD0 | OCIA doma | Q9CRD0 [1 | 0 | 1507.772 | 0 | 0.97 | 0.000682 | 11.08 | 65.9 | 134.1 | 11.66 | 7.83 | | High | High | High | High | High | High | High | High | High | High | High | 0.000204 | 0.001379 | 3.66 |
| FALSE | High | IEILDSPASK | [K].IEILDSP | 1xPhospho | Q9CX66 1x | 0.001463 | 5.66E-05 | 1 | 1 | 2 | Q9CX66 | Uncharacte | Q9CX66 [1 | 0 | 1610.881 | 0 | -0.04 | 0.419265 | 0.9 | 100.3 | 99.7 | 3.48 | 11.41 | | High | High | High | High | High | High | High | High | High | High | High | 2.58E-05 | 0.000367 | 3.29 |
| FALSE | High | IEIIQPLLDN | [K].IEIIQPL | 1xPhospho | Q6PFF0 1x | 8.4E-05 | 0 | 2 | 3 | 3 | Q6PFF0; Q | SR-related | Q6PFF0 [1 | 0 | 4209.155 | 0 | 0.15 | 0.231512 | 14.05 | 95.4 | 104.6 | 7.88 | 7.25 | Shared | High | High | High | High | High | High | High | High | High | High | High | 0 | 7.65E-05 | 5.07 |
| FALSE | High | IEIPVTPTSC | [K].IEIPVTP | 1xPhospho | Q542C3 1x | 0.000645 | 0 | 1 | 2 | 6 | Q542C3 | Putative ur | Q542C3 [1 | 0 | 3071.668 | 0 | 0.38 | 0.022397 | 12.4 | 88 | 112 | 7.66 | 4.17 | | High | High | High | High | High | High | High | High | High | High | High | 0.000585 | 0.00516 | 3 |
| FALSE | High | IEIPVTPTSC | [K].IEIPVTP | 1xDeamida | Q542C3 1x | 0.007538 | 0.00062 | 1 | 2 | 1 | Q542C3 | Putative ur | Q542C3 [1 | 0 | 3072.652 | 0 | 0.52 | 0.082 | 35.09 | 82.2 | 117.8 | 16.26 | 15.74 | | High | High | High | High | High | High | High | High | High | High | High | 0.000275 | 0.002287 | 3.65 |
| FALSE | High | LELSEDSPN | [K].LELSED | 1xPhospho | E9QAU4 1x | 0.000334 | 0 | 1 | 11 | 1 | E9QAU4 | Enhancer t | E9QAU4 [1 | 0 | 2371.132 | 0 | -0.53 | 0.000542 | 3.2 | 118 | 82 | 2.91 | 2.46 | | High | High | High | High | High | High | High | High | High | High | High | 0 | 6.93E-05 | 3.06 |
| FALSE | High | LELSEDSPN | [K].LELSED | 2xPhospho | E9QAU4 2x | 0.00195 | 5.66E-05 | 1 | 11 | 2 | E9QAU4 | Enhancer t | E9QAU4 [1 | 0 | 2451.098 | 0 | -0.05 | 0.013121 | 0.77 | 102.6 | 97.4 | 5.49 | 5.54 | | High | High | High | High | High | High | High | High | High | High | High | 2.58E-05 | 0.000505 | 3.04 |
| FALSE | High | LEISPDSSPI | [R].LEISPDS | 2xPhospho | Q3TIY5 2x | 0.002115 | 5.66E-05 | 1 | 4 | 5 | Q3TIY5 | Putative ur | Q3TIY5 [14 | 0 | 1618.696 | 0 | 0.45 | 0.003423 | 1.06 | 82.2 | 117.8 | 6.7 | 8.04 | | High | High | High | High | High | High | High | High | High | High | High | 0.000275 | 0.002061 | 3.13 |
| FALSE | High | LELVESLDS | [R].LELVES | 1xPhospho | Q8CH77 1x | 4.76E-05 | 0 | 1 | 3 | 2 | Q8CH77 | Neuron na | Q8CH77 [3 | 0 | 2242.151 | 0 | 1.01 | 0.008266 | 15.31 | 66.3 | 133.7 | 19.52 | 13.48 | | High | High | High | High | High | High | High | High | High | High | High | 0 | 7.79E-06 | 4.58 |
| FALSE | High | LEIYQSSQE | [R].LEIYQS | 1xPhospho | Q3UH85 1x | 9.74E-06 | 0 | 1 | 5 | 6 | Q3UH85 | Nucleolar p | Q3UH85 [5 | 0 | 2401.23 | 0 | -0.59 | 0.00965 | 8.91 | 122.1 | 77.9 | 3.77 | 20.3 | | High | High | High | High | High | High | High | High | High | High | High | 0 | 0.000123 | 3.63 |
| FALSE | High | LENDLFGPI | [R].LENDLF | 1xCarbami | P56960 1x | 1.28E-05 | 0 | 1 | 6 | 10 | P56960 | Exosome c | P56960 [60 | 0 | 4120.9 | 0 | -0.57 | 0.011923 | 9.93 | 120.9 | 79.1 | 13.67 | 8.26 | | High | High | High | High | High | High | High | High | High | High | High | 0 | 5.92E-06 | 4.63 |
| FALSE | High | LENEGSDE | [K].LENEGS | 1xPhospho | Q9EQQ9-3 | 5.19E-07 | 0 | 1 | 4 | 5 | Q9EQQ9-3 | Isoform 3 c | Q9EQQ9-3 | 0 | 3134.489 | 0 | 0.39 | 0.004486 | 7.75 | 86.7 | 113.3 | 3.7 | 6.92 | | High | High | High | High | High | High | High | High | High | High | High | 0 | 4.85E-08 | 5.37 |
| FALSE | High | LENEGSDE | [K].LENEGS | 1xPhospho | Q9EQQ9-3 | 8.94E-08 | 0 | 1 | 4 | 6 | Q9EQQ9-3 | Isoform 3 c | Q9EQQ9-3 | 0 | 3150.484 | 0 | 0.37 | 0.000842 | 4.15 | 86.7 | 113.3 | 6.36 | 3.98 | | High | High | High | High | High | High | High | High | High | High | High | 0 | 6.76E-09 | 4.87 |
| FALSE | High | LENGCGNV | [R].LENGC | 1xCarbami | Q3UQC2 1 | 0.005322 | 0.000599 | 1 | 6 | 1 | Q3UQC2 | Putative ur | Q3UQC2 [5 | 0 | 1941.951 | 0 | 0.07 | 0.355427 | 9.1 | 95.9 | 104.1 | 6.55 | 8.28 | | High | High | High | High | High | High | High | High | High | High | High | 0.000267 | 0.001556 | 2.97 |
| FALSE | High | LENGCGNV | [R].LENGC | 1xCarbami | Q3UQC2 1 | 0.071662 | 0.005278 | 1 | 6 | 1 | Q3UQC2 | Putative ur | Q3UQC2 [5 | 0 | 1942.935 | 0 | 0.27 | 0.061913 | 9.63 | 91.2 | 108.8 | 4.14 | 7.59 | | High | High | High | High | High | High | High | High | High | High | High | 0.002748 | 0.02934 | 2.87 |
| FALSE | High | LEILPSFEET | [R].LEILPSF | 1xPhospho | E9Q842 1x | 0.000114 | 0 | 1 | 7 | 4 | E9Q842 | Neuron na | E9Q842 [4 | 0 | 2500.206 | 0 | 1.36 | 0.004459 | 11.02 | 56.1 | 143.9 | 18.89 | 15.1 | | High | High | High | High | High | High | High | High | High | High | High | 0 | 2.08E-05 | 3.93 |
| FALSE | High | IESDEEEDF | [R].IESDEEI | 1xPhospho | E9PWW9 1 | 1.29E-05 | 0 | 1 | 1 | 6 | E9PWW9 | Remodelin | E9PWW9 [ | 0 | 2021.899 | 0 | -0.88 | 0.000466 | 3.3 | 128.7 | 71.3 | 4.28 | 8.69 | | High | High | High | High | High | High | High | High | High | High | High | 0 | 2.58E-06 | 3.9 |
| FALSE | High | LESEEDSIG | [R].LESEED | 1xCarbami | O08648 1x | 2E-08 | 0 | 1 | 5 | 2 | O08648 | Mitogen-ac | O08648 [4 | 0 | 2575.086 | 0 | -0.37 | 0.001542 | 0.37 | 112.8 | 87.2 | 5.91 | 2.36 | | High | High | High | High | High | High | High | High | High | High | High | 0 | 1.25E-09 | 4.74 |
| FALSE | High | LESFESLR | [R].LESFESI | 1xPhospho | Q60875 1x | 0.059159 | 0.004294 | 1 | 9 | 1 | Q60875 | Rho guanir | Q60875 [6 | 0 | 1289.634 | 0 | 0.89 | 0.001165 | 6.81 | 71.5 | 128.5 | 12 | 2.17 | | High | High | High | High | High | High | High | High | High | High | High | 0.002202 | 0.02341 | 2.41 |
| FALSE | High | IEYQNHLSI | [K].IEYQNH | 1xPhospho | A2A7F4 1x | 0.036731 | 0.00249 | 1 | 3 | 1 | A2A7F4 | MCG12287 | A2A7F4 [8 | 0 | 2894.38 | 0 | -0.26 | 0.110515 | 11.85 | 106.5 | 93.5 | 8.14 | 11.38 | | High | High | High | High | High | High | High | High | High | High | High | 0.001308 | 0.01354 | 2.34 |
| FALSE | High | LFASWHGL | [K].LFASWI | 1xPhospho | Q8BP00 1x | 0.097743 | 0.007804 | 1 | 2 | 1 | Q8BP00 | IQ calmodu | Q8BP00 [4 | 0 | 2053.011 | 0 | -0.4 | 0.028422 | 10.84 | 113.9 | 86.1 | 12.56 | 4.94 | | High | High | High | High | High | High | High | High | High | High | High | 0.003937 | 0.04192 | 1.89 |
| FALSE | High | LFATPLTPS | [K].LFATPL | 1xPhospho | E9Q3P4 1x | 1.52E-06 | 0 | 1 | 6 | 2 | E9Q3P4 | Centromer | E9Q3P4 [1 | 0 | 2321.208 | 0 | -0.17 | 0.060843 | 9.95 | 105.7 | 94.3 | 3.12 | 9.23 | | High | High | High | High | High | High | High | High | High | High | High | 0 | 1.62E-07 | 3.91 |
| FALSE | High | LFCPEHSPE | [K].LFCPEH | 1xCarbami | A6H5X4 1x | 1.86E-05 | 0 | 1 | 3 | 2 | A6H5X4 | PHD finger | A6H5X4 [1 | 0 | 3172.389 | 0 | 0.02 | 0.222353 | 3.63 | 95.1 | 104.9 | 7.83 | 7.07 | | High | High | High | High | High | High | High | High | High | High | High | 0 | 2.71E-06 | 2.64 |
| FALSE | High | LFCPEHSPE | [K].LFCPEH | 1xCarbami | A6H5X4 1x | 0.000656 | 0 | 1 | 3 | 3 | A6H5X4 | PHD finger | A6H5X4 [1 | 0 | 3188.384 | 0 | -0.49 | 0.003397 | 2.12 | 116.9 | 83.1 | 4.73 | 7 | | High | High | High | High | High | High | High | High | High | High | High | 0 | 0.000148 | 2.65 |
| FALSE | High | LFDDEASVI | [R].LFDDEA | 1xPhospho | A2ADL9 1x | 0.006031 | 0.00062 | 1 | 8 | 2 | A2ADL9 | InaD-like (I | A2ADL9 [6 | 0 | 1701.757 | 0 | 0.62 | 0.001227 | 9.82 | 77.9 | 122.1 | 4.69 | 4.48 | | High | High | High | High | High | High | High | High | High | High | High | 0.000275 | 0.001786 | 2.61 |
| FALSE | High | LFDDSDEK | [K].LFDDSD | 1xPhospho | Q8BGC0 1x | 0.014158 | 0.001011 | 1 | 1 | 2 | Q8BGC0 | HIV Tat-spe | Q8BGC0 [6 | 0 | 1277.55 | 0 | -0.16 | 0.718399 | 23.39 | 110.8 | 89.2 | 70 | 64.35 | | High | High | High | High | High | High | High | High | High | High | High | 0.000538 | 0.004628 | 2.13 |
| FALSE | High | LFDDSDEK | [K].LFDDSD | 1xPhospho | Q8BGC0 1x | 0.000386 | 0 | 1 | 1 | 7 | Q8BGC0 | HIV Tat-spe | Q8BGC0 [6 | 0 | 1506.713 | 0 | -0.33 | 0.007212 | 3.06 | 111.2 | 88.8 | 4.17 | 6.85 | | High | High | High | High | High | High | High | High | High | High | High | 0 | 8.21E-05 | 3.16 |
| FALSE | High | LFDDSDEKI | [K].LFDDSD | 1xPhospho | Q8BGC0 1x | 0.000822 | 0 | 1 | 1 | 1 | Q8BGC0 | HIV Tat-spe | Q8BGC0 [6 | 1 | 2754.249 | 0 | -0.42 | 0.031164 | 11.9 | 112.7 | 87.3 | 8.13 | 10.6 | | High | High | High | High | High | High | High | High | High | High | High | 0 | 0.000192 | 2.76 |
| FALSE | High | LFDDSDER | [K].LFDDSD | 1xPhospho | Q8BGC0 1x | 0.005044 | 0.000504 | 1 | 1 | 2 | Q8BGC0 | HIV Tat-spe | Q8BGC0 [7 | 0 | 1305.556 | 0 | -0.16 | 0.248272 | 4.83 | 104 | 96 | 3.89 | 9.16 | | High | High | High | High | High | High | High | High | High | High | High | 0.000246 | 0.00146 | 2.22 |
| FALSE | High | LFDDSDER( | [K].LFDDSD | 1xPhospho | Q8BGC0 1x | 0.003253 | 0.00016 | 1 | 1 | 2 | Q8BGC0 | HIV Tat-spe | Q8BGC0 [7 | 1 | 2190.084 | 0 | -0.2 | 0.073506 | 2.86 | 108.3 | 91.7 | 6.8 | 7.21 | | High | High | High | High | High | High | High | High | High | High | High | 0.001151 | 0.01247 | 1.54 |
| FALSE | High | LFDHPEVP | [K].LFDHPE | 1xPhospho | P19096 1x | 0.014033 | 0.001011 | 1 | 4 | 1 | P19096 | Fatty acid s | P19096 [9 | 0 | 2274.1 | 0 | 0.48 | 0.006564 | 12.15 | 82.7 | 117.3 | 1.71 | 8.8 | | High | High | High | High | High | High | High | High | High | High | High | 0.000538 | 0.004579 | 2.35 |
| FALSE | High | LFDSPQK | [K].LFDSPC | 1xPhospho | A6H8H2-2 | 0.033404 | 0.002195 | 1 | 3 | 2 | A6H8H2-2 | Isoform 2 c | A6H8H2-2 | 0 | 1372.728 | 0 | 0.35 | 0.003723 | 2.97 | 88 | 112 | 4.04 | 7.04 | | High | High | High | High | High | High | High | High | High | High | High | 0.001151 | 0.01214 | 2.05 |
| FALSE | High | LEVVCSDCI | [K].LEVVCS | 3xCarbami | B4XVP7 1x | 0.000496 | 0 | 1 | 11 | 1 | B4XVP7 | SET domai | B4XVP7 [4 | 0 | 2430.054 | 0 | 0.21 | 0.049442 | 6.21 | 91.7 | 108.3 | 11.48 | 13.75 | | High | High | High | High | High | High | High | High | High | High | High | 0 | 0.000109 | 2.8 |
| FALSE | High | LFDSSDDEI | [K].LFDSSD | 2xPhospho | E9QKD1 2x | 0.002058 | 5.66E-05 | 1 | 2 | 1 | E9QKD1 | Nucleolar p | E9QKD1 [8 | 0 | 2077.829 | 0 | -0.32 | 0.009589 | 3.38 | 114.1 | 85.9 | 6.56 | 9.67 | | High | High | High | High | High | High | High | High | High | High | High | 2.58E-05 | 0.000538 | 2.49 |
| FALSE | High | LFDVCGSQ | [K].LFDVCC | 1xCarbami | P70388 1x | 1.8E-05 | 0 | 1 | 4 | 2 | P70388 | DNA repair | P70388 [6 | 0 | 2119.957 | 0 | 0.29 | 0.358159 | 14.88 | 88.4 | 111.6 | 9.05 | 17.59 | | High | High | High | High | High | High | High | High | High | High | High | 0 | 2.6E-06 | 3.96 |
| FALSE | High | LFGGFNSSI | [K].LFGGFI | 1xPhospho | P05480 1x | 0.000134 | 0 | 1 | 2 | 2 | P05480 | Neuronal p | P05480 [6 | 0 | 2021.953 | 0 | 0.2 | 0.081248 | 11.74 | 93.6 | 106.4 | 1.92 | 11.02 | | High | High | High | High | High | High | High | High | High | High | High | 0 | 2.51E-05 | 3.13 |
| FALSE | High | LFGTSPAAI | [R].LFGTSP | 1xPhospho | Q5U3K5 1x | 2.47E-06 | 0 | 1 | 1 | 2 | Q5U3K5 | Rab-like pr | Q5U3K5 [3 | 0 | 2882.454 | 0 | -0.06 | 0.562369 | 36.39 | 103.3 | 96.7 | 14.26 | 15.18 | | High | High | High | High | High | High | High | High | High | High | High | 0 | 2.82E-07 | 5.33 |
| FALSE | High | IFIESSSDQI | [K].IFIESSS | 1xPhospho | E9PYW4 1x | 4.72E-05 | 0 | 1 | 2 | 2 | E9PYW4 | E3 ubiquiti | E9PYW4 [4 | 0 | 2769.327 | 0 | 0.77 | 0.00135 | 4.95 | 76.5 | 123.5 | 10.42 | 5.46 | | High | High | High | High | High | High | High | High | High | High | High | 0 | 7.72E-06 | 2.87 |
| FALSE | High | IFLGADAEI | [R].IFLGAD | 1xPhospho | Q64701 1x | 1.7E-05 | 0 | 1 | 5 | 4 | Q64701 | Retinoblast | Q64701 [3 | 0 | 1926.941 | 0 | 0.69 | 0.000838 | 8.01 | 76.5 | 123.5 | 8.75 | 1.88 | | High | High | High | High | High | High | High | High | High | High | High | 0 | 2.59E-06 | 2.94 |
| FALSE | High | IFLGADAEI | [R].IFLGAD | 1xPhospho | Q64701 1x | 1E-05 | 0 | 1 | 5 | 1 | Q64701 | Retinoblast | Q64701 [3 | 1 | 2284.199 | 0 | 0.81 | 0.001715 | 15.01 | 72.3 | 127.7 | 10.3 | 7.06 | | High | High | High | High | High | High | High | High | High | High | High | 0 | 1.35E-06 | 3.48 |
| FALSE | High | LFNPPPAST | [K].LFNPPP | 1xPhospho | Q4VBD2 1x | 0.106748 | 0.008634 | 1 | 1 | 1 | Q4VBD2 | Transmem | Q4VBD2 [4 | 0 | 2392.324 | 0 | 2.07 | 0.002006 | 18.42 | 40.1 | 159.9 | 10.92 | 17.65 | | High | High | High | High | High | High | High | High | High | High | High | 0.004327 | 0.04672 | 1.79 |
| FALSE | High | LFPACHDSI | [K].LFPACH | 1xCarbami | O88508 1x | 0.081085 | 0.006219 | 1 | 6 | 1 | O88508 | DNA (cytos | O88508 [3 | 0 | 2202.978 | 0 | -0.72 | 0.004243 | 4.77 | 125.9 | 74.1 | 9.94 | 11.14 | | High | High | High | High | High | High | High | High | High | High | High | 0.0032 | 0.03381 | 1.7 |
| FALSE | High | LFPDTPLAL | [K].LFPDTP | 1xPhospho | Q9Z1X4-3 | 0.001933 | 5.66E-05 | 1 | 6 | 1 | Q9Z1X4-3 | Isoform 3 c | Q9Z1X4-3 | 0 | 1967.066 | 0 | -0.92 | 0.000443 | 2.71 | 130.8 | 69.2 | 2.89 | 9.61 | | High | High | High | High | High | High | High | High | High | High | High | 2.58E-05 | 0.000498 | 4.02 |
| FALSE | High | LFPGSPAIY | [K].LFPGSP | 1xPhospho | Q9ERU9 1x | 0.045508 | 0.003373 | 1 | 1 | 1 | Q9ERU9 | E3 SUMO-p | Q9ERU9 [1 | 0 | 1630.901 | 0 | -0.15 | 0.041563 | 3.81 | 107.7 | 92.3 | 6.64 | 7.89 | | High | High | High | High | High | High | High | High | High | High | High | 0.001741 | 0.01736 | 2.33 |
| FALSE | High | IFPHHSISES | [R].IFPHHS | 1xPhospho | Q8CDZ5 1x | 0.001233 | 0 | 1 | 2 | 1 | Q8CDZ5 | Putative ur | Q8CDZ5 [6 | 0 | 2209.064 | 0 | 0.66 | 0.054683 | 28.53 | 73.5 | 126.5 | 9.96 | 21.89 | | High | High | High | High | High | High | High | High | High | High | High | 0 | 0.000302 | 2.18 |
| FALSE | High | LFPSPGLPT | [K].LFPSPG | 1xPhospho | Q3TRM4 1 | 0.068752 | 0.005042 | 1 | 6 | 1 | Q3TRM4 | Neuropath | Q3TRM4 [3 | 0 | 1393.744 | 0 | 0.14 | 0.045921 | 2.6 | 93.9 | 106.1 | 4.32 | 6.37 | | High | High | High | High | High | High | High | High | High | High | High | 0.002604 | 0.0279 | 1.6 |
| FALSE | High | IFQGSFGGI | [K].IFQGSF | 1xPhospho | Q9DCA5 1x | 0.000558 | 0 | 1 | 1 | 2 | Q9DCA5 | Ribosome I | Q9DCA5 [2 | 0 | 3102.413 | 0 | -0.26 | 0.09562 | 14.84 | 103.3 | 96.7 | 8.91 | 9.14 | | High | High | High | High | High | High | High | High | High | High | High | 0 | 0.00015 | 3.16 |
| FALSE | High | LFDTPHTPI | [R].LFDTPH | 1xPhospho | P47810 1x | 0.048715 | 0.00354 | 1 | 1 | 2 | P47810 | Wee1-like | P47810 [1 | 0 | 1593.844 | 0 | -1.48 | 0.000353 | 2.3 | 147.1 | 52.9 | 4.31 | 7.06 | | High | High | High | High | High | High | High | High | High | High | High | 0.001844 | 0.01866 | 1.57 |
| FALSE | High | LEVTSDSEI | [R].LEVTSD | 2xPhospho | E9Q4K7 2x | 1.1E-05 | 0 | 1 | 2 | 4 | E9Q4K7 | Kinesin fan | E9Q4K7 [1 | 0 | 2451.057 | 0 | 0.81 | 0.001295 | 4.05 | 73.5 | 126.5 | 9.04 | 6.63 | | High | High | High | High | High | High | High | High | High | High | High | 0 | 1.52E-06 | 3.64 |
| FALSE | High | LEVTSDSEI | [R].LEVTSD | 1xPhospho | E9Q4K7 1x | 1.7E-08 | 0 | 1 | 2 | 17 | E9Q4K7 | Kinesin fan | E9Q4K7 [1 | 0 | 2371.09 | 0 | 0.07 | 0.068497 | 2.88 | 97.1 | 102.9 | 2.49 | 3.61 | | High | High | High | High | High | High | High | High | High | High | High | 0 | 8.87E-08 | 4.83 |
| FALSE | High | LEVSPGDE | [K].LEVSPG | 1xPhospho | Q3TK27 1x | 0.001337 | 0 | 1 | 3 | 3 | Q3TK27 | Putative ur | Q3TK27 [9 | 1 | 3012.409 | 0 | -0.47 | 0.000869 | 1.44 | 116.3 | 83.7 | 6.06 | 4.94 | | High | High | High | High | High | High | High | High | High | High | High | 0.003178 | 0.03323 | 2.33 |
| FALSE | High | LESGEQSD | [K].LESGEC | 1xPhospho | Q3TAB2 1x | 0.000466 | 0 | 2 | 6 | 3 | Q3TAB2; H | Putative ur | Q3TAB2 [6 | 0 | 1627.798 | 0 | 0.03 | 0.598035 | 4.4 | 98.4 | 101.6 | 2.17 | 8.85 | Shared | High | High | High | High | High | High | High | High | High | High | High | 0 | 0.000101 | 3.18 |
| FALSE | High | LESGGTNP | [R].LESGGT | 1xPhospho | A6H644 1x | 8.48E-05 | 0 | 1 | 2 | 1 | A6H644 | MCG13049 | A6H644 [8 | 0 | 2081.886 | 0 | -0.63 | 0.003428 | 5.97 | 121 | 79 | 18.72 | 25.66 | | High | High | High | High | High | High | High | High | High | High | High | 0 | 1.5E-05 | 3.55 |
| FALSE | High | LESLSPK | [K].LESLSPI | 1xPhospho | Q8CJ76 1x | 0.044739 | 0.003302 | 1 | 3 | 1 | Q8CJ76 | Leukemia-a | Q8CJ76 [2 | 0 | 1311.733 | 0 | 0.55 | 0.003036 | 9.71 | 81.2 | 118.8 | 3.32 | 7.28 | | High | High | High | High | High | High | High | High | High | High | High | 0.00168 | 0.0169 | 1.37 |
| FALSE | High | LESLYSDEE | [R].LESLYSI | 1xPhospho | F8WIJ0 1x | 0.001179 | 0 | 1 | 10 | 2 | F8WIJ0 | Solute carr | F8WIJ0 [9 | 0 | 2509.164 | 0 | 0.57 | 0.123103 | 22.65 | 79 | 121 | 24.64 | 17.22 | | High | High | High | High | High | High | High | High | High | High | High | 0 | 0.000288 | 2.79 |
| FALSE | High | LESLYSDEE | [R].LESLYSI | 2xPhospho | F8WIJ0 2x | 0.000184 | 0 | 1 | 10 | 2 | F8WIJ0 | Solute carr | F8WIJ0 [9 | 0 | 2589.13 | 0 | -0.27 | 0.184555 | 16.85 | 109.9 | 90.1 | 11.62 | 22.21 | | High | High | High | High | High | High | High | High | High | High | High | 0 | 3.57E-05 | 3.35 |
| FALSE | High | LESNIDLK | [R].LESNID | 1xPhospho | G3X963 1x | 0.005228 | 0.000551 | 1 | 4 | 2 | G3X963 | ATPase fan | G3X963 [1 | 0 | 1469.802 | 0 | -0.25 | 0.025058 | 4.11 | 107.4 | 92.6 | 7.98 | 4.86 | | High | High | High | High | High | High | High | High | High | High | High | 0.000997 | 0.0103 | 2.49 |
| FALSE | High | IESPHPVDV | [K].IESPHP | 1xPhospho | Q8VHK9 1x | 0.014929 | 0.001049 | 1 | 2 | 1 | Q8VHK9 | ATP-depen | Q8VHK9 [9 | 0 | 2077.004 | 0 | 0.01 | 0.45257 | 4.75 | 100.6 | 99.4 | 8.84 | 7.23 | | High | High | High | High | High | High | High | High | High | High | High | 0.000555 | 0.004911 | 2.33 |
| FALSE | High | IESPIPNPSC | [K].IESPIPN | 1xPhospho | A2A7F4 1x | 1.7E-05 | 0 | 1 | 4 | 1 | A2A7F4 | MCG12287 | A2A7F4 [1 | 0 | 3172.544 | 0 | -1.03 | 0.0009 | 3.89 | 134.2 | 65.8 | 7.65 | 12.59 | | High | High | High | High | High | High | High | High | High | High | High | 0 | 2.45E-06 | 4.54 |
| FALSE | High | LESPSER | [R].LESPSEI | 1xPhospho | Q3TYS2 1x | 0.017192 | 0.001204 | 1 | 1 | 2 | Q3TYS2 | Uncharacte | Q3TYS2 [1 | 0 | 1126.534 | 0 | 1.74 | 0.001041 | 14.48 | 47.7 | 152.3 | 11.65 | 8.56 | | High | High | High | High | High | High | High | High | High | High | High | 0.000638 | 0.005742 | 1.61 |
| FALSE | High | LESPVAH | [K].LESPVA | 1xPhospho | A0A0R4J1F | 0.071071 | 0.005247 | 1 | 4 | 1 | A0A0R4J1F | E3 ubiquiti | A0A0R4J1F | 0 | 1061.523 | 0 | 0.98 | 0.001526 | 7.86 | 67.5 | 132.5 | 6.49 | 9.29 | | High | High | High | High | High | High | High | High | High | High | High | 0.002703 | 0.02897 | 1.93 |
| FALSE | High | IESTTGITTI | [R].IESTTG | 1xPhospho | Q9ESK9 1x | 8.18E-05 | 0 | 1 | 6 | 1 | Q9ESK9 | RB1-induci | Q9ESK9 [6 | 0 | 1975.04 | 0 | -0.49 | 0.007505 | 5.47 | 114 | 86 | 4.6 | 8.42 | | High | High | High | High | High | High | High | High | High | High | High | 0 | 1.44E-05 | 3.17 |
| FALSE | High | LESYRPDTI | [R].LESYRP | 1xPhospho | Q8BGU5 1 | 0.069903 | 0.005181 | 1 | 2 | 1 | Q8BGU5 | Cyclin-Y OS | Q8BGU5 [2 | 0 | 1760.842 | 0 | 0.1 | 0.184271 | 17.66 | 97.3 | 102.7 | 15.05 | 7.7 | | High | High | High | High | High | High | High | High | High | High | High | 0.002662 | 0.02836 | 1.2 |
| FALSE | High | IETDEESCD | [R].IETDEE | 1xCarbami | E9PWW9 1 | 0.020679 | 0.001472 | 1 | 1 | 1 | E9PWW9 | Remodelin | E9PWW9 [ | 0 | 2539.024 | 0 | -0.88 | 0.002324 | 5.77 | 130.1 | 69.9 | 12.29 | 30.69 | | High | High | High | High | High | High | High | High | High | High | High | 0.000786 | 0.007103 | 3.86 |
| FALSE | High | LETEDTSGS | [R].LETEDT | 1xPhospho | A0A0R4J0C | 2.4E-05 | 0 | 1 | 3 | 3 | A0A0R4J0C | Lamin-B2 C | A0A0R4J0C | 0 | 1587.71 | 0 | 0.05 | 0.025522 | 1.95 | 98.1 | 101.9 | 3.19 | 4.37 | | High | High | High | High | High | High | High | High | High | High | High | 0 | 3.61E-06 | 1.67 |
| FALSE | High | LETGQTVLI | [K].LETGQT | 2xPhospho | Q3UFM5 2 | 7.2E-08 | 0 | 1 | 1 | 1 | Q3UFM5 | Nucleolar I | Q3UFM5 [ | 0 | 2468.161 | 0 | 0.41 | 0.060328 | 17.74 | 80.9 | 119.1 | 12.27 | 16.35 | | High | High | High | High | High | High | High | High | High | High | High | 0 | 5.29E-09 | 4.71 |
| FALSE | High | LETPNTFPK | [K].LETPNT | 2xPhospho | E9Q5C9 2x | 0.069325 | 0.005112 | 1 | 2 | 1 | E9Q5C9 | Nucleolar a | E9Q5C9 [6 | 0 | 1435.647 | 0 | 0.14 | 0.711069 | 10.77 | 95.1 | 104.9 | 20.3 | 15.67 | | High | High | High | High | High | High | High | High | High | High | High | 0.002618 | 0.028 | 1.7 |
| FALSE | High | LETPNTFPK | [K].LETPNT | 1xPhospho | E9Q5C9 1x | 0.005768 | 0.00062 | 1 | 2 | 3 | E9Q5C9 | Nucleolar a | E9Q5C9 [6 | 0 | 1584.844 | 0 | -0.37 | 0.002669 | 4.18 | 112.9 | 87.1 | 3.63 | 9.19 | | High | High | High | High | High | High | High | High | High | High | High | 0.00196 | 0.0199 | 2.2 |
| FALSE | High | LETPNTFPK | [K].LETPNT | 2xPhospho | E9Q5C9 2x | 2.17E-05 | 0 | 1 | 2 | 6 | E9Q5C9 | Nucleolar a | E9Q5C9 [6 | 0 | 1664.81 | 0 | 0.17 | 0.083326 | 17.47 | 89.7 | 110.3 | 15 | 5.2 | | High | High | High | High | High | High | High | High | High | High | High | 0.000847 | 0.008135 | 3.3 |
| FALSE | High | LETQEILSEI | [K].LETQEI | 1xPhospho | A0A1W2P6 | 0.000359 | 0 | 1 | 12 | 1 | A0A1W2P6 | Bromodom | A0A1W2P6 | 0 | 1957.977 | 0 | -0.64 | 0.009986 | 2.71 | 120.3 | 79.7 | 13.8 | 7.26 | | High | High | High | High | High | High | High | High | High | High | High | 0 | 7.54E-05 | 2.92 |
| FALSE | High | LETSSESEA | [R].LETSSE | 1xPhospho | E9Q9R9 1x | 0.002212 | 0.000109 | 1 | 5 | 2 | E9Q9R9 | Disks large | E9Q9R9 [1 | 1 | 4668.243 | 0 | 0.21 | 0.697117 | 27.7 | 100 | 100 | 9.2 | 25.24 | | High | High | High | High | High | High | High | High | High | High | High | 4.94E-05 | 0.000583 | 3.22 |
| FALSE | High | LEVASGQG | [K].LEVASG | 1xDeamida | Q6NV66 1x | 0.069325 | 0.005112 | 1 | 3 | 1 | Q6NV66 | Zinc finger | Q6NV66 [1 | 0 | 3251.433 | 0 | -0.28 | 0.067089 | 11.66 | 109.7 | 90.3 | 14.52 | 9.58 | | High | High | High | High | High | High | High | High | High | High | High | 0.002633 | 0.02813 | 3.16 |
| FALSE | High | IEVDESNV | [K].IEVDES | 1xPhospho | Q5DTL2 1x | 0.097743 | 0.007804 | 1 | 2 | 1 | Q5DTL2 | MKIAA414 | Q5DTL2 [6 | 0 | 1801.899 | 0 | 0.95 | 0.003243 | 30.83 | 68.3 | 131.7 | 15.99 | 12.97 | | High | High | High | High | High | High | High | High | High | High | High | 0.003937 | 0.04192 | 1.93 |
| FALSE | High | IEVEEGFCS | [R].IEVEEG | 1xCarbami | Q5SWW4 1 | 2.32E-06 | 0 | 1 | 1 | 1 | Q5SWW4 | Mediator c | Q5SWW4 [ | 0 | 2616.36 | 0 | 0.58 | 0.010379 | 20.74 | 77.8 | 122.2 | 6.34 | 10.87 | | High | High | High | High | High | High | High | High | High | High | High | 0 | 2.62E-07 | 3.22 |
| FALSE | High | IEVGAPDLE | [K].IEVGAP | 1xCarbami | Q8K298 1x | 0.00172 | 5.66E-05 | 1 | 2 | 1 | Q8K298 | Anillin OS= | Q8K298 [7 | 0 | 3441.72 | 0 | 0.12 | 0.979952 | 75.68 | 94.3 | 105.7 | 24.21 | 38.05 | | High | High | High | High | High | High | High | High | High | High | High | 2.58E-05 | 0.000439 | 3.58 |
| FALSE | High | IEVGAPDLE | [K].IEVGAP | 1xCarbami | Q8K298 1x | 4.05E-05 | 0 | 1 | 2 | 1 | Q8K298 | Anillin OS= | Q8K298 [7 | 0 | 3457.715 | 0 | -0.06 | 0.47464 | 21.82 | 98.8 | 101.2 | 13.07 | 32.82 | | High | High | High | High | High | High | High | High | High | High | High | 0 | 6.5E-06 | 3.06 |
| FALSE | High | LEVLECSSS | [K].LEVLEC | 1xCarbami | E9Q166 1x | 0.00013 | 0 | 1 | 2 | 2 | E9Q166 | ATPase fan | E9Q166 [1 | 0 | 3025.504 | 0 | -0.93 | 0.004135 | 5.85 | 128.1 | 71.9 | 10.82 | 13.75 | | High | High | High | High | High | High | High | High | High | High | High | 0 | 2.39E-05 | 3.49 |
| FALSE | High | LEVPAERSF | [K].LEVPAE | 1xPhospho | E9PW37 1x | 0.111965 | 0.009053 | 1 | 4 | 1 | E9PW37 | RAS protei | E9PW37 [1 | 1 | 1462.762 | 0 | 0.71 | 0.001684 | 15.63 | 75.7 | 124.3 | 9.44 | 3.43 | | High | High | High | High | High | High | High | High | High | High | High | 0.004543 | 0.04933 | 2.26 |
| FALSE | High | LEVSPEAQI | [R].LEVSPE | 1xPhospho | E9Q8L9 1x | 0.0004 | 0 | 1 | 2 | 2 | E9Q8L9 | Rab11 fam | E9Q8L9 [5 | 0 | 1635.876 | 0 | -0.66 | 0.051831 | 4.66 | 123.7 | 76.3 | 14.73 | 36.54 | | High | High | High | High | High | High | High | High | High | High | High | 0 | 8.51E-05 | 3.68 |
| FALSE | High | LEVSPGDE | [K].LEVSPG | 1xPhospho | Q3TK27 1x | 8.43E-06 | 0 | 1 | 3 | 3 | Q3TK27 | Putative ur | Q3TK27 [9 | 0 | 1968.911 | 0 | -0.41 | 0.002364 | 2.4 | 113.8 | 86.2 | 9.26 | 5.17 | | High | High | High | High | High | High | High | High | High | High | High | 0 | 3.09E-06 | 2.98 |
| FALSE | High | LEGSPEPPE | [K].LEGSPE | 2xPhospho | Q8CFT2 2x | 0.000589 | 0 | 1 | 2 | 1 | Q8CFT2 | Histone-lys | Q8CFT2 [1 | 0 | 3346.515 | 0 | -0.22 | 0.21997 | 12.4 | 113.2 | 86.8 | 11.18 | 28.4 | | High | High | High | High | High | High | High | High | High | High | High | 0 | 0.000132 | 3.92 |
| FALSE | High | LFQQIYSD | [K].LFQQIY | 1xPhospho | Q9CX34 1x | 4.05E-05 | 0 | 1 | 1 | 1 | Q9CX34 | Protein SG | Q9CX34 [2 | 0 | 2167.072 | 0 | -1.81 | 0.001008 | 0.93 | 155 | 45 | 12.08 | 8.83 | | High | High | High | High | High | High | High | High | High | High | High | 0 | 6.5E-06 | 2.25 |
| FALSE | High | IEGSITGPS | [K].IEGSIT | 1xPhospho | E9Q616 1x | 7.98E-06 | 0 | 1 | 3 | 1 | E9Q616 | AHNAK nu | E9Q616 [4 | 0 | 2972.538 | 0 | 1.92 | 0.002067 | 2.87 | 37.8 | 162.2 | 8.36 | 16.17 | | High | High | High | High | High | High | High | High | High | High | High | 0 | 1.05E-06 | 3.24 |
| FALSE | High | LEGSGSGLI | [K].LEGSGS | 1xPhospho | F8VPN7 1x | 0.000884 | 0 | 1 | 11 | 1 | F8VPN7 | Dedicator c | F8VPN7 [3 | 0 | 2274.167 | 0 | 1.28 | 0.002806 | 17.75 | 54.9 | 145.1 | 12.59 | 15.42 | | High | High | High | High | High | High | High | High | High | High | High | 0 | 0.000207 | 2.84 |
| FALSE | High | LDSSIISEGI | [K].LDSSIIS | 1xPhospho | A2A6Q5-3 | 0.005322 | 0.000599 | 1 | 3 | 2 | A2A6Q5-3 | Isoform 3 c | A2A6Q5-3 | 0 | 1586.844 | 0 | 0.1 | 0.332097 | 9.11 | 93.4 | 106.6 | 14.39 | 11.22 | | High | High | High | High | High | High | High | High | High | High | High | 0.000267 | 0.00155 | 2.15 |
| FALSE | High | LDSSPVLSP | [K].LDSSPV | 1xPhospho | Q9JM52-3 | 2.19E-05 | 0 | 1 | 8 | 5 | Q9JM52-3 | Isoform 3 c | Q9JM52-3 | 0 | 1751.934 | 0 | 1.8 | 0.000992 | 12.99 | 43.5 | 156.5 | 8.03 | 11.44 | | High | High | High | High | High | High | High | High | High | High | High | 0 | 1.02E-05 | 3.09 |
| FALSE | High | LDSSPVLSP | [K].LDSSPV | 2xPhospho | Q9JM52-3 | 8.48E-05 | 0 | 1 | 8 | 1 | Q9JM52-3 | Isoform 3 c | Q9JM52-3 | 0 | 1831.901 | 0 | 1.69 | 0.000241 | 7.22 | 47.2 | 152.8 | 9.09 | 3.02 | | High | High | High | High | High | High | High | High | High | High | High | 0 | 1.49E-05 | 3.16 |
| FALSE | High | LDSSQLPLC | [K].LDSSQL | 1xPhospho | Q8CJF7 1x | 1.97E-05 | 0 | 1 | 1 | 2 | Q8CJF7 | Protein ELY | Q8CJF7 [1 | 0 | 2317.2 | 0 | -0.61 | 0.020177 | 3.59 | 120.7 | 79.3 | 14.85 | 9.31 | | High | High | High | High | High | High | High | High | High | High | High | 0 | 2.9E-06 | 3.77 |
| FALSE | High | LDSSQLPLC | [K].LDSSQL | 1xPhospho | Q8CJF7 1x | 0.111965 | 0.009053 | 1 | 1 | 1 | Q8CJF7 | Protein ELY | Q8CJF7 [1 | 1 | 2674.458 | 0 | -0.55 | 0.055182 | 11.47 | 122.8 | 77.2 | 18.62 | 16.14 | | High | High | High | High | High | High | High | High | High | High | High | 0.004543 | 0.0493 | 2.62 |
| FALSE | High | LDSSTCLHA | [R].LDSSTC | 1xCarbami | Q9Z2Z9 1x | 0.000133 | 0 | 1 | 2 | 2 | Q9Z2Z9 | Glutamine- | Q9Z2Z9 [2 | 0 | 1940.955 | 0 | -0.61 | 0.024177 | 6.33 | 120.9 | 79.1 | 7.27 | 17.08 | | High | High | High | High | High | High | High | High | High | High | High | 0 | 2.49E-05 | 2.56 |
| FALSE | High | IDSTEVIYQ | [R].IDSTEV | 1xPhospho | Q9WTK7 1 | 0.000107 | 0 | 1 | 2 | 2 | Q9WTK7 | Serine/thre | Q9WTK7 [ | 0 | 1629.809 | 0 | -0.08 | 0.260937 | 18.28 | 105.3 | 94.7 | 3.37 | 14.02 | | High | High | High | High | High | High | High | High | High | High | High | 0 | 1.94E-05 | 4.01 |
| FALSE | High | LDSVDGPG | [R].LDSVD( | 1xPhospho | Q5SSZ5 1x | 0.013787 | 0.001011 | 1 | 1 | 1 | Q5SSZ5 | Tensin-3 O | Q5SSZ5 [7 | 0 | 1224.582 | 0 | 0.69 | 0.002793 | 10.37 | 76.6 | 123.4 | 8.37 | 5.96 | | High | High | High | High | High | High | High | High | High | High | High | 0.000538 | 0.004474 | 1.64 |
| FALSE | High | LDSVTVNSI | [K].LDSVTV | 1xPhospho | Q9WU62 1 | 0.077188 | 0.005893 | 1 | 2 | 1 | Q9WU62 | Inner centr | Q9WU62 [ | 0 | 1613.892 | 0 | -0.68 | 0.058083 | 4.48 | 117 | 83 | 12.33 | 23.35 | | High | High | High | High | High | High | High | High | High | High | High | 0.00302 | 0.03171 | 1.98 |
| FALSE | High | LDTFCGSPI | [K].LDTFCG | 1xCarbami | Q03141 1x | 6.83E-05 | 0 | 4 | 25 | 1 | Q03141; Q | MAP/micro | Q03141 [2 | 0 | 2636.287 | 0 | -0.26 | 0.479171 | 5.98 | 107 | 93 | 11.05 | 7.63 | Shared | High | High | High | High | High | High | High | High | High | High | High | 0 | 1.17E-05 | 3.5 |
| FALSE | High | LDTFQSTSI | [K].LDTFQS | 1xPhospho | F8WJA0 1x | 0.001361 | 0 | 1 | 6 | 1 | F8WJA0 | ATP-depen | F8WJA0 [2 | 0 | 1661.855 | 0 | -0.38 | 0.105807 | 12.51 | 108.5 | 91.5 | 10.69 | 12.94 | | High | High | High | High | High | High | High | High | High | High | High | 0 | 0.000337 | 3.71 |
| FALSE | High | LDTPEEK | [R].LDTPEE | 1xPhospho | P70399 1x | 0.054844 | 0.003982 | 1 | 5 | 2 | P70399 | TP53-bindi | P70399 [7 | 0 | 1369.702 | 0 | 0.7 | 0.016017 | 12.1 | 74.3 | 125.7 | 13.45 | 10.5 | | High | High | High | High | High | High | High | High | High | High | High | 0.002159 | 0.0228 | 2.07 |
| FALSE | High | LDVFTRPAS | [R].LDVFTR | 1xPhospho | Q5I1X5 1x | 0.030104 | 0.001857 | 1 | 1 | 1 | Q5I1X5 | RelA-assoc | Q5I1X5 [2 | 0 | 2375.284 | 0 | 1.05 | 0.000605 | 8.51 | 66.3 | 133.7 | 12.33 | 6.84 | | High | High | High | High | High | High | High | High | High | High | High | 0.001011 | 0.01079 | 1.76 |
| FALSE | High | LDSSEMDH | [R].LDSSEN | 1xPhospho | F8WHU5 1 | 0.000235 | 0 | 1 | 4 | 3 | F8WHU5 | Sister chro | F8WHU5 [ | 0 | 3123.358 | 0 | -0.5 | 0.046487 | 3.65 | 118.3 | 81.7 | 8.28 | 14.78 | | High | High | High | High | High | High | High | High | High | High | High | 0 | 4.68E-05 | 3.23 |
| FALSE | High | LDVSSVASI | [K].LDVSSV | 2xPhospho | E9Q3Y4 2x | 0.017966 | 0.001282 | 1 | 7 | 1 | E9Q3Y4 | Lipopolysa | E9Q3Y4 [1 | 0 | 1667.713 | 0 | 1.24 | 0.001307 | 19.59 | 58.6 | 141.4 | 11.39 | 7.38 | | High | High | High | High | High | High | High | High | High | High | High | 0.000689 | 0.006051 | 2.24 |
| FALSE | High | LDVTILSPSI | [K].LDVTIL | 1xPhospho | Q80X90 1x | 0.006653 | 0.00062 | 1 | 1 | 2 | Q80X90 | Filamin-B C | Q80X90 [9 | 0 | 1409.76 | 0 | 0.81 | 0.000374 | 2.85 | 74.3 | 125.7 | 3.93 | 4.36 | | High | High | High | High | High | High | High | High | High | High | High | 0.00074 | 0.006443 | 1.54 |

| | | | | | | | | | | | | | | | | | | | | | | | | | | | | | | | | | | | | | |
|---|---|---|---|---|---|---|---|---|---|---|---|---|---|---|---|---|---|---|---|---|---|---|---|---|---|---|---|---|---|---|---|---|---|---|---|---|---|
| FALSE | High | LDVTTPESF | [R].LDVTTF | 1xPhospho | D3YTL8 1xI | 0.067059 | 0.004796 | 1 | 2 | 2 | D3YTL8 | Protein spi | D3YTL8 [23 | 0 | 1624.86 | 0 | 1.36 | 0.00056 | 5.51 | 57.1 | 142.9 | 7.64 | 8.22 | | High | High | High | High | High | High | High | High | High | High | High | 0.009844 | 0.1069 | 2.34 |
| FALSE | High | LDWESFK | [K].LDWES | 1xPhospho | O88271 1x | 0.103392 | 0.008249 | 1 | 1 | 1 | O88271 | Craniofacia | O88271 [2 | 0 | 1462.738 | 0 | -0.81 | 0.07606 | 12.25 | 127.4 | 72.6 | 19.63 | 18.7 | | High | High | High | High | High | High | High | High | High | High | High | 0.004138 | 0.04469 | 1.89 |
| FALSE | High | LDWESFKE | [K].LDWES | 1xPhospho | O88271 1x | 0.000822 | 0 | 1 | 1 | 1 | O88271 | Craniofacia | O88271 [2 | 1 | 2810.324 | 0 | -1.4 | 0.007 | 2.29 | 144.9 | 55.1 | 15.03 | 17.9 | | High | High | High | High | High | High | High | High | High | High | High | 0 | 0.000191 | 2.71 |
| FALSE | High | LDWESFKE | [K].LDWES | 1xPhospho | O88271 1x | 1.59E-07 | 0 | 1 | 1 | 18 | O88271 | Craniofacia | O88271 [2 | 1 | 3039.487 | 0 | -0.67 | 0.030646 | 7.44 | 121.6 | 78.4 | 11.16 | 15.56 | | High | High | High | High | High | High | High | High | High | High | High | 0 | 2.3E-05 | 4.95 |
| FALSE | High | LDYGQHVV | [K].LDYGQ | 1xPhospho | Q91VC3 1x | 0.000674 | 0 | 1 | 5 | 2 | Q91VC3 | Eukaryotic | Q91VC3 [1 | 0 | 1778.879 | 0 | -1.05 | 0.000966 | 0.94 | 129.9 | 70.1 | 6.57 | 8.12 | | High | High | High | High | High | High | High | High | High | High | High | 0.000117 | 0.001089 | 2.76 |
| FALSE | High | IDYLLPGPG | [K].IDYLLP | 1xPhospho | E9Q9R9 1x | 4.3E-07 | 0 | 1 | 5 | 1 | E9Q9R9 | Disks large | E9Q9R9 [9 | 0 | 2458.315 | 0 | 0.09 | 0.151666 | 5.24 | 94.2 | 105.8 | 7.59 | 4.8 | | High | High | High | High | High | High | High | High | High | High | High | 0 | 3.95E-08 | 3.04 |
| FALSE | High | IEAFLSHYP | [K].IEAFLSI | 1xPhospho | Q99PU8-3 | 2.11E-05 | 0 | 1 | 5 | 3 | Q99PU8-3 | Isoform 3 c | Q99PU8-3 | 0 | 2655.306 | 0 | -0.69 | 0.004111 | 6.28 | 125.7 | 74.3 | 6.7 | 11.36 | | High | High | High | High | High | High | High | High | High | High | High | 7.19E-05 | 0.000846 | 2.98 |
| FALSE | High | LEAFSPPPP | [R].LEAFSP | 1xPhospho | E9QLH9 1x | 0.000379 | 0 | 1 | 2 | 1 | E9QLH9 | PEAK1-rela | E9QLH9 [6 | 0 | 1814.986 | 0 | 1.09 | 0.000241 | 1.61 | 64.4 | 135.6 | 2.84 | 4.73 | | High | High | High | High | High | High | High | High | High | High | High | 0 | 7.99E-05 | 2.16 |
| FALSE | High | LEAGGSLTI | [R].LEAGG | 1xPhospho | A2A863-3 | 0.083101 | 0.006345 | 1 | 8 | 1 | A2A863-3 | Isoform 3 c | A2A863-3 | 0 | 1212.619 | 0 | 4.11 | 0.000154 | 15.68 | 10.6 | 189.4 | 13.98 | 4.14 | | High | High | High | High | High | High | High | High | High | High | High | 0.003265 | 0.03466 | 1.37 |
| FALSE | High | LEAKESCEI | [R].LEAKES | 1xCarbami | Q6GTE9 1x | 0.000611 | 0 | 1 | 4 | 1 | Q6GTE9 | NIMA (Nev | Q6GTE9 [6 | 1 | 2859.4 | 0 | 0.01 | 0.704157 | 14.71 | 99.8 | 100.2 | 7.85 | 10.21 | | High | High | High | High | High | High | High | High | High | High | High | 0 | 0.000137 | 3.04 |
| FALSE | High | LEAIITPPPA | [K].LEAIITP | 1xPhospho | Q9QZQ8 1 | 0.001398 | 5.66E-05 | 1 | 3 | 3 | Q9QZQ8 | Core histor | Q9QZQ8 [1 | 0 | 1687.98 | 0 | -0.05 | 0.858253 | 7.71 | 102.6 | 97.4 | 4.89 | 10.31 | | High | High | High | High | High | High | High | High | High | High | High | 2.58E-05 | 0.000349 | 2.06 |
| FALSE | High | LEASAVSDS | [K].LEASAV | 1xPhospho | P33609 1xI | 0.000712 | 0 | 1 | 3 | 1 | P33609 | DNA polym | P33609 [11 | 0 | 1863.869 | 0 | -0.34 | 0.009741 | 5.88 | 111.7 | 88.3 | 9.36 | 2.93 | | High | High | High | High | High | High | High | High | High | High | High | 0 | 0.000163 | 1.86 |
| FALSE | High | LDVTLGPVI | [R].LDVTLG | 1xPhospho | Q9D903 1x | 9E-07 | 0 | 1 | 2 | 1 | Q9D903 | Probable r | Q9D903 [7 | 0 | 3158.602 | 0 | -0.25 | 0.370227 | 12.26 | 106.2 | 93.8 | 11.13 | 9.27 | | High | High | High | High | High | High | High | High | High | High | High | 0 | 8.99E-08 | 4.92 |
| FALSE | High | LEASSVPNS | [K].LEASSV | 2xCarbami | A6H619 1x | 7.29E-06 | 0 | 1 | 2 | 2 | A6H619 | PHD and RI | A6H619 [9 | 0 | 2976.295 | 0 | -0.4 | 0.041833 | 15.72 | 116.9 | 83.1 | 12.64 | 8.33 | | High | High | High | High | High | High | High | High | High | High | High | 0 | 9.49E-07 | 4.33 |
| FALSE | High | LDSSEMDH | [R].LDSSEN | 1xPhospho | F8WHU5 1 | 0.006713 | 0.00062 | 1 | 4 | 1 | F8WHU5 | Sister chro | F8WHU5 [ | 0 | 3107.363 | 0 | 0 | 0.552232 | 20.7 | 96.2 | 103.8 | 12.43 | 21.95 | | High | High | High | High | High | High | High | High | High | High | High | 0.000275 | 0.002019 | 2.46 |
| FALSE | High | LDSSEMDH | [R].LDSSEN | 2xPhospho | F8WHU5 2 | 0.048715 | 0.00354 | 1 | 4 | 1 | F8WHU5 | Sister chro | F8WHU5 [ | 0 | 2958.166 | 0 | 0.22 | 0.160904 | 21.04 | 92.4 | 107.6 | 4.5 | 13.81 | | High | High | High | High | High | High | High | High | High | High | High | 0.001876 | 0.01875 | 3.11 |
| FALSE | High | LDNSPISSP | [R].LDNSPI | 1xPhospho | B2RT41 1xI | 0.006957 | 0.00062 | 1 | 3 | 2 | B2RT41 | Zinc finger, | B2RT41 [9 | 0 | 1394.688 | 0 | 0.05 | 0.49795 | 3.07 | 100.1 | 99.9 | 4.71 | 5.91 | | High | High | High | High | High | High | High | High | High | High | High | 0.000275 | 0.002094 | 2.43 |
| FALSE | High | LDNSPISSP | [R].LDNSPI | 2xPhospho | B2RT41 2xI | 0.054383 | 0.003951 | 1 | 3 | 1 | B2RT41 | Zinc finger, | B2RT41 [9 | 1 | 1831.912 | 0 | 0.21 | 0.276932 | 7.65 | 92.5 | 107.5 | 10.01 | 8.11 | | High | High | High | High | High | High | High | High | High | High | High | 0.002059 | 0.02127 | 1.89 |
| FALSE | High | LDNSPNMI | [K].LDNSPI | 1xPhospho | Q05BA5 1x | 6.77E-05 | 0 | 1 | 7 | 1 | Q05BA5 | Nuclear re | Q05BA5 [5 | 0 | 2083.03 | 0 | 0.42 | 0.108167 | 8.32 | 84.9 | 115.1 | 5.12 | 23.98 | | High | High | High | High | High | High | High | High | High | High | High | 0 | 1.16E-05 | 3.26 |
| FALSE | High | LDNSPNMI | [K].LDNSPI | 1xPhospho | Q05BA5 1x | 0.000171 | 0 | 1 | 7 | 2 | Q05BA5 | Nuclear re | Q05BA5 [5 | 0 | 2099.024 | 0 | 0.54 | 0.023363 | 7.03 | 83.3 | 116.7 | 6.12 | 17.3 | | High | High | High | High | High | High | High | High | High | High | High | 0 | 3.28E-05 | 3.47 |
| FALSE | High | LDNSSEEES | [R].LDNSSE | 1xCarbami | Q8BHK9 2x | 0.002398 | 0.000109 | 1 | 3 | 1 | Q8BHK9 | DNA excisi | Q8BHK9 [1 | 0 | 4870.07 | 0 | -0.31 | 0.267945 | 6.9 | 105 | 95 | 19.36 | 8.06 | | High | High | High | High | High | High | High | High | High | High | High | 4.94E-05 | 0.000636 | 3.9 |
| FALSE | High | LDPDTETP( | [R].LDPDTE | 1xPhospho | Q542Q3 1x | 0.009931 | 0.000736 | 1 | 1 | 1 | Q542Q3 | G two S ph | Q542Q3 [4 | 0 | 1908.972 | 0 | 0.93 | 0.016656 | 11.98 | 74.2 | 125.8 | 16.69 | 23.52 | | High | High | High | High | High | High | High | High | High | High | High | 0.000344 | 0.003105 | 1.98 |
| FALSE | High | IDPEGSNS( | [K].IDPEGS | 1xPhospho | E9Q9R9 1x | 9.61E-08 | 0 | 1 | 5 | 2 | E9Q9R9 | Disks large | E9Q9R9 [9 | 0 | 1982.962 | 0 | -0.11 | 0.279507 | 7.31 | 101.8 | 98.2 | 6.5 | 6.35 | | High | High | High | High | High | High | High | High | High | High | High | 0 | 7.29E-09 | 3.78 |
| FALSE | High | LDPFADGG | [R].LDPFAD | 1xPhospho | G5E866 1x | 1.42E-05 | 0 | 1 | 5 | 1 | G5E866 | Splicing fac | G5E866 [1 | 1 | 2225.182 | 0 | 0.27 | 0.209105 | 10.38 | 87.3 | 112.7 | 15.31 | 8.47 | | High | High | High | High | High | High | High | High | High | High | High | 0 | 0.000002 | 3.33 |
| FALSE | High | LDPFESSDF | [K].LDPFES | 1xPhospho | Q60902 1x | 0.008463 | 0.000736 | 1 | 3 | 2 | Q60902 | Epidermal | Q60902 [7 | 0 | 2455.168 | 0 | 0.2 | 0.065117 | 5.67 | 93.1 | 106.9 | 6.26 | 3.27 | | High | High | High | High | High | High | High | High | High | High | High | 0.000327 | 0.002611 | 2.85 |
| FALSE | High | IDPNPVDS( | [R].IDPNPV | 1xPhospho | Q8BH58 1x | 0.000389 | 0 | 1 | 1 | 3 | Q8BH58 | TIP41-like | Q8BH58 [2 | 0 | 1794.8 | 0 | -0.09 | 0.30786 | 10.11 | 102.1 | 97.9 | 12.19 | 8.36 | | High | High | High | High | High | High | High | High | High | High | High | 0.001468 | 0.01471 | 2.88 |
| FALSE | High | LDPPPSPHA | [K].LDPPPS | 1xPhospho | Q8BXK8 1x | 0.059657 | 0.004365 | 1 | 5 | 1 | Q8BXK8 | Arf-GAP wi | Q8BXK8 [5 | 0 | 1509.741 | 0 | -1.09 | 0.000447 | 0.37 | 135.2 | 64.8 | 2.42 | 12.18 | | High | High | High | High | High | High | High | High | High | High | High | 0.002233 | 0.02355 | 2.52 |
| FALSE | High | LDQLTITPE | [K].LDQLTI | 1xPhospho | Q542Q3 1x | 2.4E-05 | 0 | 1 | 1 | 3 | Q542Q3 | G two S ph | Q542Q3 [6 | 0 | 1679.857 | 0 | 0.01 | 0.797693 | 7.55 | 99.4 | 100.6 | 5.21 | 5.69 | | High | High | High | High | High | High | High | High | High | High | High | 0 | 3.6E-06 | 2.74 |
| FALSE | High | LDQPVSAP | [K].LDQPV | 1xPhospho | D3YZP9 1xI | 0.012959 | 0.000889 | 1 | 1 | 3 | D3YZP9 | Coiled-coil | D3YZP9 [2 | 0 | 1572.798 | 0 | 0.15 | 0.023421 | 3.52 | 92.8 | 107.2 | 3.79 | 6.77 | | High | High | High | High | High | High | High | High | High | High | High | 0.002187 | 0.02332 | 2.28 |
| FALSE | High | LDSSEMDH | [R].LDSSEN | 2xPhospho | F8WHU5 2 | 0.000418 | 0 | 1 | 4 | 3 | F8WHU5 | Sister chro | F8WHU5 [ | 0 | 2974.161 | 0 | 0.63 | 0.001794 | 7.57 | 77.5 | 122.5 | 8.83 | 11.6 | | High | High | High | High | High | High | High | High | High | High | High | 0 | 8.94E-05 | 2.56 |
| FALSE | High | LDQPVSAP | [K].LDQPV | 2xPhospho | D3YZP9 2xI | 8.87E-05 | 0 | 1 | 1 | 4 | D3YZP9 | Coiled-coil | D3YZP9 [2 | 0 | 1652.765 | 0 | 0.79 | 0.001887 | 11.02 | 74.4 | 125.6 | 5.48 | 6.27 | | High | High | High | High | High | High | High | High | High | High | High | 0 | 6.16E-05 | 2.69 |
| FALSE | High | IDSASLNGE | [K].IDSASLI | 1xPhospho | Q9D3S3 1x | 1.57E-05 | 0 | 1 | 2 | 1 | Q9D3S3 | Sorting ne | Q9D3S3 [3 | 0 | 2033.036 | 0 | -0.52 | 0.009526 | 8.12 | 118.2 | 81.8 | 7.4 | 9.4 | | High | High | High | High | High | High | High | High | High | High | High | 0 | 2.24E-06 | 4.26 |
| FALSE | High | IDSDDDEEI | [K].IDSDDD | 1xPhospho | Q8C1D8 1x | 0.000158 | 0 | 1 | 3 | 1 | Q8C1D8 | Protein IW | Q8C1D8 [3 | 0 | 1603.714 | 0 | -0.32 | 0.024087 | 11.97 | 110.2 | 89.8 | 9.99 | 3.91 | | High | High | High | High | High | High | High | High | High | High | High | 0 | 3.01E-05 | 2.76 |
| FALSE | High | LDSDDPSVI | [K].LDSDDI | 1xPhospho | F8WI37 1xI | 0.001233 | 0 | 1 | 2 | 1 | F8WI37 | Histone-lys | F8WI37 [2 | 0 | 1513.755 | 0 | 0.39 | 0.071622 | 8.79 | 85.4 | 114.6 | 6.91 | 18.54 | | High | High | High | High | High | High | High | High | High | High | High | 0 | 0.000302 | 2.22 |
| FALSE | High | LDSDRVSS | [R].LDSDR | 1xPhospho | E9Q9C3 1x | 0.011246 | 0.000772 | 1 | 5 | 1 | E9Q9C3 | Afadin OS= | E9Q9C3 [5 | 1 | 1889.88 | 0 | 0.35 | 0.03725 | 13.24 | 91.4 | 108.6 | 9.75 | 11.05 | | High | High | High | High | High | High | High | High | High | High | High | 0.000418 | 0.003573 | 2.06 |
| FALSE | High | IDSEISPK | [K].IDSEISP | 1xPhospho | B2RXU2 1x | 0.0026 | 0.000109 | 1 | 3 | 2 | B2RXU2 | ESF1, nucle | B2RXU2 [1 | 0 | 1426.759 | 0 | -0.77 | 0.002364 | 3.14 | 126.4 | 73.6 | 7.54 | 5.99 | | High | High | High | High | High | High | High | High | High | High | High | 0.009831 | 0.1067 | 1.65 |
| FALSE | High | LDSGTCEH | [K].LDSGT( | 1xCarbami | H3BJV2 1xI | 0.007471 | 0.00062 | 1 | 1 | 1 | H3BJV2 | A-kinase ar | H3BJV2 [4 | 0 | 2197.963 | 0 | 0.01 | 0.764886 | 14.39 | 101.9 | 98.1 | 6.26 | 10.57 | | High | High | High | High | High | High | High | High | High | High | High | 0.000275 | 0.00227 | 2.76 |
| FALSE | High | IDSHPSPSH | [K].IDSHPS | 1xPhospho | B1AZI6 1xP | 0.000132 | 0 | 1 | 3 | 2 | B1AZI6 | THO comp | B1AZI6 [14 | 0 | 2017.016 | 0 | -0.04 | 0.393776 | 19.68 | 104.4 | 95.6 | 14.09 | 13.89 | | High | High | High | High | High | High | High | High | High | High | High | 0 | 2.45E-05 | 1.79 |
| FALSE | High | LDSLAEPH( | [R].LDSLAE | 1xPhospho | E9Q7D5 1x | 5.09E-06 | 0 | 1 | 2 | 6 | E9Q7D5 | Rho guanir | E9Q7D5 [1 | 0 | 1787.889 | 0 | 2.84 | 0.000555 | 23.14 | 24.2 | 175.8 | 14.95 | 10.35 | | High | High | High | High | High | High | High | High | High | High | High | 0 | 0.000225 | 2.72 |
| FALSE | High | IDSLSAQLS | [R].IDSLSA | 1xPhospho | P48678 1xI | 0.000446 | 0 | 1 | 3 | 2 | P48678 | Prelamin-A | P48678 [2 | 0 | 1969.077 | 0 | 0.92 | 0.023825 | 35.94 | 60.1 | 139.9 | 14.45 | 23.91 | | High | High | High | High | High | High | High | High | High | High | High | 0 | 0.000194 | 3.26 |
| FALSE | High | LDSPEVNIII | [R].LDSPEV | 1xCarbami | P30415 1xI | 1.2E-05 | 0 | 1 | 1 | 2 | P30415 | NK-tumor | P30415 [11 | 0 | 2659.209 | 0 | -0.25 | 0.064607 | 4.42 | 108.6 | 91.4 | 6.05 | 20.75 | | High | High | High | High | High | High | High | High | High | High | High | 0 | 1.66E-06 | 3.83 |
| FALSE | High | LDSPEVNIII | [R].LDSPEV | 1xCarbami | P30415 1xI | 2.24E-06 | 0 | 1 | 1 | 2 | P30415 | NK-tumor | P30415 [11 | 0 | 2675.204 | 0 | 0.02 | 0.611095 | 13.67 | 96.2 | 103.8 | 4.16 | 15.56 | | High | High | High | High | High | High | High | High | High | High | High | 0 | 2.51E-07 | 4.03 |
| FALSE | High | LDSPTLSR | [K].LDSPTL | 1xPhospho | Q3U3A6 1x | 0.002844 | 0.00016 | 1 | 5 | 3 | Q3U3A6 | Sodium/hy | Q3U3A6 [6 | 0 | 1197.608 | 0 | 0.03 | 0.37746 | 5.08 | 99.3 | 100.7 | 5.24 | 2.58 | | High | High | High | High | High | High | High | High | High | High | High | 0.000571 | 0.004945 | 2.4 |
| FALSE | High | LDSSDSDS( | [K].LDSSDS | 1xPhospho | Q8K0S9 1x | 9.99E-09 | 0 | 1 | 1 | 1 | Q8K0S9 | snRNA-acti | Q8K0S9 [3 | 0 | 1903.823 | 0 | -1.4 | 0.001429 | 1.75 | 138.2 | 61.8 | 9.77 | 15.01 | | High | High | High | High | High | High | High | High | High | High | High | 0 | 5.74E-10 | 3.27 |
| FALSE | High | LDRASPDL\ | [K].LDRASI | 1xPhospho | Q8CD94 1x | 0.031989 | 0.00205 | 1 | 1 | 1 | Q8CD94 | Protein lin- | Q8CD94 [2 | 1 | 3182.633 | 0 | 0.81 | 0.028129 | 8.06 | 72.5 | 127.5 | 12.45 | 22.74 | | High | High | High | High | High | High | High | High | High | High | High | 0.001093 | 0.01154 | 2.74 |
| FALSE | High | LEASSVPNS | [K].LEASSV | 2xCarbami | A6H619 2x | 4.18E-07 | 0 | 1 | 2 | 3 | A6H619 | PHD and RI | A6H619 [9 | 0 | 3056.262 | 0 | 0.78 | 0.01791 | 15.24 | 75.4 | 124.6 | 12.87 | 18.96 | | High | High | High | High | High | High | High | High | High | High | High | 0 | 3.81E-08 | 4.92 |
| FALSE | High | LEAVSHTSI | [R].LEAVSH | 1xCarbami | Q3TC53 1x | 0.00582 | 0.00062 | 1 | 6 | 3 | Q3TC53 | Adenylyl cy | Q3TC53 [1 | 0 | 2716.215 | 0 | 1.01 | 0.00481 | 19.09 | 67.2 | 132.8 | 13.93 | 8.87 | | High | High | High | High | High | High | High | High | High | High | High | 0.000275 | 0.001706 | 2.17 |
| FALSE | High | LEAVSHTSI | [R].LEAVSH | 1xCarbami | Q3TC53 1x | 9.03E-05 | 0 | 1 | 6 | 4 | Q3TC53 | Adenylyl cy | Q3TC53 [1 | 0 | 2732.21 | 0 | 0.8 | 0.004531 | 23.97 | 75.3 | 124.7 | 9.18 | 7.16 | | High | High | High | High | High | High | High | High | High | High | High | 9.42E-05 | 0.000984 | 3.3 |
| FALSE | High | LEEDRQSLI | [R].LEEDRC | 1xPhospho | Q547N0 1x | 0.00011 | 0 | 1 | 2 | 1 | Q547N0 | BRMS1 OS | Q547N0 [1 | 1 | 2789.307 | 0 | -1.24 | 0.000401 | 4.29 | 136.7 | 63.3 | 7.33 | 24.59 | | High | High | High | High | High | High | High | High | High | High | High | 0 | 2E-05 | 4.83 |
| FALSE | High | IEEFSPR | [R].IEEFSPI | 1xPhospho | Q9EPQ8 1x | 0.011346 | 0.000816 | 1 | 2 | 1 | Q9EPQ8 | Transcripti | Q9EPQ8 [2 | 0 | 1186.571 | 0 | -0.14 | 0.479405 | 9.11 | 104.2 | 95.8 | 6.76 | 6.89 | | High | High | High | High | High | High | High | High | High | High | High | 0.000418 | 0.003615 | 1.22 |
| FALSE | High | LEELNGSQ | [K].LEELNG | 1xPhospho | E9Q4S3 1x | 0.003464 | 0.00021 | 1 | 7 | 1 | E9Q4S3 | Ninein OS= | E9Q4S3 [1 | 0 | 2241.12 | 0 | 3.05 | 0.000663 | 10.54 | 23.5 | 176.5 | 25.44 | 12.32 | | High | High | High | High | High | High | High | High | High | High | High | 9.42E-05 | 0.000963 | 2 |
| FALSE | High | LEEPEGTQI | [R].LEEPEG | 1xPhospho | Q5NCX5 1x | 0.000229 | 0 | 1 | 7 | 1 | Q5NCX5 | Neuralized | Q5NCX5 [1 | 0 | 3780.633 | 0 | 0.44 | 0.029639 | 5.34 | 78.5 | 121.5 | 15.29 | 14.55 | | High | High | High | High | High | High | High | High | High | High | High | 0 | 4.56E-05 | 3.84 |
| FALSE | High | LEEPEGTQI | [R].LEEPEG | 1xDeamida | Q5NCX5 1x | 4.56E-06 | 0 | 1 | 7 | 2 | Q5NCX5 | Neuralized | Q5NCX5 [1 | 0 | 3781.617 | 0 | 0.44 | 0.109268 | 36.58 | 85.8 | 114.2 | 18.08 | 12.49 | | High | High | High | High | High | High | High | High | High | High | High | 0 | 0.000119 | 3.96 |
| FALSE | High | LEEPEMPS( | [K].LEEPEN | 1xPhospho | P32507 1xI | 3.64E-06 | 0 | 1 | 2 | 1 | P32507 | Nectin-2 O | P32507 [4 | 0 | 2964.483 | 0 | -1.52 | 0.001965 | 2.56 | 145.3 | 54.7 | 15.57 | 22.19 | | High | High | High | High | High | High | High | High | High | High | High | 0 | 4.36E-07 | 4.36 |
| FALSE | High | LEEPEMPS( | [K].LEEPEN | 1xPhospho | P32507 1xI | 0.068184 | 0.004935 | 1 | 2 | 1 | P32507 | Nectin-2 O | P32507 [4 | 0 | 2980.478 | 0 | -1.18 | 0.033667 | 7.97 | 146 | 54 | 20.67 | 41.1 | | High | High | High | High | High | High | High | High | High | High | High | 0.002559 | 0.02757 | 2.33 |
| FALSE | High | LEESASFES | [K].LEESAS | 1xPhospho | B2RUG9 1x | 1.08E-07 | 0 | 1 | 6 | 2 | B2RUG9 | Adenomat | B2RUG9 [2 | 0 | 2588.208 | 0 | -0.67 | 0.001361 | 1.7 | 123.6 | 76.4 | 7.95 | 9.77 | | High | High | High | High | High | High | High | High | High | High | High | 0 | 8.38E-09 | 3.7 |
| FALSE | High | IEESDEEAV | [K].IEESDEI | 1xPhospho | F6YRW4 1x | 0.000279 | 0 | 1 | 4 | 3 | F6YRW4 | F-box and | F6YRW4 [6 | 0 | 1885.92 | 0 | 0.43 | 0.004129 | 4.59 | 85.3 | 114.7 | 10.52 | 5.22 | | High | High | High | High | High | High | High | High | High | High | High | 0.000555 | 0.004823 | 2.87 |
| FALSE | High | LEESENELE | [K].LEESEN | 1xPhospho | P15533 1xI | 0.000868 | 0 | 1 | 2 | 1 | P15533 | Tripartite r | P15533 [21 | 0 | 2242.033 | 0 | 0.17 | 0.410097 | 19.81 | 97.2 | 102.8 | 20.76 | 7.51 | | High | High | High | High | High | High | High | High | High | High | High | 0 | 0.000203 | 4.1 |
| FALSE | High | LEESGLPET | [K].LEESGL | 1xPhospho | Q7TML3 1 | 0.001704 | 5.66E-05 | 1 | 1 | 1 | Q7TML3 | Solute carr | Q7TML3 [3 | 0 | 1790.877 | 0 | 1.17 | 0.028841 | 5.64 | 63.3 | 136.7 | 8.33 | 29.47 | | High | High | High | High | High | High | High | High | High | High | High | 2.58E-05 | 0.000432 | 2.74 |
| FALSE | High | LEESNSIGC | [K].LEESNS | 1xPhospho | O70551 1x | 0.001117 | 0 | 1 | 2 | 3 | O70551 | SRSF prote | O70551 [3 | 0 | 2128.996 | 0 | -0.09 | 0.085631 | 4.88 | 104.4 | 95.6 | 5.7 | 3.52 | | High | High | High | High | High | High | High | High | High | High | High | 2.58E-05 | 0.000451 | 2.91 |
| FALSE | High | LEESPFVPY | [R].LEESPF | 1xPhospho | F6ZMJ4 1xI | 6.95E-08 | 0 | 1 | 7 | 2 | F6ZMJ4 | Trinucleoti | F6ZMJ4 [1 | 0 | 3633.645 | 0 | 0.86 | 0.02597 | 11.2 | 80 | 120 | 13.24 | 17.26 | | High | High | High | High | High | High | High | High | High | High | High | 0 | 5.06E-09 | 4.13 |
| FALSE | High | LEEDADSPS | [K].LEEDAI | 1xCarbami | Q3UYC8 1x | 0.005135 | 0.000551 | 1 | 5 | 2 | Q3UYC8 | Putative ur | Q3UYC8 [7 | 0 | 3710.837 | 0 | 1.42 | 0.000186 | 18.25 | 54.3 | 145.7 | 17 | 6.12 | | High | High | High | High | High | High | High | High | High | High | High | 0.000382 | 0.003346 | 3.42 |
| FALSE | High | LEESPFVPY | [R].LEESPF | 1xPhospho | F6ZMJ4 1xI | 3.72E-07 | 0 | 1 | 7 | 1 | F6ZMJ4 | Trinucleoti | F6ZMJ4 [1 | 0 | 3649.64 | 0 | 0.82 | 0.00495 | 13.48 | 72.4 | 127.6 | 16.89 | 8.24 | | High | High | High | High | High | High | High | High | High | High | High | 0 | 3.34E-08 | 3.53 |
| FALSE | High | LEFFGFEDI | [K].LEFFGF | 1xPhospho | B7ZP47 1xI | 0.000115 | 0 | 1 | 2 | 4 | B7ZP47 | Wapal prot | B7ZP47 [4 | 0 | 3219.383 | 0 | -1.46 | 0.000364 | 1.39 | 147.6 | 52.4 | 8.13 | 10.21 | | High | High | High | High | High | High | High | High | High | High | High | 7.19E-05 | 0.000791 | 3.58 |
| FALSE | High | IEGAEELEE | [R].IEGAEE | 1xCarbami | Q5BLK4 1x | 2.74E-08 | 0 | 1 | 2 | 1 | Q5BLK4 | Terminal u | Q5BLK4 [8 | 0 | 2789.29 | 0 | 1.52 | 0.000414 | 4.2 | 53.4 | 146.6 | 8.67 | 6.72 | | High | High | High | High | High | High | High | High | High | High | High | 0 | 1.79E-09 | 3.92 |
| FALSE | High | LEGDSDDL | [K].LEGDSI | 1xPhospho | A0A1S6GW | 1.81E-08 | 0 | 1 | 6 | 9 | A0A1S6GW | Uncharacte | A0A1S6GW | 0 | 2457.014 | 0 | 0.63 | 0.002019 | 6.51 | 79.4 | 120.6 | 3.75 | 5.47 | | High | High | High | High | High | High | High | High | High | High | High | 0 | 1.12E-09 | 5.38 |
| FALSE | High | LEGDSDVD | [K].LEGDSI | 1xPhospho | Q9JMH9-6 | 0.013666 | 0.001011 | 2 | 4 | 3 | Q9JMH9-6 | Isoform 6 c | Q9JMH9-6 | 1 | 2615.249 | 0 | -0.22 | 0.166849 | 16.99 | 107.7 | 92.3 | 12.97 | 8.15 | Shared | High | High | High | High | High | High | High | High | High | High | High | 0.003265 | 0.0353 | 2.03 |
| FALSE | High | LEGDSDVD | [K].LEGDSI | 2xPhospho | Q9JMH9-6 | 0.0004 | 0 | 2 | 4 | 2 | Q9JMH9-6 | Isoform 6 c | Q9JMH9-6 | 1 | 2695.215 | 0 | 0.4 | 0.002219 | 4.61 | 88.2 | 111.8 | 6.34 | 3 | Shared | High | High | High | High | High | High | High | High | High | High | High | 0.000294 | 0.002356 | 3.3 |
| FALSE | High | LEGDSPSSI | [K].LEGDSF | 1xPhospho | E0CZH3 1xI | 0.116482 | 0.009523 | 1 | 2 | 1 | E0CZH3 | Forkhead b | E0CZH3 [2 | 0 | 2946.486 | 0 | -0.42 | 0.028841 | 11.69 | 116.6 | 83.4 | 13.05 | 11 | | High | High | High | High | High | High | High | High | High | High | High | 0.004757 | 0.05174 | 2.82 |
| FALSE | High | LEGLATAV | [K].LEGLAT | 1xPhospho | G5E8A0 1x | 5.26E-05 | 0 | 1 | 2 | 2 | G5E8A0 | Oxysterol-l | G5E8A0 [2 | 0 | 1751.971 | 0 | -0.65 | 0.000362 | 2.08 | 123 | 77 | 2.13 | 4.04 | | High | High | High | High | High | High | High | High | High | High | High | 0.002471 | 0.02708 | 2.25 |
| FALSE | High | LEGNSPQG | [K].LEGNSI | 1xPhospho | Q3UHW5 1 | 0.003788 | 0.00021 | 1 | 2 | 3 | Q3UHW5 | Putative ur | Q3UHW5 [ | 0 | 1934.974 | 0 | 0.36 | 0.030888 | 20.86 | 88.7 | 111.3 | 13.99 | 9.92 | | High | High | High | High | High | High | High | High | High | High | High | 0.003646 | 0.03888 | 2.38 |
| FALSE | High | LEGPASPD\ | [K].LEGPAS | 1xPhospho | A0A0A0M( | 8.15E-07 | 0 | 1 | 2 | 2 | A0A0A0M( | Histone-lys | A0A0A0M( | 0 | 1849.971 | 0 | 0.28 | 0.012809 | 11.9 | 90.4 | 109.6 | 2.07 | 6.86 | | High | High | High | High | High | High | High | High | High | High | High | 0 | 8.11E-08 | 3.14 |
| FALSE | High | LEGPHLSLK | [K].LEGPHI | 1xPhospho | E9Q616 1x | 0.010758 | 0.000772 | 1 | 6 | 1 | E9Q616 | AHNAK nu | E9Q616 [5 | 0 | 1531.865 | 0 | 2.81 | 0.003209 | 9.76 | 23.3 | 176.7 | 9.81 | 23.42 | | High | High | High | High | High | High | High | High | High | High | High | 0.000399 | 0.00339 | 1.75 |
| FALSE | High | IEGQLDED( | [R].IEGQLD | 2xCarbami | B7ZND8 1x | 0.011146 | 0.000772 | 1 | 8 | 1 | B7ZND8 | Testis expr | B7ZND8 [1 | 0 | 3208.41 | 0 | -1.15 | 0.002007 | 3.67 | 141 | 59 | 9.1 | 19.71 | | High | High | High | High | High | High | High | High | High | High | High | 0.000399 | 0.003554 | 2.99 |
| FALSE | High | LEGQSPQC | [K].LEGQSI | 1xPhospho | Q3UKC1 1x | 0.022773 | 0.001542 | 1 | 1 | 1 | Q3UKC1 | Tax1-bindi | Q3UKC1 [6 | 0 | 1537.757 | 0 | 0.83 | 0.000275 | 4.53 | 71.1 | 128.9 | 2.33 | 4.61 | | High | High | High | High | High | High | High | High | High | High | High | 0.000832 | 0.007913 | 2.55 |
| FALSE | High | LEGSEEEK | [K].LEGSEE | 1xPhospho | P97496 1xI | 0.009416 | 0.000736 | 1 | 5 | 1 | P97496 | SWI/SNF c | P97496 [7 | 0 | 1458.713 | 0 | -0.9 | 0.000422 | 2.16 | 130.4 | 69.6 | 2.83 | 6.44 | | High | High | High | High | High | High | High | High | High | High | High | 0.000327 | 0.002934 | 2.64 |
| FALSE | High | LEESPFVPY | [R].LEESPF | 1xDeamida | F6ZMJ4 1xI | 0.012072 | 0.000889 | 1 | 7 | 1 | F6ZMJ4 | Trinucleoti | F6ZMJ4 [1 | 0 | 3650.624 | 0 | 1.01 | 0.003002 | 34.84 | 68.6 | 131.4 | 20.13 | 4.57 | | High | High | High | High | High | High | High | High | High | High | High | 0.000449 | 0.003873 | 3.67 |
| FALSE | High | LEEAQQVD | [R].LEEAQ( | 1xPhospho | Q62315 1x | 0.000558 | 0 | 1 | 5 | 1 | Q62315 | Protein Jur | Q62315 [4 | 0 | 2232.168 | 0 | -0.39 | 0.060132 | 10.2 | 116.7 | 83.3 | 8.96 | 12.58 | | High | High | High | High | High | High | High | High | High | High | High | 0 | 0.000124 | 2.75 |
| FALSE | High | IEEAPEAAP | [K].IEEAPE | 1xDeamida | Q04692 2x | 0.053473 | 0.00384 | 1 | 5 | 4 | Q04692 | SWI/SNF-r | Q04692 [1 | 0 | 4010.774 | 0 | -0.16 | 0.522376 | 16.84 | 100 | 100 | 9.78 | 10.62 | | High | High | High | High | High | High | High | High | High | High | High | 0.002015 | 0.02093 | 3.24 |
| FALSE | High | IEEAPEAAP | [K].IEEAPE | 2xPhospho | Q04692 2x | 0.0002 | 0 | 1 | 5 | 3 | Q04692 | SWI/SNF-r | Q04692 [1 | 0 | 4009.79 | 0 | -1.08 | 0.004899 | 5.35 | 133.5 | 66.5 | 8.71 | 15.12 | | High | High | High | High | High | High | High | High | High | High | High | 0 | 3.92E-05 | 3.98 |
| FALSE | High | LEAYHTQT | [R].LEAYHT | 1xPhospho | Q9WTP6 1 | 0.02933 | 0.001857 | 1 | 2 | 1 | Q9WTP6 | Adenylate | Q9WTP6 [ | 0 | 2293.11 | 0 | -0.67 | 0.007464 | 3.58 | 122.7 | 77.3 | 4.24 | 12.01 | | High | High | High | High | High | High | High | High | High | High | High | 0.000997 | 0.01054 | 2.86 |
| FALSE | High | LECLPPEPS | [K].LECLPP | 1xCarbami | Q3TDN2 1x | 0.002695 | 0.000109 | 1 | 3 | 1 | Q3TDN2 | FAS-associ | Q3TDN2 [3 | 0 | 2447.182 | 0 | 0.91 | 0.006293 | 24.23 | 68.3 | 131.7 | 10.19 | 13.7 | | High | High | High | High | High | High | High | High | High | High | High | 4.94E-05 | 0.000725 | 2.06 |
| FALSE | High | LEDAESSNI | [K].LEDAES | 1xPhospho | Q9CQE6 1x | 2.13E-07 | 0 | 1 | 2 | 4 | Q9CQE6 | Histone ch | Q9CQE6 [1 | 0 | 3125.566 | 0 | 0.08 | 0.602388 | 16.87 | 97.9 | 102.1 | 11.19 | 9.11 | | High | High | High | High | High | High | High | High | High | High | High | 0 | 1.79E-08 | 4.23 |
| FALSE | High | LEDAESSNI | [K].LEDAES | 2xPhospho | Q9CQE6 2x | 0.011146 | 0.000772 | 1 | 2 | 3 | Q9CQE6 | Histone ch | Q9CQE6 [1 | 0 | 3205.532 | 0 | 0.55 | 0.087261 | 19.19 | 84.3 | 115.7 | 20.89 | 9.96 | | High | High | High | High | High | High | High | High | High | High | High | 0.000801 | 0.007312 | 3.09 |
| FALSE | High | IEDAPAPST | [K].IEDAPA | 1xPhospho | B1AU76 1x | 1.54E-05 | 0 | 2 | 5 | 3 | B1AU76; B | Nuclear au | B1AU76 [2 | 0 | 1839.914 | 0 | -1.35 | 0.000348 | 2.93 | 146.8 | 53.2 | 3.6 | 13.36 | Shared | High | High | High | High | High | High | High | High | High | High | High | 0.000786 | 0.006824 | 2.42 |
| FALSE | High | LEDDSEESI | [K].LEDDSE | 1xPhospho | F8VPZ5 1xI | 0.000765 | 0 | 1 | 2 | 2 | F8VPZ5 | Excision re | F8VPZ5 [4 | 0 | 2500.069 | 0 | -0.45 | 0.033602 | 13.98 | 120.7 | 79.3 | 16.06 | 13.35 | | High | High | High | High | High | High | High | High | High | High | High | 0 | 0.000177 | 2.5 |
| FALSE | High | LEDDTSGSI | [K].LEDDTS | 1xPhospho | A2AIX1 1xF | 0.000244 | 0 | 1 | 2 | 3 | A2AIX1 | SEC16 hon | A2AIX1 [5 | 0 | 1783.856 | 0 | 2.34 | 0.000738 | 14.03 | 30.7 | 169.3 | 14.72 | 13.27 | | High | High | High | High | High | High | High | High | High | High | High | 0 | 4.86E-05 | 2.93 |
| FALSE | High | LEDEDDSD | [K].LEDEDI | 2xPhospho | Q09XV5 2x | 3.11E-08 | 0 | 1 | 2 | 2 | Q09XV5 | Chromodo | Q09XV5 [2 | 0 | 2327.982 | 0 | -0.41 | 0.003674 | 4.63 | 111.1 | 88.9 | 7.4 | 11.92 | | High | High | High | High | High | High | High | High | High | High | High | 0 | 2.06E-09 | 3.6 |
| FALSE | High | LEDLDTCM | [R].LEDLDT | 1xCarbami | Q6EDY6 1x | 5.81E-05 | 0 | 1 | 2 | 2 | Q6EDY6 | F-actin-unc | Q6EDY6 [9 | 0 | 2007.924 | 0 | -0.91 | 0.006301 | 1.93 | 125.7 | 74.3 | 10.19 | 11.55 | | High | High | High | High | High | High | High | High | High | High | High | 0 | 3.21E-05 | 3.05 |
| FALSE | High | LEDIFQQEF | [K].LEDIFQ | 1xPhospho | Q9EP97 1x | 7.36E-06 | 0 | 1 | 2 | 3 | Q9EP97 | Sentrin-spe | Q9EP97 [3 | 0 | 2005.947 | 0 | -0.8 | 0.005645 | 4.61 | 129.5 | 70.5 | 9.67 | 11.89 | | High | High | High | High | High | High | High | High | High | High | High | 0 | 9.56E-07 | 3.7 |
| FALSE | High | LEDNDSAT | [R].LEDND | 1xPhospho | Q99NF3 1x | 0.010853 | 0.000772 | 1 | 2 | 3 | Q99NF3 | Centrosom | Q99NF3 [9 | 0 | 2388.122 | 0 | -0.39 | 0.049524 | 14.58 | 109.5 | 90.5 | 12.34 | 17.89 | | High | High | High | High | High | High | High | High | High | High | High | 0.001308 | 0.01351 | 2.69 |
| FALSE | High | LEDNPCSG | [K].LEDNP( | 1xCarbami | Q8BG15 1x | 0.000162 | 0 | 1 | 5 | 4 | Q8BG15 | CTD small | Q8BG15 [1 | 0 | 1637.719 | 0 | -1.19 | 0.001227 | 1.92 | 137.4 | 62.6 | 5.34 | 10.57 | | High | High | High | High | High | High | High | High | High | High | High | 0 | 3.08E-05 | 1.41 |
| FALSE | High | LEDQAAAP | [R].LEDQA | 1xPhospho | Q8BX02 1x | 0.001544 | 5.66E-05 | 1 | 2 | 1 | Q8BX02 | KN motif a | Q8BX02 [1 | 0 | 3799.932 | 0 | 0.01 | 0.849682 | 37.25 | 94 | 106 | 12.55 | 18.43 | | High | High | High | High | High | High | High | High | High | High | High | 2.58E-05 | 0.000389 | 3.9 |
| FALSE | High | LEDQGAQ( | [R].LEDQG | 1xCarbami | O35245 1x | 2.31E-05 | 0 | 1 | 5 | 1 | O35245 | Polycystin- | O35245 [1 | 0 | 2371.07 | 0 | -0.08 | 0.552505 | 7.45 | 102.7 | 97.3 | 12.7 | 14.14 | | High | High | High | High | High | High | High | High | High | High | High | 0 | 3.48E-06 | 3.06 |
| FALSE | High | LEDSANAD | [K].LEDSAN | 1xPhospho | Q9DB00 1x | 0.001704 | 5.66E-05 | 2 | 6 | 1 | Q9DB00; C | GON-4-like | Q9DB00 [1 | 0 | 2811.309 | 0 | -0.67 | 0.009477 | 5.78 | 123 | 77 | 9.53 | 12.2 | Shared | High | High | High | High | High | High | High | High | High | High | High | 2.58E-05 | 0.000432 | 4.41 |
| FALSE | High | IEDSEPHIPI | [R].IEDSEPI | 1xPhospho | G5E829 1x | 3.86E-07 | 0 | 1 | 5 | 5 | G5E829 | Plasma me | G5E829 [1 | 0 | 3074.45 | 0 | 2.69 | 0.000434 | 8.82 | 25.1 | 174.9 | 5.98 | 10.31 | | High | High | High | High | High | High | High | High | High | High | High | 0 | 9.92E-08 | 4.8 |
| FALSE | High | IEDSEPHIPI | [R].IEDSEPI | 2xPhospho | G5E829 2x | 6.52E-08 | 0 | 1 | 5 | 2 | G5E829 | Plasma me | G5E829 [1 | 0 | 3154.416 | 0 | 3.01 | 0.009891 | 20.9 | 22.2 | 177.8 | 24.22 | 35.06 | | High | High | High | High | High | High | High | High | High | High | High | 0 | 7.13E-08 | 6.07 |
| FALSE | High | IEDVGSDEI | [K].IEDVGS | 1xPhospho | P11499 1xI | 8.78E-08 | 0 | 1 | 2 | 26 | P11499 | Heat shock | P11499 [2 | 0 | 2032.9 | 0 | -0.45 | 0.00554 | 9.56 | 116.7 | 83.3 | 3.36 | 9.5 | | High | High | High | High | High | High | High | High | High | High | High | 0 | 8.23E-06 | 3.58 |
| FALSE | High | IEDVGSDEI | [K].IEDVGS | 2xPhospho | P11499 2xI | 8.22E-07 | 0 | 1 | 2 | 6 | P11499 | Heat shock | P11499 [2 | 0 | 2112.866 | 0 | -0.24 | 0.012446 | 5.48 | 110 | 90 | 1.94 | 6.54 | | High | High | High | High | High | High | High | High | High | High | High | 0 | 1.13E-07 | 5.06 |
| FALSE | High | IEDVGSDEI | [K].IEDVGS | 1xPhospho | P11499 1xI | 0.007605 | 0.000664 | 1 | 2 | 2 | P11499 | Heat shock | P11499 [2 | 1 | 2505.185 | 0 | -0.56 | 0.780534 | 17.85 | 105 | 95 | 46.66 | 90.94 | | High | High | High | High | High | High | High | High | High | High | High | 0.000294 | 0.002304 | 2.92 |
| FALSE | High | IEDVTPIPSI | [R].IEDVTP | 1xPhospho | O70569 1x | 0.002719 | 0.000109 | 1 | 3 | 3 | O70569 | Ribosomal | O70569 [1 | 0 | 1738.846 | 0 | 0.09 | 0.050326 | 0.88 | 96.9 | 103.1 | 2.39 | 3.17 | | High | High | High | High | High | High | High | High | High | High | High | 4.94E-05 | 0.000731 | 2.52 |
| FALSE | High | LEDVTTHH | [R].LEDVTT | 1xCarbami | B1AQY2 1x | 0.011146 | 0.000772 | 1 | 3 | 2 | B1AQY2 | Rho GTPas | B1AQY2 [3 | 0 | 3682.658 | 0 | -0.31 | 0.120951 | 15.3 | 108.7 | 91.3 | 12.92 | 25.49 | | High | High | High | High | High | High | High | High | High | High | High | 0.000399 | 0.003529 | 3.96 |
| FALSE | High | LEEADSPPL | [K].LEEADS | 1xPhospho | Q8CFL8 1xI | 0.017498 | 0.001245 | 1 | 1 | 1 | Q8CFL8 | Zinc finger | Q8CFL8 [6 | 0 | 2048.909 | 0 | -0.66 | 0.005346 | 8.43 | 123.9 | 76.1 | 5.21 | 16.52 | | High | High | High | High | High | High | High | High | High | High | High | 0.000689 | 0.005883 | 2.64 |
| FALSE | High | LEEALSTEG | [K].LEEALS | 1xPhospho | Q61074 1x | 1.7E-05 | 0 | 1 | 2 | 2 | Q61074 | Protein ph | Q61074 [5 | 0 | 2404.117 | 0 | 0.29 | 0.072141 | 4.59 | 91.3 | 108.7 | 6.86 | 8.45 | | High | High | High | High | High | High | High | High | High | High | High | 0 | 2.46E-06 | 4.23 |
| FALSE | High | LEEALSTEG | [K].LEEALS | 2xPhospho | Q61074 2x | 0.000211 | 0 | 1 | 2 | 1 | Q61074 | Protein ph | Q61074 [5 | 0 | 2484.083 | 0 | -0.23 | 0.166985 | 19.87 | 112.4 | 87.6 | 9.09 | 15.53 | | High | High | High | High | High | High | High | High | High | High | High | 0 | 4.16E-05 | 2.94 |
| FALSE | High | LEEALSTEG | [K].LEEALS | 1xPhospho | Q61074 1x | 0.000718 | 0 | 1 | 2 | 1 | Q61074 | Protein ph | Q61074 [5 | 1 | 2761.375 | 0 | 0.22 | 0.311428 | 24.66 | 92.4 | 107.6 | 17.25 | 8.05 | | High | High | High | High | High | High | High | High | High | High | High | 0 | 0.000165 | 2.66 |
| FALSE | High | IEEAMDGS | [K].IEEAMI | 1xPhospho | Q3UAI4 1x | 3.77E-05 | 0 | 1 | 4 | 4 | Q3UAI4 | Putative ur | Q3UAI4 [7 | 0 | 2772.382 | 0 | 0.2 | 0.893242 | 21.24 | 93.8 | 106.2 | 15.66 | 17.13 | | High | High | High | High | High | High | High | High | High | High | High | 0 | 7.66E-05 | 3.24 |
| FALSE | High | IEEAMDGS | [K].IEEAMI | 1xPhospho | Q3UAI4 1x | 1.13E-06 | 0 | 1 | 4 | 1 | Q3UAI4 | Putative ur | Q3UAI4 [7 | 0 | 2788.377 | 0 | -0.03 | 0.752088 | 5.82 | 100.8 | 99.2 | 18.53 | 36.23 | | High | High | High | High | High | High | High | High | High | High | High | 0 | 1.17E-07 | 3.07 |
| FALSE | High | IEEAPEAAP | [K].IEEAPE | 1xDeamida | Q04692 1x | 6.02E-05 | 0 | 1 | 5 | 7 | Q04692 | SWI/SNF-r | Q04692 [1 | 0 | 3930.808 | 0 | -1.47 | 0.000534 | 1.51 | 149.8 | 50.2 | 9.53 | 18.71 | | High | High | High | High | High | High | High | High | High | High | High | 0 | 1.02E-05 | 5.04 |
| FALSE | High | LEGSHTPEI | [R].LEGSHT | 1xPhospho | Q6NZP2 1x | 0.021797 | 0.001507 | 1 | 1 | 1 | Q6NZP2 | Vasculin-lil | Q6NZP2 [3 | 0 | 1632.84 | 0 | 0.08 | 0.089289 | 4.4 | 94.3 | 105.7 | 4.57 | 7.29 | | High | High | High | High | High | High | High | High | High | High | High | 0.000814 | 0.00754 | 1.89 |

| | | | | | | | | | | | | | | | | | | | | | | | | | | | | | | | | | | | | | | |
|---|---|---|---|---|---|---|---|---|---|---|---|---|---|---|---|---|---|---|---|---|---|---|---|---|---|---|---|---|---|---|---|---|---|---|---|---|---|---|
| FALSE | High | LFSADPFDL | [R].LFSADP | 1xPhospho | A2ADY9 1x | 0.000194 | 0 | 1 | 1 | 1 | A2ADY9 | Protein DD | A2ADY9 [1 | 0 | 2090.061 | 0 | -0.5 | 0.009986 | 2.52 | 125.5 | 74.5 | 16.62 | 15.18 | | High | High | High | High | High | High | High | High | High | High | High | 0 | 3.79E-05 | 3.38 |
| FALSE | High | LFSLSNPSL | [R].LFSLSN | 1xPhospho | Q91X84 1x | 0.023792 | 0.001612 | 1 | 2 | 1 | Q91X84 | CREB-regu | Q91X84 [3 | 0 | 2287.153 | 0 | 2.95 | 0.000322 | 41.02 | 22.8 | 177.2 | 24.83 | 6.04 | | High | High | High | High | High | High | High | High | High | High | High | 0.000859 | 0.00828 | 1.9 |
| FALSE | High | LFSPVISR | [K].LFSPVIS | 1xPhospho | A6H8H2-2 | 0.041781 | 0.003126 | 1 | 3 | 1 | A6H8H2-2 | Isoform 2 ( | A6H8H2-2 | 0 | 1227.67 | 0 | 2.35 | 0.000227 | 7.54 | 32.5 | 167.5 | 8.65 | 6.59 | | High | High | High | High | High | High | High | High | High | High | High | 0.001609 | 0.01573 | 2.76 |
| FALSE | High | IHLPLNYPP | [R].IHLPLN | 1xPhospho | F7BJK1 1xP | 3.77E-05 | 0 | 1 | 7 | 2 | F7BJK1 | Protocadh | F7BJK1 [79 | 0 | 2055.063 | 0 | -0.45 | 0.005264 | 4.02 | 116.2 | 83.8 | 8.45 | 4.74 | | High | High | High | High | High | High | High | High | High | High | High | 0 | 6.02E-06 | 2.98 |
| FALSE | High | LHLSPPLGF | [K].LHLSPP | 1xPhospho | E9Q1Q1 1x | 0.121159 | 0.009984 | 1 | 6 | 1 | E9Q1Q1 | Microtubu | E9Q1Q1 [1 | 0 | 1298.718 | 0 | -1.16 | 0.001546 | 3.62 | 141.2 | 58.8 | 7.27 | 12.52 | | High | High | High | High | High | High | High | High | High | High | High | 0.004961 | 0.05407 | 1.42 |
| FALSE | High | IHMSGPK | [K].IHMSGI | 1xPhospho | E9Q616 1x | 0.093123 | 0.007276 | 1 | 5 | 1 | E9Q616 | AHNAK nu | E9Q616 [4 | 0 | 1323.69 | 0 | 1.88 | 0.007 | 10.38 | 35.8 | 164.2 | 30.12 | 24.97 | | High | High | High | High | High | High | High | High | High | High | High | 0.003685 | 0.03957 | 1.9 |
| FALSE | High | LHNGSASP | [R].LHNGSA | 1xPhospho | Q80U38 1x | 0.000765 | 0 | 1 | 1 | 1 | Q80U38 | Protein KH | Q80U38 [3 | 0 | 1344.662 | 0 | -0.55 | 0.006634 | 4 | 120.3 | 79.7 | 6.04 | 15.62 | | High | High | High | High | High | High | High | High | High | High | High | 0 | 0.000176 | 3.07 |
| FALSE | High | LHNGSASP | [R].LHNGSA | 1xDeamida | Q80U38 1x | 0.023792 | 0.001612 | 1 | 1 | 1 | Q80U38 | Protein KH | Q80U38 [3 | 0 | 1345.646 | 0 | -0.41 | 0.245991 | 30.7 | 106.6 | 93.4 | 35.82 | 27.34 | | High | High | High | High | High | High | High | High | High | High | High | 0.000859 | 0.008292 | 2.2 |
| FALSE | High | LHNGSASP | [R].LHNGSA | 1xCarbami | Q80U38 2x | 0.001544 | 5.66E-05 | 1 | 1 | 1 | Q80U38 | Protein KH | Q80U38 [3 | 1 | 3262.474 | 0 | -0.01 | 0.470683 | 6.68 | 102 | 98 | 6.27 | 3.18 | | High | High | High | High | High | High | High | High | High | High | High | 2.58E-05 | 0.000389 | 2.55 |
| FALSE | High | LHNQNSVF | [K].LHNQN | 1xPhospho | E9QP99 1x | 0.01861 | 0.001362 | 1 | 4 | 1 | E9QP99 | Golgin sub | E9QP99 [1 | 0 | 1373.689 | 0 | 0.47 | 0.011503 | 14.01 | 83.7 | 116.3 | 7.08 | 13.37 | | High | High | High | High | High | High | High | High | High | High | High | 0.000723 | 0.006305 | 1.59 |
| FALSE | High | IHNVGSPLI | [K].IHNVGS | 1xPhospho | P54276 1xI | 0.000319 | 0 | 1 | 2 | 2 | P54276 | DNA mism | P54276 [82 | 0 | 1502.85 | 0 | -1.16 | 0.000222 | 1.37 | 139.1 | 60.9 | 3.5 | 8.46 | | High | High | High | High | High | High | High | High | High | High | High | 0 | 6.59E-05 | 2.99 |
| FALSE | High | LHPDGSPD | [K].LHPDG | 1xPhospho | Q8BQ33 1x | 0.000124 | 0 | 1 | 4 | 1 | Q8BQ33 | Treslin OS= | Q8BQ33 [5 | 0 | 1903.957 | 0 | -0.58 | 0.000758 | 2.12 | 120.2 | 79.8 | 5.7 | 7.07 | | High | High | High | High | High | High | High | High | High | High | High | 0 | 2.28E-05 | 3.05 |
| FALSE | High | IHPEASHPA | [K].IHPEAS | 1xCarbami | Q3UW53 1 | 9.5E-07 | 0 | 1 | 1 | 2 | Q3UW53 | Protein Nib | Q3UW53 [ | 0 | 2484.118 | 0 | 0.63 | 0.002468 | 14.49 | 76.3 | 123.7 | 4.62 | 8.67 | | High | High | High | High | High | High | High | High | High | High | High | 0 | 8.42E-05 | 3.57 |
| FALSE | High | LHPPSPVPQ | [R].LHPPSP | 1xCarbami | Q99PQ2-2 | 0.004824 | 0.000456 | 1 | 3 | 1 | Q99PQ2-2 | Isoform 2 ( | Q99PQ2-2 | 0 | 2032.015 | 0 | 1.21 | 0.000647 | 0.87 | 59.3 | 140.7 | 6.08 | 6.71 | | High | High | High | High | High | High | High | High | High | High | High | 0.000204 | 0.001387 | 2.22 |
| FALSE | High | LHQLAMQ | [K].LHQLAI | 1xPhospho | Q61990 1x | 0.000725 | 0 | 1 | 4 | 3 | Q61990 | Poly(rC)-bi | Q61990 [2 | 0 | 3936.881 | 0 | -0.29 | 0.210368 | 15.7 | 109.9 | 90.1 | 23.66 | 9.34 | | High | High | High | High | High | High | High | High | High | High | High | 4.94E-05 | 0.000701 | 3.07 |
| FALSE | High | IHRASDPGI | [R].IHRASD | 1xPhospho | B2RQC6 1x | 7.86E-07 | 0 | 1 | 6 | 3 | B2RQC6 | CAD protei | B2RQC6 [1 | 1 | 2155.131 | 0 | -0.45 | 0.040221 | 21.1 | 116.6 | 83.4 | 10.19 | 22.35 | | High | High | High | High | High | High | High | High | High | High | High | 0 | 7.79E-08 | 2.54 |
| FALSE | High | LHLESPQPS | [R].LHLESP | 3xPhospho | Q8JZX9 3xI | 0.007951 | 0.000736 | 1 | 1 | 1 | Q8JZX9 | Cdc42 effe | Q8JZX9 [13 | 0 | 3065.352 | 0 | -0.31 | 0.605262 | 20.1 | 102.4 | 97.6 | 23.1 | 20.09 | | High | High | High | High | High | High | High | High | High | High | High | 0.000327 | 0.002433 | 2.57 |
| FALSE | High | LHSAPNLSI | [R].LHSAPN | 1xPhospho | Q3UFT4 1x | 0.001313 | 0 | 1 | 7 | 1 | Q3UFT4 | Putative ur | Q3UFT4 [5 | 0 | 2355.274 | 0 | -1.56 | 0.001948 | 1.84 | 152.5 | 47.5 | 11.09 | 13.78 | | High | High | High | High | High | High | High | High | High | High | High | 0 | 0.000324 | 1.53 |
| FALSE | High | LHSAQLSPV | [R].LHSAQI | 1xDeamida | Q149H7 1x | 1.86E-07 | 0 | 1 | 2 | 3 | Q149H7 | Myeloid le | Q149H7 [1 | 0 | 2689.385 | 0 | -0.29 | 0.246983 | 9.8 | 106.6 | 93.4 | 12.17 | 9.26 | | High | High | High | High | High | High | High | High | High | High | High | 0 | 5.55E-08 | 4.61 |
| FALSE | High | LHSAQLSPV | [R].LHSAQI | 2xPhospho | Q149H7 2x | 4.03E-07 | 0 | 1 | 2 | 2 | Q149H7 | Myeloid le | Q149H7 [1 | 0 | 2768.367 | 0 | -0.5 | 0.006223 | 4.48 | 114.7 | 85.3 | 5.85 | 6.17 | | High | High | High | High | High | High | High | High | High | High | High | 0 | 3.65E-08 | 4.84 |
| FALSE | High | LHSDSGISV | [K].LHSDS( | 1xPhospho | Q8CFT3 1x | 4.56E-06 | 0 | 1 | 2 | 5 | Q8CFT3 | Nerve grov | Q8CFT3 [3 | 0 | 3714.812 | 0 | 2.62 | 0.005516 | 33.21 | 28.5 | 171.5 | 19.61 | 23.87 | | High | High | High | High | High | High | High | High | High | High | High | 0 | 5.57E-07 | 4.03 |
| FALSE | High | LHSDSGISV | [K].LHSDS( | 2xPhospho | Q8CFT3 2x | 4.36E-06 | 0 | 1 | 2 | 3 | Q8CFT3 | Nerve grov | Q8CFT3 [3 | 0 | 3794.779 | 0 | 1.87 | 0.00302 | 2.49 | 42.9 | 157.1 | 13.61 | 18.11 | | High | High | High | High | High | High | High | High | High | High | High | 0 | 5.33E-07 | 4.86 |
| FALSE | High | LHSGSEGQ | [R].LHSGSE | 1xPhospho | Q5SVL6 1x | 4.27E-05 | 0 | 1 | 3 | 1 | Q5SVL6 | Rap1 GTPa | Q5SVL6 [5 | 0 | 1538.68 | 0 | 2.44 | 0.002159 | 19.75 | 27.8 | 172.2 | 12.46 | 19.43 | | High | High | High | High | High | High | High | High | High | High | High | 0 | 6.93E-06 | 3.64 |
| FALSE | High | LHSQDPLLS | [R].LHSQDI | 1xPhospho | A0A172Q4 | 0.000595 | 0 | 1 | 4 | 1 | A0A172Q4 | Cysteine-k | A0A172Q4 | 0 | 1700.857 | 0 | 2.3 | 0.000406 | 20.71 | 33.7 | 166.3 | 14.99 | 8.95 | | High | High | High | High | High | High | High | High | High | High | High | 0 | 0.000133 | 3.71 |
| FALSE | High | LHSSNPNLS | [R].LHSSNF | 1xPhospho | D3YTT6 1xI | 2.43E-07 | 0 | 1 | 7 | 6 | D3YTT6 | Oxysterol-l | D3YTT6 [3 | 0 | 2426.2 | 0 | 1.36 | 0.000608 | 5.94 | 57.3 | 142.7 | 4.27 | 8.26 | | High | High | High | High | High | High | High | High | High | High | High | 0 | 2.74E-06 | 3.96 |
| FALSE | High | IHSSQQESE | [R].IHSSQC | 1xPhospho | Q8BQ33 1x | 0.000967 | 0 | 1 | 1 | 2 | Q8BQ33 | Treslin OS= | Q8BQ33 [1 | 0 | 1637.748 | 0 | 0.51 | 0.00863 | 7.15 | 84.9 | 115.1 | 16.13 | 5.97 | | High | High | High | High | High | High | High | High | High | High | High | 0.009151 | 0.09882 | 2.01 |
| FALSE | High | LHSYSSPST | [R].LHSYSS | 1xPhospho | Q0VBL3 1x | 0.00095 | 0 | 1 | 7 | 2 | Q0VBL3 | RNA bindir | Q0VBL3 [1 | 0 | 1644.84 | 0 | -0.05 | 0.287525 | 14.57 | 101.4 | 98.6 | 9.9 | 10.05 | | High | High | High | High | High | High | High | High | High | High | High | 0 | 0.000225 | 2.61 |
| FALSE | High | IHTGDRPY | [R].IHTGDF | 2xCarbami | Q00899 1x | 0.031713 | 0.002011 | 1 | 3 | 1 | Q00899 | Transcripti | Q00899 [3 | 0 | 2574.203 | 0 | -0.5 | 0.007206 | 7.14 | 116 | 84 | 9.67 | 6.98 | | High | High | High | High | High | High | High | High | High | High | High | 0.001076 | 0.01145 | 2.57 |
| FALSE | High | LHVGNISPT | [K].LHVGN | 1xCarbami | F7BGR7 1x | 0.000589 | 0 | 1 | 10 | 1 | F7BGR7 | Uncharacte | F7BGR7 [1 | 0 | 2148.047 | 0 | -1.01 | 0.000611 | 2.07 | 134.3 | 65.7 | 3.58 | 10.14 | | High | High | High | High | High | High | High | High | High | High | High | 0 | 0.000131 | 3.2 |
| FALSE | High | IHVSDQEL( | [K].IHVSDC | 1xPhospho | A0A1S6GW | 7.27E-08 | 0 | 1 | 2 | 2 | A0A1S6GW | Uncharacte | A0A1S6GW | 0 | 2380.098 | 0 | -0.01 | 0.951381 | 23.4 | 100.4 | 99.6 | 6.88 | 15.77 | | High | High | High | High | High | High | High | High | High | High | High | 0 | 8.24E-08 | 4.67 |
| FALSE | High | LHVSTINLC | [K].LHVSTI | 1xPhospho | Q9JLN9 1xI | 3.15E-06 | 0 | 1 | 1 | 1 | Q9JLN9 | Serine/thre | Q9JLN9 [1 | 0 | 1690.966 | 0 | 0.25 | 0.060968 | 10.06 | 89.1 | 110.9 | 10 | 11.25 | | High | High | High | High | High | High | High | High | High | High | High | 0 | 3.7E-07 | 3.16 |
| FALSE | High | LHSAQLSPV | [R].LHSAQI | 1xPhospho | Q149H7 1x | 1.49E-10 | 0 | 1 | 2 | 5 | Q149H7 | Myeloid le | Q149H7 [1 | 0 | 2688.401 | 0 | -1.12 | 0.000492 | 0.78 | 136.5 | 63.5 | 6.15 | 3.81 | | High | High | High | High | High | High | High | High | High | High | High | 0 | 1.85E-08 | 4.26 |
| FALSE | High | LHVVENDF | [K].LHVVEI | 1xPhospho | Q3TIY5 1xI | 0.002793 | 0.000109 | 1 | 4 | 1 | Q3TIY5 | Putative ur | Q3TIY5 [9 | 1 | 2262.159 | 0 | 0.8 | 0.000703 | 8.1 | 73.1 | 126.9 | 8.99 | 3.2 | | High | High | High | High | High | High | High | High | High | High | High | 4.94E-05 | 0.000755 | 1.2 |
| FALSE | High | LHLESPQPS | [R].LHLESP | 1xPhospho | Q8JZX9 1xI | 0.001107 | 0 | 1 | 1 | 1 | Q8JZX9 | Cdc42 effe | Q8JZX9 [1 | 0 | 2905.42 | 0 | -0.29 | 0.687595 | 34.01 | 110.1 | 89.9 | 19.85 | 54.96 | | High | High | High | High | High | High | High | High | High | High | High | 0 | 0.000266 | 3.05 |
| FALSE | High | IHIDPESQD | [R].IHIDPES | 1xCarbami | P70271 1xI | 8.47E-08 | 0 | 1 | 3 | 3 | P70271 | PDZ and LI | P70271 [1 | 0 | 2121.947 | 0 | 0.7 | 0.005343 | 0.74 | 77.8 | 122.2 | 11.69 | 8.87 | | High | High | High | High | High | High | High | High | High | High | High | 0 | 9.01E-07 | 5.2 |
| FALSE | High | LGTHEGLSI | [R].LGTHE( | 1xPhospho | Q61286-2 | 0.000257 | 0 | 2 | 7 | 2 | Q61286-2; | Isoform AL | Q61286-2 | 0 | 2651.367 | 0 | 0.07 | 0.265018 | 3.26 | 97.8 | 102.2 | 8.08 | 7.07 | Shared | High | High | High | High | High | High | High | High | High | High | High | 0 | 5.18E-05 | 3.4 |
| FALSE | High | IGTLHGSPE | [K].IGTLHG | 1xPhospho | P59438 1xI | 0.090888 | 0.007081 | 1 | 7 | 1 | P59438 | Hermansky | P59438 [5 | 0 | 1689.934 | 0 | 1.84 | 0.000191 | 2.8 | 43.1 | 156.9 | 8.29 | 3.1 | | High | High | High | High | High | High | High | High | High | High | High | 0.003606 | 0.03847 | 2.34 |
| FALSE | High | LGTPPSLPH | [K].LGTPPS | 1xPhospho | Q8R5K2 1x | 0.00204 | 5.66E-05 | 1 | 4 | 1 | Q8R5K2 | Ubiquitin c | Q8R5K2 [9 | 0 | 1482.803 | 0 | -0.44 | 0.018923 | 4.43 | 115.1 | 84.9 | 3.17 | 17.06 | | High | High | High | High | High | High | High | High | High | High | High | 2.58E-05 | 0.000529 | 3.23 |
| FALSE | High | LGTSDGEE | [K].LGTSD( | 1xPhospho | B9EIT8 1xP | 0.030104 | 0.001857 | 2 | 6 | 2 | B9EIT8; A0 | Rg9mtd2 p | B9EIT8 [19 | 0 | 1401.61 | 0 | -0.6 | 0.018619 | 8.46 | 120.6 | 79.4 | 13.76 | 14.6 | Shared | High | High | High | High | High | High | High | High | High | High | High | 0.001011 | 0.01076 | 2.46 |
| FALSE | High | LGVAGVPE | [R].LGVAG | 1xPhospho | F8VQH0 1x | 5.09E-06 | 0 | 1 | 5 | 3 | F8VQH0 | Tyrosine-p | F8VQH0 [8 | 0 | 2535.29 | 0 | -0.89 | 0.000659 | 3.21 | 129.8 | 70.2 | 5.55 | 3.1 | | High | High | High | High | High | High | High | High | High | High | High | 0 | 6.35E-07 | 3.54 |
| FALSE | High | LGVDNTLS | [KR].LGVDI | 1xPhospho | Q8C9G3 1x | 0.000422 | 0 | 1 | 6 | 2 | Q8C9G3 | Putative ur | Q8C9G3 [3 | 0 | 1493.793 | 0 | -0.85 | 0.000486 | 3.49 | 127.5 | 72.5 | 3.92 | 5.3 | | High | High | High | High | High | High | High | High | High | High | High | 0.000327 | 0.002785 | 2.01 |
| FALSE | High | LGVDNTLS | [KR].LGVDI | 1xDeamida | Q8C9G3 1x | 0.0095 | 0.000736 | 1 | 6 | 2 | Q8C9G3 | Putative ur | Q8C9G3 [3 | 1 | 1935.026 | 0 | -0.7 | 0.002422 | 4.85 | 123.8 | 76.2 | 6.99 | 4.66 | | High | High | High | High | High | High | High | High | High | High | High | 0.000327 | 0.002949 | 2.5 |
| FALSE | High | LGVDQDGT | [R].LGVDQ | 1xPhospho | Q921C3 1x | 4.22E-07 | 0 | 1 | 6 | 3 | Q921C3 | Bromodom | Q921C3 [6 | 0 | 2250.06 | 0 | -1.07 | 0.000278 | 1.41 | 136 | 64 | 3.65 | 7.45 | | High | High | High | High | High | High | High | High | High | High | High | 0 | 3.87E-08 | 3.17 |
| FALSE | High | LGVDQDGT | [R].LGVDQ | 1xDeamida | Q921C3 1x | 1.99E-06 | 0 | 1 | 6 | 1 | Q921C3 | Bromodom | Q921C3 [6 | 0 | 2251.044 | 0 | -0.72 | 0.03466 | 2.31 | 122.6 | 77.4 | 16.44 | 11.96 | | High | High | High | High | High | High | High | High | High | High | High | 0 | 2.19E-07 | 3.73 |
| FALSE | High | LGVGETLE | [R].LGVGE | 1xPhospho | Q6PJN8 1x | 2.68E-06 | 0 | 1 | 2 | 1 | Q6PJN8 | DALR antic | Q6PJN8 [6 | 0 | 2850.521 | 0 | -0.14 | 0.429766 | 21.52 | 105.8 | 94.2 | 9.89 | 15 | | High | High | High | High | High | High | High | High | High | High | High | 0 | 3.08E-07 | 3.09 |
| FALSE | High | LGVNNISG | [K].LGVNN | 1xPhospho | Q64152 1x | 4.95E-06 | 0 | 1 | 2 | 1 | Q64152 | Transcripti | Q64152 [9 | 0 | 3624.829 | 0 | -1.34 | 0.000405 | 1.04 | 144.8 | 55.2 | 11.31 | 13.63 | | High | High | High | High | High | High | High | High | High | High | High | 0 | 6.15E-07 | 4.5 |
| FALSE | High | LGVPTEVST | [R].LGVPTE | 1xPhospho | Q9DB42 1x | 4.37E-07 | 0 | 1 | 2 | 9 | Q9DB42 | Zinc finger | Q9DB42 [1 | 0 | 2201.013 | 0 | -0.26 | 0.187292 | 12.78 | 114.2 | 85.8 | 19.58 | 42.57 | | High | High | High | High | High | High | High | High | High | High | High | 0 | 4.01E-08 | 3.13 |
| FALSE | High | LGVPTEVST | [R].LGVPTE | 1xPhospho | Q9DB42 1x | 1.49E-05 | 0 | 1 | 2 | 14 | Q9DB42 | Zinc finger | Q9DB42 [1 | 0 | 2217.008 | 0 | -0.73 | 0.767297 | 7.7 | 110.6 | 89.4 | 34.49 | 57.33 | | High | High | High | High | High | High | High | High | High | High | High | 0 | 5.13E-06 | 3.15 |
| FALSE | High | IHIDPESQD | [R].IHIDPES | 1xCarbami | P70271 2xI | 3.94E-08 | 0 | 1 | 3 | 3 | P70271 | PDZ and LI | P70271 [1 | 0 | 2201.914 | 0 | 0.16 | 0.530827 | 8.62 | 97.9 | 102.1 | 13.01 | 4.06 | | High | High | High | High | High | High | High | High | High | High | High | 0 | 2.69E-09 | 4.37 |
| FALSE | High | LGVSVSPSF | [K].LGVSVS | 1xPhospho | G3X8T2 1x | 0.003495 | 0.00021 | 1 | 3 | 1 | G3X8T2 | RIKEN cDN | G3X8T2 [5 | 0 | 1210.639 | 0 | -0.6 | 0.004623 | 5.19 | 118.5 | 81.5 | 7.26 | 7.17 | | High | High | High | High | High | High | High | High | High | High | High | 9.42E-05 | 0.000973 | 1.74 |
| FALSE | High | LHDFLAHSS | [K].LHDFLA | 1xCarbami | Q61687 1x | 7.98E-06 | 0 | 1 | 4 | 5 | Q61687 | Transcripti | Q61687 [1 | 0 | 2627.128 | 0 | 0.62 | 0.003746 | 10.4 | 76.7 | 123.3 | 6.67 | 9.43 | | High | High | High | High | High | High | High | High | High | High | High | 0 | 1.05E-06 | 3.77 |
| FALSE | High | IHDLEDDLE | [R].IHDLED | 1xPhospho | Q3THK3 1x | 0.000126 | 0 | 1 | 2 | 1 | Q3THK3 | General tra | Q3THK3 [2 | 0 | 2890.177 | 0 | 0.73 | 0.021591 | 17.26 | 75.7 | 124.3 | 28.85 | 17.22 | | High | High | High | High | High | High | High | High | High | High | High | 0 | 2.33E-05 | 2.71 |
| FALSE | High | IHDLEDDLE | [R].IHDLED | 2xPhospho | Q3THK3 2x | 0.000622 | 0 | 1 | 2 | 3 | Q3THK3 | General tra | Q3THK3 [2 | 0 | 2970.143 | 0 | -0.07 | 0.42202 | 19.4 | 99 | 101 | 4.51 | 19.15 | | High | High | High | High | High | High | High | High | High | High | High | 0 | 0.000274 | 3.98 |
| FALSE | High | LHEDGIDN | [R].LHEDGI | 1xPhospho | Q9D3U0 1x | 1.7E-05 | 0 | 1 | 2 | 3 | Q9D3U0 | Putative tR | Q9D3U0 [7 | 0 | 2080 | 0 | 0.71 | 0.00014 | 1.5 | 75.4 | 124.6 | 1.18 | 1.1 | | High | High | High | High | High | High | High | High | High | High | High | 0 | 0.000163 | 3.28 |
| FALSE | High | LHEKPPPSF | [R].LHEKPF | 1xPhospho | Q9CZX2 1x | 0.007274 | 0.00062 | 1 | 3 | 1 | Q9CZX2 | Centrosom | Q9CZX2 [2 | 0 | 2038.089 | 0 | -0.33 | 0.038186 | 9.09 | 111.4 | 88.6 | 15.4 | 2.1 | | High | High | High | High | High | High | High | High | High | High | High | 0.000275 | 0.002191 | 1.91 |
| FALSE | High | LHESEEAFI | [K].LHESEE | 1xPhospho | Q9CPS8 1x | 0.002463 | 0.000109 | 1 | 1 | 2 | Q9CPS8 | Small mem | Q9CPS8 [2 | 0 | 2332.033 | 0 | 0.01 | 0.904664 | 12.91 | 99.4 | 100.6 | 3.76 | 18.3 | | High | High | High | High | High | High | High | High | High | High | High | 4.94E-05 | 0.000654 | 2.72 |
| FALSE | High | LHFLAPSED | [K].LHFLAP | 1xPhospho | Q3U0K8 1x | 5.45E-05 | 0 | 1 | 4 | 2 | Q3U0K8 | Prolyl 3-hy | Q3U0K8 [3 | 0 | 2298.094 | 0 | -0.76 | 0.000385 | 2.75 | 125.3 | 74.7 | 5.06 | 5.43 | | High | High | High | High | High | High | High | High | High | High | High | 0 | 9.05E-06 | 3.67 |
| FALSE | High | LHFLGIPED | [K].LHFLGI | 1xPhospho | Q6P5H2-2 | 0.042141 | 0.003162 | 1 | 2 | 5 | Q6P5H2-2 | Isoform 2 ( | Q6P5H2-2 | 0 | 4937.651 | 0 | 2.98 | 0.002072 | 32.49 | 20.6 | 179.4 | 23.11 | 18.57 | | High | High | High | High | High | High | High | High | High | High | High | 0.003148 | 0.03287 | 2.88 |
| FALSE | High | IHFSPLESTI | [R].IHFSPLI | 1xPhospho | Q8CJ27 1xI | 0.000492 | 0 | 1 | 2 | 1 | Q8CJ27 | Abnormal s | Q8CJ27 [5 | 0 | 1950.989 | 0 | -0.01 | 0.933401 | 21.04 | 100.8 | 99.2 | 5.57 | 11.58 | | High | High | High | High | High | High | High | High | High | High | High | 0 | 0.000108 | 2.46 |
| FALSE | High | LHGADTED | [K].LHGAD | 1xCarbami | Q3U2W2 1 | 6.98E-10 | 0 | 1 | 2 | 2 | Q3U2W2 | MYB bindir | Q3U2W2 [ | 0 | 3001.308 | 0 | 0.02 | 0.916853 | 2.14 | 95.7 | 104.3 | 8.73 | 16.53 | | High | High | High | High | High | High | High | High | High | High | High | 0 | 3.61E-11 | 5.96 |
| FALSE | High | LHGGFDSD | [R].LHGGFI | 1xCarbami | O08582 1x | 5.4E-05 | 0 | 1 | 1 | 1 | O08582 | GTP-bindir | O08582 [3 | 0 | 3430.54 | 0 | -0.12 | 0.248435 | 21.94 | 104.2 | 95.8 | 17.29 | 13.34 | | High | High | High | High | High | High | High | High | High | High | High | 0 | 8.97E-06 | 4.89 |
| FALSE | High | LHGGFDSD | [R].LHGGFI | 1xCarbami | O08582 2x | 2.44E-05 | 0 | 1 | 1 | 7 | O08582 | GTP-bindir | O08582 [3 | 0 | 3510.506 | 0 | -0.24 | 0.049865 | 5.86 | 107.9 | 92.1 | 2.36 | 9.56 | | High | High | High | High | High | High | High | High | High | High | High | 0 | 3.69E-06 | 5.96 |
| FALSE | High | LHGSVPNL | [R].LHGSVI | 1xPhospho | A2A716 1x | 0.000941 | 0 | 1 | 3 | 2 | A2A716 | Oxysterol-l | A2A716 [3 | 0 | 1388.725 | 0 | 0.2 | 0.031546 | 6.67 | 93.2 | 106.8 | 1.67 | 6.75 | | High | High | High | High | High | High | High | High | High | High | High | 0.000449 | 0.003727 | 2.43 |
| FALSE | High | LGVSVSPSF | [K].LGVSVS | 2xPhospho | G3X8T2 2x | 0.041425 | 0.003126 | 1 | 3 | 4 | G3X8T2 | RIKEN cDN | G3X8T2 [5 | 0 | 1290.606 | 0 | -0.28 | 0.027489 | 0.85 | 109.6 | 90.4 | 5.47 | 6.37 | | High | High | High | High | High | High | High | High | High | High | High | 0.009162 | 0.09916 | 2.17 |
| FALSE | High | LHYTPPLQS | [R].LHYTPP | 1xPhospho | A0A0A6YV. | 5.28E-10 | 0 | 1 | 4 | 1 | A0A0A6YV. | Protein lin- | A0A0A6YV. | 0 | 2997.507 | 0 | 0.88 | 0.068753 | 18.34 | 75.9 | 124.1 | 26.24 | 15.69 | | High | High | High | High | High | High | High | High | High | High | High | 0 | 2.12E-11 | 4.59 |
| FALSE | High | IKDPDLTTP | [K].IKDPDL | 1xPhospho | Q3V1H1 1x | 0.055308 | 0.003982 | 1 | 1 | 2 | Q3V1H1 | Cytoskelet | Q3V1H1 [5 | 1 | 2097.145 | 0 | 0 | 0.690972 | 15.81 | 99.3 | 100.7 | 4.78 | 8.99 | | High | High | High | High | High | High | High | High | High | High | High | 0.009992 | 0.1092 | 2.89 |
| FALSE | High | IKDSPLGGV | [R].IKDSPL | 1xPhospho | Q8R3F9 1x | 0.000738 | 0 | 1 | 1 | 1 | Q8R3F9 | Speckle tar | Q8R3F9 [6 | 1 | 1895.097 | 0 | -0.68 | 0.046128 | 15.85 | 127.8 | 72.2 | 24.95 | 14.63 | | High | High | High | High | High | High | High | High | High | High | High | 0 | 0.000169 | 1.64 |
| FALSE | High | LLDFGSLSN | [K].LLDFGS | 1xPhospho | Q8BT90 1x | 6.41E-08 | 0 | 1 | 3 | 7 | Q8BT90 | Putative ur | Q8BT90 [1 | 0 | 2963.535 | 0 | -0.16 | 0.989115 | 13.14 | 95.2 | 104.8 | 24.3 | 20.29 | | High | High | High | High | High | High | High | High | High | High | High | 0 | 4.66E-09 | 4.31 |
| FALSE | High | LLDFSEEEE | [K].LLDFSE | 1xPhospho | Q8VBW5 1 | 0.000229 | 0 | 1 | 7 | 1 | Q8VBW5 | HMG box t | Q8VBW5 [ | 0 | 2880.249 | 0 | 0.13 | 0.159655 | 2.88 | 94.8 | 105.2 | 7.03 | 9.76 | | High | High | High | High | High | High | High | High | High | High | High | 0 | 4.54E-05 | 2.86 |
| FALSE | High | LLDFSPIQK | [K].LLDFSP | 1xPhospho | Q9R0X5-5 | 0.009168 | 0.000736 | 1 | 11 | 1 | Q9R0X5-5 | Isoform 5 ( | Q9R0X5-5 | 0 | 1598.896 | 0 | -0.41 | 0.005963 | 7.89 | 113.7 | 86.3 | 5.12 | 7.26 | | High | High | High | High | High | High | High | High | High | High | High | 0.000327 | 0.002859 | 2.43 |
| FALSE | High | LLDGNPLD | [K].LLDGNI | 1xPhospho | Q920Q8 1x | 1.49E-05 | 0 | 1 | 3 | 1 | Q920Q8 | Influenza v | Q920Q8 [2 | 0 | 3395.656 | 0 | -0.16 | 0.033993 | 5.48 | 104.1 | 95.9 | 6.01 | 10.38 | | High | High | High | High | High | High | High | High | High | High | High | 0 | 2.12E-06 | 4.4 |
| FALSE | High | LLDGPTGS | [K].LLDGPT | 1xPhospho | P70399 1xI | 5.12E-05 | 0 | 1 | 6 | 6 | P70399 | TP53-bindi | P70399 [17 | 0 | 3115.516 | 0 | 0.54 | 0.017418 | 20.68 | 76.1 | 123.9 | 6.78 | 13.73 | | High | High | High | High | High | High | High | High | High | High | High | 0 | 8.47E-06 | 3.19 |
| FALSE | High | LLDHMAPF | [R].LLDHM | 1xPhospho | Q9WTP2 1 | 7.41E-05 | 0 | 1 | 1 | 2 | Q9WTP2 | Protein spr | Q9WTP2 [ | 0 | 2138.067 | 0 | -0.1 | 0.056819 | 6.73 | 103.6 | 96.4 | 4.83 | 8.29 | | High | High | High | High | High | High | High | High | High | High | High | 0 | 1.29E-05 | 2.69 |
| FALSE | High | LLDHMAPF | [R].LLDHM | 1xPhospho | Q9WTP2 1 | 1.28E-05 | 0 | 1 | 1 | 6 | Q9WTP2 | Protein spr | Q9WTP2 [ | 0 | 2154.062 | 0 | -0.01 | 0.517945 | 7.12 | 98.1 | 101.9 | 7.99 | 13.12 | | High | High | High | High | High | High | High | High | High | High | High | 0 | 1.78E-06 | 2.91 |
| FALSE | High | LIDLEDSAD | [K].LIDLED | 1xPhospho | O54916 1x | 3.09E-08 | 0 | 1 | 8 | 9 | O54916 | RalBP1-ass | O54916 [3 | 0 | 3552.703 | 0 | -0.34 | 0.04254 | 6.57 | 108.3 | 91.7 | 11.99 | 9.27 | | High | High | High | High | High | High | High | High | High | High | High | 0 | 1.63E-06 | 4.67 |
| FALSE | High | LIDLEDSAD | [K].LIDLED | 1xDeamida | O54916 1x | 0.000563 | 0 | 1 | 8 | 3 | O54916 | RalBP1-ass | O54916 [3 | 0 | 3553.688 | 0 | -0.3 | 0.033799 | 8.55 | 110.2 | 89.8 | 9.54 | 6.18 | | High | High | High | High | High | High | High | High | High | High | High | 0 | 0.000125 | 4 |
| FALSE | High | LIDLESPTP | [K].LIDLESI | 1xPhospho | Q8C0T5 1x | 3.72E-07 | 0 | 1 | 3 | 4 | Q8C0T5 | Signal-indu | Q8C0T5 [1 | 0 | 1995.045 | 0 | 2.22 | 0.000354 | 4.95 | 33.6 | 166.4 | 3.51 | 8.08 | | High | High | High | High | High | High | High | High | High | High | High | 0 | 3.34E-08 | 3.04 |
| FALSE | High | LLDLSDSTS | [K].LLDLSD | 1xPhospho | P70335 1xI | 0.000248 | 0 | 1 | 3 | 2 | P70335 | Rho-associ | P70335 [1 | 0 | 3132.446 | 0 | 0.42 | 0.033145 | 8.61 | 84.8 | 115.2 | 13.64 | 5.22 | | High | High | High | High | High | High | High | High | High | High | High | 0 | 0.000135 | 3.76 |
| FALSE | High | LLDLSDSTS | [K].LLDLSD | 2xPhospho | P70335 2xI | 1.69E-05 | 0 | 1 | 3 | 5 | P70335 | Rho-associ | P70335 [1 | 0 | 3212.412 | 0 | 0.86 | 0.001006 | 3.57 | 70.1 | 129.9 | 12.33 | 10.34 | | High | High | High | High | High | High | High | High | High | High | High | 7.19E-05 | 0.000801 | 4.27 |
| FALSE | High | LLDLVVHP | [R].LLDLVV | 1xCarbami | Q8CJF7 1xI | 5.09E-06 | 0 | 1 | 1 | 1 | Q8CJF7 | Protein EL | Q8CJF7 [1 | 0 | 3210.658 | 0 | -0.59 | 0.001002 | 2.8 | 120.1 | 79.9 | 3.44 | 6.05 | | High | High | High | High | High | High | High | High | High | High | High | 0 | 6.35E-07 | 3.57 |
| FALSE | High | LLDFGSLSN | [K].LLDFGS | 1xPhospho | Q8BT90 1x | 1.47E-05 | 0 | 1 | 3 | 1 | Q8BT90 | Putative ur | Q8BT90 [1 | 0 | 2947.54 | 0 | -0.22 | 0.636469 | 27.76 | 101.8 | 98.2 | 27.04 | 28.11 | | High | High | High | High | High | High | High | High | High | High | High | 0 | 2.08E-06 | 4.23 |
| FALSE | High | ILDPDPDPF | [K].ILDPDP | 1xPhospho | Q05AH6 1x | 7.68E-05 | 0 | 1 | 2 | 3 | Q05AH6 | Uncharacte | Q05AH6 [2 | 0 | 2957.402 | 0 | 0.82 | 0.001415 | 11.19 | 72.4 | 127.6 | 20.4 | 4.98 | | High | High | High | High | High | High | High | High | High | High | High | 0 | 1.34E-05 | 2.77 |
| FALSE | High | LLDPEDVD | [R].LLDPED | 1xPhospho | S4R1P5 1xI | 1.23E-06 | 0 | 1 | 9 | 2 | S4R1P5 | Dystonin O | S4R1P5 [4 | 0 | 2196.073 | 0 | 1.01 | 0.000209 | 3.72 | 66.7 | 133.3 | 2.61 | 3.86 | | High | High | High | High | High | High | High | High | High | High | High | 0 | 1.29E-07 | 4.21 |
| FALSE | High | ILDPEGLAL | [R].ILDPEG | 1xPhospho | Q9DBE9 1x | 1.61E-07 | 0 | 1 | 1 | 2 | Q9DBE9 | pre-rRNA p | Q9DBE9 [6 | 0 | 2192.234 | 0 | -0.11 | 0.268902 | 6.67 | 98.8 | 101.2 | 10.51 | 9.29 | | High | High | High | High | High | High | High | High | High | High | High | 0 | 1.31E-08 | 4.77 |
| FALSE | High | ILDPNSGEF | [K].ILDPNS | 1xPhospho | Q6ZPH4 1x | 8.87E-05 | 0 | 1 | 7 | 4 | Q6ZPH4 | MKIAA183 | Q6ZPH4 [4 | 0 | 3878.976 | 0 | -0.38 | 0.013454 | 2.31 | 111.9 | 88.1 | 5.89 | 7.76 | | High | High | High | High | High | High | High | High | High | High | High | 0.000344 | 0.003203 | 3.91 |
| FALSE | High | ILDPNSGEF | [K].ILDPNS | 1xDeamida | Q6ZPH4 1x | 0.115566 | 0.009504 | 1 | 7 | 1 | Q6ZPH4 | MKIAA183 | Q6ZPH4 [4 | 0 | 3879.96 | 0 | 1 | 0.003103 | 4.03 | 66.7 | 133.3 | 1.85 | 12.6 | | High | High | High | High | High | High | High | High | High | High | High | 0.004747 | 0.05127 | 3.03 |
| FALSE | High | ILDQMPAT | [K].ILDQMI | 1xPhospho | Q543X6 1x | 0.000645 | 0 | 1 | 2 | 2 | Q543X6 | Putative ur | Q543X6 [3 | 0 | 2105.937 | 0 | -0.06 | 0.699093 | 14.74 | 101.6 | 98.4 | 13.8 | 9.41 | | High | High | High | High | High | High | High | High | High | High | High | 0 | 0.000145 | 2.01 |
| FALSE | High | LLDSLEPPG | [R].LLDSLE | 1xPhospho | Q9Z1A1 1x | 0.027121 | 0.001676 | 1 | 3 | 2 | Q9Z1A1 | Putative ur | Q9Z1A1 [1 | 0 | 3001.424 | 0 | 0.09 | 0.13066 | 10.36 | 89.5 | 110.5 | 10.29 | 14.01 | | High | High | High | High | High | High | High | High | High | High | High | 0.001183 | 0.01268 | 2.7 |
| FALSE | High | LLDSPGAAT | [K].LLDSPG | 1xPhospho | Q8BYH8 1x | 0.000184 | 0 | 1 | 3 | 1 | Q8BYH8 | Chromodo | Q8BYH8 [2 | 0 | 1438.714 | 0 | 0.33 | 0.001744 | 2.96 | 90.2 | 109.8 | 7.89 | 4.97 | | High | High | High | High | High | High | High | High | High | High | High | 0 | 3.58E-05 | 2.39 |
| FALSE | High | LIDSSGSAS | [R].LIDSSG | 1xPhospho | A0A1S6GW | 2.91E-06 | 0 | 1 | 9 | 3 | A0A1S6GW | Oxysterol-l | A0A1S6GW | 0 | 2585.322 | 0 | 0.39 | 0.272189 | 14.99 | 84 | 116 | 20.85 | 17.68 | | High | High | High | High | High | High | High | High | High | High | High | 0 | 3.39E-07 | 3.94 |
| FALSE | High | LLDTNPEIN | [R].LLDTNF | 1xPhospho | G5E870 1x | 0.008769 | 0.000736 | 1 | 3 | 1 | G5E870 | E3 ubiquiti | G5E870 [1 | 0 | 2241.023 | 0 | -0.02 | 0.486575 | 6.51 | 98.5 | 101.5 | 6.01 | 9.04 | | High | High | High | High | High | High | High | High | High | High | High | 0.000327 | 0.002706 | 2.09 |
| FALSE | High | ILDTSSLTQ | [R].ILDTSSI | 1xPhospho | A2ACM0 1 | 8.15E-07 | 0 | 1 | 4 | 1 | A2ACM0 | Putative ur | A2ACM0 [8 | 0 | 2369.237 | 0 | 0.54 | 0.022446 | 20.18 | 82.3 | 117.7 | 11.49 | 6.14 | | High | High | High | High | High | High | High | High | High | High | High | 0 | 8.07E-08 | 3.77 |
| FALSE | High | ILDTSSLTQ | [R].ILDTSSI | 2xPhospho | A2ACM0 2 | 1.67E-05 | 0 | 1 | 4 | 2 | A2ACM0 | Putative ur | A2ACM0 [8 | 0 | 2449.203 | 0 | 0.66 | 0.005999 | 9.73 | 78.1 | 121.9 | 3.51 | 10.72 | | High | High | High | High | High | High | High | High | High | High | High | 0 | 2.41E-06 | 3.63 |
| FALSE | High | LIEDDIVCSI | [R].LIEDDI | 1xCarbami | A2A761 1x | 0.000292 | 0 | 1 | 4 | 1 | A2A761 | Zinc finger | A2A761 [2 | 0 | 1940.022 | 0 | -0.86 | 0.013196 | 5.36 | 125.4 | 74.6 | 21.99 | 11.17 | | High | High | High | High | High | High | High | High | High | High | High | 0 | 5.96E-05 | 2.85 |
| FALSE | High | LIEDFTQVS | [R].LIEDFT | 1xCarbami | Q9DB00 1x | 1.3E-05 | 0 | 1 | 5 | 1 | Q9DB00 | GON-4-like | Q9DB00 [7 | 0 | 2427.203 | 0 | -1.63 | 0.000263 | 1.67 | 151.1 | 48.9 | 4.59 | 9.62 | | High | High | High | High | High | High | High | High | High | High | High | 0 | 1.82E-06 | 3.97 |
| FALSE | High | ILDPDPDPF | [K].ILDPDP | 2xPhospho | Q05AH6 2x | 1.23E-05 | 0 | 1 | 2 | 9 | Q05AH6 | Uncharacte | Q05AH6 [2 | 0 | 3037.368 | 0 | -0.61 | 0.003831 | 8.29 | 121.3 | 78.7 | 5.81 | 8.51 | | High | High | High | High | High | High | High | High | High | High | High | 0 | 1.71E-06 | 3.96 |
| FALSE | High | LLDEYNVTI | [R].LLDEYN | 1xCarbami | G5E8J0 1xI | 0.003495 | 0.00021 | 1 | 5 | 1 | G5E8J0 | Neurogeni | G5E8J0 [2 | 0 | 3376.706 | 0 | 0.79 | 0.006653 | 6.55 | 68.2 | 131.8 | 20.67 | 9.67 | | High | High | High | High | High | High | High | High | High | High | High | 9.42E-05 | 0.000968 | 2.53 |
| FALSE | High | ILDEDSWSI | [R].ILDEDS | 2xPhospho | Q8BQR4 2x | 3.44E-05 | 0 | 1 | 3 | 3 | Q8BQR4 | KAT8 regul | Q8BQR4 [1 | 0 | 2994.264 | 0 | -0.39 | 0.018076 | 5.88 | 113.9 | 86.1 | 9.25 | 11.61 | | High | High | High | High | High | High | High | High | High | High | High | 0 | 5.59E-06 | 4.12 |
| FALSE | High | LLDEALSPS | [K].LLDEAL | 1xPhospho | A0A140T8 | 0.002441 | 0.000109 | 1 | 1 | 5 | A0A140T8 | Chromoso | A0A140T8 | 0 | 1697.913 | 0 | -0.58 | 0.000394 | 1.71 | 121.3 | 78.7 | 2.99 | 3.2 | | High | High | High | High | High | High | High | High | High | High | High | 0.000847 | 0.007963 | 2.17 |
| FALSE | High | LKEEEEGIS | [K].LKEEEE | 2xPhospho | Q3TE85 2x | 0.001463 | 5.66E-05 | 1 | 1 | 3 | Q3TE85 | Putative ur | Q3TE85 [9 | 1 | 2598.115 | 0 | 0.02 | 0.370032 | 3.33 | 98.7 | 101.3 | 5.69 | 1.77 | | High | High | High | High | High | High | High | High | High | High | High | 2.58E-05 | 0.000364 | 3.9 |
| FALSE | High | IKEEVLSESI | [K].IKEEVL | 1xPhospho | Q3TI31 1xP | 0.00311 | 0.00016 | 1 | 3 | 8 | Q3TI31 | Putative ur | Q3TI31 [7 | 1 | 3703.929 | 0 | -0.24 | 0.046076 | 9.95 | 109.6 | 90.4 | 10.3 | 4.43 | | High | High | High | High | High | High | High | High | High | High | High | 7.19E-05 | 0.000847 | 3.79 |
| FALSE | High | LKFSDDEDI | [K].LKFSDD | 1xPhospho | Q7TPM1 1 | 0.037692 | 0.00249 | 1 | 3 | 2 | Q7TPM1 | Protein PR | Q7TPM1 [3 | 1 | 2206.057 | 0 | -0.43 | 0.636871 | 21.93 | 123 | 77 | 48.97 | 53.88 | | High | High | High | High | High | High | High | High | High | High | High | 0.001324 | 0.01401 | 3.34 |
| FALSE | High | LKFSDDEDI | [K].LKFSDD | 1xPhospho | Q7TPM1 1 | 0.000407 | 0 | 1 | 3 | 2 | Q7TPM1 | Protein PR | Q7TPM1 [3 | 1 | 2435.22 | 0 | -0.34 | 0.003576 | 3.14 | 113 | 87 | 3.21 | 7.09 | | High | High | High | High | High | High | High | High | High | High | High | 0.00186 | 0.01873 | 2.54 |
| FALSE | High | IKNSPSEAC | [K].IKNSPS | 1xPhospho | Q5XJV7-2 | 0.000262 | 0 | 1 | 2 | 1 | Q5XJV7-2 | Isoform 2 ( | Q5XJV7-2 | 1 | 3070.452 | 0 | -0.96 | 0.000401 | 2.41 | 130.4 | 69.6 | 4.24 | 6.5 | | High | High | High | High | High | High | High | High | High | High | High | 0 | 5.3E-05 | 3.82 |
| FALSE | High | IKPDEDLPS | [K].IKPDED | 1xPhospho | B7ZNV2 1x | 0.039347 | 0.002793 | 1 | 4 | 1 | B7ZNV2 | GLI-Kruppe | B7ZNV2 [4 | 0 | 1949.015 | 0 | 0.87 | 0.000549 | 13.22 | 69.5 | 130.5 | 8.68 | 2.24 | | High | High | High | High | High | High | High | High | High | High | High | 0.001453 | 0.01462 | 1.33 |
| FALSE | High | IKPSPPLLV | [K].IKPSPPI | 1xPhospho | Z4YKQ9 1x | 6.07E-05 | 0 | 1 | 4 | 1 | Z4YKQ9 | ATPase fan | Z4YKQ9 [8 | 0 | 2117.27 | 0 | -0.66 | 0.000964 | 4.1 | 124.3 | 75.7 | 7.11 | 5.84 | | High | High | High | High | High | High | High | High | High | High | High | 0 | 1.03E-05 | 2.83 |
| FALSE | High | IKPVTENLV | [K].IKPVTE | 1xCarbami | Q61687 1x | 0.001137 | 0 | 1 | 3 | 2 | Q61687 | Transcripti | Q61687 [1 | 0 | 3681.917 | 0 | -0.11 | 0.823892 | 6.74 | 103.4 | 96.6 | 9.1 | 10.42 | | High | High | High | High | High | High | High | High | High | High | High | 2.58E-05 | 0.000493 | 3.54 |
| FALSE | High | LKSDEDEPA | [K].LKSDED | 1xPhospho | Q8CIL4 1xP | 0.000434 | 0 | 1 | 3 | 1 | Q8CIL4 | Uncharacte | Q8CIL4 [12 | 1 | 1899.008 | 0 | -0.77 | 0.000505 | 1.79 | 127.1 | 72.9 | 4.51 | 3.87 | | High | High | High | High | High | High | High | High | High | High | High | 0 | 9.32E-05 | 1.25 |
| FALSE | High | LKSEDELRF | [R].LKSEDE | 1xPhospho | O54916 1x | 5.21E-05 | 0 | 1 | 6 | 2 | O54916 | RalBP1-ass | O54916 [7 | 1 | 2806.459 | 0 | 0.11 | 0.083985 | 5.86 | 97.4 | 102.6 | 8.72 | 11.19 | | High | High | High | High | High | High | High | High | High | High | High | 0 | 8.59E-06 | 2.35 |
| FALSE | High | IKSENVEGT | [K].IKSENV | 1xPhospho | Q01320 1x | 4.39E-05 | 0 | 2 | 3 | 1 | Q01320; Q | DNA topois | Q01320 [1 | 1 | 2694.339 | 0 | -1.38 | 0.001293 | 3.14 | 143.2 | 56.8 | 9.88 | 9.08 | Shared | High | High | High | High | High | High | High | High | High | High | High | 0 | 7.1E-06 | 3.51 |

| | | | | | | | | | | | | | | | | | | | | | | | | | | | | | | | | | | | | | | |
|---|---|---|---|---|---|---|---|---|---|---|---|---|---|---|---|---|---|---|---|---|---|---|---|---|---|---|---|---|---|---|---|---|---|---|---|---|---|---|
| FALSE | High | LLAAVEAFY | [R].LLAAVE | 1xPhospho | G5E8I8 1xP | 0.003001 | 0.00016 | 1 | 3 | 3 | G5E8I8 | Calcium ho | G5E8I8 [70 | 0 | 2335.18 | 0 | -1.05 | 0.002108 | 3.43 | 135.8 | 64.2 | 8.68 | 7.5 | | High | High | High | High | High | High | High | High | High | High | High | 0.00014 | 0.001118 | 2.61 |
| FALSE | High | IIAEGANGF | [K].IIAEGAI | 1xPhospho | P26443 1xP | 1.49E-05 | 0 | 1 | 3 | 1 | P26443 | Glutamate | P26443 [40 | 0 | 2122.083 | 0 | -0.75 | 0.000996 | 4.72 | 124.8 | 75.2 | 5.18 | 7.35 | | High | High | High | High | High | High | High | High | High | High | High | 0 | 2.11E-06 | 2.81 |
| FALSE | High | ILAETPDSF | [R].ILAETPI | 1xPhospho | Q8CEC0 1x | 0.000543 | 0 | 1 | 5 | 4 | Q8CEC0 | Nuclear po | Q8CEC0 [5 | 0 | 1787.923 | 0 | -0.13 | 0.079029 | 9.1 | 104.8 | 95.2 | 4.57 | 7.05 | | High | High | High | High | High | High | High | High | High | High | High | 0 | 0.00012 | 3.6 |
| FALSE | High | LIAHAGSLT | [R].LIAHAG | 1xPhospho | Q9D6Z1 1x | 0.001815 | 5.66E-05 | 1 | 3 | 1 | Q9D6Z1 | Nucleolar p | Q9D6Z1 [3 | 0 | 1847.056 | 0 | 0.7 | 0.104087 | 36.04 | 72.7 | 127.3 | 6.41 | 30.46 | | High | High | High | High | High | High | High | High | High | High | High | 2.58E-05 | 0.000465 | 1.78 |
| FALSE | High | LLAPDSLPP | [R].LLAPDS | 1xPhospho | Q8VC51 1x | 6.83E-05 | 0 | 1 | 2 | 2 | Q8VC51 | Telomerase | Q8VC51 [8 | 0 | 2918.568 | 0 | -0.47 | 0.002619 | 4.38 | 115.2 | 84.8 | 2.54 | 5.34 | | High | High | High | High | High | High | High | High | High | High | High | 0 | 1.17E-05 | 3.54 |
| FALSE | High | IIAPSPVK | [K].IIAPSPV | 1xPhospho | P52963 1xP | 0.026885 | 0.001676 | 1 | 2 | 1 | P52963 | Band 4.1-li | P52963 [38 | 0 | 1362.816 | 0 | -1.9 | 0.00026 | 0.58 | 156.1 | 43.9 | 3.62 | 9.5 | | High | High | High | High | High | High | High | High | High | High | High | 0.000903 | 0.009498 | 1.55 |
| FALSE | High | LLAQAVHN | [R].LLAQAV | 1xCarbami | P70196 1xP | 0.006834 | 0.00062 | 1 | 1 | 1 | P70196 | TNF recept | P70196 [27 | 0 | 2782.439 | 0 | -0.27 | 0.050392 | 6.08 | 110 | 90 | 4.99 | 8.05 | | High | High | High | High | High | High | High | High | High | High | High | 0.000275 | 0.002042 | 1.5 |
| FALSE | High | ILASLPK | [R].ILASLPK | 1xPhospho | Q4GX45 1x | 0.004451 | 0.000409 | 1 | 3 | 1 | Q4GX45 | Peroxisom | Q4GX45 [2 | 0 | 1279.779 | 0 | -0.12 | 0.045199 | 6.73 | 101.9 | 98.1 | 7.49 | 8.05 | | High | High | High | High | High | High | High | High | High | High | High | 0.000183 | 0.001266 | 2.13 |
| FALSE | High | ILASNSIICP | [K].ILASNSI | 1xCarbami | E9QAE4 1x | 0.075927 | 0.005826 | 1 | 2 | 1 | E9QAE4 | Histone-lys | E9QAE4 [1 | 0 | 2149.083 | 0 | -0.5 | 0.003537 | 7.7 | 120.2 | 79.8 | 5.79 | 6.1 | | High | High | High | High | High | High | High | High | High | High | High | 0.002976 | 0.03127 | 2.38 |
| FALSE | High | LLASPDAST | [K].LLASPD | 1xPhospho | Q91W06 1 | 5.13E-09 | 0 | 1 | 4 | 6 | Q91W06 | V-type pro | Q91W06 [6 | 0 | 2727.317 | 0 | 1.48 | 0.001843 | 23.75 | 52 | 148 | 5.65 | 12.65 | | High | High | High | High | High | High | High | High | High | High | High | 0 | 8.14E-10 | 5.95 |
| FALSE | High | LLASPDAST | [K].LLASPD | 2xPhospho | Q91W06 2 | 2.19E-05 | 0 | 1 | 4 | 1 | Q91W06 | V-type pro | Q91W06 [6 | 0 | 2807.283 | 0 | 1.72 | 0.005619 | 32.18 | 51.8 | 148.2 | 12.06 | 19.82 | | High | High | High | High | High | High | High | High | High | High | High | 0 | 3.28E-06 | 4.87 |
| FALSE | High | LLASPTYR | [K].LLASPT | 1xPhospho | P58468 1xP | 0.049971 | 0.003641 | 1 | 4 | 2 | P58468 | Protein FAI | P58468 [18 | 0 | 1229.649 | 0 | -0.04 | 0.26239 | 5.67 | 101.4 | 98.6 | 1.54 | 2.26 | | High | High | High | High | High | High | High | High | High | High | High | 0.001902 | 0.01935 | 1.49 |
| FALSE | High | LIASSPTLIS | [R].LIASSPT | 1xPhospho | B1ASA5 1x | 3.03E-05 | 0 | 1 | 2 | 3 | B1ASA5 | Zinc finger | B1ASA5 [1 | 0 | 2761.577 | 0 | 1.53 | 0.000415 | 14.72 | 51.1 | 148.9 | 13.1 | 3.45 | | High | High | High | High | High | High | High | High | High | High | High | 0 | 4.71E-06 | 3.44 |
| FALSE | High | ILATPPQED | [K].ILATPP | 1xPhospho | P40142 1xP | 2.21E-08 | 0 | 1 | 1 | 2 | P40142 | Transketol | P40142 [28 | 0 | 2329.2 | 0 | -0.54 | 0.001647 | 3.57 | 120.2 | 79.8 | 5.3 | 5.03 | | High | High | High | High | High | High | High | High | High | High | High | 0 | 1.41E-09 | 4.12 |
| FALSE | High | LLCELQSIM | [R].LLCELQ | 1xCarbami | Q8BP27 1x | 2.18E-07 | 0 | 1 | 3 | 2 | Q8BP27 | Swi5-depe | Q8BP27 [2 | 0 | 2493.154 | 0 | 0.18 | 0.756062 | 17.05 | 91.6 | 108.4 | 4.83 | 21.56 | | High | High | High | High | High | High | High | High | High | High | High | 0.000327 | 0.002895 | 2.93 |
| FALSE | High | LLCQEETPE | [K].LLCQEE | 1xCarbami | Q5SW15 1 | 6.85E-06 | 0 | 1 | 3 | 5 | Q5SW15 | Methyltran | Q5SW15 [4 | 1 | 2658.144 | 0 | 0.35 | 0.027625 | 8.98 | 87.7 | 112.3 | 9.88 | 4.99 | | High | High | High | High | High | High | High | High | High | High | High | 0.000571 | 0.005089 | 3.05 |
| FALSE | High | LLDAEDVD | [R].LLDAED | 1xPhospho | F7ACR9 1x | 6.89E-05 | 0 | 1 | 13 | 1 | F7ACR9 | Microtubu | F7ACR9 [1 | 0 | 2180.078 | 0 | 1.13 | 0.001302 | 12.42 | 64.9 | 135.1 | 6.92 | 7.95 | | High | High | High | High | High | High | High | High | High | High | High | 0 | 1.19E-05 | 2.45 |
| FALSE | High | LIDASRVSE | [K].LIDASR | 1xPhospho | P56485 1xP | 7.98E-06 | 0 | 1 | 1 | 1 | P56485 | Atypical ch | P56485 [34 | 1 | 2792.412 | 0 | 2.35 | 0.000158 | 4.7 | 32.1 | 167.9 | 9.15 | 4.98 | | High | High | High | High | High | High | High | High | High | High | High | 0 | 1.05E-06 | 3.94 |
| FALSE | High | LGTGGLRS | [K].LGTGG | 1xPhospho | G3UYU0 1x | 0.08447 | 0.006345 | 1 | 1 | 1 | G3UYU0 | Mex3 RNA- | G3UYU0 [4 | 1 | 1523.789 | 0 | -0.91 | 0.000964 | 1.36 | 130.5 | 69.5 | 10.39 | 7.89 | | High | High | High | High | High | High | High | High | High | High | High | 0.003265 | 0.03541 | 1.91 |
| FALSE | High | LGTGGGGS | [R].LGTGG | 1xPhospho | Q3TYJ0 1xP | 0.000146 | 0 | 1 | 2 | 2 | Q3TYJ0 | Putative ur | Q3TYJ0 [13 | 0 | 1483.756 | 0 | -1.04 | 0.000207 | 2.08 | 135.6 | 64.4 | 4.49 | 11.16 | | High | High | High | High | High | High | High | High | High | High | High | 0 | 2.74E-05 | 2.16 |
| FALSE | High | LGTGGGGS | [R].LGTGG | 1xPhospho | Q3TYJ0 1xP | 0.00051 | 0 | 1 | 2 | 2 | Q3TYJ0 | Putative ur | Q3TYJ0 [13 | 0 | 1254.593 | 0 | -0.94 | 0.436293 | 4.77 | 127.8 | 72.2 | 75.85 | 77.52 | | High | High | High | High | High | High | High | High | High | High | High | 0.008836 | 0.09443 | 1.5 |
| FALSE | High | LGTEPFVTF | [R].LGTEPF | 1xPhospho | Q14B71 1x | 0.000595 | 0 | 1 | 4 | 2 | Q14B71 | Cell divisio | Q14B71 [2 | 0 | 1754.949 | 0 | -0.93 | 0.003984 | 2.07 | 128.9 | 71.1 | 8.13 | 12.98 | | High | High | High | High | High | High | High | High | High | High | High | 0 | 0.000133 | 3.32 |
| FALSE | High | LGEQGPEP | [R].LGEQG | 2xPhospho | Q3UIA2 2x | 1.51E-05 | 0 | 1 | 6 | 5 | Q3UIA2 | Rho GTPas | Q3UIA2 [7 | 0 | 3209.594 | 0 | -0.35 | 0.004179 | 3.08 | 112.2 | 87.8 | 2.5 | 6.39 | | High | High | High | High | High | High | High | High | High | High | High | 4.94E-05 | 0.000585 | 3.81 |
| FALSE | High | LGESQTLQ | [K].LGESQT | 1xPhospho | A3KGU9 1x | 0.005044 | 0.000504 | 1 | 9 | 1 | A3KGU9 | Spectrin al | A3KGU9 [1 | 0 | 1702.836 | 0 | 1.24 | 0.011434 | 39.98 | 55.5 | 144.5 | 10.4 | 23.91 | | High | High | High | High | High | High | High | High | High | High | High | 0.000246 | 0.001461 | 3.07 |
| FALSE | High | LGETLQGT | [K].LGETLC | 1xCarbami | P97412 1xP | 3.39E-06 | 0 | 1 | 4 | 3 | P97412 | Lysosomal- | P97412 [65 | 0 | 2728.321 | 0 | 1.06 | 0.005117 | 34.36 | 63.3 | 136.7 | 19.69 | 8.4 | | High | High | High | High | High | High | High | High | High | High | High | 0 | 4E-07 | 4.14 |
| FALSE | High | IGFPETAEE | [K].IGFPET. | 1xPhospho | Q9Z1Z2 1xP | 8.12E-09 | 0 | 1 | 1 | 16 | Q9Z1Z2 | Serine-thre | Q9Z1Z2 [3 | 0 | 3838.809 | 0 | -0.35 | 0.019583 | 6.61 | 108.7 | 91.3 | 6.43 | 7.66 | | High | High | High | High | High | High | High | High | High | High | High | 0 | 4.53E-10 | 4.89 |
| FALSE | High | IGFPETAEE | [K].IGFPET. | 2xPhospho | Q9Z1Z2 2xP | 0.024424 | 0.001612 | 1 | 1 | 2 | Q9Z1Z2 | Serine-thre | Q9Z1Z2 [3 | 0 | 3918.775 | 0 | -1.01 | 0.08733 | 12.9 | 123.6 | 76.4 | 15.17 | 39.66 | | High | High | High | High | High | High | High | High | High | High | High | 0.002403 | 0.02576 | 2.86 |
| FALSE | High | IGFPSTSPA | [R].IGFPST | 1xPhospho | Q9Z0H8 1x | 0.001424 | 5.66E-05 | 1 | 2 | 3 | Q9Z0H8 | CAP-Gly dc | Q9Z0H8 [2 | 0 | 1542.833 | 0 | -1.38 | 0.000472 | 2.52 | 143.5 | 56.5 | 3.56 | 17.7 | | High | High | High | High | High | High | High | High | High | High | High | 0.000786 | 0.007119 | 2.64 |
| FALSE | High | LGFVDVQN | [R].LGFVD | 1xCarbami | P70227 1xP | 0.000393 | 0 | 1 | 3 | 3 | P70227 | Inositol 1,4 | P70227 [26 | 0 | 1750.786 | 0 | 1.79 | 0.000278 | 8.61 | 45.5 | 154.5 | 6.94 | 6.86 | | High | High | High | High | High | High | High | High | High | High | High | 0 | 8.39E-05 | 2.93 |
| FALSE | High | IGGDAATT | [K].IGGDA | 1xPhospho | Q3U0V1 1x | 2.55E-08 | 0 | 1 | 1 | 1 | Q3U0V1 | Far upstrea | Q3U0V1 [8 | 0 | 2719.313 | 0 | 0.93 | 0.000441 | 6.49 | 70.4 | 129.6 | 8.64 | 3.26 | | High | High | High | High | High | High | High | High | High | High | High | 0 | 1.65E-09 | 5.24 |
| FALSE | High | IGGDAATT | [K].IGGDA | 1xDeamida | Q3U0V1 1x | 5.15E-07 | 0 | 1 | 1 | 1 | Q3U0V1 | Far upstrea | Q3U0V1 [8 | 0 | 2720.297 | 0 | 1.11 | 0.000406 | 9.22 | 65.4 | 134.6 | 14.67 | 4.46 | | High | High | High | High | High | High | High | High | High | High | High | 0 | 4.8E-08 | 4.73 |
| FALSE | High | IGGDAGTS | [K].IGGDA | 1xPhospho | A0A0G2JG | 2.93E-06 | 0 | 1 | 11 | 3 | A0A0G2JG | Far upstrea | A0A0G2JG | 0 | 2512.176 | 0 | -0.94 | 0.001954 | 4.2 | 131.4 | 68.6 | 5.28 | 13.88 | | High | High | High | High | High | High | High | High | High | High | High | 0 | 4.61E-06 | 3.77 |
| FALSE | High | LGGDSEYS | [R].LGGDS | 1xCarbami | A2A654 1x | 1.2E-05 | 0 | 1 | 4 | 2 | A2A654 | Bromodom | A2A654 [1 | 0 | 2660.105 | 0 | -0.09 | 0.850505 | 14.89 | 102.1 | 97.9 | 6.89 | 11.14 | | High | High | High | High | High | High | High | High | High | High | High | 0 | 1.65E-06 | 3.88 |
| FALSE | High | LGGDVSPT | [K].LGGDV. | 1xPhospho | P14873 1xP | 3.24E-06 | 0 | 2 | 2 | 3 | P14873; B2 | Microtubu | P14873 [14 | 0 | 2520.279 | 0 | -0.51 | 0.010743 | 4.06 | 116 | 84 | 8.34 | 8.39 | Shared | High | High | High | High | High | High | High | High | High | High | High | 0 | 3.82E-07 | 3.64 |
| FALSE | High | IGGKSPPPF | [K].IGGKSP | 1xPhospho | E9QAU4 1x | 0.061176 | 0.004646 | 1 | 11 | 1 | E9QAU4 | Enhancer t | E9QAU4 [1 | 1 | 1932.087 | 0 | 0.03 | 0.854322 | 15.78 | 94.7 | 105.3 | 15.47 | 11.45 | | High | High | High | High | High | High | High | High | High | High | High | 0.002381 | 0.02442 | 2.15 |
| FALSE | High | LGELWEEN | [K].LGELW | 1xPhospho | Q3UGI2 1x | 0.003892 | 0.000261 | 1 | 3 | 1 | Q3UGI2 | Putative ur | Q3UGI2 [2 | 0 | 1839.891 | 0 | 0.17 | 0.529496 | 34.6 | 94 | 106 | 17.06 | 20.27 | | High | High | High | High | High | High | High | High | High | High | High | 0.00014 | 0.001095 | 2.39 |
| FALSE | High | IGGLENLNI | [R].IGGLEN | 1xCarbami | B7ZP47 1xP | 0.000611 | 0 | 1 | 2 | 1 | B7ZP47 | Wapal prot | B7ZP47 [27 | 0 | 3312.613 | 0 | -0.93 | 0.005251 | 5.99 | 131.2 | 68.8 | 12.11 | 9.51 | | High | High | High | High | High | High | High | High | High | High | High | 0 | 0.000137 | 2.25 |
| FALSE | High | LGGNGQLC | [R].LGGNG | 1xPhospho | A0A068F12 | 4.07E-07 | 0 | 1 | 11 | 3 | A0A068F12 | Glyco-gag | A0A068F12 | 0 | 2998.454 | 0 | 6.64 | | 0 | | 200 | | 51.59 | | Not Found | Not Found | Not Found | Not Found | Not Found | High | High | High | High | High | High | 0 | 3.7E-08 | 4.35 |
| FALSE | High | LGGNGQLC | [R].LGGNG | 1xDeamida | A0A068F12 | 0.000209 | 0 | 1 | 11 | 2 | A0A068F12 | Glyco-gag | A0A068F12 | 0 | 2999.438 | 0 | 6.64 | | 0 | | 200 | | 47.88 | | Not Found | Not Found | Not Found | Not Found | Not Found | High | High | High | High | High | High | 0 | 0.000103 | 3.08 |
| FALSE | High | LGGPAVGT | [K].LGGPA | 1xPhospho | Q7TSH3 1x | 1.25E-06 | 0 | 1 | 1 | 1 | Q7TSH3 | Zinc finger | Q7TSH3 [9 | 0 | 1951.045 | 0 | -0.43 | 0.006824 | 1.76 | 113.3 | 86.7 | 9.75 | 10.29 | | High | High | High | High | High | High | High | High | High | High | High | 0 | 1.29E-07 | 4.23 |
| FALSE | High | LGGPEGPG | [R].LGGPE | 1xPhospho | Q99PT3 1x | 0.003997 | 0.000311 | 1 | 3 | 2 | Q99PT3 | INO80 com | Q99PT3 [3 | 0 | 1574.803 | 0 | -0.27 | 0.001895 | 3.39 | 108.5 | 91.5 | 4.8 | 4.51 | | High | High | High | High | High | High | High | High | High | High | High | 0.000585 | 0.005234 | 3.07 |
| FALSE | High | LGGPSASLF | [R].LGGPS | 1xPhospho | Q8VBT9 1x | 0.000219 | 0 | 1 | 4 | 2 | Q8VBT9 | Tether con | Q8VBT9 [2 | 0 | 2465.317 | 0 | -1.32 | 0.000668 | 1.32 | 142.1 | 57.9 | 6.33 | 12.16 | | High | High | High | High | High | High | High | High | High | High | High | 0 | 0.000152 | 1.58 |
| FALSE | High | LGGPSASLF | [R].LGGPS | 1xDeamida | Q8VBT9 1x | 0.00824 | 0.000736 | 1 | 4 | 3 | Q8VBT9 | Tether con | Q8VBT9 [2 | 0 | 2466.301 | 0 | -1.13 | 0.000985 | 1.94 | 133.7 | 66.3 | 6.12 | 9.73 | | High | High | High | High | High | High | High | High | High | High | High | 0.000327 | 0.002533 | 2.18 |
| FALSE | High | LGGQTAIG | [R].LGGQT. | 1xPhospho | Q91VS8 1x | 2.58E-05 | 0 | 1 | 2 | 4 | Q91VS8 | FERM, Rho | Q91VS8 [1 | 0 | 2449.254 | 0 | 1.93 | 0.000263 | 11.79 | 42.2 | 157.8 | 7.06 | 4.63 | | High | High | High | High | High | High | High | High | High | High | High | 0 | 3.9E-06 | 4 |
| FALSE | High | LGGSLIVAF | [K].LGGSLI | 1xPhospho | Q9R0P5 1x | 2.77E-05 | 0 | 1 | 1 | 1 | Q9R0P5 | Destrin OS | Q9R0P5 [1 | 0 | 1654.865 | 0 | -0.91 | 0.000671 | 4.7 | 130.6 | 69.4 | 8.5 | 15.96 | | High | High | High | High | High | High | High | High | High | High | High | 0 | 4.25E-06 | 3.28 |
| FALSE | High | LGGSPTNG | [K].LGGSPT | 1xDeamida | P0C027 1xP | 0.113753 | 0.009241 | 1 | 1 | 3 | P0C027 | Diphospho | P0C027 [14 | 0 | 2175.964 | 0 | -0.55 | 0.003705 | 5.19 | 118.8 | 81.2 | 8.15 | 11.56 | | High | High | High | High | High | High | High | High | High | High | High | 0.004667 | 0.05028 | 2.64 |
| FALSE | High | IGHHSTSDI | [R].IGHHST | 1xPhospho | Q3U3J1 1x | 0.000712 | 0 | 1 | 3 | 3 | Q3U3J1 | 2-oxoisova | Q3U3J1 [3 | 0 | 1841.802 | 0 | -1.52 | 0.016702 | 1.86 | 148.4 | 51.6 | 22.07 | 30.05 | | High | High | High | High | High | High | High | High | High | High | High | 0 | 0.000162 | 2.35 |
| FALSE | High | IGHHSTSDI | [R].IGHHST | 2xPhospho | Q3U3J1 2x | 0.002252 | 0.000109 | 1 | 3 | 1 | Q3U3J1 | 2-oxoisova | Q3U3J1 [3 | 0 | 1921.768 | 0 | -1.07 | 0.015519 | 12.45 | 128.3 | 71.7 | 17.35 | 28.99 | | High | High | High | High | High | High | High | High | High | High | High | 4.94E-05 | 0.000593 | 3.44 |
| FALSE | High | IGHHSTSDI | [R].IGHHST | 2xPhospho | Q3U3J1 2x | 0.000233 | 0 | 1 | 3 | 2 | Q3U3J1 | 2-oxoisova | Q3U3J1 [3 | 1 | 3386.477 | 0 | -0.69 | 0.008997 | 1.16 | 124.8 | 75.2 | 12.09 | 10.93 | | High | High | High | High | High | High | High | High | High | High | High | 0 | 0.000167 | 3.93 |
| FALSE | High | LGHVVMG | [K].LGHVV | 1xPhospho | Q3UIG0 1x | 1.03E-08 | 0 | 1 | 2 | 1 | Q3UIG0 | Eukaryotic | Q3UIG0 [3 | 0 | 2721.42 | 1 | -0.44 | 0.000813 | 3.62 | 115.2 | 84.8 | 6.31 | 9.78 | | High | High | High | High | High | High | High | High | High | High | High | 0 | 5.95E-10 | 4.46 |
| FALSE | High | LGGLLPAG | [K].LGGLLP | 1xPhospho | G5DDB7 1x | 2.54E-09 | 0 | 1 | 3 | 4 | G5DDB7 | Phosphoin | G5DDB7 [4 | 0 | 3668.6 | 0 | -1.74 | 0.000352 | 1.18 | 155.6 | 44.4 | 5.62 | 14.92 | | High | High | High | High | High | High | High | High | High | High | High | 0 | 1.24E-10 | 5.56 |
| FALSE | High | LGELTMQL | [K].LGELTM | 1xPhospho | Q9DCT6-3 | 0.000331 | 0 | 2 | 4 | 2 | Q9DCT6-3; | Isoform 3 c | Q9DCT6-3 | 0 | 2833.457 | 0 | -0.25 | 0.287075 | 19.04 | 104.1 | 95.9 | 11.55 | 11.81 | Shared | High | High | High | High | High | High | High | High | High | High | High | 0 | 6.92E-05 | 4.05 |
| FALSE | High | LGELTMQL | [K].LGELTM | 1xPhospho | Q9DCT6-3 | 1.18E-05 | 0 | 2 | 4 | 1 | Q9DCT6-3; | Isoform 3 c | Q9DCT6-3 | 0 | 2817.462 | 0 | -0.05 | 0.552232 | 39.8 | 97.7 | 102.3 | 11.71 | 29.82 | Shared | High | High | High | High | High | High | High | High | High | High | High | 0 | 1.62E-06 | 4.54 |
| FALSE | High | IGELGAPE | [R].IGELGA | 1xPhospho | Q9D0F4 1x | 0.011755 | 0.000889 | 1 | 2 | 2 | Q9D0F4 | NF-kappa- | Q9D0F4 [1 | 0 | 2091.129 | 0 | 0.45 | 0.00759 | 6.55 | 86 | 114 | 4.66 | 7.82 | | High | High | High | High | High | High | High | High | High | High | High | 0.000449 | 0.003771 | 3.07 |
| FALSE | High | IFSQDSLCS | [R].IFSQDS | 1xCarbami | E9Q9Q2 1x | 0.029844 | 0.001857 | 1 | 5 | 2 | E9Q9Q2 | R3H domai | E9Q9Q2 [2 | 0 | 2598.256 | 0 | 0.87 | 0.000964 | 9.88 | 72.5 | 127.5 | 7.27 | 4.86 | | High | High | High | High | High | High | High | High | High | High | High | 0.001011 | 0.01075 | 2.07 |
| FALSE | High | LFSQGQDV | [R].LFSQG | 1xPhospho | E9Q9C3 1x | 3.9E-05 | 0 | 1 | 4 | 2 | E9Q9C3 | Afadin OS= | E9Q9C3 [1 | 0 | 1761.882 | 0 | 0.28 | 0.043679 | 5.64 | 90.4 | 109.6 | 9.05 | 15.45 | | High | High | High | High | High | High | High | High | High | High | High | 0 | 6.26E-06 | 3.04 |
| FALSE | High | LFSSEETK | [R].LFSSEE | 1xPhospho | F8WJA0 1x | 0.110202 | 0.008961 | 1 | 6 | 1 | F8WJA0 | ATP-depen | F8WJA0 [1 | 0 | 1478.754 | 0 | 0.06 | 0.546276 | 13.03 | 97.8 | 102.2 | 3.5 | 8.74 | | High | High | High | High | High | High | High | High | High | High | High | 0.004462 | 0.04819 | 2.73 |
| FALSE | High | LFSSSGLK | [K].LFSSSG | 1xPhospho | B2RRE0 1x | 0.020319 | 0.001472 | 1 | 3 | 2 | B2RRE0 | A kinase (P | B2RRE0 [5 | 0 | 1376.759 | 0 | -1.82 | 0.003526 | 2.95 | 155.7 | 44.3 | 12.71 | 21.73 | | High | High | High | High | High | High | High | High | High | High | High | 0.000786 | 0.006931 | 1.65 |
| FALSE | High | LFTDSWAV | [R].LFTDSV | 2xPhospho | V9H0W7 2 | 0.049549 | 0.00361 | 1 | 2 | 2 | V9H0W7 | Pro-Pol-dU | V9H0W7 [6 | 0 | 2785.319 | 0 | 0.77 | 0.090869 | 45.01 | 78.3 | 121.7 | 27.9 | 23.02 | | High | High | High | High | High | High | High | High | High | High | High | 0.002662 | 0.02848 | 2.82 |
| FALSE | High | IFVGGLSPD | [K].IFVGGL | 1xPhospho | Q60668 1x | 7.96E-05 | 0 | 1 | 9 | 2 | Q60668 | Heterogen | Q60668 [1 | 0 | 2027.05 | 0 | -1.15 | 0.000153 | 0.6 | 137.8 | 62.2 | 1.88 | 5.25 | | High | High | High | High | High | High | High | High | High | High | High | 2.58E-05 | 0.00049 | 2.42 |
| FALSE | High | LFVLLPEQS | [R].LFVLLP | 1xPhospho | A0A1L1SV7 | 0.000108 | 0 | 1 | 3 | 1 | A0A1L1SV7 | Ubiquitin c | A0A1L1SV7 | 0 | 2203.182 | 0 | -0.56 | 0.009589 | 2.56 | 117.3 | 82.7 | 5.07 | 10.28 | | High | High | High | High | High | High | High | High | High | High | High | 0 | 1.97E-05 | 2.86 |
| FALSE | High | IFYYFSNSP | [K].IFYYFS | 1xPhospho | Q64700 1x | 0.001411 | 5.66E-05 | 1 | 5 | 1 | Q64700 | Retinoblast | Q64700 [1 | 0 | 1890.944 | 0 | 2.01 | 0.000337 | 6.94 | 38.4 | 161.6 | 2.72 | 7.78 | | High | High | High | High | High | High | High | High | High | High | High | 2.58E-05 | 0.000352 | 3.84 |
| FALSE | High | LGADESEEI | [R].LGADE | 1xPhospho | A2A9Q2 1x | 0.005275 | 0.000599 | 1 | 4 | 1 | A2A9Q2 | Nardilysin, | A2A9Q2 [8 | 0 | 1500.642 | 0 | 0.83 | 0.015809 | 9.49 | 70.7 | 129.3 | 10.11 | 18.23 | | High | High | High | High | High | High | High | High | High | High | High | 0.000267 | 0.001529 | 2.33 |
| FALSE | High | LGADGAGS | [R].LGADG. | 1xPhospho | Q4QRL3-2 | 0.000211 | 0 | 1 | 3 | 1 | Q4QRL3-2 | Isoform 2 c | Q4QRL3-2 | 0 | 2622.25 | 0 | 0.68 | 0.009769 | 27.36 | 76.9 | 123.1 | 21.98 | 8.46 | | High | High | High | High | High | High | High | High | High | High | High | 0 | 4.14E-05 | 4.62 |
| FALSE | High | LGAESPSIA | [R].LGAESF | 1xPhospho | Q68FH0 1x | 3.8E-05 | 0 | 1 | 7 | 2 | Q68FH0 | Plakophilin | Q68FH0 [7 | 0 | 2103.062 | 0 | 0.04 | 0.8287 | 6.61 | 100.4 | 99.6 | 4.87 | 7.4 | | High | High | High | High | High | High | High | High | High | High | High | 0 | 6.04E-06 | 2.13 |
| FALSE | High | LGAGEQAL | [R].LGAGE | 1xPhospho | Q80WC3 1 | 0.000274 | 0 | 1 | 2 | 1 | Q80WC3 | Trinucleoti | Q80WC3 [1 | 0 | 2477.285 | 0 | -0.78 | 0.007657 | 5.74 | 127.3 | 72.7 | 11.26 | 12.52 | | High | High | High | High | High | High | High | High | High | High | High | 0 | 5.58E-05 | 2.81 |
| FALSE | High | LGAPENSG | [R].LGAPEN | 1xPhospho | F8VPU2 1x | 9.53E-08 | 0 | 1 | 3 | 1 | F8VPU2 | FERM, Rho | F8VPU2 [1 | 0 | 1752.873 | 0 | -0.27 | 0.458311 | 6.16 | 105.6 | 94.4 | 3.86 | 16.16 | | High | High | High | High | High | High | High | High | High | High | High | 0 | 7.26E-09 | 3.26 |
| FALSE | High | LGASLTPD | [R].LGASLT | 1xPhospho | S4R1P5 1xP | 0.000237 | 0 | 1 | 6 | 2 | S4R1P5 | Dystonin O | S4R1P5 [26 | 0 | 1618.794 | 0 | 0.61 | 0.023289 | 13.89 | 75.1 | 124.9 | 9.96 | 18.04 | | High | High | High | High | High | High | High | High | High | High | High | 0.000814 | 0.007497 | 1.91 |
| FALSE | High | LGASNSPG | [K].LGASN | 1xPhospho | B2RY56 1xP | 1.58E-07 | 0 | 1 | 2 | 6 | B2RY56 | RNA-bindir | B2RY56 [6 | 0 | 1893.984 | 0 | -0.32 | 0.019604 | 5.12 | 111.9 | 88.1 | 6.97 | 5.43 | | High | High | High | High | High | High | High | High | High | High | High | 0 | 1.28E-08 | 3.42 |
| FALSE | High | LGASNSPG | [K].LGASN | 1xDeamida | B2RY56 1xP | 0.000967 | 0 | 1 | 2 | 4 | B2RY56 | RNA-bindir | B2RY56 [6 | 0 | 1894.968 | 0 | -0.01 | 0.421383 | 2.75 | 99.9 | 100.1 | 1.67 | 5.26 | | High | High | High | High | High | High | High | High | High | High | High | 0.000832 | 0.007905 | 3.06 |
| FALSE | High | LGASNSPG | [K].LGASN | 2xPhospho | B2RY56 2xP | 0.026652 | 0.001676 | 1 | 2 | 2 | B2RY56 | RNA-bindir | B2RY56 [6 | 0 | 1973.95 | 0 | -0.38 | 0.001524 | 1.6 | 109.8 | 90.2 | 7.11 | 8.7 | | High | High | High | High | High | High | High | High | High | High | High | 0.004341 | 0.04696 | 2.94 |
| FALSE | High | IGCSDVPDI | [K].IGCSDV | 1xCarbami | B2RUC3 1x | 0.017192 | 0.001204 | 1 | 1 | 1 | B2RUC3 | Excision re | B2RUC3 [1 | 1 | 2189.037 | 0 | 1.62 | 0.000213 | 9.25 | 47.4 | 152.6 | 10.29 | 4.93 | | High | High | High | High | High | High | High | High | High | High | High | 0.000638 | 0.005756 | 2.06 |
| FALSE | High | LGCTTGVS | [R].LGCTT | 2xCarbami | Q8BYN3 1x | 0.000475 | 0 | 1 | 1 | 2 | Q8BYN3 | Inositol-tet | Q8BYN3 [3 | 0 | 2887.425 | 0 | -0.46 | 0.268295 | 20.3 | 115.9 | 84.1 | 22.29 | 20.63 | | High | High | High | High | High | High | High | High | High | High | High | 0 | 0.000316 | 2.62 |
| FALSE | High | LGDASDTE | [R].LGDASI | 1xPhospho | Q80U49 1x | 0.001411 | 5.66E-05 | 1 | 2 | 1 | Q80U49 | Centrosom | Q80U49 [1 | 0 | 1743.764 | 0 | 0.56 | 0.019319 | 3.4 | 77.9 | 122.1 | 12.81 | 15.95 | | High | High | High | High | High | High | High | High | High | High | High | 2.58E-05 | 0.000352 | 2.22 |
| FALSE | High | IGDGFPFDI | [R].IGDGFF | 1xPhospho | Q6NVF4 1x | 0.001489 | 5.66E-05 | 1 | 1 | 2 | Q6NVF4 | DNA helica | Q6NVF4 [1 | 0 | 1833.815 | 0 | -0.46 | 0.006609 | 7.19 | 119.9 | 80.1 | 8.66 | 5.79 | | High | High | High | High | High | High | High | High | High | High | High | 2.58E-05 | 0.000374 | 2.73 |
| FALSE | High | IGDGFPFDI | [R].IGDGFF | 1xPhospho | Q6NVF4 1x | 4.74E-07 | 0 | 1 | 1 | 6 | Q6NVF4 | DNA helica | Q6NVF4 [1 | 0 | 2062.977 | 0 | -1.02 | 0.001043 | 1.52 | 134.6 | 65.4 | 3.46 | 10.41 | | High | High | High | High | High | High | High | High | High | High | High | 0 | 4.38E-08 | 4.5 |
| FALSE | High | LGEASDSEI | [R].LGEASD | 1xPhospho | Q6A065 1x | 0.002463 | 0.000109 | 1 | 3 | 4 | Q6A065 | Centrosom | Q6A065 [1 | 0 | 1958.936 | 0 | 1.04 | 0.001401 | 13.5 | 62.5 | 137.5 | 12.48 | 10.79 | | High | High | High | High | High | High | High | High | High | High | High | 7.19E-05 | 0.000868 | 2.75 |
| FALSE | High | LGEELVAIV | [R].LGEELV | 1xPhospho | A0A140LI3 | 3.98E-05 | 0 | 1 | 1 | 1 | A0A140LI3 | Zinc finger | A0A140LI3 | 0 | 2725.437 | 0 | -0.04 | 0.576438 | 9.55 | 104 | 96 | 9.63 | 12.48 | | High | High | High | High | High | High | High | High | High | High | High | 0 | 6.35E-06 | 3.68 |
| FALSE | High | IGEEQSAEI | [K].IGEEQS | 1xPhospho | Q3UJI2 1xP | 1.88E-08 | 0 | 1 | 6 | 1 | Q3UJI2 | Putative ur | Q3UJI2 [34 | 0 | 3385.635 | 0 | 2.39 | 0.000261 | 15.23 | 31.4 | 168.6 | 8.28 | 6.11 | | High | High | High | High | High | High | High | High | High | High | High | 0 | 1.17E-09 | 4.23 |
| FALSE | High | IGEGTYGV | [K].IGEGTY | 1xPhospho | P97377 1xP | 0.06169 | 0.004646 | 2 | 7 | 1 | P97377; P1 | Cyclin-dep | P97377 [1 | 0 | 1494.744 | 0 | -1.06 | 0.365045 | 10.09 | 138.8 | 61.2 | 91.3 | 79.57 | Shared | High | High | High | High | High | High | High | High | High | High | High | 0.002381 | 0.02458 | 1.78 |
| FALSE | High | IGEGTYGV | [K].IGEGTY | 1xPhospho | P97377 1xP | 1.97E-06 | 0 | 2 | 7 | 19 | P97377; P1 | Cyclin-dep | P97377 [1 | 0 | 1723.907 | 0 | -1.07 | 0.000166 | 2.04 | 137.6 | 62.4 | 4.1 | 9.29 | Shared | High | High | High | High | High | High | High | High | High | High | High | 0 | 2.18E-07 | 3.27 |
| FALSE | High | IGEGTYGV | [K].IGEGTY | 1xPhospho | P11440 1xP | 0.107603 | 0.008634 | 1 | 3 | 1 | P11440 | Cyclin-dep | P11440 [1 | 1 | 1937.03 | 0 | -1.04 | 0.00445 | 4.89 | 136.1 | 63.9 | 14.32 | 19.36 | | High | High | High | High | High | High | High | High | High | High | High | 0.004327 | 0.04689 | 2.6 |
| FALSE | High | IGELGAPE | [R].IGELGA | 1xPhospho | Q9D0F4 1x | 3.6E-05 | 0 | 1 | 2 | 1 | Q9D0F4 | NF-kappa- | Q9D0F4 [1 | 0 | 1861.966 | 0 | 0.82 | 0.69426 | 46.68 | 79.2 | 120.8 | 73.43 | 60.3 | | High | High | High | High | High | High | High | High | High | High | High | 0 | 5.69E-06 | 2.7 |
| FALSE | High | LGHVVMG | [K].LGHVV | 1xPhospho | Q3UIG0 1x | 1.69E-08 | 0 | 1 | 2 | 1 | Q3UIG0 | Eukaryotic | Q3UIG0 [3 | 0 | 2737.415 | 1 | -0.3 | 0.006384 | 1.25 | 114 | 86 | 9.49 | 5.25 | | High | High | High | High | High | High | High | High | High | High | High | 0 | 1.03E-09 | 4.6 |
| FALSE | High | LQMEAPHI | [K].LQMEA | 1xPhospho | P60843 1xP | 0.021418 | 0.001507 | 1 | 9 | 1 | P60843 | Eukaryotic | P60843 [14 | 0 | 1943.998 | 0 | -0.42 | 0.005002 | 3.66 | 113.8 | 86.2 | 8.72 | 4.75 | | High | High | High | High | High | High | High | High | High | High | High | 0.000801 | 0.007368 | 1.48 |
| FALSE | High | LGLAVIHGI | [R].LGLAVI | 1xCarbami | B1AT82 1xP | 0.000668 | 0 | 1 | 3 | 2 | B1AT82 | MCG6846, | B1AT82 [2 | 1 | 3630.752 | 0 | -0.13 | 0.426785 | 1.94 | 104.8 | 95.2 | 2.19 | 12.63 | | High | High | High | High | High | High | High | High | High | High | High | 0 | 0.000152 | 4.22 |
| FALSE | High | LGIAVIHGE | [R].LGIAVI | 1xDeamida | Q8R574 1x | 0.047086 | 0.00354 | 1 | 2 | 1 | Q8R574 | Phosphorit | Q8R574 [2 | 1 | 3279.603 | 0 | -1.03 | 0.001802 | 5.44 | 134.8 | 65.2 | 9.22 | 8.17 | | High | High | High | High | High | High | High | High | High | High | High | 0.001812 | 0.01798 | 3.43 |
| FALSE | High | LGSIAEIDL | [R].LGSIAE | 1xPhospho | Q99MR6 1 | 0.01588 | 0.001124 | 1 | 5 | 1 | Q99MR6 | Serrate RN | Q99MR6 [1 | 0 | 2288.274 | 0 | 0.51 | 0.008105 | 14.85 | 79.2 | 120.8 | 10.85 | 14.04 | | High | High | High | High | High | High | High | High | High | High | High | 0.000585 | 0.005268 | 2.84 |
| FALSE | High | LGSIAEIDL | [R].LGSIAE | 1xPhospho | Q99MR6 1 | 0.0002 | 0 | 1 | 5 | 1 | Q99MR6 | Serrate RN | Q99MR6 [1 | 0 | 2304.269 | 0 | 1.01 | 0.000599 | 4.3 | 71 | 129 | 12.54 | 9.58 | | High | High | High | High | High | High | High | High | High | High | High | 0 | 3.91E-05 | 2.27 |
| FALSE | High | IGSLIDVNC | [K].IGSLID | 1xPhospho | A0A023T7 | 0.000237 | 0 | 1 | 4 | 1 | A0A023T7 | Mago nash | A0A023T7 | 0 | 1711.94 | 0 | -1.03 | 0.002136 | 4.68 | 136.7 | 63.3 | 9.83 | 7.38 | | High | High | High | High | High | High | High | High | High | High | High | 0 | 4.74E-05 | 2.36 |
| FALSE | High | LGSLNFSTE | [K].LGSLNF | 1xPhospho | Q76I79 1xP | 1.39E-06 | 0 | 1 | 3 | 4 | Q76I79 | Protein ph | Q76I79 [97 | 0 | 3340.572 | 0 | 0.64 | 0.014138 | 16.08 | 78.2 | 121.8 | 6.01 | 14.96 | | High | High | High | High | High | High | High | High | High | High | High | 0 | 1.46E-07 | 5.61 |
| FALSE | High | LGSLTPEIV | [K].LGSLTP | 1xPhospho | Q0KL01 1x | 7.91E-06 | 0 | 1 | 1 | 2 | Q0KL01 | UBX domai | Q0KL01 [2 | 0 | 2653.326 | 0 | -0.84 | 0.004886 | 4.79 | 129.4 | 70.6 | 10.51 | 14.25 | | High | High | High | High | High | High | High | High | High | High | High | 0 | 3.29E-05 | 3.83 |
| FALSE | High | LGSLTPEIV | [K].LGSLTP | 2xPhospho | Q0KL01 2x | 8.28E-06 | 0 | 1 | 1 | 3 | Q0KL01 | UBX domai | Q0KL01 [2 | 0 | 2733.292 | 0 | -1.16 | 0.000946 | 0.73 | 137.5 | 62.5 | 5.36 | 18.93 | | High | High | High | High | High | High | High | High | High | High | High | 0 | 1.09E-06 | 3.82 |
| FALSE | High | LGSPAGSP | [R].LGSPAG | 1xPhospho | Q6P5E8 1x | 0.096181 | 0.007535 | 1 | 2 | 1 | Q6P5E8 | Diacylglyce | Q6P5E8 [2 | 0 | 1676.893 | 0 | 0.22 | 0.133243 | 14.93 | 92.2 | 107.8 | 8.08 | 15 | | High | High | High | High | High | High | High | High | High | High | High | 0.003823 | 0.04125 | 2.07 |
| FALSE | High | LGSPAGSP | [R].LGSPAG | 2xPhospho | Q6P5E8 2x | 0.000622 | 0 | 1 | 2 | 2 | Q6P5E8 | Diacylglyce | Q6P5E8 [2 | 0 | 1756.86 | 0 | 0.68 | 0.001858 | 14.15 | 75.8 | 124.2 | 9.05 | 8.68 | | High | High | High | High | High | High | High | High | High | High | High | 0 | 0.00014 | 3.88 |
| FALSE | High | LGSPAHEA | [R].LGSPAH | 1xCarbami | A0A087WF | 1.79E-07 | 0 | 1 | 4 | 1 | A0A087WF | MyoD fami | A0A087WF | 0 | 2310.047 | 0 | 2.55 | 0.000217 | 9.39 | 30.1 | 169.9 | 13.66 | 7.98 | | High | High | High | High | High | High | High | High | High | High | High | 0 | 1.48E-08 | 4.05 |
| FALSE | High | LGSPDDGN | [R].LGSPDI | 1xPhospho | E9Q7G0 1x | 0.062209 | 0.004646 | 1 | 4 | 1 | E9Q7G0 | Nuclear mi | E9Q7G0 [1 | 0 | 2596.297 | 0 | -0.45 | 0.008488 | 2.13 | 114.3 | 85.7 | 8.56 | 5.24 | | High | High | High | High | High | High | High | High | High | High | High | 0.002381 | 0.02474 | 1.57 |
| FALSE | High | LGSPDSTSF | [K].LGSPDS | 1xPhospho | Q8BRG8 1x | 0.000687 | 0 | 1 | 5 | 7 | Q8BRG8 | Transmem | Q8BRG8 [2 | 0 | 2496.128 | 0 | -0.24 | 0.137056 | 15.02 | 108.3 | 91.7 | 10.39 | 14.22 | | High | High | High | High | High | High | High | High | High | High | High | 0 | 0.000156 | 2.29 |
| FALSE | High | LGSPIR | [R].LGSPIR. | 1xPhospho | A2ASY1 1xP | 0.096181 | 0.007535 | 1 | 6 | 2 | A2ASY1 | TAF3 RNA p | A2ASY1 [2 | 0 | 951.5226 | 0 | -0.35 | 0.002573 | 2.83 | 112.2 | 87.8 | 4.13 | 2.26 | | High | High | High | High | High | High | High | High | High | High | High | 0.003809 | 0.04109 | 1.45 |
| FALSE | High | LGSPPPHK | [K].LGSPPP | 1xPhospho | Q6A028 1x | 0.110202 | 0.008961 | 1 | 1 | 1 | Q6A028 | Switch-ass | Q6A028 [3 | 0 | 1370.76 | 0 | 1.22 | 0.000181 | 5.73 | 59.1 | 140.9 | 5.05 | 2.64 | | High | High | High | High | High | High | High | High | High | High | High | 0.00449 | 0.04859 | 1.86 |
| FALSE | High | LGSLADEFI | [R].LGSLAD | 1xPhospho | A0A0R4J18 | 0.000337 | 0 | 1 | 2 | 1 | A0A0R4J18 | X-ray repai | A0A0R4J18 | 0 | 1517.802 | 0 | 0.77 | 0.014454 | 12 | 76.4 | 123.6 | 16.14 | 7.97 | | High | High | High | High | High | High | High | High | High | High | High | 0 | 7.03E-05 | 2.12 |
| FALSE | High | LGSPVDGL | [R].LGSPVD | 1xPhospho | B0V2M3 1x | 0.052133 | 0.00384 | 1 | 2 | 1 | B0V2M3 | Zinc finger | B0V2M3 [2 | 1 | 2992.344 | 0 | -0.28 | 0.153014 | 15.82 | 109.6 | 90.4 | 16.08 | 14.26 | | High | High | High | High | High | High | High | High | High | High | High | 0.001999 | 0.02023 | 2.66 |
| FALSE | High | LGSSESLPC | [R].LGSSES | 1xCarbami | Q8BZ03 2x | 0.001544 | 5.66E-05 | 1 | 2 | 4 | Q8BZ03 | Serine/thre | Q8BZ03 [2 | 0 | 2124.912 | 0 | 0.67 | 0.004646 | 9.49 | 77.3 | 122.7 | 12.26 | 8.96 | | High | High | High | High | High | High | High | High | High | High | High | 9.42E-05 | 0.001 | 2.38 |
| FALSE | High | LGSSGLGS | [R].LGSSGL | 1xPhospho | A2AFQ0 1x | 3.12E-05 | 0 | 1 | 5 | 2 | A2AFQ0 | E3 ubiquiti | A2AFQ0 [3 | 0 | 1869.963 | 0 | -0.42 | 0.00939 | 4.26 | 113.7 | 86.3 | 3.41 | 10.42 | | High | High | High | High | High | High | High | High | High | High | High | 0 | 1.2E-05 | 2.52 |
| FALSE | High | LGSSPTSSC | [K].LGSSPT | 1xCarbami | Q5SXY1 1x | 5.88E-06 | 0 | 1 | 8 | 9 | Q5SXY1 | Cytospin-B | Q5SXY1 [1 | 0 | 2072.014 | 0 | 0.84 | 0.00251 | 2.77 | 72.3 | 127.7 | 2.42 | 10.75 | | High | High | High | High | High | High | High | High | High | High | High | 0 | 7.41E-07 | 3.62 |
| FALSE | High | LGSSSNLQI | [R].LGSSSN | 1xPhospho | E9Q9R9 1x | 0.033404 | 0.002195 | 1 | 5 | 1 | E9Q9R9 | Disks large | E9Q9R9 [1 | 0 | 1618.861 | 0 | 0.18 | 0.029324 | 5.63 | 92.6 | 107.4 | 1.9 | 9.65 | | High | High | High | High | High | High | High | High | High | High | High | 0.001151 | 0.01213 | 2.04 |
| FALSE | High | LGSTAPQV | [K].LGSTAF | 1xPhospho | Q3UVN5 1 | 1.63E-09 | 0 | 1 | 4 | 3 | Q3UVN5 | Putative ur | Q3UVN5 [2 | 0 | 2879.455 | 0 | 0.21 | 0.050066 | 11.67 | 93.8 | 106.2 | 5.75 | 4.44 | | High | High | High | High | High | High | High | High | High | High | High | 0 | 7.46E-11 | 5.05 |
| FALSE | High | LGSTEATIE | [K].LGSTEA | 1xPhospho | Q6ZQL4 1x | 0.000269 | 0 | 1 | 2 | 1 | Q6ZQL4 | WD repeat | Q6ZQL4 [4 | 0 | 1514.73 | 0 | -0.08 | 0.10711 | 4.48 | 105.9 | 94.1 | 5 | 7.62 | | High | High | High | High | High | High | High | High | High | High | High | 0 | 5.45E-05 | 2.13 |
| FALSE | High | IGSTENLK | [K].IGSTEN | 1xPhospho | A2A5Y6 1x | 0.030367 | 0.001857 | 1 | 11 | 1 | A2A5Y6 | Microtubu | A2A5Y6 [5 | 0 | 1170.597 | 0 | -0.24 | 0.846248 | 8.11 | 105.5 | 94.5 | 27.18 | 36.72 | | High | High | High | High | High | High | High | High | High | High | High | 0.001011 | 0.01088 | 1.92 |
| FALSE | High | IGSTENLK | [K].IGSTEN | 1xPhospho | A2A5Y6 1x | 0.002232 | 0.000109 | 1 | 11 | 3 | A2A5Y6 | Microtubu | A2A5Y6 [5 | 0 | 1399.76 | 0 | -0.28 | 0.223004 | 15.71 | 111.8 | 88.2 | 9.46 | 22.7 | | High | High | High | High | High | High | High | High | High | High | High | 4.94E-05 | 0.000585 | 1.71 |

| | | | | | | | | | | | | | | | | | | | | | | | | | | | | | | | | | | | | | | |
|---|---|---|---|---|---|---|---|---|---|---|---|---|---|---|---|---|---|---|---|---|---|---|---|---|---|---|---|---|---|---|---|---|---|---|---|---|---|---|
| FALSE | High | IGSTEVEKP | [R].IGSTEV | 1xPhospho | E9Q8T1 1x | 0.00172 | 5.66E-05 | 2 | 14 | 2 | E9Q8T1; Q | Transformi | E9Q8T1 [2 | 0 | 3446.834 | 0 | 1.05 | 0.013329 | 27.16 | 72.6 | 127.4 | 14.92 | 18.41 | Shared | High | High | High | High | High | High | High | High | High | High | High | 2.58E-05 | 0.000437 | 3.94 |
| FALSE | High | LGSTGGK | [R].LGSTG | 1xPhospho | E9PVB7 1x | 0.111965 | 0.009053 | 1 | 4 | 1 | E9PVB7 | DNA-bindi | E9PVB7 [4 | 0 | 1157.633 | 0 | 0.32 | 0.068652 | 19.05 | 86.9 | 113.1 | 20.34 | 13.14 | | High | High | High | High | High | High | High | High | High | High | High | 0.004543 | 0.04941 | 0.98 |
| FALSE | High | LGSTGSQP | [R].LGSTGS | 2xPhospho | Q501J7-3 2 | 0.088701 | 0.006669 | 1 | 5 | 2 | Q501J7-3 | Isoform 3 | Q501J7-3 [ | 0 | 2479.074 | 0 | 0.42 | 0.093976 | 28.61 | 83.8 | 116.2 | 20.41 | 23.77 | | High | High | High | High | High | High | High | High | High | High | High | 0.003461 | 0.03754 | 1.66 |
| FALSE | High | LGSTGSQP | [R].LGSTGS | 1xPhospho | Q501J7-3 1 | 3.35E-05 | 0 | 1 | 5 | 1 | Q501J7-3 | Isoform 3 | Q501J7-3 [ | 0 | 2628.271 | 0 | 1.27 | 0.00392 | 12.49 | 56.8 | 143.2 | 17.08 | 10.77 | | High | High | High | High | High | High | High | High | High | High | High | 0 | 5.26E-06 | 2.66 |
| FALSE | High | LGSTGSQP | [R].LGSTGS | 2xPhospho | Q501J7-3 2 | 0.002376 | 0.000109 | 1 | 5 | 1 | Q501J7-3 | Isoform 3 | Q501J7-3 [ | 0 | 2708.237 | 0 | 1.35 | 0.002137 | 18.42 | 57.3 | 142.7 | 7.7 | 10.89 | | High | High | High | High | High | High | High | High | High | High | High | 4.94E-05 | 0.000628 | 2.49 |
| FALSE | High | LGSSESLPC | [R].LGSSES | 1xCarbami | Q8BZ03 1x | 0.000337 | 0 | 1 | 2 | 2 | Q8BZ03 | Serine/thr | Q8BZ03 [2 | 0 | 2044.946 | 0 | 1.1 | 0.00021 | 2.64 | 63.6 | 136.4 | 7.56 | 4.17 | | High | High | High | High | High | High | High | High | High | High | High | 0 | 0.00013 | 2.5 |
| FALSE | High | LGSLADEF | [R].LGSLAD | 1xPhospho | A0A0R4J18 | 0.017344 | 0.001204 | 1 | 2 | 1 | A0A0R4J18 | X-ray repai | A0A0R4J18 | 0 | 1288.639 | 0 | 0.8 | 0.140717 | 24.5 | 74.9 | 125.1 | 16.61 | 29.79 | | High | High | High | High | High | High | High | High | High | High | High | 0.000656 | 0.00583 | 2.43 |
| FALSE | High | LGSGPDSE | [R].LGSGP | 1xPhospho | E9Q9M8 1 | 0.036103 | 0.002342 | 1 | 3 | 1 | E9Q9M8 | PWWP dor | E9Q9M8 [2 | 0 | 1363.62 | 0 | 2.32 | 0.000246 | 1.8 | 33.1 | 166.9 | 7.43 | 7.89 | | High | High | High | High | High | High | High | High | High | High | High | 0.001279 | 0.01333 | 1.26 |
| FALSE | High | LGSFGSITR | [R].LGSFGS | 1xPhospho | Q8VHX6 1x | 0.011246 | 0.000772 | 1 | 3 | 2 | Q8VHX6 | Filamin-C | Q8VHX6 [2 | 0 | 1246.639 | 0 | -0.97 | 0.005619 | 7.62 | 137.2 | 62.8 | 12.92 | 19.76 | | High | High | High | High | High | High | High | High | High | High | High | 0.000418 | 0.003593 | 1.18 |
| FALSE | High | LGIAVIHGE | [R].LGIAVI | 1xPhospho | Q8R574 1x | 0.002355 | 0.000109 | 1 | 2 | 2 | Q8R574 | Phosphori | Q8R574 [2 | 1 | 3294.614 | 0 | -0.66 | 0.000467 | 2.66 | 124.3 | 75.7 | 5.24 | 8.57 | | High | High | High | High | High | High | High | High | High | High | High | 4.94E-05 | 0.000622 | 3.64 |
| FALSE | High | LGLGLDDE | [K].LGLGL | 1xPhospho | B2RWS6 1 | 1.97E-05 | 0 | 1 | 1 | 1 | B2RWS6 | Histone ac | B2RWS6 [1 | 0 | 2753.258 | 0 | -0.22 | 0.099213 | 11.11 | 106.2 | 93.8 | 4.45 | 7.63 | | High | High | High | High | High | High | High | High | High | High | High | 0 | 2.88E-06 | 4.25 |
| FALSE | High | LGIHEDSTN | [K].LGIHED | 1xPhospho | P11499 1x | 0.00614 | 0.00062 | 1 | 4 | 1 | P11499 | Heat shock | P11499 [43 | 0 | 1450.689 | 0 | -0.08 | 0.270912 | 6.44 | 101.9 | 98.1 | 5.57 | 6.7 | | High | High | High | High | High | High | High | High | High | High | High | 0.000275 | 0.001811 | 1.92 |
| FALSE | High | LGLHVSPS | [K].LGLHV | 1xPhospho | G3UWZ0 1 | 4.62E-07 | 0 | 1 | 4 | 2 | G3UWZ0 | Bromodom | G3UWZ0 [ | 0 | 2558.4 | 0 | 0.49 | 0.04804 | 9.2 | 86.3 | 113.7 | 12.04 | 15.92 | | High | High | High | High | High | High | High | High | High | High | High | 0 | 4.25E-08 | 3.84 |
| FALSE | High | LGLLSQES | [K].LGLLSQ | 1xPhospho | Q9EP71 1x | 5.21E-05 | 0 | 1 | 1 | 1 | Q9EP71 | Ankycorbi | Q9EP71 [5 | 0 | 2055.011 | 0 | 0.53 | 0.006581 | 12.7 | 81.9 | 118.1 | 3.45 | 12.11 | | High | High | High | High | High | High | High | High | High | High | High | 0 | 8.64E-06 | 4.9 |
| FALSE | High | LGIPETISPF | [K].LGIPET | 1xPhospho | E9Q6J5-3 1 | 0.005978 | 0.00062 | 1 | 2 | 2 | E9Q6J5-3 | Isoform Ne | E9Q6J5-3 [ | 0 | 1391.75 | 0 | 0.62 | 0.000928 | 3.99 | 76.9 | 123.1 | 5.54 | 8.42 | | High | High | High | High | High | High | High | High | High | High | High | 0.000903 | 0.009532 | 1.92 |
| FALSE | High | IGLPHSIK | [K].IGLPHS | 1xPhospho | Q8VH51 1x | 0.042141 | 0.003162 | 1 | 5 | 1 | Q8VH51 | RNA-bindi | Q8VH51 [1 | 0 | 1402.822 | 0 | 0.36 | 0.246891 | 19.94 | 98.5 | 101.5 | 16.4 | 20.44 | | High | High | High | High | High | High | High | High | High | High | High | 0.001625 | 0.0158 | 1.85 |
| FALSE | High | LGLPLPSDI | [R].LGLPLP | 1xPhospho | Q3TWW5 | 0.007951 | 0.000736 | 1 | 2 | 2 | Q3TWW5 | Putative u | Q3TWW5 | 0 | 3769.008 | 0 | 0.33 | 0.042084 | 12.9 | 92.6 | 107.4 | 10.28 | 9.1 | | High | High | High | High | High | High | High | High | High | High | High | 0.001812 | 0.01842 | 3.2 |
| FALSE | High | LGLPLPSDI | [R].LGLPLP | 1xDeamida | Q3TWW5 | 6.88E-08 | 0 | 1 | 2 | 4 | Q3TWW5 | Putative u | Q3TWW5 | 0 | 3769.992 | 0 | 0.21 | 0.051254 | 16.88 | 92.2 | 107.8 | 10.75 | 9.46 | | High | High | High | High | High | High | High | High | High | High | High | 0 | 5.03E-09 | 5.24 |
| FALSE | High | LGLPPLTPE | [K].LGLPPL | 1xPhospho | Q3UEB3 1x | 7.47E-05 | 0 | 1 | 3 | 1 | Q3UEB3 | Poly(U)-bi | Q3UEB3 [5 | 0 | 2300.267 | 0 | 0.49 | 0.003426 | 5.95 | 85.1 | 114.9 | 8.67 | 2.91 | | High | High | High | High | High | High | High | High | High | High | High | 0 | 0.000013 | 3.05 |
| FALSE | High | LGLPTSPLS | [R].LGLPTS | 1xPhospho | Q6P5D4 1x | 0.001674 | 5.66E-05 | 1 | 1 | 2 | Q6P5D4 | Centrosom | Q6P5D4 [1 | 0 | 1852.06 | 0 | 0 | 0.754096 | 23.42 | 99.8 | 100.2 | 10.61 | 11.04 | | High | High | High | High | High | High | High | High | High | High | High | 0.00014 | 0.001098 | 1.95 |
| FALSE | High | LGLTPPPEF | [R].LGLTPP | 1xPhospho | Q3UT19 1x | 0.041781 | 0.003126 | 1 | 2 | 3 | Q3UT19 | Putative u | Q3UT19 [4 | 0 | 1586.896 | 0 | -0.77 | 0.002162 | 2 | 126 | 74 | 5.29 | 8.04 | | High | High | High | High | High | High | High | High | High | High | High | 0.002279 | 0.02373 | 2.49 |
| FALSE | High | LGNDVISQ | [K].LGNDV | 1xCarbami | Q6EDY6 1x | 4.01E-05 | 0 | 1 | 4 | 1 | Q6EDY6 | F-actin-un | Q6EDY6 [1 | 0 | 2484.128 | 0 | 1.39 | 0.000806 | 7.97 | 58.4 | 141.6 | 9.12 | 7.52 | | High | High | High | High | High | High | High | High | High | High | High | 0 | 6.44E-06 | 4.33 |
| FALSE | High | IGNEESDLE | [K].IGNEES | 1xCarbami | Q4U2R1 1x | 1.57E-08 | 0 | 1 | 3 | 3 | Q4U2R1 | E3 ubiquiti | Q4U2R1 [1 | 0 | 3103.506 | 0 | 1.01 | 0.000281 | 7.24 | 66.5 | 133.5 | 3.29 | 4.51 | | High | High | High | High | High | High | High | High | High | High | High | 0 | 1.14E-05 | 4.93 |
| FALSE | High | LGNGLSPE | [R].LGNGL | 1xPhospho | Q4VBD9 1x | 0.000738 | 0 | 1 | 3 | 3 | Q4VBD9 | GDNF-indu | Q4VBD9 [1 | 0 | 1737.919 | 0 | -0.49 | 0.00109 | 3.24 | 118.2 | 81.8 | 3.85 | 3.01 | | High | High | High | High | High | High | High | High | High | High | High | 2.58E-05 | 0.000362 | 2.94 |
| FALSE | High | LGNGLSPE | [R].LGNGL | 1xDeamida | Q4VBD9 1x | 1.31E-05 | 0 | 1 | 3 | 1 | Q4VBD9 | GDNF-indu | Q4VBD9 [1 | 0 | 1738.903 | 0 | -0.18 | 0.087893 | 8.42 | 106.1 | 93.9 | 6.69 | 9.05 | | High | High | High | High | High | High | High | High | High | High | High | 0 | 1.83E-06 | 2.78 |
| FALSE | High | LGNPYCSP | [R].LGNPY | 1xCarbami | A2AM67 1 | 3.94E-05 | 0 | 1 | 3 | 2 | A2AM67 | SH2 domai | A2AM67 [2 | 0 | 1685.828 | 0 | 2.06 | 0.000695 | 21.53 | 39.6 | 160.4 | 6.95 | 10.37 | | High | High | High | High | High | High | High | High | High | High | High | 0 | 6.32E-06 | 2.74 |
| FALSE | High | LGPGLPQD | [R].LGPGLF | 1xPhospho | O54825 1x | 1.32E-05 | 0 | 1 | 1 | 13 | O54825 | Bystin OS= | O54825 [8 | 0 | 2979.391 | 0 | -0.09 | 0.05986 | 5.48 | 103.9 | 96.1 | 1.87 | 5.76 | | High | High | High | High | High | High | High | High | High | High | High | 0 | 1.86E-06 | 4.36 |
| FALSE | High | IGPLGLSPK | [K].IGPLGL | 1xPhospho | Q3TIQ2 1x | 0.00129 | 0 | 1 | 4 | 1 | Q3TIQ2 | Putative u | Q3TIQ2 [3 | 0 | 1419.838 | 0 | -0.35 | 0.00935 | 4.98 | 116.1 | 83.9 | 8.88 | 10.17 | | High | High | High | High | High | High | High | High | High | High | High | 0 | 0.000317 | 2.04 |
| FALSE | High | IGPPSPGD | [K].IGPPSP | 1xPhospho | O70494 1x | 7.86E-07 | 0 | 1 | 1 | 1 | O70494 | Transcripti | O70494 [6 | 0 | 4135.977 | 0 | -0.89 | 0.004563 | 1.91 | 128.9 | 71.1 | 17.32 | 13.04 | | High | High | High | High | High | High | High | High | High | High | High | 0 | 7.77E-08 | 4.81 |
| FALSE | High | LGPQDSDP | [K].LGPQD | 1xCarbami | Q0GNC1 1 | 2.59E-06 | 0 | 1 | 3 | 1 | Q0GNC1 | Inverted fo | Q0GNC1 [1 | 0 | 2850.307 | 0 | 0.66 | 0.016677 | 9.26 | 82.5 | 117.5 | 12.65 | 12.12 | | High | High | High | High | High | High | High | High | High | High | High | 0 | 2.96E-07 | 3.98 |
| FALSE | High | LGPSSPAHS | [K].LGPSSP | 1xPhospho | A2AAY5 1x | 0.001437 | 5.66E-05 | 1 | 2 | 2 | A2AAY5 | SH3 and PX | A2AAY5 [2 | 0 | 2145.054 | 0 | 2.26 | 0.000679 | 6.4 | 31.8 | 168.2 | 7.52 | 10.84 | | High | High | High | High | High | High | High | High | High | High | High | 2.58E-05 | 0.000356 | 2.29 |
| FALSE | High | LGPVSPAPS | [R].LGPVSF | 1xPhospho | Q8C9B9 1x | 7.41E-05 | 0 | 1 | 3 | 1 | Q8C9B9 | Death-indu | Q8C9B9 [6 | 0 | 1957.011 | 0 | -0.11 | 0.137679 | 11.3 | 104.1 | 95.9 | 7.97 | 6.02 | | High | High | High | High | High | High | High | High | High | High | High | 0 | 1.29E-05 | 3.1 |
| FALSE | High | LGQGVLVC | [K].LGQGV | 1xPhospho | Q8VBV3 1x | 2.06E-06 | 0 | 1 | 2 | 1 | Q8VBV3 | Exosome c | Q8VBV3 [1 | 0 | 2062.208 | 0 | -0.48 | 0.004896 | 2.21 | 120.4 | 79.6 | 8.34 | 6.88 | | High | High | High | High | High | High | High | High | High | High | High | 0 | 2.29E-07 | 4.33 |
| FALSE | High | LGRGSVSD | [R].LGRGS | 1xCarbami | A0A0R4J0I | 0.027121 | 0.001676 | 1 | 2 | 1 | A0A0R4J0I | ETS domai | A0A0R4J0I | 1 | 2971.319 | 0 | -0.49 | 0.095596 | 17.57 | 115.3 | 84.7 | 23.72 | 17.49 | | High | High | High | High | High | High | High | High | High | High | High | 0.00092 | 0.009613 | 3.32 |
| FALSE | High | LGSDEFCV | [K].LGSDEF | 1xCarbami | E9QLG3 1x | 0.004256 | 0.00036 | 1 | 4 | 1 | E9QLG3 | MAX gene- | E9QLG3 [2 | 0 | 1575.708 | 0 | -0.96 | 0.001634 | 3.31 | 130.6 | 69.4 | 7.42 | 4.42 | | High | High | High | High | High | High | High | High | High | High | High | 0.000161 | 0.001208 | 1.78 |
| FALSE | High | IGSDPLAYE | [R].IGSDPL | 1xPhospho | Q3U3A6 1x | 0.002486 | 0.000109 | 1 | 5 | 2 | Q3U3A6 | Sodium/hy | Q3U3A6 [7 | 0 | 1498.739 | 0 | 0.5 | 0.367444 | 21.87 | 85.2 | 114.8 | 25.03 | 35.37 | | High | High | High | High | High | High | High | High | High | High | High | 4.94E-05 | 0.000661 | 2.4 |
| FALSE | High | IGSDPLAYE | [R].IGSDPL | 1xPhospho | Q3U3A6 1x | 0.001337 | 0 | 1 | 5 | 2 | Q3U3A6 | Sodium/hy | Q3U3A6 [7 | 0 | 1727.902 | 0 | 0.41 | 0.012867 | 3.86 | 84.2 | 115.8 | 9.19 | 6.91 | | High | High | High | High | High | High | High | High | High | High | High | 0 | 0.00033 | 3.18 |
| FALSE | High | LGSEEEVDI | [R].LGSEEE | 1xPhospho | A2A5Y6 1x | 3.15E-06 | 0 | 1 | 3 | 2 | A2A5Y6 | Microtubu | A2A5Y6 [2 | 0 | 3583.601 | 0 | 1.29 | 0.003708 | 33.38 | 61.2 | 138.8 | 17.65 | 8.9 | | High | High | High | High | High | High | High | High | High | High | High | 0 | 3.68E-07 | 5 |
| FALSE | High | LGIAVIHGE | [R].LGIAVI | 1xPhospho | Q8R574 1x | 0.009756 | 0.000736 | 1 | 2 | 1 | Q8R574 | Phosphori | Q8R574 [2 | 1 | 3278.619 | 0 | -1.13 | 0.000223 | 1.05 | 137.8 | 62.2 | 4.11 | 12.51 | | High | High | High | High | High | High | High | High | High | High | High | 0.000344 | 0.003042 | 2.99 |
| FALSE | High | LQNEDENI | [R].LQNED | 1xPhospho | A2AG58 1x | 3.01E-05 | 0 | 1 | 2 | 1 | A2AG58 | Uncharacte | A2AG58 [5 | 0 | 2340.053 | 0 | -1.29 | 0.000485 | 1.01 | 141.7 | 58.3 | 6.52 | 1.99 | | High | High | High | High | High | High | High | High | High | High | High | 0 | 4.63E-06 | 3.27 |
| FALSE | High | LQNEDENI | [R].LQNED | 1xPhospho | A2AG58 1x | 0.004867 | 0.000504 | 1 | 2 | 2 | A2AG58 | Uncharacte | A2AG58 [5 | 0 | 2356.048 | 0 | -0.67 | 0.003805 | 3.82 | 121.8 | 78.2 | 6.18 | 6.88 | | High | High | High | High | High | High | High | High | High | High | High | 0.001296 | 0.01342 | 3.18 |
| FALSE | High | LQNSDDDE | [K].LQNSD | 1xPhospho | Q5XJE5 1x | 0.006773 | 0.00062 | 1 | 2 | 4 | Q5XJE5 | RNA polym | Q5XJE5 [1 | 0 | 1730.789 | 0 | -0.67 | 0.002332 | 5.6 | 119.9 | 80.1 | 6.39 | 4.72 | | High | High | High | High | High | High | High | High | High | High | High | 0.000275 | 0.002025 | 3.42 |
| FALSE | High | LYHQPLLES | [R].LYHQP | 1xPhospho | E9PV87 1x | 1.6E-06 | 0 | 1 | 4 | 1 | E9PV87 | Protein TA | E9PV87 [7 | 0 | 2118.14 | 0 | 0.12 | 0.07728 | 6.72 | 94.7 | 105.3 | 7.49 | 7.62 | | High | High | High | High | High | High | High | High | High | High | High | 0 | 1.73E-07 | 3.64 |
| FALSE | High | LYHSLGPVT | [R].LYHSLG | 1xPhospho | Q9ERD6 1x | 0.006834 | 0.00062 | 1 | 5 | 1 | Q9ERD6 | Ras-specifi | Q9ERD6 [4 | 0 | 1451.761 | 0 | 0.12 | 0.209005 | 14.06 | 91.2 | 108.8 | 6.67 | 11.67 | | High | High | High | High | High | High | High | High | High | High | High | 0.000275 | 0.002041 | 2.14 |
| FALSE | High | LYLDELEGS | [R].LYLDEL | 1xCarbami | K3W4Q9 1 | 0.000568 | 0 | 1 | 5 | 1 | K3W4Q9 | Golgi-asso | K3W4Q9 [3 | 0 | 2338.104 | 0 | 0.56 | 0.050633 | 23.16 | 81.3 | 118.7 | 8.89 | 12.75 | | High | High | High | High | High | High | High | High | High | High | High | 0 | 0.000126 | 2.89 |
| FALSE | High | IYISNTFSPS | [R].IYISNTF | 1xPhospho | Q3TXH6 1x | 1.12E-06 | 0 | 1 | 3 | 1 | Q3TXH6 | Putative u | Q3TXH6 [1 | 0 | 1794.944 | 0 | -0.26 | 0.010849 | 6.85 | 108.9 | 91.1 | 2.84 | 7.12 | | High | High | High | High | High | High | High | High | High | High | High | 0 | 1.16E-07 | 2.67 |
| FALSE | High | LYPPAHFG | [R].LYPPAH | 1xPhospho | P70218 1x | 0.048715 | 0.00354 | 1 | 4 | 1 | P70218 | Mitogen-a | P70218 [3 | 0 | 1738.852 | 0 | -1.91 | 0.000283 | 0.5 | 160.7 | 39.3 | 9.88 | 14.49 | | High | High | High | High | High | High | High | High | High | High | High | 0.001876 | 0.01874 | 2.28 |
| FALSE | High | LYQDTRGS | [R].LYQDT | 2xPhospho | Q9Z0U1 2x | 0.025073 | 0.001645 | 1 | 1 | 2 | Q9Z0U1 | Tight junct | Q9Z0U1 [1 | 1 | 2683.08 | 0 | 0.62 | 0.004531 | 4.56 | 73.9 | 126.1 | 8.88 | 8.97 | | High | High | High | High | High | High | High | High | High | High | High | 0.000968 | 0.01003 | 3.05 |
| FALSE | High | LYQSAGGN | [K].LYQSA | 1xPhospho | P63017 1x | 0.003892 | 0.000261 | 1 | 15 | 4 | P63017 | Heat shock | P63017 [6 | 0 | 3687.618 | 0 | -0.37 | 0.359049 | 23.6 | 108.5 | 91.5 | 23.49 | 13.87 | | High | High | High | High | High | High | High | High | High | High | High | 0.000275 | 0.002174 | 3.49 |
| FALSE | High | IYQYIQSR | [R].IYQYIQ | 1xPhospho | A1L341 1x | 0.003224 | 0.00016 | 1 | 8 | 4 | A1L341 | Dual-speci | A1L341 [3 | 0 | 1379.692 | 0 | -0.29 | 0.001617 | 4.15 | 111.3 | 88.7 | 2.19 | 6.61 | | High | High | High | High | High | High | High | High | High | High | High | 7.19E-05 | 0.000883 | 2.54 |
| FALSE | High | IYSFTDNAF | [K].IYSFTD | 1xPhospho | Q8BZ47 1x | 0.000237 | 0 | 1 | 2 | 1 | Q8BZ47 | Zinc finger | Q8BZ47 [7 | 0 | 2165.011 | 0 | 0.93 | 0.000889 | 12.57 | 68 | 132 | 9.26 | 5.32 | | High | High | High | High | High | High | High | High | High | High | High | 0 | 4.75E-05 | 2.36 |
| FALSE | High | LYSLLGIDL | [K].LYSLLG | 1xPhospho | B2RRI2 1x | 0.002115 | 5.66E-05 | 1 | 8 | 2 | B2RRI2 | Protein str | B2RRI2 [67 | 0 | 2327.185 | 0 | 0.77 | 0.004822 | 29.52 | 69.1 | 130.9 | 17.49 | 9.23 | | High | High | High | High | High | High | High | High | High | High | High | 4.94E-05 | 0.000628 | 3.46 |
| FALSE | High | LYSLLGIDL | [K].LYSLLG | 1xCarbami | B2RRI2 1x | 5.5E-05 | 0 | 1 | 7 | 1 | B2RRI2 | Protein str | B2RRI2 [67 | 1 | 3143.585 | 0 | -0.87 | 0.002998 | 4.4 | 131.2 | 68.8 | 7.93 | 10.34 | | High | High | High | High | High | High | High | High | High | High | High | 0 | 9.16E-06 | 3.2 |
| FALSE | High | IYSSDSDEG | [R].IYSSDS | 1xPhospho | Q5XJE5 1x | 0.001147 | 0 | 1 | 1 | 1 | Q5XJE5 | RNA polym | Q5XJE5 [6 | 0 | 2098.911 | 0 | -0.15 | 0.30495 | 53 | 104 | 96 | 18.06 | 52.45 | | High | High | High | High | High | High | High | High | High | High | High | 0 | 0.000278 | 1.98 |
| FALSE | High | IYSSDSDEG | [R].IYSSDS | 2xPhospho | Q5XJE5 2x | 0.046687 | 0.00354 | 1 | 1 | 1 | Q5XJE5 | RNA polym | Q5XJE5 [6 | 0 | 2178.877 | 0 | 0.45 | 0.25029 | 17.49 | 84.7 | 115.3 | 22.83 | 48.5 | | High | High | High | High | High | High | High | High | High | High | High | 0.001812 | 0.01785 | 2.36 |
| FALSE | High | IYHLPDAES | [K].IYHLPD | 1xPhospho | P42208 1x | 9.19E-08 | 0 | 1 | 2 | 17 | P42208 | Septin-2 O | P42208 [2 | 0 | 2461.121 | 0 | 0.25 | 0.009145 | 5.38 | 90.6 | 109.4 | 2.02 | 6.08 | | High | High | High | High | High | High | High | High | High | High | High | 0 | 5.59E-07 | 4.71 |
| FALSE | High | LYSSGSCTF | [K].LYSSGS | 1xCarbami | G3X9J0 1x | 6.47E-05 | 0 | 2 | 10 | 1 | G3X9J0; Q | Signal-indu | G3X9J0 [1 | 0 | 2003.946 | 0 | 0.83 | 0.000704 | 4.63 | 72.5 | 127.5 | 11.58 | 9.73 | Shared | High | High | High | High | High | High | High | High | High | High | High | 0 | 1.1E-05 | 1.45 |
| FALSE | High | LYTTDQSTI | [K].LYTTDQ | 1xPhospho | E9PYV1 1x | 5.27E-06 | 0 | 1 | 12 | 1 | E9PYV1 | Glucocorti | E9PYV1 [2 | 0 | 3171.554 | 0 | 0.76 | 0.027325 | 6.27 | 80.2 | 119.8 | 22.4 | 14.44 | | High | High | High | High | High | High | High | High | High | High | High | 0 | 6.59E-07 | 3.38 |
| FALSE | High | LYTYIQSR | [K].LYTYIQ | 1xPhospho | Q922Y0 1x | 0.00824 | 0.000736 | 1 | 1 | 2 | Q922Y0 | Dual speci | Q922Y0 [3 | 0 | 1352.681 | 0 | -0.04 | 0.459709 | 1.86 | 100.6 | 99.4 | 5.58 | 15.22 | | High | High | High | High | High | High | High | High | High | High | High | 0.003782 | 0.04084 | 1.95 |
| FALSE | High | LYVAQQM | [K].LYVAQ | 1xPhospho | Q8VC70 1x | 0.000442 | 0 | 1 | 2 | 1 | Q8VC70 | RNA-bindi | Q8VC70 [2 | 0 | 1766.886 | 0 | 1.99 | 0.001312 | 18.84 | 38.3 | 161.7 | 16.67 | 13.72 | | High | High | High | High | High | High | High | High | High | High | High | 0 | 9.49E-05 | 2.92 |
| FALSE | High | LYVAQQM | [K].LYVAQ | 1xPhospho | Q8VC70 1x | 6.18E-05 | 0 | 1 | 2 | 2 | Q8VC70 | RNA-bindi | Q8VC70 [2 | 0 | 1782.881 | 0 | 0.63 | 0.001248 | 12.51 | 80.5 | 119.5 | 6.57 | 6.94 | | High | High | High | High | High | High | High | High | High | High | High | 0 | 1.05E-05 | 3.66 |
| FALSE | High | LYVLEAATI | [R].LYVLEA | 1xPhospho | B2RXS1 1x | 2.49E-05 | 0 | 1 | 7 | 3 | B2RXS1 | Nbeal2 pro | B2RXS1 [1 | 0 | 1827.909 | 0 | 1.49 | 0.000927 | 8.67 | 51.5 | 148.5 | 8.06 | 8.84 | | High | High | High | High | High | High | High | High | High | High | High | 0 | 7.29E-05 | 3.33 |
| FALSE | High | IYYSEETSS | [K].IYYSEE | 1xPhospho | Q62077 1x | 0.000109 | 0 | 1 | 4 | 4 | Q62077 | 1-phospha | Q62077 [5 | 0 | 2887.245 | 0 | -0.39 | 0.010676 | 8.01 | 113.3 | 86.7 | 9.7 | 10.91 | | High | High | High | High | High | High | High | High | High | High | High | 0 | 6.93E-05 | 2.85 |
| FALSE | High | MAEESSSS | [K].MAEES | 1xPhospho | A2A432 1x | 7.95E-11 | 0 | 1 | 4 | 6 | A2A432 | Cullin-4B | A2A432 [1 | 0 | 3466.605 | 2 | -1.56 | 0.00037 | 2.02 | 149.3 | 50.7 | 5.53 | 12.63 | | High | High | High | High | High | High | High | High | High | High | High | 0 | 1.06E-11 | 5.45 |
| FALSE | High | MAEESSSS | [K].MAEES | 2xPhospho | A2A432 2x | 1.88E-05 | 0 | 1 | 4 | 4 | A2A432 | Cullin-4B | A2A432 [1 | 0 | 3546.571 | 0 | 0.03 | 0.722859 | 13.18 | 99 | 101 | 10.85 | 6.52 | | High | High | High | High | High | High | High | High | High | High | High | 0 | 5.86E-06 | 3.88 |
| FALSE | High | MAPPIDDL | [R].MAPPI | 1xPhospho | B2RQG2 1x | 2.87E-05 | 0 | 1 | 3 | 1 | B2RQG2 | PHD finger | B2RQG2 [1 | 0 | 1721.895 | 0 | -0.86 | 0.004693 | 8.07 | 129.8 | 70.2 | 8.13 | 13.58 | | High | High | High | High | High | High | High | High | High | High | High | 0 | 4.44E-06 | 3.33 |
| FALSE | High | MAQTAVTI | [K].MAQTA | 1xDeamida | Q3UQS8 1 | 0.060159 | 0.004476 | 1 | 3 | 2 | Q3UQS8 | RNA-bindi | Q3UQS8 [1 | 0 | 2556.294 | 0 | 0.2 | 0.013655 | 6.65 | 93 | 107 | 6.39 | 8.73 | | High | High | High | High | High | High | High | High | High | High | High | 0.002293 | 0.02392 | 2.71 |
| FALSE | High | MASVVPG | [R].MASVV | 1xPhospho | Q922Q2 1x | 4.56E-06 | 0 | 1 | 1 | 2 | Q922Q2 | Serine/thr | Q922Q2 [8 | 0 | 2637.179 | 0 | -1.03 | 0.003636 | 0.41 | 130.7 | 69.3 | 8.52 | 16.58 | | High | High | High | High | High | High | High | High | High | High | High | 0 | 5.59E-07 | 3.98 |
| FALSE | High | MASVVPG | [R].MASVV | 2xPhospho | Q922Q2 2x | 0.000191 | 0 | 1 | 1 | 4 | Q922Q2 | Serine/thr | Q922Q2 [8 | 0 | 2717.145 | 0 | -0.11 | 0.12344 | 11.43 | 106.3 | 93.7 | 5.86 | 9.07 | | High | High | High | High | High | High | High | High | High | High | High | 0 | 3.76E-05 | 4.14 |
| FALSE | High | MAVESPSD | [K].MAVES | 1xPhospho | A7E215 1x | 0.017344 | 0.001204 | 1 | 2 | 3 | A7E215 | Rps6ka3 pr | A7E215 [1 | 0 | 4460.971 | 0 | -0.05 | 0.649671 | 7.77 | 99.5 | 100.5 | 3.52 | 7.92 | | High | High | High | High | High | High | High | High | High | High | High | 0.000756 | 0.006702 | 4.08 |
| FALSE | High | IYSYQMAL | [RK].IYSYQ | 1xPhospho | P30285 1x | 0.000292 | 0 | 1 | 6 | 1 | P30285 | Cyclin-dep | P30285 [1 | 0 | 2512.291 | 0 | 1.07 | 0.009969 | 21.61 | 65.3 | 134.7 | 6.07 | 18.28 | | High | High | High | High | High | High | High | High | High | High | High | 0 | 5.96E-05 | 3.01 |
| FALSE | High | MDAGDTFI | [K].MDAGI | 1xPhospho | E9Q5F9 1x | 3.48E-06 | 0 | 1 | 2 | 2 | E9Q5F9 | Histone-lys | E9Q5F9 [1 | 0 | 2318.066 | 0 | 0.31 | 0.114871 | 8.1 | 89.6 | 110.4 | 3.49 | 14.25 | | High | High | High | High | High | High | High | High | High | High | High | 0 | 4.14E-07 | 3.53 |
| FALSE | High | IYHLPDAES | [K].IYHLPD | 1xPhospho | P42208 1x | 1.11E-05 | 0 | 1 | 2 | 5 | P42208 | Septin-2 O | P42208 [2 | 0 | 2231.958 | 0 | 0.38 | 0.187121 | 9.12 | 86.8 | 113.2 | 15.09 | 28.91 | | High | High | High | High | High | High | High | High | High | High | High | 0 | 3.5E-05 | 3.49 |
| FALSE | High | LYHDSTNII | [R].LYHDST | 1xPhospho | B9EKC3 1x | 0.004294 | 0.00036 | 1 | 3 | 1 | B9EKC3 | Rho GTPas | B9EKC3 [5 | 0 | 1743.872 | 0 | -0.03 | 0.585463 | 17.24 | 107.3 | 92.7 | 9.71 | 15.98 | | High | High | High | High | High | High | High | High | High | High | High | 0.000161 | 0.001221 | 1.97 |
| FALSE | High | LVVSSPTSP | [K].LVVSSP | 1xPhospho | Q7TSI1 1x | 0.02933 | 0.001857 | 1 | 1 | 2 | Q7TSI1 | Pleckstrin | Q7TSI1 [42 | 0 | 1552.875 | 0 | 0.25 | 0.004579 | 5.76 | 92.6 | 107.4 | 4.26 | 4.35 | | High | High | High | High | High | High | High | High | High | High | High | 0.000997 | 0.01051 | 2.75 |
| FALSE | High | LWAEDSTE | [R].LWAED | 1xPhospho | Q9DBR7 1x | 0.003402 | 0.00021 | 1 | 3 | 1 | Q9DBR7 | Protein ph | Q9DBR7 [6 | 0 | 1616.797 | 0 | 0.39 | 0.217482 | 2.7 | 86.6 | 113.4 | 5.91 | 22.84 | | High | High | High | High | High | High | High | High | High | High | High | 9.42E-05 | 0.000936 | 3.29 |
| FALSE | High | LWAGPDPS | [K].LWAGP | 1xPhospho | A2SW42 1x | 1.05E-05 | 0 | 1 | 4 | 2 | A2SW42 | Zinc finger | A2SW42 [1 | 0 | 2497.245 | 0 | -1.55 | 0.000158 | 0.59 | 148.8 | 51.2 | 2.46 | 8.36 | | High | High | High | High | High | High | High | High | High | High | High | 0 | 1.42E-06 | 2.41 |
| FALSE | High | LWAGPDPS | [K].LWAGP | 1xPhospho | A2SW42 1x | 3.94E-05 | 0 | 1 | 4 | 5 | A2SW42 | Zinc finger | A2SW42 [1 | 0 | 2513.24 | 0 | -1.11 | 0.000616 | 3 | 137.1 | 62.9 | 4.95 | 11.28 | | High | High | High | High | High | High | High | High | High | High | High | 0 | 6.34E-06 | 3.96 |
| FALSE | High | IWDPTPSH | [K].IWDPTI | 1xPhospho | G5E866 1x | 0.014283 | 0.001011 | 1 | 5 | 1 | G5E866 | Splicing fac | G5E866 [2 | 0 | 2141.038 | 0 | -0.37 | 0.022501 | 1.62 | 112.9 | 87.1 | 11.56 | 8.17 | | High | High | High | High | High | High | High | High | High | High | High | 0.000538 | 0.004659 | 2.58 |
| FALSE | High | IWDPTPSH | [K].IWDPTI | 2xPhospho | G5E866 2x | 0.050826 | 0.003676 | 1 | 5 | 1 | G5E866 | Splicing fac | G5E866 [2 | 0 | 2221.004 | 0 | 0.21 | 0.808651 | 1.92 | 93.2 | 106.8 | 15.09 | 16.61 | | High | High | High | High | High | High | High | High | High | High | High | 0.00196 | 0.01972 | 1.58 |
| FALSE | High | IWDPTPSH | [K].IWDPTI | 2xPhospho | G5E866 2x | 0.051694 | 0.003745 | 1 | 5 | 2 | G5E866 | Splicing fac | G5E866 [2 | 1 | 3935.781 | 0 | 0.04 | 0.663564 | 35.95 | 100.9 | 99.1 | 11.8 | 21.32 | | High | High | High | High | High | High | High | High | High | High | High | 0.005349 | 0.05846 | 3.6 |
| FALSE | High | LWDSSQK | [R].LWDSS | 1xPhospho | Q7TSZ8 1x | 0.033116 | 0.002195 | 1 | 1 | 2 | Q7TSZ8 | Nucleus ac | Q7TSZ8 [1 | 0 | 1401.718 | 0 | -1.07 | 0.001831 | 1.45 | 135.8 | 64.2 | 8.83 | 10.62 | | High | High | High | High | High | High | High | High | High | High | High | 0.001151 | 0.01205 | 1.98 |
| FALSE | High | LWEDSAGE | [R].LWEDS | 1xPhospho | F8VQD1 1x | 0.006834 | 0.00062 | 1 | 9 | 1 | F8VQD1 | Protein pol | F8VQD1 [9 | 0 | 1572.771 | 0 | 0.33 | 0.067176 | 4.55 | 91 | 109 | 4.52 | 11.87 | | High | High | High | High | High | High | High | High | High | High | High | 0.000275 | 0.002042 | 2.14 |
| FALSE | High | LWESMERS | [R].LWESM | 1xPhospho | D3Z0X5 1x | 0.002096 | 5.66E-05 | 1 | 7 | 2 | D3Z0X5 | Pleckstrin | D3Z0X5 [6 | 1 | 2320.093 | 0 | 1.68 | 0.004737 | 27.45 | 47.7 | 152.3 | 14.44 | 16.25 | | High | High | High | High | High | High | High | High | High | High | High | 0.004206 | 0.04523 | 1.99 |
| FALSE | High | IWGPAPFII | [R].IWGPA | 1xCarbami | Q8C6G8 1x | 0.047488 | 0.00354 | 1 | 3 | 1 | Q8C6G8 | WD repeat | Q8C6G8 [6 | 0 | 2887.215 | 0 | 0.58 | 0.009937 | 16.14 | 74.8 | 125.2 | 18.51 | 11.98 | | High | High | High | High | High | High | High | High | High | High | High | 0.001812 | 0.01811 | 2.83 |
| FALSE | High | LWLISEGY | [R].LWLISE | 1xPhospho | Q5EBP3 1x | 0.000107 | 0 | 1 | 2 | 1 | Q5EBP3 | Armadillo r | Q5EBP3 [4 | 0 | 2825.338 | 0 | 0.45 | 0.04282 | 26.67 | 89.8 | 110.2 | 11.62 | 13.66 | | High | High | High | High | High | High | High | High | High | High | High | 0 | 1.95E-05 | 2.89 |
| FALSE | High | LWQGHHA | [R].LWQG | 1xPhospho | Q61122 1x | 0.017809 | 0.001282 | 1 | 3 | 1 | Q61122 | NGFI-A-bin | Q61122 [1 | 0 | 3277.604 | 0 | 0.29 | 0.41246 | 43.49 | 93.2 | 106.8 | 24.98 | 21.79 | | High | High | High | High | High | High | High | High | High | High | High | 0.000689 | 0.005958 | 2.73 |
| FALSE | High | LYHGGYRS | [R].LYHGG | 1xPhospho | V9GXX7 1x | 0.017041 | 0.001204 | 1 | 3 | 1 | V9GXX7 | DNA repair | V9GXX7 [2 | 1 | 4203.026 | 0 | -0.6 | 0.016602 | 3.63 | 118 | 82 | 13.78 | 17.01 | | High | High | High | High | High | High | High | High | High | High | High | 0.000638 | 0.005694 | 3.1 |
| FALSE | High | IWSPLVTEE | [K].IWSPLV | 1xPhospho | O70400 1x | 2.34E-05 | 0 | 1 | 3 | 2 | O70400 | PDZ and LI | O70400 [8 | 0 | 1796.96 | 0 | -0.02 | 0.928356 | 12.46 | 100.7 | 99.3 | 2.2 | 15.77 | | High | High | High | High | High | High | High | High | High | High | High | 0 | 3.52E-06 | 3.14 |
| FALSE | High | IWSTVPTD | [K].IWSTVF | 1xPhospho | Q4V9X1 1x | 0.000687 | 0 | 1 | 1 | 1 | Q4V9X1 | Ribosome | Q4V9X1 [2 | 0 | 2814.249 | 0 | -0.01 | 0.971553 | 19.5 | 99.1 | 100.9 | 2.72 | 15.73 | | High | High | High | High | High | High | High | High | High | High | High | 0 | 0.000156 | 1.83 |
| FALSE | High | IWTHPGW | [K].IWTHP | 1xPhospho | Q059T9 1x | 0.028082 | 0.001791 | 1 | 3 | 1 | Q059T9 | PRP4 pre-r | Q059T9 [4 | 0 | 1859.997 | 0 | -0.5 | 0.000479 | 3.13 | 116 | 84 | 4.01 | 4.27 | | High | High | High | High | High | High | High | High | High | High | High | 0.000968 | 0.01001 | 1.46 |
| FALSE | High | LWYLPFQN | [K].LWYLP | 1xPhospho | A2AWT3-2 | 0.106748 | 0.008634 | 1 | 2 | 1 | A2AWT3-2 | Isoform 2 | A2AWT3-2 | 0 | 1940.962 | 0 | -0.69 | 0.021493 | 2.44 | 116.2 | 83.8 | 13.62 | 16.16 | | High | High | High | High | High | High | High | High | High | High | High | 0.004327 | 0.0465 | 2.1 |
| FALSE | High | LYADSDSG | [R].LYADSD | 1xPhospho | O35071 1x | 0.00047 | 0 | 1 | 2 | 1 | O35071 | Kinesin-lik | O35071 [6 | 0 | 1939.857 | 0 | 0.47 | 0.009016 | 10.5 | 86.8 | 113.2 | 9.04 | 3.87 | | High | High | High | High | High | High | High | High | High | High | High | 0 | 0.000102 | 2.44 |
| FALSE | High | LYADSDSG | [R].LYADSD | 2xPhospho | O35071 2x | 6.56E-07 | 0 | 1 | 2 | 2 | O35071 | Kinesin-lik | O35071 [6 | 0 | 2019.824 | 0 | 0.97 | 0.000266 | 3.56 | 67.7 | 132.3 | 3.4 | 5.43 | | High | High | High | High | High | High | High | High | High | High | High | 0 | 6.33E-08 | 3.57 |
| FALSE | High | IYASSSPPD | [R].IYASSS | 1xPhospho | Q3U4D5 1x | 0.031168 | 0.001935 | 1 | 3 | 1 | Q3U4D5 | Protein kin | Q3U4D5 [1 | 0 | 1687.789 | 0 | 1.14 | 0.002736 | 17.92 | 66.1 | 133.9 | 14.98 | 10.72 | | High | High | High | High | High | High | High | High | High | High | High | 0.001076 | 0.01126 | 2.52 |
| FALSE | High | IYCVACQE | [K].IYCVAC | 2xCarbami | P56873 1x | 5.08E-09 | 0 | 1 | 4 | 2 | P56873 | Sjoegren s | P56873 [6 | 0 | 2353.086 | 0 | 0.24 | 0.290173 | 25.89 | 93.5 | 106.5 | 7.08 | 16.77 | | High | High | High | High | High | High | High | High | High | High | High | 0 | 2.69E-10 | 3.71 |
| FALSE | High | IYEFPETDD | [K].IYEFPE | 1xPhospho | E9Q1G8 1x | 2.74E-08 | 0 | 1 | 6 | 4 | E9Q1G8 | Septin-7 O | E9Q1G8 [2 | 0 | 2296.031 | 0 | 0.62 | 0.000138 | 0.29 | 78.9 | 121.1 | 3.38 | 2.93 | | High | High | High | High | High | High | High | High | High | High | High | 0 | 4.73E-09 | 4.14 |
| FALSE | High | IYESVLTPP | [K].IYESVL | 1xPhospho | F8WJ40 1x | 0.01894 | 0.001362 | 1 | 2 | 3 | F8WJ40 | Histone-lys | F8WJ40 [6 | 0 | 3932.93 | 0 | -0.31 | 0.24759 | 11.45 | 104.6 | 95.4 | 20.14 | 25.11 | | High | High | High | High | High | High | High | High | High | High | High | 0.000786 | 0.006963 | 3.31 |
| FALSE | High | LYFHSDTC | [R].LYFHSD | 1xCarbami | Q80U70 1x | 0.005418 | 0.00062 | 1 | 5 | 1 | Q80U70 | Polycomb | Q80U70 [5 | 0 | 3476.616 | 0 | -0.74 | 0.002668 | 6.98 | 121.8 | 78.2 | 4.89 | 11.42 | | High | High | High | High | High | High | High | High | High | High | High | 0.000275 | 0.001589 | 1.76 |
| FALSE | High | LYFHSDTC | [R].LYFHSD | 1xCarbami | Q80U70 1x | 0.002022 | 5.66E-05 | 1 | 5 | 1 | Q80U70 | Polycomb | Q80U70 [5 | 0 | 3477.6 | 0 | -0.52 | 0.026792 | 14.61 | 121.3 | 78.7 | 11.78 | 15.95 | | High | High | High | High | High | High | High | High | High | High | High | 2.58E-05 | 0.000527 | 3.13 |
| FALSE | High | LYFHSDTC | [R].LYFHSD | 1xCarbami | Q80U70 1x | 2.46E-08 | 0 | 1 | 5 | 10 | Q80U70 | Polycomb | Q80U70 [5 | 0 | 3492.61 | 0 | -0.48 | 0.001931 | 4.54 | 116.8 | 83.2 | 4.3 | 4.08 | | High | High | High | High | High | High | High | High | High | High | High | 0 | 2.21E-06 | 4.49 |
| FALSE | High | LYGSGGPP | [K].LYGSG | 1xPhospho | P20029 1x | 0.011966 | 0.000889 | 1 | 7 | 1 | P20029 | 78 kDa glu | P20029 [6 | 1 | 2716.264 | 0 | -0.49 | 0.014763 | 8.21 | 118.9 | 81.1 | 12.15 | 14.33 | | High | High | High | High | High | High | High | High | High | High | High | 0.000449 | 0.00383 | 2.65 |
| FALSE | High | IWSPTVGR | [R].IWSPT | 1xPhospho | O70472 1x | 0.116482 | 0.009523 | 1 | 3 | 1 | O70472 | Transmem | O70472 [1 | 0 | 1224.634 | 0 | 0.26 | 0.001333 | 4.74 | 90.7 | 109.3 | 8.97 | 5.48 | | High | High | High | High | High | High | High | High | High | High | High | 0.004757 | 0.05176 | 2.21 |
| FALSE | High | MDDISPVC | [K].MDDIS | 1xPhospho | E9QLG3 1x | 0.005717 | 0.00062 | 1 | 5 | 1 | E9QLG3 | MAX gene- | E9QLG3 [1 | 0 | 2070.962 | 0 | -0.29 | 0.005749 | 4.95 | 107.3 | 92.7 | 5.22 | 12.08 | | High | High | High | High | High | High | High | High | High | High | High | 0.000275 | 0.001684 | 2.53 |
| FALSE | High | MDIDRSPG | [R].MDIDR | 1xPhospho | P97434-2 1 | 0.010758 | 0.000772 | 1 | 11 | 1 | P97434-2 | Isoform 2 | P97434-2 [ | 1 | 2266.192 | 0 | 0.9 | 0.007147 | 2.92 | 70.1 | 129.9 | 10.55 | 14.95 | | High | High | High | High | High | High | High | High | High | High | High | 0.000399 | 0.003409 | 3.31 |
| FALSE | High | MDLVGVSS | [-].MDLVG | 1xAcetyl [N | Q3TC66 1x | 8.68E-07 | 0 | 1 | 5 | 3 | Q3TC66 | Putative u | Q3TC66 [1 | 0 | 2404.146 | 0 | -0.34 | 0.01248 | 1.83 | 113 | 87 | 3.46 | 11.62 | | High | High | High | High | High | High | High | High | High | High | High | 0 | 8.64E-08 | 3.92 |
| FALSE | High | MGLLSPSG | [R].MGLLS | 1xPhospho | G5E8A7 1x | 2.18E-06 | 0 | 1 | 3 | 2 | G5E8A7 | Protein Sm | G5E8A7 [5 | 0 | 1736.897 | 0 | 1.89 | 0.000325 | 17.43 | 42.6 | 157.4 | 8.31 | 5.65 | | High | High | High | High | High | High | High | High | High | High | High | 0 | 3.67E-06 | 2.42 |
| FALSE | High | MGNLQNS | [K].MGNL | 1xPhospho | Q61026 1x | 0.000127 | 0 | 1 | 5 | 2 | Q61026 | Nuclear re | Q61026 [5 | 0 | 2492.244 | 0 | 0.37 | 0.015951 | 6.01 | 87.2 | 112.8 | 10.43 | 14.21 | | High | High | High | High | High | High | High | High | High | High | High | 0 | 2.36E-05 | 2.69 |
| FALSE | High | MGSDGESI | [K].MGSD | 1xCarbami | Q8K0D0 1x | 4.62E-11 | 0 | 1 | 4 | 2 | Q8K0D0 | Cyclin-dep | Q8K0D0 [1 | 0 | 3066.287 | 0 | -0.21 | 0.084116 | 6.34 | 108.9 | 91.1 | 5.44 | 7.11 | | High | High | High | High | High | High | High | High | High | High | High | 0 | 1.36E-12 | 5.77 |
| FALSE | High | MGSPAPLL | [R].MGSPA | 1xPhospho | Q8BKI2 1x | 0.019966 | 0.001438 | 1 | 7 | 1 | Q8BKI2 | Trinucleoti | Q8BKI2 [1 | 0 | 2163.061 | 0 | 1.69 | 0.000734 | 14.9 | 46.9 | 153.1 | 23.97 | 15.25 | | High | High | High | High | High | High | High | High | High | High | High | 0.000786 | 0.006804 | 2.61 |

| | | | | | | | | | | | | | | | | | | | | | | | | | | | | | | | | | | | | | | |
|---|---|---|---|---|---|---|---|---|---|---|---|---|---|---|---|---|---|---|---|---|---|---|---|---|---|---|---|---|---|---|---|---|---|---|---|---|---|---|
| FALSE | High | MHDGELE( | [K].MHDGI | 1xPhosphc | A2ANY6 1x | 2.05E-07 | 0 | 1 | 2 | 2 | A2ANY6 | Midasin OS | A2ANY6 [4 | 1 | 3532.54 | 0 | -0.85 | 0.002834 | 2.68 | 128.6 | 71.4 | 6.03 | 14.66 | | High | High | High | High | High | High | High | High | High | High | High | 0 | 1.71E-08 | 4.53 |
| FALSE | High | MHLPSPTD | [R].MHLPS | 1xPhosphc | Q01279 1x | 0.071071 | 0.005247 | 1 | 3 | 1 | Q01279 | Epidermal | Q01279 [9 | 0 | 1873.851 | 0 | -0.15 | 0.092222 | 3.62 | 106.9 | 93.1 | 4.35 | 8.87 | | High | High | High | High | High | High | High | High | High | High | High | 0.002703 | 0.02904 | 2.33 |
| FALSE | High | MHRDSCPL | [-].MHRDS | 1xAcetyl [N | P84104 1x | 4.69E-06 | 0 | 1 | 3 | 1 | P84104 | Serine/argi | P84104 [1- | 1 | 1769.737 | 0 | 0.01 | 0.941718 | 29.09 | 96.1 | 103.9 | 10.96 | 23.85 | | High | High | High | High | High | High | High | High | High | High | High | 0 | 5.77E-07 | 3.77 |
| FALSE | High | MLAESDDS | [K].MLAESI | 2xPhosphc | Q3B7Z2 2x | 4.2E-05 | 0 | 1 | 1 | 2 | Q3B7Z2 | Oxysterol-I | Q3B7Z2 [1 | 0 | 2661.093 | 0 | 0.31 | 0.013404 | 13.57 | 90 | 110 | 6.78 | 11.39 | | High | High | High | High | High | High | High | High | High | High | High | 0 | 6.75E-06 | 3.13 |
| FALSE | High | MLGEDSDE | [K-].MLGEI | 1xPhosphc | E9PX68 1xI | 1.19E-06 | 0 | 1 | 8 | 1 | E9PX68 | Solute carr | E9PX68 [25 | 0 | 2522.089 | 0 | 0.11 | 0.731992 | 8.77 | 96.1 | 103.9 | 8.83 | 11.21 | | High | High | High | High | High | High | High | High | High | High | High | 0 | 1.24E-07 | 3.54 |
| FALSE | High | MLLNLGLG | [-].MLLNLG | 1xDeamida | B2X2D8 1x | 0.109329 | 0.008865 | 1 | 1 | 1 | B2X2D8 | ASL1/FUT8 | B2X2D8 [1- | 0 | 1710.87 | 0 | -0.22 | 0.016239 | 5.25 | 107.4 | 92.6 | 4.35 | 10.38 | | High | High | High | High | High | High | High | High | High | High | High | 0.004462 | 0.04815 | 2.44 |
| FALSE | High | MLPHAPG\ | [R].MLPHA | 1xPhosphc | D3YYI8 1xF | 6.13E-05 | 0 | 1 | 2 | 3 | D3YYI8 | Histone de | D3YYI8 [37 | 0 | 4027.823 | 0 | -0.73 | 0.00176 | 1.32 | 121.5 | 78.5 | 4.81 | 17.93 | | High | High | High | High | High | High | High | High | High | High | High | 0 | 1.03E-05 | 3.87 |
| FALSE | High | MLPHAPG\ | [R].MLPHA | 1xPhosphc | D3YYI8 1xF | 1.57E-05 | 0 | 1 | 2 | 21 | D3YYI8 | Histone de | D3YYI8 [37 | 0 | 4043.818 | 0 | -0.71 | 0.001207 | 3.11 | 122.5 | 77.5 | 2.57 | 10.4 | | High | High | High | High | High | High | High | High | High | High | High | 0 | 3.05E-05 | 4.6 |
| FALSE | High | MLPHAPG\ | [R].MLPHA | 1xPhosphc | D3YYI8 1xF | 0.001558 | 5.66E-05 | 1 | 2 | 1 | D3YYI8 | Histone de | D3YYI8 [37 | 1 | 4199.919 | 0 | -1 | 0.039214 | 7.04 | 130.5 | 69.5 | 7.15 | 40.93 | | High | High | High | High | High | High | High | High | High | High | High | 2.58E-05 | 0.000391 | 4.12 |
| FALSE | High | MGLEESSE | [K].MGLEE | 1xPhosphc | Q8BLH7 1x | 2.87E-05 | 0 | 1 | 2 | 2 | Q8BLH7 | HIRA-intera | Q8BLH7 [1 | 0 | 1975.852 | 0 | -1.85 | 0.000377 | 1.78 | 157.3 | 42.7 | 7.34 | 18.89 | | High | High | High | High | High | High | High | High | High | High | High | 0 | 4.42E-06 | 2.22 |
| FALSE | High | MLPHAPG\ | [R].MLPHA | 1xPhosphc | A0A0R4J0C | 2.48E-07 | 0 | 1 | 3 | 3 | A0A0R4J0C | Histone de | A0A0R4J0C | 0 | 3967.777 | 0 | 0.31 | 0.435381 | 19.45 | 90.1 | 109.9 | 4.76 | 22.24 | | High | High | High | High | High | High | High | High | High | High | High | 0 | 2.11E-08 | 4.52 |
| FALSE | High | MLPHAPG\ | [R].MLPHA | 1xPhosphc | A0A0R4J0C | 1.32E-08 | 0 | 1 | 3 | 9 | A0A0R4J0C | Histone de | A0A0R4J0C | 0 | 3983.772 | 0 | 0.29 | 0.003737 | 1.93 | 92.3 | 107.7 | 2.75 | 4.65 | | High | High | High | High | High | High | High | High | High | High | High | 0 | 7.83E-10 | 4.67 |
| FALSE | High | MLQAISPK | [R].MLQAI | 1xPhosphc | P49586 1xI | 0.004572 | 0.000456 | 2 | 3 | 2 | P49586; Q | Choline-ph | P49586 [31 | 0 | 1425.794 | 0 | 1.24 | 0.001238 | 12.6 | 58.6 | 141.4 | 5.91 | 9.75 | Shared | High | High | High | High | High | High | High | High | High | High | High | 0.000204 | 0.001307 | 1.35 |
| FALSE | High | MLQDDED | [R].MLQDD | 1xPhosphc | I1VGT4 1xI | 0.081085 | 0.006219 | 2 | 19 | 2 | I1VGT4; I1 | Anion exch | I1VGT4 [10 | 0 | 2153.978 | 0 | 1.01 | 0.026879 | 32.85 | 61.7 | 138.3 | 10.27 | 30.43 | Shared | High | High | High | High | High | High | High | High | High | High | High | 0.00673 | 0.07294 | 2.23 |
| FALSE | High | MLSGSELT | [R].MLSGS | 1xPhosphc | A0A0R4J2D | 9.91E-06 | 0 | 1 | 3 | 1 | A0A0R4J2D | Ubiquitin c | A0A0R4J2D | 0 | 2007.975 | 0 | -1.2 | 0.00032 | 2.38 | 141.2 | 58.8 | 3.04 | 11.82 | | High | High | High | High | High | High | High | High | High | High | High | 0 | 1.33E-06 | 2.74 |
| FALSE | High | MLSPAMSI | [-].MLSPAN | 1xAcetyl [N | Q923F1 1x | 0.003822 | 0.000261 | 1 | 2 | 1 | Q923F1 | Chloride ch | Q923F1 [1- | 0 | 1475.724 | 0 | -0.42 | 0.0747 | 20.3 | 114.6 | 85.4 | 10.8 | 13.75 | | High | High | High | High | High | High | High | High | High | High | High | 0.000117 | 0.00107 | 2.14 |
| FALSE | High | MNDFHISD | [K].MNDFH | 1xPhosphc | Q3TRR0 1x | 0.002869 | 0.00016 | 1 | 5 | 1 | Q3TRR0 | Microtubu | Q3TRR0 [7 | 0 | 2017.898 | 0 | 0.59 | 0.115372 | 3.48 | 81.9 | 118.1 | 3.26 | 26.1 | | High | High | High | High | High | High | High | High | High | High | High | 7.19E-05 | 0.000777 | 3.29 |
| FALSE | High | MNGVMFF | [R].MNGVI | 1xPhosphc | Q0VGB7 1x | 0.015061 | 0.001049 | 2 | 2 | 1 | Q0VGB7; A | Serine/thre | Q0VGB7 [1 | 0 | 2124.908 | 0 | -0.07 | 0.805593 | 11.69 | 106.4 | 93.6 | 15.37 | 19.83 | Shared | High | High | High | High | High | High | High | High | High | High | High | 0.000571 | 0.004958 | 1.68 |
| FALSE | High | MNIPSTSP\ | [R].MNIPST | 1xPhosphc | A0A1B0GS | 0.000146 | 0 | 1 | 6 | 1 | A0A1B0GS | Nuclear po | A0A1B0GS | 0 | 3842.062 | 0 | -0.54 | 0.010538 | 5.19 | 126.4 | 73.6 | 11.43 | 12.66 | | High | High | High | High | High | High | High | High | High | High | High | 0 | 2.73E-05 | 5.76 |
| FALSE | High | MNPNPQD | [-].MNPNP | 3xPhosphc | Q3TDX5 3x | 0.097743 | 0.007804 | 1 | 3 | 1 | Q3TDX5 | Putative ur | Q3TDX5 [1 | 1 | 2890.26 | 0 | -0.49 | 0.142439 | 13.75 | 115.1 | 84.9 | 9.63 | 23.91 | | High | High | High | High | High | High | High | High | High | High | High | 0.003922 | 0.04187 | 2.37 |
| FALSE | High | MNVSPQT( | [R].MNVSP | 1xPhosphc | Q2VPQ9-2 | 0.021231 | 0.001507 | 1 | 1 | 1 | Q2VPQ9-2 | Isoform 2 c | Q2VPQ9-2 | 0 | 1956.935 | 0 | -0.12 | 0.24176 | 3.84 | 103.8 | 96.2 | 9.22 | 11.7 | | High | High | High | High | High | High | High | High | High | High | High | 0.000801 | 0.007259 | 1.62 |
| FALSE | High | MPDEPEEP | [K].MPDEP | 1xPhosphc | Q3UH70 1x | 0.000422 | 0 | 1 | 8 | 4 | Q3UH70 | Bromodom | Q3UH70 [4 | 0 | 2841.44 | 0 | -0.57 | 0.012767 | 7.54 | 119.9 | 80.1 | 7.01 | 11.28 | | High | High | High | High | High | High | High | High | High | High | High | 0 | 9.03E-05 | 2.49 |
| FALSE | High | MPFLSISSP | [K].MPFLSI | 1xPhosphc | E9Q616 1x | 0.01861 | 0.001362 | 1 | 2 | 1 | E9Q616 | AHNAK nu | E9Q616 [3 | 0 | 1644.884 | 0 | 4.22 | 0.002546 | 26 | 9.9 | 190.1 | 18.27 | 23.44 | | High | High | High | High | High | High | High | High | High | High | High | 0.000723 | 0.006287 | 1.63 |
| FALSE | High | MPGGTDQ | [K].MPGGT | 1xPhosphc | A2AQH4 1x | 3.28E-07 | 0 | 1 | 3 | 1 | A2AQH4 | BCL-6 core | A2AQH4 [5 | 0 | 2619.278 | 0 | 0.63 | 0.015324 | 10.2 | 79.4 | 120.6 | 7.59 | 12.57 | | High | High | High | High | High | High | High | High | High | High | High | 0 | 2.89E-08 | 4.32 |
| FALSE | High | MLPHAPG\ | [R].MLPHA | 1xDeamida | A0A0R4J0C | 0.000543 | 0 | 1 | 3 | 2 | A0A0R4J0C | Histone de | A0A0R4J0C | 0 | 3968.761 | 0 | 0.32 | 0.021674 | 0.86 | 75.2 | 124.8 | 15.67 | 17.3 | | High | High | High | High | High | High | High | High | High | High | High | 0 | 0.000121 | 4.25 |
| FALSE | High | MFSTKIEIV | [-].MFSTKI | 2xPhosphc | Q3UP34 2x | 0.054383 | 0.003951 | 1 | 1 | 2 | Q3UP34 | Putative ur | Q3UP34 [1 | 1 | 1800.882 | 0 | 1.42 | 0.00041 | 16.23 | 55 | 145 | 14.22 | 2.44 | | High | High | High | High | High | High | High | High | High | High | High | 0.002059 | 0.02124 | 2.06 |
| FALSE | High | MEVQESG( | [-].MEVQE | 1xAcetyl [N | Q9D0K3 1x | 9.42E-07 | 0 | 1 | 2 | 2 | Q9D0K3 | Putative ur | Q9D0K3 [1 | 0 | 2883.226 | 0 | -0.21 | 0.239849 | 15.73 | 106.7 | 93.3 | 7.33 | 16.55 | | High | High | High | High | High | High | High | High | High | High | High | 0 | 4.24E-07 | 5.09 |
| FALSE | High | MEVLSPAS | [K].MEVLSI | 2xPhosphc | Q6NZR2 2x | 0.003857 | 0.000261 | 1 | 1 | 1 | Q6NZR2 | Myb/SANT | Q6NZR2 [2 | 0 | 3015.326 | 0 | -0.8 | 0.014134 | 11.32 | 125.3 | 74.7 | 16.57 | 11.44 | | High | High | High | High | High | High | High | High | High | High | High | 0.000117 | 0.001082 | 3.94 |
| FALSE | High | MDSASQDI | [-].MDSAS( | 1xAcetyl [N | A2AUD5 1x | 2.19E-05 | 0 | 1 | 6 | 2 | A2AUD5 | Tumor pro | A2AUD5 [1 | 0 | 1984.888 | 0 | 0.46 | 0.079561 | 16.62 | 81.1 | 118.9 | 5.95 | 20.9 | | High | High | High | High | High | High | High | High | High | High | High | 0 | 3.26E-06 | 3.31 |
| FALSE | High | MDSFDEDI | [R].MDSFD | 1xPhosphc | Q569Z6 1x | 0.080423 | 0.006219 | 1 | 3 | 1 | Q569Z6 | Thyroid ho | Q569Z6 [5 | 0 | 2459.148 | 0 | -0.12 | 0.272234 | 7.55 | 99.8 | 100.2 | 10.6 | 12.1 | | High | High | High | High | High | High | High | High | High | High | High | 0.003188 | 0.03335 | 2.37 |
| FALSE | High | MDSNFGL/ | [R].MDSNF | 1xPhosphc | O54864-2 | 1.74E-06 | 0 | 1 | 3 | 1 | O54864-2 | Isoform 2 c | O54864-2 | 0 | 2029.023 | 0 | 1.78 | 0.001114 | 1.39 | 45.2 | 154.8 | 5.21 | 12.94 | | High | High | High | High | High | High | High | High | High | High | High | 0 | 1.89E-07 | 2.67 |
| FALSE | High | MEAEGSLN | [R].MEAEG | 1xPhosphc | A0A0A0M( | 4.89E-05 | 0 | 1 | 2 | 7 | A0A0A0M( | F-box only | A0A0A0M( | 0 | 3086.449 | 0 | 1.61 | 0.000154 | 8.53 | 51.1 | 148.9 | 12.04 | 5.96 | | High | High | High | High | High | High | High | High | High | High | High | 0 | 1.69E-05 | 4.01 |
| FALSE | High | MEAEGSLN | [R].MEAEG | 2xPhosphc | A0A0A0M( | 0.000418 | 0 | 1 | 2 | 7 | A0A0A0M( | F-box only | A0A0A0M( | 0 | 3166.416 | 0 | 3.39 | 0.000261 | 20.18 | 15.5 | 184.5 | 15.7 | 7.95 | | High | High | High | High | High | High | High | High | High | High | High | 0.000382 | 0.003261 | 3.27 |
| FALSE | High | MEAEGSLN | [R].MEAEG | 1xDeamida | A0A0A0M( | 0.016743 | 0.001166 | 1 | 2 | 1 | A0A0A0M( | F-box only | A0A0A0M( | 0 | 3167.4 | 0 | 1.56 | 0.000927 | 19.22 | 49.3 | 150.7 | 11.41 | 10.11 | | High | High | High | High | High | High | High | High | High | High | High | 0.000622 | 0.005561 | 3.2 |
| FALSE | High | MEAEVDSF | [K].MEAEV | 1xPhosphc | B1AR17 1x | 0.000681 | 0 | 1 | 3 | 1 | B1AR17 | Chromodo | B1AR17 [1 | 0 | 2080.946 | 0 | 0.35 | 0.019763 | 11.49 | 83.4 | 116.6 | 11.24 | 14.46 | | High | High | High | High | High | High | High | High | High | High | High | 0 | 0.000155 | 2.55 |
| FALSE | High | MEAQELG! | [-].MEAQE | 1xAcetyl [N | Q8BGK6 1x | 0.000876 | 0 | 1 | 4 | 2 | Q8BGK6 | Y+L amino | Q8BGK6 [1 | 0 | 2363.156 | 0 | 0.78 | 0.002608 | 9.04 | 73.7 | 126.3 | 9.25 | 9.7 | | High | High | High | High | High | High | High | High | High | High | High | 0 | 0.000336 | 2.6 |
| FALSE | High | MEASAAE( | [-].MEASA/ | 1xAcetyl [N | Q8CE96 1x | 2.38E-05 | 0 | 1 | 1 | 5 | Q8CE96 | tRNA (ader | Q8CE96 [1- | 0 | 3552.638 | 0 | -0.03 | 0.807198 | 25.64 | 96.4 | 103.6 | 19.26 | 8.79 | | High | High | High | High | High | High | High | High | High | High | High | 0 | 3.59E-06 | 3.86 |
| FALSE | High | MEDIDQSS | [-].MEDID( | 1xAcetyl [N | Q8K3G7 1x | 5.89E-07 | 0 | 3 | 4 | 7 | Q8K3G7; C | Reticulon ( | Q8K3G7 [1 | 0 | 3009.411 | 0 | 0.69 | 0.068817 | 18.23 | 76.5 | 123.5 | 5.46 | 25.9 | Shared | High | High | High | High | High | High | High | High | High | High | High | 0 | 3.02E-07 | 3.64 |
| FALSE | High | MEDQELM | [R].MEDQE | 1xPhosphc | Q6P5H2-2 | 0.029076 | 0.001857 | 1 | 2 | 1 | Q6P5H2-2 | Isoform 2 c | Q6P5H2-2 | 0 | 2959.423 | 0 | 2.96 | 0.022435 | 22.17 | 23.1 | 176.9 | 20.1 | 42.69 | | High | High | High | High | High | High | High | High | High | High | High | 0.000997 | 0.01036 | 2.33 |
| FALSE | High | MEDQELM | [R].MEDQE | 1xPhosphc | Q6P5H2-2 | 3.29E-05 | 0 | 1 | 2 | 3 | Q6P5H2-2 | Isoform 2 c | Q6P5H2-2 | 0 | 2975.418 | 0 | 2.29 | 0.003952 | 9.47 | 35 | 165 | 14.8 | 20.7 | | High | High | High | High | High | High | High | High | High | High | High | 0 | 5.16E-06 | 3.21 |
| FALSE | High | MEDQGQS | [R-].MEDQ | 1xPhosphc | Q3TJP2 1xI | 0.083783 | 0.006345 | 1 | 5 | 1 | Q3TJP2 | Putative ur | Q3TJP2 [16 | 0 | 3301.604 | 0 | -0.03 | 0.981451 | 27.24 | 99 | 101 | 14.7 | 11.17 | | High | High | High | High | High | High | High | High | High | High | High | 0.003265 | 0.0352 | 2.42 |
| FALSE | High | MEDSMDN | [-].MEDSM | 1xAcetyl [N | Q5SQB7 1x | 2.23E-05 | 0 | 2 | 8 | 3 | Q5SQB7; C | MCG68069 | Q5SQB7 [1 | 0 | 3258.389 | 0 | -0.52 | 0.123449 | 22.59 | 109.7 | 90.3 | 13.8 | 33.22 | Shared | High | High | High | High | High | High | High | High | High | High | High | 0 | 6.53E-06 | 4.23 |
| FALSE | High | MEESFSSK | [R].MEESF! | 1xPhosphc | Q3U955 1x | 0.112856 | 0.009146 | 1 | 8 | 1 | Q3U955 | Putative ur | Q3U955 [3 | 0 | 1482.695 | 0 | 0.94 | 0.101001 | 16.31 | 69.2 | 130.8 | 16.85 | 41.6 | | High | High | High | High | High | High | High | High | High | High | High | 0.004583 | 0.04977 | 1.53 |
| FALSE | High | MELGTPLR | [K].MELGT | 1xPhosphc | Q8BTI8 1xF | 0.058664 | 0.004294 | 1 | 3 | 2 | Q8BTI8 | Serine/argi | Q8BTI8 [95 | 0 | 1225.621 | 0 | -0.25 | 0.149476 | 6.34 | 110.4 | 89.6 | 10.19 | 29.54 | | High | High | High | High | High | High | High | High | High | High | High | 0.002187 | 0.0232 | 1.09 |
| FALSE | High | MESEAGAI | [K].MESEA | 1xPhosphc | Q3U6P5 1x | 2.57E-11 | 0 | 2 | 6 | 23 | Q3U6P5; C | Putative ur | Q3U6P5 [2 | 1 | 4149.749 | 0 | -0.45 | 0.008588 | 1.33 | 115.5 | 84.5 | 1.87 | 16.13 | Shared | High | High | High | High | High | High | High | High | High | High | High | 0 | 7.11E-13 | 5.64 |
| FALSE | High | MESEAGAI | [K].MESEA | 1xDeamida | Q3U6P5 1x | 1.09E-06 | 0 | 2 | 6 | 15 | Q3U6P5; C | Putative ur | Q3U6P5 [2 | 1 | 4150.733 | 0 | -0.49 | 0.065117 | 19.92 | 112.5 | 87.5 | 15.41 | 20.14 | Shared | High | High | High | High | High | High | High | High | High | High | High | 0 | 1.11E-07 | 4.83 |
| FALSE | High | MESEAGAI | [K].MESEA | 2xPhosphc | Q3U6P5 2x | 9.11E-05 | 0 | 2 | 6 | 3 | Q3U6P5; C | Putative ur | Q3U6P5 [2 | 1 | 4229.715 | 0 | -0.92 | 0.000614 | 2.43 | 130.3 | 69.7 | 3.62 | 6.29 | Shared | High | High | High | High | High | High | High | High | High | High | High | 0 | 1.62E-05 | 5.28 |
| FALSE | High | MESEAGAI | [K].MESEA | 1xPhosphc | Q3U6P5 1x | 0.064862 | 0.004664 | 2 | 6 | 1 | Q3U6P5; C | Putative ur | Q3U6P5 [2 | 2 | 4866.104 | 0 | -1.29 | 0.007465 | 7.21 | 142 | 58 | 17.3 | 20.74 | Shared | High | High | High | High | High | High | High | High | High | High | High | 0.002432 | 0.02606 | 3.84 |
| FALSE | High | MESLGSPR | [K].MESLG | 1xPhosphc | P70399 1xI | 0.088701 | 0.006669 | 1 | 5 | 1 | P70399 | TP53-bindi | P70399 [52 | 0 | 1185.554 | 0 | 0.33 | 0.626553 | 8.3 | 89.2 | 110.8 | 9.72 | 27.51 | | High | High | High | High | High | High | High | High | High | High | High | 0.003447 | 0.03744 | 1.42 |
| FALSE | High | MESPIPILPI | [R].MESPIF | 1xPhosphc | B7ZNM7 1 | 0.016451 | 0.001124 | 1 | 3 | 1 | B7ZNM7 | Sept5 prot | B7ZNM7 [3 | 0 | 2616.365 | 0 | 3.18 | 0.000173 | 18.81 | 18.5 | 181.5 | 26.41 | 6.47 | | High | High | High | High | High | High | High | High | High | High | High | 0.000585 | 0.005478 | 2.85 |
| FALSE | High | MESTTVSP | [R].MESTT | 1xCarbami | B0V2M3 1 | 9.2E-05 | 0 | 1 | 3 | 2 | B0V2M3 | Zinc finger | B0V2M3 [2 | 0 | 3071.467 | 0 | -0.45 | 0.025269 | 5.73 | 124.7 | 75.3 | 9.86 | 41.35 | | High | High | High | High | High | High | High | High | High | High | High | 0 | 1.92E-05 | 4.29 |
| FALSE | High | METAPIDT | [K].METAP | 1xPhosphc | Q9JHI7 1xF | 1.75E-05 | 0 | 1 | 2 | 1 | Q9JHI7 | Exosome c | Q9JHI7 [29 | 0 | 1914.872 | 0 | -0.56 | 0.001818 | 2.49 | 116.2 | 83.8 | 6 | 8 | | High | High | High | High | High | High | High | High | High | High | High | 0 | 2.54E-06 | 3.08 |
| FALSE | High | METQALEF | [-].METQA | 1xAcetyl [N | Q9CXG9 1x | 0.027839 | 0.001755 | 1 | 2 | 2 | Q9CXG9 | PHD finger | Q9CXG9 [1 | 0 | 2543.194 | 0 | -1.29 | 0.000928 | 2.23 | 145.6 | 54.4 | 9.68 | 20.81 | | High | High | High | High | High | High | High | High | High | High | High | 0.000953 | 0.00988 | 2.53 |
| FALSE | High | METVSNAS | [R].METVS | 1xPhosphc | F8WHU5 1 | 1.57E-08 | 0 | 1 | 4 | 1 | F8WHU5 | Sister chro | F8WHU5 [ | 0 | 2103.922 | 0 | 0.51 | 0.019821 | 1.84 | 82.3 | 117.7 | 3.6 | 14.53 | | High | High | High | High | High | High | High | High | High | High | High | 0 | 9.54E-10 | 5.03 |
| FALSE | High | METVSNAS | [R].METVS | 2xPhosphc | F8WHU5 2 | 0.001107 | 0 | 1 | 4 | 1 | F8WHU5 | Sister chro | F8WHU5 [ | 0 | 2183.888 | 0 | 1.23 | 0.002078 | 8.94 | 57.4 | 142.6 | 10.61 | 12.48 | | High | High | High | High | High | High | High | High | High | High | High | 0 | 0.000267 | 2.51 |
| FALSE | High | MEVKPPPG | [-].MEVKPI | 1xAcetyl [N | Q3UZG3 1x | 0.012731 | 0.000889 | 1 | 4 | 1 | Q3UZG3 | Putative ur | Q3UZG3 [1 | 0 | 2099.031 | 0 | -1.12 | 0.000289 | 0.57 | 134.9 | 65.1 | 5.95 | 6.16 | | High | High | High | High | High | High | High | High | High | High | High | 0.000449 | 0.00412 | 0.89 |
| FALSE | High | MEVLSPAS | [K].MEVLSI | 1xPhosphc | Q6NZR2 1x | 3.32E-05 | 0 | 1 | 1 | 2 | Q6NZR2 | Myb/SANT | Q6NZR2 [2 | 0 | 2935.359 | 0 | -0.73 | 0.001471 | 3.4 | 125.5 | 74.5 | 5.34 | 8.38 | | High | High | High | High | High | High | High | High | High | High | High | 0 | 5.19E-06 | 5.02 |
| FALSE | High | LVTTASYSS | [R].LVTTAS | 1xPhosphc | Q9D666 1x | 0.002793 | 0.000109 | 1 | 10 | 2 | Q9D666 | SUN doma | Q9D666 [5 | 0 | 2692.266 | 0 | 0.46 | 0.091402 | 31.27 | 87 | 113 | 13.7 | 14.38 | | High | High | High | High | High | High | High | High | High | High | High | 0.000585 | 0.005455 | 2.85 |
| FALSE | High | MPHLDSPG | [R].MPHLD | 1xPhosphc | Q6ZWQ0 1 | 0.003055 | 0.00016 | 1 | 10 | 1 | Q6ZWQ0 | Nesprin-2 ( | Q6ZWQ0 [ | 0 | 1717.793 | 0 | 1.88 | 0.001686 | 24.7 | 43.9 | 156.1 | 24.3 | 13.33 | | High | High | High | High | High | High | High | High | High | High | High | 7.19E-05 | 0.000837 | 2.59 |
| FALSE | High | IVTSPDGLD | [R].IVTSPD | 1xPhosphc | Q0VF22 1x | 0.000231 | 0 | 1 | 1 | 2 | Q0VF22 | Coiled-coil | Q0VF22 [6 | 0 | 2193.073 | 0 | -1.47 | 0.000431 | 1.9 | 146.8 | 53.2 | 6.47 | 8.32 | | High | High | High | High | High | High | High | High | High | High | High | 0 | 4.57E-05 | 2.75 |
| FALSE | High | LVSYTNLT( | [K].LVSYTN | 1xPhosphc | F8WIJ0 1xF | 0.009756 | 0.000736 | 1 | 12 | 2 | F8WIJ0 | Solute carr | F8WIJ0 [88 | 0 | 1832.992 | 0 | 1.08 | 0.018219 | 36.66 | 66.5 | 133.5 | 7.11 | 23.38 | | High | High | High | High | High | High | High | High | High | High | High | 0.000399 | 0.003454 | 2.84 |
| FALSE | High | LTSVSSSVD | [R].LTSVSS | 3xPhosphc | Q3TWI9 3x | 0.076555 | 0.005826 | 1 | 5 | 1 | Q3TWI9 | CSC1-like p | Q3TWI9 [1 | 0 | 1909.758 | 0 | 2.03 | 0.001754 | 28.07 | 41.3 | 158.7 | 13.87 | 16.15 | | High | High | High | High | High | High | High | High | High | High | High | 0.002976 | 0.03154 | 2.57 |
| FALSE | High | ITSVSTGNL | [K].ITSVST( | 1xCarbami | E9Q9C3 1x | 0.064862 | 0.004664 | 1 | 5 | 2 | E9Q9C3 | Afadin OS= | E9Q9C3 [1 | 0 | 3899.872 | 0 | -1.07 | 0.03022 | 9.37 | 135.6 | 64.4 | 15.24 | 25.64 | | High | High | High | High | High | High | High | High | High | High | High | 0.002432 | 0.02606 | 2.85 |
| FALSE | High | ITSVSTGNL | [K].ITSVST( | 1xCarbami | E9Q9C3 2x | 0.003433 | 0.00021 | 1 | 5 | 1 | E9Q9C3 | Afadin OS= | E9Q9C3 [1 | 0 | 3979.839 | 0 | -0.78 | 0.007561 | 5.09 | 127.3 | 72.7 | 15.33 | 6.19 | | High | High | High | High | High | High | High | High | High | High | High | 9.42E-05 | 0.000951 | 4.26 |
| FALSE | High | LTTLANGE/ | [K].LTTLAN | 1xPhosphc | B2RUM8 1 | 3.47E-05 | 0 | 1 | 2 | 4 | B2RUM8 | RNA helica | B2RUM8 [ | 0 | 2546.316 | 0 | -0.97 | 0.001532 | 1.61 | 132.5 | 67.5 | 7.48 | 7.22 | | High | High | High | High | High | High | High | High | High | High | High | 0 | 5.46E-06 | 4.04 |
| FALSE | High | LTTLANGE/ | [K].LTTLAN | 1xDeamida | B2RUM8 1 | 1.15E-05 | 0 | 1 | 2 | 3 | B2RUM8 | RNA helica | B2RUM8 [ | 0 | 2547.3 | 0 | -1.02 | 0.001489 | 2.48 | 132 | 68 | 8.15 | 5.65 | | High | High | High | High | High | High | High | High | High | High | High | 0 | 1.58E-06 | 2.94 |
| FALSE | High | ITTITPGSTI | [K].ITTITPG | 1xPhosphc | E9Q7F2 1x | 0.00311 | 0.00016 | 1 | 1 | 5 | E9Q7F2 | E3 ubiquiti | E9Q7F2 [3 | 0 | 2605.416 | 0 | 0.4 | 0.08169 | 15.44 | 86.1 | 113.9 | 14.09 | 12.57 | | High | High | High | High | High | High | High | High | High | High | High | 0.000449 | 0.00421 | 3.09 |
| FALSE | High | LTTTGQVT | [K].LTTTGC | 1xPhosphc | K4DI59 1xF | 0.000173 | 0 | 1 | 8 | 4 | K4DI59 | DNA ligase | K4DI59 [20 | 0 | 1769.981 | 0 | -0.22 | 0.086994 | 11.44 | 106.8 | 93.2 | 5.25 | 10.76 | | High | High | High | High | High | High | High | High | High | High | High | 0 | 0.00012 | 2.44 |
| FALSE | High | LTTTSGVQ | [K].LTTTSG | 1xPhosphc | Q6PCM2 1 | 1.26E-07 | 0 | 1 | 3 | 1 | Q6PCM2 | Integrator | Q6PCM2 [ | 0 | 3285.738 | 0 | 0.73 | 0.004615 | 11.47 | 73.8 | 126.2 | 9.35 | 13.31 | | High | High | High | High | High | High | High | High | High | High | High | 0 | 9.94E-09 | 3.35 |
| FALSE | High | ITTVTPQIQ | [K].ITTVTP | 1xPhosphc | A2A6Q5-3 | 0.000191 | 0 | 1 | 3 | 3 | A2A6Q5-3 | Isoform 3 c | A2A6Q5-3 | 0 | 2240.246 | 0 | 0.25 | 0.049395 | 18.14 | 88.9 | 111.1 | 8.82 | 8.11 | | High | High | High | High | High | High | High | High | High | High | High | 0 | 3.7E-05 | 2.84 |
| FALSE | High | LTVASPK | [KR].LTVAS | 1xPhosphc | Q99NH0 1x | 0.093879 | 0.007306 | 1 | 7 | 1 | Q99NH0 | Ankyrin re | Q99NH0 [1 | 0 | 1253.727 | 0 | -0.37 | 0.004809 | 2.45 | 113.2 | 86.8 | 3.11 | 7.73 | | High | High | High | High | High | High | High | High | High | High | High | 0.003727 | 0.04004 | 1.06 |
| FALSE | High | LTVENSPK | [K].LTVENS | 1xPhosphc | Q9CQJ6 1x | 0.001629 | 5.66E-05 | 1 | 2 | 2 | Q9CQJ6 | Density-re | Q9CQJ6 [6 | 0 | 1425.775 | 0 | -0.12 | 0.505108 | 13.3 | 106.1 | 93.9 | 2.07 | 14.19 | | High | High | High | High | High | High | High | High | High | High | High | 0.00196 | 0.02006 | 2.09 |
| FALSE | High | ITVHSSPPL | [K].ITVHSS | 1xPhosphc | G3UZM1 1 | 0.007274 | 0.00062 | 1 | 4 | 1 | G3UZM1 | Probable J | G3UZM1 [ | 0 | 1717.965 | 0 | -0.47 | 0.000731 | 2.44 | 115.8 | 84.2 | 4.26 | 8.56 | | High | High | High | High | High | High | High | High | High | High | High | 0.000275 | 0.0022 | 2.07 |
| FALSE | High | ITVNGDSS/ | [K].ITVNGI | 1xPhosphc | G3X8Y3 1x | 1.23E-06 | 0 | 1 | 5 | 9 | G3X8Y3 | N-alpha-ac | G3X8Y3 [8 | 0 | 2201.006 | 0 | 0 | 0.383348 | 7.23 | 96.7 | 103.3 | 9.64 | 7.4 | | High | High | High | High | High | High | High | High | High | High | High | 0 | 1.29E-07 | 4.08 |
| FALSE | High | LTSVSSSVD | [R].LTSVSS | 2xPhosphc | Q3TWI9 2x | 7.15E-05 | 0 | 1 | 5 | 5 | Q3TWI9 | CSC1-like p | Q3TWI9 [1 | 0 | 1829.792 | 0 | 3.34 | 0.00159 | 13.39 | 16.7 | 183.3 | 16.54 | 17.67 | | High | High | High | High | High | High | High | High | High | High | High | 0 | 1.23E-05 | 3.62 |
| FALSE | High | ITVNGDSS/ | [K].ITVNGI | 1xDeamida | G3X8Y3 1x | 4.02E-09 | 0 | 1 | 5 | 27 | G3X8Y3 | N-alpha-ac | G3X8Y3 [8 | 0 | 2201.99 | 0 | 0 | 0.781856 | 5.25 | 99.1 | 100.9 | 7.27 | 7.49 | | High | High | High | High | High | High | High | High | High | High | High | 0 | 6.5E-10 | 4.53 |
| FALSE | High | ITVQASPGI | [R].ITVQAS | 1xPhosphc | Q80XI6 1xF | 0.009168 | 0.000736 | 1 | 1 | 1 | Q80XI6 | Mitogen-ac | Q80XI6 [50 | 0 | 1465.761 | 0 | 0.85 | 0.002155 | 5.76 | 69.3 | 130.7 | 9.67 | 9.09 | | High | High | High | High | High | High | High | High | High | High | High | 0.000327 | 0.002861 | 1.94 |
| FALSE | High | LTVVSPNR | [R].LTVVSP | 1xPhosphc | Q14AM7 1 | 0.119269 | 0.00979 | 1 | 1 | 1 | Q14AM7 | UPF0472 p | Q14AM7 [ | 0 | 1194.645 | 0 | -0.24 | 0.017728 | 5.16 | 108.4 | 91.6 | 9.2 | 8.06 | | High | High | High | High | High | High | High | High | High | High | High | 0.004894 | 0.05329 | 0.47 |
| FALSE | High | LTWHSCPE | [R].LTWHS | 1xCarbami | Q32P00 1x | 3.5E-05 | 0 | 1 | 3 | 3 | Q32P00 | Chromobo | Q32P00 [1 | 0 | 1781.74 | 0 | 0.58 | 0.031858 | 15.64 | 80.2 | 119.8 | 8.27 | 11.59 | | High | High | High | High | High | High | High | High | High | High | High | 0 | 5.51E-06 | 3.63 |
| FALSE | High | LTWHSYPS | [R].LTWHS | 1xPhosphc | P83917 1xI | 0.008614 | 0.000736 | 1 | 1 | 1 | P83917 | Chromobo | P83917 [16 | 0 | 2130.979 | 0 | -0.34 | 0.016852 | 5.16 | 111.3 | 88.7 | 8.04 | 13.45 | | High | High | High | High | High | High | High | High | High | High | High | 0.000327 | 0.002659 | 2.6 |
| FALSE | High | LTWHSYPS | [R].LTWHS | 1xPhosphc | P83917 1xI | 0.026885 | 0.001676 | 1 | 1 | 1 | P83917 | Chromobo | P83917 [16 | 1 | 2488.236 | 0 | -0.36 | 0.040799 | 10.14 | 112.4 | 87.6 | 7.91 | 14.63 | | High | High | High | High | High | High | High | High | High | High | High | 0.000903 | 0.009498 | 1.28 |
| FALSE | High | LTYIPENTP | [K].LTYIPEI | 1xDeamida | Q2PZL6 2xI | 0.024638 | 0.001612 | 1 | 1 | 1 | Q2PZL6 | Protocadh | Q2PZL6 [2 | 0 | 2240.074 | 0 | -1.59 | 0.000618 | 0.53 | 149.1 | 50.9 | 8.46 | 4.02 | | High | High | High | High | High | High | High | High | High | High | High | 0.000872 | 0.00865 | 2.86 |
| FALSE | High | LVAEQVM( | [R].LVAEQ\ | 1xPhosphc | Q80TJ7 1xI | 1.52E-06 | 0 | 1 | 3 | 3 | Q80TJ7 | Histone lys | Q80TJ7 [60 | 0 | 3106.365 | 0 | -0.03 | 0.38408 | 13.93 | 104.8 | 95.2 | 9.67 | 14.26 | | High | High | High | High | High | High | High | High | High | High | High | 0 | 1.63E-07 | 4.94 |
| FALSE | High | LVAEQVM( | [R].LVAEQ\ | 1xDeamida | Q80TJ7 1xI | 0.002553 | 0.000109 | 1 | 3 | 1 | Q80TJ7 | Histone lys | Q80TJ7 [60 | 0 | 3107.349 | 0 | -0.31 | 0.268555 | 6.02 | 108.8 | 91.2 | 17.92 | 8.23 | | High | High | High | High | High | High | High | High | High | High | High | 4.94E-05 | 0.000681 | 3.88 |
| FALSE | High | IVAHVPQTI | [K].IVAHVF | 1xPhosphc | A2ASY1 1xI | 0.000668 | 0 | 1 | 5 | 1 | A2ASY1 | TAF3 RNA | A2ASY1 [3 | 0 | 2220.185 | 0 | -0.15 | 0.199714 | 5.56 | 104.7 | 95.3 | 8.67 | 5.63 | | High | High | High | High | High | High | High | High | High | High | High | 0 | 0.000152 | 1.87 |
| FALSE | High | IVAPTVSQI | [K].IVAPTV | 1xCarbami | Q8CFQ3 1x | 5.7E-05 | 0 | 1 | 2 | 2 | Q8CFQ3 | Intron-binc | Q8CFQ3 [1 | 0 | 2727.456 | 0 | 0.63 | 0.04241 | 15.7 | 78.5 | 121.5 | 10.41 | 23.13 | | High | High | High | High | High | High | High | High | High | High | High | 0 | 9.54E-06 | 3.21 |
| FALSE | High | LVATSPGS! | [R].LVATSP | 1xPhosphc | A2RSY1 1xI | 7.08E-05 | 0 | 1 | 3 | 2 | A2RSY1 | KAT8 regul | A2RSY1 [7 | 0 | 3744.971 | 0 | -0.9 | 0.014634 | 3.17 | 129.7 | 70.3 | 17.77 | 13.36 | | High | High | High | High | High | High | High | High | High | High | High | 0 | 1.9E-05 | 3.88 |
| FALSE | High | LVDSDGSL/ | [R].LVDSD( | 1xPhosphc | E9QN92 1x | 0.000107 | 0 | 1 | 5 | 2 | E9QN92 | Metallored | E9QN92 [5 | 0 | 1867.982 | 0 | -1.4 | 0.011586 | 2.68 | 145.4 | 54.6 | 9.24 | 34.66 | | High | High | High | High | High | High | High | High | High | High | High | 0 | 1.94E-05 | 2.7 |
| FALSE | High | LVDSDGSL/ | [R].LVDSD( | 2xPhosphc | E9QN92 2x | 0.043236 | 0.003228 | 1 | 5 | 1 | E9QN92 | Metallored | E9QN92 [5 | 0 | 1947.948 | 0 | -0.22 | 0.1296 | 6.34 | 107.6 | 92.4 | 10.81 | 12.68 | | High | High | High | High | High | High | High | High | High | High | High | 0.001651 | 0.01628 | 2.78 |
| FALSE | High | ITVPPEGPL | [R].ITVPPE | 1xPhosphc | Q61037 1x | 0.006031 | 0.00062 | 1 | 14 | 2 | Q61037 | Tuberin OS | Q61037 [1 | 0 | 1842.956 | 0 | -0.33 | 0.167906 | 6.28 | 110.8 | 89.2 | 11.83 | 16.96 | | High | High | High | High | High | High | High | High | High | High | High | 0.000327 | 0.002539 | 2.02 |
| FALSE | High | LVDSPANIE | [K].LVDSPA | 1xPhosphc | Q923A2 1x | 1.54E-05 | 0 | 1 | 1 | 2 | Q923A2 | Protein Spi | Q923A2 [5 | 0 | 2783.356 | 0 | -0.58 | 0.004997 | 1.76 | 121.6 | 78.4 | 12.37 | 5.4 | | High | High | High | High | High | High | High | High | High | High | High | 0 | 2.78E-06 | 3.58 |
| FALSE | High | LTSVSSSVD | [R].LTSVSS | 1xPhosphc | Q3TWI9 1x | 2.66E-06 | 0 | 1 | 5 | 8 | Q3TWI9 | CSC1-like p | Q3TWI9 [1 | 0 | 1749.826 | 0 | 1.22 | 0.000158 | 9.2 | 60.9 | 139.1 | 7.95 | 3.38 | | High | High | High | High | High | High | High | High | High | High | High | 0 | 3.05E-07 | 3.46 |
| FALSE | High | LTSTSTSDT | [R].LTSTST | 1xPhosphc | Q8C180 1x | 7.47E-08 | 0 | 1 | 1 | 4 | Q8C180 | Fibroblast | Q8C180 [3 | 0 | 2247.045 | 0 | 1.3 | 0.00034 | 5.83 | 57.8 | 142.2 | 5.78 | 6.37 | | High | High | High | High | High | High | High | High | High | High | High | 0 | 5.51E-09 | 4.94 |
| FALSE | High | LTPVSLSNS | [K].LTPVSL | 1xDeamida | Q61687 1x | 0.021797 | 0.001507 | 1 | 4 | 1 | Q61687 | Transcripti | Q61687 [5 | 0 | 1795.002 | 0 | -0.2 | 0.013703 | 5.02 | 102.4 | 97.6 | 6.15 | 9.46 | | High | High | High | High | High | High | High | High | High | High | High | 0.000801 | 0.007477 | 1.91 |
| FALSE | High | LTPVSLSNS | [K].LTPVSL | 2xPhosphc | Q61687 2x | 9.56E-06 | 0 | 1 | 4 | 7 | Q61687 | Transcripti | Q61687 [5 | 0 | 1873.984 | 0 | 0.29 | 0.010126 | 5.22 | 89.2 | 110.8 | 6.23 | 9.18 | | High | High | High | High | High | High | High | High | High | High | High | 0 | 1.29E-06 | 3.03 |
| FALSE | High | LTPVSPVQI | [R].LTPVSP | 1xPhosphc | F8VPJ6 1xF | 6.15E-06 | 0 | 1 | 3 | 1 | F8VPJ6 | Regulatory | F8VPJ6 [11 | 0 | 2429.295 | 0 | -0.53 | 0.001375 | 1.19 | 115.5 | 84.5 | 4.75 | 3.87 | | High | High | High | High | High | High | High | High | High | High | High | 0 | 7.79E-07 | 2.84 |
| FALSE | High | LTQADISEC | [R].LTQADI | 1xPhosphc | F2Z3U3 1xI | 0.000143 | 0 | 1 | 1 | 2 | F2Z3U3 | Ras associa | F2Z3U3 [1 | 0 | 3076.604 | 0 | 1.26 | 0.003773 | 21.48 | 58 | 142 | 15.6 | 17.24 | | High | High | High | High | High | High | High | High | High | High | High | 0 | 2.7E-05 | 3.26 |
| FALSE | High | LTQADISEC | [R].LTQADI | 1xPhosphc | F2Z3U3 1xI | 1.23E-05 | 0 | 1 | 1 | 2 | F2Z3U3 | Ras associa | F2Z3U3 [1 | 0 | 3092.599 | 0 | 1.06 | 0.018772 | 31.68 | 63.5 | 136.5 | 18.84 | 22.83 | | High | High | High | High | High | High | High | High | High | High | High | 0 | 1.7E-06 | 3.88 |
| FALSE | High | LTQYHGGS | [R].LTQYH( | 1xPhosphc | Q91X84 1x | 6.59E-05 | 0 | 1 | 1 | 1 | Q91X84 | CREB-regu | Q91X84 [5 | 0 | 2079.059 | 0 | 2.75 | 0.000282 | 11.12 | 28 | 172 | 13.55 | 7.02 | | High | High | High | High | High | High | High | High | High | High | High | 0 | 1.12E-05 | 1.25 |
| FALSE | High | LTRPAASP/ | [K].LTRPAA | 1xPhosphc | Q8CFC7 1x | 1.78E-05 | 0 | 1 | 4 | 2 | Q8CFC7 | CLK4-assoc | Q8CFC7 [5 | 0 | 1835.019 | 0 | -0.04 | 0.35328 | 11.67 | 103 | 97 | 4.62 | 8.27 | | High | High | High | High | High | High | High | High | High | High | High | 0 | 0.000208 | 1.45 |
| FALSE | High | LTSDAEDLS | [K].LTSDAE | 1xCarbami | Q921C3 1x | 8.7E-08 | 0 | 1 | 4 | 8 | Q921C3 | Bromodom | Q921C3 [1 | 0 | 2104.967 | 0 | -1.12 | 0.001317 | 2.36 | 137.8 | 62.2 | 7.22 | 10.09 | | High | High | High | High | High | High | High | High | High | High | High | 0 | 7.08E-09 | 4.46 |
| FALSE | High | LTSDAEDLS | [K].LTSDAE | 1xCarbami | Q921C3 2x | 0.015329 | 0.001089 | 1 | 4 | 1 | Q921C3 | Bromodom | Q921C3 [1 | 0 | 2184.933 | 0 | -0.3 | 0.047897 | 4.27 | 112.5 | 87.5 | 13.19 | 6.98 | | High | High | High | High | High | High | High | High | High | High | High | 0.000571 | 0.005078 | 2.55 |
| FALSE | High | LTSDAEK | [RK].LTSDA | 1xPhosphc | Q497V5 1x | 0.004824 | 0.000456 | 1 | 10 | 1 | Q497V5 | S1 RNA-bir | Q497V5 [5 | 0 | 1301.675 | 0 | 2.36 | 0.000435 | 17.14 | 32 | 168 | 9.97 | 9.59 | | High | High | High | High | High | High | High | High | High | High | High | 0.000204 | 0.001386 | 2.13 |
| FALSE | High | ITSEAEDLV | [R].ITSEAEI | 1xPhosphc | P61290 1xI | 5.98E-06 | 0 | 1 | 3 | 4 | P61290 | Proteasom | P61290 [22 | 0 | 2219.14 | 0 | 1.84 | 0.000908 | 14.86 | 43.6 | 156.4 | 5.17 | 10.95 | | High | High | High | High | High | High | High | High | High | High | High | 0.000183 | 0.001259 | 2.73 |
| FALSE | High | LTSHTPGLI | [R].LTSHTP | 1xPhosphc | Q6ZWQ0 1 | 0.032831 | 0.002195 | 1 | 10 | 2 | Q6ZWQ0 | Nesprin-2 ( | Q6ZWQ0 [ | 0 | 1850.93 | 0 | 4.01 | 0.001657 | 22.14 | 11.6 | 188.4 | 19.85 | 20.04 | | High | High | High | High | High | High | High | High | High | High | High | 0.001151 | 0.01188 | 1.92 |
| FALSE | High | LTSLDGWC | [R].LTSLDG | 1xCarbami | G3UW83 1 | 0.022378 | 0.001542 | 1 | 2 | 1 | G3UW83 | Nuclear fac | G3UW83 [ | 0 | 1487.692 | 0 | 0.09 | 0.197066 | 17.26 | 96.3 | 103.7 | 14.8 | 4.97 | | High | High | High | High | High | High | High | High | High | High | High | 0.000814 | 0.00775 | 2.4 |
| FALSE | High | LTSVSGESS | [R].LTSVSG | 1xPhosphc | A0A1B0GS | 0.033404 | 0.002195 | 1 | 2 | 3 | A0A1B0GS | GRB2-asso | A0A1B0GS | 0 | 1931.968 | 0 | -1.29 | 0.000729 | 0.46 | 142.6 | 57.4 | 7.15 | 15.48 | | High | High | High | High | High | High | High | High | High | High | High | 0.001151 | 0.01216 | 2.09 |
| FALSE | High | LTSIGSDED | [R].LTSIGSI | 1xPhosphc | Q924N4 1x | 0.001011 | 0 | 2 | 4 | 3 | Q924N4; C | Solute carr | Q924N4 [1 | 0 | 2612.19 | 0 | 1.9 | 0.002235 | 25.55 | 40.7 | 159.3 | 20.26 | 18.35 | Shared | High | High | High | High | High | High | High | High | High | High | High | 0 | 0.000242 | 3.36 |
| FALSE | High | ITSISPTPDF | [R].ITSISPT | 1xPhosphc | Q8K120 1x | 1.45E-06 | 0 | 1 | 2 | 3 | Q8K120 | Nuclear fac | Q8K120 [1 | 0 | 3152.451 | 0 | -4.28 | | 0.08 | 188.3 | 11.7 | 14.95 | 69.06 | | High | High | High | High | High | High | High | High | High | Not Found | High | 0 | 2.07E-07 | 4.67 |
| FALSE | High | ITSPFQLK | [K].ITSPFQ | 1xPhosphc | Q6PGB8-2 | 0.08727 | 0.006537 | 1 | 6 | 1 | Q6PGB8-2 | Isoform 2 c | Q6PGB8-2 | 0 | 1471.833 | 0 | -1.94 | 0.00028 | 0.65 | 158.8 | 41.2 | 5.16 | 10.3 | | High | High | High | High | High | High | High | High | High | High | High | 0.003375 | 0.03685 | 1.65 |

| | | | | | | | | | | | | | | | | | | | | | | | | | | | | | | | | | | | | | | |
|---|---|---|---|---|---|---|---|---|---|---|---|---|---|---|---|---|---|---|---|---|---|---|---|---|---|---|---|---|---|---|---|---|---|---|---|---|---|---|
| FALSE | High | ITSPLMEPS | [R].ITSPLM | 1xPhosphc | Q3TUI9 1xl | 0.001629 | 5.66E-05 | 1 | 3 | 2 | Q3TUI9 | Proteasom | Q3TUI9 [54 | 0 | 1970.032 | 0 | -0.64 | 0.020226 | 3.16 | 121.7 | 78.3 | 13.45 | 9.51 | | High | High | High | High | High | High | High | High | High | High | High | 9.42E-05 | 0.001029 | 2.7 |
| FALSE | High | ITSPLMEPS | [R].ITSPLM | 1xPhosphc | Q3TUI9 1xl | 1.86E-05 | 0 | 1 | 3 | 8 | Q3TUI9 | Proteasom | Q3TUI9 [54 | 0 | 1986.027 | 0 | -0.65 | 0.000928 | 4.85 | 124 | 76 | 8.77 | 12.07 | | High | High | High | High | High | High | High | High | High | High | High | 0.001011 | 0.01094 | 3.36 |
| FALSE | High | LTSPLQENS | [K].LTSPLQ | 1xPhosphc | Q8CIV7 1xl | 2.16E-06 | 0 | 1 | 3 | 1 | Q8CIV7 | Transcripti | Q8CIV7 [25 | 0 | 2472.216 | 0 | -1.35 | 0.000928 | 1.39 | 145 | 55 | 8.4 | 10.54 | | High | High | High | High | High | High | High | High | High | High | High | 0 | 2.41E-07 | 4.31 |
| FALSE | High | ITSPVLVGE | [R].ITSPVLV | 1xPhosphc | Q8R4H2 1x | 3.58E-06 | 0 | 2 | 2 | 2 | Q8R4H2; F | Rho guanir | Q8R4H2 [1 | 0 | 2515.332 | 0 | 1.85 | 0.000266 | 7.02 | 43.1 | 156.9 | 4.26 | 5.92 | Shared | High | High | High | High | High | High | High | High | High | High | High | 0 | 4.26E-07 | 3.29 |
| FALSE | High | LTSPVTSISI | [K].LTSPVT | 1xPhosphc | Q8BMB0 1 | 6.23E-08 | 0 | 1 | 2 | 3 | Q8BMB0 | BRCA2-inte | Q8BMB0 [ | 0 | 2196.193 | 0 | -0.86 | 0.000978 | 4.21 | 129.1 | 70.9 | 4.37 | 8.27 | | High | High | High | High | High | High | High | High | High | High | High | 0 | 4.48E-09 | 3.81 |
| FALSE | High | LTSPVTSISI | [K].LTSPVT | 2xPhosphc | Q8BMB0 2 | 0.036416 | 0.002419 | 1 | 2 | 2 | Q8BMB0 | BRCA2-inte | Q8BMB0 [ | 0 | 2276.159 | 0 | -0.96 | 0.001211 | 0.92 | 134 | 66 | 10.98 | 10.31 | | High | High | High | High | High | High | High | High | High | High | High | 0.001296 | 0.01342 | 2.77 |
| FALSE | High | LTSQASVE( | [K].LTSQAS | 1xCarbami | Q64701 1x | 0.000203 | 0 | 1 | 5 | 1 | Q64701 | Retinoblast | Q64701 [34 | 0 | 2557.252 | 0 | 1.99 | 0.000191 | 6.59 | 38.8 | 161.2 | 4.8 | 4.38 | | High | High | High | High | High | High | High | High | High | High | High | 0 | 3.98E-05 | 2.2 |
| FALSE | High | LTSQDSYN | [K].LTSQDS | 1xPhosphc | Q8CCN5 1x | 2.59E-06 | 0 | 1 | 7 | 7 | Q8CCN5 | Breast carc | Q8CCN5 [4 | 0 | 2462.094 | 0 | 0.55 | 0.000964 | 3.33 | 80.1 | 119.9 | 8.36 | 3.37 | | High | High | High | High | High | High | High | High | High | High | High | 0 | 2.82E-06 | 4.29 |
| FALSE | High | LTSQDSYN | [K].LTSQDS | 1xDeamida | Q8CCN5 1x | 0.000352 | 0 | 1 | 7 | 10 | Q8CCN5 | Breast carc | Q8CCN5 [4 | 0 | 2463.078 | 0 | 0.14 | 0.437303 | 16.36 | 91.7 | 108.3 | 11.55 | 11.2 | | High | High | High | High | High | High | High | High | High | High | High | 0.000449 | 0.003913 | 3.34 |
| FALSE | High | LTSQDYEVI | [R].LTSQDY | 1xPhosphc | Q09XV5 1x | 0.014158 | 0.001011 | 1 | 2 | 5 | Q09XV5 | Chromodo | Q09XV5 [2 | 0 | 1419.672 | 0 | -0.35 | 0.051057 | 2.32 | 112.3 | 87.7 | 9.88 | 5.22 | | High | High | High | High | High | High | High | High | High | High | High | 0.000538 | 0.004623 | 3.18 |
| FALSE | High | LTSTSAITR | [K].LTSTSA | 1xPhosphc | Q8K212 1x | 0.016163 | 0.001124 | 1 | 3 | 2 | Q8K212 | Phosphofu | Q8K212 [3 | 0 | 1258.661 | 0 | 1.89 | 0.00135 | 21.71 | 37.1 | 162.9 | 11.72 | 14.05 | | High | High | High | High | High | High | High | High | High | High | High | 0.001183 | 0.01273 | 2.17 |
| FALSE | High | LTSIGSDED | [R].LTSIGSI | 2xPhosphc | Q924N4 2x | 0.000202 | 0 | 2 | 4 | 2 | Q924N4; Q | Solute carr | Q924N4 [1 | 0 | 2692.157 | 0 | 1.37 | 0.000724 | 6.37 | 54.3 | 145.7 | 16.38 | 8.29 | Shared | High | High | High | High | High | High | High | High | High | High | High | 0 | 3.96E-05 | 3.66 |
| FALSE | High | LVDSPANIE | [K].LVDSPA | 2xPhosphc | Q923A2 2x | 0.011246 | 0.000772 | 1 | 1 | 1 | Q923A2 | Protein Spi | Q923A2 [5 | 0 | 2863.323 | 0 | -0.37 | 0.019688 | 7.21 | 110.9 | 89.1 | 7.52 | 6.71 | | High | High | High | High | High | High | High | High | High | High | High | 0.000418 | 0.003589 | 3.1 |
| FALSE | High | LVELPVSSC | [K].LVELPV | 1xPhosphc | Q4U2R1 1x | 3.61E-06 | 0 | 1 | 3 | 3 | Q4U2R1 | E3 ubiquiti | Q4U2R1 [1 | 0 | 3328.443 | 0 | 0.03 | 0.590415 | 7.22 | 102.1 | 97.9 | 8.99 | 7.51 | | High | High | High | High | High | High | High | High | High | High | High | 0 | 2.53E-05 | 4.63 |
| FALSE | High | LVELPVSSC | [K].LVELPV | 2xPhosphc | Q4U2R1 2x | 0.000538 | 0 | 1 | 3 | 3 | Q4U2R1 | E3 ubiquiti | Q4U2R1 [1 | 0 | 3408.409 | 0 | -0.04 | 0.83043 | 12.58 | 101.8 | 98.2 | 8.26 | 9.32 | | High | High | High | High | High | High | High | High | High | High | High | 0 | 0.000149 | 5.39 |
| FALSE | High | IVPQSQVPI | [K].IVPQSC | 1xDeamida | Q3TUF7 1x | 0.059159 | 0.004294 | 1 | 4 | 3 | Q3TUF7 | YEATS dom | Q3TUF7 [4 | 0 | 2116.109 | 0 | 0.22 | 0.080129 | 17.26 | 92.5 | 107.5 | 8.66 | 3.64 | | High | High | High | High | High | High | High | High | High | High | High | 0.005561 | 0.06089 | 2.45 |
| FALSE | High | LVPVERNSI | [K].LVPVER | 1xPhosphc | Q8CB77 1x | 0.000223 | 0 | 1 | 1 | 2 | Q8CB77 | Elongin-A ( | Q8CB77 [7 | 1 | 2543.243 | 0 | 0.52 | 0.000648 | 1.6 | 82.8 | 117.2 | 2.01 | 4.05 | | High | High | High | High | High | High | High | High | High | High | High | 0 | 4.43E-05 | 2.6 |
| FALSE | High | LVQAIQAT/ | [R].LVQAIC | 1xPhosphc | Q6P4T0 1x | 4.08E-05 | 0 | 1 | 2 | 1 | Q6P4T0 | Autophagy | Q6P4T0 [1 | 0 | 2838.485 | 0 | 0.09 | 0.962676 | 4.27 | 96.7 | 103.3 | 25.25 | 8.9 | | High | High | High | High | High | High | High | High | High | High | High | 0 | 6.54E-06 | 3.18 |
| FALSE | High | IVQGSDVLI | [R].IVQGSD | 1xPhosphc | A2AKX3 1x | 9.15E-10 | 0 | 1 | 2 | 3 | A2AKX3 | Probable h | A2AKX3 [1 | 0 | 2406.066 | 0 | 0.02 | 0.486962 | 10.43 | 96.7 | 103.3 | 6.62 | 8.56 | | High | High | High | High | High | High | High | High | High | High | High | 0 | 3.9E-11 | 5.46 |
| FALSE | High | LVQSPNSYI | [R].LVQSPN | 1xPhosphc | Q6ZWU9 1 | 0.00034 | 0 | 1 | 6 | 1 | Q6ZWU9 | 40S riboso | Q6ZWU9 [ | 0 | 2066.043 | 0 | -0.26 | 0.04193 | 7.34 | 109.7 | 90.3 | 2.84 | 10 | | High | High | High | High | High | High | High | High | High | High | High | 0 | 7.11E-05 | 2.79 |
| FALSE | High | LVQSPNSYI | [R].LVQSPN | 1xPhosphc | Q6ZWU9 1 | 0.074071 | 0.0057 | 1 | 6 | 3 | Q6ZWU9 | 40S riboso | Q6ZWU9 [ | 0 | 2082.038 | 0 | -0.29 | 0.108361 | 2.67 | 110.1 | 89.9 | 4.08 | 15.3 | | High | High | High | High | High | High | High | High | High | High | High | 0.002924 | 0.03037 | 1.98 |
| FALSE | High | LVRPVTMV | [K].LVRPVT | 1xPhosphc | Q640L6 1xl | 0.002232 | 0.000109 | 1 | 2 | 2 | Q640L6 | Atf2 protei | Q640L6 [13 | 0 | 3408.837 | 0 | -0.46 | 0.010742 | 2.64 | 113 | 87 | 3.95 | 14.29 | | High | High | High | High | High | High | High | High | High | High | High | 4.94E-05 | 0.00059 | 3.33 |
| FALSE | High | LVSAAASPS | [R].LVSAAA | 1xPhosphc | Q61033 1x | 0.023175 | 0.001578 | 1 | 1 | 2 | Q61033 | Lamina-ass | Q61033 [3( | 0 | 1493.829 | 0 | 0.59 | 0.003074 | 7.25 | 77.8 | 122.2 | 8.78 | 6.52 | | High | High | High | High | High | High | High | High | High | High | High | 0.000847 | 0.008077 | 2.72 |
| FALSE | High | IVSAQSLAE | [R].IVSAQS | 1xPhosphc | Q4KL41 1x | 1.62E-06 | 0 | 1 | 2 | 7 | Q4KL41 | MCG10089 | Q4KL41 [1 | 0 | 1684.788 | 0 | 0.71 | 0.10419 | 18.79 | 75.7 | 124.3 | 6.91 | 26.5 | | High | High | High | High | High | High | High | High | High | High | High | 0 | 2.57E-06 | 3.9 |
| FALSE | High | LVSESSDVV | [K].LVSESS | 1xPhosphc | P11679 1xl | 4.43E-05 | 0 | 1 | 1 | 3 | P11679 | Keratin, ty | P11679 [48 | 0 | 1687.892 | 0 | 3.67 | 0.000906 | 21.2 | 13.5 | 186.5 | 14.44 | 15.27 | | High | High | High | High | High | High | High | High | High | High | High | 0 | 4.18E-05 | 3.8 |
| FALSE | High | LVSFHDDSI | [K].LVSFHD | 1xPhosphc | B7ZWC4 1x | 2.95E-05 | 0 | 1 | 8 | 6 | B7ZWC4 | Insulin-like | B7ZWC4 [2 | 0 | 2063.952 | 0 | -1.54 | 0.000251 | 1.24 | 149 | 51 | 3.5 | 10.68 | | High | High | High | High | High | High | High | High | High | High | High | 0 | 0.000195 | 3.06 |
| FALSE | High | LVSFHDDSI | [K].LVSFHD | 2xPhosphc | B7ZWC4 2x | 0.020679 | 0.001472 | 1 | 8 | 1 | B7ZWC4 | Insulin-like | B7ZWC4 [2 | 0 | 2143.919 | 0 | -0.93 | 0.005411 | 3.76 | 131.3 | 68.7 | 12.34 | 13.8 | | High | High | High | High | High | High | High | High | High | High | High | 0.000786 | 0.007095 | 3.05 |
| FALSE | High | IVSGSPISTI | [K].IVSGSP | 1xPhosphc | Q3TUF7 1x | 0.000167 | 0 | 1 | 4 | 1 | Q3TUF7 | YEATS dom | Q3TUF7 [4 | 0 | 2001.062 | 0 | -0.17 | 0.006152 | 4.04 | 106.5 | 93.5 | 3.43 | 6.28 | | High | High | High | High | High | High | High | High | High | High | High | 0 | 3.18E-05 | 2.46 |
| FALSE | High | IVPQSQVPI | [K].IVPQSC | 1xPhosphc | Q3TUF7 1x | 2.77E-05 | 0 | 1 | 4 | 2 | Q3TUF7 | YEATS dom | Q3TUF7 [4 | 0 | 2115.125 | 0 | -0.74 | 0.000207 | 1.1 | 124.6 | 75.4 | 2.07 | 1.67 | | High | High | High | High | High | High | High | High | High | High | High | 0 | 9.39E-06 | 3.97 |
| FALSE | High | IVSGSPISTI | [K].IVSGSP | 2xPhosphc | Q3TUF7 2x | 0.000543 | 0 | 1 | 4 | 2 | Q3TUF7 | YEATS dom | Q3TUF7 [4 | 0 | 2081.028 | 0 | -0.06 | 0.617773 | 12.35 | 106.9 | 93.1 | 16.74 | 7.77 | | High | High | High | High | High | High | High | High | High | High | High | 0 | 0.000121 | 2.52 |
| FALSE | High | IVSNLLGVE | [K].IVSNLL( | 1xPhosphc | G5E8P8 1x | 0.000131 | 0 | 1 | 6 | 2 | G5E8P8 | Estrogen re | G5E8P8 [2 | 0 | 1853.019 | 0 | -0.74 | 0.002467 | 3.27 | 126.7 | 73.3 | 7.13 | 5.31 | | High | High | High | High | High | High | High | High | High | High | High | 0 | 2.44E-05 | 2.79 |
| FALSE | High | LVSPEQPPI | [R].LVSPEC | 1xPhosphc | B6ZHD0 1x | 0.064323 | 0.004664 | 2 | 14 | 1 | B6ZHD0; Q | Erythrocyt | B6ZHD0 [4 | 0 | 1532.849 | 0 | -0.97 | 0.00198 | 5.92 | 132.3 | 67.7 | 5.51 | 9.76 | Shared | High | High | High | High | High | High | High | High | High | High | High | 0.002403 | 0.02574 | 2.68 |
| FALSE | High | LVSPGSANI | [R].LVSPGS | 1xPhosphc | Q80X90 1x | 2.12E-06 | 0 | 1 | 2 | 1 | Q80X90 | Filamin-B C | Q80X90 [2 | 0 | 2355.201 | 0 | 1.34 | 0.000551 | 14.42 | 57.2 | 142.8 | 22.98 | 12.18 | | High | High | High | High | High | High | High | High | High | High | High | 0 | 2.37E-07 | 4.76 |
| FALSE | High | LVSPLASPV | [K].LVSPLA | 1xPhosphc | Q8CJF7 1xl | 0.020862 | 0.001472 | 1 | 1 | 1 | Q8CJF7 | Protein ELY | Q8CJF7 [21 | 0 | 1906.07 | 0 | -0.12 | 0.114387 | 13.72 | 108.3 | 91.7 | 15.86 | 5.48 | | High | High | High | High | High | High | High | High | High | High | High | 0.000786 | 0.007149 | 2.4 |
| FALSE | High | LVSPLASPV | [K].LVSPLA | 2xPhosphc | Q8CJF7 2xl | 0.000203 | 0 | 1 | 1 | 3 | Q8CJF7 | Protein ELY | Q8CJF7 [21 | 0 | 1986.037 | 0 | -0.33 | 0.013281 | 5.51 | 113.2 | 86.8 | 4.2 | 13.6 | | High | High | High | High | High | High | High | High | High | High | High | 0.000585 | 0.005502 | 3.24 |
| FALSE | High | IVSPSGAAV | [K].IVSPSG | 1xCarbami | Q8BTM8 1 | 0.002869 | 0.00016 | 1 | 6 | 1 | Q8BTM8 | Filamin-A C | Q8BTM8 [ | 0 | 1723.922 | 0 | 0.6 | 0.003494 | 7.38 | 80.3 | 119.7 | 10.96 | 5.68 | | High | High | High | High | High | High | High | High | High | High | High | 7.19E-05 | 0.000776 | 2.78 |
| FALSE | High | LVSQDGK | [R].LVSQD( | 1xPhosphc | P09405 1xl | 0.016743 | 0.001166 | 1 | 7 | 1 | P09405 | Nucleolin ( | P09405 [42 | 0 | 1284.696 | 0 | -0.42 | 0.000486 | 1.06 | 114.6 | 85.4 | 2.5 | 2.24 | | High | High | High | High | High | High | High | High | High | High | High | 0.000638 | 0.005593 | 1.75 |
| FALSE | High | IVSQNVGD | [R].IVSQNV | 1xPhosphc | Q8K0L9 1x | 1.01E-08 | 0 | 1 | 3 | 2 | Q8K0L9 | Zinc finger | Q8K0L9 [1 | 0 | 2638.271 | 0 | 4.07 | 0.000619 | 9.78 | 12.4 | 187.6 | 46.38 | 12.62 | | High | High | High | High | High | High | High | High | High | High | High | 0 | 5.82E-10 | 3.89 |
| FALSE | High | LVSRPQDA | [R].LVSRPC | 1xPhosphc | Q925I1 1xI | 0.114657 | 0.009411 | 1 | 2 | 1 | Q925I1 | ATPase fan | Q925I1 [30 | 0 | 2559.374 | 0 | -0.06 | 0.179875 | 9.56 | 102.6 | 97.4 | 6.83 | 6.69 | | High | High | High | High | High | High | High | High | High | High | High | 0.004733 | 0.05087 | 1.27 |
| FALSE | High | LVSSESK | [R].LVSSES | 1xPhosphc | P56695 1xl | 0.029076 | 0.001857 | 1 | 4 | 1 | P56695 | Wolframin | P56695 [23 | 0 | 1287.696 | 0 | 0.83 | 0.002176 | 13.06 | 67.8 | 132.2 | 9.41 | 9.8 | | High | High | High | High | High | High | High | High | High | High | High | 0.000997 | 0.01037 | 2.21 |
| FALSE | High | LVSSPCCIV | [R].LVSSPC | 2xCarbami | P11499 1xl | 9.2E-05 | 0 | 1 | 3 | 3 | P11499 | Heat shock | P11499 [58 | 0 | 2741.234 | 0 | -0.31 | 0.049519 | 10.11 | 114.7 | 85.3 | 12.41 | 14.29 | | High | High | High | High | High | High | High | High | High | High | High | 0 | 1.64E-05 | 3.22 |
| FALSE | High | LVSSPCCIV | [R].LVSSPC | 2xCarbami | P11499 1xl | 0.000519 | 0 | 1 | 3 | 2 | P11499 | Heat shock | P11499 [58 | 0 | 2757.228 | 0 | -0.81 | 0.006187 | 1.85 | 127.3 | 72.7 | 9.94 | 9.79 | | High | High | High | High | High | High | High | High | High | High | High | 0 | 0.000166 | 3.48 |
| FALSE | High | IVSSPLQFC | [R].IVSSPL( | 1xCarbami | E9Q286 1x | 0.003558 | 0.00021 | 1 | 1 | 2 | E9Q286 | Little elong | E9Q286 [1( | 0 | 2057.091 | 0 | -1.02 | 0.000696 | 1.79 | 135.9 | 64.1 | 5.01 | 8.28 | | High | High | High | High | High | High | High | High | High | High | High | 0.000267 | 0.001571 | 2.08 |
| FALSE | High | IVSNASCTT | [K].IVSNAS | 2xCarbami | A0A0A0M( | 9.14E-06 | 0 | 1 | 6 | 2 | A0A0A0M( | Glyceralde | A0A0A0M( | 0 | 2358.196 | 0 | 0.2 | 0.355892 | 12.68 | 93.1 | 106.9 | 9.82 | 9.68 | | High | High | High | High | High | High | High | High | High | High | High | 0 | 1.22E-06 | 2.14 |
| FALSE | High | IVNSFEDDS | [R].IVNSFE | 1xPhosphc | E9QLG3 1x | 4.28E-06 | 0 | 1 | 7 | 2 | E9QLG3 | MAX gene- | E9QLG3 [3 | 0 | 3359.656 | 0 | -0.37 | 0.008232 | 7.11 | 111.4 | 88.6 | 6.77 | 3.62 | | High | High | High | High | High | High | High | High | High | High | High | 0 | 5.24E-07 | 4.11 |
| FALSE | High | IVMPDLEPS | [R].IVMPDI | 1xPhosphc | P30415 1xl | 0.102567 | 0.008249 | 1 | 2 | 2 | P30415 | NK-tumor r | P30415 [46 | 1 | 2060.02 | 0 | -0.2 | 0.928827 | 12.05 | 106.9 | 93.1 | 7.24 | 29.04 | | High | High | High | High | High | High | High | High | High | High | High | 0.004138 | 0.04466 | 2.06 |
| FALSE | High | IVMPDLEPS | [R].IVMPDI | 1xPhosphc | P30415 1xl | 0.031168 | 0.001935 | 1 | 2 | 2 | P30415 | NK-tumor r | P30415 [46 | 1 | 2044.025 | 0 | -0.19 | 0.266686 | 13.54 | 104.7 | 95.3 | 5 | 14.26 | | High | High | High | High | High | High | High | High | High | High | High | 0.002867 | 0.02972 | 1.79 |
| FALSE | High | IVEPEVVGE | [K].IVEPEV | 1xPhosphc | C0HKD9 1x | 9.38E-09 | 0 | 1 | 1 | 5 | C0HKD9 | Microfibrill | C0HKD9 [1 | 0 | 2511.149 | 0 | -0.4 | 0.032726 | 14.37 | 111.9 | 88.1 | 8.73 | 14.32 | | High | High | High | High | High | High | High | High | High | High | High | 0 | 5.35E-10 | 4.13 |
| FALSE | High | IVEPEVVGE | [K].IVEPEV | 2xPhosphc | C0HKD9 2x | 6.22E-07 | 0 | 1 | 1 | 13 | C0HKD9 | Microfibrill | C0HKD9 [1 | 0 | 2591.115 | 0 | -0.32 | 0.019567 | 7.47 | 111 | 89 | 3.58 | 13.61 | | High | High | High | High | High | High | High | High | High | High | High | 0 | 5.74E-07 | 4.83 |
| FALSE | High | LVEPHSPSF | [R].LVEPHS | 1xPhosphc | Q8BZ47 1x | 2.65E-05 | 0 | 1 | 2 | 2 | Q8BZ47 | Zinc finger | Q8BZ47 [5 | 0 | 1802.945 | 0 | -0.01 | 0.436686 | 14.95 | 102.9 | 97.1 | 8.14 | 7.38 | | High | High | High | High | High | High | High | High | High | High | High | 0 | 0.000204 | 2.26 |
| FALSE | High | LVEPHSPSF | [R].LVEPHS | 2xPhosphc | Q8BZ47 2x | 0.007951 | 0.000736 | 1 | 2 | 2 | Q8BZ47 | Zinc finger | Q8BZ47 [5 | 0 | 1882.912 | 0 | -0.33 | 0.008011 | 6.83 | 111.7 | 88.3 | 2.47 | 6.42 | | High | High | High | High | High | High | High | High | High | High | High | 0.000847 | 0.008067 | 2.07 |
| FALSE | High | LVEQANSP | [K].LVEQAI | 1xPhosphc | Q8BHJ9 1x | 1.51E-05 | 0 | 1 | 3 | 2 | Q8BHJ9 | Pre-mRNA- | Q8BHJ9 [2( | 0 | 1523.823 | 0 | -0.1 | 0.069547 | 5.89 | 103.8 | 96.2 | 2.54 | 6.04 | | High | High | High | High | High | High | High | High | High | High | High | 0 | 2.16E-06 | 3.24 |
| FALSE | High | IVESPELSK | [K].IVESPEI | 1xPhosphc | G3X928 1x | 0.070485 | 0.005214 | 1 | 4 | 2 | G3X928 | SEC23-inte | G3X928 [9 | 0 | 1539.844 | 0 | -0.02 | 0.357224 | 7.23 | 100.8 | 99.2 | 6.41 | 4.44 | | High | High | High | High | High | High | High | High | High | High | High | 0.00269 | 0.02879 | 2.25 |
| FALSE | High | IVETINSDSI | [K].IVETINS | 1xPhosphc | Q64511 1x | 1.49E-05 | 0 | 1 | 1 | 4 | Q64511 | DNA topois | Q64511 [1 | 0 | 2389.194 | 0 | 0.31 | 0.012889 | 7.66 | 85.1 | 114.9 | 6.11 | 9.83 | | High | High | High | High | High | High | High | High | High | High | High | 0 | 2.12E-06 | 3.84 |
| FALSE | High | IVETINSDSI | [K].IVETINS | 2xPhosphc | Q64511 2x | 2.23E-05 | 0 | 1 | 1 | 5 | Q64511 | DNA topois | Q64511 [1 | 0 | 2469.16 | 0 | 0.13 | 0.355427 | 10.08 | 94.4 | 105.6 | 6.06 | 11.06 | | High | High | High | High | High | High | High | High | High | High | High | 0 | 3.32E-06 | 2.8 |
| FALSE | High | IVEYSPPSA | [R].IVEYSPI | 1xPhosphc | E9PZ26 1xl | 0.011246 | 0.000772 | 1 | 5 | 4 | E9PZ26 | Bromodom | E9PZ26 [12 | 0 | 1524.766 | 0 | -0.98 | 0.000779 | 2.5 | 134.5 | 65.5 | 6.47 | 3.82 | | High | High | High | High | High | High | High | High | High | High | High | 0.000418 | 0.003571 | 2.33 |
| FALSE | High | IVFVPGCSV | [K].IVFVPG | 1xCarbami | Q9D0I9 1xl | 0.000933 | 0 | 1 | 3 | 1 | Q9D0I9 | Arginine--t | Q9D0I9 [36 | 0 | 2167.237 | 0 | -0.03 | 0.313592 | 6.8 | 108.1 | 91.9 | 15.63 | 12.93 | | High | High | High | High | High | High | High | High | High | High | High | 0 | 0.000222 | 2.64 |
| FALSE | High | LVGATATPI | [R].LVGAT/ | 1xPhosphc | Q91Z49 1x | 8.25E-05 | 0 | 1 | 2 | 1 | Q91Z49 | UAP56-inte | Q91Z49 [8 | 0 | 1784.012 | 0 | 0.52 | 0.006826 | 7.37 | 83.1 | 116.9 | 7.67 | 6.09 | | High | High | High | High | High | High | High | High | High | High | High | 0 | 1.44E-05 | 1.77 |
| FALSE | High | LVGGTTPG | [K].LVGGT | 1xPhosphc | Q9WUM5 | 0.000382 | 0 | 1 | 1 | 2 | Q9WUM5 | Succinate-- | Q9WUM5 | 0 | 1367.77 | 0 | -0.76 | 0.00082 | 1.41 | 126.6 | 73.4 | 4.93 | 4.98 | | High | High | High | High | High | High | High | High | High | High | High | 0 | 8.11E-05 | 1.9 |
| FALSE | High | LVGLNLSSS | [K].LVGLNI | 1xPhosphc | Q8C804 1x | 2.14E-06 | 0 | 1 | 1 | 2 | Q8C804 | Spindle an | Q8C804 [7 | 0 | 2260.215 | 0 | -0.21 | 0.002832 | 3.25 | 105.4 | 94.6 | 3.23 | 3.3 | | High | High | High | High | High | High | High | High | High | High | High | 0.000571 | 0.004979 | 3.54 |
| FALSE | High | LVGSQEEL/ | [R].LVGSQI | 1xPhosphc | Q3TI61 1xF | 5.32E-06 | 0 | 1 | 7 | 1 | Q3TI61 | Putative ur | Q3TI61 [14 | 0 | 2269.085 | 0 | 0.78 | 0.025615 | 27.04 | 73.8 | 126.2 | 17.58 | 17.71 | | High | High | High | High | High | High | High | High | High | High | High | 0 | 6.64E-07 | 5.03 |
| FALSE | High | LVGSSPSLG | [K].LVGSSP | 1xPhosphc | E9Q3P4 1x | 0.002134 | 5.66E-05 | 1 | 5 | 2 | E9Q3P4 | Centromer | E9Q3P4 [2 | 0 | 1482.833 | 0 | -0.08 | 0.092894 | 9.06 | 103.2 | 96.8 | 4.73 | 5.92 | | High | High | High | High | High | High | High | High | High | High | High | 0.000327 | 0.002576 | 3.4 |
| FALSE | High | LVGTPNPS | [K].LVGTPN | 1xPhosphc | Q9WVB0-2 | 0.000221 | 0 | 1 | 7 | 3 | Q9WVB0-2 | Isoform 2 c | Q9WVB0-2 | 0 | 2528.348 | 0 | -0.03 | 0.323743 | 0.93 | 103.6 | 96.4 | 13.21 | 13.75 | | High | High | High | High | High | High | High | High | High | High | High | 0 | 4.37E-05 | 4.15 |
| FALSE | High | LVHNLTSPI | [K].LVHNLT | 1xPhosphc | Q80TY5 1x | 0.01002 | 0.000736 | 1 | 2 | 1 | Q80TY5 | Vacuolar p | Q80TY5 [3( | 0 | 1546.876 | 0 | 0.26 | 0.00709 | 2.63 | 91 | 109 | 3.7 | 7.43 | | High | High | High | High | High | High | High | High | High | High | High | 0.000344 | 0.003136 | 2.33 |
| FALSE | High | LVHTGSLNI | [K].LVHTGS | 1xPhosphc | B2RTL2 1xl | 0.000479 | 0 | 1 | 6 | 1 | B2RTL2 | Phosphodi | B2RTL2 [32 | 0 | 1728.936 | 0 | 0.7 | 0.004778 | 13.56 | 75.7 | 124.3 | 12.05 | 11.08 | | High | High | High | High | High | High | High | High | High | High | High | 0 | 0.000104 | 2.53 |
| FALSE | High | LVIGSLPAH | [R].LVIGSLI | 1xPhosphc | Q71FD5 1x | 0.001674 | 5.66E-05 | 1 | 1 | 1 | Q71FD5 | E3 ubiquiti | Q71FD5 [1 | 0 | 2528.419 | 0 | 0.52 | 0.002998 | 9.24 | 82.1 | 117.9 | 7.42 | 4.11 | | High | High | High | High | High | High | High | High | High | High | High | 2.58E-05 | 0.000425 | 2.3 |
| FALSE | High | IVLLDSSLEY | [R].IVLLDSS | 1xPhosphc | Q3U4U6 1x | 4.98E-05 | 0 | 1 | 5 | 2 | Q3U4U6 | T-complex | Q3U4U6 [2 | 0 | 1818.007 | 0 | -0.58 | 0.005045 | 7.92 | 119.2 | 80.8 | 3.29 | 14.21 | | High | High | High | High | High | High | High | High | High | High | High | 0 | 8.24E-06 | 2.63 |
| FALSE | High | LVINGGNG | [K].LVINGG | 1xPhosphc | A2APM2 1: | 4.49E-07 | 0 | 1 | 22 | 2 | A2APM2 | CD44 antig | A2APM2 [7 | 0 | 1652.821 | 0 | 3.05 | 0.003294 | 11.57 | 21.5 | 178.5 | 19.08 | 25.57 | | High | High | High | High | High | High | High | High | High | High | High | 0 | 4.16E-08 | 3.41 |
| FALSE | High | LVINGGNG | [K].LVINGG | 1xDeamida | A2APM2 1: | 5.15E-07 | 0 | 1 | 22 | 19 | A2APM2 | CD44 antig | A2APM2 [7 | 0 | 1653.805 | 0 | 1.81 | 0.001799 | 22.84 | 49.2 | 150.8 | 20.4 | 16.3 | | High | High | High | High | High | High | High | High | High | High | High | 0 | 6.94E-08 | 4.18 |
| FALSE | High | LVINGGNG | [K].LVINGG | 2xDeamida | A2APM2 1: | 3.37E-07 | 0 | 1 | 22 | 11 | A2APM2 | CD44 antig | A2APM2 [7 | 0 | 1654.789 | 0 | 2.77 | 0.00226 | 17.45 | 28.5 | 171.5 | 25.33 | 21.85 | | High | High | High | High | High | High | High | High | High | High | High | 0 | 3E-08 | 4.3 |
| FALSE | High | LVLPAPQIS | [K].LVLPAP | 1xPhosphc | A0A1W2P7 | 0.019791 | 0.001438 | 1 | 7 | 1 | A0A1W2P7 | Uncharacte | A0A1W2P7 | 0 | 2487.388 | 0 | -1.23 | 0.00317 | 5.67 | 138.5 | 61.5 | 12.63 | 14.4 | | High | High | High | High | High | High | High | High | High | High | High | 0.000773 | 0.00677 | 3.2 |
| FALSE | High | LVLPMDDS | [R].LVLPMI | 1xPhosphc | A1L3P4 1xl | 2.66E-06 | 0 | 1 | 3 | 3 | A1L3P4 | Sodium/hy | A1L3P4 [68 | 1 | 2818.317 | 0 | 0.74 | 0.007339 | 20.21 | 74.4 | 125.6 | 11.43 | 14.14 | | High | High | High | High | High | High | High | High | High | High | High | 0 | 3.05E-07 | 4.86 |
| FALSE | High | LVLPSNSPN | [K].LVLPSN | 1xPhosphc | P49452 1xl | 0.000731 | 0 | 1 | 1 | 4 | P49452 | Centromer | P49452 [69 | 0 | 1504.809 | 0 | -0.95 | 0.000181 | 1.06 | 133 | 67 | 3.09 | 3.06 | | High | High | High | High | High | High | High | High | High | High | High | 0.000344 | 0.003201 | 3.05 |
| FALSE | High | LVLSGEK | [R].LVLSGE | 1xPhosphc | A0A0N4SV | 0.120211 | 0.009851 | 1 | 9 | 1 | A0A0N4SV | Eukaryotic | A0A0N4SV | 0 | 1283.738 | 0 | 1.39 | 0.000264 | 5.27 | 54.1 | 145.9 | 5.57 | 4.82 | | High | High | High | High | High | High | High | High | High | High | High | 0.004961 | 0.05386 | 0.92 |
| FALSE | High | LVLVLPDVE | [K].LVLVLP | 1xPhosphc | Q80TX5 1x | 8.87E-05 | 0 | 1 | 4 | 1 | Q80TX5 | MKIAA057 | Q80TX5 [1 | 0 | 2287.178 | 0 | 0.06 | 0.130506 | 2.66 | 97.8 | 102.2 | 5.2 | 13.68 | | High | High | High | High | High | High | High | High | High | High | High | 0 | 1.57E-05 | 4.31 |
| FALSE | High | LVMNQSTI | [R].LVMNC | 1xPhosphc | Q61687 1x | 0.025514 | 0.001645 | 1 | 4 | 1 | Q61687 | Transcripti | Q61687 [3 | 0 | 1589.801 | 0 | -0.16 | 0.203403 | 22.85 | 105.9 | 94.1 | 11.69 | 12.9 | | High | High | High | High | High | High | High | High | High | High | High | 0.000872 | 0.008961 | 2.07 |
| FALSE | High | LVTSPCCIV | [R].LVTSPC | 2xCarbami | Q3UIF3 1xl | 2.46E-07 | 0 | 1 | 4 | 1 | Q3UIF3 | Putative ur | Q3UIF3 [59 | 0 | 2755.249 | 0 | -0.89 | 0.004628 | 6.57 | 127.3 | 72.7 | 10.07 | 10.02 | | High | High | High | High | High | High | High | High | High | High | High | 0 | 2.1E-08 | 3.88 |
| FALSE | High | MPLDLSPL/ | [K].MPLDLS | 1xPhosphc | Q640L6 1xl | 0.000933 | 0 | 1 | 13 | 1 | Q640L6 | Atf2 protei | Q640L6 [8 | 0 | 1846.011 | 0 | -0.47 | 0.031162 | 8.71 | 124.5 | 75.5 | 11.95 | 26.1 | | High | High | High | High | High | High | High | High | High | High | High | 0 | 0.00022 | 1.75 |
| FALSE | High | MPIEESSPE | [R].MPIEES | 1xPhosphc | Q5RIM6 1x | 7.15E-05 | 0 | 1 | 10 | 2 | Q5RIM6 | Nuclear re | Q5RIM6 [1 | 0 | 1684.827 | 0 | -0.51 | 0.012971 | 3.27 | 118.4 | 81.6 | 5.67 | 14.8 | | High | High | High | High | High | High | High | High | High | High | High | 0 | 1.23E-05 | 2.34 |
| FALSE | High | MPSLEVSV | [K].MPSLEV | 1xPhosphc | E9Q616 1x | 0.00582 | 0.00062 | 1 | 5 | 1 | E9Q616 | AHNAK nu | E9Q616 [4( | 0 | 1624.879 | 0 | 2.58 | 0.002276 | 14.12 | 27.1 | 172.9 | 14.35 | 20.72 | | High | High | High | High | High | High | High | High | High | High | High | 0.000275 | 0.001706 | 2.43 |
| FALSE | High | NEVSLNPLI | [K].NEVSLN | 1xPhosphc | Q5U647 1x | 1.72E-08 | 0 | 1 | 4 | 1 | Q5U647 | Amino acid | Q5U647 [5 | 0 | 2686.447 | 0 | 0.04 | 0.831131 | 7.9 | 100.3 | 99.7 | 6.86 | 11.53 | | High | High | High | High | High | High | High | High | High | High | High | 0 | 1.06E-09 | 5.09 |
| FALSE | High | NFCSIELNV | [R].NFCSIE | 2xCarbami | E9Q5F9 1x | 3.54E-05 | 0 | 1 | 3 | 1 | E9Q5F9 | Histone-lys | E9Q5F9 [6( | 0 | 3428.579 | 0 | -0.19 | 0.094588 | 17.5 | 109.2 | 90.8 | 6.78 | 13.75 | | High | High | High | High | High | High | High | High | High | High | High | 0 | 5.56E-06 | 4 |
| FALSE | High | NFDDEDSV | [K].NFDDEI | 1xPhosphc | B2RY22 1xl | 5.97E-05 | 0 | 2 | 18 | 4 | B2RY22; A( | Clasp1 pro | B2RY22 [24 | 0 | 2826.247 | 0 | -0.88 | 0.000623 | 2.31 | 129.2 | 70.8 | 5.11 | 10.35 | Shared | High | High | High | High | High | High | High | High | High | High | High | 0.000275 | 0.00223 | 2.54 |
| FALSE | High | NFDDLDNF | [K].NFDDLI | 1xCarbami | Q6P5E3 1x | 0.001615 | 5.66E-05 | 1 | 3 | 2 | Q6P5E3 | Sal-like 1 (I | Q6P5E3 [8 | 0 | 3569.501 | 0 | -0.99 | 0.000209 | 1.75 | 136 | 64 | 4.84 | 7.34 | | High | High | High | High | High | High | High | High | High | High | High | 4.94E-05 | 0.000757 | 4.03 |
| FALSE | High | NFDDLDNF | [K].NFDDLI | 1xCarbami | Q6P5E3 2x | 0.000745 | 0 | 1 | 3 | 1 | Q6P5E3 | Sal-like 1 (I | Q6P5E3 [8 | 0 | 3633.473 | 0 | -1.16 | 0.00062 | 1.09 | 138.6 | 61.4 | 5.68 | 11.24 | | High | High | High | High | High | High | High | High | High | High | High | 0 | 0.000172 | 3.72 |
| FALSE | High | NFDGYRSP | [R].NFDGYI | 1xPhosphc | Q8CCN5 1x | 0.11108 | 0.008991 | 1 | 5 | 1 | Q8CCN5 | Breast carc | Q8CCN5 [9 | 1 | 3090.481 | 0 | 1.31 | 0.00584 | 21.4 | 47.7 | 152.3 | 25.36 | 18.86 | | High | High | High | High | High | High | High | High | High | High | High | 0.004503 | 0.04882 | 2.53 |
| FALSE | High | NFDTLDLPI | [R].NFDTLI | 1xPhosphc | P98203-3 1 | 0.000281 | 0 | 1 | 6 | 1 | P98203-3 | Isoform 3 c | P98203-3 [ | 0 | 1600.839 | 0 | 0.46 | 0.155686 | 43.08 | 79 | 121 | 14.33 | 24.8 | | High | High | High | High | High | High | High | High | High | High | High | 0 | 5.76E-05 | 2.11 |
| FALSE | High | NFEDSSPET | [K].NFEDSS | 1xPhosphc | Q8K2V6-2 | 0.070485 | 0.005214 | 1 | 3 | 2 | Q8K2V6-2 | Isoform 2 c | Q8K2V6-2 | 0 | 2142.016 | 0 | -0.84 | 0.002177 | 5.75 | 128.2 | 71.8 | 2.98 | 12.26 | | High | High | High | High | High | High | High | High | High | High | High | 0.002675 | 0.02862 | 2.43 |
| FALSE | High | NFHLVDSSI | [R].NFHLVI | 1xCarbami | Q8C0P0 1x | 2.25E-05 | 0 | 1 | 1 | 2 | Q8C0P0 | Serine/thre | Q8C0P0 [4 | 0 | 2558.218 | 0 | -0.43 | 0.057757 | 17.8 | 110.9 | 89.1 | 13.6 | 13.16 | | High | High | High | High | High | High | High | High | High | High | High | 0 | 3.35E-06 | 2.91 |
| FALSE | High | NFHLVDSSI | [R].NFHLVI | 1xCarbami | Q8C0P0 1x | 7.23E-06 | 0 | 1 | 1 | 2 | Q8C0P0 | Serine/thre | Q8C0P0 [4 | 0 | 2574.213 | 0 | -0.6 | 0.010885 | 4.94 | 121.9 | 78.1 | 14.94 | 6.99 | | High | High | High | High | High | High | High | High | High | High | High | 0 | 9.4E-07 | 4.06 |
| FALSE | High | NFLNALTSI | [R].NFLNAI | 1xPhosphc | Q0KL02-4 1 | 0.016306 | 0.001124 | 1 | 3 | 1 | Q0KL02-4 | Isoform 4 c | Q0KL02-4 | | 0 | 1974.989 | 0 | 0.91 | 0.003716 | 15.33 | 66.5 | 133.5 | 34.28 | 15.66 | | High | High | High | High | High | High | High | High | High | High | High | 0.000585 | 0.005417 | 3.56 |
| FALSE | High | NFMYDQSI | [R].NFMYD | 1xPhosphc | Q9CZA6 1x | 0.001222 | 0 | 1 | 5 | 2 | Q9CZA6 | Nuclear dis | Q9CZA6 [2 | 0 | 1569.661 | 0 | -0.83 | 0.000491 | 3.71 | 128.2 | 71.8 | 3.08 | 5.67 | | High | High | High | High | High | High | High | High | High | High | High | 0 | 0.000299 | 1.76 |
| FALSE | High | NFQAATES | [R].NFQAA | 1xPhosphc | M0QWZ1 1 | 1.64E-05 | 0 | 1 | 3 | 4 | M0QWZ1 | Protein FAI | M0QWZ1 [ | 0 | 4123.832 | 0 | -0.96 | 0.055546 | 14.83 | 132 | 68 | 18.6 | 50.82 | | High | High | High | High | High | High | High | High | High | High | High | 0 | 2.35E-06 | 5.63 |
| FALSE | High | NEVNGTSE | [K].NEVNG | 2xPhosphc | Q5SUF2 2x | 0.003622 | 0.00021 | 1 | 1 | 1 | Q5SUF2 | Luc7-like p | Q5SUF2 [4 | 1 | 2642.127 | 0 | -0.53 | 0.005307 | 6.15 | 119.4 | 80.6 | 4.82 | 8.57 | | High | High | High | High | High | High | High | High | High | High | High | 9.42E-05 | 0.001008 | 4.8 |
| FALSE | High | NFSCGSME | [R].NFSCGS | 1xCarbami | Q8K1N2-2 | 5.47E-06 | 0 | 1 | 3 | 3 | Q8K1N2-2 | Isoform 2 c | Q8K1N2-2 | 0 | 2383.998 | 0 | 0.43 | 0.003594 | 4.88 | 85.1 | 114.9 | 7.13 | 11.33 | | High | High | High | High | High | High | High | High | High | High | High | 0 | 1E-06 | 3.69 |
| FALSE | High | NFSCGSME | [R].NFSCGS | 1xCarbami | Q8K1N2-2 | 0.006594 | 0.00062 | 1 | 3 | 1 | Q8K1N2-2 | Isoform 2 c | Q8K1N2-2 | 0 | 2463.965 | 0 | 3.18 | 0.001841 | 26.94 | 19.8 | 180.2 | 9.32 | 19.21 | | High | High | High | High | High | High | High | High | High | High | High | 0.000275 | 0.001968 | 2.8 |
| FALSE | High | NFSDNQLC | [R].NFSDN( | 1xPhosphc | Q9WVA4 1 | 3.32E-05 | 0 | 1 | 1 | 2 | Q9WVA4 | Transgelin- | Q9WVA4 [ | 0 | 1588.721 | 0 | 0.99 | 0.041935 | 28.07 | 72.6 | 127.4 | 12.43 | 24.7 | | High | High | High | High | High | High | High | High | High | High | High | 0 | 5.18E-06 | 3.02 |
| FALSE | High | NFSDNQLC | [R].NFSDN( | 1xPhosphc | Q9WVA4 1 | 2.96E-06 | 0 | 1 | 1 | 2 | Q9WVA4 | Transgelin- | Q9WVA4 [ | 0 | 1817.884 | 0 | 2.44 | 0.001265 | 18.05 | 33 | 167 | 9.45 | 14.87 | | High | High | High | High | High | High | High | High | High | High | High | 0 | 3.44E-07 | 3.42 |
| FALSE | High | NFSNSPGP | [R].NFSNSF | 1xCarbami | Q3LR78 1x | 8.22E-07 | 0 | 1 | 6 | 2 | Q3LR78 | CtBP-intera | Q3LR78 [1 | 0 | 2350.155 | 0 | 1.21 | 0.002306 | 20.04 | 62.3 | 137.7 | 7.7 | 12.32 | | High | High | High | High | High | High | High | High | High | High | High | 0 | 7.29E-07 | 3.44 |
| FALSE | High | NFSPDQSP | [R].NFSPD( | 1xPhosphc | Q80Y19 1x | 0.000138 | 0 | 1 | 2 | 2 | Q80Y19 | Rho GTPas | Q80Y19 [6 | 0 | 1703.763 | 0 | -0.56 | 0.001082 | 3.47 | 120.5 | 79.5 | 7.2 | 3.8 | | High | High | High | High | High | High | High | High | High | High | High | 0 | 2.57E-05 | 3.14 |
| FALSE | High | NFSPDQSP | [R].NFSPD( | 2xPhosphc | Q80Y19 2x | 0.001489 | 5.66E-05 | 1 | 2 | 1 | Q80Y19 | Rho GTPas | Q80Y19 [6 | 0 | 1783.729 | 0 | -0.29 | 0.009682 | 3.23 | 110.6 | 89.4 | 2.89 | 7.16 | | High | High | High | High | High | High | High | High | High | High | High | 2.58E-05 | 0.000375 | 3.13 |
| FALSE | High | NFSSASAL( | [K].NFSSAS | 1xPhosphc | Q8BX22 1x | 0.021046 | 0.001507 | 1 | 3 | 1 | Q8BX22 | Sal-like pro | Q8BX22 [8 | 0 | 1768.858 | 0 | -1.33 | 0.000605 | 2.82 | 143.1 | 56.9 | 4.56 | 11.65 | | High | High | High | High | High | High | High | High | High | High | High | 0.000801 | 0.007222 | 1.2 |
| FALSE | High | NFSSEIEDL | [K].NFSSEI | 1xPhosphc | Q3UHF3 1x | 0.000375 | 0 | 1 | 5 | 2 | Q3UHF3 | Mesoderm | Q3UHF3 [5 | 0 | 1848.903 | 0 | -1.23 | 0.000442 | 1.47 | 140.2 | 59.8 | 4.58 | 8.68 | | High | High | High | High | High | High | High | High | High | High | High | 0 | 0.000203 | 3.3 |
| FALSE | High | NFSSEIEDL | [K].NFSSEI | 2xPhosphc | Q3UHF3 2x | 0.00145 | 5.66E-05 | 1 | 5 | 2 | Q3UHF3 | Mesoderm | Q3UHF3 [5 | 0 | 1928.87 | 0 | -1.03 | 0.00714 | 10.26 | 134.2 | 65.8 | 10.61 | 41.46 | | High | High | High | High | High | High | High | High | High | High | High | 2.58E-05 | 0.000362 | 2.74 |
| FALSE | High | NFSSTEDDI | [K].NFSSTE | 1xPhosphc | B2RY61 1xl | 0.001158 | 0 | 1 | 3 | 3 | B2RY61 | AT rich inte | B2RY61 [84 | 0 | 2197.994 | 0 | 0.38 | 0.003891 | 3.44 | 87 | 113 | 6.21 | 8.24 | | High | High | High | High | High | High | High | High | High | High | High | 0 | 0.000282 | 3.27 |
| FALSE | High | NFTDDLGL | [K].NFTDDI | 1xCarbami | P70399 1xl | 0.000167 | 0 | 1 | 5 | 1 | P70399 | TP53-bindi | P70399 [49 | 0 | 2315.034 | 0 | 0.2 | 0.587306 | 73.33 | 91.2 | 108.8 | 23.23 | 29.48 | | High | High | High | High | High | High | High | High | High | High | High | 0 | 3.2E-05 | 2.57 |
| FALSE | High | NFTQSPEQ | [K].NFTQSI | 1xPhosphc | Q9JJA9 1xF | 0.000772 | 0 | 1 | 1 | 1 | Q9JJA9 | General re | Q9JJA9 [89 | 0 | 1543.71 | 0 | 2.08 | 0.000618 | 15.89 | 36.5 | 163.5 | 22.26 | 8.04 | | High | High | High | High | High | High | High | High | High | High | High | 0 | 0.000179 | 2.66 |
| FALSE | High | NFVFHTLSI | [R].NFVFHT | 1xPhosphc | Q80YR7 1x | 0.011048 | 0.000772 | 1 | 2 | 1 | Q80YR7 | Claspin OS | Q80YR7 [1 | 0 | 1599.813 | 0 | -0.34 | 0.567196 | 21.55 | 109.2 | 90.8 | 45.72 | 47.81 | | High | High | High | High | High | High | High | High | High | High | High | 0.000399 | 0.003497 | 2.66 |
| FALSE | High | NFSCGSME | [R].NFSCGS | 1xCarbami | Q8K1N2-2 | 0.00153 | 5.66E-05 | 1 | 3 | 2 | Q8K1N2-2 | Isoform 2 c | Q8K1N2-2 | 0 | 2399.993 | 0 | 1.6 | 0.002636 | 30.68 | 47.9 | 152.1 | 12.79 | 15.89 | | High | High | High | High | High | High | High | High | High | High | High | 2.58E-05 | 0.000385 | 2.34 |
| FALSE | High | NFVFHTLSI | [R].NFVFHT | 1xPhosphc | Q80YR7 1x | 0.038346 | 0.002563 | 1 | 2 | 1 | Q80YR7 | Claspin OS | Q80YR7 [1 | 0 | 1828.976 | 0 | -0.57 | 0.018429 | 6.82 | 119.7 | 80.3 | 5.62 | 11.83 | | High | High | High | High | High | High | High | High | High | High | High | 0.001355 | 0.01421 | 1.79 |

| | | | | | | | | | | | | | | | | | | | | | | | | | | | | | | | | | | | | | | |
|---|---|---|---|---|---|---|---|---|---|---|---|---|---|---|---|---|---|---|---|---|---|---|---|---|---|---|---|---|---|---|---|---|---|---|---|---|---|---|
| FALSE | High | NEVNGTSE | [K].NEVNG | 1xPhosphc | Q5SUF2 1x | 2.56E-05 | 0 | 1 | 1 | 3 | Q5SUF2 | Luc7-like p | Q5SUF2 [4 | 1 | 2562.161 | 0 | -0.8 | 0.001286 | 2.11 | 128.9 | 71.1 | 5.41 | 7.75 | | High | High | High | High | High | High | High | High | High | High | High | 0 | 6.94E-06 | 4.27 |
| FALSE | High | NETETISSP | [R].NETETI | 1xPhosphc | J3QPN0 1x | 0.000993 | 0 | 1 | 3 | 1 | J3QPN0 | MCG14131 | J3QPN0 [1: | 0 | 1856.941 | 0 | -1.67 | 0.000225 | 0.84 | 151.3 | 48.7 | 4.21 | 6.99 | | High | High | High | High | High | High | High | High | High | High | High | 0 | 0.000236 | 2.69 |
| FALSE | High | NEEDSEGS | [K].NEEDSE | 2xPhosphc | Q8R1B4 2x | 2.96E-06 | 0 | 1 | 4 | 5 | Q8R1B4 | Eukaryotic | Q8R1B4 [1 | 0 | 3336.324 | 0 | 0.59 | 0.031006 | 12.76 | 79.9 | 120.1 | 8.21 | 23.21 | | High | High | High | High | High | High | High | High | High | High | High | 0 | 3.44E-07 | 5.19 |
| FALSE | High | NEEENIYSV | [R].NEEENI | 1xPhosphc | B2RTN5 1x | 0.066503 | 0.004796 | 1 | 4 | 2 | B2RTN5 | Glucocorti | B2RTN5 [1( | 0 | 2256.002 | 0 | -0.15 | 0.588263 | 32.74 | 97.9 | 102.1 | 22.26 | 18.95 | | High | High | High | High | High | High | High | High | High | High | High | 0.002455 | 0.02669 | 1.74 |
| FALSE | High | NEEGDDEN | [R].NEEGD | 1xPhosphc | Q6PG04 1x | 3.71E-06 | 0 | 1 | 2 | 1 | Q6PG04 | Coiled-coil | Q6PG04 [2 | 0 | 2988.347 | 0 | 1.01 | 0.00148 | 18.71 | 66.2 | 133.8 | 13.31 | 5.72 | | High | High | High | High | High | High | High | High | High | High | High | 0 | 4.43E-07 | 2.84 |
| FALSE | High | NEEQFVFS | [K].NEEQF | 1xPhosphc | Q3TU15 1x | 4.86E-06 | 0 | 2 | 4 | 1 | Q3TU15; C | Non-specif | Q3TU15 [4 | 0 | 1891.924 | 0 | -0.71 | 0.006725 | 1.13 | 123.3 | 76.7 | 4.01 | 13.99 | Shared | High | High | High | High | High | High | High | High | High | High | High | 0 | 6.01E-07 | 3.15 |
| FALSE | High | YYSVDDNC | [R].YYSVDD | 1xPhosphc | B9EIW5 1x | 0.00095 | 0 | 1 | 5 | 3 | B9EIW5 | Ncoa7 prot | B9EIW5 [9( | 0 | 1783.83 | 0 | 3.58 | 0.00538 | 51.66 | 18.8 | 181.2 | 23.86 | 27.86 | | High | High | High | High | High | High | High | High | High | High | High | 0 | 0.000226 | 3.06 |
| FALSE | High | NEGITQPLF | [K].NEGITC | 1xPhosphc | P51125-3 1 | 0.063258 | 0.004664 | 1 | 1 | 1 | P51125-3 | Isoform 3 c | P51125-3 [ | 0 | 1704.841 | 0 | 1.28 | 0.005174 | 4.95 | 57.3 | 142.7 | 10.89 | 17.53 | | High | High | High | High | High | High | High | High | High | High | High | 0.002403 | 0.02532 | 2.51 |
| FALSE | High | NEGITQPLF | [K].NEGITC | 1xPhosphc | P51125-3 1 | 0.000158 | 0 | 1 | 1 | 2 | P51125-3 | Isoform 3 c | P51125-3 [ | 0 | 1934.004 | 0 | 2.54 | 0.00849 | 40.43 | 29.4 | 170.6 | 18.4 | 29.3 | | High | High | High | High | High | High | High | High | High | High | High | 0 | 0.000185 | 2.98 |
| FALSE | High | NEGSESAP | [K].NEGSE | 1xPhosphc | Q3UBT1 1x | 1E-05 | 0 | 1 | 5 | 2 | Q3UBT1 | Putative ur | Q3UBT1 [1 | 0 | 1896.829 | 0 | 0.13 | 0.312713 | 7.68 | 95.5 | 104.5 | 10.32 | 12.77 | | High | High | High | High | High | High | High | High | High | High | High | 0 | 1.35E-06 | 2.9 |
| FALSE | High | NEKSEEEQ | [K].NEKSEE | 1xPhosphc | Q3U6P5 1x | 3.44E-05 | 0 | 2 | 6 | 6 | Q3U6P5; C | Putative ur | Q3U6P5 [2 | 1 | 2319.168 | 0 | -0.36 | 0.085865 | 10.2 | 112.6 | 87.4 | 12.62 | 11.83 | Shared | High | High | High | High | High | High | High | High | High | High | High | 0 | 0.000142 | 2.91 |
| FALSE | High | NELETLGY( | [R].NELETL | 1xPhosphc | A0PJJ1 1xP | 0.002212 | 0.000109 | 1 | 1 | 1 | A0PJJ1 | Phf3 prote | A0PJJ1 [38 | 0 | 3470.589 | 0 | -1.77 | 0.001593 | 4.16 | 154.8 | 45.2 | 6.6 | 35.58 | | High | High | High | High | High | High | High | High | High | High | High | 4.94E-05 | 0.00058 | 2.8 |
| FALSE | High | NELETLGY( | [R].NELETL | 1xPhosphc | B2RQG2 1x | 4.89E-05 | 0 | 1 | 1 | 2 | B2RQG2 | PHD finger | B2RQG2 [3 | 0 | 2321.997 | 0 | -0.08 | 0.186812 | 6.2 | 102.2 | 97.8 | 2.53 | 7.93 | | High | High | High | High | High | High | High | High | High | High | High | 0 | 8.06E-06 | 3.79 |
| FALSE | High | NEIHGDNL | [R].NEIHGD | 2xCarbami | Q6PCN7 1x | 4.91E-06 | 0 | 1 | 1 | 1 | Q6PCN7 | Helicase-lik | Q6PCN7 [7 | 0 | 3093.395 | 0 | 0.71 | 0.025 | 12.62 | 73 | 127 | 8.28 | 16.41 | | High | High | High | High | High | High | High | High | High | High | High | 0 | 6.1E-07 | 5.18 |
| FALSE | High | NELQEPCE: | [K].NELQEI | 1xCarbami | F6UFZ5 1xl | 2.51E-08 | 0 | 2 | 7 | 2 | F6UFZ5; Q( | PCF11 clea | F6UFZ5 [14 | 0 | 1868.887 | 0 | 0.13 | 0.25455 | 11.14 | 95.4 | 104.6 | 8.79 | 9.22 | Shared | High | High | High | High | High | High | High | High | High | High | High | 0 | 1.61E-09 | 3.35 |
| FALSE | High | NETPTVSG. | [R].NETPTV | 1xPhosphc | Q5SS00 1xl | 2.06E-05 | 0 | 1 | 1 | 2 | Q5SS00 | DBF4-type | Q5SS00 [4! | 0 | 2075.908 | 0 | -1.66 | 0.000986 | 2.98 | 151.9 | 48.1 | 7.68 | 20.12 | | High | High | High | High | High | High | High | High | High | High | High | 0 | 3.05E-06 | 3.54 |
| FALSE | High | NENASFPR | [R].NENASI | 1xPhosphc | Q99P72 1x | 0.015061 | 0.001049 | 1 | 2 | 1 | Q99P72 | Reticulon-4 | Q99P72 [4: | 1 | 2140.12 | 0 | 0.79 | 0.010719 | 9.57 | 72.7 | 127.3 | 9.76 | 13.16 | | High | High | High | High | High | High | High | High | High | High | High | 0.000571 | 0.004977 | 2.14 |
| FALSE | High | NENPVDYT | [K].NENPV | 1xPhosphc | Q9Z1X4-3 : | 0.000501 | 0 | 1 | 6 | 4 | Q9Z1X4-3 | Isoform 3 c | Q9Z1X4-3 | 0 | 3117.562 | 0 | -0.38 | 0.048364 | 13.73 | 106.8 | 93.2 | 10.17 | 16.02 | | High | High | High | High | High | High | High | High | High | High | High | 0 | 0.00011 | 2.49 |
| FALSE | High | NENPVDYT | [K].NENPV | 1xPhosphc | Q9Z1X4-3 : | 3.6E-08 | 0 | 1 | 6 | 6 | Q9Z1X4-3 | Isoform 3 c | Q9Z1X4-3 | 0 | 3133.557 | 0 | -0.3 | 0.044354 | 12.26 | 110.4 | 89.6 | 8.91 | 8.96 | | High | High | High | High | High | High | High | High | High | High | High | 0 | 6.87E-07 | 3.26 |
| FALSE | High | NEPCSSSSS | [K].NEPCSS | 1xCarbami | Q8BU04 1x | 0.025964 | 0.001645 | 1 | 4 | 4 | Q8BU04 | Putative E3 | Q8BU04 [2 | 0 | 2311 | 0 | -0.7 | 0.270831 | 11.51 | 122.3 | 77.7 | 29.64 | 34.94 | | High | High | High | High | High | High | High | High | High | High | High | 0.001076 | 0.01142 | 1.93 |
| FALSE | High | NEPCSSSSS | [K].NEPCSS | 1xCarbami | Q8BU04 1x | 0.000101 | 0 | 1 | 4 | 6 | Q8BU04 | Putative E3 | Q8BU04 [2 | 0 | 2540.163 | 0 | -0.83 | 0.000509 | 3.83 | 126.2 | 73.8 | 2.54 | 7.36 | | High | High | High | High | High | High | High | High | High | High | High | 0 | 0.000294 | 3.14 |
| FALSE | High | NEPCSSSSS | [K].NEPCSS | 1xCarbami | Q8BU04 2x | 0.093123 | 0.007276 | 1 | 4 | 2 | Q8BU04 | Putative E3 | Q8BU04 [2 | 0 | 2620.129 | 0 | -0.73 | 0.00045 | 0.86 | 124.7 | 75.3 | 6.9 | 6.26 | | High | High | High | High | High | High | High | High | High | High | High | 0.003685 | 0.0396 | 2.51 |
| FALSE | High | NEPLLTPIPI | [R].NEPLLT | 1xPhosphc | P33609 1xl | 0.007274 | 0.00062 | 1 | 3 | 1 | P33609 | DNA polym | P33609 [22 | 0 | 1543.87 | 0 | 0.34 | 0.739281 | 18.71 | 88.4 | 111.6 | 25.97 | 38.18 | | High | High | High | High | High | High | High | High | High | High | High | 0.000275 | 0.0022 | 2.41 |
| FALSE | High | NEPLLTPIPI | [R].NEPLLT | 1xPhosphc | P33609 1xl | 0.002134 | 5.66E-05 | 1 | 3 | 2 | P33609 | DNA polym | P33609 [22 | 0 | 1773.033 | 0 | 0.33 | 0.012721 | 9.38 | 86.6 | 113.4 | 10.37 | 9.07 | | High | High | High | High | High | High | High | High | High | High | High | 2.58E-05 | 0.000556 | 2.27 |
| FALSE | High | NEPTTPSW | [R].NEPTTF | 1xPhosphc | A2ARP8 1x | 0.000845 | 0 | 1 | 3 | 2 | A2ARP8 | Microtubu | A2ARP8 [2( | 0 | 2600.357 | 0 | -0.98 | 0.011296 | 3.15 | 147.3 | 52.7 | 18.42 | 38.42 | | High | High | High | High | High | High | High | High | High | High | High | 0 | 0.000329 | 2.26 |
| FALSE | High | NESEEEEN/ | [K].NESEEE | 1xCarbami | Q80TJ7 2xl | 4.91E-06 | 0 | 1 | 3 | 1 | Q80TJ7 | Histone lys | Q80TJ7 [78 | 0 | 2890.121 | 0 | -0.43 | 0.34879 | 23.52 | 113.9 | 86.1 | 23.38 | 18.73 | | High | High | High | High | High | High | High | High | High | High | High | 0 | 6.08E-07 | 5.39 |
| FALSE | High | NESEEEEN/ | [K].NESEEE | 1xCarbami | Q80TJ7 1xl | 1.05E-05 | 0 | 1 | 3 | 1 | Q80TJ7 | Histone lys | Q80TJ7 [78 | 0 | 3039.318 | 0 | -0.32 | 0.058087 | 18.85 | 111 | 89 | 8.75 | 15.77 | | High | High | High | High | High | High | High | High | High | High | High | 0 | 1.42E-06 | 3.37 |
| FALSE | High | NESEEEEN/ | [K].NESEEE | 1xCarbami | Q80TJ7 1xl | 0.042503 | 0.003197 | 1 | 3 | 1 | Q80TJ7 | Histone lys | Q80TJ7 [78 | 0 | 3040.302 | 0 | -0.22 | 0.289084 | 25.32 | 101.6 | 98.4 | 8.53 | 19.73 | | High | High | High | High | High | High | High | High | High | High | High | 0.001639 | 0.01601 | 3.61 |
| FALSE | High | NESEEEEN/ | [K].NESEEE | 1xCarbami | Q80TJ7 2xl | 5.35E-08 | 0 | 1 | 3 | 6 | Q80TJ7 | Histone lys | Q80TJ7 [78 | 0 | 3119.284 | 0 | -0.26 | 0.153745 | 9.07 | 110.9 | 89.1 | 7.19 | 14.22 | | High | High | High | High | High | High | High | High | High | High | High | 0 | 3.77E-09 | 4.68 |
| FALSE | High | NESLTATD! | [R].NESLTA | 1xPhosphc | O89032 1x | 0.024424 | 0.001612 | 1 | 3 | 2 | O89032 | SH3 and PX | O89032 [9! | 0 | 1515.725 | 0 | 1.44 | 0.000892 | 13.72 | 52.3 | 147.7 | 10.88 | 10.48 | | High | High | High | High | High | High | High | High | High | High | High | 0.000859 | 0.008507 | 1.84 |
| FALSE | High | NENGVNSI | [R].NENGV | 1xPhosphc | Q6ZQB8 1x | 5.57E-06 | 0 | 1 | 2 | 1 | Q6ZQB8 | MKIAA042 | Q6ZQB8 [2 | 0 | 2132.006 | 0 | -0.15 | 0.02626 | 9.1 | 105.7 | 94.3 | 4.6 | 8.79 | | High | High | High | High | High | High | High | High | High | High | High | 0 | 7E-07 | 2.85 |
| FALSE | High | NFYESEEEI | [K].NFYESE | 1xPhosphc | Q9Z1T1 1x | 1.03E-05 | 0 | 1 | 3 | 10 | Q9Z1T1 | AP-3 comp | Q9Z1T1 [2 | 0 | 2099.91 | 0 | 0.16 | 0.016938 | 3.39 | 94.3 | 105.7 | 1.74 | 5.62 | | High | High | High | High | High | High | High | High | High | High | High | 0 | 4.08E-06 | 4.46 |
| FALSE | High | NGDECVYH | [K].NGDEC | 2xCarbami | Q8BJ05 1xl | 0.004911 | 0.000504 | 1 | 4 | 1 | Q8BJ05 | Zinc finger | Q8BJ05 [6( | 0 | 2351.071 | 0 | -0.24 | 0.226904 | 14.71 | 106.9 | 93.1 | 17.55 | 19.77 | | High | High | High | High | High | High | High | High | High | High | High | 0.000225 | 0.001423 | 2.47 |
| FALSE | High | NGDICETPI | [K].NGDICI | 1xCarbami | P97801 1xl | 0.000352 | 0 | 1 | 5 | 1 | P97801 | Survival m | P97801 [53 | 0 | 1911.929 | 0 | -0.02 | 0.613033 | 39.55 | 102.6 | 97.4 | 27.75 | 16.97 | | High | High | High | High | High | High | High | High | High | High | High | 0 | 7.42E-05 | 2.58 |
| FALSE | High | NGQHVASS | [K].NGQHV | 1xDeamida | Q8BTM8 1: | 2.13E-05 | 0 | 1 | 8 | 2 | Q8BTM8 | Filamin-A C | Q8BTM8 [2 | 0 | 2758.361 | 0 | 0.39 | 0.098604 | 5.31 | 88.5 | 111.5 | 21.73 | 15.89 | | High | High | High | High | High | High | High | High | High | High | High | 0 | 3.42E-06 | 4.05 |
| FALSE | High | NGSDDDDI | [K].NGSDD | 1xPhosphc | Q80WC1-4 | 0.000202 | 0 | 1 | 5 | 1 | Q80WC1-4 | Isoform 4 c | Q80WC1-4 | 0 | 2159.011 | 0 | -0.28 | 0.051254 | 11.77 | 112.5 | 87.5 | 9.38 | 11.79 | | High | High | High | High | High | High | High | High | High | High | High | 0 | 3.94E-05 | 2.61 |
| FALSE | High | NGSDDDNI | [R].NGSDD | 1xPhosphc | Q3TYF6 1x | 0.001674 | 5.66E-05 | 1 | 3 | 2 | Q3TYF6 | DNA segme | Q3TYF6 [8: | 0 | 2059.949 | 2 | 0.09 | 0.294411 | 7.87 | 97.2 | 102.8 | 8.59 | 7.25 | | High | High | High | High | High | High | High | High | High | High | High | 0.000327 | 0.002782 | 2.6 |
| FALSE | High | NGSDDDNI | [R].NGSDD | 1xDeamida | Q3TYF6 1x | 0.01441 | 0.001011 | 1 | 3 | 3 | Q3TYF6 | DNA segme | Q3TYF6 [8: | 0 | 2060.933 | 0 | 0.17 | 0.046657 | 5.49 | 94.3 | 105.7 | 6.45 | 6.39 | | High | High | High | High | High | High | High | High | High | High | High | 0.001142 | 0.01179 | 3.04 |
| FALSE | High | NGSEADIDI | [K].NGSEAI | 1xPhosphc | A0A1S6GW | 1.32E-07 | 0 | 1 | 2 | 2 | A0A1S6GW | Uncharacte | A0A1S6GW | 0 | 1864.816 | 0 | -0.23 | 0.135636 | 8.84 | 107.7 | 92.3 | 4.73 | 8.86 | | High | High | High | High | High | High | High | High | High | High | High | 2.58E-05 | 0.000422 | 3.84 |
| FALSE | High | NGSLDSPG | [K].NGSLD! | 1xPhosphc | O35685 1x | 0.001222 | 0 | 1 | 1 | 2 | O35685 | Nuclear mi | O35685 [1: | 0 | 1183.556 | 0 | -0.97 | 0.143546 | 9.27 | 132.3 | 67.7 | 37.68 | 40.54 | | High | High | High | High | High | High | High | High | High | High | High | 0.000449 | 0.00406 | 1.99 |
| FALSE | High | NGSLDSPG | [K].NGSLD! | 1xPhosphc | O35685 1x | 0.008927 | 0.000736 | 1 | 1 | 3 | O35685 | Nuclear mi | O35685 [1: | 0 | 1412.719 | 0 | -0.73 | 0.002723 | 1.8 | 124.6 | 75.4 | 4.54 | 8.19 | | High | High | High | High | High | High | High | High | High | High | High | 0.000327 | 0.002773 | 1.52 |
| FALSE | High | NGSLDSPG | [K].NGSLD! | 1xPhosphc | O35685 1x | 0.000118 | 0 | 1 | 1 | 3 | O35685 | Nuclear mi | O35685 [1: | 1 | 3203.436 | 0 | -1.27 | 0.006502 | 3.93 | 141.3 | 58.7 | 19.52 | 8.87 | | High | High | High | High | High | High | High | High | High | High | High | 0 | 2.18E-05 | 3.19 |
| FALSE | High | NGSLQSLS\ | [R].NGSLQ: | 1xDeamida | O54826 1x | 0.000475 | 0 | 1 | 4 | 2 | O54826 | Protein AF- | O54826 [5: | 0 | 2709.264 | 0 | -0.46 | 0.662639 | 4.95 | 116.5 | 83.5 | 5.29 | 36.01 | | High | High | High | High | High | High | High | High | High | High | High | 0 | 0.000103 | 3.92 |
| FALSE | High | NGSPPPGA | [R].NGSPPI | 1xPhosphc | Q3TCX6 1x | 0.076555 | 0.005826 | 1 | 7 | 1 | Q3TCX6 | Putative ur | Q3TCX6 [6 | 0 | 1416.683 | 0 | 0.67 | 0.000687 | 4.44 | 77.9 | 122.1 | 5.56 | 7.25 | | High | High | High | High | High | High | High | High | High | High | High | 0.002976 | 0.03147 | 1.49 |
| FALSE | High | NGSPVPVP | [R].NGSPVI | 1xPhosphc | Q8BSP2-2 : | 0.073462 | 0.005631 | 1 | 4 | 1 | Q8BSP2-2 | Isoform 2 c | Q8BSP2-2 | 0 | 4646.151 | 0 | -0.41 | 0.024864 | 8.37 | 115.8 | 84.2 | 11.96 | 6.08 | | High | High | High | High | High | High | High | High | High | High | High | 0.002896 | 0.03008 | 4.24 |
| FALSE | High | NGSPVPVP | [R].NGSPVI | 1xDeamida | Q8BSP2-2 : | 0.000134 | 0 | 1 | 4 | 5 | Q8BSP2-2 | Isoform 2 c | Q8BSP2-2 | 0 | 4647.135 | 0 | -0.18 | 0.235139 | 25.67 | 109.7 | 90.3 | 13.22 | 11.1 | | High | High | High | High | High | High | High | High | High | High | High | 0 | 2.49E-05 | 4.52 |
| FALSE | High | NGSRLDIVI | [R].NGSRLI | 1xPhosphc | Q8K0D0 1x | 9.65E-06 | 0 | 1 | 4 | 1 | Q8K0D0 | Cyclin-dep | Q8K0D0 [9 | 1 | 2033.095 | 0 | 0.34 | 0.182961 | 26.85 | 88.2 | 111.8 | 9.09 | 16.2 | | High | High | High | High | High | High | High | High | High | High | High | 0 | 1.3E-06 | 2.4 |
| FALSE | High | NGQHVASS | [K].NGQHV | 1xPhosphc | Q8BTM8 1: | 1.45E-05 | 0 | 1 | 8 | 2 | Q8BTM8 | Filamin-A C | Q8BTM8 [2 | 0 | 2757.377 | 0 | 0.32 | 0.036234 | 24.52 | 89 | 111 | 25.93 | 7.14 | | High | High | High | High | High | High | High | High | High | High | High | 0 | 9.16E-06 | 3.58 |
| FALSE | High | NGSRLDIVI | [R].NGSRLI | 1xDeamida | Q8K0D0 1x | 0.065405 | 0.004734 | 1 | 4 | 1 | Q8K0D0 | Cyclin-dep | Q8K0D0 [9 | 1 | 2034.079 | 0 | 0.77 | 0.133502 | 36.47 | 73.8 | 126.2 | 16.67 | 24.31 | | High | High | High | High | High | High | High | High | High | High | High | 0.002432 | 0.02621 | 1.62 |
| FALSE | High | NGTSGSDS | [K].NGTSG | 1xDeamida | Q8BG15 1x | 1.09E-05 | 0 | 1 | 4 | 11 | Q8BG15 | CTD small ; | Q8BG15 [1 | 0 | 2514.201 | 0 | -0.19 | 0.028719 | 3.81 | 107.5 | 92.5 | 7.67 | 8.42 | | High | High | High | High | High | High | High | High | High | High | High | 0 | 1.86E-06 | 3.76 |
| FALSE | High | NGVNCEH! | [K].NGVNC | 1xCarbami | Q8JZR6 1xl | 0.007811 | 0.000664 | 1 | 2 | 1 | Q8JZR6 | Electroneu | Q8JZR6 [27 | 0 | 2094.993 | 0 | 2.61 | 0.000748 | 6.95 | 26.7 | 173.3 | 11.15 | 13.38 | | High | High | High | High | High | High | High | High | High | High | High | 0.000294 | 0.00237 | 2.34 |
| FALSE | High | NGVSLFNS | [R].NGVSLI | 1xPhosphc | Q8CJF7 1xl | 0.001659 | 5.66E-05 | 1 | 2 | 1 | Q8CJF7 | Protein ELY | Q8CJF7 [1( | 0 | 1371.687 | 0 | -0.28 | 0.358071 | 29.17 | 109.7 | 90.3 | 36.83 | 32.09 | | High | High | High | High | High | High | High | High | High | High | High | 2.58E-05 | 0.000422 | 2.08 |
| FALSE | High | NGVSLFNS | [R].NGVSLI | 1xPhosphc | Q8CJF7 1xl | 0.0006 | 0 | 1 | 2 | 1 | Q8CJF7 | Protein ELY | Q8CJF7 [1( | 0 | 1600.85 | 0 | -0.62 | 0.00131 | 3.24 | 120.8 | 79.2 | 2.25 | 7.56 | | High | High | High | High | High | High | High | High | High | High | High | 0 | 0.000135 | 2.87 |
| FALSE | High | NGVSLFNS | [R].NGVSLI | 1xDeamida | Q8CJF7 1xl | 0.022378 | 0.001542 | 1 | 2 | 2 | Q8CJF7 | Protein ELY | Q8CJF7 [1( | 0 | 1601.834 | 0 | -0.08 | 0.038553 | 1.78 | 106 | 94 | 3.73 | 6.3 | | High | High | High | High | High | High | High | High | High | High | High | 0.006969 | 0.07527 | 1.92 |
| FALSE | High | NGVSTPCC | [R].NGVSTI | 1xCarbami | Q5SXJ3 1xl | 0.000102 | 0 | 1 | 3 | 3 | Q5SXJ3 | Fanconi an | Q5SXJ3 [11 | 0 | 1957.898 | 0 | 0.19 | 0.012951 | 4.19 | 93.1 | 106.9 | 3.84 | 3.17 | | High | High | High | High | High | High | High | High | High | High | High | 0 | 0.000112 | 3.05 |
| FALSE | High | NGYGFVEF | [K].NGYGF | 1xPhosphc | Q3TWW8 : | 0.001463 | 5.66E-05 | 1 | 2 | 1 | Q3TWW8 | Serine/argi | Q3TWW8 | 0 | 1728.747 | 0 | -0.67 | 0.01184 | 7.18 | 122.8 | 77.2 | 8.63 | 8.8 | | High | High | High | High | High | High | High | High | High | High | High | 2.58E-05 | 0.000365 | 2.12 |
| FALSE | High | NGYGFVEF | [K].NGYGF | 1xPhosphc | Q3TWW8 : | 0.005135 | 0.000551 | 1 | 2 | 3 | Q3TWW8 | Serine/argi | Q3TWW8 | 1 | 3278.493 | 0 | -0.67 | 0.003069 | 2.41 | 121.9 | 78.1 | 5.81 | 7.92 | | High | High | High | High | High | High | High | High | High | High | High | 0.000246 | 0.00149 | 3.57 |
| FALSE | High | NGYGFVEF | [K].NGYGF | 1xDeamida | Q3TWW8 : | 0.066503 | 0.004796 | 1 | 2 | 2 | Q3TWW8 | Serine/argi | Q3TWW8 | 1 | 3279.477 | 0 | -0.43 | 0.003322 | 3.15 | 114.9 | 85.1 | 0.97 | 8.74 | | High | High | High | High | High | High | High | High | High | High | High | 0.002867 | 0.02987 | 2.79 |
| FALSE | High | NHCCNGEI | [R].NHCCN | 2xCarbami | F8VPV0 1xl | 0.003892 | 0.000261 | 1 | 8 | 1 | F8VPV0 | Pericentrin | F8VPV0 [1: | 0 | 3248.568 | 0 | -0.43 | 0.037319 | 7.69 | 113.1 | 86.9 | 9.9 | 15.76 | | High | High | High | High | High | High | High | High | High | High | High | 0.00014 | 0.001097 | 3.15 |
| FALSE | High | NHGPDSGI | [K].NHGPD | 1xPhosphc | F6VQ19 1x | 0.002895 | 0.00016 | 2 | 6 | 1 | F6VQ19; Q | Leucine-ric | F6VQ19 [3( | 0 | 2021.933 | 0 | -1.04 | 0.000496 | 3.74 | 131.8 | 68.2 | 4.04 | 15.07 | Shared | High | High | High | High | High | High | High | High | High | High | High | 7.19E-05 | 0.000788 | 3.56 |
| FALSE | High | NHGYASPL | [R].NHGYA | 1xPhosphc | Q9QZ59 1x | 0.00195 | 5.66E-05 | 1 | 5 | 1 | Q9QZ59 | Doublesex- | Q9QZ59 [7 | 0 | 1524.798 | 0 | -1.63 | 0.00252 | 2.33 | 151.1 | 48.9 | 11.5 | 15.5 | | High | High | High | High | High | High | High | High | High | High | High | 2.58E-05 | 0.000505 | 2.15 |
| FALSE | High | NHLSPQEG | [K].NHLSP( | 2xCarbami | Q9EP97 1x | 9.69E-11 | 0 | 1 | 1 | 6 | Q9EP97 | Sentrin-spe | Q9EP97 [1( | 0 | 2501.126 | 0 | -0.81 | 0.010953 | 10.04 | 127.4 | 72.6 | 12.06 | 11.34 | | High | High | High | High | High | High | High | High | High | High | High | 0 | 5.3E-12 | 4.31 |
| FALSE | High | NGTSGSDS | [K].NGTSG | 1xPhosphc | Q8BG15 1x | 3.63E-05 | 0 | 1 | 4 | 2 | Q8BG15 | CTD small ; | Q8BG15 [1 | 0 | 2513.217 | 0 | 0 | 0.363066 | 8.29 | 102.5 | 97.5 | 5.04 | 7.31 | | High | High | High | High | High | High | High | High | High | High | High | 0 | 5.73E-06 | 3.65 |
| FALSE | High | NGQEPVRF | [K].NGQEP | 1xDeamida | A2AI08 1xF | 5.31E-05 | 0 | 1 | 1 | 2 | A2AI08 | Taperin OS | A2AI08 [48 | 0 | 3769.825 | 0 | 1.62 | 0.000725 | 17.44 | 50.4 | 149.6 | 12.2 | 10.04 | | High | High | High | High | High | High | High | High | High | High | High | 0 | 8.8E-06 | 3.57 |
| FALSE | High | NGQEPVRF | [K].NGQEP | 1xPhosphc | A2AI08 1xF | 0.000104 | 0 | 1 | 1 | 3 | A2AI08 | Taperin OS | A2AI08 [48 | 0 | 3768.841 | 0 | 1.63 | 0.001302 | 25.91 | 51.5 | 148.5 | 20.27 | 8.58 | | High | High | High | High | High | High | High | High | High | High | High | 0 | 1.89E-05 | 3.22 |
| FALSE | High | NGQEAGP/ | [K].NGQEA | 1xDeamida | P0DOV1 1x | 0.000651 | 0 | 2 | 3 | 1 | P0DOV1; P | Interferon- | P0DOV1 [9 | 0 | 2569.144 | 0 | 0.98 | 0.102193 | 16.24 | 63.3 | 136.7 | 12.42 | 56.2 | Shared | High | High | High | High | High | High | High | High | High | High | High | 0 | 0.000147 | 3.31 |
| FALSE | High | NGEVVHTF | [K].NGEVV | 1xPhosphc | Q3TMA0 1 | 0.00582 | 0.00062 | 1 | 2 | 5 | Q3TMA0 | Putative ur | Q3TMA0 [4 | 0 | 1577.741 | 0 | 2.82 | 0.000435 | 3.86 | 25.2 | 174.8 | 8.22 | 8.93 | | High | High | High | High | High | High | High | High | High | High | High | 0.001324 | 0.01398 | 1.8 |
| FALSE | High | NGFYCHSC | [K].NGFYCI | 1xCarbami | G5E8J4 1xF | 0.106748 | 0.008634 | 1 | 5 | 1 | G5E8J4 | Tyrosine-p | G5E8J4 [36 | 0 | 1893.815 | 0 | 0.38 | 0.023156 | 14.13 | 86.9 | 113.1 | 12.33 | 9.77 | | High | High | High | High | High | High | High | High | High | High | High | 0.004327 | 0.04656 | 2 |
| FALSE | High | NGGLEHVF | [K].NGGLEI | 1xPhosphc | Q3TEW0 1: | 0.022182 | 0.001542 | 1 | 3 | 1 | Q3TEW0 | Putative ur | Q3TEW0 [! | 0 | 2649.238 | 0 | -0.62 | 0.456743 | 18.76 | 121.1 | 78.9 | 25.53 | 41.27 | | High | High | High | High | High | High | High | High | High | High | High | 0.000814 | 0.007665 | 2.19 |
| FALSE | High | NGGIQHTG | [R].NGGIQ | 1xPhosphc | A0A0R4J0K | 8.5E-06 | 0 | 1 | 2 | 1 | A0A0R4J0K | E3 ubiquiti | A0A0R4J0K | 0 | 2355.009 | 0 | 0.14 | 0.426744 | 7.4 | 97.6 | 102.4 | 6.05 | 8.75 | | High | High | High | High | High | High | High | High | High | High | High | 0 | 1.12E-06 | 3.14 |
| FALSE | High | NGGQASSS | [R].NGGQ/ | 1xPhosphc | E9Q8G4 1x | 0.017041 | 0.001204 | 1 | 21 | 1 | E9Q8G4 | Transcripti | E9Q8G4 [3 | 0 | 2495.151 | 0 | -0.56 | 0.028635 | 5.77 | 116.3 | 83.7 | 11.42 | 6.82 | | High | High | High | High | High | High | High | High | High | High | High | 0.000638 | 0.005686 | 2.16 |
| FALSE | High | NGGQASSS | [R].NGGQ/ | 1xDeamida | E9Q8G4 1x | 0.016743 | 0.001166 | 1 | 21 | 5 | E9Q8G4 | Transcripti | E9Q8G4 [3 | 0 | 2496.135 | 0 | -0.63 | 0.001325 | 0.29 | 121.4 | 78.6 | 2.78 | 7.27 | | High | High | High | High | High | High | High | High | High | High | High | 0.000801 | 0.007226 | 2.17 |
| FALSE | High | NGGQTFIC | [K].NGGQT | 2xCarbami | Q8BQ33 1x | 1.2E-07 | 0 | 1 | 1 | 1 | Q8BQ33 | Treslin OS= | Q8BQ33 [1 | 0 | 2021.877 | 0 | -0.64 | 0.00289 | 6.55 | 121.7 | 78.3 | 7.06 | 11.93 | | High | High | High | High | High | High | High | High | High | High | High | 0 | 9.46E-09 | 3.86 |
| FALSE | High | NGGQTFIC | [K].NGGQT | 2xCarbami | Q8BQ33 1x | 3.2E-05 | 0 | 1 | 1 | 2 | Q8BQ33 | Treslin OS= | Q8BQ33 [1 | 0 | 2022.861 | 0 | -0.87 | 0.000774 | 3.76 | 131.9 | 68.1 | 5.78 | 4.81 | | High | High | High | High | High | High | High | High | High | High | High | 0 | 7.37E-05 | 3.24 |
| FALSE | High | NGGSFEDD | [R].NGGSF | 1xCarbami | Q922B9 1x | 0.003822 | 0.000261 | 1 | 4 | 1 | Q922B9 | Sperm-spe | Q922B9 [1( | 0 | 4216.965 | 0 | 0.26 | 0.270929 | 22.48 | 93 | 107 | 6.27 | 13.89 | | High | High | High | High | High | High | High | High | High | High | High | 0.000117 | 0.001067 | 3.13 |
| FALSE | High | NGGSPALN | [K].NGGSP. | 1xPhosphc | Q99JY8 1xl | 0.00083 | 0 | 1 | 1 | 1 | Q99JY8 | Phospholip | Q99JY8 [16 | 0 | 1519.722 | 0 | -1.73 | 0.00037 | 1.31 | 154.4 | 45.6 | 2.72 | 28.91 | | High | High | High | High | High | High | High | High | High | High | High | 0 | 0.000193 | 2.24 |
| FALSE | High | NGILAIEGT | [R].NGILAII | 1xPhosphc | P70398 1xl | 1E-06 | 0 | 1 | 2 | 1 | P70398 | Probable u | P70398 [15 | 0 | 2547.101 | 0 | -0.21 | 0.088098 | 4.37 | 106.5 | 93.5 | 10.29 | 7.66 | | High | High | High | High | High | High | High | High | High | High | High | 0 | 1.02E-07 | 3.54 |
| FALSE | High | NGILAIEGT | [R].NGILAII | 1xPhosphc | P70398 1xl | 6.29E-08 | 0 | 1 | 2 | 9 | P70398 | Probable u | P70398 [15 | 0 | 2563.096 | 0 | 0.02 | 0.949501 | 20.45 | 103.2 | 96.8 | 18.82 | 26.02 | | High | High | High | High | High | High | High | High | High | High | High | 0 | 4.55E-06 | 4.66 |
| FALSE | High | NGILAIEGT | [R].NGILAII | 1xDeamida | P70398 1xl | 4.49E-07 | 0 | 1 | 2 | 3 | P70398 | Probable u | P70398 [15 | 0 | 2564.08 | 0 | -0.24 | 0.057129 | 16.48 | 115.3 | 84.7 | 9.74 | 16.97 | | High | High | High | High | High | High | High | High | High | High | High | 0 | 4.15E-08 | 3.43 |
| FALSE | High | NGILAIEGT | [R].NGILAII | 2xPhosphc | P70398 2xl | 1.49E-05 | 0 | 1 | 2 | 2 | P70398 | Probable u | P70398 [15 | 0 | 2643.062 | 0 | 0.35 | 0.524954 | 16.39 | 88.4 | 111.6 | 35.3 | 38.62 | | High | High | High | High | High | High | High | High | High | High | High | 0 | 2.11E-06 | 5.09 |
| FALSE | High | NGILAIEGT | [R].NGILAII | 1xPhosphc | P70398 1xl | 1.57E-09 | 0 | 1 | 2 | 2 | P70398 | Probable u | P70398 [15 | 0 | 2776.264 | 0 | -0.34 | 0.138408 | 14.84 | 111.7 | 88.3 | 11.25 | 13.46 | | High | High | High | High | High | High | High | High | High | High | High | 0 | 1.09E-07 | 3.67 |
| FALSE | High | NGILAIEGT | [R].NGILAII | 1xPhosphc | P70398 1xl | 1.99E-12 | 0 | 1 | 2 | 16 | P70398 | Probable u | P70398 [15 | 0 | 2792.259 | 0 | -0.1 | 0.802471 | 7.65 | 102.3 | 97.7 | 2.62 | 9.6 | | High | High | High | High | High | High | High | High | High | High | High | 0 | 2.36E-12 | 7.26 |
| FALSE | High | NGILASPQ! | [R].NGILAS | 1xPhosphc | I1VGT4 1xF | 1.09E-06 | 0 | 1 | 7 | 2 | I1VGT4 | Anion exch | I1VGT4 [25 | 0 | 2092.027 | 0 | 0.91 | 0.190481 | 31.99 | 64.4 | 135.6 | 37.9 | 40.85 | | High | High | High | High | High | High | High | High | High | High | High | 0 | 1.12E-07 | 4.9 |
| FALSE | High | NGILASPQ! | [R].NGILAS | 1xDeamida | I1VGT4 1xF | 4.27E-05 | 0 | 1 | 7 | 1 | I1VGT4 | Anion exch | I1VGT4 [25 | 0 | 2093.011 | 0 | 0.95 | 0.01747 | 26.63 | 66.8 | 133.2 | 13.47 | 19.77 | | High | High | High | High | High | High | High | High | High | High | High | 0 | 6.94E-06 | 3.23 |
| FALSE | High | NGILASPQ! | [R].NGILAS | 1xPhosphc | I1VGT4 1xF | 1.32E-07 | 0 | 1 | 7 | 4 | I1VGT4 | Anion exch | I1VGT4 [25 | 0 | 2321.19 | 0 | 0.9 | 0.000925 | 16.83 | 67.4 | 132.6 | 6.5 | 5.9 | | High | High | High | High | High | High | High | High | High | High | High | 0 | 1.37E-05 | 3.36 |
| FALSE | High | NGILASPQ! | [R].NGILAS | 1xDeamida | I1VGT4 1xF | 1.17E-05 | 0 | 1 | 7 | 6 | I1VGT4 | Anion exch | I1VGT4 [25 | 0 | 2322.174 | 0 | 0.94 | 0.000209 | 3.85 | 67.3 | 132.7 | 7.47 | 2.88 | | High | High | High | High | High | High | High | High | High | High | High | 0 | 3.94E-06 | 3.42 |
| FALSE | High | NGILASPQ! | [R].NGILAS | 2xPhosphc | I1VGT4 2xF | 0.029586 | 0.001857 | 1 | 7 | 1 | I1VGT4 | Anion exch | I1VGT4 [25 | 0 | 2401.157 | 0 | -0.2 | 0.001656 | 2.82 | 105.6 | 94.4 | 8.82 | 8.1 | | High | High | High | High | High | High | High | High | High | High | High | 0.000997 | 0.01064 | 2.89 |
| FALSE | High | NGLPGSRP | [K].NGLPG! | 1xPhosphc | Q8CHK4-3 | 0.104222 | 0.008348 | 1 | 5 | 1 | Q8CHK4-3 | Isoform 3 c | Q8CHK4-3 | 0 | 1632.806 | 0 | -0.59 | 0.000278 | 1.16 | 121 | 79 | 7.67 | 8.83 | | High | High | High | High | High | High | High | High | High | High | High | 0.00425 | 0.04545 | 1.27 |
| FALSE | High | NGISPTNPT | [R].NGISPT | 1xPhosphc | E9PW37 1x | 0.101749 | 0.008194 | 1 | 3 | 1 | E9PW37 | RAS protei | E9PW37 [1 | 0 | 1566.829 | 0 | 0.35 | 0.011088 | 6.79 | 86.7 | 113.3 | 4.74 | 12.62 | | High | High | High | High | High | High | High | High | High | High | High | 0.004126 | 0.04405 | 2.01 |
| FALSE | High | NGLSQPSE | [K].NGLSQ | 1xPhosphc | Q3ULC7 1x | 6.83E-05 | 0 | 1 | 3 | 1 | Q3ULC7 | Putative ur | Q3ULC7 [1 | 0 | 2606.368 | 0 | 0.24 | 0.260322 | 18.51 | 91.6 | 108.4 | 19.5 | 15.37 | | High | High | High | High | High | High | High | High | High | High | High | 0 | 1.17E-05 | 1.78 |
| FALSE | High | NGLSQPSE | [K].NGLSQ | 1xPhosphc | Q3ULC7 1x | 4.35E-05 | 0 | 1 | 5 | 1 | Q3ULC7 | Putative ur | Q3ULC7 [1 | 0 | 2634.399 | 0 | 0.05 | 0.633954 | 31.45 | 96.7 | 103.3 | 22.1 | 19.48 | | High | High | High | High | High | High | High | High | High | High | High | 0 | 7.02E-06 | 1.63 |
| FALSE | High | NGPLNESQ | [R].NGPLN | 1xPhosphc | B2RY27 1xl | 1.93E-07 | 0 | 1 | 3 | 2 | B2RY27 | MCG12578 | B2RY27 [31 | 0 | 2754.169 | 0 | 0.07 | 0.19443 | 6.37 | 92.2 | 107.8 | 14.62 | 33.49 | | High | High | High | High | High | High | High | High | High | High | High | 0 | 7.84E-08 | 4.95 |
| FALSE | High | NGQEAGP/ | [K].NGQEA | 1xPhosphc | P0DOV1 1x | 2.71E-07 | 0 | 2 | 3 | 1 | P0DOV1; P | Interferon- | P0DOV1 [9 | 0 | 2552.165 | 0 | 1.01 | 0.010398 | 23.03 | 69.5 | 130.5 | 15.86 | 19.61 | Shared | High | High | High | High | High | High | High | High | High | High | High | 0 | 2.35E-08 | 3.3 |
| FALSE | High | NGQEAGP/ | [K].NGQEA | 1xDeamida | P0DOV1 1x | 1.85E-08 | 0 | 2 | 3 | 1 | P0DOV1; P | Interferon- | P0DOV1 [9 | 0 | 2553.149 | 0 | 1.16 | 0.000547 | 3.87 | 61.8 | 138.2 | 13.8 | 12.09 | Shared | High | High | High | High | High | High | High | High | High | High | High | 0 | 1.14E-09 | 4.19 |
| FALSE | High | NGQEAGP/ | [K].NGQEA | 1xPhosphc | P0DOV1 1x | 2.99E-06 | 0 | 2 | 3 | 1 | P0DOV1; P | Interferon- | P0DOV1 [9 | 0 | 2568.16 | 0 | 1.56 | 0.001839 | 14.04 | 54.6 | 145.4 | 22.72 | 20.58 | Shared | High | High | High | High | High | High | High | High | High | High | High | 0 | 3.47E-07 | 4.73 |
| FALSE | High | NEEDSEGS | [K].NEEDSE | 1xPhosphc | Q8R1B4 1x | 6.65E-05 | 0 | 1 | 4 | 2 | Q8R1B4 | Eukaryotic | Q8R1B4 [1 | 0 | 3256.358 | 0 | -0.56 | 0.287002 | 24.83 | 109.1 | 90.9 | 30.13 | 50.68 | | High | High | High | High | High | High | High | High | High | High | High | 0 | 1.77E-05 | 4.68 |
| FALSE | High | NEEDEGHS | [K].NEEDE( | 1xPhosphc | Q60668 1x | 0.002441 | 0.000109 | 1 | 3 | 2 | Q60668 | Heterogen | Q60668 [7: | 0 | 1766.718 | 0 | -0.51 | 0.040179 | 5.2 | 122.4 | 77.6 | 22.58 | 47.08 | | High | High | High | High | High | High | High | High | High | High | High | 4.94E-05 | 0.000647 | 2.44 |
| FALSE | High | NEALISQEK | [K].NEALIS( | 1xDeamida | A2AEG2 1x | 0.064323 | 0.004664 | 1 | 2 | 1 | A2AEG2 | Oral-facial- | A2AEG2 [2 | 0 | 1341.65 | 0 | 0.12 | 0.398634 | 38.2 | 91.2 | 108.8 | 12.21 | 123.3 | | High | High | High | High | High | High | High | High | High | High | High | 0.002403 | 0.02585 | 2.5 |
| FALSE | High | NEALLGELI | [K].NEALLC | 1xPhosphc | Q8BKT8 1x | 3.06E-05 | 0 | 1 | 1 | 2 | Q8BKT8 | HAUS augn | Q8BKT8 [1 | 0 | 1968.999 | 0 | -0.83 | 0.011821 | 1.5 | 129.1 | 70.9 | 12.36 | 11.13 | | High | High | High | High | High | High | High | High | High | High | High | 0 | 4.74E-06 | 3.46 |
| FALSE | High | NAASFPLR! | [R].NAASFI | 1xCarbami | Q6KCD5 1x | 5.07E-08 | 0 | 1 | 5 | 2 | Q6KCD5 | Nipped-B-l | Q6KCD5 [2 | 1 | 2979.48 | 0 | 0.17 | 0.112662 | 11.21 | 94.2 | 105.8 | 10.46 | 5.13 | | High | High | High | High | High | High | High | High | High | High | High | 0 | 3.55E-09 | 4.48 |
| FALSE | High | NAASFPLR! | [R].NAASFI | 1xCarbami | Q6KCD5 2x | 0.004033 | 0.000311 | 1 | 5 | 1 | Q6KCD5 | Nipped-B-l | Q6KCD5 [2 | 1 | 3059.446 | 0 | 1.3 | 0.000756 | 8.57 | 57.7 | 142.3 | 9.8 | 11.45 | | High | High | High | High | High | High | High | High | High | High | High | 0.00014 | 0.001133 | 3.06 |
| FALSE | High | NACILQAS( | [K].NACILC | 1xCarbami | P41183 1xl | 0.039347 | 0.002793 | 1 | 2 | 2 | P41183 | B-cell lymp | P41183 [35 | 0 | 1952.008 | 0 | 0.85 | 0.015829 | 37.59 | 78.8 | 121.2 | 37 | 15.77 | | High | High | High | High | High | High | High | High | High | High | High | 0.001453 | 0.01459 | 1.5 |
| FALSE | High | NAEEEDSA | [R].NAEEEI | 1xPhosphc | Q8BP78 1x | 2.31E-05 | 0 | 1 | 1 | 3 | Q8BP78 | Protein FR/ | Q8BP78 [2' | 0 | 2248.006 | 0 | 0 | 0.772636 | 11.95 | 101.9 | 98.1 | 3.87 | 6.9 | | High | High | High | High | High | High | High | High | High | High | High | 0 | 3.45E-06 | 3.78 |
| FALSE | High | NAEEEDSA | [R].NAEEEI | 2xPhosphc | Q8BP78 2x | 5.94E-07 | 0 | 1 | 1 | 8 | Q8BP78 | Protein FR/ | Q8BP78 [2' | 0 | 2327.972 | 0 | 0.29 | 0.001578 | 3.45 | 88.1 | 111.9 | 4.04 | 3.42 | | High | High | High | High | High | High | High | High | High | High | High | 0 | 5.66E-08 | 3.82 |
| FALSE | High | NAEISDVAI | [R].NAEISD | 1xPhosphc | Q3TK27 1x | 0.001659 | 5.66E-05 | 1 | 2 | 2 | Q3TK27 | Putative ur | Q3TK27 [4: | 0 | 1838.873 | 0 | -0.84 | 0.00076 | 3.2 | 129.2 | 70.8 | 6.82 | 15.83 | | High | High | High | High | High | High | High | High | High | High | High | 2.58E-05 | 0.000421 | 3.5 |
| FALSE | High | NAEPPSPTI | [K].NAEPP! | 1xPhosphc | F8VQ52 1x | 3.35E-05 | 0 | 1 | 2 | 2 | F8VQ52 | Adenylate | F8VQ52 [4: | 0 | 1754.842 | 0 | 1.59 | 0.000255 | 1.1 | 47.9 | 152.1 | 6.15 | 6.79 | | High | High | High | High | High | High | High | High | High | High | High | 0 | 5.23E-06 | 1.95 |
| FALSE | High | NAFFPEVF! | [R].NAFFPE | 1xPhosphc | Q9D279 1x | 2.15E-08 | 0 | 1 | 1 | 5 | Q9D279 | Mitotic inte | Q9D279 [5 | 0 | 2943.304 | 0 | 3.41 | 0.000181 | 10.85 | 18.1 | 181.9 | 12.88 | 6.41 | | High | High | High | High | High | High | High | High | High | High | High | 0 | 3.7E-08 | 4.36 |
| FALSE | High | NAFFPEVF! | [R].NAFFPE | 2xPhosphc | Q9D279 2x | 9.89E-05 | 0 | 1 | 1 | 1 | Q9D279 | Mitotic inte | Q9D279 [5 | 0 | 3023.27 | 0 | 5.77 | 0.000847 | 74.57 | 3 | 197 | 74.89 | 16.56 | | High | High | High | High | High | High | High | High | High | High | High | 0 | 1.78E-05 | 3.3 |
| FALSE | High | NAFFPEVF! | [R].NAFFPE | 1xDeamida | Q9D279 2x | 0.003167 | 0.00016 | 1 | 1 | 1 | Q9D279 | Mitotic inte | Q9D279 [5 | 0 | 3024.254 | 0 | 4.67 | | 72.72 | 10.1 | 189.9 | 23.58 | 22.43 | | Not Found | High | High | Not Found | High | High | High | High | High | High | High | 7.19E-05 | 0.000866 | 3.87 |
| FALSE | High | NAFFPEVF! | [R].NAFFPE | 3xPhosphc | Q9D279 3x | 0.058664 | 0.004294 | 1 | 1 | 1 | Q9D279 | Mitotic inte | Q9D279 [5 | 1 | 3520.433 | 0 | 2.73 | 0.001787 | 19.87 | 27.5 | 172.5 | 19 | 19.96 | | High | High | High | High | High | High | High | High | High | High | High | 0.002187 | 0.02317 | 2.82 |

| | | | | | | | | | | | | | | | | | | | | | | | | | | | | | | | | | | | | | | |
|---|---|---|---|---|---|---|---|---|---|---|---|---|---|---|---|---|---|---|---|---|---|---|---|---|---|---|---|---|---|---|---|---|---|---|---|---|---|---|
| FALSE | High | NAGFTSAL | [K].NAGFT | 1xPhospho | Q3TYD6 1x | 1.9E-06 | 0 | 1 | 1 | 1 | Q3TYD6 | Serine/thr | Q3TYD6 [7 | 0 | 1799.889 | 0 | -1.24 | 0.006343 | 3.25 | 136.6 | 63.4 | 14.39 | 9.22 | | High | High | High | High | High | High | High | High | High | High | High | 0 | 2.1E-07 | 3.56 |
| FALSE | High | NAGPEPET | [R].NAGPE | 2xPhospho | Q3V1Z0 2x | 0.047086 | 0.00354 | 1 | 5 | 2 | Q3V1Z0 | Iporin OS= | Q3V1Z0 [7 | 0 | 2587.179 | 0 | 1.61 | 0.000526 | 15.17 | 48 | 152 | 12.35 | 8.58 | | High | High | High | High | High | High | High | High | High | High | High | 0.001812 | 0.01792 | 2.19 |
| FALSE | High | MVQASSQ | [R].MVQAS | 1xPhospho | Q8BTI8 1xP | 0.000933 | 0 | 1 | 3 | 3 | Q8BTI8 | Serine/argi | Q8BTI8 [23 | 1 | 3548.752 | 0 | -0.53 | 0.017728 | 3.38 | 121.9 | 78.1 | 10.9 | 18.35 | | High | High | High | High | High | High | High | High | High | High | High | 0 | 0.00022 | 2.53 |
| FALSE | High | NAGSASSS | [K].NAGSA | 1xPhospho | Q3UZR5 1x | 3.47E-05 | 0 | 1 | 5 | 2 | Q3UZR5 | Calcium-tra | Q3UZR5 [9 | 0 | 1477.677 | 0 | 1.01 | 0.001562 | 12.45 | 62.9 | 137.1 | 13.11 | 14.47 | | High | High | High | High | High | High | High | High | High | High | High | 0 | 5.5E-06 | 3.1 |
| FALSE | High | NAGVGSTT | [R].NAGVG | 1xPhospho | Q6A037 1x | 6.36E-09 | 0 | 1 | 2 | 3 | Q6A037 | NEDD4-bin | Q6A037 [4 | 0 | 2301.138 | 0 | -0.77 | 0.005464 | 1.25 | 126.8 | 73.2 | 7.6 | 11.79 | | High | High | High | High | High | High | High | High | High | High | High | 0 | 3.45E-10 | 4.09 |
| FALSE | High | NALGSLPK | [R].NALGSI | 1xDeamida | Q78JN3 1x | 0.083783 | 0.006345 | 1 | 2 | 1 | Q78JN3 | Enoyl-CoA | Q78JN3 [1 | 0 | 1109.581 | 0 | 0.45 | 0.523073 | 40.81 | 82.9 | 117.1 | 142.53 | 16.09 | | High | High | High | High | High | High | High | High | High | High | High | 0.003265 | 0.03513 | 1.6 |
| FALSE | High | NAIPSDSEA | [K].NAIPSD | 2xPhospho | Q5XJE5 2xI | 0.002022 | 5.66E-05 | 1 | 2 | 3 | Q5XJE5 | RNA polym | Q5XJE5 [2 | 0 | 2393.056 | 0 | -0.4 | 0.018266 | 4.3 | 115.7 | 84.3 | 9.96 | 9.37 | | High | High | High | High | High | High | High | High | High | High | High | 2.58E-05 | 0.000527 | 3.28 |
| FALSE | High | NALSLLTLG | [K].NALSLL | 1xPhospho | A0A1L1SSA | 0.000325 | 0 | 1 | 6 | 1 | A0A1L1SSA | Centrosom | A0A1L1SSA | 0 | 3230.442 | 0 | -0.35 | 0.073736 | 13.23 | 112.1 | 87.9 | 10.15 | 15.17 | | High | High | High | High | High | High | High | High | High | High | High | 0 | 6.77E-05 | 4.6 |
| FALSE | High | NAISLPQLN | [K].NAISLP | 1xPhospho | A0A0R4J0S | 1.04E-09 | 0 | 1 | 2 | 1 | A0A0R4J0S | Cdc42 effe | A0A0R4J0S | 0 | 2717.399 | 0 | 1.48 | 0.000427 | 4.88 | 54.6 | 145.4 | 9.64 | 5.33 | | High | High | High | High | High | High | High | High | High | High | High | 0 | 4.51E-11 | 5.62 |
| FALSE | High | NAISLPTIG | [K].NAISLP | 1xPhospho | Q9CQC5 1x | 0.034878 | 0.002269 | 1 | 1 | 1 | Q9CQC5 | Cdc42 effe | Q9CQC5 [1 | 0 | 3346.825 | 0 | 2.68 | 0.001316 | 32.48 | 31.4 | 168.6 | 34.51 | 16.8 | | High | High | High | High | High | High | High | High | High | High | High | 0.001183 | 0.01272 | 2.57 |
| FALSE | High | NALTTLAG | [K].NALTTL | 1xPhospho | Q8BG30 1x | 4.18E-07 | 0 | 1 | 2 | 1 | Q8BG30 | Negative e | Q8BG30 [1 | 0 | 2031.166 | 0 | 0.08 | 0.156366 | 12.9 | 96.6 | 103.4 | 10 | 3.86 | | High | High | High | High | High | High | High | High | High | High | High | 0 | 3.81E-08 | 2.56 |
| FALSE | High | NAMSLPQI | [K].NAMSL | 1xPhospho | Q3TH99 1x | 0.000845 | 0 | 1 | 3 | 2 | Q3TH99 | Putative ur | Q3TH99 [1 | 0 | 1782.923 | 0 | -1.02 | 0.004167 | 4.17 | 132.8 | 67.2 | 11.26 | 13.32 | | High | High | High | High | High | High | High | High | High | High | High | 0 | 0.000198 | 2.71 |
| FALSE | High | NAMSLPQI | [K].NAMSL | 1xPhospho | Q3TH99 1x | 0.003312 | 0.00016 | 1 | 3 | 2 | Q3TH99 | Putative ur | Q3TH99 [1 | 0 | 1798.917 | 0 | -0.94 | 0.005566 | 5.76 | 135.1 | 64.9 | 6.38 | 25.97 | | High | High | High | High | High | High | High | High | High | High | High | 0.002662 | 0.02838 | 3.27 |
| FALSE | High | NAPHGDDI | [R].NAPHG | 1xCarbami | Q9DBR4-2 | 8.47E-08 | 0 | 2 | 8 | 1 | Q9DBR4-2; | Isoform 2 c | Q9DBR4-2 | 0 | 2682.139 | 0 | 0.52 | 0.177016 | 37.52 | 82.9 | 117.1 | 21.03 | 12.78 | Shared | High | High | High | High | High | High | High | High | High | High | High | 0 | 6.34E-09 | 2.9 |
| FALSE | High | NASGVVNS | [K].NASGV | 1xPhospho | Q7TQE6 1x | 0.000101 | 0 | 1 | 3 | 4 | Q7TQE6 | Macoilin O | Q7TQE6 [3 | 0 | 1396.678 | 0 | 0.48 | 0.015691 | 10.32 | 84.7 | 115.3 | 7.92 | 9.4 | | High | High | High | High | High | High | High | High | High | High | High | 0 | 1.92E-05 | 1.66 |
| FALSE | High | NASILLEELI | [K].NASILLI | 1xPhospho | Q99NH0 1x | 6.3E-05 | 0 | 1 | 8 | 2 | Q99NH0 | Ankyrin rep | Q99NH0 [1 | 0 | 2025.092 | 0 | -0.22 | 0.263415 | 10.66 | 103 | 97 | 15.04 | 13.16 | | High | High | High | High | High | High | High | High | High | High | High | 0 | 1.07E-05 | 3.28 |
| FALSE | High | NASPQDIH | [R].NASPQ | 1xPhospho | Q8CBD1 1x | 0.001222 | 0 | 1 | 1 | 1 | Q8CBD1 | Nuclear rec | Q8CBD1 [5 | 0 | 1907.926 | 0 | 0.71 | 0.00194 | 2.14 | 76.2 | 123.8 | 7.75 | 4.68 | | High | High | High | High | High | High | High | High | High | High | High | 0 | 0.000299 | 2.6 |
| FALSE | High | NAGSSSDG | [R].NAGSS | 1xPhospho | Q5SVQ0 1x | 0.000211 | 0 | 1 | 5 | 1 | Q5SVQ0 | Histone ac | Q5SVQ0 [9 | 0 | 3502.372 | 0 | 0.74 | 0.000202 | 3.09 | 74.5 | 125.5 | 14.97 | 10.55 | | High | High | High | High | High | High | High | High | High | High | High | 0 | 4.14E-05 | 3.78 |
| FALSE | High | MVEPENA | [R].MVEPE | 1xPhospho | Q80U72-4 | 4.59E-05 | 0 | 2 | 6 | 2 | Q80U72-4; | Isoform 4 c | Q80U72-4 | 0 | 2838.358 | 0 | 1.36 | 0.002888 | 15.6 | 52.8 | 147.2 | 6.99 | 16.72 | Shared | High | High | High | High | High | High | High | High | High | High | High | 0 | 7.51E-06 | 1.9 |
| FALSE | High | MVEAPSPC | [R].MVEAP | 1xPhospho | Q6NVF4 1x | 0.00011 | 0 | 1 | 1 | 1 | Q6NVF4 | DNA helica | Q6NVF4 [1 | 0 | 2216.044 | 0 | -0.64 | 0.002399 | 5.85 | 126.1 | 73.9 | 8.17 | 11.76 | | High | High | High | High | High | High | High | High | High | High | High | 0 | 2E-05 | 2.61 |
| FALSE | High | MTSGVLSP | [K].MTSGV | 1xPhospho | Q9DCT6-3 | 0.075304 | 0.005727 | 2 | 10 | 2 | Q9DCT6-3; | Isoform 3 c | Q9DCT6-3 | 0 | 3238.683 | 0 | -1.64 | 0.000961 | 1.04 | 152.5 | 47.5 | 11.38 | 24.07 | Shared | High | High | High | High | High | High | High | High | High | High | High | 0.003782 | 0.04074 | 2.33 |
| FALSE | High | MPSPEVEE | [R].MPSPE | 1xCarbami | A0A0R4J07 | 0.009416 | 0.000736 | 1 | 5 | 1 | A0A0R4J07 | BRCA1-A c | A0A0R4J07 | 0 | 1787.754 | 0 | 0.44 | 0.014477 | 8.17 | 84.7 | 115.3 | 5.11 | 12.29 | | High | High | High | High | High | High | High | High | High | High | High | 0.000327 | 0.002946 | 1.88 |
| FALSE | High | MPSSEPELI | [K].MPSSEI | 1xPhospho | Q5U647 1x | 0.007209 | 0.00062 | 1 | 4 | 2 | Q5U647 | Amino acid | Q5U647 [5 | 0 | 1895.995 | 0 | 0.51 | 0.02848 | 17.98 | 86 | 114 | 4.32 | 14.48 | | High | High | High | High | High | High | High | High | High | High | High | 0.000275 | 0.002178 | 2.65 |
| FALSE | High | MPTQSDEE | [K].MPTQS | 2xPhospho | Q924K8 2x | 0.012508 | 0.000889 | 1 | 10 | 1 | Q924K8 | Metastasis | Q924K8 [4 | 1 | 2720.193 | 0 | -0.14 | 0.166782 | 12.42 | 107.7 | 92.3 | 8.72 | 9.84 | | High | High | High | High | High | High | High | High | High | High | High | 0.000449 | 0.004011 | 1.67 |
| FALSE | High | MQAESQSI | [R].MQAES | 1xPhospho | B9EKJ7 1xP | 7.91E-06 | 0 | 1 | 3 | 4 | B9EKJ7 | Tankyrase | B9EKJ7 [10 | 0 | 2330.099 | 0 | 0.02 | 0.754239 | 14.65 | 96.8 | 103.2 | 8.77 | 22.59 | | High | High | High | High | High | High | High | High | High | High | High | 0 | 1.04E-06 | 4.16 |
| FALSE | High | MQSPELPA | [R].MQSPE | 1xPhospho | Q80U72-4 | 0.005 | 0.000504 | 2 | 5 | 1 | Q80U72-4; | Isoform 4 c | Q80U72-4 | 0 | 1563.744 | 0 | 0.44 | 0.183589 | 14.84 | 86.5 | 113.5 | 6.61 | 20.91 | Shared | High | High | High | High | High | High | High | High | High | High | High | 0.000225 | 0.001446 | 2.97 |
| FALSE | High | MQVDQEE | [K].MQVD | 1xPhospho | Q571M2 1 | 1.07E-05 | 0 | 1 | 3 | 1 | Q571M2 | MKIAA402 | Q571M2 [6 | 0 | 3387.556 | 0 | 0.13 | 0.38387 | 20.21 | 92.9 | 107.1 | 7.52 | 16.9 | | High | High | High | High | High | High | High | High | High | High | High | 0 | 1.45E-06 | 4.75 |
| FALSE | High | MRSPPGEN | [-K].MRSPI | 1xPhospho | Q5SXI5 1xP | 3.77E-05 | 0 | 1 | 2 | 1 | Q5SXI5 | Zinc finger | Q5SXI5 [21 | 1 | 2629.192 | 0 | -0.27 | 0.042938 | 11.27 | 107.9 | 92.1 | 14.27 | 4.5 | | High | High | High | High | High | High | High | High | High | High | High | 0 | 5.98E-06 | 2.26 |
| FALSE | High | MSDSPTPC | [R].MSDSP | 1xCarbami | E9Q8T1 1x | 0.000334 | 0 | 2 | 14 | 1 | E9Q8T1; Q | Transformi | E9Q8T1 [2 | 0 | 3140.421 | 0 | 0.23 | 0.77127 | 8.69 | 94.6 | 105.4 | 11.08 | 16.59 | Shared | High | High | High | High | High | High | High | High | High | High | High | 0 | 6.96E-05 | 3.17 |
| FALSE | High | MSGFIYQG | [R].MSGFI | 1xPhospho | Q9ES28 1x | 0.000203 | 0 | 1 | 10 | 1 | Q9ES28 | Rho guanir | Q9ES28 [4 | 0 | 1568.795 | 0 | 1.02 | 0.011601 | 33.37 | 55.7 | 144.3 | 24.71 | 19 | | High | High | High | High | High | High | High | High | High | High | High | 0 | 3.97E-05 | 1.75 |
| FALSE | High | MSGGSLIP | [-R].MSGG | 1xPhospho | Q8VCF0 1x | 0.000924 | 0 | 1 | 2 | 1 | Q8VCF0 | Mitochond | Q8VCF0 [1 | 0 | 2461.211 | 0 | 1.97 | 0.002516 | 18.87 | 37.6 | 162.4 | 11.6 | 18.45 | | High | High | High | High | High | High | High | High | High | High | High | 0 | 0.000219 | 2.96 |
| FALSE | High | MSGGSTM | [K].MSGGS | 1xPhospho | P70662 1xP | 1.93E-08 | 0 | 2 | 4 | 29 | P70662; D3 | LIM domai | P70662 [30 | 0 | 2429.047 | 0 | 0.14 | 0.039469 | 8.54 | 93.7 | 106.3 | 7.14 | 2.58 | Shared | High | High | High | High | High | High | High | High | High | High | High | 0 | 1.45E-09 | 5.01 |
| FALSE | High | MSGSPGVS | [R].MSGSP | 1xPhospho | Q3USH5 1x | 0.059159 | 0.004294 | 1 | 2 | 1 | Q3USH5 | Splicing fac | Q3USH5 [8 | 0 | 1186.549 | 0 | -0.25 | 0.076264 | 18.91 | 108.1 | 91.9 | 5.64 | 13.4 | | High | High | High | High | High | High | High | High | High | High | High | 0.002217 | 0.0235 | 2.29 |
| FALSE | High | MSPHLLPE | [K].MSPHL | 1xPhospho | Q8K205 1x | 3.11E-08 | 0 | 1 | 2 | 1 | Q8K205 | Processing | Q8K205 [6 | 0 | 2523.22 | 0 | -0.27 | 0.023159 | 9.37 | 110 | 90 | 6.45 | 5.41 | | High | High | High | High | High | High | High | High | High | High | High | 0 | 2.06E-09 | 5.33 |
| FALSE | High | MSPMGTA | [R].MSPM | 1xPhospho | O88878 1x | 2.68E-06 | 0 | 1 | 3 | 1 | O88878 | AN1-type z | O88878 [4 | 0 | 2464.068 | 0 | 0.03 | 0.270069 | 7.31 | 96 | 104 | 6.14 | 6.71 | | High | High | High | High | High | High | High | High | High | High | High | 0 | 3.07E-07 | 3.57 |
| FALSE | High | MSPMGTA | [R].MSPM | 1xPhospho | O88878 1x | 3.07E-09 | 0 | 1 | 3 | 6 | O88878 | AN1-type z | O88878 [4 | 0 | 2480.063 | 0 | 0.12 | 0.087247 | 7.87 | 89.4 | 110.6 | 8.13 | 15.22 | | High | High | High | High | High | High | High | High | High | High | High | 0 | 9.89E-10 | 3.9 |
| FALSE | High | MSPPPSSFI | [K].MSPPP | 1xPhospho | Q9CW46 1 | 0.005275 | 0.000599 | 1 | 2 | 1 | Q9CW46 | Ribonuclec | Q9CW46 [ | 0 | 1654.75 | 0 | -0.07 | 0.263443 | 8.46 | 102.6 | 97.4 | 5.92 | 14.27 | | High | High | High | High | High | High | High | High | High | High | High | 0.000267 | 0.00153 | 1.79 |
| FALSE | High | MSPTDPLD | [-].MSPTDI | 1xPhospho | Q3URU4 1 | 0.007811 | 0.000664 | 1 | 6 | 3 | Q3URU4 | Meiotic rec | Q3URU4 [1 | 0 | 2148.981 | 0 | 0.21 | 0.230232 | 2.15 | 89.3 | 110.7 | 7.33 | 19.17 | | High | High | High | High | High | High | High | High | High | High | High | 0.000689 | 0.005915 | 2.77 |
| FALSE | High | MSQPGSPS | [K].MSQPG | 1xPhospho | Q6PDQ2 1 | 0.017966 | 0.001282 | 1 | 6 | 2 | Q6PDQ2 | Chromodo | Q6PDQ2 [1 | 0 | 1340.612 | 0 | -0.59 | 0.924574 | 28.56 | 114.3 | 85.7 | 49.67 | 113.05 | | High | High | High | High | High | High | High | High | High | High | High | 0.000689 | 0.006017 | 1.98 |
| FALSE | High | MSQPGSPS | [K].MSQPG | 1xPhospho | Q6PDQ2 1 | 4.35E-05 | 0 | 1 | 6 | 2 | Q6PDQ2 | Chromodo | Q6PDQ2 [1 | 0 | 1553.78 | 0 | -0.45 | 0.02287 | 3.28 | 115.9 | 84.1 | 6.7 | 16.4 | | High | High | High | High | High | High | High | High | High | High | High | 0 | 7.05E-06 | 2.17 |
| FALSE | High | MSSHTETS | [R].MSSHT | 1xPhospho | A2A9M4 1 | 0.00172 | 5.66E-05 | 1 | 7 | 2 | A2A9M4 | Dedicator | A2A9M4 [ | 0 | 2192.026 | 0 | 0.47 | 0.06747 | 26.44 | 76.7 | 123.3 | 11.43 | 25.64 | | High | High | High | High | High | High | High | High | High | High | High | 4.94E-05 | 0.000636 | 2.84 |
| FALSE | High | MSSPLADA | [K].MSSPL | 1xPhospho | E9Q3G8 1x | 0.005182 | 0.000551 | 1 | 2 | 2 | E9Q3G8 | Nucleopor | E9Q3G8 [3 | 0 | 1457.748 | 0 | -0.15 | 0.860382 | 6.96 | 101 | 99 | 6.94 | 14.49 | | High | High | High | High | High | High | High | High | High | High | High | 0.000984 | 0.01019 | 2.01 |
| FALSE | High | MSSPPPSP | [K].MSSPP | 2xPhospho | Q8BX90 2x | 0.006773 | 0.00062 | 1 | 3 | 1 | Q8BX90 | Fibronectir | Q8BX90 [2 | 0 | 1673.778 | 0 | 0.48 | 0.03861 | 18.98 | 81.5 | 118.5 | 4.73 | 15.13 | | High | High | High | High | High | High | High | High | High | High | High | 0.000275 | 0.002029 | 0.95 |
| FALSE | High | MSTSGTAN | [K].MSTSG | 1xPhospho | Q07231 1x | 0.107603 | 0.008634 | 1 | 2 | 1 | Q07231 | Zinc finger | Q07231 [2 | 0 | 3036.394 | 0 | -0.06 | 0.472929 | 15.97 | 107.6 | 92.4 | 7.05 | 36.77 | | High | High | High | High | High | High | High | High | High | High | High | 0.004341 | 0.04691 | 2.03 |
| FALSE | High | MSVDAVEI | [R].MSVDA | 1xPhospho | Q9ESZ8-5 | 3.54E-05 | 0 | 2 | 12 | 1 | Q9ESZ8-5; | Isoform 5 c | Q9ESZ8-5 | 0 | 1671.823 | 0 | 0.66 | 0.00755 | 9.23 | 74.8 | 125.2 | 13.86 | 7.85 | Shared | High | High | High | High | High | High | High | High | High | High | High | 0 | 5.59E-06 | 2.94 |
| FALSE | High | MSVQPTVS | [K].MSVQF | 1xPhospho | Q5SQB7 1x | 0.00172 | 5.66E-05 | 2 | 9 | 1 | Q5SQB7; C | MCG68069 | Q5SQB7 [8 | 0 | 2536.345 | 0 | -0.8 | 0.001548 | 2.06 | 135.7 | 64.3 | 9.64 | 17.04 | Shared | High | High | High | High | High | High | High | High | High | High | High | 2.58E-05 | 0.000439 | 3.32 |
| FALSE | High | MSYLTAM | [R].MSYLT | 1xCarbami | Q9Z0U1 1x | 5.96E-08 | 0 | 1 | 1 | 1 | Q9Z0U1 | Tight junct | Q9Z0U1 [8 | 0 | 2249.948 | 0 | -0.9 | 0.001517 | 1.2 | 128.7 | 71.3 | 10.73 | 11.36 | | High | High | High | High | High | High | High | High | High | High | High | 0 | 4.29E-09 | 4.65 |
| FALSE | High | MSYLTAM | [R].MSYLT | 1xCarbami | Q9Z0U1 1x | 7.81E-08 | 0 | 1 | 1 | 2 | Q9Z0U1 | Tight junct | Q9Z0U1 [8 | 0 | 2265.943 | 0 | 0 | 0.904565 | 11.08 | 101.1 | 98.9 | 2.62 | 9.25 | | High | High | High | High | High | High | High | High | High | High | High | 0 | 5.77E-09 | 4.01 |
| FALSE | High | MTAANVA | [R].MTAAN | 1xPhospho | A2A863-3 | 1.09E-07 | 0 | 1 | 6 | 1 | A2A863-3 | Isoform 3 c | A2A863-3 | 0 | 2372.153 | 0 | 2.63 | 0.000819 | 1.62 | 25.9 | 174.1 | 17.52 | 12.02 | | High | High | High | High | High | High | High | High | High | High | High | 0 | 8.43E-09 | 3.14 |
| FALSE | High | MTGSAPPF | [K].MTGSA | 1xPhospho | Q9CU65 1x | 0.034579 | 0.002233 | 1 | 1 | 2 | Q9CU65 | Zinc finger | Q9CU65 [8 | 0 | 1919.97 | 0 | -0.95 | 0.000301 | 0.82 | 134.3 | 65.7 | 3.28 | 7.27 | | High | High | High | High | High | High | High | High | High | High | High | 0.002455 | 0.02698 | 1.78 |
| FALSE | High | NASSNSSPI | [R].NASSN | 1xPhospho | Q6VGS5-2 | 0.005615 | 0.00062 | 1 | 3 | 2 | Q6VGS5-2 | Isoform 2 c | Q6VGS5-2 | 0 | 1742.909 | 0 | -0.28 | 0.053572 | 7.09 | 109.8 | 90.2 | 5.68 | 9.95 | | High | High | High | High | High | High | High | High | High | High | High | 0.000275 | 0.001651 | 2.43 |
| FALSE | High | LTPVSLSNS | [K].LTPVSL | 1xPhospho | Q61687 1x | 4.05E-05 | 0 | 1 | 4 | 2 | Q61687 | Transcripti | Q61687 [5 | 0 | 1794.018 | 0 | -0.48 | 0.006813 | 7.07 | 119.4 | 80.6 | 9.06 | 3.55 | | High | High | High | High | High | High | High | High | High | High | High | 0 | 1.27E-05 | 3.65 |
| FALSE | High | NASTSFQE | [R].NASTSF | 1xPhospho | P54103 1xP | 6.83E-05 | 0 | 1 | 6 | 4 | P54103 | DnaJ homo | P54103 [45 | 0 | 1677.757 | 0 | -0.23 | 0.881854 | 43.34 | 108 | 92 | 58.76 | 60.47 | | High | High | High | High | High | High | High | High | High | High | High | 0 | 0.000226 | 3.85 |
| FALSE | High | NASTSFQE | [R].NASTSF | 1xPhospho | P54103 1xP | 9.65E-06 | 0 | 1 | 6 | 9 | P54103 | DnaJ homo | P54103 [45 | 0 | 1906.92 | 0 | -0.38 | 0.059306 | 9.52 | 113.2 | 86.8 | 8.73 | 8.19 | | High | High | High | High | High | High | High | High | High | High | High | 0 | 1.3E-06 | 3.45 |
| FALSE | High | NDESSEQE | [R].NDESSE | 1xPhospho | Q69ZS0 1xI | 3.04E-06 | 0 | 1 | 4 | 3 | Q69ZS0 | E3 ubiquiti | Q69ZS0 [5 | 0 | 2913.233 | 0 | 0.76 | 0.013567 | 38.1 | 79.2 | 120.8 | 43.95 | 11.48 | | High | High | High | High | High | High | High | High | High | High | High | 0 | 3.53E-07 | 5.72 |
| FALSE | High | NDMAVPTI | [K].NDMA | 1xPhospho | Q924H2 1x | 0.003195 | 0.00016 | 1 | 9 | 1 | Q924H2 | Mediator c | Q924H2 [5 | 0 | 2096.07 | 0 | 0.25 | 0.360338 | 21.37 | 90.7 | 109.3 | 18.39 | 13 | | High | High | High | High | High | High | High | High | High | High | High | 7.19E-05 | 0.000873 | 1.97 |
| FALSE | High | NDMAVPTI | [K].NDMA | 1xPhospho | Q924H2 1x | 1.25E-05 | 0 | 1 | 9 | 1 | Q924H2 | Mediator c | Q924H2 [5 | 0 | 2309.238 | 0 | 0.6 | 0.011106 | 5.2 | 79.4 | 120.6 | 5.3 | 14.71 | | High | High | High | High | High | High | High | High | High | High | High | 0 | 1.73E-06 | 2.85 |
| FALSE | High | NDMAVPTI | [K].NDMA | 1xPhospho | Q924H2 1x | 2.82E-05 | 0 | 1 | 9 | 3 | Q924H2 | Mediator c | Q924H2 [5 | 0 | 2325.233 | 0 | 0.62 | 0.000255 | 5.18 | 81.3 | 118.7 | 6.29 | 4.8 | | High | High | High | High | High | High | High | High | High | High | High | 0.00014 | 0.001125 | 2.49 |
| FALSE | High | NDMDEPPI | [K].NDMDI | 1xPhospho | B1AR17 1x | 2E-05 | 0 | 1 | 4 | 1 | B1AR17 | Chromodo | B1AR17 [6 | 0 | 2605.105 | 0 | 1.57 | 0.00034 | 3.52 | 51.3 | 148.7 | 5.95 | 7.49 | | High | High | High | High | High | High | High | High | High | High | High | 0 | 2.95E-06 | 4.11 |
| FALSE | High | NDNTLVVS | [K].NDNTL | 1xPhospho | E9Q555 1x | 6.45E-07 | 0 | 1 | 2 | 3 | E9Q555 | E3 ubiquiti | E9Q555 [3 | 0 | 1999.015 | 0 | 2.44 | 0.000824 | 34.46 | 32.4 | 167.6 | 17.47 | 10.55 | | High | High | High | High | High | High | High | High | High | High | High | 0 | 1.1E-06 | 3.73 |
| FALSE | High | NDQDTWD | [K].NDQDT | 1xPhospho | G5E924 1x | 1.1E-07 | 0 | 1 | 4 | 3 | G5E924 | Heterogen | G5E924 [2 | 0 | 3272.453 | 0 | -0.38 | 0.003644 | 1.29 | 112.2 | 87.8 | 6.45 | 3.93 | | High | High | High | High | High | High | High | High | High | High | High | 0 | 1.48E-05 | 4.45 |
| FALSE | High | NDQDTWD | [K].NDQDT | 1xDeamida | G5E924 1x | 0.008313 | 0.000736 | 1 | 4 | 3 | G5E924 | Heterogen | G5E924 [2 | 0 | 3273.437 | 0 | -0.64 | 0.024307 | 8.56 | 120.5 | 79.5 | 6.36 | 20.49 | | High | High | High | High | High | High | High | High | High | High | High | 0.000327 | 0.002546 | 3.7 |
| FALSE | High | NDQEPPPE | [K].NDQEP | 1xPhospho | E9QJT2 1xI | 2.93E-09 | 0 | 1 | 2 | 6 | E9QJT2 | H/ACA ribo | E9QJT2 [4 | 0 | 2282.99 | 0 | -0.29 | 0.608294 | 17.51 | 109.9 | 90.1 | 44.96 | 44.72 | | High | High | High | High | High | High | High | High | High | High | High | 0 | 1.45E-10 | 4.45 |
| FALSE | High | NDQEPPPE | [K].NDQEP | 1xPhospho | E9QJT2 1xI | 4.92E-07 | 0 | 1 | 2 | 6 | E9QJT2 | H/ACA ribo | E9QJT2 [4 | 0 | 2512.153 | 0 | -0.46 | 0.026579 | 7.36 | 112.9 | 87.1 | 7.5 | 8.62 | | High | High | High | High | High | High | High | High | High | High | High | 0 | 4.56E-08 | 4.09 |
| FALSE | High | NDQEPPPE | [K].NDQEP | 1xDeamida | E9QJT2 1xI | 2.89E-10 | 0 | 1 | 2 | 4 | E9QJT2 | H/ACA ribo | E9QJT2 [4 | 0 | 2513.137 | 0 | -0.32 | 0.028045 | 7.22 | 111.8 | 88.2 | 8.75 | 4.61 | | High | High | High | High | High | High | High | High | High | High | High | 0 | 1.07E-11 | 4.89 |
| FALSE | High | NDSGDHSD | [R].NDSGD | 1xPhospho | S4R1W5 1x | 0.009585 | 0.000736 | 1 | 7 | 2 | S4R1W5 | RNA-bindir | S4R1W5 [7 | 0 | 2445.033 | 0 | -0.92 | 0.004678 | 8.19 | 131.6 | 68.4 | 7.34 | 15.54 | | High | High | High | High | High | High | High | High | High | High | High | 0.000872 | 0.00877 | 1.99 |
| FALSE | High | NDSGEENV | [R].NDSGE | 1xPhospho | Q6QI06 1xI | 9.65E-06 | 0 | 1 | 5 | 3 | Q6QI06 | Rapamycin | Q6QI06 [1 | 0 | 1867.864 | 0 | 0.78 | 0.000925 | 3.4 | 73.6 | 126.4 | 4.53 | 5.3 | | High | High | High | High | High | High | High | High | High | High | High | 0 | 1.29E-06 | 3.75 |
| FALSE | High | NDESQLSP | [R].NDESQ | 1xPhospho | Q9QXS1-3 | 0.009332 | 0.000736 | 3 | 31 | 2 | Q9QXS1-3; | Isoform PL | Q9QXS1-3 | 0 | 1526.705 | 0 | 1.68 | 0.001106 | 19 | 50.5 | 149.5 | 7.91 | 11.36 | Shared | High | High | High | High | High | High | High | High | High | High | High | 0.001999 | 0.02078 | 2.15 |
| FALSE | High | NDSIPVLR | [R].NDSIPV | 1xPhospho | Q6PCP5-2 | 0.001986 | 5.66E-05 | 1 | 2 | 2 | Q6PCP5-2 | Isoform 2 c | Q6PCP5-2 | 0 | 1222.639 | 0 | 1.15 | 0.00116 | 11.97 | 60.4 | 139.6 | 8.24 | 8.79 | | High | High | High | High | High | High | High | High | High | High | High | 2.58E-05 | 0.000514 | 1.92 |
| FALSE | High | NDSIVTPSF | [R].NDSIVT | 1xPhospho | Q6PCP5 1x | 0.040721 | 0.003058 | 1 | 3 | 1 | Q6PCP5 | Mitochond | Q6PCP5 [1 | 0 | 1690.836 | 0 | 0.94 | 0.000714 | 7.74 | 67.2 | 132.8 | 3.99 | 7.26 | | High | High | High | High | High | High | High | High | High | High | High | 0.001597 | 0.01531 | 1.71 |
| FALSE | High | NDSPQAK | [R].NDSPQ | 1xPhospho | Q8R1J3 1xI | 0.00407 | 0.00036 | 1 | 3 | 1 | Q8R1J3 | Zinc finger | Q8R1J3 [5 | 0 | 1297.655 | 0 | -0.84 | 0.000225 | 0.73 | 128.4 | 71.6 | 2.55 | 3.29 | | High | High | High | High | High | High | High | High | High | High | High | 0.000161 | 0.00115 | 2.33 |
| FALSE | High | NDSSELEDI | [R].NDSSEI | 1xPhospho | Q04692 1x | 4.56E-06 | 0 | 1 | 1 | 9 | Q04692 | SWI/SNF-r | Q04692 [1 | 0 | 2374.131 | 0 | -0.8 | 0.002675 | 3.24 | 124.2 | 75.8 | 9.37 | 6.03 | | High | High | High | High | High | High | High | High | High | High | High | 0.006181 | 0.06687 | 3.71 |
| FALSE | High | NDSVIVAD | [R].NDSVI | 1xPhospho | Q640L6 1xI | 1.43E-05 | 0 | 1 | 10 | 2 | Q640L6 | Atf2 protei | Q640L6 [4 | 0 | 1922.942 | 0 | 0.06 | 0.989115 | 7.3 | 100.1 | 99.9 | 7.15 | 5.35 | | High | High | High | High | High | High | High | High | High | High | High | 0 | 0.000153 | 3.36 |
| FALSE | High | NDSVIVAD | [R].NDSVI | 2xPhospho | Q640L6 2xI | 6.79E-06 | 0 | 1 | 10 | 3 | Q640L6 | Atf2 protei | Q640L6 [4 | 0 | 2002.909 | 0 | -0.03 | 0.478174 | 10.03 | 101.4 | 98.6 | 9.9 | 5.39 | | High | High | High | High | High | High | High | High | High | High | High | 0 | 3.75E-06 | 3.22 |
| FALSE | High | NDSWGSFI | [R].NDSW | 1xPhospho | Q5F2E7-2 | 0.000112 | 0 | 1 | 2 | 5 | Q5F2E7-2 | Isoform 2 c | Q5F2E7-2 | 0 | 1505.662 | 0 | 0.21 | 0.076811 | 12.57 | 92.6 | 107.4 | 5.64 | 7.67 | | High | High | High | High | High | High | High | High | High | High | High | 0 | 2.03E-05 | 3.01 |
| FALSE | High | NDSYTEEEI | [R].NDSYT | 1xPhospho | Q9EST3 1x | 1.61E-07 | 0 | 1 | 3 | 5 | Q9EST3 | Eukaryotic | Q9EST3 [2 | 0 | 4047.795 | 0 | 0.54 | 0.007654 | 14.4 | 85.1 | 114.9 | 9.5 | 14.98 | | High | High | High | High | High | High | High | High | High | High | High | 0 | 7.33E-07 | 4.6 |
| FALSE | High | NDSYTEEEI | [R].NDSYT | 1xDeamida | Q9EST3 1x | 2.59E-07 | 0 | 1 | 3 | 2 | Q9EST3 | Eukaryotic | Q9EST3 [2 | 0 | 4048.779 | 0 | 0.4 | 0.035886 | 32.69 | 86.6 | 113.4 | 23.61 | 9.22 | | High | High | High | High | High | High | High | High | High | High | High | 0 | 2.24E-08 | 5.45 |
| FALSE | High | NDYEEDDL | [R].NDYEEI | 1xCarbami | Q923W1 1 | 1.59E-06 | 0 | 1 | 1 | 2 | Q923W1 | Trimethylg | Q923W1 [ | 0 | 3146.273 | 0 | -0.87 | 0.00341 | 1.68 | 126 | 74 | 6.5 | 9.45 | | High | High | High | High | High | High | High | High | High | High | High | 0 | 1.72E-07 | 3.62 |
| FALSE | High | NDYEEDDL | [R].NDYEEI | 1xCarbami | Q923W1 1 | 1.75E-05 | 0 | 1 | 1 | 4 | Q923W1 | Trimethylg | Q923W1 [ | 0 | 3147.257 | 0 | -0.78 | 0.005316 | 1.69 | 126.2 | 73.8 | 8.34 | 8.22 | | High | High | High | High | High | High | High | High | High | High | High | 0 | 2.53E-06 | 3.75 |
| FALSE | High | NEAFVCKP | [R].NEAFV | 1xCarbami | Q80U72-4 | 3.98E-05 | 0 | 2 | 5 | 1 | Q80U72-4; | Isoform 4 c | Q80U72-4 | 0 | 2782.305 | 0 | 0.01 | 0.821195 | 3.79 | 99.5 | 100.5 | 7.24 | 10.9 | Shared | High | High | High | High | High | High | High | High | High | High | High | 0 | 6.39E-06 | 4.12 |
| FALSE | High | NEAFVCKP | [R].NEAFV | 1xCarbami | Q80U72-4 | 0.003312 | 0.00016 | 2 | 5 | 1 | Q80U72-4; | Isoform 4 c | Q80U72-4 | 0 | 3011.468 | 0 | -0.01 | 0.785814 | 20.83 | 100.4 | 99.6 | 13.9 | 9.32 | Shared | High | High | High | High | High | High | High | High | High | High | High | 7.19E-05 | 0.000914 | 2.99 |
| FALSE | High | NEAGGLSL | [R].NEAGG | 2xPhospho | P59222 2xI | 0.085859 | 0.006406 | 1 | 1 | 1 | P59222 | Scavenger | P59222 [6 | 0 | 1802.792 | 0 | -0.26 | 0.082811 | 12.29 | 107.1 | 92.9 | 8.28 | 16.94 | | High | High | High | High | High | High | High | High | High | High | High | 0.003303 | 0.03623 | 2.84 |
| FALSE | High | NDSLSSDQ | [R].NDSLS | 1xPhospho | D9HP81 1x | 0.001301 | 0 | 1 | 5 | 1 | D9HP81 | Regulating | D9HP81 [4 | 0 | 2868.441 | 0 | 0.73 | 0.000656 | 7.36 | 74.4 | 125.6 | 7.83 | 4.05 | | High | High | High | High | High | High | High | High | High | High | High | 0 | 0.000319 | 1.77 |
| FALSE | High | NDENGNCS | [K].NDENG | 1xCarbami | E9PV45 1xI | 3.94E-05 | 0 | 1 | 3 | 1 | E9PV45 | Ubiquitin c | E9PV45 [1 | 0 | 3154.351 | 0 | -0.58 | 0.028395 | 6.84 | 119.6 | 80.4 | 20.97 | 9.6 | | High | High | High | High | High | High | High | High | High | High | High | 0 | 6.3E-06 | 3.72 |
| FALSE | High | NDDTPEDF | [K].NDDTP | 1xPhospho | P09926 1xI | 8.13E-06 | 0 | 1 | 2 | 1 | P09926 | Surfeit locu | P09926 [1 | 0 | 3513.615 | 0 | 0.21 | 0.058666 | 11.51 | 92.6 | 107.4 | 9.21 | 6.57 | | High | High | High | High | High | High | High | High | High | High | High | 0 | 1.07E-06 | 5.42 |
| FALSE | High | NDAPTPGT | [R].NDAPT | 1xPhospho | Q80TC1 1x | 0.001349 | 0 | 1 | 13 | 6 | Q80TC1 | MKIAA154 | Q80TC1 [4 | 0 | 1793.863 | 0 | -1.05 | 0.002451 | 3.41 | 135.1 | 64.9 | 9.35 | 10.39 | | High | High | High | High | High | High | High | High | High | High | High | 0 | 0.000335 | 2.94 |
| FALSE | High | NASTSFQE | [R].NASTSF | 2xPhospho | P54103 2xI | 0.000168 | 0 | 1 | 6 | 9 | P54103 | DnaJ homo | P54103 [45 | 0 | 1986.886 | 1 | -1.09 | 0.009489 | 2.89 | 127.4 | 72.6 | 15.36 | 15.17 | | High | High | High | High | High | High | High | High | High | High | High | 0 | 3.22E-05 | 4.32 |
| FALSE | High | NASTSFQE | [R].NASTSF | 1xPhospho | P54103 1xI | 0.121159 | 0.009984 | 1 | 6 | 1 | P54103 | DnaJ homo | P54103 [45 | 1 | 2264.178 | 0 | -0.45 | 0.169096 | 23.85 | 115.6 | 84.4 | 26.22 | 15.41 | | High | High | High | High | High | High | High | High | High | High | High | 0.004974 | 0.05434 | 1.42 |
| FALSE | High | NASTVSSN | [R].NASTV | 1xPhospho | D3Z5I1 1xP | 0.003433 | 0.00021 | 1 | 2 | 3 | D3Z5I1 | Zinc finger | D3Z5I1 [80 | 0 | 3332.525 | 0 | 0.11 | 0.960786 | 42.47 | 101.7 | 98.3 | 11.97 | 27.46 | | High | High | High | High | High | High | High | High | High | High | High | 9.42E-05 | 0.000948 | 3.65 |
| FALSE | High | NASVDGEII | [R].NASVD | 1xPhospho | Q80Y83 1x | 7.04E-06 | 0 | 1 | 8 | 2 | Q80Y83 | Dixin OS=M | Q80Y83 [2 | 0 | 2144.985 | 0 | 0.74 | 0.004656 | 8.82 | 71.4 | 128.6 | 13.57 | 7.79 | | High | High | High | High | High | High | High | High | High | High | High | 0 | 9.06E-07 | 4.04 |
| FALSE | High | NASVEEVV | [R].NASVEI | 1xPhospho | A0A087WF | 0.008691 | 0.000736 | 1 | 2 | 1 | A0A087WF | Proline-ricl | A0A087WF | 0 | 1398.683 | 0 | 2 | 0.001147 | 22.94 | 40.1 | 159.9 | 10.56 | 11.59 | | High | High | High | High | High | High | High | High | High | High | High | 0.000327 | 0.00269 | 2.44 |
| FALSE | High | NASVPNLR | [R].NASVPI | 1xPhospho | Q9CWE0 1 | 0.01441 | 0.001011 | 1 | 4 | 2 | Q9CWE0 | Mitochond | Q9CWE0 [ | 0 | 1179.608 | 0 | 0.1 | 0.165643 | 2.16 | 97.1 | 102.9 | 4.99 | 4.82 | | High | High | High | High | High | High | High | High | High | High | High | 0.000538 | 0.004723 | 2.54 |
| FALSE | High | NATLAWDS | [K].NATLA | 1xPhospho | Q9R1X5 1x | 0.000434 | 0 | 1 | 2 | 1 | Q9R1X5 | Multidrug | Q9R1X5 [4 | 0 | 2252.055 | 0 | 0.9 | 0.095714 | 56.58 | 73.3 | 126.7 | 20.04 | 30.07 | | High | High | High | High | High | High | High | High | High | High | High | 0 | 9.33E-05 | 3.22 |
| FALSE | High | NATLAWDS | [K].NATLA | 2xPhospho | Q9R1X5 2x | 0.028082 | 0.001791 | 1 | 2 | 1 | Q9R1X5 | Multidrug | Q9R1X5 [4 | 0 | 2561.184 | 0 | 1.56 | 0.002921 | 17.62 | 50.7 | 149.3 | 8.49 | 19.1 | | High | High | High | High | High | High | High | High | High | High | High | 0.000968 | 0.01003 | 2.26 |
| FALSE | High | NATSPQVC | [R].NATSP | 1xPhospho | Q9JI10 1xP | 0.001233 | 0 | 1 | 3 | 1 | Q9JI10 | Serine/thr | Q9JI10 [38 | 0 | 2669.284 | 0 | 1.45 | 0.001606 | 2.03 | 54 | 146 | 4 | 11.61 | | High | High | High | High | High | High | High | High | High | High | High | 0 | 0.000302 | 1.99 |
| FALSE | High | NATSPQVC | [R].NATSP | 1xPhospho | Q9JI10 1xP | 0.009843 | 0.000736 | 1 | 3 | 1 | Q9JI10 | Serine/thr | Q9JI10 [38 | 0 | 2685.278 | 0 | 1.43 | 0.001926 | 19.81 | 57.9 | 142.1 | 9.62 | 12.72 | | High | High | High | High | High | High | High | High | High | High | High | 0.000344 | 0.003094 | 1.86 |
| FALSE | High | NATTPLK | [K].NATTPI | 1xPhospho | P97801 1xI | 0.067619 | 0.004864 | 1 | 4 | 1 | P97801 | Survival m | P97801 [81 | 0 | 1282.717 | 0 | -0.91 | 0.001029 | 3.22 | 129.6 | 70.4 | 6.44 | 6.14 | | High | High | High | High | High | High | High | High | High | High | High | 0.002485 | 0.02719 | 2.2 |
| FALSE | High | NAVASVQN | [R].NAVAS | 1xPhospho | B1AZI6 1xP | 2.12E-09 | 0 | 1 | 1 | 3 | B1AZI6 | THO comp | B1AZI6 [11 | 1 | 4062.934 | 0 | 0.33 | 0.194612 | 26.83 | 87.9 | 112.1 | 16.52 | 8.04 | | High | High | High | High | High | High | High | High | High | High | High | 0 | 1.02E-09 | 6.47 |
| FALSE | High | NAWNPSG | [R].NAWNI | 1xPhospho | Q8BLH7 1x | 2.66E-07 | 0 | 1 | 1 | 7 | Q8BLH7 | HIRA-inter | Q8BLH7 [5 | 0 | 2131.928 | 0 | -1.28 | 0.000986 | 2.99 | 142.3 | 57.7 | 4.19 | 16.39 | | High | High | High | High | High | High | High | High | High | High | High | 0 | 1.41E-05 | 2.62 |
| FALSE | High | NAWNPSG | [R].NAWNI | 1xPhospho | Q8BLH7 1x | 0.049971 | 0.003641 | 1 | 1 | 1 | Q8BLH7 | HIRA-inter | Q8BLH7 [5 | 1 | 2288.029 | 0 | -1.23 | 0.001414 | 0.69 | 141.2 | 58.8 | 9.97 | 5.35 | | High | High | High | High | High | High | High | High | High | High | High | 0.001902 | 0.0193 | 1.25 |
| FALSE | High | NCEAPYIPE | [R].NCEAP | 2xCarbami | A0A1D5RL | 5.65E-05 | 0 | 1 | 7 | 4 | A0A1D5RL | Serine/thr | A0A1D5RL | 0 | 3626.596 | 0 | 1.54 | 0.008488 | 28.49 | 52.5 | 147.5 | 21.93 | 19.7 | | High | High | High | High | High | High | High | High | High | High | High | 0 | 5.09E-05 | 4.48 |
| FALSE | High | NCEDPASP | [R].NCEDP | 1xCarbami | Q9JHI2 1xP | 0.001689 | 5.66E-05 | 1 | 4 | 1 | Q9JHI2 | tRNA-speci | Q9JHI2 [18 | 0 | 1496.665 | 0 | -0.92 | 0.003869 | 2.3 | 123.8 | 76.2 | 9.18 | 22.27 | | High | High | High | High | High | High | High | High | High | High | High | 2.58E-05 | 0.000431 | 3.08 |
| FALSE | High | NCEDPASP | [R].NCEDP | 1xCarbami | Q9JHI2 1xP | 0.000375 | 0 | 1 | 4 | 2 | Q9JHI2 | tRNA-speci | Q9JHI2 [18 | 0 | 1725.828 | 0 | -1.12 | 0.000829 | 2.95 | 138.3 | 61.7 | 5.55 | 7.65 | | High | High | High | High | High | High | High | High | High | High | High | 2.58E-05 | 0.000346 | 3.05 |
| FALSE | High | NCEEVGLF | [K].NCEEV | 1xCarbami | Q640L6 1xI | 3.88E-09 | 0 | 1 | 14 | 3 | Q640L6 | Atf2 protei | Q640L6 [6 | 0 | 2911.363 | 0 | -0.7 | 0.00267 | 4.59 | 123.9 | 76.1 | 11.15 | 5.65 | | High | High | High | High | High | High | High | High | High | High | High | 0 | 1.99E-10 | 5.2 |
| FALSE | High | NCPAVTLT | [R].NCPAV | 1xCarbami | Q9EPQ8 1x | 0.000117 | 0 | 1 | 4 | 1 | Q9EPQ8 | Transcripti | Q9EPQ8 [1 | 0 | 1567.775 | 0 | -0.21 | 0.501989 | 2.79 | 107.4 | 92.6 | 24.65 | 20.23 | | High | High | High | High | High | High | High | High | High | High | High | 0 | 2.14E-05 | 2.71 |
| FALSE | High | NCPAVTLT | [R].NCPAV | 1xCarbami | Q9EPQ8 1x | 3.27E-06 | 0 | 1 | 4 | 3 | Q9EPQ8 | Transcripti | Q9EPQ8 [1 | 0 | 1796.938 | 0 | -0.45 | 0.031121 | 8.18 | 114.9 | 85.1 | 8.99 | 8.91 | | High | High | High | High | High | High | High | High | High | High | High | 0 | 5.31E-07 | 3.7 |
| FALSE | High | NCPHIVVG | [K].NCPHI | 1xCarbami | A0A1S6GW | 0.000262 | 0 | 1 | 4 | 2 | A0A1S6GW | Uncharacte | A0A1S6GW | 0 | 1615.798 | 0 | -1.21 | 0.000275 | 0.4 | 141.1 | 58.9 | 4.31 | 4.46 | | High | High | High | High | High | High | High | High | High | High | High | 0.005414 | 0.05948 | 2.96 |
| FALSE | High | NCPHVVVG | [K].NCPHV | 1xCarbami | Q3T9L0 1xI | 1.36E-05 | 0 | 1 | 7 | 1 | Q3T9L0 | Putative ur | Q3T9L0 [1 | 0 | 1601.782 | 0 | -0.59 | 0.000862 | 2.69 | 121.8 | 78.2 | 6.45 | 4.51 | | High | High | High | High | High | High | High | High | High | High | High | 0 | 1.91E-06 | 3.18 |
| FALSE | High | NCPSPMQT | [R].NCPSPI | 1xCarbami | Q80XH9 1x | 1.02E-08 | 0 | 1 | 16 | 1 | Q80XH9 | ELAV-like p | Q80XH9 [3 | 0 | 2233.987 | 0 | 0.25 | 0.03475 | 2.84 | 91.5 | 108.5 | 10.86 | 15.03 | | High | High | High | High | High | High | High | High | High | High | High | 0 | 5.86E-10 | 4.32 |
| FALSE | High | NCPSPVLID | [R].NCPSP | 3xCarbami | Q8BZ47 1x | 1.11E-07 | 0 | 1 | 2 | 2 | Q8BZ47 | Zinc finger | Q8BZ47 [4 | 0 | 2560.191 | 0 | 0.02 | 0.813642 | 25.13 | 96.4 | 103.6 | 10.7 | 10.03 | | High | High | High | High | High | High | High | High | High | High | High | 0 | 8.63E-09 | 4.3 |

| | | | | | | | | | | | | | | | | | | | | | | | | | | | | | | | | | | | | | | |
|---|---|---|---|---|---|---|---|---|---|---|---|---|---|---|---|---|---|---|---|---|---|---|---|---|---|---|---|---|---|---|---|---|---|---|---|---|---|---|
| FALSE | High | NCPTPTRTI | [K].NCPTPT | 2xCarbami | Q8K3K8 1x | 0.005924 | 0.00062 | 1 | 1 | 1 | Q8K3K8 | Optineurin | Q8K3K8 [2 | 1 | 2714.211 | 0 | 2.53 | 0.000324 | 22.44 | 29.5 | 170.5 | 14.97 | 7 | | High | High | High | High | High | High | High | High | High | High | High | 0.000275 | 0.001754 | 3.37 |
| FALSE | High | NCSSPEFSK | [R].NCSSPE | 1xCarbami | Q3U893 1x | 0.00925 | 0.000736 | 1 | 11 | 2 | Q3U893 | Putative ur | Q3U893 [5 | 0 | 1593.738 | 0 | 0.26 | 0.029082 | 1.85 | 88.7 | 111.3 | 3.88 | 10.18 | | High | High | High | High | High | High | High | High | High | High | High | 0.000674 | 0.005878 | 2.18 |
| FALSE | High | NCSSSSHYS | [R].NCSSSS | 1xCarbami | Q9CZ70 1x | 0.103392 | 0.008249 | 1 | 1 | 1 | Q9CZ70 | MCG12980 | Q9CZ70 [6 | 0 | 2401.064 | 0 | -1.06 | 0.017204 | 11 | 141.5 | 58.5 | 21.94 | 15.75 | | High | High | High | High | High | High | High | High | High | High | High | 0.004164 | 0.04488 | 2.44 |
| FALSE | High | NCTPEVTEI | [K].NCTPEV | 1xCarbami | Q5DTT3 1x | 0.01588 | 0.001124 | 1 | 1 | 1 | Q5DTT3 | Protein FAI | Q5DTT3 [1 | 0 | 1615.78 | 0 | 0.12 | 0.024949 | 1.11 | 96.2 | 103.8 | 8.23 | 11.01 | | High | High | High | High | High | High | High | High | High | High | High | 0.000585 | 0.005282 | 2.12 |
| FALSE | High | NCWPPVLS | [R].NCWPP | 2xCarbami | Q544H6 1x | 0.006834 | 0.00062 | 1 | 3 | 1 | Q544H6 | Cyclin E2, i | Q544H6 [5 | 0 | 3013.544 | 0 | 1.69 | 0.000793 | 15.77 | 52.2 | 147.8 | 13.9 | 11.73 | | High | High | High | High | High | High | High | High | High | High | High | 0.000275 | 0.002052 | 2.64 |
| FALSE | High | NASTSFQE | [R].NASTSF | 2xPhospho | P54103 2x | 0.003083 | 0.00016 | 1 | 6 | 3 | P54103 | DnaJ homo | P54103 [45 | 0 | 1757.723 | 0 | -0.68 | 0.291149 | 16.62 | 131.3 | 68.7 | 19.18 | 60.83 | | High | High | High | High | High | High | High | High | High | High | High | 7.19E-05 | 0.000847 | 3.06 |
| FALSE | High | IDLYQQASI | [R].IDLYQC | 1xPhospho | Q7TSC1 1x | 1.46E-06 | 0 | 1 | 1 | 1 | Q7TSC1 | Protein PRI | Q7TSC1 [2 | 0 | 1895.947 | 0 | 0.14 | 0.226901 | 17.39 | 95.2 | 104.8 | 13.53 | 9.5 | | High | High | High | High | High | High | High | High | High | High | High | 0 | 1.56E-07 | 3.11 |
| FALSE | High | LTPVSLSNS | [K].LTPVSL | 1xPhospho | Q61687 1x | 0.104222 | 0.008348 | 1 | 4 | 1 | Q61687 | Transcripti | Q61687 [5 | 0 | 1564.855 | 0 | 0.15 | 0.801776 | 66.66 | 94.9 | 105.1 | 60.53 | 30.48 | | High | High | High | High | High | High | High | High | High | High | High | 0.004206 | 0.0453 | 2.3 |
| FALSE | High | LTPTTPESS | [R].LTPTTP | 2xPhospho | Q64337 2x | 0.000106 | 0 | 1 | 3 | 2 | Q64337 | Sequestoso | Q64337 [2 | 0 | 2269.029 | 0 | 1.56 | 0.000431 | 9.07 | 51 | 149 | 12.72 | 5.42 | | High | High | High | High | High | High | High | High | High | High | High | 0 | 1.92E-05 | 3.01 |
| FALSE | High | LSGCESQQ | [K].LSGCES | 1xCarbami | Q8C0P0 1x | 4.31E-05 | 0 | 1 | 1 | 1 | Q8C0P0 | Serine/thre | Q8C0P0 [4 | 0 | 2151.019 | 0 | -0.01 | 0.34586 | 3.02 | 103 | 97 | 5.27 | 8.99 | | High | High | High | High | High | High | High | High | High | High | High | 0 | 6.99E-06 | 2.82 |
| FALSE | High | LSGDLEAG. | [K].LSGDLE | 1xPhospho | H3BL37 1x | 0.000548 | 0 | 1 | 4 | 2 | H3BL37 | Treacle pro | H3BL37 [12 | 0 | 1595.845 | 0 | -0.48 | 0.007082 | 2.59 | 115.9 | 84.1 | 9.88 | 2.58 | | High | High | High | High | High | High | High | High | High | High | High | 0 | 0.000122 | 3.41 |
| FALSE | High | LSGDSEEM | [R].LSGDSE | 1xPhospho | A0A1W2P6 | 0.000221 | 0 | 1 | 8 | 1 | A0A1W2P6 | Bromodom | A0A1W2P6 | 0 | 2486.189 | 0 | -0.42 | 0.008743 | 5.62 | 116.6 | 83.4 | 4.4 | 16.86 | | High | High | High | High | High | High | High | High | High | High | High | 0 | 4.36E-05 | 3.15 |
| FALSE | High | LSGDSEEM | [R].LSGDSE | 1xPhospho | A0A1W2P6 | 2.17E-05 | 0 | 1 | 8 | 3 | A0A1W2P6 | Bromodom | A0A1W2P6 | 0 | 2502.184 | 0 | -0.48 | 0.002468 | 4.3 | 118.3 | 81.7 | 7.61 | 6.38 | | High | High | High | High | High | High | High | High | High | High | High | 0 | 3.22E-06 | 3.68 |
| FALSE | High | LSGDTSPPT | [R].LSGDTS | 1xPhospho | Q8BLB7 1x | 0.000346 | 0 | 1 | 3 | 3 | Q8BLB7 | Lethal(3)m | Q8BLB7 [6 | 0 | 1868.899 | 0 | 0.57 | 0.002304 | 5.87 | 80.2 | 119.8 | 2.84 | 7.26 | | High | High | High | High | High | High | High | High | High | High | High | 0 | 7.25E-05 | 3.03 |
| FALSE | High | LSGEGDTD | [K].LSGEGI | 1xPhospho | Q8BZ47 1x | 3.83E-05 | 0 | 1 | 2 | 2 | Q8BZ47 | Zinc finger | Q8BZ47 [5 | 0 | 4081.682 | 0 | 0.15 | 0.684935 | 14.65 | 96.7 | 103.3 | 16.86 | 16.71 | | High | High | High | High | High | High | High | High | High | High | High | 0 | 6.11E-06 | 4.82 |
| FALSE | High | LSGEGDTD | [K].LSGEGI | 1xDeamida | Q8BZ47 1x | 1.96E-07 | 0 | 1 | 2 | 10 | Q8BZ47 | Zinc finger | Q8BZ47 [5 | 0 | 4098.66 | 0 | -0.09 | 0.683354 | 29.92 | 105.8 | 94.2 | 12.34 | 19.51 | | High | High | High | High | High | High | High | High | High | High | High | 0 | 1.04E-06 | 4.79 |
| FALSE | High | LSGFSFK | [K].LSGFSF | 1xPhospho | P26645 1x | 0.053926 | 0.00384 | 1 | 2 | 1 | P26645 | Myristoyla | P26645 [15 | 0 | 1094.548 | 0 | 0.41 | 0.227792 | 32.86 | 79.3 | 120.7 | 19.27 | 60.55 | | High | High | High | High | High | High | High | High | High | High | High | 0.002045 | 0.02105 | 1.63 |
| FALSE | High | LSGFSFK | [K].LSGFSF | 1xPhospho | P26645 1x | 0.006478 | 0.00062 | 1 | 2 | 3 | P26645 | Myristoyla | P26645 [15 | 0 | 1323.711 | 0 | 1.75 | 0.001117 | 24.76 | 45.8 | 154.2 | 13.08 | 9.28 | | High | High | High | High | High | High | High | High | High | High | High | 0.000327 | 0.002964 | 2.09 |
| FALSE | High | LSGGSGSD | [R].LSGGS | 1xPhospho | A0A0A0M | 0.115566 | 0.009504 | 1 | 2 | 1 | A0A0A0M | Histone-lys | A0A0A0M | 0 | 2362.099 | 0 | 0.64 | 0.003158 | 13.72 | 75 | 125 | 11.48 | 6.63 | | High | High | High | High | High | High | High | High | High | High | High | 0.004733 | 0.05108 | 2.6 |
| FALSE | High | LSGGSHYN | [R].LSGGSI | 1xPhospho | Q922G2 1x | 0.101749 | 0.008194 | 1 | 1 | 1 | Q922G2 | Protein FAI | Q922G2 [1 | 0 | 1715.852 | 0 | 0 | 0.898721 | 5.2 | 100.8 | 99.2 | 8.01 | 7.15 | | High | High | High | High | High | High | High | High | High | High | High | 0.004126 | 0.04422 | 2.53 |
| FALSE | High | LSGKPQNA | [R].LSGKPC | 1xPhospho | Q9JJ66 1xP | 0.071071 | 0.005247 | 1 | 2 | 1 | Q9JJ66 | Cell divisio | Q9JJ66 [13 | 0 | 2197.117 | 0 | -0.5 | 0.005743 | 4.74 | 118 | 82 | 6.22 | 7.89 | | High | High | High | High | High | High | High | High | High | High | High | 0.002703 | 0.02901 | 2.35 |
| FALSE | High | LSGIMVPA | [R].LSGIMV | 1xPhospho | P28658 1x | 0.083783 | 0.006345 | 1 | 4 | 1 | P28658 | Ataxin-10 ( | P28658 [11 | 0 | 2148.134 | 0 | -0.39 | 0.169251 | 3.75 | 114.1 | 85.9 | 17.97 | 6.84 | | High | High | High | High | High | High | High | High | High | High | High | 0.003265 | 0.03507 | 2.63 |
| FALSE | High | LSFTPETPA | [R].LSFTPE | 2xPhospho | O88466 2x | 0.002232 | 0.000109 | 2 | 4 | 1 | O88466; R | Zinc finger | O88466 [1 | 0 | 1837.879 | 0 | -0.54 | 0.000711 | 2.94 | 120 | 80 | 6.33 | 6.21 | Shared | High | High | High | High | High | High | High | High | High | High | High | 4.94E-05 | 0.000589 | 2.11 |
| FALSE | High | LSGINEIPRI | [K].LSGINE | 1xPhospho | Q3UQN2 1 | 0.108463 | 0.008839 | 1 | 3 | 1 | Q3UQN2 | F-BAR dom | Q3UQN2 [ | 0 | 3322.74 | 0 | 0.77 | 0.001922 | 14.4 | 73.9 | 126.1 | 10.66 | 7.78 | | High | High | High | High | High | High | High | High | High | High | High | 0.004411 | 0.04729 | 3.01 |
| FALSE | High | LSGLSFK | [K].LSGLSF | 1xPhospho | P28667 1x | 0.012508 | 0.000889 | 1 | 1 | 2 | P28667 | MARCKS-re | P28667 [10 | 0 | 1289.727 | 0 | -1.78 | 0.000564 | 2.5 | 158.3 | 41.7 | 5.61 | 22.25 | | High | High | High | High | High | High | High | High | High | High | High | 0.009953 | 0.1088 | 1.34 |
| FALSE | High | ISGISSPIGS | [K].ISGISSP | 1xPhospho | A2A7F4 1x | 0.006536 | 0.00062 | 1 | 3 | 2 | A2A7F4 | MCG12287 | A2A7F4 [1 | 0 | 1519.783 | 0 | 0.03 | 0.583167 | 12.86 | 98.8 | 101.2 | 12.01 | 10.29 | | High | High | High | High | High | High | High | High | High | High | High | 0.000984 | 0.01024 | 2.01 |
| FALSE | High | LSGPAELR | [R].LSGPAE | 1xPhospho | Q91ZV0 1x | 0.088701 | 0.006669 | 1 | 12 | 1 | Q91ZV0 | Melanoma | Q91ZV0 [1 | 0 | 1151.602 | 0 | -0.03 | 0.59594 | 30.92 | 98.9 | 101.1 | 9.58 | 18.13 | | High | High | High | High | High | High | High | High | High | High | High | 0.003447 | 0.03742 | 2.45 |
| FALSE | High | LSGQNTMC | [K].LSGQN | 1xPhospho | Q9D3A8 1x | 0.000165 | 0 | 1 | 2 | 1 | Q9D3A8 | Carboxyl-te | Q9D3A8 [3 | 0 | 2406.157 | 0 | 1.13 | 0.003734 | 21.08 | 65.1 | 134.9 | 10.5 | 13.26 | | High | High | High | High | High | High | High | High | High | High | High | 0 | 3.17E-05 | 2.92 |
| FALSE | High | LSGQNTMC | [K].LSGQN | 1xPhospho | Q9D3A8 1x | 0.000426 | 0 | 1 | 2 | 1 | Q9D3A8 | Carboxyl-te | Q9D3A8 [3 | 0 | 2422.152 | 0 | 0.44 | 0.10194 | 14.88 | 89.9 | 110.1 | 20.47 | 12.74 | | High | High | High | High | High | High | High | High | High | High | High | 0 | 9.14E-05 | 3.29 |
| FALSE | High | LSGSCAPST | [K].LSGSCA | 1xCarbami | Q6NVF4 1x | 5.4E-08 | 0 | 1 | 1 | 4 | Q6NVF4 | DNA helica | Q6NVF4 [9 | 0 | 2203.005 | 0 | -0.7 | 0.001917 | 5.44 | 125.3 | 74.7 | 6.16 | 7.93 | | High | High | High | High | High | High | High | High | High | High | High | 0 | 3.82E-09 | 4.97 |
| FALSE | High | LSGSCAPST | [K].LSGSCA | 1xCarbami | Q6NVF4 2x | 2.06E-05 | 0 | 1 | 1 | 2 | Q6NVF4 | DNA helica | Q6NVF4 [9 | 0 | 2282.972 | 0 | -0.49 | 0.002793 | 4.65 | 115.7 | 84.3 | 4.67 | 8.05 | | High | High | High | High | High | High | High | High | High | High | High | 0 | 3.05E-06 | 4.38 |
| FALSE | High | LSGSGPAEI | [R].LSGSGF | 1xPhospho | Q1HDU4 1 | 1.97E-05 | 0 | 1 | 4 | 1 | Q1HDU4 | ArhGAP9 C | Q1HDU4 [3 | 0 | 3254.561 | 0 | -1.65 | 0.000159 | 0.62 | 152.7 | 47.3 | 3.96 | 6.39 | | High | High | High | High | High | High | High | High | High | High | High | 0 | 2.9E-06 | 4.83 |
| FALSE | High | LSGSNLCSS | [R].LSGSNL | 1xCarbami | Q9R1C7 1x | 0.010382 | 0.000772 | 1 | 4 | 2 | Q9R1C7 | Pre-mRNA- | Q9R1C7 [3 | 0 | 2352.089 | 0 | 0.14 | 0.64213 | 18.85 | 97.8 | 102.2 | 8.1 | 13.17 | | High | High | High | High | High | High | High | High | High | High | High | 0.000382 | 0.003279 | 3.58 |
| FALSE | High | ISGSPAQSS | [R].ISGSPA | 1xCarbami | Q9CV28 1x | 5.65E-08 | 0 | 1 | 2 | 2 | Q9CV28 | Ubiquitin c | Q9CV28 [1 | 0 | 3184.482 | 0 | 0.35 | 0.183581 | 17.23 | 87.9 | 112.1 | 18.19 | 8.62 | | High | High | High | High | High | High | High | High | High | High | High | 0 | 4.02E-09 | 4.58 |
| FALSE | High | ISGTTALQE | [K].ISGTTA | 1xPhospho | Q9Z0H8 1x | 4.17E-06 | 0 | 2 | 8 | 1 | Q9Z0H8; F | CAP-Gly do | Q9Z0H8 [3 | 0 | 1769.981 | 0 | 1.63 | 0.001068 | 16.74 | 50.3 | 149.7 | 15.25 | 9.36 | Shared | High | High | High | High | High | High | High | High | High | High | High | 0 | 5.06E-07 | 2.81 |
| FALSE | High | ISHELESSS | [R].ISHELE | 1xPhospho | F8WIP8 1x | 0.000168 | 0 | 1 | 7 | 3 | F8WIP8 | Osteopont | F8WIP8 [2 | 0 | 1726.773 | 0 | -0.53 | 0.054678 | 2.63 | 116.4 | 83.6 | 18.41 | 11.3 | | High | High | High | High | High | High | High | High | High | High | High | 0 | 3.24E-05 | 2.39 |
| FALSE | High | ISHELESSS | [R].ISHELE | 2xPhospho | F8WIP8 2x | 0.0016 | 5.66E-05 | 1 | 7 | 3 | F8WIP8 | Osteopont | F8WIP8 [2 | 0 | 1806.74 | 0 | 0.6 | 0.200261 | 30.45 | 75.4 | 124.6 | 33.3 | 12.47 | | High | High | High | High | High | High | High | High | High | High | High | 2.58E-05 | 0.000404 | 2.15 |
| FALSE | High | LSGINEIPRI | [K].LSGINE | 2xPhospho | Q3UQN2 2 | 0.000346 | 0 | 1 | 3 | 2 | Q3UQN2 | F-BAR dom | Q3UQN2 [ | 0 | 3402.706 | 0 | 0.56 | 0.005178 | 14.01 | 79.4 | 120.6 | 8.59 | 12.19 | | High | High | High | High | High | High | High | High | High | High | High | 0 | 7.28E-05 | 2.77 |
| FALSE | High | ISHLSGSGS | [K].ISHLSG | 1xPhospho | Q0VBL3 1x | 1.11E-05 | 0 | 1 | 7 | 2 | Q0VBL3 | RNA bindin | Q0VBL3 [2 | 0 | 1610.737 | 0 | -0.19 | 0.101437 | 7.45 | 108.9 | 91.1 | 11.52 | 13.36 | | High | High | High | High | High | High | High | High | High | High | High | 0 | 1.53E-06 | 1.99 |
| FALSE | High | LSFTPETPA | [R].LSFTPE | 1xPhospho | O88466 1x | 1.42E-05 | 0 | 2 | 4 | 2 | O88466; R | Zinc finger | O88466 [1 | 0 | 1757.913 | 0 | -0.95 | 0.000704 | 1.05 | 133 | 67 | 5.85 | 4.87 | Shared | High | High | High | High | High | High | High | High | High | High | High | 0 | 2.01E-06 | 3.03 |
| FALSE | High | ISFSTSDVS | [R].ISFSTSD | 1xPhospho | E9PZW8 1x | 0.042868 | 0.003228 | 1 | 6 | 3 | E9PZW8 | Unconvent | E9PZW8 [1 | 0 | 1608.829 | 0 | 1.78 | 0.003573 | 18.64 | 43.6 | 156.4 | 18.52 | 18.56 | | High | High | High | High | High | High | High | High | High | High | High | 0.001651 | 0.01613 | 2.52 |
| FALSE | High | LSEALEPEV | [K].LSEALE | 1xCarbami | Q6ZQF0 1x | 6.77E-05 | 0 | 1 | 1 | 1 | Q6ZQF0 | DNA topois | Q6ZQF0 [5 | 0 | 2215.042 | 0 | 0.26 | 0.128692 | 21.25 | 90.9 | 109.1 | 10.3 | 13.66 | | High | High | High | High | High | High | High | High | High | High | High | 0 | 1.16E-05 | 3.02 |
| FALSE | High | LSEASDER | [K].LSEASD | 1xPhospho | Q3UVG3 1 | 0.112856 | 0.009146 | 1 | 1 | 1 | Q3UVG3 | Protein FAI | Q3UVG3 [6 | 0 | 1215.546 | 0 | 0.75 | 0.006247 | 18.74 | 77.8 | 122.2 | 17.85 | 9.12 | | High | High | High | High | High | High | High | High | High | High | High | 0.004583 | 0.04985 | 1.77 |
| FALSE | High | LSEASGGL | [K].LSEASG | 1xPhospho | Q3TIV5 1x | 0.026652 | 0.001676 | 1 | 2 | 1 | Q3TIV5 | Zinc finger | Q3TIV5 [36 | 0 | 1698.79 | 0 | -0.5 | 0.059833 | 14.54 | 113.4 | 86.6 | 15.66 | 11.92 | | High | High | High | High | High | High | High | High | High | High | High | 0.000903 | 0.009464 | 2.79 |
| FALSE | High | ISEASWLPE | [K].ISEASW | 1xPhospho | F8WGX5 1 | 0.000462 | 0 | 1 | 2 | 1 | F8WGX5 | DmX-like p | F8WGX5 [9 | 0 | 2614.248 | 0 | 0.14 | 0.419928 | 9.3 | 95.4 | 104.6 | 8.62 | 2.21 | | High | High | High | High | High | High | High | High | High | High | High | 0 | 0.0001 | 2.98 |
| FALSE | High | LSEDAMST | [K].LSEDAM | 1xPhospho | E9QPR5 1x | 0.0003 | 0 | 1 | 5 | 1 | E9QPR5 | Serine/thre | E9QPR5 [9 | 0 | 1671.702 | 0 | -0.06 | 0.604658 | 10.52 | 106.1 | 93.9 | 8.28 | 14.43 | | High | High | High | High | High | High | High | High | High | High | High | 0 | 6.14E-05 | 1.63 |
| FALSE | High | LSEDAMST | [K].LSEDAM | 2xPhospho | E9QPR5 2x | 0.030104 | 0.001857 | 1 | 5 | 1 | E9QPR5 | Serine/thre | E9QPR5 [9 | 0 | 1767.663 | 0 | 0.02 | 0.764244 | 8.76 | 102.1 | 97.9 | 6.64 | 16.03 | | High | High | High | High | High | High | High | High | High | High | High | 0.001011 | 0.01085 | 1.76 |
| FALSE | High | ISEDETERN | [R].ISEDETI | 1xDeamida | Q9JIX8-3 1 | 0.000852 | 0 | 3 | 12 | 1 | Q9JIX8-3; F | Isoform 3 c | Q9JIX8-3 [ | 1 | 2457.082 | 0 | -0.37 | 0.004232 | 1.19 | 109.1 | 90.9 | 8.57 | 8.98 | Shared | High | High | High | High | High | High | High | High | High | High | High | 0 | 0.000199 | 2.8 |
| FALSE | High | LSEEACPGV | [K].LSEEAC | 1xCarbami | O08715-4 | 0.003755 | 0.00021 | 1 | 6 | 1 | O08715-4 | Isoform 4 c | O08715-4 | 0 | 2574.284 | 0 | 0.06 | 0.499473 | 22.98 | 96.9 | 103.1 | 5.15 | 22.12 | | High | High | High | High | High | High | High | High | High | High | High | 9.42E-05 | 0.00105 | 3.95 |
| FALSE | High | LSEEAEPRF | [K].LSEEAE | 1xPhospho | Q5DTS2 1x | 0.017344 | 0.001204 | 1 | 4 | 1 | Q5DTS2 | MKIAA402 | Q5DTS2 [9 | 0 | 2080.073 | 0 | -0.15 | 0.035429 | 6.82 | 105.8 | 94.2 | 4.57 | 7.74 | | High | High | High | High | High | High | High | High | High | High | High | 0.000656 | 0.00581 | 1.96 |
| FALSE | High | LSEEIVVTP | [K].LSEEIV | 1xPhospho | Q8BJ05 1x | 0.002058 | 5.66E-05 | 1 | 1 | 1 | Q8BJ05 | Zinc finger | Q8BJ05 [47 | 0 | 2285.15 | 0 | -0.21 | 0.127986 | 17.01 | 111.5 | 88.5 | 8.96 | 13.62 | | High | High | High | High | High | High | High | High | High | High | High | 2.58E-05 | 0.000534 | 3.16 |
| FALSE | High | LSEEIVVTP | [K].LSEEIV | 1xPhospho | Q8BJ05 1x | 5.07E-05 | 0 | 1 | 1 | 1 | Q8BJ05 | Zinc finger | Q8BJ05 [47 | 0 | 2301.145 | 0 | -0.3 | 0.226478 | 6.17 | 110.5 | 89.5 | 4.64 | 22.31 | | High | High | High | High | High | High | High | High | High | High | High | 0 | 8.39E-06 | 2.76 |
| FALSE | High | ISEEQQQL | [K].ISEEQQ | 1xPhospho | Q91VW5 1 | 0.000595 | 0 | 1 | 6 | 3 | Q91VW5 | Golgin sub | Q91VW5 [ | 0 | 3052.479 | 0 | 0.55 | 0.005667 | 14.11 | 77.3 | 122.7 | 6.47 | 8.73 | | High | High | High | High | High | High | High | High | High | High | High | 0 | 0.000134 | 3.23 |
| FALSE | High | ISEGLPTPT | [K].ISEGLP | 1xPhospho | P13405 1x | 0.002844 | 0.00016 | 1 | 4 | 2 | P13405 | Retinoblas | P13405 [8 | 0 | 1580.87 | 0 | 1.25 | 0.000375 | 4.81 | 59.4 | 140.6 | 6.03 | 4.53 | | High | High | High | High | High | High | High | High | High | High | High | 0.0032 | 0.03394 | 2.05 |
| FALSE | High | LSFTESLTG | [K].LSFTESI | 1xPhospho | P25799 1x | 2.63E-06 | 0 | 1 | 2 | 3 | P25799 | Nuclear fac | P25799 [9 | 0 | 2460.304 | 0 | -0.07 | 0.317116 | 33.39 | 109.5 | 90.5 | 14.32 | 15.4 | | High | High | High | High | High | High | High | High | High | High | High | 0 | 7.83E-07 | 4.04 |
| FALSE | High | LSEGPAALA | [R].LSEGPA | 1xPhospho | Q3V1Z0 1x | 0.000165 | 0 | 1 | 6 | 2 | Q3V1Z0 | Iporin OS= | Q3V1Z0 [5 | 0 | 1799.925 | 0 | 1.72 | 0.000239 | 10.95 | 45.2 | 154.8 | 10.58 | 5.17 | | High | High | High | High | High | High | High | High | High | High | High | 0 | 0.000281 | 2.95 |
| FALSE | High | LSEPGELR | [R].LSEPGE | 1xPhospho | E9PYW4 1x | 0.042141 | 0.003162 | 1 | 5 | 1 | E9PYW4 | E3 ubiquiti | E9PYW4 [1 | 0 | 1209.608 | 0 | 0.69 | 0.019068 | 17.47 | 73.5 | 126.5 | 8.39 | 19.09 | | High | High | High | High | High | High | High | High | High | High | High | 0.001639 | 0.01583 | 1.72 |
| FALSE | High | LSEPSLPVE | [K].LSEPSLI | 1xPhospho | Q80YR7 1x | 2.54E-06 | 0 | 1 | 2 | 1 | Q80YR7 | Claspin OS | Q80YR7 [8 | 0 | 2940.453 | 0 | 0.15 | 0.441744 | 8.63 | 94.8 | 105.2 | 11.7 | 10.5 | | High | High | High | High | High | High | High | High | High | High | High | 0 | 2.89E-07 | 4.11 |
| FALSE | High | LSEPSLPVE | [K].LSEPSLI | 1xDeamida | Q80YR7 1x | 1.31E-05 | 0 | 1 | 2 | 2 | Q80YR7 | Claspin OS | Q80YR7 [8 | 0 | 2941.437 | 0 | 0.24 | 0.172363 | 18.36 | 94.7 | 105.3 | 13.25 | 9.03 | | High | High | High | High | High | High | High | High | High | High | High | 0 | 1.19E-05 | 3.24 |
| FALSE | High | ISERVSLES | [R].ISERVSI | 1xPhospho | P59729 1x | 0.000164 | 0 | 1 | 4 | 1 | P59729 | Ras and Ra | P59729 [3 | 1 | 2287.113 | 0 | 2.42 | 0.000495 | 19.72 | 31.4 | 168.6 | 18.11 | 7.29 | | High | High | High | High | High | High | High | High | High | High | High | 0 | 3.13E-05 | 2.62 |
| FALSE | High | LSESLHSAD | [R].LSESLH | 1xPhospho | E9Q6J5-3 1 | 0.003558 | 0.00021 | 1 | 3 | 1 | E9Q6J5-3 | Isoform Ne | E9Q6J5-3 [ | 0 | 1867.92 | 0 | -0.45 | 0.008081 | 9.8 | 119.7 | 80.3 | 5.73 | 11.41 | | High | High | High | High | High | High | High | High | High | High | High | 9.42E-05 | 0.000987 | 1.91 |
| FALSE | High | ISESPEVAR | [R].ISESPE | 1xPhospho | D3Z3Y7 1x | 0.08447 | 0.006345 | 1 | 6 | 1 | D3Z3Y7 | Pleckstrin | D3Z3Y7 [1 | 0 | 1296.64 | 0 | -0.22 | 0.120676 | 14.91 | 104.9 | 95.1 | 9.79 | 7.53 | | High | High | High | High | High | High | High | High | High | High | High | 0.003279 | 0.0355 | 1.37 |
| FALSE | High | LSESSALK | [R].LSESSA | 1xPhospho | Q91XU0 1x | 0.051258 | 0.003745 | 1 | 2 | 2 | Q91XU0 | ATPase WF | Q91XU0 [7 | 0 | 1372.749 | 0 | -0.47 | 0.003767 | 3.03 | 117.4 | 82.6 | 2.76 | 10.05 | | High | High | High | High | High | High | High | High | High | High | High | 0.003064 | 0.03186 | 1.65 |
| FALSE | High | LSESSPAFR | [R].LSESSP | 1xPhospho | A0A0R4IZV | 0.003655 | 0.00021 | 1 | 7 | 1 | A0A0R4IZV | tRNA (guar | A0A0R4IZV | 0 | 1302.629 | 0 | -0.38 | 0.024907 | 5.94 | 111.3 | 88.7 | 8.98 | 8.4 | | High | High | High | High | High | High | High | High | High | High | High | 9.42E-05 | 0.00102 | 2.48 |
| FALSE | High | LSESSSSPQ | [R].LSESSS | 1xPhospho | Q9WVG9 1 | 0.013074 | 0.000889 | 1 | 2 | 2 | Q9WVG9 | Male-speci | Q9WVG9 [ | 0 | 1684.856 | 0 | 1.69 | 0.000181 | 4.31 | 49 | 151 | 8.71 | 5.18 | | High | High | High | High | High | High | High | High | High | High | High | 0.000467 | 0.004227 | 2.4 |
| FALSE | High | LSFEGAAK | [R].LSFEGA | 1xPhospho | Q9CXS4 1x | 0.000674 | 0 | 1 | 2 | 2 | Q9CXS4 | Centromer | Q9CXS4 [1 | 0 | 1360.728 | 0 | 0.2 | 0.023148 | 8.42 | 92.9 | 107.1 | 3.52 | 6.87 | | High | High | High | High | High | High | High | High | High | High | High | 0 | 0.000199 | 1.95 |
| FALSE | High | LSFHSPSAC | [R].LSFHSP | 1xPhospho | E9Q4K7 1x | 0.047894 | 0.00354 | 1 | 1 | 1 | E9Q4K7 | Kinesin fan | E9Q4K7 [6 | 0 | 1566.763 | 0 | 0.5 | 0.001993 | 5 | 80.4 | 119.6 | 7.39 | 5.98 | | High | High | High | High | High | High | High | High | High | High | High | 0.001812 | 0.01839 | 1.92 |
| FALSE | High | ISFILEK | [R].ISFILEK | 1xPhospho | Q9CPY3 1x | 0.003195 | 0.00016 | 1 | 1 | 1 | Q9CPY3 | Sororin OS | Q9CPY3 [8 | 0 | 1387.8 | 0 | -0.37 | 0.00295 | 2.62 | 111.3 | 88.7 | 6.59 | 4.73 | | High | High | High | High | High | High | High | High | High | High | High | 7.19E-05 | 0.000876 | 2.18 |
| FALSE | High | ISFPANCPD | [R].ISFPAN | 1xCarbami | Q5DTU0-2 | 0.000656 | 0 | 1 | 3 | 1 | Q5DTU0-2 | Isoform 2 c | Q5DTU0-2 | 0 | 2896.439 | 0 | 1.47 | 0.001078 | 2.26 | 46.8 | 153.2 | 10.11 | 11.12 | | High | High | High | High | High | High | High | High | High | High | High | 0 | 0.000148 | 2.68 |
| FALSE | High | LSEGPAALA | [R].LSEGPA | 1xPhospho | Q3V1Z0 1x | 0.000662 | 0 | 1 | 6 | 1 | Q3V1Z0 | Iporin OS= | Q3V1Z0 [5 | 1 | 1956.027 | 0 | -0.4 | 0.0118 | 6.37 | 112 | 88 | 5.32 | 5.99 | | High | High | High | High | High | High | High | High | High | High | High | 0 | 0.00015 | 2.71 |
| FALSE | High | LSHPGQLM | [R].LSHPG | 1xCarbami | O88351 1x | 0.000434 | 0 | 1 | 4 | 2 | O88351 | Inhibitor o | O88351 [6 | 0 | 3025.405 | 0 | 0.93 | 0.141876 | 89.12 | 68.9 | 131.1 | 31.79 | 48.44 | | High | High | High | High | High | High | High | High | High | High | High | 0 | 9.32E-05 | 3.74 |
| FALSE | High | LSHPPSPPS | [R].LSHPPS | 1xCarbami | B2RUG8 1x | 2.7E-06 | 0 | 1 | 7 | 2 | B2RUG8 | Asxl2 prote | B2RUG8 [1 | 0 | 2552.298 | 0 | -0.43 | 0.028599 | 11.98 | 114.9 | 85.1 | 9.3 | 8.57 | | High | High | High | High | High | High | High | High | High | High | High | 0 | 3.1E-07 | 3.4 |
| FALSE | High | LSHPPSPPS | [R].LSHPPS | 1xCarbami | B2RUG8 2x | 0.001267 | 0 | 1 | 7 | 1 | B2RUG8 | Asxl2 prote | B2RUG8 [1 | 0 | 2632.265 | 0 | 0.25 | 0.021129 | 9.06 | 88.3 | 111.7 | 9.5 | 14.37 | | High | High | High | High | High | High | High | High | High | High | High | 0 | 0.000312 | 2.31 |
| FALSE | High | ISMPDFDLI | [K].ISMPDI | 1xPhospho | E9Q616 1x | 0.009843 | 0.000736 | 1 | 3 | 1 | E9Q616 | AHNAK nuc | E9Q616 [2 | 0 | 1869.959 | 0 | 3.27 | 0.015299 | 42.18 | 23.3 | 176.7 | 40.39 | 40.51 | | High | High | High | High | High | High | High | High | High | High | High | 0.000344 | 0.003087 | 2.67 |
| FALSE | High | ISMPDFDLI | [K].ISMPDI | 1xPhospho | E9Q616 1x | 0.064862 | 0.004664 | 1 | 2 | 1 | E9Q616 | AHNAK nuc | E9Q616 [4 | 0 | 1830.948 | 0 | 1.95 | 0.002884 | 20.3 | 41 | 159 | 5.04 | 18.23 | | High | High | High | High | High | High | High | High | High | High | High | 0.002432 | 0.02607 | 2.59 |
| FALSE | High | ISMPDIDLN | [K].ISMPDI | 1xPhospho | E9Q616 1x | 0.001644 | 5.66E-05 | 1 | 4 | 1 | E9Q616 | AHNAK nuc | E9Q616 [2 | 0 | 1796.963 | 1 | 2.48 | 0.008085 | 25.02 | 30.4 | 169.6 | 7.12 | 29.92 | | High | High | High | High | High | High | High | High | High | High | High | 2.58E-05 | 0.000418 | 3.18 |
| FALSE | High | ISMPDIDLN | [K].ISMPDI | 1xPhospho | E9Q616 1x | 0.003892 | 0.000261 | 1 | 4 | 2 | E9Q616 | AHNAK nuc | E9Q616 [2 | 0 | 1812.958 | 0 | 1.92 | 0.00041 | 4.01 | 43 | 157 | 4.8 | 8.36 | | High | High | High | High | High | High | High | High | High | High | High | 0.00014 | 0.001096 | 2.53 |
| FALSE | High | ISMPDLHLI | [K].ISMPDI | 1xPhospho | E9Q616 1x | 9.62E-05 | 0 | 1 | 4 | 1 | E9Q616 | AHNAK nuc | E9Q616 [1 | 0 | 1591.868 | 0 | 3.87 | 0.007445 | 12.68 | 12.5 | 187.5 | 25.11 | 32.74 | | High | High | High | High | High | High | High | High | High | High | High | 0 | 1.72E-05 | 2.25 |
| FALSE | High | ISMPDLHLI | [K].ISMPDI | 1xPhospho | E9Q616 1x | 0.000745 | 0 | 1 | 4 | 2 | E9Q616 | AHNAK nuc | E9Q616 [1 | 0 | 1607.863 | 0 | 3.41 | 0.014367 | 54.11 | 16.3 | 183.7 | 32.38 | 39.48 | | High | High | High | High | High | High | High | High | High | High | High | 0 | 0.000171 | 2.15 |
| FALSE | High | ISMPDLNLI | [K].ISMPDI | 1xPhospho | E9Q616 1x | 0.017652 | 0.001282 | 1 | 2 | 1 | E9Q616 | AHNAK nuc | E9Q616 [2 | 0 | 1795.979 | 0 | 2.69 | 0.007702 | 9.94 | 26.4 | 173.6 | 13.63 | 30.52 | | High | High | High | High | High | High | High | High | High | High | High | 0.000689 | 0.005912 | 2.84 |
| FALSE | High | ISMPDLNLI | [K].ISMPDI | 1xPhospho | E9Q616 1x | 0.040373 | 0.002982 | 1 | 2 | 1 | E9Q616 | AHNAK nuc | E9Q616 [2 | 0 | 1811.974 | 0 | 2.59 | 0.000311 | 12.07 | 26.6 | 173.4 | 14.24 | 8.33 | | High | High | High | High | High | High | High | High | High | High | High | 0.001581 | 0.01516 | 2.45 |
| FALSE | High | ISMPDVDL | [K].ISMPD\ | 1xPhospho | E9Q616 1x | 0.000633 | 0 | 1 | 3 | 2 | E9Q616 | AHNAK nuc | E9Q616 [6 | 0 | 1805.964 | 0 | 2.51 | 0.007864 | 9.82 | 30.2 | 169.8 | 9.89 | 27.98 | | High | High | High | High | High | High | High | High | High | High | High | 0 | 0.000143 | 2.77 |
| FALSE | High | ISMPDVDL | [K].ISMPD\ | 1xPhospho | E9Q616 1x | 0.010853 | 0.000772 | 1 | 3 | 4 | E9Q616 | AHNAK nuc | E9Q616 [6 | 0 | 1821.959 | 0 | 1.97 | 0.000637 | 5.1 | 40.6 | 159.4 | 13.62 | 11.11 | | High | High | High | High | High | High | High | High | High | High | High | 0.000936 | 0.009772 | 3.21 |
| FALSE | High | ISMPDVSLI | [K].ISMPD\ | 1xPhospho | E9Q616 1x | 0.001704 | 5.66E-05 | 1 | 3 | 1 | E9Q616 | AHNAK nuc | E9Q616 [1 | 0 | 1754.953 | 0 | 1.4 | 0.026608 | 17.06 | 57.3 | 142.7 | 15.83 | 35.26 | | High | High | High | High | High | High | High | High | High | High | High | 2.58E-05 | 0.000433 | 2.88 |
| FALSE | High | LSMPQSAA | [R].LSMPQ | 1xPhospho | B2RRF1 1x | 0.002376 | 0.000109 | 1 | 7 | 2 | B2RRF1 | B230339M | B2RRF1 [3 | 0 | 2130.036 | 0 | 1 | 0.001025 | 4.53 | 68.6 | 131.4 | 9.04 | 8.08 | | High | High | High | High | High | High | High | High | High | High | High | 4.94E-05 | 0.000627 | 3.24 |
| FALSE | High | LSMPQSAA | [R].LSMPQ | 1xPhospho | B2RRF1 1x | 7.07E-08 | 0 | 1 | 7 | 6 | B2RRF1 | B230339M | B2RRF1 [3 | 0 | 2146.031 | 0 | 0.13 | 0.298589 | 17.58 | 96.1 | 103.9 | 4.43 | 9.43 | | High | High | High | High | High | High | High | High | High | High | High | 0 | 5.2E-09 | 4.62 |
| FALSE | High | ISMPDFDLI | [K].ISMPDI | 1xPhospho | E9Q616 1x | 0.006653 | 0.00062 | 1 | 3 | 1 | E9Q616 | AHNAK nuc | E9Q616 [2 | 0 | 1853.964 | 0 | 2.49 | 0.008472 | 19.89 | 30.8 | 169.2 | 9.07 | 30.33 | | High | High | High | High | High | High | High | High | High | High | High | 0.000275 | 0.00199 | 2.45 |
| FALSE | High | ISMQDVDL | [K].ISMQD' | 1xCarbami | E9Q616 1x | 0.097743 | 0.007804 | 1 | 3 | 1 | E9Q616 | AHNAK nuc | E9Q616 [5 | 0 | 2091.027 | 0 | 1.89 | 0.040046 | 16.63 | 44.1 | 155.9 | 15.67 | 47.9 | | High | High | High | High | High | High | High | High | High | High | High | 0.00395 | 0.04195 | 2.37 |
| FALSE | High | LSMTDDSK | [R].LSMTD | 1xPhospho | A0A1S6GW | 0.032548 | 0.002167 | 1 | 6 | 1 | A0A1S6GW | Uncharacte | A0A1S6GW | 0 | 1450.69 | 0 | 0.87 | 0.006162 | 5.73 | 71.6 | 128.4 | 6.35 | 14.31 | | High | High | High | High | High | High | High | High | High | High | High | 0.001142 | 0.01183 | 1.6 |
| FALSE | High | LSNASPNPI | [K].LSNASF | 1xPhospho | B9EKX8 1x | 0.010382 | 0.000772 | 1 | 7 | 2 | B9EKX8 | Ell2 protei | B9EKX8 [4 | 0 | 1864.957 | 0 | -1.32 | 0.000139 | 0.89 | 143.6 | 56.4 | 2.43 | 7.15 | | High | High | High | High | High | High | High | High | High | High | High | 0.000382 | 0.003258 | 1.8 |
| FALSE | High | LSNGSFSAI | [R].LSNGSF | 1xPhospho | Q5FWX6 1 | 6.89E-05 | 0 | 1 | 3 | 1 | Q5FWX6 | Serine/thre | Q5FWX6 [4 | 0 | 1846.89 | 0 | 0.57 | 0.018759 | 17.15 | 80.4 | 119.6 | 15.8 | 12.3 | | High | High | High | High | High | High | High | High | High | High | High | 0 | 1.18E-05 | 2.34 |
| FALSE | High | LSNIFVIGK | [R].LSNIFV | 1xPhospho | Q3UXQ6 1 | 0.001815 | 5.66E-05 | 1 | 5 | 1 | Q3UXQ6 | 40S riboso | Q3UXQ6 [2 | 0 | 1528.89 | 0 | 0.04 | 0.95065 | 14.59 | 98.3 | 101.7 | 14.85 | 12.39 | | High | High | High | High | High | High | High | High | High | High | High | 2.58E-05 | 0.000466 | 2.49 |
| FALSE | High | LSNLPTVDI | [R].LSNLPT | 1xCarbami | F8VQD1 1x | 3.6E-05 | 0 | 1 | 14 | 2 | F8VQD1 | Protein pol | F8VQD1 [3 | 0 | 2734.384 | 0 | 0.56 | 0.023025 | 22.28 | 76.9 | 123.1 | 10.76 | 14.05 | | High | High | High | High | High | High | High | High | High | High | High | 0 | 5.69E-06 | 3.47 |
| FALSE | High | LSNPDIFAP | [K].LSNPDI | 1xPhospho | E9Q1Q1 1x | 0.041072 | 0.003094 | 1 | 5 | 1 | E9Q1Q1 | Microtubu | E9Q1Q1 [7 | 0 | 1797.955 | 0 | -1 | 0.007175 | 4.35 | 133.3 | 66.7 | 9.93 | 17.54 | | High | High | High | High | High | High | High | High | High | High | High | 0.001597 | 0.01536 | 2.25 |
| FALSE | High | ISNPSDDTS | [R].ISNPSD | 1xCarbami | Q8BHD8 1 | 0.032548 | 0.002167 | 1 | 1 | 2 | Q8BHD8 | Protein-L-i | Q8BHD8 [3 | 1 | 2404.976 | 0 | 0.45 | 0.068262 | 16.03 | 89.9 | 110.1 | 6.32 | 18.96 | | High | High | High | High | High | High | High | High | High | High | High | 0.001142 | 0.01177 | 2.01 |
| FALSE | High | LSPALPHK | [R].LSPALP | 1xPhospho | D3YYY8 1x | 0.040028 | 0.002867 | 1 | 1 | 1 | D3YYY8 | Rho guanir | D3YYY8 [2 | 0 | 1400.807 | 0 | 2.89 | 0.000197 | 19.14 | 23.7 | 176.3 | 22.75 | 3.82 | | High | High | High | High | High | High | High | High | High | High | High | 0.001484 | 0.01489 | 1.83 |
| FALSE | High | LSPANQR | [K].LSPANC | 1xPhospho | Q62318 1x | 0.067059 | 0.004796 | 1 | 2 | 4 | Q62318 | Transcripti | Q62318 [6 | 0 | 1094.556 | 0 | -1.96 | 0.000175 | 0.71 | 160.8 | 39.2 | 4.98 | 9.41 | | High | High | High | High | High | High | High | High | High | High | High | 0.009441 | 0.1022 | 2.28 |
| FALSE | High | ISPAPSDSS | [R].ISPAPSI | 1xPhospho | Q8CHE4 1x | 1.74E-06 | 0 | 1 | 1 | 1 | Q8CHE4 | PH domain | Q8CHE4 [2 | 0 | 2489.191 | 0 | 0.72 | 0.002585 | 2.92 | 75.6 | 124.4 | 5.17 | 10.74 | | High | High | High | High | High | High | High | High | High | High | High | 0 | 1.89E-07 | 4.71 |
| FALSE | High | ISPAPSDSS | [R].ISPAPSI | 2xPhospho | Q8CHE4 2x | 0.003001 | 0.00016 | 1 | 1 | 1 | Q8CHE4 | PH domain | Q8CHE4 [2 | 0 | 2569.157 | 0 | -0.08 | 0.688871 | 12.42 | 95.7 | 104.3 | 16.33 | 12.17 | | High | High | High | High | High | High | High | High | High | High | High | 7.19E-05 | 0.00082 | 2.54 |
| FALSE | High | LSPAPSVLT | [R].LSPAPS | 1xCarbami | B2RQC2 1x | 2.77E-08 | 0 | 1 | 3 | 3 | B2RQC2 | Ubiquitin c | B2RQC2 [8 | 0 | 2197.098 | 0 | 0.16 | 0.2261 | 13.08 | 97.2 | 102.8 | 8.46 | 9.94 | | High | High | High | High | High | High | High | High | High | High | High | 0 | 2.65E-09 | 5.5 |
| FALSE | High | ISPASPAGS | [R].ISPASP | 2xPhospho | A2AIW0 2x | 7.27E-08 | 0 | 1 | 6 | 2 | A2AIW0 | Serological | A2AIW0 [2 | 0 | 2785.207 | 0 | 0.06 | 0.087828 | 3.14 | 98.7 | 101.3 | 4.64 | 9.75 | | High | High | High | High | High | High | High | High | High | High | High | 0 | 5.35E-09 | 5.26 |
| FALSE | High | ISMQDVDL | [K].ISMQD' | 1xCarbami | E9Q616 1x | 0.000337 | 0 | 1 | 3 | 2 | E9Q616 | AHNAK nuc | E9Q616 [5 | 0 | 2107.022 | 0 | 1.84 | 0.005611 | 12.1 | 40.4 | 159.6 | 16.77 | 25.64 | | High | High | High | High | High | High | High | High | High | High | High | 2.58E-05 | 0.000469 | 3.15 |
| FALSE | High | LSMPDCPN | [R].LSMPD | 2xCarbami | M0QWZ1 1 | 0.100936 | 0.008097 | 1 | 4 | 1 | M0QWZ1 | Protein FAI | M0QWZ1 [ | 0 | 1851.743 | 0 | 1.36 | 0.003101 | 9 | 57.6 | 142.4 | 6.44 | 15.39 | | High | High | High | High | High | High | High | High | High | High | High | 0.004086 | 0.04376 | 2.18 |
| FALSE | High | LSMLSQEI | [R].LSMLS | 1xPhospho | D3Z6D5 1x | 0.059657 | 0.004365 | 1 | 4 | 1 | D3Z6D5 | Fasciculati | D3Z6D5 [1 | 0 | 1945.048 | 0 | 1.27 | 0.000859 | 5.16 | 59.8 | 140.2 | 10.29 | 11.27 | | High | High | High | High | High | High | High | High | High | High | High | 0.002279 | 0.02373 | 1.69 |
| FALSE | High | ISMADVDL | [R].ISMAD | 1xPhospho | E9Q616 1x | 0.000712 | 0 | 1 | 3 | 2 | E9Q616 | AHNAK nuc | E9Q616 [5 | 0 | 1998.038 | 0 | 2.15 | 0.001256 | 19.48 | 35.4 | 164.6 | 15.49 | 13.3 | | High | High | High | High | High | High | High | High | High | High | High | 0.00607 | 0.06614 | 1.89 |
| FALSE | High | ISHSLYSGIE | [R].ISHSLY | 1xPhospho | A5D6P2 1x | 7.42E-09 | 0 | 1 | 23 | 5 | A5D6P2 | Par-3 (Part | A5D6P2 [7 | 0 | 2270.09 | 0 | -0.86 | 0.001117 | 2.19 | 128.8 | 71.2 | 6.46 | 4.04 | | High | High | High | High | High | High | High | High | High | High | High | 0 | 4.1E-10 | 4.46 |
| FALSE | High | LSHSSGYA | [R].LSHSSG | 1xPhospho | Q80W68 1 | 0.018447 | 0.001324 | 1 | 1 | 2 | Q80W68 | Kin of IRRE | Q80W68 [6 | 0 | 1992.938 | 0 | 0.22 | 0.088543 | 13.59 | 91.9 | 108.1 | 11.08 | 6.34 | | High | High | High | High | High | High | High | High | High | High | High | 0.000723 | 0.006257 | 3.15 |
| FALSE | High | ISLASSQAE | [K].ISLASS | 2xPhospho | E9PVX6 2x | 0.080423 | 0.006219 | 1 | 2 | 1 | E9PVX6 | Proliferatic | E9PVX6 [1 | 1 | 2461.25 | 0 | 0.09 | 0.215412 | 3.68 | 98.3 | 101.7 | 3.58 | 3.67 | | High | High | High | High | High | High | High | High | High | High | High | 0.003188 | 0.03341 | 2.17 |
| FALSE | High | ISICSSDK | [R].ISICSSD | 1xCarbami | D3YYI8 1xF | 0.0095 | 0.000736 | 1 | 2 | 1 | D3YYI8 | Histone de | D3YYI8 [40 | 0 | 1447.727 | 0 | -0.5 | 0.009284 | 3.98 | 118.7 | 81.3 | 4.24 | 10.17 | | High | High | High | High | High | High | High | High | High | High | High | 0.000327 | 0.002951 | 2.45 |
| FALSE | High | LSLDLSHR | [R].LSLDLSI | 1xPhospho | E9Q9R9 1x | 0.071071 | 0.005247 | 1 | 5 | 1 | E9Q9R9 | Disks large | E9Q9R9 [1 | 0 | 1249.65 | 0 | 1.16 | 0.010426 | 24.69 | 67.9 | 132.1 | 21 | 14.52 | | High | High | High | High | High | High | High | High | High | High | High | 0.002703 | 0.02883 | 1.88 |

| | | | | | | | | | | | | | | | | | | | | | | | | | | | | | | | | | | | | | | |
|---|---|---|---|---|---|---|---|---|---|---|---|---|---|---|---|---|---|---|---|---|---|---|---|---|---|---|---|---|---|---|---|---|---|---|---|---|---|---|
| FALSE | High | ISIESACEK | [R].ISIESAC | 1xCarbami | Q8VCD5 1x | 0.001831 | 5.66E-05 | 1 | 2 | 1 | Q8VCD5 | Mediator c | Q8VCD5 [9 | 0 | 1574.79 | 0 | -0.24 | 0.779137 | 11.41 | 104.3 | 95.7 | 17.62 | 9.21 | | High | High | High | High | High | High | High | High | High | High | High | 2.58E-05 | 0.00047 | 2.49 |
| FALSE | High | LSLESK | [R].LSLESK. | 1xPhospho | Q0VGR5 1x | 0.067619 | 0.004864 | 1 | 3 | 1 | Q0VGR5 | CDK5 regul | Q0VGR5 [1 | 0 | 1214.68 | 1 | -0.14 | 0.279329 | 6.13 | 103.2 | 96.8 | 9.12 | 6.37 | | High | High | High | High | High | High | High | High | High | High | High | 0.002516 | 0.02729 | 1.37 |
| FALSE | High | ISIFDNLDV | [R].ISIFDNI | 1xPhospho | A0A0N4SU | 1.16E-05 | 0 | 1 | 10 | 5 | A0A0N4SU | Peregrin O | A0A0N4SU | 0 | 3485.625 | 0 | -0.25 | 0.034179 | 5.26 | 106.1 | 93.9 | 4.04 | 10.16 | | High | High | High | High | High | High | High | High | High | High | High | 0 | 1.59E-06 | 4.6 |
| FALSE | High | ISIFDNLDV | [R].ISIFDNI | 1xDeamida | A0A0N4SU | 6.31E-06 | 0 | 1 | 10 | 6 | A0A0N4SU | Peregrin O | A0A0N4SU | 0 | 3486.609 | 0 | -0.25 | 0.018697 | 6.31 | 108.2 | 91.8 | 6.49 | 7.95 | | High | High | High | High | High | High | High | High | High | High | High | 0 | 1.45E-05 | 3.99 |
| FALSE | High | LSLGEFFAE | [R].LSLGEF | 1xPhospho | E9Q4Y4 1x | 0.000151 | 0 | 1 | 1 | 1 | E9Q4Y4 | Centrosom | E9Q4Y4 [5 | 0 | 1477.729 | 0 | -0.38 | 0.002507 | 3.36 | 110.7 | 89.3 | 6.3 | 7.35 | | High | High | High | High | High | High | High | High | High | High | High | 0 | 2.86E-05 | 2.71 |
| FALSE | High | ISIGGGSCG | [R].ISIGGG | 1xCarbami | P50446 1x | 3.17E-08 | 0 | 1 | 5 | 2 | P50446 | Keratin, typ | P50446 [68 | 0 | 1863.862 | 0 | 2.84 | 0.003357 | 27.17 | 23.9 | 176.1 | 6.94 | 23.67 | | High | High | High | High | High | High | High | High | High | High | High | 0.000585 | 0.005411 | 2.44 |
| FALSE | High | LSIGQYDNI | [K].LSIGQY | 1xPhospho | Q5SSZ5 1x | 9.31E-06 | 0 | 1 | 1 | 1 | Q5SSZ5 | Tensin-3 O | Q5SSZ5 [7 | 0 | 2482.227 | 0 | -0.42 | 0.018921 | 7.14 | 116.5 | 83.5 | 9.18 | 9.63 | | High | High | High | High | High | High | High | High | High | High | High | 0 | 1.25E-06 | 4.03 |
| FALSE | High | LSLLIGGLQ | [K].LSLLIG | 1xPhospho | P97329 1x | 0.004491 | 0.000409 | 1 | 1 | 1 | P97329 | Kinesin-like | P97329 [22 | 0 | 2454.351 | 0 | -0.32 | 0.035769 | 12.35 | 110.7 | 89.3 | 8.02 | 15.11 | | High | High | High | High | High | High | High | High | High | High | High | 0.000204 | 0.001286 | 2.23 |
| FALSE | High | LSIMTSENI | [K].LSIMTS | 1xPhospho | Q8CIB5 1x | 0.001986 | 5.66E-05 | 1 | 3 | 1 | Q8CIB5 | Fermitin fa | Q8CIB5 [32 | 0 | 2241.099 | 0 | -0.9 | 0.00125 | 2.86 | 127.3 | 72.7 | 8.41 | 18.01 | | High | High | High | High | High | High | High | High | High | High | High | 2.58E-05 | 0.000515 | 3.13 |
| FALSE | High | LSIMTSENI | [K].LSIMTS | 1xPhospho | Q8CIB5 1x | 0.038677 | 0.002563 | 1 | 3 | 1 | Q8CIB5 | Fermitin fa | Q8CIB5 [32 | 0 | 2257.094 | 0 | -0.91 | 0.005487 | 5.34 | 127.8 | 72.2 | 10.83 | 10.91 | | High | High | High | High | High | High | High | High | High | High | High | 0.001388 | 0.01443 | 1.64 |
| FALSE | High | LSLNNDIFE | [R].LSLNNI | 1xPhospho | Q3TYA6 1x | 0.033986 | 0.002195 | 1 | 5 | 2 | Q3TYA6 | M-phase p | Q3TYA6 [1 | 0 | 2880.319 | 0 | -1.42 | 0.001196 | 3.01 | 145.5 | 54.5 | 10.08 | 20.29 | | High | High | High | High | High | High | High | High | High | High | High | 0.001151 | 0.0124 | 2.49 |
| FALSE | High | LSLNNDIFE | [R].LSLNNI | 2xPhospho | Q3TYA6 2x | 0.000616 | 0 | 1 | 5 | 4 | Q3TYA6 | M-phase p | Q3TYA6 [1 | 0 | 2960.285 | 0 | -1 | 0.001512 | 2.74 | 131.1 | 68.9 | 5.37 | 11.69 | | High | High | High | High | High | High | High | High | High | High | High | 0 | 0.000138 | 3.76 |
| FALSE | High | ISLNTYTPT | [K].ISLNTY | 1xPhospho | E9Q9J4 1x | 0.013909 | 0.001011 | 1 | 6 | 3 | E9Q9J4 | Inositol he | E9Q9J4 [11 | 0 | 1675.907 | 0 | 0.07 | 0.166382 | 6.51 | 97.4 | 102.6 | 2.64 | 5.52 | | High | High | High | High | High | High | High | High | High | High | High | 0.000997 | 0.01048 | 2.12 |
| FALSE | High | LSLPGLLSQ | [R].LSLPGL | 1xPhospho | Q920R0 1x | 0.000668 | 0 | 1 | 2 | 1 | Q920R0 | Alsin OS=M | Q920R0 [4 | 0 | 1675.935 | 0 | -1.58 | 0.000295 | 1.81 | 149.9 | 50.1 | 3.09 | 12.85 | | High | High | High | High | High | High | High | High | High | High | High | 0 | 0.000152 | 3.29 |
| FALSE | High | ISLPGQMT | [K].ISLPGQ | 1xPhospho | Q3TIK1 1x | 1.12E-05 | 0 | 1 | 9 | 1 | Q3TIK1 | Putative ur | Q3TIK1 [21 | 0 | 2192.217 | 0 | 0.19 | 0.405947 | 9.7 | 95.6 | 104.4 | 8.56 | 17.87 | | High | High | High | High | High | High | High | High | High | High | High | 0 | 1.55E-06 | 2.63 |
| FALSE | High | ISLPGQMT | [K].ISLPGQ | 1xPhospho | Q3TIK1 1x | 1.83E-05 | 0 | 1 | 9 | 2 | Q3TIK1 | Putative ur | Q3TIK1 [21 | 0 | 2208.212 | 0 | 0.19 | 0.078897 | 12.45 | 91.8 | 108.2 | 5.53 | 13.25 | | High | High | High | High | High | High | High | High | High | High | High | 0 | 2.68E-06 | 2.9 |
| FALSE | High | ISLPLPTFSS | [R].ISLPLPT | 1xPhospho | P20152 1x | 2.36E-05 | 0 | 1 | 7 | 4 | P20152 | Vimentin C | P20152 [41 | 0 | 1867.029 | 0 | 2.87 | 0.010295 | 26.01 | 22.6 | 177.4 | 11.92 | 34.59 | | High | High | High | High | High | High | High | High | High | High | High | 0 | 9.85E-06 | 2.59 |
| FALSE | High | LSLPTTCSG | [R].LSLPTT | 1xCarbami | Q03160 1x | 3.27E-06 | 0 | 1 | 2 | 1 | Q03160 | Growth fac | Q03160 [4 | 0 | 2167.078 | 0 | -0.89 | 0.004314 | 6.66 | 131.9 | 68.1 | 8.93 | 14.73 | | High | High | High | High | High | High | High | High | High | High | High | 0 | 3.83E-07 | 3.77 |
| FALSE | High | LSLQHTQQ | [K].LSLQHT | 1xPhospho | Q9D3A8 1x | 0.041425 | 0.003126 | 1 | 6 | 2 | Q9D3A8 | Carboxyl-te | Q9D3A8 [1 | 1 | 3493.505 | 0 | 2.73 | 0.000762 | 25.68 | 26.2 | 173.8 | 46.48 | 12.62 | | High | High | High | High | High | High | High | High | High | High | High | 0.001609 | 0.01551 | 2.55 |
| FALSE | High | ISLSPFSASI | [K].ISLSPFS | 1xPhospho | F8VQ54 1x | 2.95E-05 | 0 | 1 | 5 | 5 | F8VQ54 | Endoribon | F8VQ54 [1 | 0 | 2326.141 | 0 | 0.24 | 0.005655 | 2.48 | 90.3 | 109.7 | 3.38 | 6.18 | | High | High | High | High | High | High | High | High | High | High | High | 0 | 6E-05 | 3.31 |
| FALSE | High | LSLSPISLAE | [K].LSLSPIS | 1xPhospho | J3QNH8 1x | 7.23E-06 | 0 | 1 | 1 | 5 | J3QNH8 | Uncharacte | J3QNH8 [2 | 0 | 2675.328 | 0 | 1.17 | 0.002946 | 25.74 | 58 | 142 | 11.23 | 14.96 | | High | High | High | High | High | High | High | High | High | High | High | 0 | 9.43E-07 | 3.26 |
| FALSE | High | LSLSTGTPC | [R].LSLSTG | 1xPhospho | Q91VY5 1x | 0.001349 | 0 | 1 | 6 | 2 | Q91VY5 | Lysine-spe | Q91VY5 [1 | 0 | 2503.237 | 0 | -1.11 | 0.001663 | 2.82 | 135.4 | 64.6 | 5.66 | 17.35 | | High | High | High | High | High | High | High | High | High | High | High | 0 | 0.000333 | 2.75 |
| FALSE | High | LSISTPDR | [K].LSISTPI | 1xPhospho | Q8CI75-2 1 | 0.020142 | 0.001472 | 1 | 3 | 2 | Q8CI75-2 | Isoform 2 c | Q8CI75-2 [ | 0 | 1197.608 | 0 | 0.22 | 0.004273 | 5.27 | 93.3 | 106.7 | 6.73 | 3.45 | | High | High | High | High | High | High | High | High | High | High | High | 0.000984 | 0.01013 | 2.09 |
| FALSE | High | LSLTPTDVK | [K].LSLTPTI | 1xPhospho | A0A0U1RP | 0.002576 | 0.000109 | 1 | 7 | 4 | A0A0U1RP | Ataxin-2-lik | A0A0U1RP | 0 | 1511.849 | 0 | 0.2 | 0.401601 | 18.53 | 93.5 | 106.5 | 20.07 | 3.61 | | High | High | High | High | High | High | High | High | High | High | High | 0.002403 | 0.02586 | 2.29 |
| FALSE | High | LSEADNR | [K].LSEADN | 1xPhospho | Q4FK79 1x | 0.020862 | 0.001472 | 1 | 3 | 1 | Q4FK79 | Putative ur | Q4FK79 [1 | 0 | 1113.514 | 0 | 0.52 | 0.126798 | 10 | 82 | 118 | 19.83 | 10.82 | | High | High | High | High | High | High | High | High | High | High | High | 0.000786 | 0.007132 | 2.11 |
| FALSE | High | LSPAYSLGS | [R].LSPAYS | 1xPhospho | D3Z0X5 1x | 2.93E-05 | 0 | 1 | 7 | 1 | D3Z0X5 | Pleckstrin h | D3Z0X5 [5 | 0 | 1885.962 | 0 | 0.96 | 0.002026 | 15.1 | 69.1 | 130.9 | 13.5 | 4.15 | | High | High | High | High | High | High | High | High | High | High | High | 0 | 4.53E-06 | 3.83 |
| FALSE | High | ISDYFEYQG | [K].ISDYFEY | 1xDeamida | Q8C0V0 1x | 8.4E-05 | 0 | 1 | 2 | 7 | Q8C0V0 | Serine/thre | Q8C0V0 [1 | 0 | 2185.964 | 0 | -0.42 | 0.003805 | 5.29 | 113.4 | 86.6 | 4.45 | 5.87 | | High | High | High | High | High | High | High | High | High | High | High | 0 | 1.47E-05 | 4.35 |
| FALSE | High | ISDYFEFAG | [K].ISDYFEI | 2xPhospho | B1ASU9 2x | 0.001278 | 0 | 1 | 3 | 3 | B1ASU9 | Serine/thre | B1ASU9 [8 | 0 | 2290.962 | 0 | 0.38 | 0.040593 | 16.53 | 85.1 | 114.9 | 11.66 | 14.75 | | High | High | High | High | High | High | High | High | High | High | High | 0.002841 | 0.02944 | 3.01 |
| FALSE | High | LQQQAALS | [R].LQQQA | 1xPhospho | Q8VHR5 1x | 0.000143 | 0 | 1 | 5 | 1 | Q8VHR5 | Transcripti | Q8VHR5 [4 | 0 | 2522.363 | 0 | 0.22 | 0.070942 | 20.72 | 89.2 | 110.8 | 8.97 | 8.84 | | High | High | High | High | High | High | High | High | High | High | High | 0 | 2.68E-05 | 2.75 |
| FALSE | High | IQQQAGTV | [R].IQQQA | 1xPhospho | Q99LG1 1x | 8.47E-08 | 0 | 1 | 1 | 6 | Q99LG1 | Transmem | Q99LG1 [9 | 0 | 3251.417 | 0 | -0.62 | 0.00131 | 3.29 | 120.9 | 79.1 | 5.78 | 4.61 | | High | High | High | High | High | High | High | High | High | High | High | 0 | 6.36E-09 | 5.82 |
| FALSE | High | IQQQAGTV | [R].IQQQA | 1xDeamida | Q99LG1 1x | 1.34E-06 | 0 | 1 | 1 | 6 | Q99LG1 | Transmem | Q99LG1 [9 | 0 | 3252.402 | 0 | -0.63 | 0.000687 | 2.93 | 120.7 | 79.3 | 2.96 | 6.31 | | High | High | High | High | High | High | High | High | High | High | High | 0 | 1.41E-07 | 5.79 |
| FALSE | High | IQQQAGTV | [R].IQQQA | 2xPhospho | Q99LG1 2x | 0.000284 | 0 | 1 | 1 | 3 | Q99LG1 | Transmem | Q99LG1 [9 | 0 | 3331.384 | 0 | -0.49 | 0.024064 | 2.2 | 113.5 | 86.5 | 11.62 | 5.27 | | High | High | High | High | High | High | High | High | High | High | High | 0 | 8.16E-05 | 3.33 |
| FALSE | High | IQQQAGTV | [R].IQQQA | 1xDeamida | Q99LG1 2x | 0.000308 | 0 | 1 | 1 | 3 | Q99LG1 | Transmem | Q99LG1 [9 | 0 | 3332.368 | 0 | -0.19 | 0.182946 | 16.74 | 103.6 | 96.4 | 13.55 | 6.55 | | High | High | High | High | High | High | High | High | High | High | High | 0 | 9.53E-05 | 4.58 |
| FALSE | High | IQQQLGEE | [R].IQQQL | 1xPhospho | S4R189 1x | 0.001189 | 0 | 1 | 6 | 1 | S4R189 | Rho guanir | S4R189 [95 | 0 | 1664.821 | 0 | -0.3 | 0.132276 | 15.31 | 105.8 | 94.2 | 12.16 | 16.48 | | High | High | High | High | High | High | High | High | High | High | High | 0 | 0.00029 | 2.22 |
| FALSE | High | LQQSGPK | [R].LQQSG | 1xDeamida | A0A0A6YY | 0.036731 | 0.00249 | 1 | 2 | 1 | A0A0A6YY | Immunoglo | A0A0A6YY | 0 | 1296.696 | 0 | -0.96 | 0.000218 | 1.01 | 135.4 | 64.6 | 3.63 | 6.61 | | High | High | High | High | High | High | High | High | High | High | High | 0.001308 | 0.0136 | 1.79 |
| FALSE | High | IQQTGNTS | [R].IQQTGI | 1xPhospho | Q9JMK2 1x | 0.071662 | 0.005278 | 1 | 3 | 2 | Q9JMK2 | Casein kina | Q9JMK2 [3 | 0 | 1410.694 | 0 | -0.47 | 0.003786 | 3.8 | 119.3 | 80.7 | 6.56 | 9.59 | | High | High | High | High | High | High | High | High | High | High | High | 0.009466 | 0.1027 | 1.36 |
| FALSE | High | LQQTVPAD | [K].LQQTV | 1xPhospho | Q80Z37 1x | 1.52E-06 | 0 | 1 | 2 | 2 | Q80Z37 | E3 ubiquiti | Q80Z37 [9 | 0 | 1995.02 | 0 | 0.23 | 0.092066 | 7.99 | 91.9 | 108.1 | 4.44 | 9.22 | | High | High | High | High | High | High | High | High | High | High | High | 0 | 1.62E-07 | 3.81 |
| FALSE | High | LQSEDSTK | [K].LQSEDS | 1xPhospho | Q9Z277 1x | 0.039011 | 0.002718 | 1 | 2 | 1 | Q9Z277 | Tyrosine-p | Q9Z277 [1 | 0 | 1445.729 | 0 | 0.63 | 0.004557 | 12.1 | 82.2 | 117.8 | 7.9 | 7.47 | | High | High | High | High | High | High | High | High | High | High | High | 0.001421 | 0.01445 | 1.84 |
| FALSE | High | LQSESASG | [K].LQSESA | 1xPhospho | Q9CZU3 1x | 0.000434 | 0 | 1 | 3 | 4 | Q9CZU3 | Superkiller | Q9CZU3 [5 | 0 | 1501.766 | 0 | -0.58 | 0.000399 | 2.9 | 120.1 | 79.9 | 3.43 | 4.14 | | High | High | High | High | High | High | High | High | High | High | High | 0 | 9.34E-05 | 1.25 |
| FALSE | High | LQSFDSPEF | [K].LQSFDS | 1xPhospho | S4R1W5 1x | 0.080423 | 0.006219 | 1 | 5 | 1 | S4R1W5 | RNA-bindir | S4R1W5 [1 | 0 | 1387.646 | 0 | -0.26 | 0.009535 | 2.41 | 108.1 | 91.9 | 5.28 | 1.97 | | High | High | High | High | High | High | High | High | High | High | High | 0.003188 | 0.03355 | 1.79 |
| FALSE | High | LQSFDSPEF | [K].LQSFDS | 1xPhospho | S4R1W5 1x | 0.085162 | 0.006376 | 1 | 5 | 2 | S4R1W5 | RNA-bindir | S4R1W5 [1 | 1 | 1744.904 | 0 | -0.34 | 0.002581 | 4.57 | 110.9 | 89.1 | 6.44 | 3.66 | | High | High | High | High | High | High | High | High | High | High | High | 0.003291 | 0.03574 | 1.81 |
| FALSE | High | LQQGFVTP | [K].LQQGF | 1xPhospho | A2APB8 1x | 0.039347 | 0.002793 | 1 | 3 | 1 | A2APB8 | Targeting p | A2APB8 [6 | 0 | 1668.949 | 0 | 0.37 | 0.02481 | 9.64 | 85.3 | 114.7 | 19.13 | 8.17 | | High | High | High | High | High | High | High | High | High | High | High | 0.001453 | 0.01461 | 2.08 |
| FALSE | High | LQSIGTENT | [R].LQSIGT | 1xPhospho | A6ZI44 1xP | 2.28E-06 | 0 | 1 | 11 | 2 | A6ZI44 | Fructose-b | A6ZI44 [98 | 0 | 1799.837 | 0 | 0.55 | 0.003183 | 4.66 | 81.1 | 118.9 | 2.99 | 7.56 | | High | High | High | High | High | High | High | High | High | High | High | 0 | 2.57E-07 | 3.64 |
| FALSE | High | LQSLLTSQE | [R].LQSLLT | 2xPhospho | Q64735 2x | 0.007605 | 0.000664 | 1 | 2 | 1 | Q64735 | Compleme | Q64735 [4 | 1 | 3167.471 | 0 | 1.35 | 0.001301 | 27.87 | 53.8 | 146.2 | 19.49 | 6.54 | | High | High | High | High | High | High | High | High | High | High | High | 0.000294 | 0.002304 | 3.41 |
| FALSE | High | LQSPLPGES | [K].LQSPLP | 1xPhospho | F8VPJ6 1xP | 0.004106 | 0.00036 | 1 | 1 | 1 | F8VPJ6 | Regulatory | F8VPJ6 [32 | 0 | 1751.934 | 0 | -0.65 | 0.000714 | 0.97 | 120.4 | 79.6 | 3.48 | 6.86 | | High | High | High | High | High | High | High | High | High | High | High | 0.000161 | 0.001167 | 2.61 |
| FALSE | High | LQSPQAVL | [K].LQSPQ/ | 1xPhospho | Q8JZY2 1x | 0.003558 | 0.00021 | 1 | 2 | 1 | Q8JZY2 | COMM dor | Q8JZY2 [15 | 0 | 2006.145 | 0 | -0.14 | 0.262446 | 1.61 | 104.7 | 95.3 | 5.3 | 9.57 | | High | High | High | High | High | High | High | High | High | High | High | 9.42E-05 | 0.000988 | 3 |
| FALSE | High | LQSQVPGT | [K].LQSQVI | 1xPhospho | Q8VDM1 1 | 1.87E-06 | 0 | 1 | 2 | 1 | Q8VDM1 | Zinc finger | Q8VDM1 [ | 1 | 2332.15 | 0 | 0.7 | 0.003322 | 7.22 | 75.9 | 124.1 | 8.83 | 7.77 | | High | High | High | High | High | High | High | High | High | High | High | 0 | 2.06E-07 | 3.88 |
| FALSE | High | IQSSLSVNN | [K].IQSSLS | 1xPhospho | A0A087WF | 0.000322 | 0 | 1 | 5 | 4 | A0A087WF | MyoD fami | A0A087WF | 0 | 1943.025 | 0 | 1.57 | 0.000154 | 3.93 | 49 | 151 | 7.82 | 3.38 | | High | High | High | High | High | High | High | High | High | High | High | 0.000997 | 0.01035 | 2.75 |
| FALSE | High | IQSSQSAE | [R].IQSSQS | 1xPhospho | P56183 1x | 0.001674 | 5.66E-05 | 1 | 5 | 3 | P56183 | Ribosomal | P56183 [48 | 0 | 1158.524 | 0 | 0.32 | 0.065703 | 13.42 | 93.7 | 106.3 | 13.02 | 16.04 | | High | High | High | High | High | High | High | High | High | High | High | 2.58E-05 | 0.000427 | 1.77 |
| FALSE | High | LQSTEFSPS | [R].LQSTEF | 1xPhospho | Q9WV69-2 | 0.033694 | 0.002195 | 1 | 2 | 2 | Q9WV69-2 | Isoform 2 c | Q9WV69-2 | 0 | 3867.917 | 0 | -0.14 | 0.034866 | 4.71 | 108.5 | 91.5 | 10.72 | 14.95 | | High | High | High | High | High | High | High | High | High | High | High | 0.001151 | 0.01231 | 2.51 |
| FALSE | High | IQSTPVK | [K].IQSTPV | 1xPhospho | Q0PD67 1x | 0.053926 | 0.00384 | 1 | 6 | 4 | Q0PD67 | RAB1, men | Q0PD67 [1 | 0 | 1310.749 | 0 | -0.54 | 0.000779 | 2.99 | 115.6 | 84.4 | 5.7 | 6.48 | | High | High | High | High | High | High | High | High | High | High | High | 0.005613 | 0.06171 | 1.68 |
| FALSE | High | LQSVQATG | [R].LQSVQ | 1xPhospho | E9PWI3 1x | 1.32E-05 | 0 | 1 | 3 | 2 | E9PWI3 | Mutated in | E9PWI3 [4 | 0 | 1693.847 | 0 | 2.21 | 0.000466 | 19.55 | 35.1 | 164.9 | 13.29 | 7.08 | | High | High | High | High | High | High | High | High | High | High | High | 0 | 1.86E-06 | 3.01 |
| FALSE | High | LQSWTPAS | [R].LQSWT | 1xPhospho | Q542Q3 1x | 0.094641 | 0.007408 | 1 | 1 | 1 | Q542Q3 | G two S ph | Q542Q3 [4 | 0 | 1354.672 | 0 | 0.7 | 0.008048 | 22.28 | 79.8 | 120.2 | 12.23 | 13.58 | | High | High | High | High | High | High | High | High | High | High | High | 0.003754 | 0.0405 | 1.4 |
| FALSE | High | LQTHDVSS | [K].LQTHD | 1xPhospho | D9HP81 1x | 0.006478 | 0.00062 | 1 | 5 | 1 | D9HP81 | Regulating | D9HP81 [8 | 0 | 2582.369 | 0 | 0.2 | 0.475963 | 15.79 | 94.5 | 105.5 | 7.21 | 13.95 | | High | High | High | High | High | High | High | High | High | High | High | 0.000275 | 0.001922 | 1.73 |
| FALSE | High | LQTHPSLG | [K].LQTHP | 1xPhospho | P70331 1x | 0.005135 | 0.000551 | 1 | 5 | 1 | P70331 | MyoD fami | P70331 [13 | 0 | 1660.837 | 0 | -1.33 | 0.001414 | 2.69 | 140.6 | 59.4 | 9.87 | 8.55 | | High | High | High | High | High | High | High | High | High | High | High | 0.000246 | 0.001488 | 2.01 |
| FALSE | High | LQTHPVCL | [K].LQTHP | 1xCarbami | Q7TS72 1x | 3.09E-08 | 0 | 1 | 1 | 2 | Q7TS72 | Inositol-tri | Q7TS72 [2 | 0 | 2565.242 | 0 | -0.67 | 0.002311 | 5.5 | 119.9 | 80.1 | 6.57 | 4.56 | | High | High | High | High | High | High | High | High | High | High | High | 0 | 2.04E-09 | 3.58 |
| FALSE | High | LQSIGTENT | [R].LQSIGT | 1xPhospho | A6ZI44 1xP | 0.02933 | 0.001857 | 1 | 11 | 1 | A6ZI44 | Fructose-b | A6ZI44 [98 | 1 | 1955.939 | 0 | 0.59 | 0.011587 | 21.66 | 80.1 | 119.9 | 7.39 | 10.98 | | High | High | High | High | High | High | High | High | High | High | High | 0.000997 | 0.01047 | 0.91 |
| FALSE | High | LQTTDNLL | [R].LQTTDI | 1xPhospho | A0A087WS | 0.000237 | 0 | 1 | 9 | 1 | A0A087WS | Signal tran | A0A087WS | 0 | 2678.193 | 0 | 0.36 | 0.151502 | 30.86 | 85.7 | 114.3 | 16.37 | 13.62 | | High | High | High | High | High | High | High | High | High | High | High | 0 | 4.73E-05 | 3.4 |
| FALSE | High | IQQFDDGG | [R].IQQFDI | 1xDeamida | G5E8R4 1x | 0.001437 | 5.66E-05 | 1 | 3 | 2 | G5E8R4 | SAPS doma | G5E8R4 [6 | 0 | 2679.175 | 0 | -0.06 | 0.978662 | 28.73 | 96 | 104 | 15.24 | 10.37 | | High | High | High | High | High | High | High | High | High | High | High | 2.58E-05 | 0.000358 | 4.67 |
| FALSE | High | IQQFDDEE | [R].IQQFDI | 1xDeamida | B2RUA5 2x | 0.000362 | 0 | 1 | 2 | 1 | B2RUA5 | SAPS doma | B2RUA5 [6 | 0 | 4205.65 | 0 | -0.88 | 0.000434 | 1.07 | 129.6 | 70.4 | 3.53 | 4.48 | | High | High | High | High | High | High | High | High | High | High | High | 0 | 7.63E-05 | 4.68 |
| FALSE | High | LQNSDDDE | [K].LQNSDI | 1xDeamida | Q5XJE5 1x | 0.003997 | 0.000311 | 1 | 2 | 2 | Q5XJE5 | RNA polym | Q5XJE5 [1 | 0 | 1731.773 | 0 | -0.47 | 0.018897 | 6.62 | 117.3 | 82.7 | 9.54 | 8.26 | | High | High | High | High | High | High | High | High | High | High | High | 0.00014 | 0.001122 | 3.07 |
| FALSE | High | LQNSDDED | [K].LQNSDI | 1xPhospho | Q5XJE5 1x | 0.065952 | 0.004766 | 1 | 2 | 1 | Q5XJE5 | RNA polym | Q5XJE5 [1 | 1 | 2518.098 | 0 | 0.05 | 0.476393 | 7.82 | 99.7 | 100.3 | 13.45 | 13.43 | | High | High | High | High | High | High | High | High | High | High | High | 0.002455 | 0.02648 | 1.88 |
| FALSE | High | LQNSNSAG | [K].LQNSN | 1xPhospho | G3UZM1 1 | 6.53E-05 | 0 | 1 | 2 | 1 | G3UZM1 | Probable Ji | G3UZM1 [ | 0 | 1451.721 | 0 | 0.06 | 0.963715 | 4.19 | 97.8 | 102.2 | 10.91 | 6.92 | | High | High | High | High | High | High | High | High | High | High | High | 0 | 1.11E-05 | 3.25 |
| FALSE | High | LQNSPTEM | [R].LQNSP | 1xPhospho | Q8BQ33 1x | 1.77E-05 | 0 | 1 | 1 | 4 | Q8BQ33 | Treslin OS= | Q8BQ33 [1 | 0 | 2746.362 | 0 | -0.75 | 0.004571 | 2.74 | 125.3 | 74.7 | 1.98 | 18.73 | | High | High | High | High | High | High | High | High | High | High | High | 0 | 4.24E-06 | 3.85 |
| FALSE | High | LQNSPTEM | [R].LQNSP | 1xPhospho | Q8BQ33 1x | 0.000138 | 0 | 1 | 1 | 4 | Q8BQ33 | Treslin OS= | Q8BQ33 [1 | 0 | 2762.357 | 0 | -0.28 | 0.141276 | 15.07 | 109.6 | 90.4 | 7.09 | 14.51 | | High | High | High | High | High | High | High | High | High | High | High | 0 | 2.59E-05 | 3.79 |
| FALSE | High | IQNTGDYY | [K].IQNTGI | 1xPhospho | P35235 1x | 0.00047 | 0 | 1 | 4 | 1 | P35235 | Tyrosine-p | P35235 [5 | 0 | 2274.073 | 0 | 0.02 | 0.36316 | 10.92 | 99.2 | 100.8 | 12.57 | 5.89 | | High | High | High | High | High | High | High | High | High | High | High | 0 | 0.000102 | 2.44 |
| FALSE | High | LQPEQGSP | [R].LQPEQ | 1xPhospho | F6RJ39 1xP | 0.001137 | 0 | 2 | 4 | 4 | F6RJ39; Q8 | Apoptotic | F6RJ39 [66 | 0 | 1521.808 | 0 | -0.05 | 0.291149 | 6.86 | 99.4 | 100.6 | 8.11 | 6.38 | Shared | High | High | High | High | High | High | High | High | High | High | High | 0.0054 | 0.05929 | 1.66 |
| FALSE | High | IQPLEPDSP | [K].IQPLEP | 1xPhospho | E9Q444 1x | 3.97E-08 | 0 | 1 | 9 | 2 | E9Q444 | Zinc finger | E9Q444 [1 | 0 | 2869.464 | 0 | 0.41 | 0.014137 | 15.89 | 85.9 | 114.1 | 5.67 | 7.02 | | High | High | High | High | High | High | High | High | High | High | High | 0 | 2.72E-09 | 4.53 |
| FALSE | High | LQPLPSCE | [K].LQPLPS | 2xCarbami | B7ZCT1 1x | 0.000174 | 0 | 1 | 5 | 1 | B7ZCT1 | Regulator c | B7ZCT1 [9 | 0 | 2348.18 | 0 | -0.08 | 0.404665 | 1.52 | 101.4 | 98.6 | 5.92 | 6.26 | | High | High | High | High | High | High | High | High | High | High | High | 0 | 3.35E-05 | 3.49 |
| FALSE | High | LQPLSPVPS | [K].LQPLSP | 1xPhospho | Q3V3R1 1x | 0.00129 | 0 | 1 | 1 | 1 | Q3V3R1 | Monofunc | Q3V3R1 [3 | 0 | 1960.035 | 0 | 0.31 | 0.003221 | 6.04 | 87.9 | 112.1 | 9.19 | 6.7 | | High | High | High | High | High | High | High | High | High | High | High | 0 | 0.000316 | 3 |
| FALSE | High | LQPPSPLGI | [R].LQPPSF | 1xPhospho | A0A087WC | 4.51E-05 | 0 | 1 | 8 | 2 | A0A087WC | Snf2-relate | A0A087WC | 0 | 3589.59 | 0 | -0.01 | 0.560401 | 8.45 | 97.3 | 102.7 | 6.57 | 5.31 | | High | High | High | High | High | High | High | High | High | High | High | 0 | 7.36E-06 | 4.36 |
| FALSE | High | LQPPSPLGI | [R].LQPPSF | 1xDeamida | A0A087WC | 3.2E-05 | 0 | 1 | 8 | 1 | A0A087WC | Snf2-relate | A0A087WC | 0 | 3590.574 | 0 | -0.3 | 0.06656 | 21.79 | 115.8 | 84.2 | 11.04 | 12.39 | | High | High | High | High | High | High | High | High | High | High | High | 0 | 5E-06 | 4.86 |
| FALSE | High | LQPQEISPF | [RK].LQPQ | 1xPhospho | P34152-4 1 | 0.003402 | 0.00021 | 1 | 6 | 1 | P34152-4 | Isoform 4 c | P34152-4 [ | 0 | 2085.058 | 0 | 1.09 | 0.00362 | 32.77 | 63.9 | 136.1 | 16.91 | 6.55 | | High | High | High | High | High | High | High | High | High | High | High | 9.42E-05 | 0.000941 | 1.76 |
| FALSE | High | IQQFDDGG | [R].IQQFDI | 1xPhospho | G5E8R4 1x | 3.96E-07 | 0 | 1 | 3 | 8 | G5E8R4 | SAPS doma | G5E8R4 [6 | 0 | 2678.191 | 0 | -0.24 | 0.057461 | 2.4 | 104.9 | 95.1 | 5.83 | 8.67 | | High | High | High | High | High | High | High | High | High | High | High | 0 | 3.59E-08 | 3.51 |
| FALSE | High | IQPQLPDEI | [K].IQPQLP | 1xPhospho | Q8K039 1x | 0.001049 | 0 | 1 | 1 | 1 | Q8K039 | Uncharacte | Q8K039 [3 | 0 | 2001.912 | 0 | 0.12 | 0.900699 | 16.24 | 97.1 | 102.9 | 54.14 | 40.17 | | High | High | High | High | High | High | High | High | High | High | High | 0 | 0.000251 | 2.5 |
| FALSE | High | IQPSFQSPV | [K].IQPSFQ | 1xPhospho | Q8JZU6 1x | 0.049549 | 0.00361 | 1 | 4 | 1 | Q8JZU6 | PX domain | Q8JZU6 [1 | 0 | 1668.913 | 0 | 3.09 | 0.000225 | 25.35 | 21.8 | 178.2 | 28.27 | 6.31 | | High | High | High | High | High | High | High | High | High | High | High | 0.001889 | 0.01903 | 2.15 |
| FALSE | High | LQPSRPAA | [R].LQPSRF | 1xPhospho | Q3TZW5 1 | 3.63E-05 | 0 | 1 | 4 | 2 | Q3TZW5 | Putative ur | Q3TZW5 [9 | 0 | 2839.492 | 0 | -0.95 | 0.002406 | 2.48 | 129.1 | 70.9 | 10.58 | 9.94 | | High | High | High | High | High | High | High | High | High | High | High | 0 | 5.76E-06 | 2.55 |
| FALSE | High | LQPSSSPET | [K].LQPSSS | 1xPhospho | A0A0U1RP | 3.48E-06 | 0 | 1 | 7 | 6 | A0A0U1RP | Ataxin-2-lik | A0A0U1RP | 0 | 2124.021 | 0 | 0.84 | 0.000677 | 0.68 | 71.8 | 128.2 | 5.06 | 5.56 | | High | High | High | High | High | High | High | High | High | High | High | 0 | 4.13E-07 | 3.44 |
| FALSE | High | LQPSTPEHI | [R].LQPSTF | 1xPhospho | Q3TQ38 1x | 0.005717 | 0.00062 | 1 | 5 | 2 | Q3TQ38 | Putative ur | Q3TQ38 [1 | 0 | 1574.834 | 0 | 0.36 | 0.01004 | 13.31 | 85.5 | 114.5 | 6.53 | 6.97 | | High | High | High | High | High | High | High | High | High | High | High | 0.000275 | 0.002059 | 1.71 |
| FALSE | High | LQQAEQSP | [R].LQQAE | 1xPhospho | Q8BH05 1x | 0.081752 | 0.006219 | 1 | 1 | 1 | Q8BH05 | Zinc finger | Q8BH05 [4 | 0 | 2366.263 | 0 | 0.92 | 0.002353 | 11.93 | 73.3 | 126.7 | 9.52 | 13.81 | | High | High | High | High | High | High | High | High | High | High | High | 0.0032 | 0.03402 | 2.09 |
| FALSE | High | IQQALTSPL | [R].IQQALT | 1xPhospho | Q501J7-3 1 | 0.000411 | 0 | 1 | 3 | 3 | Q501J7-3 | Isoform 3 c | Q501J7-3 [ | 0 | 2464.316 | 0 | 1.06 | 0.001804 | 13.57 | 64.9 | 135.1 | 7.61 | 8.65 | | High | High | High | High | High | High | High | High | High | High | High | 0 | 8.74E-05 | 2.46 |
| FALSE | High | IQQALTSPL | [R].IQQALT | 2xPhospho | Q501J7-3 2 | 0.004491 | 0.000409 | 1 | 3 | 1 | Q501J7-3 | Isoform 3 c | Q501J7-3 [ | 0 | 2544.283 | 0 | 0.58 | 0.009495 | 18.61 | 80.3 | 119.7 | 14.03 | 11.3 | | High | High | High | High | High | High | High | High | High | High | High | 0.000183 | 0.001281 | 2.38 |
| FALSE | High | IQQDSESG | [R].IQQDSE | 1xPhospho | S4R221 1x | 3.45E-06 | 0 | 1 | 9 | 2 | S4R221 | Rho GTPas | S4R221 [2 | 0 | 2523.076 | 0 | -0.27 | 0.076241 | 14.12 | 115 | 85 | 7.06 | 20.96 | | High | High | High | High | High | High | High | High | High | High | High | 0 | 4.08E-07 | 4.07 |
| FALSE | High | IQQDSESG | [R].IQQDSE | 1xPhospho | S4R221 1x | 0.002486 | 0.000109 | 1 | 9 | 6 | S4R221 | Rho GTPas | S4R221 [2 | 0 | 2539.07 | 0 | -0.75 | 0.045409 | 8.25 | 125.4 | 74.6 | 10.89 | 19.77 | | High | High | High | High | High | High | High | High | High | High | High | 9.42E-05 | 0.001014 | 3.58 |
| FALSE | High | IQQDSESG | [R].IQQDSE | 2xPhospho | S4R221 2x | 0.007274 | 0.00062 | 1 | 9 | 1 | S4R221 | Rho GTPas | S4R221 [2 | 0 | 2619.037 | 0 | -0.78 | 0.00062 | 4.81 | 126.3 | 73.7 | 7.16 | 18.31 | | High | High | High | High | High | High | High | High | High | High | High | 0.000275 | 0.002208 | 2.95 |
| FALSE | High | LQQENNH | [R].LQQEN | 1xCarbami | P97360 1x | 0.010663 | 0.000772 | 1 | 6 | 3 | P97360 | Transcripti | P97360 [22 | 0 | 4315.955 | 0 | -0.07 | 0.264764 | 18.73 | 105.4 | 94.6 | 8.02 | 20.73 | | High | High | High | High | High | High | High | High | High | High | High | 0.000538 | 0.004558 | 3.02 |
| FALSE | High | LQQENNH | [R].LQQEN | 1xCarbami | P97360 2x | 0.079766 | 0.00613 | 1 | 6 | 1 | P97360 | Transcripti | P97360 [22 | 0 | 4394.938 | 0 | 0.75 | 0.010011 | 13.93 | 78.1 | 121.9 | 6.44 | 14.1 | | High | High | High | High | High | High | High | High | High | High | High | 0.003148 | 0.03296 | 2.95 |
| FALSE | High | IQQFDDEE | [R].IQQFDI | 2xPhospho | B2RUA5 2x | 1.23E-06 | 0 | 1 | 2 | 2 | B2RUA5 | SAPS doma | B2RUA5 [6 | 0 | 4204.666 | 0 | -0.65 | 0.002491 | 3.7 | 119.3 | 80.7 | 11.34 | 14.15 | | High | High | High | High | High | High | High | High | High | High | High | 0 | 1.28E-07 | 4.77 |
| FALSE | High | IQPQLPDEI | [K].IQPQLP | 1xPhospho | Q8K039 1x | 0.006713 | 0.00062 | 1 | 1 | 3 | Q8K039 | Uncharacte | Q8K039 [3 | 1 | 3724.904 | 0 | -0.16 | 0.030876 | 7.99 | 104.7 | 95.3 | 4.87 | 6.69 | | High | High | High | High | High | High | High | High | High | High | High | 0.000872 | 0.008647 | 3.33 |
| FALSE | High | IQVQEVDG | [R].IQVQEV | 2xPhospho | Q80ZX8 2x | 0.025738 | 0.001645 | 1 | 3 | 1 | Q80ZX8 | Sperm-ass | Q80ZX8 [7 | 0 | 3315.462 | 0 | -1.25 | 0.000717 | 1.91 | 137.8 | 62.2 | 11.23 | 11.19 | | High | High | High | High | High | High | High | High | High | High | High | 0.000872 | 0.009036 | 3.25 |
| FALSE | High | LQVVSESEI | [K].LQVVSI | 1xPhospho | Q3UYD0 1x | 7.06E-07 | 0 | 1 | 2 | 7 | Q3UYD0 | Putative ur | Q3UYD0 [3 | 0 | 3679.788 | 0 | -0.33 | 0.049292 | 9.99 | 109.9 | 90.1 | 10.04 | 12.63 | | High | High | High | High | High | High | High | High | High | High | High | 0 | 6.84E-08 | 5.9 |
| FALSE | High | LQVVSESEI | [K].LQVVSI | 2xPhospho | Q3UYD0 2x | 1.35E-06 | 0 | 1 | 2 | 12 | Q3UYD0 | Putative ur | Q3UYD0 [3 | 0 | 3759.754 | 0 | -0.23 | 0.261896 | 10.7 | 108.3 | 91.7 | 0.89 | 14.6 | | High | High | High | High | High | High | High | High | High | High | High | 0 | 1.42E-07 | 5.36 |
| FALSE | High | LSAGTTVAI | [R].LSAGTT | 1xPhospho | Q8K1N2-2 | 0.047894 | 0.00354 | 1 | 3 | 1 | Q8K1N2-2 | Isoform 2 c | Q8K1N2-2 | 0 | 1727.934 | 0 | 0.82 | 0.010558 | 23.68 | 67.7 | 132.3 | 15.47 | 13.27 | | High | High | High | High | High | High | High | High | High | High | High | 0.001812 | 0.01832 | 2.07 |
| FALSE | High | LSALSQVTF | [K].LSALSQ | 1xPhospho | G3UYN0 1x | 3.61E-06 | 0 | 1 | 2 | 1 | G3UYN0 | L1TD1 OS= | G3UYN0 [5 | 0 | 2265.287 | 0 | -1.35 | 0.000368 | 0.77 | 145.3 | 54.7 | 5.85 | 5.6 | | High | High | High | High | High | High | High | High | High | High | High | 0 | 4.31E-07 | 2.21 |
| FALSE | High | LSALSTGSN | [K].LSALST | 1xPhospho | O35099 1x | 0.036731 | 0.00249 | 1 | 3 | 1 | O35099 | Mitogen-ac | O35099 [9 | 0 | 1719.852 | 0 | 1.22 | 0.000656 | 14.69 | 57.2 | 142.8 | 5.67 | 7.48 | | High | High | High | High | High | High | High | High | High | High | High | 0.001308 | 0.01355 | 2.04 |
| FALSE | High | LSAQSNSN | [R].LSAQSN | 1xPhospho | G5E870 1x | 7.02E-05 | 0 | 1 | 2 | 2 | G5E870 | E3 ubiquiti | G5E870 [1 | 0 | 1809.869 | 0 | -1.56 | 0.000986 | 2.37 | 150 | 50 | 7.89 | 13.18 | | High | High | High | High | High | High | High | High | High | High | High | 0 | 1.31E-05 | 3.29 |
| FALSE | High | LSAQSNSN | [R].LSAQSN | 1xDeamida | G5E870 1x | 0.010758 | 0.000772 | 1 | 2 | 3 | G5E870 | E3 ubiquiti | G5E870 [1 | 0 | 1810.853 | 0 | -0.69 | 0.117397 | 19.71 | 121.3 | 78.7 | 17.59 | 23.24 | | High | High | High | High | High | High | High | High | High | High | High | 0.000399 | 0.003414 | 2.84 |
| FALSE | High | LSASAAESF | [R].LSASAA | 1xPhospho | Q9WU40-2 | 1.07E-05 | 0 | 1 | 4 | 5 | Q9WU40-2 | Isoform 2 c | Q9WU40-2 | 0 | 2679.376 | 0 | 1.05 | 0.000236 | 6.57 | 65 | 135 | 10.62 | 5.73 | | High | High | High | High | High | High | High | High | High | High | High | 0 | 1.45E-06 | 3.63 |
| FALSE | High | ISASSATR | [R].ISASSAT | 1xPhospho | F8VQ28 1x | 0.001689 | 5.66E-05 | 1 | 7 | 1 | F8VQ28 | Paxillin OS= | F8VQ28 [2 | 0 | 1101.55 | 0 | 0.2 | 0.128293 | 11.6 | 97 | 103 | 10.76 | 15.73 | | High | High | High | High | High | High | High | High | High | High | High | 2.58E-05 | 0.00043 | 1.45 |
| FALSE | High | LSASSEDISI | [R].LSASSE | 1xPhospho | P53349 1x | 0.01186 | 0.000889 | 2 | 2 | 3 | P53349; F8 | Mitogen-ac | P53349 [91 | 0 | 1488.678 | 0 | -0.74 | 0.008659 | 3.28 | 124.2 | 75.8 | 4.34 | 13.89 | Shared | High | High | High | High | High | High | High | High | High | High | High | 0.000449 | 0.003796 | 2.23 |
| FALSE | High | LSAVSLR | [K].LSAVSL | 1xPhospho | Q8BVL3 1x | 0.036731 | 0.00249 | 1 | 1 | 2 | Q8BVL3 | Sorting nex | Q8BVL3 [4 | 0 | 1054.586 | 0 | 1.63 | 0.010209 | 29.84 | 47.9 | 152.1 | 14.13 | 25.65 | | High | High | High | High | High | High | High | High | High | High | High | 0.002455 | 0.02667 | 1.52 |
| FALSE | High | LSAVSSRPT | [R].LSAVSS | 1xPhospho | A2AP82 1x | 0.000136 | 0 | 1 | 2 | 3 | A2AP82 | Transcripti | A2AP82 [1 | 0 | 2896.57 | 0 | 1.02 | 0.001871 | 12.6 | 69.3 | 130.7 | 11.93 | 11.57 | | High | High | High | High | High | High | High | High | High | High | High | 0 | 2.52E-05 | 3.87 |
| FALSE | High | LSCLFAGPH | [K].LSCLFA | 2xCarbami | Q8CFD4 1x | 0.066503 | 0.004796 | 1 | 3 | 1 | Q8CFD4 | Sorting nex | Q8CFD4 [4 | 1 | 3167.501 | 0 | -0.07 | 0.934211 | 9.17 | 98 | 102 | 4.69 | 10.44 | | High | High | High | High | High | High | High | High | High | High | High | 0.002455 | 0.02667 | 2.78 |
| FALSE | High | LSCQTTPA | [R].LSCQTT | 1xCarbami | E9QK89 1x | 0.060159 | 0.004476 | 1 | 2 | 2 | E9QK89 | Mediator c | E9QK89 [9 | 0 | 1600.817 | 0 | -1.04 | 0.00125 | 2.49 | 133.9 | 66.1 | 6.57 | 7.22 | | High | High | High | High | High | High | High | High | High | High | High | 0.003838 | 0.04128 | 1.81 |
| FALSE | High | LSCSLEDLR | [K].LSCSLEI | 1xCarbami | O35711-4 | 0.001158 | 0 | 1 | 4 | 1 | O35711-4 | Isoform 4 c | O35711-4 | 0 | 1401.665 | 0 | -1.29 | 0.000248 | 0.8 | 140.3 | 59.7 | 4.17 | 5.86 | | High | High | High | High | High | High | High | High | High | High | High | 0 | 0.00028 | 2.13 |
| FALSE | High | LSAGSFAR | [R].LSAGSF | 1xPhospho | Q9D3J3 1x | 0.053473 | 0.00384 | 1 | 1 | 2 | Q9D3J3 | BCL2-like 1 | Q9D3J3 [1 | 0 | 1117.56 | 0 | 0.14 | 0.46064 | 8.14 | 96.1 | 103.9 | 3.04 | 11.32 | | High | High | High | High | High | High | High | High | High | High | High | 0.004115 | 0.04382 | 1.88 |
| FALSE | High | ISDAHLADT | [R].ISDAHL | 1xPhospho | Q3UWM7 | 0.003253 | 0.00016 | 1 | 7 | 2 | Q3UWM7 | Putative ur | Q3UWM7 | 0 | 1909.986 | 0 | -0.87 | 0.000211 | 1 | 129.3 | 70.7 | 3.37 | 5.23 | | High | High | High | High | High | High | High | High | High | High | High | 0.000449 | 0.004184 | 2.23 |
| FALSE | High | LSDASPAEI | [K].LSDASP | 1xPhospho | F8VQC7 1x | 3.22E-07 | 0 | 1 | 35 | 1 | F8VQC7 | Kinectin OS | F8VQC7 [8 | 0 | 3125.488 | 0 | 1.46 | 0.003358 | 15.23 | 50.2 | 149.8 | 9.79 | 13.53 | | High | High | High | High | High | High | High | High | High | High | High | 0 | 2.84E-08 | 4.57 |
| FALSE | High | LSDATPEH | [R].LSDATF | 1xPhospho | B7ZMW4 1 | 0.004411 | 0.000409 | 1 | 5 | 1 | B7ZMW4 | X99384 pro | B7ZMW4 [ | 0 | 1894.926 | 0 | 0.21 | 0.821032 | 8.24 | 100.6 | 99.4 | 15.22 | 14.28 | | High | High | High | High | High | High | High | High | High | High | High | 0.000183 | 0.001262 | 2.13 |
| FALSE | High | LSDDFDSP | [K].LSDDFI | 1xPhospho | E9PWW9 1 | 0.00925 | 0.000736 | 1 | 3 | 2 | E9PWW9 | Remodelin | E9PWW9 [ | 0 | 1660.824 | 0 | -0.75 | 0.000489 | 2.01 | 124.2 | 75.8 | 2.45 | 7.05 | | High | High | High | High | High | High | High | High | High | High | High | 0.001199 | 0.01286 | 2.84 |

| | | | | | | | | | | | | | | | | | | | | | | | | | | | | | | | | | | | | | | |
|---|---|---|---|---|---|---|---|---|---|---|---|---|---|---|---|---|---|---|---|---|---|---|---|---|---|---|---|---|---|---|---|---|---|---|---|---|---|---|
| FALSE | High | LSDGEYISD | [K].LSDGEY | 1xPhosphc | G5E8K3 1x | 0.000107 | 0 | 2 | 5 | 1 | G5E8K3; A | Ankyrin 3, | G5E8K3 [8 | 0 | 2954.308 | 0 | -1.56 | 0.000928 | 2.28 | 149.8 | 50.2 | 11.91 | 24.46 | Shared | High | High | High | High | High | High | High | High | High | High | High | 0 | 1.94E-05 | 3.61 |
| FALSE | High | ISDPLTSSP( | [R].ISDPLTS | 1xPhosphc | Q3UJN1 1x | 0.005717 | 0.00062 | 1 | 3 | 6 | Q3UJN1 | DNA helica | Q3UJN1 [2 | 0 | 1438.714 | 0 | -0.75 | 0.000311 | 1.5 | 124.8 | 75.2 | 3.18 | 4.61 | | High | High | High | High | High | High | High | High | High | High | High | 0.000872 | 0.008887 | 1.9 |
| FALSE | High | ISDPLTSSP( | [R].ISDPLTS | 2xPhosphc | Q3UJN1 2x | 0.04913 | 0.003576 | 1 | 3 | 4 | Q3UJN1 | DNA helica | Q3UJN1 [2 | 0 | 1518.68 | 0 | -0.85 | 0.000596 | 1.39 | 126.6 | 73.4 | 4.49 | 5.53 | | High | High | High | High | High | High | High | High | High | High | High | 0.004907 | 0.05336 | 2.77 |
| FALSE | High | ISDPLTSSP( | [R].ISDPLTS | 3xPhosphc | Q3UJN1 3x | 0.001117 | 0 | 1 | 3 | 7 | Q3UJN1 | DNA helica | Q3UJN1 [2 | 0 | 1598.647 | 0 | -1.08 | 0.001559 | 1.36 | 134.2 | 65.8 | 6.8 | 10.13 | | High | High | High | High | High | High | High | High | High | High | High | 4.94E-05 | 0.000703 | 3.37 |
| FALSE | High | ISDSESEELI | [R].ISDSESI | 2xPhosphc | Q8C1D8 2x | 0.028575 | 0.001829 | 1 | 3 | 1 | Q8C1D8 | Protein IW | Q8C1D8 [1 | 0 | 2104.997 | 0 | -0.57 | 0.015201 | 4.31 | 120.3 | 79.7 | 12.73 | 6.4 | | High | High | High | High | High | High | High | High | High | High | High | 0.000984 | 0.01018 | 1.99 |
| FALSE | High | LSDSGFK | [R].LSDSGF | 1xPhosphc | P06684 1xl | 0.086562 | 0.006435 | 1 | 2 | 1 | P06684 | Compleme | P06684 [13 | 0 | 1291.67 | 0 | -0.06 | 0.440435 | 11.78 | 102.1 | 97.9 | 12.44 | 5.09 | | High | High | High | High | High | High | High | High | High | High | High | 0.003346 | 0.03656 | 1.62 |
| FALSE | High | LSDSPSVK | [R].LSDSPS | 1xPhosphc | Q571G8 1x | 0.003083 | 0.00016 | 1 | 5 | 2 | Q571G8 | MKIAA164 | Q571G8 [3 | 0 | 1370.733 | 0 | 0.43 | 0.000648 | 4.72 | 83.6 | 116.4 | 4.92 | 4.82 | | High | High | High | High | High | High | High | High | High | High | High | 7.19E-05 | 0.00084 | 1.95 |
| FALSE | High | LSDTPAPK | [K].LSDTPA | 1xPhosphc | A0A0U1RP | 0.005978 | 0.00062 | 1 | 7 | 1 | A0A0U1RP | INO80 com | A0A0U1RP | 0 | 1366.738 | 0 | -0.04 | 0.323206 | 8.89 | 99.7 | 100.3 | 4.85 | 10.87 | | High | High | High | High | High | High | High | High | High | High | High | 0.000275 | 0.001766 | 2.18 |
| FALSE | High | LSDVLK | [R].LSDVLK | 1xPhosphc | G5E870 1x | 0.015603 | 0.001124 | 1 | 3 | 3 | G5E870 | E3 ubiquiti | G5E870 [1( | 0 | 1212.701 | 0 | -1.91 | 0.000225 | 1.14 | 160.3 | 39.7 | 4.84 | 21.74 | | High | High | High | High | High | High | High | High | High | High | High | 0.000585 | 0.005183 | 2.25 |
| FALSE | High | ISDYFEFAG | [K].ISDYFEI | 1xPhosphc | B1ASU9 1x | 9.61E-08 | 0 | 1 | 3 | 1 | B1ASU9 | Serine/thre | B1ASU9 [8 | 0 | 2210.996 | 0 | -0.12 | 0.221255 | 13.19 | 103.8 | 96.2 | 6.97 | 6.33 | | High | High | High | High | High | High | High | High | High | High | High | 0 | 7.33E-09 | 3.76 |
| FALSE | High | ISDAHLADT | [R].ISDAHL | 1xPhosphc | Q3UWM7 | 0.00311 | 0.00016 | 1 | 7 | 1 | Q3UWM7 | Putative ur | Q3UWM7 | 0 | 1925.981 | 0 | -0.7 | 0.005319 | 1.02 | 122.2 | 77.8 | 6.69 | 14.72 | | High | High | High | High | High | High | High | High | High | High | High | 7.19E-05 | 0.000855 | 2.98 |
| FALSE | High | LSAELQLTK | [R].LSAELQ | 1xPhosphc | Q9WU62 1 | 0.000941 | 0 | 1 | 2 | 2 | Q9WU62 | Inner centi | Q9WU62 [ | 0 | 1540.875 | 0 | -0.58 | 0.015875 | 3.95 | 115.3 | 84.7 | 10.46 | 10.99 | | High | High | High | High | High | High | High | High | High | High | High | 0 | 0.000223 | 2.24 |
| FALSE | High | LSAELLK | [K].LSAELLI | 1xPhosphc | E9Q284 1x | 0.007209 | 0.00062 | 1 | 3 | 1 | E9Q284 | Coilin OS=l | E9Q284 [2 | 0 | 1311.769 | 0 | -1.45 | 0.000167 | 1.46 | 145.5 | 54.5 | 1.45 | 7.43 | | High | High | High | High | High | High | High | High | High | High | High | 0.000275 | 0.002187 | 2.51 |
| FALSE | High | ISAEDRSPE | [R].ISAEDR | 1xPhosphc | Q8C9S4 1x | 0.013666 | 0.001011 | 1 | 2 | 1 | Q8C9S4 | Coiled-coil | Q8C9S4 [7 | 1 | 3231.546 | 0 | 0.65 | 0.002492 | 9.94 | 79 | 121 | 3.4 | 9.5 | | High | High | High | High | High | High | High | High | High | High | High | 0.000538 | 0.004463 | 2.64 |
| FALSE | High | LQWDGSSI | [R].LQWDC | 1xPhosphc | A2AWT3-2 | 4.01E-05 | 0 | 1 | 4 | 3 | A2AWT3-2 | Isoform 2 c | A2AWT3-2 | 0 | 2391.112 | 0 | 0.2 | 0.069694 | 15.6 | 95.1 | 104.9 | 7.35 | 4.66 | | High | High | High | High | High | High | High | High | High | High | High | 2.58E-05 | 0.000431 | 2.59 |
| FALSE | High | LQYYYSTSE | [K].LQYYYS | 2xPhosphc | Q3UZN6 2x | 0.001751 | 5.66E-05 | 1 | 5 | 1 | Q3UZN6 | Putative ur | Q3UZN6 [1 | 0 | 2842.142 | 0 | -0.39 | 0.010845 | 9.11 | 113.2 | 86.8 | 10.39 | 7.1 | | High | High | High | High | High | High | High | High | High | High | High | 2.58E-05 | 0.000449 | 3.04 |
| FALSE | High | LREELQSAI | [K].LREELQ | 1xPhosphc | Q8CIL4 1xF | 0.108463 | 0.008839 | 1 | 3 | 1 | Q8CIL4 | Uncharacte | Q8CIL4 [83 | 1 | 3083.623 | 0 | -0.43 | 0.626279 | 48.27 | 102.7 | 97.3 | 64.87 | 46.14 | | High | High | High | High | High | High | High | High | High | High | High | 0.004422 | 0.04768 | 3.37 |
| FALSE | High | LREELQSAI | [K].LREELQ | 1xPhosphc | Q8CIL4 1xF | 0.000454 | 0 | 1 | 3 | 1 | Q8CIL4 | Uncharacte | Q8CIL4 [83 | 1 | 3312.786 | 0 | -0.82 | 0.000408 | 1.71 | 127.8 | 72.2 | 5.03 | 5.79 | | High | High | High | High | High | High | High | High | High | High | High | 0 | 9.78E-05 | 2.65 |
| FALSE | High | LRLSDSGSA | [K].LRLSDS | 1xPhosphc | Q91XC0 1x | 0.056248 | 0.004124 | 1 | 1 | 1 | Q91XC0 | LIM domai | Q91XC0 [1 | 1 | 1571.856 | 0 | -1.02 | 0.001246 | 3.12 | 133.9 | 66.1 | 7.33 | 5.57 | | High | High | High | High | High | High | High | High | High | High | High | 0.002117 | 0.02215 | 2.47 |
| FALSE | High | LRLSFQGDI | [R].LRLSFQ | 1xPhosphc | E9Q4Y4 1x | 0.004824 | 0.000456 | 1 | 2 | 1 | E9Q4Y4 | Centrosom | E9Q4Y4 [9 | 1 | 1786.905 | 0 | -0.65 | 0.003313 | 5.34 | 125.6 | 74.4 | 7.7 | 10.13 | | High | High | High | High | High | High | High | High | High | High | High | 0.000204 | 0.001395 | 2.02 |
| FALSE | High | LRLSPSEEA | [K].LRLSPS | 1xPhosphc | A0A1N9NP | 0.002077 | 5.66E-05 | 1 | 8 | 1 | A0A1N9NP | Centrosom | A0A1N9NP | 1 | 2041.15 | 0 | -0.91 | 0.000987 | 1.81 | 129.5 | 70.5 | 4.17 | 8.71 | | High | High | High | High | High | High | High | High | High | High | High | 2.58E-05 | 0.00054 | 2.22 |
| FALSE | High | LRLSPSPTS | [R].LRLSPS | 1xPhosphc | P48678 1xl | 0.030898 | 0.001896 | 1 | 3 | 2 | P48678 | Prelamin-A | P48678 [38 | 1 | 1550.825 | 0 | 1.23 | 0.001363 | 21.45 | 59.7 | 140.3 | 12.08 | 8.23 | | High | High | High | High | High | High | High | High | High | High | High | 0.004164 | 0.04493 | 2.38 |
| FALSE | High | LRPLLSQLG | [R].LRPLLS | 1xPhosphc | Q61210-5 | 0.000651 | 0 | 1 | 5 | 1 | Q61210-5 | Isoform 5 c | Q61210-5 | 0 | 2512.385 | 0 | 1.01 | 0.00119 | 15.36 | 65.3 | 134.7 | 9.13 | 6.51 | | High | High | High | High | High | High | High | High | High | High | High | 0 | 0.000147 | 2.67 |
| FALSE | High | LRPLPESQF | [-].LRPLPES | 1xPhosphc | F6YAU7 1x | 0.000191 | 0 | 1 | 1 | 2 | F6YAU7 | Nuclear re | F6YAU7 [1 | 0 | 2854.584 | 0 | 1.25 | 0.000571 | 14.47 | 57.6 | 142.4 | 16.63 | 2.02 | | High | High | High | High | High | High | High | High | High | High | High | 0 | 7.36E-05 | 2.35 |
| FALSE | High | LRPPGSTGI | [R].LRPPGS | 1xPhosphc | D3YWA2 1 | 0.012508 | 0.000889 | 1 | 3 | 1 | D3YWA2 | Kinase OS= | D3YWA2 [ | 0 | 1841.947 | 0 | -0.45 | 0.008108 | 5.91 | 114.6 | 85.4 | 6.03 | 12.9 | | High | High | High | High | High | High | High | High | High | High | High | 0.000449 | 0.004018 | 1.93 |
| FALSE | High | IRPPPELET | [K].IRPPPEI | 1xPhosphc | Q3V0J1 1xl | 5.16E-05 | 0 | 1 | 5 | 2 | Q3V0J1 | Transmem | Q3V0J1 [1 | 0 | 2784.486 | 0 | -0.13 | 0.08502 | 6 | 106.9 | 93.1 | 5.29 | 7.56 | | High | High | High | High | High | High | High | High | High | High | High | 0 | 8.54E-06 | 1.92 |
| FALSE | High | LRPQSVQS | [R].LRPQSV | 1xPhosphc | Q3U3K9 1x | 9.02E-08 | 0 | 1 | 5 | 2 | Q3U3K9 | Ancient ub | Q3U3K9 [3 | 0 | 3328.689 | 0 | 1.09 | 0.002026 | 9.29 | 59.8 | 140.2 | 9.28 | 11.22 | | High | High | High | High | High | High | High | High | High | High | High | 0 | 6.8E-09 | 4.08 |
| FALSE | High | LRPSLDFDK | [R].LRPSLD | 1xPhosphc | Q8BXQ8 1x | 3.83E-05 | 0 | 1 | 2 | 2 | Q8BXQ8 | Protein FAI | Q8BXQ8 [2 | 0 | 1628.881 | 0 | -0.72 | 0.003739 | 9.22 | 126.7 | 73.3 | 3.06 | 18.88 | | High | High | High | High | High | High | High | High | High | High | High | 0 | 6.13E-06 | 1.99 |
| FALSE | High | LRPSTSVDE | [R].LRPSTS | 1xPhosphc | Q8CFT2 1x | 0.000136 | 0 | 1 | 2 | 3 | Q8CFT2 | Histone-lys | Q8CFT2 [9 | 0 | 2186.965 | 0 | -0.4 | 0.021809 | 6.3 | 111.3 | 88.7 | 9.05 | 5.02 | | High | High | High | High | High | High | High | High | High | High | High | 2.58E-05 | 0.000359 | 2.65 |
| FALSE | High | LRPVSPEEII | [K].LRPVSP | 1xPhosphc | D3YTL8 1xl | 0.031713 | 0.002011 | 1 | 5 | 1 | D3YTL8 | Protein spi | D3YTL8 [22 | 0 | 1504.809 | 0 | 0.6 | 0.003152 | 10.79 | 81.4 | 118.6 | 6.47 | 5.78 | | High | High | High | High | High | High | High | High | High | High | High | 0.001076 | 0.01142 | 1.71 |
| FALSE | High | LRQDHADS | [R].LRQDH | 1xCarbami | Q3ULB8 2x | 0.058664 | 0.004294 | 1 | 3 | 1 | Q3ULB8 | Putative ur | Q3ULB8 [8 | 1 | 2707.127 | 0 | 0.06 | 0.287125 | 6.94 | 97.8 | 102.2 | 10.24 | 11.96 | | High | High | High | High | High | High | High | High | High | High | High | 0.002187 | 0.02309 | 1.88 |
| FALSE | High | LRQSNPEF( | [K].LRQSNI | 1xCarbami | Q3TBN1 1x | 0.00407 | 0.00036 | 1 | 1 | 1 | Q3TBN1 | Transmem | Q3TBN1 [2 | 1 | 2043.013 | 0 | 0.84 | 0.000965 | 18.67 | 69.4 | 130.6 | 18.12 | 9.53 | | High | High | High | High | High | High | High | High | High | High | High | 0.000161 | 0.001149 | 2.06 |
| FALSE | High | LRSADSEN/ | [R].LRSADS | 1xPhosphc | Q3UVM5 1 | 0.000107 | 0 | 1 | 3 | 1 | Q3UVM5 | Putative ur | Q3UVM5 [ | 1 | 1997.985 | 0 | 0.28 | 0.578502 | 21.91 | 95.4 | 104.6 | 23.69 | 12.66 | | High | High | High | High | High | High | High | High | High | High | High | 0 | 1.95E-05 | 3.76 |
| FALSE | High | LRSEDGVE( | [R].LRSEDC | 1xPhosphc | E9Q616 1x | 7.95E-08 | 0 | 1 | 2 | 11 | E9Q616 | AHNAK nu | E9Q616 [1 | 1 | 2157.002 | 0 | 2.34 | 0.006315 | 6.84 | 32.7 | 167.3 | 4.33 | 27.16 | | High | High | High | High | High | High | High | High | High | High | High | 0 | 2.63E-08 | 4.38 |
| FALSE | High | IRSFPDFPIF | [R].IRSFPDI | 1xPhosphc | P08030 1xl | 0.000213 | 0 | 1 | 2 | 1 | P08030 | Adenine pl | P08030 [13 | 1 | 2070.114 | 0 | 0.55 | 0.008533 | 12.2 | 81 | 119 | 8.05 | 13.08 | | High | High | High | High | High | High | High | High | High | High | High | 0 | 4.17E-05 | 3.22 |
| FALSE | High | LRSPSNDS/ | [R].LRSPSN | 1xPhosphc | E9Q784 1x | 0.001629 | 5.66E-05 | 1 | 1 | 2 | E9Q784 | Zinc finger | E9Q784 [1 | 1 | 1548.748 | 0 | -0.44 | 0.013178 | 11.15 | 114.5 | 85.5 | 6.49 | 9.33 | | High | High | High | High | High | High | High | High | High | High | High | 0.000449 | 0.003706 | 2.5 |
| FALSE | High | LRSPSNDS/ | [R].LRSPSN | 2xPhosphc | E9Q784 2x | 0.019618 | 0.001438 | 1 | 1 | 2 | E9Q784 | Zinc finger | E9Q784 [1 | 1 | 1628.714 | 0 | -0.89 | 0.000878 | 1.76 | 130.3 | 69.7 | 7.14 | 3.24 | | High | High | High | High | High | High | High | High | High | High | High | 0.000984 | 0.01021 | 2.27 |
| FALSE | High | LRSYEDMI( | [R].LRSYED | 1xPhosphc | E9Q8Z8 1x | 0.00045 | 0 | 1 | 20 | 3 | E9Q8Z8 | Catenin de | E9Q8Z8 [3 | 1 | 3601.666 | 0 | 0.69 | 0.0034 | 10.38 | 76.5 | 123.5 | 4.56 | 11.18 | | High | High | High | High | High | High | High | High | High | High | High | 7.19E-05 | 0.000794 | 4.37 |
| FALSE | High | LRSYEDMI( | [R].LRSYED | 1xPhosphc | E9Q8Z8 1x | 0.016743 | 0.001166 | 1 | 20 | 4 | E9Q8Z8 | Catenin de | E9Q8Z8 [3 | 1 | 3617.661 | 0 | 0.8 | 0.207659 | 32.58 | 75.9 | 124.1 | 25.03 | 46.82 | | High | High | High | High | High | High | High | High | High | High | High | 0.007424 | 0.08178 | 3.48 |
| FALSE | High | LRTQDDGE | [R].LRTQDI | 1xPhosphc | A0A0R4J07 | 1.16E-05 | 0 | 1 | 4 | 2 | A0A0R4J07 | UBX domai | A0A0R4J07 | 1 | 2878.178 | 0 | 0.92 | 0.001414 | 16.64 | 68.5 | 131.5 | 15.86 | 12.03 | | High | High | High | High | High | High | High | High | High | High | High | 0 | 1.59E-06 | 3.47 |
| FALSE | High | ISAAASDSG | [K].ISAAAS | 1xPhosphc | Q3U2Z5 1x | 0.013074 | 0.000889 | 1 | 3 | 1 | Q3U2Z5 | Putative ur | Q3U2Z5 [8 | 1 | 2161.912 | 0 | 1.08 | 0.06595 | 76 | 60.9 | 139.1 | 38.89 | 28.56 | | High | High | High | High | High | High | High | High | High | High | High | 0.000467 | 0.004223 | 3.25 |
| FALSE | High | ISAAASDSG | [K].ISAAAS | 2xPhosphc | Q3U2Z5 2x | 0.000876 | 0 | 1 | 3 | 1 | Q3U2Z5 | Putative ur | Q3U2Z5 [8 | 0 | 2241.879 | 0 | 1.26 | 0.014827 | 44.88 | 59 | 141 | 10.75 | 23.52 | | High | High | High | High | High | High | High | High | High | High | High | 0 | 0.000205 | 3.13 |
| FALSE | High | LSADYEK | [R].LSADYE | 1xPhosphc | B2RRE0 1x | 0.004613 | 0.000456 | 1 | 3 | 4 | B2RRE0 | A kinase (P | B2RRE0 [3 | 0 | 1363.691 | 0 | -1.96 | 0.00034 | 1.57 | 162.9 | 37.1 | 6.78 | 22.93 | | High | High | High | High | High | High | High | High | High | High | High | 0.000467 | 0.004258 | 2.3 |
| FALSE | High | ISDYFEYQC | [K].ISDYFEY | 1xPhosphc | Q8C0V0 1x | 6.51E-07 | 0 | 1 | 2 | 4 | Q8C0V0 | Serine/thre | Q8C0V0 [1 | 0 | 2184.98 | 0 | -0.14 | 0.014217 | 2.43 | 105.4 | 94.6 | 2.46 | 4.95 | | High | High | High | High | High | High | High | High | High | High | High | 0 | 6.39E-06 | 4.29 |
| FALSE | High | LSPAYSLGS | [K].LSPAYS | 2xPhosphc | D3Z0X5 2x | 0.000131 | 0 | 1 | 7 | 2 | D3Z0X5 | Pleckstrin I | D3Z0X5 [5 | 0 | 1965.929 | 0 | 0.92 | 0.002997 | 10.27 | 68.7 | 131.3 | 17.35 | 6.94 | | High | High | High | High | High | High | High | High | High | High | High | 0 | 2.42E-05 | 3.45 |
| FALSE | High | LSPCCQSPT | [K].LSPCCC | 2xCarbami | Q3UHR0 1 | 0.031168 | 0.001935 | 1 | 2 | 1 | Q3UHR0 | BAH and cc | Q3UHR0 [1 | 0 | 1828.91 | 0 | -0.05 | 0.587938 | 9.94 | 97.2 | 102.8 | 9.62 | 16.23 | | High | High | High | High | High | High | High | High | High | High | High | 0.001076 | 0.01129 | 2.43 |
| FALSE | High | LSPCLHR | [R].LSPCLH | 1xCarbami | Q7TNF9 1x | 0.081085 | 0.006219 | 1 | 2 | 1 | Q7TNF9 | Protein FAI | Q7TNF9 [2 | 0 | 1191.591 | 0 | -1.19 | 0.000746 | 3.52 | 145.2 | 54.8 | 9.42 | 14.78 | | High | High | High | High | High | High | High | High | High | High | High | 0.0032 | 0.03366 | 1.81 |
| FALSE | High | LSTGPALDS | [R].LSTGPA | 1xPhosphc | G3UWX1 1 | 4.96E-10 | 0 | 2 | 9 | 7 | G3UWX1; I | Replication | G3UWX1 [ | 0 | 3243.451 | 0 | -0.05 | 0.603181 | 3.93 | 101.8 | 98.2 | 5.65 | 5.59 | Shared | High | High | High | High | High | High | High | High | High | High | High | 0 | 7.67E-09 | 6.23 |
| FALSE | High | LSTGPALDS | [R].LSTGPA | 2xPhosphc | G3UWX1 2 | 1.41E-05 | 0 | 2 | 9 | 7 | G3UWX1; I | Replication | G3UWX1 [ | 0 | 3323.417 | 0 | -0.1 | 0.820445 | 9.89 | 101.6 | 98.4 | 6.59 | 7.08 | Shared | High | High | High | High | High | High | High | High | High | High | High | 0 | 1.98E-06 | 5.15 |
| FALSE | High | LSTGSELSP | [R].LSTGSE | 1xPhosphc | A2AJI0 1xP | 0.000534 | 0 | 1 | 5 | 2 | A2AJI0 | MAP7 dom | A2AJI0 [45 | 0 | 1556.834 | 0 | 0.87 | 0.002264 | 5.62 | 69.6 | 130.4 | 11.26 | 7.83 | | High | High | High | High | High | High | High | High | High | High | High | 0 | 0.000118 | 2.08 |
| FALSE | High | LSTGSELSP | [R].LSTGSE | 2xPhosphc | A2AJI0 2xP | 0.036731 | 0.00249 | 1 | 5 | 1 | A2AJI0 | MAP7 dom | A2AJI0 [45 | 0 | 1636.8 | 0 | 1.49 | 0.000278 | 13.12 | 53.3 | 146.7 | 9.41 | 3.62 | | High | High | High | High | High | High | High | High | High | High | High | 0.001308 | 0.01349 | 1.61 |
| FALSE | High | LSTGSFPEC | [R].LSTGSF | 1xPhosphc | Q68FM7 1 | 3.17E-05 | 0 | 1 | 3 | 13 | Q68FM7 | Rho guanir | Q68FM7 [6 | 0 | 2148.99 | 0 | -0.31 | 0.074544 | 5.83 | 110.7 | 89.3 | 12.48 | 3.51 | | High | High | High | High | High | High | High | High | High | High | High | 0 | 1.72E-05 | 3.83 |
| FALSE | High | LSTGSFPEC | [R].LSTGSF | 2xPhosphc | Q68FM7 2 | 5.4E-05 | 0 | 1 | 3 | 3 | Q68FM7 | Rho guanir | Q68FM7 [6 | 0 | 2228.956 | 0 | 0.06 | 0.396776 | 7.47 | 98 | 102 | 12.87 | 14.7 | | High | High | High | High | High | High | High | High | High | High | High | 0.000246 | 0.001494 | 3.22 |
| FALSE | High | LSTIDESGS | [R].LSTIDES | 2xPhosphc | Q9WVM1 | 0.000113 | 0 | 1 | 3 | 2 | Q9WVM1 | Rac GTPase | Q9WVM1 | 0 | 2545.213 | 0 | -0.86 | 0.024426 | 8.97 | 137.4 | 62.6 | 19.95 | 21.27 | | High | High | High | High | High | High | High | High | High | High | High | 9.42E-05 | 0.000938 | 3.2 |
| FALSE | High | LSTLEPFED | [R].LSTLEPI | 1xPhosphc | Q6NSQ7 1 | 7.16E-06 | 0 | 1 | 1 | 2 | Q6NSQ7 | Protein LT\ | Q6NSQ7 [3 | 0 | 3087.47 | 0 | -0.59 | 0.031553 | 18.89 | 121.8 | 78.2 | 9.16 | 21.98 | | High | High | High | High | High | High | High | High | High | High | High | 0 | 9.31E-07 | 4.51 |
| FALSE | High | LSTPLQTN( | [R].LSTPLQ | 1xCarbami | Q3UZG4 1 | 3.55E-07 | 0 | 1 | 3 | 1 | Q3UZG4 | Aminoacyl | Q3UZG4 [9 | 0 | 3571.797 | 0 | 0.51 | 0.005082 | 8.26 | 83.7 | 116.3 | 4.06 | 6.8 | | High | High | High | High | High | High | High | High | High | High | High | 0 | 3.18E-08 | 3.53 |
| FALSE | High | LSTPNVSSF | [K].LSTPNV | 1xPhosphc | Q8VCB1 1x | 2.8E-05 | 0 | 1 | 2 | 4 | Q8VCB1 | Nucleopor | Q8VCB1 [4 | 0 | 2459.217 | 0 | -0.51 | 0.011195 | 4.23 | 115 | 85 | 13.99 | 7.91 | | High | High | High | High | High | High | High | High | High | High | High | 0 | 4.28E-06 | 4.68 |
| FALSE | High | LSTSPATR | [K].LSTSPA | 1xPhosphc | Q9EPQ8 1x | 0.012619 | 0.000889 | 1 | 3 | 2 | Q9EPQ8 | Transcripti | Q9EPQ8 [6 | 0 | 1141.582 | 0 | -0.15 | 0.113717 | 10.28 | 105 | 95 | 5.53 | 5.59 | | High | High | High | High | High | High | High | High | High | High | High | 0.000585 | 0.005336 | 1.87 |
| FALSE | High | LSTSPTPGS | [R].LSTSPT | 1xPhosphc | Q8C0V0 1x | 4.47E-05 | 0 | 1 | 1 | 2 | Q8C0V0 | Serine/thre | Q8C0V0 [1 | 0 | 1595.799 | 0 | -0.66 | 0.000616 | 3.05 | 123.6 | 76.4 | 4.02 | 5.4 | | High | High | High | High | High | High | High | High | High | High | High | 0 | 7.25E-06 | 3.13 |
| FALSE | High | LSTSQDAA( | [K].LSTSQD | 1xCarbami | G5E8R4 1x | 7.43E-06 | 0 | 1 | 3 | 1 | G5E8R4 | SAPS doma | G5E8R4 [8 | 0 | 1618.791 | 0 | -0.74 | 0.002167 | 2.83 | 124.9 | 75.1 | 10.2 | 2.49 | | High | High | High | High | High | High | High | High | High | High | High | 0 | 9.66E-07 | 3.28 |
| FALSE | High | LSTGPALDS | [R].LSTGPA | 1xDeamida | G3UWX1 1 | 0.031713 | 0.002011 | 2 | 9 | 1 | G3UWX1; I | Replication | G3UWX1 [ | 0 | 3015.272 | 0 | -0.62 | 0.017128 | 8.33 | 124.8 | 75.2 | 12.49 | 17.86 | Shared | High | High | High | High | High | High | High | High | High | High | High | 0.001093 | 0.01148 | 2.62 |
| FALSE | High | LSTTPSPTN | [R].LSTTPS | 2xPhosphc | Q6A006 2x | 0.00614 | 0.00062 | 1 | 3 | 3 | Q6A006 | MKIAA073 | Q6A006 [1 | 0 | 2577.111 | 0 | -0.51 | 0.002756 | 1.26 | 118.8 | 81.2 | 8.91 | 5.37 | | High | High | High | High | High | High | High | High | High | High | High | 0.000294 | 0.002299 | 2.44 |
| FALSE | High | LSTTPSPTN | [R].LSTTPS | 2xPhosphc | Q6A006 2x | 0.056724 | 0.004124 | 1 | 3 | 1 | Q6A006 | MKIAA073 | Q6A006 [1 | 1 | 2733.212 | 0 | -0.66 | 0.000672 | 3.53 | 122.6 | 77.4 | 3.36 | 10.01 | | High | High | High | High | High | High | High | High | High | High | High | 0.002117 | 0.0222 | 2 |
| FALSE | High | LSTVTPEPP | [K].LSTVTP | 1xDeamida | M0QWZ1 1 | 4.98E-05 | 0 | 1 | 4 | 1 | M0QWZ1 | Protein FAI | M0QWZ1 [ | 0 | 2331.179 | 0 | 0.46 | 0.000913 | 2.64 | 84.4 | 115.6 | 4.86 | 6.91 | | High | High | High | High | High | High | High | High | High | High | High | 0 | 8.18E-06 | 2.44 |
| FALSE | High | LSTVTPEPP | [K].LSTVTP | 2xDeamida | M0QWZ1 | 0.019618 | 0.001438 | 1 | 4 | 2 | M0QWZ1 | Protein FAI | M0QWZ1 [ | 0 | 2332.163 | 0 | 0.39 | 0.032358 | 15.17 | 84.7 | 115.3 | 12.39 | 6.96 | | High | High | High | High | High | High | High | High | High | High | High | 0.000756 | 0.006672 | 3.13 |
| FALSE | High | ISVDDYGM | [K].ISVDDY | 2xPhosphc | Q9WU62 2 | 0.014667 | 0.001049 | 1 | 2 | 1 | Q9WU62 | Inner centi | Q9WU62 [ | 0 | 2972.138 | 0 | -0.4 | 0.036557 | 12.68 | 109.2 | 90.8 | 15.99 | 10.76 | | High | High | High | High | High | High | High | High | High | High | High | 0.000555 | 0.004793 | 3.82 |
| FALSE | High | ISVDDYGM | [K].ISVDDY | 2xPhosphc | Q9WU62 2 | 0.022773 | 0.001542 | 1 | 2 | 1 | Q9WU62 | Inner centi | Q9WU62 [ | 0 | 2988.133 | 0 | -0.41 | 0.017746 | 10.29 | 115.1 | 84.9 | 5.35 | 13.84 | | High | High | High | High | High | High | High | High | High | High | High | 0.000832 | 0.00791 | 2.79 |
| FALSE | High | LSVDNNTS | [K].LSVDNI | 1xPhosphc | A0A1B0GS | 0.00582 | 0.00062 | 1 | 6 | 3 | A0A1B0GS | Supervillin | A0A1B0GS | 0 | 1965.957 | 0 | 0.63 | 0.000738 | 10.71 | 78.2 | 121.8 | 5.51 | 2.7 | | High | High | High | High | High | High | High | High | High | High | High | 0.002662 | 0.02827 | 2.64 |
| FALSE | High | LSVDSQEA | [R].LSVDSC | 1xPhosphc | Q68FM7 1 | 1.01E-06 | 0 | 1 | 4 | 2 | Q68FM7 | Rho guanir | Q68FM7 [2 | 0 | 2174.965 | 0 | 0.02 | 0.509548 | 18.76 | 102.1 | 97.9 | 8.49 | 18.81 | | High | High | High | High | High | High | High | High | High | High | High | 0 | 1.03E-07 | 4.49 |
| FALSE | High | LSVDYGK | [R].LSVDYC | 1xPhosphc | P68369 1xl | 0.004372 | 0.000409 | 3 | 8 | 2 | P68369; Q | Tubulin alp | P68369 [1 | 0 | 1319.701 | 0 | 1.92 | 0.000707 | 12.01 | 42 | 158 | 7.07 | 9.98 | Shared | High | High | High | High | High | High | High | High | High | High | High | 0.000183 | 0.001245 | 2.37 |
| FALSE | High | LSVELTSSLI | [R].LSVELT | 1xPhosphc | Q02248 1x | 0.008538 | 0.000736 | 1 | 4 | 1 | Q02248 | Catenin be | Q02248 [6 | 0 | 1560.824 | 0 | -0.26 | 0.077209 | 9.75 | 110.3 | 89.7 | 14.4 | 13.5 | | High | High | High | High | High | High | High | High | High | High | High | 0.000327 | 0.002637 | 2.23 |
| FALSE | High | LSVGAYVS | [R].LSVGAY | 1xPhosphc | G3X928 1x | 0.000255 | 0 | 1 | 4 | 1 | G3X928 | SEC23-inte | G3X928 [7 | 0 | 1446.756 | 0 | 0.97 | 0.000648 | 8.36 | 67.7 | 132.3 | 6.39 | 8.29 | | High | High | High | High | High | High | High | High | High | High | High | 0 | 5.13E-05 | 2.34 |
| FALSE | High | LSVGEQPD | [R].LSVGE( | 1xPhosphc | A0A0R4J17 | 6.37E-06 | 0 | 1 | 2 | 1 | A0A0R4J17 | Bromodom | A0A0R4J17 | 0 | 3430.697 | 0 | -0.92 | 0.006091 | 3.39 | 126.1 | 73.9 | 12.29 | 8.42 | | High | High | High | High | High | High | High | High | High | High | High | 0 | 8.15E-07 | 5.23 |
| FALSE | High | LSVGGDSD | [R].LSVGGI | 1xPhosphc | Q3TIV2 1xl | 2.02E-05 | 0 | 1 | 3 | 2 | Q3TIV2 | Putative ur | Q3TIV2 [28 | 0 | 1722.908 | 0 | -0.12 | 0.425704 | 19.74 | 104.2 | 95.8 | 15.35 | 6.66 | | High | High | High | High | High | High | High | High | High | High | High | 0 | 2.44E-05 | 2.5 |
| FALSE | High | ISVGGLPVL | [R].ISVGGL | 1xPhosphc | Q8VCF1-3 | 0.007404 | 0.00062 | 1 | 3 | 1 | Q8VCF1-3 | Isoform 3 c | Q8VCF1-3 | 0 | 1911.079 | 0 | 1.74 | 0.025094 | 49.62 | 51.9 | 148.1 | 22.32 | 40.52 | | High | High | High | High | High | High | High | High | High | High | High | 0.000275 | 0.002233 | 2.3 |
| FALSE | High | LSTTPSPTN | [R].LSTTPS | 3xPhosphc | Q6A006 3x | 0.076555 | 0.005826 | 1 | 3 | 2 | Q6A006 | MKIAA073 | Q6A006 [1 | 0 | 2657.077 | 0 | -0.22 | 0.159943 | 7.46 | 107.6 | 92.4 | 9.67 | 7.8 | | High | High | High | High | High | High | High | High | High | High | High | 0.00299 | 0.03158 | 1.73 |
| FALSE | High | ISVGSDADI | [R].ISVGSD | 1xPhosphc | O08553 1x | 4.76E-05 | 0 | 1 | 1 | 2 | O08553 | Dihydropyi | O08553 [4 | 0 | 2454.257 | 0 | 0.93 | 0.006278 | 22.18 | 69 | 131 | 14.03 | 13.56 | | High | High | High | High | High | High | High | High | High | High | High | 0 | 7.81E-06 | 2.89 |
| FALSE | High | LSTGPALDS | [R].LSTGPA | 1xPhosphc | G3UWX1 1 | 6.72E-09 | 0 | 2 | 9 | 1 | G3UWX1; I | Replication | G3UWX1 [ | 0 | 3014.288 | 0 | 0.18 | 0.904301 | 58.56 | 90.9 | 109.1 | 50.27 | 52.06 | Shared | High | High | High | High | High | High | High | High | High | High | High | 0 | 3.68E-10 | 5.93 |
| FALSE | High | ISTEDINK | [K].ISTEDIN | 1xPhosphc | Q3UPI3 1xl | 0.006713 | 0.00062 | 1 | 4 | 1 | Q3UPI3 | Putative ur | Q3UPI3 [1( | 0 | 1457.765 | 0 | 0.2 | 0.434002 | 29.28 | 96.8 | 103.2 | 13.41 | 10.3 | | High | High | High | High | High | High | High | High | High | High | High | 0.000275 | 0.002009 | 2.63 |
| FALSE | High | ISSPPGPPG | [R].ISSPPGI | 1xPhosphc | Q99MR1 1 | 2.32E-06 | 0 | 1 | 1 | 6 | Q99MR1 | GRB10-inte | Q99MR1 [4 | 0 | 2375.178 | 0 | 0.08 | 0.194478 | 4.29 | 97.7 | 102.3 | 5.58 | 3.03 | | High | High | High | High | High | High | High | High | High | High | High | 0 | 2.03E-06 | 4.07 |
| FALSE | High | LSSPPQAE/ | [K].LSSPPQ | 1xPhosphc | Q8R5A0 1x | 0.087983 | 0.006639 | 1 | 2 | 1 | Q8R5A0 | N-lysine m | Q8R5A0 [2 | 0 | 1477.761 | 0 | 0.12 | 0.407998 | 12.02 | 96.6 | 103.4 | 5.72 | 11.34 | | High | High | High | High | High | High | High | High | High | High | High | 0.003402 | 0.03703 | 2.17 |
| FALSE | High | ISSPGK | [K].ISSPGI | 1xPhosphc | E9Q616 1x | 0.103392 | 0.008249 | 1 | 3 | 1 | E9Q616 | AHNAK nu | E9Q616 [5 | 0 | 1213.659 | 0 | 3.04 | 0.000222 | 5.74 | 21.3 | 178.7 | 4.67 | 6.06 | | High | High | High | High | High | High | High | High | High | High | High | 0.004164 | 0.04502 | 1.62 |
| FALSE | High | ISSQDSPDN | [R].ISSQDS | 1xPhosphc | Q14B71 1x | 9.79E-08 | 0 | 1 | 4 | 4 | Q14B71 | Cell divisio | Q14B71 [2 | 0 | 2150.042 | 0 | -0.41 | 0.00537 | 5.12 | 117.5 | 82.5 | 6.55 | 7.52 | | High | High | High | High | High | High | High | High | High | High | High | 0 | 1.35E-07 | 3.44 |
| FALSE | High | LSSQLSEEK | [K].LSSQLS | 1xPhosphc | Q80WJ7 1x | 0.009006 | 0.000736 | 1 | 5 | 3 | Q80WJ7 | Protein LYI | Q80WJ7 [2 | 0 | 1558.813 | 0 | 1.64 | 0.000675 | 14.73 | 46.5 | 153.5 | 4.03 | 9.75 | | High | High | High | High | High | High | High | High | High | High | High | 0.000327 | 0.002784 | 1.87 |
| FALSE | High | LSSQPSTD\ | [K].LSSQPS | 1xPhosphc | E9Q6J5-3 1 | 4.27E-05 | 0 | 1 | 3 | 3 | E9Q6J5-3 | Isoform Ne | E9Q6J5-3 [ | 0 | 1902.946 | 0 | 0.43 | 0.007973 | 16.28 | 83.6 | 116.4 | 9.61 | 5.58 | | High | High | High | High | High | High | High | High | High | High | High | 0 | 6.88E-06 | 3.07 |
| FALSE | High | LSSRPSEPE | [K].LSSRPS | 1xPhosphc | Q501J7-3 1 | 0.003755 | 0.00021 | 1 | 3 | 2 | Q501J7-3 | Isoform 3 c | Q501J7-3 [ | 0 | 2279.102 | 0 | 1.22 | 0.001712 | 13.38 | 60.4 | 139.6 | 9.32 | 8.18 | | High | High | High | High | High | High | High | High | High | High | High | 0.006622 | 0.07194 | 1.66 |
| FALSE | High | LSSSESPAP | [K].LSSSESI | 1xPhosphc | E9Q0J5 1xl | 5.5E-05 | 0 | 1 | 7 | 1 | E9Q0J5 | Kinesin-like | E9Q0J5 [8 | 0 | 3444.617 | 0 | 2.7 | 0.000235 | 16.38 | 28.9 | 171.1 | 22.54 | 6.17 | | High | High | High | High | High | High | High | High | High | High | High | 0 | 9.13E-06 | 2.42 |
| FALSE | High | LSSSEWSL\ | [K].LSSSEW | 1xPhosphc | Q64512 1x | 0.005666 | 0.00062 | 1 | 4 | 1 | Q64512 | Tyrosine-p | Q64512 [9 | 0 | 2492.247 | 0 | 1.59 | 0.005696 | 33.55 | 44.7 | 155.3 | 16.72 | 21.69 | | High | High | High | High | High | High | High | High | High | High | High | 0.000275 | 0.001668 | 2.95 |
| FALSE | High | LSSSGSNCS | [R].LSSSGS | 2xCarbami | E9PV45 1x | 1.85E-06 | 0 | 1 | 2 | 10 | E9PV45 | Ubiquitin c | E9PV45 [1 | 0 | 2929.302 | 0 | 0.89 | 0.012331 | 6.7 | 72.1 | 127.9 | 9.47 | 13.84 | | High | High | High | High | High | High | High | High | High | High | High | 0 | 3.18E-07 | 4.28 |
| FALSE | High | LSSSLSSYK | [K].LSSSLS | 1xPhosphc | A0A1L1SSL | 0.007605 | 0.000664 | 1 | 4 | 2 | A0A1L1SSL | Serine/thre | A0A1L1SSL | 0 | 1509.797 | 0 | -0.3 | 0.060483 | 4.1 | 110.7 | 89.3 | 6.92 | 11.46 | | High | High | High | High | High | High | High | High | High | High | High | 0.000294 | 0.002317 | 1.8 |
| FALSE | High | LSSSPGK | [K].LSSSPG | 1xPhosphc | G3X9X7 1x | 0.085859 | 0.006406 | 1 | 8 | 1 | G3X9X7 | Activated ( | G3X9X7 [8 | 0 | 1213.659 | 0 | 3.02 | 0.000181 | 13.14 | 22.3 | 177.7 | 14.93 | 3.35 | | High | High | High | High | High | High | High | High | High | High | High | 0.003291 | 0.03599 | 1.71 |
| FALSE | High | LSSSPPSGS | [K].LSSSPP | 2xCarbami | Q9JHT5 1x | 2.15E-07 | 0 | 1 | 1 | 2 | Q9JHT5 | AMME syn | Q9JHT5 [1 | 0 | 2933.199 | 0 | 0.74 | 0.276131 | 74.86 | 75 | 125 | 32.81 | 46.81 | | High | High | High | High | High | High | High | High | High | High | High | 0 | 1.81E-08 | 5.17 |
| FALSE | High | LSTEPSPEF | [K].LSTEPS | 1xPhosphc | Q99P72 1x | 0.017041 | 0.001204 | 1 | 2 | 1 | Q99P72 | Reticulon- | Q99P72 [6 | 0 | 2437.194 | 0 | -0.11 | 0.769885 | 40.41 | 104.7 | 95.3 | 18.91 | 14.78 | | High | High | High | High | High | High | High | High | High | High | High | 0.000638 | 0.005686 | 2.85 |
| FALSE | High | ISSSSAQPC | [R].ISSSSA( | 1xCarbami | A0A0R4J0Y | 1.32E-08 | 0 | 1 | 4 | 4 | A0A0R4J0Y | Probable h | A0A0R4J0Y | 0 | 2985.428 | 0 | 0.08 | 0.117612 | 6.12 | 94.1 | 105.9 | 7.07 | 5.37 | | High | High | High | High | High | High | High | High | High | High | High | 0 | 7.88E-10 | 5.05 |
| FALSE | High | LSSSSLGAP | [R].LSSSSL( | 1xPhosphc | B0QZP6 1x | 0.026885 | 0.001676 | 1 | 4 | 1 | B0QZP6 | Transmem | B0QZP6 [6 | 0 | 1882.947 | 0 | 0.73 | 0.003869 | 4.63 | 75.1 | 124.9 | 7.14 | 10.85 | | High | High | High | High | High | High | High | High | High | High | High | 0.000903 | 0.009497 | 1.93 |
| FALSE | High | LSSSSSNDS | [R].LSSSSSI | 1xPhosphc | E9Q616 1x | 0.000159 | 0 | 1 | 4 | 3 | E9Q616 | AHNAK nu | E9Q616 [5 | 0 | 1707.82 | 0 | 3.29 | 0.000445 | 10.64 | 19.6 | 180.4 | 14.18 | 11.3 | | High | High | High | High | High | High | High | High | High | High | High | 0 | 3.04E-05 | 2.71 |
| FALSE | High | LSSSSSNDS | [R].LSSSSSI | 2xPhosphc | E9Q616 2x | 0.000223 | 0 | 1 | 4 | 3 | E9Q616 | AHNAK nu | E9Q616 [5 | 0 | 1787.787 | 0 | 2.01 | 0.000191 | 5.74 | 41.1 | 158.9 | 13.33 | 4.83 | | High | High | High | High | High | High | High | High | High | High | High | 0 | 4.4E-05 | 3.62 |
| FALSE | High | ISSSTPVVE | [R].ISSSTP\ | 1xPhosphc | A2APV2 1x | 0.010475 | 0.000772 | 1 | 1 | 2 | A2APV2 | Formin-like | A2APV2 [1 | 0 | 1658.772 | 0 | 0.13 | 0.643815 | 2.96 | 95.9 | 104.1 | 7.64 | 8.27 | | High | High | High | High | High | High | High | High | High | High | High | 0.001011 | 0.01092 | 2.7 |
| FALSE | High | LSSTGGQT | [R].LSSTGC | 1xPhosphc | Q8C1D8 1x | 0.000375 | 0 | 1 | 3 | 6 | Q8C1D8 | Protein IW | Q8C1D8 [6 | 0 | 1312.646 | 0 | -0.42 | 0.050286 | 8.6 | 112.8 | 87.2 | 8.76 | 9.41 | | High | High | High | High | High | High | High | High | High | High | High | 0 | 7.9E-05 | 2.13 |
| FALSE | High | ISSTLYQAT | [K].ISSTLY( | 1xPhosphc | Q5RL79 1x | 1.58E-07 | 0 | 1 | 2 | 1 | Q5RL79 | Keratinocy | Q5RL79 [1 | 0 | 2299.271 | 0 | 0.14 | 0.043598 | 9.01 | 96 | 104 | 9.08 | 6.36 | | High | High | High | High | High | High | High | High | High | High | High | 0 | 1.27E-08 | 3.14 |
| FALSE | High | LSSTPPLSA | [K].LSSTPP | 1xPhosphc | K3W4P2 1 | 0.003372 | 0.00016 | 1 | 5 | 2 | K3W4P2 | Integrator | K3W4P2 [1 | 0 | 1507.808 | 0 | 0.46 | 0.011835 | 12.89 | 85.6 | 114.4 | 12.14 | 9.13 | | High | High | High | High | High | High | High | High | High | High | High | 9.42E-05 | 0.000929 | 2.78 |
| FALSE | High | LSSTSSLAG | [R].LSSTSSI | 1xPhosphc | Q07139 1x | 0.00025 | 0 | 1 | 2 | 5 | Q07139 | Protein EC | Q07139 [8 | 0 | 2788.437 | 0 | -0.11 | 0.851728 | 6.73 | 100.6 | 99.4 | 11.21 | 9.72 | | High | High | High | High | High | High | High | High | High | High | High | 0 | 6.88E-05 | 3.88 |
| FALSE | High | LSSWDQAE | [K].LSSWD | 1xPhosphc | G5E866 1x | 3.55E-06 | 0 | 1 | 5 | 5 | G5E866 | Splicing fac | G5E866 [2 | 0 | 2191.038 | 0 | 0.34 | 0.387559 | 16.26 | 90.8 | 109.2 | 7.38 | 21.79 | | High | High | High | High | High | High | High | High | High | High | High | 0 | 1.09E-05 | 4.1 |
| FALSE | High | LSSWDQAE | [K].LSSWD | 2xPhosphc | G5E866 2x | 0.001127 | 0 | 1 | 5 | 4 | G5E866 | Splicing fac | G5E866 [2 | 0 | 2271.005 | 0 | 0.42 | 0.350962 | 16.09 | 85.7 | 114.3 | 11.8 | 28.63 | | High | High | High | High | High | High | High | High | High | High | High | 0.000225 | 0.001451 | 2.81 |
| FALSE | High | ISTASGDGI | [R].ISTASG | 1xPhosphc | Q6R0H7 1x | 0.039347 | 0.002793 | 1 | 6 | 1 | Q6R0H7 | Guanine nu | Q6R0H7 [1 | 0 | 1172.551 | 0 | 1.35 | 0.000805 | 8.6 | 55.8 | 144.2 | 7.9 | 7.91 | | High | High | High | High | High | High | High | High | High | High | High | 0.001453 | 0.01464 | 0.95 |
| FALSE | High | ISTCDSSM\ | [R].ISTCDS | 1xCarbami | Q6ZWQ0 1 | 2.96E-06 | 0 | 1 | 2 | 1 | Q6ZWQ0 | Nesprin-2 ( | Q6ZWQ0 [ | 0 | 3207.454 | 0 | 4.01 | | 46.21 | 11.6 | 188.4 | 30.15 | 14.56 | | High | High | High | High | Not Found | High | High | High | High | High | High | 0 | 3.44E-07 | 3.75 |
| FALSE | High | ISTCDSSM\ | [R].ISTCDS | 1xCarbami | Q6ZWQ0 1 | 0.001986 | 5.66E-05 | 1 | 2 | 2 | Q6ZWQ0 | Nesprin-2 ( | Q6ZWQ0 [ | 0 | 3223.449 | 0 | 4.28 | 0.001451 | 33.95 | 9.8 | 190.2 | 44.32 | 19.34 | | High | High | High | High | High | High | High | High | High | High | High | 2.58E-05 | 0.000515 | 3.94 |
| FALSE | High | LSSSSATLLI | [R].LSSSSA | 1xPhosphc | E9QMU3 1 | 1.45E-06 | 0 | 1 | 5 | 5 | E9QMU3 | Ensconsin | E9QMU3 [ | 0 | 1756.868 | 0 | -0.57 | 0.000401 | 1.38 | 120.6 | 79.4 | 3.2 | 4.96 | | High | High | High | High | High | High | High | High | High | High | High | 0 | 1.55E-07 | 3.02 |
| FALSE | High | ISVLEIK | [K].ISVLEIK | 1xPhosphc | Q91ZX6 1x | 0.003138 | 0.00016 | 1 | 3 | 1 | Q91ZX6 | Sentrin-spe | Q91ZX6 [3 | 0 | 1339.8 | 0 | 1.32 | 0.002603 | 19.33 | 58.6 | 141.4 | 5.09 | 12.3 | | High | High | High | High | High | High | High | High | High | High | High | 7.19E-05 | 0.000859 | 2.76 |
| FALSE | High | LSVPAQQA | [R].LSVPAC | 1xPhosphc | Q9JHC9 1x | 0.000281 | 0 | 1 | 7 | 2 | Q9JHC9 | ETS-related | Q9JHC9 [5 | 0 | 1845.942 | 0 | -0.16 | 0.31112 | 13.03 | 101.8 | 98.2 | 12.56 | 10.22 | | High | High | High | High | High | High | High | High | High | High | High | 0 | 5.76E-05 | 3.28 |

| | | | | | | | | | | | | | | | | | | | | | | | | | | | | | | | | | | | | | | |
|---|---|---|---|---|---|---|---|---|---|---|---|---|---|---|---|---|---|---|---|---|---|---|---|---|---|---|---|---|---|---|---|---|---|---|---|---|---|---|
| FALSE | High | LSVQDPVV | [R].LSVQDI | 1xPhospho | J3QNE8 1x | 1.13E-06 | 0 | 1 | 5 | 2 | J3QNE8 | Putative so | J3QNE8 [4 | 0 | 2380.241 | 0 | 1.21 | 0.001786 | 24 | 58.8 | 141.2 | 11.23 | 9.48 | | High | High | High | High | High | High | High | High | High | High | High | 0 | 1.17E-07 | 3.9 |
| FALSE | High | LTGSTSSLN | [K].LTGSTS | 1xPhospho | Q3U1G4 1x | 0.004451 | 0.000409 | 1 | 4 | 10 | Q3U1G4 | Putative ur | Q3U1G4 [4 | 0 | 1545.829 | 0 | 0.02 | 0.588415 | 8.21 | 99.5 | 100.5 | 2.7 | 12.85 | | High | High | High | High | High | High | High | High | High | High | High | 0.008705 | 0.09297 | 2.08 |
| FALSE | High | LTGSTSSLN | [K].LTGSTS | 2xPhospho | Q3U1G4 2x | 0.064323 | 0.004664 | 1 | 4 | 1 | Q3U1G4 | Putative ur | Q3U1G4 [4 | 0 | 1625.795 | 0 | -0.52 | 0.000222 | 1.49 | 118 | 82 | 4.47 | 7.16 | | High | High | High | High | High | High | High | High | High | High | High | 0.002403 | 0.02576 | 2.87 |
| FALSE | High | LTHGDSPSI | [R].LTHGD | 1xPhospho | Q3U7R1 1x | 0.00011 | 0 | 1 | 1 | 2 | Q3U7R1 | Extended s | Q3U7R1 [9 | 0 | 2427.269 | 0 | 0.34 | 0.013933 | 11.02 | 87.2 | 112.8 | 2.32 | 10.57 | | High | High | High | High | High | High | High | High | High | High | High | 2.58E-05 | 0.000451 | 4.85 |
| FALSE | High | ITHSPDSVK | [K].ITHSPD | 1xPhospho | G3UZM1 1 | 0.000188 | 0 | 1 | 4 | 3 | G3UZM1 | Probable Ji | G3UZM1 [6 | 0 | 1521.808 | 0 | -0.97 | 0.000565 | 1.7 | 132.7 | 67.3 | 6.93 | 1.32 | | High | High | High | High | High | High | High | High | High | High | High | 0 | 3.66E-05 | 2.58 |
| FALSE | High | LTHSPEGQ | [K].LTHSPE | 1xPhospho | Q2KHI9-2 1 | 2.36E-05 | 0 | 1 | 3 | 2 | Q2KHI9-2 | Isoform 2 c | Q2KHI9-2 [ | 0 | 2942.491 | 1 | -0.94 | 0.000355 | 0.81 | 132.7 | 67.3 | 4.89 | 2.97 | | High | High | High | High | High | High | High | High | High | High | High | 0 | 3.53E-06 | 4.65 |
| FALSE | High | LTLEQVVAI | [R].LTLEQV | 1xPhospho | D0VLQ4 1x | 6.79E-06 | 0 | 1 | 3 | 1 | D0VLQ4 | Tet oncoge | D0VLQ4 [8 | 0 | 3936.107 | 0 | -1.12 | 0.020729 | 11.26 | 131.5 | 68.5 | 16.96 | 31.15 | | High | High | High | High | High | High | High | High | High | High | High | 0 | 8.75E-07 | 4 |
| FALSE | High | ITIGQSPTE | [K].ITIGQSI | 1xPhospho | Q3UZH3 1x | 0.000356 | 0 | 1 | 5 | 2 | Q3UZH3 | Putative ur | Q3UZH3 [5 | 0 | 1611.876 | 0 | -0.36 | 0.078548 | 6.2 | 114.5 | 85.5 | 6.14 | 17.31 | | High | High | High | High | High | High | High | High | High | High | High | 0 | 7.51E-05 | 2.93 |
| FALSE | High | LTLPTVPTP | [R].LTLPTV | 1xCarbami | P23798 1x | 0.000162 | 0 | 1 | 2 | 2 | P23798 | Polycomb g | P23798 [23 | 0 | 3302.575 | 0 | -1.06 | 0.005876 | 7.46 | 134.4 | 65.6 | 9.82 | 14.04 | | High | High | High | High | High | High | High | High | High | High | High | 0 | 3.1E-05 | 2.99 |
| FALSE | High | LTLSGSDSV | [K].LTLSGS | 1xPhospho | Q3TWZ3 1 | 3.62E-07 | 0 | 1 | 2 | 10 | Q3TWZ3 | Putative ur | Q3TWZ3 [3 | 0 | 3070.405 | 0 | -0.28 | 0.301488 | 6.12 | 106.4 | 93.6 | 7.89 | 16.67 | | High | High | High | High | High | High | High | High | High | High | High | 0 | 3.24E-08 | 4.94 |
| FALSE | High | LTISLTQLSI | [K].LTISLTC | 1xPhospho | Q05860 1x | 9.11E-05 | 0 | 1 | 8 | 1 | Q05860 | Formin-1 C | Q05860 [8 | 0 | 1826.044 | 0 | -1.04 | 0.003291 | 6.47 | 134.5 | 65.5 | 9.09 | 13.08 | | High | High | High | High | High | High | High | High | High | High | High | 0 | 1.63E-05 | 3.52 |
| FALSE | High | ITISSLQDLT | [K].ITISSLQ | 1xPhospho | O88477 1x | 0.000113 | 0 | 1 | 2 | 2 | O88477 | Insulin-like | O88477 [3 | 0 | 2172.115 | 0 | -1.18 | 0.000877 | 0.97 | 137 | 63 | 8.68 | 10.96 | | High | High | High | High | High | High | High | High | High | High | High | 0 | 2.06E-05 | 4.16 |
| FALSE | High | LTISSPLEAI | [R].LTISSPL | 1xPhospho | B2RSV4 1x | 0.061176 | 0.004646 | 1 | 2 | 2 | B2RSV4 | Splicing fac | B2RSV4 [1 | 0 | 1733.96 | 0 | -0.6 | 0.000909 | 3.86 | 120.3 | 79.7 | 4.93 | 9.87 | | High | High | High | High | High | High | High | High | High | High | High | 0.00237 | 0.02438 | 1.3 |
| FALSE | High | LTLTSDEST | [K].LTLTSD | 1xPhospho | A9CLV6 1x | 5.62E-06 | 0 | 1 | 2 | 6 | A9CLV6 | Axotomy ir | A9CLV6 [3 | 0 | 2130.016 | 0 | 0.98 | 0.001547 | 14.31 | 67.3 | 132.7 | 8.27 | 8.96 | | High | High | High | High | High | High | High | High | High | High | High | 0 | 2.4E-06 | 3.73 |
| FALSE | High | LTGQESSL( | [K].LTGQE | 2xPhospho | O88622 2x | 6.89E-05 | 0 | 1 | 4 | 1 | O88622 | Poly(ADP-r | O88622 [2 | 0 | 2310.071 | 0 | -0.35 | 0.012625 | 4.04 | 114.9 | 85.1 | 5.83 | 13.96 | | High | High | High | High | High | High | High | High | High | High | High | 0 | 1.18E-05 | 2.9 |
| FALSE | High | LTNLSSNSI | [R].LTNLSS | 1xPhospho | Q9Z0L0 1x | 0.013787 | 0.001011 | 1 | 2 | 1 | Q9Z0L0 | Trophoblas | Q9Z0L0 [4 | 0 | 1358.64 | 0 | 0.34 | 0.04423 | 21.06 | 87.9 | 112.1 | 12.08 | 13.76 | | High | High | High | High | High | High | High | High | High | High | High | 0.000538 | 0.004492 | 1.74 |
| FALSE | High | LTPAAIGIA | [R].LTPAAI | 1xPhospho | A2AW15 1 | 0.000151 | 0 | 1 | 5 | 1 | A2AW15 | Tau-tubulii | A2AW15 [3 | 0 | 2283.268 | 0 | 0.25 | 0.044768 | 14.37 | 90.8 | 109.2 | 8.08 | 9.49 | | High | High | High | High | High | High | High | High | High | High | High | 0 | 2.86E-05 | 2.97 |
| FALSE | High | LTPAAIGIA | [R].LTPAAI | 2xPhospho | A2AW15 2 | 4.76E-05 | 0 | 1 | 5 | 1 | A2AW15 | Tau-tubulii | A2AW15 [3 | 0 | 2363.234 | 0 | 0.13 | 0.083111 | 6.21 | 96.6 | 103.4 | 20.93 | 6.62 | | High | High | High | High | High | High | High | High | High | High | High | 0 | 7.83E-06 | 3.97 |
| FALSE | High | LTPAEEESN | [K].LTPAEE | 1xPhospho | D3YTP3 1x | 0.001424 | 5.66E-05 | 1 | 1 | 1 | D3YTP3 | Metaxin 3 | D3YTP3 [2 | 0 | 2036.999 | 0 | 1.03 | 0.001448 | 6.46 | 65.9 | 134.1 | 10.11 | 11.3 | | High | High | High | High | High | High | High | High | High | High | High | 2.58E-05 | 0.000354 | 2.57 |
| FALSE | High | LTPEEEEILI | [K].LTPEEE | 1xPhospho | Q497E9 1x | 8.87E-05 | 0 | 1 | 5 | 3 | Q497E9 | 40S riboso | Q497E9 [1 | 0 | 1852.971 | 0 | 1.14 | 0.000216 | 5.97 | 60.6 | 139.4 | 4.24 | 4.16 | | High | High | High | High | High | High | High | High | High | High | High | 0 | 1.57E-05 | 3.25 |
| FALSE | High | LTPFHSPSP | [R].LTPFHS | 1xPhospho | B2RY04 1x | 0.000286 | 0 | 1 | 2 | 1 | B2RY04 | Dedicator c | B2RY04 [1 | 0 | 2951.605 | 0 | 0.31 | 0.479451 | 26.06 | 93.6 | 106.4 | 15.44 | 16.31 | | High | High | High | High | High | High | High | High | High | High | High | 0 | 5.88E-05 | 3.61 |
| FALSE | High | ITPHTSPAP | [K].ITPHTSI | 1xPhospho | H3BKF3 1x | 1.72E-05 | 0 | 1 | 3 | 2 | H3BKF3 | PAB-deper | H3BKF3 [3 | 0 | 1385.714 | 0 | -0.07 | 0.465936 | 6.14 | 102.3 | 97.7 | 0.45 | 10.12 | | High | High | High | High | High | High | High | High | High | High | High | 0 | 2.47E-06 | 2.73 |
| FALSE | High | ITPHTSPAP | [K].ITPHTSI | 2xPhospho | H3BKF3 2x | 0.005516 | 0.00062 | 1 | 3 | 1 | H3BKF3 | PAB-deper | H3BKF3 [3 | 0 | 1465.68 | 0 | 1.13 | 0.001253 | 10.99 | 64.8 | 135.2 | 4.89 | 10.81 | | High | High | High | High | High | High | High | High | High | High | High | 0.000275 | 0.001611 | 2.31 |
| FALSE | High | LTPPPADLF | [R].LTPPPA | 1xPhospho | Q61127 1x | 2.43E-06 | 0 | 1 | 3 | 1 | Q61127 | NGFI-A-bin | Q61127 [4 | 0 | 2431.31 | 0 | 0.47 | 0.004647 | 10.11 | 90.1 | 109.9 | 14.67 | 9.11 | | High | High | High | High | High | High | High | High | High | High | High | 0 | 2.76E-07 | 2.92 |
| FALSE | High | LTPSPDIIVL | [R].LTPSPD | 1xPhospho | Q8VHR5 1x | 9.21E-09 | 0 | 1 | 3 | 6 | Q8VHR5 | Transcripti | Q8VHR5 [1 | 0 | 2320.163 | 0 | -0.55 | 0.003626 | 1.41 | 119 | 81 | 3.64 | 7.9 | | High | High | High | High | High | High | High | High | High | High | High | 0 | 5.23E-10 | 4.53 |
| FALSE | High | LTPSPDIIVL | [R].LTPSPD | 2xPhospho | Q8VHR5 2x | 1.45E-05 | 0 | 1 | 3 | 7 | Q8VHR5 | Transcripti | Q8VHR5 [1 | 0 | 2400.13 | 0 | -0.38 | 0.003331 | 6.19 | 112.6 | 87.4 | 3.14 | 6.81 | | High | High | High | High | High | High | High | High | High | High | High | 0 | 3.61E-06 | 4.34 |
| FALSE | High | LTPTSVLDY | [K].LTPTSV | 1xPhospho | G3UWX1 1 | 6.61E-06 | 0 | 2 | 9 | 3 | G3UWX1; I | Replicatior | G3UWX1 [ | 0 | 2222.094 | 0 | -0.01 | 0.165394 | 1.05 | 101 | 99 | 3.61 | 3.2 | Shared | High | High | High | High | High | High | High | High | High | High | High | 0 | 1.53E-05 | 3.48 |
| FALSE | High | LTPTTPESS | [R].LTPTTP | 1xPhospho | Q64337 1x | 0.000415 | 0 | 1 | 3 | 1 | Q64337 | Sequestos | Q64337 [2 | 0 | 1959.9 | 0 | 2.11 | 0.002585 | 10.41 | 39.8 | 160.2 | 32.63 | 14.01 | | High | High | High | High | High | High | High | High | High | High | High | 0 | 8.89E-05 | 2.3 |
| FALSE | High | LTPTTPESS | [R].LTPTTP | 1xPhospho | Q64337 1x | 9.76E-07 | 0 | 1 | 3 | 5 | Q64337 | Sequestos | Q64337 [2 | 0 | 2189.063 | 0 | 1.54 | 0.000325 | 8.68 | 49.8 | 150.2 | 1.95 | 6.31 | | High | High | High | High | High | High | High | High | High | High | High | 0 | 1.68E-07 | 4.22 |
| FALSE | High | LTNTSFTEII | [K].LTNTSF | 1xPhospho | A0A0R4J0T | 2.87E-05 | 0 | 1 | 2 | 1 | A0A0R4J0T | ADP-ribosy | A0A0R4J0T | 0 | 1820.945 | 0 | 1.12 | 0.008087 | 33.66 | 63 | 137 | 9.63 | 16.44 | | High | High | High | High | High | High | High | High | High | High | High | 0 | 4.42E-06 | 3.67 |
| FALSE | High | LTGQESSL( | [K].LTGQE | 1xPhospho | O88622 1x | 6E-07 | 0 | 1 | 4 | 2 | O88622 | Poly(ADP-r | O88622 [2 | 0 | 2230.104 | 0 | -1.48 | 0.000154 | 0.34 | 147.4 | 52.6 | 5.88 | 9.82 | | High | High | High | High | High | High | High | High | High | High | High | 0 | 4.14E-07 | 3.6 |
| FALSE | High | ITGLSSGGA | [R].ITGLSS | 1xCarbami | Q80ZK9 1x | 0.008022 | 0.000736 | 1 | 3 | 1 | Q80ZK9 | WD and te | Q80ZK9 [6 | 0 | 2960.278 | 0 | -0.17 | 0.055685 | 4.06 | 110.6 | 89.4 | 11.54 | 7.3 | | High | High | High | High | High | High | High | High | High | High | High | 0.000327 | 0.002442 | 1.91 |
| FALSE | High | ITGCTSPGK | [K].ITGCTS | 1xCarbami | P80315 1x | 0.000568 | 0 | 1 | 4 | 4 | P80315 | T-complex | P80315 [37 | 0 | 1458.743 | 0 | -1.46 | 0.000364 | 0.63 | 145.3 | 54.7 | 2.72 | 11.74 | | High | High | High | High | High | High | High | High | High | High | High | 9.42E-05 | 0.001018 | 2.46 |
| FALSE | High | LSVQSNPSI | [K].LSVQSN | 1xPhospho | Q3TIV2 1x | 2.72E-05 | 0 | 1 | 4 | 2 | Q3TIV2 | Putative ur | Q3TIV2 [12 | 0 | 1634.846 | 0 | -0.62 | 0.000628 | 2.47 | 118.4 | 81.6 | 3.02 | 6.29 | | High | High | High | High | High | High | High | High | High | High | High | 0 | 0.000109 | 2.53 |
| FALSE | High | LSVSSLLAA | [R].LSVSSLI | 1xPhospho | Q69ZR2 1x | 6.47E-05 | 0 | 1 | 3 | 5 | Q69ZR2 | E3 ubiquiti | Q69ZR2 [1 | 0 | 2698.368 | 0 | 0.66 | 0.009634 | 12.14 | 73.7 | 126.3 | 14.59 | 12.28 | | High | High | High | High | High | High | High | High | High | High | High | 0 | 1.1E-05 | 3.21 |
| FALSE | High | LSVSSLLAA | [R].LSVSSLI | 1xPhospho | Q69ZR2 1x | 8.11E-05 | 0 | 1 | 3 | 2 | Q69ZR2 | E3 ubiquiti | Q69ZR2 [1 | 0 | 2714.363 | 0 | 1.4 | 0.006998 | 9.66 | 56.3 | 143.7 | 13.25 | 21.49 | | High | High | High | High | High | High | High | High | High | High | High | 0 | 1.42E-05 | 3.74 |
| FALSE | High | LSVSSNDT( | [R].LSVSSN | 1xPhospho | G3X9K3 1x | 3.37E-11 | 0 | 1 | 2 | 6 | G3X9K3 | Brefeldin A | G3X9K3 [3 | 0 | 2644.25 | 0 | 1.01 | 0.004178 | 11.06 | 69.2 | 130.8 | 4.71 | 15.67 | | High | High | High | High | High | High | High | High | High | High | High | 0 | 8.92E-12 | 6.23 |
| FALSE | High | ISVVSATK | [K].ISVVSA | 1xPhospho | Q9JIX8-3 1 | 0.004531 | 0.000456 | 3 | 10 | 1 | Q9JIX8-3; F | Isoform 3 c | Q9JIX8-3 [ | 0 | 1342.775 | 0 | -0.01 | 0.603791 | 22.4 | 98.2 | 101.8 | 18.54 | 15.81 | Shared | High | High | High | High | High | High | High | High | High | High | High | 0.000204 | 0.001298 | 2.06 |
| FALSE | High | LSWGYGEH | [R].LSWGY | 1xPhospho | Q9D6N1 1x | 0.012959 | 0.000889 | 1 | 1 | 2 | Q9D6N1 | Carbonic a | Q9D6N1 [4 | 0 | 4186.048 | 0 | 2.39 | 0.000313 | 11.32 | 29.2 | 170.8 | 18.89 | 6.57 | | High | High | High | High | High | High | High | High | High | High | High | 0.000585 | 0.005266 | 3.08 |
| FALSE | High | LSWGYGEH | [R].LSWGY | 1xDeamida | Q9D6N1 1x | 0.000438 | 0 | 1 | 1 | 1 | Q9D6N1 | Carbonic a | Q9D6N1 [4 | 0 | 4187.032 | 0 | 2.12 | 0.001262 | 49.59 | 35.7 | 164.3 | 48.52 | 10.58 | | High | High | High | High | High | High | High | High | High | High | High | 0 | 9.44E-05 | 4.08 |
| FALSE | High | ISYIPDEEVS | [K].ISYIPDE | 1xPhospho | Q5SF07 1x | 3.31E-07 | 0 | 1 | 3 | 7 | Q5SF07 | Insulin-like | Q5SF07 [1 | 0 | 2219.058 | 0 | 0.56 | 0.001821 | 0.66 | 81.6 | 118.4 | 3.6 | 4.24 | | High | High | High | High | High | High | High | High | High | High | High | 0 | 1.61E-06 | 3.89 |
| FALSE | High | ISYIPDEEVS | [K].ISYIPDE | 2xPhospho | Q5SF07 2x | 0.000845 | 0 | 1 | 3 | 2 | Q5SF07 | Insulin-like | Q5SF07 [1 | 0 | 2299.025 | 0 | -0.39 | 0.000222 | 0.96 | 112.9 | 87.1 | 3.73 | 4.39 | | High | High | High | High | High | High | High | High | High | High | High | 0.000161 | 0.001196 | 3.53 |
| FALSE | High | LSYLSAPGS | [R].LSYLSA | 1xPhospho | P39447 1x | 0.039686 | 0.00283 | 1 | 3 | 1 | P39447 | Tight junct | P39447 [8 | 0 | 2423.068 | 0 | 0.16 | 0.96892 | 22.18 | 94.4 | 105.6 | 10.29 | 22.04 | | High | High | High | High | High | High | High | High | High | High | High | 0.001468 | 0.0148 | 2.34 |
| FALSE | High | ISYPDLEEV | [R].ISYPDLI | 1xPhospho | Q8VCG1 1x | 0.02933 | 0.001857 | 1 | 3 | 1 | Q8VCG1 | Deoxyuridi | Q8VCG1 [1 | 0 | 2332.079 | 0 | -0.46 | 0.010452 | 8.45 | 117.9 | 82.1 | 6.96 | 10.34 | | High | High | High | High | High | High | High | High | High | High | High | 0.000997 | 0.0105 | 3.54 |
| FALSE | High | ISYPDLEEV | [R].ISYPDLI | 1xPhospho | Q8VCG1 1x | 0.004696 | 0.000456 | 1 | 3 | 1 | Q8VCG1 | Deoxyuridi | Q8VCG1 [1 | 1 | 3396.625 | 0 | -1.16 | 0.001573 | 4.31 | 138.2 | 61.8 | 8.95 | 14.2 | | High | High | High | High | High | High | High | High | High | High | High | 0.000204 | 0.001344 | 3.79 |
| FALSE | High | LSYSTSQEF | [K].LSYSTS | 1xPhospho | A2ATK9 1x | 0.000244 | 0 | 1 | 3 | 1 | A2ATK9 | Family with | A2ATK9 [4 | 0 | 1700.773 | 0 | 0.99 | 0.001814 | 14.59 | 66.6 | 133.4 | 3.98 | 9.34 | | High | High | High | High | High | High | High | High | High | High | High | 0 | 4.88E-05 | 2.9 |
| FALSE | High | ISYTPPESP\ | [R].ISYTPPI | 1xPhospho | P97360 1x | 1.52E-06 | 0 | 1 | 3 | 10 | P97360 | Transcripti | P97360 [1 | 0 | 1849.905 | 0 | 0 | 0.384303 | 4.2 | 101.4 | 98.6 | 2.35 | 7.45 | | High | High | High | High | High | High | High | High | High | High | High | 0 | 1.63E-07 | 3.57 |
| FALSE | High | ISYTPPESP\ | [R].ISYTPPI | 2xPhospho | P97360 2x | 0.003224 | 0.00016 | 1 | 3 | 2 | P97360 | Transcripti | P97360 [1 | 0 | 1929.871 | 0 | 0.79 | 0.000358 | 2.46 | 72.9 | 127.1 | 3.4 | 4.32 | | High | High | High | High | High | High | High | High | High | High | High | 7.19E-05 | 0.000882 | 3.57 |
| FALSE | High | ITAAQHSV | [R].ITAAQH | 1xPhospho | Q570Z8 1x | 0.000793 | 0 | 1 | 7 | 1 | Q570Z8 | MKIAA411 | Q570Z8 [2 | 0 | 1995.068 | 0 | 1.34 | 0.040347 | 56.26 | 59.1 | 140.9 | 44.78 | 22.77 | | High | High | High | High | High | High | High | High | High | High | High | 0 | 0.000185 | 1.53 |
| FALSE | High | LTATPTPLG | [K].LTATPT | 1xPhospho | G5E866 1x | 2.19E-05 | 0 | 1 | 5 | 2 | G5E866 | Splicing fac | G5E866 [4 | 0 | 2570.198 | 0 | 0.46 | 0.00251 | 8.01 | 84 | 116 | 10.19 | 7.35 | | High | High | High | High | High | High | High | High | High | High | High | 0 | 3.25E-06 | 3.77 |
| FALSE | High | LTATPTPLG | [K].LTATPT | 1xPhospho | G5E866 1x | 0.062209 | 0.004646 | 1 | 5 | 1 | G5E866 | Splicing fac | G5E866 [4 | 0 | 2602.188 | 0 | -0.33 | 0.006581 | 4.09 | 114 | 86 | 6.45 | 10.44 | | High | High | High | High | High | High | High | High | High | High | High | 0.002381 | 0.02475 | 1.82 |
| FALSE | High | LTDESPIQE | [K].LTDESP | 1xPhospho | Q6PR54-3 | 0.000529 | 0 | 2 | 4 | 4 | Q6PR54-3; | Isoform 3 c | Q6PR54-3 | 0 | 2155.094 | 0 | -1.25 | 0.000193 | 0.53 | 141.2 | 58.8 | 3.71 | 5.03 | Shared | High | High | High | High | High | High | High | High | High | High | High | 0 | 0.000116 | 3.18 |
| FALSE | High | LTDPDLTSF | [R].LTDPDL | 1xPhospho | A0A0R4J0F | 0.003282 | 0.00016 | 1 | 4 | 2 | A0A0R4J0F | Germ cell-s | A0A0R4J0F | 0 | 1681.881 | 0 | -0.15 | 0.246465 | 3.1 | 105.4 | 94.6 | 10.77 | 6.35 | | High | High | High | High | High | High | High | High | High | High | High | 7.19E-05 | 0.000903 | 3.12 |
| FALSE | High | LTEGEGSEI | [R].LTEGEG | 1xPhospho | E9QLG3 1x | 0.001463 | 5.66E-05 | 1 | 7 | 1 | E9QLG3 | MAX gene- | E9QLG3 [2 | 0 | 2367.107 | 0 | -0.51 | 0.013013 | 3.8 | 116.1 | 83.9 | 6.07 | 10.44 | | High | High | High | High | High | High | High | High | High | High | High | 2.58E-05 | 0.000366 | 2.04 |
| FALSE | High | ITEQEEDSP | [K].ITEQEE | 1xPhospho | Q5SS00 1x | 0.000117 | 0 | 1 | 1 | 3 | Q5SS00 | DBF4-type | Q5SS00 [1 | 0 | 1713.835 | 0 | -1.34 | 0.000909 | 3.12 | 140.8 | 59.2 | 7.13 | 11.19 | | High | High | High | High | High | High | High | High | High | High | High | 0 | 0.000148 | 3.27 |
| FALSE | High | ITESEEVVS | [R].ITESEE\ | 1xPhospho | P48678 1x | 0.051258 | 0.003745 | 1 | 3 | 1 | P48678 | Prelamin-A | P48678 [6 | 0 | 1457.709 | 0 | 0.49 | 0.009861 | 12.74 | 82.9 | 117.1 | 7.41 | 13.92 | | High | High | High | High | High | High | High | High | High | High | High | 0.00196 | 0.01978 | 2.33 |
| FALSE | High | LTFDSSFSP | [K].LTFDSS | 1xPhospho | Q60932 1x | 0.000211 | 0 | 1 | 6 | 3 | Q60932 | Voltage-de | Q60932 [1 | 0 | 1938.961 | 0 | -0.43 | 0.004354 | 2.65 | 113.2 | 86.8 | 4.4 | 7.85 | | High | High | High | High | High | High | High | High | High | High | High | 0 | 4.13E-05 | 3.46 |
| FALSE | High | LTFDTTFSP | [K].LTFDTT | 1xPhospho | Q60930 1x | 0.005924 | 0.00062 | 1 | 3 | 1 | Q60930 | Voltage-de | Q60930 [1 | 0 | 1966.993 | 0 | -0.98 | 0.000431 | 0.75 | 132.8 | 67.2 | 8 | 5.53 | | High | High | High | High | High | High | High | High | High | High | High | 0.000275 | 0.001743 | 2.24 |
| FALSE | High | LTFHSPEAS | [K].LTFHSP | 1xCarbami | Q8BS90 1x | 3.14E-08 | 0 | 1 | 1 | 1 | Q8BS90 | Protein aur | Q8BS90 [2 | 0 | 2725.322 | 0 | -0.33 | 0.015051 | 4.78 | 111.5 | 88.5 | 11.32 | 12.03 | | High | High | High | High | High | High | High | High | High | High | High | 0 | 2.08E-09 | 3.91 |
| FALSE | High | ITFITSFGG | [K].ITFITSF | 1xPhospho | Q8CFC7 1x | 0.001799 | 5.66E-05 | 1 | 4 | 1 | Q8CFC7 | CLK4-assoc | Q8CFC7 [3 | 0 | 4500.26 | 0 | -0.73 | 0.002237 | 2.91 | 122.5 | 77.5 | 5.72 | 6.55 | | High | High | High | High | High | High | High | High | High | High | High | 2.58E-05 | 0.000459 | 3.37 |
| FALSE | High | LTFSENRPT | [K].LTFSEN | 1xPhospho | A2AKX3 1x | 0.015061 | 0.001049 | 1 | 2 | 2 | A2AKX3 | Probable h | A2AKX3 [1 | 0 | 3832.034 | 0 | 0.05 | 0.133814 | 4.17 | 93.9 | 106.1 | 14.92 | 8.77 | | High | High | High | High | High | High | High | High | High | High | High | 0.000571 | 0.004983 | 2.76 |
| FALSE | High | LTFSPSTDK | [R].LTFSPS | 1xPhospho | Q6NS45 1x | 0.001278 | 0 | 1 | 2 | 2 | Q6NS45 | Coiled-coil | Q6NS45 [8 | 0 | 1533.797 | 0 | -0.3 | 0.01134 | 4.4 | 109.2 | 90.8 | 6.09 | 5.06 | | High | High | High | High | High | High | High | High | High | High | High | 0 | 0.000315 | 2.24 |
| FALSE | High | LSSPPGLFG | [R].LSSPPG | 1xPhospho | Q6NXI6 1x | 3.74E-06 | 0 | 1 | 3 | 1 | Q6NXI6 | Regulation | Q6NXI6 [9 | 0 | 1784.93 | 0 | 0.16 | 0.113855 | 8.41 | 94.4 | 105.6 | 6.38 | 4.02 | | High | High | High | High | High | High | High | High | High | High | High | 0 | 4.46E-07 | 2.19 |
| FALSE | High | LSSPLSEVI\ | [K].LSSPLSI | 2xPhospho | Q6ZQF0 2x | 0.066503 | 0.004796 | 1 | 2 | 1 | Q6ZQF0 | DNA topois | Q6ZQF0 [8 | 0 | 1588.795 | 0 | -0.1 | 0.476576 | 8.58 | 100.2 | 99.8 | 14.66 | 22.84 | | High | High | High | High | High | High | High | High | High | High | High | 0.002455 | 0.02687 | 2.79 |
| FALSE | High | LSSPLSEVI\ | [K].LSSPLSI | 1xPhospho | Q6ZQF0 1x | 0.001463 | 5.66E-05 | 1 | 2 | 2 | Q6ZQF0 | DNA topois | Q6ZQF0 [8 | 0 | 1508.829 | 0 | -0.37 | 0.029023 | 10.79 | 111.4 | 88.6 | 7.14 | 8.28 | | High | High | High | High | High | High | High | High | High | High | High | 2.58E-05 | 0.000364 | 3.6 |
| FALSE | High | ISSPADK | [K].ISSPAD | 1xPhospho | Q8BVE8-2 | 0.096959 | 0.007777 | 2 | 7 | 1 | Q8BVE8-2; | Isoform 2 c | Q8BVE8-2 | 0 | 1255.67 | 0 | 1.31 | 0.000154 | 1.45 | 57.7 | 142.3 | 6.59 | 4.71 | Shared | High | High | High | High | High | High | High | High | High | High | High | 0.003911 | 0.04145 | 1.57 |
| FALSE | High | LSPPLHMA | [R].LSPPLH | 1xPhospho | Q69Z36 1x | 0.001881 | 5.66E-05 | 1 | 2 | 1 | Q69Z36 | RNA-bindir | Q69Z36 [4 | 0 | 2220.095 | 0 | 0.12 | 0.592305 | 31.24 | 86.5 | 113.5 | 18.9 | 17.86 | | High | High | High | High | High | High | High | High | High | High | High | 2.58E-05 | 0.000486 | 2.74 |
| FALSE | High | LSPPPEDSP | [PR].LSPPP | 1xPhospho | A0A0A0M( | 7.23E-06 | 0 | 1 | 2 | 1 | A0A0A0M( | Histone-lys | A0A0A0M( | 0 | 2140.016 | 0 | -0.01 | 0.681732 | 13.32 | 101.1 | 98.9 | 8.98 | 12 | | High | High | High | High | High | High | High | High | High | High | High | 0 | 9.35E-07 | 3.59 |
| FALSE | High | LSPPPGDHI | [R].LSPPPG | 1xCarbami | Q8R0K9 1x | 0.00047 | 0 | 1 | 1 | 1 | Q8R0K9 | Transcripti | Q8R0K9 [3 | 0 | 3941.896 | 0 | -0.81 | 0.00106 | 4.62 | 131.1 | 68.9 | 5.05 | 12.95 | | High | High | High | High | High | High | High | High | High | High | High | 0 | 0.000103 | 2.83 |
| FALSE | High | LSPPPPVTT | [R].LSPPPP | 1xPhospho | Q3TTA7 1x | 1.75E-05 | 0 | 1 | 3 | 3 | Q3TTA7 | E3 ubiquiti | Q3TTA7 [6 | 0 | 2098.233 | 0 | 0.53 | 0.038656 | 19.83 | 81.8 | 118.2 | 10.82 | 14.92 | | High | High | High | High | High | High | High | High | High | High | High | 0 | 2.53E-06 | 3.07 |
| FALSE | High | LSPPPSLA | [K].LSPPPS | 1xPhospho | Q9D6Y4 1x | 0.027121 | 0.001676 | 1 | 2 | 1 | Q9D6Y4 | BLOC-1-rel | Q9D6Y4 [1 | 0 | 1090.575 | 0 | 1.35 | 0.045824 | 17.58 | 56.6 | 143.4 | 11.84 | 32.92 | | High | High | High | High | High | High | High | High | High | High | High | 0.00092 | 0.009578 | 0.77 |
| FALSE | High | ISPPPSVVL | [R].ISPPPS\ | 1xPhospho | B2RS11 1x | 6.31E-06 | 0 | 1 | 3 | 3 | B2RS11 | Cyclin E1 C | B2RS11 [3 | 0 | 2141.202 | 0 | 0.59 | 0.000471 | 3.32 | 79.7 | 120.3 | 4.95 | 3.16 | | High | High | High | High | High | High | High | High | High | High | High | 0 | 2.4E-06 | 3.08 |
| FALSE | High | LSPPYSSPQ | [K].LSPPYS | 1xPhospho | Q62318 1x | 8.3E-07 | 0 | 1 | 2 | 9 | Q62318 | Transcripti | Q62318 [7 | 0 | 2187.032 | 0 | -1.33 | 0.0002 | 0.59 | 144.1 | 55.9 | 3.7 | 4.6 | | High | High | High | High | High | High | High | High | High | High | High | 0 | 2.75E-07 | 4.81 |
| FALSE | High | LSPPYSSPQ | [K].LSPPYS | 2xPhospho | Q62318 2x | 0.002531 | 0.000109 | 1 | 2 | 3 | Q62318 | Transcripti | Q62318 [7 | 0 | 2266.998 | 0 | -1.78 | 0.000254 | 0.83 | 154.6 | 45.4 | 4.04 | 13.33 | | High | High | High | High | High | High | High | High | High | High | High | 4.94E-05 | 0.000674 | 3.88 |
| FALSE | High | LSPQDPPL/ | [K].LSPQDI | 1xPhospho | H9LAN6 1x | 1.59E-06 | 0 | 1 | 6 | 2 | H9LAN6 | Histone-lys | H9LAN6 [8 | 0 | 2574.347 | 0 | -0.92 | 0.000261 | 0.52 | 130.8 | 69.2 | 3.32 | 9.59 | | High | High | High | High | High | High | High | High | High | High | High | 0 | 1.71E-07 | 4.26 |
| FALSE | High | LSPQDPPL/ | [K].LSPQDI | 2xPhospho | H9LAN6 2x | 2.58E-08 | 0 | 1 | 6 | 2 | H9LAN6 | Histone-lys | H9LAN6 [8 | 0 | 2654.313 | 0 | -0.23 | 0.074848 | 7.29 | 108 | 92 | 9.42 | 9.81 | | High | High | High | High | High | High | High | High | High | High | High | 0 | 1.66E-09 | 4.77 |
| FALSE | High | ISPQSNVDI | [K].ISPQSN | 1xPhospho | Q9DCS2 1x | 1.46E-05 | 0 | 1 | 3 | 1 | Q9DCS2 | Methyltrar | Q9DCS2 [1 | 0 | 1913.957 | 0 | 1.09 | 0.000167 | 3.99 | 62.7 | 137.3 | 6.86 | 5.13 | | High | High | High | High | High | High | High | High | High | High | High | 0 | 2.07E-06 | 2.91 |
| FALSE | High | LSPSASPPR | [R].LSPSAS | 3xPhospho | Q52KI8 3x | 0.109329 | 0.008865 | 1 | 6 | 1 | Q52KI8 | Serine/argi | Q52KI8 [3 | 0 | 1380.556 | 0 | -0.17 | 0.06695 | 3.29 | 104.5 | 95.5 | 5.98 | 5.41 | | High | High | High | High | High | High | High | High | High | High | High | 0.004422 | 0.04778 | 2.35 |
| FALSE | High | LSPSAVLAC | [R].LSPSAV | 1xPhospho | Q7TME2 1 | 4.02E-06 | 0 | 1 | 2 | 3 | Q7TME2 | Sperm-ass | Q7TME2 [2 | 0 | 3120.472 | 0 | -0.01 | 0.936021 | 6.06 | 103 | 97 | 14.14 | 15.06 | | High | High | High | High | High | High | High | High | High | High | High | 0 | 4.84E-07 | 4.66 |
| FALSE | High | ISPPETQNL | [R].ISPPET( | 1xPhospho | E9QLG3 1x | 0.007339 | 0.00062 | 1 | 5 | 2 | E9QLG3 | MAX gene- | E9QLG3 [1 | 0 | 1822.972 | 0 | -1.07 | 0.000761 | 1.45 | 136.3 | 63.7 | 7.02 | 3.8 | | High | High | High | High | High | High | High | High | High | High | High | 0.000275 | 0.002217 | 2.31 |
| FALSE | High | LSPSLGSW | [R].LSPSLG | 1xPhospho | Q7TN08 1x | 2.67E-05 | 0 | 1 | 1 | 3 | Q7TN08 | Dapper ho | Q7TN08 [1 | 0 | 2281.24 | 0 | 0.76 | 0.01797 | 23.93 | 74.1 | 125.9 | 12.73 | 9.63 | | High | High | High | High | High | High | High | High | High | High | High | 0 | 5.4E-05 | 2.77 |
| FALSE | High | ISPSPELNLI | [K].ISPSPEI | 1xPhospho | E9QML5 1x | 1.69E-05 | 0 | 1 | 5 | 4 | E9QML5 | Zinc finger | E9QML5 [1 | 0 | 1635.912 | 0 | -0.76 | 0.000202 | 1.52 | 126.5 | 73.5 | 2.57 | 4.95 | | High | High | High | High | High | High | High | High | High | High | High | 0 | 0.000115 | 2.58 |
| FALSE | High | ISPSPELNLI | [K].ISPSPEI | 2xPhospho | E9QML5 2x | 0.021607 | 0.001507 | 1 | 5 | 2 | E9QML5 | Zinc finger | E9QML5 [1 | 0 | 1715.879 | 0 | -0.62 | 0.003095 | 4.97 | 119.4 | 80.6 | 5.21 | 8.15 | | High | High | High | High | High | High | High | High | High | High | High | 0.000801 | 0.007459 | 2.63 |
| FALSE | High | LSPSPSLR | [K].LSPSPSI | 2xPhospho | E9Q784 2x | 0.033694 | 0.002195 | 1 | 3 | 1 | E9Q784 | Zinc finger | E9Q784 [2 | 0 | 1245.584 | 0 | -0.5 | 0.015401 | 1.16 | 116.7 | 83.3 | 12.63 | 5.3 | | High | High | High | High | High | High | High | High | High | High | High | 0.001151 | 0.01232 | 1.64 |
| FALSE | High | LSPSPSSR | [K].LSPSPS | 1xPhospho | P14733 1x | 0.00537 | 0.000599 | 2 | 5 | 2 | P14733; A( | Lamin-B1 C | P14733 [3 | 0 | 1139.566 | 0 | -0.82 | 0.000243 | 1.28 | 127.3 | 72.7 | 3.16 | 7.4 | Shared | High | High | High | High | High | High | High | High | High | High | High | 0.000267 | 0.001564 | 1.52 |
| FALSE | High | LSPSPTAPII | [K].LSPSPT. | 1xPhospho | A0A068F12 | 4.36E-06 | 0 | 1 | 10 | 15 | A0A068F12 | Glyco-gag | A0A068F12 | 0 | 2406.263 | 0 | 2.43 | 0.000213 | 6.93 | 29.5 | 170.5 | 10.51 | 6.42 | | High | High | High | High | High | High | High | High | High | High | High | 0.000275 | 0.002173 | 4.33 |
| FALSE | High | LSPSPTAPII | [K].LSPSPT. | 2xPhospho | A0A068F12 | 0.005768 | 0.00062 | 1 | 10 | 4 | A0A068F12 | Glyco-gag | A0A068F12 | 0 | 2486.229 | 0 | 3.43 | 0.000964 | 27.65 | 20.4 | 179.6 | 34.49 | 17.46 | | High | High | High | High | High | High | High | High | High | High | High | 0.000449 | 0.003951 | 2.86 |
| FALSE | High | LSPSPTSQF | [R].LSPSPT | 1xPhospho | P48678 1x | 0.017041 | 0.001204 | 1 | 3 | 3 | P48678 | Prelamin-A | P48678 [3 | 0 | 1281.64 | 0 | 1.44 | 0.000622 | 5.24 | 55 | 145 | 5.7 | 7.7 | | High | High | High | High | High | High | High | High | High | High | High | 0.000859 | 0.008285 | 2 |
| FALSE | High | ISPSSLSNN | [K].ISPSSLS | 1xPhospho | Q8R5H1 1x | 8.6E-07 | 0 | 1 | 2 | 2 | Q8R5H1 | Ubiquitin c | Q8R5H1 [2 | 0 | 2116.002 | 0 | 0.43 | 0.006552 | 14.32 | 85.2 | 114.8 | 5.38 | 4.95 | | High | High | High | High | High | High | High | High | High | High | High | 0 | 8.59E-08 | 3.38 |
| FALSE | High | LSPSTDSSS | [R].LSPSTD | 1xPhospho | A2A5Y4 1x | 1.24E-05 | 0 | 1 | 5 | 4 | A2A5Y4 | KAT8 regul | A2A5Y4 [2 | 0 | 2088.063 | 0 | -1.07 | 0.000496 | 2.52 | 136.7 | 63.3 | 5.82 | 3.07 | | High | High | High | High | High | High | High | High | High | High | High | 0 | 0.000114 | 2.96 |
| FALSE | High | LSPTDVFC\ | [R].LSPTDV | 1xCarbami | Q9JK23 1x | 3.98E-05 | 0 | 1 | 2 | 2 | Q9JK23 | Proteasom | Q9JK23 [1 | 0 | 2255.159 | 0 | -1.07 | 0.009865 | 6.72 | 134 | 66 | 15.04 | 15.56 | | High | High | High | High | High | High | High | High | High | High | High | 0 | 6.39E-06 | 2.93 |
| FALSE | High | LSPTFPESIE | [R].LSPTFP | 1xPhospho | Q05A36 1x | 1.38E-05 | 0 | 1 | 4 | 3 | Q05A36 | RNA-bindir | Q05A36 [5 | 0 | 2003.02 | 0 | -0.68 | 0.01328 | 1.86 | 121.6 | 78.4 | 11.11 | 7.36 | | High | High | High | High | High | High | High | High | High | High | High | 0 | 1.95E-06 | 4.18 |
| FALSE | High | ISPTGQQS\ | [R].ISPTGQ | 1xCarbami | P97929 1x | 0.000718 | 0 | 1 | 3 | 4 | P97929 | Breast canc | P97929 [3 | 0 | 2670.336 | 0 | -1.46 | 0.019686 | 5.36 | 150.6 | 49.4 | 24.98 | 25.59 | | High | High | High | High | High | High | High | High | High | High | High | 0 | 0.000165 | 3 |
| FALSE | High | LSPTPSMQ | [R].LSPTPS | 1xPhospho | Q52KE7 1x | 5.35E-08 | 0 | 1 | 5 | 1 | Q52KE7 | Cyclin-L1 C | Q52KE7 [7 | 0 | 2497.173 | 0 | -0.51 | 0.006889 | 5.64 | 116.1 | 83.9 | 11.12 | 14.72 | | High | High | High | High | High | High | High | High | High | High | High | 0 | 3.79E-09 | 5.8 |
| FALSE | High | LSPSLLATA | [R].LSPSLL/ | 1xCarbami | Q80U72-4 | 9.07E-10 | 0 | 2 | 5 | 4 | Q80U72-4; | Isoform 4 c | Q80U72-4 | 0 | 2834.461 | 0 | 0.61 | 0.016084 | 19.99 | 78.6 | 121.4 | 14.07 | 10.02 | Shared | High | High | High | High | High | High | High | High | High | High | High | 0 | 3.88E-11 | 4.97 |
| FALSE | High | LSPPAGLFT | [R].LSPPAG | 1xPhospho | P42230 1x | 0.034878 | 0.002269 | 1 | 2 | 1 | P42230 | Signal trans | P42230 [7 | 0 | 1525.798 | 0 | 2.21 | 0.000335 | 3.38 | 37.5 | 162.5 | 15.05 | 9.71 | | High | High | High | High | High | High | High | High | High | High | High | 0.001183 | 0.01272 | 1.58 |
| FALSE | High | LSPNSDRN | [R].LSPNSD | 1xDeamida | Q9WUF3 1 | 0.00407 | 0.00036 | 1 | 1 | 2 | Q9WUF3 | CASP8-asso | Q9WUF3 [ | 1 | 1748.791 | 0 | -0.5 | 0.007864 | 7.98 | 115.4 | 84.6 | 13.47 | 12.25 | | High | High | High | High | High | High | High | High | High | High | High | 0.001214 | 0.0131 | 2.26 |
| FALSE | High | LSPNPPNLT | [K].LSPNPP | 1xPhospho | A0A0R4J17 | 0.00925 | 0.000736 | 1 | 6 | 3 | A0A0R4J17 | Transcripti | A0A0R4J17 | 0 | 1618.897 | 0 | -0.92 | 0.000155 | 1.01 | 131.1 | 68.9 | 0.75 | 4.03 | | High | High | High | High | High | High | High | High | High | High | High | 0.000555 | 0.004808 | 2.21 |
| FALSE | High | LSPCSPPK | [R].LSPCSP | 1xCarbami | A0A0R4J14 | 0.040028 | 0.002867 | 1 | 3 | 2 | A0A0R4J14 | Cell divisior | A0A0R4J14 | 0 | 1423.742 | 0 | -0.76 | 0.00049 | 1.92 | 123.9 | 76.1 | 4.83 | 6.81 | | High | High | High | High | High | High | High | High | High | High | High | 0.002896 | 0.03015 | 1.64 |
| FALSE | High | LSPCSPPK | [R].LSPCSP | 1xCarbami | A0A0R4J14 | 0.045898 | 0.003408 | 1 | 3 | 1 | A0A0R4J14 | Cell divisior | A0A0R4J14 | 0 | 1503.708 | 0 | -1.18 | 0.001972 | 5.52 | 138.5 | 61.5 | 8.71 | 15.26 | | High | High | High | High | High | High | High | High | High | High | High | 0.001741 | 0.01744 | 1.67 |
| FALSE | High | LSPCSSTGK | [R].LSPCSS | 1xCarbami | Q3UHC1 1 | 0.104222 | 0.008348 | 1 | 5 | 1 | Q3UHC1 | Putative ur | Q3UHC1 [3 | 0 | 1474.738 | 0 | -0.16 | 0.01883 | 4.27 | 106.5 | 93.5 | 6 | 5.47 | | High | High | High | High | High | High | High | High | High | High | High | 0.004206 | 0.04524 | 1.07 |
| FALSE | High | LSPDFVEEL | [R].LSPDFV | 1xPhospho | Q80U72-4 | 0.011146 | 0.000772 | 2 | 3 | 1 | Q80U72-4; | Isoform 4 c | Q80U72-4 | 0 | 1513.75 | 0 | -0.1 | 0.355359 | 4.37 | 101.8 | 98.2 | 7.56 | 5.21 | Shared | High | High | High | High | High | High | High | High | High | High | High | 0.000399 | 0.003544 | 2.77 |
| FALSE | High | LSPDIVAEH | [R].LSPDIV. | 1xPhospho | Q64669 1x | 0.000462 | 0 | 1 | 2 | 1 | Q64669 | NAD(P)H d | Q64669 [8 | 0 | 1646.892 | 0 | 2.09 | 0.000191 | 5.85 | 39.6 | 160.4 | 12.65 | 5.35 | | High | High | High | High | High | High | High | High | High | High | High | 0 | 0.0001 | 2.91 |
| FALSE | High | LSPENNQV | [R].LSPENN | 1xPhospho | E9QNT5 1x | 0.000193 | 0 | 1 | 3 | 2 | E9QNT5 | Transcripti | E9QNT5 [1 | 0 | 1780.961 | 0 | -0.59 | 0.006795 | 5.82 | 118 | 82 | 8.64 | 8.8 | | High | High | High | High | High | High | High | High | High | High | High | 7.19E-05 | 0.000858 | 3.11 |
| FALSE | High | LSPEPVAHI | [R].LSPEPV | 1xPhospho | Q8BIZ6 1x | 0.002096 | 5.66E-05 | 1 | 1 | 4 | Q8BIZ6 | Smad nucle | Q8BIZ6 [3 | 0 | 1314.677 | 0 | -0.44 | 0.000317 | 0.4 | 115.3 | 84.7 | 3.94 | 3.6 | | High | High | High | High | High | High | High | High | High | High | High | 2.58E-05 | 0.000549 | 2.37 |
| FALSE | High | LSPEPWTP | [R].LSPEPW | 1xPhospho | Q9QUJ7 1x | 0.000589 | 0 | 1 | 3 | 1 | Q9QUJ7 | Long-chain | Q9QUJ7 [6 | 0 | 2526.293 | 0 | 1.22 | 0.000748 | 5.03 | 53.8 | 146.2 | 21.24 | 7.79 | | High | High | High | High | High | High | High | High | High | High | High | 0 | 0.000132 | 2.61 |
| FALSE | High | LSPEQEGQ | [K].LSPEQE | 1xPhospho | K3W4P2 1x | 0.000305 | 0 | 1 | 1 | 2 | K3W4P2 | Integrator | K3W4P2 [3 | 0 | 1693.818 | 0 | 0.06 | 0.97851 | 8.4 | 99.4 | 100.6 | 10.89 | 2.44 | | High | High | High | High | High | High | High | High | High | High | High | 0 | 9.06E-05 | 2.62 |
| FALSE | High | LSPEQEGQ | [K].LSPEQE | 1xPhospho | K3W4P2 1x | 0.003195 | 0.00016 | 1 | 1 | 2 | K3W4P2 | Integrator | K3W4P2 [3 | 0 | 1709.813 | 0 | -0.09 | 0.631967 | 6.25 | 101.6 | 98.4 | 8 | 8.72 | | High | High | High | High | High | High | High | High | High | High | High | 7.19E-05 | 0.000874 | 2.6 |
| FALSE | High | ISPEQHWD | [K].ISPEQH | 1xPhospho | Q543X6 1x | 0.000372 | 0 | 1 | 2 | 1 | Q543X6 | Putative ur | Q543X6 [8 | 0 | 2354.147 | 0 | 0.61 | 0.011324 | 15.41 | 74.4 | 125.6 | 21.68 | 11.31 | | High | High | High | High | High | High | High | High | High | High | High | 0 | 7.84E-05 | 3.23 |
| FALSE | High | ISPETVNLS | [K].ISPETVI | 1xPhospho | D3Z7B5 1x | 0.00017 | 0 | 1 | 2 | 4 | D3Z7B5 | RIKEN cDN | D3Z7B5 [8 | 0 | 1381.718 | 0 | 0.56 | 0.004776 | 14.69 | 80 | 120 | 6.3 | 6.88 | | High | High | High | High | High | High | High | High | High | High | High | 0 | 3.25E-05 | 2.32 |

| | | | | | | | | | | | | | | | | | | | | | | | | | | | | | | | | | | | | | | |
|---|---|---|---|---|---|---|---|---|---|---|---|---|---|---|---|---|---|---|---|---|---|---|---|---|---|---|---|---|---|---|---|---|---|---|---|---|---|---|
| FALSE | High | LSPEVQSVA | [R].LSPEVC | 1xPhospho | B7ZCJ1 1xP | 0.001011 | 0 | 1 | 5 | 2 | B7ZCJ1 | Rho GTPas | B7ZCJ1 [16 | 0 | 1811.956 | 0 | 0.9 | 0.000265 | 2.35 | 70.9 | 129.1 | 4.51 | 4.13 | | High | High | High | High | High | High | High | High | High | High | High | 2.58E-05 | 0.000391 | 2.15 |
| FALSE | High | LSPFFTLDL | [K].LSPFFTI | 1xPhospho | Q811P8 1x | 0.000106 | 0 | 1 | 3 | 1 | Q811P8 | Rho GTPas | Q811P8 [8 | 0 | 2234.14 | 0 | -1.55 | 0.002836 | 2.43 | 149.3 | 50.7 | 16.12 | 10.53 | | High | High | High | High | High | High | High | High | High | High | High | 0 | 1.92E-05 | 3.21 |
| FALSE | High | LSPFSTPVN | [K].LSPFSTI | 1xPhospho | E9Q5G3 1x | 0.006834 | 0.00062 | 1 | 1 | 1 | E9Q5G3 | Kinesin-like | E9Q5G3 [7 | 0 | 1897.999 | 0 | -0.45 | 0.000401 | 0.55 | 114.7 | 85.3 | 3.84 | 2.24 | | High | High | High | High | High | High | High | High | High | High | High | 0.000275 | 0.002046 | 2.47 |
| FALSE | High | LSPGAYPTV | [R].LSPGAY | 1xPhospho | Q60591 1x | 8.03E-05 | 0 | 1 | 10 | 1 | Q60591 | Nuclear fac | Q60591 [8 | 0 | 2351.196 | 0 | 2.62 | 0.000549 | 28.62 | 28.4 | 171.6 | 12.99 | 8.79 | | High | High | High | High | High | High | High | High | High | High | High | 0 | 1.41E-05 | 3.18 |
| FALSE | High | LSPGEALPS | [R].LSPGEA | 1xPhospho | Q8CGB6-3 | 1.88E-05 | 0 | 1 | 4 | 1 | Q8CGB6-3 | Isoform 3 c | Q8CGB6-3 | 0 | 2219.209 | 0 | -0.59 | 0.004585 | 3.74 | 116.4 | 83.6 | 7.51 | 8.73 | | High | High | High | High | High | High | High | High | High | High | High | 0 | 2.75E-06 | 3.07 |
| FALSE | High | LSPGETPEE | [R].LSPGET | 1xPhospho | Q8BQ30-2 | 0.000563 | 0 | 1 | 4 | 2 | Q8BQ30-2 | Isoform 2 c | Q8BQ30-2 | 0 | 1623.783 | 0 | 0.87 | 0.024667 | 19.85 | 76 | 124 | 8.6 | 27.92 | | High | High | High | High | High | High | High | High | High | High | High | 0 | 0.000125 | 2.33 |
| FALSE | High | LSPGSDIYV | [R].LSPGSD | 1xPhospho | Q8C0T5 1x | 3.29E-05 | 0 | 1 | 3 | 1 | Q8C0T5 | Signal-indu | Q8C0T5 [1 | 0 | 2176.092 | 0 | 0.69 | 0.009442 | 16.16 | 75.6 | 124.4 | 14.35 | 14.94 | | High | High | High | High | High | High | High | High | High | High | High | 0 | 5.14E-06 | 3.78 |
| FALSE | High | LSPLLPGGC | [R].LSPLLP | 1xPhospho | Q91ZV0 1x | 0.021797 | 0.001507 | 1 | 9 | 1 | Q91ZV0 | Melanoma | Q91ZV0 [1 | 0 | 1332.724 | 0 | 0.36 | 0.063936 | 13.08 | 87.5 | 112.5 | 10.57 | 8.55 | | High | High | High | High | High | High | High | High | High | High | High | 0.000814 | 0.007547 | 1.62 |
| FALSE | High | LSPLISSLPS | [K].LSPLISS | 1xPhospho | E9Q921 1x | 2.85E-05 | 0 | 1 | 8 | 1 | E9Q921 | AF4/FMR2 | E9Q921 [1 | 0 | 2489.373 | 0 | -0.11 | 0.404404 | 37.67 | 95.8 | 104.2 | 46.32 | 17.69 | | High | High | High | High | High | High | High | High | High | High | High | 0 | 4.37E-06 | 3.33 |
| FALSE | High | LSPIMTEQI | [R].LSPIMT | 1xPhospho | Q9R1E0 1x | 0.012959 | 0.000889 | 1 | 3 | 2 | Q9R1E0 | Forkhead b | Q9R1E0 [3 | 0 | 3393.669 | 0 | 0.2 | 0.180726 | 19.84 | 95.9 | 104.1 | 11.1 | 11.02 | | High | High | High | High | High | High | High | High | High | High | High | 0.000449 | 0.004206 | 2.91 |
| FALSE | High | LSPIMTEQI | [R].LSPIMT | 1xPhospho | Q9R1E0 1x | 0.000438 | 0 | 1 | 3 | 2 | Q9R1E0 | Forkhead b | Q9R1E0 [3 | 0 | 3409.664 | 0 | 0.04 | 0.831979 | 3.83 | 99.8 | 100.2 | 13.42 | 23.88 | | High | High | High | High | High | High | High | High | High | High | High | 0 | 0.000094 | 3.68 |
| FALSE | High | LSPLPGGPC | [K].LSPLPG | 1xPhospho | Q61127 1x | 3.9E-05 | 0 | 1 | 4 | 2 | Q61127 | NGFI-A-bin | Q61127 [1 | 0 | 1599.809 | 0 | 0.21 | 0.171387 | 16.98 | 92.6 | 107.4 | 6.24 | 7.48 | | High | High | High | High | High | High | High | High | High | High | High | 0.001453 | 0.01451 | 2.81 |
| FALSE | High | LSPLPSLMV | [R].LSPLPSI | 1xPhospho | Q8CCN5 1x | 3.87E-05 | 0 | 1 | 7 | 2 | Q8CCN5 | Breast carc | Q8CCN5 [5 | 0 | 2345.368 | 0 | -0.79 | 0.004939 | 6.67 | 130.4 | 69.6 | 10.96 | 11.64 | | High | High | High | High | High | High | High | High | High | High | High | 0 | 6.17E-06 | 4.25 |
| FALSE | High | LSPLPSLMV | [R].LSPLPSI | 1xPhospho | Q8CCN5 1x | 0.001301 | 0 | 1 | 7 | 1 | Q8CCN5 | Breast carc | Q8CCN5 [5 | 0 | 2361.363 | 0 | -0.67 | 0.057117 | 6.86 | 124.8 | 75.2 | 16.88 | 15.16 | | High | High | High | High | High | High | High | High | High | High | High | 0 | 0.000321 | 2.86 |
| FALSE | High | LSPLSPGSP | [R].LSPLSP | 1xPhospho | Q9Z248 1x | 0.028082 | 0.001791 | 1 | 4 | 4 | Q9Z248 | Zinc finger | Q9Z248 [1 | 0 | 1615.841 | 0 | -0.26 | 0.003001 | 3.94 | 110.4 | 89.6 | 4.52 | 3.16 | | High | High | High | High | High | High | High | High | High | High | High | 0.007201 | 0.07789 | 2.21 |
| FALSE | High | LSPLSPGSP | [R].LSPLSP | 2xPhospho | Q9Z248 2x | 0.014797 | 0.001049 | 1 | 4 | 1 | Q9Z248 | Zinc finger | Q9Z248 [1 | 0 | 1695.807 | 0 | 0.77 | 0.002525 | 16.72 | 74.7 | 125.3 | 8.4 | 7.31 | | High | High | High | High | High | High | High | High | High | High | High | 0.000555 | 0.00487 | 1.81 |
| FALSE | High | ISPLVLFSNI | [R].ISPLVLF | 1xPhospho | Q922M7 1 | 2.34E-06 | 0 | 1 | 1 | 1 | Q922M7 | Ashwin OS | Q922M7 [1 | 0 | 2216.261 | 0 | -0.32 | 0.038164 | 8 | 111.7 | 88.3 | 11.54 | 2.39 | | High | High | High | High | High | High | High | High | High | High | High | 0 | 2.64E-07 | 3.39 |
| FALSE | High | LSPVPPVPC | [R].LSPVPP | 1xPhospho | Q80VP1-2 | 0.038017 | 0.002528 | 1 | 2 | 2 | Q80VP1-2 | Isoform 2 c | Q80VP1-2 | 0 | 4434.288 | 0 | -0.42 | 0.004472 | 7.25 | 112.7 | 87.3 | 8.31 | 13.71 | | High | High | High | High | High | High | High | High | High | High | High | 0.001324 | 0.01404 | 4.19 |
| FALSE | High | LTPVRPAAA | [R].LTPVRP | 2xPhospho | F6YRW4 2x | 0.000687 | 0 | 1 | 4 | 1 | F6YRW4 | F-box and l | F6YRW4 [5 | 0 | 2052.061 | 0 | 0.72 | 0.000518 | 1.83 | 74.6 | 125.4 | 8.47 | 5.79 | | High | High | High | High | High | High | High | High | High | High | High | 0 | 0.000157 | 1.79 |
| FALSE | High | LSPVPVPR | [R].LSPVPV | 1xPhospho | Q64012 1x | 0.031713 | 0.002011 | 1 | 6 | 2 | Q64012 | RNA-bindir | Q64012 [1 | 0 | 1173.659 | 0 | -0.15 | 0.053698 | 6.63 | 105.1 | 94.9 | 7.38 | 4.29 | | High | High | High | High | High | High | High | High | High | High | High | 0.005216 | 0.05692 | 1.67 |
| FALSE | High | ISPVSLPQC | [K].ISPVSLF | 1xCarbami | E9PVX6 1x | 0.003055 | 0.00016 | 1 | 2 | 2 | E9PVX6 | Proliferatic | E9PVX6 [1 | 0 | 3540.77 | 0 | 0.01 | 0.83934 | 8.56 | 99.6 | 100.4 | 10.31 | 14.53 | | High | High | High | High | High | High | High | High | High | High | High | 7.19E-05 | 0.000833 | 3.93 |
| FALSE | High | LSSDDETE( | [R].LSSDDE | 2xPhospho | B2RUC3 2x | 0.000207 | 0 | 1 | 8 | 2 | B2RUC3 | Excision re | B2RUC3 [5 | 0 | 2262.993 | 0 | -0.34 | 0.005251 | 0.88 | 112.1 | 87.9 | 8.65 | 3.34 | | High | High | High | High | High | High | High | High | High | High | High | 0 | 4.06E-05 | 2.96 |
| FALSE | High | LSSDDGDS | [R].LSSDDG | 1xPhospho | Q6KCD5 1x | 2.75E-05 | 0 | 1 | 4 | 3 | Q6KCD5 | Nipped-B-l | Q6KCD5 [2 | 0 | 1595.646 | 0 | 0.65 | 0.08547 | 66.1 | 75.3 | 124.7 | 15.69 | 28.83 | | High | High | High | High | High | High | High | High | High | High | High | 0 | 4.22E-06 | 2.64 |
| FALSE | High | LSSDENSNI | [K].LSSDEN | 1xPhospho | Q921E6-2 | 2.02E-05 | 0 | 1 | 3 | 2 | Q921E6-2 | Isoform 2 c | Q921E6-2 | 0 | 4795.093 | 0 | -1.39 | 0.001189 | 2.5 | 144.9 | 55.1 | 5.12 | 17.64 | | High | High | High | High | High | High | High | High | High | High | High | 0 | 2.99E-06 | 5.64 |
| FALSE | High | LSSDENSNI | [K].LSSDEN | 1xDeamida | Q921E6-2 | 1.38E-10 | 0 | 1 | 3 | 9 | Q921E6-2 | Isoform 2 c | Q921E6-2 | 0 | 4796.077 | 0 | -1.38 | 0.000495 | 2.11 | 144.2 | 55.8 | 4.19 | 14.49 | | High | High | High | High | High | High | High | High | High | High | High | 0 | 4.67E-12 | 6.19 |
| FALSE | High | LSSDENSNI | [K].LSSDEN | 3xPhospho | Q921E6-2 | 0.000106 | 0 | 1 | 3 | 4 | Q921E6-2 | Isoform 2 c | Q921E6-2 | 0 | 4955.025 | 0 | -1.3 | 0.001988 | 3.1 | 136.9 | 63.1 | 9.09 | 15.56 | | High | High | High | High | High | High | High | High | High | High | High | 0 | 1.92E-05 | 4.6 |
| FALSE | High | LSSDLSGSII | [R].LSSDLS | 1xPhospho | F8VPJ6 1xP | 1.12E-05 | 0 | 1 | 3 | 3 | F8VPJ6 | Regulatory | F8VPJ6 [13 | 0 | 2982.466 | 0 | -1.02 | 0.005481 | 2.23 | 134 | 66 | 11.15 | 19.54 | | High | High | High | High | High | High | High | High | High | High | High | 0 | 1.54E-06 | 4.54 |
| FALSE | High | LSSDLTR | [R].LSSDLT | 1xPhospho | P97868 1x | 0.093123 | 0.007276 | 1 | 3 | 2 | P97868 | E3 ubiquiti | P97868 [16 | 0 | 1100.555 | 0 | 0.7 | 0.001603 | 8.36 | 76 | 124 | 3.5 | 8.76 | | High | High | High | High | High | High | High | High | High | High | High | 0.003685 | 0.03952 | 2.02 |
| FALSE | High | LSSEMNTS | [R].LSSEMI | 1xPhospho | P14733 1x | 0.020319 | 0.001472 | 1 | 2 | 2 | P14733 | Lamin-B1 ( | P14733 [27 | 0 | 1805.83 | 0 | -0.8 | 0.000662 | 2.16 | 126.2 | 73.8 | 3.66 | 8.73 | | High | High | High | High | High | High | High | High | High | High | High | 0.000786 | 0.006971 | 1.95 |
| FALSE | High | LSSERPSSD | [K].LSSERP | 1xCarbami | Q80YS6 1x | 0.002022 | 5.66E-05 | 1 | 2 | 2 | Q80YS6 | Actin filam | Q80YS6 [2 | 0 | 2975.382 | 0 | 0.14 | 0.330864 | 11.58 | 92.3 | 107.7 | 10.89 | 11.3 | | High | High | High | High | High | High | High | High | High | High | High | 0.00014 | 0.001127 | 2.32 |
| FALSE | High | LSSERPSSD | [K].LSSERP | 1xCarbami | Q80YS6 1x | 9.06E-06 | 0 | 1 | 2 | 4 | Q80YS6 | Actin filam | Q80YS6 [2 | 0 | 2976.366 | 0 | 0.15 | 0.372143 | 12.39 | 95.1 | 104.9 | 6.15 | 10.61 | | High | High | High | High | High | High | High | High | High | High | High | 0 | 1.21E-06 | 3.73 |
| FALSE | High | LSSESHHG( | [R].LSSESH | 1xPhospho | E9Q5K9 1x | 0.000404 | 0 | 1 | 4 | 1 | E9Q5K9 | YTH domai | E9Q5K9 [4 | 0 | 2939.464 | 0 | -0.18 | 0.427917 | 42.81 | 106.4 | 93.6 | 28.57 | 31.18 | | High | High | High | High | High | High | High | High | High | High | High | 0 | 8.62E-05 | 2.83 |
| FALSE | High | LSSETGGM | [R].LSSETG | 1xPhospho | Q61990 1x | 2.63E-05 | 0 | 2 | 6 | 8 | Q61990; Q | Poly(rC)-bi | Q61990 [3 | 0 | 1337.549 | 0 | 0.01 | 0.917905 | 7.3 | 99.8 | 100.2 | 2.94 | 5.07 | Shared | High | High | High | High | High | High | High | High | High | High | High | 0 | 1.58E-05 | 2.99 |
| FALSE | High | LSSEVAETD | [R].LSSEVA | 1xPhospho | Q9DB00 1x | 0.036416 | 0.002419 | 1 | 5 | 3 | Q9DB00 | GON-4-like | Q9DB00 [4 | 0 | 2707.267 | 0 | -0.25 | 0.380791 | 15.49 | 116.4 | 83.6 | 21.6 | 26.17 | | High | High | High | High | High | High | High | High | High | High | High | 0.001279 | 0.01339 | 2.95 |
| FALSE | High | LSSDDETE( | [R].LSSDDE | 1xPhospho | B2RUC3 1x | 1.62E-06 | 0 | 1 | 8 | 3 | B2RUC3 | Excision re | B2RUC3 [5 | 0 | 2183.027 | 0 | -0.7 | 0.004301 | 2.02 | 122.6 | 77.4 | 10.16 | 3.58 | | High | High | High | High | High | High | High | High | High | High | High | 0 | 1.75E-07 | 3.77 |
| FALSE | High | LSSEVAETD | [R].LSSEVA | 1xPhospho | Q9DB00 1x | 2.32E-06 | 0 | 1 | 5 | 2 | Q9DB00 | GON-4-like | Q9DB00 [4 | 0 | 2723.262 | 0 | -0.57 | 0.222397 | 7.66 | 120.8 | 79.2 | 13.67 | 25.69 | | High | High | High | High | High | High | High | High | High | High | High | 0 | 2.62E-07 | 4.53 |
| FALSE | High | LSSGFDDID | [R].LSSGFD | 1xPhospho | Q9QY23-2 | 4.56E-06 | 0 | 1 | 2 | 1 | Q9QY23-2 | Isoform 2 c | Q9QY23-2 | 0 | 2102.082 | 0 | 1.37 | 0.000895 | 8.11 | 56 | 144 | 4.81 | 9.42 | | High | High | High | High | High | High | High | High | High | High | High | 0 | 5.59E-07 | 3.18 |
| FALSE | High | LSSGLVSPA | [R].LSSGLV | 1xPhospho | E9QMD3 1 | 0.000352 | 0 | 1 | 5 | 1 | E9QMD3 | Zinc finger | E9QMD3 [ | 0 | 2078.098 | 0 | 2.85 | 0.000289 | 8.7 | 23.2 | 176.8 | 14.61 | 7.58 | | High | High | High | High | High | High | High | High | High | High | High | 0 | 7.43E-05 | 3.22 |
| FALSE | High | LSSGPPAQI | [R].LSSGPP | 1xPhospho | A6H5X4 1x | 0.014538 | 0.001011 | 2 | 6 | 2 | A6H5X4; B | PHD finger | A6H5X4 [1 | 0 | 1519.829 | 0 | -1.43 | 0.001617 | 2.63 | 145.8 | 54.2 | 9.21 | 15.12 | Shared | High | High | High | High | High | High | High | High | High | High | High | 0.000555 | 0.004791 | 3.16 |
| FALSE | High | LSSGQISGF | [K].LSSGQI | 1xPhospho | E9Q616 1x | 0.000217 | 0 | 1 | 6 | 1 | E9Q616 | AHNAK nuc | E9Q616 [5 | 0 | 1753.95 | 0 | 2.33 | 0.002461 | 12.48 | 34.4 | 165.6 | 12.5 | 19.68 | | High | High | High | High | High | High | High | High | High | High | High | 0 | 4.28E-05 | 2.79 |
| FALSE | High | LSSGSEEDK | [K].LSSGSE | 1xPhospho | Q9D1P4 1x | 0.052576 | 0.00384 | 1 | 3 | 1 | Q9D1P4 | Cysteine ar | Q9D1P4 [1 | 1 | 1846.977 | 0 | 0.63 | 0.000701 | 4.27 | 78.6 | 121.4 | 3.86 | 5.77 | | High | High | High | High | High | High | High | High | High | High | High | 0.001999 | 0.02033 | 1.3 |
| FALSE | High | LSSIGIQVD | [K].LSSIGIC | 1xCarbami | B1AZP2 1x | 0.000127 | 0 | 1 | 4 | 2 | B1AZP2 | Disks large | B1AZP2 [6 | 0 | 2292.244 | 0 | 0.42 | 0.020384 | 13.62 | 81.8 | 118.2 | 11.29 | 11.72 | | High | High | High | High | High | High | High | High | High | High | High | 0 | 2.37E-05 | 3.52 |
| FALSE | High | ISSLLEEQF( | [K].ISSLLEE | 1xPhospho | Q497E9 1x | 0.001039 | 0 | 1 | 2 | 2 | Q497E9 | 40S ribosoi | Q497E9 [1 | 0 | 2045.072 | 0 | 0.47 | 0.00868 | 11.05 | 83.3 | 116.7 | 10.25 | 6.5 | | High | High | High | High | High | High | High | High | High | High | High | 0 | 0.00025 | 2.78 |
| FALSE | High | LSSLPVSR | [R].LSSLPV | 1xPhospho | E9QAU4 1x | 0.019791 | 0.001438 | 1 | 13 | 2 | E9QAU4 | Enhancer t | E9QAU4 [4 | 0 | 1167.634 | 0 | 1.93 | 0.003397 | 13.21 | 41.6 | 158.4 | 15.87 | 19.19 | | High | High | High | High | High | High | High | High | High | High | High | 0.002031 | 0.02099 | 1.95 |
| FALSE | High | LSSLTAPVG | [K].LSSLTA | 1xPhospho | Q3UGE1 1x | 0.007951 | 0.000736 | 1 | 5 | 2 | Q3UGE1 | Putative ur | Q3UGE1 [2 | 0 | 1639.907 | 0 | 1.26 | 0.002429 | 16.41 | 56.1 | 143.9 | 12.4 | 14.48 | | High | High | High | High | High | High | High | High | High | High | High | 0.000327 | 0.002434 | 1.92 |
| FALSE | High | LSSNCSGVI | [K].LSSNCS | 1xCarbami | Q8K394 1x | 5.27E-09 | 0 | 1 | 1 | 1 | Q8K394 | Inactive ph | Q8K394 [5 | 0 | 2897.165 | 0 | -0.81 | 0.019288 | 10.56 | 126.6 | 73.4 | 8.92 | 18.07 | | High | High | High | High | High | High | High | High | High | High | High | 0 | 2.79E-10 | 6.65 |
| FALSE | High | LSSNGLQS | [R].LSSNGL | 2xPhospho | Q9Z2F6 2x | 0.036416 | 0.002419 | 1 | 2 | 1 | Q9Z2F6 | B-cell lymp | Q9Z2F6 [3 | 0 | 2575.246 | 0 | -0.75 | 0.000486 | 3.55 | 125.6 | 74.4 | 8.75 | 15.49 | | High | High | High | High | High | High | High | High | High | High | High | 0.001279 | 0.01336 | 2.27 |
| FALSE | High | LSSPAAFLP | [K].LSSPAA | 1xCarbami | Q7TPS5 1x | 3.09E-05 | 0 | 2 | 6 | 3 | Q7TPS5; A | C2 domain | Q7TPS5 [2 | 0 | 1956.945 | 0 | -0.05 | 0.738899 | 7.64 | 99.4 | 100.6 | 8.5 | 7.14 | Shared | High | High | High | High | High | High | High | High | High | High | High | 0 | 4.12E-05 | 3.46 |
| FALSE | High | LSSPAAFLP | [K].LSSPAA | 1xCarbami | Q7TPS5 2x | 0.000745 | 0 | 2 | 6 | 1 | Q7TPS5; A | C2 domain | Q7TPS5 [2 | 0 | 2036.912 | 0 | 0.43 | 0.18195 | 38.77 | 78.7 | 121.3 | 34.7 | 19.53 | Shared | High | High | High | High | High | High | High | High | High | High | High | 0 | 0.000171 | 2.42 |
| FALSE | High | LSSGDLR | [K].LSSGDL | 1xPhospho | D3Z0X5 1x | 0.024855 | 0.001645 | 1 | 7 | 1 | D3Z0X5 | Pleckstrin h | D3Z0X5 [5 | 0 | 1056.529 | 1 | 1.01 | 0.004541 | 4.42 | 65.8 | 134.2 | 9.25 | 13.56 | | High | High | High | High | High | High | High | High | High | High | High | 0.000872 | 0.008733 | 1.68 |
| FALSE | High | LSSDAITQI | [K].LSSDAI | 1xPhospho | A0A1D5RM | 1.91E-05 | 0 | 1 | 2 | 1 | A0A1D5RM | Alstrom sy | A0A1D5RM | 0 | 2258.157 | 0 | -0.44 | 0.00251 | 5.4 | 116.5 | 83.5 | 3.23 | 5.98 | | High | High | High | High | High | High | High | High | High | High | High | 0 | 2.81E-06 | 4.61 |
| FALSE | High | ISSCSDESSI | [R].ISSCSDI | 1xCarbami | Q99NH0 1x | 0.067059 | 0.004796 | 1 | 6 | 1 | Q99NH0 | Ankyrin rep | Q99NH0 [1 | 1 | 2283.032 | 0 | -0.38 | 0.014287 | 6.54 | 113.1 | 86.9 | 8.7 | 3.74 | | High | High | High | High | High | High | High | High | High | High | High | 0.002455 | 0.02693 | 1.69 |
| FALSE | High | ISSCSDESSI | [R].ISSCSDI | 1xCarbami | Q99NH0 1x | 0.019276 | 0.001402 | 1 | 6 | 3 | Q99NH0 | Ankyrin rep | Q99NH0 [1 | 0 | 1925.775 | 0 | 0.17 | 0.450973 | 40.47 | 94.7 | 105.3 | 10.86 | 21.52 | | High | High | High | High | High | High | High | High | High | High | High | 0.000756 | 0.006546 | 2.74 |
| FALSE | High | LSPVTPTPE | [K].LSPVTP | 1xPhospho | Q9D2D7 1x | 0.001659 | 5.66E-05 | 1 | 3 | 2 | Q9D2D7 | Zinc finger | Q9D2D7 [3 | 0 | 1760.96 | 0 | -0.41 | 0.006276 | 4.47 | 115.7 | 84.3 | 4.09 | 11.09 | | High | High | High | High | High | High | High | High | High | High | High | 0.000275 | 0.001926 | 3.26 |
| FALSE | High | LSPYPTPSP | [K].LSPYPT | 1xPhospho | I7H459 1xP | 2.42E-05 | 0 | 1 | 2 | 3 | I7H459 | Protein inc | I7H459 [30 | 0 | 2185.104 | 0 | 0.97 | 0.00055 | 8.84 | 67.8 | 132.2 | 8.74 | 3.65 | | High | High | High | High | High | High | High | High | High | High | High | 0 | 3.63E-06 | 3.38 |
| FALSE | High | ISQEHLADH | [R].ISQEHL | 1xPhospho | Q62077 1x | 1.52E-06 | 0 | 1 | 5 | 1 | Q62077 | 1-phospha | Q62077 [1 | 0 | 1863.859 | 0 | -0.61 | 0.094797 | 6.64 | 122 | 78 | 4.18 | 26.74 | | High | High | High | High | High | High | High | High | High | High | High | 0 | 1.63E-07 | 3.57 |
| FALSE | High | LSQELDFVS | [K].LSQELD | 1xPhospho | Q02819 1x | 0.005044 | 0.000504 | 1 | 7 | 1 | Q02819 | Nucleobinc | Q02819 [8 | 0 | 1852.916 | 0 | -0.42 | 0.000758 | 1.12 | 117.2 | 82.8 | 3.6 | 5.18 | | High | High | High | High | High | High | High | High | High | High | High | 0.000246 | 0.001462 | 2.5 |
| FALSE | High | ISQHGGSST | [R].ISQHGC | 1xPhospho | Q8CHG1 1x | 0.005467 | 0.00062 | 1 | 13 | 1 | Q8CHG1 | MKIAA036 | Q8CHG1 [3 | 0 | 2015.021 | 0 | 0.82 | 0.034691 | 8.95 | 74.2 | 125.8 | 6.61 | 29.59 | | High | High | High | High | High | High | High | High | High | High | High | 0.000275 | 0.001602 | 2.66 |
| FALSE | High | LSQPAPPIS | [R].LSQPAF | 1xPhospho | A2AI08 1xP | 0.000127 | 0 | 1 | 1 | 1 | A2AI08 | Taperin OS | A2AI08 [18 | 0 | 1966.072 | 0 | 1.9 | 0.000466 | 13.99 | 42.3 | 157.7 | 13.86 | 5.96 | | High | High | High | High | High | High | High | High | High | High | High | 0 | 2.36E-05 | 2.74 |
| FALSE | High | ISQQHSLYV | [R].ISQQHS | 1xPhospho | Q64701 1x | 0.021046 | 0.001507 | 1 | 3 | 1 | Q64701 | Retinoblast | Q64701 [9 | 0 | 2062.089 | 0 | 0.59 | 0.088063 | 32.54 | 81.3 | 118.7 | 16.62 | 13.61 | | High | High | High | High | High | High | High | High | High | High | High | 0.000801 | 0.007189 | 2.98 |
| FALSE | High | LSQQHSQA | [K].LSQQH | 1xPhospho | E9PZ26 1xP | 0.007404 | 0.00062 | 1 | 2 | 1 | E9PZ26 | Bromodom | E9PZ26 [75 | 0 | 4369.066 | 0 | 0.2 | 0.318338 | 45.91 | 91.3 | 108.7 | 29.76 | 17.22 | | High | High | High | High | High | High | High | High | High | High | High | 0.000275 | 0.002247 | 3.46 |
| FALSE | High | LSQQLDK | [R].LSQQLI | 1xPhospho | Q8R0A0 1x | 0.103392 | 0.008249 | 1 | 1 | 1 | Q8R0A0 | General tra | Q8R0A0 [1 | 0 | 1369.749 | 0 | -0.35 | 0.04905 | 11.97 | 109.5 | 90.5 | 11.69 | 13.71 | | High | High | High | High | High | High | High | High | High | High | High | 0.004164 | 0.04493 | 1.33 |
| FALSE | High | ISQSALPPL | [K].ISQSAL | 1xPhospho | S4R221 1xP | 0.014283 | 0.001011 | 1 | 8 | 1 | S4R221 | Rho GTPas | S4R221 [24 | 0 | 3194.665 | 0 | -0.18 | 0.072975 | 5.39 | 109.1 | 90.9 | 17.64 | 16.82 | | High | High | High | High | High | High | High | High | High | High | High | 0.000538 | 0.004669 | 2.59 |
| FALSE | High | ISQSDSEEV | [K].ISQSDS | 2xPhospho | Q9CZ92 2x | 0.064323 | 0.004664 | 1 | 2 | 1 | Q9CZ92 | Centromer | Q9CZ92 [4 | 0 | 2155.96 | 0 | -0.44 | 0.000474 | 1.35 | 115.2 | 84.8 | 7.5 | 6.67 | | High | High | High | High | High | High | High | High | High | High | High | 0.002403 | 0.02579 | 1.87 |
| FALSE | High | LSQSIPFVP | [R].LSQSIPI | 1xPhospho | B9EK83 1x | 0.016451 | 0.001124 | 2 | 3 | 2 | B9EK83; A | Transient r | B9EK83 [1 | 0 | 1745.955 | 0 | 2.15 | 0.000222 | 8.89 | 36.7 | 163.3 | 19.65 | 9.59 | Shared | High | High | High | High | High | High | High | High | High | High | High | 0.000585 | 0.005457 | 2.66 |
| FALSE | High | ISQSQPVR | [R].ISQSQP | 1xPhospho | E9Q2H1 1x | 0.049971 | 0.003641 | 1 | 2 | 2 | E9Q2H1 | E3 ubiquiti | E9Q2H1 [1 | 0 | 1223.635 | 0 | 0.34 | 0.001827 | 2.13 | 88.5 | 111.5 | 1.71 | 4.14 | | High | High | High | High | High | High | High | High | High | High | High | 0.001902 | 0.01919 | 2.03 |
| FALSE | High | LSQSSQDS | [R].LSQSSC | 1xPhospho | Q5SVQ0 1x | 0.000488 | 0 | 1 | 5 | 3 | Q5SVQ0 | Histone ace | Q5SVQ0 [5 | 0 | 1599.758 | 0 | 0.08 | 0.562682 | 17.61 | 97.3 | 102.7 | 5.43 | 6.48 | | High | High | High | High | High | High | High | High | High | High | High | 0 | 0.000118 | 2.22 |
| FALSE | High | LSQSSQDS | [R].LSQSSC | 2xPhospho | Q5SVQ0 2x | 0.024211 | 0.001612 | 1 | 5 | 3 | Q5SVQ0 | Histone ace | Q5SVQ0 [5 | 0 | 1679.724 | 0 | -0.38 | 0.049145 | 12.67 | 111.2 | 88.8 | 11.07 | 10.53 | | High | High | High | High | High | High | High | High | High | High | High | 0.000859 | 0.008436 | 3.23 |
| FALSE | High | LSQSSVNLS | [R].LSQSSV | 1xPhospho | B2KF52 1x | 0.000438 | 0 | 1 | 2 | 3 | B2KF52 | Uncharacte | B2KF52 [2 | 0 | 3017.56 | 0 | -0.07 | 0.150616 | 4.14 | 104.1 | 95.9 | 6.76 | 7.15 | | High | High | High | High | High | High | High | High | High | High | High | 0 | 0.000148 | 3.31 |
| FALSE | High | LSQSSYQD | [R].LSQSSY | 1xPhospho | Q61033 1x | 1.83E-05 | 0 | 1 | 1 | 7 | Q61033 | Lamina-ass | Q61033 [4 | 0 | 2089.964 | 0 | 0.65 | 0.024813 | 26.35 | 78 | 122 | 6.17 | 14.43 | | High | High | High | High | High | High | High | High | High | High | High | 0 | 4.39E-06 | 3.93 |
| FALSE | High | LSQSSYQD | [R].LSQSSY | 2xPhospho | Q61033 2x | 0.013909 | 0.001011 | 1 | 1 | 1 | Q61033 | Lamina-ass | Q61033 [4 | 0 | 2169.93 | 0 | 0.31 | 0.313549 | 20.79 | 90.9 | 109.1 | 21.71 | 14.89 | | High | High | High | High | High | High | High | High | High | High | High | 0.000538 | 0.004516 | 2.73 |
| FALSE | High | LSQTDEGA | [R].LSQTDE | 1xPhospho | Q6ZWQ0 1 | 3.44E-08 | 0 | 1 | 2 | 11 | Q6ZWQ0 | Nesprin-2 ( | Q6ZWQ0 [ | 0 | 2550.269 | 0 | 4.61 | 0.000436 | 5.32 | 8.2 | 191.8 | 28.39 | 12.08 | | High | High | High | High | High | High | High | High | High | High | High | 0 | 2.22E-06 | 4.34 |
| FALSE | High | LSQTPK | [K].LSQTPK | 1xPhospho | Q80TR8-2 | 0.089425 | 0.006807 | 1 | 4 | 1 | Q80TR8-2 | Isoform 2 c | Q80TR8-2 | 0 | 1211.68 | 0 | 0.14 | 0.199224 | 9.12 | 95.8 | 104.2 | 10.9 | 8.2 | | High | High | High | High | High | High | High | High | High | High | High | 0.003521 | 0.03773 | 1.46 |
| FALSE | High | LSQVNGAT | [K].LSQVN | 1xPhospho | Q3U2W2 1 | 4.99E-09 | 0 | 1 | 2 | 5 | Q3U2W2 | MYB bindir | Q3U2W2 [ | 0 | 2391.257 | 0 | 0.3 | 0.130434 | 5.38 | 89.6 | 110.4 | 1.94 | 15.25 | | High | High | High | High | High | High | High | High | High | High | High | 0 | 2.64E-10 | 4.18 |
| FALSE | High | LSQVNGAT | [K].LSQVN | 1xDeamida | Q3U2W2 1 | 6.23E-08 | 0 | 1 | 2 | 10 | Q3U2W2 | MYB bindir | Q3U2W2 [ | 0 | 2392.241 | 0 | 0.45 | 0.0997 | 7.28 | 86 | 114 | 4.82 | 17.13 | | High | High | High | High | High | High | High | High | High | High | High | 0 | 4.51E-09 | 4.08 |
| FALSE | High | LSQVNGAT | [K].LSQVN | 2xPhospho | Q3U2W2 2 | 6.47E-05 | 0 | 1 | 2 | 2 | Q3U2W2 | MYB bindir | Q3U2W2 [ | 0 | 2471.224 | 0 | -0.1 | 0.359563 | 3.17 | 100.1 | 99.9 | 6.48 | 5.41 | | High | High | High | High | High | High | High | High | High | High | High | 0 | 1.1E-05 | 3.01 |
| FALSE | High | LSRSDEQLS | [K].LSRSDE | 1xPhospho | Q3UHC1 1x | 9.54E-05 | 0 | 1 | 5 | 1 | Q3UHC1 | Putative ur | Q3UHC1 [3 | 1 | 1814.885 | 0 | -0.14 | 0.105874 | 12.98 | 102.8 | 97.2 | 2.9 | 10.66 | | High | High | High | High | High | High | High | High | High | High | High | 0 | 1.7E-05 | 2.35 |
| FALSE | High | LSSADSPDS | [K].LSSADS | 1xPhospho | E9QAU4 1x | 1.17E-05 | 0 | 1 | 3 | 2 | E9QAU4 | Enhancer t | E9QAU4 [1 | 0 | 1715.825 | 0 | -0.42 | 0.001919 | 2.15 | 114.7 | 85.3 | 3.64 | 5.7 | | High | High | High | High | High | High | High | High | High | High | High | 0 | 1.61E-06 | 3.25 |
| FALSE | High | LSSAEETAF | [K].LSSAEE | 1xPhospho | Q8VCB1 1x | 4.76E-05 | 0 | 1 | 2 | 1 | Q8VCB1 | Nucleopor | Q8VCB1 [4 | 0 | 3406.848 | 0 | -0.41 | 0.01382 | 7.28 | 114.1 | 85.9 | 4.93 | 13.76 | | High | High | High | High | High | High | High | High | High | High | High | 0 | 7.78E-06 | 3.13 |
| FALSE | High | LSSAMSAA | [K].LSSAMS | 1xPhospho | P14152 1x | 0.017192 | 0.001204 | 1 | 1 | 1 | P14152 | Malate del | P14152 [24 | 0 | 1419.732 | 0 | 1.23 | 0.002405 | 22.09 | 62.5 | 137.5 | 14.34 | 9.58 | | High | High | High | High | High | High | High | High | High | High | High | 0.000638 | 0.005769 | 2.17 |
| FALSE | High | ISSAPSSPVI | [K].ISSAPSS | 1xPhospho | Q58FA4 1x | 0.000442 | 0 | 1 | 2 | 1 | Q58FA4 | Transcripti | Q58FA4 [4 | 0 | 1510.828 | 0 | -1.24 | 0.00043 | 1.84 | 143.9 | 56.1 | 5.98 | 7.38 | | High | High | High | High | High | High | High | High | High | High | High | 0 | 9.5E-05 | 1.74 |
| FALSE | High | LSSASTGKF | [R].LSSAST | 1xPhospho | Q3TIV2 1x | 0.071662 | 0.005278 | 1 | 3 | 2 | Q3TIV2 | Putative ur | Q3TIV2 [73 | 0 | 2774.447 | 0 | -0.56 | 0.727393 | 25.15 | 111.7 | 88.3 | 40.91 | 71.69 | | High | High | High | High | High | High | High | High | High | High | High | 0.002703 | 0.02911 | 2.37 |
| FALSE | High | LSPVSHGN | [R].LSPVSH | 1xPhospho | Q8R3L2 1x | 0.001411 | 5.66E-05 | 1 | 9 | 1 | Q8R3L2 | Transcripti | Q8R3L2 [6 | 0 | 1968.031 | 0 | -0.21 | 0.043673 | 10.69 | 105.2 | 94.8 | 9.83 | 9.89 | | High | High | High | High | High | High | High | High | High | High | High | 2.58E-05 | 0.000351 | 3.3 |
| FALSE | High | FLESDSEDE | [R].FLESDS | 2xPhospho | E9QKD1 2x | 1.35E-05 | 0 | 1 | 3 | 8 | E9QKD1 | Nucleolar p | E9QKD1 [8 | 0 | 1816.77 | 0 | -0.44 | 0.000912 | 2.07 | 115.4 | 84.6 | 4.24 | 6.1 | | High | High | High | High | High | High | High | High | High | High | High | 0 | 1.89E-06 | 3.47 |
| FALSE | High | LDISSLNRP | [K].LDISSLN | 1xPhospho | E9PXP7 1x | 0.062731 | 0.004664 | 1 | 2 | 1 | E9PXP7 | Stonin-2 O | E9PXP7 [28 | 0 | 2456.29 | 0 | -0.36 | 0.002004 | 2.04 | 115.7 | 84.3 | 4.37 | 9.01 | | High | High | High | High | High | High | High | High | High | High | High | 0.002403 | 0.02497 | 1.48 |
| FALSE | High | IDISPSALR | [R].IDISPSA | 1xPhospho | Q8K019 1x | 0.001325 | 0 | 1 | 9 | 8 | Q8K019 | Bcl-2-assoc | Q8K019 [6 | 0 | 1280.681 | 0 | -0.42 | 0.003513 | 3.01 | 114.9 | 85.1 | 4.54 | 4.63 | | High | High | High | High | High | High | High | High | High | High | High | 0 | 0.000328 | 2.6 |
| FALSE | High | GLQLTPGI( | [R].GLQLTI | 1xPhospho | F6ZDS4 1x | 1.56E-05 | 0 | 1 | 5 | 2 | F6ZDS4 | Nucleopro | F6ZDS4 [21 | 0 | 2686.217 | 0 | 1.63 | 0.011789 | 27.34 | 48.9 | 151.1 | 16.02 | 22.68 | | High | High | High | High | High | High | High | High | High | High | High | 0 | 2.23E-06 | 2.84 |
| FALSE | High | GIQLYDTPY | [K].GIQLYD | 1xPhospho | Q6PD21 1x | 0.000633 | 0 | 1 | 3 | 2 | Q6PD21 | SH2 domai | Q6PD21 [2 | 0 | 3281.494 | 0 | 1.7 | 0.00043 | 16.33 | 46.7 | 153.3 | 18.04 | 8.96 | | High | High | High | High | High | High | High | High | High | High | High | 0 | 0.000143 | 3.97 |
| FALSE | High | GLQPPPPT | [R].GLQPPI | 2xCarbami | E9PYL2 1xP | 0.007209 | 0.00062 | 1 | 2 | 1 | E9PYL2 | Proline-ricl | E9PYL2 [13 | 0 | 4839.429 | 0 | -0.15 | 0.244161 | 24.96 | 101.4 | 98.6 | 9.22 | 17.84 | | High | High | High | High | High | High | High | High | High | High | High | 0.000275 | 0.002176 | 3.54 |
| FALSE | High | GLQVDLQS | [K].GLQVD | 1xPhospho | Q3UA65 1x | 3.61E-09 | 0 | 1 | 5 | 2 | Q3UA65 | DNA helica | Q3UA65 [1 | 0 | 3193.603 | 0 | -0.6 | 0.009552 | 5.7 | 118.1 | 81.9 | 7.61 | 10.99 | | High | High | High | High | High | High | High | High | High | High | High | 0 | 6.44E-10 | 4.2 |
| FALSE | High | GLSADSPA | [K].GLSADS | 1xPhospho | E9PVX6 1x | 9.82E-06 | 0 | 1 | 2 | 2 | E9PVX6 | Proliferatic | E9PVX6 [1 | 0 | 1729.841 | 0 | 0.49 | 0.002467 | 7.2 | 83.1 | 116.9 | 4.86 | 4.94 | | High | High | High | High | High | High | High | High | High | High | High | 0 | 5.28E-05 | 2.32 |
| FALSE | High | GLSASLPDI | [K].GLSASL | 1xPhospho | Q6ZQ58 1x | 0.001572 | 5.66E-05 | 1 | 3 | 8 | Q6ZQ58 | La-related | Q6ZQ58 [5 | 0 | 2484.268 | 0 | 0.54 | 0.002237 | 8.55 | 79.7 | 120.3 | 5.44 | 5.04 | | High | High | High | High | High | High | High | High | High | High | High | 2.58E-05 | 0.000397 | 3.36 |
| FALSE | High | GLSASTVDI | [R].GLSAST | 1xPhospho | Q9DBS5 1x | 0.0004 | 0 | 1 | 1 | 6 | Q9DBS5 | Kinesin ligh | Q9DBS5 [6 | 0 | 1519.709 | 0 | 0.31 | 0.043319 | 16.01 | 87.6 | 112.4 | 10.56 | 5.53 | | High | High | High | High | High | High | High | High | High | High | High | 2.58E-05 | 0.000409 | 2.44 |
| FALSE | High | GLSDHASLI | [R].GLSDHA | 1xPhospho | Q99LH9 1x | 0.003997 | 0.000311 | 1 | 2 | 1 | Q99LH9 | SH3 domai | Q99LH9 [3 | 0 | 2150.008 | 0 | 0.01 | 0.659361 | 4.94 | 98.9 | 101.1 | 11.57 | 6.08 | | High | High | High | High | High | High | High | High | High | High | High | 0.00014 | 0.001124 | 3.08 |
| FALSE | High | GLSDHASLI | [R].GLSDHA | 2xPhospho | Q99LH9 2x | 2.06E-05 | 0 | 1 | 2 | 1 | Q99LH9 | SH3 domai | Q99LH9 [3 | 0 | 2229.974 | 0 | 0.28 | 0.002097 | 5.68 | 90.8 | 109.2 | 3.59 | 5.03 | | High | High | High | High | High | High | High | High | High | High | High | 0 | 3.05E-06 | 3.99 |
| FALSE | High | GLSEDTTEE | [K].GLSEDT | 1xPhospho | P09405 1x | 0.000133 | 0 | 1 | 5 | 2 | P09405 | Nucleolin ( | P09405 [57 | 0 | 1860.924 | 2 | -0.39 | 0.022074 | 8.96 | 114.5 | 85.5 | 8.92 | 21.2 | | High | High | High | High | High | High | High | High | High | High | High | 0.007335 | 0.08049 | 2.29 |
| FALSE | High | GISEGDDVI | [R].GISEGD | 1xPhospho | Q0VBD2 1x | 4.89E-05 | 0 | 1 | 1 | 4 | Q0VBD2 | Protein MC | Q0VBD2 [6 | 0 | 2726.348 | 0 | -0.09 | 0.663633 | 9.8 | 102.2 | 97.8 | 8.18 | 3.35 | | High | High | High | High | High | High | High | High | High | High | High | 0 | 8.08E-06 | 3.57 |
| FALSE | High | GLSHSPWA | [R].GLSHSF | 1xPhospho | Q9JJ78 1xP | 0.009931 | 0.000736 | 1 | 2 | 1 | Q9JJ78 | Lymphokin | Q9JJ78 [54 | 0 | 1619.871 | 0 | 0.55 | 0.000596 | 2.83 | 80.2 | 119.8 | 5.45 | 6.77 | | High | High | High | High | High | High | High | High | High | High | High | 0.000344 | 0.003103 | 2.12 |
| FALSE | High | GLSHVSDLI | [R].GLSHVS | 1xCarbami | G5E893 1x | 0.000209 | 0 | 1 | 2 | 3 | G5E893 | Ankyrin rep | G5E893 [1 | 0 | 2244.043 | 0 | 0.73 | 0.024821 | 33.09 | 76.8 | 123.2 | 10.95 | 12.51 | | High | High | High | High | High | High | High | High | High | High | High | 0 | 4.12E-05 | 2.9 |
| FALSE | High | GLQLTPGI( | [R].GLQLTI | 1xPhospho | F6ZDS4 1x | 7.15E-05 | 0 | 1 | 5 | 1 | F6ZDS4 | Nucleopro | F6ZDS4 [21 | 0 | 2670.222 | 0 | 1.67 | 0.032971 | 35.25 | 48.6 | 151.4 | 14.22 | 38.52 | | High | High | High | High | High | High | High | High | High | High | High | 0 | 1.23E-05 | 3.62 |
| FALSE | High | GLSLGSAVI | [R].GLSLGS | 1xPhospho | Q8CDZ5 1x | 3.48E-09 | 0 | 1 | 2 | 3 | Q8CDZ5 | Putative ur | Q8CDZ5 [3 | 0 | 2138.121 | 0 | -0.27 | 0.044682 | 11.43 | 110.5 | 89.5 | 8.44 | 8.19 | | High | High | High | High | High | High | High | High | High | High | High | 0 | 1.75E-10 | 4.69 |
| FALSE | High | GLSLTETEL | [R].GLSLTE | 1xPhospho | E9QP94 1x | 0.000154 | 0 | 1 | 9 | 1 | E9QP94 | Coiled-coil | E9QP94 [9 | 0 | 1798.904 | 0 | -0.56 | 0.00462 | 2.49 | 119.1 | 80.9 | 7.15 | 11.4 | | High | High | High | High | High | High | High | High | High | High | High | 0 | 2.9E-05 | 3.14 |
| FALSE | High | GISNGPNPI | [R].GISNGF | 1xCarbami | Q5SSZ5 1x | 0.000152 | 0 | 1 | 1 | 1 | Q5SSZ5 | Tensin-3 O | Q5SSZ5 [6 | 0 | 2418.189 | 0 | 1.51 | 0.001921 | 17.94 | 48.3 | 151.7 | 6.86 | 14.06 | | High | High | High | High | High | High | High | High | High | High | High | 0 | 2.89E-05 | 2.52 |
| FALSE | High | GISNGPNPI | [R].GISNGF | 1xCarbami | Q5SSZ5 1x | 2.53E-05 | 0 | 1 | 1 | 2 | Q5SSZ5 | Tensin-3 O | Q5SSZ5 [6 | 0 | 2419.173 | 0 | 0.95 | 0.003929 | 25.75 | 68.1 | 131.9 | 10.3 | 9.7 | | High | High | High | High | High | High | High | High | High | High | High | 0 | 6.76E-06 | 3.57 |
| FALSE | High | GLSPADVA | [R].GLSPAD | 1xPhospho | Q0VGT2 1x | 0.090153 | 0.00698 | 1 | 3 | 1 | Q0VGT2 | Zinc finger | Q0VGT2 [1 | 0 | 1912.009 | 0 | -1.66 | 0.001105 | 0.85 | 151.8 | 48.2 | 6.9 | 16.84 | | High | High | High | High | High | High | High | High | High | High | High | 0.003564 | 0.03804 | 1.88 |
| FALSE | High | GLSPAMSP | [K].GLSPAN | 2xPhospho | Q3UFM6 2 | 0.008847 | 0.000736 | 1 | 3 | 1 | Q3UFM6 | Expressed | Q3UFM6 [ | 0 | 1616.747 | 0 | -0.07 | 0.196082 | 6.68 | 104.5 | 95.5 | 4.34 | 14.3 | | High | High | High | High | High | High | High | High | High | High | High | 0.000327 | 0.002747 | 2.52 |
| FALSE | High | GISPIVFDR | [R].GISPIVF | 1xPhospho | E9Q5K9 1x | 0.037048 | 0.00249 | 1 | 4 | 2 | E9Q5K9 | YTH domai | E9Q5K9 [3 | 0 | 1312.686 | 0 | -0.18 | 0.04844 | 8.39 | 107.3 | 92.7 | 5.66 | 5.96 | | High | High | High | High | High | High | High | High | High | High | High | 0.00468 | 0.0505 | 2.06 |
| FALSE | High | GISPMNRP | [K].GISPMI | 1xPhospho | Q91Z49 1x | 0.057203 | 0.004227 | 1 | 2 | 1 | Q91Z49 | UAP56-inte | Q91Z49 [1 | 0 | 1966.023 | 0 | -0.19 | 0.039225 | 12.43 | 107.2 | 92.8 | 5.97 | 5.18 | | High | High | High | High | High | High | High | High | High | High | High | 0.002159 | 0.02258 | 1.59 |
| FALSE | High | GLSPSYLG( | [R].GLSPSY | 1xPhospho | Q8C9B9 1x | 0.002115 | 5.66E-05 | 1 | 1 | 1 | Q8C9B9 | Death-indu | Q8C9B9 [1 | 0 | 1412.714 | 0 | 0.66 | 0.005175 | 21.8 | 80.2 | 119.8 | 10.11 | 8.28 | | High | High | High | High | High | High | High | High | High | High | High | 2.58E-05 | 0.000555 | 1.76 |
| FALSE | High | GISQTNLIT | [K].GISQTN | 1xPhospho | Q9WV92-7 | 1.34E-06 | 0 | 1 | 13 | 4 | Q9WV92-7 | Isoform 7 c | Q9WV92-7 | 0 | 2140.167 | 0 | -1.24 | 0.005725 | 6.24 | 135.8 | 64.2 | 11.14 | 19.96 | | High | High | High | High | High | High | High | High | High | High | High | 0 | 1.41E-07 | 4.65 |

| | | | | | | | | | | | | | | | | | | | | | | | | | | | | | | | | | | | | | | |
|---|---|---|---|---|---|---|---|---|---|---|---|---|---|---|---|---|---|---|---|---|---|---|---|---|---|---|---|---|---|---|---|---|---|---|---|---|---|---|
| FALSE | High | GISSDSESV | [K].GISSDS | 1xCarbami | E9Q842 1x | 8.74E-06 | 0 | 1 | 4 | 4 | E9Q842 | Neuron na | E9Q842 [1 | 0 | 2165.02 | 0 | 1.53 | 0.000429 | 11.48 | 51.4 | 148.6 | 6.93 | 5.81 | | High | High | High | High | High | High | High | High | High | High | High | 0 | 0.000191 | 2.84 |
| FALSE | High | GLSSPNEL( | [R].GLSSPN | 1xPhospho | Q62431 1x | 2.75E-06 | 0 | 1 | 2 | 2 | Q62431 | AT-rich int | Q62431 [3 | 0 | 1980.959 | 0 | -1.23 | 0.002995 | 5.43 | 141.6 | 58.4 | 12.11 | 20.61 | | High | High | High | High | High | High | High | High | High | High | High | 0 | 3.17E-07 | 4.01 |
| FALSE | High | GISSSNEG\ | [R].GISSSN | 1xPhospho | Q8BI72 1xP | 0.002334 | 0.000109 | 1 | 1 | 2 | Q8BI72 | CDKN2A-in | Q8BI72 [12 | 0 | 1957.952 | 0 | 1.3 | 0.001247 | 22.86 | 59.9 | 140.1 | 18.71 | 11.77 | | High | High | High | High | High | High | High | High | High | High | High | 4.94E-05 | 0.00062 | 3.16 |
| FALSE | High | GLSVDSAQ | [R].GLSVDS | 1xPhospho | A0A1D5RLI | 0.001337 | 0 | 1 | 7 | 2 | A0A1D5RLI | Transform | A0A1D5RLI | 0 | 1441.714 | 0 | 0.94 | 0.000816 | 7.53 | 69.8 | 130.2 | 9.28 | 6.04 | | High | High | High | High | High | High | High | High | High | High | High | 0.001774 | 0.01749 | 2.59 |
| FALSE | High | GLSLGSAVI | [R].GLSLGS | 2xPhospho | Q8CDZ5 2x | 0.000434 | 0 | 1 | 2 | 1 | Q8CDZ5 | Putative ur | Q8CDZ5 [3 | 0 | 2218.087 | 0 | -0.36 | 0.017728 | 9.4 | 113.1 | 86.9 | 5.45 | 9.7 | | High | High | High | High | High | High | High | High | High | High | High | 0 | 9.35E-05 | 2.95 |
| FALSE | High | GLSVDSAQ | [R].GLSVDS | 1xPhospho | A0A1D5RLI | 4.02E-06 | 0 | 1 | 7 | 1 | A0A1D5RLI | Transform | A0A1D5RLI | 0 | 1670.877 | 0 | 1.19 | 0.004084 | 31.34 | 61 | 139 | 13.52 | 9.54 | | High | High | High | High | High | High | High | High | High | High | High | 0 | 4.85E-07 | 2.58 |
| FALSE | High | GLQGFSPK | [K].GLQGF | 1xPhospho | Q6P9L6 1x | 0.015465 | 0.001089 | 1 | 2 | 2 | Q6P9L6 | Kinesin-like | Q6P9L6 [5 | 0 | 1371.744 | 0 | -0.93 | 0.000533 | 0.96 | 130.7 | 69.3 | 3.45 | 10.62 | | High | High | High | High | High | High | High | High | High | High | High | 0.001076 | 0.01128 | 1.89 |
| FALSE | High | GLQASPGC | [K].GLQASI | 2xPhospho | Q8BHK9 2x | 0.0102 | 0.000736 | 1 | 1 | 2 | Q8BHK9 | DNA excisi | Q8BHK9 [8 | 0 | 2844.296 | 0 | 0.3 | 0.022046 | 16.76 | 85.6 | 114.4 | 6.12 | 9.56 | | High | High | High | High | High | High | High | High | High | High | High | 0.000382 | 0.003219 | 3.6 |
| FALSE | High | GLLSQNGS | [R].GLLSQN | 1xCarbami | Q8BI84 1xP | 5.37E-06 | 0 | 1 | 4 | 4 | Q8BI84 | Melanoma | Q8BI84 [16 | 0 | 4084.005 | 0 | 0.14 | 0.178116 | 5.16 | 95.1 | 104.9 | 11.92 | 12.77 | | High | High | High | High | High | High | High | High | High | High | High | 0 | 8.74E-07 | 4.1 |
| FALSE | High | GIITDSFGR | [R].GIITDSI | 1xPhospho | Q8K2Y9 1x | 0.014158 | 0.001011 | 1 | 4 | 1 | Q8K2Y9 | Cerebral ca | Q8K2Y9 [3 | 0 | 1274.634 | 0 | 0.75 | 0.000313 | 3.7 | 74.5 | 125.5 | 2.26 | 4.17 | | High | High | High | High | High | High | High | High | High | High | High | 0.000538 | 0.004623 | 0.77 |
| FALSE | High | GLLTDSPA/ | [R].GLLTDS | 1xPhospho | D3Z0X5 1xI | 0.000407 | 0 | 1 | 6 | 2 | D3Z0X5 | Pleckstrin I | D3Z0X5 [3 | 0 | 1893.988 | 0 | 1.07 | 0.000408 | 1.54 | 64.8 | 135.2 | 11.26 | 7.23 | | High | High | High | High | High | High | High | High | High | High | High | 0 | 0.000184 | 3.4 |
| FALSE | High | GLLTLPSPT | [R].GLLTLP | 1xPhospho | Q9QWF0 1 | 0.037048 | 0.00249 | 1 | 1 | 1 | Q9QWF0 | Chromatin | Q9QWF0 [ | 0 | 3263.752 | 0 | -0.13 | 0.044682 | 6.32 | 104.9 | 95.1 | 5.79 | 6.1 | | High | High | High | High | High | High | High | High | High | High | High | 0.001308 | 0.01369 | 2.6 |
| FALSE | High | GLIVYCVTS | [K].GLIVYC | 1xCarbami | B2RUJ6 1xI | 0.002553 | 0.000109 | 1 | 6 | 1 | B2RUJ6 | Rap guanin | B2RUJ6 [14 | 0 | 1774.958 | 0 | 0.29 | 0.238501 | 21.24 | 93.6 | 106.4 | 12.81 | 7.41 | | High | High | High | High | High | High | High | High | High | High | High | 4.94E-05 | 0.000685 | 3.24 |
| FALSE | High | GLLYDSSEE | [R].GLLYDS | 1xPhospho | Q3UJN1 1x | 0.000205 | 0 | 1 | 3 | 16 | Q3UJN1 | DNA helica | Q3UJN1 [1 | 0 | 2174.98 | 0 | -0.69 | 0.000223 | 0.62 | 123.4 | 76.6 | 1.82 | 2.83 | | High | High | High | High | High | High | High | High | High | High | High | 0.000786 | 0.006846 | 2.02 |
| FALSE | High | GLLYDSSEE | [R].GLLYDS | 2xPhospho | Q3UJN1 2x | 7.41E-05 | 0 | 1 | 3 | 14 | Q3UJN1 | DNA helica | Q3UJN1 [1 | 0 | 2254.947 | 0 | -0.3 | 0.002809 | 1.79 | 110.2 | 89.8 | 3.41 | 3.69 | | High | High | High | High | High | High | High | High | High | High | High | 0.000275 | 0.002088 | 2.67 |
| FALSE | High | GLMAGGR | [R].GLMAG | 1xPhospho | Q3TIR3 1xP | 0.000984 | 0 | 1 | 2 | 2 | Q3TIR3 | Synembryr | Q3TIR3 [42 | 0 | 3014.256 | 0 | 0.16 | 0.022081 | 5.96 | 92.5 | 107.5 | 8.52 | 1.94 | | High | High | High | High | High | High | High | High | High | High | High | 0 | 0.000235 | 2.79 |
| FALSE | High | GLMAGGR | [R].GLMAG | 1xPhospho | Q3TIR3 1xP | 0.000179 | 0 | 1 | 2 | 3 | Q3TIR3 | Synembryr | Q3TIR3 [42 | 0 | 3030.251 | 0 | 0.1 | 0.115794 | 8.81 | 97 | 103 | 5.69 | 13.85 | | High | High | High | High | High | High | High | High | High | High | High | 0.000275 | 0.001775 | 3.22 |
| FALSE | High | GLMAGGR | [R].GLMAG | 2xPhospho | Q3TIR3 2xP | 0.000488 | 0 | 1 | 2 | 3 | Q3TIR3 | Synembryr | Q3TIR3 [42 | 0 | 3094.222 | 0 | 0.05 | 0.980556 | 4.28 | 99.7 | 100.3 | 5.71 | 6.27 | | High | High | High | High | High | High | High | High | High | High | High | 0 | 0.000107 | 3.12 |
| FALSE | High | GLMAGGR | [R].GLMAG | 2xPhospho | Q3TIR3 2xP | 8.37E-07 | 0 | 1 | 2 | 17 | Q3TIR3 | Synembryr | Q3TIR3 [42 | 0 | 3110.217 | 0 | -0.05 | 0.083542 | 4.85 | 102.5 | 97.5 | 1.26 | 3.91 | | High | High | High | High | High | High | High | High | High | High | High | 0 | 1.41E-05 | 3.32 |
| FALSE | High | GLNGCESP | [K].GLNGC | 1xCarbami | Q60929 2x | 0.092372 | 0.007211 | 1 | 3 | 1 | Q60929 | Myocyte-s | Q60929 [9 | 0 | 3000.148 | 0 | 0.88 | 0.002475 | 13.82 | 70.8 | 129.2 | 6.83 | 9.83 | | High | High | High | High | High | High | High | High | High | High | High | 0.003672 | 0.03938 | 3.35 |
| FALSE | High | GLNGGITPI | [R].GLNGG | 1xPhospho | Q7TN98 1x | 8.25E-05 | 0 | 1 | 7 | 1 | Q7TN98 | Cytoplasmi | Q7TN98 [3 | 0 | 2119.193 | 0 | 1.53 | 0.001291 | 27.04 | 45.3 | 154.7 | 19.76 | 12.01 | | High | High | High | High | High | High | High | High | High | High | High | 0 | 1.45E-05 | 3.79 |
| FALSE | High | GLQDHSAS | [R].GLQDH | 1xPhospho | Q60949 1x | 2.74E-07 | 0 | 1 | 5 | 2 | Q60949 | TBC1 doma | Q60949 [5 | 0 | 2975.425 | 0 | 0.37 | 0.210806 | 47.87 | 86.9 | 113.1 | 18.12 | 21.92 | | High | High | High | High | High | High | High | High | High | High | High | 0 | 2.36E-08 | 5.25 |
| FALSE | High | GLNLDGTP | [K].GLNLD( | 2xPhospho | Q52KE7 2x | 0.002355 | 0.000109 | 1 | 1 | 1 | Q52KE7 | Cyclin-L1 C | Q52KE7 [3 | 0 | 3145.574 | 0 | -0.89 | 0.005489 | 1.4 | 130 | 70 | 13.27 | 5.02 | | High | High | High | High | High | High | High | High | High | High | High | 4.94E-05 | 0.000622 | 2.8 |
| FALSE | High | GIPCGSDTI | [K].GIPCGS | 2xCarbami | Q3UZ18 1x | 2.03E-06 | 0 | 1 | 1 | 4 | Q3UZ18 | Little elong | Q3UZ18 [5 | 0 | 2562.094 | 0 | -0.39 | 0.010157 | 8.04 | 114.3 | 85.7 | 6.03 | 14.69 | | High | High | High | High | High | High | High | High | High | High | High | 0 | 3.24E-07 | 3.9 |
| FALSE | High | GLPFYAGL! | [K].GLPFYA | 1xPhospho | E9QLG3 1x | 0.00204 | 5.66E-05 | 1 | 5 | 1 | E9QLG3 | MAX gene- | E9QLG3 [1 | 0 | 1831.976 | 0 | -0.68 | 0.003776 | 6.51 | 123.3 | 76.7 | 7.63 | 6.53 | | High | High | High | High | High | High | High | High | High | High | High | 2.58E-05 | 0.000529 | 2.4 |
| FALSE | High | GIPLPTGDT | [K].GIPLPT( | 1xPhospho | Q9Z1D1 1x | 0.063258 | 0.004664 | 1 | 2 | 4 | Q9Z1D1 | Eukaryotic | Q9Z1D1 [3 | 0 | 2843.468 | 0 | -0.28 | 0.70507 | 9.67 | 108.8 | 91.2 | 20.75 | 85.72 | | High | High | High | High | High | High | High | High | High | High | High | 0.006234 | 0.06724 | 2.72 |
| FALSE | High | GIPLPTGDT | [K].GIPLPT( | 1xPhospho | Q9Z1D1 1x | 2.72E-05 | 0 | 1 | 2 | 5 | Q9Z1D1 | Eukaryotic | Q9Z1D1 [3 | 0 | 3072.631 | 0 | -0.19 | 0.039745 | 6.4 | 109.6 | 90.4 | 6.66 | 7.76 | | High | High | High | High | High | High | High | High | High | High | High | 2.58E-05 | 0.000359 | 3.63 |
| FALSE | High | GLPPEAW( | [K].GLPPEA | 1xCarbami | P23188 2xI | 0.002022 | 5.66E-05 | 1 | 2 | 1 | P23188 | Furin OS=N | P23188 [76 | 1 | 3148.244 | 0 | 0.16 | 0.085479 | 11.06 | 92.4 | 107.6 | 6.76 | 12.56 | | High | High | High | High | High | High | High | High | High | High | High | 2.58E-05 | 0.000524 | 2.77 |
| FALSE | High | GLPPSMER | [R].GLPPSM | 1xPhospho | Q91VM5 1 | 0.032267 | 0.002129 | 1 | 3 | 10 | Q91VM5 | RNA bindir | Q91VM5 [3 | 0 | 1211.569 | 0 | -0.41 | 0.011653 | 2.32 | 111.8 | 88.2 | 5.02 | 10.7 | | High | High | High | High | High | High | High | High | High | High | High | 0.004072 | 0.04352 | 1.11 |
| FALSE | High | GLPSPPQL\ | [R].GLPSPP | 1xPhospho | B7ZNH9 1x | 0.01894 | 0.001362 | 1 | 3 | 1 | B7ZNH9 | Phosphoin | B7ZNH9 [2 | 0 | 2262.215 | 0 | 0.56 | 0.01808 | 17.25 | 80.8 | 119.2 | 9.52 | 12.65 | | High | High | High | High | High | High | High | High | High | High | High | 0.000723 | 0.006413 | 2.64 |
| FALSE | High | GLPSPYNM | [K].GLPSPY | 1xPhospho | E9Q557 1x | 0.00824 | 0.000736 | 1 | 2 | 4 | E9Q557 | Desmoplak | E9Q557 [2 | 0 | 1758.808 | 0 | 2.32 | 0.002051 | 7.44 | 33.3 | 166.7 | 6.53 | 17.32 | | High | High | High | High | High | High | High | High | High | High | High | 0.005561 | 0.06091 | 2.4 |
| FALSE | High | GLPSPYNM | [K].GLPSPY | 2xPhospho | E9Q557 2x | 0.014929 | 0.001049 | 1 | 2 | 2 | E9Q557 | Desmoplak | E9Q557 [2 | 0 | 1838.775 | 0 | 2.39 | 0.001775 | 9.56 | 30.9 | 169.1 | 13.25 | 16.06 | | High | High | High | High | High | High | High | High | High | High | High | 0.000555 | 0.004917 | 2.93 |
| FALSE | High | GLPSTASGI | [R].GLPSTA | 1xPhospho | Q3USK2 1x | 0.102567 | 0.008249 | 1 | 3 | 1 | Q3USK2 | Casein kina | Q3USK2 [3 | 0 | 1154.577 | 0 | 0.35 | 0.008827 | 8.78 | 88.1 | 111.9 | 2.63 | 7.07 | | High | High | High | High | High | High | High | High | High | High | High | 0.004138 | 0.04436 | 0.97 |
| FALSE | High | GLPWSCSII | [R].GLPWS | 2xCarbami | Q9Z2X1 1x | 0.000892 | 0 | 1 | 4 | 2 | Q9Z2X1 | Heterogen | Q9Z2X1 [1 | 0 | 4102.903 | 0 | 0.36 | 0.197115 | 17.22 | 89.1 | 110.9 | 7.68 | 14.43 | | High | High | High | High | High | High | High | High | High | High | High | 0 | 0.000211 | 3.13 |
| FALSE | High | GLQAGTA/ | [K].GLQAG | 1xPhospho | Q3UQJ0 1x | 0.01002 | 0.000736 | 1 | 4 | 1 | Q3UQJ0 | Putative ur | Q3UQJ0 [1 | 0 | 2307.138 | 0 | -1.2 | 0.000445 | 0.57 | 139.4 | 60.6 | 2.67 | 9.54 | | High | High | High | High | High | High | High | High | High | High | High | 0.000344 | 0.003159 | 1.84 |
| FALSE | High | GLQASPGC | [K].GLQASI | 1xPhospho | Q8BHK9 1x | 0.000186 | 0 | 1 | 1 | 1 | Q8BHK9 | DNA excisi | Q8BHK9 [8 | 0 | 2764.329 | 0 | 0.29 | 0.086112 | 19.08 | 90.6 | 109.4 | 10.39 | 7.68 | | High | High | High | High | High | High | High | High | High | High | High | 0 | 3.61E-05 | 3.39 |
| FALSE | High | GLNSGSEA | [R].GLNSG! | 1xPhospho | Q8K298 1x | 0.053023 | 0.00384 | 1 | 2 | 1 | Q8K298 | Anillin OS= | Q8K298 [1 | 0 | 1932.968 | 0 | -0.07 | 0.690441 | 27.92 | 97 | 103 | 19.5 | 27.11 | | High | High | High | High | High | High | High | High | High | High | High | 0.001999 | 0.02055 | 1.86 |
| FALSE | High | GLSVEK | [R].GLSVEK | 1xPhospho | Q80WT5 1 | 0.113753 | 0.009241 | 1 | 6 | 1 | Q80WT5 | Aftiphilin C | Q80WT5 [3 | 0 | 1170.654 | 0 | 0.59 | 0.016714 | 32.01 | 78.6 | 121.4 | 14.48 | 9.41 | | High | High | High | High | High | High | High | High | High | High | High | 0.004625 | 0.05004 | 1.08 |
| FALSE | High | GLSVSGPC | [K].GLSVSG | 1xCarbami | Q99LL5 1xI | 0.000365 | 0 | 1 | 3 | 1 | Q99LL5 | Periodic try | Q99LL5 [48 | 1 | 2387.057 | 0 | -0.33 | 0.014316 | 10.52 | 116.3 | 83.7 | 6.35 | 15.15 | | High | High | High | High | High | High | High | High | High | High | High | 0 | 7.68E-05 | 2.04 |
| FALSE | High | GLSVTAEQ | [R].GLSVTA | 1xPhospho | E9Q166 1x | 6.91E-06 | 0 | 1 | 2 | 3 | E9Q166 | ATPase fan | E9Q166 [1 | 0 | 1974.001 | 0 | -0.31 | 0.022733 | 6.29 | 109.6 | 90.4 | 5.46 | 10.28 | | High | High | High | High | High | High | High | High | High | High | High | 0 | 8.93E-07 | 2.74 |
| FALSE | High | GNHSGSPG | [R].GNHSG | 1xDeamida | P30658 1xI | 0.001476 | 5.66E-05 | 1 | 2 | 3 | P30658 | Chromobo | P30658 [32 | 0 | 3413.772 | 0 | -0.63 | 0.009654 | 9.18 | 120.5 | 79.5 | 7.15 | 10.34 | | High | High | High | High | High | High | High | High | High | High | High | 2.58E-05 | 0.000371 | 3.13 |
| FALSE | High | GNKSPSPPI | [K].GNKSP! | 2xPhospho | O08539 2x | 0.000876 | 0 | 1 | 3 | 1 | O08539 | Myc box-d | O08539 [2 | 1 | 2593.246 | 0 | 0.69 | 0.009741 | 14.44 | 76.9 | 123.1 | 9.99 | 11.64 | | High | High | High | High | High | High | High | High | High | High | High | 0 | 0.000206 | 2.86 |
| FALSE | High | GNLETHED | [R].GNLETI | 1xPhospho | E9Q784 1x | 0.000573 | 0 | 1 | 1 | 2 | E9Q784 | Zinc finger | E9Q784 [1 | 0 | 2226.084 | 0 | -0.82 | 0.000386 | 2.3 | 128.7 | 71.3 | 2.66 | 7.44 | | High | High | High | High | High | High | High | High | High | High | High | 0 | 0.000128 | 2.28 |
| FALSE | High | GNIIISTPEK | [K].GNIIIST | 1xPhospho | Q6P4T2 1x | 0.033404 | 0.002195 | 1 | 1 | 1 | Q6P4T2 | U5 small nu | Q6P4T2 [1 | 0 | 1609.897 | 0 | -0.56 | 0.003389 | 3.49 | 120.9 | 79.1 | 6.92 | 9.75 | | High | High | High | High | High | High | High | High | High | High | High | 0.001151 | 0.01221 | 2.1 |
| FALSE | High | GNLQPAPA | [R].GNLQP. | 1xDeamida | Q78JW9 1x | 0.005044 | 0.000504 | 1 | 2 | 2 | Q78JW9 | Ubiquitin d | Q78JW9 [1 | 0 | 3453.669 | 0 | -0.94 | 0.055707 | 7.25 | 127.8 | 72.2 | 12.89 | 23.81 | | High | High | High | High | High | High | High | High | High | High | High | 0.000246 | 0.001457 | 3.76 |
| FALSE | High | GNISLVAFF | [K].GNISLV | 1xPhospho | A0A087WF | 1.02E-07 | 0 | 1 | 2 | 1 | A0A087WF | Transcripti | A0A087WF | 0 | 2357.252 | 0 | -1.45 | 0.000154 | 1.22 | 143.7 | 56.3 | 4.16 | 9.6 | | High | High | High | High | High | High | High | High | High | High | High | 0 | 7.85E-09 | 4.15 |
| FALSE | High | GNLTPHPA | [R].GNLTPI | 1xCarbami | A0A0N4SU | 0.001349 | 0 | 1 | 11 | 2 | A0A0N4SU | Peregrin O | A0A0N4SU | 0 | 1818.897 | 0 | -0.44 | 0.001402 | 1.83 | 115.1 | 84.9 | 4.42 | 3.17 | | High | High | High | High | High | High | High | High | High | High | High | 0 | 0.000333 | 2.14 |
| FALSE | High | GNLWSAEI | [K].GNLWS | 1xPhospho | Q8K298 1x | 0.005924 | 0.00062 | 1 | 1 | 2 | Q8K298 | Anillin OS= | Q8K298 [4 | 0 | 1442.745 | 0 | 0.11 | 0.325483 | 4.81 | 94.1 | 105.9 | 3.41 | 11.93 | | High | High | High | High | High | High | High | High | High | High | High | 0.000382 | 0.003271 | 2.09 |
| FALSE | High | GNPTVEVD | [R].GNPTV | 1xPhospho | Q5FW97 1 | 0.000519 | 0 | 1 | 14 | 1 | Q5FW97 | Enolase 1, | Q5FW97 [1 | 0 | 1945.008 | 0 | -0.41 | 0.030712 | 16.88 | 117.4 | 82.6 | 11.31 | 10 | | High | High | High | High | High | High | High | High | High | High | High | 0 | 0.000114 | 2.22 |
| FALSE | High | GNSDGYIIF | [R].GNSDG | 1xPhospho | Q8JZZ7-2 1 | 0.081085 | 0.006219 | 1 | 11 | 2 | Q8JZZ7-2 | Isoform 2 c | Q8JZZ7-2 [ | 0 | 1828.961 | 0 | -0.49 | 0.006382 | 3.29 | 115.9 | 84.1 | 8.14 | 7.73 | | High | High | High | High | High | High | High | High | High | High | High | 0.003188 | 0.0336 | 1.85 |
| FALSE | High | GNSELSEAI | [K].GNSEL! | 1xPhospho | Q9Z1X4-3 | 0.022773 | 0.001542 | 1 | 8 | 1 | Q9Z1X4-3 | Isoform 3 c | Q9Z1X4-3 | 0 | 2703.174 | 0 | -0.25 | 0.006348 | 5.68 | 108.7 | 91.3 | 3.11 | 5.82 | | High | High | High | High | High | High | High | High | High | High | High | 0.000832 | 0.007925 | 3.1 |
| FALSE | High | GNSELSEAI | [K].GNSEL! | 1xPhospho | Q9Z1X4-3 | 1.51E-06 | 0 | 1 | 8 | 8 | Q9Z1X4-3 | Isoform 3 c | Q9Z1X4-3 | 0 | 2719.169 | 0 | -0.14 | 0.366054 | 18.25 | 101.2 | 98.8 | 4.35 | 12.54 | | High | High | High | High | High | High | High | High | High | High | High | 0 | 2.82E-07 | 4.2 |
| FALSE | High | GNSESVST | [R].GNSES\ | 1xPhospho | Q9DBY8 1x | 0.00359 | 0.00021 | 1 | 2 | 1 | Q9DBY8 | Nuclear va | Q9DBY8 [1 | 0 | 1644.825 | 0 | -0.64 | 0.007777 | 7.26 | 121.4 | 78.6 | 9.09 | 11.28 | | High | High | High | High | High | High | High | High | High | High | High | 9.42E-05 | 0.001003 | 1.56 |
| FALSE | High | GNGSGGSI | [K].GNGSG | 2xPhospho | I1VGT4 2xI | 0.029076 | 0.001857 | 2 | 20 | 1 | I1VGT4; I1\ | Anion exch | I1VGT4 [27 | 1 | 2317.026 | 0 | 0.83 | 0.003287 | 11.66 | 74.6 | 125.4 | 9.92 | 7.29 | Shared | High | High | High | High | High | High | High | High | High | High | High | 0.000997 | 0.01044 | 1.87 |
| FALSE | High | GNSPSCQS | [K].GNSPS( | 1xCarbami | O88466 1x | 2.93E-05 | 0 | 2 | 4 | 4 | O88466; R | Zinc finger | O88466 [1 | 0 | 2007.916 | 0 | -1.07 | 0.000865 | 2.07 | 132.5 | 67.5 | 6.05 | 6.48 | Shared | High | High | High | High | High | High | High | High | High | High | High | 0 | 4.53E-06 | 3.7 |
| FALSE | High | GNSPVTAK | [K].GNSPV | 1xPhospho | Q9DCE5 1x | 0.018285 | 0.001282 | 1 | 2 | 6 | Q9DCE5 | p21-activat | Q9DCE5 [3 | 0 | 1311.707 | 0 | -0.64 | 0.000263 | 1.52 | 121.8 | 78.2 | 4.11 | 5.62 | | High | High | High | High | High | High | High | High | High | High | High | 0.003035 | 0.03181 | 1.38 |
| FALSE | High | GNSQPLLS | [K].GNSQP | 1xPhospho | E9QN31 1x | 0.051694 | 0.003745 | 2 | 3 | 1 | E9QN31; C | Probable 2 | E9QN31 [7 | 0 | 1124.555 | 0 | -0.11 | 0.080205 | 9.93 | 104.5 | 95.5 | 2.96 | 8.71 | Shared | High | High | High | High | High | High | High | High | High | High | High | 0.00196 | 0.01995 | 1.36 |
| FALSE | High | GNSSDHEG | [K].GNSSDI | 1xCarbami | Q8VBW5 1 | 4.73E-06 | 0 | 1 | 5 | 4 | Q8VBW5 | HMG box t | Q8VBW5 [ | 0 | 3399.426 | 0 | 0.08 | 0.608621 | 14.95 | 93.9 | 106.1 | 10.1 | 15.71 | | High | High | High | High | High | High | High | High | High | High | High | 0 | 1.89E-06 | 4.67 |
| FALSE | High | GNSSDHEG | [K].GNSSDI | 1xCarbami | Q8VBW5 1 | 0.001349 | 0 | 1 | 5 | 3 | Q8VBW5 | HMG box t | Q8VBW5 [ | 0 | 3400.41 | 0 | -0.3 | 0.180029 | 8.59 | 108.7 | 91.3 | 14.74 | 9.89 | | High | High | High | High | High | High | High | High | High | High | High | 0 | 0.000334 | 3.79 |
| FALSE | High | GNSSPTVP | [K].GNSSPT | 1xPhospho | E9Q1M1 1 | 0.001815 | 5.66E-05 | 1 | 3 | 1 | E9Q1M1 | PDZ domai | E9Q1M1 [2 | 0 | 1468.699 | 0 | 2.24 | 0.000252 | 14.09 | 35.4 | 164.6 | 16.9 | 3.74 | | High | High | High | High | High | High | High | High | High | High | High | 2.58E-05 | 0.000467 | 1.39 |
| FALSE | High | GNVAGDTI | [K].GNVAG | 1xPhospho | B9EKB5 1xI | 8.73E-09 | 0 | 1 | 2 | 2 | B9EKB5 | Baz2b prot | B9EKB5 [1 | 0 | 2508.139 | 0 | -0.72 | 0.000246 | 1.53 | 124.1 | 75.9 | 2.82 | 3.52 | | High | High | High | High | High | High | High | High | High | High | High | 0 | 4.92E-10 | 3.24 |
| FALSE | High | GNVAGDTI | [K].GNVAG | 2xPhospho | B9EKB5 2xI | 0.004033 | 0.000311 | 1 | 2 | 1 | B9EKB5 | Baz2b prot | B9EKB5 [1 | 0 | 2588.105 | 0 | -0.22 | 0.133211 | 4.96 | 107.8 | 92.2 | 10.57 | 7.08 | | High | High | High | High | High | High | High | High | High | High | High | 0.000161 | 0.001135 | 2.19 |
| FALSE | High | GNVVPSPL | [R].GNVVP | 1xPhospho | Q3U434 1x | 0.001087 | 0 | 1 | 3 | 3 | Q3U434 | Putative ur | Q3U434 [3 | 0 | 1445.772 | 0 | 1.1 | 0.000704 | 7.09 | 64.1 | 135.9 | 5.54 | 9.73 | | High | High | High | High | High | High | High | High | High | High | High | 0.002589 | 0.02765 | 1.4 |
| FALSE | High | GNYPSPSSI | [R].GNYPSI | 1xPhospho | D3Z3A8 1x | 0.00824 | 0.000736 | 1 | 3 | 4 | D3Z3A8 | Unconvent | D3Z3A8 [2 | 0 | 1681.851 | 0 | 0.24 | 0.000825 | 1.74 | 91.9 | 108.1 | 2.35 | 2.64 | | High | High | High | High | High | High | High | High | High | High | High | 0.000859 | 0.008191 | 2.87 |
| FALSE | High | GNYPSPSSI | [R].GNYPSI | 2xPhospho | D3Z3A8 2x | 0.01588 | 0.001124 | 1 | 3 | 1 | D3Z3A8 | Unconvent | D3Z3A8 [2 | 0 | 1761.818 | 0 | -0.04 | 0.302729 | 2.77 | 101.4 | 98.6 | 6.06 | 2.56 | | High | High | High | High | High | High | High | High | High | High | High | 0.000585 | 0.005244 | 3.37 |
| FALSE | High | GPAAAPGA | [R].GPAAA | 1xPhospho | Q9QZM0 1 | 0.000122 | 0 | 1 | 1 | 1 | Q9QZM0 | Ubiquilin-2 | Q9QZM0 [ | 0 | 1927.009 | 0 | 0.36 | 0.002161 | 1.96 | 88.1 | 111.9 | 2.87 | 7.28 | | High | High | High | High | High | High | High | High | High | High | High | 0 | 2.24E-05 | 1.63 |
| FALSE | High | GPAAPLTP( | [K].GPAAPI | 1xPhospho | A0A1B0GR | 2.04E-05 | 0 | 1 | 8 | 2 | A0A1B0GR | Deformed | A0A1B0GR | 0 | 2746.458 | 0 | 0.49 | 0.023498 | 17.08 | 83.2 | 116.8 | 16.06 | 12.16 | | High | High | High | High | High | High | High | High | High | High | High | 0 | 3.01E-06 | 3.78 |
| FALSE | High | GPAFQSDV | [R].GPAFQ! | 1xPhospho | Q80XK6 1x | 0.000379 | 0 | 1 | 2 | 1 | Q80XK6 | Autophagy | Q80XK6 [4 | 0 | 2219.163 | 0 | 0.81 | 0.000181 | 2.77 | 72.6 | 127.4 | 3.1 | 3.62 | | High | High | High | High | High | High | High | High | High | High | High | 0 | 8.01E-05 | 3.06 |
| FALSE | High | GNSPSCQS | [K].GNSPS( | 1xCarbami | O88466 2x | 6.04E-06 | 0 | 2 | 4 | 3 | O88466; R | Zinc finger | O88466 [1 | 0 | 2087.882 | 0 | -1.31 | 0.000241 | 0.93 | 143.9 | 56.1 | 4.28 | 8.7 | Shared | High | High | High | High | High | High | High | High | High | High | High | 0 | 7.65E-07 | 4.25 |
| FALSE | High | GNFGGSFA | [R].GNFGG | 1xPhospho | A0A1S6GW | 0.000108 | 0 | 1 | 9 | 1 | A0A1S6GW | Uncharacte | A0A1S6GW | 0 | 2344.082 | 0 | 0.45 | 0.160085 | 16.61 | 80.4 | 119.6 | 15.23 | 16.76 | | High | High | High | High | High | High | High | High | High | High | High | 0 | 1.96E-05 | 2.16 |
| FALSE | High | GNDSDGE/ | [K].GNDSD | 1xPhospho | Q3TN07 1x | 0.045898 | 0.003408 | 1 | 2 | 1 | Q3TN07 | Putative ur | Q3TN07 [9 | 1 | 2460.138 | 0 | -0.53 | 0.011653 | 1.43 | 113.8 | 86.2 | 9.4 | 10.91 | | High | High | High | High | High | High | High | High | High | High | High | 0.001774 | 0.01749 | 2.35 |
| FALSE | High | GNDISSGT\ | [K].GNDISS | 1xPhospho | Q3TGC5 1x | 0.016306 | 0.001124 | 1 | 2 | 2 | Q3TGC5 | Putative ur | Q3TGC5 [9 | 0 | 2839.346 | 0 | 0.32 | 0.078753 | 9.13 | 92 | 108 | 7.46 | 10.7 | | High | High | High | High | High | High | High | High | High | High | High | 0.000585 | 0.005413 | 3.58 |
| FALSE | High | GLSVTAEQ | [R].GLSVTA | 2xPhospho | E9Q166 2x | 0.003312 | 0.00016 | 1 | 2 | 1 | E9Q166 | ATPase fan | E9Q166 [1 | 0 | 2053.967 | 0 | -0.5 | 0.000488 | 2.23 | 118.8 | 81.2 | 5.81 | 4.95 | | High | High | High | High | High | High | High | High | High | High | High | 7.19E-05 | 0.000916 | 2.98 |
| FALSE | High | GLTHPSSSF | [R].GLTHP! | 1xPhospho | F6ZBR8 1xI | 0.001767 | 5.66E-05 | 2 | 5 | 2 | F6ZBR8; O | Protein Wi | F6ZBR8 [16 | 0 | 1774.987 | 0 | -0.05 | 0.940589 | 36.29 | 99.2 | 100.8 | 18.12 | 20.86 | Shared | High | High | High | High | High | High | High | High | High | High | High | 2.58E-05 | 0.000452 | 1.36 |
| FALSE | High | GLTLSDDLE | [K].GLTLSD | 1xPhospho | P59325 1xI | 0.016451 | 0.001124 | 1 | 5 | 1 | P59325 | Eukaryotic | P59325 [22 | 0 | 1427.698 | 0 | -0.84 | 0.000406 | 2.04 | 128.7 | 71.3 | 5.5 | 5.91 | | High | High | High | High | High | High | High | High | High | High | High | 0.000585 | 0.005459 | 1.75 |
| FALSE | High | GLTPGAAA | [K].GLTPG/ | 1xPhospho | A2A654 1x | 2.23E-08 | 0 | 1 | 7 | 1 | A2A654 | Bromodom | A2A654 [2 | 0 | 3185.699 | 0 | 0.43 | 0.072893 | 18.49 | 87 | 113 | 14.42 | 11.61 | | High | High | High | High | High | High | High | High | High | High | High | 0 | 1.42E-09 | 3.58 |
| FALSE | High | GITQGPNE | [K].GITQGI | 1xPhospho | A0A068F1 | 0.000181 | 0 | 2 | 9 | 2 | A0A068F1 | Glyco-gag | A0A068F1 | 0 | 2011.969 | 0 | 2.82 | 0.000606 | 25 | 24.6 | 175.4 | 27.68 | 8.06 | Shared | High | High | High | High | High | High | High | High | High | High | High | 0 | 3.51E-05 | 3.7 |
| FALSE | High | GLTSSIDYS | [K].GLTSSII | 1xPhospho | B9EKT8 1xI | 0.002463 | 0.000109 | 1 | 3 | 2 | B9EKT8 | Uncharacte | B9EKT8 [1 | 0 | 1608.829 | 0 | -0.33 | 0.001695 | 1.2 | 110.2 | 89.8 | 2.01 | 5.07 | | High | High | High | High | High | High | High | High | High | High | High | 0.00196 | 0.01993 | 1.76 |
| FALSE | High | GLVAAYSG | [R].GLVAA\ | 1xPhospho | Q91YE7 1x | 1.75E-09 | 0 | 1 | 2 | 4 | Q91YE7 | RNA-bindir | Q91YE7 [6 | 0 | 2362.065 | 0 | -0.19 | 0.069242 | 7.34 | 106.6 | 93.4 | 4.6 | 6.32 | | High | High | High | High | High | High | High | High | High | High | High | 0 | 8.13E-11 | 4.81 |
| FALSE | High | GLVAAYSG | [R].GLVAA\ | 1xDeamida | Q91YE7 1x | 0.001704 | 5.66E-05 | 1 | 2 | 3 | Q91YE7 | RNA-bindir | Q91YE7 [6 | 0 | 2363.049 | 0 | 0.01 | 0.934341 | 12.81 | 99.7 | 100.3 | 9.42 | 13.24 | | High | High | High | High | High | High | High | High | High | High | High | 2.58E-05 | 0.000435 | 3.46 |
| FALSE | High | GLVAAYSG | [R].GLVAA\ | 2xPhospho | Q91YE7 2x | 4.25E-06 | 0 | 1 | 2 | 4 | Q91YE7 | RNA-bindir | Q91YE7 [6 | 0 | 2442.031 | 0 | 0.32 | 0.01107 | 11.05 | 85.6 | 114.4 | 7.26 | 9.04 | | High | High | High | High | High | High | High | High | High | High | High | 0 | 5.17E-07 | 4.19 |
| FALSE | High | GLVAAYSG | [R].GLVAA\ | 1xPhospho | Q99KG3 1x | 3.69E-07 | 0 | 2 | 3 | 3 | Q99KG3; C | RNA-bindir | Q99KG3 [7 | 0 | 2264.976 | 0 | -0.45 | 0.008619 | 0.72 | 115.5 | 84.5 | 5.83 | 6.22 | Shared | High | High | High | High | High | High | High | High | High | High | High | 0 | 3.36E-08 | 3.67 |
| FALSE | High | GLVAAYSG | [R].GLVAA\ | 2xPhospho | Q99KG3 2x | 8.78E-08 | 0 | 2 | 3 | 9 | Q99KG3; C | RNA-bindir | Q99KG3 [7 | 0 | 2344.942 | 0 | -0.08 | 0.117687 | 8.14 | 106.7 | 93.3 | 6.67 | 7.41 | Shared | High | High | High | High | High | High | High | High | High | High | High | 0 | 3.7E-05 | 3.95 |
| FALSE | High | GLVAAYSG | [R].GLVAA\ | 3xPhospho | Q99KG3 3x | 0.000346 | 0 | 2 | 3 | 14 | Q99KG3; C | RNA-bindir | Q99KG3 [7 | 0 | 2424.908 | 0 | -0.08 | 0.900537 | 7.87 | 99.9 | 100.1 | 9.53 | 14.5 | Shared | High | High | High | High | High | High | High | High | High | High | High | 0 | 7.23E-05 | 3.2 |
| FALSE | High | GLVLPAEPI | [R].GLVLPA | 1xPhospho | I7H459 1xP | 7.47E-05 | 0 | 1 | 2 | 3 | I7H459 | Protein inc | I7H459 [45 | 0 | 3777.729 | 0 | -0.01 | 0.52028 | 18.89 | 96.6 | 103.4 | 9.78 | 55.88 | | High | High | High | High | High | High | High | High | High | High | High | 0 | 1.3E-05 | 3.65 |
| FALSE | High | GLVSPQSP( | [K].GLVSP( | 1xPhospho | P41183 1xI | 0.001049 | 0 | 1 | 2 | 1 | P41183 | B-cell lymp | P41183 [32 | 0 | 1578.866 | 0 | 3.27 | 0.000196 | 10.87 | 18.8 | 181.2 | 15.2 | 6.61 | | High | High | High | High | High | High | High | High | High | High | High | 0 | 0.000253 | 2.3 |
| FALSE | High | GIVVYTGDI | [R].GIVVYT | 1xPhospho | Q3TXF9 1x | 0.000924 | 0 | 1 | 2 | 1 | Q3TXF9 | Sodium/pc | Q3TXF9 [2 | 0 | 1288.65 | 0 | -0.72 | 0.024537 | 6.84 | 120.1 | 79.9 | 17.02 | 12.5 | | High | High | High | High | High | High | High | High | High | High | High | 0 | 0.000219 | 2.3 |
| FALSE | High | GLYAAATV | [R].GLYAA/ | 1xPhospho | A0A0A6YW | 4.47E-08 | 0 | 1 | 4 | 5 | A0A0A6YW | Rap guanin | A0A0A6YW | 0 | 2813.319 | 0 | 0.14 | 0.205521 | 20.68 | 95.2 | 104.8 | 9.7 | 17.77 | | High | High | High | High | High | High | High | High | High | High | High | 0 | 3.09E-09 | 3.69 |
| FALSE | High | GLYDGPVC | [R].GLYDGI | 1xCarbami | O08553 1x | 9.14E-06 | 0 | 1 | 1 | 2 | O08553 | Dihydropy | O08553 [4 | 0 | 2159.086 | 0 | 1.33 | 0.001503 | 8.43 | 57.1 | 142.9 | 7.34 | 11.93 | | High | High | High | High | High | High | High | High | High | High | High | 0 | 8.9E-06 | 3.6 |
| FALSE | High | GMEGLISP | [R].GMEGL | 2xCarbami | Q69Z43 1x | 0.000186 | 0 | 1 | 2 | 1 | Q69Z43 | MKIAA199 | Q69Z43 [8 | 0 | 3089.373 | 0 | 1.28 | 0.018948 | 33.1 | 57.5 | 142.5 | 21.82 | 20.26 | | High | High | High | High | High | High | High | High | High | High | High | 0 | 3.59E-05 | 4.9 |
| FALSE | High | GMEGLISP | [R].GMEGL | 2xCarbami | Q69Z43 1x | 1.28E-06 | 0 | 1 | 2 | 3 | Q69Z43 | MKIAA199 | Q69Z43 [8 | 0 | 3105.368 | 0 | 1.38 | 0.006823 | 17.31 | 60.4 | 139.6 | 15.99 | 20.89 | | High | High | High | High | High | High | High | High | High | High | High | 0 | 1.34E-07 | 4.32 |
| FALSE | High | GMGPGTP/ | [R].GMGP( | 1xPhospho | Q8VIJ6 1xP | 0.000563 | 0 | 1 | 3 | 1 | Q8VIJ6 | Splicing fac | Q8VIJ6 [67 | 0 | 1445.645 | 0 | 1.61 | 0.022482 | 37.38 | 45 | 155 | 32.66 | 29.79 | | High | High | High | High | High | High | High | High | High | High | High | 0 | 0.000125 | 2.37 |
| FALSE | High | GMPPGLQ( | [R].GMPPG | 1xPhospho | E9Q8G4 1x | 0.011652 | 0.000889 | 1 | 17 | 3 | E9Q8G4 | Transcripti | E9Q8G4 [5 | 1 | 4059.931 | 0 | -0.07 | 0.532074 | 34.04 | 103.4 | 96.6 | 14.1 | 17.69 | | High | High | High | High | High | High | High | High | High | High | High | 0.000555 | 0.004784 | 3.46 |
| FALSE | High | GMSPVVSE | [K].GMSPV | 1xPhospho | E9QM77 1 | 0.000438 | 0 | 1 | 9 | 3 | E9QM77 | Ataxin-2 O | E9QM77 [7 | 0 | 1423.66 | 0 | 0.2 | 0.13367 | 8.22 | 92.7 | 107.3 | 3.08 | 23.81 | | High | High | High | High | High | High | High | High | High | High | High | 0 | 9.44E-05 | 3.16 |
| FALSE | High | GMVSLSEII | [K].GMVSL | 1xPhospho | P27546 1xI | 0.023379 | 0.001578 | 1 | 3 | 2 | P27546 | Microtubu | P27546 [37 | 0 | 2031.049 | 0 | 2.11 | 0.000855 | 9.07 | 36.2 | 163.8 | 9.39 | 14.08 | | High | High | High | High | High | High | High | High | High | High | High | 0.000847 | 0.008101 | 3.17 |
| FALSE | High | GMYDGPV | [R].GMYD( | 1xPhospho | E9PWE8 1x | 0.000492 | 0 | 1 | 4 | 2 | E9PWE8 | Dihydropy | E9PWE8 [6 | 0 | 2180.075 | 0 | 1.19 | 0.001062 | 8.41 | 61.1 | 138.9 | 14.69 | 9.8 | | High | High | High | High | High | High | High | High | High | High | High | 0 | 0.000108 | 2.24 |
| FALSE | High | GMYDGPV | [R].GMYD( | 1xPhospho | E9PWE8 1x | 3.44E-05 | 0 | 1 | 4 | 6 | E9PWE8 | Dihydropy | E9PWE8 [6 | 0 | 2196.07 | 0 | 1.11 | 0.00332 | 22.6 | 64.4 | 135.6 | 10.42 | 14.37 | | High | High | High | High | High | High | High | High | High | High | High | 0 | 5.44E-06 | 3.06 |
| FALSE | High | GMYLSPEG | [K].GMYLS | 1xPhospho | Q8C796 1x | 0.000198 | 0 | 1 | 2 | 2 | Q8C796 | REST corep | Q8C796 [2 | 0 | 2787.384 | 0 | -1.86 | 0.001195 | 2.81 | 156.6 | 43.4 | 10.4 | 13.15 | | High | High | High | High | High | High | High | High | High | High | High | 0 | 4.79E-05 | 4.17 |
| FALSE | High | GNAGGGG | [R].GNAGG | 1xPhospho | Q8BXQ8 1x | 0.000139 | 0 | 1 | 2 | 2 | Q8BXQ8 | Protein FAI | Q8BXQ8 [1 | 0 | 2114.08 | 0 | -1.11 | 0.001648 | 1.92 | 133.7 | 66.3 | 9.51 | 9.34 | | High | High | High | High | High | High | High | High | High | High | High | 0 | 2.62E-05 | 2.85 |
| FALSE | High | GNASPGA/ | [R].GNASP( | 1xPhospho | Q9EPQ8 1x | 4.08E-05 | 0 | 1 | 4 | 1 | Q9EPQ8 | Transcripti | Q9EPQ8 [9 | 0 | 2482.083 | 0 | 0.62 | 0.015363 | 5.45 | 79.6 | 120.4 | 14.4 | 7.34 | | High | High | High | High | High | High | High | High | High | High | High | 0 | 6.58E-06 | 2.77 |
| FALSE | High | GNASPVLD | [K].GNASP' | 1xPhospho | Q4VAB2 1x | 0.000186 | 0 | 1 | 1 | 4 | Q4VAB2 | GPN-loop ( | Q4VAB2 [3 | 0 | 1977.026 | 0 | 1 | 0.000275 | 5.05 | 68.3 | 131.7 | 2.84 | 3.47 | | High | High | High | High | High | High | High | High | High | High | High | 0 | 0.000144 | 2.62 |
| FALSE | High | GLLSQNGS | [R].GLLSQN | 1xCarbami | Q8BI84 1xP | 0.024001 | 0.001612 | 1 | 4 | 2 | Q8BI84 | Melanoma | Q8BI84 [16 | 0 | 4083.021 | 0 | -0.06 | 0.51574 | 8.8 | 104.7 | 95.3 | 14.22 | 14.97 | | High | High | High | High | High | High | High | High | High | High | High | 0.000859 | 0.008332 | 3.9 |
| FALSE | High | GPAGEASA | [R].GPAGE. | 1xPhospho | B7ZNU9 1x | 2.85E-05 | 0 | 1 | 3 | 3 | B7ZNU9 | Beta-2-syn | B7ZNU9 [8 | 0 | 1504.736 | 0 | -0.42 | 0.002468 | 2.41 | 115.4 | 84.6 | 5.18 | 3.26 | | High | High | High | High | High | High | High | High | High | High | High | 0.005231 | 0.05702 | 1.95 |
| FALSE | High | GILSLPHQ/ | [K].GILSLPI | 1xPhospho | Q2M4G9 1 | 0.000141 | 0 | 1 | 2 | 1 | Q2M4G9 | Protein inh | Q2M4G9 [ | 0 | 1770.947 | 0 | 0.56 | 0.00252 | 8.35 | 81.4 | 118.6 | 8.2 | 7.42 | | High | High | High | High | High | High | High | High | High | High | High | 0 | 2.62E-05 | 2.37 |
| FALSE | High | GLILVSPPIT | [K].GLILVSI | 1xPhospho | Q80WE4 1 | 5.05E-07 | 0 | 1 | 4 | 2 | Q80WE4 | Kinesin-like | Q80WE4 [8 | 0 | 2176.203 | 0 | -0.1 | 0.4707 | 18.05 | 103.3 | 96.7 | 3.99 | 8.78 | | High | High | High | High | High | High | High | High | High | High | High | 0 | 2.87E-06 | 3.83 |
| FALSE | High | GGSPYNQF | [R].GGSPYI | 1xPhospho | Q8BKI2 1xI | 5.81E-05 | 0 | 1 | 5 | 2 | Q8BKI2 | Trinucleoti | Q8BKI2 [14 | 0 | 3663.789 | 0 | 0.92 | 0.010291 | 25.12 | 69.3 | 130.7 | 9.09 | 13.81 | | High | High | High | High | High | High | High | High | High | High | High | 0 | 9.77E-06 | 4.24 |
| FALSE | High | GGSSEELHI | [R].GGSSEI | 1xPhospho | Q3UMU9-3 | 0.002232 | 0.000109 | 1 | 2 | 7 | Q3UMU9-3 | Isoform 3 c | Q3UMU9-3 | 0 | 1579.695 | 0 | 0.03 | 0.229152 | 4.55 | 97.5 | 102.5 | 6.68 | 4.59 | | High | High | High | High | High | High | High | High | High | High | High | 4.94E-05 | 0.000589 | 2.74 |
| FALSE | High | GGSSENGV | [K].GGSSEI | 1xPhospho | Q9Z2U2 1x | 3.85E-06 | 0 | 1 | 3 | 3 | Q9Z2U2 | Zinc finger | Q9Z2U2 [1 | 0 | 3496.616 | 0 | -0.88 | 0.000222 | 1.27 | 128.9 | 71.1 | 3.45 | 5.44 | | High | High | High | High | High | High | High | High | High | High | High | 0 | 4.61E-07 | 3.73 |
| FALSE | High | GGSSENGV | [K].GGSSEI | 1xDeamida | Q9Z2U2 1x | 2.38E-05 | 0 | 1 | 3 | 2 | Q9Z2U2 | Zinc finger | Q9Z2U2 [1 | 0 | 3497.6 | 0 | -0.65 | 0.003376 | 5.97 | 126 | 74 | 9.98 | 5.82 | | High | High | High | High | High | High | High | High | High | High | High | 0 | 3.56E-06 | 4.29 |
| FALSE | High | GGSSGATG | [R].GGSSG. | 1xPhospho | A0A1N9PT | 0.017966 | 0.001282 | 1 | 1 | 2 | A0A1N9PT | Probable fi | A0A1N9PT | 1 | 2950.324 | 0 | -0.2 | 0.03449 | 6.25 | 106.3 | 93.7 | 6.83 | 6.88 | | High | High | High | High | High | High | High | High | High | High | High | 0.000689 | 0.006036 | 1.74 |

| | | | | | | | | | | | | | | | | | | | | | | | | | | | | | | | | | | | | | | |
|---|---|---|---|---|---|---|---|---|---|---|---|---|---|---|---|---|---|---|---|---|---|---|---|---|---|---|---|---|---|---|---|---|---|---|---|---|---|---|
| FALSE | High | GGSSGATC | [R].GGSSG | 2xPhospho | A0A1N9PT | 0.003857 | 0.000261 | 1 | 1 | 3 | A0A1N9PT | Probable fi | A0A1N9PT | 1 | 3030.29 | 0 | -0.23 | 0.309329 | 14.26 | 104.3 | 95.7 | 9.02 | 10.56 | | High | High | High | High | High | High | High | High | High | High | High | 0.000585 | 0.005309 | 2.83 |
| FALSE | High | GGSSGEEL | [R].GGSSG | 2xPhospho | Q3UMU9-3 | 5.5E-05 | 0 | 1 | 4 | 3 | Q3UMU9-3 | Isoform 3 c | Q3UMU9-3 | 0 | 2179.945 | 0 | -0.09 | 0.60052 | 6.26 | 104.5 | 95.5 | 7.24 | 5.75 | | High | High | High | High | High | High | High | High | High | High | High | 0 | 9.16E-06 | 3.39 |
| FALSE | High | GGSSPSQS | [K].GGSSPS | 1xCarbami | Q8CAF4-3 | 0.000178 | 0 | 1 | 4 | 2 | Q8CAF4-3 | Isoform 3 c | Q8CAF4-3 | 0 | 2609.062 | 0 | 0.58 | 0.10719 | 26.45 | 81.1 | 118.9 | 15.84 | 15.34 | | High | High | High | High | High | High | High | High | High | High | High | 0 | 0.000219 | 3.69 |
| FALSE | High | GGSTGGG | [R].GGSTG | 1xPhospho | E9PV45 1x | 0.000248 | 0 | 1 | 3 | 4 | E9PV45 | Ubiquitin c | E9PV45 [8 | 0 | 2839.334 | 0 | 0.59 | 0.003054 | 14.21 | 81.3 | 118.7 | 9.21 | 3.51 | | High | High | High | High | High | High | High | High | High | High | High | 0.004152 | 0.04485 | 2.95 |
| FALSE | High | GGTTSWG | [R].GGTTS | 1xPhospho | Q61286-2 | 4.27E-05 | 0 | 2 | 7 | 4 | Q61286-2; | Isoform AL | Q61286-2 | 0 | 2324.003 | 0 | 0.17 | 0.007099 | 5.39 | 93.4 | 106.6 | 9.97 | 8.6 | Shared | High | High | High | High | High | High | High | High | High | High | High | 4.94E-05 | 0.000591 | 3.2 |
| FALSE | High | GGTVADLD | [K].GGTVA | 1xPhospho | P97496 1x | 7.33E-08 | 0 | 1 | 6 | 2 | P97496 | SWI/SNF c | P97496 [4 | 0 | 2530.196 | 0 | -1.16 | 0.001213 | 3.27 | 138.2 | 61.8 | 7.83 | 10 | | High | High | High | High | High | High | High | High | High | High | High | 0 | 5.41E-09 | 4.8 |
| FALSE | High | GGVTGSPE | [K].GGVTG | 1xPhospho | E9Q616 1x | 3.87E-08 | 0 | 1 | 4 | 7 | E9Q616 | AHNAK nu | E9Q616 [5 | 0 | 1871.952 | 0 | 2.27 | 0.000209 | 7.35 | 36.4 | 163.6 | 12.6 | 7.45 | | High | High | High | High | High | High | High | High | High | High | High | 0 | 1.97E-05 | 3.83 |
| FALSE | High | GGVTGSPE | [K].GGVTG | 2xPhospho | E9Q616 2x | 3.98E-05 | 0 | 1 | 4 | 1 | E9Q616 | AHNAK nu | E9Q616 [5 | 0 | 1951.918 | 0 | 1.63 | 0.000412 | 18.63 | 48.2 | 151.8 | 14.87 | 8.52 | | High | High | High | High | High | High | High | High | High | High | High | 0 | 6.34E-06 | 3.74 |
| FALSE | High | GGSPQNLF | [R].GGSPQ | 1xCarbami | G3X904 1x | 0.081752 | 0.006219 | 1 | 5 | 1 | G3X904 | Zinc finger | G3X904 [1 | 0 | 2330.152 | 0 | 0.69 | 0.012188 | 28.99 | 77.2 | 122.8 | 12.76 | 15.54 | | High | High | High | High | High | High | High | High | High | High | High | 0.0032 | 0.03397 | 2.42 |
| FALSE | High | GGWDHGD | [K].GGWD | 1xPhospho | A2A7S8-2 | 1.13E-05 | 0 | 1 | 3 | 8 | A2A7S8-2 | Isoform 2 c | A2A7S8-2 | 0 | 3675.685 | 0 | -0.82 | 0.002046 | 1.92 | 128.4 | 71.6 | 6.79 | 14.96 | | High | High | High | High | High | High | High | High | High | High | High | 0 | 2.55E-06 | 6.1 |
| FALSE | High | GGYHGSLE | [R].GGYHG | 1xPhospho | B2RWU2 1 | 7.28E-05 | 0 | 1 | 3 | 1 | B2RWU2 | Immunode | B2RWU2 [ | 0 | 2173.04 | 0 | 2.92 | 0.000891 | 17.34 | 21 | 179 | 17.84 | 13.97 | | High | High | High | High | High | High | High | High | High | High | High | 0 | 1.26E-05 | 3 |
| FALSE | High | GGYLQGNI | [K].GGYLQ | 1xPhospho | Q3TN05 1x | 0.010853 | 0.000772 | 1 | 4 | 1 | Q3TN05 | Putative ur | Q3TN05 [1 | 0 | 1464.65 | 0 | 2.77 | 0.006018 | 23.61 | 26.5 | 173.5 | 11.42 | 29.43 | | High | High | High | High | High | High | High | High | High | High | High | 0.000399 | 0.003445 | 1.28 |
| FALSE | High | GGYLSDGD | [K].GGYLS | 2xPhospho | E9PUP1 2x | 5.91E-05 | 0 | 1 | 4 | 2 | E9PUP1 | Synapse de | E9PUP1 [7 | 0 | 1754.724 | 0 | -1.83 | 0.00034 | 1.94 | 157.7 | 42.3 | 3.32 | 21.56 | | High | High | High | High | High | High | High | High | High | High | High | 0 | 9.9E-06 | 2.61 |
| FALSE | High | GHAGGQR | [R].GHAGG | 1xPhospho | E9Q4D5 1x | 0.001489 | 5.66E-05 | 1 | 3 | 2 | E9Q4D5 | GTP-bindin | E9Q4D5 [2 | 0 | 2377.125 | 0 | -0.39 | 0.043884 | 12.89 | 112.7 | 87.3 | 7.18 | 14.32 | | High | High | High | High | High | High | High | High | High | High | High | 2.58E-05 | 0.000375 | 2.09 |
| FALSE | High | GHFQPLEN | [R].GHFQP | 1xCarbami | Q5DTG8 1x | 1.46E-09 | 0 | 1 | 8 | 2 | Q5DTG8 | MKIAA422 | Q5DTG8 [1 | 0 | 2450.137 | 0 | -0.23 | 0.030379 | 9.98 | 109.2 | 90.8 | 3.68 | 6.87 | | High | High | High | High | High | High | High | High | High | High | High | 0 | 6.63E-11 | 5.26 |
| FALSE | High | GHFQPLEN | [R].GHFQP | 1xCarbami | Q5DTG8 1x | 0.000255 | 0 | 1 | 8 | 2 | Q5DTG8 | MKIAA422 | Q5DTG8 [1 | 0 | 2451.121 | 0 | -0.35 | 0.198433 | 2.89 | 112.2 | 87.8 | 9.63 | 18.39 | | High | High | High | High | High | High | High | High | High | High | High | 0 | 5.14E-05 | 4.77 |
| FALSE | High | GHGLEPTA | [R].GHGLE | 1xPhospho | E9PYL2 1x | 0.003927 | 0.000311 | 1 | 2 | 1 | E9PYL2 | Proline-ricl | E9PYL2 [8 | 0 | 1527.752 | 0 | 0.54 | 0.00109 | 7.06 | 82.1 | 117.9 | 5.6 | 4.62 | | High | High | High | High | High | High | High | High | High | High | High | 0.00014 | 0.001106 | 3.3 |
| FALSE | High | GHHGSLGT | [R].GHHGS | 1xPhospho | Q8R149 1x | 0.000681 | 0 | 1 | 2 | 6 | Q8R149 | BUD13 hor | Q8R149 [2 | 0 | 1501.711 | 0 | -0.02 | 0.322602 | 10.81 | 100.8 | 99.2 | 8.95 | 14.42 | | High | High | High | High | High | High | High | High | High | High | High | 4.94E-05 | 0.000716 | 2.75 |
| FALSE | High | GHLDAELD | [K].GHLDA | 1xPhospho | Q9CY57 1x | 0.000207 | 0 | 1 | 7 | 2 | Q9CY57 | Chromatin | Q9CY57 [2 | 0 | 2531.048 | 0 | 0.58 | 0.020901 | 5.34 | 80.1 | 119.9 | 12.48 | 16.15 | | High | High | High | High | High | High | High | High | High | High | High | 0 | 4.08E-05 | 3.15 |
| FALSE | High | GHLSEGLV | [R].GHLSE | 1xPhospho | Q61532 1x | 0.003055 | 0.00016 | 1 | 2 | 2 | Q61532 | Mitogen-a | Q61532 [1 | 0 | 1578.866 | 0 | 1.15 | 0.002321 | 15.33 | 58.1 | 141.9 | 19.4 | 12.67 | | High | High | High | High | High | High | High | High | High | High | High | 0.000968 | 0.01007 | 2.51 |
| FALSE | High | GHLSNEEA | [K].GHLSN | 1xCarbami | Q8C9B9 1x | 0.022973 | 0.001578 | 1 | 3 | 1 | Q8C9B9 | Death-indu | Q8C9B9 [5 | 0 | 1619.819 | 0 | -0.22 | 0.029171 | 8.92 | 106.5 | 93.5 | 6.93 | 8.27 | | High | High | High | High | High | High | High | High | High | High | High | 0.000847 | 0.007997 | 2.37 |
| FALSE | High | GHPSAGAE | [R].GHPSA | 1xPhospho | Q8CIJ3 1xP | 2.23E-08 | 0 | 1 | 2 | 15 | Q8CIJ3 | Eukaryotic | Q8CIJ3 [10 | 0 | 2477.042 | 0 | 0.37 | 0.019821 | 11.03 | 87.2 | 112.8 | 2.86 | 13.11 | | High | High | High | High | High | High | High | High | High | High | High | 0 | 1.42E-06 | 5.31 |
| FALSE | High | GHPSAGAE | [R].GHPSA | 2xPhospho | Q8CIJ3 2xP | 6.62E-10 | 0 | 1 | 2 | 6 | Q8CIJ3 | Eukaryotic | Q8CIJ3 [10 | 0 | 2557.008 | 0 | 0.47 | 0.012418 | 12.18 | 84.9 | 115.1 | 6.04 | 8.16 | | High | High | High | High | High | High | High | High | High | High | High | 0 | 2.73E-11 | 4.8 |
| FALSE | High | GGWLAPSI | [R].GGWLA | 1xPhospho | F6RGI3 1x | 0.005872 | 0.00062 | 1 | 1 | 1 | F6RGI3 | Uncharacte | F6RGI3 [11 | 0 | 1784.905 | 0 | -0.08 | 0.07742 | 3.12 | 102.3 | 97.7 | 3.74 | 10.28 | | High | High | High | High | High | High | High | High | High | High | High | 0.000275 | 0.001727 | 2.61 |
| FALSE | High | GHPTQSPG | [R].GHPTQ | 1xPhospho | Q5XJV7-2 | 0.004143 | 0.00036 | 1 | 3 | 1 | Q5XJV7-2 | Isoform 2 c | Q5XJV7-2 | 0 | 1658.753 | 0 | -0.32 | 0.181413 | 0.91 | 106.8 | 93.2 | 14.55 | 11.28 | | High | High | High | High | High | High | High | High | High | High | High | 0.000161 | 0.001178 | 2.63 |
| FALSE | High | GGSPNVES | [K].GGSPN | 1xPhospho | Q3TK27 1x | 0.000573 | 0 | 1 | 3 | 2 | Q3TK27 | Putative ur | Q3TK27 [3 | 0 | 1554.793 | 0 | -1.24 | 0.000463 | 1.56 | 142.2 | 57.8 | 6.03 | 4.91 | | High | High | High | High | High | High | High | High | High | High | High | 0 | 0.000196 | 1.53 |
| FALSE | High | GGSPGLES | [K].GGSPG | 1xPhospho | Q8CHP6 1x | 0.037369 | 0.00249 | 1 | 6 | 1 | Q8CHP6 | Polyhomeo | Q8CHP6 [2 | 0 | 1168.556 | 0 | 1.93 | 0.000487 | 14.72 | 40.4 | 159.6 | 4.4 | 8.41 | | High | High | High | High | High | High | High | High | High | High | High | 0.001308 | 0.01376 | 1.51 |
| FALSE | High | GGNVFEAL | [K].GGNVF | 1xPhospho | Q6P542 1x | 2.25E-08 | 0 | 2 | 2 | 31 | Q6P542; Q | ATP-bindin | Q6P542 [1 | 0 | 2976.247 | 0 | -0.36 | 0.001432 | 2.8 | 111.2 | 88.8 | 2.25 | 3.1 | Shared | High | High | High | High | High | High | High | High | High | High | High | 0 | 3.08E-09 | 5.47 |
| FALSE | High | GGPAEAPS | [R].GGPAE | 1xPhospho | D3YWD3 1 | 0.024855 | 0.001645 | 1 | 3 | 2 | D3YWD3 | Transmem | D3YWD3 [ | 0 | 1247.598 | 0 | 1.68 | 0.00044 | 12.42 | 48.9 | 151.1 | 5.59 | 7.33 | | High | High | High | High | High | High | High | High | High | High | High | 0.000872 | 0.008731 | 1.06 |
| FALSE | High | GGPASVPS | [R].GGPAS | 1xPhospho | Q6PDL0 1x | 7.91E-06 | 0 | 1 | 3 | 1 | Q6PDL0 | Cytoplasmi | Q6PDL0 [3 | 0 | 1937.015 | 0 | 1.1 | 0.000971 | 12.08 | 63.5 | 136.5 | 8.4 | 7.33 | | High | High | High | High | High | High | High | High | High | High | High | 0 | 1.04E-06 | 3.33 |
| FALSE | High | GGPCSPPS | [R].GGPCS | 1xCarbami | Q80U72-4 | 0.000693 | 0 | 1 | 4 | 2 | Q80U72-4 | Isoform 4 c | Q80U72-4 | 0 | 2260.109 | 0 | 2.17 | 0.000899 | 7.62 | 42 | 158 | 13.19 | 13.72 | | High | High | High | High | High | High | High | High | High | High | High | 0 | 0.000158 | 2.91 |
| FALSE | High | GGPCSPTK | [R].GGPCS | 1xCarbami | G5E8T6 1x | 0.011652 | 0.000889 | 1 | 3 | 1 | G5E8T6 | Membrane | G5E8T6 [6 | 0 | 1341.664 | 0 | -1.2 | 0.000265 | 1.2 | 141.1 | 58.9 | 3.99 | 6.24 | | High | High | High | High | High | High | High | High | High | High | High | 0.000449 | 0.003721 | 1.3 |
| FALSE | High | GGPEGGA | [K].GGPEG | 1xCarbami | Q91ZW3 1 | 0.007145 | 0.00062 | 1 | 1 | 2 | Q91ZW3 | SWI/SNF-r | Q91ZW3 [ | 0 | 3590.54 | 0 | -0.49 | 0.466803 | 25.41 | 102.5 | 97.5 | 66.68 | 35.86 | | High | High | High | High | High | High | High | High | High | High | High | 0.000344 | 0.003074 | 3.17 |
| FALSE | High | GGPEGGA | [K].GGPEG | 1xCarbami | Q91ZW3 1 | 1.96E-07 | 0 | 1 | 1 | 5 | Q91ZW3 | SWI/SNF-r | Q91ZW3 [ | 0 | 3819.703 | 0 | -0.11 | 0.357413 | 6.87 | 103.8 | 96.2 | 5.88 | 22.86 | | High | High | High | High | High | High | High | High | High | High | High | 0 | 1.63E-08 | 4.96 |
| FALSE | High | GGPEGGA | [K].GGPEG | 1xCarbami | Q91ZW3 1 | 2.68E-06 | 0 | 1 | 1 | 21 | Q91ZW3 | SWI/SNF-r | Q91ZW3 [ | 0 | 3835.698 | 0 | -0.12 | 0.113282 | 8.13 | 104.2 | 95.8 | 3.59 | 11.23 | | High | High | High | High | High | High | High | High | High | High | High | 0 | 2.36E-05 | 4.69 |
| FALSE | High | GGPGSAVS | [K].GGPGS | 1xPhospho | B7STB7 1x | 4.26E-07 | 0 | 1 | 5 | 3 | B7STB7 | Annexin O | B7STB7 [3 | 0 | 2882.449 | 0 | 3.91 | 0.000162 | 27.76 | 12.7 | 187.3 | 22.14 | 5.64 | | High | High | High | High | High | High | High | High | High | High | High | 0 | 3.88E-08 | 3.79 |
| FALSE | High | GGPPSHPS | [K].GGPPS | 1xCarbami | Q62311 1x | 0.003167 | 0.00016 | 1 | 1 | 1 | Q62311 | Transcripti | Q62311 [6 | 0 | 3223.586 | 0 | -0.31 | 0.940589 | 7.16 | 110.2 | 89.8 | 12.54 | 26.77 | | High | High | High | High | High | High | High | High | High | High | High | 7.19E-05 | 0.000871 | 2.97 |
| FALSE | High | GGPPSHPS | [K].GGPPS | 1xCarbami | Q62311 2x | 0.020319 | 0.001472 | 1 | 1 | 5 | Q62311 | Transcripti | Q62311 [6 | 0 | 3303.552 | 0 | 0.15 | 0.869621 | 38.21 | 98.8 | 101.2 | 19.49 | 6.08 | | High | High | High | High | High | High | High | High | High | High | High | 0.000786 | 0.006967 | 3.43 |
| FALSE | High | GGPVVATA | [R].GGPVV | 1xPhospho | P58334 1x | 2.02E-09 | 0 | 1 | 2 | 5 | P58334 | Krueppel-li | P58334 [8 | 0 | 2957.405 | 0 | 0.2 | 0.021664 | 10.06 | 90.8 | 109.2 | 4.3 | 9.93 | | High | High | High | High | High | High | High | High | High | High | High | 0 | 9.54E-11 | 4.74 |
| FALSE | High | GGQEVGN | [R].GGQEV | 1xPhospho | A0A0R4J1 | 4.09E-09 | 0 | 1 | 6 | 2 | A0A0R4J1 | Menin OS= | A0A0R4J1 | 0 | 3540.778 | 0 | -0.27 | 0.088526 | 3.76 | 109.4 | 90.6 | 3.71 | 13.19 | | High | High | High | High | High | High | High | High | High | High | High | 0 | 1.08E-07 | 5.52 |
| FALSE | High | GGSPHHSD | [R].GGSPH | 1xPhospho | F6M3K1 1x | 0.121159 | 0.009984 | 1 | 1 | 1 | F6M3K1 | MCG2244 | F6M3K1 [2 | 0 | 1619.74 | 0 | 1.44 | 0.004587 | 24.98 | 53.7 | 146.3 | 18.11 | 12.51 | | High | High | High | High | High | High | High | High | High | High | High | 0.004961 | 0.05404 | 2.11 |
| FALSE | High | GGQNLPTS | [R].GGQNL | 1xCarbami | Q99NF3 1x | 5.05E-07 | 0 | 1 | 4 | 4 | Q99NF3 | Centrosom | Q99NF3 [3 | 0 | 2061.937 | 0 | -0.76 | 0.001421 | 2.09 | 126.5 | 73.5 | 6.04 | 5.63 | | High | High | High | High | High | High | High | High | High | High | High | 0 | 4.71E-08 | 4.18 |
| FALSE | High | GGRSPDEV | [K].GGRSP | 1xPhospho | Q543N7 1x | 0.001489 | 5.66E-05 | 1 | 2 | 2 | Q543N7 | Protein kin | Q543N7 [3 | 1 | 2094.079 | 0 | -0.01 | 0.401904 | 8.22 | 99.7 | 100.3 | 10.95 | 3.08 | | High | High | High | High | High | High | High | High | High | High | High | 2.58E-05 | 0.000375 | 3.12 |
| FALSE | High | GGSDTNLN | [R].GGSDT | 1xPhospho | Q8C440 1x | 1.56E-07 | 0 | 1 | 6 | 2 | Q8C440 | Putative ur | Q8C440 [4 | 0 | 2699.312 | 0 | 2.1 | 0.000729 | 16.32 | 41.1 | 158.9 | 13.66 | 8.84 | | High | High | High | High | High | High | High | High | High | High | High | 0 | 1.27E-08 | 4.62 |
| FALSE | High | GGSEGQPE | [K].GGSEG | 1xPhospho | A2A7S8-2 | 8.68E-07 | 0 | 1 | 4 | 2 | A2A7S8-2 | Isoform 2 c | A2A7S8-2 | 0 | 2671.231 | 0 | -0.47 | 0.009436 | 8.16 | 118.2 | 81.8 | 8.3 | 8.12 | | High | High | High | High | High | High | High | High | High | High | High | 0 | 8.69E-08 | 5.15 |
| FALSE | High | GGSFEMTD | [R].GGSFE | 1xPhospho | Q99PU8-3 | 0.003342 | 0.00016 | 1 | 5 | 1 | Q99PU8-3 | Isoform 3 c | Q99PU8-3 | 0 | 1825.751 | 0 | 0.37 | 0.007966 | 12.23 | 89.9 | 110.1 | 13.09 | 8.68 | | High | High | High | High | High | High | High | High | High | High | High | 7.19E-05 | 0.000925 | 2.68 |
| FALSE | High | GGSFSGTL | [R].GGSFS | 1xPhospho | P19001 1x | 7.97E-09 | 0 | 1 | 1 | 9 | P19001 | Keratin, typ | P19001 [6 | 0 | 2434.227 | 0 | 3.82 | 0.001001 | 9.53 | 11.3 | 188.7 | 18.03 | 16.27 | | High | High | High | High | High | High | High | High | High | High | High | 0 | 4.48E-10 | 3.81 |
| FALSE | High | GGSGDSAV | [K].GGSGD | 1xPhospho | Q3TIK5 1x | 0.089425 | 0.006807 | 1 | 6 | 1 | Q3TIK5 | Eukaryotic | Q3TIK5 [2 | 0 | 1315.666 | 0 | -0.32 | 0.072934 | 7.24 | 109 | 91 | 7.96 | 14.72 | | High | High | High | High | High | High | High | High | High | High | High | 0.003535 | 0.03783 | 1.31 |
| FALSE | High | GGSGSQIN | [K].GGSGS | 1xPhospho | P23949 1x | 0.112856 | 0.009146 | 1 | 1 | 1 | P23949 | mRNA dec | P23949 [1 | 0 | 1372.642 | 0 | 1.08 | 0.004951 | 13.71 | 64.3 | 135.7 | 11.4 | 14.84 | | High | High | High | High | High | High | High | High | High | High | High | 0.004569 | 0.04961 | 1.62 |
| FALSE | High | GGSLGENC | [K].GGSLG | 1xPhospho | E9QA16 1x | 0.001137 | 0 | 1 | 7 | 1 | E9QA16 | Caldesmon | E9QA16 [1 | 0 | 1540.814 | 0 | -0.07 | 0.222265 | 12.61 | 108.5 | 91.5 | 14.66 | 14.79 | | High | High | High | High | High | High | High | High | High | High | High | 0 | 0.000274 | 1.46 |
| FALSE | High | GGSLPPVK | [R].GGSLP | 1xPhospho | M1VMF6 1 | 0.009585 | 0.000736 | 1 | 7 | 2 | M1VMF6 | Metastasis | M1VMF6 [ | 0 | 1292.738 | 0 | 0.89 | 0.000489 | 5.26 | 67.1 | 132.9 | 10.67 | 4.19 | | High | High | High | High | High | High | High | High | High | High | High | 0.000344 | 0.002991 | 1.48 |
| FALSE | High | GGSISVQV | [R].GGSISV | 1xPhospho | E9Q5C9 1x | 0.000699 | 0 | 1 | 1 | 2 | E9Q5C9 | Nucleolar a | E9Q5C9 [6 | 0 | 1712.935 | 0 | 0.75 | 0.002266 | 10.06 | 72.8 | 127.2 | 11.53 | 12.32 | | High | High | High | High | High | High | High | High | High | High | High | 0 | 0.000253 | 2.47 |
| FALSE | High | GGSISVQV | [R].GGSISV | 1xPhospho | E9Q5C9 1x | 0.000616 | 0 | 1 | 1 | 1 | E9Q5C9 | Nucleolar a | E9Q5C9 [6 | 1 | 2191.105 | 0 | -0.16 | 0.679473 | 33.07 | 94.9 | 105.1 | 29.15 | 59.62 | | High | High | High | High | High | High | High | High | High | High | High | 0 | 0.000138 | 3.17 |
| FALSE | High | GGSISVQV | [R].GGSISV | 1xDeamida | E9Q5C9 1x | 0.046687 | 0.00354 | 1 | 1 | 2 | E9Q5C9 | Nucleolar a | E9Q5C9 [6 | 1 | 2192.089 | 0 | -1.19 | 0.000765 | 3.82 | 138.4 | 61.6 | 10.23 | 14.6 | | High | High | High | High | High | High | High | High | High | High | High | 0.009953 | 0.1088 | 2.63 |
| FALSE | High | GGSPAPAC | [R].GGSPA | 3xCarbami | Q5NCQ5 1 | 5.84E-07 | 0 | 1 | 1 | 3 | Q5NCQ5 | 2-(3-amino | Q5NCQ5 [4 | 0 | 2346.992 | 0 | 0.76 | 0.003427 | 8.28 | 75.3 | 124.7 | 3.58 | 11.07 | | High | High | High | High | High | High | High | High | High | High | High | 0 | 5.64E-07 | 4.69 |
| FALSE | High | GGREDMD | [K].GGRED | 1xPhospho | B1AU75 1x | 0.000422 | 0 | 1 | 2 | 6 | B1AU75 | Nuclear au | B1AU75 [1 | 1 | 2145.978 | 0 | -1.96 | 0.00044 | 1.04 | 159.8 | 40.2 | 7.85 | 5.82 | | High | High | High | High | High | High | High | High | High | High | High | 0.001609 | 0.01573 | 2 |
| FALSE | High | GHRPASPA | [R].GHRPA | 1xCarbami | A2A7F4 1x | 0.000718 | 0 | 1 | 1 | 1 | A2A7F4 | MCG12287 | A2A7F4 [3 | 0 | 3165.598 | 0 | 0.27 | 0.141158 | 22.14 | 88.6 | 111.4 | 10.83 | 13.21 | | High | High | High | High | High | High | High | High | High | High | High | 0 | 0.000165 | 2.37 |
| FALSE | High | GHSDSSAS | [K].GHSDS | 1xPhospho | Q0VGB7 1x | 1.34E-07 | 0 | 2 | 3 | 4 | Q0VGB7; A | Serine/thre | Q0VGB7 [2 | 0 | 2454.217 | 0 | -0.66 | 0.023225 | 2.18 | 122.5 | 77.5 | 14.18 | 14.1 | Shared | High | High | High | High | High | High | High | High | High | High | High | 0 | 1.06E-08 | 4.19 |
| FALSE | High | GHSEGDYP | [R].GHSEG | 1xPhospho | Q8CAS9 1x | 0.000104 | 0 | 1 | 3 | 1 | Q8CAS9 | Poly [ADP- | Q8CAS9 [4 | 0 | 1536.705 | 0 | 1.84 | 0.00409 | 30.69 | 42.9 | 157.1 | 19.51 | 16.48 | | High | High | High | High | High | High | High | High | High | High | High | 0 | 1.89E-05 | 2.66 |
| FALSE | High | GLCSGPGA | [R].GLCSG | 1xCarbami | Q80UU9 1 | 6.31E-09 | 0 | 1 | 1 | 1 | Q80UU9 | Membrane | Q80UU9 [7 | 0 | 2135.999 | 0 | 0.2 | 0.189702 | 13.21 | 92 | 108 | 19.05 | 8.63 | | High | High | High | High | High | High | High | High | High | High | High | 0 | 3.43E-10 | 3.81 |
| FALSE | High | GLEDSNISY | [R].GLEDS | 1xPhospho | D3YZK4 1x | 0.000246 | 0 | 1 | 5 | 1 | D3YZK4 | Transcripti | D3YZK4 [1 | 0 | 2538.169 | 0 | -1.01 | 0.003874 | 7.02 | 136.1 | 63.9 | 12.15 | 16.71 | | High | High | High | High | High | High | High | High | High | High | High | 0 | 4.93E-05 | 2.86 |
| FALSE | High | GLEDSNISY | [R].GLEDS | 2xPhospho | D3YZK4 2x | 0.000189 | 0 | 1 | 5 | 4 | D3YZK4 | Transcripti | D3YZK4 [1 | 0 | 2618.135 | 0 | -0.73 | 0.006832 | 0.97 | 122.5 | 77.5 | 9.38 | 5.87 | | High | High | High | High | High | High | High | High | High | High | High | 0 | 3.69E-05 | 3.42 |
| FALSE | High | GLEDVRPS | [R].GLEDV | 1xPhospho | F8VQ28 1x | 0.02933 | 0.001857 | 1 | 8 | 1 | F8VQ28 | Paxillin OS | F8VQ28 [2 | 0 | 4458.195 | 0 | 0.97 | 0.00346 | 12.15 | 71.1 | 128.9 | 8.48 | 17.39 | | High | High | High | High | High | High | High | High | High | High | High | 0.000997 | 0.01052 | 3.07 |
| FALSE | High | GLEDVRPS | [R].GLEDV | 1xPhospho | F8VQ28 1x | 1.95E-05 | 0 | 1 | 8 | 2 | F8VQ28 | Paxillin OS | F8VQ28 [2 | 0 | 4474.19 | 0 | 0.75 | 0.015748 | 11.1 | 77.1 | 122.9 | 12.77 | 14.24 | | High | High | High | High | High | High | High | High | High | High | High | 0 | 3.02E-06 | 6.04 |
| FALSE | High | GIEFPMAD | [R].GIEFPM | 1xPhospho | Q5SRX1 1x | 0.000466 | 0 | 1 | 6 | 1 | Q5SRX1 | TOM1-like | Q5SRX1 [1 | 0 | 2533.236 | 0 | 1.35 | 0.003799 | 18.41 | 57.2 | 142.8 | 26.14 | 5.87 | | High | High | High | High | High | High | High | High | High | High | High | 0 | 0.000101 | 2.58 |
| FALSE | High | GLEFPMTD | [K].GLEFPM | 1xPhospho | Q3UDC3 1 | 0.030104 | 0.001857 | 1 | 3 | 1 | Q3UDC3 | Putative ur | Q3UDC3 [1 | 0 | 2623.25 | 0 | 0.72 | 0.003738 | 3.14 | 77.1 | 122.9 | 8.46 | 8.73 | | High | High | High | High | High | High | High | High | High | High | High | 0.001011 | 0.01083 | 3.27 |
| FALSE | High | GLEPSPAA | [R].GLEPSP | 1xPhospho | Q3UH60 1 | 0.0026 | 0.000109 | 1 | 1 | 1 | Q3UH60 | Disco-inter | Q3UH60 [5 | 0 | 1983.051 | 0 | 0.15 | 0.324605 | 22.51 | 96.9 | 103.1 | 11.19 | 14.77 | | High | High | High | High | High | High | High | High | High | High | High | 4.94E-05 | 0.000696 | 2.56 |
| FALSE | High | GLESPLK | [K].GLESPL | 1xPhospho | Q8CDA1 1 | 0.092372 | 0.007211 | 1 | 3 | 1 | Q8CDA1 | Phosphatic | Q8CDA1 [9 | 0 | 1281.722 | 0 | 1.36 | 0.000473 | 7.49 | 55.8 | 144.2 | 8.57 | 4.71 | | High | High | High | High | High | High | High | High | High | High | High | 0.003659 | 0.03927 | 1.93 |
| FALSE | High | GIFTSPTNS | [R].GIFTSP | 1xPhospho | Q8BM75 1 | 0.004294 | 0.00036 | 1 | 2 | 2 | Q8BM75 | AT-rich inte | Q8BM75 [ | 0 | 1920.942 | 0 | 0.54 | 0.003257 | 11.79 | 75.4 | 124.6 | 23.21 | 12.42 | | High | High | High | High | High | High | High | High | High | High | High | 0.000161 | 0.001224 | 2.2 |
| FALSE | High | GLGESDAL | [K].GLGES | 1xPhospho | Q3SYK5 1x | 4.39E-08 | 0 | 1 | 6 | 2 | Q3SYK5 | Tyrosine-p | Q3SYK5 [5 | 0 | 2332.163 | 0 | -0.3 | 0.529255 | 4.22 | 110.1 | 89.9 | 7.75 | 19.04 | | High | High | High | High | High | High | High | High | High | High | High | 0 | 2.06E-08 | 3.44 |
| FALSE | High | GLGGPDGE | [K].GLGGP | 1xPhospho | Q8VI59 1x | 0.000125 | 0 | 1 | 4 | 2 | Q8VI59 | Pecanex-lik | Q8VI59 [1 | 0 | 1863.925 | 0 | 1.5 | 0.002013 | 14.44 | 56 | 144 | 14.27 | 14.18 | | High | High | High | High | High | High | High | High | High | High | High | 0 | 2.32E-05 | 2.38 |
| FALSE | High | GLGGVGG | [K].GLGGV | 1xPhospho | A2A9C3 1x | 0.000141 | 0 | 1 | 2 | 2 | A2A9C3 | KICSTOR co | A2A9C3 [7 | 0 | 1540.814 | 0 | -0.23 | 0.016958 | 6.91 | 105.9 | 94.1 | 6.08 | 8.33 | | High | High | High | High | High | High | High | High | High | High | High | 0 | 0.000254 | 1.8 |
| FALSE | High | GLAYGHID | [R].GLAYG | 1xPhospho | O55176 1x | 0.000352 | 0 | 1 | 4 | 2 | O55176 | E3 ubiquiti | O55176 [7 | 1 | 2922.383 | 0 | 0.75 | 0.00199 | 11.7 | 74.5 | 125.5 | 5.62 | 9.21 | | High | High | High | High | High | High | High | High | High | High | High | 0 | 7.39E-05 | 3.3 |
| FALSE | High | GLGPNMS | [R].GLGPN | 1xPhospho | A0A0N4SU | 0.100129 | 0.008067 | 1 | 9 | 1 | A0A0N4SU | Peregrin O | A0A0N4SU | 0 | 1934.899 | 0 | 0.33 | 0.129956 | 24.45 | 89.2 | 110.8 | 8.82 | 11.74 | | High | High | High | High | High | High | High | High | High | High | High | 0.004046 | 0.0432 | 2.69 |
| FALSE | High | GLGSNEDG | [R].GLGSN | 1xPhospho | Q8C0E3-2 | 0.000173 | 0 | 1 | 7 | 1 | Q8C0E3-2 | Isoform 2 c | Q8C0E3-2 | 0 | 1655.841 | 0 | 1.6 | 0.000753 | 14.7 | 46.8 | 153.2 | 11.75 | 11.83 | | High | High | High | High | High | High | High | High | High | High | High | 0 | 3.32E-05 | 2.25 |
| FALSE | High | GLGSPEPG | [R].GLGSP | 1xPhospho | Q8VDZ4 1x | 0.001168 | 0 | 1 | 2 | 2 | Q8VDZ4 | Palmitoyltr | Q8VDZ4 [6 | 0 | 1881.931 | 0 | 0.95 | 0.000291 | 5.82 | 68.2 | 131.8 | 2.4 | 3.56 | | High | High | High | High | High | High | High | High | High | High | High | 0 | 0.000284 | 2.17 |
| FALSE | High | GLGSSTDD | [R].GLGSS | 1xPhospho | B1AQY2 1x | 0.000184 | 0 | 1 | 4 | 3 | B1AQY2 | Rho GTPas | B1AQY2 [1 | 0 | 2213.996 | 0 | 0.04 | 0.892576 | 14.98 | 99.1 | 100.9 | 5.93 | 6.11 | | High | High | High | High | High | High | High | High | High | High | High | 0 | 3.58E-05 | 3.42 |
| FALSE | High | GLGSSTDD | [R].GLGSS | 2xPhospho | B1AQY2 2x | 0.026421 | 0.001676 | 1 | 4 | 1 | B1AQY2 | Rho GTPas | B1AQY2 [1 | 0 | 2293.963 | 0 | 0.49 | 0.006742 | 4.56 | 79 | 121 | 12.35 | 5.54 | | High | High | High | High | High | High | High | High | High | High | High | 0.000903 | 0.009342 | 2.64 |
| FALSE | High | GLGTGLPE | [R].GLGTG | 1xPhospho | Q9ET78 1x | 0.023792 | 0.001612 | 1 | 1 | 1 | Q9ET78 | Junctophili | Q9ET78 [4 | 1 | 2438.25 | 0 | 0.08 | 0.912288 | 13.18 | 102 | 98 | 19.48 | 8.77 | | High | High | High | High | High | High | High | High | High | High | High | 0.000859 | 0.008277 | 1.7 |
| FALSE | High | GIGTPPNT | [K].GIGTPP | 1xPhospho | Q8BVE8-2 | 0.002293 | 0.000109 | 2 | 7 | 2 | Q8BVE8-2; | Isoform 2 c | Q8BVE8-2 | 0 | 1733.96 | 0 | 0.37 | 0.003058 | 8.11 | 85.6 | 114.4 | 3.15 | 5.29 | Shared | High | High | High | High | High | High | High | High | High | High | High | 0.000786 | 0.006975 | 2.58 |
| FALSE | High | GIGTPPNT | [K].GIGTPP | 2xPhospho | Q8BVE8-2 | 0.012508 | 0.000889 | 2 | 7 | 3 | Q8BVE8-2; | Isoform 2 c | Q8BVE8-2 | 0 | 1813.927 | 0 | 1.67 | 0.000158 | 1.98 | 48.2 | 151.8 | 8.55 | 5.89 | Shared | High | High | High | High | High | High | High | High | High | High | High | 0.000449 | 0.004032 | 2.82 |
| FALSE | High | GLHSSDSE | [K].GLHSS | 1xPhospho | Q8C1D8 1x | 4.45E-07 | 0 | 1 | 3 | 2 | Q8C1D8 | Protein IW | Q8C1D8 [3 | 0 | 1981.916 | 0 | 0.36 | 0.282479 | 33.28 | 88.7 | 111.3 | 6.57 | 30.81 | | High | High | High | High | High | High | High | High | High | High | High | 0 | 4.09E-08 | 2.81 |
| FALSE | High | GLHSSDSE | [K].GLHSS | 2xPhospho | Q8C1D8 2x | 0.007082 | 0.00062 | 1 | 3 | 3 | Q8C1D8 | Protein IW | Q8C1D8 [3 | 0 | 2061.882 | 0 | -0.1 | 0.25479 | 13.66 | 101.9 | 98.1 | 6.36 | 14.26 | | High | High | High | High | High | High | High | High | High | High | High | 0.000275 | 0.002124 | 2.83 |
| FALSE | High | GLKEGINP | [R].GLKEG | 1xPhospho | Q08943-2 | 0.000404 | 0 | 1 | 4 | 1 | Q08943-2 | Isoform 2 c | Q08943-2 | 1 | 3394.459 | 0 | -0.44 | 0.042861 | 16.3 | 125.7 | 74.3 | 10.33 | 32.22 | | High | High | High | High | High | High | High | High | High | High | High | 0 | 8.6E-05 | 3.19 |
| FALSE | High | GILAADES | [R].GILAAD | 1xPhospho | A6ZI44 1xP | 0.019276 | 0.001402 | 1 | 9 | 1 | A6ZI44 | Fructose-b | A6ZI44 [83 | 0 | 1870.993 | 0 | 0.15 | 0.073646 | 5.1 | 93.7 | 106.3 | 5.27 | 4.36 | | High | High | High | High | High | High | High | High | High | High | High | 0.000756 | 0.006513 | 1.75 |
| FALSE | High | GILLDDGS | [R].GILLDD | 1xPhospho | Q64700 1x | 0.000259 | 0 | 1 | 5 | 1 | Q64700 | Retinoblas | Q64700 [1 | 0 | 1839.951 | 0 | 2.53 | 0.001387 | 17.85 | 29.1 | 170.9 | 27.43 | 16.19 | | High | High | High | High | High | High | High | High | High | High | High | 0 | 5.25E-05 | 3.52 |
| FALSE | High | GILISPAK | [R].GILISPA | 1xPhospho | Q6P1D7 1x | 0.00537 | 0.000599 | 1 | 2 | 1 | Q6P1D7 | Structure-s | Q6P1D7 [9 | 0 | 1336.801 | 0 | -0.58 | 0.000829 | 2.97 | 120.4 | 79.6 | 6.33 | 2.21 | | High | High | High | High | High | High | High | High | High | High | High | 0.000267 | 0.00156 | 1.69 |
| FALSE | High | GLGPPSPP | [R].GLGPP | 1xPhospho | B7ZNU9 1x | 0.000125 | 0 | 1 | 3 | 2 | B7ZNU9 | Beta-2-syn | B7ZNU9 [7 | 0 | 1451.761 | 0 | -0.42 | 0.000566 | 1.86 | 112.6 | 87.4 | 3.3 | 5.16 | | High | High | High | High | High | High | High | High | High | High | High | 0 | 2.3E-05 | 2.28 |
| FALSE | High | GLAVAPAS | [K].GLAVA | 1xPhospho | A2A7S8-2 | 2.95E-05 | 0 | 1 | 4 | 3 | A2A7S8-2 | Isoform 2 c | A2A7S8-2 | 0 | 1505.849 | 0 | -0.62 | 0.001351 | 1.33 | 122.9 | 77.1 | 7.63 | 5.35 | | High | High | High | High | High | High | High | High | High | High | High | 0 | 1.24E-05 | 2.27 |
| FALSE | High | GIASTSDP | [R].GIASTS | 1xPhospho | F6ZDS4 1x | 0.000311 | 0 | 1 | 5 | 1 | F6ZDS4 | Nucleopro | F6ZDS4 [1 | 0 | 3132.716 | 0 | -1.04 | 0.013281 | 7.16 | 134.5 | 65.5 | 16.08 | 14.44 | | High | High | High | High | High | High | High | High | High | High | High | 0 | 6.39E-05 | 3.11 |
| FALSE | High | GLASPTSIT | [K].GLASPT | 1xCarbami | A0A1B0GS | 4.49E-07 | 0 | 1 | 8 | 2 | A0A1B0GS | Supervillin | A0A1B0GS | 0 | 2354.244 | 0 | 1.83 | 0.000908 | 10.7 | 42.5 | 157.5 | 4.97 | 11.34 | | High | High | High | High | High | High | High | High | High | High | High | 0 | 4.12E-08 | 4.75 |
| FALSE | High | GHSKPPHS | [R].GHSKP | 1xPhospho | B2RSN1 1x | 0.015195 | 0.001089 | 1 | 8 | 1 | B2RSN1 | Myc proto- | B2RSN1 [3 | 0 | 2164.261 | 0 | 1.45 | 0.001525 | 3.34 | 58.1 | 141.9 | 11.37 | 8.62 | | High | High | High | High | High | High | High | High | High | High | High | 0.000571 | 0.004987 | 1.3 |
| FALSE | High | GHSITEYLY | [R].GHSITE | 1xPhospho | Q3UI33 1x | 0.007082 | 0.00062 | 1 | 1 | 3 | Q3UI33 | Methionin | Q3UI33 [1 | 0 | 2034.071 | 0 | 0.72 | 0.002578 | 11.14 | 74.3 | 125.7 | 10.69 | 8 | | High | High | High | High | High | High | High | High | High | High | High | 0.000275 | 0.002133 | 2.65 |
| FALSE | High | GHSSLTNS | [R].GHSSLT | 1xCarbami | Q05BA5 1x | 2.01E-07 | 0 | 1 | 6 | 2 | Q05BA5 | Nuclear re | Q05BA5 [6 | 0 | 2165.046 | 0 | 0.56 | 0.004615 | 12.65 | 83.4 | 116.6 | 5.73 | 6.93 | | High | High | High | High | High | High | High | High | High | High | High | 0 | 1.68E-08 | 3.78 |
| FALSE | High | GHTEENLS | [K].GHTEE | 1xPhospho | Q5XG69 1x | 5.4E-05 | 0 | 1 | 1 | 2 | Q5XG69 | Soluble lan | Q5XG69 [5 | 0 | 1835.93 | 0 | -1.35 | 0.001004 | 1.43 | 142.5 | 57.5 | 6.94 | 13.38 | | High | High | High | High | High | High | High | High | High | High | High | 0 | 8.95E-06 | 2.81 |
| FALSE | High | GHTPPFTT | [K].GHTPP | 1xPhospho | B2RT41 1x | 0.007082 | 0.00062 | 1 | 3 | 1 | B2RT41 | Zinc finger, | B2RT41 [1 | 0 | 1686.866 | 0 | -0.75 | 0.004022 | 5.24 | 129.2 | 70.8 | 7.43 | 13.96 | | High | High | High | High | High | High | High | High | High | High | High | 0.000275 | 0.002135 | 2.31 |
| FALSE | High | GHTQTWP | [K].GHTQT | 1xPhospho | Q8BTI8 1x | 4.43E-05 | 0 | 1 | 3 | 2 | Q8BTI8 | Serine/arg | Q8BTI8 [1 | 0 | 3549.734 | 0 | -0.2 | 0.172112 | 14.85 | 109.8 | 90.2 | 12.43 | 37.71 | | High | High | High | High | High | High | High | High | High | High | High | 0 | 7.21E-06 | 5.57 |
| FALSE | High | GHTQTWP | [K].GHTQT | 1xDeamida | Q8BTI8 1x | 3.29E-08 | 0 | 1 | 3 | 2 | Q8BTI8 | Serine/arg | Q8BTI8 [1 | 0 | 3550.718 | 0 | -0.56 | 0.078697 | 20.58 | 125.8 | 74.2 | 19.16 | 32.08 | | High | High | High | High | High | High | High | High | High | High | High | 0 | 2.18E-09 | 6.49 |
| FALSE | High | GHTQTWP | [K].GHTQT | 1xPhospho | Q8BTI8 1x | 1.09E-08 | 0 | 1 | 3 | 8 | Q8BTI8 | Serine/arg | Q8BTI8 [1 | 0 | 3565.729 | 0 | -0.43 | 0.005756 | 6.67 | 121 | 79 | 11.54 | 13.62 | | High | High | High | High | High | High | High | High | High | High | High | 0 | 6.61E-09 | 5.21 |
| FALSE | High | GHTQTWP | [K].GHTQT | 2xPhospho | Q8BTI8 2x | 5.86E-05 | 0 | 1 | 3 | 1 | Q8BTI8 | Serine/arg | Q8BTI8 [1 | 0 | 3629.7 | 0 | 0.18 | 0.802186 | 14.93 | 104.2 | 95.8 | 13.6 | 37.46 | | High | High | High | High | High | High | High | High | High | High | High | 0 | 9.81E-06 | 3.99 |
| FALSE | High | GHVQSAQ | [K].GHVQS | 1xPhospho | B9EIW5 1x | 8.47E-08 | 0 | 1 | 3 | 2 | B9EIW5 | Ncoa7 prot | B9EIW5 [4 | 0 | 2580.242 | 0 | 0.91 | 0.002164 | 10.26 | 69.5 | 130.5 | 2.34 | 11.79 | | High | High | High | High | High | High | High | High | High | High | High | 0 | 6.35E-09 | 3.68 |
| FALSE | High | GHVTWSL | [K].GHVTW | 1xPhospho | P35831 1x | 0.027358 | 0.001676 | 1 | 1 | 1 | P35831 | Tyrosine-p | P35831 [5 | 0 | 2936.434 | 0 | 0.97 | 0.034575 | 5.45 | 66.4 | 133.6 | 21.81 | 18.89 | | High | High | High | High | High | High | High | High | High | High | High | 0.000936 | 0.009705 | 1.55 |
| FALSE | High | GHYEVTGS | [K].GHYEV | 1xPhospho | E9Q616 1x | 8.66E-06 | 0 | 1 | 4 | 2 | E9Q616 | AHNAK nu | E9Q616 [5 | 0 | 1773.753 | 0 | 2.1 | 0.188452 | 49.53 | 37.9 | 162.1 | 50.58 | 98.67 | | High | High | High | High | High | High | High | High | High | High | High | 0 | 1.95E-06 | 4.84 |
| FALSE | High | GHYEVTGS | [K].GHYEV | 1xPhospho | E9Q616 1x | 8.05E-09 | 0 | 1 | 4 | 14 | E9Q616 | AHNAK nu | E9Q616 [5 | 0 | 2002.916 | 0 | 2.93 | 0.000172 | 4.35 | 23.1 | 176.9 | 7.15 | 5.13 | | High | High | High | High | High | High | High | High | High | High | High | 0 | 4.52E-10 | 3.79 |
| FALSE | High | GKASPFEE | [K].GKASP | 1xPhospho | Q80U58 1x | 0.034579 | 0.002233 | 1 | 7 | 1 | Q80U58 | Pumilio ho | Q80U58 [1 | 1 | 1915.932 | 0 | 0.75 | 0.008509 | 18.28 | 74.7 | 125.3 | 17.84 | 5.3 | | High | High | High | High | High | High | High | High | High | High | High | 0.001183 | 0.01267 | 1.41 |
| FALSE | High | GKLEAIITP | [K].GKLEA | 1xPhospho | Q9QZQ8 1 | 0.016596 | 0.001124 | 1 | 3 | 1 | Q9QZQ8 | Core histo | Q9QZQ8 [1 | 1 | 2102.259 | 0 | -0.33 | 0.197159 | 33.85 | 114.4 | 85.6 | 27.92 | 25.08 | | High | High | High | High | High | High | High | High | High | High | High | 0.000585 | 0.005515 | 2.52 |
| FALSE | High | GKNSDEEA | [K].GKNSD | 1xPhospho | Q8C0D7 1x | 0.004333 | 0.00036 | 1 | 6 | 2 | Q8C0D7 | Inhibitor o | Q8C0D7 [1 | 1 | 1841.961 | 0 | -1.03 | 0.000246 | 2.33 | 132.8 | 67.2 | 3.96 | 7.6 | | High | High | High | High | High | High | High | High | High | High | High | 0.001151 | 0.01233 | 1.41 |
| FALSE | High | GKPETDAV | [K].GKPET | 1xPhospho | P39447 1x | 0.000183 | 0 | 1 | 3 | 3 | P39447 | Tight junct | P39447 [1 | 1 | 2433.263 | 0 | -0.7 | 0.003434 | 4.82 | 122.3 | 77.7 | 9.75 | 7.18 | | High | High | High | High | High | High | High | High | High | High | High | 0 | 3.55E-05 | 1.42 |
| FALSE | High | GKPGPDTC | [K].GKPGP | 1xPhospho | Q6ZQH0 1 | 5.5E-05 | 0 | 1 | 5 | 4 | Q6ZQH0 | MKIAA018 | Q6ZQH0 [8 | 0 | 2754.369 | 0 | -0.43 | 0.004684 | 5.08 | 114.6 | 85.4 | 9.35 | 6.08 | | High | High | High | High | High | High | High | High | High | High | High | 0 | 1.68E-05 | 2.76 |
| FALSE | High | GKPQPLLP | [R].GKPQP | 1xPhospho | Q9CW46 1 | 0.012072 | 0.000889 | 1 | 2 | 1 | Q9CW46 | Ribonuclec | Q9CW46 [ | 1 | 4514.476 | 0 | -0.61 | 0.034123 | 13.77 | 121.1 | 78.9 | 9.95 | 17.08 | | High | High | High | High | High | High | High | High | High | High | High | 0.000449 | 0.003857 | 1.83 |

| | | | | | | | | | | | | | | | | | | | | | | | | | | | | | | | | | | | | | | |
|---|---|---|---|---|---|---|---|---|---|---|---|---|---|---|---|---|---|---|---|---|---|---|---|---|---|---|---|---|---|---|---|---|---|---|---|---|---|---|
| FALSE | High | GKPQPLLPI | [R].GKPQP | 1xPhospho | Q9CW46 1 | 0.003433 | 0.00021 | 1 | 2 | 1 | Q9CW46 | Ribonuclec | Q9CW46 [ | 1 | 4530.471 | 0 | -0.38 | 0.024805 | 7.6 | 119.2 | 80.8 | 8.69 | 14.81 | | High | High | High | High | High | High | High | High | High | High | High | 9.42E-05 | 0.000952 | 2.45 |
| FALSE | High | GKYSPDQL | [R].GKYSPD | 1xCarbami | Q3USC0 1x | 7.43E-06 | 0 | 1 | 6 | 2 | Q3USC0 | Putative ur | Q3USC0 [6 | 1 | 2503.323 | 0 | -1.58 | 0.002706 | 2.02 | 150.3 | 49.7 | 17.51 | 21.22 | | High | High | High | High | High | High | High | High | High | High | High | 0 | 9.67E-07 | 2.98 |
| FALSE | High | GLAAEDSG | [K].GLAAEI | 1xPhospho | Q540H0 1x | 7.84E-06 | 0 | 1 | 4 | 2 | Q540H0 | N-acetyltra | Q540H0 [1 | 0 | 1644.788 | 0 | -0.15 | 0.122336 | 7.08 | 104.1 | 95.9 | 2.26 | 7.62 | | High | High | High | High | High | High | High | High | High | High | High | 0 | 1.03E-06 | 2.06 |
| FALSE | High | GLAAEDSG | [K].GLAAEI | 2xPhospho | Q540H0 2x | 0.000462 | 0 | 1 | 4 | 1 | Q540H0 | N-acetyltra | Q540H0 [1 | 0 | 1724.755 | 0 | 0.76 | 0.001745 | 14.36 | 71.2 | 128.8 | 6.36 | 6.44 | | High | High | High | High | High | High | High | High | High | High | High | 0 | 9.99E-05 | 1.8 |
| FALSE | High | GIAIAYEDE | [R].GIAIAYI | 1xPhospho | Q9CR25 1x | 1.38E-05 | 0 | 1 | 1 | 6 | Q9CR25 | 2-(3-amino | Q9CR25 [4 | 0 | 1520.672 | 0 | 1.21 | 0.001432 | 5.98 | 58.1 | 141.9 | 6.68 | 9.95 | | High | High | High | High | High | High | High | High | High | High | High | 0 | 1.94E-06 | 3.31 |
| FALSE | High | GIAPASPM | [K].GIAPAS | 1xPhospho | Q03141 1x | 3.41E-08 | 0 | 1 | 5 | 1 | Q03141 | MAP/micro | Q03141 [4 | 0 | 2147.108 | 0 | -0.65 | 0.001008 | 1.34 | 117.6 | 82.4 | 7.5 | 8.56 | | High | High | High | High | High | High | High | High | High | High | High | 0 | 2.27E-09 | 3.8 |
| FALSE | High | GIAPASPM | [K].GIAPAS | 1xPhospho | Q03141 1x | 1.06E-06 | 0 | 1 | 5 | 2 | Q03141 | MAP/micro | Q03141 [4 | 0 | 2163.103 | 0 | -0.55 | 0.233921 | 8.87 | 118.7 | 81.3 | 3.99 | 32.46 | | High | High | High | High | High | High | High | High | High | High | High | 0 | 1.09E-07 | 3.84 |
| FALSE | High | GLAQWVD | [R].GLAQW | 1xPhospho | O08715-4 | 0.008094 | 0.000736 | 1 | 3 | 1 | O08715-4 | Isoform 4 c | O08715-4 | 0 | 1781.835 | 0 | 0.36 | 0.033918 | 1.23 | 87.9 | 112.1 | 8.65 | 9.11 | | High | High | High | High | High | High | High | High | High | High | High | 0.000327 | 0.002481 | 1.99 |
| FALSE | High | GIASEEGSF | [K].GIASEE | 1xPhospho | E9Q309 1x | 0.022378 | 0.001542 | 1 | 1 | 1 | E9Q309 | Centrosom | E9Q309 [1 | 0 | 1989.883 | 0 | -0.14 | 0.78576 | 22.26 | 99.6 | 100.4 | 13.97 | 13.07 | | High | High | High | High | High | High | High | High | High | High | High | 0.000814 | 0.007718 | 2.94 |
| FALSE | High | GLASPSLK | [K].GLASPS | 1xPhospho | B2RWU2 1 | 0.019966 | 0.001438 | 1 | 2 | 1 | B2RWU2 | Immunode | B2RWU2 [ | 0 | 1310.749 | 0 | 3.96 | 0.000158 | 31.26 | 11.4 | 188.6 | 19.63 | 3.95 | | High | High | High | High | High | High | High | High | High | High | High | 0.000773 | 0.006796 | 1.93 |
| FALSE | High | GIIRPGSLD | [R].GIIRPG | 1xPhospho | Q9Z1Y4 1x | 0.002212 | 0.000109 | 1 | 2 | 1 | Q9Z1Y4 | Thyroid rec | Q9Z1Y4 [8 | 0 | 2981.485 | 0 | 0.88 | 0.007798 | 32.08 | 65.1 | 134.9 | 13.89 | 13.34 | | High | High | High | High | High | High | High | High | High | High | High | 4.94E-05 | 0.000583 | 3.63 |
| FALSE | High | GGLVSDAY | [K].GGLVSI | 1xPhospho | Q02614 1x | 0.00086 | 0 | 1 | 1 | 1 | Q02614 | SAP30-binc | Q02614 [3 | 0 | 4005.671 | 0 | 0.85 | 0.1618 | 43.02 | 71.3 | 128.7 | 13.74 | 37.33 | | High | High | High | High | High | High | High | High | High | High | High | 0 | 0.000201 | 3.94 |
| FALSE | High | GPAGEASA | [R].GPAGE | 1xPhospho | B7ZNU9 1x | 0.079114 | 0.00613 | 1 | 3 | 1 | B7ZNU9 | Beta-2-syn | B7ZNU9 [8 | 1 | 1660.837 | 0 | -0.28 | 0.005963 | 3.45 | 109.6 | 90.4 | 6.61 | 8.15 | | High | High | High | High | High | High | High | High | High | High | High | 0.003148 | 0.03286 | 1.5 |
| FALSE | High | GPAGEESW | [K].GPAGE | 1xPhospho | B2RQC2 1x | 2.26E-06 | 0 | 1 | 1 | 4 | B2RQC2 | Ubiquitin c | B2RQC2 [7 | 0 | 2187.037 | 0 | 0.16 | 0.927554 | 2.61 | 96.2 | 103.8 | 9.7 | 17.04 | | High | High | High | High | High | High | High | High | High | High | High | 0 | 6.55E-06 | 3.16 |
| FALSE | High | GQSSQGLF | [R].GQSSQ | 1xPhospho | A0A0R4J07 | 0.000297 | 0 | 1 | 3 | 3 | A0A0R4J07 | BRCA1-A c | A0A0R4J07 | 0 | 2463.206 | 0 | 0.77 | 0.005179 | 9.19 | 70 | 130 | 9.39 | 11.6 | | High | High | High | High | High | High | High | High | High | High | High | 0 | 0.000106 | 3.12 |
| FALSE | High | GQSSWENS | [K].GQSSW | 2xPhospho | P49452 2x | 0.00129 | 0 | 1 | 3 | 1 | P49452 | Centromer | P49452 [4 | 0 | 2456.053 | 0 | -0.54 | 0.002566 | 2.06 | 119.8 | 80.2 | 5.73 | 3.78 | | High | High | High | High | High | High | High | High | High | High | High | 0 | 0.000317 | 2.32 |
| FALSE | High | GQSTDSLP | [R].GQSTD | 1xCarbami | Q8C0V9 1x | 0.000967 | 0 | 1 | 2 | 2 | Q8C0V9 | FERM dom | Q8C0V9 [5 | 0 | 1771.825 | 0 | 0.38 | 0.011006 | 12.71 | 83.7 | 116.3 | 6.44 | 7.61 | | High | High | High | High | High | High | High | High | High | High | High | 0 | 0.000231 | 2.89 |
| FALSE | High | GQSTDSLP | [R].GQSTD | 1xCarbami | Q8C0V9 2x | 0.002134 | 5.66E-05 | 1 | 2 | 2 | Q8C0V9 | FERM dom | Q8C0V9 [5 | 0 | 1851.791 | 0 | 0.57 | 0.015903 | 14.43 | 80.6 | 119.4 | 4.35 | 14.27 | | High | High | High | High | High | High | High | High | High | High | High | 2.58E-05 | 0.000559 | 2.77 |
| FALSE | High | GQSTVPPC | [K].GQSTV | 1xCarbami | P27546 1x | 7.06E-07 | 0 | 1 | 3 | 10 | P27546 | Microtubu | P27546 [5 | 0 | 2193.103 | 0 | 0.69 | 0.012372 | 8.54 | 79.7 | 120.3 | 10.21 | 10.9 | | High | High | High | High | High | High | High | High | High | High | High | 0 | 2.19E-05 | 3.26 |
| FALSE | High | GQTLGLGN | [K].GQTLG | 1xPhospho | P27612 1x | 0.001735 | 5.66E-05 | 1 | 1 | 2 | P27612 | Phospholip | P27612 [4 | 0 | 2278.07 | 0 | -0.59 | 0.001262 | 2.84 | 118.3 | 81.7 | 9.86 | 7.98 | | High | High | High | High | High | High | High | High | High | High | High | 2.58E-05 | 0.000443 | 2.5 |
| FALSE | High | GQTLQSSIF | [R-].GQTLC | 1xPhospho | A2BIE1 1xF | 0.000718 | 0 | 1 | 2 | 1 | A2BIE1 | Glutamine | A2BIE1 [49 | 0 | 1993.041 | 0 | -0.05 | 0.404482 | 12.67 | 102.8 | 97.2 | 7.44 | 5.93 | | High | High | High | High | High | High | High | High | High | High | High | 0 | 0.000165 | 2.17 |
| FALSE | High | GQTPLTEG | [R].GQTPL | 1xPhospho | Q9CQV4 1x | 8.25E-05 | 0 | 1 | 2 | 4 | Q9CQV4 | Reticuloph | Q9CQV4 [3 | 0 | 2883.252 | 0 | 0.01 | 0.870475 | 26.32 | 101.2 | 98.8 | 9.54 | 10.43 | | High | High | High | High | High | High | High | High | High | High | High | 2.58E-05 | 0.000441 | 4.77 |
| FALSE | High | GQTPLTEG | [R].GQTPL | 1xDeamida | Q9CQV4 1x | 0.000346 | 0 | 1 | 2 | 1 | Q9CQV4 | Reticuloph | Q9CQV4 [3 | 0 | 2884.236 | 0 | -0.01 | 0.840277 | 16.49 | 101.1 | 98.9 | 22.1 | 13.36 | | High | High | High | High | High | High | High | High | High | High | High | 0 | 7.28E-05 | 5.23 |
| FALSE | High | GQTPLTEG | [R].GQTPL | 2xPhospho | Q9CQV4 2x | 0.000422 | 0 | 1 | 2 | 4 | Q9CQV4 | Reticuloph | Q9CQV4 [3 | 0 | 2963.218 | 0 | 0.53 | 0.010307 | 6.76 | 81.5 | 118.5 | 3.48 | 10.39 | | High | High | High | High | High | High | High | High | High | High | High | 0 | 9.01E-05 | 4.62 |
| FALSE | High | GRASPGGV | [R].GRASP | 1xPhospho | O35126 1x | 4.13E-06 | 0 | 1 | 2 | 1 | O35126 | Atrophin-1 | O35126 [3 | 1 | 1987.985 | 0 | 1.14 | 0.000707 | 8.65 | 63.2 | 136.8 | 8.71 | 5.37 | | High | High | High | High | High | High | High | High | High | High | High | 0 | 5.01E-07 | 3.11 |
| FALSE | High | GRASPLLLS | [K].GRASPI | 1xPhospho | E9QAU9 1x | 5.37E-06 | 0 | 1 | 5 | 1 | E9QAU9 | Leucine-ric | E9QAU9 [6 | 1 | 2519.307 | 0 | 1.14 | 0.001293 | 6.56 | 62.4 | 137.6 | 12.09 | 8.98 | | High | High | High | High | High | High | High | High | High | High | High | 0 | 6.71E-07 | 3.86 |
| FALSE | High | GRASSHSS | [R].GRASSH | 1xPhospho | P48678 1x | 0.004106 | 0.00036 | 1 | 3 | 2 | P48678 | Prelamin-A | P48678 [4 | 1 | 2256.113 | 0 | 1.82 | 0.003531 | 32.33 | 47.2 | 152.8 | 20.61 | 15.35 | | High | High | High | High | High | High | High | High | High | High | High | 0.000275 | 0.001945 | 2.63 |
| FALSE | High | GQSQLSNF | [R].GQSQL | 2xPhospho | A2A5R2 2x | 0.013426 | 0.000976 | 1 | 2 | 1 | A2A5R2 | Brefeldin A | A2A5R2 [1 | 0 | 2180.967 | 0 | -0.35 | 0.031818 | 11.65 | 112 | 88 | 7.16 | 9.27 | | High | High | High | High | High | High | High | High | High | High | High | 0.000505 | 0.004373 | 1.84 |
| FALSE | High | GRDSYGGF | [R].GRDSY | 1xPhospho | Q91VM5 1 | 0.063788 | 0.004664 | 1 | 1 | 1 | Q91VM5 | RNA bindir | Q91VM5 [ | 1 | 1370.642 | 0 | -0.34 | 0.007224 | 2.49 | 110.8 | 89.2 | 4 | 5.61 | | High | High | High | High | High | High | High | High | High | High | High | 0.002403 | 0.02559 | 2.1 |
| FALSE | High | GREDPGQ | [K].GREDP | 1xPhospho | Q69ZC8 1x | 0.012845 | 0.000889 | 1 | 1 | 1 | Q69ZC8 | GPALPP mc | Q69ZC8 [1 | 1 | 3964.784 | 0 | -0.3 | 0.450172 | 14.13 | 105.7 | 94.3 | 23.15 | 37.28 | | High | High | High | High | High | High | High | High | High | High | High | 0.000449 | 0.00413 | 2.58 |
| FALSE | High | GREDVSNF | [R].GREDV | 1xPhospho | Q8BWW9 | 0.004572 | 0.000456 | 1 | 5 | 1 | Q8BWW9 | Serine/thre | Q8BWW9 | 1 | 2901.351 | 0 | 0.15 | 0.129956 | 10.96 | 93.3 | 106.7 | 4.8 | 11.05 | | High | High | High | High | High | High | High | High | High | High | High | 0.000204 | 0.00131 | 2.75 |
| FALSE | High | GREPSSSQ | [R].GREPSS | 1xPhospho | Q3TVI8 1x | 3.39E-06 | 0 | 1 | 2 | 1 | Q3TVI8 | Pre-B-cell l | Q3TVI8 [16 | 1 | 2662.349 | 0 | 0.09 | 0.407564 | 7.89 | 99.7 | 100.3 | 8.25 | 7.56 | | High | High | High | High | High | High | High | High | High | High | High | 0 | 3.99E-07 | 3.83 |
| FALSE | High | GRGSLASG | [K].GRGSL | 1xPhospho | Q6PAJ1 1x | 0.000389 | 0 | 1 | 2 | 1 | Q6PAJ1 | Breakpoint | Q6PAJ1 [4 | 1 | 1868.004 | 0 | 0.93 | 0.017448 | 22.36 | 73.8 | 126.2 | 11.27 | 17.97 | | High | High | High | High | High | High | High | High | High | High | High | 0 | 8.3E-05 | 3.49 |
| FALSE | High | GRHSSPLAI | [R].GRHSS | 2xPhospho | E9Q4K7 2x | 0.004333 | 0.00036 | 1 | 2 | 1 | E9Q4K7 | Kinesin fan | E9Q4K7 [1 | 1 | 1927.925 | 0 | 0 | 0.982202 | 3.47 | 101.1 | 98.9 | 11.98 | 8.4 | | High | High | High | High | High | High | High | High | High | High | High | 0.000161 | 0.001237 | 2.36 |
| FALSE | High | GRLSLEQTI | [R].GRLSLE | 1xPhospho | Q3UYY2 1x | 0.027839 | 0.001755 | 1 | 1 | 1 | Q3UYY2 | Putative ur | Q3UYY2 [6 | 1 | 1993.007 | 0 | 0.43 | 0.173291 | 24.02 | 89.3 | 110.7 | 13.62 | 15.22 | | High | High | High | High | High | High | High | High | High | High | High | 0.000953 | 0.009895 | 2.11 |
| FALSE | High | GRLSPVPVI | [R].GRLSPV | 1xPhospho | Q64012 1x | 0.038677 | 0.002563 | 1 | 6 | 3 | Q64012 | RNA-bindir | Q64012 [1 | 1 | 1386.782 | 0 | -0.13 | 0.102883 | 3.79 | 103.8 | 96.2 | 7.18 | 3.48 | | High | High | High | High | High | High | High | High | High | High | High | 0.005414 | 0.05941 | 1.58 |
| FALSE | High | GRISVGSDA | [K].GRISVG | 1xPhospho | O08553 1x | 0.000933 | 0 | 1 | 1 | 1 | O08553 | Dihydropy | O08553 [3 | 1 | 2667.38 | 0 | 1.08 | 0.004221 | 10.5 | 65.6 | 134.4 | 10.88 | 11.63 | | High | High | High | High | High | High | High | High | High | High | High | 0 | 0.00022 | 3.26 |
| FALSE | High | GRLTPSPDI | [R].GRLTPS | 1xPhospho | Q8VHR5 1x | 9.91E-06 | 0 | 1 | 3 | 2 | Q8VHR5 | Transcripti | Q8VHR5 [1 | 1 | 2533.286 | 0 | -0.08 | 0.378034 | 10.33 | 103.7 | 96.3 | 5.21 | 5.7 | | High | High | High | High | High | High | High | High | High | High | High | 0 | 1.34E-06 | 3.35 |
| FALSE | High | GRLTPSPDI | [R].GRLTPS | 2xPhospho | Q8VHR5 2x | 0.003655 | 0.00021 | 1 | 3 | 2 | Q8VHR5 | Transcripti | Q8VHR5 [1 | 1 | 2613.252 | 0 | 0.27 | 0.010538 | 10.34 | 92.4 | 107.6 | 4.18 | 4.92 | | High | High | High | High | High | High | High | High | High | High | High | 0.000689 | 0.006124 | 4.5 |
| FALSE | High | GRLTPSPDI | [R].GRLTPS | 3xPhospho | Q8VHR5 3x | 0.000146 | 0 | 1 | 3 | 2 | Q8VHR5 | Transcripti | Q8VHR5 [1 | 1 | 2693.219 | 0 | 0.15 | 0.147736 | 10.49 | 97.9 | 102.1 | 4.16 | 8.67 | | High | High | High | High | High | High | High | High | High | High | High | 0.000161 | 0.001168 | 4.28 |
| FALSE | High | GRNSATSA | [R].GRNSA | 1xPhospho | Q571J8 1x | 5.77E-06 | 0 | 1 | 17 | 3 | Q571J8 | Non-specif | Q571J8 [4 | 1 | 2282.051 | 0 | 1.03 | 0.009308 | 15.95 | 66.5 | 133.5 | 21.15 | 12.5 | | High | High | High | High | High | High | High | High | High | High | High | 0 | 7.32E-07 | 3.11 |
| FALSE | High | GRNSATSA | [R].GRNSA | 2xPhospho | Q571J8 2x | 0.035486 | 0.002304 | 1 | 17 | 1 | Q571J8 | Non-specif | Q571J8 [4 | 1 | 2362.018 | 0 | 0.52 | 0.050049 | 25.96 | 82.2 | 117.8 | 14.83 | 16.86 | | High | High | High | High | High | High | High | High | High | High | High | 0.001214 | 0.01305 | 1.58 |
| FALSE | High | GREDAVAE | [K].GREDA | 2xCarbami | P70399 1x | 0.000616 | 0 | 1 | 4 | 4 | P70399 | TP53-bindi | P70399 [5 | 1 | 3385.524 | 0 | 0.39 | 0.025039 | 4.85 | 84.9 | 115.1 | 9.36 | 13.18 | | High | High | High | High | High | High | High | High | High | High | High | 0 | 0.000138 | 3.63 |
| FALSE | High | GRPAHPSP | [R].GRPAH | 2xPhospho | Q3UJV1 2x | 0.024638 | 0.001612 | 1 | 2 | 2 | Q3UJV1 | Coiled-coil | Q3UJV1 [3 | 0 | 1792.809 | 0 | 1.33 | 0.001881 | 19.52 | 55.6 | 144.4 | 4.97 | 12.01 | | High | High | High | High | High | High | High | High | High | High | High | 0.000872 | 0.008649 | 1.68 |
| FALSE | High | GQSQLSNF | [R].GQSQL | 1xPhospho | A2A5R2 1x | 2.44E-05 | 0 | 1 | 2 | 1 | A2A5R2 | Brefeldin A | A2A5R2 [1 | 0 | 2101 | 0 | 0.57 | 0.009717 | 10.79 | 82 | 118 | 10.63 | 7.41 | | High | High | High | High | High | High | High | High | High | High | High | 0 | 3.69E-06 | 2.29 |
| FALSE | High | GQRSPEDS | [K].GQRSP | 1xPhospho | Q9QXY1 1x | 0.002004 | 5.66E-05 | 1 | 3 | 1 | Q9QXY1 | Tight junct | Q9QXY1 [3 | 1 | 2520.157 | 0 | -1.58 | 0.000356 | 1.39 | 149.9 | 50.1 | 6.83 | 19.08 | | High | High | High | High | High | High | High | High | High | High | High | 2.58E-05 | 0.000519 | 3.59 |
| FALSE | High | GQENPGEI | [K].GQENP | 1xPhospho | Q921N6 1x | 0.012845 | 0.000889 | 1 | 3 | 1 | Q921N6 | Probable A | Q921N6 [1 | 0 | 1855.848 | 0 | 0.15 | 0.079201 | 8.29 | 92.2 | 107.8 | 9.52 | 8.46 | | High | High | High | High | High | High | High | High | High | High | High | 0.000449 | 0.004156 | 2.07 |
| FALSE | High | GQESSSDQ | [K].GQESS | 1xPhospho | A0A140T8 | 9.36E-08 | 0 | 1 | 1 | 6 | A0A140T8 | Chromoso | A0A140T8 | 0 | 2752.26 | 0 | -0.53 | 0.00388 | 4.93 | 116.5 | 83.5 | 4.01 | 12.21 | | High | High | High | High | High | High | High | High | High | High | High | 0 | 7.06E-09 | 4.31 |
| FALSE | High | GQESSSDQ | [K].GQESS | 2xPhospho | A0A140T8 | 3.42E-06 | 0 | 1 | 1 | 10 | A0A140T8 | Chromoso | A0A140T8 | 0 | 2832.227 | 0 | -0.52 | 0.000618 | 2.62 | 119.7 | 80.3 | 3.64 | 2.15 | | High | High | High | High | High | High | High | High | High | High | High | 0 | 8.35E-06 | 4.47 |
| FALSE | High | GQGASPVA | [R].GQGAS | 1xPhospho | Q64727 1x | 0.069325 | 0.005112 | 1 | 1 | 1 | Q64727 | Vinculin OS | Q64727 [3 | 0 | 1627.828 | 0 | 0.19 | 0.118714 | 14.57 | 93.5 | 106.5 | 5.31 | 20.18 | | High | High | High | High | High | High | High | High | High | High | High | 0.002633 | 0.0281 | 1.41 |
| FALSE | High | GQGATFVE | [K].GQGAT | 1xPhospho | G3UVV3 1x | 5.46E-09 | 0 | 1 | 3 | 1 | G3UVV3 | MCG1556, | G3UVV3 [1 | 0 | 3091.528 | 0 | -0.89 | 0.00024 | 1.99 | 130.2 | 69.8 | 3.8 | 5.31 | | High | High | High | High | High | High | High | High | High | High | High | 0 | 2.93E-10 | 4.53 |
| FALSE | High | GQGSEPAA | [R].GQGSE | 1xPhospho | E9QP49 1x | 1.85E-08 | 0 | 1 | 4 | 4 | E9QP49 | EH domain | E9QP49 [2 | 0 | 3149.499 | 0 | 2.03 | 0.000294 | 31.55 | 37.5 | 162.5 | 21.5 | 5.32 | | High | High | High | High | High | High | High | High | High | High | High | 0 | 7.85E-09 | 5.41 |
| FALSE | High | GQGSEPAA | [R].GQGSE | 2xPhospho | E9QP49 2x | 0.000269 | 0 | 1 | 4 | 1 | E9QP49 | EH domain | E9QP49 [2 | 0 | 3229.465 | 0 | 0.09 | 0.275412 | 11.51 | 98.5 | 101.5 | 6.77 | 13.26 | | High | High | High | High | High | High | High | High | High | High | High | 0 | 5.47E-05 | 2.81 |
| FALSE | High | GQGSVSAS | [K].GQGSV | 1xPhospho | Q9D1M0 1 | 9.06E-06 | 0 | 1 | 1 | 6 | Q9D1M0 | Protein SE | Q9D1M0 [ | 0 | 2028.907 | 0 | 0.17 | 0.228867 | 15.06 | 96.5 | 103.5 | 4.26 | 15.97 | | High | High | High | High | High | High | High | High | High | High | High | 0 | 3.9E-05 | 4.14 |
| FALSE | High | GQGTASPC | [R].GQGTA | 1xPhospho | P51830 1x | 0.033986 | 0.002195 | 1 | 2 | 2 | P51830 | Adenylate | P51830 [6 | 0 | 2241.153 | 0 | 1.05 | 0.001622 | 21.46 | 64.1 | 135.9 | 9.41 | 7.31 | | High | High | High | High | High | High | High | High | High | High | High | 0.001665 | 0.01672 | 2.52 |
| FALSE | High | GQGTFDGI | [K].GQGTF | 1xPhospho | D3YYY8 1x | 3.88E-06 | 0 | 1 | 1 | 1 | D3YYY8 | Rho guanir | D3YYY8 [3 | 0 | 2571.217 | 0 | 2.73 | 0.007624 | 27.66 | 26.1 | 173.9 | 21.77 | 29.7 | | High | High | High | High | High | High | High | High | High | High | High | 0 | 4.65E-07 | 3.45 |
| FALSE | High | GQGTLQHS | [R].GQGTL | 1xPhospho | B3V097 1x | 0.000104 | 0 | 1 | 4 | 2 | B3V097 | BetaCstF-6 | B3V097 [2 | 0 | 2268.134 | 0 | -0.66 | 0.001265 | 4.29 | 120.2 | 79.8 | 5.09 | 5.24 | | High | High | High | High | High | High | High | High | High | High | High | 0 | 1.88E-05 | 1.86 |
| FALSE | High | GQGTPPSC | [R].GQGTP | 1xPhospho | F8WJ40 1x | 0.001767 | 5.66E-05 | 1 | 5 | 3 | F8WJ40 | Histone-lys | F8WJ40 [2 | 0 | 1475.721 | 0 | -1.03 | 0.000217 | 1.18 | 133.5 | 66.5 | 3.22 | 3.72 | | High | High | High | High | High | High | High | High | High | High | High | 2.58E-05 | 0.000454 | 1.91 |
| FALSE | High | GQLEDSAV | [K].GQLED | 1xPhospho | Q8C080 1x | 0.002077 | 5.66E-05 | 1 | 3 | 1 | Q8C080 | Sorting ne | Q8C080 [4 | 0 | 1741.914 | 0 | -1.09 | 0.012008 | 4.8 | 133.2 | 66.8 | 18.29 | 7.77 | | High | High | High | High | High | High | High | High | High | High | High | 2.58E-05 | 0.000544 | 2.07 |
| FALSE | High | GQSPPESV | [K].GQSPP | 1xPhospho | Q3TYD6 1x | 0.000152 | 0 | 1 | 1 | 1 | Q3TYD6 | Serine/thre | Q3TYD6 [1 | 0 | 2107.975 | 0 | -0.43 | 0.000222 | 1.33 | 115.4 | 84.6 | 11.22 | 15.47 | | High | High | High | High | High | High | High | High | High | High | High | 0 | 2.88E-05 | 2.87 |
| FALSE | High | GQLHGSSD | [R].GQLHG | 2xPhospho | Q3U3Q3 2 | 5.1E-07 | 0 | 1 | 2 | 5 | Q3U3Q3 | Putative ur | Q3U3Q3 [1 | 0 | 2434.058 | 0 | -0.57 | 0.017099 | 11 | 122.5 | 77.5 | 7.64 | 16.38 | | High | High | High | High | High | High | High | High | High | High | High | 0 | 4.77E-08 | 5.02 |
| FALSE | High | GQISEDDC | [K].GQISED | 1xCarbami | Q8K3J5 1x | 1E-06 | 0 | 1 | 3 | 1 | Q8K3J5 | Zinc finger | Q8K3J5 [2 | 0 | 2159.993 | 0 | -0.9 | 0.000281 | 0.72 | 131 | 69 | 5.06 | 1.94 | | High | High | High | High | High | High | High | High | High | High | High | 0 | 1.02E-07 | 3.56 |
| FALSE | High | GQLTNIVSI | [K].GQLTN | 1xPhospho | Q8CH77 1x | 3.33E-06 | 0 | 1 | 4 | 2 | Q8CH77 | Neuron na | Q8CH77 [9 | 0 | 1936.01 | 0 | 1.16 | 0.000167 | 2.48 | 61.7 | 138.3 | 5.97 | 4.8 | | High | High | High | High | High | High | High | High | High | High | High | 0 | 3.92E-07 | 3.75 |
| FALSE | High | GQNGEDLS | [K].GQNGE | 1xPhospho | Q8R409 1x | 5.6E-05 | 0 | 1 | 1 | 3 | Q8R409 | Protein HE | Q8R409 [9 | 0 | 4094.854 | 0 | -0.12 | 0.262039 | 27.47 | 104.2 | 95.8 | 10.62 | 25.74 | | High | High | High | High | High | High | High | High | High | High | High | 0.000275 | 0.002053 | 2.6 |
| FALSE | High | GQNGEDLS | [K].GQNGE | 1xPhospho | Q8R409 1x | 0.007471 | 0.00062 | 1 | 1 | 4 | Q8R409 | Protein HE | Q8R409 [9 | 0 | 4110.849 | 0 | -0.51 | 0.013862 | 6.96 | 118.8 | 81.2 | 5.22 | 15.47 | | High | High | High | High | High | High | High | High | High | High | High | 0.001011 | 0.0108 | 3.79 |
| FALSE | High | GQPGSAYD | [R].GQPGS | 1xPhospho | Q8C2Q3 1x | 5.63E-07 | 0 | 1 | 4 | 13 | Q8C2Q3 | RNA-bindir | Q8C2Q3 [5 | 0 | 4022.807 | 0 | -0.45 | 0.016212 | 6.5 | 114.9 | 85.1 | 7.21 | 7.36 | | High | High | High | High | High | High | High | High | High | High | High | 0 | 5.33E-08 | 5.72 |
| FALSE | High | GQPGSAYD | [R].GQPGS | 1xDeamida | Q8C2Q3 1x | 6.18E-08 | 0 | 1 | 4 | 19 | Q8C2Q3 | RNA-bindir | Q8C2Q3 [5 | 0 | 4023.791 | 0 | -0.24 | 0.11941 | 19.19 | 106 | 94 | 12.17 | 13.41 | | High | High | High | High | High | High | High | High | High | High | High | 0 | 4.46E-09 | 5.67 |
| FALSE | High | GQPSPLAQ | [R].GQPSP | 1xPhospho | A6P3E4 1x | 0.012288 | 0.000889 | 1 | 3 | 1 | A6P3E4 | Mitogen-a | A6P3E4 [3 | 0 | 1461.73 | 0 | -0.37 | 0.000445 | 1.19 | 113.3 | 86.7 | 4.87 | 7.68 | | High | High | High | High | High | High | High | High | High | High | High | 0.000449 | 0.003963 | 3.2 |
| FALSE | High | GQPVLTPP | [R].GQPVL | 1xPhospho | Q4VA93 1x | 4.72E-05 | 0 | 1 | 3 | 1 | Q4VA93 | Protein kin | Q4VA93 [6 | 0 | 4689.348 | 0 | 2.09 | 0.000378 | 5.2 | 37.1 | 162.9 | 22.43 | 10.88 | | High | High | High | High | High | High | High | High | High | High | High | 0 | 7.76E-06 | 5.5 |
| FALSE | High | GQPVLTPP | [R].GQPVL | 1xDeamida | Q4VA93 1x | 3.61E-06 | 0 | 1 | 3 | 1 | Q4VA93 | Protein kin | Q4VA93 [6 | 0 | 4690.332 | 0 | 2.68 | 0.000864 | 15.9 | 25.4 | 174.6 | 17.83 | 13.75 | | High | High | High | High | High | High | High | High | High | High | High | 0 | 4.3E-07 | 5.37 |
| FALSE | High | GQQASPFV | [K].GQQAS | 1xPhospho | Q9CZ82 1x | 0.062731 | 0.004664 | 1 | 2 | 1 | Q9CZ82 | Mediator c | Q9CZ82 [6 | 0 | 1411.73 | 0 | -0.13 | 0.081763 | 10.74 | 105.8 | 94.2 | 5.64 | 6.72 | | High | High | High | High | High | High | High | High | High | High | High | 0.002403 | 0.02497 | 1.68 |
| FALSE | High | GQQDVLSS | [R].GQQD | 1xPhospho | Q8C156 1x | 0.00017 | 0 | 1 | 1 | 2 | Q8C156 | Condensin | Q8C156 [1 | 0 | 1637.81 | 0 | -0.14 | 0.05342 | 10.54 | 104.8 | 95.2 | 5.32 | 6.52 | | High | High | High | High | High | High | High | High | High | High | High | 0.000161 | 0.001156 | 2.59 |
| FALSE | High | GQQSAEED | [R].GQQSA | 1xPhospho | Q3TXN7 1x | 0.063258 | 0.004664 | 1 | 10 | 1 | Q3TXN7 | Putative ur | Q3TXN7 [1 | 0 | 1790.836 | 0 | -0.64 | 0.001287 | 3.7 | 119.6 | 80.4 | 6.27 | 10.88 | | High | High | High | High | High | High | High | High | High | High | High | 0.002403 | 0.02519 | 2.39 |
| FALSE | High | GQQSSPIA | [R].GQQSS | 1xPhospho | Q6P1D7 1x | 2.58E-05 | 0 | 1 | 2 | 2 | Q6P1D7 | Structure-s | Q6P1D7 [9 | 0 | 1446.705 | 0 | -0.21 | 0.023374 | 3.05 | 106.3 | 93.7 | 3.84 | 4.88 | | High | High | High | High | High | High | High | High | High | High | High | 0 | 3.94E-06 | 1.76 |
| FALSE | High | GQLISSPTF | [K].GQLISS | 1xPhospho | A2AEV7 1x | 2.73E-09 | 0 | 1 | 2 | 4 | A2AEV7 | Coiled-coil | A2AEV7 [5 | 0 | 2924.594 | 0 | 3.07 | 0.000246 | 1.89 | 22.6 | 177.4 | 6.89 | 7.3 | | High | High | High | High | High | High | High | High | High | High | High | 0 | 1.17E-08 | 4.62 |
| FALSE | High | GRPSDVVS | [K].GRPSD | 1xPhospho | Q9CXH7 1x | 0.000179 | 0 | 1 | 2 | 1 | Q9CXH7 | Shugoshin | Q9CXH7 [1 | 0 | 1912.933 | 0 | -0.01 | 0.395807 | 11.35 | 107.5 | 92.5 | 8.96 | 9.18 | | High | High | High | High | High | High | High | High | High | High | High | 0 | 3.46E-05 | 2.73 |
| FALSE | High | GRPSLYVQ | [R].GRPSLY | 1xPhospho | Q9D279 1x | 0.012397 | 0.000889 | 1 | 1 | 1 | Q9D279 | Mitotic inte | Q9D279 [3 | 0 | 1384.73 | 0 | 3.95 | 0.015861 | 44.06 | 11.9 | 188.1 | 17.86 | 42.74 | | High | High | High | High | High | High | High | High | High | High | High | 0.000449 | 0.003986 | 2.03 |
| FALSE | High | GRSPAQPA | [R].GRSPA | 1xPhospho | Q6RUT7 1x | 8.4E-05 | 0 | 1 | 1 | 1 | Q6RUT7 | Protein CC | Q6RUT7 [3 | 2 | 3493.664 | 0 | -0.08 | 0.082368 | 2.08 | 108.7 | 91.3 | 7.45 | 11.53 | | High | High | High | High | High | High | High | High | High | High | High | 0 | 1.47E-05 | 4.27 |
| FALSE | High | GSDLNISSH | [K].GSDLNI | 1xPhospho | Q9DBS8 1x | 0.020499 | 0.001472 | 1 | 2 | 1 | Q9DBS8 | Centrosom | Q9DBS8 [8 | 0 | 1682.851 | 0 | 0.48 | 0.035613 | 26.73 | 83.4 | 116.6 | 3.83 | 15.57 | | High | High | High | High | High | High | High | High | High | High | High | 0.000786 | 0.007004 | 1.89 |
| FALSE | High | GSDNSSYD | [R].GSDNS | 1xPhospho | Q64028 1x | 1.76E-07 | 0 | 1 | 7 | 4 | Q64028 | Polyhomec | Q64028 [8 | 0 | 2545.166 | 0 | -0.79 | 0.000992 | 3.41 | 127.6 | 72.4 | 3.23 | 11.14 | | High | High | High | High | High | High | High | High | High | High | High | 0 | 1.45E-08 | 4.28 |
| FALSE | High | GSDNSSYD | [R].GSDNS | 2xPhospho | Q64028 2x | 3.7E-05 | 0 | 1 | 7 | 1 | Q64028 | Polyhomec | Q64028 [8 | 0 | 2625.132 | 0 | -1.13 | 0.008216 | 6.62 | 132.8 | 67.2 | 13.22 | 17.3 | | High | High | High | High | High | High | High | High | High | High | High | 0 | 5.9E-06 | 4.09 |
| FALSE | High | GSDSLIK | [R].GSDSLI | 1xPhospho | Q3V235 1x | 0.075304 | 0.005727 | 1 | 2 | 1 | Q3V235 | Prohibitin | Q3V235 [2 | 0 | 1257.686 | 0 | 0.34 | 0.047537 | 15.8 | 88.4 | 111.6 | 4.09 | 13.79 | | High | High | High | High | High | High | High | High | High | High | High | 0.002967 | 0.03093 | 1.05 |
| FALSE | High | GSDSPEDL | [K].GSDSPE | 1xPhospho | D0VLQ4 1x | 0.118334 | 0.009621 | 1 | 3 | 1 | D0VLQ4 | Tet oncoge | D0VLQ4 [3 | 0 | 3149.368 | 0 | -0.34 | 0.155541 | 6.11 | 110.2 | 89.8 | 4.57 | 19.39 | | High | High | High | High | High | High | High | High | High | High | High | 0.004855 | 0.05261 | 2.34 |
| FALSE | High | GSDYSWSY | [K].GSDYS | 2xPhospho | Q6ZQB7 2x | 0.001848 | 5.66E-05 | 1 | 4 | 1 | Q6ZQB7 | MKIAA042 | Q6ZQB7 [2 | 0 | 2711.094 | 0 | -0.43 | 0.207887 | 9.98 | 114.8 | 85.2 | 18.16 | 14.86 | | High | High | High | High | High | High | High | High | High | High | High | 2.58E-05 | 0.000474 | 2.9 |
| FALSE | High | GSDYSWSY | [K].GSDYS | 2xPhospho | Q6ZQB7 2x | 8.45E-07 | 0 | 1 | 4 | 2 | Q6ZQB7 | MKIAA042 | Q6ZQB7 [2 | 0 | 2727.088 | 0 | -0.02 | 0.947588 | 3.58 | 100.7 | 99.3 | 9.22 | 10.38 | | High | High | High | High | High | High | High | High | High | High | High | 0 | 8.4E-08 | 4.23 |
| FALSE | High | GSEAESNT | [K].GSEAES | 1xPhospho | B7ZNA0 1x | 6.26E-06 | 0 | 2 | 8 | 1 | B7ZNA0; Q | Shroom3 p | B7ZNA0 [1 | 0 | 2839.351 | 0 | -0.47 | 0.011027 | 4.62 | 114.6 | 85.4 | 9.23 | 5.12 | Shared | High | High | High | High | High | High | High | High | High | High | High | 0 | 7.96E-07 | 3.39 |
| FALSE | High | GSEAESNT | [K].GSEAES | 1xDeamida | B7ZNA0 1x | 6.58E-08 | 0 | 2 | 8 | 1 | B7ZNA0; Q | Shroom3 p | B7ZNA0 [1 | 0 | 2840.335 | 0 | -0.47 | 0.024411 | 11.39 | 112.5 | 87.5 | 4.57 | 14.86 | Shared | High | High | High | High | High | High | High | High | High | High | High | 0 | 4.76E-09 | 4.37 |
| FALSE | High | GSEDEWE | [K].GSEDE | 1xPhospho | P52479-2 1 | 0.001222 | 0 | 1 | 2 | 2 | P52479-2 | Isoform 2 c | P52479-2 [ | 0 | 1898.894 | 0 | -0.71 | 0.015673 | 7.4 | 122.9 | 77.1 | 11.21 | 11.25 | | High | High | High | High | High | High | High | High | High | High | High | 7.19E-05 | 0.000792 | 3.14 |
| FALSE | High | GSEEDPLLS | [R].GSEEDI | 1xPhospho | B9EKC3 1x | 2.35E-08 | 0 | 1 | 3 | 2 | B9EKC3 | Rho GTPas | B9EKC3 [1 | 0 | 2225.114 | 0 | 0.19 | 0.006836 | 6.06 | 91.8 | 108.2 | 5.82 | 3.11 | | High | High | High | High | High | High | High | High | High | High | High | 0 | 1.5E-09 | 3.56 |
| FALSE | High | GSEGYLAA | [K].GSEGYI | 1xPhospho | Q04690 1x | 1.04E-08 | 0 | 1 | 6 | 3 | Q04690 | Neurofibro | Q04690 [2 | 0 | 2234.069 | 0 | 1.55 | 0.000405 | 3.99 | 51.3 | 148.7 | 9.7 | 3.68 | | High | High | High | High | High | High | High | High | High | High | High | 0 | 1.65E-08 | 3.54 |
| FALSE | High | GSEIDQVV | [R].GSEIDC | 1xCarbami | Q8BJQ2 1x | 5.68E-10 | 0 | 1 | 1 | 2 | Q8BJQ2 | Ubiquitin c | Q8BJQ2 [5 | 0 | 2626.284 | 0 | 0.14 | 0.124977 | 11.51 | 97.6 | 102.4 | 7.97 | 4.24 | | High | High | High | High | High | High | High | High | High | High | High | 0 | 2.3E-11 | 4.22 |
| FALSE | High | GSDLDTQE | [K].GSDLD | 1xCarbami | Q3U0S4 1x | 0.007673 | 0.000664 | 1 | 2 | 2 | Q3U0S4 | Putative ur | Q3U0S4 [5 | 1 | 2975.289 | 0 | -0.85 | 0.002071 | 5.88 | 127.4 | 72.6 | 4.71 | 15.06 | | High | High | High | High | High | High | High | High | High | High | High | 0.000399 | 0.00343 | 2.36 |
| FALSE | High | GSEQDSVA | [K].GSEQD | 1xPhospho | Q8BRN9 1x | 0.032267 | 0.002129 | 1 | 3 | 1 | Q8BRN9 | Coiled-coil | Q8BRN9 [4 | 0 | 2115.074 | 0 | 0.7 | 0.029719 | 27.42 | 77 | 123 | 10.5 | 15.41 | | High | High | High | High | High | High | High | High | High | High | High | 0.001142 | 0.01174 | 2.11 |
| FALSE | High | GSESMVFE | [R].GSESM | 1xPhospho | Q8CDZ5 1x | 0.006478 | 0.00062 | 1 | 2 | 4 | Q8CDZ5 | Putative ur | Q8CDZ5 [4 | 0 | 1870.891 | 0 | -0.09 | 0.771516 | 5.53 | 102.8 | 97.2 | 2.96 | 8.14 | | High | High | High | High | High | High | High | High | High | High | High | 0.000275 | 0.00194 | 2.13 |
| FALSE | High | GSESPNSFL | [K].GSESPN | 1xPhospho | Q8BMA3 1 | 0.000189 | 0 | 1 | 2 | 2 | Q8BMA3 | Connector | Q8BMA3 [ | 0 | 1989.875 | 0 | -0.42 | 0.005591 | 8.76 | 114 | 86 | 8.05 | 6.58 | | High | High | High | High | High | High | High | High | High | High | High | 0 | 3.7E-05 | 3.71 |
| FALSE | High | GSETGSETI | [R].GSETG | 1xPhospho | Q7TSC1 1x | 0.0003 | 0 | 1 | 3 | 3 | Q7TSC1 | Protein PRI | Q7TSC1 [1 | 0 | 2385.086 | 0 | 0.06 | 0.368454 | 16.2 | 97.8 | 102.2 | 7.05 | 5.05 | | High | High | High | High | High | High | High | High | High | High | High | 0 | 6.17E-05 | 3.02 |
| FALSE | High | GSETPGAT | [K].GSETP | 1xPhospho | G5E866 1x | 3.65E-07 | 0 | 1 | 5 | 6 | G5E866 | Splicing fac | G5E866 [2 | 0 | 1626.814 | 0 | 0.13 | 0.273488 | 8.02 | 96.6 | 103.4 | 9.17 | 4.63 | | High | High | High | High | High | High | High | High | High | High | High | 0 | 3.29E-08 | 2.44 |
| FALSE | High | GSETPGAT | [K].GSETP | 2xPhospho | G5E866 2x | 0.062209 | 0.004646 | 1 | 5 | 1 | G5E866 | Splicing fac | G5E866 [2 | 0 | 1706.78 | 0 | 0.52 | 0.096113 | 16.37 | 77.4 | 122.6 | 13.94 | 22.47 | | High | High | High | High | High | High | High | High | High | High | High | 0.002397 | 0.02487 | 2.48 |
| FALSE | High | GSETSAVV | [K].GSETSA | 1xPhospho | Q8CIN4 1x | 0.000227 | 0 | 1 | 1 | 1 | Q8CIN4 | Serine/thre | Q8CIN4 [1 | 0 | 3801.774 | 0 | 0.7 | 0.000429 | 2.98 | 75.3 | 124.7 | 3.72 | 5.04 | | High | High | High | High | High | High | High | High | High | High | High | 0 | 4.5E-05 | 3.79 |
| FALSE | High | GSFEAGR | [R].GSFEA | 1xPhospho | Q99M08 1 | 0.029586 | 0.001857 | 1 | 1 | 1 | Q99M08 | Uncharacte | Q99M08 [1 | 0 | 1032.471 | 0 | -0.23 | 0.467453 | 27.39 | 107.8 | 92.2 | 7.65 | 28.04 | | High | High | High | High | High | High | High | High | High | High | High | 0.000997 | 0.01063 | 1.67 |
| FALSE | High | GSFEAQR | [R].GSFEA | 1xPhospho | Q91XC0 1x | 0.014033 | 0.001011 | 1 | 1 | 2 | Q91XC0 | LIM domai | Q91XC0 [8 | 0 | 1103.508 | 0 | -1.22 | 0.001787 | 4.07 | 137.9 | 62.1 | 7.82 | 13.17 | | High | High | High | High | High | High | High | High | High | High | High | 0.000538 | 0.004561 | 2.39 |
| FALSE | High | GSFGLHAQ | [K].GSFGLH | 1xPhospho | Q3UHE1 1x | 0.017041 | 0.001204 | 1 | 2 | 1 | Q3UHE1 | Membrane | Q3UHE1 [9 | 0 | 1767.878 | 0 | 1.63 | 0.000596 | 10.53 | 49.6 | 150.4 | 17.14 | 10.07 | | High | High | High | High | High | High | High | High | High | High | High | 0.000638 | 0.005715 | 1.94 |
| FALSE | High | GSFNPGDA | [K].GSFNP | 1xPhospho | D3Z7R7 1x | 0.000207 | 0 | 1 | 6 | 1 | D3Z7R7 | Rho GTPas | D3Z7R7 [5 | 0 | 3323.511 | 0 | 0.29 | 0.071785 | 6.15 | 90 | 110 | 13.04 | 9.42 | | High | High | High | High | High | High | High | High | High | High | High | 0 | 4.07E-05 | 3.72 |
| FALSE | High | GSFSLVPDS | [K].GSFSLV | 1xPhospho | E9Q7D5 1x | 0.000352 | 0 | 1 | 2 | 2 | E9Q7D5 | Rho guanir | E9Q7D5 [7 | 0 | 1644.82 | 0 | 2.52 | 0.000209 | 16.83 | 27.5 | 172.5 | 10.9 | 5.28 | | High | High | High | High | High | High | High | High | High | High | High | 0 | 7.38E-05 | 2.86 |
| FALSE | High | GSFSSENY | [R].GSFSSE | 1xPhospho | Q3TSG4 1x | 0.020499 | 0.001472 | 1 | 1 | 1 | Q3TSG4 | RNA deme | Q3TSG4 [3 | 0 | 1541.662 | 0 | -0.05 | 0.485823 | 3.94 | 103.2 | 96.8 | 8.96 | 8.25 | | High | High | High | High | High | High | High | High | High | High | High | 0.000786 | 0.006989 | 2.91 |
| FALSE | High | GSGAAEEG | [R].GSGAA | 1xPhospho | Q8CC12 1x | 9.3E-09 | 0 | 1 | 3 | 1 | Q8CC12 | Codanin-1 | Q8CC12 [1 | 0 | 2138.934 | 0 | -1.33 | 0.000243 | 1.22 | 141.7 | 58.3 | 1.74 | 11.46 | | High | High | High | High | High | High | High | High | High | High | High | 0 | 5.32E-10 | 4.26 |
| FALSE | High | GSERPSSSS | [R].GSERPS | 1xPhospho | Q91W18 1 | 0.002272 | 0.000109 | 1 | 3 | 3 | Q91W18 | Tudor dom | Q91W18 [4 | 0 | 2231.977 | 0 | 0.38 | 0.010517 | 11.04 | 90.8 | 109.2 | 9.69 | 5.1 | | High | High | High | High | High | High | High | High | High | High | High | 4.94E-05 | 0.000601 | 2.29 |
| FALSE | High | GSDGEDSA | [R].GSDGE | 1xPhospho | A6H6E1 1x | 0.101749 | 0.008194 | 1 | 4 | 1 | A6H6E1 | Espl1 prote | A6H6E1 [1 | 0 | 1676.742 | 0 | -0.93 | 0.056772 | 8.24 | 129.9 | 70.1 | 15.67 | 33.95 | | High | High | High | High | High | High | High | High | High | High | High | 0.004126 | 0.04401 | 1.73 |
| FALSE | High | GSDGDPGA | [R].GSDGD | 1xCarbami | Q80T69 1x | 6.13E-05 | 0 | 1 | 2 | 1 | Q80T69 | Round spe | Q80T69 [1 | 0 | 1982.929 | 0 | -0.17 | 0.077334 | 9.56 | 105.1 | 94.9 | 5.52 | 14.55 | | High | High | High | High | High | High | High | High | High | High | High | 0 | 1.03E-05 | 3.64 |
| FALSE | High | GSDFDCELI | [K].GSDFD | 1xCarbami | Q5FWJ5 1x | 0.103392 | 0.008249 | 2 | 15 | 1 | Q5FWJ5; C | Hnrpk prot | Q5FWJ5 [1 | 0 | 1407.581 | 0 | 0.35 | 0.057993 | 10.27 | 91.8 | 108.2 | 10.15 | 11.26 | Shared | High | High | High | High | High | High | High | High | High | High | High | 0.004193 | 0.0451 | 2.13 |

| | | | | | | | | | | | | | | | | | | | | | | | | | | | | | | | | | | | | | | |
|---|---|---|---|---|---|---|---|---|---|---|---|---|---|---|---|---|---|---|---|---|---|---|---|---|---|---|---|---|---|---|---|---|---|---|---|---|---|---|
| FALSE | High | GRSPDELP | [K].GRSPDI | 1xPhospho | Q5DTQ4 1x | 0.00013 | 0 | 1 | 6 | 2 | Q5DTQ4 | MKIAA406 | Q5DTQ4 [4 | 1 | 2052.016 | 0 | 0.08 | 0.218922 | 5.23 | 96.6 | 103.4 | 6.39 | 4.48 | | High | High | High | High | High | High | High | High | High | High | High | 0.000538 | 0.004429 | 2.96 |
| FALSE | High | GRSPQPPA | [R].GRSPQ | 1xCarbami | Q8VDM6 1 | 1.25E-06 | 0 | 1 | 3 | 6 | Q8VDM6 | Heterogen | Q8VDM6 [ | 1 | 4247.916 | 0 | 0.28 | 0.087597 | 7.92 | 92.1 | 107.9 | 12.74 | 8.3 | | High | High | High | High | High | High | High | High | High | High | High | 0 | 2.24E-07 | 5.74 |
| FALSE | High | GRSPSPLG | [R].GRSPSF | 2xPhospho | Q3UHU5 2 | 0.117405 | 0.009523 | 1 | 5 | 1 | Q3UHU5 | Microtubu | Q3UHU5 [ | 1 | 1935.893 | 0 | -1.71 | 0.000687 | 2.56 | 153.1 | 46.9 | 5.78 | 14.19 | | High | High | High | High | High | High | High | High | High | High | High | 0.004769 | 0.052 | 2.18 |
| FALSE | High | GRTSVDGF | [K].GRTSVI | 2xPhospho | Q9WTK2 2 | 0.024638 | 0.001612 | 1 | 2 | 3 | Q9WTK2 | Chromodo | Q9WTK2 [ | 1 | 2212.009 | 0 | 1.05 | 0.002364 | 28.35 | 62.2 | 137.8 | 10.85 | 8.37 | | High | High | High | High | High | High | High | High | High | High | High | 0.003633 | 0.03878 | 2.58 |
| FALSE | High | GRVTPTQS | [R].GRVTP | 1xPhospho | Q3UA65 1x | 0.090153 | 0.00698 | 1 | 6 | 1 | Q3UA65 | DNA helica | Q3UA65 [1 | 1 | 1423.762 | 0 | -0.1 | 0.968258 | 15.48 | 98.6 | 101.4 | 11.98 | 14.06 | | High | High | High | High | High | High | High | High | High | High | High | 0.003564 | 0.03804 | 1.73 |
| FALSE | High | GRWESQQ | [K].GRWES | 1xPhospho | E9Q4K7 1x | 0.011549 | 0.00086 | 1 | 2 | 2 | E9Q4K7 | Kinesin fan | E9Q4K7 [1 | 1 | 2185.06 | 0 | 0.24 | 0.005811 | 6.4 | 89.5 | 110.5 | 3.92 | 3.84 | | High | High | High | High | High | High | High | High | High | High | High | 0.000437 | 0.003677 | 1.98 |
| FALSE | High | GSAEGSSD | [K].GSAEG | 1xPhospho | P51859 1x | 3.13E-07 | 0 | 1 | 2 | 4 | P51859 | Hepatoma- | P51859 [12 | 0 | 1690.757 | 0 | -0.83 | 0.001426 | 1.72 | 126.4 | 73.6 | 5.03 | 10.4 | | High | High | High | High | High | High | High | High | High | High | High | 0 | 2.76E-08 | 2.77 |
| FALSE | High | GSAEGSSD | [K].GSAEG | 2xPhospho | P51859 2x | 4.45E-07 | 0 | 1 | 2 | 1 | P51859 | Hepatoma- | P51859 [12 | 0 | 1770.724 | 0 | -0.01 | 0.503685 | 4.67 | 96.5 | 103.5 | 4.83 | 6.72 | | High | High | High | High | High | High | High | High | High | High | High | 0 | 4.09E-08 | 2.75 |
| FALSE | High | GSAEGTSD | [K].GSAEG | 1xPhospho | A6H619 1x | 2.2E-06 | 0 | 1 | 2 | 8 | A6H619 | PHD and RI | A6H619 [9 | 0 | 2726.261 | 0 | -0.32 | 0.046516 | 1.04 | 109.4 | 90.6 | 9.28 | 5.1 | | High | High | High | High | High | High | High | High | High | High | High | 0 | 3.03E-07 | 4.06 |
| FALSE | High | GSAGNEVI | [K].GSAGN | 1xPhospho | Q6A037 1x | 2.81E-07 | 0 | 1 | 2 | 2 | Q6A037 | NEDD4-bin | Q6A037 [3 | 0 | 2983.43 | 0 | -0.83 | 0.001001 | 5.2 | 128 | 72 | 4.23 | 7.45 | | High | High | High | High | High | High | High | High | High | High | High | 0 | 2.45E-08 | 4.88 |
| FALSE | High | GSAGNEVI | [K].GSAGN | 1xDeamida | Q6A037 1x | 0.000276 | 0 | 1 | 2 | 2 | Q6A037 | NEDD4-bin | Q6A037 [3 | 0 | 2984.414 | 0 | -0.98 | 0.004227 | 4.1 | 128.4 | 71.6 | 9.67 | 18.48 | | High | High | High | High | High | High | High | High | High | High | High | 0 | 0.000138 | 3.66 |
| FALSE | High | GSALDAAV | [R].GSALD | 1xCarbami | Q6PAR7 1x | 4.25E-06 | 0 | 1 | 3 | 1 | Q6PAR7 | Phf20 prot | Q6PAR7 [8 | 0 | 2540.154 | 0 | -1.25 | 0.00034 | 0.3 | 139.7 | 60.3 | 8.11 | 6.08 | | High | High | High | High | High | High | High | High | High | High | High | 0 | 5.18E-07 | 5.13 |
| FALSE | High | GSALDAAV | [R].GSALD | 1xCarbami | Q6PAR7 2x | 5.88E-06 | 0 | 1 | 3 | 2 | Q6PAR7 | Phf20 prot | Q6PAR7 [8 | 0 | 2620.12 | 0 | -0.95 | 0.000218 | 0.57 | 131.6 | 68.4 | 3.43 | 6.59 | | High | High | High | High | High | High | High | High | High | High | High | 0 | 7.45E-07 | 4.54 |
| FALSE | High | GSALSQLS | [R].GSALS | 1xPhospho | A2SW42 1x | 1.43E-06 | 0 | 1 | 2 | 2 | A2SW42 | Zinc finger | A2SW42 [1 | 0 | 2360.197 | 0 | -1.17 | 0.001191 | 2.32 | 138 | 62 | 6.32 | 9.2 | | High | High | High | High | High | High | High | High | High | High | High | 0 | 1.51E-07 | 3.99 |
| FALSE | High | GSAPHSESI | [R].GSAPH | 1xCarbami | O70551 1x | 6.74E-07 | 0 | 1 | 4 | 10 | O70551 | SRSF prote | O70551 [3 | 0 | 4272.796 | 0 | -0.83 | 0.001462 | 4.14 | 127.9 | 72.1 | 4.64 | 8.52 | | High | High | High | High | High | High | High | High | High | High | High | 0 | 3.06E-06 | 4.38 |
| FALSE | High | GSAPHSESI | [R].GSAPH | 1xCarbami | O70551 1x | 1.8E-08 | 0 | 1 | 4 | 6 | O70551 | SRSF prote | O70551 [3 | 0 | 4273.78 | 0 | -0.85 | 0.000662 | 2.09 | 128.8 | 71.2 | 3.07 | 8.11 | | High | High | High | High | High | High | High | High | High | High | High | 0 | 1.11E-09 | 5.17 |
| FALSE | High | GSAPHSESI | [R].GSAPH | 1xCarbami | O70551 2x | 8.47E-08 | 0 | 1 | 4 | 4 | O70551 | SRSF prote | O70551 [3 | 0 | 4352.763 | 0 | -0.37 | 0.008909 | 5.83 | 113 | 87 | 4.27 | 7.42 | | High | High | High | High | High | High | High | High | High | High | High | 0 | 6.36E-09 | 5.59 |
| FALSE | High | GSASESPS | [K].GSASES | 1xPhospho | Q3UL54 1x | 5.29E-07 | 0 | 1 | 6 | 24 | Q3UL54 | Putative ur | Q3UL54 [1 | 0 | 3411.673 | 0 | 0.02 | 0.552232 | 5.18 | 99.5 | 100.5 | 6.27 | 9.76 | | High | High | High | High | High | High | High | High | High | High | High | 0 | 4.99E-08 | 5.89 |
| FALSE | High | GSASPGLSI | [R].GSASP | 1xPhospho | Q6PR54-3 | 4.03E-07 | 0 | 2 | 4 | 4 | Q6PR54-3; | Isoform 3 c | Q6PR54-3 | 0 | 1944.036 | 0 | -1.02 | 0.002834 | 4.1 | 133.2 | 66.8 | 8.19 | 12.15 | Shared | High | High | High | High | High | High | High | High | High | High | High | 0 | 3.66E-08 | 3.93 |
| FALSE | High | GSASPGLSI | [R].GSASP | 2xPhospho | Q6PR54-3 | 4.43E-05 | 0 | 2 | 4 | 3 | Q6PR54-3; | Isoform 3 c | Q6PR54-3 | 0 | 2024.002 | 0 | -0.86 | 0.001287 | 3.13 | 129.9 | 70.1 | 9.45 | 2.7 | Shared | High | High | High | High | High | High | High | High | High | High | High | 2.58E-05 | 0.000483 | 3.62 |
| FALSE | High | GSATDYQE | [K].GSATD | 1xPhospho | Q3UHZ2 1x | 0.000568 | 0 | 1 | 4 | 2 | Q3UHZ2 | Aryl hydro | Q3UHZ2 [4 | 0 | 2101.904 | 0 | 0.93 | 0.001678 | 16.03 | 68.8 | 131.2 | 13.92 | 6.09 | | High | High | High | High | High | High | High | High | High | High | High | 0 | 0.000127 | 3.34 |
| FALSE | High | GSATSSSN | [R].GSATS | 1xPhospho | Q7TPM6 1 | 9.2E-05 | 0 | 1 | 2 | 2 | Q7TPM6 | Fibronectir | Q7TPM6 [4 | 0 | 1421.636 | 0 | -1.3 | 0.000193 | 2.11 | 142.3 | 57.7 | 4.4 | 10.49 | | High | High | High | High | High | High | High | High | High | High | High | 0 | 1.64E-05 | 2.86 |
| FALSE | High | GSDASDFD | [R].GSDASI | 1xCarbami | B1ARU1 1x | 2.8E-05 | 0 | 2 | 19 | 1 | B1ARU1; F | Microtubu | B1ARU1 [5 | 0 | 3489.443 | 0 | -0.25 | 0.162875 | 4.32 | 106.5 | 93.5 | 12.81 | 11.11 | Shared | High | High | High | High | High | High | High | High | High | High | High | 0 | 4.27E-06 | 4.15 |
| FALSE | High | GSDASDFD | [R].GSDASI | 1xCarbami | B1ARU1 2x | 0.032831 | 0.002195 | 2 | 19 | 1 | B1ARU1; F | Microtubu | B1ARU1 [5 | 0 | 3570.394 | 0 | 0.55 | 0.031294 | 10.51 | 85 | 115 | 19.74 | 13.12 | Shared | High | High | High | High | High | High | High | High | High | High | High | 0.001151 | 0.01189 | 2.88 |
| FALSE | High | GSDAVSET | [R].GSDAV | 1xPhospho | Q99K01-4 | 0.000181 | 0 | 1 | 4 | 7 | Q99K01-4 | Isoform 4 c | Q99K01-4 | 0 | 2528.217 | 0 | 0.35 | 0.080129 | 25.54 | 87.6 | 112.4 | 15.78 | 10.01 | | High | High | High | High | High | High | High | High | High | High | High | 0.000204 | 0.001312 | 3.18 |
| FALSE | High | GSDAVSET | [R].GSDAV | 2xPhospho | Q99K01-4 | 0.003282 | 0.00016 | 1 | 4 | 1 | Q99K01-4 | Isoform 4 c | Q99K01-4 | 0 | 2608.183 | 0 | 0.43 | 0.002834 | 4.81 | 87.8 | 112.2 | 2.68 | 7.11 | | High | High | High | High | High | High | High | High | High | High | High | 7.19E-05 | 0.000902 | 2.15 |
| FALSE | High | GSDDVLVS | [K].GSDDV | 1xCarbami | E9Q6J5-3 1 | 4.59E-08 | 0 | 1 | 3 | 2 | E9Q6J5-3 | Isoform Ne | E9Q6J5-3 [ | 0 | 2665.184 | 0 | 0.02 | 0.588428 | 16.96 | 98.3 | 101.7 | 9.59 | 23.52 | | High | High | High | High | High | High | High | High | High | High | High | 0 | 3.18E-09 | 5.29 |
| FALSE | High | GSDDVLVS | [K].GSDDV | 1xCarbami | E9Q6J5-3 1 | 1.08E-08 | 0 | 1 | 3 | 8 | E9Q6J5-3 | Isoform Ne | E9Q6J5-3 [ | 0 | 2681.179 | 0 | -0.03 | 0.755634 | 0.52 | 99.6 | 100.4 | 6.52 | 6.52 | | High | High | High | High | High | High | High | High | High | High | High | 0 | 6.3E-10 | 5.68 |
| FALSE | High | GSDEENLD | [R].GSDEEI | 1xPhospho | Q9QY06-2 | 3.27E-10 | 0 | 2 | 9 | 5 | Q9QY06-2; | Isoform 2 c | Q9QY06-2 | 0 | 2707.167 | 0 | 0.17 | 0.061812 | 7.63 | 96.7 | 103.3 | 12.22 | 6.35 | Shared | High | High | High | High | High | High | High | High | High | High | High | 0 | 1.24E-11 | 4.14 |
| FALSE | High | GQELAFPL | [R].GQELA | 1xPhospho | Q9D8N0 1x | 0.003655 | 0.00021 | 1 | 2 | 1 | Q9D8N0 | Elongation | Q9D8N0 [3 | 0 | 2996.371 | 0 | 0.48 | 0.014296 | 9.56 | 85.7 | 114.3 | 11.87 | 9.96 | | High | High | High | High | High | High | High | High | High | High | High | 9.42E-05 | 0.001019 | 3.68 |
| FALSE | High | GPAGEESW | [K].GPAGE | 1xPhospho | B2RQC2 1x | 0.000529 | 0 | 1 | 1 | 1 | B2RQC2 | Ubiquitin c | B2RQC2 [7 | 0 | 1957.874 | 0 | -0.3 | 0.215072 | 12.61 | 109.5 | 90.5 | 10.74 | 14.49 | | High | High | High | High | High | High | High | High | High | High | High | 0 | 0.000117 | 2.41 |
| FALSE | High | GQEETSGA | [K].GQEET | 1xPhospho | Q8VBT6 1x | 3.77E-05 | 0 | 1 | 1 | 1 | Q8VBT6 | Apolipopro | Q8VBT6 [4 | 0 | 1881.843 | 0 | 2.96 | 0.000587 | 9.69 | 20.8 | 179.2 | 9.12 | 12.46 | | High | High | High | High | High | High | High | High | High | High | High | 0 | 6E-06 | 2.86 |
| FALSE | High | GQDTVAIE | [K].GQDTV | 2xPhospho | A2BIE1 2xP | 9.98E-05 | 0 | 1 | 1 | 6 | A2BIE1 | Glutamine | A2BIE1 [12 | 0 | 3090.282 | 0 | -0.19 | 0.05713 | 6.63 | 104.6 | 95.4 | 8.14 | 7.67 | | High | High | High | High | High | High | High | High | High | High | High | 0.000723 | 0.006278 | 3.86 |
| FALSE | High | GPLQTPQN | [R].GPLQTI | 1xPhospho | E9PYU8 1x | 0.014158 | 0.001011 | 1 | 4 | 2 | E9PYU8 | E3 ubiquiti | E9PYU8 [2 | 0 | 3161.512 | 0 | 3.02 | 0.000406 | 31.72 | 24.5 | 175.5 | 28.75 | 9.26 | | High | High | High | High | High | High | High | High | High | High | High | 0.000538 | 0.004617 | 2.35 |
| FALSE | High | GPLSPFNG | [K].GPLSPF | 1xDeamida | Q4VAC9 1x | 0.033404 | 0.002195 | 1 | 5 | 1 | Q4VAC9 | Pleckstrin I | Q4VAC9 [7 | 0 | 1254.608 | 0 | 1.81 | 0.000669 | 11 | 42.9 | 157.1 | 11.3 | 8.2 | | High | High | High | High | High | High | High | High | High | High | High | 0.001151 | 0.01221 | 1.57 |
| FALSE | High | GPLSQLCD | [K].GPLSQI | 1xCarbami | Q1W617 1 | 0.007605 | 0.000664 | 1 | 2 | 3 | Q1W617 | Protein Shi | Q1W617 [6 | 0 | 4494.168 | 0 | 2.79 | 0.000886 | 8.37 | 25.8 | 174.2 | 13.44 | 14.79 | | High | High | High | High | High | High | High | High | High | High | High | 0.000294 | 0.002303 | 3.99 |
| FALSE | High | GPLSQLCD | [K].GPLSQI | 1xCarbami | Q1W617 1 | 9.76E-07 | 0 | 1 | 2 | 2 | Q1W617 | Protein Shi | Q1W617 [6 | 0 | 4495.152 | 0 | 2.57 | 0.001876 | 22 | 25.9 | 174.1 | 30.55 | 16.04 | | High | High | High | High | High | High | High | High | High | High | High | 0 | 9.88E-08 | 4.5 |
| FALSE | High | GPLVGIGP | [R].GPLVGI | 1xPhospho | Q8CDZ5 1x | 0.010758 | 0.000772 | 1 | 2 | 2 | Q8CDZ5 | Putative ur | Q8CDZ5 [1 | 0 | 1560.835 | 0 | -0.03 | 0.977432 | 22.44 | 101 | 99 | 9.13 | 5.68 | | High | High | High | High | High | High | High | High | High | High | High | 0.000399 | 0.003409 | 2.9 |
| FALSE | High | GPNSPPPG | [R].GPNSPI | 1xPhospho | Q9D2Q3 1x | 0.047488 | 0.00354 | 1 | 1 | 1 | Q9D2Q3 | Uncharacte | Q9D2Q3 [4 | 0 | 1528.754 | 0 | 0.35 | 0.003799 | 5.46 | 88 | 112 | 3.33 | 8.04 | | High | High | High | High | High | High | High | High | High | High | High | 0.001812 | 0.01823 | 1.75 |
| FALSE | High | GPNYTSGY | [R].GPNYT | 1xPhospho | Q61584-3 | 1.88E-09 | 0 | 2 | 9 | 3 | Q61584-3; | Isoform B c | Q61584-3 | 0 | 2885.231 | 0 | -0.55 | 0.018799 | 4.45 | 119 | 81 | 4.06 | 14.19 | Shared | High | High | High | High | High | High | High | High | High | High | High | 0 | 8.79E-11 | 5.38 |
| FALSE | High | GPNYTSGY | [R].GPNYT | 2xPhospho | Q61584-3 | 5.71E-09 | 0 | 2 | 9 | 3 | Q61584-3; | Isoform B c | Q61584-3 | 0 | 2965.198 | 0 | -0.79 | 0.002019 | 5.54 | 123.5 | 76.5 | 6.06 | 11.72 | Shared | High | High | High | High | High | High | High | High | High | High | High | 0 | 3.06E-10 | 5.99 |
| FALSE | High | GPNYTSGY | [R].GPNYT | 3xPhospho | Q61584-3 | 0.038677 | 0.002563 | 2 | 9 | 1 | Q61584-3; | Isoform B c | Q61584-3 | 0 | 3045.164 | 0 | -0.66 | 0.007824 | 2.39 | 122.2 | 77.8 | 10.74 | 8.63 | Shared | High | High | High | High | High | High | High | High | High | High | High | 0.001421 | 0.01444 | 3.21 |
| FALSE | High | GPNYTSGY | [R].GPNYT | 1xPhospho | Q61584-3 | 0.001244 | 0 | 2 | 9 | 1 | Q61584-3; | Isoform B c | Q61584-3 | 1 | 3013.326 | 0 | -0.58 | 0.038994 | 4.28 | 117.8 | 82.2 | 14.81 | 11.67 | Shared | High | High | High | High | High | High | High | High | High | High | High | 0 | 0.000306 | 3.45 |
| FALSE | High | GPNYTSGY | [R].GPNYT | 2xPhospho | Q61584-3 | 0.11108 | 0.008991 | 2 | 9 | 1 | Q61584-3; | Isoform B c | Q61584-3 | 1 | 3093.292 | 0 | -0.86 | 0.010517 | 2.4 | 123.8 | 76.2 | 16.17 | 11.28 | Shared | High | High | High | High | High | High | High | High | High | High | High | 0.004503 | 0.04877 | 3.24 |
| FALSE | High | GPNYTSGY | [R].GPNYT | 1xPhospho | Q61584-3 | 1.1E-06 | 0 | 2 | 9 | 3 | Q61584-3; | Isoform B c | Q61584-3 | 1 | 3242.489 | 0 | -0.01 | 0.887425 | 4.68 | 98.6 | 101.4 | 5.03 | 5.91 | Shared | High | High | High | High | High | High | High | High | High | High | High | 0 | 1.13E-07 | 3.01 |
| FALSE | High | GPNYTSGY | [R].GPNYT | 2xPhospho | Q61584-3 | 0.004451 | 0.000409 | 2 | 9 | 2 | Q61584-3; | Isoform B c | Q61584-3 | 1 | 3322.455 | 0 | -0.58 | 0.001646 | 3.32 | 121.8 | 78.2 | 6.23 | 2.59 | Shared | High | High | High | High | High | High | High | High | High | High | High | 0.005041 | 0.05512 | 2.72 |
| FALSE | High | GPLNSESSI | [K].GPLNSI | 1xCarbami | B1ASU9 1x | 7.74E-08 | 0 | 1 | 7 | 2 | B1ASU9 | Serine/thre | B1ASU9 [2 | 0 | 2704.29 | 0 | -0.71 | 0.005314 | 7.78 | 121.3 | 78.7 | 8.31 | 10.74 | | High | High | High | High | High | High | High | High | High | High | High | 0 | 5.73E-09 | 4.1 |
| FALSE | High | GPPDFSSDI | [K].GPPDF | 1xPhospho | Q3U955 1x | 0.000967 | 0 | 2 | 8 | 7 | Q3U955; C | Putative ur | Q3U955 [6 | 1 | 2852.33 | 0 | 1.74 | 0.002161 | 21.93 | 45.6 | 154.4 | 9.57 | 14.11 | Shared | High | High | High | High | High | High | High | High | High | High | High | 0 | 0.000229 | 2.76 |
| FALSE | High | GPPEGSETI | [K].GPPEG | 1xPhospho | Q5SSL4-2 1 | 0.026652 | 0.001676 | 1 | 2 | 1 | Q5SSL4-2 | Isoform 2 c | Q5SSL4-2 [ | 0 | 2986.341 | 0 | 0.56 | 0.058529 | 22.55 | 88.9 | 111.1 | 24.45 | 15.45 | | High | High | High | High | High | High | High | High | High | High | High | 0.000903 | 0.009422 | 3.15 |
| FALSE | High | GPPEGSETI | [K].GPPEG | 1xPhospho | Q5SSL4-2 1 | 2.85E-05 | 0 | 1 | 2 | 5 | Q5SSL4-2 | Isoform 2 c | Q5SSL4-2 [ | 0 | 3002.336 | 0 | 0.46 | 0.056578 | 23.01 | 87.4 | 112.6 | 12.31 | 11.56 | | High | High | High | High | High | High | High | High | High | High | High | 0 | 7.03E-06 | 4.26 |
| FALSE | High | GPPEGSETI | [K].GPPEG | 1xPhospho | Q5SSL4-2 1 | 0.00204 | 5.66E-05 | 1 | 2 | 1 | Q5SSL4-2 | Isoform 2 c | Q5SSL4-2 [ | 0 | 3018.331 | 0 | 0.84 | 0.006795 | 15.93 | 74.9 | 125.1 | 15.67 | 19.8 | | High | High | High | High | High | High | High | High | High | High | High | 2.58E-05 | 0.000529 | 2.88 |
| FALSE | High | GPPEGSETI | [K].GPPEG | 2xPhospho | Q5SSL4-2 2 | 1.68E-06 | 0 | 1 | 2 | 1 | Q5SSL4-2 | Isoform 2 c | Q5SSL4-2 [ | 0 | 3066.308 | 0 | 0.64 | 0.024998 | 11.76 | 78.2 | 121.8 | 12.44 | 14.46 | | High | High | High | High | High | High | High | High | High | High | High | 0 | 1.81E-07 | 4.5 |
| FALSE | High | GPPEGSETI | [K].GPPEG | 2xPhospho | Q5SSL4-2 2 | 3.28E-07 | 0 | 1 | 2 | 1 | Q5SSL4-2 | Isoform 2 c | Q5SSL4-2 [ | 0 | 3082.303 | 0 | 0.71 | 0.000209 | 4.06 | 75.7 | 124.3 | 8.91 | 3.97 | | High | High | High | High | High | High | High | High | High | High | High | 0 | 2.89E-08 | 4.14 |
| FALSE | High | GPPGDDVS | [K].GPPGD | 1xPhospho | Q69ZS7 1x | 0.0008 | 0 | 1 | 4 | 3 | Q69ZS7 | HBS1-like p | Q69ZS7 [1 | 0 | 2573.29 | 0 | 0.85 | 0.000158 | 3.53 | 72.2 | 127.8 | 4.09 | 3.27 | | High | High | High | High | High | High | High | High | High | High | High | 0 | 0.000204 | 3.29 |
| FALSE | High | GPPGSADK | [K].GPPGS | 1xPhospho | Q9CZU3 1x | 0.017498 | 0.001245 | 1 | 3 | 2 | Q9CZU3 | Superkiller | Q9CZU3 [3 | 0 | 1266.65 | 0 | -0.58 | 0.000855 | 2.31 | 120.4 | 79.6 | 2.22 | 6.77 | | High | High | High | High | High | High | High | High | High | High | High | 0.000674 | 0.005879 | 1.52 |
| FALSE | High | GPPQSPVF | [K].GPPQSI | 1xPhospho | A0A0U1RP | 3.79E-07 | 0 | 1 | 6 | 7 | A0A0U1RP | Ataxin-2-lik | A0A0U1RP | 0 | 2056.969 | 0 | 0.26 | 0.052762 | 9.29 | 91.2 | 108.8 | 5.63 | 4.31 | | High | High | High | High | High | High | High | High | High | High | High | 0 | 3.4E-08 | 4.38 |
| FALSE | High | GPPQSPVF | [K].GPPQSI | 1xDeamida | A0A0U1RP | 7.53E-08 | 0 | 1 | 6 | 8 | A0A0U1RP | Ataxin-2-lik | A0A0U1RP | 0 | 2057.953 | 0 | 0.37 | 0.029572 | 15.55 | 87.1 | 112.9 | 11.19 | 6.85 | | High | High | High | High | High | High | High | High | High | High | High | 0 | 5.57E-09 | 4.58 |
| FALSE | High | GPPRNYQC | [R].GPPRN | 1xDeamida | Q3UBT1 1x | 1.7E-05 | 0 | 1 | 6 | 1 | Q3UBT1 | Putative ur | Q3UBT1 [1 | 1 | 2635.219 | 0 | -0.35 | 0.176079 | 3.36 | 106.7 | 93.3 | 9.4 | 9.92 | | High | High | High | High | High | High | High | High | High | High | High | 0 | 2.46E-06 | 2.92 |
| FALSE | High | GPPSPPAP | [R].GPPSPF | 1xPhospho | A0A0B4J1E | 8.03E-05 | 0 | 1 | 4 | 1 | A0A0B4J1E | SNW doma | A0A0B4J1E | 0 | 1822.887 | 0 | -0.37 | 0.074436 | 15.88 | 112.7 | 87.3 | 8.63 | 13.29 | | High | High | High | High | High | High | High | High | High | High | High | 0 | 1.4E-05 | 2.66 |
| FALSE | High | GPPSPPAP | [R].GPPSPF | 1xPhospho | A0A0B4J1E | 0.002058 | 5.66E-05 | 1 | 4 | 4 | A0A0B4J1E | SNW doma | A0A0B4J1E | 0 | 1838.882 | 0 | -0.23 | 0.046056 | 2.8 | 107.9 | 92.1 | 7.42 | 8.57 | | High | High | High | High | High | High | High | High | High | High | High | 0.002867 | 0.02993 | 2.08 |
| FALSE | High | GPPSPPAP | [R].GPPSPF | 2xPhospho | A0A0B4J1E | 1.2E-05 | 0 | 1 | 4 | 3 | A0A0B4J1E | SNW doma | A0A0B4J1E | 0 | 1902.854 | 0 | 0.39 | 0.049004 | 5.16 | 85.5 | 114.5 | 3.8 | 11.17 | | High | High | High | High | High | High | High | High | High | High | High | 0.003698 | 0.03978 | 2.62 |
| FALSE | High | GPPDFSSDI | [K].GPPDF | 2xPhospho | Q3U955 2x | 0.031439 | 0.002011 | 2 | 8 | 2 | Q3U955; C | Putative ur | Q3U955 [6 | 1 | 2932.296 | 0 | 1.56 | 0.00085 | 17.79 | 51.8 | 148.2 | 7.06 | 8.56 | Shared | High | High | High | High | High | High | High | High | High | High | High | 0.001076 | 0.01137 | 2.99 |
| FALSE | High | GPPSPPAP | [R].GPPSPF | 2xPhospho | A0A0B4J1E | 0.008022 | 0.000736 | 1 | 4 | 7 | A0A0B4J1E | SNW doma | A0A0B4J1E | 0 | 1918.849 | 0 | -0.04 | 0.284827 | 2.08 | 101.4 | 98.6 | 1.95 | 3.85 | | High | High | High | High | High | High | High | High | High | High | High | 0.002675 | 0.02857 | 2.09 |
| FALSE | High | GPLELSPAY | [R].GPLELS | 1xPhospho | A2AIX1 1xP | 0.007339 | 0.00062 | 1 | 7 | 1 | A2AIX1 | SEC16 hom | A2AIX1 [19 | 0 | 2595.269 | 0 | 0.2 | 0.005687 | 4.87 | 96 | 104 | 10.17 | 9.28 | | High | High | High | High | High | High | High | High | High | High | High | 0.000275 | 0.002225 | 2.44 |
| FALSE | High | GPHHQQQ | [R].GPHHQ | 1xPhospho | Q8K3A9 1x | 0.025964 | 0.001645 | 1 | 1 | 2 | Q8K3A9 | 7SK snRNA | Q8K3A9 [2 | 0 | 4709.204 | 0 | -0.97 | 0.037249 | 3.31 | 132.1 | 67.9 | 14.01 | 25.18 | | High | High | High | High | High | High | High | High | High | High | High | 0.000872 | 0.009149 | 2.91 |
| FALSE | High | GPAGSPCE | [R].GPAGSI | 1xCarbami | G3X8T2 1x | 2.49E-06 | 0 | 1 | 4 | 8 | G3X8T2 | RIKEN cDN | G3X8T2 [8 | 1 | 3446.438 | 0 | -0.15 | 0.019706 | 1.95 | 103.6 | 96.4 | 3.24 | 5.9 | | High | High | High | High | High | High | High | High | High | High | High | 0.006429 | 0.0695 | 3.16 |
| FALSE | High | GPAGSPCE | [R].GPAGSI | 1xCarbami | G3X8T2 2x | 0.115566 | 0.009504 | 1 | 4 | 1 | G3X8T2 | RIKEN cDN | G3X8T2 [8 | 1 | 3526.404 | 0 | -0.24 | 0.078763 | 10.93 | 111.1 | 88.9 | 6.17 | 10.65 | | High | High | High | High | High | High | High | High | High | High | High | 0.004733 | 0.05114 | 2.33 |
| FALSE | High | GPAHPGTP | [K].GPAHP | 1xPhospho | Q5PRE5 1x | 0.00642 | 0.00062 | 1 | 2 | 2 | Q5PRE5 | Proline anc | Q5PRE5 [6 | 0 | 1297.662 | 0 | -1.32 | 0.00016 | 0.33 | 142.3 | 57.7 | 4.22 | 4.61 | | High | High | High | High | High | High | High | High | High | High | High | 0.001308 | 0.01347 | 2.78 |
| FALSE | High | GPALELVD | [K].GPALEL | 1xPhospho | A6H6E1 1x | 9.28E-05 | 0 | 1 | 4 | 1 | A6H6E1 | Espl1 prote | A6H6E1 [1 | 0 | 3398.822 | 0 | -1.03 | 0.000909 | 0.47 | 133.3 | 66.7 | 2.09 | 11.3 | | High | High | High | High | High | High | High | High | High | High | High | 0 | 1.66E-05 | 2.97 |
| FALSE | High | GPALELVD | [K].GPALEL | 1xDeamida | A6H6E1 1x | 0.000102 | 0 | 1 | 4 | 2 | A6H6E1 | Espl1 prote | A6H6E1 [1 | 0 | 3399.806 | 0 | -0.69 | 0.006832 | 7.14 | 127.3 | 72.7 | 7.21 | 17.71 | | High | High | High | High | High | High | High | High | High | High | High | 0 | 1.85E-05 | 3.83 |
| FALSE | High | GPASQFYI | [R].GPASQ | 1xPhospho | D3YWX2 1 | 8.79E-05 | 0 | 1 | 2 | 1 | D3YWX2 | YLP motif-c | D3YWX2 [8 | 0 | 2145.058 | 0 | 0.62 | 0.092344 | 21.67 | 79.6 | 120.4 | 14.42 | 13.49 | | High | High | High | High | High | High | High | High | High | High | High | 0 | 1.56E-05 | 3.48 |
| FALSE | High | GPATPDVP | [K].GPATPI | 1xPhospho | Q3TW36 1 | 0.000123 | 0 | 1 | 6 | 1 | Q3TW36 | Putative ur | Q3TW36 [8 | 0 | 2314.188 | 0 | 0.94 | 0.00033 | 0.29 | 68.4 | 131.6 | 7.04 | 5.29 | | High | High | High | High | High | High | High | High | High | High | High | 0 | 2.26E-05 | 2.23 |
| FALSE | High | GPDEAIED | [R].GPDEA | 1xPhospho | Q8BH64 1x | 9.48E-06 | 0 | 1 | 2 | 3 | Q8BH64 | EH domain | Q8BH64 [4 | 0 | 3015.34 | 0 | 0.96 | 0.010731 | 12.06 | 69.7 | 130.3 | 14.79 | 13.06 | | High | High | High | High | High | High | High | High | High | High | High | 0 | 1.27E-06 | 4.09 |
| FALSE | High | GPDHQPVI | [R].GPDHQ | 1xPhospho | D3Z5I1 1xP | 0.016163 | 0.001124 | 1 | 2 | 2 | D3Z5I1 | Zinc finger | D3Z5I1 [78 | 0 | 2717.296 | 0 | 0.32 | 0.044936 | 15.43 | 88.1 | 111.9 | 4.54 | 16.41 | | High | High | High | High | High | High | High | High | High | High | High | 0.000585 | 0.005393 | 3.05 |
| FALSE | High | GPDTGLVP | [K].GPDTG | 1xPhospho | Q3V1Z0 1x | 0.015603 | 0.001124 | 1 | 2 | 2 | Q3V1Z0 | Iporin OS= | Q3V1Z0 [1 | 0 | 2962.465 | 0 | 1.88 | 0.000438 | 3.05 | 42.9 | 157.1 | 9.5 | 8.53 | | High | High | High | High | High | High | High | High | High | High | High | 0.001061 | 0.01124 | 3.1 |
| FALSE | High | GPEASTQD | [K].GPEAST | 1xPhospho | Q8K2Z4-2 | 0.000674 | 0 | 1 | 2 | 1 | Q8K2Z4-2 | Isoform 2 c | Q8K2Z4-2 | 0 | 2595.198 | 0 | 0.34 | 0.000957 | 3.62 | 88.3 | 111.7 | 6.23 | 2.42 | | High | High | High | High | High | High | High | High | High | High | High | 0 | 0.000153 | 4.09 |
| FALSE | High | GPEASTQD | [K].GPEAST | 1xPhospho | Q8K2Z4-2 | 0.000302 | 0 | 1 | 2 | 4 | Q8K2Z4-2 | Isoform 2 c | Q8K2Z4-2 | 1 | 4819.267 | 0 | 0.39 | 0.079015 | 15.98 | 83.9 | 116.1 | 10.09 | 13.22 | | High | High | High | High | High | High | High | High | High | High | High | 0 | 6.22E-05 | 4.1 |
| FALSE | High | GPEEESTG | [K].GPEEES | 1xPhospho | Q3V300 1x | 0.000515 | 0 | 1 | 1 | 6 | Q3V300 | Kinesin-like | Q3V300 [3 | 0 | 2656.239 | 0 | -0.48 | 0.00443 | 2.5 | 117 | 83 | 5.07 | 8.44 | | High | High | High | High | High | High | High | High | High | High | High | 0 | 0.000113 | 3.71 |
| FALSE | High | GPHTPPPP | [R].GPHTPI | 1xPhospho | Q9D281 1x | 0.017041 | 0.001204 | 1 | 1 | 2 | Q9D281 | Protein No | Q9D281 [1 | 0 | 1607.789 | 0 | -0.97 | 0.014367 | 6.59 | 127.5 | 72.5 | 15.94 | 18.47 | | High | High | High | High | High | High | High | High | High | High | High | 0.004399 | 0.04716 | 2.31 |
| FALSE | High | GPEGSSPE | [R].GPEGS | 1xPhospho | Q9CPV5 1x | 7.98E-06 | 0 | 1 | 4 | 1 | Q9CPV5 | Polyamine- | Q9CPV5 [1 | 0 | 2045.974 | 0 | -0.4 | 0.023028 | 10.72 | 111.7 | 88.3 | 5.16 | 10.33 | | High | High | High | High | High | High | High | High | High | High | High | 0 | 1.05E-06 | 2.73 |
| FALSE | High | GPEQHRYS | [K].GPEQH | 1xPhospho | Q7TSH3 1x | 0.006085 | 0.00062 | 1 | 1 | 2 | Q7TSH3 | Zinc finger | Q7TSH3 [6 | 1 | 1981.031 | 0 | -0.07 | 0.671652 | 17.64 | 101.5 | 98.5 | 13.61 | 9.4 | | High | High | High | High | High | High | High | High | High | High | High | 0.000275 | 0.001795 | 4.1 |
| FALSE | High | GPESPLRSF | [K].GPESPL | 1xPhospho | A0A0R4J0V | 0.074685 | 0.005727 | 1 | 4 | 1 | A0A0R4J0V | Junction-m | A0A0R4J0V | 1 | 1475.757 | 0 | 0.16 | 0.276212 | 19.69 | 94.5 | 105.5 | 4.15 | 11.07 | | High | High | High | High | High | High | High | High | High | High | High | 0.002937 | 0.03064 | 1.66 |
| FALSE | High | GPEVEGSP | [R].GPEVE | 1xPhospho | Q921C3 1x | 2.24E-06 | 0 | 1 | 5 | 1 | Q921C3 | Bromodom | Q921C3 [1 | 0 | 1735.847 | 0 | -0.96 | 0.000489 | 1.3 | 133.9 | 66.1 | 3.58 | 10.99 | | High | High | High | High | High | High | High | High | High | High | High | 0 | 2.52E-07 | 4.18 |
| FALSE | High | GPEVTSQG | [R].GPEVTS | 1xCarbami | E9QM77 1 | 8.6E-07 | 0 | 1 | 8 | 1 | E9QM77 | Ataxin-2 O | E9QM77 [8 | 0 | 2271.109 | 0 | 0.73 | 0.004596 | 12.95 | 74.4 | 125.6 | 12.42 | 6.28 | | High | High | High | High | High | High | High | High | High | High | High | 0 | 8.58E-08 | 4.58 |
| FALSE | High | GPGEPDSP | [K].GPGEPI | 2xPhospho | P55200 2x | 0.014929 | 0.001049 | 1 | 3 | 1 | P55200 | Histone-lys | P55200 [18 | 0 | 2767.294 | 0 | 0.09 | 0.521295 | 27.67 | 96.9 | 103.1 | 10.34 | 7.66 | | High | High | High | High | High | High | High | High | High | High | High | 0.000555 | 0.004887 | 2.69 |
| FALSE | High | GPGLLESPS | [R].GPGLLE | 1xPhospho | Q6A0A2 1x | 6.31E-06 | 0 | 1 | 2 | 1 | Q6A0A2 | La-related | Q6A0A2 [4 | 0 | 2130.047 | 0 | 1.51 | 0.014034 | 31.73 | 52 | 148 | 25.68 | 25 | | High | High | High | High | High | High | High | High | High | High | High | 0 | 8.05E-07 | 2.98 |
| FALSE | High | GPGIYEDL | [R].GPGIYE | 2xDeamida | Q8C0L0 1x | 0.056248 | 0.004124 | 1 | 2 | 1 | Q8C0L0 | Thioredoxi | Q8C0L0 [1 | 0 | 2062.982 | 0 | 0.37 | 0.031596 | 17.91 | 92 | 108 | 18.73 | 9.21 | | High | High | High | High | High | High | High | High | High | High | High | 0.002117 | 0.02205 | 2.57 |
| FALSE | High | GPGQVSTA | [K].GPGQV | 1xPhospho | Q3UJB9 1x | 1.44E-07 | 0 | 1 | 6 | 3 | Q3UJB9 | Enhancer c | Q3UJB9 [7 | 0 | 3384.725 | 0 | -0.1 | 0.315725 | 6.98 | 100.9 | 99.1 | 2.95 | 7.95 | | High | High | High | High | High | High | High | High | High | High | High | 0 | 2.84E-08 | 4.57 |
| FALSE | High | GPGSHDAV | [K].GPGSH | 3xPhospho | O54916 3x | 0.112856 | 0.009146 | 1 | 7 | 1 | O54916 | RalBP1-ass | O54916 [1 | 0 | 4524.037 | 0 | -1.91 | 0.001456 | 3.44 | 155.3 | 44.7 | 12.03 | 40.26 | | High | High | High | High | High | High | High | High | High | High | High | 0.004596 | 0.04992 | 3.61 |
| FALSE | High | GPGSSPET | [R].GPGSSI | 1xPhospho | Q9D2D7 1x | 7.63E-06 | 0 | 1 | 3 | 4 | Q9D2D7 | Zinc finger | Q9D2D7 [2 | 0 | 2962.533 | 0 | -0.52 | 0.012515 | 9.63 | 117.8 | 82.2 | 8.39 | 9.85 | | High | High | High | High | High | High | High | High | High | High | High | 0 | 2.01E-06 | 3.98 |
| FALSE | High | GPGSWWT | [R].GPGSW | 1xCarbami | Q8BX02 1x | 1.05E-09 | 0 | 1 | 2 | 11 | Q8BX02 | KN motif a | Q8BX02 [7 | 0 | 2552.06 | 0 | -0.09 | 0.231147 | 17.26 | 104.9 | 95.1 | 8.64 | 5.73 | | High | High | High | High | High | High | High | High | High | High | High | 0 | 1.62E-08 | 5.09 |
| FALSE | High | GPGVGVH | [R].GPGVG | 1xPhospho | Q8C5W3-2 | 0.003721 | 0.00021 | 1 | 3 | 1 | Q8C5W3-2 | Isoform 2 c | Q8C5W3-2 | 0 | 2254.182 | 0 | -0.07 | 0.40308 | 5.12 | 104.5 | 95.5 | 5.26 | 7.76 | | High | High | High | High | High | High | High | High | High | High | High | 9.42E-05 | 0.001037 | 2.43 |
| FALSE | High | GPHHLDNS | [R].GPHHLI | 1xPhospho | A0A0U1RP | 2.38E-05 | 0 | 1 | 7 | 14 | A0A0U1RP | Ataxin-2-lik | A0A0U1RP | 0 | 2023.918 | 0 | 0.16 | 0.636243 | 21.64 | 96 | 104 | 8.87 | 10.63 | | High | High | High | High | High | High | High | High | High | High | High | 0 | 4.26E-05 | 4.07 |
| FALSE | High | GPENPQVE | [K].GPENP | 1xPhospho | F6WJB7 1x | 0.001107 | 0 | 1 | 6 | 1 | F6WJB7 | Ubiquitin c | F6WJB7 [2 | 0 | 4052.852 | 0 | 0.4 | 0.02649 | 15.96 | 82.2 | 117.8 | 11.49 | 10.86 | | High | High | High | High | High | High | High | High | High | High | High | 0 | 0.000268 | 3.88 |
| FALSE | High | GPPTQMSI | [K].GPPTQI | 1xPhospho | Q6ZQF7 1x | 3.07E-06 | 0 | 1 | 3 | 2 | Q6ZQF7 | E3 ubiquiti | Q6ZQF7 [1 | 0 | 2956.313 | 0 | 1.38 | 0.000202 | 4.78 | 55.3 | 144.7 | 14.59 | 6.79 | | High | High | High | High | High | High | High | High | High | High | High | 0 | 3.6E-07 | 3.44 |
| FALSE | High | GPPTQMSI | [K].GPPTQI | 1xDeamida | Q6ZQF7 1x | 3.67E-05 | 0 | 1 | 3 | 1 | Q6ZQF7 | E3 ubiquiti | Q6ZQF7 [1 | 0 | 2957.297 | 0 | 1.31 | 0.007167 | 24.98 | 54.5 | 145.5 | 9.89 | 16.96 | | High | High | High | High | High | High | High | High | High | High | High | 0 | 5.81E-06 | 4.45 |
| FALSE | High | GPPVTPSSE | [R].GPPVTI | 1xPhospho | B2RY22 1x | 0.014797 | 0.001049 | 2 | 18 | 2 | B2RY22; A | Clasp1 pro | B2RY22 [6 | 0 | 1536.808 | 0 | -0.78 | 0.001117 | 6 | 133.6 | 66.4 | 11.64 | 21.06 | Shared | High | High | High | High | High | High | High | High | High | High | High | 0.000555 | 0.004841 | 1.95 |
| FALSE | High | GPSPSPVG | [R].GPSPSF | 1xPhospho | E9QAQ7 1x | 1.04E-06 | 0 | 1 | 5 | 6 | E9QAQ7 | AT-rich inte | E9QAQ7 [6 | 0 | 1889.932 | 0 | -0.85 | 0.000359 | 1.3 | 126.5 | 73.5 | 4.89 | 5.51 | | High | High | High | High | High | High | High | High | High | High | High | 0 | 1.06E-07 | 3.87 |
| FALSE | High | GPSPSPVG | [R].GPSPSF | 2xPhospho | E9QAQ7 2x | 0.000205 | 0 | 1 | 5 | 3 | E9QAQ7 | AT-rich inte | E9QAQ7 [6 | 0 | 1969.898 | 0 | 0.19 | 0.129992 | 2.45 | 94.4 | 105.6 | 2.75 | 9.5 | | High | High | High | High | High | High | High | High | High | High | High | 0 | 4.04E-05 | 2.98 |
| FALSE | High | GPSPSPVG | [R].GPSPSF | 3xPhospho | E9QAQ7 3x | 7.7E-06 | 0 | 1 | 5 | 3 | E9QAQ7 | AT-rich inte | E9QAQ7 [6 | 0 | 2049.865 | 0 | 0.98 | 0.001089 | 2 | 59.4 | 140.6 | 10.72 | 11.79 | | High | High | High | High | High | High | High | High | High | High | High | 0 | 1.28E-06 | 3.33 |
| FALSE | High | GPSPSSPTF | [R].GPSPSS | 2xPhospho | E9QKK4 2x | 0.001049 | 0 | 1 | 2 | 2 | E9QKK4 | Glucocortic | E9QKK4 [9 | 0 | 2372.089 | 0 | 0.74 | 0.008372 | 9.14 | 75.3 | 124.7 | 8.62 | 11.83 | | High | High | High | High | High | High | High | High | High | High | High | 0 | 0.000251 | 2.8 |
| FALSE | High | GPSPVSLGI | [R].GPSPVS | 1xPhospho | Q3UQN2 1 | 5.6E-05 | 0 | 1 | 3 | 3 | Q3UQN2 | F-BAR dom | Q3UQN2 [ | 0 | 3426.796 | 0 | 0.16 | 0.062805 | 7.38 | 98.2 | 101.8 | 6.68 | 11.01 | | High | High | High | High | High | High | High | High | High | High | High | 0 | 4E-05 | 3.61 |
| FALSE | High | GPSPVVLG | [R].GPSPV | 1xPhospho | Q8K285 1x | 3.29E-05 | 0 | 1 | 2 | 1 | Q8K285 | F-BAR dom | Q8K285 [5 | 0 | 3301.65 | 0 | -0.29 | 0.107565 | 1.71 | 115.8 | 84.2 | 13.83 | 33.98 | | High | High | High | High | High | High | High | High | High | High | High | 0 | 5.15E-06 | 4.49 |
| FALSE | High | GPSSTGAS | [K].GPSSTC | 1xPhospho | Q8CH25-2 | 0.005275 | 0.000599 | 1 | 4 | 3 | Q8CH25-2 | Isoform 2 c | Q8CH25-2 | 0 | 1585.799 | 0 | -0.32 | 0.019769 | 4.05 | 108.4 | 91.6 | 8.61 | 6.32 | | High | High | High | High | High | High | High | High | High | High | High | 0.000267 | 0.001539 | 1.98 |
| FALSE | High | GPSSVEDIK | [K].GPSSVE | 1xPhospho | Q5SQB7 1x | 0.064323 | 0.004664 | 1 | 6 | 2 | Q5SQB7 | MCG68069 | Q5SQB7 [2 | 0 | 1469.765 | 0 | -0.49 | 0.009131 | 8.29 | 114.5 | 85.5 | 5.83 | 9.6 | | High | High | High | High | High | High | High | High | High | High | High | 0.002403 | 0.02573 | 2.31 |
| FALSE | High | GPSTPNVK | [K].GPSTPN | 1xPhospho | Q8R5H1-2 | 0.021989 | 0.001542 | 1 | 2 | 2 | Q8R5H1-2 | Isoform 2 c | Q8R5H1-2 | 0 | 1337.723 | 0 | -0.12 | 0.425127 | 7.65 | 101 | 99 | 5.15 | 7.21 | | High | High | High | High | High | High | High | High | High | High | High | 0.002896 | 0.03005 | 2.05 |
| FALSE | High | GPSTPYTK | [K].GPSTPY | 1xPhospho | Q6PDG5 1x | 0.086562 | 0.006435 | 1 | 3 | 1 | Q6PDG5 | SWI/SNF c | Q6PDG5 [3 | 0 | 1388.723 | 0 | 1.79 | 0.000181 | 8.14 | 44.9 | 155.1 | 9.07 | 3.79 | | High | High | High | High | High | High | High | High | High | High | High | 0.003317 | 0.03646 | 1.42 |
| FALSE | High | GPSVDVEV | [K].GPSVD | 1xCarbami | E9Q616 1x | 1.44E-07 | 0 | 1 | 1 | 2 | E9Q616 | AHNAK nuc | E9Q616 [1 | 0 | 2579.235 | 0 | 2.47 | 0.006604 | 64.3 | 30 | 170 | 18.67 | 26.18 | | High | High | High | High | High | High | High | High | High | High | High | 0 | 1.15E-08 | 3.96 |
| FALSE | High | GPTPTATG | [R].GPTPTA | 1xPhospho | A2AJI0 1xP | 0.004106 | 0.00036 | 1 | 4 | 1 | A2AJI0 | MAP7 dom | A2AJI0 [97 | 0 | 1263.63 | 0 | 1.04 | 0.000707 | 5.92 | 66.1 | 133.9 | 5.91 | 5.02 | | High | High | High | High | High | High | High | High | High | High | High | 0.000161 | 0.001162 | 1.84 |
| FALSE | High | GPTQMPCI | [K].GPTQM | 1xCarbami | E9PVX6 1x | 0.002313 | 0.000109 | 1 | 2 | 2 | E9PVX6 | Proliferatic | E9PVX6 [2 | 0 | 2899.295 | 0 | 0.45 | 0.074853 | 7.58 | 93.8 | 106.2 | 13.13 | 11.45 | | High | High | High | High | High | High | High | High | High | High | High | 0.000204 | 0.001363 | 3.24 |

| | | | | | | | | | | | | | | | | | | | | | | | | | | | | | | | | | | | | | | |
|---|---|---|---|---|---|---|---|---|---|---|---|---|---|---|---|---|---|---|---|---|---|---|---|---|---|---|---|---|---|---|---|---|---|---|---|---|---|---|
| FALSE | High | GPSPSLLST | [R].GPSPSL | 1xPhospho | E9Q8S5 1x | 0.000239 | 0 | 1 | 3 | 1 | E9Q8S5 | Microtubu | E9Q8S5 [7 | 0 | 2296.228 | 0 | 0.44 | 0.009302 | 12.21 | 88.3 | 111.7 | 3.61 | 10.37 | | High | High | High | High | High | High | High | High | High | High | High | 0 | 4.8E-05 | 2.27 |
| FALSE | High | GPTQMPCI | [K].GPTQM | 1xCarbami | E9PVX6 1x | 0.000311 | 0 | 1 | 2 | 5 | E9PVX6 | Proliferatic | E9PVX6 [2 | 0 | 2900.279 | 0 | 0.21 | 0.416362 | 20.22 | 87.2 | 112.8 | 17.33 | 20.95 | | High | High | High | High | High | High | High | High | High | High | High | 0 | 0.000167 | 3.57 |
| FALSE | High | GPTSTSVD | [R].GPTSTS | 1xPhospho | Q6NXI6 1x | 1.28E-07 | 0 | 1 | 3 | 2 | Q6NXI6 | Regulation | Q6NXI6 [7 | 1 | 2298.081 | 0 | 0.33 | 0.000424 | 4.12 | 88.1 | 111.9 | 3 | 1.31 | | High | High | High | High | High | High | High | High | High | High | High | 0 | 1.01E-08 | 3.35 |
| FALSE | High | GPVAGTG | [K].GPVAG | 1xPhospho | Q9DB90 1x | 4.08E-05 | 0 | 1 | 3 | 2 | Q9DB90 | Protein SM | Q9DB90 [1 | 0 | 2861.485 | 0 | 0.31 | 0.012947 | 12.57 | 89.5 | 110.5 | 17.03 | 6.4 | | High | High | High | High | High | High | High | High | High | High | High | 0 | 8.05E-06 | 3.97 |
| FALSE | High | GPVAQLSS | [R].GPVAQ | 1xPhospho | Q6PGL7 1x | 9.91E-06 | 0 | 1 | 3 | 3 | Q6PGL7 | WASH com | Q6PGL7 [1 | 0 | 2701.439 | 0 | 0.18 | 0.086473 | 19.29 | 90.9 | 109.1 | 6.95 | 7.52 | | High | High | High | High | High | High | High | High | High | High | High | 0 | 1.34E-06 | 2.88 |
| FALSE | High | GPVAQLSS | [R].GPVAQ | 1xDeamida | Q6PGL7 1x | 0.000483 | 0 | 1 | 3 | 6 | Q6PGL7 | WASH com | Q6PGL7 [1 | 0 | 2702.423 | 0 | 0.2 | 0.075012 | 0.62 | 97.7 | 102.3 | 5.62 | 4.88 | | High | High | High | High | High | High | High | High | High | High | High | 0.008618 | 0.09211 | 3.23 |
| FALSE | High | GPVFGEPP | [R].GPVFG | 1xPhospho | Q3UKS3 1x | 0.000319 | 0 | 1 | 2 | 1 | Q3UKS3 | Putative ur | Q3UKS3 [3 | 0 | 1933.962 | 0 | 3.58 | 0.006124 | 37.16 | 14 | 186 | 10.66 | 30.73 | | High | High | High | High | High | High | High | High | High | High | High | 0 | 6.62E-05 | 2.89 |
| FALSE | High | GPVSGSPD | [R].GPVSG | 1xPhospho | O88351 1x | 5.45E-05 | 0 | 1 | 4 | 1 | O88351 | Inhibitor of | O88351 [6 | 0 | 1698.772 | 0 | 0.75 | 0.000583 | 1.65 | 74.1 | 125.9 | 3.13 | 6.63 | | High | High | High | High | High | High | High | High | High | High | High | 0 | 9.12E-06 | 2.35 |
| FALSE | High | GPVSGSPD | [R].GPVSG | 1xPhospho | O88351 1x | 4.76E-05 | 0 | 1 | 4 | 2 | O88351 | Inhibitor of | O88351 [6 | 0 | 1714.767 | 0 | 0.78 | 0.017126 | 7.33 | 71.1 | 128.9 | 5.68 | 23.44 | | High | High | High | High | High | High | High | High | High | High | High | 0 | 4.03E-05 | 3.33 |
| FALSE | High | GPVSPSYD | [K].GPVSPS | 1xPhospho | Q8K2W9 1 | 3.49E-07 | 0 | 1 | 1 | 1 | Q8K2W9 | Uncharacte | Q8K2W9 [5 | 0 | 1849.925 | 0 | -0.81 | 0.00039 | 0.27 | 128.5 | 71.5 | 5.31 | 3.34 | | High | High | High | High | High | High | High | High | High | High | High | 0 | 3.13E-08 | 2.88 |
| FALSE | High | GQAPLETS | [R].GQAPL | 1xPhospho | E9Q7G0 1x | 2.82E-05 | 0 | 1 | 3 | 2 | E9Q7G0 | Nuclear mi | E9Q7G0 [1 | 0 | 2442.139 | 0 | 0.17 | 0.1347 | 21.2 | 97.9 | 102.1 | 13.36 | 11.99 | | High | High | High | High | High | High | High | High | High | High | High | 0 | 4.34E-06 | 3.2 |
| FALSE | High | GQAPLETS | [R].GQAPL | 2xPhospho | E9Q7G0 2x | 0.049971 | 0.003641 | 1 | 3 | 1 | E9Q7G0 | Nuclear mi | E9Q7G0 [1 | 1 | 2879.363 | 0 | -0.1 | 0.30131 | 14.74 | 103.1 | 96.9 | 10.63 | 9.46 | | High | High | High | High | High | High | High | High | High | High | High | 0.001902 | 0.0193 | 3.01 |
| FALSE | High | GQASLYGK | [K].GQASL | 1xPhospho | P97496 1x | 0.108463 | 0.008839 | 1 | 6 | 1 | P97496 | SWI/SNF c | P97496 [3 | 0 | 1361.723 | 0 | -1.21 | 0.017237 | 5.45 | 136.9 | 63.1 | 26.11 | 10.02 | | High | High | High | High | High | High | High | High | High | High | High | 0.004411 | 0.04757 | 1.4 |
| FALSE | High | GQCCDSTE | [K].GQCCD | 2xCarbami | Q61687 1x | 0.007404 | 0.00062 | 1 | 4 | 1 | Q61687 | Transcripti | Q61687 [1 | 0 | 1622.696 | 0 | -0.31 | 0.006784 | 5.68 | 109.3 | 90.7 | 3.37 | 7.02 | | High | High | High | High | High | High | High | High | High | High | High | 0.000275 | 0.002247 | 1.55 |
| FALSE | High | GQDTVAIE | [K].GQDTV | 1xPhospho | A2BIE1 1xP | 5.6E-08 | 0 | 1 | 1 | 1 | A2BIE1 | Glutamine | A2BIE1 [12 | 0 | 3010.315 | 0 | -0.39 | 0.014173 | 3.46 | 116.1 | 83.9 | 7.13 | 12.42 | | High | High | High | High | High | High | High | High | High | High | High | 0 | 4E-09 | 4.42 |
| FALSE | High | GPTQMPCI | [K].GPTQM | 1xCarbami | E9PVX6 1x | 7.56E-06 | 0 | 1 | 2 | 1 | E9PVX6 | Proliferatic | E9PVX6 [2 | 0 | 2915.29 | 0 | -0.15 | 0.908077 | 31.36 | 105.3 | 94.7 | 18.09 | 21.29 | | High | High | High | High | High | High | High | High | High | High | High | 0 | 9.86E-07 | 3.92 |
| FALSE | High | GPSPPGAK | [K].GPSPP | 1xPhospho | Q91XU0 1x | 0.085162 | 0.006376 | 1 | 2 | 1 | Q91XU0 | ATPase WF | Q91XU0 [6 | 0 | 1248.675 | 0 | -0.97 | 0.000181 | 0.74 | 132.4 | 67.6 | 4.05 | 3.66 | | High | High | High | High | High | High | High | High | High | High | High | 0.003291 | 0.03569 | 1.38 |
| FALSE | High | GPSPLVTM | [R].GPSPLV | 1xPhospho | E9PU87 1x | 4.12E-05 | 0 | 1 | 5 | 4 | E9PU87 | Serine/thre | E9PU87 [5 | 0 | 3830.788 | 0 | 0.25 | 0.123121 | 6.18 | 91.3 | 108.7 | 10.08 | 7.88 | | High | High | High | High | High | High | High | High | High | High | High | 0 | 6.61E-06 | 5.1 |
| FALSE | High | GPSPLVTM | [R].GPSPLV | 1xPhospho | E9PU87 1x | 0.001615 | 5.66E-05 | 1 | 5 | 3 | E9PU87 | Serine/thre | E9PU87 [5 | 0 | 3814.793 | 0 | -0.04 | 0.587689 | 34.09 | 102.2 | 97.8 | 14.41 | 19.32 | | High | High | High | High | High | High | High | High | High | High | High | 2.58E-05 | 0.000407 | 3.95 |
| FALSE | High | GPPVTPSS | [R].GPPVT | 2xPhospho | B2RY22 2x | 0.042141 | 0.003162 | 2 | 18 | 1 | B2RY22; A | Clasp1 pro | B2RY22 [6 | 0 | 1616.774 | 0 | -0.83 | 0.000641 | 1.39 | 125.7 | 74.3 | 4.17 | 8.24 | Shared | High | High | High | High | High | High | High | High | High | High | High | 0.001625 | 0.01579 | 3.1 |
| FALSE | High | GPQFGSEV | [R].GPQFG | 1xPhospho | P09470 1x | 0.021607 | 0.001507 | 1 | 4 | 1 | P09470 | Angiotensi | P09470 [1 | 0 | 1527.741 | 0 | 3.1 | 0.000779 | 4.09 | 22.1 | 177.9 | 13.54 | 14.27 | | High | High | High | High | High | High | High | High | High | High | High | 0.000801 | 0.007437 | 2.23 |
| FALSE | High | GPQLPPLS | [R].GPQLP | 1xDeamida | Q68ED7 1x | 0.0003 | 0 | 1 | 1 | 6 | Q68ED7 | CREB-regu | Q68ED7 [4 | 0 | 4510.241 | 0 | -0.08 | 0.663449 | 37.5 | 100.4 | 99.6 | 16.72 | 20.22 | | High | High | High | High | High | High | High | High | High | High | High | 0 | 0.00016 | 3.97 |
| FALSE | High | GPQLPPLS | [R].GPQLP | 1xPhospho | Q68ED7 1x | 0.002948 | 0.00016 | 1 | 1 | 2 | Q68ED7 | CREB-regu | Q68ED7 [4 | 0 | 4525.252 | 0 | -0.1 | 0.827399 | 13.92 | 104.3 | 95.7 | 9.41 | 20.99 | | High | High | High | High | High | High | High | High | High | High | High | 7.19E-05 | 0.000803 | 2.86 |
| FALSE | High | GPQISAPS | [K].GPQISA | 1xPhospho | E9Q616 1x | 0.002895 | 0.00016 | 1 | 6 | 2 | E9Q616 | AHNAK nu | E9Q616 [5 | 0 | 2326.155 | 0 | 1.78 | 0.004236 | 14.45 | 38.1 | 161.9 | 17.8 | 21.73 | | High | High | High | High | High | High | High | High | High | High | High | 7.19E-05 | 0.000786 | 2.97 |
| FALSE | High | GPQISAPS | [K].GPQISA | 1xPhospho | E9Q616 1x | 0.008927 | 0.000736 | 1 | 6 | 2 | E9Q616 | AHNAK nu | E9Q616 [5 | 0 | 2342.15 | 0 | 2.25 | 0.001353 | 9.09 | 33.2 | 166.8 | 16.78 | 17.89 | | High | High | High | High | High | High | High | High | High | High | High | 0.000418 | 0.003583 | 2.07 |
| FALSE | High | GPQITGPS | [K].GPQITG | 1xPhospho | E9Q616 1x | 0.000492 | 0 | 1 | 3 | 1 | E9Q616 | AHNAK nu | E9Q616 [3 | 0 | 2120.14 | 0 | 1.98 | 0.004357 | 41.63 | 39.9 | 160.1 | 24.38 | 17.89 | | High | High | High | High | High | High | High | High | High | High | High | 0 | 0.000108 | 2.58 |
| FALSE | High | GPQQDSD | [K].GPQQD | 2xPhospho | Q5U3K5 2x | 2.1E-05 | 0 | 1 | 1 | 3 | Q5U3K5 | Rab-like pr | Q5U3K5 [4 | 0 | 4362.842 | 0 | 0.06 | 0.755242 | 4.8 | 99.3 | 100.7 | 9.25 | 16.01 | | High | High | High | High | High | High | High | High | High | High | High | 0.000246 | 0.001505 | 4.68 |
| FALSE | High | GPQVSSSL | [K].GPQVS | 1xPhospho | E9Q616 1x | 0.000434 | 0 | 1 | 6 | 3 | E9Q616 | AHNAK nu | E9Q616 [5 | 0 | 1971.02 | 0 | 2.43 | 0.006057 | 18.73 | 33.8 | 166.2 | 11.02 | 24.62 | | High | High | High | High | High | High | High | High | High | High | High | 0 | 9.33E-05 | 2.29 |
| FALSE | High | GPRPQTSL | [R].GPRPQ | 1xPhospho | A0A068F1 | 0.030104 | 0.001857 | 2 | 8 | 2 | A0A068F1 | Glyco-gag | A0A068F1 | 0 | 1721.867 | 0 | 2.96 | 0.020745 | 25.08 | 23.8 | 176.2 | 28.24 | 42.58 | Shared | High | High | High | High | High | High | High | High | High | High | High | 0.004961 | 0.0539 | 1.85 |
| FALSE | High | GPRTPPGP | [R].GPRTP | 1xPhospho | E9PZY8 1x | 0.106748 | 0.008634 | 1 | 4 | 1 | E9PZY8 | RIKEN cDN | E9PZY8 [1 | 1 | 3323.554 | 0 | -0.22 | 0.034415 | 5.65 | 107.2 | 92.8 | 13.92 | 14.81 | | High | High | High | High | High | High | High | High | High | High | High | 0.004327 | 0.04678 | 2.49 |
| FALSE | High | GPRTPPGP | [R].GPRTP | 1xPhospho | E9PZY8 1x | 0.054383 | 0.003951 | 1 | 4 | 1 | E9PZY8 | RIKEN cDN | E9PZY8 [1 | 1 | 3339.549 | 0 | -0.4 | 0.637056 | 19.5 | 101.6 | 98.4 | 15.87 | 32.26 | | High | High | High | High | High | High | High | High | High | High | High | 0.002059 | 0.0213 | 3.19 |
| FALSE | High | GPRTPPLA | [R].GPRTP | 1xPhospho | E9Q784 1x | 0.096181 | 0.007535 | 1 | 1 | 1 | E9Q784 | Zinc finger | E9Q784 [1 | 1 | 1358.776 | 0 | -0.83 | 0.021493 | 6.97 | 122.4 | 77.6 | 13.48 | 16.11 | | High | High | High | High | High | High | High | High | High | High | High | 0.003823 | 0.04119 | 3.21 |
| FALSE | High | GPRTPSPP | [R].GPRTPS | 2xPhospho | E9Q784 2x | 0.000186 | 0 | 1 | 3 | 5 | E9Q784 | Zinc finger | E9Q784 [2 | 1 | 2615.402 | 0 | -0.5 | 0.001617 | 4.65 | 119.9 | 80.1 | 7.27 | 5.68 | | High | High | High | High | High | High | High | High | High | High | High | 0 | 3.61E-05 | 3.64 |
| FALSE | High | GPSATTGG | [R].GPSAT | 1xPhospho | Q80U72-4 | 0.117405 | 0.009523 | 2 | 5 | 1 | Q80U72-4; | Isoform 4 c | Q80U72-4 | 0 | 1313.687 | 0 | 0.95 | 0.002799 | 14.96 | 68.2 | 131.8 | 6.84 | 11.69 | Shared | High | High | High | High | High | High | High | High | High | High | High | 0.004797 | 0.05241 | 0.71 |
| FALSE | High | GPSDFSGL | [K].GPSDFS | 1xDeamida | Q9CX00 1x | 0.003402 | 0.00021 | 1 | 1 | 7 | Q9CX00 | IST1 homo | Q9CX00 [2 | 0 | 4864.344 | 0 | 0.64 | 0.022501 | 27.84 | 66.7 | 133.3 | 18.5 | 19.25 | | High | High | High | High | High | High | High | High | High | High | High | 0.000399 | 0.003376 | 4.54 |
| FALSE | High | GPSEAPQE | [K].GPSEA | 1xPhospho | B2RRE0 1x | 4.71E-08 | 0 | 1 | 3 | 9 | B2RRE0 | A kinase (P | B2RRE0 [5 | 0 | 2627.176 | 0 | -0.77 | 0.001818 | 3.79 | 127.7 | 72.3 | 7.28 | 6.34 | | High | High | High | High | High | High | High | High | High | High | High | 0 | 2.41E-05 | 3.85 |
| FALSE | High | GPSEFVSS | [K].GPSEFV | 1xPhospho | A2AIX1 1x | 0.047488 | 0.00354 | 1 | 2 | 2 | A2AIX1 | SEC16 hon | A2AIX1 [9 | 0 | 3625.823 | 0 | 0.96 | 0.017402 | 37.5 | 62.7 | 137.3 | 27.13 | 20.62 | | High | High | High | High | High | High | High | High | High | High | High | 0.001812 | 0.0181 | 2.79 |
| FALSE | High | GPSEYPTK | [R].GPSEY | 1xPhospho | Q0GH64 1x | 0.007811 | 0.000664 | 1 | 1 | 1 | Q0GH64 | Claudin OS | Q0GH64 [2 | 0 | 1416.718 | 0 | 3.03 | 0.007864 | 19.69 | 19.3 | 180.7 | 22.53 | 33.07 | | High | High | High | High | High | High | High | High | High | High | High | 0.000327 | 0.00239 | 1.57 |
| FALSE | High | GPSFNVAS | [K].GPSFN | 1xPhospho | E9Q616 1x | 1.17E-07 | 0 | 1 | 6 | 2 | E9Q616 | AHNAK nu | E9Q616 [5 | 0 | 2376.189 | 0 | 3.23 | 0.014206 | 21.27 | 20.2 | 179.8 | 21.02 | 40.84 | | High | High | High | High | High | High | High | High | High | High | High | 0 | 1.66E-07 | 4.15 |
| FALSE | High | GPSGTVSE | [K].GPSGT | 1xPhospho | B9EK95 1x | 0.000984 | 0 | 1 | 6 | 3 | B9EK95 | Myof prote | B9EK95 [1 | 0 | 1581.784 | 0 | 2.42 | 0.025992 | 50.35 | 30.5 | 169.5 | 9.16 | 43.82 | | High | High | High | High | High | High | High | High | High | High | High | 0 | 0.000234 | 1.21 |
| FALSE | High | GPSLNPVL | [R].GPSLN | 1xPhospho | Q6ZQ61 1x | 0.000993 | 0 | 1 | 6 | 1 | Q6ZQ61 | MCG12197 | Q6ZQ61 [1 | 0 | 1935.916 | 0 | 0.46 | 0.027215 | 23.16 | 77.2 | 122.8 | 14.02 | 19.16 | | High | High | High | High | High | High | High | High | High | High | High | 0 | 0.000237 | 1.71 |
| FALSE | High | GPSPAPAD | [R].GPSPA | 1xPhospho | Q6ZQ70 1x | 0.000573 | 0 | 1 | 1 | 1 | Q6ZQ70 | MKIAA068 | Q6ZQ70 [3 | 0 | 2387.161 | 0 | 0.12 | 0.083663 | 9.35 | 94.8 | 105.2 | 5.65 | 7.98 | | High | High | High | High | High | High | High | High | High | High | High | 0 | 0.000128 | 2.47 |
| FALSE | High | GPSPAPAD | [R].GPSPA | 1xPhospho | Q6ZQ70 1x | 0.000122 | 0 | 1 | 1 | 1 | Q6ZQ70 | MKIAA068 | Q6ZQ70 [3 | 0 | 2403.156 | 0 | -0.11 | 0.768029 | 15.2 | 101.3 | 98.7 | 3.64 | 14.44 | | High | High | High | High | High | High | High | High | High | High | High | 0 | 2.24E-05 | 2.83 |
| FALSE | High | GPSPAPLH | [K].GPSPA | 1xPhospho | Q9QXD8 1 | 0.003433 | 0.00021 | 1 | 1 | 1 | Q9QXD8 | LIM domai | Q9QXD8 [6 | 0 | 1633.784 | 0 | 1.15 | 0.000434 | 8.4 | 62.7 | 137.3 | 2.64 | 5.28 | | High | High | High | High | High | High | High | High | High | High | High | 9.42E-05 | 0.000947 | 1.98 |
| FALSE | High | GPSPAPVD | [R].GPSPA | 1xPhospho | Q9DC40 1x | 4.19E-08 | 0 | 1 | 2 | 1 | Q9DC40 | Telomere l | Q9DC40 [4 | 0 | 2405.171 | 0 | -0.6 | 0.00125 | 2.38 | 120.4 | 79.6 | 6.98 | 2.5 | | High | High | High | High | High | High | High | High | High | High | High | 0 | 2.87E-09 | 3.92 |
| FALSE | High | GPSPEGSS | [R].GPSPE | 1xPhospho | Q8BTI8 1x | 0.001222 | 0 | 1 | 3 | 1 | Q8BTI8 | Serine/argi | Q8BTI8 [1 | 0 | 2162.944 | 0 | -0.64 | 0.005189 | 1.5 | 119.4 | 80.6 | 7.74 | 14.58 | | High | High | High | High | High | High | High | High | High | High | High | 0 | 0.000297 | 2.71 |
| FALSE | High | GPSPESVK | [R].GPSPES | 2xPhospho | A0A140T8 | 0.016892 | 0.001204 | 1 | 1 | 1 | A0A140T8 | Chromoso | A0A140T8 | 0 | 1418.673 | 0 | -1.01 | 0.000609 | 1.5 | 133.6 | 66.4 | 4.8 | 6.37 | | High | High | High | High | High | High | High | High | High | High | High | 0.000638 | 0.005618 | 1.75 |
| FALSE | High | GPSPGPEL | [R].GPSPG | 1xPhospho | Q8JZP9 1x | 1.45E-05 | 0 | 1 | 1 | 1 | Q8JZP9 | GAS2-like | Q8JZP9 [4 | 0 | 1977.004 | 0 | 0.74 | 0.000748 | 10.56 | 73.5 | 126.5 | 5.73 | 5.51 | | High | High | High | High | High | High | High | High | High | High | High | 0 | 2.05E-06 | 2.21 |
| FALSE | High | GQEDSLAS | [K].GQEDS | 1xCarbami | Q8R344 1x | 1.35E-10 | 0 | 1 | 1 | 10 | Q8R344 | Coiled-coil | Q8R344 [1 | 0 | 2536.023 | 0 | -0.28 | 0.001787 | 2.07 | 110.4 | 89.6 | 5.19 | 4.11 | | High | High | High | High | High | High | High | High | High | High | High | 0 | 4.57E-12 | 6.86 |
| FALSE | High | GGLTQPSI | [K].GGLTQ | 1xPhospho | A2A6Q5-3 | 0.001337 | 0 | 1 | 3 | 2 | A2A6Q5-3 | Isoform 3 c | A2A6Q5-3 | 0 | 2211.167 | 0 | 0.19 | 0.002031 | 3.77 | 93.8 | 106.2 | 12.15 | 8.74 | | High | High | High | High | High | High | High | High | High | High | High | 0 | 0.000329 | 2.98 |
| FALSE | High | GGLNTPLH | [K].GGLNT | 2xPhospho | A0A1W2P7 | 4.07E-07 | 0 | 1 | 6 | 2 | A0A1W2P7 | Uncharacte | A0A1W2P7 | 0 | 2401.079 | 0 | -0.64 | 0.006424 | 1.46 | 121.6 | 78.4 | 6.5 | 12.97 | | High | High | High | High | High | High | High | High | High | High | High | 0 | 3.69E-08 | 2.84 |
| FALSE | High | GGLNTPLH | [K].GGLNT | 1xPhospho | A0A1W2P7 | 2.93E-05 | 0 | 1 | 6 | 3 | A0A1W2P7 | Uncharacte | A0A1W2P7 | 0 | 2321.112 | 0 | -0.4 | 0.0535 | 0.82 | 114.1 | 85.9 | 8.05 | 10.21 | | High | High | High | High | High | High | High | High | High | High | High | 0 | 4.52E-06 | 3.13 |
| FALSE | High | FSVGSASP | [R].FSVGS | 1xPhospho | P98192 1x | 0.00172 | 5.66E-05 | 1 | 5 | 1 | P98192 | Dihydroxya | P98192 [1 | 0 | 2151.15 | 0 | -1.85 | 0.000209 | 0.42 | 156 | 44 | 4.52 | 16.16 | | High | High | High | High | High | High | High | High | High | High | High | 2.58E-05 | 0.000439 | 2.32 |
| FALSE | High | FSVQEQDV | [R].FSVQE | 1xPhospho | Q542P8 1x | 8.22E-07 | 0 | 1 | 2 | 7 | Q542P8 | EMG1 nucl | Q542P8 [1 | 0 | 2230.083 | 0 | -0.31 | 0.006444 | 6.47 | 115 | 85 | 5.91 | 7.67 | | High | High | High | High | High | High | High | High | High | High | High | 0 | 9.99E-07 | 3.27 |
| FALSE | High | FSVQEQDV | [R].FSVQE | 2xPhospho | Q542P8 2x | 6.67E-06 | 0 | 1 | 2 | 1 | Q542P8 | EMG1 nucl | Q542P8 [1 | 0 | 2310.05 | 0 | -0.46 | 0.003647 | 2.09 | 116.6 | 83.4 | 3.52 | 6.13 | | High | High | High | High | High | High | High | High | High | High | High | 0 | 8.56E-07 | 3.31 |
| FALSE | High | FSVQEQDV | [R].FSVQE | 1xPhospho | Q542P8 1x | 0.007605 | 0.000664 | 1 | 2 | 2 | Q542P8 | EMG1 nucl | Q542P8 [1 | 1 | 2587.341 | 0 | -0.84 | 0.004087 | 6.07 | 129 | 71 | 7.71 | 11.19 | | High | High | High | High | High | High | High | High | High | High | High | 0.000294 | 0.002307 | 2.01 |
| FALSE | High | FSVSPASPS | [R].FSVSPA | 1xPhospho | Q8BUR4 1x | 0.004411 | 0.000409 | 1 | 1 | 2 | Q8BUR4 | Dedicator | Q8BUR4 [1 | 0 | 2376.18 | 0 | 0.06 | 0.499929 | 22.15 | 92.2 | 107.8 | 16.84 | 9.57 | | High | High | High | High | High | High | High | High | High | High | High | 0.000183 | 0.001253 | 1.6 |
| FALSE | High | FSVSPVVR | [K].FSVSPV | 1xPhospho | P58252 1x | 0.012508 | 0.000889 | 1 | 16 | 1 | P58252 | Elongation | P58252 [4 | 0 | 1199.639 | 0 | -0.47 | 0.005517 | 6.01 | 116.4 | 83.6 | 4.93 | 9.65 | | High | High | High | High | High | High | High | High | High | High | High | 0.000449 | 0.004041 | 1.36 |
| FALSE | High | FSVSTPSQ | [K].FSVSTP | 1xCarbami | B9EK83 1x | 0.009416 | 0.000736 | 2 | 3 | 5 | B9EK83; A | Transient r | B9EK83 [1 | 0 | 1862.912 | 0 | 2.43 | 0.000291 | 10.84 | 30.1 | 169.9 | 8.67 | 7.43 | Shared | High | High | High | High | High | High | High | High | High | High | High | 0.000449 | 0.003844 | 2.33 |
| FALSE | High | FSWGQQR | [K].FSWGC | 1xPhospho | Q8C6M1 1 | 0.005924 | 0.00062 | 1 | 1 | 1 | Q8C6M1 | Ubiquitin c | Q8C6M1 [3 | 0 | 1217.567 | 0 | 0.74 | 0.003021 | 9.07 | 74.9 | 125.1 | 11.71 | 12 | | High | High | High | High | High | High | High | High | High | High | High | 0.000275 | 0.001747 | 2.21 |
| FALSE | High | FTASPLEEE | [R].FTASPL | 1xPhospho | Q9CY64 1x | 0.063788 | 0.004664 | 1 | 1 | 1 | Q9CY64 | Biliverdin r | Q9CY64 [1 | 0 | 1688.855 | 0 | 0.94 | 0.001864 | 2.57 | 68.5 | 131.5 | 8.96 | 8.14 | | High | High | High | High | High | High | High | High | High | High | High | 0.002403 | 0.02551 | 2.15 |
| FALSE | High | FTFPDPPPL | [R].FTFPDP | 1xPhospho | Q9JL47 1x | 0.000479 | 0 | 1 | 1 | 1 | Q9JL47 | ATP-bindin | Q9JL47 [35 | 0 | 3435.779 | 0 | 0.31 | 0.012326 | 10.76 | 88.8 | 111.2 | 9.16 | 4.26 | | High | High | High | High | High | High | High | High | High | High | High | 0 | 0.000104 | 3.95 |
| FALSE | High | FTGSFDDD | [R].FTGSFD | 1xPhospho | A2AIX1 1x | 4.36E-06 | 0 | 1 | 2 | 2 | A2AIX1 | SEC16 hon | A2AIX1 [1 | 0 | 1818.79 | 0 | 0.52 | 0.011615 | 17.54 | 82.8 | 117.2 | 6.44 | 9.38 | | High | High | High | High | High | High | High | High | High | High | High | 0 | 5.33E-07 | 3.2 |
| FALSE | High | FTGVATGS | [R].FTGVA | 1xCarbami | Q8C0V0 1x | 0.000605 | 0 | 1 | 1 | 2 | Q8C0V0 | Serine/thre | Q8C0V0 [6 | 0 | 2370.141 | 0 | -0.21 | 0.205777 | 8.28 | 105 | 95 | 8.83 | 11.15 | | High | High | High | High | High | High | High | High | High | High | High | 0 | 0.000135 | 2.8 |
| FALSE | High | FTGYQTIQ | [K].FTGYQ | 2xPhospho | Q9JMG7-2 | 0.000259 | 0 | 1 | 3 | 3 | Q9JMG7-2 | Isoform 2 c | Q9JMG7-2 | 1 | 4641.967 | 0 | 0.53 | 0.001633 | 9.14 | 81.3 | 118.7 | 6.16 | 8.19 | | High | High | High | High | High | High | High | High | High | High | High | 0 | 5.24E-05 | 4.77 |
| FALSE | High | FSVASSVV | [R].FSVASS | 1xPhospho | E9Q4F8 1x | 0.005978 | 0.00062 | 1 | 3 | 1 | E9Q4F8 | Ankyrin re | E9Q4F8 [1 | 0 | 3161.583 | 0 | -0.33 | 0.047966 | 9.33 | 111.5 | 88.5 | 10.3 | 7.54 | | High | High | High | High | High | High | High | High | High | High | High | 0.000275 | 0.001764 | 1.91 |
| FALSE | High | FTLPGDIDI | [R].FTLPGD | 1xPhospho | E9Q411 1x | 0.000548 | 0 | 1 | 2 | 1 | E9Q411 | Neuroblast | E9Q411 [6 | 0 | 2947.463 | 0 | 0.63 | 0.069853 | 20.52 | 81.8 | 118.2 | 15.71 | 12.05 | | High | High | High | High | High | High | High | High | High | High | High | 0 | 0.000122 | 3.17 |
| FALSE | High | FTLSGVDG | [K].FTLSGV | 1xPhospho | Q6P5H2-2 | 2.37E-07 | 0 | 1 | 3 | 4 | Q6P5H2-2 | Isoform 2 c | Q6P5H2-2 | 0 | 1967.811 | 0 | 2.39 | 0.039899 | 40.1 | 28.3 | 171.7 | 16.11 | 50.95 | | High | High | High | High | High | High | High | High | High | High | High | 0 | 2.01E-08 | 3.47 |
| FALSE | High | FTLSGVDG | [K].FTLSGV | 2xPhospho | Q6P5H2-2 | 1.5E-08 | 0 | 1 | 3 | 22 | Q6P5H2-2 | Isoform 2 c | Q6P5H2-2 | 0 | 2047.777 | 0 | 2.69 | 0.009857 | 35.99 | 23.7 | 176.3 | 18.22 | 33.1 | | High | High | High | High | High | High | High | High | High | High | High | 0 | 9.03E-10 | 4.05 |
| FALSE | High | FTPPSTALS | [R].FTPPST | 1xPhospho | Q03347-4 | 0.000568 | 0 | 1 | 10 | 1 | Q03347-4 | Isoform 4 c | Q03347-4 | 0 | 1740.934 | 0 | 3.44 | 0.000177 | 12.58 | 15.8 | 184.2 | 16.73 | 5.32 | | High | High | High | High | High | High | High | High | High | High | High | 0 | 0.000127 | 2.97 |
| FALSE | High | FTSSPGLST | [K].FTSSPG | 1xPhospho | B1AUH5 1x | 0.000725 | 0 | 1 | 1 | 5 | B1AUH5 | Peripheral | B1AUH5 [4 | 0 | 1748.887 | 0 | 3.78 | 0.000513 | 44.98 | 15.9 | 184.1 | 24.17 | 10.8 | | High | High | High | High | High | High | High | High | High | High | High | 0.004422 | 0.04778 | 2.48 |
| FALSE | High | FTVDLDSD | [K].FTVDLD | 2xPhospho | Q01320 2x | 0.057203 | 0.004227 | 2 | 4 | 1 | Q01320; Q | DNA topois | Q01320 [1 | 0 | 2320.935 | 0 | -0.37 | 0.175454 | 12.84 | 113.6 | 86.4 | 27.07 | 23.47 | Shared | High | High | High | High | High | High | High | High | High | High | High | 0.002159 | 0.02257 | 2.83 |
| FALSE | High | FTVDLDSD | [K].FTVDLD | 1xPhospho | Q01320 1x | 3.95E-09 | 0 | 2 | 4 | 12 | Q01320; Q | DNA topois | Q01320 [1 | 0 | 2470.131 | 0 | -0.91 | 0.000764 | 3.49 | 130.5 | 69.5 | 6.65 | 12.69 | Shared | High | High | High | High | High | High | High | High | High | High | High | 0 | 4E-09 | 4.09 |
| FALSE | High | FTVDLDSD | [K].FTVDLD | 2xPhospho | Q01320 2x | 2E-08 | 0 | 2 | 4 | 25 | Q01320; Q | DNA topois | Q01320 [1 | 0 | 2550.098 | 0 | -0.92 | 0.000261 | 1.22 | 127.7 | 72.3 | 4.2 | 5.07 | Shared | High | High | High | High | High | High | High | High | High | High | High | 0 | 8.05E-09 | 4.12 |
| FALSE | High | FTVQMPAS | [R].FTVQM | 1xPhospho | Q8VI33 1x | 6.55E-06 | 0 | 1 | 1 | 1 | Q8VI33 | Transcripti | Q8VI33 [1 | 0 | 2045.054 | 0 | 0.01 | 0.810501 | 19.74 | 99.9 | 100.1 | 16.43 | 7.47 | | High | High | High | High | High | High | High | High | High | High | High | 0 | 8.36E-07 | 4.01 |
| FALSE | High | FVETSPSGL | [K].FVETSP | 1xPhospho | Q7TQE6 1x | 2.8E-06 | 0 | 1 | 3 | 2 | Q7TQE6 | Macoilin O | Q7TQE6 [6 | 0 | 2687.373 | 0 | 0.06 | 0.451407 | 2.6 | 101 | 99 | 3.77 | 7.32 | | High | High | High | High | High | High | High | High | High | High | High | 0 | 3.23E-07 | 3.4 |
| FALSE | High | FVQDHFDH | [R].FVQDH | 1xPhospho | Q3TXV4 1x | 0.069903 | 0.005181 | 1 | 2 | 1 | Q3TXV4 | Putative ur | Q3TXV4 [2 | 0 | 2930.431 | 0 | 2.18 | 0.001183 | 15.86 | 36.6 | 163.4 | 21.12 | 15.83 | | High | High | High | High | High | High | High | High | High | High | High | 0.002662 | 0.02829 | 1.8 |
| FALSE | High | FVQDHFDH | [R].FVQDH | 1xPhospho | Q3TXV4 1x | 0.008614 | 0.000736 | 1 | 2 | 1 | Q3TXV4 | Putative ur | Q3TXV4 [2 | 0 | 2946.426 | 0 | 2.07 | 0.001644 | 22.42 | 36.9 | 163.1 | 12.73 | 16.25 | | High | High | High | High | High | High | High | High | High | High | High | 0.000327 | 0.002662 | 3.58 |
| FALSE | High | FVQWQSSI | [R].FVQWC | 1xPhospho | Q9D3A9 1x | 0.007209 | 0.00062 | 1 | 3 | 1 | Q9D3A9 | Protein tw | Q9D3A9 [4 | 0 | 1303.629 | 0 | -1 | 0.006049 | 5.34 | 135 | 65 | 14.78 | 8.29 | | High | High | High | High | High | High | High | High | High | High | High | 0.000275 | 0.002185 | 2.39 |
| FALSE | High | FVSEGDGG | [R].FVSEGD | 1xPhospho | Q61823 1x | 0.001049 | 0 | 1 | 1 | 4 | Q61823 | Programm | Q61823 [4 | 0 | 1232.551 | 0 | -1.77 | 0.000749 | 0.56 | 153.6 | 46.4 | 5.33 | 16.52 | | High | High | High | High | High | High | High | High | High | High | High | 0.000275 | 0.001582 | 1.36 |
| FALSE | High | FTIQLQLSC | [K].FTIQLQ | 1xPhospho | Q6A065 1x | 0.000699 | 0 | 1 | 3 | 1 | Q6A065 | Centrosom | Q6A065 [1 | 0 | 1743.981 | 0 | 1.58 | 0.000591 | 10.52 | 49.3 | 150.7 | 11.67 | 10.73 | | High | High | High | High | High | High | High | High | High | High | High | 0 | 0.00016 | 2.12 |
| FALSE | High | FVSVVPQY | [K].FVSVVF | 1xPhospho | Q8C4D5 1x | 0.008847 | 0.000736 | 1 | 2 | 2 | Q8C4D5 | Putative ur | Q8C4D5 [1 | 0 | 2204.095 | 0 | 0.1 | 0.124689 | 5.61 | 93 | 107 | 11.86 | 11.08 | | High | High | High | High | High | High | High | High | High | High | High | 0.000467 | 0.004232 | 2.95 |
| FALSE | High | FSTYSQSPF | [R].FSTYSQ | 1xPhospho | Q68FE6 1x | 0.000253 | 0 | 1 | 1 | 2 | Q68FE6 | Rho family | Q68FE6 [3 | 0 | 1991.931 | 0 | 0.44 | 0.004161 | 8.81 | 83.4 | 116.6 | 10.17 | 4.01 | | High | High | High | High | High | High | High | High | High | High | High | 0 | 5.06E-05 | 2.57 |
| FALSE | High | FSTPDLALN | [K].FSTPDL | 1xPhospho | Q7TSZ8 1x | 0.001011 | 0 | 1 | 1 | 4 | Q7TSZ8 | Nucleus ac | Q7TSZ8 [2 | 0 | 1442.724 | 0 | -0.95 | 0.000697 | 2.48 | 131.9 | 68.1 | 5.87 | 11.29 | | High | High | High | High | High | High | High | High | High | High | High | 0.000275 | 0.0018 | 2.51 |
| FALSE | High | FSLGPLSPE | [R].FSLGPL | 2xPhospho | Q9D0P7 2x | 0.009585 | 0.000736 | 1 | 3 | 1 | Q9D0P7 | Proline/ser | Q9D0P7 [6 | 0 | 1692.842 | 0 | -0.69 | 0.011261 | 3.17 | 120.8 | 79.2 | 15.11 | 4.46 | | High | High | High | High | High | High | High | High | High | High | High | 0.000344 | 0.002988 | 2.6 |
| FALSE | High | FSLSDDQQ | [K].FSLSDD | 1xPhospho | E9Q4F8 1x | 0.005 | 0.000504 | 1 | 2 | 3 | E9Q4F8 | Ankyrin re | E9Q4F8 [8 | 0 | 1533.678 | 0 | -0.78 | 0.000619 | 2.53 | 126.7 | 73.3 | 5.42 | 3.14 | | High | High | High | High | High | High | High | High | High | High | High | 0.000327 | 0.002418 | 2.59 |
| FALSE | High | FSPADWEF | [K].FSPADV | 1xPhospho | Q8CGN4 1 | 0.110202 | 0.008961 | 1 | 5 | 1 | Q8CGN4 | BCL-6 core | Q8CGN4 [1 | 0 | 1316.587 | 0 | 0.77 | 0.003806 | 6.42 | 77.1 | 122.9 | 12.75 | 7.15 | | High | High | High | High | High | High | High | High | High | High | High | 0.004462 | 0.04837 | 2.02 |
| FALSE | High | FSPAGLQH | [R].FSPAGL | 1xPhospho | E9Q309 1x | 0.04913 | 0.003576 | 1 | 1 | 1 | E9Q309 | Centrosom | E9Q309 [1 | 0 | 1321.662 | 0 | -0.54 | 0.004498 | 5.5 | 114.1 | 85.9 | 16.14 | 19.03 | | High | High | High | High | High | High | High | High | High | High | High | 0.001889 | 0.01881 | 2.35 |
| FALSE | High | FSPDDK | [R].FSPDDK | 1xPhospho | Q9CQS2 1x | 0.083101 | 0.006345 | 1 | 1 | 1 | Q9CQS2 | H/ACA ribc | Q9CQS2 [3 | 0 | 1246.612 | 0 | -1.31 | 0.000429 | 1.11 | 142.2 | 57.8 | 4.53 | 7.2 | | High | High | High | High | High | High | High | High | High | High | High | 0.003265 | 0.03477 | 1.77 |
| FALSE | High | FSPDSHCV | [R].FSPDSH | 1xCarbami | A5D6P2 1x | 9.98E-05 | 0 | 1 | 23 | 1 | A5D6P2 | Par-3 (Part | A5D6P2 [3 | 0 | 1598.698 | 0 | -0.41 | 0.008293 | 3.32 | 114 | 86 | 8.75 | 1.15 | | High | High | High | High | High | High | High | High | High | High | High | 0 | 1.79E-05 | 2.85 |
| FALSE | High | FSPNSLVAL | [R].FSPNSL | 1xPhospho | Q6ZPF1 1x | 0.000337 | 0 | 1 | 4 | 1 | Q6ZPF1 | MKIAA301 | Q6ZPF1 [4 | 0 | 2100.099 | 0 | -0.12 | 0.352892 | 14.67 | 108.4 | 91.6 | 15.14 | 6.72 | | High | High | High | High | High | High | High | High | High | High | High | 0 | 7.06E-05 | 3.73 |
| FALSE | High | FSPNSSNPI | [R].FSPNSS | 1xCarbami | P68040 1x | 6.61E-06 | 0 | 1 | 1 | 2 | P68040 | Receptor o | P68040 [1 | 0 | 2446.188 | 0 | -0.09 | 0.961905 | 9.17 | 99 | 101 | 10.04 | 10.07 | | High | High | High | High | High | High | High | High | High | High | High | 0 | 1.61E-06 | 3.43 |
| FALSE | High | FSPPPVGG | [R].FSPPPV | 1xPhospho | A2A690-2 | 5.81E-05 | 0 | 1 | 3 | 1 | A2A690-2 | Isoform 2 c | A2A690-2 | 0 | 1638.866 | 0 | 1.55 | 0.000154 | 7.95 | 51.1 | 148.9 | 17.95 | 4.18 | | High | High | High | High | High | High | High | High | High | High | High | 0 | 9.75E-06 | 3.03 |
| FALSE | High | FSPPSPLPQ | [K].FSPPSP | 1xPhospho | Q8VCA7 1x | 0.013545 | 0.000976 | 1 | 5 | 2 | Q8VCA7 | Tead3 prot | Q8VCA7 [1 | 0 | 2100.036 | 0 | 0.82 | 0.001325 | 13.36 | 71.7 | 128.3 | 8.84 | 9.51 | | High | High | High | High | High | High | High | High | High | High | High | 0.000505 | 0.004391 | 2.1 |
| FALSE | High | FSPQNYTR | [K].FSPQN | 1xPhospho | Q8CBD1 1x | 0.032267 | 0.002129 | 1 | 1 | 1 | Q8CBD1 | Nuclear re | Q8CBD1 [5 | 0 | 1321.614 | 0 | 0.97 | 0.005028 | 19.83 | 66.6 | 133.4 | 9.68 | 13.94 | | High | High | High | High | High | High | High | High | High | High | High | 0.001142 | 0.01172 | 1.83 |
| FALSE | High | FSPQPQGC | [K].FSPQP | 1xPhospho | Q7TSH3 1x | 0.004955 | 0.000504 | 1 | 1 | 1 | Q7TSH3 | Zinc finger | Q7TSH3 [9 | 0 | 1861.998 | 0 | 0.16 | 0.79128 | 23.22 | 93.6 | 106.4 | 19.17 | 5.22 | | High | High | High | High | High | High | High | High | High | High | High | 0.000225 | 0.001437 | 1.72 |
| FALSE | High | FSPTMGR | [R].FSPTM | 1xPhospho | A0JNY7 1x | 0.022773 | 0.001542 | 1 | 7 | 4 | A0JNY7 | Eukaryotic | A0JNY7 [3 | 0 | 1120.506 | 0 | 0.31 | 0.073896 | 5.96 | 89.3 | 110.7 | 7.25 | 23.78 | | High | High | High | High | High | High | High | High | High | High | High | 0.008139 | 0.08784 | 2.04 |
| FALSE | High | FSTPEQAA | [R].FSTPEC | 1xPhospho | G5E924 1x | 0.011447 | 0.000816 | 1 | 4 | 2 | G5E924 | Heterogen | G5E924 [5 | 0 | 1516.781 | 0 | -0.67 | 0.000235 | 1.23 | 123.5 | 76.5 | 6.32 | 7.41 | | High | High | High | High | High | High | High | High | High | High | High | 0.000437 | 0.003658 | 1.98 |
| FALSE | High | FSPTQDRP | [K].FSPTQD | 1xPhospho | Q8BTI8 1x | 0.041425 | 0.003126 | 1 | 3 | 1 | Q8BTI8 | Serine/argi | Q8BTI8 [1 | 0 | 2000.989 | 0 | -0.23 | 0.599727 | 31.1 | 117 | 83 | 77.28 | 60.22 | | High | High | High | High | High | High | High | High | High | High | High | 0.001609 | 0.01552 | 1.75 |
| FALSE | High | FSQEEPVP | [R].FSQEEF | 1xPhospho | Q571H2 1x | 0.025738 | 0.001645 | 1 | 5 | 1 | Q571H2 | MKIAA021 | Q571H2 [3 | 0 | 1711.907 | 0 | -0.24 | 0.576405 | 18.61 | 105.5 | 94.5 | 6.65 | 21.44 | | High | High | High | High | High | High | High | High | High | High | High | 0.000872 | 0.009049 | 2.07 |
| FALSE | High | FSQGPNNI | [K].FSQGPI | 1xPhospho | E9PYK3 1x | 8.74E-06 | 0 | 1 | 2 | 1 | E9PYK3 | Poly [ADP- | E9PYK3 [1 | 0 | 1960.993 | 0 | 2.94 | 0.00022 | 13.97 | 22.9 | 177.1 | 13.13 | 7.02 | | High | High | High | High | High | High | High | High | High | High | High | 0 | 1.17E-06 | 3.13 |
| FALSE | High | FSREEFPTL | [R].FSREEF | 1xPhospho | Q7TSC1 1x | 1.4E-07 | 0 | 1 | 3 | 1 | Q7TSC1 | Protein PRI | Q7TSC1 [1 | 1 | 2477.211 | 0 | 1.98 | 0.001353 | 11.73 | 36.5 | 163.5 | 10.17 | 14.47 | | High | High | High | High | High | High | High | High | High | High | High | 0 | 1.12E-08 | 4.35 |
| FALSE | High | FSSASSPRP | [R].FSSASS | 1xPhospho | A2A7S8-2 | 0.001002 | 0 | 1 | 4 | 2 | A2A7S8-2 | Isoform 2 c | A2A7S8-2 | 0 | 1400.688 | 0 | -0.65 | 0.005078 | 4.05 | 119.3 | 80.7 | 8.2 | 16.27 | | High | High | High | High | High | High | High | High | High | High | High | 0 | 0.000239 | 2.16 |
| FALSE | High | FSSDPPVSF | [K].FSSDPP | 1xPhospho | Q0VBM1 1 | 0.062731 | 0.004664 | 1 | 5 | 1 | Q0VBM1 | AI894139 | Q0VBM1 [ | 0 | 1798.939 | 0 | -0.2 | 0.274268 | 8.84 | 106.4 | 93.6 | 13.36 | 7.1 | | High | High | High | High | High | High | High | High | High | High | High | 0.002403 | 0.02497 | 2.21 |
| FALSE | High | FSSGGACG | [R].FSSGG | 1xCarbami | Q61781 1x | 2.13E-12 | 0 | 1 | 1 | 3 | Q61781 | Keratin, ty | Q61781 [5 | 0 | 3244.399 | 0 | 3.99 | | 89.12 | 18.6 | 181.4 | 17.98 | 25.63 | | High | High | High | High | High | High | High | High | High | High | High | 0 | 2.07E-12 | 6.11 |
| FALSE | High | FSSLDLEED | [R].FSSLDL | 1xPhospho | Q8R1G6 1x | 0.009168 | 0.000736 | 1 | 1 | 2 | Q8R1G6 | PDZ and LI | Q8R1G6 [2 | 0 | 2183.056 | 0 | 0.39 | 0.095958 | 36.82 | 85.8 | 114.2 | 23.22 | 25.77 | | High | High | High | High | High | High | High | High | High | High | High | 0.000327 | 0.002843 | 2.4 |
| FALSE | High | FSSLTETLH | [R].FSSLTE | 1xPhospho | Q99LE3 1x | 0.066503 | 0.004796 | 1 | 1 | 2 | Q99LE3 | LysM and | Q99LE3 [1 | 0 | 1911.039 | 0 | 0.8 | 0.00584 | 8.45 | 70.4 | 129.6 | 13.32 | 17.53 | | High | High | High | High | High | High | High | High | High | High | High | 0.003754 | 0.04049 | 1.59 |
| FALSE | High | FSSPGLENI | [K].FSSPGL | 1xPhospho | Q6A027 1x | 2.98E-05 | 0 | 1 | 5 | 2 | Q6A027 | MKIAA064 | Q6A027 [1 | 0 | 2185.038 | 0 | -0.29 | 0.105287 | 18.78 | 110.1 | 89.9 | 5.46 | 16.97 | | High | High | High | High | High | High | High | High | High | High | High | 0 | 1.39E-05 | 2.99 |
| FALSE | High | FSSSQPEPF | [K].FSSSQP | 1xPhospho | O35280 1x | 0.014929 | 0.001049 | 1 | 4 | 1 | O35280 | Serine/thre | O35280 [3 | 0 | 1343.619 | 0 | -1.65 | 0.010571 | 3.28 | 151.5 | 48.5 | 20.75 | 17.39 | | High | High | High | High | High | High | High | High | High | High | High | 0.000555 | 0.004899 | 1.36 |
| FALSE | High | FSSSTEQSS | [R].FSSSTE | 1xPhospho | I6L9H1 1xP | 0.053023 | 0.00384 | 1 | 6 | 2 | I6L9H1 | Dishevelle | I6L9H1 [22 | 0 | 1582.695 | 0 | -0.35 | 0.011966 | 8.46 | 111.6 | 88.4 | 3.81 | 7.89 | | High | High | High | High | High | High | High | High | High | High | High | 0.001999 | 0.0206 | 1.66 |

| | | | | | | | | | | | | | | | | | | | | | | | | | | | | | | | | | | | | | | |
|---|---|---|---|---|---|---|---|---|---|---|---|---|---|---|---|---|---|---|---|---|---|---|---|---|---|---|---|---|---|---|---|---|---|---|---|---|---|---|
| FALSE | High | FSSTSSDK | [K].FSSTSSI | 1xPhosphc | V9GX40 1x | 0.051258 | 0.003745 | 1 | 4 | 1 | V9GX40 | Synergin g | V9GX40 [8 | 0 | 1396.676 | 0 | 0.75 | 0.009984 | 18.03 | 73.4 | 126.6 | 4.87 | 16.73 | | High | High | High | High | High | High | High | High | High | High | High | 0.00196 | 0.01993 | 1.42 |
| FALSE | High | FSTLTDEPS | [R].FSTLTD | 1xPhosphc | Q6PAR5 1x | 0.011549 | 0.00086 | 1 | 6 | 1 | Q6PAR5 | GTPase-act | Q6PAR5 [1 | 0 | 1558.735 | 0 | 0.73 | 0.000207 | 4.58 | 74.5 | 125.5 | 1.47 | 2.64 | | High | High | High | High | High | High | High | High | High | High | High | 0.000437 | 0.003688 | 1.57 |
| FALSE | High | FSPTQDRPI | [K].FSPTQD | 1xPhosphc | Q8BTI8 1xP | 0.014283 | 0.001011 | 1 | 3 | 4 | Q8BTI8 | Serine/argi | Q8BTI8 [11 | 0 | 2230.152 | 0 | -0.67 | 0.002692 | 5.43 | 120.4 | 79.6 | 8.62 | 4.73 | | High | High | High | High | High | High | High | High | High | High | High | 0.000859 | 0.008306 | 1.8 |
| FALSE | High | FVTSSSGSY | [R].FVTSSS | 1xPhosphc | P19001 1xP | 0.000738 | 0 | 1 | 1 | 10 | P19001 | Keratin, typ | P19001 [52 | 0 | 1612.757 | 0 | 3.58 | 0.000953 | 5.19 | 14.5 | 185.5 | 15.2 | 15.72 | | High | High | High | High | High | High | High | High | High | High | High | 0 | 0.00017 | 2.32 |
| FALSE | High | FVVIQNED | [R].FVVIQN | 1xPhosphc | P70424 1xP | 0.000257 | 0 | 1 | 3 | 2 | P70424 | Receptor t | P70424 [98 | 0 | 2711.263 | 0 | 0.49 | 0.007885 | 13.36 | 82 | 118 | 18.87 | 12.9 | | High | High | High | High | High | High | High | High | High | High | High | 0 | 0.000118 | 3.45 |
| FALSE | High | FYCDYCDT | [K].FYCDYC | 2xCarbami | Q569X3 1x | 6E-07 | 0 | 1 | 1 | 4 | Q569X3 | U1 small nu | Q569X3 [4 | 0 | 2608.072 | 0 | 0.08 | 0.634499 | 17.73 | 97.3 | 102.7 | 5.64 | 11.63 | | High | High | High | High | High | High | High | High | High | High | High | 0 | 5.7E-08 | 4.39 |
| FALSE | High | GAGGVESC | [K].GAGGV | 1xCarbami | Q3TYU7 1x | 0.002398 | 0.000109 | 1 | 3 | 1 | Q3TYU7 | Putative ur | Q3TYU7 [7 | 1 | 2463.122 | 0 | -0.27 | 0.109367 | 11.41 | 109.8 | 90.2 | 13.35 | 5.77 | | High | High | High | High | High | High | High | High | High | High | High | 4.94E-05 | 0.000635 | 2.59 |
| FALSE | High | GAGGYTCC | [K].GAGGY | 1xCarbami | E9Q557 1x | 2.25E-11 | 0 | 1 | 2 | 1 | E9Q557 | Desmoplak | E9Q557 [1 | 0 | 2446.079 | 0 | 3.03 | 0.001318 | 28.89 | 19.9 | 180.1 | 12.51 | 17.11 | | High | High | High | High | High | High | High | High | High | High | High | 0 | 6.1E-13 | 5.57 |
| FALSE | High | GAGLGAQT | [K].GAGLG | 1xPhosphc | F8VPZ9 1xP | 0.003622 | 0.00021 | 1 | 1 | 1 | F8VPZ9 | BRD4-inter | F8VPZ9 [74 | 0 | 1438.689 | 0 | 0.28 | 0.084024 | 19.35 | 90.8 | 109.2 | 12.31 | 5.64 | | High | High | High | High | High | High | High | High | High | High | High | 9.42E-05 | 0.00101 | 1.47 |
| FALSE | High | GAGSPPPA | [R].GAGSPI | 1xPhosphc | Q8CHQ0 1 | 0.000808 | 0 | 1 | 1 | 1 | Q8CHQ0 | F-box only | Q8CHQ0 [8 | 0 | 1563.715 | 0 | 2.38 | 0.000222 | 8.19 | 33.4 | 166.6 | 5.76 | 5.92 | | High | High | High | High | High | High | High | High | High | High | High | 0 | 0.000187 | 1.71 |
| FALSE | High | GAGSPTPLI | [R].GAGSPT | 1xPhosphc | F8VPZ9 1xP | 0.012845 | 0.000889 | 1 | 2 | 2 | F8VPZ9 | BRD4-inter | F8VPZ9 [14 | 0 | 1563.855 | 0 | -0.09 | 0.211805 | 5 | 101.7 | 98.3 | 2.15 | 4.84 | | High | High | High | High | High | High | High | High | High | High | High | 0.000449 | 0.004151 | 1.89 |
| FALSE | High | GAGWLSEI | [R].GAGWI | 1xPhosphc | A0A0N4SU | 9.14E-06 | 0 | 1 | 11 | 1 | A0A0N4SU | Peregrin O | A0A0N4SU | 0 | 2925.432 | 0 | -0.29 | 0.157647 | 7.04 | 108.7 | 91.3 | 7.25 | 9.42 | | High | High | High | High | High | High | High | High | High | High | High | 0 | 1.22E-06 | 3.59 |
| FALSE | High | GAHPGTPG | [K].GAHPG | 1xPhosphc | H3BL37 1xP | 0.01588 | 0.001124 | 1 | 7 | 3 | H3BL37 | Treacle pro | H3BL37 [79 | 0 | 1359.719 | 0 | -0.25 | 0.050995 | 4.76 | 108.6 | 91.4 | 6.68 | 6.7 | | High | High | High | High | High | High | High | High | High | High | High | 0.000585 | 0.005255 | 2.23 |
| FALSE | High | GAHPSGGA | [K].GAHPS | 1xPhosphc | Q8BMK4 1 | 2.85E-05 | 0 | 1 | 1 | 2 | Q8BMK4 | Cytoskelet | Q8BMK4 [2 | 0 | 1719.847 | 0 | -1.72 | 0.000483 | 1.04 | 151.2 | 48.8 | 6.51 | 9.63 | | High | High | High | High | High | High | High | High | High | High | High | 0 | 4.39E-06 | 2.75 |
| FALSE | High | GAHSPQVF | [K].GAHSPQ | 1xPhosphc | Q8C0T5 1x | 0.002793 | 0.000109 | 1 | 3 | 3 | Q8C0T5 | Signal-indu | Q8C0T5 [1 | 0 | 2233.063 | 0 | 2 | 0.002796 | 35.92 | 42.3 | 157.7 | 53.5 | 18.05 | | High | High | High | High | High | High | High | High | High | High | High | 4.94E-05 | 0.000756 | 3.25 |
| FALSE | High | GAHTTVSC | [R].GAHTT | 1xPhosphc | Q9ESJ4 1xP | 1.08E-05 | 0 | 1 | 1 | 1 | Q9ESJ4 | NCK-intera | Q9ESJ4 [24 | 0 | 2060.058 | 0 | -0.7 | 0.087214 | 12.92 | 128.5 | 71.5 | 19 | 21.33 | | High | High | High | High | High | High | High | High | High | High | High | 0 | 1.48E-06 | 2.63 |
| FALSE | High | GAILSEEEL | [K].GAILSE | 1xPhosphc | F6ZDS4 1xP | 1.22E-05 | 0 | 1 | 8 | 3 | F6ZDS4 | Nucleopro | F6ZDS4 [44 | 0 | 2568.34 | 0 | -0.24 | 0.030586 | 6.05 | 108.8 | 91.2 | 3.56 | 12.19 | | High | High | High | High | High | High | High | High | High | High | High | 0 | 1.69E-06 | 4.28 |
| FALSE | High | GAILSEEEL | [K].GAILSE | 1xPhosphc | F6ZDS4 1xP | 2.06E-06 | 0 | 1 | 8 | 14 | F6ZDS4 | Nucleopro | F6ZDS4 [44 | 0 | 2584.335 | 0 | 0.01 | 0.239874 | 2.74 | 98.4 | 101.6 | 2.46 | 3.59 | | High | High | High | High | High | High | High | High | High | High | High | 0 | 1.62E-06 | 4.5 |
| FALSE | High | GALSPITDT | [K].GALSPI | 1xPhosphc | H7BX08 1x | 0.040028 | 0.002867 | 1 | 5 | 3 | H7BX08 | Calmodulin | H7BX08 [5 | 0 | 3936.815 | 0 | -0.45 | 0.063249 | 9.88 | 116.1 | 83.9 | 16.41 | 13.91 | | High | High | High | High | High | High | High | High | High | High | High | 0.004437 | 0.04793 | 3.85 |
| FALSE | High | GAGGPASA | [R].GAGGP | 2xPhosphc | Q5SWW4 2 | 0.01588 | 0.001124 | 1 | 1 | 1 | Q5SWW4 | Mediator c | Q5SWW4 [ | 0 | 1704.812 | 0 | 1.47 | 0.001434 | 21.47 | 51.5 | 148.5 | 17.14 | 14 | | High | High | High | High | High | High | High | High | High | High | High | 0.000585 | 0.005267 | 1.76 |
| FALSE | High | GAMPPAPV | [R].GAMPF | 1xPhosphc | Q3TMM5 1 | 0.006195 | 0.00062 | 2 | 6 | 4 | Q3TMM5; | Putative ur | Q3TMM5 [ | 0 | 3802.846 | 0 | 0.2 | 0.097469 | 11.87 | 89.9 | 110.1 | 13.85 | 13.52 | Shared | High | High | High | High | High | High | High | High | High | High | High | 0.000327 | 0.002432 | 2.88 |
| FALSE | High | GAPAQDPV | [R].GAPAQ | 1xPhosphc | Q9D7E4 1x | 5.5E-05 | 0 | 1 | 1 | 2 | Q9D7E4 | UPF0449 p | Q9D7E4 [2 | 0 | 2654.27 | 0 | 0.49 | 0.032136 | 9.06 | 69.9 | 130.1 | 24.98 | 25.67 | | High | High | High | High | High | High | High | High | High | High | High | 0 | 9.18E-06 | 3.92 |
| FALSE | High | GAPAVTPG | [K].GAPAV | 1xPhosphc | H3BL37 1xP | 0.000492 | 0 | 1 | 7 | 1 | H3BL37 | Treacle pro | H3BL37 [39 | 0 | 1335.744 | 0 | -0.84 | 0.000879 | 2.42 | 129.5 | 70.5 | 4.26 | 6.61 | | High | High | High | High | High | High | High | High | High | High | High | 0 | 0.000108 | 1.53 |
| FALSE | High | GAPEDTEE | [R].GAPED | 1xPhosphc | E9PZ97 1xP | 3.56E-08 | 0 | 1 | 3 | 3 | E9PZ97 | Bloom synd | E9PZ97 [13 | 0 | 3131.332 | 0 | -1.13 | 0.000245 | 0.81 | 136.5 | 63.5 | 5.88 | 4.27 | | High | High | High | High | High | High | High | High | High | High | High | 0 | 2.4E-09 | 4.74 |
| FALSE | High | GAPHTVSH | [K].GAPHT | 1xPhosphc | B8QI33 1xP | 5.03E-05 | 0 | 1 | 4 | 2 | B8QI33 | Liprin-alph | B8QI33 [79 | 0 | 1627.779 | 0 | -0.68 | 0.001543 | 6.91 | 123.3 | 76.7 | 8.02 | 20.15 | | High | High | High | High | High | High | High | High | High | High | High | 0 | 8.31E-06 | 2.73 |
| FALSE | High | GAPQLSPE | [R].GAPQL | 1xPhosphc | A2BFB0 1xP | 8.48E-05 | 0 | 1 | 4 | 3 | A2BFB0 | MAX-like p | A2BFB0 [4 | 0 | 1888.973 | 0 | 0 | 0.985067 | 12.26 | 101.5 | 98.5 | 4.73 | 7.97 | | High | High | High | High | High | High | High | High | High | High | High | 0.006741 | 0.07301 | 3.01 |
| FALSE | High | GAPQLSPE | [R].GAPQL | 2xPhosphc | A2BFB0 2xP | 6.59E-05 | 0 | 1 | 4 | 1 | A2BFB0 | MAX-like p | A2BFB0 [4 | 0 | 1968.939 | 0 | 0.66 | 0.004921 | 16.76 | 77.6 | 122.4 | 7.69 | 7.96 | | High | High | High | High | High | High | High | High | High | High | High | 0 | 1.13E-05 | 3.92 |
| FALSE | High | GAPSPGVL | [R].GAPSP | 1xPhosphc | Q8BTW9 1 | 0.000297 | 0 | 1 | 2 | 3 | Q8BTW9 | Serine/thre | Q8BTW9 [2 | 0 | 2415.238 | 0 | 0.23 | 0.021136 | 7.31 | 92.1 | 107.9 | 4.34 | 6.12 | | High | High | High | High | High | High | High | High | High | High | High | 0 | 6.12E-05 | 2.51 |
| FALSE | High | GAPSSPAA | [R].GAPSSF | 1xPhosphc | Q6ZQH8 1x | 3.27E-06 | 0 | 1 | 1 | 3 | Q6ZQH8 | Nucleopor | Q6ZQH8 [1 | 0 | 2059.099 | 0 | -0.12 | 0.102983 | 8.97 | 104.7 | 95.3 | 2.21 | 5.96 | | High | High | High | High | High | High | High | High | High | High | High | 0 | 3.83E-07 | 3.9 |
| FALSE | High | GAPSSPAA | [R].GAPSSF | 2xPhosphc | Q6ZQH8 2x | 2.7E-06 | 0 | 1 | 1 | 1 | Q6ZQH8 | Nucleopor | Q6ZQH8 [1 | 0 | 2139.065 | 0 | -0.33 | 0.025014 | 6.57 | 109 | 91 | 4.88 | 10.57 | | High | High | High | High | High | High | High | High | High | High | High | 0 | 3.11E-07 | 3.09 |
| FALSE | High | GAPVNVSS | [R].GAPVN | 1xPhosphc | Q3USK2 1x | 0.000136 | 0 | 1 | 3 | 4 | Q3USK2 | Casein kina | Q3USK2 [3 | 0 | 1668.816 | 0 | 0.06 | 0.266401 | 8.69 | 94.8 | 105.2 | 10.14 | 5.83 | | High | High | High | High | High | High | High | High | High | High | High | 0 | 2.56E-05 | 2.24 |
| FALSE | High | GAQADHFT | [R].GAQAD | 1xPhosphc | A2A4P4 1x | 2.14E-06 | 0 | 1 | 3 | 4 | A2A4P4 | Uncharacte | A2A4P4 [2 | 0 | 2520.208 | 0 | 0.46 | 0.057517 | 9.68 | 83.5 | 116.5 | 10.44 | 17 | | High | High | High | High | High | High | High | High | High | High | High | 0 | 2.39E-07 | 5.7 |
| FALSE | High | GASAASSP | [R].GASAA | 1xPhosphc | Q80ZM5 1 | 0.000573 | 0 | 1 | 1 | 1 | Q80ZM5 | H1 histone | Q80ZM5 [1 | 0 | 1481.777 | 0 | -0.07 | 0.518829 | 11.02 | 102.3 | 97.7 | 8.97 | 2.1 | | High | High | High | High | High | High | High | High | High | High | High | 0 | 0.000127 | 1.72 |
| FALSE | High | GASAATGII | [K].GASAA | 1xCarbami | Q8R5H1 1x | 0.000411 | 0 | 2 | 4 | 3 | Q8R5H1; C | Ubiquitin c | Q8R5H1 [9 | 0 | 3758.49 | 0 | 0.03 | 0.710222 | 14.81 | 99.1 | 100.9 | 4.01 | 9.81 | Shared | High | High | High | High | High | High | High | High | High | High | High | 0 | 8.8E-05 | 4.72 |
| FALSE | High | GAPANVSS | [R].GAPAN | 1xPhosphc | Q9JMK2 1x | 0.002192 | 0.000109 | 1 | 3 | 2 | Q9JMK2 | Casein kina | Q9JMK2 [3 | 0 | 1640.784 | 0 | 0.45 | 0.093713 | 27.78 | 84.7 | 115.3 | 10.95 | 19.39 | | High | High | High | High | High | High | High | High | High | High | High | 4.94E-05 | 0.000574 | 1.31 |
| FALSE | High | GAGGPASA | [R].GAGGP | 1xPhosphc | Q5SWW4 1 | 0.024211 | 0.001612 | 1 | 1 | 1 | Q5SWW4 | Mediator c | Q5SWW4 [ | 0 | 1624.846 | 0 | 1.4 | 0.000969 | 2.04 | 54.3 | 145.7 | 10.95 | 12.88 | | High | High | High | High | High | High | High | High | High | High | High | 0.000859 | 0.008464 | 1.49 |
| FALSE | High | GAGFPSTP | [K].GAGFP | 1xPhosphc | Q8BM41 1 | 0.007082 | 0.00062 | 1 | 1 | 1 | Q8BM41 | Protein LSM | Q8BM41 [6 | 0 | 1555.829 | 0 | 0.69 | 0.01097 | 20.34 | 80.3 | 119.7 | 6.91 | 14.56 | | High | High | High | High | High | High | High | High | High | High | High | 0.000275 | 0.002128 | 1.97 |
| FALSE | High | GAGAGASS | [R].GAGAG | 1xCarbami | A0A068BF | 7.41E-12 | 0 | 1 | 3 | 1 | A0A068BF | Ring finger | A0A068BF | 0 | 2790.158 | 0 | 0.12 | 0.05 | 5.87 | 94.9 | 105.1 | 2.02 | 8.29 | | High | High | High | High | High | High | High | High | High | High | High | 0 | 1.74E-13 | 7.02 |
| FALSE | High | FYDEIYFDS | [K].FYDEIY | 2xPhosphc | Q3UM97 2 | 7.36E-06 | 0 | 1 | 5 | 4 | Q3UM97 | Putative ur | Q3UM97 [ | 0 | 2635.045 | 0 | 0.3 | 0.099445 | 3.63 | 90.4 | 109.6 | 7.82 | 9.49 | | High | High | High | High | High | High | High | High | High | High | High | 0 | 9.57E-07 | 4.15 |
| FALSE | High | FYDSNTVK | [K].FYDSNT | 1xPhosphc | Q8CIN4 1x | 0.002419 | 0.000109 | 1 | 1 | 2 | Q8CIN4 | Serine/thre | Q8CIN4 [1 | 0 | 1511.755 | 0 | 1.22 | 0.006089 | 23.2 | 53.6 | 146.4 | 13.4 | 19.19 | | High | High | High | High | High | High | High | High | High | High | High | 4.94E-05 | 0.00064 | 2.3 |
| FALSE | High | FYQGLPW | [R].FYQGLI | 1xPhosphc | Q7TS75 1x | 0.007605 | 0.000664 | 1 | 1 | 2 | Q7TS75 | APC memb | Q7TS75 [8 | 0 | 1915.967 | 0 | 0.67 | 0.116265 | 28.08 | 80.7 | 119.3 | 12.55 | 18.14 | | High | High | High | High | High | High | High | High | High | High | High | 0.000294 | 0.00231 | 2.35 |
| FALSE | High | GAAAADLL | [R].GAAAA | 1xPhosphc | Q6PB44 1x | 0.005135 | 0.000551 | 1 | 2 | 1 | Q6PB44 | Tyrosine-p | Q6PB44 [1 | 0 | 3692.869 | 0 | 0.36 | 0.040822 | 6.22 | 88.1 | 111.9 | 5.19 | 12.6 | | High | High | High | High | High | High | High | High | High | High | High | 0.000246 | 0.00149 | 2.54 |
| FALSE | High | GAAATTTP | [R].GAAAT | 1xPhosphc | P83741-3 1 | 0.028575 | 0.001829 | 1 | 8 | 1 | P83741-3 | Isoform 3 c | P83741-3 [ | 0 | 1521.726 | 0 | 1.42 | 0.002675 | 29.9 | 53.9 | 146.1 | 18.77 | 10.21 | | High | High | High | High | High | High | High | High | High | High | High | 0.000984 | 0.01015 | 2.56 |
| FALSE | High | GAADSDLL | [K].GAADS | 1xPhosphc | Q6P1H6 1x | 0.000147 | 0 | 1 | 3 | 1 | Q6P1H6 | Ankyrin re | Q6P1H6 [7 | 0 | 2292.225 | 0 | 0.12 | 0.012918 | 5.11 | 95.7 | 104.3 | 5.75 | 4.66 | | High | High | High | High | High | High | High | High | High | High | High | 0 | 2.77E-05 | 2.36 |
| FALSE | High | GAAGPLDN | [K].GAAGP | 2xPhosphc | Q3TZR9 2x | 0.023175 | 0.001578 | 1 | 5 | 1 | Q3TZR9 | Activating | Q3TZR9 [8 | 0 | 2288.105 | 0 | 1.16 | 0.023215 | 5.73 | 58.8 | 141.2 | 10.17 | 28.7 | | High | High | High | High | High | High | High | High | High | High | High | 0.000847 | 0.008048 | 2.35 |
| FALSE | High | GAALGDLV | [R].GAALG | 1xCarbami | Q921G6 1x | 0.014667 | 0.001049 | 1 | 5 | 1 | Q921G6 | Leucine-ric | Q921G6 [2 | 0 | 3119.522 | 0 | 0.08 | 0.909944 | 10.73 | 107.7 | 92.3 | 13.62 | 19.46 | | High | High | High | High | High | High | High | High | High | High | High | 0.000555 | 0.004837 | 2.48 |
| FALSE | High | GAASPAQK | [R].GAASP | 1xPhosphc | A2A5Y6 1x | 0.008927 | 0.000736 | 1 | 11 | 2 | A2A5Y6 | Microtubu | A2A5Y6 [4 | 0 | 1267.681 | 0 | 0.24 | 0.011984 | 11.1 | 90.7 | 109.3 | 2.76 | 7.02 | | High | High | High | High | High | High | High | High | High | High | High | 0.000327 | 0.00277 | 1.73 |
| FALSE | High | GAASSSVSI | [R].GAASS | 1xCarbami | Q6NZR2 1x | 2.31E-11 | 0 | 1 | 1 | 2 | Q6NZR2 | Myb/SANT | Q6NZR2 [7 | 0 | 2562.161 | 0 | -1.06 | 0.000476 | 1.26 | 136.8 | 63.2 | 4.61 | 7.31 | | High | High | High | High | High | High | High | High | High | High | High | 0 | 6.3E-13 | 5.47 |
| FALSE | High | GAATPAK | [K].GAATP | 1xPhosphc | P09405 1xP | 0.011346 | 0.000816 | 1 | 6 | 5 | P09405 | Nucleolin ( | P09405 [12 | 0 | 1153.638 | 0 | -1.1 | 0.00108 | 2.12 | 138.7 | 61.3 | 13.13 | 16.44 | | High | High | High | High | High | High | High | High | High | High | High | 0.000449 | 0.004201 | 1.92 |
| FALSE | High | GACSTPEM | [R].GACSTI | 1xCarbami | A2AUK5 1x | 0.000305 | 0 | 2 | 5 | 2 | A2AUK5; C | Band 4.1-li | A2AUK5 [6 | 0 | 2222.027 | 0 | -0.22 | 0.181963 | 5.92 | 108.5 | 91.5 | 6.04 | 13.01 | Shared | High | High | High | High | High | High | High | High | High | High | High | 0 | 0.000139 | 2.83 |
| FALSE | High | GADAASPP | [R].GADAA | 1xPhosphc | A2AKB9 1x | 0.000162 | 0 | 1 | 3 | 2 | A2AKB9 | DDB1- and | A2AKB9 [4 | 0 | 1658.81 | 0 | -0.77 | 0.000163 | 0.43 | 127.5 | 72.5 | 4.39 | 5.57 | | High | High | High | High | High | High | High | High | High | High | High | 0 | 3.11E-05 | 2.54 |
| FALSE | High | GADHTPEH | [K].GADHT | 1xPhosphc | G3UWZ0 1 | 0.000126 | 0 | 1 | 4 | 1 | G3UWZ0 | Bromodom | G3UWZ0 [ | 0 | 2108.939 | 0 | 1.38 | 0.001288 | 7.81 | 57.6 | 142.4 | 19.55 | 7.1 | | High | High | High | High | High | High | High | High | High | High | High | 0 | 2.32E-05 | 2.83 |
| FALSE | High | GADHTPEH | [K].GADHT | 2xPhosphc | G3UWZ0 2 | 0.004867 | 0.000504 | 1 | 4 | 2 | G3UWZ0 | Bromodom | G3UWZ0 [ | 0 | 2188.905 | 0 | 1.49 | 0.000551 | 4.26 | 48.8 | 151.2 | 10.96 | 7.83 | | High | High | High | High | High | High | High | High | High | High | High | 0.003911 | 0.04147 | 2.29 |
| FALSE | High | GADNDGS | [K].GADND | 1xPhosphc | Q5F2E7-2 1 | 1.74E-06 | 0 | 1 | 2 | 16 | Q5F2E7-2 | Isoform 2 c | Q5F2E7-2 [ | 0 | 2180.975 | 0 | 0.19 | 0.007369 | 5.58 | 93.7 | 106.3 | 1.73 | 5.75 | | High | High | High | High | High | High | High | High | High | High | High | 0 | 1.89E-07 | 3.09 |
| FALSE | High | GADNDGS | [K].GADND | 1xDeamida | Q5F2E7-2 1 | 6.91E-06 | 0 | 1 | 2 | 3 | Q5F2E7-2 | Isoform 2 c | Q5F2E7-2 [ | 0 | 2181.959 | 0 | 0.51 | 0.005517 | 18.37 | 82.4 | 117.6 | 5.17 | 7.29 | | High | High | High | High | High | High | High | High | High | High | High | 0 | 8.93E-07 | 3.49 |
| FALSE | High | GADNDGS | [K].GADND | 2xPhosphc | Q5F2E7-2 2 | 4.28E-06 | 0 | 1 | 2 | 6 | Q5F2E7-2 | Isoform 2 c | Q5F2E7-2 [ | 0 | 2260.941 | 0 | 0.77 | 0.000816 | 6.81 | 74 | 126 | 4.93 | 6.37 | | High | High | High | High | High | High | High | High | High | High | High | 0 | 1.06E-05 | 3.76 |
| FALSE | High | GADNDGS | [K].GADND | 1xPhosphc | Q5F2E7-2 1 | 0.000284 | 0 | 1 | 2 | 1 | Q5F2E7-2 | Isoform 2 c | Q5F2E7-2 [ | 1 | 2538.233 | 0 | -0.51 | 0.003991 | 2.48 | 113.5 | 86.5 | 8.77 | 9.09 | | High | High | High | High | High | High | High | High | High | High | High | 0 | 5.81E-05 | 2.76 |
| FALSE | High | GADPGSPG | [R].GADPG | 1xPhosphc | Q6PB97 1x | 0.004256 | 0.00036 | 1 | 1 | 1 | Q6PB97 | F-box/LRR- | Q6PB97 [2 | 0 | 1542.788 | 0 | -0.3 | 0.096355 | 14.65 | 107.1 | 92.9 | 17.58 | 6.94 | | High | High | High | High | High | High | High | High | High | High | High | 0.000161 | 0.001203 | 2.51 |
| FALSE | High | GADSGGEK | [K].GADSG | 1xPhosphc | Q9D0B6 1x | 0.01186 | 0.000889 | 1 | 1 | 1 | Q9D0B6 | Protein PBI | Q9D0B6 [1 | 0 | 1258.608 | 0 | -0.36 | 0.005406 | 5.27 | 114.7 | 85.3 | 4.93 | 12.52 | | High | High | High | High | High | High | High | High | High | High | High | 0.000449 | 0.003804 | 1.74 |
| FALSE | High | GAEAGSQA | [K].GAEAG | 1xPhosphc | B1AS46 1xP | 7.69E-09 | 0 | 1 | 4 | 1 | B1AS46 | Zinc finger | B1AS46 [4 | 0 | 1872.88 | 0 | -0.7 | 0.001404 | 3.03 | 126.9 | 73.1 | 4.92 | 12.18 | | High | High | High | High | High | High | High | High | High | High | High | 0 | 4.27E-10 | 4.32 |
| FALSE | High | GAEDYPDP | [K].GAEDY | 1xPhosphc | Q3UTJ2-2 1 | 0.003312 | 0.00016 | 1 | 14 | 1 | Q3UTJ2-2 | Isoform 2 c | Q3UTJ2-2 [ | 0 | 2312.007 | 0 | 0.77 | 0.009366 | 10.71 | 73.8 | 126.2 | 17.28 | 11.94 | | High | High | High | High | High | High | High | High | High | High | High | 7.19E-05 | 0.000914 | 1.83 |
| FALSE | High | GAELGPEP | [R].GAELGI | 2xPhosphc | Q922H1 2x | 3.99E-06 | 0 | 1 | 5 | 6 | Q922H1 | Protein arg | Q922H1 [1 | 0 | 3826.548 | 0 | -0.45 | 0.002101 | 5.61 | 115.1 | 84.9 | 2.58 | 6.96 | | High | High | High | High | High | High | High | High | High | High | High | 0 | 4.79E-07 | 4.79 |
| FALSE | High | GAEPSPGG | [K].GAEPSF | 1xPhosphc | A0A0U1RN | 0.054383 | 0.003951 | 1 | 3 | 3 | A0A0U1RN | B-cell CLL/l | A0A0U1RN | 0 | 4157.998 | 0 | -0.66 | 0.002719 | 3.69 | 119.8 | 80.2 | 10.73 | 9.2 | | High | High | High | High | High | High | High | High | High | High | High | 0.002059 | 0.02126 | 3.94 |
| FALSE | High | GAEPSPGG | [K].GAEPSF | 3xPhosphc | A0A0U1RN | 0.049971 | 0.003641 | 1 | 3 | 2 | A0A0U1RN | B-cell CLL/l | A0A0U1RN | 0 | 4317.93 | 0 | -0.21 | 0.046805 | 12.36 | 109 | 91 | 7.55 | 14.86 | | High | High | High | High | High | High | High | High | High | High | High | 0.001902 | 0.01935 | 3.93 |
| FALSE | High | GAEVLLEEI | [R].GAEVLI | 1xPhosphc | Q7TMI6 1x | 8.06E-06 | 0 | 1 | 3 | 2 | Q7TMI6 | Zinc finger | Q7TMI6 [1 | 0 | 2851.478 | 0 | 0.26 | 0.085001 | 17.7 | 94.9 | 105.1 | 5.86 | 9.81 | | High | High | High | High | High | High | High | High | High | High | High | 0 | 1.06E-06 | 3.86 |
| FALSE | High | GAFDGSVP | [K].GAFDG | 1xPhosphc | E9Q616 1x | 0.000328 | 0 | 1 | 5 | 1 | E9Q616 | AHNAK nuc | E9Q616 [4 | 0 | 1415.734 | 0 | 4.54 | 0.007751 | 10.29 | 7.4 | 192.6 | 31.1 | 35.65 | | High | High | High | High | High | High | High | High | High | High | High | 0 | 6.84E-05 | 0.98 |
| FALSE | High | GAFPESYSI | [K].GAFPES | 1xPhosphc | P97348 1xP | 0.053023 | 0.00384 | 1 | 2 | 3 | P97348 | Rho-relate | P97348 [4 | 0 | 1895.878 | 0 | 0.57 | 0.004899 | 13.95 | 78.5 | 121.5 | 5.12 | 9.96 | | High | High | High | High | High | High | High | High | High | High | High | 0.004782 | 0.0524 | 2.02 |
| FALSE | High | FSLGPLSPE | [R].FSLGPL | 1xPhosphc | Q9D0P7 1x | 0.08727 | 0.006537 | 1 | 3 | 1 | Q9D0P7 | Proline/ser | Q9D0P7 [6 | 0 | 1612.875 | 0 | -0.66 | 0.005788 | 4.22 | 120.8 | 79.2 | 6.63 | 11.06 | | High | High | High | High | High | High | High | High | High | High | High | 0.003361 | 0.03672 | 1.92 |
| FALSE | High | GASAATGII | [K].GASAA | 1xCarbami | Q8R5H1 1x | 5.37E-06 | 0 | 2 | 4 | 5 | Q8R5H1; C | Ubiquitin c | Q8R5H1 [9 | 0 | 3759.474 | 0 | 0.05 | 0.808304 | 2.91 | 99.3 | 100.7 | 7.04 | 16.15 | Shared | High | High | High | High | High | High | High | High | High | High | High | 0 | 1.15E-06 | 5.12 |
| FALSE | High | FSLDGEAW | [R].FSLDGE | 1xPhosphc | Q9Z2B9 1x | 0.000765 | 0 | 1 | 1 | 1 | Q9Z2B9 | Ribosomal | Q9Z2B9 [6 | 0 | 2291.1 | 0 | 1.15 | 0.001999 | 3.6 | 60.1 | 139.9 | 34.08 | 7.8 | | High | High | High | High | High | High | High | High | High | High | High | 0 | 0.000177 | 3 |
| FALSE | High | FSHSYLSDS | [R].FSHSYL | 2xPhosphc | Q9JMH9-6 | 0.000107 | 0 | 1 | 14 | 7 | Q9JMH9-6 | Isoform 6 c | Q9JMH9-6 | 0 | 2204.955 | 0 | 0.05 | 0.219975 | 8.52 | 95.1 | 104.9 | 7.9 | 5.35 | | High | High | High | High | High | High | High | High | High | High | High | 0.000275 | 0.002099 | 3.5 |
| FALSE | High | FNSESESSS | [R].FNSESE | 1xPhosphc | Q8BLH7 1x | 1.91E-11 | 0 | 1 | 2 | 6 | Q8BLH7 | HIRA-intera | Q8BLH7 [9 | 0 | 2696.198 | 0 | -1.1 | 0.000435 | 2.16 | 132.9 | 67.1 | 5.54 | 10.31 | | High | High | High | High | High | High | High | High | High | High | High | 0 | 5.09E-13 | 4.86 |
| FALSE | High | FNSLPKPDS | [R].FNSLPK | 1xPhosphc | Q4VAC9 1x | 0.001503 | 5.66E-05 | 1 | 3 | 1 | Q4VAC9 | Pleckstrin h | Q4VAC9 [8 | 0 | 2684.43 | 0 | 2.51 | 0.000441 | 14.93 | 30.2 | 169.8 | 9.49 | 9.05 | | High | High | High | High | High | High | High | High | High | High | High | 2.58E-05 | 0.000378 | 3.05 |
| FALSE | High | FNSLPQYSI | [K].FNSLPC | 1xPhosphc | Q8VBW5 1 | 0.027121 | 0.001676 | 1 | 5 | 2 | Q8VBW5 | HMG box t | Q8VBW5 [ | 0 | 1993.962 | 0 | 0.94 | 0.001684 | 15.56 | 67.9 | 132.1 | 14.98 | 9.44 | | High | High | High | High | High | High | High | High | High | High | High | 0.00092 | 0.009587 | 2.84 |
| FALSE | High | FNSITSAYY | [R].FNSITS | 1xPhosphc | A2CG35 1x | 0.050826 | 0.003676 | 1 | 2 | 1 | A2CG35 | Ras-relatec | A2CG35 [1 | 0 | 1530.719 | 0 | 0.61 | 0.001293 | 6.64 | 81.8 | 118.2 | 11.56 | 8.9 | | High | High | High | High | High | High | High | High | High | High | High | 0.001916 | 0.01963 | 2.07 |
| FALSE | High | FNSPSEELD | [K].FNSPSE | 1xPhosphc | Q9JJG0 1xP | 0.006594 | 0.00062 | 1 | 10 | 2 | Q9JJG0 | Transformi | Q9JJG0 [71 | 0 | 1665.736 | 0 | -0.48 | 0.015545 | 8.79 | 113.4 | 86.6 | 7.85 | 7.88 | | High | High | High | High | High | High | High | High | High | High | High | 0.001651 | 0.01623 | 2.37 |
| FALSE | High | FNSPSEGK | [K].FNSPSE | 1xPhosphc | E9Q8T1 1x | 0.004572 | 0.000456 | 1 | 5 | 1 | E9Q8T1 | Transformi | E9Q8T1 [2 | 0 | 1403.697 | 0 | -0.6 | 0.01706 | 10.59 | 120.6 | 79.4 | 7.79 | 12.99 | | High | High | High | High | High | High | High | High | High | High | High | 0.000204 | 0.00131 | 2.24 |
| FALSE | High | FPANEDSIL | [K].FPANEI | 1xCarbami | E9Q4Y4 1x | 2.79E-08 | 0 | 1 | 2 | 1 | E9Q4Y4 | Centrosom | E9Q4Y4 [2 | 0 | 2777.208 | 0 | -0.05 | 0.159187 | 2 | 105.2 | 94.8 | 5.19 | 10.99 | | High | High | High | High | High | High | High | High | High | High | High | 0 | 1.82E-09 | 4.03 |
| FALSE | High | FPDLDYSEY | [R].FPDLDY | 1xCarbami | D2CGM4 1 | 1.13E-07 | 0 | 1 | 4 | 8 | D2CGM4 | Melanoma | D2CGM4 [ | 0 | 2339.914 | 0 | 0.98 | 0.00409 | 14.94 | 63.4 | 136.6 | 7.81 | 17.42 | | High | High | High | High | High | High | High | High | High | High | High | 0 | 8.79E-09 | 4.04 |
| FALSE | High | FPEFCSSPS | [K].FPEFCS | 1xCarbami | Q7TPD0 1x | 0.001644 | 5.66E-05 | 1 | 3 | 2 | Q7TPD0 | Integrator | Q7TPD0 [4 | 0 | 2245.102 | 0 | -0.64 | 0.001846 | 2.63 | 122 | 78 | 6.68 | 3.62 | | High | High | High | High | High | High | High | High | High | High | High | 0.000275 | 0.001969 | 2.86 |
| FALSE | High | FPEFCSSPS | [K].FPEFCS | 1xCarbami | Q7TPD0 2x | 0.003755 | 0.00021 | 1 | 3 | 2 | Q7TPD0 | Integrator | Q7TPD0 [4 | 0 | 2325.068 | 0 | -0.96 | 0.002578 | 6.76 | 135.6 | 64.4 | 12.05 | 25.34 | | High | High | High | High | High | High | High | High | High | High | High | 0.000786 | 0.007139 | 3.13 |
| FALSE | High | FPFGISPAC | [K].FPFGISI | 1xPhosphc | E9QAQ7 1x | 0.00153 | 5.66E-05 | 1 | 5 | 1 | E9QAQ7 | AT-rich inte | E9QAQ7 [1 | 0 | 1652.815 | 0 | -1.96 | 0.000196 | 1.47 | 157.6 | 42.4 | 3.93 | 14.62 | | High | High | High | High | High | High | High | High | High | High | High | 2.58E-05 | 0.000385 | 1.43 |
| FALSE | High | FPPPPQPQ | [K].FPPPPC | 1xPhosphc | Q5NCY0 1x | 7.47E-05 | 0 | 1 | 1 | 2 | Q5NCY0 | Lysine-spe | Q5NCY0 [7 | 0 | 2795.53 | 0 | -0.59 | 0.002004 | 8.03 | 117.2 | 82.8 | 6.46 | 17.12 | | High | High | High | High | High | High | High | High | High | High | High | 4.94E-05 | 0.000714 | 2.23 |
| FALSE | High | FPPPQELSC | [R].FPPPQE | 1xPhosphc | E9QAQ7 1x | 3.81E-06 | 0 | 1 | 5 | 1 | E9QAQ7 | AT-rich inte | E9QAQ7 [5 | 0 | 3236.522 | 0 | 0.64 | 0.263552 | 38.29 | 77.7 | 122.3 | 24.13 | 23.63 | | High | High | High | High | High | High | High | High | High | High | High | 0 | 4.55E-07 | 2.86 |
| FALSE | High | FNSDSHSPI | [K].FNSDSH | 1xPhosphc | S4R221 1xP | 0.041072 | 0.003094 | 1 | 9 | 2 | S4R221 | Rho GTPas | S4R221 [15 | 0 | 1556.751 | 0 | -0.56 | 0.00378 | 4.26 | 121.9 | 78.1 | 10.27 | 8.23 | | High | High | High | High | High | High | High | High | High | High | High | 0.002432 | 0.02611 | 1.79 |
| FALSE | High | FPPPQELSC | [R].FPPPQE | 1xDeamida | E9QAQ7 1x | 0.000171 | 0 | 1 | 5 | 3 | E9QAQ7 | AT-rich inte | E9QAQ7 [5 | 0 | 3237.506 | 0 | 0.7 | 0.294792 | 16.36 | 72.4 | 127.6 | 25.88 | 21.4 | | High | High | High | High | High | High | High | High | High | High | High | 0 | 0.000109 | 4.63 |
| FALSE | High | FPPSTPSEV | [R].FPPSTP | 1xPhosphc | A0A0R4J0I | 0.025514 | 0.001645 | 1 | 2 | 4 | A0A0R4J0I | ETS domain | A0A0R4J0I | 0 | 2165.037 | 0 | -0.59 | 0.054717 | 12.32 | 119.7 | 80.3 | 12.29 | 15.2 | | High | High | High | High | High | High | High | High | High | High | High | 0.003188 | 0.03358 | 3.28 |
| FALSE | High | FPQALTCN | [R].FPQALT | 1xCarbami | Q9WU40-2 | 0.006536 | 0.00062 | 1 | 4 | 1 | Q9WU40-2 | Isoform 2 c | Q9WU40-2 | 0 | 2567.402 | 0 | 0.28 | 0.443327 | 9.41 | 94.9 | 105.1 | 13.16 | 15.09 | | High | High | High | High | High | High | High | High | High | High | High | 0.000275 | 0.001956 | 2.97 |
| FALSE | High | FPQGPPSQ | [R].FPQGPI | 1xPhosphc | Q8R326 1x | 8.78E-08 | 0 | 1 | 2 | 13 | Q8R326 | Paraspeckl | Q8R326 [4 | 0 | 2028.887 | 0 | 0.21 | 0.022964 | 9.64 | 91 | 109 | 5.64 | 7.39 | | High | High | High | High | High | High | High | High | High | High | High | 0 | 6.61E-09 | 4.15 |
| FALSE | High | FPRSPEEDC | [R].FPRSPE | 1xPhosphc | Q6P5H2-2 | 0.021607 | 0.001507 | 1 | 2 | 1 | Q6P5H2-2 | Isoform 2 c | Q6P5H2-2 | 1 | 2612.327 | 0 | 2.88 | 0.001759 | 15.55 | 25.7 | 174.3 | 25.74 | 16.67 | | High | High | High | High | High | High | High | High | High | High | High | 0.000801 | 0.007452 | 1.67 |
| FALSE | High | FPSGNQGC | [R].FPSGN | 1xPhosphc | Q01853 1x | 1.84E-07 | 0 | 1 | 2 | 9 | Q01853 | Transitiona | Q01853 [7 | 0 | 3529.45 | 0 | 0.01 | 0.704126 | 4.86 | 102.7 | 97.3 | 6.8 | 7.36 | | High | High | High | High | High | High | High | High | High | High | High | 0 | 1.52E-08 | 6.14 |
| FALSE | High | FPSGNQGC | [R].FPSGN | 1xDeamida | Q01853 1x | 0.00103 | 0 | 1 | 2 | 10 | Q01853 | Transitiona | Q01853 [7 | 0 | 3530.434 | 0 | 0.33 | 0.099798 | 19.47 | 90.4 | 109.6 | 9.92 | 7.62 | | High | High | High | High | High | High | High | High | High | High | High | 0 | 0.000246 | 4.98 |
| FALSE | High | FPSPHPSPA | [K].FPSPHP | 1xPhosphc | A0A0R4J25 | 0.026191 | 0.001676 | 1 | 3 | 2 | A0A0R4J25 | Etoposide | A0A0R4J25 | 0 | 1602.845 | 0 | 0.18 | 0.226037 | 12.49 | 93 | 107 | 7.92 | 10.45 | | High | High | High | High | High | High | High | High | High | High | High | 0.000903 | 0.009243 | 2.03 |
| FALSE | High | FPSPHPSPA | [K].FPSPHP | 2xPhosphc | A0A0R4J25 | 0.001689 | 5.66E-05 | 1 | 3 | 2 | A0A0R4J25 | Etoposide | A0A0R4J25 | 0 | 1682.811 | 0 | 0.55 | 0.001591 | 9.35 | 78.9 | 121.1 | 7.75 | 3.75 | | High | High | High | High | High | High | High | High | High | High | High | 2.58E-05 | 0.000431 | 2.86 |
| FALSE | High | FPSSAESCE | [R].FPSSAE | 2xCarbami | O09053 1x | 0.00153 | 5.66E-05 | 1 | 3 | 2 | O09053 | Werner syn | O09053 [1 | 0 | 1926.838 | 0 | 0.25 | 0.126788 | 27.52 | 86.1 | 113.9 | 13.95 | 11.72 | | High | High | High | High | High | High | High | High | High | High | High | 4.94E-05 | 0.000715 | 2.59 |
| FALSE | High | FPSSGLATF | [K].FPSSGL | 1xPhosphc | Q3U3K9 1x | 0.000225 | 0 | 1 | 4 | 1 | Q3U3K9 | Ancient ub | Q3U3K9 [3 | 0 | 2372.267 | 0 | 2.33 | 0.000965 | 10.45 | 33.4 | 166.6 | 14.6 | 11.45 | | High | High | High | High | High | High | High | High | High | High | High | 0 | 4.47E-05 | 2.71 |
| FALSE | High | FPSTGSCTE | [R].FPSTGS | 1xCarbami | Q8BIE6-2 1 | 0.001544 | 5.66E-05 | 1 | 4 | 3 | Q8BIE6-2 | Isoform 2 c | Q8BIE6-2 [ | 0 | 2591.201 | 0 | 0.78 | 0.003272 | 11.78 | 72 | 128 | 17.82 | 6.23 | | High | High | High | High | High | High | High | High | High | High | High | 2.58E-05 | 0.000389 | 3.71 |
| FALSE | High | FPSWVTFE | [R].FPSWV | 1xCarbami | E9PXP7 1xP | 2.77E-05 | 0 | 1 | 2 | 1 | E9PXP7 | Stonin-2 O | E9PXP7 [2 | 0 | 3111.494 | 0 | -1.19 | 0.002816 | 1.59 | 136 | 64 | 12.22 | 7.28 | | High | High | High | High | High | High | High | High | High | High | High | 0 | 4.23E-06 | 4.27 |
| FALSE | High | FPSWVTFE | [R].FPSWV | 1xCarbami | E9PXP7 1xP | 0.001097 | 0 | 1 | 2 | 1 | E9PXP7 | Stonin-2 O | E9PXP7 [2 | 0 | 3112.478 | 0 | -0.53 | 0.013742 | 4.8 | 118.3 | 81.7 | 9.61 | 12.67 | | High | High | High | High | High | High | High | High | High | High | High | 0 | 0.000265 | 4.34 |
| FALSE | High | FPPPQELSC | [R].FPPPQE | 1xPhosphc | E9QAQ7 1x | 1.23E-05 | 0 | 1 | 5 | 2 | E9QAQ7 | AT-rich inte | E9QAQ7 [5 | 0 | 3252.517 | 0 | 0.03 | 0.818443 | 20.18 | 103 | 97 | 8.5 | 16.95 | | High | High | High | High | High | High | High | High | High | High | High | 0 | 1.7E-06 | 3.81 |
| FALSE | High | FPTDYASTS | [R].FPTDYA | 1xPhosphc | Q6A065 1x | 2.01E-07 | 0 | 1 | 3 | 1 | Q6A065 | Centrosom | Q6A065 [1 | 0 | 2231.933 | 0 | 0.41 | 0.042126 | 20.72 | 86 | 114 | 10.62 | 13.71 | | High | High | High | High | High | High | High | High | High | High | High | 0 | 1.68E-08 | 3.11 |
| FALSE | High | FNLSSHSQS | [R].FNLSSH | 1xPhosphc | B7ZNX2 1x | 0.006536 | 0.00062 | 1 | 5 | 1 | B7ZNX2 | Anaphase p | B7ZNX2 [3 | 0 | 1769.899 | 0 | -0.4 | 0.003576 | 6.08 | 112.5 | 87.5 | 5.19 | 6.04 | | High | High | High | High | High | High | High | High | High | High | High | 0.000275 | 0.001942 | 2.25 |
| FALSE | High | FNLSFPPSE | [K].FNLSFP | 1xPhosphc | A2AT37 1x | 0.006957 | 0.00062 | 1 | 1 | 1 | A2AT37 | UPF2 regul | A2AT37 [3 | 0 | 2243.177 | 0 | -0.48 | 0.000487 | 2.29 | 117.3 | 82.7 | 3.18 | 3.27 | | High | High | High | High | High | High | High | High | High | High | High | 0.000275 | 0.002096 | 2.86 |
| FALSE | High | FIIGSVSEDI | [R].FIIGSVS | 1xPhosphc | Q5SWU9-2 | 6.3E-05 | 0 | 1 | 6 | 5 | Q5SWU9-2 | Isoform 2 c | Q5SWU9-2 | 0 | 2733.364 | 0 | 0.76 | 0.001223 | 9.33 | 74.5 | 125.5 | 7.42 | 6.67 | | High | High | High | High | High | High | High | High | High | High | High | 2.58E-05 | 0.000348 | 4.06 |
| FALSE | High | FIIGSVSEDI | [R].FIIGSVS | 2xPhosphc | Q5SWU9-2 | 0.000362 | 0 | 1 | 6 | 3 | Q5SWU9-2 | Isoform 2 c | Q5SWU9-2 | 0 | 2813.33 | 0 | 1 | 0.005504 | 23.47 | 63.1 | 136.9 | 23.14 | 18.01 | | High | High | High | High | High | High | High | High | High | High | High | 0 | 7.63E-05 | 3.51 |
| FALSE | High | FLMPEAYP | [R].FLMPEA | 1xPhosphc | Q8C0T5 1x | 0.030104 | 0.001857 | 1 | 4 | 3 | Q8C0T5 | Signal-indu | Q8C0T5 [1 | 0 | 1719.801 | 0 | 0.87 | 0.003612 | 17.02 | 65.9 | 134.1 | 10.64 | 12.8 | | High | High | High | High | High | High | High | High | High | High | High | 0.001011 | 0.01082 | 3.28 |
| FALSE | High | FLNTMSVY | [K].FLNTM | 1xPhosphc | Q8BP40 1x | 0.000262 | 0 | 1 | 1 | 1 | Q8BP40 | Lysophosp | Q8BP40 [3 | 0 | 2156.075 | 0 | -0.88 | 0.003922 | 3.1 | 130.8 | 69.2 | 12.74 | 15.67 | | High | High | High | High | High | High | High | High | High | High | High | 0 | 5.28E-05 | 4.27 |
| FALSE | High | FIPCSPFSD | [R].FIPCSPI | 1xCarbami | E9PWE8 1x | 0.000255 | 0 | 1 | 4 | 1 | E9PWE8 | Dihydropy | E9PWE8 [5 | 0 | 2161.048 | 0 | -0.12 | 0.602936 | 5.99 | 104.7 | 95.3 | 6.48 | 7.7 | | High | High | High | High | High | High | High | High | High | High | High | 0 | 5.16E-05 | 2.24 |
| FALSE | High | FLPLPVLSH | [R].FLPLPV | 1xPhosphc | A0A0A0M( | 4.33E-07 | 0 | 1 | 2 | 1 | A0A0A0M( | Histone-lys | A0A0A0M( | 0 | 2310.282 | 0 | 0.63 | 0.003259 | 7.79 | 79.9 | 120.1 | 4.6 | 8.6 | | High | High | High | High | High | High | High | High | High | High | High | 0 | 3.96E-08 | 4.08 |
| FALSE | High | FLPISSTPQ | [K].FLPISST | 1xPhosphc | D3YTL8 1xP | 0.113753 | 0.009241 | 1 | 5 | 2 | D3YTL8 | Protein spi | D3YTL8 [3 | 0 | 1680.856 | 0 | 0.2 | 0.604658 | 13.95 | 93 | 107 | 13.31 | 16.68 | | High | High | High | High | High | High | High | High | High | High | High | 0.004667 | 0.05036 | 2.14 |

| | | | | | | | | | | | | | | | | | | | | | | | | | | | | | | | | | | | | | | |
|---|---|---|---|---|---|---|---|---|---|---|---|---|---|---|---|---|---|---|---|---|---|---|---|---|---|---|---|---|---|---|---|---|---|---|---|---|---|---|
| FALSE | High | FLQDHGSD | [K].FLQDH | 1xPhospho | Q62446 1x | 0.021418 | 0.001507 | 1 | 1 | 1 | Q62446 | Peptidyl-pr | Q62446 [2 | 0 | 2269.105 | 0 | -0.8 | 0.012366 | 7.49 | 127 | 73 | 11.54 | 10.48 | | High | High | High | High | High | High | High | High | High | High | High | 0.000801 | 0.007382 | 2.12 |
| FALSE | High | FIQELSGSS | [R].FIQELS | 1xPhospho | Q9JIZ5 1xP | 3.1E-06 | 0 | 1 | 2 | 2 | Q9JIZ5 | Activator p | Q9JIZ5 [11 | 0 | 1730.913 | 0 | 0.27 | 0.017627 | 8.43 | 90.6 | 109.4 | 5.52 | 8.7 | | High | High | High | High | High | High | High | High | High | High | High | 0 | 3.63E-07 | 3.91 |
| FALSE | High | FLSHSTDSL | [K].FLSHST | 1xPhospho | A0A140LJJ | 0.003464 | 0.00021 | 1 | 5 | 2 | A0A140LJJ | A-kinase ar | A0A140LJJ | 0 | 1786.914 | 0 | -0.5 | 0.132869 | 5.63 | 117.1 | 82.9 | 13.97 | 24.43 | | High | High | High | High | High | High | High | High | High | High | High | 0.002233 | 0.02355 | 2.51 |
| FALSE | High | FLSHSTDSL | [K].FLSHST | 2xPhospho | A0A140LJJ | 0.010568 | 0.000772 | 1 | 5 | 1 | A0A140LJJ | A-kinase ar | A0A140LJJ | 0 | 1866.88 | 0 | -1.07 | 0.002461 | 2.79 | 134.4 | 65.6 | 12.03 | 10.99 | | High | High | High | High | High | High | High | High | High | High | High | 0.000382 | 0.003322 | 2.23 |
| FALSE | High | FLSLEYSPV | [R].FLSLEY | 1xPhospho | G3X932 1x | 0.026652 | 0.001676 | 1 | 4 | 1 | G3X932 | DNA segme | G3X932 [3 | 0 | 1777.954 | 0 | -0.64 | 0.006132 | 9.05 | 120.3 | 79.7 | 7.46 | 8.3 | | High | High | High | High | High | High | High | High | High | High | High | 0.000903 | 0.009415 | 2.11 |
| FALSE | High | FLSNSWYIF | [K].FLSNSV | 1xPhospho | Q8BJL1 1x | 0.000584 | 0 | 1 | 1 | 2 | Q8BJL1 | F-box only | Q8BJL1 [39 | 0 | 1792.944 | 0 | 0.19 | 0.088621 | 2.96 | 93.4 | 106.6 | 8.23 | 4.09 | | High | High | High | High | High | High | High | High | High | High | High | 0 | 0.000131 | 2.03 |
| FALSE | High | FNISPDEDS | [R].FNISPD | 1xPhospho | Q8CFE6 1x | 0.000668 | 0 | 1 | 2 | 4 | Q8CFE6 | Sodium-co | Q8CFE6 [9 | 0 | 3389.478 | 0 | 2.22 | 0.004967 | 56.38 | 42.7 | 157.3 | 48.88 | 21.93 | | High | High | High | High | High | High | High | High | High | High | High | 0 | 0.000151 | 3.49 |
| FALSE | High | FLSSPQFSH | [R].FLSSPQ | 1xPhospho | Q8BI22 1x | 1.19E-05 | 0 | 1 | 2 | 2 | Q8BI22 | Centrosom | Q8BI22 [10 | 0 | 2360.246 | 0 | -0.04 | 0.561785 | 29.22 | 104.6 | 95.4 | 8.79 | 12.47 | | High | High | High | High | High | High | High | High | High | High | High | 0 | 1.64E-06 | 3.29 |
| FALSE | High | FIVSPVPES | [R].FIVSPV | 1xPhospho | P83741-3 1 | 0.119269 | 0.00979 | 1 | 6 | 1 | P83741-3 | Isoform 3 | P83741-3 [ | 0 | 1439.75 | 0 | 1.3 | 0.000912 | 10.46 | 55.8 | 144.2 | 9.14 | 8.06 | | High | High | High | High | High | High | High | High | High | High | High | 0.004894 | 0.05331 | 2 |
| FALSE | High | FMSSPSLSD | [K].FMSSPS | 1xPhospho | Q812A5 1x | 0.026191 | 0.001676 | 1 | 1 | 1 | Q812A5 | Proline-ricl | Q812A5 [9 | 0 | 1822.906 | 0 | -0.72 | 0.005436 | 7.65 | 125.7 | 74.3 | 7.48 | 17.41 | | High | High | High | High | High | High | High | High | High | High | High | 0.000903 | 0.009238 | 2.39 |
| FALSE | High | FMTGADSF | [K].FMTGA | 1xPhospho | Q9D0F1 1x | 0.000124 | 0 | 1 | 1 | 2 | Q9D0F1 | Kinetochor | Q9D0F1 [2 | 0 | 2556.162 | 0 | 0.69 | 0.011541 | 31.62 | 76.4 | 123.6 | 13.91 | 8.69 | | High | High | High | High | High | High | High | High | High | High | High | 2.58E-05 | 0.000389 | 3.63 |
| FALSE | High | FNDPSGCA | [R].FNDPS | 1xCarbami | Q8VDR2 1x | 0.000127 | 0 | 1 | 6 | 1 | Q8VDR2 | Hmmr prot | Q8VDR2 [1 | 0 | 2320.072 | 0 | 0.29 | 0.139823 | 21.31 | 89.4 | 110.6 | 17.5 | 9.42 | | High | High | High | High | High | High | High | High | High | High | High | 0 | 2.37E-05 | 3.26 |
| FALSE | High | FNEEHIPDS | [K].FNEEHI | 1xPhospho | Q8BTM8 1 | 2.4E-09 | 0 | 1 | 6 | 3 | Q8BTM8 | Filamin-A | Q8BTM8 [2 | 0 | 2776.318 | 0 | 0.38 | 0.058666 | 11.52 | 88.6 | 111.4 | 7.48 | 9.78 | | High | High | High | High | High | High | High | High | High | High | High | 0 | 1.16E-10 | 4.97 |
| FALSE | High | FNFESVPES | [K].FNFESV | 1xPhospho | Q3U0M1 1 | 0.000884 | 0 | 1 | 3 | 2 | Q3U0M1 | Trafficking | Q3U0M1 [ | 0 | 2004.972 | 0 | 1.97 | 0.000401 | 18.81 | 39.3 | 160.7 | 8.03 | 7.19 | | High | High | High | High | High | High | High | High | High | High | High | 0.001011 | 0.01091 | 2.42 |
| FALSE | High | FNFSGSK | [K].FNFSGS | 1xPhospho | E9Q616 1x | 0.052576 | 0.00384 | 1 | 2 | 1 | E9Q616 | AHNAK nu | E9Q616 [3 | 0 | 1324.67 | 0 | 2.06 | 0.004077 | 15.55 | 38.5 | 161.5 | 18.93 | 23.96 | | High | High | High | High | High | High | High | High | High | High | High | 0.001999 | 0.02045 | 1.54 |
| FALSE | High | FNGEDVDT | [K].FNGED | 1xPhospho | Q9D0J4 1x | 6.53E-05 | 0 | 1 | 1 | 1 | Q9D0J4 | ADP-ribosy | Q9D0J4 [3 | 0 | 2505.268 | 0 | -0.27 | 0.135977 | 4.83 | 109.1 | 90.9 | 4.35 | 12.27 | | High | High | High | High | High | High | High | High | High | High | High | 0 | 1.11E-05 | 3.09 |
| FALSE | High | FNGGHSPT | [R].FNGGH | 1xPhospho | B2RRE8 1x | 8.63E-05 | 0 | 1 | 2 | 3 | B2RRE8 | CD2-associ | B2RRE8 [5 | 0 | 1923.937 | 0 | -0.03 | 0.480645 | 5.99 | 96.5 | 103.5 | 6.93 | 11.4 | | High | High | High | High | High | High | High | High | High | High | High | 0 | 1.53E-05 | 2.87 |
| FALSE | High | FNGGHSPT | [R].FNGGH | 1xDeamida | B2RRE8 1x | 3.32E-05 | 0 | 1 | 2 | 3 | B2RRE8 | CD2-associ | B2RRE8 [5 | 0 | 1924.921 | 0 | 0.01 | 0.521741 | 4.88 | 98.3 | 101.7 | 2.82 | 9.27 | | High | High | High | High | High | High | High | High | High | High | High | 0 | 5.21E-06 | 2.99 |
| FALSE | High | FNGGHSPT | [R].FNGGH | 2xPhospho | B2RRE8 2x | 6.55E-06 | 0 | 1 | 2 | 2 | B2RRE8 | CD2-associ | B2RRE8 [5 | 0 | 2003.903 | 0 | -0.56 | 0.006518 | 5.36 | 119.8 | 80.2 | 4.43 | 12.08 | | High | High | High | High | High | High | High | High | High | High | High | 0 | 9.6E-05 | 2.44 |
| FALSE | High | FNGSTEALI | [R].FNGSTI | 1xPhospho | Q6R891 1x | 0.003001 | 0.00016 | 1 | 1 | 2 | Q6R891 | Neurabin-2 | Q6R891 [1 | 0 | 1619.808 | 0 | 1.07 | 0.001035 | 6.21 | 67.8 | 132.2 | 5.07 | 10.84 | | High | High | High | High | High | High | High | High | High | High | High | 7.19E-05 | 0.000819 | 1.98 |
| FALSE | High | FNGSTEALI | [R].FNGSTI | 1xDeamida | Q6R891 1x | 0.003526 | 0.00021 | 1 | 1 | 5 | Q6R891 | Neurabin-2 | Q6R891 [1 | 0 | 1620.792 | 0 | 1.02 | 0.00205 | 7.91 | 65.6 | 134.4 | 6.73 | 10.44 | | High | High | High | High | High | High | High | High | High | High | High | 0.000859 | 0.008317 | 2.17 |
| FALSE | High | FLTVPVPES | [R].FLTVPV | 1xPhospho | G5E8L6 1x | 2.31E-05 | 0 | 1 | 3 | 1 | G5E8L6 | Killer cell le | G5E8L6 [13 | 0 | 1839.961 | 0 | 6.64 | | 0 | 4.2 | 195.8 | | 16.82 | | Not Found | Not Found | Not Found | High | Not Found | High | High | High | High | High | High | 0 | 3.46E-06 | 2.96 |
| FALSE | High | FPTDYASTS | [R].FPTDYA | 2xPhospho | Q6A065 2x | 0.001011 | 0 | 1 | 3 | 1 | Q6A065 | Centrosom | Q6A065 [1 | 0 | 2311.899 | 0 | 0.54 | 0.408193 | 21.8 | 87.3 | 112.7 | 37.84 | 16.45 | | High | High | High | High | High | High | High | High | High | High | High | 0 | 0.000241 | 3.05 |
| FALSE | High | FPTDYASTS | [R].FPTDYA | 1xPhospho | Q6A065 1x | 0.043236 | 0.003228 | 1 | 3 | 1 | Q6A065 | Centrosom | Q6A065 [1 | 1 | 2887.319 | 0 | 1.35 | 0.000459 | 15.32 | 56.4 | 143.6 | 17.19 | 11.09 | | High | High | High | High | High | High | High | High | High | High | High | 0.001651 | 0.01639 | 3.15 |
| FALSE | High | FPTQDASP | [K].FPTQD | 1xPhospho | Q80YR7 1x | 4.8E-05 | 0 | 1 | 1 | 1 | Q80YR7 | Claspin OS | Q80YR7 [9 | 0 | 1877.972 | 0 | -0.35 | 0.005206 | 3.23 | 115.7 | 84.3 | 5 | 8.33 | | High | High | High | High | High | High | High | High | High | High | High | 0 | 7.89E-06 | 2.46 |
| FALSE | High | FSAVLVEPF | [K].FSAVLV | 1xPhospho | Q62318 1x | 3.38E-05 | 0 | 1 | 1 | 6 | Q62318 | Transcripti | Q62318 [8 | 0 | 3241.623 | 0 | -1.81 | 0.001021 | 1.6 | 155.7 | 44.3 | 10.53 | 10.05 | | High | High | High | High | High | High | High | High | High | High | High | 2.58E-05 | 0.000412 | 5.61 |
| FALSE | High | FSDDEDEEI | [K].FSDDED | 1xPhospho | Q7TPM1 1 | 1.03E-06 | 0 | 1 | 3 | 2 | Q7TPM1 | Protein PRI | Q7TPM1 [3 | 0 | 1964.878 | 0 | -0.01 | 0.375116 | 4.13 | 102.2 | 97.8 | 15 | 6.23 | | High | High | High | High | High | High | High | High | High | High | High | 0 | 5.86E-06 | 3.6 |
| FALSE | High | FSDDSDDD | [K].FSDDSD | 1xPhospho | Q64127 1x | 1.26E-06 | 0 | 1 | 5 | 3 | Q64127 | Transcripti | Q64127 [1 | 0 | 1851.764 | 0 | -0.79 | 0.000772 | 3.01 | 129.4 | 70.6 | 8.88 | 8.77 | | High | High | High | High | High | High | High | High | High | High | High | 0 | 1.32E-07 | 4.51 |
| FALSE | High | FSDDSDDD | [K].FSDDSD | 2xPhospho | Q64127 2x | 3.04E-06 | 0 | 1 | 5 | 2 | Q64127 | Transcripti | Q64127 [1 | 0 | 1931.73 | 0 | -1.31 | 0.001128 | 4.1 | 142.5 | 57.5 | 10.98 | 21.86 | | High | High | High | High | High | High | High | High | High | High | High | 0 | 3.56E-07 | 4.47 |
| FALSE | High | FSDEFSPSH | [R].FSDEFS | 1xPhospho | Q99MJ9 1x | 0.011447 | 0.000816 | 1 | 3 | 2 | Q99MJ9 | ATP-depen | Q99MJ9 [8 | 0 | 1718.819 | 0 | -0.48 | 0.000774 | 2.08 | 117.2 | 82.8 | 3.52 | 3.24 | | High | High | High | High | High | High | High | High | High | High | High | 0.003809 | 0.04105 | 2 |
| FALSE | High | FSDFLGLHS | [R].FSDFLG | 1xPhospho | Q3U5L3 1x | 0.000322 | 0 | 1 | 3 | 1 | Q3U5L3 | Putative ur | Q3U5L3 [1 | 0 | 1688.881 | 0 | 1.75 | 0.000791 | 2.44 | 44.9 | 155.1 | 16.82 | 9.89 | | High | High | High | High | High | High | High | High | High | High | High | 0 | 6.69E-05 | 2.39 |
| FALSE | High | FSDSEDDV | [R].FSDSED | 2xPhospho | Q7TT28 2x | 8.58E-06 | 0 | 1 | 4 | 2 | Q7TT28 | RNA exonu | Q7TT28 [2 | 0 | 2041.881 | 0 | -0.65 | 0.015314 | 3.88 | 121.9 | 78.1 | 11.8 | 8.09 | | High | High | High | High | High | High | High | High | High | High | High | 0 | 1.14E-06 | 4.08 |
| FALSE | High | FSDTSTPQI | [R].FSDTST | 1xCarbami | S4R1P5 1x | 3.57E-09 | 0 | 1 | 6 | 4 | S4R1P5 | Dystonin | S4R1P5 [24 | 0 | 3071.35 | 0 | 0.7 | 0.00196 | 6.14 | 76.3 | 123.7 | 9.33 | 7.03 | | High | High | High | High | High | High | High | High | High | High | High | 0 | 1.81E-10 | 4.72 |
| FALSE | High | FSEMMDH | [R].FSEMM | 1xPhospho | Q9R0Q7 1x | 6.22E-07 | 0 | 1 | 2 | 3 | Q9R0Q7 | Prostaglan | Q9R0Q7 [1 | 0 | 4183.65 | 0 | -0.48 | 0.033416 | 3.09 | 118.5 | 81.5 | 6.79 | 20.71 | | High | High | High | High | High | High | High | High | High | High | High | 0 | 5.98E-08 | 5.59 |
| FALSE | High | FSEMMDH | [R].FSEMM | 1xDeamida | Q9R0Q7 1x | 3.77E-05 | 0 | 1 | 2 | 4 | Q9R0Q7 | Prostaglan | Q9R0Q7 [1 | 0 | 4184.634 | 0 | -0.24 | 0.293624 | 11.07 | 106.6 | 93.4 | 9.17 | 40.16 | | High | High | High | High | High | High | High | High | High | High | High | 0.000161 | 0.001236 | 4.33 |
| FALSE | High | FSEMMDH | [R].FSEMM | 1xPhospho | Q9R0Q7 1x | 8.2E-06 | 0 | 1 | 2 | 30 | Q9R0Q7 | Prostaglan | Q9R0Q7 [1 | 0 | 4199.645 | 0 | -0.48 | 0.005262 | 6.76 | 113.6 | 86.4 | 4.14 | 14.28 | | High | High | High | High | High | High | High | High | High | High | High | 0 | 2.06E-06 | 5.41 |
| FALSE | High | FSEMMDH | [R].FSEMM | 2xPhospho | Q9R0Q7 2x | 7.98E-06 | 0 | 1 | 2 | 16 | Q9R0Q7 | Prostaglan | Q9R0Q7 [1 | 0 | 4263.617 | 0 | -0.24 | 0.059055 | 14.59 | 111 | 89 | 4.63 | 16.4 | | High | High | High | High | High | High | High | High | High | High | High | 0 | 1.09E-05 | 4.58 |
| FALSE | High | FSEPNTYID | [K].FSEPNT | 1xPhospho | Q5DTU0-2 | 0.003721 | 0.00021 | 1 | 3 | 1 | Q5DTU0-2 | Isoform 2 | Q5DTU0-2 | 0 | 1904.899 | 0 | 2.94 | 0.000527 | 32.81 | 25.4 | 174.6 | 19.7 | 9.59 | | High | High | High | High | High | High | High | High | High | High | High | 9.42E-05 | 0.001043 | 3.07 |
| FALSE | High | FSAFGGSPI | [R].FSAFGC | 1xPhospho | Q76KJ5 1x | 0.000271 | 0 | 1 | 2 | 2 | Q76KJ5 | DNA-direct | Q76KJ5 [15 | 1 | 2958.513 | 0 | -0.22 | 0.046453 | 10.48 | 104.5 | 95.5 | 5.17 | 12.17 | | High | High | High | High | High | High | High | High | High | High | High | 0 | 5.49E-05 | 3.23 |
| FALSE | High | FSFGLLDSP | [K].FSFGLL | 1xPhospho | Q6P5D8 1x | 0.034281 | 0.002195 | 1 | 1 | 1 | Q6P5D8 | Structural | Q6P5D8 [8 | 0 | 1594.787 | 0 | 0.16 | 0.155644 | 9.34 | 96.9 | 103.1 | 5.69 | 10.9 | | High | High | High | High | High | High | High | High | High | High | High | 0.001168 | 0.01256 | 1.93 |
| FALSE | High | FSFLASDTE | [K].FSFLAS | 2xPhospho | E9PYH6 2x | 0.000155 | 0 | 1 | 1 | 7 | E9PYH6 | SET domai | E9PYH6 [5 | 0 | 2719.101 | 0 | -0.13 | 0.033021 | 1.69 | 105.1 | 94.9 | 4.36 | 2.56 | | High | High | High | High | High | High | High | High | High | High | High | 0 | 2.94E-05 | 4.31 |
| FALSE | High | FSFRSQEDI | [K].FSFRSQ | 1xPhospho | B2RY22 1x | 0.001586 | 5.66E-05 | 2 | 18 | 2 | B2RY22; A | Clasp1 pro | B2RY22 [1 | 1 | 2248.142 | 0 | -0.42 | 0.011632 | 6.11 | 114.5 | 85.5 | 14.35 | 10.61 | Shared | High | High | High | High | High | High | High | High | High | High | High | 7.19E-05 | 0.000784 | 2.24 |
| FALSE | High | FSFSPSAVS | [R].FSFSPS | 1xPhospho | Q4VAC9 1x | 0.01218 | 0.000889 | 1 | 3 | 4 | Q4VAC9 | Pleckstrin | Q4VAC9 [1 | 0 | 1490.724 | 0 | 1.71 | 0.000242 | 5.49 | 45.3 | 154.7 | 8 | 5.48 | | High | High | High | High | High | High | High | High | High | High | High | 0.007586 | 0.08283 | 2.22 |
| FALSE | High | FSFSPSAVS | [R].FSFSPS | 2xPhospho | Q4VAC9 2x | 0.110202 | 0.008961 | 1 | 3 | 1 | Q4VAC9 | Pleckstrin | Q4VAC9 [1 | 0 | 1570.691 | 0 | 2.57 | 0.000181 | 15.86 | 29.4 | 170.6 | 7.55 | 3.75 | | High | High | High | High | High | High | High | High | High | High | High | 0.004462 | 0.04824 | 2.42 |
| FALSE | High | FSFYGNVSI | [K].FSFYGN | 1xPhospho | B9EK76 1x | 0.000662 | 0 | 1 | 5 | 1 | B9EK76 | Signal-indu | B9EK76 [14 | 0 | 1482.698 | 0 | -0.98 | 0.000466 | 2.47 | 131.6 | 68.4 | 3.28 | 8.87 | | High | High | High | High | High | High | High | High | High | High | High | 0 | 0.00015 | 1.18 |
| FALSE | High | FSGDVVLA | [K].FSGDV | 1xPhospho | B2RXY7 1x | 0.000712 | 0 | 1 | 1 | 1 | B2RXY7 | Carbonyl r | B2RXY7 [2 | 0 | 1343.692 | 0 | 1.77 | 0.00054 | 10.76 | 46.9 | 153.1 | 6.97 | 7.55 | | High | High | High | High | High | High | High | High | High | High | High | 0 | 0.000162 | 2.25 |
| FALSE | High | FSGEEGEIE | [K].FSGEEG | 1xPhospho | Q569Z6 1x | 0.000227 | 0 | 1 | 3 | 7 | Q569Z6 | Thyroid ho | Q569Z6 [9 | 1 | 2924.272 | 0 | -0.04 | 0.651806 | 5.69 | 101.5 | 98.5 | 4 | 5.37 | | High | High | High | High | High | High | High | High | High | High | High | 7.19E-05 | 0.000777 | 3.16 |
| FALSE | High | FSGEEGEIE | [K].FSGEEG | 1xDeamida | Q569Z6 1x | 0.025073 | 0.001645 | 1 | 3 | 1 | Q569Z6 | Thyroid ho | Q569Z6 [9 | 1 | 2925.256 | 0 | 0.01 | 0.476197 | 13.39 | 100.4 | 99.6 | 9.09 | 8.81 | | High | High | High | High | High | High | High | High | High | High | High | 0.000872 | 0.008822 | 2.49 |
| FALSE | High | FSGETSASA | [K].FSGETS | 1xPhospho | Q8CI12 1x | 0.007471 | 0.00062 | 1 | 2 | 1 | Q8CI12 | Smoothelin | Q8CI12 [2 | 0 | 4388.163 | 0 | -0.77 | 0.024714 | 9.85 | 126.2 | 73.8 | 11.27 | 22.19 | | High | High | High | High | High | High | High | High | High | High | High | 0.000275 | 0.002255 | 3.03 |
| FALSE | High | FSGGFGAR | [R].FSGGF | 1xPhospho | Q3TQX5 1x | 0.000681 | 0 | 1 | 7 | 3 | Q3TQX5 | Putative ur | Q3TQX5 [5 | 0 | 1107.519 | 0 | -0.52 | 0.000363 | 1.9 | 117.2 | 82.8 | 2.51 | 6.16 | | High | High | High | High | High | High | High | High | High | High | High | 0 | 0.000155 | 2.52 |
| FALSE | High | FSGISGCSD | [K].FSGISG | 1xCarbami | Q6QI06 1x | 2.55E-07 | 0 | 1 | 2 | 5 | Q6QI06 | Rapamycin | Q6QI06 [1 | 0 | 2687.191 | 0 | 0.59 | 0.000304 | 5.25 | 79.9 | 120.1 | 5.07 | 3.05 | | High | High | High | High | High | High | High | High | High | High | High | 0 | 3.55E-05 | 3.9 |
| FALSE | High | FSGWYDAI | [R].FSGWY | 1xPhospho | Q3U7Z6 1x | 4.22E-07 | 0 | 1 | 1 | 4 | Q3U7Z6 | Phosphogl | Q3U7Z6 [2 | 0 | 2518.169 | 0 | 0.36 | 0.049377 | 2.13 | 87.3 | 112.7 | 12.78 | 4.08 | | High | High | High | High | High | High | High | High | High | High | High | 0 | 3.87E-08 | 4.34 |
| FALSE | High | FSHSYLSDS | [R].FSHSYL | 1xPhospho | Q9JMH9-6 | 2.05E-06 | 0 | 1 | 14 | 3 | Q9JMH9-6 | Isoform 6 | Q9JMH9-6 | 0 | 2124.989 | 0 | -0.57 | 0.04158 | 6.17 | 117.8 | 82.2 | 15.17 | 10.44 | | High | High | High | High | High | High | High | High | High | High | High | 0 | 3.89E-06 | 4.42 |
| FALSE | High | FSFLASDTE | [K].FSFLAS | 1xPhospho | E9PYH6 1x | 1.02E-05 | 0 | 1 | 1 | 4 | E9PYH6 | SET domai | E9PYH6 [5 | 0 | 2639.135 | 0 | -0.11 | 0.221664 | 11.43 | 107.7 | 92.3 | 11.79 | 8.53 | | High | High | High | High | High | High | High | High | High | High | High | 0 | 1.38E-06 | 4.14 |
| FALSE | High | FSAFGGSPI | [R].FSAFGC | 1xPhospho | Q76KJ5 1x | 0.016451 | 0.001124 | 1 | 2 | 1 | Q76KJ5 | DNA-direct | Q76KJ5 [15 | 0 | 2172.069 | 0 | 0.19 | 0.042366 | 8.89 | 93.8 | 106.2 | 5.65 | 4.86 | | High | High | High | High | High | High | High | High | High | High | High | 0.000585 | 0.005487 | 2.9 |
| FALSE | High | FRSPAAPSF | [K].FRSPAA | 2xPhospho | Q8CI08-2 2 | 0.041781 | 0.003126 | 1 | 2 | 1 | Q8CI08-2 | Isoform 2 | Q8CI08-2 [ | 1 | 1771.886 | 0 | 0.33 | 0.196073 | 8.73 | 89.1 | 110.9 | 9.07 | 11.12 | | High | High | High | High | High | High | High | High | High | High | High | 0.001625 | 0.01575 | 2.5 |
| FALSE | High | FRNSVEGLI | [R].FRNSVE | 1xPhospho | Q3U3E2 1x | 0.000188 | 0 | 1 | 1 | 1 | Q3U3E2 | Protein FAI | Q3U3E2 [3 | 1 | 2540.389 | 0 | 1 | 0.001779 | 16.62 | 66.9 | 133.1 | 4.36 | 8.94 | | High | High | High | High | High | High | High | High | High | High | High | 0 | 3.63E-05 | 3.26 |
| FALSE | High | FPVREDLSD | [R].FPVRED | 2xPhospho | Q5U3K5 2x | 0.00204 | 5.66E-05 | 1 | 1 | 2 | Q5U3K5 | Rab-like pr | Q5U3K5 [5 | 1 | 3312.548 | 0 | 0.01 | 0.71419 | 16.18 | 103.1 | 96.9 | 7.59 | 10.13 | | High | High | High | High | High | High | High | High | High | High | High | 0.003064 | 0.03193 | 2.77 |
| FALSE | High | FPVSEHRN | [K].FPVSEH | 1xPhospho | Q80WE4 1 | 0.042141 | 0.003162 | 1 | 4 | 1 | Q80WE4 | Kinesin-like | Q80WE4 [ | 1 | 1935.025 | 0 | -0.44 | 0.030813 | 11.13 | 116.4 | 83.6 | 11.85 | 7.36 | | High | High | High | High | High | High | High | High | High | High | High | 0.001639 | 0.01581 | 1.46 |
| FALSE | High | FQALSMQS | [K].FQALSM | 1xPhospho | Q9CU65 1x | 0.001087 | 0 | 1 | 1 | 2 | Q9CU65 | Zinc finger | Q9CU65 [6 | 0 | 3047.531 | 0 | -0.85 | 0.001028 | 3.13 | 125.4 | 74.6 | 9.1 | 12.76 | | High | High | High | High | High | High | High | High | High | High | High | 0 | 0.000262 | 3.02 |
| FALSE | High | FQEGAEMI | [K].FQEGA | 1xPhospho | Q8R0T2 1x | 0.101749 | 0.008194 | 1 | 2 | 1 | Q8R0T2 | Zinc finger | Q8R0T2 [2 | 0 | 2130.023 | 0 | 2.86 | 0.007347 | 16.02 | 24.2 | 175.8 | 9.16 | 28.74 | | High | High | High | High | High | High | High | High | High | High | High | 0.004126 | 0.04404 | 2.64 |
| FALSE | High | FQEGAEMI | [K].FQEGA | 1xPhospho | Q8R0T2 1x | 9.54E-05 | 0 | 1 | 2 | 1 | Q8R0T2 | Zinc finger | Q8R0T2 [2 | 0 | 2146.018 | 0 | 2.74 | 0.04343 | 25.52 | 25.8 | 174.2 | 18.23 | 57.24 | | High | High | High | High | High | High | High | High | High | High | High | 0 | 1.69E-05 | 3.21 |
| FALSE | High | FQEQECPP | [K].FQEQE | 1xCarbami | P62071 1x | 0.000233 | 0 | 1 | 2 | 2 | P62071 | Ras-related | P62071 [17 | 0 | 2010.883 | 0 | 0.05 | 0.184962 | 6.97 | 97.7 | 102.3 | 3.1 | 6.61 | | High | High | High | High | High | High | High | High | High | High | High | 2.58E-05 | 0.000398 | 1.43 |
| FALSE | High | FQEQECPP | [K].FQEQE | 1xCarbami | P62071 1x | 4.61E-06 | 0 | 1 | 2 | 2 | P62071 | Ras-related | P62071 [17 | 1 | 2368.141 | 0 | -0.02 | 0.816475 | 1.5 | 100.7 | 99.3 | 6.99 | 5.21 | | High | High | High | High | High | High | High | High | High | High | High | 0 | 5.66E-07 | 2.05 |
| FALSE | High | FQLAHPQS | [K].FQLAHI | 1xPhospho | Q8BHE0 1x | 0.005275 | 0.000599 | 1 | 1 | 2 | Q8BHE0 | Proline-ricl | Q8BHE0 [3 | 0 | 2968.393 | 0 | 1.75 | 0.003564 | 26.22 | 45.2 | 154.8 | 8.85 | 19.99 | | High | High | High | High | High | High | High | High | High | High | High | 0.000267 | 0.001543 | 2.81 |
| FALSE | High | FQLAPPSP | [R].FQLAPF | 1xPhospho | Q9D8C8 1x | 0.030632 | 0.001857 | 1 | 2 | 2 | Q9D8C8 | Protein ph | Q9D8C8 [7 | 0 | 1420.755 | 0 | 1.39 | 0.000372 | 10.67 | 54.8 | 145.2 | 6.67 | 4.16 | | High | High | High | High | High | High | High | High | High | High | High | 0.001011 | 0.01101 | 1.41 |
| FALSE | High | FQLPSELQ | [R].FQLPSE | 1xPhospho | Q8K4K6 1x | 1.97E-05 | 0 | 1 | 2 | 1 | Q8K4K6 | Pantothen | Q8K4K6 [1 | 0 | 2916.506 | 0 | -0.15 | 0.213721 | 6.22 | 107.5 | 92.5 | 6.99 | 8.03 | | High | High | High | High | High | High | High | High | High | High | High | 0 | 2.88E-06 | 3.97 |
| FALSE | High | FQLQHLEG | [K].FQLQH | 1xPhospho | P48754 1x | 0.000758 | 0 | 1 | 3 | 1 | P48754 | Breast can | P48754 [14 | 0 | 3255.555 | 0 | -0.44 | 0.066589 | 14.02 | 115.3 | 84.7 | 16.63 | 15.57 | | High | High | High | High | High | High | High | High | High | High | High | 0 | 0.000176 | 2.37 |
| FALSE | High | FQLSSVVSF | [R].FQLSSV | 1xPhospho | Q8C2A2 1x | 0.014797 | 0.001049 | 1 | 2 | 1 | Q8C2A2 | tRNA-splici | Q8C2A2 [1 | 0 | 1720.851 | 0 | 0.32 | 0.004874 | 7.76 | 88.4 | 111.6 | 6.11 | 3.55 | | High | High | High | High | High | High | High | High | High | High | High | 0.000555 | 0.004877 | 1.73 |
| FALSE | High | FQNYSPPP | [R].FQNYSI | 1xPhospho | Q8BZT2 1x | 0.112856 | 0.009146 | 1 | 2 | 1 | Q8BZT2 | Putative E3 | Q8BZT2 [6 | 0 | 1716.876 | 0 | 4.14 | 0.000181 | 41.74 | 10.5 | 189.5 | 35.28 | 5.61 | | High | High | High | High | High | High | High | High | High | High | High | 0.004569 | 0.04967 | 1.36 |
| FALSE | High | FQPDGESD | [K].FQPDG | 1xPhospho | A0A0N4SU | 2.75E-06 | 0 | 1 | 2 | 2 | A0A0N4SU | Pumilio ho | A0A0N4SU | 0 | 1904.868 | 0 | 0.21 | 0.013231 | 3.17 | 91.7 | 108.3 | 2.46 | 7.52 | | High | High | High | High | High | High | High | High | High | High | High | 0 | 4.62E-05 | 3.67 |
| FALSE | High | FQPDGESD | [K].FQPDG | 2xPhospho | A0A0N4SU | 0.007742 | 0.000664 | 1 | 2 | 1 | A0A0N4SU | Pumilio ho | A0A0N4SU | 0 | 1984.834 | 0 | 0.32 | 0.007591 | 8.73 | 90.6 | 109.4 | 10.94 | 10.19 | | High | High | High | High | High | High | High | High | High | High | High | 0.000294 | 0.002354 | 3.33 |
| FALSE | High | FQPHFGQ | [K].FQPHF | 1xCarbami | B2RUG9 1x | 0.065952 | 0.004766 | 1 | 7 | 1 | B2RUG9 | Adenomat | B2RUG9 [1 | 0 | 2188.984 | 0 | -0.31 | 0.059075 | 13.66 | 110.1 | 89.9 | 10.47 | 10.78 | | High | High | High | High | High | High | High | High | High | High | High | 0.002446 | 0.02641 | 1.89 |
| FALSE | High | FQQPPQPE | [R].FQQPP | 1xPhospho | A0A1S6GW | 3.07E-06 | 0 | 1 | 3 | 5 | A0A1S6GW | Uncharacte | A0A1S6GW | 0 | 2252.966 | 0 | 0.05 | 0.497915 | 4.48 | 98.1 | 101.9 | 1.34 | 7.22 | | High | High | High | High | High | High | High | High | High | High | High | 0 | 3.59E-07 | 3.76 |
| FALSE | High | FQQPPQPE | [R].FQQPP | 1xDeamida | A0A1S6GW | 0.060159 | 0.004476 | 1 | 3 | 1 | A0A1S6GW | Uncharacte | A0A1S6GW | 0 | 2253.95 | 0 | 0.1 | 0.35489 | 8 | 98.1 | 101.9 | 9.01 | 8.39 | | High | High | High | High | High | High | High | High | High | High | High | 0.002293 | 0.02384 | 2.78 |
| FALSE | High | FQQPPQPE | [R].FQQPP | 1xPhospho | A0A1S6GW | 9.54E-05 | 0 | 1 | 3 | 4 | A0A1S6GW | Uncharacte | A0A1S6GW | 1 | 2610.224 | 0 | 0.58 | 0.000726 | 3.73 | 81.3 | 118.7 | 4.66 | 5.23 | | High | High | High | High | High | High | High | High | High | High | High | 7.19E-05 | 0.000871 | 2.66 |
| FALSE | High | FQSPVPAP | [K].FQSPVF | 1xPhospho | D3Z0X5 1x | 0.083783 | 0.006345 | 1 | 6 | 1 | D3Z0X5 | Pleckstrin | D3Z0X5 [3 | 0 | 1307.671 | 0 | 0.75 | 0.000482 | 5.79 | 75.8 | 124.2 | 5.34 | 4.4 | | High | High | High | High | High | High | High | High | High | High | High | 0.003265 | 0.03518 | 1.57 |
| FALSE | High | FQSQADQ | [K].FQSQA | 1xPhospho | Q99LJ0 1x | 5.13E-09 | 0 | 1 | 1 | 3 | Q99LJ0 | CTTNBP2 N | Q99LJ0 [46 | 0 | 2484.135 | 0 | 0.57 | 0.000676 | 2.85 | 81.8 | 118.2 | 2.19 | 4.38 | | High | High | High | High | High | High | High | High | High | High | High | 0 | 2.72E-10 | 4.85 |
| FALSE | High | FQSQADQ | [K].FQSQA | 2xPhospho | Q99LJ0 2x | 0.000102 | 0 | 1 | 1 | 1 | Q99LJ0 | CTTNBP2 N | Q99LJ0 [46 | 0 | 2564.102 | 0 | -0.06 | 0.254772 | 9.29 | 103.7 | 96.3 | 10.33 | 9.65 | | High | High | High | High | High | High | High | High | High | High | High | 0 | 1.83E-05 | 4.38 |
| FALSE | High | FQSVESGA | [R].FQSVES | 1xPhospho | Q99J36 1x | 6.87E-07 | 0 | 1 | 1 | 1 | Q99J36 | THUMP do | Q99J36 [11 | 0 | 1975.984 | 0 | -0.18 | 0.6153 | 9.42 | 104.5 | 95.5 | 7.94 | 13.55 | | High | High | High | High | High | High | High | High | High | High | High | 0 | 6.64E-08 | 3.28 |
| FALSE | High | FQSVPAQP | [R].FQSVP | 1xPhospho | Q80U89 1x | 2.29E-05 | 0 | 1 | 5 | 3 | Q80U89 | MKIAA003 | Q80U89 [4 | 0 | 3725.971 | 0 | 0.17 | 0.137252 | 4.42 | 95.1 | 104.9 | 6.73 | 7.49 | | High | High | High | High | High | High | High | High | High | High | High | 0 | 3.67E-06 | 3.53 |
| FALSE | High | FQSVQIQA | [K].FQSVQ | 1xPhospho | Q6PGC1 1x | 2.62E-07 | 0 | 1 | 2 | 1 | Q6PGC1 | ATP-depen | Q6PGC1 [1 | 0 | 2439.305 | 0 | -0.23 | 0.009536 | 0.54 | 108.1 | 91.9 | 3.54 | 5.55 | | High | High | High | High | High | High | High | High | High | High | High | 0 | 2.26E-08 | 3.21 |
| FALSE | High | FQTSPAVK | [R].FQTSPA | 1xPhospho | Q6NS45 1x | 0.039011 | 0.002718 | 1 | 3 | 2 | Q6NS45 | Coiled-coil | Q6NS45 [7 | 0 | 1415.77 | 0 | -0.32 | 0.135317 | 17.46 | 112.2 | 87.8 | 7.69 | 16.19 | | High | High | High | High | High | High | High | High | High | High | High | 0.001665 | 0.01673 | 1.99 |
| FALSE | High | FRGSQEDL | [R].FRGSQ | 1xCarbami | Q3TZ34 1x | 1.66E-05 | 0 | 1 | 3 | 2 | Q3TZ34 | Putative ur | Q3TZ34 [1 | 1 | 2331.075 | 0 | -0.08 | 0.634657 | 6.35 | 104.1 | 95.9 | 6.11 | 5.84 | | High | High | High | High | High | High | High | High | High | High | High | 0 | 2.39E-06 | 3.74 |
| FALSE | High | FRGSQEDL | [R].FRGSQ | 1xCarbami | Q3TZ34 1x | 0.000128 | 0 | 1 | 3 | 2 | Q3TZ34 | Putative ur | Q3TZ34 [1 | 1 | 2332.059 | 0 | 0.58 | 0.003277 | 15.15 | 77.7 | 122.3 | 7.59 | 9.62 | | High | High | High | High | High | High | High | High | High | High | High | 0 | 2.38E-05 | 3.99 |
| FALSE | High | FRNGTLILS | [K].FRNGTI | 1xDeamida | Q8CIL4 1x | 0.004256 | 0.00036 | 1 | 1 | 1 | Q8CIL4 | Uncharacte | Q8CIL4 [25 | 1 | 2114.166 | 0 | -0.27 | 0.007798 | 5.81 | 109.8 | 90.2 | 6.91 | 3.23 | | High | High | High | High | High | High | High | High | High | High | High | 0.000161 | 0.001203 | 3.02 |
| FALSE | High | FSLCSDNLE | [R].FSLCSD | 1xCarbami | Q6PAR5 1x | 3.23E-05 | 0 | 1 | 4 | 2 | Q6PAR5 | GTPase-act | Q6PAR5 [5 | 0 | 2291.021 | 0 | 0.11 | 0.986826 | 22.6 | 101.8 | 98.2 | 17.85 | 10.42 | | High | High | High | High | High | High | High | High | High | High | High | 0 | 5.04E-06 | 3.34 |
| FALSE | High | GASAATGII | [K].GASAA | 1xCarbami | Q8R5H1 1x | 9.11E-08 | 0 | 2 | 4 | 8 | Q8R5H1; Q | Ubiquitin c | Q8R5H1 [9 | 0 | 3774.485 | 0 | -0.06 | 0.961025 | 11.87 | 101.9 | 98.1 | 2.72 | 13.97 | Shared | High | High | High | High | High | High | High | High | High | High | High | 0 | 6.89E-09 | 5.36 |
| FALSE | High | GASAATGII | [K].GASAA | 1xCarbami | Q8R5H1 1x | 4.91E-06 | 0 | 2 | 4 | 13 | Q8R5H1; Q | Ubiquitin c | Q8R5H1 [9 | 0 | 3775.469 | 0 | -0.01 | 0.734569 | 1.43 | 100.4 | 99.6 | 19.06 | 18.26 | Shared | High | High | High | High | High | High | High | High | High | High | High | 0 | 7.66E-07 | 4.72 |
| FALSE | High | GASAATGII | [K].GASAA | 1xCarbami | Q8R5H1 2x | 8.63E-05 | 0 | 2 | 4 | 2 | Q8R5H1; Q | Ubiquitin c | Q8R5H1 [9 | 0 | 3838.457 | 0 | 0.27 | 0.023525 | 9.48 | 90.7 | 109.3 | 4.06 | 11.77 | Shared | High | High | High | High | High | High | High | High | High | High | High | 0 | 1.52E-05 | 4.45 |
| FALSE | High | GEVQTVAS | [K].GEVQT | 1xPhospho | G5E8B9 1x | 7.21E-05 | 0 | 1 | 2 | 1 | G5E8B9 | MCG13089 | G5E8B9 [3 | 0 | 2818.357 | 0 | -0.56 | 0.005809 | 8.36 | 124 | 76 | 7.19 | 18.12 | | High | High | High | High | High | High | High | High | High | High | High | 0 | 1.24E-05 | 3.56 |
| FALSE | High | GEVVSPLA | [K].GEVVSI | 1xPhospho | P30415 1x | 0.004613 | 0.000456 | 1 | 1 | 1 | P30415 | NK-tumor | P30415 [11 | 0 | 1487.757 | 0 | -0.31 | 0.005952 | 1.39 | 111.3 | 88.7 | 5.61 | 4.2 | | High | High | High | High | High | High | High | High | High | High | High | 0.000204 | 0.001325 | 1.23 |
| FALSE | High | GEWFLLGS | [R].GEWFL | 1xPhospho | Q9D071 1x | 0.002743 | 0.000109 | 1 | 3 | 2 | Q9D071 | MMS19 nu | Q9D071 [1 | 0 | 1458.687 | 0 | -0.37 | 0.005655 | 3.98 | 112.5 | 87.5 | 4.83 | 4.45 | | High | High | High | High | High | High | High | High | High | High | High | 0.000275 | 0.001977 | 2.25 |
| FALSE | High | GEYEVTPPI | [K].GEYEV | 1xPhospho | E9Q616 1x | 0.016892 | 0.001204 | 1 | 1 | 2 | E9Q616 | AHNAK nu | E9Q616 [7 | 0 | 1557.797 | 0 | 3.28 | 9.77E-05 | 14.84 | 18.7 | 181.3 | 19.17 | 2.42 | | High | High | High | High | High | High | High | High | High | High | High | 0.000638 | 0.005634 | 2.44 |
| FALSE | High | GEYEVTPPI | [K].GEYEV | 1xPhospho | E9Q616 1x | 0.001767 | 5.66E-05 | 1 | 1 | 1 | E9Q616 | AHNAK nu | E9Q616 [7 | 1 | 2456.329 | 0 | 3.35 | 0.000475 | 6.6 | 17.3 | 182.7 | 18.93 | 12.58 | | High | High | High | High | High | High | High | High | High | High | High | 2.58E-05 | 0.000454 | 2.78 |
| FALSE | High | GFAFVTFES | [R].GFAFVT | 1xPhospho | Q91VM5 1 | 0.022574 | 0.001542 | 1 | 5 | 1 | Q91VM5 | RNA bindir | Q91VM5 [ | 0 | 2025.013 | 0 | -0.39 | 0.033078 | 4.59 | 113.3 | 86.7 | 4.91 | 9.92 | | High | High | High | High | High | High | High | High | High | High | High | 0.000814 | 0.007845 | 2.87 |
| FALSE | High | GFDTSPEH | [K].GFDTSF | 1xPhospho | Q8CHI8 1x | 0.000289 | 0 | 1 | 5 | 4 | Q8CHI8 | E1A-bindin | Q8CHI8 [9 | 0 | 2096.985 | 0 | -0.87 | 0.00131 | 1.26 | 127.7 | 72.3 | 6.28 | 8.49 | | High | High | High | High | High | High | High | High | High | High | High | 0 | 0.000178 | 2.06 |
| FALSE | High | GFGDGYN | [R].GFGDG | 1xPhospho | O88569 1x | 1.27E-09 | 0 | 1 | 3 | 11 | O88569 | Heterogen | O88569 [2 | 0 | 2804.169 | 0 | 0.77 | 0.002272 | 6.24 | 69.7 | 130.3 | 6.61 | 12.24 | | High | High | High | High | High | High | High | High | High | High | High | 0 | 1.06E-09 | 6.39 |
| FALSE | High | GFGDGYN | [R].GFGDG | 1xDeamida | O88569 1x | 2.44E-11 | 0 | 1 | 3 | 23 | O88569 | Heterogen | O88569 [2 | 0 | 2805.153 | 0 | 0.85 | 0.000278 | 2.18 | 69.6 | 130.4 | 7.22 | 6.51 | | High | High | High | High | High | High | High | High | High | High | High | 0 | 1.17E-11 | 5.29 |
| FALSE | High | GFGDGYN | [R].GFGDG | 2xDeamida | O88569 1x | 0.021607 | 0.001507 | 1 | 3 | 3 | O88569 | Heterogen | O88569 [2 | 0 | 2806.137 | 0 | 0.55 | 0.004532 | 6.13 | 84.7 | 115.3 | 7.11 | 6.46 | | High | High | High | High | High | High | High | High | High | High | High | 0.001999 | 0.0205 | 3.43 |
| FALSE | High | GFGDSFGR | [R].GFGDS | 1xPhospho | Q8C854 1x | 0.006834 | 0.00062 | 1 | 7 | 1 | Q8C854 | Myelin exp | Q8C854 [4 | 0 | 1151.508 | 0 | 1.18 | 0.046805 | 12.53 | 60.1 | 139.9 | 16.7 | 29.19 | | High | High | High | High | High | High | High | High | High | High | High | 0.000275 | 0.002056 | 0.96 |
| FALSE | High | GFGFGQG | [K].GFGFG | 1xPhospho | Q4FJX4 1x | 4.19E-08 | 0 | 1 | 1 | 2 | Q4FJX4 | Csrp1 prot | Q4FJX4 [17 | 0 | 1742.81 | 0 | 0.19 | 0.066436 | 11.36 | 93.5 | 106.5 | 5.86 | 8.51 | | High | High | High | High | High | High | High | High | High | High | High | 0 | 2.89E-09 | 4.76 |
| FALSE | High | GFGFVCFS | [K].GFGFV | 1xCarbami | P29341 1x | 0.0009 | 0 | 1 | 16 | 1 | P29341 | Polyadenyl | P29341 [33 | 0 | 2201.039 | 0 | -0.41 | 0.020251 | 4.54 | 113.5 | 86.5 | 4.97 | 7.4 | | High | High | High | High | High | High | High | High | High | High | High | 0 | 0.000211 | 2.5 |
| FALSE | High | GETVQEICS | [K].GETVQ | 1xCarbami | A0A1B0GS | 0.000446 | 0 | 1 | 6 | 3 | A0A1B0GS | Nuclear po | A0A1B0GS | 0 | 1584.729 | 0 | -0.26 | 0.019611 | 6.58 | 110 | 90 | 3.34 | 7.65 | | High | High | High | High | High | High | High | High | High | High | High | 0 | 9.63E-05 | 2.44 |
| FALSE | High | GFGFVDFN | [K].GFGFVI | 1xPhospho | P09405 1x | 1.03E-05 | 0 | 1 | 5 | 2 | P09405 | Nucleolin | P09405 [6 | 0 | 2099.973 | 0 | -0.32 | 0.001788 | 0.51 | 113.7 | 86.3 | 3.78 | 4.4 | | High | High | High | High | High | High | High | High | High | High | High | 0 | 1.39E-06 | 3.57 |
| FALSE | High | GFGSLPCIL | [R].GFGSLF | 2xCarbami | Q8K1S6 1x | 3.64E-06 | 0 | 1 | 2 | 1 | Q8K1S6 | Protein spi | Q8K1S6 [3 | 0 | 2347.159 | 0 | -1.17 | 0.001654 | 5.03 | 141.3 | 58.7 | 7.45 | 16.22 | | High | High | High | High | High | High | High | High | High | High | High | 0 | 4.33E-07 | 3.58 |
| FALSE | High | GFGSSQNF | [R].GFGSS | 1xPhospho | E9QL43 1x | 6.47E-05 | 0 | 1 | 4 | 2 | E9QL43 | Nucleopor | E9QL43 [9 | 0 | 2434.169 | 0 | -0.13 | 0.02355 | 0.59 | 104.8 | 95.2 | 5.45 | 1.74 | | High | High | High | High | High | High | High | High | High | High | High | 0 | 1.1E-05 | 3.45 |
| FALSE | High | GFGSSVPE | [R].GFGSSV | 1xDeamida | Q3UJB9 1x | 0.001674 | 5.66E-05 | 1 | 2 | 1 | Q3UJB9 | Enhancer | Q3UJB9 [7 | 0 | 3474.729 | 1 | -0.01 | 0.703297 | 36.32 | 100.2 | 99.8 | 18.84 | 40.38 | | High | High | High | High | High | High | High | High | High | High | High | 2.58E-05 | 0.000424 | 2.41 |
| FALSE | High | GFLDGVYS | [K].GFLDG | 1xPhospho | Q80X90 1x | 0.001058 | 0 | 1 | 1 | 2 | Q80X90 | Filamin-B | Q80X90 [5 | 0 | 2565.235 | 0 | 0.46 | 0.029397 | 13.32 | 85.4 | 114.6 | 5.98 | 13.51 | | High | High | High | High | High | High | High | High | High | High | High | 0 | 0.000255 | 3.41 |
| FALSE | High | GFIDQNFSI | [R].GFIDQN | 1xPhospho | A0A087WF | 0.009843 | 0.000736 | 1 | 1 | 1 | A0A087WF | Transcripti | A0A087WF | 0 | 1791.908 | 0 | -1.18 | 0.001599 | 4.61 | 139 | 61 | 6.84 | 13.56 | | High | High | High | High | High | High | High | High | High | High | High | 0.000344 | 0.003097 | 1.63 |

| | | | | | | | | | | | | | | | | | | | | | | | | | | | | | | | | | | | | | | |
|---|---|---|---|---|---|---|---|---|---|---|---|---|---|---|---|---|---|---|---|---|---|---|---|---|---|---|---|---|---|---|---|---|---|---|---|---|---|---|
| FALSE | High | GFLSPGSQ | [K].GFLSPG | 1xPhospho | Q6PR54-3 1 | 0.009756 | 0.000736 | 2 | 3 | 2 | Q6PR54-3; | Isoform 3 c | Q6PR54-3 | 0 | 1632.84 | 0 | -1.62 | 0.000281 | 0.3 | 151.1 | 48.9 | 3.7 | 12.96 | Shared | High | High | High | High | High | High | High | High | High | High | High | 0.000344 | 0.003061 | 1.96 |
| FALSE | High | GFNGCDSF | [K].GFNGC | 1xCarbami | Q63943 1x | 2.51E-08 | 0 | 1 | 4 | 2 | Q63943 | Myocyte-s | Q63943 [9 | 0 | 3273.455 | 0 | 0.31 | 0.100254 | 15.43 | 88.7 | 111.3 | 10.18 | 8.62 | | High | High | High | High | High | High | High | High | High | High | High | 0 | 1.61E-09 | 5.15 |
| FALSE | High | GFNGCDSF | [K].GFNGC | 1xCarbami | Q63943 1x | 2.08E-06 | 0 | 1 | 4 | 4 | Q63943 | Myocyte-s | Q63943 [9 | 0 | 3274.439 | 0 | 0.38 | 0.005931 | 5.91 | 86.5 | 113.5 | 4.64 | 7.87 | | High | High | High | High | High | High | High | High | High | High | High | 0 | 2.31E-07 | 5.33 |
| FALSE | High | GFNGCDSF | [K].GFNGC | 1xCarbami | Q63943 2x | 0.004033 | 0.000311 | 1 | 4 | 2 | Q63943 | Myocyte-s | Q63943 [9 | 0 | 3353.421 | 0 | 1.5 | 0.004293 | 10.19 | 53.7 | 146.3 | 11.44 | 21.67 | | High | High | High | High | High | High | High | High | High | High | High | 0.001651 | 0.01652 | 3.19 |
| FALSE | High | GFQPSRPT | [K].GFQPSI | 1xPhospho | O55029 1x | 2.08E-05 | 0 | 1 | 1 | 1 | O55029 | Coatomer | O55029 [8 | 0 | 4046.077 | 0 | 0.12 | 0.148461 | 9.35 | 92.9 | 107.1 | 17.66 | 9.08 | | High | High | High | High | High | High | High | High | High | High | High | 0 | 3.08E-06 | 2.98 |
| FALSE | High | GFSDSGGG | [R].GFSDS | 1xPhospho | Q3TUQ5 1 | 0.000284 | 0 | 1 | 3 | 1 | Q3TUQ5 | Pinin OS=M | Q3TUQ5 [6 | 0 | 1385.63 | 0 | 1.18 | 0.008616 | 24.89 | 61.2 | 138.8 | 11.96 | 14.21 | | High | High | High | High | High | High | High | High | High | High | High | 0 | 5.82E-05 | 2.46 |
| FALSE | High | GFSDSGGG | [R].GFSDS | 1xPhospho | Q3TUQ5 1 | 4.55E-05 | 0 | 1 | 3 | 2 | Q3TUQ5 | Pinin OS=M | Q3TUQ5 [6 | 0 | 1614.793 | 0 | 1.32 | 0.007755 | 29.26 | 58.8 | 141.2 | 19.11 | 12.03 | | High | High | High | High | High | High | High | High | High | High | High | 0 | 5.98E-05 | 3.85 |
| FALSE | High | GFSFVATSI | [K].GFSFVA | 1xPhospho | E9QK39 1x | 1.1E-06 | 0 | 1 | 8 | 1 | E9QK39 | Ribosomal | E9QK39 [5 | 0 | 2087.05 | 0 | -1.29 | 0.009994 | 4.87 | 142.3 | 57.7 | 22.49 | 10.03 | | High | High | High | High | High | High | High | High | High | High | High | 0 | 1.13E-07 | 3.21 |
| FALSE | High | GFGLVAYA | [K].GFGLV | 1xPhospho | Q3TCX3 1x | 5.5E-08 | 0 | 1 | 1 | 2 | Q3TCX3 | UPF0469 p | Q3TCX3 [5 | 0 | 2843.293 | 0 | -1.45 | 0.003135 | 1.3 | 149.1 | 50.9 | 14.76 | 30.14 | | High | High | High | High | High | High | High | High | High | High | High | 0 | 3.89E-09 | 3.46 |
| FALSE | High | GFSQYGVS | [R].GFSQY | 1xPhospho | Q8VE19 1x | 0.000305 | 0 | 1 | 2 | 1 | Q8VE19 | GATOR cor | Q8VE19 [7 | 0 | 1852.925 | 0 | -1.21 | 0.000511 | 0.58 | 141 | 59 | 6.85 | 10.67 | | High | High | High | High | High | High | High | High | High | High | High | 0 | 6.29E-05 | 1.42 |
| FALSE | High | GETSGNES | [K].GETSGI | 2xPhospho | E9Q4E4 2x | 0.002895 | 0.00016 | 1 | 4 | 3 | E9Q4E4 | DENN dom | E9Q4E4 [4 | 0 | 2214.961 | 0 | -0.54 | 0.001095 | 2.86 | 122.3 | 77.7 | 4.96 | 7.09 | | High | High | High | High | High | High | High | High | High | High | High | 9.42E-05 | 0.000998 | 2.35 |
| FALSE | High | GETPPSGT | [K].GETPPS | 1xPhospho | A0A0R4J1C | 1.02E-06 | 0 | 1 | 11 | 2 | A0A0R4J1C | Disabled h | A0A0R4J1C | 0 | 2516.175 | 0 | 1.78 | 0.000384 | 15.82 | 43.1 | 156.9 | 22.26 | 3.62 | | High | High | High | High | High | High | High | High | High | High | High | 0 | 1.04E-07 | 4.41 |
| FALSE | High | GEGAPGEE | [K].GEGAP | 1xPhospho | G3X8T2 1x | 0.003342 | 0.00016 | 1 | 4 | 2 | G3X8T2 | RIKEN cDN | G3X8T2 [1 | 0 | 3363.656 | 0 | -0.67 | 0.000396 | 1.01 | 124.5 | 75.5 | 8.36 | 8.39 | | High | High | High | High | High | High | High | High | High | High | High | 9.42E-05 | 0.00103 | 2.91 |
| FALSE | High | GEGESPPV | [K].GEGESI | 1xDeamida | P28667 1x | 0.00582 | 0.00062 | 1 | 1 | 2 | P28667 | MARCKS-re | P28667 [44 | 0 | 3788.743 | 0 | -0.28 | 0.110284 | 12.6 | 105.8 | 94.2 | 19.93 | 10.97 | | High | High | High | High | High | High | High | High | High | High | High | 0.000327 | 0.002744 | 3.66 |
| FALSE | High | GEGLEPQA | [R].GEGLEI | 1xPhospho | B7ZCJ1 1xF | 0.022378 | 0.001542 | 1 | 5 | 3 | B7ZCJ1 | Rho GTPas | B7ZCJ1 [18 | 0 | 3492.685 | 0 | 0.78 | 0.00162 | 6.88 | 72.7 | 127.3 | 28.48 | 11.87 | | High | High | High | High | High | High | High | High | High | High | High | 0.000814 | 0.007721 | 2.53 |
| FALSE | High | GEGPPTPP | [R].GEGPP | 2xPhospho | Q9ES52 2x | 0.002623 | 0.000109 | 1 | 3 | 1 | Q9ES52 | Phosphatic | Q9ES52 [9 | 0 | 2204.081 | 0 | -1.03 | 0.002067 | 2.1 | 134.5 | 65.5 | 5.98 | 11.7 | | High | High | High | High | High | High | High | High | High | High | High | 4.94E-05 | 0.000706 | 2.69 |
| FALSE | High | GEGVSQV | [R].GEGVS | 1xPhospho | B9EKJ7 1xF | 0.020499 | 0.001472 | 1 | 2 | 1 | B9EKJ7 | Tankyrase | B9EKJ7 [52 | 0 | 2128.027 | 0 | -0.55 | 0.00112 | 3.28 | 119.5 | 80.5 | 4.5 | 6.96 | | High | High | High | High | High | High | High | High | High | High | High | 0.000786 | 0.007032 | 3.06 |
| FALSE | High | GEGVSQV | [R].GEGVS | 2xPhospho | B9EKJ7 2xF | 7.74E-08 | 0 | 1 | 2 | 2 | B9EKJ7 | Tankyrase | B9EKJ7 [52 | 0 | 2207.994 | 0 | -0.57 | 0.017093 | 5.22 | 118.9 | 81.1 | 12.04 | 6.41 | | High | High | High | High | High | High | High | High | High | High | High | 0 | 5.75E-09 | 4.21 |
| FALSE | High | GEGVSQV | [R].GEGVS | 2xPhospho | B9EKJ7 2xF | 0.077188 | 0.005893 | 1 | 2 | 1 | B9EKJ7 | Tankyrase | B9EKJ7 [52 | 1 | 4281.147 | 0 | -0.99 | 0.009192 | 7.18 | 134.5 | 65.5 | 23 | 20.25 | | High | High | High | High | High | High | High | High | High | High | High | 0.003035 | 0.03179 | 3.04 |
| FALSE | High | GELDLPLEN | [K].GELDLF | 1xPhospho | B2RRE7 1x | 8.74E-06 | 0 | 1 | 2 | 2 | B2RRE7 | OTU doma | B2RRE7 [8 | 0 | 2094.114 | 0 | 0.41 | 0.048207 | 19.9 | 87.3 | 112.7 | 12.21 | 11.09 | | High | High | High | High | High | High | High | High | High | High | High | 0.001453 | 0.0146 | 3.05 |
| FALSE | High | GELIPISPST | [R].GELIPIS | 1xPhospho | P48972 1x | 0.000157 | 0 | 1 | 1 | 4 | P48972 | Myb-relate | P48972 [39 | 0 | 2978.589 | 0 | -1.04 | 0.000474 | 1.35 | 134.8 | 65.2 | 6.22 | 3.68 | | High | High | High | High | High | High | High | High | High | High | High | 0.003035 | 0.03181 | 2.61 |
| FALSE | High | GELPSDPAI | [K].GELPSD | 1xPhospho | H9LAN6 1x | 6.27E-07 | 0 | 1 | 7 | 1 | H9LAN6 | Histone-lys | H9LAN6 [1 | 0 | 2314.089 | 0 | -0.19 | 0.019369 | 5.82 | 106.6 | 93.4 | 4.54 | 4.35 | | High | High | High | High | High | High | High | High | High | High | High | 0 | 6.04E-08 | 3.41 |
| FALSE | High | GENKPVLF | [K].GENKP | 1xPhospho | Q9ERU9 1x | 8.5E-06 | 0 | 1 | 1 | 1 | Q9ERU9 | E3 SUMO- | Q9ERU9 [2 | 0 | 4136.062 | 0 | 0.15 | 0.536322 | 81.43 | 85.2 | 114.8 | 17.91 | 47.44 | | High | High | High | High | High | High | High | High | High | High | High | 0 | 1.12E-06 | 4.82 |
| FALSE | High | GEPGLDSE | [R].GEPGLI | 1xPhospho | Q99MU3 1 | 1.3E-05 | 0 | 1 | 4 | 3 | Q99MU3 | Double-str | Q99MU3 [ | 0 | 3508.597 | 0 | 0.46 | 0.007032 | 4.99 | 81.6 | 118.4 | 7.03 | 8.49 | | High | High | High | High | High | High | High | High | High | High | High | 0 | 1.81E-06 | 4.47 |
| FALSE | High | GEPNVSYI | [R].GEPNV | 1xCarbami | Q2NL51 1x | 6.37E-06 | 0 | 2 | 4 | 11 | Q2NL51; E | Glycogen s | Q2NL51 [2 | 0 | 1590.718 | 0 | 0.03 | 0.269767 | 2.95 | 97.6 | 102.4 | 2.55 | 5.82 | Shared | High | High | High | High | High | High | High | High | High | High | High | 0 | 6.87E-06 | 3.48 |
| FALSE | High | GETSGNES | [K].GETSGI | 1xPhospho | E9Q4E4 1x | 0.000772 | 0 | 1 | 4 | 3 | E9Q4E4 | DENN dom | E9Q4E4 [4 | 0 | 2134.995 | 0 | -0.76 | 0.000904 | 5.08 | 126.5 | 73.5 | 1.24 | 11.7 | | High | High | High | High | High | High | High | High | High | High | High | 0 | 0.000178 | 2.68 |
| FALSE | High | GEPPTPPSI | [R].GEPPTF | 1xPhospho | Q924A2 1x | 5.65E-05 | 0 | 1 | 3 | 4 | Q924A2 | Protein cap | Q924A2 [2 | 0 | 2891.289 | 0 | 0.22 | 0.034505 | 10.6 | 88.5 | 111.5 | 8.02 | 10.94 | | High | High | High | High | High | High | High | High | High | High | High | 0 | 3.31E-05 | 3.37 |
| FALSE | High | GEPQTPTQ | [R].GEPQTI | 1xPhospho | Q62187 1x | 1.97E-05 | 0 | 1 | 1 | 3 | Q62187 | Transcripti | Q62187 [1 | 0 | 2989.517 | 0 | 0.89 | 0.002238 | 1.5 | 73 | 127 | 12.17 | 7.04 | | High | High | High | High | High | High | High | High | High | High | High | 0 | 2.9E-06 | 4.19 |
| FALSE | High | GEPSGIEPS | [R].GEPSGI | 1xPhospho | Q91XC0 1x | 3.87E-08 | 0 | 1 | 1 | 1 | Q91XC0 | LIM domai | Q91XC0 [3 | 0 | 2734.281 | 0 | -1.53 | 0.000765 | 2.76 | 148.7 | 51.3 | 7.85 | 12.48 | | High | High | High | High | High | High | High | High | High | High | High | 0 | 2.64E-09 | 3.91 |
| FALSE | High | GEPSPTWE | [K].GEPSPT | 1xPhospho | Q5XJV7-2 1 | 6.02E-05 | 0 | 1 | 3 | 2 | Q5XJV7-2 | Isoform 2 c | Q5XJV7-2 [ | 0 | 2113.025 | 0 | -0.73 | 0.003994 | 7.57 | 125.3 | 74.7 | 3.32 | 16.87 | | High | High | High | High | High | High | High | High | High | High | High | 0 | 7.3E-05 | 3.4 |
| FALSE | High | GERDSQTC | [R].GERDS | 1xPhospho | G3UXW9 1 | 0.118334 | 0.009621 | 1 | 6 | 1 | G3UXW9 | COP9 signa | G3UXW9 [ | 1 | 1985.047 | 0 | 0.74 | 0.056061 | 50.01 | 65.5 | 134.5 | 45.06 | 9.6 | | High | High | High | High | High | High | High | High | High | High | High | 0.004866 | 0.05279 | 3.02 |
| FALSE | High | GESAADSD | [R].GESAAI | 1xPhospho | Q8BSN5 1x | 0.000274 | 0 | 1 | 2 | 4 | Q8BSN5 | PHD finger | Q8BSN5 [6 | 0 | 2144.897 | 0 | -1.05 | 0.00074 | 3.18 | 136.6 | 63.4 | 6.55 | 5.91 | | High | High | High | High | High | High | High | High | High | High | High | 0 | 5.58E-05 | 3.82 |
| FALSE | High | GESENEIED | [K].GESENI | 1xPhospho | O09053 1x | 0.00095 | 0 | 1 | 4 | 1 | O09053 | Werner syı | O09053 [3 | 0 | 1826.837 | 0 | 0.49 | 0.054444 | 9.51 | 76.2 | 123.8 | 19.77 | 24.99 | | High | High | High | High | High | High | High | High | High | High | High | 0 | 0.000224 | 2.96 |
| FALSE | High | GESGTSTA | [K].GESGTS | 1xPhospho | A3KGF9 1x | 0.100129 | 0.008067 | 1 | 1 | 1 | A3KGF9 | Uncharacte | A3KGF9 [4 | 0 | 3476.638 | 0 | 2.37 | 0.000191 | 45.03 | 33.3 | 166.7 | 49.87 | 11.09 | | High | High | High | High | High | High | High | High | High | High | High | 0.004032 | 0.0431 | 2.57 |
| FALSE | High | GESGTSTA | [K].GESGTS | 1xDeamida | A3KGF9 1x | 0.019107 | 0.001402 | 1 | 1 | 1 | A3KGF9 | Uncharacte | A3KGF9 [4 | 0 | 3477.622 | 0 | 3.06 | 0.001964 | 11.7 | 25.6 | 174.4 | 44.43 | 13.13 | | High | High | High | High | High | High | High | High | High | High | High | 0.00074 | 0.006447 | 2.72 |
| FALSE | High | GESGVPEE | [K].GESGVI | 1xPhospho | A6H624 1x | 2.88E-06 | 0 | 1 | 1 | 4 | A6H624 | LysM, puta | A6H624 [1 | 0 | 2185.985 | 0 | 0.37 | 0.031329 | 12.8 | 86.3 | 113.7 | 3.53 | 12.36 | | High | High | High | High | High | High | High | High | High | High | High | 2.58E-05 | 0.000369 | 4.19 |
| FALSE | High | GESIEPLDP | [R].GESIEP | 1xPhospho | Q61526 1x | 0.001437 | 5.66E-05 | 1 | 1 | 2 | Q61526 | Receptor t | Q61526 [6 | 0 | 1838.919 | 0 | 3.16 | 0.000832 | 37.61 | 20.4 | 179.6 | 13.69 | 12.39 | | High | High | High | High | High | High | High | High | High | High | High | 0.000161 | 0.001141 | 2.99 |
| FALSE | High | GESQTPVC | [R].GESQTI | 1xPhospho | A0A1W2P6 | 0.085859 | 0.006406 | 1 | 8 | 1 | A0A1W2P6 | Bromodom | A0A1W2P6 | 0 | 1566.747 | 0 | -0.06 | 0.484159 | 23.07 | 102.2 | 97.8 | 13.7 | 25.82 | | High | High | High | High | High | High | High | High | High | High | High | 0.003303 | 0.03622 | 1.84 |
| FALSE | High | GESSASSPE | [K].GESSAS | 1xCarbami | B2RRE0 1x | 2.13E-05 | 0 | 1 | 3 | 17 | B2RRE0 | A kinase (P | B2RRE0 [5 | 0 | 2587.189 | 0 | -1.78 | 0.000642 | 0.65 | 153.2 | 46.8 | 10.51 | 13.21 | | High | High | High | High | High | High | High | High | High | High | High | 0 | 3.87E-06 | 4.02 |
| FALSE | High | GESSASSPE | [K].GESSAS | 1xCarbami | B2RRE0 2x | 9.27E-08 | 0 | 1 | 3 | 11 | B2RRE0 | A kinase (P | B2RRE0 [5 | 0 | 2667.155 | 0 | -0.98 | 0.001711 | 3.08 | 130.6 | 69.4 | 7.62 | 6.09 | | High | High | High | High | High | High | High | High | High | High | High | 0 | 6.99E-09 | 5.44 |
| FALSE | High | GEPPTPPSI | [R].GEPPTF | 2xPhospho | Q924A2 2x | 0.018774 | 0.001362 | 1 | 3 | 2 | Q924A2 | Protein cap | Q924A2 [2 | 0 | 2971.256 | 0 | 0.05 | 0.983921 | 49.06 | 98.3 | 101.7 | 31.52 | 7.83 | | High | High | High | High | High | High | High | High | High | High | High | 0.008451 | 0.09026 | 2.71 |
| FALSE | High | GFSVVADT | [K].GFSVV | 1xPhospho | Q543N3 1x | 0.000633 | 0 | 1 | 7 | 3 | Q543N3 | LIM and SH | Q543N3 [9 | 0 | 1727.857 | 0 | 0.4 | 0.003702 | 5.6 | 85.6 | 114.4 | 8.64 | 8.03 | | High | High | High | High | High | High | High | High | High | High | High | 0 | 0.000143 | 3.34 |
| FALSE | High | GFWSDICS | [K].GFWSD | 2xCarbami | Q3URL4 1x | 5.09E-06 | 0 | 1 | 2 | 1 | Q3URL4 | BTB and CN | Q3URL4 [4 | 0 | 3153.45 | 0 | 0.34 | 0.217835 | 34.46 | 86.6 | 113.4 | 14.2 | 29.8 | | High | High | High | High | High | High | High | High | High | High | High | 0 | 6.32E-07 | 3.18 |
| FALSE | High | GGAGTSQI | [R].GGAGT | 1xPhospho | Q8BP60 1x | 0.000191 | 0 | 1 | 3 | 1 | Q8BP60 | Putative ur | Q8BP60 [1 | 0 | 1617.789 | 0 | 0.18 | 0.024573 | 4.21 | 92.1 | 107.9 | 4.1 | 7.01 | | High | High | High | High | High | High | High | High | High | High | High | 0 | 3.73E-05 | 2.24 |
| FALSE | High | GGGGSGQ | [R].GGGGS | 1xPhospho | A2RS68 1x | 1.01E-07 | 0 | 1 | 2 | 7 | A2RS68 | Small nucle | A2RS68 [4 | 0 | 1998.872 | 0 | -0.37 | 0.010241 | 1.57 | 112.9 | 87.1 | 3.69 | 6.77 | | High | High | High | High | High | High | High | High | High | High | High | 0 | 1.87E-06 | 2.8 |
| FALSE | High | GGGGSGQ | [R].GGGGS | 1xDeamida | A2RS68 1x | 0.000165 | 0 | 1 | 2 | 3 | A2RS68 | Small nucle | A2RS68 [4 | 0 | 1999.856 | 0 | -0.27 | 0.014586 | 7.09 | 109.9 | 90.1 | 4.1 | 6.57 | | High | High | High | High | High | High | High | High | High | High | High | 0 | 3.16E-05 | 3.05 |
| FALSE | High | GGGGYGG | [R].GGGGY | 1xPhospho | Q5EBP8 1x | 4.51E-08 | 0 | 1 | 1 | 14 | Q5EBP8 | Heterogen | Q5EBP8 [2 | 0 | 3213.277 | 0 | 0.76 | 0.002238 | 2.59 | 70.6 | 129.4 | 4.65 | 11.73 | | High | High | High | High | High | High | High | High | High | High | High | 0 | 3.12E-09 | 5.94 |
| FALSE | High | GGGGYGG | [R].GGGGY | 1xDeamida | Q5EBP8 1x | 1.44E-10 | 0 | 1 | 1 | 25 | Q5EBP8 | Heterogen | Q5EBP8 [2 | 0 | 3214.261 | 0 | 0.03 | 0.916572 | 10.85 | 98.8 | 101.2 | 2.02 | 9.99 | | High | High | High | High | High | High | High | High | High | High | High | 0 | 7.22E-09 | 5.55 |
| FALSE | High | GGGGYGG | [R].GGGGY | 2xDeamida | Q5EBP8 1x | 2.9E-08 | 0 | 1 | 1 | 3 | Q5EBP8 | Heterogen | Q5EBP8 [2 | 0 | 3215.245 | 0 | 0.47 | 0.004506 | 10.82 | 80.5 | 119.5 | 7.36 | 12.33 | | High | High | High | High | High | High | High | High | High | High | High | 0 | 1.9E-09 | 5.96 |
| FALSE | High | GGGLHSSS | [K].GGGLH | 1xCarbami | P70399 1x | 0.109329 | 0.008865 | 1 | 4 | 1 | P70399 | TP53-bindi | P70399 [47 | 0 | 2057.014 | 0 | -0.27 | 0.076017 | 11.82 | 112.1 | 87.9 | 9.68 | 16.18 | | High | High | High | High | High | High | High | High | High | High | High | 0.004462 | 0.04802 | 2.08 |
| FALSE | High | GGGQGQP | [R].GGGQ | 1xPhospho | A2AIH5 1x | 0.063258 | 0.004664 | 1 | 5 | 4 | A2AIH5 | Phosphatic | A2AIH5 [6- | 0 | 2549.147 | 0 | -0.12 | 0.41947 | 4.95 | 103.9 | 96.1 | 7.08 | 3.81 | | High | High | High | High | High | High | High | High | High | High | High | 0.002403 | 0.02532 | 3.36 |
| FALSE | High | GGGQSSP | [R].GGGQS | 1xPhospho | Q8C5N3 1x | 0.001489 | 5.66E-05 | 1 | 6 | 5 | Q8C5N3 | Pre-mRNA- | Q8C5N3 [1 | 0 | 2025.968 | 0 | -1.1 | 0.002786 | 1.45 | 135.9 | 64.1 | 11.84 | 2.91 | | High | High | High | High | High | High | High | High | High | High | High | 0.000327 | 0.002495 | 2.72 |
| FALSE | High | GGGSAAA | [K].GGGSA | 1xPhospho | Q3TWM8 1 | 4.35E-05 | 0 | 1 | 7 | 2 | Q3TWM8 | Putative ur | Q3TWM8 [ | 0 | 2983.338 | 0 | 0.49 | 0.005807 | 6.79 | 82.3 | 117.7 | 12.22 | 10.57 | | High | High | High | High | High | High | High | High | High | High | High | 0 | 7.08E-06 | 4.9 |
| FALSE | High | GGGSCSGS | [R].GGGSC | 1xCarbami | Q8K3W3 1 | 0.000183 | 0 | 1 | 1 | 2 | Q8K3W3 | Protein CA | Q8K3W3 [ | 0 | 2116.928 | 0 | -0.49 | 0.008111 | 8.67 | 116.3 | 83.7 | 7.58 | 12.52 | | High | High | High | High | High | High | High | High | High | High | High | 0 | 3.54E-05 | 2.87 |
| FALSE | High | GGGSCSGS | [R].GGGSC | 1xCarbami | Q8K3W3 2 | 0.000157 | 0 | 1 | 1 | 1 | Q8K3W3 | Protein CA | Q8K3W3 [ | 0 | 2196.894 | 0 | -0.55 | 0.041946 | 5.62 | 118.2 | 81.8 | 4.54 | 18.29 | | High | High | High | High | High | High | High | High | High | High | High | 0 | 2.98E-05 | 3.61 |
| FALSE | High | GGGSDSAS | [K].GGGSD | 1xPhospho | Q3TTA7 1x | 0.013545 | 0.000976 | 1 | 2 | 1 | Q3TTA7 | E3 ubiquiti | Q3TTA7 [7 | 1 | 2992.529 | 0 | -0.12 | 0.102262 | 11.6 | 107.1 | 92.9 | 8.72 | 10.03 | | High | High | High | High | High | High | High | High | High | High | High | 0.000505 | 0.004388 | 3.23 |
| FALSE | High | GGGSGGG | [R].GGGSG | 1xPhospho | P09450 1x | 7.6E-08 | 0 | 1 | 4 | 2 | P09450 | Transcripti | P09450 [11 | 0 | 2795.293 | 0 | 0.76 | 0.005934 | 14.25 | 74.8 | 125.2 | 4.33 | 11.24 | | High | High | High | High | High | High | High | High | High | High | High | 0 | 5.6E-09 | 3.92 |
| FALSE | High | GGGGPGG | [R].GGGGF | 1xPhospho | Q3U0V1 1x | 1.82E-07 | 0 | 1 | 1 | 1 | Q3U0V1 | Far upstrea | Q3U0V1 [4 | 0 | 2636.216 | 0 | 0.31 | 0.100269 | 28.35 | 88.3 | 111.7 | 29.14 | 10.44 | | High | High | High | High | High | High | High | High | High | High | High | 0 | 1.5E-08 | 5.23 |
| FALSE | High | GGGTPDAI | [R].GGGTP | 1xPhospho | Q80V76 1x | 0.015465 | 0.001089 | 1 | 2 | 1 | Q80V76 | Pygopus 2 | Q80V76 [2 | 0 | 1876.957 | 0 | 1.08 | 0.000433 | 12.41 | 65.6 | 134.4 | 7.69 | 3.53 | | High | High | High | High | High | High | High | High | High | High | High | 0.000571 | 0.005122 | 2.75 |
| FALSE | High | GGHDDAPI | [R].GGHDD | 1xPhospho | Q9JLY0 1xF | 2.43E-06 | 0 | 1 | 2 | 1 | Q9JLY0 | Suppressor | Q9JLY0 [30 | 0 | 2773.369 | 0 | 2.32 | 0.000523 | 22.27 | 34.8 | 165.2 | 11.79 | 8.85 | | High | High | High | High | High | High | High | High | High | High | High | 0 | 2.75E-07 | 3.25 |
| FALSE | High | GGHDDAPI | [R].GGHDD | 1xPhospho | Q9JLY0 1xF | 0.000718 | 0 | 1 | 2 | 1 | Q9JLY0 | Suppressor | Q9JLY0 [30 | 0 | 2789.364 | 0 | 1.44 | 0.000746 | 22.31 | 55 | 145 | 33.4 | 14.04 | | High | High | High | High | High | High | High | High | High | High | High | 0 | 0.000164 | 2.1 |
| FALSE | High | GGHMIAVS | [K].GGHMI | 1xPhospho | Q3TUF7 1x | 0.000106 | 0 | 1 | 4 | 2 | Q3TUF7 | YEATS dom | Q3TUF7 [6 | 0 | 1678.875 | 0 | -0.64 | 0.00185 | 3.99 | 123.9 | 76.1 | 5.29 | 7 | | High | High | High | High | High | High | High | High | High | High | High | 0 | 1.91E-05 | 2.11 |
| FALSE | High | GGHTEEND | [R].GGHTE | 1xCarbami | Q8C0L3 1x | 2.7E-05 | 0 | 1 | 2 | 2 | Q8C0L3 | Putative ur | Q8C0L3 [1 | 0 | 2328.98 | 0 | -0.28 | 0.049686 | 9.27 | 112.2 | 87.8 | 9.74 | 13.53 | | High | High | High | High | High | High | High | High | High | High | High | 0 | 4.13E-06 | 3.94 |
| FALSE | High | GGHTNVVS | [K].GGHTN | 1xPhospho | E9PUR0 1x | 7.08E-05 | 0 | 1 | 4 | 4 | E9PUR0 | Ankyrin rep | E9PUR0 [6 | 0 | 4623.245 | 0 | -0.9 | 0.000391 | 1.19 | 131.3 | 68.7 | 5.98 | 8.4 | | High | High | High | High | High | High | High | High | High | High | High | 0 | 1.22E-05 | 4.05 |
| FALSE | High | GGKESDPG | [K].GGKESI | 1xPhospho | E9PWG6 1 | 1.26E-06 | 0 | 1 | 2 | 2 | E9PWG6 | Non-SMC c | E9PWG6 [8 | 1 | 4003.936 | 0 | -0.79 | 0.005802 | 2.93 | 127.7 | 72.3 | 22.91 | 25.4 | | High | High | High | High | High | High | High | High | High | High | High | 0 | 1.32E-07 | 4.05 |
| FALSE | High | GGKPSPPS | [K].GGKPSI | 1xPhospho | Q9JJA7 1xF | 0.004294 | 0.00036 | 1 | 1 | 2 | Q9JJA7 | Cyclin-L2 C | Q9JJA7 [34 | 0 | 1678.95 | 0 | -1.55 | 0.000167 | 0.93 | 149.1 | 50.9 | 3.4 | 6.33 | | High | High | High | High | High | High | High | High | High | High | High | 0.005122 | 0.05567 | 1.62 |
| FALSE | High | GGIDNPAI | [K].GGIDNI | 1xPhospho | B9EKC3 1x | 9.93E-10 | 0 | 1 | 3 | 3 | B9EKC3 | Rho GTPas | B9EKC3 [1 | 0 | 2312.106 | 0 | 0.03 | 0.821304 | 19.63 | 100.2 | 99.8 | 6.71 | 8.8 | | High | High | High | High | High | High | High | High | High | High | High | 0 | 4.31E-11 | 4.55 |
| FALSE | High | GGIDNPAI | [K].GGIDNI | 1xPhospho | B9EKC3 1x | 2.04E-05 | 0 | 1 | 3 | 2 | B9EKC3 | Rho GTPas | B9EKC3 [1 | 1 | 2669.364 | 0 | -0.13 | 0.087 | 8.64 | 108 | 92 | 9.8 | 11.05 | | High | High | High | High | High | High | High | High | High | High | High | 0 | 3.02E-06 | 2.15 |
| FALSE | High | GGIHSNLC | [R].GGIHSN | 1xCarbami | Q62210 1x | 4.53E-07 | 0 | 1 | 1 | 2 | Q62210 | Baculoviral | Q62210 [1 | 0 | 1907.9 | 0 | -0.32 | 0.090605 | 3.79 | 111.2 | 88.8 | 6.82 | 8.49 | | High | High | High | High | High | High | High | High | High | High | High | 0 | 4.19E-08 | 3.37 |
| FALSE | High | GGLLTSEEI | [K].GGLLTS | 1xPhospho | Q3UJ69 1x | 9.2E-05 | 0 | 1 | 4 | 2 | Q3UJ69 | Putative ur | Q3UJ69 [2 | 0 | 2020.931 | 0 | 1.21 | 0.177316 | 28.99 | 70 | 130 | 33.35 | 52.41 | | High | High | High | High | High | High | High | High | High | High | High | 0 | 1.64E-05 | 2.25 |
| FALSE | High | GGLLTSEEI | [K].GGLLTS | 1xPhospho | Q3UJ69 1x | 3.07E-09 | 0 | 1 | 4 | 5 | Q3UJ69 | Putative ur | Q3UJ69 [2 | 0 | 2250.094 | 0 | 1.62 | 0.000796 | 14.66 | 49.5 | 150.5 | 2.33 | 10.89 | | High | High | High | High | High | High | High | High | High | High | High | 0 | 1.2E-07 | 3.84 |
| FALSE | High | GGLLTSEEI | [K].GGLLTS | 2xPhospho | Q3UJ69 2x | 2.56E-06 | 0 | 1 | 4 | 3 | Q3UJ69 | Putative ur | Q3UJ69 [2 | 0 | 2330.061 | 0 | 1.48 | 0.001354 | 8.53 | 50.5 | 149.5 | 10.15 | 12.03 | | High | High | High | High | High | High | High | High | High | High | High | 0 | 2.94E-07 | 3.2 |
| FALSE | High | GGGVPFDT | [R].GGGVP | 1xPhospho | B2RUC3 1x | 1.54E-05 | 0 | 1 | 8 | 1 | B2RUC3 | Excision re | B2RUC3 [4 | 0 | 2410.267 | 0 | 0.5 | 0.007659 | 5.2 | 82.9 | 117.1 | 4.47 | 7.59 | | High | High | High | High | High | High | High | High | High | High | High | 0 | 2.21E-06 | 3.64 |
| FALSE | High | GGGGNFG | [R].GGGGN | 1xPhospho | O88569-3 1 | 0.009006 | 0.000736 | 2 | 4 | 2 | O88569-3; | Isoform 3 c | O88569-3 | 0 | 1686.759 | 0 | -0.01 | 0.940579 | 18.54 | 96.3 | 103.7 | 12.53 | 4.28 | Shared | High | High | High | High | High | High | High | High | High | High | High | 0.000327 | 0.00279 | 1.68 |
| FALSE | High | GGGGGGG | [R].GGGGG | 1xPhospho | A2RS68 1x | 2.31E-08 | 0 | 1 | 1 | 6 | A2RS68 | Small nucle | A2RS68 [3 | 0 | 4615.932 | 0 | -0.73 | 0.005269 | 3.59 | 124.2 | 75.8 | 10.43 | 9.81 | | High | High | High | High | High | High | High | High | High | High | High | 0 | 1.48E-09 | 5.37 |
| FALSE | High | GGGGGGG | [R].GGGGG | 1xPhospho | A2RS68 1x | 9.74E-06 | 0 | 1 | 1 | 2 | A2RS68 | Small nucle | A2RS68 [3 | 0 | 4599.937 | 0 | -0.61 | 0.033773 | 9.6 | 120.7 | 79.3 | 14.67 | 22.22 | | High | High | High | High | High | High | High | High | High | High | High | 0 | 1E-05 | 4.65 |
| FALSE | High | GGCGSPSL | [K].GGCGS | 2xCarbami | E9QLH9 1x | 0.002272 | 0.000109 | 1 | 2 | 1 | E9QLH9 | PEAK1-rela | E9QLH9 [8 | 0 | 1487.633 | 0 | 0.65 | 0.005219 | 11.78 | 76.6 | 123.4 | 7.32 | 12.52 | | High | High | High | High | High | High | High | High | High | High | High | 4.94E-05 | 0.000597 | 1.19 |
| FALSE | High | GGDAAWS | [R].GGDAA | 1xPhospho | P54276 1x | 0.025073 | 0.001645 | 1 | 2 | 1 | P54276 | DNA misma | P54276 [42 | 0 | 1698.732 | 0 | -0.67 | 0.027933 | 7.57 | 118.3 | 81.7 | 8.76 | 15.4 | | High | High | High | High | High | High | High | High | High | High | High | 0.000872 | 0.008814 | 2.02 |
| FALSE | High | GGDGATHS | [R].GGDGA | 1xPhospho | E9QMU3 1 | 0.003195 | 0.00016 | 1 | 5 | 1 | E9QMU3 | Ensconsin | E9QMU3 [ | 0 | 2072.933 | 0 | -0.08 | 0.73252 | 5.97 | 105.1 | 94.9 | 8.33 | 12.14 | | High | High | High | High | High | High | High | High | High | High | High | 7.19E-05 | 0.000877 | 3.35 |
| FALSE | High | GGDPAASP | [R].GGDPA | 1xPhospho | Z4YJP0 1xP | 1.14E-06 | 0 | 1 | 2 | 1 | Z4YJP0 | Hypermeth | Z4YJP0 [52 | 0 | 2118.095 | 0 | 0.72 | 0.005153 | 8.78 | 75.6 | 124.4 | 10.82 | 10.59 | | High | High | High | High | High | High | High | High | High | High | High | 0 | 1.18E-07 | 2.3 |
| FALSE | High | GGDPGDEI | [R].GGDPG | 1xPhospho | Q8BYJ6 1x | 1.08E-05 | 0 | 1 | 7 | 1 | Q8BYJ6 | TBC1 doma | Q8BYJ6 [24 | 0 | 3193.49 | 0 | 0.97 | 0.012808 | 23.42 | 66.7 | 133.3 | 10.92 | 19.37 | | High | High | High | High | High | High | High | High | High | High | High | 0 | 1.47E-06 | 4.37 |
| FALSE | High | GGDSGSCV | [R].GGDSG | 1xCarbami | Q61033 1x | 6.95E-08 | 0 | 1 | 1 | 3 | Q61033 | Lamina-ass | Q61033 [4 | 0 | 2271.965 | 0 | 0.48 | 0.017098 | 10.34 | 84.4 | 115.6 | 5.1 | 10.8 | | High | High | High | High | High | High | High | High | High | High | High | 0 | 5.08E-09 | 5.95 |
| FALSE | High | GGDSGSCV | [R].GGDSG | 1xCarbami | Q61033 2x | 0.000198 | 0 | 1 | 1 | 2 | Q61033 | Lamina-ass | Q61033 [4 | 0 | 2351.931 | 0 | 0.6 | 0.008699 | 16.91 | 78.3 | 121.7 | 12.62 | 7.03 | | High | High | High | High | High | High | High | High | High | High | High | 0 | 0.000195 | 4.04 |
| FALSE | High | GGDSIGETI | [R].GGDSIC | 1xPhospho | G5E866 1x | 0.003526 | 0.00021 | 1 | 5 | 1 | G5E866 | Splicing fac | G5E866 [3 | 0 | 1911.947 | 0 | -0.15 | 0.284952 | 11.03 | 106.3 | 93.7 | 4.29 | 9.88 | | High | High | High | High | High | High | High | High | High | High | High | 9.42E-05 | 0.000981 | 1.73 |
| FALSE | High | GGDSIGETI | [R].GGDSIC | 2xPhospho | G5E866 2x | 0.005228 | 0.000551 | 1 | 5 | 6 | G5E866 | Splicing fac | G5E866 [3 | 0 | 1991.913 | 0 | 0.39 | 0.083347 | 8.87 | 87.5 | 112.5 | 11.21 | 8.3 | | High | High | High | High | High | High | High | High | High | High | High | 0.000505 | 0.004368 | 3 |
| FALSE | High | GGDTSDTE | [R].GGDTS | 1xPhospho | Q8C405 1x | 0.000158 | 0 | 1 | 8 | 2 | Q8C405 | Nuclear au | Q8C405 [3 | 0 | 1759.831 | 0 | 2.79 | 0.002147 | 26.7 | 26.8 | 173.2 | 29.46 | 20.27 | | High | High | High | High | High | High | High | High | High | High | High | 0 | 2.99E-05 | 2.64 |
| FALSE | High | GGDVSPSP | [R].GGDVS | 1xPhospho | Q69ZA1 1x | 8.84E-07 | 0 | 1 | 2 | 2 | Q69ZA1 | Cyclin-dep | Q69ZA1 [3 | 0 | 1877.827 | 0 | -0.17 | 0.057872 | 6.86 | 107.9 | 92.1 | 6.56 | 8.73 | | High | High | High | High | High | High | High | High | High | High | High | 0 | 8.81E-08 | 2.15 |
| FALSE | High | GGDVSPSP | [R].GGDVS | 2xPhospho | Q69ZA1 2x | 0.004613 | 0.000456 | 1 | 2 | 1 | Q69ZA1 | Cyclin-dep | Q69ZA1 [3 | 0 | 1957.793 | 0 | 0.12 | 0.055522 | 9.02 | 94.8 | 105.2 | 5.4 | 8.32 | | High | High | High | High | High | High | High | High | High | High | High | 0.000204 | 0.001327 | 2.35 |
| FALSE | High | GGDYALAP | [K].GGDYA | 2xPhospho | Q61892 2x | 0.000967 | 0 | 1 | 16 | 2 | Q61892 | mRNA OS= | Q61892 [3 | 0 | 2230.929 | 0 | 0.2 | 0.083697 | 8.72 | 92.9 | 107.1 | 15.82 | 15.97 | | High | High | High | High | High | High | High | High | High | High | High | 0 | 0.000229 | 3.83 |
| FALSE | High | GGDYALAP | [K].GGDYA | 1xCarbami | Q31173 1x | 0.000529 | 0 | 1 | 22 | 1 | Q31173 | MHC H-2K- | Q31173 [3 | 0 | 2690.278 | 0 | 2.02 | 0.000416 | 9.56 | 39.3 | 160.7 | 6.73 | 9.33 | | High | High | High | High | High | High | High | High | High | High | High | 0 | 0.000116 | 4.23 |
| FALSE | High | GGEATWS | [R].GGEAT | 1xPhospho | Q3TCN5 1x | 0.00204 | 5.66E-05 | 1 | 1 | 1 | Q3TCN5 | Putative ur | Q3TCN5 [3 | 0 | 3453.559 | 0 | -0.11 | 0.729754 | 25.58 | 103.8 | 96.2 | 14.05 | 7.35 | | High | High | High | High | High | High | High | High | High | High | High | 2.58E-05 | 0.000529 | 2.74 |
| FALSE | High | GGEDEDGS | [K].GGEDE | 1xPhospho | Q9ERU9 1x | 1.72E-05 | 0 | 1 | 1 | 6 | Q9ERU9 | E3 SUMO- | Q9ERU9 [2 | 0 | 4072.895 | 0 | -0.4 | 0.000588 | 1.05 | 113.6 | 86.4 | 3.3 | 4.01 | | High | High | High | High | High | High | High | High | High | High | High | 0 | 2.48E-06 | 4.79 |
| FALSE | High | GGEFDEFV | [K].GGEFD | 1xPhospho | Q62018 1x | 6.99E-07 | 0 | 1 | 4 | 5 | Q62018 | RNA polym | Q62018 [9 | 0 | 2752.228 | 0 | -0.62 | 0.000416 | 1.13 | 121.1 | 78.9 | 2.23 | 2.49 | | High | High | High | High | High | High | High | High | High | High | High | 0 | 1.04E-05 | 3.85 |
| FALSE | High | GGELPLRPI | [R].GGELPI | 1xPhospho | Q9WVK0 1 | 0.00043 | 0 | 1 | 1 | 1 | Q9WVK0 | Type-1 ang | Q9WVK0 [ | 0 | 4790.266 | 0 | 1.84 | 0.007364 | 48.15 | 46.7 | 153.3 | 33.11 | 23.54 | | High | High | High | High | High | High | High | High | High | High | High | 0 | 9.25E-05 | 3.39 |
| FALSE | High | GGFDSPFY | [R].GGFDSI | 1xPhospho | Q9EP53 1x | 0.006834 | 0.00062 | 1 | 4 | 1 | Q9EP53 | Hamartin C | Q9EP53 [4 | 0 | 1354.603 | 0 | -0.04 | 0.318624 | 8.82 | 104.4 | 95.6 | 3.98 | 6.41 | | High | High | High | High | High | High | High | High | High | High | High | 0.000275 | 0.002051 | 2.54 |
| FALSE | High | GGFSDADV | [K].GGFSD | 1xPhospho | Q569Z6 1x | 0.000479 | 0 | 1 | 4 | 3 | Q569Z6 | Thyroid ho | Q569Z6 [3 | 0 | 1433.708 | 0 | -0.22 | 0.007564 | 4.26 | 108.6 | 91.4 | 1.79 | 6.13 | | High | High | High | High | High | High | High | High | High | High | High | 7.19E-05 | 0.000878 | 1.57 |
| FALSE | High | GGFSPFGN | [K].GGFSPI | 1xPhospho | E9QKE4 1x | 0.000136 | 0 | 1 | 3 | 1 | E9QKE4 | Rab3 GTPa | E9QKE4 [4 | 0 | 1717.79 | 0 | -0.88 | 0.00502 | 3.03 | 129.6 | 70.4 | 16.42 | 7.88 | | High | High | High | High | High | High | High | High | High | High | High | 0 | 2.53E-05 | 1.82 |
| FALSE | High | GGFTEDDL | [K].GGFTEI | 1xPhospho | Q3TRR0 1x | 1.26E-07 | 0 | 1 | 5 | 2 | Q3TRR0 | Microtubu | Q3TRR0 [3 | 0 | 3724.85 | 0 | 1.3 | 0.00101 | 7.69 | 60.2 | 139.8 | 4.2 | 10.19 | | High | High | High | High | High | High | High | High | High | High | High | 0 | 9.87E-09 | 4.1 |
| FALSE | High | GGGAGGN | [R].GGGAG | 1xCarbami | E9PUR0 1x | 5.36E-09 | 0 | 1 | 4 | 1 | E9PUR0 | Ankyrin rep | E9PUR0 [1 | 0 | 3070.258 | 0 | -0.88 | 0.001059 | 3.22 | 127.5 | 72.5 | 7.81 | 10.7 | | High | High | High | High | High | High | High | High | High | High | High | 0 | 2.86E-10 | 5.22 |
| FALSE | High | GGGASSPA | [R].GGGAS | 1xPhospho | Q3UFT4 1x | 1.13E-05 | 0 | 1 | 8 | 2 | Q3UFT4 | Putative ur | Q3UFT4 [7 | 0 | 2875.419 | 0 | -1.76 | 0.000427 | 0.19 | 155.5 | 44.5 | 6.26 | 20.4 | | High | High | High | High | High | High | High | High | High | High | High | 0 | 1.55E-06 | 3.99 |
| FALSE | High | GGGASSPA | [R].GGGAS | 2xPhospho | Q3UFT4 2x | 8.56E-05 | 0 | 1 | 8 | 3 | Q3UFT4 | Putative ur | Q3UFT4 [7 | 0 | 2955.385 | 0 | -0.94 | 0.005316 | 2.07 | 131.3 | 68.7 | 11.11 | 14.36 | | High | High | High | High | High | High | High | High | High | High | High | 0 | 1.51E-05 | 4.41 |
| FALSE | High | GGGDGNV | [R].GGGDG | 1xPhospho | Q61687 1x | 4.26E-07 | 0 | 1 | 3 | 2 | Q61687 | Transcripti | Q61687 [1 | 0 | 2341.107 | 0 | -0.14 | 0.658957 | 3.26 | 101.9 | 98.1 | 11.99 | 6.9 | | High | High | High | High | High | High | High | High | High | High | High | 0 | 2.16E-06 | 3.8 |
| FALSE | High | GGGGGDE | [R].GGGGG | 1xPhospho | B2RRE0 1x | 5.91E-05 | 0 | 1 | 3 | 3 | B2RRE0 | A kinase (P | B2RRE0 [5 | 0 | 3140.317 | 0 | -0.65 | 0.011498 | 4.32 | 122.7 | 77.3 | 9.2 | 17.23 | | High | High | High | High | High | High | High | High | High | High | High | 0 | 9.99E-06 | 3.31 |
| FALSE | High | GGGGGDE | [R].GGGGG | 1xPhospho | B2RRE0 1x | 1.43E-07 | 0 | 1 | 3 | 11 | B2RRE0 | A kinase (P | B2RRE0 [5 | 0 | 3369.48 | 0 | -1.43 | 0.000587 | 1.8 | 143.1 | 56.9 | 4.62 | 14.58 | | High | High | High | High | High | High | High | High | High | High | High | 0 | 8.37E-05 | 4.7 |
| FALSE | High | GGGGGDE | [R].GGGGG | 2xPhospho | B2RRE0 2x | 0.000382 | 0 | 1 | 3 | 1 | B2RRE0 | A kinase (P | B2RRE0 [5 | 0 | 3449.446 | 0 | -1.19 | 0.003531 | 3.52 | 137.8 | 62.2 | 12.34 | 10.58 | | High | High | High | High | High | High | High | High | High | High | High | 0 | 8.08E-05 | 3.44 |
| FALSE | High | GEGAGQPS | [R].GEGAG | 1xPhospho | Q6NSP9 1x | 8.18E-05 | 0 | 1 | 2 | 2 | Q6NSP9 | High mobil | Q6NSP9 [5 | 0 | 2601.307 | 0 | 1.97 | 0.000371 | 22.41 | 42.5 | 157.5 | 24.99 | 10.41 | | High | High | High | High | High | High | High | High | High | High | High | 0 | 2.55E-05 | 3.52 |
| FALSE | High | GEESNESA | [R].GEESNI | 2xPhospho | Q04692 2x | 0.000101 | 0 | 1 | 2 | 3 | Q04692 | SWI/SNF-r | Q04692 [2 | 0 | 2371.973 | 0 | -1 | 0.000693 | 2.41 | 133.3 | 66.7 | 4.44 | 8.34 | | High | High | High | High | High | High | High | High | High | High | High | 0 | 1.81E-05 | 3.71 |
| FALSE | High | GEESNESA | [R].GEESNI | 1xPhospho | Q04692 1x | 6.72E-09 | 0 | 1 | 2 | 2 | Q04692 | SWI/SNF-r | Q04692 [2 | 0 | 2292.007 | 0 | -0.93 | 0.000946 | 2.92 | 130.9 | 69.1 | 5.09 | 6.34 | | High | High | High | High | High | High | High | High | High | High | High | 0 | 1.11E-08 | 5.23 |
| FALSE | High | GEEGASQE | [R].GEEGA | 1xCarbami | D3Z5N5 1x | 0.007881 | 0.000736 | 2 | 3 | 1 | D3Z5N5; Q | 1-phospha | D3Z5N5 [8 | 0 | 2753.192 | 0 | -0.06 | 0.208005 | 6.86 | 102.1 | 97.9 | 14.9 | 22.2 | Shared | High | High | High | High | High | High | High | High | High | High | High | 0.000327 | 0.002412 | 2.76 |
| FALSE | High | GCTSEVGL | [K].GCTSEV | 1xCarbami | Q60774 1x | 0.007019 | 0.00062 | 1 | 2 | 3 | Q60774 | Transmem | Q60774 [2 | 0 | 1601.837 | 0 | 2.92 | 0.001003 | 43.92 | 25.6 | 174.4 | 41.2 | 16.89 | | High | High | High | High | High | High | High | High | High | High | High | 0.000275 | 0.002115 | 1.88 |

| | | | | | | | | | | | | | | | | | | | | | | | | | | | | | | | | | | | | | | |
|---|---|---|---|---|---|---|---|---|---|---|---|---|---|---|---|---|---|---|---|---|---|---|---|---|---|---|---|---|---|---|---|---|---|---|---|---|---|---|
| FALSE | High | GCVRSEDE | [R].GCVRSI | 1xCarbami | F8WJ40 1x | 0.000693 | 0 | 1 | 4 | 1 | F8WJ40 | Histone-lys | F8WJ40 [8 | 1 | 2035.923 | 0 | -1.01 | 0.000844 | 2.01 | 133.4 | 66.6 | 3.68 | 13 | | High | High | High | High | High | High | High | High | High | High | High | 0 | 0.000157 | 1.85 |
| FALSE | High | GCVRSEDE | [R].GCVRSI | 1xCarbami | F8WJ40 1x | 0.000137 | 0 | 1 | 4 | 1 | F8WJ40 | Histone-lys | F8WJ40 [8 | 1 | 2051.918 | 0 | -0.82 | 0.006456 | 4.09 | 128.7 | 71.3 | 7.99 | 15.36 | | High | High | High | High | High | High | High | High | High | High | High | 0 | 2.54E-05 | 3.38 |
| FALSE | High | GCVTTPNS | [K].GCVTTI | 1xCarbami | G5E8P4 1x | 4.44E-06 | 0 | 1 | 3 | 2 | G5E8P4 | Protein cor | G5E8P4 [2 | 0 | 1821.852 | 0 | -0.17 | 0.588415 | 5.04 | 106 | 94 | 6.51 | 13.61 | | High | High | High | High | High | High | High | High | High | High | High | 0 | 5.44E-07 | 3.71 |
| FALSE | High | GDAETDEE | [K].GDAETI | 1xPhospho | Q922Q2 1x | 5.32E-06 | 0 | 1 | 1 | 11 | Q922Q2 | Serine/thre | Q922Q2 [5 | 0 | 3421.332 | 0 | -0.61 | 0.012561 | 8.17 | 123.8 | 76.2 | 6.04 | 15.27 | | High | High | High | High | High | High | High | High | High | High | High | 0 | 6.65E-07 | 5.26 |
| FALSE | High | GDASASQV | [R].GDASA | 1xPhospho | G3X963 1x | 1.4E-05 | 0 | 1 | 4 | 7 | G3X963 | ATPase fan | G3X963 [1 | 0 | 2374.106 | 0 | 0.11 | 0.432372 | 13.57 | 93.7 | 106.3 | 7.53 | 10.48 | | High | High | High | High | High | High | High | High | High | High | High | 0 | 1.82E-05 | 3.55 |
| FALSE | High | GDASASQV | [R].GDASA | 2xPhospho | G3X963 2x | 0.01218 | 0.000889 | 1 | 4 | 1 | G3X963 | ATPase fan | G3X963 [1 | 0 | 2454.073 | 0 | -0.07 | 0.512102 | 15.9 | 103.9 | 96.1 | 12.35 | 16.63 | | High | High | High | High | High | High | High | High | High | High | High | 0.000449 | 0.003917 | 2.45 |
| FALSE | High | GDCQVLAE | [K].GDCQV | 1xCarbami | Q6PCX9 1x | 1.06E-07 | 0 | 1 | 1 | 1 | Q6PCX9 | E3 ubiquiti | Q6PCX9 [7 | 0 | 2177.933 | 0 | 0.49 | 0.003741 | 2.01 | 81.6 | 118.4 | 7.28 | 12.7 | | High | High | High | High | High | High | High | High | High | High | High | 0 | 8.18E-09 | 3.49 |
| FALSE | High | GDCSSGSP | [R].GDCSS( | 1xCarbami | E9QAE4 1x | 7.25E-07 | 0 | 1 | 2 | 3 | E9QAE4 | Histone-lys | E9QAE4 [9 | 0 | 1776.824 | 0 | -0.73 | 0.000675 | 2.95 | 124.7 | 75.3 | 4.05 | 6.93 | | High | High | High | High | High | High | High | High | High | High | High | 0 | 7.1E-08 | 2.7 |
| FALSE | High | GDDASEEG | [R].GDDAS | 1xPhospho | Q8K232 1x | 0.000237 | 0 | 2 | 7 | 4 | Q8K232; E | Adducin 1 | Q8K232 [4 | 0 | 2015.896 | 0 | 1.06 | 0.001375 | 15.22 | 61.1 | 138.9 | 6.71 | 12.35 | Shared | High | High | High | High | High | High | High | High | High | High | High | 0 | 4.73E-05 | 3.51 |
| FALSE | High | GDDASEEG | [R].GDDAS | 1xDeamida | Q8K232 1x | 8.06E-06 | 0 | 2 | 7 | 1 | Q8K232; E | Adducin 1 | Q8K232 [4 | 0 | 2016.88 | 0 | 1.35 | 0.000926 | 20.52 | 52.3 | 147.7 | 13.93 | 7.19 | Shared | High | High | High | High | High | High | High | High | High | High | High | 0 | 1.06E-06 | 3.69 |
| FALSE | High | GDDDWGF | [R].GDDDV | 1xPhospho | Q9JLF6 1xF | 0.000311 | 0 | 2 | 2 | 2 | Q9JLF6; A0 | Protein-glu | Q9JLF6 [60 | 0 | 1683.685 | 0 | 2.99 | 0.007706 | 63.16 | 23.2 | 176.8 | 35.47 | 27.49 | Shared | High | High | High | High | High | High | High | High | High | High | High | 0 | 6.45E-05 | 2.4 |
| FALSE | High | GDDEGIFIS | [K].GDDEG | 1xPhospho | Q80U72-4 | 0.039347 | 0.002793 | 2 | 6 | 1 | Q80U72-4; | Isoform 4 c | Q80U72-4 | 1 | 2314.091 | 0 | -0.51 | 0.00488 | 5.16 | 117.3 | 82.7 | 6.95 | 4.4 | Shared | High | High | High | High | High | High | High | High | High | High | High | 0.001468 | 0.01467 | 2.52 |
| FALSE | High | GCSPSSLD( | [R].GCSPSS | 1xCarbami | Q5DTT3 2x | 0.062731 | 0.004664 | 1 | 2 | 1 | Q5DTT3 | Protein FAI | Q5DTT3 [2 | 0 | 2219.949 | 0 | -0.14 | 0.567297 | 7.15 | 100.6 | 99.4 | 7.94 | 15.46 | | High | High | High | High | High | High | High | High | High | High | High | 0.002403 | 0.02505 | 1.8 |
| FALSE | High | GDDGPDV/ | [K].GDDGP | 1xPhospho | A0A0A0MC | 0.008094 | 0.000736 | 1 | 3 | 2 | A0A0A0MC | Histone-lys | A0A0A0MC | 0 | 3473.563 | 0 | 1 | 0.00438 | 8.54 | 67.4 | 132.6 | 7.81 | 16.92 | | High | High | High | High | High | High | High | High | High | High | High | 0.000327 | 0.002871 | 2.82 |
| FALSE | High | GDEDFVDL | [K].GDEDF | 1xPhospho | B9EIW5 1x | 2.08E-06 | 0 | 1 | 3 | 1 | B9EIW5 | Ncoa7 prot | B9EIW5 [4 | 0 | 2846.339 | 0 | 1.44 | 0.000534 | 20.61 | 52.9 | 147.1 | 17.23 | 5.53 | | High | High | High | High | High | High | High | High | High | High | High | 0 | 2.32E-07 | 3.3 |
| FALSE | High | GDEPQASC | [K].GDEPQ | 1xPhospho | Q8BL06 1x | 0.005565 | 0.00062 | 1 | 2 | 2 | Q8BL06 | Inactive ub | Q8BL06 [4 | 0 | 2343.091 | 0 | -0.77 | 0.000633 | 3.92 | 126.2 | 73.8 | 3.3 | 7.43 | | High | High | High | High | High | High | High | High | High | High | High | 0.000275 | 0.001627 | 2.22 |
| FALSE | High | GDGGSTPT | [R].GDGGS | 1xPhospho | Q8BK03 1x | 2.6E-05 | 0 | 1 | 2 | 6 | Q8BK03 | Mitoguard | Q8BK03 [2 | 0 | 3309.459 | 0 | 1.49 | 0.000443 | 15.34 | 55.3 | 144.7 | 23.89 | 6.54 | | High | High | High | High | High | High | High | High | High | High | High | 0 | 3.96E-06 | 3.93 |
| FALSE | High | GDGGSTPT | [R].GDGGS | 1xDeamida | Q8BK03 1x | 0.069903 | 0.005181 | 1 | 2 | 1 | Q8BK03 | Mitoguard | Q8BK03 [2 | 0 | 3310.443 | 0 | 0.87 | 0.027625 | 15.42 | 81.2 | 118.8 | 15.48 | 15.9 | | High | High | High | High | High | High | High | High | High | High | High | 0.002662 | 0.02831 | 3.5 |
| FALSE | High | GDGGSTPT | [R].GDGGS | 3xPhospho | Q8BK03 3x | 0.072858 | 0.005599 | 1 | 2 | 1 | Q8BK03 | Mitoguard | Q8BK03 [2 | 1 | 3625.493 | 0 | 0.12 | 0.153911 | 9.5 | 94.3 | 105.7 | 6.23 | 11.38 | | High | High | High | High | High | High | High | High | High | High | High | 0.002867 | 0.02972 | 3.36 |
| FALSE | High | GDGGSTTC | [R].GDGGS | 1xPhospho | F6SZ47 1xF | 1.93E-05 | 0 | 1 | 2 | 1 | F6SZ47 | Serine/thre | F6SZ47 [37 | 0 | 2823.454 | 0 | -0.06 | 0.917119 | 7.78 | 101.6 | 98.4 | 11.48 | 11.2 | | High | High | High | High | High | High | High | High | High | High | High | 0 | 2.83E-06 | 3.57 |
| FALSE | High | GDGGTPPI | [R].GDGGT | 1xPhospho | A2CG76 1x | 2.5E-10 | 0 | 2 | 6 | 3 | A2CG76; A | Histone-lys | A2CG76 [5 | 0 | 2615.318 | 0 | -0.48 | 0.008384 | 6.28 | 116.4 | 83.6 | 4.72 | 11.66 | Shared | High | High | High | High | High | High | High | High | High | High | High | 0 | 9.07E-12 | 4.3 |
| FALSE | High | GDGRPETP | [K].GDGRP | 1xPhospho | Q6KCD5 1x | 0.047894 | 0.00354 | 1 | 3 | 2 | Q6KCD5 | Nipped-B-l | Q6KCD5 [7 | 0 | 1494.772 | 0 | 0.42 | 0.000983 | 2.86 | 84.6 | 115.4 | 3.12 | 4.67 | | High | High | High | High | High | High | High | High | High | High | High | 0.001812 | 0.01844 | 1.8 |
| FALSE | High | GDHSLTPG | [R].GDHSL | 1xPhospho | A3KCG1 1x | 0.096181 | 0.007535 | 1 | 1 | 1 | A3KCG1 | Calcium rel | A3KCG1 [2 | 0 | 1564.699 | 0 | 1.17 | 0.001514 | 13.47 | 62 | 138 | 8 | 8.15 | | High | High | High | High | High | High | High | High | High | High | High | 0.003883 | 0.04135 | 2.18 |
| FALSE | High | GDLAFGAP | [R].GDLAF( | 1xPhospho | A2AGH6 1x | 0.000411 | 0 | 1 | 4 | 2 | A2AGH6 | Mediator c | A2AGH6 [6 | 0 | 2932.335 | 0 | 0.39 | 0.007281 | 6.51 | 86.4 | 113.6 | 2.71 | 8.94 | | High | High | High | High | High | High | High | High | High | High | High | 0.002059 | 0.02143 | 2.89 |
| FALSE | High | GDLGASSP | [K].GDLGA | 1xPhospho | E9Q616 1x | 2.1E-05 | 0 | 1 | 6 | 4 | E9Q616 | AHNAK nu | E9Q616 [5 | 0 | 1587.785 | 0 | 2.94 | 0.003457 | 8.81 | 26.2 | 173.8 | 9.74 | 23.2 | | High | High | High | High | High | High | High | High | High | High | High | 0 | 3.1E-06 | 2.7 |
| FALSE | High | GDLGASSP | [K].GDLGA | 1xPhospho | E9Q616 1x | 1.9E-05 | 0 | 1 | 6 | 31 | E9Q616 | AHNAK nu | E9Q616 [5 | 0 | 1603.78 | 0 | 2.24 | 0.000361 | 7.22 | 35.6 | 164.4 | 8.48 | 8.9 | | High | High | High | High | High | High | High | High | High | High | High | 0 | 7.43E-06 | 3.01 |
| FALSE | High | GDILSNPCI | [K].GDILSN | 1xCarbami | Q5SW75 1 | 0.00537 | 0.000599 | 1 | 4 | 1 | Q5SW75 | Protein ph | Q5SW75 [7 | 0 | 1876.939 | 0 | 1.61 | 0.000915 | 17.91 | 48.9 | 151.1 | 11.52 | 10.92 | | High | High | High | High | High | High | High | High | High | High | High | 0.000267 | 0.001572 | 2.02 |
| FALSE | High | GDDLEEGV | [K].GDDLEI | 1xPhospho | Q5SRX1 1x | 0.00043 | 0 | 1 | 6 | 1 | Q5SRX1 | TOM1-like | Q5SRX1 [4 | 0 | 2208 | 0 | 1.34 | 0.00024 | 13.35 | 56 | 144 | 10.88 | 5.21 | | High | High | High | High | High | High | High | High | High | High | High | 0 | 9.28E-05 | 3.16 |
| FALSE | High | GCSPSSLD( | [R].GCSPSS | 1xCarbami | Q5DTT3 1x | 2.31E-07 | 0 | 1 | 2 | 3 | Q5DTT3 | Protein FAI | Q5DTT3 [2 | 0 | 2139.982 | 0 | -0.41 | 0.00222 | 6.47 | 116.2 | 83.8 | 3.16 | 10.55 | | High | High | High | High | High | High | High | High | High | High | High | 0 | 2.08E-07 | 2.84 |
| FALSE | High | GCSPPAPA | [R].GCSPPA | 1xCarbami | Q7TNF9 1x | 0.001278 | 0 | 1 | 2 | 2 | Q7TNF9 | Protein FAI | Q7TNF9 [2 | 0 | 1462.671 | 0 | -1.14 | 0.000267 | 2.35 | 137.7 | 62.3 | 2.3 | 13.49 | | High | High | High | High | High | High | High | High | High | High | High | 0 | 0.000314 | 2.88 |
| FALSE | High | GCISPQVEI | [K].GCISPQ | 1xCarbami | Q80ZK9 1x | 0.000104 | 0 | 1 | 3 | 1 | Q80ZK9 | WD and te | Q80ZK9 [3 | 0 | 2067.018 | 0 | -1.14 | 0.000348 | 1.03 | 136 | 64 | 6.52 | 5.67 | | High | High | High | High | High | High | High | High | High | High | High | 0 | 1.89E-05 | 3.42 |
| FALSE | High | GASAATGII | [K].GASAA | 1xCarbami | Q8R5H1 2x | 1.91E-05 | 0 | 2 | 4 | 7 | Q8R5H1; C | Ubiquitin c | Q8R5H1 [9 | 0 | 3854.452 | 0 | 0.2 | 0.03064 | 5.3 | 88.4 | 111.6 | 5.99 | 8.25 | Shared | High | High | High | High | High | High | High | High | High | High | High | 0 | 2.8E-06 | 4.32 |
| FALSE | High | GASAEELL/ | [R].GASAEI | 1xPhospho | Q80TX5 1x | 0.014283 | 0.001011 | 1 | 3 | 2 | Q80TX5 | MKIAA057 | Q80TX5 [1 | 0 | 2600.347 | 0 | -0.29 | 0.016845 | 6.83 | 110.6 | 89.4 | 6.65 | 9.78 | | High | High | High | High | High | High | High | High | High | High | High | 0.000585 | 0.005172 | 2.73 |
| FALSE | High | GASFGSAG | [R].GASFG | 1xPhospho | E9PXP7 1xI | 0.001659 | 5.66E-05 | 1 | 2 | 1 | E9PXP7 | Stonin-2 O | E9PXP7 [7 | 0 | 1876.846 | 0 | -0.76 | 0.020999 | 8.6 | 124.2 | 75.8 | 14.02 | 12.42 | | High | High | High | High | High | High | High | High | High | High | High | 2.58E-05 | 0.000422 | 2.8 |
| FALSE | High | GASFGSAG | [R].GASFG | 1xPhospho | E9PXP7 1xI | 0.000731 | 0 | 1 | 2 | 1 | E9PXP7 | Stonin-2 O | E9PXP7 [7 | 0 | 1892.841 | 0 | -0.75 | 0.003296 | 2.48 | 124.9 | 75.1 | 13.39 | 8.27 | | High | High | High | High | High | High | High | High | High | High | High | 0 | 0.000169 | 2.39 |
| FALSE | High | GASPLGPG | [R].GASPL( | 1xPhospho | Q6NZR2 1x | 1.07E-07 | 0 | 1 | 1 | 2 | Q6NZR2 | Myb/SANT | Q6NZR2 [4 | 0 | 2532.277 | 0 | 0.03 | 0.621094 | 4.65 | 102.8 | 97.2 | 7.65 | 13.1 | | High | High | High | High | High | High | High | High | High | High | High | 0 | 8.29E-09 | 4.02 |
| FALSE | High | GASPVTPS | [R].GASPV | 1xPhospho | Q80U49 1x | 8.5E-06 | 0 | 1 | 2 | 3 | Q80U49 | Centrosom | Q80U49 [5 | 0 | 2627.373 | 0 | 0.68 | 0.00028 | 5.82 | 76 | 124 | 5.33 | 3.53 | | High | High | High | High | High | High | High | High | High | High | High | 0 | 1.12E-06 | 3.24 |
| FALSE | High | GASPYGSP | [K].GASPY( | 1xPhospho | Q6PR54-3 | 0.004294 | 0.00036 | 2 | 4 | 3 | Q6PR54-3; | Isoform 3 c | Q6PR54-3 | 0 | 1200.561 | 0 | -0.85 | 0.000236 | 0.55 | 128.3 | 71.7 | 3.91 | 3.68 | Shared | High | High | High | High | High | High | High | High | High | High | High | 0.007983 | 0.08625 | 1.43 |
| FALSE | High | GASPYGSP | [K].GASPY( | 2xPhospho | Q6PR54-3 | 0.053473 | 0.00384 | 2 | 4 | 1 | Q6PR54-3; | Isoform 3 c | Q6PR54-3 | 0 | 1280.528 | 0 | -1.06 | 0.002397 | 1.85 | 134 | 66 | 14.6 | 16.43 | Shared | High | High | High | High | High | High | High | High | High | High | High | 0.001999 | 0.02083 | 1.42 |
| FALSE | High | GASYSSIPK | [R].GASYSS | 1xPhospho | Q8VHX6 1x | 0.006594 | 0.00062 | 1 | 3 | 2 | Q8VHX6 | Filamin-C C | Q8VHX6 [2 | 0 | 1447.76 | 0 | -1.01 | 0.000479 | 1.29 | 133.5 | 66.5 | 12.19 | 16.13 | | High | High | High | High | High | High | High | High | High | High | High | 0.000555 | 0.004809 | 1.4 |
| FALSE | High | GATADDSC | [R].GATAD | 1xPhospho | Q9JL04 1xF | 0.005322 | 0.000599 | 1 | 1 | 1 | Q9JL04 | Formin-2 C | Q9JL04 [48 | 0 | 2012.01 | 0 | 0.53 | 0.001241 | 8.64 | 82.7 | 117.3 | 15.74 | 13.81 | | High | High | High | High | High | High | High | High | High | High | High | 0.000267 | 0.001548 | 2.05 |
| FALSE | High | GATDTSPL | [R].GATDT | 1xPhospho | Q9QXL1 1x | 0.00642 | 0.00062 | 1 | 3 | 3 | Q9QXL1 | Kinesin-like | Q9QXL1 [1 | 0 | 1327.646 | 0 | -0.71 | 0.002987 | 4.5 | 127.3 | 72.7 | 7.07 | 13.93 | | High | High | High | High | High | High | High | High | High | High | High | 0.000275 | 0.001908 | 1.71 |
| FALSE | High | GATPAEDD | [K].GATPAI | 1xPhospho | F6ZFU0 1xI | 0.031439 | 0.002011 | 2 | 12 | 1 | F6ZFU0; A( | Elongation | F6ZFU0 [4 | 0 | 1685.767 | 0 | 0.48 | 0.198739 | 18.65 | 88.5 | 111.5 | 11.27 | 24.12 | Shared | High | High | High | High | High | High | High | High | High | High | High | 0.001076 | 0.01134 | 2.04 |
| FALSE | High | GATPAEDD | [K].GATPAI | 1xPhospho | F6ZFU0 1xI | 0.001398 | 5.66E-05 | 2 | 12 | 2 | F6ZFU0; A( | Elongation | F6ZFU0 [4 | 1 | 3536.593 | 0 | 0.4 | 0.196708 | 17.6 | 92.1 | 107.9 | 11.74 | 47.5 | Shared | High | High | High | High | High | High | High | High | High | High | High | 2.58E-05 | 0.000348 | 2.42 |
| FALSE | High | GATPPAPP | [R].GATPPA | 1xPhospho | D3YXK1 1x | 0.005 | 0.000504 | 1 | 1 | 1 | D3YXK1 | Atherin OS | D3YXK1 [1 | 0 | 1172.603 | 0 | 1.05 | 0.001041 | 16.64 | 58.4 | 141.6 | 13.79 | 7.32 | | High | High | High | High | High | High | High | High | High | High | High | 0.000225 | 0.001445 | 1.25 |
| FALSE | High | GATSLQQE | [R].GATSL( | 1xCarbami | B9EKC1 1x | 0.000622 | 0 | 1 | 2 | 2 | B9EKC1 | PAS domai | B9EKC1 [9 | 0 | 2693.223 | 0 | -0.14 | 0.184779 | 17.34 | 105 | 95 | 8.75 | 14.86 | | High | High | High | High | High | High | High | High | High | High | High | 0 | 0.00014 | 3.08 |
| FALSE | High | GATVLPAN | [R].GATVLI | 1xPhospho | Q3B7Z2 1x | 0.00642 | 0.00062 | 1 | 2 | 2 | Q3B7Z2 | Oxysterol-l | Q3B7Z2 [3 | 0 | 2019.068 | 0 | 0.5 | 0.011835 | 17.83 | 77.9 | 122.1 | 13.79 | 8.34 | | High | High | High | High | High | High | High | High | High | High | High | 0.000275 | 0.001915 | 2.41 |
| FALSE | High | GAVEAAHC | [R].GAVEA | 1xPhospho | A0A1B0GS | 0.000168 | 0 | 1 | 3 | 1 | A0A1B0GS | Aldehyde c | A0A1B0GS | 0 | 2170.039 | 0 | -0.54 | 0.056774 | 8.64 | 116.6 | 83.4 | 9.17 | 16.68 | | High | High | High | High | High | High | High | High | High | High | High | 0 | 3.21E-05 | 3.19 |
| FALSE | High | GAVEGPPA | [R].GAVEG | 1xCarbami | Q9QY06-2 | 1.12E-05 | 0 | 2 | 3 | 1 | Q9QY06-2; | Isoform 2 c | Q9QY06-2 | 0 | 3239.674 | 0 | 1.5 | 0.001433 | 25.34 | 52.3 | 147.7 | 15.59 | 7.89 | Shared | High | High | High | High | High | High | High | High | High | High | High | 0 | 1.54E-06 | 5.28 |
| FALSE | High | GAVGVPDT | [R].GAVGV | 1xPhospho | A2A863-3 | 0.024424 | 0.001612 | 1 | 6 | 1 | A2A863-3 | Isoform 3 c | A2A863-3 | 0 | 1378.693 | 0 | 1.47 | 0.001266 | 20.84 | 50.5 | 149.5 | 5.29 | 11.52 | | High | High | High | High | High | High | High | High | High | High | High | 0.000859 | 0.008503 | 1.11 |
| FALSE | High | GAVSEGEP | [R].GAVSE( | 1xPhospho | Q8C796 1x | 1.43E-06 | 0 | 1 | 2 | 4 | Q8C796 | REST corep | Q8C796 [2 | 0 | 1896.899 | 0 | -1.87 | 0.00014 | 0.92 | 155.7 | 44.3 | 3.16 | 13.02 | | High | High | High | High | High | High | High | High | High | High | High | 0 | 1.33E-05 | 2.55 |
| FALSE | High | GAYCDLSSI | [K].GAYCD | 1xCarbami | Q9D7A3 1x | 2.18E-06 | 0 | 1 | 2 | 3 | Q9D7A3 | Putative ur | Q9D7A3 [7 | 0 | 2054.903 | 0 | 0 | 0.802157 | 16.3 | 99.6 | 100.4 | 6.45 | 8.26 | | High | High | High | High | High | High | High | High | High | High | High | 0 | 2.44E-07 | 3.39 |
| FALSE | High | GAYCDLSSI | [R].GAYCD | 1xCarbami | Q9D7A3 2x | 0.000579 | 0 | 1 | 2 | 7 | Q9D7A3 | Putative ur | Q9D7A3 [7 | 0 | 2134.869 | 0 | 0.29 | 0.002817 | 2.26 | 86.4 | 113.6 | 5.07 | 3.89 | | High | High | High | High | High | High | High | High | High | High | High | 9.42E-05 | 0.000994 | 2.93 |
| FALSE | High | GAYSLSIR | [K].GAYSLS | 1xPhospho | Q04736 1x | 0.046687 | 0.00354 | 2 | 9 | 1 | Q04736; P | Tyrosine-p | Q04736 [1 | 0 | 1175.602 | 0 | -0.24 | 0.042678 | 3.71 | 105.8 | 94.2 | 12.87 | 7.36 | Shared | High | High | High | High | High | High | High | High | High | High | High | 0.001812 | 0.01784 | 2.22 |
| FALSE | High | GAYVPSSP | [K].GAYVP | 1xPhospho | Q80W47 1 | 0.025293 | 0.001645 | 1 | 1 | 2 | Q80W47 | WD repeat | Q80W47 [3 | 0 | 1343.656 | 0 | -0.14 | 0.288824 | 9.11 | 105.3 | 94.7 | 3.2 | 11.06 | | High | High | High | High | High | High | High | High | High | High | High | 0.000872 | 0.008875 | 1.34 |
| FALSE | High | GCALQCAII | [R].GCALQ | 2xCarbami | Q571M2 1 | 0.037369 | 0.00249 | 2 | 10 | 1 | Q571M2; ( | MKIAA402 | Q571M2 [4 | 0 | 2074.063 | 0 | -0.57 | 0.007206 | 4.64 | 120.4 | 79.6 | 7.62 | 6.09 | Shared | High | High | High | High | High | High | High | High | High | High | High | 0.001308 | 0.01384 | 3.06 |
| FALSE | High | GCDSPDPD | [R].GCDSPI | 1xCarbami | Q60929-3 | 6.73E-06 | 0 | 1 | 1 | 2 | Q60929-3 | Isoform 3 c | Q60929-3 | 0 | 2786.263 | 0 | 1.61 | 0.000917 | 5.36 | 51.6 | 148.4 | 6.87 | 10.4 | | High | High | High | High | High | High | High | High | High | High | High | 0 | 8.64E-07 | 4.58 |
| FALSE | High | GCDSPDPD | [R].GCDSPI | 1xCarbami | Q60929-3 | 0.096181 | 0.007535 | 1 | 1 | 1 | Q60929-3 | Isoform 3 c | Q60929-3 | 0 | 2866.23 | 0 | 2.02 | 0.000704 | 10.97 | 40.4 | 159.6 | 18.7 | 14.14 | | High | High | High | High | High | High | High | High | High | High | High | 0.003809 | 0.04109 | 2.77 |
| FALSE | High | GCEGGTTS | [R].GCEGG | 1xCarbami | A0A140LJ | 0.000553 | 0 | 1 | 3 | 2 | A0A140LJ | A-kinase ar | A0A140LJ | 0 | 2907.325 | 0 | 0.41 | 0.038661 | 5.79 | 83.8 | 116.2 | 12.31 | 15.07 | | High | High | High | High | High | High | High | High | High | High | High | 0 | 0.000211 | 3.16 |
| FALSE | High | GCHLLVAT | [R].GCHLLV | 1xCarbami | Q3TQX5 1x | 0.001137 | 0 | 1 | 6 | 1 | Q3TQX5 | Putative ur | Q3TQX5 [3 | 0 | 1489.755 | 0 | -0.38 | 0.054381 | 13.84 | 113.2 | 86.8 | 11.06 | 7.41 | | High | High | High | High | High | High | High | High | High | High | High | 0 | 0.000275 | 2.91 |
| FALSE | High | GDLLVNDN | [K].GDLLVI | 1xPhospho | V9GX74 1x | 5.63E-07 | 0 | 1 | 9 | 2 | V9GX74 | Zinc finger | V9GX74 [2 | 0 | 2932.475 | 0 | -0.37 | 0.033477 | 5.61 | 115.7 | 84.3 | 14.75 | 9.59 | | High | High | High | High | High | High | High | High | High | High | High | 0 | 5.32E-08 | 4.35 |
| FALSE | High | GSGDTSISN | [R].GSGDT | 1xPhospho | Q3UZ39 1x | 1.99E-06 | 0 | 1 | 10 | 3 | Q3UZ39 | Leucine-ric | Q3UZ39 [8 | 0 | 1951.852 | 0 | 1 | 0.047977 | 4.66 | 63.5 | 136.5 | 9.52 | 40.54 | | High | High | High | High | High | High | High | High | High | High | High | 0 | 2.61E-06 | 3.05 |
| FALSE | High | GDINIEGPS | [K].GDINIE | 1xCarbami | E9Q616 1x | 0.000808 | 0 | 1 | 2 | 1 | E9Q616 | AHNAK nu | E9Q616 [3 | 0 | 3436.642 | 0 | 1.64 | 0.024506 | 57.37 | 51.4 | 148.6 | 37.51 | 35.37 | | High | High | High | High | High | High | High | High | High | High | High | 0 | 0.000188 | 4.43 |
| FALSE | High | GDLSDVEE | [R].GDLSD | 1xPhospho | Q60520-1 | 0.000908 | 0 | 1 | 2 | 2 | Q60520-1 | Isoform 2 c | Q60520-1 | 0 | 3161.365 | 0 | -0.99 | 0.00022 | 2.7 | 135.1 | 64.9 | 4 | 8.54 | | High | High | High | High | High | High | High | High | High | High | High | 0 | 0.000213 | 3.39 |
| FALSE | High | GDSIGSSK | [R].GDSIGS | 1xPhospho | Q80TM6 1 | 0.025738 | 0.001645 | 1 | 8 | 2 | Q80TM6 | R3H domai | Q80TM6 [3 | 0 | 1288.655 | 0 | 1.08 | 0.000261 | 4.34 | 66 | 134 | 7.02 | 5.93 | | High | High | High | High | High | High | High | High | High | High | High | 0.000968 | 0.01009 | 1.51 |
| FALSE | High | GDSLILEHC | [R].GDSLIL | 1xPhospho | Q60575 1x | 0.001933 | 5.66E-05 | 1 | 3 | 3 | Q60575 | Kinesin-like | Q60575 [1 | 0 | 2235.146 | 0 | 1.26 | 0.000459 | 14.18 | 59.6 | 140.4 | 8.17 | 3.91 | | High | High | High | High | High | High | High | High | High | High | High | 0.001453 | 0.01452 | 3.38 |
| FALSE | High | GDSPIIEK | [R].GDSPIII | 1xPhospho | A6H5X4 1x | 0.002844 | 0.00016 | 1 | 3 | 1 | A6H5X4 | PHD finger | A6H5X4 [1 | 0 | 1396.749 | 0 | 3.84 | 0.000199 | 5.79 | 14.1 | 185.9 | 20.76 | 6.38 | | High | High | High | High | High | High | High | High | High | High | High | 7.19E-05 | 0.00077 | 1.77 |
| FALSE | High | GDSPTQEP | [K].GDSPT( | 1xCarbami | A3KFM7 1 | 2.22E-07 | 0 | 1 | 3 | 1 | A3KFM7 | Chromodo | A3KFM7 [2 | 0 | 2042.951 | 0 | -0.28 | 0.014555 | 7.1 | 109.4 | 90.6 | 6.27 | 7 | | High | High | High | High | High | High | High | High | High | High | High | 0 | 1.88E-08 | 3.56 |
| FALSE | High | GDSSSSSPE | [K].GDSSSS | 1xPhospho | B1AQJ2 1x | 0.001489 | 5.66E-05 | 1 | 2 | 5 | B1AQJ2 | Ubiquitin c | B1AQJ2 [6 | 0 | 2271.018 | 0 | -1.21 | 0.00057 | 1.79 | 139.8 | 60.2 | 4.98 | 8.61 | | High | High | High | High | High | High | High | High | High | High | High | 0.000505 | 0.004365 | 2.44 |
| FALSE | High | GDSSSSSPE | [K].GDSSSS | 1xPhospho | B1AQJ2 2x | 0.00642 | 0.00062 | 1 | 2 | 3 | B1AQJ2 | Ubiquitin c | B1AQJ2 [6 | 0 | 2350.984 | 0 | -0.71 | 0.008715 | 2.99 | 121.1 | 78.9 | 12.28 | 12.14 | | High | High | High | High | High | High | High | High | High | High | High | 0.003148 | 0.03286 | 2.34 |
| FALSE | High | GDTPGHAT | [R].GDTPG | 2xPhospho | G5E866 2x | 0.006713 | 0.00062 | 1 | 5 | 1 | G5E866 | Splicing fac | G5E866 [2 | 0 | 2122.891 | 0 | 0.7 | 0.027704 | 32.09 | 76.1 | 123.9 | 18.72 | 13.57 | | High | High | High | High | High | High | High | High | High | High | High | 0.000275 | 0.002019 | 2.19 |
| FALSE | High | GDTSHSPT | [R].GDTSH | 1xPhospho | Q99LJ0 1xF | 0.005418 | 0.00062 | 1 | 1 | 2 | Q99LJ0 | CTTNBP2 N | Q99LJ0 [54 | 0 | 1621.799 | 0 | 1.45 | 0.000285 | 5.54 | 54.4 | 145.6 | 5.08 | 4.83 | | High | High | High | High | High | High | High | High | High | High | High | 0.000555 | 0.004841 | 2.05 |
| FALSE | High | GDVDVSLP | [K].GDVDV | 1xPhospho | E9Q616 1x | 0.022574 | 0.001542 | 1 | 5 | 2 | E9Q616 | AHNAK nu | E9Q616 [1 | 0 | 1467.786 | 0 | 3.12 | 0.000717 | 26.59 | 19.4 | 180.6 | 15.47 | 13.75 | | High | High | High | High | High | High | High | High | High | High | High | 0.001142 | 0.01169 | 2.62 |
| FALSE | High | GDVGPGES | [R].GDVGP | 1xPhospho | A0A0R4J0I | 1.91E-08 | 0 | 1 | 2 | 2 | A0A0R4J0I | ETS domai | A0A0R4J0I | 0 | 1704.816 | 0 | 0.26 | 0.192783 | 16.63 | 96.8 | 103.2 | 7.87 | 12.54 | | High | High | High | High | High | High | High | High | High | High | High | 0 | 1.2E-09 | 3.23 |
| FALSE | High | GDVPSVGL | [K].GDVPS | 1xPhospho | E9Q616 1x | 0.000845 | 0 | 1 | 5 | 1 | E9Q616 | AHNAK nu | E9Q616 [4 | 0 | 2616.332 | 0 | 0.94 | 0.005132 | 23.35 | 68.7 | 131.3 | 24.97 | 17.62 | | High | High | High | High | High | High | High | High | High | High | High | 0 | 0.000197 | 2.47 |
| FALSE | High | GDVSEDEP | [R].GDVSE | 1xPhospho | Q9D0L8 1x | 0.002793 | 0.000109 | 1 | 6 | 2 | Q9D0L8 | mRNA cap | Q9D0L8 [9 | 0 | 1569.7 | 0 | -0.4 | 0.183589 | 7.77 | 116 | 84 | 7.07 | 18.44 | | High | High | High | High | High | High | High | High | High | High | High | 4.94E-05 | 0.000752 | 2.54 |
| FALSE | High | GDVVHCW | [K].GDVVH | 1xCarbami | Q62446 1x | 0.001865 | 5.66E-05 | 1 | 1 | 2 | Q62446 | Peptidyl-pr | Q62446 [1 | 0 | 3536.681 | 0 | -0.86 | 0.009082 | 7.59 | 135 | 65 | 11.06 | 33.33 | | High | High | High | High | High | High | High | High | High | High | High | 2.58E-05 | 0.000481 | 3.79 |
| FALSE | High | GDSIEEILAI | [K].GDSIEE | 1xPhospho | Q6P5B0 1x | 0.000362 | 0 | 1 | 1 | 2 | Q6P5B0 | RRP12-like | Q6P5B0 [1 | 1 | 2831.194 | 0 | -0.42 | 0.005307 | 5.81 | 113.8 | 86.2 | 3.9 | 10.16 | | High | High | High | High | High | High | High | High | High | High | High | 0 | 7.62E-05 | 2.88 |
| FALSE | High | GEADDEDI | [K].GEADD | 1xPhospho | F8VQD1 1x | 1.46E-08 | 0 | 1 | 13 | 11 | F8VQD1 | Protein pol | F8VQD1 [1 | 0 | 3551.486 | 0 | -0.35 | 0.007376 | 7.03 | 113.3 | 86.7 | 4.85 | 7.59 | | High | High | High | High | High | High | High | High | High | High | High | 0 | 7.46E-08 | 5.48 |
| FALSE | High | GEAGQELL | [K].GEAGQ | 1xPhospho | Q8BYU6 1x | 0.008313 | 0.000736 | 1 | 1 | 1 | Q8BYU6 | Torsin-1A-i | Q8BYU6 [1 | 1 | 3705.694 | 0 | 0.01 | 0.584031 | 10.38 | 100.1 | 99.9 | 3.55 | 5.94 | | High | High | High | High | High | High | High | High | High | High | High | 0.000327 | 0.002545 | 3.89 |
| FALSE | High | GEAGVLGS | [K].GEAGV | 1xPhospho | D3Z3A8 1x | 8.87E-05 | 0 | 1 | 3 | 2 | D3Z3A8 | Unconvent | D3Z3A8 [1 | 0 | 1810.972 | 0 | 0.09 | 0.196402 | 5.33 | 96.4 | 103.6 | 11.14 | 8.03 | | High | High | High | High | High | High | High | High | High | High | High | 0 | 1.56E-05 | 2.49 |
| FALSE | High | GEAMATV | [K].GEAMA | 1xPhospho | Q69ZP8 1x | 3.29E-05 | 0 | 1 | 1 | 2 | Q69ZP8 | MKIAA117 | Q69ZP8 [1 | 0 | 3278.683 | 0 | 1.3 | 0.03411 | 24.67 | 58.5 | 141.5 | 14.21 | 27.02 | | High | High | High | High | High | High | High | High | High | High | High | 0 | 6.98E-06 | 4.36 |
| FALSE | High | GEAMATV | [K].GEAMA | 1xPhospho | Q6PFF0 1x | 0.001586 | 5.66E-05 | 1 | 3 | 4 | Q6PFF0 | SR-related | Q6PFF0 [1 | 0 | 3306.715 | 0 | -0.25 | 0.057461 | 7.27 | 111.2 | 88.8 | 5.37 | 18.69 | | High | High | High | High | High | High | High | High | High | High | High | 2.58E-05 | 0.0004 | 2.67 |
| FALSE | High | GEAYYDTD | [R].GEAYYI | 1xPhospho | B2M0S2 1x | 0.060666 | 0.00451 | 1 | 8 | 2 | B2M0S2 | Secretory ( | B2M0S2 [8 | 1 | 2509.066 | 0 | -0.6 | 0.006943 | 3.23 | 118 | 82 | 8.29 | 7.98 | | High | High | High | High | High | High | High | High | High | High | High | 0.002293 | 0.024 | 2.13 |
| FALSE | High | GEDLDFWL | [K].GEDLDI | 1xPhospho | O54774 1x | 0.025964 | 0.001645 | 1 | 1 | 3 | O54774 | AP-3 comp | O54774 [8 | 0 | 4217.135 | 0 | 0.03 | 0.277367 | 3.84 | 98.9 | 101.1 | 13.74 | 13.54 | | High | High | High | High | High | High | High | High | High | High | High | 0.001876 | 0.01879 | 2.89 |
| FALSE | High | GEDPFVNE | [K].GEDPF | 1xPhospho | Q059U9 1x | 1.65E-07 | 0 | 1 | 1 | 3 | Q059U9 | A kinase (P | Q059U9 [5 | 0 | 3161.409 | 0 | 0.59 | 0.045602 | 22.14 | 76.4 | 123.6 | 10.79 | 22.25 | | High | High | High | High | High | High | High | High | High | High | High | 0 | 1.34E-08 | 4.92 |
| FALSE | High | GEDSWDS | [K].GEDSW | 2xPhospho | E9Q4F8 2x | 0.005228 | 0.000551 | 1 | 3 | 2 | E9Q4F8 | Ankyrin re | E9Q4F8 [8 | 0 | 1818.682 | 0 | -0.15 | 0.312405 | 6.64 | 105.3 | 94.7 | 9.52 | 12.02 | | High | High | High | High | High | High | High | High | High | High | High | 0.000267 | 0.001527 | 3.38 |
| FALSE | High | GEEAPEELS | [K].GEEAPI | 1xPhospho | Q6P9R1 1x | 1.35E-06 | 0 | 1 | 1 | 2 | Q6P9R1 | ATP-depen | Q6P9R1 [1 | 0 | 3479.591 | 0 | -0.05 | 0.403566 | 13.83 | 104.1 | 95.9 | 11.74 | 10.72 | | High | High | High | High | High | High | High | High | High | High | High | 0 | 1.43E-07 | 3.55 |
| FALSE | High | GEEDGPEE | [K].GEEDG | 1xPhospho | Q00PI9 1xF | 0.002818 | 0.000109 | 1 | 3 | 1 | Q00PI9 | Heterogen | Q00PI9 [15 | 1 | 2883.257 | 0 | 0.55 | 0.050145 | 11.49 | 83.6 | 116.4 | 12.87 | 23.02 | | High | High | High | High | High | High | High | High | High | High | High | 7.19E-05 | 0.000763 | 3.44 |
| FALSE | High | GEEDSDVS | [R].GEEDSI | 1xPhospho | Q3UQS8 1 | 7.72E-07 | 0 | 1 | 3 | 4 | Q3UQS8 | RNA-bindir | Q3UQS8 [1 | 0 | 2952.313 | 0 | 2.4 | 0.007475 | 41.57 | 30.3 | 169.7 | 10.31 | 27.63 | | High | High | High | High | High | High | High | High | High | High | High | 0 | 7.6E-08 | 3.75 |
| FALSE | High | GEEDSDVS | [R].GEEDSI | 1xPhospho | Q3UQS8 1 | 1.81E-10 | 0 | 1 | 3 | 6 | Q3UQS8 | RNA-bindir | Q3UQS8 [1 | 0 | 2968.308 | 0 | 3.69 | 0.00122 | 61.94 | 12.1 | 187.9 | 44.89 | 16.33 | | High | High | High | High | High | High | High | High | High | High | High | 0 | 7.79E-11 | 5.21 |
| FALSE | High | GEEEPLVG | [K].GEEEPL | 1xPhospho | Q544C5 1x | 2.86E-07 | 0 | 1 | 2 | 4 | Q544C5 | MCG14526 | Q544C5 [3 | 0 | 3109.45 | 0 | -0.65 | 0.003293 | 3.83 | 123.2 | 76.8 | 2.25 | 12.13 | | High | High | High | High | High | High | High | High | High | High | High | 0 | 2.5E-08 | 4.62 |
| FALSE | High | GEAGPDW | [R].GEAGP | 1xCarbami | Q2MHN3 1 | 9.3E-09 | 0 | 1 | 1 | 7 | Q2MHN3 | ZFH-5 OS=I | Q2MHN3 [ | 0 | 3740.719 | 0 | -0.69 | 0.012012 | 5.82 | 125 | 75 | 11.96 | 12.2 | | High | High | High | High | High | High | High | High | High | High | High | 0 | 9.72E-06 | 5.33 |
| FALSE | High | GDSIEEILAI | [K].GDSIEE | 1xPhospho | Q6P5B0 1x | 1.63E-08 | 0 | 1 | 1 | 15 | Q6P5B0 | RRP12-like | Q6P5B0 [1 | 0 | 2618.072 | 0 | -0.37 | 0.001684 | 3.34 | 112.5 | 87.5 | 4.08 | 7.12 | | High | High | High | High | High | High | High | High | High | High | High | 0 | 4.2E-09 | 5.26 |
| FALSE | High | GDSLDVVL | [K].GDSLD | 1xPhospho | A2ASY1 1x | 0.003342 | 0.00016 | 1 | 5 | 1 | A2ASY1 | TAF3 RNA | A2ASY1 [2 | 0 | 1595.824 | 0 | 0.14 | 0.138879 | 14.55 | 94.7 | 105.3 | 3.88 | 7.84 | | High | High | High | High | High | High | High | High | High | High | High | 7.19E-05 | 0.000924 | 3.09 |
| FALSE | High | GDSHSPGH | [R].GDSHS | 2xPhospho | Q8BTI8 2x | 0.000668 | 0 | 1 | 3 | 4 | Q8BTI8 | Serine/arg | Q8BTI8 [2 | 0 | 1539.676 | 0 | -0.52 | 0.001968 | 2.82 | 118 | 82 | 2.63 | 7.04 | | High | High | High | High | High | High | High | High | High | High | High | 0.000275 | 0.002181 | 1.84 |
| FALSE | High | GDLSDVEE | [R].GDLSD | 1xPhospho | Q60520-1 | 2.31E-05 | 0 | 1 | 2 | 8 | Q60520-1 | Isoform 2 c | Q60520-1 | 0 | 3177.359 | 0 | -0.7 | 0.005006 | 5.18 | 125.5 | 74.5 | 8.81 | 11.54 | | High | High | High | High | High | High | High | High | High | High | High | 0 | 0.000131 | 4.2 |
| FALSE | High | GDMSAVN | [K].GDMSA | 1xPhospho | Q8CCI5 1x | 0.000352 | 0 | 1 | 1 | 1 | Q8CCI5 | RING1 and | Q8CCI5 [2 | 0 | 1480.586 | 0 | -1.62 | 0.00049 | 1.45 | 150 | 50 | 12.11 | 16.42 | | High | High | High | High | High | High | High | High | High | High | High | 0 | 7.39E-05 | 3.09 |
| FALSE | High | GDMSAVN | [K].GDMSA | 1xPhospho | Q8CCI5 1x | 1.25E-05 | 0 | 1 | 1 | 18 | Q8CCI5 | RING1 and | Q8CCI5 [2 | 0 | 1496.581 | 0 | -1.45 | 0.000158 | 0.42 | 146.5 | 53.5 | 3.86 | 8.35 | | High | High | High | High | High | High | High | High | High | High | High | 0 | 1.75E-06 | 3.15 |
| FALSE | High | GDMSDED | [K].GDMSI | 1xPhospho | Q3B7Z2 1x | 0.000259 | 0 | 1 | 2 | 3 | Q3B7Z2 | Oxysterol-l | Q3B7Z2 [3 | 0 | 3733.633 | 0 | 0.49 | 0.054444 | 14.33 | 84.2 | 115.8 | 12 | 19.28 | | High | High | High | High | High | High | High | High | High | High | High | 0 | 0.000203 | 3.53 |
| FALSE | High | GDMSDED | [K].GDMSI | 1xDeamida | Q3B7Z2 1x | 0.026885 | 0.001676 | 1 | 2 | 2 | Q3B7Z2 | Oxysterol-l | Q3B7Z2 [3 | 0 | 3734.617 | 0 | 0.02 | 0.538694 | 40.6 | 94 | 106 | 26.5 | 22.88 | | High | High | High | High | High | High | High | High | High | High | High | 0.000903 | 0.009528 | 3.32 |
| FALSE | High | GDMSDED | [K].GDMSI | 1xPhospho | Q3B7Z2 1x | 3.65E-07 | 0 | 1 | 2 | 11 | Q3B7Z2 | Oxysterol-l | Q3B7Z2 [3 | 0 | 3749.628 | 0 | 0.56 | 0.00044 | 5.6 | 81.7 | 118.3 | 4.85 | 4.86 | | High | High | High | High | High | High | High | High | High | High | High | 0 | 1.13E-06 | 5.28 |
| FALSE | High | GDNASPSP | [R].GDNAS | 1xPhospho | Q3TW36 1 | 3.34E-07 | 0 | 1 | 5 | 4 | Q3TW36 | Putative ur | Q3TW36 [ | 0 | 1763.853 | 0 | 0.53 | 0.001931 | 9.64 | 81.1 | 118.9 | 4.91 | 5.8 | | High | High | High | High | High | High | High | High | High | High | High | 0 | 2.24E-06 | 2.17 |
| FALSE | High | GDNASPSP | [R].GDNAS | 2xPhospho | Q3TW36 2 | 0.000538 | 0 | 1 | 5 | 2 | Q3TW36 | Putative ur | Q3TW36 [ | 0 | 1843.819 | 0 | 0.18 | 0.094315 | 8.5 | 96.1 | 103.9 | 7.61 | 7.54 | | High | High | High | High | High | High | High | High | High | High | High | 0 | 0.000119 | 2.69 |
| FALSE | High | GDNWYSEI | [R].GDNW | 2xPhospho | Q80XE1-1 | 1.12E-05 | 0 | 1 | 2 | 12 | Q80XE1-1 | Isoform 1 c | Q80XE1-1 | 0 | 2714.047 | 0 | -0.07 | 0.099193 | 4.02 | 101.9 | 98.1 | 2.35 | 8.87 | | High | High | High | High | High | High | High | High | High | High | High | 0 | 4.84E-05 | 3.86 |
| FALSE | High | GDPFTDLC | [R].GDPFTI | 1xCarbami | Q9CXH7 1x | 3.68E-06 | 0 | 1 | 1 | 1 | Q9CXH7 | Shugoshin | Q9CXH7 [4 | 0 | 2409.197 | 0 | -0.18 | 0.655982 | 11.88 | 105.9 | 94.1 | 8.11 | 12.59 | | High | High | High | High | High | High | High | High | High | High | High | 0 | 4.4E-07 | 3.36 |
| FALSE | High | GDPLSPGR | [K].GDPLSI | 1xPhospho | E9QK89 1x | 0.002134 | 5.66E-05 | 1 | 2 | 1 | E9QK89 | Mediator c | E9QK89 [9 | 0 | 1107.54 | 0 | -0.79 | 0.001421 | 4.02 | 124.4 | 75.6 | 6.74 | 4.07 | | High | High | High | High | High | High | High | High | High | High | High | 2.58E-05 | 0.000556 | 1.81 |
| FALSE | High | GDQEEELD | [R].GDQEE | 1xPhospho | A0A140LJ0 | 0.000605 | 0 | 1 | 4 | 2 | A0A140LJ0 | Zinc finger | A0A140LJ0 | 0 | 2738.292 | 0 | 1.31 | 0.001147 | 14.39 | 60.5 | 139.5 | 6.76 | 10.65 | | High | High | High | High | High | High | High | High | High | High | High | 0 | 0.000221 | 2.49 |
| FALSE | High | GDQPLHSC | [R].GDQPL | 1xCarbami | Q7TN98 1x | 0.001572 | 5.66E-05 | 1 | 9 | 1 | Q7TN98 | Cytoplasmi | Q7TN98 [4 | 0 | 2740.225 | 0 | 1.7 | 0.000734 | 10.36 | 44.9 | 155.1 | 9.55 | 8.86 | | High | High | High | High | High | High | High | High | High | High | High | 2.58E-05 | 0.000396 | 1.76 |
| FALSE | High | GDQPLHSC | [R].GDQPL | 1xCarbami | Q7TN98 1x | 0.001021 | 0 | 1 | 9 | 2 | Q7TN98 | Cytoplasmi | Q7TN98 [4 | 0 | 2741.209 | 0 | 0.04 | 0.723234 | 17.27 | 96.8 | 103.2 | 8.64 | 8.14 | | High | High | High | High | High | High | High | High | High | High | High | 0 | 0.000244 | 4.33 |

| | | | | | | | | | | | | | | | | | | | | | | | | | | | | | | | | | | | | | | |
|---|---|---|---|---|---|---|---|---|---|---|---|---|---|---|---|---|---|---|---|---|---|---|---|---|---|---|---|---|---|---|---|---|---|---|---|---|---|---|
| FALSE | High | GDQSPEDL | [K].GDQSP | 1xPhosphc | Q6PCQ0 1x | 0.081752 | 0.006219 | 1 | 3 | 1 | Q6PCQ0 | IQ domain- | Q6PCQ0 [3 | 0 | 1623.803 | 0 | 1.35 | 0.000704 | 16.37 | 56.3 | 143.7 | 10.46 | 5.31 | | High | High | High | High | High | High | High | High | High | High | High | 0.003229 | 0.03417 | 2.47 |
| FALSE | High | GDQVLNFS | [K].GDQVL | 1xPhosphc | Q3TYU7 1x | 7.74E-08 | 0 | 1 | 3 | 5 | Q3TYU7 | Putative ur | Q3TYU7 [2 | 0 | 2519.195 | 0 | -0.75 | 0.000527 | 0.56 | 122.7 | 77.3 | 2.89 | 5.49 | | High | High | High | High | High | High | High | High | High | High | High | 0 | 3.56E-08 | 3.87 |
| FALSE | High | GDQVLNFS | [K].GDQVL | 1xDeamida | Q3TYU7 1x | 0.000178 | 0 | 1 | 3 | 1 | Q3TYU7 | Putative ur | Q3TYU7 [2 | 0 | 2520.179 | 0 | -0.6 | 0.070061 | 14.79 | 120.4 | 79.6 | 14.48 | 26.81 | | High | High | High | High | High | High | High | High | High | High | High | 0 | 3.42E-05 | 2.57 |
| FALSE | High | GDQVSQN | [K].GDQVS | 1xPhosphc | Q62261 1x | 1.68E-10 | 0 | 2 | 6 | 8 | Q62261; Q | Spectrin be | Q62261 [2 | 0 | 2048.96 | 0 | 1.56 | 0.000285 | 5.54 | 51.1 | 148.9 | 3.27 | 5.18 | Shared | High | High | High | High | High | High | High | High | High | High | High | 0 | 5.8E-12 | 4.56 |
| FALSE | High | GDQVSQN | [K].GDQVS | 1xDeamida | Q62261 1x | 5.36E-09 | 0 | 2 | 6 | 10 | Q62261; Q | Spectrin be | Q62261 [2 | 0 | 2049.944 | 0 | 1.54 | 0.000236 | 10.2 | 51.4 | 148.6 | 7.01 | 5.49 | Shared | High | High | High | High | High | High | High | High | High | High | High | 0 | 3.19E-09 | 4.25 |
| FALSE | High | GDQVSQN | [K].GDQVS | 2xPhosphc | Q62261 2x | 0.001058 | 0 | 2 | 6 | 2 | Q62261; Q | Spectrin be | Q62261 [2 | 0 | 2128.926 | 0 | 0.55 | 0.019805 | 19.2 | 81 | 119 | 8.31 | 13.68 | Shared | High | High | High | High | High | High | High | High | High | High | High | 0 | 0.000254 | 3.35 |
| FALSE | High | GDRGEESN | [R].GDRGE | 1xPhosphc | Q04692 1x | 0.001325 | 0 | 1 | 2 | 3 | Q04692 | SWI/SNF-r | Q04692 [2 | 1 | 2620.156 | 0 | -1.03 | 0.001059 | 5.02 | 134.2 | 65.8 | 3.61 | 11.66 | | High | High | High | High | High | High | High | High | High | High | High | 0 | 0.000326 | 3.72 |
| FALSE | High | GDRGEESN | [R].GDRGE | 1xDeamida | Q04692 1x | 0.095408 | 0.007437 | 1 | 2 | 1 | Q04692 | SWI/SNF-r | Q04692 [2 | 1 | 2621.14 | 0 | -0.71 | 0.003045 | 3.71 | 120.5 | 79.5 | 6.51 | 9.77 | | High | High | High | High | High | High | High | High | High | High | High | 0.003782 | 0.04072 | 2.31 |
| FALSE | High | GDRGEESN | [R].GDRGE | 2xPhosphc | Q04692 2x | 0.004294 | 0.00036 | 1 | 2 | 3 | Q04692 | SWI/SNF-r | Q04692 [2 | 1 | 2700.123 | 0 | -1.11 | 0.000389 | 0.63 | 135.8 | 64.2 | 3.55 | 5.68 | | High | High | High | High | High | High | High | High | High | High | High | 0.000161 | 0.001225 | 4.17 |
| FALSE | High | GDRSPEPG | [K].GDRSPI | 1xPhosphc | E9Q616 1x | 1.9E-06 | 0 | 1 | 5 | 4 | E9Q616 | AHNAK nu | E9Q616 [9 | 1 | 2482.135 | 0 | 2.64 | 0.001179 | 9.43 | 27.7 | 172.3 | 6.11 | 15.54 | | High | High | High | High | High | High | High | High | High | High | High | 0 | 2.09E-07 | 2.95 |
| FALSE | High | GDSEEVSG | [R].GDSEE\ | 1xPhosphc | Q8CHG2 1x | 3.82E-07 | 0 | 1 | 3 | 1 | Q8CHG2 | MKIAA036 | Q8CHG2 [8 | 0 | 2901.402 | 0 | -0.24 | 0.029224 | 8.99 | 110.3 | 89.7 | 10.92 | 3.95 | | High | High | High | High | High | High | High | High | High | High | High | 0 | 3.44E-08 | 4.21 |
| FALSE | High | GDSETDLE/ | [R].GDSETI | 1xPhosphc | P46938 1xI | 2.98E-05 | 0 | 1 | 6 | 10 | P46938 | Transcripti | P46938 [44 | 0 | 2489.204 | 0 | 1.12 | 0.005649 | 6.31 | 64.1 | 135.9 | 10.17 | 21.72 | | High | High | High | High | High | High | High | High | High | High | High | 0 | 1.21E-05 | 3.6 |
| FALSE | High | GDSETDLE/ | [R].GDSETI | 1xPhosphc | P46938 1xI | 3.72E-07 | 0 | 1 | 6 | 19 | P46938 | Transcripti | P46938 [44 | 0 | 2505.198 | 0 | 0.52 | 0.021399 | 18.64 | 82.2 | 117.8 | 11.41 | 11.3 | | High | High | High | High | High | High | High | High | High | High | High | 0 | 3.98E-08 | 3.91 |
| FALSE | High | GDSGVDTC | [K].GDSGV | 1xCarbami | Q8K2F8 1x | 1.4E-09 | 0 | 1 | 1 | 5 | Q8K2F8 | Protein LSI | Q8K2F8 [3 | 0 | 3584.505 | 3 | 0 | 0.978662 | 21.4 | 95.5 | 104.5 | 7.81 | 11.29 | | High | High | High | High | High | High | High | High | High | High | High | 0 | 1.32E-10 | 6.83 |
| FALSE | High | GDSHSPGH | [R].GDSHSI | 1xPhosphc | Q8BTI8 1xI | 0.070485 | 0.005214 | 1 | 3 | 2 | Q8BTI8 | Serine/argi | Q8BTI8 [26 | 0 | 1459.71 | 0 | -0.35 | 0.129748 | 7.49 | 114.1 | 85.9 | 12.31 | 15.57 | | High | High | High | High | High | High | High | High | High | High | High | 0.004757 | 0.0514 | 1.59 |
| FALSE | High | GDINIEGPS | [K].GDINIE | 1xCarbami | E9Q616 1x | 3.35E-08 | 0 | 1 | 2 | 2 | E9Q616 | AHNAK nu | E9Q616 [3 | 0 | 3452.637 | 0 | 2.17 | 0.000724 | 17.39 | 36.3 | 163.7 | 21.32 | 12.38 | | High | High | High | High | High | High | High | High | High | High | High | 0 | 2.23E-09 | 6.6 |
| FALSE | High | GSGDTSSLI | [R].GSGDT | 1xPhosphc | A0A1L1SSC | 0.016892 | 0.001204 | 1 | 12 | 1 | A0A1L1SSC | Leucine-ric | A0A1L1SSC | 0 | 2259.059 | 0 | 0.33 | 0.042886 | 15.05 | 90.8 | 109.2 | 6.47 | 10.23 | | High | High | High | High | High | High | High | High | High | High | High | 0.000638 | 0.005649 | 2.74 |
| FALSE | High | GSGEASSD | [K].GSGEA | 1xPhosphc | A0A0R4J07 | 7.98E-06 | 0 | 1 | 5 | 2 | A0A0R4J07 | Histone-lys | A0A0R4J07 | 0 | 2082.975 | 0 | -0.63 | 0.009137 | 8.11 | 120.3 | 79.7 | 8.51 | 9.8 | | High | High | High | High | High | High | High | High | High | High | High | 0 | 1.05E-06 | 2.87 |
| FALSE | High | GSGEEPAT | [R].GSGEEI | 1xPhosphc | Q6ZPH4 1x | 0.003927 | 0.000311 | 1 | 3 | 1 | Q6ZPH4 | MKIAA183 | Q6ZPH4 [4 | 0 | 1422.683 | 0 | -1.61 | 0.000207 | 1.68 | 150.5 | 49.5 | 2.87 | 12.17 | | High | High | High | High | High | High | High | High | High | High | High | 0.00014 | 0.001101 | 1.88 |
| FALSE | High | HSLSGSSP( | [R].HSLSGS | 1xPhosphc | Q8BTI8 1xI | 7.04E-06 | 0 | 1 | 3 | 8 | Q8BTI8 | Serine/argi | Q8BTI8 [14 | 0 | 1641.807 | 0 | -0.4 | 0.014183 | 3.14 | 112.2 | 87.8 | 4.27 | 6.28 | | High | High | High | High | High | High | High | High | High | High | High | 0.000571 | 0.004959 | 1.88 |
| FALSE | High | HSLSLDDIR | [R].HSLSLD | 1xPhosphc | Q8C0V9 1x | 0.040028 | 0.002867 | 1 | 2 | 2 | Q8C0V9 | FERM dom | Q8C0V9 [5 | 0 | 1364.677 | 0 | 1.11 | 0.000446 | 2.45 | 61.5 | 138.5 | 6.36 | 8 | | High | High | High | High | High | High | High | High | High | High | High | 0.001484 | 0.0149 | 2.03 |
| FALSE | High | HSLSSESQ/ | [K].HSLSSE | 1xPhosphc | Q99K30 1x | 0.002232 | 0.000109 | 1 | 1 | 2 | Q99K30 | Epidermal | Q99K30 [4 | 0 | 2780.284 | 0 | 0.57 | 0.02249 | 10.12 | 81.2 | 118.8 | 16.52 | 12.17 | | High | High | High | High | High | High | High | High | High | High | High | 4.94E-05 | 0.000587 | 2.78 |
| FALSE | High | HSLSSESQ/ | [K].HSLSSE | 2xPhosphc | Q99K30 2x | 0.009006 | 0.000736 | 1 | 1 | 1 | Q99K30 | Epidermal | Q99K30 [4 | 0 | 2860.25 | 0 | 1.18 | 0.006214 | 14.67 | 60.2 | 139.8 | 13.98 | 22.44 | | High | High | High | High | High | High | High | High | High | High | High | 0.000327 | 0.002804 | 3.13 |
| FALSE | High | HSMGPGS\ | [R].HSMGF | 1xPhosphc | Q3TWM8 1 | 1.87E-09 | 0 | 1 | 6 | 2 | Q3TWM8 | Putative ur | Q3TWM8 [ | 0 | 2475.096 | 0 | 0.89 | 0.004068 | 14.38 | 71.3 | 128.7 | 13.53 | 8.66 | | High | High | High | High | High | High | High | High | High | High | High | 0 | 8.72E-11 | 4.36 |
| FALSE | High | HSNTLEQT. | [R].HSNTLE | 1xPhosphc | E9Q7M2 1: | 0.000822 | 0 | 1 | 3 | 1 | E9Q7M2 | TSC22 dom | E9Q7M2 [1 | 0 | 1594.742 | 0 | 1.88 | 0.000816 | 8.67 | 44.8 | 155.2 | 14.53 | 8.93 | | High | High | High | High | High | High | High | High | High | High | High | 0 | 0.000191 | 3.18 |
| FALSE | High | HSPATISSS | [R].HSPATI | 1xPhosphc | P70324 1xI | 0.029076 | 0.001857 | 1 | 4 | 1 | P70324 | T-box trans | P70324 [43 | 0 | 1452.705 | 0 | -1.7 | 0.000596 | 1.5 | 156.6 | 43.4 | 9.09 | 10.81 | | High | High | High | High | High | High | High | High | High | High | High | 0.000997 | 0.01038 | 1.25 |
| FALSE | High | HSPDHPGN | [R].HSPDHI | 1xPhosphc | Q8C1S0 1x | 0.001767 | 5.66E-05 | 1 | 1 | 1 | Q8C1S0 | Mediator c | Q8C1S0 [2 | 0 | 2337.013 | 0 | -0.66 | 0.139521 | 33.42 | 118.8 | 81.2 | 18.07 | 60.87 | | High | High | High | High | High | High | High | High | High | High | High | 2.58E-05 | 0.000451 | 2.42 |
| FALSE | High | HSPDRQDG | [K].HSPDR( | 1xPhosphc | Q56A08 1x | 0.045122 | 0.003373 | 1 | 2 | 1 | Q56A08 | G patch do | Q56A08 [3 | 1 | 1641.795 | 0 | 1.3 | 0.246496 | 56.75 | 61.6 | 138.4 | 50.43 | 68.7 | | High | High | High | High | High | High | High | High | High | High | High | 0.001727 | 0.01715 | 2.73 |
| FALSE | High | HSPDRQDG | [K].HSPDR( | 1xPhosphc | Q56A08 1x | 0.000246 | 0 | 1 | 2 | 2 | Q56A08 | G patch do | Q56A08 [3 | 1 | 1870.958 | 0 | 1.38 | 0.001684 | 13.68 | 55.5 | 144.5 | 6.77 | 12.12 | | High | High | High | High | High | High | High | High | High | High | High | 0 | 4.93E-05 | 1.36 |
| FALSE | High | HSPDSACS\ | [K].HSPDS/ | 1xCarbami | Q921Q9 1x | 0.000967 | 0 | 2 | 4 | 1 | Q921Q9; C | Mapkbp1 p | Q921Q9 [1 | 0 | 1963.805 | 0 | -0.46 | 0.026294 | 11.56 | 110.9 | 89.1 | 8.25 | 16.83 | Shared | High | High | High | High | High | High | High | High | High | High | High | 0 | 0.000229 | 2.92 |
| FALSE | High | HSPPELSQE | [R].HSPPEL | 1xDeamida | A2AIX1 1xI | 0.043607 | 0.003228 | 1 | 7 | 1 | A2AIX1 | SEC16 hon | A2AIX1 [20 | 0 | 2700.166 | 0 | -0.52 | 0.006959 | 7.25 | 117.7 | 82.3 | 5.81 | 14.25 | | High | High | High | High | High | High | High | High | High | High | High | 0.001651 | 0.01646 | 2.38 |
| FALSE | High | HSPSALQD | [R].HSPSAL | 1xPhosphc | G3UWZ0 1 | 7.02E-05 | 0 | 1 | 4 | 3 | G3UWZ0 | Bromodom | G3UWZ0 [ | 0 | 2629.379 | 0 | 0.92 | 0.014383 | 26.85 | 70.5 | 129.5 | 16.94 | 17 | | High | High | High | High | High | High | High | High | High | High | High | 0 | 1.21E-05 | 2.91 |
| FALSE | High | HSLPSTFTN | [R].HSLPST | 1xPhosphc | A2A9F4 1xI | 0.007673 | 0.000664 | 1 | 3 | 2 | A2A9F4 | Keratinocy | A2A9F4 [2( | 0 | 1652.8 | 0 | -0.1 | 0.113595 | 11.76 | 103.7 | 96.3 | 22.51 | 27.04 | | High | High | High | High | High | High | High | High | High | High | High | 0.000294 | 0.002325 | 3.12 |
| FALSE | High | HSPSALQD | [R].HSPSAL | 2xPhosphc | G3UWZ0 2 | 0.002508 | 0.000109 | 1 | 4 | 1 | G3UWZ0 | Bromodom | G3UWZ0 [ | 0 | 2709.345 | 0 | 1.32 | 0.003307 | 21.22 | 55.4 | 144.6 | 9.93 | 15.71 | | High | High | High | High | High | High | High | High | High | High | High | 4.94E-05 | 0.000669 | 2.89 |
| FALSE | High | HSPTGPPG | [R].HSPTGF | 1xPhosphc | Q99KG3 1x | 0.006773 | 0.00062 | 1 | 2 | 2 | Q99KG3 | RNA-bindir | Q99KG3 [8 | 0 | 1458.709 | 0 | -0.23 | 0.517082 | 12.7 | 108.4 | 91.6 | 12.66 | 26.97 | | High | High | High | High | High | High | High | High | High | High | High | 0.001214 | 0.01297 | 2.44 |
| FALSE | High | HSPTLPEPG | [R].HSPTLP | 1xPhosphc | D3YTR3 1x | 1.09E-05 | 0 | 1 | 4 | 2 | D3YTR3 | RNA-bindir | D3YTR3 [5( | 0 | 2293.215 | 0 | -0.15 | 0.081184 | 5.26 | 103.8 | 96.2 | 5.12 | 9.34 | | High | High | High | High | High | High | High | High | High | High | High | 0 | 1.5E-06 | 3.15 |
| FALSE | High | HSPTLVPT/ | [K].HSPTLV | 1xPhosphc | Q9DB91 1x | 0.003495 | 0.00021 | 1 | 1 | 3 | Q9DB91 | Mediator c | Q9DB91 [1 | 0 | 4014.196 | 0 | -0.2 | 0.351835 | 17.89 | 106.7 | 93.3 | 16.45 | 38.87 | | High | High | High | High | High | High | High | High | High | High | High | 9.42E-05 | 0.000973 | 3.59 |
| FALSE | High | HSPVQMN | [R].HSPVQI | 1xPhosphc | Q3UQN2 1 | 0.040721 | 0.003058 | 1 | 3 | 2 | Q3UQN2 | F-BAR dom | Q3UQN2 [ | 0 | 1277.602 | 0 | 0.74 | 0.079135 | 14.93 | 86.8 | 113.2 | 18.77 | 29.48 | | High | High | High | High | High | High | High | High | High | High | High | 0.009735 | 0.1052 | 1.52 |
| FALSE | High | HSQPATPT | [R].HSQPAT | 1xPhosphc | Q3TJD7 1xI | 1.23E-05 | 0 | 1 | 1 | 3 | Q3TJD7 | PDZ and LII | Q3TJD7 [2 | 0 | 1755.874 | 0 | 0.75 | 0.000658 | 3.8 | 73.2 | 126.8 | 9.4 | 3.99 | | High | High | High | High | High | High | High | High | High | High | High | 0 | 1.71E-06 | 2.98 |
| FALSE | High | HSQQLFPE | [R].HSQQLI | 1xPhosphc | Q3U671 1x | 1.58E-05 | 0 | 1 | 2 | 1 | Q3U671 | Putative ur | Q3U671 [2 | 0 | 2508.232 | 0 | -0.64 | 0.000256 | 0.49 | 121.4 | 78.6 | 4.27 | 11.84 | | High | High | High | High | High | High | High | High | High | High | High | 0 | 2.27E-06 | 4.76 |
| FALSE | High | HSSAAADA | [K].HSSAA/ | 1xPhosphc | E9Q0A7 1x | 0.031713 | 0.002011 | 2 | 5 | 2 | E9Q0A7; Q | Oxidation r | E9Q0A7 [5 | 0 | 2176.08 | 0 | -1.55 | 0.006834 | 4.67 | 149.1 | 50.9 | 18.34 | 16.22 | Shared | High | High | High | High | High | High | High | High | High | High | High | 0.001093 | 0.01148 | 2.34 |
| FALSE | High | HSSAPPK | [R].HSSAPF | 1xPhosphc | G3X956 1x | 0.032548 | 0.002167 | 1 | 2 | 1 | G3X956 | Suppressor | G3X956 [1( | 0 | 1261.671 | 0 | -1.83 | 0.001383 | 0.4 | 154.3 | 45.7 | 8.5 | 17.09 | | High | High | High | High | High | High | High | High | High | High | High | 0.001142 | 0.01178 | 1.92 |
| FALSE | High | HSSDINHL\ | [R].HSSDIN | 1xPhosphc | E9PUI5 1xI | 0.000294 | 0 | 1 | 5 | 5 | E9PUI5 | Formin-bin | E9PUI5 [48 | 1 | 3550.603 | 0 | -0.06 | 0.252626 | 15.28 | 101.4 | 98.6 | 9.41 | 6.91 | | High | High | High | High | High | High | High | High | High | High | High | 0.00014 | 0.001122 | 3.4 |
| FALSE | High | HSSDINHL\ | [R].HSSDIN | 2xPhosphc | E9PUI5 2xI | 0.110202 | 0.008961 | 1 | 5 | 1 | E9PUI5 | Formin-bin | E9PUI5 [48 | 1 | 3630.57 | 0 | -0.84 | 0.000618 | 2.47 | 128.8 | 71.2 | 5.65 | 10.83 | | High | High | High | High | High | High | High | High | High | High | High | 0.004462 | 0.04838 | 3.1 |
| FALSE | High | HSSEDSDIT | [R].HSSEDS | 1xPhosphc | O70472 1x | 0.000496 | 0 | 1 | 2 | 3 | O70472 | Transmem | O70472 [1 | 0 | 2432.13 | 0 | -0.3 | 0.796267 | 4.51 | 103.6 | 96.4 | 9.55 | 21.3 | | High | High | High | High | High | High | High | High | High | High | High | 0 | 0.000206 | 2.83 |
| FALSE | High | HSSEDSDIT | [R].HSSEDS | 2xPhosphc | O70472 2x | 0.001751 | 5.66E-05 | 1 | 2 | 1 | O70472 | Transmem | O70472 [1 | 0 | 2496.102 | 0 | -0.91 | 0.010879 | 8.06 | 122.2 | 77.8 | 12.58 | 25.45 | | High | High | High | High | High | High | High | High | High | High | High | 2.58E-05 | 0.000448 | 2.19 |
| FALSE | High | HSSGEQET | [K].HSSGE( | 2xPhosphc | Q6PAL5 2x | 0.000104 | 0 | 1 | 6 | 3 | Q6PAL5 | Uncharacte | Q6PAL5 [9 | 0 | 3641.703 | 3 | 0.79 | 0.023274 | 15.8 | 73.2 | 126.8 | 3.13 | 20.66 | | High | High | High | High | High | High | High | High | High | High | High | 0 | 1.88E-05 | 3.97 |
| FALSE | High | HSPSSPPAK | [K].HSPSSP | 1xPhosphc | B2RY19 1xI | 0.041781 | 0.003126 | 1 | 7 | 1 | B2RY19 | Cgnl1 prot | B2RY19 [43 | 0 | 1445.755 | 0 | 1.19 | 0.012794 | 35.11 | 55.5 | 144.5 | 10.05 | 23.94 | | High | High | High | High | High | High | High | High | High | High | High | 0.001609 | 0.01571 | 0.67 |
| FALSE | High | HSSGEQET | [K].HSSGE( | 1xPhosphc | Q6PAL5 1x | 0.067619 | 0.004864 | 1 | 6 | 2 | Q6PAL5 | Uncharacte | Q6PAL5 [9 | 1 | 3561.716 | 1 | 0.37 | 0.103725 | 5.21 | 86 | 114 | 5.48 | 18.3 | | High | High | High | High | High | High | High | High | High | High | High | 0.002911 | 0.03021 | 3.4 |
| FALSE | High | HSLPSPSRP | [K].HSLPSP | 1xPhosphc | Q3TJM4 1x | 0.032267 | 0.002129 | 1 | 1 | 3 | Q3TJM4 | Centromer | Q3TJM4 [2 | 0 | 2071.017 | 0 | -0.5 | 0.021533 | 6.31 | 116.6 | 83.4 | 4.83 | 17.89 | | High | High | High | High | High | High | High | High | High | High | High | 0.001142 | 0.01174 | 1.6 |
| FALSE | High | HSLPFSLPS | [R].HSLPFS | 1xPhosphc | P22682 1xI | 0.000745 | 0 | 1 | 5 | 1 | P22682 | E3 ubiquiti | P22682 [61 | 0 | 1950.935 | 0 | 0.84 | 0.02533 | 25.81 | 72.9 | 127.1 | 15.19 | 19.4 | | High | High | High | High | High | High | High | High | High | High | High | 0 | 0.000171 | 2.78 |
| FALSE | High | HSFCDNLPI | [K].HSFCDI | 1xCarbami | A2SW42 1x | 0.00407 | 0.00036 | 1 | 2 | 1 | A2SW42 | Zinc finger | A2SW42 [6 | 0 | 1655.802 | 0 | -1.66 | 0.00064 | 0.35 | 148.9 | 51.1 | 10.31 | 10.94 | | High | High | High | High | High | High | High | High | High | High | High | 0.000161 | 0.001151 | 1.94 |
| FALSE | High | HSFSAGPEI | [R].HSFSAG | 1xPhosphc | Q3UGY8 1x | 0.001337 | 0 | 1 | 1 | 1 | Q3UGY8 | Brefeldin A | Q3UGY8 [2 | 0 | 1522.762 | 0 | 2.35 | 0.001345 | 18.2 | 32.8 | 167.2 | 11.81 | 13.71 | | High | High | High | High | High | High | High | High | High | High | High | 0 | 0.000332 | 3.16 |
| FALSE | High | HSFSGLPE( | [R].HSFSGL | 1xCarbami | A2AEG2 1x | 0.004218 | 0.00036 | 1 | 4 | 2 | A2AEG2 | Oral-facial- | A2AEG2 [7 | 0 | 2216.016 | 0 | 0.19 | 0.532404 | 23.28 | 93.3 | 106.7 | 15.13 | 9.88 | | High | High | High | High | High | High | High | High | High | High | High | 0.000161 | 0.001193 | 2.48 |
| FALSE | High | HSGAESGY | [K].HSGAE | 1xPhosphc | Q08288 1x | 0.007209 | 0.00062 | 1 | 1 | 2 | Q08288 | Cell growth | Q08288 [2 | 0 | 1473.714 | 0 | -0.64 | 0.004621 | 3.74 | 121.4 | 78.6 | 1.89 | 18.56 | | High | High | High | High | High | High | High | High | High | High | High | 0.001076 | 0.01143 | 2.59 |
| FALSE | High | HSGGHFAA | [R].HSGGH | 1xCarbami | A1L341 1xI | 0.01218 | 0.000889 | 1 | 5 | 2 | A1L341 | Dual-specit | A1L341 [52 | 0 | 2802.244 | 0 | 0.18 | 0.560099 | 12.25 | 94.1 | 105.9 | 11.78 | 10.03 | | High | High | High | High | High | High | High | High | High | High | High | 0.009311 | 0.101 | 2.1 |
| FALSE | High | HSGGHFAA | [R].HSGGH | 1xCarbami | A1L341 1xI | 0.026421 | 0.001676 | 1 | 5 | 1 | A1L341 | Dual-specit | A1L341 [52 | 0 | 2818.239 | 0 | 0.81 | 0.07722 | 28.89 | 72.7 | 127.3 | 25.99 | 24.33 | | High | High | High | High | High | High | High | High | High | High | High | 0.000903 | 0.009343 | 1.87 |
| FALSE | High | HSGPNSAD | [K].HSGPN | 1xPhosphc | Q9Z2X1 1x | 0.004143 | 0.00036 | 1 | 4 | 1 | Q9Z2X1 | Heterogen | Q9Z2X1 [9 | 0 | 1939.85 | 0 | 0.7 | 0.017003 | 28.54 | 76.3 | 123.7 | 11.69 | 12.87 | | High | High | High | High | High | High | High | High | High | High | High | 0.000161 | 0.001171 | 3.21 |
| FALSE | High | HSGQDVH\ | [R].HSGQD | 1xPhosphc | Q3UVN5 1: | 0.00095 | 0 | 1 | 4 | 1 | Q3UVN5 | Putative ur | Q3UVN5 [1 | 0 | 1527.788 | 0 | 0.22 | 0.157647 | 16.16 | 92.4 | 107.6 | 7.76 | 21.46 | | High | High | High | High | High | High | High | High | High | High | High | 0 | 0.000226 | 2.73 |
| FALSE | High | HSGQDVH\ | [R].HSGQD | 1xPhosphc | Q3UVN5 1: | 0.001267 | 0 | 1 | 4 | 2 | Q3UVN5 | Putative ur | Q3UVN5 [1 | 0 | 1756.951 | 0 | 0.48 | 0.004058 | 4.69 | 82.9 | 117.1 | 6.44 | 5.08 | | High | High | High | High | High | High | High | High | High | High | High | 2.58E-05 | 0.000479 | 3.52 |
| FALSE | High | HSGSEASG | [R].HSGSE/ | 1xPhosphc | A0A1L1SSA | 0.002671 | 0.000109 | 1 | 7 | 1 | A0A1L1SSA | Centrosom | A0A1L1SSA | 0 | 1494.735 | 0 | -0.48 | 0.022836 | 14 | 113.7 | 86.3 | 8.77 | 14.09 | | High | High | High | High | High | High | High | High | High | High | High | 4.94E-05 | 0.00072 | 2.88 |
| FALSE | High | HSGSFLSAF | [R].HSGSFL | 1xPhosphc | B9EKS2 1xI | 5.4E-05 | 0 | 1 | 3 | 1 | B9EKS2 | Jumonji do | B9EKS2 [77 | 0 | 2358.153 | 0 | -0.35 | 0.021746 | 1.4 | 108.1 | 91.9 | 13.77 | 6.29 | | High | High | High | High | High | High | High | High | High | High | High | 0 | 8.97E-06 | 3.03 |
| FALSE | High | HSGSPENL( | [K].HSGSPE | 1xPhosphc | B9EIW5 1x | 1.05E-07 | 0 | 1 | 3 | 3 | B9EIW5 | Ncoa7 prot | B9EIW5 [4 | 0 | 2830.366 | 0 | 1.7 | 0.022774 | 18.21 | 45.9 | 154.1 | 5.97 | 33.41 | | High | High | High | High | High | High | High | High | High | High | High | 0 | 8.09E-09 | 3.7 |
| FALSE | High | HSGSTSPYI | [R].HSGSTS | 1xPhosphc | Q8BTI8 1xI | 0.007404 | 0.00062 | 1 | 3 | 1 | Q8BTI8 | Serine/argi | Q8BTI8 [95 | 0 | 1385.666 | 0 | -0.07 | 0.870112 | 19.99 | 104.7 | 95.3 | 28.03 | 29.87 | | High | High | High | High | High | High | High | High | High | High | High | 0.000275 | 0.002252 | 2.02 |
| FALSE | High | HSLPSGLGI | [K].HSLPSG | 1xPhosphc | G5E8V9 1x | 0.011346 | 0.000816 | 1 | 3 | 1 | G5E8V9 | ADP-ribosy | G5E8V9 [3 | 0 | 2937.476 | 0 | 2.5 | 0.022604 | 59.55 | 28.4 | 171.6 | 13.48 | 43.8 | | High | High | High | High | High | High | High | High | High | High | High | 0.000418 | 0.003615 | 3.24 |
| FALSE | High | HSGSTSPYI | [R].HSGSTS | 2xPhosphc | Q8BTI8 2xI | 0.00625 | 0.00062 | 1 | 3 | 2 | Q8BTI8 | Serine/argi | Q8BTI8 [95 | 0 | 1465.633 | 0 | -1 | 0.008876 | 6.64 | 138.7 | 61.3 | 14.57 | 20.28 | | High | High | High | High | High | High | High | High | High | High | High | 0.000859 | 0.008187 | 3.52 |
| FALSE | High | HSGSTSPYI | [R].HSGSTS | 2xPhosphc | Q8BTI8 2xI | 0.056248 | 0.004124 | 1 | 3 | 2 | Q8BTI8 | Serine/argi | Q8BTI8 [95 | 0 | 1694.796 | 0 | -1.15 | 0.00237 | 3.84 | 137.9 | 62.1 | 11.41 | 5.24 | | High | High | High | High | High | High | High | High | High | High | High | 0.002381 | 0.02464 | 2.46 |
| FALSE | High | HSGTGATP | [R].HSGTG. | 1xPhosphc | Q60520-1 | 0.000176 | 0 | 1 | 2 | 3 | Q60520-1 | Isoform 2 c | Q60520-1 [ | 0 | 1589.845 | 0 | -0.8 | 0.001282 | 5.68 | 127 | 73 | 3.77 | 10.76 | | High | High | High | High | High | High | High | High | High | High | High | 0.001651 | 0.01628 | 2.06 |
| FALSE | High | HSHSHSPN | [R].HSHSH | 2xPhosphc | P62996 2xI | 0.020679 | 0.001472 | 1 | 2 | 2 | P62996 | Transform | P62996 [94 | 0 | 1652.66 | 0 | -0.07 | 0.536028 | 26.22 | 102.4 | 97.6 | 21.05 | 34.9 | | High | High | High | High | High | High | High | High | High | High | High | 0.001999 | 0.02076 | 1.59 |
| FALSE | High | HSLASDSA/ | [R].HSLASD | 1xPhosphc | Q6ZQH8 1x | 3.55E-06 | 0 | 1 | 1 | 1 | Q6ZQH8 | Nucleopor | Q6ZQH8 [1 | 0 | 1768.852 | 0 | 0.55 | 0.045199 | 13.9 | 82.5 | 117.5 | 5.72 | 14.91 | | High | High | High | High | High | High | High | High | High | High | High | 0 | 4.22E-07 | 2.97 |
| FALSE | High | HSLASTDEI | [R].HSLAST | 1xPhosphc | A0A0R4J0I | 0.001189 | 0 | 1 | 4 | 3 | A0A0R4J0I | Low densit | A0A0R4J0I | 0 | 1525.766 | 0 | 1.39 | 0.00146 | 10.73 | 55.3 | 144.7 | 7.98 | 11.57 | | High | High | High | High | High | High | High | High | High | High | High | 0 | 0.000291 | 2.31 |
| FALSE | High | HSLASTDEI | [R].HSLAST | 2xPhosphc | A0A0R4J0I | 0.002134 | 5.66E-05 | 1 | 4 | 2 | A0A0R4J0I | Low densit | A0A0R4J0I | 0 | 1605.733 | 0 | 1.57 | 0.000567 | 4.64 | 51.3 | 148.7 | 6.22 | 7.45 | | High | High | High | High | High | High | High | High | High | High | High | 0.002105 | 0.02177 | 2.59 |
| FALSE | High | HSLDLPHG | [K].HSLDLP | 1xPhosphc | E9Q1Q1 1x | 0.000479 | 0 | 1 | 6 | 2 | E9Q1Q1 | Microtubu | E9Q1Q1 [1 | 0 | 1783.951 | 0 | -0.69 | 0.05654 | 5.49 | 119.6 | 80.4 | 17.59 | 13.56 | | High | High | High | High | High | High | High | High | High | High | High | 0.0032 | 0.03372 | 1.93 |
| FALSE | High | HSLDSDEEI | [K].HSLDSD | 1xPhosphc | Q69ZP6 1x | 2.27E-08 | 0 | 1 | 6 | 9 | Q69ZP6 | MKIAA117 | Q69ZP6 [4 | 0 | 2432.003 | 0 | 0.36 | 0.137036 | 42.05 | 87.7 | 112.3 | 8.72 | 21.41 | | High | High | High | High | High | High | High | High | High | High | High | 0 | 1.44E-09 | 3.76 |
| FALSE | High | HSLDSDEEI | [K].HSLDSD | 1xDeamida | Q69ZP6 1x | 0.044358 | 0.003264 | 1 | 5 | 1 | Q69ZP6 | MKIAA117 | Q69ZP6 [4 | 1 | 4920.163 | 0 | 0.19 | 0.865701 | 30.15 | 97.1 | 102.9 | 19.78 | 9.93 | | High | High | High | High | High | High | High | High | High | High | High | 0.00168 | 0.01685 | 2.98 |
| FALSE | High | HSLECIQAN | [R].HSLECI( | 1xCarbami | Q8K296 1x | 1.47E-05 | 0 | 1 | 7 | 2 | Q8K296 | Myotubula | Q8K296 [7 | 0 | 2021.983 | 0 | 0.99 | 0.000737 | 5.59 | 68.8 | 131.2 | 12.91 | 12.92 | | High | High | High | High | High | High | High | High | High | High | High | 0 | 2.08E-06 | 4.1 |
| FALSE | High | HSLEVTQEI | [K].HSLEVT | 1xPhosphc | Q811L6 1xI | 0.000264 | 0 | 1 | 4 | 1 | Q811L6 | Microtubu | Q811L6 [14 | 0 | 1763.853 | 0 | 0.54 | 0.010221 | 14.25 | 76.6 | 123.4 | 16.63 | 11.6 | | High | High | High | High | High | High | High | High | High | High | High | 0 | 5.35E-05 | 3.81 |
| FALSE | High | HSLFDSCK | [K].HSLFDS | 1xCarbami | Q91W98 1: | 0.011246 | 0.000772 | 1 | 2 | 2 | Q91W98 | Solute carr | Q91W98 [2 | 0 | 1531.738 | 0 | -0.04 | 0.485023 | 19.21 | 105.4 | 94.6 | 9.27 | 21.9 | | High | High | High | High | High | High | High | High | High | High | High | 0.000418 | 0.003585 | 3.08 |
| FALSE | High | HSLGQEVI( | [R].HSLGQI | 1xPhosphc | B9EKJ7 1xF | 3.32E-08 | 0 | 1 | 3 | 3 | B9EKJ7 | Tankyrase | B9EKJ7 [79 | 0 | 2602.271 | 0 | 0.25 | 0.287181 | 34.12 | 90.4 | 109.6 | 14.84 | 17.03 | | High | High | High | High | High | High | High | High | High | High | High | 0 | 2.21E-09 | 4.55 |
| FALSE | High | HSGSTSPYI | [R].HSGSTS | 1xPhosphc | Q8BTI8 1xF | 0.003655 | 0.00021 | 1 | 3 | 2 | Q8BTI8 | Serine/argi | Q8BTI8 [95 | 0 | 1614.829 | 0 | -0.7 | 0.001433 | 0.18 | 124.3 | 75.7 | 7.36 | 5.08 | | High | High | High | High | High | High | High | High | High | High | High | 0.000847 | 0.007977 | 2.12 |
| FALSE | High | HSSGTPAS\ | [R].HSSGTF | 1xPhosphc | Q9QXV1 1x | 1.37E-05 | 0 | 1 | 2 | 5 | Q9QXV1 | Chromobo | Q9QXV1 [2 | 0 | 1727.832 | 0 | 0.49 | 0.001744 | 10.03 | 83.1 | 116.9 | 8.15 | 3.72 | | High | High | High | High | High | High | High | High | High | High | High | 4.94E-05 | 0.000599 | 2.32 |
| FALSE | High | HSSHDMLS | [R].HSSHDI | 1xPhosphc | Q9Z0U1 1x | 0.01588 | 0.001124 | 1 | 1 | 1 | Q9Z0U1 | Tight junct | Q9Z0U1 [1 | 0 | 2088.891 | 0 | -0.22 | 0.123374 | 11.59 | 105 | 95 | 9.9 | 15.48 | | High | High | High | High | High | High | High | High | High | High | High | 0.000585 | 0.005247 | 3.09 |
| FALSE | High | HSSHDMLS | [R].HSSHDI | 1xPhosphc | Q9Z0U1 1x | 0.00129 | 0 | 1 | 1 | 8 | Q9Z0U1 | Tight junct | Q9Z0U1 [1 | 0 | 2104.886 | 0 | -0.26 | 0.667492 | 15.56 | 104.5 | 95.5 | 6.93 | 26.21 | | High | High | High | High | High | High | High | High | High | High | High | 0.000275 | 0.00225 | 2.65 |
| FALSE | High | HSYPPALGS | [R].HSYPPA | 1xPhosphc | Q91XC0 1x | 1.02E-08 | 0 | 1 | 1 | 3 | Q91XC0 | LIM domai | Q91XC0 [2 | 0 | 2784.428 | 0 | -1.96 | 0.000374 | 0.64 | 158.9 | 41.1 | 5.08 | 14.55 | | High | High | High | High | High | High | High | High | High | High | High | 0 | 5.85E-10 | 3.31 |
| FALSE | High | HTAAVPFSI | [R].HTAAVI | 1xPhosphc | P49452 1xI | 0.001644 | 5.66E-05 | 1 | 3 | 1 | P49452 | Centromer | P49452 [25 | 0 | 2233.113 | 0 | -0.27 | 0.030853 | 6.16 | 109.9 | 90.1 | 7.28 | 9.1 | | High | High | High | High | High | High | High | High | High | High | High | 2.58E-05 | 0.000419 | 1.98 |
| FALSE | High | HTADGMPI | [K].HTADG | 1xPhosphc | Q3U832 1x | 0.011966 | 0.000889 | 1 | 3 | 1 | Q3U832 | Putative ur | Q3U832 [5 | 1 | 3142.555 | 0 | -0.62 | 0.017137 | 12.46 | 116.9 | 83.1 | 8.23 | 19.1 | | High | High | High | High | High | High | High | High | High | High | High | 0.000449 | 0.003827 | 2.78 |
| FALSE | High | HTADGMPI | [K].HTADG | 1xPhosphc | Q3U832 1x | 9.79E-08 | 0 | 1 | 3 | 2 | Q3U832 | Putative ur | Q3U832 [5 | 1 | 3158.55 | 0 | -1.03 | 0.002941 | 4.24 | 133 | 67 | 12.13 | 11.41 | | High | High | High | High | High | High | High | High | High | High | High | 0 | 7.48E-09 | 3.32 |
| FALSE | High | HTAPSSPSF | [R].HTAPSS | 1xPhosphc | Q3UR70 1x | 0.000573 | 0 | 1 | 1 | 4 | Q3UR70 | Transformi | Q3UR70 [8 | 0 | 1503.715 | 0 | 0.64 | 0.001448 | 3.78 | 76.3 | 123.7 | 6.41 | 9.92 | | High | High | High | High | High | High | High | High | High | High | High | 0 | 0.000127 | 2.74 |
| FALSE | High | HTASGSSE | [K].HTASG | 1xPhosphc | A0A140T8 | 4.79E-07 | 0 | 1 | 1 | 5 | A0A140T8 | Chromosor | A0A140T8 | 0 | 1625.794 | 0 | 0.34 | 0.272702 | 23.85 | 83.9 | 116.1 | 6.61 | 25.69 | | High | High | High | High | High | High | High | High | High | High | High | 0 | 4.45E-08 | 2.98 |
| FALSE | High | HTAVETSP | [R].HTAVE | 1xPhosphc | Q7TSJ0 1xF | 0.013426 | 0.000976 | 1 | 1 | 2 | Q7TSJ0 | DNA (cytos | Q7TSJ0 [15 | 0 | 1751.853 | 0 | 0.86 | 0.001347 | 18.42 | 74.3 | 125.7 | 10.88 | 8.26 | | High | High | High | High | High | High | High | High | High | High | High | 0.000505 | 0.004346 | 2.73 |
| FALSE | High | HTCAEPDA | [R].HTCAEI | 1xCarbami | Q8K337 1x | 3.63E-05 | 0 | 1 | 4 | 2 | Q8K337 | Type II ino: | Q8K337 [1 | 0 | 2133.02 | 0 | 0.8 | 0.00148 | 14.63 | 73.1 | 126.9 | 4.99 | 5.39 | | High | High | High | High | High | High | High | High | High | High | High | 0 | 5.74E-06 | 2.78 |
| FALSE | High | HTDDEMT( | [R].HTDDEI | 1xPhosphc | P47811 1xI | 0.000139 | 0 | 1 | 6 | 5 | P47811 | Mitogen-a( | P47811 [17 | 0 | 1804.777 | 0 | 0.26 | 0.031185 | 9.56 | 95.8 | 104.2 | 11.33 | 6.01 | | High | High | High | High | High | High | High | High | High | High | High | 0.000638 | 0.005777 | 3.05 |
| FALSE | High | HTDDEMT( | [R].HTDDEI | 1xPhosphc | P47811 1xI | 2.43E-06 | 0 | 1 | 6 | 18 | P47811 | Mitogen-a( | P47811 [17 | 0 | 1820.772 | 0 | 0.79 | 0.002177 | 16.77 | 74.4 | 125.6 | 10.8 | 4.16 | | High | High | High | High | High | High | High | High | High | High | High | 0 | 0.000183 | 3.44 |
| FALSE | High | HTDTEMT( | [R].HTDTEI | 1xPhosphc | Q3UIB2 1xI | 6.05E-07 | 0 | 1 | 2 | 2 | Q3UIB2 | Mitogen-a( | Q3UIB2 [1 | 0 | 1834.824 | 0 | 2.06 | 0.000975 | 41.64 | 37.3 | 162.7 | 23.44 | 9.51 | | High | High | High | High | High | High | High | High | High | High | High | 0 | 9.69E-08 | 3.33 |
| FALSE | High | HTGHRPFC | [K].HTGHR | 1xCarbami | Q60793 1x | 0.042141 | 0.003162 | 1 | 1 | 1 | Q60793 | Krueppel-li | Q60793 [4 | 0 | 1933.962 | 0 | 1.17 | 0.049333 | 41.03 | 62.9 | 137.1 | 15.37 | 29.07 | | High | High | High | High | High | High | High | High | High | High | High | 0.001625 | 0.01579 | 2.22 |
| FALSE | High | HTGMASID | [K].HTGM/ | 1xPhosphc | Q3UHH1 1: | 2.72E-08 | 0 | 1 | 5 | 3 | Q3UHH1 | Zinc finger | Q3UHH1 [1 | 0 | 2744.228 | 0 | -0.74 | 0.002152 | 2.1 | 125.5 | 74.5 | 14.22 | 10.13 | | High | High | High | High | High | High | High | High | High | High | High | 0 | 1.77E-09 | 4.78 |
| FALSE | High | HSYENDGG | [R].HSYENI | 1xPhosphc | Q3UYV9 1x | 3.1E-06 | 0 | 1 | 2 | 4 | Q3UYV9 | Nuclear ca | Q3UYV9 [6 | 0 | 1906.885 | 0 | -0.31 | 0.050332 | 4.72 | 109.5 | 90.5 | 17.64 | 4.62 | | High | High | High | High | High | High | High | High | High | High | High | 0 | 3.62E-07 | 3.77 |
| FALSE | High | HTGMASID | [K].HTGM/ | 1xPhosphc | Q3UHH1 1: | 1.31E-09 | 0 | 1 | 5 | 3 | Q3UHH1 | Zinc finger | Q3UHH1 [1 | 0 | 2760.223 | 0 | -0.31 | 0.020246 | 2 | 110.4 | 89.6 | 15.15 | 8.01 | | High | High | High | High | High | High | High | High | High | High | High | 0 | 5.85E-11 | 3.82 |
| FALSE | High | HTGPNSPD | [K].HTGPN. | 1xDeamida | Q8C2Q7 1x | 9.25E-07 | 0 | 2 | 4 | 1 | Q8C2Q7; P | Heterogen | Q8C2Q7 [9 | 0 | 1994.881 | 0 | 0.12 | 0.740174 | 4.25 | 97.4 | 102.6 | 10.73 | 24.74 | Shared | High | High | High | High | High | High | High | High | High | High | High | 0 | 9.33E-08 | 3.83 |
| FALSE | High | HTLQQHQE | [K].HTLQQ | 1xPhosphc | Q5F2E7-2 | 0.000346 | 0 | 1 | 2 | 2 | Q5F2E7-2 | Isoform 2 c | Q5F2E7-2 [ | 0 | 1884.973 | 0 | 0.34 | 0.14524 | 24.34 | 85.1 | 114.9 | 8.69 | 25.28 | | High | High | High | High | High | High | High | High | High | High | High | 0 | 0.000309 | 3.11 |
| FALSE | High | HTNSVICK | [K].HTNSVI | 1xCarbami | Q9WTP2 1: | 0.00614 | 0.00062 | 1 | 1 | 2 | Q9WTP2 | Protein spr | Q9WTP2 [2 | 0 | 1496.77 | 0 | 0.52 | 0.157647 | 25.11 | 83.2 | 116.8 | 13.6 | 34.58 | | High | High | High | High | High | High | High | High | High | High | High | 0.000275 | 0.001819 | 2.67 |
| FALSE | High | HTPLYER | [K].HTPLYE | 1xPhosphc | Q6A0A9 1x | 0.058664 | 0.004294 | 1 | 1 | 1 | Q6A0A9 | Constitutiv | Q6A0A9 [4 | 0 | 1224.598 | 0 | 2.49 | 0.001706 | 29.21 | 27.2 | 172.8 | 31.02 | 16.38 | | High | High | High | High | High | High | High | High | High | High | High | 0.002187 | 0.02325 | 2.24 |
| FALSE | High | HTPNTSDN | [K].HTPNT | 1xCarbami | B9EJ80 1xF | 1.45E-06 | 0 | 1 | 2 | 2 | B9EJ80 | PDZ domai | B9EJ80 [96 | 0 | 2958.282 | 0 | -0.26 | 0.444105 | 22.05 | 104.4 | 95.6 | 17.47 | 20.2 | | High | High | High | High | High | High | High | High | High | High | High | 0 | 1.54E-07 | 3.2 |
| FALSE | High | HTPNTSDN | [K].HTPNT | 1xCarbami | B9EJ80 2xF | 5.73E-10 | 0 | 1 | 2 | 2 | B9EJ80 | PDZ domai | B9EJ80 [96 | 0 | 3038.249 | 0 | 0.69 | 0.060345 | 26.08 | 72.6 | 127.4 | 8.2 | 25.99 | | High | High | High | High | High | High | High | High | High | High | High | 0 | 2.32E-11 | 5.58 |
| FALSE | High | HTPPTIGGS | [R].HTPPTI( | 1xPhosphc | Q6AXD2 1x | 0.000181 | 0 | 1 | 9 | 3 | Q6AXD2 | Abi2 prote | Q6AXD2 [3 | 0 | 1704.867 | 0 | -0.22 | 0.122518 | 8.31 | 106.3 | 93.7 | 13.33 | 10.67 | | High | High | High | High | High | High | High | High | High | High | High | 0 | 8.29E-05 | 3.49 |
| FALSE | High | HTPSLALTF | [R].HTPSLA | 1xPhosphc | G5E8J0 1xF | 6.32E-10 | 0 | 1 | 2 | 1 | G5E8J0 | Neurogeni | G5E8J0 [18 | 0 | 2937.492 | 0 | 0.82 | 0.010376 | 26.76 | 72.4 | 127.6 | 12.13 | 13.48 | | High | High | High | High | High | High | High | High | High | High | High | 0 | 2.58E-11 | 4.27 |
| FALSE | High | HTPYGNQT | [R].HTPYGI | 1xPhosphc | P70662 1xI | 0.000725 | 0 | 2 | 4 | 2 | P70662; D3 | LIM domai | P70662 [6( | 0 | 1660.732 | 0 | 1.7 | 0.000587 | 10.88 | 43.4 | 156.6 | 13 | 10.43 | Shared | High | High | High | High | High | High | High | High | High | High | High | 0 | 0.000166 | 2.92 |
| FALSE | High | HTSADTLD' | [R].HTSAD | 1xCarbami | S4R1P5 1xI | 2.47E-06 | 0 | 1 | 6 | 3 | S4R1P5 | Dystonin O | S4R1P5 [32 | 0 | 2282.032 | 0 | -0.15 | 0.20868 | 4.93 | 101 | 99 | 14.95 | 12.8 | | High | High | High | High | High | High | High | High | High | High | High | 0 | 2.8E-07 | 4.21 |
| FALSE | High | HTSCTGDF | [R].HTSCT( | 1xCarbami | B2RSV7 1xI | 6.53E-05 | 0 | 1 | 2 | 3 | B2RSV7 | Fgd6 prote | B2RSV7 [7( | 0 | 2177.916 | 0 | 0.34 | 0.022501 | 3.1 | 89.9 | 110.1 | 9.13 | 7.01 | | High | High | High | High | High | High | High | High | High | High | High | 0 | 1.12E-05 | 2.82 |
| FALSE | High | HTSDYEDT | [R].HTSDYI | 1xPhosphc | P39447 1xI | 5.16E-05 | 0 | 1 | 3 | 10 | P39447 | Tight junct | P39447 [83 | 0 | 4480.915 | 0 | -0.64 | 0.034617 | 4.99 | 111.1 | 88.9 | 20.06 | 22.29 | | High | High | High | High | High | High | High | High | High | High | High | 0 | 8.57E-06 | 5.02 |

| | | | | | | | | | | | | | | | | | | | | | | | | | | | | | | | | | | | | | | |
|---|---|---|---|---|---|---|---|---|---|---|---|---|---|---|---|---|---|---|---|---|---|---|---|---|---|---|---|---|---|---|---|---|---|---|---|---|---|---|
| FALSE | High | HTSLEEEEF | [R].HTSLEE | 1xPhospho | D3Z357 1xl | 0.000475 | 0 | 1 | 6 | 3 | D3Z357 | Histone-lys | D3Z357 [46 | 0 | 2030.025 | 0 | 1.16 | 0.001622 | 10.41 | 61.8 | 138.2 | 8.42 | 11.47 | | High | High | High | High | High | High | High | High | High | High | High | 0.006863 | 0.07412 | 2.72 |
| FALSE | High | HTGPNSPD | [K].HTGPN | 1xPhospho | Q8C2Q7 1x | 1.33E-06 | 0 | 2 | 4 | 6 | Q8C2Q7; P | Heterogen | Q8C2Q7 [9 | 0 | 1993.897 | 0 | 0.03 | 0.632225 | 2.27 | 99.7 | 100.3 | 5.18 | 11.38 | Shared | High | High | High | High | High | High | High | High | High | High | High | 0 | 1.6E-07 | 3.52 |
| FALSE | High | HSYENDGG | [R].HSYENI | 1xPhospho | Q3UYV9 1x | 6.65E-05 | 0 | 1 | 2 | 1 | Q3UYV9 | Nuclear ca | Q3UYV9 [6 | 0 | 1677.722 | 0 | 0.17 | 0.430608 | 56.9 | 88.4 | 111.6 | 15.41 | 20.61 | | High | High | High | High | High | High | High | High | High | High | High | 0 | 1.14E-05 | 3.4 |
| FALSE | High | HSVTNEVII | [R].HSVTNI | 1xPhospho | Q3ULS5 1x | 0.000356 | 0 | 1 | 3 | 1 | Q3ULS5 | Putative ur | Q3ULS5 [1 | 0 | 1862.978 | 0 | -0.46 | 0.001066 | 1.97 | 116.7 | 83.3 | 3.85 | 6.1 | | High | High | High | High | High | High | High | High | High | High | High | 0 | 7.46E-05 | 3.48 |
| FALSE | High | HSVTAATP | [K].HSVTA | 2xPhospho | Q924Z6 2x | 0.018447 | 0.001324 | 1 | 2 | 1 | Q924Z6 | Exportin-6 | Q924Z6 [1 | 0 | 3751.823 | 0 | 0.45 | 0.168586 | 23.65 | 84.2 | 115.8 | 12.27 | 15.87 | | High | High | High | High | High | High | High | High | High | High | High | 0.000706 | 0.006228 | 3.02 |
| FALSE | High | HSSHDMLS | [R].HSSHDI | 2xPhospho | Q9Z0U1 2x | 0.115566 | 0.009504 | 1 | 1 | 1 | Q9Z0U1 | Tight junct | Q9Z0U1 [1 | 0 | 2168.857 | 0 | -0.34 | 0.350088 | 13.86 | 106.6 | 93.4 | 6.76 | 29.9 | | High | High | High | High | High | High | High | High | High | High | High | 0.004733 | 0.05108 | 2.4 |
| FALSE | High | HSSLPTESD | [R].HSSLPT | 1xPhospho | O54774 1x | 5.23E-10 | 0 | 1 | 2 | 1 | O54774 | AP-3 comp | O54774 [7 | 0 | 2161.996 | 0 | 0.49 | 0.003574 | 8.92 | 79.9 | 120.1 | 6.52 | 8.82 | | High | High | High | High | High | High | High | High | High | High | High | 0 | 2.09E-11 | 3.29 |
| FALSE | High | HSSLPTESD | [R].HSSLPT | 2xPhospho | O54774 2x | 2.09E-08 | 0 | 1 | 2 | 11 | O54774 | AP-3 comp | O54774 [7 | 0 | 2241.963 | 0 | -0.03 | 0.949356 | 8.92 | 100.3 | 99.7 | 4.38 | 3.35 | | High | High | High | High | High | High | High | High | High | High | High | 0 | 5.28E-06 | 3.47 |
| FALSE | High | HSSISPSTLT | [R].HSSISPS | 2xPhospho | Q69ZA1 2x | 4.31E-05 | 0 | 1 | 3 | 1 | Q69ZA1 | Cyclin-dep | Q69ZA1 [4 | 0 | 1888.959 | 0 | -0.15 | 0.152984 | 12.15 | 108.3 | 91.7 | 6.94 | 16.5 | | High | High | High | High | High | High | High | High | High | High | High | 0 | 6.95E-06 | 3.26 |
| FALSE | High | HSSISPVR | [R].HSSISP\ | 2xPhospho | Q14AX6-2 | 0.005135 | 0.000551 | 3 | 5 | 1 | Q14AX6-2; | Isoform 2 c | Q14AX6-2 | 0 | 1271.575 | 0 | -0.45 | 0.225053 | 3.36 | 115.5 | 84.5 | 16.66 | 17.56 | Shared | High | High | High | High | High | High | High | High | High | High | High | 0.000246 | 0.001495 | 2.91 |
| FALSE | High | HSSNLASPI | [K].HSSNLA | 1xPhospho | Q8C263 1x | 1.37E-05 | 0 | 1 | 1 | 1 | Q8C263 | Spindle an | Q8C263 [3 | 0 | 1805.956 | 0 | -0.57 | 0.012827 | 4.2 | 118 | 82 | 12.79 | 18.42 | | High | High | High | High | High | High | High | High | High | High | High | 0 | 1.92E-06 | 2.3 |
| FALSE | High | HSSNPPLES | [R].HSSNPI | 1xPhospho | Q6ZQ58 1x | 3.35E-05 | 0 | 1 | 2 | 2 | Q6ZQ58 | La-related | Q6ZQ58 [7 | 0 | 2344.074 | 0 | -0.53 | 0.001425 | 0.92 | 118.2 | 81.8 | 2.18 | 8.42 | | High | High | High | High | High | High | High | High | High | High | High | 0 | 5.26E-06 | 3.37 |
| FALSE | High | HSSNPPLES | [R].HSSNPI | 1xPhospho | Q6ZQ58 1x | 1.81E-08 | 0 | 1 | 2 | 32 | Q6ZQ58 | La-related | Q6ZQ58 [7 | 0 | 2360.069 | 0 | -0.58 | 0.007474 | 2.15 | 118.9 | 81.1 | 3.87 | 9.29 | | High | High | High | High | High | High | High | High | High | High | High | 0.000997 | 0.01062 | 3.78 |
| FALSE | High | HSSPPLTVK | [R].HSSPPL | 1xPhospho | Q3UMZ4 1 | 0.007951 | 0.000736 | 1 | 2 | 1 | Q3UMZ4 | Putative ur | Q3UMZ4 [ | 0 | 1274.671 | 0 | -0.03 | 0.993752 | 32.54 | 100.5 | 99.5 | 22.4 | 32.39 | | High | High | High | High | High | High | High | High | High | High | High | 0.000327 | 0.00242 | 1.91 |
| FALSE | High | HSSPPLTVK | [R].HSSPPL | 1xPhospho | Q3UMZ4 1 | 0.000161 | 0 | 1 | 2 | 6 | Q3UMZ4 | Putative ur | Q3UMZ4 [ | 0 | 1503.834 | 0 | -1.15 | 0.000293 | 1.89 | 137.1 | 62.9 | 4.41 | 5.52 | | High | High | High | High | High | High | High | High | High | High | High | 0 | 0.000231 | 3.5 |
| FALSE | High | HSSPSGTV | [K].HSSPSG | 1xPhospho | B2RUG9 1x | 0.016743 | 0.001166 | 1 | 6 | 1 | B2RUG9 | Adenomat | B2RUG9 [2 | 0 | 1378.668 | 0 | 0.01 | 0.399444 | 14.02 | 99.3 | 100.7 | 13.6 | 19.6 | | High | High | High | High | High | High | High | High | High | High | High | 0.000638 | 0.005599 | 1.53 |
| FALSE | High | HSSPTEERI | [R].HSSPTE | 1xPhospho | O08648 1x | 0.00509 | 0.000551 | 1 | 4 | 2 | O08648 | Mitogen-ac | O08648 [1 | 1 | 2079.933 | 0 | -0.16 | 0.038891 | 9.41 | 107.6 | 92.4 | 4.53 | 6.15 | | High | High | High | High | High | High | High | High | High | High | High | 0.000246 | 0.001476 | 2.51 |
| FALSE | High | HSSSSGPGI | [K].HSSSSG | 1xPhospho | Q810L3 1xl | 1.59E-06 | 0 | 1 | 4 | 3 | Q810L3 | E3 ubiquiti | Q810L3 [2 | 0 | 1980.979 | 0 | -0.64 | 0.021592 | 7.14 | 119.4 | 80.6 | 16.51 | 13.12 | | High | High | High | High | High | High | High | High | High | High | High | 0 | 1.71E-07 | 4.05 |
| FALSE | High | HSSSSPVSG | [K].HSSSSP | 1xPhospho | Q9ERU9 1x | 0.062209 | 0.004646 | 1 | 1 | 3 | Q9ERU9 | E3 SUMO-p | Q9ERU9 [2 | 0 | 2642.29 | 0 | -0.2 | 0.667492 | 66.7 | 104.5 | 95.5 | 27.88 | 43.22 | | High | High | High | High | High | High | High | High | High | High | High | 0.008188 | 0.08847 | 2.36 |
| FALSE | High | HSSSSSQSS | [R].HSSSSS | 1xPhospho | O35671 1x | 0.051258 | 0.003745 | 1 | 2 | 1 | O35671 | Integrin be | O35671 [9 | 0 | 1989.953 | 0 | 0.06 | 0.999212 | 30.57 | 91.9 | 108.1 | 30.19 | 15.39 | | High | High | High | High | High | High | High | High | High | High | High | 0.00196 | 0.01989 | 1.78 |
| FALSE | High | HSSTGDLLS | [K].HSSTGI | 1xPhospho | P0C090 1xl | 0.001986 | 5.66E-05 | 1 | 4 | 3 | P0C090 | Roquin-2 C | P0C090 [97 | 0 | 2265.189 | 0 | 1.32 | 0.009166 | 14.92 | 61.3 | 138.7 | 17.46 | 20.12 | | High | High | High | High | High | High | High | High | High | High | High | 2.58E-05 | 0.000517 | 2.65 |
| FALSE | High | HSSYPAGTI | [R].HSSYPA | 1xPhospho | Q61337 1x | 2.18E-07 | 0 | 1 | 5 | 3 | Q61337 | Bcl2-associ | Q61337 [1 | 0 | 2806.175 | 0 | 0.77 | 0.16872 | 9.32 | 78.3 | 121.7 | 13.29 | 28.23 | | High | High | High | High | High | High | High | High | High | High | High | 0 | 1.83E-08 | 4.23 |
| FALSE | High | HSSYPAGTI | [R].HSSYPA | 1xPhospho | Q61337 1x | 1.54E-07 | 0 | 1 | 5 | 6 | Q61337 | Bcl2-associ | Q61337 [1 | 0 | 2822.17 | 0 | 0.06 | 0.433912 | 12.86 | 94.8 | 105.2 | 15.65 | 12.69 | | High | High | High | High | High | High | High | High | High | High | High | 0 | 1.23E-08 | 4.61 |
| FALSE | High | HSTPSNSSI | [K].HSTPSN | 1xPhospho | B2RQQ7 1x | 2.47E-09 | 0 | 1 | 2 | 3 | B2RQQ7 | CDC42 bin | B2RQQ7 [1 | 0 | 2450.105 | 0 | 0.35 | 0.616073 | 29.1 | 90 | 110 | 30.05 | 15.75 | | High | High | High | High | High | High | High | High | High | High | High | 0 | 4.8E-09 | 5.55 |
| FALSE | High | HSTPSNSSI | [K].HSTPSN | 2xPhospho | B2RQQ7 2x | 4.84E-08 | 0 | 1 | 2 | 3 | B2RQQ7 | CDC42 bin | B2RQQ7 [1 | 0 | 2530.071 | 0 | 0.22 | 0.440727 | 10.5 | 89.4 | 110.6 | 8.55 | 22.56 | | High | High | High | High | High | High | High | High | High | High | High | 0 | 2.62E-08 | 4.75 |
| FALSE | High | HSTPSNSSI | [K].HSTPSN | 1xDeamida | B2RQQ7 2x | 7.54E-05 | 0 | 1 | 2 | 3 | B2RQQ7 | CDC42 bin | B2RQQ7 [1 | 0 | 2531.055 | 0 | 0.59 | 0.394196 | 28.37 | 79.9 | 120.1 | 22.77 | 21.36 | | High | High | High | High | High | High | High | High | High | High | High | 0 | 3.88E-05 | 4.05 |
| FALSE | High | HSTPSNSSI | [K].HSTPSN | 3xPhospho | B2RQQ7 3x | 1.59E-06 | 0 | 1 | 2 | 4 | B2RQQ7 | CDC42 bin | B2RQQ7 [1 | 0 | 2610.037 | 0 | 0.62 | 0.375801 | 54.26 | 79.7 | 120.3 | 51.84 | 7.73 | | High | High | High | High | High | High | High | High | High | High | High | 0 | 1.71E-07 | 4.22 |
| FALSE | High | HSTPSPTK | [K].HSTPSP | 1xPhospho | Q9ERU9 1x | 0.061176 | 0.004646 | 1 | 1 | 1 | Q9ERU9 | E3 SUMO-p | Q9ERU9 [7 | 0 | 1392.729 | 0 | -1.43 | 0.008105 | 5.45 | 145.4 | 54.6 | 18.28 | 19.42 | | High | High | High | High | High | High | High | High | High | High | High | 0.002354 | 0.02428 | 2.08 |
| FALSE | High | HSTPSPTK | [K].HSTPSP | 2xPhospho | Q9ERU9 2x | 0.115566 | 0.009504 | 1 | 1 | 1 | Q9ERU9 | E3 SUMO-p | Q9ERU9 [7 | 0 | 1472.695 | 0 | -0.81 | 0.003831 | 2.57 | 126.8 | 73.2 | 5.06 | 20.24 | | High | High | High | High | High | High | High | High | High | High | High | 0.004757 | 0.05145 | 1.16 |
| FALSE | High | HSVDLIGRI | [R].HSVDLI | 1xPhospho | Q80XC2 1x | 0.006195 | 0.00062 | 1 | 1 | 1 | Q80XC2 | tRNA (ader | Q80XC2 [4 | 0 | 1951.057 | 0 | 1.11 | 0.000793 | 15.08 | 64.2 | 135.8 | 7.57 | 7.99 | | High | High | High | High | High | High | High | High | High | High | High | 0.000275 | 0.001835 | 1.15 |
| FALSE | High | HSVNLDEG | [R].HSVNLI | 1xPhospho | E9PVX6 1xl | 7.89E-05 | 0 | 1 | 2 | 1 | E9PVX6 | Proliferatic | E9PVX6 [3 | 0 | 2259.117 | 0 | 0.02 | 0.984015 | 9.2 | 98.8 | 101.2 | 8.13 | 10.31 | | High | High | High | High | High | High | High | High | High | High | High | 0 | 1.37E-05 | 1.81 |
| FALSE | High | HSVSPPPPY | [K].HSVSPI | 1xPhospho | Q3TLN5 1x | 0.000219 | 0 | 1 | 2 | 2 | Q3TLN5 | Putative ur | Q3TLN5 [1 | 0 | 1767.878 | 0 | 2.01 | 0.003216 | 22.09 | 41.1 | 158.9 | 9.19 | 18.58 | | High | High | High | High | High | High | High | High | High | High | High | 0 | 4.31E-05 | 3.06 |
| FALSE | High | HSVSPPPPY | [K].HSVSPI | 2xPhospho | Q3TLN5 2x | 6.13E-05 | 0 | 1 | 2 | 1 | Q3TLN5 | Putative ur | Q3TLN5 [1 | 0 | 1847.844 | 0 | -0.15 | 0.63775 | 23.99 | 105.2 | 94.8 | 17.15 | 21.73 | | High | High | High | High | High | High | High | High | High | High | High | 0 | 1.04E-05 | 2.6 |
| FALSE | High | HSVTAATP | [K].HSVTA | 1xPhospho | Q924Z6 1x | 6.25E-09 | 0 | 1 | 2 | 1 | Q924Z6 | Exportin-6 | Q924Z6 [1 | 0 | 3671.857 | 0 | -0.59 | 0.014164 | 5.88 | 120.1 | 79.9 | 12.71 | 17 | | High | High | High | High | High | High | High | High | High | High | High | 0 | 3.4E-10 | 6.06 |
| FALSE | High | HSETTLVNI | [R].HSETTL | 1xPhospho | Q3UQI9 1x | 5.8E-08 | 0 | 1 | 4 | 1 | Q3UQI9 | Probable u | Q3UQI9 [1 | 0 | 2474.129 | 0 | 0.37 | 0.010813 | 8.68 | 91.4 | 108.6 | 9.45 | 8.72 | | High | High | High | High | High | High | High | High | High | High | High | 0 | 4.14E-09 | 3.7 |
| FALSE | High | HTSPLSVAI | [K].HTSPLS | 1xPhospho | B2RXS2 1xl | 0.041072 | 0.003094 | 1 | 5 | 2 | B2RXS2 | Fam179b p | B2RXS2 [8 | 0 | 1574.871 | 0 | 0.14 | 0.098705 | 11.58 | 95.4 | 104.6 | 12.46 | 4.54 | | High | High | High | High | High | High | High | High | High | High | High | 0.001597 | 0.01539 | 1.4 |
| FALSE | High | HSEEANFES | [K].HSEEAI | 1xPhospho | B2RT77 1xl | 2.87E-05 | 0 | 1 | 1 | 1 | B2RT77 | TWIST neig | B2RT77 [3 | 1 | 2170.115 | 0 | -0.23 | 0.207074 | 14.64 | 108.1 | 91.9 | 6.16 | 26.23 | | High | High | High | High | High | High | High | High | High | High | High | 0 | 4.43E-06 | 3.35 |
| FALSE | High | HSEEANFES | [K].HSEEAI | 1xPhospho | B2RT77 1xl | 8.22E-07 | 0 | 1 | 1 | 2 | B2RT77 | TWIST neig | B2RT77 [3 | 0 | 1812.857 | 0 | -0.02 | 0.759311 | 11.56 | 96.7 | 103.3 | 7.56 | 24.92 | | High | High | High | High | High | High | High | High | High | High | High | 0 | 1.13E-07 | 4.51 |
| FALSE | High | HNSWSSSS | [R].HNSWS | 1xPhospho | Q8K2D3 1x | 0.079766 | 0.00613 | 1 | 1 | 2 | Q8K2D3 | Enhancer c | Q8K2D3 [1 | 0 | 1356.59 | 0 | -0.93 | 0.006027 | 2.7 | 131 | 69 | 11.98 | 9.61 | | High | High | High | High | High | High | High | High | High | High | High | 0.003163 | 0.03314 | 2.13 |
| FALSE | High | HNVHSPYK | [K].HNVHS | 1xPhospho | A0A140T8 | 0.002974 | 0.00016 | 1 | 1 | 2 | A0A140T8 | Chromoso | A0A140T8 | 0 | 1519.782 | 0 | -1.09 | 0.001127 | 2.8 | 135.9 | 64.1 | 5.27 | 19.29 | | High | High | High | High | High | High | High | High | High | High | High | 7.19E-05 | 0.000813 | 2.32 |
| FALSE | High | HPAGTSCP | [R].HPAGT | 1xCarbami | P41274 1xl | 0.000158 | 0 | 1 | 2 | 1 | P41274 | Tumor nec | P41274 [16 | 0 | 1861.883 | 0 | 0.78 | 0.006776 | 21.31 | 70.9 | 129.1 | 19.15 | 9.15 | | High | High | High | High | High | High | High | High | High | High | High | 0 | 3.02E-05 | 3.21 |
| FALSE | High | HPASAQST | [K].HPASA | 2xPhospho | Q9DBN4 2x | 8.56E-05 | 0 | 1 | 3 | 2 | Q9DBN4 | Putative m | Q9DBN4 [1 | 0 | 2406.126 | 0 | 0.28 | 0.028747 | 2.13 | 88.8 | 111.2 | 10.75 | 11.01 | | High | High | High | High | High | High | High | High | High | High | High | 0 | 1.51E-05 | 3.07 |
| FALSE | High | HPASLSSSG | [R].HPASLS | 1xPhospho | Q0VGY8 1x | 2.47E-06 | 0 | 1 | 5 | 3 | Q0VGY8 | Protein TA | Q0VGY8 [1 | 0 | 2298.138 | 0 | 0.18 | 0.555991 | 21.84 | 93.7 | 106.3 | 12.38 | 17.61 | | High | High | High | High | High | High | High | High | High | High | High | 0 | 2.81E-07 | 4.25 |
| FALSE | High | HPDASVNF | [K].HPDAS\ | 1xPhospho | Q3UBK2 1x | 2.38E-05 | 0 | 1 | 13 | 1 | Q3UBK2 | Putative ur | Q3UBK2 [3 | 0 | 2002.968 | 0 | 0.43 | 0.131076 | 45.41 | 72.5 | 127.5 | 20.45 | 27.5 | | High | High | High | High | High | High | High | High | High | High | High | 0 | 3.57E-06 | 3.54 |
| FALSE | High | HPDISYILG | [R].HPDISY | 1xPhospho | Q569X8 1x | 2.87E-05 | 0 | 1 | 6 | 1 | Q569X8 | Nucleopor | Q569X8 [7 | 1 | 2107.053 | 0 | -0.4 | 0.01984 | 1.8 | 112.2 | 87.8 | 11.18 | 5.43 | | High | High | High | High | High | High | High | High | High | High | High | 0 | 4.44E-06 | 2.79 |
| FALSE | High | HPEGIVSEV | [K].HPEGIV | 1xPhospho | B2RRE0 1x | 2.71E-07 | 0 | 1 | 3 | 2 | B2RRE0 | A kinase (P | B2RRE0 [4 | 0 | 2236.052 | 0 | -0.7 | 0.001142 | 1.65 | 123.8 | 76.2 | 4.23 | 8.66 | | High | High | High | High | High | High | High | High | High | High | High | 0 | 2.34E-08 | 4.2 |
| FALSE | High | HPEGIVSEV | [K].HPEGIV | 1xPhospho | B2RRE0 1x | 2.99E-06 | 0 | 1 | 3 | 2 | B2RRE0 | A kinase (P | B2RRE0 [4 | 0 | 2252.047 | 0 | -1.11 | 0.002986 | 5.99 | 136.6 | 63.4 | 7.28 | 14.69 | | High | High | High | High | High | High | High | High | High | High | High | 0 | 3.49E-07 | 3.89 |
| FALSE | High | HPEPSPSPF | [K].HPEPSF | 2xPhospho | Q80XC3-2 | 0.0009 | 0 | 1 | 2 | 1 | Q80XC3-2 | Isoform 2 c | Q80XC3-2 | 0 | 1690.801 | 0 | 0.22 | 0.342881 | 10.66 | 91.3 | 108.7 | 7.33 | 22.45 | | High | High | High | High | High | High | High | High | High | High | High | 0 | 0.000212 | 2.35 |
| FALSE | High | HPEPVPEE | [R].HPEPVI | 1xPhospho | A0A087WI | 8.5E-06 | 0 | 1 | 3 | 1 | A0A087WI | Autophagy | A0A087WI | 0 | 2788.36 | 0 | -0.15 | 0.057085 | 7.24 | 110.6 | 89.4 | 7.92 | 7.8 | | High | High | High | High | High | High | High | High | High | High | High | 0 | 1.12E-06 | 2.11 |
| FALSE | High | HPGAHQGI | [R].HPGAH | 1xPhospho | E9Q0S6 1x | 6.68E-07 | 0 | 1 | 9 | 1 | E9Q0S6 | Tensin 1 O | E9Q0S6 [1 | 0 | 3057.522 | 0 | -0.32 | 0.18909 | 30.1 | 115.2 | 84.8 | 31.57 | 18.23 | | High | High | High | High | High | High | High | High | High | High | High | 0 | 6.48E-08 | 3.94 |
| FALSE | High | HPGCPTPV | [K].HPGCP | 1xCarbami | Q80WE4 1 | 0.081085 | 0.006219 | 1 | 4 | 1 | Q80WE4 | Kinesin-like | Q80WE4 [ | 0 | 1744.922 | 0 | -0.15 | 0.476758 | 21.6 | 100 | 100 | 13.9 | 18.56 | | High | High | High | High | High | High | High | High | High | High | High | 0.0032 | 0.03382 | 2.43 |
| FALSE | High | HNSWDTV | [R].HNSWI | 1xPhospho | Q3U8L9 1x | 0.007811 | 0.000664 | 1 | 2 | 2 | Q3U8L9 | Putative ur | Q3U8L9 [6 | 0 | 2950.382 | 0 | -0.91 | 0.002739 | 5.3 | 133.8 | 66.2 | 14 | 17.6 | | High | High | High | High | High | High | High | High | High | High | High | 0.000327 | 0.002387 | 3.06 |
| FALSE | High | HPGDAPVL | [K].HPGDA | 1xPhospho | Q68FM7 1 | 2.31E-08 | 0 | 1 | 5 | 2 | Q68FM7 | Rho guanir | Q68FM7 [1 | 0 | 3179.474 | 0 | -0.41 | 0.098651 | 9.45 | 114 | 86 | 9.55 | 14.17 | | High | High | High | High | High | High | High | High | High | High | High | 0 | 1.47E-09 | 4.37 |
| FALSE | High | HPLLSPGGI | [R].HPLLSP | 1xPhospho | U5LK19 1xl | 1.03E-06 | 0 | 2 | 17 | 2 | U5LK19; E | Marshalin- | U5LK19 [3 | 0 | 1764.936 | 0 | -1.69 | 0.000399 | 1.18 | 152.5 | 47.5 | 6.07 | 10.88 | Shared | High | High | High | High | High | High | High | High | High | High | High | 0 | 1.05E-07 | 3.81 |
| FALSE | High | HPLLSPGGI | [R].HPLLSP | 1xPhospho | U5LK19 1xl | 0.001325 | 0 | 2 | 17 | 3 | U5LK19; E | Marshalin- | U5LK19 [3 | 1 | 2624.433 | 0 | -0.38 | 0.068412 | 4.76 | 110.4 | 89.6 | 8.04 | 22.24 | Shared | High | High | High | High | High | High | High | High | High | High | High | 2.58E-05 | 0.000517 | 3.5 |
| FALSE | High | HPNQATPK | [R].HPNQA | 1xPhospho | Q8K2D3 1x | 0.069325 | 0.005112 | 1 | 1 | 3 | Q8K2D3 | Enhancer c | Q8K2D3 [1 | 0 | 1430.756 | 0 | -1.24 | 0.000518 | 3.31 | 143.1 | 56.9 | 5.35 | 16.41 | | High | High | High | High | High | High | High | High | High | High | High | 0.002618 | 0.02806 | 2.73 |
| FALSE | High | HPNSSPR | [K].HPNSSI | 1xPhospho | B2RY61 1xl | 0.038677 | 0.002563 | 1 | 4 | 3 | B2RY61 | AT rich inte | B2RY61 [1 | 0 | 1103.52 | 0 | 0.15 | 0.768029 | 18.06 | 100.9 | 99.1 | 7.87 | 37.17 | | High | High | High | High | High | High | High | High | High | High | High | 0.003258 | 0.03436 | 1.18 |
| FALSE | High | HPNYSPPG | [K].HPNYSI | 2xPhospho | V9GX74 2x | 0.014033 | 0.001011 | 1 | 10 | 2 | V9GX74 | Zinc finger | V9GX74 [1 | 0 | 2040.96 | 0 | -0.28 | 0.183161 | 4.27 | 111.3 | 88.7 | 12.83 | 33.22 | | High | High | High | High | High | High | High | High | High | High | High | 0.000538 | 0.004569 | 2.72 |
| FALSE | High | HPPEQTNG | [R].HPPEQ | 2xPhospho | Q3TDH0 2x | 0.001586 | 5.66E-05 | 1 | 7 | 1 | Q3TDH0 | Putative ur | Q3TDH0 [5 | 0 | 3257.582 | 0 | -0.38 | 0.082689 | 10.93 | 111.3 | 88.7 | 14.47 | 10.53 | | High | High | High | High | High | High | High | High | High | High | High | 2.58E-05 | 0.000401 | 3.29 |
| FALSE | High | HPPEQTNG | [R].HPPEQ | 1xDeamida | Q3TDH0 2x | 1.9E-06 | 0 | 1 | 7 | 5 | Q3TDH0 | Putative ur | Q3TDH0 [5 | 0 | 3258.566 | 0 | -0.16 | 0.108958 | 8.26 | 107.6 | 92.4 | 9.51 | 6.07 | | High | High | High | High | High | High | High | High | High | High | High | 0 | 0.000136 | 4.26 |
| FALSE | High | HPPGSSITN | [R].HPPGS | 1xPhospho | A6H8H2-2 | 0.000246 | 0 | 1 | 3 | 1 | A6H8H2-2 | Isoform 2 c | A6H8H2-2 | 0 | 2146.113 | 0 | 0.3 | 0.037136 | 18.48 | 91.9 | 108.1 | 13.87 | 6.34 | | High | High | High | High | High | High | High | High | High | High | High | 0 | 4.92E-05 | 3.22 |
| FALSE | High | HPPSDIGLL | [R].HPPSDI | 1xPhospho | Q99KW3-2 | 1.54E-05 | 0 | 1 | 5 | 1 | Q99KW3-2 | Isoform 2 c | Q99KW3-2 | 0 | 2335.164 | 0 | -0.35 | 0.088375 | 10.98 | 115.1 | 84.9 | 13.6 | 24.39 | | High | High | High | High | High | High | High | High | High | High | High | 0 | 2.21E-06 | 3.22 |
| FALSE | High | HPPSLGTPV | [K].HPPSLG | 1xPhospho | Q924A2 1x | 0.006536 | 0.00062 | 1 | 1 | 1 | Q924A2 | Protein cap | Q924A2 [4 | 0 | 3030.756 | 0 | 0.39 | 0.151519 | 37.56 | 81.5 | 118.5 | 9.06 | 24.73 | | High | High | High | High | High | High | High | High | High | High | High | 0.000275 | 0.001947 | 3.06 |
| FALSE | High | HPPSLGTPV | [K].HPPSLG | 2xPhospho | Q924A2 2x | 5.57E-06 | 0 | 1 | 1 | 2 | Q924A2 | Protein cap | Q924A2 [4 | 0 | 3110.722 | 0 | -0.13 | 0.200838 | 8.62 | 107.9 | 92.1 | 12.38 | 17.31 | | High | High | High | High | High | High | High | High | High | High | High | 0 | 1.14E-06 | 2.78 |
| FALSE | High | HPPSYTLPL | [K].HPPSYT | 1xPhospho | Q91YD3 1x | 2.38E-05 | 0 | 1 | 2 | 6 | Q91YD3 | mRNA-dec | Q91YD3 [3 | 0 | 3622.924 | 0 | -0.5 | 0.002502 | 5.2 | 116.3 | 83.7 | 7.74 | 12.24 | | High | High | High | High | High | High | High | High | High | High | High | 0.000327 | 0.002549 | 3.62 |
| FALSE | High | HPPSYTLPL | [K].HPPSYT | 2xPhospho | Q91YD3 2x | 0.079114 | 0.00613 | 1 | 2 | 1 | Q91YD3 | mRNA-dec | Q91YD3 [3 | 0 | 3702.89 | 0 | -0.42 | 0.021779 | 6.59 | 120.3 | 79.7 | 10.18 | 19.17 | | High | High | High | High | High | High | High | High | High | High | High | 0.003148 | 0.03279 | 3.72 |
| FALSE | High | HPLGSVPS | [K].HPLGS\ | 1xPhospho | B9EKB5 1xl | 4.05E-09 | 0 | 1 | 2 | 1 | B9EKB5 | Baz2b prot | B9EKB5 [1 | 0 | 2272.21 | 0 | -0.93 | 0.004946 | 7.43 | 131.1 | 68.9 | 11.24 | 17.43 | | High | High | High | High | High | High | High | High | High | High | High | 0 | 2.09E-10 | 4.45 |
| FALSE | High | HPPSYTLPL | [K].HPPSYT | 1xDeamida | Q91YD3 2x | 0.026652 | 0.001676 | 1 | 2 | 1 | Q91YD3 | mRNA-dec | Q91YD3 [3 | 0 | 3703.874 | 0 | -0.27 | 0.11161 | 17.17 | 109.6 | 90.4 | 11.29 | 18.46 | | High | High | High | High | High | High | High | High | High | High | High | 0.000903 | 0.009441 | 2.96 |
| FALSE | High | HNSTDLPM | [R].HNSTDI | 1xPhospho | Q8C180 1x | 0.054844 | 0.003982 | 1 | 1 | 1 | Q8C180 | Fibroblast | Q8C180 [5 | 0 | 1239.528 | 0 | 1.24 | 0.001758 | 13.46 | 59.4 | 140.6 | 8.9 | 9.82 | | High | High | High | High | High | High | High | High | High | High | High | 0.002059 | 0.02144 | 1.48 |
| FALSE | High | HNSSDGFF | [R].HNSSD | 1xPhospho | Q6NZP2 1x | 1.05E-08 | 0 | 1 | 1 | 4 | Q6NZP2 | Vasculin-lik | Q6NZP2 [4 | 0 | 1870.843 | 0 | -0.09 | 0.510837 | 6.59 | 103.6 | 96.4 | 2.55 | 7.52 | | High | High | High | High | High | High | High | High | High | High | High | 0 | 6.03E-10 | 4.51 |
| FALSE | High | HLSSTSDDI | [R].HLSSTS | 2xPhospho | Q9DBC3 2x | 0.000155 | 0 | 1 | 2 | 2 | Q9DBC3 | Cap-specifi | Q9DBC3 [2 | 0 | 2613.2 | 0 | 0.89 | 0.007117 | 18.01 | 70.7 | 129.3 | 12.96 | 8.31 | | High | High | High | High | High | High | High | High | High | High | High | 0 | 7.58E-05 | 3.83 |
| FALSE | High | HLSTPSSVS | [K].HLSTPS | 1xPhospho | Q8R4H2 1x | 0.012397 | 0.000889 | 2 | 2 | 1 | Q8R4H2; F | Rho guanir | Q8R4H2 [6 | 0 | 2315.168 | 0 | 1.26 | 0.010229 | 7.79 | 61.6 | 138.4 | 10.16 | 20.84 | Shared | High | High | High | High | High | High | High | High | High | High | High | 0.000449 | 0.003977 | 3 |
| FALSE | High | HLSVSSQN | [R].HLSVSS | 1xPhospho | Q3UVM5 1 | 1.52E-06 | 0 | 1 | 3 | 2 | Q3UVM5 | Putative ur | Q3UVM5 [ | 0 | 1490.731 | 0 | 0.77 | 0.001722 | 10.86 | 73.9 | 126.1 | 4.86 | 9.5 | | High | High | High | High | High | High | High | High | High | High | High | 0 | 1.63E-07 | 2.24 |
| FALSE | High | HLTPEPDIV | [R].HLTPEP | 1xPhospho | Q6NVE8 1x | 0.000297 | 0 | 1 | 1 | 2 | Q6NVE8 | WD repeat | Q6NVE8 [2 | 0 | 1946.04 | 0 | 0.9 | 0.000945 | 4.72 | 69.7 | 130.3 | 2.61 | 8.1 | | High | High | High | High | High | High | High | High | High | High | High | 0 | 6.08E-05 | 2.89 |
| FALSE | High | HLTTVEGTI | [K].HLTTVE | 1xPhospho | E9QAQ7 1x | 9.16E-07 | 0 | 1 | 5 | 4 | E9QAQ7 | AT-rich inte | E9QAQ7 [1 | 0 | 3125.544 | 0 | -1.48 | 0.002619 | 4.21 | 150.9 | 49.1 | 12.39 | 20.01 | | High | High | High | High | High | High | High | High | High | High | High | 0 | 9.22E-08 | 4.25 |
| FALSE | High | HIVSNDSSI | [K].HIVSND | 1xPhospho | P47713 1xl | 0.0004 | 0 | 1 | 2 | 1 | P47713 | Cytosolic p | P47713 [42 | 0 | 2209.945 | 0 | -0.98 | 0.042151 | 12.18 | 127.7 | 72.3 | 31.57 | 13.93 | | High | High | High | High | High | High | High | High | High | High | High | 0 | 8.52E-05 | 2.76 |
| FALSE | High | HIVSNDSSI | [K].HIVSND | 1xPhospho | P47713 1xl | 4.02E-09 | 0 | 1 | 2 | 3 | P47713 | Cytosolic p | P47713 [42 | 0 | 2439.108 | 0 | -0.74 | 0.037307 | 16.45 | 128.8 | 71.2 | 11.33 | 29.13 | | High | High | High | High | High | High | High | High | High | High | High | 0 | 2.07E-10 | 4.69 |
| FALSE | High | HIVSNDSSI | [K].HIVSND | 2xPhospho | P47713 2xl | 0.000181 | 0 | 1 | 2 | 1 | P47713 | Cytosolic p | P47713 [42 | 0 | 2519.074 | 0 | -0.62 | 0.007528 | 8.71 | 122.4 | 77.6 | 6.77 | 11.7 | | High | High | High | High | High | High | High | High | High | High | High | 0 | 3.49E-05 | 3.21 |
| FALSE | High | HLYISSSNP | [R].HLYISS | 1xPhospho | G5E8J4 1xl | 2.65E-08 | 0 | 1 | 5 | 3 | G5E8J4 | Tyrosine-p | G5E8J4 [58 | 0 | 1924.973 | 0 | 0.4 | 0.005072 | 3.9 | 85.4 | 114.6 | 3.91 | 7.53 | | High | High | High | High | High | High | High | High | High | High | High | 0 | 6.89E-05 | 3.96 |
| FALSE | High | HMSESPNF | [R].HMSES | 1xPhospho | A2AR02 1x | 0.08727 | 0.006537 | 1 | 3 | 1 | A2AR02 | Peptidyl-pr | A2AR02 [4 | 0 | 1282.545 | 0 | -0.86 | 0.0105 | 5.89 | 118 | 82 | 14.21 | 16.75 | | High | High | High | High | High | High | High | High | High | High | High | 0.003346 | 0.0366 | 0.89 |
| FALSE | High | HNDIVDSD | [K].HNDIVI | 2xPhospho | Q8R3N1 2x | 0.001386 | 5.66E-05 | 1 | 3 | 3 | Q8R3N1 | Nucleolar | Q8R3N1 [1 | 1 | 3955.79 | 0 | -0.2 | 0.553814 | 14.59 | 105.2 | 94.8 | 11.08 | 10.81 | | High | High | High | High | High | High | High | High | High | High | High | 0.000399 | 0.003429 | 4.59 |
| FALSE | High | HNELGTTT | [R].HNELG | 1xPhospho | A0A087WC | 4.22E-07 | 0 | 1 | 6 | 5 | A0A087WC | Snf2-relate | A0A087WC | 0 | 3315.651 | 0 | 0.08 | 0.82376 | 15.73 | 97.3 | 102.7 | 8.54 | 11.05 | | High | High | High | High | High | High | High | High | High | High | High | 0 | 6.54E-08 | 4.02 |
| FALSE | High | HNELGTTT | [R].HNELG | 1xDeamida | A0A087WC | 0.000203 | 0 | 1 | 6 | 3 | A0A087WC | Snf2-relate | A0A087WC | 0 | 3316.635 | 0 | -0.05 | 0.221759 | 7.53 | 103.7 | 96.3 | 19.82 | 17.08 | | High | High | High | High | High | High | High | High | High | High | High | 0 | 4E-05 | 3.53 |
| FALSE | High | HNSSDGFF | [R].HNSSD | 1xDeamida | Q6NZP2 1x | 2.28E-07 | 0 | 1 | 1 | 2 | Q6NZP2 | Vasculin-lik | Q6NZP2 [4 | 0 | 1871.828 | 0 | -0.41 | 0.37379 | 3.19 | 109.8 | 90.2 | 20.49 | 11.77 | | High | High | High | High | High | High | High | High | High | High | High | 0 | 1.94E-08 | 4.25 |
| FALSE | High | HNGVGGSI | [K].HNGVG | 1xPhospho | Q60520-1 | 0.101749 | 0.008194 | 1 | 3 | 1 | Q60520-1 | Isoform 2 c | Q60520-1 | 0 | 1258.614 | 0 | -0.75 | 0.13115 | 16.62 | 135.1 | 64.9 | 39.33 | 39.38 | | High | High | High | High | High | High | High | High | High | High | High | 0.004115 | 0.04385 | 2.02 |
| FALSE | High | HNGVGGSI | [K].HNGVG | 1xDeamida | Q60520-1 | 0.000118 | 0 | 1 | 3 | 5 | Q60520-1 | Isoform 2 c | Q60520-1 | 0 | 1488.761 | 0 | -0.96 | 0.002005 | 1.69 | 131.8 | 68.2 | 4.35 | 15.67 | | High | High | High | High | High | High | High | High | High | High | High | 4.94E-05 | 0.000594 | 2.14 |
| FALSE | High | HNLDNSQS | [K].HNLDN | 1xPhospho | E9PZM7 1x | 7.77E-06 | 0 | 1 | 3 | 4 | E9PZM7 | SR-related | E9PZM7 [6 | 0 | 1695.81 | 0 | 0.2 | 0.974051 | 23.9 | 95.7 | 104.3 | 8.34 | 26.15 | | High | High | High | High | High | High | High | High | High | High | High | 0 | 1.01E-06 | 2.98 |
| FALSE | High | HNLDNSQS | [K].HNLDN | 1xDeamida | E9PZM7 1x | 8.33E-05 | 0 | 1 | 3 | 1 | E9PZM7 | SR-related | E9PZM7 [6 | 0 | 1696.794 | 0 | -0.02 | 0.52028 | 5.57 | 99.1 | 100.9 | 6.03 | 24.51 | | High | High | High | High | High | High | High | High | High | High | High | 0 | 1.46E-05 | 2.8 |
| FALSE | High | HNLDVVSP | [K].HNLDV | 1xPhospho | B9EKC3 1xl | 2.49E-05 | 0 | 1 | 3 | 2 | B9EKC3 | Rho GTPas | B9EKC3 [7 | 0 | 1970.088 | 0 | 0.33 | 0.379089 | 9.54 | 90.7 | 109.3 | 6.77 | 21.38 | | High | High | High | High | High | High | High | High | High | High | High | 0 | 3.77E-06 | 3.43 |
| FALSE | High | HNLFEDNN | [K].HNLFEI | 1xPhospho | Q3UW53 1 | 8.29E-10 | 0 | 1 | 1 | 3 | Q3UW53 | Protein Nib | Q3UW53 [ | 0 | 2985.468 | 0 | 0.55 | 0.193705 | 6.42 | 85.1 | 114.9 | 5.66 | 27.71 | | High | High | High | High | High | High | High | High | High | High | High | 0 | 8.23E-09 | 4.94 |
| FALSE | High | HNLFEDNN | [K].HNLFEI | 1xPhospho | Q3UW53 1 | 7.64E-10 | 0 | 1 | 1 | 13 | Q3UW53 | Protein Nib | Q3UW53 [ | 0 | 3001.463 | 0 | 0.49 | 0.008175 | 15.63 | 82.9 | 117.1 | 7.71 | 9.5 | | High | High | High | High | High | High | High | High | High | High | High | 0 | 4.55E-08 | 3.99 |
| FALSE | High | HNPQSPLQ | [R].HNPQS | 1xPhospho | Q8CGC3 1x | 8.91E-10 | 0 | 1 | 6 | 3 | Q8CGC3 | Histone de | Q8CGC3 [1 | 0 | 2175.121 | 0 | -0.85 | 0.00026 | 0.94 | 129.5 | 70.5 | 3.62 | 5.85 | | High | High | High | High | High | High | High | High | High | High | High | 0 | 3.81E-11 | 4.98 |
| FALSE | High | HNQDSQH | [R].HNQDS | 1xCarbami | Q6P9Q6-2 | 4.3E-07 | 0 | 1 | 3 | 1 | Q6P9Q6-2 | Isoform B c | Q6P9Q6-2 | 0 | 2684.113 | 0 | 0.1 | 0.673493 | 32.51 | 99.3 | 100.7 | 13.94 | 22.45 | | High | High | High | High | High | High | High | High | High | High | High | 0 | 3.93E-08 | 4.35 |
| FALSE | High | HNQDSQH | [R].HNQDS | 1xCarbami | Q6P9Q6-2 | 0.000584 | 0 | 1 | 3 | 2 | Q6P9Q6-2 | Isoform B c | Q6P9Q6-2 | 0 | 2993.243 | 0 | 0.6 | 0.00146 | 5.39 | 80.6 | 119.4 | 8.05 | 6.98 | | High | High | High | High | High | High | High | High | High | High | High | 0 | 0.00013 | 3.94 |
| FALSE | High | HNRPSNGN | [K].HNRPSI | 1xPhospho | J3QJU5 1xl | 0.007881 | 0.000736 | 1 | 4 | 1 | J3QJU5 | Lethal(2) gi | J3QJU5 [96 | 0 | 1889.975 | 0 | -0.1 | 0.539824 | 28.93 | 102.5 | 97.5 | 20.19 | 22.95 | | High | High | High | High | High | High | High | High | High | High | High | 0.000327 | 0.002392 | 1.77 |
| FALSE | High | HNSFDTVN | [R].HNSFD | 2xCarbami | Q6GQW0 1 | 0.003253 | 0.00016 | 1 | 2 | 1 | Q6GQW0 | Ankyrin re | Q6GQW0 [ | 0 | 3328.42 | 0 | -0.16 | 0.009736 | 3.28 | 104.6 | 95.4 | 6.32 | 7.43 | | High | High | High | High | High | High | High | High | High | High | High | 7.19E-05 | 0.000892 | 3.34 |
| FALSE | High | HNSGSSSV | [R].HNSGS | 1xPhospho | P50636 1xl | 1.87E-06 | 0 | 1 | 1 | 1 | P50636 | E3 ubiquiti | P50636 [62 | 0 | 1785.76 | 0 | 0.25 | 0.237536 | 26.6 | 90.9 | 109.1 | 13.32 | 12.79 | | High | High | High | High | High | High | High | High | High | High | High | 0 | 2.04E-07 | 3.24 |
| FALSE | High | HNSSDGFD | [R].HNSSD | 1xPhospho | Q6NXH3-2 | 0.007019 | 0.00062 | 1 | 2 | 2 | Q6NXH3-2 | Isoform 2 c | Q6NXH3-2 | 0 | 2457.089 | 0 | -1.07 | 0.004102 | 6.62 | 135.6 | 64.4 | 10.78 | 12.82 | | High | High | High | High | High | High | High | High | High | High | High | 0.000382 | 0.003256 | 1.57 |
| FALSE | High | HNGVGGSI | [K].HNGVG | 1xPhospho | Q60520-1 | 1.21E-05 | 0 | 1 | 3 | 3 | Q60520-1 | Isoform 2 c | Q60520-1 | 0 | 1487.777 | 0 | -1.08 | 0.003058 | 1.8 | 135 | 65 | 6.86 | 20.42 | | High | High | High | High | High | High | High | High | High | High | High | 0 | 1.68E-06 | 3.15 |
| FALSE | High | HPPVLTPPI | [R].HPPVLT | 1xPhospho | P68404-2 1 | 0.002623 | 0.000109 | 1 | 1 | 1 | P68404-2 | Isoform Be | P68404-2 [ | 0 | 1906.999 | 0 | -1.08 | 0.002895 | 2.19 | 137.4 | 62.6 | 11.35 | 10.45 | | High | High | High | High | High | High | High | High | High | High | High | 4.94E-05 | 0.000705 | 2.22 |
| FALSE | High | HPQTSSPG | [KR].HPQT | 2xCarbami | Q80TH2-2 | 0.047488 | 0.00354 | 1 | 4 | 2 | Q80TH2-2 | Isoform 2 c | Q80TH2-2 | 0 | 2082.821 | 0 | 1.17 | 0.02923 | 69.69 | 55.7 | 144.3 | 48.35 | 29.07 | | High | High | High | High | High | High | High | High | High | High | High | 0.001812 | 0.01809 | 2.74 |
| FALSE | High | HPSGTSFQ | [K].HPSGT | 1xCarbami | Q8C0D9 1x | 3.03E-05 | 0 | 1 | 5 | 2 | Q8C0D9 | Centrosom | Q8C0D9 [5 | 0 | 2282.038 | 0 | -0.08 | 0.387768 | 15.29 | 100.9 | 99.1 | 10.69 | 5.99 | | High | High | High | High | High | High | High | High | High | High | High | 0 | 4.69E-06 | 4.12 |
| FALSE | High | HQSPALGC | [R].HQSPA | 1xCarbami | Q99LF7 1xl | 9.13E-09 | 0 | 1 | 1 | 2 | Q99LF7 | DDB1 and | Q99LF7 [1 | 0 | 1880.816 | 0 | -0.56 | 0.002003 | 5.06 | 119.3 | 80.7 | 4.25 | 8.19 | | High | High | High | High | High | High | High | High | High | High | High | 0 | 5.2E-10 | 2.94 |
| FALSE | High | HQVSSDAP | [K].HQVSS | 1xPhospho | Q6NZQ2 1x | 0.053023 | 0.00384 | 1 | 1 | 2 | Q6NZQ2 | Probable A | Q6NZQ2 [1 | 0 | 1674.862 | 0 | 0.57 | 0.000868 | 5.11 | 83.6 | 116.4 | 6.68 | 6.05 | | High | High | High | High | High | High | High | High | High | High | High | 0.001999 | 0.0207 | 2.33 |
| FALSE | High | HREDSDVE | [R].HREDSI | 1xPhospho | Q8VDI4 1x | 6.93E-07 | 0 | 1 | 4 | 1 | Q8VDI4 | MutL homo | Q8VDI4 [4 | 1 | 2357.084 | 0 | -0.47 | 0.011079 | 7.11 | 116.1 | 83.9 | 7.34 | 10.29 | | High | High | High | High | High | High | High | High | High | High | High | 0 | 6.71E-08 | 3.49 |
| FALSE | High | HRESWSST | [R].HRESW | 1xPhospho | F6UFZ5 1xl | 0.019107 | 0.001402 | 2 | 7 | 2 | F6UFZ5; Q | PCF11 clea | F6UFZ5 [4 | 1 | 1655.831 | 0 | 0.38 | 0.009244 | 6 | 87 | 113 | 7.13 | 14.08 | Shared | High | High | High | High | High | High | High | High | High | High | High | 0.005717 | 0.06318 | 0.89 |
| FALSE | High | HRGSADYS | [K].HRGSA | 1xPhospho | Q80TC1 1x | 0.002553 | 0.000109 | 1 | 15 | 2 | Q80TC1 | MKIAA154 | Q80TC1 [3 | 1 | 1905.893 | 0 | -1.36 | 0.001246 | 2.22 | 143.1 | 56.9 | 8.53 | 12.74 | | High | High | High | High | High | High | High | High | High | High | High | 4.94E-05 | 0.000685 | 3.12 |
| FALSE | High | HRGSEEDP | [R].HRGSEI | 1xPhospho | B9EKC3 1xl | 1.47E-07 | 0 | 1 | 3 | 3 | B9EKC3 | Rho GTPas | B9EKC3 [1 | 1 | 2518.274 | 0 | -0.7 | 0.000357 | 2.3 | 125.8 | 74.2 | 6.76 | 12.32 | | High | High | High | High | High | High | High | High | High | High | High | 0 | 1.18E-08 | 3.82 |
| FALSE | High | HRGSEEDP | [R].HRGSEI | 2xPhospho | B9EKC3 2xl | 0.000264 | 0 | 1 | 3 | 2 | B9EKC3 | Rho GTPas | B9EKC3 [1 | 1 | 2598.241 | 0 | 0.34 | 0.152311 | 20.22 | 85.8 | 114.2 | 4.16 | 17.99 | | High | High | High | High | High | High | High | High | High | High | High | 0 | 5.36E-05 | 3.83 |
| FALSE | High | HRHSPTGP | [R].HRHSPT | 1xPhospho | Q99KG3 1x | 0.000793 | 0 | 1 | 2 | 3 | Q99KG3 | RNA-bindir | Q99KG3 [8 | 1 | 1751.869 | 0 | -0.72 | 0.010331 | 8.06 | 125.8 | 74.2 | 8.49 | 21.79 | | High | High | High | High | High | High | High | High | High | High | High | 0 | 0.000183 | 2.82 |

| | | | | | | | | | | | | | | | | | | | | | | | | | | | | | | | | | | | | | | |
|---|---|---|---|---|---|---|---|---|---|---|---|---|---|---|---|---|---|---|---|---|---|---|---|---|---|---|---|---|---|---|---|---|---|---|---|---|---|---|
| FALSE | High | HRNSLSDSA | [R].HRNSLS | 1xPhosphc | Q9WVB3 1 | 0.002334 | 0.000109 | 1 | 3 | 1 | Q9WVB3 | Transducin | Q9WVB3 [ | 1 | 2238.168 | 0 | 0.75 | 0.011841 | 12.71 | 73.6 | 126.4 | 2.39 | 16.87 | | High | High | High | High | High | High | High | High | High | High | High | 4.94E-05 | 0.000616 | 3.24 |
| FALSE | High | HRNSTDYS | [K].HRNSTI | 1xPhosphc | Q3UY41 1x | 6.77E-05 | 0 | 1 | 4 | 2 | Q3UY41 | Putative ur | Q3UY41 [2 | 1 | 1948.917 | 0 | 0.75 | 0.116436 | 8.2 | 76.9 | 123.1 | 10.1 | 33.93 | | High | High | High | High | High | High | High | High | High | High | High | 0 | 1.16E-05 | 2.58 |
| FALSE | High | HRNSTDYS | [K].HRNSTI | 1xDeamida | Q3UY41 1x | 0.003892 | 0.000261 | 1 | 4 | 1 | Q3UY41 | Putative ur | Q3UY41 [2 | 1 | 1949.901 | 0 | 0.63 | 0.096351 | 19.76 | 72.7 | 127.3 | 12.43 | 31.78 | | High | High | High | High | High | High | High | High | High | High | High | 0.00014 | 0.001094 | 2.07 |
| FALSE | High | HRPSSPATI | [R].HRPSSP | 2xPhosphc | Q52KI8 2xP | 4.56E-06 | 0 | 1 | 6 | 6 | Q52KI8 | Serine/argi | Q52KI8 [39 | 0 | 1660.781 | 0 | -0.22 | 0.307446 | 45.68 | 112.6 | 87.4 | 26.32 | 26.64 | | High | High | High | High | High | High | High | High | High | High | High | 0 | 3.98E-06 | 3.54 |
| FALSE | High | HRPSSPATI | [R].HRPSSP | 1xPhosphc | Q52KI8 1xP | 0.000286 | 0 | 1 | 6 | 5 | Q52KI8 | Serine/argi | Q52KI8 [39 | 0 | 1809.978 | 0 | -1.24 | 0.003299 | 2.07 | 142.1 | 57.9 | 11.02 | 11.07 | | High | High | High | High | High | High | High | High | High | High | High | 0 | 0.000103 | 2.26 |
| FALSE | High | HQSDGNEI | [R].HQSDG | 1xPhosphc | Q8C0T5 1x | 7.47E-08 | 0 | 1 | 3 | 2 | Q8C0T5 | Signal-indu | Q8C0T5 [1 | 0 | 1673.759 | 0 | 1 | 0.000506 | 10.51 | 66.5 | 133.5 | 5.83 | 4.06 | | High | High | High | High | High | High | High | High | High | High | High | 0 | 6.35E-06 | 2.38 |
| FALSE | High | HRPSSPATI | [R].HRPSSP | 2xPhosphc | Q52KI8 2xP | 5.07E-05 | 0 | 1 | 6 | 10 | Q52KI8 | Serine/argi | Q52KI8 [39 | 0 | 1889.944 | 0 | -0.29 | 0.018972 | 8.19 | 110.2 | 89.8 | 4.35 | 9.34 | | High | High | High | High | High | High | High | High | High | High | High | 0 | 0.00019 | 2.73 |
| FALSE | High | HSAGSPGIA | [R].HSAGSI | 1xPhosphc | E9PXF8 1xP | 3.14E-05 | 0 | 1 | 6 | 2 | E9PXF8 | SET-bindin | E9PXF8 [17 | 0 | 2339.18 | 0 | 1.08 | 0.008729 | 23.52 | 66.3 | 133.7 | 7.43 | 22.62 | | High | High | High | High | High | High | High | High | High | High | High | 0 | 5.36E-06 | 3.38 |
| FALSE | High | HSAPEQLLA | [R].HSAPEC | 1xPhosphc | Q1W617 1 | 0.000124 | 0 | 1 | 2 | 1 | Q1W617 | Protein Shi | Q1W617 [4 | 0 | 2575.298 | 0 | 2.05 | 0.001559 | 9.53 | 39.7 | 160.3 | 10.11 | 14.9 | | High | High | High | High | High | High | High | High | High | High | High | 0 | 2.29E-05 | 2.44 |
| FALSE | High | HSAPGSTP | [R].HSAPGS | 1xPhosphc | Q80XH7 1x | 0.002376 | 0.000109 | 1 | 7 | 3 | Q80XH7 | Polymerase | Q80XH7 [2 | 0 | 1518.808 | 0 | -1.26 | 0.000494 | 1.02 | 139.5 | 60.5 | 6.07 | 4.97 | | High | High | High | High | High | High | High | High | High | High | High | 4.94E-05 | 0.000628 | 2.41 |
| FALSE | High | HSASDPGP | [K].HSASDI | 1xPhosphc | Q8BI84 1xP | 1.2E-10 | 0 | 1 | 4 | 10 | Q8BI84 | Melanoma | Q8BI84 [17 | 0 | 1973.928 | 0 | 0.29 | 0.632225 | 13.48 | 90.7 | 109.3 | 8.06 | 20.8 | | High | High | High | High | High | High | High | High | High | High | High | 0 | 4.01E-12 | 3.73 |
| FALSE | High | HSASDPGP | [K].HSASDI | 1xDeamida | Q8BI84 1xP | 0.000852 | 0 | 1 | 4 | 2 | Q8BI84 | Melanoma | Q8BI84 [17 | 0 | 1974.912 | 0 | -0.03 | 0.650913 | 4.19 | 100.6 | 99.4 | 10.55 | 3.56 | | High | High | High | High | High | High | High | High | High | High | High | 0 | 0.0002 | 2.89 |
| FALSE | High | HSASPVVFS | [R].HSASPV | 1xPhosphc | Q8C0T5 1x | 0.00045 | 0 | 1 | 3 | 2 | Q8C0T5 | Signal-indu | Q8C0T5 [1 | 0 | 1553.767 | 0 | 2.41 | 0.001361 | 5.12 | 31.6 | 168.4 | 6.32 | 15.9 | | High | High | High | High | High | High | High | High | High | High | High | 0.000275 | 0.001987 | 2.96 |
| FALSE | High | HSASPVVFS | [R].HSASPV | 2xPhosphc | Q8C0T5 2x | 0.00083 | 0 | 1 | 3 | 1 | Q8C0T5 | Signal-indu | Q8C0T5 [1 | 0 | 1633.734 | 0 | 1.16 | 0.00098 | 7.09 | 64.4 | 135.6 | 17.98 | 16.59 | | High | High | High | High | High | High | High | High | High | High | High | 0 | 0.000194 | 2.87 |
| FALSE | High | HSASSPK | [R].HSASSF | 1xPhosphc | E9QKL0 1x | 0.019276 | 0.001402 | 1 | 2 | 2 | E9QKL0 | Helicase AF | E9QKL0 [1 | 0 | 1251.65 | 0 | -0.43 | 0.020679 | 11.74 | 117 | 83 | 7.78 | 13.93 | | High | High | High | High | High | High | High | High | High | High | High | 0.001421 | 0.01446 | 2.25 |
| FALSE | High | HSASSPTSC | [K].HSASSF | 1xPhosphc | Q8BZT2 1x | 0.000845 | 0 | 1 | 2 | 2 | Q8BZT2 | Putative E3 | Q8BZT2 [6 | 0 | 1496.751 | 0 | 5.12 | | 12.06 | 5.8 | 194.2 | 6.08 | 28.9 | | High | Not Found | High | High | High | High | High | High | High | High | High | 0 | 0.000197 | 2.58 |
| FALSE | High | HSCAEALV | [R].HSCAEA | 1xCarbami | Q60591 1x | 1.35E-06 | 0 | 1 | 14 | 2 | Q60591 | Nuclear fac | Q60591 [2 | 0 | 2184.083 | 0 | 1.95 | 0.000447 | 5.42 | 38.8 | 161.2 | 7.58 | 9.59 | | High | High | High | High | High | High | High | High | High | High | High | 0 | 1.43E-07 | 3.05 |
| FALSE | High | HSDCDSSG | [R].HSDCD | 1xCarbami | Q8K3G5 1x | 1.45E-10 | 0 | 1 | 5 | 2 | Q8K3G5 | Inactive se | Q8K3G5 [7 | 0 | 2259.902 | 0 | 0.01 | 0.448961 | 20.43 | 101 | 99 | 8.14 | 15.29 | | High | High | High | High | High | High | High | High | High | High | High | 0 | 4.95E-12 | 4.98 |
| FALSE | High | HSDDRPYS | [K].HSDDR | 1xPhosphc | Q80YR7 1x | 0.068184 | 0.004935 | 1 | 2 | 1 | Q80YR7 | Claspin OS | Q80YR7 [4 | 0 | 1710.743 | 0 | -0.6 | 0.03839 | 17.92 | 120.3 | 79.7 | 8.47 | 17.28 | | High | High | High | High | High | High | High | High | High | High | High | 0.002559 | 0.02754 | 2.18 |
| FALSE | High | HSDFSPCSI | [R].HSDFSF | 1xCarbami | Q99LX5 1x | 0.011549 | 0.00086 | 1 | 1 | 2 | Q99LX5 | Multiple m | Q99LX5 [2 | 0 | 1602.739 | 0 | 0.3 | 0.094114 | 14.97 | 89.1 | 110.9 | 5.07 | 11.6 | | High | High | High | High | High | High | High | High | High | High | High | 0.002455 | 0.027 | 1.45 |
| FALSE | High | HRSGDALT | [R].HRSGD | 1xPhosphc | Q8BIZ6 1xP | 0.000155 | 0 | 1 | 1 | 1 | Q8BIZ6 | Smad nucle | Q8BIZ6 [16 | 1 | 1921.067 | 0 | -0.7 | 0.02321 | 10.06 | 119.9 | 80.1 | 15.2 | 12.47 | | High | High | High | High | High | High | High | High | High | High | High | 0 | 2.93E-05 | 3.16 |
| FALSE | High | HQQQFLAS | [R].HQQQF | 1xPhosphc | A0A0R4J25 | 6.16E-10 | 0 | 1 | 2 | 4 | A0A0R4J25 | Exportin-4 | A0A0R4J25 | 0 | 2396.201 | 0 | -1.48 | 0.000357 | 2.37 | 148.5 | 51.5 | 4.05 | 9.93 | | High | High | High | High | High | High | High | High | High | High | High | 0 | 8.2E-07 | 4.34 |
| FALSE | High | HQQQFLAS | [R].HQQQF | 1xPhosphc | A0A0R4J25 | 0.0002 | 0 | 1 | 2 | 1 | A0A0R4J25 | Exportin-4 | A0A0R4J25 | 0 | 2167.038 | 0 | -1.25 | 0.288036 | 7.95 | 137.1 | 62.9 | 26.76 | 69.9 | | High | High | High | High | High | High | High | High | High | High | High | 0 | 3.9E-05 | 2.53 |
| FALSE | High | HQQIPLHSI | [K].HQQIPI | 1xPhosphc | Q8CHP6 1x | 0.000466 | 0 | 1 | 6 | 1 | Q8CHP6 | Polyhomec | Q8CHP6 [3 | 0 | 1917.051 | 0 | 1.23 | 0.02265 | 29.84 | 57.1 | 142.9 | 6.4 | 27.68 | | High | High | High | High | High | High | High | High | High | High | High | 0 | 0.000101 | 2.14 |
| FALSE | High | HPSHSTSPS | [R].HPSHST | 1xPhosphc | Q8K221 1x | 3.13E-07 | 0 | 1 | 6 | 2 | Q8K221 | Arfaptin-2 | Q8K221 [7 | 0 | 2026.918 | 0 | 0.17 | 0.914434 | 23.9 | 97 | 103 | 18.58 | 14.76 | | High | High | High | High | High | High | High | High | High | High | High | 0 | 2.75E-08 | 3.52 |
| FALSE | High | HPSIETLDS | [R].HPSIETI | 1xCarbami | I7HJQ9 1xP | 0.000316 | 0 | 1 | 4 | 1 | I7HJQ9 | Myotubula | I7HJQ9 [45 | 0 | 2718.3 | 0 | 0.62 | 0.005148 | 10.16 | 78.7 | 121.3 | 10.01 | 7.45 | | High | High | High | High | High | High | High | High | High | High | High | 0 | 6.57E-05 | 2.16 |
| FALSE | High | HPSIETLDS | [R].HPSIETI | 1xCarbami | I7HJQ9 2xP | 0.001373 | 5.66E-05 | 1 | 4 | 1 | I7HJQ9 | Myotubula | I7HJQ9 [45 | 0 | 2798.266 | 0 | 1.88 | 0.00058 | 13.78 | 42.2 | 157.8 | 11.94 | 8.02 | | High | High | High | High | High | High | High | High | High | High | High | 2.58E-05 | 0.000341 | 2.88 |
| FALSE | High | HPSPLLTQT | [K].HPSPLL | 1xPhosphc | Q7TSH4 1x | 0.00582 | 0.00062 | 1 | 3 | 2 | Q7TSH4 | Centriolar | Q7TSH4 [5 | 0 | 1774.905 | 0 | -0.34 | 0.118947 | 9.82 | 107.3 | 92.7 | 15.25 | 8.76 | | High | High | High | High | High | High | High | High | High | High | High | 0.000275 | 0.001708 | 2.49 |
| FALSE | High | HPSPQSPSI | [R].HPSPQS | 1xPhosphc | E9Q3E2 1x | 0.00034 | 0 | 1 | 3 | 1 | E9Q3E2 | Synaptopo | E9Q3E2 [2 | 0 | 1301.62 | 0 | -0.34 | 0.103359 | 18.66 | 124.7 | 75.3 | 15.94 | 48.03 | | High | High | High | High | High | High | High | High | High | High | High | 0 | 7.11E-05 | 2.92 |
| FALSE | High | HPSVESVV | [K].HPSVES | 1xCarbami | Q3UJL7 1xl | 0.00045 | 0 | 1 | 3 | 1 | Q3UJL7 | Putative ur | Q3UJL7 [1 | 0 | 2936.436 | 0 | -0.31 | 0.026542 | 12.44 | 115.9 | 84.1 | 14.09 | 20.95 | | High | High | High | High | High | High | High | High | High | High | High | 0 | 9.74E-05 | 4.27 |
| FALSE | High | HPSVGSAR | [R].HPSVGS | 1xPhosphc | Q8C180 1x | 0.065405 | 0.004734 | 1 | 1 | 1 | Q8C180 | Fibroblast | Q8C180 [1 | 0 | 1119.551 | 0 | 0.83 | 0.001084 | 11.57 | 72 | 128 | 6.99 | 5.99 | | High | High | High | High | High | High | High | High | High | High | High | 0.002432 | 0.02619 | 0.61 |
| FALSE | High | HPSVGSAR | [R].HPSVGS | 2xPhosphc | Q8C180 2x | 0.001704 | 5.66E-05 | 1 | 1 | 1 | Q8C180 | Fibroblast | Q8C180 [1 | 0 | 1199.517 | 0 | 1.55 | 0.000377 | 8.81 | 50.1 | 149.9 | 8.55 | 6.68 | | High | High | High | High | High | High | High | High | High | High | High | 2.58E-05 | 0.000434 | 1.44 |
| FALSE | High | HPSWEEED | [R].HPSWE | 1xPhosphc | Q8BKI2 1xl | 2.9E-05 | 0 | 1 | 4 | 2 | Q8BKI2 | Trinucleoti | Q8BKI2 [1 | 0 | 4167.811 | 0 | 0.63 | 0.145551 | 29.89 | 91.8 | 108.2 | 18.7 | 32.57 | | High | High | High | High | High | High | High | High | High | High | High | 0 | 4.45E-06 | 5.39 |
| FALSE | High | HPVGSHQ | [R].HPVGS | 1xPhosphc | E9Q0S6 1x | 1.3E-05 | 0 | 1 | 11 | 2 | E9Q0S6 | Tensin 1 O | E9Q0S6 [1 | 0 | 2899.478 | 0 | 0.01 | 0.702368 | 17.01 | 98.5 | 101.5 | 8.65 | 12.09 | | High | High | High | High | High | High | High | High | High | High | High | 0 | 1.82E-06 | 3.42 |
| FALSE | High | HPVLDESLI | [R].HPVLDI | 1xPhosphc | Q9D666 1x | 4.59E-05 | 0 | 1 | 9 | 1 | Q9D666 | SUN doma | Q9D666 [1 | 0 | 1487.782 | 0 | 0.58 | 0.040415 | 25.16 | 72.9 | 127.1 | 15.29 | 29.32 | | High | High | High | High | High | High | High | High | High | High | High | 0 | 7.48E-06 | 2.88 |
| FALSE | High | HPVVAGGS | [R].HPVVA | 1xPhosphc | Q6ZQ58 1x | 3.2E-05 | 0 | 1 | 3 | 1 | Q6ZQ58 | La-related | Q6ZQ58 [1 | 0 | 1431.694 | 0 | 0.06 | 0.74443 | 8.7 | 98.2 | 101.8 | 5.89 | 3.6 | | High | High | High | High | High | High | High | High | High | High | High | 0 | 4.99E-06 | 2.86 |
| FALSE | High | HQASINELI | [R].HQASIN | 1xPhosphc | A2AUK5 1x | 4.4E-06 | 0 | 2 | 10 | 2 | A2AUK5; C | Band 4.1-li | A2AUK5 [5 | 0 | 1577.845 | 0 | 0.47 | 0.399037 | 16.37 | 86.6 | 113.4 | 13.69 | 37.15 | Shared | High | High | High | High | High | High | High | High | High | High | High | 0 | 5.39E-07 | 2.79 |
| FALSE | High | HQASISELK | [K].HQASIS | 1xPhosphc | B6ZHD0 1x | 1.8E-05 | 0 | 1 | 5 | 1 | B6ZHD0 | Erythrocyt | B6ZHD0 [6 | 0 | 1550.834 | 0 | 2.72 | 0.035831 | 21.22 | 25.7 | 174.3 | 13.98 | 52.28 | | High | High | High | High | High | High | High | High | High | High | High | 0 | 2.61E-06 | 2.54 |
| FALSE | High | HQDGLPYII | [R].HQDGL | 2xPhosphc | Q6PAJ1 2xl | 0.001107 | 0 | 1 | 2 | 1 | Q6PAJ1 | Breakpoint | Q6PAJ1 [4 | 0 | 2656.153 | 0 | 1.07 | 0.005343 | 30.06 | 68.5 | 131.5 | 10.57 | 13.56 | | High | High | High | High | High | High | High | High | High | High | High | 0 | 0.000268 | 2.74 |
| FALSE | High | HQEDQDPI | [R].HQEDC | 1xPhosphc | B2RT77 1xl | 1.09E-05 | 0 | 1 | 1 | 4 | B2RT77 | TWIST neig | B2RT77 [2 | 0 | 2942.243 | 0 | -0.11 | 0.194194 | 13.2 | 107.3 | 92.7 | 12.32 | 10.32 | | High | High | High | High | High | High | High | High | High | High | High | 0 | 1.5E-06 | 5.33 |
| FALSE | High | HQEDQDPI | [R].HQEDC | 1xDeamida | B2RT77 1xl | 0.049549 | 0.00361 | 1 | 1 | 2 | B2RT77 | TWIST neig | B2RT77 [2 | 0 | 2943.227 | 0 | -0.25 | 0.05264 | 11.08 | 104 | 96 | 10.29 | 10.93 | | High | High | High | High | High | High | High | High | High | High | High | 0.003621 | 0.03851 | 2.65 |
| FALSE | High | HQEDQDPI | [R].HQEDC | 1xPhosphc | B2RT77 1xl | 0.003083 | 0.00016 | 1 | 1 | 1 | B2RT77 | TWIST neig | B2RT77 [2 | 0 | 3171.406 | 0 | 0.26 | 0.360527 | 11.65 | 93.7 | 106.3 | 11.87 | 78.77 | | High | High | High | High | High | High | High | High | High | High | High | 7.19E-05 | 0.000844 | 2.21 |
| FALSE | High | HQEESDSD | [K].HQEESI | 1xPhosphc | Q80WQ8 1 | 0.000102 | 0 | 1 | 2 | 2 | Q80WQ8 | Mis18-binc | Q80WQ8 [ | 0 | 2480.134 | 0 | -0.84 | 0.029722 | 10.18 | 126.5 | 73.5 | 18.74 | 15.62 | | High | High | High | High | High | High | High | High | High | High | High | 0 | 1.85E-05 | 3.26 |
| FALSE | High | HQEESDSD | [K].HQEESI | 2xPhosphc | Q80WQ8 2 | 0.005978 | 0.00062 | 1 | 2 | 2 | Q80WQ8 | Mis18-binc | Q80WQ8 [ | 0 | 2560.101 | 0 | -0.71 | 0.050844 | 10.84 | 121.4 | 78.6 | 14.75 | 24.33 | | High | High | High | High | High | High | High | High | High | High | High | 0.000859 | 0.008531 | 3.02 |
| FALSE | High | HQEETAHA | [K].HQEET/ | 1xCarbami | E9QAU4 1x | 6.13E-05 | 0 | 1 | 12 | 6 | E9QAU4 | Enhancer t | E9QAU4 [9 | 0 | 3078.45 | 0 | -0.2 | 0.079585 | 2.59 | 107.3 | 92.7 | 2.47 | 16.82 | | High | High | High | High | High | High | High | High | High | High | High | 0 | 1.04E-05 | 2.07 |
| FALSE | High | HQIVCINNI | [R].HQIVCI | 1xCarbami | Q3TAH3 3x | 0.000325 | 0 | 1 | 2 | 3 | Q3TAH3 | Putative ur | Q3TAH3 [1 | 0 | 3445.315 | 0 | -0.57 | 0.20352 | 48.82 | 119.4 | 80.6 | 35.09 | 51.76 | | High | High | High | High | High | High | High | High | High | High | High | 0 | 6.78E-05 | 4.48 |
| FALSE | High | HQIVCINNI | [R].HQIVCI | 1xCarbami | Q3TAH3 2x | 1.01E-05 | 0 | 1 | 2 | 4 | Q3TAH3 | Putative ur | Q3TAH3 [1 | 0 | 3594.512 | 0 | -0.41 | 0.004387 | 2.95 | 109.3 | 90.7 | 9.17 | 13.46 | | High | High | High | High | High | High | High | High | High | High | High | 0 | 1.52E-05 | 4.26 |
| FALSE | High | HQMLEISLS | [K].HQMLE | 1xPhosphc | O09130 1x | 1.03E-05 | 0 | 1 | 1 | 1 | O09130 | NFATC2-int | O09130 [3 | 0 | 2249.165 | 0 | -1.45 | 0.000398 | 0.44 | 146.5 | 53.5 | 5.73 | 7.02 | | High | High | High | High | High | High | High | High | High | High | High | 0 | 1.39E-06 | 3.77 |
| FALSE | High | HQNQDNS | [R].HQNQ | 1xPhosphc | P33174 1xl | 2.94E-07 | 0 | 1 | 2 | 1 | P33174 | Chromoso | P33174 [11 | 0 | 2971.347 | 0 | -0.09 | 0.159941 | 11.42 | 105 | 95 | 5.83 | 10.79 | | High | High | High | High | High | High | High | High | High | High | High | 0 | 2.57E-08 | 4.48 |
| FALSE | High | HQPGQEST | [R].HQPGC | 1xPhosphc | A0A0R4J22 | 4.2E-05 | 0 | 1 | 2 | 1 | A0A0R4J22 | Bernardine | A0A0R4J22 | 0 | 4893.231 | 0 | 2.04 | 0.000708 | 37.93 | 44.1 | 155.9 | 31.88 | 8.05 | | High | High | High | High | High | High | High | High | High | High | High | 0 | 6.76E-06 | 7.02 |
| FALSE | High | HQQESDFS | [R].HQQES | 1xPhosphc | B2RQC2 1x | 7.98E-06 | 0 | 1 | 2 | 2 | B2RQC2 | Ubiquitin c | B2RQC2 [1 | 0 | 2372.014 | 0 | 0.05 | 0.837663 | 36.34 | 98.1 | 101.9 | 42.36 | 22.95 | | High | High | High | High | High | High | High | High | High | High | High | 0 | 1.05E-06 | 4.08 |
| FALSE | High | HQQESDFS | [R].HQQES | 2xPhosphc | B2RQC2 2x | 1.9E-05 | 0 | 1 | 2 | 2 | B2RQC2 | Ubiquitin c | B2RQC2 [1 | 0 | 2451.981 | 0 | 0.23 | 0.311557 | 5.5 | 95 | 105 | 6.14 | 13.95 | | High | High | High | High | High | High | High | High | High | High | High | 0 | 2.78E-06 | 3.19 |
| FALSE | High | HQQESDFS | [R].HQQES | 3xPhosphc | B2RQC2 3x | 1.33E-05 | 0 | 1 | 2 | 2 | B2RQC2 | Ubiquitin c | B2RQC2 [1 | 0 | 2531.947 | 0 | 0.09 | 0.642554 | 10.11 | 98.1 | 101.9 | 4.28 | 11.69 | | High | High | High | High | High | High | High | High | High | High | High | 0 | 1.86E-06 | 4.41 |
| FALSE | High | HSEEANFE | [K].HSEEAN | 1xPhosphc | B2RT77 1xl | 0.087983 | 0.006639 | 1 | 1 | 1 | B2RT77 | TWIST neig | B2RT77 [3 | 1 | 1940.952 | 0 | -0.32 | 0.519339 | 13.96 | 105.4 | 94.6 | 12.87 | 26.6 | | High | High | High | High | High | High | High | High | High | High | High | 0.003402 | 0.03705 | 2.11 |
| FALSE | High | HTSSPEVV | [R].HTSSPE | 1xPhosphc | Q68FE6 1x | 0.000141 | 0 | 1 | 1 | 4 | Q68FE6 | Rho family | Q68FE6 [8 | 0 | 1635.758 | 0 | 0.13 | 0.809082 | 7.38 | 96 | 104 | 11.57 | 13.42 | | High | High | High | High | High | High | High | High | High | High | High | 0.000449 | 0.003986 | 3.09 |
| FALSE | High | HTSVVSSG | [R].HTSVVS | 1xPhosphc | Q5RIM6 1x | 2.44E-05 | 0 | 1 | 10 | 2 | Q5RIM6 | Nuclear re | Q5RIM6 [1 | 0 | 1634.846 | 0 | -0.73 | 0.005498 | 3.72 | 125.3 | 74.7 | 14.25 | 9.26 | | High | High | High | High | High | High | High | High | High | High | High | 0 | 3.7E-06 | 3.4 |
| FALSE | High | HTVLYISPP | [R].HTVLYI | 1xPhosphc | Q9WV30-2 | 0.003167 | 0.00016 | 1 | 9 | 1 | Q9WV30-2 | Isoform 2 c | Q9WV30-2 | 0 | 2375.185 | 0 | 0.92 | 0.011655 | 7.88 | 69.9 | 130.1 | 4.34 | 15.66 | | High | High | High | High | High | High | High | High | High | High | High | 7.19E-05 | 0.000868 | 2.16 |
| FALSE | High | LAYLSDEM | [R].LAYLSD | 1xPhosphc | Q9CSH3 1x | 0.000563 | 0 | 1 | 2 | 4 | Q9CSH3 | Exosome c | Q9CSH3 [2 | 0 | 2253.076 | 0 | -0.92 | 0.000445 | 0.64 | 132.3 | 67.7 | 4.13 | 8.4 | | High | High | High | High | High | High | High | High | High | High | High | 7.19E-05 | 0.000778 | 3.3 |
| FALSE | High | IAYQSSASP | [R].IAYQSS | 1xPhosphc | Q91VX2 1x | 1.65E-09 | 0 | 1 | 2 | 10 | Q91VX2 | Ubiquitin-a | Q91VX2 [6 | 0 | 2649.226 | 0 | -0.06 | 0.227824 | 8 | 109.9 | 90.1 | 11.93 | 12.38 | | High | High | High | High | High | High | High | High | High | High | High | 0 | 4.6E-10 | 3.43 |
| FALSE | High | IAYQSSASP | [R].IAYQSS | 1xDeamida | Q91VX2 1x | 1.29E-09 | 0 | 1 | 2 | 11 | Q91VX2 | Ubiquitin-a | Q91VX2 [6 | 0 | 2650.21 | 0 | -0.36 | 0.007281 | 5.07 | 110.4 | 89.6 | 5.68 | 6.28 | | High | High | High | High | High | High | High | High | High | High | High | 0 | 3.15E-09 | 4.17 |
| FALSE | High | LAYVTPTIP | [R].LAYVTF | 1xPhosphc | Q9DBR7 1x | 0.013666 | 0.001011 | 1 | 3 | 1 | Q9DBR7 | Protein ph | Q9DBR7 [4 | 0 | 1439.786 | 0 | -1.13 | 0.001797 | 0.9 | 138.7 | 61.3 | 17.62 | 23.91 | | High | High | High | High | High | High | High | High | High | High | High | 0.000538 | 0.004462 | 2.14 |
| FALSE | High | LCAGIMITA | [K].LCAGIN | 1xCarbami | Q7TSV4 1x | 9.98E-05 | 0 | 1 | 2 | 1 | Q7TSV4 | Phosphogl | Q7TSV4 [1 | 0 | 2051.058 | 0 | -0.27 | 0.590667 | 17.91 | 109.5 | 90.5 | 21.03 | 9.89 | | High | High | High | High | High | High | High | High | High | High | High | 0 | 1.79E-05 | 2.52 |
| FALSE | High | LCAGIMITA | [K].LCAGIN | 1xCarbami | Q7TSV4 1x | 0.000349 | 0 | 1 | 2 | 1 | Q7TSV4 | Phosphogl | Q7TSV4 [1 | 0 | 2067.053 | 0 | -0.23 | 0.671967 | 30.02 | 101.6 | 98.4 | 21.2 | 19.01 | | High | High | High | High | High | High | High | High | High | High | High | 0 | 7.34E-05 | 2.02 |
| FALSE | High | LCDFGSASI | [K].LCDFGS | 1xCarbami | C7G3P2 1x | 1.12E-06 | 0 | 1 | 3 | 3 | C7G3P2 | MKIAA053 | C7G3P2 [8 | 0 | 2746.275 | 0 | -0.04 | 0.469404 | 26.39 | 100.4 | 99.6 | 10.66 | 16.62 | | High | High | High | High | High | High | High | High | High | High | High | 0 | 1.16E-07 | 4.65 |
| FALSE | High | LCDFGSASI | [K].LCDFGS | 1xCarbami | C7G3P2 1x | 0.000203 | 0 | 1 | 3 | 3 | C7G3P2 | MKIAA053 | C7G3P2 [8 | 0 | 2747.259 | 0 | -0.01 | 0.33427 | 2.47 | 101.9 | 98.1 | 6.08 | 12.27 | | High | High | High | High | High | High | High | High | High | High | High | 0 | 4E-05 | 4.92 |
| FALSE | High | LCDFGVSG | [K].LCDFGV | 1xCarbami | P31938 1xl | 0.000411 | 0 | 1 | 7 | 1 | P31938 | Dual specif | P31938 [2 | 0 | 2683.246 | 0 | 1.53 | 0.062644 | 63.6 | 48.6 | 151.4 | 11.96 | 48.62 | | High | High | High | High | High | High | High | High | High | High | High | 0 | 8.77E-05 | 3.44 |
| FALSE | High | LCEPPVLSS | [R].LCEPPV | 1xCarbami | Q8BJ05 1xl | 0.000125 | 0 | 1 | 3 | 2 | Q8BJ05 | Zinc finger | Q8BJ05 [2 | 0 | 2552.198 | 0 | 0.6 | 0.050085 | 7.36 | 82.2 | 117.8 | 13.14 | 16.66 | | High | High | High | High | High | High | High | High | High | High | High | 0 | 2.32E-05 | 3.52 |
| FALSE | High | LCIDGYPSP | [R].LCIDGY | 1xCarbami | Q3TJB1 2xl | 0.026885 | 0.001676 | 1 | 5 | 2 | Q3TJB1 | Putative ur | Q3TJB1 [4 | 0 | 2403.184 | 0 | -0.66 | 0.027488 | 6.22 | 132.1 | 67.9 | 17.04 | 23.5 | | High | High | High | High | High | High | High | High | High | High | High | 0.000903 | 0.009541 | 2.53 |
| FALSE | High | LCIEVTPQS | [K].LCIEVTI | 1xCarbami | P61222 1xl | 0.016451 | 0.001124 | 1 | 4 | 1 | P61222 | ATP-bindin | P61222 [3 | 0 | 1712.906 | 0 | -0.94 | 0.000317 | 1.06 | 133.4 | 66.6 | 5.19 | 5.81 | | High | High | High | High | High | High | High | High | High | High | High | 0.000585 | 0.005502 | 2.75 |
| FALSE | High | ICIGESHGS | [R].ICIGESH | 1xCarbami | Q3UKS3 1x | 9.2E-05 | 0 | 1 | 2 | 1 | Q3UKS3 | Putative ur | Q3UKS3 [3 | 0 | 2358.03 | 0 | 1.54 | 0.013787 | 46.98 | 47.7 | 152.3 | 6.33 | 28.56 | | High | High | High | High | High | High | High | High | High | High | High | 0 | 1.64E-05 | 2.92 |
| FALSE | High | LAYLSDEM | [R].LAYLSD | 1xPhosphc | Q9CSH3 1x | 0.000125 | 0 | 1 | 2 | 2 | Q9CSH3 | Exosome c | Q9CSH3 [2 | 0 | 2237.081 | 0 | -1.21 | 0.000748 | 3.12 | 140.3 | 59.7 | 7.77 | 13 | | High | High | High | High | High | High | High | High | High | High | High | 0 | 3.25E-05 | 3.14 |
| FALSE | High | ICIGESHGS | [R].ICIGESH | 1xCarbami | Q3UKS3 1x | 0.000181 | 0 | 1 | 2 | 2 | Q3UKS3 | Putative ur | Q3UKS3 [3 | 0 | 2374.025 | 0 | 2.12 | 0.001888 | 10.81 | 34.4 | 165.6 | 10.02 | 18.63 | | High | High | High | High | High | High | High | High | High | High | High | 0 | 0.00022 | 2.96 |
| FALSE | High | LCLSPTMSI | [K].LCLSPT | 1xCarbami | Q7TT79 1x | 2.14E-06 | 0 | 1 | 5 | 2 | Q7TT79 | Microceph | Q7TT79 [3 | 0 | 2222.091 | 0 | -0.84 | 0.001455 | 1.98 | 127.3 | 72.7 | 7.62 | 9.29 | | High | High | High | High | High | High | High | High | High | High | High | 0 | 3.66E-07 | 4.93 |
| FALSE | High | LCLSPTMSI | [K].LCLSPT | 1xCarbami | Q7TT79 1x | 1.9E-05 | 0 | 1 | 5 | 2 | Q7TT79 | Microceph | Q7TT79 [3 | 0 | 2238.086 | 0 | -0.65 | 0.001079 | 0.1 | 122.3 | 77.7 | 4.1 | 4.25 | | High | High | High | High | High | High | High | High | High | High | High | 0 | 2.79E-06 | 3.18 |
| FALSE | High | LCLSTVDLE | [K].LCLSTV | 1xCarbami | Q8CGA3 1 | 9.08E-07 | 0 | 1 | 1 | 1 | Q8CGA3 | Large neut | Q8CGA3 [2 | 0 | 1814.974 | 0 | 0.39 | 0.19041 | 25.32 | 79 | 121 | 13.45 | 21.29 | | High | High | High | High | High | High | High | High | High | High | High | 0 | 9.1E-08 | 4 |
| FALSE | High | LCPADDSIK | [R].LCPADI | 1xCarbami | Q3ULL6 1x | 0.00614 | 0.00062 | 1 | 3 | 2 | Q3ULL6 | Putative ur | Q3ULL6 [4 | 0 | 2581.319 | 0 | 0.22 | 0.011109 | 4.57 | 92.4 | 107.6 | 4.35 | 3.96 | | High | High | High | High | High | High | High | High | High | High | High | 0.000449 | 0.003933 | 2.1 |
| FALSE | High | ICPDPGSDI | [K].ICPDPG | 1xCarbami | Q3UHL0 1x | 0.000229 | 0 | 1 | 6 | 1 | Q3UHL0 | Putative ur | Q3UHL0 [1 | 0 | 1808.809 | 0 | 0.49 | 0.023737 | 17.87 | 82.5 | 117.5 | 8.4 | 10.81 | | High | High | High | High | High | High | High | High | High | High | High | 0 | 4.57E-05 | 2.64 |
| FALSE | High | LCPQPEEP | [R].LCPQPE | 1xCarbami | A0A0A0MC | 1.46E-06 | 0 | 1 | 3 | 2 | A0A0A0MC | Histone-lys | A0A0A0MC | 0 | 2475.146 | 0 | 0.4 | 0.204146 | 25.8 | 86.3 | 113.7 | 18.11 | 18.46 | | High | High | High | High | High | High | High | High | High | High | High | 0 | 1.69E-07 | 4.8 |
| FALSE | High | LCPQPEEP | [R].LCPQPE | 1xCarbami | A0A0A0MC | 0.019966 | 0.001438 | 1 | 3 | 2 | A0A0A0MC | Histone-lys | A0A0A0MC | 0 | 2476.13 | 0 | 0.7 | 0.086759 | 56.71 | 75.5 | 124.5 | 24.97 | 27.95 | | High | High | High | High | High | High | High | High | High | High | High | 0.002403 | 0.0257 | 2.43 |
| FALSE | High | LCQPQNAT | [K].LCQPQ | 1xCarbami | O08856 1x | 0.0008 | 0 | 1 | 1 | 3 | O08856 | RNA polym | O08856 [2 | 0 | 1980.905 | 0 | -0.17 | 0.105965 | 9.49 | 103.5 | 96.5 | 7.53 | 17.21 | | High | High | High | High | High | High | High | High | High | High | High | 0 | 0.000187 | 2.75 |
| FALSE | High | LCQPQNAT | [K].LCQPQ | 1xCarbami | O08856 1x | 0.020862 | 0.001472 | 1 | 1 | 2 | O08856 | RNA polym | O08856 [2 | 1 | 2460.129 | 0 | -0.47 | 0.017258 | 7.94 | 114.2 | 85.8 | 8.72 | 11.47 | | High | High | High | High | High | High | High | High | High | High | High | 0.000786 | 0.007121 | 1.92 |
| FALSE | High | LCSDSPK | [K].LCSDSP | 1xCarbami | Q8K4J0 1xl | 0.033694 | 0.002195 | 1 | 1 | 1 | Q8K4J0 | Protein art | Q8K4J0 [5 | 0 | 1344.663 | 0 | -0.54 | 0.025445 | 8.43 | 118.4 | 81.6 | 8.51 | 16.84 | | High | High | High | High | High | High | High | High | High | High | High | 0.001151 | 0.0123 | 1.73 |
| FALSE | High | ICSFEEAK | [R].ICSFEE/ | 1xCarbami | P48453 1xl | 0.012288 | 0.000889 | 1 | 4 | 1 | P48453 | Serine/thre | P48453 [4 | 0 | 1521.742 | 0 | 0.33 | 0.052108 | 23.05 | 88.7 | 111.3 | 6.45 | 12.61 | | High | High | High | High | High | High | High | High | High | High | High | 0.000449 | 0.003945 | 2.34 |
| FALSE | High | LCSSSSSDT | [R].LCSSSS | 1xCarbami | Q80YV2 1x | 8.03E-05 | 0 | 1 | 4 | 8 | Q80YV2 | Nuclear-int | Q80YV2 [4 | 0 | 1592.682 | 0 | -0.54 | 0.003309 | 2.37 | 117.2 | 82.8 | 2.07 | 8.57 | | High | High | High | High | High | High | High | High | High | High | High | 0 | 1.41E-05 | 2.41 |
| FALSE | High | LCSSSSSDT | [R].LCSSSS | 1xCarbami | Q80YV2 2x | 0.030104 | 0.001857 | 1 | 4 | 3 | Q80YV2 | Nuclear-int | Q80YV2 [4 | 0 | 1672.649 | 0 | -0.32 | 0.008328 | 9.52 | 110.9 | 89.1 | 3.66 | 8.1 | | High | High | High | High | High | High | High | High | High | High | High | 0.001011 | 0.01078 | 2.18 |
| FALSE | High | LCLPENESF | [K].LCLPEN | 1xCarbami | S4R221 1xl | 1.75E-05 | 0 | 1 | 4 | 2 | S4R221 | Rho GTPas | S4R221 [4 | 0 | 2097.034 | 0 | -0.47 | 0.004171 | 4.04 | 114.7 | 85.3 | 3.6 | 6.68 | | High | High | High | High | High | High | High | High | High | High | High | 0 | 2.53E-06 | 2.47 |
| FALSE | High | LCTDSLPET | [R].LCTDSL | 2xCarbami | A0A0A0MC | 0.002486 | 0.000109 | 1 | 3 | 3 | A0A0A0MC | Histone-lys | A0A0A0MC | 0 | 3152.442 | 0 | 0.22 | 0.022417 | 8.5 | 95.3 | 104.7 | 23.37 | 18.17 | | High | High | High | High | High | High | High | High | High | High | High | 4.94E-05 | 0.000662 | 3.14 |
| FALSE | High | LAWVGDV | [K].LAWVG | 2xPhosphc | Q7TSC1 2x | 0.00824 | 0.000736 | 1 | 3 | 1 | Q7TSC1 | Protein PRI | Q7TSC1 [7 | 0 | 2929.446 | 0 | 1.62 | 0.010878 | 47.47 | 43.3 | 156.7 | 12.52 | 28.35 | | High | High | High | High | High | High | High | High | High | High | High | 0.000327 | 0.002534 | 2.59 |
| FALSE | High | LAWTLTGN | [R].LAWTL | 1xPhosphc | P58854 1xl | 6.18E-05 | 0 | 1 | 5 | 1 | P58854 | Gamma-tu | P58854 [2 | 0 | 2455.263 | 0 | 0.13 | 0.44178 | 20.37 | 97.7 | 102.3 | 8.42 | 10.85 | | High | High | High | High | High | High | High | High | High | High | High | 0 | 1.05E-05 | 3.37 |
| FALSE | High | LASEYLSPE | [R].LASEYL | 1xPhosphc | Q9Z315 1x | 0.004451 | 0.000409 | 1 | 1 | 2 | Q9Z315 | U4/U6.U5 | Q9Z315 [3 | 0 | 2298.138 | 0 | -0.27 | 0.373708 | 11.94 | 109.5 | 90.5 | 1.81 | 22.86 | | High | High | High | High | High | High | High | High | High | High | High | 0.000183 | 0.001271 | 2.94 |
| FALSE | High | LASGAQSIF | [R].LASGA | 1xPhosphc | Q9CQ13 1 | 0.0008 | 0 | 1 | 1 | 2 | Q9CQ13 | Coordinat | Q9CQ13 [4 | 0 | 3532.725 | 0 | -0.81 | 0.001041 | 3.07 | 127.9 | 72.1 | 7.04 | 14.71 | | High | High | High | High | High | High | High | High | High | High | High | 0 | 0.000185 | 3.21 |
| FALSE | High | LASGDGDE | [R].LASGD | 1xPhosphc | Q80Y44 1x | 1.44E-07 | 0 | 1 | 6 | 10 | Q80Y44 | Probable A | Q80Y44 [5 | 0 | 3315.395 | 0 | -0.56 | 0.001464 | 4.48 | 119.2 | 80.8 | 2.4 | 8.1 | | High | High | High | High | High | High | High | High | High | High | High | 0 | 7.06E-07 | 5.19 |
| FALSE | High | LASGDGDE | [R].LASGD | 1xDeamida | Q80Y44 1x | 0.000219 | 0 | 1 | 6 | 3 | Q80Y44 | Probable A | Q80Y44 [5 | 0 | 3316.379 | 0 | -0.68 | 0.000222 | 2.27 | 123.1 | 76.9 | 3.07 | 8.88 | | High | High | High | High | High | High | High | High | High | High | High | 0 | 4.34E-05 | 4.85 |
| FALSE | High | LASGSSELA | [R].LASGSS | 1xPhosphc | F8WI30 1x | 3.82E-07 | 0 | 1 | 1 | 3 | F8WI30 | Sorting ne | F8WI30 [6 | 0 | 1768.868 | 0 | 0.85 | 0.001084 | 6.33 | 72.2 | 127.8 | 7.74 | 5.57 | | High | High | High | High | High | High | High | High | High | High | High | 0 | 4.13E-07 | 3.26 |
| FALSE | High | LASGSSELA | [R].LASGSS | 2xPhosphc | F8WI30 2x | 0.000369 | 0 | 1 | 1 | 4 | F8WI30 | Sorting ne | F8WI30 [6 | 0 | 1848.834 | 0 | 1.39 | 0.000363 | 4.53 | 55.2 | 144.8 | 8.39 | 5.75 | | High | High | High | High | High | High | High | High | High | High | High | 0 | 7.82E-05 | 3.88 |
| FALSE | High | LASLNAEAI | [R].LASLNA | 1xPhosphc | Q497V6 1x | 0.000168 | 0 | 1 | 4 | 1 | Q497V6 | Bromo adj | Q497V6 [1 | 0 | 2063.11 | 0 | 1.35 | 0.007307 | 36.4 | 60 | 140 | 17.42 | 15.41 | | High | High | High | High | High | High | High | High | High | High | High | 0 | 3.23E-05 | 3.79 |
| FALSE | High | LASNSPALF | [R].LASNSF | 1xPhosphc | Q3THC1 1x | 0.00047 | 0 | 1 | 1 | 1 | Q3THC1 | Proteasom | Q3THC1 [1 | 0 | 1751.904 | 0 | 1.23 | 0.00235 | 10.23 | 64.2 | 135.8 | 7.43 | 11.49 | | High | High | High | High | High | High | High | High | High | High | High | 0 | 0.000103 | 2.95 |
| FALSE | High | LASPQPSYA | [K].LASPQF | 1xPhosphc | Q6PAL7 1x | 7.1E-06 | 0 | 1 | 1 | 2 | Q6PAL7 | AT-hook D | Q6PAL7 [5 | 0 | 2173.058 | 0 | -1.27 | 0.000181 | 0.85 | 140.9 | 59.1 | 2.89 | 9.71 | | High | High | High | High | High | High | High | High | High | High | High | 0 | 9.19E-07 | 3.8 |
| FALSE | High | LASPSGSTS | [R].LASPSG | 1xCarbami | Q62318 1x | 5.55E-05 | 0 | 1 | 2 | 18 | Q62318 | Transcripti | Q62318 [5 | 0 | 3993.957 | 0 | -1.82 | 0.020955 | 3.85 | 151.9 | 48.1 | 32.12 | 21.42 | | High | High | High | High | High | High | High | High | High | High | High | 0 | 1.01E-05 | 4.81 |
| FALSE | High | LASPSGSTS | [R].LASPSG | 1xCarbami | Q62318 2x | 3.32E-05 | 0 | 1 | 2 | 29 | Q62318 | Transcripti | Q62318 [5 | 0 | 4073.923 | 0 | -1.62 | 0.026644 | 9.43 | 147.2 | 52.8 | 31.89 | 44.66 | | High | High | High | High | High | High | High | High | High | High | High | 2.58E-05 | 0.000488 | 5.08 |
| FALSE | High | LASPVAPG | [R].LASPVA | 1xPhosphc | Q8CHI8 1xl | 0.000418 | 0 | 1 | 5 | 1 | Q8CHI8 | E1A-bindin | Q8CHI8 [2 | 0 | 2940.514 | 0 | -0.87 | 0.002826 | 2.47 | 128.5 | 71.5 | 8.86 | 10.69 | | High | High | High | High | High | High | High | High | High | High | High | 0 | 9.01E-05 | 3.61 |
| FALSE | High | LASSLGNLI | [R].LASSLG | 1xPhosphc | Q6GQT0 1 | 8.68E-07 | 0 | 1 | 2 | 2 | Q6GQT0 | Dual specif | Q6GQT0 [4 | 0 | 2369.2 | 0 | 1.61 | 0.001466 | 19.82 | 45.1 | 154.9 | 9.01 | 12.76 | | High | High | High | High | High | High | High | High | High | High | High | 0 | 1.63E-07 | 4.29 |
| FALSE | High | LAWVGDV | [K].LAWVG | 1xPhosphc | Q7TSC1 1x | 7.18E-07 | 0 | 1 | 3 | 5 | Q7TSC1 | Protein PRI | Q7TSC1 [7 | 0 | 2849.48 | 0 | 1.3 | 0.018556 | 26.48 | 57.8 | 142.2 | 12.22 | 27.1 | | High | High | High | High | High | High | High | High | High | High | High | 0 | 2.32E-07 | 4.02 |
| FALSE | High | IASSTTPAK | [K].IASSTI | 1xPhosphc | P97868 1xl | 0.001386 | 5.66E-05 | 1 | 3 | 2 | P97868 | E3 ubiquiti | P97868 [1 | 0 | 1413.775 | 0 | -0.76 | 0.00085 | 1.83 | 127.3 | 72.7 | 3.7 | 7.53 | | High | High | High | High | High | High | High | High | High | High | High | 2.58E-05 | 0.000345 | 2.47 |
| FALSE | High | LASTDDGY | [R].LASTDI | 1xPhosphc | Q8CIB6 1xl | 0.000908 | 0 | 1 | 1 | 3 | Q8CIB6 | Transmem | Q8CIB6 [2 | 0 | 2124.067 | 0 | 1.76 | 0.000377 | 18.08 | 45.5 | 154.5 | 10.13 | 5.67 | | High | High | High | High | High | High | High | High | High | High | High | 0.000275 | 0.001887 | 2.66 |
| FALSE | High | LASTSDIEE | [R].LASTSD | 1xPhosphc | Q9DBR7 1x | 0.003464 | 0.00021 | 1 | 3 | 1 | Q9DBR7 | Protein ph | Q9DBR7 [5 | 0 | 1401.671 | 0 | -0.16 | 0.266795 | 16.87 | 108.3 | 91.7 | 11.78 | 12.09 | | High | High | High | High | High | High | High | High | High | High | High | 9.42E-05 | 0.000963 | 3.53 |
| FALSE | High | LASTSDIEE | [R].LASTSD | 1xPhosphc | Q9DBR7 1x | 0.000131 | 0 | 1 | 3 | 5 | Q9DBR7 | Protein ph | Q9DBR7 [5 | 0 | 1630.834 | 0 | -0.21 | 0.009539 | 5.51 | 110.5 | 89.5 | 5.85 | 4.71 | | High | High | High | High | High | High | High | High | High | High | High | 0 | 9.86E-05 | 3.15 |
| FALSE | High | LASTSDIEE | [R].LASTSD | 2xPhosphc | Q9DBR7 2x | 0.001783 | 5.66E-05 | 1 | 3 | 1 | Q9DBR7 | Protein ph | Q9DBR7 [5 | 0 | 1710.8 | 0 | 0.54 | 0.015798 | 11.16 | 83.8 | 116.2 | 7.71 | 13.03 | | High | High | High | High | High | High | High | High | High | High | High | 2.58E-05 | 0.000456 | 3.24 |
| FALSE | High | LASVPAGG | [K].LASVPA | 1xPhosphc | P99027 1xl | 3.46E-07 | 0 | 1 | 1 | 1 | P99027 | 60S acidic | P99027 [6 | 0 | 3312.724 | 0 | 0.59 | 0.265943 | 35.8 | 82.9 | 117.1 | 15.48 | 27.59 | | High | High | High | High | High | High | High | High | High | High | High | 0 | 3.08E-08 | 4.68 |
| FALSE | High | LASWTPYP | [R].LASWT | 1xPhosphc | B9EKK8 1xl | 0.006306 | 0.00062 | 1 | 4 | 1 | B9EKK8 | Uncharacte | B9EKK8 [6 | 0 | 2542.294 | 0 | 0.43 | 0.002501 | 12.06 | 84.4 | 115.6 | 12.61 | 6.3 | | High | High | High | High | High | High | High | High | High | High | High | 0.000275 | 0.001878 | 3.43 |

| | | | | | | | | | | | | | | | | | | | | | | | | | | | | | | | | | | | | | | |
|---|---|---|---|---|---|---|---|---|---|---|---|---|---|---|---|---|---|---|---|---|---|---|---|---|---|---|---|---|---|---|---|---|---|---|---|---|---|---|
| FALSE | High | LATAAEQT | [K].LATAAE | 1xCarbami | E9PVX6 1x | 0.035181 | 0.002269 | 1 | 3 | 3 | E9PVX6 | Proliferatic | E9PVX6 [42 | 0 | 3490.718 | 0 | -0.45 | 0.055133 | 14.16 | 118 | 82 | 12.43 | 14.76 | | High | High | High | High | High | High | High | High | High | High | High | 0.003521 | 0.03773 | 2.61 |
| FALSE | High | LATLFGDVI | [R].LATLFG | 1xPhospho | Q0VBD2 1x | 0.000141 | 0 | 1 | 1 | 9 | Q0VBD2 | Protein MC | Q0VBD2 [7 | 0 | 2677.326 | 0 | -0.35 | 0.003249 | 5.36 | 112.9 | 87.1 | 2.4 | 5.39 | | High | High | High | High | High | High | High | High | High | High | High | 0 | 0.000111 | 3.57 |
| FALSE | High | LATSSPEQS | [R].LATSSP | 1xPhospho | Q60953 1x | 0.000196 | 0 | 2 | 6 | 4 | Q60953; Q | Protein PM | Q60953 [5 | 0 | 2204.104 | 0 | -0.23 | 0.052355 | 10.68 | 109.3 | 90.7 | 4.53 | 10.22 | Shared | High | High | High | High | High | High | High | High | High | High | High | 0 | 3.84E-05 | 2.65 |
| FALSE | High | LATSSPEQS | [R].LATSSP | 2xPhospho | Q60953 2x | 0.001751 | 5.66E-05 | 2 | 6 | 13 | Q60953; Q | Protein PM | Q60953 [5 | 0 | 2284.07 | 0 | -0.02 | 0.333089 | 0.84 | 102 | 98 | 4.58 | 9.84 | Shared | High | High | High | High | High | High | High | High | High | High | High | 0.000859 | 0.008626 | 3.32 |
| FALSE | High | IATTTTGTA | [R].IATTTT | 1xPhospho | E9Q7G0 1x | 0.002463 | 0.000109 | 1 | 5 | 1 | E9Q7G0 | Nuclear mi | E9Q7G0 [2 | 0 | 1871.968 | 0 | 0.94 | 0.006486 | 33.64 | 67.8 | 132.2 | 22.41 | 12.46 | | High | High | High | High | High | High | High | High | High | High | High | 4.94E-05 | 0.000655 | 2.23 |
| FALSE | High | LATTVSAPI | [R].LATTVS | 1xPhospho | P27546 1x | 0.000215 | 0 | 1 | 12 | 2 | P27546 | Microtubu | P27546 [89 | 0 | 1653.923 | 0 | 1.86 | 0.007622 | 22.11 | 40.7 | 159.3 | 9.74 | 26.54 | | High | High | High | High | High | High | High | High | High | High | High | 0 | 4.23E-05 | 2.53 |
| FALSE | High | IAVDLSDQI | [K].IAVDLS | 1xPhospho | A0A0R4IZV | 4.96E-07 | 0 | 1 | 11 | 7 | A0A0R4IZV | tRNA (guar | A0A0R4IZV | 0 | 2143.057 | 0 | -0.36 | 0.006216 | 5.3 | 113.3 | 86.7 | 4.18 | 6.28 | | High | High | High | High | High | High | High | High | High | High | High | 0 | 4.62E-08 | 3.36 |
| FALSE | High | IASSYHSEG | [R].IASSYH | 1xPhospho | E9Q9R9 1x | 7.41E-05 | 0 | 1 | 5 | 1 | E9Q9R9 | Disks large | E9Q9R9 [1 | 0 | 2536.201 | 0 | 0.31 | 0.121868 | 18.9 | 90.4 | 109.6 | 12.33 | 21.04 | | High | High | High | High | High | High | High | High | High | High | High | 0 | 1.28E-05 | 2.2 |
| FALSE | High | ICTEQSNSF | [K].ICTEQS | 1xCarbami | Q62130 1x | 8.63E-08 | 0 | 1 | 6 | 2 | Q62130 | Tyrosine-p | Q62130 [3 | 0 | 1807.861 | 0 | 0.91 | 0.00238 | 12.92 | 69.5 | 130.5 | 8.98 | 5.47 | | High | High | High | High | High | High | High | High | High | High | High | 0 | 6.45E-09 | 4.53 |
| FALSE | High | ICTPSSPSE | [K].ICTPSS | 1xCarbami | O08736 1x | 0.025073 | 0.001645 | 1 | 5 | 2 | O08736 | Caspase-12 | O08736 [9 | 0 | 1730.844 | 0 | 0.22 | 0.448094 | 21.4 | 96.6 | 103.4 | 12.39 | 22.23 | | High | High | High | High | High | High | High | High | High | High | High | 0.000872 | 0.008804 | 2.12 |
| FALSE | High | LCYVALDFE | [K].LCYVAL | 1xCarbami | P60710 1x | 9.31E-06 | 0 | 1 | 20 | 3 | P60710 | Actin, cyto | P60710 [21 | 0 | 3088.466 | 0 | 1.11 | 0.022732 | 21.67 | 63.2 | 136.8 | 9.22 | 21.28 | | High | High | High | High | High | High | High | High | High | High | High | 0 | 1.25E-06 | 3.36 |
| FALSE | High | IDFIPVSPAI | [R].IDFIPVS | 1xPhospho | Q9DB52 1x | 0.106748 | 0.008634 | 1 | 1 | 1 | Q9DB52 | Protein FAI | Q9DB52 [1 | 0 | 1805.94 | 0 | 1.24 | 0.000975 | 9.99 | 61.4 | 138.6 | 17.17 | 3.79 | | High | High | High | High | High | High | High | High | High | High | High | 0.004327 | 0.04667 | 2.07 |
| FALSE | High | LDFPELQSF | [R].LDFPEL | 1xPhospho | Q3TGQ4 1x | 0.025738 | 0.001645 | 1 | 3 | 1 | Q3TGQ4 | Putative ur | Q3TGQ4 [2 | 0 | 1711.907 | 0 | 1.24 | 0.000218 | 5.24 | 60.3 | 139.7 | 9.32 | 6.42 | | High | High | High | High | High | High | High | High | High | High | High | 0.000872 | 0.009079 | 2.92 |
| FALSE | High | IDFSQGSGS | [R].IDFSQG | 1xPhospho | Q8VCD5 1x | 0.004372 | 0.000409 | 1 | 1 | 3 | Q8VCD5 | Mediator c | Q8VCD5 [4 | 0 | 4445.909 | 0 | 0.15 | 0.790035 | 25.02 | 94.7 | 105.3 | 10.3 | 16.98 | | High | High | High | High | High | High | High | High | High | High | High | 0.000183 | 0.001246 | 3.14 |
| FALSE | High | LDFTGNSS | [K].LDFTGN | 1xPhospho | E9PVX6 1x | 0.01186 | 0.000889 | 1 | 2 | 2 | E9PVX6 | Proliferatic | E9PVX6 [20 | 0 | 1700.841 | 0 | 0.02 | 0.818613 | 21.33 | 101 | 99 | 15.03 | 6.06 | | High | High | High | High | High | High | High | High | High | High | High | 0.000449 | 0.003787 | 2.31 |
| FALSE | High | LDFTGNST | [K].LDFTGN | 1xPhospho | E9PVX6 1x | 0.018125 | 0.001282 | 1 | 1 | 1 | E9PVX6 | Proliferatic | E9PVX6 [13 | 0 | 1714.857 | 0 | -0.08 | 0.419138 | 9.77 | 102.1 | 97.9 | 9.68 | 5.66 | | High | High | High | High | High | High | High | High | High | High | High | 0.000689 | 0.006079 | 2.29 |
| FALSE | High | IDFTPVSPA | [R].IDFTPV | 1xPhospho | Q6NZE7-2 | 0.014283 | 0.001011 | 1 | 4 | 1 | Q6NZE7-2 | Isoform 2 c | Q6NZE7-2 | 0 | 1793.904 | 0 | -0.12 | 0.008699 | 1.55 | 105.7 | 94.3 | 4.31 | 5.05 | | High | High | High | High | High | High | High | High | High | High | High | 0.000538 | 0.004671 | 1.87 |
| FALSE | High | IDFTPVSPA | [R].IDFTPV | 2xPhospho | Q6NZE7-2 | 0.000605 | 0 | 1 | 4 | 2 | Q6NZE7-2 | Isoform 2 c | Q6NZE7-2 | 0 | 1873.87 | 0 | -0.96 | 0.003378 | 0.87 | 132 | 68 | 9.26 | 11.3 | | High | High | High | High | High | High | High | High | High | High | High | 0 | 0.000136 | 2.93 |
| FALSE | High | IDGAVISES | [R].IDGAVI | 1xPhospho | Q3U955 1x | 3.87E-08 | 0 | 1 | 6 | 5 | Q3U955 | Putative ur | Q3U955 [2 | 0 | 2282.23 | 0 | 1.85 | 0.001663 | 31.42 | 39.1 | 160.9 | 21.77 | 11.69 | | High | High | High | High | High | High | High | High | High | High | High | 0 | 1.24E-05 | 3.18 |
| FALSE | High | IDGIAEAAP | [K].IDGIAE | 1xPhospho | Q149F1 1x | 1.5E-07 | 0 | 1 | 1 | 2 | Q149F1 | RNA pseud | Q149F1 [4 | 0 | 2343.2 | 0 | -0.84 | 0.001139 | 2.09 | 126 | 74 | 5.27 | 6.41 | | High | High | High | High | High | High | High | High | High | High | High | 0 | 6.74E-08 | 3.06 |
| FALSE | High | LDGLPNSPI | [R].LDGLPN | 1xPhospho | Q8BMG1 1 | 0.006478 | 0.00062 | 1 | 4 | 1 | Q8BMG1 | ATR-intera | Q8BMG1 [ | 0 | 1390.729 | 0 | -0.03 | 0.427114 | 6.57 | 100.3 | 99.7 | 9 | 2.9 | | High | High | High | High | High | High | High | High | High | High | High | 0.000275 | 0.00194 | 1.12 |
| FALSE | High | LDGLPNSPI | [R].LDGLPN | 1xPhospho | Q8BMG1 1 | 0.000674 | 0 | 1 | 4 | 1 | Q8BMG1 | ATR-intera | Q8BMG1 [ | 1 | 1747.987 | 0 | -0.2 | 0.030165 | 1.47 | 106.9 | 93.1 | 2.92 | 5.59 | | High | High | High | High | High | High | High | High | High | High | High | 0 | 0.000154 | 1.3 |
| FALSE | High | LDGLVDTP | [R].LDGLVD | 1xPhospho | E9PW66 1x | 0.000133 | 0 | 1 | 4 | 1 | E9PW66 | Nucleosom | E9PW66 [8 | 0 | 2356.245 | 0 | 0.86 | 0.000576 | 8 | 67.1 | 132.9 | 9.54 | 6.17 | | High | High | High | High | High | High | High | High | High | High | High | 0 | 2.47E-05 | 3.47 |
| FALSE | High | LDGNELDL | [K].LDGNE | 1xPhospho | Q922Q8 1x | 0.001244 | 0 | 1 | 2 | 1 | Q922Q8 | Leucine-ric | Q922Q8 [1 | 0 | 2608.352 | 0 | 0.3 | 0.013263 | 4.71 | 88 | 112 | 13.16 | 11.16 | | High | High | High | High | High | High | High | High | High | High | High | 0 | 0.000305 | 2.26 |
| FALSE | High | LDFLPEMM | [K].LDFLPE | 1xCarbami | P28659-4 1 | 0.015465 | 0.001089 | 1 | 1 | 3 | P28659-4 | Isoform 4 c | P28659-4 [ | 0 | 2960.401 | 0 | -0.37 | 0.038718 | 5.65 | 111.8 | 88.2 | 10.86 | 3.5 | | High | High | High | High | High | High | High | High | High | High | High | 0.000953 | 0.009822 | 3.36 |
| FALSE | High | IDGPPTPGS | [R].IDGPPT | 1xPhospho | F6UFZ5 1x | 0.022574 | 0.001542 | 2 | 7 | 2 | F6UFZ5; Q | PCF11 clea | F6UFZ5 [70 | 0 | 1418.724 | 0 | 0.39 | 0.023697 | 21.63 | 85.6 | 114.4 | 7.58 | 9.81 | Shared | High | High | High | High | High | High | High | High | High | High | High | 0.000814 | 0.007845 | 2.47 |
| FALSE | High | LDHINFPVF | [K].LDHINF | 1xPhospho | Q8VE18 1x | 3.47E-05 | 0 | 1 | 2 | 1 | Q8VE18 | Protein SM | Q8VE18 [6 | 0 | 2730.394 | 0 | 0.26 | 0.090219 | 12.11 | 95.2 | 104.8 | 7.23 | 8.15 | | High | High | High | High | High | High | High | High | High | High | High | 0 | 5.48E-06 | 2.84 |
| FALSE | High | LDHLSSSAF | [K].LDHLSS | 2xPhospho | Q3UT88 2x | 0.004613 | 0.000456 | 1 | 3 | 2 | Q3UT88 | Putative ur | Q3UT88 [5 | 0 | 2736.33 | 0 | 0.14 | 0.025498 | 1.14 | 97.4 | 102.6 | 6.63 | 7.53 | | High | High | High | High | High | High | High | High | High | High | High | 0.000204 | 0.001327 | 2.74 |
| FALSE | High | LDLDLTSDS | [R].LDLDLT | 1xPhospho | Q62318 1x | 0.000815 | 0 | 1 | 2 | 3 | Q62318 | Transcripti | Q62318 [4 | 0 | 2213.151 | 0 | -0.24 | 0.41778 | 11.36 | 110.1 | 89.9 | 17.32 | 9.29 | | High | High | High | High | High | High | High | High | High | High | High | 0 | 0.00019 | 2.89 |
| FALSE | High | LDIDTPDID | [K].LDIDTP | 1xPhospho | E9Q616 1x | 3.95E-09 | 0 | 1 | 2 | 2 | E9Q616 | AHNAK nuc | E9Q616 [4 | 0 | 2273.147 | 0 | 3.66 | 0.000544 | 31.51 | 14.7 | 185.3 | 14.46 | 11.24 | | High | High | High | High | High | High | High | High | High | High | High | 0 | 2.02E-10 | 4.38 |
| FALSE | High | LDIETSDVS | [K].LDIETS | 2xPhospho | E9Q616 2x | 0.009168 | 0.000736 | 1 | 3 | 1 | E9Q616 | AHNAK nuc | E9Q616 [1 | 0 | 2307.081 | 0 | 2.77 | 0.000446 | 30.45 | 25.8 | 174.2 | 41.94 | 14.59 | | High | High | High | High | High | High | High | High | High | High | High | 0.000327 | 0.002842 | 2.62 |
| FALSE | High | LDLGEDYP | [R].LDLGED | 1xPhospho | Q3TFE5 1x | 0.065952 | 0.004766 | 1 | 7 | 1 | Q3TFE5 | Putative ur | Q3TFE5 [7 | 0 | 1731.861 | 0 | 0.48 | 0.005325 | 7.33 | 87.7 | 112.3 | 5.9 | 11.51 | | High | High | High | High | High | High | High | High | High | High | High | 0.002455 | 0.0265 | 2.65 |
| FALSE | High | LDIPTGMT | [K].LDIPTG | 1xPhospho | Q6P9P6 1x | 0.02933 | 0.001857 | 1 | 1 | 1 | Q6P9P6 | Kinesin-like | Q6P9P6 [9 | 0 | 1538.749 | 0 | -0.78 | 0.003847 | 4.02 | 126.6 | 73.4 | 10.64 | 6.48 | | High | High | High | High | High | High | High | High | High | High | High | 0.000997 | 0.01049 | 2.28 |
| FALSE | High | LDIPTGMT | [K].LDIPTG | 1xPhospho | Q6P9P6 1x | 0.023584 | 0.001578 | 1 | 1 | 2 | Q6P9P6 | Kinesin-like | Q6P9P6 [9 | 0 | 1554.744 | 0 | -0.31 | 0.000818 | 2.89 | 109.6 | 90.4 | 3.04 | 6.63 | | High | High | High | High | High | High | High | High | High | High | High | 0.000859 | 0.008205 | 3.06 |
| FALSE | High | LDIPTGMT | [K].LDIPTG | 1xPhospho | Q6P9P6 1x | 0.023792 | 0.001612 | 1 | 1 | 1 | Q6P9P6 | Kinesin-like | Q6P9P6 [9 | 1 | 1896.007 | 0 | -0.44 | 0.046805 | 4.8 | 116.1 | 83.9 | 8.16 | 20.02 | | High | High | High | High | High | High | High | High | High | High | High | 0.000859 | 0.008292 | 1.98 |
| FALSE | High | LDISAPDLN | [K].LDISAP | 1xPhospho | E9Q616 1x | 1.33E-05 | 0 | 1 | 3 | 1 | E9Q616 | AHNAK nuc | E9Q616 [1 | 0 | 2206.141 | 0 | 3.56 | 0.000987 | 9.09 | 16.4 | 183.6 | 20.07 | 15.3 | | High | High | High | High | High | High | High | High | High | High | High | 0 | 1.88E-06 | 3.16 |
| FALSE | High | LDLSAYEM | [R].LDLSAY | 1xPhospho | K4DI77 1xF | 0.000884 | 0 | 1 | 4 | 1 | K4DI77 | WD repeat | K4DI77 [29 | 0 | 2954.29 | 0 | 0.78 | 0.000988 | 6.05 | 74.3 | 125.7 | 9.09 | 9.65 | | High | High | High | High | High | High | High | High | High | High | High | 0 | 0.000209 | 2.85 |
| FALSE | High | LDLSLEDR | [K].LDLSLE | 1xPhospho | Q3TN28 1x | 0.008313 | 0.000736 | 1 | 4 | 1 | Q3TN28 | Putative ur | Q3TN28 [2 | 0 | 1269.629 | 0 | -0.87 | 0.011181 | 0.21 | 128.2 | 71.8 | 14.04 | 6.35 | | High | High | High | High | High | High | High | High | High | High | High | 0.000327 | 0.002539 | 2.09 |
| FALSE | High | IDLSPESSD | [K].IDLSPES | 1xPhospho | Q7TQG0 1x | 0.036416 | 0.002419 | 1 | 1 | 1 | Q7TQG0 | Zinc finger | Q7TQG0 [3 | 0 | 1427.662 | 0 | -0.23 | 0.093151 | 3.24 | 107 | 93 | 6.44 | 9.12 | | High | High | High | High | High | High | High | High | High | High | High | 0.001279 | 0.01339 | 1.88 |
| FALSE | High | LDGPTENIS | [K].LDGPTE | 1xPhospho | A0A0A6YX | 1.83E-05 | 0 | 1 | 4 | 2 | A0A0A6YX | La ribonucl | A0A0A6YX | 0 | 2372.045 | 0 | 0.69 | 0.000729 | 2.37 | 73.5 | 126.5 | 6.49 | 4.3 | | High | High | High | High | High | High | High | High | High | High | High | 0 | 2.67E-06 | 3.58 |
| FALSE | High | LDFLPEMM | [K].LDFLPE | 1xCarbami | P28659-4 1 | 0.048715 | 0.00354 | 1 | 1 | 2 | P28659-4 | Isoform 4 c | P28659-4 [ | 0 | 2944.406 | 0 | -0.36 | 0.006199 | 4.67 | 113.3 | 86.7 | 5.78 | 12.19 | | High | High | High | High | High | High | High | High | High | High | High | 0.001876 | 0.0188 | 3.38 |
| FALSE | High | LDFIGNSTG | [K].LDFIGN | 1xPhospho | E9PVX6 1x | 0.005666 | 0.00062 | 1 | 2 | 1 | E9PVX6 | Proliferatic | E9PVX6 [18 | 0 | 1726.893 | 0 | -0.14 | 0.130887 | 9.88 | 105.5 | 94.5 | 11.44 | 7.7 | | High | High | High | High | High | High | High | High | High | High | High | 0.000275 | 0.001661 | 2.89 |
| FALSE | High | IDFDSVSSII | [K].IDFDSV | 1xPhospho | Q9DBZ5 1x | 1.06E-05 | 0 | 1 | 2 | 4 | Q9DBZ5 | Eukaryotic | Q9DBZ5 [2 | 0 | 1811.797 | 0 | -0.75 | 0.003373 | 6.74 | 125.3 | 74.7 | 5.37 | 9.98 | | High | High | High | High | High | High | High | High | High | High | High | 0 | 1.44E-06 | 3.5 |
| FALSE | High | LCYVALDFE | [K].LCYVAL | 1xCarbami | P60710 1x | 6.65E-05 | 0 | 1 | 20 | 4 | P60710 | Actin, cyto | P60710 [21 | 0 | 3089.45 | 0 | 1.37 | 0.013915 | 21.79 | 55.9 | 144.1 | 12.48 | 22.12 | | High | High | High | High | High | High | High | High | High | High | High | 0 | 1.14E-05 | 4.47 |
| FALSE | High | LCYVALDFE | [K].LCYVAL | 1xCarbami | P60710 1x | 2.8E-05 | 0 | 1 | 20 | 2 | P60710 | Actin, cyto | P60710 [21 | 0 | 3104.461 | 0 | 1.26 | 0.00333 | 5.91 | 61.9 | 138.1 | 6.51 | 13.21 | | High | High | High | High | High | High | High | High | High | High | High | 0 | 4.28E-06 | 3.96 |
| FALSE | High | LDADAVSP | [K].LDADA | 1xPhospho | Q6R891 1x | 0.000203 | 0 | 1 | 1 | 2 | Q6R891 | Neurabin-2 | Q6R891 [1 | 0 | 2515.31 | 0 | 3 | 0.000704 | 21.93 | 23.8 | 176.2 | 12.07 | 12.55 | | High | High | High | High | High | High | High | High | High | High | High | 0 | 6.15E-05 | 3.43 |
| FALSE | High | IDASKNEED | [K].IDASKN | 1xDeamida | Q60668 1x | 0.096181 | 0.007535 | 1 | 3 | 2 | Q60668 | Heterogen | Q60668 [6 | 1 | 2511.14 | 0 | -0.96 | 0.000945 | 2 | 130.8 | 69.2 | 5.01 | 7.93 | | High | High | High | High | High | High | High | High | High | High | High | 0.007212 | 0.07827 | 1.85 |
| FALSE | High | LDDDSEED | [K].LDDDS | 1xCarbami | Q3USH5 1x | 0.000127 | 0 | 1 | 2 | 8 | Q3USH5 | Splicing fac | Q3USH5 [5 | 1 | 4600.045 | 0 | -0.5 | 0.009203 | 7.93 | 117.4 | 82.6 | 8.26 | 8.93 | | High | High | High | High | High | High | High | High | High | High | High | 0 | 2.36E-05 | 4.12 |
| FALSE | High | LDDDSEED | [K].LDDDS | 1xCarbami | Q3USH5 2x | 0.012072 | 0.000889 | 1 | 2 | 1 | Q3USH5 | Splicing fac | Q3USH5 [5 | 1 | 4680.011 | 0 | -0.28 | 0.029249 | 8.15 | 113.3 | 86.7 | 10.5 | 12.34 | | High | High | High | High | High | High | High | High | High | High | High | 0.000449 | 0.003873 | 3.11 |
| FALSE | High | LDDGHLNN | [K].LDDGH | 1xPhospho | E9QAC6 1x | 4.61E-10 | 0 | 1 | 3 | 2 | E9QAC6 | Galectin O | E9QAC6 [3 | 0 | 2723.303 | 0 | 0.78 | 0.000613 | 7.51 | 74.2 | 125.8 | 2.96 | 4.2 | | High | High | High | High | High | High | High | High | High | High | High | 0 | 1.82E-11 | 5.29 |
| FALSE | High | LDDGHLNN | [K].LDDGH | 1xDeamida | E9QAC6 1x | 3.19E-10 | 0 | 1 | 3 | 2 | E9QAC6 | Galectin O | E9QAC6 [3 | 0 | 2724.287 | 0 | 0.36 | 0.025616 | 16.9 | 83.6 | 116.4 | 10.78 | 7.66 | | High | High | High | High | High | High | High | High | High | High | High | 0 | 1.2E-11 | 3.85 |
| FALSE | High | LDDQESPV | [R].LDDQE | 1xPhospho | Q60520-1 | 4.76E-08 | 0 | 1 | 4 | 2 | Q60520-1 | Isoform 2 c | Q60520-1 | 0 | 1928.895 | 0 | -0.37 | 0.001182 | 2.85 | 112.9 | 87.1 | 4.99 | 3.14 | | High | High | High | High | High | High | High | High | High | High | High | 0 | 1.13E-06 | 2.88 |
| FALSE | High | IDDRDSEEE | [K].IDDRDS | 1xPhospho | Q8CD14 1x | 0.001313 | 0 | 1 | 6 | 2 | Q8CD14 | Putative ur | Q8CD14 [7 | 1 | 2055.882 | 0 | 0.18 | 0.003624 | 0.17 | 93.9 | 106.1 | 4.02 | 5.59 | | High | High | High | High | High | High | High | High | High | High | High | 0 | 0.000324 | 3.04 |
| FALSE | High | LDDTWPVS | [R].LDDTW | 1xPhospho | O35387 1x | 0.007082 | 0.00062 | 1 | 4 | 1 | O35387 | HCLS1-asso | O35387 [1 | 0 | 1718.81 | 0 | -1.41 | 0.000758 | 1.86 | 145 | 55 | 9.71 | 9.03 | | High | High | High | High | High | High | High | High | High | High | High | 0.000275 | 0.002123 | 2.47 |
| FALSE | High | LDEAEPEDS | [K].LDEAEP | 1xPhospho | Q9CXL3 1x | 3.5E-05 | 0 | 1 | 1 | 3 | Q9CXL3 | Uncharacte | Q9CXL3 [1 | 0 | 2025.942 | 0 | -0.61 | 0.000738 | 3.77 | 118.6 | 81.4 | 2.2 | 7.62 | | High | High | High | High | High | High | High | High | High | High | High | 0 | 5.51E-06 | 3.1 |
| FALSE | High | LDEDEDED | [K].LDEDED | 1xPhospho | Q9R020 1x | 1.38E-05 | 0 | 1 | 5 | 1 | Q9R020 | Zinc finger | Q9R020 [1 | 0 | 2146.932 | 0 | -0.37 | 0.084914 | 18.43 | 117.2 | 82.8 | 19.6 | 9.87 | | High | High | High | High | High | High | High | High | High | High | High | 0 | 1.94E-06 | 3.37 |
| FALSE | High | LDEEDTDFI | [R].LDEEDT | 1xPhospho | Q9R0A5 1x | 1.63E-06 | 0 | 1 | 2 | 3 | Q9R0A5 | Serine/thre | Q9R0A5 [4 | 0 | 3206.398 | 0 | -0.95 | 0.000309 | 0.4 | 133.9 | 66.1 | 7.42 | 7.47 | | High | High | High | High | High | High | High | High | High | High | High | 0 | 5.3E-06 | 4.78 |
| FALSE | High | IDEFESM | [R].IDEFES | 1xPhospho | P26041 1x | 0.069903 | 0.005181 | 1 | 3 | 1 | P26041 | Moesin OS | P26041 [57 | 0 | 1195.479 | 0 | 2.38 | 0.000311 | 4.27 | 31.1 | 168.9 | 19.2 | 5.07 | | High | High | High | High | High | High | High | High | High | High | High | 0.002662 | 0.02848 | 1.26 |
| FALSE | High | LDEGTPPEI | [K].LDEGTI | 1xPhospho | A0A1L1SQ | 0.000492 | 0 | 1 | 8 | 2 | A0A1L1SQ | Talin-2 OS= | A0A1L1SQ | 0 | 1620.829 | 0 | -1.54 | 0.000261 | 1.26 | 146.9 | 53.1 | 6.81 | 12.03 | | High | High | High | High | High | High | High | High | High | High | High | 0.001844 | 0.01868 | 2.73 |
| FALSE | High | IDEISDK | [K].IDEISD | 1xPhospho | A0A0R4IZZ | 0.050397 | 0.003641 | 1 | 4 | 3 | A0A0R4IZZ | Formin bin | A0A0R4IZZ | 0 | 1357.702 | 0 | -0.43 | 0.001566 | 2.19 | 114.6 | 85.4 | 5.89 | 4.6 | | High | High | High | High | High | High | High | High | High | High | High | 0.002937 | 0.03074 | 1.83 |
| FALSE | High | IDEISDKETI | [K].IDEISD | 1xPhospho | A0A0R4IZZ | 0.010475 | 0.000772 | 1 | 4 | 1 | A0A0R4IZZ | Formin bin | A0A0R4IZZ | 1 | 2505.19 | 0 | -0.21 | 0.502244 | 41.37 | 118.3 | 81.7 | 57.66 | 49.72 | | High | High | High | High | High | High | High | High | High | High | High | 0.000382 | 0.003291 | 2.49 |
| FALSE | High | IDEISDKETI | [K].IDEISD | 1xPhospho | A0A0R4IZZ | 0.000993 | 0 | 1 | 4 | 2 | A0A0R4IZZ | Formin bin | A0A0R4IZZ | 1 | 2734.353 | 0 | -0.75 | 0.008645 | 2.46 | 126.5 | 73.5 | 13.31 | 3.61 | | High | High | High | High | High | High | High | High | High | High | High | 7.19E-05 | 0.000827 | 2.64 |
| FALSE | High | IDEPNTPYH | [K].IDEPNT | 1xPhospho | Q9DCL8 1x | 7.18E-07 | 0 | 1 | 4 | 7 | Q9DCL8 | Protein phi | Q9DCL8 [6 | 0 | 4433.972 | 0 | 0.53 | 0.01263 | 6.36 | 83.9 | 116.1 | 21.37 | 17.49 | | High | High | High | High | High | High | High | High | High | High | High | 0 | 6.98E-08 | 4.63 |
| FALSE | High | IDEPNTPYH | [K].IDEPNT | 1xPhospho | Q9DCL8 1x | 7.56E-06 | 0 | 1 | 4 | 16 | Q9DCL8 | Protein phi | Q9DCL8 [6 | 0 | 4449.967 | 0 | 0.49 | 0.08861 | 14.74 | 82.8 | 117.2 | 10.47 | 19.58 | | High | High | High | High | High | High | High | High | High | High | High | 0 | 1.11E-06 | 6.01 |
| FALSE | High | LDERPSSPI | [R].LDERPS | 1xPhospho | E9PYH6 1x | 0.009168 | 0.000736 | 1 | 3 | 2 | E9PYH6 | SET domai | E9PYH6 [1 | 0 | 2294.293 | 0 | -0.54 | 0.000154 | 1.33 | 118.8 | 81.2 | 1.89 | 4.09 | | High | High | High | High | High | High | High | High | High | High | High | 0.000327 | 0.002851 | 2.93 |
| FALSE | High | LDESPVLKF | [K].LDESPV | 1xPhospho | E9Q5F9 1x | 0.0026 | 0.000109 | 1 | 3 | 1 | E9Q5F9 | Histone-lys | E9Q5F9 [6 | 0 | 2447.274 | 0 | -1.76 | 0.000707 | 2.36 | 154.4 | 45.6 | 6.83 | 21.14 | | High | High | High | High | High | High | High | High | High | High | High | 4.94E-05 | 0.000693 | 2.79 |
| FALSE | High | LDFAENSS | [K].LDFAEN | 1xPhospho | E9PVX6 1x | 0.001337 | 0 | 1 | 2 | 2 | E9PVX6 | Proliferatic | E9PVX6 [2 | 0 | 1692.825 | 0 | 0.11 | 0.168742 | 12.27 | 95.1 | 104.9 | 9.88 | 12.31 | | High | High | High | High | High | High | High | High | High | High | High | 0 | 0.000331 | 3.08 |
| FALSE | High | LDFAGNSS | [K].LDFAGI | 1xPhospho | E9PVX6 1x | 0.000852 | 0 | 1 | 1 | 2 | E9PVX6 | Proliferatic | E9PVX6 [1 | 0 | 1620.803 | 0 | -0.01 | 0.37423 | 5.12 | 97.6 | 102.4 | 10 | 7 | | High | High | High | High | High | High | High | High | High | High | High | 0 | 0.000199 | 2.3 |
| FALSE | High | LDFAGNSS | [K].LDFAGI | 2xPhospho | E9PVX6 2x | 0.022973 | 0.001578 | 1 | 1 | 1 | E9PVX6 | Proliferatic | E9PVX6 [1 | 0 | 1700.77 | 0 | -0.12 | 0.168773 | 5.68 | 101.5 | 98.5 | 5.47 | 7.29 | | High | High | High | High | High | High | High | High | High | High | High | 0.000832 | 0.007951 | 2.45 |
| FALSE | High | LDFCSALSS | [K].LDFCSA | 1xCarbami | Q5CZY4 1x | 0.000211 | 0 | 1 | 5 | 1 | Q5CZY4 | Nuclear fac | Q5CZY4 [3 | 0 | 1920.872 | 0 | 1.52 | 0.001585 | 28.74 | 60.4 | 139.6 | 18.6 | 13.2 | | High | High | High | High | High | High | High | High | High | High | High | 0 | 4.16E-05 | 2.9 |
| FALSE | High | IDFDDERSV | [K].IDFDDE | 1xPhospho | Q569L8 1x | 0.000976 | 0 | 1 | 1 | 2 | Q569L8 | Centromer | Q569L8 [7 | 1 | 3281.474 | 0 | -0.26 | 0.635918 | 9.33 | 104 | 96 | 16.27 | 14.98 | | High | High | High | High | High | High | High | High | High | High | High | 4.94E-05 | 0.000582 | 2.83 |
| FALSE | High | IDFDSVSSII | [K].IDFDSV | 1xPhospho | Q9DBZ5 1x | 4.86E-06 | 0 | 1 | 2 | 2 | Q9DBZ5 | Eukaryotic | Q9DBZ5 [2 | 0 | 1795.802 | 0 | -0.72 | 0.004606 | 5.2 | 127.5 | 72.5 | 8.9 | 10.15 | | High | High | High | High | High | High | High | High | High | High | High | 0 | 6E-07 | 3.28 |
| FALSE | High | LASEYLSPE | [R].LASEYL | 1xPhospho | Q9Z315 1x | 4.08E-05 | 0 | 1 | 1 | 1 | Q9Z315 | U4/U6.U5 | Q9Z315 [3 | 0 | 2282.143 | 0 | -0.2 | 0.090117 | 11.73 | 110.9 | 89.1 | 7.68 | 15.28 | | High | High | High | High | High | High | High | High | High | High | High | 0 | 6.54E-06 | 2.59 |
| FALSE | High | LASEDAAL | [K].LASEDA | 1xPhospho | Q8BRN9 1x | 8.08E-07 | 0 | 1 | 3 | 4 | Q8BRN9 | Coiled-coil | Q8BRN9 [4 | 0 | 3109.487 | 0 | 0.02 | 0.621829 | 4.33 | 99.2 | 100.8 | 3.45 | 5.9 | | High | High | High | High | High | High | High | High | High | High | High | 0 | 2.41E-05 | 3.63 |
| FALSE | High | IASDSLSFLI | [R].IASDSL | 1xPhospho | G3UYN0 1x | 0.027839 | 0.001755 | 1 | 2 | 2 | G3UYN0 | L1TD1 OS= | G3UYN0 [3 | 0 | 1879.038 | 0 | -1.1 | 0.005068 | 4.98 | 133.6 | 66.4 | 11.64 | 21.21 | | High | High | High | High | High | High | High | High | High | High | High | 0.000953 | 0.009892 | 2.11 |
| FALSE | High | LASDFPLDL | [K].LASDFP | 1xPhospho | A2SW42 1x | 7.7E-06 | 0 | 1 | 2 | 2 | A2SW42 | Zinc finger | A2SW42 [6 | 0 | 1940.055 | 0 | -1.77 | 0.000154 | 1.01 | 155.5 | 44.5 | 1.88 | 10.73 | | High | High | High | High | High | High | High | High | High | High | High | 0 | 1.11E-05 | 4.17 |
| FALSE | High | KDTPPPLVI | [R].KDTPPI | 1xPhospho | Q99J09 1x | 0.000164 | 0 | 1 | 2 | 2 | Q99J09 | Methyloso | Q99J09 [3- | 1 | 1667.908 | 0 | -1.39 | 0.043443 | 6.54 | 144.7 | 55.3 | 20.87 | 31.58 | | High | High | High | High | High | High | High | High | High | High | High | 0 | 3.13E-05 | 4.09 |
| FALSE | High | KEDSDEDE | [K].KEDSDI | 2xPhospho | P09405 2x | 0.004106 | 0.00036 | 1 | 6 | 12 | P09405 | Nucleolin ( | P09405 [14 | 1 | 4678.734 | 0 | -0.63 | 0.4569 | 17.09 | 116.8 | 83.2 | 58.82 | 73.98 | | High | High | High | High | High | High | High | High | High | High | High | 0.001625 | 0.01576 | 4.33 |
| FALSE | High | KEESEESDI | [K].KEESEE | 2xPhospho | P99027 2x | 0.00509 | 0.000551 | 1 | 1 | 8 | P99027 | 60S acidic r | P99027 [9 | 1 | 2354.85 | 0 | -0.47 | 0.661072 | 12.38 | 116.4 | 83.6 | 54.22 | 66.88 | | High | High | High | High | High | High | High | High | High | High | High | 0.001844 | 0.01859 | 3.71 |
| FALSE | High | KEESEESED | [K].KEESEE | 2xPhospho | E9Q3T0 2x | 9.71E-05 | 0 | 1 | 2 | 30 | E9Q3T0 | Uncharacte | E9Q3T0 [9 | 1 | 2368.865 | 0 | -0.71 | 0.833673 | 25.37 | 124.2 | 75.8 | 84.44 | 105.61 | | High | High | High | High | High | High | High | High | High | High | High | 7.19E-05 | 0.00077 | 4.59 |
| FALSE | High | KFSEEPEVA | [R].KFSEEP | 1xPhospho | Q9D6Z1 1x | 0.000404 | 0 | 1 | 3 | 2 | Q9D6Z1 | Nucleolar p | Q9D6Z1 [5 | 1 | 2135.083 | 0 | -0.31 | 0.566568 | 31.93 | 122.1 | 77.9 | 54.86 | 61.04 | | High | High | High | High | High | High | High | High | High | High | High | 0 | 8.65E-05 | 3.5 |
| FALSE | High | KHSEEANF | [R].KHSEEA | 1xPhospho | B2RT77 1x | 0.001278 | 0 | 1 | 1 | 2 | B2RT77 | TWIST neig | B2RT77 [3 | 1 | 1940.952 | 0 | 0.22 | 0.452911 | 12.06 | 94.3 | 105.7 | 11.79 | 6.68 | | High | High | High | High | High | High | High | High | High | High | High | 0 | 0.000314 | 3.65 |
| FALSE | High | KHSPDRQD | [R].KHSPDI | 1xPhospho | Q56A08 1x | 0.021231 | 0.001507 | 1 | 2 | 2 | Q56A08 | G patch do | Q56A08 [3 | 2 | 1999.053 | 0 | 1.18 | 0.002662 | 25.75 | 66.2 | 133.8 | 9.29 | 11.98 | | High | High | High | High | High | High | High | High | High | High | High | 0.000801 | 0.007305 | 2.48 |
| FALSE | High | KHSQTDLV | [K].KHSQTI | 1xPhospho | Q4QQM4 1 | 0.008538 | 0.000736 | 1 | 2 | 2 | Q4QQM4 | Tumor pro | Q4QQM4 [ | 1 | 1479.752 | 0 | -1.73 | 0.097122 | 2.99 | 149.2 | 50.8 | 63.21 | 37.12 | | High | High | High | High | High | High | High | High | High | High | High | 0.000327 | 0.002635 | 3.02 |
| FALSE | High | KLEEALSTE | [R].KLEEAL | 1xPhospho | Q61074 1x | 4.78E-10 | 0 | 1 | 2 | 4 | Q61074 | Protein phi | Q61074 [5 | 1 | 2532.212 | 0 | 0.09 | 0.58159 | 25.07 | 95 | 105 | 15.05 | 13.66 | | High | High | High | High | High | High | High | High | High | High | High | 0 | 1.89E-11 | 4.64 |
| FALSE | High | KIIEDCSNSI | [K].KIIEDCS | 1xCarbami | P70398 1x | 0.106748 | 0.008634 | 1 | 4 | 2 | P70398 | Probable u | P70398 [22 | 1 | 2177.045 | 0 | 1.97 | 0.001284 | 31.43 | 36.7 | 163.3 | 25.04 | 14.35 | | High | High | High | High | High | High | High | High | High | High | High | 0.004327 | 0.04667 | 2.16 |
| FALSE | High | KLSGDLEA | [R].KLSGDL | 1xPhospho | H3BL37 1x | 0.002272 | 0.000109 | 1 | 3 | 2 | H3BL37 | Treacle prc | H3BL37 [1 | 1 | 1723.94 | 0 | -0.66 | 0.009668 | 5.56 | 122 | 78 | 5.99 | 14.95 | | High | High | High | High | High | High | High | High | High | High | High | 4.94E-05 | 0.000597 | 2.71 |
| FALSE | High | KLSVGGDS | [R].KLSVGC | 1xPhospho | Q3TIV2 1x | 0.000123 | 0 | 1 | 3 | 2 | Q3TIV2 | Putative ur | Q3TIV2 [28 | 1 | 1851.003 | 0 | -0.35 | 0.139666 | 6.48 | 108.8 | 91.2 | 16.85 | 17.46 | | High | High | High | High | High | High | High | High | High | High | High | 0 | 2.27E-05 | 3.35 |
| FALSE | High | KNSTDLDS | [R].KNSTDI | 2xPhospho | Q924A2 2x | 3.06E-05 | 0 | 1 | 3 | 4 | Q924A2 | Protein cap | Q924A2 [2 | 1 | 2420.104 | 0 | 0.11 | 0.806714 | 5.52 | 98.7 | 101.3 | 10.86 | 6.3 | | High | High | High | High | High | High | High | High | High | High | High | 0 | 4.77E-06 | 3.99 |
| FALSE | High | KDSEEGEE | [R].KDSEEC | 1xPhospho | G3UWX1 1 | 5.12E-05 | 0 | 2 | 9 | 2 | G3UWX1; | Replication | G3UWX1 [ | 1 | 2654.212 | 0 | -0.32 | 0.125906 | 7.3 | 112 | 88 | 13.83 | 13.82 | Shared | High | High | High | High | High | High | High | High | High | High | High | 0 | 8.5E-06 | 3.73 |
| FALSE | High | KPGAGGSF | [R].KPGAG | 1xPhospho | A2AJI0 1xP | 3.65E-07 | 0 | 1 | 5 | 4 | A2AJI0 | MAP7 dom | A2AJI0 [39 | 0 | 1390.741 | 0 | 0.39 | 0.021575 | 1.75 | 87.1 | 112.9 | 12.15 | 8.86 | | High | High | High | High | High | High | High | High | High | High | High | 0 | 3.26E-08 | 3.99 |
| FALSE | High | KQSGLLLEE | [R].KQSGLI | 1xPhospho | O88271 1x | 1.29E-06 | 0 | 1 | 1 | 2 | O88271 | Craniofacia | O88271 [6 | 1 | 2113.083 | 0 | -0.56 | 0.004945 | 0.74 | 117.5 | 82.5 | 6.4 | 12.58 | | High | High | High | High | High | High | High | High | High | High | High | 0 | 1.35E-07 | 3.4 |
| FALSE | High | LAAAEETIT | [R].LAAAEE | 1xPhospho | Q8K0Y7 1x | 0.000178 | 0 | 1 | 1 | 1 | Q8K0Y7 | Protein PA | Q8K0Y7 [1 | 0 | 1768.95 | 0 | 0.23 | 0.054888 | 8.71 | 92 | 108 | 5.17 | 5.63 | | High | High | High | High | High | High | High | High | High | High | High | 0 | 3.42E-05 | 1.98 |
| FALSE | High | LAAAQGQT | [K].LAAAQ | 1xPhospho | Q8C0D5 1x | 3.22E-07 | 0 | 1 | 1 | 2 | Q8C0D5 | Elongation | Q8C0D5 [4 | 0 | 3257.491 | 0 | 0.26 | 0.113006 | 5.23 | 92.3 | 107.7 | 6.71 | 12.2 | | High | High | High | High | High | High | High | High | High | High | High | 0 | 2.85E-08 | 5.62 |
| FALSE | High | IAAEEEEEN | [R].IAAEEE | 1xPhospho | Q80XK6 1x | 0.000223 | 0 | 1 | 1 | 1 | Q80XK6 | Autophagy | Q80XK6 [8 | 0 | 3324.458 | 0 | 0.9 | 0.001435 | 13.19 | 70.3 | 129.7 | 8.6 | 5.38 | | High | High | High | High | High | High | High | High | High | High | High | 0 | 4.41E-05 | 3.54 |
| FALSE | High | IAAEEEEEN | [R].IAAEEE | 1xDeamida | Q80XK6 1x | 0.000259 | 0 | 1 | 1 | 1 | Q80XK6 | Autophagy | Q80XK6 [8 | 0 | 3325.442 | 0 | 0.73 | 0.001529 | 5.52 | 77.9 | 122.1 | 13.44 | 14.52 | | High | High | High | High | High | High | High | High | High | High | High | 0 | 5.23E-05 | 3.46 |
| FALSE | High | LAAGAESP | [K].LAAGAI | 1xPhospho | P16054 1x | 2.11E-07 | 0 | 1 | 1 | 1 | P16054 | Protein kin | P16054 [32 | 0 | 2365.051 | 0 | 1.15 | 0.000877 | 5.65 | 63.4 | 136.6 | 5.76 | 8.12 | | High | High | High | High | High | High | High | High | High | High | High | 0 | 1.78E-08 | 3.26 |
| FALSE | High | LAAGRPFN | [K].LAAGRI | 1xPhospho | Q8BTW9 1 | 0.023584 | 0.001578 | 1 | 2 | 1 | Q8BTW9 | Serine/thre | Q8BTW9 [ | 0 | 1671.857 | 0 | 0.92 | 0.00147 | 16.04 | 69.1 | 130.9 | 9.56 | 10.33 | | High | High | High | High | High | High | High | High | High | High | High | 0.000847 | 0.00817 | 1.29 |
| FALSE | High | LAALALASS | [R].LAALAL | 1xCarbami | Q9D6Z1 1x | 4.52E-09 | 0 | 1 | 4 | 10 | Q9D6Z1 | Nucleolar p | Q9D6Z1 [4 | 0 | 3116.475 | 0 | -0.32 | 0.032983 | 7.19 | 111.2 | 88.8 | 6.41 | 8.04 | | High | High | High | High | High | High | High | High | High | High | High | 0 | 2.36E-10 | 4.38 |
| FALSE | High | LAALALASS | [R].LAALAL | 1xCarbami | Q9D6Z1 2x | 0.001058 | 0 | 1 | 4 | 4 | Q9D6Z1 | Nucleolar p | Q9D6Z1 [4 | 0 | 3196.441 | 0 | 0.24 | 0.103033 | 15.42 | 96 | 104 | 13.87 | 5.47 | | High | High | High | High | High | High | High | High | High | High | High | 0.002976 | 0.03119 | 3.27 |
| FALSE | High | LAAQESSEA | [R].LAAQE | 1xPhospho | Q6PGL7 1x | 2.36E-06 | 0 | 1 | 3 | 2 | Q6PGL7 | WASH com | Q6PGL7 [1 | 0 | 2028.932 | 0 | 0.51 | 0.029906 | 18.54 | 84.1 | 115.9 | 12.42 | 11.3 | | High | High | High | High | High | High | High | High | High | High | High | 0 | 2.68E-07 | 3.75 |
| FALSE | High | LAAQESSEA | [R].LAAQE | 2xPhospho | Q6PGL7 2x | 0.044358 | 0.003264 | 1 | 3 | 1 | Q6PGL7 | WASH com | Q6PGL7 [1 | 0 | 2108.899 | 0 | 0.25 | 0.010269 | 11.69 | 90.2 | 109.8 | 18.81 | 11.46 | | High | High | High | High | High | High | High | High | High | High | High | 0.001665 | 0.01673 | 3.22 |
| FALSE | High | LAARPPSPY | [K].LAARPI | 1xCarbami | E9Q9T0 1x | 0.093123 | 0.007276 | 1 | 7 | 1 | E9Q9T0 | Fibrosin-lik | E9Q9T0 [8 | 0 | 1797.949 | 0 | -0.36 | 0.002972 | 4.42 | 112.5 | 87.5 | 2.97 | 4.41 | | High | High | High | High | High | High | High | High | High | High | High | 0.003698 | 0.03979 | 1.56 |
| FALSE | High | LAASYSSDS | [R].LAASYS | 1xPhospho | O54916 1x | 4.51E-05 | 0 | 1 | 7 | 8 | O54916 | RalBP1-ass | O54916 [1 | 0 | 2845.324 | 0 | 0.41 | 0.043771 | 14.47 | 87.3 | 112.7 | 8.56 | 7.34 | | High | High | High | High | High | High | High | High | High | High | High | 0 | 1.18E-05 | 4.22 |
| FALSE | High | KPSPSPPPP | [R].KPSPSP | 2xPhospho | P97496 2x | 0.000786 | 0 | 1 | 6 | 6 | P97496 | SWI/SNF c | P97496 [32 | 0 | 1937.897 | 0 | -0.78 | 0.000447 | 1.71 | 127.3 | 72.7 | 4.1 | 3.26 | | High | High | High | High | High | High | High | High | High | High | High | 0 | 0.000276 | 2.38 |
| FALSE | High | KDSEEEVSI | [R].KDSEEE | 1xPhospho | P15920 1x | 2E-07 | 0 | 1 | 2 | 4 | P15920 | V-type pro | P15920 [69 | 1 | 2657.214 | 0 | 0.26 | 0.445746 | 18.42 | 94.2 | 105.8 | 18.24 | 22.78 | | High | High | High | High | High | High | High | High | High | High | High | 0 | 1.65E-08 | 5.31 |
| FALSE | High | HYSQDCSS | [R].HYSQD | 1xCarbami | B1AQY2 1x | 0.000779 | 0 | 1 | 4 | 4 | B1AQY2 | Rho GTPas | B1AQY2 [5 | 0 | 1762.824 | 0 | 0.94 | 0.001981 | 10.67 | 70.6 | 129.4 | 5.45 | 7.92 | | High | High | High | High | High | High | High | High | High | High | High | 9.42E-05 | 0.000994 | 1.88 |
| FALSE | High | HYSPEDELS | [R].HYSPED | 1xPhospho | Q6PIU9 1x | 2.22E-07 | 0 | 1 | 1 | 4 | Q6PIU9 | Uncharacte | Q6PIU9 [2 | 0 | 2657.29 | 0 | -0.62 | 0.00171 | 1.05 | 121.9 | 78.1 | 5.09 | 6.53 | | High | High | High | High | High | High | High | High | High | High | High | 0 | 3.78E-08 | 4.04 |
| FALSE | High | HTYGTINS | [R].HTYGTI | 1xPhospho | Q8CBX0 1x | 5.54E-08 | 0 | 1 | 1 | 2 | Q8CBX0 | Calcium pe | Q8CBX0 [7 | 0 | 2542.13 | 0 | 2.38 | 0.00128 | 25.88 | 30.8 | 169.2 | 9.85 | 14.87 | | High | High | High | High | High | High | High | High | High | High | High | 0 | 3.96E-09 | 5 |
| FALSE | High | HVDEFSPR | [K].HVDEF | 1xPhospho | Q3UC10 1x | 0.086562 | 0.006435 | 1 | 4 | 1 | Q3UC10 | DNA helica | Q3UC10 [4 | 0 | 1295.598 | 0 | -0.75 | 0.001274 | 3.3 | 129 | 71 | 5.24 | 12.17 | | High | High | High | High | High | High | High | High | High | High | High | 0.003303 | 0.0363 | 1.06 |

| | | | | | | | | | | | | | | | | | | | | | | | | | | | | | | | | | | | | | | |
|---|---|---|---|---|---|---|---|---|---|---|---|---|---|---|---|---|---|---|---|---|---|---|---|---|---|---|---|---|---|---|---|---|---|---|---|---|---|---|
| FALSE | High | HVDPPSPR | [R].HVDPP | 1xPhospho | Q3U6K5 1x | 0.014797 | 0.001049 | 1 | 4 | 2 | Q3U6K5 | Spermatog | Q3U6K5 [2 | 0 | 1213.593 | 0 | 1.95 | 0.000726 | 17.29 | 40.8 | 159.2 | 9.42 | 10.01 | | High | High | High | High | High | High | High | High | High | High | High | 0.000555 | 0.004857 | 2.87 |
| FALSE | High | HVEGQLAH | [R].HVEGQ | 1xPhospho | Q52KB6 1x | 0.001615 | 5.66E-05 | 1 | 6 | 2 | Q52KB6 | C2 domain | Q52KB6 [1 | 0 | 1841.968 | 0 | -0.03 | 0.484157 | 15.8 | 100.9 | 99.1 | 4.28 | 10.42 | | High | High | High | High | High | High | High | High | High | High | High | 2.58E-05 | 0.00041 | 2.26 |
| FALSE | High | HVEGQSAA | [K].HVEGQ | 1xCarbami | Q6A065 1x | 0.000203 | 0 | 1 | 3 | 1 | Q6A065 | Centrosom | Q6A065 [2 | 0 | 2415.1 | 0 | 1.11 | 0.008241 | 27 | 58 | 142 | 12.94 | 19.95 | | High | High | High | High | High | High | High | High | High | High | High | 0 | 4.01E-05 | 2.78 |
| FALSE | High | HVIAEQEVI | [K].HVIAEC | 1xPhospho | Q3TUQ5 1: | 1.87E-07 | 0 | 1 | 2 | 1 | Q3TUQ5 | Pinin OS=M | Q3TUQ5 [3 | 0 | 3495.66 | 0 | -0.45 | 0.036278 | 1.27 | 121.2 | 78.8 | 6.84 | 36.74 | | High | High | High | High | High | High | High | High | High | High | High | 0 | 1.55E-08 | 4.42 |
| FALSE | High | HVIAEQEVI | [K].HVIAEC | 1xPhospho | Q3TUQ5 1: | 0.003372 | 0.00016 | 1 | 2 | 2 | Q3TUQ5 | Pinin OS=M | Q3TUQ5 [3 | 0 | 3511.655 | 0 | -0.53 | 0.011706 | 3.65 | 120.5 | 79.5 | 16.58 | 14.65 | | High | High | High | High | High | High | High | High | High | High | High | 9.42E-05 | 0.000929 | 3.44 |
| FALSE | High | HVIGLQMC | [K].HVIGLC | 1xPhospho | P37804 1xF | 0.012288 | 0.000889 | 1 | 1 | 2 | P37804 | Transgelin | P37804 [17 | 0 | 1536.756 | 0 | -0.53 | 0.036293 | 13.45 | 116.9 | 83.1 | 11.96 | 8.81 | | High | High | High | High | High | High | High | High | High | High | High | 0.000505 | 0.004377 | 3.41 |
| FALSE | High | HVILSGSAE | [K].HVILSG | 1xPhospho | Q69Z38 1x | 1.25E-05 | 0 | 1 | 1 | 1 | Q69Z38 | Pseudopoc | Q69Z38 [2C | 0 | 2034.022 | 0 | -0.48 | 0.010914 | 8.17 | 117.9 | 82.1 | 5.32 | 16.09 | | High | High | High | High | High | High | High | High | High | High | High | 0 | 1.74E-06 | 3.45 |
| FALSE | High | HVLPASAPI | [K].HVLPAS | 1xPhospho | P97434-2 1 | 0.000331 | 0 | 1 | 10 | 1 | P97434-2 | Isoform 2 c | P97434-2 [ | 0 | 2343.236 | 0 | 2.35 | 0.004931 | 8.27 | 33.2 | 166.8 | 11.34 | 22.52 | | High | High | High | High | High | High | High | High | High | High | High | 0 | 6.91E-05 | 2.68 |
| FALSE | High | HVIQSPSTD | [R].HVIQSF | 1xPhospho | F8VQ70 1x | 0.003433 | 0.00021 | 1 | 8 | 2 | F8VQ70 | S phase cy | F8VQ70 [1 | 0 | 1448.71 | 0 | 0.6 | 0.000951 | 7.44 | 81.7 | 118.3 | 8.09 | 5.68 | | High | High | High | High | High | High | High | High | High | High | High | 0.000294 | 0.002358 | 2.69 |
| FALSE | High | HVSDANIK | [R].HVSDAI | 1xPhospho | G5E8P0 1x | 0.008847 | 0.000736 | 1 | 2 | 2 | G5E8P0 | Gamma-tu | G5E8P0 [1 | 0 | 1421.755 | 0 | 1.16 | 0.003422 | 26.6 | 59.9 | 140.1 | 9.4 | 13.61 | | High | High | High | High | High | High | High | High | High | High | High | 0.000327 | 0.002724 | 2.38 |
| FALSE | High | HVSFQDED | [R].HVSFQI | 1xPhospho | Q924S8 1x | 4.17E-06 | 0 | 1 | 1 | 2 | Q924S8 | Sprouty-re | Q924S8 [2 | 0 | 1782.826 | 0 | 0.06 | 0.541777 | 23.45 | 101.4 | 98.6 | 6.83 | 17.52 | | High | High | High | High | High | High | High | High | High | High | High | 0 | 5.05E-07 | 2.86 |
| FALSE | High | HVSISYDIPI | [R].HVSISY | 1xPhospho | A0A1B0GS | 0.016743 | 0.001166 | 1 | 2 | 2 | A0A1B0GS | GRB2-asso | A0A1B0GS | 0 | 2564.275 | 0 | -1 | 0.013241 | 2.99 | 128.1 | 71.9 | 12.07 | 33.66 | | High | High | High | High | High | High | High | High | High | High | High | 0.000622 | 0.005563 | 2.51 |
| FALSE | High | HVSSPDVT | [K].HVSSPD | 1xPhospho | B8JJC1 1xP | 5.5E-05 | 0 | 1 | 2 | 8 | B8JJC1 | Tyrosyl-DN | B8JJC1 [11 | 0 | 1807.936 | 0 | -0.05 | 0.296934 | 9.89 | 105.8 | 94.2 | 4.56 | 17.52 | | High | High | High | High | High | High | High | High | High | High | High | 0 | 9.17E-06 | 3.16 |
| FALSE | High | HVSSTVVK | [R].HVSSTV | 1xPhospho | Q99M31-2 | 0.003224 | 0.00016 | 1 | 2 | 2 | Q99M31-2 | Isoform 2 c | Q99M31-2 | 0 | 1394.781 | 0 | -1.3 | 0.000689 | 3.39 | 142.7 | 57.3 | 5.62 | 11.43 | | High | High | High | High | High | High | High | High | High | High | High | 0.009089 | 0.09781 | 2.33 |
| FALSE | High | HVTLPSSPF | [R].HVTLPS | 1xPhospho | A2AN08-5 | 0.000267 | 0 | 1 | 4 | 7 | A2AN08-5 | Isoform 5 c | A2AN08-5 | 0 | 1302.677 | 0 | -0.16 | 0.278265 | 2.14 | 105.7 | 94.3 | 10.22 | 9.63 | | High | High | High | High | High | High | High | High | High | High | High | 2.58E-05 | 0.000379 | 3.06 |
| FALSE | High | HVTLPSSPF | [R].HVTLPS | 2xPhospho | A2AN08-5 | 0.091627 | 0.007146 | 1 | 4 | 1 | A2AN08-5 | Isoform 5 c | A2AN08-5 | 0 | 1382.643 | 0 | 0.12 | 0.035165 | 8.71 | 96.8 | 103.2 | 6.87 | 4.03 | | High | High | High | High | High | High | High | High | High | High | High | 0.003633 | 0.03878 | 2.08 |
| FALSE | High | HVTSNASD | [K].HVTSN/ | 1xPhospho | H7BWY4 1: | 4.23E-05 | 0 | 1 | 3 | 2 | H7BWY4 | Disks large | H7BWY4 [6 | 0 | 1848.796 | 0 | 0.39 | 0.157049 | 47.33 | 81.7 | 118.3 | 13.25 | 22.77 | | High | High | High | High | High | High | High | High | High | High | High | 0 | 6.85E-06 | 3.07 |
| FALSE | High | HVVRPPPR | [R].HVVRPI | 1xPhospho | Q6PB97 1x | 0.000128 | 0 | 1 | 4 | 1 | Q6PB97 | F-box/LRR- | Q6PB97 [3 | 1 | 3217.683 | 0 | -0.1 | 0.101199 | 6.02 | 104.6 | 95.4 | 7.53 | 9.01 | | High | High | High | High | High | High | High | High | High | High | High | 0 | 2.38E-05 | 2.06 |
| FALSE | High | HWLQLSPT | [R].HWLQL | 1xPhospho | A0A0A0MC | 2.58E-05 | 0 | 1 | 4 | 1 | A0A0A0MC | Ral GTPase | A0A0A0MC | 0 | 2250.075 | 0 | 0.33 | 0.07084 | 4.62 | 88.5 | 111.5 | 11.68 | 10.77 | | High | High | High | High | High | High | High | High | High | High | High | 0 | 3.92E-06 | 2.92 |
| FALSE | High | HWVAPGG | [R].HWVAF | 1xPhospho | Q8R3Y8 1x | 1.19E-07 | 0 | 1 | 2 | 1 | Q8R3Y8 | Interferon | Q8R3Y8 [4 | 0 | 2956.537 | 0 | 1.87 | 0.003241 | 13.19 | 43.6 | 156.4 | 10.34 | 17.7 | | High | High | High | High | High | High | High | High | High | High | High | 0 | 9.31E-09 | 4.31 |
| FALSE | High | HYEDGYPG | [R].HYEDG | 1xPhospho | E9Q8Z8 1x | 6.54E-09 | 0 | 1 | 19 | 3 | E9Q8Z8 | Catenin de | E9Q8Z8 [2 | 0 | 2282.955 | 0 | 0.21 | 0.244753 | 18.46 | 92.9 | 107.1 | 10.81 | 10.85 | | High | High | High | High | High | High | High | High | High | High | High | 0 | 9.22E-08 | 4.27 |
| FALSE | High | HYGGLTGL | [R].HYGGL | 1xPhospho | Q3U7Z6 1x | 2.51E-05 | 0 | 1 | 3 | 1 | Q3U7Z6 | Phosphogl | Q3U7Z6 [9 | 0 | 1597.85 | 0 | 0.06 | 0.047229 | 3.19 | 98.1 | 101.9 | 2.95 | 4.23 | | High | High | High | High | High | High | High | High | High | High | High | 0 | 3.79E-06 | 2.96 |
| FALSE | High | HYGITSPISI | [R].HYGITS | 1xPhospho | Q3V141 1x | 5.88E-06 | 0 | 1 | 12 | 2 | Q3V141 | Putative ur | Q3V141 [1 | 0 | 1993.092 | 0 | -0.22 | 0.158267 | 15.91 | 107.3 | 92.7 | 5.07 | 14.44 | | High | High | High | High | High | High | High | High | High | High | High | 0 | 7.42E-07 | 2.93 |
| FALSE | High | HYMACYST | [R].HYMAC | 1xCarbami | Q0GH64 1x | 0.010382 | 0.000772 | 1 | 2 | 2 | Q0GH64 | Claudin OS | Q0GH64 [1 | 0 | 2004.866 | 0 | 0.89 | 0.001928 | 16.81 | 72.3 | 127.7 | 21.11 | 7.57 | | High | High | High | High | High | High | High | High | High | High | High | 0.000382 | 0.00328 | 2.56 |
| FALSE | High | HYMACYST | [R].HYMAC | 1xCarbami | Q0GH64 1x | 1.13E-05 | 0 | 1 | 2 | 10 | Q0GH64 | Claudin OS | Q0GH64 [1 | 0 | 2020.861 | 0 | 0.69 | 0.003612 | 20.02 | 77.4 | 122.6 | 17.03 | 9.71 | | High | High | High | High | High | High | High | High | High | High | High | 0 | 4.54E-06 | 4.32 |
| FALSE | High | HYMACYST | [R].HYMAC | 1xCarbami | Q0GH64 2x | 0.003622 | 0.00021 | 1 | 2 | 1 | Q0GH64 | Claudin OS | Q0GH64 [1 | 0 | 2100.827 | 0 | -0.29 | 0.273309 | 24.43 | 109.6 | 90.4 | 15.35 | 13.31 | | High | High | High | High | High | High | High | High | High | High | High | 9.42E-05 | 0.00101 | 2.36 |
| FALSE | High | HYQTEEKS | [R].HYQTEI | 1xPhospho | Q8R0A0 1x | 0.071071 | 0.005247 | 1 | 2 | 2 | Q8R0A0 | General tra | Q8R0A0 [2 | 1 | 1674.778 | 0 | -0.09 | 0.373017 | 13.16 | 103 | 97 | 15.21 | 22.7 | | High | High | High | High | High | High | High | High | High | High | High | 0.002703 | 0.02903 | 1.73 |
| FALSE | High | IAATILTSPD | [R].IAATILT | 1xPhospho | P05202 1xF | 0.023792 | 0.001612 | 1 | 1 | 2 | P05202 | Aspartate a | P05202 [32 | 0 | 1579.866 | 0 | 0.02 | 0.417037 | 3.15 | 99.4 | 100.6 | 5.22 | 4.71 | | High | High | High | High | High | High | High | High | High | High | High | 0.000859 | 0.008283 | 2.5 |
| FALSE | High | HLSSTSDDI | [R].HLSSTS | 1xPhospho | Q9DBC3 1x | 9.48E-06 | 0 | 1 | 2 | 12 | Q9DBC3 | Cap-specifi | Q9DBC3 [2 | 0 | 2533.234 | 0 | 0.29 | 0.274348 | 31.27 | 89.9 | 110.1 | 19.62 | 11.31 | | High | High | High | High | High | High | High | High | High | High | High | 2.58E-05 | 0.000534 | 3.43 |
| FALSE | High | LACCVVGT | [R].LACCV\ | 2xCarbami | O88592 2x | 0.099328 | 0.008003 | 1 | 4 | 1 | O88592 | Superoxide | O88592 [2 | 0 | 2660.19 | 0 | -0.29 | 0.173749 | 11.67 | 110.1 | 89.9 | 13.76 | 7.94 | | High | High | High | High | High | High | High | High | High | High | High | 0.004032 | 0.04288 | 2.52 |
| FALSE | High | IACEEEFSD | [R].IACEEE | 1xCarbami | D3YYI8 2xP | 1.43E-05 | 0 | 1 | 2 | 9 | D3YYI8 | Histone de | D3YYI8 [41 | 1 | 2762.131 | 0 | -0.7 | 0.000222 | 1.69 | 123.4 | 76.6 | 1.99 | 2.83 | | High | High | High | High | High | High | High | High | High | High | High | 0 | 3.51E-05 | 3.3 |
| FALSE | High | IALESVGQF | [K].IALESV | 1xCarbami | Q64337 1x | 7.7E-06 | 0 | 1 | 2 | 2 | Q64337 | Sequestos | Q64337 [3 | 0 | 3901.73 | 0 | 1.76 | 0.012493 | 24.64 | 45.6 | 154.4 | 8.93 | 28.98 | | High | High | High | High | High | High | High | High | High | High | High | 0 | 3.76E-06 | 5.05 |
| FALSE | High | IALESVGQF | [K].IALESV | 1xCarbami | Q64337 1x | 2.33E-07 | 0 | 1 | 2 | 6 | Q64337 | Sequestos | Q64337 [3 | 0 | 3917.725 | 0 | 1.88 | 0.000344 | 11.69 | 43.8 | 156.2 | 6.25 | 7.25 | | High | High | High | High | High | High | High | High | High | High | High | 0 | 1.97E-08 | 4.29 |
| FALSE | High | LAIFGDAAS | [K].LAIFGD | 1xPhospho | Q80Y19 1x | 0.001674 | 5.66E-05 | 1 | 2 | 1 | Q80Y19 | Rho GTPas | Q80Y19 [8 | 0 | 1840.039 | 0 | 0 | 0.855482 | 2.45 | 100.1 | 99.9 | 9.34 | 8.42 | | High | High | High | High | High | High | High | High | High | High | High | 2.58E-05 | 0.000426 | 1.92 |
| FALSE | High | LAIHNPNLF | [K].LAIHNP | 1xPhospho | Q9Z0Z7 1xF | 3.1E-06 | 0 | 1 | 2 | 2 | Q9Z0Z7 | Krueppel-li | Q9Z0Z7 [3 | 0 | 2740.511 | 0 | -1.5 | 0.012524 | 0.14 | 147.8 | 52.2 | 18.83 | 21.61 | | High | High | High | High | High | High | High | High | High | High | High | 0 | 3.63E-07 | 4.22 |
| FALSE | High | LALLIHNDS | [R].LALLIH | 1xPhospho | Q8CBX0 1x | 0.00129 | 0 | 1 | 3 | 2 | Q8CBX0 | Calcium pe | Q8CBX0 [6 | 0 | 2882.532 | 0 | 2.04 | 0.00192 | 8.35 | 39.3 | 160.7 | 14.3 | 13.05 | | High | High | High | High | High | High | High | High | High | High | High | 4.94E-05 | 0.00068 | 2.68 |
| FALSE | High | IALNLEGCA | [K].IALNLE | 1xCarbami | Q80UU6 1x | 6.58E-08 | 0 | 1 | 5 | 4 | Q80UU6 | Opioid gro | Q80UU6 [3 | 0 | 2265.115 | 0 | 0.85 | 0.001361 | 18.12 | 71.1 | 128.9 | 11.64 | 6.28 | | High | High | High | High | High | High | High | High | High | High | High | 0 | 4.79E-09 | 5.08 |
| FALSE | High | LAIQGPEDS | [R].LAIQGF | 1xPhospho | Q3URS8 1x | 2.49E-05 | 0 | 1 | 2 | 2 | Q3URS8 | Putative ur | Q3URS8 [2 | 0 | 1578.773 | 0 | -0.37 | 0.084137 | 5.8 | 111.4 | 88.6 | 7.78 | 10.55 | | High | High | High | High | High | High | High | High | High | High | High | 0 | 3.76E-06 | 2.76 |
| FALSE | High | LALSQNQC | [R].LALSQN | 1xPhospho | Q5SUF2 1x | 0.000118 | 0 | 1 | 3 | 1 | Q5SUF2 | Luc7-like p | Q5SUF2 [1 | 0 | 2253.164 | 0 | 0.46 | 0.009519 | 10.91 | 88.2 | 111.8 | 9.3 | 6.16 | | High | High | High | High | High | High | High | High | High | High | High | 0 | 2.17E-05 | 2.55 |
| FALSE | High | LAISTPDAP | [R].LAISTPI | 1xPhospho | A0A0R4J04 | 0.000924 | 0 | 1 | 6 | 2 | A0A0R4J04 | N6-adenos | A0A0R4J04 | 0 | 2692.421 | 0 | -1.72 | 0.00074 | 0.49 | 156.7 | 43.3 | 14.24 | 14.92 | | High | High | High | High | High | High | High | High | High | High | High | 0 | 0.000219 | 3.42 |
| FALSE | High | LANALSEDT | [R].LANALS | 1xPhospho | P97468 1xF | 4.36E-06 | 0 | 1 | 1 | 1 | P97468 | Chemokine | P97468 [3 | 0 | 2211.028 | 0 | -1.39 | 0.008043 | 5.59 | 144.8 | 55.2 | 17.83 | 16.78 | | High | High | High | High | High | High | High | High | High | High | High | 0 | 5.31E-07 | 2.94 |
| FALSE | High | LANHDEGS | [R].LANHD | 1xPhospho | Q8K1A6 1x | 3.98E-05 | 0 | 1 | 2 | 6 | Q8K1A6 | Coiled-coil | Q8K1A6 [4 | 0 | 2453.076 | 0 | -0.03 | 0.506923 | 11.19 | 101.1 | 98.9 | 1.7 | 7.49 | | High | High | High | High | High | High | High | High | High | High | High | 0 | 1.1E-05 | 3.52 |
| FALSE | High | LANHDEGS | [R].LANHD | 1xPhospho | Q8K1A6 1x | 0.000993 | 0 | 1 | 2 | 2 | Q8K1A6 | Coiled-coil | Q8K1A6 [4 | 1 | 2810.334 | 0 | -0.25 | 0.061461 | 7.33 | 108.1 | 91.9 | 10.21 | 6.23 | | High | High | High | High | High | High | High | High | High | High | High | 0 | 0.000238 | 2.31 |
| FALSE | High | LANISVPAS | [R].LANISV | 1xPhospho | B9EKJ7 1xF | 0.002058 | 5.66E-05 | 1 | 2 | 3 | B9EKJ7 | Tankyrase | B9EKJ7 [29 | 0 | 1649.846 | 0 | 0.55 | 0.003121 | 10.14 | 80 | 120 | 4.38 | 6.88 | | High | High | High | High | High | High | High | High | High | High | High | 2.58E-05 | 0.000536 | 2.27 |
| FALSE | High | LALAPDPE( | [K].LALAPD | 1xCarbami | Q3UY28 1x | 0.003342 | 0.00016 | 1 | 1 | 1 | Q3UY28 | Putative ur | Q3UY28 [1 | 1 | 2678.362 | 0 | -1.06 | 0.000868 | 3.97 | 135.6 | 64.4 | 3.94 | 11.58 | | High | High | High | High | High | High | High | High | High | High | High | 7.19E-05 | 0.000921 | 2.9 |
| FALSE | High | LANYTNLS( | [K].LANYTN | 1xPhospho | Q9WVL3 1: | 0.108463 | 0.008839 | 1 | 3 | 1 | Q9WVL3 | Solute carr | Q9WVL3 [8 | 2 | 3026.417 | 1 | -0.31 | 0.143719 | 9.68 | 109 | 91 | 13.06 | 6.27 | | High | High | High | High | High | High | High | High | High | High | High | 0.004411 | 0.04736 | 3.01 |
| FALSE | High | LAPPPPRTF | [R].LAPPPP | 1xPhospho | Q9D880 1x | 0.024855 | 0.001645 | 1 | 1 | 1 | Q9D880 | Mitochond | Q9D880 [2 | 1 | 2295.206 | 0 | -0.38 | 0.020841 | 7.08 | 110.9 | 89.1 | 7.93 | 9.7 | | High | High | High | High | High | High | High | High | High | High | High | 0.000872 | 0.00872 | 2.28 |
| FALSE | High | LAPSFPSPP | [K].LAPSFP | 1xPhospho | B9EIG6 1xF | 0.026652 | 0.001676 | 1 | 3 | 1 | B9EIG6 | Pogo trans | B9EIG6 [28 | 0 | 2553.413 | 0 | -0.65 | 0.005472 | 9.95 | 124.5 | 75.5 | 6.63 | 13.44 | | High | High | High | High | High | High | High | High | High | High | High | 0.000903 | 0.009454 | 2.28 |
| FALSE | High | LAPSPGSGS | [R].LAPSPG | 1xPhospho | Q7TSH3 1x | 0.003028 | 0.00016 | 1 | 1 | 4 | Q7TSH3 | Zinc finger | Q7TSH3 [9 | 0 | 1437.73 | 0 | -0.06 | 0.504304 | 3.36 | 101.2 | 98.8 | 4.58 | 3.68 | | High | High | High | High | High | High | High | High | High | High | High | 7.19E-05 | 0.000822 | 1.73 |
| FALSE | High | LAPSPSEEP | [R].LAPSPS | 1xPhospho | Q6ZQ32 1x | 0.004143 | 0.00036 | 2 | 7 | 2 | Q6ZQ32; F | MKIAA085 | Q6ZQ32 [7 | 0 | 1391.677 | 0 | 2.11 | 0.000395 | 16.25 | 36.9 | 163.1 | 4.78 | 7.66 | Shared | High | High | High | High | High | High | High | High | High | High | High | 0.000161 | 0.001171 | 2.74 |
| FALSE | High | LAPSSNWP | [R].LAPSSN | 1xPhospho | Q3URR7 1x | 1.04E-06 | 0 | 1 | 3 | 6 | Q3URR7 | Zinc finger | Q3URR7 [1 | 0 | 2717.284 | 0 | -1.59 | 0.000892 | 2.57 | 149.6 | 50.4 | 8.26 | 8.34 | | High | High | High | High | High | High | High | High | High | High | High | 0 | 1.07E-05 | 4.21 |
| FALSE | High | IAQEIASLSI | [K].IAQEIA | 1xPhospho | A6H6B3 1x | 0.000194 | 0 | 1 | 5 | 2 | A6H6B3 | RalA bindir | A6H6B3 [4 | 0 | 1597.897 | 0 | -0.92 | 0.001132 | 6 | 130.8 | 69.2 | 5.37 | 19.4 | | High | High | High | High | High | High | High | High | High | High | High | 0 | 5.58E-05 | 2.83 |
| FALSE | High | LAQTLSCDY | [K].LAQTLS | 1xCarbami | Q6ZWQ0 1 | 0.015195 | 0.001089 | 1 | 4 | 3 | Q6ZWQ0 | Nesprin-2 | Q6ZWQ0 [ | 0 | 3756.844 | 0 | 4.37 | 0.000706 | 6.35 | 7.9 | 192.1 | 17.54 | 14.36 | | High | High | High | High | High | High | High | High | High | High | High | 0.000571 | 0.004993 | 3.42 |
| FALSE | High | LAQTLSCDY | [K].LAQTLS | 1xCarbami | Q6ZWQ0 2 | 0.000837 | 0 | 1 | 4 | 3 | Q6ZWQ0 | Nesprin-2 | Q6ZWQ0 [ | 0 | 3836.811 | 0 | 2.94 | 0.000213 | 37.78 | 23 | 177 | 35.37 | 8.31 | | High | High | High | High | High | High | High | High | High | High | High | 4.94E-05 | 0.000742 | 2.76 |
| FALSE | High | LAQTTPVD | [R].LAQTTF | 1xPhospho | F8VPJ6 1xF | 5.6E-05 | 0 | 1 | 2 | 4 | F8VPJ6 | Regulatory | F8VPJ6 [98 | 0 | 1811.91 | 0 | -0.61 | 0.002332 | 6.95 | 123.5 | 76.5 | 6.54 | 13.2 | | High | High | High | High | High | High | High | High | High | High | High | 0 | 0.000265 | 3.1 |
| FALSE | High | LASATANA | [R].LASATA | 1xPhospho | Q148V8 1x | 4.63E-05 | 0 | 1 | 1 | 2 | Q148V8 | Protein FAI | Q148V8 [1 | 0 | 1860.942 | 0 | 1.04 | 0.020729 | 44.42 | 61.3 | 138.7 | 15.22 | 22.76 | | High | High | High | High | High | High | High | High | High | High | High | 0 | 7.59E-06 | 2.89 |
| FALSE | High | LASAVLSQS | [K].LASAVL | 1xPhospho | Q9QY06-2 | 0.000422 | 0 | 2 | 10 | 1 | Q9QY06-2; | Isoform 2 c | Q9QY06-2 | 0 | 2099.14 | 0 | 0.88 | 0.001473 | 9.95 | 67.9 | 132.1 | 4.16 | 10.4 | Shared | High | High | High | High | High | High | High | High | High | High | High | 0 | 9.07E-05 | 2.28 |
| FALSE | High | LASDDRPSI | [K].LASDDF | 1xPhospho | Q99KG3 1x | 9.28E-05 | 0 | 2 | 3 | 4 | Q99KG3; C | RNA-bindir | Q99KG3 [7 | 0 | 1519.747 | 0 | -0.07 | 0.270831 | 9 | 103.5 | 96.5 | 1.74 | 6.39 | Shared | High | High | High | High | High | High | High | High | High | High | High | 0.000689 | 0.006135 | 2 |
| FALSE | High | IASDEEIQG | [R].IASDEE | 1xPhospho | Q9ET54 1x | 0.004613 | 0.000456 | 1 | 9 | 3 | Q9ET54 | Palladin OS | Q9ET54 [8 | 0 | 1728.882 | 0 | -1.48 | 0.001723 | 1.09 | 141.3 | 58.7 | 13.32 | 21.89 | | High | High | High | High | High | High | High | High | High | High | High | 0.000327 | 0.002414 | 3.21 |
| FALSE | High | IAPPETPDS | [K].IAPPETI | 1xPhospho | O88477 1x | 0.030367 | 0.001857 | 1 | 2 | 1 | O88477 | Insulin-like | O88477 [4 | 0 | 1592.834 | 0 | -1.55 | 0.000911 | 1.77 | 149.1 | 50.9 | 4.52 | 19.48 | | High | High | High | High | High | High | High | High | High | High | High | 0.001011 | 0.0109 | 2.01 |
| FALSE | High | LAHDSLEDI | [K].LAHDSL | 1xPhospho | Q5XF89-2 | 1.31E-06 | 0 | 1 | 2 | 2 | Q5XF89-2 | Isoform 2 c | Q5XF89-2 | 0 | 1806.884 | 0 | 0.5 | 0.025097 | 26.52 | 88.8 | 111.2 | 13.77 | 10.56 | | High | High | High | High | High | High | High | High | High | High | High | 0 | 1.39E-07 | 3.57 |
| FALSE | High | IAGTPGTG( | [K].IAGTPG | 1xDeamida | E9QNT5 1x | 0.051258 | 0.003745 | 1 | 3 | 3 | E9QNT5 | Transcripti | E9QNT5 [9 | 1 | 2649.401 | 0 | -0.47 | 0.001021 | 2.25 | 115.3 | 84.7 | 8.8 | 6.62 | | High | High | High | High | High | High | High | High | High | High | High | 0.00196 | 0.01974 | 2.06 |
| FALSE | High | IAGTPGTG( | [K].IAGTPG | 1xPhospho | E9QNT5 1x | 0.087983 | 0.006639 | 1 | 3 | 1 | E9QNT5 | Transcripti | E9QNT5 [9 | 1 | 2648.417 | 0 | -0.44 | 0.008538 | 2.98 | 115 | 85 | 5.78 | 7.71 | | High | High | High | High | High | High | High | High | High | High | High | 0.003389 | 0.03699 | 2.25 |
| FALSE | High | LADAMAES | [R].LADAM | 1xPhospho | Q8CCN5 1x | 0.005666 | 0.00062 | 1 | 7 | 2 | Q8CCN5 | Breast carc | Q8CCN5 [8 | 0 | 1472.665 | 0 | -0.33 | 0.028915 | 10.93 | 113.7 | 86.3 | 10.67 | 20.28 | | High | High | High | High | High | High | High | High | High | High | High | 0.000872 | 0.009092 | 1.86 |
| FALSE | High | LADDEGDS | [R].LADDE( | 1xPhospho | Q9Z277 1x | 2.09E-08 | 0 | 1 | 2 | 2 | Q9Z277 | Tyrosine-p | Q9Z277 [1 | 0 | 2075.86 | 0 | 0.29 | 0.062209 | 15.96 | 90 | 110 | 23.56 | 9.99 | | High | High | High | High | High | High | High | High | High | High | High | 0 | 1.32E-09 | 3.33 |
| FALSE | High | LADFGLSTF | [K].LADFGI | 1xPhospho | Q9QZL0 1x | 1.25E-05 | 0 | 1 | 2 | 1 | Q9QZL0 | Receptor-ii | Q9QZL0 [1 | 0 | 2556.157 | 0 | -0.89 | 0.006523 | 4.78 | 128.5 | 71.5 | 8.65 | 12.94 | | High | High | High | High | High | High | High | High | High | High | High | 0 | 1.75E-06 | 4.83 |
| FALSE | High | LADLEEESE | [K].LADLEE | 1xPhospho | Q8QZS3 1x | 9.76E-07 | 0 | 1 | 1 | 1 | Q8QZS3 | Folliculin O | Q8QZS3 [2 | 0 | 3006.303 | 0 | 0.1 | 0.813937 | 19.56 | 97 | 103 | 5.14 | 12.29 | | High | High | High | High | High | High | High | High | High | High | High | 0 | 9.88E-08 | 4.78 |
| FALSE | High | LADLYGSK | [K].LADLYG | 1xPhospho | P55288 1xF | 0.081752 | 0.006219 | 1 | 2 | 1 | P55288 | Cadherin-1 | P55288 [78 | 0 | 1404.754 | 0 | 0.23 | 0.124689 | 1.46 | 88.2 | 111.8 | 14.67 | 20.76 | | High | High | High | High | High | High | High | High | High | High | High | 0.003229 | 0.03412 | 2.12 |
| FALSE | High | IADPEHDH | [R].IADPEH | 1xPhospho | D3Z3G6 1x | 0.000213 | 0 | 1 | 5 | 1 | D3Z3G6 | Mitogen-ac | D3Z3G6 [1 | 0 | 2481.165 | 0 | 1.27 | 0.126426 | 65.48 | 52 | 148 | 21.08 | 46.77 | | High | High | High | High | High | High | High | High | High | High | High | 0 | 4.2E-05 | 3.75 |
| FALSE | High | LADVADAD | [K].LADVAI | 1xPhospho | B2RUJ6 1xF | 7.7E-06 | 0 | 1 | 6 | 2 | B2RUJ6 | Rap guanin | B2RUJ6 [15 | 0 | 2357.023 | 0 | 1.17 | 0.004935 | 13.02 | 59.7 | 140.3 | 28.21 | 7 | | High | High | High | High | High | High | High | High | High | High | High | 0 | 1.01E-06 | 4.21 |
| FALSE | High | LAEAPSPAF | [R].LAEAPS | 1xDeamida | Q80U72-4 | 0.000822 | 0 | 2 | 5 | 2 | Q80U72-4; | Isoform 4 c | Q80U72-4 | 0 | 3201.555 | 0 | 0.7 | 0.010948 | 4.67 | 81.5 | 118.5 | 18.14 | 15.14 | Shared | High | High | High | High | High | High | High | High | High | High | High | 0 | 0.000269 | 4.16 |
| FALSE | High | LAEAYLSPG | [K].LAEAYL | 1xPhospho | Q9D2X8 1x | 0.007605 | 0.000664 | 1 | 2 | 1 | Q9D2X8 | Uncharacte | Q9D2X8 [1 | 0 | 2675.421 | 0 | -0.22 | 0.249087 | 6.67 | 106.2 | 93.8 | 11.48 | 14.49 | | High | High | High | High | High | High | High | High | High | High | High | 0.000294 | 0.002316 | 3.06 |
| FALSE | High | LAEDDWT( | [R].LAEDD\ | 1xPhospho | Q80W37 1: | 5.5E-05 | 0 | 1 | 1 | 2 | Q80W37 | Snurportin | Q80W37 [6 | 0 | 2478.078 | 0 | 0.57 | 0.014672 | 1.89 | 76.6 | 123.4 | 9.44 | 14.09 | | High | High | High | High | High | High | High | High | High | High | High | 0.000275 | 0.002278 | 3.03 |
| FALSE | High | LAEDDWT( | [R].LAEDD\ | 1xPhospho | Q80W37 1: | 0.000845 | 0 | 1 | 1 | 2 | Q80W37 | Snurportin | Q80W37 [6 | 0 | 2494.073 | 0 | 0.01 | 0.232167 | 3.23 | 91.8 | 108.2 | 9.63 | 21.66 | | High | High | High | High | High | High | High | High | High | High | High | 0 | 0.000198 | 3.26 |
| FALSE | High | IAEEIGIEGS | [K].IAEEIGI | 1xPhospho | A0A1S6GW | 1.18E-06 | 0 | 1 | 4 | 2 | A0A1S6GW | Uncharacte | A0A1S6GW | 0 | 3099.409 | 0 | -0.51 | 0.021592 | 11.56 | 119 | 81 | 7.38 | 14.35 | | High | High | High | High | High | High | High | High | High | High | High | 0 | 1.22E-07 | 5.2 |
| FALSE | High | LAEPAHCF | [K].LAEPAH | 1xCarbami | Q8K1J5 1xF | 0.000144 | 0 | 1 | 1 | 3 | Q8K1J5 | Protein SDI | Q8K1J5 [14 | 0 | 2100.941 | 0 | -1.58 | 0.000181 | 1.62 | 152.1 | 47.9 | 4.34 | 15.26 | | High | High | High | High | High | High | High | High | High | High | High | 0 | 4.29E-05 | 3.09 |
| FALSE | High | LAESPQLSK | [R].LAESPC | 1xPhospho | B2RTE5 1xF | 0.007082 | 0.00062 | 1 | 2 | 1 | B2RTE5 | Coiled-coil | B2RTE5 [9- | 0 | 1510.828 | 0 | -0.93 | 0.003574 | 4.18 | 129.1 | 70.9 | 10.25 | 13.72 | | High | High | High | High | High | High | High | High | High | High | High | 0.000275 | 0.002139 | 2.63 |
| FALSE | High | IAFDAESEP | [R].IAFDAE | 1xPhospho | Q571D6 1x | 0.01002 | 0.000736 | 1 | 3 | 2 | Q571D6 | MKIAA028 | Q571D6 [3 | 0 | 2467.11 | 0 | 1.64 | 0.000879 | 14.32 | 48.4 | 151.6 | 5.67 | 9.64 | | High | High | High | High | High | High | High | High | High | High | High | 0.000344 | 0.00315 | 2.91 |
| FALSE | High | LAGASPAT | [R].LAGASF | 1xPhospho | Q9JJ28 1xP | 0.015465 | 0.001089 | 1 | 1 | 4 | Q9JJ28 | Protein flig | Q9JJ28 [40 | 0 | 2195.184 | 0 | 0.44 | 0.016906 | 6.26 | 89.1 | 110.9 | 8.96 | 9.55 | | High | High | High | High | High | High | High | High | High | High | High | 0.000571 | 0.005087 | 3.12 |
| FALSE | High | LAGGGVLG | [R].LAGGG | 1xPhospho | E9PYL2 1xF | 0.002553 | 0.000109 | 1 | 1 | 1 | E9PYL2 | Proline-rich | E9PYL2 [20 | 0 | 3362.761 | 0 | 0.15 | 0.265411 | 9.12 | 92.8 | 107.2 | 19.91 | 7.67 | | High | High | High | High | High | High | High | High | High | High | High | 4.94E-05 | 0.00068 | 1.98 |
| FALSE | High | LAGGQTSC | [R].LAGGQ | 1xPhospho | Q80TU3 1x | 5.65E-08 | 0 | 1 | 6 | 4 | Q80TU3 | MKIAA068 | Q80TU3 [3 | 0 | 2149.085 | 0 | -1.07 | 0.000805 | 3.92 | 135.5 | 64.5 | 5.93 | 9.53 | | High | High | High | High | High | High | High | High | High | High | High | 0 | 4.04E-09 | 3.79 |
| FALSE | High | LAGGQTSC | [R].LAGGQ | 2xPhospho | Q80TU3 2x | 0.026885 | 0.001676 | 1 | 6 | 3 | Q80TU3 | MKIAA068 | Q80TU3 [3 | 0 | 2229.052 | 0 | -0.19 | 0.021965 | 5.92 | 105.4 | 94.6 | 5.55 | 1.97 | | High | High | High | High | High | High | High | High | High | High | High | 0.000903 | 0.009547 | 3.4 |
| FALSE | High | LAGLEQAV | [R].LAGLEC | 1xPhospho | Q9EST3 1x | 0.000524 | 0 | 1 | 6 | 1 | Q9EST3 | Eukaryotic | Q9EST3 [3 | 0 | 3791.868 | 0 | 0.62 | 0.003461 | 13.72 | 72.1 | 127.9 | 10.7 | 11.02 | | High | High | High | High | High | High | High | High | High | High | High | 0 | 0.000115 | 3.59 |
| FALSE | High | LAGLTVSSF | [R].LAGLTV | 1xPhospho | Q76I79 1xF | 0.000308 | 0 | 1 | 3 | 2 | Q76I79 | Protein ph | Q76I79 [95 | 0 | 1623.949 | 0 | 0.28 | 0.012168 | 4.19 | 91.2 | 108.8 | 6.57 | 5.36 | | High | High | High | High | High | High | High | High | High | High | High | 0 | 6.34E-05 | 2.99 |
| FALSE | High | LAGNEALSI | [K].LAGNE/ | 1xPhospho | Q8C5R2 1x | 0.0004 | 0 | 1 | 1 | 2 | Q8C5R2 | Proline anc | Q8C5R2 [2 | 0 | 1910.004 | 0 | 0.52 | 0.006837 | 17.1 | 82 | 118 | 16.39 | 7.72 | | High | High | High | High | High | High | High | High | High | High | High | 0.000275 | 0.001895 | 2.96 |
| FALSE | High | LAGNEALSI | [K].LAGNE/ | 2xPhospho | Q8C5R2 2x | 0.054844 | 0.003982 | 1 | 1 | 1 | Q8C5R2 | Proline anc | Q8C5R2 [2 | 0 | 1989.97 | 0 | 1.58 | 0.000362 | 9.93 | 48.3 | 151.7 | 10.57 | 7.62 | | High | High | High | High | High | High | High | High | High | High | High | 0.002059 | 0.02151 | 2 |
| FALSE | High | LAGNLHFS( | [K].LAGNLH | 1xPhospho | E9Q616 1x | 0.000519 | 0 | 1 | 6 | 1 | E9Q616 | AHNAK nuc | E9Q616 [5 | 0 | 1678.908 | 0 | 2.8 | 0.000829 | 5.09 | 24.8 | 175.2 | 7.94 | 13.84 | | High | High | High | High | High | High | High | High | High | High | High | 0 | 0.000114 | 1.22 |
| FALSE | High | LAGSGDDS | [R].LAGSGI | 1xPhospho | Q8BQ30-2 | 0.102567 | 0.008249 | 1 | 4 | 1 | Q8BQ30-2 | Isoform 2 c | Q8BQ30-2 | 0 | 1255.577 | 0 | 0.87 | 0.130977 | 29.43 | 68.5 | 131.5 | 22.07 | 41.06 | | High | High | High | High | High | High | High | High | High | High | High | 0.004138 | 0.04434 | 1.64 |
| FALSE | High | LAGSGDDS | [R].LAGSGI | 1xPhospho | Q8BQ30-2 | 9.36E-05 | 0 | 1 | 4 | 3 | Q8BQ30-2 | Isoform 2 c | Q8BQ30-2 | 0 | 1484.74 | 0 | 0.65 | 0.00291 | 3.39 | 76.1 | 123.9 | 6.9 | 6.63 | | High | High | High | High | High | High | High | High | High | High | High | 0 | 1.73E-05 | 2.57 |
| FALSE | High | LAGSGDDS | [R].LAGSGI | 2xPhospho | Q8BQ30-2 | 0.060666 | 0.00451 | 1 | 4 | 1 | Q8BQ30-2 | Isoform 2 c | Q8BQ30-2 | 0 | 1564.706 | 0 | 1.21 | 0.00251 | 10.04 | 58.2 | 141.8 | 15.24 | 11.62 | | High | High | High | High | High | High | High | High | High | High | High | 0.002341 | 0.02411 | 1.78 |
| FALSE | High | LAGSSAASS | [K].LAGSSA | 1xPhospho | Q80TS5 1x | 9.42E-07 | 0 | 1 | 4 | 2 | Q80TS5 | Zinc finger | Q80TS5 [1 | 0 | 2111.09 | 0 | -1.92 | 0.000193 | 1.04 | 158.3 | 41.7 | 4.2 | 15.34 | | High | High | High | High | High | High | High | High | High | High | High | 0 | 5.51E-05 | 2.37 |
| FALSE | High | LAGSSAASS | [K].LAGSSA | 1xDeamida | Q80TS5 1x | 0.00582 | 0.00062 | 1 | 4 | 5 | Q80TS5 | Zinc finger | Q80TS5 [1 | 0 | 2112.074 | 0 | -1.48 | 0.002069 | 4.83 | 147.3 | 52.7 | 7.97 | 21.77 | | High | High | High | High | High | High | High | High | High | High | High | 0.000275 | 0.001713 | 2.49 |
| FALSE | High | IACDEEFSD | [R].IACDEE | 1xCarbami | A0A0R4J0C | 0.103392 | 0.008249 | 1 | 4 | 1 | A0A0R4J0C | Histone de | A0A0R4J0C | 1 | 2546.958 | 0 | 0.36 | 0.086759 | 4.23 | 86.8 | 113.2 | 22.1 | 7.51 | | High | High | High | High | High | High | High | High | High | High | High | 0.004164 | 0.04501 | 2.38 |
| FALSE | High | IDISPSTFR | [R].IDISPST | 1xPhospho | Q569Z6 1x | 0.035793 | 0.002304 | 1 | 4 | 3 | Q569Z6 | Thyroid ho | Q569Z6 [6 | 0 | 1344.676 | 0 | -0.22 | 0.008068 | 1 | 106.6 | 93.4 | 4.67 | 3.8 | | High | High | High | High | High | High | High | High | High | High | High | 0.00639 | 0.06892 | 1.99 |
| FALSE | High | HLSSTSDDI | [R].HLSSTS | 2xPhospho | Q9DBC3 2x | 0.002192 | 0.000109 | 1 | 2 | 1 | Q9DBC3 | Cap-specifi | Q9DBC3 [2 | 0 | 2384.037 | 0 | 1.37 | 0.008132 | 32.29 | 59.6 | 140.4 | 19.68 | 13.66 | | High | High | High | High | High | High | High | High | High | High | High | 4.94E-05 | 0.000577 | 2.57 |
| FALSE | High | HLSQDHTG | [R].HLSQDI | 1xPhospho | H9LAN6 1x | 0.097743 | 0.007804 | 1 | 6 | 1 | H9LAN6 | Histone-lys | H9LAN6 [7 | 0 | 1718.863 | 0 | -0.78 | 0.000225 | 1.76 | 127.8 | 72.2 | 3.24 | 6.03 | | High | High | High | High | High | High | High | High | High | High | High | 0.00395 | 0.042 | 1.17 |
| FALSE | High | GSSPTSPCS | [K].GSSPTS | 1xCarbami | Q8CJ67 1xF | 4.07E-07 | 0 | 1 | 6 | 2 | Q8CJ67 | Double-str | Q8CJ67 [48 | 0 | 2043.982 | 0 | 1.19 | 0.000515 | 3.95 | 60.9 | 139.1 | 3.64 | 5.96 | | High | High | High | High | High | High | High | High | High | High | High | 0 | 3.72E-08 | 2.94 |
| FALSE | High | GSSPTSPCS | [K].GSSPTS | 1xCarbami | Q8CJ67 2xF | 0.00359 | 0.00021 | 1 | 6 | 1 | Q8CJ67 | Double-str | Q8CJ67 [48 | 0 | 2123.949 | 0 | 0.84 | 0.001151 | 11 | 68.6 | 131.4 | 10.36 | 7.11 | | High | High | High | High | High | High | High | High | High | High | High | 9.42E-05 | 0.001001 | 3.17 |
| FALSE | High | GSSQLDVN | [K].GSSQLI | 1xPhospho | O35379 1x | 0.000786 | 0 | 1 | 3 | 13 | O35379 | Multidrug r | O35379 [2 | 0 | 2368.241 | 0 | 1.16 | 0.000729 | 8.02 | 62.4 | 137.6 | 3.57 | 6.95 | | High | High | High | High | High | High | High | High | High | High | High | 4.94E-05 | 0.000587 | 3.16 |
| FALSE | High | GSSQPNLST | [R].GSSQPI | 1xPhospho | Q8CHU3 1x | 5.31E-09 | 0 | 1 | 7 | 7 | Q8CHU3 | Epsin-2 OS | Q8CHU3 [1 | 0 | 2500.165 | 0 | 1.15 | 0.000308 | 3.63 | 62 | 138 | 6.03 | 5.57 | | High | High | High | High | High | High | High | High | High | High | High | 0 | 2.82E-10 | 4.85 |
| FALSE | High | GSSQPNLST | [R].GSSQPI | 2xPhospho | Q8CHU3 2x | 0.029844 | 0.001857 | 1 | 7 | 4 | Q8CHU3 | Epsin-2 OS | Q8CHU3 [1 | 0 | 2580.131 | 0 | 0.25 | 0.020522 | 5.78 | 88 | 112 | 5.61 | 8.6 | | High | High | High | High | High | High | High | High | High | High | High | 0.004422 | 0.04759 | 2.67 |
| FALSE | High | GSSQVGSC | [K].GSSQV( | 1xPhospho | Q8BUM9-2 | 0.006713 | 0.00062 | 1 | 3 | 6 | Q8BUM9-2 | Isoform 2 c | Q8BUM9-2 | 0 | 2238.096 | 0 | 0.55 | 0.005069 | 2.94 | 78.3 | 121.7 | 6.13 | 10.63 | | High | High | High | High | High | High | High | High | High | High | High | 0.000275 | 0.002006 | 4.06 |
| FALSE | High | GSSSASVTI | [K].GSSSAS | 1xPhospho | O55111 1x | 0.003721 | 0.00021 | 1 | 2 | 3 | O55111 | Desmoglein | O55111 [7 | 0 | 1361.708 | 0 | 0.64 | 0.004363 | 15.75 | 79.2 | 120.8 | 5.13 | 10.88 | | High | High | High | High | High | High | High | High | High | High | High | 9.42E-05 | 0.001042 | 1.41 |
| FALSE | High | GSSSEVFSD | [K].GSSSEV | 1xPhospho | B7ZCJ1 1xF | 0.009087 | 0.000736 | 1 | 7 | 1 | B7ZCJ1 | Rho GTPas | B7ZCJ1 [92 | 0 | 1521.678 | 0 | 0.55 | 0.00302 | 4.92 | 82.9 | 117.1 | 7.89 | 5.14 | | High | High | High | High | High | High | High | High | High | High | High | 0.000327 | 0.002827 | 3.33 |
| FALSE | High | GSSSPLGVT | [K].GSSSPL | 1xPhospho | Q7TSH3 1x | 0.000147 | 0 | 1 | 1 | 3 | Q7TSH3 | Zinc finger | Q7TSH3 [8 | 0 | 1571.845 | 0 | -0.25 | 0.015402 | 4.37 | 106 | 94 | 6.6 | 7.35 | | High | High | High | High | High | High | High | High | High | High | High | 0 | 2.77E-05 | 1.64 |
| FALSE | High | GSSSQGPS | [R].GSSSQ( | 1xPhospho | Q3TSW4 1: | 2.54E-06 | 0 | 1 | 3 | 2 | Q3TSW4 | MCG15576 | Q3TSW4 [2 | 0 | 2380.123 | 0 | -0.35 | 0.150403 | 6.99 | 111.9 | 88.1 | 13.31 | 13.08 | | High | High | High | High | High | High | High | High | High | High | High | 0 | 2.89E-07 | 2.77 |
| FALSE | High | GSSSQGPS | [R].GSSSQ( | 2xPhospho | Q3TSW4 2: | 0.006594 | 0.00062 | 1 | 3 | 1 | Q3TSW4 | MCG15576 | Q3TSW4 [2 | 0 | 2460.089 | 0 | 0.68 | 0.002422 | 9.01 | 74.2 | 125.8 | 11.17 | 8.57 | | High | High | High | High | High | High | High | High | High | High | High | 0.000275 | 0.001966 | 2.71 |
| FALSE | High | GSSSSVCS\ | [K].GSSSSV | 1xCarbami | Q69ZR2 1x | 0.000237 | 0 | 1 | 3 | 6 | Q69ZR2 | E3 ubiquiti | Q69ZR2 [1 | 0 | 2555.231 | 0 | 0.8 | 0.000787 | 8.58 | 71.7 | 128.3 | 7.13 | 3.89 | | High | High | High | High | High | High | High | High | High | High | High | 0 | 6.12E-05 | 2.71 |
| FALSE | High | GSSSTPNSI | [K].GSSSTP | 1xPhospho | Q69Z38 1x | 5.75E-05 | 0 | 1 | 1 | 3 | Q69Z38 | Pseudopoc | Q69Z38 [4 | 0 | 2999.56 | 0 | 0.08 | 0.938323 | 12.57 | 99.7 | 100.3 | 9.38 | 23.89 | | High | High | High | High | High | High | High | High | High | High | High | 0 | 3.13E-05 | 4.78 |
| FALSE | High | GSSPTHSA | [R].GSSPTH | 1xPhospho | I7HJQ9 1xF | 0.021607 | 0.001507 | 1 | 4 | 2 | I7HJQ9 | Myotubula | I7HJQ9 [65 | 0 | 1773.837 | 0 | 1.13 | 0.008588 | 21.34 | 63.9 | 136.1 | 8.32 | 15.79 | | High | High | High | High | High | High | High | High | High | High | High | 0.000801 | 0.007474 | 3.25 |

| | | | | | | | | | | | | | | | | | | | | | | | | | | | | | | | | | | | | | | |
|---|---|---|---|---|---|---|---|---|---|---|---|---|---|---|---|---|---|---|---|---|---|---|---|---|---|---|---|---|---|---|---|---|---|---|---|---|---|---|
| FALSE | High | GSSTDISED | [K].GSSTDI | 1xPhospho | Q80Y55 1x | 0.012619 | 0.000889 | 1 | 1 | 2 | Q80Y55 | BSD domai | Q80Y55 [3 | 0 | 1891.873 | 0 | -0.22 | 0.063511 | 12.12 | 106.7 | 93.3 | 8.81 | 15.83 | | High | High | High | High | High | High | High | High | High | High | High | 0.000859 | 0.008608 | 2.03 |
| FALSE | High | GSSYSESSE | [K].GSSYSE | 1xPhospho | B7ZP47 1xl | 4.77E-06 | 0 | 1 | 2 | 12 | B7ZP47 | Wapal prot | B7ZP47 [94 | 0 | 3174.477 | 0 | -1.09 | 0.006349 | 8.86 | 141.7 | 58.3 | 14.41 | 42.45 | | High | High | High | High | High | High | High | High | High | High | High | 0 | 1.63E-05 | 4.51 |
| FALSE | High | GSTAVLAA | [K].GSTAVI | 1xPhospho | Q9WUF3 1 | 0.000276 | 0 | 1 | 1 | 1 | Q9WUF3 | CASP8-assc | Q9WUF3 [ | 0 | 1969.077 | 0 | -0.43 | 0.006891 | 3.31 | 114.7 | 85.3 | 2.87 | 8.92 | | High | High | High | High | High | High | High | High | High | High | High | 0 | 5.63E-05 | 2.72 |
| FALSE | High | GSTDNLMI | [R].GSTDN | 1xPhospho | Q6A0F1 1x | 0.018447 | 0.001324 | 1 | 9 | 1 | Q6A0F1 | MKIAA000 | Q6A0F1 [3 | 0 | 1690.719 | 0 | -0.83 | 0.001562 | 5.48 | 128 | 72 | 2.45 | 12.36 | | High | High | High | High | High | High | High | High | High | High | High | 0.000706 | 0.006227 | 2.68 |
| FALSE | High | GSTEALSG | [K].GSTEAL | 1xCarbami | E9Q6J5-3 1 | 0.00017 | 0 | 1 | 3 | 3 | E9Q6J5-3 | Isoform Ne | E9Q6J5-3 [ | 0 | 3019.338 | 0 | 0.51 | 0.109669 | 12.97 | 79 | 121 | 17.16 | 21.49 | | High | High | High | High | High | High | High | High | High | High | High | 0 | 3.27E-05 | 4.33 |
| FALSE | High | GSTESPQW | [R].GSTESP | 1xPhospho | E9Q921 1x | 0.017498 | 0.001245 | 1 | 7 | 2 | E9Q921 | AF4/FMR2 | E9Q921 [5 | 0 | 1356.615 | 0 | 0.05 | 0.063731 | 1.9 | 94.6 | 105.4 | 4.16 | 4.75 | | High | High | High | High | High | High | High | High | High | High | High | 0.004072 | 0.04357 | 2.18 |
| FALSE | High | GSTFSPDLK | [K].GSTFSP | 1xPhospho | E9QML5 1x | 0.008769 | 0.000736 | 1 | 10 | 2 | E9QML5 | Zinc finger | E9QML5 [1 | 0 | 1489.77 | 0 | -0.46 | 0.002933 | 4.15 | 117 | 83 | 4.69 | 5.8 | | High | High | High | High | High | High | High | High | High | High | High | 0.000903 | 0.009344 | 2.09 |
| FALSE | High | GSTGSAGG | [K].GSTGS | 1xPhospho | Q8CH25-2 | 0.000633 | 0 | 1 | 4 | 1 | Q8CH25-2 | Isoform 2 c | Q8CH25-2 | 0 | 1765.837 | 0 | -0.85 | 0.008588 | 7.46 | 126.9 | 73.1 | 7.04 | 14.24 | | High | High | High | High | High | High | High | High | High | High | High | 0 | 0.000143 | 2.29 |
| FALSE | High | GSTIETEQK | [K].GSTIET | 1xPhospho | E9QLZ9 1xl | 0.015329 | 0.001089 | 1 | 12 | 2 | E9QLZ9 | Protein ena | E9QLZ9 [69 | 0 | 1530.782 | 0 | -0.19 | 0.657541 | 15.27 | 109.3 | 90.7 | 15.85 | 34.64 | | High | High | High | High | High | High | High | High | High | High | High | 0.000571 | 0.005064 | 1.68 |
| FALSE | High | GSTNLFSK | [R].GSTNLF | 1xPhospho | Q03141 1x | 0.08727 | 0.006537 | 1 | 5 | 2 | Q03141 | MAP/micrc | Q03141 [6 | 0 | 1162.571 | 0 | -1.04 | 0.002349 | 3.28 | 134.6 | 65.4 | 11.22 | 17.35 | | High | High | High | High | High | High | High | High | High | High | High | 0.006342 | 0.06813 | 1.94 |
| FALSE | High | GSTNLFSK | [R].GSTNLF | 1xPhospho | Q03141 1x | 0.014158 | 0.001011 | 1 | 5 | 1 | Q03141 | MAP/micrc | Q03141 [6 | 0 | 1391.734 | 0 | -0.93 | 0.00194 | 2.21 | 131 | 69 | 10.27 | 9.55 | | High | High | High | High | High | High | High | High | High | High | High | 0.000538 | 0.004614 | 1.39 |
| FALSE | High | GSTSPDLL | [K].GSTSPD | 1xPhospho | Q9CQX8 1x | 3.38E-05 | 0 | 1 | 2 | 3 | Q9CQX8 | 28S ribosо | Q9CQX8 [5 | 0 | 2761.388 | 0 | -0.88 | 0.001817 | 2.22 | 127.3 | 72.7 | 7.97 | 10 | | High | High | High | High | High | High | High | High | High | High | High | 0 | 2.02E-05 | 3.65 |
| FALSE | High | GSTSQSTS | [R].GSTSQ | 1xCarbami | Q3UW53 1 | 4.52E-06 | 0 | 1 | 1 | 4 | Q3UW53 | Protein Nit | Q3UW53 [ | 0 | 3960.81 | 0 | 0.11 | 0.373511 | 13.96 | 93.8 | 106.2 | 18.73 | 8.77 | | High | High | High | High | High | High | High | High | High | High | High | 0 | 5.57E-07 | 5.37 |
| FALSE | High | GSTSQSTS | [R].GSTSQ | 1xCarbami | Q3UW53 1 | 0.003857 | 0.000261 | 1 | 1 | 2 | Q3UW53 | Protein Nit | Q3UW53 [ | 0 | 3961.794 | 0 | 0.31 | 0.311557 | 17.07 | 89.5 | 110.5 | 29.23 | 17.39 | | High | High | High | High | High | High | High | High | High | High | High | 0.000117 | 0.001079 | 4.13 |
| FALSE | High | GSSWSPPP | [K].GSSWS | 1xPhospho | Q8CJF7 1xl | 6.41E-05 | 0 | 1 | 1 | 4 | Q8CJF7 | Protein EL | Q8CJF7 [21 | 0 | 1821.955 | 0 | 0.25 | 0.438206 | 31.26 | 90.9 | 109.1 | 15.8 | 17.97 | | High | High | High | High | High | High | High | High | High | High | High | 0 | 1.09E-05 | 2.18 |
| FALSE | High | GSTSSSPYV | [R].GSTSSS | 1xDeamida | Q61286-2 | 0.111965 | 0.009053 | 2 | 11 | 1 | Q61286-2; | Isoform AL | Q61286-2 | 0 | 2969.409 | 0 | 0.11 | 0.973475 | 44.37 | 96.2 | 103.8 | 24.5 | 9.5 | Shared | High | High | High | High | High | High | High | High | High | High | High | 0.004543 | 0.04934 | 2.4 |
| FALSE | High | GSSPPSAM | [R].GSSPPS | 1xPhospho | Q05BA5 1x | 0.023584 | 0.001578 | 1 | 6 | 2 | Q05BA5 | Nuclear re | Q05BA5 [1 | 0 | 1535.643 | 0 | -0.22 | 0.511932 | 5.57 | 107.8 | 92.2 | 8.62 | 16.57 | | High | High | High | High | High | High | High | High | High | High | High | 0.004317 | 0.04607 | 2.06 |
| FALSE | High | GSSLSSESS | [R].GSSLSS | 1xPhospho | Q3U3C9-2 | 0.015195 | 0.001089 | 1 | 4 | 2 | Q3U3C9-2 | Isoform 2 c | Q3U3C9-2 | 0 | 3024.456 | 0 | -1.25 | 0.033496 | 17.91 | 141.4 | 58.6 | 13.53 | 57.11 | | High | High | High | High | High | High | High | High | High | High | High | 0.000571 | 0.005028 | 2.86 |
| FALSE | High | GSQGTSPF | [K].GSQGT | 2xPhospho | F6YZ95 2xF | 0.001476 | 5.66E-05 | 2 | 7 | 1 | F6YZ95; Q8 | Liprin-beta | F6YZ95 [12 | 0 | 2236.918 | 0 | 0.23 | 0.095945 | 17.33 | 93.7 | 106.3 | 12.43 | 15.82 | Shared | High | High | High | High | High | High | High | High | High | High | High | 2.58E-05 | 0.000369 | 2.83 |
| FALSE | High | GSQITQQS | [K].GSQITC | 1xPhospho | Q99J95 1xl | 9.56E-06 | 0 | 1 | 3 | 2 | Q99J95 | Cyclin-dep | Q99J95 [34 | 0 | 1743.822 | 0 | -0.27 | 0.01126 | 5.84 | 109.9 | 90.1 | 2.45 | 7.94 | | High | High | High | High | High | High | High | High | High | High | High | 0 | 5.71E-06 | 4.27 |
| FALSE | High | GSQNSSEH | [R].GSQNS | 1xPhospho | Q8BZ47 1x | 0.001898 | 5.66E-05 | 1 | 2 | 2 | Q8BZ47 | Zinc finger | Q8BZ47 [4 | 0 | 2638.251 | 0 | 0 | 0.779137 | 18.4 | 103.9 | 96.1 | 14.35 | 12.84 | | High | High | High | High | High | High | High | High | High | High | High | 0.000161 | 0.001145 | 2.42 |
| FALSE | High | GSQSPK | [R].GSQSPI | 2xPhospho | Q569Z6 2x | 0.096959 | 0.007777 | 1 | 5 | 1 | Q569Z6 | Thyroid ho | Q569Z6 [4 | 0 | 1221.568 | 0 | -0.16 | 0.410585 | 36.81 | 104.8 | 95.2 | 24.9 | 28.84 | | High | High | High | High | High | High | High | High | High | High | High | 0.003922 | 0.04179 | 1.89 |
| FALSE | High | GSQVTHSS | [R].GSQVT | 1xCarbami | Q8CH77 1x | 0.025293 | 0.001645 | 1 | 3 | 2 | Q8CH77 | Neuron na | Q8CH77 [2 | 0 | 2775.159 | 0 | 0.13 | 0.1347 | 16.36 | 97.6 | 102.4 | 17.22 | 6.64 | | High | High | High | High | High | High | High | High | High | High | High | 0.000872 | 0.008848 | 3.06 |
| FALSE | High | GSQVTHSS | [R].GSQVT | 1xCarbami | Q8CH77 1x | 1.91E-08 | 0 | 1 | 3 | 6 | Q8CH77 | Neuron na | Q8CH77 [2 | 0 | 2791.154 | 0 | 1.09 | 0.001172 | 14.5 | 65.1 | 134.9 | 14.14 | 14.01 | | High | High | High | High | High | High | High | High | High | High | High | 0 | 1.19E-09 | 5.27 |
| FALSE | High | GSRGPENL | [R].GSRGP | 1xPhospho | Q91ZV0 1x | 0.001179 | 0 | 1 | 12 | 2 | Q91ZV0 | Melanoma | Q91ZV0 [1 | 1 | 2279.044 | 0 | 0.18 | 0.030215 | 6.78 | 93.7 | 106.3 | 4.71 | 12.11 | | High | High | High | High | High | High | High | High | High | High | High | 2.58E-05 | 0.000487 | 2.33 |
| FALSE | High | GSRSDELL | [R].GSRSD | 1xPhospho | Q8K1N2-2 | 0.087983 | 0.006639 | 1 | 3 | 3 | Q8K1N2-2 | Isoform 2 c | Q8K1N2-2 | 1 | 1727.853 | 0 | 1.64 | 0.000364 | 9.15 | 49 | 151 | 9.17 | 8.31 | | High | High | High | High | High | High | High | High | High | High | High | 0.004667 | 0.05037 | 2.37 |
| FALSE | High | GSRSEDSE | [K].GSRSED | 1xPhospho | O89032 1x | 1.17E-05 | 0 | 1 | 3 | 2 | O89032 | SH3 and PX | O89032 [7 | 1 | 2329.033 | 0 | 0.6 | 0.004907 | 19.53 | 76.5 | 123.5 | 11.22 | 6.63 | | High | High | High | High | High | High | High | High | High | High | High | 0 | 6.6E-06 | 3.31 |
| FALSE | High | GSRSEDSE | [K].GSRSED | 1xPhospho | O89032 1x | 0.000438 | 0 | 1 | 3 | 6 | O89032 | SH3 and PX | O89032 [7 | 1 | 2345.028 | 0 | 0.73 | 0.00292 | 11.74 | 75.6 | 124.4 | 5.45 | 11.97 | | High | High | High | High | High | High | High | High | High | High | High | 0 | 9.47E-05 | 3.69 |
| FALSE | High | GSSDDEEN | [K].GSSDD | 1xCarbami | Z4YKQ9 1x | 0.028082 | 0.001791 | 1 | 3 | 2 | Z4YKQ9 | ATPase fan | Z4YKQ9 [1 | 1 | 3058.48 | 0 | 0.48 | 0.015988 | 10.84 | 79.6 | 120.4 | 11.79 | 12.93 | | High | High | High | High | High | High | High | High | High | High | High | 0.000968 | 0.01003 | 2.41 |
| FALSE | High | GSSDHLHS | [K].GSSDH | 1xPhospho | Q6VGS5-2 | 0.075304 | 0.005727 | 1 | 3 | 1 | Q6VGS5-2 | Isoform 2 c | Q6VGS5-2 | 0 | 1304.595 | 0 | -0.87 | 0.005966 | 6.07 | 131.9 | 68.1 | 10.04 | 16.43 | | High | High | High | High | High | High | High | High | High | High | High | 0.002951 | 0.03086 | 1.22 |
| FALSE | High | GSSEGSES | [K].GSSEG | 1xPhospho | Q3UC10 1x | 3.85E-06 | 0 | 1 | 2 | 20 | Q3UC10 | DNA helica | Q3UC10 [7 | 0 | 3114.355 | 0 | -0.16 | 0.196312 | 19.79 | 106.1 | 93.9 | 12.45 | 10.98 | | High | High | High | High | High | High | High | High | High | High | High | 0 | 4.6E-07 | 4.85 |
| FALSE | High | GSSPDVHT | [R].GSSPD | 1xPhospho | O54724 1x | 2.76E-07 | 0 | 1 | 1 | 12 | O54724 | Caveolae-a | O54724 [3 | 0 | 2992.517 | 0 | 0.94 | 0.00521 | 14.42 | 68.7 | 131.3 | 11.66 | 9.43 | | High | High | High | High | High | High | High | High | High | High | High | 0 | 2.38E-08 | 4.3 |
| FALSE | High | GSSESQSL | [R].GSSESC | 1xPhospho | E9PWW9 1 | 3.6E-05 | 0 | 1 | 3 | 2 | E9PWW9 | Remodelin | E9PWW9 [ | 0 | 2314.158 | 0 | -0.94 | 0.000143 | 0.48 | 130.9 | 69.1 | 3.27 | 4.08 | | High | High | High | High | High | High | High | High | High | High | High | 0 | 5.83E-05 | 2.73 |
| FALSE | High | GSSFQSGR | [R].GSSFQ | 1xPhospho | Q9D4H1 1x | 0.023584 | 0.001578 | 1 | 2 | 3 | Q9D4H1 | Exocyst co | Q9D4H1 [4 | 0 | 1134.514 | 0 | 0.18 | 0.033138 | 9.34 | 93.7 | 106.3 | 4.2 | 6.81 | | High | High | High | High | High | High | High | High | High | High | High | 0.000872 | 0.00911 | 1.11 |
| FALSE | High | GSSFTFLTP | [R].GSSFTF | 1xPhospho | Q9Z206 1x | 0.000289 | 0 | 1 | 1 | 2 | Q9Z206 | Neuroepith | Q9Z206 [4 | 0 | 2576.299 | 0 | -1.02 | 0.000373 | 2.37 | 135.2 | 64.8 | 4.2 | 7.73 | | High | High | High | High | High | High | High | High | High | High | High | 0 | 0.000138 | 3.64 |
| FALSE | High | GSSGDSDS | [R].GSSGD | 1xPhospho | O89032 1x | 0.000134 | 0 | 1 | 3 | 9 | O89032 | SH3 and PX | O89032 [6 | 0 | 2276.106 | 0 | 1.54 | 0.000556 | 14.87 | 51.2 | 148.8 | 13.44 | 4.52 | | High | High | High | High | High | High | High | High | High | High | High | 0 | 9.39E-05 | 3.45 |
| FALSE | High | GSSGEGLP | [R].GSSGE | 1xPhospho | Q8VHK1 1x | 0.001179 | 0 | 1 | 1 | 2 | Q8VHK1 | Caskin-2 O | Q8VHK1 [9 | 0 | 2344.184 | 0 | -0.02 | 0.816363 | 6.81 | 100.2 | 99.8 | 5.53 | 11.02 | | High | High | High | High | High | High | High | High | High | High | High | 2.58E-05 | 0.000357 | 2.21 |
| FALSE | High | GSSGLNLG | [K].GSSGLN | 1xPhospho | P97467 1xl | 1.19E-06 | 0 | 1 | 5 | 2 | P97467 | Peptidyl-gl | P97467 [92 | 0 | 1735.837 | 0 | 3.23 | 0.000167 | 1.77 | 19.3 | 180.7 | 16.89 | 4.68 | | High | High | High | High | High | High | High | High | High | High | High | 0 | 1.24E-07 | 2.15 |
| FALSE | High | GSSGNASE | [K].GSSGN | 1xCarbami | A0A0R4J0J | 4.37E-07 | 0 | 1 | 4 | 4 | A0A0R4J0J | Cytospin-A | A0A0R4J0J | 0 | 2032.921 | 0 | 1.19 | 0.000459 | 7.45 | 62.1 | 137.9 | 4.78 | 5.39 | | High | High | High | High | High | High | High | High | High | High | High | 0 | 4.02E-08 | 3.73 |
| FALSE | High | GSSGNASE | [K].GSSGN | 1xCarbami | A0A0R4J0J | 2.13E-08 | 0 | 1 | 4 | 1 | A0A0R4J0J | Cytospin-A | A0A0R4J0J | 0 | 2112.887 | 0 | 1.43 | 0.001615 | 6.8 | 52.5 | 147.5 | 15.93 | 9.58 | | High | High | High | High | High | High | High | High | High | High | High | 0 | 1.35E-09 | 3.77 |
| FALSE | High | GSSGSEAC | [K].GSSGSE | 1xCarbami | O88466 1x | 1.43E-05 | 0 | 2 | 4 | 3 | O88466; R | Zinc finger | O88466 [1 | 0 | 1740.746 | 0 | -1.19 | 0.000897 | 0.66 | 139.4 | 60.6 | 7.23 | 4.34 | Shared | High | High | High | High | High | High | High | High | High | High | High | 0 | 2.04E-06 | 1.83 |
| FALSE | High | GSSGSPAH | [R].GSSGSI | 1xPhospho | P47930 1xl | 3.53E-08 | 0 | 1 | 1 | 3 | P47930 | Fos-relatec | P47930 [15 | 0 | 2474.135 | 0 | 1.17 | 0.023293 | 21.79 | 63.1 | 136.9 | 13.66 | 19.5 | | High | High | High | High | High | High | High | High | High | High | High | 0 | 2.37E-09 | 4.49 |
| FALSE | High | GSSLADQR | [R].GSSLAD | 1xPhospho | Q3UDC3 1x | 0.022773 | 0.001542 | 1 | 3 | 1 | Q3UDC3 | Putative ur | Q3UDC3 [3 | 0 | 1142.54 | 0 | 0.44 | 0.002525 | 8.41 | 82.9 | 117.1 | 5.82 | 9.79 | | High | High | High | High | High | High | High | High | High | High | High | 0.000814 | 0.007875 | 1.47 |
| FALSE | High | GSSICHVD | [R].GSSICH | 1xCarbami | E9Q842 1x | 0.012397 | 0.000889 | 1 | 4 | 2 | E9Q842 | Neuron na | E9Q842 [8 | 0 | 1841.887 | 0 | 1.75 | 0.000261 | 3.56 | 46.5 | 153.5 | 8.13 | 5.47 | | High | High | High | High | High | High | High | High | High | High | High | 0.000555 | 0.004831 | 2.3 |
| FALSE | High | GSSLQSSP | [K].GSSLQS | 1xPhospho | B1AR09 1x | 0.01095 | 0.000772 | 1 | 3 | 2 | B1AR09 | Myeloid/ly | B1AR09 [3 | 0 | 2077.041 | 0 | 0.06 | 0.171498 | 5.83 | 95.4 | 104.6 | 5.92 | 8.36 | | High | High | High | High | High | High | High | High | High | High | High | 0.000399 | 0.003468 | 2.09 |
| FALSE | High | GSSLQSSP | [K].GSSLQS | 2xPhospho | B1AR09 2x | 0.051694 | 0.003745 | 1 | 3 | 2 | B1AR09 | Myeloid/ly | B1AR09 [3 | 0 | 2157.007 | 0 | 0.33 | 0.071794 | 22.87 | 84.8 | 115.2 | 13.94 | 16.01 | | High | High | High | High | High | High | High | High | High | High | High | 0.00196 | 0.01999 | 2.48 |
| FALSE | High | GSSETETEF | [K].GSSETE | 1xPhospho | O08580 1x | 3.57E-05 | 0 | 1 | 2 | 6 | O08580 | Steroid ho | O08580 [2 | 0 | 2363.133 | 0 | 1.58 | 0.000432 | 9.49 | 50.2 | 149.8 | 9.08 | 7.89 | | High | High | High | High | High | High | High | High | High | High | High | 0 | 1.16E-05 | 3.07 |
| FALSE | High | GSTYSPTSF | [K].GSTYSP | 1xPhospho | P08775 1xl | 8.4E-05 | 0 | 1 | 2 | 7 | P08775 | DNA-direct | P08775 [1 | 0 | 3816.637 | 0 | -0.15 | 0.312427 | 13.88 | 107 | 93 | 8.64 | 19.76 | | High | High | High | High | High | High | High | High | High | High | High | 2.58E-05 | 0.000539 | 4.52 |
| FALSE | High | GSVAANPD | [K].GSVAA | 1xPhospho | Q7TQI8 1xl | 1.15E-05 | 0 | 1 | 1 | 1 | Q7TQI8 | Testis-spec | Q7TQI8 [4 | 0 | 2298.018 | 0 | -0.43 | 0.001357 | 4.21 | 115.4 | 84.6 | 3.57 | 3.76 | | High | High | High | High | High | High | High | High | High | High | High | 0 | 1.58E-06 | 4.25 |
| FALSE | High | GSVDEDAV | [K].GSVDE | 1xPhospho | B2RUA6 1x | 0.004867 | 0.000504 | 1 | 4 | 1 | B2RUA6 | tRNA-yW s | B2RUA6 [2 | 0 | 2140.037 | 0 | 0.4 | 0.029451 | 1.41 | 86.2 | 113.8 | 5.86 | 8.63 | | High | High | High | High | High | High | High | High | High | High | High | 0.000225 | 0.001405 | 3.35 |
| FALSE | High | GTDNQPEE | [R].GTDNC | 1xPhospho | Q3KNJ2 1x | 0.003927 | 0.000311 | 1 | 3 | 1 | Q3KNJ2 | Non-homo | Q3KNJ2 [2 | 0 | 3736.689 | 0 | -1.18 | 0.00911 | 1.62 | 136.5 | 63.5 | 10.11 | 20.81 | | High | High | High | High | High | High | High | High | High | High | High | 0.00014 | 0.0011 | 2.87 |
| FALSE | High | GTEDGSGS | [R].GTEDG | 1xCarbami | P83741-3 1 | 1.28E-07 | 0 | 1 | 8 | 2 | P83741-3 | Isoform 3 c | P83741-3 [ | 0 | 2388.106 | 0 | -0.01 | 0.934397 | 25.55 | 107.5 | 92.5 | 13.69 | 33.21 | | High | High | High | High | High | High | High | High | High | High | High | 0 | 1.01E-08 | 4.52 |
| FALSE | High | GTEDGSGS | [R].GTEDG | 1xCarbami | P83741-3 2 | 0.026421 | 0.001676 | 1 | 8 | 2 | P83741-3 | Isoform 3 c | P83741-3 [ | 0 | 2468.072 | 0 | 0.95 | 0.00057 | 4.42 | 69.5 | 130.5 | 4.6 | 8.17 | | High | High | High | High | High | High | High | High | High | High | High | 0.003265 | 0.03467 | 2.36 |
| FALSE | High | GTEETSWS | [K].GTEETS | 1xPhospho | Q5U4C3 1x | 0.001933 | 5.66E-05 | 1 | 1 | 3 | Q5U4C3 | Splicing fac | Q5U4C3 [9 | 0 | 1676.7 | 0 | -0.03 | 0.519659 | 2.99 | 99.2 | 100.8 | 9.39 | 5.27 | | High | High | High | High | High | High | High | High | High | High | High | 2.58E-05 | 0.000502 | 1.65 |
| FALSE | High | GTELQIAV | [K].GTELQI | 1xPhospho | B2KFD2 1x | 0.00051 | 0 | 1 | 4 | 1 | B2KFD2 | 28104740 | B2KFD2 [5 | 0 | 2890.594 | 0 | -1.45 | 0.000484 | 1.5 | 147.2 | 52.8 | 6.57 | 6.78 | | High | High | High | High | High | High | High | High | High | High | High | 0 | 0.000112 | 2.82 |
| FALSE | High | GTENGVN | [K].GTENG | 1xPhospho | Q3UAT2 1x | 1.22E-07 | 0 | 1 | 5 | 3 | Q3UAT2 | Translocati | Q3UAT2 [3 | 0 | 2141.966 | 0 | 0.83 | 0.002082 | 13.66 | 69 | 131 | 7.43 | 7.73 | | High | High | High | High | High | High | High | High | High | High | High | 0 | 9.49E-09 | 4.44 |
| FALSE | High | GTENGVN | [K].GTENG | 1xDeamida | Q3UAT2 1x | 1.34E-07 | 0 | 1 | 5 | 15 | Q3UAT2 | Translocati | Q3UAT2 [3 | 0 | 2142.95 | 0 | 0.75 | 0.001957 | 12.01 | 74.2 | 125.8 | 4.59 | 8.34 | | High | High | High | High | High | High | High | High | High | High | High | 0 | 1.16E-07 | 4.48 |
| FALSE | High | GTENGVN | [K].GTENG | 2xDeamida | Q3UAT2 1x | 1.93E-08 | 0 | 1 | 5 | 7 | Q3UAT2 | Translocati | Q3UAT2 [3 | 0 | 2143.934 | 0 | 0.77 | 0.00043 | 7.16 | 71.9 | 128.1 | 6.25 | 6.43 | | High | High | High | High | High | High | High | High | High | High | High | 0 | 2.88E-06 | 3.32 |
| FALSE | High | GTFSDQEL | [R].GTFSD | 2xPhospho | Q8CI08-2 2 | 0.039011 | 0.002718 | 1 | 2 | 1 | Q8CI08-2 | Isoform 2 c | Q8CI08-2 [ | 0 | 3712.6 | 0 | 2.53 | 0.000965 | 21.8 | 28.4 | 171.6 | 19.33 | 13.92 | | High | High | High | High | High | High | High | High | High | High | High | 0.001421 | 0.01447 | 3.01 |
| FALSE | High | GTFTDDLH | [K].GTFTD | 1xPhospho | E9QM73 1 | 0.01186 | 0.000889 | 2 | 25 | 2 | E9QM73; F | Serine/thre | E9QM73 [2 | 0 | 1571.787 | 0 | 2.52 | 0.000755 | 21.13 | 31.8 | 168.2 | 20.6 | 12.35 | Shared | High | High | High | High | High | High | High | High | High | High | High | 0.000449 | 0.003803 | 2.57 |
| FALSE | High | GTGDCSDE | [K].GTGDC | 1xCarbami | Q8VDD5 1 | 2.43E-06 | 0 | 1 | 5 | 9 | Q8VDD5 | Myosin-9 C | Q8VDD5 [1 | 0 | 1906.814 | 0 | 0.97 | 0.000985 | 12.74 | 68.5 | 131.5 | 7.12 | 6.94 | | High | High | High | High | High | High | High | High | High | High | High | 0 | 2.74E-07 | 2.72 |
| FALSE | High | GTGDCSDE | [K].GTGDC | 1xCarbami | Q8VDD5 1 | 0.000884 | 0 | 1 | 5 | 2 | Q8VDD5 | Myosin-9 C | Q8VDD5 [1 | 1 | 2764.259 | 0 | -0.34 | 0.588864 | 63.37 | 103.9 | 96.1 | 86.78 | 157.59 | | High | High | High | High | High | High | High | High | High | High | High | 0 | 0.000209 | 3.16 |
| FALSE | High | GTGFAHPS | [K].GTGFA | 1xPhospho | Q3U1H6 1x | 0.000253 | 0 | 1 | 2 | 1 | Q3U1H6 | Putative ur | Q3U1H6 [4 | 0 | 3794.691 | 0 | 0.42 | 0.068714 | 19.67 | 84.9 | 115.1 | 8.26 | 20.55 | | High | High | High | High | High | High | High | High | High | High | High | 0 | 5.1E-05 | 3.89 |
| FALSE | High | GTCSEPSLF | [K].GTCSEF | 2xCarbami | E9PZ97 2xl | 0.059657 | 0.004365 | 1 | 2 | 1 | E9PZ97 | Bloom sync | E9PZ97 [1 | 0 | 3181.46 | 0 | 0.3 | 0.020975 | 19.12 | 89.5 | 110.5 | 10.56 | 6.41 | | High | High | High | High | High | High | High | High | High | High | High | 0.002264 | 0.02367 | 2.47 |
| FALSE | High | GTGFNQNI | [R].GTGFN | 1xPhospho | Q8BYK6-3 | 0.000154 | 0 | 1 | 5 | 2 | Q8BYK6-3 | Isoform 3 c | Q8BYK6-3 | 0 | 4250.117 | 0 | 0.88 | 0.00198 | 21.39 | 75.5 | 124.5 | 9.41 | 7.17 | | High | High | High | High | High | High | High | High | High | High | High | 0 | 2.91E-05 | 2.92 |
| FALSE | High | GTGQSDDS | [R].GTGQS | 1xPhospho | P97801 1xl | 2.06E-06 | 0 | 1 | 3 | 12 | P97801 | Survival mc | P97801 [21 | 0 | 2475.169 | 0 | -0.68 | 0.001084 | 2.37 | 124.4 | 75.6 | 3.02 | 9.65 | | High | High | High | High | High | High | High | High | High | High | High | 0 | 3.97E-07 | 4.06 |
| FALSE | High | GTGQSDDS | [R].GTGQS | 2xPhospho | P97801 2xl | 2.8E-06 | 0 | 1 | 3 | 5 | P97801 | Survival mc | P97801 [21 | 0 | 2555.136 | 0 | -0.31 | 0.080088 | 6.6 | 113.7 | 86.3 | 10.87 | 9.59 | | High | High | High | High | High | High | High | High | High | High | High | 0 | 3.23E-07 | 4.31 |
| FALSE | High | GTKPPSPY | [K].GTKPPS | 1xPhospho | Q3U7R1 1x | 0.00017 | 0 | 1 | 3 | 2 | Q3U7R1 | Extended s | Q3U7R1 [8 | 0 | 2738.473 | 0 | 0.62 | 0.003131 | 6.47 | 76.9 | 123.1 | 4.73 | 8.99 | | High | High | High | High | High | High | High | High | High | High | High | 0 | 7.32E-05 | 1.96 |
| FALSE | High | GTLSTTGH | [R].GTLSTT | 2xPhospho | L7N1Y0 2xl | 0.074685 | 0.005727 | 1 | 2 | 2 | L7N1Y0 | Dynein, axo | L7N1Y0 [4 | 0 | 2920.377 | 0 | 0.53 | 0.011608 | 7.17 | 79.6 | 120.4 | 7.53 | 8.88 | | High | High | High | High | High | High | High | High | High | High | High | 0.002937 | 0.03079 | 3.07 |
| FALSE | High | GTMDDISC | [K].GTMDD | 1xPhospho | Q9CU62 1x | 2.24E-06 | 0 | 1 | 2 | 5 | Q9CU62 | Structural | Q9CU62 [9 | 0 | 2825.162 | 0 | -0.31 | 0.33546 | 18.64 | 109 | 91 | 11.41 | 16.29 | | High | High | High | High | High | High | High | High | High | High | High | 0 | 2.52E-07 | 4.33 |
| FALSE | High | GTMYSMIC | [R].GTMYS | 1xCarbami | Q9ESE5 2x | 0.074071 | 0.0057 | 1 | 4 | 1 | Q9ESE5 | NK cell rec | Q9ESE5 [3 | 0 | 2628.067 | 0 | -0.53 | 0.017796 | 5.79 | 118.2 | 81.8 | 11.9 | 30.2 | | High | High | High | High | High | High | High | High | High | High | High | 0.002924 | 0.03037 | 3.3 |
| FALSE | High | GTNSSDSE | [K].GTNSSI | 1xPhospho | A2CG63 1x | 0.000519 | 0 | 1 | 4 | 2 | A2CG63 | AT-rich inte | A2CG63 [1 | 0 | 2336.091 | 0 | 1.57 | 0.000675 | 12.53 | 49.1 | 150.9 | 13.45 | 6.92 | | High | High | High | High | High | High | High | High | High | High | High | 0 | 0.000124 | 3.25 |
| FALSE | High | GTNVFGSL | [K].GTNVF | 1xPhospho | Q3TU15 1x | 0.065952 | 0.004766 | 1 | 3 | 1 | Q3TU15 | Non-specif | Q3TU15 [5 | 0 | 1388.677 | 0 | -1.13 | 0.000783 | 3.35 | 139.3 | 60.7 | 5.87 | 16.28 | | High | High | High | High | High | High | High | High | High | High | High | 0.002455 | 0.02647 | 1.16 |
| FALSE | High | GTPALTSE | [K].GTPALT | 1xPhospho | Q69Z89 1x | 6.27E-07 | 0 | 1 | 6 | 2 | Q69Z89 | Ras-associa | Q69Z89 [2 | 0 | 1979.964 | 0 | 1.01 | 0.000591 | 11.43 | 67.2 | 132.8 | 8.59 | 4.92 | | High | High | High | High | High | High | High | High | High | High | High | 0 | 6E-08 | 3.69 |
| FALSE | High | GTPASATG | [K].GTPAS | 1xCarbami | H3BL37 1xl | 2.45E-06 | 0 | 1 | 7 | 2 | H3BL37 | Treacle prc | H3BL37 [5 | 0 | 2057.973 | 0 | 0.58 | 0.027704 | 22.84 | 82 | 118 | 13.19 | 9.76 | | High | High | High | High | High | High | High | High | High | High | High | 0 | 2.77E-07 | 2.67 |
| FALSE | High | GTPATAGS | [R].GTPAT | 1xPhospho | Q3UI84 1xl | 3.26E-05 | 0 | 1 | 4 | 2 | Q3UI84 | Putative ur | Q3UI84 [2 | 0 | 1701.846 | 0 | 0.37 | 0.022733 | 16.14 | 87.8 | 112.2 | 3.51 | 9.3 | | High | High | High | High | High | High | High | High | High | High | High | 0 | 5.11E-06 | 2.53 |
| FALSE | High | GTPESGTS | [R].GTPESC | 1xPhospho | Q8K0L9 1x | 7.29E-06 | 0 | 1 | 3 | 1 | Q8K0L9 | Zinc finger | Q8K0L9 [2 | 0 | 3340.455 | 0 | 1.95 | 0.008104 | 43.35 | 30.4 | 169.6 | 59.32 | 25.93 | | High | High | High | High | High | High | High | High | High | High | High | 0 | 9.44E-07 | 2.79 |
| FALSE | High | GTPKPPGP | [R].GTPKPI | 1xPhospho | Q9JM52-3 | 2.87E-05 | 0 | 1 | 6 | 2 | Q9JM52-3 | Isoform 3 c | Q9JM52-3 | 0 | 2984.54 | 0 | 1.44 | 0.005547 | 33.22 | 59.3 | 140.7 | 9.98 | 18.98 | | High | High | High | High | High | High | High | High | High | High | High | 0 | 4.42E-06 | 3.62 |
| FALSE | High | GTGLQPGE | [R].GTGLQ | 1xPhospho | P56399 1xl | 6.33E-07 | 0 | 1 | 2 | 2 | P56399 | Ubiquitin c | P56399 [6 | 0 | 3137.606 | 0 | -0.55 | 0.024529 | 8.08 | 124.2 | 75.8 | 25.27 | 12.07 | | High | High | High | High | High | High | High | High | High | High | High | 0 | 6.11E-08 | 3.8 |
| FALSE | High | GTAVASAC | [R].GTAVA | 1xPhospho | E9QMN5 1 | 1.5E-08 | 0 | 1 | 3 | 1 | E9QMN5 | Transcripti | E9QMN5 [ | 0 | 2983.457 | 0 | -0.37 | 0.015617 | 8.78 | 108.7 | 91.3 | 4.53 | 11.52 | | High | High | High | High | High | High | High | High | High | High | High | 0 | 9.09E-10 | 5.65 |
| FALSE | High | GTADWFG | [R].GTADW | 1xPhospho | Q6PIX5 1xl | 0.1059 | 0.008582 | 1 | 4 | 1 | Q6PIX5 | Inactive rh | Q6PIX5 [84 | 0 | 1376.645 | 0 | 0.18 | 0.029736 | 7.97 | 96.1 | 103.9 | 7.06 | 4.9 | | High | High | High | High | High | High | High | High | High | High | High | 0.004327 | 0.04636 | 1.81 |
| FALSE | High | GSYVTYEP | [K].GSYVTY | 1xPhospho | Q99KC7 1x | 9.49E-10 | 0 | 1 | 1 | 3 | Q99KC7 | Small cell a | Q99KC7 [6 | 0 | 3227.511 | 0 | 0.3 | 0.805593 | 12.92 | 91.6 | 108.4 | 5.03 | 32.05 | | High | High | High | High | High | High | High | High | High | High | High | 0 | 4.09E-11 | 6.33 |
| FALSE | High | GSVDQYLL | [K].GSVDQ | 1xPhospho | E9Q7F2 1x | 0.001137 | 0 | 1 | 1 | 2 | E9Q7F2 | E3 ubiquiti | E9Q7F2 [6 | 0 | 1359.687 | 0 | 0.04 | 0.914068 | 8.72 | 99.5 | 100.5 | 12.09 | 7.32 | | High | High | High | High | High | High | High | High | High | High | High | 7.19E-05 | 0.000903 | 2.22 |
| FALSE | High | GSVELEK | [R].GSVELE | 1xPhospho | E9Q0N0 1x | 0.002671 | 0.000109 | 1 | 10 | 1 | E9Q0N0 | Intersectin | E9Q0N0 [3 | 0 | 1299.696 | 0 | 1.07 | 0.00058 | 14.99 | 64.5 | 135.5 | 13.86 | 5.59 | | High | High | High | High | High | High | High | High | High | High | High | 4.94E-05 | 0.000719 | 2.06 |
| FALSE | High | GSVFSAPS | [K].GSVFS | 1xPhospho | E9QA16 1x | 0.010853 | 0.000772 | 1 | 7 | 2 | E9QA16 | Caldesmon | E9QA16 [4 | 0 | 1944.983 | 0 | 1.49 | 0.007108 | 18.33 | 57.9 | 142.1 | 11.81 | 21.87 | | High | High | High | High | High | High | High | High | High | High | High | 0.000399 | 0.003423 | 1.58 |
| FALSE | High | GSVGCCSF | [R].GSVGC | 2xCarbami | O55111 1x | 2.41E-06 | 0 | 1 | 2 | 1 | O55111 | Desmoglei | O55111 [8 | 0 | 3197.519 | 0 | -0.21 | 0.186791 | 23.41 | 107.3 | 92.7 | 17.35 | 21.83 | | High | High | High | High | High | High | High | High | High | High | High | 0 | 2.73E-07 | 4.33 |
| FALSE | High | GSVISEGSF | [-].GSVISE | 1xPhospho | Q8CBI1 1xl | 0.001127 | 0 | 1 | 1 | 2 | Q8CBI1 | Putative ur | Q8CBI1 [1- | 0 | 2637.188 | 0 | -0.11 | 0.795358 | 9.46 | 97.8 | 102.2 | 15.96 | 15.31 | | High | High | High | High | High | High | High | High | High | High | High | 0 | 0.000272 | 2.08 |
| FALSE | High | GSVSALMG | [K].GSVSAL | 1xPhospho | Q3TCH7 1x | 0.028575 | 0.001829 | 1 | 5 | 1 | Q3TCH7 | Cullin-4A C | Q3TCH7 [9 | 0 | 1202.58 | 0 | 0.81 | 0.003875 | 17.9 | 67.1 | 132.9 | 14.54 | 9.63 | | High | High | High | High | High | High | High | High | High | High | High | 0.000984 | 0.01018 | 1.01 |
| FALSE | High | GSVSDCSD | [R].GSVSD | 1xCarbami | A0A0R4J0I | 1.56E-09 | 0 | 1 | 2 | 13 | A0A0R4J0I | ETS domain | A0A0R4J0I | 0 | 2645.112 | 0 | -0.18 | 0.012884 | 0.22 | 111.4 | 88.6 | 5.04 | 6.65 | | High | High | High | High | High | High | High | High | High | High | High | 0 | 7.09E-11 | 5.93 |
| FALSE | High | GSVSETIVII | [K].GSVSET | 1xPhospho | Q9CU65 1x | 6.23E-08 | 0 | 1 | 2 | 4 | Q9CU65 | Zinc finger | Q9CU65 [1 | 0 | 3107.402 | 0 | -0.85 | 0.150281 | 12.36 | 132.6 | 67.4 | 22.32 | 45.23 | | High | High | High | High | High | High | High | High | High | High | High | 0 | 1.96E-08 | 5.55 |
| FALSE | High | GSVSLNSSF | [R].GSVSLN | 1xPhospho | Q9ESL4 1xl | 0.003822 | 0.000261 | 1 | 1 | 5 | Q9ESL4 | Mitogen-ac | Q9ESL4 [6 | 0 | 1513.803 | 0 | 0.79 | 0.000442 | 6.01 | 73.2 | 126.8 | 4.82 | 5.34 | | High | High | High | High | High | High | High | High | High | High | High | 0.001609 | 0.01562 | 1.82 |
| FALSE | High | GSVSLNSSF | [R].GSVSLN | 2xPhospho | Q9ESL4 2xl | 2.58E-05 | 0 | 1 | 1 | 1 | Q9ESL4 | Mitogen-ac | Q9ESL4 [6 | 0 | 1593.769 | 0 | 1.41 | 0.001134 | 25.33 | 54.2 | 145.8 | 10.43 | 9.31 | | High | High | High | High | High | High | High | High | High | High | High | 0 | 3.91E-06 | 1.98 |
| FALSE | High | GSVSSTSEV | [R].GSVSST | 1xPhospho | F8VQ93 1x | 1.08E-07 | 0 | 1 | 5 | 5 | F8VQ93 | PH-interac | F8VQ93 [6 | 0 | 2133.054 | 0 | 0.58 | 0.052865 | 28.89 | 80.6 | 119.4 | 15.94 | 15.01 | | High | High | High | High | High | High | High | High | High | High | High | 0 | 8.36E-09 | 3.72 |
| FALSE | High | GSVSSTSEV | [R].GSVSST | 2xPhospho | F8VQ93 2x | 0.000725 | 0 | 1 | 5 | 2 | F8VQ93 | PH-interac | F8VQ93 [6 | 0 | 2213.02 | 0 | 0.09 | 0.342957 | 1.81 | 97.2 | 102.8 | 15.42 | 16.66 | | High | High | High | High | High | High | High | High | High | High | High | 0 | 0.000166 | 3.03 |
| FALSE | High | GSVVNVNF | [K].GSVVN | 1xPhospho | A0A0A6YW | 0.000884 | 0 | 2 | 10 | 4 | A0A0A6YW | Mitogen-ac | A0A0A6YW | 0 | 2590.282 | 0 | -0.79 | 0.001246 | 2.36 | 126.4 | 73.6 | 5.14 | 7.3 | Shared | High | High | High | High | High | High | High | High | High | High | High | 2.58E-05 | 0.000355 | 1.64 |
| FALSE | High | GSWGHESI | [K].GSWGH | 1xPhospho | A2A6A1 1x | 0.019618 | 0.001438 | 1 | 1 | 1 | A2A6A1 | G patch do | A2A6A1 [1 | 0 | 1579.674 | 0 | 0 | 0.671967 | 34.52 | 96.7 | 103.3 | 8.75 | 18.49 | | High | High | High | High | High | High | High | High | High | High | High | 0.000756 | 0.006679 | 2.71 |
| FALSE | High | GSWGHESI | [K].GSWGH | 2xPhospho | A2A6A1 2x | 0.006653 | 0.00062 | 1 | 1 | 2 | A2A6A1 | G patch do | A2A6A1 [1 | 0 | 1659.64 | 0 | 0.14 | 0.47736 | 12.81 | 98.6 | 101.4 | 17.15 | 21.73 | | High | High | High | High | High | High | High | High | High | High | High | 0.002403 | 0.02539 | 2.66 |
| FALSE | High | GSWSQASC | [K].GSWSC | 1xPhospho | B9EKS2 1xl | 0.065952 | 0.004766 | 1 | 1 | 2 | B9EKS2 | Jumonji do | B9EKS2 [4 | 0 | 1574.68 | 0 | 0.51 | 0.048532 | 27.97 | 82.4 | 117.6 | 12.36 | 11.84 | | High | High | High | High | High | High | High | High | High | High | High | 0.004462 | 0.04824 | 1.15 |
| FALSE | High | GSWTPDLE | [R].GSWTP | 1xPhospho | A0A0A6YY | 0.008847 | 0.000736 | 1 | 1 | 1 | A0A0A6YY | BTB (POZ) | A0A0A6YY | 0 | 2783.354 | 0 | -1.77 | 0.001504 | 2.08 | 149.9 | 50.1 | 13.69 | 13.38 | | High | High | High | High | High | High | High | High | High | High | High | 0.000327 | 0.002739 | 2.97 |
| FALSE | High | GSWTSCSS | [R].GSWTS | 1xCarbami | A0A0A6YW | 0.000239 | 0 | 1 | 4 | 1 | A0A0A6YW | Rap guanin | A0A0A6YW | 0 | 2695.188 | 0 | 0.3 | 0.366555 | 27.11 | 88.5 | 111.5 | 11.61 | 19.25 | | High | High | High | High | High | High | High | High | High | High | High | 0 | 4.79E-05 | 3.41 |
| FALSE | High | GSYGDLGG | [R].GSYGD | 1xPhospho | Q5FWJ5 1x | 0.003688 | 0.00021 | 2 | 12 | 2 | Q5FWJ5; C | Hnrpk prot | Q5FWJ5 [3 | 0 | 2455.325 | 0 | -0.18 | 0.015643 | 0.86 | 103.4 | 96.6 | 3.17 | 3.29 | Shared | High | High | High | High | High | High | High | High | High | High | High | 9.42E-05 | 0.001026 | 1.47 |
| FALSE | High | GSYGSDPE | [R].GSYGSI | 1xPhospho | Q9Z0U1 1x | 3.69E-07 | 0 | 1 | 1 | 5 | Q9Z0U1 | Tight junct | Q9Z0U1 [1 | 0 | 1826.732 | 0 | 0.5 | 0.029273 | 19.2 | 84.2 | 115.8 | 12.06 | 7.23 | | High | High | High | High | High | High | High | High | High | High | High | 0 | 0.000211 | 3.23 |
| FALSE | High | GSYGSDPE | [R].GSYGSI | 1xPhospho | Q9Z0U1 1x | 0.000808 | 0 | 1 | 1 | 3 | Q9Z0U1 | Tight junct | Q9Z0U1 [1 | 1 | 2919.356 | 0 | 0.16 | 0.96896 | 20.5 | 97 | 103 | 22.17 | 10.74 | | High | High | High | High | High | High | High | High | High | High | High | 2.58E-05 | 0.000469 | 1.75 |
| FALSE | High | GSYGSDPE | [R].GSYGSI | 2xPhospho | Q9Z0U1 2x | 0.014538 | 0.001011 | 1 | 1 | 1 | Q9Z0U1 | Tight junct | Q9Z0U1 [1 | 1 | 2999.323 | 0 | 0.8 | 0.048151 | 26.68 | 75.7 | 124.3 | 25.64 | 14.76 | | High | High | High | High | High | High | High | High | High | High | High | 0.000555 | 0.004785 | 3.18 |
| FALSE | High | GSYHEHYQ | [R].GSYHEI | 1xPhospho | B2M0S2 1x | 0.000438 | 0 | 1 | 8 | 2 | B2M0S2 | Secretory c | B2M0S2 [1 | 0 | 1572.679 | 0 | -0.22 | 0.022388 | 1.13 | 107.8 | 92.2 | 4.71 | 6.12 | | High | High | High | High | High | High | High | High | High | High | High | 0 | 9.43E-05 | 2.81 |
| FALSE | High | GSYLTHEAS | [R].GSYLTH | 1xPhospho | Q8R464 1x | 0.00051 | 0 | 1 | 1 | 1 | Q8R464 | Cell adhesi | Q8R464 [3 | 0 | 2229.002 | 0 | -1.12 | 0.005954 | 3.67 | 136.4 | 63.6 | 18.42 | 7.9 | | High | High | High | High | High | High | High | High | High | High | High | 0 | 0.000112 | 3.71 |
| FALSE | High | GSYNGQIG | [R].GSYNG | 1xDeamida | Q99M51 1 | 0.000573 | 0 | 1 | 3 | 2 | Q99M51 | Cytoplasmi | Q99M51 [1 | 0 | 4065.88 | 0 | -0.13 | 0.817362 | 17.62 | 107.1 | 92.9 | 8.54 | 16.86 | | High | High | High | High | High | High | High | High | High | High | High | 0 | 0.000127 | 6.52 |
| FALSE | High | GSYNTVLS | [R].GSYNTY | 1xPhospho | A2ATK9 1x | 0.002695 | 0.000109 | 1 | 3 | 1 | A2ATK9 | Family with | A2ATK9 [4 | 0 | 2111.083 | 0 | 1.31 | 0.002642 | 16.89 | 57.4 | 142.6 | 9.15 | 12.67 | | High | High | High | High | High | High | High | High | High | High | High | 4.94E-05 | 0.000726 | 3.2 |
| FALSE | High | GSYSALNTI | [K].GSYSAL | 1xPhospho | Q6NS46 1x | 0.006085 | 0.00062 | 1 | 1 | 1 | Q6NS46 | Protein RR | Q6NS46 [1 | 0 | 2244.095 | 0 | -0.06 | 0.459287 | 32.12 | 103.2 | 96.8 | 2.99 | 21.88 | | High | High | High | High | High | High | High | High | High | High | High | 0.000275 | 0.001794 | 2.49 |
| FALSE | High | GSYSSVVN | [R].GSYSSV | 1xPhospho | O70472 1x | 4.43E-05 | 0 | 1 | 2 | 3 | O70472 | Transmem | O70472 [1 | 0 | 2012.958 | 0 | 0.38 | 0.021612 | 9.4 | 87.8 | 112.2 | 9.93 | 18.26 | | High | High | High | High | High | High | High | High | High | High | High | 0 | 7.17E-06 | 3.55 |

| | | | | | | | | | | | | | | | | | | | | | | | | | | | | | | | | | | | | | | |
|---|---|---|---|---|---|---|---|---|---|---|---|---|---|---|---|---|---|---|---|---|---|---|---|---|---|---|---|---|---|---|---|---|---|---|---|---|---|---|
| FALSE | High | GSYVTYEPA | [K].GSYVTY | 1xPhospho | Q99KC7 1x | 1.43E-07 | 0 | 1 | 1 | 1 | Q99KC7 | Small cell a | Q99KC7 [6 | 0 | 3211.516 | 0 | 0 | 0.85475 | 38.13 | 99.9 | 100.1 | 6.79 | 36.26 | | High | High | High | High | High | High | High | High | High | High | High | 0 | 1.14E-08 | 5.54 |
| FALSE | High | GSQGTSPF | [K].GSQGT | 1xPhospho | F6YZ95 1xP | 7.41E-05 | 0 | 2 | 7 | 3 | F6YZ95; Q8 | Liprin-beta | F6YZ95 [12 | 0 | 2156.952 | 0 | 0.53 | 0.0115 | 15.46 | 81.9 | 118.1 | 4.24 | 14.88 | Shared | High | High | High | High | High | High | High | High | High | High | High | 0 | 1.29E-05 | 3.81 |
| FALSE | High | GTPIPDSSS | [R].GTPIPD | 1xPhospho | Q60902 1x | 1.47E-05 | 0 | 1 | 5 | 8 | Q60902 | Epidermal | Q60902 [3 | 0 | 2489.258 | 0 | -0.35 | 0.005354 | 2.58 | 112.3 | 87.7 | 5.46 | 4.58 | | High | High | High | High | High | High | High | High | High | High | High | 0 | 5.1E-06 | 3.21 |
| FALSE | High | GSQGTSPF | [K].GSQGT | 1xPhospho | F6YZ95 1xP | 0.025738 | 0.001645 | 2 | 7 | 3 | F6YZ95; Q8 | Liprin-beta | F6YZ95 [12 | 0 | 2140.957 | 0 | -0.02 | 0.550185 | 19.02 | 102.4 | 97.6 | 9.48 | 8.97 | Shared | High | High | High | High | High | High | High | High | High | High | High | 0.000872 | 0.009039 | 3.08 |
| FALSE | High | GSQELTK | [R].GSQELT | 1xPhospho | Q5DTX6 1x | 0.027597 | 0.001716 | 1 | 1 | 1 | Q5DTX6 | Junctional | Q5DTX6 [9 | 0 | 1300.691 | 0 | 2.11 | 0.000479 | 3.44 | 37.6 | 162.4 | 10.41 | 11.09 | | High | High | High | High | High | High | High | High | High | High | High | 0.000936 | 0.009766 | 1.25 |
| FALSE | High | GSLASLDSL | [R].GSLASL | 1xPhospho | E9Q8Z8 1x | 0.046687 | 0.00354 | 1 | 21 | 1 | E9Q8Z8 | Catenin de | E9Q8Z8 [3 | 0 | 1327.682 | 0 | -0.47 | 0.058129 | 7.95 | 116.2 | 83.8 | 8.93 | 12.38 | | High | High | High | High | High | High | High | High | High | High | High | 0.001812 | 0.01783 | 1 |
| FALSE | High | GSLASLDSL | [R].GSLASL | 2xPhospho | E9Q8Z8 2x | 0.001039 | 0 | 1 | 21 | 3 | E9Q8Z8 | Catenin de | E9Q8Z8 [3 | 0 | 1407.648 | 0 | -0.32 | 0.678183 | 13.02 | 111.7 | 88.3 | 9.42 | 28.59 | | High | High | High | High | High | High | High | High | High | High | High | 0.001093 | 0.01152 | 2.45 |
| FALSE | High | GSLDESSLG | [R].GSLDES | 1xPhospho | Q61382 1x | 5.72E-06 | 0 | 1 | 1 | 7 | Q61382 | TNF recept | Q61382 [4 | 0 | 1994.988 | 0 | -1.49 | 0.000328 | 0.73 | 145.9 | 54.1 | 7.3 | 7.6 | | High | High | High | High | High | High | High | High | High | High | High | 0 | 9.93E-06 | 3.42 |
| FALSE | High | GSLDWPEE | [R].GSLDW | 1xPhospho | Q80U49 1x | 8.18E-05 | 0 | 1 | 2 | 2 | Q80U49 | Centrosom | Q80U49 [1 | 0 | 1526.673 | 0 | 0.06 | 0.82105 | 16.48 | 103.8 | 96.2 | 16.12 | 17.79 | | High | High | High | High | High | High | High | High | High | High | High | 0 | 3.6E-05 | 2.72 |
| FALSE | High | GSLDYLK | [K].GSLDYL | 1xPhospho | Q8R4U7 1x | 0.008538 | 0.000736 | 2 | 3 | 1 | Q8R4U7; B | Leucine zip | Q8R4U7 [2 | 0 | 1333.717 | 0 | -0.12 | 0.982226 | 10.84 | 101.9 | 98.1 | 7.38 | 16.26 | Shared | High | High | High | High | High | High | High | High | High | High | High | 0.000327 | 0.002632 | 2.03 |
| FALSE | High | GSLEGQFS | [R].GSLEG | 1xPhospho | Q8BS90 1x | 1.82E-06 | 0 | 1 | 2 | 1 | Q8BS90 | Protein au | Q8BS90 [2 | 0 | 2303.168 | 0 | -1.45 | 0.00043 | 1 | 144 | 56 | 3.91 | 14.06 | | High | High | High | High | High | High | High | High | High | High | High | 0 | 1.99E-07 | 3.51 |
| FALSE | High | GSLEGQFS | [R].GSLEG | 2xPhospho | Q8BS90 2x | 0.01319 | 0.000889 | 1 | 2 | 1 | Q8BS90 | Protein au | Q8BS90 [2 | 0 | 2383.135 | 0 | -0.66 | 0.030877 | 13.75 | 122.6 | 77.4 | 15.24 | 15.37 | | High | High | High | High | High | High | High | High | High | High | High | 0.000467 | 0.004287 | 3.56 |
| FALSE | High | GSLELGNP | [R].GSLELG | 1xPhospho | A0A1B0GS | 0.002463 | 0.000109 | 1 | 6 | 1 | A0A1B0GS | Supervillin | A0A1B0GS | 0 | 2346.211 | 0 | 0.66 | 0.005271 | 14.01 | 79.2 | 120.8 | 11.22 | 6.91 | | High | High | High | High | High | High | High | High | High | High | High | 4.94E-05 | 0.000655 | 2.4 |
| FALSE | High | GSLEPEATI | [K].GSLEPE | 1xPhospho | Q3TV65 1x | 0.001137 | 0 | 1 | 6 | 1 | Q3TV65 | MPN doma | Q3TV65 [2 | 0 | 1720.892 | 0 | -0.17 | 0.044575 | 5.95 | 107.6 | 92.4 | 5.07 | 7.39 | | High | High | High | High | High | High | High | High | High | High | High | 0 | 0.000276 | 3.36 |
| FALSE | High | GSIGFLLGK | [K].GSIGFL | 1xPhospho | Q9D4H4 1x | 8.71E-05 | 0 | 1 | 2 | 1 | Q9D4H4 | Angiomotir | Q9D4H4 [8 | 0 | 1429.822 | 0 | 2.04 | 0.001458 | 28.15 | 36.2 | 163.8 | 15.49 | 15.21 | | High | High | High | High | High | High | High | High | High | High | High | 0 | 1.54E-05 | 3.04 |
| FALSE | High | GSLLPTSPR | [K].GSLLPT | 1xPhospho | D3YZC8 1x | 0.045508 | 0.003373 | 1 | 3 | 3 | D3YZC8 | Bromodom | D3YZC8 [2 | 0 | 1236.655 | 0 | -0.19 | 0.003738 | 2.56 | 108.1 | 91.9 | 5.01 | 4.97 | | High | High | High | High | High | High | High | High | High | High | High | 0.004757 | 0.05163 | 1.22 |
| FALSE | High | GSLLQIPVK | [R].GSLLQI | 1xPhospho | I1VGT4 1xP | 0.000496 | 0 | 2 | 19 | 1 | I1VGT4; I1 | Anion exch | I1VGT4 [10 | 0 | 1492.89 | 0 | 1.12 | 0.002897 | 6.5 | 66.7 | 133.3 | 7.48 | 12.63 | Shared | High | High | High | High | High | High | High | High | High | High | High | 0 | 0.000108 | 2.72 |
| FALSE | High | GSLNEQIAI | [R].GSLNE | 1xPhospho | Q5CZX6 1x | 0.101749 | 0.008194 | 1 | 5 | 1 | Q5CZX6 | Acbd5 prot | Q5CZX6 [4 | 0 | 1734.972 | 0 | 2.29 | 0.000429 | 5.72 | 38.1 | 161.9 | 9.78 | 9.85 | | High | High | High | High | High | High | High | High | High | High | High | 0.004126 | 0.0442 | 2.34 |
| FALSE | High | GSLAALYDI | [K].GSLAAL | 1xPhospho | Q6PIJ4 1xP | 0.038346 | 0.002563 | 2 | 2 | 1 | Q6PIJ4; A0 | Nuclear fac | Q6PIJ4 [29 | 0 | 1872.065 | 0 | -0.09 | 0.877983 | 29.62 | 103.1 | 96.9 | 11.94 | 18.51 | Shared | High | High | High | High | High | High | High | High | High | High | High | 0.001355 | 0.01426 | 2.29 |
| FALSE | High | GSLNINEFK | [K].GSLNIN | 1xPhospho | A0A0A0M | 0.046687 | 0.00354 | 1 | 2 | 1 | A0A0A0M | Protein S1 | A0A0A0M | 0 | 1559.823 | 0 | 0.82 | 0.088375 | 20.73 | 68.6 | 131.4 | 22 | 19.27 | | High | High | High | High | High | High | High | High | High | High | High | 0.001812 | 0.0179 | 1.86 |
| FALSE | High | GSLPPSPVS | [K].GSLPPS | 2xPhospho | Q9WU62 2 | 0.000223 | 0 | 1 | 2 | 10 | Q9WU62 | Inner centr | Q9WU62 [ | 0 | 2872.403 | 0 | -0.66 | 0.000492 | 2.98 | 122.4 | 77.6 | 3.71 | 3.32 | | High | High | High | High | High | High | High | High | High | High | High | 0 | 4.4E-05 | 3.59 |
| FALSE | High | GSLPPSPVS | [K].GSLPPS | 1xPhospho | Q9WU62 1 | 0.001411 | 5.66E-05 | 1 | 2 | 1 | Q9WU62 | Inner centr | Q9WU62 [ | 1 | 3149.695 | 0 | -0.56 | 0.01422 | 7.37 | 119.1 | 80.9 | 10.86 | 4.94 | | High | High | High | High | High | High | High | High | High | High | High | 2.58E-05 | 0.000352 | 2.98 |
| FALSE | High | GSLPPVSSF | [R].GSLPPV | 1xPhospho | B2RQR2 1x | 0.069903 | 0.005181 | 1 | 5 | 2 | B2RQR2 | Mpdz prot | B2RQR2 [2 | 0 | 1305.677 | 0 | -1.13 | 0.001395 | 2.77 | 136.8 | 63.2 | 6.18 | 11.75 | | High | High | High | High | High | High | High | High | High | High | High | 0.002662 | 0.02838 | 1.26 |
| FALSE | High | GSLQDVDV | [K].GSLQD | 1xPhospho | Q8K1N2-2 | 7.64E-13 | 0 | 1 | 3 | 1 | Q8K1N2-2 | Isoform 2 c | Q8K1N2-2 | 0 | 2724.334 | 0 | 2.34 | 0.000223 | 19.62 | 35.9 | 164.1 | 13.97 | 5.37 | | High | High | High | High | High | High | High | High | High | High | High | 0 | 1.36E-14 | 7.22 |
| FALSE | High | GSISSGVSY | [R].GSISSG | 1xPhospho | O35245 1x | 0.000146 | 0 | 1 | 2 | 3 | O35245 | Polycystin- | O35245 [8 | 0 | 2166.068 | 0 | -0.14 | 0.375093 | 14 | 104.7 | 95.3 | 11.26 | 5.32 | | High | High | High | High | High | High | High | High | High | High | High | 0 | 2.75E-05 | 2.91 |
| FALSE | High | GSISSVDGE | [R].GSISSV | 1xCarbami | Q8K0F1 1x | 0.000563 | 0 | 1 | 1 | 1 | Q8K0F1 | TBC1 doma | Q8K0F1 [4 | 0 | 2049.838 | 0 | 0.97 | 0.00799 | 14.96 | 67.1 | 132.9 | 18.29 | 13.79 | | High | High | High | High | High | High | High | High | High | High | High | 0 | 0.000125 | 2.86 |
| FALSE | High | GSLSSVTPS | [K].GSLSSV | 1xPhospho | Q9DBR4-2 | 0.002869 | 0.00016 | 1 | 5 | 2 | Q9DBR4-2 | Isoform 2 c | Q9DBR4-2 | 0 | 3498.741 | 0 | -1.59 | 0.008259 | 2.12 | 144.7 | 55.3 | 15.35 | 25.28 | | High | High | High | High | High | High | High | High | High | High | High | 7.19E-05 | 0.000778 | 3.12 |
| FALSE | High | GSLSSVTPS | [K].GSLSSV | 1xPhospho | E9QPX0 1x | 8.11E-05 | 0 | 1 | 4 | 4 | E9QPX0 | Amyloid be | E9QPX0 [3 | 0 | 2168.089 | 0 | 0.84 | 0.000975 | 11.4 | 69.1 | 130.9 | 8 | 7.58 | | High | High | High | High | High | High | High | High | High | High | High | 0.006366 | 0.06848 | 2.62 |
| FALSE | High | GSLSSVTPS | [K].GSLSSV | 2xPhospho | E9QPX0 2x | 0.002818 | 0.000109 | 1 | 4 | 1 | E9QPX0 | Amyloid be | E9QPX0 [3 | 0 | 2248.055 | 0 | 2.05 | 0.00024 | 15.83 | 36.3 | 163.7 | 15.99 | 4.98 | | High | High | High | High | High | High | High | High | High | High | High | 7.19E-05 | 0.000765 | 1.74 |
| FALSE | High | GSITEYTAT | [K].GSITEY | 1xPhospho | Q3TWC8 1 | 2.97E-07 | 0 | 1 | 6 | 3 | Q3TWC8 | Putative ur | Q3TWC8 [ | 0 | 1866.914 | 0 | 2.29 | 0.000179 | 9.41 | 33 | 167 | 8.6 | 4.18 | | High | High | High | High | High | High | High | High | High | High | High | 0 | 2.61E-08 | 3.13 |
| FALSE | High | GSLTFAGES | [R].GSLTFA | 1xPhospho | Q148V8 1x | 0.000515 | 0 | 1 | 1 | 1 | Q148V8 | Protein FAI | Q148V8 [9 | 0 | 1621.824 | 0 | 1.76 | 0.000278 | 20.44 | 46.5 | 153.5 | 14.7 | 4.87 | | High | High | High | High | High | High | High | High | High | High | High | 0 | 0.000113 | 3.03 |
| FALSE | High | GSLTFEPLT | [R].GSLTFE | 1xPhospho | Q3UE92 1x | 0.000352 | 0 | 1 | 7 | 2 | Q3UE92 | Putative ur | Q3UE92 [5 | 0 | 2282.281 | 0 | 0.8 | 0.000191 | 1.59 | 71.3 | 128.7 | 6.83 | 4.14 | | High | High | High | High | High | High | High | High | High | High | High | 0 | 7.36E-05 | 2.87 |
| FALSE | High | GSMGTGV | [R].GSMGT | 1xPhospho | P11679 1xP | 5.01E-07 | 0 | 1 | 1 | 1 | P11679 | Keratin, typ | P11679 [5 | 0 | 3597.912 | 0 | 5.47 | 0.001214 | 28.91 | 4.3 | 195.7 | 56.85 | 17.94 | | High | High | High | High | High | High | High | High | High | High | High | 0 | 4.68E-08 | 5.2 |
| FALSE | High | GSLPPSPVS | [K].GSLPPS | 1xPhospho | Q9WU62 1 | 1.21E-06 | 0 | 1 | 2 | 6 | Q9WU62 | Inner centr | Q9WU62 [ | 0 | 2792.437 | 0 | -0.74 | 0.001132 | 1.7 | 123.5 | 76.5 | 3.65 | 6.53 | | High | High | High | High | High | High | High | High | High | High | High | 0 | 1.26E-07 | 4.17 |
| FALSE | High | GSMGTGV | [R].GSMGT | 1xPhospho | P11679 1xP | 6.04E-06 | 0 | 1 | 1 | 3 | P11679 | Keratin, typ | P11679 [5 | 0 | 3613.907 | 0 | 3.57 | 0.000513 | 22.35 | 16.9 | 183.1 | 21.81 | 11.65 | | High | High | High | High | High | High | High | High | High | High | High | 0 | 1.67E-06 | 5.37 |
| FALSE | High | GSHTNSINI | [K].GSHTN | 2xPhospho | Q8BHK9 2x | 0.000568 | 0 | 1 | 3 | 1 | Q8BHK9 | DNA excisi | Q8BHK9 [1 | 0 | 2537.224 | 0 | -0.23 | 0.04282 | 4.64 | 107.7 | 92.3 | 11.05 | 4.5 | | High | High | High | High | High | High | High | High | High | High | High | 0 | 0.000127 | 3.61 |
| FALSE | High | GSHQMSLI | [K].GSHQM | 1xPhospho | Q01279 1x | 8.32E-08 | 0 | 1 | 3 | 1 | Q01279 | Epidermal | Q01279 [1 | 0 | 2808.274 | 0 | 1.97 | 0.027332 | 50.36 | 37.7 | 162.3 | 22.21 | 43.21 | | High | High | High | High | High | High | High | High | High | High | High | 0 | 6.21E-09 | 3.61 |
| FALSE | High | GSGFSLDVI | [K].GSGFSL | 1xPhospho | Q8C0T5 1x | 1.63E-05 | 0 | 1 | 4 | 8 | Q8C0T5 | Signal-indu | Q8C0T5 [2 | 0 | 1956.963 | 0 | 1.69 | 0.000368 | 10.2 | 48.7 | 151.3 | 7.44 | 5.78 | | High | High | High | High | High | High | High | High | High | High | High | 0 | 1.65E-05 | 4.33 |
| FALSE | High | GSGGFGG | [R].GSGGF | 1xPhospho | O08583 1x | 0.000305 | 0 | 1 | 2 | 1 | O08583 | THO comp | O08583 [2 | 0 | 1218.547 | 0 | 0.94 | 0.002086 | 13.98 | 69 | 131 | 10.83 | 5.81 | | High | High | High | High | High | High | High | High | High | High | High | 0 | 6.28E-05 | 1.28 |
| FALSE | High | GSGGFGST | [R].GSGGF | 1xPhospho | Q8VCG1 1x | 0.000289 | 0 | 1 | 3 | 3 | Q8VCG1 | Deoxyuridi | Q8VCG1 [1 | 0 | 1392.692 | 0 | -1.03 | 0.00333 | 1.58 | 131.1 | 68.9 | 6.4 | 15.09 | | High | High | High | High | High | High | High | High | High | High | High | 0 | 5.94E-05 | 1.85 |
| FALSE | High | GSGGGQG | [R].GSGGG | 1xPhospho | Q99020 1x | 8.3E-07 | 0 | 1 | 1 | 5 | Q99020 | Heterogen | Q99020 [2 | 0 | 1707.81 | 0 | 0.04 | 0.644453 | 6.66 | 99.9 | 100.1 | 4.2 | 3.74 | | High | High | High | High | High | High | High | High | High | High | High | 0 | 5.35E-06 | 3.08 |
| FALSE | High | GSGGSSAN | [R].GSGGS | 1xCarbami | P50446 1xP | 0.062209 | 0.004646 | 1 | 5 | 1 | P50446 | Keratin, typ | P50446 [41 | 0 | 1827.735 | 0 | 3.37 | 0.001303 | 60.68 | 13.1 | 186.9 | 49.13 | 17.3 | | High | High | High | High | High | High | High | High | High | High | High | 0.002381 | 0.02471 | 2.07 |
| FALSE | High | GSGHPAYA | [R].GSGHP | 1xPhospho | Q9DC11 1x | 1.48E-09 | 0 | 1 | 3 | 1 | Q9DC11 | Plexin dom | Q9DC11 [5 | 0 | 2165.068 | 0 | -0.4 | 0.066809 | 10.08 | 113.5 | 86.5 | 4.67 | 17.62 | | High | High | High | High | High | High | High | High | High | High | High | 0 | 6.69E-11 | 4.59 |
| FALSE | High | GSGHSQEC | [R].GSGHS | 1xPhospho | Q8VHK1 1x | 0.002463 | 0.000109 | 1 | 1 | 1 | Q8VHK1 | Caskin-2 O | Q8VHK1 [7 | 0 | 2780.284 | 0 | -0.02 | 0.304236 | 10.77 | 106.3 | 93.7 | 8.93 | 9.3 | | High | High | High | High | High | High | High | High | High | High | High | 4.94E-05 | 0.000658 | 3.34 |
| FALSE | High | GSGLGLHG | [K].GSGLG | 1xPhospho | E9Q616 1x | 0.000202 | 0 | 1 | 5 | 1 | E9Q616 | AHNAK nuc | E9Q616 [5 | 0 | 2015.109 | 0 | 3.73 | 0.004648 | 19.55 | 13.7 | 186.3 | 14.12 | 28.02 | | High | High | High | High | High | High | High | High | High | High | High | 0 | 3.96E-05 | 2.86 |
| FALSE | High | GSGRSSPA | [R].GSGRS | 3xPhospho | A0A068BIR | 0.000993 | 0 | 1 | 3 | 1 | A0A068BIR | KN motif a | A0A068BIR | 1 | 2897.367 | 0 | -1.37 | 0.001949 | 2.49 | 142.7 | 57.3 | 4.7 | 24.33 | | High | High | High | High | High | High | High | High | High | High | High | 0 | 0.000237 | 3.5 |
| FALSE | High | GSGSQSSV | [R].GSGSQ | 1xPhospho | Q1HL32 1x | 0.000276 | 0 | 1 | 4 | 2 | Q1HL32 | Structural | Q1HL32 [1 | 0 | 2274.096 | 0 | -0.49 | 0.034177 | 13.81 | 117.1 | 82.9 | 8.21 | 16.72 | | High | High | High | High | High | High | High | High | High | High | High | 0 | 5.6E-05 | 2.54 |
| FALSE | High | GSGSSPQL | [K].GSGSS | 1xPhospho | A2ADB0 1x | 0.041425 | 0.003126 | 1 | 5 | 3 | A2ADB0 | Msx2-inter | A2ADB0 [2 | 0 | 1991.902 | 0 | -0.32 | 0.663505 | 8.09 | 109.9 | 90.1 | 24 | 40 | | High | High | High | High | High | High | High | High | High | High | High | 0.001609 | 0.01554 | 1.98 |
| FALSE | High | GSGSSPQL | [K].GSGSS | 1xPhospho | A2ADB0 1x | 0.009416 | 0.000736 | 1 | 5 | 3 | A2ADB0 | Msx2-inter | A2ADB0 [2 | 1 | 4103.863 | 0 | -0.6 | 0.004221 | 4.75 | 125.6 | 74.4 | 10.07 | 13.14 | | High | High | High | High | High | High | High | High | High | High | High | 0.000571 | 0.00501 | 3.18 |
| FALSE | High | GSGSVDET | [R].GSGSV | 1xPhospho | Q99P72 1x | 0.000191 | 0 | 1 | 1 | 6 | Q99P72 | Reticulon- | Q99P72 [1 | 0 | 2868.412 | 0 | -0.59 | 0.032599 | 10.61 | 117 | 83 | 24.89 | 15.18 | | High | High | High | High | High | High | High | High | High | High | High | 0 | 3.71E-05 | 3.5 |
| FALSE | High | GSHTNSINI | [K].GSHTN | 1xPhospho | Q8BHK9 1x | 3.29E-05 | 0 | 1 | 3 | 1 | Q8BHK9 | DNA excisi | Q8BHK9 [1 | 0 | 2457.258 | 0 | 0.6 | 0.053128 | 21.72 | 83.6 | 116.4 | 18.23 | 14.32 | | High | High | High | High | High | High | High | High | High | High | High | 0 | 5.17E-06 | 2.8 |
| FALSE | High | GSGSVDET | [R].GSGSV | 1xPhospho | Q99P72 1x | 0.000656 | 0 | 1 | 1 | 4 | Q99P72 | Reticulon- | Q99P72 [1 | 0 | 3097.575 | 0 | -0.36 | 0.007243 | 2.9 | 118.6 | 81.4 | 7.63 | 10.3 | | High | High | High | High | High | High | High | High | High | High | High | 9.42E-05 | 0.000944 | 2.89 |
| FALSE | High | GSGSVVVD | [R].GSGSV | 1xPhospho | Q8K3G7 1x | 6.15E-06 | 0 | 1 | 2 | 1 | Q8K3G7 | Reticulon | Q8K3G7 [1 | 0 | 1759.898 | 0 | 2.75 | 0.000513 | 5.76 | 25.2 | 174.8 | 14.42 | 10.67 | | High | High | High | High | High | High | High | High | High | High | High | 0 | 7.84E-07 | 3.2 |
| FALSE | High | GSGTASDD | [R].GSGTA | 1xPhospho | A2AFQ0 1x | 1.11E-07 | 0 | 1 | 6 | 10 | A2AFQ0 | E3 ubiquiti | A2AFQ0 [1 | 0 | 1806.774 | 0 | -0.25 | 0.006381 | 4.23 | 110.6 | 89.4 | 3.39 | 4.63 | | High | High | High | High | High | High | High | High | High | High | High | 0 | 8.59E-09 | 2.78 |
| FALSE | High | GSGTGHPV | [R].GSGTG | 1xPhospho | Q99L88 1x | 0.001586 | 5.66E-05 | 1 | 2 | 1 | Q99L88 | Beta-1-syn | Q99L88 [7 | 0 | 2323.139 | 0 | -0.49 | 0.127698 | 12.59 | 112 | 88 | 12.14 | 14.96 | | High | High | High | High | High | High | High | High | High | High | High | 2.58E-05 | 0.0004 | 4.03 |
| FALSE | High | GSGTHMN | [K].GSGTH | 1xPhospho | P18572 1xP | 0.01861 | 0.001362 | 1 | 5 | 2 | P18572 | Basigin OS- | P18572 [37 | 0 | 1484.697 | 0 | 1.35 | 0.009792 | 13.04 | 57.1 | 142.9 | 6.23 | 22.94 | | High | High | High | High | High | High | High | High | High | High | High | 0.000723 | 0.006296 | 1.9 |
| FALSE | High | GSGTHMN | [K].GSGTH | 1xPhospho | P18572 1xP | 0.083783 | 0.006345 | 1 | 5 | 2 | P18572 | Basigin OS- | P18572 [37 | 0 | 1500.692 | 0 | 1.31 | 0.009304 | 17.45 | 55.9 | 144.1 | 5.44 | 23.05 | | High | High | High | High | High | High | High | High | High | High | High | 0.006622 | 0.07122 | 1.65 |
| FALSE | High | GSGVAQSF | [R].GSGVA | 1xPhospho | Q8K212 1x | 6.67E-06 | 0 | 1 | 2 | 2 | Q8K212 | Phosphofu | Q8K212 [2 | 0 | 3301.673 | 0 | 3.71 | 0.000242 | 72.03 | 14.1 | 185.9 | 67.62 | 12.32 | | High | High | High | High | High | High | High | High | High | High | High | 0 | 8.56E-07 | 3.81 |
| FALSE | High | GSGVAQSF | [R].GSGVA | 2xPhospho | Q8K212 2x | 1.21E-05 | 0 | 1 | 2 | 2 | Q8K212 | Phosphofu | Q8K212 [2 | 0 | 3381.639 | 0 | 2.12 | 0.000362 | 13.48 | 35.2 | 164.8 | 14.01 | 7.57 | | High | High | High | High | High | High | High | High | High | High | High | 0 | 1.67E-06 | 4.18 |
| FALSE | High | GSGVNAAC | [R].GSGVN | 1xPhospho | Q9JLF6 1xP | 4.74E-07 | 0 | 2 | 2 | 2 | Q9JLF6; A0 | Protein-glu | Q9JLF6 [93 | 0 | 1483.71 | 0 | 0.29 | 0.01751 | 11.64 | 89.1 | 110.9 | 4.42 | 7.17 | Shared | High | High | High | High | High | High | High | High | High | High | High | 0 | 4.39E-08 | 3.29 |
| FALSE | High | GSHCSGSG | [R].GSHCS | 1xCarbami | P48678 1xP | 2.59E-07 | 0 | 1 | 2 | 6 | P48678 | Prelamin-A | P48678 [5 | 0 | 2015.848 | 0 | 1.35 | 0.002198 | 25.97 | 54.6 | 145.4 | 6.09 | 13.22 | | High | High | High | High | High | High | High | High | High | High | High | 0 | 2.23E-08 | 4.13 |
| FALSE | High | GSHCSGSG | [R].GSHCS | 1xCarbami | P48678 2xP | 0.000434 | 0 | 1 | 2 | 2 | P48678 | Prelamin-A | P48678 [5 | 0 | 2095.814 | 0 | 1.05 | 0.004552 | 16.4 | 68 | 132 | 17.09 | 15.94 | | High | High | High | High | High | High | High | High | High | High | High | 2.58E-05 | 0.000366 | 4.02 |
| FALSE | High | GSHNGQV | [R].GSHNG | 1xCarbami | F6YRW4 1x | 2.35E-08 | 0 | 1 | 5 | 2 | F6YRW4 | F-box and | F6YRW4 [4 | 0 | 2235.042 | 0 | 0.86 | 0.002388 | 3.54 | 71.1 | 128.9 | 4.65 | 8.93 | | High | High | High | High | High | High | High | High | High | High | High | 0 | 5.82E-09 | 4.45 |
| FALSE | High | GSHNGQV | [R].GSHNG | 1xCarbami | F6YRW4 1x | 1.49E-08 | 0 | 1 | 5 | 8 | F6YRW4 | F-box and | F6YRW4 [4 | 0 | 2236.026 | 0 | 0.68 | 0.001037 | 14.84 | 77 | 123 | 11.82 | 7.7 | | High | High | High | High | High | High | High | High | High | High | High | 0 | 1.39E-08 | 4.08 |
| FALSE | High | GSHPDFPD | [R].GSHPD | 1xPhospho | Q8K4P0 1x | 0.003138 | 0.00016 | 1 | 2 | 1 | Q8K4P0 | pre-mRNA | Q8K4P0 [9 | 0 | 2779.255 | 0 | 0.67 | 0.005642 | 6.65 | 78.2 | 121.8 | 16.16 | 9.75 | | High | High | High | High | High | High | High | High | High | High | High | 7.19E-05 | 0.000863 | 2.78 |
| FALSE | High | GSGSVDET | [R].GSGSV | 2xPhospho | Q99P72 2x | 6.59E-05 | 0 | 1 | 1 | 3 | Q99P72 | Reticulon- | Q99P72 [1 | 0 | 3177.541 | 0 | -0.46 | 0.028395 | 6.04 | 121.5 | 78.5 | 16.07 | 8.61 | | High | High | High | High | High | High | High | High | High | High | High | 0 | 1.12E-05 | 3.93 |
| FALSE | High | GSMSNGH | [K].GSMSN | 1xPhospho | Q61127 1x | 0.031713 | 0.002011 | 1 | 5 | 1 | Q61127 | NGFI-A-bin | Q61127 [1 | 0 | 1798.82 | 0 | 0.06 | 0.316599 | 14.32 | 90.3 | 109.7 | 11.13 | 35.77 | | High | High | High | High | High | High | High | High | High | High | High | 0.001076 | 0.01145 | 1.58 |
| FALSE | High | GSMSPGGI | [R].GSMSP | 1xPhospho | Q5XJV7-2 | 0.000758 | 0 | 1 | 3 | 1 | Q5XJV7-2 | Isoform 2 c | Q5XJV7-2 [ | 0 | 1202.507 | 0 | -0.81 | 0.000997 | 4.57 | 127.5 | 72.5 | 5.63 | 11.76 | | High | High | High | High | High | High | High | High | High | High | High | 0 | 0.000174 | 2.36 |
| FALSE | High | GSNLPSPV | [R].GSNLPS | 1xPhospho | A2APV2 1x | 0.032548 | 0.002167 | 1 | 5 | 1 | A2APV2 | Formin-like | A2APV2 [1 | 0 | 1679.832 | 0 | 1.2 | 0.000574 | 14.97 | 59.1 | 140.9 | 14.59 | 7.65 | | High | High | High | High | High | High | High | High | High | High | High | 0.001142 | 0.01186 | 1.82 |
| FALSE | High | GSPPVPSG | [R].GSPPVF | 1xPhospho | Q9EP97 1x | 0.000194 | 0 | 1 | 1 | 2 | Q9EP97 | Sentrin-spe | Q9EP97 [2 | 0 | 2030.946 | 0 | 0 | 0.671948 | 5.38 | 99.9 | 100.1 | 2.81 | 7.54 | | High | High | High | High | High | High | High | High | High | High | High | 0 | 0.000154 | 3.28 |
| FALSE | High | GSPPVPSG | [R].GSPPVF | 1xPhospho | Q9EP97 1x | 1.67E-07 | 0 | 1 | 1 | 2 | Q9EP97 | Sentrin-spe | Q9EP97 [2 | 0 | 2046.94 | 0 | -0.29 | 0.156553 | 1.38 | 105.9 | 94.1 | 5.47 | 9.72 | | High | High | High | High | High | High | High | High | High | High | High | 0 | 1.36E-08 | 2.79 |
| FALSE | High | GSPQEEDA | [K].GSPQEI | 1xPhospho | Q07230 1x | 5.91E-05 | 0 | 1 | 4 | 1 | Q07230 | Zinc finger | Q07230 [6 | 0 | 2119.986 | 0 | -0.15 | 0.131011 | 6.92 | 107.4 | 92.6 | 10.64 | 10.46 | | High | High | High | High | High | High | High | High | High | High | High | 0 | 9.91E-06 | 3.66 |
| FALSE | High | GSPQGSAA | [R].GSPQG | 1xPhospho | Q61572 1x | 4.8E-05 | 0 | 1 | 1 | 1 | Q61572 | Forkhead b | Q61572 [3 | 0 | 2354.155 | 0 | 1.59 | 0.000693 | 17.07 | 49.4 | 150.6 | 11.59 | 7.57 | | High | High | High | High | High | High | High | High | High | High | High | 0 | 7.91E-06 | 3.15 |
| FALSE | High | GSPQQIDH | [R].GSPQQ | 1xPhospho | Q3U0V1 1x | 0.020499 | 0.001472 | 1 | 1 | 2 | Q3U0V1 | Far upstrea | Q3U0V1 [4 | 0 | 1618.835 | 0 | -0.12 | 0.023225 | 4.78 | 102.6 | 97.4 | 2.94 | 2.56 | | High | High | High | High | High | High | High | High | High | High | High | 0.000786 | 0.007028 | 3.44 |
| FALSE | High | GSPSGGST | [R].GSPSG | 1xPhospho | A2A7S8-2 | 2.41E-07 | 0 | 1 | 4 | 4 | A2A7S8-2 | Isoform 2 c | A2A7S8-2 [ | 0 | 1989.896 | 0 | -0.2 | 0.154898 | 3.33 | 107.5 | 92.5 | 9.72 | 5.65 | | High | High | High | High | High | High | High | High | High | High | High | 0 | 2.44E-05 | 3.44 |
| FALSE | High | GSPSSPVG | [K].GSPSSP | 1xPhospho | S4R2I9 1xP | 3.39E-06 | 0 | 1 | 4 | 5 | S4R2I9 | Zinc finger | S4R2I9 [34 | 0 | 1920.024 | 0 | -0.87 | 0.000179 | 1.91 | 128.4 | 71.6 | 3.25 | 8.36 | | High | High | High | High | High | High | High | High | High | High | High | 0 | 4.02E-07 | 3.28 |
| FALSE | High | GSPSSPVG | [K].GSPSSP | 2xPhospho | S4R2I9 2xP | 0.004572 | 0.000456 | 1 | 4 | 3 | S4R2I9 | Zinc finger | S4R2I9 [34 | 0 | 1999.991 | 0 | -0.41 | 0.035723 | 9.6 | 111.9 | 88.1 | 6.06 | 11.39 | | High | High | High | High | High | High | High | High | High | High | High | 0.000585 | 0.005252 | 2.68 |
| FALSE | High | GSPSVAAS | [K].GSPSVA | 1xPhospho | Q8BHZ4 1x | 1.41E-06 | 0 | 1 | 4 | 4 | Q8BHZ4 | Zinc finger | Q8BHZ4 [3 | 0 | 1904.029 | 0 | -0.21 | 0.045143 | 5.32 | 107.4 | 92.6 | 2.02 | 10.58 | | High | High | High | High | High | High | High | High | High | High | High | 0 | 1.51E-07 | 2.49 |
| FALSE | High | GSPSVAAS | [K].GSPSVA | 2xPhospho | Q8BHZ4 2x | 0.012845 | 0.000889 | 1 | 4 | 2 | Q8BHZ4 | Zinc finger | Q8BHZ4 [3 | 0 | 1983.996 | 0 | 0.27 | 0.0039 | 6.96 | 90 | 110 | 3.97 | 3.5 | | High | High | High | High | High | High | High | High | High | High | High | 0.001011 | 0.01079 | 2.09 |
| FALSE | High | GSPTGSSPI | [K].GSPTGS | 1xPhospho | Q5SXY1 1x | 1.39E-06 | 0 | 1 | 8 | 15 | Q5SXY1 | Cytospin-B | Q5SXY1 [3 | 0 | 2685.338 | 0 | 1.04 | 0.00471 | 14.56 | 64.7 | 135.3 | 7.52 | 12.35 | | High | High | High | High | High | High | High | High | High | High | High | 0 | 6.76E-07 | 3.82 |
| FALSE | High | GSPTGSSPI | [K].GSPTGS | 1xDeamida | Q5SXY1 1x | 0.001267 | 0 | 1 | 8 | 4 | Q5SXY1 | Cytospin-B | Q5SXY1 [3 | 0 | 2686.322 | 0 | 0.44 | 0.030095 | 15.05 | 80.5 | 119.5 | 17.7 | 14.58 | | High | High | High | High | High | High | High | High | High | High | High | 0.000161 | 0.001189 | 2.39 |
| FALSE | High | GSPTGSSPI | [K].GSPTGS | 2xPhospho | Q5SXY1 2x | 0.000941 | 0 | 1 | 8 | 4 | Q5SXY1 | Cytospin-B | Q5SXY1 [3 | 0 | 2765.305 | 0 | 0.68 | 0.011372 | 18.82 | 78.9 | 121.1 | 17.35 | 7.88 | | High | High | High | High | High | High | High | High | High | High | High | 0 | 0.000223 | 2.83 |
| FALSE | High | GSPPSSGP | [R].GSPPSS | 1xPhospho | Q8BGG6 1x | 7.21E-05 | 0 | 1 | 3 | 1 | Q8BGG6 | Motile spe | Q8BGG6 [2 | 0 | 2572.378 | 0 | 1.49 | 0.003574 | 28.97 | 54.3 | 145.7 | 35.44 | 8 | | High | High | High | High | High | High | High | High | High | High | High | 0 | 1.24E-05 | 3.24 |
| FALSE | High | GSPTPAYPI | [K].GSPTPA | 1xPhospho | Q148V8 1x | 0.08727 | 0.006537 | 1 | 1 | 2 | Q148V8 | Protein FAI | Q148V8 [8 | 0 | 1383.651 | 0 | 1.36 | 0.00128 | 2.18 | 55.1 | 144.9 | 4.5 | 12.09 | | High | High | High | High | High | High | High | High | High | High | High | 0.003346 | 0.03667 | 2.23 |
| FALSE | High | GSPTRPNP | [R].GSPTRF | 1xPhospho | E9PWE8 1x | 0.029076 | 0.001857 | 1 | 4 | 2 | E9PWE8 | Dihydropyr | E9PWE8 [6 | 0 | 1486.773 | 0 | 0.09 | 0.076734 | 5.26 | 95.1 | 104.9 | 5.61 | 8.86 | | High | High | High | High | High | High | High | High | High | High | High | 0.001151 | 0.01224 | 2.47 |
| FALSE | High | GSPTSGFPI | [K].GSPTSG | 1xPhospho | Q148V8 1x | 0.025073 | 0.001645 | 1 | 1 | 1 | Q148V8 | Protein FAI | Q148V8 [9 | 0 | 1328.62 | 0 | 1.22 | 0.000181 | 4.79 | 60 | 140 | 3.59 | 3.31 | | High | High | High | High | High | High | High | High | High | High | High | 0.000872 | 0.008835 | 2.49 |
| FALSE | High | GSPTSTYPI | [K].GSPTST | 1xPhospho | Q148V8 1x | 0.001011 | 0 | 1 | 1 | 2 | Q148V8 | Protein FAI | Q148V8 [9 | 0 | 1389.625 | 0 | 1.6 | 0.00024 | 5.94 | 48.9 | 151.1 | 3.5 | 5.2 | | High | High | High | High | High | High | High | High | High | High | High | 0 | 0.000241 | 1.96 |
| FALSE | High | GSPTTGLM | [R].GSPTTC | 1xPhospho | Q148V8 1x | 2.34E-05 | 0 | 1 | 1 | 3 | Q148V8 | Protein FAI | Q148V8 [9 | 0 | 1702.849 | 0 | 1.07 | 0.009671 | 23.21 | 61.4 | 138.6 | 9.48 | 19.39 | | High | High | High | High | High | High | High | High | High | High | High | 0 | 3.52E-06 | 4.3 |
| FALSE | High | GSPVDAGS | [K].GSPVD | 1xPhospho | A0A0A0M | 1.82E-06 | 0 | 1 | 6 | 1 | A0A0A0M | Anoctamin | A0A0A0M | 0 | 2683.2 | 0 | 3.69 | 0.000947 | 3.27 | 14.4 | 185.6 | 15.31 | 16.17 | | High | High | High | High | High | High | High | High | High | High | High | 0 | 1.99E-07 | 3.96 |
| FALSE | High | GSPVDAGS | [K].GSPVD | 1xPhospho | A0A0A0M | 2.06E-06 | 0 | 1 | 6 | 5 | A0A0A0M | Anoctamin | A0A0A0M | 0 | 2699.195 | 0 | 2.89 | 0.000303 | 8.24 | 23.8 | 176.2 | 15.91 | 8.45 | | High | High | High | High | High | High | High | High | High | High | High | 0 | 3.02E-07 | 3.31 |
| FALSE | High | GSPVLLDPI | [K].GSPVLL | 1xCarbami | A0A0A0M | 0.000207 | 0 | 1 | 3 | 3 | A0A0A0M | Histone-lys | A0A0A0M | 0 | 3295.628 | 0 | 0.35 | 0.759017 | 19.58 | 87.2 | 112.8 | 16.43 | 37.63 | | High | High | High | High | High | High | High | High | High | High | High | 0.000275 | 0.002127 | 3.94 |
| FALSE | High | GSPVLLDPI | [K].GSPVLL | 1xCarbami | A0A0A0M | 3.77E-05 | 0 | 1 | 3 | 2 | A0A0A0M | Histone-lys | A0A0A0M | 0 | 3311.623 | 0 | 0.26 | 0.142646 | 4.44 | 86.2 | 113.8 | 12.9 | 30.25 | | High | High | High | High | High | High | High | High | High | High | High | 0 | 6.94E-06 | 4.3 |
| FALSE | High | GSPVRHSP | [R].GSPVRI | 2xPhospho | A2AIX1 2xP | 0.055308 | 0.003982 | 1 | 7 | 1 | A2AIX1 | SEC16 hom | A2AIX1 [2 | 1 | 3275.425 | 0 | 0.03 | 0.415173 | 3.31 | 99.5 | 100.5 | 7.01 | 7.9 | | High | High | High | High | High | High | High | High | High | High | High | 0.002105 | 0.02163 | 2.94 |
| FALSE | High | GSPVSQGS | [K].GSPVS | 1xPhospho | Q8C0T5 1x | 6.83E-05 | 0 | 1 | 4 | 3 | Q8C0T5 | Signal-indu | Q8C0T5 [1 | 0 | 2603.279 | 0 | 1.51 | 0.00246 | 15.06 | 54.1 | 145.9 | 15.78 | 15.48 | | High | High | High | High | High | High | High | High | High | High | High | 0 | 0.000131 | 3.09 |
| FALSE | High | GSPVSQGS | [K].GSPVS | 1xPhospho | Q8C0T5 1x | 3.01E-09 | 0 | 1 | 4 | 9 | Q8C0T5 | Signal-indu | Q8C0T5 [1 | 0 | 2619.274 | 0 | 2.01 | 0.000965 | 23.35 | 38.4 | 161.6 | 11.12 | 10.14 | | High | High | High | High | High | High | High | High | High | High | High | 0 | 1.49E-10 | 4.81 |
| FALSE | High | GSPYHTGC | [K].GSPYH | 1xPhospho | B1AUH1 1x | 0.003857 | 0.000261 | 1 | 3 | 1 | B1AUH1 | Receptor-t | B1AUH1 [8 | 0 | 1828.906 | 0 | 0.07 | 0.092684 | 4.89 | 94 | 106 | 10.6 | 9.63 | | High | High | High | High | High | High | High | High | High | High | High | 0.000117 | 0.001087 | 2.86 |
| FALSE | High | GSQEDDPA | [K].GSQED | 1xPhospho | Q9EPQ8 1x | 4.35E-08 | 0 | 1 | 3 | 1 | Q9EPQ8 | Transcripti | Q9EPQ8 [6 | 0 | 2565.235 | 0 | 0.58 | 0.011507 | 15.29 | 78.7 | 121.3 | 2.09 | 13.1 | | High | High | High | High | High | High | High | High | High | High | High | 0 | 2.98E-09 | 2.27 |
| FALSE | High | GSPTQAYP | [K].GSPTQ | 1xPhospho | Q148V8 1x | 0.039347 | 0.002793 | 1 | 1 | 1 | Q148V8 | Protein FAI | Q148V8 [9 | 0 | 1414.657 | 0 | 1.24 | 0.000286 | 7.8 | 59.4 | 140.6 | 6.42 | 2.64 | | High | High | High | High | High | High | High | High | High | High | High | 0.001468 | 0.01473 | 1.84 |
| FALSE | High | GSPPDCTC | [R].GSPPD | 2xCarbami | Q69ZZ9 1x | 9.14E-06 | 0 | 1 | 1 | 2 | Q69ZZ9 | Uncharacte | Q69ZZ9 [2 | 0 | 2627.25 | 0 | 0.38 | 0.013931 | 18.65 | 85.7 | 114.3 | 14.3 | 13.36 | | High | High | High | High | High | High | High | High | High | High | High | 0 | 4.55E-05 | 3.17 |
| FALSE | High | GSPPAAGG | [R].GSPPA | 2xPhospho | Q545J5 2x | 0.000276 | 0 | 1 | 2 | 1 | Q545J5 | Basic Krup | Q545J5 [7 | 0 | 1830.88 | 0 | 0.57 | 0.003356 | 5.57 | 78.7 | 121.3 | 6.1 | 10.88 | | High | High | High | High | High | High | High | High | High | High | High | 0 | 5.62E-05 | 2.58 |
| FALSE | High | GSPPAAGG | [R].GSPPA | 1xPhospho | Q545J5 1x | 4.21E-06 | 0 | 1 | 2 | 2 | Q545J5 | Basic Krup | Q545J5 [7 | 0 | 1750.914 | 0 | 0.65 | 0.014379 | 25.24 | 82.5 | 117.5 | 12.51 | 8.68 | | High | High | High | High | High | High | High | High | High | High | High | 0 | 1.68E-06 | 2.82 |
| FALSE | High | GSNPFEDE | [K].GSNPFI | 1xPhospho | Q6PGF7 1x | 2.98E-09 | 0 | 1 | 1 | 2 | Q6PGF7 | Exocyst co | Q6PGF7 [2 | 0 | 3018.303 | 0 | 0.98 | 0.000361 | 5.24 | 66.7 | 133.3 | 5.56 | 6.28 | | High | High | High | High | High | High | High | High | High | High | High | 0 | 4.96E-09 | 4.79 |
| FALSE | High | GSPAEELR | [K].GSPAEI | 1xPhospho | Q8C0P7 1x | 0.002022 | 5.66E-05 | 1 | 6 | 2 | Q8C0P7 | E3 SUMO- | Q8C0P7 [1 | 0 | 1167.561 | 0 | -1.05 | 0.000154 | 0.12 | 134.8 | 65.2 | 2.14 | 6.18 | | High | High | High | High | High | High | High | High | High | High | High | 0.001142 | 0.0118 | 2.2 |
| FALSE | High | GSPALLPST | [R].GSPALL | 1xPhospho | F8VPP8 1x | 0.008022 | 0.000736 | 1 | 4 | 2 | F8VPP8 | Zinc finger | F8VPP8 [2 | 0 | 3408.784 | 0 | 1.81 | 0.002679 | 34.4 | 47.8 | 152.2 | 23.16 | 12.68 | | High | High | High | High | High | High | High | High | High | High | High | 0.000327 | 0.00245 | 2.49 |
| FALSE | High | GSPALLPST | [R].GSPALL | 2xPhospho | F8VPP8 2x | 0.000645 | 0 | 1 | 4 | 1 | F8VPP8 | Zinc finger | F8VPP8 [2 | 0 | 3488.75 | 0 | 1.71 | 0.00207 | 28.98 | 47.6 | 152.4 | 37.69 | 16.12 | | High | High | High | High | High | High | High | High | High | High | High | 0 | 0.000146 | 4.48 |
| FALSE | High | GSPAPSSAS | [R].GSPAPS | 1xPhospho | G3UXW9 1 | 5.94E-07 | 0 | 1 | 1 | 6 | G3UXW9 | COP9 signa | G3UXW9 [ | 0 | 1872.854 | 0 | 1.08 | 0.000181 | 1.75 | 64.3 | 135.7 | 7.36 | 6.15 | | High | High | High | High | High | High | High | High | High | High | High | 0 | 4.77E-07 | 3.61 |
| FALSE | High | GSPAPSSAS | [R].GSPAPS | 2xPhospho | G3UXW9 2 | 0.014538 | 0.001011 | 1 | 1 | 4 | G3UXW9 | COP9 signa | G3UXW9 [ | 0 | 1952.82 | 0 | 1.73 | 0.008493 | 19.17 | 42 | 158 | 23.59 | 31.14 | | High | High | High | High | High | High | High | High | High | High | High | 0.000538 | 0.004758 | 2.84 |
| FALSE | High | GSPCAEEEI | [K].GSPCAI | 1xCarbami | O54839 1x | 2.35E-08 | 0 | 1 | 3 | 2 | O54839 | Eomesode | O54839 [1 | 0 | 2269.037 | 0 | -0.8 | 0.023554 | 7.12 | 125.2 | 74.8 | 5.74 | 26.1 | | High | High | High | High | High | High | High | High | High | High | High | 0 | 2.13E-06 | 3.69 |
| FALSE | High | GSPCDTLAS | [K].GSPCD | 1xCarbami | P70365 1xP | 0.002376 | 0.000109 | 1 | 5 | 1 | P70365 | Nuclear rec | P70365 [21 | 0 | 1890.892 | 0 | -0.26 | 0.011453 | 5.77 | 109.2 | 90.8 | 6.45 | 6.54 | | High | High | High | High | High | High | High | High | High | High | High | 4.94E-05 | 0.000633 | 2.13 |
| FALSE | High | GSPDGSHP | [K].GSPDG | 1xPhospho | Q60793 1x | 6.56E-07 | 0 | 1 | 1 | 4 | Q60793 | Krueppel-li | Q60793 [2 | 0 | 2242.086 | 0 | 0.51 | 0.00378 | 14.04 | 79.7 | 120.3 | 4.19 | 9.22 | | High | High | High | High | High | High | High | High | High | High | High | 0 | 6.36E-08 | 4.55 |

| | | | | | | | | | | | | | | | | | | | | | | | | | | | | | | | | | | | | | | |
|---|---|---|---|---|---|---|---|---|---|---|---|---|---|---|---|---|---|---|---|---|---|---|---|---|---|---|---|---|---|---|---|---|---|---|---|---|---|---|
| FALSE | High | GSPDGSHP | [K].GSPDG | 2xPhospho | Q60793 2x | 0.001674 | 5.66E-05 | 1 | 1 | 1 | Q60793 | Krueppel-li | Q60793 [2 | 0 | 2322.052 | 0 | 0.43 | 0.000986 | 2.96 | 84.5 | 115.5 | 7.53 | 6.25 | | High | High | High | High | High | High | High | High | High | High | High | 2.58E-05 | 0.000425 | 3.46 |
| FALSE | High | GSPDQYQL | [K].GSPDQ | 1xPhospho | E9PYW4 1x | 0.00509 | 0.000551 | 1 | 2 | 1 | E9PYW4 | E3 ubiquiti | E9PYW4 [4 | 0 | 1372.646 | 0 | 0.72 | 0.000986 | 9.53 | 75 | 125 | 5.39 | 6.13 | | High | High | High | High | High | High | High | High | High | High | High | 0.000246 | 0.00148 | 1.77 |
| FALSE | High | GSPECVGL | [R].GSPEC | 1xCarbami | B2RQE3 1x | 0.007339 | 0.00062 | 1 | 6 | 1 | B2RQE3 | Stim2 prot | B2RQE3 [7 | 0 | 1815.896 | 0 | 1.23 | 0.000762 | 2.72 | 60 | 140 | 10.84 | 6.82 | | High | High | High | High | High | High | High | High | High | High | High | 0.000275 | 0.002211 | 2.64 |
| FALSE | High | GSPEDGSH | [R].GSPED | 1xPhospho | Q3UQS8 1x | 0.001373 | 5.66E-05 | 1 | 3 | 1 | Q3UQS8 | RNA-bindir | Q3UQS8 [1 | 0 | 1905.843 | 0 | 1.36 | 0.000242 | 5.24 | 55.4 | 144.6 | 7.25 | 6.49 | | High | High | High | High | High | High | High | High | High | High | High | 2.58E-05 | 0.000342 | 2.97 |
| FALSE | High | GSPEDGSH | [R].GSPED | 1xPhospho | Q3UQS8 1x | 8.81E-09 | 0 | 1 | 3 | 2 | Q3UQS8 | RNA-bindir | Q3UQS8 [1 | 0 | 2135.006 | 0 | 2.88 | 0.001501 | 25.55 | 24.5 | 175.5 | 13.73 | 17.35 | | High | High | High | High | High | High | High | High | High | High | High | 0 | 4.98E-10 | 4.98 |
| FALSE | High | GSPETNCLI | [R].GSPETN | 1xCarbami | Q14B71 1x | 0.081752 | 0.006219 | 1 | 4 | 1 | Q14B71 | Cell divisio | Q14B71 [9 | 0 | 1455.686 | 0 | -0.31 | 0.001066 | 0.75 | 109.6 | 90.4 | 6.37 | 5.97 | | High | High | High | High | High | High | High | High | High | High | High | 0.003229 | 0.03414 | 1.68 |
| FALSE | High | GSPGALSSI | [R].GSPGAI | 1xPhospho | D3YUW7 1 | 5.23E-06 | 0 | 1 | 8 | 3 | D3YUW7 | Cingulin OS | D3YUW7 [ | 0 | 2700.317 | 0 | -0.48 | 0.009984 | 7.87 | 117 | 83 | 4.76 | 12.75 | | High | High | High | High | High | High | High | High | High | High | High | 0 | 6.5E-07 | 3.33 |
| FALSE | High | GSPGALSSI | [R].GSPGAI | 2xPhospho | D3YUW7 2 | 1.01E-05 | 0 | 1 | 8 | 1 | D3YUW7 | Cingulin OS | D3YUW7 [ | 0 | 2780.283 | 0 | -1.27 | 0.000899 | 1.13 | 139.6 | 60.4 | 7.44 | 4.93 | | High | High | High | High | High | High | High | High | High | High | High | 0 | 1.37E-06 | 4.37 |
| FALSE | High | GSPGGDLG | [K].GSPGG | 1xPhospho | O70311 1x | 0.000415 | 0 | 1 | 2 | 5 | O70311 | Glycylpept | O70311 [3 | 0 | 1396.724 | 0 | -0.66 | 0.005504 | 1.28 | 121.6 | 78.4 | 2.33 | 11.74 | | High | High | High | High | High | High | High | High | High | High | High | 0 | 0.000279 | 1.93 |
| FALSE | High | GSPGLEEK | [K].GSPGLE | 1xPhospho | O70469 1x | 0.102567 | 0.008249 | 1 | 2 | 1 | O70469 | Docking pr | O70469 [1 | 0 | 1354.702 | 0 | 1.98 | 0.000323 | 7.37 | 40.8 | 159.2 | 3.55 | 6.73 | | High | High | High | High | High | High | High | High | High | High | High | 0.004138 | 0.04459 | 1.5 |
| FALSE | High | GSPGPQGA | [K].GSPGP | 1xPhospho | Q3TAB2 1x | 0.000396 | 0 | 2 | 6 | 4 | Q3TAB2; H | Putative ur | Q3TAB2 [6 | 0 | 1336.703 | 0 | -0.62 | 0.001035 | 2 | 121.2 | 78.8 | 4.15 | 3.81 | Shared | High | High | High | High | High | High | High | High | High | High | High | 0.00607 | 0.06605 | 1.87 |
| FALSE | High | GSPGSLSSK | [K].GSPGSI | 1xPhospho | Q5BLK4 1x | 0.022973 | 0.001578 | 1 | 1 | 1 | Q5BLK4 | Terminal u | Q5BLK4 [1 | 0 | 1357.713 | 0 | 0.16 | 0.007693 | 4.1 | 94.4 | 105.6 | 3.7 | 3.24 | | High | High | High | High | High | High | High | High | High | High | High | 0.000847 | 0.00796 | 1.25 |
| FALSE | High | GSPHPGVG | [K].GSPHP | 1xPhospho | E9Q2I4 1x | 4.31E-05 | 0 | 1 | 3 | 1 | E9Q2I4 | ELM2 and | E9Q2I4 [14 | 0 | 2654.353 | 0 | 1.18 | 0.004533 | 26.52 | 60.4 | 139.6 | 20.38 | 10.58 | | High | High | High | High | High | High | High | High | High | High | High | 0 | 6.98E-06 | 3.07 |
| FALSE | High | GSPHYFSPI | [R].GSPHYI | 1xPhospho | Q3TXM9 1 | 0.095408 | 0.007437 | 1 | 3 | 2 | Q3TXM9 | Putative ur | Q3TXM9 [ | 0 | 1763.814 | 0 | 0.45 | 0.147754 | 10.62 | 85.9 | 114.1 | 8.32 | 15.6 | | High | High | High | High | High | High | High | High | High | High | High | 0.004517 | 0.04898 | 1.97 |
| FALSE | High | GSPHYFSPI | [R].GSPHYI | 2xPhospho | Q3TXM9 2 | 0.003857 | 0.000261 | 1 | 3 | 6 | Q3TXM9 | Putative ur | Q3TXM9 [ | 0 | 1843.781 | 0 | 0.36 | 0.004522 | 5.97 | 86.7 | 113.3 | 6.38 | 6.26 | | High | High | High | High | High | High | High | High | High | High | High | 0.000275 | 0.002059 | 3.18 |
| FALSE | High | GSPLAANA | [R].GSPLAA | 2xCarbami | G5E8L6 1x | 3.69E-07 | 0 | 1 | 3 | 2 | G5E8L6 | Killer cell le | G5E8L6 [1 | 0 | 3095.383 | 0 | 0.6 | 0.001999 | 2.99 | 79.4 | 120.6 | 4.01 | 7.23 | | High | High | High | High | High | High | High | High | High | High | High | 0 | 5.98E-07 | 3.84 |
| FALSE | High | GSPLAEGPI | [R].GSPLAE | 1xPhospho | Q5I1X5 1x | 0.053023 | 0.00384 | 1 | 1 | 1 | Q5I1X5 | RelA-assoc | Q5I1X5 [18 | 0 | 1911.92 | 0 | 0.48 | 0.003622 | 6.18 | 83.2 | 116.8 | 11.19 | 8.43 | | High | High | High | High | High | High | High | High | High | High | High | 0.001999 | 0.02055 | 1.45 |
| FALSE | High | GSPIDVLHC | [K].GSPIDV | 1xPhospho | F6ZBR8 1x | 5.86E-05 | 0 | 1 | 1 | 1 | F6ZBR8 | Protein Wi | F6ZBR8 [27 | 0 | 1585.866 | 0 | 0.16 | 0.269502 | 20.85 | 94.4 | 105.6 | 8.35 | 16.24 | | High | High | High | High | High | High | High | High | High | High | High | 0 | 9.81E-06 | 2.81 |
| FALSE | High | GSPLHPAH | [K].GSPLHF | 1xPhospho | Q571H2 1x | 0.000725 | 0 | 1 | 5 | 1 | Q571H2 | MKIAA021 | Q571H2 [1 | 0 | 2244.068 | 0 | -0.45 | 0.023088 | 5.92 | 117.6 | 82.4 | 12.83 | 11.11 | | High | High | High | High | High | High | High | High | High | High | High | 0 | 0.000165 | 3.3 |
| FALSE | High | GSPLSNRQ | [K].GSPLSN | 2xPhospho | Q8K3G5 2x | 0.1059 | 0.008582 | 1 | 3 | 1 | Q8K3G5 | Inactive se | Q8K3G5 [1 | 1 | 2131.071 | 0 | 0.51 | 0.012889 | 9.62 | 82 | 118 | 10.24 | 7.05 | | High | High | High | High | High | High | High | High | High | High | High | 0.004317 | 0.0462 | 1.07 |
| FALSE | High | GSQGDIK | [R].GSQGD | 1xPhospho | Q8CI51 1x | 0.043981 | 0.003228 | 2 | 11 | 2 | Q8CI51; Q | PDZ and LI | Q8CI51 [22 | 0 | 1242.65 | 0 | 0.45 | 0.020109 | 9.86 | 84.5 | 115.5 | 4.98 | 12.85 | Shared | High | High | High | High | High | High | High | High | High | High | High | 0.003258 | 0.03439 | 1.86 |
| FALSE | High | GTPNSASK | [K].GTPNS | 1xPhospho | E9Q4F8 1x | 0.019446 | 0.001438 | 1 | 6 | 2 | E9Q4F8 | Ankyrin re | E9Q4F8 [1 | 0 | 1299.671 | 0 | -0.7 | 0.000338 | 2.02 | 125.7 | 74.3 | 4.83 | 4.17 | | High | High | High | High | High | High | High | High | High | High | High | 0.001812 | 0.01842 | 1.51 |
| FALSE | High | GTPPWSDC | [R].GTPPW | 1xPhospho | Q5U3K5 1x | 0.000837 | 0 | 1 | 1 | 1 | Q5U3K5 | Rab-like pr | Q5U3K5 [5 | 0 | 2811.324 | 0 | -0.2 | 0.027774 | 2.49 | 109.1 | 90.9 | 6.01 | 8.48 | | High | High | High | High | High | High | High | High | High | High | High | 0 | 0.000195 | 1.61 |
| FALSE | High | GTPSQSPV | [K].GTPSQ | 1xPhospho | A0A087WF | 0.002441 | 0.000109 | 1 | 8 | 5 | A0A087WF | Arf-GAP dc | A0A087WF | 0 | 1393.704 | 0 | -0.2 | 0.411041 | 7.46 | 105.9 | 94.1 | 1.34 | 14.06 | | High | High | High | High | High | High | High | High | High | High | High | 0.001061 | 0.01123 | 2.18 |
| FALSE | High | HEVSGETV | [K].HEVSGI | 1xPhospho | Q6PDI5 1x | 5.88E-06 | 0 | 1 | 3 | 1 | Q6PDI5 | Proteasom | Q6PDI5 [10 | 0 | 2239.152 | 0 | 0.81 | 0.00444 | 7.14 | 72.4 | 127.6 | 10.66 | 17.08 | | High | High | High | High | High | High | High | High | High | High | High | 0 | 7.45E-07 | 3.22 |
| FALSE | High | HFGSGDLM | [K].HFGSGI | 1xPhospho | B9EK76 1x | 0.000334 | 0 | 1 | 5 | 1 | B9EK76 | Signal-indu | B9EK76 [1 | 0 | 2644.35 | 0 | -0.44 | 0.04581 | 4.69 | 116.3 | 83.7 | 10.02 | 14.9 | | High | High | High | High | High | High | High | High | High | High | High | 0 | 6.95E-05 | 2.65 |
| FALSE | High | HFGSGDLM | [K].HFGSGI | 1xPhospho | B9EK76 1x | 0.000142 | 0 | 1 | 5 | 1 | B9EK76 | Signal-indu | B9EK76 [1 | 0 | 2660.345 | 0 | -0.44 | 0.025785 | 5.2 | 115.9 | 84.1 | 7.17 | 30.79 | | High | High | High | High | High | High | High | High | High | High | High | 0 | 2.66E-05 | 2.97 |
| FALSE | High | HFKDEEED | [K].HFKDEI | 1xPhospho | B7ZWA9 1x | 0.003001 | 0.00016 | 1 | 2 | 1 | B7ZWA9 | Zbtb7a pro | B7ZWA9 [5 | 1 | 2701.167 | 0 | 2.39 | 0.026421 | 54.02 | 31 | 169 | 13.7 | 41.77 | | High | High | High | High | High | High | High | High | High | High | High | 7.19E-05 | 0.000821 | 2.2 |
| FALSE | High | HFLPLDFST | [K].HFLPLD | 1xDeamida | P28700 1x | 0.080423 | 0.006219 | 1 | 2 | 2 | P28700 | Retinoic ac | P28700 [5- | 0 | 2714.302 | 0 | -0.44 | 0.347474 | 8.91 | 104 | 96 | 17.63 | 15.99 | | High | High | High | High | High | High | High | High | High | High | High | 0.0032 | 0.03396 | 4.02 |
| FALSE | High | HFPGEDGD | [K].HFPGEI | 1xPhospho | E9Q2Z1 1x | 0.064323 | 0.004664 | 1 | 3 | 1 | E9Q2Z1 | CECR2, hist | E9Q2Z1 [5 | 0 | 2230.953 | 0 | -1.78 | 0.001039 | 1.6 | 151.8 | 48.2 | 7.27 | 18.67 | | High | High | High | High | High | High | High | High | High | High | High | 0.002403 | 0.02589 | 2.7 |
| FALSE | High | HFSISGCPL | [R].HFSISG | 2xCarbami | Q5SVQ0 1x | 0.000772 | 0 | 1 | 7 | 2 | Q5SVQ0 | Histone ac | Q5SVQ0 [2 | 0 | 2773.288 | 0 | -0.63 | 0.013135 | 13.84 | 129.2 | 70.8 | 12.7 | 21.34 | | High | High | High | High | High | High | High | High | High | High | High | 0 | 0.000178 | 2.15 |
| FALSE | High | HFSQSEDT | [R].HFSQSE | 1xPhospho | A0A1W2P7 | 1.8E-05 | 0 | 1 | 7 | 4 | A0A1W2P7 | Ral GTPase | A0A1W2P7 | 0 | 3033.394 | 0 | 0.7 | 0.001282 | 8.79 | 77.3 | 122.7 | 2.28 | 6.25 | | High | High | High | High | High | High | High | High | High | High | High | 0 | 2.62E-06 | 4.45 |
| FALSE | High | HFSQSEDT | [R].HFSQSE | 1xDeamida | A0A1W2P7 | 0.045122 | 0.003373 | 1 | 7 | 1 | A0A1W2P7 | Ral GTPase | A0A1W2P7 | 0 | 3034.378 | 0 | 0.31 | 0.016638 | 8.07 | 91.4 | 108.6 | 13.43 | 6.4 | | High | High | High | High | High | High | High | High | High | High | High | 0.001712 | 0.0171 | 3.14 |
| FALSE | High | HGAPAAPS | [R].HGAPA | 1xPhospho | Q8BHL3 1x | 1.91E-05 | 0 | 1 | 1 | 4 | Q8BHL3 | TBC1 doma | Q8BHL3 [1 | 0 | 1463.736 | 0 | 0.13 | 0.136702 | 4.65 | 95.3 | 104.7 | 7.67 | 8.24 | | High | High | High | High | High | High | High | High | High | High | High | 0 | 2.79E-06 | 2.53 |
| FALSE | High | HGASSGCL | [R].HGASS | 1xCarbami | Q9JK83 1x | 1.38E-06 | 0 | 1 | 2 | 8 | Q9JK83 | Partitionin | Q9JK83 [7- | 0 | 1987.938 | 0 | -0.99 | 0.001401 | 3.36 | 133 | 67 | 9.5 | 4.68 | | High | High | High | High | High | High | High | High | High | High | High | 0 | 2.94E-07 | 4.64 |
| FALSE | High | HGATVLGS | [R].HGATV | 1xPhospho | E9PZP8 1x | 0.007339 | 0.00062 | 1 | 3 | 1 | E9PZP8 | HECT and | E9PZP8 [28 | 0 | 1521.819 | 0 | 0.46 | 0.006763 | 7.49 | 85.5 | 114.5 | 8.26 | 5.72 | | High | High | High | High | High | High | High | High | High | High | High | 0.000275 | 0.002218 | 2.77 |
| FALSE | High | HGCLSSPK | [R].HGCLS | 1xCarbami | E9Q6H8 1x | 0.002293 | 0.000109 | 1 | 12 | 1 | E9Q6H8 | Pleckstrin | E9Q6H8 [5 | 0 | 1423.717 | 0 | -0.47 | 0.031664 | 3.88 | 112.7 | 87.3 | 5.49 | 13.35 | | High | High | High | High | High | High | High | High | High | High | High | 4.94E-05 | 0.000607 | 2.31 |
| FALSE | High | HETEESSV | [K].HETEES | 1xPhospho | Q6NS59 1x | 1.89E-07 | 0 | 1 | 2 | 1 | Q6NS59 | Protein FA | Q6NS59 [4 | 0 | 2366.189 | 0 | 0.49 | 0.026756 | 10.89 | 83.1 | 116.9 | 9.51 | 9.47 | | High | High | High | High | High | High | High | High | High | High | High | 0 | 1.57E-08 | 3.64 |
| FALSE | High | HGFASEPT | [K].HGFASI | 1xPhospho | Q9QZ41 1x | 0.017498 | 0.001245 | 1 | 2 | 1 | Q9QZ41 | Protein DB | Q9QZ41 [4 | 0 | 2191.084 | 0 | -0.44 | 0.046737 | 15.42 | 113.5 | 86.5 | 5.82 | 17.24 | | High | High | High | High | High | High | High | High | High | High | High | 0.000674 | 0.00587 | 1.75 |
| FALSE | High | HGGGGIVA | [K].HGGG | 2xPhospho | Q3UM45 2 | 0.002576 | 0.000109 | 1 | 2 | 1 | Q3UM45 | Protein ph | Q3UM45 [ | 0 | 1955.919 | 0 | -0.56 | 0.170952 | 32.18 | 121.5 | 78.5 | 38.57 | 26.8 | | High | High | High | High | High | High | High | High | High | High | High | 4.94E-05 | 0.000691 | 1.92 |
| FALSE | High | HGGGGIVA | [K].HGGG | 1xPhospho | Q3UM45 1 | 0.000331 | 0 | 1 | 2 | 3 | Q3UM45 | Protein ph | Q3UM45 [ | 0 | 2105.116 | 0 | 0.09 | 0.963397 | 17.54 | 93.8 | 106.2 | 13.33 | 24.79 | | High | High | High | High | High | High | High | High | High | High | High | 0.000275 | 0.002261 | 2.93 |
| FALSE | High | HGGGGIVA | [K].HGGG | 2xPhospho | Q3UM45 2 | 2.29E-05 | 0 | 1 | 2 | 5 | Q3UM45 | Protein ph | Q3UM45 [ | 0 | 2185.082 | 0 | -0.22 | 0.123284 | 9.77 | 108.1 | 91.9 | 2.59 | 24.04 | | High | High | High | High | High | High | High | High | High | High | High | 0 | 3.42E-06 | 3.09 |
| FALSE | High | HGHFGWP | [K].HGHFG | 1xPhospho | Q64337 1x | 0.000375 | 0 | 1 | 3 | 1 | Q64337 | Sequestos | Q64337 [1 | 0 | 2964.35 | 0 | 1.06 | 0.00146 | 5.09 | 61.9 | 138.1 | 11.9 | 14.58 | | High | High | High | High | High | High | High | High | High | High | High | 0 | 7.97E-05 | 3.37 |
| FALSE | High | HGLLLPASF | [R].HGLLLP | 1xPhospho | Q9DB52 1x | 0.003028 | 0.00016 | 1 | 1 | 1 | Q9DB52 | Protein FA | Q9DB52 [6 | 0 | 1468.824 | 0 | 0.77 | 0.000983 | 3.27 | 74.7 | 125.3 | 5.73 | 7.69 | | High | High | High | High | High | High | High | High | High | High | High | 7.19E-05 | 0.000825 | 3.05 |
| FALSE | High | HGILSSVNY | [K].HGILSS | 1xPhospho | O88413 1x | 6.67E-06 | 0 | 1 | 1 | 3 | O88413 | Tubby-rela | O88413 [1 | 1 | 3669.696 | 0 | -0.37 | 0.254588 | 11.61 | 106.9 | 93.1 | 11.74 | 18.52 | | High | High | High | High | High | High | High | High | High | High | High | 0 | 8.57E-07 | 4.96 |
| FALSE | High | HGPAQAVT | [K].HGPAQ | 1xPhospho | Q3U832 1x | 0.001516 | 5.66E-05 | 1 | 3 | 1 | Q3U832 | Putative ur | Q3U832 [4 | 0 | 1960.01 | 0 | -1.72 | 0.150519 | 6.73 | 148.6 | 51.4 | 52.96 | 54.65 | | High | High | High | High | High | High | High | High | High | High | High | 2.58E-05 | 0.000379 | 3.12 |
| FALSE | High | HGPAQAVT | [K].HGPAQ | 1xPhospho | Q3U832 1x | 1.03E-08 | 0 | 1 | 3 | 3 | Q3U832 | Putative ur | Q3U832 [4 | 0 | 2189.173 | 0 | -1.54 | 0.000476 | 0.95 | 149.9 | 50.1 | 4.35 | 15.99 | | High | High | High | High | High | High | High | High | High | High | High | 0 | 5.92E-10 | 5.44 |
| FALSE | High | HGSASQVC | [R].HGSAS | 1xPhospho | Q04690 1x | 0.010853 | 0.000772 | 1 | 5 | 1 | Q04690 | Neurofibrc | Q04690 [2 | 0 | 1479.772 | 0 | 1.21 | 0.000624 | 4.9 | 60.6 | 139.4 | 1.66 | 7.04 | | High | High | High | High | High | High | High | High | High | High | High | 0.000399 | 0.003431 | 2.85 |
| FALSE | High | HGSDPAFG | [R].HGSDP | 1xPhospho | Q148V8 1x | 0.00043 | 0 | 1 | 1 | 2 | Q148V8 | Protein FA | Q148V8 [5 | 0 | 1533.705 | 0 | 0.59 | 0.000236 | 3.88 | 78.8 | 121.2 | 3.98 | 4.03 | | High | High | High | High | High | High | High | High | High | High | High | 0 | 9.2E-05 | 3.52 |
| FALSE | High | HGSGADSD | [R].HGSGA | 1xPhospho | Q68FF6 1x | 8.33E-05 | 0 | 1 | 3 | 1 | Q68FF6 | ARF GTPas | Q68FF6 [5 | 0 | 2756.225 | 0 | -0.06 | 0.925932 | 40.27 | 106.2 | 93.8 | 24.09 | 46.99 | | High | High | High | High | High | High | High | High | High | High | High | 0 | 1.47E-05 | 3.64 |
| FALSE | High | HGSGADSD | [R].HGSGA | 2xPhospho | Q68FF6 2x | 1.02E-07 | 0 | 1 | 3 | 3 | Q68FF6 | ARF GTPas | Q68FF6 [5 | 0 | 2836.191 | 0 | 0.71 | 0.011092 | 25.51 | 72.7 | 127.3 | 11.88 | 13.28 | | High | High | High | High | High | High | High | High | High | High | High | 0 | 7.88E-09 | 4.98 |
| FALSE | High | HGSGADSD | [R].HGSGA | 1xPhospho | Q68FF6 1x | 1.15E-07 | 0 | 1 | 3 | 4 | Q68FF6 | ARF GTPas | Q68FF6 [5 | 0 | 2985.388 | 0 | 0.17 | 0.976492 | 13.21 | 99.2 | 100.8 | 9.5 | 16.16 | | High | High | High | High | High | High | High | High | High | High | High | 0 | 8.94E-09 | 4.66 |
| FALSE | High | HGGAGGSI | [R].HGGAG | 1xPhospho | F8VPZ9 1x | 0.002508 | 0.000109 | 1 | 2 | 1 | F8VPZ9 | BRD4-inter | F8VPZ9 [12 | 0 | 1648.779 | 0 | 0.33 | 0.074732 | 25.45 | 88.6 | 111.4 | 9.39 | 10.02 | | High | High | High | High | High | High | High | High | High | High | High | 4.94E-05 | 0.000672 | 1.46 |
| FALSE | High | HGSGADSD | [R].HGSGA | 2xPhospho | Q68FF6 2x | 0.008167 | 0.000736 | 1 | 3 | 1 | Q68FF6 | ARF GTPas | Q68FF6 [5 | 0 | 3065.354 | 0 | 1.1 | 0.004261 | 23.07 | 66.9 | 133.1 | 10.85 | 11.29 | | High | High | High | High | High | High | High | High | High | High | High | 0.000327 | 0.002512 | 1.97 |
| FALSE | High | HERSPTGEI | [R].HERSPT | 1xCarbami | Q6PEE4 1x | 1.69E-06 | 0 | 1 | 2 | 1 | Q6PEE4 | Zinc finger | Q6PEE4 [6 | 1 | 2371.079 | 0 | -0.23 | 0.164026 | 12.06 | 108.7 | 91.3 | 4.1 | 10.74 | | High | High | High | High | High | High | High | High | High | High | High | 0 | 1.84E-07 | 3.48 |
| FALSE | High | HERPAVPG | [R].HERPA | 2xCarbami | A2AUM0 1 | 0.001767 | 5.66E-05 | 1 | 7 | 1 | A2AUM0 | eIF-2-alpha | A2AUM0 [ | 0 | 3409.607 | 0 | 0.2 | 0.193806 | 21.85 | 90.9 | 109.1 | 5.73 | 12.86 | | High | High | High | High | High | High | High | High | High | High | High | 2.58E-05 | 0.00045 | 3.92 |
| FALSE | High | HCSTYQLSI | [R].HCSTY | 1xCarbami | B7ZWN9 1 | 8.56E-05 | 0 | 2 | 5 | 1 | B7ZWN9; F | SUMO/sen | B7ZWN9 [2 | 0 | 2214.103 | 0 | -0.42 | 0.015681 | 4.66 | 111.9 | 88.1 | 11.87 | 10.76 | Shared | High | High | High | High | High | High | High | High | High | High | High | 0 | 1.51E-05 | 3.18 |
| FALSE | High | HDAGGIHT | [K].HDAGG | 1xPhospho | P35831 1x | 2.89E-10 | 0 | 1 | 1 | 2 | P35831 | Tyrosine-p | P35831 [72 | 0 | 2419.076 | 0 | 0.34 | 0.09649 | 39.03 | 82 | 118 | 12.63 | 20.65 | | High | High | High | High | High | High | High | High | High | High | High | 0 | 1.1E-10 | 5.09 |
| FALSE | High | HDDGSLSP | [K].HDDGS | 1xPhospho | Q2MHN3 1 | 1.61E-05 | 0 | 1 | 1 | 5 | Q2MHN3 | ZFH-5 OS= | Q2MHN3 [ | 0 | 2676.219 | 0 | -0.52 | 0.019496 | 8.09 | 116.7 | 83.3 | 8.74 | 13.43 | | High | High | High | High | High | High | High | High | High | High | High | 0 | 1.55E-05 | 4.01 |
| FALSE | High | HDFPSENSI | [K].HDFPSE | 1xPhospho | Q8CIB9 1x | 3.44E-05 | 0 | 1 | 2 | 2 | Q8CIB9 | N-acetyltra | Q8CIB9 [3 | 0 | 2069.958 | 0 | -1.26 | 0.000699 | 1.88 | 141 | 59 | 2.53 | 17.83 | | High | High | High | High | High | High | High | High | High | High | High | 0 | 5.43E-06 | 3 |
| FALSE | High | HDGGSPSII | [K].HDGGS | 1xPhospho | Q925J9 1x | 0.061176 | 0.004646 | 1 | 2 | 1 | Q925J9 | Mediator c | Q925J9 [1 | 0 | 1435.735 | 0 | 1.56 | 0.02073 | 7.25 | 50.5 | 149.5 | 6.54 | 33.41 | | High | High | High | High | High | High | High | High | High | High | High | 0.00237 | 0.02435 | 1.66 |
| FALSE | High | HDLDASPP | [R].HDLDA | 1xPhospho | Q8R149 1x | 0.004411 | 0.000409 | 1 | 2 | 1 | Q8R149 | BUD13 hor | Q8R149 [2 | 0 | 1316.62 | 0 | -0.49 | 0.008908 | 8.22 | 117.1 | 82.9 | 7.91 | 8.45 | | High | High | High | High | High | High | High | High | High | High | High | 0.000183 | 0.001263 | 3.26 |
| FALSE | High | HDSEQDNS | [R].HDSEQ | 1xPhospho | Q3USY4 1x | 0.057203 | 0.004227 | 1 | 3 | 2 | Q3USY4 | Putative ur | Q3USY4 [2 | 0 | 4123.917 | 0 | -0.88 | 0.002273 | 0.51 | 128.6 | 71.4 | 8.66 | 8.23 | | High | High | High | High | High | High | High | High | High | High | High | 0.002159 | 0.02246 | 3.36 |
| FALSE | High | HDSLLHGT | [K].HDSLLH | 1xPhospho | P25206 1x | 0.088701 | 0.006669 | 2 | 3 | 1 | P25206; Q | DNA replic | P25206 [54 | 0 | 1545.819 | 0 | -0.58 | 0.003195 | 7.72 | 118.9 | 81.1 | 7.6 | 16.7 | Shared | High | High | High | High | High | High | High | High | High | High | High | 0.003432 | 0.03738 | 1.2 |
| FALSE | High | HDSLPDPG | [R].HDSLPD | 1xPhospho | Q3UHP6 1x | 9.45E-05 | 0 | 1 | 5 | 1 | Q3UHP6 | Putative ur | Q3UHP6 [9 | 0 | 1561.721 | 0 | 0.13 | 0.142554 | 10.18 | 95.7 | 104.3 | 10.25 | 10.19 | | High | High | High | High | High | High | High | High | High | High | High | 0 | 1.68E-05 | 2.73 |
| FALSE | High | HDSPEDVK | [R].HDSPEI | 1xPhospho | Q3THQ5 1x | 0.001021 | 0 | 1 | 2 | 5 | Q3THQ5 | Putative ur | Q3THQ5 [4 | 0 | 1464.714 | 0 | -0.28 | 0.028841 | 3.54 | 110.8 | 89.2 | 3.99 | 9.44 | | High | High | High | High | High | High | High | High | High | High | High | 0.000555 | 0.004779 | 2.5 |
| FALSE | High | HDSSPPNA | [R].HDSSPF | 1xPhospho | Q8C9B9 1x | 2.8E-05 | 0 | 1 | 3 | 5 | Q8C9B9 | Death-indu | Q8C9B9 [8 | 0 | 4498 | 0 | -0.26 | 0.27402 | 7.25 | 112.4 | 87.6 | 19.6 | 53.3 | | High | High | High | High | High | High | High | High | High | High | High | 0 | 4.29E-06 | 3.88 |
| FALSE | High | HDSVDSGV | [R].HDSVD | 1xPhospho | Q6NZP2 1x | 2.87E-05 | 0 | 1 | 1 | 2 | Q6NZP2 | Vasculin-lik | Q6NZP2 [7 | 0 | 1568.772 | 0 | 0.61 | 0.001021 | 3.32 | 79.3 | 120.7 | 6.09 | 4.21 | | High | High | High | High | High | High | High | High | High | High | High | 0 | 4.41E-06 | 3.58 |
| FALSE | High | HDSYLAVEI | [K].HDSYLA | 1xPhospho | E9Q3P4 1x | 5.5E-05 | 0 | 1 | 3 | 3 | E9Q3P4 | Centromer | E9Q3P4 [1 | 0 | 2225.126 | 0 | -0.05 | 0.334301 | 9.26 | 101.6 | 98.4 | 4.6 | 14.19 | | High | High | High | High | High | High | High | High | High | High | High | 0 | 9.17E-06 | 3.65 |
| FALSE | High | HERSPAGE | [R].HERSPA | 1xCarbami | E9Q6S4 1x | 0.000611 | 0 | 1 | 1 | 2 | E9Q6S4 | Uncharacte | E9Q6S4 [6 | 1 | 2710.208 | 0 | -1.46 | 0.0044 | 2.5 | 150.2 | 49.8 | 14.77 | 34.03 | | High | High | High | High | High | High | High | High | High | High | High | 0 | 0.000138 | 3.18 |
| FALSE | High | HDTPDLSPI | [R].HDTPDI | 1xPhospho | Q8R149 1x | 0.000176 | 0 | 1 | 2 | 1 | Q8R149 | BUD13 hor | Q8R149 [1 | 0 | 1443.683 | 0 | -0.92 | 0.014049 | 5.94 | 133.4 | 66.6 | 19.94 | 8.69 | | High | High | High | High | High | High | High | High | High | High | High | 0 | 3.4E-05 | 3.52 |
| FALSE | High | HDTPDPSP | [R].HDTPDI | 1xPhospho | Q8R149 1x | 0.002921 | 0.00016 | 1 | 2 | 1 | Q8R149 | BUD13 hor | Q8R149 [1 | 0 | 1427.652 | 0 | -1.34 | 0.000658 | 4.1 | 143.4 | 56.6 | 3.81 | 15.64 | | High | High | High | High | High | High | High | High | High | High | High | 7.19E-05 | 0.000795 | 3.09 |
| FALSE | High | HDTPDTSP | [R].HDTPD | 1xPhospho | Q8R149 1x | 0.022182 | 0.001542 | 1 | 2 | 2 | Q8R149 | BUD13 hor | Q8R149 [1 | 0 | 1431.647 | 0 | -0.56 | 0.011749 | 6.81 | 125.1 | 74.9 | 12.79 | 8.35 | | High | High | High | High | High | High | High | High | High | High | High | 0.004072 | 0.04348 | 3.36 |
| FALSE | High | HDTSASTQ | [K].HDTSA | 1xPhospho | Q62261 1x | 2.74E-07 | 0 | 1 | 3 | 5 | Q62261 | Spectrin be | Q62261 [2 | 0 | 1841.823 | 0 | 0.69 | 0.03698 | 21.66 | 75.4 | 124.6 | 22.15 | 28.36 | | High | High | High | High | High | High | High | High | High | High | High | 0 | 2.37E-08 | 3.68 |
| FALSE | High | HEAEDCKF | [R].HEAED | 1xCarbami | Q64127 2x | 0.005615 | 0.00062 | 1 | 5 | 2 | Q64127 | Transcripti | Q64127 [1 | 1 | 2801.064 | 0 | -0.88 | 0.030391 | 15.97 | 131.6 | 68.4 | 22.14 | 18.34 | | High | High | High | High | High | High | High | High | High | High | High | 0.000275 | 0.001646 | 3.97 |
| FALSE | High | HEAEDCKF | [R].HEAED | 1xCarbami | Q64127 2x | 0.000173 | 0 | 1 | 5 | 7 | Q64127 | Transcripti | Q64127 [1 | 1 | 3030.227 | 0 | -1.03 | 0.000671 | 2.53 | 133.6 | 66.4 | 3.69 | 7.64 | | High | High | High | High | High | High | High | High | High | High | High | 0 | 3.31E-05 | 5.49 |
| FALSE | High | HEAPSSPLA | [K].HEAPSS | 1xCarbami | Q5RIM6 1x | 1.15E-06 | 0 | 1 | 8 | 5 | Q5RIM6 | Nuclear re | Q5RIM6 [1 | 0 | 2888.329 | 0 | 0.1 | 0.883214 | 9.36 | 99.4 | 100.6 | 5.31 | 18.36 | | High | High | High | High | High | High | High | High | High | High | High | 0 | 1.51E-07 | 4.21 |
| FALSE | High | HEAPSSPLA | [K].HEAPSS | 1xCarbami | Q5RIM6 2x | 1.23E-09 | 0 | 1 | 8 | 3 | Q5RIM6 | Nuclear re | Q5RIM6 [1 | 0 | 2968.295 | 0 | -0.33 | 0.057144 | 6.21 | 108.8 | 91.2 | 5.91 | 17.58 | | High | High | High | High | High | High | High | High | High | High | High | 0 | 5.48E-11 | 5.44 |
| FALSE | High | HEDGTQSD | [K].HEDGT | 1xPhospho | Q80U49 1x | 1.9E-06 | 0 | 1 | 2 | 3 | Q80U49 | Centrosom | Q80U49 [3 | 0 | 2166.996 | 0 | 0.83 | 0.022669 | 5.58 | 71.1 | 128.9 | 4.2 | 20.09 | | High | High | High | High | High | High | High | High | High | High | High | 0 | 2.09E-07 | 3.03 |
| FALSE | High | HEDWSAD | [K].HEDWS | 1xPhospho | A2ADB0 1x | 7.02E-05 | 0 | 1 | 3 | 1 | A2ADB0 | Msx2-inter | A2ADB0 [6 | 0 | 2045.948 | 0 | 0.35 | 0.3263 | 12.51 | 87.5 | 112.5 | 5.48 | 22.81 | | High | High | High | High | High | High | High | High | High | High | High | 0 | 1.21E-05 | 2.28 |
| FALSE | High | HEGGPGQ | [K].HEGGP | 1xPhospho | A0A0R4J0T | 3.01E-09 | 0 | 1 | 2 | 1 | A0A0R4J0T | ADP-ribosy | A0A0R4J0T | 0 | 2356.17 | 0 | 0.73 | 0.007633 | 6.28 | 74 | 126 | 3.89 | 14.78 | | High | High | High | High | High | High | High | High | High | High | High | 0 | 1.49E-10 | 4.33 |
| FALSE | High | HEIANSDG | [K].HEIANS | 1xPhospho | Q9R207 1x | 0.000198 | 0 | 1 | 1 | 2 | Q9R207 | Nibrin OS= | Q9R207 [6 | 0 | 2306.05 | 0 | 0.19 | 0.027917 | 2.21 | 93.6 | 106.4 | 8.01 | 12.86 | | High | High | High | High | High | High | High | High | High | High | High | 0 | 3.89E-05 | 3 |
| FALSE | High | HELSPPQK | [R].HELSPP | 1xPhospho | Q99MR6 1 | 0.0095 | 0.000736 | 1 | 5 | 2 | Q99MR6 | Serrate RN | Q99MR6 [ | 0 | 1473.787 | 0 | -0.21 | 0.048856 | 6.21 | 107.6 | 92.4 | 2.95 | 12.1 | | High | High | High | High | High | High | High | High | High | High | High | 0.003243 | 0.03423 | 1.69 |
| FALSE | High | HENTVQTF | [K].HENTV | 1xPhospho | E9PVX6 1x | 7.54E-05 | 0 | 1 | 2 | 2 | E9PVX6 | Proliferatic | E9PVX6 [7 | 0 | 1591.825 | 0 | 0.19 | 0.509753 | 2 | 93.8 | 106.2 | 4.16 | 16.45 | | High | High | High | High | High | High | High | High | High | High | High | 0 | 6.62E-05 | 3.66 |
| FALSE | High | HDTPDLSPI | [R].HDTPDI | 2xPhospho | Q8R149 2x | 0.000186 | 0 | 1 | 2 | 3 | Q8R149 | BUD13 hor | Q8R149 [1 | 0 | 1523.649 | 0 | 0.42 | 0.034255 | 6 | 86.1 | 113.9 | 9.81 | 7.6 | | High | High | High | High | High | High | High | High | High | High | High | 0 | 3.59E-05 | 2.79 |
| FALSE | High | HGSGPNIIL | [R].HGSGPI | 1xPhospho | Q3U182 1x | 5.02E-08 | 0 | 2 | 4 | 4 | Q3U182; C | CREB-regu | Q3U182 [6 | 0 | 2409.222 | 0 | 0.65 | 0.008965 | 11.83 | 77.1 | 122.9 | 4.33 | 14.28 | Shared | High | High | High | High | High | High | High | High | High | High | High | 0 | 4.37E-09 | 4.71 |
| FALSE | High | HGSGPNIIL | [R].HGSGPI | 2xPhospho | Q3U182 2x | 1.44E-06 | 0 | 2 | 4 | 1 | Q3U182; C | CREB-regu | Q3U182 [6 | 0 | 2489.188 | 0 | 0.96 | 0.012892 | 7.48 | 67.9 | 132.1 | 10.54 | 15.06 | Shared | High | High | High | High | High | High | High | High | High | High | High | 0 | 1.53E-07 | 4.53 |
| FALSE | High | HGSLSADD | [R].HGSLSA | 1xPhospho | Q5HZI2 1x | 0.01588 | 0.001124 | 1 | 1 | 2 | Q5HZI2 | C2 calcium | Q5HZI2 [2 | 0 | 2107.949 | 0 | -0.63 | 0.006424 | 8.31 | 122 | 78 | 6.11 | 19.7 | | High | High | High | High | High | High | High | High | High | High | High | 0.000585 | 0.005258 | 2.34 |
| FALSE | High | HLGGSGSV | [R].HLGGS | 1xPhospho | E9Q0S6 1x | 0.000908 | 0 | 1 | 10 | 3 | E9Q0S6 | Tensin 1 O | E9Q0S6 [1 | 0 | 1930.959 | 0 | -0.27 | 0.011643 | 8.15 | 111.5 | 88.5 | 9.94 | 8.26 | | High | High | High | High | High | High | High | High | High | High | High | 0 | 0.000214 | 3.82 |
| FALSE | High | HLGNAATS | [K].HLGNA | 1xPhospho | Q9JJ48 1xP | 1.54E-05 | 0 | 1 | 3 | 2 | Q9JJ48 | Zinc finger | Q9JJ48 [60 | 0 | 1533.819 | 0 | -1.04 | 0.002756 | 1.68 | 134.7 | 65.3 | 5.29 | 22.59 | | High | High | High | High | High | High | High | High | High | High | High | 0 | 2.2E-06 | 3.15 |
| FALSE | High | HLGNAATS | [K].HLGNA | 1xDeamida | Q9JJ48 1xP | 4.16E-05 | 0 | 1 | 3 | 2 | Q9JJ48 | Zinc finger | Q9JJ48 [60 | 0 | 1534.803 | 0 | -0.64 | 0.001311 | 2.33 | 122.3 | 77.7 | 2.97 | 10.81 | | High | High | High | High | High | High | High | High | High | High | High | 0 | 6.71E-06 | 2.96 |
| FALSE | High | HIHSNLDFS | [K].HIHSNL | 1xPhospho | O35280 1x | 1.45E-05 | 0 | 1 | 4 | 1 | O35280 | Serine/thre | O35280 [2 | 0 | 2849.387 | 0 | -0.9 | 0.000992 | 1.89 | 127 | 73 | 10.35 | 19.06 | | High | High | High | High | High | High | High | High | High | High | High | 0 | 2.04E-06 | 4.69 |
| FALSE | High | HIHSNLDFS | [K].HIHSNL | 1xDeamida | O35280 1x | 5.81E-05 | 0 | 1 | 4 | 1 | O35280 | Serine/thre | O35280 [2 | 0 | 2850.371 | 0 | -0.78 | 0.01523 | 8.33 | 124.1 | 75.9 | 11.01 | 18.1 | | High | High | High | High | High | High | High | High | High | High | High | 0 | 9.8E-06 | 3.2 |
| FALSE | High | HLLANSPK | [R].HLLANS | 1xPhospho | Q5HZJ0 1x | 0.020142 | 0.001472 | 1 | 8 | 1 | Q5HZJ0 | Ribonuclea | Q5HZJ0 [8 | 0 | 1417.797 | 0 | -0.68 | 0.011451 | 6.4 | 121.3 | 78.7 | 6.61 | 16.04 | | High | High | High | High | High | High | High | High | High | High | High | 0.000786 | 0.006878 | 1.67 |
| FALSE | High | HLLETHSSF | [K].HLLETH | 1xPhospho | Q80WT5 1 | 0.001068 | 0 | 1 | 6 | 1 | Q80WT5 | Aftiphilin C | Q80WT5 [ | 0 | 1757.935 | 0 | -0.18 | 0.073509 | 3.41 | 113.9 | 86.1 | 8.64 | 17.28 | | High | High | High | High | High | High | High | High | High | High | High | 0 | 0.000257 | 2.79 |
| FALSE | High | HLLNSPTDI | [R].HLLNSF | 1xPhospho | Q80TU3 1x | 1.56E-05 | 0 | 1 | 4 | 2 | Q80TU3 | MKIAA068 | Q80TU3 [1 | 0 | 1719.842 | 0 | -0.93 | 0.000158 | 2.09 | 131.2 | 68.8 | 3.04 | 7.84 | | High | High | High | High | High | High | High | High | High | High | High | 0.000638 | 0.005716 | 2.5 |
| FALSE | High | HLLPISTSDI | [R].HLLPIST | 1xPhospho | Q8CD26 1x | 0.008388 | 0.000736 | 1 | 2 | 1 | Q8CD26 | Solute carr | Q8CD26 [3 | 0 | 1734.899 | 0 | 0.42 | 0.018742 | 14.02 | 82.6 | 117.4 | 11.2 | 19.51 | | High | High | High | High | High | High | High | High | High | High | High | 0.000327 | 0.002576 | 1.94 |
| FALSE | High | HLLSPLSIQI | [R].HLLSPL | 1xCarbami | Q9D3R3 1x | 7.08E-05 | 0 | 1 | 1 | 1 | Q9D3R3 | Centrosom | Q9D3R3 [2 | 0 | 2228.084 | 0 | -1.24 | 0.003543 | 4.98 | 146.7 | 53.3 | 16.07 | 10.54 | | High | High | High | High | High | High | High | High | High | High | High | 0 | 1.21E-05 | 2.52 |
| FALSE | High | HLLTDLPLP | [R].HLLTDL | 1xPhospho | Q14AX6-2 | 3.81E-06 | 0 | 3 | 4 | 2 | Q14AX6-2; | Isoform 2 c | Q14AX6-2 | 0 | 3253.68 | 0 | -0.38 | 0.042531 | 8.33 | 109.3 | 90.7 | 6.73 | 10.73 | Shared | High | High | High | High | High | High | High | High | High | High | High | 0 | 4.55E-07 | 4.21 |
| FALSE | High | HLMDVGSI | [R].HLMDV | 1xPhospho | Q62431 1x | 5.19E-07 | 0 | 1 | 2 | 3 | Q62431 | AT-rich int | Q62431 [1 | 0 | 2001.888 | 0 | -1.1 | 0.000263 | 2.14 | 137.2 | 62.8 | 4.73 | 8.22 | | High | High | High | High | High | High | High | High | High | High | High | 0 | 5.3E-08 | 3.88 |
| FALSE | High | HLPGPSNP | [K].HLPGPS | 1xPhospho | D3YTR4 1x | 0.005418 | 0.00062 | 1 | 1 | 1 | D3YTR4 | Methyl-Cp | D3YTR4 [6 | 0 | 2063.043 | 0 | -1.87 | 0.000158 | 0.5 | 156.5 | 43.5 | 4.16 | 7.44 | | High | High | High | High | High | High | High | High | High | High | High | 0.000275 | 0.001575 | 2.03 |
| FALSE | High | HLFSSTENL | [K].HLFSST | 1xPhospho | E9Q8L9 1x | 6.26E-06 | 0 | 1 | 3 | 4 | E9Q8L9 | Rab11 fam | E9Q8L9 [3 | 0 | 1654.815 | 0 | -0.28 | 0.065992 | 4.32 | 110.4 | 89.6 | 6.94 | 16.88 | | High | High | High | High | High | High | High | High | High | High | High | 0 | 7.98E-07 | 4.11 |
| FALSE | High | HLPGSGQC | [R].HLPGS | 1xPhospho | Q8CGB6-3 | 0.011652 | 0.000889 | 1 | 4 | 1 | Q8CGB6-3 | Isoform 3 c | Q8CGB6-3 | 0 | 1737.864 | 0 | 0.31 | 0.028136 | 11.55 | 85.4 | 114.6 | 15.07 | 7.95 | | High | High | High | High | High | High | High | High | High | High | High | 0.000449 | 0.003713 | 2.69 |
| FALSE | High | HLPSPPLTC | [R].HLPSPP | 1xPhospho | Q62245 1x | 0.000281 | 0 | 1 | 2 | 1 | Q62245 | Son of seve | Q62245 [1 | 0 | 2911.474 | 0 | -0.35 | 0.39588 | 1.24 | 107 | 93 | 10.07 | 22.2 | | High | High | High | High | High | High | High | High | High | High | High | 0 | 5.73E-05 | 2.77 |
| FALSE | High | HLPSPPTLD | [R].HLPSPP | 1xPhospho | Q80TR8-2 | 1.16E-05 | 0 | 1 | 3 | 2 | Q80TR8-2 | Isoform 2 c | Q80TR8-2 | 0 | 2261.178 | 0 | -0.57 | 0.010701 | 2.35 | 121.2 | 78.8 | 6.94 | 10.9 | | High | High | High | High | High | High | High | High | High | High | High | 0 | 1.58E-06 | 4.27 |
| FALSE | High | HLQNTSPL | [R].HLQNT | 1xPhospho | Q5DTT3 1x | 4.56E-06 | 0 | 1 | 1 | 2 | Q5DTT3 | Protein FA | Q5DTT3 [6 | 0 | 1703.925 | 0 | -0.96 | 0.000781 | 3.83 | 131.6 | 68.4 | 3.17 | 10 | | High | High | High | High | High | High | High | High | High | High | High | 0 | 5.6E-07 | 2.31 |
| FALSE | High | HLQPTPEP | [K].HLQPTI | 1xPhospho | A6H8H2-2 | 0.019276 | 0.001402 | 1 | 3 | 2 | A6H8H2-2 | Isoform 2 c | A6H8H2-2 | 0 | 3021.549 | 0 | 1.12 | 0.021607 | 3.9 | 64 | 136 | 11.82 | 19.11 | | High | High | High | High | High | High | High | High | High | High | High | 0.000756 | 0.00654 | 1.82 |
| FALSE | High | HLQPTPEP | [K].HLQPTI | 2xPhospho | A6H8H2-2 | 0.001107 | 0 | 1 | 3 | 1 | A6H8H2-2 | Isoform 2 c | A6H8H2-2 | 0 | 3101.515 | 0 | 1.58 | 0.018486 | 24.34 | 49.5 | 150.5 | 12.27 | 31.37 | | High | High | High | High | High | High | High | High | High | High | High | 0 | 0.000266 | 2.92 |
| FALSE | High | HLQQGSES | [R].HLQQG | 1xPhospho | E9PV45 1x | 8.06E-06 | 0 | 1 | 2 | 2 | E9PV45 | Ubiquitin c | E9PV45 [2 | 0 | 2153.992 | 0 | 0.16 | 0.060388 | 7.6 | 96.7 | 103.3 | 8.32 | 6.82 | | High | High | High | High | High | High | High | High | High | High | High | 0 | 5.02E-05 | 3.94 |
| FALSE | High | HLQVVLLD | [R].HLQVV | 1xCarbami | Q921G6 1x | 0.033404 | 0.002195 | 1 | 5 | 1 | Q921G6 | Leucine-ric | Q921G6 [2 | 0 | 3008.641 | 0 | -0.49 | 0.082148 | 8.6 | 117.4 | 82.6 | 10.33 | 27.92 | | High | High | High | High | High | High | High | High | High | High | High | 0.001151 | 0.01212 | 2.73 |

| | | | | | | | | | | | | | | | | | | | | | | | | | | | | | | | | | | | | | | |
|---|---|---|---|---|---|---|---|---|---|---|---|---|---|---|---|---|---|---|---|---|---|---|---|---|---|---|---|---|---|---|---|---|---|---|---|---|---|---|
| FALSE | High | HLSAHKPSI | [R].HLSAHK | 1xPhospho | P49452 1xP | 0.086562 | 0.006435 | 1 | 3 | 1 | P49452 | Centromer | P49452 [29 | 0 | 2695.49 | 0 | -0.53 | 0.027013 | 14.87 | 116.3 | 83.7 | 8.23 | 15.14 | | High | High | High | High | High | High | High | High | High | High | High | 0.003317 | 0.0365 | 1.96 |
| FALSE | High | HLSASTPK | [R].HLSAST | 1xPhospho | Q924A2 1x | 0.009087 | 0.000736 | 1 | 1 | 2 | Q924A2 | Protein cap | Q924A2 [6 | 0 | 1378.75 | 0 | 0.27 | 0.004838 | 5.81 | 90.5 | 109.5 | 3.45 | 2.86 | | High | High | High | High | High | High | High | High | High | High | High | 0.000327 | 0.002805 | 2.74 |
| FALSE | High | HLSEDSQA | [R].HLSEDS | 1xPhospho | E9Q6J5-3 1 | 0.026652 | 0.001676 | 1 | 4 | 1 | E9Q6J5-3 | Isoform Ne | E9Q6J5-3 [ | 0 | 2130.088 | 0 | -0.07 | 0.731877 | 24.49 | 99.7 | 100.3 | 17.89 | 15.57 | | High | High | High | High | High | High | High | High | High | High | High | 0.000903 | 0.009383 | 2.05 |
| FALSE | High | HLSGSSSPF | [R].HLSGSS | 1xPhospho | Q3UIL6-2 1 | 1.83E-05 | 0 | 1 | 8 | 2 | Q3UIL6-2 | Isoform 2 c | Q3UIL6-2 [ | 0 | 1430.699 | 0 | -0.95 | 0.002357 | 5.27 | 133.2 | 66.8 | 9.16 | 11.27 | | High | High | High | High | High | High | High | High | High | High | High | 0 | 2.66E-06 | 3.87 |
| FALSE | High | HISHQTSVI | [R].HISHQT | 1xPhospho | Q6ZPR5-4 1 | 0.000656 | 0 | 1 | 7 | 2 | Q6ZPR5-4 | Isoform 4 c | Q6ZPR5-4 [ | 0 | 2494.183 | 0 | 1.13 | 0.005418 | 19.19 | 66.8 | 133.2 | 19.46 | 10.79 | | High | High | High | High | High | High | High | High | High | High | High | 0 | 0.000149 | 1.85 |
| FALSE | High | HLSMQSFD | [R].HLSMQ | 1xPhospho | E9Q4S3 1x | 0.01861 | 0.001362 | 1 | 7 | 1 | E9Q4S3 | Ninein OS= | E9Q4S3 [2 | 0 | 1702.746 | 0 | 1.25 | 0.000192 | 4.56 | 56.6 | 143.4 | 9.44 | 5.43 | | High | High | High | High | High | High | High | High | High | High | High | 0.000723 | 0.00629 | 2.03 |
| FALSE | High | HLPLSFLSS | [R].HLPLSF | 1xPhospho | Q3U1N2 1x | 0.026421 | 0.001676 | 1 | 2 | 2 | Q3U1N2 | Sterol regu | Q3U1N2 [1 | 0 | 1747.909 | 0 | 0.03 | 0.558712 | 14.38 | 99.4 | 100.6 | 13.8 | 15.76 | | High | High | High | High | High | High | High | High | High | High | High | 0.000903 | 0.009366 | 2.51 |
| FALSE | High | HLESSPVSC | [R].HLESSP | 1xPhospho | A0A1B0GR | 0.030104 | 0.001857 | 1 | 1 | 1 | A0A1B0GR | RIKEN cDN | A0A1B0GR | 0 | 1448.71 | 0 | 0.92 | 0.000313 | 3.22 | 68.5 | 131.5 | 8.28 | 8.01 | | High | High | High | High | High | High | High | High | High | High | High | 0.001011 | 0.01081 | 2.4 |
| FALSE | High | HIEPELAGR | [R].HIEPEL | 1xPhospho | Q8C1Y8 1x | 0.000712 | 0 | 1 | 1 | 2 | Q8C1Y8 | Vacuolar fu | Q8C1Y8 [2 | 1 | 1884.953 | 0 | 0.36 | 0.001788 | 4.23 | 86.3 | 113.7 | 6.46 | 6.59 | | High | High | High | High | High | High | High | High | High | High | High | 0 | 0.000163 | 1.98 |
| FALSE | High | HLELSPKPK | [K].HLELSP | 1xPhospho | Q8BGX7 1x | 0.031439 | 0.002011 | 1 | 2 | 1 | Q8BGX7 | DNA repair | Q8BGX7 [7 | 0 | 1816.07 | 0 | -0.32 | 0.100547 | 8.64 | 113.1 | 86.9 | 10.63 | 21.27 | | High | High | High | High | High | High | High | High | High | High | High | 0.001076 | 0.01141 | 2.25 |
| FALSE | High | HGSLSSSR | [R].HGSLSS | 1xPhospho | F2Z3U3 1xP | 0.033116 | 0.002195 | 1 | 1 | 1 | F2Z3U3 | Ras associa | F2Z3U3 [1 | 0 | 1139.541 | 0 | 1.21 | 0.003284 | 17.83 | 60.7 | 139.3 | 8.94 | 16 | | High | High | High | High | High | High | High | High | High | High | High | 0.001151 | 0.01203 | 1.02 |
| FALSE | High | HGSPEFCG | [K].HGSPEF | 1xCarbami | Q6PD21 1x | 0.000793 | 0 | 1 | 2 | 2 | Q6PD21 | SH2 domai | Q6PD21 [3 | 0 | 1767.809 | 0 | 0.45 | 0.00863 | 10.12 | 83 | 117 | 10.06 | 7.66 | | High | High | High | High | High | High | High | High | High | High | High | 0 | 0.000184 | 2.61 |
| FALSE | High | HGSPLPAG | [R].HGSPLP | 1xCarbami | Q91XC0 1x | 0.001011 | 0 | 1 | 1 | 1 | Q91XC0 | LIM domai | Q91XC0 [1 | 0 | 4306.136 | 0 | -1.13 | 0.002021 | 3.75 | 138.3 | 61.7 | 14.38 | 18.32 | | High | High | High | High | High | High | High | High | High | High | High | 0 | 0.000242 | 4.18 |
| FALSE | High | HGSPTAPIC | [R].HGSPTA | 1xPhospho | Q3UIL6-2 1 | 1.37E-08 | 0 | 1 | 8 | 1 | Q3UIL6-2 | Isoform 2 c | Q3UIL6-2 [ | 0 | 2305.081 | 0 | 0.16 | 0.012749 | 7.44 | 93.5 | 106.5 | 4.79 | 5.29 | | High | High | High | High | High | High | High | High | High | High | High | 0 | 8.17E-10 | 3.95 |
| FALSE | High | HGSSDDYL | [R].HGSSDI | 1xPhospho | Q6KCD5 1x | 0.001267 | 0 | 1 | 4 | 1 | Q6KCD5 | Nipped-B-l | Q6KCD5 [2 | 0 | 1878.816 | 0 | 0.06 | 0.677847 | 20.73 | 96 | 104 | 13.49 | 14.04 | | High | High | High | High | High | High | High | High | High | High | High | 0 | 0.000311 | 2.52 |
| FALSE | High | HGSSEDEE | [R].HGSSED | 1xPhospho | P70398 1xP | 9.39E-06 | 0 | 1 | 4 | 3 | P70398 | Probable u | P70398 [37 | 0 | 1954.838 | 0 | -0.62 | 0.002921 | 1.83 | 120.6 | 79.4 | 9.68 | 8.01 | | High | High | High | High | High | High | High | High | High | High | High | 0 | 1.35E-06 | 3.77 |
| FALSE | High | HGSYENAV | [K].HGSYEN | 1xPhospho | P11983 1xP | 1.25E-05 | 0 | 1 | 2 | 5 | P11983 | T-complex | P11983 [54 | 0 | 1880.801 | 0 | -0.25 | 0.060565 | 4.46 | 109.7 | 90.3 | 4.13 | 16.08 | | High | High | High | High | High | High | High | High | High | High | High | 0 | 2.98E-05 | 3.11 |
| FALSE | High | HGTDLWLD | [R].HGTDLV | 1xCarbami | A0A0R4J0T | 0.000105 | 0 | 1 | 2 | 1 | A0A0R4J0T | ADP-ribosy | A0A0R4J0T | 0 | 2683.336 | 0 | 0.95 | 0.004854 | 7.2 | 70.6 | 129.4 | 5.72 | 15.48 | | High | High | High | High | High | High | High | High | High | High | High | 0 | 1.91E-05 | 3.13 |
| FALSE | High | HGTDLWID | [R].HGTDLV | 1xPhospho | Q99K28-2 1 | 0.001244 | 0 | 1 | 2 | 2 | Q99K28-2 | Isoform 2 c | Q99K28-2 | 0 | 2747.29 | 0 | -0.03 | 0.471011 | 12.21 | 100.4 | 99.6 | 5.97 | 6.64 | | High | High | High | High | High | High | High | High | High | High | High | 0 | 0.000305 | 2.64 |
| FALSE | High | HGTDLWID | [R].HGTDLV | 1xDeamida | Q99K28-2 1 | 0.00582 | 0.00062 | 1 | 2 | 2 | Q99K28-2 | Isoform 2 c | Q99K28-2 | 0 | 2748.274 | 0 | -0.02 | 0.973452 | 6.33 | 99.7 | 100.3 | 22.8 | 23.69 | | High | High | High | High | High | High | High | High | High | High | High | 0.000275 | 0.001715 | 3.03 |
| FALSE | High | HGTDLWID | [R].HGTDLV | 1xPhospho | Q99K28-2 1 | 6.07E-05 | 0 | 1 | 2 | 3 | Q99K28-2 | Isoform 2 c | Q99K28-2 | 0 | 2763.285 | 0 | 0 | 0.857387 | 5 | 101.6 | 98.4 | 17.69 | 24.76 | | High | High | High | High | High | High | High | High | High | High | High | 0 | 2.47E-05 | 3.2 |
| FALSE | High | HHGLNVLA | [R].HHGLN | 1xCarbami | Q9D8T2 1x | 0.018774 | 0.001362 | 1 | 1 | 1 | Q9D8T2 | Gasdermin | Q9D8T2 [2 | 0 | 2128.15 | 0 | 0.17 | 0.187947 | 27.41 | 92.2 | 107.8 | 14.63 | 23.97 | | High | High | High | High | High | High | High | High | High | High | High | 0.000723 | 0.006349 | 1.58 |
| FALSE | High | HHGTPTSA | [R].HHGTP | 1xPhospho | Q91Z96 1x | 0.040373 | 0.002982 | 1 | 1 | 2 | Q91Z96 | BMP-2-ind | Q91Z96 [9 | 0 | 1473.762 | 0 | -0.84 | 0.008895 | 6.79 | 130.5 | 69.5 | 14.36 | 11.25 | | High | High | High | High | High | High | High | High | High | High | High | 0.00155 | 0.01506 | 2.34 |
| FALSE | High | HHNHGSPH | [R].HHNHG | 1xPhospho | Q52KE7 1x | 0.000668 | 0 | 1 | 1 | 8 | Q52KE7 | Cyclin-L1 C | Q52KE7 [4 | 0 | 1701.874 | 0 | -0.28 | 0.052076 | 5.7 | 110 | 90 | 8.04 | 10.14 | | High | High | High | High | High | High | High | High | High | High | High | 0.000327 | 0.002557 | 2.62 |
| FALSE | High | HHNLSLNG | [R].HHNLSI | 1xPhospho | O70374 1x | 5.26E-05 | 0 | 1 | 4 | 1 | O70374 | Protein CB | O70374 [3 | 0 | 2413.125 | 0 | 0.11 | 0.154658 | 8.21 | 91.9 | 108.1 | 12.79 | 7.43 | | High | High | High | High | High | High | High | High | High | High | High | 0 | 8.75E-06 | 4.1 |
| FALSE | High | HHSDTNFG | [R].HHSDTI | 1xPhospho | E9Q0V6 1x | 0.002212 | 0.000109 | 1 | 2 | 1 | E9Q0V6 | NHS-like pr | E9Q0V6 [7 | 0 | 1679.831 | 0 | 0.35 | 0.000756 | 5.01 | 87.9 | 112.1 | 3.14 | 4.69 | | High | High | High | High | High | High | High | High | High | High | High | 4.94E-05 | 0.000583 | 2.55 |
| FALSE | High | HHSVEAAP | [R].HHSVEA | 1xPhospho | Q8C8T7 1x | 0.100129 | 0.008067 | 1 | 2 | 1 | Q8C8T7 | Protein ELF | Q8C8T7 [6 | 0 | 1563.763 | 0 | 0.29 | 0.289827 | 21.28 | 84.8 | 115.2 | 13.19 | 26.49 | | High | High | High | High | High | High | High | High | High | High | High | 0.004046 | 0.04322 | 1.76 |
| FALSE | High | HLACSPER | [R].HLACSF | 1xCarbami | Q9CZ62 1x | 0.003822 | 0.000261 | 1 | 2 | 2 | Q9CZ62 | Centrosom | Q9CZ62 [7 | 0 | 1278.586 | 0 | 0 | 0.545001 | 12.79 | 100.4 | 99.6 | 6.33 | 10.97 | | High | High | High | High | High | High | High | High | High | High | High | 0.001812 | 0.01846 | 3.18 |
| FALSE | High | HLAESESLL | [K].HLAESE | 1xPhospho | G5E870 1x | 1.84E-06 | 0 | 1 | 3 | 5 | G5E870 | E3 ubiquiti | G5E870 [9 | 0 | 2047.088 | 0 | -1.3 | 0.000154 | 0.82 | 142.8 | 57.2 | 3.98 | 11.08 | | High | High | High | High | High | High | High | High | High | High | High | 0 | 2.01E-07 | 4.22 |
| FALSE | High | HIAFTPESC | [K].HIAFTP | 1xPhospho | G5E8R4 1x | 0.000553 | 0 | 1 | 3 | 2 | G5E8R4 | SAPS doma | G5E8R4 [6 | 0 | 1494.73 | 0 | 0.73 | 0.022417 | 3.03 | 72.1 | 127.9 | 4.46 | 20.46 | | High | High | High | High | High | High | High | High | High | High | High | 0 | 0.000123 | 2.6 |
| FALSE | High | HLALESSVF | [K].HLALES | 1xCarbami | Q5DTT3 1x | 1.65E-06 | 0 | 1 | 2 | 3 | Q5DTT3 | Protein FA | Q5DTT3 [1 | 0 | 1848.888 | 0 | -0.74 | 0.009604 | 8.9 | 123.5 | 76.5 | 6.34 | 18.2 | | High | High | High | High | High | High | High | High | High | High | High | 0 | 2.5E-07 | 3.2 |
| FALSE | High | HLASSGGW | [R].HLASSG | 1xPhospho | Q3UJV1 1x | 0.007019 | 0.00062 | 1 | 2 | 2 | Q3UJV1 | Coiled-coil | Q3UJV1 [4 | 0 | 1789.977 | 0 | 0.15 | 0.10275 | 15.18 | 90.9 | 109.1 | 5.12 | 15.62 | | High | High | High | High | High | High | High | High | High | High | High | 0.000275 | 0.002119 | 2.17 |
| FALSE | High | HLATEPTAI | [R].HLATEF | 1xPhospho | H3BLC5 1x | 0.067619 | 0.004864 | 1 | 2 | 1 | H3BLC5 | Histone de | H3BLC5 [6 | 1 | 2154.127 | 0 | -1.32 | 0.000992 | 1.73 | 141 | 59 | 12.47 | 8.07 | | High | High | High | High | High | High | High | High | High | High | High | 0.00253 | 0.0274 | 2.3 |
| FALSE | High | HLDRSPESE | [R].HLDRSF | 1xPhospho | Q0VBL3 1x | 0.007605 | 0.000664 | 1 | 7 | 1 | Q0VBL3 | RNA bindir | Q0VBL3 [6 | 1 | 1787.875 | 0 | -0.09 | 0.135762 | 4.82 | 104 | 96 | 4.27 | 7.87 | | High | High | High | High | High | High | High | High | High | High | High | 0.000294 | 0.002318 | 2.01 |
| FALSE | High | HLDSPPAIP | [K].HLDSPF | 1xPhospho | Q62245 1x | 1.22E-05 | 0 | 1 | 2 | 1 | Q62245 | Son of seve | Q62245 [1 | 0 | 1508.782 | 0 | 0.65 | 0.133808 | 6.05 | 77 | 123 | 7.86 | 53.14 | | High | High | High | High | High | High | High | High | High | High | High | 0 | 1.69E-06 | 2.28 |
| FALSE | High | HIDSSPYSP | [R].HIDSSP | 1xPhospho | Q3U182 1x | 1.19E-05 | 0 | 1 | 2 | 1 | Q3U182 | CREB-regu | Q3U182 [1 | 0 | 2652.233 | 0 | 0.62 | 0.002241 | 5.61 | 83.8 | 116.2 | 7.24 | 10.68 | | High | High | High | High | High | High | High | High | High | High | High | 0 | 1.64E-06 | 3.71 |
| FALSE | High | HLDVDLDR | [R].HLDVDI | 1xPhospho | A2A5R2 1x | 0.002232 | 0.000109 | 1 | 2 | 1 | A2A5R2 | Brefeldin A | A2A5R2 [1 | 1 | 2178.075 | 0 | -0.03 | 0.258306 | 5.4 | 105.2 | 94.8 | 9.25 | 10.51 | | High | High | High | High | High | High | High | High | High | High | High | 4.94E-05 | 0.000586 | 2.46 |
| FALSE | High | HLEEVNSD | [K].HLEEVN | 1xPhospho | Q5SS00 1xP | 1.37E-08 | 0 | 1 | 1 | 2 | Q5SS00 | DBF4-type | Q5SS00 [9 | 0 | 2690.239 | 2 | -1.36 | 0.000834 | 3.65 | 141.2 | 58.8 | 7.34 | 11.58 | | High | High | High | High | High | High | High | High | High | High | High | 0 | 8.23E-10 | 5.32 |
| FALSE | High | HLEHAAST | [K].HLEHA | 1xPhospho | Q6PCM1 1 | 3.38E-05 | 0 | 1 | 5 | 3 | Q6PCM1 | Lysine-spe | Q6PCM1 [4 | 0 | 2846.454 | 0 | -0.94 | 0.004501 | 2.77 | 131.3 | 68.7 | 4.49 | 18.68 | | High | High | High | High | High | High | High | High | High | High | High | 0 | 4.38E-05 | 3.31 |
| FALSE | High | HCSEEK | [R].HCSEEK | 1xCarbami | Q9CXG3 1x | 0.11108 | 0.008991 | 1 | 3 | 1 | Q9CXG3 | Peptidyl-pr | Q9CXG3 [3 | 0 | 1327.612 | 0 | -0.75 | 0.001797 | 1.87 | 124.6 | 75.4 | 6.86 | 8.54 | | High | High | High | High | High | High | High | High | High | High | High | 0.004517 | 0.04894 | 0.99 |
| FALSE | High | HCASPVLN | [K].HCASPV | 1xCarbami | Q3U6K5 1x | 0.052133 | 0.00384 | 1 | 4 | 1 | Q3U6K5 | Spermatog | Q3U6K5 [3 | 0 | 1362.655 | 0 | 1.18 | 0.008266 | 15.33 | 59.1 | 140.9 | 6.71 | 21.61 | | High | High | High | High | High | High | High | High | High | High | High | 0.001999 | 0.02026 | 1.74 |
| FALSE | High | HAVSEGTK | [K].HAVSE | 1xPhospho | Q8CGP2-2 | 0.086562 | 0.006435 | 1 | 17 | 2 | Q8CGP2-2 | Isoform 2 c | Q8CGP2-2 | 0 | 1366.713 | 0 | -0.35 | 0.217316 | 17.7 | 112.5 | 87.5 | 14.11 | 32.69 | | High | High | High | High | High | High | High | High | High | High | High | 0.008559 | 0.09077 | 3.31 |
| FALSE | High | HAVSDPSIL | [K].HAVSD | 1xPhospho | Q6ZWX6 1 | 2.28E-06 | 0 | 1 | 1 | 1 | Q6ZWX6 | Eukaryotic | Q6ZWX6 [1 | 0 | 2434.134 | 0 | -0.21 | 0.191639 | 9.04 | 104 | 96 | 11.62 | 30.66 | | High | High | High | High | High | High | High | High | High | High | High | 0 | 2.57E-07 | 4.49 |
| FALSE | High | GVDFESDE | [K].GVDFE | 1xCarbami | Q3URU4 1 | 4.23E-08 | 0 | 1 | 4 | 5 | Q3URU4 | Meiotic rec | Q3URU4 [6 | 0 | 2831.053 | 0 | -0.38 | 0.00557 | 2.41 | 113.8 | 86.2 | 7.16 | 7.99 | | High | High | High | High | High | High | High | High | High | High | High | 0 | 1.2E-08 | 4.91 |
| FALSE | High | GVDFESDE | [K].GVDFE | 1xCarbami | Q3URU4 1 | 9.24E-10 | 0 | 1 | 4 | 13 | Q3URU4 | Meiotic rec | Q3URU4 [6 | 0 | 2847.048 | 0 | -0.25 | 0.065746 | 10.85 | 112.3 | 87.7 | 4.21 | 14.29 | | High | High | High | High | High | High | High | High | High | High | High | 0 | 3.97E-11 | 5.72 |
| FALSE | High | GVDLLLEG | [K].GVDLLI | 1xPhospho | E9QMX4 1 | 7.93E-07 | 0 | 1 | 4 | 6 | E9QMX4 | Ribosomal | E9QMX4 [2 | 0 | 2066.037 | 0 | 0.21 | 0.011363 | 5.35 | 91 | 109 | 4.25 | 4.23 | | High | High | High | High | High | High | High | High | High | High | High | 0 | 6.71E-05 | 3.99 |
| FALSE | High | GVDLLLEG | [K].GVDLLI | 1xPhospho | E9QMX4 1 | 0.000668 | 0 | 1 | 4 | 1 | E9QMX4 | Ribosomal | E9QMX4 [2 | 1 | 2222.138 | 0 | 0.28 | 0.010252 | 9.55 | 90.6 | 109.4 | 2.21 | 8.61 | | High | High | High | High | High | High | High | High | High | High | High | 0 | 0.000152 | 3.29 |
| FALSE | High | GVEGSADF | [K].GVEGS | 1xPhospho | A2AJK6 1xP | 0.015465 | 0.001089 | 1 | 2 | 1 | A2AJK6 | Chromodo | A2AJK6 [22 | 0 | 1541.761 | 0 | -0.68 | 0.004946 | 4.22 | 121 | 79 | 7.5 | 11 | | High | High | High | High | High | High | High | High | High | High | High | 0.000571 | 0.005113 | 1.99 |
| FALSE | High | GVEPSPGIF | [R].GVEPSF | 1xPhospho | Q5NCY0 1x | 0.000553 | 0 | 1 | 1 | 1 | Q5NCY0 | Lysine-spe | Q5NCY0 [4 | 0 | 3146.526 | 0 | -0.82 | 0.00106 | 2.74 | 128.1 | 71.9 | 6.62 | 7.97 | | High | High | High | High | High | High | High | High | High | High | High | 0 | 0.000123 | 3.37 |
| FALSE | High | GVEPSPSPI | [K].GVEPSF | 1xPhospho | P70362 1xP | 0.023175 | 0.001578 | 1 | 3 | 1 | P70362 | Ubiquitin r | P70362 [24 | 0 | 2288.287 | 0 | 0.21 | 0.088923 | 13.37 | 91.6 | 108.4 | 10.91 | 5.43 | | High | High | High | High | High | High | High | High | High | High | High | 0.000847 | 0.008023 | 1.38 |
| FALSE | High | GVGDDQL | [R].GVGDD | 1xPhospho | P24668 1xP | 0.067619 | 0.004864 | 1 | 2 | 1 | P24668 | Cation-dep | P24668 [25 | 1 | 2666.149 | 0 | 0.84 | 0.013718 | 6.67 | 72 | 128 | 9.89 | 17.23 | | High | High | High | High | High | High | High | High | High | High | High | 0.0025 | 0.02724 | 1.56 |
| FALSE | High | GVGDVLLC | [R].GVGDV | 1xCarbami | Q7TSV9 1x | 3.94E-08 | 0 | 1 | 2 | 1 | Q7TSV9 | Enkurin do | Q7TSV9 [7 | 0 | 2594.403 | 0 | 0.39 | 0.020745 | 0.91 | 86.6 | 113.4 | 9.98 | 6.73 | | High | High | High | High | High | High | High | High | High | High | High | 0 | 2.69E-09 | 3.77 |
| FALSE | High | GVGSPAGF | [R].GVGSP | 1xPhospho | Q8BVE8-2 | 0.018125 | 0.001282 | 2 | 6 | 1 | Q8BVE8-2; | Isoform 2 c | Q8BVE8-2 | 0 | 1009.503 | 0 | 1.38 | 0.000843 | 9.07 | 55.5 | 144.5 | 6.42 | 10.9 | Shared | High | High | High | High | High | High | High | High | High | High | High | 0.000689 | 0.006128 | 1.83 |
| FALSE | High | GVLDLNNP | [R].GVLDLI | 1xPhospho | F8WJ40 1x | 0.00013 | 0 | 1 | 5 | 2 | F8WJ40 | Histone-lys | F8WJ40 [2 | 0 | 2842.321 | 0 | -0.83 | 0.00059 | 2.91 | 127.9 | 72.1 | 4.07 | 3.66 | | High | High | High | High | High | High | High | High | High | High | High | 0.000327 | 0.002518 | 2.28 |
| FALSE | High | GVLDLNNP | [R].GVLDLI | 1xDeamida | F8WJ40 1x | 0.113753 | 0.009241 | 1 | 5 | 1 | F8WJ40 | Histone-lys | F8WJ40 [2 | 0 | 2843.305 | 0 | -0.71 | 0.002382 | 5.11 | 124.6 | 75.4 | 6.56 | 6.37 | | High | High | High | High | High | High | High | High | High | High | High | 0.00468 | 0.0504 | 2.21 |
| FALSE | High | GVLHTFSQ | [R].GVLHTI | 1xPhospho | E9QMN5 1 | 0.003055 | 0.00016 | 1 | 3 | 1 | E9QMN5 | Transcripti | E9QMN5 [ | 0 | 1738.929 | 0 | -0.49 | 0.011102 | 1.5 | 116.3 | 83.7 | 7.69 | 7.55 | | High | High | High | High | High | High | High | High | High | High | High | 7.19E-05 | 0.000837 | 1.9 |
| FALSE | High | GVAPADSP | [R].GVAPA | 1xPhospho | B2RX15 1x | 1.05E-05 | 0 | 1 | 3 | 1 | B2RX15 | Ankib1 pro | B2RX15 [7 | 0 | 1461.694 | 0 | 1.71 | 0.000162 | 5.44 | 47.5 | 152.5 | 11.65 | 2.66 | | High | High | High | High | High | High | High | High | High | High | High | 0 | 1.42E-06 | 2.81 |
| FALSE | High | GVLSSPSLA | [R].GVLSSF | 1xPhospho | Q8R4R6 1x | 2.63E-06 | 0 | 1 | 1 | 3 | Q8R4R6 | Nucleopor | Q8R4R6 [2 | 0 | 1854.993 | 0 | 0.3 | 0.012442 | 9.74 | 93.4 | 106.6 | 7.31 | 6.25 | | High | High | High | High | High | High | High | High | High | High | High | 0 | 6.72E-07 | 4.15 |
| FALSE | High | GVNFAEEP | [K].GVNFA | 2xPhospho | P31324 2xP | 0.000712 | 0 | 1 | 1 | 1 | P31324 | cAMP-dep | P31324 [7 | 1 | 4041.705 | 0 | 0.19 | 0.497332 | 24.59 | 94.9 | 105.1 | 9.61 | 17 | | High | High | High | High | High | High | High | High | High | High | High | 0 | 0.000163 | 3.31 |
| FALSE | High | GVNTFSPE | [R].GVNTF | 1xPhospho | Q3TUI9 1xP | 0.052576 | 0.00384 | 1 | 3 | 1 | Q3TUI9 | Proteasom | Q3TUI9 [1 | 0 | 1372.646 | 0 | 0.02 | 0.894495 | 8.46 | 98.7 | 101.3 | 6.52 | 3.78 | | High | High | High | High | High | High | High | High | High | High | High | 0.001999 | 0.02049 | 1.8 |
| FALSE | High | GVPHSASP | [R].GVPHS | 2xPhospho | Q80WQ6 2 | 0.001002 | 0 | 1 | 2 | 2 | Q80WQ6 | Inactive rh | Q80WQ6 [ | 0 | 2512.277 | 0 | 0.3 | 0.009137 | 10.13 | 87.4 | 112.6 | 4.63 | 5.55 | | High | High | High | High | High | High | High | High | High | High | High | 0 | 0.00024 | 2.97 |
| FALSE | High | GVQAGNSI | [K].GVQAG | 1xPhospho | Q9JIX8-3 1 | 2.18E-07 | 0 | 3 | 12 | 3 | Q9JIX8-3; F | Isoform 3 c | Q9JIX8-3 [ | 0 | 1838.823 | 0 | 0.12 | 0.116693 | 7.59 | 96.1 | 103.9 | 4.71 | 6.27 | Shared | High | High | High | High | High | High | High | High | High | High | High | 0 | 1.84E-08 | 3.68 |
| FALSE | High | GVQAGNSI | [K].GVQAG | 1xDeamida | Q9JIX8-3 1 | 0.000181 | 0 | 3 | 12 | 3 | Q9JIX8-3; F | Isoform 3 c | Q9JIX8-3 [ | 0 | 1839.807 | 0 | 0.09 | 0.364447 | 10.57 | 92.5 | 107.5 | 28.44 | 30.25 | Shared | High | High | High | High | High | High | High | High | High | High | High | 0 | 3.51E-05 | 3.34 |
| FALSE | High | GVQAGNSI | [K].GVQAG | 1xPhospho | Q9JIX8-3 1 | 0.000154 | 0 | 3 | 12 | 2 | Q9JIX8-3; F | Isoform 3 c | Q9JIX8-3 [ | 1 | 2196.081 | 0 | -0.03 | 0.433168 | 10.1 | 99.7 | 100.3 | 8.13 | 7.07 | Shared | High | High | High | High | High | High | High | High | High | High | High | 0 | 2.92E-05 | 2.48 |
| FALSE | High | GVSASSVPI | [K].GVSAS | 1xCarbami | Q68FF6 1xP | 3.9E-05 | 0 | 1 | 3 | 2 | Q68FF6 | ARF GTPas | Q68FF6 [5 | 0 | 2746.296 | 0 | 0.62 | 0.002607 | 8.58 | 79.3 | 120.7 | 9.36 | 9.17 | | High | High | High | High | High | High | High | High | High | High | High | 0 | 6.24E-06 | 2.94 |
| FALSE | High | GVSDEDTD | [K].GVSDEI | 1xPhospho | A2AWA9 1 | 0.025964 | 0.001645 | 1 | 1 | 1 | A2AWA9 | Rab GTPas | A2AWA9 [ | 0 | 1761.783 | 0 | 0.63 | 0.029164 | 28.91 | 73.5 | 126.5 | 14.56 | 19.77 | | High | High | High | High | High | High | High | High | High | High | High | 0.000872 | 0.009169 | 1.72 |
| FALSE | High | GVSFDQAN | [K].GVSFD | 1xPhospho | Q80U72 1x | 0.023379 | 0.001578 | 1 | 4 | 1 | Q80U72 | Protein scr | Q80U72 [6 | 0 | 2053.032 | 0 | 0.39 | 0.077437 | 13.03 | 91.7 | 108.3 | 7.11 | 14.63 | | High | High | High | High | High | High | High | High | High | High | High | 0.000847 | 0.008108 | 2 |
| FALSE | High | GVSFGSGC | [R].GVSFG | 1xPhospho | Q9EP53 1x | 0.034579 | 0.002233 | 1 | 4 | 2 | Q9EP53 | Hamartin C | Q9EP53 [5 | 0 | 2896.505 | 0 | 0.62 | 0.000725 | 1.92 | 79.8 | 120.2 | 10.64 | 6.71 | | High | High | High | High | High | High | High | High | High | High | High | 0.001183 | 0.01263 | 2.85 |
| FALSE | High | GVSGDREE | [R].GVSGD | 1xPhospho | P20357 1xP | 0.079766 | 0.00613 | 1 | 8 | 2 | P20357 | Microtubu | P20357 [15 | 1 | 2508.157 | 0 | 0.92 | 0.001361 | 17.83 | 67.9 | 132.1 | 14.18 | 5.46 | | High | High | High | High | High | High | High | High | High | High | High | 0.003148 | 0.03299 | 1.6 |
| FALSE | High | GVSGQCTC | [R].GVSGQ | 1xCarbami | G3UY09 1x | 5.86E-05 | 0 | 1 | 1 | 1 | G3UY09 | MCG61508 | G3UY09 [2 | 0 | 2841.317 | 0 | -1.03 | 0.005984 | 2.23 | 134.9 | 65.1 | 8.6 | 19.99 | | High | High | High | High | High | High | High | High | High | High | High | 0 | 9.82E-06 | 3.85 |
| FALSE | High | GVSMSLPS | [K].GVSMS | 1xPhospho | Q0VGY8 1x | 0.000639 | 0 | 1 | 3 | 2 | Q0VGY8 | Protein TA | Q0VGY8 [5 | 0 | 1749.917 | 0 | -0.03 | 0.939248 | 11.58 | 101.7 | 98.3 | 7.96 | 13.5 | | High | High | High | High | High | High | High | High | High | High | High | 0 | 0.000144 | 2.25 |
| FALSE | High | GVLSSPSLA | [R].GVLSSF | 2xPhospho | Q8R4R6 2x | 0.041781 | 0.003126 | 1 | 1 | 3 | Q8R4R6 | Nucleopor | Q8R4R6 [2 | 0 | 1934.959 | 0 | -0.29 | 0.020727 | 9.3 | 110.8 | 89.2 | 5.7 | 6.62 | | High | High | High | High | High | High | High | High | High | High | High | 0.001609 | 0.01564 | 2.07 |
| FALSE | High | GTYFPAILN | [K].GTYFPA | 1xPhospho | A0A0R4J0A | 0.004738 | 0.000456 | 1 | 4 | 2 | A0A0R4J0A | Low-densit | A0A0R4J0A | 0 | 2237.121 | 0 | -0.52 | 0.006006 | 6.46 | 122.1 | 77.9 | 11.49 | 8.93 | | High | High | High | High | High | High | High | High | High | High | High | 0.000801 | 0.007468 | 2.72 |
| FALSE | High | GTVTGERC | [K].GTVTG | 1xPhospho | Q8K3W3 1 | 1.16E-05 | 0 | 1 | 1 | 3 | Q8K3W3 | Protein CA | Q8K3W3 [1 | 1 | 2843.346 | 0 | -0.44 | 0.000222 | 0.39 | 115 | 85 | 3.59 | 2.69 | | High | High | High | High | High | High | High | High | High | High | High | 0 | 1.58E-06 | 3.71 |
| FALSE | High | GTVAVGDS | [K].GTVAV | 1xPhospho | Q3TYA6 1x | 3.14E-08 | 0 | 1 | 5 | 6 | Q3TYA6 | M-phase p | Q3TYA6 [4 | 0 | 2448.065 | 0 | -1.09 | 0.000568 | 1.74 | 134.7 | 65.3 | 5.73 | 5.46 | | High | High | High | High | High | High | High | High | High | High | High | 0 | 4.34E-08 | 4.29 |
| FALSE | High | GTPSQSPV | [K].GTPSQ | 2xPhospho | A0A087WF | 0.047488 | 0.00354 | 1 | 8 | 1 | A0A087WF | Arf-GAP dc | A0A087WF | 0 | 1473.67 | 0 | 0.77 | 0.005567 | 11.05 | 78.4 | 121.6 | 6.73 | 14.93 | | High | High | High | High | High | High | High | High | High | High | High | 0.001812 | 0.01808 | 2.64 |
| FALSE | High | GTPSSSPVS | [R].GTPSSS | 1xPhospho | A0A1L7NR | 8.63E-05 | 0 | 2 | 12 | 2 | A0A1L7NR | C-Cbl assoc | A0A1L7NR | 0 | 2023.015 | 0 | 0.92 | 0.000222 | 4.57 | 67.9 | 132.1 | 10.14 | 4.46 | Shared | High | High | High | High | High | High | High | High | High | High | High | 0 | 1.52E-05 | 1.77 |
| FALSE | High | GTPSSSPVS | [R].GTPSSS | 2xPhospho | A0A1L7NR | 3.54E-05 | 0 | 2 | 12 | 2 | A0A1L7NR | C-Cbl assoc | A0A1L7NR | 0 | 2102.981 | 0 | 1.2 | 0.000616 | 10.09 | 58.8 | 141.2 | 9.56 | 4.68 | Shared | High | High | High | High | High | High | High | High | High | High | High | 0 | 5.58E-06 | 2.9 |
| FALSE | High | GTQCPNLE | [R].GTQCP | 1xCarbami | Q1WG82 1 | 1.5E-05 | 0 | 1 | 1 | 3 | Q1WG82 | GATA-type | Q1WG82 [ | 0 | 2426.158 | 0 | -0.47 | 0.044442 | 8.72 | 116.1 | 83.9 | 11.71 | 7.76 | | High | High | High | High | High | High | High | High | High | High | High | 0 | 2.13E-06 | 4 |
| FALSE | High | GTQEVAPP | [R].GTQEV | 1xPhospho | Q3USK2 1x | 0.039686 | 0.00283 | 1 | 3 | 1 | Q3USK2 | Casein kina | Q3USK2 [3 | 0 | 3425.672 | 0 | 0.73 | 0.001573 | 9.88 | 73.7 | 126.3 | 14.48 | 8.56 | | High | High | High | High | High | High | High | High | High | High | High | 0.001468 | 0.0148 | 2.34 |
| FALSE | High | GTQEVAPP | [R].GTQEV | 1xPhospho | Q3USK2 1x | 0.000108 | 0 | 1 | 3 | 6 | Q3USK2 | Casein kina | Q3USK2 [3 | 0 | 3441.667 | 0 | 0.65 | 0.004574 | 10.53 | 83.4 | 116.6 | 13.36 | 11.32 | | High | High | High | High | High | High | High | High | High | High | High | 0 | 3.63E-05 | 3.81 |
| FALSE | High | GTQEVAPP | [R].GTQEV | 1xDeamida | Q3USK2 1x | 0.000984 | 0 | 1 | 3 | 2 | Q3USK2 | Casein kina | Q3USK2 [3 | 0 | 3442.651 | 0 | 0.99 | 0.00333 | 18.13 | 68.9 | 131.1 | 11.94 | 11.88 | | High | High | High | High | High | High | High | High | High | High | High | 4.94E-05 | 0.00058 | 3.04 |
| FALSE | High | GTQEVAPP | [R].GTQEV | 2xPhospho | Q3USK2 2x | 0.012731 | 0.000889 | 1 | 3 | 3 | Q3USK2 | Casein kina | Q3USK2 [3 | 0 | 3521.633 | 0 | 0.8 | 0.007173 | 24.2 | 66.4 | 133.6 | 14.63 | 12.29 | | High | High | High | High | High | High | High | High | High | High | High | 0.000449 | 0.004101 | 3.43 |
| FALSE | High | GTQLPTVS | [K].GTQLPT | 1xPhospho | Q7TSJ2 1xP | 0.074685 | 0.005727 | 1 | 2 | 1 | Q7TSJ2 | Microtubu | Q7TSJ2 [8 | 0 | 3256.605 | 0 | 0.79 | 0.004584 | 10.12 | 73.1 | 126.9 | 20.34 | 22.94 | | High | High | High | High | High | High | High | High | High | High | High | 0.002937 | 0.03079 | 3.54 |
| FALSE | High | GTQPGLQE | [K].GTQPG | 1xPhospho | B1ATU5 1x | 0.026885 | 0.001676 | 1 | 3 | 1 | B1ATU5 | tRNA-dihy | B1ATU5 [4 | 0 | 3219.625 | 0 | -1.01 | 0.000945 | 1.51 | 133 | 67 | 7.32 | 12.22 | | High | High | High | High | High | High | High | High | High | High | High | 0.000903 | 0.009497 | 2.9 |
| FALSE | High | GTSDSSSGI | [K].GTSDSS | 1xPhospho | K4DI61 1xP | 9.5E-07 | 0 | 1 | 3 | 7 | K4DI61 | PHD finger | K4DI61 [31 | 0 | 3061.249 | 0 | -0.17 | 0.093976 | 12.61 | 102.9 | 97.1 | 6.54 | 11.26 | | High | High | High | High | High | High | High | High | High | High | High | 0 | 9.58E-08 | 4.33 |
| FALSE | High | GTSPGSELI | [K].GTSPGS | 1xPhospho | Q7TSC1 1x | 0.045898 | 0.003408 | 1 | 5 | 2 | Q7TSC1 | Protein PR | Q7TSC1 [3 | 0 | 1704.897 | 0 | 0.08 | 0.180738 | 15.59 | 95.5 | 104.5 | 6.21 | 4.88 | | High | High | High | High | High | High | High | High | High | High | High | 0.001741 | 0.01743 | 2.85 |
| FALSE | High | GTSPPYGV | [K].GTSPPY | 1xCarbami | P47806 1xP | 4.91E-06 | 0 | 1 | 2 | 2 | P47806 | Zinc finger | P47806 [14 | 0 | 2264.037 | 0 | -1.02 | 0.004623 | 5.93 | 130.8 | 69.2 | 12.05 | 21.13 | | High | High | High | High | High | High | High | High | High | High | High | 0 | 6.06E-07 | 3.27 |
| FALSE | High | GTSPRPPE | [R].GTSPRI | 1xPhospho | P24788 1xP | 2.07E-07 | 0 | 1 | 4 | 6 | P24788 | Cyclin-dep | P24788 [73 | 0 | 2568.218 | 0 | -0.51 | 0.000602 | 2.23 | 117.7 | 82.3 | 3.46 | 3.97 | | High | High | High | High | High | High | High | High | High | High | High | 0 | 1.74E-08 | 3.92 |
| FALSE | High | GTSPRPPE | [R].GTSPRI | 1xPhospho | P24788 1xP | 1.35E-06 | 0 | 1 | 4 | 2 | P24788 | Cyclin-dep | P24788 [73 | 0 | 2797.381 | 0 | -0.57 | 0.000334 | 1.45 | 119.3 | 80.7 | 4.57 | 4.97 | | High | High | High | High | High | High | High | High | High | High | High | 0 | 1.43E-07 | 2.91 |
| FALSE | High | GTSPTQIW | [K].GTSPTC | 1xCarbami | F8WHL2 1x | 0.002252 | 0.000109 | 1 | 6 | 2 | F8WHL2 | Coatomer s | F8WHL2 [9 | 0 | 3539.665 | 0 | -1.31 | 0.002197 | 3.76 | 145.2 | 54.8 | 13.25 | 5.2 | | High | High | High | High | High | High | High | High | High | High | High | 4.94E-05 | 0.000595 | 4.65 |
| FALSE | High | GTSPTQIW | [K].GTSPTC | 1xCarbami | F8WHL2 1x | 0.086562 | 0.006435 | 1 | 6 | 1 | F8WHL2 | Coatomer s | F8WHL2 [9 | 0 | 3555.66 | 0 | -0.72 | 0.000905 | 2.61 | 128.4 | 71.6 | 8.28 | 13.11 | | High | High | High | High | High | High | High | High | High | High | High | 0.003346 | 0.03655 | 2.12 |
| FALSE | High | GTSQVSSP | [R].GTSQV | 1xPhospho | Q8CJ19 1xP | 2.08E-05 | 0 | 1 | 3 | 2 | Q8CJ19 | [F-actin]-m | Q8CJ19 [1 | 0 | 1966.989 | 0 | -0.79 | 0.003064 | 5.02 | 120.7 | 79.3 | 11.32 | 13.52 | | High | High | High | High | High | High | High | High | High | High | High | 0 | 3.09E-06 | 2.65 |
| FALSE | High | GTSRPGTP | [K].GTSRPC | 1xPhospho | Q3THK3 1x | 0.001049 | 0 | 1 | 1 | 5 | Q3THK3 | General tra | Q3THK3 [3 | 0 | 2243.087 | 0 | -0.31 | 0.029912 | 5.49 | 109.3 | 90.7 | 7.93 | 4.76 | | High | High | High | High | High | High | High | High | High | High | High | 0.001639 | 0.01594 | 2.69 |
| FALSE | High | GTSRPGTP | [K].GTSRPC | 2xPhospho | Q3THK3 2x | 0.00034 | 0 | 1 | 1 | 7 | Q3THK3 | General tra | Q3THK3 [3 | 0 | 2323.053 | 0 | -0.08 | 0.20201 | 6.44 | 103.8 | 96.2 | 3.95 | 6.96 | | High | High | High | High | High | High | High | High | High | High | High | 0 | 0.000131 | 3.03 |
| FALSE | High | GTSSSPAAS | [R].GTSSSP | 1xPhospho | Q3UIR3 1x | 0.002153 | 5.66E-05 | 1 | 3 | 2 | Q3UIR3 | E3 ubiquiti | Q3UIR3 [5 | 0 | 1430.729 | 0 | 1.48 | 0.000378 | 5.88 | 50.9 | 149.1 | 3.41 | 6.99 | | High | High | High | High | High | High | High | High | High | High | High | 2.58E-05 | 0.000564 | 2.89 |
| FALSE | High | GTSVMVTH | [K].GTSVM | 1xPhospho | Q61191 1x | 0.015741 | 0.001124 | 1 | 3 | 2 | Q61191 | Host cell fa | Q61191 [1 | 0 | 4149.94 | 0 | 0.15 | 0.872375 | 4.22 | 98.5 | 101.5 | 4.55 | 17.14 | | High | High | High | High | High | High | High | High | High | High | High | 0.000756 | 0.006636 | 3.3 |
| FALSE | High | GTSVMVTH | [K].GTSVM | 1xPhospho | Q61191 1x | 0.002115 | 5.66E-05 | 1 | 3 | 3 | Q61191 | Host cell fa | Q61191 [1 | 0 | 4165.935 | 0 | 0.06 | 0.797656 | 7.7 | 96 | 104 | 8.36 | 9.06 | | High | High | High | High | High | High | High | High | High | High | High | 9.42E-05 | 0.000958 | 3.85 |
| FALSE | High | GTTASAAS | [K].GTTASA | 1xCarbami | Q6PAL7 1x | 4.37E-10 | 0 | 1 | 1 | 2 | Q6PAL7 | AT-hook D | Q6PAL7 [1 | 0 | 2782.219 | 0 | -1.25 | 0.000361 | 0.8 | 139.2 | 60.8 | 5.56 | 12.18 | | High | High | High | High | High | High | High | High | High | High | High | 0 | 1.7E-11 | 6.44 |
| FALSE | High | GTTASPK | [R].GTTASF | 1xPhospho | A2AJI0 1xP | 0.079766 | 0.00613 | 1 | 5 | 4 | A2AJI0 | MAP7 dom | A2AJI0 [49 | 0 | 1199.644 | 0 | 1.49 | 0.000486 | 3.98 | 51.4 | 148.6 | 8.04 | 5.99 | | High | High | High | High | High | High | High | High | High | High | High | 0.003291 | 0.03584 | 0.89 |
| FALSE | High | GTTPPPPS | [R].GTTPPF | 1xPhospho | Q8BYR2 1x | 3.44E-05 | 0 | 1 | 1 | 2 | Q8BYR2 | Serine/thre | Q8BYR2 [2 | 0 | 2235.11 | 0 | 1.75 | 0.000285 | 2.28 | 46.1 | 153.9 | 7.87 | 5.69 | | High | High | High | High | High | High | High | High | High | High | High | 0 | 5.43E-06 | 3.43 |
| FALSE | High | GTTQEVDG | [K].GTTQE | 1xPhospho | F6Q8D3 1x | 2.8E-06 | 0 | 1 | 4 | 1 | F6Q8D3 | Phospholip | F6Q8D3 [4 | 0 | 2780.343 | 0 | 1.47 | 0.000959 | 29.7 | 54.8 | 145.2 | 21.18 | 10.24 | | High | High | High | High | High | High | High | High | High | High | High | 0 | 3.23E-07 | 3.19 |
| FALSE | High | GTTSAQLP | [R].GTTSA | 1xPhospho | Q3TDD9 1x | 0.000524 | 0 | 1 | 1 | 2 | Q3TDD9 | Protein ph | Q3TDD9 [6 | 0 | 3378.719 | 0 | 1.04 | 0.004701 | 24.68 | 65 | 135 | 10.28 | 10.67 | | High | High | High | High | High | High | High | High | High | High | High | 0 | 0.000116 | 3.51 |
| FALSE | High | GTVASGEE | [K].GTVAS | 1xPhospho | E9PZM4 1x | 0.104222 | 0.008348 | 1 | 2 | 1 | E9PZM4 | Chromodo | E9PZM4 [1 | 0 | 1486.755 | 0 | -0.17 | 0.692878 | 15.45 | 100.8 | 99.2 | 17.3 | 16.86 | | High | High | High | High | High | High | High | High | High | High | High | 0.004221 | 0.04537 | 1.3 |
| FALSE | High | GVSNDTSS | [R].GVSND | 1xPhospho | A2CG76 1x | 0.045122 | 0.003373 | 1 | 3 | 3 | A2CG76 | Histone-lys | A2CG76 [4 | 0 | 1703.769 | 0 | 0.09 | 0.560237 | 19.84 | 102.4 | 97.6 | 12.78 | 13.41 | | High | High | High | High | High | High | High | High | High | High | High | 0.001727 | 0.01714 | 1.97 |
| FALSE | High | HLSSSTVAI | [K].HLSSST | 1xPhospho | A2AG50 1x | 0.028082 | 0.001791 | 1 | 5 | 1 | A2AG50 | MAP7 dom | A2AG50 [2 | 0 | 1928.932 | 0 | 0.24 | 0.017252 | 8.61 | 89.5 | 110.5 | 12.4 | 7.53 | | High | High | High | High | High | High | High | High | High | High | High | 0.000968 | 0.009956 | 3.44 |
| FALSE | High | GVSNESVP | [K].GVSNE | 1xPhospho | A0A0R4J1A | 0.001373 | 5.66E-05 | 1 | 3 | 2 | A0A0R4J1A | GAS2-like p | A0A0R4J1A | 0 | 1623.758 | 0 | 1.06 | 0.000484 | 8.46 | 64.4 | 135.6 | 9.31 | 2.61 | | High | High | High | High | High | High | High | High | High | High | High | 0.000294 | 0.002315 | 2.39 |
| FALSE | High | GVSSVVGP | [R].GVSSV | 1xPhospho | Q8CI75-2 1 | 5.36E-09 | 0 | 1 | 3 | 1 | Q8CI75-2 | Isoform 2 c | Q8CI75-2 [ | 0 | 2109.099 | 0 | 0.02 | 0.485791 | 11.39 | 96.3 | 103.7 | 13.16 | 7.52 | | High | High | High | High | High | High | High | High | High | High | High | 0 | 2.85E-10 | 4.29 |
| FALSE | High | GYYSPYSVS | [K].GYYSPY | 1xPhospho | Q9QXS1-3 | 8.82E-06 | 0 | 3 | 23 | 14 | Q9QXS1-3; | Isoform PL | Q9QXS1-3 | 0 | 2091.922 | 0 | 2.52 | 0.002109 | 0.47 | 24.1 | 175.9 | 14.15 | 18.73 | Shared | High | High | High | High | High | High | High | High | High | High | High | 0 | 3.55E-06 | 3.02 |
| FALSE | High | GYYSPYSVS | [K].GYYSPY | 2xPhospho | Q9QXS1-3 | 0.001168 | 0 | 3 | 23 | 8 | Q9QXS1-3; | Isoform PL | Q9QXS1-3 | 0 | 2171.889 | 0 | 1.72 | 0.002178 | 18.95 | 44.2 | 155.8 | 10.27 | 16.17 | Shared | High | High | High | High | High | High | High | High | High | High | High | 0.000859 | 0.008528 | 2.85 |

| | | | | | | | | | | | | | | | | | | | | | | | | | | | | | | | | | | | | | | | |
|---|---|---|---|---|---|---|---|---|---|---|---|---|---|---|---|---|---|---|---|---|---|---|---|---|---|---|---|---|---|---|---|---|---|---|---|---|---|---|---|
| FALSE | High | GYYSPYSVS | [K].GYYSPY | 3xPhospho | Q9QXS1-3 | 0.031168 | 0.001935 | 3 | 23 | 1 | Q9QXS1-3; | Isoform PL | Q9QXS1-3 | 0 | 2251.855 | 0 | 0.04 | 0.384923 | 16.13 | 93.6 | 106.4 | 7.41 | 16 | Shared | High | High | High | High | High | High | High | High | High | High | High | High | 0.001076 | 0.0113 | 3.16 |
| FALSE | High | GYYYDLDD | [K].GYYYDI | 1xPhospho | Q3U3C9-2 | 2.08E-05 | 0 | 1 | 4 | 4 | Q3U3C9-2 | Isoform 2 c | Q3U3C9-2 | 0 | 2671.045 | 0 | 0 | 0.593847 | 18.47 | 103.2 | 96.8 | 12.26 | 10.69 | | High | High | High | High | High | High | High | High | High | High | High | High | 0 | 3.41E-06 | 4.14 |
| FALSE | High | GYYYDLDD | [K].GYYYDI | 2xPhospho | Q3U3C9-2 | 4.76E-05 | 0 | 1 | 4 | 2 | Q3U3C9-2 | Isoform 2 c | Q3U3C9-2 | 0 | 2751.011 | 0 | -0.76 | 0.003746 | 5.41 | 126.5 | 73.5 | 2.53 | 16.58 | | High | High | High | High | High | High | High | High | High | High | High | High | 0.000847 | 0.007983 | 3.26 |
| FALSE | High | HAADSAEK | [K].HAADS. | 1xPhospho | Q3U0X6 1x | 0.02933 | 0.001857 | 1 | 1 | 1 | Q3U0X6 | Putative ur | Q3U0X6 [6 | 0 | 1366.677 | 0 | -0.37 | 0.196121 | 31.5 | 107.4 | 92.6 | 15.52 | 20.95 | | High | High | High | High | High | High | High | High | High | High | High | High | 0.000997 | 0.01049 | 2.95 |
| FALSE | High | HAASDVDL | [K].HAASD' | 1xPhospho | A0A0R4J04 | 0.013074 | 0.000889 | 1 | 7 | 1 | A0A0R4J04 | N6-adenos | A0A0R4J04 | 0 | 2736.386 | 0 | -0.35 | 0.17026 | 10.1 | 107.5 | 92.5 | 10.86 | 10.53 | | High | High | High | High | High | High | High | High | High | High | High | High | 0.000467 | 0.004253 | 2.78 |
| FALSE | High | HAATHTPT | [R].HAATH | 1xCarbami | Q8K2R5 1x | 1.3E-05 | 0 | 1 | 2 | 2 | Q8K2R5 | Zinc finger | Q8K2R5 [4 | 0 | 2033.895 | 0 | 0.39 | 0.003925 | 5.78 | 87 | 113 | 9.65 | 6.64 | | High | High | High | High | High | High | High | High | High | High | High | High | 0 | 1.81E-06 | 3.21 |
| FALSE | High | HACNNQG | [K].HACNN | 2xCarbami | B2RQG2 1x | 0.000184 | 0 | 2 | 2 | 4 | B2RQG2; A | PHD finger | B2RQG2 [2 | 0 | 3948.84 | 0 | -0.75 | 0.019187 | 2.39 | 127 | 73 | 11.9 | 26.02 | Shared | High | High | High | High | High | High | High | High | High | High | High | High | 0 | 0.000108 | 3.42 |
| FALSE | High | HADGESGS | [R].HADGE | 1xPhospho | Q24JP2 1xl | 0.000316 | 0 | 1 | 1 | 2 | Q24JP2 | Additional | Q24JP2 [19 | 0 | 2640.21 | 0 | 0.12 | 0.907358 | 15.59 | 99 | 101 | 24.26 | 16.93 | | High | High | High | High | High | High | High | High | High | High | High | High | 0 | 6.58E-05 | 4.24 |
| FALSE | High | HADHSSLTI | [R].HADHS | 1xPhospho | A2AFQ0 1x | 0.005467 | 0.00062 | 1 | 6 | 3 | A2AFQ0 | E3 ubiquiti | A2AFQ0 [2 | 0 | 2022.944 | 0 | 0.14 | 0.060514 | 9.83 | 98.8 | 101.2 | 9.59 | 6.21 | | High | High | High | High | High | High | High | High | High | High | High | High | 0.000275 | 0.001592 | 2.98 |
| FALSE | High | HAELSGSPI | [R].HAELSC | 1xPhospho | Q924K8 1x | 1.2E-05 | 0 | 1 | 5 | 2 | Q924K8 | Metastasis | Q924K8 [5 | 0 | 1576.85 | 0 | -0.15 | 0.034383 | 5.17 | 105 | 95 | 7.55 | 9.72 | | High | High | High | High | High | High | High | High | High | High | High | High | 0 | 1.61E-05 | 2.69 |
| FALSE | High | HAEQNGPV | [K].HAEQN | 1xPhospho | Q571M2 1: | 3.67E-09 | 0 | 1 | 4 | 4 | Q571M2 | MKIAA402 | Q571M2 [8 | 0 | 3866.726 | 0 | -0.58 | 0.005966 | 2.56 | 118.1 | 81.9 | 4.74 | 15.85 | | High | High | High | High | High | High | High | High | High | High | High | High | 0 | 4.33E-08 | 5.17 |
| FALSE | High | GYVTDDGC | [K].GYVTDI | 1xPhospho | E9QAS8 1x | 0.002134 | 5.66E-05 | 1 | 12 | 1 | E9QAS8 | Synaptic fu | E9QAS8 [4 | 0 | 1580.661 | 0 | 0.98 | 0.001351 | 16.43 | 67.2 | 132.8 | 5.17 | 7.52 | | High | High | High | High | High | High | High | High | High | High | High | High | 2.58E-05 | 0.000561 | 1.12 |
| FALSE | High | HAEQNGPV | [K].HAEQN | 1xDeamida | Q571M2 1: | 6.14E-09 | 0 | 1 | 4 | 8 | Q571M2 | MKIAA402 | Q571M2 [8 | 0 | 3867.71 | 0 | -0.2 | 0.086112 | 3.87 | 107.1 | 92.9 | 3.36 | 20.83 | | High | High | High | High | High | High | High | High | High | High | High | High | 0 | 6.49E-10 | 5.64 |
| FALSE | High | HALNLSQE | [R].HALNLS | 1xPhospho | P48754 1xl | 0.001097 | 0 | 1 | 7 | 2 | P48754 | Breast canc | P48754 [82 | 0 | 1577.845 | 0 | -0.79 | 0.000333 | 1.57 | 127.8 | 72.2 | 3.95 | 6.25 | | High | High | High | High | High | High | High | High | High | High | High | High | 0 | 0.000265 | 2.49 |
| FALSE | High | HALTSPSLG | [R].HALTSP | 1xPhospho | G5E8A7 1x | 1.66E-05 | 0 | 1 | 3 | 2 | G5E8A7 | Protein Sm | G5E8A7 [5 | 0 | 1589.8 | 0 | 1.46 | 0.000729 | 10.17 | 51.8 | 148.2 | 12.2 | 8.28 | | High | High | High | High | High | High | High | High | High | High | High | High | 0 | 5.6E-06 | 2.48 |
| FALSE | High | HASAAASK | [K].HASAA | 1xPhospho | Q5F2E7-2 | 0.001107 | 0 | 1 | 2 | 1 | Q5F2E7-2 | Isoform 2 c | Q5F2E7-2 | 0 | 1280.676 | 0 | -0.6 | 0.028179 | 14.8 | 107.6 | 92.4 | 16.14 | 28.65 | | High | High | High | High | High | High | High | High | High | High | High | High | 0 | 0.000269 | 1.45 |
| FALSE | High | HASAPSHV | [R].HASAPS | 1xPhospho | Q8BYJ6 1xl | 0.004451 | 0.000409 | 1 | 8 | 3 | Q8BYJ6 | TBC1 doma | Q8BYJ6 [34 | 0 | 2115.027 | 0 | -0.4 | 0.023228 | 9.91 | 113.9 | 86.1 | 4.44 | 14.37 | | High | High | High | High | High | High | High | High | High | High | High | High | 0.000183 | 0.001274 | 2.41 |
| FALSE | High | HASEESPQ | [R].HASEES | 1xPhospho | Q6P1D7 1x | 0.022574 | 0.001542 | 1 | 2 | 1 | Q6P1D7 | Structure-s | Q6P1D7 [5 | 0 | 1550.762 | 0 | -0.87 | 0.007067 | 5.43 | 130.4 | 69.6 | 16.34 | 14.65 | | High | High | High | High | High | High | High | High | High | High | High | High | 0.000814 | 0.007798 | 2.26 |
| FALSE | High | HASLDGAS | [R].HASLDC | 1xPhospho | E9PW07 1x | 0.005228 | 0.000551 | 1 | 2 | 1 | E9PW07 | Voltage-de | E9PW07 [2 | 0 | 1830.919 | 0 | 0.66 | 0.017213 | 22.67 | 75.6 | 124.4 | 13.27 | 8.74 | | High | High | High | High | High | High | High | High | High | High | High | High | 0.000246 | 0.001517 | 1.97 |
| FALSE | High | HASPEDIK | [R].HASPED | 1xPhospho | O54946 1x | 0.000967 | 0 | 1 | 7 | 2 | O54946 | DnaJ homc | O54946 [1 | 0 | 1434.739 | 0 | -0.12 | 0.110147 | 7.25 | 104.2 | 95.8 | 15.96 | 6.86 | | High | High | High | High | High | High | High | High | High | High | High | High | 0 | 0.000231 | 2.09 |
| FALSE | High | HASSANQY | [R].HASSAN | 1xPhospho | Q8BZI0 1xP | 0.090888 | 0.007081 | 1 | 1 | 2 | Q8BZI0 | Actin filam | Q8BZI0 [60 | 0 | 1543.767 | 0 | -1.34 | 0.002683 | 0.55 | 146.1 | 53.9 | 13.73 | 19.26 | | High | High | High | High | High | High | High | High | High | High | High | High | 0.003633 | 0.03859 | 2.61 |
| FALSE | High | HASSSPESL | [R].HASSSP | 3xPhospho | Q8BTI8 3xP | 0.00509 | 0.000551 | 1 | 3 | 1 | Q8BTI8 | Serine/argi | Q8BTI8 [43 | 0 | 2274.976 | 0 | -0.51 | 0.040041 | 15.33 | 126 | 74 | 17.57 | 16.55 | | High | High | High | High | High | High | High | High | High | High | High | High | 0.000246 | 0.001476 | 2.04 |
| FALSE | High | HASSSPESL | [R].HASSSP | 2xPhospho | Q8BTI8 2xP | 0.067619 | 0.004864 | 1 | 3 | 3 | Q8BTI8 | Serine/argi | Q8BTI8 [43 | 0 | 2424.173 | 0 | -0.97 | 0.001359 | 2.34 | 132.3 | 67.7 | 6.54 | 8.24 | | High | High | High | High | High | High | High | High | High | High | High | High | 0.003659 | 0.03928 | 2.39 |
| FALSE | High | HASTSPAD | [R].HASTSP | 1xPhospho | A2AN08-5 | 0.001313 | 0 | 1 | 4 | 3 | A2AN08-5 | Isoform 5 c | A2AN08-5 | 0 | 1222.567 | 0 | -0.63 | 0.997403 | 47.7 | 121.5 | 78.5 | 69.51 | 99.46 | | High | High | High | High | High | High | High | High | High | High | High | High | 0 | 0.000325 | 2.19 |
| FALSE | High | HASTSPAD | [R].HASTSP | 1xPhospho | A2AN08-5 | 0.001158 | 0 | 1 | 4 | 2 | A2AN08-5 | Isoform 5 c | A2AN08-5 | 0 | 1451.73 | 0 | -0.01 | 0.998251 | 3.45 | 100.6 | 99.4 | 4.26 | 4.41 | | High | High | High | High | High | High | High | High | High | High | High | High | 0 | 0.000282 | 2.27 |
| FALSE | High | HASTSPAD | [R].HASTSP | 2xPhospho | A2AN08-5 | 0.002419 | 0.000109 | 1 | 4 | 1 | A2AN08-5 | Isoform 5 c | A2AN08-5 | 0 | 1531.696 | 0 | 0.06 | 0.756194 | 9.65 | 102.1 | 97.9 | 11.63 | 11.02 | | High | High | High | High | High | High | High | High | High | High | High | High | 4.94E-05 | 0.000643 | 2.03 |
| FALSE | High | HAGSAGAL | [K].HAGSA | 1xPhospho | Q4VAC9 1x | 2.7E-06 | 0 | 1 | 2 | 1 | Q4VAC9 | Pleckstrin I | Q4VAC9 [4 | 0 | 2414.238 | 0 | 1.98 | 0.045591 | 16.82 | 43.5 | 156.5 | 15.18 | 45.81 | | High | High | High | High | High | High | High | High | High | High | High | High | 0 | 3.1E-07 | 3.66 |
| FALSE | High | GYTSEDDT | [K].GYTSED | 1xCarbami | Q3TYA6 1x | 3.61E-06 | 0 | 1 | 5 | 14 | Q3TYA6 | M-phase p | Q3TYA6 [8 | 0 | 2848.242 | 0 | -1.07 | 0.000569 | 1.11 | 135.4 | 64.6 | 3.68 | 7.12 | | High | High | High | High | High | High | High | High | High | High | High | High | 0 | 6.02E-05 | 3.98 |
| FALSE | High | GYTALSQSI | [K].GYTALS | 2xPhospho | Q14DQ1 2x | 0.000492 | 0 | 1 | 1 | 1 | Q14DQ1 | Protein FAI | Q14DQ1 [1 | 0 | 3273.392 | 0 | 0.33 | 0.061943 | 24.6 | 91.7 | 108.3 | 7.81 | 14.54 | | High | High | High | High | High | High | High | High | High | High | High | High | 0 | 0.000108 | 3.47 |
| FALSE | High | GYSYDDSM | [R].GYSYDI | 1xPhospho | Q8C5N3 1x | 0.000438 | 0 | 1 | 14 | 17 | Q8C5N3 | Pre-mRNA- | Q8C5N3 [5 | 0 | 1634.624 | 0 | -0.47 | 0.00085 | 3.15 | 115 | 85 | 2.86 | 2.77 | | High | High | High | High | High | High | High | High | High | High | High | High | 0 | 0.000127 | 2.4 |
| FALSE | High | GVTASSSSI | [K].GVTAS! | 1xPhospho | P13595 1xl | 7.34E-05 | 0 | 1 | 2 | 1 | P13595 | Neural cell | P13595 [88 | 0 | 1784.92 | 0 | -0.19 | 0.223567 | 15.63 | 106.5 | 93.5 | 7.05 | 12.37 | | High | High | High | High | High | High | High | High | High | High | High | High | 0 | 1.27E-05 | 3.49 |
| FALSE | High | GVVAVDST | [K].GVVAV | 1xPhospho | Q5U647 1x | 8.92E-07 | 0 | 1 | 3 | 1 | Q5U647 | Amino acid | Q5U647 [1 | 1 | 2824.424 | 0 | 0.38 | 0.067257 | 15.52 | 90.3 | 109.7 | 13.12 | 6.02 | | High | High | High | High | High | High | High | High | High | High | High | High | 0 | 8.94E-08 | 4.1 |
| FALSE | High | GVVAVDST | [K].GVVAV | 1xDeamida | Q5U647 1x | 0.049549 | 0.00361 | 1 | 3 | 1 | Q5U647 | Amino acid | Q5U647 [1 | 1 | 2825.408 | 0 | 0.33 | 0.06008 | 20.56 | 88.3 | 111.7 | 8.59 | 9.98 | | High | High | High | High | High | High | High | High | High | High | High | High | 0.001889 | 0.01904 | 2.07 |
| FALSE | High | GVVDSEDII | [R].GVVDS | 1xPhospho | Q3UIF3 1xl | 0.0004 | 0 | 3 | 18 | 4 | Q3UIF3; P1 | Putative ur | Q3UIF3 [38 | 0 | 1822.915 | 0 | -0.67 | 0.000161 | 0.05 | 123 | 77 | 3.29 | 2.26 | Shared | High | High | High | High | High | High | High | High | High | High | High | High | 0 | 8.5E-05 | 1.88 |
| FALSE | High | GVVIPTWN | [K].GVVIPT | 1xPhospho | Q9DB00 1x | 0.000152 | 0 | 1 | 3 | 1 | Q9DB00 | GON-4-like | Q9DB00 [3 | 0 | 1962.123 | 0 | -1.57 | 0.000986 | 1.99 | 150.3 | 49.7 | 9.72 | 10.46 | | High | High | High | High | High | High | High | High | High | High | High | High | 0 | 2.88E-05 | 3.61 |
| FALSE | High | GVVPEDAV | [K].GVVPEI | 1xPhospho | P51125-3 1 | 0.000276 | 0 | 1 | 12 | 2 | P51125-3 | Isoform 3 c | P51125-3 [ | 0 | 2080.052 | 0 | 2.66 | 0.000801 | 21.89 | 27.3 | 172.7 | 47.5 | 7.18 | | High | High | High | High | High | High | High | High | High | High | High | High | 0 | 5.63E-05 | 3.73 |
| FALSE | High | GWAFGSN | [K].GWAFC | 1xPhospho | G5E8A7 1x | 6.09E-06 | 0 | 1 | 2 | 2 | G5E8A7 | Protein Sm | G5E8A7 [5 | 0 | 2343.152 | 0 | 0.96 | 0.003287 | 15.83 | 67.8 | 132.2 | 9.3 | 9.28 | | High | High | High | High | High | High | High | High | High | High | High | High | 0 | 4.35E-06 | 4.21 |
| FALSE | High | GWAFGSN | [K].GWAFC | 1xPhospho | G5E8A7 1x | 7.82E-05 | 0 | 1 | 2 | 2 | G5E8A7 | Protein Sm | G5E8A7 [5 | 0 | 2359.147 | 0 | 1.32 | 0.000744 | 1.11 | 59 | 141 | 18.93 | 14.37 | | High | High | High | High | High | High | High | High | High | High | High | High | 0 | 1.36E-05 | 3.52 |
| FALSE | High | GWAPPGPS | [R].GWAPF | 1xPhospho | Q9JLM4 1x | 2.63E-06 | 0 | 1 | 7 | 5 | Q9JLM4 | Zinc finger | Q9JLM4 [4 | 0 | 2987.517 | 0 | -0.19 | 0.246496 | 16.92 | 106.6 | 93.4 | 8.05 | 18.33 | | High | High | High | High | High | High | High | High | High | High | High | High | 0 | 2.86E-05 | 3.73 |
| FALSE | High | GWDAEGS | [K].GWDAI | 1xPhospho | B0FTY3 1xl | 0.003253 | 0.00016 | 1 | 4 | 1 | B0FTY3 | NudC dom | B0FTY3 [33 | 0 | 1430.63 | 0 | 1.32 | 0.000504 | 10.15 | 54.7 | 145.3 | 17.05 | 9.63 | | High | High | High | High | High | High | High | High | High | High | High | High | 7.19E-05 | 0.000894 | 1.96 |
| FALSE | High | GWEESGEC | [K].GWEES | 1xDeamida | Q6P5H2-2 | 0.038017 | 0.002528 | 1 | 2 | 3 | Q6P5H2-2 | Isoform 2 c | Q6P5H2-2 | 0 | 4141.845 | 0 | 0.67 | 0.002071 | 9.35 | 73 | 127 | 4.86 | 10 | | High | High | High | High | High | High | High | High | High | High | High | High | 0.001609 | 0.01564 | 3.68 |
| FALSE | High | GWPSPYAA | [R].GWPSP | 1xPhospho | O08715-4 | 0.000107 | 0 | 1 | 6 | 1 | O08715-4 | Isoform 4 c | O08715-4 | 0 | 1787.877 | 0 | 0.32 | 0.146654 | 4.59 | 90 | 110 | 6 | 15.67 | | High | High | High | High | High | High | High | High | High | High | High | High | 0 | 1.95E-05 | 2.5 |
| FALSE | High | GWSESEQS | [K].GWSES | 1xPhospho | Q8C7S2 1x | 0.000128 | 0 | 1 | 5 | 4 | Q8C7S2 | Putative ur | Q8C7S2 [6 | 0 | 2496.059 | 0 | 0.77 | 0.006465 | 11.06 | 80.6 | 119.4 | 15.15 | 13.88 | | High | High | High | High | High | High | High | High | High | High | High | High | 0 | 2.42E-05 | 3.56 |
| FALSE | High | GWSESEQS | [K].GWSES | 1xPhospho | Q8C7S2 1x | 5.16E-08 | 0 | 1 | 5 | 6 | Q8C7S2 | Putative ur | Q8C7S2 [6 | 0 | 2512.054 | 0 | 0.95 | 0.010548 | 27.52 | 63.9 | 136.1 | 10.96 | 17.98 | | High | High | High | High | High | High | High | High | High | High | High | High | 0 | 1.01E-07 | 4.08 |
| FALSE | High | GWSGPDV | [K].GWSGF | 1xPhospho | Q8CI75-2 1 | 1E-06 | 0 | 1 | 3 | 7 | Q8CI75-2 | Isoform 2 c | Q8CI75-2 [ | 0 | 2647.14 | 0 | 0.19 | 0.022934 | 5.23 | 94.4 | 105.6 | 5.67 | 6.78 | | High | High | High | High | High | High | High | High | High | High | High | High | 0 | 1.02E-07 | 4.47 |
| FALSE | High | GWSGPDV | [K].GWSGF | 1xPhospho | Q8CI75-2 1 | 0.010109 | 0.000736 | 1 | 3 | 1 | Q8CI75-2 | Isoform 2 c | Q8CI75-2 [ | 1 | 4040.892 | 0 | 0.39 | 0.090722 | 12.92 | 88.5 | 111.5 | 5.97 | 18.29 | | High | High | High | High | High | High | High | High | High | High | High | High | 0.000344 | 0.003178 | 4.23 |
| FALSE | High | GWSPPPEV | [R].GWSPP | 1xPhospho | A2AQN4 1x | 0.030104 | 0.001857 | 1 | 9 | 1 | A2AQN4 | Acetyl-coe | A2AQN4 [2 | 0 | 1333.65 | 0 | 1.35 | 0.000154 | 2.92 | 55.5 | 144.5 | 6.57 | 3.98 | | High | High | High | High | High | High | High | High | High | High | High | High | 0.001011 | 0.01078 | 1.36 |
| FALSE | High | GYALVPIIV | [K].GYALVF | 1xPhospho | B9EKI3 1xP | 0.000191 | 0 | 1 | 2 | 2 | B9EKI3 | TATA elem | B9EKI3 [34 | 0 | 2080.186 | 0 | 0.41 | 0.074657 | 4.23 | 86.1 | 113.9 | 10.22 | 13.15 | | High | High | High | High | High | High | High | High | High | High | High | High | 0 | 3.72E-05 | 3.22 |
| FALSE | High | GYATDEST | [K].GYATDI | 1xPhospho | Q61584-3 | 0.000595 | 0 | 2 | 5 | 3 | Q61584-3; | Isoform B c | Q61584-3 | 0 | 1952.88 | 0 | -0.86 | 0.000486 | 4.19 | 132.1 | 67.9 | 4.57 | 13 | Shared | High | High | High | High | High | High | High | High | High | High | High | High | 0 | 0.000133 | 2.83 |
| FALSE | High | GYDSEDDT | [K].GYDSEI | 1xCarbami | Q9WTK2 1 | 4.13E-06 | 0 | 1 | 2 | 10 | Q9WTK2 | Chromodo | Q9WTK2 [8 | 0 | 3706.527 | 0 | 0.28 | 0.029862 | 13.69 | 89.3 | 110.7 | 7.57 | 7.96 | | High | High | High | High | High | High | High | High | High | High | High | High | 0 | 8.83E-05 | 4.34 |
| FALSE | High | GYEDYLQS | [K].GYEDYI | 1xPhospho | A0A0R4J04 | 8.06E-06 | 0 | 1 | 3 | 1 | A0A0R4J04 | Protein arg | A0A0R4J04 | 0 | 3774.79 | 0 | -0.95 | 0.00194 | 6.72 | 124 | 76 | 16.47 | 23.61 | | High | High | High | High | High | High | High | High | High | High | High | High | 0 | 1.06E-06 | 4.11 |
| FALSE | High | GYESPTLSK | [R].GYESPT | 1xPhospho | Q80W49-2 | 0.000779 | 0 | 1 | 1 | 1 | Q80W49-2 | Isoform 2 c | Q80W49-2 | 0 | 1519.781 | 0 | 3.22 | 0.000188 | 27.39 | 18.9 | 181.1 | 15.78 | 4.31 | | High | High | High | High | High | High | High | High | High | High | High | High | 0 | 0.00018 | 1.88 |
| FALSE | High | GYGFVDFD | [K].GYGFVI | 1xPhospho | Q91W59 1: | 0.000529 | 0 | 1 | 17 | 1 | Q91W59 | RNA-bindir | Q91W59 [1 | 0 | 2111.025 | 0 | 0.71 | 0.005759 | 10.11 | 76.4 | 123.6 | 11.14 | 4.37 | | High | High | High | High | High | High | High | High | High | High | High | High | 0 | 0.000117 | 2.6 |
| FALSE | High | GYGFVDFD | [K].GYGFVI | 1xPhospho | Q8VC70 1x | 4E-07 | 0 | 1 | 4 | 2 | Q8VC70 | RNA-bindir | Q8VC70 [1 | 0 | 2143.015 | 0 | 0.73 | 0.006859 | 8.15 | 75.3 | 124.7 | 10.04 | 5.88 | | High | High | High | High | High | High | High | High | High | High | High | High | 0 | 3.62E-08 | 2.06 |
| FALSE | High | GYGHGQG | [K].GYGHG | 1xPhospho | Q9Z1X4-3 | 1.02E-05 | 0 | 1 | 3 | 3 | Q9Z1X4-3 | Isoform 3 c | Q9Z1X4-3 | 0 | 3780.576 | 0 | -0.61 | 0.0747 | 20.63 | 119.9 | 80.1 | 17.75 | 17.86 | | High | High | High | High | High | High | High | High | High | High | High | High | 0 | 1.38E-06 | 3.87 |
| FALSE | High | GYNYDEDF | [R].GYNYD | 1xCarbami | L7N209 1xl | 0.000274 | 0 | 2 | 7 | 7 | L7N209; Q | HBS1-like p | L7N209 [9- | 0 | 4556.875 | 0 | 0.35 | 0.028578 | 7.21 | 80.2 | 119.8 | 14.96 | 10.33 | Shared | High | High | High | High | High | High | High | High | High | High | High | High | 0 | 8.21E-05 | 4.23 |
| FALSE | High | GYQALSDS | [R].GYQAL! | 1xPhospho | Q62179 1x | 3.38E-05 | 0 | 1 | 2 | 2 | Q62179 | Semaphori | Q62179 [7 | 0 | 1643.763 | 0 | 0.13 | 0.179118 | 10.46 | 93.8 | 106.2 | 11.25 | 6.04 | | High | High | High | High | High | High | High | High | High | High | High | High | 0 | 5.33E-06 | 2.33 |
| FALSE | High | GYSFSLTTF | [R].GYSFSL | 1xPhospho | Q3UWT6 1 | 0.000699 | 0 | 1 | 3 | 1 | Q3UWT6 | Proteasom | Q3UWT6 [ | 0 | 2017.008 | 0 | -0.84 | 0.002286 | 2.53 | 129 | 71 | 6 | 12.76 | | High | High | High | High | High | High | High | High | High | High | High | High | 0 | 0.000159 | 3.68 |
| FALSE | High | GYSYDDSM | [R].GYSYDI | 1xPhospho | Q8C5N3 1x | 0.002172 | 5.66E-05 | 1 | 14 | 1 | Q8C5N3 | Pre-mRNA- | Q8C5N3 [5 | 0 | 1618.629 | 0 | -0.57 | 0.003124 | 1.07 | 118.8 | 81.2 | 8.06 | 5.03 | | High | High | High | High | High | High | High | High | High | High | High | High | 4.94E-05 | 0.000572 | 2.26 |
| FALSE | High | GVSQEK | [R].GVSQEI | 1xPhospho | Q3USH5 1x | 0.094641 | 0.007408 | 1 | 2 | 1 | Q3USH5 | Splicing fac | Q3USH5 [9 | 0 | 1185.628 | 0 | -0.36 | 0.001627 | 1.44 | 112.4 | 87.6 | 2.87 | 2.39 | | High | High | High | High | High | High | High | High | High | High | High | High | 0.003768 | 0.04058 | 1.11 |
| FALSE | High | YYTSASGDI | [R].YYTSAS | 1xPhospho | Q3UIF3 1xl | 0.01002 | 0.000736 | 1 | 8 | 2 | Q3UIF3 | Putative ur | Q3UIF3 [46 | 0 | 2104.991 | 0 | -0.76 | 0.000518 | 3.97 | 125.7 | 74.3 | 3.03 | 9.74 | | High | High | High | High | High | High | High | High | High | High | High | High | 0.000344 | 0.003141 | 2.07 |
