## Supplementary_Table10 for "Aging-dependent dysregulation of EXOSC2 is maintained in cancer as a dependency"

CLIP Summary of annotations

Summary of gene annotation

------------------------------

| Region | No. | % |
|---|---|---|
| Intervals overlapping with UCSC/RefSeq genes | 64230 | 80.6 |
| Other | 15468 | 19.4 |

Summary of region annotation

------------------------------

| Region | No. | % |
|---|---|---|
| Total | 79698 | 100 |
| Genic | 64914 | 81.4 |
| Exon | 29951 | 37.6 |
| CDS exon | 12494 | 15.7 |
| 5' UTR exon | 1526 | 1.9 |
| 3' UTR exon | 7085 | 8.9 |
| Intron | 34963 | 43.9 |
| Upstream 10K | 1545 | 1.9 |
| Downstream 10K | 2944 | 3.7 |
| Genic+ext10K | 68810 | 86.3 |
| Deep intergeic | 10888 | 13.7 |

Percentage of intervals in each region (adjusted):

| Region | % |
|---|---|
| CDS exon | 22.2 |
| 5' UTR exon | 2.7 |
| Upstream 10K | 1.7 |
| 3' UTR exon | 12.6 |
| Downstream 10K | 3.2 |
| Intron | 43.9 |
| Deep intergenic | 13.7 |
